# Cultivated Robusta coffee meets wild *Coffea canephora*: Evidence of cultivated-wild hybridisation in the Democratic Republic of the Congo

**DOI:** 10.1101/2023.10.30.564669

**Authors:** Verleysen Lauren, Depecker Jonas, Bollen Robrecht, Asimonyio Justin, Hatangi Yves, Kambale Jean-Léon, Mwanga Mwanga Ithe, Ebele Tshimi, Dhed’a Benoit, Stoffelen Piet, Ruttink Tom, Vandelook Filip, Honnay Olivier

**Affiliations:** Division of Ecology, Evolution and Biodiversity Conservation, KU Leuven, Leuven, Belgium; Plant Sciences Unit, Flanders Research Institute for Agriculture, Fisheries and Food (ILVO), Melle, Belgium; Meise Botanic Garden, Meise, Belgium; KU Leuven Plant Institute, Leuven, Belgium; Centre de Surveillance de la Biodiversité et Université de Kisangani, Kisangani, DR Congo; Université de Kisangani, Kisangani, DR Congo; Liège University, Gembloux Agro-Bio Tech, Gembloux, Belgium; Centre de Recherche en Science Naturelles, Lwiro, DR Congo; Institut National des Etudes et Recherches Agronomique, Yangambi, DR Congo; Department of Plant Biotechnology and Bioinformatics, Ghent University, Ghent, Belgium

**Keywords:** *Coffea canephora*, crop wild relatives (CWR), cultivated-wild hybridisation, Congo Basin, field gene bank, gene flow, introgression

## Abstract

**Background and aims:** Plant breeders are increasingly turning to crop wild relatives (CWRs) to ensure global food security amidst a rapidly changing environment. However, CWR populations are confronted with various human-induced threats, including hybridisation with their nearby cultivated crops, posing a significant threat to the wild gene pool. Here, we focussed on Robusta coffee in the Yangambi region in the DR Congo, where cultivated Robusta coffee coexists with its wild relative (Coffea canephora), creating ample opportunity for genetic exchange. We examined the geographical distribution of cultivated Robusta coffee plants and the incidence of hybridisation between cultivated and wild individuals within the rainforest.

**Methods:** We used Genotyping-by-Sequencing data from 471 C. canephora individuals across various locations, including natural rainforests, home gardens, and the INERA Coffee Collection, a Robusta coffee field genebank. Our analyses involved establishing robust diagnostic fingerprints that differentiate cultivated from wild coffee, identifying cultivated-wild hybrids and mapping their geographical position.

**Key results:** We identified cultivated genotypes and cultivated-wild hybrids in zones with clear anthropogenic activity, and where Robusta coffee cultivated in the home gardens may serve as a source for crop-to-wild gene flow. We found relatively few hybrids and backcrosses in the rainforests, which is so far likely not sufficient for the stable incorporation of alleles from the cultivated gene pool into the CWR gene pool.

**Conclusions:** The cultivation of Robusta coffee in close proximity to its wild gene pool has led to cultivated genotypes and cultivated-wild hybrids appearing within the natural habitats of C. canephora. Yet, given the high overlap between the cultivated and wild gene pool, together with the relatively low frequency of hybridisation, our results indicate that the overall impact in terms of risk of introgression remains limited so far. Nevertheless, it is important to continue monitoring crop-to-wild gene flow to safeguard the wild gene pool of C. canephora.

## Introduction

A growing human population in an increasingly warmer world may jeopardize the food security of many households (FAO, 2018; Bohra *et al*., 2022). To cope with this future challenge, innovations in plant breeding and agricultural systems are an important part of the solution (Zhang *et al*., 2017; Bohra *et al*., 2022). Crop improvement should focus on increasing yields, enhancing crop climate change resilience, and providing nutritional security (Brozynska *et al*., 2016). To do so, plant breeders can fall back on crop wild relatives (CWRs), which harbour genetic diversity that is not present in their cultivated relatives, and may underlie desirable traits, including resistance to pests and diseases, and tolerance to abiotic stresses (Heywood *et al*., 2007; Zhang *et al*., 2017; Saeed and Fatima, 2021). The conservation of the genetic resources in populations of CWRs is thus of utmost importance and can be realised both *in-situ* and *ex-situ*. Solely relying on *ex-situ* conservation of CWRs is, however, problematic for several reasons (Meilleur and Hodgkin, 2004). First, it remains challenging to cover all extant CWR species and their genetic diversity, and place duplicates in conservation repositories (Castañeda-Álvarez *et al*., 2016; Wambugu and Henry, 2022). Second, evolutionary adaptive processes are strongly impeded in *ex-situ* germplasm collections. More so, artificial selection during *ex-situ* conservation can drive populations away from their phenotypic optimum in nature (Ensslin and Godefroid, 2020). As a consequence, there is a considerable risk of maladaptation when the CWRs are reintroduced into their original environments (Schoen and Brown, 2001; Heywood, 2015; Ensslin *et al*., 2023). Finally, for some species *ex-situ* conservation can be a very costly option due to seed recalcitrance (Mertens *et al*., 2022).

*In-situ* conservation of CWRs on the other hand is increasingly compromised by multiple anthropogenic processes. One conspicuous global threat is habitat degradation and habitat loss, the latter mainly caused by the conversion of natural habitat into agricultural land (Balvanera *et al*., 2019; Jaureguiberry *et al*., 2022). Agricultural encroachment has furthermore resulted in the introduction of cultivated crops in or near the natural habitats of their wild relatives, increasing the chance of genetic exchange (Ellstrand *et al*., 1999; Kareiva *et al*., 2007; Hufford *et al*., 2013). Such genetic exchange has already been observed in multiple CWR species, including wild apple (*Malus sieversii*) in Kazakhstan (Ha *et al*., 2021), Macademia trees in Australia (O’Connor *et al*., 2015), and Japanese chestnut (*Castanea crenata*) (Nishio *et al*., 2021). The transfer of cultivated genetic material can lead to cultivated-wild hybrids and subsequent backcrosses (Kwit *et al*., 2011). Hybridisation can be detrimental to the wild populations through genetic swamping if it happens at a large scale (Todesco *et al*., 2016; Macková *et al*., 2018). Ultimately, crop-to-wild gene flow can result in introgression, *i.e.* the stable incorporation of alleles of the cultivar genepool into the CWR genepool at a relatively high frequency, which can cause the loss of genetic variation and even local extinction of the original populations (Anderson and Hubricht 1938; Ellstrand, 2003; Laikre *et al*., 2010). The extent of these processes is, however, expected to vary among species, populations, pollinating agents, and distances between the crop and wild populations, and thus needs to be carefully studied across systems (Arriola and Ellstrand, 1996; Ellstrand *et al*., 1999; O’Connor *et al*., 2015).

A CWR where crop-to-wild gene flow has already been observed is *Coffea arabica* (Arabica coffee) (Aerts *et al*., 2013), of which the crop accounts for ± 56% of the global coffee market share (ICO, 2023). The remaining share of the global coffee market is attributed to Robusta coffee (*Coffea canephora*) (ICO, 2023). The importance of Robusta coffee is growing in the coffee sector presumably due to its high disease resistance and broad climatic range fit for cultivation (Craparo *et al*., 2015; Davis *et al*., 2019). The genetic diversity still present in wild populations of *C. canephora* is key for future coffee breeding efforts, not only to improve the cultivated Robusta gene pool, but also the Arabica gene pool. For example, Castro Caicedo *et al*. (2013) derived Arabica genotypes with high levels of resistance to pathogens by crossing Arabica cultivars with Robusta. Many of the Robusta coffee genotypes that are currently cultivated worldwide originate from the Congo Basin (Cubry *et al*., 2013), a region that harbours genetically highly diverse wild *C. canephora* populations (Ferrão *et al*., 2019; Merot-L’anthoene *et al*., 2019; Depecker *et al*., 2023). Furthermore, previous studies have shown that the genetic diversity of these wild populations is significantly higher than material that is currently available for cultivation (Krishnan, 2013; Leroy *et al*., 2014; Vanden Abeele *et al*., 2021).

Anthropogenic factors that may compromise the genetic integrity of the wild *C. canephora* populations include rainforest disturbance, agricultural encroachment, and cultivated-wild introgression. In Yangambi, in the heart of the Congo Basin in the Democratic Republic of the Congo (DR Congo), cultivated coffee is grown in home gardens and in the field genebank of the INERA (*Institut National des Etudes et Recherches Agronomiques*) Coffee Collection, whereas wild populations of *C. canephora* grow in the understory of the surrounding rainforests. The Yangambi Research Station has played a very significant role in the history of Robusta coffee breeding and commercialisation, but has lost its prominent position (Coste 1955; Capot 1962; Vanden Abeele *et al*., 2021; Verleysen *et al*., 2023). Verleysen *et al*. (2023) recently catalogued genetic fingerprints of the accessions of the INERA Coffee Collection and showed that most accessions of the germplasm collection were highly similar to “Lula” cultivar – material originating from the INERA Coffee Research Station in Lula in the DR Congo –, whereas some accessions were more similar to the Congolese subgroup A (Dussert *et al*., 2003) or to local wild genotypes. In the local villages, Robusta coffee is predominantly cultivated in home garden systems, which are often located very closely (sometimes < 1 km) to wild *C. canephora* populations (Vanden Abeele *et al*., 2021). This close proximity of the cultivated Robusta coffee in the small-scale home garden systems to the wild *C. canephora* populations may provide ample opportunities for cultivated-wild hybridisation. In that regard, using SSR markers, two putative cultivated-wild hybrids were already identified in the rainforests surrounding the Yangambi region by Vanden Abeele *et al*. (2021). Yet a more comprehensive assessment of the occurrence of cultivated-wild hybrids and the potential impact on the genetic composition of wild populations of *C. canephora* with more high resolution genetic markers remains to be done.

Here, we aimed to assess the extent of the threat to the genetic integrity of wild Robusta coffee through investigating the overlap in geographical distribution of cultivated Robusta coffee and wild coffee, and the occurrence of cultivated-wild hybridisation in the Yangambi region in the DR Congo. We used Genotyping-by-Sequencing data of 471 *C. canephora* individuals sampled from natural rainforests, home gardens, and the INERA Coffee Collection in Yangambi. Our specific objectives were to: i) compare the genetic diversity within and between cultivated Robusta coffee and local wild populations, and identify genetic fingerprints that can discriminate between both groups; ii) assign individuals to a cultivated or wild origin; identify clonal material; and identify individuals derived from cultivated-wild hybridisation events; iii) map the geographic position of the cultivated-wild hybrids in the landscape; and iv) discuss scenarios that may affect the genetic integrity of wild *C. canephora* wild populations.

## Materials and methods

### Sampling

We collected leaf material of 71 cultivated *C. canephora* individuals from 21 home gardens spread across villages in the Yangambi region. We complemented this with samples previously obtained by Depecker *et al*. (2023) and Verleysen *et al*. (2023). Depecker *et al*. (2023) established 24 survey plots in the understory of the rainforests in the Yangambi region, covering an area of ca. 50 km-by-20 km. From this study, the corresponding GBS data of 249 putatively wild *C. canephora* individuals was retrieved from Sequence Read Archive (SRA, see below) and twelve additional individuals, including individuals from one additional survey plot were included to create novel GBS data. Additionally, GBS data from 139 individuals from Verleysen *et* al. (2023), covering cultivated and local wild material now maintained in the field genebank of the INERA Coffee Collection. Metadata on the sampling site and classification based on the genetic analysis of all 471 individuals are available in Supplementary **Table S1**. The leaf material of the 71 home garden individuals and the 12 additional rainforest individuals was dried with silica gel and genomic DNA was extracted from 20-30 mg dried leaf material using an optimised cetyltrimethylammonium bromide (CTAB) protocol adapted from Doyle and Doyle (1987). DNA quantities were measured with the Quantifluor dsDNA system on a Promega Quantus Fluorometer (Promega, Madison, USA).

### Genotyping-by-Sequencing (GBS) and read data processing

DNA extracts of the 83 individuals were subjected to GBS. Following Depecker *et al*. (2023), GBS libraries were prepared using a double-enzyme GBS protocol adapted from Elshire (2011) and Poland *et al*. (2012). In short, 100 ng of genomic DNA was digested with PstI and MseI restriction enzymes (New England Biolabs, Ipswich, USA), and barcoded and common adapter constructs were ligated with T4 ligase (New England Biolabs, Ipswich, USA) in a final volume of 35 µL. Ligation products were purified with 1.6x MagNA magnetic beads (GE Healthcare Europe, Machelen, BE) and eluted in 30 µL TE. Of the purified DNA eluate, 3 µL was used for amplification with Taq 2x Master Mix (New England Biolabs, Ipswitch, USA) using an 18 cycles PCR protocol. PCR products were bead-purified with 1.6x MagNA, and their DNA concentrations were quantified using a Quantus Fluorometer. The library quality and fragment size distributions were assessed using a QIAxcel system (Qiagen, Venlo, NL). Equimolar amounts of the GBS libraries were pooled, bead-purified, and 150 bp paired-end sequenced on an Illumina HiSeq-X instrument by Admera Health (South Plainfield, USA).

Reads were processed with a customised script available on Gitlab (https://gitlab.com/ilvo/GBprocesS). The quality of sequence data was validated with FastQC 0.11 (Andrews, 2010) and reads were demultiplexed using Cutadapt 2.10 (Martin, 2011), allowing zero mismatches in barcodes or barcode-restriction site remnant combination. The 3’ restriction site remnant and the common adapter sequence of forward reads and the 3’ restriction site remnant, the barcode, and the barcode adapter sequence of reverse reads were removed based on sequence-specific pattern recognition and positional trimming using Cutadapt. After trimming the 5’ restriction site remnant of forward and reverse reads using positional trimming in Cutadapt, forward and reverse reads with a minimum read length of 60 bp and a minimum overlap of 10 bp were merged using PEAR 0.9.11 (Zhang *et al*., 2014). Merged reads with a mean base quality below 25 or with more than 5% of the nucleotides uncalled and reads containing internal restriction sites were discarded using GBprocesS.

The GBS data of the 83 newly sampled individuals was complemented with GBS data derived from SRA (Bioproject PRJNA901681) of the 249 *C. canephora* individuals studied by Depecker *et al*. (2023), and the 139 *C. canephora* individuals studied by Verleysen *et al*. (2023). GBS data of all 471 *C. canephora* individuals were mapped onto the *C. canephora* reference genome sequence (Denoeud *et al*., 2014) with the BWA-mem algorithm in BWA 0.7.17 with default parameters (Li and Durbin, 2009). Alignments were sorted, indexed, and filtered on mapping quality above 20 with SAMtools 1.10 (Li *et al*., 2009). Next, high-quality GBS loci and Stack Mapping Anchor Points (SMAPs) were identified in the mapped reads using the SMAP *delineate* module within the SMAP package v4.4.0 (Schaumont *et al*., 2022; https://ngs-smap.readthedocs.io/en/latest/home.html) with parameters: *mapping_orientation* ignore, *min_stack_depth* 4, *max_stack_depth* 400, *min_cluster_depth* 8, *max_cluster_depth* 400, *completeness* 80, and *min_mapping_quality* 20.

### SNP and haplotype calling

Single nucleotide polymorphisms (SNPs) within the high-quality GBS loci were called with GATK (Genome Analysis Toolkit) Unified Genotyper 3.7.0. (McKenna *et al*., 2010). Multi-allelic SNPs were removed with GATK, and the remaining SNPs were filtered using the following parameters: *min-meanDP* 30, *mac* 4, and *minQ* 20. The remaining SNPs were then subjected to further filtering with the following parameters: *minDP10*, *minGQ* 30, *max-missing* 0.7, *mac* 3, *minQ* 30, *min-alleles* 2, *max-alleles* 2, and *maf* 0.05 using VCFtools 0.1.16 (Danecek *et al*., 2011).

Read-backed haplotyping was conducted based on the combined variation in SMAPs and SNPs using the SMAP *haplotype-sites* module within the SMAP package v4.4.0 with parameters: *mapping_orientation* ignore, *partial* include, *no_indels*, *min_read_count* 10, *min_distinct_haplotypes* 2, *min_haplotype_frequency* 5, *discrete_calls* dosage, *frequency_interval_bounds* 10 10 90 90, and *dosage_filter* 2.

### Genetic structure

To investigate the genetic structure among all 471 individuals, a principal component analysis (PCA) was performed using the R package ADEGENET (Jombart, 2008; Rstudio Team, 2016). Additionally, a Bayesian clustering implemented in fastSTRUCTURE v1.0 (Raj *et al*., 2014) was run. Hundred iterations were run for each expected cluster setting K, ranging from 2 to 9. The StructureSelector software (Li and Liu, 2018) was used to determine the optimum number of K, by first plotting the mean log probability of each successive K and then using the Delta K method following Evanno *et al*. (2005).

### Genetic similarity

The genetic similarity between all 471 individuals was quantified with the SMAP *grm* module within the SMAP package v4.4.0, using the Jaccard Inversed Distance (Jaccard, 1912) that was calculated based on the discrete dosage haplotype calls in high-quality GBS loci. SMAP *grm* was run with parameters: *locus_completeness* 0.1, *similarity_coefficient* Jaccard, *distance_method* Euclidean, *locus_information_content* shared, and *partial* FALSE creating a pairwise Jaccard Inversed Distance (JID) matrix. The procedure of Verleysen *et al*. (2023) was used to calculate the minimal JID as a threshold to identify all pairs of genetically identical individuals (*i.e.*, clones).

### Genetic diversity

Cluster setting K = 2, separated the cultivated from the wild genotypes. The estimated admixture proportions, Bayesian Q-value (Pritchard *et al*., 2000), for each individual for cluster setting K = 2 were used to establish a “wild reference group”, containing all individuals collected from the Yangambi rainforest with a proportion of wild genotype higher than 0.9 (n = 249), and a “cultivated reference group”, containing all individuals collected from the INERA Coffee Collection and home gardens with a proportion of cultivated genotype higher than 0.9 (n = 152). These reference groups were used to calculate genetic diversity. Allelic richness (Ar) was calculated according to El Mousadik and Petit (1996) using the *allelic.richness* function of R package hierfstat (Goudet, 2013). The observed and expected heterozygosity (*H_o_* and *H_e_*, respectively) and inbreeding coefficient (*F_IS_*) were calculated using the *gl.report.heterozygosity* function of R package dartR (Gruber *et al*., 2017). All genetic diversity indices were compared between the wild and cultivated reference group using a Mann-Whitney U test. Genetic differentiation between the two reference groups was further quantified with a pairwise *F_ST_* according to Weir and Cockerham (1984).

### Hybridisation analysis

For the hybridisation analysis, we only retained SNPs with large differences in allele frequency (*F_ST_* > 0.8) between the “wild” and “cultivated” reference groups. Hybridisation levels were tested for all individuals not included in the two established reference groups (n = 70). First, heterozygosity (H) within the individuals was compared to their proportion of ancestry (S) from either reference group using the r package *Hiest* (Fitzpatrick, 2012). Next, likelihoods for six early generation hybrid classes (wild genotype, cultivated genotype, F_1_ hybrid, F_2_ hybrid, hybrid-wild backcross, hybrid-cultivated backcross) were calculated using the *Hiclass* function. The best fit of these hybrid classes was compared to the maximum likelihood genotype described by ancestry (S) and individual heterozygosity (H). We accepted a putative classification as credible if the log-likelihood of the best fit was within 2 units of the maximum log-likelihood.

## Results

The 471 coffee individuals collected from rainforests, the INERA Coffee Collection, and home gardens (Supplementary **Fig. S1**) yielded a total of 19 678 bi-allelic SNPs within 14 251 high-quality GBS loci with a completeness of at least 80% across all 471 individuals. Of these, 8 131 SNPs with a minimum minor allele count of 3 and a minimum minor allele frequency of 0.05 were used for all further analyses.

### Genetic identities

The PCA was performed on the 8 131 SNPs showed that individuals collected from the rainforests and the INERA Coffee Collection were separated along the PC1-axis (**Fig. 1A**). Along the PC2-axis, 32 rainforest individuals were separated from all other individuals collected in the rainforest. Likewise, 15 individuals belonging to the Congolese subgroup A were separated from all the other individuals collected from the INERA Coffee Collection along the PC2-axis.

**Figure 1:**
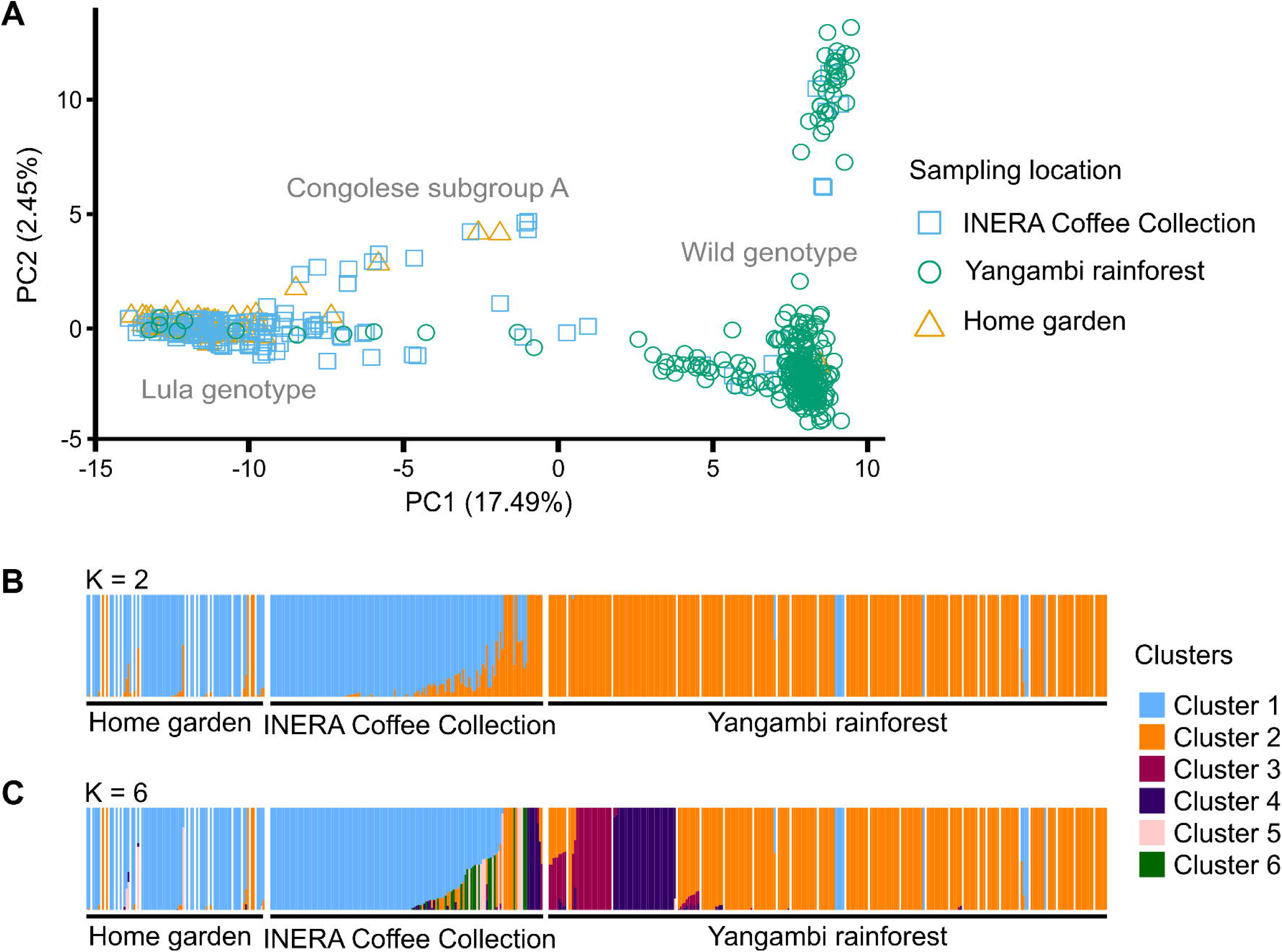
Population genetic structure within the *Coffea canephora* sample set. **A)** Principal component analysis using 8 131 SNPs indicating individuals collected from the INERA Coffee Collection, the Yangambi rainforest and the home gardens in the Yangambi region. **B)** fastSTRUCTURE bar plot representing two (K= 2) and six clusters (K=6). Colours define subpopulations: blue (”Lula” cultivars), orange (wild genotype), dark pink (wild genotypes from), purple (wild genotypes from), light pink (Congolese subgroup A), and green (unknown origin). Individuals are shown by thin vertical lines, which are divided into K coloured segments representing the estimated membership probabilities (Q) of each individual.

FastSTRUCTURE revealed that the cultivated individuals collected in the home gardens and INERA Coffee Collection (Cluster 1) were separated from wild individuals collected in the rainforest (Cluster 2) for a minimal number of clusters (K=2) (**Fig. 1B**). The optimum number of clusters for wild and cultivated individuals together was six (K=6) (**Fig. 1C**). Three genetic clusters were present in the Yangambi rainforest (Clusters 2, 3, and 4). All three wild clusters were also present in the INERA Coffee Collection, along with three cultivated clusters. Based on Verleysen *et al*. (2023), Cluster 1 represents the “Lula” cultivar genotype and Cluster 5 represents the “Congolese subgroup A”. The origin of Cluster 6 remained unknown in the frame of this analysis. Most of the individuals (n=61) cultivated in home gardens belonged to Cluster 1 and were classified as Lula cultivars, but also wild (n=5) and one Congolese subgroup A genotype were present in the home gardens. From the individuals collected in the rainforest, three were positioned between the cultivated and the wild group in the PCA and were assigned an admixed wild-Lula genotype by fastSTRUCTURE, and nine individuals were positioned on the negative PC1-axis and were assigned a Lula genotype by fastSTRUCTURE.

The pairwise JID calculated for all 471 individuals based on 41 522 haplotypes within 10 045 polymorphic high-quality GBS loci, showed a group of replicates with pairwise JID values greater than 0.979 (**Supplementary Table S2, Fig. S2**). Using the minimal JID values of 0.979 to identify pairs of clones, one rainforest individual was genetically identical to a Lula genotype from the INERA Coffee Collection. Within the home gardens, one individual collected was genetically identical to an individual collected from another home garden, one individual was genetically identical to a Lula genotype collected in the INERA Coffee Collection and another was genetically identical to an individual collected from the rainforest.

### Genetic diversity and differentiation within and between reference groups

To calculate the genetic diversity and differentiation within and between wild and cultivated groups, 249 individuals collected from the Yangambi rainforest with a proportion of wild genotype higher than 0.9 for K=2 were used as a “wild reference group”, which includes all three wild genetic clusters, and 152 individuals collected from the INERA Coffee Collection and home gardens with a proportion of cultivated genotype higher than 0.9 were used as a “cultivated reference group”, which includes the Lula genetic group, Congolese subgroup A, and Cluster 6.

No significant differences were found in allelic richness (A_r_) (p=0.5) and observed heterozygosity (*H_o_*) (p=0.35) between wild and cultivated reference groups (**Table 1**). Expected heterozygosity (*H_e_*) (p=0.00074) and inbreeding coefficient (*F_IS_*) (p < 2.2e^-16)^ were significantly lower in the cultivated reference group than in the wild reference group. Genetic differentiation as measured by overall *F_ST_* between both groups was 0.142.

**Table 1:** Genetic diversity estimates for the wild and cultivated reference group.

| | N | $A_r$ | $H_o$ | $H_e$ | $F_{IS}$ |
| --- | --- | --- | --- | --- | --- |
| wild reference group | 249 | 1.90 | 0.38 | 0.30 | -0.28 |
| cutlivated reference group | 152 | 1.90 | 0.38 | 0.28 | 0.38 |

### Admixture and hybridisation

To explore putative cultivated-wild hybridisation, the wild and cultivated reference groups were used to identify SNPs with large differences in allele frequency between wild and cultivated material, resulting in 24 SNPs with *F_ST_* > 0.8, here considered as discriminatory SNPs (**Fig. 3A**). The proportion of ancestry (S) and individual heterozygosity (H) were calculated on the 24 SNPs for all 70 individuals not included in the two reference groups (**Fig. 3C**). Using the ancestry-heterozygosity ratio (**Fig. 3B**), 40 individuals were assigned to six different hybrid classes based on statistical support (**Fig. 3D**): two individuals were assigned as F_1_ hybrids (one in the INERA collection and one in a home garden), four as F_2_ hybrids (one in the rainforest, one in a home garden and two in the INERA collection), 16 as hybrid-cultivated backcrosses (one in the rainforest, one in a home garden and 14 in the INERA collection), one as a hybrid-wild backcross in the INERA collection, ten as cultivated genotype (nine in the rainforest and one in the INERA collection) and seven as wild genotypes (two in the home gardens and five in the INERA collection) (**Supplementary Table S1**).

### Wild, cultivated, and cultivated-wild genotypes in the Yangambi rainforest

To assess the risk of cultivated-wild hybridisation, the fastSTRUCTURE results (K=6) were placed on the landscape map, revealing four different situations (**Fig. 3**).

Situation I: Four plots (Plots 16-19) from undisturbed old-growth rainforest more than 16 km away from home gardens contained only wild genotypes (Cluster 2).

Situation II: Plot 3 established in disturbed old-growth rainforest was located in close vicinity (< 1-4 km) of 14 home gardens. All individuals collected from this plot were assigned a wild (Cluster 4) genotype and we did not detect any cultivated or cultivated-wild genotypes. The home gardens in this locality contained individuals assigned to a Lula genotype and individuals with an admixed Lula-Congolese subgroup A genotype, of which one individual was identified as an F_1_ hybrid and one individual as a hybrid-cultivated backcross.

Situation III: Five plots located in disturbed old-growth and four plots in regrowth rainforest, which were surrounded by home gardens. Almost all individuals collected in these nine rainforest plots had a wild genotype, except for one individual from Plot 13 that was assigned a Lula genotype and one individual from Plot 7 that showed indications of an admixed wild-Lula genotype, although this was not statistically supported. Four individuals collected from three home gardens were assigned a wild genotype. In one home garden, all four individuals were identified as a Lula genotype, and another one contained one wild genotype, one Lula genotype, and one admixed wild-Lula genotype that was statistically assigned as a F_2_ hybrid.

Situation IV: one plot in regrowth rainforest, two plots in disturbed old-growth rainforest and six plots in undisturbed old-growth rainforest (Depecker *et al*., 2022;2023) with no home gardens nearby (> 7 km). Plot 11, Plot 12, Plot 22, Plot 23, Plot 24, and Plot 25 contained only individuals with wild genotypes. Plot 10 in disturbed old-growth rainforest on the other hand, contained nine individuals with a wild genotype and five individuals with a Lula genotype but no admixed wild-Lula genotypes. Plot 21 in undisturbed old-growth rainforest had seven individuals with a wild genotype and one individual with a Lula genotype, but no admixed wild-Lula genotypes. Plot 20 in undisturbed old-growth rainforest contained no individuals with a wild genotype, two individuals with a Lula genotype, one F_2_ hybrid, and one hybrid-cultivated backcross.

## Discussion

We combined novel GBS genotyping data of coffee trees from home gardens and rainforests with GBS data from previous studies on the genetic diversity in wild populations (Depecker et al., 2023), and cultivated source material of the INERA Coffee Collection (Verleysen et al., 2023) to investigate whether planting of selected cultivated materials in home gardens in close vicinity to the wild populations may lead to crop-to-wild gene flow; and whether cultivated-wild hybridisation could lead to changes in the genetic composition of the wild populations.

### INERA Coffee Collection germplasm has a broad genetic base

Our results suggest that cultivated materials found in the area were most likely created and distributed by the Yangambi INERA station. Notably, the cultivated materials present in the INERA Coffee Collection and distributed in the Yangambi area should be considered as one or more heterogeneous “genetic groups”, and not as ‘pure’ cultivars with a narrow genetic base. The INERA Coffee Collection that is used for the breeding and distribution of germplasm for cultivation encompasses genetic material from several different origins, mostly Lula material, Congolese subgroup A, and wild material from local wild populations, as previously described in detail in Verleysen *et al*. (2023). In addition to Verleysen *et al*. (2023), a novel group with unknown origin was found. The creation of new genetic diversity by crossings between materials of different genetic origin and between historically cultivated materials and local wild materials further adds to the broad genetic base (Verleysen *et al*., 2023). Furthermore, most of this material is distributed after seed-based propagation, produced by open-pollination, and it appears that only a fraction of the germplasm is multiplied clonally for distribution, as illustrated by a relatively low portion of clonal pairs between the INERA Coffee Collection, the wild populations, and the home gardens. Our data also showed that most cultivated coffees in the home gardens belong to the “Lula” cultivar type, confirming an earlier study (Vanden Abeele *et al*., 2021).

We found a relatively high allelic richness and observed heterozygosity in the cultivated reference group that was similar to that observed in the wild reference group, which was in contrast to our expectations. Typically, CWRs are considered to exhibit greater genetic diversity than their cultivated counterparts (Cubry *et al*., 2013; Zhang *et al*; 2017). For instance, Vanden Abeele *et al*. (2021) previously reported higher levels of observed heterozygosity in cultivated materials in the Yangambi region, as compared to the wild populations of *C. canephora* in the region, and equivalent or higher levels of observed heterozygosity were found in cultivated Robusta coffee elsewhere in the Afrotropical region compared to wild *C. canephora* populations (Musoli *et al*., 2009; Kiwuka *et al*., 2021). However, such comparisons strongly rely on the composition and structure of the cultivated germplasm (*i.e.*, the breeding history), the geographical area of wild populations covered and their population structure (previously described for the Yangambi region in Depecker *et al*., 2023), the number of individuals sampled per group, and the type and number of molecular markers. For instance, Vanden Abeele *et al*. (2021) analysed a substantially lower number of individuals with 18 multi-allelic SSRs compared to our study with hundreds of individuals and thousands of bi-allelic SNP markers. Not only was *within*-genetic group allelic richness and observed heterozygosity similar in the cultivar and wild reference groups, but also *between*-reference group genetic diversity comparisons showed a relatively high overlap in allele composition, albeit at different allele frequency per group (no strictly private alleles and only 24 out of 8131 SNPs with an *F_ST_* > 0.8 in our study). The relatively close genetic relatedness between cultivated and wild *C. canephora* was also highlighted by Cubry *et al*. (2013); and Kiwuka *et al*. (2021) found a low number of private alleles in cultivated Robusta coffee populations. This is likely related to the long generation time of coffee, limited breeding selection, and a broad genetic base (Stoffelen, 1998; Cubry *et al*., 2013; Gomez *et al*., 2013), and because “Lula” cultivars came from sources relatively close to the Yangambi region (Verleysen *et al*., 2023).

### Differentiation of cultivated, wild, and cultivated-wild hybrids, and reconstruction of spreading patterns

Despite the relatively high proportion of common alleles in cultivated materials and local wild populations, genetic analyses could delineate ‘diagnostic’ molecular markers that distinguish between the groups of cultivated and wild genotypes. Our analyses are thus consistent with previous studies that genetically separated wild and cultivated *C. canephora* in the Congo Basin (Vanden Abeele *et al*., 2021; Verleysen *et al*., 2023) and other studies in Africa comparing wild with cultivated *C. canephora* (Musoli *et al*., 2009; Kiwuka *et al*., 2021). The genetic distinction between cultivated and wild was instrumental in our study to assign genotypes to cultivar or wild origins and identify cultivated-wild hybridisation events, and placing them onto the landscape map revealed their distribution pattern. Genetic analyses also help to disentangle the different ways cultivated materials spread in the area, for instance by distinguishing between the different cultivated genetic groups and first or second-generation cultivated-wild hybrids and/or backcrosses. As expected, the first and most prominent route of spreading cultivated materials is the intentional planting of cultivars in home gardens. The second type of location where cultivars were found was the regrowth rainforest, disturbed old-growth rainforest and even presumed undisturbed old-growth rainforest. Cultivar individuals may end up there via different routes, either being planted or as remnants of abandoned plantations (historical land-use maps), which would be expected to yield clusters of cultivars in a given small area (like Plot 20). Alternatively, seed flow may partly account for the dispersal of berries or seeds from within-cultivar-group pollinated trees, which may be expected to yield more sporadic instances of cultivar individuals in otherwise predominantly wild populations, like Plot 7 or Plot 10. Seeds of *Coffea* species are predominantly dispersed by birds and mammals that can cover long distances (Stoffelen, 1998; Noirot et al., 2016). Conversely, cultivated-wild hybrids may be derived from pollination between cultivated and wild materials. Such materials are known to be generated in the INERA Coffee Collection (Verleysen *et al*., 2023), and may be distributed and planted, just like the cultivated material. Alternatively, cultivated-wild hybrids may result from the natural process of cross-pollination by pollinators in the rainforest. Crops grown in the vicinity of the rainforest have been shown to be visited by a rich pollinator community, as compared to more isolated crop fields (Klein *et al*., 2008;2009). Such spread of cultivar genetic material would then also lead to more sporadic patterns of F_1_, F_2_, or backcross materials. The frequency of cultivated-wild hybrid occurrence in nature would be determined by a combination of the density of cultivars as source material, pollen transport over distance in that area, followed by seed dispersal from the mother plant. The relatively short geographical distance between cultivated Robusta coffee genotypes and the wild populations creates opportunities for gene flow, while Isolation-by-Distance (IBD) may limit gene flow. Interestingly, IBD may not be homogeneous across the landscape in the study system (Depecker *et al*., 2023), with Plot 3 genetically distinct from the closest neighbouring Plot 1 and Plot 2, while the population in the northern part displayed more genetic similarity (*i.e.*, connectivity) between plots at a comparatively longer distance. Taking the human and natural factors that potentially facilitate cultivar spread and cultivated-wild gene flow together with the locations at which cultivars and cultivated-wild hybrids were found in the landscape, we could distinguish four different situations that revealed heterogeneity of potential cultivated-wild interactions at the landscape level within the relatively small study area.

First, in the most northern part of the Yangambi region (**Fig. 2**, Situation I), where rainforests have been classified as undisturbed old-growth (Depecker *et al*., 2022;2023), genetic analysis confirmed the presence of exclusively wild individuals. This remote area contained no cultivated materials, thus representing a pristine rainforest fit for *in*-*situ* conservation as genetic exchange between cultivated and wild material is still absent.

**Figure 2:**
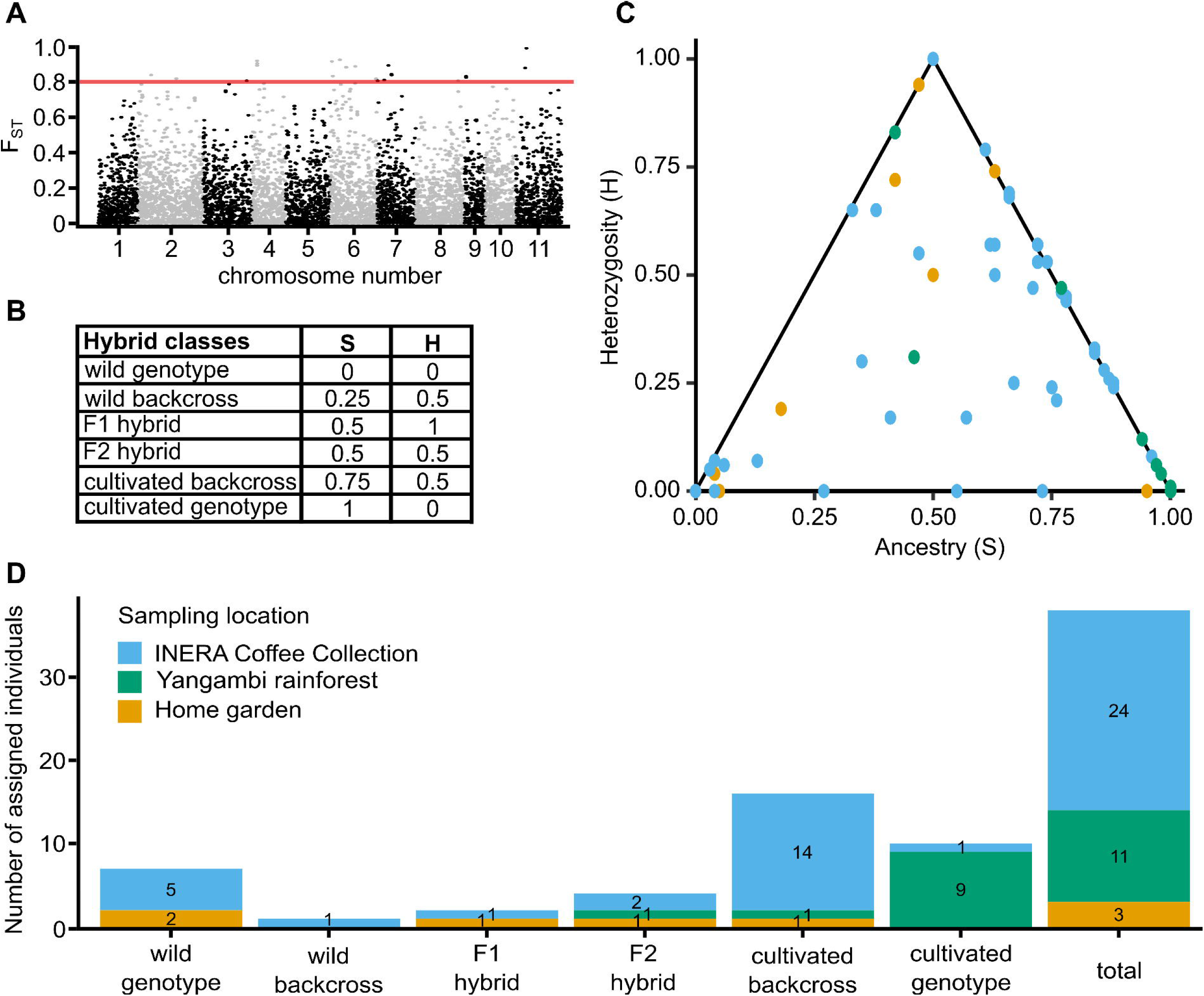
Estimation of hybridisation within all 70 individuals, not included in the wild and cultivated reference groups. **A)** *F_ST_* of all 8 131 SNPs calculated between the wild and cultivated reference group. Red line indicating the threshold to identify differentiating SNPs (*F_ST_* = 0.8). **B)** Theoretical hybrid assignments based on the ratio between the proportion of ancestry (S) and individual heterozygosity (H). **C)** The proportion of ancestry (S) and individual heterozygosity (H) calculated on the 24 SNPs for all 70 individuals coloured by their sampling location. **D)** All 38 statistically supported hybrid assignments into six hybrid classes coloured by their sampling location.

**Figure 3:**
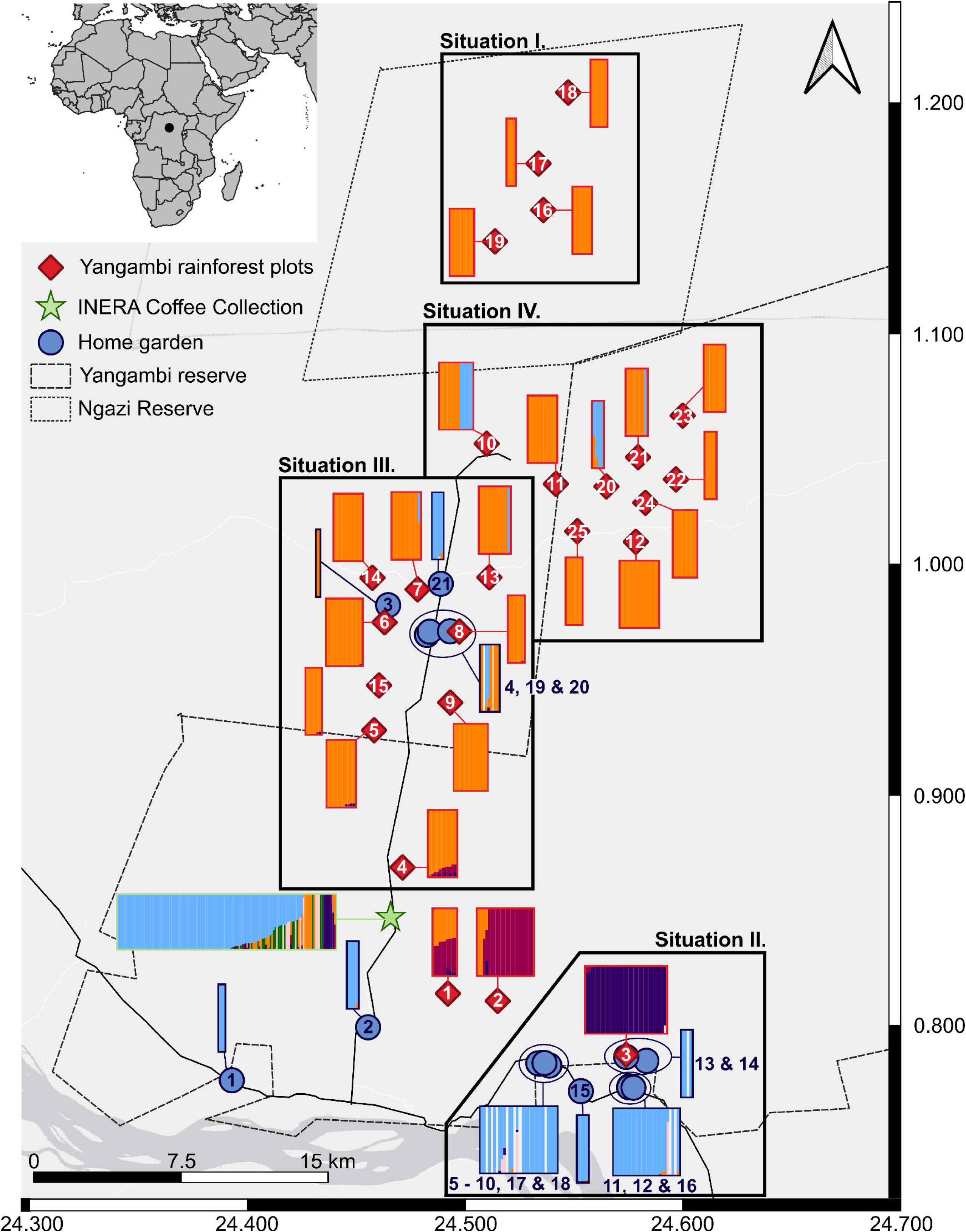
Map of the Yangambi region showing the location of each Yangambi rainforest plot (red rhombuses), each home garden (blue rhombuses) and the INERA Coffee Collection (green star), as well as the K coloured segments (K=6) of each individual within each location. Individuals are shown by thin vertical lines and grouped together based on sample location. All individuals collected in the Yangambi rainforest are red-bordered and all individuals collected in the home gardens are bleu-bordered.

Second, in the south-eastern part (**Fig. 2**, Situation II), wild individuals were collected in disturbed old-growth rainforest (Depecker *et al*., 2022;2023) in the close vicinity of several home gardens containing cultivated coffee genotypes. Despite the substantial level of anthropogenic activity, both in terms of disturbance and agricultural encroachment, no signs of admixture and hybridisation between cultivated Robusta coffee and wild *C. canephora* were detected. As reported by Kearsley *et al*. (2017), a monodominant *Gilbertiodendron dewevrei* forest isolates this part of the rainforest from the surrounding environment, and Depecker *et al*. (2023) coined this to be a natural barrier explaining the high genetic differentiation between the *C. canephora* population in this area and the other populations in the Yangambi region, observing IBD even at relatively small distances.

Third, in the centre of the Yangambi region (**Fig. 2**, Situation III), disturbed old-growth and regrowth rainforests are patched together (Depecker *et al*., 2022;2023), with a strip of village development with home gardens along the road that crosses these rainforests. Here, wild *C. canephora* populations and cultivated Robusta coffee can be found in very close proximity, in some occasions even less than one km, without an apparent natural barrier as observed in situation II. A cultivated genotype and a cultivated-wild hybrid were present within the wild population, indicating a low level of spreading of cultivated material and crop-to-wild gene flow. Such spurious individuals are consistent with several scenarios of spreading; in regrowth rainforest, they could be founders originating from neighbouring populations (Depecker *et al*., 2023), in disturbed old-growth or regrowth rainforest cultivars might be occasionally planted, or cultivated-wild hybridisation may arise from cross-pollination and seed flow from neighbouring fields or gardens.

Finally, in the fourth situation, just north-east of the centre of the region (**Fig. 2**, Situation IV), wild genotypes of *C. canephora* were found in disturbed and undisturbed old-growth rainforests, without any home gardens in close proximity (Depecker *et al*., 2022;2023). Nevertheless, multiple cultivar genotypes and cultivated-wild hybrid genotypes were also identified in three different plots. The high level of grouping of the cultivated genotypes raises questions about whether these individuals have emerged naturally following pollen and/or seed dispersal. For instance, the cultivated coffees in Plot 10 might have been a remnant of a former coffee plantation based on the historical land-use maps (see Depecker *et al*., 2022;2023). In addition to the home gardens, these cultivated genotypes should also be considered as sources for crop-to-wild gene flow, and we indeed found hybrids in close proximity to these cultivated genotypes. Notably, in Plot 20, in an area of the rainforest that was previously classified as undisturbed old-growth rainforest based on canopy structure, previous land-use maps, and absence of logging, an individual was found that was genetically identical to a cultivated genotype of the INERA Coffee Collection, suggesting that this individual is derived from clonal propagation and was dispersed by human intervention, rather than by seed dispersal.

In conclusion, we identified cultivated genotypes and cultivated-wild hybrids in zones with clear anthropogenic activity, and where the Robusta coffee cultivated in the home gardens may serve as a source for crop-to-wild gene flow. Our data further show that the current distribution area of the cultivated genotypes is more extensive than previously believed. The presence of clonally propagated cultivated genetic material in presumed undisturbed locations is a sign of past or present human activity and expands the region of the Yangambi rainforest that is subject to disturbance with possible consequences for habitat integrity. Our study further illustrates that genetic analyses may uncover sites of human activities in parallel to field observations of forest canopy structure, vegetation composition, or logging, and therefore may be of complementary value in landscape-level monitoring of anthropogenic activities and habitat disturbance. In other African regions, hybridisation between wild *C. canephora* and Robusta coffee in cultivation has also been reported, for example in Uganda (Musoli *et al*., 2009). In *C. arabica*, hybridisation and introgression were reported in montane rainforests in south-western Ethiopia (Aerts *et al*., 2013). Introgression, however, will only occur when cultivated-wild hybrids form backcrosses with the wild population for many generations and at a large scale (Ridley, 2004; Verónica *et al*., 2017). In our study, we found only relatively few F_1_ and F_2_ hybrids and backcrosses in the wild, which is likely not sufficient for the stable incorporation of alleles of the cultivated gene pool into the CWR gene pool (Anderson and Hubricht 1938; Ellstrand, 2003; Laikre *et al*., 2010). Furthermore, the cultivated material distributed in the Yangambi region generally has a broad genetic base, is derived from several origins, and is genetically closely related to the local wild populations. Therefore, if hybridisation occurs, mostly genetic material is exchanged that was already part of the wild gene pool. Nevertheless, it is important to continue monitoring habitat integrity and cultivated-wild gene flow to safeguard the wild gene pool of *C. canephora*, as the *in-situ* conservation of CWR is important to guarantee future food security.

## Acknowledgments

We are grateful for the field assistance by the Institut National pour l’Étude et la Recherche Agronomiques (INERA) and the FORETS project, which is financed by the 11^th^ European Development Fund. We would also like to sincerely thank the Ministère de L’Environnement et Développement Durable (MEDD) for their indispensable help with obtaining research and export permits in accordance with the Nagoya regulations of the DR Congo (N°003/ANCCB-RDC/SG-EDD/BTB/02/2020, N°008/ANCCB-RDC/SG-EDD/BTB/11/2020, N°001/ANCCB-RDC/SG-EDD/BTB/01/2021, N°004/ANCCB-RDC/SG-EDD/BTB/2021, N°014/ANCCB-RDC/SG-EDD/BTB/11/2021, N°025/ANCCB-RDC/SG-EDD/BTB/11/2022)

## Data accessibility

The data that support the findings of this study are available on request. In accordance with DR Congo and international regulations, restrictions apply on the availability of these data, which were used under license for this study.

## Author contributions

OH, FV, TR, JD, and LV designed this study. JD, JA, YH, J-LK, IMM, TE, and RB participated in fieldwork. LV executed the lab work. LV and JD analysed the data. JD, LV, FV, TR, and OH wrote the manuscript. All authors contributed to finalising the manuscript.

## Supplementary data

**Table S1:** Source data of all 471 individuals collected from the Yangambi rainforest, the home gardens and the INERA Coffee Collection in Yangambi.

**Table S2:** Pairwise Jaccard Inversed Distance (JID) calculated for all 471 individuals based on 41 522 haplotypes within 10 045 polymorphic high-quality GBS loci. The JID matric was arranged based on the sample location. Red-shaded JID values are JID values that were above the minimal JID and by this indicating genetically identical individuals.

**Figure S1:** Distribution of all pairwise Jaccard Inversed Distance (JID) was calculated for all 471 individuals based on 41 522 haplotypes within 10 045 polymorphic high-quality GBS loci.

