## Supplementary figures and images for "Cultivated Robusta coffee meets wild *Coffea canephora*: Evidence of cultivated-wild hybridisation in the Democratic Republic of the Congo"

### Supplemental Figure S1

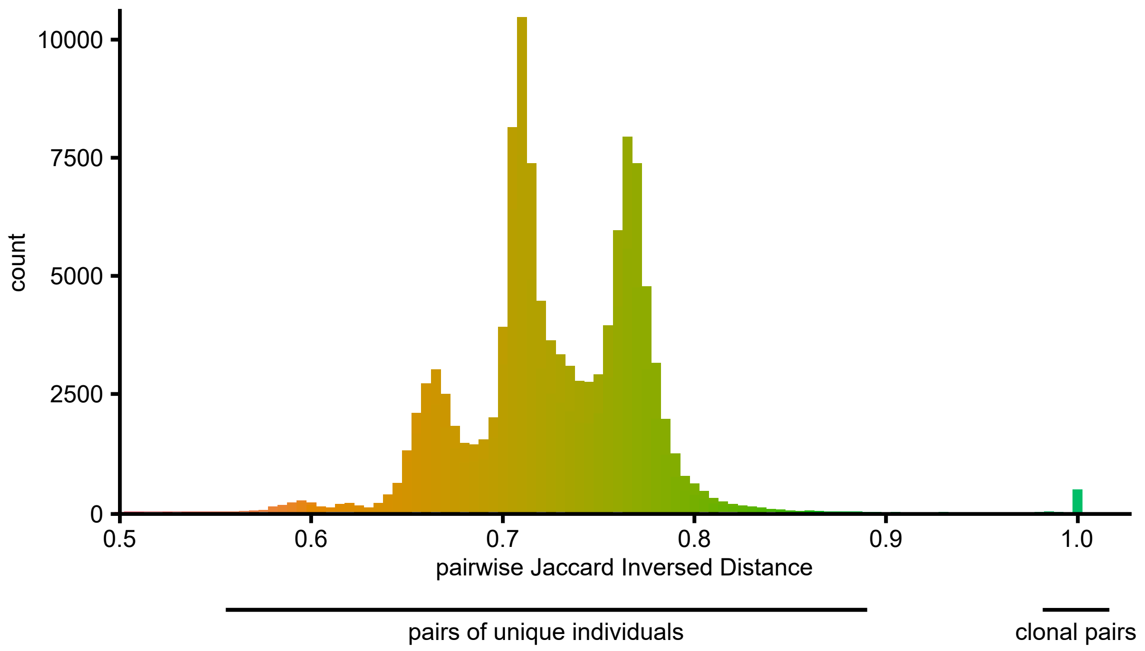
