## Supplemental Table S1 for "Cultivated Robusta coffee meets wild *Coffea canephora*: Evidence of cultivated-wild hybridisation in the Democratic Republic of the Congo"

**Supplemental Table S1**: Overview of the sample ID, sample location and plot number for all 471 *C. canep*
the home gardens with an indication of which individuals belong to cultivated or wild reference group, w

| Sample info | | | |
|---|---|---|---|
| Sample_ID | Sample location | Plot | Cultivated reference group |
| Home_garden_1347 | Home garden | HG_01 | x |
| Home_garden_1348 | Home garden | HG_01 | x |
| Home_garden_1330 | Home garden | HG_02 | x |
| Home_garden_1331 | Home garden | HG_02 | x |
| Home_garden_1332 | Home garden | HG_02 | x |
| Home_garden_1328 | Home garden | HG_02 | x |
| Home_garden_1350 | Home garden | HG_03 | |
| Home_garden_1358 | Home garden | HG_04 | |
| Home_garden_1397 | Home garden | HG_05 | x |
| Home_garden_1396 | Home garden | HG_05 | x |
| Home_garden_1398 | Home garden | HG_06 | x |
| Home_garden_1399 | Home garden | HG_07 | x |
| Home_garden_1402 | Home garden | HG_08 | x |
| Home_garden_1401 | Home garden | HG_08 | |
| Home_garden_1400 | Home garden | HG_08 | |
| Home_garden_1403 | Home garden | HG_09 | x |
| Home_garden_1404 | Home garden | HG_09 | x |
| Home_garden_1405 | Home garden | HG_10 | |
| Home_garden_1421 | Home garden | HG_11 | x |
| Home_garden_1424 | Home garden | HG_11 | x |
| Home_garden_1425 | Home garden | HG_11 | x |
| Home_garden_1426 | Home garden | HG_11 | x |
| Home_garden_1422 | Home garden | HG_11 | x |
| Home_garden_1410 | Home garden | HG_11 | x |
| Home_garden_1407 | Home garden | HG_11 | x |
| Home_garden_1416 | Home garden | HG_11 | x |
| Home_garden_1413 | Home garden | HG_11 | x |
| Home_garden_1427 | Home garden | HG_11 | x |
| Home_garden_1414 | Home garden | HG_11 | x |
| Home_garden_1411 | Home garden | HG_11 | x |
| Home_garden_1409 | Home garden | HG_11 | x |
| Home_garden_1408 | Home garden | HG_11 | x |
| Home_garden_1423 | Home garden | HG_11 | x |
| Home_garden_1415 | Home garden | HG_11 | x |
| Home_garden_1417 | Home garden | HG_11 | x |
| Home_garden_1419 | Home garden | HG_11 | x |
| Home_garden_1420 | Home garden | HG_11 | x |
| Home_garden_1418 | Home garden | HG_11 | x |
| Home_garden_1412 | Home garden | HG_11 | x |
| Home_garden_1406 | Home garden | HG_11 | |
| Home_garden_1428 | Home garden | HG_12 | x |
| Home_garden_1441 | Home garden | HG_13 | x |
| Home_garden_1438 | Home garden | HG_13 | x |

| | | | |
|---|---|---|---|
| Home_garden_1442 | Home garden | HG_14 | x |
| Home_garden_1456 | Home garden | HG_15 | x |
| Home_garden_1453 | Home garden | HG_15 | x |
| Home_garden_1455 | Home garden | HG_15 | x |
| Home_garden_1454 | Home garden | HG_15 | x |
| Home_garden_1462 | Home garden | HG_16 | x |
| Home_garden_1485 | Home garden | HG_17 | x |
| Home_garden_1489 | Home garden | HG_17 | x |
| Home_garden_1481 | Home garden | HG_17 | x |
| Home_garden_1484 | Home garden | HG_17 | x |
| Home_garden_1487 | Home garden | HG_17 | x |
| Home_garden_1488 | Home garden | HG_17 | x |
| Home_garden_1482 | Home garden | HG_17 | x |
| Home_garden_1483 | Home garden | HG_17 | x |
| Home_garden_1486 | Home garden | HG_17 | x |
| Home_garden_1490 | Home garden | HG_18 | x |
| Home_garden_1492 | Home garden | HG_18 | x |
| Home_garden_1493 | Home garden | HG_18 | x |
| Home_garden_1491 | Home garden | HG_18 | x |
| Home_garden_1601 | Home garden | HG_19 | x |
| Home_garden_1600 | Home garden | HG_19 | |
| Home_garden_1585 | Home garden | HG_19 | |
| Home_garden_1613 | Home garden | HG_20 | |
| Home_garden_1614 | Home garden | HG_20 | |
| Home_garden_1698 | Home garden | HG_21 | x |
| Home_garden_1699 | Home garden | HG_21 | x |
| Home_garden_1701 | Home garden | HG_21 | x |
| Home_garden_1700 | Home garden | HG_21 | x |
| Collection_G0095_2067 | Collection | INERA collection | x |
| Collection_G0119_1391 | Collection | INERA collection | x |
| Collection_G0004_1968 | Collection | INERA collection | x |
| Collection_G0106_2083 | Collection | INERA collection | x |
| Collection_G0034_2001 | Collection | INERA collection | x |
| Collection_G0040_2007 | Collection | INERA collection | x |
| Collection_G0089FOG_2060 | Collection | INERA collection | x |
| Collection_G0093_2064 | Collection | INERA collection | x |
| Collection_G0096_2068 | Collection | INERA collection | x |
| Collection_G0250FOG_1470 | Collection | INERA collection | x |
| Collection_G0007_1971 | Collection | INERA collection | x |
| Collection_G0014_1387 | Collection | INERA collection | x |
| Collection_G0022_1386 | Collection | INERA collection | x |
| Collection_G0037_2004 | Collection | INERA collection | x |
| Collection_G0075_2044 | Collection | INERA collection | x |
| Collection_G0198_1463 | Collection | INERA collection | x |
| Collection_G0001_1965 | Collection | INERA collection | x |
| Collection_G0084_2055 | Collection | INERA collection | x |
| Collection_G0046_2013 | Collection | INERA collection | x |
| Collection_G0015_1980 | Collection | INERA collection | x |
| Collection_G0047_2014 | Collection | INERA collection | x |
| Collection_G0025_1383 | Collection | INERA collection | x |

| | | | |
|---|---|---|---|
| Collection_G0024_1374 | Collection | INERA collection | x |
| Collection_G0027_1377 | Collection | INERA collection | x |
| Collection_G0029_1995 | Collection | INERA collection | x |
| Collection_G0044_2011 | Collection | INERA collection | x |
| Collection_G0055_1392 | Collection | INERA collection | x |
| Collection_G0077_2046 | Collection | INERA collection | x |
| Collection_G0017_1376 | Collection | INERA collection | x |
| Collection_G0008_1476 | Collection | INERA collection | x |
| Collection_G0230_1394 | Collection | INERA collection | x |
| Collection_G0107_2084 | Collection | INERA collection | x |
| Collection_G0080_2050 | Collection | INERA collection | x |
| Collection_G0021_1375 | Collection | INERA collection | x |
| Collection_G0036_2003 | Collection | INERA collection | x |
| Collection_G0013_1978 | Collection | INERA collection | x |
| Collection_G0032_1999 | Collection | INERA collection | x |
| Collection_G0104_2080 | Collection | INERA collection | x |
| Collection_G0091_2062 | Collection | INERA collection | x |
| Collection_G0235_1497 | Collection | INERA collection | x |
| Collection_G0090_2061 | Collection | INERA collection | x |
| Collection_G0232_1393 | Collection | INERA collection | x |
| Collection_G0019_1321 | Collection | INERA collection | x |
| Collection_G0098_2070 | Collection | INERA collection | x |
| Collection_G0071_2040 | Collection | INERA collection | x |
| Collection_G0033_2000 | Collection | INERA collection | x |
| Collection_G0018_2071 | Collection | INERA collection | x |
| Collection_G0072_2041 | Collection | INERA collection | x |
| Collection_G0099_2074 | Collection | INERA collection | x |
| Collection_G0068_2037 | Collection | INERA collection | x |
| Collection_G0056_2025 | Collection | INERA collection | x |
| Collection_G0094_2112 | Collection | INERA collection | x |
| Collection_G0105_2133 | Collection | INERA collection | x |
| Collection_G0113_2110 | Collection | INERA collection | x |
| Collection_G0248_2137 | Collection | INERA collection | x |
| Collection_G0006_2143 | Collection | INERA collection | x |
| Collection_G0009_2160 | Collection | INERA collection | x |
| Collection_G0031_2108 | Collection | INERA collection | x |
| Collection_G0234_2145 | Collection | INERA collection | x |
| Collection_G0197_2151 | Collection | INERA collection | x |
| Collection_G0245_2162 | Collection | INERA collection | x |
| Collection_G0127_2142 | Collection | INERA collection | x |
| Collection_G0005_2157 | Collection | INERA collection | x |
| Collection_G0114_2111 | Collection | INERA collection | x |
| Collection_G0020_2161 | Collection | INERA collection | x |
| Collection_G0246_2163 | Collection | INERA collection | x |
| Collection_G0233_2115 | Collection | INERA collection | x |
| Collection_G0138_2139 | Collection | INERA collection | x |
| Collection_G0121_2155 | Collection | INERA collection | x |
| Collection_G0002_2141 | Collection | INERA collection | x |
| Collection_G0003_2148 | Collection | INERA collection | x |
| Collection_G0135_2131 | Collection | INERA collection | x |

| | | | |
|---|---|---|---|
| Collection_G0012_1976 | Collection | INERA collection | x |
| Collection_G0030_1997 | Collection | INERA collection | x |
| Collection_G0134_2150 | Collection | INERA collection | x |
| Collection_G0035_2002 | Collection | INERA collection | x |
| Collection_G0079_2049 | Collection | INERA collection | x |
| Collection_G0057_1395 | Collection | INERA collection | x |
| Collection_G0016_1390 | Collection | INERA collection | x |
| Collection_G0010_1974 | Collection | INERA collection | x |
| Collection_G0101_2077 | Collection | INERA collection | |
| Collection_G0247_2136 | Collection | INERA collection | |
| Collection_G0026_1385 | Collection | INERA collection | |
| Collection_G0175_1477 | Collection | INERA collection | |
| Collection_G0115_2134 | Collection | INERA collection | |
| Collection_G0092FOG_2063 | Collection | INERA collection | x |
| Collection_G0112_2109 | Collection | INERA collection | |
| Collection_G0256FOG_1498 | Collection | INERA collection | |
| Collection_G0255FOG_1494 | Collection | INERA collection | |
| Collection_G0039_2006 | Collection | INERA collection | x |
| Collection_G0011_1975 | Collection | INERA collection | |
| Collection_G0023_1389 | Collection | INERA collection | |
| Collection_G0085_2056 | Collection | INERA collection | x |
| Collection_G0083_2054 | Collection | INERA collection | x |
| Collection_G0103_2079 | Collection | INERA collection | |
| Collection_G0097_2069 | Collection | INERA collection | |
| Collection_G0078_2048 | Collection | INERA collection | |
| Collection_G0100_2076 | Collection | INERA collection | |
| Collection_G0038_2005 | Collection | INERA collection | |
| Collection_G0088_2059 | Collection | INERA collection | |
| Collection_G0251FOG_1472 | Collection | INERA collection | x |
| Collection_G0063_2032 | Collection | INERA collection | |
| Collection_G0074_2043 | Collection | INERA collection | x |
| Collection_G0102_2078 | Collection | INERA collection | x |
| Collection_G0086_2057 | Collection | INERA collection | x |
| Collection_G0064_2033 | Collection | INERA collection | |
| Collection_G0253FOG_1474 | Collection | INERA collection | x |
| Collection_G0061_2030 | Collection | INERA collection | |
| Collection_G0069_2038 | Collection | INERA collection | |
| Collection_G0067_2036 | Collection | INERA collection | |
| Collection_G0041_2008 | Collection | INERA collection | |
| Collection_G0252FOG_1473 | Collection | INERA collection | x |
| Collection_G0073_2042 | Collection | INERA collection | x |
| Collection_G0066_2035 | Collection | INERA collection | |
| Collection_G0043_2073 | Collection | INERA collection | |
| Collection_G0042_1496 | Collection | INERA collection | |
| Collection_G0062_2031 | Collection | INERA collection | |
| Collection_G0028_1388 | Collection | INERA collection | |
| Collection_G0059_2028 | Collection | INERA collection | |
| Collection_G0109_2105 | Collection | INERA collection | |
| Collection_G0110_2106 | Collection | INERA collection | |
| Collection_G0076_2102 | Collection | INERA collection | |

| | | |
|---|---|---|
| Collection_G0050_2101 | Collection | INERA collection |
| Collection_G0111_2107 | Collection | INERA collection |
| Collection_G0081_2120 | Collection | INERA collection |
| Collection_G0108_2104 | Collection | INERA collection |
| Collection_G0065_2034 | Collection | INERA collection |
| Collection_G0060_2029 | Collection | INERA collection |
| Collection_G0058_2027 | Collection | INERA collection |
| Collection_G0254FOG_1475 | Collection | INERA collection |
| Collection_G0087_2058 | Collection | INERA collection |
| Collection_G0048_2016 | Collection | INERA collection |
| Collection_G0049_2017 | Collection | INERA collection |
| Collection_G0054_2022 | Collection | INERA collection |
| Collection_G0052_2020 | Collection | INERA collection |
| Collection_G0053_2021 | Collection | INERA collection |
| Collection_G0070_2039 | Collection | INERA collection |
| Collection_G0082_2053 | Collection | INERA collection |
| Collection_G0051_2019 | Collection | INERA collection |
| Wild_plot_01_1501 | Forest | plot_01 |
| Wild_plot_01_1502 | Forest | plot_01 |
| Wild_plot_01_1562 | Forest | plot_01 |
| Wild_plot_01_1504 | Forest | plot_01 |
| Wild_plot_01_1505 | Forest | plot_01 |
| Wild_plot_01_1503 | Forest | plot_01 |
| Wild_plot_01_1507 | Forest | plot_01 |
| Wild_plot_01_1506 | Forest | plot_01 |
| Wild_plot_01_1561 | Forest | plot_01 |
| Wild_plot_02_2244 | Forest | plot_02 |
| Wild_plot_02_2245 | Forest | plot_02 |
| Wild_plot_02_1517 | Forest | plot_02 |
| Wild_plot_02_1509 | Forest | plot_02 |
| Wild_plot_02_2246 | Forest | plot_02 |
| Wild_plot_02_1520 | Forest | plot_02 |
| Wild_plot_02_1512 | Forest | plot_02 |
| Wild_plot_02_1519 | Forest | plot_02 |
| Wild_plot_02_1525 | Forest | plot_02 |
| Wild_plot_02_2088 | Forest | plot_02 |
| Wild_plot_02_1528 | Forest | plot_02 |
| Wild_plot_02_1822 | Forest | plot_02 |
| Wild_plot_02_1527 | Forest | plot_02 |
| Wild_plot_02_1521 | Forest | plot_02 |
| Wild_plot_02_1518 | Forest | plot_02 |
| Wild_plot_02_1513 | Forest | plot_02 |
| Wild_plot_02_1523 | Forest | plot_02 |
| Wild_plot_02_1510 | Forest | plot_02 |
| Wild_plot_02_1511 | Forest | plot_02 |
| Wild_plot_02_1524 | Forest | plot_02 |
| Wild_plot_02_1508 | Forest | plot_02 |
| Wild_plot_02_2089 | Forest | plot_02 |
| Wild_plot_03_2091 | Forest | plot_03 |
| Wild_plot_03_1549 | Forest | plot_03 |

| | | |
|---|---|---|
| Wild_plot_03_1541 | Forest | plot_03 |
| Wild_plot_03_1534 | Forest | plot_03 |
| Wild_plot_03_1557 | Forest | plot_03 |
| Wild_plot_03_1539 | Forest | plot_03 |
| Wild_plot_03_1558 | Forest | plot_03 |
| Wild_plot_03_1552 | Forest | plot_03 |
| Wild_plot_03_1540 | Forest | plot_03 |
| Wild_plot_03_1544 | Forest | plot_03 |
| Wild_plot_03_1551 | Forest | plot_03 |
| Wild_plot_03_1542 | Forest | plot_03 |
| Wild_plot_03_1536 | Forest | plot_03 |
| Wild_plot_03_1560 | Forest | plot_03 |
| Wild_plot_03_1535 | Forest | plot_03 |
| Wild_plot_03_1538 | Forest | plot_03 |
| Wild_plot_03_1545 | Forest | plot_03 |
| Wild_plot_03_2092 | Forest | plot_03 |
| Wild_plot_03_2090 | Forest | plot_03 |
| Wild_plot_03_1533 | Forest | plot_03 |
| Wild_plot_03_1537 | Forest | plot_03 |
| Wild_plot_03_1547 | Forest | plot_03 |
| Wild_plot_03_1559 | Forest | plot_03 |
| Wild_plot_03_1548 | Forest | plot_03 |
| Wild_plot_03_2093 | Forest | plot_03 |
| Wild_plot_03_1550 | Forest | plot_03 |
| Wild_plot_03_1556 | Forest | plot_03 |
| Wild_plot_03_1546 | Forest | plot_03 |
| Wild_plot_03_1554 | Forest | plot_03 |
| Wild_plot_03_1555 | Forest | plot_03 |
| Wild_plot_03_1543 | Forest | plot_03 |
| Wild_plot_03_1532 | Forest | plot_03 |
| Wild_plot_04_1567 | Forest | plot_04 |
| Wild_plot_04_1566 | Forest | plot_04 |
| Wild_plot_04_1564 | Forest | plot_04 |
| Wild_plot_04_1571 | Forest | plot_04 |
| Wild_plot_04_1573 | Forest | plot_04 |
| Wild_plot_04_1570 | Forest | plot_04 |
| Wild_plot_04_1569 | Forest | plot_04 |
| Wild_plot_04_1568 | Forest | plot_04 |
| Wild_plot_04_1563 | Forest | plot_04 |
| Wild_plot_04_1572 | Forest | plot_04 |
| Wild_plot_04_1565 | Forest | plot_04 |
| Wild_plot_05_1580 | Forest | plot_05 |
| Wild_plot_05_1578 | Forest | plot_05 |
| Wild_plot_05_1584 | Forest | plot_05 |
| Wild_plot_05_1581 | Forest | plot_05 |
| Wild_plot_05_1575 | Forest | plot_05 |
| Wild_plot_05_1574 | Forest | plot_05 |
| Wild_plot_05_1577 | Forest | plot_05 |
| Wild_plot_05_1576 | Forest | plot_05 |
| Wild_plot_05_1579 | Forest | plot_05 |

| | | |
|---|---|---|
| Wild_plot_05_1583 | Forest | plot_05 |
| Wild_plot_05_1582 | Forest | plot_05 |
| Wild_plot_06_1588 | Forest | plot_06 |
| Wild_plot_06_1589 | Forest | plot_06 |
| Wild_plot_06_1592 | Forest | plot_06 |
| Wild_plot_06_1590 | Forest | plot_06 |
| Wild_plot_06_1591 | Forest | plot_06 |
| Wild_plot_06_1593 | Forest | plot_06 |
| Wild_plot_06_1597 | Forest | plot_06 |
| Wild_plot_06_1599 | Forest | plot_06 |
| Wild_plot_06_1595 | Forest | plot_06 |
| Wild_plot_06_1598 | Forest | plot_06 |
| Wild_plot_06_1594 | Forest | plot_06 |
| Wild_plot_06_1596 | Forest | plot_06 |
| Wild_plot_06_1587 | Forest | plot_06 |
| Wild_plot_06_1586 | Forest | plot_06 |
| Wild_plot_07_1607 | Forest | plot_07 |
| Wild_plot_07_1611 | Forest | plot_07 |
| Wild_plot_07_1605 | Forest | plot_07 |
| Wild_plot_07_1606 | Forest | plot_07 |
| Wild_plot_07_1608 | Forest | plot_07 |
| Wild_plot_07_1609 | Forest | plot_07 |
| Wild_plot_07_1604 | Forest | plot_07 |
| Wild_plot_07_1602 | Forest | plot_07 |
| Wild_plot_07_1603 | Forest | plot_07 |
| Wild_plot_07_1610 | Forest | plot_07 |
| Wild_plot_07_1612 | Forest | plot_07 |
| Wild_plot_08_1620 | Forest | plot_08 |
| Wild_plot_08_1616 | Forest | plot_08 |
| Wild_plot_08_1619 | Forest | plot_08 |
| Wild_plot_08_1618 | Forest | plot_08 |
| Wild_plot_08_1615 | Forest | plot_08 |
| Wild_plot_08_1617 | Forest | plot_08 |
| Wild_plot_09_1625 | Forest | plot_09 |
| Wild_plot_09_1630 | Forest | plot_09 |
| Wild_plot_09_1629 | Forest | plot_09 |
| Wild_plot_09_1623 | Forest | plot_09 |
| Wild_plot_09_1633 | Forest | plot_09 |
| Wild_plot_09_1624 | Forest | plot_09 |
| Wild_plot_09_1631 | Forest | plot_09 |
| Wild_plot_09_2095 | Forest | plot_09 |
| Wild_plot_09_1621 | Forest | plot_09 |
| Wild_plot_09_1632 | Forest | plot_09 |
| Wild_plot_09_2094 | Forest | plot_09 |
| Wild_plot_09_1626 | Forest | plot_09 |
| Wild_plot_09_1628 | Forest | plot_09 |
| Wild_plot_10_1643 | Forest | plot_10 |
| Wild_plot_10_1634 | Forest | plot_10 |
| Wild_plot_10_1636 | Forest | plot_10 |
| Wild_plot_10_1638 | Forest | plot_10 |

| | | |
|---|---|---|
| Wild_plot_10_1635 | Forest | plot_10 |
| Wild_plot_10_1637 | Forest | plot_10 |
| Wild_plot_10_1645 | Forest | plot_10 |
| Wild_plot_10_1639 | Forest | plot_10 |
| Wild_plot_10_1646 | Forest | plot_10 |
| Wild_plot_10_1642 | Forest | plot_10 |
| Wild_plot_10_1641 | Forest | plot_10 |
| Wild_plot_10_1644 | Forest | plot_10 |
| Wild_plot_10_1640 | Forest | plot_10 |
| Wild_plot_11_1653 | Forest | plot_11 |
| Wild_plot_11_1650 | Forest | plot_11 |
| Wild_plot_11_1656 | Forest | plot_11 |
| Wild_plot_11_1647 | Forest | plot_11 |
| Wild_plot_11_2098 | Forest | plot_11 |
| Wild_plot_11_2096 | Forest | plot_11 |
| Wild_plot_11_1658 | Forest | plot_11 |
| Wild_plot_11_1648 | Forest | plot_11 |
| Wild_plot_11_1651 | Forest | plot_11 |
| Wild_plot_11_1652 | Forest | plot_11 |
| Wild_plot_11_1657 | Forest | plot_11 |
| Wild_plot_12_1661 | Forest | plot_12 |
| Wild_plot_12_1674 | Forest | plot_12 |
| Wild_plot_12_1664 | Forest | plot_12 |
| Wild_plot_12_1666 | Forest | plot_12 |
| Wild_plot_12_1669 | Forest | plot_12 |
| Wild_plot_12_1673 | Forest | plot_12 |
| Wild_plot_12_1659 | Forest | plot_12 |
| Wild_plot_12_1668 | Forest | plot_12 |
| Wild_plot_12_1663 | Forest | plot_12 |
| Wild_plot_12_1665 | Forest | plot_12 |
| Wild_plot_12_1662 | Forest | plot_12 |
| Wild_plot_12_1667 | Forest | plot_12 |
| Wild_plot_12_1671 | Forest | plot_12 |
| Wild_plot_12_1670 | Forest | plot_12 |
| Wild_plot_12_1660 | Forest | plot_12 |
| Wild_plot_13_1678 | Forest | plot_13 |
| Wild_plot_13_1683 | Forest | plot_13 |
| Wild_plot_13_1681 | Forest | plot_13 |
| Wild_plot_13_1676 | Forest | plot_13 |
| Wild_plot_13_1677 | Forest | plot_13 |
| Wild_plot_13_1680 | Forest | plot_13 |
| Wild_plot_13_1685 | Forest | plot_13 |
| Wild_plot_13_1679 | Forest | plot_13 |
| Wild_plot_13_1682 | Forest | plot_13 |
| Wild_plot_13_1675 | Forest | plot_13 |
| Wild_plot_13_1684 | Forest | plot_13 |
| Wild_plot_13_1686 | Forest | plot_13 |
| Wild_plot_14_1696 | Forest | plot_14 |
| Wild_plot_14_1690 | Forest | plot_14 |
| Wild_plot_14_1689 | Forest | plot_14 |

| | | |
|---|---|---|
| Wild_plot_14_1687 | Forest | plot_14 |
| Wild_plot_14_1695 | Forest | plot_14 |
| Wild_plot_14_1693 | Forest | plot_14 |
| Wild_plot_14_1694 | Forest | plot_14 |
| Wild_plot_14_1692 | Forest | plot_14 |
| Wild_plot_14_1688 | Forest | plot_14 |
| Wild_plot_14_1691 | Forest | plot_14 |
| Wild_plot_14_1697 | Forest | plot_14 |
| Wild_plot_15_1705 | Forest | plot_15 |
| Wild_plot_15_1702 | Forest | plot_15 |
| Wild_plot_15_1707 | Forest | plot_15 |
| Wild_plot_15_1704 | Forest | plot_15 |
| Wild_plot_15_1706 | Forest | plot_15 |
| Wild_plot_15_1703 | Forest | plot_15 |
| Wild_plot_16_1827 | Forest | plot_16 |
| Wild_plot_16_1830 | Forest | plot_16 |
| Wild_plot_16_1824 | Forest | plot_16 |
| Wild_plot_16_1832 | Forest | plot_16 |
| Wild_plot_16_1829 | Forest | plot_16 |
| Wild_plot_16_1828 | Forest | plot_16 |
| Wild_plot_16_1826 | Forest | plot_16 |
| Wild_plot_17_1833 | Forest | plot_17 |
| Wild_plot_17_1835 | Forest | plot_17 |
| Wild_plot_17_1834 | Forest | plot_17 |
| Wild_plot_18_1836 | Forest | plot_18 |
| Wild_plot_18_1840 | Forest | plot_18 |
| Wild_plot_18_1841 | Forest | plot_18 |
| Wild_plot_18_1838 | Forest | plot_18 |
| Wild_plot_18_1839 | Forest | plot_18 |
| Wild_plot_18_1837 | Forest | plot_18 |
| Wild_plot_19_1851 | Forest | plot_19 |
| Wild_plot_19_1850 | Forest | plot_19 |
| Wild_plot_19_1842 | Forest | plot_19 |
| Wild_plot_19_1848 | Forest | plot_19 |
| Wild_plot_19_1846 | Forest | plot_19 |
| Wild_plot_19_1844 | Forest | plot_19 |
| Wild_plot_19_1852 | Forest | plot_19 |
| Wild_plot_19_1854 | Forest | plot_19 |
| Wild_plot_19_1843 | Forest | plot_19 |
| Wild_plot_20_2168 | Forest | plot_20 |
| Wild_plot_20_2170 | Forest | plot_20 |
| Wild_plot_20_2166 | Forest | plot_20 |
| Wild_plot_20_2167 | Forest | plot_20 |
| Wild_plot_21_2174 | Forest | plot_21 |
| Wild_plot_21_2176 | Forest | plot_21 |
| Wild_plot_21_2172 | Forest | plot_21 |
| Wild_plot_21_2185 | Forest | plot_21 |
| Wild_plot_21_2173 | Forest | plot_21 |
| Wild_plot_21_2186 | Forest | plot_21 |
| Wild_plot_21_2175 | Forest | plot_21 |

| | | |
|---|---|---|
| Wild_plot_21_2171 | Forest | plot_21 |
| Wild_plot_22_2190 | Forest | plot_22 |
| Wild_plot_22_2196 | Forest | plot_22 |
| Wild_plot_22_2200 | Forest | plot_22 |
| Wild_plot_22_2197 | Forest | plot_22 |
| Wild_plot_23_2203 | Forest | plot_23 |
| Wild_plot_23_2202 | Forest | plot_23 |
| Wild_plot_23_2201 | Forest | plot_23 |
| Wild_plot_23_2204 | Forest | plot_23 |
| Wild_plot_23_2208 | Forest | plot_23 |
| Wild_plot_23_2205 | Forest | plot_23 |
| Wild_plot_23_2206 | Forest | plot_23 |
| Wild_plot_23_2207 | Forest | plot_23 |
| Wild_plot_24_2213 | Forest | plot_24 |
| Wild_plot_24_2218 | Forest | plot_24 |
| Wild_plot_24_2214 | Forest | plot_24 |
| Wild_plot_24_2224 | Forest | plot_24 |
| Wild_plot_24_2215 | Forest | plot_24 |
| Wild_plot_24_2212 | Forest | plot_24 |
| Wild_plot_24_2217 | Forest | plot_24 |
| Wild_plot_24_2211 | Forest | plot_24 |
| Wild_plot_24_2219 | Forest | plot_24 |
| Wild_plot_25_2233 | Forest | plot_25 |
| Wild_plot_25_2231 | Forest | plot_25 |
| Wild_plot_25_2234 | Forest | plot_25 |
| Wild_plot_25_2232 | Forest | plot_25 |
| Wild_plot_25_2235 | Forest | plot_25 |
| Wild_plot_25_2230 | Forest | plot_25 |

*phora* individuals collected from the Yangambi rainforest, the INERA Coffee Colle
hich indiviudals were used in the hybridisation analysis and the hybrid class assig

| Sample sets | | PCA | |
|---|---|---|---|
| Wild reference group | Hybrid analysis | PC1 | PC2 |
| | | -12.17340502 | 0.092892396 |
| | | -11.27631773 | -0.363773028 |
| | | -12.51906694 | 0.269506106 |
| | | -11.43591293 | 0.362639161 |
| | | -10.91992129 | -0.05104781 |
| | | -9.712126285 | 0.220356246 |
| | x | 7.818818599 | -1.787622153 |
| | x | 8.465548355 | -1.542781626 |
| | | -11.47140239 | -0.108309955 |
| | | -11.75061629 | -0.240512145 |
| | | -10.98804705 | 0.292110711 |
| | | -13.15390517 | 0.513756071 |
| | | -10.18713992 | 0.185821065 |
| | x | -8.473087705 | 1.754928688 |
| | x | -5.803636769 | 2.826945106 |
| | | -11.0955558 | 0.341678059 |
| | | -11.32041035 | 0.387529533 |
| | x | -2.575893495 | 4.174425299 |
| | | -12.85402669 | -0.038181624 |
| | | -12.5706442 | -0.262204727 |
| | | -12.28958635 | 0.010947672 |
| | | -12.81466524 | 0.027378463 |
| | | -12.11516482 | 0.107931563 |
| | | -12.823668 | 0.117532762 |
| | | -11.59153911 | 0.11570417 |
| | | -11.94272896 | 0.386122419 |
| | | -11.85865893 | -0.24812705 |
| | | -11.483895 | -0.307055368 |
| | | -11.10976075 | -0.300116086 |
| | | -11.29668375 | -0.669034919 |
| | | -11.63110145 | 0.509405377 |
| | | -11.73169377 | 0.239340215 |
| | | -11.46488506 | 0.013562424 |
| | | -11.01778507 | -0.052215278 |
| | | -11.31416094 | 0.018647323 |
| | | -11.87548846 | -0.311602408 |
| | | -10.82895663 | -0.283252627 |
| | | -10.22228043 | -0.321208073 |
| | | -10.03786521 | 0.483037144 |
| | x | -1.878608349 | 4.13845235 |
| | | -13.042715 | -0.047669158 |
| | | -13.4285479 | 0.543223817 |
| | | -12.33077284 | -0.181722916 |

| | | | |
|---|---|---|---|
| | | -12.32813628 | -0.074295011 |
| | | -11.41271706 | 0.364015849 |
| | | -11.16028134 | 0.290842453 |
| | | -10.51452215 | 0.504622055 |
| | | -10.56506928 | -0.251861746 |
| | | -10.14163148 | 0.012371039 |
| | | -13.79032883 | 0.546921592 |
| | | -13.40956774 | 0.170093845 |
| | | -13.08302191 | 0.073291912 |
| | | -12.85984837 | -0.04035837 |
| | | -12.6740901 | 0.483004063 |
| | | -12.29732051 | 0.714122821 |
| | | -12.81241248 | 0.243735213 |
| | | -12.96460231 | 0.00146035 |
| | | -11.53376244 | 0.110260337 |
| | | -13.31052565 | 0.501309844 |
| | | -13.0657533 | 0.268185204 |
| | | -12.25596761 | -0.270775707 |
| | | -11.92533399 | -0.229743903 |
| | | -9.759171936 | 0.622659858 |
| | x | -7.329038723 | 0.541711557 |
| | x | 8.416836065 | -1.613931354 |
| | x | 8.06290354 | -2.327419693 |
| | x | 7.870759799 | -1.915091925 |
| | | -11.10626891 | -0.151725782 |
| | | -11.0922435 | -0.17523854 |
| | | -10.14463659 | -0.268760476 |
| | | -9.500912343 | -0.343833926 |
| | | -13.86933522 | 0.446432878 |
| | | -13.45755761 | 0.041710948 |
| | | -13.62507441 | -0.177827804 |
| | | -13.33701269 | 0.196138572 |
| | | -12.73835024 | 0.163736268 |
| | | -13.23697999 | -0.034094899 |
| | | -13.01764571 | 0.254918223 |
| | | -12.94750625 | -0.167752407 |
| | | -13.10846273 | 0.309316302 |
| | | -12.97180645 | 0.001760259 |
| | | -12.58403215 | -0.186066112 |
| | | -12.86361411 | 0.483282312 |
| | | -12.26492147 | 0.065800917 |
| | | -12.33691612 | -0.40031208 |
| | | -12.48146135 | -0.097938695 |
| | | -11.77476278 | 0.265138591 |
| | | -12.1531181 | -0.208984185 |
| | | -12.49565969 | 0.234939856 |
| | | -11.83549069 | -0.305533514 |
| | | -12.03187307 | 0.404420876 |
| | | -12.35771405 | 0.169396147 |
| | | -11.52550477 | -0.437523588 |

| | | | |
|---|---|---|---|
| | | -11.81793153 | -0.03523024 |
| | | -11.5923469 | 0.441616738 |
| | | -11.79106587 | 0.093295738 |
| | | -11.50662201 | -0.150378166 |
| | | -11.59454253 | 0.038574221 |
| | | -11.64190094 | -0.242447138 |
| | | -11.37738465 | -0.453782967 |
| | | -11.5393114 | -0.427828011 |
| | | -11.15377015 | 0.181536161 |
| | | -12.20152176 | -0.136679774 |
| | | -11.13723124 | -0.534509049 |
| | | -11.05307754 | 0.265241721 |
| | | -11.19952975 | -0.168302653 |
| | | -11.61299754 | -0.144863517 |
| | | -11.50349953 | 0.14828583 |
| | | -11.70591187 | -0.234023289 |
| | | -11.82278035 | 0.126997365 |
| | | -11.88882022 | 0.095976469 |
| | | -11.68897395 | 0.374851044 |
| | | -10.61218328 | 0.060698266 |
| | | -11.11628083 | 0.386080285 |
| | | -11.25356964 | 0.35613375 |
| | | -10.95165929 | 0.383428366 |
| | | -10.56360561 | 0.196480221 |
| | | -11.50248356 | -0.524023919 |
| | | -12.32651688 | -0.2586799 |
| | | -11.74890421 | 0.399847449 |
| | | -11.56220881 | 0.117534771 |
| | | -10.3865835 | 0.235053834 |
| | | -9.980244483 | -0.574950826 |
| | | -11.16576038 | 0.01985272 |
| | | -10.42361749 | 0.020272253 |
| | | -11.66620273 | -0.064272148 |
| | | -11.14176304 | -0.187647525 |
| | | -11.20257348 | 0.246968213 |
| | | -10.55001199 | -0.100195548 |
| | | -9.401342312 | 0.223560485 |
| | | -10.93165367 | 0.342607851 |
| | | -10.18750352 | 0.069928674 |
| | | -8.991487386 | -0.659793344 |
| | | -10.88128995 | 0.210210366 |
| | | -7.866120819 | 0.104888982 |
| | | -10.24442235 | -0.010359482 |
| | | -11.0394324 | 0.072180993 |
| | | -9.530437201 | -0.212587145 |
| | | -9.378255086 | 0.297794476 |
| | | -10.12103734 | 0.291912037 |
| | | -7.955389975 | -0.077562776 |
| | | -9.744853317 | -0.149692449 |
| | | -8.332923355 | 0.048611287 |

| | | | |
|---|---|---|---|
| | | -12.5193793 | 0.343101912 |
| | | -10.89310704 | -0.732003107 |
| | | -8.860134731 | 0.374121359 |
| | | -9.657486985 | 0.326280155 |
| | | -10.6287205 | -0.325265445 |
| | | -9.925106106 | -0.383275675 |
| | | -9.483755909 | 0.201839054 |
| | | -10.40418047 | -0.612773955 |
| | x | -9.271220234 | 0.22819788 |
| | x | -7.723351183 | 0.18941162 |
| | x | -9.406597595 | 0.95690147 |
| | x | -8.815086481 | 0.665240641 |
| | x | -7.959944089 | 0.360139495 |
| | | -11.03096384 | -0.632922677 |
| | x | -6.299248623 | -0.222775397 |
| | x | -9.42934909 | 0.08723215 |
| | x | -8.908422006 | 0.09895966 |
| | | -10.29646311 | -0.598936367 |
| | x | -7.919415896 | -0.027868276 |
| | x | -7.582110467 | -0.091409921 |
| | | -10.96524723 | -0.398217241 |
| | | -10.93989606 | -0.708420356 |
| | x | -7.200125936 | 0.383715764 |
| | x | -7.025875223 | -0.329945304 |
| | x | -6.9938588 | -0.385820683 |
| | x | -8.652708679 | -0.229511544 |
| | x | -6.206691712 | -0.154608857 |
| | x | -9.42147427 | -1.045295599 |
| | | -9.543439061 | -0.797189338 |
| | x | -8.332121106 | 2.368486469 |
| | | -9.377246655 | -0.623480732 |
| | | -10.82333312 | -0.773034945 |
| | | -10.00719204 | -0.69613711 |
| | x | -6.803725647 | 1.972357645 |
| | | -9.589674203 | -1.162568875 |
| | x | -7.772170133 | 2.678389793 |
| | x | -5.792127462 | 3.252901713 |
| | x | -6.772872108 | 2.596365964 |
| | x | -1.874759986 | 1.097965977 |
| | | -9.995182602 | -0.760791431 |
| | | -9.098545332 | -0.996595206 |
| | x | -5.984600238 | 2.926072589 |
| | x | -7.457323123 | -1.422473639 |
| | x | -1.131996976 | -0.368463342 |
| | x | -4.647560935 | 3.075104648 |
| | x | 0.964863392 | 0.098805238 |
| | x | -2.829057689 | 4.227125861 |
| | x | 0.286266684 | -0.182633932 |
| | x | 4.64738537 | -1.572927461 |
| | x | 5.649025312 | -2.064769183 |

| | | | |
|---|---|---|---|
| | x | 5.86400132 | -2.491961926 |
| | x | 6.667798431 | -2.313068805 |
| | x | -4.544879724 | -1.185528189 |
| | x | 6.929392342 | -1.529717971 |
| | x | -0.964578211 | 4.685925064 |
| | x | -1.062419797 | 4.612841134 |
| | x | -0.974453305 | 4.323823401 |
| | x | -6.033255336 | -1.25467785 |
| | x | -4.694801748 | -1.145252276 |
| | x | 8.341594819 | 10.47941361 |
| | x | 8.668824873 | 9.502074501 |
| | x | 8.766304386 | 11.14550413 |
| | x | 9.114872509 | 9.792220485 |
| | x | 9.005600058 | 11.82259844 |
| | x | 8.57680214 | 6.1627343 |
| | x | 8.537571956 | 6.219282417 |
| | x | 7.206959271 | -0.488595835 |
| x | | 7.571464081 | -0.73856178 |
| x | | 7.072504004 | -0.501210322 |
| x | | 7.137413282 | -0.477480346 |
| x | | 8.571789068 | -0.848522389 |
| x | | 7.465392528 | -0.191008624 |
| x | | 7.975775978 | 0.682318581 |
| x | | 7.417992865 | 0.703371912 |
| x | | 7.839017321 | -0.137921301 |
| x | | 7.761818899 | 0.31715891 |
| x | | 4.8542007 | -1.729039924 |
| x | | 4.494637378 | -1.260215659 |
| x | | 6.99223562 | -0.429363551 |
| x | | 7.81403246 | 2.064297827 |
| x | | 5.631586758 | -0.060316856 |
| x | | 8.451151244 | -0.529085579 |
| x | | 8.67507992 | -0.436651 |
| x | | 8.494362655 | 0.628030163 |
| x | | 8.315196655 | 0.104864243 |
| x | | 7.983416969 | -0.473635768 |
| x | | 8.036147638 | 0.263678633 |
| x | | 7.721968035 | -0.23567551 |
| x | | 8.11804264 | 0.00825418 |
| x | | 7.955019665 | 0.201572555 |
| x | | 7.793170084 | 0.219304065 |
| x | | 8.333070381 | 0.208977696 |
| x | | 7.677516603 | -0.533388399 |
| x | | 7.467256618 | -0.24864203 |
| x | | 8.466746451 | 0.532524057 |
| x | | 7.999872583 | -0.863409843 |
| x | | 8.120048787 | -0.140117838 |
| x | | 7.286135261 | 0.409732807 |
| x | | 8.516060595 | 8.524140966 |
| x | | 9.255178256 | 7.259350993 |

| | | | |
|---|---|---|---|
| x | | 9.016880239 | 12.12934984 |
| x | | 9.46536408 | 11.92805879 |
| x | | 9.277562575 | 12.0176217 |
| x | | 9.297233357 | 11.20106668 |
| x | | 9.029975235 | 11.65037952 |
| x | | 8.941036353 | 11.41604216 |
| x | | 8.728680367 | 9.368478187 |
| x | | 9.314597796 | 9.858819023 |
| x | | 9.061033111 | 10.99191278 |
| x | | 8.636094831 | 11.92557824 |
| x | | 8.633171947 | 9.455204398 |
| x | | 8.91759316 | 10.94444475 |
| x | | 8.901615548 | 10.19146469 |
| x | | 9.057828282 | 10.86362084 |
| x | | 8.509708393 | 10.9427533 |
| x | | 8.86966996 | 10.62827923 |
| x | | 8.637891166 | 8.808912253 |
| x | | 8.507174645 | 10.68826187 |
| x | | 8.668816273 | 10.33038242 |
| x | | 8.475208265 | 9.729823403 |
| x | | 8.103770021 | 9.047710486 |
| x | | 9.466010006 | 13.14710699 |
| x | | 8.800433116 | 9.523063657 |
| x | | 9.018883475 | 11.22215815 |
| x | | 8.871943448 | 11.4257932 |
| x | | 8.704985885 | 12.92643786 |
| x | | 8.935341224 | 11.68803631 |
| x | | 8.411617627 | 9.163170648 |
| x | | 7.853321566 | 7.709395434 |
| x | | 8.486191145 | 9.707963013 |
| x | | 8.282672864 | -1.056021037 |
| x | | 7.952696066 | -1.139423902 |
| x | | 7.648506245 | -1.417021597 |
| x | | 7.713889742 | -1.037621027 |
| x | | 8.005205946 | -0.768145135 |
| x | | 7.981031507 | -0.529014124 |
| x | | 7.60228979 | -0.856325174 |
| x | | 7.391249137 | -1.011572594 |
| x | | 7.310836295 | -0.40040346 |
| x | | 7.674174282 | -0.73759173 |
| x | | 7.85819274 | -0.373018385 |
| x | | 8.308406537 | -1.749942575 |
| x | | 7.699065867 | -1.704256652 |
| x | | 8.085447108 | -2.371027473 |
| x | | 7.782272885 | -1.508168912 |
| x | | 7.766310374 | -1.771436193 |
| x | | 7.703845836 | -2.286714785 |
| x | | 7.743657528 | -1.938695374 |
| x | | 8.91542123 | -1.420644185 |
| x | | 7.340328474 | -1.440268634 |

| | | | |
|---|---|---|---|
| x | | 7.914568562 | -1.178488336 |
| x | | 7.998912283 | -0.67319264 |
| x | | 8.121235782 | -2.454742625 |
| x | | 8.767995196 | -1.957880509 |
| x | | 8.172800516 | -1.751539794 |
| x | | 8.583656002 | -2.228503804 |
| x | | 8.047319669 | -2.477996362 |
| x | | 8.035895482 | -2.116795132 |
| x | | 7.86756379 | -2.396776599 |
| x | | 7.794440805 | -1.818722638 |
| x | | 7.681318408 | -1.476894402 |
| x | | 7.899465781 | -1.38023736 |
| x | | 7.918815671 | -1.623114303 |
| x | | 7.816502343 | -2.869750116 |
| x | | 7.893534696 | -2.062855277 |
| x | | 7.580422143 | -1.505020196 |
| x | | 8.249880289 | -1.886877085 |
| x | | 7.973554199 | -1.901393024 |
| x | | 7.925542303 | -2.130536121 |
| x | | 7.629567451 | -1.921747035 |
| x | | 8.070121111 | -2.832507649 |
| x | | 8.26291966 | -2.352048602 |
| x | | 7.729222834 | -1.858689464 |
| x | | 8.199078606 | -1.760109329 |
| x | | 8.535050552 | -1.924503837 |
| x | | 7.677547845 | -1.603946643 |
| | x | -0.776156296 | -0.832605835 |
| x | | 8.157481504 | -2.645406717 |
| x | | 7.492460041 | -1.843761665 |
| x | | 8.045488904 | -1.700800641 |
| x | | 7.957488904 | -1.917875274 |
| x | | 8.237641241 | -1.962871396 |
| x | | 7.832537723 | -2.061169401 |
| x | | 8.825621857 | -2.024174735 |
| x | | 8.0412972 | -2.287457229 |
| x | | 8.166210837 | -2.265379562 |
| x | | 8.036168598 | -2.3194879 |
| x | | 7.982358822 | -1.741853772 |
| x | | 8.144024331 | -2.031386777 |
| x | | 8.347050821 | -1.924406785 |
| x | | 7.606134884 | -2.170719092 |
| x | | 7.700644877 | -1.56351723 |
| x | | 7.782425693 | -1.779046536 |
| x | | 8.06898358 | -1.889287863 |
| x | | 7.953969157 | -1.349970495 |
| x | | 7.642365649 | -2.619151673 |
| x | | 8.202033857 | -3.468791749 |
| x | | 7.935919916 | -3.529584585 |
| x | | 8.09052366 | -3.043408083 |
| x | | 8.026325357 | -2.69554749 |

| | | | |
|---|---|---|---|
| x | | 7.680585748 | -3.27987842 |
| x | | 8.163668643 | -2.639515751 |
| x | | 8.011952161 | -1.749346527 |
| x | | 7.798214044 | -3.205480726 |
| | x | -10.40935617 | -0.09938866 |
| | x | -12.30275327 | -0.091051092 |
| | x | -12.07157431 | 0.339036932 |
| | x | -12.86293922 | 0.477874794 |
| | x | -13.20696707 | -0.068845951 |
| x | | 9.149112737 | -4.026656823 |
| x | | 8.620634299 | -3.144650351 |
| x | | 8.368497075 | -3.479925403 |
| x | | 8.469148604 | -4.039995 |
| x | | 8.407142321 | -3.041292443 |
| x | | 8.493323956 | -3.333215754 |
| x | | 8.360886102 | -3.210989275 |
| x | | 7.844360329 | -3.400693852 |
| x | | 8.671130177 | -3.699181856 |
| x | | 8.555308722 | -3.974792292 |
| x | | 8.082335611 | -4.075829176 |
| x | | 8.44959653 | -3.844814846 |
| x | | 7.940413246 | -3.037943499 |
| x | | 8.198968032 | -3.138679891 |
| x | | 8.159787774 | -3.154619778 |
| x | | 7.620692194 | -3.113901559 |
| x | | 8.780474417 | -3.028007148 |
| x | | 8.403905544 | -2.858960849 |
| x | | 7.910789181 | -2.737890737 |
| x | | 7.927546322 | -2.929506766 |
| x | | 8.183137238 | -3.02933906 |
| x | | 7.734506464 | -3.217105585 |
| x | | 8.080819826 | -2.888910578 |
| x | | 7.986010787 | -1.899425876 |
| x | | 8.455536293 | -2.616676689 |
| x | | 7.768901942 | -2.342876235 |
| x | | 7.391650874 | -1.421074436 |
| x | | 7.75391641 | -1.969960309 |
| x | | 7.748444158 | -1.867532548 |
| x | | 7.493985332 | -2.118253121 |
| x | | 8.406529924 | -2.154040823 |
| x | | 8.066450736 | -1.80465971 |
| x | | 7.989938786 | -1.689303152 |
| x | | 7.483514578 | -1.739350198 |
| x | | 7.697139059 | -1.716588833 |
| x | | 7.691812125 | -3.110360762 |
| x | | 7.44968025 | -1.599424978 |
| | x | -12.88085109 | 0.169095949 |
| x | | 8.460816645 | -2.334460886 |
| x | | 8.333544234 | -3.077028751 |
| x | | 8.166809795 | -2.345585674 |

| | | | |
|---|---|---|---|
| x | | 8.36610988 | -2.785788836 |
| x | | 7.816926331 | -2.32209201 |
| x | | 8.224843307 | -1.907099555 |
| x | | 7.797292632 | -1.798886265 |
| x | | 7.965094333 | -1.966473794 |
| x | | 7.693523329 | -2.55781855 |
| x | | 8.059452251 | -3.199858451 |
| x | | 7.481824618 | -1.974547239 |
| x | | 7.675164758 | -1.6451739 |
| x | | 8.222746781 | -1.559746501 |
| x | | 8.190422942 | -1.130395349 |
| x | | 7.970852826 | -1.575571657 |
| x | | 8.265392536 | -1.454304339 |
| x | | 7.710884927 | -0.494252579 |
| x | | 7.638584312 | -2.360501001 |
| x | | 8.62115955 | -3.004782474 |
| x | | 8.287284157 | -2.18157886 |
| x | | 8.119187197 | -2.759802187 |
| x | | 8.62769699 | -2.314080094 |
| x | | 8.056118134 | -2.426998205 |
| x | | 2.593648177 | -0.467483712 |
| x | | 7.846454143 | -2.735121679 |
| x | | 7.428896033 | -2.269591649 |
| x | | 8.071167283 | -1.156616468 |
| x | | 7.950318813 | -2.672823113 |
| x | | 7.589299525 | -2.476575046 |
| x | | 8.091720744 | -3.242182597 |
| x | | 7.881242607 | -2.454002855 |
| x | | 7.529750739 | -2.711049014 |
| x | | 7.568467353 | -2.347184324 |
| x | | 8.667917397 | -2.837034084 |
| x | | 8.6353521 | -3.010685421 |
| x | | 8.720191416 | -2.123005757 |
| x | | 8.244957516 | -3.149493084 |
| x | | 8.413596829 | -3.147671325 |
| x | | 8.117334145 | -2.629572146 |
| x | | 8.455215252 | -2.473611591 |
| x | | 8.438688564 | -2.577296896 |
| x | | 8.307639209 | -2.8648636 |
| | x | -1.308336235 | -0.157375689 |
| | x | -4.26339215 | -0.172660673 |
| | x | -5.953497402 | -0.132755737 |
| | x | -8.430016926 | -0.289815868 |
| x | | 6.160814484 | -2.582547295 |
| x | | 4.648212902 | -1.685558133 |
| x | | 5.309247998 | -1.361174377 |
| x | | 4.561136908 | -1.588476751 |
| x | | 4.310108656 | -1.568754619 |
| x | | 3.503823276 | -1.680158935 |
| x | | 3.464718532 | -1.446539153 |

| | | | |
|---|---|---|---|
| | x | -6.934804282 | -0.236169023 |
| x | | 7.845825036 | -3.100119149 |
| x | | 7.161509975 | -3.10895132 |
| x | | 6.412791477 | -2.378597599 |
| x | | 4.939275658 | -1.89591417 |
| x | | 7.174778668 | -2.794058002 |
| x | | 6.109022225 | -1.597715466 |
| x | | 5.838460573 | -1.477094866 |
| x | | 6.032337124 | -1.809988589 |
| x | | 5.42222675 | -1.792351316 |
| x | | 4.763707025 | -2.236050935 |
| x | | 3.348538414 | -1.860830235 |
| x | | 3.067601746 | -1.148073582 |
| x | | 7.695552716 | -3.151188257 |
| x | | 7.009246724 | -2.424692983 |
| x | | 6.076096404 | -2.252659212 |
| x | | 5.712903627 | -2.503825014 |
| x | | 4.464498822 | -1.719652438 |
| x | | 4.03411109 | -1.573655276 |
| x | | 3.77055551 | -1.949264007 |
| x | | 4.08881926 | -1.086716117 |
| x | | 3.780909576 | -1.520276907 |
| x | | 7.358370981 | -1.939812455 |
| x | | 6.923399975 | -2.26759405 |
| x | | 6.757632872 | -2.625441273 |
| x | | 6.065101755 | -2.250388874 |
| x | | 5.882614709 | -2.145474554 |
| x | | 6.555044537 | -2.340637077 |

ction in Yangambi and
nment for all 471 individuals.

| | | fastSTRUCTURE results | | |
|---|---|---|---|---|
| **Cluster assignment** | **Cluster 1** | **Cluster 2** | **Cluster 3** | **Cluster 4** |
| Cluster 1 | 0.999947 | 0.000011 | 0.000011 | 0.000011 |
| Cluster 1 | 0.984927 | 0.015031 | 0.000011 | 0.000011 |
| Cluster 1 | 0.999947 | 0.000011 | 0.000011 | 0.000011 |
| Cluster 1 | 0.999947 | 0.000011 | 0.000011 | 0.000011 |
| Cluster 1 | 0.999947 | 0.000011 | 0.000011 | 0.000011 |
| Cluster 1 | 0.950257 | 0.049701 | 0.000011 | 0.000011 |
| Cluster 1 | 0.000011 | 0.999947 | 0.000011 | 0.000011 |
| Cluster 1 | 0.000011 | 0.999947 | 0.000011 | 0.000011 |
| Cluster 1 | 0.999947 | 0.000011 | 0.000011 | 0.000011 |
| Cluster 1 | 0.999947 | 0.000011 | 0.000011 | 0.000011 |
| Cluster 1 | 0.999947 | 0.000011 | 0.000011 | 0.000011 |
| Cluster 1 | 0.999947 | 0.000011 | 0.000011 | 0.000011 |
| Cluster 1 | 0.971396 | 0.000011 | 0.000011 | 0.028562 |
| Cluster 1 - Cluster 5 | 0.7951 | 0.000011 | 0.000011 | 0.000011 |
| Cluster 1 - Cluster 5 | 0.628452 | 0.000011 | 0.000011 | 0.101707 |
| Cluster 1 | 0.999947 | 0.000011 | 0.000011 | 0.000011 |
| Cluster 1 | 0.943248 | 0.05671 | 0.000011 | 0.000011 |
| Cluster 5 | 0.347863 | 0.000011 | 0.000011 | 0.035931 |
| Cluster 1 | 0.999947 | 0.000011 | 0.000011 | 0.000011 |
| Cluster 1 | 0.999947 | 0.000011 | 0.000011 | 0.000011 |
| Cluster 1 | 0.999947 | 0.000011 | 0.000011 | 0.000011 |
| Cluster 1 | 0.999947 | 0.000011 | 0.000011 | 0.000011 |
| Cluster 1 | 0.999947 | 0.000011 | 0.000011 | 0.000011 |
| Cluster 1 | 0.999947 | 0.000011 | 0.000011 | 0.000011 |
| Cluster 1 | 0.999947 | 0.000011 | 0.000011 | 0.000011 |
| Cluster 1 | 0.999947 | 0.000011 | 0.000011 | 0.000011 |
| Cluster 1 | 0.999947 | 0.000011 | 0.000011 | 0.000011 |
| Cluster 1 | 0.999947 | 0.000011 | 0.000011 | 0.000011 |
| Cluster 1 | 0.999947 | 0.000011 | 0.000011 | 0.000011 |
| Cluster 1 | 0.999947 | 0.000011 | 0.000011 | 0.000011 |
| Cluster 1 | 0.999947 | 0.000011 | 0.000011 | 0.000011 |
| Cluster 1 | 0.999947 | 0.000011 | 0.000011 | 0.000011 |
| Cluster 1 | 0.999947 | 0.000011 | 0.000011 | 0.000011 |
| Cluster 1 | 0.999947 | 0.000011 | 0.000011 | 0.000011 |
| Cluster 1 | 0.999947 | 0.000011 | 0.000011 | 0.000011 |
| Cluster 1 | 0.999947 | 0.000011 | 0.000011 | 0.000011 |
| Cluster 1 | 0.992532 | 0.000011 | 0.000011 | 0.007426 |
| Cluster 1 | 0.95501 | 0.044948 | 0.000011 | 0.000011 |
| Cluster 1 | 0.951368 | 0.048589 | 0.000011 | 0.000011 |
| Cluster 5 | 0.19538 | 0.000011 | 0.000011 | 0.000011 |
| Cluster 1 | 0.999947 | 0.000011 | 0.000011 | 0.000011 |
| Cluster 1 | 0.999947 | 0.000011 | 0.000011 | 0.000011 |
| Cluster 1 | 0.999947 | 0.000011 | 0.000011 | 0.000011 |

| | | | | |
|---|---|---|---|---|
| Cluster 1 | 0.999947 | 0.000011 | 0.000011 | 0.000011 |
| Cluster 1 | 0.999947 | 0.000011 | 0.000011 | 0.000011 |
| Cluster 1 | 0.999947 | 0.000011 | 0.000011 | 0.000011 |
| Cluster 1 | 0.999946 | 0.000011 | 0.000011 | 0.000011 |
| Cluster 1 | 0.96936 | 0.030598 | 0.000011 | 0.000011 |
| Cluster 1 | 0.955346 | 0.044611 | 0.000011 | 0.000011 |
| Cluster 1 | 0.999947 | 0.000011 | 0.000011 | 0.000011 |
| Cluster 1 | 0.999947 | 0.000011 | 0.000011 | 0.000011 |
| Cluster 1 | 0.999947 | 0.000011 | 0.000011 | 0.000011 |
| Cluster 1 | 0.999947 | 0.000011 | 0.000011 | 0.000011 |
| Cluster 1 | 0.999947 | 0.000011 | 0.000011 | 0.000011 |
| Cluster 1 | 0.999947 | 0.000011 | 0.000011 | 0.000011 |
| Cluster 1 | 0.999947 | 0.000011 | 0.000011 | 0.000011 |
| Cluster 1 | 0.999947 | 0.000011 | 0.000011 | 0.000011 |
| Cluster 1 | 0.999947 | 0.000011 | 0.000011 | 0.000011 |
| Cluster 1 | 0.999947 | 0.000011 | 0.000011 | 0.000011 |
| Cluster 1 | 0.999947 | 0.000011 | 0.000011 | 0.000011 |
| Cluster 1 | 0.999947 | 0.000011 | 0.000011 | 0.000011 |
| Cluster 1 | 0.999947 | 0.000011 | 0.000011 | 0.000011 |
| Cluster 1 | 0.93089 | 0.069068 | 0.000011 | 0.000011 |
| Cluster 1 | 0.797187 | 0.150945 | 0.000011 | 0.051836 |
| Cluster 2 | 0.000011 | 0.999947 | 0.000011 | 0.000011 |
| Cluster 2 | 0.000011 | 0.999947 | 0.00001 | 0.000011 |
| Cluster 2 | 0.000011 | 0.999947 | 0.000011 | 0.000011 |
| Cluster 1 | 0.999947 | 0.000011 | 0.000011 | 0.000011 |
| Cluster 1 | 0.999946 | 0.000011 | 0.000011 | 0.000011 |
| Cluster 1 | 0.961332 | 0.000011 | 0.000011 | 0.000011 |
| Cluster 1 | 0.906777 | 0.09318 | 0.000011 | 0.000011 |
| Cluster 1 | 0.999947 | 0.000011 | 0.000011 | 0.000011 |
| Cluster 1 | 0.999947 | 0.000011 | 0.000011 | 0.000011 |
| Cluster 1 | 0.999947 | 0.000011 | 0.000011 | 0.000011 |
| Cluster 1 | 0.999947 | 0.000011 | 0.000011 | 0.000011 |
| Cluster 1 | 0.999947 | 0.000011 | 0.000011 | 0.000011 |
| Cluster 1 | 0.999947 | 0.000011 | 0.000011 | 0.000011 |
| Cluster 1 | 0.999947 | 0.000011 | 0.000011 | 0.000011 |
| Cluster 1 | 0.999947 | 0.000011 | 0.000011 | 0.000011 |
| Cluster 1 | 0.999947 | 0.000011 | 0.000011 | 0.000011 |
| Cluster 1 | 0.999947 | 0.000011 | 0.000011 | 0.000011 |
| Cluster 1 | 0.999947 | 0.000011 | 0.000011 | 0.000011 |
| Cluster 1 | 0.999947 | 0.000011 | 0.000011 | 0.000011 |
| Cluster 1 | 0.999947 | 0.000011 | 0.000011 | 0.000011 |
| Cluster 1 | 0.999947 | 0.000011 | 0.000011 | 0.000011 |
| Cluster 1 | 0.999947 | 0.000011 | 0.000011 | 0.000011 |
| Cluster 1 | 0.999947 | 0.000011 | 0.000011 | 0.000011 |
| Cluster 1 | 0.999947 | 0.000011 | 0.000011 | 0.000011 |
| Cluster 1 | 0.999947 | 0.000011 | 0.000011 | 0.000011 |
| Cluster 1 | 0.999947 | 0.000011 | 0.000011 | 0.000011 |
| Cluster 1 | 0.999947 | 0.000011 | 0.000011 | 0.000011 |
| Cluster 1 | 0.999947 | 0.000011 | 0.000011 | 0.000011 |
| Cluster 1 | 0.999947 | 0.000011 | 0.000011 | 0.000011 |

| | | | | |
|---|---|---|---|---|
| Cluster 1 | 0.999947 | 0.000011 | 0.000011 | 0.000011 |
| Cluster 1 | 0.999947 | 0.000011 | 0.000011 | 0.000011 |
| Cluster 1 | 0.999947 | 0.000011 | 0.000011 | 0.000011 |
| Cluster 1 | 0.999947 | 0.000011 | 0.000011 | 0.000011 |
| Cluster 1 | 0.999947 | 0.000011 | 0.000011 | 0.000011 |
| Cluster 1 | 0.999947 | 0.000011 | 0.000011 | 0.000011 |
| Cluster 1 | 0.999947 | 0.000011 | 0.000011 | 0.000011 |
| Cluster 1 | 0.999947 | 0.000011 | 0.000011 | 0.000011 |
| Cluster 1 | 0.999947 | 0.000011 | 0.000011 | 0.000011 |
| Cluster 1 | 0.999947 | 0.000011 | 0.000011 | 0.000011 |
| Cluster 1 | 0.999947 | 0.000011 | 0.000011 | 0.000011 |
| Cluster 1 | 0.999947 | 0.000011 | 0.000011 | 0.000011 |
| Cluster 1 | 0.999947 | 0.000011 | 0.000011 | 0.000011 |
| Cluster 1 | 0.999947 | 0.000011 | 0.000011 | 0.000011 |
| Cluster 1 | 0.999947 | 0.000011 | 0.000011 | 0.000011 |
| Cluster 1 | 0.999947 | 0.000011 | 0.000011 | 0.000011 |
| Cluster 1 | 0.999947 | 0.000011 | 0.000011 | 0.000011 |
| Cluster 1 | 0.999947 | 0.000011 | 0.000011 | 0.000011 |
| Cluster 1 | 0.999947 | 0.000011 | 0.000011 | 0.000011 |
| Cluster 1 | 0.999947 | 0.000011 | 0.000011 | 0.000011 |
| Cluster 1 | 0.999947 | 0.000011 | 0.000011 | 0.000011 |
| Cluster 1 | 0.999947 | 0.000011 | 0.000011 | 0.000011 |
| Cluster 1 | 0.999947 | 0.000011 | 0.000011 | 0.000011 |
| Cluster 1 | 0.999947 | 0.000011 | 0.000011 | 0.000011 |
| Cluster 1 | 0.999946 | 0.000011 | 0.000011 | 0.000011 |
| Cluster 1 | 0.999946 | 0.000011 | 0.000011 | 0.000011 |
| Cluster 1 | 0.999946 | 0.000011 | 0.000011 | 0.000011 |
| Cluster 1 | 0.999946 | 0.000011 | 0.000011 | 0.000011 |
| Cluster 1 | 0.999943 | 0.000011 | 0.000011 | 0.000011 |
| Cluster 1 | 0.999943 | 0.000011 | 0.000011 | 0.000011 |
| Cluster 1 | 0.999942 | 0.000012 | 0.000012 | 0.000012 |
| Cluster 1 | 0.999941 | 0.000012 | 0.000012 | 0.000012 |
| Cluster 1 | 0.999939 | 0.000012 | 0.000012 | 0.000012 |
| Cluster 1 | 0.999936 | 0.000013 | 0.000013 | 0.000013 |
| Cluster 1 | 0.999935 | 0.000013 | 0.000013 | 0.000013 |
| Cluster 1 | 0.999935 | 0.000013 | 0.000013 | 0.000013 |
| Cluster 1 | 0.999935 | 0.000013 | 0.000013 | 0.000013 |
| Cluster 1 | 0.999934 | 0.000013 | 0.000013 | 0.000013 |
| Cluster 1 | 0.999934 | 0.000013 | 0.000013 | 0.000013 |
| Cluster 1 | 0.999934 | 0.000013 | 0.000013 | 0.000013 |
| Cluster 1 | 0.999933 | 0.000013 | 0.000013 | 0.000013 |
| Cluster 1 | 0.999933 | 0.000013 | 0.000013 | 0.000013 |
| Cluster 1 | 0.999932 | 0.000014 | 0.000014 | 0.000014 |
| Cluster 1 | 0.999931 | 0.000014 | 0.000014 | 0.000014 |
| Cluster 1 | 0.99993 | 0.000014 | 0.000014 | 0.000014 |
| Cluster 1 | 0.99993 | 0.000014 | 0.000014 | 0.000014 |
| Cluster 1 | 0.999927 | 0.000015 | 0.000015 | 0.000015 |
| Cluster 1 | 0.999927 | 0.000015 | 0.000015 | 0.000015 |
| Cluster 1 | 0.999926 | 0.000015 | 0.000015 | 0.000015 |
| Cluster 1 | 0.999921 | 0.000016 | 0.000016 | 0.000016 |

| | | | | |
|---|---|---|---|---|
| Cluster 1 | 0.980999 | 0.000011 | 0.000011 | 0.000011 |
| Cluster 1 | 0.975271 | 0.000011 | 0.000011 | 0.024687 |
| Cluster 1 | 0.960633 | 0.000013 | 0.000013 | 0.039317 |
| Cluster 1 | 0.954476 | 0.000011 | 0.000011 | 0.045482 |
| Cluster 1 | 0.950842 | 0.000011 | 0.000011 | 0.000011 |
| Cluster 1 | 0.950724 | 0.049233 | 0.000011 | 0.000011 |
| Cluster 1 | 0.939895 | 0.000011 | 0.000011 | 0.060062 |
| Cluster 1 | 0.91739 | 0.000011 | 0.000011 | 0.000011 |
| Cluster 1 | 0.913956 | 0.000011 | 0.000011 | 0.086001 |
| Cluster 1 | 0.903005 | 0.000013 | 0.000013 | 0.096944 |
| Cluster 1 | 0.901951 | 0.081845 | 0.000011 | 0.016172 |
| Cluster 1 | 0.901868 | 0.000011 | 0.000011 | 0.09809 |
| Cluster 1 | 0.899269 | 0.10068 | 0.000013 | 0.000013 |
| Cluster 1 | 0.886445 | 0.000011 | 0.000011 | 0.000011 |
| Cluster 1 | 0.88496 | 0.114983 | 0.000014 | 0.000014 |
| Cluster 1 | 0.883855 | 0.116102 | 0.000011 | 0.000011 |
| Cluster 1 | 0.877622 | 0.122336 | 0.000011 | 0.000011 |
| Cluster 1 | 0.862076 | 0.000011 | 0.000011 | 0.000011 |
| Cluster 1 | 0.834111 | 0.12403 | 0.000011 | 0.041827 |
| Cluster 1 | 0.82198 | 0.177978 | 0.000011 | 0.000011 |
| Cluster 1 | 0.816889 | 0.000011 | 0.000011 | 0.000011 |
| Cluster 1 | 0.801662 | 0.000011 | 0.000011 | 0.000011 |
| Cluster 1 | 0.796905 | 0.128581 | 0.000011 | 0.074481 |
| Cluster 1 | 0.794643 | 0.205315 | 0.00001 | 0.00001 |
| Cluster 1 | 0.774605 | 0.217986 | 0.000011 | 0.007377 |
| Cluster 1 | 0.74996 | 0.000011 | 0.000011 | 0.000011 |
| Cluster 1 - Cluster 2 | 0.727228 | 0.27273 | 0.000011 | 0.000011 |
| Cluster 1 - Cluster 6 | 0.656813 | 0.000011 | 0.000011 | 0.000011 |
| Cluster 1 - Cluster 6 | 0.600001 | 0.000011 | 0.000011 | 0.000011 |
| Cluster 1 - Cluster 5 | 0.598397 | 0.000011 | 0.000011 | 0.000011 |
| Cluster 1 - Cluster 6 | 0.593339 | 0.000011 | 0.000011 | 0.000011 |
| Cluster 1 - Cluster 6 | 0.575345 | 0.000011 | 0.000011 | 0.000011 |
| Cluster 1 - Cluster 6 | 0.56108 | 0.000011 | 0.000011 | 0.000011 |
| Cluster 1 - Cluster 5 | 0.556743 | 0.000011 | 0.000011 | 0.000011 |
| Cluster 1 - Cluster 6 | 0.556697 | 0.000011 | 0.000011 | 0.000011 |
| Cluster 1 - Cluster 5 | 0.5549 | 0.000011 | 0.000011 | 0.000011 |
| Cluster 1 - Cluster 5 | 0.50637 | 0.000011 | 0.000011 | 0.000011 |
| Cluster 1 - Cluster 5 | 0.503972 | 0.000011 | 0.000011 | 0.000011 |
| Cluster 1 - Cluster 5 | 0.499529 | 0.247207 | 0.164245 | 0.088997 |
| Cluster 1 - Cluster 6 | 0.497804 | 0.000011 | 0.000011 | 0.000011 |
| Cluster 1 - Cluster 6 | 0.489136 | 0.000011 | 0.000011 | 0.000011 |
| Cluster 1 - Cluster 5 | 0.486856 | 0.000011 | 0.000011 | 0.000011 |
| Cluster 1 - Cluster 6 | 0.468689 | 0.000011 | 0.000011 | 0.000011 |
| Cluster 1 - Cluster 2 | 0.454742 | 0.519042 | 0.026184 | 0.000011 |
| Cluster 1 - Cluster 5 | 0.423146 | 0.000011 | 0.000011 | 0.000011 |
| Cluster 1 - Cluster 2 | 0.353671 | 0.586504 | 0.052272 | 0.000011 |
| Cluster 5 | 0.058737 | 0.000011 | 0.000011 | 0.000011 |
| Cluster 2 | 0.000221 | 0.998893 | 0.000221 | 0.000221 |
| Cluster 2 | 0.000018 | 0.999909 | 0.000018 | 0.000018 |
| Cluster 2 | 0.000015 | 0.999925 | 0.000015 | 0.000015 |

| | | | | |
|---|---|---|---|---|
| Cluster 2 | 0.000015 | 0.999927 | 0.000015 | 0.000015 |
| Cluster 2 | 0.000014 | 0.999931 | 0.000014 | 0.000014 |
| Cluster 6 | 0.000013 | 0.000013 | 0.000013 | 0.000013 |
| Cluster 2 | 0.000013 | 0.999934 | 0.000013 | 0.000013 |
| Cluster 5 | 0.000011 | 0.000011 | 0.000011 | 0.000011 |
| Cluster 5 | 0.000011 | 0.000011 | 0.000011 | 0.000011 |
| Cluster 5 | 0.000011 | 0.000011 | 0.000011 | 0.000011 |
| Cluster 6 | 0.000011 | 0.000011 | 0.000011 | 0.000011 |
| Cluster 6 | 0.000011 | 0.000011 | 0.000011 | 0.000011 |
| Cluster 4 | 0.000011 | 0.000011 | 0.000011 | 0.999947 |
| Cluster 4 | 0.000011 | 0.000011 | 0.000011 | 0.999947 |
| Cluster 4 | 0.000011 | 0.000011 | 0.000011 | 0.999947 |
| Cluster 4 | 0.000011 | 0.000011 | 0.000011 | 0.999947 |
| Cluster 4 | 0.000011 | 0.000011 | 0.000011 | 0.999947 |
| Cluster 4 | 0.000011 | 0.000011 | 0.151347 | 0.84861 |
| Cluster 2 - Cluster 4 | 0.000011 | 0.330181 | 0.000011 | 0.669777 |
| Cluster 2 - Cluster 3 | 0.000011 | 0.553152 | 0.446805 | 0.000011 |
| Cluster 2 - Cluster 3 | 0.000011 | 0.557305 | 0.442652 | 0.000011 |
| Cluster 2 - Cluster 3 | 0.000011 | 0.556235 | 0.443722 | 0.000011 |
| Cluster 2 - Cluster 3 | 0.000011 | 0.492204 | 0.507754 | 0.000011 |
| Cluster 2 - Cluster 3 | 0.000011 | 0.489348 | 0.51061 | 0.000011 |
| Cluster 2 - Cluster 3 | 0.000011 | 0.487574 | 0.512383 | 0.000011 |
| Cluster 2 - Cluster 3 | 0.000011 | 0.454933 | 0.434397 | 0.110638 |
| Cluster 2 - Cluster 3 | 0.000011 | 0.449815 | 0.47747 | 0.072683 |
| Cluster 2 - Cluster 3 | 0.000011 | 0.437806 | 0.535239 | 0.026923 |
| Cluster 2 - Cluster 3 | 0.000011 | 0.430674 | 0.525115 | 0.044179 |
| Cluster 2 | 0.000016 | 0.999918 | 0.000016 | 0.000016 |
| Cluster 2 | 0.000017 | 0.999917 | 0.000017 | 0.000017 |
| Cluster 2 - Cluster 3 | 0.000011 | 0.450462 | 0.549496 | 0.000011 |
| Cluster 2 - Cluster 3 - Cluster 4 | 0.000011 | 0.317173 | 0.46609 | 0.216705 |
| Cluster 3 | 0.000018 | 0.000018 | 0.999912 | 0.000018 |
| Cluster 3 | 0.000011 | 0.000011 | 0.999947 | 0.000011 |
| Cluster 3 | 0.000011 | 0.000011 | 0.999947 | 0.000011 |
| Cluster 3 | 0.000011 | 0.000011 | 0.999947 | 0.000011 |
| Cluster 3 | 0.000011 | 0.000011 | 0.999947 | 0.000011 |
| Cluster 3 | 0.000011 | 0.000011 | 0.999947 | 0.000011 |
| Cluster 3 | 0.000011 | 0.000011 | 0.999947 | 0.000011 |
| Cluster 3 | 0.000011 | 0.000011 | 0.999947 | 0.000011 |
| Cluster 3 | 0.000011 | 0.000011 | 0.999947 | 0.000011 |
| Cluster 3 | 0.000011 | 0.000011 | 0.999947 | 0.000011 |
| Cluster 3 | 0.000011 | 0.000011 | 0.999947 | 0.000011 |
| Cluster 3 | 0.000011 | 0.000011 | 0.999947 | 0.000011 |
| Cluster 3 | 0.000011 | 0.000011 | 0.999947 | 0.000011 |
| Cluster 3 | 0.000011 | 0.000011 | 0.999947 | 0.000011 |
| Cluster 3 | 0.000011 | 0.000011 | 0.999946 | 0.000011 |
| Cluster 3 | 0.000011 | 0.000011 | 0.999946 | 0.000011 |
| Cluster 3 | 0.000011 | 0.000011 | 0.999946 | 0.000011 |
| Cluster 3 | 0.00001 | 0.00001 | 0.999948 | 0.00001 |
| Cluster 4 | 0.000011 | 0.000011 | 0.089514 | 0.910444 |
| Cluster 4 | 0.000011 | 0.000011 | 0.067875 | 0.932083 |

| | | | | |
|---|---|---|---|---|
| Cluster 4 | 0.000011 | 0.000011 | 0.000011 | 0.999947 |
| Cluster 4 | 0.000011 | 0.000011 | 0.000011 | 0.999947 |
| Cluster 4 | 0.000011 | 0.000011 | 0.000011 | 0.999947 |
| Cluster 4 | 0.000011 | 0.000011 | 0.000011 | 0.999947 |
| Cluster 4 | 0.000011 | 0.000011 | 0.000011 | 0.999947 |
| Cluster 4 | 0.000011 | 0.000011 | 0.000011 | 0.999947 |
| Cluster 4 | 0.000011 | 0.000011 | 0.000011 | 0.999947 |
| Cluster 4 | 0.000011 | 0.000011 | 0.000011 | 0.999947 |
| Cluster 4 | 0.000011 | 0.000011 | 0.000011 | 0.999947 |
| Cluster 4 | 0.000011 | 0.000011 | 0.000011 | 0.999947 |
| Cluster 4 | 0.000011 | 0.000011 | 0.000011 | 0.999947 |
| Cluster 4 | 0.000011 | 0.000011 | 0.000011 | 0.999947 |
| Cluster 4 | 0.000011 | 0.000011 | 0.000011 | 0.999947 |
| Cluster 4 | 0.00001 | 0.000011 | 0.000011 | 0.999947 |
| Cluster 4 | 0.000011 | 0.000011 | 0.000011 | 0.999947 |
| Cluster 4 | 0.000011 | 0.000011 | 0.000011 | 0.999947 |
| Cluster 4 | 0.000011 | 0.000011 | 0.000011 | 0.999947 |
| Cluster 4 | 0.000011 | 0.000011 | 0.000011 | 0.999947 |
| Cluster 4 | 0.000011 | 0.000011 | 0.000011 | 0.999947 |
| Cluster 4 | 0.000011 | 0.000011 | 0.000011 | 0.999947 |
| Cluster 4 | 0.000011 | 0.000011 | 0.000011 | 0.999947 |
| Cluster 4 | 0.000011 | 0.000011 | 0.000011 | 0.999946 |
| Cluster 4 | 0.000011 | 0.000011 | 0.000011 | 0.999946 |
| Cluster 4 | 0.000011 | 0.000011 | 0.000011 | 0.999946 |
| Cluster 4 | 0.000011 | 0.000011 | 0.000011 | 0.999946 |
| Cluster 4 | 0.000011 | 0.000011 | 0.000011 | 0.999946 |
| Cluster 4 | 0.000011 | 0.000011 | 0.000011 | 0.999946 |
| Cluster 4 | 0.000011 | 0.000011 | 0.000011 | 0.999946 |
| Cluster 4 | 0.000011 | 0.000011 | 0.000011 | 0.999944 |
| Cluster 4 | 0.000011 | 0.000011 | 0.000011 | 0.88829 |
| Cluster 2 | 0.000011 | 0.992123 | 0.000011 | 0.007834 |
| Cluster 2 | 0.000011 | 0.965708 | 0.034249 | 0.000011 |
| Cluster 2 | 0.000011 | 0.936807 | 0.06315 | 0.000011 |
| Cluster 2 | 0.000011 | 0.932596 | 0.046019 | 0.021352 |
| Cluster 2 | 0.000011 | 0.89626 | 0.065596 | 0.038112 |
| Cluster 2 | 0.000011 | 0.871603 | 0.128354 | 0.000011 |
| Cluster 2 | 0.000011 | 0.859679 | 0.078072 | 0.062217 |
| Cluster 2 | 0.000011 | 0.855419 | 0.144537 | 0.000011 |
| Cluster 2 | 0.000011 | 0.849597 | 0.10456 | 0.045811 |
| Cluster 2 | 0.000011 | 0.821674 | 0.178284 | 0.000011 |
| Cluster 2 | 0.000011 | 0.815651 | 0.122594 | 0.061723 |
| Cluster 2 | 0.000011 | 0.999947 | 0.000011 | 0.000011 |
| Cluster 2 | 0.000011 | 0.999947 | 0.000011 | 0.000011 |
| Cluster 2 | 0.000011 | 0.999947 | 0.000011 | 0.000011 |
| Cluster 2 | 0.000011 | 0.999947 | 0.000011 | 0.000011 |
| Cluster 2 | 0.000011 | 0.999947 | 0.000011 | 0.000011 |
| Cluster 2 | 0.000011 | 0.999947 | 0.000011 | 0.000011 |
| Cluster 2 | 0.000011 | 0.999947 | 0.000011 | 0.000011 |
| Cluster 2 | 0.000011 | 0.966471 | 0.000011 | 0.033487 |
| Cluster 2 | 0.000011 | 0.964811 | 0.017021 | 0.018136 |

| | | | | |
|---|---|---|---|---|
| Cluster 2 | 0.000011 | 0.952744 | 0.000011 | 0.047213 |
| Cluster 2 | 0.000011 | 0.95194 | 0.000011 | 0.048018 |
| Cluster 2 | 0.00001 | 0.999948 | 0.00001 | 0.00001 |
| Cluster 2 | 0.000011 | 0.999947 | 0.000011 | 0.000011 |
| Cluster 2 | 0.000011 | 0.999947 | 0.000011 | 0.000011 |
| Cluster 2 | 0.000011 | 0.999947 | 0.000011 | 0.000011 |
| Cluster 2 | 0.000011 | 0.999947 | 0.000011 | 0.000011 |
| Cluster 2 | 0.000011 | 0.999947 | 0.000011 | 0.000011 |
| Cluster 2 | 0.000011 | 0.999947 | 0.000011 | 0.000011 |
| Cluster 2 | 0.000011 | 0.999947 | 0.000011 | 0.000011 |
| Cluster 2 | 0.000011 | 0.999947 | 0.000011 | 0.000011 |
| Cluster 2 | 0.000011 | 0.999947 | 0.000011 | 0.000011 |
| Cluster 2 | 0.000011 | 0.999946 | 0.000011 | 0.000011 |
| Cluster 2 | 0.000011 | 0.999946 | 0.000011 | 0.000011 |
| Cluster 2 | 0.000011 | 0.999946 | 0.000011 | 0.000011 |
| Cluster 2 | 0.000011 | 0.982008 | 0.017949 | 0.000011 |
| Cluster 2 | 0.000011 | 0.999947 | 0.000011 | 0.000011 |
| Cluster 2 | 0.000011 | 0.999947 | 0.000011 | 0.000011 |
| Cluster 2 | 0.000011 | 0.999947 | 0.000011 | 0.000011 |
| Cluster 2 | 0.000011 | 0.999947 | 0.000011 | 0.000011 |
| Cluster 2 | 0.000011 | 0.999947 | 0.000011 | 0.000011 |
| Cluster 2 | 0.000011 | 0.999947 | 0.000011 | 0.000011 |
| Cluster 2 | 0.000011 | 0.999947 | 0.000011 | 0.000011 |
| Cluster 2 | 0.000011 | 0.999947 | 0.000011 | 0.000011 |
| Cluster 2 | 0.000011 | 0.999946 | 0.000011 | 0.000011 |
| Cluster 2 | 0.000011 | 0.999944 | 0.000011 | 0.000011 |
| Cluster 1 - Cluster 2 | 0.443198 | 0.556759 | 0.000011 | 0.000011 |
| Cluster 2 | 0.000011 | 0.999947 | 0.000011 | 0.000011 |
| Cluster 2 | 0.000011 | 0.999947 | 0.000011 | 0.000011 |
| Cluster 2 | 0.000011 | 0.999946 | 0.000011 | 0.000011 |
| Cluster 2 | 0.000011 | 0.999946 | 0.000011 | 0.000011 |
| Cluster 2 | 0.000011 | 0.999945 | 0.000011 | 0.000011 |
| Cluster 2 | 0.000011 | 0.98503 | 0.000011 | 0.014926 |
| Cluster 2 | 0.000011 | 0.999947 | 0.000011 | 0.000011 |
| Cluster 2 | 0.000011 | 0.999947 | 0.000011 | 0.000011 |
| Cluster 2 | 0.000011 | 0.999947 | 0.000011 | 0.000011 |
| Cluster 2 | 0.000011 | 0.999947 | 0.000011 | 0.000011 |
| Cluster 2 | 0.000011 | 0.999947 | 0.000011 | 0.000011 |
| Cluster 2 | 0.000011 | 0.999947 | 0.000011 | 0.000011 |
| Cluster 2 | 0.000011 | 0.999947 | 0.000011 | 0.000011 |
| Cluster 2 | 0.000011 | 0.999947 | 0.000011 | 0.000011 |
| Cluster 2 | 0.000011 | 0.999947 | 0.000011 | 0.000011 |
| Cluster 2 | 0.000011 | 0.999947 | 0.000011 | 0.000011 |
| Cluster 2 | 0.000011 | 0.999946 | 0.000011 | 0.000011 |
| Cluster 2 | 0.000011 | 0.999946 | 0.000011 | 0.000011 |
| Cluster 2 | 0.000011 | 0.999946 | 0.000011 | 0.000011 |
| Cluster 2 | 0.000011 | 0.999947 | 0.000011 | 0.000011 |
| Cluster 2 | 0.000011 | 0.999947 | 0.000011 | 0.000011 |
| Cluster 2 | 0.000011 | 0.999947 | 0.000011 | 0.000011 |
| Cluster 2 | 0.000011 | 0.999947 | 0.000011 | 0.000011 |

| | | | | |
|---|---|---|---|---|
| Cluster 2 | 0.000011 | 0.999947 | 0.000011 | 0.000011 |
| Cluster 2 | 0.000011 | 0.999946 | 0.000011 | 0.000011 |
| Cluster 2 | 0.000011 | 0.999946 | 0.000011 | 0.000011 |
| Cluster 2 | 0.000011 | 0.999946 | 0.000011 | 0.000011 |
| Cluster 1 | 0.999944 | 0.000011 | 0.000011 | 0.000011 |
| Cluster 1 | 0.999947 | 0.000011 | 0.000011 | 0.000011 |
| Cluster 1 | 0.999947 | 0.000011 | 0.000011 | 0.000011 |
| Cluster 1 | 0.999945 | 0.000011 | 0.000011 | 0.000011 |
| Cluster 1 | 0.999948 | 0.00001 | 0.00001 | 0.00001 |
| Cluster 2 | 0.000011 | 0.999947 | 0.000011 | 0.000011 |
| Cluster 2 | 0.000011 | 0.999947 | 0.000011 | 0.000011 |
| Cluster 2 | 0.000011 | 0.999947 | 0.000011 | 0.000011 |
| Cluster 2 | 0.000011 | 0.999947 | 0.000011 | 0.000011 |
| Cluster 2 | 0.000011 | 0.999947 | 0.000011 | 0.000011 |
| Cluster 2 | 0.000011 | 0.999947 | 0.000011 | 0.000011 |
| Cluster 2 | 0.000011 | 0.999947 | 0.000011 | 0.000011 |
| Cluster 2 | 0.000011 | 0.999947 | 0.000011 | 0.000011 |
| Cluster 2 | 0.000011 | 0.999946 | 0.000011 | 0.000011 |
| Cluster 2 | 0.000011 | 0.999946 | 0.000011 | 0.000011 |
| Cluster 2 | 0.000011 | 0.999946 | 0.000011 | 0.000011 |
| Cluster 2 | 0.000011 | 0.999947 | 0.000011 | 0.000011 |
| Cluster 2 | 0.000011 | 0.999947 | 0.000011 | 0.000011 |
| Cluster 2 | 0.000011 | 0.999947 | 0.000011 | 0.000011 |
| Cluster 2 | 0.000011 | 0.999947 | 0.000011 | 0.000011 |
| Cluster 2 | 0.000011 | 0.999947 | 0.000011 | 0.000011 |
| Cluster 2 | 0.000011 | 0.999946 | 0.000011 | 0.000011 |
| Cluster 2 | 0.000011 | 0.999946 | 0.000011 | 0.000011 |
| Cluster 2 | 0.000011 | 0.999946 | 0.000011 | 0.000011 |
| Cluster 2 | 0.000011 | 0.999946 | 0.000011 | 0.000011 |
| Cluster 2 | 0.000011 | 0.999946 | 0.000011 | 0.000011 |
| Cluster 2 | 0.000011 | 0.999946 | 0.000011 | 0.000011 |
| Cluster 2 | 0.000011 | 0.999946 | 0.000011 | 0.000011 |
| Cluster 2 | 0.000011 | 0.999946 | 0.000011 | 0.000011 |
| Cluster 2 | 0.000011 | 0.999945 | 0.000011 | 0.000011 |
| Cluster 2 | 0.000011 | 0.999944 | 0.000011 | 0.000011 |
| Cluster 2 | 0.000011 | 0.999947 | 0.000011 | 0.000011 |
| Cluster 2 | 0.000011 | 0.999947 | 0.000011 | 0.000011 |
| Cluster 2 | 0.000011 | 0.999947 | 0.000011 | 0.000011 |
| Cluster 2 | 0.000011 | 0.999947 | 0.000011 | 0.000011 |
| Cluster 2 | 0.000011 | 0.999946 | 0.000011 | 0.000011 |
| Cluster 2 | 0.000011 | 0.999946 | 0.000011 | 0.000011 |
| Cluster 2 | 0.000011 | 0.999946 | 0.000011 | 0.000011 |
| Cluster 2 | 0.000011 | 0.999946 | 0.000011 | 0.000011 |
| Cluster 2 | 0.000011 | 0.999946 | 0.000011 | 0.000011 |
| Cluster 2 | 0.000011 | 0.999945 | 0.000011 | 0.000011 |
| Cluster 2 | 0.000011 | 0.999945 | 0.000011 | 0.000011 |
| Cluster 1 | 0.999946 | 0.000011 | 0.000011 | 0.000011 |
| Cluster 2 | 0.000011 | 0.999947 | 0.000011 | 0.000011 |
| Cluster 2 | 0.000011 | 0.999947 | 0.000011 | 0.000011 |
| Cluster 2 | 0.000011 | 0.999947 | 0.000011 | 0.000011 |

| | | | | |
|---|---|---|---|---|
| Cluster 2 | 0.000011 | 0.999947 | 0.000011 | 0.000011 |
| Cluster 2 | 0.000011 | 0.999946 | 0.000011 | 0.000011 |
| Cluster 2 | 0.000011 | 0.999946 | 0.000011 | 0.000011 |
| Cluster 2 | 0.000011 | 0.999946 | 0.000011 | 0.000011 |
| Cluster 2 | 0.000011 | 0.999946 | 0.000011 | 0.000011 |
| Cluster 2 | 0.000011 | 0.999946 | 0.000011 | 0.000011 |
| Cluster 2 | 0.000011 | 0.999946 | 0.000011 | 0.000011 |
| Cluster 2 | 0.000011 | 0.999009 | 0.000948 | 0.000011 |
| Cluster 2 | 0.000011 | 0.999947 | 0.000011 | 0.000011 |
| Cluster 2 | 0.000011 | 0.999946 | 0.000011 | 0.000011 |
| Cluster 2 | 0.000011 | 0.999946 | 0.000011 | 0.000011 |
| Cluster 2 | 0.000011 | 0.999946 | 0.000011 | 0.000011 |
| Cluster 2 | 0.000011 | 0.973873 | 0.026084 | 0.000011 |
| Cluster 2 | 0.000011 | 0.961946 | 0.000011 | 0.038011 |
| Cluster 2 | 0.000011 | 0.999948 | 0.00001 | 0.000011 |
| Cluster 2 | 0.000011 | 0.999947 | 0.000011 | 0.000011 |
| Cluster 2 | 0.000011 | 0.999947 | 0.000011 | 0.000011 |
| Cluster 2 | 0.000011 | 0.999947 | 0.000011 | 0.000011 |
| Cluster 2 | 0.000011 | 0.999947 | 0.000011 | 0.000011 |
| Cluster 2 | 0.000011 | 0.999946 | 0.000011 | 0.000011 |
| Cluster 2 | 0.000024 | 0.999882 | 0.000024 | 0.000024 |
| Cluster 2 | 0.000011 | 0.999947 | 0.000011 | 0.000011 |
| Cluster 2 | 0.000011 | 0.999947 | 0.000011 | 0.000011 |
| Cluster 2 | 0.000011 | 0.999947 | 0.000011 | 0.000011 |
| Cluster 2 | 0.000011 | 0.999947 | 0.000011 | 0.000011 |
| Cluster 2 | 0.000011 | 0.999947 | 0.000011 | 0.000011 |
| Cluster 2 | 0.000011 | 0.999947 | 0.000011 | 0.000011 |
| Cluster 2 | 0.000011 | 0.999947 | 0.000011 | 0.000011 |
| Cluster 2 | 0.000011 | 0.999947 | 0.000011 | 0.000011 |
| Cluster 2 | 0.000011 | 0.999947 | 0.000011 | 0.000011 |
| Cluster 2 | 0.000011 | 0.999947 | 0.000011 | 0.000011 |
| Cluster 2 | 0.000011 | 0.999947 | 0.000011 | 0.000011 |
| Cluster 2 | 0.000011 | 0.999947 | 0.000011 | 0.000011 |
| Cluster 2 | 0.000011 | 0.999947 | 0.000011 | 0.000011 |
| Cluster 2 | 0.000011 | 0.999947 | 0.000011 | 0.000011 |
| Cluster 2 | 0.000011 | 0.999947 | 0.000011 | 0.000011 |
| Cluster 2 | 0.000011 | 0.999947 | 0.000011 | 0.000011 |
| Cluster 2 | 0.000011 | 0.999947 | 0.000011 | 0.000011 |
| Cluster 2 | 0.000011 | 0.999947 | 0.000011 | 0.000011 |
| Cluster 1 - Cluster 2 | 0.522945 | 0.476989 | 0.000017 | 0.000017 |
| Cluster 1 | 0.854189 | 0.14573 | 0.00002 | 0.00002 |
| Cluster 1 | 0.999902 | 0.00002 | 0.00002 | 0.00002 |
| Cluster 1 | 0.99992 | 0.000016 | 0.000016 | 0.000016 |
| Cluster 2 | 0.000015 | 0.999927 | 0.000015 | 0.000015 |
| Cluster 2 | 0.000017 | 0.999917 | 0.000017 | 0.000017 |
| Cluster 2 | 0.000017 | 0.999915 | 0.000017 | 0.000017 |
| Cluster 2 | 0.000017 | 0.999914 | 0.000017 | 0.000017 |
| Cluster 2 | 0.000019 | 0.999904 | 0.000019 | 0.000019 |
| Cluster 2 | 0.000022 | 0.999891 | 0.000022 | 0.000022 |
| Cluster 2 | 0.000023 | 0.999883 | 0.000023 | 0.000023 |

| | | | | |
|---|---|---|---|---|
| Cluster 1 | 0.999916 | 0.000017 | 0.000017 | 0.000017 |
| Cluster 2 | 0.000011 | 0.999945 | 0.000011 | 0.000011 |
| Cluster 2 | 0.000012 | 0.999938 | 0.000012 | 0.000012 |
| Cluster 2 | 0.000014 | 0.999931 | 0.000014 | 0.000014 |
| Cluster 2 | 0.000017 | 0.999914 | 0.000017 | 0.000017 |
| Cluster 2 | 0.000013 | 0.999937 | 0.000013 | 0.000013 |
| Cluster 2 | 0.000013 | 0.999934 | 0.000013 | 0.000013 |
| Cluster 2 | 0.000015 | 0.999926 | 0.000015 | 0.000015 |
| Cluster 2 | 0.000015 | 0.999926 | 0.000015 | 0.000015 |
| Cluster 2 | 0.000017 | 0.999916 | 0.000017 | 0.000017 |
| Cluster 2 | 0.000017 | 0.999913 | 0.000017 | 0.000017 |
| Cluster 2 | 0.000022 | 0.999891 | 0.000022 | 0.000022 |
| Cluster 2 | 0.000027 | 0.999864 | 0.000027 | 0.000027 |
| Cluster 2 | 0.000012 | 0.999941 | 0.000012 | 0.000012 |
| Cluster 2 | 0.000014 | 0.999929 | 0.000014 | 0.000014 |
| Cluster 2 | 0.000015 | 0.999926 | 0.000015 | 0.000015 |
| Cluster 2 | 0.000015 | 0.999926 | 0.000015 | 0.000015 |
| Cluster 2 | 0.000019 | 0.999906 | 0.000019 | 0.000019 |
| Cluster 2 | 0.000021 | 0.999896 | 0.000021 | 0.000021 |
| Cluster 2 | 0.000022 | 0.999889 | 0.000022 | 0.000022 |
| Cluster 2 | 0.000023 | 0.999884 | 0.000023 | 0.000023 |
| Cluster 2 | 0.000023 | 0.999884 | 0.000023 | 0.000023 |
| Cluster 2 | 0.000012 | 0.999942 | 0.000012 | 0.000012 |
| Cluster 2 | 0.000012 | 0.999942 | 0.000012 | 0.000012 |
| Cluster 2 | 0.000012 | 0.999939 | 0.000012 | 0.000012 |
| Cluster 2 | 0.000013 | 0.999936 | 0.000013 | 0.000013 |
| Cluster 2 | 0.000013 | 0.999936 | 0.000013 | 0.000013 |
| Cluster 2 | 0.000013 | 0.999935 | 0.000013 | 0.000013 |

| | | Hybridizatior | |
|---|---|---|---|
| Cluster 5 | Cluster 6 | S | H |
| 0.000011 | 0.000011 | 1 | 0 |
| 0.000011 | 0.000011 | 0.975 | 0.05 |
| 0.000011 | 0.000011 | 1 | 0 |
| 0.000011 | 0.000011 | 0.955 | 0.09 |
| 0.000011 | 0.000011 | 0.88 | 0.2 |
| 0.000011 | 0.000011 | 0.78 | 0.28 |
| 0.000011 | 0.000011 | 0.04 | 0 |
| 0.000011 | 0.000011 | 0 | 0 |
| 0.000011 | 0.000011 | 0.94 | 0.12 |
| 0.000011 | 0.000011 | 0.96 | 0.08 |
| 0.000011 | 0.000011 | 0.885 | 0.23 |
| 0.000011 | 0.000011 | 1 | 0 |
| 0.000011 | 0.000011 | 0.88 | 0.24 |
| 0.204858 | 0.000011 | 0.95 | 0 |
| 0.269809 | 0.000011 | 0.63 | 0.74 |
| 0.000011 | 0.000011 | 0.935 | 0.13 |
| 0.000011 | 0.000011 | 1 | 0 |
| 0.616174 | 0.000011 | 0.47 | 0.94 |
| 0.000011 | 0.000011 | 1 | 0 |
| 0.000011 | 0.000011 | 0.99 | 0.02 |
| 0.000011 | 0.000011 | 0.93 | 0.14 |
| 0.000011 | 0.000011 | 1 | 0 |
| 0.000011 | 0.000011 | 1 | 0 |
| 0.000011 | 0.000011 | 0.78 | 0.12 |
| 0.000011 | 0.000011 | 1 | 0 |
| 0.000011 | 0.000011 | 0.985 | 0.03 |
| 0.000011 | 0.000011 | 1 | 0 |
| 0.000011 | 0.000011 | 0.925 | 0.15 |
| 0.000011 | 0.000011 | 0.97 | 0.06 |
| 0.000011 | 0.000011 | 0.855 | 0.29 |
| 0.000011 | 0.000011 | 1 | 0 |
| 0.000011 | 0.000011 | 0.895 | 0.21 |
| 0.000011 | 0.000011 | 1 | 0 |
| 0.000011 | 0.000011 | 1 | 0 |
| 0.000011 | 0.000011 | 1 | 0 |
| 0.000011 | 0.000011 | 0.93 | 0.14 |
| 0.000011 | 0.000011 | 1 | 0 |
| 0.000011 | 0.000011 | 0.99 | 0.02 |
| 0.000011 | 0.000011 | 1 | 0 |
| 0.804577 | 0.000011 | 0.42 | 0.72 |
| 0.000011 | 0.000011 | 1 | 0 |
| 0.000011 | 0.000011 | 1 | 0 |
| 0.000011 | 0.000011 | 0.955 | 0.09 |

| | | | |
|---|---|---|---|
| 0.000011 | 0.000011 | 1 | 0 |
| 0.000011 | 0.000011 | 0.93 | 0 |
| 0.000011 | 0.000011 | 0.895 | 0.21 |
| 0.000011 | 0.000011 | 0.92 | 0.16 |
| 0.000011 | 0.000011 | 1 | 0 |
| 0.000011 | 0.000011 | 0.9 | 0.2 |
| 0.000011 | 0.000011 | 0.99 | 0.02 |
| 0.000011 | 0.000011 | 0.98 | 0.04 |
| 0.000011 | 0.000011 | 1 | 0 |
| 0.000011 | 0.000011 | 1 | 0 |
| 0.000011 | 0.000011 | 1 | 0 |
| 0.000011 | 0.000011 | 0.96 | 0.08 |
| 0.000011 | 0.000011 | 0.98 | 0 |
| 0.000011 | 0.000011 | 0.97 | 0 |
| 0.000011 | 0.000011 | 1 | 0 |
| 0.000011 | 0.000011 | 0.955 | 0.09 |
| 0.000011 | 0.000011 | 1 | 0 |
| 0.000011 | 0.000011 | 0.975 | 0.05 |
| 0.000011 | 0.000011 | 0.87 | 0.2 |
| 0.000011 | 0.000011 | 0.91 | 0.08 |
| 0.000011 | 0.000011 | 0.5 | 0.5 |
| 0.000011 | 0.000011 | 0.04 | 0.04 |
| 0.000011 | 0.000011 | 0.175 | 0.19 |
| 0.000011 | 0.000011 | 0.05 | 0 |
| 0.000011 | 0.000011 | 1 | 0 |
| 0.000011 | 0.000011 | 0.96 | 0.08 |
| 0.038625 | 0.000011 | 0.925 | 0.15 |
| 0.000011 | 0.000011 | 0.82 | 0.36 |
| 0.000011 | 0.000011 | 0.99 | 0.02 |
| 0.000011 | 0.000011 | 1 | 0 |
| 0.000011 | 0.000011 | 0.87 | 0.14 |
| 0.000011 | 0.000011 | 1 | 0 |
| 0.000011 | 0.000011 | 0.985 | 0.03 |
| 0.000011 | 0.000011 | 0.98 | 0.04 |
| 0.000011 | 0.000011 | 0.915 | 0.17 |
| 0.000011 | 0.000011 | 1 | 0 |
| 0.000011 | 0.000011 | 1 | 0 |
| 0.000011 | 0.000011 | 0.965 | 0.07 |
| 0.000011 | 0.000011 | 1 | 0 |
| 0.000011 | 0.000011 | 0.965 | 0.07 |
| 0.000011 | 0.000011 | 1 | 0 |
| 0.000011 | 0.000011 | 0.965 | 0.07 |
| 0.000011 | 0.000011 | 1 | 0 |
| 0.000011 | 0.000011 | 0.91 | 0.18 |
| 0.000011 | 0.000011 | 1 | 0 |
| 0.000011 | 0.000011 | 0.88 | 0.24 |
| 0.000011 | 0.000011 | 1 | 0 |
| 0.000011 | 0.000011 | 0.995 | 0.01 |
| 0.000011 | 0.000011 | 0.965 | 0.07 |
| 0.000011 | 0.000011 | 1 | 0 |

| | | | |
|---|---|---|---|
| 0.000011 | 0.000011 | 1 | 0 |
| 0.000011 | 0.000011 | 0.88 | 0.24 |
| 0.000011 | 0.000011 | 0.985 | 0.03 |
| 0.000011 | 0.000011 | 1 | 0 |
| 0.000011 | 0.000011 | 0.905 | 0.19 |
| 0.000011 | 0.000011 | 0.73 | 0.3 |
| 0.000011 | 0.000011 | 0.965 | 0.07 |
| 0.000011 | 0.000011 | 1 | 0 |
| 0.000011 | 0.000011 | 1 | 0 |
| 0.000011 | 0.000011 | 1 | 0 |
| 0.000011 | 0.000011 | 0.895 | 0.21 |
| 0.000011 | 0.000011 | 0.765 | 0.37 |
| 0.000011 | 0.000011 | 0.88 | 0 |
| 0.000011 | 0.000011 | 0.945 | 0.11 |
| 0.000011 | 0.000011 | 0.955 | 0.09 |
| 0.000011 | 0.000011 | 0.965 | 0.07 |
| 0.000011 | 0.000011 | 1 | 0 |
| 0.000011 | 0.000011 | 0.835 | 0.13 |
| 0.000011 | 0.000011 | 1 | 0 |
| 0.000011 | 0.000011 | 0.855 | 0.29 |
| 0.000011 | 0.000011 | 0.815 | 0.37 |
| 0.000011 | 0.000011 | 1 | 0 |
| 0.000011 | 0.000011 | 0.945 | 0.11 |
| 0.000011 | 0.000011 | 0.83 | 0.22 |
| 0.000011 | 0.000011 | 0.895 | 0.21 |
| 0.000011 | 0.000011 | 0.945 | 0.07 |
| 0.000011 | 0.000011 | 1 | 0 |
| 0.000011 | 0.000011 | 1 | 0 |
| 0.000011 | 0.000011 | 0.92 | 0 |
| 0.000011 | 0.000011 | 1 | 0 |
| 0.000012 | 0.000012 | 1 | 0 |
| 0.000012 | 0.000012 | 0.925 | 0.15 |
| 0.000012 | 0.000012 | 0.97 | 0.06 |
| 0.000013 | 0.000013 | 0.98 | 0.04 |
| 0.000013 | 0.000013 | 1 | 0 |
| 0.000013 | 0.000013 | 0.975 | 0.05 |
| 0.000013 | 0.000013 | 1 | 0 |
| 0.000013 | 0.000013 | 0.955 | 0.09 |
| 0.000013 | 0.000013 | 0.935 | 0.07 |
| 0.000013 | 0.000013 | 0.9 | 0 |
| 0.000013 | 0.000013 | 0.915 | 0.17 |
| 0.000013 | 0.000013 | 0.73 | 0.3 |
| 0.000014 | 0.000014 | 0.96 | 0.08 |
| 0.000014 | 0.000014 | 0.97 | 0.06 |
| 0.000014 | 0.000014 | 1 | 0 |
| 0.000014 | 0.000014 | 1 | 0 |
| 0.000015 | 0.000015 | 1 | 0 |
| 0.000015 | 0.000015 | 0.98 | 0.04 |
| 0.000015 | 0.000015 | 0.88 | 0.06 |
| 0.000016 | 0.000016 | 0.95 | 0.1 |

| | | | |
|---|---|---|---|
| 0.000011 | 0.018959 | 0.93 | 0.14 |
| 0.000011 | 0.000011 | 0.965 | 0.07 |
| 0.000013 | 0.000013 | 0.75 | 0.4 |
| 0.000011 | 0.000011 | 1 | 0 |
| 0.000011 | 0.049116 | 0.89 | 0.12 |
| 0.000011 | 0.000011 | 0.81 | 0.38 |
| 0.000011 | 0.000011 | 0.75 | 0.5 |
| 0.000011 | 0.082568 | 0.885 | 0.23 |
| 0.000011 | 0.000011 | 0.88 | 0.24 |
| 0.000013 | 0.000013 | 0.86 | 0.28 |
| 0.000011 | 0.000011 | 0.835 | 0.33 |
| 0.000011 | 0.000011 | 0.565 | 0.17 |
| 0.000013 | 0.000013 | 0.715 | 0.57 |
| 0.000011 | 0.113512 | 1 | 0 |
| 0.000014 | 0.000014 | 0.775 | 0.45 |
| 0.000011 | 0.000011 | 0.96 | 0.08 |
| 0.000011 | 0.000011 | 0.66 | 0.68 |
| 0.000011 | 0.137881 | 0.88 | 0.24 |
| 0.000011 | 0.000011 | 0.63 | 0.5 |
| 0.000011 | 0.000011 | 0.715 | 0.53 |
| 0.000011 | 0.183069 | 0.925 | 0.15 |
| 0.000011 | 0.198296 | 1 | 0 |
| 0.000011 | 0.000011 | 0.705 | 0.47 |
| 0.00001 | 0.00001 | 0.755 | 0.21 |
| 0.000011 | 0.000011 | 0.625 | 0.57 |
| 0.000011 | 0.249997 | 0.875 | 0.25 |
| 0.000011 | 0.000011 | 0.75 | 0.24 |
| 0.000011 | 0.343144 | 0.87 | 0.26 |
| 0.000011 | 0.399957 | 0.92 | 0 |
| 0.401561 | 0.000011 | 0.715 | 0.57 |
| 0.000011 | 0.406619 | 0.685 | 0.29 |
| 0.000011 | 0.424612 | 0.855 | 0.29 |
| 0.000011 | 0.438877 | 0.825 | 0.23 |
| 0.443214 | 0.000011 | 0.665 | 0.25 |
| 0.000011 | 0.44326 | 0.7 | 0.4 |
| 0.445057 | 0.000011 | 0.78 | 0.44 |
| 0.493587 | 0.000011 | 0.605 | 0.79 |
| 0.495986 | 0.000011 | 0.735 | 0.53 |
| 0.000011 | 0.000011 | 0.5 | 1 |
| 0.000011 | 0.502153 | 0.87 | 0.26 |
| 0.000011 | 0.51082 | 0.675 | 0.61 |
| 0.513102 | 0.000011 | 0.655 | 0.69 |
| 0.000011 | 0.531268 | 0.84 | 0.32 |
| 0.000011 | 0.000011 | 0.35 | 0.3 |
| 0.576812 | 0.000011 | 0.615 | 0.57 |
| 0.000011 | 0.007532 | 0.27 | 0 |
| 0.941221 | 0.000011 | 0.325 | 0.65 |
| 0.000221 | 0.000221 | 0 | 0 |
| 0.000018 | 0.000018 | 0.035 | 0.07 |
| 0.000015 | 0.000015 | 0.05 | 0 |

| | | | |
|---|---|---|---|
| 0.000015 | 0.000015 | 0 | 0 |
| 0.000014 | 0.000014 | 0.06 | 0.06 |
| 0.000013 | 0.999935 | 0.77 | 0.46 |
| 0.000013 | 0.000013 | 0.035 | 0.07 |
| 0.999947 | 0.000011 | 0.465 | 0.55 |
| 0.999946 | 0.000011 | 0.375 | 0.65 |
| 0.999946 | 0.000011 | 0.405 | 0.17 |
| 0.000011 | 0.999946 | 0.73 | 0 |
| 0.000011 | 0.999944 | 0.55 | 0 |
| 0.000011 | 0.000011 | 0.025 | 0.05 |
| 0.000011 | 0.000011 | 0.025 | 0.05 |
| 0.000011 | 0.000011 | 0 | 0 |
| 0.000011 | 0.000011 | 0.04 | 0 |
| 0.000011 | 0.000011 | 0 | 0 |
| 0.000011 | 0.000011 | 0 | 0 |
| 0.000011 | 0.000011 | 0.125 | 0.07 |
| 0.000011 | 0.000011 | 0 | 0 |
| 0.000011 | 0.000011 | 0.075 | 0.15 |
| 0.000011 | 0.000011 | 0.085 | 0.07 |
| 0.000011 | 0.000011 | 0.03 | 0.06 |
| 0.000011 | 0.000011 | 0.025 | 0.05 |
| 0.000011 | 0.000011 | 0 | 0 |
| 0.000011 | 0.000011 | 0.05 | 0.1 |
| 0.000011 | 0.000011 | 0.025 | 0.05 |
| 0.000011 | 0.000011 | 0.035 | 0.07 |
| 0.000011 | 0.000011 | 0.095 | 0.19 |
| 0.000016 | 0.000016 | 0.06 | 0 |
| 0.000017 | 0.000017 | 0.48 | 0.54 |
| 0.000011 | 0.000011 | 0.15 | 0.3 |
| 0.000011 | 0.000011 | 0.035 | 0.07 |
| 0.000018 | 0.000018 | 0 | 0 |
| 0.000011 | 0.000011 | 0 | 0 |
| 0.000011 | 0.000011 | 0 | 0 |
| 0.000011 | 0.000011 | 0.025 | 0.05 |
| 0.000011 | 0.000011 | 0 | 0 |
| 0.000011 | 0.000011 | 0 | 0 |
| 0.000011 | 0.000011 | 0 | 0 |
| 0.000011 | 0.000011 | 0 | 0 |
| 0.000011 | 0.000011 | 0 | 0 |
| 0.000011 | 0.000011 | 0 | 0 |
| 0.000011 | 0.000011 | 0.06 | 0.12 |
| 0.000011 | 0.000011 | 0.04 | 0.02 |
| 0.000011 | 0.000011 | 0.1 | 0.2 |
| 0.000011 | 0.000011 | 0 | 0 |
| 0.000011 | 0.000011 | 0 | 0 |
| 0.000011 | 0.000011 | 0 | 0 |
| 0.000011 | 0.000011 | 0.04 | 0.08 |
| 0.00001 | 0.00001 | 0.075 | 0.07 |
| 0.000011 | 0.000011 | 0 | 0 |
| 0.000011 | 0.000011 | 0.025 | 0.05 |

| | | | |
|---|---|---|---|
| 0.000011 | 0.000011 | 0.04 | 0 |
| 0.000011 | 0.000011 | 0 | 0 |
| 0.000011 | 0.000011 | 0 | 0 |
| 0.000011 | 0.000011 | 0.025 | 0.05 |
| 0.000011 | 0.000011 | 0.04 | 0 |
| 0.000011 | 0.000011 | 0.5 | 0.5 |
| 0.000011 | 0.000011 | 0.04 | 0 |
| 0.000011 | 0.000011 | 0.025 | 0.05 |
| 0.000011 | 0.000011 | 0 | 0 |
| 0.000011 | 0.000011 | 0.025 | 0.05 |
| 0.000011 | 0.000011 | 0 | 0 |
| 0.000011 | 0.000011 | 0 | 0 |
| 0.000011 | 0.000011 | 0.04 | 0.08 |
| 0.00001 | 0.00001 | 0.025 | 0.05 |
| 0.000011 | 0.000011 | 0.025 | 0.05 |
| 0.000011 | 0.000011 | 0.075 | 0.15 |
| 0.000011 | 0.000011 | 0.08 | 0.06 |
| 0.000011 | 0.000011 | 0 | 0 |
| 0.000011 | 0.000011 | 0.025 | 0.05 |
| 0.000011 | 0.000011 | 0 | 0 |
| 0.000011 | 0.000011 | 0.04 | 0.08 |
| 0.000011 | 0.000011 | 0.025 | 0.05 |
| 0.000011 | 0.000011 | 0 | 0 |
| 0.000011 | 0.000011 | 0 | 0 |
| 0.000011 | 0.000011 | 0 | 0 |
| 0.000011 | 0.000011 | 0 | 0 |
| 0.000011 | 0.000011 | 0 | 0 |
| 0.000011 | 0.000011 | 0.065 | 0.13 |
| 0.000011 | 0.000011 | 0.025 | 0.05 |
| 0.111667 | 0.000011 | 0 | 0 |
| 0.000011 | 0.000011 | 0 | 0 |
| 0.000011 | 0.000011 | 0.03 | 0.06 |
| 0.000011 | 0.000011 | 0 | 0 |
| 0.000011 | 0.000011 | 0 | 0 |
| 0.000011 | 0.000011 | 0 | 0 |
| 0.000011 | 0.000011 | 0 | 0 |
| 0.000011 | 0.000011 | 0.075 | 0.15 |
| 0.000011 | 0.000011 | 0.075 | 0.15 |
| 0.000011 | 0.000011 | 0 | 0 |
| 0.000011 | 0.000011 | 0 | 0 |
| 0.000011 | 0.000011 | 0 | 0 |
| 0.000011 | 0.000011 | 0.025 | 0.05 |
| 0.000011 | 0.000011 | 0 | 0 |
| 0.000011 | 0.000011 | 0 | 0 |
| 0.000011 | 0.000011 | 0 | 0 |
| 0.000011 | 0.000011 | 0.04 | 0 |
| 0.000011 | 0.000011 | 0.04 | 0 |
| 0.000011 | 0.000011 | 0.025 | 0.05 |
| 0.000011 | 0.000011 | 0.005 | 0.01 |
| 0.000011 | 0.000011 | 0 | 0 |

| | | | |
|---|---|---|---|
| 0.000011 | 0.000011 | 0 | 0 |
| 0.000011 | 0.000011 | 0.025 | 0.05 |
| 0.00001 | 0.00001 | 0 | 0 |
| 0.000011 | 0.000011 | 0 | 0 |
| 0.000011 | 0.000011 | 0 | 0 |
| 0.000011 | 0.000011 | 0.11 | 0.1 |
| 0.000011 | 0.000011 | 0.05 | 0.1 |
| 0.000011 | 0.000011 | 0 | 0 |
| 0.000011 | 0.000011 | 0.105 | 0.21 |
| 0.000011 | 0.000011 | 0 | 0 |
| 0.000011 | 0.000011 | 0.5 | 0.5 |
| 0.000011 | 0.000011 | 0 | 0 |
| 0.000011 | 0.000011 | 0.025 | 0.05 |
| 0.000011 | 0.000011 | 0.025 | 0.05 |
| 0.000011 | 0.000011 | 0.04 | 0 |
| 0.000011 | 0.000011 | 0.05 | 0.1 |
| 0.000011 | 0.000011 | 0 | 0 |
| 0.000011 | 0.000011 | 0 | 0 |
| 0.000011 | 0.000011 | 0.045 | 0.03 |
| 0.000011 | 0.000011 | 0.025 | 0.05 |
| 0.000011 | 0.000011 | 0.05 | 0 |
| 0.000011 | 0.000011 | 0.05 | 0.1 |
| 0.000011 | 0.000011 | 0.04 | 0.08 |
| 0.000011 | 0.000011 | 0 | 0 |
| 0.000011 | 0.000011 | 0 | 0 |
| 0.000011 | 0.000011 | 0.07 | 0.14 |
| 0.000011 | 0.000011 | 0.415 | 0.83 |
| 0.000011 | 0.000011 | 0.03 | 0.06 |
| 0.000011 | 0.000011 | 0 | 0 |
| 0.000011 | 0.000011 | 0.03 | 0.06 |
| 0.000011 | 0.000011 | 0.025 | 0.05 |
| 0.000011 | 0.000011 | 0 | 0 |
| 0.000011 | 0.000011 | 0 | 0 |
| 0.000011 | 0.000011 | 0.1 | 0 |
| 0.000011 | 0.000011 | 0.06 | 0.12 |
| 0.000011 | 0.000011 | 0.04 | 0 |
| 0.000011 | 0.000011 | 0 | 0 |
| 0.000011 | 0.000011 | 0 | 0 |
| 0.000011 | 0.000011 | 0.065 | 0.13 |
| 0.000011 | 0.000011 | 0.15 | 0.26 |
| 0.000011 | 0.000011 | 0.12 | 0.06 |
| 0.000011 | 0.000011 | 0.09 | 0.18 |
| 0.000011 | 0.000011 | 0.11 | 0.22 |
| 0.000011 | 0.000011 | 0 | 0 |
| 0.000011 | 0.000011 | 0.05 | 0 |
| 0.000011 | 0.000011 | 0.05 | 0.04 |
| 0.000011 | 0.000011 | 0 | 0 |
| 0.000011 | 0.000011 | 0 | 0 |
| 0.000011 | 0.000011 | 0.005 | 0.01 |
| 0.000011 | 0.000011 | 0 | 0 |

| | | | |
|---|---|---|---|
| 0.000011 | 0.000011 | 0.005 | 0.01 |
| 0.000011 | 0.000011 | 0 | 0 |
| 0.000011 | 0.000011 | 0.005 | 0.01 |
| 0.000011 | 0.000011 | 0 | 0 |
| 0.000011 | 0.000011 | 1 | 0 |
| 0.000011 | 0.000011 | 0.995 | 0.01 |
| 0.000011 | 0.000011 | 1 | 0 |
| 0.000011 | 0.000011 | 1 | 0 |
| 0.00001 | 0.00001 | 0.98 | 0.04 |
| 0.000011 | 0.000011 | 0.025 | 0.05 |
| 0.000011 | 0.000011 | 0 | 0 |
| 0.000011 | 0.000011 | 0.01 | 0.02 |
| 0.000011 | 0.000011 | 0 | 0 |
| 0.000011 | 0.000011 | 0 | 0 |
| 0.000011 | 0.000011 | 0.025 | 0.05 |
| 0.000011 | 0.000011 | 0.025 | 0.05 |
| 0.000011 | 0.000011 | 0.025 | 0.05 |
| 0.000011 | 0.000011 | 0.025 | 0.05 |
| 0.000011 | 0.000011 | 0 | 0 |
| 0.000011 | 0.000011 | 0.025 | 0.05 |
| 0.000011 | 0.000011 | 0.05 | 0.1 |
| 0.000011 | 0.000011 | 0 | 0 |
| 0.000011 | 0.000011 | 0.04 | 0 |
| 0.000011 | 0.000011 | 0 | 0 |
| 0.000011 | 0.000011 | 0.05 | 0.1 |
| 0.000011 | 0.000011 | 0 | 0 |
| 0.000011 | 0.000011 | 0 | 0 |
| 0.000011 | 0.000011 | 0 | 0 |
| 0.000011 | 0.000011 | 0 | 0 |
| 0.000011 | 0.000011 | 0.025 | 0.05 |
| 0.000011 | 0.000011 | 0.065 | 0.13 |
| 0.000011 | 0.000011 | 0 | 0 |
| 0.000011 | 0.000011 | 0 | 0 |
| 0.000011 | 0.000011 | 0 | 0 |
| 0.000011 | 0.000011 | 0 | 0 |
| 0.000011 | 0.000011 | 0 | 0 |
| 0.000011 | 0.000011 | 0 | 0 |
| 0.000011 | 0.000011 | 0 | 0 |
| 0.000011 | 0.000011 | 0.025 | 0.05 |
| 0.000011 | 0.000011 | 0.04 | 0 |
| 0.000011 | 0.000011 | 0 | 0 |
| 0.000011 | 0.000011 | 0 | 0 |
| 0.000011 | 0.000011 | 0.04 | 0 |
| 0.000011 | 0.000011 | 0 | 0 |
| 0.000011 | 0.000011 | 0 | 0 |
| 0.000011 | 0.000011 | 0 | 0 |
| 0.000011 | 0.000011 | 0.94 | 0.12 |
| 0.000011 | 0.000011 | 0 | 0 |
| 0.000011 | 0.000011 | 0 | 0 |
| 0.000011 | 0.000011 | 0 | 0 |

| | | | |
|---|---|---|---|
| 0.000011 | 0.000011 | 0 | 0 |
| 0.000011 | 0.000011 | 0.025 | 0.05 |
| 0.000011 | 0.000011 | 0.025 | 0.05 |
| 0.000011 | 0.000011 | 0.025 | 0.05 |
| 0.000011 | 0.000011 | 0.185 | 0.17 |
| 0.000011 | 0.000011 | 0.05 | 0 |
| 0.000011 | 0.000011 | 0.005 | 0.01 |
| 0.000011 | 0.000011 | 0 | 0 |
| 0.000011 | 0.000011 | 0.035 | 0.07 |
| 0.000011 | 0.000011 | 0.065 | 0.13 |
| 0.000011 | 0.000011 | 0.09 | 0 |
| 0.000011 | 0.000011 | 0.055 | 0.11 |
| 0.000011 | 0.000011 | 0.025 | 0.05 |
| 0.000011 | 0.000011 | 0 | 0 |
| 0.00001 | 0.00001 | 0.04 | 0 |
| 0.000011 | 0.000011 | 0.04 | 0 |
| 0.000011 | 0.000011 | 0 | 0 |
| 0.000011 | 0.000011 | 0 | 0 |
| 0.000011 | 0.000011 | 0 | 0 |
| 0.000011 | 0.000011 | 0.025 | 0.05 |
| 0.000024 | 0.000024 | 0 | 0 |
| 0.000011 | 0.000011 | 0.025 | 0.05 |
| 0.000011 | 0.000011 | 0 | 0 |
| 0.000011 | 0.000011 | 0 | 0 |
| 0.000011 | 0.000011 | 0.05 | 0.1 |
| 0.000011 | 0.000011 | 0.04 | 0 |
| 0.000011 | 0.000011 | 0 | 0 |
| 0.000011 | 0.000011 | 0 | 0 |
| 0.000011 | 0.000011 | 0.05 | 0 |
| 0.000011 | 0.000011 | 0 | 0 |
| 0.000011 | 0.000011 | 0 | 0 |
| 0.000011 | 0.000011 | 0 | 0 |
| 0.000011 | 0.000011 | 0 | 0 |
| 0.000011 | 0.000011 | 0.09 | 0 |
| 0.000011 | 0.000011 | 0 | 0 |
| 0.000011 | 0.000011 | 0.05 | 0.1 |
| 0.000011 | 0.000011 | 0 | 0 |
| 0.000011 | 0.000011 | 0 | 0 |
| 0.000011 | 0.000011 | 0.025 | 0.05 |
| 0.000017 | 0.000017 | 0.455 | 0.31 |
| 0.00002 | 0.00002 | 0.765 | 0.47 |
| 0.00002 | 0.00002 | 1 | 0 |
| 0.000016 | 0.000016 | 0.97 | 0.06 |
| 0.000015 | 0.000015 | 0 | 0 |
| 0.000017 | 0.000017 | 0.08 | 0.16 |
| 0.000017 | 0.000017 | 0 | 0 |
| 0.000017 | 0.000017 | 0.06 | 0.12 |
| 0.000019 | 0.000019 | 0.045 | 0.09 |
| 0.000022 | 0.000022 | 0 | 0 |
| 0.000023 | 0.000023 | 0 | 0 |

| | | | |
|---|---|---|---|
| 0.000017 | 0.000017 | 0.97 | 0.06 |
| 0.000011 | 0.000011 | 0 | 0 |
| 0.000012 | 0.000012 | 0.03 | 0.06 |
| 0.000014 | 0.000014 | 0.445 | 0.57 |
| 0.000017 | 0.000017 | 0 | 0 |
| 0.000013 | 0.000013 | 0.05 | 0 |
| 0.000013 | 0.000013 | 0.195 | 0.25 |
| 0.000015 | 0.000015 | 0.075 | 0.15 |
| 0.000015 | 0.000015 | 0 | 0 |
| 0.000017 | 0.000017 | 0 | 0 |
| 0.000017 | 0.000017 | 0.1 | 0.2 |
| 0.000022 | 0.000022 | 0 | 0 |
| 0.000027 | 0.000027 | 0 | 0 |
| 0.000012 | 0.000012 | 0.055 | 0.11 |
| 0.000014 | 0.000014 | 0 | 0 |
| 0.000015 | 0.000015 | 0.025 | 0.05 |
| 0.000015 | 0.000015 | 0.04 | 0.02 |
| 0.000019 | 0.000019 | 0 | 0 |
| 0.000021 | 0.000021 | 0 | 0 |
| 0.000022 | 0.000022 | 0 | 0 |
| 0.000023 | 0.000023 | 0 | 0 |
| 0.000023 | 0.000023 | 0 | 0 |
| 0.000012 | 0.000012 | 0 | 0 |
| 0.000012 | 0.000012 | 0 | 0 |
| 0.000012 | 0.000012 | 0.115 | 0.11 |
| 0.000013 | 0.000013 | 0.03 | 0.06 |
| 0.000013 | 0.000013 | 0 | 0 |
| 0.000013 | 0.000013 | 0 | 0 |

| ı analysis |
| --- |
| **Hybrid assignment** |
| Cultivated genotype |
| Cultivated genotype |
| Cultivated genotype |
| Cultivated genotype |
| Cultivated genotype |
| Cultivated genotype |
| No statistical supported assignment |
| Wild genotype |
| Cultivated genotype |
| Cultivated genotype |
| Cultivated genotype |
| Cultivated genotype |
| Cultivated genotype |
| No statistical supported assignment |
| hybrid-cultivated backcross |
| Cultivated genotype |
| Cultivated genotype |
| F1 hybrid |
| Cultivated genotype |
| Cultivated genotype |
| Cultivated genotype |
| Cultivated genotype |
| Cultivated genotype |
| Cultivated genotype |
| Cultivated genotype |
| Cultivated genotype |
| Cultivated genotype |
| Cultivated genotype |
| Cultivated genotype |
| Cultivated genotype |
| Cultivated genotype |
| Cultivated genotype |
| Cultivated genotype |
| Cultivated genotype |
| Cultivated genotype |
| Cultivated genotype |
| Cultivated genotype |
| Cultivated genotype |
| Cultivated genotype |
| No statistical supported assignment |
| Cultivated genotype |
| Cultivated genotype |
| Cultivated genotype |

| |
|---|
| Cultivated genotype |
| Cultivated genotype |
| Cultivated genotype |
| Cultivated genotype |
| Cultivated genotype |
| Cultivated genotype |
| Cultivated genotype |
| Cultivated genotype |
| Cultivated genotype |
| Cultivated genotype |
| Cultivated genotype |
| Cultivated genotype |
| Cultivated genotype |
| Cultivated genotype |
| Cultivated genotype |
| Cultivated genotype |
| Cultivated genotype |
| Cultivated genotype |
| Cultivated genotype |
| Cultivated genotype |
| F2 hybrid |
| Wild genotype |
| No statistical supported assignment |
| No statistical supported assignment |
| Cultivated genotype |
| Cultivated genotype |
| Cultivated genotype |
| Cultivated genotype |
| Cultivated genotype |
| Cultivated genotype |
| Cultivated genotype |
| Cultivated genotype |
| Cultivated genotype |
| Cultivated genotype |
| Cultivated genotype |
| Cultivated genotype |
| Cultivated genotype |
| Cultivated genotype |
| Cultivated genotype |
| Cultivated genotype |
| Cultivated genotype |
| Cultivated genotype |
| Cultivated genotype |
| Cultivated genotype |
| Cultivated genotype |
| Cultivated genotype |
| Cultivated genotype |
| Cultivated genotype |
| Cultivated genotype |
| Cultivated genotype |

Cultivated genotype
Cultivated genotype
Cultivated genotype
Cultivated genotype
Cultivated genotype
Cultivated genotype
Cultivated genotype
Cultivated genotype
Cultivated genotype
Cultivated genotype
Cultivated genotype
Cultivated genotype
Cultivated genotype
Cultivated genotype
Cultivated genotype
Cultivated genotype
Cultivated genotype
Cultivated genotype
Cultivated genotype
Cultivated genotype
Cultivated genotype
Cultivated genotype
Cultivated genotype
Cultivated genotype
Cultivated genotype
Cultivated genotype
Cultivated genotype
Cultivated genotype
Cultivated genotype
Cultivated genotype
Cultivated genotype
Cultivated genotype
Cultivated genotype
Cultivated genotype
Cultivated genotype
Cultivated genotype
Cultivated genotype
Cultivated genotype
Cultivated genotype
Cultivated genotype
Cultivated genotype
Cultivated genotype
Cultivated genotype
Cultivated genotype
Cultivated genotype
Cultivated genotype
Cultivated genotype
Cultivated genotype
Cultivated genotype
Cultivated genotype

| |
|---|
| Cultivated genotype |
| Cultivated genotype |
| Cultivated genotype |
| Cultivated genotype |
| Cultivated genotype |
| Cultivated genotype |
| Cultivated genotype |
| Cultivated genotype |
| No statistical supported assignment |
| hybrid-cultivated backcross |
| Cultivated genotype |
| No statistical supported assignment |
| hybrid-cultivated backcross |
| Cultivated genotype |
| hybrid-cultivated backcross |
| Cultivated genotype |
| hybrid-cultivated backcross |
| Cultivated genotype |
| F2 hybrid |
| hybrid-cultivated backcross |
| Cultivated genotype |
| Cultivated genotype |
| hybrid-cultivated backcross |
| No statistical supported assignment |
| No statistical supported assignment |
| No statistical supported assignment |
| No statistical supported assignment |
| hybrid-cultivated backcross |
| Cultivated genotype |
| hybrid-cultivated backcross |
| Cultivated genotype |
| Cultivated genotype |
| Cultivated genotype |
| No statistical supported assignment |
| Cultivated genotype |
| hybrid-cultivated backcross |
| No statistical supported assignment |
| hybrid-cultivated backcross |
| F1 hybrid |
| Cultivated genotype |
| Cultivated genotype |
| hybrid-cultivated backcross |
| hybrid-cultivated backcross |
| No statistical supported assignment |
| No statistical supported assignment |
| No statistical supported assignment |
| No statistical supported assignment |
| No statistical supported assignment |
| No statistical supported assignment |
| No statistical supported assignment |

| |
|---|
| Wild genotype |
| No statistical supported assignment |
| hybrid-cultivated backcross |
| No statistical supported assignment |
| F2 hybrid |
| No statistical supported assignment |
| No statistical supported assignment |
| No statistical supported assignment |
| No statistical supported assignment |
| No statistical supported assignment |
| No statistical supported assignment |
| Wild genotype |
| No statistical supported assignment |
| Wild genotype |
| Wild genotype |
| hybrid-wild backcross |
| Wild genotype |
| Wild genotype |
| Wild genotype |
| Wild genotype |
| Wild genotype |
| Wild genotype |
| Wild genotype |
| Wild genotype |
| Wild genotype |
| Wild genotype |
| Wild genotype |
| Wild genotype |
| Wild genotype |
| Wild genotype |
| Wild genotype |
| Wild genotype |
| Wild genotype |
| Wild genotype |
| Wild genotype |
| Wild genotype |
| Wild genotype |
| Wild genotype |
| Wild genotype |
| Wild genotype |
| Wild genotype |
| Wild genotype |
| Wild genotype |
| Wild genotype |
| Wild genotype |
| Wild genotype |
| Wild genotype |
| Wild genotype |
| Wild genotype |

Wild genotype
Wild genotype
Wild genotype
Wild genotype
Wild genotype
Wild genotype
Wild genotype
Wild genotype
Wild genotype
Wild genotype
Wild genotype
Wild genotype
Wild genotype
Wild genotype
Wild genotype
Wild genotype
Wild genotype
Wild genotype
Wild genotype
Wild genotype
Wild genotype
Wild genotype
Wild genotype
Wild genotype
Wild genotype
Wild genotype
Wild genotype
Wild genotype
Wild genotype
Wild genotype
Wild genotype
Wild genotype
Wild genotype
Wild genotype
Wild genotype
Wild genotype
Wild genotype
Wild genotype
Wild genotype
Wild genotype
Wild genotype
Wild genotype
Wild genotype
Wild genotype
Wild genotype
Wild genotype
Wild genotype
Wild genotype
Wild genotype
Wild genotype

| |
|---|
| Wild genotype |
| Wild genotype |
| Wild genotype |
| Wild genotype |
| Wild genotype |
| Wild genotype |
| Wild genotype |
| Wild genotype |
| Wild genotype |
| Wild genotype |
| Wild genotype |
| Wild genotype |
| Wild genotype |
| Wild genotype |
| Wild genotype |
| Wild genotype |
| Wild genotype |
| Wild genotype |
| Wild genotype |
| Wild genotype |
| Wild genotype |
| Wild genotype |
| Wild genotype |
| Wild genotype |
| Wild genotype |
| Wild genotype |
| No statistical supported assignment |
| Wild genotype |
| Wild genotype |
| Wild genotype |
| Wild genotype |
| Wild genotype |
| Wild genotype |
| Wild genotype |
| Wild genotype |
| Wild genotype |
| Wild genotype |
| Wild genotype |
| Wild genotype |
| Wild genotype |
| Wild genotype |
| Wild genotype |
| Wild genotype |
| Wild genotype |
| Wild genotype |
| Wild genotype |
| Wild genotype |
| Wild genotype |
| Wild genotype |
| Wild genotype |

Wild genotype
Wild genotype
Wild genotype
Wild genotype
Cultivated genotype
Cultivated genotype
Cultivated genotype
Cultivated genotype
Cultivated genotype
Wild genotype
Wild genotype
Wild genotype
Wild genotype
Wild genotype
Wild genotype
Wild genotype
Wild genotype
Wild genotype
Wild genotype
Wild genotype
Wild genotype
Wild genotype
Wild genotype
Wild genotype
Wild genotype
Wild genotype
Wild genotype
Wild genotype
Wild genotype
Wild genotype
Wild genotype
Wild genotype
Wild genotype
Wild genotype
Wild genotype
Wild genotype
Wild genotype
Wild genotype
Wild genotype
Wild genotype
Wild genotype
Wild genotype
Wild genotype
Wild genotype
Wild genotype
Wild genotype
Cultivated genotype
Wild genotype
Wild genotype
Wild genotype

| |
|---|
| Wild genotype |
| Wild genotype |
| Wild genotype |
| Wild genotype |
| Wild genotype |
| Wild genotype |
| Wild genotype |
| Wild genotype |
| Wild genotype |
| Wild genotype |
| Wild genotype |
| Wild genotype |
| Wild genotype |
| Wild genotype |
| Wild genotype |
| Wild genotype |
| Wild genotype |
| Wild genotype |
| Wild genotype |
| Wild genotype |
| Wild genotype |
| Wild genotype |
| Wild genotype |
| Wild genotype |
| Wild genotype |
| Wild genotype |
| Wild genotype |
| Wild genotype |
| Wild genotype |
| Wild genotype |
| Wild genotype |
| Wild genotype |
| Wild genotype |
| Wild genotype |
| Wild genotype |
| Wild genotype |
| Wild genotype |
| Wild genotype |
| Wild genotype |
| F2 hybrid |
| hybrid-cultivated backcross |
| Cultivated genotype |
| Cultivated genotype |
| Wild genotype |
| Wild genotype |
| Wild genotype |
| Wild genotype |
| Wild genotype |
| Wild genotype |
| Wild genotype |

Cultivated genotype
Wild genotype
Wild genotype
Wild genotype
Wild genotype
Wild genotype
Wild genotype
Wild genotype
Wild genotype
Wild genotype
Wild genotype
Wild genotype
Wild genotype
Wild genotype
Wild genotype
Wild genotype
Wild genotype
Wild genotype
Wild genotype
Wild genotype
Wild genotype
Wild genotype
Wild genotype
Wild genotype
Wild genotype
Wild genotype
Wild genotype
Wild genotype
