## Supplemental Table S2 for "Cultivated Robusta coffee meets wild *Coffea canephora*: Evidence of cultivated-wild hybridisation in the Democratic Republic of the Congo"

**Supplemental Table S2:** Pairwise Jaccard Inversed Distance (JID) calculated for all 481 indiv
The JID matric was arranged based on the sample location. Red-shaded JID values are JID v

| | | Sample location | | |
|---|---|---|---|---|
| | | Plot number | HG_01 | |
| Sample location | Plot number | Sample ID | Home_garc | Home_garc |
| Home gard | HG_01 | Home_garden_1347 | 1 | 0.7841 |
| | | Home_garden_1348 | 0.7841 | 1 |
| | HG_02 | Home_garden_1330 | 0.783621 | 0.794251 |
| | | Home_garden_1331 | 0.785562 | 0.78633 |
| | | Home_garden_1332 | 0.782962 | 0.771092 |
| | | Home_garden_1328 | 0.779546 | 0.775609 |
| | HG_03 | Home_garden_1350 | 0.710025 | 0.718979 |
| | HG_04 | Home_garden_1358 | 0.705975 | 0.71748 |
| | HG_05 | Home_garden_1397 | 0.784427 | 0.784264 |
| | | Home_garden_1396 | 0.788511 | 0.780227 |
| | HG_06 | Home_garden_1398 | 0.783585 | 0.776086 |
| | HG_07 | Home_garden_1399 | 0.796432 | 0.787801 |
| | HG_08 | Home_garden_1402 | 0.772298 | 0.787167 |
| | | Home_garden_1401 | 0.78424 | 0.784833 |
| | | Home_garden_1400 | 0.763446 | 0.777532 |
| | HG_09 | Home_garden_1403 | 0.792252 | 0.781655 |
| | | Home_garden_1404 | 0.794434 | 0.779911 |
| | HG_10 | Home_garden_1405 | 0.759537 | 0.765304 |
| | HG_11 | Home_garden_1421 | 0.790135 | 0.792483 |
| | | Home_garden_1424 | 0.783487 | 0.789404 |
| | | Home_garden_1425 | 0.790272 | 0.794583 |
| | | Home_garden_1426 | 0.802914 | 0.789552 |
| | | Home_garden_1422 | 0.796602 | 0.799468 |
| | | Home_garden_1410 | 0.798603 | 0.796676 |
| | | Home_garden_1407 | 0.778906 | 0.787614 |
| | | Home_garden_1416 | 0.784356 | 0.794304 |
| | | Home_garden_1413 | 0.777072 | 0.786442 |
| | | Home_garden_1427 | 0.779532 | 0.790021 |
| | | Home_garden_1414 | 0.782859 | 0.792937 |
| | | Home_garden_1411 | 0.788007 | 0.788866 |
| | | Home_garden_1409 | 0.773644 | 0.77797 |
| | | Home_garden_1408 | 0.775597 | 0.777259 |
| | | Home_garden_1423 | 0.776425 | 0.787273 |
| | | Home_garden_1415 | 0.767413 | 0.779349 |
| | | Home_garden_1417 | 0.767082 | 0.779141 |
| | | Home_garden_1419 | 0.780094 | 0.792031 |
| | | Home_garden_1420 | 0.77622 | 0.786481 |
| | | Home_garden_1418 | 0.762943 | 0.778693 |
| | | Home_garden_1412 | 0.771999 | 0.780761 |

| | | | | |
|---|---|---|---|---|
| e<br>n | | Home_garden_1406 | 0.753684 | 0.75928 |
| | HG_12 | Home_garden_1428 | 0.799752 | 0.785131 |
| | HG_13 | Home_garden_1441 | 0.796671 | 0.790443 |
| | | Home_garden_1438 | 0.800016 | 0.793876 |
| | HG_14 | Home_garden_1442 | 0.792796 | 0.781757 |
| | HG_15 | Home_garden_1456 | 0.779465 | 0.779241 |
| | | Home_garden_1453 | 0.777168 | 0.769784 |
| | | Home_garden_1455 | 0.790021 | 0.792724 |
| | | Home_garden_1454 | 0.772206 | 0.775849 |
| | HG_16 | Home_garden_1462 | 0.778379 | 0.775049 |
| | HG_17 | Home_garden_1485 | 0.804176 | 0.792135 |
| | | Home_garden_1489 | 0.7971 | 0.789766 |
| | | Home_garden_1481 | 0.793921 | 0.7895 |
| | | Home_garden_1484 | 0.785027 | 0.787521 |
| | | Home_garden_1487 | 0.790865 | 0.795905 |
| | | Home_garden_1488 | 0.791362 | 0.788525 |
| | | Home_garden_1482 | 0.796791 | 0.777648 |
| | | Home_garden_1483 | 0.804504 | 0.800183 |
| | | Home_garden_1486 | 0.789504 | 0.7848 |
| | HG_18 | Home_garden_1490 | 0.79014 | 0.792648 |
| | | Home_garden_1492 | 0.804271 | 0.795062 |
| | | Home_garden_1493 | 0.804453 | 0.791882 |
| | | Home_garden_1491 | 0.775789 | 0.789973 |
| | HG_19 | Home_garden_1601 | 0.766156 | 0.781961 |
| | | Home_garden_1600 | 0.75913 | 0.769734 |
| | | Home_garden_1585 | 0.706493 | 0.717701 |
| | HG_20 | Home_garden_1613 | 0.711378 | 0.718254 |
| | | Home_garden_1614 | 0.710343 | 0.718946 |
| | HG_21 | Home_garden_1698 | 0.773037 | 0.784285 |
| | | Home_garden_1699 | 0.774284 | 0.794259 |
| | | Home_garden_1701 | 0.784586 | 0.775798 |
| | | Home_garden_1700 | 0.773513 | 0.780447 |
| | | Collection_G0095_2067 | 0.805373 | 0.817086 |
| | | Collection_G0119_1391 | 0.796923 | 0.812721 |
| | | Collection_G0004_1968 | 0.806179 | 0.809775 |
| | | Collection_G0106_2083 | 0.811555 | 0.820751 |
| | | Collection_G0034_2001 | 0.783508 | 0.790085 |
| | | Collection_G0040_2007 | 0.834365 | 0.816015 |
| | | Collection_G0089FOG_2060 | 0.783856 | 0.793689 |
| | | Collection_G0093_2064 | 0.781955 | 0.789988 |
| | | Collection_G0096_2068 | 0.794268 | 0.796371 |
| | | Collection_G0250FOG_1470 | 0.827677 | 0.815475 |
| | | Collection_G0007_1971 | 0.798382 | 0.779293 |
| | | Collection_G0014_1387 | 0.794701 | 0.781604 |
| | | Collection_G0022_1386 | 0.782076 | 0.781509 |
| | | Collection_G0037_2004 | 0.77956 | 0.797444 |

| | | |
|---|---|---|
| Collection_G0075_2044 | 0.804619 | 0.778659 |
| Collection_G0198_1463 | 0.781011 | 0.778863 |
| Collection_G0001_1965 | 0.788391 | 0.777167 |
| Collection_G0084_2055 | 0.81178 | 0.803348 |
| Collection_G0046_2013 | 0.781457 | 0.785108 |
| Collection_G0015_1980 | 0.77752 | 0.79189 |
| Collection_G0047_2014 | 0.786212 | 0.794455 |
| Collection_G0025_1383 | 0.777087 | 0.784141 |
| Collection_G0024_1374 | 0.776922 | 0.776555 |
| Collection_G0027_1377 | 0.774612 | 0.774787 |
| Collection_G0029_1995 | 0.779413 | 0.776564 |
| Collection_G0044_2011 | 0.773007 | 0.780109 |
| Collection_G0055_1392 | 0.784812 | 0.785554 |
| Collection_G0077_2046 | 0.771536 | 0.773205 |
| Collection_G0017_1376 | 0.778651 | 0.779606 |
| Collection_G0008_1476 | 0.77541 | 0.775693 |
| Collection_G0230_1394 | 0.774623 | 0.776234 |
| Collection_G0107_2084 | 0.798268 | 0.79511 |
| Collection_G0080_2050 | 0.7761 | 0.779265 |
| Collection_G0021_1375 | 0.773549 | 0.784048 |
| Collection_G0036_2003 | 0.777585 | 0.775429 |
| Collection_G0013_1978 | 0.767556 | 0.774613 |
| Collection_G0032_1999 | 0.785524 | 0.789333 |
| Collection_G0104_2080 | 0.775051 | 0.780267 |
| Collection_G0091_2062 | 0.778492 | 0.784139 |
| Collection_G0235_1497 | 0.783301 | 0.802756 |
| Collection_G0090_2061 | 0.782534 | 0.784835 |
| Collection_G0232_1393 | 0.775803 | 0.774312 |
| Collection_G0019_1321 | 0.776248 | 0.783797 |
| Collection_G0098_2070 | 0.771917 | 0.782102 |
| Collection_G0071_2040 | 0.768615 | 0.779114 |
| Collection_G0033_2000 | 0.767668 | 0.772405 |
| Collection_G0018_2071 | 0.790493 | 0.790133 |
| Collection_G0072_2041 | 0.796355 | 0.790607 |
| Collection_G0099_2074 | 0.789645 | 0.781159 |
| Collection_G0068_2037 | 0.791997 | 0.790486 |
| Collection_G0056_2025 | 0.78077 | 0.776486 |
| Collection_G0094_2112 | 0.753568 | 0.749239 |
| Collection_G0105_2133 | 0.760426 | 0.772847 |
| Collection_G0113_2110 | 0.748402 | 0.749787 |
| Collection_G0248_2137 | 0.775774 | 0.787534 |
| Collection_G0006_2143 | 0.743677 | 0.742227 |
| Collection_G0009_2160 | 0.778048 | 0.753526 |
| Collection_G0031_2108 | 0.761082 | 0.758502 |
| I Collection_G0234_2145 | 0.740003 | 0.744171 |
| N Collection_G0197_2151 | 0.758796 | 0.772082 |
| E Collection_G0245_2162 | 0.76604 | 0.779146 |

| | | | |
|---|---|---|---|
| - RA Coffee Collection | Collection_G0127_2142 | 0.740472 | 0.742242 |
| | Collection_G0005_2157 | 0.783051 | 0.791765 |
| | Collection_G0114_2111 | 0.729373 | 0.738709 |
| | Collection_G0020_2161 | 0.764256 | 0.740358 |
| | Collection_G0246_2163 | 0.746852 | 0.760255 |
| | Collection_G0233_2115 | 0.74043 | 0.745941 |
| | Collection_G0138_2139 | 0.743088 | 0.758024 |
| | Collection_G0121_2155 | 0.762999 | 0.765324 |
| | Collection_G0002_2141 | 0.718359 | 0.726138 |
| | Collection_G0003_2148 | 0.766893 | 0.784185 |
| | Collection_G0135_2131 | 0.744848 | 0.742201 |
| | Collection_G0012_1976 | 0.788517 | 0.765131 |
| | Collection_G0030_1997 | 0.772775 | 0.769444 |
| | Collection_G0134_2150 | 0.75 | 0.758744 |
| | Collection_G0035_2002 | 0.778472 | 0.772304 |
| | Collection_G0079_2049 | 0.767888 | 0.768668 |
| | Collection_G0057_1395 | 0.772192 | 0.779765 |
| | Collection_G0016_1390 | 0.769358 | 0.772287 |
| | Collection_G0010_1974 | 0.749577 | 0.750564 |
| | Collection_G0101_2077 | 0.774262 | 0.770647 |
| | Collection_G0247_2136 | 0.747209 | 0.754274 |
| | Collection_G0026_1385 | 0.771377 | 0.772462 |
| | Collection_G0175_1477 | 0.774379 | 0.769041 |
| | Collection_G0115_2134 | 0.743994 | 0.759067 |
| | Collection_G0092FOG_2063 | 0.753383 | 0.752511 |
| | Collection_G0112_2109 | 0.761899 | 0.768445 |
| | Collection_G0256FOG_1498 | 0.761654 | 0.76147 |
| | Collection_G0255FOG_1494 | 0.774423 | 0.778937 |
| | Collection_G0039_2006 | 0.743575 | 0.743753 |
| | Collection_G0011_1975 | 0.787844 | 0.779455 |
| | Collection_G0023_1389 | 0.765532 | 0.769392 |
| | Collection_G0085_2056 | 0.773952 | 0.766193 |
| | Collection_G0083_2054 | 0.774461 | 0.767759 |
| | Collection_G0103_2079 | 0.769861 | 0.769533 |
| | Collection_G0097_2069 | 0.769807 | 0.761748 |
| | Collection_G0078_2048 | 0.766434 | 0.769693 |
| | Collection_G0100_2076 | 0.721715 | 0.721542 |
| | Collection_G0038_2005 | 0.753372 | 0.758498 |
| | Collection_G0088_2059 | 0.726586 | 0.723951 |
| | Collection_G0251FOG_1472 | 0.721017 | 0.7215 |
| | Collection_G0063_2032 | 0.769193 | 0.768999 |
| | Collection_G0074_2043 | 0.716822 | 0.719526 |
| | Collection_G0102_2078 | 0.715575 | 0.725375 |
| | Collection_G0086_2057 | 0.707707 | 0.721651 |
| | Collection_G0064_2033 | 0.766545 | 0.765791 |
| | Collection_G0253FOG_1474 | 0.716247 | 0.724117 |
| | Collection_G0061_2030 | 0.764927 | 0.770724 |

| | | | | |
|---|---|---|---|---|
| | | Collection_G0069_2038 | 0.761635 | 0.765688 |
| | | Collection_G0067_2036 | 0.767854 | 0.762373 |
| | | Collection_G0041_2008 | 0.746113 | 0.747757 |
| | | Collection_G0252FOG_1473 | 0.704312 | 0.711549 |
| | | Collection_G0073_2042 | 0.703276 | 0.714853 |
| | | Collection_G0066_2035 | 0.77745 | 0.777268 |
| | | Collection_G0043_2073 | 0.687514 | 0.693687 |
| | | Collection_G0042_1496 | 0.72996 | 0.734498 |
| | | Collection_G0062_2031 | 0.763575 | 0.75935 |
| | | Collection_G0028_1388 | 0.718164 | 0.72293 |
| | | Collection_G0059_2028 | 0.749654 | 0.750285 |
| | | Collection_G0109_2105 | 0.611111 | 0.659722 |
| | | Collection_G0110_2106 | 0.644146 | 0.65848 |
| | | Collection_G0076_2102 | 0.654254 | 0.665391 |
| | | Collection_G0050_2101 | 0.664536 | 0.674231 |
| | | Collection_G0111_2107 | 0.658055 | 0.666449 |
| | | Collection_G0081_2120 | 0.619723 | 0.62072 |
| | | Collection_G0108_2104 | 0.657767 | 0.676982 |
| | | Collection_G0065_2034 | 0.737777 | 0.744483 |
| | | Collection_G0060_2029 | 0.732755 | 0.740667 |
| | | Collection_G0058_2027 | 0.742359 | 0.744366 |
| | | Collection_G0254FOG_1475 | 0.618364 | 0.617729 |
| | | Collection_G0087_2058 | 0.601559 | 0.601051 |
| | | Collection_G0048_2016 | 0.705206 | 0.711419 |
| | | Collection_G0049_2017 | 0.708304 | 0.711851 |
| | | Collection_G0054_2022 | 0.698548 | 0.706275 |
| | | Collection_G0052_2020 | 0.701006 | 0.707441 |
| | | Collection_G0053_2021 | 0.706302 | 0.710779 |
| | | Collection_G0070_2039 | 0.704051 | 0.711244 |
| | | Collection_G0082_2053 | 0.706927 | 0.709896 |
| | | Collection_G0051_2019 | 0.706181 | 0.710665 |
| | plot 01 | Wild_plot_01_1501 | 0.711827 | 0.717772 |
| | | Wild_plot_01_1502 | 0.709002 | 0.717605 |
| | | Wild_plot_01_1562 | 0.709644 | 0.715268 |
| | | Wild_plot_01_1504 | 0.716041 | 0.719437 |
| | | Wild_plot_01_1505 | 0.706549 | 0.713679 |
| | | Wild_plot_01_1503 | 0.712652 | 0.719434 |
| | | Wild_plot_01_1507 | 0.709455 | 0.716587 |
| | | Wild_plot_01_1506 | 0.704872 | 0.710339 |
| | | Wild_plot_01_1561 | 0.709534 | 0.715172 |
| | | Wild_plot_02_2244 | 0.65758 | 0.663723 |
| | | Wild_plot_02_2245 | 0.653972 | 0.66489 |
| | | Wild_plot_02_1517 | 0.710722 | 0.719656 |
| | | Wild_plot_02_1509 | 0.712104 | 0.718223 |
| | | Wild_plot_02_2246 | 0.642305 | 0.649805 |
| | | Wild_plot_02_1520 | 0.7088 | 0.715795 |
| | | Wild_plot_02_1512 | 0.700284 | 0.706582 |

| | | | | |
|---|---|---|---|---|
| | plot 02 | Wild_plot_02_1519 | 0.705238 | 0.712051 |
| | | Wild_plot_02_1525 | 0.707361 | 0.716439 |
| | | Wild_plot_02_2088 | 0.710586 | 0.713202 |
| | | Wild_plot_02_1528 | 0.706615 | 0.716042 |
| | | Wild_plot_02_1822 | 0.715263 | 0.716882 |
| | | Wild_plot_02_1527 | 0.704402 | 0.710915 |
| | | Wild_plot_02_1521 | 0.705339 | 0.714774 |
| | | Wild_plot_02_1518 | 0.699992 | 0.709993 |
| | | Wild_plot_02_1513 | 0.700514 | 0.711362 |
| | | Wild_plot_02_1523 | 0.705658 | 0.711715 |
| | | Wild_plot_02_1510 | 0.706602 | 0.715454 |
| | | Wild_plot_02_1511 | 0.700767 | 0.712182 |
| | | Wild_plot_02_1524 | 0.700538 | 0.708902 |
| | | Wild_plot_02_1508 | 0.697828 | 0.706866 |
| | | Wild_plot_02_2089 | 0.713099 | 0.717711 |
| | plot 03 | Wild_plot_03_2091 | 0.706152 | 0.711455 |
| | | Wild_plot_03_1549 | 0.703034 | 0.708121 |
| | | Wild_plot_03_1541 | 0.705231 | 0.710783 |
| | | Wild_plot_03_1534 | 0.704281 | 0.709913 |
| | | Wild_plot_03_1557 | 0.705676 | 0.713244 |
| | | Wild_plot_03_1539 | 0.699782 | 0.705635 |
| | | Wild_plot_03_1558 | 0.703832 | 0.709162 |
| | | Wild_plot_03_1552 | 0.707174 | 0.715158 |
| | | Wild_plot_03_1540 | 0.70736 | 0.712646 |
| | | Wild_plot_03_1544 | 0.705523 | 0.710713 |
| | | Wild_plot_03_1551 | 0.701726 | 0.706619 |
| | | Wild_plot_03_1542 | 0.702709 | 0.708243 |
| | | Wild_plot_03_1536 | 0.700476 | 0.708554 |
| | | Wild_plot_03_1560 | 0.701639 | 0.708877 |
| | | Wild_plot_03_1535 | 0.708615 | 0.71489 |
| | | Wild_plot_03_1538 | 0.707561 | 0.713351 |
| | | Wild_plot_03_1545 | 0.708209 | 0.714931 |
| | | Wild_plot_03_2092 | 0.710833 | 0.717197 |
| | | Wild_plot_03_2090 | 0.710608 | 0.718346 |
| | | Wild_plot_03_1533 | 0.705101 | 0.708407 |
| | | Wild_plot_03_1537 | 0.704646 | 0.707027 |
| | | Wild_plot_03_1547 | 0.70843 | 0.712542 |
| | | Wild_plot_03_1559 | 0.703656 | 0.70831 |
| | | Wild_plot_03_1548 | 0.697155 | 0.703758 |
| | | Wild_plot_03_2093 | 0.697502 | 0.698892 |
| | | Wild_plot_03_1550 | 0.69244 | 0.701688 |
| | | Wild_plot_03_1556 | 0.696967 | 0.704383 |
| | | Wild_plot_03_1546 | 0.69748 | 0.708393 |
| | | Wild_plot_03_1554 | 0.690807 | 0.698889 |
| | | Wild_plot_03_1555 | 0.701435 | 0.706469 |
| | | Wild_plot_03_1543 | 0.698077 | 0.704933 |

| | | | | |
|---|---|---|---|---|
| | | Wild_plot_03_1532 | 0.708131 | 0.7114 |
| | plot 04 | Wild_plot_04_1567 | 0.704108 | 0.712498 |
| | | Wild_plot_04_1566 | 0.703246 | 0.711138 |
| | | Wild_plot_04_1564 | 0.709227 | 0.716372 |
| | | Wild_plot_04_1571 | 0.71341 | 0.719862 |
| | | Wild_plot_04_1573 | 0.710311 | 0.714434 |
| | | Wild_plot_04_1570 | 0.704957 | 0.709664 |
| | | Wild_plot_04_1569 | 0.707338 | 0.712007 |
| | | Wild_plot_04_1568 | 0.700948 | 0.707566 |
| | | Wild_plot_04_1563 | 0.710615 | 0.715314 |
| | | Wild_plot_04_1572 | 0.709578 | 0.720291 |
| | | Wild_plot_04_1565 | 0.708849 | 0.716176 |
| | plot 05 | Wild_plot_05_1580 | 0.707136 | 0.712174 |
| | | Wild_plot_05_1578 | 0.708108 | 0.715129 |
| | | Wild_plot_05_1584 | 0.707723 | 0.717945 |
| | | Wild_plot_05_1581 | 0.708201 | 0.713003 |
| | | Wild_plot_05_1575 | 0.711563 | 0.720687 |
| | | Wild_plot_05_1574 | 0.709 | 0.717484 |
| | | Wild_plot_05_1577 | 0.712676 | 0.718713 |
| | | Wild_plot_05_1576 | 0.708478 | 0.716742 |
| | | Wild_plot_05_1579 | 0.708866 | 0.712147 |
| | | Wild_plot_05_1583 | 0.706541 | 0.713953 |
| | | Wild_plot_05_1582 | 0.708906 | 0.715871 |
| | plot 06 | Wild_plot_06_1588 | 0.71146 | 0.717989 |
| | | Wild_plot_06_1589 | 0.708164 | 0.713679 |
| | | Wild_plot_06_1592 | 0.709156 | 0.715878 |
| | | Wild_plot_06_1590 | 0.706827 | 0.713369 |
| | | Wild_plot_06_1591 | 0.705535 | 0.709693 |
| | | Wild_plot_06_1593 | 0.711749 | 0.718095 |
| | | Wild_plot_06_1597 | 0.709931 | 0.717559 |
| | | Wild_plot_06_1599 | 0.705295 | 0.714714 |
| | | Wild_plot_06_1595 | 0.711202 | 0.717479 |
| | | Wild_plot_06_1598 | 0.710047 | 0.719421 |
| | | Wild_plot_06_1594 | 0.704508 | 0.709349 |
| | | Wild_plot_06_1596 | 0.706897 | 0.715225 |
| | | Wild_plot_06_1587 | 0.702761 | 0.710866 |
| | | Wild_plot_06_1586 | 0.714728 | 0.72137 |
| | plot 07 | Wild_plot_07_1607 | 0.706216 | 0.716178 |
| | | Wild_plot_07_1611 | 0.709175 | 0.715665 |
| | | Wild_plot_07_1605 | 0.709453 | 0.714867 |
| | | Wild_plot_07_1606 | 0.713336 | 0.719276 |
| | | Wild_plot_07_1608 | 0.712435 | 0.721987 |
| | | Wild_plot_07_1609 | 0.711059 | 0.722081 |
| | | Wild_plot_07_1604 | 0.710028 | 0.713137 |
| | | Wild_plot_07_1602 | 0.709994 | 0.714492 |
| | | Wild_plot_07_1603 | 0.703331 | 0.712272 |

| | | | | |
|---|---|---|---|---|
| Yangambi rainforest | | Wild_plot_07_1610 | 0.693725 | 0.698322 |
| | | Wild_plot_07_1612 | 0.750041 | 0.753497 |
| | plot 08 | Wild_plot_08_1620 | 0.706934 | 0.709496 |
| | | Wild_plot_08_1616 | 0.711975 | 0.712268 |
| | | Wild_plot_08_1619 | 0.704796 | 0.708233 |
| | | Wild_plot_08_1618 | 0.708394 | 0.711458 |
| | | Wild_plot_08_1615 | 0.704187 | 0.710104 |
| | | Wild_plot_08_1617 | 0.701897 | 0.710189 |
| | plot 09 | Wild_plot_09_1625 | 0.705404 | 0.712025 |
| | | Wild_plot_09_1630 | 0.714058 | 0.72116 |
| | | Wild_plot_09_1629 | 0.710774 | 0.717545 |
| | | Wild_plot_09_1623 | 0.710133 | 0.718123 |
| | | Wild_plot_09_1633 | 0.712274 | 0.720566 |
| | | Wild_plot_09_1624 | 0.709379 | 0.719733 |
| | | Wild_plot_09_1631 | 0.706689 | 0.71468 |
| | | Wild_plot_09_2095 | 0.708768 | 0.713489 |
| | | Wild_plot_09_1621 | 0.710838 | 0.716542 |
| | | Wild_plot_09_1632 | 0.70921 | 0.714 |
| | | Wild_plot_09_2094 | 0.701576 | 0.70956 |
| | | Wild_plot_09_1626 | 0.705086 | 0.710298 |
| | | Wild_plot_09_1628 | 0.709044 | 0.714367 |
| | plot 10 | Wild_plot_10_1643 | 0.694501 | 0.703646 |
| | | Wild_plot_10_1634 | 0.706615 | 0.710265 |
| | | Wild_plot_10_1636 | 0.708981 | 0.71456 |
| | | Wild_plot_10_1638 | 0.710528 | 0.719476 |
| | | Wild_plot_10_1635 | 0.708034 | 0.714912 |
| | | Wild_plot_10_1637 | 0.702705 | 0.715018 |
| | | Wild_plot_10_1645 | 0.702983 | 0.709172 |
| | | Wild_plot_10_1639 | 0.70041 | 0.711677 |
| | | Wild_plot_10_1646 | 0.76263 | 0.769481 |
| | | Wild_plot_10_1642 | 0.788616 | 0.797597 |
| | | Wild_plot_10_1641 | 0.776487 | 0.779137 |
| | | Wild_plot_10_1644 | 0.793909 | 0.780721 |
| | | Wild_plot_10_1640 | 0.791423 | 0.792473 |
| | plot 11 | Wild_plot_11_1653 | 0.702667 | 0.707608 |
| | | Wild_plot_11_1650 | 0.703135 | 0.71179 |
| | | Wild_plot_11_1656 | 0.705156 | 0.712481 |
| | | Wild_plot_11_1647 | 0.710352 | 0.711301 |
| | | Wild_plot_11_2098 | 0.703124 | 0.709693 |
| | | Wild_plot_11_2096 | 0.709546 | 0.715078 |
| | | Wild_plot_11_1658 | 0.704918 | 0.711874 |
| | | Wild_plot_11_1648 | 0.70653 | 0.71101 |
| | | Wild_plot_11_1651 | 0.698374 | 0.705803 |
| | | Wild_plot_11_1652 | 0.700615 | 0.704047 |
| | | Wild_plot_11_1657 | 0.704072 | 0.71364 |
| | | Wild_plot_12_1661 | 0.706588 | 0.713835 |

| | | | |
|---|---|---|---|
| plot 12 | Wild_plot_12_1674 | 0.705514 | 0.71542 |
| | Wild_plot_12_1664 | 0.705699 | 0.713368 |
| | Wild_plot_12_1666 | 0.707476 | 0.720191 |
| | Wild_plot_12_1669 | 0.705831 | 0.714332 |
| | Wild_plot_12_1673 | 0.705695 | 0.711491 |
| | Wild_plot_12_1659 | 0.702894 | 0.711758 |
| | Wild_plot_12_1668 | 0.705892 | 0.716794 |
| | Wild_plot_12_1663 | 0.70594 | 0.715658 |
| | Wild_plot_12_1665 | 0.704597 | 0.712627 |
| | Wild_plot_12_1662 | 0.707331 | 0.713537 |
| | Wild_plot_12_1667 | 0.70267 | 0.714007 |
| | Wild_plot_12_1671 | 0.701655 | 0.713424 |
| | Wild_plot_12_1670 | 0.698264 | 0.709831 |
| | Wild_plot_12_1660 | 0.699085 | 0.7054 |
| plot 13 | Wild_plot_13_1678 | 0.7077 | 0.714484 |
| | Wild_plot_13_1683 | 0.71256 | 0.722884 |
| | Wild_plot_13_1681 | 0.710091 | 0.715822 |
| | Wild_plot_13_1676 | 0.71082 | 0.71807 |
| | Wild_plot_13_1677 | 0.702367 | 0.712855 |
| | Wild_plot_13_1680 | 0.705351 | 0.715965 |
| | Wild_plot_13_1685 | 0.701718 | 0.711264 |
| | Wild_plot_13_1679 | 0.705387 | 0.717818 |
| | Wild_plot_13_1682 | 0.708559 | 0.719506 |
| | Wild_plot_13_1675 | 0.705562 | 0.714004 |
| | Wild_plot_13_1684 | 0.70553 | 0.715419 |
| | Wild_plot_13_1686 | 0.80424 | 0.785882 |
| plot 14 | Wild_plot_14_1696 | 0.709797 | 0.718026 |
| | Wild_plot_14_1690 | 0.711925 | 0.715149 |
| | Wild_plot_14_1689 | 0.711913 | 0.715389 |
| | Wild_plot_14_1687 | 0.710077 | 0.714046 |
| | Wild_plot_14_1695 | 0.705828 | 0.714549 |
| | Wild_plot_14_1693 | 0.704661 | 0.712471 |
| | Wild_plot_14_1694 | 0.707313 | 0.717015 |
| | Wild_plot_14_1692 | 0.704337 | 0.7131 |
| | Wild_plot_14_1688 | 0.704744 | 0.708797 |
| | Wild_plot_14_1691 | 0.7064 | 0.710089 |
| | Wild_plot_14_1697 | 0.708006 | 0.717052 |
| plot 15 | Wild_plot_15_1705 | 0.712114 | 0.717903 |
| | Wild_plot_15_1702 | 0.709139 | 0.712904 |
| | Wild_plot_15_1707 | 0.708773 | 0.712071 |
| | Wild_plot_15_1704 | 0.70335 | 0.708091 |
| | Wild_plot_15_1706 | 0.711246 | 0.715156 |
| | Wild_plot_15_1703 | 0.703912 | 0.711942 |
| | Wild_plot_16_1827 | 0.704183 | 0.711758 |
| | Wild_plot_16_1830 | 0.707469 | 0.71495 |
| | Wild_plot_16_1824 | 0.710992 | 0.711782 |

| | | | | |
|---|---|---|---|---|
| | plot 16 | Wild_plot_16_1832 | 0.70406 | 0.712744 |
| | | Wild_plot_16_1829 | 0.705538 | 0.712062 |
| | | Wild_plot_16_1828 | 0.699912 | 0.712271 |
| | | Wild_plot_16_1826 | 0.65652 | 0.67428 |
| | plot 17 | Wild_plot_17_1833 | 0.701393 | 0.711277 |
| | | Wild_plot_17_1835 | 0.704867 | 0.709935 |
| | | Wild_plot_17_1834 | 0.70425 | 0.712354 |
| | plot 18 | Wild_plot_18_1836 | 0.704687 | 0.709882 |
| | | Wild_plot_18_1840 | 0.709808 | 0.715764 |
| | | Wild_plot_18_1841 | 0.705035 | 0.714252 |
| | | Wild_plot_18_1838 | 0.708217 | 0.713098 |
| | | Wild_plot_18_1839 | 0.708294 | 0.71519 |
| | | Wild_plot_18_1837 | 0.710037 | 0.716157 |
| | plot 19 | Wild_plot_19_1851 | 0.704912 | 0.712936 |
| | | Wild_plot_19_1850 | 0.711297 | 0.719133 |
| | | Wild_plot_19_1842 | 0.700832 | 0.710719 |
| | | Wild_plot_19_1848 | 0.706354 | 0.715009 |
| | | Wild_plot_19_1846 | 0.708248 | 0.713638 |
| | | Wild_plot_19_1844 | 0.698535 | 0.707966 |
| | | Wild_plot_19_1852 | 0.707271 | 0.712443 |
| | | Wild_plot_19_1854 | 0.705579 | 0.714856 |
| | | Wild_plot_19_1843 | 0.706511 | 0.716084 |
| | plot 20 | Wild_plot_20_2168 | 0.690922 | 0.701016 |
| | | Wild_plot_20_2170 | 0.72468 | 0.715479 |
| | | Wild_plot_20_2166 | 0.742266 | 0.756092 |
| | | Wild_plot_20_2167 | 0.760727 | 0.734634 |
| | plot 21 | Wild_plot_21_2174 | 0.663535 | 0.669143 |
| | | Wild_plot_21_2176 | 0.65636 | 0.66979 |
| | | Wild_plot_21_2172 | 0.661223 | 0.664452 |
| | | Wild_plot_21_2185 | 0.669754 | 0.684971 |
| | | Wild_plot_21_2173 | 0.648356 | 0.661477 |
| | | Wild_plot_21_2186 | 0.645135 | 0.649671 |
| | | Wild_plot_21_2175 | 0.637809 | 0.650115 |
| | | Wild_plot_21_2171 | 0.722929 | 0.724215 |
| | plot 22 | Wild_plot_22_2190 | 0.698673 | 0.707002 |
| | | Wild_plot_22_2196 | 0.674874 | 0.679079 |
| | | Wild_plot_22_2200 | 0.663464 | 0.674849 |
| | | Wild_plot_22_2197 | 0.653929 | 0.661799 |
| | plot 23 | Wild_plot_23_2203 | 0.673183 | 0.681699 |
| | | Wild_plot_23_2202 | 0.666956 | 0.676199 |
| | | Wild_plot_23_2201 | 0.656331 | 0.668732 |
| | | Wild_plot_23_2204 | 0.656891 | 0.666063 |
| | | Wild_plot_23_2208 | 0.655442 | 0.662009 |
| | | Wild_plot_23_2205 | 0.655153 | 0.660302 |
| | | Wild_plot_23_2206 | 0.653799 | 0.654949 |
| | | Wild_plot_23_2207 | 0.646665 | 0.651065 |

| | | | | |
|---|---|---|---|---|
| | plot 24 | Wild_plot_24_2213 | 0.685917 | 0.696809 |
| | | Wild_plot_24_2218 | 0.652572 | 0.662091 |
| | | Wild_plot_24_2214 | 0.658273 | 0.662453 |
| | | Wild_plot_24_2224 | 0.659211 | 0.668853 |
| | | Wild_plot_24_2215 | 0.65345 | 0.659961 |
| | | Wild_plot_24_2212 | 0.65873 | 0.669915 |
| | | Wild_plot_24_2217 | 0.646552 | 0.661546 |
| | | Wild_plot_24_2211 | 0.645425 | 0.659434 |
| | | Wild_plot_24_2219 | 0.641791 | 0.650659 |
| | plot 25 | Wild_plot_25_2233 | 0.686674 | 0.694474 |
| | | Wild_plot_25_2231 | 0.68705 | 0.69158 |
| | | Wild_plot_25_2234 | 0.686272 | 0.689214 |
| | | Wild_plot_25_2232 | 0.669625 | 0.680254 |
| | | Wild_plot_25_2235 | 0.676783 | 0.689262 |
| | | Wild_plot_25_2230 | 0.671975 | 0.678061 |

ˈiduals based on 41 522 haplotypes within 10 045 polymorphic high-quality loci.
alues that were above the minimal JID and by this indicating genetically identical individuals.

| HG_02 | | | | HG_03 | HG_04 | HG_05 | | HG_06 |
|---|---|---|---|---|---|---|---|---|
| Home_garc | Home_garc | Home_garc | Home_gard | Home_gard | Home_gard | Home_garc | Home_gard | Home_garc |
| 0.783621 | 0.785562 | 0.782962 | 0.779546 | 0.710025 | 0.705975 | 0.784427 | 0.788511 | 0.783585 |
| 0.794251 | 0.78633 | 0.771092 | 0.775609 | 0.718979 | 0.71748 | 0.784264 | 0.780227 | 0.776086 |
| 1 | 0.793707 | 0.775782 | 0.766988 | 0.705868 | 0.701352 | 0.783321 | 0.780824 | 0.777705 |
| 0.793707 | 1 | 0.780892 | 0.793067 | 0.715585 | 0.710481 | 0.784822 | 0.77726 | 0.786551 |
| 0.775782 | 0.780892 | 1 | 0.778503 | 0.709147 | 0.706873 | 0.77577 | 0.779464 | 0.780637 |
| 0.766988 | 0.793067 | 0.778503 | 1 | 0.718787 | 0.718752 | 0.783726 | 0.765873 | 0.767258 |
| 0.705868 | 0.715585 | 0.709147 | 0.718787 | 1 | 0.785679 | 0.716458 | 0.709754 | 0.715173 |
| 0.701352 | 0.710481 | 0.706873 | 0.718752 | 0.785679 | 1 | 0.709529 | 0.70867 | 0.705962 |
| 0.783321 | 0.784822 | 0.77577 | 0.783726 | 0.716458 | 0.709529 | 1 | 0.777705 | 0.779356 |
| 0.780824 | 0.77726 | 0.779464 | 0.765873 | 0.709754 | 0.70867 | 0.777705 | 1 | 0.79176 |
| 0.777705 | 0.786551 | 0.780637 | 0.767258 | 0.715173 | 0.705962 | 0.779356 | 0.79176 | 1 |
| 0.788953 | 0.784488 | 0.783944 | 0.772408 | 0.704459 | 0.699338 | 0.781472 | 0.784309 | 0.783008 |
| 0.778106 | 0.776262 | 0.767971 | 0.770939 | 0.713514 | 0.70729 | 0.777998 | 0.769363 | 0.772843 |
| 0.782787 | 0.786807 | 0.77677 | 0.771233 | 0.723701 | 0.720526 | 0.774582 | 0.785285 | 0.78045 |
| 0.764106 | 0.760793 | 0.767133 | 0.768752 | 0.73394 | 0.730576 | 0.765337 | 0.766561 | 0.76228 |
| 0.793209 | 0.787241 | 0.783886 | 0.782598 | 0.716645 | 0.715875 | 0.783336 | 0.786407 | 0.785939 |
| 0.791373 | 0.788759 | 0.767451 | 0.767813 | 0.715718 | 0.712505 | 0.773337 | 0.79638 | 0.786748 |
| 0.750997 | 0.760291 | 0.755902 | 0.753808 | 0.742343 | 0.741286 | 0.758587 | 0.753281 | 0.755194 |
| 0.789883 | 0.783229 | 0.770563 | 0.767872 | 0.707002 | 0.701668 | 0.78572 | 0.774584 | 0.775405 |
| 0.781743 | 0.791832 | 0.781039 | 0.781065 | 0.715013 | 0.708227 | 0.78637 | 0.773384 | 0.780218 |
| 0.775902 | 0.788158 | 0.778709 | 0.784281 | 0.71407 | 0.706194 | 0.788937 | 0.779502 | 0.780546 |
| 0.790884 | 0.793872 | 0.786774 | 0.776714 | 0.711723 | 0.709143 | 0.782727 | 0.798806 | 0.790964 |
| 0.787919 | 0.798035 | 0.78436 | 0.7885 | 0.718077 | 0.712722 | 0.792732 | 0.785797 | 0.791068 |
| 0.787225 | 0.784141 | 0.779081 | 0.78521 | 0.717979 | 0.711971 | 0.786696 | 0.790313 | 0.790875 |
| 0.779061 | 0.782333 | 0.783008 | 0.780379 | 0.716685 | 0.71187 | 0.787536 | 0.773583 | 0.776194 |
| 0.787002 | 0.787187 | 0.778304 | 0.768284 | 0.712157 | 0.705577 | 0.784768 | 0.778766 | 0.775967 |
| 0.771597 | 0.781101 | 0.771648 | 0.766195 | 0.703982 | 0.701938 | 0.775919 | 0.771424 | 0.772873 |
| 0.785363 | 0.784438 | 0.780573 | 0.776965 | 0.711974 | 0.70984 | 0.783631 | 0.773925 | 0.777186 |
| 0.781847 | 0.787558 | 0.780251 | 0.782313 | 0.716518 | 0.714502 | 0.780041 | 0.77622 | 0.787291 |
| 0.778776 | 0.774678 | 0.77443 | 0.775441 | 0.713419 | 0.709058 | 0.778902 | 0.788492 | 0.778911 |
| 0.77297 | 0.779697 | 0.782856 | 0.781098 | 0.710973 | 0.708244 | 0.783918 | 0.772671 | 0.778008 |
| 0.775041 | 0.788155 | 0.776228 | 0.770894 | 0.706852 | 0.705033 | 0.783379 | 0.771012 | 0.768979 |
| 0.78232 | 0.785025 | 0.776062 | 0.779102 | 0.714421 | 0.704115 | 0.780965 | 0.776796 | 0.775619 |
| 0.773912 | 0.784745 | 0.778214 | 0.77698 | 0.709514 | 0.708241 | 0.777395 | 0.770059 | 0.770953 |
| 0.778736 | 0.77961 | 0.765436 | 0.767321 | 0.705599 | 0.703305 | 0.769717 | 0.766209 | 0.762583 |
| 0.782903 | 0.789748 | 0.773793 | 0.772768 | 0.711482 | 0.708972 | 0.782212 | 0.774887 | 0.782493 |
| 0.781602 | 0.780323 | 0.77381 | 0.780805 | 0.716522 | 0.711859 | 0.784249 | 0.777524 | 0.779044 |
| 0.763844 | 0.773259 | 0.769663 | 0.778246 | 0.717703 | 0.711988 | 0.772252 | 0.762775 | 0.774234 |
| 0.76955 | 0.78422 | 0.774898 | 0.780877 | 0.71882 | 0.713345 | 0.778417 | 0.77498 | 0.774776 |

| | | | | | | | | |
|---|---|---|---|---|---|---|---|---|
| 0.760978 | 0.760443 | 0.754564 | 0.751417 | 0.748219 | 0.744701 | 0.759948 | 0.766394 | 0.765436 |
| 0.78358 | 0.780157 | 0.780478 | 0.771747 | 0.704099 | 0.698338 | 0.773325 | 0.795212 | 0.782469 |
| 0.7899 | 0.784863 | 0.784241 | 0.776911 | 0.709025 | 0.704787 | 0.784099 | 0.788847 | 0.792203 |
| 0.793886 | 0.799458 | 0.787886 | 0.773044 | 0.719458 | 0.708232 | 0.783123 | 0.810822 | 0.802321 |
| 0.786497 | 0.784627 | 0.781371 | 0.779153 | 0.71313 | 0.706505 | 0.784424 | 0.778534 | 0.785691 |
| 0.776497 | 0.78953 | 0.778502 | 0.781949 | 0.71627 | 0.71195 | 0.786693 | 0.784289 | 0.782861 |
| 0.767402 | 0.775037 | 0.780354 | 0.780685 | 0.709586 | 0.7026 | 0.782079 | 0.774207 | 0.767754 |
| 0.776927 | 0.787871 | 0.777613 | 0.781461 | 0.716184 | 0.709809 | 0.793129 | 0.775723 | 0.783721 |
| 0.774433 | 0.775567 | 0.774807 | 0.769597 | 0.714834 | 0.708377 | 0.782442 | 0.776538 | 0.777593 |
| 0.77014 | 0.789569 | 0.766445 | 0.776381 | 0.711136 | 0.711018 | 0.787741 | 0.770292 | 0.769912 |
| 0.783625 | 0.7869 | 0.790025 | 0.780502 | 0.705397 | 0.700731 | 0.781165 | 0.788893 | 0.783111 |
| 0.782573 | 0.787824 | 0.784463 | 0.783984 | 0.70902 | 0.705138 | 0.788895 | 0.785768 | 0.787372 |
| 0.803286 | 0.787026 | 0.783544 | 0.768777 | 0.712457 | 0.706517 | 0.782678 | 0.793636 | 0.788223 |
| 0.785945 | 0.789713 | 0.78085 | 0.783133 | 0.707891 | 0.706943 | 0.7905 | 0.787949 | 0.785756 |
| 0.786 | 0.782125 | 0.778238 | 0.768813 | 0.707906 | 0.703324 | 0.778639 | 0.782349 | 0.77966 |
| 0.789695 | 0.772237 | 0.77998 | 0.77252 | 0.707484 | 0.701515 | 0.782269 | 0.779363 | 0.774204 |
| 0.789395 | 0.791029 | 0.774007 | 0.766211 | 0.709491 | 0.701537 | 0.772554 | 0.784762 | 0.778975 |
| 0.799039 | 0.792353 | 0.788094 | 0.780406 | 0.714263 | 0.707841 | 0.786931 | 0.795675 | 0.793221 |
| 0.78037 | 0.784964 | 0.785485 | 0.775395 | 0.714024 | 0.714047 | 0.783405 | 0.809759 | 0.787751 |
| 0.789496 | 0.784609 | 0.777594 | 0.767366 | 0.708626 | 0.705272 | 0.782065 | 0.792569 | 0.773157 |
| 0.801925 | 0.795279 | 0.78312 | 0.770488 | 0.710671 | 0.706281 | 0.780066 | 0.788897 | 0.788678 |
| 0.794118 | 0.776639 | 0.780627 | 0.767239 | 0.709844 | 0.702977 | 0.779949 | 0.794066 | 0.796176 |
| 0.781848 | 0.783007 | 0.769793 | 0.771063 | 0.712978 | 0.705799 | 0.821989 | 0.785 | 0.77482 |
| 0.775309 | 0.788166 | 0.770012 | 0.784519 | 0.718395 | 0.711586 | 0.789601 | 0.771321 | 0.778421 |
| 0.758476 | 0.7761 | 0.766713 | 0.778267 | 0.727019 | 0.719469 | 0.777641 | 0.759153 | 0.763458 |
| 0.702829 | 0.710698 | 0.707393 | 0.718401 | 0.786387 | 0.987885 | 0.711828 | 0.708297 | 0.706396 |
| 0.707711 | 0.71504 | 0.708919 | 0.719221 | 0.776718 | 0.77208 | 0.715011 | 0.709739 | 0.711687 |
| 0.709045 | 0.718928 | 0.71166 | 0.719923 | 0.776327 | 0.773126 | 0.717344 | 0.709703 | 0.712562 |
| 0.772623 | 0.778087 | 0.767968 | 0.77266 | 0.708727 | 0.705337 | 0.766948 | 0.767664 | 0.769041 |
| 0.770035 | 0.770804 | 0.771457 | 0.772382 | 0.706799 | 0.704297 | 0.773808 | 0.769879 | 0.762969 |
| 0.766168 | 0.777003 | 0.777942 | 0.77295 | 0.715014 | 0.712679 | 0.773274 | 0.7806 | 0.767778 |
| 0.772392 | 0.786321 | 0.772761 | 0.77397 | 0.720352 | 0.714055 | 0.778028 | 0.769091 | 0.766175 |
| 0.816223 | 0.795795 | 0.774579 | 0.773841 | 0.709406 | 0.705373 | 0.79132 | 0.795647 | 0.789602 |
| 0.794775 | 0.792258 | 0.781628 | 0.772697 | 0.710974 | 0.706702 | 0.782001 | 0.780908 | 0.791355 |
| 0.803073 | 0.79013 | 0.772958 | 0.777371 | 0.707484 | 0.705551 | 0.787057 | 0.785936 | 0.776298 |
| 0.824656 | 0.798158 | 0.77487 | 0.775247 | 0.713032 | 0.70848 | 0.786889 | 0.786201 | 0.792032 |
| 0.789538 | 0.783665 | 0.77568 | 0.782258 | 0.710684 | 0.706035 | 0.790947 | 0.782168 | 0.769143 |
| 0.810675 | 0.813048 | 0.803671 | 0.793494 | 0.723956 | 0.718286 | 0.803094 | 0.820614 | 0.810288 |
| 0.792148 | 0.782092 | 0.766218 | 0.762669 | 0.702892 | 0.702228 | 0.770639 | 0.782889 | 0.771778 |
| 0.776529 | 0.781156 | 0.781163 | 0.777687 | 0.710107 | 0.706067 | 0.782213 | 0.773737 | 0.781163 |
| 0.793608 | 0.789702 | 0.774432 | 0.770828 | 0.707294 | 0.702546 | 0.773885 | 0.786241 | 0.778362 |
| 0.815346 | 0.820049 | 0.811303 | 0.797974 | 0.727702 | 0.721898 | 0.805632 | 0.824637 | 0.811875 |
| 0.788012 | 0.796439 | 0.789817 | 0.774709 | 0.710075 | 0.708661 | 0.782152 | 0.803755 | 0.792297 |
| 0.784164 | 0.797987 | 0.801452 | 0.774117 | 0.710508 | 0.703217 | 0.779108 | 0.797527 | 0.794307 |
| 0.782655 | 0.783029 | 0.78258 | 0.784983 | 0.713168 | 0.705671 | 0.784604 | 0.782461 | 0.780778 |
| 0.801292 | 0.783271 | 0.773502 | 0.768952 | 0.709257 | 0.705095 | 0.78113 | 0.781625 | 0.781655 |

| | | | | | | | | |
|---|---|---|---|---|---|---|---|---|
| 0.787648 | 0.787959 | 0.789104 | 0.781227 | 0.710244 | 0.704919 | 0.780968 | 0.80637 | 0.781102 |
| 0.772244 | 0.776834 | 0.770502 | 0.778849 | 0.713347 | 0.711173 | 0.79816 | 0.775059 | 0.778914 |
| 0.796984 | 0.78306 | 0.779022 | 0.786327 | 0.710153 | 0.708026 | 0.791499 | 0.779609 | 0.781083 |
| 0.796146 | 0.801108 | 0.799361 | 0.792178 | 0.720087 | 0.713653 | 0.791239 | 0.801399 | 0.788934 |
| 0.785292 | 0.788063 | 0.780366 | 0.790682 | 0.715486 | 0.711921 | 0.797249 | 0.780123 | 0.778711 |
| 0.777842 | 0.773423 | 0.777163 | 0.772806 | 0.707678 | 0.709708 | 0.786508 | 0.76878 | 0.768169 |
| 0.781467 | 0.786216 | 0.772602 | 0.781517 | 0.710978 | 0.710983 | 0.795949 | 0.783516 | 0.773924 |
| 0.781833 | 0.777714 | 0.778577 | 0.788579 | 0.713931 | 0.709139 | 0.800854 | 0.775109 | 0.777105 |
| 0.778521 | 0.78665 | 0.781911 | 0.773591 | 0.711614 | 0.711397 | 0.777455 | 0.779048 | 0.77631 |
| 0.775309 | 0.774863 | 0.775974 | 0.768515 | 0.707383 | 0.705794 | 0.778786 | 0.772117 | 0.770513 |
| 0.781025 | 0.777453 | 0.778984 | 0.775026 | 0.707281 | 0.707691 | 0.78027 | 0.772188 | 0.768723 |
| 0.769709 | 0.775199 | 0.777537 | 0.770735 | 0.708046 | 0.704381 | 0.77367 | 0.769804 | 0.774536 |
| 0.776474 | 0.773899 | 0.781201 | 0.770091 | 0.717139 | 0.709595 | 0.780502 | 0.778743 | 0.773873 |
| 0.765423 | 0.776246 | 0.775167 | 0.772625 | 0.707199 | 0.701938 | 0.774339 | 0.764169 | 0.772028 |
| 0.783533 | 0.778564 | 0.774632 | 0.771937 | 0.713658 | 0.708244 | 0.784286 | 0.775303 | 0.773267 |
| 0.782823 | 0.778095 | 0.777887 | 0.776581 | 0.709127 | 0.707523 | 0.779254 | 0.768911 | 0.769218 |
| 0.76954 | 0.7853 | 0.773941 | 0.776871 | 0.711981 | 0.710541 | 0.783209 | 0.776105 | 0.777649 |
| 0.793875 | 0.780376 | 0.77539 | 0.778542 | 0.710395 | 0.710993 | 0.777924 | 0.771756 | 0.78022 |
| 0.773869 | 0.776811 | 0.777869 | 0.775626 | 0.712013 | 0.709677 | 0.78289 | 0.772768 | 0.769015 |
| 0.775689 | 0.779119 | 0.777732 | 0.780885 | 0.711095 | 0.709288 | 0.776659 | 0.774879 | 0.77537 |
| 0.774008 | 0.774501 | 0.767236 | 0.773007 | 0.708184 | 0.707725 | 0.772847 | 0.779889 | 0.76676 |
| 0.77342 | 0.776559 | 0.773452 | 0.770774 | 0.705417 | 0.703165 | 0.775517 | 0.771114 | 0.770315 |
| 0.778275 | 0.775002 | 0.773557 | 0.781333 | 0.71016 | 0.705676 | 0.784906 | 0.769193 | 0.778435 |
| 0.770947 | 0.77532 | 0.773526 | 0.779521 | 0.711882 | 0.708779 | 0.779761 | 0.771466 | 0.769989 |
| 0.775904 | 0.771946 | 0.765737 | 0.769835 | 0.714228 | 0.710754 | 0.77723 | 0.781016 | 0.764765 |
| 0.781989 | 0.792315 | 0.77319 | 0.78768 | 0.718003 | 0.714049 | 0.804963 | 0.793996 | 0.782881 |
| 0.77717 | 0.774388 | 0.764482 | 0.766624 | 0.71187 | 0.708886 | 0.77961 | 0.785896 | 0.766171 |
| 0.774528 | 0.781365 | 0.771207 | 0.776581 | 0.715577 | 0.708416 | 0.777313 | 0.774054 | 0.769136 |
| 0.779118 | 0.781199 | 0.778424 | 0.778077 | 0.709657 | 0.71003 | 0.780488 | 0.773798 | 0.765398 |
| 0.777722 | 0.774049 | 0.772254 | 0.774996 | 0.713732 | 0.708897 | 0.785294 | 0.773804 | 0.762273 |
| 0.770677 | 0.776074 | 0.772806 | 0.778474 | 0.712878 | 0.709842 | 0.776118 | 0.770384 | 0.76622 |
| 0.775608 | 0.784054 | 0.766656 | 0.775817 | 0.711436 | 0.701713 | 0.777288 | 0.771719 | 0.768911 |
| 0.781765 | 0.785137 | 0.77506 | 0.771351 | 0.713527 | 0.709094 | 0.790287 | 0.78014 | 0.774875 |
| 0.79098 | 0.79637 | 0.784478 | 0.768806 | 0.714927 | 0.70459 | 0.779171 | 0.807975 | 0.800892 |
| 0.778017 | 0.777448 | 0.774911 | 0.766997 | 0.7095 | 0.706897 | 0.779581 | 0.772851 | 0.773632 |
| 0.784894 | 0.783262 | 0.777648 | 0.778676 | 0.713305 | 0.710611 | 0.783516 | 0.78742 | 0.774898 |
| 0.782339 | 0.778318 | 0.776251 | 0.776949 | 0.706993 | 0.704035 | 0.777433 | 0.769669 | 0.773882 |
| 0.744535 | 0.750443 | 0.750418 | 0.761351 | 0.685948 | 0.675216 | 0.75273 | 0.752285 | 0.757474 |
| 0.761298 | 0.757588 | 0.76181 | 0.758316 | 0.691678 | 0.686996 | 0.767168 | 0.756396 | 0.76192 |
| 0.744778 | 0.757472 | 0.750826 | 0.75754 | 0.682247 | 0.674242 | 0.76048 | 0.741597 | 0.742154 |
| 0.779041 | 0.766916 | 0.755212 | 0.748775 | 0.683199 | 0.677344 | 0.767523 | 0.757261 | 0.759114 |
| 0.733809 | 0.737792 | 0.740217 | 0.730599 | 0.665428 | 0.659904 | 0.736747 | 0.737167 | 0.745437 |
| 0.758001 | 0.760369 | 0.765957 | 0.744545 | 0.669239 | 0.659853 | 0.755063 | 0.777622 | 0.765661 |
| 0.780216 | 0.748422 | 0.742456 | 0.740422 | 0.668838 | 0.665134 | 0.760478 | 0.753265 | 0.753541 |
| 0.741186 | 0.741497 | 0.745337 | 0.752781 | 0.668779 | 0.659787 | 0.75345 | 0.731762 | 0.741327 |
| 0.769493 | 0.756553 | 0.75068 | 0.746848 | 0.666419 | 0.665424 | 0.763782 | 0.758531 | 0.754199 |
| 0.768832 | 0.74984 | 0.740306 | 0.745411 | 0.676836 | 0.670571 | 0.760031 | 0.752031 | 0.752344 |

| | | | | | | | | |
|---|---|---|---|---|---|---|---|---|
| 0.737177 | 0.739862 | 0.740237 | 0.741294 | 0.673129 | 0.666544 | 0.742171 | 0.731562 | 0.734955 |
| 0.7877 | 0.767945 | 0.736068 | 0.745103 | 0.674105 | 0.670211 | 0.758466 | 0.766175 | 0.759767 |
| 0.734087 | 0.748922 | 0.744051 | 0.740419 | 0.675762 | 0.667242 | 0.748613 | 0.73164 | 0.738322 |
| 0.753753 | 0.746769 | 0.760182 | 0.732828 | 0.663562 | 0.658245 | 0.743301 | 0.780966 | 0.751236 |
| 0.743658 | 0.750568 | 0.746866 | 0.737985 | 0.667608 | 0.654499 | 0.744111 | 0.752881 | 0.7499 |
| 0.760478 | 0.73818 | 0.735008 | 0.760074 | 0.666621 | 0.663384 | 0.772767 | 0.735428 | 0.741163 |
| 0.748555 | 0.747157 | 0.751296 | 0.738565 | 0.664413 | 0.661302 | 0.746036 | 0.75159 | 0.742711 |
| 0.762135 | 0.754584 | 0.743177 | 0.74275 | 0.662765 | 0.663777 | 0.748615 | 0.752565 | 0.754324 |
| 0.719867 | 0.723063 | 0.727438 | 0.7307 | 0.660992 | 0.659877 | 0.733597 | 0.721607 | 0.723779 |
| 0.761231 | 0.758975 | 0.738218 | 0.740517 | 0.671437 | 0.657987 | 0.753092 | 0.764986 | 0.745513 |
| 0.749033 | 0.743163 | 0.740451 | 0.750913 | 0.655098 | 0.650907 | 0.768687 | 0.736392 | 0.745772 |
| 0.765587 | 0.767893 | 0.778719 | 0.761136 | 0.697263 | 0.695221 | 0.773299 | 0.79044 | 0.786624 |
| 0.766161 | 0.778257 | 0.768032 | 0.77167 | 0.70856 | 0.706831 | 0.77836 | 0.769731 | 0.770361 |
| 0.750763 | 0.763133 | 0.744273 | 0.759268 | 0.687309 | 0.679416 | 0.767558 | 0.748912 | 0.748566 |
| 0.768848 | 0.78001 | 0.775675 | 0.774948 | 0.716624 | 0.714412 | 0.776301 | 0.778723 | 0.783661 |
| 0.757991 | 0.768428 | 0.770622 | 0.761808 | 0.703422 | 0.705831 | 0.769212 | 0.754991 | 0.758643 |
| 0.77737 | 0.772809 | 0.77844 | 0.769866 | 0.714511 | 0.714184 | 0.777523 | 0.775006 | 0.771288 |
| 0.774036 | 0.785868 | 0.772218 | 0.77929 | 0.721923 | 0.716808 | 0.782316 | 0.775271 | 0.774338 |
| 0.756877 | 0.754926 | 0.757058 | 0.7625 | 0.695428 | 0.692752 | 0.763955 | 0.753607 | 0.757731 |
| 0.764078 | 0.783832 | 0.769162 | 0.780091 | 0.71832 | 0.714793 | 0.774183 | 0.769381 | 0.783555 |
| 0.750279 | 0.759174 | 0.736752 | 0.766016 | 0.693482 | 0.687988 | 0.776269 | 0.741428 | 0.752944 |
| 0.764143 | 0.773058 | 0.775443 | 0.778124 | 0.717334 | 0.718287 | 0.774066 | 0.774055 | 0.76819 |
| 0.767585 | 0.775006 | 0.772761 | 0.777038 | 0.720757 | 0.719151 | 0.776108 | 0.775807 | 0.768351 |
| 0.7358 | 0.757088 | 0.734624 | 0.742646 | 0.688029 | 0.68492 | 0.762118 | 0.734869 | 0.75087 |
| 0.756391 | 0.75214 | 0.742802 | 0.742908 | 0.68574 | 0.685886 | 0.745669 | 0.749342 | 0.748173 |
| 0.762209 | 0.760767 | 0.748514 | 0.759168 | 0.69287 | 0.688543 | 0.763913 | 0.744984 | 0.752106 |
| 0.76713 | 0.777694 | 0.760113 | 0.775755 | 0.70834 | 0.708255 | 0.779506 | 0.755391 | 0.762796 |
| 0.770908 | 0.78237 | 0.770725 | 0.777932 | 0.723552 | 0.715102 | 0.791801 | 0.764797 | 0.772824 |
| 0.740829 | 0.749741 | 0.749261 | 0.742494 | 0.685381 | 0.683396 | 0.750921 | 0.74401 | 0.737807 |
| 0.766853 | 0.780706 | 0.764141 | 0.770112 | 0.723445 | 0.720917 | 0.770782 | 0.790816 | 0.770297 |
| 0.769324 | 0.764241 | 0.763858 | 0.76513 | 0.727416 | 0.725603 | 0.771666 | 0.763801 | 0.76215 |
| 0.765947 | 0.76448 | 0.755335 | 0.74812 | 0.694945 | 0.693865 | 0.75388 | 0.771727 | 0.763563 |
| 0.765425 | 0.7615 | 0.765419 | 0.755457 | 0.696583 | 0.693666 | 0.755233 | 0.773565 | 0.759259 |
| 0.758872 | 0.78603 | 0.765985 | 0.778122 | 0.725736 | 0.721311 | 0.784555 | 0.766501 | 0.775224 |
| 0.756651 | 0.763982 | 0.763814 | 0.766208 | 0.723125 | 0.717274 | 0.765192 | 0.769669 | 0.772239 |
| 0.759464 | 0.770516 | 0.765062 | 0.769803 | 0.724528 | 0.720205 | 0.769675 | 0.767545 | 0.765868 |
| 0.723224 | 0.722442 | 0.719604 | 0.724097 | 0.677499 | 0.670559 | 0.736155 | 0.716491 | 0.725216 |
| 0.751892 | 0.760333 | 0.751391 | 0.763982 | 0.718824 | 0.719197 | 0.757134 | 0.752747 | 0.749555 |
| 0.733449 | 0.731903 | 0.72977 | 0.733863 | 0.673611 | 0.673134 | 0.732175 | 0.722109 | 0.728954 |
| 0.727152 | 0.717344 | 0.726319 | 0.724475 | 0.671175 | 0.669037 | 0.72996 | 0.711593 | 0.71962 |
| 0.763385 | 0.776307 | 0.763296 | 0.765552 | 0.72173 | 0.717467 | 0.770521 | 0.766089 | 0.766018 |
| 0.713346 | 0.714927 | 0.720692 | 0.718547 | 0.672853 | 0.667322 | 0.722456 | 0.709735 | 0.710496 |
| 0.727153 | 0.719285 | 0.716209 | 0.705671 | 0.662794 | 0.658052 | 0.715576 | 0.727118 | 0.716397 |
| 0.709529 | 0.709087 | 0.709554 | 0.702949 | 0.659275 | 0.655456 | 0.706463 | 0.715251 | 0.703956 |
| 0.767334 | 0.770731 | 0.765196 | 0.75595 | 0.729125 | 0.724877 | 0.771291 | 0.793123 | 0.791488 |
| 0.715177 | 0.709455 | 0.708359 | 0.702727 | 0.664143 | 0.659099 | 0.710075 | 0.714286 | 0.710549 |
| 0.76147 | 0.770575 | 0.759288 | 0.761029 | 0.720418 | 0.718301 | 0.769667 | 0.777778 | 0.762763 |

| | | | | | | | | |
|---|---|---|---|---|---|---|---|---|
| 0.763994 | 0.772536 | 0.756655 | 0.763251 | 0.731329 | 0.727121 | 0.76422 | 0.769618 | 0.756512 |
| 0.763962 | 0.765715 | 0.757791 | 0.758957 | 0.724494 | 0.723309 | 0.768534 | 0.776571 | 0.768841 |
| 0.74588 | 0.749838 | 0.750081 | 0.755592 | 0.748577 | 0.742069 | 0.747291 | 0.751061 | 0.748367 |
| 0.714918 | 0.702909 | 0.697548 | 0.697257 | 0.654261 | 0.64979 | 0.702584 | 0.708029 | 0.702034 |
| 0.704195 | 0.696282 | 0.692008 | 0.696289 | 0.655393 | 0.651428 | 0.702424 | 0.698092 | 0.69795 |
| 0.76833 | 0.773356 | 0.762767 | 0.766669 | 0.735021 | 0.726953 | 0.771667 | 0.768849 | 0.768739 |
| 0.689092 | 0.693175 | 0.694524 | 0.688937 | 0.651955 | 0.652232 | 0.688781 | 0.69119 | 0.692495 |
| 0.728298 | 0.745996 | 0.734046 | 0.74108 | 0.740229 | 0.732624 | 0.737775 | 0.729773 | 0.730994 |
| 0.761368 | 0.762738 | 0.760877 | 0.755662 | 0.728046 | 0.730927 | 0.765058 | 0.763611 | 0.760592 |
| 0.71843 | 0.729266 | 0.722308 | 0.734047 | 0.738352 | 0.732285 | 0.725382 | 0.720635 | 0.71573 |
| 0.750611 | 0.753826 | 0.745359 | 0.75136 | 0.733603 | 0.730999 | 0.756659 | 0.763571 | 0.753196 |
| 0.643836 | 0.655844 | 0.623529 | 0.625 | 0.716216 | 0.671053 | 0.6125 | 0.610778 | 0.621118 |
| 0.64387 | 0.661449 | 0.657945 | 0.663793 | 0.724378 | 0.722543 | 0.661836 | 0.656455 | 0.656851 |
| 0.650784 | 0.656694 | 0.652265 | 0.661919 | 0.725582 | 0.7199 | 0.659146 | 0.647572 | 0.654746 |
| 0.657629 | 0.667828 | 0.662912 | 0.668243 | 0.729242 | 0.722897 | 0.663048 | 0.663945 | 0.667344 |
| 0.657117 | 0.668922 | 0.658094 | 0.669416 | 0.728614 | 0.733683 | 0.665845 | 0.660336 | 0.668479 |
| 0.624986 | 0.623583 | 0.626071 | 0.618902 | 0.590337 | 0.587097 | 0.617818 | 0.616643 | 0.626661 |
| 0.659935 | 0.668672 | 0.663661 | 0.670411 | 0.728607 | 0.728597 | 0.672927 | 0.667002 | 0.668378 |
| 0.732903 | 0.743031 | 0.734432 | 0.745924 | 0.737657 | 0.732988 | 0.739187 | 0.749206 | 0.740883 |
| 0.735684 | 0.745706 | 0.734701 | 0.741579 | 0.736501 | 0.735998 | 0.740152 | 0.738816 | 0.744933 |
| 0.741497 | 0.749206 | 0.741377 | 0.746586 | 0.74249 | 0.738364 | 0.75382 | 0.747528 | 0.749365 |
| 0.619287 | 0.618461 | 0.624053 | 0.620868 | 0.591931 | 0.594366 | 0.625262 | 0.618412 | 0.6201 |
| 0.599508 | 0.600364 | 0.605117 | 0.59891 | 0.580863 | 0.580954 | 0.601224 | 0.599603 | 0.599555 |
| 0.69817 | 0.709654 | 0.705533 | 0.713333 | 0.763402 | 0.759407 | 0.707813 | 0.704606 | 0.706604 |
| 0.700039 | 0.708551 | 0.703777 | 0.717407 | 0.765694 | 0.762551 | 0.70831 | 0.705445 | 0.708327 |
| 0.693772 | 0.702574 | 0.697726 | 0.707109 | 0.760254 | 0.752527 | 0.701704 | 0.700128 | 0.702022 |
| 0.702008 | 0.706946 | 0.701968 | 0.714478 | 0.761088 | 0.756651 | 0.706732 | 0.704281 | 0.704281 |
| 0.703543 | 0.712379 | 0.707681 | 0.718142 | 0.762452 | 0.758879 | 0.708142 | 0.704535 | 0.709036 |
| 0.701582 | 0.710623 | 0.70592 | 0.715134 | 0.765334 | 0.767054 | 0.707644 | 0.704022 | 0.708185 |
| 0.70019 | 0.707868 | 0.704322 | 0.714661 | 0.766382 | 0.763644 | 0.711912 | 0.70434 | 0.707389 |
| 0.702524 | 0.710487 | 0.703927 | 0.713622 | 0.76053 | 0.752258 | 0.709239 | 0.704103 | 0.706714 |
| 0.708258 | 0.72017 | 0.711887 | 0.724943 | 0.771559 | 0.767917 | 0.721084 | 0.714894 | 0.714637 |
| 0.703349 | 0.714766 | 0.711984 | 0.716644 | 0.766755 | 0.756761 | 0.713295 | 0.713242 | 0.711475 |
| 0.705062 | 0.715117 | 0.707246 | 0.717953 | 0.765742 | 0.7644 | 0.717764 | 0.711595 | 0.710674 |
| 0.706393 | 0.717357 | 0.709999 | 0.721776 | 0.768003 | 0.769212 | 0.719225 | 0.71466 | 0.713795 |
| 0.703192 | 0.711806 | 0.706149 | 0.716564 | 0.759649 | 0.760612 | 0.715032 | 0.705488 | 0.709245 |
| 0.704861 | 0.714889 | 0.710145 | 0.716105 | 0.768356 | 0.760272 | 0.712823 | 0.708719 | 0.711063 |
| 0.70806 | 0.717471 | 0.710042 | 0.720631 | 0.766531 | 0.762883 | 0.713877 | 0.711423 | 0.711932 |
| 0.701137 | 0.709438 | 0.703887 | 0.713576 | 0.759071 | 0.763551 | 0.708412 | 0.704555 | 0.706499 |
| 0.704748 | 0.713271 | 0.710963 | 0.716144 | 0.761544 | 0.759329 | 0.710266 | 0.71172 | 0.709636 |
| 0.650548 | 0.659317 | 0.657128 | 0.667507 | 0.72538 | 0.716467 | 0.664467 | 0.652489 | 0.658849 |
| 0.657914 | 0.666059 | 0.666849 | 0.666667 | 0.727136 | 0.724864 | 0.667031 | 0.659675 | 0.669408 |
| 0.708639 | 0.716121 | 0.712308 | 0.722147 | 0.766686 | 0.760673 | 0.717694 | 0.709504 | 0.710938 |
| 0.708686 | 0.720353 | 0.717881 | 0.724703 | 0.772623 | 0.768594 | 0.714457 | 0.712609 | 0.716687 |
| 0.636081 | 0.654903 | 0.645351 | 0.653988 | 0.713438 | 0.715881 | 0.651257 | 0.645914 | 0.647331 |
| 0.703698 | 0.716433 | 0.712785 | 0.721275 | 0.763569 | 0.7633 | 0.715578 | 0.711023 | 0.714377 |
| 0.696668 | 0.705234 | 0.699064 | 0.709403 | 0.761532 | 0.756965 | 0.705719 | 0.703119 | 0.702468 |

| | | | | | | | | |
|---|---|---|---|---|---|---|---|---|
| 0.702658 | 0.710462 | 0.707375 | 0.714172 | 0.765544 | 0.760408 | 0.712803 | 0.709908 | 0.710358 |
| 0.70391 | 0.713029 | 0.712507 | 0.718457 | 0.766713 | 0.766134 | 0.714455 | 0.713586 | 0.713526 |
| 0.705378 | 0.712546 | 0.708671 | 0.718717 | 0.762 | 0.758681 | 0.715079 | 0.710824 | 0.710094 |
| 0.701703 | 0.714297 | 0.708756 | 0.722784 | 0.76689 | 0.764441 | 0.716568 | 0.712195 | 0.714069 |
| 0.708103 | 0.719122 | 0.713754 | 0.721591 | 0.76918 | 0.764773 | 0.722031 | 0.715876 | 0.714628 |
| 0.699548 | 0.712268 | 0.707055 | 0.716126 | 0.766551 | 0.763292 | 0.715767 | 0.708656 | 0.706546 |
| 0.701256 | 0.711519 | 0.706151 | 0.716138 | 0.768271 | 0.7604 | 0.711644 | 0.711519 | 0.714604 |
| 0.69838 | 0.706663 | 0.70307 | 0.709763 | 0.756636 | 0.752514 | 0.707954 | 0.701528 | 0.703742 |
| 0.698014 | 0.710334 | 0.706409 | 0.714955 | 0.765173 | 0.757132 | 0.70895 | 0.709811 | 0.709004 |
| 0.702092 | 0.709183 | 0.70585 | 0.715265 | 0.75879 | 0.756615 | 0.713702 | 0.703998 | 0.707963 |
| 0.706731 | 0.713441 | 0.70855 | 0.718192 | 0.763786 | 0.764202 | 0.713299 | 0.711745 | 0.712922 |
| 0.701911 | 0.709727 | 0.706776 | 0.711548 | 0.766672 | 0.762826 | 0.71014 | 0.70631 | 0.70783 |
| 0.697292 | 0.70087 | 0.696402 | 0.709362 | 0.756835 | 0.752974 | 0.707183 | 0.700698 | 0.698567 |
| 0.69547 | 0.707275 | 0.700322 | 0.70932 | 0.758267 | 0.75234 | 0.705951 | 0.70192 | 0.699743 |
| 0.70622 | 0.718782 | 0.71298 | 0.721389 | 0.764438 | 0.76132 | 0.720714 | 0.707377 | 0.714638 |
| 0.707248 | 0.711963 | 0.70943 | 0.716726 | 0.768729 | 0.763349 | 0.714013 | 0.709887 | 0.710646 |
| 0.700204 | 0.706311 | 0.704411 | 0.713407 | 0.764786 | 0.763136 | 0.707849 | 0.702548 | 0.70499 |
| 0.701341 | 0.712455 | 0.707972 | 0.719464 | 0.761678 | 0.760272 | 0.711475 | 0.708187 | 0.706805 |
| 0.703739 | 0.710406 | 0.707767 | 0.717072 | 0.764639 | 0.758983 | 0.709772 | 0.70484 | 0.706466 |
| 0.702274 | 0.710098 | 0.705093 | 0.715866 | 0.76319 | 0.760216 | 0.706758 | 0.704554 | 0.706715 |
| 0.694957 | 0.707055 | 0.700395 | 0.713131 | 0.756705 | 0.757487 | 0.704566 | 0.703584 | 0.702953 |
| 0.703354 | 0.707467 | 0.704863 | 0.714713 | 0.761476 | 0.756122 | 0.70706 | 0.701951 | 0.704921 |
| 0.707235 | 0.714511 | 0.706003 | 0.718052 | 0.768442 | 0.763291 | 0.709626 | 0.706753 | 0.709808 |
| 0.703595 | 0.713232 | 0.710116 | 0.720653 | 0.764908 | 0.760964 | 0.71205 | 0.708933 | 0.711565 |
| 0.700809 | 0.712013 | 0.70711 | 0.715526 | 0.764153 | 0.762256 | 0.710107 | 0.706374 | 0.703994 |
| 0.700375 | 0.708759 | 0.702845 | 0.712333 | 0.763794 | 0.76161 | 0.707892 | 0.701876 | 0.706751 |
| 0.697479 | 0.709755 | 0.707272 | 0.716668 | 0.757034 | 0.752791 | 0.706614 | 0.704279 | 0.704269 |
| 0.698776 | 0.706994 | 0.700742 | 0.711963 | 0.755336 | 0.755131 | 0.704082 | 0.699927 | 0.704558 |
| 0.699953 | 0.707872 | 0.70412 | 0.717001 | 0.761573 | 0.761211 | 0.706753 | 0.701825 | 0.706585 |
| 0.700426 | 0.712366 | 0.708314 | 0.716433 | 0.762029 | 0.762497 | 0.710819 | 0.704542 | 0.707955 |
| 0.703598 | 0.712575 | 0.709381 | 0.720013 | 0.762598 | 0.760203 | 0.713989 | 0.705929 | 0.71051 |
| 0.705351 | 0.714721 | 0.710752 | 0.720502 | 0.763321 | 0.758925 | 0.712133 | 0.708771 | 0.709292 |
| 0.707509 | 0.715602 | 0.709594 | 0.723377 | 0.767444 | 0.767784 | 0.715474 | 0.708875 | 0.710835 |
| 0.707184 | 0.715158 | 0.711046 | 0.722452 | 0.768538 | 0.766855 | 0.715357 | 0.708839 | 0.712698 |
| 0.698043 | 0.704884 | 0.702889 | 0.713273 | 0.762222 | 0.760691 | 0.707605 | 0.705097 | 0.706877 |
| 0.700714 | 0.710147 | 0.707246 | 0.715137 | 0.761308 | 0.760375 | 0.706713 | 0.705441 | 0.7065 |
| 0.7046 | 0.712298 | 0.707454 | 0.716264 | 0.7661 | 0.759364 | 0.710799 | 0.707611 | 0.7099 |
| 0.700665 | 0.710942 | 0.704567 | 0.713013 | 0.760882 | 0.756927 | 0.708159 | 0.702586 | 0.702507 |
| 0.696016 | 0.704421 | 0.700121 | 0.711061 | 0.7548 | 0.755284 | 0.702819 | 0.698631 | 0.698148 |
| 0.693734 | 0.700707 | 0.695472 | 0.70292 | 0.75999 | 0.7524 | 0.699434 | 0.696371 | 0.700367 |
| 0.689276 | 0.702102 | 0.695503 | 0.705897 | 0.755821 | 0.748483 | 0.700025 | 0.695969 | 0.699153 |
| 0.696875 | 0.69979 | 0.697084 | 0.705134 | 0.753753 | 0.750977 | 0.700692 | 0.696681 | 0.700763 |
| 0.69722 | 0.707538 | 0.699545 | 0.712017 | 0.752971 | 0.75316 | 0.70334 | 0.698099 | 0.702167 |
| 0.690838 | 0.697561 | 0.693162 | 0.703292 | 0.750202 | 0.748722 | 0.696978 | 0.691091 | 0.692716 |
| 0.698769 | 0.710234 | 0.701463 | 0.714799 | 0.76087 | 0.753882 | 0.707211 | 0.701357 | 0.705414 |
| 0.692162 | 0.698945 | 0.6967 | 0.705947 | 0.751754 | 0.74476 | 0.700119 | 0.694687 | 0.700092 |

| | | | | | | | | |
|---|---|---|---|---|---|---|---|---|
| 0.705026 | 0.712866 | 0.709549 | 0.718104 | 0.765107 | 0.761212 | 0.715947 | 0.706475 | 0.709757 |
| 0.703244 | 0.715378 | 0.710709 | 0.716999 | 0.775609 | 0.771626 | 0.710427 | 0.706623 | 0.708918 |
| 0.70315 | 0.708427 | 0.704893 | 0.712223 | 0.772595 | 0.772193 | 0.710202 | 0.705727 | 0.702764 |
| 0.704792 | 0.715242 | 0.710852 | 0.718577 | 0.765887 | 0.766408 | 0.711945 | 0.709281 | 0.707958 |
| 0.708541 | 0.722059 | 0.714683 | 0.724765 | 0.775232 | 0.77972 | 0.71867 | 0.714593 | 0.715769 |
| 0.702275 | 0.714737 | 0.711271 | 0.717837 | 0.770318 | 0.764658 | 0.715161 | 0.707972 | 0.710392 |
| 0.700214 | 0.711093 | 0.705276 | 0.712776 | 0.767825 | 0.761317 | 0.713313 | 0.707618 | 0.707548 |
| 0.705139 | 0.7151 | 0.711099 | 0.72149 | 0.763312 | 0.76171 | 0.710503 | 0.710014 | 0.711275 |
| 0.695521 | 0.707942 | 0.699359 | 0.710487 | 0.768389 | 0.761014 | 0.705944 | 0.702362 | 0.704213 |
| 0.706541 | 0.715606 | 0.711508 | 0.716695 | 0.767037 | 0.766839 | 0.711532 | 0.708588 | 0.707074 |
| 0.704179 | 0.714106 | 0.713297 | 0.717937 | 0.772513 | 0.767967 | 0.716174 | 0.712497 | 0.713382 |
| 0.70658 | 0.718333 | 0.714695 | 0.721239 | 0.77234 | 0.766585 | 0.71252 | 0.710409 | 0.710748 |
| 0.699717 | 0.710899 | 0.708742 | 0.718138 | 0.787113 | 0.779727 | 0.711077 | 0.708099 | 0.70833 |
| 0.705654 | 0.711324 | 0.710189 | 0.718172 | 0.779964 | 0.778061 | 0.712055 | 0.708551 | 0.710952 |
| 0.706662 | 0.715928 | 0.710978 | 0.720875 | 0.777841 | 0.784484 | 0.712579 | 0.709423 | 0.711918 |
| 0.703607 | 0.714546 | 0.709848 | 0.717412 | 0.77796 | 0.779196 | 0.712103 | 0.713242 | 0.712682 |
| 0.708291 | 0.717666 | 0.711277 | 0.721192 | 0.986099 | 0.787737 | 0.717197 | 0.713082 | 0.71657 |
| 0.70474 | 0.711375 | 0.708708 | 0.717328 | 0.781456 | 0.780697 | 0.714387 | 0.710906 | 0.712442 |
| 0.70942 | 0.719451 | 0.715553 | 0.721452 | 0.782641 | 0.780877 | 0.718429 | 0.714819 | 0.713944 |
| 0.708736 | 0.717137 | 0.710002 | 0.717739 | 0.781273 | 0.785843 | 0.712553 | 0.710759 | 0.709197 |
| 0.702748 | 0.713273 | 0.709621 | 0.717525 | 0.76899 | 0.774622 | 0.713062 | 0.710818 | 0.709452 |
| 0.702258 | 0.709654 | 0.708752 | 0.718043 | 0.774401 | 0.773329 | 0.710159 | 0.709242 | 0.706716 |
| 0.704285 | 0.718074 | 0.711113 | 0.720363 | 0.778925 | 0.779053 | 0.715182 | 0.712932 | 0.713763 |
| 0.706963 | 0.716232 | 0.711711 | 0.718199 | 0.777062 | 0.7756 | 0.718658 | 0.71315 | 0.714082 |
| 0.704946 | 0.714934 | 0.709839 | 0.716981 | 0.77457 | 0.77433 | 0.712378 | 0.707468 | 0.708769 |
| 0.705279 | 0.713924 | 0.706403 | 0.716392 | 0.774241 | 0.769256 | 0.710024 | 0.71046 | 0.710759 |
| 0.70198 | 0.710027 | 0.707704 | 0.715734 | 0.769493 | 0.766992 | 0.711088 | 0.709223 | 0.708442 |
| 0.699329 | 0.713971 | 0.706611 | 0.71542 | 0.773699 | 0.773985 | 0.710458 | 0.707379 | 0.711805 |
| 0.704209 | 0.717176 | 0.712048 | 0.717444 | 0.780773 | 0.775053 | 0.719574 | 0.709908 | 0.711523 |
| 0.707444 | 0.714553 | 0.708272 | 0.720955 | 0.778662 | 0.766933 | 0.716373 | 0.712152 | 0.715837 |
| 0.701885 | 0.712365 | 0.708857 | 0.714929 | 0.771398 | 0.764681 | 0.71276 | 0.70974 | 0.708333 |
| 0.708529 | 0.719284 | 0.71384 | 0.721211 | 0.776226 | 0.768768 | 0.713588 | 0.711786 | 0.711382 |
| 0.706796 | 0.717142 | 0.712918 | 0.718866 | 0.775933 | 0.770728 | 0.715788 | 0.712272 | 0.713934 |
| 0.699172 | 0.710475 | 0.706853 | 0.713388 | 0.776611 | 0.771409 | 0.708937 | 0.706748 | 0.705458 |
| 0.707265 | 0.715535 | 0.70918 | 0.717194 | 0.769711 | 0.768964 | 0.714896 | 0.709522 | 0.708525 |
| 0.697563 | 0.709135 | 0.706502 | 0.710025 | 0.772857 | 0.770598 | 0.708139 | 0.704121 | 0.707264 |
| 0.709976 | 0.722165 | 0.712432 | 0.725314 | 0.78585 | 0.77958 | 0.722435 | 0.716542 | 0.718573 |
| 0.701807 | 0.713444 | 0.706816 | 0.718364 | 0.777183 | 0.770195 | 0.710906 | 0.708179 | 0.709222 |
| 0.701982 | 0.718261 | 0.706221 | 0.720149 | 0.769704 | 0.768529 | 0.714743 | 0.708489 | 0.710808 |
| 0.706205 | 0.713743 | 0.710063 | 0.714931 | 0.778383 | 0.768562 | 0.712906 | 0.708431 | 0.711506 |
| 0.710096 | 0.720534 | 0.713948 | 0.719779 | 0.779223 | 0.76815 | 0.716043 | 0.713755 | 0.71618 |
| 0.704829 | 0.716469 | 0.709688 | 0.718785 | 0.777183 | 0.771074 | 0.717081 | 0.709922 | 0.714503 |
| 0.707905 | 0.716813 | 0.711394 | 0.71863 | 0.776806 | 0.774523 | 0.71909 | 0.711151 | 0.714988 |
| 0.707667 | 0.711602 | 0.70787 | 0.712867 | 0.769745 | 0.773482 | 0.711848 | 0.709458 | 0.709926 |
| 0.708824 | 0.714297 | 0.711871 | 0.717922 | 0.776303 | 0.776666 | 0.715825 | 0.707531 | 0.710239 |
| 0.7039 | 0.711869 | 0.7074 | 0.715218 | 0.772177 | 0.771281 | 0.711822 | 0.702117 | 0.707972 |

| | | | | | | | | |
|---|---|---|---|---|---|---|---|---|
| 0.690294 | 0.700191 | 0.695057 | 0.70647 | 0.764684 | 0.767822 | 0.700623 | 0.697396 | 0.699377 |
| 0.745167 | 0.760332 | 0.743996 | 0.763728 | 0.755422 | 0.749014 | 0.756911 | 0.742787 | 0.750577 |
| 0.701661 | 0.710849 | 0.708676 | 0.716582 | 0.768423 | 0.761845 | 0.709159 | 0.705543 | 0.709293 |
| 0.706976 | 0.714263 | 0.71252 | 0.722292 | 0.775434 | 0.772764 | 0.717965 | 0.706552 | 0.709642 |
| 0.699109 | 0.705091 | 0.706057 | 0.713862 | 0.76678 | 0.762663 | 0.705167 | 0.70454 | 0.705702 |
| 0.701498 | 0.710484 | 0.709978 | 0.718278 | 0.770338 | 0.767061 | 0.709234 | 0.706953 | 0.7091 |
| 0.701375 | 0.711499 | 0.706543 | 0.71608 | 0.776294 | 0.767086 | 0.710862 | 0.704306 | 0.704326 |
| 0.697949 | 0.704606 | 0.704786 | 0.710367 | 0.765635 | 0.762503 | 0.706475 | 0.700239 | 0.698692 |
| 0.704225 | 0.711267 | 0.712489 | 0.718881 | 0.776986 | 0.769795 | 0.709315 | 0.704245 | 0.706801 |
| 0.708049 | 0.717148 | 0.716269 | 0.725573 | 0.776386 | 0.772634 | 0.716038 | 0.713865 | 0.710421 |
| 0.704824 | 0.716372 | 0.711824 | 0.716652 | 0.771382 | 0.774088 | 0.714422 | 0.70873 | 0.710958 |
| 0.707216 | 0.714343 | 0.711941 | 0.721664 | 0.772705 | 0.769738 | 0.71867 | 0.713614 | 0.712599 |
| 0.708729 | 0.718983 | 0.714399 | 0.721772 | 0.776126 | 0.772143 | 0.718785 | 0.709501 | 0.7103 |
| 0.705836 | 0.717432 | 0.711884 | 0.721708 | 0.775464 | 0.771795 | 0.71669 | 0.713665 | 0.713605 |
| 0.704677 | 0.713215 | 0.708374 | 0.715724 | 0.77379 | 0.770816 | 0.710998 | 0.708384 | 0.70863 |
| 0.704188 | 0.714163 | 0.70766 | 0.714174 | 0.773945 | 0.76999 | 0.710912 | 0.707985 | 0.710634 |
| 0.706949 | 0.715512 | 0.714183 | 0.719465 | 0.770899 | 0.77003 | 0.714663 | 0.712133 | 0.713289 |
| 0.704918 | 0.714286 | 0.712415 | 0.71718 | 0.76804 | 0.764029 | 0.714792 | 0.711744 | 0.709763 |
| 0.698667 | 0.704637 | 0.701653 | 0.706818 | 0.764455 | 0.764669 | 0.703777 | 0.698934 | 0.701329 |
| 0.698821 | 0.708844 | 0.706567 | 0.712144 | 0.771488 | 0.769846 | 0.710382 | 0.701488 | 0.703305 |
| 0.705628 | 0.715651 | 0.710084 | 0.713621 | 0.771886 | 0.767322 | 0.7108 | 0.710245 | 0.71175 |
| 0.696642 | 0.703195 | 0.699366 | 0.706055 | 0.764017 | 0.758548 | 0.701013 | 0.704212 | 0.701934 |
| 0.702609 | 0.714184 | 0.706678 | 0.714704 | 0.77002 | 0.758972 | 0.714948 | 0.706902 | 0.708197 |
| 0.707797 | 0.715175 | 0.708704 | 0.71915 | 0.77058 | 0.761149 | 0.715593 | 0.711272 | 0.709806 |
| 0.710238 | 0.714448 | 0.71146 | 0.721083 | 0.769882 | 0.770149 | 0.715582 | 0.712868 | 0.711051 |
| 0.703885 | 0.716377 | 0.708851 | 0.715104 | 0.767399 | 0.760864 | 0.71337 | 0.711219 | 0.710183 |
| 0.703231 | 0.709966 | 0.703667 | 0.709253 | 0.763229 | 0.761252 | 0.707997 | 0.703334 | 0.706494 |
| 0.700443 | 0.707726 | 0.701363 | 0.711978 | 0.766851 | 0.759864 | 0.70297 | 0.705829 | 0.706661 |
| 0.699357 | 0.708082 | 0.70316 | 0.710707 | 0.763144 | 0.759969 | 0.70617 | 0.703729 | 0.705783 |
| 0.758043 | 0.760242 | 0.769868 | 0.759079 | 0.699807 | 0.693145 | 0.766396 | 0.75702 | 0.7596 |
| 0.785994 | 0.782425 | 0.78262 | 0.779947 | 0.709135 | 0.706565 | 0.792041 | 0.785804 | 0.776248 |
| 0.772276 | 0.776143 | 0.778385 | 0.774239 | 0.706291 | 0.703106 | 0.791077 | 0.775971 | 0.775833 |
| 0.790293 | 0.780702 | 0.787628 | 0.773341 | 0.697521 | 0.693677 | 0.783137 | 0.802225 | 0.786518 |
| 0.789863 | 0.789707 | 0.781194 | 0.788164 | 0.708991 | 0.703336 | 0.804143 | 0.785433 | 0.77682 |
| 0.696091 | 0.706027 | 0.702291 | 0.709733 | 0.765866 | 0.763987 | 0.706672 | 0.698386 | 0.701633 |
| 0.699784 | 0.709062 | 0.704871 | 0.712851 | 0.770993 | 0.770592 | 0.709863 | 0.705193 | 0.704152 |
| 0.703345 | 0.711444 | 0.706622 | 0.714962 | 0.771341 | 0.769942 | 0.712012 | 0.703189 | 0.707183 |
| 0.701989 | 0.71249 | 0.708675 | 0.719821 | 0.77165 | 0.770081 | 0.714446 | 0.708244 | 0.707799 |
| 0.700231 | 0.706121 | 0.703916 | 0.715671 | 0.770211 | 0.766334 | 0.707822 | 0.702263 | 0.70324 |
| 0.70143 | 0.711816 | 0.706773 | 0.716438 | 0.769903 | 0.767987 | 0.713711 | 0.707492 | 0.708996 |
| 0.699369 | 0.711001 | 0.706553 | 0.713689 | 0.768796 | 0.767628 | 0.710675 | 0.704535 | 0.707081 |
| 0.700289 | 0.70921 | 0.705086 | 0.715233 | 0.763353 | 0.759563 | 0.708384 | 0.705308 | 0.707206 |
| 0.695235 | 0.706213 | 0.700939 | 0.707758 | 0.764038 | 0.760138 | 0.705247 | 0.69736 | 0.703238 |
| 0.696368 | 0.706891 | 0.701238 | 0.706088 | 0.76569 | 0.760664 | 0.702916 | 0.701597 | 0.701723 |
| 0.698994 | 0.709672 | 0.704174 | 0.715348 | 0.768733 | 0.765945 | 0.707832 | 0.705351 | 0.705805 |
| 0.702222 | 0.71158 | 0.708608 | 0.717589 | 0.768975 | 0.767246 | 0.711167 | 0.705944 | 0.707349 |

| | | | | | | | | |
|---|---|---|---|---|---|---|---|---|
| 0.702085 | 0.714506 | 0.706274 | 0.719456 | 0.767667 | 0.768106 | 0.711556 | 0.707119 | 0.70676 |
| 0.703209 | 0.712858 | 0.702984 | 0.715494 | 0.763011 | 0.765041 | 0.708223 | 0.706253 | 0.712449 |
| 0.706822 | 0.71547 | 0.710395 | 0.716816 | 0.770681 | 0.772447 | 0.713188 | 0.709924 | 0.712158 |
| 0.697956 | 0.710392 | 0.708307 | 0.714936 | 0.763283 | 0.767194 | 0.707793 | 0.707391 | 0.710068 |
| 0.699052 | 0.708455 | 0.704861 | 0.71225 | 0.764232 | 0.762121 | 0.703773 | 0.702841 | 0.707146 |
| 0.703807 | 0.710989 | 0.705222 | 0.710799 | 0.766154 | 0.764156 | 0.704439 | 0.706467 | 0.705955 |
| 0.706318 | 0.71544 | 0.708818 | 0.717581 | 0.764316 | 0.765052 | 0.710895 | 0.710312 | 0.709736 |
| 0.703338 | 0.712796 | 0.709063 | 0.713774 | 0.769929 | 0.766978 | 0.711978 | 0.707722 | 0.710916 |
| 0.696916 | 0.708354 | 0.705787 | 0.712893 | 0.763922 | 0.768764 | 0.705743 | 0.705456 | 0.704547 |
| 0.703624 | 0.717493 | 0.707221 | 0.716613 | 0.76 | 0.766788 | 0.710904 | 0.708434 | 0.710984 |
| 0.699863 | 0.711375 | 0.705092 | 0.71296 | 0.765015 | 0.76421 | 0.708621 | 0.70391 | 0.710708 |
| 0.700572 | 0.711097 | 0.706591 | 0.712496 | 0.76252 | 0.76419 | 0.709961 | 0.704345 | 0.705261 |
| 0.693472 | 0.704739 | 0.699846 | 0.703545 | 0.762234 | 0.757013 | 0.7005 | 0.699031 | 0.697032 |
| 0.693075 | 0.704807 | 0.700351 | 0.707825 | 0.759299 | 0.756608 | 0.703994 | 0.700754 | 0.701136 |
| 0.705963 | 0.717763 | 0.705749 | 0.719381 | 0.768708 | 0.76646 | 0.709888 | 0.707268 | 0.711718 |
| 0.711085 | 0.719811 | 0.713326 | 0.722093 | 0.778467 | 0.780749 | 0.718149 | 0.712052 | 0.713919 |
| 0.706709 | 0.716081 | 0.709655 | 0.720602 | 0.778004 | 0.770509 | 0.712289 | 0.711155 | 0.713257 |
| 0.711713 | 0.718872 | 0.714559 | 0.724416 | 0.772241 | 0.775694 | 0.711674 | 0.714536 | 0.714618 |
| 0.70207 | 0.714509 | 0.703646 | 0.712782 | 0.768786 | 0.768784 | 0.705654 | 0.702063 | 0.705202 |
| 0.701242 | 0.709622 | 0.706945 | 0.713631 | 0.771927 | 0.76834 | 0.711032 | 0.707067 | 0.707317 |
| 0.701657 | 0.708519 | 0.704782 | 0.709895 | 0.766986 | 0.76649 | 0.706292 | 0.706433 | 0.704636 |
| 0.704809 | 0.713885 | 0.7077 | 0.719345 | 0.773127 | 0.769563 | 0.713604 | 0.708243 | 0.710168 |
| 0.711288 | 0.714391 | 0.70849 | 0.719651 | 0.777683 | 0.774199 | 0.715338 | 0.712065 | 0.712845 |
| 0.703169 | 0.709834 | 0.705723 | 0.71352 | 0.76239 | 0.759068 | 0.708822 | 0.703707 | 0.70723 |
| 0.702224 | 0.710421 | 0.703461 | 0.719283 | 0.772239 | 0.771188 | 0.711646 | 0.706696 | 0.70665 |
| 0.788381 | 0.794108 | 0.790731 | 0.777224 | 0.706491 | 0.703591 | 0.782857 | 0.784686 | 0.788192 |
| 0.704506 | 0.711134 | 0.707909 | 0.716023 | 0.775517 | 0.766438 | 0.710453 | 0.710684 | 0.710194 |
| 0.703222 | 0.713857 | 0.709446 | 0.715131 | 0.773781 | 0.766982 | 0.715482 | 0.709249 | 0.70918 |
| 0.707455 | 0.716159 | 0.710522 | 0.722548 | 0.774095 | 0.773349 | 0.714182 | 0.714366 | 0.716481 |
| 0.703127 | 0.714138 | 0.709469 | 0.721587 | 0.77373 | 0.766716 | 0.712671 | 0.707305 | 0.712794 |
| 0.701842 | 0.712083 | 0.70515 | 0.712031 | 0.76348 | 0.760319 | 0.709942 | 0.707539 | 0.710575 |
| 0.699517 | 0.710447 | 0.70425 | 0.714415 | 0.773141 | 0.771436 | 0.709268 | 0.705926 | 0.710852 |
| 0.703345 | 0.713615 | 0.710825 | 0.717076 | 0.77204 | 0.765553 | 0.708394 | 0.707205 | 0.71146 |
| 0.700648 | 0.711915 | 0.706861 | 0.714262 | 0.774414 | 0.762812 | 0.712095 | 0.705089 | 0.71096 |
| 0.700008 | 0.709391 | 0.703673 | 0.711795 | 0.771722 | 0.756561 | 0.708555 | 0.701391 | 0.710244 |
| 0.70017 | 0.708594 | 0.703163 | 0.712568 | 0.768067 | 0.75877 | 0.710935 | 0.705701 | 0.706719 |
| 0.707234 | 0.715085 | 0.709913 | 0.71728 | 0.780005 | 0.770346 | 0.715406 | 0.711028 | 0.711646 |
| 0.703621 | 0.717456 | 0.712786 | 0.722605 | 0.788306 | 0.779478 | 0.715956 | 0.71037 | 0.714746 |
| 0.706118 | 0.714618 | 0.710649 | 0.715054 | 0.782482 | 0.773277 | 0.714782 | 0.70543 | 0.711459 |
| 0.703364 | 0.714357 | 0.708838 | 0.716631 | 0.781871 | 0.777471 | 0.711272 | 0.709889 | 0.707613 |
| 0.698098 | 0.708654 | 0.706951 | 0.712781 | 0.774482 | 0.778448 | 0.710065 | 0.702073 | 0.703556 |
| 0.706924 | 0.715623 | 0.713443 | 0.720229 | 0.786495 | 0.779207 | 0.716264 | 0.707127 | 0.71332 |
| 0.699324 | 0.712544 | 0.704344 | 0.717761 | 0.781255 | 0.772794 | 0.709123 | 0.708032 | 0.710147 |
| 0.702094 | 0.7113 | 0.705326 | 0.714342 | 0.766176 | 0.761245 | 0.708492 | 0.704131 | 0.706918 |
| 0.705261 | 0.716074 | 0.707707 | 0.721574 | 0.768433 | 0.769713 | 0.711124 | 0.708124 | 0.708764 |
| 0.708221 | 0.713063 | 0.706992 | 0.720181 | 0.767834 | 0.766267 | 0.712701 | 0.710541 | 0.709474 |

| | | | | | | | | |
|---|---|---|---|---|---|---|---|---|
| 0.703456 | 0.713351 | 0.706095 | 0.714365 | 0.766912 | 0.766585 | 0.709823 | 0.706291 | 0.707536 |
| 0.705061 | 0.71255 | 0.71145 | 0.717579 | 0.768679 | 0.765786 | 0.711238 | 0.707304 | 0.708191 |
| 0.704293 | 0.710265 | 0.703461 | 0.712699 | 0.766146 | 0.76185 | 0.706841 | 0.704053 | 0.704904 |
| 0.658511 | 0.669705 | 0.663212 | 0.673043 | 0.724534 | 0.718706 | 0.664348 | 0.656423 | 0.668635 |
| 0.702363 | 0.711304 | 0.704633 | 0.715334 | 0.766243 | 0.76671 | 0.708993 | 0.706229 | 0.707564 |
| 0.700815 | 0.710535 | 0.705327 | 0.715555 | 0.76472 | 0.767518 | 0.708483 | 0.706158 | 0.706847 |
| 0.706952 | 0.713449 | 0.705093 | 0.718081 | 0.767037 | 0.765602 | 0.713716 | 0.707278 | 0.709079 |
| 0.702263 | 0.71398 | 0.71042 | 0.718852 | 0.764514 | 0.761653 | 0.710503 | 0.704619 | 0.70828 |
| 0.707623 | 0.714974 | 0.707785 | 0.719594 | 0.76898 | 0.773642 | 0.718748 | 0.711863 | 0.711198 |
| 0.702 | 0.712656 | 0.708238 | 0.717083 | 0.772887 | 0.766475 | 0.715676 | 0.706381 | 0.708078 |
| 0.704328 | 0.712505 | 0.708926 | 0.713707 | 0.767577 | 0.765054 | 0.71207 | 0.707429 | 0.70817 |
| 0.705257 | 0.714843 | 0.711844 | 0.720613 | 0.768643 | 0.766804 | 0.715178 | 0.710338 | 0.709198 |
| 0.706115 | 0.714422 | 0.713957 | 0.716411 | 0.771743 | 0.767137 | 0.714195 | 0.712128 | 0.712757 |
| 0.70447 | 0.712886 | 0.708067 | 0.715346 | 0.768697 | 0.772422 | 0.709782 | 0.704844 | 0.706394 |
| 0.708076 | 0.716575 | 0.709529 | 0.718844 | 0.768859 | 0.768839 | 0.715368 | 0.710522 | 0.714343 |
| 0.702372 | 0.711196 | 0.702799 | 0.709726 | 0.765452 | 0.768883 | 0.705047 | 0.704421 | 0.708337 |
| 0.704809 | 0.715284 | 0.706971 | 0.715632 | 0.770168 | 0.767712 | 0.713822 | 0.706541 | 0.712162 |
| 0.704667 | 0.716242 | 0.709412 | 0.719499 | 0.770462 | 0.774759 | 0.712331 | 0.708491 | 0.711681 |
| 0.70089 | 0.710024 | 0.701535 | 0.715803 | 0.764365 | 0.763803 | 0.707083 | 0.706805 | 0.702107 |
| 0.705349 | 0.712027 | 0.705961 | 0.713088 | 0.765869 | 0.768197 | 0.712061 | 0.706813 | 0.709754 |
| 0.708347 | 0.713811 | 0.707267 | 0.717924 | 0.767872 | 0.772073 | 0.713568 | 0.710189 | 0.709981 |
| 0.707259 | 0.716854 | 0.708732 | 0.717908 | 0.770847 | 0.769546 | 0.714513 | 0.712789 | 0.711689 |
| 0.684652 | 0.709071 | 0.680014 | 0.694263 | 0.681802 | 0.67417 | 0.697619 | 0.69543 | 0.681039 |
| 0.713237 | 0.715753 | 0.715505 | 0.715741 | 0.65962 | 0.658583 | 0.71537 | 0.723749 | 0.720624 |
| 0.748885 | 0.759883 | 0.755923 | 0.746673 | 0.672879 | 0.670334 | 0.755331 | 0.743304 | 0.751741 |
| 0.746498 | 0.742789 | 0.757373 | 0.726325 | 0.653827 | 0.647997 | 0.741913 | 0.77182 | 0.74471 |
| 0.659317 | 0.669358 | 0.659822 | 0.671864 | 0.733992 | 0.733263 | 0.670052 | 0.664399 | 0.665703 |
| 0.653938 | 0.670009 | 0.662635 | 0.666113 | 0.725791 | 0.721819 | 0.669015 | 0.665057 | 0.66537 |
| 0.654238 | 0.662843 | 0.662277 | 0.67036 | 0.71767 | 0.728553 | 0.660829 | 0.653398 | 0.665974 |
| 0.669747 | 0.680886 | 0.6736 | 0.677562 | 0.73988 | 0.734592 | 0.679032 | 0.681468 | 0.669954 |
| 0.643956 | 0.657604 | 0.646724 | 0.655346 | 0.711753 | 0.71751 | 0.655821 | 0.655494 | 0.658256 |
| 0.635259 | 0.647814 | 0.641422 | 0.64958 | 0.718043 | 0.7136 | 0.652233 | 0.647768 | 0.650946 |
| 0.64212 | 0.649343 | 0.64043 | 0.647875 | 0.714707 | 0.715205 | 0.644684 | 0.643444 | 0.643182 |
| 0.724068 | 0.7299 | 0.726422 | 0.723012 | 0.659893 | 0.655662 | 0.727894 | 0.728619 | 0.719501 |
| 0.697979 | 0.705707 | 0.70253 | 0.706405 | 0.760836 | 0.760058 | 0.701937 | 0.701252 | 0.702016 |
| 0.668005 | 0.681864 | 0.672452 | 0.681405 | 0.738908 | 0.74246 | 0.678354 | 0.673435 | 0.67601 |
| 0.664069 | 0.668835 | 0.670754 | 0.669169 | 0.731458 | 0.722563 | 0.665585 | 0.668881 | 0.671176 |
| 0.653832 | 0.662473 | 0.651159 | 0.656189 | 0.723239 | 0.716596 | 0.652182 | 0.652158 | 0.656949 |
| 0.671499 | 0.676246 | 0.67421 | 0.683382 | 0.735202 | 0.736493 | 0.677525 | 0.674242 | 0.677525 |
| 0.662611 | 0.672397 | 0.662541 | 0.673472 | 0.724481 | 0.724129 | 0.672784 | 0.665387 | 0.671013 |
| 0.651102 | 0.661236 | 0.655361 | 0.663413 | 0.726969 | 0.729054 | 0.665693 | 0.656771 | 0.661508 |
| 0.657586 | 0.669384 | 0.660468 | 0.668814 | 0.724213 | 0.72962 | 0.667472 | 0.659424 | 0.664361 |
| 0.649371 | 0.661386 | 0.652498 | 0.667071 | 0.724838 | 0.720511 | 0.661711 | 0.65806 | 0.657789 |
| 0.653073 | 0.664377 | 0.661162 | 0.670588 | 0.72704 | 0.715208 | 0.667813 | 0.656041 | 0.662162 |
| 0.649028 | 0.662759 | 0.662952 | 0.665925 | 0.721282 | 0.71052 | 0.658591 | 0.660284 | 0.659349 |
| 0.64181 | 0.647507 | 0.633417 | 0.646966 | 0.716368 | 0.712927 | 0.652951 | 0.653303 | 0.645707 |

| | | | | | | | | |
|---|---|---|---|---|---|---|---|---|
| 0.686123 | 0.691689 | 0.685058 | 0.695868 | 0.750843 | 0.753136 | 0.691738 | 0.68813 | 0.691169 |
| 0.654435 | 0.663781 | 0.651994 | 0.667117 | 0.727935 | 0.723762 | 0.662464 | 0.654146 | 0.658681 |
| 0.656317 | 0.671362 | 0.662534 | 0.67066 | 0.728469 | 0.719141 | 0.670893 | 0.66285 | 0.66426 |
| 0.660667 | 0.664847 | 0.660985 | 0.668039 | 0.735083 | 0.730108 | 0.664697 | 0.66287 | 0.665051 |
| 0.650137 | 0.658141 | 0.660359 | 0.664846 | 0.736391 | 0.725082 | 0.65835 | 0.665674 | 0.65736 |
| 0.653975 | 0.664734 | 0.660757 | 0.666906 | 0.722469 | 0.72112 | 0.658396 | 0.655537 | 0.659661 |
| 0.645229 | 0.656762 | 0.652408 | 0.657276 | 0.718426 | 0.719572 | 0.652204 | 0.647209 | 0.656856 |
| 0.638769 | 0.648325 | 0.644648 | 0.657267 | 0.723068 | 0.713083 | 0.649605 | 0.647331 | 0.646615 |
| 0.644228 | 0.651283 | 0.645214 | 0.658696 | 0.713926 | 0.717525 | 0.651972 | 0.645998 | 0.652399 |
| 0.685262 | 0.692919 | 0.68986 | 0.697388 | 0.75541 | 0.754092 | 0.691045 | 0.686226 | 0.692797 |
| 0.682771 | 0.692465 | 0.686271 | 0.695825 | 0.750452 | 0.749496 | 0.690978 | 0.686752 | 0.688684 |
| 0.676214 | 0.6855 | 0.682177 | 0.689387 | 0.745892 | 0.747892 | 0.685114 | 0.682436 | 0.681949 |
| 0.674874 | 0.676936 | 0.677132 | 0.680327 | 0.737905 | 0.738512 | 0.677612 | 0.675932 | 0.677246 |
| 0.674503 | 0.681872 | 0.680836 | 0.68452 | 0.747741 | 0.73991 | 0.684335 | 0.681226 | 0.681523 |
| 0.666706 | 0.673313 | 0.669496 | 0.682056 | 0.73984 | 0.739793 | 0.678699 | 0.673252 | 0.670203 |

| HG_07 | HG_08 | | | HG_09 | | HG_10 | | |
|---|---|---|---|---|---|---|---|---|
| Home_garc | Home_garc | Home_garc | Home_garc | Home_garc | Home_garc | Home_garc | Home_garc | Home_garc |
| 0.796432 | 0.772298 | 0.78424 | 0.763446 | 0.792252 | 0.794434 | 0.759537 | 0.790135 | 0.783487 |
| 0.787801 | 0.787167 | 0.784833 | 0.777532 | 0.781655 | 0.779911 | 0.765304 | 0.792483 | 0.789404 |
| 0.788953 | 0.778106 | 0.782787 | 0.764106 | 0.793209 | 0.791373 | 0.750997 | 0.789883 | 0.781743 |
| 0.784488 | 0.776262 | 0.786807 | 0.760793 | 0.787241 | 0.788759 | 0.760291 | 0.783229 | 0.791832 |
| 0.783944 | 0.767971 | 0.77677 | 0.767133 | 0.783886 | 0.767451 | 0.755902 | 0.770563 | 0.781039 |
| 0.772408 | 0.770939 | 0.771233 | 0.768752 | 0.782598 | 0.767813 | 0.753808 | 0.767872 | 0.781065 |
| 0.704459 | 0.713514 | 0.723701 | 0.73394 | 0.716645 | 0.715718 | 0.742343 | 0.707002 | 0.715013 |
| 0.699338 | 0.70729 | 0.720526 | 0.730576 | 0.715875 | 0.712505 | 0.741286 | 0.701668 | 0.708227 |
| 0.781472 | 0.777998 | 0.774582 | 0.765337 | 0.783336 | 0.773337 | 0.758587 | 0.78572 | 0.78637 |
| 0.784309 | 0.769363 | 0.785285 | 0.766561 | 0.786407 | 0.79638 | 0.753281 | 0.774584 | 0.773384 |
| 0.783008 | 0.772843 | 0.78045 | 0.76228 | 0.785939 | 0.786748 | 0.755194 | 0.775405 | 0.780218 |
| 1 | 0.780498 | 0.785048 | 0.781135 | 0.781409 | 0.787461 | 0.764863 | 0.794581 | 0.786594 |
| 0.780498 | 1 | 0.772701 | 0.766096 | 0.777577 | 0.774077 | 0.75685 | 0.787411 | 0.780166 |
| 0.785048 | 0.772701 | 1 | 0.778194 | 0.786994 | 0.781762 | 0.805152 | 0.777704 | 0.776463 |
| 0.781135 | 0.766096 | 0.778194 | 1 | 0.775816 | 0.770994 | 0.807777 | 0.769701 | 0.765838 |
| 0.781409 | 0.777577 | 0.786994 | 0.775816 | 1 | 0.841698 | 0.765241 | 0.787789 | 0.781312 |
| 0.787461 | 0.774077 | 0.781762 | 0.770994 | 0.841698 | 1 | 0.762572 | 0.77269 | 0.782721 |
| 0.764863 | 0.75685 | 0.805152 | 0.807777 | 0.765241 | 0.762572 | 1 | 0.754791 | 0.760696 |
| 0.794581 | 0.787411 | 0.777704 | 0.769701 | 0.787789 | 0.77269 | 0.754791 | 1 | 0.785844 |
| 0.786594 | 0.780166 | 0.776463 | 0.765838 | 0.781312 | 0.782721 | 0.760696 | 0.785844 | 1 |
| 0.789151 | 0.770883 | 0.778929 | 0.765852 | 0.78238 | 0.780877 | 0.761744 | 0.783232 | 0.824253 |
| 0.803104 | 0.774681 | 0.786279 | 0.771077 | 0.791257 | 0.793664 | 0.765713 | 0.787929 | 0.799517 |
| 0.797625 | 0.782029 | 0.78297 | 0.781201 | 0.782502 | 0.784791 | 0.767408 | 0.79302 | 0.810538 |
| 0.815631 | 0.786923 | 0.788257 | 0.780432 | 0.801758 | 0.800701 | 0.770468 | 0.804113 | 0.790376 |
| 0.791674 | 0.781659 | 0.776229 | 0.767459 | 0.778289 | 0.770204 | 0.753482 | 0.790527 | 0.808157 |
| 0.791111 | 0.781258 | 0.785216 | 0.769623 | 0.78482 | 0.77151 | 0.761722 | 0.803881 | 0.788063 |
| 0.792334 | 0.775438 | 0.764951 | 0.765404 | 0.777085 | 0.774752 | 0.752089 | 0.804388 | 0.780198 |
| 0.788382 | 0.781678 | 0.778142 | 0.7704 | 0.773274 | 0.77347 | 0.759313 | 0.77671 | 0.79055 |
| 0.787972 | 0.772966 | 0.784608 | 0.769531 | 0.775562 | 0.775529 | 0.756342 | 0.781309 | 0.793656 |
| 0.795608 | 0.784524 | 0.786691 | 0.770966 | 0.799833 | 0.803755 | 0.758526 | 0.783658 | 0.777897 |
| 0.789172 | 0.766762 | 0.775838 | 0.77162 | 0.781343 | 0.769656 | 0.76022 | 0.774439 | 0.782259 |
| 0.776876 | 0.771994 | 0.774228 | 0.76702 | 0.781452 | 0.767434 | 0.754053 | 0.773939 | 0.78152 |
| 0.784986 | 0.769382 | 0.780606 | 0.764749 | 0.782847 | 0.769288 | 0.750717 | 0.806956 | 0.783148 |
| 0.776023 | 0.771153 | 0.770818 | 0.762398 | 0.773934 | 0.760081 | 0.750419 | 0.772508 | 0.779971 |
| 0.775676 | 0.768325 | 0.763626 | 0.762512 | 0.778097 | 0.768035 | 0.743682 | 0.78154 | 0.773231 |
| 0.792392 | 0.78736 | 0.780591 | 0.777924 | 0.783003 | 0.778266 | 0.761077 | 0.811566 | 0.781729 |
| 0.787428 | 0.781774 | 0.778313 | 0.774457 | 0.782364 | 0.782548 | 0.767584 | 0.785337 | 0.790479 |
| 0.768435 | 0.770315 | 0.768888 | 0.762018 | 0.767683 | 0.760619 | 0.745365 | 0.770948 | 0.776835 |
| 0.780265 | 0.78048 | 0.770631 | 0.765473 | 0.782128 | 0.772765 | 0.755556 | 0.772306 | 0.783976 |

| | | | | | | | | |
|---|---|---|---|---|---|---|---|---|
| 0.76222 | 0.761897 | 0.79651 | 0.801061 | 0.773718 | 0.786578 | 0.83406 | 0.754603 | 0.758446 |
| 0.793361 | 0.771239 | 0.780105 | 0.763286 | 0.785091 | 0.793304 | 0.764815 | 0.787537 | 0.782122 |
| 0.808619 | 0.774308 | 0.788573 | 0.774434 | 0.796913 | 0.791288 | 0.761082 | 0.790795 | 0.788715 |
| 0.805533 | 0.791427 | 0.791585 | 0.769573 | 0.79476 | 0.806347 | 0.766859 | 0.789014 | 0.795915 |
| 0.798057 | 0.777312 | 0.779915 | 0.769587 | 0.788223 | 0.785122 | 0.759235 | 0.795179 | 0.784071 |
| 0.786319 | 0.774247 | 0.776165 | 0.770142 | 0.783147 | 0.775539 | 0.764758 | 0.783126 | 0.78307 |
| 0.783955 | 0.763432 | 0.768051 | 0.769912 | 0.775365 | 0.765657 | 0.74998 | 0.775362 | 0.777111 |
| 0.790066 | 0.782677 | 0.779257 | 0.766081 | 0.786158 | 0.769969 | 0.760794 | 0.796534 | 0.79923 |
| 0.774777 | 0.76785 | 0.772083 | 0.762999 | 0.773531 | 0.773365 | 0.751067 | 0.777057 | 0.780522 |
| 0.768459 | 0.767668 | 0.771711 | 0.766602 | 0.771577 | 0.766517 | 0.750996 | 0.773209 | 0.778854 |
| 0.818166 | 0.78024 | 0.784809 | 0.769634 | 0.792574 | 0.799731 | 0.767657 | 0.787772 | 0.795954 |
| 0.816445 | 0.786787 | 0.777759 | 0.774515 | 0.78657 | 0.793106 | 0.763797 | 0.792827 | 0.791154 |
| 0.808434 | 0.780627 | 0.781001 | 0.781438 | 0.80025 | 0.79774 | 0.762605 | 0.799395 | 0.797758 |
| 0.809155 | 0.782908 | 0.778972 | 0.772787 | 0.792169 | 0.786119 | 0.757745 | 0.791025 | 0.794471 |
| 0.795831 | 0.774143 | 0.78556 | 0.77383 | 0.78796 | 0.801998 | 0.765205 | 0.793234 | 0.796338 |
| 0.801473 | 0.770333 | 0.781054 | 0.770458 | 0.785684 | 0.794476 | 0.763311 | 0.787963 | 0.790241 |
| 0.791457 | 0.780315 | 0.781151 | 0.768625 | 0.797353 | 0.794229 | 0.756952 | 0.791298 | 0.782652 |
| 0.804754 | 0.793477 | 0.792761 | 0.776202 | 0.805828 | 0.812253 | 0.772628 | 0.791899 | 0.795587 |
| 0.798836 | 0.783648 | 0.786623 | 0.7756 | 0.798923 | 0.795794 | 0.766949 | 0.788796 | 0.789347 |
| 0.797402 | 0.787234 | 0.787386 | 0.768897 | 0.806747 | 0.786409 | 0.763271 | 0.799865 | 0.79794 |
| 0.791687 | 0.774215 | 0.78311 | 0.77156 | 0.811326 | 0.805298 | 0.756637 | 0.785446 | 0.784456 |
| 0.793098 | 0.798548 | 0.784403 | 0.764682 | 0.799967 | 0.80188 | 0.759455 | 0.780328 | 0.782663 |
| 0.785491 | 0.771 | 0.775773 | 0.768436 | 0.792871 | 0.775577 | 0.759764 | 0.785726 | 0.792728 |
| 0.773708 | 0.77787 | 0.76409 | 0.772274 | 0.770183 | 0.763102 | 0.753199 | 0.769548 | 0.784655 |
| 0.762788 | 0.772386 | 0.764917 | 0.769853 | 0.773365 | 0.763502 | 0.75494 | 0.761194 | 0.771293 |
| 0.70169 | 0.706277 | 0.721664 | 0.731501 | 0.717163 | 0.714621 | 0.741987 | 0.702518 | 0.708744 |
| 0.704142 | 0.713413 | 0.724174 | 0.737225 | 0.717074 | 0.719364 | 0.738949 | 0.711427 | 0.713366 |
| 0.706395 | 0.715145 | 0.726307 | 0.740367 | 0.717494 | 0.716479 | 0.74347 | 0.711986 | 0.717961 |
| 0.781492 | 0.790561 | 0.778537 | 0.758788 | 0.766093 | 0.760367 | 0.759382 | 0.7801 | 0.786384 |
| 0.77742 | 0.784858 | 0.776435 | 0.763057 | 0.768241 | 0.760429 | 0.760468 | 0.778773 | 0.78586 |
| 0.782389 | 0.780036 | 0.777458 | 0.766257 | 0.782132 | 0.778955 | 0.767604 | 0.762851 | 0.78771 |
| 0.777546 | 0.786522 | 0.776504 | 0.760804 | 0.775502 | 0.770478 | 0.769451 | 0.782594 | 0.783135 |
| 0.812638 | 0.797338 | 0.79412 | 0.787193 | 0.797767 | 0.79934 | 0.767863 | 0.807123 | 0.800068 |
| 0.821589 | 0.801093 | 0.792879 | 0.783639 | 0.802698 | 0.797872 | 0.757399 | 0.837106 | 0.793649 |
| 0.802369 | 0.785288 | 0.791145 | 0.77538 | 0.803344 | 0.794668 | 0.764379 | 0.828657 | 0.804543 |
| 0.806143 | 0.784206 | 0.800417 | 0.780536 | 0.815966 | 0.826065 | 0.758269 | 0.818205 | 0.796927 |
| 0.790365 | 0.796105 | 0.777035 | 0.769505 | 0.784073 | 0.770685 | 0.768491 | 0.79206 | 0.783892 |
| 0.813744 | 0.811944 | 0.805711 | 0.795107 | 0.818174 | 0.823659 | 0.783429 | 0.811276 | 0.811561 |
| 0.789806 | 0.774097 | 0.780856 | 0.765732 | 0.782266 | 0.78139 | 0.75523 | 0.801431 | 0.789606 |
| 0.800149 | 0.782505 | 0.775281 | 0.770234 | 0.779224 | 0.773698 | 0.767377 | 0.777041 | 0.802402 |
| 0.791536 | 0.778159 | 0.78685 | 0.76537 | 0.793697 | 0.803632 | 0.755043 | 0.786463 | 0.792375 |
| 0.814631 | 0.816378 | 0.803655 | 0.796862 | 0.823693 | 0.821811 | 0.78257 | 0.818428 | 0.813266 |
| 0.800284 | 0.774119 | 0.788573 | 0.767294 | 0.796475 | 0.806628 | 0.766872 | 0.774175 | 0.788546 |
| 0.814426 | 0.77312 | 0.789482 | 0.769896 | 0.791766 | 0.789378 | 0.761874 | 0.78051 | 0.781455 |
| 0.782544 | 0.777447 | 0.780379 | 0.771996 | 0.785915 | 0.774156 | 0.753485 | 0.784809 | 0.785934 |
| 0.795388 | 0.78463 | 0.779014 | 0.772413 | 0.792193 | 0.795564 | 0.752916 | 0.799646 | 0.789562 |

| | | | | | | | | |
|---|---|---|---|---|---|---|---|---|
| 0.798897 | 0.772858 | 0.785325 | 0.772344 | 0.798749 | 0.805404 | 0.755233 | 0.7768 | 0.781949 |
| 0.786038 | 0.776856 | 0.772902 | 0.759628 | 0.782387 | 0.769092 | 0.766167 | 0.777668 | 0.79789 |
| 0.781648 | 0.78022 | 0.779414 | 0.771134 | 0.785976 | 0.773908 | 0.750808 | 0.778679 | 0.779537 |
| 0.795292 | 0.796582 | 0.78961 | 0.778127 | 0.799428 | 0.798425 | 0.771281 | 0.798713 | 0.79936 |
| 0.788438 | 0.788101 | 0.782049 | 0.768351 | 0.789422 | 0.775163 | 0.76158 | 0.782551 | 0.78718 |
| 0.79174 | 0.779569 | 0.774164 | 0.760105 | 0.774159 | 0.771936 | 0.766241 | 0.781281 | 0.792458 |
| 0.7831 | 0.787573 | 0.77631 | 0.769399 | 0.785557 | 0.777175 | 0.76538 | 0.780865 | 0.794552 |
| 0.773343 | 0.774164 | 0.774857 | 0.772899 | 0.791785 | 0.769384 | 0.75515 | 0.775643 | 0.782695 |
| 0.782248 | 0.769668 | 0.774127 | 0.765878 | 0.784525 | 0.77238 | 0.751571 | 0.775808 | 0.783714 |
| 0.790559 | 0.77303 | 0.771503 | 0.758548 | 0.775268 | 0.767121 | 0.74966 | 0.784831 | 0.772187 |
| 0.775875 | 0.773819 | 0.769187 | 0.766349 | 0.783297 | 0.769407 | 0.755043 | 0.78476 | 0.780206 |
| 0.779457 | 0.770741 | 0.765115 | 0.768539 | 0.774591 | 0.768916 | 0.751565 | 0.787403 | 0.783847 |
| 0.791034 | 0.776515 | 0.776645 | 0.768504 | 0.780344 | 0.774981 | 0.755331 | 0.783194 | 0.786626 |
| 0.798256 | 0.776781 | 0.769391 | 0.759252 | 0.7737 | 0.764558 | 0.745939 | 0.789807 | 0.775595 |
| 0.778926 | 0.776785 | 0.780007 | 0.762681 | 0.780452 | 0.769409 | 0.75866 | 0.774677 | 0.782926 |
| 0.780903 | 0.771106 | 0.77483 | 0.764134 | 0.779592 | 0.772384 | 0.749102 | 0.778651 | 0.783239 |
| 0.781886 | 0.77418 | 0.769846 | 0.76906 | 0.778612 | 0.767257 | 0.755865 | 0.780512 | 0.785328 |
| 0.788169 | 0.776068 | 0.784296 | 0.77715 | 0.801463 | 0.800402 | 0.756274 | 0.79141 | 0.78598 |
| 0.785288 | 0.773034 | 0.775297 | 0.770503 | 0.781707 | 0.769854 | 0.753037 | 0.779311 | 0.786966 |
| 0.775519 | 0.774281 | 0.769212 | 0.767156 | 0.781217 | 0.774071 | 0.746622 | 0.778118 | 0.782303 |
| 0.777162 | 0.763647 | 0.768181 | 0.755857 | 0.773933 | 0.772059 | 0.748411 | 0.77915 | 0.787796 |
| 0.771813 | 0.766986 | 0.765832 | 0.759271 | 0.778608 | 0.768673 | 0.7503 | 0.772851 | 0.776116 |
| 0.781805 | 0.778581 | 0.772047 | 0.768329 | 0.777133 | 0.772693 | 0.754874 | 0.780881 | 0.783599 |
| 0.791366 | 0.783115 | 0.775336 | 0.762815 | 0.776084 | 0.770981 | 0.760784 | 0.792351 | 0.782975 |
| 0.777824 | 0.774145 | 0.770175 | 0.771374 | 0.777676 | 0.765759 | 0.761178 | 0.785912 | 0.777129 |
| 0.786782 | 0.790098 | 0.783832 | 0.780228 | 0.787272 | 0.770441 | 0.767861 | 0.799153 | 0.792193 |
| 0.77393 | 0.776622 | 0.769513 | 0.773758 | 0.781961 | 0.768152 | 0.759241 | 0.78621 | 0.781178 |
| 0.783012 | 0.770779 | 0.775029 | 0.766242 | 0.781368 | 0.771621 | 0.756614 | 0.773097 | 0.780446 |
| 0.785632 | 0.770762 | 0.771699 | 0.767969 | 0.77969 | 0.776459 | 0.758184 | 0.783997 | 0.779895 |
| 0.77814 | 0.777796 | 0.770726 | 0.771488 | 0.782365 | 0.766527 | 0.761394 | 0.779995 | 0.780596 |
| 0.786012 | 0.767177 | 0.77706 | 0.767872 | 0.774061 | 0.766067 | 0.752316 | 0.782453 | 0.780158 |
| 0.771155 | 0.77499 | 0.763158 | 0.768215 | 0.77516 | 0.770456 | 0.752617 | 0.772256 | 0.78268 |
| 0.782805 | 0.790801 | 0.779678 | 0.769472 | 0.777824 | 0.765725 | 0.758202 | 0.7978 | 0.792828 |
| 0.801676 | 0.78749 | 0.788881 | 0.767702 | 0.79008 | 0.803495 | 0.762903 | 0.785976 | 0.793587 |
| 0.789908 | 0.776783 | 0.776877 | 0.76243 | 0.778412 | 0.769148 | 0.755865 | 0.781029 | 0.792232 |
| 0.792194 | 0.770059 | 0.778437 | 0.767038 | 0.782201 | 0.775284 | 0.757427 | 0.78104 | 0.793159 |
| 0.784642 | 0.7742 | 0.769869 | 0.764711 | 0.778007 | 0.77102 | 0.749156 | 0.784787 | 0.783351 |
| 0.75549 | 0.759324 | 0.752064 | 0.736316 | 0.75748 | 0.746773 | 0.725696 | 0.759632 | 0.756495 |
| 0.772281 | 0.763442 | 0.765334 | 0.74549 | 0.759753 | 0.756632 | 0.748156 | 0.767623 | 0.778165 |
| 0.744651 | 0.755342 | 0.741414 | 0.738605 | 0.747211 | 0.73962 | 0.727603 | 0.753003 | 0.759751 |
| 0.776986 | 0.764366 | 0.758905 | 0.75447 | 0.768537 | 0.77451 | 0.733371 | 0.781144 | 0.77072 |
| 0.799925 | 0.759107 | 0.745609 | 0.736354 | 0.741087 | 0.731425 | 0.707432 | 0.788903 | 0.73667 |
| 0.781005 | 0.747108 | 0.752112 | 0.735217 | 0.76743 | 0.780225 | 0.724951 | 0.761281 | 0.756503 |
| 0.763834 | 0.744756 | 0.758717 | 0.732821 | 0.766955 | 0.749685 | 0.719907 | 0.761162 | 0.752781 |
| 0.74633 | 0.742921 | 0.744961 | 0.730428 | 0.756504 | 0.73621 | 0.720708 | 0.752632 | 0.749303 |
| 0.757003 | 0.749903 | 0.762672 | 0.741935 | 0.761433 | 0.760727 | 0.7313 | 0.765023 | 0.764197 |
| 0.769561 | 0.749388 | 0.756292 | 0.754881 | 0.775712 | 0.769432 | 0.725935 | 0.778998 | 0.755912 |

| | | | | | | | | |
|---|---|---|---|---|---|---|---|---|
| 0.734153 | 0.741543 | 0.730725 | 0.723289 | 0.737707 | 0.725349 | 0.71585 | 0.744725 | 0.746986 |
| 0.778956 | 0.768496 | 0.759946 | 0.749597 | 0.776475 | 0.762243 | 0.735329 | 0.80064 | 0.772055 |
| 0.729622 | 0.739836 | 0.728353 | 0.728834 | 0.739546 | 0.722931 | 0.708188 | 0.741689 | 0.748955 |
| 0.759536 | 0.742094 | 0.75203 | 0.735542 | 0.774216 | 0.760885 | 0.722772 | 0.737637 | 0.748756 |
| 0.787551 | 0.770292 | 0.751734 | 0.730567 | 0.741793 | 0.726122 | 0.720135 | 0.793607 | 0.760394 |
| 0.73252 | 0.752736 | 0.742906 | 0.725661 | 0.757689 | 0.732869 | 0.71486 | 0.741829 | 0.744346 |
| 0.76337 | 0.752443 | 0.74585 | 0.738293 | 0.735182 | 0.734205 | 0.734343 | 0.752334 | 0.761096 |
| 0.782131 | 0.746824 | 0.75911 | 0.738861 | 0.769469 | 0.758631 | 0.721939 | 0.786226 | 0.758357 |
| 0.748965 | 0.737647 | 0.720042 | 0.713069 | 0.736633 | 0.719227 | 0.698689 | 0.752092 | 0.730863 |
| 0.757171 | 0.785403 | 0.745573 | 0.73786 | 0.745959 | 0.739243 | 0.736999 | 0.787247 | 0.776725 |
| 0.743206 | 0.74154 | 0.744315 | 0.720619 | 0.752639 | 0.736695 | 0.710535 | 0.737469 | 0.742777 |
| 0.781572 | 0.758131 | 0.773267 | 0.75724 | 0.784022 | 0.793183 | 0.748353 | 0.764485 | 0.770881 |
| 0.77039 | 0.767472 | 0.765064 | 0.757184 | 0.771259 | 0.76328 | 0.747778 | 0.77056 | 0.77886 |
| 0.747325 | 0.753664 | 0.747567 | 0.740728 | 0.744643 | 0.743113 | 0.721577 | 0.749823 | 0.758861 |
| 0.773662 | 0.792987 | 0.779548 | 0.764459 | 0.781427 | 0.777387 | 0.752966 | 0.782525 | 0.774 |
| 0.774691 | 0.774948 | 0.758166 | 0.753072 | 0.765957 | 0.756216 | 0.740862 | 0.772269 | 0.770679 |
| 0.781863 | 0.781147 | 0.769519 | 0.770727 | 0.778886 | 0.772828 | 0.758521 | 0.776718 | 0.782677 |
| 0.776509 | 0.782674 | 0.776216 | 0.773976 | 0.779166 | 0.775289 | 0.754142 | 0.775424 | 0.781076 |
| 0.755145 | 0.751673 | 0.749577 | 0.74686 | 0.761878 | 0.745882 | 0.734884 | 0.753867 | 0.757157 |
| 0.790231 | 0.795969 | 0.776437 | 0.766895 | 0.771802 | 0.770608 | 0.751807 | 0.791268 | 0.768718 |
| 0.743013 | 0.752395 | 0.75204 | 0.743978 | 0.754902 | 0.743485 | 0.728742 | 0.738225 | 0.743091 |
| 0.774672 | 0.771003 | 0.768051 | 0.764299 | 0.772272 | 0.76555 | 0.757593 | 0.770394 | 0.775949 |
| 0.767904 | 0.767408 | 0.77235 | 0.767598 | 0.780405 | 0.772284 | 0.748296 | 0.777925 | 0.773864 |
| 0.757993 | 0.761098 | 0.73465 | 0.73668 | 0.741839 | 0.732122 | 0.745998 | 0.746721 | 0.765674 |
| 0.74934 | 0.757389 | 0.744477 | 0.736094 | 0.754818 | 0.748656 | 0.729355 | 0.755788 | 0.756193 |
| 0.760324 | 0.752666 | 0.759859 | 0.746971 | 0.773498 | 0.758986 | 0.733698 | 0.758897 | 0.757903 |
| 0.760955 | 0.762491 | 0.763055 | 0.755038 | 0.764804 | 0.758563 | 0.73946 | 0.749311 | 0.763972 |
| 0.777705 | 0.772079 | 0.766212 | 0.761289 | 0.775454 | 0.765487 | 0.764739 | 0.77667 | 0.78424 |
| 0.740785 | 0.741683 | 0.746232 | 0.732362 | 0.75118 | 0.738524 | 0.724149 | 0.74803 | 0.74914 |
| 0.763973 | 0.797576 | 0.777914 | 0.769014 | 0.775114 | 0.768492 | 0.763257 | 0.774117 | 0.774729 |
| 0.765534 | 0.765672 | 0.771943 | 0.764894 | 0.764654 | 0.764093 | 0.75712 | 0.765947 | 0.776732 |
| 0.763527 | 0.758414 | 0.761149 | 0.754316 | 0.767686 | 0.76932 | 0.742642 | 0.761477 | 0.76272 |
| 0.766217 | 0.757428 | 0.760429 | 0.750221 | 0.765304 | 0.767092 | 0.746037 | 0.766486 | 0.767195 |
| 0.764639 | 0.767468 | 0.76908 | 0.768182 | 0.770908 | 0.767556 | 0.754372 | 0.759138 | 0.771157 |
| 0.756337 | 0.771297 | 0.760964 | 0.757884 | 0.766718 | 0.765552 | 0.751279 | 0.759398 | 0.76316 |
| 0.773245 | 0.76762 | 0.769756 | 0.760822 | 0.764123 | 0.75772 | 0.764768 | 0.76252 | 0.767042 |
| 0.721176 | 0.714716 | 0.718987 | 0.716265 | 0.730266 | 0.719122 | 0.706184 | 0.720579 | 0.725692 |
| 0.755365 | 0.745709 | 0.76013 | 0.755099 | 0.760975 | 0.753949 | 0.751931 | 0.75684 | 0.759172 |
| 0.721843 | 0.726407 | 0.725433 | 0.720097 | 0.732274 | 0.725004 | 0.710427 | 0.725472 | 0.734699 |
| 0.714186 | 0.715116 | 0.721705 | 0.711354 | 0.727422 | 0.709133 | 0.714973 | 0.714018 | 0.724464 |
| 0.78979 | 0.783618 | 0.776265 | 0.766895 | 0.769992 | 0.765919 | 0.789499 | 0.782469 | 0.771242 |
| 0.719305 | 0.713155 | 0.713678 | 0.70824 | 0.718667 | 0.711395 | 0.710327 | 0.709034 | 0.721595 |
| 0.723849 | 0.720077 | 0.713351 | 0.715897 | 0.726463 | 0.718908 | 0.708137 | 0.720782 | 0.726411 |
| 0.708765 | 0.715399 | 0.713086 | 0.705219 | 0.709229 | 0.70856 | 0.701302 | 0.719588 | 0.715838 |
| 0.770473 | 0.771868 | 0.780043 | 0.782765 | 0.774951 | 0.781309 | 0.797365 | 0.76798 | 0.770874 |
| 0.713789 | 0.713802 | 0.711762 | 0.70805 | 0.719639 | 0.710157 | 0.702474 | 0.727315 | 0.709767 |
| 0.778116 | 0.777347 | 0.776013 | 0.779077 | 0.772348 | 0.770404 | 0.788933 | 0.782551 | 0.771823 |

| | | | | | | | | |
|---|---|---|---|---|---|---|---|---|
| 0.774729 | 0.773046 | 0.783686 | 0.779363 | 0.772133 | 0.767907 | 0.811145 | 0.762435 | 0.762783 |
| 0.773388 | 0.767499 | 0.777732 | 0.793016 | 0.776135 | 0.769635 | 0.804846 | 0.771492 | 0.767594 |
| 0.748226 | 0.745763 | 0.756026 | 0.760075 | 0.754346 | 0.751335 | 0.762167 | 0.742974 | 0.754321 |
| 0.70966 | 0.706365 | 0.700455 | 0.703975 | 0.715255 | 0.711084 | 0.695834 | 0.718158 | 0.705528 |
| 0.695656 | 0.713613 | 0.699869 | 0.694913 | 0.699193 | 0.688419 | 0.687831 | 0.70792 | 0.701103 |
| 0.77113 | 0.771246 | 0.785802 | 0.790479 | 0.799983 | 0.783905 | 0.806951 | 0.770837 | 0.771141 |
| 0.686583 | 0.685902 | 0.690345 | 0.683174 | 0.692115 | 0.688887 | 0.683147 | 0.691489 | 0.694726 |
| 0.728023 | 0.73183 | 0.74248 | 0.741509 | 0.739865 | 0.730474 | 0.741142 | 0.731453 | 0.737581 |
| 0.763078 | 0.762817 | 0.772269 | 0.793943 | 0.82552 | 0.784677 | 0.802549 | 0.77254 | 0.759001 |
| 0.717182 | 0.720981 | 0.725032 | 0.734841 | 0.724679 | 0.722816 | 0.73792 | 0.713591 | 0.719925 |
| 0.750326 | 0.757509 | 0.773596 | 0.788943 | 0.757318 | 0.762538 | 0.805148 | 0.756657 | 0.75096 |
| 0.597561 | 0.595092 | 0.649007 | 0.689189 | 0.624204 | 0.684211 | 0.643312 | 0.652482 | 0.602484 |
| 0.64637 | 0.659459 | 0.672565 | 0.689305 | 0.668093 | 0.658937 | 0.691196 | 0.650658 | 0.661014 |
| 0.644391 | 0.65895 | 0.666616 | 0.687882 | 0.666869 | 0.656155 | 0.682831 | 0.646785 | 0.649481 |
| 0.657673 | 0.673843 | 0.675417 | 0.69472 | 0.670438 | 0.666085 | 0.694513 | 0.660026 | 0.672595 |
| 0.65486 | 0.665282 | 0.678669 | 0.696863 | 0.672349 | 0.66472 | 0.704002 | 0.658075 | 0.661965 |
| 0.616124 | 0.621839 | 0.624973 | 0.617782 | 0.620493 | 0.616197 | 0.612855 | 0.614912 | 0.628051 |
| 0.657522 | 0.66847 | 0.677383 | 0.698635 | 0.673261 | 0.669322 | 0.706653 | 0.663061 | 0.667085 |
| 0.747691 | 0.748815 | 0.759039 | 0.785785 | 0.746301 | 0.741988 | 0.814324 | 0.74246 | 0.742636 |
| 0.740475 | 0.745352 | 0.764275 | 0.784347 | 0.749773 | 0.748422 | 0.810113 | 0.740195 | 0.738725 |
| 0.739765 | 0.756071 | 0.768456 | 0.789692 | 0.756334 | 0.753837 | 0.810003 | 0.748479 | 0.750205 |
| 0.614625 | 0.619169 | 0.62196 | 0.61439 | 0.619753 | 0.61676 | 0.616673 | 0.615928 | 0.621928 |
| 0.598746 | 0.597538 | 0.602526 | 0.596848 | 0.602846 | 0.598499 | 0.595125 | 0.597145 | 0.602802 |
| 0.699149 | 0.709456 | 0.731497 | 0.743726 | 0.71454 | 0.713702 | 0.761905 | 0.699661 | 0.707205 |
| 0.70087 | 0.710239 | 0.728644 | 0.748971 | 0.71366 | 0.712942 | 0.762012 | 0.702498 | 0.708151 |
| 0.692791 | 0.703062 | 0.720427 | 0.732589 | 0.702437 | 0.703647 | 0.741385 | 0.6942 | 0.701347 |
| 0.696132 | 0.705583 | 0.721395 | 0.735734 | 0.711348 | 0.709344 | 0.74634 | 0.700032 | 0.707375 |
| 0.699921 | 0.711468 | 0.7296 | 0.744263 | 0.712687 | 0.710958 | 0.75843 | 0.702151 | 0.707099 |
| 0.700465 | 0.712571 | 0.726073 | 0.741616 | 0.713871 | 0.713915 | 0.7527 | 0.703128 | 0.709135 |
| 0.698107 | 0.715476 | 0.726184 | 0.745935 | 0.713375 | 0.713057 | 0.756574 | 0.703323 | 0.707589 |
| 0.704866 | 0.711193 | 0.718528 | 0.731239 | 0.71519 | 0.714958 | 0.736097 | 0.703321 | 0.710035 |
| 0.709817 | 0.718222 | 0.724359 | 0.736508 | 0.723033 | 0.721499 | 0.743111 | 0.710038 | 0.717309 |
| 0.708039 | 0.712769 | 0.719751 | 0.737856 | 0.715743 | 0.713628 | 0.740786 | 0.707697 | 0.710273 |
| 0.703004 | 0.715912 | 0.724431 | 0.73421 | 0.71819 | 0.720437 | 0.7391 | 0.708475 | 0.712548 |
| 0.709624 | 0.714843 | 0.725606 | 0.738783 | 0.719407 | 0.719274 | 0.746441 | 0.709191 | 0.716823 |
| 0.701879 | 0.709564 | 0.72145 | 0.732974 | 0.712836 | 0.711583 | 0.736872 | 0.70643 | 0.709326 |
| 0.706998 | 0.711924 | 0.724833 | 0.733264 | 0.716535 | 0.718427 | 0.744532 | 0.705553 | 0.711452 |
| 0.71001 | 0.714036 | 0.72294 | 0.737818 | 0.719459 | 0.715098 | 0.741425 | 0.707548 | 0.713206 |
| 0.700147 | 0.710231 | 0.721066 | 0.733366 | 0.712286 | 0.712401 | 0.737932 | 0.703328 | 0.708163 |
| 0.703821 | 0.715785 | 0.722368 | 0.732639 | 0.717965 | 0.716448 | 0.738872 | 0.708768 | 0.713593 |
| 0.650708 | 0.661877 | 0.675199 | 0.686118 | 0.664014 | 0.659963 | 0.69552 | 0.649874 | 0.6549 |
| 0.655574 | 0.665147 | 0.67441 | 0.687192 | 0.670088 | 0.665814 | 0.696533 | 0.658862 | 0.665754 |
| 0.708109 | 0.717215 | 0.7251 | 0.741956 | 0.720267 | 0.719123 | 0.748087 | 0.712025 | 0.714592 |
| 0.708598 | 0.716769 | 0.728782 | 0.743066 | 0.718524 | 0.719585 | 0.754237 | 0.707858 | 0.714035 |
| 0.63789 | 0.654204 | 0.66263 | 0.681852 | 0.65994 | 0.657207 | 0.687952 | 0.63851 | 0.64379 |
| 0.707995 | 0.716055 | 0.721005 | 0.735535 | 0.718358 | 0.720436 | 0.741048 | 0.70834 | 0.711164 |
| 0.696528 | 0.706591 | 0.715988 | 0.72854 | 0.7086 | 0.710453 | 0.736909 | 0.700175 | 0.704769 |

| | | | | | | | | |
|---|---|---|---|---|---|---|---|---|
| 0.702892 | 0.714058 | 0.718436 | 0.734664 | 0.712763 | 0.717437 | 0.74311 | 0.706093 | 0.708628 |
| 0.70891 | 0.715487 | 0.723617 | 0.738251 | 0.716385 | 0.718558 | 0.747693 | 0.709944 | 0.711619 |
| 0.703962 | 0.711772 | 0.725047 | 0.736091 | 0.716927 | 0.719865 | 0.739481 | 0.70944 | 0.712038 |
| 0.705178 | 0.715574 | 0.724182 | 0.736088 | 0.716706 | 0.717204 | 0.744448 | 0.709049 | 0.712057 |
| 0.710299 | 0.716789 | 0.729334 | 0.738006 | 0.721434 | 0.725316 | 0.747266 | 0.7115 | 0.716761 |
| 0.70253 | 0.713002 | 0.716427 | 0.734983 | 0.712563 | 0.714008 | 0.742758 | 0.705693 | 0.708484 |
| 0.705283 | 0.712198 | 0.722487 | 0.732299 | 0.712851 | 0.717405 | 0.742943 | 0.703921 | 0.70768 |
| 0.700569 | 0.709553 | 0.715382 | 0.72948 | 0.711534 | 0.705758 | 0.731094 | 0.699139 | 0.704023 |
| 0.701951 | 0.711646 | 0.718972 | 0.732826 | 0.712094 | 0.71424 | 0.739068 | 0.702647 | 0.70969 |
| 0.704182 | 0.713954 | 0.718181 | 0.734743 | 0.716777 | 0.713192 | 0.739472 | 0.705736 | 0.710451 |
| 0.704739 | 0.714558 | 0.725745 | 0.738596 | 0.718363 | 0.718107 | 0.746345 | 0.708234 | 0.714342 |
| 0.702049 | 0.707934 | 0.718229 | 0.733086 | 0.713302 | 0.715043 | 0.744347 | 0.702636 | 0.706949 |
| 0.697473 | 0.705484 | 0.71282 | 0.725992 | 0.706275 | 0.706612 | 0.734011 | 0.697397 | 0.703635 |
| 0.696091 | 0.707412 | 0.714536 | 0.725327 | 0.702399 | 0.707148 | 0.730683 | 0.69774 | 0.702025 |
| 0.709466 | 0.716314 | 0.725813 | 0.737553 | 0.720035 | 0.718475 | 0.750219 | 0.709192 | 0.713209 |
| 0.704395 | 0.714513 | 0.7238 | 0.748332 | 0.715623 | 0.713761 | 0.763794 | 0.707552 | 0.710817 |
| 0.70233 | 0.709988 | 0.720486 | 0.738174 | 0.710882 | 0.711959 | 0.745283 | 0.700915 | 0.704361 |
| 0.702004 | 0.709076 | 0.720122 | 0.746421 | 0.712701 | 0.712917 | 0.752398 | 0.702124 | 0.705915 |
| 0.701455 | 0.709867 | 0.728256 | 0.747304 | 0.714331 | 0.710144 | 0.765171 | 0.699944 | 0.709154 |
| 0.699905 | 0.708074 | 0.722811 | 0.745195 | 0.710621 | 0.710141 | 0.758966 | 0.70216 | 0.706337 |
| 0.6984 | 0.70743 | 0.72033 | 0.752331 | 0.711062 | 0.707275 | 0.762592 | 0.70087 | 0.703803 |
| 0.699339 | 0.708019 | 0.728217 | 0.742346 | 0.713242 | 0.711231 | 0.758822 | 0.699716 | 0.707691 |
| 0.703296 | 0.710626 | 0.727962 | 0.743336 | 0.714716 | 0.714217 | 0.75974 | 0.70599 | 0.711363 |
| 0.701427 | 0.714229 | 0.725096 | 0.73955 | 0.715242 | 0.714688 | 0.750281 | 0.705099 | 0.710205 |
| 0.702522 | 0.710367 | 0.725325 | 0.745309 | 0.715778 | 0.714111 | 0.764537 | 0.702138 | 0.706499 |
| 0.696883 | 0.707708 | 0.733295 | 0.744141 | 0.709597 | 0.706196 | 0.766283 | 0.702614 | 0.705528 |
| 0.700121 | 0.709029 | 0.720689 | 0.740635 | 0.711829 | 0.711263 | 0.74817 | 0.702676 | 0.704077 |
| 0.695954 | 0.707335 | 0.719053 | 0.733529 | 0.706725 | 0.710986 | 0.748835 | 0.696501 | 0.705615 |
| 0.698735 | 0.71211 | 0.725922 | 0.752754 | 0.711629 | 0.710358 | 0.773289 | 0.702316 | 0.706013 |
| 0.703146 | 0.708039 | 0.726894 | 0.745333 | 0.713445 | 0.713255 | 0.761878 | 0.702885 | 0.71083 |
| 0.704592 | 0.710642 | 0.725136 | 0.750464 | 0.717822 | 0.714791 | 0.762574 | 0.707378 | 0.709834 |
| 0.707025 | 0.714655 | 0.723739 | 0.747233 | 0.716173 | 0.715688 | 0.752653 | 0.707063 | 0.711684 |
| 0.706109 | 0.716938 | 0.729799 | 0.751584 | 0.719642 | 0.717957 | 0.76087 | 0.710314 | 0.713731 |
| 0.706579 | 0.716327 | 0.72833 | 0.74518 | 0.717402 | 0.719099 | 0.752885 | 0.708151 | 0.712567 |
| 0.69903 | 0.710297 | 0.729728 | 0.74482 | 0.711969 | 0.710314 | 0.764639 | 0.699984 | 0.704206 |
| 0.703107 | 0.709044 | 0.725931 | 0.74809 | 0.714818 | 0.710039 | 0.764064 | 0.704516 | 0.70829 |
| 0.704642 | 0.715796 | 0.726258 | 0.739137 | 0.712509 | 0.712138 | 0.755148 | 0.704325 | 0.711251 |
| 0.701291 | 0.708343 | 0.722452 | 0.739212 | 0.711849 | 0.709769 | 0.748926 | 0.699638 | 0.704445 |
| 0.698669 | 0.701811 | 0.715462 | 0.743452 | 0.706853 | 0.706569 | 0.751025 | 0.697139 | 0.700712 |
| 0.691196 | 0.700584 | 0.713221 | 0.728104 | 0.703645 | 0.702914 | 0.744356 | 0.691942 | 0.698932 |
| 0.690336 | 0.701069 | 0.721403 | 0.743916 | 0.705268 | 0.702112 | 0.762276 | 0.694911 | 0.696895 |
| 0.692612 | 0.702337 | 0.720918 | 0.736766 | 0.705604 | 0.703811 | 0.753455 | 0.694762 | 0.699619 |
| 0.70119 | 0.701963 | 0.716079 | 0.74147 | 0.707614 | 0.709181 | 0.74958 | 0.698475 | 0.701617 |
| 0.687283 | 0.695645 | 0.709955 | 0.729069 | 0.700551 | 0.697781 | 0.739031 | 0.688683 | 0.695478 |
| 0.699317 | 0.703728 | 0.722213 | 0.733861 | 0.70908 | 0.708826 | 0.743165 | 0.6972 | 0.706164 |
| 0.694609 | 0.703439 | 0.711111 | 0.724883 | 0.704764 | 0.702847 | 0.73015 | 0.696131 | 0.701017 |

| | | | | | | | | |
|---|---|---|---|---|---|---|---|---|
| 0.704099 | 0.715958 | 0.741614 | 0.761571 | 0.717365 | 0.716695 | 0.789452 | 0.706226 | 0.709381 |
| 0.706128 | 0.710486 | 0.723127 | 0.734106 | 0.715356 | 0.717181 | 0.743389 | 0.704173 | 0.710167 |
| 0.698798 | 0.709482 | 0.72106 | 0.733268 | 0.714217 | 0.712041 | 0.741113 | 0.702655 | 0.70611 |
| 0.703147 | 0.71206 | 0.723839 | 0.73423 | 0.718597 | 0.714115 | 0.746836 | 0.705612 | 0.710745 |
| 0.712013 | 0.718264 | 0.728278 | 0.743264 | 0.719426 | 0.719475 | 0.751563 | 0.709698 | 0.717249 |
| 0.710245 | 0.711543 | 0.722603 | 0.734237 | 0.718952 | 0.715543 | 0.744545 | 0.706322 | 0.71172 |
| 0.701049 | 0.708053 | 0.722741 | 0.730074 | 0.711966 | 0.710594 | 0.742633 | 0.705589 | 0.704497 |
| 0.702564 | 0.711166 | 0.723176 | 0.735174 | 0.714104 | 0.713624 | 0.741248 | 0.705396 | 0.711529 |
| 0.698654 | 0.708745 | 0.717416 | 0.726947 | 0.70771 | 0.708187 | 0.738245 | 0.702078 | 0.705017 |
| 0.703548 | 0.713051 | 0.720368 | 0.7339 | 0.717763 | 0.714263 | 0.740978 | 0.703911 | 0.707593 |
| 0.705392 | 0.717589 | 0.722152 | 0.736746 | 0.71691 | 0.717339 | 0.743343 | 0.709912 | 0.715279 |
| 0.705239 | 0.716371 | 0.720713 | 0.733381 | 0.719547 | 0.717821 | 0.741493 | 0.712524 | 0.713694 |
| 0.701753 | 0.713411 | 0.720051 | 0.734244 | 0.714843 | 0.713632 | 0.742342 | 0.704891 | 0.709785 |
| 0.702375 | 0.71342 | 0.720688 | 0.733732 | 0.71432 | 0.716387 | 0.741382 | 0.702505 | 0.709536 |
| 0.705855 | 0.716097 | 0.722985 | 0.73797 | 0.720301 | 0.71506 | 0.744919 | 0.708845 | 0.715304 |
| 0.707099 | 0.715628 | 0.721845 | 0.735177 | 0.715873 | 0.717782 | 0.744091 | 0.7065 | 0.713104 |
| 0.705818 | 0.71657 | 0.726782 | 0.737553 | 0.717926 | 0.717402 | 0.744149 | 0.709537 | 0.716104 |
| 0.705574 | 0.71709 | 0.720454 | 0.736492 | 0.714768 | 0.710587 | 0.742625 | 0.706123 | 0.713858 |
| 0.709485 | 0.716563 | 0.727615 | 0.739089 | 0.720013 | 0.721847 | 0.744323 | 0.711038 | 0.715253 |
| 0.704115 | 0.714342 | 0.722496 | 0.735804 | 0.717257 | 0.716598 | 0.744444 | 0.707818 | 0.713459 |
| 0.700659 | 0.714938 | 0.723103 | 0.733955 | 0.71408 | 0.715588 | 0.739909 | 0.703618 | 0.71023 |
| 0.699165 | 0.710533 | 0.719524 | 0.736028 | 0.714297 | 0.710932 | 0.740247 | 0.703829 | 0.70738 |
| 0.70787 | 0.715021 | 0.726193 | 0.739813 | 0.716611 | 0.71677 | 0.752196 | 0.708747 | 0.71424 |
| 0.704572 | 0.712705 | 0.722447 | 0.737876 | 0.717888 | 0.715093 | 0.744153 | 0.713696 | 0.712974 |
| 0.703226 | 0.711245 | 0.721789 | 0.734175 | 0.713715 | 0.714116 | 0.740764 | 0.706779 | 0.712993 |
| 0.706241 | 0.711624 | 0.723799 | 0.739426 | 0.714569 | 0.712444 | 0.742384 | 0.704287 | 0.710591 |
| 0.703564 | 0.711476 | 0.720941 | 0.731779 | 0.712542 | 0.708975 | 0.738626 | 0.702626 | 0.710558 |
| 0.703132 | 0.710184 | 0.720915 | 0.734708 | 0.713745 | 0.710234 | 0.73668 | 0.702408 | 0.711099 |
| 0.704922 | 0.713011 | 0.724102 | 0.738553 | 0.714567 | 0.716855 | 0.746349 | 0.706867 | 0.712954 |
| 0.708871 | 0.717436 | 0.724416 | 0.734965 | 0.717254 | 0.715064 | 0.751072 | 0.709675 | 0.714896 |
| 0.703894 | 0.711964 | 0.722466 | 0.736307 | 0.71499 | 0.711742 | 0.739296 | 0.707173 | 0.710805 |
| 0.705971 | 0.716374 | 0.725926 | 0.740624 | 0.71654 | 0.715704 | 0.746739 | 0.710757 | 0.713781 |
| 0.705989 | 0.716218 | 0.723835 | 0.739694 | 0.71544 | 0.71464 | 0.74526 | 0.707459 | 0.715673 |
| 0.705117 | 0.70936 | 0.720161 | 0.729937 | 0.710397 | 0.714061 | 0.738709 | 0.703928 | 0.70847 |
| 0.702593 | 0.713922 | 0.72499 | 0.737691 | 0.714895 | 0.714675 | 0.742341 | 0.708748 | 0.714297 |
| 0.700422 | 0.7109 | 0.717718 | 0.734646 | 0.713106 | 0.70928 | 0.739875 | 0.705794 | 0.708903 |
| 0.709608 | 0.721318 | 0.729814 | 0.739294 | 0.721476 | 0.721229 | 0.751642 | 0.715439 | 0.718882 |
| 0.704522 | 0.713301 | 0.720013 | 0.736975 | 0.711495 | 0.710736 | 0.739874 | 0.704516 | 0.708975 |
| 0.707092 | 0.715699 | 0.724912 | 0.736369 | 0.714728 | 0.712173 | 0.740425 | 0.707971 | 0.714765 |
| 0.705141 | 0.712998 | 0.720785 | 0.73572 | 0.710582 | 0.714075 | 0.743269 | 0.707079 | 0.713484 |
| 0.710067 | 0.719657 | 0.725264 | 0.739748 | 0.717316 | 0.717154 | 0.749005 | 0.709829 | 0.715647 |
| 0.709196 | 0.713072 | 0.725884 | 0.739047 | 0.71669 | 0.712459 | 0.745333 | 0.714377 | 0.71473 |
| 0.708054 | 0.717503 | 0.723875 | 0.740223 | 0.717182 | 0.714354 | 0.748372 | 0.714252 | 0.716062 |
| 0.704599 | 0.70962 | 0.720316 | 0.73778 | 0.715456 | 0.71298 | 0.741836 | 0.709309 | 0.710046 |
| 0.708004 | 0.714882 | 0.723129 | 0.737318 | 0.719349 | 0.714055 | 0.746039 | 0.708856 | 0.71519 |
| 0.704435 | 0.711819 | 0.71911 | 0.736967 | 0.710662 | 0.709866 | 0.742198 | 0.704684 | 0.708769 |

| | | | | | | | | |
|---|---|---|---|---|---|---|---|---|
| 0.689463 | 0.700679 | 0.705148 | 0.721823 | 0.706309 | 0.701274 | 0.727256 | 0.693021 | 0.696903 |
| 0.742927 | 0.748521 | 0.758666 | 0.764009 | 0.753532 | 0.74423 | 0.765865 | 0.749277 | 0.752182 |
| 0.700162 | 0.715411 | 0.71998 | 0.733542 | 0.714951 | 0.712875 | 0.739778 | 0.706774 | 0.710068 |
| 0.704308 | 0.717739 | 0.723571 | 0.738764 | 0.718253 | 0.714445 | 0.744429 | 0.708756 | 0.715497 |
| 0.7 | 0.706888 | 0.718804 | 0.730138 | 0.710444 | 0.710629 | 0.732756 | 0.69944 | 0.70783 |
| 0.703903 | 0.712226 | 0.720815 | 0.731381 | 0.713391 | 0.711277 | 0.736084 | 0.704765 | 0.71149 |
| 0.69779 | 0.707632 | 0.720971 | 0.728999 | 0.710662 | 0.710842 | 0.738528 | 0.701779 | 0.708775 |
| 0.694888 | 0.703945 | 0.712214 | 0.726405 | 0.707208 | 0.707887 | 0.731674 | 0.69697 | 0.704514 |
| 0.703689 | 0.717428 | 0.721117 | 0.736649 | 0.713642 | 0.711042 | 0.74578 | 0.707946 | 0.70833 |
| 0.709641 | 0.720173 | 0.729257 | 0.740851 | 0.723826 | 0.720624 | 0.74421 | 0.715241 | 0.719224 |
| 0.706138 | 0.719132 | 0.726535 | 0.735807 | 0.716013 | 0.714821 | 0.745149 | 0.710162 | 0.717112 |
| 0.709907 | 0.719393 | 0.724096 | 0.737681 | 0.718212 | 0.718056 | 0.743505 | 0.714113 | 0.712684 |
| 0.710671 | 0.721609 | 0.729519 | 0.740319 | 0.718631 | 0.719646 | 0.74762 | 0.711306 | 0.714627 |
| 0.706793 | 0.715713 | 0.727316 | 0.736917 | 0.720626 | 0.71882 | 0.74326 | 0.711747 | 0.718745 |
| 0.70468 | 0.713927 | 0.721983 | 0.73284 | 0.715963 | 0.718456 | 0.737611 | 0.709468 | 0.710622 |
| 0.70557 | 0.712044 | 0.721225 | 0.73519 | 0.716437 | 0.71509 | 0.740508 | 0.709983 | 0.711726 |
| 0.706248 | 0.720116 | 0.723087 | 0.740272 | 0.720878 | 0.71943 | 0.740975 | 0.712343 | 0.7162 |
| 0.704839 | 0.71794 | 0.719184 | 0.736918 | 0.716554 | 0.717928 | 0.739695 | 0.7098 | 0.715193 |
| 0.696235 | 0.707153 | 0.714038 | 0.7322 | 0.706055 | 0.709632 | 0.736058 | 0.701463 | 0.705923 |
| 0.700444 | 0.708606 | 0.716929 | 0.730558 | 0.715396 | 0.711211 | 0.736598 | 0.699882 | 0.706402 |
| 0.705107 | 0.716516 | 0.724269 | 0.734294 | 0.717121 | 0.714844 | 0.742209 | 0.708481 | 0.707193 |
| 0.695564 | 0.70513 | 0.710116 | 0.72698 | 0.705561 | 0.701373 | 0.733229 | 0.700968 | 0.701372 |
| 0.699297 | 0.708714 | 0.718354 | 0.731411 | 0.71253 | 0.709512 | 0.740322 | 0.707247 | 0.707474 |
| 0.706248 | 0.711448 | 0.727214 | 0.73545 | 0.714388 | 0.713538 | 0.74524 | 0.706386 | 0.711729 |
| 0.710128 | 0.713819 | 0.723441 | 0.736855 | 0.715066 | 0.714133 | 0.745315 | 0.709532 | 0.714998 |
| 0.704749 | 0.712213 | 0.72409 | 0.736063 | 0.715938 | 0.714776 | 0.742296 | 0.707925 | 0.713486 |
| 0.699206 | 0.707181 | 0.715499 | 0.731877 | 0.707975 | 0.708579 | 0.737873 | 0.701221 | 0.705553 |
| 0.699745 | 0.706617 | 0.715979 | 0.733139 | 0.709424 | 0.70908 | 0.740679 | 0.700224 | 0.704953 |
| 0.702149 | 0.708928 | 0.717736 | 0.727265 | 0.70726 | 0.705055 | 0.737764 | 0.702395 | 0.709108 |
| 0.774539 | 0.763982 | 0.760762 | 0.751759 | 0.761589 | 0.753378 | 0.754953 | 0.768929 | 0.784049 |
| 0.789535 | 0.801684 | 0.778543 | 0.771134 | 0.778938 | 0.773088 | 0.77003 | 0.792897 | 0.795254 |
| 0.793121 | 0.781349 | 0.771223 | 0.760779 | 0.773764 | 0.764848 | 0.764069 | 0.775454 | 0.795129 |
| 0.790276 | 0.762204 | 0.786462 | 0.757407 | 0.799741 | 0.793305 | 0.756631 | 0.765585 | 0.776312 |
| 0.792713 | 0.799834 | 0.783839 | 0.76905 | 0.78704 | 0.773318 | 0.767434 | 0.792187 | 0.793588 |
| 0.694712 | 0.705152 | 0.714886 | 0.72656 | 0.710247 | 0.708464 | 0.732686 | 0.69949 | 0.70628 |
| 0.698877 | 0.708307 | 0.720697 | 0.726862 | 0.708096 | 0.710067 | 0.740178 | 0.703234 | 0.705915 |
| 0.70201 | 0.711937 | 0.720915 | 0.731903 | 0.712655 | 0.710908 | 0.742068 | 0.703047 | 0.708625 |
| 0.701605 | 0.714045 | 0.721504 | 0.734617 | 0.714994 | 0.713445 | 0.740753 | 0.703591 | 0.713577 |
| 0.69639 | 0.70954 | 0.719752 | 0.731963 | 0.70983 | 0.708905 | 0.738868 | 0.702119 | 0.70729 |
| 0.70302 | 0.712303 | 0.720251 | 0.732496 | 0.713623 | 0.714182 | 0.746172 | 0.705102 | 0.713106 |
| 0.698032 | 0.71387 | 0.720642 | 0.732397 | 0.710075 | 0.71121 | 0.741316 | 0.70091 | 0.709493 |
| 0.69946 | 0.709764 | 0.720667 | 0.733388 | 0.713846 | 0.713772 | 0.740407 | 0.70141 | 0.707121 |
| 0.695361 | 0.702716 | 0.713852 | 0.726127 | 0.706853 | 0.705504 | 0.736567 | 0.699078 | 0.700008 |
| 0.694544 | 0.706054 | 0.715381 | 0.723604 | 0.705197 | 0.70395 | 0.730614 | 0.696618 | 0.701616 |
| 0.69678 | 0.707781 | 0.716451 | 0.728414 | 0.709251 | 0.713318 | 0.736786 | 0.702729 | 0.707577 |
| 0.701189 | 0.711677 | 0.721394 | 0.730685 | 0.712443 | 0.712798 | 0.740687 | 0.704027 | 0.70937 |

| | | | | | | | | |
|---|---|---|---|---|---|---|---|---|
| 0.701548 | 0.711733 | 0.721218 | 0.737603 | 0.720163 | 0.718735 | 0.742434 | 0.704436 | 0.711815 |
| 0.702472 | 0.714113 | 0.719312 | 0.733317 | 0.714057 | 0.712579 | 0.739941 | 0.705278 | 0.709938 |
| 0.704881 | 0.715429 | 0.723699 | 0.737124 | 0.717458 | 0.715833 | 0.742573 | 0.706206 | 0.71491 |
| 0.701654 | 0.712694 | 0.719836 | 0.735169 | 0.717296 | 0.711844 | 0.740435 | 0.703393 | 0.711779 |
| 0.697922 | 0.70684 | 0.71632 | 0.728823 | 0.709935 | 0.708601 | 0.738041 | 0.700065 | 0.706654 |
| 0.697291 | 0.707624 | 0.718509 | 0.729911 | 0.710487 | 0.710352 | 0.735524 | 0.703125 | 0.706178 |
| 0.703975 | 0.714344 | 0.722838 | 0.733153 | 0.717061 | 0.714592 | 0.740738 | 0.706253 | 0.712403 |
| 0.70152 | 0.715622 | 0.723327 | 0.732846 | 0.716532 | 0.713426 | 0.742351 | 0.701987 | 0.710255 |
| 0.701277 | 0.709989 | 0.718671 | 0.733344 | 0.712245 | 0.706808 | 0.739696 | 0.69991 | 0.70618 |
| 0.704334 | 0.709955 | 0.720384 | 0.733646 | 0.712316 | 0.714123 | 0.740514 | 0.705203 | 0.710304 |
| 0.700823 | 0.711028 | 0.718816 | 0.732149 | 0.715666 | 0.710391 | 0.742255 | 0.702633 | 0.711256 |
| 0.701457 | 0.712436 | 0.717788 | 0.730509 | 0.711759 | 0.711073 | 0.739254 | 0.701706 | 0.710173 |
| 0.690391 | 0.702691 | 0.710495 | 0.724684 | 0.70673 | 0.704301 | 0.729776 | 0.694145 | 0.705369 |
| 0.692955 | 0.704241 | 0.712164 | 0.728665 | 0.706307 | 0.708485 | 0.733944 | 0.696342 | 0.699956 |
| 0.7038 | 0.715442 | 0.722927 | 0.734186 | 0.717166 | 0.716436 | 0.746234 | 0.709949 | 0.711906 |
| 0.711047 | 0.721489 | 0.731083 | 0.742827 | 0.719263 | 0.715865 | 0.75139 | 0.714194 | 0.715214 |
| 0.708238 | 0.713496 | 0.722561 | 0.737645 | 0.712418 | 0.717044 | 0.747936 | 0.708347 | 0.712335 |
| 0.708781 | 0.716321 | 0.725327 | 0.740818 | 0.717457 | 0.718159 | 0.748855 | 0.705051 | 0.716183 |
| 0.700247 | 0.709952 | 0.718183 | 0.732658 | 0.713483 | 0.712668 | 0.741935 | 0.704675 | 0.710957 |
| 0.698576 | 0.712371 | 0.722034 | 0.734266 | 0.71331 | 0.711089 | 0.740814 | 0.705839 | 0.711007 |
| 0.697732 | 0.708451 | 0.718149 | 0.727551 | 0.710654 | 0.710945 | 0.73591 | 0.698686 | 0.708427 |
| 0.704064 | 0.715544 | 0.720808 | 0.736411 | 0.714309 | 0.714541 | 0.739877 | 0.707574 | 0.711627 |
| 0.704312 | 0.717595 | 0.728494 | 0.741739 | 0.716371 | 0.717522 | 0.75079 | 0.70905 | 0.713353 |
| 0.701201 | 0.711043 | 0.72313 | 0.731605 | 0.712506 | 0.710332 | 0.741362 | 0.703871 | 0.708159 |
| 0.70231 | 0.711212 | 0.720762 | 0.734298 | 0.713305 | 0.712235 | 0.743992 | 0.705746 | 0.708205 |
| 0.796139 | 0.774723 | 0.789738 | 0.759031 | 0.797846 | 0.796862 | 0.771234 | 0.797647 | 0.787739 |
| 0.705696 | 0.711907 | 0.721687 | 0.731444 | 0.714783 | 0.715085 | 0.742292 | 0.705453 | 0.712904 |
| 0.701943 | 0.714775 | 0.720226 | 0.73788 | 0.715898 | 0.71495 | 0.743404 | 0.70556 | 0.713207 |
| 0.706216 | 0.714998 | 0.726405 | 0.740959 | 0.719939 | 0.717837 | 0.74659 | 0.712889 | 0.717566 |
| 0.702207 | 0.71416 | 0.720906 | 0.736787 | 0.716883 | 0.713606 | 0.740281 | 0.704383 | 0.709092 |
| 0.701991 | 0.707046 | 0.717725 | 0.731883 | 0.709906 | 0.710013 | 0.741517 | 0.701911 | 0.70982 |
| 0.699459 | 0.710635 | 0.718211 | 0.731158 | 0.709606 | 0.712341 | 0.738736 | 0.702285 | 0.708388 |
| 0.700315 | 0.714833 | 0.723451 | 0.738099 | 0.712605 | 0.711682 | 0.742521 | 0.707383 | 0.711659 |
| 0.701282 | 0.712383 | 0.714644 | 0.733003 | 0.71264 | 0.715046 | 0.738854 | 0.70235 | 0.707382 |
| 0.696227 | 0.708411 | 0.717743 | 0.733905 | 0.708872 | 0.710176 | 0.735401 | 0.700825 | 0.707519 |
| 0.700545 | 0.709198 | 0.713983 | 0.731709 | 0.712619 | 0.710784 | 0.73553 | 0.703613 | 0.707383 |
| 0.705597 | 0.711021 | 0.722277 | 0.738117 | 0.713694 | 0.717641 | 0.74254 | 0.708347 | 0.712726 |
| 0.70824 | 0.71563 | 0.723696 | 0.738522 | 0.715488 | 0.715003 | 0.743393 | 0.710586 | 0.715748 |
| 0.701746 | 0.712476 | 0.720155 | 0.739166 | 0.716946 | 0.711746 | 0.743218 | 0.707933 | 0.710554 |
| 0.702716 | 0.710088 | 0.719807 | 0.734542 | 0.715 | 0.714741 | 0.742057 | 0.708629 | 0.711733 |
| 0.695943 | 0.71021 | 0.719155 | 0.730442 | 0.711356 | 0.711923 | 0.739562 | 0.698663 | 0.704073 |
| 0.704162 | 0.718475 | 0.725078 | 0.741244 | 0.717739 | 0.714239 | 0.747374 | 0.709649 | 0.709343 |
| 0.70348 | 0.707281 | 0.717002 | 0.734168 | 0.714107 | 0.710934 | 0.73987 | 0.702914 | 0.712009 |
| 0.699126 | 0.711813 | 0.716968 | 0.72921 | 0.71058 | 0.71001 | 0.732837 | 0.700856 | 0.709657 |
| 0.702914 | 0.712705 | 0.721767 | 0.734161 | 0.716482 | 0.716411 | 0.738549 | 0.70398 | 0.71393 |
| 0.701603 | 0.712543 | 0.720157 | 0.731378 | 0.716473 | 0.714206 | 0.741111 | 0.70428 | 0.714286 |

| | | | | | | | | |
|---|---|---|---|---|---|---|---|---|
| 0.703014 | 0.711226 | 0.718126 | 0.729138 | 0.714444 | 0.713691 | 0.73665 | 0.703106 | 0.713185 |
| 0.701444 | 0.714683 | 0.723827 | 0.737301 | 0.717554 | 0.715316 | 0.743612 | 0.705593 | 0.712967 |
| 0.703158 | 0.710982 | 0.720673 | 0.731219 | 0.715724 | 0.713405 | 0.737915 | 0.702544 | 0.712973 |
| 0.659052 | 0.676419 | 0.686473 | 0.696422 | 0.680042 | 0.664921 | 0.696641 | 0.662583 | 0.667482 |
| 0.698587 | 0.707747 | 0.719346 | 0.730409 | 0.709286 | 0.710468 | 0.738254 | 0.70187 | 0.709997 |
| 0.699446 | 0.704962 | 0.716966 | 0.730609 | 0.712453 | 0.711708 | 0.737924 | 0.702912 | 0.710405 |
| 0.700815 | 0.711629 | 0.717786 | 0.732224 | 0.716424 | 0.715652 | 0.737385 | 0.706032 | 0.713773 |
| 0.70093 | 0.711688 | 0.717041 | 0.733062 | 0.714455 | 0.711581 | 0.737055 | 0.704345 | 0.709143 |
| 0.702718 | 0.714615 | 0.725812 | 0.732446 | 0.719241 | 0.718956 | 0.744784 | 0.70839 | 0.715378 |
| 0.699109 | 0.708049 | 0.723802 | 0.732251 | 0.713775 | 0.713128 | 0.735229 | 0.706792 | 0.708608 |
| 0.703186 | 0.710831 | 0.71884 | 0.730766 | 0.71749 | 0.713768 | 0.73904 | 0.707059 | 0.710724 |
| 0.706098 | 0.712458 | 0.722138 | 0.738134 | 0.714947 | 0.712172 | 0.740847 | 0.706978 | 0.713576 |
| 0.707218 | 0.71083 | 0.724036 | 0.734842 | 0.716492 | 0.71566 | 0.740054 | 0.709448 | 0.7127 |
| 0.702833 | 0.710835 | 0.718395 | 0.729302 | 0.716681 | 0.717579 | 0.737406 | 0.705663 | 0.710898 |
| 0.705472 | 0.713353 | 0.72472 | 0.734736 | 0.7173 | 0.715293 | 0.742791 | 0.709922 | 0.714707 |
| 0.700079 | 0.712475 | 0.716758 | 0.732815 | 0.710721 | 0.710946 | 0.73502 | 0.701767 | 0.709644 |
| 0.700894 | 0.714083 | 0.724195 | 0.734338 | 0.715086 | 0.711916 | 0.745632 | 0.707773 | 0.710283 |
| 0.703958 | 0.71094 | 0.721386 | 0.734062 | 0.714694 | 0.714686 | 0.741802 | 0.705864 | 0.712957 |
| 0.697445 | 0.709777 | 0.716127 | 0.730754 | 0.710426 | 0.710348 | 0.736139 | 0.702486 | 0.707373 |
| 0.703519 | 0.715833 | 0.71933 | 0.733669 | 0.712929 | 0.714366 | 0.740558 | 0.702349 | 0.709483 |
| 0.701543 | 0.715938 | 0.722787 | 0.738796 | 0.718503 | 0.717353 | 0.743033 | 0.708317 | 0.709436 |
| 0.702692 | 0.714354 | 0.727033 | 0.737941 | 0.714399 | 0.714984 | 0.743144 | 0.707577 | 0.712228 |
| 0.69356 | 0.691489 | 0.710628 | 0.704313 | 0.699502 | 0.69514 | 0.698933 | 0.692789 | 0.694103 |
| 0.724951 | 0.725185 | 0.731875 | 0.715677 | 0.734467 | 0.726737 | 0.705798 | 0.723232 | 0.718551 |
| 0.758119 | 0.75524 | 0.750728 | 0.741528 | 0.764825 | 0.745243 | 0.729034 | 0.766735 | 0.758155 |
| 0.756547 | 0.736609 | 0.744884 | 0.72553 | 0.765954 | 0.748714 | 0.715211 | 0.735274 | 0.74351 |
| 0.654839 | 0.671395 | 0.683236 | 0.693708 | 0.670231 | 0.66982 | 0.705573 | 0.656232 | 0.66831 |
| 0.653141 | 0.664189 | 0.678292 | 0.687 | 0.668026 | 0.667723 | 0.697635 | 0.65808 | 0.671279 |
| 0.65465 | 0.66829 | 0.672208 | 0.693942 | 0.665102 | 0.666959 | 0.697975 | 0.653292 | 0.66423 |
| 0.66485 | 0.686803 | 0.681882 | 0.700851 | 0.686358 | 0.676979 | 0.698231 | 0.669825 | 0.680243 |
| 0.638639 | 0.661611 | 0.660607 | 0.678253 | 0.659207 | 0.655196 | 0.683882 | 0.647215 | 0.648678 |
| 0.632421 | 0.654641 | 0.664103 | 0.675979 | 0.650082 | 0.648819 | 0.681532 | 0.632973 | 0.649089 |
| 0.638214 | 0.649332 | 0.660679 | 0.678706 | 0.656849 | 0.648997 | 0.678348 | 0.640011 | 0.650771 |
| 0.731213 | 0.73206 | 0.734562 | 0.718761 | 0.739866 | 0.72449 | 0.699928 | 0.733445 | 0.730891 |
| 0.697387 | 0.708511 | 0.717002 | 0.726686 | 0.707453 | 0.709935 | 0.73638 | 0.697141 | 0.705063 |
| 0.669727 | 0.682356 | 0.688515 | 0.704383 | 0.680894 | 0.674752 | 0.711326 | 0.675395 | 0.680018 |
| 0.662328 | 0.674385 | 0.678338 | 0.702018 | 0.674873 | 0.675269 | 0.704231 | 0.662848 | 0.668321 |
| 0.647373 | 0.657348 | 0.667985 | 0.679245 | 0.657285 | 0.654764 | 0.68267 | 0.64178 | 0.653412 |
| 0.66341 | 0.682148 | 0.691976 | 0.700304 | 0.684628 | 0.679792 | 0.709285 | 0.669589 | 0.675073 |
| 0.654111 | 0.670053 | 0.680174 | 0.697645 | 0.675217 | 0.674384 | 0.704246 | 0.664516 | 0.670015 |
| 0.649402 | 0.6623 | 0.676421 | 0.689331 | 0.667537 | 0.657673 | 0.69829 | 0.652955 | 0.657729 |
| 0.646089 | 0.667624 | 0.68057 | 0.690859 | 0.670955 | 0.668585 | 0.695606 | 0.656113 | 0.664498 |
| 0.64658 | 0.660864 | 0.66916 | 0.680744 | 0.662615 | 0.658056 | 0.695978 | 0.650321 | 0.658952 |
| 0.650625 | 0.658754 | 0.669275 | 0.689205 | 0.660975 | 0.659078 | 0.692718 | 0.652562 | 0.658857 |
| 0.650174 | 0.661783 | 0.674286 | 0.683791 | 0.658483 | 0.656475 | 0.685636 | 0.656335 | 0.664188 |
| 0.634695 | 0.641635 | 0.648119 | 0.676631 | 0.660462 | 0.640215 | 0.669398 | 0.639208 | 0.651714 |

| | | | | | | | | |
|---|---|---|---|---|---|---|---|---|
| 0.681101 | 0.696815 | 0.699548 | 0.71699 | 0.6966 | 0.693399 | 0.720115 | 0.685153 | 0.691871 |
| 0.644949 | 0.670558 | 0.671133 | 0.689683 | 0.666802 | 0.660421 | 0.696005 | 0.652827 | 0.661645 |
| 0.650423 | 0.66546 | 0.677541 | 0.691007 | 0.669617 | 0.666375 | 0.696149 | 0.660717 | 0.669288 |
| 0.647369 | 0.665143 | 0.680918 | 0.686587 | 0.668885 | 0.661391 | 0.695864 | 0.657421 | 0.66411 |
| 0.656268 | 0.66588 | 0.675212 | 0.680487 | 0.665818 | 0.659207 | 0.691766 | 0.649892 | 0.660106 |
| 0.655572 | 0.671973 | 0.682018 | 0.689571 | 0.67157 | 0.669494 | 0.700652 | 0.655785 | 0.664906 |
| 0.647183 | 0.666197 | 0.664401 | 0.679027 | 0.660576 | 0.651462 | 0.68235 | 0.641457 | 0.65309 |
| 0.640443 | 0.65159 | 0.663366 | 0.676164 | 0.653919 | 0.651436 | 0.684584 | 0.643226 | 0.643516 |
| 0.636242 | 0.654947 | 0.660336 | 0.677784 | 0.656532 | 0.646755 | 0.681203 | 0.647752 | 0.649699 |
| 0.688227 | 0.696759 | 0.70413 | 0.720839 | 0.69545 | 0.69649 | 0.725302 | 0.685048 | 0.689225 |
| 0.685009 | 0.692405 | 0.702224 | 0.713281 | 0.69104 | 0.69258 | 0.721702 | 0.684539 | 0.688072 |
| 0.675918 | 0.685358 | 0.695335 | 0.71056 | 0.690942 | 0.688491 | 0.715522 | 0.680645 | 0.682859 |
| 0.670256 | 0.68449 | 0.687125 | 0.703865 | 0.679112 | 0.679495 | 0.711231 | 0.677673 | 0.676412 |
| 0.671418 | 0.688152 | 0.695621 | 0.702409 | 0.687951 | 0.685986 | 0.709967 | 0.670977 | 0.682583 |
| 0.662702 | 0.677603 | 0.6828 | 0.701037 | 0.6819 | 0.674488 | 0.702268 | 0.665486 | 0.671398 |

| | | | | | | | | |
|---|---|---|---|---|---|---|---|---|
| | | | | | | | | HG_ |
| Home_garc | Home_garc | Home_garc | Home_garc | Home_garc | Home_garc | Home_garc | Home_garc | Home_garc |
| 0.790272 | 0.802914 | 0.796602 | 0.798603 | 0.778906 | 0.784356 | 0.777072 | 0.779532 | 0.782859 |
| 0.794583 | 0.789552 | 0.799468 | 0.796676 | 0.787614 | 0.794304 | 0.786442 | 0.790021 | 0.792937 |
| 0.775902 | 0.790884 | 0.787919 | 0.787225 | 0.779061 | 0.787002 | 0.771597 | 0.785363 | 0.781847 |
| 0.788158 | 0.793872 | 0.798035 | 0.784141 | 0.782333 | 0.787187 | 0.781101 | 0.784438 | 0.787558 |
| 0.778709 | 0.786774 | 0.78436 | 0.779081 | 0.783008 | 0.778304 | 0.771648 | 0.780573 | 0.780251 |
| 0.784281 | 0.776714 | 0.7885 | 0.78521 | 0.780379 | 0.768284 | 0.766195 | 0.776965 | 0.782313 |
| 0.71407 | 0.711723 | 0.718077 | 0.717979 | 0.716685 | 0.712157 | 0.703982 | 0.711974 | 0.716518 |
| 0.706194 | 0.709143 | 0.712722 | 0.711971 | 0.71187 | 0.705577 | 0.701938 | 0.70984 | 0.714502 |
| 0.788937 | 0.782727 | 0.792732 | 0.786696 | 0.787536 | 0.784768 | 0.775919 | 0.783631 | 0.780041 |
| 0.779502 | 0.798806 | 0.785797 | 0.790313 | 0.773583 | 0.778766 | 0.771424 | 0.773925 | 0.77622 |
| 0.780546 | 0.790964 | 0.791068 | 0.790875 | 0.776194 | 0.775967 | 0.772873 | 0.777186 | 0.787291 |
| 0.789151 | 0.803104 | 0.797625 | 0.815631 | 0.791674 | 0.791111 | 0.792334 | 0.788382 | 0.787972 |
| 0.770883 | 0.774681 | 0.782029 | 0.786923 | 0.781659 | 0.781258 | 0.775438 | 0.781678 | 0.772966 |
| 0.778929 | 0.786279 | 0.78297 | 0.788257 | 0.776229 | 0.785216 | 0.764951 | 0.778142 | 0.784608 |
| 0.765852 | 0.771077 | 0.781201 | 0.780432 | 0.767459 | 0.769623 | 0.765404 | 0.7704 | 0.769531 |
| 0.78238 | 0.791257 | 0.782502 | 0.801758 | 0.778289 | 0.78482 | 0.777085 | 0.773274 | 0.775562 |
| 0.780877 | 0.793664 | 0.784791 | 0.800701 | 0.770204 | 0.77151 | 0.774752 | 0.77347 | 0.775529 |
| 0.761744 | 0.765713 | 0.767408 | 0.770468 | 0.753482 | 0.761722 | 0.752089 | 0.759313 | 0.756342 |
| 0.783232 | 0.787929 | 0.79302 | 0.804113 | 0.790527 | 0.803881 | 0.804388 | 0.77671 | 0.781309 |
| 0.824253 | 0.799517 | 0.810538 | 0.790376 | 0.808157 | 0.788063 | 0.780198 | 0.79055 | 0.793656 |
| 1 | 0.813202 | 0.800033 | 0.797657 | 0.796541 | 0.785539 | 0.778428 | 0.787707 | 0.79069 |
| 0.813202 | 1 | 0.808925 | 0.809209 | 0.791042 | 0.794683 | 0.784649 | 0.78491 | 0.792866 |
| 0.800033 | 0.808925 | 1 | 0.802077 | 0.854042 | 0.791013 | 0.782361 | 0.803197 | 0.829328 |
| 0.797657 | 0.809209 | 0.802077 | 1 | 0.790721 | 0.793701 | 0.790938 | 0.791928 | 0.791674 |
| 0.796541 | 0.791042 | 0.854042 | 0.790721 | 1 | 0.782841 | 0.778771 | 0.786192 | 0.819646 |
| 0.785539 | 0.794683 | 0.791013 | 0.793701 | 0.782841 | 1 | 0.794931 | 0.784198 | 0.783325 |
| 0.778428 | 0.784649 | 0.782361 | 0.790938 | 0.778771 | 0.794931 | 1 | 0.775524 | 0.782659 |
| 0.787707 | 0.78491 | 0.803197 | 0.791928 | 0.786192 | 0.784198 | 0.775524 | 1 | 0.798298 |
| 0.79069 | 0.792866 | 0.829328 | 0.791674 | 0.819646 | 0.783325 | 0.782659 | 0.798298 | 1 |
| 0.777392 | 0.783217 | 0.784631 | 0.820595 | 0.785756 | 0.784049 | 0.776252 | 0.772835 | 0.78102 |
| 0.781315 | 0.786129 | 0.791455 | 0.788958 | 0.786982 | 0.829494 | 0.773827 | 0.781513 | 0.781547 |
| 0.781194 | 0.792559 | 0.785908 | 0.78663 | 0.78046 | 0.830321 | 0.772958 | 0.785293 | 0.780085 |
| 0.786309 | 0.783617 | 0.790154 | 0.789734 | 0.781036 | 0.78835 | 0.79385 | 0.792118 | 0.784709 |
| 0.778925 | 0.78065 | 0.810227 | 0.779768 | 0.801196 | 0.802243 | 0.775646 | 0.788551 | 0.807101 |
| 0.773594 | 0.775973 | 0.775074 | 0.779042 | 0.7729 | 0.802168 | 0.820349 | 0.772479 | 0.7753 |
| 0.779762 | 0.788113 | 0.786146 | 0.798599 | 0.784354 | 0.799418 | 0.80727 | 0.779128 | 0.783228 |
| 0.787258 | 0.795957 | 0.798091 | 0.794106 | 0.777605 | 0.794151 | 0.792913 | 0.786996 | 0.790382 |
| 0.771018 | 0.773884 | 0.827134 | 0.781599 | 0.814859 | 0.772038 | 0.770556 | 0.783692 | 0.844745 |
| 0.783993 | 0.777158 | 0.793853 | 0.778982 | 0.791681 | 0.778866 | 0.772384 | 0.786451 | 0.788266 |

| | | | | | | | | |
|---|---|---|---|---|---|---|---|---|
| 0.753695 | 0.763798 | 0.760698 | 0.778013 | 0.758289 | 0.758748 | 0.749598 | 0.755357 | 0.758135 |
| 0.777823 | 0.793867 | 0.790669 | 0.796405 | 0.776961 | 0.783844 | 0.77786 | 0.785106 | 0.780404 |
| 0.789626 | 0.806382 | 0.799817 | 0.819252 | 0.791402 | 0.785767 | 0.784998 | 0.781655 | 0.79325 |
| 0.791626 | 0.808535 | 0.798813 | 0.808933 | 0.789921 | 0.782888 | 0.777705 | 0.781387 | 0.7851 |
| 0.789374 | 0.799137 | 0.79578 | 0.807849 | 0.792702 | 0.777361 | 0.777741 | 0.776944 | 0.793417 |
| 0.788015 | 0.79716 | 0.794382 | 0.797814 | 0.785982 | 0.780309 | 0.774839 | 0.784496 | 0.785156 |
| 0.782766 | 0.786433 | 0.785826 | 0.774085 | 0.78004 | 0.781107 | 0.76891 | 0.78127 | 0.782362 |
| 0.78911 | 0.807094 | 0.797581 | 0.794135 | 0.792895 | 0.793238 | 0.786427 | 0.778829 | 0.785322 |
| 0.779019 | 0.779258 | 0.782486 | 0.779313 | 0.779006 | 0.778656 | 0.769999 | 0.778409 | 0.781385 |
| 0.784162 | 0.782066 | 0.781353 | 0.778353 | 0.777924 | 0.785018 | 0.769457 | 0.786842 | 0.7795 |
| 0.796335 | 0.803621 | 0.791106 | 0.804561 | 0.783257 | 0.7819 | 0.780164 | 0.784323 | 0.780548 |
| 0.798 | 0.799197 | 0.796219 | 0.806303 | 0.797266 | 0.784478 | 0.790043 | 0.785855 | 0.782627 |
| 0.793147 | 0.797739 | 0.800251 | 0.808647 | 0.78246 | 0.789461 | 0.791259 | 0.787549 | 0.782573 |
| 0.793385 | 0.796052 | 0.790369 | 0.803736 | 0.791191 | 0.786441 | 0.785833 | 0.7875 | 0.781779 |
| 0.792341 | 0.80616 | 0.790433 | 0.809238 | 0.782423 | 0.791833 | 0.781551 | 0.780562 | 0.783312 |
| 0.797787 | 0.79978 | 0.788431 | 0.791849 | 0.781884 | 0.791193 | 0.780022 | 0.784564 | 0.779485 |
| 0.788269 | 0.791978 | 0.791688 | 0.808274 | 0.773846 | 0.785251 | 0.781574 | 0.780161 | 0.777335 |
| 0.788117 | 0.808628 | 0.795744 | 0.799054 | 0.787555 | 0.799123 | 0.795396 | 0.789569 | 0.782017 |
| 0.78859 | 0.802934 | 0.791849 | 0.799618 | 0.782429 | 0.78535 | 0.78249 | 0.779998 | 0.786287 |
| 0.787395 | 0.793311 | 0.796109 | 0.810037 | 0.783888 | 0.799883 | 0.778552 | 0.791691 | 0.779896 |
| 0.793627 | 0.801056 | 0.788695 | 0.799432 | 0.784469 | 0.78469 | 0.774752 | 0.783105 | 0.785922 |
| 0.782468 | 0.790474 | 0.786165 | 0.79223 | 0.769783 | 0.785032 | 0.769941 | 0.785714 | 0.775953 |
| 0.786659 | 0.790982 | 0.79168 | 0.794265 | 0.788024 | 0.785277 | 0.775969 | 0.781158 | 0.785697 |
| 0.796802 | 0.781222 | 0.79347 | 0.779754 | 0.7875 | 0.78387 | 0.774183 | 0.788527 | 0.788351 |
| 0.77622 | 0.762908 | 0.784343 | 0.76405 | 0.783382 | 0.769137 | 0.768465 | 0.774711 | 0.780233 |
| 0.706169 | 0.709063 | 0.714561 | 0.713438 | 0.712641 | 0.706083 | 0.702353 | 0.709595 | 0.714769 |
| 0.711329 | 0.713872 | 0.718508 | 0.713861 | 0.716247 | 0.71179 | 0.707586 | 0.713589 | 0.713172 |
| 0.712973 | 0.714182 | 0.722535 | 0.71517 | 0.717162 | 0.716746 | 0.708013 | 0.717035 | 0.715273 |
| 0.779272 | 0.779909 | 0.783061 | 0.781875 | 0.780944 | 0.77786 | 0.770744 | 0.776431 | 0.770698 |
| 0.784503 | 0.777117 | 0.785471 | 0.784232 | 0.779038 | 0.775087 | 0.770084 | 0.778971 | 0.774371 |
| 0.785821 | 0.786224 | 0.782292 | 0.788811 | 0.768545 | 0.770638 | 0.760659 | 0.774967 | 0.770344 |
| 0.781068 | 0.778083 | 0.781521 | 0.781555 | 0.777054 | 0.773797 | 0.777435 | 0.780928 | 0.774183 |
| 0.794349 | 0.808158 | 0.810589 | 0.815075 | 0.788773 | 0.793594 | 0.797889 | 0.796374 | 0.78443 |
| 0.792015 | 0.808532 | 0.806072 | 0.819175 | 0.804637 | 0.79872 | 0.806791 | 0.788037 | 0.794403 |
| 0.788126 | 0.797665 | 0.803473 | 0.80878 | 0.791494 | 0.79669 | 0.795518 | 0.784568 | 0.782948 |
| 0.795746 | 0.809376 | 0.795419 | 0.815334 | 0.789926 | 0.812442 | 0.805834 | 0.788652 | 0.791528 |
| 0.775069 | 0.784433 | 0.790018 | 0.791245 | 0.780402 | 0.782205 | 0.786102 | 0.787765 | 0.777769 |
| 0.804813 | 0.821576 | 0.816534 | 0.821534 | 0.800301 | 0.810417 | 0.812106 | 0.802776 | 0.805525 |
| 0.779466 | 0.799202 | 0.784745 | 0.78438 | 0.778294 | 0.788167 | 0.776652 | 0.77698 | 0.776816 |
| 0.799488 | 0.794096 | 0.792715 | 0.801208 | 0.793044 | 0.78844 | 0.778768 | 0.785418 | 0.785995 |
| 0.782791 | 0.800348 | 0.790916 | 0.793972 | 0.778901 | 0.787532 | 0.780721 | 0.778004 | 0.784515 |
| 0.805795 | 0.822952 | 0.821377 | 0.820256 | 0.803429 | 0.815633 | 0.814376 | 0.805847 | 0.805432 |
| 0.79023 | 0.80294 | 0.797627 | 0.798314 | 0.782104 | 0.782477 | 0.772383 | 0.782245 | 0.784492 |
| 0.795443 | 0.806339 | 0.794963 | 0.798167 | 0.784503 | 0.781928 | 0.773426 | 0.787198 | 0.795742 |
| 0.789552 | 0.796193 | 0.788524 | 0.793262 | 0.785619 | 0.790613 | 0.777345 | 0.786251 | 0.783165 |
| 0.785047 | 0.790219 | 0.781489 | 0.794721 | 0.787057 | 0.794152 | 0.788097 | 0.778542 | 0.782674 |

| | | | | | | | | |
|---|---|---|---|---|---|---|---|---|
| 0.786449 | 0.804729 | 0.794565 | 0.804776 | 0.779412 | 0.782698 | 0.773321 | 0.786121 | 0.785702 |
| 0.79874 | 0.789487 | 0.793388 | 0.792011 | 0.800233 | 0.791913 | 0.781517 | 0.784865 | 0.783564 |
| 0.781075 | 0.784556 | 0.787664 | 0.785511 | 0.783368 | 0.774172 | 0.763936 | 0.78049 | 0.780656 |
| 0.795862 | 0.802281 | 0.803779 | 0.801181 | 0.793959 | 0.798426 | 0.793865 | 0.79526 | 0.795644 |
| 0.788178 | 0.795421 | 0.789984 | 0.79927 | 0.788267 | 0.783337 | 0.776611 | 0.78857 | 0.783562 |
| 0.799143 | 0.798621 | 0.791238 | 0.796033 | 0.79337 | 0.782627 | 0.77369 | 0.786904 | 0.778931 |
| 0.782175 | 0.787764 | 0.791341 | 0.7924 | 0.789235 | 0.787569 | 0.775082 | 0.786076 | 0.779624 |
| 0.779429 | 0.781991 | 0.784296 | 0.782569 | 0.780773 | 0.788065 | 0.769447 | 0.77596 | 0.78034 |
| 0.784397 | 0.789491 | 0.785494 | 0.784284 | 0.787476 | 0.776771 | 0.771296 | 0.782935 | 0.778302 |
| 0.778454 | 0.774745 | 0.787771 | 0.788535 | 0.783987 | 0.779258 | 0.779647 | 0.774312 | 0.782397 |
| 0.777433 | 0.779923 | 0.78032 | 0.778441 | 0.778781 | 0.776642 | 0.773627 | 0.780347 | 0.779449 |
| 0.779674 | 0.783005 | 0.789085 | 0.787573 | 0.781737 | 0.773646 | 0.775923 | 0.779153 | 0.774505 |
| 0.783856 | 0.791217 | 0.788474 | 0.794314 | 0.792138 | 0.786187 | 0.779275 | 0.783161 | 0.778394 |
| 0.777806 | 0.775008 | 0.788521 | 0.79001 | 0.796045 | 0.778249 | 0.778876 | 0.772282 | 0.778911 |
| 0.793479 | 0.782569 | 0.78454 | 0.783383 | 0.782583 | 0.781222 | 0.775711 | 0.779872 | 0.779856 |
| 0.778981 | 0.775974 | 0.783369 | 0.782773 | 0.782035 | 0.78146 | 0.769369 | 0.78103 | 0.78023 |
| 0.776978 | 0.779565 | 0.792965 | 0.78553 | 0.782765 | 0.778244 | 0.774781 | 0.781165 | 0.783306 |
| 0.783154 | 0.793903 | 0.793696 | 0.797259 | 0.781315 | 0.794673 | 0.789399 | 0.778605 | 0.785278 |
| 0.779929 | 0.777181 | 0.786731 | 0.785957 | 0.787131 | 0.781745 | 0.777024 | 0.780373 | 0.781931 |
| 0.785378 | 0.780859 | 0.786666 | 0.782103 | 0.786945 | 0.775545 | 0.774422 | 0.778944 | 0.781449 |
| 0.77654 | 0.784338 | 0.779874 | 0.783993 | 0.779395 | 0.7725 | 0.771341 | 0.771756 | 0.783454 |
| 0.780645 | 0.775882 | 0.780872 | 0.782935 | 0.778348 | 0.776352 | 0.773277 | 0.782502 | 0.779344 |
| 0.78063 | 0.782547 | 0.789848 | 0.786293 | 0.786366 | 0.783676 | 0.768548 | 0.780341 | 0.784577 |
| 0.791288 | 0.786003 | 0.78046 | 0.792087 | 0.791217 | 0.786799 | 0.774 | 0.778287 | 0.775736 |
| 0.778167 | 0.78879 | 0.788335 | 0.783842 | 0.776344 | 0.782131 | 0.786098 | 0.778387 | 0.774443 |
| 0.786611 | 0.793528 | 0.803419 | 0.794006 | 0.789796 | 0.78831 | 0.789912 | 0.784991 | 0.783788 |
| 0.777441 | 0.787816 | 0.790918 | 0.78095 | 0.776408 | 0.779743 | 0.779949 | 0.779245 | 0.772014 |
| 0.777077 | 0.780138 | 0.785256 | 0.785673 | 0.783893 | 0.779113 | 0.77112 | 0.779646 | 0.779915 |
| 0.780418 | 0.782576 | 0.787309 | 0.788635 | 0.784603 | 0.781398 | 0.776872 | 0.780966 | 0.779915 |
| 0.780357 | 0.777861 | 0.787965 | 0.789283 | 0.777451 | 0.77571 | 0.774496 | 0.77848 | 0.77356 |
| 0.779374 | 0.7907 | 0.78801 | 0.785167 | 0.785362 | 0.771281 | 0.775234 | 0.777457 | 0.782815 |
| 0.780747 | 0.770376 | 0.78125 | 0.779545 | 0.772234 | 0.778237 | 0.770032 | 0.777913 | 0.778004 |
| 0.782478 | 0.789995 | 0.79909 | 0.788021 | 0.784054 | 0.779368 | 0.774923 | 0.789394 | 0.781765 |
| 0.787969 | 0.803507 | 0.794053 | 0.804734 | 0.785156 | 0.781271 | 0.773268 | 0.778798 | 0.781551 |
| 0.791228 | 0.797277 | 0.789294 | 0.797742 | 0.786706 | 0.792868 | 0.776044 | 0.78263 | 0.777897 |
| 0.796218 | 0.859214 | 0.802658 | 0.792825 | 0.790688 | 0.78759 | 0.780289 | 0.793639 | 0.79787 |
| 0.781232 | 0.781866 | 0.786145 | 0.78222 | 0.784429 | 0.773895 | 0.768503 | 0.775784 | 0.782908 |
| 0.755149 | 0.758417 | 0.764129 | 0.762623 | 0.758795 | 0.766067 | 0.752212 | 0.758215 | 0.756912 |
| 0.786798 | 0.780866 | 0.772947 | 0.776565 | 0.783643 | 0.772651 | 0.760721 | 0.761771 | 0.766315 |
| 0.763981 | 0.753502 | 0.76537 | 0.75765 | 0.750241 | 0.754037 | 0.74425 | 0.761327 | 0.750266 |
| 0.774805 | 0.769936 | 0.772938 | 0.779388 | 0.762532 | 0.775678 | 0.766628 | 0.758732 | 0.76034 |
| 0.740709 | 0.755987 | 0.757257 | 0.783431 | 0.762662 | 0.74534 | 0.758971 | 0.736988 | 0.764778 |
| 0.755561 | 0.775422 | 0.778132 | 0.779272 | 0.757637 | 0.755595 | 0.744098 | 0.760631 | 0.760177 |
| 0.74959 | 0.770628 | 0.753665 | 0.761105 | 0.752726 | 0.759146 | 0.745294 | 0.752527 | 0.746866 |
| 0.752292 | 0.7492 | 0.755717 | 0.750631 | 0.752291 | 0.745026 | 0.738975 | 0.745562 | 0.750829 |
| 0.755856 | 0.769898 | 0.768475 | 0.768676 | 0.754702 | 0.758858 | 0.756225 | 0.759652 | 0.759329 |
| 0.760682 | 0.767212 | 0.766469 | 0.770584 | 0.750161 | 0.781116 | 0.757427 | 0.750895 | 0.754874 |

| | | | | | | | | |
|---|---|---|---|---|---|---|---|---|
| 0.752703 | 0.743586 | 0.745722 | 0.743518 | 0.748811 | 0.762853 | 0.737577 | 0.742436 | 0.746682 |
| 0.765402 | 0.780313 | 0.779854 | 0.788944 | 0.763033 | 0.774484 | 0.778683 | 0.760445 | 0.759588 |
| 0.744254 | 0.741671 | 0.741191 | 0.742655 | 0.736791 | 0.744796 | 0.735063 | 0.742198 | 0.745505 |
| 0.743881 | 0.757727 | 0.751923 | 0.759851 | 0.742908 | 0.746438 | 0.739796 | 0.744483 | 0.749211 |
| 0.752222 | 0.768532 | 0.769168 | 0.773457 | 0.768761 | 0.755075 | 0.781084 | 0.746891 | 0.760543 |
| 0.735541 | 0.744766 | 0.751036 | 0.753277 | 0.742935 | 0.742715 | 0.73417 | 0.747336 | 0.748614 |
| 0.754123 | 0.760776 | 0.757791 | 0.760802 | 0.756137 | 0.749411 | 0.746531 | 0.755457 | 0.759805 |
| 0.764226 | 0.773112 | 0.763081 | 0.769549 | 0.761743 | 0.769129 | 0.761602 | 0.753068 | 0.757953 |
| 0.726333 | 0.726077 | 0.730886 | 0.748 | 0.7443 | 0.729705 | 0.7304 | 0.728261 | 0.727437 |
| 0.75498 | 0.76092 | 0.778103 | 0.765944 | 0.759877 | 0.757191 | 0.769883 | 0.756344 | 0.745933 |
| 0.741025 | 0.743742 | 0.757708 | 0.747337 | 0.746895 | 0.749913 | 0.737263 | 0.744448 | 0.745742 |
| 0.772557 | 0.791211 | 0.784219 | 0.783927 | 0.768277 | 0.765849 | 0.75774 | 0.769679 | 0.769181 |
| 0.781171 | 0.774867 | 0.781756 | 0.775557 | 0.779383 | 0.775363 | 0.771536 | 0.774578 | 0.785144 |
| 0.760303 | 0.747883 | 0.769231 | 0.761967 | 0.755656 | 0.757829 | 0.742158 | 0.769077 | 0.765656 |
| 0.771034 | 0.784583 | 0.77824 | 0.778946 | 0.786676 | 0.765486 | 0.760945 | 0.776892 | 0.776187 |
| 0.76775 | 0.763834 | 0.775436 | 0.769414 | 0.771846 | 0.769884 | 0.761803 | 0.761579 | 0.776395 |
| 0.776232 | 0.775041 | 0.781483 | 0.788594 | 0.783509 | 0.775335 | 0.772976 | 0.779173 | 0.785896 |
| 0.779988 | 0.780271 | 0.787672 | 0.777529 | 0.785881 | 0.779098 | 0.771183 | 0.786514 | 0.781071 |
| 0.757946 | 0.761739 | 0.759648 | 0.758139 | 0.760198 | 0.762024 | 0.748165 | 0.76273 | 0.759156 |
| 0.769824 | 0.772876 | 0.790493 | 0.799273 | 0.789899 | 0.777553 | 0.776589 | 0.772797 | 0.795827 |
| 0.747085 | 0.748265 | 0.760065 | 0.744831 | 0.749127 | 0.755641 | 0.735439 | 0.755427 | 0.759011 |
| 0.775146 | 0.773826 | 0.780637 | 0.778934 | 0.783782 | 0.770551 | 0.76937 | 0.773871 | 0.779116 |
| 0.776469 | 0.767826 | 0.775839 | 0.776047 | 0.778708 | 0.773356 | 0.766579 | 0.774865 | 0.772862 |
| 0.777453 | 0.761295 | 0.750808 | 0.765818 | 0.75778 | 0.771095 | 0.738585 | 0.754609 | 0.754803 |
| 0.74983 | 0.753172 | 0.755574 | 0.758217 | 0.750619 | 0.755474 | 0.754014 | 0.751443 | 0.744874 |
| 0.768483 | 0.767174 | 0.768171 | 0.769423 | 0.762328 | 0.76206 | 0.752248 | 0.760867 | 0.764821 |
| 0.766697 | 0.761789 | 0.77146 | 0.762988 | 0.764745 | 0.765934 | 0.751728 | 0.769609 | 0.766195 |
| 0.788506 | 0.777213 | 0.783717 | 0.787659 | 0.785974 | 0.782094 | 0.7684 | 0.781079 | 0.778625 |
| 0.749319 | 0.750581 | 0.753993 | 0.750962 | 0.749417 | 0.742212 | 0.744783 | 0.751716 | 0.749359 |
| 0.770075 | 0.77221 | 0.773122 | 0.775469 | 0.772297 | 0.768685 | 0.765607 | 0.768739 | 0.769343 |
| 0.779445 | 0.76956 | 0.770301 | 0.772166 | 0.772458 | 0.765463 | 0.760289 | 0.773374 | 0.774939 |
| 0.758037 | 0.770644 | 0.769205 | 0.766932 | 0.757753 | 0.764495 | 0.761193 | 0.763128 | 0.764092 |
| 0.759964 | 0.769867 | 0.765684 | 0.770238 | 0.757114 | 0.76442 | 0.767136 | 0.762657 | 0.759031 |
| 0.77113 | 0.766858 | 0.784022 | 0.768217 | 0.775694 | 0.773865 | 0.760854 | 0.779104 | 0.783935 |
| 0.766693 | 0.759423 | 0.763636 | 0.771979 | 0.764929 | 0.7706 | 0.752539 | 0.762414 | 0.764512 |
| 0.772948 | 0.770232 | 0.76882 | 0.775943 | 0.769523 | 0.767083 | 0.766889 | 0.766808 | 0.766746 |
| 0.730484 | 0.730926 | 0.73034 | 0.732129 | 0.728101 | 0.720585 | 0.71475 | 0.724113 | 0.730052 |
| 0.755064 | 0.763094 | 0.763633 | 0.756847 | 0.759521 | 0.755393 | 0.751964 | 0.755899 | 0.759659 |
| 0.728582 | 0.727809 | 0.732714 | 0.734355 | 0.730398 | 0.724267 | 0.727222 | 0.731814 | 0.7288 |
| 0.724243 | 0.727949 | 0.729105 | 0.727237 | 0.726261 | 0.720452 | 0.718692 | 0.729555 | 0.723939 |
| 0.774249 | 0.779475 | 0.77766 | 0.777868 | 0.776985 | 0.780198 | 0.77103 | 0.765043 | 0.770601 |
| 0.726697 | 0.724151 | 0.725481 | 0.729618 | 0.727244 | 0.714308 | 0.711979 | 0.725689 | 0.723769 |
| 0.719737 | 0.727138 | 0.723022 | 0.726286 | 0.722564 | 0.714516 | 0.722089 | 0.716106 | 0.716692 |
| 0.710052 | 0.71113 | 0.718066 | 0.714555 | 0.716011 | 0.711798 | 0.714612 | 0.712743 | 0.705136 |
| 0.765769 | 0.774154 | 0.781577 | 0.773783 | 0.768407 | 0.766107 | 0.755815 | 0.767711 | 0.766496 |
| 0.709095 | 0.716186 | 0.720813 | 0.727824 | 0.715692 | 0.71271 | 0.709178 | 0.715942 | 0.710132 |
| 0.770868 | 0.778365 | 0.777458 | 0.778409 | 0.772589 | 0.775297 | 0.765631 | 0.757826 | 0.767768 |

| | | | | | | | | |
|---|---|---|---|---|---|---|---|---|
| 0.761374 | 0.772076 | 0.771459 | 0.777126 | 0.762067 | 0.766591 | 0.756283 | 0.760752 | 0.774015 |
| 0.771538 | 0.783023 | 0.780566 | 0.781319 | 0.769785 | 0.774149 | 0.767659 | 0.765649 | 0.774209 |
| 0.749307 | 0.751452 | 0.75737 | 0.755503 | 0.760423 | 0.748237 | 0.743964 | 0.758891 | 0.756856 |
| 0.709253 | 0.71243 | 0.715171 | 0.716383 | 0.707028 | 0.701327 | 0.707163 | 0.705037 | 0.702409 |
| 0.695098 | 0.694608 | 0.701055 | 0.711554 | 0.701632 | 0.704638 | 0.705752 | 0.703834 | 0.699746 |
| 0.769702 | 0.771135 | 0.775353 | 0.788296 | 0.773 | 0.76753 | 0.757894 | 0.766537 | 0.772114 |
| 0.694265 | 0.69302 | 0.697264 | 0.695418 | 0.694529 | 0.688834 | 0.688196 | 0.696811 | 0.695649 |
| 0.732134 | 0.736324 | 0.741284 | 0.74015 | 0.738804 | 0.732839 | 0.72787 | 0.738286 | 0.733652 |
| 0.755368 | 0.767602 | 0.764691 | 0.766759 | 0.765117 | 0.768014 | 0.754831 | 0.754285 | 0.758241 |
| 0.720622 | 0.729314 | 0.733165 | 0.726741 | 0.729483 | 0.721488 | 0.716216 | 0.724865 | 0.726306 |
| 0.748494 | 0.758235 | 0.762841 | 0.762276 | 0.748755 | 0.747591 | 0.744846 | 0.745085 | 0.752265 |
| 0.586826 | 0.606061 | 0.633987 | 0.568862 | 0.594771 | 0.624204 | 0.570552 | 0.641892 | 0.605263 |
| 0.653824 | 0.6543 | 0.656238 | 0.651913 | 0.652805 | 0.654849 | 0.648628 | 0.654948 | 0.661343 |
| 0.654677 | 0.654855 | 0.661183 | 0.656524 | 0.661695 | 0.655922 | 0.649376 | 0.652911 | 0.660466 |
| 0.662276 | 0.660887 | 0.672627 | 0.660683 | 0.670455 | 0.663792 | 0.660255 | 0.671637 | 0.671016 |
| 0.659948 | 0.660931 | 0.665894 | 0.663293 | 0.668835 | 0.662632 | 0.660143 | 0.66028 | 0.669048 |
| 0.621851 | 0.620606 | 0.623558 | 0.620773 | 0.623347 | 0.61793 | 0.621315 | 0.622596 | 0.619982 |
| 0.662316 | 0.664371 | 0.671377 | 0.66629 | 0.671966 | 0.668169 | 0.659812 | 0.661854 | 0.669553 |
| 0.739636 | 0.74336 | 0.752939 | 0.7527 | 0.748135 | 0.740471 | 0.739368 | 0.741894 | 0.74462 |
| 0.731965 | 0.743069 | 0.746641 | 0.746908 | 0.745024 | 0.734208 | 0.731747 | 0.735311 | 0.742664 |
| 0.745919 | 0.750348 | 0.758445 | 0.75244 | 0.753269 | 0.747116 | 0.74049 | 0.746123 | 0.747486 |
| 0.618011 | 0.617734 | 0.619084 | 0.623196 | 0.622493 | 0.619264 | 0.618896 | 0.618502 | 0.620738 |
| 0.596806 | 0.599889 | 0.602921 | 0.603999 | 0.603389 | 0.596482 | 0.600541 | 0.600965 | 0.600953 |
| 0.704878 | 0.708121 | 0.711545 | 0.708837 | 0.710772 | 0.711769 | 0.702381 | 0.707405 | 0.708145 |
| 0.705845 | 0.709356 | 0.714115 | 0.707917 | 0.714675 | 0.706598 | 0.701772 | 0.707675 | 0.708429 |
| 0.699522 | 0.70108 | 0.703275 | 0.699129 | 0.703185 | 0.701916 | 0.694176 | 0.700978 | 0.704491 |
| 0.703434 | 0.705317 | 0.709222 | 0.704006 | 0.707735 | 0.706499 | 0.69871 | 0.705771 | 0.708784 |
| 0.707058 | 0.706587 | 0.711636 | 0.709823 | 0.708589 | 0.710024 | 0.704787 | 0.709527 | 0.712103 |
| 0.704828 | 0.710312 | 0.714562 | 0.711473 | 0.711752 | 0.710933 | 0.703278 | 0.709938 | 0.712269 |
| 0.701974 | 0.709045 | 0.714834 | 0.707441 | 0.710444 | 0.709288 | 0.702378 | 0.708931 | 0.71064 |
| 0.703103 | 0.706928 | 0.713191 | 0.710255 | 0.710454 | 0.707351 | 0.700573 | 0.710008 | 0.712775 |
| 0.709876 | 0.713552 | 0.719744 | 0.719099 | 0.717377 | 0.716717 | 0.710955 | 0.715837 | 0.719882 |
| 0.710446 | 0.712791 | 0.714861 | 0.712355 | 0.717073 | 0.714366 | 0.706585 | 0.711902 | 0.712518 |
| 0.710175 | 0.710613 | 0.719061 | 0.712519 | 0.714456 | 0.713155 | 0.706756 | 0.715598 | 0.716729 |
| 0.713103 | 0.712647 | 0.721038 | 0.71792 | 0.718632 | 0.716755 | 0.707334 | 0.714694 | 0.718436 |
| 0.704314 | 0.709353 | 0.715237 | 0.709649 | 0.714274 | 0.709131 | 0.704413 | 0.714115 | 0.712957 |
| 0.711262 | 0.707387 | 0.71798 | 0.715189 | 0.716641 | 0.71473 | 0.707687 | 0.713786 | 0.713384 |
| 0.71094 | 0.710643 | 0.717398 | 0.719081 | 0.715564 | 0.711387 | 0.705412 | 0.714173 | 0.711719 |
| 0.702524 | 0.706243 | 0.712568 | 0.706594 | 0.709058 | 0.704844 | 0.701772 | 0.710962 | 0.710653 |
| 0.711844 | 0.711392 | 0.716915 | 0.715215 | 0.71555 | 0.71217 | 0.705534 | 0.717448 | 0.714865 |
| 0.65175 | 0.657278 | 0.664763 | 0.661479 | 0.664974 | 0.657555 | 0.64636 | 0.664027 | 0.666441 |
| 0.658847 | 0.662406 | 0.668852 | 0.667155 | 0.669844 | 0.658651 | 0.659455 | 0.659378 | 0.664552 |
| 0.710593 | 0.713651 | 0.723241 | 0.715907 | 0.719559 | 0.716848 | 0.709274 | 0.716329 | 0.71626 |
| 0.711572 | 0.715949 | 0.726567 | 0.719185 | 0.719158 | 0.714274 | 0.706606 | 0.71768 | 0.719866 |
| 0.6432 | 0.643425 | 0.649853 | 0.648694 | 0.650298 | 0.648514 | 0.64381 | 0.650902 | 0.648838 |
| 0.709898 | 0.711159 | 0.720013 | 0.716181 | 0.714962 | 0.714115 | 0.709464 | 0.715716 | 0.713875 |
| 0.698811 | 0.70049 | 0.707599 | 0.703258 | 0.707498 | 0.705559 | 0.696102 | 0.704596 | 0.703179 |

| | | | | | | | | |
|---|---|---|---|---|---|---|---|---|
| 0.704867 | 0.707239 | 0.715459 | 0.713181 | 0.715111 | 0.710505 | 0.703521 | 0.711672 | 0.709942 |
| 0.707924 | 0.709769 | 0.718421 | 0.71501 | 0.715523 | 0.714241 | 0.707261 | 0.71459 | 0.713608 |
| 0.708915 | 0.711006 | 0.715403 | 0.713424 | 0.713293 | 0.710199 | 0.701099 | 0.711809 | 0.714659 |
| 0.706977 | 0.710505 | 0.717787 | 0.715675 | 0.715384 | 0.714207 | 0.707102 | 0.714602 | 0.713557 |
| 0.714661 | 0.715868 | 0.719857 | 0.721548 | 0.71837 | 0.718311 | 0.709865 | 0.717117 | 0.716492 |
| 0.704902 | 0.708701 | 0.714297 | 0.711939 | 0.712309 | 0.710762 | 0.699411 | 0.71078 | 0.71014 |
| 0.705628 | 0.708997 | 0.717464 | 0.715659 | 0.714685 | 0.708002 | 0.70344 | 0.710941 | 0.713457 |
| 0.701219 | 0.704251 | 0.710319 | 0.70609 | 0.708975 | 0.706639 | 0.697661 | 0.704527 | 0.707537 |
| 0.700822 | 0.704071 | 0.711652 | 0.712202 | 0.712007 | 0.711446 | 0.703713 | 0.710364 | 0.710598 |
| 0.704142 | 0.704814 | 0.717682 | 0.711807 | 0.717274 | 0.712018 | 0.703101 | 0.710424 | 0.712798 |
| 0.707304 | 0.710799 | 0.71972 | 0.713889 | 0.716462 | 0.715233 | 0.707097 | 0.714985 | 0.71568 |
| 0.703622 | 0.70541 | 0.712366 | 0.713003 | 0.709752 | 0.707434 | 0.702831 | 0.707373 | 0.708798 |
| 0.699039 | 0.700129 | 0.709729 | 0.70053 | 0.708232 | 0.704106 | 0.695914 | 0.701822 | 0.706195 |
| 0.698026 | 0.702586 | 0.71012 | 0.702806 | 0.706726 | 0.7007 | 0.693763 | 0.703658 | 0.707536 |
| 0.710225 | 0.711722 | 0.718054 | 0.718678 | 0.718177 | 0.717307 | 0.706128 | 0.716392 | 0.715792 |
| 0.706552 | 0.709391 | 0.714172 | 0.713411 | 0.715667 | 0.708494 | 0.707021 | 0.714625 | 0.715105 |
| 0.697889 | 0.704938 | 0.7105 | 0.710952 | 0.707339 | 0.706908 | 0.70276 | 0.707764 | 0.708165 |
| 0.702079 | 0.707239 | 0.710518 | 0.710996 | 0.709639 | 0.708479 | 0.702397 | 0.711846 | 0.708526 |
| 0.704949 | 0.7098 | 0.711042 | 0.706674 | 0.707498 | 0.707972 | 0.70221 | 0.710443 | 0.710485 |
| 0.704994 | 0.705502 | 0.713395 | 0.710507 | 0.712691 | 0.707024 | 0.701233 | 0.70736 | 0.709544 |
| 0.698789 | 0.702821 | 0.709999 | 0.708912 | 0.707706 | 0.703552 | 0.70097 | 0.704789 | 0.707971 |
| 0.703455 | 0.707904 | 0.710286 | 0.707248 | 0.70625 | 0.705221 | 0.700663 | 0.70513 | 0.710848 |
| 0.705248 | 0.709438 | 0.715891 | 0.712626 | 0.710838 | 0.712948 | 0.705265 | 0.712934 | 0.714218 |
| 0.703962 | 0.708111 | 0.715004 | 0.713441 | 0.712201 | 0.709123 | 0.703021 | 0.711817 | 0.714823 |
| 0.70393 | 0.709706 | 0.713974 | 0.711134 | 0.710752 | 0.709889 | 0.705541 | 0.709714 | 0.710673 |
| 0.703384 | 0.703766 | 0.708935 | 0.706819 | 0.704547 | 0.706215 | 0.701928 | 0.706028 | 0.709732 |
| 0.700972 | 0.702707 | 0.711632 | 0.708676 | 0.708982 | 0.707565 | 0.699839 | 0.708102 | 0.710237 |
| 0.701179 | 0.704286 | 0.710294 | 0.703887 | 0.70753 | 0.702816 | 0.696367 | 0.703313 | 0.706092 |
| 0.703011 | 0.704172 | 0.713263 | 0.708668 | 0.711702 | 0.707086 | 0.700571 | 0.707641 | 0.709521 |
| 0.70516 | 0.710514 | 0.716003 | 0.712059 | 0.711889 | 0.707675 | 0.701995 | 0.709367 | 0.712464 |
| 0.707247 | 0.710327 | 0.71619 | 0.713568 | 0.715523 | 0.712849 | 0.705293 | 0.715536 | 0.712748 |
| 0.710446 | 0.711825 | 0.715668 | 0.714251 | 0.71513 | 0.712466 | 0.705703 | 0.71274 | 0.71371 |
| 0.709494 | 0.714795 | 0.71792 | 0.716539 | 0.714172 | 0.713668 | 0.710116 | 0.71358 | 0.714681 |
| 0.707466 | 0.713603 | 0.7182 | 0.716292 | 0.716461 | 0.713539 | 0.708363 | 0.714874 | 0.715905 |
| 0.703155 | 0.706555 | 0.710766 | 0.710057 | 0.709952 | 0.704913 | 0.698173 | 0.706405 | 0.70789 |
| 0.705665 | 0.707451 | 0.711827 | 0.708555 | 0.71325 | 0.709849 | 0.70399 | 0.710414 | 0.712078 |
| 0.704103 | 0.707382 | 0.71479 | 0.713288 | 0.710439 | 0.709261 | 0.705194 | 0.710535 | 0.711118 |
| 0.69881 | 0.704724 | 0.710877 | 0.708876 | 0.705119 | 0.709017 | 0.700721 | 0.706414 | 0.704662 |
| 0.696706 | 0.701938 | 0.708144 | 0.702827 | 0.7042 | 0.701617 | 0.696017 | 0.704489 | 0.705104 |
| 0.692881 | 0.700477 | 0.705637 | 0.702312 | 0.698413 | 0.701476 | 0.6928 | 0.702465 | 0.703608 |
| 0.69727 | 0.696097 | 0.703761 | 0.701355 | 0.70364 | 0.700311 | 0.694907 | 0.700196 | 0.701323 |
| 0.696656 | 0.69741 | 0.703801 | 0.700651 | 0.701095 | 0.701368 | 0.692592 | 0.702138 | 0.70397 |
| 0.698581 | 0.702 | 0.706892 | 0.705319 | 0.703376 | 0.70157 | 0.698452 | 0.703774 | 0.70309 |
| 0.691699 | 0.696438 | 0.698981 | 0.695133 | 0.695804 | 0.694118 | 0.690329 | 0.697043 | 0.699564 |
| 0.701268 | 0.704492 | 0.711987 | 0.705247 | 0.707311 | 0.702479 | 0.699026 | 0.705159 | 0.705116 |
| 0.69355 | 0.700366 | 0.703029 | 0.700137 | 0.702954 | 0.699463 | 0.696381 | 0.700282 | 0.703164 |

| | | | | | | | | |
|---|---|---|---|---|---|---|---|---|
| 0.709577 | 0.709443 | 0.71537 | 0.716194 | 0.713301 | 0.712698 | 0.709819 | 0.710338 | 0.714331 |
| 0.708139 | 0.708096 | 0.716226 | 0.715509 | 0.713868 | 0.712109 | 0.706081 | 0.709321 | 0.714343 |
| 0.70566 | 0.70752 | 0.712902 | 0.709125 | 0.709931 | 0.706432 | 0.701117 | 0.70867 | 0.711481 |
| 0.709955 | 0.711705 | 0.717051 | 0.714987 | 0.713945 | 0.710723 | 0.706482 | 0.7136 | 0.71329 |
| 0.717109 | 0.719262 | 0.724016 | 0.720964 | 0.718572 | 0.718134 | 0.712668 | 0.717469 | 0.719981 |
| 0.712993 | 0.711065 | 0.720134 | 0.716524 | 0.716969 | 0.711717 | 0.709288 | 0.714229 | 0.716846 |
| 0.70367 | 0.705318 | 0.710677 | 0.709426 | 0.70815 | 0.706976 | 0.701495 | 0.708842 | 0.709302 |
| 0.708587 | 0.706498 | 0.717124 | 0.712583 | 0.714845 | 0.709462 | 0.706274 | 0.714681 | 0.713706 |
| 0.702771 | 0.702191 | 0.709088 | 0.706128 | 0.70974 | 0.702911 | 0.701325 | 0.704628 | 0.708261 |
| 0.709672 | 0.709307 | 0.71497 | 0.713238 | 0.710227 | 0.710151 | 0.705396 | 0.713355 | 0.712755 |
| 0.711381 | 0.711974 | 0.719343 | 0.716045 | 0.71475 | 0.712147 | 0.705642 | 0.712223 | 0.718131 |
| 0.710579 | 0.711525 | 0.720726 | 0.716451 | 0.715601 | 0.710243 | 0.70628 | 0.716099 | 0.716918 |
| 0.706376 | 0.709795 | 0.715217 | 0.709417 | 0.710327 | 0.705929 | 0.703 | 0.709721 | 0.712376 |
| 0.709398 | 0.708977 | 0.714059 | 0.713477 | 0.713829 | 0.708574 | 0.704813 | 0.708893 | 0.712475 |
| 0.711143 | 0.710339 | 0.721284 | 0.714873 | 0.712871 | 0.71105 | 0.708511 | 0.713522 | 0.716995 |
| 0.706152 | 0.714411 | 0.718096 | 0.709634 | 0.715565 | 0.707392 | 0.706798 | 0.711997 | 0.714468 |
| 0.714815 | 0.714557 | 0.721228 | 0.720978 | 0.718263 | 0.715149 | 0.707025 | 0.713681 | 0.718611 |
| 0.707189 | 0.712176 | 0.720404 | 0.714398 | 0.71477 | 0.706029 | 0.705601 | 0.715002 | 0.716834 |
| 0.711108 | 0.717685 | 0.721051 | 0.717784 | 0.71783 | 0.715824 | 0.707454 | 0.719446 | 0.717838 |
| 0.70444 | 0.709532 | 0.718728 | 0.712874 | 0.714331 | 0.711602 | 0.705762 | 0.713016 | 0.714785 |
| 0.707802 | 0.709476 | 0.71859 | 0.711674 | 0.71501 | 0.713398 | 0.7063 | 0.713044 | 0.714069 |
| 0.703742 | 0.709111 | 0.71504 | 0.712316 | 0.710975 | 0.709978 | 0.703855 | 0.708867 | 0.712393 |
| 0.711781 | 0.714444 | 0.722372 | 0.715315 | 0.717435 | 0.713466 | 0.707668 | 0.713319 | 0.715656 |
| 0.711783 | 0.716945 | 0.719731 | 0.712843 | 0.717839 | 0.7135 | 0.708396 | 0.714949 | 0.715598 |
| 0.707143 | 0.707026 | 0.716627 | 0.71132 | 0.712653 | 0.711201 | 0.7087 | 0.710742 | 0.712631 |
| 0.704228 | 0.708502 | 0.713593 | 0.712879 | 0.710459 | 0.709652 | 0.703041 | 0.708646 | 0.710236 |
| 0.710769 | 0.711101 | 0.713981 | 0.709168 | 0.710798 | 0.710528 | 0.706022 | 0.708565 | 0.712501 |
| 0.706192 | 0.707963 | 0.713767 | 0.709352 | 0.711963 | 0.707922 | 0.701464 | 0.711132 | 0.712217 |
| 0.709028 | 0.712222 | 0.719858 | 0.713464 | 0.716634 | 0.711685 | 0.70831 | 0.713287 | 0.719038 |
| 0.711298 | 0.713268 | 0.720365 | 0.717432 | 0.721791 | 0.713983 | 0.709034 | 0.714495 | 0.717522 |
| 0.706417 | 0.70923 | 0.711744 | 0.711278 | 0.712489 | 0.70828 | 0.701776 | 0.708263 | 0.711233 |
| 0.712377 | 0.714206 | 0.721267 | 0.715916 | 0.717791 | 0.714468 | 0.70663 | 0.716856 | 0.71541 |
| 0.710096 | 0.712572 | 0.718107 | 0.71363 | 0.718825 | 0.709891 | 0.706478 | 0.71415 | 0.718072 |
| 0.707269 | 0.710932 | 0.713723 | 0.710768 | 0.711475 | 0.705433 | 0.701153 | 0.708507 | 0.711894 |
| 0.707439 | 0.712222 | 0.714075 | 0.7106 | 0.714615 | 0.708401 | 0.701648 | 0.715359 | 0.713617 |
| 0.705589 | 0.709325 | 0.713892 | 0.71 | 0.709008 | 0.705464 | 0.700498 | 0.709942 | 0.713008 |
| 0.71373 | 0.716912 | 0.722617 | 0.72043 | 0.721929 | 0.716774 | 0.708776 | 0.718949 | 0.72091 |
| 0.705764 | 0.709649 | 0.716153 | 0.711318 | 0.713495 | 0.711095 | 0.704788 | 0.710617 | 0.716909 |
| 0.711819 | 0.711612 | 0.717227 | 0.713468 | 0.714137 | 0.712118 | 0.708476 | 0.712296 | 0.714491 |
| 0.708084 | 0.707699 | 0.719541 | 0.713032 | 0.715172 | 0.709329 | 0.706278 | 0.71275 | 0.71475 |
| 0.709161 | 0.715658 | 0.723626 | 0.71657 | 0.720373 | 0.714398 | 0.708534 | 0.716336 | 0.718499 |
| 0.713411 | 0.713614 | 0.719505 | 0.716129 | 0.715634 | 0.713451 | 0.710121 | 0.714037 | 0.718623 |
| 0.709733 | 0.714173 | 0.719234 | 0.715561 | 0.720271 | 0.716897 | 0.70923 | 0.714241 | 0.720638 |
| 0.705312 | 0.709455 | 0.715505 | 0.712884 | 0.712459 | 0.70838 | 0.70324 | 0.712246 | 0.711521 |
| 0.71184 | 0.710709 | 0.718249 | 0.715705 | 0.715654 | 0.711639 | 0.710406 | 0.712239 | 0.715477 |
| 0.705042 | 0.706842 | 0.714829 | 0.712192 | 0.711001 | 0.705548 | 0.704219 | 0.707791 | 0.711694 |

| | | | | | | | | |
|---|---|---|---|---|---|---|---|---|
| 0.695898 | 0.695947 | 0.701855 | 0.698251 | 0.699057 | 0.697986 | 0.695086 | 0.698845 | 0.698326 |
| 0.755055 | 0.747887 | 0.760531 | 0.755416 | 0.760335 | 0.752289 | 0.749588 | 0.760118 | 0.754976 |
| 0.703851 | 0.71064 | 0.71785 | 0.711069 | 0.712762 | 0.710443 | 0.702775 | 0.708455 | 0.712791 |
| 0.709256 | 0.712921 | 0.721056 | 0.716975 | 0.718315 | 0.713166 | 0.708517 | 0.712951 | 0.717806 |
| 0.702122 | 0.706082 | 0.713043 | 0.710722 | 0.711154 | 0.706274 | 0.699374 | 0.705578 | 0.712556 |
| 0.708266 | 0.710249 | 0.71478 | 0.71455 | 0.71402 | 0.708912 | 0.705764 | 0.709482 | 0.712822 |
| 0.702449 | 0.702407 | 0.713273 | 0.708764 | 0.711651 | 0.706165 | 0.701807 | 0.706685 | 0.713018 |
| 0.699932 | 0.701549 | 0.710766 | 0.706266 | 0.705017 | 0.701216 | 0.699171 | 0.702364 | 0.708616 |
| 0.70389 | 0.707876 | 0.715881 | 0.712061 | 0.715049 | 0.712114 | 0.704883 | 0.709388 | 0.71161 |
| 0.715141 | 0.715979 | 0.722584 | 0.718029 | 0.719955 | 0.718026 | 0.711816 | 0.716195 | 0.718169 |
| 0.710578 | 0.716326 | 0.719965 | 0.717493 | 0.715208 | 0.715073 | 0.707807 | 0.711354 | 0.714274 |
| 0.709657 | 0.713965 | 0.719564 | 0.717881 | 0.719862 | 0.716038 | 0.71248 | 0.712888 | 0.71684 |
| 0.714365 | 0.714718 | 0.721468 | 0.721188 | 0.720308 | 0.71754 | 0.708972 | 0.716395 | 0.717893 |
| 0.713008 | 0.71337 | 0.718814 | 0.717672 | 0.720979 | 0.714601 | 0.71106 | 0.71351 | 0.716566 |
| 0.709401 | 0.711109 | 0.718003 | 0.715062 | 0.71909 | 0.71302 | 0.709461 | 0.710384 | 0.713949 |
| 0.70878 | 0.71032 | 0.715318 | 0.712361 | 0.71459 | 0.710789 | 0.706246 | 0.711507 | 0.713163 |
| 0.710971 | 0.71093 | 0.718302 | 0.715691 | 0.720606 | 0.714252 | 0.710436 | 0.715082 | 0.715845 |
| 0.711415 | 0.712509 | 0.717428 | 0.715328 | 0.718077 | 0.711591 | 0.708487 | 0.711923 | 0.712041 |
| 0.702467 | 0.701863 | 0.711826 | 0.702607 | 0.708866 | 0.703244 | 0.697988 | 0.704076 | 0.705872 |
| 0.70159 | 0.70604 | 0.71121 | 0.707473 | 0.7109 | 0.704621 | 0.699539 | 0.709864 | 0.706432 |
| 0.705472 | 0.714447 | 0.722533 | 0.716051 | 0.71779 | 0.712832 | 0.707005 | 0.71556 | 0.712776 |
| 0.697234 | 0.704944 | 0.706041 | 0.70493 | 0.709401 | 0.698962 | 0.695652 | 0.702058 | 0.704982 |
| 0.706167 | 0.709324 | 0.714161 | 0.711831 | 0.711718 | 0.711036 | 0.704545 | 0.709358 | 0.713261 |
| 0.710071 | 0.714023 | 0.718878 | 0.714651 | 0.717404 | 0.714706 | 0.707146 | 0.710067 | 0.718002 |
| 0.712496 | 0.714344 | 0.721301 | 0.715346 | 0.719546 | 0.713106 | 0.707795 | 0.714738 | 0.717712 |
| 0.707987 | 0.713407 | 0.717599 | 0.713997 | 0.715927 | 0.712193 | 0.706158 | 0.711113 | 0.715468 |
| 0.70479 | 0.710672 | 0.710676 | 0.710217 | 0.713113 | 0.706412 | 0.704078 | 0.707415 | 0.711799 |
| 0.704003 | 0.707166 | 0.709001 | 0.70916 | 0.711259 | 0.705385 | 0.700767 | 0.705969 | 0.707903 |
| 0.70251 | 0.707385 | 0.711448 | 0.706526 | 0.713149 | 0.705582 | 0.701563 | 0.706001 | 0.711057 |
| 0.777203 | 0.776041 | 0.780663 | 0.780953 | 0.781042 | 0.776742 | 0.765515 | 0.768931 | 0.772831 |
| 0.792243 | 0.793141 | 0.79921 | 0.798065 | 0.802491 | 0.795049 | 0.784053 | 0.780366 | 0.781573 |
| 0.789703 | 0.785118 | 0.791872 | 0.789588 | 0.794576 | 0.783463 | 0.773875 | 0.78332 | 0.783315 |
| 0.7779 | 0.789537 | 0.784079 | 0.791963 | 0.769323 | 0.775758 | 0.7619 | 0.782657 | 0.774927 |
| 0.785121 | 0.796619 | 0.801159 | 0.792065 | 0.787333 | 0.786885 | 0.781917 | 0.789331 | 0.781245 |
| 0.703124 | 0.703479 | 0.706882 | 0.707223 | 0.708547 | 0.702128 | 0.700197 | 0.702857 | 0.705467 |
| 0.703481 | 0.708565 | 0.708846 | 0.712521 | 0.709125 | 0.708233 | 0.703554 | 0.707221 | 0.707315 |
| 0.707715 | 0.712719 | 0.714816 | 0.712614 | 0.710261 | 0.709206 | 0.704998 | 0.708243 | 0.710249 |
| 0.710997 | 0.713919 | 0.715051 | 0.713473 | 0.712693 | 0.708588 | 0.706821 | 0.709264 | 0.713714 |
| 0.70339 | 0.705717 | 0.71083 | 0.707962 | 0.709488 | 0.70358 | 0.702354 | 0.707977 | 0.708654 |
| 0.705709 | 0.712582 | 0.715499 | 0.712031 | 0.715173 | 0.712056 | 0.705262 | 0.708864 | 0.715996 |
| 0.704195 | 0.708384 | 0.711971 | 0.710369 | 0.710732 | 0.708687 | 0.703784 | 0.70838 | 0.714286 |
| 0.705095 | 0.707274 | 0.709427 | 0.712711 | 0.709007 | 0.709363 | 0.703227 | 0.70886 | 0.711145 |
| 0.69872 | 0.701824 | 0.706625 | 0.706756 | 0.70337 | 0.701009 | 0.6977 | 0.702757 | 0.705724 |
| 0.697502 | 0.705266 | 0.704124 | 0.704224 | 0.708441 | 0.699041 | 0.698512 | 0.698572 | 0.703431 |
| 0.70924 | 0.70864 | 0.71314 | 0.713337 | 0.709762 | 0.706083 | 0.704018 | 0.709814 | 0.710345 |
| 0.709073 | 0.710094 | 0.71371 | 0.713743 | 0.71381 | 0.709776 | 0.707859 | 0.711441 | 0.717539 |

| | | | | | | | | |
|---|---|---|---|---|---|---|---|---|
| 0.707287 | 0.712398 | 0.717631 | 0.713411 | 0.712943 | 0.712292 | 0.704327 | 0.707278 | 0.713352 |
| 0.704743 | 0.710758 | 0.715788 | 0.713828 | 0.711791 | 0.708221 | 0.704774 | 0.711131 | 0.711035 |
| 0.710334 | 0.716587 | 0.717707 | 0.715349 | 0.715596 | 0.713039 | 0.707725 | 0.713368 | 0.720705 |
| 0.706365 | 0.708882 | 0.717229 | 0.708879 | 0.714895 | 0.706279 | 0.701068 | 0.709421 | 0.711711 |
| 0.702522 | 0.707462 | 0.711999 | 0.7057 | 0.709461 | 0.706945 | 0.698529 | 0.706945 | 0.709762 |
| 0.705088 | 0.70876 | 0.714676 | 0.709637 | 0.712981 | 0.706104 | 0.698823 | 0.706128 | 0.711394 |
| 0.705217 | 0.712168 | 0.717831 | 0.714695 | 0.71691 | 0.710475 | 0.705028 | 0.71325 | 0.71856 |
| 0.708823 | 0.713793 | 0.714531 | 0.71126 | 0.714733 | 0.711183 | 0.704878 | 0.71361 | 0.712964 |
| 0.702654 | 0.707208 | 0.714087 | 0.70849 | 0.713558 | 0.705625 | 0.702982 | 0.711545 | 0.714403 |
| 0.705557 | 0.708853 | 0.715321 | 0.71233 | 0.71216 | 0.708007 | 0.704802 | 0.712554 | 0.710677 |
| 0.703698 | 0.709215 | 0.715215 | 0.710242 | 0.713156 | 0.711172 | 0.702078 | 0.7095 | 0.713368 |
| 0.706559 | 0.708387 | 0.714599 | 0.707904 | 0.713223 | 0.709789 | 0.702703 | 0.707392 | 0.713505 |
| 0.697441 | 0.701406 | 0.707813 | 0.702542 | 0.705358 | 0.701463 | 0.693482 | 0.701759 | 0.704394 |
| 0.698054 | 0.701882 | 0.706644 | 0.702453 | 0.70594 | 0.701014 | 0.699216 | 0.703512 | 0.707306 |
| 0.707004 | 0.708642 | 0.717695 | 0.715044 | 0.714438 | 0.707167 | 0.703267 | 0.711833 | 0.720726 |
| 0.711259 | 0.717815 | 0.722222 | 0.717414 | 0.724185 | 0.716231 | 0.710419 | 0.718495 | 0.723046 |
| 0.706196 | 0.712683 | 0.716837 | 0.712155 | 0.713583 | 0.710953 | 0.706364 | 0.715072 | 0.715688 |
| 0.711209 | 0.71326 | 0.718445 | 0.71878 | 0.716227 | 0.716217 | 0.708602 | 0.716742 | 0.720986 |
| 0.70562 | 0.708837 | 0.714179 | 0.711159 | 0.714393 | 0.708686 | 0.70238 | 0.707143 | 0.712001 |
| 0.70553 | 0.708443 | 0.714498 | 0.712637 | 0.714499 | 0.710887 | 0.702581 | 0.709799 | 0.715149 |
| 0.702187 | 0.708105 | 0.710147 | 0.706936 | 0.710678 | 0.703704 | 0.698822 | 0.707687 | 0.712217 |
| 0.70835 | 0.709145 | 0.717698 | 0.714032 | 0.71527 | 0.712616 | 0.705613 | 0.714538 | 0.716676 |
| 0.708886 | 0.71366 | 0.717781 | 0.714085 | 0.714842 | 0.713383 | 0.710277 | 0.716024 | 0.718377 |
| 0.70353 | 0.711273 | 0.713919 | 0.707067 | 0.711628 | 0.708773 | 0.699498 | 0.707773 | 0.709575 |
| 0.708351 | 0.708398 | 0.713403 | 0.713057 | 0.713459 | 0.707587 | 0.705999 | 0.71133 | 0.714052 |
| 0.798127 | 0.8 | 0.78969 | 0.804353 | 0.782105 | 0.793034 | 0.778588 | 0.791001 | 0.791912 |
| 0.708725 | 0.711718 | 0.714161 | 0.712803 | 0.715402 | 0.70812 | 0.707957 | 0.710673 | 0.713175 |
| 0.710626 | 0.714661 | 0.72044 | 0.715081 | 0.716641 | 0.714829 | 0.705705 | 0.715656 | 0.717397 |
| 0.714205 | 0.716716 | 0.722827 | 0.714079 | 0.715888 | 0.714604 | 0.704584 | 0.714297 | 0.717792 |
| 0.709724 | 0.712958 | 0.71764 | 0.712927 | 0.714711 | 0.714479 | 0.706221 | 0.716159 | 0.712376 |
| 0.703316 | 0.707804 | 0.713868 | 0.708278 | 0.707842 | 0.706175 | 0.702347 | 0.711742 | 0.71006 |
| 0.705482 | 0.711162 | 0.714486 | 0.708577 | 0.712441 | 0.708967 | 0.702166 | 0.71131 | 0.709378 |
| 0.706587 | 0.708858 | 0.718156 | 0.712093 | 0.716675 | 0.712862 | 0.701624 | 0.715063 | 0.714344 |
| 0.706422 | 0.707268 | 0.714095 | 0.709753 | 0.711897 | 0.70825 | 0.700067 | 0.710776 | 0.712753 |
| 0.699674 | 0.707436 | 0.709265 | 0.707688 | 0.71048 | 0.707023 | 0.699085 | 0.708207 | 0.709835 |
| 0.70717 | 0.708868 | 0.714286 | 0.710209 | 0.712351 | 0.707357 | 0.703568 | 0.709743 | 0.713046 |
| 0.706996 | 0.711093 | 0.715956 | 0.715378 | 0.715315 | 0.715431 | 0.707062 | 0.712643 | 0.715276 |
| 0.710999 | 0.712757 | 0.721272 | 0.715801 | 0.717479 | 0.715429 | 0.705812 | 0.714045 | 0.717405 |
| 0.708712 | 0.712162 | 0.716382 | 0.711494 | 0.712979 | 0.709786 | 0.704919 | 0.710697 | 0.713963 |
| 0.707315 | 0.710174 | 0.71669 | 0.711222 | 0.716046 | 0.706862 | 0.702274 | 0.711554 | 0.716123 |
| 0.702286 | 0.704276 | 0.708506 | 0.70597 | 0.704682 | 0.70469 | 0.700732 | 0.7045 | 0.708368 |
| 0.709154 | 0.715644 | 0.718205 | 0.714668 | 0.716152 | 0.714182 | 0.707902 | 0.713985 | 0.715969 |
| 0.708949 | 0.708191 | 0.716261 | 0.712154 | 0.712329 | 0.706382 | 0.69995 | 0.711257 | 0.711595 |
| 0.703304 | 0.709017 | 0.711799 | 0.708904 | 0.711528 | 0.713687 | 0.702281 | 0.711343 | 0.71256 |
| 0.706613 | 0.710044 | 0.716508 | 0.712468 | 0.714642 | 0.709427 | 0.703547 | 0.712009 | 0.71597 |
| 0.7078 | 0.709495 | 0.715712 | 0.712885 | 0.715158 | 0.710629 | 0.706782 | 0.711577 | 0.716218 |

| | | | | | | | | |
|---|---|---|---|---|---|---|---|---|
| 0.708508 | 0.70551 | 0.711595 | 0.711465 | 0.711927 | 0.710636 | 0.704214 | 0.711025 | 0.711097 |
| 0.703683 | 0.70935 | 0.713788 | 0.712665 | 0.711906 | 0.707704 | 0.703967 | 0.71108 | 0.713479 |
| 0.704738 | 0.706175 | 0.709398 | 0.711823 | 0.711428 | 0.711134 | 0.7006 | 0.712865 | 0.709307 |
| 0.658766 | 0.660384 | 0.672963 | 0.673043 | 0.67507 | 0.66204 | 0.656153 | 0.672504 | 0.670744 |
| 0.705209 | 0.705984 | 0.711991 | 0.709458 | 0.711037 | 0.707469 | 0.702148 | 0.708396 | 0.713276 |
| 0.706326 | 0.710701 | 0.711251 | 0.709464 | 0.712867 | 0.708762 | 0.70406 | 0.709856 | 0.710415 |
| 0.707508 | 0.710853 | 0.712085 | 0.712628 | 0.710772 | 0.710508 | 0.70344 | 0.710821 | 0.713443 |
| 0.706561 | 0.708396 | 0.713538 | 0.710067 | 0.712579 | 0.708993 | 0.70592 | 0.710829 | 0.71205 |
| 0.71165 | 0.712923 | 0.720354 | 0.716947 | 0.717744 | 0.715653 | 0.708148 | 0.718708 | 0.717395 |
| 0.708856 | 0.712191 | 0.717528 | 0.707433 | 0.713064 | 0.711754 | 0.706724 | 0.712082 | 0.712886 |
| 0.706051 | 0.712324 | 0.714787 | 0.714594 | 0.71338 | 0.710222 | 0.706028 | 0.713832 | 0.716653 |
| 0.708287 | 0.711671 | 0.718068 | 0.715602 | 0.715741 | 0.710822 | 0.705671 | 0.714694 | 0.716552 |
| 0.710167 | 0.714001 | 0.7189 | 0.714957 | 0.717595 | 0.714546 | 0.706603 | 0.715964 | 0.716326 |
| 0.704066 | 0.711173 | 0.714399 | 0.712307 | 0.714388 | 0.708481 | 0.703698 | 0.711232 | 0.710925 |
| 0.711836 | 0.713489 | 0.718683 | 0.715266 | 0.719554 | 0.712726 | 0.708184 | 0.716382 | 0.717353 |
| 0.703777 | 0.707094 | 0.713716 | 0.708168 | 0.714183 | 0.70744 | 0.701841 | 0.706426 | 0.71129 |
| 0.704899 | 0.708809 | 0.716204 | 0.711698 | 0.714841 | 0.707737 | 0.704102 | 0.714062 | 0.714908 |
| 0.710516 | 0.709189 | 0.719402 | 0.712185 | 0.718068 | 0.708333 | 0.707292 | 0.714816 | 0.716389 |
| 0.704169 | 0.70515 | 0.711554 | 0.707422 | 0.710696 | 0.703599 | 0.700918 | 0.707907 | 0.710708 |
| 0.704768 | 0.710374 | 0.713221 | 0.711742 | 0.710296 | 0.706262 | 0.70508 | 0.710342 | 0.712716 |
| 0.708842 | 0.71199 | 0.715843 | 0.711844 | 0.715361 | 0.709461 | 0.708559 | 0.71398 | 0.71595 |
| 0.7093 | 0.711418 | 0.718188 | 0.711416 | 0.715518 | 0.711478 | 0.70668 | 0.714829 | 0.719288 |
| 0.693614 | 0.695064 | 0.695636 | 0.694031 | 0.693309 | 0.69855 | 0.699262 | 0.691233 | 0.693307 |
| 0.721963 | 0.722685 | 0.724079 | 0.714495 | 0.72001 | 0.723655 | 0.7139 | 0.714563 | 0.722562 |
| 0.755735 | 0.756537 | 0.76578 | 0.765158 | 0.758108 | 0.761104 | 0.75134 | 0.760054 | 0.763371 |
| 0.741536 | 0.755176 | 0.749316 | 0.757158 | 0.73768 | 0.742057 | 0.727133 | 0.741156 | 0.740224 |
| 0.661614 | 0.670001 | 0.672682 | 0.668605 | 0.674236 | 0.664418 | 0.66044 | 0.66638 | 0.675923 |
| 0.661977 | 0.663648 | 0.671808 | 0.6603 | 0.67075 | 0.660081 | 0.660185 | 0.66193 | 0.668837 |
| 0.656017 | 0.658082 | 0.668634 | 0.661603 | 0.663174 | 0.657414 | 0.661327 | 0.657532 | 0.672381 |
| 0.667536 | 0.680984 | 0.680718 | 0.674848 | 0.679211 | 0.680648 | 0.67163 | 0.676065 | 0.679976 |
| 0.651569 | 0.65754 | 0.658035 | 0.641834 | 0.659537 | 0.654228 | 0.650376 | 0.651239 | 0.655051 |
| 0.639228 | 0.647811 | 0.645388 | 0.646738 | 0.647802 | 0.640389 | 0.640391 | 0.635827 | 0.652884 |
| 0.636236 | 0.65261 | 0.653451 | 0.643718 | 0.644647 | 0.645721 | 0.636003 | 0.63758 | 0.64798 |
| 0.738936 | 0.740748 | 0.739026 | 0.744117 | 0.732898 | 0.736725 | 0.722048 | 0.730621 | 0.733837 |
| 0.702855 | 0.705293 | 0.708711 | 0.705083 | 0.704671 | 0.703538 | 0.699384 | 0.702745 | 0.707795 |
| 0.671964 | 0.678962 | 0.684898 | 0.674798 | 0.68037 | 0.676313 | 0.674592 | 0.673561 | 0.682018 |
| 0.66306 | 0.670916 | 0.682014 | 0.672427 | 0.67318 | 0.668959 | 0.670367 | 0.671258 | 0.675503 |
| 0.648118 | 0.654993 | 0.660953 | 0.654496 | 0.660804 | 0.649503 | 0.64643 | 0.650853 | 0.662903 |
| 0.672365 | 0.678584 | 0.679326 | 0.678483 | 0.681283 | 0.676217 | 0.670902 | 0.679152 | 0.683125 |
| 0.666956 | 0.666295 | 0.675133 | 0.670764 | 0.67127 | 0.670523 | 0.661901 | 0.668842 | 0.677524 |
| 0.659283 | 0.657911 | 0.663909 | 0.660838 | 0.663842 | 0.661092 | 0.661434 | 0.664673 | 0.666769 |
| 0.662174 | 0.662236 | 0.669746 | 0.659873 | 0.662887 | 0.665165 | 0.659732 | 0.663423 | 0.672221 |
| 0.653813 | 0.656864 | 0.665273 | 0.656038 | 0.656761 | 0.654721 | 0.656315 | 0.650567 | 0.666376 |
| 0.651901 | 0.660486 | 0.664131 | 0.658 | 0.658448 | 0.657495 | 0.654455 | 0.656185 | 0.667751 |
| 0.65555 | 0.651358 | 0.667992 | 0.653061 | 0.65377 | 0.658651 | 0.654865 | 0.659459 | 0.659458 |
| 0.634463 | 0.647473 | 0.64665 | 0.64832 | 0.641551 | 0.646928 | 0.639785 | 0.634192 | 0.648113 |

| | | | | | | | | |
|---|---|---|---|---|---|---|---|---|
| 0.690604 | 0.692138 | 0.696964 | 0.692751 | 0.693383 | 0.69051 | 0.682619 | 0.693273 | 0.696271 |
| 0.656575 | 0.661823 | 0.667614 | 0.660382 | 0.660249 | 0.65793 | 0.651975 | 0.656024 | 0.666622 |
| 0.664339 | 0.661556 | 0.669658 | 0.666377 | 0.666909 | 0.664692 | 0.659015 | 0.665901 | 0.671499 |
| 0.657189 | 0.662983 | 0.664654 | 0.66213 | 0.669087 | 0.667307 | 0.661934 | 0.661667 | 0.671209 |
| 0.653975 | 0.65942 | 0.664374 | 0.65978 | 0.663583 | 0.657199 | 0.65524 | 0.65214 | 0.664118 |
| 0.657742 | 0.664399 | 0.670926 | 0.658877 | 0.6687 | 0.662506 | 0.65006 | 0.662919 | 0.672543 |
| 0.645576 | 0.652409 | 0.657063 | 0.647733 | 0.663534 | 0.64993 | 0.649193 | 0.655452 | 0.667136 |
| 0.643678 | 0.648033 | 0.655809 | 0.647059 | 0.653878 | 0.64398 | 0.649032 | 0.647807 | 0.662657 |
| 0.642157 | 0.654947 | 0.652657 | 0.644336 | 0.652519 | 0.650068 | 0.646272 | 0.648714 | 0.662195 |
| 0.684602 | 0.688447 | 0.697476 | 0.693994 | 0.695532 | 0.691437 | 0.687831 | 0.691851 | 0.697407 |
| 0.686493 | 0.688555 | 0.695054 | 0.693021 | 0.688398 | 0.686628 | 0.686486 | 0.687488 | 0.696294 |
| 0.68101 | 0.684453 | 0.691921 | 0.685163 | 0.689035 | 0.680697 | 0.680557 | 0.687713 | 0.69078 |
| 0.671993 | 0.676919 | 0.677548 | 0.681818 | 0.681867 | 0.675443 | 0.674623 | 0.680486 | 0.67909 |
| 0.678398 | 0.677982 | 0.686701 | 0.67974 | 0.685989 | 0.679498 | 0.674317 | 0.684725 | 0.689139 |
| 0.669621 | 0.671552 | 0.678288 | 0.674132 | 0.6794 | 0.671607 | 0.668997 | 0.669848 | 0.678974 |

| | | | | | | | | |
|---|---|---|---|---|---|---|---|---|
| | | | | | Home garden | | | |
| _11 | | | | | | | | |
| Home_garc | Home_garc | Home_garc | Home_garc | Home_garc | Home_garc | Home_garc | Home_garc | Home_garc |
| 0.788007 | 0.773644 | 0.775597 | 0.776425 | 0.767413 | 0.767082 | 0.780094 | 0.77622 | 0.762943 |
| 0.788866 | 0.77797 | 0.777259 | 0.787273 | 0.779349 | 0.779141 | 0.792031 | 0.786481 | 0.778693 |
| 0.778776 | 0.77297 | 0.775041 | 0.78232 | 0.773912 | 0.778736 | 0.782903 | 0.781602 | 0.763844 |
| 0.774678 | 0.779697 | 0.788155 | 0.785025 | 0.784745 | 0.77961 | 0.789748 | 0.780323 | 0.773259 |
| 0.77443 | 0.782856 | 0.776228 | 0.776062 | 0.778214 | 0.765436 | 0.773793 | 0.77381 | 0.769663 |
| 0.775441 | 0.781098 | 0.770894 | 0.779102 | 0.77698 | 0.767321 | 0.772768 | 0.780805 | 0.778246 |
| 0.713419 | 0.710973 | 0.706852 | 0.714421 | 0.709514 | 0.705599 | 0.711482 | 0.716522 | 0.717703 |
| 0.709058 | 0.708244 | 0.705033 | 0.704115 | 0.708241 | 0.703305 | 0.708972 | 0.711859 | 0.711988 |
| 0.778902 | 0.783918 | 0.783379 | 0.780965 | 0.777395 | 0.769717 | 0.782212 | 0.784249 | 0.772252 |
| 0.788492 | 0.772671 | 0.771012 | 0.776796 | 0.770059 | 0.766209 | 0.774887 | 0.777524 | 0.762775 |
| 0.778911 | 0.778008 | 0.768979 | 0.775619 | 0.770953 | 0.762583 | 0.782493 | 0.779044 | 0.774234 |
| 0.795608 | 0.789172 | 0.776876 | 0.784986 | 0.776023 | 0.775676 | 0.792392 | 0.787428 | 0.768435 |
| 0.784524 | 0.766762 | 0.771994 | 0.769382 | 0.771153 | 0.768325 | 0.78736 | 0.781774 | 0.770315 |
| 0.786691 | 0.775838 | 0.774228 | 0.780606 | 0.770818 | 0.763626 | 0.780591 | 0.778313 | 0.768888 |
| 0.770966 | 0.77162 | 0.76702 | 0.764749 | 0.762398 | 0.762512 | 0.777924 | 0.774457 | 0.762018 |
| 0.799833 | 0.781343 | 0.781452 | 0.782847 | 0.773934 | 0.778097 | 0.783003 | 0.782364 | 0.767683 |
| 0.803755 | 0.769656 | 0.767434 | 0.769288 | 0.760081 | 0.768035 | 0.778266 | 0.782548 | 0.760619 |
| 0.758526 | 0.76022 | 0.754053 | 0.750717 | 0.750419 | 0.743682 | 0.761077 | 0.767584 | 0.745365 |
| 0.783658 | 0.774439 | 0.773939 | 0.806956 | 0.772508 | 0.78154 | 0.811566 | 0.785337 | 0.770948 |
| 0.777897 | 0.782259 | 0.78152 | 0.783148 | 0.779971 | 0.773231 | 0.781729 | 0.790479 | 0.776835 |
| 0.777392 | 0.781315 | 0.781194 | 0.786309 | 0.778925 | 0.773594 | 0.779762 | 0.787258 | 0.771018 |
| 0.783217 | 0.786129 | 0.792559 | 0.783617 | 0.78065 | 0.775973 | 0.788113 | 0.795957 | 0.773884 |
| 0.784631 | 0.791455 | 0.785908 | 0.790154 | 0.810227 | 0.775074 | 0.786146 | 0.798091 | 0.827134 |
| 0.820595 | 0.788958 | 0.78663 | 0.789734 | 0.779768 | 0.779042 | 0.798599 | 0.794106 | 0.781599 |
| 0.785756 | 0.786982 | 0.78046 | 0.781036 | 0.801196 | 0.7729 | 0.784354 | 0.777605 | 0.814859 |
| 0.784049 | 0.829494 | 0.830321 | 0.78835 | 0.802243 | 0.802168 | 0.799418 | 0.794151 | 0.772038 |
| 0.776252 | 0.773827 | 0.772958 | 0.79385 | 0.775646 | 0.820349 | 0.80727 | 0.792913 | 0.770556 |
| 0.772835 | 0.781513 | 0.785293 | 0.792118 | 0.788551 | 0.772479 | 0.779128 | 0.786996 | 0.783692 |
| 0.78102 | 0.781547 | 0.780085 | 0.784709 | 0.807101 | 0.7753 | 0.783228 | 0.790382 | 0.844745 |
| 1 | 0.770025 | 0.761423 | 0.779585 | 0.769086 | 0.767725 | 0.788515 | 0.784185 | 0.768151 |
| 0.770025 | 1 | 0.818121 | 0.781044 | 0.80474 | 0.779413 | 0.785423 | 0.783775 | 0.779199 |
| 0.761423 | 0.818121 | 1 | 0.777787 | 0.819827 | 0.797648 | 0.777485 | 0.777416 | 0.772127 |
| 0.779585 | 0.781044 | 0.777787 | 1 | 0.782299 | 0.772746 | 0.798952 | 0.780158 | 0.773339 |
| 0.769086 | 0.80474 | 0.819827 | 0.782299 | 1 | 0.796427 | 0.784745 | 0.781128 | 0.800452 |
| 0.767725 | 0.779413 | 0.797648 | 0.772746 | 0.796427 | 1 | 0.789316 | 0.783511 | 0.760373 |
| 0.788515 | 0.785423 | 0.777485 | 0.798952 | 0.784745 | 0.789316 | 1 | 0.771779 | 0.774357 |
| 0.784185 | 0.783775 | 0.777416 | 0.780158 | 0.781128 | 0.783511 | 0.771779 | 1 | 0.781542 |
| 0.768151 | 0.779199 | 0.772127 | 0.773339 | 0.800452 | 0.760373 | 0.774357 | 0.781542 | 1 |
| 0.774669 | 0.790513 | 0.77372 | 0.77877 | 0.789296 | 0.765758 | 0.778263 | 0.777877 | 0.798524 |

| | | | | | | | | |
|---|---|---|---|---|---|---|---|---|
| 0.770089 | 0.749718 | 0.749121 | 0.75234 | 0.75008 | 0.750422 | 0.761785 | 0.765754 | 0.752009 |
| 0.778659 | 0.77616 | 0.780943 | 0.77988 | 0.776777 | 0.770029 | 0.781374 | 0.778032 | 0.766711 |
| 0.786718 | 0.786996 | 0.776064 | 0.785508 | 0.777823 | 0.775398 | 0.788343 | 0.786466 | 0.777352 |
| 0.799602 | 0.785591 | 0.774282 | 0.784205 | 0.77466 | 0.773662 | 0.796286 | 0.781225 | 0.774794 |
| 0.779106 | 0.78413 | 0.780868 | 0.786067 | 0.776046 | 0.777039 | 0.78533 | 0.778922 | 0.777289 |
| 0.784466 | 0.787752 | 0.806187 | 0.780956 | 0.777997 | 0.778247 | 0.784701 | 0.784861 | 0.770896 |
| 0.772298 | 0.784887 | 0.776768 | 0.784869 | 0.782124 | 0.764116 | 0.776945 | 0.774614 | 0.767211 |
| 0.780963 | 0.78326 | 0.779418 | 0.793224 | 0.777796 | 0.7743 | 0.790522 | 0.784171 | 0.77906 |
| 0.775634 | 0.777778 | 0.775544 | 0.778767 | 0.776369 | 0.766842 | 0.774113 | 0.773197 | 0.771645 |
| 0.769844 | 0.779023 | 0.779322 | 0.778786 | 0.768774 | 0.765966 | 0.77275 | 0.770361 | 0.768483 |
| 0.786115 | 0.786865 | 0.776252 | 0.784374 | 0.767745 | 0.769003 | 0.778147 | 0.78071 | 0.768956 |
| 0.786101 | 0.784495 | 0.778051 | 0.789073 | 0.777385 | 0.774036 | 0.787046 | 0.789005 | 0.770766 |
| 0.796788 | 0.785076 | 0.784967 | 0.78315 | 0.774852 | 0.780508 | 0.791653 | 0.794936 | 0.768187 |
| 0.799109 | 0.777676 | 0.778824 | 0.785809 | 0.777016 | 0.77375 | 0.783786 | 0.795856 | 0.765189 |
| 0.782114 | 0.779995 | 0.777447 | 0.780952 | 0.76904 | 0.767006 | 0.78481 | 0.787333 | 0.761774 |
| 0.78049 | 0.774029 | 0.77618 | 0.77837 | 0.773309 | 0.771405 | 0.781416 | 0.784483 | 0.759806 |
| 0.782328 | 0.775517 | 0.777263 | 0.785559 | 0.768869 | 0.766198 | 0.790879 | 0.78094 | 0.76011 |
| 0.794164 | 0.78524 | 0.783351 | 0.78459 | 0.777069 | 0.782297 | 0.792125 | 0.792993 | 0.771857 |
| 0.794275 | 0.785004 | 0.779857 | 0.786985 | 0.77725 | 0.770133 | 0.787627 | 0.791872 | 0.769326 |
| 0.803508 | 0.778629 | 0.776521 | 0.780311 | 0.767005 | 0.762726 | 0.783536 | 0.789404 | 0.768519 |
| 0.79061 | 0.77078 | 0.781534 | 0.783355 | 0.775389 | 0.775738 | 0.794389 | 0.783708 | 0.762678 |
| 0.790228 | 0.775607 | 0.776792 | 0.78141 | 0.774257 | 0.768952 | 0.781673 | 0.774059 | 0.763247 |
| 0.786482 | 0.775226 | 0.780806 | 0.781173 | 0.77785 | 0.772564 | 0.783674 | 0.782261 | 0.763346 |
| 0.765534 | 0.785096 | 0.780838 | 0.783407 | 0.78185 | 0.772645 | 0.777289 | 0.780402 | 0.780718 |
| 0.765318 | 0.772877 | 0.770521 | 0.77735 | 0.77498 | 0.761475 | 0.764566 | 0.778484 | 0.777667 |
| 0.709921 | 0.707909 | 0.704732 | 0.705845 | 0.708934 | 0.703488 | 0.710325 | 0.712727 | 0.713173 |
| 0.715243 | 0.711424 | 0.709391 | 0.711331 | 0.711466 | 0.709468 | 0.714021 | 0.716571 | 0.713706 |
| 0.714988 | 0.71668 | 0.712837 | 0.716322 | 0.716868 | 0.710764 | 0.716177 | 0.717067 | 0.715813 |
| 0.770911 | 0.772031 | 0.772283 | 0.770499 | 0.771763 | 0.768115 | 0.773125 | 0.767878 | 0.761491 |
| 0.771171 | 0.770096 | 0.772877 | 0.767003 | 0.766322 | 0.762118 | 0.767108 | 0.782903 | 0.763008 |
| 0.779186 | 0.771221 | 0.768224 | 0.766519 | 0.767546 | 0.761366 | 0.764386 | 0.783733 | 0.758143 |
| 0.772321 | 0.772038 | 0.777418 | 0.772385 | 0.779105 | 0.771322 | 0.771193 | 0.773765 | 0.768137 |
| 0.799307 | 0.777954 | 0.782591 | 0.794689 | 0.775104 | 0.778639 | 0.790095 | 0.797924 | 0.777872 |
| 0.798298 | 0.78643 | 0.783246 | 0.801306 | 0.78214 | 0.799446 | 0.8249 | 0.786051 | 0.783245 |
| 0.786914 | 0.775187 | 0.780335 | 0.800876 | 0.768677 | 0.784432 | 0.799422 | 0.781794 | 0.766961 |
| 0.797165 | 0.779423 | 0.784544 | 0.802201 | 0.779838 | 0.793931 | 0.815756 | 0.800764 | 0.774089 |
| 0.779734 | 0.772668 | 0.772207 | 0.784734 | 0.766901 | 0.766764 | 0.791339 | 0.783335 | 0.769839 |
| 0.807317 | 0.802738 | 0.795573 | 0.810053 | 0.794266 | 0.792035 | 0.818682 | 0.807606 | 0.789686 |
| 0.78111 | 0.766181 | 0.766743 | 0.779774 | 0.761341 | 0.772673 | 0.780245 | 0.784779 | 0.762412 |
| 0.780159 | 0.790241 | 0.784613 | 0.774194 | 0.783662 | 0.769838 | 0.777796 | 0.780971 | 0.770835 |
| 0.785661 | 0.771192 | 0.775057 | 0.786736 | 0.775893 | 0.781219 | 0.788047 | 0.787446 | 0.770716 |
| 0.81038 | 0.802302 | 0.800133 | 0.818082 | 0.797024 | 0.794957 | 0.826756 | 0.811226 | 0.789019 |
| 0.791931 | 0.786963 | 0.782086 | 0.772092 | 0.772407 | 0.767375 | 0.781159 | 0.780284 | 0.765784 |
| 0.790074 | 0.785349 | 0.780047 | 0.781668 | 0.777243 | 0.762671 | 0.78539 | 0.783724 | 0.773943 |
| 0.781752 | 0.791618 | 0.781859 | 0.788814 | 0.783683 | 0.782627 | 0.78187 | 0.780237 | 0.769627 |
| 0.793632 | 0.774118 | 0.775856 | 0.7861 | 0.775122 | 0.781698 | 0.79752 | 0.78562 | 0.765638 |

| | | | | | | | | |
|---|---|---|---|---|---|---|---|---|
| 0.79849 | 0.782883 | 0.781214 | 0.781304 | 0.773571 | 0.769694 | 0.778831 | 0.781427 | 0.76628 |
| 0.775196 | 0.787206 | 0.779872 | 0.774655 | 0.774486 | 0.770206 | 0.776578 | 0.789227 | 0.768069 |
| 0.775466 | 0.776841 | 0.777115 | 0.771247 | 0.771817 | 0.766598 | 0.780365 | 0.77666 | 0.76284 |
| 0.792208 | 0.793124 | 0.785714 | 0.795729 | 0.789126 | 0.784269 | 0.8 | 0.795842 | 0.779594 |
| 0.781108 | 0.792109 | 0.781836 | 0.784989 | 0.776571 | 0.769419 | 0.786998 | 0.786762 | 0.780553 |
| 0.777361 | 0.781153 | 0.777521 | 0.775023 | 0.7738 | 0.765808 | 0.771167 | 0.783482 | 0.764877 |
| 0.790584 | 0.789252 | 0.779262 | 0.784373 | 0.775441 | 0.77225 | 0.773735 | 0.776949 | 0.77922 |
| 0.771557 | 0.778731 | 0.785584 | 0.781821 | 0.780403 | 0.774747 | 0.783336 | 0.782347 | 0.772382 |
| 0.772504 | 0.780035 | 0.778264 | 0.784501 | 0.776629 | 0.771178 | 0.781216 | 0.776919 | 0.771715 |
| 0.774053 | 0.773644 | 0.774961 | 0.774041 | 0.775803 | 0.771588 | 0.778779 | 0.76606 | 0.771061 |
| 0.771681 | 0.778836 | 0.775465 | 0.777298 | 0.780931 | 0.768968 | 0.774635 | 0.782584 | 0.769875 |
| 0.777666 | 0.770795 | 0.77467 | 0.78345 | 0.774946 | 0.775165 | 0.782346 | 0.770444 | 0.773079 |
| 0.778696 | 0.781073 | 0.779126 | 0.778365 | 0.779982 | 0.771263 | 0.789752 | 0.77818 | 0.770795 |
| 0.782462 | 0.777516 | 0.765612 | 0.782362 | 0.76902 | 0.765604 | 0.778095 | 0.772584 | 0.771174 |
| 0.769428 | 0.785126 | 0.776034 | 0.77775 | 0.777147 | 0.768048 | 0.776663 | 0.780929 | 0.774975 |
| 0.770292 | 0.779426 | 0.775924 | 0.782512 | 0.773272 | 0.769388 | 0.77918 | 0.7801 | 0.771561 |
| 0.773079 | 0.777251 | 0.778824 | 0.778621 | 0.776284 | 0.770164 | 0.779857 | 0.781716 | 0.773539 |
| 0.780833 | 0.779798 | 0.776382 | 0.783633 | 0.774724 | 0.783545 | 0.797041 | 0.788444 | 0.771658 |
| 0.780618 | 0.775686 | 0.777659 | 0.77797 | 0.771705 | 0.772784 | 0.776485 | 0.77477 | 0.776924 |
| 0.779696 | 0.780106 | 0.773766 | 0.783122 | 0.782745 | 0.767205 | 0.779158 | 0.781998 | 0.777324 |
| 0.771398 | 0.777741 | 0.767897 | 0.785342 | 0.771041 | 0.763428 | 0.778239 | 0.778656 | 0.768757 |
| 0.769826 | 0.776166 | 0.776075 | 0.780202 | 0.776292 | 0.766669 | 0.779794 | 0.772542 | 0.764822 |
| 0.776417 | 0.778454 | 0.779591 | 0.774738 | 0.777106 | 0.761818 | 0.781307 | 0.772394 | 0.768017 |
| 0.780769 | 0.784791 | 0.776752 | 0.780963 | 0.770315 | 0.771588 | 0.781053 | 0.782296 | 0.768532 |
| 0.77099 | 0.775706 | 0.774023 | 0.776514 | 0.76739 | 0.767231 | 0.779596 | 0.782544 | 0.76934 |
| 0.783964 | 0.787654 | 0.781401 | 0.792053 | 0.774799 | 0.77836 | 0.791467 | 0.784456 | 0.782014 |
| 0.77732 | 0.777358 | 0.773999 | 0.781079 | 0.768312 | 0.764792 | 0.783172 | 0.779354 | 0.771709 |
| 0.774951 | 0.781944 | 0.776315 | 0.773496 | 0.773668 | 0.772802 | 0.778402 | 0.77411 | 0.767872 |
| 0.774751 | 0.782217 | 0.779153 | 0.781942 | 0.776099 | 0.773514 | 0.779302 | 0.784436 | 0.769949 |
| 0.77596 | 0.775276 | 0.771459 | 0.772006 | 0.76654 | 0.768281 | 0.77746 | 0.778369 | 0.764676 |
| 0.770098 | 0.779337 | 0.780803 | 0.784448 | 0.774758 | 0.772607 | 0.782091 | 0.77742 | 0.773875 |
| 0.765471 | 0.773072 | 0.770926 | 0.776786 | 0.771609 | 0.766123 | 0.780294 | 0.776825 | 0.770292 |
| 0.788076 | 0.778252 | 0.774884 | 0.78899 | 0.777196 | 0.771747 | 0.781583 | 0.785054 | 0.773944 |
| 0.797618 | 0.781929 | 0.771027 | 0.780558 | 0.771728 | 0.770496 | 0.792572 | 0.77775 | 0.771518 |
| 0.768746 | 0.781803 | 0.779524 | 0.776584 | 0.777704 | 0.772095 | 0.779892 | 0.782986 | 0.764203 |
| 0.778862 | 0.785937 | 0.778841 | 0.778314 | 0.791184 | 0.768 | 0.781899 | 0.790505 | 0.786582 |
| 0.764827 | 0.779416 | 0.770793 | 0.782535 | 0.772396 | 0.764223 | 0.77696 | 0.779667 | 0.767477 |
| 0.756886 | 0.766653 | 0.75865 | 0.760756 | 0.75633 | 0.752228 | 0.755786 | 0.754305 | 0.743992 |
| 0.766924 | 0.769077 | 0.76182 | 0.757806 | 0.755507 | 0.751251 | 0.755621 | 0.779234 | 0.744612 |
| 0.743161 | 0.753205 | 0.747537 | 0.745472 | 0.742625 | 0.73251 | 0.754614 | 0.746306 | 0.742056 |
| 0.759977 | 0.756336 | 0.763972 | 0.765044 | 0.756368 | 0.760236 | 0.776011 | 0.768815 | 0.744428 |
| 0.759763 | 0.74717 | 0.731431 | 0.756717 | 0.732716 | 0.749054 | 0.771273 | 0.732437 | 0.753114 |
| 0.767677 | 0.76043 | 0.749311 | 0.753142 | 0.746904 | 0.739932 | 0.750761 | 0.757015 | 0.742683 |
| 0.751338 | 0.74492 | 0.742914 | 0.74843 | 0.741524 | 0.742489 | 0.757553 | 0.757889 | 0.733863 |
| 0.743508 | 0.746852 | 0.74581 | 0.755877 | 0.745433 | 0.734474 | 0.74359 | 0.744073 | 0.733883 |
| 0.759543 | 0.751392 | 0.747016 | 0.750968 | 0.738288 | 0.749707 | 0.759519 | 0.760109 | 0.739267 |
| 0.747805 | 0.759995 | 0.765298 | 0.759205 | 0.743961 | 0.757274 | 0.774063 | 0.765206 | 0.740186 |

| | | | | | | | | |
|---|---|---|---|---|---|---|---|---|
| 0.741674 | 0.75589 | 0.753172 | 0.738038 | 0.758585 | 0.733097 | 0.741443 | 0.731848 | 0.730946 |
| 0.772427 | 0.756471 | 0.750947 | 0.769147 | 0.743387 | 0.747363 | 0.764891 | 0.766354 | 0.749074 |
| 0.729299 | 0.746746 | 0.748544 | 0.741078 | 0.739749 | 0.731596 | 0.742291 | 0.739896 | 0.730652 |
| 0.761958 | 0.744298 | 0.737416 | 0.743779 | 0.736418 | 0.732398 | 0.749029 | 0.737743 | 0.73159 |
| 0.763104 | 0.754259 | 0.74277 | 0.767107 | 0.741267 | 0.753189 | 0.773161 | 0.747929 | 0.746122 |
| 0.734609 | 0.744129 | 0.741945 | 0.741035 | 0.740709 | 0.736759 | 0.748777 | 0.741611 | 0.734379 |
| 0.755721 | 0.741973 | 0.739924 | 0.740616 | 0.735165 | 0.731764 | 0.744028 | 0.745115 | 0.732467 |
| 0.756533 | 0.747445 | 0.74894 | 0.758116 | 0.742587 | 0.744332 | 0.768056 | 0.761266 | 0.743382 |
| 0.732606 | 0.727562 | 0.725393 | 0.730166 | 0.720174 | 0.73373 | 0.738259 | 0.722757 | 0.73054 |
| 0.769862 | 0.745714 | 0.744119 | 0.758084 | 0.737384 | 0.740701 | 0.767923 | 0.751292 | 0.739257 |
| 0.736962 | 0.746862 | 0.74 | 0.735787 | 0.728913 | 0.734892 | 0.746704 | 0.738062 | 0.732713 |
| 0.78659 | 0.767286 | 0.767961 | 0.76197 | 0.755146 | 0.753933 | 0.766147 | 0.765951 | 0.759013 |
| 0.76668 | 0.775248 | 0.775006 | 0.776475 | 0.772864 | 0.766105 | 0.773823 | 0.768628 | 0.771889 |
| 0.744304 | 0.761826 | 0.756976 | 0.754772 | 0.749036 | 0.749498 | 0.742891 | 0.748013 | 0.754634 |
| 0.779325 | 0.770884 | 0.768618 | 0.774515 | 0.76995 | 0.764023 | 0.774483 | 0.776795 | 0.765169 |
| 0.762427 | 0.760937 | 0.765446 | 0.771818 | 0.768966 | 0.767728 | 0.767651 | 0.761716 | 0.759993 |
| 0.780647 | 0.774408 | 0.7739 | 0.77609 | 0.77756 | 0.766949 | 0.777038 | 0.777353 | 0.766271 |
| 0.772289 | 0.781586 | 0.770078 | 0.780851 | 0.779368 | 0.770866 | 0.775245 | 0.782777 | 0.775435 |
| 0.746466 | 0.756048 | 0.759964 | 0.751888 | 0.756879 | 0.75111 | 0.758929 | 0.758266 | 0.748475 |
| 0.778498 | 0.778717 | 0.77313 | 0.78702 | 0.774846 | 0.775614 | 0.785462 | 0.773551 | 0.787352 |
| 0.741783 | 0.754996 | 0.746388 | 0.745091 | 0.750802 | 0.743796 | 0.744319 | 0.743398 | 0.753227 |
| 0.773408 | 0.775644 | 0.765086 | 0.779461 | 0.776079 | 0.76446 | 0.76601 | 0.777214 | 0.769244 |
| 0.770813 | 0.77082 | 0.767902 | 0.770908 | 0.770647 | 0.770098 | 0.773204 | 0.77101 | 0.772425 |
| 0.742706 | 0.766997 | 0.752698 | 0.738903 | 0.740732 | 0.735147 | 0.73839 | 0.765619 | 0.741134 |
| 0.749018 | 0.743119 | 0.742303 | 0.751992 | 0.744467 | 0.744591 | 0.753484 | 0.751882 | 0.734258 |
| 0.759011 | 0.750974 | 0.751521 | 0.754345 | 0.75582 | 0.749675 | 0.756289 | 0.761595 | 0.747872 |
| 0.75496 | 0.773092 | 0.759427 | 0.765157 | 0.75377 | 0.758445 | 0.756804 | 0.758824 | 0.765632 |
| 0.771215 | 0.782281 | 0.771715 | 0.778723 | 0.77138 | 0.76764 | 0.777604 | 0.78337 | 0.769752 |
| 0.746201 | 0.743182 | 0.744116 | 0.749621 | 0.745185 | 0.742848 | 0.745425 | 0.744899 | 0.737797 |
| 0.77714 | 0.764677 | 0.767087 | 0.769231 | 0.763135 | 0.760434 | 0.781956 | 0.766597 | 0.759357 |
| 0.764353 | 0.767205 | 0.765766 | 0.769905 | 0.767526 | 0.762798 | 0.762685 | 0.773081 | 0.76494 |
| 0.759657 | 0.757325 | 0.751808 | 0.76395 | 0.760683 | 0.75239 | 0.770868 | 0.760345 | 0.746858 |
| 0.759951 | 0.75608 | 0.751008 | 0.76844 | 0.75778 | 0.755572 | 0.769952 | 0.759935 | 0.746938 |
| 0.762841 | 0.781743 | 0.775557 | 0.78155 | 0.776171 | 0.76647 | 0.765562 | 0.770112 | 0.784074 |
| 0.777236 | 0.777435 | 0.772702 | 0.765036 | 0.775209 | 0.754114 | 0.770656 | 0.759987 | 0.759165 |
| 0.766956 | 0.769796 | 0.768216 | 0.76486 | 0.764658 | 0.759029 | 0.768165 | 0.774597 | 0.761795 |
| 0.720272 | 0.726436 | 0.720491 | 0.721738 | 0.720978 | 0.715293 | 0.720006 | 0.725617 | 0.721385 |
| 0.748745 | 0.764076 | 0.756072 | 0.757573 | 0.755217 | 0.749091 | 0.75529 | 0.756709 | 0.757437 |
| 0.719402 | 0.723054 | 0.725229 | 0.724443 | 0.72426 | 0.726915 | 0.728231 | 0.72516 | 0.715683 |
| 0.710663 | 0.718726 | 0.718211 | 0.71217 | 0.721697 | 0.715816 | 0.721965 | 0.715982 | 0.711111 |
| 0.76766 | 0.76708 | 0.764872 | 0.766648 | 0.764844 | 0.766111 | 0.772139 | 0.773739 | 0.759199 |
| 0.706093 | 0.716613 | 0.713821 | 0.712194 | 0.719026 | 0.714852 | 0.717251 | 0.71541 | 0.710467 |
| 0.721802 | 0.711589 | 0.715122 | 0.714593 | 0.714637 | 0.717794 | 0.720675 | 0.719975 | 0.708048 |
| 0.719987 | 0.705688 | 0.701431 | 0.712288 | 0.708789 | 0.708523 | 0.717204 | 0.70834 | 0.697303 |
| 0.767105 | 0.76472 | 0.758265 | 0.764844 | 0.755452 | 0.754751 | 0.76737 | 0.766527 | 0.760157 |
| 0.721409 | 0.703456 | 0.708151 | 0.712553 | 0.706617 | 0.709183 | 0.719216 | 0.718702 | 0.705417 |
| 0.769421 | 0.76001 | 0.756801 | 0.761932 | 0.75541 | 0.75928 | 0.771843 | 0.766352 | 0.755406 |

| | | | | | | | | |
|---|---|---|---|---|---|---|---|---|
| 0.760455 | 0.759708 | 0.760364 | 0.763404 | 0.758394 | 0.754939 | 0.765421 | 0.759244 | 0.7581 |
| 0.774936 | 0.758789 | 0.755974 | 0.765459 | 0.756145 | 0.75452 | 0.769231 | 0.766624 | 0.761455 |
| 0.746059 | 0.750777 | 0.749491 | 0.750386 | 0.75596 | 0.739615 | 0.747218 | 0.747653 | 0.749526 |
| 0.711643 | 0.697306 | 0.700423 | 0.70932 | 0.702893 | 0.711272 | 0.711252 | 0.701032 | 0.694453 |
| 0.70934 | 0.694943 | 0.693416 | 0.699707 | 0.696256 | 0.695827 | 0.704907 | 0.697671 | 0.692674 |
| 0.771177 | 0.758041 | 0.75902 | 0.767344 | 0.75395 | 0.762847 | 0.764513 | 0.768127 | 0.761944 |
| 0.683356 | 0.687842 | 0.687458 | 0.695004 | 0.692092 | 0.691899 | 0.696476 | 0.689655 | 0.685147 |
| 0.729363 | 0.734795 | 0.736701 | 0.73445 | 0.737881 | 0.72999 | 0.733377 | 0.741025 | 0.73621 |
| 0.774673 | 0.757789 | 0.75641 | 0.760451 | 0.753069 | 0.760226 | 0.765676 | 0.754196 | 0.747829 |
| 0.718938 | 0.719606 | 0.723219 | 0.721506 | 0.726022 | 0.717652 | 0.721463 | 0.724216 | 0.727825 |
| 0.760662 | 0.740313 | 0.740965 | 0.746168 | 0.739215 | 0.742624 | 0.762593 | 0.750486 | 0.745586 |
| 0.653061 | 0.6375 | 0.632911 | 0.628931 | 0.608696 | 0.616438 | 0.646259 | 0.590909 | 0.601266 |
| 0.659144 | 0.660151 | 0.652614 | 0.655939 | 0.649571 | 0.646463 | 0.651896 | 0.659812 | 0.656384 |
| 0.662755 | 0.656588 | 0.650022 | 0.658727 | 0.656306 | 0.649655 | 0.652357 | 0.661356 | 0.654568 |
| 0.667543 | 0.66768 | 0.65498 | 0.660196 | 0.665803 | 0.660026 | 0.662495 | 0.671154 | 0.67097 |
| 0.665319 | 0.661761 | 0.654206 | 0.660385 | 0.661446 | 0.657276 | 0.667801 | 0.669897 | 0.666405 |
| 0.615885 | 0.618142 | 0.617786 | 0.618106 | 0.623263 | 0.623709 | 0.619815 | 0.618037 | 0.610457 |
| 0.667425 | 0.666792 | 0.660067 | 0.663757 | 0.6665 | 0.66365 | 0.665365 | 0.669082 | 0.670391 |
| 0.753244 | 0.741678 | 0.735928 | 0.738007 | 0.735099 | 0.733668 | 0.745592 | 0.745343 | 0.742079 |
| 0.740243 | 0.733416 | 0.733895 | 0.736575 | 0.735128 | 0.728912 | 0.741395 | 0.743668 | 0.741267 |
| 0.754081 | 0.735932 | 0.738668 | 0.741071 | 0.737243 | 0.744256 | 0.75446 | 0.743831 | 0.743839 |
| 0.619189 | 0.615963 | 0.617334 | 0.616841 | 0.620153 | 0.627375 | 0.620716 | 0.620568 | 0.614155 |
| 0.599264 | 0.598949 | 0.596934 | 0.596919 | 0.599462 | 0.603745 | 0.602587 | 0.6 | 0.59923 |
| 0.706239 | 0.709132 | 0.707223 | 0.706468 | 0.712052 | 0.70316 | 0.708837 | 0.712702 | 0.708999 |
| 0.707792 | 0.707888 | 0.705501 | 0.704405 | 0.70956 | 0.702761 | 0.706796 | 0.714286 | 0.707896 |
| 0.69826 | 0.70209 | 0.697939 | 0.696338 | 0.704467 | 0.695132 | 0.697684 | 0.708094 | 0.700379 |
| 0.703506 | 0.705166 | 0.701633 | 0.700631 | 0.707555 | 0.699945 | 0.70502 | 0.710909 | 0.709623 |
| 0.704414 | 0.70914 | 0.705743 | 0.70521 | 0.711975 | 0.704556 | 0.708217 | 0.715924 | 0.711764 |
| 0.706668 | 0.708805 | 0.708159 | 0.705506 | 0.71097 | 0.703537 | 0.704577 | 0.709592 | 0.713052 |
| 0.705225 | 0.705248 | 0.706045 | 0.704191 | 0.710832 | 0.701146 | 0.71022 | 0.714979 | 0.709309 |
| 0.708709 | 0.703658 | 0.702808 | 0.703348 | 0.708752 | 0.700033 | 0.708825 | 0.714203 | 0.709884 |
| 0.714796 | 0.715558 | 0.714342 | 0.71316 | 0.718454 | 0.711519 | 0.715181 | 0.720483 | 0.71836 |
| 0.712542 | 0.711888 | 0.707545 | 0.709804 | 0.709827 | 0.705953 | 0.708333 | 0.71152 | 0.710287 |
| 0.713176 | 0.711044 | 0.708997 | 0.710031 | 0.713298 | 0.708363 | 0.711468 | 0.717067 | 0.714081 |
| 0.715578 | 0.711764 | 0.712654 | 0.709981 | 0.712871 | 0.711095 | 0.716663 | 0.718847 | 0.715699 |
| 0.712031 | 0.705849 | 0.70915 | 0.704677 | 0.707554 | 0.7077 | 0.706363 | 0.715355 | 0.710139 |
| 0.712209 | 0.708967 | 0.71016 | 0.711798 | 0.714217 | 0.706299 | 0.709677 | 0.716977 | 0.713385 |
| 0.711253 | 0.710019 | 0.706731 | 0.709234 | 0.714694 | 0.706722 | 0.709523 | 0.715369 | 0.712574 |
| 0.708234 | 0.709013 | 0.700405 | 0.704594 | 0.707967 | 0.703997 | 0.706545 | 0.709698 | 0.709883 |
| 0.711681 | 0.710144 | 0.708462 | 0.713054 | 0.712501 | 0.705845 | 0.71424 | 0.713746 | 0.712178 |
| 0.667294 | 0.658693 | 0.65287 | 0.655743 | 0.655276 | 0.650479 | 0.664289 | 0.662855 | 0.663712 |
| 0.667402 | 0.665577 | 0.658138 | 0.666062 | 0.664078 | 0.657995 | 0.663312 | 0.662157 | 0.668452 |
| 0.713512 | 0.716236 | 0.712793 | 0.711529 | 0.71502 | 0.710789 | 0.716259 | 0.717945 | 0.716877 |
| 0.712082 | 0.716125 | 0.71086 | 0.712588 | 0.718208 | 0.708806 | 0.713097 | 0.719291 | 0.719981 |
| 0.645845 | 0.656297 | 0.647417 | 0.650267 | 0.648962 | 0.647681 | 0.647092 | 0.654294 | 0.648432 |
| 0.715905 | 0.716121 | 0.713627 | 0.709326 | 0.716996 | 0.708778 | 0.712932 | 0.714874 | 0.716297 |
| 0.703242 | 0.704551 | 0.700196 | 0.701517 | 0.704585 | 0.695741 | 0.702293 | 0.704529 | 0.705892 |

| | | | | | | | | |
|---|---|---|---|---|---|---|---|---|
| 0.712477 | 0.710425 | 0.706809 | 0.708333 | 0.71315 | 0.701397 | 0.709159 | 0.711599 | 0.711299 |
| 0.714479 | 0.714047 | 0.71184 | 0.710877 | 0.715209 | 0.706429 | 0.711279 | 0.717343 | 0.71456 |
| 0.713043 | 0.711122 | 0.709116 | 0.709029 | 0.715004 | 0.70572 | 0.711131 | 0.71246 | 0.71155 |
| 0.711469 | 0.713371 | 0.71203 | 0.709549 | 0.715877 | 0.706274 | 0.712527 | 0.716414 | 0.713422 |
| 0.715759 | 0.716869 | 0.716065 | 0.715544 | 0.720712 | 0.70928 | 0.715419 | 0.718243 | 0.715559 |
| 0.707884 | 0.710325 | 0.703966 | 0.708725 | 0.711028 | 0.701619 | 0.705255 | 0.712063 | 0.709826 |
| 0.71253 | 0.710133 | 0.70999 | 0.709766 | 0.712975 | 0.702592 | 0.709438 | 0.713925 | 0.713885 |
| 0.707362 | 0.704905 | 0.702021 | 0.702761 | 0.706184 | 0.699526 | 0.70463 | 0.703078 | 0.709838 |
| 0.709639 | 0.705568 | 0.70686 | 0.707894 | 0.710128 | 0.702548 | 0.710349 | 0.709774 | 0.709402 |
| 0.7082 | 0.711738 | 0.707975 | 0.70878 | 0.710235 | 0.706601 | 0.706782 | 0.710785 | 0.713812 |
| 0.712502 | 0.714695 | 0.71284 | 0.712017 | 0.715551 | 0.708941 | 0.710376 | 0.716717 | 0.713669 |
| 0.707257 | 0.704868 | 0.704481 | 0.705815 | 0.709519 | 0.701232 | 0.705354 | 0.710226 | 0.709883 |
| 0.703212 | 0.702855 | 0.696929 | 0.699704 | 0.704593 | 0.696943 | 0.701621 | 0.70474 | 0.703559 |
| 0.701933 | 0.70249 | 0.696124 | 0.702493 | 0.704812 | 0.698319 | 0.701556 | 0.703576 | 0.703741 |
| 0.71246 | 0.716034 | 0.71162 | 0.712167 | 0.712281 | 0.706997 | 0.713616 | 0.718698 | 0.715028 |
| 0.709657 | 0.707565 | 0.706383 | 0.707567 | 0.709504 | 0.704465 | 0.714594 | 0.714951 | 0.711339 |
| 0.706466 | 0.706334 | 0.701169 | 0.705318 | 0.706003 | 0.696408 | 0.707214 | 0.705474 | 0.707202 |
| 0.708006 | 0.711509 | 0.703325 | 0.704085 | 0.707288 | 0.703166 | 0.706047 | 0.712674 | 0.70961 |
| 0.704042 | 0.707587 | 0.706505 | 0.703975 | 0.710631 | 0.704183 | 0.708383 | 0.711646 | 0.71144 |
| 0.706551 | 0.707159 | 0.705667 | 0.706186 | 0.709623 | 0.702353 | 0.706382 | 0.713172 | 0.708858 |
| 0.704605 | 0.703008 | 0.698931 | 0.699236 | 0.705322 | 0.700517 | 0.703966 | 0.708286 | 0.704095 |
| 0.706016 | 0.705023 | 0.703552 | 0.701076 | 0.707421 | 0.701441 | 0.706313 | 0.709166 | 0.708719 |
| 0.707795 | 0.709163 | 0.707085 | 0.708865 | 0.713124 | 0.70632 | 0.709075 | 0.716242 | 0.712333 |
| 0.711287 | 0.711703 | 0.706237 | 0.709082 | 0.710635 | 0.702853 | 0.710375 | 0.71338 | 0.713698 |
| 0.705407 | 0.707242 | 0.706585 | 0.700008 | 0.711607 | 0.703355 | 0.706353 | 0.710396 | 0.707991 |
| 0.702158 | 0.704177 | 0.701936 | 0.704582 | 0.708403 | 0.701057 | 0.704926 | 0.711767 | 0.706524 |
| 0.707679 | 0.708495 | 0.702599 | 0.70317 | 0.70893 | 0.699019 | 0.70915 | 0.710133 | 0.70914 |
| 0.701186 | 0.704782 | 0.70497 | 0.696527 | 0.706601 | 0.69841 | 0.697782 | 0.705916 | 0.705416 |
| 0.705756 | 0.706521 | 0.703991 | 0.70256 | 0.70584 | 0.702003 | 0.707943 | 0.711006 | 0.706822 |
| 0.708959 | 0.710833 | 0.707119 | 0.705957 | 0.709213 | 0.701642 | 0.705653 | 0.712781 | 0.71184 |
| 0.70875 | 0.710966 | 0.706585 | 0.706334 | 0.710903 | 0.70434 | 0.7122 | 0.713199 | 0.712515 |
| 0.713071 | 0.711766 | 0.708547 | 0.707011 | 0.711969 | 0.70646 | 0.71439 | 0.715604 | 0.711012 |
| 0.712429 | 0.714308 | 0.706224 | 0.710985 | 0.713353 | 0.707525 | 0.715253 | 0.718285 | 0.71344 |
| 0.715305 | 0.713075 | 0.709897 | 0.709279 | 0.714467 | 0.7057 | 0.711844 | 0.715564 | 0.716811 |
| 0.705279 | 0.706259 | 0.702799 | 0.703821 | 0.703789 | 0.699411 | 0.706161 | 0.707798 | 0.707008 |
| 0.707236 | 0.70613 | 0.706469 | 0.707211 | 0.708454 | 0.702565 | 0.709898 | 0.711108 | 0.710262 |
| 0.705104 | 0.711819 | 0.705864 | 0.706135 | 0.710745 | 0.70176 | 0.710265 | 0.713032 | 0.708973 |
| 0.70408 | 0.70547 | 0.703105 | 0.703177 | 0.706967 | 0.69891 | 0.701819 | 0.709014 | 0.704976 |
| 0.698581 | 0.701991 | 0.696585 | 0.696145 | 0.699122 | 0.697738 | 0.70126 | 0.703989 | 0.702952 |
| 0.698777 | 0.700527 | 0.695749 | 0.692829 | 0.69956 | 0.694239 | 0.696929 | 0.704672 | 0.701247 |
| 0.696636 | 0.698502 | 0.695482 | 0.695261 | 0.698308 | 0.695656 | 0.699389 | 0.701273 | 0.702424 |
| 0.699339 | 0.696567 | 0.693527 | 0.69995 | 0.69968 | 0.696371 | 0.697822 | 0.704433 | 0.701512 |
| 0.700375 | 0.702106 | 0.699693 | 0.7015 | 0.700207 | 0.697476 | 0.705389 | 0.705025 | 0.7025 |
| 0.694158 | 0.693146 | 0.691526 | 0.691921 | 0.697412 | 0.691905 | 0.694955 | 0.698404 | 0.697421 |
| 0.705793 | 0.705912 | 0.696834 | 0.701512 | 0.706103 | 0.702362 | 0.702748 | 0.709156 | 0.708094 |
| 0.699917 | 0.699478 | 0.69448 | 0.701194 | 0.700456 | 0.692546 | 0.698156 | 0.701574 | 0.701719 |

| | | | | | | | | |
|---|---|---|---|---|---|---|---|---|
| 0.705586 | 0.707861 | 0.710042 | 0.708635 | 0.709685 | 0.705826 | 0.712885 | 0.713378 | 0.709958 |
| 0.713926 | 0.708609 | 0.707088 | 0.709584 | 0.714493 | 0.706787 | 0.714297 | 0.71589 | 0.71073 |
| 0.707828 | 0.705977 | 0.700855 | 0.705892 | 0.707781 | 0.703224 | 0.709424 | 0.71219 | 0.710269 |
| 0.715878 | 0.710765 | 0.708494 | 0.709998 | 0.712878 | 0.707392 | 0.713022 | 0.717121 | 0.712233 |
| 0.717152 | 0.716528 | 0.712312 | 0.715886 | 0.720769 | 0.711612 | 0.716256 | 0.722213 | 0.719179 |
| 0.715335 | 0.712635 | 0.709978 | 0.715304 | 0.714105 | 0.707052 | 0.709927 | 0.720472 | 0.714584 |
| 0.711065 | 0.705375 | 0.70142 | 0.706188 | 0.708388 | 0.701743 | 0.705026 | 0.710509 | 0.707668 |
| 0.713384 | 0.710734 | 0.706216 | 0.710391 | 0.712591 | 0.704932 | 0.711374 | 0.711934 | 0.71647 |
| 0.709867 | 0.705096 | 0.699792 | 0.70412 | 0.70673 | 0.700244 | 0.706011 | 0.710156 | 0.708363 |
| 0.71275 | 0.708673 | 0.708366 | 0.710744 | 0.711502 | 0.705329 | 0.712452 | 0.711872 | 0.710905 |
| 0.715634 | 0.709308 | 0.711151 | 0.713769 | 0.71745 | 0.707559 | 0.710858 | 0.714857 | 0.711416 |
| 0.716577 | 0.70914 | 0.712416 | 0.713707 | 0.715227 | 0.70619 | 0.710974 | 0.715358 | 0.714045 |
| 0.708493 | 0.707257 | 0.703797 | 0.706779 | 0.709005 | 0.698484 | 0.710022 | 0.713078 | 0.710393 |
| 0.710863 | 0.709458 | 0.707257 | 0.70506 | 0.712768 | 0.705962 | 0.708337 | 0.71746 | 0.710076 |
| 0.715693 | 0.711227 | 0.708938 | 0.711992 | 0.713915 | 0.705748 | 0.713514 | 0.715587 | 0.715678 |
| 0.71139 | 0.712518 | 0.709953 | 0.707452 | 0.712961 | 0.705509 | 0.714892 | 0.716092 | 0.713942 |
| 0.716908 | 0.713799 | 0.708379 | 0.716364 | 0.712569 | 0.708694 | 0.714501 | 0.7174 | 0.718322 |
| 0.711181 | 0.710043 | 0.70742 | 0.709488 | 0.713156 | 0.704312 | 0.710657 | 0.715245 | 0.714036 |
| 0.71318 | 0.715221 | 0.716257 | 0.719142 | 0.71731 | 0.705226 | 0.712848 | 0.717523 | 0.716238 |
| 0.71042 | 0.708373 | 0.707846 | 0.706627 | 0.713148 | 0.709417 | 0.712059 | 0.713848 | 0.713453 |
| 0.712555 | 0.712507 | 0.707519 | 0.710967 | 0.716601 | 0.705028 | 0.710907 | 0.716847 | 0.713963 |
| 0.713364 | 0.70817 | 0.705995 | 0.708257 | 0.712836 | 0.703256 | 0.711082 | 0.714422 | 0.712976 |
| 0.715014 | 0.714683 | 0.708929 | 0.709614 | 0.717321 | 0.708406 | 0.714559 | 0.715376 | 0.718895 |
| 0.713231 | 0.712939 | 0.712328 | 0.714027 | 0.713859 | 0.710215 | 0.71716 | 0.714745 | 0.714616 |
| 0.712978 | 0.708183 | 0.705557 | 0.707569 | 0.712341 | 0.708422 | 0.711637 | 0.710661 | 0.710923 |
| 0.707965 | 0.709094 | 0.707728 | 0.70996 | 0.711377 | 0.707185 | 0.709178 | 0.712842 | 0.710419 |
| 0.709025 | 0.70631 | 0.706298 | 0.708036 | 0.713026 | 0.704855 | 0.70897 | 0.712393 | 0.710281 |
| 0.711132 | 0.708271 | 0.707045 | 0.706247 | 0.713284 | 0.705089 | 0.709552 | 0.707727 | 0.709098 |
| 0.713697 | 0.711872 | 0.709931 | 0.708487 | 0.714241 | 0.708123 | 0.713866 | 0.713345 | 0.714514 |
| 0.718071 | 0.711179 | 0.707953 | 0.71205 | 0.712955 | 0.710652 | 0.710662 | 0.719271 | 0.717774 |
| 0.710567 | 0.707075 | 0.707768 | 0.710253 | 0.712157 | 0.704683 | 0.711048 | 0.712051 | 0.711323 |
| 0.715853 | 0.714791 | 0.712565 | 0.713624 | 0.715963 | 0.707375 | 0.716154 | 0.718777 | 0.71542 |
| 0.713216 | 0.711569 | 0.710853 | 0.712501 | 0.716232 | 0.707908 | 0.71408 | 0.717596 | 0.713919 |
| 0.708059 | 0.706813 | 0.702641 | 0.703887 | 0.705213 | 0.703952 | 0.706677 | 0.70942 | 0.708371 |
| 0.709877 | 0.710343 | 0.708632 | 0.712302 | 0.712105 | 0.702326 | 0.713826 | 0.713484 | 0.713065 |
| 0.710026 | 0.704411 | 0.704427 | 0.702238 | 0.711089 | 0.701603 | 0.707182 | 0.709976 | 0.708505 |
| 0.718345 | 0.717494 | 0.711776 | 0.715633 | 0.718864 | 0.711855 | 0.718119 | 0.722323 | 0.719727 |
| 0.713599 | 0.70859 | 0.708145 | 0.711154 | 0.713156 | 0.708772 | 0.712304 | 0.715322 | 0.713105 |
| 0.71052 | 0.71106 | 0.705868 | 0.711406 | 0.713674 | 0.707416 | 0.714618 | 0.715066 | 0.711791 |
| 0.712362 | 0.709044 | 0.707134 | 0.710507 | 0.712923 | 0.70485 | 0.713982 | 0.713134 | 0.713254 |
| 0.713502 | 0.715364 | 0.710628 | 0.713395 | 0.715923 | 0.711134 | 0.715413 | 0.720183 | 0.719759 |
| 0.713295 | 0.71456 | 0.709183 | 0.715238 | 0.71704 | 0.710796 | 0.71629 | 0.71626 | 0.717746 |
| 0.717839 | 0.716701 | 0.710764 | 0.714555 | 0.720372 | 0.711545 | 0.714819 | 0.717651 | 0.717848 |
| 0.711156 | 0.71076 | 0.705721 | 0.709321 | 0.708925 | 0.706426 | 0.706083 | 0.711215 | 0.71019 |
| 0.711316 | 0.710864 | 0.708227 | 0.713498 | 0.712212 | 0.70645 | 0.71107 | 0.720457 | 0.714563 |
| 0.710871 | 0.704781 | 0.703608 | 0.710151 | 0.710756 | 0.701032 | 0.709112 | 0.715045 | 0.711796 |

| | | | | | | | | |
|---|---|---|---|---|---|---|---|---|
| 0.6983 | 0.699395 | 0.696183 | 0.697772 | 0.699563 | 0.69783 | 0.697546 | 0.70134 | 0.697423 |
| 0.746422 | 0.758487 | 0.752255 | 0.757795 | 0.750921 | 0.749712 | 0.755692 | 0.753761 | 0.756467 |
| 0.710895 | 0.706219 | 0.705768 | 0.707373 | 0.710629 | 0.706472 | 0.708866 | 0.712941 | 0.711869 |
| 0.713923 | 0.712756 | 0.711478 | 0.710522 | 0.712937 | 0.711339 | 0.71295 | 0.71786 | 0.718712 |
| 0.706541 | 0.707026 | 0.70579 | 0.703244 | 0.708165 | 0.705221 | 0.702647 | 0.708724 | 0.711931 |
| 0.711253 | 0.708119 | 0.707171 | 0.705699 | 0.709881 | 0.7076 | 0.70961 | 0.713085 | 0.713425 |
| 0.70513 | 0.703602 | 0.706141 | 0.707113 | 0.704572 | 0.703821 | 0.709028 | 0.710596 | 0.711209 |
| 0.702635 | 0.703218 | 0.696638 | 0.700469 | 0.701639 | 0.702483 | 0.702989 | 0.707344 | 0.707895 |
| 0.712864 | 0.706339 | 0.705365 | 0.708415 | 0.709881 | 0.705534 | 0.707478 | 0.714668 | 0.710713 |
| 0.717728 | 0.714492 | 0.713205 | 0.713081 | 0.718847 | 0.710579 | 0.715962 | 0.721539 | 0.716791 |
| 0.714502 | 0.71136 | 0.709738 | 0.709594 | 0.716535 | 0.707995 | 0.712297 | 0.714905 | 0.714686 |
| 0.718112 | 0.713873 | 0.712759 | 0.715435 | 0.71539 | 0.70765 | 0.713918 | 0.718675 | 0.719008 |
| 0.721364 | 0.714331 | 0.713198 | 0.712408 | 0.716512 | 0.71299 | 0.717484 | 0.717512 | 0.719022 |
| 0.714922 | 0.713866 | 0.709944 | 0.71397 | 0.715855 | 0.708244 | 0.71588 | 0.715645 | 0.715925 |
| 0.715322 | 0.71092 | 0.707907 | 0.708562 | 0.712935 | 0.706101 | 0.713738 | 0.717256 | 0.71541 |
| 0.708461 | 0.709827 | 0.709733 | 0.707889 | 0.713069 | 0.707978 | 0.708835 | 0.714063 | 0.710316 |
| 0.713184 | 0.714837 | 0.709824 | 0.708883 | 0.715072 | 0.710381 | 0.713368 | 0.720656 | 0.716065 |
| 0.715196 | 0.715012 | 0.709034 | 0.711913 | 0.712666 | 0.709062 | 0.711732 | 0.715152 | 0.712399 |
| 0.70353 | 0.703572 | 0.697079 | 0.700523 | 0.703748 | 0.702521 | 0.704161 | 0.705368 | 0.708945 |
| 0.70692 | 0.702632 | 0.703335 | 0.70164 | 0.708253 | 0.700729 | 0.700833 | 0.70864 | 0.707826 |
| 0.712456 | 0.71029 | 0.708504 | 0.712116 | 0.715424 | 0.706824 | 0.71171 | 0.716363 | 0.714774 |
| 0.70437 | 0.700008 | 0.695642 | 0.697614 | 0.701675 | 0.69938 | 0.699484 | 0.706691 | 0.705483 |
| 0.707759 | 0.708633 | 0.705403 | 0.707516 | 0.706826 | 0.705602 | 0.707835 | 0.71554 | 0.711252 |
| 0.714217 | 0.714883 | 0.708519 | 0.711644 | 0.711841 | 0.709603 | 0.711134 | 0.719305 | 0.71402 |
| 0.713957 | 0.715067 | 0.708482 | 0.713173 | 0.714367 | 0.709833 | 0.710225 | 0.717934 | 0.717201 |
| 0.713879 | 0.71214 | 0.709088 | 0.7088 | 0.708956 | 0.709322 | 0.70726 | 0.713998 | 0.714613 |
| 0.705907 | 0.711683 | 0.702756 | 0.706935 | 0.705926 | 0.705094 | 0.703853 | 0.709854 | 0.704655 |
| 0.706517 | 0.70289 | 0.701603 | 0.706086 | 0.706294 | 0.704512 | 0.705843 | 0.708128 | 0.705107 |
| 0.705351 | 0.706775 | 0.703638 | 0.700852 | 0.707063 | 0.700134 | 0.701669 | 0.709715 | 0.708362 |
| 0.765279 | 0.775038 | 0.767784 | 0.76381 | 0.771453 | 0.756259 | 0.762437 | 0.777334 | 0.751959 |
| 0.788054 | 0.781786 | 0.777483 | 0.783307 | 0.774263 | 0.76839 | 0.787107 | 0.788778 | 0.774166 |
| 0.773307 | 0.787115 | 0.778625 | 0.778091 | 0.775782 | 0.766892 | 0.775733 | 0.778575 | 0.765114 |
| 0.781689 | 0.771439 | 0.765112 | 0.770389 | 0.766312 | 0.760889 | 0.771909 | 0.77193 | 0.759022 |
| 0.782267 | 0.781044 | 0.774772 | 0.788544 | 0.778485 | 0.770801 | 0.792835 | 0.781449 | 0.768546 |
| 0.704574 | 0.700049 | 0.698573 | 0.705337 | 0.704052 | 0.697545 | 0.702366 | 0.70818 | 0.70555 |
| 0.707453 | 0.703611 | 0.703582 | 0.705223 | 0.706325 | 0.699968 | 0.704123 | 0.71086 | 0.710528 |
| 0.706961 | 0.704889 | 0.703612 | 0.705685 | 0.707916 | 0.704198 | 0.708192 | 0.713178 | 0.709586 |
| 0.710414 | 0.706056 | 0.706611 | 0.711382 | 0.709252 | 0.704593 | 0.708136 | 0.717977 | 0.712233 |
| 0.706763 | 0.70387 | 0.70247 | 0.70624 | 0.706 | 0.700496 | 0.705338 | 0.713123 | 0.711337 |
| 0.70924 | 0.707206 | 0.706285 | 0.709654 | 0.710596 | 0.705868 | 0.7091 | 0.716268 | 0.713429 |
| 0.707471 | 0.705233 | 0.705854 | 0.707621 | 0.707698 | 0.701437 | 0.708087 | 0.713241 | 0.710803 |
| 0.710473 | 0.708597 | 0.705427 | 0.707715 | 0.708642 | 0.699088 | 0.70972 | 0.715579 | 0.709839 |
| 0.704896 | 0.699294 | 0.696825 | 0.699407 | 0.703601 | 0.695331 | 0.703018 | 0.707235 | 0.705952 |
| 0.705209 | 0.700086 | 0.698563 | 0.701434 | 0.701632 | 0.697078 | 0.699159 | 0.705852 | 0.70466 |
| 0.705285 | 0.704306 | 0.701305 | 0.707506 | 0.705667 | 0.701753 | 0.708059 | 0.712495 | 0.707546 |
| 0.708448 | 0.709206 | 0.703458 | 0.710988 | 0.711436 | 0.705963 | 0.710516 | 0.714274 | 0.714146 |

| | | | | | | | | |
|---|---|---|---|---|---|---|---|---|
| 0.712144 | 0.70763 | 0.708465 | 0.710131 | 0.710088 | 0.701604 | 0.710369 | 0.714089 | 0.712942 |
| 0.708598 | 0.707874 | 0.704603 | 0.708323 | 0.712141 | 0.703147 | 0.71172 | 0.714468 | 0.712099 |
| 0.713845 | 0.712455 | 0.710493 | 0.712607 | 0.717525 | 0.707594 | 0.713403 | 0.716537 | 0.716458 |
| 0.711599 | 0.708594 | 0.706299 | 0.708121 | 0.71269 | 0.700773 | 0.713354 | 0.712244 | 0.713026 |
| 0.706987 | 0.706094 | 0.701743 | 0.704923 | 0.706527 | 0.700203 | 0.706147 | 0.708208 | 0.70964 |
| 0.707826 | 0.705605 | 0.70292 | 0.701899 | 0.709563 | 0.703008 | 0.706778 | 0.709907 | 0.709121 |
| 0.713499 | 0.708921 | 0.707454 | 0.711465 | 0.712113 | 0.707499 | 0.712721 | 0.715772 | 0.713576 |
| 0.710611 | 0.708566 | 0.706016 | 0.708323 | 0.711719 | 0.703948 | 0.709987 | 0.714274 | 0.710446 |
| 0.709006 | 0.70787 | 0.705327 | 0.708486 | 0.708255 | 0.703441 | 0.708039 | 0.711053 | 0.712691 |
| 0.709521 | 0.707453 | 0.705948 | 0.708343 | 0.710138 | 0.703234 | 0.709101 | 0.715407 | 0.711936 |
| 0.70988 | 0.710715 | 0.707467 | 0.7083 | 0.712978 | 0.701464 | 0.712258 | 0.712465 | 0.711007 |
| 0.709738 | 0.708011 | 0.707639 | 0.706457 | 0.711311 | 0.703599 | 0.708959 | 0.712522 | 0.71237 |
| 0.703145 | 0.699285 | 0.698158 | 0.697818 | 0.700103 | 0.695562 | 0.700606 | 0.705282 | 0.700557 |
| 0.702662 | 0.701579 | 0.697221 | 0.69894 | 0.702558 | 0.701444 | 0.701387 | 0.702552 | 0.700027 |
| 0.711899 | 0.705297 | 0.704959 | 0.707719 | 0.710161 | 0.705581 | 0.713242 | 0.713568 | 0.711923 |
| 0.718443 | 0.714446 | 0.709942 | 0.715347 | 0.715503 | 0.712444 | 0.715634 | 0.720984 | 0.718958 |
| 0.71209 | 0.708044 | 0.708742 | 0.710269 | 0.711628 | 0.706081 | 0.710483 | 0.716496 | 0.711963 |
| 0.715594 | 0.713645 | 0.711607 | 0.712603 | 0.713119 | 0.708234 | 0.718496 | 0.719787 | 0.71678 |
| 0.709121 | 0.706889 | 0.703261 | 0.705264 | 0.707494 | 0.701935 | 0.709375 | 0.710664 | 0.711195 |
| 0.711966 | 0.708175 | 0.705391 | 0.708576 | 0.707941 | 0.703473 | 0.711542 | 0.712936 | 0.713081 |
| 0.708212 | 0.704026 | 0.701547 | 0.703914 | 0.707824 | 0.703456 | 0.709976 | 0.711978 | 0.710202 |
| 0.71151 | 0.709696 | 0.707456 | 0.709191 | 0.711775 | 0.705367 | 0.713069 | 0.71257 | 0.714623 |
| 0.715235 | 0.706963 | 0.707637 | 0.71263 | 0.71351 | 0.707751 | 0.712177 | 0.719243 | 0.715737 |
| 0.7103 | 0.707956 | 0.702187 | 0.704626 | 0.709834 | 0.70193 | 0.708144 | 0.713229 | 0.709246 |
| 0.710908 | 0.707206 | 0.707067 | 0.70843 | 0.707219 | 0.704466 | 0.70815 | 0.714237 | 0.711645 |
| 0.78601 | 0.778141 | 0.777704 | 0.791555 | 0.777075 | 0.774831 | 0.788591 | 0.78669 | 0.771736 |
| 0.710016 | 0.708942 | 0.707223 | 0.709708 | 0.712293 | 0.708356 | 0.710681 | 0.711747 | 0.710827 |
| 0.712917 | 0.7096 | 0.704118 | 0.71067 | 0.711224 | 0.707884 | 0.71189 | 0.715539 | 0.714171 |
| 0.717395 | 0.714389 | 0.708047 | 0.712378 | 0.71319 | 0.70848 | 0.713397 | 0.717029 | 0.7169 |
| 0.71162 | 0.711239 | 0.708562 | 0.707313 | 0.709243 | 0.706866 | 0.710292 | 0.715696 | 0.711447 |
| 0.707166 | 0.703973 | 0.703731 | 0.70712 | 0.706224 | 0.704466 | 0.706616 | 0.709886 | 0.707001 |
| 0.706669 | 0.706943 | 0.70361 | 0.702524 | 0.708624 | 0.700796 | 0.706106 | 0.714789 | 0.709249 |
| 0.712265 | 0.710689 | 0.704263 | 0.708084 | 0.713845 | 0.707911 | 0.709559 | 0.717252 | 0.715887 |
| 0.710323 | 0.70461 | 0.702451 | 0.704827 | 0.710307 | 0.701192 | 0.706331 | 0.713157 | 0.710266 |
| 0.706646 | 0.704306 | 0.701119 | 0.703784 | 0.705873 | 0.697721 | 0.706991 | 0.707101 | 0.705822 |
| 0.708751 | 0.704523 | 0.700956 | 0.708841 | 0.70603 | 0.703825 | 0.709823 | 0.710339 | 0.710537 |
| 0.711617 | 0.70974 | 0.709633 | 0.711738 | 0.709385 | 0.710458 | 0.712699 | 0.714898 | 0.712762 |
| 0.713709 | 0.714228 | 0.708989 | 0.710129 | 0.710539 | 0.706836 | 0.713974 | 0.718187 | 0.71845 |
| 0.708347 | 0.709925 | 0.707022 | 0.710963 | 0.712346 | 0.703971 | 0.713927 | 0.714629 | 0.713004 |
| 0.708104 | 0.707841 | 0.706881 | 0.708337 | 0.710805 | 0.705482 | 0.705745 | 0.716562 | 0.712566 |
| 0.706187 | 0.704598 | 0.700594 | 0.703169 | 0.707557 | 0.702555 | 0.706402 | 0.710448 | 0.709924 |
| 0.712634 | 0.710595 | 0.710618 | 0.707101 | 0.713695 | 0.709772 | 0.715009 | 0.717676 | 0.715765 |
| 0.708897 | 0.710603 | 0.704346 | 0.707992 | 0.708796 | 0.703106 | 0.70891 | 0.713931 | 0.712506 |
| 0.70782 | 0.709888 | 0.707646 | 0.703783 | 0.709685 | 0.703258 | 0.708828 | 0.711673 | 0.71165 |
| 0.70836 | 0.709644 | 0.704612 | 0.710688 | 0.709444 | 0.704061 | 0.711874 | 0.717016 | 0.718968 |
| 0.712727 | 0.70884 | 0.705887 | 0.71084 | 0.712957 | 0.70366 | 0.710978 | 0.714988 | 0.717505 |

| | | | | | | | | |
|---|---|---|---|---|---|---|---|---|
| 0.709128 | 0.706246 | 0.704039 | 0.706865 | 0.705401 | 0.702598 | 0.708499 | 0.711735 | 0.70936 |
| 0.710495 | 0.712352 | 0.707837 | 0.710111 | 0.710368 | 0.700868 | 0.71285 | 0.718594 | 0.71106 |
| 0.708578 | 0.708625 | 0.704061 | 0.706888 | 0.706211 | 0.704502 | 0.706409 | 0.710717 | 0.708962 |
| 0.678282 | 0.675157 | 0.65884 | 0.66368 | 0.654898 | 0.660417 | 0.664806 | 0.671384 | 0.669346 |
| 0.706723 | 0.704844 | 0.702395 | 0.705425 | 0.708389 | 0.701478 | 0.707969 | 0.711224 | 0.70837 |
| 0.704406 | 0.705409 | 0.703013 | 0.707894 | 0.708327 | 0.70613 | 0.709696 | 0.711642 | 0.709722 |
| 0.708633 | 0.709081 | 0.706789 | 0.711317 | 0.706505 | 0.70349 | 0.709659 | 0.714624 | 0.711489 |
| 0.710049 | 0.70728 | 0.703266 | 0.707521 | 0.711013 | 0.704301 | 0.710665 | 0.711237 | 0.710185 |
| 0.716377 | 0.712054 | 0.712841 | 0.71187 | 0.714252 | 0.710873 | 0.712253 | 0.717345 | 0.717334 |
| 0.709167 | 0.709268 | 0.706045 | 0.709902 | 0.710814 | 0.705826 | 0.709275 | 0.715632 | 0.712138 |
| 0.708603 | 0.71016 | 0.705524 | 0.708905 | 0.712502 | 0.705251 | 0.709215 | 0.71174 | 0.71095 |
| 0.716719 | 0.715723 | 0.711265 | 0.708641 | 0.713251 | 0.706014 | 0.711605 | 0.715657 | 0.714689 |
| 0.711618 | 0.714172 | 0.709723 | 0.7109 | 0.716099 | 0.707816 | 0.715317 | 0.715279 | 0.714999 |
| 0.70918 | 0.711134 | 0.701984 | 0.701562 | 0.70831 | 0.703023 | 0.70906 | 0.715606 | 0.708833 |
| 0.716059 | 0.712609 | 0.708241 | 0.710954 | 0.715488 | 0.706274 | 0.713897 | 0.719851 | 0.716208 |
| 0.707387 | 0.705967 | 0.703136 | 0.702905 | 0.708849 | 0.700807 | 0.706897 | 0.713822 | 0.709312 |
| 0.712091 | 0.710101 | 0.706081 | 0.709272 | 0.710752 | 0.703669 | 0.711432 | 0.713714 | 0.713567 |
| 0.71309 | 0.711123 | 0.705492 | 0.709731 | 0.713019 | 0.70354 | 0.713395 | 0.719477 | 0.716594 |
| 0.702604 | 0.706973 | 0.700819 | 0.703859 | 0.704649 | 0.701677 | 0.705934 | 0.711622 | 0.709947 |
| 0.710163 | 0.704805 | 0.703386 | 0.703718 | 0.709874 | 0.70285 | 0.711572 | 0.715469 | 0.712049 |
| 0.710556 | 0.712289 | 0.705395 | 0.707622 | 0.712054 | 0.70892 | 0.709876 | 0.714444 | 0.715737 |
| 0.712089 | 0.711951 | 0.70702 | 0.710318 | 0.713439 | 0.704871 | 0.712486 | 0.717928 | 0.714905 |
| 0.689699 | 0.690707 | 0.690701 | 0.700604 | 0.693248 | 0.691039 | 0.692364 | 0.697318 | 0.686652 |
| 0.719961 | 0.716064 | 0.710061 | 0.71737 | 0.702102 | 0.705596 | 0.7195 | 0.724096 | 0.71516 |
| 0.751807 | 0.753597 | 0.759131 | 0.75663 | 0.749059 | 0.741503 | 0.760417 | 0.745446 | 0.757465 |
| 0.756794 | 0.739443 | 0.733696 | 0.734942 | 0.730554 | 0.731282 | 0.74208 | 0.732871 | 0.72592 |
| 0.667835 | 0.664691 | 0.662635 | 0.660924 | 0.663522 | 0.663669 | 0.663821 | 0.668926 | 0.671197 |
| 0.667659 | 0.67103 | 0.661036 | 0.661876 | 0.665133 | 0.656932 | 0.660037 | 0.667465 | 0.668035 |
| 0.668308 | 0.661153 | 0.655953 | 0.658617 | 0.657433 | 0.660745 | 0.664041 | 0.672585 | 0.664269 |
| 0.680456 | 0.673569 | 0.673363 | 0.673188 | 0.6796 | 0.67179 | 0.680782 | 0.678119 | 0.677723 |
| 0.647797 | 0.647952 | 0.648964 | 0.647136 | 0.64458 | 0.646838 | 0.652184 | 0.652499 | 0.650475 |
| 0.647991 | 0.641026 | 0.626298 | 0.641046 | 0.641835 | 0.635428 | 0.643806 | 0.647297 | 0.643289 |
| 0.646604 | 0.64818 | 0.638944 | 0.642336 | 0.644897 | 0.637916 | 0.639561 | 0.651865 | 0.653468 |
| 0.719402 | 0.733037 | 0.729535 | 0.733715 | 0.730246 | 0.732001 | 0.739831 | 0.726722 | 0.71744 |
| 0.703955 | 0.702709 | 0.701747 | 0.7012 | 0.705082 | 0.699593 | 0.703481 | 0.706038 | 0.705302 |
| 0.676102 | 0.675059 | 0.672003 | 0.672916 | 0.673292 | 0.67169 | 0.672538 | 0.67891 | 0.67623 |
| 0.669975 | 0.67258 | 0.666319 | 0.669695 | 0.667313 | 0.66671 | 0.669324 | 0.673459 | 0.67311 |
| 0.651933 | 0.648783 | 0.650446 | 0.648057 | 0.65296 | 0.647101 | 0.65355 | 0.653307 | 0.657519 |
| 0.679538 | 0.676649 | 0.671491 | 0.672245 | 0.675416 | 0.668277 | 0.674673 | 0.678085 | 0.680853 |
| 0.672258 | 0.666255 | 0.663224 | 0.668719 | 0.669747 | 0.663494 | 0.667949 | 0.67317 | 0.667497 |
| 0.662087 | 0.654521 | 0.660294 | 0.656925 | 0.662389 | 0.653564 | 0.663473 | 0.664324 | 0.66646 |
| 0.664143 | 0.666515 | 0.659271 | 0.66164 | 0.662562 | 0.656851 | 0.662491 | 0.668274 | 0.667377 |
| 0.657886 | 0.653012 | 0.647858 | 0.656228 | 0.657296 | 0.650805 | 0.655809 | 0.662094 | 0.658793 |
| 0.662263 | 0.660377 | 0.657825 | 0.655478 | 0.65473 | 0.65089 | 0.656304 | 0.662945 | 0.663456 |
| 0.662648 | 0.657895 | 0.642355 | 0.652961 | 0.652547 | 0.650198 | 0.658215 | 0.664183 | 0.655026 |
| 0.651598 | 0.639029 | 0.636819 | 0.645694 | 0.6399 | 0.642926 | 0.64736 | 0.650829 | 0.646188 |

| | | | | | | | | |
|---|---|---|---|---|---|---|---|---|
| 0.692122 | 0.692352 | 0.684005 | 0.685872 | 0.688334 | 0.684554 | 0.690323 | 0.692792 | 0.695246 |
| 0.660503 | 0.659517 | 0.658901 | 0.65776 | 0.658667 | 0.654609 | 0.662551 | 0.664149 | 0.658271 |
| 0.668031 | 0.66363 | 0.658996 | 0.663828 | 0.662744 | 0.663759 | 0.6656 | 0.670406 | 0.671788 |
| 0.664982 | 0.660027 | 0.659518 | 0.66134 | 0.665394 | 0.659138 | 0.660894 | 0.666227 | 0.666667 |
| 0.668918 | 0.661768 | 0.649273 | 0.658101 | 0.65401 | 0.649065 | 0.664423 | 0.656274 | 0.664878 |
| 0.663366 | 0.654031 | 0.66001 | 0.65359 | 0.665462 | 0.657768 | 0.664491 | 0.668589 | 0.663833 |
| 0.656243 | 0.656769 | 0.652768 | 0.648324 | 0.663097 | 0.649287 | 0.648465 | 0.655981 | 0.661332 |
| 0.655941 | 0.642896 | 0.640785 | 0.648355 | 0.646361 | 0.641396 | 0.654585 | 0.657794 | 0.656741 |
| 0.651756 | 0.650527 | 0.644961 | 0.647792 | 0.652901 | 0.641275 | 0.654375 | 0.644875 | 0.658794 |
| 0.695267 | 0.693257 | 0.685378 | 0.694035 | 0.695188 | 0.687004 | 0.693662 | 0.695039 | 0.694051 |
| 0.692541 | 0.689733 | 0.685507 | 0.685918 | 0.692367 | 0.684062 | 0.688489 | 0.691243 | 0.689367 |
| 0.685185 | 0.684948 | 0.676902 | 0.683426 | 0.690664 | 0.68 | 0.684244 | 0.686387 | 0.683182 |
| 0.680406 | 0.67656 | 0.673731 | 0.674779 | 0.677855 | 0.674754 | 0.678588 | 0.680496 | 0.675219 |
| 0.681091 | 0.680708 | 0.675565 | 0.681033 | 0.681333 | 0.677808 | 0.678318 | 0.68559 | 0.682517 |
| 0.677755 | 0.674664 | 0.665728 | 0.667371 | 0.673604 | 0.669873 | 0.671628 | 0.678369 | 0.679361 |

| | | HG_12 | HG_13 | | HG_14 | HG_15 | | |
|---|---|---|---|---|---|---|---|---|
| Home_garc | Home_garc | Home_garc | Home_garc | Home_garc | Home_garc | Home_garc | Home_garc | Home_garc |
| 0.771999 | 0.753684 | 0.799752 | 0.796671 | 0.800016 | 0.792796 | 0.779465 | 0.777168 | 0.790021 |
| 0.780761 | 0.75928 | 0.785131 | 0.790443 | 0.793876 | 0.781757 | 0.779241 | 0.769784 | 0.792724 |
| 0.76955 | 0.760978 | 0.78358 | 0.7899 | 0.793886 | 0.786497 | 0.776497 | 0.767402 | 0.776927 |
| 0.78422 | 0.760443 | 0.780157 | 0.784863 | 0.799458 | 0.784627 | 0.78953 | 0.775037 | 0.787871 |
| 0.774898 | 0.754564 | 0.780478 | 0.784241 | 0.787886 | 0.781371 | 0.778502 | 0.780354 | 0.777613 |
| 0.780877 | 0.751417 | 0.771747 | 0.776911 | 0.773044 | 0.779153 | 0.781949 | 0.780685 | 0.781461 |
| 0.71882 | 0.748219 | 0.704099 | 0.709025 | 0.719458 | 0.71313 | 0.71627 | 0.709586 | 0.716184 |
| 0.713345 | 0.744701 | 0.698338 | 0.704787 | 0.708232 | 0.706505 | 0.71195 | 0.7026 | 0.709809 |
| 0.778417 | 0.759948 | 0.773325 | 0.784099 | 0.783123 | 0.784424 | 0.786693 | 0.782079 | 0.793129 |
| 0.77498 | 0.766394 | 0.795212 | 0.788847 | 0.810822 | 0.778534 | 0.784289 | 0.774207 | 0.775723 |
| 0.774776 | 0.765436 | 0.782469 | 0.792203 | 0.802321 | 0.785691 | 0.782861 | 0.767754 | 0.783721 |
| 0.780265 | 0.76222 | 0.793361 | 0.808619 | 0.805533 | 0.798057 | 0.786319 | 0.783955 | 0.790066 |
| 0.78048 | 0.761897 | 0.771239 | 0.774308 | 0.791427 | 0.777312 | 0.774247 | 0.763432 | 0.782677 |
| 0.770631 | 0.79651 | 0.780105 | 0.788573 | 0.791585 | 0.779915 | 0.776165 | 0.768051 | 0.779257 |
| 0.765473 | 0.801061 | 0.763286 | 0.774434 | 0.769573 | 0.769587 | 0.770142 | 0.769912 | 0.766081 |
| 0.782128 | 0.773718 | 0.785091 | 0.796913 | 0.79476 | 0.788223 | 0.783147 | 0.775365 | 0.786158 |
| 0.772765 | 0.786578 | 0.793304 | 0.791288 | 0.806347 | 0.785122 | 0.775539 | 0.765657 | 0.769969 |
| 0.755556 | 0.83406 | 0.764815 | 0.761082 | 0.766859 | 0.759235 | 0.764758 | 0.74998 | 0.760794 |
| 0.772306 | 0.754603 | 0.787537 | 0.790795 | 0.789014 | 0.795179 | 0.783126 | 0.775362 | 0.796534 |
| 0.783976 | 0.758446 | 0.782122 | 0.788715 | 0.795915 | 0.784071 | 0.78307 | 0.777111 | 0.79923 |
| 0.783993 | 0.753695 | 0.777823 | 0.789626 | 0.791626 | 0.789374 | 0.788015 | 0.782766 | 0.78911 |
| 0.777158 | 0.763798 | 0.793867 | 0.806382 | 0.808535 | 0.799137 | 0.79716 | 0.786433 | 0.807094 |
| 0.793853 | 0.760698 | 0.790669 | 0.799817 | 0.798813 | 0.79578 | 0.794382 | 0.785826 | 0.797581 |
| 0.778982 | 0.778013 | 0.796405 | 0.819252 | 0.808933 | 0.807849 | 0.797814 | 0.774085 | 0.794135 |
| 0.791681 | 0.758289 | 0.776961 | 0.791402 | 0.789921 | 0.792702 | 0.785982 | 0.78004 | 0.792895 |
| 0.778866 | 0.758748 | 0.783844 | 0.785767 | 0.782888 | 0.777361 | 0.780309 | 0.781107 | 0.793238 |
| 0.772384 | 0.749598 | 0.77786 | 0.784998 | 0.777705 | 0.777741 | 0.774839 | 0.76891 | 0.786427 |
| 0.786451 | 0.755357 | 0.785106 | 0.781655 | 0.781387 | 0.776944 | 0.784496 | 0.78127 | 0.778829 |
| 0.788266 | 0.758135 | 0.780404 | 0.79325 | 0.7851 | 0.793417 | 0.785156 | 0.782362 | 0.785322 |
| 0.774669 | 0.770089 | 0.778659 | 0.786718 | 0.799602 | 0.779106 | 0.784466 | 0.772298 | 0.780963 |
| 0.790513 | 0.749718 | 0.77616 | 0.786996 | 0.785591 | 0.78413 | 0.787752 | 0.784887 | 0.78326 |
| 0.77372 | 0.749121 | 0.780943 | 0.776064 | 0.774282 | 0.780868 | 0.806187 | 0.776768 | 0.779418 |
| 0.77877 | 0.75234 | 0.77988 | 0.785508 | 0.784205 | 0.786067 | 0.780956 | 0.784869 | 0.793224 |
| 0.789296 | 0.75008 | 0.776777 | 0.777823 | 0.77466 | 0.776046 | 0.777997 | 0.782124 | 0.777796 |
| 0.765758 | 0.750422 | 0.770029 | 0.775398 | 0.773662 | 0.777039 | 0.778247 | 0.764116 | 0.7743 |
| 0.778263 | 0.761785 | 0.781374 | 0.788343 | 0.796286 | 0.78533 | 0.784701 | 0.776945 | 0.790522 |
| 0.777877 | 0.765754 | 0.778032 | 0.786466 | 0.781225 | 0.778922 | 0.784861 | 0.774614 | 0.784171 |
| 0.798524 | 0.752009 | 0.766711 | 0.777352 | 0.774794 | 0.777289 | 0.770896 | 0.767211 | 0.77906 |
| 1 | 0.758637 | 0.773977 | 0.775203 | 0.782825 | 0.774257 | 0.785407 | 0.779569 | 0.777567 |

| | | | | | | | | |
|---|---|---|---|---|---|---|---|---|
| 0.758637 | 1 | 0.76388 | 0.765182 | 0.780561 | 0.761537 | 0.762628 | 0.75048 | 0.758383 |
| 0.773977 | 0.76388 | 1 | 0.80353 | 0.792267 | 0.792344 | 0.790538 | 0.7751 | 0.792728 |
| 0.775203 | 0.765182 | 0.80353 | 1 | 0.819839 | 0.838387 | 0.788131 | 0.775633 | 0.78553 |
| 0.782825 | 0.780561 | 0.792267 | 0.819839 | 1 | 0.799405 | 0.789343 | 0.777506 | 0.791732 |
| 0.774257 | 0.761537 | 0.792344 | 0.838387 | 0.799405 | 1 | 0.788052 | 0.779575 | 0.796032 |
| 0.785407 | 0.762628 | 0.790538 | 0.788131 | 0.789343 | 0.788052 | 1 | 0.778364 | 0.784673 |
| 0.779569 | 0.75048 | 0.7751 | 0.775633 | 0.777506 | 0.779575 | 0.778364 | 1 | 0.777238 |
| 0.777567 | 0.758383 | 0.792728 | 0.78553 | 0.791732 | 0.796032 | 0.784673 | 0.777238 | 1 |
| 0.77566 | 0.756643 | 0.770097 | 0.774295 | 0.777978 | 0.771671 | 0.779582 | 0.771685 | 0.776484 |
| 0.782869 | 0.75212 | 0.767844 | 0.772791 | 0.770529 | 0.776628 | 0.780876 | 0.784167 | 0.781085 |
| 0.776065 | 0.75686 | 0.803175 | 0.801325 | 0.800763 | 0.789229 | 0.788625 | 0.77477 | 0.794495 |
| 0.77808 | 0.755147 | 0.804094 | 0.803738 | 0.802221 | 0.790592 | 0.784397 | 0.780755 | 0.801142 |
| 0.77854 | 0.768383 | 0.793428 | 0.800922 | 0.810221 | 0.793574 | 0.791041 | 0.778531 | 0.790731 |
| 0.777219 | 0.763904 | 0.796562 | 0.803556 | 0.800931 | 0.793006 | 0.787836 | 0.777686 | 0.787582 |
| 0.770864 | 0.755695 | 0.785631 | 0.79543 | 0.804768 | 0.785827 | 0.778149 | 0.771802 | 0.791873 |
| 0.768078 | 0.758742 | 0.785995 | 0.787614 | 0.79704 | 0.784753 | 0.781636 | 0.77526 | 0.78938 |
| 0.772358 | 0.764538 | 0.793141 | 0.793853 | 0.802856 | 0.78833 | 0.779004 | 0.768766 | 0.780163 |
| 0.781558 | 0.773475 | 0.805722 | 0.798884 | 0.809938 | 0.796983 | 0.793146 | 0.776731 | 0.794069 |
| 0.783846 | 0.76406 | 0.78469 | 0.797404 | 0.80369 | 0.782312 | 0.79099 | 0.781649 | 0.797494 |
| 0.773887 | 0.762622 | 0.796233 | 0.805303 | 0.808317 | 0.794763 | 0.785244 | 0.769997 | 0.795208 |
| 0.771736 | 0.761193 | 0.801861 | 0.806281 | 0.798524 | 0.799332 | 0.788482 | 0.773699 | 0.779545 |
| 0.7752 | 0.7644 | 0.7989 | 0.797024 | 0.81522 | 0.789513 | 0.777815 | 0.768676 | 0.779291 |
| 0.781918 | 0.755385 | 0.780366 | 0.782051 | 0.7878 | 0.783493 | 0.778079 | 0.777524 | 0.779428 |
| 0.780709 | 0.754941 | 0.767247 | 0.776558 | 0.776396 | 0.773676 | 0.779619 | 0.778318 | 0.783127 |
| 0.776597 | 0.751519 | 0.763015 | 0.765296 | 0.766594 | 0.773917 | 0.774281 | 0.773076 | 0.771208 |
| 0.714984 | 0.745698 | 0.699172 | 0.706099 | 0.7095 | 0.707399 | 0.713617 | 0.702527 | 0.711605 |
| 0.717201 | 0.74834 | 0.704749 | 0.705557 | 0.713624 | 0.712545 | 0.714355 | 0.710644 | 0.713316 |
| 0.715862 | 0.750767 | 0.706695 | 0.708043 | 0.71873 | 0.711131 | 0.71639 | 0.712335 | 0.714493 |
| 0.770189 | 0.745393 | 0.766609 | 0.782587 | 0.776424 | 0.778427 | 0.77128 | 0.773616 | 0.783075 |
| 0.764666 | 0.751635 | 0.773566 | 0.782052 | 0.776032 | 0.776713 | 0.776067 | 0.770055 | 0.782862 |
| 0.768135 | 0.763332 | 0.78892 | 0.798598 | 0.801923 | 0.781175 | 0.777142 | 0.774578 | 0.776544 |
| 0.769492 | 0.754383 | 0.781977 | 0.780897 | 0.778118 | 0.77333 | 0.772378 | 0.774301 | 0.779174 |
| 0.784945 | 0.774082 | 0.799681 | 0.805676 | 0.802142 | 0.799479 | 0.789905 | 0.775802 | 0.801292 |
| 0.779484 | 0.771384 | 0.786677 | 0.82065 | 0.81917 | 0.813631 | 0.790977 | 0.776253 | 0.803986 |
| 0.777564 | 0.770617 | 0.802938 | 0.800459 | 0.809901 | 0.800846 | 0.788603 | 0.77566 | 0.804723 |
| 0.777163 | 0.785744 | 0.792811 | 0.803254 | 0.802212 | 0.804375 | 0.78368 | 0.772506 | 0.792973 |
| 0.780245 | 0.751076 | 0.797004 | 0.786202 | 0.789066 | 0.779173 | 0.782873 | 0.776283 | 0.802892 |
| 0.796375 | 0.782538 | 0.833701 | 0.81512 | 0.83161 | 0.809603 | 0.805278 | 0.79613 | 0.823041 |
| 0.772964 | 0.762474 | 0.783027 | 0.784155 | 0.789907 | 0.783206 | 0.775987 | 0.765157 | 0.791132 |
| 0.782179 | 0.75704 | 0.777207 | 0.804878 | 0.790546 | 0.800858 | 0.787123 | 0.783965 | 0.789465 |
| 0.772276 | 0.768104 | 0.783395 | 0.791378 | 0.798146 | 0.787374 | 0.77531 | 0.769062 | 0.780564 |
| 0.800317 | 0.789893 | 0.830978 | 0.822938 | 0.82902 | 0.82154 | 0.811372 | 0.79955 | 0.826113 |
| 0.776836 | 0.768918 | 0.808043 | 0.806124 | 0.813963 | 0.795083 | 0.797904 | 0.777037 | 0.78071 |
| 0.78034 | 0.765459 | 0.796174 | 0.818828 | 0.821618 | 0.796631 | 0.793149 | 0.777207 | 0.783534 |
| 0.784323 | 0.758476 | 0.776624 | 0.784397 | 0.789831 | 0.795687 | 0.787067 | 0.783084 | 0.790634 |
| 0.778783 | 0.766724 | 0.775964 | 0.785947 | 0.788115 | 0.783131 | 0.78394 | 0.768932 | 0.78632 |

| | | | | | | | | |
|---|---|---|---|---|---|---|---|---|
| 0.772686 | 0.763365 | 0.800785 | 0.797943 | 0.807073 | 0.787294 | 0.787497 | 0.781691 | 0.7822 |
| 0.775996 | 0.755743 | 0.778389 | 0.794427 | 0.781217 | 0.788043 | 0.787359 | 0.784086 | 0.791314 |
| 0.781484 | 0.755776 | 0.786383 | 0.790981 | 0.789688 | 0.782515 | 0.783809 | 0.770172 | 0.777945 |
| 0.787149 | 0.76865 | 0.804932 | 0.797368 | 0.812269 | 0.795697 | 0.793211 | 0.786767 | 0.809962 |
| 0.787152 | 0.761289 | 0.780528 | 0.787769 | 0.788983 | 0.79447 | 0.790276 | 0.781212 | 0.790792 |
| 0.772506 | 0.756597 | 0.774364 | 0.785834 | 0.785507 | 0.78822 | 0.783027 | 0.788606 | 0.788793 |
| 0.791899 | 0.753212 | 0.794014 | 0.781455 | 0.792515 | 0.786962 | 0.79196 | 0.77873 | 0.794859 |
| 0.775852 | 0.754108 | 0.77418 | 0.780711 | 0.778061 | 0.78397 | 0.781081 | 0.779396 | 0.785577 |
| 0.781343 | 0.756625 | 0.775426 | 0.778425 | 0.782691 | 0.783932 | 0.781591 | 0.783699 | 0.787996 |
| 0.776528 | 0.751524 | 0.767885 | 0.78388 | 0.7837 | 0.781446 | 0.777741 | 0.778736 | 0.77764 |
| 0.777732 | 0.753191 | 0.76965 | 0.775854 | 0.783656 | 0.774417 | 0.778471 | 0.773949 | 0.779909 |
| 0.775964 | 0.751872 | 0.771794 | 0.788601 | 0.785756 | 0.784849 | 0.78757 | 0.771483 | 0.77747 |
| 0.779497 | 0.755426 | 0.774791 | 0.787223 | 0.794929 | 0.784706 | 0.784961 | 0.777079 | 0.788492 |
| 0.772137 | 0.751796 | 0.766326 | 0.780676 | 0.781696 | 0.780105 | 0.775174 | 0.771336 | 0.776862 |
| 0.776762 | 0.757351 | 0.777448 | 0.786202 | 0.784393 | 0.783602 | 0.780421 | 0.775078 | 0.780699 |
| 0.780074 | 0.757753 | 0.78041 | 0.78552 | 0.776169 | 0.784067 | 0.780637 | 0.777896 | 0.783683 |
| 0.777851 | 0.754635 | 0.772825 | 0.779452 | 0.782619 | 0.786652 | 0.782627 | 0.776363 | 0.786683 |
| 0.775008 | 0.767966 | 0.770798 | 0.796518 | 0.789141 | 0.786907 | 0.779829 | 0.772605 | 0.783373 |
| 0.780377 | 0.755335 | 0.774183 | 0.788918 | 0.788034 | 0.785436 | 0.783898 | 0.773562 | 0.785506 |
| 0.782877 | 0.751667 | 0.771931 | 0.778623 | 0.777896 | 0.774818 | 0.77799 | 0.772198 | 0.788237 |
| 0.775258 | 0.752684 | 0.771956 | 0.776609 | 0.778161 | 0.778503 | 0.775709 | 0.779734 | 0.7805 |
| 0.771295 | 0.748277 | 0.772593 | 0.774825 | 0.775865 | 0.773525 | 0.778219 | 0.772776 | 0.781006 |
| 0.776315 | 0.753599 | 0.77433 | 0.784351 | 0.784746 | 0.78348 | 0.782918 | 0.774837 | 0.778195 |
| 0.77431 | 0.752868 | 0.771534 | 0.787245 | 0.782516 | 0.789712 | 0.784927 | 0.775418 | 0.78191 |
| 0.77275 | 0.752277 | 0.779379 | 0.778205 | 0.783304 | 0.78048 | 0.773073 | 0.773018 | 0.790715 |
| 0.787032 | 0.757901 | 0.798859 | 0.78542 | 0.793256 | 0.795265 | 0.795475 | 0.77776 | 0.803284 |
| 0.77649 | 0.751897 | 0.78748 | 0.781389 | 0.7934 | 0.782948 | 0.777329 | 0.771557 | 0.793595 |
| 0.772479 | 0.75136 | 0.774307 | 0.784519 | 0.77939 | 0.780496 | 0.775794 | 0.775338 | 0.785288 |
| 0.771461 | 0.760605 | 0.773248 | 0.779926 | 0.781652 | 0.783815 | 0.781602 | 0.77827 | 0.788095 |
| 0.775063 | 0.755029 | 0.772913 | 0.778048 | 0.779403 | 0.780518 | 0.772175 | 0.77192 | 0.784182 |
| 0.77088 | 0.752289 | 0.772866 | 0.780894 | 0.784264 | 0.786017 | 0.781782 | 0.778266 | 0.786403 |
| 0.771028 | 0.750834 | 0.770751 | 0.772898 | 0.777454 | 0.775422 | 0.774159 | 0.773443 | 0.780783 |
| 0.780909 | 0.755612 | 0.791981 | 0.785456 | 0.796674 | 0.784975 | 0.78382 | 0.779186 | 0.801152 |
| 0.778358 | 0.777388 | 0.789532 | 0.814709 | 0.985718 | 0.795302 | 0.784826 | 0.773653 | 0.78747 |
| 0.7717 | 0.758915 | 0.774971 | 0.788975 | 0.789443 | 0.785145 | 0.783477 | 0.776803 | 0.790975 |
| 0.79539 | 0.757375 | 0.773394 | 0.786942 | 0.792717 | 0.782558 | 0.781576 | 0.778724 | 0.790512 |
| 0.77232 | 0.754441 | 0.773194 | 0.781515 | 0.782983 | 0.781297 | 0.779057 | 0.777382 | 0.789088 |
| 0.757372 | 0.729496 | 0.746233 | 0.754771 | 0.756164 | 0.760623 | 0.75409 | 0.750105 | 0.75627 |
| 0.758305 | 0.736002 | 0.756484 | 0.773288 | 0.769255 | 0.773401 | 0.763822 | 0.755283 | 0.777013 |
| 0.756717 | 0.723468 | 0.749149 | 0.752509 | 0.749367 | 0.752664 | 0.75342 | 0.752848 | 0.76191 |
| 0.747614 | 0.751335 | 0.76018 | 0.770314 | 0.777534 | 0.770596 | 0.762219 | 0.746892 | 0.763533 |
| 0.737784 | 0.715676 | 0.729834 | 0.777406 | 0.774724 | 0.770184 | 0.748506 | 0.731646 | 0.758778 |
| 0.750031 | 0.734555 | 0.79126 | 0.786552 | 0.798152 | 0.771585 | 0.766066 | 0.75173 | 0.754326 |
| 0.743879 | 0.728728 | 0.755035 | 0.765729 | 0.760394 | 0.764766 | 0.747596 | 0.734931 | 0.760372 |
| 0.741855 | 0.721246 | 0.734554 | 0.744106 | 0.74855 | 0.755677 | 0.748472 | 0.746619 | 0.759222 |
| 0.746159 | 0.728923 | 0.760517 | 0.763803 | 0.76769 | 0.762537 | 0.750195 | 0.745184 | 0.759242 |
| 0.745506 | 0.736822 | 0.752823 | 0.765684 | 0.7674 | 0.758994 | 0.754164 | 0.743224 | 0.763483 |

| | | | | | | | | |
|---|---|---|---|---|---|---|---|---|
| 0.741099 | 0.715106 | 0.736395 | 0.738104 | 0.738866 | 0.744183 | 0.733911 | 0.73228 | 0.751191 |
| 0.751552 | 0.749095 | 0.777444 | 0.778146 | 0.780895 | 0.772231 | 0.751663 | 0.744715 | 0.778298 |
| 0.73128 | 0.715152 | 0.733729 | 0.746171 | 0.738495 | 0.742663 | 0.738389 | 0.74136 | 0.749232 |
| 0.734349 | 0.722686 | 0.773551 | 0.768844 | 0.778365 | 0.753515 | 0.747141 | 0.744902 | 0.744394 |
| 0.747461 | 0.714975 | 0.755045 | 0.7639 | 0.786122 | 0.771939 | 0.756042 | 0.741716 | 0.779855 |
| 0.736349 | 0.7125 | 0.740788 | 0.748745 | 0.742772 | 0.747535 | 0.747227 | 0.742523 | 0.745126 |
| 0.738608 | 0.713183 | 0.753065 | 0.756549 | 0.75337 | 0.752656 | 0.751733 | 0.745992 | 0.75637 |
| 0.741514 | 0.73253 | 0.750255 | 0.776948 | 0.768825 | 0.768289 | 0.757393 | 0.736636 | 0.760018 |
| 0.727561 | 0.706897 | 0.713233 | 0.73989 | 0.742653 | 0.737378 | 0.734339 | 0.724336 | 0.729533 |
| 0.747946 | 0.70853 | 0.779616 | 0.754106 | 0.776839 | 0.758953 | 0.755576 | 0.742715 | 0.790265 |
| 0.737864 | 0.709546 | 0.744186 | 0.749035 | 0.747696 | 0.748209 | 0.742747 | 0.741964 | 0.751535 |
| 0.765869 | 0.757552 | 0.795587 | 0.798464 | 0.790527 | 0.787452 | 0.781827 | 0.763592 | 0.767389 |
| 0.775984 | 0.748308 | 0.76382 | 0.771233 | 0.774899 | 0.775683 | 0.773443 | 0.772687 | 0.782734 |
| 0.753756 | 0.724874 | 0.74672 | 0.749088 | 0.752079 | 0.75285 | 0.761341 | 0.757972 | 0.759479 |
| 0.771991 | 0.756113 | 0.769435 | 0.775188 | 0.784231 | 0.778388 | 0.780998 | 0.769607 | 0.778263 |
| 0.760601 | 0.743074 | 0.755094 | 0.773981 | 0.775253 | 0.76831 | 0.771495 | 0.767276 | 0.770223 |
| 0.7693 | 0.759682 | 0.77855 | 0.786092 | 0.779232 | 0.778295 | 0.779799 | 0.770764 | 0.780405 |
| 0.780833 | 0.757605 | 0.771157 | 0.784553 | 0.778875 | 0.779712 | 0.775699 | 0.776896 | 0.783318 |
| 0.754919 | 0.73326 | 0.754537 | 0.752421 | 0.753961 | 0.75884 | 0.755907 | 0.755529 | 0.756375 |
| 0.7792 | 0.761821 | 0.767953 | 0.78751 | 0.787382 | 0.785289 | 0.788631 | 0.767361 | 0.782476 |
| 0.755517 | 0.73076 | 0.745767 | 0.751028 | 0.745513 | 0.753111 | 0.753237 | 0.753999 | 0.746765 |
| 0.775604 | 0.752515 | 0.76434 | 0.773275 | 0.771007 | 0.775719 | 0.779699 | 0.779826 | 0.777023 |
| 0.77334 | 0.756107 | 0.765872 | 0.769737 | 0.775143 | 0.769477 | 0.777676 | 0.77439 | 0.778312 |
| 0.755489 | 0.725387 | 0.74192 | 0.750481 | 0.741299 | 0.75635 | 0.752739 | 0.741805 | 0.756917 |
| 0.74585 | 0.733023 | 0.754592 | 0.750298 | 0.751901 | 0.74818 | 0.748655 | 0.737661 | 0.749317 |
| 0.758706 | 0.7374 | 0.756454 | 0.77002 | 0.7574 | 0.762869 | 0.761661 | 0.747387 | 0.760912 |
| 0.771237 | 0.743588 | 0.760084 | 0.763251 | 0.760664 | 0.762096 | 0.772109 | 0.758113 | 0.758875 |
| 0.776262 | 0.7551 | 0.7723 | 0.774569 | 0.775816 | 0.791347 | 0.782967 | 0.771799 | 0.782767 |
| 0.74828 | 0.726298 | 0.739638 | 0.744842 | 0.747659 | 0.744423 | 0.752192 | 0.744455 | 0.7543 |
| 0.774456 | 0.766453 | 0.779941 | 0.76461 | 0.787057 | 0.773059 | 0.772412 | 0.76885 | 0.786889 |
| 0.772958 | 0.756697 | 0.764971 | 0.765308 | 0.771426 | 0.768495 | 0.779778 | 0.763628 | 0.77488 |
| 0.755904 | 0.747005 | 0.777137 | 0.766368 | 0.776648 | 0.764566 | 0.761113 | 0.751915 | 0.767922 |
| 0.752794 | 0.74874 | 0.781712 | 0.763946 | 0.775532 | 0.76211 | 0.762704 | 0.758055 | 0.773886 |
| 0.786943 | 0.757168 | 0.767045 | 0.759951 | 0.762578 | 0.769162 | 0.786413 | 0.779739 | 0.769199 |
| 0.764376 | 0.759753 | 0.758433 | 0.763474 | 0.778564 | 0.761056 | 0.763578 | 0.76805 | 0.76566 |
| 0.767175 | 0.752594 | 0.758722 | 0.76628 | 0.77197 | 0.767459 | 0.770547 | 0.761691 | 0.768996 |
| 0.724163 | 0.711507 | 0.716153 | 0.72866 | 0.724536 | 0.726673 | 0.730309 | 0.731021 | 0.723346 |
| 0.765322 | 0.74994 | 0.75006 | 0.754198 | 0.758266 | 0.753589 | 0.759421 | 0.755412 | 0.762867 |
| 0.722401 | 0.708766 | 0.731846 | 0.727731 | 0.727124 | 0.724284 | 0.724629 | 0.728404 | 0.732461 |
| 0.714308 | 0.704753 | 0.715669 | 0.724612 | 0.723047 | 0.722865 | 0.720979 | 0.719929 | 0.720491 |
| 0.766294 | 0.791023 | 0.766175 | 0.784518 | 0.783619 | 0.785369 | 0.774571 | 0.763423 | 0.773895 |
| 0.712718 | 0.705043 | 0.715514 | 0.723384 | 0.720887 | 0.717412 | 0.719019 | 0.718898 | 0.719236 |
| 0.715342 | 0.706052 | 0.715916 | 0.718828 | 0.734294 | 0.722897 | 0.718451 | 0.715543 | 0.725513 |
| 0.706563 | 0.701523 | 0.715609 | 0.707797 | 0.723534 | 0.711168 | 0.710632 | 0.705238 | 0.713553 |
| 0.771901 | 0.799804 | 0.772168 | 0.768334 | 0.79493 | 0.766277 | 0.764663 | 0.755235 | 0.763158 |
| 0.709886 | 0.70548 | 0.710828 | 0.713877 | 0.720769 | 0.714737 | 0.712871 | 0.706401 | 0.721751 |
| 0.770985 | 0.782609 | 0.767282 | 0.770703 | 0.788026 | 0.774478 | 0.770832 | 0.764663 | 0.76918 |

| | | | | | | | | |
|---|---|---|---|---|---|---|---|---|
| 0.760817 | 0.791606 | 0.762213 | 0.766903 | 0.777463 | 0.766033 | 0.770367 | 0.756959 | 0.763272 |
| 0.767084 | 0.797285 | 0.768717 | 0.787627 | 0.782233 | 0.778096 | 0.77285 | 0.761654 | 0.766598 |
| 0.751711 | 0.756905 | 0.754148 | 0.75418 | 0.755832 | 0.753728 | 0.753598 | 0.747112 | 0.744966 |
| 0.702834 | 0.688653 | 0.704962 | 0.708614 | 0.717795 | 0.707789 | 0.707696 | 0.701425 | 0.714823 |
| 0.69691 | 0.694563 | 0.714074 | 0.701602 | 0.701913 | 0.699199 | 0.694971 | 0.695716 | 0.71237 |
| 0.768836 | 0.813252 | 0.771569 | 0.794174 | 0.799951 | 0.769281 | 0.766134 | 0.754674 | 0.771584 |
| 0.68845 | 0.684678 | 0.691871 | 0.692963 | 0.690541 | 0.690145 | 0.689527 | 0.686298 | 0.700069 |
| 0.740635 | 0.742724 | 0.72717 | 0.731828 | 0.738255 | 0.734247 | 0.735922 | 0.730064 | 0.737032 |
| 0.762503 | 0.805748 | 0.756902 | 0.765281 | 0.773172 | 0.766475 | 0.768876 | 0.759365 | 0.765393 |
| 0.727295 | 0.737774 | 0.714954 | 0.717876 | 0.723242 | 0.721844 | 0.725742 | 0.721469 | 0.723058 |
| 0.750571 | 0.858201 | 0.752428 | 0.751988 | 0.767818 | 0.747741 | 0.754067 | 0.742727 | 0.758183 |
| 0.6125 | 0.660256 | 0.690141 | 0.569697 | 0.6 | 0.625 | 0.642384 | 0.623377 | 0.616766 |
| 0.666925 | 0.707647 | 0.646946 | 0.651989 | 0.659778 | 0.653742 | 0.663295 | 0.650019 | 0.650732 |
| 0.65795 | 0.695135 | 0.649647 | 0.655073 | 0.661208 | 0.660118 | 0.659778 | 0.648786 | 0.65972 |
| 0.670571 | 0.7079 | 0.660944 | 0.660484 | 0.671334 | 0.663915 | 0.668949 | 0.66225 | 0.666181 |
| 0.663952 | 0.707755 | 0.653708 | 0.659494 | 0.667655 | 0.662063 | 0.666321 | 0.654672 | 0.669437 |
| 0.614838 | 0.614826 | 0.621898 | 0.619873 | 0.617474 | 0.620584 | 0.616274 | 0.616968 | 0.627468 |
| 0.670133 | 0.711477 | 0.660928 | 0.660236 | 0.675016 | 0.667797 | 0.665829 | 0.655001 | 0.671376 |
| 0.74512 | 0.809631 | 0.739092 | 0.745321 | 0.750932 | 0.745825 | 0.741416 | 0.737729 | 0.749712 |
| 0.747108 | 0.861809 | 0.741058 | 0.74131 | 0.757318 | 0.7415 | 0.738739 | 0.731869 | 0.742497 |
| 0.745143 | 0.85801 | 0.747301 | 0.745818 | 0.759072 | 0.740046 | 0.745708 | 0.739766 | 0.749116 |
| 0.614205 | 0.616095 | 0.618269 | 0.620214 | 0.61863 | 0.616131 | 0.619185 | 0.619816 | 0.620988 |
| 0.599555 | 0.59785 | 0.599714 | 0.597901 | 0.598959 | 0.60046 | 0.601727 | 0.598416 | 0.603703 |
| 0.713628 | 0.769301 | 0.698493 | 0.700692 | 0.711992 | 0.704125 | 0.709821 | 0.705363 | 0.71033 |
| 0.712353 | 0.764115 | 0.696358 | 0.703098 | 0.712012 | 0.708 | 0.712026 | 0.703036 | 0.712453 |
| 0.706796 | 0.746517 | 0.691032 | 0.693886 | 0.704281 | 0.698247 | 0.699438 | 0.695815 | 0.701312 |
| 0.710796 | 0.750483 | 0.697425 | 0.697544 | 0.709143 | 0.704473 | 0.705915 | 0.70109 | 0.703973 |
| 0.715807 | 0.760151 | 0.701887 | 0.702193 | 0.710035 | 0.705859 | 0.708473 | 0.70578 | 0.71156 |
| 0.714977 | 0.757801 | 0.697615 | 0.704196 | 0.713001 | 0.704116 | 0.711994 | 0.703349 | 0.71092 |
| 0.713284 | 0.761399 | 0.697474 | 0.70275 | 0.711366 | 0.703962 | 0.708643 | 0.704394 | 0.708534 |
| 0.713737 | 0.742208 | 0.69764 | 0.703522 | 0.714321 | 0.702473 | 0.711232 | 0.706756 | 0.707378 |
| 0.720612 | 0.748667 | 0.706368 | 0.712502 | 0.719965 | 0.717359 | 0.716157 | 0.714714 | 0.714774 |
| 0.719027 | 0.750976 | 0.704145 | 0.706043 | 0.714686 | 0.711991 | 0.714851 | 0.710708 | 0.712527 |
| 0.718992 | 0.746054 | 0.705527 | 0.706857 | 0.717979 | 0.711902 | 0.715713 | 0.708579 | 0.712994 |
| 0.721708 | 0.749639 | 0.708621 | 0.707922 | 0.715021 | 0.712881 | 0.717348 | 0.712798 | 0.714663 |
| 0.714822 | 0.744061 | 0.701535 | 0.705596 | 0.714502 | 0.705564 | 0.712666 | 0.705466 | 0.708729 |
| 0.720054 | 0.746881 | 0.70473 | 0.705535 | 0.714024 | 0.711324 | 0.713183 | 0.706657 | 0.70947 |
| 0.718238 | 0.746627 | 0.70359 | 0.70783 | 0.714669 | 0.713549 | 0.714868 | 0.712758 | 0.711659 |
| 0.714251 | 0.744535 | 0.699537 | 0.703822 | 0.709141 | 0.706516 | 0.706519 | 0.704732 | 0.70796 |
| 0.719936 | 0.746365 | 0.704897 | 0.707039 | 0.71432 | 0.712813 | 0.715678 | 0.709624 | 0.713498 |
| 0.660969 | 0.697892 | 0.641707 | 0.656165 | 0.664764 | 0.654072 | 0.663669 | 0.647784 | 0.659333 |
| 0.667882 | 0.700714 | 0.648795 | 0.659074 | 0.670663 | 0.662668 | 0.670398 | 0.654076 | 0.667642 |
| 0.72091 | 0.747405 | 0.711569 | 0.708673 | 0.721427 | 0.714229 | 0.716281 | 0.713743 | 0.715049 |
| 0.721872 | 0.756198 | 0.707362 | 0.710315 | 0.719711 | 0.712709 | 0.718785 | 0.712517 | 0.715385 |
| 0.658129 | 0.696362 | 0.634897 | 0.641764 | 0.65223 | 0.644531 | 0.64755 | 0.643699 | 0.650847 |
| 0.722712 | 0.74593 | 0.706835 | 0.709226 | 0.713878 | 0.712531 | 0.716236 | 0.708919 | 0.713987 |
| 0.714558 | 0.743097 | 0.695974 | 0.694215 | 0.705457 | 0.700228 | 0.7029 | 0.698334 | 0.701697 |

| | | | | | | | | |
|---|---|---|---|---|---|---|---|---|
| 0.716986 | 0.750886 | 0.701282 | 0.703018 | 0.711941 | 0.708085 | 0.708227 | 0.708948 | 0.708144 |
| 0.720307 | 0.751815 | 0.706624 | 0.707462 | 0.714623 | 0.710323 | 0.714786 | 0.70986 | 0.713911 |
| 0.716669 | 0.746944 | 0.704959 | 0.705935 | 0.715526 | 0.711907 | 0.711504 | 0.707742 | 0.711268 |
| 0.7182 | 0.748195 | 0.705487 | 0.708085 | 0.715136 | 0.713207 | 0.713829 | 0.708548 | 0.715134 |
| 0.721355 | 0.751825 | 0.708604 | 0.712829 | 0.723145 | 0.715295 | 0.71416 | 0.710947 | 0.717139 |
| 0.716439 | 0.746646 | 0.70233 | 0.704493 | 0.7131 | 0.707821 | 0.711469 | 0.704397 | 0.708739 |
| 0.716333 | 0.749259 | 0.700944 | 0.704883 | 0.71459 | 0.708893 | 0.71283 | 0.70956 | 0.708429 |
| 0.713076 | 0.741365 | 0.699064 | 0.699534 | 0.709264 | 0.702508 | 0.706566 | 0.703327 | 0.705312 |
| 0.715504 | 0.742075 | 0.700531 | 0.703977 | 0.711831 | 0.704419 | 0.710707 | 0.706675 | 0.708517 |
| 0.715725 | 0.743559 | 0.702554 | 0.705939 | 0.714206 | 0.708041 | 0.706942 | 0.704172 | 0.709157 |
| 0.720295 | 0.7503 | 0.70533 | 0.707746 | 0.713914 | 0.711213 | 0.71391 | 0.709158 | 0.710296 |
| 0.713806 | 0.750849 | 0.699788 | 0.704019 | 0.710973 | 0.706233 | 0.708299 | 0.70799 | 0.708069 |
| 0.710259 | 0.738866 | 0.696351 | 0.695635 | 0.70663 | 0.699656 | 0.701364 | 0.700537 | 0.701932 |
| 0.709026 | 0.741765 | 0.693233 | 0.695069 | 0.705887 | 0.698588 | 0.703463 | 0.697381 | 0.701223 |
| 0.720543 | 0.750299 | 0.706068 | 0.710947 | 0.71987 | 0.716614 | 0.71894 | 0.711763 | 0.714639 |
| 0.716194 | 0.768055 | 0.703242 | 0.704132 | 0.715121 | 0.709029 | 0.710309 | 0.707001 | 0.710713 |
| 0.712285 | 0.749626 | 0.697996 | 0.701306 | 0.711135 | 0.705287 | 0.708295 | 0.701759 | 0.703667 |
| 0.715829 | 0.761387 | 0.699126 | 0.703166 | 0.713014 | 0.707789 | 0.709295 | 0.704018 | 0.709325 |
| 0.713934 | 0.770402 | 0.701534 | 0.701574 | 0.710056 | 0.707964 | 0.709436 | 0.701602 | 0.712447 |
| 0.713989 | 0.758903 | 0.700389 | 0.700621 | 0.70913 | 0.707708 | 0.710858 | 0.701672 | 0.707581 |
| 0.706977 | 0.764322 | 0.694806 | 0.699733 | 0.705949 | 0.702035 | 0.708286 | 0.70191 | 0.703948 |
| 0.711428 | 0.762022 | 0.699612 | 0.699968 | 0.707784 | 0.7045 | 0.707536 | 0.701239 | 0.709598 |
| 0.713913 | 0.761691 | 0.70296 | 0.705158 | 0.713319 | 0.710601 | 0.711159 | 0.707 | 0.712818 |
| 0.716876 | 0.758002 | 0.702332 | 0.706463 | 0.714263 | 0.709962 | 0.711897 | 0.705934 | 0.712769 |
| 0.712341 | 0.766103 | 0.699254 | 0.701883 | 0.709605 | 0.706498 | 0.71213 | 0.703386 | 0.707226 |
| 0.710727 | 0.771159 | 0.697366 | 0.698579 | 0.708107 | 0.700956 | 0.707967 | 0.701577 | 0.707746 |
| 0.71207 | 0.754748 | 0.696933 | 0.701596 | 0.709414 | 0.704417 | 0.70935 | 0.702874 | 0.707085 |
| 0.707608 | 0.755594 | 0.696339 | 0.698682 | 0.706617 | 0.699847 | 0.705442 | 0.698792 | 0.703187 |
| 0.714422 | 0.774798 | 0.696212 | 0.700254 | 0.708805 | 0.706302 | 0.713565 | 0.703466 | 0.708667 |
| 0.713593 | 0.772772 | 0.70148 | 0.704869 | 0.71301 | 0.709396 | 0.710264 | 0.703523 | 0.71352 |
| 0.715469 | 0.763982 | 0.700673 | 0.704695 | 0.713493 | 0.709293 | 0.715168 | 0.706315 | 0.710461 |
| 0.718553 | 0.759768 | 0.701544 | 0.707056 | 0.716585 | 0.709341 | 0.71395 | 0.709843 | 0.710384 |
| 0.717813 | 0.762709 | 0.703861 | 0.708007 | 0.717234 | 0.714274 | 0.716162 | 0.709581 | 0.715707 |
| 0.718368 | 0.757768 | 0.705322 | 0.708755 | 0.718941 | 0.712617 | 0.712951 | 0.708934 | 0.715476 |
| 0.711875 | 0.771765 | 0.694669 | 0.700734 | 0.710813 | 0.706337 | 0.708394 | 0.701326 | 0.706973 |
| 0.71257 | 0.767392 | 0.7 | 0.705963 | 0.709714 | 0.707439 | 0.713821 | 0.705196 | 0.705615 |
| 0.716834 | 0.760815 | 0.701519 | 0.706496 | 0.712483 | 0.707204 | 0.712705 | 0.703063 | 0.711837 |
| 0.712742 | 0.752421 | 0.696556 | 0.701326 | 0.710533 | 0.704758 | 0.705935 | 0.702473 | 0.705455 |
| 0.707556 | 0.757631 | 0.692023 | 0.696358 | 0.702831 | 0.700915 | 0.704449 | 0.697714 | 0.701033 |
| 0.703122 | 0.752293 | 0.688913 | 0.693575 | 0.702582 | 0.696519 | 0.701224 | 0.696756 | 0.70047 |
| 0.707305 | 0.764374 | 0.689224 | 0.693646 | 0.702518 | 0.696893 | 0.704356 | 0.697619 | 0.699752 |
| 0.701734 | 0.756687 | 0.692003 | 0.695612 | 0.702491 | 0.699451 | 0.701248 | 0.691491 | 0.698708 |
| 0.707532 | 0.756967 | 0.693382 | 0.699509 | 0.703716 | 0.700806 | 0.704481 | 0.698485 | 0.702883 |
| 0.698382 | 0.750887 | 0.685615 | 0.688891 | 0.697574 | 0.692936 | 0.694391 | 0.690487 | 0.69753 |
| 0.710498 | 0.755114 | 0.693978 | 0.699599 | 0.707341 | 0.701851 | 0.706175 | 0.700465 | 0.709072 |
| 0.706702 | 0.739587 | 0.69195 | 0.696826 | 0.70337 | 0.697304 | 0.696836 | 0.692119 | 0.700166 |

| | | | | | | | | |
|---|---|---|---|---|---|---|---|---|
| 0.717017 | 0.796154 | 0.701502 | 0.707 | 0.714683 | 0.708469 | 0.718974 | 0.709754 | 0.712161 |
| 0.712504 | 0.748882 | 0.703338 | 0.708515 | 0.713988 | 0.709031 | 0.713081 | 0.706631 | 0.708276 |
| 0.712296 | 0.743867 | 0.6981 | 0.702218 | 0.707557 | 0.708122 | 0.710603 | 0.701344 | 0.708654 |
| 0.715675 | 0.748068 | 0.704281 | 0.705971 | 0.714218 | 0.711884 | 0.716029 | 0.709446 | 0.711697 |
| 0.721131 | 0.754767 | 0.710215 | 0.713909 | 0.723039 | 0.717548 | 0.717757 | 0.714763 | 0.72019 |
| 0.715578 | 0.75004 | 0.704078 | 0.707913 | 0.718001 | 0.714433 | 0.714365 | 0.711834 | 0.712859 |
| 0.710837 | 0.748199 | 0.698287 | 0.702783 | 0.710364 | 0.704741 | 0.710581 | 0.704425 | 0.705203 |
| 0.718312 | 0.74616 | 0.706148 | 0.707608 | 0.714003 | 0.713072 | 0.711436 | 0.708686 | 0.711264 |
| 0.709388 | 0.739237 | 0.694435 | 0.695509 | 0.711528 | 0.704334 | 0.706242 | 0.702665 | 0.705003 |
| 0.714149 | 0.74497 | 0.702741 | 0.703904 | 0.71558 | 0.710619 | 0.713896 | 0.707181 | 0.711967 |
| 0.716878 | 0.745769 | 0.70446 | 0.709204 | 0.719635 | 0.712162 | 0.717122 | 0.710802 | 0.716923 |
| 0.717433 | 0.744056 | 0.699653 | 0.70829 | 0.717709 | 0.713364 | 0.717486 | 0.707236 | 0.714434 |
| 0.71441 | 0.745723 | 0.70165 | 0.702568 | 0.712774 | 0.709206 | 0.717906 | 0.703029 | 0.714343 |
| 0.714479 | 0.744516 | 0.700981 | 0.707344 | 0.717063 | 0.713774 | 0.712446 | 0.705366 | 0.71002 |
| 0.715141 | 0.746082 | 0.703152 | 0.708258 | 0.7146 | 0.711003 | 0.715274 | 0.708373 | 0.713184 |
| 0.718616 | 0.748439 | 0.70279 | 0.705468 | 0.717689 | 0.712439 | 0.717327 | 0.709435 | 0.715796 |
| 0.721427 | 0.748947 | 0.705435 | 0.71223 | 0.7221 | 0.714274 | 0.717255 | 0.711806 | 0.718355 |
| 0.718263 | 0.747901 | 0.704165 | 0.706937 | 0.717584 | 0.711814 | 0.715741 | 0.709609 | 0.715291 |
| 0.722355 | 0.74716 | 0.707238 | 0.708998 | 0.718673 | 0.716987 | 0.718587 | 0.71381 | 0.712074 |
| 0.718678 | 0.750399 | 0.701618 | 0.7081 | 0.711763 | 0.709728 | 0.713717 | 0.709167 | 0.713057 |
| 0.717906 | 0.750525 | 0.700215 | 0.703677 | 0.712686 | 0.708967 | 0.713998 | 0.708824 | 0.70911 |
| 0.711933 | 0.746744 | 0.700295 | 0.705355 | 0.71359 | 0.711134 | 0.710664 | 0.707958 | 0.709138 |
| 0.721613 | 0.750999 | 0.703075 | 0.709297 | 0.718604 | 0.716618 | 0.713047 | 0.710458 | 0.712991 |
| 0.718427 | 0.743689 | 0.708179 | 0.706963 | 0.71845 | 0.710976 | 0.715895 | 0.712013 | 0.712303 |
| 0.717675 | 0.752817 | 0.704219 | 0.705177 | 0.716376 | 0.708871 | 0.716464 | 0.707176 | 0.709609 |
| 0.71289 | 0.748299 | 0.703686 | 0.708005 | 0.717286 | 0.711735 | 0.718432 | 0.707244 | 0.709999 |
| 0.717324 | 0.744225 | 0.700581 | 0.704282 | 0.712703 | 0.706155 | 0.71262 | 0.707006 | 0.708955 |
| 0.712731 | 0.744422 | 0.701608 | 0.704447 | 0.713759 | 0.708674 | 0.712911 | 0.710147 | 0.712956 |
| 0.719427 | 0.752033 | 0.706743 | 0.70831 | 0.720032 | 0.711484 | 0.716917 | 0.710228 | 0.711465 |
| 0.719139 | 0.751466 | 0.707773 | 0.705806 | 0.719957 | 0.710604 | 0.719775 | 0.710939 | 0.712286 |
| 0.716562 | 0.747149 | 0.702247 | 0.706795 | 0.716091 | 0.707529 | 0.71518 | 0.70593 | 0.708084 |
| 0.721512 | 0.749737 | 0.707247 | 0.710463 | 0.718513 | 0.71287 | 0.719345 | 0.712516 | 0.717173 |
| 0.720035 | 0.752886 | 0.705316 | 0.708525 | 0.718036 | 0.712949 | 0.719162 | 0.709525 | 0.713579 |
| 0.711783 | 0.745697 | 0.699153 | 0.703073 | 0.714064 | 0.708286 | 0.710728 | 0.706871 | 0.70846 |
| 0.713981 | 0.745315 | 0.703015 | 0.705791 | 0.715722 | 0.707307 | 0.712535 | 0.710234 | 0.711675 |
| 0.714975 | 0.749264 | 0.700224 | 0.70521 | 0.707359 | 0.708427 | 0.710312 | 0.707473 | 0.711917 |
| 0.723343 | 0.756005 | 0.710649 | 0.713876 | 0.722161 | 0.716282 | 0.720067 | 0.714818 | 0.719414 |
| 0.718009 | 0.747558 | 0.701021 | 0.704404 | 0.712801 | 0.7087 | 0.713496 | 0.706171 | 0.713252 |
| 0.716628 | 0.747774 | 0.704223 | 0.707637 | 0.716256 | 0.710439 | 0.713817 | 0.710338 | 0.714205 |
| 0.718166 | 0.749448 | 0.705682 | 0.704273 | 0.714147 | 0.709137 | 0.709788 | 0.706267 | 0.711656 |
| 0.720441 | 0.756889 | 0.710458 | 0.707853 | 0.718863 | 0.714071 | 0.718077 | 0.709546 | 0.715847 |
| 0.720395 | 0.7494 | 0.702188 | 0.70833 | 0.72047 | 0.712553 | 0.715919 | 0.710053 | 0.716297 |
| 0.725014 | 0.750279 | 0.703663 | 0.710778 | 0.71861 | 0.712745 | 0.717181 | 0.711844 | 0.716412 |
| 0.714529 | 0.749241 | 0.702447 | 0.706278 | 0.712461 | 0.711626 | 0.711867 | 0.706224 | 0.71073 |
| 0.717296 | 0.750869 | 0.706239 | 0.709087 | 0.714479 | 0.712 | 0.712199 | 0.71187 | 0.714988 |
| 0.7141 | 0.745716 | 0.700386 | 0.702241 | 0.710243 | 0.707073 | 0.710952 | 0.706181 | 0.710492 |

| | | | | | | | | |
|---|---|---|---|---|---|---|---|---|
| 0.706222 | 0.735503 | 0.688075 | 0.688348 | 0.696843 | 0.695814 | 0.698513 | 0.693909 | 0.699411 |
| 0.759799 | 0.761496 | 0.742088 | 0.747885 | 0.747881 | 0.754619 | 0.763757 | 0.748181 | 0.751159 |
| 0.713097 | 0.744559 | 0.701537 | 0.704401 | 0.710853 | 0.705452 | 0.711647 | 0.707014 | 0.710466 |
| 0.720114 | 0.749801 | 0.703011 | 0.71077 | 0.716466 | 0.713982 | 0.715979 | 0.711517 | 0.71508 |
| 0.708671 | 0.740449 | 0.694857 | 0.698839 | 0.707435 | 0.70745 | 0.708898 | 0.70207 | 0.706569 |
| 0.713103 | 0.742462 | 0.700912 | 0.702926 | 0.715778 | 0.710159 | 0.712635 | 0.705901 | 0.710237 |
| 0.711598 | 0.744547 | 0.699229 | 0.699252 | 0.710884 | 0.706621 | 0.711691 | 0.702242 | 0.707593 |
| 0.705359 | 0.740114 | 0.690767 | 0.695723 | 0.705421 | 0.702481 | 0.705953 | 0.697305 | 0.705462 |
| 0.713856 | 0.747567 | 0.700016 | 0.704432 | 0.711892 | 0.711834 | 0.711978 | 0.708084 | 0.711332 |
| 0.722967 | 0.7523 | 0.708615 | 0.713451 | 0.722763 | 0.718088 | 0.719144 | 0.713546 | 0.720532 |
| 0.719316 | 0.749641 | 0.704549 | 0.706166 | 0.716111 | 0.711616 | 0.714695 | 0.710999 | 0.716152 |
| 0.720071 | 0.750605 | 0.70284 | 0.709899 | 0.719263 | 0.716664 | 0.716196 | 0.70939 | 0.718114 |
| 0.717941 | 0.751446 | 0.709875 | 0.708887 | 0.722611 | 0.715104 | 0.720292 | 0.714002 | 0.718936 |
| 0.72167 | 0.748563 | 0.705451 | 0.710337 | 0.71838 | 0.71511 | 0.718765 | 0.712006 | 0.716566 |
| 0.716854 | 0.746648 | 0.700443 | 0.70885 | 0.714217 | 0.710637 | 0.711265 | 0.705343 | 0.712865 |
| 0.7159 | 0.745417 | 0.702823 | 0.704732 | 0.714509 | 0.71616 | 0.714692 | 0.709114 | 0.714173 |
| 0.718312 | 0.752019 | 0.705985 | 0.707857 | 0.72162 | 0.711848 | 0.716369 | 0.710598 | 0.716045 |
| 0.71361 | 0.747275 | 0.704113 | 0.707472 | 0.72101 | 0.714755 | 0.711147 | 0.709088 | 0.715001 |
| 0.704857 | 0.743592 | 0.69267 | 0.697592 | 0.708567 | 0.700207 | 0.70505 | 0.700851 | 0.707533 |
| 0.709037 | 0.741796 | 0.69845 | 0.700395 | 0.712452 | 0.70564 | 0.709798 | 0.704342 | 0.709778 |
| 0.71977 | 0.746409 | 0.705542 | 0.707769 | 0.714698 | 0.712756 | 0.714529 | 0.707912 | 0.711719 |
| 0.709847 | 0.732316 | 0.693475 | 0.69578 | 0.706978 | 0.701839 | 0.708931 | 0.697735 | 0.70343 |
| 0.713015 | 0.740859 | 0.706148 | 0.706182 | 0.713427 | 0.710234 | 0.714641 | 0.707317 | 0.713268 |
| 0.714845 | 0.744154 | 0.705953 | 0.705868 | 0.714569 | 0.712312 | 0.7151 | 0.711669 | 0.71494 |
| 0.716355 | 0.748256 | 0.701183 | 0.707069 | 0.716745 | 0.712602 | 0.716713 | 0.710652 | 0.71293 |
| 0.715324 | 0.742973 | 0.705697 | 0.706487 | 0.717491 | 0.711725 | 0.715205 | 0.711031 | 0.713159 |
| 0.709025 | 0.743466 | 0.699787 | 0.701593 | 0.710705 | 0.707315 | 0.708216 | 0.703115 | 0.706106 |
| 0.706712 | 0.743588 | 0.696713 | 0.701569 | 0.711635 | 0.706605 | 0.708247 | 0.702213 | 0.70335 |
| 0.707694 | 0.738851 | 0.696476 | 0.702341 | 0.708887 | 0.705098 | 0.712217 | 0.704212 | 0.708147 |
| 0.763476 | 0.743046 | 0.761109 | 0.781767 | 0.765692 | 0.775941 | 0.768751 | 0.764082 | 0.777949 |
| 0.777096 | 0.751561 | 0.793514 | 0.791313 | 0.793101 | 0.79705 | 0.784685 | 0.77237 | 0.809026 |
| 0.773257 | 0.751236 | 0.774944 | 0.79763 | 0.786388 | 0.788251 | 0.785822 | 0.783608 | 0.785882 |
| 0.767416 | 0.755544 | 0.795945 | 0.795706 | 0.815493 | 0.778562 | 0.777102 | 0.766906 | 0.771497 |
| 0.779297 | 0.751299 | 0.790884 | 0.79275 | 0.805281 | 0.788906 | 0.782752 | 0.775996 | 0.805862 |
| 0.710515 | 0.736038 | 0.696504 | 0.700213 | 0.707391 | 0.707648 | 0.706309 | 0.702053 | 0.703511 |
| 0.709698 | 0.744475 | 0.70196 | 0.703486 | 0.711434 | 0.706356 | 0.708681 | 0.702932 | 0.710161 |
| 0.708965 | 0.745138 | 0.701986 | 0.706138 | 0.714023 | 0.708658 | 0.710714 | 0.706203 | 0.706959 |
| 0.713909 | 0.743013 | 0.702598 | 0.708247 | 0.716471 | 0.71035 | 0.713332 | 0.708052 | 0.709887 |
| 0.71028 | 0.742391 | 0.69559 | 0.70352 | 0.710572 | 0.707623 | 0.706972 | 0.703562 | 0.7068 |
| 0.713553 | 0.747175 | 0.703034 | 0.70824 | 0.715276 | 0.712087 | 0.714366 | 0.707676 | 0.712075 |
| 0.71031 | 0.75053 | 0.701829 | 0.704157 | 0.711387 | 0.70754 | 0.711318 | 0.706062 | 0.708908 |
| 0.708468 | 0.74306 | 0.700701 | 0.702222 | 0.713917 | 0.707623 | 0.711316 | 0.705001 | 0.707256 |
| 0.706586 | 0.741338 | 0.694827 | 0.699782 | 0.704019 | 0.704099 | 0.705634 | 0.699448 | 0.704933 |
| 0.705571 | 0.733602 | 0.692563 | 0.699082 | 0.705071 | 0.700513 | 0.705659 | 0.699464 | 0.703091 |
| 0.710473 | 0.743269 | 0.698725 | 0.702864 | 0.708841 | 0.703436 | 0.713684 | 0.702962 | 0.707689 |
| 0.714793 | 0.74379 | 0.701196 | 0.707116 | 0.714894 | 0.707894 | 0.712986 | 0.705222 | 0.708938 |

| | | | | | | | | |
|---|---|---|---|---|---|---|---|---|
| 0.711743 | 0.748274 | 0.703091 | 0.705049 | 0.714796 | 0.708797 | 0.713537 | 0.705549 | 0.710058 |
| 0.71573 | 0.743416 | 0.702421 | 0.702077 | 0.712533 | 0.708107 | 0.710677 | 0.704738 | 0.710658 |
| 0.716878 | 0.749206 | 0.704013 | 0.705792 | 0.718287 | 0.712121 | 0.713475 | 0.708089 | 0.714971 |
| 0.713908 | 0.741684 | 0.701042 | 0.706108 | 0.716518 | 0.709024 | 0.711954 | 0.705142 | 0.70845 |
| 0.708442 | 0.739163 | 0.700154 | 0.700423 | 0.709476 | 0.701199 | 0.707699 | 0.699515 | 0.707556 |
| 0.712647 | 0.740897 | 0.699245 | 0.701039 | 0.709696 | 0.70229 | 0.709712 | 0.701146 | 0.706004 |
| 0.719341 | 0.744167 | 0.703828 | 0.706012 | 0.714646 | 0.711779 | 0.716764 | 0.704794 | 0.711396 |
| 0.718079 | 0.744198 | 0.700853 | 0.704198 | 0.713601 | 0.706892 | 0.713486 | 0.703791 | 0.710027 |
| 0.712409 | 0.743164 | 0.697124 | 0.699748 | 0.70633 | 0.706371 | 0.712527 | 0.705386 | 0.708412 |
| 0.715783 | 0.746109 | 0.702878 | 0.701228 | 0.715432 | 0.706881 | 0.712121 | 0.705171 | 0.710922 |
| 0.709904 | 0.741999 | 0.698776 | 0.704389 | 0.711245 | 0.704164 | 0.711584 | 0.703567 | 0.708174 |
| 0.713194 | 0.743133 | 0.697858 | 0.702083 | 0.711176 | 0.706606 | 0.712929 | 0.705198 | 0.706567 |
| 0.702998 | 0.733345 | 0.692499 | 0.694444 | 0.703561 | 0.697529 | 0.703688 | 0.694998 | 0.70083 |
| 0.705566 | 0.736642 | 0.691125 | 0.695427 | 0.70758 | 0.702216 | 0.704592 | 0.697589 | 0.701503 |
| 0.713902 | 0.748295 | 0.704116 | 0.708119 | 0.716013 | 0.715021 | 0.716563 | 0.705406 | 0.713253 |
| 0.719865 | 0.756827 | 0.707809 | 0.712921 | 0.721378 | 0.717025 | 0.719761 | 0.71416 | 0.718838 |
| 0.717039 | 0.749363 | 0.705356 | 0.708962 | 0.718249 | 0.709969 | 0.716636 | 0.711814 | 0.714501 |
| 0.722774 | 0.75262 | 0.704984 | 0.709925 | 0.71879 | 0.713967 | 0.717582 | 0.712537 | 0.714435 |
| 0.710691 | 0.746437 | 0.701767 | 0.701532 | 0.710981 | 0.706705 | 0.711032 | 0.702829 | 0.711942 |
| 0.711288 | 0.745554 | 0.699264 | 0.702994 | 0.710632 | 0.70654 | 0.71342 | 0.703209 | 0.711583 |
| 0.712533 | 0.742672 | 0.698609 | 0.698157 | 0.708125 | 0.705892 | 0.710274 | 0.701061 | 0.705267 |
| 0.714366 | 0.74165 | 0.701944 | 0.707695 | 0.714812 | 0.707208 | 0.712931 | 0.710869 | 0.713288 |
| 0.721362 | 0.755238 | 0.707726 | 0.710031 | 0.719223 | 0.710617 | 0.715469 | 0.708922 | 0.71457 |
| 0.711396 | 0.745101 | 0.700959 | 0.703833 | 0.711205 | 0.705021 | 0.709824 | 0.701002 | 0.711924 |
| 0.714765 | 0.750325 | 0.703475 | 0.703113 | 0.711345 | 0.706824 | 0.712724 | 0.708255 | 0.70953 |
| 0.77377 | 0.762362 | 0.798651 | 0.807493 | 0.797894 | 0.792439 | 0.790321 | 0.782348 | 0.788457 |
| 0.712805 | 0.746177 | 0.702093 | 0.705719 | 0.712545 | 0.708804 | 0.710604 | 0.706879 | 0.709154 |
| 0.716387 | 0.748957 | 0.704748 | 0.706923 | 0.715842 | 0.70863 | 0.710629 | 0.709448 | 0.713326 |
| 0.719537 | 0.754035 | 0.70848 | 0.70997 | 0.72158 | 0.711348 | 0.719258 | 0.712592 | 0.717934 |
| 0.714766 | 0.748105 | 0.701174 | 0.708807 | 0.712244 | 0.710711 | 0.711896 | 0.710329 | 0.712216 |
| 0.711376 | 0.742155 | 0.700374 | 0.701289 | 0.709677 | 0.705275 | 0.707064 | 0.703194 | 0.708347 |
| 0.715569 | 0.748336 | 0.697999 | 0.702871 | 0.712294 | 0.707604 | 0.709077 | 0.707069 | 0.709288 |
| 0.71726 | 0.746769 | 0.705173 | 0.704496 | 0.712566 | 0.711804 | 0.712931 | 0.708394 | 0.713268 |
| 0.71413 | 0.747406 | 0.699444 | 0.703663 | 0.71318 | 0.703173 | 0.708239 | 0.707215 | 0.707485 |
| 0.712133 | 0.741299 | 0.698005 | 0.701843 | 0.707999 | 0.705344 | 0.710274 | 0.705147 | 0.704473 |
| 0.708771 | 0.740154 | 0.700869 | 0.703359 | 0.709787 | 0.705532 | 0.707705 | 0.704154 | 0.705767 |
| 0.717053 | 0.750369 | 0.706898 | 0.70978 | 0.715418 | 0.714018 | 0.71383 | 0.708671 | 0.712979 |
| 0.719278 | 0.751174 | 0.706743 | 0.709745 | 0.71898 | 0.711051 | 0.716839 | 0.710263 | 0.712269 |
| 0.714654 | 0.749004 | 0.704711 | 0.70684 | 0.712305 | 0.709439 | 0.71366 | 0.70627 | 0.712121 |
| 0.717532 | 0.745529 | 0.70425 | 0.707398 | 0.713931 | 0.7096 | 0.711921 | 0.705235 | 0.712228 |
| 0.711845 | 0.741239 | 0.69676 | 0.702398 | 0.709565 | 0.707141 | 0.706815 | 0.703453 | 0.706118 |
| 0.716373 | 0.747676 | 0.703877 | 0.708168 | 0.714654 | 0.714935 | 0.718272 | 0.70834 | 0.713737 |
| 0.715792 | 0.74424 | 0.702084 | 0.705516 | 0.712785 | 0.708029 | 0.714095 | 0.707984 | 0.707236 |
| 0.716099 | 0.742463 | 0.699677 | 0.70098 | 0.712217 | 0.706287 | 0.708098 | 0.704411 | 0.70952 |
| 0.714251 | 0.746895 | 0.703624 | 0.705652 | 0.714992 | 0.709227 | 0.708494 | 0.704396 | 0.712825 |
| 0.715606 | 0.747371 | 0.700864 | 0.704349 | 0.716839 | 0.710018 | 0.710514 | 0.708015 | 0.716169 |

| | | | | | | | | |
|---|---|---|---|---|---|---|---|---|
| 0.714048 | 0.744984 | 0.699119 | 0.703648 | 0.713509 | 0.70635 | 0.71109 | 0.703298 | 0.70979 |
| 0.717141 | 0.750862 | 0.701645 | 0.706717 | 0.714229 | 0.709813 | 0.712678 | 0.704549 | 0.710757 |
| 0.715607 | 0.745586 | 0.699107 | 0.703371 | 0.714788 | 0.704575 | 0.709828 | 0.702923 | 0.70979 |
| 0.66852 | 0.69972 | 0.655817 | 0.66655 | 0.664932 | 0.668772 | 0.66713 | 0.667821 | 0.671016 |
| 0.713573 | 0.745501 | 0.698443 | 0.703677 | 0.708768 | 0.705743 | 0.709079 | 0.704476 | 0.706154 |
| 0.711931 | 0.744464 | 0.700443 | 0.702492 | 0.711414 | 0.707251 | 0.7108 | 0.708085 | 0.709194 |
| 0.712947 | 0.744798 | 0.699203 | 0.705952 | 0.714715 | 0.707811 | 0.71148 | 0.707838 | 0.710295 |
| 0.715556 | 0.744957 | 0.699172 | 0.703554 | 0.712519 | 0.706918 | 0.707338 | 0.705252 | 0.709554 |
| 0.717149 | 0.74996 | 0.703982 | 0.707402 | 0.713655 | 0.712486 | 0.717084 | 0.709598 | 0.714605 |
| 0.714933 | 0.740081 | 0.702003 | 0.707369 | 0.713313 | 0.71231 | 0.711237 | 0.706175 | 0.7132 |
| 0.712192 | 0.742158 | 0.704295 | 0.707195 | 0.71115 | 0.708956 | 0.710758 | 0.707532 | 0.710008 |
| 0.716008 | 0.747464 | 0.704324 | 0.710667 | 0.714569 | 0.715884 | 0.714171 | 0.711716 | 0.712289 |
| 0.716722 | 0.746736 | 0.709275 | 0.712442 | 0.716425 | 0.716029 | 0.716218 | 0.710189 | 0.712777 |
| 0.711774 | 0.74551 | 0.700677 | 0.707093 | 0.710148 | 0.709764 | 0.712418 | 0.703421 | 0.707298 |
| 0.716685 | 0.749236 | 0.70488 | 0.708884 | 0.714104 | 0.713661 | 0.715292 | 0.709172 | 0.711292 |
| 0.710917 | 0.741894 | 0.697978 | 0.703031 | 0.71006 | 0.704988 | 0.710964 | 0.705836 | 0.705681 |
| 0.716132 | 0.749502 | 0.703183 | 0.705841 | 0.713098 | 0.712427 | 0.71207 | 0.705767 | 0.712585 |
| 0.717765 | 0.749379 | 0.702393 | 0.705766 | 0.716827 | 0.713991 | 0.712601 | 0.711266 | 0.711251 |
| 0.710775 | 0.742039 | 0.696929 | 0.700609 | 0.709677 | 0.705985 | 0.70776 | 0.701395 | 0.704146 |
| 0.713708 | 0.747738 | 0.699007 | 0.703806 | 0.710122 | 0.70982 | 0.710927 | 0.702905 | 0.709505 |
| 0.715125 | 0.752071 | 0.702506 | 0.708211 | 0.714173 | 0.712109 | 0.711901 | 0.705752 | 0.710823 |
| 0.718385 | 0.750742 | 0.699109 | 0.707758 | 0.715562 | 0.713651 | 0.717774 | 0.708838 | 0.710648 |
| 0.697258 | 0.704512 | 0.686953 | 0.686428 | 0.700329 | 0.685041 | 0.70203 | 0.686221 | 0.696702 |
| 0.711431 | 0.699041 | 0.724691 | 0.714042 | 0.735872 | 0.729776 | 0.72396 | 0.716411 | 0.717841 |
| 0.739516 | 0.718285 | 0.749106 | 0.772129 | 0.760179 | 0.759118 | 0.759568 | 0.751738 | 0.753829 |
| 0.732339 | 0.712787 | 0.765315 | 0.765598 | 0.776316 | 0.747304 | 0.742416 | 0.740918 | 0.737446 |
| 0.670423 | 0.71175 | 0.659599 | 0.660317 | 0.67534 | 0.661707 | 0.663384 | 0.657534 | 0.662522 |
| 0.671013 | 0.699608 | 0.65211 | 0.655853 | 0.669309 | 0.661237 | 0.666729 | 0.659496 | 0.661964 |
| 0.661867 | 0.706973 | 0.66196 | 0.662026 | 0.670091 | 0.665279 | 0.662581 | 0.661299 | 0.663929 |
| 0.678078 | 0.70813 | 0.670076 | 0.671705 | 0.67541 | 0.674157 | 0.676715 | 0.671282 | 0.681198 |
| 0.652775 | 0.689033 | 0.639308 | 0.650232 | 0.667108 | 0.651685 | 0.650993 | 0.643731 | 0.660667 |
| 0.64831 | 0.688619 | 0.630691 | 0.638258 | 0.652529 | 0.641622 | 0.638453 | 0.637389 | 0.653668 |
| 0.649093 | 0.683832 | 0.63285 | 0.639228 | 0.647943 | 0.63796 | 0.64622 | 0.634222 | 0.649624 |
| 0.726188 | 0.704562 | 0.719076 | 0.730504 | 0.728091 | 0.734875 | 0.739017 | 0.729695 | 0.734477 |
| 0.708955 | 0.742139 | 0.696178 | 0.699593 | 0.711034 | 0.702824 | 0.708487 | 0.700091 | 0.705185 |
| 0.679502 | 0.715406 | 0.666592 | 0.667414 | 0.677624 | 0.678543 | 0.675879 | 0.671277 | 0.680208 |
| 0.675257 | 0.70742 | 0.659061 | 0.66658 | 0.676195 | 0.670875 | 0.66449 | 0.662693 | 0.671294 |
| 0.659692 | 0.695817 | 0.641503 | 0.648673 | 0.659339 | 0.651275 | 0.651534 | 0.645042 | 0.656407 |
| 0.68468 | 0.713571 | 0.665587 | 0.670267 | 0.681706 | 0.671299 | 0.6752 | 0.672273 | 0.675715 |
| 0.676379 | 0.711064 | 0.657716 | 0.660971 | 0.672999 | 0.66317 | 0.670149 | 0.660736 | 0.671308 |
| 0.668714 | 0.70295 | 0.64629 | 0.65659 | 0.66442 | 0.662669 | 0.659552 | 0.655567 | 0.66309 |
| 0.671536 | 0.705371 | 0.650388 | 0.654302 | 0.664761 | 0.663199 | 0.660636 | 0.656428 | 0.668532 |
| 0.659434 | 0.69884 | 0.650474 | 0.653266 | 0.664873 | 0.651283 | 0.659589 | 0.655018 | 0.655908 |
| 0.66584 | 0.700712 | 0.649331 | 0.652215 | 0.670292 | 0.658049 | 0.663482 | 0.64835 | 0.666284 |
| 0.66067 | 0.705436 | 0.649901 | 0.659796 | 0.66131 | 0.652475 | 0.654046 | 0.647189 | 0.659664 |
| 0.639126 | 0.678526 | 0.629782 | 0.639516 | 0.652615 | 0.639339 | 0.647811 | 0.63375 | 0.650783 |

| | | | | | | | | |
|---|---|---|---|---|---|---|---|---|
| 0.697636 | 0.726033 | 0.681398 | 0.685684 | 0.697332 | 0.690104 | 0.694182 | 0.684998 | 0.696228 |
| 0.662935 | 0.704427 | 0.64687 | 0.648794 | 0.662095 | 0.654685 | 0.663268 | 0.655786 | 0.664948 |
| 0.665223 | 0.707253 | 0.653301 | 0.655098 | 0.668401 | 0.659206 | 0.663242 | 0.658091 | 0.662619 |
| 0.669821 | 0.703421 | 0.654996 | 0.66224 | 0.668676 | 0.666716 | 0.663473 | 0.65569 | 0.663951 |
| 0.663143 | 0.704315 | 0.645444 | 0.653808 | 0.670789 | 0.654004 | 0.661411 | 0.652986 | 0.658024 |
| 0.666827 | 0.703713 | 0.652538 | 0.660972 | 0.671822 | 0.65961 | 0.665381 | 0.658355 | 0.661928 |
| 0.664637 | 0.689103 | 0.643817 | 0.648364 | 0.658231 | 0.654206 | 0.655026 | 0.64665 | 0.655261 |
| 0.644951 | 0.691314 | 0.636217 | 0.639318 | 0.660593 | 0.644192 | 0.653184 | 0.643109 | 0.644155 |
| 0.652517 | 0.686135 | 0.643379 | 0.646418 | 0.659771 | 0.654277 | 0.653286 | 0.646788 | 0.652399 |
| 0.700123 | 0.731962 | 0.680612 | 0.682863 | 0.694502 | 0.687818 | 0.690099 | 0.688296 | 0.691686 |
| 0.692935 | 0.725228 | 0.679389 | 0.680882 | 0.692501 | 0.688149 | 0.693876 | 0.681823 | 0.69212 |
| 0.69041 | 0.726584 | 0.672008 | 0.676558 | 0.690919 | 0.678781 | 0.686037 | 0.681426 | 0.68455 |
| 0.679322 | 0.716402 | 0.663765 | 0.671787 | 0.679118 | 0.675252 | 0.681385 | 0.67062 | 0.675942 |
| 0.686585 | 0.716014 | 0.665227 | 0.675386 | 0.686597 | 0.675948 | 0.687619 | 0.676088 | 0.681 |
| 0.684398 | 0.717151 | 0.657768 | 0.662663 | 0.676678 | 0.669819 | 0.676033 | 0.668983 | 0.677419 |

| | HG_16 | HG_17 | | | | | | |
|---|---|---|---|---|---|---|---|---|
| Home_gard | Home_gard | Home_garc | Home_garc | Home_garc | Home_garc | Home_garc | Home_garc | Home_garc |
| 0.772206 | 0.778379 | 0.804176 | 0.7971 | 0.793921 | 0.785027 | 0.790865 | 0.791362 | 0.796791 |
| 0.775849 | 0.775049 | 0.792135 | 0.789766 | 0.7895 | 0.787521 | 0.795905 | 0.788525 | 0.777648 |
| 0.774433 | 0.77014 | 0.783625 | 0.782573 | 0.803286 | 0.785945 | 0.786 | 0.789695 | 0.789395 |
| 0.775567 | 0.789569 | 0.7869 | 0.787824 | 0.787026 | 0.789713 | 0.782125 | 0.772237 | 0.791029 |
| 0.774807 | 0.766445 | 0.790025 | 0.784463 | 0.783544 | 0.78085 | 0.778238 | 0.77998 | 0.774007 |
| 0.769597 | 0.776381 | 0.780502 | 0.783984 | 0.768777 | 0.783133 | 0.768813 | 0.77252 | 0.766211 |
| 0.714834 | 0.711136 | 0.705397 | 0.70902 | 0.712457 | 0.707891 | 0.707906 | 0.707484 | 0.709491 |
| 0.708377 | 0.711018 | 0.700731 | 0.705138 | 0.706517 | 0.706943 | 0.703324 | 0.701515 | 0.701537 |
| 0.782442 | 0.787741 | 0.781165 | 0.788895 | 0.782678 | 0.7905 | 0.778639 | 0.782269 | 0.772554 |
| 0.776538 | 0.770292 | 0.788893 | 0.785768 | 0.793636 | 0.787949 | 0.782349 | 0.779363 | 0.784762 |
| 0.777593 | 0.769912 | 0.783111 | 0.787372 | 0.788223 | 0.785756 | 0.77966 | 0.774204 | 0.778975 |
| 0.774777 | 0.768459 | 0.818166 | 0.816445 | 0.808434 | 0.809155 | 0.795831 | 0.801473 | 0.791457 |
| 0.76785 | 0.767668 | 0.78024 | 0.786787 | 0.780627 | 0.782908 | 0.774143 | 0.770333 | 0.780315 |
| 0.772083 | 0.771711 | 0.784809 | 0.777759 | 0.781001 | 0.778972 | 0.78556 | 0.781054 | 0.781151 |
| 0.762999 | 0.766602 | 0.769634 | 0.774515 | 0.781438 | 0.772787 | 0.77383 | 0.770458 | 0.768625 |
| 0.773531 | 0.771577 | 0.792574 | 0.78657 | 0.80025 | 0.792169 | 0.78796 | 0.785684 | 0.797353 |
| 0.773365 | 0.766517 | 0.799731 | 0.793106 | 0.79774 | 0.786119 | 0.801998 | 0.794476 | 0.794229 |
| 0.751067 | 0.750996 | 0.767657 | 0.763797 | 0.762605 | 0.757745 | 0.765205 | 0.763311 | 0.756952 |
| 0.777057 | 0.773209 | 0.787772 | 0.792827 | 0.799395 | 0.791025 | 0.793234 | 0.787963 | 0.791298 |
| 0.780522 | 0.778854 | 0.795954 | 0.791154 | 0.797758 | 0.794471 | 0.796338 | 0.790241 | 0.782652 |
| 0.779019 | 0.784162 | 0.796335 | 0.798 | 0.793147 | 0.793385 | 0.792341 | 0.797787 | 0.788269 |
| 0.779258 | 0.782066 | 0.803621 | 0.799197 | 0.797739 | 0.796052 | 0.80616 | 0.79978 | 0.791978 |
| 0.782486 | 0.781353 | 0.791106 | 0.796219 | 0.800251 | 0.790369 | 0.790433 | 0.788431 | 0.791688 |
| 0.779313 | 0.778353 | 0.804561 | 0.806303 | 0.808647 | 0.803736 | 0.809238 | 0.791849 | 0.808274 |
| 0.779006 | 0.777924 | 0.783257 | 0.797266 | 0.78246 | 0.791191 | 0.782423 | 0.781884 | 0.773846 |
| 0.778656 | 0.785018 | 0.7819 | 0.784478 | 0.789461 | 0.786441 | 0.791833 | 0.791193 | 0.785251 |
| 0.769999 | 0.769457 | 0.780164 | 0.790043 | 0.791259 | 0.785833 | 0.781551 | 0.780022 | 0.781574 |
| 0.778409 | 0.786842 | 0.784323 | 0.785855 | 0.787549 | 0.7875 | 0.780562 | 0.784564 | 0.780161 |
| 0.781385 | 0.7795 | 0.780548 | 0.782627 | 0.782573 | 0.781779 | 0.783312 | 0.779485 | 0.777335 |
| 0.775634 | 0.769844 | 0.786115 | 0.786101 | 0.796788 | 0.799109 | 0.782114 | 0.78049 | 0.782328 |
| 0.777778 | 0.779023 | 0.786865 | 0.784495 | 0.785076 | 0.777676 | 0.779995 | 0.774029 | 0.775517 |
| 0.775544 | 0.779322 | 0.776252 | 0.778051 | 0.784967 | 0.778824 | 0.777447 | 0.77618 | 0.777263 |
| 0.778767 | 0.778786 | 0.784374 | 0.789073 | 0.78315 | 0.785809 | 0.780952 | 0.77837 | 0.785559 |
| 0.776369 | 0.768774 | 0.767745 | 0.777385 | 0.774852 | 0.777016 | 0.76904 | 0.773309 | 0.768869 |
| 0.766842 | 0.765966 | 0.769003 | 0.774036 | 0.780508 | 0.77375 | 0.767006 | 0.771405 | 0.766198 |
| 0.774113 | 0.77275 | 0.778147 | 0.787046 | 0.791653 | 0.783786 | 0.78481 | 0.781416 | 0.790879 |
| 0.773197 | 0.770361 | 0.78071 | 0.789005 | 0.794936 | 0.795856 | 0.787333 | 0.784483 | 0.78094 |
| 0.771645 | 0.768483 | 0.768956 | 0.770766 | 0.768187 | 0.765189 | 0.761774 | 0.759806 | 0.76011 |
| 0.77566 | 0.782869 | 0.776065 | 0.77808 | 0.77854 | 0.777219 | 0.770864 | 0.768078 | 0.772358 |

| | | | | | | | | |
|---|---|---|---|---|---|---|---|---|
| 0.756643 | 0.75212 | 0.75686 | 0.755147 | 0.768383 | 0.763904 | 0.755695 | 0.758742 | 0.764538 |
| 0.770097 | 0.767844 | 0.803175 | 0.804094 | 0.793428 | 0.796562 | 0.785631 | 0.785995 | 0.793141 |
| 0.774295 | 0.772791 | 0.801325 | 0.803738 | 0.800922 | 0.803556 | 0.79543 | 0.787614 | 0.793853 |
| 0.777978 | 0.770529 | 0.800763 | 0.802221 | 0.810221 | 0.800931 | 0.804768 | 0.79704 | 0.802856 |
| 0.771671 | 0.776628 | 0.789229 | 0.790592 | 0.793574 | 0.793006 | 0.785827 | 0.784753 | 0.78833 |
| 0.779582 | 0.780876 | 0.788625 | 0.784397 | 0.791041 | 0.787836 | 0.778149 | 0.781636 | 0.779004 |
| 0.771685 | 0.784167 | 0.77477 | 0.780755 | 0.778531 | 0.777686 | 0.771802 | 0.77526 | 0.768766 |
| 0.776484 | 0.781085 | 0.794495 | 0.801142 | 0.790731 | 0.787582 | 0.791873 | 0.78938 | 0.780163 |
| 1 | 0.787603 | 0.775235 | 0.773583 | 0.778009 | 0.774979 | 0.783058 | 0.776496 | 0.766705 |
| 0.787603 | 1 | 0.77334 | 0.771885 | 0.777769 | 0.772873 | 0.774025 | 0.767511 | 0.770294 |
| 0.775235 | 0.77334 | 1 | 0.855825 | 0.802904 | 0.797376 | 0.806142 | 0.824688 | 0.798629 |
| 0.773583 | 0.771885 | 0.855825 | 1 | 0.801499 | 0.835855 | 0.799612 | 0.824101 | 0.794619 |
| 0.778009 | 0.777769 | 0.802904 | 0.801499 | 1 | 0.810792 | 0.799796 | 0.797926 | 0.865401 |
| 0.774979 | 0.772873 | 0.797376 | 0.835855 | 0.810792 | 1 | 0.803086 | 0.811677 | 0.811637 |
| 0.783058 | 0.774025 | 0.806142 | 0.799612 | 0.799796 | 0.803086 | 1 | 0.855325 | 0.795617 |
| 0.776496 | 0.767511 | 0.824688 | 0.824101 | 0.797926 | 0.811677 | 0.855325 | 1 | 0.796771 |
| 0.766705 | 0.770294 | 0.798629 | 0.794619 | 0.865401 | 0.811637 | 0.795617 | 0.796771 | 1 |
| 0.780734 | 0.776805 | 0.818251 | 0.817807 | 0.816076 | 0.806313 | 0.801752 | 0.807924 | 0.818878 |
| 0.774506 | 0.77992 | 0.799014 | 0.792731 | 0.806283 | 0.79348 | 0.787636 | 0.795378 | 0.790918 |
| 0.772334 | 0.774704 | 0.80664 | 0.80172 | 0.835452 | 0.809649 | 0.793743 | 0.789621 | 0.813412 |
| 0.767913 | 0.770089 | 0.798318 | 0.791576 | 0.809024 | 0.804981 | 0.792882 | 0.801941 | 0.81191 |
| 0.770596 | 0.775138 | 0.802213 | 0.7944 | 0.809327 | 0.792085 | 0.790013 | 0.79046 | 0.810242 |
| 0.781131 | 0.778485 | 0.781456 | 0.786853 | 0.787622 | 0.791072 | 0.790026 | 0.790144 | 0.782652 |
| 0.774533 | 0.795001 | 0.777123 | 0.779022 | 0.772075 | 0.776653 | 0.773491 | 0.771178 | 0.775451 |
| 0.777137 | 0.78257 | 0.763072 | 0.765244 | 0.765931 | 0.762178 | 0.764338 | 0.758672 | 0.75442 |
| 0.708974 | 0.713633 | 0.70121 | 0.70575 | 0.707561 | 0.707726 | 0.704075 | 0.703454 | 0.70312 |
| 0.713754 | 0.713509 | 0.704109 | 0.70905 | 0.711917 | 0.71001 | 0.708134 | 0.707671 | 0.70813 |
| 0.714909 | 0.71666 | 0.706034 | 0.712584 | 0.714436 | 0.710452 | 0.705792 | 0.708947 | 0.708853 |
| 0.765427 | 0.767901 | 0.776318 | 0.781496 | 0.776495 | 0.771762 | 0.775256 | 0.772167 | 0.765524 |
| 0.767369 | 0.771269 | 0.774092 | 0.784144 | 0.77909 | 0.782258 | 0.783253 | 0.776802 | 0.769369 |
| 0.770132 | 0.766914 | 0.793066 | 0.780718 | 0.792039 | 0.783887 | 0.780898 | 0.778728 | 0.775928 |
| 0.772484 | 0.77871 | 0.774261 | 0.787018 | 0.77495 | 0.777564 | 0.777216 | 0.771844 | 0.769386 |
| 0.781914 | 0.779887 | 0.808281 | 0.807454 | 0.813694 | 0.795343 | 0.811164 | 0.806391 | 0.802821 |
| 0.778214 | 0.775219 | 0.800927 | 0.798152 | 0.80899 | 0.799746 | 0.795412 | 0.789327 | 0.808424 |
| 0.784844 | 0.769966 | 0.807512 | 0.807282 | 0.800649 | 0.793695 | 0.807441 | 0.805505 | 0.796844 |
| 0.784789 | 0.778254 | 0.809898 | 0.805796 | 0.816365 | 0.805715 | 0.821987 | 0.821198 | 0.829732 |
| 0.774639 | 0.77241 | 0.796011 | 0.802308 | 0.789849 | 0.789774 | 0.782843 | 0.778821 | 0.77018 |
| 0.795441 | 0.792254 | 0.826541 | 0.819773 | 0.823054 | 0.808677 | 0.81747 | 0.814701 | 0.819857 |
| 0.774241 | 0.758901 | 0.783807 | 0.77642 | 0.791997 | 0.773996 | 0.785664 | 0.783394 | 0.782287 |
| 0.784714 | 0.790047 | 0.792664 | 0.794887 | 0.790931 | 0.790031 | 0.806541 | 0.792538 | 0.781485 |
| 0.779106 | 0.773892 | 0.802364 | 0.785138 | 0.791442 | 0.790162 | 0.792629 | 0.793294 | 0.785364 |
| 0.800117 | 0.796722 | 0.82861 | 0.820205 | 0.823514 | 0.811318 | 0.816016 | 0.810776 | 0.81946 |
| 0.780271 | 0.775555 | 0.802068 | 0.79649 | 0.807913 | 0.804086 | 0.791415 | 0.793214 | 0.801842 |
| 0.776901 | 0.775583 | 0.799467 | 0.797231 | 0.801551 | 0.805428 | 0.783099 | 0.781459 | 0.794322 |
| 0.787611 | 0.789875 | 0.791782 | 0.790455 | 0.797 | 0.788808 | 0.789491 | 0.781639 | 0.78694 |
| 0.779584 | 0.766126 | 0.783575 | 0.783555 | 0.79956 | 0.790093 | 0.795944 | 0.79045 | 0.798384 |

| | | | | | | | | |
|---|---|---|---|---|---|---|---|---|
| 0.779035 | 0.77851 | 0.809351 | 0.794578 | 0.803965 | 0.812119 | 0.800557 | 0.797808 | 0.791277 |
| 0.778374 | 0.775267 | 0.784894 | 0.803843 | 0.787468 | 0.805804 | 0.788301 | 0.789363 | 0.772059 |
| 0.776376 | 0.778128 | 0.783114 | 0.786588 | 0.795262 | 0.801186 | 0.778702 | 0.779472 | 0.780362 |
| 0.789144 | 0.782862 | 0.808072 | 0.804748 | 0.800968 | 0.801917 | 0.801345 | 0.800395 | 0.795714 |
| 0.777686 | 0.785288 | 0.793565 | 0.797991 | 0.787836 | 0.79043 | 0.786237 | 0.785882 | 0.780023 |
| 0.773954 | 0.777115 | 0.794095 | 0.802497 | 0.786685 | 0.798968 | 0.803983 | 0.804859 | 0.778132 |
| 0.771773 | 0.779107 | 0.79218 | 0.790859 | 0.788102 | 0.780995 | 0.77979 | 0.778637 | 0.774308 |
| 0.781519 | 0.77775 | 0.78204 | 0.78828 | 0.780482 | 0.792884 | 0.777973 | 0.777059 | 0.774828 |
| 0.777565 | 0.775342 | 0.781626 | 0.777501 | 0.78578 | 0.786522 | 0.778692 | 0.775594 | 0.773172 |
| 0.775159 | 0.768095 | 0.77537 | 0.778752 | 0.780933 | 0.78201 | 0.776162 | 0.77462 | 0.773813 |
| 0.782233 | 0.768626 | 0.779073 | 0.780297 | 0.779032 | 0.78305 | 0.776245 | 0.781732 | 0.767945 |
| 0.777135 | 0.771872 | 0.770329 | 0.779137 | 0.782085 | 0.776398 | 0.771112 | 0.770941 | 0.769295 |
| 0.773306 | 0.769686 | 0.776026 | 0.784597 | 0.78515 | 0.789514 | 0.784483 | 0.782113 | 0.772476 |
| 0.769102 | 0.772176 | 0.784178 | 0.779729 | 0.77546 | 0.783645 | 0.772559 | 0.775263 | 0.769905 |
| 0.779448 | 0.777329 | 0.776672 | 0.785595 | 0.781503 | 0.78417 | 0.78141 | 0.777283 | 0.772367 |
| 0.774582 | 0.775868 | 0.779576 | 0.781227 | 0.784577 | 0.78594 | 0.779789 | 0.773239 | 0.774768 |
| 0.77499 | 0.777814 | 0.781795 | 0.774678 | 0.784577 | 0.778806 | 0.772138 | 0.768065 | 0.777768 |
| 0.782468 | 0.778607 | 0.790934 | 0.786638 | 0.79455 | 0.783371 | 0.804487 | 0.800169 | 0.784745 |
| 0.780313 | 0.773459 | 0.776587 | 0.780241 | 0.777084 | 0.786419 | 0.779358 | 0.776182 | 0.773213 |
| 0.779781 | 0.770654 | 0.778349 | 0.782685 | 0.785131 | 0.782931 | 0.780478 | 0.780439 | 0.775585 |
| 0.776456 | 0.775828 | 0.774247 | 0.776496 | 0.775611 | 0.774963 | 0.771505 | 0.769591 | 0.767364 |
| 0.77523 | 0.761255 | 0.772409 | 0.778365 | 0.772168 | 0.780111 | 0.770557 | 0.772955 | 0.76932 |
| 0.775759 | 0.772465 | 0.781587 | 0.78328 | 0.779908 | 0.789137 | 0.782287 | 0.775307 | 0.775872 |
| 0.774866 | 0.766056 | 0.786914 | 0.79925 | 0.787321 | 0.791004 | 0.792806 | 0.792504 | 0.775081 |
| 0.769263 | 0.765513 | 0.777824 | 0.782298 | 0.780599 | 0.780259 | 0.780869 | 0.774623 | 0.773064 |
| 0.784492 | 0.779791 | 0.793303 | 0.796768 | 0.79131 | 0.787042 | 0.788833 | 0.78266 | 0.77418 |
| 0.771396 | 0.770457 | 0.781941 | 0.785832 | 0.779962 | 0.780689 | 0.773844 | 0.770015 | 0.77266 |
| 0.779588 | 0.771857 | 0.778072 | 0.778499 | 0.779019 | 0.779219 | 0.775632 | 0.776454 | 0.768944 |
| 0.774367 | 0.772148 | 0.779759 | 0.781994 | 0.78051 | 0.783347 | 0.785506 | 0.783791 | 0.776341 |
| 0.771137 | 0.765334 | 0.771077 | 0.780223 | 0.780791 | 0.788205 | 0.776603 | 0.77514 | 0.776387 |
| 0.775509 | 0.767903 | 0.774161 | 0.777898 | 0.781559 | 0.779783 | 0.772952 | 0.773336 | 0.768961 |
| 0.7761 | 0.771237 | 0.771053 | 0.775217 | 0.781168 | 0.788809 | 0.778721 | 0.773579 | 0.775976 |
| 0.776119 | 0.772708 | 0.782974 | 0.790809 | 0.794078 | 0.785244 | 0.780894 | 0.781521 | 0.774653 |
| 0.774582 | 0.765617 | 0.798646 | 0.798647 | 0.805647 | 0.79832 | 0.800553 | 0.792806 | 0.800104 |
| 0.771239 | 0.764205 | 0.77812 | 0.783034 | 0.782802 | 0.781234 | 0.78432 | 0.783747 | 0.775165 |
| 0.777152 | 0.776731 | 0.790911 | 0.784558 | 0.787362 | 0.787369 | 0.788802 | 0.793714 | 0.781595 |
| 0.770294 | 0.764049 | 0.779554 | 0.786289 | 0.787052 | 0.780236 | 0.774184 | 0.775877 | 0.774692 |
| 0.748604 | 0.749011 | 0.748214 | 0.760216 | 0.759405 | 0.759128 | 0.755442 | 0.753756 | 0.749867 |
| 0.765551 | 0.758452 | 0.780268 | 0.789479 | 0.769247 | 0.787979 | 0.790835 | 0.79374 | 0.75773 |
| 0.753868 | 0.751462 | 0.749198 | 0.752491 | 0.750509 | 0.747889 | 0.74721 | 0.744541 | 0.738241 |
| 0.760046 | 0.755191 | 0.775119 | 0.773277 | 0.780316 | 0.776919 | 0.777227 | 0.777317 | 0.783286 |
| 0.734137 | 0.731097 | 0.760602 | 0.754108 | 0.755962 | 0.749631 | 0.73998 | 0.737925 | 0.750588 |
| 0.745466 | 0.746642 | 0.775308 | 0.763875 | 0.785863 | 0.783164 | 0.761764 | 0.772198 | 0.770473 |
| 0.751497 | 0.746974 | 0.759852 | 0.764834 | 0.762823 | 0.769339 | 0.76036 | 0.75718 | 0.758448 |
| 0.747111 | 0.744336 | 0.749301 | 0.751337 | 0.750704 | 0.747766 | 0.746023 | 0.747941 | 0.742156 |
| 0.752205 | 0.740903 | 0.764498 | 0.76312 | 0.77213 | 0.761593 | 0.761911 | 0.757568 | 0.762535 |
| 0.752528 | 0.75251 | 0.771048 | 0.768508 | 0.775257 | 0.766176 | 0.77967 | 0.780935 | 0.764235 |

| | | | | | | | | |
|---|---|---|---|---|---|---|---|---|
| 0.738135 | 0.739693 | 0.739275 | 0.741253 | 0.740459 | 0.743521 | 0.74606 | 0.743616 | 0.734938 |
| 0.755501 | 0.747743 | 0.782555 | 0.782482 | 0.774609 | 0.763908 | 0.765784 | 0.771564 | 0.78044 |
| 0.745808 | 0.737429 | 0.736879 | 0.74022 | 0.740263 | 0.746089 | 0.739994 | 0.739963 | 0.733474 |
| 0.744389 | 0.739719 | 0.782431 | 0.75971 | 0.7731 | 0.760284 | 0.760398 | 0.763424 | 0.768253 |
| 0.739218 | 0.733723 | 0.762402 | 0.760967 | 0.764077 | 0.753113 | 0.749932 | 0.742919 | 0.745857 |
| 0.743925 | 0.739495 | 0.748814 | 0.749387 | 0.746856 | 0.7667 | 0.734368 | 0.735629 | 0.73757 |
| 0.744877 | 0.742487 | 0.754025 | 0.760785 | 0.752662 | 0.764205 | 0.758543 | 0.75304 | 0.737978 |
| 0.753953 | 0.741344 | 0.776866 | 0.769345 | 0.771967 | 0.762173 | 0.764294 | 0.772495 | 0.773718 |
| 0.727579 | 0.71923 | 0.735299 | 0.732508 | 0.731399 | 0.72801 | 0.726158 | 0.726937 | 0.720566 |
| 0.743637 | 0.737456 | 0.76714 | 0.774545 | 0.771625 | 0.753965 | 0.75884 | 0.753418 | 0.74666 |
| 0.746429 | 0.739667 | 0.742687 | 0.746305 | 0.746501 | 0.758316 | 0.731464 | 0.735445 | 0.732959 |
| 0.767695 | 0.765758 | 0.793146 | 0.773225 | 0.786589 | 0.78292 | 0.769401 | 0.770414 | 0.776758 |
| 0.772328 | 0.777959 | 0.765695 | 0.77101 | 0.771915 | 0.765605 | 0.770629 | 0.766854 | 0.763559 |
| 0.760779 | 0.776366 | 0.745437 | 0.743802 | 0.743917 | 0.746959 | 0.748186 | 0.738662 | 0.736637 |
| 0.771974 | 0.769791 | 0.78319 | 0.775751 | 0.780621 | 0.789909 | 0.77524 | 0.772704 | 0.776615 |
| 0.76522 | 0.757522 | 0.764623 | 0.76792 | 0.761815 | 0.766796 | 0.767236 | 0.765444 | 0.755971 |
| 0.775665 | 0.77355 | 0.778794 | 0.781881 | 0.787627 | 0.776789 | 0.776345 | 0.779943 | 0.775993 |
| 0.778474 | 0.771252 | 0.770003 | 0.779947 | 0.78411 | 0.779262 | 0.771835 | 0.773302 | 0.769969 |
| 0.7561 | 0.752128 | 0.75866 | 0.75998 | 0.757966 | 0.763577 | 0.762341 | 0.760873 | 0.75394 |
| 0.77218 | 0.780518 | 0.776193 | 0.771573 | 0.772686 | 0.775517 | 0.764971 | 0.760471 | 0.768548 |
| 0.747847 | 0.766766 | 0.747574 | 0.747794 | 0.750063 | 0.754406 | 0.736587 | 0.73735 | 0.739114 |
| 0.776683 | 0.77215 | 0.772423 | 0.778561 | 0.776431 | 0.772212 | 0.767286 | 0.767156 | 0.759264 |
| 0.782457 | 0.769867 | 0.771946 | 0.771785 | 0.773361 | 0.769638 | 0.765942 | 0.767708 | 0.762696 |
| 0.741169 | 0.773063 | 0.752751 | 0.753627 | 0.746915 | 0.745216 | 0.751971 | 0.758096 | 0.735958 |
| 0.746461 | 0.73372 | 0.749829 | 0.755914 | 0.760566 | 0.749487 | 0.759026 | 0.75458 | 0.75125 |
| 0.758784 | 0.762016 | 0.769231 | 0.763348 | 0.762213 | 0.76643 | 0.755985 | 0.7544 | 0.76367 |
| 0.764765 | 0.784536 | 0.764311 | 0.766027 | 0.758192 | 0.769672 | 0.754741 | 0.756155 | 0.75543 |
| 0.77499 | 0.781009 | 0.779776 | 0.785514 | 0.777566 | 0.775021 | 0.774311 | 0.773264 | 0.766066 |
| 0.751534 | 0.740514 | 0.744154 | 0.74677 | 0.746472 | 0.749738 | 0.741213 | 0.743571 | 0.737432 |
| 0.767051 | 0.771847 | 0.770687 | 0.774244 | 0.784865 | 0.760299 | 0.768014 | 0.758621 | 0.769035 |
| 0.772642 | 0.770868 | 0.77119 | 0.76661 | 0.769068 | 0.765602 | 0.763139 | 0.761249 | 0.761507 |
| 0.752849 | 0.750427 | 0.767661 | 0.764022 | 0.768327 | 0.75615 | 0.765705 | 0.764789 | 0.761873 |
| 0.75715 | 0.751859 | 0.772095 | 0.768612 | 0.770063 | 0.765398 | 0.765604 | 0.764149 | 0.765778 |
| 0.778191 | 0.795372 | 0.764412 | 0.762968 | 0.765117 | 0.766319 | 0.760424 | 0.763025 | 0.761042 |
| 0.765468 | 0.76151 | 0.759034 | 0.760793 | 0.758866 | 0.760194 | 0.76307 | 0.760091 | 0.761754 |
| 0.766595 | 0.762733 | 0.76789 | 0.774717 | 0.767348 | 0.770894 | 0.756444 | 0.759721 | 0.764823 |
| 0.732116 | 0.724474 | 0.720208 | 0.723647 | 0.723229 | 0.724707 | 0.71837 | 0.719106 | 0.715857 |
| 0.75924 | 0.75171 | 0.753876 | 0.754591 | 0.757957 | 0.752014 | 0.750897 | 0.750964 | 0.741676 |
| 0.730833 | 0.723906 | 0.720186 | 0.727539 | 0.730458 | 0.729308 | 0.728797 | 0.727862 | 0.723957 |
| 0.712124 | 0.714407 | 0.716079 | 0.719338 | 0.718985 | 0.721209 | 0.718588 | 0.717717 | 0.712797 |
| 0.767701 | 0.760235 | 0.776575 | 0.783933 | 0.768815 | 0.774588 | 0.771229 | 0.76758 | 0.764045 |
| 0.712582 | 0.712785 | 0.717042 | 0.721201 | 0.712243 | 0.719743 | 0.716701 | 0.716259 | 0.710242 |
| 0.712213 | 0.706959 | 0.72051 | 0.725629 | 0.727016 | 0.717963 | 0.723414 | 0.713593 | 0.714542 |
| 0.704696 | 0.695848 | 0.71017 | 0.713982 | 0.719914 | 0.710618 | 0.713763 | 0.706932 | 0.706139 |
| 0.760928 | 0.757524 | 0.774573 | 0.771515 | 0.776049 | 0.769578 | 0.76868 | 0.764473 | 0.7678 |
| 0.712612 | 0.69808 | 0.71097 | 0.712309 | 0.724842 | 0.708362 | 0.715099 | 0.707191 | 0.707321 |
| 0.759054 | 0.761034 | 0.772153 | 0.775488 | 0.773752 | 0.771837 | 0.777722 | 0.763503 | 0.766854 |

| | | | | | | | | |
|---|---|---|---|---|---|---|---|---|
| 0.759271 | 0.755015 | 0.768772 | 0.763438 | 0.763918 | 0.753139 | 0.76781 | 0.76268 | 0.759314 |
| 0.765067 | 0.760664 | 0.767491 | 0.77418 | 0.764292 | 0.772448 | 0.771203 | 0.762365 | 0.762433 |
| 0.756899 | 0.755031 | 0.747482 | 0.746809 | 0.749466 | 0.748728 | 0.749423 | 0.751327 | 0.745303 |
| 0.698664 | 0.693557 | 0.7079 | 0.719324 | 0.71378 | 0.704301 | 0.711541 | 0.706418 | 0.700307 |
| 0.699333 | 0.690784 | 0.697352 | 0.703889 | 0.71217 | 0.698688 | 0.698561 | 0.690837 | 0.692136 |
| 0.759355 | 0.757559 | 0.773452 | 0.781899 | 0.775804 | 0.780369 | 0.774535 | 0.772527 | 0.771433 |
| 0.692049 | 0.6905 | 0.685326 | 0.690511 | 0.691033 | 0.688252 | 0.688615 | 0.685626 | 0.68508 |
| 0.735962 | 0.73397 | 0.729662 | 0.732681 | 0.735106 | 0.731061 | 0.730325 | 0.728365 | 0.728346 |
| 0.756936 | 0.756005 | 0.771773 | 0.764233 | 0.769193 | 0.76459 | 0.756997 | 0.755859 | 0.765799 |
| 0.725883 | 0.723286 | 0.716995 | 0.722344 | 0.720862 | 0.72058 | 0.717588 | 0.719279 | 0.71474 |
| 0.745684 | 0.745603 | 0.75115 | 0.756242 | 0.757491 | 0.755351 | 0.747753 | 0.743844 | 0.749381 |
| 0.640523 | 0.613924 | 0.651007 | 0.602564 | 0.631206 | 0.627329 | 0.587097 | 0.633987 | 0.591195 |
| 0.659401 | 0.671442 | 0.652772 | 0.644351 | 0.653013 | 0.654362 | 0.65 | 0.650828 | 0.645575 |
| 0.658544 | 0.665404 | 0.653018 | 0.649758 | 0.653343 | 0.65342 | 0.65279 | 0.650307 | 0.649595 |
| 0.677267 | 0.673135 | 0.660977 | 0.663268 | 0.662121 | 0.662438 | 0.66155 | 0.665341 | 0.655001 |
| 0.66436 | 0.666581 | 0.656891 | 0.658546 | 0.658314 | 0.659212 | 0.65995 | 0.660919 | 0.657038 |
| 0.625277 | 0.618985 | 0.612101 | 0.617559 | 0.621481 | 0.619766 | 0.618262 | 0.618788 | 0.615444 |
| 0.669876 | 0.675091 | 0.659411 | 0.656906 | 0.66358 | 0.6644 | 0.665309 | 0.664286 | 0.6588 |
| 0.739961 | 0.736591 | 0.746391 | 0.741176 | 0.739801 | 0.744463 | 0.743795 | 0.740349 | 0.737223 |
| 0.736698 | 0.734876 | 0.7357 | 0.737111 | 0.746978 | 0.743792 | 0.73387 | 0.7337 | 0.740954 |
| 0.744266 | 0.7454 | 0.745195 | 0.748849 | 0.748885 | 0.751795 | 0.746075 | 0.743771 | 0.740292 |
| 0.61957 | 0.616006 | 0.611515 | 0.616731 | 0.619734 | 0.618302 | 0.615612 | 0.614692 | 0.609747 |
| 0.602465 | 0.594416 | 0.594726 | 0.596925 | 0.601136 | 0.594674 | 0.598267 | 0.595165 | 0.589072 |
| 0.707052 | 0.710776 | 0.69895 | 0.702358 | 0.703551 | 0.705706 | 0.702872 | 0.703667 | 0.698265 |
| 0.70771 | 0.708188 | 0.697265 | 0.700833 | 0.707563 | 0.704651 | 0.702404 | 0.701867 | 0.697621 |
| 0.699301 | 0.699976 | 0.692652 | 0.696411 | 0.697426 | 0.696979 | 0.698618 | 0.696388 | 0.691558 |
| 0.705474 | 0.703748 | 0.697661 | 0.700397 | 0.702075 | 0.700955 | 0.704518 | 0.700056 | 0.698002 |
| 0.70815 | 0.705794 | 0.699682 | 0.702506 | 0.705193 | 0.702776 | 0.706137 | 0.709906 | 0.701935 |
| 0.710697 | 0.709734 | 0.700992 | 0.704762 | 0.705701 | 0.702777 | 0.701524 | 0.701344 | 0.700495 |
| 0.706693 | 0.7108 | 0.700279 | 0.703373 | 0.704779 | 0.702346 | 0.70208 | 0.701089 | 0.698916 |
| 0.706483 | 0.708237 | 0.701627 | 0.700857 | 0.706104 | 0.70366 | 0.702306 | 0.705306 | 0.703109 |
| 0.716154 | 0.71772 | 0.710353 | 0.708251 | 0.714263 | 0.712014 | 0.711798 | 0.713368 | 0.706126 |
| 0.714205 | 0.713016 | 0.706333 | 0.70654 | 0.712389 | 0.708152 | 0.702764 | 0.70442 | 0.706499 |
| 0.715921 | 0.716008 | 0.709156 | 0.708337 | 0.715242 | 0.707651 | 0.707936 | 0.711292 | 0.709281 |
| 0.717918 | 0.719309 | 0.707379 | 0.708765 | 0.715926 | 0.712239 | 0.709341 | 0.708653 | 0.71132 |
| 0.713559 | 0.711063 | 0.700625 | 0.700678 | 0.707474 | 0.704679 | 0.70274 | 0.704961 | 0.702694 |
| 0.716731 | 0.711462 | 0.706524 | 0.704455 | 0.709265 | 0.707835 | 0.708693 | 0.710479 | 0.707337 |
| 0.715407 | 0.715736 | 0.705545 | 0.708131 | 0.709056 | 0.707177 | 0.707423 | 0.707754 | 0.704222 |
| 0.707067 | 0.710133 | 0.700884 | 0.704202 | 0.703251 | 0.702323 | 0.699441 | 0.698751 | 0.700832 |
| 0.714343 | 0.718222 | 0.700761 | 0.704012 | 0.712994 | 0.70641 | 0.708347 | 0.702575 | 0.70608 |
| 0.663655 | 0.664315 | 0.652005 | 0.645253 | 0.651746 | 0.652299 | 0.655785 | 0.655308 | 0.650179 |
| 0.665564 | 0.664305 | 0.659216 | 0.657504 | 0.659908 | 0.655066 | 0.658505 | 0.653655 | 0.654995 |
| 0.716565 | 0.712334 | 0.707876 | 0.709357 | 0.714686 | 0.710617 | 0.708004 | 0.710378 | 0.710123 |
| 0.718356 | 0.714965 | 0.708027 | 0.711177 | 0.713804 | 0.708902 | 0.708206 | 0.709113 | 0.70893 |
| 0.650037 | 0.6565 | 0.633717 | 0.635041 | 0.639993 | 0.636953 | 0.64064 | 0.63905 | 0.636584 |
| 0.715644 | 0.715492 | 0.705634 | 0.707635 | 0.711454 | 0.707916 | 0.708656 | 0.703393 | 0.708921 |
| 0.703654 | 0.70055 | 0.695319 | 0.695216 | 0.701224 | 0.697875 | 0.698025 | 0.697566 | 0.69786 |

| | | | | | | | | |
|---|---|---|---|---|---|---|---|---|
| 0.712302 | 0.711676 | 0.701696 | 0.70263 | 0.70418 | 0.706616 | 0.704259 | 0.699236 | 0.701754 |
| 0.71521 | 0.711992 | 0.702096 | 0.705836 | 0.709952 | 0.708 | 0.707722 | 0.706189 | 0.70812 |
| 0.713864 | 0.71032 | 0.703655 | 0.704075 | 0.707333 | 0.705882 | 0.706734 | 0.7078 | 0.70346 |
| 0.715501 | 0.713448 | 0.703813 | 0.709796 | 0.707315 | 0.70652 | 0.709151 | 0.707421 | 0.705377 |
| 0.72145 | 0.718765 | 0.711692 | 0.709683 | 0.710463 | 0.710946 | 0.712388 | 0.710881 | 0.708939 |
| 0.71319 | 0.709605 | 0.702757 | 0.703772 | 0.706541 | 0.705291 | 0.703357 | 0.701826 | 0.701348 |
| 0.711932 | 0.709807 | 0.702448 | 0.702242 | 0.70636 | 0.703604 | 0.705093 | 0.701952 | 0.703893 |
| 0.70573 | 0.706057 | 0.698046 | 0.700548 | 0.705096 | 0.697588 | 0.69698 | 0.695995 | 0.699489 |
| 0.710151 | 0.710543 | 0.699921 | 0.703503 | 0.703225 | 0.703429 | 0.705462 | 0.701756 | 0.701314 |
| 0.713192 | 0.710706 | 0.700886 | 0.702129 | 0.707649 | 0.70748 | 0.704214 | 0.70284 | 0.705783 |
| 0.714617 | 0.712064 | 0.704428 | 0.706223 | 0.710045 | 0.707564 | 0.706606 | 0.705722 | 0.706451 |
| 0.707935 | 0.709042 | 0.700016 | 0.699951 | 0.705344 | 0.70325 | 0.702324 | 0.698526 | 0.703152 |
| 0.707172 | 0.701248 | 0.695127 | 0.695747 | 0.700874 | 0.696994 | 0.69867 | 0.697229 | 0.69785 |
| 0.70955 | 0.703909 | 0.692977 | 0.695778 | 0.698743 | 0.697835 | 0.697117 | 0.695087 | 0.698482 |
| 0.715667 | 0.715908 | 0.70746 | 0.709294 | 0.70987 | 0.710775 | 0.710082 | 0.710543 | 0.705478 |
| 0.711969 | 0.711071 | 0.705135 | 0.707191 | 0.710789 | 0.707117 | 0.705078 | 0.706531 | 0.705704 |
| 0.707095 | 0.706885 | 0.698494 | 0.699786 | 0.70128 | 0.70066 | 0.701898 | 0.701592 | 0.696023 |
| 0.709019 | 0.708575 | 0.699736 | 0.704183 | 0.706005 | 0.705339 | 0.703725 | 0.702071 | 0.700587 |
| 0.709368 | 0.709392 | 0.700302 | 0.700293 | 0.70508 | 0.703282 | 0.703351 | 0.705395 | 0.698905 |
| 0.705379 | 0.707106 | 0.697626 | 0.697804 | 0.705821 | 0.699753 | 0.704906 | 0.703312 | 0.701636 |
| 0.703354 | 0.703808 | 0.697493 | 0.697639 | 0.707221 | 0.698871 | 0.703725 | 0.699082 | 0.698281 |
| 0.709788 | 0.704778 | 0.698346 | 0.699814 | 0.701413 | 0.699627 | 0.704737 | 0.704601 | 0.69746 |
| 0.709978 | 0.708589 | 0.701106 | 0.703791 | 0.708025 | 0.701541 | 0.708526 | 0.706906 | 0.702752 |
| 0.7116 | 0.709708 | 0.702571 | 0.706643 | 0.707728 | 0.705943 | 0.707401 | 0.704191 | 0.70479 |
| 0.708182 | 0.704862 | 0.702417 | 0.704226 | 0.706939 | 0.705773 | 0.704557 | 0.705459 | 0.702813 |
| 0.706352 | 0.706056 | 0.696394 | 0.697706 | 0.702328 | 0.698742 | 0.70121 | 0.700957 | 0.696508 |
| 0.705348 | 0.704206 | 0.699055 | 0.701041 | 0.707859 | 0.701869 | 0.70214 | 0.702149 | 0.700691 |
| 0.705844 | 0.706244 | 0.697459 | 0.699635 | 0.69978 | 0.701049 | 0.703002 | 0.697486 | 0.697401 |
| 0.707676 | 0.709337 | 0.699769 | 0.699309 | 0.705195 | 0.703323 | 0.704825 | 0.702071 | 0.699047 |
| 0.709206 | 0.709075 | 0.702232 | 0.703548 | 0.706432 | 0.708243 | 0.70593 | 0.701697 | 0.703084 |
| 0.713152 | 0.711099 | 0.705422 | 0.706009 | 0.711795 | 0.707943 | 0.708008 | 0.70471 | 0.704206 |
| 0.713371 | 0.711881 | 0.705925 | 0.706527 | 0.713973 | 0.708856 | 0.710255 | 0.709063 | 0.704953 |
| 0.713432 | 0.713634 | 0.703551 | 0.706594 | 0.713535 | 0.708327 | 0.707709 | 0.708223 | 0.704823 |
| 0.714309 | 0.71491 | 0.707165 | 0.710102 | 0.710758 | 0.709275 | 0.711295 | 0.709602 | 0.706208 |
| 0.706964 | 0.709068 | 0.698345 | 0.69913 | 0.703055 | 0.700953 | 0.701569 | 0.700461 | 0.697316 |
| 0.70814 | 0.706577 | 0.703464 | 0.702989 | 0.706577 | 0.705204 | 0.705187 | 0.703341 | 0.702178 |
| 0.711258 | 0.710785 | 0.704472 | 0.706176 | 0.704283 | 0.70739 | 0.706249 | 0.705489 | 0.701825 |
| 0.709866 | 0.704744 | 0.697893 | 0.697337 | 0.702602 | 0.700895 | 0.703911 | 0.699846 | 0.696634 |
| 0.703758 | 0.698238 | 0.693846 | 0.695459 | 0.700603 | 0.696188 | 0.700033 | 0.698016 | 0.691687 |
| 0.700931 | 0.698869 | 0.691962 | 0.689787 | 0.697379 | 0.695103 | 0.695835 | 0.693306 | 0.687385 |
| 0.701266 | 0.701073 | 0.692985 | 0.692528 | 0.697653 | 0.694424 | 0.69758 | 0.693196 | 0.69211 |
| 0.700136 | 0.700455 | 0.690818 | 0.692139 | 0.697101 | 0.691748 | 0.695745 | 0.69351 | 0.693895 |
| 0.703266 | 0.702723 | 0.69558 | 0.698972 | 0.703943 | 0.697935 | 0.6992 | 0.696389 | 0.695088 |
| 0.698456 | 0.693441 | 0.686835 | 0.686175 | 0.690472 | 0.689923 | 0.693845 | 0.690895 | 0.687445 |
| 0.705591 | 0.703682 | 0.696845 | 0.699547 | 0.701657 | 0.698072 | 0.700447 | 0.700229 | 0.698018 |
| 0.70023 | 0.69585 | 0.69247 | 0.695026 | 0.697235 | 0.693006 | 0.693363 | 0.694712 | 0.692599 |

| | | | | | | | | |
|---|---|---|---|---|---|---|---|---|
| 0.710947 | 0.711392 | 0.704753 | 0.702425 | 0.710433 | 0.706354 | 0.708007 | 0.708023 | 0.704958 |
| 0.713612 | 0.711084 | 0.705122 | 0.70405 | 0.710479 | 0.70833 | 0.705255 | 0.705911 | 0.703845 |
| 0.710783 | 0.706297 | 0.700194 | 0.702047 | 0.706064 | 0.702281 | 0.702451 | 0.704009 | 0.702747 |
| 0.70967 | 0.712505 | 0.70689 | 0.705757 | 0.710761 | 0.705216 | 0.702651 | 0.705326 | 0.704129 |
| 0.721301 | 0.718004 | 0.707858 | 0.712229 | 0.718058 | 0.713301 | 0.714193 | 0.712214 | 0.710755 |
| 0.715235 | 0.714014 | 0.707164 | 0.707092 | 0.70959 | 0.70629 | 0.709908 | 0.705267 | 0.707022 |
| 0.710173 | 0.708498 | 0.702651 | 0.700265 | 0.704988 | 0.70077 | 0.701067 | 0.702998 | 0.702386 |
| 0.715784 | 0.713122 | 0.70038 | 0.705612 | 0.707282 | 0.706037 | 0.705995 | 0.703448 | 0.705024 |
| 0.709578 | 0.705698 | 0.700655 | 0.699321 | 0.701256 | 0.700166 | 0.698126 | 0.698201 | 0.693772 |
| 0.715795 | 0.713785 | 0.706019 | 0.704344 | 0.707948 | 0.706278 | 0.706511 | 0.703557 | 0.70803 |
| 0.713049 | 0.712569 | 0.704131 | 0.705553 | 0.710556 | 0.708732 | 0.707509 | 0.706094 | 0.708327 |
| 0.713576 | 0.712213 | 0.706375 | 0.705513 | 0.709566 | 0.70802 | 0.707272 | 0.704339 | 0.707047 |
| 0.710535 | 0.708073 | 0.701447 | 0.707739 | 0.708937 | 0.707617 | 0.705048 | 0.705678 | 0.703487 |
| 0.712547 | 0.711012 | 0.70438 | 0.705011 | 0.711796 | 0.705859 | 0.706732 | 0.70874 | 0.703805 |
| 0.71782 | 0.716291 | 0.706778 | 0.70605 | 0.710405 | 0.709659 | 0.706973 | 0.7082 | 0.705836 |
| 0.715842 | 0.709915 | 0.707364 | 0.710182 | 0.709422 | 0.7084 | 0.707982 | 0.70968 | 0.704398 |
| 0.716058 | 0.712513 | 0.706142 | 0.710593 | 0.714047 | 0.70864 | 0.708333 | 0.708573 | 0.711891 |
| 0.715859 | 0.712554 | 0.705947 | 0.708242 | 0.7094 | 0.705933 | 0.708069 | 0.707175 | 0.70598 |
| 0.716412 | 0.716597 | 0.711388 | 0.711384 | 0.715592 | 0.711129 | 0.710961 | 0.711821 | 0.710607 |
| 0.712534 | 0.715652 | 0.703223 | 0.705785 | 0.706265 | 0.704074 | 0.70835 | 0.707732 | 0.701818 |
| 0.71394 | 0.714628 | 0.702675 | 0.703991 | 0.709404 | 0.703505 | 0.708596 | 0.707647 | 0.702733 |
| 0.712282 | 0.712082 | 0.700953 | 0.703203 | 0.707641 | 0.702742 | 0.704476 | 0.707097 | 0.702026 |
| 0.720925 | 0.718377 | 0.708363 | 0.70945 | 0.713738 | 0.710158 | 0.707897 | 0.708541 | 0.707321 |
| 0.717379 | 0.716776 | 0.706297 | 0.711426 | 0.711603 | 0.70734 | 0.709253 | 0.705403 | 0.705188 |
| 0.714749 | 0.714902 | 0.703634 | 0.704175 | 0.711438 | 0.708094 | 0.702359 | 0.702863 | 0.706498 |
| 0.712605 | 0.709575 | 0.706126 | 0.70905 | 0.711369 | 0.705544 | 0.703301 | 0.703704 | 0.700998 |
| 0.711143 | 0.710907 | 0.701745 | 0.705375 | 0.70876 | 0.704932 | 0.701591 | 0.700917 | 0.702265 |
| 0.711393 | 0.712677 | 0.701753 | 0.70359 | 0.707253 | 0.707375 | 0.702892 | 0.703763 | 0.703612 |
| 0.716787 | 0.714848 | 0.706431 | 0.709565 | 0.709875 | 0.708241 | 0.706757 | 0.7087 | 0.707729 |
| 0.720315 | 0.714892 | 0.704191 | 0.710686 | 0.712964 | 0.707692 | 0.708237 | 0.708824 | 0.709122 |
| 0.715265 | 0.713078 | 0.703105 | 0.706573 | 0.71143 | 0.703922 | 0.704766 | 0.702888 | 0.706094 |
| 0.721347 | 0.717748 | 0.706798 | 0.711281 | 0.714343 | 0.707664 | 0.709774 | 0.708073 | 0.711563 |
| 0.720308 | 0.716926 | 0.707296 | 0.710526 | 0.712617 | 0.709939 | 0.708547 | 0.708904 | 0.710788 |
| 0.710324 | 0.713841 | 0.698673 | 0.702393 | 0.709608 | 0.701802 | 0.703587 | 0.700514 | 0.700711 |
| 0.716593 | 0.714368 | 0.70249 | 0.707964 | 0.710696 | 0.705839 | 0.708816 | 0.704845 | 0.703337 |
| 0.711948 | 0.713823 | 0.699658 | 0.70161 | 0.707106 | 0.701624 | 0.699774 | 0.703161 | 0.705506 |
| 0.721673 | 0.721335 | 0.710004 | 0.711298 | 0.715053 | 0.711106 | 0.712007 | 0.710812 | 0.70822 |
| 0.71333 | 0.715257 | 0.703538 | 0.703293 | 0.708938 | 0.705959 | 0.706718 | 0.705501 | 0.704451 |
| 0.713826 | 0.714184 | 0.703819 | 0.705915 | 0.70952 | 0.709525 | 0.706562 | 0.705868 | 0.70653 |
| 0.715068 | 0.71637 | 0.707567 | 0.706998 | 0.710862 | 0.702538 | 0.707385 | 0.705592 | 0.704158 |
| 0.720768 | 0.720127 | 0.708905 | 0.712945 | 0.714548 | 0.710211 | 0.711466 | 0.709374 | 0.70846 |
| 0.717173 | 0.717079 | 0.708889 | 0.707658 | 0.714859 | 0.711317 | 0.707606 | 0.708676 | 0.70729 |
| 0.719116 | 0.72112 | 0.706516 | 0.708712 | 0.714877 | 0.711113 | 0.705195 | 0.709011 | 0.708058 |
| 0.712328 | 0.713696 | 0.704898 | 0.707192 | 0.7123 | 0.70693 | 0.705469 | 0.703746 | 0.704067 |
| 0.713502 | 0.711316 | 0.70832 | 0.711566 | 0.711222 | 0.711728 | 0.708956 | 0.706975 | 0.705773 |
| 0.710646 | 0.707742 | 0.703863 | 0.703665 | 0.706361 | 0.706948 | 0.703824 | 0.702906 | 0.701068 |

| | | | | | | | | |
|---|---|---|---|---|---|---|---|---|
| 0.697988 | 0.699918 | 0.689113 | 0.693473 | 0.692555 | 0.693262 | 0.689585 | 0.690793 | 0.690542 |
| 0.744972 | 0.763434 | 0.742458 | 0.746869 | 0.749132 | 0.74666 | 0.741154 | 0.741153 | 0.742379 |
| 0.713568 | 0.712282 | 0.702658 | 0.702588 | 0.706872 | 0.702838 | 0.704493 | 0.703888 | 0.701431 |
| 0.716027 | 0.71478 | 0.706899 | 0.707342 | 0.714081 | 0.708168 | 0.706637 | 0.704763 | 0.70546 |
| 0.707208 | 0.707016 | 0.699711 | 0.700182 | 0.705159 | 0.699561 | 0.70231 | 0.700619 | 0.698496 |
| 0.71381 | 0.712033 | 0.701862 | 0.705925 | 0.712244 | 0.705085 | 0.705539 | 0.702514 | 0.706657 |
| 0.709813 | 0.710064 | 0.698757 | 0.702399 | 0.70604 | 0.703118 | 0.702555 | 0.702418 | 0.700151 |
| 0.704893 | 0.702405 | 0.697331 | 0.696367 | 0.704059 | 0.697988 | 0.69753 | 0.698197 | 0.6973 |
| 0.713283 | 0.714139 | 0.704262 | 0.707429 | 0.708652 | 0.705163 | 0.704096 | 0.706365 | 0.702077 |
| 0.723452 | 0.719221 | 0.711056 | 0.713358 | 0.716131 | 0.712002 | 0.713039 | 0.71287 | 0.711388 |
| 0.71424 | 0.716959 | 0.704961 | 0.709179 | 0.708967 | 0.708056 | 0.70828 | 0.705755 | 0.705606 |
| 0.717246 | 0.716966 | 0.709747 | 0.712303 | 0.711213 | 0.710087 | 0.710762 | 0.712002 | 0.70574 |
| 0.719283 | 0.718934 | 0.709958 | 0.709842 | 0.714091 | 0.711099 | 0.71202 | 0.710097 | 0.709535 |
| 0.716545 | 0.717823 | 0.707864 | 0.709979 | 0.71187 | 0.707956 | 0.71069 | 0.709119 | 0.705252 |
| 0.713784 | 0.712339 | 0.705361 | 0.709481 | 0.708557 | 0.706508 | 0.707774 | 0.705179 | 0.703202 |
| 0.713099 | 0.712933 | 0.702486 | 0.705471 | 0.712082 | 0.704149 | 0.706557 | 0.707849 | 0.704623 |
| 0.712966 | 0.713772 | 0.705496 | 0.711213 | 0.713618 | 0.708347 | 0.710424 | 0.709884 | 0.709266 |
| 0.715151 | 0.712554 | 0.705523 | 0.710653 | 0.711917 | 0.709101 | 0.709163 | 0.709384 | 0.710234 |
| 0.709059 | 0.707139 | 0.694334 | 0.700329 | 0.698351 | 0.697904 | 0.702893 | 0.697469 | 0.698084 |
| 0.710917 | 0.709954 | 0.700438 | 0.702974 | 0.704407 | 0.703722 | 0.703316 | 0.700145 | 0.696546 |
| 0.71568 | 0.716179 | 0.705536 | 0.707738 | 0.711715 | 0.70835 | 0.706002 | 0.703333 | 0.704849 |
| 0.702222 | 0.701767 | 0.694213 | 0.696762 | 0.698488 | 0.696404 | 0.702725 | 0.698517 | 0.694678 |
| 0.706831 | 0.710428 | 0.702617 | 0.70447 | 0.706423 | 0.704073 | 0.7055 | 0.705133 | 0.704286 |
| 0.712634 | 0.715345 | 0.706165 | 0.702849 | 0.70898 | 0.705505 | 0.707851 | 0.707143 | 0.707772 |
| 0.714262 | 0.712211 | 0.70606 | 0.706559 | 0.711122 | 0.709152 | 0.709871 | 0.706848 | 0.706899 |
| 0.712308 | 0.711168 | 0.702517 | 0.704327 | 0.707561 | 0.707115 | 0.709124 | 0.709895 | 0.705806 |
| 0.708813 | 0.711463 | 0.703026 | 0.701137 | 0.704369 | 0.701154 | 0.706323 | 0.703722 | 0.698706 |
| 0.70817 | 0.706845 | 0.696942 | 0.697055 | 0.704493 | 0.701036 | 0.701633 | 0.70112 | 0.698859 |
| 0.704271 | 0.705384 | 0.697283 | 0.701187 | 0.702522 | 0.698569 | 0.700466 | 0.699157 | 0.696736 |
| 0.764289 | 0.759258 | 0.772043 | 0.783259 | 0.775613 | 0.777178 | 0.784765 | 0.774712 | 0.755664 |
| 0.776254 | 0.778423 | 0.794125 | 0.80891 | 0.797898 | 0.783918 | 0.797791 | 0.793691 | 0.779294 |
| 0.778782 | 0.775294 | 0.791786 | 0.796398 | 0.789159 | 0.791575 | 0.78691 | 0.77878 | 0.772033 |
| 0.77219 | 0.766112 | 0.803136 | 0.795332 | 0.795896 | 0.804987 | 0.781683 | 0.793213 | 0.788314 |
| 0.781178 | 0.779474 | 0.788973 | 0.796333 | 0.785815 | 0.787821 | 0.779861 | 0.774086 | 0.773615 |
| 0.700882 | 0.706167 | 0.698388 | 0.699268 | 0.703642 | 0.699424 | 0.700613 | 0.699369 | 0.699203 |
| 0.705759 | 0.710913 | 0.701227 | 0.700346 | 0.70646 | 0.70075 | 0.702959 | 0.704312 | 0.70026 |
| 0.712325 | 0.713014 | 0.701352 | 0.702414 | 0.704902 | 0.702858 | 0.70739 | 0.704918 | 0.702426 |
| 0.70844 | 0.711357 | 0.704751 | 0.706915 | 0.707898 | 0.70828 | 0.705338 | 0.70603 | 0.705778 |
| 0.705428 | 0.704061 | 0.698362 | 0.7002 | 0.705579 | 0.701245 | 0.703758 | 0.702389 | 0.701471 |
| 0.711785 | 0.712929 | 0.704135 | 0.703401 | 0.705097 | 0.704859 | 0.70916 | 0.706843 | 0.700483 |
| 0.710371 | 0.71013 | 0.701556 | 0.701303 | 0.706397 | 0.705492 | 0.703912 | 0.705815 | 0.698435 |
| 0.711557 | 0.710182 | 0.701321 | 0.704289 | 0.703471 | 0.700917 | 0.705061 | 0.70199 | 0.700662 |
| 0.704011 | 0.705631 | 0.694634 | 0.69749 | 0.700745 | 0.696676 | 0.70079 | 0.699123 | 0.697317 |
| 0.700611 | 0.702829 | 0.696913 | 0.697431 | 0.701545 | 0.696447 | 0.698188 | 0.696112 | 0.697282 |
| 0.707309 | 0.711631 | 0.700147 | 0.704769 | 0.700805 | 0.705482 | 0.706857 | 0.705428 | 0.698632 |
| 0.714413 | 0.714816 | 0.701234 | 0.703838 | 0.706958 | 0.703854 | 0.705772 | 0.70515 | 0.702885 |

| | | | | | | | | |
|---|---|---|---|---|---|---|---|---|
| 0.714684 | 0.71273 | 0.703383 | 0.70705 | 0.708611 | 0.704266 | 0.707431 | 0.70511 | 0.705553 |
| 0.711423 | 0.712674 | 0.703549 | 0.706657 | 0.709032 | 0.704752 | 0.703695 | 0.700534 | 0.702325 |
| 0.715726 | 0.717744 | 0.705821 | 0.70864 | 0.709048 | 0.7083 | 0.709851 | 0.707656 | 0.705926 |
| 0.710715 | 0.710922 | 0.706133 | 0.704837 | 0.707158 | 0.706303 | 0.705773 | 0.707028 | 0.70124 |
| 0.711534 | 0.709423 | 0.698549 | 0.698606 | 0.706332 | 0.70482 | 0.700484 | 0.702024 | 0.70084 |
| 0.709696 | 0.709508 | 0.699876 | 0.701673 | 0.70596 | 0.704504 | 0.701702 | 0.703846 | 0.700607 |
| 0.714392 | 0.713192 | 0.704232 | 0.705911 | 0.708131 | 0.704605 | 0.707976 | 0.707567 | 0.703795 |
| 0.711453 | 0.711942 | 0.70223 | 0.703365 | 0.709058 | 0.704229 | 0.702133 | 0.703183 | 0.702436 |
| 0.709489 | 0.712221 | 0.699584 | 0.704088 | 0.705684 | 0.702795 | 0.703557 | 0.702618 | 0.698644 |
| 0.712847 | 0.711464 | 0.705375 | 0.705745 | 0.706941 | 0.704403 | 0.704085 | 0.704736 | 0.701272 |
| 0.712662 | 0.71131 | 0.702422 | 0.7021 | 0.705239 | 0.705523 | 0.705402 | 0.702067 | 0.703154 |
| 0.710769 | 0.712362 | 0.699545 | 0.701844 | 0.704378 | 0.70535 | 0.702667 | 0.701116 | 0.700402 |
| 0.703974 | 0.703898 | 0.692321 | 0.69364 | 0.698586 | 0.696894 | 0.697438 | 0.692214 | 0.694338 |
| 0.706822 | 0.705453 | 0.698726 | 0.699154 | 0.698725 | 0.696128 | 0.695156 | 0.697965 | 0.691599 |
| 0.712489 | 0.711342 | 0.702039 | 0.70546 | 0.709097 | 0.707764 | 0.705433 | 0.705494 | 0.707373 |
| 0.720492 | 0.716154 | 0.710628 | 0.71277 | 0.715984 | 0.708618 | 0.716872 | 0.714997 | 0.710656 |
| 0.713524 | 0.714218 | 0.705664 | 0.706801 | 0.712501 | 0.708916 | 0.710189 | 0.708377 | 0.707218 |
| 0.71493 | 0.717914 | 0.70746 | 0.707947 | 0.713171 | 0.709409 | 0.708642 | 0.708313 | 0.710015 |
| 0.709932 | 0.70916 | 0.702172 | 0.701917 | 0.706567 | 0.703376 | 0.706093 | 0.704107 | 0.699775 |
| 0.711709 | 0.709683 | 0.699447 | 0.702345 | 0.707248 | 0.706198 | 0.702862 | 0.703515 | 0.70214 |
| 0.707077 | 0.705689 | 0.69695 | 0.697448 | 0.703494 | 0.702599 | 0.704069 | 0.70156 | 0.699567 |
| 0.715492 | 0.714366 | 0.704663 | 0.707897 | 0.709594 | 0.706934 | 0.709767 | 0.705219 | 0.703986 |
| 0.716783 | 0.713992 | 0.707517 | 0.706602 | 0.715085 | 0.708133 | 0.709852 | 0.710434 | 0.708462 |
| 0.710676 | 0.709931 | 0.701351 | 0.702304 | 0.706044 | 0.7041 | 0.703054 | 0.70477 | 0.701514 |
| 0.715004 | 0.710475 | 0.701476 | 0.702533 | 0.709781 | 0.702047 | 0.705887 | 0.705488 | 0.702567 |
| 0.777414 | 0.776282 | 0.806988 | 0.793112 | 0.796177 | 0.799252 | 0.80434 | 0.795662 | 0.792398 |
| 0.713032 | 0.710771 | 0.702872 | 0.705766 | 0.707958 | 0.705747 | 0.703949 | 0.703894 | 0.700605 |
| 0.714514 | 0.714716 | 0.703816 | 0.705775 | 0.711403 | 0.705003 | 0.707999 | 0.705129 | 0.706704 |
| 0.718621 | 0.717001 | 0.710388 | 0.710624 | 0.71077 | 0.707392 | 0.708849 | 0.707547 | 0.710031 |
| 0.714758 | 0.712269 | 0.702577 | 0.706184 | 0.711726 | 0.70601 | 0.706372 | 0.704505 | 0.702296 |
| 0.709808 | 0.708257 | 0.700451 | 0.702099 | 0.704098 | 0.702094 | 0.70084 | 0.703738 | 0.698774 |
| 0.7122 | 0.709553 | 0.699481 | 0.701987 | 0.705643 | 0.701361 | 0.704636 | 0.703682 | 0.698584 |
| 0.71384 | 0.714228 | 0.700146 | 0.708146 | 0.708678 | 0.703854 | 0.707467 | 0.70232 | 0.704601 |
| 0.708064 | 0.712496 | 0.701089 | 0.700753 | 0.706333 | 0.702477 | 0.702737 | 0.700778 | 0.702061 |
| 0.707867 | 0.708442 | 0.698112 | 0.700968 | 0.702829 | 0.699217 | 0.702216 | 0.700085 | 0.698195 |
| 0.71129 | 0.711098 | 0.699281 | 0.700497 | 0.707439 | 0.701323 | 0.701094 | 0.697249 | 0.702224 |
| 0.716053 | 0.718862 | 0.707247 | 0.705973 | 0.712857 | 0.705039 | 0.708634 | 0.709576 | 0.705617 |
| 0.718217 | 0.717107 | 0.708988 | 0.70637 | 0.714043 | 0.707923 | 0.710353 | 0.706772 | 0.707711 |
| 0.715797 | 0.710299 | 0.705916 | 0.704778 | 0.711431 | 0.704078 | 0.705629 | 0.704338 | 0.705457 |
| 0.712204 | 0.710758 | 0.702763 | 0.705691 | 0.710458 | 0.70778 | 0.708715 | 0.704494 | 0.69969 |
| 0.709996 | 0.705625 | 0.699086 | 0.699138 | 0.702915 | 0.702479 | 0.69808 | 0.699471 | 0.700931 |
| 0.717259 | 0.715246 | 0.706479 | 0.709069 | 0.712394 | 0.708771 | 0.709767 | 0.708525 | 0.707337 |
| 0.71063 | 0.712687 | 0.704201 | 0.702527 | 0.710487 | 0.703241 | 0.704863 | 0.702002 | 0.702103 |
| 0.712723 | 0.711236 | 0.701263 | 0.698249 | 0.70893 | 0.700643 | 0.700869 | 0.703052 | 0.700503 |
| 0.712918 | 0.712558 | 0.704256 | 0.705171 | 0.70875 | 0.708086 | 0.705526 | 0.705348 | 0.703151 |
| 0.712237 | 0.713888 | 0.705365 | 0.703426 | 0.709796 | 0.707696 | 0.706696 | 0.70425 | 0.706412 |

| | | | | | | | | |
|---|---|---|---|---|---|---|---|---|
| 0.713111 | 0.708764 | 0.702377 | 0.703888 | 0.706262 | 0.703201 | 0.700668 | 0.701256 | 0.699268 |
| 0.708489 | 0.712265 | 0.700412 | 0.704109 | 0.708018 | 0.707358 | 0.705727 | 0.703267 | 0.704186 |
| 0.70955 | 0.710922 | 0.699743 | 0.700577 | 0.708249 | 0.703083 | 0.703245 | 0.704847 | 0.70226 |
| 0.667489 | 0.666435 | 0.649686 | 0.66156 | 0.671353 | 0.660695 | 0.65979 | 0.652189 | 0.659067 |
| 0.709829 | 0.710294 | 0.700768 | 0.699258 | 0.705541 | 0.70152 | 0.702927 | 0.698906 | 0.697451 |
| 0.706089 | 0.710627 | 0.697348 | 0.703063 | 0.708243 | 0.706255 | 0.703793 | 0.705061 | 0.703859 |
| 0.711069 | 0.713492 | 0.701676 | 0.703866 | 0.710082 | 0.703978 | 0.70837 | 0.705485 | 0.705349 |
| 0.712528 | 0.71257 | 0.700515 | 0.69801 | 0.705643 | 0.702763 | 0.70569 | 0.703356 | 0.702126 |
| 0.715567 | 0.718695 | 0.704578 | 0.706965 | 0.711326 | 0.709107 | 0.70866 | 0.710444 | 0.705529 |
| 0.713598 | 0.712475 | 0.703314 | 0.703222 | 0.707337 | 0.705006 | 0.706345 | 0.70631 | 0.702722 |
| 0.712219 | 0.711755 | 0.702985 | 0.701194 | 0.70819 | 0.705234 | 0.706261 | 0.707619 | 0.701052 |
| 0.716691 | 0.71358 | 0.701276 | 0.704129 | 0.713527 | 0.710872 | 0.708176 | 0.709008 | 0.708112 |
| 0.714079 | 0.715308 | 0.705104 | 0.705667 | 0.71269 | 0.711289 | 0.706571 | 0.70998 | 0.7076 |
| 0.71335 | 0.710913 | 0.700641 | 0.702816 | 0.707965 | 0.704331 | 0.703997 | 0.705401 | 0.70277 |
| 0.718499 | 0.713515 | 0.705249 | 0.70754 | 0.710938 | 0.706488 | 0.704747 | 0.703141 | 0.703849 |
| 0.70661 | 0.70686 | 0.699101 | 0.700755 | 0.704509 | 0.701602 | 0.701483 | 0.703486 | 0.700048 |
| 0.716149 | 0.713511 | 0.701207 | 0.705381 | 0.709818 | 0.701918 | 0.708323 | 0.704482 | 0.706472 |
| 0.714343 | 0.714387 | 0.700928 | 0.704206 | 0.709224 | 0.708184 | 0.705403 | 0.705996 | 0.705119 |
| 0.705573 | 0.708442 | 0.697595 | 0.70046 | 0.70584 | 0.700199 | 0.701522 | 0.697268 | 0.699177 |
| 0.710621 | 0.710002 | 0.70085 | 0.703812 | 0.707249 | 0.701869 | 0.700614 | 0.703186 | 0.701245 |
| 0.713703 | 0.710751 | 0.701805 | 0.705102 | 0.709411 | 0.703288 | 0.70728 | 0.70678 | 0.705585 |
| 0.717079 | 0.714535 | 0.702538 | 0.704621 | 0.708686 | 0.705456 | 0.705534 | 0.705428 | 0.705544 |
| 0.690353 | 0.697882 | 0.696077 | 0.691558 | 0.6897 | 0.692195 | 0.689292 | 0.693653 | 0.689841 |
| 0.707197 | 0.705385 | 0.727878 | 0.727048 | 0.727681 | 0.733449 | 0.721685 | 0.728907 | 0.722775 |
| 0.749101 | 0.750112 | 0.763624 | 0.759074 | 0.748933 | 0.755134 | 0.750396 | 0.747171 | 0.743192 |
| 0.741355 | 0.731546 | 0.780839 | 0.751293 | 0.767948 | 0.757723 | 0.756959 | 0.765304 | 0.760603 |
| 0.667251 | 0.671203 | 0.658636 | 0.656436 | 0.655187 | 0.656941 | 0.657853 | 0.661871 | 0.654842 |
| 0.664314 | 0.665868 | 0.652399 | 0.656459 | 0.656513 | 0.656899 | 0.654833 | 0.653471 | 0.649869 |
| 0.66353 | 0.670028 | 0.654776 | 0.653772 | 0.657328 | 0.655635 | 0.655046 | 0.659593 | 0.649694 |
| 0.68118 | 0.683109 | 0.666937 | 0.66998 | 0.677005 | 0.676125 | 0.670672 | 0.671292 | 0.666127 |
| 0.649402 | 0.656986 | 0.644612 | 0.644337 | 0.64032 | 0.650323 | 0.652603 | 0.64439 | 0.638501 |
| 0.643033 | 0.656011 | 0.634847 | 0.634279 | 0.63287 | 0.640851 | 0.636314 | 0.637653 | 0.624763 |
| 0.649288 | 0.647743 | 0.640104 | 0.636467 | 0.642426 | 0.63934 | 0.641217 | 0.638139 | 0.632513 |
| 0.722925 | 0.726401 | 0.729379 | 0.730784 | 0.740343 | 0.728455 | 0.724022 | 0.725546 | 0.725487 |
| 0.706386 | 0.704872 | 0.696061 | 0.697315 | 0.701759 | 0.697973 | 0.698616 | 0.698145 | 0.698713 |
| 0.678543 | 0.683676 | 0.669184 | 0.670234 | 0.675105 | 0.674242 | 0.671979 | 0.67599 | 0.669244 |
| 0.673459 | 0.669389 | 0.661192 | 0.661751 | 0.668907 | 0.662842 | 0.662855 | 0.667108 | 0.663015 |
| 0.657606 | 0.656479 | 0.639161 | 0.636622 | 0.649855 | 0.648774 | 0.652529 | 0.646697 | 0.644894 |
| 0.681642 | 0.682839 | 0.670401 | 0.66825 | 0.670474 | 0.675516 | 0.677112 | 0.673891 | 0.666818 |
| 0.669444 | 0.673013 | 0.658214 | 0.657914 | 0.663386 | 0.667331 | 0.664753 | 0.66312 | 0.659848 |
| 0.659719 | 0.676135 | 0.650916 | 0.650146 | 0.656352 | 0.657077 | 0.657396 | 0.657928 | 0.650558 |
| 0.661531 | 0.670023 | 0.651945 | 0.64772 | 0.6543 | 0.657151 | 0.658043 | 0.661402 | 0.654438 |
| 0.660658 | 0.655647 | 0.646977 | 0.649065 | 0.651542 | 0.653799 | 0.647469 | 0.651534 | 0.649842 |
| 0.666731 | 0.666161 | 0.652839 | 0.658621 | 0.65663 | 0.657925 | 0.655491 | 0.656728 | 0.651261 |
| 0.658234 | 0.668567 | 0.646562 | 0.649344 | 0.648272 | 0.648823 | 0.647192 | 0.656949 | 0.646708 |
| 0.654405 | 0.653274 | 0.637909 | 0.635959 | 0.638181 | 0.637876 | 0.640351 | 0.634374 | 0.636511 |

| | | | | | | | | |
|---|---|---|---|---|---|---|---|---|
| 0.69416 | 0.693207 | 0.686485 | 0.687928 | 0.687198 | 0.685317 | 0.68756 | 0.688472 | 0.68262 |
| 0.663353 | 0.668112 | 0.648051 | 0.64931 | 0.659414 | 0.65333 | 0.653778 | 0.652364 | 0.650826 |
| 0.674535 | 0.671009 | 0.657409 | 0.655422 | 0.656831 | 0.661822 | 0.653337 | 0.656616 | 0.65429 |
| 0.667757 | 0.673545 | 0.659182 | 0.658136 | 0.661504 | 0.658374 | 0.659257 | 0.659047 | 0.660544 |
| 0.66166 | 0.663861 | 0.653459 | 0.645772 | 0.656027 | 0.650294 | 0.64794 | 0.65509 | 0.646896 |
| 0.666667 | 0.664661 | 0.659559 | 0.658182 | 0.665527 | 0.6627 | 0.655742 | 0.663812 | 0.657108 |
| 0.657634 | 0.657123 | 0.641421 | 0.65113 | 0.646503 | 0.644398 | 0.64645 | 0.645085 | 0.638713 |
| 0.650041 | 0.646025 | 0.644142 | 0.643013 | 0.643094 | 0.641975 | 0.630494 | 0.63863 | 0.63609 |
| 0.65 | 0.656011 | 0.640885 | 0.643798 | 0.645954 | 0.645981 | 0.640969 | 0.639353 | 0.641992 |
| 0.693092 | 0.695068 | 0.688029 | 0.687453 | 0.690612 | 0.688462 | 0.689401 | 0.686342 | 0.685215 |
| 0.691704 | 0.695077 | 0.68539 | 0.682799 | 0.68739 | 0.683067 | 0.687666 | 0.684195 | 0.686348 |
| 0.6906 | 0.691739 | 0.677623 | 0.676552 | 0.679449 | 0.679352 | 0.680505 | 0.677918 | 0.673941 |
| 0.680265 | 0.685734 | 0.669487 | 0.668584 | 0.671689 | 0.673642 | 0.674941 | 0.676551 | 0.671445 |
| 0.688134 | 0.686903 | 0.674882 | 0.67505 | 0.675464 | 0.675272 | 0.681726 | 0.677924 | 0.673277 |
| 0.676973 | 0.67902 | 0.66568 | 0.663277 | 0.670335 | 0.667696 | 0.669405 | 0.671113 | 0.663995 |

| | | HG_18 | | | | HG_19 | | |
|---|---|---|---|---|---|---|---|---|
| Home_garc | Home_garc | Home_garc | Home_garc | Home_garc | Home_garc | Home_garc | Home_garc | Home_garc |
| 0.804504 | 0.789504 | 0.79014 | 0.804271 | 0.804453 | 0.775789 | 0.766156 | 0.75913 | 0.706493 |
| 0.800183 | 0.7848 | 0.792648 | 0.795062 | 0.791882 | 0.789973 | 0.781961 | 0.769734 | 0.717701 |
| 0.799039 | 0.78037 | 0.789496 | 0.801925 | 0.794118 | 0.781848 | 0.775309 | 0.758476 | 0.702829 |
| 0.792353 | 0.784964 | 0.784609 | 0.795279 | 0.776639 | 0.783007 | 0.788166 | 0.7761 | 0.710698 |
| 0.788094 | 0.785485 | 0.777594 | 0.78312 | 0.780627 | 0.769793 | 0.770012 | 0.766713 | 0.707393 |
| 0.780406 | 0.775395 | 0.767366 | 0.770488 | 0.767239 | 0.771063 | 0.784519 | 0.778267 | 0.718401 |
| 0.714263 | 0.714024 | 0.708626 | 0.710671 | 0.709844 | 0.712978 | 0.718395 | 0.727019 | 0.786387 |
| 0.707841 | 0.714047 | 0.705272 | 0.706281 | 0.702977 | 0.705799 | 0.711586 | 0.719469 | 0.987885 |
| 0.786931 | 0.783405 | 0.782065 | 0.780066 | 0.779949 | 0.821989 | 0.789601 | 0.777641 | 0.711828 |
| 0.795675 | 0.809759 | 0.792569 | 0.788897 | 0.794066 | 0.785 | 0.771321 | 0.759153 | 0.708297 |
| 0.793221 | 0.787751 | 0.773157 | 0.788678 | 0.796176 | 0.77482 | 0.778421 | 0.763458 | 0.706396 |
| 0.804754 | 0.798836 | 0.797402 | 0.791687 | 0.793098 | 0.785491 | 0.773708 | 0.762788 | 0.70169 |
| 0.793477 | 0.783648 | 0.787234 | 0.774215 | 0.798548 | 0.771 | 0.77787 | 0.772386 | 0.706277 |
| 0.792761 | 0.786623 | 0.787386 | 0.78311 | 0.784403 | 0.775773 | 0.76409 | 0.764917 | 0.721664 |
| 0.776202 | 0.7756 | 0.768897 | 0.77156 | 0.764682 | 0.768436 | 0.772274 | 0.769853 | 0.731501 |
| 0.805828 | 0.798923 | 0.806747 | 0.811326 | 0.799967 | 0.792871 | 0.770183 | 0.773365 | 0.717163 |
| 0.812253 | 0.795794 | 0.786409 | 0.805298 | 0.80188 | 0.775577 | 0.763102 | 0.763502 | 0.714621 |
| 0.772628 | 0.766949 | 0.763271 | 0.756637 | 0.759455 | 0.759764 | 0.753199 | 0.75494 | 0.741987 |
| 0.791899 | 0.788796 | 0.799865 | 0.785446 | 0.780328 | 0.785726 | 0.769548 | 0.761194 | 0.702518 |
| 0.795587 | 0.789347 | 0.79794 | 0.784456 | 0.782663 | 0.792728 | 0.784655 | 0.771293 | 0.708744 |
| 0.788117 | 0.78859 | 0.787395 | 0.793627 | 0.782468 | 0.786659 | 0.796802 | 0.77622 | 0.706169 |
| 0.808628 | 0.802934 | 0.793311 | 0.801056 | 0.790474 | 0.790982 | 0.781222 | 0.762908 | 0.709063 |
| 0.795744 | 0.791849 | 0.796109 | 0.788695 | 0.786165 | 0.79168 | 0.79347 | 0.784343 | 0.714561 |
| 0.799054 | 0.799618 | 0.810037 | 0.799432 | 0.79223 | 0.794265 | 0.779754 | 0.76405 | 0.713438 |
| 0.787555 | 0.782429 | 0.783888 | 0.784469 | 0.769783 | 0.788024 | 0.7875 | 0.783382 | 0.712641 |
| 0.799123 | 0.78535 | 0.799883 | 0.78469 | 0.785032 | 0.785277 | 0.78387 | 0.769137 | 0.706083 |
| 0.795396 | 0.78249 | 0.778552 | 0.774752 | 0.769941 | 0.775969 | 0.774183 | 0.768465 | 0.702353 |
| 0.789569 | 0.779998 | 0.791691 | 0.783105 | 0.785714 | 0.781158 | 0.788527 | 0.774711 | 0.709595 |
| 0.782017 | 0.786287 | 0.779896 | 0.785922 | 0.775953 | 0.785697 | 0.788351 | 0.780233 | 0.714769 |
| 0.794164 | 0.794275 | 0.803508 | 0.79061 | 0.790228 | 0.786482 | 0.765534 | 0.765318 | 0.709921 |
| 0.78524 | 0.785004 | 0.778629 | 0.77078 | 0.775607 | 0.775226 | 0.785096 | 0.772877 | 0.707909 |
| 0.783351 | 0.779857 | 0.776521 | 0.781534 | 0.776792 | 0.780806 | 0.780838 | 0.770521 | 0.704732 |
| 0.78459 | 0.786985 | 0.780311 | 0.783355 | 0.78141 | 0.781173 | 0.783407 | 0.77735 | 0.705845 |
| 0.777069 | 0.77725 | 0.767005 | 0.775389 | 0.774257 | 0.77785 | 0.78185 | 0.77498 | 0.708934 |
| 0.782297 | 0.770133 | 0.762726 | 0.775738 | 0.768952 | 0.772564 | 0.772645 | 0.761475 | 0.703488 |
| 0.792125 | 0.787627 | 0.783536 | 0.794389 | 0.781673 | 0.783674 | 0.777289 | 0.764566 | 0.710325 |
| 0.792993 | 0.791872 | 0.789404 | 0.783708 | 0.774059 | 0.782261 | 0.780402 | 0.778484 | 0.712727 |
| 0.771857 | 0.769326 | 0.768519 | 0.762678 | 0.763247 | 0.763346 | 0.780718 | 0.777667 | 0.713173 |
| 0.781558 | 0.783846 | 0.773887 | 0.771736 | 0.7752 | 0.781918 | 0.780709 | 0.776597 | 0.714984 |

| | | | | | | | | |
|---|---|---|---|---|---|---|---|---|
| 0.773475 | 0.76406 | 0.762622 | 0.761193 | 0.7644 | 0.755385 | 0.754941 | 0.751519 | 0.745698 |
| 0.805722 | 0.78469 | 0.796233 | 0.801861 | 0.7989 | 0.780366 | 0.767247 | 0.763015 | 0.699172 |
| 0.798884 | 0.797404 | 0.805303 | 0.806281 | 0.797024 | 0.782051 | 0.776558 | 0.765296 | 0.706099 |
| 0.809938 | 0.80369 | 0.808317 | 0.798524 | 0.81522 | 0.7878 | 0.776396 | 0.766594 | 0.7095 |
| 0.796983 | 0.782312 | 0.794763 | 0.799332 | 0.789513 | 0.783493 | 0.773676 | 0.773917 | 0.707399 |
| 0.793146 | 0.79099 | 0.785244 | 0.788482 | 0.777815 | 0.778079 | 0.779619 | 0.774281 | 0.713617 |
| 0.776731 | 0.781649 | 0.769997 | 0.773699 | 0.768676 | 0.777524 | 0.778318 | 0.773076 | 0.702527 |
| 0.794069 | 0.797494 | 0.795208 | 0.779545 | 0.779291 | 0.779428 | 0.783127 | 0.771208 | 0.711605 |
| 0.780734 | 0.774506 | 0.772334 | 0.767913 | 0.770596 | 0.781131 | 0.774533 | 0.777137 | 0.708974 |
| 0.776805 | 0.77992 | 0.774704 | 0.770089 | 0.775138 | 0.778485 | 0.795001 | 0.78257 | 0.713633 |
| 0.818251 | 0.799014 | 0.80664 | 0.798318 | 0.802213 | 0.781456 | 0.777123 | 0.763072 | 0.70121 |
| 0.817807 | 0.792731 | 0.80172 | 0.791576 | 0.7944 | 0.786853 | 0.779022 | 0.765244 | 0.70575 |
| 0.816076 | 0.806283 | 0.835452 | 0.809024 | 0.809327 | 0.787622 | 0.772075 | 0.765931 | 0.707561 |
| 0.806313 | 0.79348 | 0.809649 | 0.804981 | 0.792085 | 0.791072 | 0.776653 | 0.762178 | 0.707726 |
| 0.801752 | 0.787636 | 0.793743 | 0.792882 | 0.790013 | 0.790026 | 0.773491 | 0.764338 | 0.704075 |
| 0.807924 | 0.795378 | 0.789621 | 0.801941 | 0.79046 | 0.790144 | 0.771178 | 0.758672 | 0.703454 |
| 0.818878 | 0.790918 | 0.813412 | 0.81191 | 0.810242 | 0.782652 | 0.775451 | 0.75442 | 0.70312 |
| 1 | 0.796896 | 0.808568 | 0.813871 | 0.819008 | 0.789461 | 0.783735 | 0.769909 | 0.709071 |
| 0.796896 | 1 | 0.795415 | 0.789417 | 0.7854 | 0.784417 | 0.772836 | 0.769162 | 0.714022 |
| 0.808568 | 0.795415 | 1 | 0.793613 | 0.808536 | 0.790622 | 0.77048 | 0.764138 | 0.707614 |
| 0.813871 | 0.789417 | 0.793613 | 1 | 0.809661 | 0.78588 | 0.769504 | 0.757138 | 0.707857 |
| 0.819008 | 0.7854 | 0.808536 | 0.809661 | 1 | 0.787239 | 0.770285 | 0.760059 | 0.703271 |
| 0.789461 | 0.784417 | 0.790622 | 0.78588 | 0.787239 | 1 | 0.779118 | 0.761681 | 0.707112 |
| 0.783735 | 0.772836 | 0.77048 | 0.769504 | 0.770285 | 0.779118 | 1 | 0.786091 | 0.713457 |
| 0.769909 | 0.769162 | 0.764138 | 0.757138 | 0.760059 | 0.761681 | 0.786091 | 1 | 0.722361 |
| 0.709071 | 0.714022 | 0.707614 | 0.707857 | 0.703271 | 0.707112 | 0.713457 | 0.722361 | 1 |
| 0.714641 | 0.716437 | 0.710367 | 0.704449 | 0.706899 | 0.714012 | 0.71649 | 0.727397 | 0.773252 |
| 0.716725 | 0.716268 | 0.707985 | 0.706355 | 0.708895 | 0.713295 | 0.719574 | 0.728093 | 0.773771 |
| 0.782429 | 0.783073 | 0.773138 | 0.76443 | 0.774159 | 0.77336 | 0.771137 | 0.763745 | 0.707104 |
| 0.786767 | 0.78138 | 0.781389 | 0.765871 | 0.777494 | 0.774996 | 0.771303 | 0.769017 | 0.705442 |
| 0.786963 | 0.795472 | 0.788276 | 0.78575 | 0.784345 | 0.776037 | 0.768377 | 0.768482 | 0.713894 |
| 0.779774 | 0.777186 | 0.774788 | 0.769244 | 0.77787 | 0.780403 | 0.774457 | 0.777879 | 0.714587 |
| 0.819032 | 0.795541 | 0.806919 | 0.810654 | 0.805523 | 0.798641 | 0.783861 | 0.768575 | 0.705625 |
| 0.810774 | 0.795325 | 0.805417 | 0.815357 | 0.78917 | 0.785336 | 0.775756 | 0.768635 | 0.707993 |
| 0.822076 | 0.788837 | 0.801415 | 0.803477 | 0.800883 | 0.798572 | 0.772135 | 0.762511 | 0.705706 |
| 0.83731 | 0.785542 | 0.800051 | 0.852426 | 0.807226 | 0.796906 | 0.777243 | 0.763941 | 0.708521 |
| 0.793384 | 0.790122 | 0.7884 | 0.774791 | 0.785967 | 0.78835 | 0.769739 | 0.763249 | 0.707302 |
| 0.842989 | 0.813629 | 0.825006 | 0.827116 | 0.818819 | 0.799619 | 0.794096 | 0.783093 | 0.719075 |
| 0.798509 | 0.790164 | 0.793362 | 0.790031 | 0.774626 | 0.786622 | 0.765283 | 0.75545 | 0.702847 |
| 0.791989 | 0.795068 | 0.793418 | 0.774558 | 0.786644 | 0.793219 | 0.784952 | 0.765763 | 0.706227 |
| 0.810343 | 0.788175 | 0.785484 | 0.807029 | 0.795949 | 0.78139 | 0.771277 | 0.757071 | 0.702182 |
| 0.837146 | 0.816672 | 0.826384 | 0.833077 | 0.825295 | 0.804697 | 0.795898 | 0.787716 | 0.722869 |
| 0.811786 | 0.800501 | 0.797832 | 0.80247 | 0.801948 | 0.782587 | 0.77887 | 0.768288 | 0.710827 |
| 0.798246 | 0.793584 | 0.805999 | 0.805419 | 0.791267 | 0.779967 | 0.776949 | 0.76452 | 0.704443 |
| 0.791604 | 0.783616 | 0.790641 | 0.78584 | 0.785405 | 0.788606 | 0.785792 | 0.766844 | 0.706686 |
| 0.807391 | 0.785028 | 0.787943 | 0.805603 | 0.79061 | 0.79149 | 0.77292 | 0.762331 | 0.704861 |

| | | | | | | | | |
|---|---|---|---|---|---|---|---|---|
| 0.810048 | 0.805023 | 0.803121 | 0.806247 | 0.806039 | 0.780175 | 0.771033 | 0.765332 | 0.703858 |
| 0.783733 | 0.796143 | 0.791521 | 0.77384 | 0.772298 | 0.79339 | 0.777083 | 0.763381 | 0.710772 |
| 0.788938 | 0.788299 | 0.78911 | 0.794229 | 0.78447 | 0.793941 | 0.77699 | 0.763522 | 0.708192 |
| 0.812548 | 0.795455 | 0.801326 | 0.804311 | 0.799189 | 0.792697 | 0.786392 | 0.773597 | 0.714054 |
| 0.791055 | 0.791133 | 0.794462 | 0.787595 | 0.782863 | 0.78461 | 0.78045 | 0.767861 | 0.713256 |
| 0.788336 | 0.788291 | 0.797195 | 0.773046 | 0.774014 | 0.782583 | 0.772512 | 0.764039 | 0.709605 |
| 0.799736 | 0.78895 | 0.796992 | 0.771436 | 0.785544 | 0.786999 | 0.770349 | 0.769717 | 0.712077 |
| 0.789263 | 0.783152 | 0.779885 | 0.784005 | 0.775899 | 0.789234 | 0.777408 | 0.76377 | 0.710458 |
| 0.791632 | 0.784402 | 0.781033 | 0.776108 | 0.778295 | 0.78332 | 0.777178 | 0.769231 | 0.711604 |
| 0.783623 | 0.777668 | 0.779962 | 0.77809 | 0.769111 | 0.772312 | 0.773866 | 0.764787 | 0.707146 |
| 0.784772 | 0.775893 | 0.778117 | 0.779402 | 0.771396 | 0.777669 | 0.769972 | 0.767204 | 0.707656 |
| 0.776424 | 0.776555 | 0.780389 | 0.774838 | 0.771038 | 0.775762 | 0.775827 | 0.765177 | 0.705382 |
| 0.78752 | 0.784273 | 0.784762 | 0.779579 | 0.778886 | 0.77896 | 0.767413 | 0.763826 | 0.71123 |
| 0.773557 | 0.777555 | 0.783141 | 0.768071 | 0.7671 | 0.772557 | 0.769411 | 0.759327 | 0.702066 |
| 0.780526 | 0.77512 | 0.779491 | 0.783565 | 0.782154 | 0.780741 | 0.779877 | 0.762239 | 0.708916 |
| 0.787023 | 0.782791 | 0.779644 | 0.774734 | 0.774517 | 0.776825 | 0.776518 | 0.763162 | 0.707745 |
| 0.782223 | 0.785803 | 0.780604 | 0.776416 | 0.778768 | 0.784101 | 0.775259 | 0.768164 | 0.710839 |
| 0.800833 | 0.776068 | 0.785619 | 0.791663 | 0.789417 | 0.782052 | 0.778401 | 0.767832 | 0.710942 |
| 0.783585 | 0.779964 | 0.780823 | 0.773342 | 0.775434 | 0.776217 | 0.780306 | 0.767332 | 0.709654 |
| 0.786135 | 0.779651 | 0.782128 | 0.778009 | 0.775025 | 0.783657 | 0.780063 | 0.770509 | 0.710687 |
| 0.782767 | 0.780182 | 0.779381 | 0.770359 | 0.771405 | 0.777019 | 0.77184 | 0.759532 | 0.707817 |
| 0.779674 | 0.774636 | 0.772686 | 0.777217 | 0.765376 | 0.774215 | 0.769914 | 0.759389 | 0.703514 |
| 0.787085 | 0.776468 | 0.778122 | 0.782944 | 0.774387 | 0.781939 | 0.775017 | 0.761545 | 0.707111 |
| 0.783577 | 0.786133 | 0.792934 | 0.778192 | 0.772435 | 0.785327 | 0.772921 | 0.757861 | 0.709136 |
| 0.787897 | 0.781616 | 0.782704 | 0.775144 | 0.774328 | 0.784715 | 0.770083 | 0.762172 | 0.712952 |
| 0.797749 | 0.795308 | 0.795557 | 0.778518 | 0.78701 | 0.802158 | 0.77786 | 0.780448 | 0.715518 |
| 0.791862 | 0.782369 | 0.782569 | 0.771304 | 0.782061 | 0.785317 | 0.768646 | 0.75968 | 0.709611 |
| 0.782004 | 0.77498 | 0.775215 | 0.776265 | 0.780051 | 0.776643 | 0.770068 | 0.765012 | 0.709608 |
| 0.787584 | 0.785767 | 0.777061 | 0.776441 | 0.772966 | 0.776046 | 0.778536 | 0.766303 | 0.710987 |
| 0.791225 | 0.78098 | 0.78506 | 0.77243 | 0.776361 | 0.7838 | 0.767572 | 0.762654 | 0.710197 |
| 0.777234 | 0.774845 | 0.775129 | 0.772655 | 0.768085 | 0.779006 | 0.778759 | 0.768034 | 0.711712 |
| 0.778212 | 0.776743 | 0.774524 | 0.777131 | 0.766482 | 0.778868 | 0.781821 | 0.764559 | 0.702639 |
| 0.800236 | 0.789982 | 0.787559 | 0.775522 | 0.788764 | 0.792895 | 0.777731 | 0.765293 | 0.70882 |
| 0.80657 | 0.800154 | 0.805294 | 0.795591 | 0.811256 | 0.782261 | 0.772497 | 0.762171 | 0.706013 |
| 0.789614 | 0.779667 | 0.784019 | 0.777129 | 0.778221 | 0.77831 | 0.773516 | 0.763787 | 0.709362 |
| 0.786736 | 0.793862 | 0.787402 | 0.782121 | 0.777461 | 0.784533 | 0.783113 | 0.777092 | 0.711665 |
| 0.785714 | 0.786206 | 0.780267 | 0.778586 | 0.774027 | 0.781016 | 0.774668 | 0.761772 | 0.706236 |
| 0.762911 | 0.755231 | 0.754581 | 0.753776 | 0.761769 | 0.758103 | 0.749817 | 0.740733 | 0.677701 |
| 0.767073 | 0.77369 | 0.784104 | 0.758443 | 0.758914 | 0.775489 | 0.763371 | 0.746321 | 0.688821 |
| 0.756658 | 0.755159 | 0.756751 | 0.744313 | 0.747522 | 0.752179 | 0.75253 | 0.74836 | 0.675765 |
| 0.786166 | 0.758692 | 0.768047 | 0.792381 | 0.776829 | 0.769701 | 0.756199 | 0.743056 | 0.677394 |
| 0.740389 | 0.76084 | 0.759936 | 0.736643 | 0.736491 | 0.739251 | 0.736969 | 0.729969 | 0.660965 |
| 0.778476 | 0.776217 | 0.774127 | 0.778897 | 0.778876 | 0.752167 | 0.743719 | 0.735896 | 0.662881 |
| 0.776922 | 0.751327 | 0.755954 | 0.774434 | 0.76291 | 0.762001 | 0.751326 | 0.732529 | 0.66541 |
| 0.757738 | 0.747119 | 0.752634 | 0.748163 | 0.743688 | 0.750561 | 0.746286 | 0.735709 | 0.661032 |
| 0.772446 | 0.758994 | 0.76269 | 0.758281 | 0.759138 | 0.763291 | 0.751915 | 0.732414 | 0.665128 |
| 0.780779 | 0.753906 | 0.770136 | 0.775898 | 0.757631 | 0.762236 | 0.74568 | 0.739208 | 0.671865 |

| | | | | | | | | |
|---|---|---|---|---|---|---|---|---|
| 0.757442 | 0.733037 | 0.739019 | 0.740988 | 0.737634 | 0.743988 | 0.738311 | 0.729794 | 0.668269 |
| 0.786692 | 0.763889 | 0.785299 | 0.781086 | 0.765531 | 0.776381 | 0.757469 | 0.738839 | 0.670053 |
| 0.739336 | 0.738462 | 0.742657 | 0.738245 | 0.743529 | 0.739807 | 0.753694 | 0.740135 | 0.667335 |
| 0.779071 | 0.769713 | 0.7752 | 0.772785 | 0.773619 | 0.745621 | 0.739452 | 0.722853 | 0.658872 |
| 0.759555 | 0.76764 | 0.772827 | 0.73574 | 0.746134 | 0.757625 | 0.744444 | 0.736404 | 0.655677 |
| 0.74903 | 0.744902 | 0.745968 | 0.757725 | 0.743062 | 0.758245 | 0.740256 | 0.725587 | 0.6663 |
| 0.75799 | 0.768967 | 0.75716 | 0.743151 | 0.748329 | 0.763612 | 0.740229 | 0.730529 | 0.664719 |
| 0.783167 | 0.758793 | 0.769516 | 0.79598 | 0.757 | 0.756268 | 0.743381 | 0.728719 | 0.664605 |
| 0.728074 | 0.727508 | 0.73202 | 0.726715 | 0.725118 | 0.728492 | 0.723388 | 0.71618 | 0.66015 |
| 0.781581 | 0.766815 | 0.782949 | 0.736834 | 0.774595 | 0.76682 | 0.743104 | 0.733634 | 0.65755 |
| 0.747272 | 0.740527 | 0.749691 | 0.745772 | 0.742562 | 0.751225 | 0.743201 | 0.723801 | 0.65232 |
| 0.788015 | 0.779802 | 0.784467 | 0.797641 | 0.785667 | 0.766583 | 0.756938 | 0.760694 | 0.696781 |
| 0.773831 | 0.773276 | 0.76954 | 0.770052 | 0.766776 | 0.777579 | 0.78126 | 0.764283 | 0.706153 |
| 0.745653 | 0.751149 | 0.750266 | 0.740627 | 0.742076 | 0.743114 | 0.772198 | 0.756026 | 0.680446 |
| 0.788856 | 0.777833 | 0.776173 | 0.778925 | 0.773043 | 0.772065 | 0.77192 | 0.769046 | 0.716216 |
| 0.768522 | 0.766533 | 0.766223 | 0.761167 | 0.762295 | 0.763303 | 0.758094 | 0.752536 | 0.706457 |
| 0.781176 | 0.781596 | 0.780669 | 0.779825 | 0.772899 | 0.775691 | 0.770366 | 0.76554 | 0.715005 |
| 0.780899 | 0.774263 | 0.77573 | 0.770651 | 0.770628 | 0.774798 | 0.788038 | 0.790682 | 0.71839 |
| 0.765364 | 0.754443 | 0.756609 | 0.753897 | 0.751152 | 0.770513 | 0.763443 | 0.742676 | 0.692653 |
| 0.774236 | 0.784805 | 0.777211 | 0.764994 | 0.764318 | 0.76482 | 0.789365 | 0.785586 | 0.716781 |
| 0.746802 | 0.750309 | 0.741763 | 0.754793 | 0.746577 | 0.748336 | 0.765498 | 0.757328 | 0.691497 |
| 0.779024 | 0.772311 | 0.770428 | 0.771218 | 0.762585 | 0.773848 | 0.771862 | 0.790085 | 0.718679 |
| 0.776305 | 0.770528 | 0.772071 | 0.766606 | 0.765634 | 0.770085 | 0.772908 | 0.791252 | 0.719376 |
| 0.748497 | 0.754363 | 0.754591 | 0.742634 | 0.744479 | 0.759793 | 0.764309 | 0.758277 | 0.685995 |
| 0.770367 | 0.742591 | 0.754197 | 0.750963 | 0.752346 | 0.757661 | 0.743387 | 0.73292 | 0.688086 |
| 0.762854 | 0.755483 | 0.762212 | 0.775645 | 0.755099 | 0.761322 | 0.755799 | 0.750647 | 0.688173 |
| 0.767695 | 0.760793 | 0.755338 | 0.763293 | 0.763491 | 0.757241 | 0.774335 | 0.761948 | 0.708084 |
| 0.776221 | 0.773826 | 0.777952 | 0.774839 | 0.773429 | 0.778195 | 0.774721 | 0.780224 | 0.717252 |
| 0.752265 | 0.748378 | 0.746554 | 0.744525 | 0.739141 | 0.744075 | 0.740187 | 0.735062 | 0.684009 |
| 0.788241 | 0.776729 | 0.776465 | 0.761266 | 0.782138 | 0.781258 | 0.774172 | 0.770611 | 0.721552 |
| 0.774507 | 0.767899 | 0.765959 | 0.761689 | 0.766963 | 0.765262 | 0.776002 | 0.772438 | 0.725362 |
| 0.783005 | 0.763711 | 0.76491 | 0.771912 | 0.767394 | 0.759697 | 0.753396 | 0.744098 | 0.693406 |
| 0.781466 | 0.767187 | 0.765821 | 0.770416 | 0.763404 | 0.758794 | 0.753827 | 0.745798 | 0.693471 |
| 0.771522 | 0.770338 | 0.760348 | 0.760574 | 0.761948 | 0.764319 | 0.790844 | 0.796917 | 0.721901 |
| 0.777509 | 0.764115 | 0.758079 | 0.757075 | 0.764 | 0.766486 | 0.776102 | 0.762152 | 0.718686 |
| 0.770874 | 0.773473 | 0.767039 | 0.758524 | 0.761193 | 0.778271 | 0.773501 | 0.768966 | 0.720763 |
| 0.727019 | 0.722853 | 0.719785 | 0.722663 | 0.718772 | 0.729329 | 0.726349 | 0.717916 | 0.670146 |
| 0.76163 | 0.75887 | 0.755054 | 0.748883 | 0.748787 | 0.755911 | 0.7552 | 0.757972 | 0.720205 |
| 0.728561 | 0.72892 | 0.725016 | 0.727929 | 0.721541 | 0.729147 | 0.727711 | 0.72217 | 0.672748 |
| 0.726506 | 0.724407 | 0.720352 | 0.720022 | 0.714663 | 0.730685 | 0.714464 | 0.71264 | 0.669055 |
| 0.782214 | 0.772472 | 0.772429 | 0.767026 | 0.767713 | 0.774348 | 0.770367 | 0.758479 | 0.717229 |
| 0.721764 | 0.722395 | 0.716903 | 0.716656 | 0.712066 | 0.723424 | 0.710083 | 0.707691 | 0.66743 |
| 0.73701 | 0.726885 | 0.728548 | 0.729542 | 0.718392 | 0.72896 | 0.715072 | 0.702201 | 0.658432 |
| 0.724514 | 0.714916 | 0.718869 | 0.714263 | 0.712014 | 0.723117 | 0.703599 | 0.701238 | 0.65397 |
| 0.780634 | 0.777869 | 0.770599 | 0.774011 | 0.787334 | 0.772388 | 0.763094 | 0.757231 | 0.725913 |
| 0.72471 | 0.718694 | 0.727358 | 0.714896 | 0.717692 | 0.724865 | 0.699908 | 0.69853 | 0.658715 |
| 0.781469 | 0.768108 | 0.782417 | 0.765635 | 0.763674 | 0.774867 | 0.768902 | 0.754417 | 0.718878 |

| | | | | | | | | |
|---|---|---|---|---|---|---|---|---|
| 0.765954 | 0.774312 | 0.769375 | 0.768621 | 0.767552 | 0.765556 | 0.756173 | 0.755774 | 0.7279 |
| 0.774483 | 0.769086 | 0.774257 | 0.768052 | 0.768218 | 0.775951 | 0.767412 | 0.756001 | 0.723792 |
| 0.756732 | 0.751122 | 0.750288 | 0.743665 | 0.749242 | 0.748033 | 0.753337 | 0.765978 | 0.743296 |
| 0.724651 | 0.705583 | 0.712221 | 0.709129 | 0.70577 | 0.709589 | 0.700213 | 0.691221 | 0.649242 |
| 0.71132 | 0.704777 | 0.712684 | 0.695174 | 0.70597 | 0.703864 | 0.693478 | 0.688691 | 0.65173 |
| 0.78032 | 0.779757 | 0.76868 | 0.779412 | 0.777137 | 0.767764 | 0.762527 | 0.762664 | 0.728678 |
| 0.694506 | 0.692577 | 0.690376 | 0.6899 | 0.685201 | 0.690782 | 0.688506 | 0.682456 | 0.652541 |
| 0.735918 | 0.738398 | 0.73329 | 0.72594 | 0.731352 | 0.731126 | 0.738697 | 0.740211 | 0.732549 |
| 0.776636 | 0.767011 | 0.777123 | 0.776517 | 0.766716 | 0.764802 | 0.763566 | 0.755987 | 0.73217 |
| 0.72077 | 0.72175 | 0.722231 | 0.714089 | 0.715627 | 0.720039 | 0.725985 | 0.733274 | 0.732156 |
| 0.758663 | 0.762054 | 0.754015 | 0.750821 | 0.756675 | 0.75555 | 0.753014 | 0.741002 | 0.731126 |
| 0.612903 | 0.607595 | 0.607595 | 0.613924 | 0.580247 | 0.634731 | 0.608696 | 0.647799 | 0.675497 |
| 0.657885 | 0.653234 | 0.644573 | 0.64508 | 0.653876 | 0.661272 | 0.662217 | 0.67485 | 0.722078 |
| 0.652365 | 0.659263 | 0.654121 | 0.651771 | 0.652517 | 0.652655 | 0.662306 | 0.675713 | 0.71968 |
| 0.666084 | 0.668212 | 0.662782 | 0.660309 | 0.661574 | 0.660632 | 0.673348 | 0.682346 | 0.724102 |
| 0.663204 | 0.66395 | 0.655704 | 0.658355 | 0.663082 | 0.658577 | 0.670899 | 0.675978 | 0.733252 |
| 0.622401 | 0.621642 | 0.617254 | 0.613697 | 0.614254 | 0.618605 | 0.615545 | 0.618892 | 0.589844 |
| 0.664484 | 0.665746 | 0.660692 | 0.659987 | 0.661761 | 0.665297 | 0.673341 | 0.686198 | 0.72974 |
| 0.745666 | 0.749673 | 0.746326 | 0.739706 | 0.742988 | 0.743505 | 0.744981 | 0.740015 | 0.734701 |
| 0.755342 | 0.74515 | 0.737652 | 0.74281 | 0.743899 | 0.739013 | 0.74124 | 0.737649 | 0.73655 |
| 0.753586 | 0.752032 | 0.74868 | 0.743737 | 0.75247 | 0.754611 | 0.752986 | 0.744927 | 0.739527 |
| 0.623914 | 0.618827 | 0.618632 | 0.615905 | 0.618193 | 0.619888 | 0.614386 | 0.611883 | 0.593591 |
| 0.604142 | 0.601064 | 0.59984 | 0.596455 | 0.597769 | 0.600348 | 0.59497 | 0.596304 | 0.579581 |
| 0.708491 | 0.714297 | 0.706417 | 0.699984 | 0.702849 | 0.704707 | 0.712423 | 0.721912 | 0.76007 |
| 0.709554 | 0.712785 | 0.706974 | 0.700119 | 0.70358 | 0.704533 | 0.712233 | 0.719551 | 0.76324 |
| 0.704751 | 0.705562 | 0.698367 | 0.696549 | 0.698709 | 0.696027 | 0.707777 | 0.711653 | 0.753191 |
| 0.705724 | 0.70949 | 0.704032 | 0.69865 | 0.701303 | 0.703996 | 0.712219 | 0.715818 | 0.756497 |
| 0.70948 | 0.713376 | 0.704573 | 0.702617 | 0.703833 | 0.708422 | 0.714331 | 0.721112 | 0.759717 |
| 0.706345 | 0.710927 | 0.705578 | 0.699068 | 0.703208 | 0.705953 | 0.716544 | 0.724806 | 0.767942 |
| 0.708496 | 0.707123 | 0.704873 | 0.698547 | 0.705487 | 0.707435 | 0.713279 | 0.722432 | 0.765154 |
| 0.709224 | 0.712117 | 0.705129 | 0.702126 | 0.705294 | 0.705693 | 0.709877 | 0.719746 | 0.751457 |
| 0.716242 | 0.720659 | 0.712501 | 0.709375 | 0.710861 | 0.715804 | 0.724673 | 0.72684 | 0.769794 |
| 0.711358 | 0.71582 | 0.709672 | 0.703722 | 0.706611 | 0.710806 | 0.718558 | 0.722854 | 0.757959 |
| 0.71486 | 0.716501 | 0.710261 | 0.70831 | 0.709193 | 0.714128 | 0.720603 | 0.726675 | 0.764586 |
| 0.716935 | 0.719329 | 0.712385 | 0.705671 | 0.708519 | 0.713268 | 0.723078 | 0.72662 | 0.76932 |
| 0.708283 | 0.714079 | 0.70418 | 0.705009 | 0.703736 | 0.709752 | 0.71486 | 0.722672 | 0.760828 |
| 0.712166 | 0.714583 | 0.708115 | 0.707006 | 0.704762 | 0.708654 | 0.716236 | 0.723713 | 0.760046 |
| 0.713605 | 0.712664 | 0.707122 | 0.705812 | 0.711549 | 0.71178 | 0.718995 | 0.720481 | 0.762571 |
| 0.704621 | 0.713757 | 0.706018 | 0.698249 | 0.701325 | 0.703935 | 0.712783 | 0.721658 | 0.763698 |
| 0.712645 | 0.718571 | 0.715865 | 0.702662 | 0.707864 | 0.710244 | 0.71908 | 0.721931 | 0.758745 |
| 0.655045 | 0.663055 | 0.650397 | 0.654809 | 0.654146 | 0.649269 | 0.6652 | 0.673281 | 0.716693 |
| 0.659629 | 0.66169 | 0.656619 | 0.651677 | 0.657842 | 0.658545 | 0.669641 | 0.680743 | 0.725649 |
| 0.713766 | 0.71432 | 0.712587 | 0.707872 | 0.710114 | 0.712888 | 0.721779 | 0.727352 | 0.761698 |
| 0.714809 | 0.715946 | 0.709396 | 0.708506 | 0.708606 | 0.712625 | 0.723693 | 0.72832 | 0.76951 |
| 0.64238 | 0.648124 | 0.644118 | 0.632105 | 0.636464 | 0.639678 | 0.653667 | 0.66333 | 0.717949 |
| 0.714741 | 0.718292 | 0.711463 | 0.706216 | 0.708536 | 0.710361 | 0.722634 | 0.725859 | 0.763783 |
| 0.702839 | 0.705664 | 0.698149 | 0.694273 | 0.698307 | 0.699623 | 0.710011 | 0.713178 | 0.756175 |

| | | | | | | | | |
|---|---|---|---|---|---|---|---|---|
| 0.710748 | 0.7095 | 0.70616 | 0.700717 | 0.703801 | 0.708412 | 0.717936 | 0.724358 | 0.761885 |
| 0.712627 | 0.716107 | 0.709665 | 0.703413 | 0.707174 | 0.711349 | 0.719605 | 0.725167 | 0.765561 |
| 0.708435 | 0.716294 | 0.705099 | 0.70439 | 0.707391 | 0.710258 | 0.718012 | 0.720143 | 0.758862 |
| 0.713194 | 0.714183 | 0.708253 | 0.702731 | 0.705031 | 0.711842 | 0.722294 | 0.727897 | 0.76565 |
| 0.716184 | 0.720645 | 0.710341 | 0.707507 | 0.713408 | 0.716371 | 0.72515 | 0.727527 | 0.765229 |
| 0.708103 | 0.710463 | 0.706689 | 0.701202 | 0.703555 | 0.707067 | 0.715526 | 0.723245 | 0.764321 |
| 0.710493 | 0.715023 | 0.703203 | 0.700728 | 0.705188 | 0.70676 | 0.715514 | 0.723367 | 0.761039 |
| 0.705193 | 0.706441 | 0.706107 | 0.696065 | 0.701241 | 0.702905 | 0.712607 | 0.71668 | 0.751038 |
| 0.710794 | 0.711955 | 0.703269 | 0.700851 | 0.703125 | 0.708779 | 0.718747 | 0.721355 | 0.758425 |
| 0.709194 | 0.711259 | 0.706005 | 0.703051 | 0.708263 | 0.707013 | 0.714618 | 0.719158 | 0.756218 |
| 0.713549 | 0.715829 | 0.710007 | 0.703724 | 0.705477 | 0.71086 | 0.721769 | 0.723374 | 0.763203 |
| 0.707163 | 0.709273 | 0.702262 | 0.701371 | 0.70214 | 0.704185 | 0.713463 | 0.720108 | 0.760563 |
| 0.699664 | 0.705674 | 0.699199 | 0.693506 | 0.70046 | 0.698755 | 0.70799 | 0.712757 | 0.751997 |
| 0.700828 | 0.701763 | 0.697529 | 0.694523 | 0.696457 | 0.699597 | 0.710092 | 0.716631 | 0.753907 |
| 0.713755 | 0.719758 | 0.711845 | 0.706386 | 0.710714 | 0.716696 | 0.718521 | 0.728161 | 0.760275 |
| 0.71245 | 0.714706 | 0.706692 | 0.704272 | 0.70407 | 0.708593 | 0.715391 | 0.722505 | 0.764295 |
| 0.703868 | 0.707847 | 0.703056 | 0.698648 | 0.700412 | 0.703235 | 0.712964 | 0.71779 | 0.765148 |
| 0.707269 | 0.709842 | 0.705807 | 0.700224 | 0.702724 | 0.704342 | 0.714744 | 0.720118 | 0.761727 |
| 0.708581 | 0.711528 | 0.702341 | 0.698508 | 0.703945 | 0.705957 | 0.71367 | 0.719831 | 0.760988 |
| 0.706466 | 0.710865 | 0.70152 | 0.700631 | 0.702043 | 0.704346 | 0.715032 | 0.720772 | 0.762165 |
| 0.706712 | 0.710173 | 0.702317 | 0.699017 | 0.697999 | 0.701675 | 0.710597 | 0.716365 | 0.758656 |
| 0.708816 | 0.70935 | 0.703079 | 0.699895 | 0.705806 | 0.70237 | 0.709394 | 0.720625 | 0.758543 |
| 0.712885 | 0.717315 | 0.708018 | 0.702414 | 0.705091 | 0.708301 | 0.716684 | 0.723257 | 0.763803 |
| 0.71006 | 0.713258 | 0.708153 | 0.700686 | 0.707385 | 0.705985 | 0.717536 | 0.722453 | 0.762313 |
| 0.709831 | 0.714598 | 0.704949 | 0.701623 | 0.70083 | 0.706429 | 0.715514 | 0.72394 | 0.762407 |
| 0.703556 | 0.708919 | 0.700684 | 0.698941 | 0.700875 | 0.704187 | 0.713572 | 0.717965 | 0.762523 |
| 0.704567 | 0.709651 | 0.70304 | 0.700494 | 0.700654 | 0.700048 | 0.712298 | 0.720058 | 0.755163 |
| 0.705582 | 0.706516 | 0.703392 | 0.697327 | 0.697999 | 0.703003 | 0.712595 | 0.716909 | 0.755755 |
| 0.70606 | 0.711839 | 0.70504 | 0.701958 | 0.699849 | 0.705245 | 0.713077 | 0.722728 | 0.761629 |
| 0.710432 | 0.711638 | 0.705831 | 0.702569 | 0.703598 | 0.706903 | 0.716547 | 0.721319 | 0.763293 |
| 0.712535 | 0.714914 | 0.708878 | 0.701761 | 0.704625 | 0.708627 | 0.716595 | 0.722817 | 0.760769 |
| 0.712174 | 0.713607 | 0.712881 | 0.707569 | 0.707054 | 0.707226 | 0.716572 | 0.72298 | 0.760734 |
| 0.714906 | 0.716288 | 0.711433 | 0.704006 | 0.709317 | 0.70867 | 0.718891 | 0.724943 | 0.768156 |
| 0.714388 | 0.717398 | 0.714205 | 0.70599 | 0.710717 | 0.711109 | 0.720766 | 0.72408 | 0.767096 |
| 0.706727 | 0.710169 | 0.705363 | 0.698378 | 0.701596 | 0.704852 | 0.711449 | 0.716959 | 0.761001 |
| 0.708068 | 0.711777 | 0.704049 | 0.701514 | 0.70176 | 0.705552 | 0.71417 | 0.720829 | 0.762688 |
| 0.71238 | 0.711049 | 0.707599 | 0.701904 | 0.704642 | 0.708104 | 0.713331 | 0.722312 | 0.76162 |
| 0.707125 | 0.711048 | 0.701008 | 0.698728 | 0.700177 | 0.704103 | 0.712207 | 0.720926 | 0.757947 |
| 0.701113 | 0.703575 | 0.699439 | 0.695522 | 0.694133 | 0.698106 | 0.708299 | 0.71645 | 0.756516 |
| 0.700667 | 0.703911 | 0.696484 | 0.691307 | 0.694741 | 0.697807 | 0.705011 | 0.712302 | 0.752634 |
| 0.699877 | 0.700279 | 0.693997 | 0.693037 | 0.691652 | 0.697925 | 0.706679 | 0.71218 | 0.749596 |
| 0.699189 | 0.702149 | 0.698248 | 0.690185 | 0.696171 | 0.695539 | 0.708231 | 0.709571 | 0.750852 |
| 0.703497 | 0.706767 | 0.699322 | 0.696008 | 0.697607 | 0.699992 | 0.707303 | 0.715252 | 0.754826 |
| 0.695379 | 0.699764 | 0.688353 | 0.688237 | 0.69294 | 0.692141 | 0.700218 | 0.711468 | 0.750904 |
| 0.707362 | 0.708445 | 0.702054 | 0.697704 | 0.700588 | 0.701045 | 0.709942 | 0.718412 | 0.754321 |
| 0.700238 | 0.701289 | 0.698426 | 0.691674 | 0.695449 | 0.693861 | 0.704908 | 0.712466 | 0.746204 |

| | | | | | | | | |
|---|---|---|---|---|---|---|---|---|
| 0.711017 | 0.716334 | 0.706038 | 0.704837 | 0.70553 | 0.709629 | 0.720463 | 0.72504 | 0.762379 |
| 0.711365 | 0.715199 | 0.707709 | 0.703259 | 0.703984 | 0.704576 | 0.718057 | 0.725154 | 0.772422 |
| 0.707613 | 0.709994 | 0.706417 | 0.698445 | 0.703226 | 0.705572 | 0.71344 | 0.720071 | 0.770482 |
| 0.711 | 0.712606 | 0.70921 | 0.703344 | 0.705351 | 0.708546 | 0.717761 | 0.724647 | 0.766615 |
| 0.718349 | 0.720125 | 0.715879 | 0.714377 | 0.712883 | 0.713278 | 0.724614 | 0.731748 | 0.780514 |
| 0.712698 | 0.716709 | 0.713999 | 0.70255 | 0.706797 | 0.709515 | 0.717687 | 0.729481 | 0.766186 |
| 0.707509 | 0.711642 | 0.704814 | 0.702105 | 0.703172 | 0.706337 | 0.715797 | 0.720123 | 0.762268 |
| 0.706538 | 0.715002 | 0.711203 | 0.702593 | 0.704672 | 0.709968 | 0.717388 | 0.725449 | 0.763114 |
| 0.701677 | 0.710377 | 0.701109 | 0.696744 | 0.699564 | 0.69756 | 0.708253 | 0.720084 | 0.761085 |
| 0.71235 | 0.71295 | 0.707006 | 0.704649 | 0.706536 | 0.70713 | 0.718612 | 0.724729 | 0.768745 |
| 0.712564 | 0.716772 | 0.712842 | 0.706422 | 0.710958 | 0.711118 | 0.719219 | 0.725499 | 0.770127 |
| 0.713706 | 0.716258 | 0.713208 | 0.704562 | 0.70934 | 0.709833 | 0.717988 | 0.727811 | 0.76796 |
| 0.711383 | 0.714888 | 0.703695 | 0.702916 | 0.704629 | 0.705526 | 0.7169 | 0.722718 | 0.780443 |
| 0.714331 | 0.71316 | 0.708517 | 0.704466 | 0.705967 | 0.709877 | 0.715084 | 0.724493 | 0.778257 |
| 0.715088 | 0.716573 | 0.714889 | 0.705133 | 0.708729 | 0.711127 | 0.7168 | 0.725022 | 0.785549 |
| 0.712686 | 0.716015 | 0.709087 | 0.708111 | 0.709536 | 0.707622 | 0.718293 | 0.723399 | 0.78017 |
| 0.716679 | 0.715606 | 0.710489 | 0.712088 | 0.712409 | 0.714767 | 0.720149 | 0.729091 | 0.788907 |
| 0.713882 | 0.71331 | 0.714422 | 0.707552 | 0.709888 | 0.710934 | 0.716585 | 0.724429 | 0.782348 |
| 0.716456 | 0.715525 | 0.712137 | 0.710957 | 0.709958 | 0.716636 | 0.721862 | 0.727403 | 0.782461 |
| 0.712671 | 0.714807 | 0.707674 | 0.704536 | 0.705789 | 0.708193 | 0.716537 | 0.724833 | 0.786241 |
| 0.710547 | 0.713954 | 0.703612 | 0.705387 | 0.706028 | 0.711805 | 0.715904 | 0.725885 | 0.775843 |
| 0.712386 | 0.713369 | 0.704067 | 0.702196 | 0.707141 | 0.706579 | 0.714274 | 0.724799 | 0.774301 |
| 0.711942 | 0.718926 | 0.711258 | 0.705262 | 0.707891 | 0.711434 | 0.718969 | 0.72754 | 0.778999 |
| 0.713812 | 0.714173 | 0.711684 | 0.705505 | 0.709121 | 0.71469 | 0.722408 | 0.726912 | 0.776422 |
| 0.711988 | 0.713309 | 0.710514 | 0.701052 | 0.706854 | 0.710719 | 0.716327 | 0.725107 | 0.776066 |
| 0.711053 | 0.714002 | 0.708627 | 0.706371 | 0.705639 | 0.706763 | 0.715495 | 0.725493 | 0.771095 |
| 0.713551 | 0.711552 | 0.705966 | 0.701589 | 0.706133 | 0.704691 | 0.717431 | 0.723156 | 0.768573 |
| 0.710394 | 0.710869 | 0.707013 | 0.701161 | 0.703946 | 0.70533 | 0.717175 | 0.723484 | 0.773728 |
| 0.713677 | 0.716989 | 0.708595 | 0.705153 | 0.705645 | 0.713926 | 0.720827 | 0.72217 | 0.776559 |
| 0.716769 | 0.719569 | 0.711223 | 0.707442 | 0.70995 | 0.714193 | 0.718451 | 0.725883 | 0.769852 |
| 0.708646 | 0.713535 | 0.70721 | 0.703395 | 0.703644 | 0.705977 | 0.71439 | 0.722915 | 0.764474 |
| 0.717827 | 0.717638 | 0.71143 | 0.707437 | 0.71193 | 0.712442 | 0.721657 | 0.726431 | 0.769928 |
| 0.714989 | 0.712894 | 0.713167 | 0.704704 | 0.708569 | 0.711006 | 0.72068 | 0.727345 | 0.773434 |
| 0.707958 | 0.711466 | 0.702881 | 0.697619 | 0.70045 | 0.70517 | 0.717084 | 0.720869 | 0.772588 |
| 0.711721 | 0.712271 | 0.709675 | 0.70457 | 0.705477 | 0.711893 | 0.718484 | 0.724634 | 0.768701 |
| 0.709197 | 0.710891 | 0.707603 | 0.698286 | 0.703392 | 0.703538 | 0.712529 | 0.725831 | 0.771297 |
| 0.718601 | 0.719873 | 0.717384 | 0.710002 | 0.71424 | 0.716295 | 0.722382 | 0.732507 | 0.779041 |
| 0.710364 | 0.716281 | 0.708866 | 0.699733 | 0.704372 | 0.708343 | 0.717738 | 0.72487 | 0.770546 |
| 0.715948 | 0.713693 | 0.706343 | 0.705117 | 0.708659 | 0.708551 | 0.722883 | 0.725681 | 0.768652 |
| 0.711946 | 0.711968 | 0.707558 | 0.705495 | 0.708615 | 0.709542 | 0.719902 | 0.724902 | 0.769633 |
| 0.718069 | 0.714184 | 0.712384 | 0.708456 | 0.7129 | 0.713197 | 0.720854 | 0.73061 | 0.77043 |
| 0.716936 | 0.718501 | 0.713438 | 0.705457 | 0.709731 | 0.711925 | 0.722155 | 0.725503 | 0.771689 |
| 0.718202 | 0.71709 | 0.711389 | 0.706124 | 0.709963 | 0.712583 | 0.724259 | 0.729685 | 0.775699 |
| 0.710247 | 0.712802 | 0.710425 | 0.701632 | 0.708188 | 0.707833 | 0.715498 | 0.721667 | 0.773877 |
| 0.715121 | 0.711623 | 0.71134 | 0.705099 | 0.707038 | 0.714252 | 0.719452 | 0.724777 | 0.776666 |
| 0.710737 | 0.710168 | 0.70815 | 0.699587 | 0.70323 | 0.705656 | 0.718465 | 0.722173 | 0.770388 |

| | | | | | | | | |
|---|---|---|---|---|---|---|---|---|
| 0.692644 | 0.700046 | 0.691764 | 0.688678 | 0.692082 | 0.695921 | 0.706185 | 0.715047 | 0.769165 |
| 0.749836 | 0.74881 | 0.74604 | 0.743393 | 0.744927 | 0.747302 | 0.76537 | 0.766562 | 0.749959 |
| 0.714204 | 0.712196 | 0.707522 | 0.699257 | 0.704696 | 0.705944 | 0.715342 | 0.721731 | 0.764063 |
| 0.714511 | 0.713315 | 0.711366 | 0.704439 | 0.710099 | 0.709688 | 0.719376 | 0.721679 | 0.773702 |
| 0.707333 | 0.70768 | 0.699561 | 0.69803 | 0.699637 | 0.704193 | 0.712777 | 0.717649 | 0.763666 |
| 0.712701 | 0.709184 | 0.709361 | 0.701351 | 0.703781 | 0.706235 | 0.717419 | 0.724915 | 0.769334 |
| 0.709747 | 0.708354 | 0.702913 | 0.698458 | 0.702092 | 0.703743 | 0.715916 | 0.720204 | 0.767815 |
| 0.705524 | 0.706963 | 0.69818 | 0.694168 | 0.697868 | 0.699474 | 0.705508 | 0.715329 | 0.763306 |
| 0.710682 | 0.713618 | 0.70768 | 0.699019 | 0.707095 | 0.706903 | 0.716677 | 0.725177 | 0.772173 |
| 0.720477 | 0.717754 | 0.715102 | 0.712886 | 0.71518 | 0.715342 | 0.721972 | 0.729634 | 0.772689 |
| 0.716561 | 0.713008 | 0.712249 | 0.704502 | 0.709434 | 0.71195 | 0.716864 | 0.72385 | 0.77595 |
| 0.716263 | 0.716909 | 0.7106 | 0.704757 | 0.709375 | 0.712209 | 0.719955 | 0.727967 | 0.770762 |
| 0.718082 | 0.717091 | 0.712922 | 0.71051 | 0.713166 | 0.71187 | 0.721937 | 0.731974 | 0.774515 |
| 0.716857 | 0.713969 | 0.711293 | 0.706004 | 0.70794 | 0.716067 | 0.720808 | 0.729444 | 0.771986 |
| 0.712258 | 0.711612 | 0.708785 | 0.703325 | 0.706336 | 0.705963 | 0.716355 | 0.723505 | 0.77303 |
| 0.713031 | 0.711743 | 0.707683 | 0.703002 | 0.705249 | 0.710688 | 0.716324 | 0.722388 | 0.769795 |
| 0.719331 | 0.713418 | 0.711111 | 0.704696 | 0.71036 | 0.714081 | 0.721475 | 0.727755 | 0.769702 |
| 0.716995 | 0.713872 | 0.708886 | 0.705604 | 0.710952 | 0.710124 | 0.716625 | 0.725501 | 0.766537 |
| 0.703138 | 0.705171 | 0.698838 | 0.694379 | 0.697904 | 0.698107 | 0.710972 | 0.716733 | 0.76578 |
| 0.706303 | 0.705739 | 0.704278 | 0.69984 | 0.703092 | 0.702671 | 0.710582 | 0.721659 | 0.771631 |
| 0.711308 | 0.712066 | 0.708728 | 0.704775 | 0.707862 | 0.70911 | 0.716019 | 0.723907 | 0.768349 |
| 0.701232 | 0.707543 | 0.69638 | 0.694124 | 0.694347 | 0.700244 | 0.710035 | 0.714145 | 0.760367 |
| 0.710729 | 0.712557 | 0.706709 | 0.702821 | 0.703356 | 0.708248 | 0.714913 | 0.720348 | 0.761187 |
| 0.712607 | 0.716794 | 0.708701 | 0.707251 | 0.708377 | 0.710061 | 0.719116 | 0.725056 | 0.764501 |
| 0.71368 | 0.719392 | 0.711142 | 0.703991 | 0.706563 | 0.71316 | 0.722894 | 0.727801 | 0.770877 |
| 0.710788 | 0.714518 | 0.708493 | 0.704957 | 0.703276 | 0.709012 | 0.721258 | 0.723942 | 0.763187 |
| 0.708998 | 0.706844 | 0.701702 | 0.7007 | 0.700675 | 0.705147 | 0.716012 | 0.718351 | 0.762185 |
| 0.706334 | 0.7067 | 0.701861 | 0.699917 | 0.700488 | 0.69776 | 0.713977 | 0.717348 | 0.761815 |
| 0.708488 | 0.704064 | 0.70236 | 0.697767 | 0.698293 | 0.7034 | 0.711873 | 0.717248 | 0.761087 |
| 0.772675 | 0.77863 | 0.783025 | 0.761577 | 0.763414 | 0.770064 | 0.758565 | 0.749365 | 0.693263 |
| 0.803048 | 0.800504 | 0.792243 | 0.771862 | 0.790791 | 0.792601 | 0.780455 | 0.767246 | 0.706774 |
| 0.781001 | 0.787062 | 0.793639 | 0.769801 | 0.777546 | 0.788857 | 0.773604 | 0.760481 | 0.703364 |
| 0.797813 | 0.78721 | 0.787865 | 0.801235 | 0.793333 | 0.779835 | 0.76024 | 0.748558 | 0.694813 |
| 0.799967 | 0.79614 | 0.793 | 0.777064 | 0.793272 | 0.792495 | 0.78062 | 0.764944 | 0.705107 |
| 0.704378 | 0.706037 | 0.701048 | 0.697169 | 0.700806 | 0.701214 | 0.712152 | 0.719528 | 0.765098 |
| 0.708199 | 0.709154 | 0.704872 | 0.701477 | 0.703047 | 0.70434 | 0.71147 | 0.722523 | 0.7729 |
| 0.706563 | 0.709532 | 0.704604 | 0.704074 | 0.706186 | 0.706164 | 0.716133 | 0.721133 | 0.770276 |
| 0.711483 | 0.713703 | 0.708943 | 0.703051 | 0.707802 | 0.708462 | 0.7201 | 0.725493 | 0.771902 |
| 0.707243 | 0.71026 | 0.701305 | 0.700313 | 0.702158 | 0.704096 | 0.713814 | 0.719589 | 0.764417 |
| 0.707438 | 0.71235 | 0.705703 | 0.703727 | 0.705543 | 0.70779 | 0.719368 | 0.723065 | 0.76868 |
| 0.707182 | 0.711845 | 0.707559 | 0.700185 | 0.70342 | 0.705892 | 0.712473 | 0.721533 | 0.768011 |
| 0.709544 | 0.712973 | 0.705164 | 0.701135 | 0.704118 | 0.706873 | 0.713936 | 0.722516 | 0.762245 |
| 0.703128 | 0.705392 | 0.699467 | 0.696903 | 0.697168 | 0.699232 | 0.707173 | 0.715527 | 0.761726 |
| 0.702765 | 0.701754 | 0.698392 | 0.694114 | 0.700575 | 0.700196 | 0.710116 | 0.716346 | 0.76278 |
| 0.706973 | 0.710989 | 0.701747 | 0.700647 | 0.705165 | 0.707093 | 0.715954 | 0.721674 | 0.767689 |
| 0.711335 | 0.714077 | 0.707281 | 0.704051 | 0.70836 | 0.705045 | 0.716465 | 0.723146 | 0.770123 |

| | | | | | | | | |
|---|---|---|---|---|---|---|---|---|
| 0.709381 | 0.712421 | 0.712099 | 0.705729 | 0.707162 | 0.707901 | 0.715925 | 0.725268 | 0.768267 |
| 0.711154 | 0.710967 | 0.707058 | 0.703299 | 0.704092 | 0.706466 | 0.713849 | 0.719311 | 0.766575 |
| 0.713282 | 0.71417 | 0.709478 | 0.706462 | 0.709028 | 0.711981 | 0.715926 | 0.723435 | 0.77251 |
| 0.707924 | 0.711711 | 0.708139 | 0.703855 | 0.705566 | 0.706845 | 0.713713 | 0.716598 | 0.767393 |
| 0.706596 | 0.70617 | 0.704793 | 0.70242 | 0.703988 | 0.698799 | 0.714333 | 0.71797 | 0.762937 |
| 0.70489 | 0.708804 | 0.701828 | 0.701363 | 0.699769 | 0.704068 | 0.712689 | 0.716465 | 0.764907 |
| 0.708901 | 0.714742 | 0.710182 | 0.705098 | 0.710485 | 0.70782 | 0.714239 | 0.724728 | 0.767292 |
| 0.711834 | 0.709622 | 0.708546 | 0.706929 | 0.707347 | 0.708732 | 0.719772 | 0.723985 | 0.769486 |
| 0.708388 | 0.709303 | 0.70241 | 0.700172 | 0.701412 | 0.708252 | 0.717035 | 0.720894 | 0.770225 |
| 0.712093 | 0.709787 | 0.709123 | 0.701328 | 0.704461 | 0.705723 | 0.714274 | 0.721441 | 0.767802 |
| 0.709612 | 0.708769 | 0.707341 | 0.700796 | 0.704049 | 0.705612 | 0.712925 | 0.718196 | 0.765057 |
| 0.706044 | 0.710652 | 0.705926 | 0.700162 | 0.706026 | 0.704617 | 0.712727 | 0.720504 | 0.766388 |
| 0.701567 | 0.702854 | 0.698881 | 0.693291 | 0.694653 | 0.698333 | 0.704927 | 0.712637 | 0.758627 |
| 0.700114 | 0.703143 | 0.699239 | 0.695553 | 0.695128 | 0.698313 | 0.708592 | 0.72056 | 0.759898 |
| 0.710839 | 0.71326 | 0.707731 | 0.706696 | 0.708129 | 0.708256 | 0.716803 | 0.720985 | 0.767452 |
| 0.713464 | 0.718793 | 0.714101 | 0.711356 | 0.715299 | 0.714877 | 0.726655 | 0.726388 | 0.780371 |
| 0.712438 | 0.717077 | 0.70936 | 0.706469 | 0.709484 | 0.705717 | 0.721115 | 0.722961 | 0.773763 |
| 0.715243 | 0.716214 | 0.710979 | 0.709251 | 0.713073 | 0.7123 | 0.720308 | 0.725809 | 0.777963 |
| 0.706139 | 0.708443 | 0.70693 | 0.703115 | 0.704822 | 0.707447 | 0.71348 | 0.720698 | 0.768173 |
| 0.707977 | 0.713341 | 0.709375 | 0.703391 | 0.704055 | 0.707706 | 0.715339 | 0.72066 | 0.770882 |
| 0.706267 | 0.708561 | 0.70602 | 0.701801 | 0.702853 | 0.702643 | 0.713573 | 0.720288 | 0.767322 |
| 0.712985 | 0.711836 | 0.707564 | 0.704454 | 0.704584 | 0.709193 | 0.71891 | 0.723397 | 0.76954 |
| 0.713435 | 0.717193 | 0.709977 | 0.705337 | 0.712593 | 0.706522 | 0.72231 | 0.721938 | 0.775307 |
| 0.708914 | 0.712158 | 0.705781 | 0.70233 | 0.706573 | 0.706781 | 0.711214 | 0.724704 | 0.762671 |
| 0.709293 | 0.711654 | 0.706217 | 0.701695 | 0.706632 | 0.70667 | 0.717307 | 0.720155 | 0.771112 |
| 0.804944 | 0.792392 | 0.80467 | 0.80471 | 0.789815 | 0.786442 | 0.771583 | 0.760244 | 0.703914 |
| 0.709644 | 0.708847 | 0.704677 | 0.702555 | 0.705482 | 0.709591 | 0.716574 | 0.72179 | 0.767934 |
| 0.710814 | 0.714683 | 0.709435 | 0.707971 | 0.70641 | 0.711444 | 0.719 | 0.726859 | 0.767419 |
| 0.715973 | 0.719619 | 0.712818 | 0.70831 | 0.707784 | 0.709798 | 0.721378 | 0.727419 | 0.77272 |
| 0.712346 | 0.712618 | 0.706412 | 0.702796 | 0.705661 | 0.712363 | 0.71708 | 0.726285 | 0.76767 |
| 0.707817 | 0.710192 | 0.703276 | 0.701089 | 0.701428 | 0.706131 | 0.714513 | 0.718411 | 0.760999 |
| 0.711082 | 0.708292 | 0.705118 | 0.700157 | 0.702819 | 0.705155 | 0.716093 | 0.720727 | 0.772583 |
| 0.710099 | 0.71225 | 0.708841 | 0.702588 | 0.704008 | 0.709471 | 0.71716 | 0.725192 | 0.766633 |
| 0.706702 | 0.713339 | 0.705205 | 0.702154 | 0.704413 | 0.704832 | 0.715848 | 0.724337 | 0.764284 |
| 0.704801 | 0.708295 | 0.700338 | 0.697204 | 0.702241 | 0.705311 | 0.710738 | 0.720932 | 0.757494 |
| 0.706246 | 0.707144 | 0.704512 | 0.699869 | 0.702147 | 0.704801 | 0.715714 | 0.721227 | 0.760051 |
| 0.713368 | 0.715332 | 0.711009 | 0.707303 | 0.705303 | 0.709873 | 0.720023 | 0.726433 | 0.772446 |
| 0.712488 | 0.715839 | 0.713568 | 0.704005 | 0.709542 | 0.711822 | 0.717683 | 0.725151 | 0.779997 |
| 0.712816 | 0.712847 | 0.711151 | 0.702369 | 0.706257 | 0.705995 | 0.714713 | 0.727052 | 0.774775 |
| 0.710275 | 0.718442 | 0.705273 | 0.704686 | 0.704095 | 0.705417 | 0.713414 | 0.720073 | 0.778347 |
| 0.705619 | 0.709675 | 0.704722 | 0.699325 | 0.705318 | 0.704944 | 0.709395 | 0.719773 | 0.778289 |
| 0.715086 | 0.715772 | 0.710359 | 0.70748 | 0.707776 | 0.709548 | 0.718604 | 0.725778 | 0.780759 |
| 0.70636 | 0.714763 | 0.70786 | 0.698678 | 0.70672 | 0.707525 | 0.714047 | 0.718885 | 0.775549 |
| 0.707991 | 0.713186 | 0.705292 | 0.699905 | 0.705901 | 0.703088 | 0.712004 | 0.72108 | 0.761924 |
| 0.712092 | 0.711769 | 0.7093 | 0.704137 | 0.70844 | 0.708655 | 0.71836 | 0.724243 | 0.769474 |
| 0.711534 | 0.714388 | 0.7107 | 0.704571 | 0.70827 | 0.709376 | 0.716918 | 0.722698 | 0.767897 |

| | | | | | | | | |
|---|---|---|---|---|---|---|---|---|
| 0.704554 | 0.711051 | 0.702924 | 0.700736 | 0.703071 | 0.704918 | 0.714172 | 0.718237 | 0.766623 |
| 0.711241 | 0.712847 | 0.709888 | 0.701754 | 0.706958 | 0.707884 | 0.719287 | 0.723257 | 0.76699 |
| 0.709652 | 0.713021 | 0.705811 | 0.703119 | 0.702012 | 0.705148 | 0.715611 | 0.720848 | 0.762504 |
| 0.672498 | 0.668171 | 0.66016 | 0.661982 | 0.661437 | 0.655483 | 0.67993 | 0.685806 | 0.719929 |
| 0.706012 | 0.71037 | 0.704089 | 0.697003 | 0.70205 | 0.70364 | 0.714104 | 0.720559 | 0.768554 |
| 0.709162 | 0.712224 | 0.702567 | 0.700487 | 0.705464 | 0.706175 | 0.715236 | 0.718401 | 0.768823 |
| 0.709718 | 0.712287 | 0.707557 | 0.703356 | 0.705592 | 0.70446 | 0.719763 | 0.725803 | 0.766314 |
| 0.710257 | 0.708858 | 0.707229 | 0.701272 | 0.704257 | 0.706267 | 0.714968 | 0.723452 | 0.761858 |
| 0.714682 | 0.715797 | 0.711595 | 0.70591 | 0.708131 | 0.712108 | 0.720746 | 0.726561 | 0.773017 |
| 0.713604 | 0.711702 | 0.709205 | 0.705203 | 0.707938 | 0.70914 | 0.71521 | 0.724646 | 0.767842 |
| 0.711283 | 0.711471 | 0.709677 | 0.707676 | 0.705882 | 0.711881 | 0.715248 | 0.722586 | 0.766098 |
| 0.715587 | 0.712779 | 0.710235 | 0.711885 | 0.707216 | 0.709844 | 0.71881 | 0.724584 | 0.767695 |
| 0.713945 | 0.717204 | 0.710404 | 0.71193 | 0.709742 | 0.713448 | 0.717327 | 0.725464 | 0.767773 |
| 0.710343 | 0.713006 | 0.705162 | 0.702949 | 0.703625 | 0.704774 | 0.713376 | 0.724896 | 0.772494 |
| 0.713865 | 0.71506 | 0.709858 | 0.707565 | 0.708975 | 0.7094 | 0.720365 | 0.729725 | 0.7689 |
| 0.706706 | 0.712746 | 0.702796 | 0.703125 | 0.704719 | 0.703295 | 0.713429 | 0.721064 | 0.768969 |
| 0.711186 | 0.709771 | 0.710516 | 0.703894 | 0.708936 | 0.710423 | 0.720768 | 0.725676 | 0.767668 |
| 0.709956 | 0.715924 | 0.707705 | 0.704354 | 0.705854 | 0.706688 | 0.71867 | 0.725056 | 0.775498 |
| 0.707358 | 0.708356 | 0.702649 | 0.69973 | 0.699612 | 0.703497 | 0.715246 | 0.721304 | 0.763325 |
| 0.709578 | 0.714399 | 0.704683 | 0.701525 | 0.70703 | 0.705873 | 0.714696 | 0.721748 | 0.767651 |
| 0.713389 | 0.713603 | 0.704854 | 0.706154 | 0.707527 | 0.708041 | 0.720571 | 0.725421 | 0.772905 |
| 0.712032 | 0.714036 | 0.707605 | 0.705037 | 0.70721 | 0.709532 | 0.721431 | 0.729438 | 0.770198 |
| 0.698089 | 0.699872 | 0.694922 | 0.690006 | 0.68294 | 0.695779 | 0.694561 | 0.695676 | 0.674791 |
| 0.732986 | 0.719242 | 0.719303 | 0.72004 | 0.721159 | 0.72175 | 0.713902 | 0.728589 | 0.657476 |
| 0.762742 | 0.751344 | 0.760722 | 0.748083 | 0.749269 | 0.76153 | 0.744857 | 0.738301 | 0.669913 |
| 0.775201 | 0.76087 | 0.766829 | 0.770888 | 0.767985 | 0.743301 | 0.727319 | 0.716675 | 0.650484 |
| 0.665896 | 0.66287 | 0.655854 | 0.66079 | 0.663665 | 0.665995 | 0.675632 | 0.684614 | 0.734601 |
| 0.659041 | 0.662528 | 0.658405 | 0.651366 | 0.653363 | 0.65431 | 0.669941 | 0.676247 | 0.721594 |
| 0.659012 | 0.662385 | 0.657307 | 0.655106 | 0.660456 | 0.658726 | 0.671312 | 0.675182 | 0.729019 |
| 0.6739 | 0.674203 | 0.667201 | 0.671675 | 0.665462 | 0.67482 | 0.689366 | 0.695095 | 0.735539 |
| 0.656174 | 0.65483 | 0.648971 | 0.643 | 0.647862 | 0.642142 | 0.658365 | 0.672456 | 0.7168 |
| 0.641979 | 0.640845 | 0.635501 | 0.635767 | 0.635208 | 0.642395 | 0.654098 | 0.665027 | 0.714326 |
| 0.64306 | 0.649631 | 0.631817 | 0.632486 | 0.633644 | 0.639201 | 0.657012 | 0.655309 | 0.718137 |
| 0.728271 | 0.735092 | 0.727087 | 0.730123 | 0.717283 | 0.729969 | 0.737632 | 0.714624 | 0.658593 |
| 0.70499 | 0.705153 | 0.701114 | 0.700434 | 0.69844 | 0.698364 | 0.707863 | 0.716874 | 0.759762 |
| 0.67982 | 0.673937 | 0.673064 | 0.669334 | 0.674185 | 0.671419 | 0.68726 | 0.691243 | 0.743082 |
| 0.666493 | 0.671013 | 0.666754 | 0.665049 | 0.664321 | 0.663871 | 0.678132 | 0.683709 | 0.72385 |
| 0.646074 | 0.651864 | 0.64833 | 0.648619 | 0.643138 | 0.652652 | 0.662657 | 0.666904 | 0.719985 |
| 0.676946 | 0.678326 | 0.667641 | 0.664972 | 0.67235 | 0.66957 | 0.684828 | 0.694209 | 0.736479 |
| 0.668196 | 0.670847 | 0.66236 | 0.659582 | 0.661145 | 0.666175 | 0.677659 | 0.682427 | 0.7254 |
| 0.660169 | 0.663638 | 0.654018 | 0.649371 | 0.652388 | 0.6547 | 0.671495 | 0.678368 | 0.730874 |
| 0.66305 | 0.663803 | 0.651187 | 0.654675 | 0.660257 | 0.661213 | 0.6708 | 0.674436 | 0.730485 |
| 0.657163 | 0.65681 | 0.651963 | 0.645724 | 0.657312 | 0.646928 | 0.663123 | 0.665514 | 0.719863 |
| 0.65798 | 0.655896 | 0.654088 | 0.647227 | 0.653715 | 0.658657 | 0.666348 | 0.67804 | 0.715938 |
| 0.654074 | 0.658391 | 0.657292 | 0.641975 | 0.647232 | 0.64524 | 0.666998 | 0.673827 | 0.711982 |
| 0.642789 | 0.65619 | 0.63486 | 0.63648 | 0.636827 | 0.628241 | 0.647397 | 0.654188 | 0.712969 |

| | | | | | | | | |
|---|---|---|---|---|---|---|---|---|
| 0.690577 | 0.69396 | 0.685249 | 0.685321 | 0.685231 | 0.692764 | 0.701224 | 0.708538 | 0.753819 |
| 0.660664 | 0.661978 | 0.653603 | 0.650434 | 0.652637 | 0.656006 | 0.66961 | 0.677778 | 0.724268 |
| 0.658537 | 0.662866 | 0.656544 | 0.655585 | 0.659242 | 0.660133 | 0.675983 | 0.680318 | 0.718881 |
| 0.663155 | 0.663366 | 0.660296 | 0.655551 | 0.661993 | 0.662269 | 0.678497 | 0.677443 | 0.72973 |
| 0.66088 | 0.663237 | 0.656657 | 0.651034 | 0.650508 | 0.650291 | 0.667059 | 0.667383 | 0.725655 |
| 0.66287 | 0.667392 | 0.661434 | 0.657607 | 0.65961 | 0.650651 | 0.668438 | 0.671898 | 0.722126 |
| 0.651786 | 0.657746 | 0.647677 | 0.644684 | 0.649895 | 0.652507 | 0.663545 | 0.674071 | 0.721395 |
| 0.638738 | 0.64958 | 0.64828 | 0.635194 | 0.636091 | 0.641033 | 0.650815 | 0.662677 | 0.711722 |
| 0.658739 | 0.650698 | 0.643948 | 0.644365 | 0.636758 | 0.639106 | 0.65521 | 0.671331 | 0.720236 |
| 0.695462 | 0.6983 | 0.687298 | 0.68528 | 0.685939 | 0.690353 | 0.70146 | 0.705654 | 0.75431 |
| 0.693615 | 0.692338 | 0.68457 | 0.683845 | 0.683437 | 0.68715 | 0.698534 | 0.704883 | 0.749823 |
| 0.68124 | 0.689899 | 0.678387 | 0.678318 | 0.676219 | 0.685748 | 0.690002 | 0.695136 | 0.748441 |
| 0.679354 | 0.680724 | 0.672509 | 0.67122 | 0.673072 | 0.672708 | 0.681712 | 0.692407 | 0.739003 |
| 0.681276 | 0.683807 | 0.674793 | 0.675544 | 0.677251 | 0.677537 | 0.690993 | 0.69833 | 0.742885 |
| 0.671836 | 0.682458 | 0.667897 | 0.668491 | 0.66892 | 0.666549 | 0.682521 | 0.692895 | 0.740654 |

| HG_20 | | HG_21 | | | | | | |
|---|---|---|---|---|---|---|---|---|
| Home_garc | Home_garc | Home_garc | Home_garc | Home_garc | Home_gar | Collection_ | Collection_ | Collection_ |
| 0.711378 | 0.710343 | 0.773037 | 0.774284 | 0.784586 | 0.773513 | 0.805373 | 0.796923 | 0.806179 |
| 0.718254 | 0.718946 | 0.784285 | 0.794259 | 0.775798 | 0.780447 | 0.817086 | 0.812721 | 0.809775 |
| 0.707711 | 0.709045 | 0.772623 | 0.770035 | 0.766168 | 0.772392 | 0.816223 | 0.794775 | 0.803073 |
| 0.71504 | 0.718928 | 0.778087 | 0.770804 | 0.777003 | 0.786321 | 0.795795 | 0.792258 | 0.79013 |
| 0.708919 | 0.71166 | 0.767968 | 0.771457 | 0.777942 | 0.772761 | 0.774579 | 0.781628 | 0.772958 |
| 0.719221 | 0.719923 | 0.77266 | 0.772382 | 0.77295 | 0.77397 | 0.773841 | 0.772697 | 0.777371 |
| 0.776718 | 0.776327 | 0.708727 | 0.706799 | 0.715014 | 0.720352 | 0.709406 | 0.710974 | 0.707484 |
| 0.77208 | 0.773126 | 0.705337 | 0.704297 | 0.712679 | 0.714055 | 0.705373 | 0.706702 | 0.705551 |
| 0.715011 | 0.717344 | 0.766948 | 0.773808 | 0.773274 | 0.778028 | 0.79132 | 0.782001 | 0.787057 |
| 0.709739 | 0.709703 | 0.767664 | 0.769879 | 0.7806 | 0.769091 | 0.795647 | 0.780908 | 0.785936 |
| 0.711687 | 0.712562 | 0.769041 | 0.762969 | 0.767778 | 0.766175 | 0.789602 | 0.791355 | 0.776298 |
| 0.704142 | 0.706395 | 0.781492 | 0.77742 | 0.782389 | 0.777546 | 0.812638 | 0.821589 | 0.802369 |
| 0.713413 | 0.715145 | 0.790561 | 0.784858 | 0.780036 | 0.786522 | 0.797338 | 0.801093 | 0.785288 |
| 0.724174 | 0.726307 | 0.778537 | 0.776435 | 0.777458 | 0.776504 | 0.79412 | 0.792879 | 0.791145 |
| 0.737225 | 0.740367 | 0.758788 | 0.763057 | 0.766257 | 0.760804 | 0.787193 | 0.783639 | 0.77538 |
| 0.717074 | 0.717494 | 0.766093 | 0.768241 | 0.782132 | 0.775502 | 0.797767 | 0.802698 | 0.803344 |
| 0.719364 | 0.716479 | 0.760367 | 0.760429 | 0.778955 | 0.770478 | 0.79934 | 0.797872 | 0.794668 |
| 0.738949 | 0.74347 | 0.759382 | 0.760468 | 0.767604 | 0.769451 | 0.767863 | 0.757399 | 0.764379 |
| 0.711427 | 0.711986 | 0.7801 | 0.778773 | 0.762851 | 0.782594 | 0.807123 | 0.837106 | 0.828657 |
| 0.713366 | 0.717961 | 0.786384 | 0.78586 | 0.78771 | 0.783135 | 0.800068 | 0.793649 | 0.804543 |
| 0.711329 | 0.712973 | 0.779272 | 0.784503 | 0.785821 | 0.781068 | 0.794349 | 0.792015 | 0.788126 |
| 0.713872 | 0.714182 | 0.779909 | 0.777117 | 0.786224 | 0.778083 | 0.808158 | 0.808532 | 0.797665 |
| 0.718508 | 0.722535 | 0.783061 | 0.785471 | 0.782292 | 0.781521 | 0.810589 | 0.806072 | 0.803473 |
| 0.713861 | 0.71517 | 0.781875 | 0.784232 | 0.788811 | 0.781555 | 0.815075 | 0.819175 | 0.80878 |
| 0.716247 | 0.717162 | 0.780944 | 0.779038 | 0.768545 | 0.777054 | 0.788773 | 0.804637 | 0.791494 |
| 0.71179 | 0.716746 | 0.77786 | 0.775087 | 0.770638 | 0.773797 | 0.793594 | 0.79872 | 0.79669 |
| 0.707586 | 0.708013 | 0.770744 | 0.770084 | 0.760659 | 0.777435 | 0.797889 | 0.806791 | 0.795518 |
| 0.713589 | 0.717035 | 0.776431 | 0.778971 | 0.774967 | 0.780928 | 0.796374 | 0.788037 | 0.784568 |
| 0.713172 | 0.715273 | 0.770698 | 0.774371 | 0.770344 | 0.774183 | 0.78443 | 0.794403 | 0.782948 |
| 0.715243 | 0.714988 | 0.770911 | 0.771171 | 0.779186 | 0.772321 | 0.799307 | 0.798298 | 0.786914 |
| 0.711424 | 0.71668 | 0.772031 | 0.770096 | 0.771221 | 0.772038 | 0.777954 | 0.78643 | 0.775187 |
| 0.709391 | 0.712837 | 0.772283 | 0.772877 | 0.768224 | 0.777418 | 0.782591 | 0.783246 | 0.780335 |
| 0.711331 | 0.716322 | 0.770499 | 0.767003 | 0.766519 | 0.772385 | 0.794689 | 0.801306 | 0.800876 |
| 0.711466 | 0.716868 | 0.771763 | 0.766322 | 0.767546 | 0.779105 | 0.775104 | 0.78214 | 0.768677 |
| 0.709468 | 0.710764 | 0.768115 | 0.762118 | 0.761366 | 0.771322 | 0.778639 | 0.799446 | 0.784432 |
| 0.714021 | 0.716177 | 0.773125 | 0.767108 | 0.764386 | 0.771193 | 0.790095 | 0.8249 | 0.799422 |
| 0.716571 | 0.717067 | 0.767878 | 0.782903 | 0.783733 | 0.773765 | 0.797924 | 0.786051 | 0.781794 |
| 0.713706 | 0.715813 | 0.761491 | 0.763008 | 0.758143 | 0.768137 | 0.777872 | 0.783245 | 0.766961 |
| 0.717201 | 0.715862 | 0.770189 | 0.764666 | 0.768135 | 0.769492 | 0.784945 | 0.779484 | 0.777564 |

| | | | | | | | | |
|---|---|---|---|---|---|---|---|---|
| 0.74834 | 0.750767 | 0.745393 | 0.751635 | 0.763332 | 0.754383 | 0.774082 | 0.771384 | 0.770617 |
| 0.704749 | 0.706695 | 0.766609 | 0.773566 | 0.78892 | 0.781977 | 0.799681 | 0.786677 | 0.802938 |
| 0.705557 | 0.708043 | 0.782587 | 0.782052 | 0.798598 | 0.780897 | 0.805676 | 0.82065 | 0.800459 |
| 0.713624 | 0.71873 | 0.776424 | 0.776032 | 0.801923 | 0.778118 | 0.802142 | 0.81917 | 0.809901 |
| 0.712545 | 0.711131 | 0.778427 | 0.776713 | 0.781175 | 0.77333 | 0.799479 | 0.813631 | 0.800846 |
| 0.714355 | 0.71639 | 0.77128 | 0.776067 | 0.777142 | 0.772378 | 0.789905 | 0.790977 | 0.788603 |
| 0.710644 | 0.712335 | 0.773616 | 0.770055 | 0.774578 | 0.774301 | 0.775802 | 0.776253 | 0.77566 |
| 0.713316 | 0.714493 | 0.783075 | 0.782862 | 0.776544 | 0.779174 | 0.801292 | 0.803986 | 0.804723 |
| 0.713754 | 0.714909 | 0.765427 | 0.767369 | 0.770132 | 0.772484 | 0.781914 | 0.778214 | 0.784844 |
| 0.713509 | 0.71666 | 0.767901 | 0.771269 | 0.766914 | 0.77871 | 0.779887 | 0.775219 | 0.769966 |
| 0.704109 | 0.706034 | 0.776318 | 0.774092 | 0.793066 | 0.774261 | 0.808281 | 0.800927 | 0.807512 |
| 0.70905 | 0.712584 | 0.781496 | 0.784144 | 0.780718 | 0.787018 | 0.807454 | 0.798152 | 0.807282 |
| 0.711917 | 0.714436 | 0.776495 | 0.77909 | 0.792039 | 0.77495 | 0.813694 | 0.80899 | 0.800649 |
| 0.71001 | 0.710452 | 0.771762 | 0.782258 | 0.783887 | 0.777564 | 0.795343 | 0.799746 | 0.793695 |
| 0.708134 | 0.705792 | 0.775256 | 0.783253 | 0.780898 | 0.777216 | 0.811164 | 0.795412 | 0.807441 |
| 0.707671 | 0.708947 | 0.772167 | 0.776802 | 0.778728 | 0.771844 | 0.806391 | 0.789327 | 0.805505 |
| 0.70813 | 0.708853 | 0.765524 | 0.769369 | 0.775928 | 0.769386 | 0.802821 | 0.808424 | 0.796844 |
| 0.714641 | 0.716725 | 0.782429 | 0.786767 | 0.786963 | 0.779774 | 0.819032 | 0.810774 | 0.822076 |
| 0.716437 | 0.716268 | 0.783073 | 0.78138 | 0.795472 | 0.777186 | 0.795541 | 0.795325 | 0.788837 |
| 0.710367 | 0.707985 | 0.773138 | 0.781389 | 0.788276 | 0.774788 | 0.806919 | 0.805417 | 0.801415 |
| 0.704449 | 0.706355 | 0.76443 | 0.765871 | 0.78575 | 0.769244 | 0.810654 | 0.815357 | 0.803477 |
| 0.706899 | 0.708895 | 0.774159 | 0.777494 | 0.784345 | 0.77787 | 0.805523 | 0.78917 | 0.800883 |
| 0.714012 | 0.713295 | 0.77336 | 0.774996 | 0.776037 | 0.780403 | 0.798641 | 0.785336 | 0.798572 |
| 0.71649 | 0.719574 | 0.771137 | 0.771303 | 0.768377 | 0.774457 | 0.783861 | 0.775756 | 0.772135 |
| 0.727397 | 0.728093 | 0.763745 | 0.769017 | 0.768482 | 0.777879 | 0.768575 | 0.768635 | 0.762511 |
| 0.773252 | 0.773771 | 0.707104 | 0.705442 | 0.713894 | 0.714587 | 0.705625 | 0.707993 | 0.705706 |
| 1 | 0.87923 | 0.707866 | 0.70947 | 0.715254 | 0.714088 | 0.706121 | 0.709141 | 0.711639 |
| 0.87923 | 1 | 0.711206 | 0.711456 | 0.71824 | 0.717218 | 0.71107 | 0.711332 | 0.711945 |
| 0.707866 | 0.711206 | 1 | 0.816217 | 0.785833 | 0.819886 | 0.794984 | 0.781221 | 0.791069 |
| 0.70947 | 0.711456 | 0.816217 | 1 | 0.795667 | 0.81402 | 0.802136 | 0.777749 | 0.790184 |
| 0.715254 | 0.71824 | 0.785833 | 0.795667 | 1 | 0.78517 | 0.775708 | 0.769027 | 0.770233 |
| 0.714088 | 0.717218 | 0.819886 | 0.81402 | 0.78517 | 1 | 0.795296 | 0.771283 | 0.786247 |
| 0.706121 | 0.71107 | 0.794984 | 0.802136 | 0.775708 | 0.795296 | 1 | 0.831405 | 0.87058 |
| 0.709141 | 0.711332 | 0.781221 | 0.777749 | 0.769027 | 0.771283 | 0.831405 | 1 | 0.831757 |
| 0.711639 | 0.711945 | 0.791069 | 0.790184 | 0.770233 | 0.786247 | 0.87058 | 0.831757 | 1 |
| 0.71063 | 0.712306 | 0.768558 | 0.767541 | 0.764159 | 0.766556 | 0.863118 | 0.863881 | 0.86812 |
| 0.710214 | 0.711392 | 0.787442 | 0.789487 | 0.776986 | 0.792793 | 0.825844 | 0.784533 | 0.81165 |
| 0.724814 | 0.725693 | 0.799783 | 0.799054 | 0.808207 | 0.805033 | 0.839402 | 0.824226 | 0.847215 |
| 0.703719 | 0.704209 | 0.782699 | 0.778552 | 0.774157 | 0.776894 | 0.866974 | 0.809173 | 0.841125 |
| 0.708688 | 0.711584 | 0.786546 | 0.783887 | 0.792674 | 0.789934 | 0.787813 | 0.784092 | 0.785845 |
| 0.704531 | 0.704484 | 0.763784 | 0.765857 | 0.762052 | 0.764135 | 0.824377 | 0.830974 | 0.823044 |
| 0.726343 | 0.727655 | 0.805146 | 0.799966 | 0.806595 | 0.803659 | 0.84146 | 0.829066 | 0.844059 |
| 0.711746 | 0.712655 | 0.772139 | 0.776372 | 0.796063 | 0.771483 | 0.792024 | 0.787914 | 0.787792 |
| 0.708068 | 0.709016 | 0.771034 | 0.778021 | 0.797731 | 0.78003 | 0.786787 | 0.804264 | 0.77654 |
| 0.71186 | 0.712452 | 0.775979 | 0.778166 | 0.778884 | 0.778846 | 0.790815 | 0.795822 | 0.784025 |
| 0.711585 | 0.712769 | 0.774954 | 0.77249 | 0.766788 | 0.767648 | 0.834732 | 0.81558 | 0.818276 |

| | | | | | | | | |
|---|---|---|---|---|---|---|---|---|
| 0.70907 | 0.710385 | 0.770675 | 0.772229 | 0.798259 | 0.773015 | 0.786146 | 0.786437 | 0.785101 |
| 0.712708 | 0.713029 | 0.777586 | 0.775619 | 0.795198 | 0.776303 | 0.781065 | 0.780666 | 0.7744 |
| 0.715432 | 0.711853 | 0.769148 | 0.773038 | 0.775677 | 0.769644 | 0.784617 | 0.787485 | 0.775762 |
| 0.718172 | 0.719811 | 0.786158 | 0.784868 | 0.799527 | 0.786161 | 0.809915 | 0.805023 | 0.824414 |
| 0.717181 | 0.717649 | 0.77695 | 0.781716 | 0.774183 | 0.782605 | 0.793476 | 0.796145 | 0.786581 |
| 0.711727 | 0.713661 | 0.778841 | 0.777188 | 0.777962 | 0.77916 | 0.78741 | 0.783934 | 0.788161 |
| 0.711338 | 0.710729 | 0.790276 | 0.786828 | 0.776205 | 0.784861 | 0.825288 | 0.780721 | 0.823819 |
| 0.717893 | 0.719997 | 0.774581 | 0.772572 | 0.773749 | 0.770105 | 0.780812 | 0.781884 | 0.77615 |
| 0.714424 | 0.713674 | 0.774822 | 0.776736 | 0.774979 | 0.77482 | 0.788567 | 0.787709 | 0.778395 |
| 0.70862 | 0.709985 | 0.775221 | 0.766726 | 0.768118 | 0.768618 | 0.778093 | 0.791774 | 0.774948 |
| 0.709708 | 0.712377 | 0.766179 | 0.766974 | 0.76802 | 0.771332 | 0.783286 | 0.786662 | 0.778248 |
| 0.70923 | 0.709086 | 0.769577 | 0.768521 | 0.764472 | 0.771909 | 0.783213 | 0.797997 | 0.782852 |
| 0.714239 | 0.714517 | 0.769371 | 0.774522 | 0.773145 | 0.775199 | 0.787685 | 0.803502 | 0.782098 |
| 0.706898 | 0.710126 | 0.763337 | 0.764776 | 0.764387 | 0.767543 | 0.777515 | 0.811028 | 0.775611 |
| 0.714818 | 0.714504 | 0.772161 | 0.775552 | 0.772454 | 0.778719 | 0.781934 | 0.785434 | 0.782444 |
| 0.711976 | 0.711718 | 0.768168 | 0.77585 | 0.768551 | 0.772358 | 0.787856 | 0.789667 | 0.780015 |
| 0.713384 | 0.719475 | 0.768614 | 0.774091 | 0.773089 | 0.775543 | 0.784818 | 0.78409 | 0.778769 |
| 0.709452 | 0.709902 | 0.778547 | 0.78533 | 0.778254 | 0.774869 | 0.818282 | 0.815227 | 0.814575 |
| 0.712663 | 0.714343 | 0.770835 | 0.775583 | 0.775607 | 0.775033 | 0.786278 | 0.792718 | 0.787435 |
| 0.711927 | 0.711545 | 0.779232 | 0.776739 | 0.768287 | 0.776136 | 0.787046 | 0.785786 | 0.780506 |
| 0.713893 | 0.714067 | 0.773088 | 0.769231 | 0.76397 | 0.768879 | 0.780039 | 0.786277 | 0.788869 |
| 0.7048 | 0.7091 | 0.772044 | 0.768193 | 0.767001 | 0.768759 | 0.78702 | 0.779809 | 0.783326 |
| 0.710335 | 0.710318 | 0.770754 | 0.769316 | 0.772697 | 0.770293 | 0.787731 | 0.791021 | 0.788506 |
| 0.713368 | 0.712754 | 0.781886 | 0.775469 | 0.771771 | 0.780455 | 0.785379 | 0.799514 | 0.784672 |
| 0.713159 | 0.712558 | 0.778976 | 0.774844 | 0.77458 | 0.781838 | 0.803779 | 0.777871 | 0.800753 |
| 0.717497 | 0.71854 | 0.787864 | 0.791649 | 0.780526 | 0.797976 | 0.830655 | 0.786389 | 0.828806 |
| 0.710027 | 0.711473 | 0.776501 | 0.770403 | 0.778216 | 0.775004 | 0.806359 | 0.779268 | 0.80209 |
| 0.716225 | 0.714 | 0.77964 | 0.782668 | 0.768562 | 0.777096 | 0.782239 | 0.788724 | 0.780219 |
| 0.713393 | 0.715133 | 0.773062 | 0.775446 | 0.770693 | 0.775959 | 0.786743 | 0.786077 | 0.787996 |
| 0.713214 | 0.714088 | 0.777304 | 0.783625 | 0.771402 | 0.783637 | 0.805563 | 0.780636 | 0.798955 |
| 0.715675 | 0.714736 | 0.765478 | 0.774912 | 0.765741 | 0.774615 | 0.785937 | 0.798729 | 0.784489 |
| 0.70912 | 0.713072 | 0.77236 | 0.764362 | 0.760104 | 0.768972 | 0.786726 | 0.777038 | 0.782897 |
| 0.713414 | 0.714541 | 0.784871 | 0.788717 | 0.780949 | 0.787059 | 0.840891 | 0.785461 | 0.818997 |
| 0.709905 | 0.713121 | 0.773057 | 0.773912 | 0.798388 | 0.774752 | 0.802644 | 0.816107 | 0.811874 |
| 0.709342 | 0.707945 | 0.77799 | 0.778357 | 0.772829 | 0.775784 | 0.793197 | 0.795667 | 0.790235 |
| 0.71171 | 0.712305 | 0.773659 | 0.77489 | 0.770215 | 0.771883 | 0.797332 | 0.796522 | 0.791784 |
| 0.706625 | 0.705025 | 0.770607 | 0.774271 | 0.775119 | 0.771385 | 0.790575 | 0.791329 | 0.78175 |
| 0.691588 | 0.686914 | 0.753036 | 0.744868 | 0.747899 | 0.745276 | 0.761052 | 0.76584 | 0.756642 |
| 0.692989 | 0.689711 | 0.769658 | 0.770042 | 0.765244 | 0.77163 | 0.769239 | 0.762591 | 0.763725 |
| 0.684004 | 0.683392 | 0.75 | 0.756047 | 0.747264 | 0.743413 | 0.75868 | 0.752839 | 0.761817 |
| 0.679809 | 0.679679 | 0.751955 | 0.757392 | 0.744773 | 0.750774 | 0.81877 | 0.816948 | 0.811534 |
| 0.666862 | 0.667098 | 0.735876 | 0.73813 | 0.725657 | 0.730099 | 0.746513 | 0.846401 | 0.749022 |
| 0.670823 | 0.668977 | 0.738935 | 0.744323 | 0.768316 | 0.745034 | 0.764496 | 0.767593 | 0.755094 |
| 0.666788 | 0.666302 | 0.749181 | 0.74324 | 0.732635 | 0.741494 | 0.792996 | 0.779493 | 0.796513 |
| 0.672843 | 0.664194 | 0.736642 | 0.743784 | 0.734828 | 0.74097 | 0.758358 | 0.76015 | 0.765843 |
| 0.667537 | 0.666959 | 0.766989 | 0.770197 | 0.749707 | 0.764071 | 0.852385 | 0.77354 | 0.794636 |
| 0.67496 | 0.673845 | 0.744787 | 0.751234 | 0.740041 | 0.749521 | 0.800424 | 0.804991 | 0.816424 |

| | | | | | | | | |
|---|---|---|---|---|---|---|---|---|
| 0.678453 | 0.676723 | 0.734614 | 0.737119 | 0.729389 | 0.737163 | 0.750386 | 0.750998 | 0.747722 |
| 0.674818 | 0.677754 | 0.761963 | 0.762108 | 0.752439 | 0.767565 | 0.870238 | 0.805877 | 0.878242 |
| 0.672718 | 0.671862 | 0.744543 | 0.748074 | 0.73706 | 0.744546 | 0.746491 | 0.74548 | 0.741293 |
| 0.669936 | 0.666533 | 0.729799 | 0.734983 | 0.759029 | 0.733388 | 0.750207 | 0.751001 | 0.752626 |
| 0.667052 | 0.666407 | 0.767247 | 0.755932 | 0.74318 | 0.762662 | 0.79527 | 0.806247 | 0.808667 |
| 0.671858 | 0.666897 | 0.732785 | 0.736487 | 0.728018 | 0.728606 | 0.743652 | 0.749892 | 0.74291 |
| 0.670775 | 0.671906 | 0.753982 | 0.762902 | 0.741893 | 0.758417 | 0.784615 | 0.750815 | 0.777026 |
| 0.66388 | 0.666667 | 0.74164 | 0.74683 | 0.729091 | 0.747281 | 0.800452 | 0.817217 | 0.812102 |
| 0.666374 | 0.66568 | 0.728711 | 0.724212 | 0.711829 | 0.727027 | 0.728793 | 0.766739 | 0.736794 |
| 0.668221 | 0.669072 | 0.78703 | 0.782718 | 0.758647 | 0.789661 | 0.844245 | 0.757688 | 0.841676 |
| 0.660227 | 0.656008 | 0.741017 | 0.739291 | 0.731661 | 0.740356 | 0.746543 | 0.743127 | 0.741039 |
| 0.699273 | 0.699835 | 0.757093 | 0.756035 | 0.786114 | 0.760314 | 0.776659 | 0.771822 | 0.770763 |
| 0.710073 | 0.710144 | 0.76785 | 0.771046 | 0.761085 | 0.764246 | 0.77625 | 0.777349 | 0.775876 |
| 0.687131 | 0.690209 | 0.747749 | 0.75 | 0.738237 | 0.753906 | 0.760862 | 0.745912 | 0.75 |
| 0.716415 | 0.717298 | 0.776029 | 0.779966 | 0.781122 | 0.76956 | 0.783235 | 0.788767 | 0.785241 |
| 0.706284 | 0.707673 | 0.769983 | 0.765799 | 0.761542 | 0.761507 | 0.767018 | 0.788442 | 0.776811 |
| 0.715452 | 0.716683 | 0.766786 | 0.771179 | 0.772579 | 0.771998 | 0.779671 | 0.782238 | 0.77404 |
| 0.721892 | 0.722308 | 0.77985 | 0.783986 | 0.785316 | 0.773653 | 0.786094 | 0.781617 | 0.774514 |
| 0.699454 | 0.698421 | 0.75514 | 0.749294 | 0.743408 | 0.757059 | 0.756359 | 0.75599 | 0.754117 |
| 0.719291 | 0.72335 | 0.773659 | 0.770314 | 0.766895 | 0.770962 | 0.774676 | 0.820858 | 0.773306 |
| 0.689655 | 0.695018 | 0.735017 | 0.740232 | 0.737753 | 0.747588 | 0.752128 | 0.745052 | 0.744438 |
| 0.719189 | 0.719236 | 0.770746 | 0.774043 | 0.772937 | 0.772277 | 0.774818 | 0.772185 | 0.768667 |
| 0.722627 | 0.724199 | 0.773132 | 0.774705 | 0.779584 | 0.769122 | 0.773505 | 0.774191 | 0.773298 |
| 0.686086 | 0.688484 | 0.751346 | 0.761138 | 0.766024 | 0.766719 | 0.756819 | 0.743279 | 0.748675 |
| 0.690765 | 0.693716 | 0.750787 | 0.743322 | 0.743029 | 0.74556 | 0.790218 | 0.766392 | 0.777206 |
| 0.686543 | 0.687516 | 0.747485 | 0.752529 | 0.749903 | 0.754845 | 0.781518 | 0.777484 | 0.790797 |
| 0.709383 | 0.709219 | 0.759876 | 0.755365 | 0.753043 | 0.757924 | 0.7656 | 0.758885 | 0.755168 |
| 0.720193 | 0.718076 | 0.772977 | 0.769367 | 0.772201 | 0.775653 | 0.78009 | 0.782475 | 0.774778 |
| 0.689061 | 0.690872 | 0.745051 | 0.739248 | 0.743419 | 0.737524 | 0.754124 | 0.753805 | 0.755012 |
| 0.724996 | 0.724755 | 0.786616 | 0.777927 | 0.787869 | 0.780762 | 0.804087 | 0.771457 | 0.80521 |
| 0.728078 | 0.72809 | 0.764048 | 0.767185 | 0.778654 | 0.769577 | 0.769658 | 0.770319 | 0.768834 |
| 0.700665 | 0.701466 | 0.762331 | 0.755833 | 0.764522 | 0.757318 | 0.786567 | 0.774967 | 0.77898 |
| 0.700335 | 0.701597 | 0.758138 | 0.754745 | 0.766607 | 0.759427 | 0.784257 | 0.77649 | 0.783397 |
| 0.727625 | 0.730142 | 0.762731 | 0.762718 | 0.7606 | 0.76862 | 0.777049 | 0.762669 | 0.771497 |
| 0.724135 | 0.722121 | 0.75965 | 0.761369 | 0.76575 | 0.760815 | 0.769626 | 0.765178 | 0.760553 |
| 0.723313 | 0.725449 | 0.765625 | 0.765984 | 0.770619 | 0.768034 | 0.775642 | 0.767565 | 0.7712 |
| 0.683354 | 0.682639 | 0.722737 | 0.71863 | 0.71618 | 0.721676 | 0.732669 | 0.726675 | 0.730523 |
| 0.723214 | 0.723684 | 0.751459 | 0.749958 | 0.746834 | 0.75645 | 0.766711 | 0.76228 | 0.764779 |
| 0.682857 | 0.681442 | 0.735948 | 0.723984 | 0.722432 | 0.726896 | 0.731963 | 0.725852 | 0.731848 |
| 0.676161 | 0.673142 | 0.7277 | 0.719319 | 0.714119 | 0.719438 | 0.717118 | 0.726264 | 0.71142 |
| 0.723696 | 0.725578 | 0.770747 | 0.768508 | 0.765012 | 0.772825 | 0.781705 | 0.791574 | 0.791663 |
| 0.67541 | 0.675163 | 0.721793 | 0.717226 | 0.719152 | 0.717162 | 0.718566 | 0.723486 | 0.716492 |
| 0.663939 | 0.663395 | 0.72439 | 0.715991 | 0.712619 | 0.717051 | 0.761539 | 0.735276 | 0.754417 |
| 0.663454 | 0.659705 | 0.715365 | 0.71491 | 0.707692 | 0.713877 | 0.741613 | 0.715383 | 0.752843 |
| 0.726411 | 0.72906 | 0.763917 | 0.759395 | 0.764931 | 0.770183 | 0.780567 | 0.767611 | 0.780622 |
| 0.665808 | 0.664779 | 0.71611 | 0.717745 | 0.710242 | 0.716414 | 0.748905 | 0.729098 | 0.745929 |
| 0.72081 | 0.720801 | 0.762151 | 0.763204 | 0.76849 | 0.763123 | 0.776981 | 0.788081 | 0.784918 |

| | | | | | | | | |
|---|---|---|---|---|---|---|---|---|
| 0.731264 | 0.736182 | 0.763923 | 0.75877 | 0.760535 | 0.769736 | 0.781165 | 0.765547 | 0.780391 |
| 0.723825 | 0.727039 | 0.762037 | 0.761279 | 0.770735 | 0.764386 | 0.787622 | 0.777338 | 0.786196 |
| 0.743869 | 0.744529 | 0.751062 | 0.746536 | 0.753258 | 0.755453 | 0.756243 | 0.748825 | 0.750767 |
| 0.656336 | 0.654719 | 0.708054 | 0.705237 | 0.694337 | 0.70692 | 0.744468 | 0.720096 | 0.740939 |
| 0.65469 | 0.654582 | 0.709287 | 0.698708 | 0.693521 | 0.703578 | 0.72927 | 0.704761 | 0.72093 |
| 0.730806 | 0.732726 | 0.760371 | 0.762706 | 0.77659 | 0.764405 | 0.785964 | 0.781597 | 0.791317 |
| 0.658639 | 0.658325 | 0.69958 | 0.68676 | 0.685063 | 0.689352 | 0.694686 | 0.694131 | 0.695609 |
| 0.733186 | 0.733989 | 0.730186 | 0.729714 | 0.735512 | 0.749834 | 0.736059 | 0.733765 | 0.732552 |
| 0.727295 | 0.733485 | 0.752891 | 0.747519 | 0.760052 | 0.753847 | 0.781878 | 0.781091 | 0.784103 |
| 0.731373 | 0.732238 | 0.717617 | 0.718374 | 0.718639 | 0.735304 | 0.720758 | 0.719043 | 0.716373 |
| 0.730245 | 0.734033 | 0.744104 | 0.74295 | 0.75217 | 0.747486 | 0.769006 | 0.760711 | 0.779953 |
| 0.709459 | 0.702703 | 0.634615 | 0.626667 | 0.679245 | 0.637584 | 0.635762 | 0.601307 | 0.632258 |
| 0.728766 | 0.724664 | 0.653735 | 0.651208 | 0.664414 | 0.670402 | 0.653586 | 0.650482 | 0.653081 |
| 0.731772 | 0.725134 | 0.656077 | 0.654308 | 0.665452 | 0.66697 | 0.655919 | 0.654933 | 0.653957 |
| 0.738874 | 0.736394 | 0.663724 | 0.662131 | 0.667438 | 0.675774 | 0.665455 | 0.664287 | 0.666909 |
| 0.736792 | 0.73056 | 0.658301 | 0.660333 | 0.663776 | 0.675091 | 0.666797 | 0.665365 | 0.666103 |
| 0.597788 | 0.599891 | 0.631249 | 0.620867 | 0.616395 | 0.623983 | 0.621293 | 0.620185 | 0.623329 |
| 0.733412 | 0.736038 | 0.664454 | 0.662109 | 0.669848 | 0.673431 | 0.66873 | 0.665577 | 0.667254 |
| 0.736192 | 0.738739 | 0.738483 | 0.738207 | 0.747665 | 0.7446 | 0.760534 | 0.754614 | 0.76225 |
| 0.734062 | 0.739011 | 0.732972 | 0.733654 | 0.740454 | 0.739848 | 0.755294 | 0.753302 | 0.764264 |
| 0.737326 | 0.744707 | 0.737353 | 0.736749 | 0.7457 | 0.7403 | 0.77146 | 0.759161 | 0.771258 |
| 0.600201 | 0.601388 | 0.621351 | 0.61563 | 0.617576 | 0.613205 | 0.618118 | 0.621413 | 0.616254 |
| 0.58465 | 0.585923 | 0.604256 | 0.596735 | 0.594687 | 0.599984 | 0.602881 | 0.600352 | 0.600581 |
| 0.758399 | 0.766232 | 0.702979 | 0.702738 | 0.711634 | 0.712079 | 0.706746 | 0.704577 | 0.703474 |
| 0.755267 | 0.764148 | 0.702716 | 0.702612 | 0.71351 | 0.71371 | 0.705448 | 0.705379 | 0.705289 |
| 0.75301 | 0.75871 | 0.696277 | 0.697521 | 0.702136 | 0.705224 | 0.700332 | 0.695148 | 0.697686 |
| 0.7635 | 0.769364 | 0.702385 | 0.701635 | 0.707067 | 0.711613 | 0.704682 | 0.699538 | 0.700698 |
| 0.763851 | 0.770045 | 0.70669 | 0.70805 | 0.712567 | 0.712055 | 0.706912 | 0.701074 | 0.702842 |
| 0.766139 | 0.765745 | 0.706019 | 0.70494 | 0.709059 | 0.713411 | 0.706605 | 0.704882 | 0.705503 |
| 0.763193 | 0.764361 | 0.705315 | 0.700164 | 0.710036 | 0.716058 | 0.707641 | 0.706038 | 0.707613 |
| 0.762937 | 0.76396 | 0.701612 | 0.705958 | 0.711783 | 0.713371 | 0.707073 | 0.706292 | 0.703738 |
| 0.77533 | 0.772399 | 0.714274 | 0.71376 | 0.721001 | 0.722817 | 0.7115 | 0.712383 | 0.71165 |
| 0.770087 | 0.770296 | 0.707154 | 0.709376 | 0.716096 | 0.717012 | 0.708587 | 0.710155 | 0.705902 |
| 0.765357 | 0.767474 | 0.70695 | 0.70815 | 0.719485 | 0.717193 | 0.710539 | 0.71187 | 0.710018 |
| 0.776051 | 0.775554 | 0.708211 | 0.711846 | 0.719562 | 0.717066 | 0.714021 | 0.71252 | 0.709597 |
| 0.766179 | 0.768975 | 0.705069 | 0.706313 | 0.715286 | 0.716981 | 0.707335 | 0.705216 | 0.708584 |
| 0.76948 | 0.76863 | 0.706982 | 0.708089 | 0.718401 | 0.715804 | 0.707183 | 0.708635 | 0.706439 |
| 0.770443 | 0.767545 | 0.707516 | 0.7108 | 0.71652 | 0.717299 | 0.710723 | 0.711697 | 0.708817 |
| 0.767396 | 0.765595 | 0.703531 | 0.704438 | 0.714603 | 0.715829 | 0.702787 | 0.702794 | 0.70521 |
| 0.764851 | 0.764566 | 0.70837 | 0.709965 | 0.716003 | 0.717563 | 0.711418 | 0.711491 | 0.706956 |
| 0.728206 | 0.730578 | 0.658405 | 0.65529 | 0.662504 | 0.666106 | 0.658073 | 0.659603 | 0.656888 |
| 0.727632 | 0.731758 | 0.66134 | 0.657856 | 0.662921 | 0.669575 | 0.659683 | 0.662392 | 0.663495 |
| 0.76817 | 0.770447 | 0.712544 | 0.712428 | 0.722409 | 0.721157 | 0.709964 | 0.711558 | 0.709256 |
| 0.768855 | 0.772897 | 0.710344 | 0.710708 | 0.719686 | 0.720742 | 0.711258 | 0.711767 | 0.709758 |
| 0.730313 | 0.725352 | 0.644751 | 0.642964 | 0.653065 | 0.650656 | 0.646504 | 0.645383 | 0.645518 |
| 0.775042 | 0.77218 | 0.70757 | 0.708104 | 0.720794 | 0.71572 | 0.70619 | 0.706377 | 0.70805 |
| 0.767017 | 0.766875 | 0.69947 | 0.699821 | 0.708069 | 0.706118 | 0.70048 | 0.701244 | 0.695683 |

| | | | | | | | | |
|---|---|---|---|---|---|---|---|---|
| 0.76644 | 0.771578 | 0.703051 | 0.706471 | 0.716872 | 0.712741 | 0.706826 | 0.704407 | 0.705394 |
| 0.772218 | 0.772447 | 0.708686 | 0.709706 | 0.722581 | 0.716383 | 0.707438 | 0.707561 | 0.706076 |
| 0.768046 | 0.769108 | 0.705265 | 0.708571 | 0.718245 | 0.717504 | 0.710994 | 0.708101 | 0.707846 |
| 0.771047 | 0.774867 | 0.707519 | 0.70835 | 0.716238 | 0.716986 | 0.708484 | 0.706823 | 0.705906 |
| 0.770625 | 0.774319 | 0.712019 | 0.715108 | 0.725317 | 0.719535 | 0.717642 | 0.715145 | 0.716192 |
| 0.770783 | 0.768879 | 0.706666 | 0.707893 | 0.715551 | 0.712431 | 0.706539 | 0.704996 | 0.705372 |
| 0.768792 | 0.773339 | 0.706377 | 0.708177 | 0.717312 | 0.712484 | 0.707743 | 0.706965 | 0.707464 |
| 0.760782 | 0.76334 | 0.699205 | 0.700207 | 0.713256 | 0.709593 | 0.699594 | 0.701945 | 0.699902 |
| 0.76809 | 0.769613 | 0.704187 | 0.704949 | 0.715121 | 0.709553 | 0.703585 | 0.70501 | 0.704208 |
| 0.765464 | 0.766769 | 0.705338 | 0.704944 | 0.717031 | 0.714019 | 0.702648 | 0.709508 | 0.704954 |
| 0.772356 | 0.771499 | 0.707796 | 0.708361 | 0.718795 | 0.715674 | 0.707282 | 0.707473 | 0.707374 |
| 0.771707 | 0.770306 | 0.702362 | 0.705253 | 0.715923 | 0.709284 | 0.705736 | 0.700927 | 0.703253 |
| 0.764193 | 0.768366 | 0.700434 | 0.699121 | 0.708269 | 0.709293 | 0.69935 | 0.699838 | 0.699389 |
| 0.764025 | 0.763619 | 0.699329 | 0.698748 | 0.706858 | 0.706653 | 0.700855 | 0.700483 | 0.697754 |
| 0.768997 | 0.768738 | 0.713832 | 0.71173 | 0.719866 | 0.718348 | 0.714091 | 0.716011 | 0.71355 |
| 0.766488 | 0.766782 | 0.705868 | 0.706257 | 0.715836 | 0.716083 | 0.709478 | 0.70829 | 0.709459 |
| 0.762962 | 0.769356 | 0.702119 | 0.701078 | 0.710498 | 0.715536 | 0.703661 | 0.703259 | 0.700848 |
| 0.760507 | 0.763367 | 0.70514 | 0.706691 | 0.710876 | 0.713962 | 0.707325 | 0.705339 | 0.702183 |
| 0.762247 | 0.768774 | 0.705752 | 0.705902 | 0.711672 | 0.714597 | 0.704179 | 0.702079 | 0.701973 |
| 0.761621 | 0.76699 | 0.701393 | 0.701869 | 0.710613 | 0.712379 | 0.706646 | 0.704697 | 0.701275 |
| 0.755085 | 0.756807 | 0.699386 | 0.69811 | 0.711491 | 0.705068 | 0.703 | 0.705435 | 0.700607 |
| 0.764899 | 0.76475 | 0.702362 | 0.704111 | 0.709835 | 0.715087 | 0.704281 | 0.699642 | 0.702998 |
| 0.7697 | 0.773962 | 0.707481 | 0.706371 | 0.71212 | 0.716388 | 0.708985 | 0.707662 | 0.705887 |
| 0.766299 | 0.766284 | 0.708094 | 0.708227 | 0.715833 | 0.716783 | 0.709047 | 0.707758 | 0.705236 |
| 0.761905 | 0.765423 | 0.707264 | 0.70281 | 0.711615 | 0.716004 | 0.708521 | 0.706011 | 0.703456 |
| 0.762531 | 0.768358 | 0.70208 | 0.702406 | 0.708961 | 0.711287 | 0.703353 | 0.700886 | 0.700374 |
| 0.757182 | 0.756883 | 0.704328 | 0.700356 | 0.711506 | 0.707436 | 0.703758 | 0.702698 | 0.700367 |
| 0.759445 | 0.762461 | 0.698237 | 0.697564 | 0.707081 | 0.709709 | 0.701071 | 0.698918 | 0.69697 |
| 0.755589 | 0.763171 | 0.701635 | 0.6992 | 0.711831 | 0.709516 | 0.701849 | 0.706647 | 0.698247 |
| 0.764433 | 0.769256 | 0.706191 | 0.704328 | 0.713955 | 0.717812 | 0.705467 | 0.704115 | 0.703287 |
| 0.761995 | 0.763773 | 0.706018 | 0.70555 | 0.713475 | 0.716006 | 0.707237 | 0.709219 | 0.706738 |
| 0.762592 | 0.764995 | 0.707021 | 0.706145 | 0.717741 | 0.714881 | 0.713469 | 0.70951 | 0.707377 |
| 0.764754 | 0.769042 | 0.710791 | 0.709683 | 0.71966 | 0.718825 | 0.713278 | 0.712157 | 0.71183 |
| 0.765799 | 0.770031 | 0.708466 | 0.7089 | 0.716548 | 0.719311 | 0.714666 | 0.709752 | 0.712358 |
| 0.751982 | 0.760704 | 0.701514 | 0.699885 | 0.711236 | 0.711203 | 0.701662 | 0.703933 | 0.699217 |
| 0.763979 | 0.761949 | 0.707343 | 0.703891 | 0.710686 | 0.713431 | 0.705517 | 0.70697 | 0.704616 |
| 0.761294 | 0.765024 | 0.703463 | 0.705455 | 0.71605 | 0.715659 | 0.704389 | 0.708606 | 0.704791 |
| 0.763767 | 0.76844 | 0.700353 | 0.700198 | 0.70943 | 0.713072 | 0.704901 | 0.701196 | 0.703495 |
| 0.753422 | 0.756222 | 0.696017 | 0.697083 | 0.704151 | 0.705199 | 0.701483 | 0.700033 | 0.698819 |
| 0.756841 | 0.760939 | 0.690561 | 0.694456 | 0.703524 | 0.704768 | 0.697782 | 0.693389 | 0.695587 |
| 0.750873 | 0.755869 | 0.693919 | 0.692094 | 0.701457 | 0.699884 | 0.698886 | 0.698463 | 0.693353 |
| 0.75272 | 0.757562 | 0.694912 | 0.69633 | 0.703148 | 0.705711 | 0.696819 | 0.694553 | 0.695563 |
| 0.750689 | 0.752601 | 0.696731 | 0.694601 | 0.704687 | 0.704224 | 0.699253 | 0.698645 | 0.695904 |
| 0.752458 | 0.758013 | 0.688877 | 0.692294 | 0.697658 | 0.703111 | 0.693575 | 0.689266 | 0.689482 |
| 0.760277 | 0.766423 | 0.699816 | 0.701116 | 0.708603 | 0.712808 | 0.702271 | 0.702251 | 0.699366 |
| 0.751912 | 0.756264 | 0.694912 | 0.696856 | 0.703944 | 0.702698 | 0.697436 | 0.700212 | 0.697096 |

| | | | | | | | | |
|---|---|---|---|---|---|---|---|---|
| 0.760781 | 0.766722 | 0.707002 | 0.705873 | 0.713977 | 0.714679 | 0.711345 | 0.710543 | 0.706711 |
| 0.77512 | 0.778021 | 0.704677 | 0.706462 | 0.715107 | 0.715453 | 0.705074 | 0.709586 | 0.703448 |
| 0.769794 | 0.771683 | 0.703266 | 0.702662 | 0.711836 | 0.71251 | 0.70499 | 0.704448 | 0.703943 |
| 0.768566 | 0.768171 | 0.708422 | 0.706821 | 0.719095 | 0.718243 | 0.705699 | 0.709575 | 0.705991 |
| 0.774896 | 0.775863 | 0.713887 | 0.711371 | 0.721915 | 0.724144 | 0.711453 | 0.716699 | 0.710624 |
| 0.771509 | 0.772874 | 0.707798 | 0.704922 | 0.716974 | 0.715356 | 0.708286 | 0.710928 | 0.707398 |
| 0.771343 | 0.771843 | 0.701659 | 0.698206 | 0.709629 | 0.713421 | 0.70415 | 0.706054 | 0.704095 |
| 0.770651 | 0.773159 | 0.70672 | 0.705484 | 0.71539 | 0.71424 | 0.705897 | 0.70848 | 0.704138 |
| 0.764238 | 0.767366 | 0.700991 | 0.701373 | 0.707602 | 0.710644 | 0.698604 | 0.705201 | 0.701798 |
| 0.769711 | 0.768931 | 0.705957 | 0.707686 | 0.714331 | 0.714159 | 0.705745 | 0.710902 | 0.703793 |
| 0.769206 | 0.76973 | 0.709285 | 0.711368 | 0.717059 | 0.719512 | 0.708895 | 0.713808 | 0.711997 |
| 0.765847 | 0.769747 | 0.709159 | 0.709292 | 0.718094 | 0.713332 | 0.709316 | 0.711626 | 0.709173 |
| 0.775941 | 0.777732 | 0.706135 | 0.706743 | 0.713306 | 0.716602 | 0.703051 | 0.708196 | 0.704845 |
| 0.777611 | 0.776017 | 0.706051 | 0.70648 | 0.717969 | 0.716457 | 0.70669 | 0.71167 | 0.706771 |
| 0.778825 | 0.777164 | 0.709522 | 0.709167 | 0.717053 | 0.720122 | 0.70815 | 0.710715 | 0.7086 |
| 0.771947 | 0.772359 | 0.709065 | 0.70545 | 0.717055 | 0.717525 | 0.705242 | 0.711662 | 0.70435 |
| 0.778474 | 0.778024 | 0.709565 | 0.708225 | 0.7164 | 0.722931 | 0.711904 | 0.714775 | 0.708836 |
| 0.773163 | 0.777769 | 0.709665 | 0.706834 | 0.719294 | 0.715922 | 0.710152 | 0.71089 | 0.709623 |
| 0.779746 | 0.781418 | 0.708396 | 0.709144 | 0.71737 | 0.71941 | 0.713354 | 0.713999 | 0.71066 |
| 0.777227 | 0.77938 | 0.708277 | 0.705734 | 0.713333 | 0.715985 | 0.708483 | 0.707822 | 0.707949 |
| 0.774417 | 0.773965 | 0.706876 | 0.705921 | 0.718184 | 0.713603 | 0.705445 | 0.707321 | 0.706699 |
| 0.775004 | 0.774877 | 0.70569 | 0.705722 | 0.715968 | 0.714634 | 0.705801 | 0.707297 | 0.704073 |
| 0.777916 | 0.780766 | 0.711828 | 0.709152 | 0.722067 | 0.720312 | 0.709404 | 0.712322 | 0.70762 |
| 0.782059 | 0.777915 | 0.709331 | 0.711249 | 0.715974 | 0.718858 | 0.709852 | 0.710871 | 0.712992 |
| 0.780648 | 0.776479 | 0.707267 | 0.703987 | 0.719662 | 0.714706 | 0.706732 | 0.708488 | 0.707694 |
| 0.780685 | 0.778027 | 0.706153 | 0.703214 | 0.715369 | 0.716774 | 0.705344 | 0.70811 | 0.707733 |
| 0.781623 | 0.782003 | 0.701046 | 0.70605 | 0.711528 | 0.712353 | 0.701926 | 0.70606 | 0.70325 |
| 0.778946 | 0.779207 | 0.706537 | 0.703192 | 0.710903 | 0.717304 | 0.705892 | 0.707635 | 0.706985 |
| 0.782137 | 0.78244 | 0.708724 | 0.710205 | 0.719234 | 0.718785 | 0.707682 | 0.71028 | 0.706136 |
| 0.781199 | 0.78102 | 0.710267 | 0.709333 | 0.719141 | 0.720581 | 0.708695 | 0.709146 | 0.7088 |
| 0.773451 | 0.768721 | 0.704943 | 0.704959 | 0.710294 | 0.713084 | 0.705983 | 0.70852 | 0.705156 |
| 0.774695 | 0.775764 | 0.709187 | 0.711124 | 0.720607 | 0.717575 | 0.712897 | 0.712935 | 0.713308 |
| 0.773423 | 0.776264 | 0.70911 | 0.711066 | 0.71605 | 0.719445 | 0.709437 | 0.713587 | 0.705991 |
| 0.77286 | 0.776445 | 0.703846 | 0.70313 | 0.710831 | 0.71277 | 0.700709 | 0.705654 | 0.701936 |
| 0.780267 | 0.779573 | 0.706744 | 0.704841 | 0.714603 | 0.715768 | 0.708999 | 0.711748 | 0.709466 |
| 0.77804 | 0.778 | 0.704252 | 0.702926 | 0.71141 | 0.712818 | 0.704314 | 0.708183 | 0.703526 |
| 0.783042 | 0.782926 | 0.715685 | 0.710473 | 0.725208 | 0.726101 | 0.715519 | 0.714549 | 0.713536 |
| 0.778549 | 0.776132 | 0.710266 | 0.705599 | 0.712073 | 0.715036 | 0.705541 | 0.708843 | 0.702477 |
| 0.777002 | 0.783368 | 0.706597 | 0.709431 | 0.715543 | 0.718998 | 0.707433 | 0.708958 | 0.710773 |
| 0.780231 | 0.785411 | 0.709869 | 0.706815 | 0.716706 | 0.717166 | 0.709174 | 0.712367 | 0.707455 |
| 0.783092 | 0.784981 | 0.71286 | 0.7139 | 0.719898 | 0.723701 | 0.714057 | 0.714137 | 0.712538 |
| 0.778074 | 0.782396 | 0.709251 | 0.709825 | 0.717457 | 0.718228 | 0.709561 | 0.715341 | 0.711425 |
| 0.779657 | 0.785555 | 0.707906 | 0.708628 | 0.717212 | 0.71862 | 0.707871 | 0.711751 | 0.710919 |
| 0.780339 | 0.77882 | 0.706035 | 0.705799 | 0.717613 | 0.714052 | 0.708511 | 0.711315 | 0.707709 |
| 0.787945 | 0.793815 | 0.709453 | 0.710911 | 0.716375 | 0.716862 | 0.711356 | 0.711197 | 0.709816 |
| 0.785216 | 0.785502 | 0.704857 | 0.704816 | 0.712419 | 0.709812 | 0.704167 | 0.706794 | 0.7058 |

| | | | | | | | | |
|---|---|---|---|---|---|---|---|---|
| 0.768569 | 0.770616 | 0.689087 | 0.693632 | 0.701027 | 0.704189 | 0.695829 | 0.694623 | 0.694255 |
| 0.752943 | 0.754505 | 0.747288 | 0.74458 | 0.747941 | 0.757447 | 0.756849 | 0.74971 | 0.752299 |
| 0.785048 | 0.785452 | 0.708701 | 0.707022 | 0.712426 | 0.719744 | 0.706237 | 0.707501 | 0.707186 |
| 0.776377 | 0.784595 | 0.711109 | 0.712015 | 0.717778 | 0.722679 | 0.711944 | 0.713853 | 0.710455 |
| 0.778775 | 0.781982 | 0.70421 | 0.703224 | 0.709076 | 0.71207 | 0.704699 | 0.706879 | 0.701009 |
| 0.783683 | 0.786339 | 0.707299 | 0.710087 | 0.713581 | 0.712473 | 0.705261 | 0.709806 | 0.707259 |
| 0.778425 | 0.780839 | 0.703239 | 0.705113 | 0.710577 | 0.715494 | 0.703139 | 0.705724 | 0.706022 |
| 0.777489 | 0.779133 | 0.698102 | 0.699974 | 0.704101 | 0.704506 | 0.697171 | 0.701265 | 0.698892 |
| 0.776089 | 0.774151 | 0.705761 | 0.707351 | 0.713208 | 0.714882 | 0.705023 | 0.706888 | 0.702945 |
| 0.776508 | 0.780212 | 0.717757 | 0.713402 | 0.723615 | 0.724427 | 0.714448 | 0.715968 | 0.714855 |
| 0.778081 | 0.782702 | 0.711002 | 0.710832 | 0.715749 | 0.720537 | 0.711255 | 0.71253 | 0.710207 |
| 0.775573 | 0.77264 | 0.7084 | 0.707728 | 0.717938 | 0.719422 | 0.712505 | 0.713766 | 0.709499 |
| 0.778064 | 0.779042 | 0.714422 | 0.712853 | 0.722494 | 0.721395 | 0.712869 | 0.713758 | 0.712851 |
| 0.780419 | 0.779959 | 0.710216 | 0.711488 | 0.718937 | 0.720373 | 0.711206 | 0.714103 | 0.711582 |
| 0.772585 | 0.771114 | 0.708995 | 0.705462 | 0.713438 | 0.71904 | 0.706627 | 0.708018 | 0.704052 |
| 0.773441 | 0.77529 | 0.708825 | 0.707972 | 0.712712 | 0.715857 | 0.710556 | 0.711674 | 0.710904 |
| 0.773189 | 0.774177 | 0.711562 | 0.710411 | 0.71766 | 0.720757 | 0.710473 | 0.713905 | 0.711848 |
| 0.77129 | 0.773289 | 0.708901 | 0.71083 | 0.717033 | 0.719844 | 0.709604 | 0.714274 | 0.708814 |
| 0.774182 | 0.774147 | 0.700517 | 0.701483 | 0.709719 | 0.709832 | 0.70451 | 0.703282 | 0.705004 |
| 0.772121 | 0.774866 | 0.704595 | 0.702861 | 0.708965 | 0.714895 | 0.702932 | 0.70299 | 0.703814 |
| 0.776974 | 0.774788 | 0.707641 | 0.710559 | 0.715709 | 0.718666 | 0.71202 | 0.709063 | 0.706462 |
| 0.761167 | 0.761953 | 0.699461 | 0.696499 | 0.705655 | 0.708302 | 0.697801 | 0.702098 | 0.696318 |
| 0.767373 | 0.768538 | 0.704177 | 0.703023 | 0.71152 | 0.7176 | 0.705443 | 0.70854 | 0.702983 |
| 0.768713 | 0.77112 | 0.710459 | 0.70797 | 0.716417 | 0.720306 | 0.709956 | 0.708812 | 0.706891 |
| 0.769002 | 0.772623 | 0.713527 | 0.708469 | 0.719081 | 0.721928 | 0.714734 | 0.713652 | 0.710548 |
| 0.766709 | 0.76903 | 0.708263 | 0.709731 | 0.716141 | 0.721064 | 0.707279 | 0.706786 | 0.706436 |
| 0.764283 | 0.764887 | 0.703451 | 0.701595 | 0.710079 | 0.71273 | 0.706034 | 0.706765 | 0.703034 |
| 0.770054 | 0.769045 | 0.704457 | 0.700195 | 0.70934 | 0.710722 | 0.702209 | 0.705305 | 0.700242 |
| 0.765859 | 0.767189 | 0.70795 | 0.702306 | 0.712096 | 0.717592 | 0.707993 | 0.708853 | 0.705531 |
| 0.700361 | 0.70038 | 0.764745 | 0.76727 | 0.779265 | 0.77049 | 0.766939 | 0.768088 | 0.759613 |
| 0.712234 | 0.711041 | 0.798132 | 0.786764 | 0.785314 | 0.797893 | 0.817971 | 0.784517 | 0.804532 |
| 0.704791 | 0.708953 | 0.778065 | 0.776227 | 0.796179 | 0.77889 | 0.777264 | 0.782052 | 0.77316 |
| 0.695725 | 0.697267 | 0.767338 | 0.764952 | 0.796514 | 0.768594 | 0.781413 | 0.771053 | 0.774921 |
| 0.709713 | 0.713021 | 0.796815 | 0.792487 | 0.780145 | 0.790705 | 0.807349 | 0.787859 | 0.808235 |
| 0.769112 | 0.76858 | 0.702035 | 0.699571 | 0.708045 | 0.712146 | 0.702835 | 0.703621 | 0.700939 |
| 0.772161 | 0.773442 | 0.704827 | 0.703217 | 0.712088 | 0.714344 | 0.703912 | 0.705587 | 0.70235 |
| 0.771591 | 0.773764 | 0.70689 | 0.70404 | 0.718186 | 0.717196 | 0.70352 | 0.707432 | 0.701203 |
| 0.772478 | 0.775726 | 0.706723 | 0.705776 | 0.718793 | 0.715631 | 0.708559 | 0.705689 | 0.70365 |
| 0.773135 | 0.77027 | 0.703944 | 0.700986 | 0.71208 | 0.712764 | 0.707353 | 0.705764 | 0.702471 |
| 0.769686 | 0.772981 | 0.707979 | 0.707341 | 0.715893 | 0.715097 | 0.709779 | 0.709677 | 0.709407 |
| 0.774637 | 0.774088 | 0.706898 | 0.703682 | 0.713869 | 0.717987 | 0.704233 | 0.706707 | 0.70352 |
| 0.766133 | 0.768123 | 0.70495 | 0.701536 | 0.713112 | 0.715767 | 0.705112 | 0.705359 | 0.703106 |
| 0.770181 | 0.770503 | 0.695151 | 0.697696 | 0.704404 | 0.707417 | 0.70251 | 0.701642 | 0.700077 |
| 0.763847 | 0.765335 | 0.699402 | 0.699055 | 0.706348 | 0.709166 | 0.701677 | 0.70031 | 0.696435 |
| 0.762817 | 0.766757 | 0.703523 | 0.703844 | 0.711115 | 0.716564 | 0.705333 | 0.705519 | 0.704014 |
| 0.768837 | 0.773251 | 0.706837 | 0.703468 | 0.713151 | 0.717086 | 0.705112 | 0.709683 | 0.702848 |

| | | | | | | | | |
|---|---|---|---|---|---|---|---|---|
| 0.773697 | 0.771579 | 0.708894 | 0.709748 | 0.71202 | 0.716269 | 0.706245 | 0.710658 | 0.705034 |
| 0.768954 | 0.773038 | 0.706523 | 0.703999 | 0.715148 | 0.712847 | 0.707588 | 0.709125 | 0.707212 |
| 0.766616 | 0.770373 | 0.705088 | 0.709726 | 0.716297 | 0.720354 | 0.710799 | 0.709785 | 0.710625 |
| 0.76675 | 0.764769 | 0.707664 | 0.704849 | 0.716132 | 0.715107 | 0.704621 | 0.70783 | 0.704809 |
| 0.765131 | 0.764861 | 0.703477 | 0.702857 | 0.712417 | 0.711948 | 0.703143 | 0.70292 | 0.702136 |
| 0.76933 | 0.771996 | 0.706421 | 0.702364 | 0.709733 | 0.712233 | 0.703065 | 0.707358 | 0.70219 |
| 0.767416 | 0.768937 | 0.708616 | 0.705942 | 0.712914 | 0.714109 | 0.710598 | 0.712507 | 0.711719 |
| 0.767083 | 0.771069 | 0.707114 | 0.708817 | 0.71492 | 0.71618 | 0.706004 | 0.709023 | 0.70744 |
| 0.764123 | 0.767458 | 0.704659 | 0.703626 | 0.712398 | 0.712951 | 0.702534 | 0.706661 | 0.700831 |
| 0.769315 | 0.772167 | 0.706294 | 0.707532 | 0.716054 | 0.715952 | 0.705152 | 0.705297 | 0.70762 |
| 0.767228 | 0.769185 | 0.704558 | 0.705661 | 0.713948 | 0.71533 | 0.704906 | 0.707677 | 0.704975 |
| 0.760793 | 0.766818 | 0.706121 | 0.706049 | 0.712702 | 0.715685 | 0.705766 | 0.706787 | 0.7058 |
| 0.761619 | 0.7635 | 0.69607 | 0.69737 | 0.705512 | 0.707271 | 0.699521 | 0.695972 | 0.697869 |
| 0.759291 | 0.766817 | 0.697142 | 0.699531 | 0.707479 | 0.708866 | 0.699006 | 0.701897 | 0.700695 |
| 0.777258 | 0.779507 | 0.709998 | 0.710352 | 0.717441 | 0.716507 | 0.709491 | 0.709192 | 0.707841 |
| 0.785091 | 0.786685 | 0.717594 | 0.710578 | 0.721153 | 0.721847 | 0.713718 | 0.716798 | 0.714437 |
| 0.784744 | 0.785744 | 0.710223 | 0.707493 | 0.71999 | 0.713167 | 0.708094 | 0.709495 | 0.70577 |
| 0.783639 | 0.779715 | 0.712502 | 0.710833 | 0.718522 | 0.720494 | 0.710169 | 0.711689 | 0.710294 |
| 0.785374 | 0.780071 | 0.70628 | 0.704418 | 0.711421 | 0.712804 | 0.706349 | 0.707384 | 0.704295 |
| 0.780509 | 0.777481 | 0.703256 | 0.70672 | 0.712101 | 0.714823 | 0.707287 | 0.708943 | 0.705053 |
| 0.768083 | 0.766638 | 0.703593 | 0.701943 | 0.708153 | 0.70965 | 0.706246 | 0.706604 | 0.70367 |
| 0.773483 | 0.772525 | 0.710182 | 0.708416 | 0.71567 | 0.721693 | 0.709659 | 0.7103 | 0.707945 |
| 0.77884 | 0.780938 | 0.714985 | 0.710721 | 0.718108 | 0.718551 | 0.712711 | 0.713012 | 0.709613 |
| 0.766325 | 0.76166 | 0.704019 | 0.706382 | 0.715637 | 0.710812 | 0.706531 | 0.706842 | 0.708456 |
| 0.775098 | 0.777599 | 0.710235 | 0.705846 | 0.714397 | 0.71539 | 0.707675 | 0.707819 | 0.708196 |
| 0.708381 | 0.706364 | 0.77762 | 0.777692 | 0.790285 | 0.778265 | 0.796285 | 0.798929 | 0.80362 |
| 0.776709 | 0.77686 | 0.706776 | 0.70742 | 0.713957 | 0.715157 | 0.704803 | 0.70829 | 0.706085 |
| 0.778119 | 0.779584 | 0.710562 | 0.707072 | 0.722462 | 0.716776 | 0.707975 | 0.711268 | 0.705683 |
| 0.782664 | 0.781981 | 0.710452 | 0.706135 | 0.717959 | 0.715366 | 0.710847 | 0.710763 | 0.71214 |
| 0.777276 | 0.77534 | 0.710564 | 0.705708 | 0.717269 | 0.715744 | 0.70601 | 0.707576 | 0.705111 |
| 0.778514 | 0.780991 | 0.703124 | 0.702151 | 0.709982 | 0.714213 | 0.705016 | 0.707947 | 0.70425 |
| 0.777787 | 0.774243 | 0.704946 | 0.703203 | 0.714109 | 0.714763 | 0.703654 | 0.703829 | 0.701281 |
| 0.777542 | 0.774625 | 0.707543 | 0.706476 | 0.714858 | 0.716639 | 0.70637 | 0.708996 | 0.706033 |
| 0.776682 | 0.775668 | 0.701449 | 0.702398 | 0.71455 | 0.714117 | 0.701988 | 0.705261 | 0.699933 |
| 0.771476 | 0.773233 | 0.703084 | 0.701621 | 0.71477 | 0.712218 | 0.699687 | 0.702538 | 0.700374 |
| 0.772642 | 0.772299 | 0.705081 | 0.701038 | 0.714567 | 0.710943 | 0.703088 | 0.707078 | 0.701441 |
| 0.783286 | 0.781261 | 0.708286 | 0.706402 | 0.717871 | 0.715486 | 0.710734 | 0.70977 | 0.710838 |
| 0.777209 | 0.778364 | 0.708481 | 0.706105 | 0.715922 | 0.716844 | 0.712612 | 0.712607 | 0.711191 |
| 0.780691 | 0.780995 | 0.708652 | 0.705464 | 0.714794 | 0.718678 | 0.708795 | 0.711364 | 0.706084 |
| 0.776696 | 0.777884 | 0.707425 | 0.705006 | 0.71561 | 0.713554 | 0.705498 | 0.709356 | 0.705917 |
| 0.777007 | 0.779661 | 0.702226 | 0.703057 | 0.711941 | 0.713949 | 0.7029 | 0.70418 | 0.701377 |
| 0.781563 | 0.784194 | 0.712907 | 0.708692 | 0.718074 | 0.718503 | 0.711658 | 0.711927 | 0.708005 |
| 0.771725 | 0.777181 | 0.706872 | 0.703912 | 0.715659 | 0.715273 | 0.704384 | 0.704642 | 0.70338 |
| 0.760414 | 0.761576 | 0.703633 | 0.704954 | 0.711037 | 0.712929 | 0.709592 | 0.704681 | 0.705395 |
| 0.771697 | 0.771158 | 0.707241 | 0.70457 | 0.714355 | 0.720381 | 0.715479 | 0.70899 | 0.713182 |
| 0.770222 | 0.768116 | 0.707441 | 0.704948 | 0.7144 | 0.717552 | 0.711545 | 0.709768 | 0.711046 |

| | | | | | | | | |
|---|---|---|---|---|---|---|---|---|
| 0.765922 | 0.770482 | 0.703168 | 0.701832 | 0.713421 | 0.711864 | 0.709315 | 0.708337 | 0.709787 |
| 0.772351 | 0.773602 | 0.706853 | 0.705128 | 0.715117 | 0.714953 | 0.709566 | 0.708786 | 0.709631 |
| 0.766817 | 0.766805 | 0.703273 | 0.704179 | 0.712259 | 0.713925 | 0.709504 | 0.707287 | 0.708415 |
| 0.71832 | 0.7307 | 0.659508 | 0.660174 | 0.66307 | 0.671384 | 0.667597 | 0.666899 | 0.665977 |
| 0.766615 | 0.768238 | 0.703426 | 0.70721 | 0.712584 | 0.712132 | 0.708074 | 0.703374 | 0.705246 |
| 0.765578 | 0.764944 | 0.702969 | 0.70446 | 0.712026 | 0.711787 | 0.706944 | 0.706827 | 0.707288 |
| 0.76777 | 0.766969 | 0.708138 | 0.705623 | 0.713863 | 0.715115 | 0.711249 | 0.70797 | 0.710596 |
| 0.763968 | 0.76297 | 0.703944 | 0.705406 | 0.712626 | 0.715264 | 0.709424 | 0.708896 | 0.708072 |
| 0.7731 | 0.774431 | 0.712388 | 0.709009 | 0.715118 | 0.717657 | 0.714343 | 0.712456 | 0.712775 |
| 0.770492 | 0.770373 | 0.708615 | 0.707469 | 0.71472 | 0.715415 | 0.710522 | 0.7113 | 0.709636 |
| 0.770655 | 0.76748 | 0.706821 | 0.707092 | 0.713326 | 0.71678 | 0.708615 | 0.706426 | 0.70926 |
| 0.768559 | 0.770861 | 0.708825 | 0.707726 | 0.717625 | 0.72253 | 0.712255 | 0.710412 | 0.711237 |
| 0.770747 | 0.771686 | 0.707426 | 0.707303 | 0.714868 | 0.719686 | 0.711365 | 0.713333 | 0.712621 |
| 0.769822 | 0.774207 | 0.705798 | 0.70581 | 0.712943 | 0.714596 | 0.71119 | 0.710699 | 0.70961 |
| 0.769912 | 0.770802 | 0.708638 | 0.709461 | 0.713966 | 0.717942 | 0.71515 | 0.711343 | 0.711904 |
| 0.766213 | 0.767346 | 0.703023 | 0.704611 | 0.708855 | 0.710423 | 0.707647 | 0.70704 | 0.708639 |
| 0.768852 | 0.768299 | 0.70713 | 0.705725 | 0.714727 | 0.716046 | 0.7111 | 0.711384 | 0.712681 |
| 0.768124 | 0.774007 | 0.707251 | 0.70581 | 0.715869 | 0.717815 | 0.711819 | 0.710141 | 0.710798 |
| 0.761826 | 0.765002 | 0.701322 | 0.702329 | 0.7144 | 0.710383 | 0.704717 | 0.705854 | 0.70628 |
| 0.766303 | 0.76777 | 0.706247 | 0.702291 | 0.709403 | 0.715633 | 0.710571 | 0.709543 | 0.707804 |
| 0.771367 | 0.772848 | 0.708562 | 0.707451 | 0.715782 | 0.719835 | 0.714136 | 0.710343 | 0.710609 |
| 0.766513 | 0.768602 | 0.708734 | 0.709979 | 0.715197 | 0.71538 | 0.71175 | 0.712273 | 0.7123 |
| 0.68022 | 0.680973 | 0.689567 | 0.688398 | 0.686684 | 0.704771 | 0.698211 | 0.699336 | 0.703307 |
| 0.664147 | 0.659752 | 0.726054 | 0.727544 | 0.724822 | 0.722874 | 0.727273 | 0.724316 | 0.740208 |
| 0.67588 | 0.684833 | 0.750339 | 0.757169 | 0.749777 | 0.755097 | 0.762966 | 0.786685 | 0.764198 |
| 0.660703 | 0.65491 | 0.719772 | 0.731078 | 0.747599 | 0.731567 | 0.746034 | 0.747506 | 0.749225 |
| 0.734213 | 0.73685 | 0.665032 | 0.6622 | 0.670971 | 0.674395 | 0.667832 | 0.663182 | 0.665942 |
| 0.728733 | 0.731314 | 0.66463 | 0.6555 | 0.666667 | 0.670495 | 0.662069 | 0.656129 | 0.666729 |
| 0.735236 | 0.723864 | 0.662574 | 0.657188 | 0.671252 | 0.666435 | 0.661762 | 0.662945 | 0.661132 |
| 0.736897 | 0.73852 | 0.676465 | 0.672846 | 0.684807 | 0.683074 | 0.67378 | 0.670423 | 0.67538 |
| 0.714123 | 0.717541 | 0.648286 | 0.650524 | 0.66049 | 0.661009 | 0.654332 | 0.65072 | 0.659858 |
| 0.716615 | 0.708829 | 0.642993 | 0.643618 | 0.653428 | 0.655823 | 0.649329 | 0.647187 | 0.638791 |
| 0.715416 | 0.708198 | 0.640497 | 0.641048 | 0.654023 | 0.653454 | 0.637138 | 0.632174 | 0.641185 |
| 0.654963 | 0.653542 | 0.731797 | 0.735503 | 0.717224 | 0.720322 | 0.73269 | 0.744147 | 0.736085 |
| 0.764497 | 0.766898 | 0.697143 | 0.699709 | 0.705341 | 0.707736 | 0.701701 | 0.703629 | 0.701548 |
| 0.744021 | 0.743569 | 0.670428 | 0.678939 | 0.680451 | 0.682007 | 0.674717 | 0.672627 | 0.676361 |
| 0.73784 | 0.737193 | 0.665243 | 0.669506 | 0.67395 | 0.680654 | 0.670419 | 0.674227 | 0.67048 |
| 0.720569 | 0.723186 | 0.647622 | 0.654392 | 0.657559 | 0.662972 | 0.65433 | 0.654969 | 0.655926 |
| 0.747706 | 0.743433 | 0.672764 | 0.673453 | 0.683207 | 0.687415 | 0.674384 | 0.673107 | 0.673896 |
| 0.72882 | 0.73273 | 0.666171 | 0.663511 | 0.668971 | 0.675615 | 0.670039 | 0.662185 | 0.670771 |
| 0.72447 | 0.725345 | 0.664402 | 0.661567 | 0.661812 | 0.669684 | 0.658567 | 0.656154 | 0.662751 |
| 0.731023 | 0.731623 | 0.665412 | 0.658441 | 0.668281 | 0.674604 | 0.664492 | 0.662027 | 0.66783 |
| 0.721467 | 0.72357 | 0.654791 | 0.651082 | 0.671459 | 0.665741 | 0.656114 | 0.656146 | 0.655538 |
| 0.729081 | 0.729687 | 0.648445 | 0.654413 | 0.666476 | 0.67072 | 0.65662 | 0.656339 | 0.65946 |
| 0.724209 | 0.724254 | 0.654215 | 0.655104 | 0.664505 | 0.671797 | 0.660777 | 0.657856 | 0.660415 |
| 0.713815 | 0.713194 | 0.643401 | 0.642426 | 0.647208 | 0.662388 | 0.650983 | 0.635182 | 0.652951 |

| | | | | | | | | |
|---|---|---|---|---|---|---|---|---|
| 0.757836 | 0.757174 | 0.691024 | 0.688544 | 0.693542 | 0.694493 | 0.690276 | 0.689011 | 0.694176 |
| 0.737772 | 0.733314 | 0.658086 | 0.660194 | 0.665357 | 0.665854 | 0.657694 | 0.658048 | 0.659929 |
| 0.733802 | 0.729277 | 0.659841 | 0.661955 | 0.665025 | 0.669171 | 0.661898 | 0.661402 | 0.663718 |
| 0.726994 | 0.727766 | 0.66269 | 0.661747 | 0.663961 | 0.67095 | 0.66163 | 0.661488 | 0.665924 |
| 0.743368 | 0.739681 | 0.658732 | 0.653633 | 0.663601 | 0.662639 | 0.65214 | 0.660444 | 0.661955 |
| 0.728177 | 0.727683 | 0.659172 | 0.659927 | 0.670843 | 0.666347 | 0.660895 | 0.666425 | 0.669409 |
| 0.727977 | 0.723107 | 0.652529 | 0.645101 | 0.65924 | 0.66146 | 0.64791 | 0.650059 | 0.652629 |
| 0.727787 | 0.718322 | 0.640389 | 0.649003 | 0.655941 | 0.656242 | 0.646915 | 0.65708 | 0.650514 |
| 0.719495 | 0.717466 | 0.650273 | 0.648796 | 0.661017 | 0.663838 | 0.639935 | 0.653195 | 0.647746 |
| 0.752036 | 0.753826 | 0.68658 | 0.686203 | 0.694627 | 0.697291 | 0.689345 | 0.691076 | 0.689797 |
| 0.75329 | 0.758023 | 0.683485 | 0.686327 | 0.691585 | 0.694325 | 0.688976 | 0.690977 | 0.690647 |
| 0.746873 | 0.749972 | 0.6813 | 0.680758 | 0.687547 | 0.690809 | 0.677984 | 0.683459 | 0.682832 |
| 0.738075 | 0.738807 | 0.676172 | 0.676502 | 0.681577 | 0.687214 | 0.675711 | 0.676474 | 0.680033 |
| 0.739474 | 0.747778 | 0.679316 | 0.677052 | 0.683682 | 0.685876 | 0.678712 | 0.68343 | 0.680993 |
| 0.741626 | 0.741723 | 0.672112 | 0.67242 | 0.681737 | 0.679809 | 0.669769 | 0.670039 | 0.670198 |

| Collection_ | Collection_ | Collection_ | Collection_ | Collection_ | Collection_ | Collection_ | Collection_ | Collection_ |
|---|---|---|---|---|---|---|---|---|
| 0.811555 | 0.783508 | 0.834365 | 0.783856 | 0.781955 | 0.794268 | 0.827677 | 0.798382 | 0.794701 |
| 0.820751 | 0.790085 | 0.816015 | 0.793689 | 0.789988 | 0.796371 | 0.815475 | 0.779293 | 0.781604 |
| 0.824656 | 0.789538 | 0.810675 | 0.792148 | 0.776529 | 0.793608 | 0.815346 | 0.788012 | 0.784164 |
| 0.798158 | 0.783665 | 0.813048 | 0.782092 | 0.781156 | 0.789702 | 0.820049 | 0.796439 | 0.797987 |
| 0.77487 | 0.77568 | 0.803671 | 0.766218 | 0.781163 | 0.774432 | 0.811303 | 0.789817 | 0.801452 |
| 0.775247 | 0.782258 | 0.793494 | 0.762669 | 0.777687 | 0.770828 | 0.797974 | 0.774709 | 0.774117 |
| 0.713032 | 0.710684 | 0.723956 | 0.702892 | 0.710107 | 0.707294 | 0.727702 | 0.710075 | 0.710508 |
| 0.70848 | 0.706035 | 0.718286 | 0.702228 | 0.706067 | 0.702546 | 0.721898 | 0.708661 | 0.703217 |
| 0.786889 | 0.790947 | 0.803094 | 0.770639 | 0.782213 | 0.773885 | 0.805632 | 0.782152 | 0.779108 |
| 0.786201 | 0.782168 | 0.820614 | 0.782889 | 0.773737 | 0.786241 | 0.824637 | 0.803755 | 0.797527 |
| 0.792032 | 0.769143 | 0.810288 | 0.771778 | 0.781163 | 0.778362 | 0.811875 | 0.792297 | 0.794307 |
| 0.806143 | 0.790365 | 0.813744 | 0.789806 | 0.800149 | 0.791536 | 0.814631 | 0.800284 | 0.814426 |
| 0.784206 | 0.796105 | 0.811944 | 0.774097 | 0.782505 | 0.778159 | 0.816378 | 0.774119 | 0.77312 |
| 0.800417 | 0.777035 | 0.805711 | 0.780856 | 0.775281 | 0.78685 | 0.803655 | 0.788573 | 0.789482 |
| 0.780536 | 0.769505 | 0.795107 | 0.765732 | 0.770234 | 0.76537 | 0.796862 | 0.767294 | 0.769896 |
| 0.815966 | 0.784073 | 0.818174 | 0.782266 | 0.779224 | 0.793697 | 0.823693 | 0.796475 | 0.791766 |
| 0.826065 | 0.770685 | 0.823659 | 0.78139 | 0.773698 | 0.803632 | 0.821811 | 0.806628 | 0.789378 |
| 0.758269 | 0.768491 | 0.783429 | 0.75523 | 0.767377 | 0.755043 | 0.78257 | 0.766872 | 0.761874 |
| 0.818205 | 0.79206 | 0.811276 | 0.801431 | 0.777041 | 0.786463 | 0.818428 | 0.774175 | 0.78051 |
| 0.796927 | 0.783892 | 0.811561 | 0.789606 | 0.802402 | 0.792375 | 0.813266 | 0.788546 | 0.781455 |
| 0.795746 | 0.775069 | 0.804813 | 0.779466 | 0.799488 | 0.782791 | 0.805795 | 0.79023 | 0.795443 |
| 0.809376 | 0.784433 | 0.821576 | 0.799202 | 0.794096 | 0.800348 | 0.822952 | 0.80294 | 0.806339 |
| 0.795419 | 0.790018 | 0.816534 | 0.784745 | 0.792715 | 0.790916 | 0.821377 | 0.797627 | 0.794963 |
| 0.815334 | 0.791245 | 0.821534 | 0.78438 | 0.801208 | 0.793972 | 0.820256 | 0.798314 | 0.798167 |
| 0.789926 | 0.780402 | 0.800301 | 0.778294 | 0.793044 | 0.778901 | 0.803429 | 0.782104 | 0.784503 |
| 0.812442 | 0.782205 | 0.810417 | 0.788167 | 0.78844 | 0.787532 | 0.815633 | 0.782477 | 0.781928 |
| 0.805834 | 0.786102 | 0.812106 | 0.776652 | 0.778768 | 0.780721 | 0.814376 | 0.772383 | 0.773426 |
| 0.788652 | 0.787765 | 0.802776 | 0.77698 | 0.785418 | 0.778004 | 0.805847 | 0.782245 | 0.787198 |
| 0.791528 | 0.777769 | 0.805525 | 0.776816 | 0.785995 | 0.784515 | 0.805432 | 0.784492 | 0.795742 |
| 0.797165 | 0.779734 | 0.807317 | 0.78111 | 0.780159 | 0.785661 | 0.81038 | 0.791931 | 0.790074 |
| 0.779423 | 0.772668 | 0.802738 | 0.766181 | 0.790241 | 0.771192 | 0.802302 | 0.786963 | 0.785349 |
| 0.784544 | 0.772207 | 0.795573 | 0.766743 | 0.784613 | 0.775057 | 0.800133 | 0.782086 | 0.780047 |
| 0.802201 | 0.784734 | 0.810053 | 0.779774 | 0.774194 | 0.786736 | 0.818082 | 0.772092 | 0.781668 |
| 0.779838 | 0.766901 | 0.794266 | 0.761341 | 0.783662 | 0.775893 | 0.797024 | 0.772407 | 0.777243 |
| 0.793931 | 0.766764 | 0.792035 | 0.772673 | 0.769838 | 0.781219 | 0.794957 | 0.767375 | 0.762671 |
| 0.815756 | 0.791339 | 0.818682 | 0.780245 | 0.777796 | 0.788047 | 0.826756 | 0.781159 | 0.78539 |
| 0.800764 | 0.783335 | 0.807606 | 0.784779 | 0.780971 | 0.787446 | 0.811226 | 0.780284 | 0.783724 |
| 0.774089 | 0.769839 | 0.789686 | 0.762412 | 0.770835 | 0.770716 | 0.789019 | 0.765784 | 0.773943 |
| 0.777163 | 0.780245 | 0.796375 | 0.772964 | 0.782179 | 0.772276 | 0.800317 | 0.776836 | 0.78034 |

| | | | | | | | | |
|---|---|---|---|---|---|---|---|---|
| 0.785744 | 0.751076 | 0.782538 | 0.762474 | 0.75704 | 0.768104 | 0.789893 | 0.768918 | 0.765459 |
| 0.792811 | 0.797004 | 0.833701 | 0.783027 | 0.777207 | 0.783395 | 0.830978 | 0.808043 | 0.796174 |
| 0.803254 | 0.786202 | 0.81512 | 0.784155 | 0.804878 | 0.791378 | 0.822938 | 0.806124 | 0.818828 |
| 0.802212 | 0.789066 | 0.83161 | 0.789907 | 0.790546 | 0.798146 | 0.82902 | 0.813963 | 0.821618 |
| 0.804375 | 0.779173 | 0.809603 | 0.783206 | 0.800858 | 0.787374 | 0.82154 | 0.795083 | 0.796631 |
| 0.78368 | 0.782873 | 0.805278 | 0.775987 | 0.787123 | 0.77531 | 0.811372 | 0.797904 | 0.793149 |
| 0.772506 | 0.776283 | 0.79613 | 0.765157 | 0.783965 | 0.769062 | 0.79955 | 0.777037 | 0.777207 |
| 0.792973 | 0.802892 | 0.823041 | 0.791132 | 0.789465 | 0.780564 | 0.826113 | 0.78071 | 0.783534 |
| 0.784789 | 0.774639 | 0.795441 | 0.774241 | 0.784714 | 0.779106 | 0.800117 | 0.780271 | 0.776901 |
| 0.778254 | 0.77241 | 0.792254 | 0.758901 | 0.790047 | 0.773892 | 0.796722 | 0.775555 | 0.775583 |
| 0.809898 | 0.796011 | 0.826541 | 0.783807 | 0.792664 | 0.802364 | 0.82861 | 0.802068 | 0.799467 |
| 0.805796 | 0.802308 | 0.819773 | 0.77642 | 0.794887 | 0.785138 | 0.820205 | 0.79649 | 0.797231 |
| 0.816365 | 0.789849 | 0.823054 | 0.791997 | 0.790931 | 0.791442 | 0.823514 | 0.807913 | 0.801551 |
| 0.805715 | 0.789774 | 0.808677 | 0.773996 | 0.790031 | 0.790162 | 0.811318 | 0.804086 | 0.805428 |
| 0.821987 | 0.782843 | 0.81747 | 0.785664 | 0.806541 | 0.792629 | 0.816016 | 0.791415 | 0.783099 |
| 0.821198 | 0.778821 | 0.814701 | 0.783394 | 0.792538 | 0.793294 | 0.810776 | 0.793214 | 0.781459 |
| 0.829732 | 0.77018 | 0.819857 | 0.782287 | 0.781485 | 0.785364 | 0.81946 | 0.801842 | 0.794322 |
| 0.83731 | 0.793384 | 0.842989 | 0.798509 | 0.791989 | 0.810343 | 0.837146 | 0.811786 | 0.798246 |
| 0.785542 | 0.790122 | 0.813629 | 0.790164 | 0.795068 | 0.788175 | 0.816672 | 0.800501 | 0.793584 |
| 0.800051 | 0.7884 | 0.825006 | 0.793362 | 0.793418 | 0.785484 | 0.826384 | 0.797832 | 0.805999 |
| 0.852426 | 0.774791 | 0.827116 | 0.790031 | 0.774558 | 0.807029 | 0.833077 | 0.80247 | 0.805419 |
| 0.807226 | 0.785967 | 0.818819 | 0.774626 | 0.786644 | 0.795949 | 0.825295 | 0.801948 | 0.791267 |
| 0.796906 | 0.78835 | 0.799619 | 0.786622 | 0.793219 | 0.78139 | 0.804697 | 0.782587 | 0.779967 |
| 0.777243 | 0.769739 | 0.794096 | 0.765283 | 0.784952 | 0.771277 | 0.795898 | 0.77887 | 0.776949 |
| 0.763941 | 0.763249 | 0.783093 | 0.75545 | 0.765763 | 0.757071 | 0.787716 | 0.768288 | 0.76452 |
| 0.708521 | 0.707302 | 0.719075 | 0.702847 | 0.706227 | 0.702182 | 0.722869 | 0.710827 | 0.704443 |
| 0.71063 | 0.710214 | 0.724814 | 0.703719 | 0.708688 | 0.704531 | 0.726343 | 0.711746 | 0.708068 |
| 0.712306 | 0.711392 | 0.725693 | 0.704209 | 0.711584 | 0.704484 | 0.727655 | 0.712655 | 0.709016 |
| 0.768558 | 0.787442 | 0.799783 | 0.782699 | 0.786546 | 0.763784 | 0.805146 | 0.772139 | 0.771034 |
| 0.767541 | 0.789487 | 0.799054 | 0.778552 | 0.783887 | 0.765857 | 0.799966 | 0.776372 | 0.778021 |
| 0.764159 | 0.776986 | 0.808207 | 0.774157 | 0.792674 | 0.762052 | 0.806595 | 0.796063 | 0.797731 |
| 0.766556 | 0.792793 | 0.805033 | 0.776894 | 0.789934 | 0.764135 | 0.803659 | 0.771483 | 0.78003 |
| 0.863118 | 0.825844 | 0.839402 | 0.866974 | 0.787813 | 0.824377 | 0.84146 | 0.792024 | 0.786787 |
| 0.863881 | 0.784533 | 0.824226 | 0.809173 | 0.784092 | 0.830974 | 0.829066 | 0.787914 | 0.804264 |
| 0.86812 | 0.81165 | 0.847215 | 0.841125 | 0.785845 | 0.823044 | 0.844059 | 0.787792 | 0.77654 |
| 1 | 0.780962 | 0.844745 | 0.833672 | 0.783435 | 0.8652 | 0.844179 | 0.790574 | 0.777897 |
| 0.780962 | 1 | 0.834177 | 0.80714 | 0.785807 | 0.781476 | 0.830062 | 0.78007 | 0.779115 |
| 0.844745 | 0.834177 | 1 | 0.824914 | 0.804635 | 0.825002 | 0.931177 | 0.827945 | 0.81921 |
| 0.833672 | 0.80714 | 0.824914 | 1 | 0.778735 | 0.805089 | 0.823022 | 0.777044 | 0.772603 |
| 0.783435 | 0.785807 | 0.804635 | 0.778735 | 1 | 0.780621 | 0.803061 | 0.789691 | 0.787842 |
| 0.8652 | 0.781476 | 0.825002 | 0.805089 | 0.780621 | 1 | 0.826039 | 0.788503 | 0.785067 |
| 0.844179 | 0.830062 | 0.931177 | 0.823022 | 0.803061 | 0.826039 | 1 | 0.825755 | 0.82292 |
| 0.790574 | 0.78007 | 0.827945 | 0.777044 | 0.789691 | 0.788503 | 0.825755 | 1 | 0.823258 |
| 0.777897 | 0.779115 | 0.81921 | 0.772603 | 0.787842 | 0.785067 | 0.82292 | 0.823258 | 1 |
| 0.787927 | 0.788633 | 0.804053 | 0.777462 | 0.831854 | 0.778454 | 0.806392 | 0.788543 | 0.788223 |
| 0.861427 | 0.779884 | 0.812003 | 0.87848 | 0.780212 | 0.814968 | 0.817347 | 0.780852 | 0.779196 |

| | | | | | | | | |
|---|---|---|---|---|---|---|---|---|
| 0.787727 | 0.786961 | 0.829511 | 0.771652 | 0.785292 | 0.818845 | 0.823814 | 0.830174 | 0.822635 |
| 0.771058 | 0.790397 | 0.794555 | 0.770951 | 0.810057 | 0.770087 | 0.800167 | 0.780928 | 0.782157 |
| 0.784828 | 0.824002 | 0.807053 | 0.774849 | 0.795731 | 0.789924 | 0.805805 | 0.796368 | 0.794255 |
| 0.819066 | 0.816505 | 0.903283 | 0.801506 | 0.79527 | 0.807191 | 0.873639 | 0.809439 | 0.80545 |
| 0.78845 | 0.816012 | 0.815283 | 0.777029 | 0.805684 | 0.787587 | 0.812212 | 0.786221 | 0.790846 |
| 0.783833 | 0.779756 | 0.799344 | 0.774002 | 0.829916 | 0.785484 | 0.79726 | 0.787241 | 0.786704 |
| 0.783468 | 0.842608 | 0.831036 | 0.805563 | 0.785807 | 0.782095 | 0.821246 | 0.789214 | 0.781352 |
| 0.781279 | 0.81262 | 0.800924 | 0.772265 | 0.77938 | 0.777265 | 0.80406 | 0.783599 | 0.775122 |
| 0.787359 | 0.787329 | 0.802519 | 0.780859 | 0.783042 | 0.779089 | 0.807339 | 0.778333 | 0.782939 |
| 0.777603 | 0.770905 | 0.79049 | 0.771204 | 0.784725 | 0.77213 | 0.795986 | 0.779723 | 0.786418 |
| 0.789022 | 0.78326 | 0.805277 | 0.779274 | 0.784333 | 0.783218 | 0.80219 | 0.780034 | 0.781889 |
| 0.779275 | 0.778432 | 0.798871 | 0.774202 | 0.786877 | 0.773774 | 0.795149 | 0.783111 | 0.784995 |
| 0.785631 | 0.783033 | 0.811234 | 0.770656 | 0.793844 | 0.783058 | 0.81138 | 0.781632 | 0.789775 |
| 0.780613 | 0.775507 | 0.793197 | 0.766557 | 0.792839 | 0.778943 | 0.791455 | 0.774715 | 0.801029 |
| 0.787316 | 0.775595 | 0.796868 | 0.767039 | 0.787427 | 0.780921 | 0.80084 | 0.789478 | 0.789056 |
| 0.78887 | 0.782687 | 0.797692 | 0.772591 | 0.780581 | 0.780342 | 0.803801 | 0.783218 | 0.779575 |
| 0.778435 | 0.786128 | 0.798758 | 0.774349 | 0.791446 | 0.782192 | 0.804366 | 0.785267 | 0.784379 |
| 0.864379 | 0.773409 | 0.819911 | 0.801908 | 0.779021 | 0.821715 | 0.818436 | 0.781201 | 0.774929 |
| 0.78118 | 0.783296 | 0.802209 | 0.776372 | 0.784785 | 0.777613 | 0.804134 | 0.783297 | 0.791294 |
| 0.783415 | 0.776387 | 0.798632 | 0.770225 | 0.783846 | 0.776238 | 0.803134 | 0.784458 | 0.781301 |
| 0.780253 | 0.790396 | 0.803422 | 0.776506 | 0.782305 | 0.779247 | 0.798993 | 0.783981 | 0.783259 |
| 0.781722 | 0.779669 | 0.801771 | 0.80431 | 0.778694 | 0.77207 | 0.80594 | 0.778089 | 0.779992 |
| 0.795378 | 0.7964 | 0.808147 | 0.782669 | 0.792102 | 0.787449 | 0.801856 | 0.788107 | 0.782828 |
| 0.784672 | 0.782548 | 0.802537 | 0.772545 | 0.809807 | 0.778367 | 0.801806 | 0.781622 | 0.791979 |
| 0.779726 | 0.828171 | 0.814362 | 0.794092 | 0.780348 | 0.776231 | 0.807278 | 0.774677 | 0.775539 |
| 0.783893 | 0.839383 | 0.82822 | 0.807352 | 0.782499 | 0.773263 | 0.83069 | 0.78076 | 0.784379 |
| 0.779526 | 0.834925 | 0.824439 | 0.796726 | 0.784643 | 0.780942 | 0.817663 | 0.779786 | 0.780057 |
| 0.777255 | 0.78257 | 0.8003 | 0.769303 | 0.78637 | 0.771333 | 0.803701 | 0.778226 | 0.784953 |
| 0.785596 | 0.777153 | 0.801935 | 0.773427 | 0.78213 | 0.781022 | 0.80972 | 0.786323 | 0.787297 |
| 0.778412 | 0.825095 | 0.813904 | 0.782363 | 0.785488 | 0.773522 | 0.808531 | 0.77404 | 0.774788 |
| 0.781263 | 0.786445 | 0.798321 | 0.772387 | 0.778563 | 0.778382 | 0.797609 | 0.788587 | 0.786229 |
| 0.782841 | 0.790703 | 0.799471 | 0.771264 | 0.782743 | 0.774703 | 0.799619 | 0.779948 | 0.779268 |
| 0.788773 | 0.821072 | 0.827274 | 0.814966 | 0.782432 | 0.786906 | 0.822804 | 0.784273 | 0.782175 |
| 0.804053 | 0.791413 | 0.833276 | 0.792806 | 0.791754 | 0.799949 | 0.826121 | 0.815629 | 0.818158 |
| 0.792404 | 0.787906 | 0.813198 | 0.778621 | 0.799683 | 0.787711 | 0.808601 | 0.787073 | 0.783786 |
| 0.802673 | 0.782471 | 0.809747 | 0.788088 | 0.78795 | 0.788028 | 0.807109 | 0.786165 | 0.790007 |
| 0.786595 | 0.782435 | 0.80019 | 0.778574 | 0.789884 | 0.779645 | 0.8 | 0.785539 | 0.781083 |
| 0.762507 | 0.765138 | 0.778938 | 0.753643 | 0.763299 | 0.755149 | 0.780712 | 0.759131 | 0.75361 |
| 0.765125 | 0.762366 | 0.791844 | 0.757859 | 0.848031 | 0.76038 | 0.787995 | 0.768641 | 0.766688 |
| 0.748536 | 0.765308 | 0.777248 | 0.753611 | 0.758965 | 0.752414 | 0.777524 | 0.754582 | 0.753244 |
| 0.844223 | 0.763185 | 0.808473 | 0.779988 | 0.763954 | 0.79659 | 0.803563 | 0.767477 | 0.764421 |
| 0.741869 | 0.74564 | 0.759975 | 0.741114 | 0.751213 | 0.741756 | 0.75762 | 0.744508 | 0.781612 |
| 0.755713 | 0.757416 | 0.807271 | 0.751976 | 0.758786 | 0.762675 | 0.800729 | 0.819423 | 0.796506 |
| 0.81415 | 0.77193 | 0.792546 | 0.786218 | 0.765003 | 0.786181 | 0.791678 | 0.758022 | 0.752707 |
| 0.753548 | 0.755576 | 0.777613 | 0.746219 | 0.751418 | 0.751623 | 0.775281 | 0.757435 | 0.749868 |
| 0.785951 | 0.805393 | 0.803937 | 0.804629 | 0.757297 | 0.772709 | 0.804111 | 0.764721 | 0.756613 |
| 0.848878 | 0.756913 | 0.811449 | 0.783193 | 0.75361 | 0.797977 | 0.800798 | 0.760636 | 0.74888 |

| | | | | | | | | |
|---|---|---|---|---|---|---|---|---|
| 0.750064 | 0.742361 | 0.770149 | 0.748737 | 0.755321 | 0.746942 | 0.768069 | 0.748432 | 0.736268 |
| 0.845383 | 0.802079 | 0.842075 | 0.836278 | 0.753245 | 0.785479 | 0.828081 | 0.760245 | 0.754702 |
| 0.743661 | 0.750725 | 0.768473 | 0.745297 | 0.750485 | 0.73959 | 0.768329 | 0.748096 | 0.747996 |
| 0.749379 | 0.748049 | 0.808813 | 0.739358 | 0.747085 | 0.839588 | 0.808352 | 0.790483 | 0.782064 |
| 0.746783 | 0.797234 | 0.808785 | 0.785724 | 0.764049 | 0.747093 | 0.802215 | 0.750267 | 0.767062 |
| 0.743732 | 0.84566 | 0.781853 | 0.735665 | 0.740998 | 0.746551 | 0.777827 | 0.742418 | 0.741455 |
| 0.741232 | 0.795061 | 0.797615 | 0.764168 | 0.772383 | 0.746684 | 0.788666 | 0.755211 | 0.758174 |
| 0.837735 | 0.756037 | 0.802132 | 0.793213 | 0.756895 | 0.799225 | 0.802083 | 0.754758 | 0.794468 |
| 0.736834 | 0.733617 | 0.754778 | 0.725179 | 0.734907 | 0.72822 | 0.752323 | 0.732424 | 0.745967 |
| 0.75459 | 0.843581 | 0.84478 | 0.819422 | 0.752355 | 0.749051 | 0.83499 | 0.753009 | 0.748937 |
| 0.738478 | 0.793583 | 0.778355 | 0.74336 | 0.74856 | 0.736659 | 0.782232 | 0.748519 | 0.746051 |
| 0.77194 | 0.772016 | 0.814232 | 0.762976 | 0.772393 | 0.770032 | 0.816916 | 0.817863 | 0.807045 |
| 0.775755 | 0.776253 | 0.795678 | 0.774606 | 0.78696 | 0.77655 | 0.793814 | 0.776813 | 0.771862 |
| 0.745395 | 0.76497 | 0.777261 | 0.754445 | 0.751432 | 0.74177 | 0.776793 | 0.752433 | 0.751609 |
| 0.793287 | 0.773669 | 0.80736 | 0.777181 | 0.782064 | 0.784961 | 0.805402 | 0.785397 | 0.778321 |
| 0.767482 | 0.771025 | 0.785495 | 0.762858 | 0.780562 | 0.764902 | 0.784742 | 0.771125 | 0.784884 |
| 0.778181 | 0.779803 | 0.804487 | 0.765934 | 0.783951 | 0.773596 | 0.811691 | 0.782348 | 0.787398 |
| 0.778214 | 0.777384 | 0.797054 | 0.776771 | 0.78017 | 0.771104 | 0.801958 | 0.780065 | 0.777494 |
| 0.760442 | 0.772757 | 0.780636 | 0.752732 | 0.781703 | 0.752764 | 0.776156 | 0.763284 | 0.750801 |
| 0.771111 | 0.778192 | 0.796907 | 0.767902 | 0.772005 | 0.766489 | 0.796238 | 0.776615 | 0.787421 |
| 0.748569 | 0.77892 | 0.779477 | 0.740759 | 0.744358 | 0.741641 | 0.771883 | 0.749691 | 0.747684 |
| 0.767143 | 0.770084 | 0.7871 | 0.766584 | 0.77683 | 0.768904 | 0.794341 | 0.776144 | 0.776748 |
| 0.7749 | 0.771781 | 0.789351 | 0.7712 | 0.773416 | 0.764149 | 0.795133 | 0.770899 | 0.772604 |
| 0.745339 | 0.746965 | 0.771491 | 0.744504 | 0.772382 | 0.737531 | 0.770484 | 0.740957 | 0.747131 |
| 0.779143 | 0.761313 | 0.794472 | 0.759884 | 0.746275 | 0.752621 | 0.791674 | 0.754153 | 0.748541 |
| 0.801199 | 0.75863 | 0.790094 | 0.769844 | 0.764683 | 0.776407 | 0.792584 | 0.756877 | 0.756161 |
| 0.758094 | 0.783913 | 0.784029 | 0.752492 | 0.756728 | 0.756564 | 0.785047 | 0.762904 | 0.760302 |
| 0.775443 | 0.773333 | 0.795349 | 0.763366 | 0.779099 | 0.773357 | 0.798751 | 0.77343 | 0.773997 |
| 0.753063 | 0.753681 | 0.774327 | 0.778653 | 0.750457 | 0.750658 | 0.772986 | 0.74836 | 0.749043 |
| 0.772163 | 0.811677 | 0.82202 | 0.79526 | 0.773542 | 0.762984 | 0.814846 | 0.771738 | 0.763446 |
| 0.762197 | 0.768603 | 0.788005 | 0.763665 | 0.769995 | 0.756735 | 0.78904 | 0.766021 | 0.770454 |
| 0.788776 | 0.767948 | 0.857069 | 0.771208 | 0.760345 | 0.775869 | 0.835302 | 0.779111 | 0.76374 |
| 0.786152 | 0.76835 | 0.850745 | 0.773033 | 0.763982 | 0.773902 | 0.830782 | 0.778162 | 0.770063 |
| 0.770688 | 0.766448 | 0.794672 | 0.769408 | 0.763403 | 0.758587 | 0.791107 | 0.772164 | 0.767985 |
| 0.766109 | 0.764532 | 0.7837 | 0.759672 | 0.7657 | 0.761814 | 0.784986 | 0.765995 | 0.756129 |
| 0.771336 | 0.76994 | 0.789342 | 0.766194 | 0.777933 | 0.765215 | 0.785637 | 0.77752 | 0.77177 |
| 0.728379 | 0.731202 | 0.744375 | 0.725176 | 0.726432 | 0.727193 | 0.747567 | 0.727338 | 0.725174 |
| 0.759863 | 0.76854 | 0.781808 | 0.755411 | 0.760528 | 0.758495 | 0.778106 | 0.759131 | 0.75235 |
| 0.730425 | 0.741254 | 0.754738 | 0.736764 | 0.733841 | 0.723726 | 0.753765 | 0.736205 | 0.726533 |
| 0.713829 | 0.736879 | 0.736855 | 0.709188 | 0.73103 | 0.712084 | 0.736759 | 0.71865 | 0.720666 |
| 0.779601 | 0.775809 | 0.801306 | 0.77086 | 0.778021 | 0.775722 | 0.794669 | 0.772843 | 0.773726 |
| 0.719647 | 0.736789 | 0.736714 | 0.712835 | 0.732898 | 0.717628 | 0.73477 | 0.719418 | 0.71671 |
| 0.751006 | 0.733499 | 0.750473 | 0.743924 | 0.721424 | 0.732364 | 0.752846 | 0.720323 | 0.71377 |
| 0.728365 | 0.731009 | 0.743286 | 0.724625 | 0.711699 | 0.715827 | 0.738936 | 0.709688 | 0.70746 |
| 0.770497 | 0.776515 | 0.808226 | 0.771031 | 0.769231 | 0.774476 | 0.805861 | 0.782827 | 0.777651 |
| 0.735814 | 0.72583 | 0.739431 | 0.730719 | 0.708782 | 0.717087 | 0.743192 | 0.708929 | 0.709972 |
| 0.779494 | 0.772093 | 0.8014 | 0.772289 | 0.770927 | 0.772301 | 0.795668 | 0.775445 | 0.776084 |

| | | | | | | | | |
|---|---|---|---|---|---|---|---|---|
| 0.777132 | 0.774632 | 0.792462 | 0.761151 | 0.766892 | 0.771087 | 0.792799 | 0.770221 | 0.763177 |
| 0.775907 | 0.776722 | 0.799204 | 0.775234 | 0.773063 | 0.77182 | 0.798604 | 0.772411 | 0.777299 |
| 0.752688 | 0.752617 | 0.773895 | 0.744117 | 0.754058 | 0.747752 | 0.776864 | 0.762285 | 0.753127 |
| 0.737067 | 0.716773 | 0.733514 | 0.726615 | 0.700053 | 0.713516 | 0.732878 | 0.698935 | 0.695537 |
| 0.711785 | 0.73921 | 0.740512 | 0.717565 | 0.703239 | 0.704625 | 0.732847 | 0.699428 | 0.693376 |
| 0.78885 | 0.779062 | 0.797479 | 0.772503 | 0.772473 | 0.785913 | 0.792937 | 0.782856 | 0.777134 |
| 0.696248 | 0.690577 | 0.710363 | 0.697444 | 0.695846 | 0.692267 | 0.713955 | 0.692518 | 0.690097 |
| 0.732833 | 0.732108 | 0.749365 | 0.729963 | 0.73285 | 0.72919 | 0.755953 | 0.739542 | 0.734773 |
| 0.791118 | 0.763258 | 0.790731 | 0.770691 | 0.763524 | 0.772107 | 0.796473 | 0.772552 | 0.764452 |
| 0.718024 | 0.721355 | 0.734079 | 0.709824 | 0.723899 | 0.723069 | 0.737498 | 0.725917 | 0.722459 |
| 0.769365 | 0.753161 | 0.782464 | 0.764907 | 0.749273 | 0.758076 | 0.781666 | 0.760279 | 0.755067 |
| 0.629139 | 0.670968 | 0.651007 | 0.675676 | 0.607595 | 0.636364 | 0.675676 | 0.645161 | 0.611111 |
| 0.652575 | 0.658162 | 0.678858 | 0.651627 | 0.655113 | 0.647708 | 0.673883 | 0.654053 | 0.648432 |
| 0.654895 | 0.658496 | 0.675507 | 0.653465 | 0.652239 | 0.649723 | 0.675651 | 0.662477 | 0.649492 |
| 0.665989 | 0.667293 | 0.687739 | 0.663988 | 0.665802 | 0.66044 | 0.684932 | 0.669432 | 0.663004 |
| 0.666753 | 0.666882 | 0.684211 | 0.659982 | 0.656774 | 0.661334 | 0.681222 | 0.667401 | 0.658678 |
| 0.623693 | 0.617946 | 0.641505 | 0.617938 | 0.626176 | 0.617689 | 0.641212 | 0.624738 | 0.621171 |
| 0.671432 | 0.672067 | 0.684104 | 0.663412 | 0.66107 | 0.664586 | 0.68362 | 0.662592 | 0.660192 |
| 0.75259 | 0.74767 | 0.762363 | 0.752769 | 0.742608 | 0.747772 | 0.76207 | 0.743548 | 0.742811 |
| 0.763292 | 0.742782 | 0.766036 | 0.750366 | 0.741101 | 0.752283 | 0.764902 | 0.746527 | 0.744697 |
| 0.764696 | 0.75379 | 0.769581 | 0.757013 | 0.746323 | 0.753526 | 0.7705 | 0.749815 | 0.747475 |
| 0.618629 | 0.631118 | 0.630348 | 0.614237 | 0.615237 | 0.618085 | 0.634213 | 0.620127 | 0.61818 |
| 0.603085 | 0.605888 | 0.611652 | 0.60109 | 0.602145 | 0.60098 | 0.614457 | 0.60183 | 0.598283 |
| 0.708171 | 0.706164 | 0.718872 | 0.699439 | 0.70934 | 0.70831 | 0.718194 | 0.703169 | 0.701504 |
| 0.707309 | 0.709316 | 0.72329 | 0.700553 | 0.710238 | 0.705901 | 0.719395 | 0.709356 | 0.704651 |
| 0.701381 | 0.701199 | 0.713378 | 0.691541 | 0.701809 | 0.701328 | 0.711938 | 0.701122 | 0.695635 |
| 0.704969 | 0.702654 | 0.71716 | 0.698574 | 0.705517 | 0.703099 | 0.71967 | 0.704737 | 0.699757 |
| 0.709559 | 0.708868 | 0.721513 | 0.700033 | 0.707457 | 0.706963 | 0.722778 | 0.710135 | 0.701611 |
| 0.708364 | 0.708011 | 0.723634 | 0.699226 | 0.708719 | 0.702079 | 0.721442 | 0.707303 | 0.704756 |
| 0.708283 | 0.709187 | 0.723498 | 0.69905 | 0.709394 | 0.70544 | 0.722985 | 0.710318 | 0.703476 |
| 0.707403 | 0.706649 | 0.722464 | 0.702458 | 0.708271 | 0.702692 | 0.721979 | 0.709182 | 0.706388 |
| 0.712073 | 0.711813 | 0.725935 | 0.704969 | 0.714611 | 0.709473 | 0.729218 | 0.71651 | 0.712288 |
| 0.708838 | 0.707735 | 0.722685 | 0.702085 | 0.71092 | 0.70336 | 0.726564 | 0.71222 | 0.709404 |
| 0.711715 | 0.710148 | 0.723418 | 0.706037 | 0.708668 | 0.705864 | 0.729114 | 0.710574 | 0.710708 |
| 0.711429 | 0.713147 | 0.724401 | 0.704773 | 0.715066 | 0.707877 | 0.726893 | 0.713261 | 0.709084 |
| 0.708491 | 0.705728 | 0.719182 | 0.705574 | 0.706884 | 0.705579 | 0.722814 | 0.708922 | 0.709231 |
| 0.709778 | 0.70912 | 0.719453 | 0.699495 | 0.708961 | 0.706498 | 0.724683 | 0.710339 | 0.706237 |
| 0.710291 | 0.707511 | 0.721021 | 0.700948 | 0.71058 | 0.708932 | 0.722794 | 0.71158 | 0.706692 |
| 0.701095 | 0.706887 | 0.717177 | 0.698083 | 0.7016 | 0.701733 | 0.718421 | 0.708135 | 0.703445 |
| 0.707806 | 0.710238 | 0.720581 | 0.703695 | 0.707754 | 0.705411 | 0.725786 | 0.710029 | 0.708258 |
| 0.658941 | 0.657868 | 0.678828 | 0.655172 | 0.661582 | 0.65696 | 0.677995 | 0.663502 | 0.654946 |
| 0.663131 | 0.661483 | 0.683977 | 0.654177 | 0.658955 | 0.659942 | 0.679919 | 0.663275 | 0.662543 |
| 0.70938 | 0.710681 | 0.726016 | 0.703816 | 0.711132 | 0.70812 | 0.732118 | 0.715469 | 0.711646 |
| 0.709366 | 0.714297 | 0.728646 | 0.703685 | 0.71211 | 0.709996 | 0.73197 | 0.716102 | 0.711057 |
| 0.642897 | 0.644936 | 0.667354 | 0.643269 | 0.641565 | 0.636996 | 0.662655 | 0.645929 | 0.642792 |
| 0.708938 | 0.709473 | 0.723744 | 0.700455 | 0.709851 | 0.70538 | 0.728343 | 0.716592 | 0.709998 |
| 0.699857 | 0.697008 | 0.715744 | 0.693299 | 0.7004 | 0.693766 | 0.71944 | 0.701663 | 0.697869 |

| | | | | | | | | |
|---|---|---|---|---|---|---|---|---|
| 0.705552 | 0.706978 | 0.721718 | 0.702475 | 0.709996 | 0.700923 | 0.727647 | 0.710831 | 0.703718 |
| 0.706517 | 0.707661 | 0.723364 | 0.7011 | 0.71143 | 0.699571 | 0.724176 | 0.711217 | 0.705601 |
| 0.709348 | 0.708162 | 0.725833 | 0.706519 | 0.711183 | 0.705128 | 0.726703 | 0.710776 | 0.706877 |
| 0.707097 | 0.709448 | 0.722441 | 0.699637 | 0.711127 | 0.705567 | 0.725476 | 0.713316 | 0.707421 |
| 0.715704 | 0.715221 | 0.731947 | 0.709811 | 0.71591 | 0.712391 | 0.732499 | 0.718268 | 0.712493 |
| 0.702536 | 0.707358 | 0.721635 | 0.698425 | 0.704787 | 0.69977 | 0.724169 | 0.709945 | 0.706126 |
| 0.705434 | 0.707872 | 0.723685 | 0.700559 | 0.707954 | 0.701655 | 0.723476 | 0.713273 | 0.707145 |
| 0.700105 | 0.698911 | 0.716157 | 0.695761 | 0.701154 | 0.698925 | 0.71728 | 0.702248 | 0.700455 |
| 0.705906 | 0.707142 | 0.718094 | 0.69658 | 0.707893 | 0.698946 | 0.7211 | 0.708636 | 0.702309 |
| 0.707393 | 0.70194 | 0.719537 | 0.699171 | 0.708419 | 0.702998 | 0.724734 | 0.708731 | 0.704 |
| 0.709893 | 0.706963 | 0.723074 | 0.700214 | 0.711413 | 0.703506 | 0.726559 | 0.712262 | 0.707733 |
| 0.705306 | 0.704243 | 0.718804 | 0.696083 | 0.705062 | 0.699131 | 0.721446 | 0.710821 | 0.703671 |
| 0.698284 | 0.699045 | 0.715756 | 0.693347 | 0.700518 | 0.693899 | 0.719678 | 0.702918 | 0.697011 |
| 0.697431 | 0.695688 | 0.713403 | 0.693 | 0.698907 | 0.696546 | 0.717923 | 0.702867 | 0.697929 |
| 0.711879 | 0.71623 | 0.728491 | 0.706483 | 0.711531 | 0.709602 | 0.729587 | 0.713765 | 0.711123 |
| 0.70976 | 0.714387 | 0.726597 | 0.703215 | 0.709234 | 0.704038 | 0.726342 | 0.71089 | 0.707231 |
| 0.704185 | 0.703997 | 0.718592 | 0.694791 | 0.701603 | 0.699421 | 0.719463 | 0.705176 | 0.70293 |
| 0.706713 | 0.705333 | 0.719544 | 0.698796 | 0.704879 | 0.70351 | 0.72435 | 0.707219 | 0.706965 |
| 0.705887 | 0.70516 | 0.719782 | 0.697826 | 0.704724 | 0.704821 | 0.721729 | 0.703992 | 0.70228 |
| 0.704077 | 0.704655 | 0.719119 | 0.697341 | 0.706138 | 0.703768 | 0.721819 | 0.70646 | 0.703821 |
| 0.703764 | 0.702449 | 0.715042 | 0.695214 | 0.700056 | 0.697885 | 0.718997 | 0.704212 | 0.699695 |
| 0.704839 | 0.701918 | 0.717377 | 0.696484 | 0.703834 | 0.702757 | 0.720611 | 0.701011 | 0.70221 |
| 0.710952 | 0.708147 | 0.723184 | 0.701813 | 0.70785 | 0.707658 | 0.726214 | 0.709759 | 0.703762 |
| 0.70844 | 0.70859 | 0.7206 | 0.701448 | 0.708228 | 0.705794 | 0.725867 | 0.707961 | 0.708567 |
| 0.706996 | 0.70783 | 0.720925 | 0.697688 | 0.704469 | 0.704555 | 0.723989 | 0.705119 | 0.703455 |
| 0.702218 | 0.70419 | 0.716756 | 0.694027 | 0.700673 | 0.701728 | 0.716876 | 0.70092 | 0.70105 |
| 0.704016 | 0.704338 | 0.714598 | 0.694927 | 0.705562 | 0.704442 | 0.719721 | 0.705003 | 0.703784 |
| 0.702221 | 0.702104 | 0.713215 | 0.692804 | 0.701099 | 0.699133 | 0.715619 | 0.700762 | 0.698339 |
| 0.70433 | 0.706314 | 0.717026 | 0.695111 | 0.704683 | 0.702 | 0.718795 | 0.703165 | 0.702311 |
| 0.707678 | 0.707461 | 0.72001 | 0.697307 | 0.705595 | 0.705813 | 0.723558 | 0.70686 | 0.706105 |
| 0.708993 | 0.709994 | 0.719904 | 0.701159 | 0.706258 | 0.70494 | 0.724066 | 0.71125 | 0.706747 |
| 0.709425 | 0.708882 | 0.720932 | 0.702601 | 0.70978 | 0.709639 | 0.724826 | 0.713039 | 0.710369 |
| 0.713038 | 0.712662 | 0.730751 | 0.706728 | 0.714745 | 0.713668 | 0.728881 | 0.715477 | 0.709308 |
| 0.714297 | 0.713553 | 0.728141 | 0.706457 | 0.713609 | 0.711759 | 0.728434 | 0.717614 | 0.711171 |
| 0.702672 | 0.703525 | 0.715236 | 0.694555 | 0.703315 | 0.700443 | 0.717708 | 0.701645 | 0.702504 |
| 0.706646 | 0.704795 | 0.719548 | 0.696742 | 0.706717 | 0.704835 | 0.723845 | 0.708252 | 0.706307 |
| 0.707237 | 0.707406 | 0.720669 | 0.696139 | 0.7092 | 0.705976 | 0.722375 | 0.705618 | 0.704974 |
| 0.707147 | 0.7036 | 0.719229 | 0.695967 | 0.703305 | 0.702413 | 0.720789 | 0.703611 | 0.702511 |
| 0.699667 | 0.700225 | 0.712879 | 0.692475 | 0.697637 | 0.697723 | 0.714588 | 0.703131 | 0.697777 |
| 0.697797 | 0.698519 | 0.71256 | 0.691126 | 0.69749 | 0.696223 | 0.711462 | 0.698823 | 0.696217 |
| 0.698599 | 0.698793 | 0.708014 | 0.692839 | 0.695543 | 0.696747 | 0.710424 | 0.694935 | 0.694107 |
| 0.697631 | 0.69727 | 0.710171 | 0.688632 | 0.696698 | 0.696101 | 0.714921 | 0.693859 | 0.695087 |
| 0.697407 | 0.702173 | 0.712195 | 0.691647 | 0.699281 | 0.69862 | 0.714453 | 0.704509 | 0.699092 |
| 0.692979 | 0.689096 | 0.701234 | 0.686571 | 0.691897 | 0.691296 | 0.705919 | 0.690547 | 0.687667 |
| 0.705254 | 0.700058 | 0.71473 | 0.697483 | 0.701531 | 0.698102 | 0.719743 | 0.701991 | 0.698096 |
| 0.696814 | 0.697594 | 0.708173 | 0.689804 | 0.694517 | 0.694314 | 0.712889 | 0.696548 | 0.697968 |

| | | | | | | | | |
|---|---|---|---|---|---|---|---|---|
| 0.710814 | 0.711058 | 0.724227 | 0.700083 | 0.707622 | 0.707844 | 0.725793 | 0.706357 | 0.704495 |
| 0.709035 | 0.705845 | 0.720798 | 0.69844 | 0.707571 | 0.702459 | 0.723731 | 0.714239 | 0.706597 |
| 0.705416 | 0.709273 | 0.71609 | 0.697883 | 0.706493 | 0.700614 | 0.718798 | 0.704799 | 0.703325 |
| 0.709539 | 0.709921 | 0.724014 | 0.697602 | 0.71012 | 0.70413 | 0.724683 | 0.712889 | 0.709195 |
| 0.717685 | 0.71248 | 0.729312 | 0.705912 | 0.719546 | 0.713618 | 0.730145 | 0.717821 | 0.712701 |
| 0.708804 | 0.708075 | 0.72137 | 0.701697 | 0.710444 | 0.707582 | 0.724713 | 0.710719 | 0.707325 |
| 0.705418 | 0.704988 | 0.717573 | 0.696558 | 0.704396 | 0.701682 | 0.718088 | 0.707635 | 0.701916 |
| 0.705256 | 0.705596 | 0.721555 | 0.699316 | 0.708861 | 0.703008 | 0.721281 | 0.710673 | 0.710416 |
| 0.699747 | 0.703425 | 0.717008 | 0.69425 | 0.701261 | 0.697972 | 0.719922 | 0.70225 | 0.698328 |
| 0.709244 | 0.702253 | 0.719756 | 0.698999 | 0.71053 | 0.704091 | 0.723596 | 0.709855 | 0.709238 |
| 0.708689 | 0.708884 | 0.723965 | 0.702913 | 0.712028 | 0.708617 | 0.726911 | 0.71337 | 0.706472 |
| 0.71036 | 0.7071 | 0.722903 | 0.701246 | 0.711143 | 0.706907 | 0.724996 | 0.710936 | 0.705627 |
| 0.705774 | 0.706369 | 0.721133 | 0.698639 | 0.709137 | 0.702051 | 0.721622 | 0.708625 | 0.708231 |
| 0.710106 | 0.70734 | 0.72212 | 0.700693 | 0.710391 | 0.704124 | 0.722703 | 0.70841 | 0.708656 |
| 0.709998 | 0.708486 | 0.723629 | 0.703716 | 0.70969 | 0.704722 | 0.726441 | 0.71226 | 0.709915 |
| 0.708513 | 0.706511 | 0.721566 | 0.700833 | 0.710769 | 0.705679 | 0.721663 | 0.712657 | 0.712774 |
| 0.714897 | 0.712752 | 0.725727 | 0.70507 | 0.712528 | 0.710225 | 0.731042 | 0.71268 | 0.712313 |
| 0.708146 | 0.708998 | 0.720451 | 0.703979 | 0.711368 | 0.701125 | 0.724739 | 0.708088 | 0.708639 |
| 0.715931 | 0.713442 | 0.725002 | 0.701679 | 0.709477 | 0.711604 | 0.728324 | 0.71216 | 0.714557 |
| 0.710809 | 0.707537 | 0.719695 | 0.700912 | 0.70454 | 0.704339 | 0.723709 | 0.707485 | 0.707486 |
| 0.708822 | 0.704445 | 0.722581 | 0.702202 | 0.710166 | 0.704106 | 0.725541 | 0.709737 | 0.709731 |
| 0.707537 | 0.705299 | 0.720113 | 0.701015 | 0.708264 | 0.704704 | 0.724227 | 0.708253 | 0.70469 |
| 0.711468 | 0.711326 | 0.72433 | 0.70032 | 0.710904 | 0.708238 | 0.727425 | 0.713855 | 0.711969 |
| 0.709667 | 0.710594 | 0.727251 | 0.705694 | 0.708852 | 0.706546 | 0.729006 | 0.711675 | 0.708634 |
| 0.708287 | 0.708013 | 0.725631 | 0.702013 | 0.706234 | 0.702518 | 0.725316 | 0.709119 | 0.708457 |
| 0.707951 | 0.707792 | 0.723787 | 0.70014 | 0.705341 | 0.704208 | 0.723169 | 0.709939 | 0.705002 |
| 0.704936 | 0.703629 | 0.721559 | 0.695638 | 0.706789 | 0.700586 | 0.722744 | 0.708828 | 0.706928 |
| 0.706139 | 0.706665 | 0.719765 | 0.697711 | 0.703964 | 0.70318 | 0.721344 | 0.709634 | 0.7055 |
| 0.709557 | 0.713793 | 0.724201 | 0.701073 | 0.710897 | 0.706085 | 0.72605 | 0.713414 | 0.711964 |
| 0.710585 | 0.712928 | 0.726411 | 0.705927 | 0.713646 | 0.705052 | 0.72659 | 0.713476 | 0.710001 |
| 0.708059 | 0.70536 | 0.719197 | 0.701823 | 0.706354 | 0.701271 | 0.721969 | 0.708539 | 0.705219 |
| 0.714113 | 0.713683 | 0.72606 | 0.705579 | 0.710629 | 0.707128 | 0.730414 | 0.709269 | 0.711453 |
| 0.713601 | 0.710355 | 0.723216 | 0.702663 | 0.714094 | 0.709134 | 0.724662 | 0.710489 | 0.711132 |
| 0.700008 | 0.704837 | 0.71925 | 0.69638 | 0.706862 | 0.697985 | 0.721015 | 0.706259 | 0.703677 |
| 0.711027 | 0.709032 | 0.725805 | 0.70254 | 0.704737 | 0.704927 | 0.727145 | 0.708973 | 0.705887 |
| 0.702885 | 0.707283 | 0.718952 | 0.696637 | 0.705746 | 0.700652 | 0.721466 | 0.707617 | 0.704267 |
| 0.713442 | 0.717663 | 0.728395 | 0.708975 | 0.716022 | 0.712076 | 0.731335 | 0.718311 | 0.713936 |
| 0.706246 | 0.708447 | 0.721797 | 0.70005 | 0.70845 | 0.704212 | 0.726967 | 0.707098 | 0.705108 |
| 0.71117 | 0.706911 | 0.725062 | 0.701982 | 0.7121 | 0.70604 | 0.727024 | 0.711431 | 0.706664 |
| 0.709226 | 0.709174 | 0.72229 | 0.701676 | 0.710518 | 0.704617 | 0.726609 | 0.707619 | 0.706332 |
| 0.716975 | 0.715095 | 0.729423 | 0.707287 | 0.712067 | 0.709475 | 0.733488 | 0.714047 | 0.711655 |
| 0.714503 | 0.711176 | 0.72576 | 0.705029 | 0.711787 | 0.707736 | 0.727834 | 0.709256 | 0.706921 |
| 0.714308 | 0.710833 | 0.724847 | 0.703203 | 0.7128 | 0.708897 | 0.729704 | 0.7098 | 0.708275 |
| 0.708374 | 0.706818 | 0.721433 | 0.701328 | 0.711466 | 0.702327 | 0.724451 | 0.7076 | 0.708883 |
| 0.70918 | 0.7083 | 0.724589 | 0.703633 | 0.71115 | 0.702833 | 0.727597 | 0.711512 | 0.710964 |
| 0.706322 | 0.703896 | 0.72054 | 0.698199 | 0.704858 | 0.699712 | 0.722644 | 0.70523 | 0.704214 |

| | | | | | | | | |
|---|---|---|---|---|---|---|---|---|
| 0.698978 | 0.696526 | 0.707019 | 0.687195 | 0.695242 | 0.690141 | 0.710176 | 0.691429 | 0.691748 |
| 0.752807 | 0.744605 | 0.767241 | 0.741675 | 0.746383 | 0.747571 | 0.770604 | 0.747262 | 0.751247 |
| 0.702824 | 0.707637 | 0.718157 | 0.697565 | 0.706653 | 0.702281 | 0.721074 | 0.703619 | 0.703902 |
| 0.709457 | 0.71236 | 0.721261 | 0.701466 | 0.712214 | 0.705206 | 0.727802 | 0.708409 | 0.709326 |
| 0.70319 | 0.706143 | 0.714084 | 0.696119 | 0.706307 | 0.698511 | 0.717761 | 0.702788 | 0.699256 |
| 0.706874 | 0.708792 | 0.721124 | 0.698518 | 0.705069 | 0.700835 | 0.721745 | 0.707352 | 0.703565 |
| 0.703868 | 0.703481 | 0.715588 | 0.695499 | 0.704934 | 0.698961 | 0.719614 | 0.707527 | 0.703271 |
| 0.699803 | 0.703142 | 0.711197 | 0.6943 | 0.699406 | 0.692529 | 0.715668 | 0.6977 | 0.698088 |
| 0.705527 | 0.708232 | 0.72066 | 0.695471 | 0.707327 | 0.701441 | 0.722027 | 0.707924 | 0.705799 |
| 0.716473 | 0.715512 | 0.730168 | 0.707465 | 0.714614 | 0.710702 | 0.731944 | 0.713634 | 0.712266 |
| 0.710396 | 0.708076 | 0.723014 | 0.703372 | 0.709124 | 0.705044 | 0.7267 | 0.712789 | 0.706523 |
| 0.70997 | 0.712789 | 0.725587 | 0.701354 | 0.71296 | 0.707155 | 0.729894 | 0.711068 | 0.712073 |
| 0.7144 | 0.708769 | 0.728477 | 0.704876 | 0.712157 | 0.713008 | 0.728573 | 0.713373 | 0.714897 |
| 0.712126 | 0.710584 | 0.726373 | 0.704364 | 0.710919 | 0.708949 | 0.729198 | 0.711692 | 0.711328 |
| 0.706595 | 0.710579 | 0.723614 | 0.699288 | 0.710499 | 0.70473 | 0.725405 | 0.707307 | 0.709789 |
| 0.711692 | 0.709116 | 0.723871 | 0.702549 | 0.709735 | 0.706969 | 0.726344 | 0.712127 | 0.707414 |
| 0.711402 | 0.711034 | 0.724432 | 0.703369 | 0.712023 | 0.708462 | 0.727824 | 0.711061 | 0.708796 |
| 0.711442 | 0.710145 | 0.722871 | 0.699809 | 0.707861 | 0.710076 | 0.726883 | 0.713354 | 0.709878 |
| 0.704599 | 0.704616 | 0.718027 | 0.699087 | 0.704314 | 0.699152 | 0.716874 | 0.700673 | 0.696198 |
| 0.703435 | 0.705494 | 0.718607 | 0.693632 | 0.700841 | 0.699085 | 0.72017 | 0.702083 | 0.7002 |
| 0.708458 | 0.710886 | 0.722475 | 0.703055 | 0.706883 | 0.703311 | 0.724691 | 0.708364 | 0.706739 |
| 0.698558 | 0.698503 | 0.708873 | 0.692301 | 0.697398 | 0.69232 | 0.712782 | 0.702663 | 0.700268 |
| 0.705554 | 0.7053 | 0.718902 | 0.697617 | 0.706077 | 0.701022 | 0.721474 | 0.706861 | 0.706086 |
| 0.706372 | 0.708934 | 0.723199 | 0.701642 | 0.709887 | 0.70297 | 0.72819 | 0.710871 | 0.708528 |
| 0.711217 | 0.711915 | 0.723465 | 0.706111 | 0.712507 | 0.707353 | 0.728253 | 0.713421 | 0.709931 |
| 0.708117 | 0.708253 | 0.720088 | 0.70156 | 0.710178 | 0.703203 | 0.722856 | 0.710326 | 0.709371 |
| 0.705425 | 0.702379 | 0.716802 | 0.697599 | 0.704068 | 0.702568 | 0.720861 | 0.70364 | 0.701078 |
| 0.703314 | 0.699681 | 0.716293 | 0.694551 | 0.704488 | 0.700328 | 0.720287 | 0.704179 | 0.698414 |
| 0.701744 | 0.703212 | 0.719245 | 0.69762 | 0.703148 | 0.700584 | 0.720626 | 0.704741 | 0.705065 |
| 0.764082 | 0.766819 | 0.782943 | 0.762646 | 0.804245 | 0.76509 | 0.783677 | 0.768996 | 0.764912 |
| 0.780535 | 0.821483 | 0.829824 | 0.791012 | 0.821842 | 0.779598 | 0.836517 | 0.783259 | 0.779932 |
| 0.767235 | 0.789478 | 0.791614 | 0.770607 | 0.821386 | 0.769155 | 0.797539 | 0.778963 | 0.781848 |
| 0.772598 | 0.801163 | 0.825191 | 0.761932 | 0.771085 | 0.78192 | 0.823346 | 0.821751 | 0.823804 |
| 0.778374 | 0.859101 | 0.830616 | 0.797132 | 0.785819 | 0.780839 | 0.83442 | 0.783304 | 0.78113 |
| 0.700658 | 0.698603 | 0.715301 | 0.693351 | 0.701821 | 0.695585 | 0.717851 | 0.703206 | 0.701597 |
| 0.704884 | 0.701698 | 0.71739 | 0.698357 | 0.705709 | 0.699537 | 0.72056 | 0.705497 | 0.703639 |
| 0.705175 | 0.705498 | 0.720376 | 0.696289 | 0.706684 | 0.700804 | 0.723283 | 0.707016 | 0.706056 |
| 0.706562 | 0.705938 | 0.721426 | 0.699038 | 0.708881 | 0.705262 | 0.724524 | 0.709639 | 0.707443 |
| 0.707103 | 0.704621 | 0.720213 | 0.699701 | 0.706817 | 0.701458 | 0.721989 | 0.704146 | 0.702795 |
| 0.708842 | 0.71249 | 0.724408 | 0.702947 | 0.713264 | 0.70788 | 0.72579 | 0.711889 | 0.707701 |
| 0.704278 | 0.708067 | 0.719435 | 0.695739 | 0.704892 | 0.703033 | 0.723094 | 0.705555 | 0.704842 |
| 0.7054 | 0.701714 | 0.71968 | 0.698365 | 0.708711 | 0.70349 | 0.723291 | 0.707181 | 0.707162 |
| 0.701443 | 0.699042 | 0.71338 | 0.694889 | 0.698013 | 0.696083 | 0.715689 | 0.701553 | 0.696558 |
| 0.699458 | 0.698421 | 0.71351 | 0.690659 | 0.699839 | 0.694987 | 0.713978 | 0.700523 | 0.698394 |
| 0.706463 | 0.700785 | 0.718475 | 0.695999 | 0.707993 | 0.70315 | 0.720727 | 0.707039 | 0.704269 |
| 0.708013 | 0.703222 | 0.720169 | 0.6977 | 0.709073 | 0.70478 | 0.7253 | 0.709117 | 0.708587 |

| | | | | | | | | |
|---|---|---|---|---|---|---|---|---|
| 0.708787 | 0.70666 | 0.723114 | 0.700742 | 0.706475 | 0.703274 | 0.72662 | 0.708799 | 0.706449 |
| 0.708048 | 0.70741 | 0.718572 | 0.705569 | 0.705635 | 0.701331 | 0.724118 | 0.70816 | 0.703818 |
| 0.711655 | 0.709584 | 0.725369 | 0.703911 | 0.708722 | 0.706592 | 0.727829 | 0.712968 | 0.7083 |
| 0.703772 | 0.707418 | 0.720818 | 0.696026 | 0.708651 | 0.702442 | 0.725222 | 0.709189 | 0.707579 |
| 0.702625 | 0.705454 | 0.717957 | 0.697753 | 0.702487 | 0.698509 | 0.72186 | 0.70569 | 0.701666 |
| 0.705839 | 0.705387 | 0.716262 | 0.699102 | 0.70111 | 0.701134 | 0.721741 | 0.70783 | 0.70421 |
| 0.710567 | 0.709557 | 0.723569 | 0.703236 | 0.706225 | 0.704902 | 0.72368 | 0.708982 | 0.706164 |
| 0.708357 | 0.707533 | 0.718781 | 0.701289 | 0.705315 | 0.705406 | 0.723594 | 0.707944 | 0.706269 |
| 0.702039 | 0.707335 | 0.718496 | 0.697885 | 0.703647 | 0.699387 | 0.721445 | 0.704673 | 0.70374 |
| 0.70602 | 0.709654 | 0.718978 | 0.702903 | 0.704592 | 0.7035 | 0.722339 | 0.707009 | 0.705497 |
| 0.706303 | 0.70619 | 0.721763 | 0.699111 | 0.705179 | 0.70122 | 0.722845 | 0.707734 | 0.705518 |
| 0.706961 | 0.70412 | 0.717342 | 0.69821 | 0.707033 | 0.70267 | 0.720982 | 0.705558 | 0.701969 |
| 0.697694 | 0.697933 | 0.71379 | 0.695278 | 0.69405 | 0.693757 | 0.717081 | 0.697548 | 0.69452 |
| 0.698747 | 0.700858 | 0.71249 | 0.693373 | 0.702215 | 0.697597 | 0.714526 | 0.698775 | 0.699372 |
| 0.709693 | 0.706599 | 0.721834 | 0.700371 | 0.711324 | 0.702419 | 0.722008 | 0.712078 | 0.70443 |
| 0.71447 | 0.714854 | 0.729662 | 0.708129 | 0.714116 | 0.710436 | 0.732771 | 0.713231 | 0.710786 |
| 0.708337 | 0.708672 | 0.725517 | 0.703946 | 0.710664 | 0.705754 | 0.725386 | 0.712139 | 0.708836 |
| 0.715216 | 0.708389 | 0.726644 | 0.705658 | 0.713324 | 0.710309 | 0.730107 | 0.71217 | 0.711723 |
| 0.706576 | 0.706827 | 0.720173 | 0.696298 | 0.708022 | 0.701515 | 0.723585 | 0.704904 | 0.702858 |
| 0.706189 | 0.705176 | 0.718222 | 0.699634 | 0.709202 | 0.701059 | 0.721548 | 0.702005 | 0.70465 |
| 0.705398 | 0.702652 | 0.715735 | 0.697573 | 0.701016 | 0.702367 | 0.718028 | 0.703228 | 0.697638 |
| 0.708114 | 0.709353 | 0.726393 | 0.70067 | 0.71122 | 0.704921 | 0.728013 | 0.7107 | 0.7086 |
| 0.711087 | 0.713478 | 0.723471 | 0.705491 | 0.711723 | 0.704323 | 0.725087 | 0.710013 | 0.70817 |
| 0.707135 | 0.704205 | 0.722667 | 0.703628 | 0.703169 | 0.701829 | 0.723685 | 0.708686 | 0.705538 |
| 0.706156 | 0.71001 | 0.720747 | 0.699664 | 0.707502 | 0.701044 | 0.724631 | 0.705312 | 0.704766 |
| 0.814451 | 0.78384 | 0.822571 | 0.78756 | 0.798105 | 0.805858 | 0.828004 | 0.802254 | 0.814068 |
| 0.706823 | 0.708134 | 0.723827 | 0.700427 | 0.708081 | 0.701087 | 0.726779 | 0.708152 | 0.704619 |
| 0.710404 | 0.709472 | 0.725122 | 0.701905 | 0.706834 | 0.706197 | 0.728486 | 0.710088 | 0.708606 |
| 0.712301 | 0.713943 | 0.728599 | 0.707115 | 0.711557 | 0.707942 | 0.730157 | 0.714862 | 0.711932 |
| 0.70898 | 0.708033 | 0.722889 | 0.700737 | 0.704849 | 0.703121 | 0.729653 | 0.705911 | 0.704655 |
| 0.704135 | 0.708551 | 0.720104 | 0.699699 | 0.703701 | 0.7013 | 0.722255 | 0.704914 | 0.705028 |
| 0.701919 | 0.704703 | 0.717359 | 0.700374 | 0.705993 | 0.69895 | 0.720023 | 0.705819 | 0.702897 |
| 0.706608 | 0.706678 | 0.72089 | 0.701287 | 0.71 | 0.703477 | 0.723101 | 0.708245 | 0.706015 |
| 0.705119 | 0.704743 | 0.720313 | 0.693955 | 0.706019 | 0.701011 | 0.721427 | 0.705547 | 0.703108 |
| 0.703211 | 0.704669 | 0.717171 | 0.692381 | 0.702021 | 0.697433 | 0.719997 | 0.701886 | 0.699799 |
| 0.706365 | 0.701257 | 0.720142 | 0.695528 | 0.701869 | 0.70085 | 0.724411 | 0.702117 | 0.704707 |
| 0.70977 | 0.711383 | 0.723881 | 0.705188 | 0.713582 | 0.706775 | 0.727153 | 0.708882 | 0.708667 |
| 0.711433 | 0.709775 | 0.724893 | 0.703568 | 0.708911 | 0.703972 | 0.727465 | 0.710399 | 0.708463 |
| 0.707842 | 0.707257 | 0.722402 | 0.703614 | 0.709526 | 0.703718 | 0.728615 | 0.707584 | 0.708894 |
| 0.706128 | 0.707567 | 0.720318 | 0.70161 | 0.704603 | 0.704422 | 0.723099 | 0.709141 | 0.707228 |
| 0.70366 | 0.704362 | 0.718133 | 0.69531 | 0.699836 | 0.699803 | 0.72079 | 0.705441 | 0.703334 |
| 0.710917 | 0.713274 | 0.726143 | 0.704365 | 0.707708 | 0.706719 | 0.72705 | 0.712495 | 0.712279 |
| 0.704984 | 0.704097 | 0.722679 | 0.700567 | 0.704691 | 0.699294 | 0.723984 | 0.709285 | 0.705329 |
| 0.70685 | 0.707983 | 0.724132 | 0.701806 | 0.707319 | 0.703164 | 0.724966 | 0.706694 | 0.702361 |
| 0.713917 | 0.712979 | 0.727832 | 0.706366 | 0.71016 | 0.709114 | 0.725216 | 0.708193 | 0.706831 |
| 0.711232 | 0.713007 | 0.728224 | 0.707147 | 0.709408 | 0.705664 | 0.727675 | 0.710774 | 0.705282 |

| | | | | | | | | |
|---|---|---|---|---|---|---|---|---|
| 0.708784 | 0.709138 | 0.72513 | 0.703279 | 0.71014 | 0.702862 | 0.726523 | 0.709798 | 0.70591 |
| 0.712097 | 0.70853 | 0.724944 | 0.704946 | 0.710514 | 0.701879 | 0.727032 | 0.711433 | 0.706387 |
| 0.707873 | 0.709011 | 0.725235 | 0.703895 | 0.711074 | 0.704665 | 0.723917 | 0.710029 | 0.704493 |
| 0.676896 | 0.660384 | 0.694159 | 0.669831 | 0.671652 | 0.65804 | 0.689864 | 0.671223 | 0.666667 |
| 0.707807 | 0.70818 | 0.722156 | 0.702732 | 0.709933 | 0.70186 | 0.723493 | 0.70716 | 0.703448 |
| 0.709737 | 0.706239 | 0.722195 | 0.700943 | 0.710238 | 0.702784 | 0.72207 | 0.707352 | 0.7022 |
| 0.711509 | 0.706975 | 0.72476 | 0.704435 | 0.71101 | 0.707952 | 0.725459 | 0.711904 | 0.705492 |
| 0.709436 | 0.710319 | 0.727906 | 0.704701 | 0.707608 | 0.707426 | 0.724688 | 0.708495 | 0.706169 |
| 0.715417 | 0.712384 | 0.729424 | 0.710182 | 0.714196 | 0.710137 | 0.728381 | 0.712067 | 0.707801 |
| 0.713085 | 0.711274 | 0.726485 | 0.704895 | 0.709888 | 0.708442 | 0.724813 | 0.708961 | 0.706667 |
| 0.710375 | 0.712068 | 0.726696 | 0.706467 | 0.709267 | 0.70401 | 0.724766 | 0.712061 | 0.708482 |
| 0.71339 | 0.715294 | 0.730496 | 0.70876 | 0.712758 | 0.709363 | 0.728985 | 0.716002 | 0.710166 |
| 0.714457 | 0.713529 | 0.729736 | 0.707071 | 0.714331 | 0.71169 | 0.727836 | 0.714297 | 0.709935 |
| 0.711551 | 0.706892 | 0.723443 | 0.7058 | 0.708598 | 0.702569 | 0.723465 | 0.71291 | 0.705171 |
| 0.713954 | 0.708287 | 0.730744 | 0.705702 | 0.713159 | 0.707064 | 0.728647 | 0.715062 | 0.708785 |
| 0.706211 | 0.704869 | 0.722267 | 0.701853 | 0.707943 | 0.702526 | 0.720677 | 0.70837 | 0.703061 |
| 0.711465 | 0.708327 | 0.725085 | 0.703643 | 0.71395 | 0.706461 | 0.725107 | 0.71053 | 0.705652 |
| 0.712406 | 0.710155 | 0.728218 | 0.703414 | 0.714049 | 0.707841 | 0.726274 | 0.714821 | 0.70699 |
| 0.705391 | 0.704335 | 0.721556 | 0.696631 | 0.705758 | 0.699835 | 0.718743 | 0.704896 | 0.701359 |
| 0.708373 | 0.707938 | 0.725136 | 0.700354 | 0.70871 | 0.704062 | 0.724838 | 0.711523 | 0.704921 |
| 0.710398 | 0.709553 | 0.728499 | 0.707843 | 0.71201 | 0.706322 | 0.728223 | 0.712859 | 0.705215 |
| 0.711813 | 0.708045 | 0.727367 | 0.707627 | 0.711834 | 0.707039 | 0.727477 | 0.713819 | 0.708685 |
| 0.695364 | 0.694171 | 0.718715 | 0.69136 | 0.691818 | 0.695811 | 0.718494 | 0.699872 | 0.691217 |
| 0.73483 | 0.72105 | 0.760151 | 0.724138 | 0.71803 | 0.723707 | 0.754506 | 0.749191 | 0.71917 |
| 0.756087 | 0.758343 | 0.780265 | 0.745214 | 0.768608 | 0.756201 | 0.783329 | 0.764226 | 0.789007 |
| 0.744612 | 0.741523 | 0.805546 | 0.731236 | 0.737889 | 0.835208 | 0.805052 | 0.785166 | 0.776526 |
| 0.662755 | 0.669551 | 0.681487 | 0.659962 | 0.660202 | 0.661332 | 0.678884 | 0.669507 | 0.661249 |
| 0.660923 | 0.670189 | 0.685882 | 0.663801 | 0.657803 | 0.65379 | 0.681329 | 0.670268 | 0.654552 |
| 0.657817 | 0.662892 | 0.683451 | 0.652773 | 0.665568 | 0.654125 | 0.681514 | 0.661859 | 0.656266 |
| 0.675396 | 0.678888 | 0.700329 | 0.666463 | 0.671291 | 0.668202 | 0.695456 | 0.676387 | 0.669214 |
| 0.648034 | 0.655907 | 0.684891 | 0.653564 | 0.650558 | 0.645669 | 0.677055 | 0.652318 | 0.647188 |
| 0.644105 | 0.641888 | 0.665469 | 0.644621 | 0.648554 | 0.635996 | 0.666481 | 0.643167 | 0.634527 |
| 0.641033 | 0.638143 | 0.664439 | 0.63706 | 0.646015 | 0.639298 | 0.661346 | 0.647344 | 0.645061 |
| 0.733918 | 0.740262 | 0.759751 | 0.732303 | 0.744894 | 0.732642 | 0.757121 | 0.742847 | 0.732179 |
| 0.704747 | 0.704423 | 0.720064 | 0.697788 | 0.702774 | 0.69874 | 0.719653 | 0.703288 | 0.700215 |
| 0.675445 | 0.681166 | 0.691907 | 0.669273 | 0.675219 | 0.67195 | 0.688227 | 0.678346 | 0.666407 |
| 0.668757 | 0.670047 | 0.691423 | 0.67021 | 0.666581 | 0.665108 | 0.687111 | 0.670496 | 0.66658 |
| 0.653105 | 0.655831 | 0.671372 | 0.646146 | 0.649151 | 0.650089 | 0.668305 | 0.658589 | 0.649494 |
| 0.673231 | 0.677728 | 0.697395 | 0.668068 | 0.674793 | 0.67301 | 0.693044 | 0.678624 | 0.672149 |
| 0.668155 | 0.67087 | 0.687594 | 0.667629 | 0.667731 | 0.667822 | 0.682434 | 0.674534 | 0.658133 |
| 0.659196 | 0.659643 | 0.675282 | 0.656555 | 0.660562 | 0.6556 | 0.674534 | 0.661449 | 0.653729 |
| 0.663645 | 0.665367 | 0.684995 | 0.663362 | 0.660304 | 0.660006 | 0.681853 | 0.664904 | 0.657045 |
| 0.658625 | 0.653336 | 0.683903 | 0.650504 | 0.654111 | 0.650577 | 0.676958 | 0.66025 | 0.649543 |
| 0.660601 | 0.661027 | 0.677209 | 0.657626 | 0.662802 | 0.656422 | 0.675953 | 0.658718 | 0.651838 |
| 0.658193 | 0.657524 | 0.676889 | 0.65985 | 0.653317 | 0.650821 | 0.678074 | 0.663027 | 0.658344 |
| 0.643634 | 0.649714 | 0.661146 | 0.644095 | 0.642274 | 0.637805 | 0.662472 | 0.648502 | 0.633365 |

| | | | | | | | | |
|---|---|---|---|---|---|---|---|---|
| 0.692552 | 0.692028 | 0.704837 | 0.687972 | 0.690784 | 0.686047 | 0.707502 | 0.694198 | 0.686093 |
| 0.657969 | 0.662712 | 0.680977 | 0.658931 | 0.658032 | 0.651592 | 0.677769 | 0.661131 | 0.651755 |
| 0.663813 | 0.668547 | 0.684087 | 0.6615 | 0.656857 | 0.661193 | 0.685347 | 0.666715 | 0.660622 |
| 0.664146 | 0.663719 | 0.686587 | 0.65922 | 0.659907 | 0.65742 | 0.681825 | 0.669082 | 0.659154 |
| 0.660067 | 0.661773 | 0.680586 | 0.654261 | 0.654273 | 0.657126 | 0.674409 | 0.665029 | 0.650572 |
| 0.666667 | 0.665308 | 0.683771 | 0.668615 | 0.663797 | 0.663872 | 0.6836 | 0.669919 | 0.654941 |
| 0.649045 | 0.651201 | 0.674496 | 0.646356 | 0.652981 | 0.648316 | 0.678081 | 0.654554 | 0.650783 |
| 0.653028 | 0.648113 | 0.672869 | 0.641648 | 0.641178 | 0.649418 | 0.66713 | 0.655549 | 0.643338 |
| 0.64778 | 0.648143 | 0.67053 | 0.638108 | 0.64484 | 0.64353 | 0.666115 | 0.651972 | 0.645946 |
| 0.691563 | 0.691619 | 0.70884 | 0.683147 | 0.691997 | 0.686619 | 0.707039 | 0.693777 | 0.686169 |
| 0.689837 | 0.686483 | 0.707046 | 0.682382 | 0.687874 | 0.684598 | 0.705546 | 0.690145 | 0.684704 |
| 0.683584 | 0.682812 | 0.699535 | 0.677321 | 0.683397 | 0.677683 | 0.700814 | 0.687822 | 0.682186 |
| 0.674792 | 0.676781 | 0.693778 | 0.672357 | 0.675978 | 0.668569 | 0.696676 | 0.680216 | 0.669225 |
| 0.682135 | 0.680399 | 0.700441 | 0.673752 | 0.680445 | 0.676619 | 0.701244 | 0.681556 | 0.679039 |
| 0.669224 | 0.670243 | 0.689222 | 0.665588 | 0.670965 | 0.667724 | 0.689639 | 0.674692 | 0.670635 |

| Collection_ | Collection_ | Collection_ | Collection_ | Collection_ | Collection_ | Collection_ | Collection_ | Collection_ |
|---|---|---|---|---|---|---|---|---|
| 0.782076 | 0.77956 | 0.804619 | 0.781011 | 0.788391 | 0.81178 | 0.781457 | 0.77752 | 0.786212 |
| 0.781509 | 0.797444 | 0.778659 | 0.778863 | 0.777167 | 0.803348 | 0.785108 | 0.79189 | 0.794455 |
| 0.782655 | 0.801292 | 0.787648 | 0.772244 | 0.796984 | 0.796146 | 0.785292 | 0.777842 | 0.781467 |
| 0.783029 | 0.783271 | 0.787959 | 0.776834 | 0.78306 | 0.801108 | 0.788063 | 0.773423 | 0.786216 |
| 0.78258 | 0.773502 | 0.789104 | 0.770502 | 0.779022 | 0.799361 | 0.780366 | 0.777163 | 0.772602 |
| 0.784983 | 0.768952 | 0.781227 | 0.778849 | 0.786327 | 0.792178 | 0.790682 | 0.772806 | 0.781517 |
| 0.713168 | 0.709257 | 0.710244 | 0.713347 | 0.710153 | 0.720087 | 0.715486 | 0.707678 | 0.710978 |
| 0.705671 | 0.705095 | 0.704919 | 0.711173 | 0.708026 | 0.713653 | 0.711921 | 0.709708 | 0.710983 |
| 0.784604 | 0.78113 | 0.780968 | 0.79816 | 0.791499 | 0.791239 | 0.797249 | 0.786508 | 0.795949 |
| 0.782461 | 0.781625 | 0.80637 | 0.775059 | 0.779609 | 0.801399 | 0.780123 | 0.76878 | 0.783516 |
| 0.780778 | 0.781655 | 0.781102 | 0.778914 | 0.781083 | 0.788934 | 0.778711 | 0.768169 | 0.773924 |
| 0.782544 | 0.795388 | 0.798897 | 0.786038 | 0.781648 | 0.795292 | 0.788438 | 0.79174 | 0.7831 |
| 0.777447 | 0.78463 | 0.772858 | 0.776856 | 0.78022 | 0.796582 | 0.788101 | 0.779569 | 0.787573 |
| 0.780379 | 0.779014 | 0.785325 | 0.772902 | 0.779414 | 0.78961 | 0.782049 | 0.774164 | 0.77631 |
| 0.771996 | 0.772413 | 0.772344 | 0.759628 | 0.771134 | 0.778127 | 0.768351 | 0.760105 | 0.769399 |
| 0.785915 | 0.792193 | 0.798749 | 0.782387 | 0.785976 | 0.799428 | 0.789422 | 0.774159 | 0.785557 |
| 0.774156 | 0.795564 | 0.805404 | 0.769092 | 0.773908 | 0.798425 | 0.775163 | 0.771936 | 0.777175 |
| 0.753485 | 0.752916 | 0.755233 | 0.766167 | 0.750808 | 0.771281 | 0.76158 | 0.766241 | 0.76538 |
| 0.784809 | 0.799646 | 0.7768 | 0.777668 | 0.778679 | 0.798713 | 0.782551 | 0.781281 | 0.780865 |
| 0.785934 | 0.789562 | 0.781949 | 0.79789 | 0.779537 | 0.79936 | 0.78718 | 0.792458 | 0.794552 |
| 0.789552 | 0.785047 | 0.786449 | 0.79874 | 0.781075 | 0.795862 | 0.788178 | 0.799143 | 0.782175 |
| 0.796193 | 0.790219 | 0.804729 | 0.789487 | 0.784556 | 0.802281 | 0.795421 | 0.798621 | 0.787764 |
| 0.788524 | 0.781489 | 0.794565 | 0.793388 | 0.787664 | 0.803779 | 0.789984 | 0.791238 | 0.791341 |
| 0.793262 | 0.794721 | 0.804776 | 0.792011 | 0.785511 | 0.801181 | 0.79927 | 0.796033 | 0.7924 |
| 0.785619 | 0.787057 | 0.779412 | 0.800233 | 0.783368 | 0.793959 | 0.788267 | 0.79337 | 0.789235 |
| 0.790613 | 0.794152 | 0.782698 | 0.791913 | 0.774172 | 0.798426 | 0.783337 | 0.782627 | 0.787569 |
| 0.777345 | 0.788097 | 0.773321 | 0.781517 | 0.763936 | 0.793865 | 0.776611 | 0.77369 | 0.775082 |
| 0.786251 | 0.778542 | 0.786121 | 0.784865 | 0.78049 | 0.79526 | 0.78857 | 0.786904 | 0.786076 |
| 0.783165 | 0.782674 | 0.785702 | 0.783564 | 0.780656 | 0.795644 | 0.783562 | 0.778931 | 0.779624 |
| 0.781752 | 0.793632 | 0.79849 | 0.775196 | 0.775466 | 0.792208 | 0.781108 | 0.777361 | 0.790584 |
| 0.791618 | 0.774118 | 0.782883 | 0.787206 | 0.776841 | 0.793124 | 0.792109 | 0.781153 | 0.789252 |
| 0.781859 | 0.775856 | 0.781214 | 0.779872 | 0.777115 | 0.785714 | 0.781836 | 0.777521 | 0.779262 |
| 0.788814 | 0.7861 | 0.781304 | 0.774655 | 0.771247 | 0.795729 | 0.784989 | 0.775023 | 0.784373 |
| 0.783683 | 0.775122 | 0.773571 | 0.774486 | 0.771817 | 0.789126 | 0.776571 | 0.7738 | 0.775441 |
| 0.782627 | 0.781698 | 0.769694 | 0.770206 | 0.766598 | 0.784269 | 0.769419 | 0.765808 | 0.77225 |
| 0.78187 | 0.79752 | 0.778831 | 0.776578 | 0.780365 | 0.8 | 0.786998 | 0.771167 | 0.773735 |
| 0.780237 | 0.78562 | 0.781427 | 0.789227 | 0.77666 | 0.795842 | 0.786762 | 0.783482 | 0.776949 |
| 0.769627 | 0.765638 | 0.76628 | 0.768069 | 0.76284 | 0.779594 | 0.780553 | 0.764877 | 0.77922 |
| 0.784323 | 0.778783 | 0.772686 | 0.775996 | 0.781484 | 0.787149 | 0.787152 | 0.772506 | 0.791899 |

| | | | | | | | | |
|---|---|---|---|---|---|---|---|---|
| 0.758476 | 0.766724 | 0.763365 | 0.755743 | 0.755776 | 0.76865 | 0.761289 | 0.756597 | 0.753212 |
| 0.776624 | 0.775964 | 0.800785 | 0.778389 | 0.786383 | 0.804932 | 0.780528 | 0.774364 | 0.794014 |
| 0.784397 | 0.785947 | 0.797943 | 0.794427 | 0.790981 | 0.797368 | 0.787769 | 0.785834 | 0.781455 |
| 0.789831 | 0.788115 | 0.807073 | 0.781217 | 0.789688 | 0.812269 | 0.788983 | 0.785507 | 0.792515 |
| 0.795687 | 0.783131 | 0.787294 | 0.788043 | 0.782515 | 0.795697 | 0.79447 | 0.78822 | 0.786962 |
| 0.787067 | 0.78394 | 0.787497 | 0.787359 | 0.783809 | 0.793211 | 0.790276 | 0.783027 | 0.79196 |
| 0.783084 | 0.768932 | 0.781691 | 0.784086 | 0.770172 | 0.786767 | 0.781212 | 0.788606 | 0.77873 |
| 0.790634 | 0.78632 | 0.7822 | 0.791314 | 0.777945 | 0.809962 | 0.790792 | 0.788793 | 0.794859 |
| 0.787611 | 0.779584 | 0.779035 | 0.778374 | 0.776376 | 0.789144 | 0.777686 | 0.773954 | 0.771773 |
| 0.789875 | 0.766126 | 0.77851 | 0.775267 | 0.778128 | 0.782862 | 0.785288 | 0.777115 | 0.779107 |
| 0.791782 | 0.783575 | 0.809351 | 0.784894 | 0.783114 | 0.808072 | 0.793565 | 0.794095 | 0.79218 |
| 0.790455 | 0.783555 | 0.794578 | 0.803843 | 0.786588 | 0.804748 | 0.797991 | 0.802497 | 0.790859 |
| 0.797 | 0.79956 | 0.803965 | 0.787468 | 0.795262 | 0.800968 | 0.787836 | 0.786685 | 0.788102 |
| 0.788808 | 0.790093 | 0.812119 | 0.805804 | 0.801186 | 0.801917 | 0.79043 | 0.798968 | 0.780995 |
| 0.789491 | 0.795944 | 0.800557 | 0.788301 | 0.778702 | 0.801345 | 0.786237 | 0.803983 | 0.77979 |
| 0.781639 | 0.79045 | 0.797808 | 0.789363 | 0.779472 | 0.800395 | 0.785882 | 0.804859 | 0.778637 |
| 0.78694 | 0.798384 | 0.791277 | 0.772059 | 0.780362 | 0.795714 | 0.780023 | 0.778132 | 0.774308 |
| 0.791604 | 0.807391 | 0.810048 | 0.783733 | 0.788938 | 0.812548 | 0.791055 | 0.788336 | 0.799736 |
| 0.783616 | 0.785028 | 0.805023 | 0.796143 | 0.788299 | 0.795455 | 0.791133 | 0.788291 | 0.78895 |
| 0.790641 | 0.787943 | 0.803121 | 0.791521 | 0.78911 | 0.801326 | 0.794462 | 0.797195 | 0.796992 |
| 0.78584 | 0.805603 | 0.806247 | 0.77384 | 0.794229 | 0.804311 | 0.787595 | 0.773046 | 0.771436 |
| 0.785405 | 0.79061 | 0.806039 | 0.772298 | 0.78447 | 0.799189 | 0.782863 | 0.774014 | 0.785544 |
| 0.788606 | 0.79149 | 0.780175 | 0.79339 | 0.793941 | 0.792697 | 0.78461 | 0.782583 | 0.786999 |
| 0.785792 | 0.77292 | 0.771033 | 0.777083 | 0.77699 | 0.786392 | 0.78045 | 0.772512 | 0.770349 |
| 0.766844 | 0.762331 | 0.765332 | 0.763381 | 0.763522 | 0.773597 | 0.767861 | 0.764039 | 0.769717 |
| 0.706686 | 0.704861 | 0.703858 | 0.710772 | 0.708192 | 0.714054 | 0.713256 | 0.709605 | 0.712077 |
| 0.71186 | 0.711585 | 0.70907 | 0.712708 | 0.715432 | 0.718172 | 0.717181 | 0.711727 | 0.711338 |
| 0.712452 | 0.712769 | 0.710385 | 0.713029 | 0.711853 | 0.719811 | 0.717649 | 0.713661 | 0.710729 |
| 0.775979 | 0.774954 | 0.770675 | 0.777586 | 0.769148 | 0.786158 | 0.77695 | 0.778841 | 0.790276 |
| 0.778166 | 0.77249 | 0.772229 | 0.775619 | 0.773038 | 0.784868 | 0.781716 | 0.777188 | 0.786828 |
| 0.778884 | 0.766788 | 0.798259 | 0.795198 | 0.775677 | 0.799527 | 0.774183 | 0.777962 | 0.776205 |
| 0.778846 | 0.767648 | 0.773015 | 0.776303 | 0.769644 | 0.786161 | 0.782605 | 0.77916 | 0.784861 |
| 0.790815 | 0.834732 | 0.786146 | 0.781065 | 0.784617 | 0.809915 | 0.793476 | 0.78741 | 0.825288 |
| 0.795822 | 0.81558 | 0.786437 | 0.780666 | 0.787485 | 0.805023 | 0.796145 | 0.783934 | 0.780721 |
| 0.784025 | 0.818276 | 0.785101 | 0.7744 | 0.775762 | 0.824414 | 0.786581 | 0.788161 | 0.823819 |
| 0.787927 | 0.861427 | 0.787727 | 0.771058 | 0.784828 | 0.819066 | 0.78845 | 0.783833 | 0.783468 |
| 0.788633 | 0.779884 | 0.786961 | 0.790397 | 0.824002 | 0.816505 | 0.816012 | 0.779756 | 0.842608 |
| 0.804053 | 0.812003 | 0.829511 | 0.794555 | 0.807053 | 0.903283 | 0.815283 | 0.799344 | 0.831036 |
| 0.777462 | 0.87848 | 0.771652 | 0.770951 | 0.774849 | 0.801506 | 0.777029 | 0.774002 | 0.805563 |
| 0.831854 | 0.780212 | 0.785292 | 0.810057 | 0.795731 | 0.79527 | 0.805684 | 0.829916 | 0.785807 |
| 0.778454 | 0.814968 | 0.818845 | 0.770087 | 0.789924 | 0.807191 | 0.787587 | 0.785484 | 0.782095 |
| 0.806392 | 0.817347 | 0.823814 | 0.800167 | 0.805805 | 0.873639 | 0.812212 | 0.79726 | 0.821246 |
| 0.788543 | 0.780852 | 0.830174 | 0.780928 | 0.796368 | 0.809439 | 0.786221 | 0.787241 | 0.789214 |
| 0.788223 | 0.779196 | 0.822635 | 0.782157 | 0.794255 | 0.80545 | 0.790846 | 0.786704 | 0.781352 |
| 1 | 0.779042 | 0.785482 | 0.786721 | 0.792781 | 0.794893 | 0.805045 | 0.788462 | 0.78647 |
| 0.779042 | 1 | 0.778873 | 0.773672 | 0.784043 | 0.794778 | 0.786011 | 0.778763 | 0.786437 |

| | | | | | | | | |
|---|---|---|---|---|---|---|---|---|
| 0.785482 | 0.778873 | 1 | 0.780242 | 0.800927 | 0.815379 | 0.796266 | 0.791604 | 0.789865 |
| 0.786721 | 0.773672 | 0.780242 | 1 | 0.799447 | 0.792037 | 0.804089 | 0.821907 | 0.780697 |
| 0.792781 | 0.784043 | 0.800927 | 0.799447 | 1 | 0.796977 | 0.812142 | 0.780201 | 0.786206 |
| 0.794893 | 0.794778 | 0.815379 | 0.792037 | 0.796977 | 1 | 0.807029 | 0.793293 | 0.815208 |
| 0.805045 | 0.786011 | 0.796266 | 0.804089 | 0.812142 | 0.807029 | 1 | 0.800584 | 0.827491 |
| 0.788462 | 0.778763 | 0.791604 | 0.821907 | 0.780201 | 0.793293 | 0.800584 | 1 | 0.783806 |
| 0.78647 | 0.786437 | 0.789865 | 0.780697 | 0.786206 | 0.815208 | 0.827491 | 0.783806 | 1 |
| 0.782765 | 0.778178 | 0.781031 | 0.808571 | 0.804619 | 0.798912 | 0.808745 | 0.775377 | 0.799168 |
| 0.788887 | 0.781413 | 0.783102 | 0.787524 | 0.784904 | 0.797442 | 0.78371 | 0.779841 | 0.786039 |
| 0.778403 | 0.776394 | 0.77809 | 0.776838 | 0.777861 | 0.783904 | 0.781635 | 0.786291 | 0.771691 |
| 0.783357 | 0.787569 | 0.783864 | 0.773392 | 0.790974 | 0.801106 | 0.789 | 0.781644 | 0.782934 |
| 0.782207 | 0.777068 | 0.780451 | 0.771976 | 0.778217 | 0.793703 | 0.784519 | 0.777311 | 0.783353 |
| 0.793552 | 0.781043 | 0.790405 | 0.790069 | 0.782182 | 0.797335 | 0.788623 | 0.787393 | 0.775659 |
| 0.78403 | 0.772436 | 0.782503 | 0.779412 | 0.782906 | 0.784307 | 0.795204 | 0.78514 | 0.775761 |
| 0.784524 | 0.781855 | 0.789619 | 0.777769 | 0.786247 | 0.793506 | 0.787851 | 0.78615 | 0.778581 |
| 0.780205 | 0.783804 | 0.786004 | 0.785057 | 0.791966 | 0.796964 | 0.788894 | 0.78098 | 0.782924 |
| 0.78924 | 0.776541 | 0.783603 | 0.776851 | 0.780972 | 0.79059 | 0.789028 | 0.772758 | 0.789164 |
| 0.774967 | 0.813169 | 0.784649 | 0.773752 | 0.777935 | 0.797817 | 0.780598 | 0.779132 | 0.773123 |
| 0.783953 | 0.785648 | 0.784306 | 0.781811 | 0.786925 | 0.796879 | 0.793103 | 0.784342 | 0.783564 |
| 0.788202 | 0.779056 | 0.776997 | 0.782494 | 0.777371 | 0.792203 | 0.782391 | 0.775337 | 0.78 |
| 0.787403 | 0.773608 | 0.786386 | 0.78179 | 0.786385 | 0.797094 | 0.793058 | 0.78461 | 0.785555 |
| 0.778891 | 0.827572 | 0.779273 | 0.77607 | 0.776364 | 0.795544 | 0.77998 | 0.772982 | 0.776591 |
| 0.782594 | 0.786045 | 0.789447 | 0.796143 | 0.79566 | 0.796367 | 0.786069 | 0.786613 | 0.780345 |
| 0.800234 | 0.774965 | 0.785667 | 0.805964 | 0.788116 | 0.798028 | 0.862722 | 0.804796 | 0.777034 |
| 0.782039 | 0.776762 | 0.78207 | 0.786676 | 0.790217 | 0.802352 | 0.793754 | 0.77774 | 0.803486 |
| 0.789966 | 0.783456 | 0.786841 | 0.787272 | 0.788186 | 0.812811 | 0.809536 | 0.778743 | 0.856624 |
| 0.78281 | 0.783815 | 0.782319 | 0.78908 | 0.791177 | 0.814107 | 0.799654 | 0.779889 | 0.812278 |
| 0.787695 | 0.780255 | 0.778611 | 0.776389 | 0.775879 | 0.795742 | 0.782438 | 0.776536 | 0.780036 |
| 0.782152 | 0.780676 | 0.784469 | 0.779319 | 0.775733 | 0.794682 | 0.786037 | 0.860744 | 0.779658 |
| 0.791526 | 0.774758 | 0.783249 | 0.782984 | 0.797015 | 0.805339 | 0.800673 | 0.773124 | 0.810699 |
| 0.788958 | 0.776756 | 0.78163 | 0.779498 | 0.785942 | 0.794406 | 0.788729 | 0.777704 | 0.782064 |
| 0.782047 | 0.782031 | 0.777496 | 0.774986 | 0.7882 | 0.793212 | 0.787928 | 0.780184 | 0.77918 |
| 0.788287 | 0.783777 | 0.787701 | 0.778946 | 0.790914 | 0.813096 | 0.791677 | 0.780958 | 0.827881 |
| 0.78456 | 0.791638 | 0.808947 | 0.776001 | 0.789361 | 0.814083 | 0.789314 | 0.786226 | 0.795119 |
| 0.793083 | 0.783484 | 0.788391 | 0.787139 | 0.787449 | 0.79866 | 0.788377 | 0.79686 | 0.784888 |
| 0.787869 | 0.784563 | 0.79166 | 0.786922 | 0.780183 | 0.79836 | 0.787927 | 0.793545 | 0.787401 |
| 0.785518 | 0.782857 | 0.781424 | 0.775129 | 0.78229 | 0.788157 | 0.784764 | 0.775234 | 0.776237 |
| 0.765441 | 0.757812 | 0.758639 | 0.7757 | 0.762781 | 0.771982 | 0.771878 | 0.75845 | 0.753425 |
| 0.809248 | 0.75874 | 0.771832 | 0.811112 | 0.77267 | 0.782965 | 0.8119 | 0.809953 | 0.758922 |
| 0.751846 | 0.756392 | 0.759184 | 0.756069 | 0.758462 | 0.769538 | 0.76561 | 0.762308 | 0.76458 |
| 0.76087 | 0.806516 | 0.767523 | 0.757809 | 0.765641 | 0.787925 | 0.766095 | 0.766524 | 0.757753 |
| 0.750031 | 0.745419 | 0.744614 | 0.7419 | 0.747749 | 0.751038 | 0.759365 | 0.741747 | 0.741436 |
| 0.7569 | 0.751728 | 0.806155 | 0.757453 | 0.774714 | 0.782575 | 0.760237 | 0.75912 | 0.757141 |
| 0.764721 | 0.79003 | 0.754617 | 0.7649 | 0.776766 | 0.775719 | 0.763873 | 0.755412 | 0.755966 |
| 0.810463 | 0.745264 | 0.756218 | 0.739827 | 0.756172 | 0.770075 | 0.760021 | 0.751862 | 0.758233 |
| 0.756609 | 0.777209 | 0.758558 | 0.785009 | 0.771944 | 0.782356 | 0.768859 | 0.759138 | 0.785302 |
| 0.758175 | 0.793933 | 0.761431 | 0.751636 | 0.756414 | 0.787392 | 0.753044 | 0.756844 | 0.755667 |

| | | | | | | | | |
|---|---|---|---|---|---|---|---|---|
| 0.755991 | 0.75308 | 0.747475 | 0.749809 | 0.746899 | 0.762236 | 0.75124 | 0.750675 | 0.737713 |
| 0.763376 | 0.799254 | 0.75441 | 0.746164 | 0.747066 | 0.807999 | 0.765513 | 0.753118 | 0.808652 |
| 0.755764 | 0.74705 | 0.742007 | 0.743816 | 0.750208 | 0.757487 | 0.749793 | 0.743682 | 0.747231 |
| 0.747171 | 0.7427 | 0.851344 | 0.743417 | 0.769359 | 0.789429 | 0.757103 | 0.748792 | 0.749245 |
| 0.762155 | 0.75514 | 0.748328 | 0.746961 | 0.753144 | 0.785705 | 0.765244 | 0.755795 | 0.802326 |
| 0.75126 | 0.740662 | 0.761699 | 0.785071 | 0.84402 | 0.775451 | 0.819522 | 0.741069 | 0.782155 |
| 0.759507 | 0.746387 | 0.767023 | 0.757223 | 0.773134 | 0.777478 | 0.772491 | 0.75579 | 0.780182 |
| 0.755222 | 0.789918 | 0.752193 | 0.746243 | 0.757833 | 0.774889 | 0.7591 | 0.76262 | 0.750581 |
| 0.739044 | 0.730839 | 0.733829 | 0.726545 | 0.736938 | 0.747056 | 0.772454 | 0.730043 | 0.726435 |
| 0.763966 | 0.757887 | 0.75038 | 0.74787 | 0.749427 | 0.812352 | 0.759384 | 0.751298 | 0.838357 |
| 0.747106 | 0.747776 | 0.755552 | 0.838522 | 0.803812 | 0.765028 | 0.778719 | 0.750571 | 0.753484 |
| 0.773107 | 0.765665 | 0.82378 | 0.768502 | 0.784286 | 0.797562 | 0.776146 | 0.7722 | 0.770224 |
| 0.779298 | 0.776704 | 0.778033 | 0.772117 | 0.779264 | 0.78559 | 0.77729 | 0.772041 | 0.777886 |
| 0.750323 | 0.758381 | 0.757601 | 0.789133 | 0.764775 | 0.768856 | 0.769403 | 0.752773 | 0.768858 |
| 0.779524 | 0.777321 | 0.780996 | 0.778564 | 0.776187 | 0.797943 | 0.782913 | 0.781479 | 0.774148 |
| 0.771221 | 0.767053 | 0.771633 | 0.768941 | 0.774494 | 0.778881 | 0.779528 | 0.778156 | 0.772186 |
| 0.782655 | 0.771759 | 0.780289 | 0.778086 | 0.776093 | 0.79166 | 0.782022 | 0.780599 | 0.775781 |
| 0.77787 | 0.777508 | 0.772166 | 0.776749 | 0.778814 | 0.788561 | 0.776731 | 0.770154 | 0.780693 |
| 0.784309 | 0.760592 | 0.760631 | 0.773544 | 0.782393 | 0.770789 | 0.779696 | 0.761351 | 0.765587 |
| 0.773073 | 0.772009 | 0.772245 | 0.768931 | 0.778824 | 0.787086 | 0.791994 | 0.771337 | 0.790753 |
| 0.750652 | 0.743358 | 0.753699 | 0.759027 | 0.783845 | 0.774528 | 0.792569 | 0.735974 | 0.799116 |
| 0.775392 | 0.766526 | 0.77287 | 0.778647 | 0.771952 | 0.783213 | 0.783115 | 0.771433 | 0.769601 |
| 0.774681 | 0.776432 | 0.766103 | 0.775954 | 0.77093 | 0.781326 | 0.771996 | 0.764458 | 0.767611 |
| 0.74715 | 0.745119 | 0.747486 | 0.784278 | 0.737979 | 0.762112 | 0.764275 | 0.770907 | 0.762759 |
| 0.751345 | 0.757245 | 0.749893 | 0.743978 | 0.749209 | 0.775886 | 0.753968 | 0.743982 | 0.771004 |
| 0.759843 | 0.791422 | 0.762284 | 0.764813 | 0.767463 | 0.779663 | 0.770866 | 0.765362 | 0.759439 |
| 0.765819 | 0.759089 | 0.768155 | 0.781558 | 0.792382 | 0.777522 | 0.799796 | 0.754999 | 0.801967 |
| 0.783113 | 0.769823 | 0.776935 | 0.792198 | 0.768252 | 0.787135 | 0.786852 | 0.786999 | 0.779072 |
| 0.753405 | 0.80163 | 0.753482 | 0.745555 | 0.752797 | 0.765726 | 0.754955 | 0.74996 | 0.762614 |
| 0.776259 | 0.771607 | 0.766805 | 0.765286 | 0.77091 | 0.805453 | 0.782025 | 0.766865 | 0.817169 |
| 0.780524 | 0.766879 | 0.764255 | 0.769775 | 0.765189 | 0.781019 | 0.770745 | 0.765351 | 0.774411 |
| 0.758326 | 0.771041 | 0.77549 | 0.752972 | 0.76003 | 0.825738 | 0.761815 | 0.753019 | 0.771064 |
| 0.762123 | 0.777152 | 0.772608 | 0.755613 | 0.757225 | 0.817407 | 0.766848 | 0.760505 | 0.770049 |
| 0.765453 | 0.776222 | 0.771384 | 0.760065 | 0.766645 | 0.784707 | 0.791321 | 0.765789 | 0.819242 |
| 0.765172 | 0.7669 | 0.762776 | 0.764009 | 0.767194 | 0.779024 | 0.76973 | 0.763545 | 0.764004 |
| 0.775556 | 0.772428 | 0.767735 | 0.771264 | 0.767446 | 0.784384 | 0.774035 | 0.770845 | 0.767921 |
| 0.726208 | 0.728715 | 0.729312 | 0.733983 | 0.737723 | 0.734219 | 0.74085 | 0.729728 | 0.737782 |
| 0.763459 | 0.759613 | 0.754869 | 0.757975 | 0.75502 | 0.77233 | 0.766073 | 0.763173 | 0.766074 |
| 0.72649 | 0.753796 | 0.733897 | 0.744096 | 0.740933 | 0.744975 | 0.728592 | 0.727699 | 0.733632 |
| 0.720846 | 0.714879 | 0.723391 | 0.746596 | 0.7499 | 0.73354 | 0.736171 | 0.728148 | 0.72362 |
| 0.765839 | 0.775801 | 0.769619 | 0.778466 | 0.769035 | 0.790369 | 0.782881 | 0.774093 | 0.774978 |
| 0.718542 | 0.717219 | 0.725041 | 0.741584 | 0.737426 | 0.731291 | 0.737175 | 0.726379 | 0.721638 |
| 0.717873 | 0.746199 | 0.719643 | 0.714417 | 0.717744 | 0.738766 | 0.72185 | 0.716799 | 0.727609 |
| 0.710177 | 0.725989 | 0.711867 | 0.706507 | 0.71126 | 0.728628 | 0.712508 | 0.708561 | 0.731978 |
| 0.766743 | 0.769465 | 0.78338 | 0.766469 | 0.766814 | 0.791871 | 0.77069 | 0.768341 | 0.778968 |
| 0.712931 | 0.725327 | 0.710752 | 0.706576 | 0.702615 | 0.724656 | 0.708758 | 0.704979 | 0.722812 |
| 0.768977 | 0.776052 | 0.774462 | 0.773257 | 0.766239 | 0.788857 | 0.776392 | 0.771377 | 0.774636 |

| | | | | | | | | |
|---|---|---|---|---|---|---|---|---|
| 0.765257 | 0.766525 | 0.76858 | 0.758284 | 0.765601 | 0.781232 | 0.769479 | 0.772418 | 0.770109 |
| 0.759739 | 0.769717 | 0.773397 | 0.775755 | 0.768453 | 0.784278 | 0.772052 | 0.772503 | 0.772536 |
| 0.753407 | 0.749117 | 0.754528 | 0.745965 | 0.751666 | 0.769616 | 0.753176 | 0.750822 | 0.752617 |
| 0.699397 | 0.726331 | 0.700684 | 0.698371 | 0.699648 | 0.722382 | 0.703878 | 0.698884 | 0.713192 |
| 0.705451 | 0.707706 | 0.700698 | 0.705461 | 0.714155 | 0.72563 | 0.708705 | 0.696539 | 0.728082 |
| 0.770532 | 0.769104 | 0.776928 | 0.767529 | 0.771947 | 0.783862 | 0.77968 | 0.768724 | 0.779932 |
| 0.69181 | 0.715939 | 0.691096 | 0.696327 | 0.6913 | 0.713264 | 0.693361 | 0.694521 | 0.690659 |
| 0.736347 | 0.731665 | 0.73143 | 0.739431 | 0.730492 | 0.739116 | 0.738669 | 0.738123 | 0.734043 |
| 0.765455 | 0.777473 | 0.769092 | 0.759684 | 0.764322 | 0.776676 | 0.768151 | 0.759681 | 0.762452 |
| 0.726262 | 0.717739 | 0.720826 | 0.727126 | 0.718549 | 0.730195 | 0.723617 | 0.72737 | 0.721757 |
| 0.744954 | 0.761716 | 0.751185 | 0.749023 | 0.75138 | 0.768103 | 0.757363 | 0.749918 | 0.759002 |
| 0.647436 | 0.615385 | 0.639241 | 0.589744 | 0.625767 | 0.68 | 0.652695 | 0.627451 | 0.625 |
| 0.65613 | 0.656731 | 0.653573 | 0.653323 | 0.650517 | 0.671653 | 0.667568 | 0.655026 | 0.661907 |
| 0.65345 | 0.656962 | 0.658312 | 0.656409 | 0.656499 | 0.66885 | 0.664706 | 0.655967 | 0.662898 |
| 0.665654 | 0.665314 | 0.662609 | 0.667053 | 0.667921 | 0.676505 | 0.676275 | 0.66861 | 0.672899 |
| 0.662143 | 0.663868 | 0.660044 | 0.663524 | 0.662758 | 0.677005 | 0.67348 | 0.6589 | 0.668561 |
| 0.621526 | 0.620868 | 0.621087 | 0.628166 | 0.618531 | 0.638451 | 0.620389 | 0.617805 | 0.618316 |
| 0.662649 | 0.668848 | 0.664828 | 0.67114 | 0.665749 | 0.678196 | 0.670429 | 0.666248 | 0.673897 |
| 0.743143 | 0.749155 | 0.74117 | 0.743567 | 0.741443 | 0.752499 | 0.748904 | 0.744884 | 0.75441 |
| 0.738768 | 0.747583 | 0.743391 | 0.736747 | 0.741898 | 0.754646 | 0.745926 | 0.737186 | 0.745206 |
| 0.744257 | 0.759004 | 0.746374 | 0.745515 | 0.747813 | 0.75869 | 0.751018 | 0.747838 | 0.75542 |
| 0.621167 | 0.618739 | 0.620341 | 0.623027 | 0.623698 | 0.630989 | 0.625087 | 0.614475 | 0.619652 |
| 0.603172 | 0.604319 | 0.59992 | 0.601823 | 0.600048 | 0.612407 | 0.605906 | 0.601298 | 0.600473 |
| 0.709486 | 0.705318 | 0.70678 | 0.709532 | 0.709249 | 0.712124 | 0.710695 | 0.707765 | 0.711387 |
| 0.707261 | 0.706524 | 0.702862 | 0.711737 | 0.712596 | 0.714424 | 0.716331 | 0.710699 | 0.711733 |
| 0.70082 | 0.7005 | 0.699872 | 0.703296 | 0.701137 | 0.707404 | 0.705986 | 0.70133 | 0.706652 |
| 0.702912 | 0.7056 | 0.704569 | 0.70767 | 0.705948 | 0.710846 | 0.712392 | 0.706598 | 0.707965 |
| 0.706715 | 0.709442 | 0.70626 | 0.709708 | 0.707036 | 0.716153 | 0.714896 | 0.706835 | 0.711855 |
| 0.708855 | 0.706207 | 0.70422 | 0.70823 | 0.709377 | 0.714554 | 0.713861 | 0.707446 | 0.712212 |
| 0.705939 | 0.704254 | 0.702847 | 0.705691 | 0.707507 | 0.715812 | 0.713627 | 0.706961 | 0.710622 |
| 0.708066 | 0.704926 | 0.710013 | 0.707165 | 0.706177 | 0.718864 | 0.71652 | 0.707887 | 0.711139 |
| 0.716378 | 0.715006 | 0.712768 | 0.719368 | 0.716012 | 0.719445 | 0.72066 | 0.715359 | 0.716057 |
| 0.712 | 0.706025 | 0.70999 | 0.713072 | 0.711515 | 0.719217 | 0.713897 | 0.710629 | 0.710572 |
| 0.714411 | 0.710598 | 0.709529 | 0.712702 | 0.713192 | 0.716881 | 0.718385 | 0.708978 | 0.712666 |
| 0.716969 | 0.708828 | 0.71464 | 0.716115 | 0.713998 | 0.720711 | 0.718929 | 0.710632 | 0.717815 |
| 0.70828 | 0.70829 | 0.708006 | 0.706388 | 0.704628 | 0.715073 | 0.712133 | 0.70936 | 0.709035 |
| 0.714263 | 0.705953 | 0.708697 | 0.709508 | 0.707801 | 0.713094 | 0.713101 | 0.708166 | 0.7099 |
| 0.716024 | 0.709952 | 0.709154 | 0.712078 | 0.712302 | 0.717176 | 0.715607 | 0.709344 | 0.709533 |
| 0.704687 | 0.699606 | 0.707049 | 0.708082 | 0.703402 | 0.712367 | 0.712733 | 0.70431 | 0.707765 |
| 0.712128 | 0.707266 | 0.7094 | 0.714037 | 0.708363 | 0.716954 | 0.718718 | 0.710324 | 0.714489 |
| 0.660512 | 0.662561 | 0.658865 | 0.661471 | 0.660789 | 0.672393 | 0.66953 | 0.65875 | 0.664076 |
| 0.656363 | 0.663937 | 0.662844 | 0.657757 | 0.663404 | 0.674355 | 0.673544 | 0.667767 | 0.668005 |
| 0.715851 | 0.710408 | 0.712286 | 0.713812 | 0.71502 | 0.719052 | 0.717539 | 0.711535 | 0.712601 |
| 0.715879 | 0.709558 | 0.712492 | 0.715306 | 0.71479 | 0.722582 | 0.722943 | 0.715272 | 0.714602 |
| 0.649528 | 0.647937 | 0.645209 | 0.641447 | 0.640659 | 0.659777 | 0.650137 | 0.647233 | 0.655364 |
| 0.714562 | 0.708793 | 0.711978 | 0.713079 | 0.71318 | 0.721361 | 0.718355 | 0.708444 | 0.713538 |
| 0.703109 | 0.700128 | 0.69834 | 0.701786 | 0.699308 | 0.709998 | 0.706638 | 0.696977 | 0.70345 |

| | | | | | | | | |
|---|---|---|---|---|---|---|---|---|
| 0.711663 | 0.706861 | 0.706393 | 0.712072 | 0.705373 | 0.718007 | 0.714445 | 0.706005 | 0.710412 |
| 0.714366 | 0.707325 | 0.707793 | 0.712157 | 0.711195 | 0.718231 | 0.715479 | 0.709004 | 0.711787 |
| 0.712475 | 0.709265 | 0.713035 | 0.711122 | 0.709234 | 0.720794 | 0.716279 | 0.707866 | 0.712529 |
| 0.713908 | 0.707895 | 0.709376 | 0.712382 | 0.712514 | 0.718596 | 0.716123 | 0.708491 | 0.71424 |
| 0.718727 | 0.714814 | 0.715744 | 0.716189 | 0.716354 | 0.724707 | 0.725058 | 0.715512 | 0.718963 |
| 0.709506 | 0.703038 | 0.708558 | 0.707772 | 0.706702 | 0.71793 | 0.713783 | 0.704196 | 0.709257 |
| 0.710017 | 0.702823 | 0.706745 | 0.709596 | 0.709361 | 0.71463 | 0.716282 | 0.708353 | 0.712193 |
| 0.704279 | 0.700908 | 0.702966 | 0.702763 | 0.701791 | 0.708903 | 0.706462 | 0.69874 | 0.704342 |
| 0.712361 | 0.706506 | 0.704974 | 0.706544 | 0.707415 | 0.710929 | 0.71382 | 0.706005 | 0.708963 |
| 0.70735 | 0.707107 | 0.706631 | 0.708588 | 0.705783 | 0.716071 | 0.710173 | 0.705097 | 0.70829 |
| 0.71513 | 0.707707 | 0.708555 | 0.713562 | 0.711462 | 0.717951 | 0.715425 | 0.708018 | 0.710597 |
| 0.710353 | 0.701317 | 0.705998 | 0.706835 | 0.706538 | 0.712449 | 0.712057 | 0.704986 | 0.70703 |
| 0.699346 | 0.69893 | 0.700966 | 0.702276 | 0.699636 | 0.710552 | 0.706297 | 0.697109 | 0.703739 |
| 0.700983 | 0.69697 | 0.702586 | 0.700935 | 0.6996 | 0.708071 | 0.707753 | 0.697477 | 0.701515 |
| 0.716417 | 0.712549 | 0.711717 | 0.713464 | 0.716486 | 0.722361 | 0.721239 | 0.713978 | 0.719113 |
| 0.708904 | 0.709105 | 0.707559 | 0.71475 | 0.713016 | 0.721594 | 0.715155 | 0.709104 | 0.715559 |
| 0.706815 | 0.700935 | 0.704354 | 0.70634 | 0.704968 | 0.713797 | 0.712453 | 0.702196 | 0.706546 |
| 0.709683 | 0.705138 | 0.705097 | 0.708893 | 0.706603 | 0.713938 | 0.712273 | 0.705212 | 0.708621 |
| 0.710192 | 0.705392 | 0.702559 | 0.707776 | 0.704027 | 0.713548 | 0.711982 | 0.704839 | 0.707271 |
| 0.710203 | 0.707511 | 0.701826 | 0.710124 | 0.704042 | 0.712012 | 0.710656 | 0.705433 | 0.708535 |
| 0.703803 | 0.702246 | 0.6972 | 0.704946 | 0.701143 | 0.709322 | 0.707191 | 0.698844 | 0.705173 |
| 0.705202 | 0.704088 | 0.701668 | 0.706794 | 0.703121 | 0.710809 | 0.709831 | 0.705 | 0.706213 |
| 0.707685 | 0.707914 | 0.705822 | 0.711852 | 0.70467 | 0.717625 | 0.712653 | 0.708782 | 0.712087 |
| 0.71028 | 0.706373 | 0.706754 | 0.712159 | 0.708478 | 0.712693 | 0.715729 | 0.71063 | 0.710114 |
| 0.711088 | 0.705591 | 0.702249 | 0.710023 | 0.703491 | 0.712591 | 0.710799 | 0.706608 | 0.709133 |
| 0.70422 | 0.701162 | 0.700745 | 0.706399 | 0.701079 | 0.709882 | 0.70886 | 0.704137 | 0.706177 |
| 0.704273 | 0.704011 | 0.700435 | 0.708296 | 0.70334 | 0.708255 | 0.707443 | 0.702657 | 0.705802 |
| 0.703432 | 0.700948 | 0.699109 | 0.705498 | 0.707193 | 0.706703 | 0.709962 | 0.702877 | 0.706319 |
| 0.706854 | 0.70409 | 0.700739 | 0.711753 | 0.710535 | 0.709763 | 0.711566 | 0.707097 | 0.706988 |
| 0.706853 | 0.705835 | 0.704044 | 0.710407 | 0.708033 | 0.712065 | 0.711586 | 0.708063 | 0.708888 |
| 0.713586 | 0.708022 | 0.707864 | 0.713308 | 0.711941 | 0.71521 | 0.715545 | 0.708668 | 0.711705 |
| 0.713129 | 0.709585 | 0.709508 | 0.712509 | 0.710498 | 0.715265 | 0.714767 | 0.710759 | 0.713027 |
| 0.714718 | 0.711063 | 0.710828 | 0.716483 | 0.712849 | 0.723577 | 0.720945 | 0.714013 | 0.716925 |
| 0.71297 | 0.712356 | 0.712588 | 0.714898 | 0.712679 | 0.721701 | 0.7219 | 0.714584 | 0.720463 |
| 0.706099 | 0.700836 | 0.700048 | 0.709166 | 0.707641 | 0.707591 | 0.708383 | 0.708317 | 0.704091 |
| 0.706392 | 0.703384 | 0.704282 | 0.712813 | 0.709793 | 0.717719 | 0.715024 | 0.706864 | 0.711261 |
| 0.710458 | 0.706453 | 0.705594 | 0.714343 | 0.711196 | 0.712203 | 0.714766 | 0.710127 | 0.71044 |
| 0.702755 | 0.704555 | 0.702186 | 0.708581 | 0.702076 | 0.710997 | 0.709974 | 0.703332 | 0.707272 |
| 0.704105 | 0.700188 | 0.698574 | 0.703988 | 0.699886 | 0.709149 | 0.703268 | 0.697502 | 0.702822 |
| 0.697081 | 0.697501 | 0.699967 | 0.703883 | 0.696293 | 0.703978 | 0.705589 | 0.700075 | 0.705457 |
| 0.702084 | 0.696543 | 0.690921 | 0.701543 | 0.698098 | 0.700398 | 0.702269 | 0.696011 | 0.699021 |
| 0.697696 | 0.696477 | 0.695523 | 0.700379 | 0.694816 | 0.702904 | 0.703804 | 0.698656 | 0.702907 |
| 0.704355 | 0.69734 | 0.698768 | 0.705188 | 0.703019 | 0.707134 | 0.70375 | 0.69923 | 0.702886 |
| 0.693974 | 0.692477 | 0.691624 | 0.696354 | 0.689782 | 0.702417 | 0.6978 | 0.691209 | 0.692946 |
| 0.704145 | 0.700742 | 0.698867 | 0.706456 | 0.702748 | 0.710825 | 0.710017 | 0.703881 | 0.706861 |
| 0.699559 | 0.695812 | 0.694982 | 0.699927 | 0.696752 | 0.702795 | 0.702597 | 0.694646 | 0.700791 |

| | | | | | | | | |
|---|---|---|---|---|---|---|---|---|
| 0.714068 | 0.70749 | 0.704858 | 0.714696 | 0.713824 | 0.716966 | 0.71895 | 0.710306 | 0.71106 |
| 0.711517 | 0.705983 | 0.710484 | 0.710861 | 0.710226 | 0.713436 | 0.714791 | 0.709966 | 0.710946 |
| 0.70612 | 0.707003 | 0.704164 | 0.706071 | 0.704767 | 0.710073 | 0.714816 | 0.708788 | 0.71094 |
| 0.711003 | 0.705939 | 0.706898 | 0.715133 | 0.710113 | 0.717161 | 0.716622 | 0.708062 | 0.713904 |
| 0.719713 | 0.715046 | 0.716173 | 0.717626 | 0.713769 | 0.722636 | 0.72482 | 0.719336 | 0.717886 |
| 0.715727 | 0.70817 | 0.712306 | 0.712165 | 0.708746 | 0.71806 | 0.719017 | 0.710873 | 0.714252 |
| 0.709885 | 0.702321 | 0.703473 | 0.705287 | 0.70679 | 0.712265 | 0.709704 | 0.707433 | 0.708165 |
| 0.712058 | 0.706844 | 0.708303 | 0.711347 | 0.706561 | 0.712701 | 0.714794 | 0.709869 | 0.708979 |
| 0.705517 | 0.698478 | 0.702315 | 0.70751 | 0.700473 | 0.710721 | 0.709773 | 0.702367 | 0.705327 |
| 0.711641 | 0.706715 | 0.708043 | 0.710694 | 0.708286 | 0.713904 | 0.716618 | 0.709151 | 0.710399 |
| 0.71507 | 0.710405 | 0.708596 | 0.712399 | 0.707573 | 0.716966 | 0.714421 | 0.709939 | 0.713368 |
| 0.716522 | 0.709548 | 0.709886 | 0.716875 | 0.710324 | 0.719083 | 0.71875 | 0.711236 | 0.711872 |
| 0.71118 | 0.70454 | 0.705854 | 0.710107 | 0.709478 | 0.713066 | 0.714297 | 0.704005 | 0.708422 |
| 0.711209 | 0.710222 | 0.705578 | 0.708241 | 0.710258 | 0.717636 | 0.712539 | 0.709266 | 0.709897 |
| 0.712266 | 0.709174 | 0.708297 | 0.711844 | 0.710195 | 0.718517 | 0.717173 | 0.707437 | 0.712975 |
| 0.711385 | 0.70961 | 0.710152 | 0.710903 | 0.710862 | 0.714907 | 0.716179 | 0.708543 | 0.709517 |
| 0.715035 | 0.71202 | 0.711848 | 0.713388 | 0.711885 | 0.722089 | 0.717444 | 0.710121 | 0.712979 |
| 0.712964 | 0.709406 | 0.703837 | 0.710967 | 0.709657 | 0.717638 | 0.715762 | 0.710351 | 0.713651 |
| 0.71408 | 0.712349 | 0.709464 | 0.71389 | 0.713473 | 0.716694 | 0.718705 | 0.713026 | 0.713621 |
| 0.708314 | 0.709188 | 0.701504 | 0.713248 | 0.708752 | 0.712172 | 0.711644 | 0.70932 | 0.707143 |
| 0.709797 | 0.709547 | 0.709662 | 0.710143 | 0.710295 | 0.715387 | 0.713955 | 0.711009 | 0.712102 |
| 0.711509 | 0.706306 | 0.70844 | 0.707795 | 0.706751 | 0.715065 | 0.713555 | 0.706585 | 0.707383 |
| 0.712524 | 0.708972 | 0.711969 | 0.716125 | 0.710665 | 0.718241 | 0.716293 | 0.710734 | 0.716977 |
| 0.715957 | 0.708967 | 0.709277 | 0.714759 | 0.713226 | 0.72108 | 0.71665 | 0.710212 | 0.714387 |
| 0.710957 | 0.709258 | 0.707587 | 0.713158 | 0.712633 | 0.716619 | 0.717856 | 0.709058 | 0.710945 |
| 0.711897 | 0.706052 | 0.707554 | 0.711407 | 0.710081 | 0.7163 | 0.714093 | 0.708784 | 0.709266 |
| 0.711258 | 0.705208 | 0.704729 | 0.708713 | 0.708396 | 0.714927 | 0.715187 | 0.70762 | 0.712549 |
| 0.712706 | 0.70681 | 0.705957 | 0.707237 | 0.710315 | 0.714629 | 0.71315 | 0.70535 | 0.71032 |
| 0.710724 | 0.710551 | 0.709088 | 0.713904 | 0.712975 | 0.719646 | 0.715892 | 0.711955 | 0.712028 |
| 0.712231 | 0.711628 | 0.70907 | 0.715919 | 0.714274 | 0.721127 | 0.71851 | 0.714853 | 0.71637 |
| 0.707837 | 0.709241 | 0.706196 | 0.71318 | 0.708126 | 0.713747 | 0.714689 | 0.708995 | 0.7093 |
| 0.716016 | 0.712308 | 0.711571 | 0.716665 | 0.713639 | 0.718806 | 0.719872 | 0.711425 | 0.715071 |
| 0.714469 | 0.711093 | 0.710721 | 0.716902 | 0.714974 | 0.718929 | 0.719025 | 0.717386 | 0.713115 |
| 0.709035 | 0.702077 | 0.702302 | 0.707174 | 0.708227 | 0.710884 | 0.711697 | 0.705005 | 0.707012 |
| 0.708875 | 0.707933 | 0.70662 | 0.711069 | 0.70674 | 0.717983 | 0.715385 | 0.708454 | 0.710599 |
| 0.707327 | 0.705157 | 0.703929 | 0.707779 | 0.708481 | 0.711806 | 0.713538 | 0.707186 | 0.705338 |
| 0.717212 | 0.716955 | 0.711964 | 0.717737 | 0.71625 | 0.723576 | 0.722845 | 0.714171 | 0.717469 |
| 0.714938 | 0.707608 | 0.706951 | 0.712786 | 0.70868 | 0.717553 | 0.712685 | 0.708235 | 0.708009 |
| 0.71556 | 0.711093 | 0.707841 | 0.713006 | 0.706932 | 0.719369 | 0.716545 | 0.707352 | 0.710395 |
| 0.711287 | 0.7095 | 0.706503 | 0.713845 | 0.709954 | 0.716177 | 0.718334 | 0.708996 | 0.712973 |
| 0.718238 | 0.715533 | 0.712958 | 0.71459 | 0.714069 | 0.72152 | 0.71987 | 0.711299 | 0.717189 |
| 0.716529 | 0.713057 | 0.705719 | 0.715818 | 0.709274 | 0.718939 | 0.717588 | 0.710168 | 0.711595 |
| 0.715557 | 0.710363 | 0.707793 | 0.71222 | 0.711191 | 0.720087 | 0.717698 | 0.710899 | 0.713155 |
| 0.710819 | 0.708279 | 0.705682 | 0.710903 | 0.709039 | 0.714333 | 0.713858 | 0.708773 | 0.711145 |
| 0.708746 | 0.712413 | 0.70791 | 0.716728 | 0.711091 | 0.719415 | 0.714343 | 0.712074 | 0.712659 |
| 0.70646 | 0.705767 | 0.701894 | 0.709192 | 0.70716 | 0.714885 | 0.713951 | 0.707749 | 0.706302 |

| | | | | | | | | |
|---|---|---|---|---|---|---|---|---|
| 0.699339 | 0.695792 | 0.692237 | 0.696523 | 0.693245 | 0.702693 | 0.70296 | 0.694365 | 0.700757 |
| 0.748493 | 0.751183 | 0.745943 | 0.743025 | 0.7432 | 0.757811 | 0.757646 | 0.747081 | 0.757874 |
| 0.708149 | 0.70698 | 0.705421 | 0.709423 | 0.705713 | 0.711356 | 0.712935 | 0.705607 | 0.710054 |
| 0.714104 | 0.706009 | 0.707517 | 0.714782 | 0.713759 | 0.716392 | 0.717807 | 0.71152 | 0.712192 |
| 0.70833 | 0.704824 | 0.703888 | 0.706973 | 0.705284 | 0.709318 | 0.709651 | 0.704461 | 0.702479 |
| 0.711383 | 0.705267 | 0.704463 | 0.710674 | 0.70947 | 0.715415 | 0.713851 | 0.708114 | 0.711508 |
| 0.706729 | 0.705291 | 0.703722 | 0.706606 | 0.700886 | 0.712528 | 0.708596 | 0.704919 | 0.705688 |
| 0.705051 | 0.70163 | 0.697234 | 0.703738 | 0.701753 | 0.706153 | 0.707617 | 0.700465 | 0.705913 |
| 0.709588 | 0.705817 | 0.705179 | 0.707678 | 0.70844 | 0.713667 | 0.713496 | 0.708626 | 0.712016 |
| 0.715896 | 0.715048 | 0.714343 | 0.713556 | 0.713065 | 0.722479 | 0.720105 | 0.714401 | 0.717626 |
| 0.714001 | 0.712031 | 0.709846 | 0.711919 | 0.709464 | 0.718559 | 0.71742 | 0.711858 | 0.715017 |
| 0.715426 | 0.707994 | 0.709066 | 0.715711 | 0.713132 | 0.720728 | 0.719347 | 0.714609 | 0.717504 |
| 0.715523 | 0.712205 | 0.713007 | 0.715068 | 0.71485 | 0.720582 | 0.720487 | 0.714228 | 0.716673 |
| 0.714707 | 0.712308 | 0.708086 | 0.715394 | 0.713726 | 0.721191 | 0.71968 | 0.71221 | 0.715253 |
| 0.711055 | 0.707299 | 0.706827 | 0.715269 | 0.71035 | 0.716411 | 0.714794 | 0.710264 | 0.713894 |
| 0.711185 | 0.711203 | 0.709794 | 0.709148 | 0.711136 | 0.719291 | 0.712769 | 0.709502 | 0.714408 |
| 0.714033 | 0.71011 | 0.709962 | 0.713398 | 0.711627 | 0.718773 | 0.716578 | 0.708588 | 0.716663 |
| 0.712476 | 0.707866 | 0.711631 | 0.710043 | 0.712133 | 0.714993 | 0.718129 | 0.708005 | 0.713103 |
| 0.702426 | 0.704737 | 0.702289 | 0.701367 | 0.706826 | 0.710896 | 0.709939 | 0.704942 | 0.705377 |
| 0.702128 | 0.700618 | 0.703046 | 0.702314 | 0.703501 | 0.710605 | 0.707974 | 0.703219 | 0.704667 |
| 0.714077 | 0.708756 | 0.708189 | 0.70813 | 0.708097 | 0.71537 | 0.714722 | 0.703794 | 0.715327 |
| 0.698777 | 0.701676 | 0.69638 | 0.704027 | 0.699325 | 0.702743 | 0.704114 | 0.702275 | 0.702536 |
| 0.708927 | 0.705897 | 0.703739 | 0.713028 | 0.708817 | 0.712721 | 0.711866 | 0.709022 | 0.707098 |
| 0.710533 | 0.708518 | 0.709771 | 0.713819 | 0.708964 | 0.715569 | 0.716701 | 0.711362 | 0.710608 |
| 0.71452 | 0.711616 | 0.710567 | 0.714414 | 0.711235 | 0.716998 | 0.71539 | 0.715273 | 0.712239 |
| 0.708726 | 0.706693 | 0.707921 | 0.711681 | 0.707487 | 0.714075 | 0.712742 | 0.712523 | 0.70825 |
| 0.70803 | 0.705257 | 0.702252 | 0.706812 | 0.703701 | 0.70938 | 0.713263 | 0.707655 | 0.705729 |
| 0.706186 | 0.705083 | 0.703223 | 0.705669 | 0.702053 | 0.710009 | 0.711111 | 0.701144 | 0.703334 |
| 0.705843 | 0.705043 | 0.700345 | 0.708935 | 0.701673 | 0.710733 | 0.709236 | 0.708185 | 0.705813 |
| 0.779137 | 0.767941 | 0.7671 | 0.824007 | 0.768034 | 0.776083 | 0.779708 | 0.80197 | 0.75636 |
| 0.803658 | 0.774954 | 0.784845 | 0.805806 | 0.784955 | 0.81294 | 0.808842 | 0.800119 | 0.815293 |
| 0.783119 | 0.775399 | 0.780323 | 0.889375 | 0.790859 | 0.786862 | 0.788396 | 0.820166 | 0.777366 |
| 0.776489 | 0.766079 | 0.823143 | 0.816808 | 0.812746 | 0.81113 | 0.792055 | 0.766262 | 0.783588 |
| 0.791722 | 0.77996 | 0.78987 | 0.808753 | 0.812258 | 0.813462 | 0.798995 | 0.777015 | 0.82008 |
| 0.702486 | 0.70359 | 0.700658 | 0.706022 | 0.69874 | 0.706282 | 0.710358 | 0.70201 | 0.700602 |
| 0.705539 | 0.705118 | 0.704445 | 0.707292 | 0.704573 | 0.711556 | 0.710659 | 0.703461 | 0.705003 |
| 0.709059 | 0.707569 | 0.703898 | 0.711694 | 0.709698 | 0.714507 | 0.712049 | 0.706216 | 0.706431 |
| 0.708183 | 0.706797 | 0.708878 | 0.714537 | 0.709956 | 0.715697 | 0.716236 | 0.711152 | 0.707864 |
| 0.703665 | 0.706172 | 0.707068 | 0.706705 | 0.708138 | 0.713786 | 0.711437 | 0.705669 | 0.705442 |
| 0.712293 | 0.707692 | 0.707323 | 0.71408 | 0.710605 | 0.718851 | 0.71743 | 0.715094 | 0.711788 |
| 0.708134 | 0.704466 | 0.704663 | 0.708976 | 0.710925 | 0.713608 | 0.715254 | 0.706947 | 0.706005 |
| 0.71234 | 0.705167 | 0.705692 | 0.708884 | 0.705777 | 0.712572 | 0.712222 | 0.7078 | 0.704266 |
| 0.699713 | 0.700874 | 0.695422 | 0.704984 | 0.699272 | 0.70693 | 0.70834 | 0.69866 | 0.700827 |
| 0.700112 | 0.703512 | 0.701613 | 0.703672 | 0.698908 | 0.703761 | 0.706516 | 0.701915 | 0.698649 |
| 0.705105 | 0.705979 | 0.705237 | 0.710348 | 0.701429 | 0.713409 | 0.713095 | 0.706638 | 0.705177 |
| 0.707594 | 0.706787 | 0.70583 | 0.710463 | 0.707204 | 0.713003 | 0.713883 | 0.707815 | 0.707472 |

| | | | | | | | | |
|---|---|---|---|---|---|---|---|---|
| 0.710953 | 0.70868 | 0.706953 | 0.710355 | 0.708916 | 0.715785 | 0.713844 | 0.709603 | 0.711284 |
| 0.709467 | 0.709917 | 0.703203 | 0.709976 | 0.708502 | 0.710753 | 0.712269 | 0.706723 | 0.709613 |
| 0.711873 | 0.70999 | 0.713754 | 0.711852 | 0.712098 | 0.718553 | 0.717028 | 0.710454 | 0.712968 |
| 0.708611 | 0.704457 | 0.708106 | 0.710572 | 0.708747 | 0.713881 | 0.713204 | 0.703739 | 0.710146 |
| 0.706713 | 0.701419 | 0.703971 | 0.705414 | 0.706414 | 0.710944 | 0.70738 | 0.702614 | 0.70664 |
| 0.704497 | 0.706662 | 0.70374 | 0.708156 | 0.704391 | 0.710915 | 0.71133 | 0.703071 | 0.704191 |
| 0.709977 | 0.710972 | 0.704902 | 0.713399 | 0.709106 | 0.714534 | 0.716247 | 0.704849 | 0.712347 |
| 0.71049 | 0.709712 | 0.705267 | 0.710533 | 0.711421 | 0.71327 | 0.713808 | 0.704544 | 0.707368 |
| 0.708542 | 0.705863 | 0.701654 | 0.705696 | 0.709116 | 0.712986 | 0.712067 | 0.704232 | 0.708178 |
| 0.70697 | 0.707572 | 0.702555 | 0.711715 | 0.708455 | 0.711706 | 0.711367 | 0.70811 | 0.708808 |
| 0.706102 | 0.706345 | 0.704077 | 0.70728 | 0.706232 | 0.713065 | 0.710484 | 0.703209 | 0.711408 |
| 0.708194 | 0.703089 | 0.704549 | 0.711147 | 0.705868 | 0.713488 | 0.712076 | 0.704097 | 0.708555 |
| 0.699628 | 0.699064 | 0.697238 | 0.70129 | 0.698936 | 0.70483 | 0.702775 | 0.696084 | 0.703173 |
| 0.701757 | 0.697755 | 0.696383 | 0.702216 | 0.70328 | 0.707289 | 0.707326 | 0.699389 | 0.701469 |
| 0.712878 | 0.708385 | 0.708476 | 0.714996 | 0.709391 | 0.717319 | 0.714425 | 0.712746 | 0.70894 |
| 0.715126 | 0.714101 | 0.714539 | 0.716467 | 0.71617 | 0.721906 | 0.719328 | 0.719148 | 0.717983 |
| 0.714206 | 0.711618 | 0.708034 | 0.71459 | 0.712602 | 0.716421 | 0.718972 | 0.711989 | 0.7132 |
| 0.712267 | 0.712087 | 0.713209 | 0.713876 | 0.714205 | 0.721015 | 0.719592 | 0.712606 | 0.712655 |
| 0.706536 | 0.704076 | 0.70743 | 0.710604 | 0.708121 | 0.71431 | 0.710732 | 0.705936 | 0.705993 |
| 0.707951 | 0.705541 | 0.704082 | 0.710193 | 0.707495 | 0.712489 | 0.711357 | 0.708084 | 0.707724 |
| 0.704551 | 0.704041 | 0.705219 | 0.709296 | 0.703349 | 0.7113 | 0.710381 | 0.704049 | 0.706936 |
| 0.710759 | 0.706766 | 0.707225 | 0.711063 | 0.70855 | 0.718433 | 0.715182 | 0.714635 | 0.712434 |
| 0.71122 | 0.71189 | 0.709576 | 0.712805 | 0.711862 | 0.717213 | 0.714156 | 0.711144 | 0.717985 |
| 0.708012 | 0.708481 | 0.704281 | 0.708834 | 0.708606 | 0.713435 | 0.71065 | 0.705716 | 0.709011 |
| 0.709224 | 0.705201 | 0.707957 | 0.707999 | 0.708852 | 0.714323 | 0.711937 | 0.710096 | 0.710922 |
| 0.788007 | 0.786864 | 0.806803 | 0.798128 | 0.782399 | 0.805029 | 0.790281 | 0.794996 | 0.779439 |
| 0.711154 | 0.708617 | 0.70414 | 0.71159 | 0.711355 | 0.71625 | 0.713914 | 0.709262 | 0.711634 |
| 0.713509 | 0.712297 | 0.709564 | 0.715815 | 0.70997 | 0.72054 | 0.717393 | 0.708612 | 0.713733 |
| 0.713721 | 0.712051 | 0.709716 | 0.709763 | 0.713868 | 0.719267 | 0.718545 | 0.71045 | 0.714685 |
| 0.707343 | 0.706461 | 0.70808 | 0.712484 | 0.708306 | 0.716977 | 0.713307 | 0.7087 | 0.708092 |
| 0.706794 | 0.705625 | 0.701864 | 0.707021 | 0.709514 | 0.713936 | 0.709782 | 0.704906 | 0.708371 |
| 0.709672 | 0.704152 | 0.702392 | 0.711668 | 0.706185 | 0.712228 | 0.711586 | 0.706218 | 0.709783 |
| 0.712557 | 0.709064 | 0.705347 | 0.710121 | 0.708976 | 0.71586 | 0.715839 | 0.709918 | 0.712011 |
| 0.705498 | 0.70131 | 0.703351 | 0.70673 | 0.703666 | 0.714552 | 0.710011 | 0.70371 | 0.705785 |
| 0.705196 | 0.700448 | 0.702853 | 0.707217 | 0.70509 | 0.711323 | 0.710617 | 0.703381 | 0.707356 |
| 0.708066 | 0.705389 | 0.702674 | 0.709567 | 0.706672 | 0.714534 | 0.712018 | 0.706588 | 0.705335 |
| 0.71561 | 0.711647 | 0.708581 | 0.71668 | 0.709155 | 0.717894 | 0.719086 | 0.713384 | 0.716115 |
| 0.713848 | 0.712395 | 0.708551 | 0.712079 | 0.713787 | 0.717455 | 0.717095 | 0.710015 | 0.713647 |
| 0.714483 | 0.707856 | 0.706375 | 0.70866 | 0.710977 | 0.717553 | 0.713758 | 0.708039 | 0.71267 |
| 0.707681 | 0.706163 | 0.70645 | 0.712805 | 0.708072 | 0.714346 | 0.714844 | 0.70439 | 0.711118 |
| 0.705406 | 0.70268 | 0.699636 | 0.70673 | 0.704285 | 0.708824 | 0.71009 | 0.703948 | 0.707438 |
| 0.715349 | 0.709015 | 0.711703 | 0.714378 | 0.712971 | 0.718316 | 0.717546 | 0.708868 | 0.71218 |
| 0.708901 | 0.705863 | 0.707656 | 0.704202 | 0.705238 | 0.713529 | 0.712204 | 0.705302 | 0.708313 |
| 0.709791 | 0.709328 | 0.706457 | 0.708113 | 0.707793 | 0.715894 | 0.716877 | 0.708223 | 0.712586 |
| 0.711147 | 0.711377 | 0.709905 | 0.711738 | 0.712728 | 0.717936 | 0.715732 | 0.710181 | 0.712391 |
| 0.710211 | 0.714851 | 0.710754 | 0.709852 | 0.710539 | 0.719064 | 0.718349 | 0.709015 | 0.712659 |

| | | | | | | | | |
|---|---|---|---|---|---|---|---|---|
| 0.705041 | 0.709396 | 0.706652 | 0.710647 | 0.70751 | 0.715065 | 0.715822 | 0.707008 | 0.708144 |
| 0.706574 | 0.711003 | 0.708095 | 0.710481 | 0.708247 | 0.717901 | 0.714873 | 0.71134 | 0.714984 |
| 0.709317 | 0.707173 | 0.702843 | 0.70927 | 0.709855 | 0.717026 | 0.714183 | 0.709086 | 0.710017 |
| 0.663205 | 0.678534 | 0.665162 | 0.661485 | 0.666783 | 0.692775 | 0.677793 | 0.669346 | 0.673754 |
| 0.707174 | 0.706014 | 0.704515 | 0.708589 | 0.708846 | 0.715375 | 0.713113 | 0.707434 | 0.710978 |
| 0.706169 | 0.708407 | 0.704983 | 0.711662 | 0.704977 | 0.717451 | 0.716026 | 0.707214 | 0.711317 |
| 0.705099 | 0.708137 | 0.70932 | 0.708383 | 0.711822 | 0.715391 | 0.714966 | 0.709136 | 0.709766 |
| 0.707467 | 0.70852 | 0.708118 | 0.704874 | 0.707138 | 0.718521 | 0.717496 | 0.709639 | 0.711923 |
| 0.712084 | 0.714894 | 0.709537 | 0.714908 | 0.711997 | 0.719894 | 0.720228 | 0.714755 | 0.719304 |
| 0.710537 | 0.708932 | 0.707624 | 0.712413 | 0.710778 | 0.717591 | 0.719085 | 0.711719 | 0.710501 |
| 0.707059 | 0.708679 | 0.707738 | 0.710489 | 0.710868 | 0.718871 | 0.71499 | 0.709116 | 0.71252 |
| 0.710596 | 0.715431 | 0.710072 | 0.711731 | 0.712857 | 0.720097 | 0.722529 | 0.712756 | 0.714887 |
| 0.711431 | 0.714746 | 0.709055 | 0.71365 | 0.715618 | 0.721696 | 0.719344 | 0.713598 | 0.710955 |
| 0.708124 | 0.70929 | 0.706573 | 0.712592 | 0.711874 | 0.714653 | 0.717178 | 0.708842 | 0.710435 |
| 0.710558 | 0.712275 | 0.709307 | 0.717421 | 0.713437 | 0.721764 | 0.718934 | 0.714297 | 0.71327 |
| 0.70537 | 0.707064 | 0.703822 | 0.710472 | 0.708615 | 0.711927 | 0.71549 | 0.709657 | 0.707886 |
| 0.709396 | 0.711148 | 0.706854 | 0.710504 | 0.711272 | 0.719436 | 0.715892 | 0.714297 | 0.714521 |
| 0.70906 | 0.711143 | 0.706587 | 0.715295 | 0.711776 | 0.719253 | 0.71927 | 0.712878 | 0.712665 |
| 0.706174 | 0.704396 | 0.705159 | 0.706115 | 0.711259 | 0.714758 | 0.712613 | 0.704442 | 0.70941 |
| 0.707282 | 0.706263 | 0.70398 | 0.709744 | 0.709881 | 0.714596 | 0.715666 | 0.712161 | 0.711239 |
| 0.707927 | 0.712127 | 0.712142 | 0.714411 | 0.710911 | 0.721701 | 0.716824 | 0.710303 | 0.711538 |
| 0.710861 | 0.713643 | 0.70978 | 0.712329 | 0.710295 | 0.720229 | 0.718482 | 0.711973 | 0.71406 |
| 0.698186 | 0.694556 | 0.693672 | 0.697888 | 0.694038 | 0.709522 | 0.706713 | 0.70711 | 0.699196 |
| 0.721187 | 0.718591 | 0.725846 | 0.723994 | 0.719238 | 0.747439 | 0.736132 | 0.730104 | 0.725567 |
| 0.756763 | 0.753666 | 0.756318 | 0.757996 | 0.753604 | 0.777396 | 0.760494 | 0.762896 | 0.762205 |
| 0.743647 | 0.734227 | 0.850693 | 0.738401 | 0.758045 | 0.790694 | 0.756181 | 0.744018 | 0.74093 |
| 0.668749 | 0.664355 | 0.666619 | 0.666427 | 0.670322 | 0.677971 | 0.672006 | 0.667875 | 0.669356 |
| 0.665369 | 0.662297 | 0.659125 | 0.660291 | 0.657909 | 0.676745 | 0.667713 | 0.661281 | 0.669813 |
| 0.662552 | 0.657271 | 0.659615 | 0.665854 | 0.659079 | 0.682442 | 0.667535 | 0.663636 | 0.664646 |
| 0.680572 | 0.681559 | 0.670372 | 0.671747 | 0.678478 | 0.695019 | 0.682671 | 0.673147 | 0.683471 |
| 0.65 | 0.651495 | 0.652935 | 0.65735 | 0.653557 | 0.682679 | 0.662786 | 0.652088 | 0.66209 |
| 0.6469 | 0.642954 | 0.638005 | 0.643535 | 0.64884 | 0.663251 | 0.651404 | 0.639858 | 0.651585 |
| 0.645426 | 0.646421 | 0.645537 | 0.636109 | 0.646259 | 0.657028 | 0.648725 | 0.648308 | 0.645189 |
| 0.761034 | 0.73258 | 0.733506 | 0.746471 | 0.746842 | 0.753561 | 0.771659 | 0.734792 | 0.740488 |
| 0.700836 | 0.702995 | 0.702128 | 0.703028 | 0.704918 | 0.711096 | 0.709219 | 0.702048 | 0.70404 |
| 0.674865 | 0.672433 | 0.678363 | 0.67874 | 0.679275 | 0.688095 | 0.687549 | 0.679426 | 0.67709 |
| 0.666537 | 0.670657 | 0.671063 | 0.67208 | 0.67048 | 0.68167 | 0.677003 | 0.671199 | 0.672427 |
| 0.651466 | 0.654059 | 0.65607 | 0.659825 | 0.655989 | 0.663677 | 0.664407 | 0.657215 | 0.651811 |
| 0.675764 | 0.677714 | 0.673559 | 0.67655 | 0.676249 | 0.688664 | 0.687296 | 0.678616 | 0.679775 |
| 0.668402 | 0.668524 | 0.668609 | 0.665562 | 0.668887 | 0.683462 | 0.679896 | 0.667535 | 0.671157 |
| 0.660197 | 0.656398 | 0.6556 | 0.660968 | 0.658983 | 0.67466 | 0.669517 | 0.663586 | 0.663456 |
| 0.664858 | 0.663101 | 0.660938 | 0.665565 | 0.663159 | 0.678175 | 0.67615 | 0.658786 | 0.668119 |
| 0.662026 | 0.655758 | 0.652513 | 0.661352 | 0.651239 | 0.668819 | 0.659272 | 0.653287 | 0.653356 |
| 0.666092 | 0.6626 | 0.658245 | 0.660156 | 0.662819 | 0.673959 | 0.670353 | 0.664188 | 0.664948 |
| 0.653304 | 0.659896 | 0.660237 | 0.660428 | 0.65694 | 0.666667 | 0.665437 | 0.660359 | 0.660912 |
| 0.643901 | 0.645151 | 0.64148 | 0.646066 | 0.645437 | 0.654633 | 0.655227 | 0.638871 | 0.648313 |

| | | | | | | | | |
|---|---|---|---|---|---|---|---|---|
| 0.690123 | 0.691758 | 0.689301 | 0.689797 | 0.691967 | 0.700203 | 0.699596 | 0.692694 | 0.694057 |
| 0.660306 | 0.663058 | 0.659643 | 0.655848 | 0.664818 | 0.673925 | 0.666846 | 0.652847 | 0.667655 |
| 0.662614 | 0.663475 | 0.658149 | 0.663215 | 0.666234 | 0.675908 | 0.671751 | 0.658864 | 0.6697 |
| 0.667259 | 0.665532 | 0.662531 | 0.66848 | 0.662702 | 0.678291 | 0.668184 | 0.666667 | 0.665737 |
| 0.656631 | 0.657751 | 0.662615 | 0.658038 | 0.66359 | 0.677765 | 0.663933 | 0.66123 | 0.658183 |
| 0.661982 | 0.667469 | 0.665053 | 0.663072 | 0.665867 | 0.674407 | 0.670907 | 0.668765 | 0.666106 |
| 0.652661 | 0.653156 | 0.652804 | 0.653541 | 0.654779 | 0.668938 | 0.662679 | 0.649217 | 0.663008 |
| 0.642005 | 0.652316 | 0.647202 | 0.643784 | 0.652304 | 0.659265 | 0.6567 | 0.650493 | 0.654704 |
| 0.647107 | 0.646454 | 0.654432 | 0.642973 | 0.652517 | 0.663734 | 0.662288 | 0.650954 | 0.656761 |
| 0.693708 | 0.692059 | 0.692928 | 0.691473 | 0.693841 | 0.701786 | 0.699991 | 0.694122 | 0.69658 |
| 0.689803 | 0.689739 | 0.689221 | 0.687579 | 0.688967 | 0.699155 | 0.695678 | 0.689528 | 0.691749 |
| 0.687446 | 0.683483 | 0.67896 | 0.687113 | 0.686314 | 0.694928 | 0.691214 | 0.6867 | 0.685586 |
| 0.679428 | 0.675228 | 0.673893 | 0.681792 | 0.674907 | 0.687759 | 0.679459 | 0.677203 | 0.681845 |
| 0.681527 | 0.68623 | 0.680823 | 0.682947 | 0.67792 | 0.694097 | 0.689407 | 0.684086 | 0.685617 |
| 0.678576 | 0.672792 | 0.669659 | 0.674267 | 0.670485 | 0.686152 | 0.685242 | 0.670134 | 0.676839 |

| Collection_ | Collection_ | Collection_ | Collection_ | Collection_ | Collection_ | Collection_ | Collection_ | Collection_ |
|---|---|---|---|---|---|---|---|---|
| 0.777087 | 0.776922 | 0.774612 | 0.779413 | 0.773007 | 0.784812 | 0.771536 | 0.778651 | 0.77541 |
| 0.784141 | 0.776555 | 0.774787 | 0.776564 | 0.780109 | 0.785554 | 0.773205 | 0.779606 | 0.775693 |
| 0.781833 | 0.778521 | 0.775309 | 0.781025 | 0.769709 | 0.776474 | 0.765423 | 0.783533 | 0.782823 |
| 0.777714 | 0.78665 | 0.774863 | 0.777453 | 0.775199 | 0.773899 | 0.776246 | 0.778564 | 0.778095 |
| 0.778577 | 0.781911 | 0.775974 | 0.778984 | 0.777537 | 0.781201 | 0.775167 | 0.774632 | 0.777887 |
| 0.788579 | 0.773591 | 0.768515 | 0.775026 | 0.770735 | 0.770091 | 0.772625 | 0.771937 | 0.776581 |
| 0.713931 | 0.711614 | 0.707383 | 0.707281 | 0.708046 | 0.717139 | 0.707199 | 0.713658 | 0.709127 |
| 0.709139 | 0.711397 | 0.705794 | 0.707691 | 0.704381 | 0.709595 | 0.701938 | 0.708244 | 0.707523 |
| 0.800854 | 0.777455 | 0.778786 | 0.78027 | 0.77367 | 0.780502 | 0.774339 | 0.784286 | 0.779254 |
| 0.775109 | 0.779048 | 0.772117 | 0.772188 | 0.769804 | 0.778743 | 0.764169 | 0.775303 | 0.768911 |
| 0.777105 | 0.77631 | 0.770513 | 0.768723 | 0.774536 | 0.773873 | 0.772028 | 0.773267 | 0.769218 |
| 0.773343 | 0.782248 | 0.790559 | 0.775875 | 0.779457 | 0.791034 | 0.798256 | 0.778926 | 0.780903 |
| 0.774164 | 0.769668 | 0.77303 | 0.773819 | 0.770741 | 0.776515 | 0.776781 | 0.776785 | 0.771106 |
| 0.774857 | 0.774127 | 0.771503 | 0.769187 | 0.765115 | 0.776645 | 0.769391 | 0.780007 | 0.77483 |
| 0.772899 | 0.765878 | 0.758548 | 0.766349 | 0.768539 | 0.768504 | 0.759252 | 0.762681 | 0.764134 |
| 0.791785 | 0.784525 | 0.775268 | 0.783297 | 0.774591 | 0.780344 | 0.7737 | 0.780452 | 0.779592 |
| 0.769384 | 0.77238 | 0.767121 | 0.769407 | 0.768916 | 0.774981 | 0.764558 | 0.769409 | 0.772384 |
| 0.75515 | 0.751571 | 0.74966 | 0.755043 | 0.751565 | 0.755331 | 0.745939 | 0.75866 | 0.749102 |
| 0.775643 | 0.775808 | 0.784831 | 0.78476 | 0.787403 | 0.783194 | 0.789807 | 0.774677 | 0.778651 |
| 0.782695 | 0.783714 | 0.772187 | 0.780206 | 0.783847 | 0.786626 | 0.775595 | 0.782926 | 0.783239 |
| 0.779429 | 0.784397 | 0.778454 | 0.777433 | 0.779674 | 0.783856 | 0.777806 | 0.793479 | 0.778981 |
| 0.781991 | 0.789491 | 0.774745 | 0.779923 | 0.783005 | 0.791217 | 0.775008 | 0.782569 | 0.775974 |
| 0.784296 | 0.785494 | 0.787771 | 0.78032 | 0.789085 | 0.788474 | 0.788521 | 0.78454 | 0.783369 |
| 0.782569 | 0.784284 | 0.788535 | 0.778441 | 0.787573 | 0.794314 | 0.79001 | 0.783383 | 0.782773 |
| 0.780773 | 0.787476 | 0.783987 | 0.778781 | 0.781737 | 0.792138 | 0.796045 | 0.782583 | 0.782035 |
| 0.788065 | 0.776771 | 0.779258 | 0.776642 | 0.773646 | 0.786187 | 0.778249 | 0.781222 | 0.78146 |
| 0.769447 | 0.771296 | 0.779647 | 0.773627 | 0.775923 | 0.779275 | 0.778876 | 0.775711 | 0.769369 |
| 0.77596 | 0.782935 | 0.774312 | 0.780347 | 0.779153 | 0.783161 | 0.772282 | 0.779872 | 0.78103 |
| 0.78034 | 0.778302 | 0.782397 | 0.779449 | 0.774505 | 0.778394 | 0.778911 | 0.779856 | 0.78023 |
| 0.771557 | 0.772504 | 0.774053 | 0.771681 | 0.777666 | 0.778696 | 0.782462 | 0.769428 | 0.770292 |
| 0.778731 | 0.780035 | 0.773644 | 0.778836 | 0.770795 | 0.781073 | 0.777516 | 0.785126 | 0.779426 |
| 0.785584 | 0.778264 | 0.774961 | 0.775465 | 0.77467 | 0.779126 | 0.765612 | 0.776034 | 0.775924 |
| 0.781821 | 0.784501 | 0.774041 | 0.777298 | 0.78345 | 0.778365 | 0.782362 | 0.77775 | 0.782512 |
| 0.780403 | 0.776629 | 0.775803 | 0.780931 | 0.774946 | 0.779982 | 0.76902 | 0.777147 | 0.773272 |
| 0.774747 | 0.771178 | 0.771588 | 0.768968 | 0.775165 | 0.771263 | 0.765604 | 0.768048 | 0.769388 |
| 0.783336 | 0.781216 | 0.778779 | 0.774635 | 0.782346 | 0.789752 | 0.778095 | 0.776663 | 0.77918 |
| 0.782347 | 0.776919 | 0.76606 | 0.782584 | 0.770444 | 0.77818 | 0.772584 | 0.780929 | 0.7801 |
| 0.772382 | 0.771715 | 0.771061 | 0.769875 | 0.773079 | 0.770795 | 0.771174 | 0.774975 | 0.771561 |
| 0.775852 | 0.781343 | 0.776528 | 0.777732 | 0.775964 | 0.779497 | 0.772137 | 0.776762 | 0.780074 |

| | | | | | | | | |
|---|---|---|---|---|---|---|---|---|
| 0.754108 | 0.756625 | 0.751524 | 0.753191 | 0.751872 | 0.755426 | 0.751796 | 0.757351 | 0.757753 |
| 0.77418 | 0.775426 | 0.767885 | 0.76965 | 0.771794 | 0.774791 | 0.766326 | 0.777448 | 0.78041 |
| 0.780711 | 0.778425 | 0.78388 | 0.775854 | 0.788601 | 0.787223 | 0.780676 | 0.786202 | 0.78552 |
| 0.778061 | 0.782691 | 0.7837 | 0.783656 | 0.785756 | 0.794929 | 0.781696 | 0.784393 | 0.776169 |
| 0.78397 | 0.783932 | 0.781446 | 0.774417 | 0.784849 | 0.784706 | 0.780105 | 0.783602 | 0.784067 |
| 0.781081 | 0.781591 | 0.777741 | 0.778471 | 0.78757 | 0.784961 | 0.775174 | 0.780421 | 0.780637 |
| 0.779396 | 0.783699 | 0.778736 | 0.773949 | 0.771483 | 0.777079 | 0.771336 | 0.775078 | 0.777896 |
| 0.785577 | 0.787996 | 0.77764 | 0.779909 | 0.77747 | 0.788492 | 0.776862 | 0.780699 | 0.783683 |
| 0.781519 | 0.777565 | 0.775159 | 0.782233 | 0.777135 | 0.773306 | 0.769102 | 0.779448 | 0.774582 |
| 0.77775 | 0.775342 | 0.768095 | 0.768626 | 0.771872 | 0.769686 | 0.772176 | 0.777329 | 0.775868 |
| 0.78204 | 0.781626 | 0.77537 | 0.779073 | 0.770329 | 0.776026 | 0.784178 | 0.776672 | 0.779576 |
| 0.78828 | 0.777501 | 0.778752 | 0.780297 | 0.779137 | 0.784597 | 0.779729 | 0.785595 | 0.781227 |
| 0.780482 | 0.78578 | 0.780933 | 0.779032 | 0.782085 | 0.78515 | 0.77546 | 0.781503 | 0.784577 |
| 0.792884 | 0.786522 | 0.78201 | 0.78305 | 0.776398 | 0.789514 | 0.783645 | 0.78417 | 0.78594 |
| 0.777973 | 0.778692 | 0.776162 | 0.776245 | 0.771112 | 0.784483 | 0.772559 | 0.78141 | 0.779789 |
| 0.777059 | 0.775594 | 0.77462 | 0.781732 | 0.770941 | 0.782113 | 0.775263 | 0.777283 | 0.773239 |
| 0.774828 | 0.773172 | 0.773813 | 0.767945 | 0.769295 | 0.772476 | 0.769905 | 0.772367 | 0.774768 |
| 0.789263 | 0.791632 | 0.783623 | 0.784772 | 0.776424 | 0.78752 | 0.773557 | 0.780526 | 0.787023 |
| 0.783152 | 0.784402 | 0.777668 | 0.775893 | 0.776555 | 0.784273 | 0.777555 | 0.77512 | 0.782791 |
| 0.779885 | 0.781033 | 0.779962 | 0.778117 | 0.780389 | 0.784762 | 0.783141 | 0.779491 | 0.779644 |
| 0.784005 | 0.776108 | 0.77809 | 0.779402 | 0.774838 | 0.779579 | 0.768071 | 0.783565 | 0.774734 |
| 0.775899 | 0.778295 | 0.769111 | 0.771396 | 0.771038 | 0.778886 | 0.7671 | 0.782154 | 0.774517 |
| 0.789234 | 0.78332 | 0.772312 | 0.777669 | 0.775762 | 0.77896 | 0.772557 | 0.780741 | 0.776825 |
| 0.777408 | 0.777178 | 0.773866 | 0.769972 | 0.775827 | 0.767413 | 0.769411 | 0.779877 | 0.776518 |
| 0.76377 | 0.769231 | 0.764787 | 0.767204 | 0.765177 | 0.763826 | 0.759327 | 0.762239 | 0.763162 |
| 0.710458 | 0.711604 | 0.707146 | 0.707656 | 0.705382 | 0.71123 | 0.702066 | 0.708916 | 0.707745 |
| 0.717893 | 0.714424 | 0.70862 | 0.709708 | 0.70923 | 0.714239 | 0.706898 | 0.714818 | 0.711976 |
| 0.719997 | 0.713674 | 0.709985 | 0.712377 | 0.709086 | 0.714517 | 0.710126 | 0.714504 | 0.711718 |
| 0.774581 | 0.774822 | 0.775221 | 0.766179 | 0.769577 | 0.769371 | 0.763337 | 0.772161 | 0.768168 |
| 0.772572 | 0.776736 | 0.766726 | 0.766974 | 0.768521 | 0.774522 | 0.764776 | 0.775552 | 0.77585 |
| 0.773749 | 0.774979 | 0.768118 | 0.76802 | 0.764472 | 0.773145 | 0.764387 | 0.772454 | 0.768551 |
| 0.770105 | 0.77482 | 0.768618 | 0.771332 | 0.771909 | 0.775199 | 0.767543 | 0.778719 | 0.772358 |
| 0.780812 | 0.788567 | 0.778093 | 0.783286 | 0.783213 | 0.787685 | 0.777515 | 0.781934 | 0.787856 |
| 0.781884 | 0.787709 | 0.791774 | 0.786662 | 0.797997 | 0.803502 | 0.811028 | 0.785434 | 0.789667 |
| 0.77615 | 0.778395 | 0.774948 | 0.778248 | 0.782852 | 0.782098 | 0.775611 | 0.782444 | 0.780015 |
| 0.781279 | 0.787359 | 0.777603 | 0.789022 | 0.779275 | 0.785631 | 0.780613 | 0.787316 | 0.78887 |
| 0.81262 | 0.787329 | 0.770905 | 0.78326 | 0.778432 | 0.783033 | 0.775507 | 0.775595 | 0.782687 |
| 0.800924 | 0.802519 | 0.79049 | 0.805277 | 0.798871 | 0.811234 | 0.793197 | 0.796868 | 0.797692 |
| 0.772265 | 0.780859 | 0.771204 | 0.779274 | 0.774202 | 0.770656 | 0.766557 | 0.767039 | 0.772591 |
| 0.77938 | 0.783042 | 0.784725 | 0.784333 | 0.786877 | 0.793844 | 0.792839 | 0.787427 | 0.780581 |
| 0.777265 | 0.779089 | 0.77213 | 0.783218 | 0.773774 | 0.783058 | 0.778943 | 0.780921 | 0.780342 |
| 0.80406 | 0.807339 | 0.795986 | 0.80219 | 0.795149 | 0.81138 | 0.791455 | 0.80084 | 0.803801 |
| 0.783599 | 0.778333 | 0.779723 | 0.780034 | 0.783111 | 0.781632 | 0.774715 | 0.789478 | 0.783218 |
| 0.775122 | 0.782939 | 0.786418 | 0.781889 | 0.784995 | 0.789775 | 0.801029 | 0.789056 | 0.779575 |
| 0.782765 | 0.788887 | 0.778403 | 0.783357 | 0.782207 | 0.793552 | 0.78403 | 0.784524 | 0.780205 |
| 0.778178 | 0.781413 | 0.776394 | 0.787569 | 0.777068 | 0.781043 | 0.772436 | 0.781855 | 0.783804 |

| | | | | | | | | |
|---|---|---|---|---|---|---|---|---|
| 0.781031 | 0.783102 | 0.77809 | 0.783864 | 0.780451 | 0.790405 | 0.782503 | 0.789619 | 0.786004 |
| 0.808571 | 0.787524 | 0.776838 | 0.773392 | 0.771976 | 0.790069 | 0.779412 | 0.777769 | 0.785057 |
| 0.804619 | 0.784904 | 0.777861 | 0.790974 | 0.778217 | 0.782182 | 0.782906 | 0.786247 | 0.791966 |
| 0.798912 | 0.797442 | 0.783904 | 0.801106 | 0.793703 | 0.797335 | 0.784307 | 0.793506 | 0.796964 |
| 0.808745 | 0.78371 | 0.781635 | 0.789 | 0.784519 | 0.788623 | 0.795204 | 0.787851 | 0.788894 |
| 0.775377 | 0.779841 | 0.786291 | 0.781644 | 0.777311 | 0.787393 | 0.78514 | 0.78615 | 0.78098 |
| 0.799168 | 0.786039 | 0.771691 | 0.782934 | 0.783353 | 0.775659 | 0.775761 | 0.778581 | 0.782924 |
| 1 | 0.784935 | 0.77649 | 0.783594 | 0.77948 | 0.778546 | 0.775926 | 0.779705 | 0.783304 |
| 0.784935 | 1 | 0.781406 | 0.788064 | 0.774881 | 0.787347 | 0.77555 | 0.781102 | 0.782313 |
| 0.77649 | 0.781406 | 1 | 0.778713 | 0.782478 | 0.784182 | 0.789199 | 0.77923 | 0.775519 |
| 0.783594 | 0.788064 | 0.778713 | 1 | 0.78026 | 0.780381 | 0.77763 | 0.783005 | 0.785205 |
| 0.77948 | 0.774881 | 0.782478 | 0.78026 | 1 | 0.785998 | 0.790356 | 0.779212 | 0.780616 |
| 0.778546 | 0.787347 | 0.784182 | 0.780381 | 0.785998 | 1 | 0.783075 | 0.782884 | 0.780508 |
| 0.775926 | 0.77555 | 0.789199 | 0.77763 | 0.790356 | 0.783075 | 1 | 0.774334 | 0.772792 |
| 0.779705 | 0.781102 | 0.77923 | 0.783005 | 0.779212 | 0.782884 | 0.774334 | 1 | 0.789653 |
| 0.783304 | 0.782313 | 0.775519 | 0.785205 | 0.780616 | 0.780508 | 0.772792 | 0.789653 | 1 |
| 0.781718 | 0.776886 | 0.773535 | 0.77551 | 0.780742 | 0.782001 | 0.781271 | 0.779129 | 0.779904 |
| 0.77131 | 0.785613 | 0.774493 | 0.781539 | 0.776892 | 0.786441 | 0.770037 | 0.782346 | 0.780189 |
| 0.784391 | 0.783204 | 0.799967 | 0.783155 | 0.790999 | 0.781553 | 0.80289 | 0.785524 | 0.786843 |
| 0.78359 | 0.785738 | 0.78022 | 0.790596 | 0.783276 | 0.776846 | 0.771588 | 0.779059 | 0.778819 |
| 0.785237 | 0.782547 | 0.768126 | 0.781966 | 0.779826 | 0.778027 | 0.781829 | 0.779165 | 0.786941 |
| 0.780591 | 0.781107 | 0.769558 | 0.79483 | 0.779177 | 0.774506 | 0.770401 | 0.780098 | 0.779406 |
| 0.792259 | 0.778856 | 0.775798 | 0.781667 | 0.781331 | 0.781148 | 0.78111 | 0.778657 | 0.787151 |
| 0.774236 | 0.776876 | 0.786393 | 0.78555 | 0.783171 | 0.783179 | 0.807608 | 0.778605 | 0.777022 |
| 0.786689 | 0.780822 | 0.771471 | 0.780647 | 0.777225 | 0.770156 | 0.767487 | 0.777388 | 0.776587 |
| 0.798353 | 0.786541 | 0.777042 | 0.778066 | 0.783568 | 0.779686 | 0.773669 | 0.784961 | 0.783181 |
| 0.790282 | 0.783717 | 0.771538 | 0.780458 | 0.782128 | 0.77567 | 0.768683 | 0.777248 | 0.780654 |
| 0.779147 | 0.787464 | 0.778408 | 0.779028 | 0.78005 | 0.784807 | 0.771833 | 0.777759 | 0.775615 |
| 0.774718 | 0.779738 | 0.784194 | 0.780842 | 0.781563 | 0.790092 | 0.777008 | 0.788095 | 0.781294 |
| 0.799386 | 0.783339 | 0.773582 | 0.777317 | 0.781057 | 0.778538 | 0.7751 | 0.778533 | 0.777833 |
| 0.784432 | 0.782085 | 0.773988 | 0.780369 | 0.789123 | 0.781592 | 0.777843 | 0.78087 | 0.781847 |
| 0.789957 | 0.776737 | 0.774701 | 0.77919 | 0.782216 | 0.777038 | 0.775923 | 0.791845 | 0.775897 |
| 0.780478 | 0.808394 | 0.77686 | 0.786261 | 0.77927 | 0.777199 | 0.778455 | 0.780062 | 0.775089 |
| 0.774561 | 0.779944 | 0.779674 | 0.782762 | 0.78632 | 0.791049 | 0.783335 | 0.779419 | 0.772613 |
| 0.77515 | 0.788978 | 0.780106 | 0.78313 | 0.778646 | 0.832146 | 0.784912 | 0.782101 | 0.782471 |
| 0.776739 | 0.779204 | 0.769872 | 0.778266 | 0.774466 | 0.788582 | 0.777976 | 0.776295 | 0.775766 |
| 0.775393 | 0.78303 | 0.77831 | 0.778875 | 0.781259 | 0.786399 | 0.789681 | 0.777339 | 0.781847 |
| 0.784251 | 0.757451 | 0.753039 | 0.756734 | 0.754303 | 0.755798 | 0.766663 | 0.750762 | 0.754057 |
| 0.757858 | 0.761444 | 0.759242 | 0.764614 | 0.755639 | 0.775175 | 0.773065 | 0.76218 | 0.7602 |
| 0.754049 | 0.748609 | 0.747685 | 0.754317 | 0.748552 | 0.749866 | 0.745256 | 0.751976 | 0.753327 |
| 0.760009 | 0.756427 | 0.750029 | 0.758005 | 0.757478 | 0.759236 | 0.749019 | 0.767399 | 0.764339 |
| 0.742511 | 0.740085 | 0.771506 | 0.743269 | 0.777308 | 0.764117 | 0.797966 | 0.733261 | 0.741467 |
| 0.756197 | 0.75038 | 0.758083 | 0.762969 | 0.768661 | 0.760459 | 0.773817 | 0.74258 | 0.747738 |
| 0.758844 | 0.754602 | 0.739185 | 0.755089 | 0.744254 | 0.748415 | 0.747716 | 0.75098 | 0.749275 |
| 0.753432 | 0.75204 | 0.741765 | 0.752136 | 0.750366 | 0.747463 | 0.750565 | 0.746621 | 0.745062 |
| 0.779507 | 0.754333 | 0.740917 | 0.748099 | 0.750389 | 0.752094 | 0.747209 | 0.746371 | 0.757119 |
| 0.752062 | 0.756079 | 0.755833 | 0.759985 | 0.755134 | 0.756669 | 0.750901 | 0.753102 | 0.758674 |

| | | | | | | | | |
|---|---|---|---|---|---|---|---|---|
| 0.765392 | 0.75029 | 0.742453 | 0.747679 | 0.739242 | 0.749064 | 0.746912 | 0.750258 | 0.736929 |
| 0.74755 | 0.754365 | 0.743065 | 0.749197 | 0.754874 | 0.746372 | 0.746006 | 0.755898 | 0.756296 |
| 0.744793 | 0.74005 | 0.74028 | 0.743867 | 0.742702 | 0.743751 | 0.73558 | 0.752056 | 0.74141 |
| 0.745538 | 0.744938 | 0.74638 | 0.749385 | 0.74425 | 0.748852 | 0.743034 | 0.747447 | 0.751521 |
| 0.745961 | 0.755607 | 0.772505 | 0.748874 | 0.767925 | 0.768613 | 0.780215 | 0.744638 | 0.744276 |
| 0.841282 | 0.750827 | 0.732312 | 0.747944 | 0.741133 | 0.748847 | 0.748275 | 0.744166 | 0.7612 |
| 0.756439 | 0.747428 | 0.732096 | 0.744566 | 0.753015 | 0.756313 | 0.749741 | 0.741826 | 0.750663 |
| 0.744466 | 0.752019 | 0.751981 | 0.755717 | 0.765496 | 0.753322 | 0.767235 | 0.743091 | 0.750876 |
| 0.727314 | 0.729833 | 0.757207 | 0.741677 | 0.747769 | 0.731984 | 0.766972 | 0.732137 | 0.729976 |
| 0.75269 | 0.757464 | 0.738121 | 0.749585 | 0.750496 | 0.749385 | 0.741553 | 0.749542 | 0.742023 |
| 0.803765 | 0.748372 | 0.727115 | 0.74541 | 0.746177 | 0.753047 | 0.745873 | 0.739138 | 0.755245 |
| 0.769904 | 0.765539 | 0.762962 | 0.773643 | 0.766393 | 0.768215 | 0.769282 | 0.767793 | 0.764253 |
| 0.775729 | 0.775107 | 0.771149 | 0.779094 | 0.773509 | 0.774585 | 0.772821 | 0.779075 | 0.770344 |
| 0.765154 | 0.746313 | 0.741761 | 0.747351 | 0.757619 | 0.753177 | 0.749144 | 0.74331 | 0.750706 |
| 0.776097 | 0.779862 | 0.772481 | 0.778155 | 0.777104 | 0.774191 | 0.777797 | 0.767116 | 0.777667 |
| 0.76848 | 0.770492 | 0.779695 | 0.775188 | 0.789615 | 0.778765 | 0.794764 | 0.768648 | 0.762517 |
| 0.774568 | 0.777906 | 0.774739 | 0.775621 | 0.771183 | 0.781433 | 0.777473 | 0.780472 | 0.7821 |
| 0.779236 | 0.784259 | 0.777354 | 0.776635 | 0.772541 | 0.772742 | 0.770013 | 0.778314 | 0.776666 |
| 0.839102 | 0.76382 | 0.750262 | 0.76359 | 0.756564 | 0.761616 | 0.75377 | 0.755273 | 0.755525 |
| 0.774135 | 0.770644 | 0.779032 | 0.769753 | 0.786047 | 0.782336 | 0.795643 | 0.763325 | 0.778086 |
| 0.802738 | 0.744818 | 0.728919 | 0.747881 | 0.745502 | 0.74344 | 0.738543 | 0.746063 | 0.746515 |
| 0.776283 | 0.78375 | 0.773277 | 0.776676 | 0.774062 | 0.778092 | 0.765534 | 0.772535 | 0.771989 |
| 0.775909 | 0.784415 | 0.774574 | 0.772701 | 0.769519 | 0.767559 | 0.769901 | 0.771728 | 0.775896 |
| 0.743633 | 0.748735 | 0.731556 | 0.742134 | 0.735554 | 0.744578 | 0.74199 | 0.737942 | 0.73767 |
| 0.744645 | 0.748273 | 0.745686 | 0.748487 | 0.746462 | 0.750342 | 0.752093 | 0.750299 | 0.744625 |
| 0.761917 | 0.760881 | 0.752249 | 0.763777 | 0.756905 | 0.752455 | 0.758834 | 0.763224 | 0.758167 |
| 0.783604 | 0.765977 | 0.751289 | 0.760561 | 0.755728 | 0.759161 | 0.755349 | 0.757662 | 0.767007 |
| 0.779964 | 0.77426 | 0.768854 | 0.771033 | 0.767862 | 0.77662 | 0.767004 | 0.778199 | 0.77199 |
| 0.751151 | 0.755186 | 0.74408 | 0.758703 | 0.74755 | 0.752205 | 0.743369 | 0.752823 | 0.747368 |
| 0.769212 | 0.773521 | 0.76383 | 0.772052 | 0.768023 | 0.762763 | 0.765035 | 0.764687 | 0.765281 |
| 0.7697 | 0.767081 | 0.770318 | 0.769052 | 0.765436 | 0.773643 | 0.759419 | 0.772403 | 0.770513 |
| 0.759795 | 0.759726 | 0.74949 | 0.767923 | 0.754739 | 0.763173 | 0.750145 | 0.754616 | 0.761012 |
| 0.757675 | 0.758226 | 0.755409 | 0.763226 | 0.754635 | 0.766922 | 0.757806 | 0.755802 | 0.756113 |
| 0.767371 | 0.766417 | 0.760365 | 0.766602 | 0.768759 | 0.765466 | 0.755216 | 0.761951 | 0.76831 |
| 0.778782 | 0.765417 | 0.76437 | 0.771193 | 0.76598 | 0.763804 | 0.763861 | 0.763411 | 0.761676 |
| 0.765359 | 0.767791 | 0.762957 | 0.768051 | 0.769968 | 0.762491 | 0.760615 | 0.764997 | 0.764282 |
| 0.740498 | 0.727675 | 0.717507 | 0.734463 | 0.725604 | 0.723793 | 0.727457 | 0.730464 | 0.729283 |
| 0.760825 | 0.763229 | 0.751049 | 0.768348 | 0.758975 | 0.761737 | 0.753015 | 0.761055 | 0.758079 |
| 0.738004 | 0.728044 | 0.725145 | 0.732524 | 0.729004 | 0.731764 | 0.723921 | 0.727129 | 0.72848 |
| 0.745877 | 0.717945 | 0.717809 | 0.728894 | 0.720699 | 0.722057 | 0.720218 | 0.718428 | 0.727656 |
| 0.77514 | 0.764289 | 0.768437 | 0.771659 | 0.764315 | 0.761307 | 0.767842 | 0.76755 | 0.763345 |
| 0.729493 | 0.715582 | 0.713066 | 0.723983 | 0.716999 | 0.717123 | 0.718202 | 0.721497 | 0.720411 |
| 0.725203 | 0.716474 | 0.714286 | 0.720884 | 0.715817 | 0.715395 | 0.71108 | 0.716011 | 0.71478 |
| 0.707369 | 0.704021 | 0.707413 | 0.709881 | 0.70977 | 0.710874 | 0.706635 | 0.708573 | 0.707906 |
| 0.766847 | 0.768878 | 0.761341 | 0.769823 | 0.763831 | 0.763115 | 0.758961 | 0.766454 | 0.759237 |
| 0.710209 | 0.706655 | 0.709777 | 0.71282 | 0.707992 | 0.717119 | 0.697877 | 0.713512 | 0.707857 |
| 0.772517 | 0.764537 | 0.771342 | 0.768223 | 0.76357 | 0.762588 | 0.760641 | 0.762288 | 0.763792 |

| | | | | | | | | |
|---|---|---|---|---|---|---|---|---|
| 0.756956 | 0.760714 | 0.754344 | 0.759494 | 0.756285 | 0.759632 | 0.755796 | 0.762679 | 0.755895 |
| 0.770027 | 0.76732 | 0.762995 | 0.768007 | 0.762128 | 0.760838 | 0.761336 | 0.768374 | 0.760353 |
| 0.751355 | 0.748687 | 0.748407 | 0.753584 | 0.761655 | 0.750555 | 0.746591 | 0.74957 | 0.752938 |
| 0.705166 | 0.703505 | 0.702588 | 0.702958 | 0.697726 | 0.702994 | 0.69044 | 0.701204 | 0.701429 |
| 0.710017 | 0.700272 | 0.697242 | 0.704838 | 0.701091 | 0.701882 | 0.697401 | 0.700098 | 0.699584 |
| 0.769985 | 0.768782 | 0.764165 | 0.767017 | 0.763542 | 0.762283 | 0.763482 | 0.764423 | 0.760078 |
| 0.695233 | 0.68749 | 0.693476 | 0.69972 | 0.694182 | 0.694258 | 0.691555 | 0.69116 | 0.693828 |
| 0.733524 | 0.73642 | 0.733138 | 0.730618 | 0.734201 | 0.73607 | 0.729122 | 0.730488 | 0.732561 |
| 0.764526 | 0.763286 | 0.759302 | 0.766483 | 0.760803 | 0.760921 | 0.758469 | 0.759511 | 0.760896 |
| 0.729026 | 0.724382 | 0.718798 | 0.724471 | 0.721994 | 0.720881 | 0.715691 | 0.723409 | 0.722919 |
| 0.747084 | 0.751027 | 0.745199 | 0.752937 | 0.745736 | 0.747823 | 0.743372 | 0.751391 | 0.745173 |
| 0.62963 | 0.596026 | 0.636986 | 0.666667 | 0.664516 | 0.605096 | 0.62 | 0.623377 | 0.655629 |
| 0.662388 | 0.6555 | 0.650693 | 0.65433 | 0.657436 | 0.655977 | 0.651486 | 0.655485 | 0.652266 |
| 0.658481 | 0.6586 | 0.648571 | 0.656236 | 0.659545 | 0.654353 | 0.654895 | 0.656651 | 0.649236 |
| 0.67383 | 0.660621 | 0.658878 | 0.661182 | 0.669886 | 0.665152 | 0.66735 | 0.664535 | 0.662279 |
| 0.666884 | 0.660519 | 0.657513 | 0.663076 | 0.66671 | 0.664621 | 0.665103 | 0.667622 | 0.655062 |
| 0.62021 | 0.61541 | 0.622658 | 0.623212 | 0.625733 | 0.621666 | 0.61743 | 0.620238 | 0.619292 |
| 0.673038 | 0.670759 | 0.65999 | 0.661757 | 0.667592 | 0.6684 | 0.671877 | 0.666625 | 0.658291 |
| 0.739565 | 0.741199 | 0.737261 | 0.741007 | 0.74476 | 0.740401 | 0.745426 | 0.740904 | 0.734956 |
| 0.741369 | 0.741446 | 0.734234 | 0.74049 | 0.738468 | 0.739152 | 0.732158 | 0.740401 | 0.736652 |
| 0.745618 | 0.747365 | 0.738423 | 0.74591 | 0.744704 | 0.74613 | 0.740885 | 0.745265 | 0.740614 |
| 0.632996 | 0.617043 | 0.619446 | 0.623477 | 0.620576 | 0.622659 | 0.61937 | 0.622935 | 0.620588 |
| 0.60478 | 0.600288 | 0.603035 | 0.605929 | 0.608225 | 0.60573 | 0.596361 | 0.603322 | 0.602428 |
| 0.706354 | 0.706838 | 0.704926 | 0.705911 | 0.706651 | 0.708408 | 0.704968 | 0.704987 | 0.705268 |
| 0.710537 | 0.706695 | 0.703378 | 0.708409 | 0.709493 | 0.708253 | 0.705109 | 0.70846 | 0.703204 |
| 0.703345 | 0.697038 | 0.694535 | 0.700681 | 0.701402 | 0.701039 | 0.700716 | 0.6998 | 0.697956 |
| 0.705506 | 0.701841 | 0.697748 | 0.705508 | 0.704001 | 0.704607 | 0.704367 | 0.703857 | 0.701255 |
| 0.708566 | 0.709543 | 0.703071 | 0.709573 | 0.708002 | 0.713552 | 0.70619 | 0.708698 | 0.706718 |
| 0.7068 | 0.706924 | 0.704622 | 0.706754 | 0.708045 | 0.706665 | 0.706286 | 0.706029 | 0.699545 |
| 0.705934 | 0.705035 | 0.702172 | 0.704658 | 0.709644 | 0.705953 | 0.703148 | 0.705991 | 0.702429 |
| 0.708916 | 0.71074 | 0.702864 | 0.711122 | 0.705833 | 0.70958 | 0.707766 | 0.708821 | 0.705063 |
| 0.722658 | 0.712856 | 0.712186 | 0.714834 | 0.713071 | 0.718313 | 0.711307 | 0.716487 | 0.712892 |
| 0.714748 | 0.714714 | 0.705717 | 0.711439 | 0.711772 | 0.713013 | 0.708153 | 0.71055 | 0.709264 |
| 0.715778 | 0.711557 | 0.705961 | 0.70835 | 0.71036 | 0.713728 | 0.705554 | 0.710447 | 0.710547 |
| 0.720406 | 0.716693 | 0.711069 | 0.71221 | 0.714586 | 0.716834 | 0.711845 | 0.715842 | 0.71287 |
| 0.707192 | 0.710088 | 0.706591 | 0.709064 | 0.707491 | 0.707974 | 0.703212 | 0.710137 | 0.706135 |
| 0.711263 | 0.710695 | 0.707716 | 0.711792 | 0.70668 | 0.713203 | 0.707234 | 0.710438 | 0.705943 |
| 0.711348 | 0.713908 | 0.707947 | 0.707482 | 0.709085 | 0.714939 | 0.709698 | 0.714673 | 0.710241 |
| 0.708721 | 0.705989 | 0.70333 | 0.702353 | 0.706297 | 0.704704 | 0.703292 | 0.706591 | 0.703122 |
| 0.711625 | 0.711122 | 0.708762 | 0.706515 | 0.711297 | 0.712745 | 0.708005 | 0.711044 | 0.712498 |
| 0.663277 | 0.655574 | 0.652824 | 0.653122 | 0.661695 | 0.658028 | 0.654724 | 0.653342 | 0.652422 |
| 0.668748 | 0.659536 | 0.659625 | 0.655364 | 0.666605 | 0.662011 | 0.663183 | 0.661936 | 0.658311 |
| 0.716769 | 0.710719 | 0.713504 | 0.714128 | 0.712964 | 0.714446 | 0.707915 | 0.71299 | 0.712161 |
| 0.713382 | 0.713806 | 0.707702 | 0.713136 | 0.713778 | 0.715169 | 0.711185 | 0.711993 | 0.712722 |
| 0.642278 | 0.643612 | 0.639396 | 0.649936 | 0.649394 | 0.651623 | 0.644486 | 0.648035 | 0.642018 |
| 0.712163 | 0.712805 | 0.708798 | 0.709588 | 0.710922 | 0.713008 | 0.709586 | 0.713586 | 0.710908 |
| 0.705028 | 0.703766 | 0.697656 | 0.702637 | 0.697962 | 0.703716 | 0.698451 | 0.70222 | 0.701267 |

| | | | | | | | | |
|---|---|---|---|---|---|---|---|---|
| 0.70999 | 0.706988 | 0.703519 | 0.704948 | 0.702993 | 0.710683 | 0.705977 | 0.707356 | 0.704298 |
| 0.715232 | 0.710088 | 0.708438 | 0.710376 | 0.709564 | 0.713271 | 0.708631 | 0.711867 | 0.707691 |
| 0.71206 | 0.710716 | 0.706208 | 0.710955 | 0.710236 | 0.712914 | 0.707791 | 0.710619 | 0.709704 |
| 0.712646 | 0.710067 | 0.707207 | 0.708287 | 0.708246 | 0.714389 | 0.70804 | 0.710354 | 0.708446 |
| 0.718552 | 0.716338 | 0.71179 | 0.714139 | 0.715715 | 0.721131 | 0.717758 | 0.71601 | 0.710655 |
| 0.710136 | 0.706516 | 0.702679 | 0.707462 | 0.705606 | 0.709586 | 0.706713 | 0.707155 | 0.701284 |
| 0.711163 | 0.708287 | 0.703739 | 0.705482 | 0.70665 | 0.712745 | 0.707474 | 0.706689 | 0.705966 |
| 0.704213 | 0.702585 | 0.702837 | 0.701925 | 0.705954 | 0.705696 | 0.700245 | 0.705873 | 0.703558 |
| 0.710081 | 0.706833 | 0.704536 | 0.704141 | 0.702746 | 0.71074 | 0.703417 | 0.705253 | 0.706554 |
| 0.71288 | 0.707372 | 0.701174 | 0.709187 | 0.704667 | 0.708478 | 0.705283 | 0.709419 | 0.708914 |
| 0.715029 | 0.709531 | 0.709124 | 0.709677 | 0.710936 | 0.712995 | 0.707241 | 0.712124 | 0.709853 |
| 0.707793 | 0.705572 | 0.701215 | 0.704125 | 0.702841 | 0.70884 | 0.704949 | 0.705641 | 0.704885 |
| 0.705262 | 0.700388 | 0.698821 | 0.703916 | 0.702575 | 0.704223 | 0.696716 | 0.702539 | 0.701981 |
| 0.701601 | 0.699451 | 0.695986 | 0.701547 | 0.698191 | 0.701227 | 0.697799 | 0.699344 | 0.699184 |
| 0.718297 | 0.71257 | 0.710139 | 0.712269 | 0.715031 | 0.717811 | 0.714022 | 0.713197 | 0.709528 |
| 0.710969 | 0.711157 | 0.706846 | 0.711919 | 0.713042 | 0.711514 | 0.706592 | 0.710996 | 0.70922 |
| 0.705464 | 0.704321 | 0.703147 | 0.705008 | 0.703306 | 0.706375 | 0.700108 | 0.706871 | 0.703634 |
| 0.709054 | 0.707652 | 0.702888 | 0.705585 | 0.70728 | 0.71007 | 0.699282 | 0.708974 | 0.710021 |
| 0.709309 | 0.707457 | 0.701907 | 0.706068 | 0.705556 | 0.708786 | 0.700232 | 0.707929 | 0.705394 |
| 0.707768 | 0.70913 | 0.703707 | 0.707836 | 0.70603 | 0.708782 | 0.701803 | 0.708153 | 0.704899 |
| 0.705065 | 0.70333 | 0.698934 | 0.70032 | 0.702685 | 0.707377 | 0.696791 | 0.706912 | 0.702781 |
| 0.707154 | 0.702947 | 0.700709 | 0.703603 | 0.703716 | 0.70831 | 0.700827 | 0.704588 | 0.701159 |
| 0.708641 | 0.7073 | 0.70437 | 0.709131 | 0.704088 | 0.709089 | 0.705001 | 0.708079 | 0.707331 |
| 0.710006 | 0.709468 | 0.704729 | 0.70862 | 0.707067 | 0.711941 | 0.706034 | 0.710031 | 0.709847 |
| 0.70906 | 0.706582 | 0.705764 | 0.707512 | 0.704033 | 0.711332 | 0.701166 | 0.708587 | 0.707266 |
| 0.704796 | 0.704375 | 0.697083 | 0.705191 | 0.702552 | 0.705617 | 0.697571 | 0.705664 | 0.700876 |
| 0.70833 | 0.705143 | 0.70503 | 0.70458 | 0.704188 | 0.706786 | 0.700065 | 0.708398 | 0.705641 |
| 0.703927 | 0.701764 | 0.702948 | 0.702818 | 0.70159 | 0.700204 | 0.702996 | 0.704605 | 0.700807 |
| 0.709111 | 0.706377 | 0.705733 | 0.707246 | 0.704032 | 0.707093 | 0.704832 | 0.705488 | 0.704751 |
| 0.711424 | 0.707354 | 0.70385 | 0.704924 | 0.70592 | 0.708377 | 0.70371 | 0.70958 | 0.706907 |
| 0.712096 | 0.709773 | 0.705649 | 0.710248 | 0.709094 | 0.709574 | 0.702402 | 0.710606 | 0.70835 |
| 0.713207 | 0.711023 | 0.70816 | 0.709556 | 0.710196 | 0.713532 | 0.706094 | 0.713731 | 0.709685 |
| 0.71506 | 0.713033 | 0.706914 | 0.71414 | 0.714664 | 0.714991 | 0.709623 | 0.715799 | 0.70966 |
| 0.713588 | 0.713622 | 0.70997 | 0.713756 | 0.712627 | 0.711961 | 0.711314 | 0.713645 | 0.711406 |
| 0.705655 | 0.703196 | 0.702675 | 0.703145 | 0.703542 | 0.705254 | 0.701434 | 0.706988 | 0.702874 |
| 0.71037 | 0.709211 | 0.703081 | 0.706043 | 0.706036 | 0.70979 | 0.702444 | 0.710018 | 0.706095 |
| 0.709318 | 0.707187 | 0.708567 | 0.708108 | 0.706126 | 0.708162 | 0.706328 | 0.71062 | 0.706201 |
| 0.70463 | 0.706091 | 0.700328 | 0.703159 | 0.699141 | 0.702508 | 0.699325 | 0.703316 | 0.701766 |
| 0.702306 | 0.70228 | 0.697581 | 0.701124 | 0.697823 | 0.707177 | 0.692113 | 0.703092 | 0.701718 |
| 0.70036 | 0.695244 | 0.697576 | 0.693112 | 0.696438 | 0.697097 | 0.699046 | 0.696593 | 0.695526 |
| 0.699678 | 0.698976 | 0.695684 | 0.695875 | 0.697531 | 0.701542 | 0.69596 | 0.701026 | 0.696198 |
| 0.698703 | 0.697526 | 0.694393 | 0.694096 | 0.694596 | 0.697821 | 0.693661 | 0.699873 | 0.697945 |
| 0.704232 | 0.701554 | 0.698631 | 0.701709 | 0.70118 | 0.707955 | 0.695245 | 0.703409 | 0.700969 |
| 0.694982 | 0.694427 | 0.690669 | 0.693012 | 0.692906 | 0.696267 | 0.689707 | 0.694335 | 0.689428 |
| 0.704965 | 0.70376 | 0.700725 | 0.703085 | 0.69989 | 0.700008 | 0.699568 | 0.704772 | 0.702113 |
| 0.699761 | 0.698148 | 0.694855 | 0.697725 | 0.695043 | 0.697463 | 0.692436 | 0.699034 | 0.698478 |

| | | | | | | | | |
|---|---|---|---|---|---|---|---|---|
| 0.712813 | 0.710979 | 0.707327 | 0.709826 | 0.708897 | 0.711892 | 0.704444 | 0.71094 | 0.705985 |
| 0.713821 | 0.710028 | 0.706762 | 0.70834 | 0.710004 | 0.713286 | 0.709247 | 0.712374 | 0.709457 |
| 0.709363 | 0.711493 | 0.704704 | 0.704785 | 0.705786 | 0.714274 | 0.70049 | 0.707528 | 0.704294 |
| 0.71926 | 0.709403 | 0.708515 | 0.710322 | 0.707259 | 0.712487 | 0.708637 | 0.708294 | 0.710821 |
| 0.716705 | 0.715593 | 0.713774 | 0.715258 | 0.716343 | 0.713748 | 0.713742 | 0.718206 | 0.715364 |
| 0.715212 | 0.712357 | 0.710174 | 0.71136 | 0.711057 | 0.713018 | 0.709007 | 0.714012 | 0.711718 |
| 0.711195 | 0.709677 | 0.704545 | 0.704287 | 0.704218 | 0.710038 | 0.706249 | 0.706709 | 0.705024 |
| 0.713007 | 0.709284 | 0.706325 | 0.706354 | 0.707486 | 0.712099 | 0.708024 | 0.71081 | 0.710606 |
| 0.709136 | 0.704018 | 0.701442 | 0.70326 | 0.705025 | 0.706043 | 0.704867 | 0.70082 | 0.700892 |
| 0.713113 | 0.712221 | 0.707459 | 0.706449 | 0.707872 | 0.712788 | 0.708769 | 0.713415 | 0.706965 |
| 0.711384 | 0.714229 | 0.70573 | 0.710288 | 0.713232 | 0.715366 | 0.708343 | 0.712213 | 0.710638 |
| 0.714824 | 0.710791 | 0.710062 | 0.710261 | 0.709192 | 0.714503 | 0.705631 | 0.71104 | 0.709864 |
| 0.713288 | 0.712203 | 0.705715 | 0.706882 | 0.709615 | 0.711984 | 0.704553 | 0.709462 | 0.704996 |
| 0.714148 | 0.714458 | 0.7066 | 0.708524 | 0.706491 | 0.712796 | 0.706628 | 0.709791 | 0.705264 |
| 0.71341 | 0.708687 | 0.711724 | 0.70749 | 0.711965 | 0.712544 | 0.708056 | 0.710357 | 0.710061 |
| 0.714137 | 0.714491 | 0.710861 | 0.709571 | 0.711903 | 0.714938 | 0.708051 | 0.710035 | 0.707517 |
| 0.715093 | 0.71407 | 0.708546 | 0.710724 | 0.7096 | 0.718743 | 0.70894 | 0.714625 | 0.70948 |
| 0.715192 | 0.711734 | 0.709134 | 0.707618 | 0.711828 | 0.713833 | 0.70494 | 0.713349 | 0.708738 |
| 0.714823 | 0.714583 | 0.712297 | 0.712125 | 0.713652 | 0.712309 | 0.710943 | 0.71367 | 0.716054 |
| 0.713921 | 0.710286 | 0.710229 | 0.711392 | 0.706923 | 0.710623 | 0.705326 | 0.711002 | 0.711629 |
| 0.713421 | 0.71289 | 0.709412 | 0.710105 | 0.709443 | 0.712317 | 0.710422 | 0.708206 | 0.707461 |
| 0.712475 | 0.711591 | 0.70848 | 0.706531 | 0.705711 | 0.709235 | 0.706494 | 0.707491 | 0.702709 |
| 0.720737 | 0.716089 | 0.713913 | 0.711148 | 0.71323 | 0.714309 | 0.705039 | 0.7144 | 0.711742 |
| 0.718041 | 0.714161 | 0.709408 | 0.712236 | 0.712606 | 0.714161 | 0.710214 | 0.713697 | 0.71151 |
| 0.713257 | 0.713857 | 0.708242 | 0.707289 | 0.708446 | 0.712555 | 0.705611 | 0.710308 | 0.711733 |
| 0.714962 | 0.710121 | 0.708657 | 0.706373 | 0.707129 | 0.708867 | 0.703531 | 0.711436 | 0.70806 |
| 0.709215 | 0.709496 | 0.703564 | 0.705689 | 0.706051 | 0.709523 | 0.707196 | 0.707377 | 0.707127 |
| 0.714343 | 0.708996 | 0.705368 | 0.706094 | 0.706861 | 0.711636 | 0.703677 | 0.710086 | 0.705859 |
| 0.718096 | 0.715344 | 0.708425 | 0.711233 | 0.710674 | 0.71563 | 0.708884 | 0.714229 | 0.713802 |
| 0.715948 | 0.71318 | 0.712471 | 0.708945 | 0.711443 | 0.716202 | 0.709648 | 0.715841 | 0.712398 |
| 0.711671 | 0.711167 | 0.702392 | 0.70807 | 0.710978 | 0.708896 | 0.706167 | 0.710052 | 0.71119 |
| 0.720156 | 0.714332 | 0.709634 | 0.716025 | 0.71112 | 0.715916 | 0.710939 | 0.71664 | 0.713486 |
| 0.715496 | 0.716554 | 0.708699 | 0.711046 | 0.713813 | 0.713817 | 0.711231 | 0.715635 | 0.712229 |
| 0.709118 | 0.706663 | 0.704737 | 0.702939 | 0.70541 | 0.708474 | 0.706459 | 0.703619 | 0.704455 |
| 0.713283 | 0.710615 | 0.710692 | 0.709154 | 0.708834 | 0.708911 | 0.706107 | 0.713333 | 0.711553 |
| 0.709141 | 0.709144 | 0.704381 | 0.706198 | 0.70562 | 0.708519 | 0.702047 | 0.706355 | 0.705487 |
| 0.72105 | 0.717238 | 0.711737 | 0.714714 | 0.717593 | 0.718326 | 0.715395 | 0.718867 | 0.713099 |
| 0.712907 | 0.708659 | 0.70556 | 0.70617 | 0.706926 | 0.711305 | 0.70327 | 0.711321 | 0.708122 |
| 0.712772 | 0.71258 | 0.706209 | 0.708952 | 0.710399 | 0.71087 | 0.706148 | 0.709376 | 0.70545 |
| 0.713923 | 0.709135 | 0.707773 | 0.70697 | 0.708217 | 0.709078 | 0.70716 | 0.711559 | 0.709625 |
| 0.718246 | 0.714627 | 0.71303 | 0.714589 | 0.715078 | 0.713329 | 0.712161 | 0.71382 | 0.714274 |
| 0.717299 | 0.715626 | 0.709693 | 0.711982 | 0.710698 | 0.719897 | 0.712059 | 0.712811 | 0.707879 |
| 0.717372 | 0.713196 | 0.712449 | 0.712785 | 0.712683 | 0.717367 | 0.714056 | 0.71582 | 0.711425 |
| 0.71221 | 0.708364 | 0.706505 | 0.706119 | 0.713144 | 0.712497 | 0.705689 | 0.71128 | 0.708195 |
| 0.716016 | 0.711545 | 0.709879 | 0.710412 | 0.710548 | 0.71189 | 0.708645 | 0.715115 | 0.710631 |
| 0.712583 | 0.705285 | 0.702997 | 0.704202 | 0.707777 | 0.710029 | 0.70554 | 0.70484 | 0.706579 |

| | | | | | | | | |
|---|---|---|---|---|---|---|---|---|
| 0.700508 | 0.694911 | 0.695465 | 0.695353 | 0.695957 | 0.697868 | 0.692308 | 0.701482 | 0.6938 |
| 0.747048 | 0.74767 | 0.742624 | 0.743861 | 0.745269 | 0.744928 | 0.741399 | 0.746992 | 0.750533 |
| 0.710736 | 0.710241 | 0.709306 | 0.708151 | 0.708457 | 0.709324 | 0.706155 | 0.706836 | 0.706771 |
| 0.718433 | 0.715355 | 0.713394 | 0.710217 | 0.711001 | 0.712807 | 0.711716 | 0.713447 | 0.711598 |
| 0.709747 | 0.710861 | 0.701744 | 0.703446 | 0.706241 | 0.70825 | 0.703601 | 0.706072 | 0.704246 |
| 0.714507 | 0.712711 | 0.708692 | 0.707412 | 0.708483 | 0.710177 | 0.70706 | 0.708691 | 0.707485 |
| 0.712832 | 0.704696 | 0.707679 | 0.702019 | 0.706046 | 0.709254 | 0.702205 | 0.708368 | 0.703777 |
| 0.706512 | 0.70301 | 0.701011 | 0.699788 | 0.702656 | 0.701974 | 0.699618 | 0.702954 | 0.701006 |
| 0.712245 | 0.708443 | 0.707984 | 0.70568 | 0.708995 | 0.711584 | 0.70755 | 0.70801 | 0.71067 |
| 0.718962 | 0.717237 | 0.710405 | 0.713302 | 0.712087 | 0.715991 | 0.711293 | 0.713357 | 0.714411 |
| 0.715244 | 0.713841 | 0.709257 | 0.709175 | 0.711097 | 0.711322 | 0.707618 | 0.711007 | 0.711535 |
| 0.720424 | 0.712765 | 0.712165 | 0.709315 | 0.711517 | 0.715357 | 0.712499 | 0.711579 | 0.714024 |
| 0.716671 | 0.713473 | 0.715117 | 0.71068 | 0.714736 | 0.717767 | 0.712495 | 0.714092 | 0.713571 |
| 0.717038 | 0.71432 | 0.710892 | 0.711106 | 0.713586 | 0.712919 | 0.709836 | 0.715448 | 0.715912 |
| 0.710552 | 0.706776 | 0.708869 | 0.705355 | 0.707423 | 0.709851 | 0.707858 | 0.711249 | 0.708014 |
| 0.71584 | 0.709181 | 0.707065 | 0.708608 | 0.711346 | 0.714331 | 0.708977 | 0.711867 | 0.710118 |
| 0.718614 | 0.712265 | 0.708752 | 0.71263 | 0.711063 | 0.714989 | 0.710084 | 0.715655 | 0.712078 |
| 0.71622 | 0.712386 | 0.707105 | 0.709677 | 0.711684 | 0.713374 | 0.709811 | 0.711236 | 0.707862 |
| 0.708065 | 0.701269 | 0.697564 | 0.702137 | 0.706519 | 0.702813 | 0.701984 | 0.703813 | 0.700705 |
| 0.707842 | 0.704703 | 0.700402 | 0.700416 | 0.703234 | 0.704718 | 0.702102 | 0.706528 | 0.703514 |
| 0.710064 | 0.710155 | 0.710342 | 0.707811 | 0.706793 | 0.713219 | 0.705183 | 0.71359 | 0.711333 |
| 0.703463 | 0.701949 | 0.698401 | 0.697266 | 0.697613 | 0.70136 | 0.699461 | 0.700565 | 0.701783 |
| 0.709845 | 0.709765 | 0.705761 | 0.703418 | 0.706242 | 0.710176 | 0.705127 | 0.7076 | 0.709134 |
| 0.713424 | 0.711894 | 0.706567 | 0.709016 | 0.70799 | 0.712918 | 0.707103 | 0.71101 | 0.711278 |
| 0.715425 | 0.712692 | 0.710803 | 0.709399 | 0.708364 | 0.711095 | 0.708199 | 0.715783 | 0.708607 |
| 0.712958 | 0.712213 | 0.707654 | 0.707524 | 0.707433 | 0.708405 | 0.706881 | 0.711596 | 0.709797 |
| 0.710884 | 0.703633 | 0.70308 | 0.705192 | 0.702885 | 0.709101 | 0.7003 | 0.706412 | 0.703798 |
| 0.708961 | 0.702492 | 0.702406 | 0.702997 | 0.7057 | 0.705124 | 0.70103 | 0.70457 | 0.702767 |
| 0.706181 | 0.705668 | 0.702462 | 0.702298 | 0.705897 | 0.706706 | 0.699983 | 0.709175 | 0.702648 |
| 0.768619 | 0.770363 | 0.764521 | 0.762448 | 0.759774 | 0.769423 | 0.768052 | 0.766971 | 0.764025 |
| 0.784909 | 0.78581 | 0.775977 | 0.777429 | 0.776411 | 0.789718 | 0.781787 | 0.78168 | 0.777437 |
| 0.786026 | 0.780077 | 0.776855 | 0.777282 | 0.776889 | 0.776597 | 0.783951 | 0.77838 | 0.782108 |
| 0.805761 | 0.772061 | 0.761414 | 0.768499 | 0.766399 | 0.772367 | 0.762809 | 0.772673 | 0.775073 |
| 0.820212 | 0.789465 | 0.77198 | 0.779808 | 0.773746 | 0.784263 | 0.781767 | 0.779847 | 0.784413 |
| 0.705464 | 0.699383 | 0.702603 | 0.700993 | 0.703098 | 0.701453 | 0.698274 | 0.705421 | 0.700834 |
| 0.710139 | 0.705468 | 0.704228 | 0.703565 | 0.703317 | 0.704568 | 0.701591 | 0.706343 | 0.705651 |
| 0.712687 | 0.70861 | 0.7068 | 0.702976 | 0.707295 | 0.705025 | 0.704547 | 0.707098 | 0.705566 |
| 0.713065 | 0.70825 | 0.706998 | 0.707768 | 0.712376 | 0.708869 | 0.706955 | 0.711717 | 0.706641 |
| 0.70869 | 0.704848 | 0.703662 | 0.703721 | 0.707382 | 0.704914 | 0.703595 | 0.707024 | 0.70228 |
| 0.71074 | 0.706237 | 0.704252 | 0.709601 | 0.710096 | 0.706176 | 0.708249 | 0.708641 | 0.706965 |
| 0.711986 | 0.71001 | 0.703344 | 0.703313 | 0.706866 | 0.705406 | 0.703837 | 0.706802 | 0.707731 |
| 0.707142 | 0.70884 | 0.705935 | 0.705038 | 0.704823 | 0.707277 | 0.70282 | 0.708631 | 0.703674 |
| 0.705618 | 0.700743 | 0.701824 | 0.699142 | 0.700449 | 0.698024 | 0.699267 | 0.703419 | 0.699288 |
| 0.706167 | 0.69969 | 0.702687 | 0.700093 | 0.700623 | 0.697722 | 0.697727 | 0.698755 | 0.696381 |
| 0.708906 | 0.707179 | 0.706531 | 0.703074 | 0.706068 | 0.707115 | 0.702676 | 0.707844 | 0.704723 |
| 0.715345 | 0.705868 | 0.708754 | 0.70639 | 0.70849 | 0.709848 | 0.705502 | 0.70967 | 0.705362 |

| | | | | | | | | |
|---|---|---|---|---|---|---|---|---|
| 0.710565 | 0.711812 | 0.703752 | 0.707782 | 0.711534 | 0.708104 | 0.70771 | 0.710132 | 0.709877 |
| 0.712739 | 0.70691 | 0.70836 | 0.705859 | 0.706598 | 0.708162 | 0.701604 | 0.706902 | 0.704423 |
| 0.717731 | 0.713311 | 0.708732 | 0.711791 | 0.712625 | 0.714647 | 0.70594 | 0.715411 | 0.710459 |
| 0.709948 | 0.708374 | 0.705329 | 0.70871 | 0.712754 | 0.706847 | 0.703538 | 0.709996 | 0.705494 |
| 0.707742 | 0.707826 | 0.702771 | 0.702605 | 0.706257 | 0.703953 | 0.701412 | 0.705892 | 0.703149 |
| 0.710176 | 0.708219 | 0.706197 | 0.703117 | 0.705214 | 0.706614 | 0.699967 | 0.704566 | 0.706332 |
| 0.713685 | 0.712307 | 0.705782 | 0.706763 | 0.710635 | 0.711931 | 0.704692 | 0.711729 | 0.707722 |
| 0.713861 | 0.710959 | 0.70862 | 0.710738 | 0.707537 | 0.708429 | 0.702696 | 0.710492 | 0.706404 |
| 0.708244 | 0.70782 | 0.704223 | 0.707457 | 0.709029 | 0.712635 | 0.70321 | 0.711179 | 0.703217 |
| 0.711875 | 0.707993 | 0.705689 | 0.705939 | 0.709409 | 0.708036 | 0.70619 | 0.708859 | 0.709292 |
| 0.710073 | 0.708686 | 0.703023 | 0.707863 | 0.709547 | 0.706832 | 0.701871 | 0.707257 | 0.703068 |
| 0.710103 | 0.706776 | 0.704966 | 0.706219 | 0.708649 | 0.707036 | 0.705649 | 0.705546 | 0.704797 |
| 0.702426 | 0.701575 | 0.695853 | 0.697926 | 0.69946 | 0.698061 | 0.691559 | 0.699861 | 0.698519 |
| 0.704509 | 0.701485 | 0.695751 | 0.697918 | 0.704738 | 0.702607 | 0.698563 | 0.700898 | 0.69887 |
| 0.712301 | 0.709585 | 0.707107 | 0.711197 | 0.708457 | 0.708425 | 0.70805 | 0.710647 | 0.706377 |
| 0.718247 | 0.713929 | 0.713049 | 0.711456 | 0.713832 | 0.716466 | 0.712457 | 0.71539 | 0.71552 |
| 0.716141 | 0.71019 | 0.707595 | 0.710807 | 0.709175 | 0.711657 | 0.708033 | 0.712214 | 0.706988 |
| 0.718977 | 0.712921 | 0.711882 | 0.712283 | 0.712701 | 0.716577 | 0.71169 | 0.716685 | 0.713071 |
| 0.710693 | 0.7078 | 0.70382 | 0.706619 | 0.706325 | 0.708216 | 0.706326 | 0.709476 | 0.703603 |
| 0.709002 | 0.708275 | 0.702971 | 0.706531 | 0.708007 | 0.712112 | 0.70383 | 0.70825 | 0.706383 |
| 0.708243 | 0.706485 | 0.699325 | 0.703849 | 0.70494 | 0.709284 | 0.700377 | 0.70694 | 0.703789 |
| 0.716041 | 0.712097 | 0.710878 | 0.707668 | 0.708701 | 0.713392 | 0.705421 | 0.708556 | 0.712006 |
| 0.716115 | 0.712828 | 0.708416 | 0.709333 | 0.709314 | 0.712184 | 0.707915 | 0.712673 | 0.70661 |
| 0.712427 | 0.706559 | 0.70491 | 0.704672 | 0.706543 | 0.706308 | 0.703976 | 0.705923 | 0.70454 |
| 0.712381 | 0.707629 | 0.704376 | 0.704613 | 0.708993 | 0.705908 | 0.707081 | 0.705654 | 0.707246 |
| 0.78557 | 0.784653 | 0.777973 | 0.782259 | 0.775201 | 0.779001 | 0.776404 | 0.779995 | 0.780168 |
| 0.712323 | 0.712242 | 0.707406 | 0.708752 | 0.706727 | 0.711916 | 0.706502 | 0.710535 | 0.708034 |
| 0.716109 | 0.714974 | 0.706254 | 0.712945 | 0.712131 | 0.711003 | 0.711852 | 0.712701 | 0.710693 |
| 0.716153 | 0.714551 | 0.70876 | 0.710329 | 0.712021 | 0.712447 | 0.71055 | 0.712996 | 0.711841 |
| 0.716582 | 0.711261 | 0.708157 | 0.711187 | 0.705749 | 0.709896 | 0.707458 | 0.709525 | 0.711801 |
| 0.712068 | 0.707976 | 0.704341 | 0.705999 | 0.707406 | 0.708249 | 0.704981 | 0.707446 | 0.705355 |
| 0.712219 | 0.7088 | 0.70286 | 0.705254 | 0.703704 | 0.707649 | 0.704195 | 0.709016 | 0.704711 |
| 0.718056 | 0.711556 | 0.705577 | 0.705367 | 0.708608 | 0.713442 | 0.707269 | 0.710655 | 0.707035 |
| 0.709378 | 0.704374 | 0.701997 | 0.703928 | 0.703595 | 0.705476 | 0.705837 | 0.707374 | 0.703137 |
| 0.709705 | 0.70407 | 0.700921 | 0.70359 | 0.704671 | 0.708673 | 0.702666 | 0.704954 | 0.70198 |
| 0.712261 | 0.707834 | 0.702014 | 0.706544 | 0.707652 | 0.706752 | 0.707024 | 0.707961 | 0.706226 |
| 0.716916 | 0.711004 | 0.707359 | 0.709917 | 0.71257 | 0.712003 | 0.709803 | 0.709672 | 0.709889 |
| 0.715247 | 0.715686 | 0.709804 | 0.709441 | 0.711446 | 0.712156 | 0.707051 | 0.712959 | 0.711343 |
| 0.714588 | 0.70896 | 0.708625 | 0.710077 | 0.709777 | 0.710136 | 0.704799 | 0.710095 | 0.710292 |
| 0.711845 | 0.712827 | 0.709511 | 0.710146 | 0.708421 | 0.709458 | 0.707136 | 0.710566 | 0.709506 |
| 0.710588 | 0.706378 | 0.699545 | 0.703457 | 0.70494 | 0.706065 | 0.700982 | 0.704511 | 0.705522 |
| 0.718235 | 0.711281 | 0.710827 | 0.712409 | 0.711926 | 0.71306 | 0.71045 | 0.715864 | 0.711374 |
| 0.71194 | 0.713362 | 0.705897 | 0.705486 | 0.707661 | 0.709826 | 0.702453 | 0.706643 | 0.706609 |
| 0.7102 | 0.70858 | 0.703739 | 0.708271 | 0.707117 | 0.710387 | 0.704095 | 0.705251 | 0.706736 |
| 0.711899 | 0.71138 | 0.706407 | 0.71066 | 0.709578 | 0.713191 | 0.706635 | 0.707966 | 0.70978 |
| 0.712965 | 0.714217 | 0.710738 | 0.712907 | 0.711683 | 0.711749 | 0.711523 | 0.708007 | 0.708495 |

| | | | | | | | | |
|---|---|---|---|---|---|---|---|---|
| 0.710873 | 0.709716 | 0.709212 | 0.709455 | 0.708815 | 0.711466 | 0.711714 | 0.708446 | 0.706829 |
| 0.710991 | 0.711219 | 0.704011 | 0.707348 | 0.707555 | 0.708529 | 0.707533 | 0.709126 | 0.708317 |
| 0.709503 | 0.709166 | 0.703609 | 0.707929 | 0.710619 | 0.711381 | 0.705681 | 0.706115 | 0.705995 |
| 0.665502 | 0.667831 | 0.658377 | 0.662795 | 0.667487 | 0.673303 | 0.664916 | 0.660515 | 0.667129 |
| 0.710294 | 0.70773 | 0.702839 | 0.705979 | 0.706641 | 0.706462 | 0.704863 | 0.707121 | 0.706082 |
| 0.711177 | 0.707642 | 0.705307 | 0.706314 | 0.708649 | 0.710613 | 0.703719 | 0.707294 | 0.707762 |
| 0.715204 | 0.709353 | 0.705771 | 0.71041 | 0.709161 | 0.709597 | 0.707258 | 0.70763 | 0.709146 |
| 0.713782 | 0.708506 | 0.706711 | 0.709037 | 0.711025 | 0.708313 | 0.708273 | 0.708661 | 0.706393 |
| 0.718274 | 0.713749 | 0.710773 | 0.710303 | 0.715426 | 0.714743 | 0.711261 | 0.712882 | 0.71217 |
| 0.71594 | 0.71039 | 0.706897 | 0.712177 | 0.712198 | 0.711381 | 0.708236 | 0.707821 | 0.707408 |
| 0.715771 | 0.713275 | 0.70434 | 0.706936 | 0.71196 | 0.711515 | 0.707625 | 0.707483 | 0.708135 |
| 0.71904 | 0.717363 | 0.713188 | 0.715023 | 0.711971 | 0.71355 | 0.709281 | 0.712922 | 0.71234 |
| 0.716663 | 0.715606 | 0.710436 | 0.71041 | 0.713824 | 0.713427 | 0.71352 | 0.710665 | 0.711856 |
| 0.7115 | 0.709353 | 0.704379 | 0.708775 | 0.707395 | 0.708078 | 0.705174 | 0.707707 | 0.710006 |
| 0.716341 | 0.711858 | 0.709042 | 0.710744 | 0.714401 | 0.713129 | 0.71121 | 0.713783 | 0.710232 |
| 0.708571 | 0.70847 | 0.702033 | 0.705919 | 0.712082 | 0.709202 | 0.705574 | 0.707103 | 0.708749 |
| 0.715378 | 0.710579 | 0.707081 | 0.709148 | 0.712264 | 0.710984 | 0.711475 | 0.710666 | 0.708565 |
| 0.712343 | 0.71051 | 0.708703 | 0.710914 | 0.712851 | 0.7124 | 0.711192 | 0.712368 | 0.708607 |
| 0.711494 | 0.707734 | 0.700963 | 0.706086 | 0.708666 | 0.709463 | 0.70439 | 0.707473 | 0.703756 |
| 0.710629 | 0.709335 | 0.705603 | 0.706585 | 0.707607 | 0.711337 | 0.706573 | 0.709289 | 0.707651 |
| 0.710451 | 0.712957 | 0.707606 | 0.710673 | 0.711478 | 0.711335 | 0.708505 | 0.713337 | 0.710176 |
| 0.712797 | 0.71384 | 0.708023 | 0.712013 | 0.712929 | 0.715443 | 0.7114 | 0.715381 | 0.708515 |
| 0.692237 | 0.693923 | 0.690198 | 0.69778 | 0.695957 | 0.691682 | 0.697153 | 0.70397 | 0.695144 |
| 0.722456 | 0.71275 | 0.724028 | 0.718704 | 0.722031 | 0.708864 | 0.711716 | 0.709184 | 0.714216 |
| 0.76147 | 0.756026 | 0.766742 | 0.760412 | 0.781739 | 0.756787 | 0.779743 | 0.758824 | 0.748936 |
| 0.738787 | 0.740499 | 0.739693 | 0.742061 | 0.738788 | 0.745546 | 0.739361 | 0.752632 | 0.741803 |
| 0.670753 | 0.664299 | 0.663399 | 0.66123 | 0.666618 | 0.665309 | 0.659682 | 0.666909 | 0.659019 |
| 0.661891 | 0.665482 | 0.663311 | 0.668689 | 0.663324 | 0.665418 | 0.663005 | 0.662715 | 0.660155 |
| 0.672414 | 0.651759 | 0.657372 | 0.662707 | 0.66064 | 0.66415 | 0.662531 | 0.659549 | 0.660597 |
| 0.683428 | 0.67226 | 0.67567 | 0.677542 | 0.675444 | 0.675501 | 0.676644 | 0.679161 | 0.669263 |
| 0.662564 | 0.652145 | 0.652953 | 0.656736 | 0.654 | 0.649666 | 0.648763 | 0.654117 | 0.647598 |
| 0.645969 | 0.633117 | 0.638292 | 0.64485 | 0.643368 | 0.64171 | 0.643013 | 0.646305 | 0.639024 |
| 0.650243 | 0.649928 | 0.64 | 0.651669 | 0.636932 | 0.646007 | 0.637941 | 0.647259 | 0.64375 |
| 0.752012 | 0.741863 | 0.735409 | 0.730769 | 0.744569 | 0.740146 | 0.733172 | 0.728537 | 0.737321 |
| 0.704658 | 0.702845 | 0.69941 | 0.703343 | 0.705758 | 0.704803 | 0.702251 | 0.701076 | 0.702826 |
| 0.68108 | 0.676321 | 0.673297 | 0.674111 | 0.676314 | 0.674785 | 0.677518 | 0.678462 | 0.676939 |
| 0.671384 | 0.669027 | 0.666884 | 0.666108 | 0.677645 | 0.671125 | 0.669027 | 0.668836 | 0.663944 |
| 0.659946 | 0.647372 | 0.646357 | 0.652888 | 0.661488 | 0.653098 | 0.652111 | 0.663152 | 0.657336 |
| 0.680667 | 0.675807 | 0.669058 | 0.675757 | 0.675335 | 0.674276 | 0.673458 | 0.674884 | 0.673659 |
| 0.67352 | 0.666294 | 0.663741 | 0.665645 | 0.669782 | 0.667414 | 0.661759 | 0.669481 | 0.665722 |
| 0.666511 | 0.661477 | 0.657709 | 0.658274 | 0.659522 | 0.662388 | 0.656597 | 0.662619 | 0.65573 |
| 0.666515 | 0.669105 | 0.657725 | 0.658871 | 0.666011 | 0.665599 | 0.660466 | 0.663101 | 0.664863 |
| 0.660024 | 0.657433 | 0.650433 | 0.654699 | 0.660325 | 0.656709 | 0.655232 | 0.653488 | 0.652835 |
| 0.663845 | 0.655324 | 0.659372 | 0.659943 | 0.664168 | 0.66129 | 0.665899 | 0.661447 | 0.654872 |
| 0.663922 | 0.65156 | 0.658379 | 0.653245 | 0.664343 | 0.659639 | 0.66034 | 0.661881 | 0.656583 |
| 0.653995 | 0.637349 | 0.63798 | 0.653456 | 0.646419 | 0.645927 | 0.644809 | 0.644041 | 0.650032 |

| | | | | | | | | |
|---|---|---|---|---|---|---|---|---|
| 0.693065 | 0.691062 | 0.687284 | 0.689223 | 0.693858 | 0.687561 | 0.688771 | 0.691067 | 0.684778 |
| 0.661192 | 0.653596 | 0.653359 | 0.655393 | 0.662524 | 0.655813 | 0.658331 | 0.655545 | 0.656086 |
| 0.668612 | 0.655392 | 0.663126 | 0.661233 | 0.665793 | 0.663553 | 0.661545 | 0.661274 | 0.660758 |
| 0.667062 | 0.660881 | 0.666271 | 0.656714 | 0.671798 | 0.667759 | 0.662522 | 0.663902 | 0.661767 |
| 0.6641 | 0.65174 | 0.660299 | 0.661865 | 0.664554 | 0.660102 | 0.656238 | 0.656232 | 0.656593 |
| 0.667881 | 0.653883 | 0.658249 | 0.66223 | 0.659096 | 0.664239 | 0.654132 | 0.664823 | 0.662572 |
| 0.664455 | 0.65073 | 0.654669 | 0.652224 | 0.657512 | 0.65966 | 0.655514 | 0.657605 | 0.653185 |
| 0.654121 | 0.643267 | 0.640608 | 0.646174 | 0.654974 | 0.640638 | 0.649699 | 0.650668 | 0.65023 |
| 0.658651 | 0.644608 | 0.650206 | 0.638628 | 0.65375 | 0.642234 | 0.65374 | 0.645788 | 0.640706 |
| 0.696746 | 0.694505 | 0.687447 | 0.688895 | 0.690144 | 0.690958 | 0.691242 | 0.692447 | 0.68696 |
| 0.692905 | 0.687445 | 0.684154 | 0.685912 | 0.687555 | 0.685541 | 0.688252 | 0.6856 | 0.684516 |
| 0.687843 | 0.680913 | 0.682514 | 0.683312 | 0.683518 | 0.682627 | 0.684921 | 0.682199 | 0.680767 |
| 0.68112 | 0.680846 | 0.6728 | 0.676905 | 0.679953 | 0.673695 | 0.674682 | 0.677358 | 0.669107 |
| 0.686405 | 0.682234 | 0.676332 | 0.679131 | 0.682301 | 0.684236 | 0.683407 | 0.684473 | 0.675349 |
| 0.673761 | 0.674923 | 0.666943 | 0.670895 | 0.674141 | 0.673038 | 0.673121 | 0.671447 | 0.668398 |

| Collection_ | Collection_ | Collection_ | Collection_ | Collection_ | Collection_ | Collection_ | Collection_ | Collection_ |
|---|---|---|---|---|---|---|---|---|
| 0.774623 | 0.798268 | 0.7761 | 0.773549 | 0.777585 | 0.767556 | 0.785524 | 0.775051 | 0.778492 |
| 0.776234 | 0.79511 | 0.779265 | 0.784048 | 0.775429 | 0.774613 | 0.789333 | 0.780267 | 0.784139 |
| 0.76954 | 0.793875 | 0.773869 | 0.775689 | 0.774008 | 0.77342 | 0.778275 | 0.770947 | 0.775904 |
| 0.7853 | 0.780376 | 0.776811 | 0.779119 | 0.774501 | 0.776559 | 0.775002 | 0.77532 | 0.771946 |
| 0.773941 | 0.77539 | 0.777869 | 0.777732 | 0.767236 | 0.773452 | 0.773557 | 0.773526 | 0.765737 |
| 0.776871 | 0.778542 | 0.775626 | 0.780885 | 0.773007 | 0.770774 | 0.781333 | 0.779521 | 0.769835 |
| 0.711981 | 0.710395 | 0.712013 | 0.711095 | 0.708184 | 0.705417 | 0.71016 | 0.711882 | 0.714228 |
| 0.710541 | 0.710993 | 0.709677 | 0.709288 | 0.707725 | 0.703165 | 0.705676 | 0.708779 | 0.710754 |
| 0.783209 | 0.777924 | 0.78289 | 0.776659 | 0.772847 | 0.775517 | 0.784906 | 0.779761 | 0.77723 |
| 0.776105 | 0.771756 | 0.772768 | 0.774879 | 0.779889 | 0.771114 | 0.769193 | 0.771466 | 0.781016 |
| 0.777649 | 0.78022 | 0.769015 | 0.77537 | 0.76676 | 0.770315 | 0.778435 | 0.769989 | 0.764765 |
| 0.781886 | 0.788169 | 0.785288 | 0.775519 | 0.777162 | 0.771813 | 0.781805 | 0.791366 | 0.777824 |
| 0.77418 | 0.776068 | 0.773034 | 0.774281 | 0.763647 | 0.766986 | 0.778581 | 0.783115 | 0.774145 |
| 0.769846 | 0.784296 | 0.775297 | 0.769212 | 0.768181 | 0.765832 | 0.772047 | 0.775336 | 0.770175 |
| 0.76906 | 0.77715 | 0.770503 | 0.767156 | 0.755857 | 0.759271 | 0.768329 | 0.762815 | 0.771374 |
| 0.778612 | 0.801463 | 0.781707 | 0.781217 | 0.773933 | 0.778608 | 0.777133 | 0.776084 | 0.777676 |
| 0.767257 | 0.800402 | 0.769854 | 0.774071 | 0.772059 | 0.768673 | 0.772693 | 0.770981 | 0.765759 |
| 0.755865 | 0.756274 | 0.753037 | 0.746622 | 0.748411 | 0.7503 | 0.754874 | 0.760784 | 0.761178 |
| 0.780512 | 0.79141 | 0.779311 | 0.778118 | 0.77915 | 0.772851 | 0.780881 | 0.792351 | 0.785912 |
| 0.785328 | 0.78598 | 0.786966 | 0.782303 | 0.787796 | 0.776116 | 0.783599 | 0.782975 | 0.777129 |
| 0.776978 | 0.783154 | 0.779929 | 0.785378 | 0.77654 | 0.780645 | 0.78063 | 0.791288 | 0.778167 |
| 0.779565 | 0.793903 | 0.777181 | 0.780859 | 0.784338 | 0.775882 | 0.782547 | 0.786003 | 0.78879 |
| 0.792965 | 0.793696 | 0.786731 | 0.786666 | 0.779874 | 0.780872 | 0.789848 | 0.78046 | 0.788335 |
| 0.78553 | 0.797259 | 0.785957 | 0.782103 | 0.783993 | 0.782935 | 0.786293 | 0.792087 | 0.783842 |
| 0.782765 | 0.781315 | 0.787131 | 0.786945 | 0.779395 | 0.778348 | 0.786366 | 0.791217 | 0.776344 |
| 0.778244 | 0.794673 | 0.781745 | 0.775545 | 0.7725 | 0.776352 | 0.783676 | 0.786799 | 0.782131 |
| 0.774781 | 0.789399 | 0.777024 | 0.774422 | 0.771341 | 0.773277 | 0.768548 | 0.774 | 0.786098 |
| 0.781165 | 0.778605 | 0.780373 | 0.778944 | 0.771756 | 0.782502 | 0.780341 | 0.778287 | 0.778387 |
| 0.783306 | 0.785278 | 0.781931 | 0.781449 | 0.783454 | 0.779344 | 0.784577 | 0.775736 | 0.774443 |
| 0.773079 | 0.780833 | 0.780618 | 0.779696 | 0.771398 | 0.769826 | 0.776417 | 0.780769 | 0.77099 |
| 0.777251 | 0.779798 | 0.775686 | 0.780106 | 0.777741 | 0.776166 | 0.778454 | 0.784791 | 0.775706 |
| 0.778824 | 0.776382 | 0.777659 | 0.773766 | 0.767897 | 0.776075 | 0.779591 | 0.776752 | 0.774023 |
| 0.778621 | 0.783633 | 0.77797 | 0.783122 | 0.785342 | 0.780202 | 0.774738 | 0.780963 | 0.776514 |
| 0.776284 | 0.774724 | 0.771705 | 0.782745 | 0.771041 | 0.776292 | 0.777106 | 0.770315 | 0.76739 |
| 0.770164 | 0.783545 | 0.772784 | 0.767205 | 0.763428 | 0.766669 | 0.761818 | 0.771588 | 0.767231 |
| 0.779857 | 0.797041 | 0.776485 | 0.779158 | 0.778239 | 0.779794 | 0.781307 | 0.781053 | 0.779596 |
| 0.781716 | 0.788444 | 0.77477 | 0.781998 | 0.778656 | 0.772542 | 0.772394 | 0.782296 | 0.782544 |
| 0.773539 | 0.771658 | 0.776924 | 0.777324 | 0.768757 | 0.764822 | 0.768017 | 0.768532 | 0.76934 |
| 0.777851 | 0.775008 | 0.780377 | 0.782877 | 0.775258 | 0.771295 | 0.776315 | 0.77431 | 0.77275 |

| | | | | | | | | |
|---|---|---|---|---|---|---|---|---|
| 0.754635 | 0.767966 | 0.755335 | 0.751667 | 0.752684 | 0.748277 | 0.753599 | 0.752868 | 0.752277 |
| 0.772825 | 0.770798 | 0.774183 | 0.771931 | 0.771956 | 0.772593 | 0.77433 | 0.771534 | 0.779379 |
| 0.779452 | 0.796518 | 0.788918 | 0.778623 | 0.776609 | 0.774825 | 0.784351 | 0.787245 | 0.778205 |
| 0.782619 | 0.789141 | 0.788034 | 0.777896 | 0.778161 | 0.775865 | 0.784746 | 0.782516 | 0.783304 |
| 0.786652 | 0.786907 | 0.785436 | 0.774818 | 0.778503 | 0.773525 | 0.78348 | 0.789712 | 0.78048 |
| 0.782627 | 0.779829 | 0.783898 | 0.77799 | 0.775709 | 0.778219 | 0.782918 | 0.784927 | 0.773073 |
| 0.776363 | 0.772605 | 0.773562 | 0.772198 | 0.779734 | 0.772776 | 0.774837 | 0.775418 | 0.773018 |
| 0.786683 | 0.783373 | 0.785506 | 0.788237 | 0.7805 | 0.781006 | 0.778195 | 0.78191 | 0.790715 |
| 0.77499 | 0.782468 | 0.780313 | 0.779781 | 0.776456 | 0.77523 | 0.775759 | 0.774866 | 0.769263 |
| 0.777814 | 0.778607 | 0.773459 | 0.770654 | 0.775828 | 0.761255 | 0.772465 | 0.766056 | 0.765513 |
| 0.781795 | 0.790934 | 0.776587 | 0.778349 | 0.774247 | 0.772409 | 0.781587 | 0.786914 | 0.777824 |
| 0.774678 | 0.786638 | 0.780241 | 0.782685 | 0.776496 | 0.778365 | 0.78328 | 0.79925 | 0.782298 |
| 0.784577 | 0.79455 | 0.777084 | 0.785131 | 0.775611 | 0.772168 | 0.779908 | 0.787321 | 0.780599 |
| 0.778806 | 0.783371 | 0.786419 | 0.782931 | 0.774963 | 0.780111 | 0.789137 | 0.791004 | 0.780259 |
| 0.772138 | 0.804487 | 0.779358 | 0.780478 | 0.771505 | 0.770557 | 0.782287 | 0.792806 | 0.780869 |
| 0.768065 | 0.800169 | 0.776182 | 0.780439 | 0.769591 | 0.772955 | 0.775307 | 0.792504 | 0.774623 |
| 0.777768 | 0.784745 | 0.773213 | 0.775585 | 0.767364 | 0.76932 | 0.775872 | 0.775081 | 0.773064 |
| 0.782223 | 0.800833 | 0.783585 | 0.786135 | 0.782767 | 0.779674 | 0.787085 | 0.783577 | 0.787897 |
| 0.785803 | 0.776068 | 0.779964 | 0.779651 | 0.780182 | 0.774636 | 0.776468 | 0.786133 | 0.781616 |
| 0.780604 | 0.785619 | 0.780823 | 0.782128 | 0.779381 | 0.772686 | 0.778122 | 0.792934 | 0.782704 |
| 0.776416 | 0.791663 | 0.773342 | 0.778009 | 0.770359 | 0.777217 | 0.782944 | 0.778192 | 0.775144 |
| 0.778768 | 0.789417 | 0.775434 | 0.775025 | 0.771405 | 0.765376 | 0.774387 | 0.772435 | 0.774328 |
| 0.784101 | 0.782052 | 0.776217 | 0.783657 | 0.777019 | 0.774215 | 0.781939 | 0.785327 | 0.784715 |
| 0.775259 | 0.778401 | 0.780306 | 0.780063 | 0.77184 | 0.769914 | 0.775017 | 0.772921 | 0.770083 |
| 0.768164 | 0.767832 | 0.767332 | 0.770509 | 0.759532 | 0.759389 | 0.761545 | 0.757861 | 0.762172 |
| 0.710839 | 0.710942 | 0.709654 | 0.710687 | 0.707817 | 0.703514 | 0.707111 | 0.709136 | 0.712952 |
| 0.713384 | 0.709452 | 0.712663 | 0.711927 | 0.713893 | 0.7048 | 0.710335 | 0.713368 | 0.713159 |
| 0.719475 | 0.709902 | 0.714343 | 0.711545 | 0.714067 | 0.7091 | 0.710318 | 0.712754 | 0.712558 |
| 0.768614 | 0.778547 | 0.770835 | 0.779232 | 0.773088 | 0.772044 | 0.770754 | 0.781886 | 0.778976 |
| 0.774091 | 0.78533 | 0.775583 | 0.776739 | 0.769231 | 0.768193 | 0.769316 | 0.775469 | 0.774844 |
| 0.773089 | 0.778254 | 0.775607 | 0.768287 | 0.76397 | 0.767001 | 0.772697 | 0.771771 | 0.77458 |
| 0.775543 | 0.774869 | 0.775033 | 0.776136 | 0.768879 | 0.768759 | 0.770293 | 0.780455 | 0.781838 |
| 0.784818 | 0.818282 | 0.786278 | 0.787046 | 0.780039 | 0.78702 | 0.787731 | 0.785379 | 0.803779 |
| 0.78409 | 0.815227 | 0.792718 | 0.785786 | 0.786277 | 0.779809 | 0.791021 | 0.799514 | 0.777871 |
| 0.778769 | 0.814575 | 0.787435 | 0.780506 | 0.788869 | 0.783326 | 0.788506 | 0.784672 | 0.800753 |
| 0.778435 | 0.864379 | 0.78118 | 0.783415 | 0.780253 | 0.781722 | 0.795378 | 0.784672 | 0.779726 |
| 0.786128 | 0.773409 | 0.783296 | 0.776387 | 0.790396 | 0.779669 | 0.7964 | 0.782548 | 0.828171 |
| 0.798758 | 0.819911 | 0.802209 | 0.798632 | 0.803422 | 0.801771 | 0.808147 | 0.802537 | 0.814362 |
| 0.774349 | 0.801908 | 0.776372 | 0.770225 | 0.776506 | 0.80431 | 0.782669 | 0.772545 | 0.794092 |
| 0.791446 | 0.779021 | 0.784785 | 0.783846 | 0.782305 | 0.778694 | 0.792102 | 0.809807 | 0.780348 |
| 0.782192 | 0.821715 | 0.777613 | 0.776238 | 0.779247 | 0.77207 | 0.787449 | 0.778367 | 0.776231 |
| 0.804366 | 0.818436 | 0.804134 | 0.803134 | 0.798993 | 0.80594 | 0.801856 | 0.801806 | 0.807278 |
| 0.785267 | 0.781201 | 0.783297 | 0.784458 | 0.783981 | 0.778089 | 0.788107 | 0.781622 | 0.774677 |
| 0.784379 | 0.774929 | 0.791294 | 0.781301 | 0.783259 | 0.779992 | 0.782828 | 0.791979 | 0.775539 |
| 0.78924 | 0.774967 | 0.783953 | 0.788202 | 0.787403 | 0.778891 | 0.782594 | 0.800234 | 0.782039 |
| 0.776541 | 0.813169 | 0.785648 | 0.779056 | 0.773608 | 0.827572 | 0.786045 | 0.774965 | 0.776762 |

| | | | | | | | | |
|---|---|---|---|---|---|---|---|---|
| 0.783603 | 0.784649 | 0.784306 | 0.776997 | 0.786386 | 0.779273 | 0.789447 | 0.785667 | 0.78207 |
| 0.776851 | 0.773752 | 0.781811 | 0.782494 | 0.78179 | 0.77607 | 0.796143 | 0.805964 | 0.786676 |
| 0.780972 | 0.777935 | 0.786925 | 0.777371 | 0.786385 | 0.776364 | 0.79566 | 0.788116 | 0.790217 |
| 0.79059 | 0.797817 | 0.796879 | 0.792203 | 0.797094 | 0.795544 | 0.796367 | 0.798028 | 0.802352 |
| 0.789028 | 0.780598 | 0.793103 | 0.782391 | 0.793058 | 0.77998 | 0.786069 | 0.862722 | 0.793754 |
| 0.772758 | 0.779132 | 0.784342 | 0.775337 | 0.78461 | 0.772982 | 0.786613 | 0.804796 | 0.77774 |
| 0.789164 | 0.773123 | 0.783564 | 0.78 | 0.785555 | 0.776591 | 0.780345 | 0.777034 | 0.803486 |
| 0.781718 | 0.77131 | 0.784391 | 0.78359 | 0.785237 | 0.780591 | 0.792259 | 0.774236 | 0.786689 |
| 0.776886 | 0.785613 | 0.783204 | 0.785738 | 0.782547 | 0.781107 | 0.778856 | 0.776876 | 0.780822 |
| 0.773535 | 0.774493 | 0.799967 | 0.78022 | 0.768126 | 0.769558 | 0.775798 | 0.786393 | 0.771471 |
| 0.77551 | 0.781539 | 0.783155 | 0.790596 | 0.781966 | 0.79483 | 0.781667 | 0.78555 | 0.780647 |
| 0.780742 | 0.776892 | 0.790999 | 0.783276 | 0.779826 | 0.779177 | 0.781331 | 0.783171 | 0.777225 |
| 0.782001 | 0.786441 | 0.781553 | 0.776846 | 0.778027 | 0.774506 | 0.781148 | 0.783179 | 0.770156 |
| 0.781271 | 0.770037 | 0.80289 | 0.771588 | 0.781829 | 0.770401 | 0.78111 | 0.807608 | 0.767487 |
| 0.779129 | 0.782346 | 0.785524 | 0.779059 | 0.779165 | 0.780098 | 0.778657 | 0.778605 | 0.777388 |
| 0.779904 | 0.780189 | 0.786843 | 0.778819 | 0.786941 | 0.779406 | 0.787151 | 0.777022 | 0.776587 |
| 1 | 0.771426 | 0.783662 | 0.772599 | 0.777037 | 0.776168 | 0.780617 | 0.783378 | 0.776389 |
| 0.771426 | 1 | 0.775753 | 0.775268 | 0.774439 | 0.775403 | 0.786518 | 0.77765 | 0.779993 |
| 0.783662 | 0.775753 | 1 | 0.777153 | 0.778858 | 0.7802 | 0.781721 | 0.786146 | 0.781043 |
| 0.772599 | 0.775268 | 0.777153 | 1 | 0.775908 | 0.783589 | 0.774303 | 0.7771 | 0.774021 |
| 0.777037 | 0.774439 | 0.778858 | 0.775908 | 1 | 0.769901 | 0.776983 | 0.773512 | 0.78266 |
| 0.776168 | 0.775403 | 0.7802 | 0.783589 | 0.769901 | 1 | 0.790018 | 0.772731 | 0.769384 |
| 0.780617 | 0.786518 | 0.781721 | 0.774303 | 0.776983 | 0.790018 | 1 | 0.787975 | 0.780576 |
| 0.783378 | 0.77765 | 0.786146 | 0.7771 | 0.773512 | 0.772731 | 0.787975 | 1 | 0.775868 |
| 0.776389 | 0.779993 | 0.781043 | 0.774021 | 0.78266 | 0.769384 | 0.780576 | 0.775868 | 1 |
| 0.794501 | 0.774328 | 0.790088 | 0.784562 | 0.782322 | 0.784455 | 0.78109 | 0.776542 | 0.82182 |
| 0.781529 | 0.77482 | 0.781757 | 0.7731 | 0.785207 | 0.777732 | 0.785655 | 0.780142 | 0.893781 |
| 0.778771 | 0.776629 | 0.778592 | 0.781557 | 0.774959 | 0.775293 | 0.76835 | 0.772973 | 0.772048 |
| 0.778045 | 0.783092 | 0.789177 | 0.775713 | 0.778428 | 0.781736 | 0.77605 | 0.779027 | 0.775339 |
| 0.77496 | 0.773844 | 0.778989 | 0.778794 | 0.780504 | 0.771388 | 0.777946 | 0.779633 | 0.849024 |
| 0.775587 | 0.776516 | 0.78757 | 0.777612 | 0.809311 | 0.77763 | 0.779597 | 0.785768 | 0.777666 |
| 0.779165 | 0.774317 | 0.778834 | 0.777405 | 0.781209 | 0.781768 | 0.784199 | 0.784349 | 0.775332 |
| 0.781389 | 0.780768 | 0.781946 | 0.789761 | 0.78194 | 0.781524 | 0.78628 | 0.776528 | 0.8 |
| 0.77897 | 0.791773 | 0.788416 | 0.773918 | 0.780643 | 0.778069 | 0.785355 | 0.78185 | 0.784523 |
| 0.7771 | 0.790189 | 0.789799 | 0.778111 | 0.776899 | 0.782746 | 0.784857 | 0.787778 | 0.777396 |
| 0.775553 | 0.793365 | 0.776558 | 0.775656 | 0.779904 | 0.774958 | 0.778508 | 0.782143 | 0.781086 |
| 0.777578 | 0.77732 | 0.790815 | 0.780407 | 0.773006 | 0.7824 | 0.778421 | 0.783156 | 0.77969 |
| 0.756077 | 0.756361 | 0.758136 | 0.758533 | 0.756811 | 0.755066 | 0.764433 | 0.772094 | 0.761558 |
| 0.764575 | 0.761949 | 0.764478 | 0.760311 | 0.759391 | 0.754063 | 0.768453 | 0.848228 | 0.762286 |
| 0.758397 | 0.751246 | 0.757256 | 0.751098 | 0.75373 | 0.761164 | 0.758824 | 0.752977 | 0.752329 |
| 0.754555 | 0.800305 | 0.762637 | 0.755884 | 0.758172 | 0.755132 | 0.770341 | 0.760294 | 0.778751 |
| 0.751881 | 0.739771 | 0.765973 | 0.740966 | 0.748995 | 0.741986 | 0.748476 | 0.772755 | 0.743146 |
| 0.761022 | 0.752734 | 0.763261 | 0.752276 | 0.758974 | 0.754933 | 0.763803 | 0.768228 | 0.753381 |
| 0.748791 | 0.768197 | 0.748553 | 0.748483 | 0.755079 | 0.759135 | 0.768642 | 0.748715 | 0.754437 |
| 0.7498 | 0.75221 | 0.753604 | 0.747307 | 0.760757 | 0.743023 | 0.747052 | 0.747267 | 0.747049 |
| 0.759055 | 0.770207 | 0.759615 | 0.74987 | 0.751813 | 0.75754 | 0.771346 | 0.752298 | 0.776005 |
| 0.742097 | 0.817509 | 0.758514 | 0.750226 | 0.744296 | 0.750287 | 0.765415 | 0.751631 | 0.77711 |

| | | | | | | | | |
|---|---|---|---|---|---|---|---|---|
| 0.744192 | 0.746631 | 0.739086 | 0.751543 | 0.743243 | 0.748981 | 0.753824 | 0.752845 | 0.747754 |
| 0.755286 | 0.787551 | 0.758003 | 0.756573 | 0.754801 | 0.760277 | 0.752364 | 0.750808 | 0.788435 |
| 0.747033 | 0.745269 | 0.746504 | 0.744096 | 0.748272 | 0.742967 | 0.74397 | 0.740041 | 0.745098 |
| 0.752084 | 0.747085 | 0.744176 | 0.744543 | 0.746881 | 0.743764 | 0.75031 | 0.745651 | 0.743956 |
| 0.758827 | 0.741272 | 0.766869 | 0.752525 | 0.756964 | 0.756677 | 0.74896 | 0.773261 | 0.791243 |
| 0.750613 | 0.740961 | 0.748895 | 0.742546 | 0.760773 | 0.745729 | 0.775247 | 0.743447 | 0.769623 |
| 0.751894 | 0.740638 | 0.752952 | 0.749074 | 0.754979 | 0.743913 | 0.750627 | 0.747245 | 0.776019 |
| 0.747 | 0.790206 | 0.765754 | 0.743951 | 0.756717 | 0.75276 | 0.759082 | 0.763162 | 0.749963 |
| 0.735523 | 0.729881 | 0.750191 | 0.726154 | 0.73201 | 0.730132 | 0.736649 | 0.842474 | 0.72629 |
| 0.759242 | 0.745003 | 0.760441 | 0.751073 | 0.751713 | 0.752432 | 0.74741 | 0.749013 | 0.811241 |
| 0.749558 | 0.742322 | 0.752401 | 0.745573 | 0.759888 | 0.750786 | 0.780267 | 0.744077 | 0.755521 |
| 0.766934 | 0.763683 | 0.774476 | 0.76483 | 0.764068 | 0.766694 | 0.774122 | 0.768134 | 0.764021 |
| 0.771227 | 0.774817 | 0.777113 | 0.781106 | 0.771517 | 0.777059 | 0.773841 | 0.771873 | 0.774231 |
| 0.755063 | 0.754483 | 0.760061 | 0.752723 | 0.750879 | 0.762604 | 0.777592 | 0.744259 | 0.749237 |
| 0.769327 | 0.784898 | 0.77966 | 0.77375 | 0.774552 | 0.768014 | 0.774899 | 0.778258 | 0.770283 |
| 0.771819 | 0.771276 | 0.786621 | 0.764706 | 0.772056 | 0.768997 | 0.767363 | 0.789561 | 0.768355 |
| 0.790202 | 0.770656 | 0.779114 | 0.77601 | 0.779198 | 0.772779 | 0.77673 | 0.776942 | 0.771965 |
| 0.781425 | 0.785589 | 0.787998 | 0.7872 | 0.772172 | 0.787884 | 0.771367 | 0.771242 | 0.771098 |
| 0.767257 | 0.755003 | 0.765195 | 0.761258 | 0.760544 | 0.75785 | 0.76192 | 0.764806 | 0.764962 |
| 0.7766 | 0.771459 | 0.790978 | 0.772019 | 0.773869 | 0.773427 | 0.779609 | 0.781944 | 0.771925 |
| 0.75285 | 0.746489 | 0.749105 | 0.740089 | 0.756137 | 0.741212 | 0.753728 | 0.738986 | 0.755511 |
| 0.776643 | 0.773375 | 0.775843 | 0.780031 | 0.774661 | 0.76834 | 0.768779 | 0.773015 | 0.772382 |
| 0.769803 | 0.779892 | 0.775534 | 0.782334 | 0.768945 | 0.78009 | 0.775196 | 0.76576 | 0.770057 |
| 0.742981 | 0.744729 | 0.738722 | 0.741611 | 0.742374 | 0.743237 | 0.745785 | 0.762482 | 0.736509 |
| 0.74953 | 0.76325 | 0.74919 | 0.745419 | 0.740063 | 0.746971 | 0.745458 | 0.77474 | 0.764604 |
| 0.757954 | 0.771481 | 0.759719 | 0.758951 | 0.756192 | 0.775713 | 0.772178 | 0.757466 | 0.75 |
| 0.767827 | 0.758264 | 0.764498 | 0.756542 | 0.768887 | 0.760988 | 0.773522 | 0.75481 | 0.765394 |
| 0.776428 | 0.768784 | 0.772649 | 0.776415 | 0.773667 | 0.771029 | 0.772025 | 0.782099 | 0.765075 |
| 0.74843 | 0.748883 | 0.757334 | 0.753478 | 0.744473 | 0.79236 | 0.753962 | 0.745608 | 0.744225 |
| 0.770355 | 0.773826 | 0.774794 | 0.778697 | 0.77048 | 0.767804 | 0.769874 | 0.768738 | 0.797269 |
| 0.769873 | 0.762169 | 0.770066 | 0.771119 | 0.77279 | 0.764105 | 0.763405 | 0.761432 | 0.769174 |
| 0.753883 | 0.774606 | 0.755572 | 0.763835 | 0.755162 | 0.768085 | 0.762681 | 0.755857 | 0.7594 |
| 0.754706 | 0.768401 | 0.759113 | 0.757731 | 0.754712 | 0.773559 | 0.767101 | 0.763915 | 0.757738 |
| 0.768725 | 0.770733 | 0.772578 | 0.766006 | 0.76377 | 0.776702 | 0.773516 | 0.758151 | 0.759297 |
| 0.766567 | 0.773722 | 0.761786 | 0.775568 | 0.768857 | 0.772724 | 0.76807 | 0.765688 | 0.765659 |
| 0.769958 | 0.76887 | 0.769199 | 0.760187 | 0.768831 | 0.767068 | 0.769231 | 0.768411 | 0.76387 |
| 0.727595 | 0.725999 | 0.725902 | 0.720952 | 0.73519 | 0.726417 | 0.726223 | 0.727541 | 0.728483 |
| 0.756876 | 0.759877 | 0.757241 | 0.757952 | 0.757221 | 0.754368 | 0.758261 | 0.757752 | 0.764658 |
| 0.731353 | 0.728999 | 0.731707 | 0.724752 | 0.727215 | 0.751257 | 0.74291 | 0.725738 | 0.722519 |
| 0.723494 | 0.712547 | 0.722417 | 0.720143 | 0.719276 | 0.717063 | 0.731297 | 0.721186 | 0.72069 |
| 0.766422 | 0.771043 | 0.774465 | 0.765395 | 0.766126 | 0.767587 | 0.771112 | 0.7845 | 0.769104 |
| 0.721359 | 0.713866 | 0.718791 | 0.718297 | 0.717608 | 0.715096 | 0.727573 | 0.724122 | 0.716688 |
| 0.713055 | 0.726786 | 0.720824 | 0.716387 | 0.71484 | 0.716009 | 0.718602 | 0.719653 | 0.728503 |
| 0.705378 | 0.71173 | 0.71022 | 0.71351 | 0.70391 | 0.707873 | 0.712458 | 0.710522 | 0.724986 |
| 0.770406 | 0.758827 | 0.763601 | 0.761994 | 0.767679 | 0.763328 | 0.767959 | 0.762371 | 0.769751 |
| 0.711222 | 0.720031 | 0.708869 | 0.712045 | 0.700578 | 0.70464 | 0.70834 | 0.707413 | 0.714787 |
| 0.764342 | 0.769413 | 0.768465 | 0.765628 | 0.763378 | 0.762473 | 0.768939 | 0.770813 | 0.767511 |

| | | | | | | | | |
|---|---|---|---|---|---|---|---|---|
| 0.761622 | 0.767012 | 0.760211 | 0.762233 | 0.756479 | 0.756094 | 0.766217 | 0.764977 | 0.766305 |
| 0.767522 | 0.768537 | 0.766074 | 0.757381 | 0.765283 | 0.765479 | 0.770597 | 0.773817 | 0.76998 |
| 0.752655 | 0.748837 | 0.755395 | 0.750551 | 0.750041 | 0.752511 | 0.753902 | 0.745639 | 0.749403 |
| 0.696214 | 0.7218 | 0.701203 | 0.702452 | 0.694273 | 0.697955 | 0.700634 | 0.701908 | 0.708948 |
| 0.706393 | 0.70067 | 0.702288 | 0.699441 | 0.697072 | 0.694019 | 0.700807 | 0.699175 | 0.71781 |
| 0.768996 | 0.778124 | 0.766119 | 0.763506 | 0.760349 | 0.764816 | 0.765398 | 0.77187 | 0.771682 |
| 0.692607 | 0.694045 | 0.692905 | 0.697414 | 0.690787 | 0.72452 | 0.697657 | 0.694118 | 0.688274 |
| 0.734717 | 0.732117 | 0.735445 | 0.734765 | 0.730669 | 0.729042 | 0.729668 | 0.738173 | 0.728973 |
| 0.756963 | 0.777339 | 0.759797 | 0.759374 | 0.759561 | 0.757662 | 0.759221 | 0.762165 | 0.757728 |
| 0.722272 | 0.719356 | 0.720055 | 0.7221 | 0.724163 | 0.716903 | 0.717808 | 0.720281 | 0.718801 |
| 0.750267 | 0.755196 | 0.749039 | 0.750409 | 0.747306 | 0.746482 | 0.751728 | 0.747015 | 0.753518 |
| 0.61039 | 0.633333 | 0.625 | 0.607843 | 0.675862 | 0.660377 | 0.655629 | 0.62963 | 0.616352 |
| 0.654436 | 0.653987 | 0.663123 | 0.658154 | 0.660294 | 0.653186 | 0.655564 | 0.654134 | 0.655352 |
| 0.654775 | 0.656241 | 0.661562 | 0.664285 | 0.659607 | 0.656133 | 0.662775 | 0.659587 | 0.65885 |
| 0.665886 | 0.662873 | 0.671462 | 0.668807 | 0.669678 | 0.659349 | 0.66856 | 0.672131 | 0.665453 |
| 0.665661 | 0.663968 | 0.668221 | 0.661303 | 0.668829 | 0.654696 | 0.665674 | 0.665032 | 0.669886 |
| 0.619389 | 0.622563 | 0.62511 | 0.620807 | 0.619053 | 0.617188 | 0.62683 | 0.618813 | 0.619861 |
| 0.669286 | 0.667084 | 0.671389 | 0.668392 | 0.667501 | 0.659762 | 0.666124 | 0.665958 | 0.668887 |
| 0.746978 | 0.744924 | 0.744365 | 0.740705 | 0.743143 | 0.740181 | 0.744602 | 0.744057 | 0.751633 |
| 0.738295 | 0.74493 | 0.739163 | 0.739658 | 0.738843 | 0.736146 | 0.739264 | 0.737831 | 0.739098 |
| 0.746481 | 0.751518 | 0.747704 | 0.744287 | 0.743569 | 0.745103 | 0.744665 | 0.743993 | 0.750166 |
| 0.623168 | 0.616402 | 0.618744 | 0.622759 | 0.613121 | 0.616794 | 0.624619 | 0.621689 | 0.616613 |
| 0.603639 | 0.600238 | 0.602952 | 0.601387 | 0.597735 | 0.600318 | 0.60325 | 0.604296 | 0.598815 |
| 0.712516 | 0.706865 | 0.706449 | 0.706044 | 0.707511 | 0.702184 | 0.709329 | 0.707509 | 0.707546 |
| 0.712852 | 0.704191 | 0.70741 | 0.70841 | 0.709984 | 0.702921 | 0.710115 | 0.710777 | 0.71122 |
| 0.703426 | 0.698437 | 0.698889 | 0.701402 | 0.701277 | 0.698404 | 0.703772 | 0.704829 | 0.700598 |
| 0.704138 | 0.702786 | 0.703942 | 0.707793 | 0.706174 | 0.700427 | 0.703597 | 0.708426 | 0.704429 |
| 0.711463 | 0.711549 | 0.705948 | 0.707966 | 0.710094 | 0.708978 | 0.713931 | 0.709833 | 0.712096 |
| 0.712058 | 0.709511 | 0.709261 | 0.706446 | 0.70753 | 0.703947 | 0.707723 | 0.713779 | 0.707781 |
| 0.708895 | 0.705873 | 0.706286 | 0.704578 | 0.707009 | 0.704239 | 0.712354 | 0.707777 | 0.70969 |
| 0.709289 | 0.705536 | 0.708066 | 0.707532 | 0.706943 | 0.703811 | 0.710641 | 0.711294 | 0.709577 |
| 0.717079 | 0.712999 | 0.71261 | 0.716019 | 0.715502 | 0.708913 | 0.717023 | 0.717395 | 0.714069 |
| 0.715085 | 0.70989 | 0.709849 | 0.715001 | 0.711276 | 0.708132 | 0.70891 | 0.707686 | 0.709593 |
| 0.717142 | 0.710327 | 0.709922 | 0.711157 | 0.712643 | 0.708107 | 0.708799 | 0.714207 | 0.713623 |
| 0.719034 | 0.711866 | 0.715966 | 0.71522 | 0.716821 | 0.710001 | 0.715652 | 0.712752 | 0.7147 |
| 0.713363 | 0.707598 | 0.707649 | 0.706588 | 0.704642 | 0.703633 | 0.709409 | 0.709443 | 0.708644 |
| 0.712227 | 0.707626 | 0.708477 | 0.711904 | 0.71129 | 0.704939 | 0.710467 | 0.708928 | 0.710718 |
| 0.718507 | 0.710777 | 0.710656 | 0.713034 | 0.709729 | 0.706666 | 0.712076 | 0.711659 | 0.710499 |
| 0.712081 | 0.702478 | 0.705285 | 0.70608 | 0.707695 | 0.699592 | 0.703408 | 0.705733 | 0.707525 |
| 0.71432 | 0.708729 | 0.710365 | 0.710779 | 0.71144 | 0.704803 | 0.711042 | 0.712668 | 0.709812 |
| 0.657483 | 0.65523 | 0.658094 | 0.656076 | 0.659542 | 0.652224 | 0.664192 | 0.663071 | 0.653419 |
| 0.661151 | 0.664609 | 0.662194 | 0.666178 | 0.668185 | 0.655682 | 0.663024 | 0.664367 | 0.665209 |
| 0.71615 | 0.711352 | 0.713139 | 0.717201 | 0.713761 | 0.710301 | 0.712485 | 0.715668 | 0.715734 |
| 0.71914 | 0.712331 | 0.712735 | 0.713511 | 0.714343 | 0.713205 | 0.71377 | 0.716493 | 0.71705 |
| 0.650233 | 0.644616 | 0.645876 | 0.65346 | 0.6509 | 0.644167 | 0.645209 | 0.642779 | 0.648613 |
| 0.715136 | 0.709531 | 0.709313 | 0.713452 | 0.71013 | 0.707507 | 0.710097 | 0.714525 | 0.714621 |
| 0.7045 | 0.698211 | 0.698845 | 0.705402 | 0.699374 | 0.696364 | 0.699595 | 0.701743 | 0.70326 |

| | | | | | | | | |
|---|---|---|---|---|---|---|---|---|
| 0.712222 | 0.70436 | 0.700667 | 0.708105 | 0.708153 | 0.703674 | 0.708814 | 0.708386 | 0.71199 |
| 0.715129 | 0.708873 | 0.709001 | 0.710974 | 0.708032 | 0.70553 | 0.711974 | 0.712214 | 0.714194 |
| 0.715575 | 0.709559 | 0.711443 | 0.709478 | 0.711964 | 0.70881 | 0.712423 | 0.710637 | 0.716685 |
| 0.717362 | 0.710088 | 0.706997 | 0.712475 | 0.708134 | 0.706405 | 0.709719 | 0.711991 | 0.711799 |
| 0.720174 | 0.71424 | 0.717436 | 0.715711 | 0.71865 | 0.710754 | 0.714938 | 0.719516 | 0.718306 |
| 0.708999 | 0.701044 | 0.704808 | 0.710084 | 0.709296 | 0.70384 | 0.70654 | 0.711529 | 0.707919 |
| 0.712085 | 0.705079 | 0.705417 | 0.70898 | 0.706275 | 0.700961 | 0.708688 | 0.71027 | 0.711879 |
| 0.704827 | 0.700819 | 0.699582 | 0.701593 | 0.703722 | 0.698341 | 0.701601 | 0.701331 | 0.703081 |
| 0.713274 | 0.703274 | 0.703078 | 0.708469 | 0.70592 | 0.701927 | 0.706567 | 0.711054 | 0.709334 |
| 0.710859 | 0.706135 | 0.705666 | 0.705807 | 0.707577 | 0.703651 | 0.707102 | 0.7084 | 0.711109 |
| 0.714799 | 0.709547 | 0.707222 | 0.71323 | 0.709937 | 0.70532 | 0.710531 | 0.712759 | 0.713 |
| 0.707917 | 0.702392 | 0.699738 | 0.70635 | 0.703296 | 0.700196 | 0.708106 | 0.707435 | 0.709932 |
| 0.703776 | 0.698325 | 0.700662 | 0.700563 | 0.69904 | 0.696334 | 0.702091 | 0.703117 | 0.706108 |
| 0.702199 | 0.697122 | 0.697362 | 0.700464 | 0.698317 | 0.696355 | 0.701459 | 0.702779 | 0.704954 |
| 0.719385 | 0.713024 | 0.710743 | 0.713041 | 0.712432 | 0.713471 | 0.713227 | 0.716713 | 0.714103 |
| 0.715696 | 0.708749 | 0.708741 | 0.70924 | 0.710126 | 0.706287 | 0.71368 | 0.712122 | 0.715101 |
| 0.708008 | 0.700508 | 0.703121 | 0.704261 | 0.707398 | 0.699492 | 0.701463 | 0.704437 | 0.701068 |
| 0.707979 | 0.706089 | 0.704607 | 0.7067 | 0.708858 | 0.700842 | 0.707666 | 0.705279 | 0.706087 |
| 0.711471 | 0.707273 | 0.702709 | 0.708028 | 0.705082 | 0.702525 | 0.707073 | 0.705817 | 0.707284 |
| 0.710162 | 0.702858 | 0.705967 | 0.706925 | 0.70721 | 0.703521 | 0.707031 | 0.706262 | 0.708418 |
| 0.70787 | 0.703428 | 0.702727 | 0.704444 | 0.702259 | 0.700048 | 0.703326 | 0.701814 | 0.707067 |
| 0.707854 | 0.70465 | 0.701554 | 0.704031 | 0.703445 | 0.697787 | 0.706281 | 0.707206 | 0.705993 |
| 0.714388 | 0.705873 | 0.705934 | 0.710158 | 0.708655 | 0.704495 | 0.70969 | 0.70941 | 0.70815 |
| 0.711358 | 0.706894 | 0.704549 | 0.707793 | 0.708273 | 0.702308 | 0.711059 | 0.710434 | 0.708219 |
| 0.713216 | 0.705508 | 0.708306 | 0.707475 | 0.704248 | 0.70295 | 0.706793 | 0.706057 | 0.707078 |
| 0.70752 | 0.705354 | 0.701807 | 0.706265 | 0.701895 | 0.699537 | 0.70584 | 0.703716 | 0.706073 |
| 0.70853 | 0.702701 | 0.704979 | 0.705583 | 0.706383 | 0.698949 | 0.706739 | 0.703846 | 0.705301 |
| 0.707999 | 0.698934 | 0.702615 | 0.704455 | 0.700008 | 0.700381 | 0.705252 | 0.704503 | 0.703112 |
| 0.711807 | 0.702866 | 0.709136 | 0.707858 | 0.703337 | 0.700451 | 0.708012 | 0.706568 | 0.708491 |
| 0.711241 | 0.708599 | 0.705289 | 0.711242 | 0.707144 | 0.701821 | 0.708587 | 0.708932 | 0.709929 |
| 0.713746 | 0.709701 | 0.70978 | 0.711108 | 0.709499 | 0.70619 | 0.713251 | 0.707846 | 0.711264 |
| 0.712685 | 0.709836 | 0.709852 | 0.71018 | 0.710795 | 0.705802 | 0.711381 | 0.70846 | 0.709843 |
| 0.716392 | 0.712231 | 0.715399 | 0.709871 | 0.716103 | 0.709708 | 0.715163 | 0.716464 | 0.713554 |
| 0.716135 | 0.711173 | 0.712441 | 0.712999 | 0.714606 | 0.711 | 0.715468 | 0.716216 | 0.714205 |
| 0.710953 | 0.703677 | 0.701446 | 0.705469 | 0.700648 | 0.69717 | 0.707345 | 0.706557 | 0.705654 |
| 0.711661 | 0.707665 | 0.705559 | 0.706948 | 0.706813 | 0.701239 | 0.707535 | 0.709581 | 0.709764 |
| 0.71306 | 0.706566 | 0.706019 | 0.70827 | 0.707729 | 0.701934 | 0.709855 | 0.712943 | 0.705302 |
| 0.709384 | 0.702909 | 0.701076 | 0.705072 | 0.704379 | 0.700441 | 0.703278 | 0.70759 | 0.70561 |
| 0.70368 | 0.702849 | 0.698164 | 0.701998 | 0.700439 | 0.694773 | 0.701312 | 0.698492 | 0.702435 |
| 0.699303 | 0.698561 | 0.696808 | 0.700008 | 0.698923 | 0.692939 | 0.701281 | 0.700615 | 0.698797 |
| 0.701532 | 0.699458 | 0.699185 | 0.698054 | 0.693924 | 0.693342 | 0.697549 | 0.697844 | 0.696769 |
| 0.699141 | 0.695199 | 0.698342 | 0.699246 | 0.694858 | 0.691358 | 0.703786 | 0.699705 | 0.699581 |
| 0.707741 | 0.697194 | 0.70005 | 0.698178 | 0.703577 | 0.694246 | 0.703458 | 0.69917 | 0.706648 |
| 0.694766 | 0.691205 | 0.689076 | 0.699342 | 0.691678 | 0.69003 | 0.690896 | 0.692154 | 0.694701 |
| 0.707523 | 0.701429 | 0.701547 | 0.702624 | 0.703626 | 0.700067 | 0.703878 | 0.704322 | 0.703047 |
| 0.700359 | 0.69408 | 0.696057 | 0.697391 | 0.696928 | 0.69186 | 0.69879 | 0.696931 | 0.699796 |

| | | | | | | | | |
|---|---|---|---|---|---|---|---|---|
| 0.715902 | 0.711882 | 0.709197 | 0.709922 | 0.707012 | 0.704222 | 0.710484 | 0.712646 | 0.711469 |
| 0.712637 | 0.706829 | 0.712216 | 0.711364 | 0.709604 | 0.70656 | 0.71021 | 0.713548 | 0.709283 |
| 0.70781 | 0.706204 | 0.707643 | 0.707376 | 0.702509 | 0.703742 | 0.707194 | 0.705443 | 0.708867 |
| 0.713112 | 0.708041 | 0.709741 | 0.717262 | 0.709754 | 0.706454 | 0.709885 | 0.711581 | 0.711323 |
| 0.718017 | 0.7161 | 0.717778 | 0.717914 | 0.715305 | 0.708847 | 0.714068 | 0.721592 | 0.716373 |
| 0.712776 | 0.708656 | 0.714149 | 0.714434 | 0.709549 | 0.704198 | 0.71123 | 0.713038 | 0.712015 |
| 0.709338 | 0.701121 | 0.706302 | 0.710032 | 0.704214 | 0.700619 | 0.702842 | 0.705625 | 0.708679 |
| 0.713725 | 0.706116 | 0.71259 | 0.712682 | 0.709088 | 0.705962 | 0.707867 | 0.712526 | 0.708755 |
| 0.70748 | 0.701642 | 0.70413 | 0.708819 | 0.702549 | 0.699254 | 0.698942 | 0.705065 | 0.705598 |
| 0.710162 | 0.708755 | 0.712112 | 0.71 | 0.710157 | 0.704473 | 0.705402 | 0.70898 | 0.707233 |
| 0.712538 | 0.708415 | 0.71137 | 0.714263 | 0.710109 | 0.705799 | 0.709736 | 0.712902 | 0.712335 |
| 0.714722 | 0.70881 | 0.711554 | 0.713282 | 0.711543 | 0.705663 | 0.709445 | 0.71399 | 0.712205 |
| 0.71369 | 0.708254 | 0.712573 | 0.710407 | 0.707513 | 0.703334 | 0.707284 | 0.709064 | 0.707601 |
| 0.712979 | 0.709649 | 0.711329 | 0.71432 | 0.707247 | 0.702773 | 0.711177 | 0.709798 | 0.708293 |
| 0.71358 | 0.709011 | 0.712305 | 0.713719 | 0.709961 | 0.707761 | 0.710009 | 0.712457 | 0.713897 |
| 0.713829 | 0.710179 | 0.713738 | 0.710901 | 0.712643 | 0.705915 | 0.708566 | 0.712155 | 0.710277 |
| 0.713425 | 0.711484 | 0.71424 | 0.712683 | 0.709484 | 0.706979 | 0.712194 | 0.714184 | 0.715348 |
| 0.712375 | 0.708582 | 0.710878 | 0.709437 | 0.710296 | 0.706952 | 0.709609 | 0.708562 | 0.712085 |
| 0.715385 | 0.711584 | 0.713921 | 0.714308 | 0.709919 | 0.708901 | 0.712252 | 0.716286 | 0.712839 |
| 0.712511 | 0.709502 | 0.711102 | 0.710946 | 0.707725 | 0.706741 | 0.710063 | 0.707801 | 0.708027 |
| 0.714562 | 0.708902 | 0.709892 | 0.711001 | 0.708658 | 0.708267 | 0.709703 | 0.710863 | 0.711129 |
| 0.712622 | 0.708463 | 0.71063 | 0.709507 | 0.706989 | 0.703793 | 0.708411 | 0.708497 | 0.710571 |
| 0.715632 | 0.711642 | 0.713968 | 0.714637 | 0.709821 | 0.707814 | 0.717294 | 0.71204 | 0.715591 |
| 0.7149 | 0.709045 | 0.711391 | 0.709971 | 0.713229 | 0.705623 | 0.709972 | 0.714195 | 0.713302 |
| 0.712489 | 0.708947 | 0.710576 | 0.708723 | 0.710327 | 0.706113 | 0.707369 | 0.711226 | 0.710995 |
| 0.711757 | 0.70689 | 0.708752 | 0.70829 | 0.710114 | 0.705617 | 0.709592 | 0.708069 | 0.709354 |
| 0.709465 | 0.705341 | 0.707051 | 0.708604 | 0.709329 | 0.703584 | 0.704686 | 0.704504 | 0.708217 |
| 0.706679 | 0.707839 | 0.708208 | 0.705437 | 0.706797 | 0.703147 | 0.709385 | 0.706442 | 0.710148 |
| 0.714422 | 0.709052 | 0.711236 | 0.711511 | 0.715184 | 0.708064 | 0.714274 | 0.709581 | 0.712905 |
| 0.713519 | 0.708561 | 0.715812 | 0.714485 | 0.714992 | 0.707826 | 0.712907 | 0.716772 | 0.713814 |
| 0.713344 | 0.707012 | 0.709547 | 0.71072 | 0.709011 | 0.704853 | 0.711347 | 0.710654 | 0.708221 |
| 0.719031 | 0.715135 | 0.714998 | 0.716414 | 0.712343 | 0.708486 | 0.712404 | 0.711395 | 0.713602 |
| 0.716909 | 0.712255 | 0.714047 | 0.712937 | 0.713932 | 0.706834 | 0.710188 | 0.713527 | 0.711109 |
| 0.707367 | 0.704042 | 0.707265 | 0.706573 | 0.706222 | 0.698981 | 0.70436 | 0.709173 | 0.706287 |
| 0.712447 | 0.709993 | 0.710239 | 0.710377 | 0.712462 | 0.705638 | 0.708856 | 0.712343 | 0.712391 |
| 0.707293 | 0.705549 | 0.707234 | 0.706687 | 0.708093 | 0.703039 | 0.706674 | 0.704802 | 0.707407 |
| 0.722401 | 0.715648 | 0.719082 | 0.717294 | 0.717339 | 0.712256 | 0.718818 | 0.718308 | 0.716159 |
| 0.714988 | 0.706053 | 0.711071 | 0.70945 | 0.712135 | 0.707589 | 0.711022 | 0.710356 | 0.710045 |
| 0.715341 | 0.709416 | 0.710203 | 0.707346 | 0.708533 | 0.705537 | 0.708504 | 0.713341 | 0.708186 |
| 0.713911 | 0.708842 | 0.711333 | 0.714251 | 0.710812 | 0.709193 | 0.710468 | 0.71114 | 0.711778 |
| 0.717348 | 0.711274 | 0.715309 | 0.718999 | 0.716568 | 0.712938 | 0.716376 | 0.712502 | 0.71557 |
| 0.717519 | 0.712547 | 0.712816 | 0.712998 | 0.713268 | 0.708419 | 0.712936 | 0.711659 | 0.711645 |
| 0.720317 | 0.710495 | 0.716316 | 0.713403 | 0.715002 | 0.710016 | 0.712409 | 0.711249 | 0.711072 |
| 0.715421 | 0.710015 | 0.705816 | 0.710336 | 0.707761 | 0.704869 | 0.707908 | 0.707925 | 0.710095 |
| 0.716133 | 0.707492 | 0.712196 | 0.715458 | 0.71455 | 0.707851 | 0.711002 | 0.710866 | 0.71306 |
| 0.712079 | 0.706515 | 0.706516 | 0.71006 | 0.707636 | 0.704026 | 0.705863 | 0.708525 | 0.706595 |

| | | | | | | | | |
|---|---|---|---|---|---|---|---|---|
| 0.701182 | 0.695867 | 0.699432 | 0.697992 | 0.696709 | 0.692055 | 0.693647 | 0.693809 | 0.701091 |
| 0.752671 | 0.748567 | 0.751587 | 0.746422 | 0.744349 | 0.739028 | 0.747581 | 0.744951 | 0.746332 |
| 0.709831 | 0.70781 | 0.709477 | 0.707668 | 0.709865 | 0.701693 | 0.707213 | 0.707536 | 0.709915 |
| 0.717327 | 0.712267 | 0.714524 | 0.715306 | 0.713888 | 0.706875 | 0.706447 | 0.715844 | 0.713167 |
| 0.7089 | 0.704845 | 0.70469 | 0.706286 | 0.707412 | 0.701118 | 0.707581 | 0.704936 | 0.708595 |
| 0.710364 | 0.706765 | 0.708501 | 0.707458 | 0.708051 | 0.703416 | 0.711234 | 0.709088 | 0.711028 |
| 0.710201 | 0.70428 | 0.706 | 0.707602 | 0.70716 | 0.701588 | 0.705029 | 0.705677 | 0.704414 |
| 0.706259 | 0.699795 | 0.702389 | 0.705426 | 0.705066 | 0.69543 | 0.700974 | 0.699991 | 0.702242 |
| 0.712942 | 0.70519 | 0.711115 | 0.711069 | 0.707149 | 0.703721 | 0.708044 | 0.711145 | 0.706479 |
| 0.720757 | 0.714537 | 0.714469 | 0.716029 | 0.716791 | 0.711752 | 0.713998 | 0.71408 | 0.716218 |
| 0.715084 | 0.70755 | 0.710733 | 0.714195 | 0.713137 | 0.706793 | 0.708602 | 0.710385 | 0.710444 |
| 0.718177 | 0.711216 | 0.713071 | 0.713186 | 0.715483 | 0.709732 | 0.711283 | 0.714229 | 0.711131 |
| 0.716439 | 0.715139 | 0.717388 | 0.713943 | 0.711794 | 0.71257 | 0.716113 | 0.717904 | 0.714286 |
| 0.720324 | 0.710558 | 0.713286 | 0.715522 | 0.715662 | 0.708458 | 0.710682 | 0.714206 | 0.711892 |
| 0.714684 | 0.705963 | 0.709565 | 0.710612 | 0.709221 | 0.704872 | 0.708245 | 0.709432 | 0.711288 |
| 0.720267 | 0.709655 | 0.711365 | 0.709078 | 0.712686 | 0.706633 | 0.708353 | 0.712058 | 0.708591 |
| 0.717584 | 0.714149 | 0.712854 | 0.713552 | 0.712174 | 0.706629 | 0.710605 | 0.713227 | 0.713626 |
| 0.712623 | 0.71083 | 0.71279 | 0.711599 | 0.710594 | 0.706136 | 0.709735 | 0.712923 | 0.709926 |
| 0.703488 | 0.704823 | 0.706025 | 0.701876 | 0.707046 | 0.699218 | 0.704593 | 0.709624 | 0.708999 |
| 0.709094 | 0.703273 | 0.702825 | 0.705793 | 0.702666 | 0.696311 | 0.702253 | 0.70394 | 0.705528 |
| 0.716729 | 0.707733 | 0.711893 | 0.706545 | 0.708306 | 0.705901 | 0.70503 | 0.709633 | 0.710102 |
| 0.699318 | 0.700734 | 0.701889 | 0.702431 | 0.701799 | 0.691481 | 0.69748 | 0.699292 | 0.705131 |
| 0.711893 | 0.708383 | 0.706769 | 0.709089 | 0.708629 | 0.697485 | 0.705722 | 0.709074 | 0.712479 |
| 0.713906 | 0.708669 | 0.70821 | 0.712924 | 0.710229 | 0.704455 | 0.709488 | 0.713315 | 0.713222 |
| 0.713546 | 0.71213 | 0.711612 | 0.7134 | 0.708643 | 0.713095 | 0.709463 | 0.713075 | 0.710262 |
| 0.713212 | 0.706595 | 0.710864 | 0.709278 | 0.709725 | 0.704105 | 0.704663 | 0.710698 | 0.709767 |
| 0.706992 | 0.707778 | 0.704231 | 0.703445 | 0.707571 | 0.701059 | 0.707422 | 0.708813 | 0.706753 |
| 0.705687 | 0.704156 | 0.704854 | 0.705941 | 0.705022 | 0.698831 | 0.703405 | 0.705664 | 0.703571 |
| 0.704742 | 0.706209 | 0.708756 | 0.706483 | 0.703863 | 0.699648 | 0.704172 | 0.704563 | 0.70663 |
| 0.764176 | 0.762474 | 0.761025 | 0.767939 | 0.760286 | 0.764913 | 0.773645 | 0.79204 | 0.765362 |
| 0.784145 | 0.774421 | 0.785349 | 0.785332 | 0.771338 | 0.773228 | 0.777927 | 0.816477 | 0.798381 |
| 0.779127 | 0.769454 | 0.784663 | 0.781472 | 0.780151 | 0.778886 | 0.798272 | 0.79467 | 0.782106 |
| 0.76967 | 0.768183 | 0.771216 | 0.76802 | 0.774661 | 0.767782 | 0.785998 | 0.766292 | 0.77888 |
| 0.791563 | 0.774386 | 0.783549 | 0.784689 | 0.782048 | 0.781805 | 0.797294 | 0.777614 | 0.812083 |
| 0.701546 | 0.701113 | 0.703238 | 0.702469 | 0.704168 | 0.696483 | 0.701512 | 0.704749 | 0.702075 |
| 0.709174 | 0.705944 | 0.705759 | 0.705664 | 0.707112 | 0.701302 | 0.702305 | 0.707258 | 0.706711 |
| 0.706653 | 0.705166 | 0.708283 | 0.706048 | 0.710064 | 0.702425 | 0.705645 | 0.709812 | 0.711448 |
| 0.707755 | 0.708536 | 0.712953 | 0.710385 | 0.706279 | 0.703255 | 0.706181 | 0.711833 | 0.710705 |
| 0.707362 | 0.708661 | 0.708772 | 0.704381 | 0.708827 | 0.703271 | 0.705617 | 0.709623 | 0.710507 |
| 0.710953 | 0.71249 | 0.710885 | 0.709864 | 0.712103 | 0.704614 | 0.709579 | 0.712617 | 0.711774 |
| 0.709095 | 0.704761 | 0.708961 | 0.709029 | 0.710059 | 0.699188 | 0.707301 | 0.709449 | 0.705705 |
| 0.702345 | 0.707132 | 0.704862 | 0.707181 | 0.706939 | 0.702569 | 0.706616 | 0.708455 | 0.706757 |
| 0.701398 | 0.699446 | 0.701496 | 0.701955 | 0.698779 | 0.694454 | 0.700711 | 0.699239 | 0.700449 |
| 0.698993 | 0.699906 | 0.698144 | 0.702582 | 0.703309 | 0.69618 | 0.697531 | 0.703196 | 0.698915 |
| 0.707235 | 0.709714 | 0.706422 | 0.706041 | 0.706442 | 0.701694 | 0.706807 | 0.707888 | 0.705378 |
| 0.712148 | 0.708276 | 0.708926 | 0.709907 | 0.708992 | 0.705176 | 0.70679 | 0.710944 | 0.707692 |

| | | | | | | | | |
|---|---|---|---|---|---|---|---|---|
| 0.710431 | 0.709627 | 0.711414 | 0.708806 | 0.710836 | 0.706759 | 0.707879 | 0.710483 | 0.710526 |
| 0.708776 | 0.706985 | 0.709654 | 0.709652 | 0.708839 | 0.705652 | 0.705282 | 0.710954 | 0.706834 |
| 0.715276 | 0.709667 | 0.716107 | 0.712272 | 0.712361 | 0.707783 | 0.711462 | 0.711616 | 0.713492 |
| 0.708521 | 0.705033 | 0.708771 | 0.712451 | 0.706558 | 0.70316 | 0.709005 | 0.709389 | 0.70659 |
| 0.704903 | 0.700357 | 0.705767 | 0.706594 | 0.703764 | 0.698133 | 0.70415 | 0.701746 | 0.705611 |
| 0.705502 | 0.704792 | 0.707204 | 0.706675 | 0.70507 | 0.701324 | 0.707374 | 0.70642 | 0.709064 |
| 0.712498 | 0.710189 | 0.710918 | 0.711321 | 0.711387 | 0.703526 | 0.70758 | 0.710834 | 0.713185 |
| 0.710126 | 0.70619 | 0.708565 | 0.71139 | 0.707617 | 0.704588 | 0.707054 | 0.709403 | 0.7066 |
| 0.707485 | 0.704294 | 0.706479 | 0.705092 | 0.704911 | 0.701032 | 0.703755 | 0.707893 | 0.708881 |
| 0.71088 | 0.705514 | 0.710779 | 0.712127 | 0.709912 | 0.70635 | 0.707042 | 0.709103 | 0.710252 |
| 0.70796 | 0.705777 | 0.70926 | 0.708249 | 0.705744 | 0.701929 | 0.707066 | 0.708644 | 0.709934 |
| 0.708082 | 0.708878 | 0.706769 | 0.708971 | 0.705815 | 0.701359 | 0.708047 | 0.7076 | 0.706771 |
| 0.698522 | 0.69823 | 0.701493 | 0.698354 | 0.699422 | 0.695552 | 0.702215 | 0.699191 | 0.702252 |
| 0.702152 | 0.702188 | 0.70164 | 0.701965 | 0.701058 | 0.695047 | 0.698526 | 0.700272 | 0.703034 |
| 0.712689 | 0.706299 | 0.711691 | 0.709299 | 0.710468 | 0.704038 | 0.711583 | 0.710481 | 0.712587 |
| 0.719644 | 0.714733 | 0.714228 | 0.712454 | 0.712617 | 0.705366 | 0.717307 | 0.717897 | 0.716257 |
| 0.716313 | 0.709639 | 0.710306 | 0.709398 | 0.70966 | 0.705517 | 0.712512 | 0.714974 | 0.711502 |
| 0.717442 | 0.714206 | 0.715764 | 0.715693 | 0.715494 | 0.708915 | 0.713284 | 0.713636 | 0.710772 |
| 0.708825 | 0.704716 | 0.705382 | 0.704395 | 0.707119 | 0.698146 | 0.706294 | 0.70611 | 0.707906 |
| 0.710823 | 0.705293 | 0.708533 | 0.709222 | 0.70475 | 0.701541 | 0.709055 | 0.712436 | 0.70618 |
| 0.70499 | 0.703884 | 0.706053 | 0.70579 | 0.706152 | 0.70123 | 0.702018 | 0.707854 | 0.703627 |
| 0.713139 | 0.705826 | 0.711733 | 0.711871 | 0.708735 | 0.703258 | 0.710757 | 0.708698 | 0.713597 |
| 0.71412 | 0.712515 | 0.711933 | 0.710796 | 0.710539 | 0.703139 | 0.713799 | 0.711236 | 0.71671 |
| 0.708229 | 0.704391 | 0.707104 | 0.708383 | 0.707781 | 0.702719 | 0.703196 | 0.707542 | 0.704711 |
| 0.711073 | 0.707847 | 0.708247 | 0.708426 | 0.711245 | 0.701102 | 0.708542 | 0.705295 | 0.711161 |
| 0.783012 | 0.793719 | 0.777461 | 0.779837 | 0.779671 | 0.779468 | 0.788917 | 0.788392 | 0.778927 |
| 0.711881 | 0.705022 | 0.708677 | 0.706856 | 0.708519 | 0.703809 | 0.709551 | 0.711146 | 0.71276 |
| 0.712321 | 0.70948 | 0.71109 | 0.710166 | 0.712836 | 0.70667 | 0.708639 | 0.711407 | 0.712929 |
| 0.715593 | 0.713368 | 0.715197 | 0.710275 | 0.712637 | 0.707853 | 0.710014 | 0.714469 | 0.717326 |
| 0.714067 | 0.708407 | 0.708651 | 0.710493 | 0.709623 | 0.705319 | 0.708692 | 0.710552 | 0.710027 |
| 0.710599 | 0.701769 | 0.705317 | 0.702588 | 0.707528 | 0.704589 | 0.705069 | 0.705056 | 0.708358 |
| 0.709126 | 0.703543 | 0.70833 | 0.707285 | 0.7058 | 0.700819 | 0.707165 | 0.703858 | 0.710113 |
| 0.713677 | 0.707539 | 0.710003 | 0.709858 | 0.708357 | 0.703812 | 0.710488 | 0.706645 | 0.712958 |
| 0.709729 | 0.704154 | 0.704291 | 0.705318 | 0.706582 | 0.698986 | 0.707622 | 0.705677 | 0.705436 |
| 0.709382 | 0.703297 | 0.701345 | 0.705049 | 0.704572 | 0.700033 | 0.702776 | 0.703598 | 0.704407 |
| 0.708303 | 0.705518 | 0.709213 | 0.705024 | 0.707025 | 0.702206 | 0.706153 | 0.706791 | 0.704713 |
| 0.71848 | 0.708709 | 0.710494 | 0.714997 | 0.708686 | 0.706513 | 0.712797 | 0.711682 | 0.712607 |
| 0.712205 | 0.708852 | 0.713167 | 0.714182 | 0.709082 | 0.704812 | 0.710605 | 0.713381 | 0.715542 |
| 0.712491 | 0.708511 | 0.710741 | 0.712738 | 0.70957 | 0.704197 | 0.708117 | 0.709395 | 0.712163 |
| 0.709937 | 0.703249 | 0.711703 | 0.711903 | 0.705843 | 0.704585 | 0.707321 | 0.712017 | 0.708071 |
| 0.709796 | 0.703781 | 0.705725 | 0.70652 | 0.704273 | 0.699313 | 0.704394 | 0.701718 | 0.703202 |
| 0.716228 | 0.712917 | 0.716465 | 0.715681 | 0.70927 | 0.709343 | 0.71015 | 0.710277 | 0.712587 |
| 0.712295 | 0.705917 | 0.705056 | 0.705677 | 0.705657 | 0.705692 | 0.707289 | 0.707012 | 0.707683 |
| 0.710265 | 0.706613 | 0.711791 | 0.708794 | 0.709947 | 0.706805 | 0.705114 | 0.714569 | 0.70779 |
| 0.711607 | 0.713189 | 0.713804 | 0.709065 | 0.71279 | 0.706182 | 0.712293 | 0.710636 | 0.715258 |
| 0.714769 | 0.712201 | 0.71448 | 0.710394 | 0.711973 | 0.710539 | 0.713137 | 0.715494 | 0.715025 |

| | | | | | | | | |
|---|---|---|---|---|---|---|---|---|
| 0.710948 | 0.709396 | 0.711288 | 0.707296 | 0.707846 | 0.70745 | 0.711372 | 0.711911 | 0.707796 |
| 0.710002 | 0.711726 | 0.713569 | 0.710597 | 0.708977 | 0.707759 | 0.711532 | 0.710766 | 0.711909 |
| 0.710514 | 0.707724 | 0.710828 | 0.705953 | 0.706806 | 0.705246 | 0.708643 | 0.711477 | 0.708215 |
| 0.669741 | 0.667017 | 0.677351 | 0.668763 | 0.668531 | 0.665059 | 0.673305 | 0.669583 | 0.651826 |
| 0.710188 | 0.710152 | 0.709429 | 0.705976 | 0.706694 | 0.703914 | 0.710636 | 0.711231 | 0.709422 |
| 0.710389 | 0.708327 | 0.707928 | 0.711408 | 0.711697 | 0.704491 | 0.707919 | 0.712466 | 0.710635 |
| 0.712912 | 0.712045 | 0.7106 | 0.708516 | 0.711784 | 0.706511 | 0.709367 | 0.712742 | 0.709097 |
| 0.710722 | 0.712801 | 0.710489 | 0.709228 | 0.713351 | 0.703955 | 0.712628 | 0.710331 | 0.713231 |
| 0.718013 | 0.715569 | 0.715642 | 0.714263 | 0.714445 | 0.711836 | 0.714469 | 0.716369 | 0.714792 |
| 0.716696 | 0.711279 | 0.71368 | 0.711106 | 0.710967 | 0.707369 | 0.711028 | 0.710579 | 0.712824 |
| 0.713516 | 0.711056 | 0.709935 | 0.7112 | 0.71211 | 0.707736 | 0.708357 | 0.711352 | 0.713191 |
| 0.714965 | 0.713521 | 0.715317 | 0.714114 | 0.716817 | 0.71194 | 0.71643 | 0.717977 | 0.715997 |
| 0.715308 | 0.714661 | 0.716482 | 0.715601 | 0.71496 | 0.70914 | 0.714538 | 0.717691 | 0.714401 |
| 0.710295 | 0.710155 | 0.709775 | 0.708095 | 0.713066 | 0.705632 | 0.708756 | 0.709349 | 0.712007 |
| 0.715905 | 0.714035 | 0.713043 | 0.712415 | 0.715157 | 0.70939 | 0.710668 | 0.715116 | 0.711665 |
| 0.708718 | 0.710152 | 0.708589 | 0.707513 | 0.710079 | 0.706527 | 0.706165 | 0.710984 | 0.709446 |
| 0.712292 | 0.713103 | 0.713358 | 0.71122 | 0.710193 | 0.711227 | 0.710583 | 0.712643 | 0.71112 |
| 0.712573 | 0.711183 | 0.714548 | 0.713523 | 0.714457 | 0.705757 | 0.708739 | 0.715979 | 0.710782 |
| 0.707019 | 0.709362 | 0.709204 | 0.703827 | 0.709325 | 0.703317 | 0.701704 | 0.70631 | 0.708621 |
| 0.709592 | 0.707582 | 0.710619 | 0.708264 | 0.710831 | 0.702669 | 0.706854 | 0.711116 | 0.709193 |
| 0.713335 | 0.711143 | 0.713134 | 0.712255 | 0.708712 | 0.707944 | 0.712201 | 0.714286 | 0.713377 |
| 0.716049 | 0.710829 | 0.715038 | 0.71199 | 0.714252 | 0.711413 | 0.713667 | 0.713413 | 0.712484 |
| 0.692606 | 0.689285 | 0.702593 | 0.70549 | 0.693747 | 0.696341 | 0.690524 | 0.696246 | 0.692026 |
| 0.713866 | 0.732315 | 0.719363 | 0.719665 | 0.70952 | 0.721629 | 0.730523 | 0.729478 | 0.721497 |
| 0.753646 | 0.757242 | 0.7808 | 0.755006 | 0.758753 | 0.759015 | 0.754518 | 0.764851 | 0.749265 |
| 0.741958 | 0.736288 | 0.73502 | 0.736292 | 0.738805 | 0.735543 | 0.747889 | 0.746113 | 0.732119 |
| 0.666618 | 0.664937 | 0.670325 | 0.669474 | 0.669132 | 0.660753 | 0.666137 | 0.666185 | 0.672303 |
| 0.663931 | 0.657316 | 0.662362 | 0.663803 | 0.67026 | 0.658214 | 0.660447 | 0.66138 | 0.662797 |
| 0.659962 | 0.65884 | 0.668583 | 0.663226 | 0.658718 | 0.65376 | 0.668757 | 0.660446 | 0.668244 |
| 0.670043 | 0.67408 | 0.68612 | 0.68308 | 0.678968 | 0.673809 | 0.677491 | 0.670861 | 0.685499 |
| 0.653265 | 0.651619 | 0.659593 | 0.657684 | 0.651819 | 0.648725 | 0.656534 | 0.649072 | 0.661333 |
| 0.645223 | 0.644637 | 0.651581 | 0.642469 | 0.652316 | 0.640675 | 0.646883 | 0.642644 | 0.645504 |
| 0.644355 | 0.646623 | 0.65037 | 0.651263 | 0.644915 | 0.63672 | 0.650825 | 0.642028 | 0.639943 |
| 0.736557 | 0.735126 | 0.739251 | 0.72914 | 0.778452 | 0.722547 | 0.732591 | 0.748374 | 0.739881 |
| 0.70294 | 0.703824 | 0.705936 | 0.704344 | 0.705838 | 0.701329 | 0.701703 | 0.703414 | 0.705586 |
| 0.67728 | 0.67586 | 0.677173 | 0.681448 | 0.68367 | 0.669264 | 0.679986 | 0.678308 | 0.68033 |
| 0.668019 | 0.6726 | 0.668621 | 0.672194 | 0.671087 | 0.663018 | 0.666493 | 0.67157 | 0.671579 |
| 0.65529 | 0.658962 | 0.658891 | 0.66558 | 0.65601 | 0.647965 | 0.661453 | 0.659331 | 0.65666 |
| 0.680549 | 0.675824 | 0.680452 | 0.678802 | 0.682328 | 0.673672 | 0.678713 | 0.681117 | 0.68034 |
| 0.669661 | 0.668105 | 0.666378 | 0.672813 | 0.670629 | 0.665152 | 0.672768 | 0.669256 | 0.674065 |
| 0.658856 | 0.661656 | 0.662088 | 0.666151 | 0.664252 | 0.655466 | 0.66333 | 0.663809 | 0.660983 |
| 0.664945 | 0.666767 | 0.665812 | 0.668644 | 0.669177 | 0.661428 | 0.661698 | 0.667674 | 0.671714 |
| 0.657043 | 0.657548 | 0.656347 | 0.65687 | 0.659327 | 0.655285 | 0.656093 | 0.660024 | 0.654366 |
| 0.65694 | 0.657596 | 0.660819 | 0.661127 | 0.66102 | 0.657731 | 0.666032 | 0.662859 | 0.663029 |
| 0.654414 | 0.661051 | 0.662556 | 0.660345 | 0.664356 | 0.658907 | 0.654245 | 0.662713 | 0.652024 |
| 0.640845 | 0.648811 | 0.653048 | 0.648095 | 0.648967 | 0.642386 | 0.645008 | 0.641677 | 0.652077 |

| | | | | | | | | |
|---|---|---|---|---|---|---|---|---|
| 0.692048 | 0.695778 | 0.689956 | 0.692174 | 0.692953 | 0.687757 | 0.693249 | 0.694735 | 0.695707 |
| 0.660454 | 0.659131 | 0.662836 | 0.661825 | 0.66306 | 0.654611 | 0.660194 | 0.661484 | 0.662517 |
| 0.670025 | 0.661392 | 0.669022 | 0.665696 | 0.666086 | 0.662984 | 0.669241 | 0.665849 | 0.66565 |
| 0.65931 | 0.662162 | 0.670016 | 0.667608 | 0.669978 | 0.657306 | 0.668239 | 0.664501 | 0.668993 |
| 0.653362 | 0.660885 | 0.654078 | 0.661037 | 0.657967 | 0.65866 | 0.663069 | 0.658704 | 0.660841 |
| 0.66335 | 0.665065 | 0.663938 | 0.662715 | 0.660183 | 0.659929 | 0.666187 | 0.665388 | 0.660101 |
| 0.647695 | 0.658151 | 0.658048 | 0.661786 | 0.658697 | 0.648755 | 0.656594 | 0.652002 | 0.65049 |
| 0.654172 | 0.648568 | 0.651068 | 0.649575 | 0.649375 | 0.644885 | 0.653898 | 0.646964 | 0.644898 |
| 0.644542 | 0.646216 | 0.655323 | 0.648893 | 0.65851 | 0.645963 | 0.650815 | 0.656516 | 0.648501 |
| 0.692652 | 0.690995 | 0.695805 | 0.690025 | 0.690725 | 0.687435 | 0.693266 | 0.69321 | 0.689865 |
| 0.688874 | 0.687852 | 0.69156 | 0.689952 | 0.691623 | 0.683263 | 0.688232 | 0.691471 | 0.689193 |
| 0.682254 | 0.681605 | 0.685916 | 0.684742 | 0.687821 | 0.681084 | 0.684272 | 0.683633 | 0.685813 |
| 0.675873 | 0.669993 | 0.676226 | 0.673515 | 0.679709 | 0.6741 | 0.676914 | 0.678434 | 0.67673 |
| 0.677717 | 0.68185 | 0.68589 | 0.681538 | 0.684279 | 0.681375 | 0.681154 | 0.681818 | 0.68034 |
| 0.673038 | 0.669966 | 0.675323 | 0.671622 | 0.672338 | 0.66874 | 0.670412 | 0.674013 | 0.674432 |

| Collection_ | Collection_ | Collection_ | Collection_ | Collection_ | Collection_ | Collection_ | Collection_ | Collection_ |
|---|---|---|---|---|---|---|---|---|
| 0.783301 | 0.782534 | 0.775803 | 0.776248 | 0.771917 | 0.768615 | 0.767668 | 0.790493 | 0.796355 |
| 0.802756 | 0.784835 | 0.774312 | 0.783797 | 0.782102 | 0.779114 | 0.772405 | 0.790133 | 0.790607 |
| 0.781989 | 0.77717 | 0.774528 | 0.779118 | 0.777722 | 0.770677 | 0.775608 | 0.781765 | 0.79098 |
| 0.792315 | 0.774388 | 0.781365 | 0.781199 | 0.774049 | 0.776074 | 0.784054 | 0.785137 | 0.79637 |
| 0.77319 | 0.764482 | 0.771207 | 0.778424 | 0.772254 | 0.772806 | 0.766656 | 0.77506 | 0.784478 |
| 0.78768 | 0.766624 | 0.776581 | 0.778077 | 0.774996 | 0.778474 | 0.775817 | 0.771351 | 0.768806 |
| 0.718003 | 0.71187 | 0.715577 | 0.709657 | 0.713732 | 0.712878 | 0.711436 | 0.713527 | 0.714927 |
| 0.714049 | 0.708886 | 0.708416 | 0.71003 | 0.708897 | 0.709842 | 0.701713 | 0.709094 | 0.70459 |
| 0.804963 | 0.77961 | 0.777313 | 0.780488 | 0.785294 | 0.776118 | 0.777288 | 0.790287 | 0.779171 |
| 0.793996 | 0.785896 | 0.774054 | 0.773798 | 0.773804 | 0.770384 | 0.771719 | 0.78014 | 0.807975 |
| 0.782881 | 0.766171 | 0.769136 | 0.765398 | 0.762273 | 0.76622 | 0.768911 | 0.774875 | 0.800892 |
| 0.786782 | 0.77393 | 0.783012 | 0.785632 | 0.77814 | 0.786012 | 0.771155 | 0.782805 | 0.801676 |
| 0.790098 | 0.776622 | 0.770779 | 0.770762 | 0.777796 | 0.767177 | 0.77499 | 0.790801 | 0.78749 |
| 0.783832 | 0.769513 | 0.775029 | 0.771699 | 0.770726 | 0.77706 | 0.763158 | 0.779678 | 0.788881 |
| 0.780228 | 0.773758 | 0.766242 | 0.767969 | 0.771488 | 0.767872 | 0.768215 | 0.769472 | 0.767702 |
| 0.787272 | 0.781961 | 0.781368 | 0.77969 | 0.782365 | 0.774061 | 0.77516 | 0.777824 | 0.79008 |
| 0.770441 | 0.768152 | 0.771621 | 0.776459 | 0.766527 | 0.766067 | 0.770456 | 0.765725 | 0.803495 |
| 0.767861 | 0.759241 | 0.756614 | 0.758184 | 0.761394 | 0.752316 | 0.752617 | 0.758202 | 0.762903 |
| 0.799153 | 0.78621 | 0.773097 | 0.783997 | 0.779995 | 0.782453 | 0.772256 | 0.7978 | 0.785976 |
| 0.792193 | 0.781178 | 0.780446 | 0.779895 | 0.780596 | 0.780158 | 0.78268 | 0.792828 | 0.793587 |
| 0.786611 | 0.777441 | 0.777077 | 0.780418 | 0.780357 | 0.779374 | 0.780747 | 0.782478 | 0.787969 |
| 0.793528 | 0.787816 | 0.780138 | 0.782576 | 0.777861 | 0.7907 | 0.770376 | 0.789995 | 0.803507 |
| 0.803419 | 0.790918 | 0.785256 | 0.787309 | 0.787965 | 0.78801 | 0.78125 | 0.79909 | 0.794053 |
| 0.794006 | 0.78095 | 0.785673 | 0.788635 | 0.789283 | 0.785167 | 0.779545 | 0.788021 | 0.804734 |
| 0.789796 | 0.776408 | 0.783893 | 0.784603 | 0.777451 | 0.785362 | 0.772234 | 0.784054 | 0.785156 |
| 0.78831 | 0.779743 | 0.779113 | 0.781398 | 0.77571 | 0.771281 | 0.778237 | 0.779368 | 0.781271 |
| 0.789912 | 0.779949 | 0.77112 | 0.776872 | 0.774496 | 0.775234 | 0.770032 | 0.774923 | 0.773268 |
| 0.784991 | 0.779245 | 0.779646 | 0.780966 | 0.77848 | 0.777457 | 0.777913 | 0.789394 | 0.778798 |
| 0.783788 | 0.772014 | 0.779915 | 0.779915 | 0.77356 | 0.782815 | 0.778004 | 0.781765 | 0.781551 |
| 0.783964 | 0.77732 | 0.774951 | 0.774751 | 0.77596 | 0.770098 | 0.765471 | 0.788076 | 0.797618 |
| 0.787654 | 0.777358 | 0.781944 | 0.782217 | 0.775276 | 0.779337 | 0.773072 | 0.778252 | 0.781929 |
| 0.781401 | 0.773999 | 0.776315 | 0.779153 | 0.771459 | 0.780803 | 0.770926 | 0.774884 | 0.771027 |
| 0.792053 | 0.781079 | 0.773496 | 0.781942 | 0.772006 | 0.784448 | 0.776786 | 0.78899 | 0.780558 |
| 0.774799 | 0.768312 | 0.773668 | 0.776099 | 0.76654 | 0.774758 | 0.771609 | 0.777196 | 0.771728 |
| 0.77836 | 0.764792 | 0.772802 | 0.773514 | 0.768281 | 0.772607 | 0.766123 | 0.771747 | 0.770496 |
| 0.791467 | 0.783172 | 0.778402 | 0.779302 | 0.77746 | 0.782091 | 0.780294 | 0.781583 | 0.792572 |
| 0.784456 | 0.779354 | 0.77411 | 0.784436 | 0.778369 | 0.77742 | 0.776825 | 0.785054 | 0.77775 |
| 0.782014 | 0.771709 | 0.767872 | 0.769949 | 0.764676 | 0.773875 | 0.770292 | 0.773944 | 0.771518 |
| 0.787032 | 0.77649 | 0.772479 | 0.771461 | 0.775063 | 0.77088 | 0.771028 | 0.780909 | 0.778358 |

| | | | | | | | | |
|---|---|---|---|---|---|---|---|---|
| 0.757901 | 0.751897 | 0.75136 | 0.760605 | 0.755029 | 0.752289 | 0.750834 | 0.755612 | 0.777388 |
| 0.798859 | 0.78748 | 0.774307 | 0.773248 | 0.772913 | 0.772866 | 0.770751 | 0.791981 | 0.789532 |
| 0.78542 | 0.781389 | 0.784519 | 0.779926 | 0.778048 | 0.780894 | 0.772898 | 0.785456 | 0.814709 |
| 0.793256 | 0.7934 | 0.77939 | 0.781652 | 0.779403 | 0.784264 | 0.777454 | 0.796674 | 0.985718 |
| 0.795265 | 0.782948 | 0.780496 | 0.783815 | 0.780518 | 0.786017 | 0.775422 | 0.784975 | 0.795302 |
| 0.795475 | 0.777329 | 0.775794 | 0.781602 | 0.772175 | 0.781782 | 0.774159 | 0.78382 | 0.784826 |
| 0.77776 | 0.771557 | 0.775338 | 0.77827 | 0.77192 | 0.778266 | 0.773443 | 0.779186 | 0.773653 |
| 0.803284 | 0.793595 | 0.785288 | 0.788095 | 0.784182 | 0.786403 | 0.780783 | 0.801152 | 0.78747 |
| 0.784492 | 0.771396 | 0.779588 | 0.774367 | 0.771137 | 0.775509 | 0.7761 | 0.776119 | 0.774582 |
| 0.779791 | 0.770457 | 0.771857 | 0.772148 | 0.765334 | 0.767903 | 0.771237 | 0.772708 | 0.765617 |
| 0.793303 | 0.781941 | 0.778072 | 0.779759 | 0.771077 | 0.774161 | 0.771053 | 0.782974 | 0.798646 |
| 0.796768 | 0.785832 | 0.778499 | 0.781994 | 0.780223 | 0.777898 | 0.775217 | 0.790809 | 0.798647 |
| 0.79131 | 0.779962 | 0.779019 | 0.78051 | 0.780791 | 0.781559 | 0.781168 | 0.794078 | 0.805647 |
| 0.787042 | 0.780689 | 0.779219 | 0.783347 | 0.788205 | 0.779783 | 0.788809 | 0.785244 | 0.79832 |
| 0.788833 | 0.773844 | 0.775632 | 0.785506 | 0.776603 | 0.772952 | 0.778721 | 0.780894 | 0.800553 |
| 0.78266 | 0.770015 | 0.776454 | 0.783791 | 0.77514 | 0.773336 | 0.773579 | 0.781521 | 0.792806 |
| 0.77418 | 0.77266 | 0.768944 | 0.776341 | 0.776387 | 0.768961 | 0.775976 | 0.774653 | 0.800104 |
| 0.797749 | 0.791862 | 0.782004 | 0.787584 | 0.791225 | 0.777234 | 0.778212 | 0.800236 | 0.80657 |
| 0.795308 | 0.782369 | 0.77498 | 0.785767 | 0.78098 | 0.774845 | 0.776743 | 0.789982 | 0.800154 |
| 0.795557 | 0.782569 | 0.775215 | 0.777061 | 0.78506 | 0.775129 | 0.774524 | 0.787559 | 0.805294 |
| 0.778518 | 0.771304 | 0.776265 | 0.776441 | 0.77243 | 0.772655 | 0.777131 | 0.775522 | 0.795591 |
| 0.78701 | 0.782061 | 0.780051 | 0.772966 | 0.776361 | 0.768085 | 0.766482 | 0.788764 | 0.811256 |
| 0.802158 | 0.785317 | 0.776643 | 0.776046 | 0.7838 | 0.779006 | 0.778868 | 0.792895 | 0.782261 |
| 0.77786 | 0.768646 | 0.770068 | 0.778536 | 0.767572 | 0.778759 | 0.781821 | 0.777731 | 0.772497 |
| 0.780448 | 0.75968 | 0.765012 | 0.766303 | 0.762654 | 0.768034 | 0.764559 | 0.765293 | 0.762171 |
| 0.715518 | 0.709611 | 0.709608 | 0.710987 | 0.710197 | 0.711712 | 0.702639 | 0.70882 | 0.706013 |
| 0.717497 | 0.710027 | 0.716225 | 0.713393 | 0.713214 | 0.715675 | 0.70912 | 0.713414 | 0.709905 |
| 0.71854 | 0.711473 | 0.714 | 0.715133 | 0.714088 | 0.714736 | 0.713072 | 0.714541 | 0.713121 |
| 0.787864 | 0.776501 | 0.77964 | 0.773062 | 0.777304 | 0.765478 | 0.77236 | 0.784871 | 0.773057 |
| 0.791649 | 0.770403 | 0.782668 | 0.775446 | 0.783625 | 0.774912 | 0.764362 | 0.788717 | 0.773912 |
| 0.780526 | 0.778216 | 0.768562 | 0.770693 | 0.771402 | 0.765741 | 0.760104 | 0.780949 | 0.798388 |
| 0.797976 | 0.775004 | 0.777096 | 0.775959 | 0.783637 | 0.774615 | 0.768972 | 0.787059 | 0.774752 |
| 0.830655 | 0.806359 | 0.782239 | 0.786743 | 0.805563 | 0.785937 | 0.786726 | 0.840891 | 0.802644 |
| 0.786389 | 0.779268 | 0.788724 | 0.786077 | 0.780636 | 0.798729 | 0.777038 | 0.785461 | 0.816107 |
| 0.828806 | 0.80209 | 0.780219 | 0.787996 | 0.798955 | 0.784489 | 0.782897 | 0.818997 | 0.811874 |
| 0.783893 | 0.779526 | 0.777255 | 0.785596 | 0.778412 | 0.781263 | 0.782841 | 0.788773 | 0.804053 |
| 0.839383 | 0.834925 | 0.78257 | 0.777153 | 0.825095 | 0.786445 | 0.790703 | 0.821072 | 0.791413 |
| 0.82822 | 0.824439 | 0.8003 | 0.801935 | 0.813904 | 0.798321 | 0.799471 | 0.827274 | 0.833276 |
| 0.807352 | 0.796726 | 0.769303 | 0.773427 | 0.782363 | 0.772387 | 0.771264 | 0.814966 | 0.792806 |
| 0.782499 | 0.784643 | 0.78637 | 0.78213 | 0.785488 | 0.778563 | 0.782743 | 0.782432 | 0.791754 |
| 0.773263 | 0.780942 | 0.771333 | 0.781022 | 0.773522 | 0.778382 | 0.774703 | 0.786906 | 0.799949 |
| 0.83069 | 0.817663 | 0.803701 | 0.80972 | 0.808531 | 0.797609 | 0.799619 | 0.822804 | 0.826121 |
| 0.78076 | 0.779786 | 0.778226 | 0.786323 | 0.77404 | 0.788587 | 0.779948 | 0.784273 | 0.815629 |
| 0.784379 | 0.780057 | 0.784953 | 0.787297 | 0.774788 | 0.786229 | 0.779268 | 0.782175 | 0.818158 |
| 0.789966 | 0.78281 | 0.787695 | 0.782152 | 0.791526 | 0.788958 | 0.782047 | 0.788287 | 0.78456 |
| 0.783456 | 0.783815 | 0.780255 | 0.780676 | 0.774758 | 0.776756 | 0.782031 | 0.783777 | 0.791638 |

| | | | | | | | | |
|---|---|---|---|---|---|---|---|---|
| 0.786841 | 0.782319 | 0.778611 | 0.784469 | 0.783249 | 0.78163 | 0.777496 | 0.787701 | 0.808947 |
| 0.787272 | 0.78908 | 0.776389 | 0.779319 | 0.782984 | 0.779498 | 0.774986 | 0.778946 | 0.776001 |
| 0.788186 | 0.791177 | 0.775879 | 0.775733 | 0.797015 | 0.785942 | 0.7882 | 0.790914 | 0.789361 |
| 0.812811 | 0.814107 | 0.795742 | 0.794682 | 0.805339 | 0.794406 | 0.793212 | 0.813096 | 0.814083 |
| 0.809536 | 0.799654 | 0.782438 | 0.786037 | 0.800673 | 0.788729 | 0.787928 | 0.791677 | 0.789314 |
| 0.778743 | 0.779889 | 0.776536 | 0.860744 | 0.773124 | 0.777704 | 0.780184 | 0.780958 | 0.786226 |
| 0.856624 | 0.812278 | 0.780036 | 0.779658 | 0.810699 | 0.782064 | 0.77918 | 0.827881 | 0.795119 |
| 0.798353 | 0.790282 | 0.779147 | 0.774718 | 0.799386 | 0.784432 | 0.789957 | 0.780478 | 0.774561 |
| 0.786541 | 0.783717 | 0.787464 | 0.779738 | 0.783339 | 0.782085 | 0.776737 | 0.808394 | 0.779944 |
| 0.777042 | 0.771538 | 0.778408 | 0.784194 | 0.773582 | 0.773988 | 0.774701 | 0.77686 | 0.779674 |
| 0.778066 | 0.780458 | 0.779028 | 0.780842 | 0.777317 | 0.780369 | 0.77919 | 0.786261 | 0.782762 |
| 0.783568 | 0.782128 | 0.78005 | 0.781563 | 0.781057 | 0.789123 | 0.782216 | 0.77927 | 0.78632 |
| 0.779686 | 0.77567 | 0.784807 | 0.790092 | 0.778538 | 0.781592 | 0.777038 | 0.777199 | 0.791049 |
| 0.773669 | 0.768683 | 0.771833 | 0.777008 | 0.7751 | 0.777843 | 0.775923 | 0.778455 | 0.783335 |
| 0.784961 | 0.777248 | 0.777759 | 0.788095 | 0.778533 | 0.78087 | 0.791845 | 0.780062 | 0.779419 |
| 0.783181 | 0.780654 | 0.775615 | 0.781294 | 0.777833 | 0.781847 | 0.775897 | 0.775089 | 0.772613 |
| 0.794501 | 0.781529 | 0.778771 | 0.778045 | 0.77496 | 0.775587 | 0.779165 | 0.781389 | 0.77897 |
| 0.774328 | 0.77482 | 0.776629 | 0.783092 | 0.773844 | 0.776516 | 0.774317 | 0.780768 | 0.791773 |
| 0.790088 | 0.781757 | 0.778592 | 0.789177 | 0.778989 | 0.78757 | 0.778834 | 0.781946 | 0.788416 |
| 0.784562 | 0.7731 | 0.781557 | 0.775713 | 0.778794 | 0.777612 | 0.777405 | 0.789761 | 0.773918 |
| 0.782322 | 0.785207 | 0.774959 | 0.778428 | 0.780504 | 0.809311 | 0.781209 | 0.78194 | 0.780643 |
| 0.784455 | 0.777732 | 0.775293 | 0.781736 | 0.771388 | 0.77763 | 0.781768 | 0.781524 | 0.778069 |
| 0.78109 | 0.785655 | 0.76835 | 0.77605 | 0.777946 | 0.779597 | 0.784199 | 0.78628 | 0.785355 |
| 0.776542 | 0.780142 | 0.772973 | 0.779027 | 0.779633 | 0.785768 | 0.784349 | 0.776528 | 0.78185 |
| 0.82182 | 0.893781 | 0.772048 | 0.775339 | 0.849024 | 0.777666 | 0.775332 | 0.8 | 0.784523 |
| 1 | 0.823987 | 0.787335 | 0.781242 | 0.81167 | 0.783155 | 0.782527 | 0.815594 | 0.788762 |
| 0.823987 | 1 | 0.772446 | 0.780346 | 0.831015 | 0.781488 | 0.780765 | 0.8026 | 0.795476 |
| 0.787335 | 0.772446 | 1 | 0.774376 | 0.775154 | 0.774092 | 0.776586 | 0.781146 | 0.775076 |
| 0.781242 | 0.780346 | 0.774376 | 1 | 0.774767 | 0.77937 | 0.782747 | 0.782061 | 0.780494 |
| 0.81167 | 0.831015 | 0.775154 | 0.774767 | 1 | 0.783499 | 0.780781 | 0.807235 | 0.780383 |
| 0.783155 | 0.781488 | 0.774092 | 0.77937 | 0.783499 | 1 | 0.784446 | 0.777871 | 0.781561 |
| 0.782527 | 0.780765 | 0.776586 | 0.782747 | 0.780781 | 0.784446 | 1 | 0.779385 | 0.77721 |
| 0.815594 | 0.8026 | 0.781146 | 0.782061 | 0.807235 | 0.777871 | 0.779385 | 1 | 0.798534 |
| 0.788762 | 0.795476 | 0.775076 | 0.780494 | 0.780383 | 0.781561 | 0.77721 | 0.798534 | 1 |
| 0.779375 | 0.780933 | 0.777006 | 0.784935 | 0.784929 | 0.787652 | 0.778171 | 0.782346 | 0.791396 |
| 0.784003 | 0.782442 | 0.77165 | 0.779982 | 0.779655 | 0.78495 | 0.774506 | 0.782169 | 0.793002 |
| 0.776897 | 0.781502 | 0.778507 | 0.778633 | 0.778724 | 0.783525 | 0.780522 | 0.784404 | 0.782449 |
| 0.751251 | 0.761466 | 0.746717 | 0.751203 | 0.76041 | 0.758418 | 0.754322 | 0.762107 | 0.759724 |
| 0.759775 | 0.757994 | 0.762109 | 0.759242 | 0.76058 | 0.764694 | 0.761519 | 0.766103 | 0.770504 |
| 0.765214 | 0.749921 | 0.750744 | 0.761027 | 0.755803 | 0.745209 | 0.75103 | 0.764668 | 0.752095 |
| 0.758865 | 0.765767 | 0.758756 | 0.76279 | 0.764487 | 0.765923 | 0.753673 | 0.763558 | 0.781445 |
| 0.741348 | 0.743911 | 0.743338 | 0.744355 | 0.746163 | 0.771197 | 0.737713 | 0.742972 | 0.77608 |
| 0.751409 | 0.755751 | 0.748366 | 0.756842 | 0.75101 | 0.756501 | 0.749469 | 0.761626 | 0.799171 |
| 0.759975 | 0.762241 | 0.749466 | 0.751889 | 0.753708 | 0.752017 | 0.754641 | 0.763302 | 0.762712 |
| 0.760939 | 0.745767 | 0.737701 | 0.749107 | 0.746034 | 0.753504 | 0.752971 | 0.762051 | 0.753105 |
| 0.796387 | 0.777778 | 0.751813 | 0.75502 | 0.771875 | 0.750453 | 0.757025 | 0.78216 | 0.772398 |
| 0.759687 | 0.777433 | 0.755841 | 0.758984 | 0.749807 | 0.753523 | 0.749395 | 0.758594 | 0.769948 |

| | | | | | | | | |
|---|---|---|---|---|---|---|---|---|
| 0.743837 | 0.745899 | 0.738195 | 0.739726 | 0.743174 | 0.738866 | 0.737825 | 0.749616 | 0.742758 |
| 0.811219 | 0.79633 | 0.751558 | 0.758672 | 0.780003 | 0.750979 | 0.756324 | 0.803631 | 0.784666 |
| 0.746273 | 0.74449 | 0.739089 | 0.749931 | 0.745746 | 0.742493 | 0.740609 | 0.747632 | 0.742129 |
| 0.742476 | 0.753053 | 0.735867 | 0.741375 | 0.748694 | 0.742408 | 0.746058 | 0.751384 | 0.781075 |
| 0.805468 | 0.795402 | 0.746432 | 0.751609 | 0.781675 | 0.769303 | 0.742993 | 0.801292 | 0.790078 |
| 0.770092 | 0.774338 | 0.74052 | 0.737402 | 0.795074 | 0.759376 | 0.762299 | 0.755572 | 0.747352 |
| 0.791842 | 0.767794 | 0.73775 | 0.746321 | 0.817767 | 0.73934 | 0.747587 | 0.776529 | 0.756681 |
| 0.752924 | 0.756883 | 0.746802 | 0.754132 | 0.747152 | 0.757886 | 0.745217 | 0.754739 | 0.770932 |
| 0.725275 | 0.733162 | 0.71898 | 0.731533 | 0.733273 | 0.739763 | 0.726673 | 0.727148 | 0.744378 |
| 0.851686 | 0.818752 | 0.746434 | 0.749733 | 0.809262 | 0.747579 | 0.749093 | 0.847712 | 0.780216 |
| 0.75729 | 0.758303 | 0.743126 | 0.742372 | 0.759972 | 0.744951 | 0.763176 | 0.743729 | 0.75092 |
| 0.775055 | 0.772201 | 0.767934 | 0.768919 | 0.76878 | 0.763795 | 0.761183 | 0.773821 | 0.790206 |
| 0.770307 | 0.773838 | 0.772406 | 0.774309 | 0.772916 | 0.772183 | 0.775702 | 0.781629 | 0.774334 |
| 0.786105 | 0.75632 | 0.752237 | 0.755911 | 0.744208 | 0.752292 | 0.757778 | 0.758175 | 0.756568 |
| 0.77153 | 0.770364 | 0.772746 | 0.771483 | 0.768933 | 0.776594 | 0.76968 | 0.779642 | 0.786315 |
| 0.770832 | 0.765656 | 0.772161 | 0.769969 | 0.76452 | 0.769243 | 0.77345 | 0.770594 | 0.774804 |
| 0.775364 | 0.775083 | 0.769306 | 0.779679 | 0.773071 | 0.772226 | 0.773677 | 0.780652 | 0.774981 |
| 0.79092 | 0.773675 | 0.781899 | 0.777161 | 0.772421 | 0.776682 | 0.774713 | 0.777675 | 0.773548 |
| 0.767925 | 0.765561 | 0.757632 | 0.754967 | 0.76626 | 0.762308 | 0.767421 | 0.760745 | 0.753841 |
| 0.788688 | 0.771965 | 0.773525 | 0.775061 | 0.769396 | 0.783601 | 0.770687 | 0.773614 | 0.789336 |
| 0.78732 | 0.757258 | 0.745383 | 0.742043 | 0.772875 | 0.757066 | 0.754894 | 0.755617 | 0.746269 |
| 0.782612 | 0.768841 | 0.779959 | 0.776898 | 0.768698 | 0.776576 | 0.769062 | 0.773396 | 0.767118 |
| 0.782805 | 0.773033 | 0.778958 | 0.771112 | 0.768628 | 0.768072 | 0.77363 | 0.774404 | 0.770941 |
| 0.762419 | 0.737123 | 0.738599 | 0.742385 | 0.739615 | 0.742314 | 0.739711 | 0.749849 | 0.741363 |
| 0.768398 | 0.760088 | 0.74001 | 0.741535 | 0.762567 | 0.741092 | 0.74666 | 0.77829 | 0.751974 |
| 0.755708 | 0.751613 | 0.753437 | 0.756746 | 0.756835 | 0.756856 | 0.758237 | 0.762123 | 0.760477 |
| 0.802137 | 0.765874 | 0.762275 | 0.763292 | 0.766905 | 0.75962 | 0.766994 | 0.762433 | 0.757863 |
| 0.787077 | 0.769983 | 0.772482 | 0.774025 | 0.769691 | 0.793995 | 0.786942 | 0.776596 | 0.771513 |
| 0.756035 | 0.753999 | 0.752025 | 0.751981 | 0.743532 | 0.745595 | 0.752321 | 0.759814 | 0.748279 |
| 0.81932 | 0.795367 | 0.768653 | 0.768238 | 0.785815 | 0.769426 | 0.769607 | 0.815951 | 0.789181 |
| 0.776255 | 0.765526 | 0.76928 | 0.771493 | 0.76617 | 0.772194 | 0.772054 | 0.772511 | 0.768008 |
| 0.769685 | 0.764595 | 0.759791 | 0.759676 | 0.76232 | 0.756755 | 0.759057 | 0.775616 | 0.778548 |
| 0.771059 | 0.766615 | 0.754475 | 0.760778 | 0.76299 | 0.760538 | 0.757096 | 0.773613 | 0.77803 |
| 0.820342 | 0.766253 | 0.766482 | 0.767514 | 0.761401 | 0.766232 | 0.769486 | 0.772449 | 0.765409 |
| 0.769434 | 0.767766 | 0.761764 | 0.763881 | 0.759417 | 0.763711 | 0.767007 | 0.770957 | 0.780504 |
| 0.772273 | 0.764178 | 0.763149 | 0.771097 | 0.762827 | 0.773082 | 0.765862 | 0.768585 | 0.772624 |
| 0.730644 | 0.732628 | 0.72359 | 0.729868 | 0.730953 | 0.731629 | 0.722468 | 0.730477 | 0.723555 |
| 0.761329 | 0.765253 | 0.758228 | 0.764496 | 0.760382 | 0.764932 | 0.759843 | 0.769682 | 0.756992 |
| 0.734298 | 0.725743 | 0.722987 | 0.731936 | 0.724504 | 0.725537 | 0.738711 | 0.729471 | 0.727324 |
| 0.732136 | 0.720084 | 0.72303 | 0.722145 | 0.724487 | 0.719279 | 0.720758 | 0.71626 | 0.719039 |
| 0.777283 | 0.770015 | 0.768346 | 0.768537 | 0.774706 | 0.7724 | 0.769714 | 0.772934 | 0.786051 |
| 0.72248 | 0.714903 | 0.719584 | 0.716947 | 0.724751 | 0.722853 | 0.718204 | 0.718082 | 0.720233 |
| 0.737701 | 0.729531 | 0.714165 | 0.714902 | 0.724369 | 0.717388 | 0.717534 | 0.734983 | 0.734052 |
| 0.74283 | 0.720757 | 0.70647 | 0.707181 | 0.725896 | 0.709472 | 0.709156 | 0.73344 | 0.721904 |
| 0.778491 | 0.772267 | 0.761693 | 0.761766 | 0.774437 | 0.765745 | 0.76393 | 0.776253 | 0.795401 |
| 0.736736 | 0.709119 | 0.712924 | 0.707051 | 0.715306 | 0.70769 | 0.701958 | 0.729992 | 0.717597 |
| 0.776105 | 0.769469 | 0.766778 | 0.764408 | 0.770082 | 0.767346 | 0.768008 | 0.77403 | 0.788333 |

| | | | | | | | | |
|---|---|---|---|---|---|---|---|---|
| 0.771313 | 0.762853 | 0.765396 | 0.76031 | 0.76443 | 0.759538 | 0.755351 | 0.768958 | 0.777044 |
| 0.777136 | 0.773525 | 0.763448 | 0.766201 | 0.770747 | 0.767017 | 0.764399 | 0.771452 | 0.782092 |
| 0.754656 | 0.753314 | 0.751753 | 0.75244 | 0.752922 | 0.754753 | 0.751981 | 0.751137 | 0.756576 |
| 0.720866 | 0.706114 | 0.701241 | 0.698925 | 0.704693 | 0.699763 | 0.701443 | 0.720398 | 0.711725 |
| 0.728669 | 0.717037 | 0.699135 | 0.693028 | 0.710932 | 0.697663 | 0.699716 | 0.729247 | 0.701737 |
| 0.7801 | 0.772047 | 0.76873 | 0.766553 | 0.771058 | 0.768725 | 0.76327 | 0.777916 | 0.800357 |
| 0.694166 | 0.690589 | 0.695942 | 0.696484 | 0.691353 | 0.695423 | 0.695983 | 0.694442 | 0.690239 |
| 0.742103 | 0.727273 | 0.731567 | 0.745044 | 0.731279 | 0.737896 | 0.730185 | 0.734936 | 0.735994 |
| 0.766849 | 0.76027 | 0.758432 | 0.761218 | 0.759152 | 0.758793 | 0.759977 | 0.767316 | 0.773267 |
| 0.722012 | 0.718511 | 0.723008 | 0.731819 | 0.724351 | 0.729023 | 0.728897 | 0.724572 | 0.719082 |
| 0.764643 | 0.755725 | 0.75059 | 0.753257 | 0.752424 | 0.74438 | 0.747463 | 0.759213 | 0.769049 |
| 0.625806 | 0.664384 | 0.623457 | 0.613497 | 0.620482 | 0.641975 | 0.613924 | 0.627451 | 0.612121 |
| 0.663071 | 0.656363 | 0.649378 | 0.65504 | 0.662408 | 0.654089 | 0.655534 | 0.660341 | 0.659423 |
| 0.662414 | 0.660513 | 0.650675 | 0.655898 | 0.662822 | 0.663362 | 0.654328 | 0.662616 | 0.659834 |
| 0.666619 | 0.664507 | 0.668655 | 0.670435 | 0.667343 | 0.671998 | 0.665712 | 0.674466 | 0.674385 |
| 0.671049 | 0.665722 | 0.66071 | 0.669214 | 0.671044 | 0.669776 | 0.663247 | 0.669837 | 0.669397 |
| 0.621857 | 0.618778 | 0.620391 | 0.618694 | 0.618353 | 0.615561 | 0.616556 | 0.624018 | 0.620181 |
| 0.670948 | 0.670659 | 0.663458 | 0.665291 | 0.671114 | 0.667293 | 0.664513 | 0.675621 | 0.674304 |
| 0.75986 | 0.752665 | 0.740548 | 0.743533 | 0.752468 | 0.742672 | 0.743342 | 0.755812 | 0.751423 |
| 0.749199 | 0.746161 | 0.738248 | 0.741951 | 0.742329 | 0.738374 | 0.738812 | 0.745913 | 0.757447 |
| 0.756155 | 0.750326 | 0.742259 | 0.748754 | 0.753018 | 0.743326 | 0.746947 | 0.760837 | 0.758929 |
| 0.624701 | 0.616564 | 0.622755 | 0.617443 | 0.620248 | 0.616149 | 0.618409 | 0.620391 | 0.613113 |
| 0.599779 | 0.59951 | 0.605299 | 0.599826 | 0.599324 | 0.60024 | 0.599843 | 0.601591 | 0.596145 |
| 0.710817 | 0.707599 | 0.703597 | 0.708552 | 0.711723 | 0.710586 | 0.707977 | 0.708214 | 0.710104 |
| 0.709792 | 0.711097 | 0.70572 | 0.709913 | 0.715202 | 0.71307 | 0.706382 | 0.711299 | 0.71108 |
| 0.705246 | 0.704414 | 0.702578 | 0.702284 | 0.703276 | 0.702788 | 0.702392 | 0.702841 | 0.703743 |
| 0.708995 | 0.704553 | 0.706479 | 0.705397 | 0.705726 | 0.707451 | 0.704963 | 0.704108 | 0.709116 |
| 0.71458 | 0.711393 | 0.709575 | 0.709634 | 0.714263 | 0.71072 | 0.709538 | 0.710372 | 0.709704 |
| 0.709886 | 0.708883 | 0.704442 | 0.707888 | 0.712754 | 0.710242 | 0.705826 | 0.707299 | 0.711089 |
| 0.714002 | 0.709516 | 0.708546 | 0.706986 | 0.71248 | 0.710729 | 0.709155 | 0.711413 | 0.711189 |
| 0.710345 | 0.710051 | 0.707248 | 0.708579 | 0.712678 | 0.711687 | 0.7075 | 0.711902 | 0.712735 |
| 0.720328 | 0.71586 | 0.714871 | 0.717939 | 0.715445 | 0.71856 | 0.715827 | 0.714915 | 0.716551 |
| 0.716521 | 0.708662 | 0.711574 | 0.71362 | 0.710164 | 0.715978 | 0.708746 | 0.710704 | 0.711313 |
| 0.716216 | 0.711569 | 0.713544 | 0.710949 | 0.711628 | 0.716816 | 0.711738 | 0.713315 | 0.713209 |
| 0.722969 | 0.716839 | 0.717355 | 0.716413 | 0.716389 | 0.721142 | 0.716857 | 0.716107 | 0.712368 |
| 0.713798 | 0.706821 | 0.70912 | 0.710165 | 0.707931 | 0.708675 | 0.709989 | 0.709876 | 0.711407 |
| 0.714797 | 0.711774 | 0.710029 | 0.712142 | 0.712786 | 0.714228 | 0.710364 | 0.710398 | 0.710692 |
| 0.717364 | 0.710159 | 0.715966 | 0.715377 | 0.714182 | 0.713323 | 0.710376 | 0.711848 | 0.710584 |
| 0.710728 | 0.706465 | 0.705354 | 0.708742 | 0.707404 | 0.709295 | 0.705174 | 0.707553 | 0.705868 |
| 0.715638 | 0.708955 | 0.712262 | 0.710749 | 0.710934 | 0.715006 | 0.710196 | 0.712939 | 0.709962 |
| 0.657431 | 0.658516 | 0.659596 | 0.657582 | 0.656377 | 0.658754 | 0.658675 | 0.658277 | 0.668191 |
| 0.667756 | 0.661436 | 0.661757 | 0.663152 | 0.668129 | 0.663575 | 0.659414 | 0.673462 | 0.670255 |
| 0.718871 | 0.714906 | 0.713495 | 0.717372 | 0.715853 | 0.72009 | 0.714634 | 0.715712 | 0.717421 |
| 0.718299 | 0.718036 | 0.718061 | 0.718247 | 0.720562 | 0.721272 | 0.712303 | 0.712949 | 0.714801 |
| 0.652038 | 0.648926 | 0.645451 | 0.645791 | 0.653001 | 0.652254 | 0.640531 | 0.650491 | 0.655172 |
| 0.715276 | 0.713378 | 0.714012 | 0.711885 | 0.715259 | 0.714332 | 0.711648 | 0.709373 | 0.712016 |
| 0.705026 | 0.702953 | 0.702901 | 0.703873 | 0.704019 | 0.705939 | 0.700094 | 0.700857 | 0.701764 |

| | | | | | | | | |
|---|---|---|---|---|---|---|---|---|
| 0.713934 | 0.712595 | 0.711189 | 0.708773 | 0.715189 | 0.712551 | 0.708448 | 0.708196 | 0.707476 |
| 0.716843 | 0.71157 | 0.715314 | 0.714399 | 0.712655 | 0.712055 | 0.710083 | 0.70889 | 0.711349 |
| 0.712469 | 0.712599 | 0.711176 | 0.711811 | 0.713107 | 0.713595 | 0.712259 | 0.712011 | 0.715494 |
| 0.718388 | 0.708402 | 0.711349 | 0.711653 | 0.717296 | 0.713139 | 0.712002 | 0.712958 | 0.711224 |
| 0.717788 | 0.717334 | 0.716527 | 0.717845 | 0.721511 | 0.719453 | 0.717937 | 0.72179 | 0.723346 |
| 0.716772 | 0.708702 | 0.711029 | 0.710299 | 0.712891 | 0.7152 | 0.706585 | 0.707295 | 0.708617 |
| 0.715075 | 0.709492 | 0.711643 | 0.709529 | 0.714366 | 0.714229 | 0.708226 | 0.709328 | 0.710134 |
| 0.708033 | 0.705821 | 0.702277 | 0.703641 | 0.705887 | 0.706178 | 0.702104 | 0.699366 | 0.705805 |
| 0.714887 | 0.707773 | 0.70874 | 0.709629 | 0.710973 | 0.708792 | 0.709426 | 0.707953 | 0.707578 |
| 0.710071 | 0.709654 | 0.709757 | 0.708978 | 0.707681 | 0.71222 | 0.709002 | 0.705958 | 0.710898 |
| 0.714951 | 0.708733 | 0.712477 | 0.71047 | 0.711018 | 0.715086 | 0.711139 | 0.70851 | 0.710398 |
| 0.713948 | 0.71069 | 0.70532 | 0.709463 | 0.71039 | 0.708684 | 0.708205 | 0.705775 | 0.70643 |
| 0.704154 | 0.706718 | 0.703942 | 0.701594 | 0.70555 | 0.705214 | 0.701487 | 0.699341 | 0.704018 |
| 0.70547 | 0.704974 | 0.701737 | 0.703109 | 0.704511 | 0.707309 | 0.700057 | 0.700629 | 0.703414 |
| 0.720518 | 0.715184 | 0.714726 | 0.714851 | 0.718521 | 0.719744 | 0.716984 | 0.716019 | 0.718421 |
| 0.713801 | 0.713778 | 0.712148 | 0.71122 | 0.716835 | 0.71271 | 0.710469 | 0.712791 | 0.713087 |
| 0.710569 | 0.705589 | 0.703355 | 0.703828 | 0.704443 | 0.708592 | 0.702965 | 0.702797 | 0.707515 |
| 0.712801 | 0.707854 | 0.70748 | 0.711482 | 0.711005 | 0.711193 | 0.706361 | 0.705569 | 0.710678 |
| 0.71151 | 0.706921 | 0.710875 | 0.712085 | 0.709389 | 0.707802 | 0.705011 | 0.707519 | 0.707506 |
| 0.713207 | 0.708796 | 0.70901 | 0.711129 | 0.710146 | 0.708812 | 0.703108 | 0.70602 | 0.705065 |
| 0.709163 | 0.704353 | 0.705982 | 0.707258 | 0.705602 | 0.708085 | 0.702068 | 0.703022 | 0.702225 |
| 0.709173 | 0.703308 | 0.707909 | 0.708104 | 0.707518 | 0.705202 | 0.702908 | 0.704305 | 0.705887 |
| 0.714691 | 0.709966 | 0.712921 | 0.7114 | 0.709646 | 0.712484 | 0.70682 | 0.710189 | 0.7094 |
| 0.715294 | 0.70898 | 0.709491 | 0.713183 | 0.710624 | 0.712977 | 0.709178 | 0.708794 | 0.708834 |
| 0.709954 | 0.706807 | 0.709091 | 0.711396 | 0.708862 | 0.707796 | 0.705544 | 0.703942 | 0.706028 |
| 0.70822 | 0.70708 | 0.704591 | 0.708554 | 0.706624 | 0.707852 | 0.706279 | 0.704262 | 0.704383 |
| 0.709957 | 0.70436 | 0.703769 | 0.706398 | 0.704289 | 0.708476 | 0.705901 | 0.702072 | 0.705407 |
| 0.709133 | 0.700899 | 0.702598 | 0.707319 | 0.700049 | 0.704976 | 0.702859 | 0.702105 | 0.704257 |
| 0.71071 | 0.706068 | 0.707276 | 0.708879 | 0.708151 | 0.710811 | 0.703085 | 0.707272 | 0.705033 |
| 0.71241 | 0.705952 | 0.710257 | 0.711798 | 0.708599 | 0.711549 | 0.707143 | 0.70575 | 0.709245 |
| 0.715487 | 0.711106 | 0.710928 | 0.714685 | 0.712003 | 0.71401 | 0.707409 | 0.706727 | 0.711605 |
| 0.715935 | 0.710179 | 0.71154 | 0.712267 | 0.71126 | 0.717945 | 0.709475 | 0.710374 | 0.713623 |
| 0.718545 | 0.715038 | 0.712607 | 0.715538 | 0.714997 | 0.717779 | 0.712645 | 0.715868 | 0.715879 |
| 0.717904 | 0.71605 | 0.711377 | 0.713752 | 0.717247 | 0.720354 | 0.712811 | 0.714459 | 0.717634 |
| 0.708115 | 0.704385 | 0.702782 | 0.706899 | 0.707058 | 0.706544 | 0.704102 | 0.703731 | 0.707766 |
| 0.714573 | 0.708548 | 0.710645 | 0.710499 | 0.71082 | 0.713388 | 0.706269 | 0.706508 | 0.706958 |
| 0.710258 | 0.706079 | 0.708243 | 0.711824 | 0.707605 | 0.712194 | 0.707818 | 0.707967 | 0.709247 |
| 0.708966 | 0.705281 | 0.706679 | 0.708199 | 0.705335 | 0.704617 | 0.703207 | 0.703399 | 0.707959 |
| 0.706959 | 0.700628 | 0.704033 | 0.702941 | 0.701723 | 0.706767 | 0.698438 | 0.699828 | 0.698963 |
| 0.702436 | 0.699436 | 0.698377 | 0.698361 | 0.698259 | 0.700025 | 0.700182 | 0.701027 | 0.70185 |
| 0.702751 | 0.699584 | 0.699681 | 0.699811 | 0.700959 | 0.701284 | 0.6986 | 0.699118 | 0.700463 |
| 0.704798 | 0.701001 | 0.697856 | 0.700034 | 0.701517 | 0.701163 | 0.695445 | 0.696482 | 0.699888 |
| 0.707366 | 0.704775 | 0.703082 | 0.703107 | 0.705251 | 0.70793 | 0.700551 | 0.700622 | 0.698327 |
| 0.697164 | 0.69595 | 0.696034 | 0.696685 | 0.697457 | 0.695801 | 0.692778 | 0.694518 | 0.695629 |
| 0.708861 | 0.702602 | 0.70761 | 0.707007 | 0.704001 | 0.704683 | 0.699134 | 0.703309 | 0.704608 |
| 0.702033 | 0.696453 | 0.697636 | 0.699245 | 0.698771 | 0.701472 | 0.697853 | 0.697031 | 0.698284 |

| | | | | | | | | |
|---|---|---|---|---|---|---|---|---|
| 0.713583 | 0.712742 | 0.710088 | 0.71187 | 0.714077 | 0.712457 | 0.710543 | 0.708192 | 0.711222 |
| 0.713702 | 0.706508 | 0.710839 | 0.715508 | 0.709701 | 0.713588 | 0.707199 | 0.708847 | 0.710827 |
| 0.713116 | 0.708096 | 0.705161 | 0.709422 | 0.708047 | 0.710119 | 0.705324 | 0.708442 | 0.704841 |
| 0.716178 | 0.710189 | 0.713642 | 0.712045 | 0.711636 | 0.714799 | 0.711508 | 0.70787 | 0.711163 |
| 0.72164 | 0.716749 | 0.716319 | 0.718743 | 0.715686 | 0.721509 | 0.717292 | 0.715151 | 0.720355 |
| 0.718379 | 0.712434 | 0.712138 | 0.714558 | 0.714286 | 0.713131 | 0.709662 | 0.711968 | 0.714718 |
| 0.710273 | 0.707253 | 0.71011 | 0.710755 | 0.706365 | 0.707037 | 0.706152 | 0.705572 | 0.706231 |
| 0.717045 | 0.707972 | 0.711285 | 0.713776 | 0.707578 | 0.711475 | 0.707692 | 0.71141 | 0.709712 |
| 0.709092 | 0.70439 | 0.708668 | 0.709222 | 0.706808 | 0.700044 | 0.701428 | 0.70432 | 0.708508 |
| 0.714206 | 0.706944 | 0.709762 | 0.711058 | 0.708624 | 0.710706 | 0.708738 | 0.707614 | 0.712273 |
| 0.718229 | 0.710157 | 0.711041 | 0.714893 | 0.711312 | 0.712197 | 0.709203 | 0.713372 | 0.716318 |
| 0.71554 | 0.710075 | 0.712177 | 0.712732 | 0.712151 | 0.713827 | 0.710835 | 0.712728 | 0.715199 |
| 0.713101 | 0.705363 | 0.711076 | 0.709844 | 0.710414 | 0.708383 | 0.709258 | 0.710043 | 0.709312 |
| 0.714705 | 0.707286 | 0.711746 | 0.710668 | 0.710741 | 0.712507 | 0.708435 | 0.711317 | 0.712882 |
| 0.715993 | 0.71127 | 0.709865 | 0.71353 | 0.712396 | 0.716076 | 0.710058 | 0.713302 | 0.711535 |
| 0.715957 | 0.707486 | 0.712316 | 0.713018 | 0.711357 | 0.712088 | 0.708677 | 0.711893 | 0.713596 |
| 0.720274 | 0.712752 | 0.717757 | 0.712321 | 0.715077 | 0.715059 | 0.714118 | 0.715737 | 0.717711 |
| 0.718161 | 0.711016 | 0.71217 | 0.712783 | 0.716879 | 0.717136 | 0.708453 | 0.717562 | 0.713859 |
| 0.716307 | 0.713879 | 0.717396 | 0.715761 | 0.713617 | 0.717811 | 0.71213 | 0.714735 | 0.71508 |
| 0.714004 | 0.708758 | 0.709573 | 0.713281 | 0.70926 | 0.711019 | 0.705139 | 0.710054 | 0.709869 |
| 0.716542 | 0.709464 | 0.711902 | 0.713098 | 0.709437 | 0.713676 | 0.71227 | 0.710665 | 0.70973 |
| 0.711681 | 0.709675 | 0.710331 | 0.709616 | 0.710458 | 0.711756 | 0.708971 | 0.714518 | 0.710613 |
| 0.71952 | 0.713847 | 0.718411 | 0.713011 | 0.713746 | 0.715677 | 0.712049 | 0.713519 | 0.712613 |
| 0.721093 | 0.711635 | 0.713057 | 0.712796 | 0.711585 | 0.716566 | 0.713436 | 0.713805 | 0.713392 |
| 0.715112 | 0.710473 | 0.711709 | 0.710965 | 0.711621 | 0.711878 | 0.708353 | 0.710918 | 0.713269 |
| 0.716912 | 0.71023 | 0.708205 | 0.713934 | 0.710955 | 0.715615 | 0.708026 | 0.710124 | 0.713701 |
| 0.714645 | 0.705633 | 0.708684 | 0.714704 | 0.710353 | 0.707993 | 0.706327 | 0.71141 | 0.7081 |
| 0.713612 | 0.707877 | 0.707898 | 0.710804 | 0.713057 | 0.710742 | 0.705428 | 0.708733 | 0.710929 |
| 0.718961 | 0.710994 | 0.71213 | 0.717113 | 0.710602 | 0.71538 | 0.709982 | 0.71248 | 0.715575 |
| 0.719869 | 0.713123 | 0.714473 | 0.716639 | 0.713836 | 0.715348 | 0.708953 | 0.714877 | 0.715293 |
| 0.715781 | 0.707601 | 0.709353 | 0.713755 | 0.705979 | 0.71046 | 0.709484 | 0.706968 | 0.711023 |
| 0.721339 | 0.7135 | 0.71432 | 0.717486 | 0.715077 | 0.715558 | 0.71296 | 0.714867 | 0.713344 |
| 0.717845 | 0.709456 | 0.714773 | 0.714897 | 0.711229 | 0.712975 | 0.715253 | 0.711034 | 0.714788 |
| 0.71023 | 0.706567 | 0.70848 | 0.711042 | 0.707161 | 0.710474 | 0.703177 | 0.707331 | 0.709721 |
| 0.715568 | 0.712014 | 0.709448 | 0.711304 | 0.714869 | 0.713672 | 0.711638 | 0.71228 | 0.712651 |
| 0.71329 | 0.706515 | 0.707226 | 0.709471 | 0.709026 | 0.706656 | 0.705679 | 0.70925 | 0.703817 |
| 0.72246 | 0.714998 | 0.721502 | 0.717352 | 0.720817 | 0.722249 | 0.715457 | 0.7178 | 0.717661 |
| 0.7169 | 0.710158 | 0.709711 | 0.710573 | 0.710788 | 0.71515 | 0.712132 | 0.709651 | 0.710188 |
| 0.712083 | 0.708156 | 0.712357 | 0.711019 | 0.710662 | 0.713048 | 0.707551 | 0.712417 | 0.71493 |
| 0.713558 | 0.71039 | 0.712481 | 0.712954 | 0.709975 | 0.716837 | 0.708935 | 0.713838 | 0.708486 |
| 0.722165 | 0.714241 | 0.717199 | 0.715404 | 0.716092 | 0.71829 | 0.7147 | 0.717291 | 0.715308 |
| 0.717902 | 0.711101 | 0.713664 | 0.715853 | 0.716906 | 0.71674 | 0.714049 | 0.714955 | 0.717953 |
| 0.715889 | 0.710347 | 0.714432 | 0.716452 | 0.716578 | 0.718862 | 0.712488 | 0.712859 | 0.715932 |
| 0.71816 | 0.713628 | 0.710488 | 0.714611 | 0.712144 | 0.709159 | 0.705477 | 0.71033 | 0.710051 |
| 0.717159 | 0.709818 | 0.7144 | 0.714092 | 0.714599 | 0.713536 | 0.71068 | 0.714797 | 0.710038 |
| 0.712755 | 0.708333 | 0.705353 | 0.71039 | 0.708511 | 0.708136 | 0.71112 | 0.705372 | 0.707032 |

| | | | | | | | | |
|---|---|---|---|---|---|---|---|---|
| 0.705593 | 0.697182 | 0.69842 | 0.697872 | 0.702074 | 0.699265 | 0.693966 | 0.697855 | 0.693923 |
| 0.762394 | 0.744728 | 0.747443 | 0.75304 | 0.745734 | 0.750021 | 0.749796 | 0.750519 | 0.74464 |
| 0.716017 | 0.709379 | 0.711365 | 0.709283 | 0.708409 | 0.710444 | 0.706714 | 0.709587 | 0.706859 |
| 0.721714 | 0.709296 | 0.716281 | 0.713723 | 0.713988 | 0.714491 | 0.710364 | 0.712893 | 0.712211 |
| 0.710277 | 0.703971 | 0.709086 | 0.706595 | 0.707811 | 0.710163 | 0.703341 | 0.706345 | 0.704731 |
| 0.714206 | 0.707851 | 0.711058 | 0.713619 | 0.713117 | 0.712742 | 0.708771 | 0.70926 | 0.710237 |
| 0.710289 | 0.701662 | 0.706429 | 0.709519 | 0.703763 | 0.708783 | 0.704143 | 0.707175 | 0.707911 |
| 0.707242 | 0.702991 | 0.706724 | 0.707426 | 0.704887 | 0.705127 | 0.698723 | 0.703607 | 0.702239 |
| 0.714038 | 0.706972 | 0.71101 | 0.713935 | 0.711848 | 0.71086 | 0.708709 | 0.710547 | 0.706494 |
| 0.723465 | 0.715798 | 0.718473 | 0.715084 | 0.718038 | 0.719013 | 0.7138 | 0.716189 | 0.717561 |
| 0.718688 | 0.709741 | 0.713462 | 0.715124 | 0.712586 | 0.716078 | 0.709907 | 0.714849 | 0.711853 |
| 0.719528 | 0.710876 | 0.716342 | 0.714708 | 0.713438 | 0.717274 | 0.711507 | 0.715993 | 0.714286 |
| 0.721371 | 0.714739 | 0.714331 | 0.720473 | 0.717419 | 0.715855 | 0.714398 | 0.715371 | 0.718235 |
| 0.719946 | 0.710566 | 0.71468 | 0.717446 | 0.711761 | 0.716664 | 0.71151 | 0.715938 | 0.716269 |
| 0.716637 | 0.709125 | 0.712575 | 0.710971 | 0.711316 | 0.712405 | 0.702853 | 0.710827 | 0.711122 |
| 0.715 | 0.706605 | 0.710708 | 0.709935 | 0.710378 | 0.71589 | 0.70919 | 0.712765 | 0.713203 |
| 0.722009 | 0.710297 | 0.716948 | 0.71195 | 0.714669 | 0.715798 | 0.712296 | 0.715212 | 0.718195 |
| 0.717002 | 0.709621 | 0.714709 | 0.711943 | 0.713273 | 0.71342 | 0.712039 | 0.710601 | 0.716285 |
| 0.706155 | 0.70795 | 0.704585 | 0.7081 | 0.708876 | 0.7097 | 0.702938 | 0.708789 | 0.708988 |
| 0.710456 | 0.703763 | 0.706291 | 0.707327 | 0.708245 | 0.708132 | 0.701445 | 0.705687 | 0.70658 |
| 0.719874 | 0.708383 | 0.713103 | 0.711907 | 0.712343 | 0.710671 | 0.708463 | 0.714601 | 0.710788 |
| 0.703499 | 0.701362 | 0.702943 | 0.704672 | 0.701674 | 0.704482 | 0.699992 | 0.70133 | 0.702872 |
| 0.712874 | 0.708511 | 0.710819 | 0.710248 | 0.71018 | 0.71179 | 0.705855 | 0.709906 | 0.709496 |
| 0.718392 | 0.709708 | 0.713068 | 0.713784 | 0.714425 | 0.712276 | 0.71159 | 0.710724 | 0.711602 |
| 0.721639 | 0.706789 | 0.709599 | 0.718523 | 0.711702 | 0.714815 | 0.712382 | 0.715905 | 0.713009 |
| 0.714366 | 0.708353 | 0.710752 | 0.717961 | 0.713276 | 0.712366 | 0.709922 | 0.710995 | 0.714012 |
| 0.710375 | 0.704503 | 0.703461 | 0.710366 | 0.707871 | 0.708714 | 0.702114 | 0.706157 | 0.708608 |
| 0.709717 | 0.701638 | 0.706511 | 0.705049 | 0.705592 | 0.708392 | 0.704512 | 0.704373 | 0.708901 |
| 0.710467 | 0.702209 | 0.705793 | 0.709713 | 0.707363 | 0.703629 | 0.703633 | 0.708769 | 0.705614 |
| 0.766803 | 0.761193 | 0.764092 | 0.75975 | 0.758142 | 0.758929 | 0.755926 | 0.770473 | 0.762024 |
| 0.819941 | 0.802384 | 0.781234 | 0.783149 | 0.789729 | 0.776819 | 0.777192 | 0.822915 | 0.790238 |
| 0.783185 | 0.782681 | 0.778027 | 0.777815 | 0.776079 | 0.774407 | 0.773157 | 0.776561 | 0.782082 |
| 0.785531 | 0.782692 | 0.76774 | 0.770289 | 0.779446 | 0.771415 | 0.779278 | 0.775321 | 0.812073 |
| 0.828727 | 0.810009 | 0.779595 | 0.782951 | 0.814705 | 0.78002 | 0.787716 | 0.819754 | 0.80109 |
| 0.708555 | 0.701183 | 0.708652 | 0.705844 | 0.703177 | 0.702956 | 0.701559 | 0.702792 | 0.7029 |
| 0.712132 | 0.702698 | 0.70729 | 0.707229 | 0.708313 | 0.712924 | 0.703018 | 0.708306 | 0.707394 |
| 0.711598 | 0.707161 | 0.710037 | 0.705661 | 0.709293 | 0.709319 | 0.704407 | 0.709341 | 0.710485 |
| 0.716498 | 0.709772 | 0.713098 | 0.715923 | 0.711881 | 0.710857 | 0.70918 | 0.708984 | 0.712586 |
| 0.709001 | 0.708663 | 0.70789 | 0.708 | 0.711795 | 0.707984 | 0.707877 | 0.710667 | 0.708731 |
| 0.714559 | 0.712429 | 0.709886 | 0.710699 | 0.714669 | 0.715797 | 0.710655 | 0.71446 | 0.71391 |
| 0.714768 | 0.70355 | 0.71063 | 0.709579 | 0.709002 | 0.710616 | 0.706568 | 0.708272 | 0.708306 |
| 0.711097 | 0.705155 | 0.708168 | 0.707785 | 0.706412 | 0.70948 | 0.704609 | 0.709929 | 0.71076 |
| 0.706581 | 0.700217 | 0.704154 | 0.703284 | 0.703606 | 0.701701 | 0.695706 | 0.705185 | 0.700627 |
| 0.705837 | 0.698766 | 0.701119 | 0.706442 | 0.701961 | 0.702395 | 0.698418 | 0.702745 | 0.702445 |
| 0.70935 | 0.702582 | 0.70744 | 0.707909 | 0.706609 | 0.710878 | 0.703425 | 0.707228 | 0.706642 |
| 0.714688 | 0.705145 | 0.710391 | 0.71042 | 0.709006 | 0.713995 | 0.705896 | 0.709836 | 0.711656 |

| | | | | | | | | |
|---|---|---|---|---|---|---|---|---|
| 0.719234 | 0.706788 | 0.713715 | 0.710782 | 0.710931 | 0.713229 | 0.705986 | 0.711464 | 0.710022 |
| 0.713909 | 0.707538 | 0.706576 | 0.709835 | 0.707424 | 0.712324 | 0.705478 | 0.707972 | 0.708306 |
| 0.717907 | 0.710029 | 0.714044 | 0.715266 | 0.711935 | 0.714646 | 0.707888 | 0.712967 | 0.713813 |
| 0.714103 | 0.707145 | 0.70828 | 0.708769 | 0.709523 | 0.712717 | 0.70572 | 0.712025 | 0.71074 |
| 0.708316 | 0.70239 | 0.705705 | 0.704414 | 0.705882 | 0.706128 | 0.702368 | 0.709269 | 0.706064 |
| 0.713298 | 0.704177 | 0.702982 | 0.708175 | 0.706378 | 0.711392 | 0.702458 | 0.705157 | 0.705063 |
| 0.717606 | 0.708222 | 0.714111 | 0.709162 | 0.71285 | 0.714498 | 0.708994 | 0.71393 | 0.711663 |
| 0.7123 | 0.70645 | 0.711729 | 0.709567 | 0.708558 | 0.714686 | 0.705056 | 0.707769 | 0.711225 |
| 0.712323 | 0.705068 | 0.709349 | 0.710591 | 0.71015 | 0.712172 | 0.705678 | 0.710067 | 0.70205 |
| 0.713575 | 0.710269 | 0.711678 | 0.71276 | 0.710036 | 0.712119 | 0.706838 | 0.707498 | 0.711651 |
| 0.712164 | 0.705026 | 0.708788 | 0.708414 | 0.710095 | 0.712196 | 0.706392 | 0.709109 | 0.708784 |
| 0.710344 | 0.703948 | 0.709411 | 0.708523 | 0.707939 | 0.711167 | 0.703968 | 0.707451 | 0.707311 |
| 0.702619 | 0.701019 | 0.700688 | 0.701521 | 0.700427 | 0.705698 | 0.697581 | 0.700915 | 0.701683 |
| 0.704889 | 0.70042 | 0.705396 | 0.699553 | 0.704133 | 0.70463 | 0.700523 | 0.70415 | 0.702577 |
| 0.716283 | 0.711117 | 0.71211 | 0.711717 | 0.712075 | 0.712256 | 0.708216 | 0.709187 | 0.712733 |
| 0.723782 | 0.712534 | 0.718961 | 0.71801 | 0.71582 | 0.716943 | 0.711007 | 0.715923 | 0.717885 |
| 0.717912 | 0.709192 | 0.714083 | 0.714162 | 0.713015 | 0.714934 | 0.710861 | 0.710921 | 0.714657 |
| 0.716168 | 0.710616 | 0.714456 | 0.714024 | 0.714205 | 0.715321 | 0.714263 | 0.71171 | 0.715119 |
| 0.709826 | 0.707055 | 0.706623 | 0.706442 | 0.710537 | 0.71174 | 0.708221 | 0.709737 | 0.706716 |
| 0.712518 | 0.705757 | 0.708751 | 0.708354 | 0.710632 | 0.711452 | 0.706045 | 0.707796 | 0.706679 |
| 0.710401 | 0.70274 | 0.705897 | 0.707394 | 0.706914 | 0.709001 | 0.700628 | 0.703118 | 0.706012 |
| 0.717786 | 0.714148 | 0.711344 | 0.713745 | 0.71404 | 0.713624 | 0.708931 | 0.713982 | 0.711047 |
| 0.722937 | 0.714075 | 0.713803 | 0.714427 | 0.716454 | 0.712839 | 0.710786 | 0.715278 | 0.714867 |
| 0.714554 | 0.705658 | 0.706952 | 0.708213 | 0.707397 | 0.71004 | 0.703904 | 0.711542 | 0.70907 |
| 0.716913 | 0.706183 | 0.71143 | 0.706144 | 0.712842 | 0.710125 | 0.708457 | 0.710978 | 0.708202 |
| 0.782366 | 0.77971 | 0.783698 | 0.783687 | 0.774869 | 0.77145 | 0.779747 | 0.780731 | 0.79338 |
| 0.714466 | 0.710168 | 0.71213 | 0.713099 | 0.711676 | 0.7115 | 0.707099 | 0.710558 | 0.7081 |
| 0.713564 | 0.709853 | 0.712632 | 0.714445 | 0.711955 | 0.712959 | 0.710326 | 0.714067 | 0.711167 |
| 0.71975 | 0.712323 | 0.714057 | 0.718423 | 0.718334 | 0.717344 | 0.708839 | 0.713995 | 0.714427 |
| 0.715629 | 0.707424 | 0.710989 | 0.710953 | 0.709654 | 0.711841 | 0.708577 | 0.708546 | 0.707563 |
| 0.710778 | 0.707374 | 0.707599 | 0.708854 | 0.705654 | 0.709317 | 0.70597 | 0.707626 | 0.706564 |
| 0.712214 | 0.706613 | 0.706022 | 0.709643 | 0.70954 | 0.709873 | 0.706265 | 0.707688 | 0.710031 |
| 0.715292 | 0.709784 | 0.710475 | 0.712903 | 0.710657 | 0.711362 | 0.708849 | 0.709797 | 0.709138 |
| 0.710112 | 0.70227 | 0.704962 | 0.705926 | 0.704599 | 0.705848 | 0.705834 | 0.70623 | 0.708412 |
| 0.710449 | 0.702466 | 0.705094 | 0.704761 | 0.705542 | 0.702534 | 0.704052 | 0.706628 | 0.70396 |
| 0.708023 | 0.702779 | 0.708781 | 0.709175 | 0.70494 | 0.709948 | 0.705427 | 0.704804 | 0.706767 |
| 0.718717 | 0.712945 | 0.714019 | 0.714553 | 0.711904 | 0.713432 | 0.711892 | 0.71232 | 0.711276 |
| 0.720147 | 0.71125 | 0.716516 | 0.713758 | 0.712515 | 0.712217 | 0.708459 | 0.717063 | 0.71543 |
| 0.714813 | 0.70843 | 0.709152 | 0.713031 | 0.712986 | 0.713601 | 0.706983 | 0.711006 | 0.708003 |
| 0.713117 | 0.706889 | 0.710039 | 0.708471 | 0.710449 | 0.711251 | 0.706681 | 0.70999 | 0.711373 |
| 0.712936 | 0.704308 | 0.707625 | 0.709806 | 0.705221 | 0.705735 | 0.701264 | 0.706904 | 0.704772 |
| 0.718052 | 0.711702 | 0.715466 | 0.713811 | 0.715123 | 0.717648 | 0.710193 | 0.714592 | 0.712344 |
| 0.714854 | 0.704326 | 0.708942 | 0.708316 | 0.708337 | 0.708232 | 0.703661 | 0.712068 | 0.709213 |
| 0.712742 | 0.707402 | 0.709968 | 0.709289 | 0.712201 | 0.710234 | 0.708716 | 0.71394 | 0.712254 |
| 0.716862 | 0.712717 | 0.711649 | 0.711797 | 0.711216 | 0.71618 | 0.708759 | 0.712976 | 0.714133 |
| 0.715409 | 0.710108 | 0.714513 | 0.710959 | 0.711248 | 0.7119 | 0.712139 | 0.711599 | 0.715551 |

| | | | | | | | | |
|---|---|---|---|---|---|---|---|---|
| 0.711843 | 0.708915 | 0.713515 | 0.711812 | 0.705458 | 0.710768 | 0.709435 | 0.712584 | 0.712502 |
| 0.713451 | 0.712146 | 0.709323 | 0.712624 | 0.71146 | 0.712288 | 0.709857 | 0.713297 | 0.713517 |
| 0.712676 | 0.706862 | 0.711052 | 0.710094 | 0.712151 | 0.711063 | 0.705943 | 0.714565 | 0.713589 |
| 0.660281 | 0.656541 | 0.6597 | 0.665977 | 0.664011 | 0.675061 | 0.669077 | 0.666667 | 0.665503 |
| 0.71182 | 0.709317 | 0.708071 | 0.708639 | 0.707434 | 0.709069 | 0.706857 | 0.709282 | 0.708462 |
| 0.710101 | 0.712493 | 0.707702 | 0.707296 | 0.709223 | 0.71292 | 0.710522 | 0.708253 | 0.711965 |
| 0.711181 | 0.709425 | 0.710372 | 0.711134 | 0.709469 | 0.713713 | 0.712758 | 0.713121 | 0.714753 |
| 0.714365 | 0.71127 | 0.70929 | 0.711984 | 0.711066 | 0.717137 | 0.714117 | 0.710115 | 0.711831 |
| 0.718634 | 0.71512 | 0.712468 | 0.718255 | 0.713835 | 0.718056 | 0.71276 | 0.716675 | 0.713411 |
| 0.71327 | 0.711185 | 0.709997 | 0.71148 | 0.711882 | 0.712751 | 0.712521 | 0.714978 | 0.712744 |
| 0.712066 | 0.710884 | 0.711242 | 0.709311 | 0.710752 | 0.711263 | 0.711027 | 0.71267 | 0.710755 |
| 0.718053 | 0.716208 | 0.716442 | 0.714936 | 0.714274 | 0.716866 | 0.717524 | 0.718271 | 0.714122 |
| 0.715587 | 0.715384 | 0.713083 | 0.712787 | 0.713501 | 0.717671 | 0.711787 | 0.713225 | 0.717051 |
| 0.711824 | 0.710054 | 0.707177 | 0.710124 | 0.71264 | 0.711363 | 0.706928 | 0.710934 | 0.708245 |
| 0.71432 | 0.712118 | 0.714422 | 0.713026 | 0.712831 | 0.713931 | 0.711786 | 0.71379 | 0.711543 |
| 0.707709 | 0.70836 | 0.707781 | 0.709042 | 0.707531 | 0.708196 | 0.705411 | 0.710584 | 0.70765 |
| 0.714118 | 0.711242 | 0.710334 | 0.714477 | 0.714 | 0.715673 | 0.710438 | 0.71416 | 0.711495 |
| 0.714026 | 0.712076 | 0.710354 | 0.714547 | 0.713894 | 0.714834 | 0.710543 | 0.714712 | 0.714309 |
| 0.708415 | 0.705984 | 0.708558 | 0.712821 | 0.709992 | 0.708973 | 0.705638 | 0.706941 | 0.708265 |
| 0.712899 | 0.708616 | 0.709935 | 0.713911 | 0.70988 | 0.707788 | 0.708735 | 0.710484 | 0.708676 |
| 0.714851 | 0.712795 | 0.713605 | 0.713831 | 0.714899 | 0.712819 | 0.709842 | 0.717521 | 0.712734 |
| 0.715663 | 0.711269 | 0.713775 | 0.713152 | 0.7111 | 0.714366 | 0.713892 | 0.712608 | 0.715197 |
| 0.702287 | 0.691944 | 0.68926 | 0.703303 | 0.695429 | 0.705364 | 0.711634 | 0.704181 | 0.701964 |
| 0.723227 | 0.718948 | 0.708313 | 0.710194 | 0.714843 | 0.725893 | 0.717181 | 0.725711 | 0.739506 |
| 0.760582 | 0.754797 | 0.751339 | 0.75156 | 0.751817 | 0.763027 | 0.751839 | 0.758204 | 0.758769 |
| 0.731716 | 0.74591 | 0.727472 | 0.735955 | 0.744484 | 0.736555 | 0.744182 | 0.746182 | 0.778048 |
| 0.669824 | 0.667528 | 0.664398 | 0.669603 | 0.671402 | 0.668399 | 0.666905 | 0.670556 | 0.675026 |
| 0.667158 | 0.664404 | 0.666543 | 0.657798 | 0.667596 | 0.668695 | 0.659909 | 0.664564 | 0.669266 |
| 0.665747 | 0.668985 | 0.665159 | 0.664187 | 0.668589 | 0.663939 | 0.65439 | 0.668941 | 0.668825 |
| 0.685967 | 0.679101 | 0.682617 | 0.674522 | 0.684264 | 0.686239 | 0.678336 | 0.686974 | 0.675228 |
| 0.661173 | 0.666373 | 0.652499 | 0.652949 | 0.663938 | 0.65837 | 0.65245 | 0.658233 | 0.669715 |
| 0.646348 | 0.643145 | 0.647876 | 0.637416 | 0.654843 | 0.645022 | 0.646649 | 0.655675 | 0.654755 |
| 0.641632 | 0.636466 | 0.651083 | 0.651163 | 0.64539 | 0.645734 | 0.649167 | 0.646638 | 0.648417 |
| 0.735006 | 0.736065 | 0.728848 | 0.717292 | 0.739163 | 0.768812 | 0.734611 | 0.742396 | 0.72965 |
| 0.70439 | 0.704395 | 0.702031 | 0.702501 | 0.70614 | 0.707695 | 0.701832 | 0.707543 | 0.710238 |
| 0.682547 | 0.677758 | 0.676623 | 0.676151 | 0.683084 | 0.682474 | 0.676667 | 0.682004 | 0.678907 |
| 0.674156 | 0.671877 | 0.669861 | 0.671581 | 0.675715 | 0.67594 | 0.67499 | 0.674352 | 0.676667 |
| 0.653314 | 0.649735 | 0.654031 | 0.653536 | 0.652151 | 0.661276 | 0.656769 | 0.654832 | 0.658353 |
| 0.680264 | 0.678755 | 0.6795 | 0.67655 | 0.682832 | 0.682121 | 0.676827 | 0.678888 | 0.684355 |
| 0.671053 | 0.669824 | 0.669463 | 0.668597 | 0.677254 | 0.674557 | 0.674453 | 0.677411 | 0.674809 |
| 0.663621 | 0.660268 | 0.655765 | 0.65898 | 0.669651 | 0.665391 | 0.665043 | 0.668064 | 0.6626 |
| 0.67082 | 0.666567 | 0.663008 | 0.663462 | 0.6727 | 0.670934 | 0.662291 | 0.673144 | 0.666464 |
| 0.658797 | 0.658688 | 0.655554 | 0.658961 | 0.660584 | 0.659154 | 0.652716 | 0.660098 | 0.66273 |
| 0.667238 | 0.66654 | 0.659655 | 0.657959 | 0.670636 | 0.667048 | 0.65399 | 0.665391 | 0.672881 |
| 0.662891 | 0.655762 | 0.654909 | 0.660228 | 0.660833 | 0.664602 | 0.65803 | 0.670175 | 0.663857 |
| 0.652809 | 0.654649 | 0.634965 | 0.649396 | 0.653405 | 0.655921 | 0.645446 | 0.648267 | 0.653761 |

| | | | | | | | | |
|---|---|---|---|---|---|---|---|---|
| 0.693108 | 0.69194 | 0.687548 | 0.69157 | 0.694989 | 0.695526 | 0.689721 | 0.696095 | 0.698163 |
| 0.665406 | 0.659286 | 0.656078 | 0.65477 | 0.661787 | 0.664909 | 0.661221 | 0.661284 | 0.663366 |
| 0.665853 | 0.666762 | 0.662496 | 0.66388 | 0.670057 | 0.671165 | 0.667525 | 0.667541 | 0.668511 |
| 0.665934 | 0.663639 | 0.660006 | 0.666422 | 0.669732 | 0.663883 | 0.663501 | 0.668506 | 0.667111 |
| 0.658778 | 0.6591 | 0.651836 | 0.660196 | 0.667708 | 0.665817 | 0.659319 | 0.66266 | 0.670408 |
| 0.661972 | 0.661667 | 0.656175 | 0.665785 | 0.668982 | 0.667623 | 0.659816 | 0.670634 | 0.674763 |
| 0.659986 | 0.654114 | 0.651738 | 0.65072 | 0.665256 | 0.660417 | 0.64653 | 0.65991 | 0.660448 |
| 0.64891 | 0.6428 | 0.635756 | 0.644769 | 0.654694 | 0.65698 | 0.652244 | 0.656336 | 0.6614 |
| 0.652068 | 0.643836 | 0.652293 | 0.651156 | 0.649714 | 0.655958 | 0.647027 | 0.651429 | 0.661366 |
| 0.692829 | 0.691402 | 0.689337 | 0.692315 | 0.695095 | 0.694791 | 0.69225 | 0.694774 | 0.695484 |
| 0.693494 | 0.686029 | 0.686537 | 0.689357 | 0.692504 | 0.695754 | 0.690855 | 0.695336 | 0.693111 |
| 0.687819 | 0.680229 | 0.685791 | 0.686486 | 0.687783 | 0.689559 | 0.685289 | 0.688487 | 0.691089 |
| 0.680518 | 0.676196 | 0.678647 | 0.676374 | 0.684198 | 0.677484 | 0.675272 | 0.686084 | 0.678525 |
| 0.685926 | 0.679841 | 0.680924 | 0.685087 | 0.680725 | 0.685205 | 0.681117 | 0.683573 | 0.686606 |
| 0.680574 | 0.674832 | 0.673339 | 0.67588 | 0.674338 | 0.67599 | 0.671935 | 0.679378 | 0.676949 |

| Collection_ | Collection_ | Collection_ | Collection_ | Collection_ | Collection_ | Collection_ | Collection_ | Collection_ |
|---|---|---|---|---|---|---|---|---|
| 0.789645 | 0.791997 | 0.78077 | 0.753568 | 0.760426 | 0.748402 | 0.775774 | 0.743677 | 0.778048 |
| 0.781159 | 0.790486 | 0.776486 | 0.749239 | 0.772847 | 0.749787 | 0.787534 | 0.742227 | 0.753526 |
| 0.778017 | 0.784894 | 0.782339 | 0.744535 | 0.761298 | 0.744778 | 0.779041 | 0.733809 | 0.758001 |
| 0.777448 | 0.783262 | 0.778318 | 0.750443 | 0.757588 | 0.757472 | 0.766916 | 0.737792 | 0.760369 |
| 0.774911 | 0.777648 | 0.776251 | 0.750418 | 0.76181 | 0.750826 | 0.755212 | 0.740217 | 0.765957 |
| 0.766997 | 0.778676 | 0.776949 | 0.761351 | 0.758316 | 0.75754 | 0.748775 | 0.730599 | 0.744545 |
| 0.7095 | 0.713305 | 0.706993 | 0.685948 | 0.691678 | 0.682247 | 0.683199 | 0.665428 | 0.669239 |
| 0.706897 | 0.710611 | 0.704035 | 0.675216 | 0.686996 | 0.674242 | 0.677344 | 0.659904 | 0.659853 |
| 0.779581 | 0.783516 | 0.777433 | 0.75273 | 0.767168 | 0.76048 | 0.767523 | 0.736747 | 0.755063 |
| 0.772851 | 0.78742 | 0.769669 | 0.752285 | 0.756396 | 0.741597 | 0.757261 | 0.737167 | 0.777622 |
| 0.773632 | 0.774898 | 0.773882 | 0.757474 | 0.76192 | 0.742154 | 0.759114 | 0.745437 | 0.765661 |
| 0.789908 | 0.792194 | 0.784642 | 0.75549 | 0.772281 | 0.744651 | 0.776986 | 0.799925 | 0.781005 |
| 0.776783 | 0.770059 | 0.7742 | 0.759324 | 0.763442 | 0.755342 | 0.764366 | 0.759107 | 0.747108 |
| 0.776877 | 0.778437 | 0.769869 | 0.752064 | 0.765334 | 0.741414 | 0.758905 | 0.745609 | 0.752112 |
| 0.76243 | 0.767038 | 0.764711 | 0.736316 | 0.74549 | 0.738605 | 0.75447 | 0.736354 | 0.735217 |
| 0.778412 | 0.782201 | 0.778007 | 0.75748 | 0.759753 | 0.747211 | 0.768537 | 0.741087 | 0.76743 |
| 0.769148 | 0.775284 | 0.77102 | 0.746773 | 0.756632 | 0.73962 | 0.77451 | 0.731425 | 0.780225 |
| 0.755865 | 0.757427 | 0.749156 | 0.725696 | 0.748156 | 0.727603 | 0.733371 | 0.707432 | 0.724951 |
| 0.781029 | 0.78104 | 0.784787 | 0.759632 | 0.767623 | 0.753003 | 0.781144 | 0.788903 | 0.761281 |
| 0.792232 | 0.793159 | 0.783351 | 0.756495 | 0.778165 | 0.759751 | 0.77072 | 0.73667 | 0.756503 |
| 0.791228 | 0.796218 | 0.781232 | 0.755149 | 0.786798 | 0.763981 | 0.774805 | 0.740709 | 0.755561 |
| 0.797277 | 0.859214 | 0.781866 | 0.758417 | 0.780866 | 0.753502 | 0.769936 | 0.755987 | 0.775422 |
| 0.789294 | 0.802658 | 0.786145 | 0.764129 | 0.772947 | 0.76537 | 0.772938 | 0.757257 | 0.778132 |
| 0.797742 | 0.792825 | 0.78222 | 0.762623 | 0.776565 | 0.75765 | 0.779388 | 0.783431 | 0.779272 |
| 0.786706 | 0.790688 | 0.784429 | 0.758795 | 0.783643 | 0.750241 | 0.762532 | 0.762662 | 0.757637 |
| 0.792868 | 0.78759 | 0.773895 | 0.766067 | 0.772651 | 0.754037 | 0.775678 | 0.74534 | 0.755595 |
| 0.776044 | 0.780289 | 0.768503 | 0.752212 | 0.760721 | 0.74425 | 0.766628 | 0.758971 | 0.744098 |
| 0.78263 | 0.793639 | 0.775784 | 0.758215 | 0.761771 | 0.761327 | 0.758732 | 0.736988 | 0.760631 |
| 0.777897 | 0.79787 | 0.782908 | 0.756912 | 0.766315 | 0.750266 | 0.76034 | 0.764778 | 0.760177 |
| 0.768746 | 0.778862 | 0.764827 | 0.756886 | 0.766924 | 0.743161 | 0.759977 | 0.759763 | 0.767677 |
| 0.781803 | 0.785937 | 0.779416 | 0.766653 | 0.769077 | 0.753205 | 0.756336 | 0.74717 | 0.76043 |
| 0.779524 | 0.778841 | 0.770793 | 0.75865 | 0.76182 | 0.747537 | 0.763972 | 0.731431 | 0.749311 |
| 0.776584 | 0.778314 | 0.782535 | 0.760756 | 0.757806 | 0.745472 | 0.765044 | 0.756717 | 0.753142 |
| 0.777704 | 0.791184 | 0.772396 | 0.75633 | 0.755507 | 0.742625 | 0.756368 | 0.732716 | 0.746904 |
| 0.772095 | 0.768 | 0.764223 | 0.752228 | 0.751251 | 0.73251 | 0.760236 | 0.749054 | 0.739932 |
| 0.779892 | 0.781899 | 0.77696 | 0.755786 | 0.755621 | 0.754614 | 0.776011 | 0.771273 | 0.750761 |
| 0.782986 | 0.790505 | 0.779667 | 0.754305 | 0.779234 | 0.746306 | 0.768815 | 0.732437 | 0.757015 |
| 0.764203 | 0.786582 | 0.767477 | 0.743992 | 0.744612 | 0.742056 | 0.744428 | 0.753114 | 0.742683 |
| 0.7717 | 0.79539 | 0.77232 | 0.757372 | 0.758305 | 0.756717 | 0.747614 | 0.737784 | 0.750031 |

| | | | | | | | | |
|---|---|---|---|---|---|---|---|---|
| 0.758915 | 0.757375 | 0.754441 | 0.729496 | 0.736002 | 0.723468 | 0.751335 | 0.715676 | 0.734555 |
| 0.774971 | 0.773394 | 0.773194 | 0.746233 | 0.756484 | 0.749149 | 0.76018 | 0.729834 | 0.79126 |
| 0.788975 | 0.786942 | 0.781515 | 0.754771 | 0.773288 | 0.752509 | 0.770314 | 0.777406 | 0.786552 |
| 0.789443 | 0.792717 | 0.782983 | 0.756164 | 0.769255 | 0.749367 | 0.777534 | 0.774724 | 0.798152 |
| 0.785145 | 0.782558 | 0.781297 | 0.760623 | 0.773401 | 0.752664 | 0.770596 | 0.770184 | 0.771585 |
| 0.783477 | 0.781576 | 0.779057 | 0.75409 | 0.763822 | 0.75342 | 0.762219 | 0.748506 | 0.766066 |
| 0.776803 | 0.778724 | 0.777382 | 0.750105 | 0.755283 | 0.752848 | 0.746892 | 0.731646 | 0.75173 |
| 0.790975 | 0.790512 | 0.789088 | 0.75627 | 0.777013 | 0.76191 | 0.763533 | 0.758778 | 0.754326 |
| 0.771239 | 0.777152 | 0.770294 | 0.748604 | 0.765551 | 0.753868 | 0.760046 | 0.734137 | 0.745466 |
| 0.764205 | 0.776731 | 0.764049 | 0.749011 | 0.758452 | 0.751462 | 0.755191 | 0.731097 | 0.746642 |
| 0.77812 | 0.790911 | 0.779554 | 0.748214 | 0.780268 | 0.749198 | 0.775119 | 0.760602 | 0.775308 |
| 0.783034 | 0.784558 | 0.786289 | 0.760216 | 0.789479 | 0.752491 | 0.773277 | 0.754108 | 0.763875 |
| 0.782802 | 0.787362 | 0.787052 | 0.759405 | 0.769247 | 0.750509 | 0.780316 | 0.755962 | 0.785863 |
| 0.781234 | 0.787369 | 0.780236 | 0.759128 | 0.787979 | 0.747889 | 0.776919 | 0.749631 | 0.783164 |
| 0.78432 | 0.788802 | 0.774184 | 0.755442 | 0.790835 | 0.74721 | 0.777227 | 0.73998 | 0.761764 |
| 0.783747 | 0.793714 | 0.775877 | 0.753756 | 0.79374 | 0.744541 | 0.777317 | 0.737925 | 0.772198 |
| 0.775165 | 0.781595 | 0.774692 | 0.749867 | 0.75773 | 0.738241 | 0.783286 | 0.750588 | 0.770473 |
| 0.789614 | 0.786736 | 0.785714 | 0.762911 | 0.767073 | 0.756658 | 0.786166 | 0.740389 | 0.778476 |
| 0.779667 | 0.793862 | 0.786206 | 0.755231 | 0.77369 | 0.755159 | 0.758692 | 0.76084 | 0.776217 |
| 0.784019 | 0.787402 | 0.780267 | 0.754581 | 0.784104 | 0.756751 | 0.768047 | 0.759936 | 0.774127 |
| 0.777129 | 0.782121 | 0.778586 | 0.753776 | 0.758443 | 0.744313 | 0.792381 | 0.736643 | 0.778897 |
| 0.778221 | 0.777461 | 0.774027 | 0.761769 | 0.758914 | 0.747522 | 0.776829 | 0.736491 | 0.778876 |
| 0.77831 | 0.784533 | 0.781016 | 0.758103 | 0.775489 | 0.752179 | 0.769701 | 0.739251 | 0.752167 |
| 0.773516 | 0.783113 | 0.774668 | 0.749817 | 0.763371 | 0.75253 | 0.756199 | 0.736969 | 0.743719 |
| 0.763787 | 0.777092 | 0.761772 | 0.740733 | 0.746321 | 0.74836 | 0.743056 | 0.729969 | 0.735896 |
| 0.709362 | 0.711665 | 0.706236 | 0.677701 | 0.688821 | 0.675765 | 0.677394 | 0.660965 | 0.662881 |
| 0.709342 | 0.71171 | 0.706625 | 0.691588 | 0.692989 | 0.684004 | 0.679809 | 0.666862 | 0.670823 |
| 0.707945 | 0.712305 | 0.705025 | 0.686914 | 0.689711 | 0.683392 | 0.679679 | 0.667098 | 0.668977 |
| 0.77799 | 0.773659 | 0.770607 | 0.753036 | 0.769658 | 0.75 | 0.751955 | 0.735876 | 0.738935 |
| 0.778357 | 0.77489 | 0.774271 | 0.744868 | 0.770042 | 0.756047 | 0.757392 | 0.73813 | 0.744323 |
| 0.772829 | 0.770215 | 0.775119 | 0.747899 | 0.765244 | 0.747264 | 0.744773 | 0.725657 | 0.768316 |
| 0.775784 | 0.771883 | 0.771385 | 0.745276 | 0.77163 | 0.743413 | 0.750774 | 0.730099 | 0.745034 |
| 0.793197 | 0.797332 | 0.790575 | 0.761052 | 0.769239 | 0.75868 | 0.81877 | 0.746513 | 0.764496 |
| 0.795667 | 0.796522 | 0.791329 | 0.76584 | 0.762591 | 0.752839 | 0.816948 | 0.846401 | 0.767593 |
| 0.790235 | 0.791784 | 0.78175 | 0.756642 | 0.763725 | 0.761817 | 0.811534 | 0.749022 | 0.755094 |
| 0.792404 | 0.802673 | 0.786595 | 0.762507 | 0.765125 | 0.748536 | 0.844223 | 0.741869 | 0.755713 |
| 0.787906 | 0.782471 | 0.782435 | 0.765138 | 0.762366 | 0.765308 | 0.763185 | 0.74564 | 0.757416 |
| 0.813198 | 0.809747 | 0.80019 | 0.778938 | 0.791844 | 0.777248 | 0.808473 | 0.759975 | 0.807271 |
| 0.778621 | 0.788088 | 0.778574 | 0.753643 | 0.757859 | 0.753611 | 0.779988 | 0.741114 | 0.751976 |
| 0.799683 | 0.78795 | 0.789884 | 0.763299 | 0.848031 | 0.758965 | 0.763954 | 0.751213 | 0.758786 |
| 0.787711 | 0.788028 | 0.779645 | 0.755149 | 0.76038 | 0.752414 | 0.79659 | 0.741756 | 0.762675 |
| 0.808601 | 0.807109 | 0.8 | 0.780712 | 0.787995 | 0.777524 | 0.803563 | 0.75762 | 0.800729 |
| 0.787073 | 0.786165 | 0.785539 | 0.759131 | 0.768641 | 0.754582 | 0.767477 | 0.744508 | 0.819423 |
| 0.783786 | 0.790007 | 0.781083 | 0.75361 | 0.766688 | 0.753244 | 0.764421 | 0.781612 | 0.796506 |
| 0.793083 | 0.787869 | 0.785518 | 0.765441 | 0.809248 | 0.751846 | 0.76087 | 0.750031 | 0.7569 |
| 0.783484 | 0.784563 | 0.782857 | 0.757812 | 0.75874 | 0.756392 | 0.806516 | 0.745419 | 0.751728 |

| | | | | | | | | |
|---|---|---|---|---|---|---|---|---|
| 0.788391 | 0.79166 | 0.781424 | 0.758639 | 0.771832 | 0.759184 | 0.767523 | 0.744614 | 0.806155 |
| 0.787139 | 0.786922 | 0.775129 | 0.7757 | 0.811112 | 0.756069 | 0.757809 | 0.7419 | 0.757453 |
| 0.787449 | 0.780183 | 0.78229 | 0.762781 | 0.77267 | 0.758462 | 0.765641 | 0.747749 | 0.774714 |
| 0.79866 | 0.79836 | 0.788157 | 0.771982 | 0.782965 | 0.769538 | 0.787925 | 0.751038 | 0.782575 |
| 0.788377 | 0.787927 | 0.784764 | 0.771878 | 0.8119 | 0.76561 | 0.766095 | 0.759365 | 0.760237 |
| 0.79686 | 0.793545 | 0.775234 | 0.75845 | 0.809953 | 0.762308 | 0.766524 | 0.741747 | 0.75912 |
| 0.784888 | 0.787401 | 0.776237 | 0.753425 | 0.758922 | 0.76458 | 0.757753 | 0.741436 | 0.757141 |
| 0.77515 | 0.776739 | 0.775393 | 0.784251 | 0.757858 | 0.754049 | 0.760009 | 0.742511 | 0.756197 |
| 0.788978 | 0.779204 | 0.78303 | 0.757451 | 0.761444 | 0.748609 | 0.756427 | 0.740085 | 0.75038 |
| 0.780106 | 0.769872 | 0.77831 | 0.753039 | 0.759242 | 0.747685 | 0.750029 | 0.771506 | 0.758083 |
| 0.78313 | 0.778266 | 0.778875 | 0.756734 | 0.764614 | 0.754317 | 0.758005 | 0.743269 | 0.762969 |
| 0.778646 | 0.774466 | 0.781259 | 0.754303 | 0.755639 | 0.748552 | 0.757478 | 0.777308 | 0.768661 |
| 0.832146 | 0.788582 | 0.786399 | 0.755798 | 0.775175 | 0.749866 | 0.759236 | 0.764117 | 0.760459 |
| 0.784912 | 0.777976 | 0.789681 | 0.766663 | 0.773065 | 0.745256 | 0.749019 | 0.797966 | 0.773817 |
| 0.782101 | 0.776295 | 0.777339 | 0.750762 | 0.76218 | 0.751976 | 0.767399 | 0.733261 | 0.74258 |
| 0.782471 | 0.775766 | 0.781847 | 0.754057 | 0.7602 | 0.753327 | 0.764339 | 0.741467 | 0.747738 |
| 0.7771 | 0.775553 | 0.777578 | 0.756077 | 0.764575 | 0.758397 | 0.754555 | 0.751881 | 0.761022 |
| 0.790189 | 0.793365 | 0.77732 | 0.756361 | 0.761949 | 0.751246 | 0.800305 | 0.739771 | 0.752734 |
| 0.789799 | 0.776558 | 0.790815 | 0.758136 | 0.764478 | 0.757256 | 0.762637 | 0.765973 | 0.763261 |
| 0.778111 | 0.775656 | 0.780407 | 0.758533 | 0.760311 | 0.751098 | 0.755884 | 0.740966 | 0.752276 |
| 0.776899 | 0.779904 | 0.773006 | 0.756811 | 0.759391 | 0.75373 | 0.758172 | 0.748995 | 0.758974 |
| 0.782746 | 0.774958 | 0.7824 | 0.755066 | 0.754063 | 0.761164 | 0.755132 | 0.741986 | 0.754933 |
| 0.784857 | 0.778508 | 0.778421 | 0.764433 | 0.768453 | 0.758824 | 0.770341 | 0.748476 | 0.763803 |
| 0.787778 | 0.782143 | 0.783156 | 0.772094 | 0.848228 | 0.752977 | 0.760294 | 0.772755 | 0.768228 |
| 0.777396 | 0.781086 | 0.77969 | 0.761558 | 0.762286 | 0.752329 | 0.778751 | 0.743146 | 0.753381 |
| 0.779375 | 0.784003 | 0.776897 | 0.751251 | 0.759775 | 0.765214 | 0.758865 | 0.741348 | 0.751409 |
| 0.780933 | 0.782442 | 0.781502 | 0.761466 | 0.757994 | 0.749921 | 0.765767 | 0.743911 | 0.755751 |
| 0.777006 | 0.77165 | 0.778507 | 0.746717 | 0.762109 | 0.750744 | 0.758756 | 0.743338 | 0.748366 |
| 0.784935 | 0.779982 | 0.778633 | 0.751203 | 0.759242 | 0.761027 | 0.76279 | 0.744355 | 0.756842 |
| 0.784929 | 0.779655 | 0.778724 | 0.76041 | 0.76058 | 0.755803 | 0.764487 | 0.746163 | 0.75101 |
| 0.787652 | 0.78495 | 0.783525 | 0.758418 | 0.764694 | 0.745209 | 0.765923 | 0.771197 | 0.756501 |
| 0.778171 | 0.774506 | 0.780522 | 0.754322 | 0.761519 | 0.75103 | 0.753673 | 0.737713 | 0.749469 |
| 0.782346 | 0.782169 | 0.784404 | 0.762107 | 0.766103 | 0.764668 | 0.763558 | 0.742972 | 0.761626 |
| 0.791396 | 0.793002 | 0.782449 | 0.759724 | 0.770504 | 0.752095 | 0.781445 | 0.77608 | 0.799171 |
| 1 | 0.800816 | 0.791052 | 0.763711 | 0.781968 | 0.754924 | 0.768707 | 0.761695 | 0.762969 |
| 0.800816 | 1 | 0.784237 | 0.75965 | 0.773232 | 0.749331 | 0.7665 | 0.759995 | 0.761814 |
| 0.791052 | 0.784237 | 1 | 0.755634 | 0.760191 | 0.739801 | 0.762989 | 0.7585 | 0.758222 |
| 0.763711 | 0.75965 | 0.755634 | 1 | 0.75602 | 0.74931 | 0.753482 | 0.749566 | 0.757426 |
| 0.781968 | 0.773232 | 0.760191 | 0.75602 | 1 | 0.749276 | 0.759414 | 0.736961 | 0.761124 |
| 0.754924 | 0.749331 | 0.739801 | 0.74931 | 0.749276 | 1 | 0.744322 | 0.722955 | 0.74111 |
| 0.768707 | 0.7665 | 0.762989 | 0.753482 | 0.759414 | 0.744322 | 1 | 0.739472 | 0.756946 |
| 0.761695 | 0.759995 | 0.7585 | 0.749566 | 0.736961 | 0.722955 | 0.739472 | 1 | 0.770833 |
| 0.762969 | 0.761814 | 0.758222 | 0.757426 | 0.761124 | 0.74111 | 0.756946 | 0.770833 | 1 |
| 0.759241 | 0.758093 | 0.754292 | 0.755418 | 0.764917 | 0.745442 | 0.776263 | 0.731932 | 0.748913 |
| 0.755785 | 0.748476 | 0.745074 | 0.744064 | 0.749786 | 0.738633 | 0.749237 | 0.735581 | 0.745481 |
| 0.76196 | 0.761587 | 0.758724 | 0.780073 | 0.756225 | 0.751843 | 0.773599 | 0.735195 | 0.763741 |
| 0.764531 | 0.762743 | 0.756444 | 0.751898 | 0.754112 | 0.738735 | 0.846249 | 0.73593 | 0.750854 |

| | | | | | | | | |
|---|---|---|---|---|---|---|---|---|
| 0.748023 | 0.747209 | 0.742418 | 0.807351 | 0.750314 | 0.740364 | 0.744403 | 0.729953 | 0.740758 |
| 0.758831 | 0.781164 | 0.751241 | 0.741334 | 0.750697 | 0.747583 | 0.800413 | 0.734112 | 0.751649 |
| 0.745171 | 0.744031 | 0.742871 | 0.740653 | 0.74851 | 0.739804 | 0.743976 | 0.721641 | 0.739338 |
| 0.750068 | 0.747582 | 0.737887 | 0.735988 | 0.74539 | 0.74412 | 0.742875 | 0.733849 | 0.79084 |
| 0.765402 | 0.762625 | 0.757386 | 0.750036 | 0.75535 | 0.741856 | 0.749768 | 0.843037 | 0.765251 |
| 0.752929 | 0.745006 | 0.746398 | 0.754596 | 0.743527 | 0.749883 | 0.743243 | 0.735657 | 0.751102 |
| 0.756529 | 0.748008 | 0.749831 | 0.740153 | 0.76057 | 0.751724 | 0.747819 | 0.727861 | 0.751084 |
| 0.75495 | 0.775063 | 0.759607 | 0.743077 | 0.753011 | 0.737431 | 0.790741 | 0.775622 | 0.765935 |
| 0.740049 | 0.725182 | 0.744609 | 0.774035 | 0.731042 | 0.718374 | 0.734286 | 0.777698 | 0.747559 |
| 0.754749 | 0.754206 | 0.739802 | 0.735256 | 0.754894 | 0.753314 | 0.748473 | 0.726641 | 0.750589 |
| 0.75856 | 0.748209 | 0.753591 | 0.782841 | 0.747182 | 0.745434 | 0.747631 | 0.734839 | 0.750654 |
| 0.770124 | 0.778064 | 0.765684 | 0.7435 | 0.754705 | 0.738761 | 0.747816 | 0.736515 | 0.789883 |
| 0.786033 | 0.776949 | 0.777457 | 0.76079 | 0.752773 | 0.75908 | 0.751114 | 0.733822 | 0.742286 |
| 0.755073 | 0.754266 | 0.743065 | 0.766314 | 0.742081 | 0.750588 | 0.738854 | 0.734241 | 0.740768 |
| 0.78163 | 0.78204 | 0.777863 | 0.767908 | 0.76559 | 0.750133 | 0.764692 | 0.756704 | 0.759318 |
| 0.77489 | 0.769006 | 0.774118 | 0.748309 | 0.755227 | 0.739801 | 0.748022 | 0.774784 | 0.76408 |
| 0.775144 | 0.771732 | 0.776647 | 0.750052 | 0.758004 | 0.747304 | 0.748257 | 0.742785 | 0.754388 |
| 0.776925 | 0.776703 | 0.780504 | 0.752498 | 0.755719 | 0.754818 | 0.751791 | 0.741186 | 0.743835 |
| 0.760535 | 0.758972 | 0.757002 | 0.749741 | 0.767261 | 0.732098 | 0.738254 | 0.717003 | 0.731689 |
| 0.783254 | 0.787816 | 0.778554 | 0.765524 | 0.750181 | 0.750053 | 0.744967 | 0.839879 | 0.761899 |
| 0.748552 | 0.758556 | 0.737568 | 0.756632 | 0.739795 | 0.751589 | 0.736326 | 0.720363 | 0.743961 |
| 0.774573 | 0.772085 | 0.775672 | 0.750996 | 0.757221 | 0.753792 | 0.744093 | 0.731531 | 0.743303 |
| 0.772471 | 0.775132 | 0.774145 | 0.748872 | 0.749817 | 0.751834 | 0.74287 | 0.736804 | 0.734873 |
| 0.761669 | 0.768262 | 0.735489 | 0.737461 | 0.78599 | 0.734058 | 0.738703 | 0.710368 | 0.73568 |
| 0.751173 | 0.745637 | 0.750941 | 0.732244 | 0.731805 | 0.713419 | 0.751022 | 0.717159 | 0.738291 |
| 0.762152 | 0.768968 | 0.756234 | 0.758326 | 0.760547 | 0.744776 | 0.780379 | 0.738522 | 0.75309 |
| 0.760981 | 0.763466 | 0.753834 | 0.751181 | 0.734651 | 0.748506 | 0.737472 | 0.719244 | 0.730136 |
| 0.782576 | 0.773654 | 0.769202 | 0.749948 | 0.767206 | 0.745659 | 0.75539 | 0.728277 | 0.740453 |
| 0.75372 | 0.745093 | 0.754052 | 0.725051 | 0.726494 | 0.727451 | 0.725075 | 0.711158 | 0.718796 |
| 0.768045 | 0.767551 | 0.767012 | 0.762514 | 0.753916 | 0.746067 | 0.744943 | 0.724342 | 0.736628 |
| 0.770225 | 0.764068 | 0.77156 | 0.744744 | 0.751894 | 0.745894 | 0.742469 | 0.726584 | 0.732834 |
| 0.766904 | 0.761905 | 0.760904 | 0.731652 | 0.743924 | 0.734428 | 0.751292 | 0.717123 | 0.744463 |
| 0.765795 | 0.763003 | 0.763691 | 0.735233 | 0.747362 | 0.734687 | 0.754019 | 0.726563 | 0.748555 |
| 0.76346 | 0.779657 | 0.762333 | 0.746835 | 0.740607 | 0.74992 | 0.740471 | 0.717179 | 0.734414 |
| 0.763377 | 0.758559 | 0.761579 | 0.796508 | 0.743265 | 0.740985 | 0.741108 | 0.728386 | 0.735628 |
| 0.770293 | 0.775836 | 0.762395 | 0.749817 | 0.763051 | 0.74261 | 0.750143 | 0.72639 | 0.736849 |
| 0.728001 | 0.721389 | 0.727083 | 0.706715 | 0.706076 | 0.702651 | 0.712068 | 0.684459 | 0.703306 |
| 0.756909 | 0.760837 | 0.754083 | 0.731313 | 0.739184 | 0.731295 | 0.73549 | 0.717839 | 0.723452 |
| 0.733307 | 0.727862 | 0.728409 | 0.721904 | 0.707856 | 0.712117 | 0.705045 | 0.686131 | 0.698375 |
| 0.72907 | 0.723765 | 0.72104 | 0.701598 | 0.700443 | 0.706991 | 0.687493 | 0.685809 | 0.686701 |
| 0.773077 | 0.769561 | 0.76988 | 0.74552 | 0.764645 | 0.744688 | 0.757185 | 0.754758 | 0.744151 |
| 0.726809 | 0.719273 | 0.716693 | 0.697491 | 0.706529 | 0.695274 | 0.69711 | 0.678818 | 0.687882 |
| 0.721211 | 0.717605 | 0.719048 | 0.688741 | 0.696916 | 0.690978 | 0.705482 | 0.676791 | 0.683144 |
| 0.709204 | 0.707189 | 0.708561 | 0.679898 | 0.689176 | 0.682815 | 0.689443 | 0.664417 | 0.676905 |
| 0.763927 | 0.763546 | 0.757914 | 0.745112 | 0.751365 | 0.738621 | 0.744659 | 0.717827 | 0.750467 |
| 0.711289 | 0.710223 | 0.706578 | 0.678939 | 0.687281 | 0.686195 | 0.694334 | 0.665061 | 0.672742 |
| 0.765809 | 0.7656 | 0.764497 | 0.74302 | 0.755971 | 0.742214 | 0.753192 | 0.741178 | 0.743398 |

| | | | | | | | | |
|---|---|---|---|---|---|---|---|---|
| 0.763029 | 0.762559 | 0.757423 | 0.737767 | 0.744076 | 0.735654 | 0.743963 | 0.718631 | 0.738522 |
| 0.766013 | 0.771109 | 0.761989 | 0.736913 | 0.748407 | 0.738105 | 0.747789 | 0.736685 | 0.742697 |
| 0.754031 | 0.752776 | 0.753391 | 0.724177 | 0.725843 | 0.732684 | 0.728246 | 0.710828 | 0.727127 |
| 0.69955 | 0.697815 | 0.70026 | 0.66689 | 0.678174 | 0.677697 | 0.69158 | 0.651468 | 0.660002 |
| 0.699327 | 0.69328 | 0.696114 | 0.67482 | 0.680832 | 0.675297 | 0.678415 | 0.661437 | 0.666112 |
| 0.76556 | 0.767291 | 0.762652 | 0.74587 | 0.752297 | 0.736842 | 0.753127 | 0.73586 | 0.75395 |
| 0.694767 | 0.689719 | 0.692175 | 0.668524 | 0.676414 | 0.672123 | 0.666282 | 0.656357 | 0.6581 |
| 0.737947 | 0.735549 | 0.727928 | 0.710271 | 0.716009 | 0.703894 | 0.707736 | 0.691164 | 0.692157 |
| 0.754892 | 0.760046 | 0.754261 | 0.736395 | 0.742138 | 0.735861 | 0.751601 | 0.71878 | 0.734635 |
| 0.716528 | 0.723632 | 0.718439 | 0.693871 | 0.70273 | 0.693339 | 0.69095 | 0.67616 | 0.68027 |
| 0.751561 | 0.748113 | 0.748304 | 0.723444 | 0.724508 | 0.722158 | 0.739876 | 0.712138 | 0.72456 |
| 0.635762 | 0.617284 | 0.605634 | 0.61875 | 0.62 | 0.625 | 0.630137 | 0.59375 | 0.655629 |
| 0.652589 | 0.661568 | 0.641133 | 0.661517 | 0.663366 | 0.653145 | 0.657485 | 0.644763 | 0.651228 |
| 0.660512 | 0.661513 | 0.650026 | 0.658935 | 0.658216 | 0.651934 | 0.654764 | 0.644239 | 0.649249 |
| 0.662156 | 0.669334 | 0.665448 | 0.668107 | 0.670896 | 0.66331 | 0.662537 | 0.656826 | 0.659348 |
| 0.662321 | 0.661481 | 0.655561 | 0.663282 | 0.661005 | 0.657273 | 0.659699 | 0.644029 | 0.651555 |
| 0.617824 | 0.619121 | 0.619982 | 0.613532 | 0.619094 | 0.617716 | 0.615607 | 0.596312 | 0.600028 |
| 0.663668 | 0.67116 | 0.657351 | 0.661022 | 0.66789 | 0.657515 | 0.660968 | 0.651862 | 0.653724 |
| 0.744932 | 0.746795 | 0.742211 | 0.713873 | 0.723968 | 0.719245 | 0.723009 | 0.711828 | 0.708544 |
| 0.740064 | 0.742568 | 0.735602 | 0.714123 | 0.719464 | 0.713923 | 0.726316 | 0.693807 | 0.709327 |
| 0.746875 | 0.745613 | 0.743666 | 0.718015 | 0.722573 | 0.719476 | 0.737 | 0.704055 | 0.713984 |
| 0.618973 | 0.609483 | 0.615066 | 0.584548 | 0.590191 | 0.587836 | 0.584278 | 0.576099 | 0.571678 |
| 0.601435 | 0.595297 | 0.600798 | 0.569967 | 0.573782 | 0.573957 | 0.567423 | 0.560822 | 0.553841 |
| 0.711371 | 0.709371 | 0.706618 | 0.680432 | 0.68907 | 0.680664 | 0.676208 | 0.661794 | 0.666301 |
| 0.70794 | 0.708054 | 0.706839 | 0.680896 | 0.690187 | 0.679716 | 0.677849 | 0.663468 | 0.666828 |
| 0.701571 | 0.701194 | 0.699133 | 0.67302 | 0.680524 | 0.671337 | 0.668741 | 0.652804 | 0.661893 |
| 0.701633 | 0.70962 | 0.701128 | 0.677276 | 0.684402 | 0.678027 | 0.672641 | 0.656152 | 0.663621 |
| 0.709994 | 0.709745 | 0.70862 | 0.682971 | 0.686719 | 0.681073 | 0.677987 | 0.66007 | 0.66973 |
| 0.707903 | 0.712086 | 0.708406 | 0.682382 | 0.691994 | 0.676382 | 0.680652 | 0.664787 | 0.666344 |
| 0.707926 | 0.712168 | 0.707448 | 0.679723 | 0.688053 | 0.680033 | 0.680164 | 0.65629 | 0.666027 |
| 0.704779 | 0.709588 | 0.7058 | 0.683039 | 0.694331 | 0.68136 | 0.678231 | 0.668663 | 0.671631 |
| 0.71316 | 0.716513 | 0.710443 | 0.689673 | 0.697248 | 0.687083 | 0.682814 | 0.67027 | 0.676065 |
| 0.708835 | 0.710821 | 0.713223 | 0.686312 | 0.691534 | 0.682369 | 0.678211 | 0.664639 | 0.667952 |
| 0.708106 | 0.709484 | 0.707956 | 0.681805 | 0.691466 | 0.676542 | 0.678862 | 0.662302 | 0.670357 |
| 0.711686 | 0.715653 | 0.713792 | 0.69161 | 0.693167 | 0.683603 | 0.678758 | 0.66729 | 0.672253 |
| 0.703043 | 0.709834 | 0.702325 | 0.681148 | 0.68818 | 0.67465 | 0.677186 | 0.658027 | 0.66566 |
| 0.708993 | 0.712126 | 0.706636 | 0.687557 | 0.68757 | 0.680987 | 0.680196 | 0.664599 | 0.670109 |
| 0.710423 | 0.708827 | 0.709023 | 0.684599 | 0.688685 | 0.685531 | 0.682938 | 0.662739 | 0.669657 |
| 0.703414 | 0.705005 | 0.705712 | 0.679922 | 0.682538 | 0.680092 | 0.674062 | 0.661484 | 0.664555 |
| 0.708473 | 0.711556 | 0.70848 | 0.688163 | 0.69216 | 0.680796 | 0.680093 | 0.669778 | 0.668399 |
| 0.661733 | 0.662887 | 0.649714 | 0.662725 | 0.66181 | 0.656854 | 0.652886 | 0.648119 | 0.651346 |
| 0.657576 | 0.665452 | 0.65827 | 0.665141 | 0.662874 | 0.666211 | 0.654447 | 0.656327 | 0.655087 |
| 0.710423 | 0.711138 | 0.710748 | 0.692206 | 0.693767 | 0.681139 | 0.682725 | 0.668448 | 0.674598 |
| 0.711129 | 0.712457 | 0.710827 | 0.689932 | 0.693935 | 0.686959 | 0.685426 | 0.66593 | 0.675705 |
| 0.64695 | 0.648059 | 0.63944 | 0.647906 | 0.641367 | 0.642395 | 0.642726 | 0.637082 | 0.637657 |
| 0.708901 | 0.71147 | 0.708326 | 0.685897 | 0.689221 | 0.683518 | 0.678686 | 0.66866 | 0.675911 |
| 0.698387 | 0.70056 | 0.700764 | 0.672903 | 0.67622 | 0.669497 | 0.673451 | 0.656388 | 0.658678 |

| | | | | | | | | |
|---|---|---|---|---|---|---|---|---|
| 0.707591 | 0.707165 | 0.702378 | 0.682984 | 0.687343 | 0.681183 | 0.676451 | 0.661172 | 0.665825 |
| 0.708764 | 0.707317 | 0.706692 | 0.684057 | 0.689448 | 0.682809 | 0.677979 | 0.663546 | 0.668902 |
| 0.70818 | 0.710568 | 0.708936 | 0.688095 | 0.692268 | 0.681861 | 0.68043 | 0.667884 | 0.668856 |
| 0.710307 | 0.710452 | 0.704759 | 0.687399 | 0.68877 | 0.68345 | 0.680451 | 0.666279 | 0.669888 |
| 0.715272 | 0.715151 | 0.712136 | 0.692105 | 0.697098 | 0.686011 | 0.688687 | 0.678083 | 0.674651 |
| 0.703951 | 0.706011 | 0.70177 | 0.683158 | 0.689512 | 0.680541 | 0.67648 | 0.664302 | 0.667149 |
| 0.705069 | 0.70776 | 0.701895 | 0.686273 | 0.687669 | 0.682269 | 0.676138 | 0.664307 | 0.668022 |
| 0.699814 | 0.703036 | 0.700475 | 0.67173 | 0.673374 | 0.671522 | 0.671144 | 0.657416 | 0.660988 |
| 0.70501 | 0.703826 | 0.702348 | 0.685578 | 0.685116 | 0.678406 | 0.673447 | 0.662551 | 0.664376 |
| 0.70346 | 0.705925 | 0.704306 | 0.680823 | 0.688072 | 0.675793 | 0.674839 | 0.662096 | 0.664037 |
| 0.706478 | 0.711754 | 0.70895 | 0.684625 | 0.689294 | 0.675662 | 0.678935 | 0.662024 | 0.666506 |
| 0.705373 | 0.703648 | 0.70323 | 0.680322 | 0.682837 | 0.676301 | 0.675171 | 0.659451 | 0.669575 |
| 0.698829 | 0.700277 | 0.697772 | 0.67195 | 0.679355 | 0.67043 | 0.672875 | 0.657176 | 0.661172 |
| 0.696718 | 0.701253 | 0.693895 | 0.677453 | 0.681785 | 0.673444 | 0.669148 | 0.657405 | 0.662102 |
| 0.713671 | 0.71165 | 0.715047 | 0.6915 | 0.69624 | 0.68734 | 0.684942 | 0.675196 | 0.67221 |
| 0.710818 | 0.714643 | 0.707942 | 0.682757 | 0.688104 | 0.683251 | 0.683721 | 0.665271 | 0.673065 |
| 0.703449 | 0.705218 | 0.698343 | 0.678046 | 0.684671 | 0.675342 | 0.677769 | 0.659662 | 0.665529 |
| 0.707343 | 0.706044 | 0.704267 | 0.680122 | 0.682274 | 0.679212 | 0.675778 | 0.659535 | 0.666423 |
| 0.706906 | 0.706636 | 0.705701 | 0.677299 | 0.685027 | 0.677184 | 0.676343 | 0.659748 | 0.6653 |
| 0.705181 | 0.705058 | 0.703053 | 0.677364 | 0.681291 | 0.678568 | 0.676233 | 0.662338 | 0.666586 |
| 0.702166 | 0.702958 | 0.696087 | 0.673132 | 0.678334 | 0.679486 | 0.668386 | 0.661332 | 0.65903 |
| 0.703484 | 0.70536 | 0.7022 | 0.679437 | 0.684803 | 0.677251 | 0.675293 | 0.660206 | 0.664059 |
| 0.706485 | 0.709475 | 0.706179 | 0.685359 | 0.68706 | 0.679886 | 0.680835 | 0.66291 | 0.669545 |
| 0.708598 | 0.709338 | 0.703996 | 0.681612 | 0.688473 | 0.678593 | 0.679386 | 0.662738 | 0.668599 |
| 0.704066 | 0.712936 | 0.697439 | 0.67871 | 0.682243 | 0.677359 | 0.672887 | 0.66176 | 0.661224 |
| 0.70511 | 0.704472 | 0.703169 | 0.67469 | 0.680529 | 0.675062 | 0.671953 | 0.659517 | 0.662658 |
| 0.704319 | 0.703664 | 0.702177 | 0.677175 | 0.681711 | 0.67486 | 0.673412 | 0.655992 | 0.665604 |
| 0.699691 | 0.703039 | 0.697692 | 0.674409 | 0.684788 | 0.67158 | 0.672333 | 0.653629 | 0.664996 |
| 0.704442 | 0.704248 | 0.700777 | 0.678387 | 0.684487 | 0.675146 | 0.671513 | 0.663539 | 0.66376 |
| 0.705831 | 0.706541 | 0.703614 | 0.68185 | 0.685902 | 0.672463 | 0.678627 | 0.660593 | 0.666344 |
| 0.70838 | 0.709195 | 0.706731 | 0.681224 | 0.685882 | 0.681299 | 0.678919 | 0.66123 | 0.66856 |
| 0.71257 | 0.712976 | 0.70578 | 0.682762 | 0.689316 | 0.68684 | 0.682247 | 0.664307 | 0.67378 |
| 0.713228 | 0.715171 | 0.713771 | 0.688824 | 0.693995 | 0.686379 | 0.683554 | 0.668101 | 0.670834 |
| 0.712648 | 0.716474 | 0.708378 | 0.686478 | 0.694464 | 0.686398 | 0.684515 | 0.666667 | 0.674815 |
| 0.701908 | 0.705115 | 0.700426 | 0.673447 | 0.685332 | 0.676206 | 0.67066 | 0.660207 | 0.661204 |
| 0.705037 | 0.707483 | 0.702564 | 0.677839 | 0.687786 | 0.678974 | 0.675887 | 0.664316 | 0.666991 |
| 0.708098 | 0.708537 | 0.70493 | 0.677852 | 0.691728 | 0.675542 | 0.676903 | 0.667805 | 0.667113 |
| 0.702834 | 0.709905 | 0.700055 | 0.679261 | 0.68583 | 0.674704 | 0.675598 | 0.661355 | 0.663089 |
| 0.70166 | 0.699918 | 0.694447 | 0.668911 | 0.676378 | 0.674035 | 0.671044 | 0.656239 | 0.659515 |
| 0.700192 | 0.701418 | 0.69501 | 0.676546 | 0.683359 | 0.670874 | 0.670333 | 0.657 | 0.663813 |
| 0.695993 | 0.700234 | 0.691826 | 0.670605 | 0.676428 | 0.670069 | 0.668675 | 0.660588 | 0.656223 |
| 0.698826 | 0.699057 | 0.697535 | 0.677812 | 0.67682 | 0.673202 | 0.671669 | 0.658598 | 0.660045 |
| 0.698982 | 0.699141 | 0.699754 | 0.672956 | 0.675005 | 0.675699 | 0.670509 | 0.658704 | 0.661879 |
| 0.689342 | 0.695502 | 0.688884 | 0.671084 | 0.674077 | 0.669596 | 0.668032 | 0.654233 | 0.663973 |
| 0.700109 | 0.707537 | 0.696187 | 0.67823 | 0.685759 | 0.676807 | 0.677386 | 0.659974 | 0.667119 |
| 0.695253 | 0.699501 | 0.693201 | 0.670933 | 0.680158 | 0.671609 | 0.671218 | 0.657607 | 0.661425 |

| | | | | | | | | |
|---|---|---|---|---|---|---|---|---|
| 0.710605 | 0.708743 | 0.70361 | 0.679688 | 0.689403 | 0.680302 | 0.67951 | 0.665102 | 0.665536 |
| 0.707362 | 0.709372 | 0.700292 | 0.687615 | 0.689112 | 0.678948 | 0.681117 | 0.66698 | 0.673288 |
| 0.705949 | 0.707678 | 0.703619 | 0.677635 | 0.684351 | 0.676124 | 0.67754 | 0.656084 | 0.660465 |
| 0.709963 | 0.709457 | 0.704384 | 0.68633 | 0.686811 | 0.678013 | 0.681003 | 0.665003 | 0.670768 |
| 0.715031 | 0.717749 | 0.711951 | 0.690817 | 0.697264 | 0.683517 | 0.691244 | 0.67115 | 0.676104 |
| 0.708134 | 0.712061 | 0.709625 | 0.691108 | 0.688533 | 0.686659 | 0.680945 | 0.670742 | 0.663317 |
| 0.704532 | 0.705561 | 0.701205 | 0.68225 | 0.687403 | 0.680547 | 0.674521 | 0.664315 | 0.666012 |
| 0.705174 | 0.705688 | 0.702262 | 0.680795 | 0.686051 | 0.680799 | 0.680009 | 0.665732 | 0.668033 |
| 0.702797 | 0.702415 | 0.700884 | 0.687727 | 0.690002 | 0.683041 | 0.679463 | 0.671611 | 0.673186 |
| 0.709398 | 0.711691 | 0.702147 | 0.682131 | 0.687802 | 0.677002 | 0.681869 | 0.665539 | 0.666506 |
| 0.706306 | 0.710042 | 0.707892 | 0.685795 | 0.690964 | 0.682737 | 0.68283 | 0.668485 | 0.663528 |
| 0.706848 | 0.712624 | 0.704404 | 0.689121 | 0.691618 | 0.682689 | 0.680964 | 0.670876 | 0.667148 |
| 0.704555 | 0.708024 | 0.703513 | 0.680351 | 0.685441 | 0.675983 | 0.677676 | 0.664687 | 0.665501 |
| 0.70711 | 0.709912 | 0.703276 | 0.684423 | 0.689003 | 0.682197 | 0.680448 | 0.662599 | 0.666747 |
| 0.708081 | 0.708567 | 0.705044 | 0.68365 | 0.692416 | 0.682069 | 0.67959 | 0.66628 | 0.666707 |
| 0.706239 | 0.713974 | 0.708805 | 0.684707 | 0.68954 | 0.678447 | 0.679876 | 0.671088 | 0.669048 |
| 0.712062 | 0.714503 | 0.709643 | 0.688544 | 0.693012 | 0.682787 | 0.68355 | 0.666705 | 0.668743 |
| 0.710612 | 0.709526 | 0.704598 | 0.683788 | 0.689714 | 0.677573 | 0.683667 | 0.666821 | 0.667861 |
| 0.706459 | 0.714147 | 0.708085 | 0.687273 | 0.689481 | 0.680221 | 0.686418 | 0.666043 | 0.671905 |
| 0.704315 | 0.710945 | 0.702771 | 0.684163 | 0.686477 | 0.678659 | 0.68094 | 0.662602 | 0.66056 |
| 0.705325 | 0.712786 | 0.704958 | 0.688425 | 0.687424 | 0.680394 | 0.677039 | 0.663727 | 0.667829 |
| 0.70611 | 0.707095 | 0.707957 | 0.684733 | 0.687759 | 0.678157 | 0.679548 | 0.663517 | 0.666024 |
| 0.712008 | 0.714297 | 0.705322 | 0.688857 | 0.692261 | 0.684848 | 0.683256 | 0.665504 | 0.664672 |
| 0.71106 | 0.713023 | 0.707895 | 0.686271 | 0.690694 | 0.684243 | 0.679141 | 0.669144 | 0.670602 |
| 0.707023 | 0.707941 | 0.706576 | 0.682563 | 0.687319 | 0.679437 | 0.676641 | 0.662522 | 0.66834 |
| 0.70842 | 0.705404 | 0.707813 | 0.68307 | 0.68583 | 0.680814 | 0.68136 | 0.667174 | 0.672408 |
| 0.704655 | 0.704964 | 0.701534 | 0.678654 | 0.685867 | 0.680057 | 0.674664 | 0.660371 | 0.666023 |
| 0.705304 | 0.705022 | 0.703126 | 0.683064 | 0.68298 | 0.679191 | 0.676374 | 0.664234 | 0.668825 |
| 0.70869 | 0.7094 | 0.706284 | 0.681199 | 0.689399 | 0.68274 | 0.681083 | 0.668252 | 0.663791 |
| 0.710214 | 0.71287 | 0.702163 | 0.690896 | 0.691358 | 0.686367 | 0.684051 | 0.664894 | 0.672989 |
| 0.704542 | 0.707604 | 0.705584 | 0.683748 | 0.687468 | 0.683418 | 0.679686 | 0.66757 | 0.66752 |
| 0.709836 | 0.71382 | 0.710142 | 0.686841 | 0.692988 | 0.689326 | 0.683936 | 0.669478 | 0.672569 |
| 0.70967 | 0.715568 | 0.710729 | 0.688821 | 0.694051 | 0.687526 | 0.687054 | 0.665968 | 0.668919 |
| 0.706143 | 0.707599 | 0.701202 | 0.683282 | 0.683388 | 0.676299 | 0.674455 | 0.67006 | 0.670662 |
| 0.704699 | 0.706414 | 0.703214 | 0.685269 | 0.689659 | 0.681363 | 0.685528 | 0.664424 | 0.668293 |
| 0.706888 | 0.706469 | 0.700683 | 0.682886 | 0.691429 | 0.684177 | 0.679215 | 0.669063 | 0.672275 |
| 0.715052 | 0.717994 | 0.715752 | 0.692777 | 0.69105 | 0.689591 | 0.681698 | 0.667367 | 0.672631 |
| 0.709312 | 0.711356 | 0.705428 | 0.685685 | 0.690029 | 0.678997 | 0.682711 | 0.666627 | 0.667555 |
| 0.708924 | 0.70916 | 0.706581 | 0.683998 | 0.691733 | 0.681267 | 0.680929 | 0.667134 | 0.671933 |
| 0.70718 | 0.710976 | 0.707531 | 0.682737 | 0.691475 | 0.680931 | 0.679292 | 0.667058 | 0.669299 |
| 0.710145 | 0.714539 | 0.718948 | 0.686275 | 0.693717 | 0.68289 | 0.68584 | 0.66934 | 0.672756 |
| 0.713358 | 0.71432 | 0.709371 | 0.682361 | 0.692688 | 0.682112 | 0.685756 | 0.670994 | 0.670138 |
| 0.712026 | 0.71562 | 0.707113 | 0.686391 | 0.694232 | 0.682041 | 0.685496 | 0.66806 | 0.67133 |
| 0.707249 | 0.710324 | 0.701131 | 0.681027 | 0.689068 | 0.677503 | 0.679915 | 0.666275 | 0.664564 |
| 0.710448 | 0.710293 | 0.708193 | 0.688665 | 0.6918 | 0.685602 | 0.684445 | 0.668661 | 0.666425 |
| 0.708567 | 0.705661 | 0.70415 | 0.679336 | 0.686824 | 0.677666 | 0.675358 | 0.660647 | 0.658744 |

| | | | | | | | | |
|---|---|---|---|---|---|---|---|---|
| 0.695254 | 0.696732 | 0.690122 | 0.675759 | 0.680568 | 0.67518 | 0.670828 | 0.65846 | 0.658018 |
| 0.74383 | 0.74896 | 0.743877 | 0.722463 | 0.727732 | 0.728202 | 0.727696 | 0.703933 | 0.712683 |
| 0.707788 | 0.707079 | 0.702155 | 0.683012 | 0.688546 | 0.678166 | 0.682381 | 0.666272 | 0.667929 |
| 0.713422 | 0.712909 | 0.712263 | 0.689395 | 0.687682 | 0.685574 | 0.678199 | 0.672101 | 0.670398 |
| 0.705624 | 0.704413 | 0.70647 | 0.690578 | 0.683807 | 0.67478 | 0.674151 | 0.669698 | 0.668315 |
| 0.704771 | 0.706635 | 0.708891 | 0.681869 | 0.683461 | 0.681515 | 0.678379 | 0.666627 | 0.665974 |
| 0.703028 | 0.70622 | 0.703429 | 0.686338 | 0.684046 | 0.680122 | 0.680438 | 0.664948 | 0.673389 |
| 0.699837 | 0.706594 | 0.696209 | 0.683732 | 0.681958 | 0.675597 | 0.676692 | 0.664684 | 0.666417 |
| 0.707375 | 0.706095 | 0.708773 | 0.682615 | 0.68669 | 0.680442 | 0.677473 | 0.662255 | 0.660395 |
| 0.713643 | 0.713472 | 0.716426 | 0.692457 | 0.694004 | 0.688582 | 0.686748 | 0.668268 | 0.675617 |
| 0.708148 | 0.710172 | 0.707166 | 0.684706 | 0.691087 | 0.685667 | 0.685342 | 0.670675 | 0.667832 |
| 0.71234 | 0.710191 | 0.713776 | 0.691374 | 0.695714 | 0.68088 | 0.683027 | 0.671487 | 0.672588 |
| 0.713759 | 0.71483 | 0.711333 | 0.686453 | 0.692401 | 0.685673 | 0.685673 | 0.673178 | 0.676738 |
| 0.710323 | 0.709799 | 0.711656 | 0.68961 | 0.692254 | 0.68233 | 0.687708 | 0.668139 | 0.670077 |
| 0.705897 | 0.708217 | 0.706212 | 0.686901 | 0.693426 | 0.68163 | 0.680491 | 0.673327 | 0.670717 |
| 0.714467 | 0.712566 | 0.707298 | 0.686255 | 0.690217 | 0.679984 | 0.684673 | 0.668707 | 0.665591 |
| 0.710734 | 0.710641 | 0.709483 | 0.685503 | 0.689945 | 0.685886 | 0.681843 | 0.671259 | 0.673028 |
| 0.708925 | 0.70832 | 0.710333 | 0.685153 | 0.691752 | 0.685179 | 0.683988 | 0.67078 | 0.672301 |
| 0.700156 | 0.705846 | 0.703142 | 0.684521 | 0.690748 | 0.683478 | 0.680959 | 0.668597 | 0.667286 |
| 0.704194 | 0.702073 | 0.70311 | 0.680698 | 0.685518 | 0.680089 | 0.679194 | 0.660592 | 0.667284 |
| 0.709223 | 0.709941 | 0.709253 | 0.685302 | 0.688033 | 0.680033 | 0.683077 | 0.667446 | 0.667715 |
| 0.697007 | 0.702629 | 0.698606 | 0.673583 | 0.676085 | 0.671122 | 0.669945 | 0.658876 | 0.656755 |
| 0.705023 | 0.70797 | 0.703519 | 0.681583 | 0.687399 | 0.676093 | 0.678195 | 0.666159 | 0.6645 |
| 0.708859 | 0.711503 | 0.700415 | 0.684952 | 0.689543 | 0.680561 | 0.67835 | 0.661537 | 0.665579 |
| 0.709564 | 0.714463 | 0.705588 | 0.687628 | 0.696203 | 0.685866 | 0.684944 | 0.667414 | 0.670604 |
| 0.70665 | 0.713087 | 0.703292 | 0.683499 | 0.693664 | 0.680771 | 0.678432 | 0.664938 | 0.667234 |
| 0.70378 | 0.706167 | 0.702936 | 0.682143 | 0.687864 | 0.679096 | 0.675497 | 0.663149 | 0.660877 |
| 0.703421 | 0.704969 | 0.701819 | 0.681931 | 0.688257 | 0.679451 | 0.677846 | 0.664721 | 0.666135 |
| 0.699402 | 0.704609 | 0.698819 | 0.680693 | 0.689601 | 0.681124 | 0.679853 | 0.661689 | 0.665838 |
| 0.768551 | 0.769216 | 0.763484 | 0.778349 | 0.798162 | 0.74542 | 0.747219 | 0.730592 | 0.749693 |
| 0.7884 | 0.782031 | 0.778105 | 0.753835 | 0.853269 | 0.752793 | 0.762846 | 0.738458 | 0.763599 |
| 0.782492 | 0.774435 | 0.780247 | 0.786817 | 0.780202 | 0.756111 | 0.757344 | 0.745862 | 0.756163 |
| 0.77077 | 0.777623 | 0.763342 | 0.766355 | 0.755002 | 0.7498 | 0.760413 | 0.743273 | 0.801164 |
| 0.786325 | 0.78569 | 0.778566 | 0.772435 | 0.755872 | 0.769504 | 0.75983 | 0.749181 | 0.755987 |
| 0.698814 | 0.701607 | 0.694156 | 0.679125 | 0.68259 | 0.671733 | 0.67585 | 0.659728 | 0.660108 |
| 0.70458 | 0.706663 | 0.698764 | 0.68265 | 0.686019 | 0.678256 | 0.676251 | 0.664623 | 0.662505 |
| 0.706206 | 0.708858 | 0.698852 | 0.680847 | 0.6915 | 0.680099 | 0.679786 | 0.663384 | 0.660839 |
| 0.704164 | 0.709265 | 0.702506 | 0.68174 | 0.688824 | 0.682179 | 0.678096 | 0.662304 | 0.665378 |
| 0.70474 | 0.706643 | 0.698213 | 0.680069 | 0.68533 | 0.67722 | 0.672458 | 0.663354 | 0.662687 |
| 0.707807 | 0.713785 | 0.701724 | 0.686554 | 0.695295 | 0.682005 | 0.680827 | 0.664915 | 0.668354 |
| 0.705073 | 0.704748 | 0.699086 | 0.680982 | 0.685912 | 0.680485 | 0.679786 | 0.665568 | 0.66772 |
| 0.707347 | 0.706482 | 0.702907 | 0.678299 | 0.689888 | 0.680357 | 0.679431 | 0.662284 | 0.661505 |
| 0.695004 | 0.699369 | 0.689511 | 0.678112 | 0.680976 | 0.674126 | 0.672642 | 0.66152 | 0.657104 |
| 0.696527 | 0.700859 | 0.690783 | 0.681389 | 0.682034 | 0.676005 | 0.676912 | 0.662888 | 0.665374 |
| 0.704221 | 0.707197 | 0.701688 | 0.680174 | 0.687906 | 0.675546 | 0.681412 | 0.660772 | 0.661637 |
| 0.70482 | 0.708832 | 0.701144 | 0.684065 | 0.688239 | 0.678616 | 0.679423 | 0.664029 | 0.664152 |

| | | | | | | | | |
|---|---|---|---|---|---|---|---|---|
| 0.705565 | 0.714132 | 0.704775 | 0.681963 | 0.688439 | 0.678553 | 0.680811 | 0.663152 | 0.668215 |
| 0.705181 | 0.708208 | 0.701339 | 0.682082 | 0.684504 | 0.674387 | 0.676769 | 0.664941 | 0.667071 |
| 0.71108 | 0.712088 | 0.710748 | 0.68794 | 0.693713 | 0.678542 | 0.683859 | 0.669254 | 0.669778 |
| 0.703867 | 0.707236 | 0.704589 | 0.679772 | 0.688174 | 0.676258 | 0.676143 | 0.668632 | 0.668253 |
| 0.703361 | 0.70397 | 0.701738 | 0.678825 | 0.681753 | 0.67169 | 0.674096 | 0.658473 | 0.667319 |
| 0.702108 | 0.703018 | 0.702969 | 0.682062 | 0.684314 | 0.673766 | 0.67754 | 0.661562 | 0.664261 |
| 0.707385 | 0.708879 | 0.707185 | 0.68624 | 0.690969 | 0.679125 | 0.683273 | 0.668449 | 0.66402 |
| 0.705593 | 0.711622 | 0.703659 | 0.685319 | 0.690094 | 0.679051 | 0.680618 | 0.669181 | 0.673467 |
| 0.706632 | 0.70649 | 0.702826 | 0.681015 | 0.684599 | 0.674711 | 0.675745 | 0.660522 | 0.661137 |
| 0.707657 | 0.707949 | 0.707695 | 0.681684 | 0.686997 | 0.677166 | 0.678051 | 0.66205 | 0.66833 |
| 0.707257 | 0.707167 | 0.705497 | 0.683693 | 0.687117 | 0.675098 | 0.67914 | 0.662506 | 0.667685 |
| 0.710831 | 0.706972 | 0.705485 | 0.67927 | 0.689836 | 0.67533 | 0.678507 | 0.665049 | 0.664106 |
| 0.6967 | 0.700105 | 0.692705 | 0.676234 | 0.6797 | 0.671187 | 0.673701 | 0.659717 | 0.661898 |
| 0.697436 | 0.70511 | 0.696014 | 0.681603 | 0.686516 | 0.677652 | 0.676554 | 0.664106 | 0.661688 |
| 0.709225 | 0.707077 | 0.707919 | 0.686408 | 0.695769 | 0.681135 | 0.687444 | 0.667771 | 0.67168 |
| 0.711942 | 0.715101 | 0.712707 | 0.692221 | 0.699021 | 0.688586 | 0.689221 | 0.674832 | 0.67582 |
| 0.709329 | 0.713323 | 0.709608 | 0.686428 | 0.69146 | 0.679399 | 0.683308 | 0.664779 | 0.67162 |
| 0.712631 | 0.713973 | 0.709812 | 0.694853 | 0.691558 | 0.684162 | 0.689482 | 0.671355 | 0.671752 |
| 0.705228 | 0.708882 | 0.702264 | 0.686549 | 0.690632 | 0.680739 | 0.680041 | 0.667262 | 0.669861 |
| 0.704566 | 0.710244 | 0.705246 | 0.682577 | 0.69202 | 0.676973 | 0.678648 | 0.670418 | 0.671941 |
| 0.702301 | 0.705038 | 0.697821 | 0.683628 | 0.689172 | 0.679962 | 0.680335 | 0.671533 | 0.671636 |
| 0.709134 | 0.711235 | 0.708821 | 0.685577 | 0.69159 | 0.683176 | 0.67904 | 0.668955 | 0.671697 |
| 0.708164 | 0.714011 | 0.708228 | 0.684254 | 0.693398 | 0.683879 | 0.685313 | 0.669448 | 0.67516 |
| 0.707219 | 0.707631 | 0.703474 | 0.684776 | 0.68979 | 0.680295 | 0.680317 | 0.665821 | 0.675144 |
| 0.704223 | 0.707493 | 0.704944 | 0.68756 | 0.69285 | 0.678367 | 0.684058 | 0.667318 | 0.673333 |
| 0.78193 | 0.788693 | 0.779304 | 0.755753 | 0.776432 | 0.756171 | 0.766787 | 0.743084 | 0.773551 |
| 0.708609 | 0.707724 | 0.709209 | 0.68695 | 0.689248 | 0.680916 | 0.681044 | 0.664336 | 0.666145 |
| 0.706629 | 0.711473 | 0.705479 | 0.68514 | 0.688976 | 0.683333 | 0.681833 | 0.666939 | 0.669728 |
| 0.712298 | 0.712211 | 0.712284 | 0.685703 | 0.691391 | 0.684205 | 0.680156 | 0.667531 | 0.672557 |
| 0.705022 | 0.706368 | 0.703281 | 0.687214 | 0.688437 | 0.683499 | 0.681003 | 0.663164 | 0.667071 |
| 0.707577 | 0.706914 | 0.702356 | 0.682228 | 0.687935 | 0.681948 | 0.683547 | 0.672139 | 0.6705 |
| 0.701837 | 0.709845 | 0.704005 | 0.684118 | 0.692101 | 0.683172 | 0.677869 | 0.664885 | 0.667319 |
| 0.707779 | 0.708014 | 0.708603 | 0.687087 | 0.690239 | 0.68074 | 0.682842 | 0.665136 | 0.667194 |
| 0.70041 | 0.707061 | 0.703482 | 0.683622 | 0.687826 | 0.681032 | 0.678007 | 0.664632 | 0.671728 |
| 0.702691 | 0.703351 | 0.695528 | 0.684089 | 0.684645 | 0.676708 | 0.673965 | 0.665025 | 0.668495 |
| 0.700229 | 0.705085 | 0.701738 | 0.681599 | 0.683518 | 0.676825 | 0.677256 | 0.666587 | 0.667646 |
| 0.708962 | 0.71194 | 0.707572 | 0.687948 | 0.689458 | 0.678186 | 0.681491 | 0.666863 | 0.673133 |
| 0.708787 | 0.711625 | 0.708489 | 0.687157 | 0.691979 | 0.683345 | 0.680701 | 0.668704 | 0.669259 |
| 0.708988 | 0.709962 | 0.706764 | 0.686582 | 0.688042 | 0.685459 | 0.682006 | 0.67102 | 0.666747 |
| 0.703833 | 0.707801 | 0.704331 | 0.685249 | 0.688135 | 0.682476 | 0.679314 | 0.670389 | 0.674635 |
| 0.701239 | 0.702453 | 0.698879 | 0.679245 | 0.6847 | 0.680286 | 0.675682 | 0.664098 | 0.667077 |
| 0.712344 | 0.711302 | 0.709144 | 0.68894 | 0.693116 | 0.685177 | 0.681066 | 0.669766 | 0.667775 |
| 0.707692 | 0.710189 | 0.706238 | 0.688658 | 0.688255 | 0.679277 | 0.679814 | 0.666227 | 0.672463 |
| 0.709809 | 0.709802 | 0.705162 | 0.683933 | 0.691255 | 0.68145 | 0.678378 | 0.662815 | 0.665662 |
| 0.709644 | 0.711884 | 0.705476 | 0.687201 | 0.690292 | 0.683277 | 0.68148 | 0.667135 | 0.67205 |
| 0.71229 | 0.713196 | 0.709938 | 0.687475 | 0.68911 | 0.685048 | 0.678825 | 0.669328 | 0.666505 |

| | | | | | | | | |
|---|---|---|---|---|---|---|---|---|
| 0.706658 | 0.70901 | 0.705623 | 0.684088 | 0.686164 | 0.677396 | 0.677878 | 0.665965 | 0.667149 |
| 0.70906 | 0.712869 | 0.709071 | 0.688156 | 0.685511 | 0.684427 | 0.682612 | 0.665778 | 0.665392 |
| 0.710518 | 0.706985 | 0.702883 | 0.683845 | 0.687946 | 0.680397 | 0.678898 | 0.662078 | 0.667356 |
| 0.669212 | 0.660912 | 0.669361 | 0.666033 | 0.664618 | 0.653488 | 0.659367 | 0.64472 | 0.650987 |
| 0.705508 | 0.708404 | 0.704114 | 0.681178 | 0.691348 | 0.676606 | 0.678646 | 0.662642 | 0.662795 |
| 0.710674 | 0.707855 | 0.703035 | 0.684766 | 0.687588 | 0.679274 | 0.678277 | 0.665377 | 0.664889 |
| 0.711858 | 0.711655 | 0.712234 | 0.687063 | 0.68921 | 0.685263 | 0.681591 | 0.66608 | 0.670097 |
| 0.710373 | 0.712949 | 0.707194 | 0.681512 | 0.689964 | 0.678637 | 0.68015 | 0.664808 | 0.666426 |
| 0.715268 | 0.717642 | 0.711429 | 0.689914 | 0.693532 | 0.685427 | 0.687417 | 0.666279 | 0.675355 |
| 0.710602 | 0.713615 | 0.707609 | 0.687145 | 0.689732 | 0.678505 | 0.68066 | 0.666823 | 0.670661 |
| 0.709149 | 0.713534 | 0.707932 | 0.686916 | 0.689694 | 0.677823 | 0.680585 | 0.666199 | 0.669727 |
| 0.714056 | 0.714007 | 0.713037 | 0.688934 | 0.690813 | 0.681935 | 0.683847 | 0.670813 | 0.673703 |
| 0.712044 | 0.714274 | 0.712962 | 0.69501 | 0.693789 | 0.682163 | 0.688827 | 0.668337 | 0.673037 |
| 0.707071 | 0.709524 | 0.704337 | 0.683833 | 0.686609 | 0.677538 | 0.679872 | 0.663098 | 0.662509 |
| 0.711669 | 0.713904 | 0.710359 | 0.689313 | 0.695349 | 0.678648 | 0.682902 | 0.669155 | 0.672714 |
| 0.707102 | 0.704745 | 0.701715 | 0.684724 | 0.689254 | 0.675192 | 0.67677 | 0.663202 | 0.660014 |
| 0.712708 | 0.71533 | 0.706553 | 0.686387 | 0.692879 | 0.681619 | 0.680598 | 0.667597 | 0.674144 |
| 0.712056 | 0.710541 | 0.711801 | 0.688133 | 0.688889 | 0.682232 | 0.678867 | 0.667365 | 0.670574 |
| 0.705929 | 0.705982 | 0.701988 | 0.685292 | 0.684988 | 0.679257 | 0.676909 | 0.664172 | 0.665781 |
| 0.707356 | 0.708551 | 0.704801 | 0.684788 | 0.686928 | 0.677085 | 0.675851 | 0.666939 | 0.666185 |
| 0.709879 | 0.711425 | 0.70724 | 0.688272 | 0.689613 | 0.683496 | 0.684695 | 0.666082 | 0.668354 |
| 0.713041 | 0.717696 | 0.709339 | 0.688589 | 0.693501 | 0.682631 | 0.68328 | 0.670187 | 0.674232 |
| 0.697315 | 0.696307 | 0.699495 | 0.69256 | 0.695105 | 0.690448 | 0.693388 | 0.675373 | 0.688083 |
| 0.723133 | 0.728801 | 0.706543 | 0.72122 | 0.730195 | 0.717705 | 0.723797 | 0.715024 | 0.73531 |
| 0.76009 | 0.754547 | 0.770869 | 0.736921 | 0.752988 | 0.747692 | 0.754377 | 0.77371 | 0.758776 |
| 0.743485 | 0.74532 | 0.735497 | 0.732327 | 0.744735 | 0.735273 | 0.73649 | 0.728002 | 0.786771 |
| 0.665942 | 0.66739 | 0.658141 | 0.658807 | 0.665417 | 0.660826 | 0.666556 | 0.651147 | 0.661063 |
| 0.668405 | 0.665677 | 0.651148 | 0.654894 | 0.663116 | 0.661017 | 0.659261 | 0.646984 | 0.654279 |
| 0.659862 | 0.655633 | 0.65998 | 0.668207 | 0.66468 | 0.65431 | 0.653425 | 0.652434 | 0.657603 |
| 0.676204 | 0.678687 | 0.678263 | 0.667546 | 0.669598 | 0.672529 | 0.670035 | 0.655205 | 0.663391 |
| 0.651168 | 0.653838 | 0.645802 | 0.655268 | 0.658968 | 0.646165 | 0.6543 | 0.639871 | 0.655849 |
| 0.645126 | 0.648832 | 0.641832 | 0.644733 | 0.644274 | 0.645431 | 0.647253 | 0.628201 | 0.641469 |
| 0.651839 | 0.655457 | 0.64224 | 0.648889 | 0.645759 | 0.639737 | 0.642614 | 0.62313 | 0.64443 |
| 0.732928 | 0.729755 | 0.734547 | 0.744217 | 0.753948 | 0.723421 | 0.734835 | 0.730353 | 0.740346 |
| 0.705996 | 0.706728 | 0.699381 | 0.682174 | 0.686845 | 0.681045 | 0.680312 | 0.663503 | 0.666344 |
| 0.674975 | 0.679638 | 0.670822 | 0.671129 | 0.676246 | 0.666625 | 0.668348 | 0.656861 | 0.66019 |
| 0.67324 | 0.673657 | 0.666368 | 0.66597 | 0.668345 | 0.65938 | 0.671747 | 0.65907 | 0.66734 |
| 0.654155 | 0.661145 | 0.645551 | 0.65118 | 0.656916 | 0.655553 | 0.654538 | 0.64558 | 0.660501 |
| 0.677372 | 0.678044 | 0.674929 | 0.671846 | 0.67378 | 0.669754 | 0.668319 | 0.653311 | 0.661688 |
| 0.669581 | 0.66708 | 0.67127 | 0.667023 | 0.673687 | 0.66298 | 0.666905 | 0.647921 | 0.658335 |
| 0.664417 | 0.660409 | 0.654914 | 0.656738 | 0.668048 | 0.659019 | 0.651389 | 0.641049 | 0.647461 |
| 0.664211 | 0.665408 | 0.661844 | 0.666027 | 0.665476 | 0.65868 | 0.659028 | 0.651261 | 0.652044 |
| 0.661049 | 0.656071 | 0.651911 | 0.658338 | 0.661039 | 0.648464 | 0.651221 | 0.643914 | 0.65476 |
| 0.65668 | 0.662154 | 0.657923 | 0.662394 | 0.665792 | 0.653501 | 0.660202 | 0.649502 | 0.659464 |
| 0.66155 | 0.662722 | 0.655672 | 0.657948 | 0.654545 | 0.657463 | 0.656596 | 0.654706 | 0.665106 |
| 0.646004 | 0.641521 | 0.633393 | 0.651862 | 0.648063 | 0.646174 | 0.646858 | 0.637306 | 0.643869 |

| | | | | | | | | |
|---|---|---|---|---|---|---|---|---|
| 0.690797 | 0.694404 | 0.684583 | 0.680119 | 0.68465 | 0.676273 | 0.676749 | 0.66633 | 0.665565 |
| 0.6554 | 0.659951 | 0.654655 | 0.659222 | 0.66205 | 0.656025 | 0.655756 | 0.646658 | 0.649439 |
| 0.665754 | 0.663471 | 0.653119 | 0.664412 | 0.66439 | 0.65458 | 0.649608 | 0.649932 | 0.656271 |
| 0.668586 | 0.666815 | 0.654036 | 0.656684 | 0.668315 | 0.658292 | 0.656261 | 0.650087 | 0.657316 |
| 0.658527 | 0.663462 | 0.657435 | 0.662869 | 0.662738 | 0.653322 | 0.660101 | 0.647449 | 0.659598 |
| 0.662605 | 0.664181 | 0.652486 | 0.660891 | 0.6718 | 0.659327 | 0.665787 | 0.649435 | 0.665423 |
| 0.65739 | 0.651174 | 0.648816 | 0.653413 | 0.654594 | 0.65179 | 0.645955 | 0.650802 | 0.657158 |
| 0.646851 | 0.653261 | 0.634561 | 0.64958 | 0.651814 | 0.639739 | 0.650468 | 0.646135 | 0.652044 |
| 0.649132 | 0.653616 | 0.63967 | 0.651195 | 0.653846 | 0.651057 | 0.646077 | 0.639493 | 0.646172 |
| 0.694775 | 0.692388 | 0.688371 | 0.684551 | 0.684811 | 0.675305 | 0.676963 | 0.668366 | 0.668705 |
| 0.687811 | 0.688891 | 0.684057 | 0.68192 | 0.681176 | 0.674677 | 0.679133 | 0.669968 | 0.665336 |
| 0.682745 | 0.685432 | 0.678383 | 0.683617 | 0.681294 | 0.673368 | 0.670946 | 0.662791 | 0.665987 |
| 0.677268 | 0.676391 | 0.665361 | 0.678018 | 0.679165 | 0.671706 | 0.668791 | 0.659311 | 0.668497 |
| 0.684839 | 0.679185 | 0.677757 | 0.675966 | 0.677765 | 0.67615 | 0.670254 | 0.660995 | 0.665922 |
| 0.674311 | 0.671093 | 0.666258 | 0.677823 | 0.66901 | 0.667193 | 0.664625 | 0.653818 | 0.658126 |

| Collection_ | Collection_ | Collection_ | Collection_ | Collection_ | Collection_ | Collection_ | Collection_ | Collection_ |
|---|---|---|---|---|---|---|---|---|
| 0.761082 | 0.740003 | 0.758796 | 0.76604 | 0.740472 | 0.783051 | 0.729373 | 0.764256 | 0.746852 |
| 0.758502 | 0.744171 | 0.772082 | 0.779146 | 0.742242 | 0.791765 | 0.738709 | 0.740358 | 0.760255 |
| 0.780216 | 0.741186 | 0.769493 | 0.768832 | 0.737177 | 0.7877 | 0.734087 | 0.753753 | 0.743658 |
| 0.748422 | 0.741497 | 0.756553 | 0.74984 | 0.739862 | 0.767945 | 0.748922 | 0.746769 | 0.750568 |
| 0.742456 | 0.745337 | 0.75068 | 0.740306 | 0.740237 | 0.736068 | 0.744051 | 0.760182 | 0.746866 |
| 0.740422 | 0.752781 | 0.746848 | 0.745411 | 0.741294 | 0.745103 | 0.740419 | 0.732828 | 0.737985 |
| 0.668838 | 0.668779 | 0.666419 | 0.676836 | 0.673129 | 0.674105 | 0.675762 | 0.663562 | 0.667608 |
| 0.665134 | 0.659787 | 0.665424 | 0.670571 | 0.666544 | 0.670211 | 0.667242 | 0.658245 | 0.654499 |
| 0.760478 | 0.75345 | 0.763782 | 0.760031 | 0.742171 | 0.758466 | 0.748613 | 0.743301 | 0.744111 |
| 0.753265 | 0.731762 | 0.758531 | 0.752031 | 0.731562 | 0.766175 | 0.73164 | 0.780966 | 0.752881 |
| 0.753541 | 0.741327 | 0.754199 | 0.752344 | 0.734955 | 0.759767 | 0.738322 | 0.751236 | 0.7499 |
| 0.763834 | 0.74633 | 0.757003 | 0.769561 | 0.734153 | 0.778956 | 0.729622 | 0.759536 | 0.787551 |
| 0.744756 | 0.742921 | 0.749903 | 0.749388 | 0.741543 | 0.768496 | 0.739836 | 0.742094 | 0.770292 |
| 0.758717 | 0.744961 | 0.762672 | 0.756292 | 0.730725 | 0.759946 | 0.728353 | 0.75203 | 0.751734 |
| 0.732821 | 0.730428 | 0.741935 | 0.754881 | 0.723289 | 0.749597 | 0.728834 | 0.735542 | 0.730567 |
| 0.766955 | 0.756504 | 0.761433 | 0.775712 | 0.737707 | 0.776475 | 0.739546 | 0.774216 | 0.741793 |
| 0.749685 | 0.73621 | 0.760727 | 0.769432 | 0.725349 | 0.762243 | 0.722931 | 0.760885 | 0.726122 |
| 0.719907 | 0.720708 | 0.7313 | 0.725935 | 0.71585 | 0.735329 | 0.708188 | 0.722772 | 0.720135 |
| 0.761162 | 0.752632 | 0.765023 | 0.778998 | 0.744725 | 0.80064 | 0.741689 | 0.737637 | 0.793607 |
| 0.752781 | 0.749303 | 0.764197 | 0.755912 | 0.746986 | 0.772055 | 0.748955 | 0.748756 | 0.760394 |
| 0.74959 | 0.752292 | 0.755856 | 0.760682 | 0.752703 | 0.765402 | 0.744254 | 0.743881 | 0.752222 |
| 0.770628 | 0.7492 | 0.769898 | 0.767212 | 0.743586 | 0.780313 | 0.741671 | 0.757727 | 0.768532 |
| 0.753665 | 0.755717 | 0.768475 | 0.766469 | 0.745722 | 0.779854 | 0.741191 | 0.751923 | 0.769168 |
| 0.761105 | 0.750631 | 0.768676 | 0.770584 | 0.743518 | 0.788944 | 0.742655 | 0.759851 | 0.773457 |
| 0.752726 | 0.752291 | 0.754702 | 0.750161 | 0.748811 | 0.763033 | 0.736791 | 0.742908 | 0.768761 |
| 0.759146 | 0.745026 | 0.758858 | 0.781116 | 0.762853 | 0.774484 | 0.744796 | 0.746438 | 0.755075 |
| 0.745294 | 0.738975 | 0.756225 | 0.757427 | 0.737577 | 0.778683 | 0.735063 | 0.739796 | 0.781084 |
| 0.752527 | 0.745562 | 0.759652 | 0.750895 | 0.742436 | 0.760445 | 0.742198 | 0.744483 | 0.746891 |
| 0.746866 | 0.750829 | 0.759329 | 0.754874 | 0.746682 | 0.759588 | 0.745505 | 0.749211 | 0.760543 |
| 0.751338 | 0.743508 | 0.759543 | 0.747805 | 0.741674 | 0.772427 | 0.729299 | 0.761958 | 0.763104 |
| 0.74492 | 0.746852 | 0.751392 | 0.759995 | 0.75589 | 0.756471 | 0.746746 | 0.744298 | 0.754259 |
| 0.742914 | 0.74581 | 0.747016 | 0.765298 | 0.753172 | 0.750947 | 0.748544 | 0.737416 | 0.74277 |
| 0.74843 | 0.755877 | 0.750968 | 0.759205 | 0.738038 | 0.769147 | 0.741078 | 0.743779 | 0.767107 |
| 0.741524 | 0.745433 | 0.738288 | 0.743961 | 0.758585 | 0.743387 | 0.739749 | 0.736418 | 0.741267 |
| 0.742489 | 0.734474 | 0.749707 | 0.757274 | 0.733097 | 0.747363 | 0.731596 | 0.732398 | 0.753189 |
| 0.757553 | 0.74359 | 0.759519 | 0.774063 | 0.741443 | 0.764891 | 0.742291 | 0.749029 | 0.773161 |
| 0.757889 | 0.744073 | 0.760109 | 0.765206 | 0.731848 | 0.766354 | 0.739896 | 0.737743 | 0.747929 |
| 0.733863 | 0.733883 | 0.739267 | 0.740186 | 0.730946 | 0.749074 | 0.730652 | 0.73159 | 0.746122 |
| 0.743879 | 0.741855 | 0.746159 | 0.745506 | 0.741099 | 0.751552 | 0.73128 | 0.734349 | 0.747461 |

| | | | | | | | | |
|---|---|---|---|---|---|---|---|---|
| 0.728728 | 0.721246 | 0.728923 | 0.736822 | 0.715106 | 0.749095 | 0.715152 | 0.722686 | 0.714975 |
| 0.755035 | 0.734554 | 0.760517 | 0.752823 | 0.736395 | 0.777444 | 0.733729 | 0.773551 | 0.755045 |
| 0.765729 | 0.744106 | 0.763803 | 0.765684 | 0.738104 | 0.778146 | 0.746171 | 0.768844 | 0.7639 |
| 0.760394 | 0.74855 | 0.76769 | 0.7674 | 0.738866 | 0.780895 | 0.738495 | 0.778365 | 0.786122 |
| 0.764766 | 0.755677 | 0.762537 | 0.758994 | 0.744183 | 0.772231 | 0.742663 | 0.753515 | 0.771939 |
| 0.747596 | 0.748472 | 0.750195 | 0.754164 | 0.733911 | 0.751663 | 0.738389 | 0.747141 | 0.756042 |
| 0.734931 | 0.746619 | 0.745184 | 0.743224 | 0.73228 | 0.744715 | 0.74136 | 0.744902 | 0.741716 |
| 0.760372 | 0.759222 | 0.759242 | 0.763483 | 0.751191 | 0.778298 | 0.749232 | 0.744394 | 0.779855 |
| 0.751497 | 0.747111 | 0.752205 | 0.752528 | 0.738135 | 0.755501 | 0.745808 | 0.744389 | 0.739218 |
| 0.746974 | 0.744336 | 0.740903 | 0.75251 | 0.739693 | 0.747743 | 0.737429 | 0.739719 | 0.733723 |
| 0.759852 | 0.749301 | 0.764498 | 0.771048 | 0.739275 | 0.782555 | 0.736879 | 0.782431 | 0.762402 |
| 0.764834 | 0.751337 | 0.76312 | 0.768508 | 0.741253 | 0.782482 | 0.74022 | 0.75971 | 0.760967 |
| 0.762823 | 0.750704 | 0.77213 | 0.775257 | 0.740459 | 0.774609 | 0.740263 | 0.7731 | 0.764077 |
| 0.769339 | 0.747766 | 0.761593 | 0.766176 | 0.743521 | 0.763908 | 0.746089 | 0.760284 | 0.753113 |
| 0.76036 | 0.746023 | 0.761911 | 0.77967 | 0.74606 | 0.765784 | 0.739994 | 0.760398 | 0.749932 |
| 0.75718 | 0.747941 | 0.757568 | 0.780935 | 0.743616 | 0.771564 | 0.739963 | 0.763424 | 0.742919 |
| 0.758448 | 0.742156 | 0.762535 | 0.764235 | 0.734938 | 0.78044 | 0.733474 | 0.768253 | 0.745857 |
| 0.776922 | 0.757738 | 0.772446 | 0.780779 | 0.757442 | 0.786692 | 0.739336 | 0.779071 | 0.759555 |
| 0.751327 | 0.747119 | 0.758994 | 0.753906 | 0.733037 | 0.763889 | 0.738462 | 0.769713 | 0.76764 |
| 0.755954 | 0.752634 | 0.76269 | 0.770136 | 0.739019 | 0.785299 | 0.742657 | 0.7752 | 0.772827 |
| 0.774434 | 0.748163 | 0.758281 | 0.775898 | 0.740988 | 0.781086 | 0.738245 | 0.772785 | 0.73574 |
| 0.76291 | 0.743688 | 0.759138 | 0.757631 | 0.737634 | 0.765531 | 0.743529 | 0.773619 | 0.746134 |
| 0.762001 | 0.750561 | 0.763291 | 0.762236 | 0.743988 | 0.776381 | 0.739807 | 0.745621 | 0.757625 |
| 0.751326 | 0.746286 | 0.751915 | 0.74568 | 0.738311 | 0.757469 | 0.753694 | 0.739452 | 0.744444 |
| 0.732529 | 0.735709 | 0.732414 | 0.739208 | 0.729794 | 0.738839 | 0.740135 | 0.722853 | 0.736404 |
| 0.66541 | 0.661032 | 0.665128 | 0.671865 | 0.668269 | 0.670053 | 0.667335 | 0.658872 | 0.655677 |
| 0.666788 | 0.672843 | 0.667537 | 0.67496 | 0.678453 | 0.674818 | 0.672718 | 0.669936 | 0.667052 |
| 0.666302 | 0.664194 | 0.666959 | 0.673845 | 0.676723 | 0.677754 | 0.671862 | 0.666533 | 0.666407 |
| 0.749181 | 0.736642 | 0.766989 | 0.744787 | 0.734614 | 0.761963 | 0.744543 | 0.729799 | 0.767247 |
| 0.74324 | 0.743784 | 0.770197 | 0.751234 | 0.737119 | 0.762108 | 0.748074 | 0.734983 | 0.755932 |
| 0.732635 | 0.734828 | 0.749707 | 0.740041 | 0.729389 | 0.752439 | 0.73706 | 0.759029 | 0.74318 |
| 0.741494 | 0.74097 | 0.764071 | 0.749521 | 0.737163 | 0.767565 | 0.744546 | 0.733388 | 0.762662 |
| 0.792996 | 0.758358 | 0.852385 | 0.800424 | 0.750386 | 0.870238 | 0.746491 | 0.750207 | 0.79527 |
| 0.779493 | 0.76015 | 0.77354 | 0.804991 | 0.750998 | 0.805877 | 0.74548 | 0.751001 | 0.806247 |
| 0.796513 | 0.765843 | 0.794636 | 0.816424 | 0.747722 | 0.878242 | 0.741293 | 0.752626 | 0.808667 |
| 0.81415 | 0.753548 | 0.785951 | 0.848878 | 0.750064 | 0.845383 | 0.743661 | 0.749379 | 0.746783 |
| 0.77193 | 0.755576 | 0.805393 | 0.756913 | 0.742361 | 0.802079 | 0.750725 | 0.748049 | 0.797234 |
| 0.792546 | 0.777613 | 0.803937 | 0.811449 | 0.770149 | 0.842075 | 0.768473 | 0.808813 | 0.808785 |
| 0.786218 | 0.746219 | 0.804629 | 0.783193 | 0.748737 | 0.836278 | 0.745297 | 0.739358 | 0.785724 |
| 0.765003 | 0.751418 | 0.757297 | 0.75361 | 0.755321 | 0.753245 | 0.750485 | 0.747085 | 0.764049 |
| 0.786181 | 0.751623 | 0.772709 | 0.797977 | 0.746942 | 0.785479 | 0.73959 | 0.839588 | 0.747093 |
| 0.791678 | 0.775281 | 0.804111 | 0.800798 | 0.768069 | 0.828081 | 0.768329 | 0.808352 | 0.802215 |
| 0.758022 | 0.757435 | 0.764721 | 0.760636 | 0.748432 | 0.760245 | 0.748096 | 0.790483 | 0.750267 |
| 0.752707 | 0.749868 | 0.756613 | 0.74888 | 0.736268 | 0.754702 | 0.747996 | 0.782064 | 0.767062 |
| 0.764721 | 0.810463 | 0.756609 | 0.758175 | 0.755991 | 0.763376 | 0.755764 | 0.747171 | 0.762155 |
| 0.79003 | 0.745264 | 0.777209 | 0.793933 | 0.75308 | 0.799254 | 0.74705 | 0.7427 | 0.75514 |

| | | | | | | | | |
|---|---|---|---|---|---|---|---|---|
| 0.754617 | 0.756218 | 0.758558 | 0.761431 | 0.747475 | 0.75441 | 0.742007 | 0.851344 | 0.748328 |
| 0.7649 | 0.739827 | 0.785009 | 0.751636 | 0.749809 | 0.746164 | 0.743816 | 0.743417 | 0.746961 |
| 0.776766 | 0.756172 | 0.771944 | 0.756414 | 0.746899 | 0.747066 | 0.750208 | 0.769359 | 0.753144 |
| 0.775719 | 0.770075 | 0.782356 | 0.787392 | 0.762236 | 0.807999 | 0.757487 | 0.789429 | 0.785705 |
| 0.763873 | 0.760021 | 0.768859 | 0.753044 | 0.75124 | 0.765513 | 0.749793 | 0.757103 | 0.765244 |
| 0.755412 | 0.751862 | 0.759138 | 0.756844 | 0.750675 | 0.753118 | 0.743682 | 0.748792 | 0.755795 |
| 0.755966 | 0.758233 | 0.785302 | 0.755667 | 0.737713 | 0.808652 | 0.747231 | 0.749245 | 0.802326 |
| 0.758844 | 0.753432 | 0.779507 | 0.752062 | 0.765392 | 0.74755 | 0.744793 | 0.745538 | 0.745961 |
| 0.754602 | 0.75204 | 0.754333 | 0.756079 | 0.75029 | 0.754365 | 0.74005 | 0.744938 | 0.755607 |
| 0.739185 | 0.741765 | 0.740917 | 0.755833 | 0.742453 | 0.743065 | 0.74028 | 0.74638 | 0.772505 |
| 0.755089 | 0.752136 | 0.748099 | 0.759985 | 0.747679 | 0.749197 | 0.743867 | 0.749385 | 0.748874 |
| 0.744254 | 0.750366 | 0.750389 | 0.755134 | 0.739242 | 0.754874 | 0.742702 | 0.74425 | 0.767925 |
| 0.748415 | 0.747463 | 0.752094 | 0.756669 | 0.749064 | 0.746372 | 0.743751 | 0.748852 | 0.768613 |
| 0.747716 | 0.750565 | 0.747209 | 0.750901 | 0.746912 | 0.746006 | 0.73558 | 0.743034 | 0.780215 |
| 0.75098 | 0.746621 | 0.746371 | 0.753102 | 0.750258 | 0.755898 | 0.752056 | 0.747447 | 0.744638 |
| 0.749275 | 0.745062 | 0.757119 | 0.758674 | 0.736929 | 0.756296 | 0.74141 | 0.751521 | 0.744276 |
| 0.748791 | 0.7498 | 0.759055 | 0.742097 | 0.744192 | 0.755286 | 0.747033 | 0.752084 | 0.758827 |
| 0.768197 | 0.75221 | 0.770207 | 0.817509 | 0.746631 | 0.787551 | 0.745269 | 0.747085 | 0.741272 |
| 0.748553 | 0.753604 | 0.759615 | 0.758514 | 0.739086 | 0.758003 | 0.746504 | 0.744176 | 0.766869 |
| 0.748483 | 0.747307 | 0.74987 | 0.750226 | 0.751543 | 0.756573 | 0.744096 | 0.744543 | 0.752525 |
| 0.755079 | 0.760757 | 0.751813 | 0.744296 | 0.743243 | 0.754801 | 0.748272 | 0.746881 | 0.756964 |
| 0.759135 | 0.743023 | 0.75754 | 0.750287 | 0.748981 | 0.760277 | 0.742967 | 0.743764 | 0.756677 |
| 0.768642 | 0.747052 | 0.771346 | 0.765415 | 0.753824 | 0.752364 | 0.74397 | 0.75031 | 0.74896 |
| 0.748715 | 0.747267 | 0.752298 | 0.751631 | 0.752845 | 0.750808 | 0.740041 | 0.745651 | 0.773261 |
| 0.754437 | 0.747049 | 0.776005 | 0.77711 | 0.747754 | 0.788435 | 0.745098 | 0.743956 | 0.791243 |
| 0.759975 | 0.760939 | 0.796387 | 0.759687 | 0.743837 | 0.811219 | 0.746273 | 0.742476 | 0.805468 |
| 0.762241 | 0.745767 | 0.777778 | 0.777433 | 0.745899 | 0.79633 | 0.74449 | 0.753053 | 0.795402 |
| 0.749466 | 0.737701 | 0.751813 | 0.755841 | 0.738195 | 0.751558 | 0.739089 | 0.735867 | 0.746432 |
| 0.751889 | 0.749107 | 0.75502 | 0.758984 | 0.739726 | 0.758672 | 0.749931 | 0.741375 | 0.751609 |
| 0.753708 | 0.746034 | 0.771875 | 0.749807 | 0.743174 | 0.780003 | 0.745746 | 0.748694 | 0.781675 |
| 0.752017 | 0.753504 | 0.750453 | 0.753523 | 0.738866 | 0.750979 | 0.742493 | 0.742408 | 0.769303 |
| 0.754641 | 0.752971 | 0.757025 | 0.749395 | 0.737825 | 0.756324 | 0.740609 | 0.746058 | 0.742993 |
| 0.763302 | 0.762051 | 0.78216 | 0.758594 | 0.749616 | 0.803631 | 0.747632 | 0.751384 | 0.801292 |
| 0.762712 | 0.753105 | 0.772398 | 0.769948 | 0.742758 | 0.784666 | 0.742129 | 0.781075 | 0.790078 |
| 0.759241 | 0.755785 | 0.76196 | 0.764531 | 0.748023 | 0.758831 | 0.745171 | 0.750068 | 0.765402 |
| 0.758093 | 0.748476 | 0.761587 | 0.762743 | 0.747209 | 0.781164 | 0.744031 | 0.747582 | 0.762625 |
| 0.754292 | 0.745074 | 0.758724 | 0.756444 | 0.742418 | 0.751241 | 0.742871 | 0.737887 | 0.757386 |
| 0.755418 | 0.744064 | 0.780073 | 0.751898 | 0.807351 | 0.741334 | 0.740653 | 0.735988 | 0.750036 |
| 0.764917 | 0.749786 | 0.756225 | 0.754112 | 0.750314 | 0.750697 | 0.74851 | 0.74539 | 0.75535 |
| 0.745442 | 0.738633 | 0.751843 | 0.738735 | 0.740364 | 0.747583 | 0.739804 | 0.74412 | 0.741856 |
| 0.776263 | 0.749237 | 0.773599 | 0.846249 | 0.744403 | 0.800413 | 0.743976 | 0.742875 | 0.749768 |
| 0.731932 | 0.735581 | 0.735195 | 0.73593 | 0.729953 | 0.734112 | 0.721641 | 0.733849 | 0.843037 |
| 0.748913 | 0.745481 | 0.763741 | 0.750854 | 0.740758 | 0.751649 | 0.739338 | 0.79084 | 0.765251 |
| 1 | 0.7457 | 0.786167 | 0.770962 | 0.728934 | 0.794488 | 0.736564 | 0.736248 | 0.746568 |
| 0.7457 | 1 | 0.750445 | 0.744264 | 0.73591 | 0.753791 | 0.737607 | 0.742784 | 0.751942 |
| 0.786167 | 0.750445 | 1 | 0.767826 | 0.744671 | 0.803401 | 0.746477 | 0.747486 | 0.777017 |
| 0.770962 | 0.744264 | 0.767826 | 1 | 0.748148 | 0.797871 | 0.738407 | 0.747285 | 0.748822 |

| | | | | | | | | |
|---|---|---|---|---|---|---|---|---|
| 0.728934 | 0.73591 | 0.744671 | 0.748148 | 1 | 0.737634 | 0.745503 | 0.732485 | 0.737853 |
| 0.794488 | 0.753791 | 0.803401 | 0.797871 | 0.737634 | 1 | 0.742674 | 0.744659 | 0.801711 |
| 0.736564 | 0.737607 | 0.746477 | 0.738407 | 0.745503 | 0.742674 | 1 | 0.73038 | 0.746479 |
| 0.736248 | 0.742784 | 0.747486 | 0.747285 | 0.732485 | 0.744659 | 0.73038 | 1 | 0.74248 |
| 0.746568 | 0.751942 | 0.777017 | 0.748822 | 0.737853 | 0.801711 | 0.746479 | 0.74248 | 1 |
| 0.769284 | 0.752463 | 0.784073 | 0.742013 | 0.740868 | 0.740269 | 0.747003 | 0.7473 | 0.745194 |
| 0.753134 | 0.749323 | 0.774754 | 0.740331 | 0.73144 | 0.771209 | 0.73511 | 0.756373 | 0.77363 |
| 0.789483 | 0.751023 | 0.77001 | 0.799433 | 0.727942 | 0.810684 | 0.740938 | 0.740814 | 0.774756 |
| 0.729504 | 0.727857 | 0.731351 | 0.725147 | 0.728193 | 0.733044 | 0.719667 | 0.722781 | 0.765616 |
| 0.756182 | 0.76 | 0.805075 | 0.749817 | 0.733491 | 0.840877 | 0.738979 | 0.737724 | 0.842744 |
| 0.796144 | 0.74359 | 0.84122 | 0.743751 | 0.736008 | 0.742399 | 0.745086 | 0.742064 | 0.744611 |
| 0.743785 | 0.737287 | 0.74201 | 0.738351 | 0.725235 | 0.743891 | 0.72484 | 0.781813 | 0.74005 |
| 0.738642 | 0.749536 | 0.744974 | 0.747446 | 0.767733 | 0.748151 | 0.740006 | 0.739915 | 0.743293 |
| 0.756574 | 0.743336 | 0.793819 | 0.736293 | 0.736633 | 0.745003 | 0.749406 | 0.742844 | 0.750114 |
| 0.754399 | 0.748509 | 0.750195 | 0.760724 | 0.739607 | 0.753393 | 0.739401 | 0.752589 | 0.754464 |
| 0.734985 | 0.744886 | 0.738503 | 0.741837 | 0.736702 | 0.746923 | 0.738122 | 0.730533 | 0.764571 |
| 0.744396 | 0.744348 | 0.742101 | 0.739878 | 0.739919 | 0.740484 | 0.7339 | 0.753007 | 0.752812 |
| 0.747917 | 0.743535 | 0.754322 | 0.746473 | 0.742923 | 0.751873 | 0.749025 | 0.738092 | 0.743891 |
| 0.732501 | 0.732918 | 0.740094 | 0.734642 | 0.735228 | 0.728451 | 0.721803 | 0.721158 | 0.723598 |
| 0.739591 | 0.743489 | 0.745123 | 0.738801 | 0.736571 | 0.751238 | 0.743425 | 0.735589 | 0.804312 |
| 0.745763 | 0.738118 | 0.75941 | 0.74081 | 0.727175 | 0.742871 | 0.746013 | 0.741492 | 0.731836 |
| 0.738531 | 0.743254 | 0.741564 | 0.737077 | 0.753707 | 0.742745 | 0.738968 | 0.741798 | 0.743321 |
| 0.738966 | 0.73453 | 0.745558 | 0.741642 | 0.733817 | 0.744312 | 0.74355 | 0.730595 | 0.74433 |
| 0.729529 | 0.729042 | 0.739846 | 0.737547 | 0.722793 | 0.744406 | 0.735656 | 0.730431 | 0.733965 |
| 0.727124 | 0.712733 | 0.738154 | 0.738745 | 0.714594 | 0.771405 | 0.712439 | 0.720044 | 0.748151 |
| 0.799869 | 0.75412 | 0.772436 | 0.771933 | 0.739784 | 0.784378 | 0.751456 | 0.74038 | 0.753856 |
| 0.745239 | 0.728188 | 0.749768 | 0.73046 | 0.711908 | 0.732373 | 0.721911 | 0.730545 | 0.727557 |
| 0.740569 | 0.738242 | 0.741495 | 0.74696 | 0.740074 | 0.744174 | 0.73552 | 0.736354 | 0.732266 |
| 0.72328 | 0.717572 | 0.721009 | 0.720688 | 0.713599 | 0.722986 | 0.716044 | 0.711419 | 0.723355 |
| 0.74212 | 0.742827 | 0.760984 | 0.742623 | 0.736598 | 0.782335 | 0.742998 | 0.727029 | 0.782971 |
| 0.73147 | 0.750588 | 0.738672 | 0.73597 | 0.73554 | 0.740225 | 0.74137 | 0.731694 | 0.739136 |
| 0.735915 | 0.724138 | 0.745103 | 0.749009 | 0.724665 | 0.759538 | 0.727709 | 0.747263 | 0.741123 |
| 0.736203 | 0.724574 | 0.748344 | 0.751631 | 0.727284 | 0.761464 | 0.723893 | 0.746991 | 0.743029 |
| 0.732389 | 0.735125 | 0.742259 | 0.739247 | 0.723466 | 0.75223 | 0.746516 | 0.730441 | 0.733254 |
| 0.730727 | 0.725467 | 0.743531 | 0.742396 | 0.835205 | 0.728919 | 0.743919 | 0.725956 | 0.731101 |
| 0.741448 | 0.732752 | 0.741202 | 0.743398 | 0.733451 | 0.7524 | 0.73137 | 0.727493 | 0.737995 |
| 0.701327 | 0.690685 | 0.704914 | 0.699195 | 0.687228 | 0.699277 | 0.692105 | 0.690327 | 0.696893 |
| 0.725267 | 0.730729 | 0.726705 | 0.735551 | 0.721301 | 0.731347 | 0.726963 | 0.717189 | 0.72652 |
| 0.713819 | 0.693034 | 0.714197 | 0.700268 | 0.690301 | 0.69847 | 0.688522 | 0.689079 | 0.68983 |
| 0.692088 | 0.685889 | 0.719068 | 0.676748 | 0.682807 | 0.684197 | 0.67852 | 0.678622 | 0.680176 |
| 0.747407 | 0.738596 | 0.745448 | 0.74714 | 0.729883 | 0.752382 | 0.734711 | 0.736577 | 0.758487 |
| 0.691003 | 0.682071 | 0.701022 | 0.682956 | 0.678954 | 0.686426 | 0.673818 | 0.678841 | 0.681116 |
| 0.698013 | 0.690576 | 0.711013 | 0.698263 | 0.680158 | 0.720715 | 0.667988 | 0.674448 | 0.704393 |
| 0.675466 | 0.676984 | 0.696911 | 0.68127 | 0.669902 | 0.71885 | 0.662795 | 0.665737 | 0.7054 |
| 0.73181 | 0.732614 | 0.743167 | 0.736162 | 0.724047 | 0.752013 | 0.726197 | 0.761709 | 0.741829 |
| 0.680318 | 0.681404 | 0.696407 | 0.688809 | 0.665014 | 0.710031 | 0.663221 | 0.656002 | 0.693662 |
| 0.735167 | 0.733697 | 0.741882 | 0.749936 | 0.725262 | 0.752887 | 0.724382 | 0.742369 | 0.747315 |

| | | | | | | | | |
|---|---|---|---|---|---|---|---|---|
| 0.73269 | 0.72449 | 0.732505 | 0.743073 | 0.725714 | 0.742595 | 0.723315 | 0.735468 | 0.73364 |
| 0.737129 | 0.727627 | 0.744882 | 0.744773 | 0.726428 | 0.757149 | 0.728224 | 0.744186 | 0.752351 |
| 0.712537 | 0.726178 | 0.72331 | 0.725919 | 0.717356 | 0.730376 | 0.719092 | 0.723296 | 0.712933 |
| 0.680181 | 0.666061 | 0.677776 | 0.687426 | 0.655992 | 0.704835 | 0.651546 | 0.655125 | 0.674081 |
| 0.679035 | 0.673411 | 0.701818 | 0.673608 | 0.66497 | 0.716083 | 0.659862 | 0.658411 | 0.696524 |
| 0.745145 | 0.731404 | 0.75226 | 0.749936 | 0.727857 | 0.756968 | 0.734358 | 0.745318 | 0.752602 |
| 0.666667 | 0.652758 | 0.666351 | 0.656621 | 0.657502 | 0.66044 | 0.65329 | 0.648615 | 0.657738 |
| 0.697435 | 0.700651 | 0.695928 | 0.698895 | 0.693298 | 0.699205 | 0.697813 | 0.687819 | 0.692982 |
| 0.741964 | 0.73431 | 0.739532 | 0.745315 | 0.725545 | 0.752465 | 0.719112 | 0.742517 | 0.726126 |
| 0.682638 | 0.68519 | 0.682989 | 0.683589 | 0.682034 | 0.680354 | 0.692088 | 0.684218 | 0.672914 |
| 0.727056 | 0.71451 | 0.72575 | 0.727145 | 0.711614 | 0.743805 | 0.716644 | 0.715156 | 0.729089 |
| 0.61875 | 0.673469 | 0.715328 | 0.654135 | 0.646667 | 0.673913 | 0.597315 | 0.643357 | 0.648649 |
| 0.653956 | 0.657781 | 0.659663 | 0.659321 | 0.647239 | 0.663433 | 0.654892 | 0.65799 | 0.653915 |
| 0.64354 | 0.649139 | 0.655939 | 0.65494 | 0.649318 | 0.655904 | 0.648131 | 0.650994 | 0.65171 |
| 0.65596 | 0.659411 | 0.660156 | 0.662052 | 0.661977 | 0.665757 | 0.657569 | 0.656256 | 0.659315 |
| 0.656329 | 0.655105 | 0.657377 | 0.658174 | 0.658164 | 0.660825 | 0.658533 | 0.653184 | 0.653285 |
| 0.607396 | 0.603717 | 0.608274 | 0.611282 | 0.602473 | 0.609999 | 0.599404 | 0.597132 | 0.601733 |
| 0.657534 | 0.65177 | 0.663072 | 0.664515 | 0.653039 | 0.662971 | 0.658921 | 0.655806 | 0.657066 |
| 0.715261 | 0.708252 | 0.718638 | 0.716864 | 0.701346 | 0.736566 | 0.704082 | 0.706547 | 0.725056 |
| 0.721404 | 0.704498 | 0.714801 | 0.717544 | 0.702522 | 0.733821 | 0.702236 | 0.704363 | 0.7061 |
| 0.717734 | 0.71453 | 0.723404 | 0.722796 | 0.70826 | 0.744909 | 0.711673 | 0.706248 | 0.719457 |
| 0.568831 | 0.573477 | 0.575798 | 0.572784 | 0.568292 | 0.574906 | 0.562556 | 0.565383 | 0.565755 |
| 0.555727 | 0.558012 | 0.557549 | 0.554313 | 0.551204 | 0.560423 | 0.550389 | 0.548039 | 0.55352 |
| 0.666218 | 0.666283 | 0.661171 | 0.670484 | 0.67047 | 0.671474 | 0.665595 | 0.6618 | 0.660411 |
| 0.665207 | 0.667857 | 0.663086 | 0.670301 | 0.673497 | 0.670493 | 0.668098 | 0.660377 | 0.667096 |
| 0.658269 | 0.656713 | 0.656551 | 0.659175 | 0.662828 | 0.663438 | 0.666487 | 0.65676 | 0.653772 |
| 0.660589 | 0.661161 | 0.66108 | 0.665517 | 0.666093 | 0.667186 | 0.666355 | 0.657162 | 0.656027 |
| 0.664128 | 0.666624 | 0.669865 | 0.673916 | 0.671682 | 0.669267 | 0.669955 | 0.663807 | 0.662434 |
| 0.665408 | 0.669766 | 0.668082 | 0.675904 | 0.67302 | 0.674642 | 0.673239 | 0.659316 | 0.660586 |
| 0.663383 | 0.669541 | 0.666749 | 0.675104 | 0.671795 | 0.673702 | 0.667107 | 0.66322 | 0.664416 |
| 0.669404 | 0.672225 | 0.670507 | 0.674489 | 0.674981 | 0.675398 | 0.6697 | 0.673895 | 0.66469 |
| 0.672751 | 0.675772 | 0.677947 | 0.679822 | 0.675898 | 0.679382 | 0.678238 | 0.673556 | 0.666497 |
| 0.667923 | 0.670739 | 0.673149 | 0.674021 | 0.677387 | 0.67558 | 0.675295 | 0.663283 | 0.661785 |
| 0.664545 | 0.670076 | 0.671796 | 0.67543 | 0.673357 | 0.677155 | 0.670834 | 0.665791 | 0.663567 |
| 0.668041 | 0.668577 | 0.673405 | 0.674597 | 0.677172 | 0.680136 | 0.672827 | 0.671815 | 0.664778 |
| 0.666101 | 0.665433 | 0.666916 | 0.67248 | 0.667648 | 0.675768 | 0.664798 | 0.661235 | 0.656374 |
| 0.669583 | 0.671642 | 0.668374 | 0.675856 | 0.674935 | 0.677436 | 0.672375 | 0.670209 | 0.665426 |
| 0.671759 | 0.672862 | 0.675238 | 0.679553 | 0.669524 | 0.673814 | 0.674624 | 0.670876 | 0.661757 |
| 0.664101 | 0.668122 | 0.668009 | 0.668738 | 0.672684 | 0.673999 | 0.670441 | 0.669741 | 0.659827 |
| 0.668845 | 0.668742 | 0.671318 | 0.673985 | 0.676959 | 0.674422 | 0.669405 | 0.668561 | 0.667739 |
| 0.658766 | 0.658972 | 0.659914 | 0.660871 | 0.661089 | 0.658833 | 0.65684 | 0.650813 | 0.656675 |
| 0.660158 | 0.661054 | 0.663855 | 0.666101 | 0.664553 | 0.66933 | 0.66077 | 0.666161 | 0.663671 |
| 0.672323 | 0.675683 | 0.675286 | 0.679245 | 0.679396 | 0.676627 | 0.678961 | 0.675354 | 0.668455 |
| 0.670961 | 0.674116 | 0.671822 | 0.674763 | 0.677133 | 0.677344 | 0.677678 | 0.671695 | 0.670434 |
| 0.639351 | 0.647488 | 0.647491 | 0.647911 | 0.643552 | 0.652001 | 0.651634 | 0.640898 | 0.64245 |
| 0.668242 | 0.673308 | 0.672037 | 0.673991 | 0.674754 | 0.673811 | 0.671457 | 0.671612 | 0.665471 |
| 0.656657 | 0.659631 | 0.660743 | 0.660071 | 0.666051 | 0.667056 | 0.654509 | 0.658148 | 0.654285 |

| | | | | | | | | |
|---|---|---|---|---|---|---|---|---|
| 0.66881 | 0.672519 | 0.667577 | 0.670919 | 0.670747 | 0.67636 | 0.67164 | 0.662066 | 0.663988 |
| 0.667712 | 0.674095 | 0.66716 | 0.672129 | 0.676489 | 0.675655 | 0.668738 | 0.666667 | 0.663952 |
| 0.666097 | 0.676312 | 0.672312 | 0.673642 | 0.675468 | 0.675697 | 0.664928 | 0.672621 | 0.667183 |
| 0.670303 | 0.675229 | 0.671269 | 0.673037 | 0.676055 | 0.674023 | 0.673317 | 0.668122 | 0.666667 |
| 0.674824 | 0.684036 | 0.678286 | 0.684002 | 0.680597 | 0.688024 | 0.675568 | 0.677189 | 0.673938 |
| 0.666384 | 0.671193 | 0.667581 | 0.672448 | 0.671129 | 0.673735 | 0.668263 | 0.664515 | 0.66607 |
| 0.667267 | 0.670077 | 0.669315 | 0.674262 | 0.675679 | 0.673443 | 0.67245 | 0.665048 | 0.666709 |
| 0.659593 | 0.663817 | 0.655099 | 0.661184 | 0.658873 | 0.665306 | 0.660416 | 0.66059 | 0.655706 |
| 0.66594 | 0.665145 | 0.660994 | 0.669585 | 0.668911 | 0.665933 | 0.666223 | 0.664998 | 0.664912 |
| 0.664798 | 0.667434 | 0.663953 | 0.671931 | 0.673421 | 0.67401 | 0.661058 | 0.659317 | 0.655581 |
| 0.666747 | 0.672558 | 0.670229 | 0.675925 | 0.678171 | 0.673953 | 0.672492 | 0.665482 | 0.661656 |
| 0.666955 | 0.669809 | 0.67018 | 0.669204 | 0.671122 | 0.674625 | 0.665631 | 0.665326 | 0.662771 |
| 0.65701 | 0.662075 | 0.663186 | 0.662606 | 0.66291 | 0.667498 | 0.655705 | 0.656923 | 0.65408 |
| 0.661785 | 0.659814 | 0.659623 | 0.663567 | 0.662997 | 0.667508 | 0.662819 | 0.661399 | 0.65994 |
| 0.673425 | 0.681911 | 0.673543 | 0.679719 | 0.678413 | 0.684313 | 0.679728 | 0.671379 | 0.670075 |
| 0.672881 | 0.669418 | 0.671388 | 0.671341 | 0.676438 | 0.675247 | 0.673711 | 0.662727 | 0.668679 |
| 0.666174 | 0.664999 | 0.665243 | 0.673587 | 0.666749 | 0.67222 | 0.674331 | 0.662006 | 0.66307 |
| 0.663943 | 0.671825 | 0.665329 | 0.666501 | 0.667079 | 0.669421 | 0.673954 | 0.658795 | 0.657939 |
| 0.664446 | 0.665989 | 0.662443 | 0.672982 | 0.666093 | 0.668055 | 0.667779 | 0.661209 | 0.658218 |
| 0.661878 | 0.666837 | 0.663267 | 0.670696 | 0.668721 | 0.668447 | 0.66885 | 0.664595 | 0.664948 |
| 0.659211 | 0.667446 | 0.656664 | 0.666416 | 0.662889 | 0.667855 | 0.663151 | 0.651861 | 0.657816 |
| 0.663189 | 0.66512 | 0.664524 | 0.668417 | 0.666459 | 0.668825 | 0.664817 | 0.663765 | 0.661167 |
| 0.667068 | 0.663215 | 0.668068 | 0.675819 | 0.670711 | 0.674413 | 0.675938 | 0.663342 | 0.665216 |
| 0.666343 | 0.668792 | 0.666252 | 0.6688 | 0.665359 | 0.675683 | 0.676779 | 0.664596 | 0.664009 |
| 0.659177 | 0.667094 | 0.667548 | 0.67153 | 0.673589 | 0.673598 | 0.667788 | 0.659546 | 0.660636 |
| 0.660407 | 0.661768 | 0.663041 | 0.664977 | 0.66576 | 0.666317 | 0.666264 | 0.658248 | 0.660333 |
| 0.658126 | 0.665506 | 0.66168 | 0.665168 | 0.666086 | 0.666228 | 0.667208 | 0.659458 | 0.660984 |
| 0.659988 | 0.659126 | 0.660505 | 0.665462 | 0.667125 | 0.667811 | 0.670722 | 0.654313 | 0.651865 |
| 0.661572 | 0.662212 | 0.66061 | 0.664157 | 0.668352 | 0.66532 | 0.663863 | 0.655513 | 0.660409 |
| 0.664082 | 0.665733 | 0.663888 | 0.669595 | 0.670243 | 0.673384 | 0.666089 | 0.661295 | 0.658026 |
| 0.669622 | 0.669442 | 0.669834 | 0.669416 | 0.673972 | 0.674294 | 0.672773 | 0.660037 | 0.663578 |
| 0.666218 | 0.67214 | 0.66755 | 0.671002 | 0.673495 | 0.673416 | 0.67474 | 0.666488 | 0.66502 |
| 0.670573 | 0.676684 | 0.669605 | 0.676953 | 0.67759 | 0.679886 | 0.677493 | 0.665835 | 0.669744 |
| 0.668972 | 0.675163 | 0.671296 | 0.67114 | 0.675716 | 0.677782 | 0.675781 | 0.667197 | 0.664865 |
| 0.660139 | 0.661907 | 0.659489 | 0.665384 | 0.669308 | 0.667453 | 0.663663 | 0.6554 | 0.663001 |
| 0.662248 | 0.66833 | 0.664418 | 0.673113 | 0.671258 | 0.670548 | 0.670467 | 0.664363 | 0.662021 |
| 0.668861 | 0.664035 | 0.661533 | 0.670759 | 0.664946 | 0.671936 | 0.67497 | 0.663926 | 0.666969 |
| 0.665529 | 0.660234 | 0.665082 | 0.669556 | 0.670334 | 0.669405 | 0.670989 | 0.659929 | 0.660309 |
| 0.657824 | 0.663526 | 0.659879 | 0.663465 | 0.663645 | 0.667108 | 0.665853 | 0.654441 | 0.657243 |
| 0.663796 | 0.659362 | 0.656836 | 0.663968 | 0.664174 | 0.669399 | 0.665319 | 0.658653 | 0.659886 |
| 0.66114 | 0.664365 | 0.661095 | 0.663681 | 0.659515 | 0.666042 | 0.660263 | 0.651922 | 0.655448 |
| 0.661558 | 0.657481 | 0.660894 | 0.662714 | 0.661252 | 0.667516 | 0.664791 | 0.656718 | 0.658296 |
| 0.659652 | 0.65965 | 0.65999 | 0.658524 | 0.658015 | 0.664475 | 0.665522 | 0.658537 | 0.657396 |
| 0.658249 | 0.65525 | 0.655829 | 0.66465 | 0.660377 | 0.661436 | 0.663372 | 0.656947 | 0.650558 |
| 0.663639 | 0.664072 | 0.665477 | 0.67129 | 0.669481 | 0.670587 | 0.674199 | 0.658092 | 0.661208 |
| 0.66036 | 0.659075 | 0.657225 | 0.65782 | 0.661826 | 0.665662 | 0.666473 | 0.657332 | 0.658437 |

| | | | | | | | | |
|---|---|---|---|---|---|---|---|---|
| 0.668661 | 0.674502 | 0.667622 | 0.675323 | 0.66774 | 0.674087 | 0.670253 | 0.664989 | 0.667011 |
| 0.667967 | 0.671113 | 0.667419 | 0.671675 | 0.674349 | 0.672193 | 0.67155 | 0.667465 | 0.665721 |
| 0.659431 | 0.663163 | 0.664256 | 0.664502 | 0.664717 | 0.665299 | 0.663634 | 0.656794 | 0.660544 |
| 0.668111 | 0.662845 | 0.663782 | 0.671986 | 0.669721 | 0.671119 | 0.670432 | 0.662157 | 0.662258 |
| 0.67511 | 0.678754 | 0.672058 | 0.681762 | 0.679457 | 0.674772 | 0.67783 | 0.669402 | 0.667052 |
| 0.668438 | 0.672374 | 0.668034 | 0.676681 | 0.674932 | 0.676765 | 0.670785 | 0.668377 | 0.670004 |
| 0.668484 | 0.662921 | 0.662217 | 0.669407 | 0.667912 | 0.670011 | 0.669973 | 0.661239 | 0.662449 |
| 0.662165 | 0.670062 | 0.664175 | 0.670358 | 0.666993 | 0.670173 | 0.671923 | 0.668741 | 0.665215 |
| 0.667928 | 0.670369 | 0.669373 | 0.678232 | 0.676721 | 0.673098 | 0.675125 | 0.661262 | 0.668742 |
| 0.667675 | 0.666328 | 0.66505 | 0.671763 | 0.668966 | 0.670166 | 0.668266 | 0.660266 | 0.660298 |
| 0.669634 | 0.676392 | 0.671095 | 0.676084 | 0.67704 | 0.676764 | 0.67312 | 0.663792 | 0.668877 |
| 0.668721 | 0.673269 | 0.667492 | 0.674631 | 0.675018 | 0.674639 | 0.67427 | 0.667458 | 0.666326 |
| 0.666343 | 0.665696 | 0.660639 | 0.666054 | 0.669982 | 0.669863 | 0.667997 | 0.660827 | 0.661693 |
| 0.668887 | 0.666371 | 0.669025 | 0.670266 | 0.675919 | 0.671302 | 0.665341 | 0.665259 | 0.665043 |
| 0.668716 | 0.671268 | 0.664027 | 0.66977 | 0.673387 | 0.677792 | 0.669006 | 0.661872 | 0.669017 |
| 0.666019 | 0.67023 | 0.66596 | 0.669179 | 0.667775 | 0.67083 | 0.671453 | 0.666534 | 0.667482 |
| 0.670277 | 0.669871 | 0.66831 | 0.677688 | 0.674908 | 0.675742 | 0.677826 | 0.665357 | 0.668541 |
| 0.665988 | 0.668638 | 0.667283 | 0.673253 | 0.664404 | 0.672376 | 0.667325 | 0.66037 | 0.669135 |
| 0.67588 | 0.670881 | 0.67053 | 0.680079 | 0.674212 | 0.680267 | 0.670921 | 0.670863 | 0.667568 |
| 0.66526 | 0.667514 | 0.66675 | 0.6706 | 0.668582 | 0.676164 | 0.666002 | 0.65994 | 0.662998 |
| 0.668038 | 0.666033 | 0.666832 | 0.672224 | 0.672023 | 0.675707 | 0.667952 | 0.664342 | 0.668768 |
| 0.667513 | 0.668234 | 0.668241 | 0.675592 | 0.672042 | 0.670909 | 0.673311 | 0.66434 | 0.66564 |
| 0.669566 | 0.665614 | 0.664199 | 0.671117 | 0.674116 | 0.674732 | 0.672831 | 0.664346 | 0.667947 |
| 0.667749 | 0.676042 | 0.667695 | 0.678087 | 0.678493 | 0.674919 | 0.671296 | 0.667983 | 0.665605 |
| 0.664468 | 0.665368 | 0.662744 | 0.669431 | 0.668977 | 0.671331 | 0.667896 | 0.662108 | 0.662145 |
| 0.665859 | 0.66743 | 0.665461 | 0.676656 | 0.672224 | 0.673873 | 0.669209 | 0.66543 | 0.664998 |
| 0.664182 | 0.659515 | 0.658618 | 0.6663 | 0.66923 | 0.664174 | 0.666091 | 0.66128 | 0.660628 |
| 0.666347 | 0.667931 | 0.663997 | 0.669728 | 0.672269 | 0.669459 | 0.666711 | 0.660358 | 0.660109 |
| 0.668873 | 0.670249 | 0.669472 | 0.673886 | 0.675506 | 0.671173 | 0.677346 | 0.664354 | 0.66374 |
| 0.674387 | 0.672294 | 0.675508 | 0.68031 | 0.674404 | 0.679046 | 0.673036 | 0.671083 | 0.671776 |
| 0.668623 | 0.661609 | 0.667503 | 0.672705 | 0.67577 | 0.671254 | 0.676077 | 0.661365 | 0.662115 |
| 0.673792 | 0.671345 | 0.672188 | 0.684289 | 0.678479 | 0.679132 | 0.676822 | 0.666931 | 0.670785 |
| 0.670664 | 0.670822 | 0.669324 | 0.67957 | 0.675009 | 0.672463 | 0.678638 | 0.665172 | 0.662701 |
| 0.662759 | 0.66799 | 0.668678 | 0.672691 | 0.668983 | 0.668896 | 0.669001 | 0.659735 | 0.66887 |
| 0.671082 | 0.671379 | 0.668508 | 0.681279 | 0.674089 | 0.678977 | 0.674431 | 0.66751 | 0.667785 |
| 0.671126 | 0.674458 | 0.667634 | 0.673282 | 0.670547 | 0.673062 | 0.670187 | 0.664839 | 0.667841 |
| 0.672476 | 0.672018 | 0.671313 | 0.677304 | 0.680841 | 0.675363 | 0.675928 | 0.670363 | 0.66769 |
| 0.668979 | 0.672239 | 0.669248 | 0.675723 | 0.675662 | 0.673132 | 0.675141 | 0.66649 | 0.663238 |
| 0.667954 | 0.663894 | 0.666625 | 0.675596 | 0.6723 | 0.674159 | 0.66844 | 0.663861 | 0.665475 |
| 0.666991 | 0.668545 | 0.670839 | 0.674975 | 0.671409 | 0.674437 | 0.674879 | 0.661811 | 0.66942 |
| 0.674475 | 0.67363 | 0.677043 | 0.684431 | 0.671188 | 0.683418 | 0.676964 | 0.668641 | 0.668364 |
| 0.666425 | 0.668406 | 0.669979 | 0.674387 | 0.674141 | 0.676941 | 0.672846 | 0.668346 | 0.664107 |
| 0.666065 | 0.668944 | 0.665431 | 0.674598 | 0.675851 | 0.674395 | 0.675611 | 0.669486 | 0.667392 |
| 0.666424 | 0.667347 | 0.669758 | 0.677095 | 0.667283 | 0.680084 | 0.663866 | 0.667197 | 0.664735 |
| 0.673643 | 0.669309 | 0.670957 | 0.67607 | 0.674071 | 0.676925 | 0.673095 | 0.663761 | 0.66916 |
| 0.663711 | 0.666667 | 0.661333 | 0.674308 | 0.662849 | 0.66885 | 0.668275 | 0.659501 | 0.660749 |

| | | | | | | | | |
|---|---|---|---|---|---|---|---|---|
| 0.661402 | 0.663116 | 0.660707 | 0.670328 | 0.664329 | 0.670925 | 0.668484 | 0.658098 | 0.661631 |
| 0.714639 | 0.719745 | 0.723152 | 0.71743 | 0.705824 | 0.721758 | 0.714518 | 0.707981 | 0.709703 |
| 0.662087 | 0.671151 | 0.67012 | 0.674404 | 0.667286 | 0.671222 | 0.668102 | 0.66041 | 0.669 |
| 0.668394 | 0.668519 | 0.67167 | 0.674911 | 0.676215 | 0.678052 | 0.672055 | 0.663056 | 0.670206 |
| 0.662842 | 0.663556 | 0.670268 | 0.673208 | 0.668467 | 0.668253 | 0.667711 | 0.6596 | 0.665227 |
| 0.667074 | 0.65989 | 0.665913 | 0.674349 | 0.667533 | 0.672935 | 0.670085 | 0.656491 | 0.666494 |
| 0.666584 | 0.664249 | 0.666709 | 0.674154 | 0.667335 | 0.670133 | 0.669616 | 0.65917 | 0.668858 |
| 0.661396 | 0.666316 | 0.665939 | 0.666582 | 0.666582 | 0.667432 | 0.660447 | 0.656313 | 0.661809 |
| 0.659906 | 0.659585 | 0.661792 | 0.668262 | 0.667687 | 0.667703 | 0.664896 | 0.659661 | 0.659466 |
| 0.67529 | 0.675596 | 0.677176 | 0.678222 | 0.678876 | 0.683187 | 0.678221 | 0.677015 | 0.671268 |
| 0.667474 | 0.67378 | 0.667702 | 0.679009 | 0.670189 | 0.677142 | 0.673221 | 0.669316 | 0.666923 |
| 0.673823 | 0.669694 | 0.671761 | 0.674456 | 0.67812 | 0.678557 | 0.673374 | 0.668929 | 0.666924 |
| 0.673855 | 0.673305 | 0.670777 | 0.679411 | 0.675825 | 0.676574 | 0.67128 | 0.668032 | 0.66829 |
| 0.670705 | 0.673783 | 0.673352 | 0.676781 | 0.677281 | 0.675704 | 0.678367 | 0.672316 | 0.666667 |
| 0.671603 | 0.6688 | 0.669122 | 0.674213 | 0.673275 | 0.672024 | 0.672886 | 0.666446 | 0.666237 |
| 0.668262 | 0.669018 | 0.665806 | 0.677218 | 0.671206 | 0.672239 | 0.667938 | 0.658562 | 0.664559 |
| 0.671046 | 0.675475 | 0.673728 | 0.676555 | 0.673628 | 0.678361 | 0.679062 | 0.670477 | 0.668605 |
| 0.674138 | 0.67117 | 0.674746 | 0.673487 | 0.672367 | 0.679775 | 0.679606 | 0.667728 | 0.668039 |
| 0.670957 | 0.66928 | 0.669556 | 0.67232 | 0.676609 | 0.677691 | 0.671353 | 0.662665 | 0.668943 |
| 0.666048 | 0.663775 | 0.663913 | 0.671337 | 0.67085 | 0.670083 | 0.671866 | 0.660142 | 0.666054 |
| 0.669747 | 0.669943 | 0.671207 | 0.674755 | 0.672646 | 0.676371 | 0.673286 | 0.662171 | 0.668426 |
| 0.658552 | 0.657628 | 0.664028 | 0.663355 | 0.660183 | 0.665467 | 0.66248 | 0.650821 | 0.65506 |
| 0.667193 | 0.667391 | 0.664504 | 0.668558 | 0.671758 | 0.671796 | 0.669168 | 0.659892 | 0.660912 |
| 0.665779 | 0.669439 | 0.671059 | 0.669137 | 0.667647 | 0.671867 | 0.670631 | 0.663672 | 0.66671 |
| 0.672631 | 0.670727 | 0.674808 | 0.676291 | 0.676245 | 0.67835 | 0.674074 | 0.665514 | 0.671982 |
| 0.669889 | 0.670276 | 0.668505 | 0.674744 | 0.671832 | 0.674734 | 0.667249 | 0.660695 | 0.661681 |
| 0.660679 | 0.665105 | 0.66314 | 0.666204 | 0.665541 | 0.673619 | 0.664991 | 0.658664 | 0.66213 |
| 0.660521 | 0.665331 | 0.660755 | 0.671466 | 0.672422 | 0.675672 | 0.664597 | 0.664262 | 0.661959 |
| 0.666876 | 0.668365 | 0.668112 | 0.672658 | 0.668605 | 0.678798 | 0.670333 | 0.657919 | 0.667585 |
| 0.735482 | 0.73777 | 0.740813 | 0.743965 | 0.832447 | 0.735976 | 0.73649 | 0.735614 | 0.742051 |
| 0.768431 | 0.749867 | 0.77956 | 0.760021 | 0.745224 | 0.777349 | 0.748388 | 0.743877 | 0.786215 |
| 0.767445 | 0.738284 | 0.800053 | 0.745078 | 0.743125 | 0.745343 | 0.739922 | 0.743657 | 0.751691 |
| 0.773793 | 0.74616 | 0.803656 | 0.756038 | 0.737997 | 0.753609 | 0.737992 | 0.802788 | 0.741375 |
| 0.775233 | 0.755181 | 0.815354 | 0.747954 | 0.747202 | 0.794354 | 0.74469 | 0.752271 | 0.796759 |
| 0.663351 | 0.6646 | 0.661302 | 0.667957 | 0.664142 | 0.667982 | 0.666936 | 0.659776 | 0.660304 |
| 0.660792 | 0.665511 | 0.6629 | 0.669396 | 0.671336 | 0.669679 | 0.663137 | 0.665649 | 0.663177 |
| 0.662671 | 0.667306 | 0.662928 | 0.669342 | 0.669588 | 0.668968 | 0.665017 | 0.65853 | 0.662344 |
| 0.669013 | 0.662548 | 0.66621 | 0.667609 | 0.668716 | 0.66671 | 0.670371 | 0.663672 | 0.663801 |
| 0.661076 | 0.663155 | 0.66252 | 0.663349 | 0.663888 | 0.666494 | 0.670645 | 0.658241 | 0.658636 |
| 0.665738 | 0.668445 | 0.668534 | 0.674174 | 0.673442 | 0.675796 | 0.670136 | 0.667282 | 0.66581 |
| 0.663586 | 0.66641 | 0.66579 | 0.669145 | 0.669845 | 0.66541 | 0.666087 | 0.663218 | 0.665289 |
| 0.665002 | 0.670255 | 0.660307 | 0.669646 | 0.66972 | 0.672996 | 0.667068 | 0.66272 | 0.6601 |
| 0.660047 | 0.665219 | 0.663159 | 0.667092 | 0.661552 | 0.667964 | 0.660643 | 0.655691 | 0.660521 |
| 0.662506 | 0.662622 | 0.664517 | 0.665558 | 0.668371 | 0.66888 | 0.663859 | 0.658619 | 0.657989 |
| 0.664039 | 0.666753 | 0.664257 | 0.672373 | 0.667499 | 0.666182 | 0.657547 | 0.662766 | 0.660086 |
| 0.659342 | 0.666966 | 0.665954 | 0.67236 | 0.669102 | 0.665097 | 0.664025 | 0.664891 | 0.663476 |

| | | | | | | | | |
|---|---|---|---|---|---|---|---|---|
| 0.67027 | 0.666495 | 0.667002 | 0.673 | 0.669151 | 0.675069 | 0.666265 | 0.663659 | 0.664033 |
| 0.663474 | 0.667219 | 0.668252 | 0.671871 | 0.672965 | 0.670898 | 0.666444 | 0.660977 | 0.665291 |
| 0.670729 | 0.671805 | 0.667748 | 0.677147 | 0.679224 | 0.675386 | 0.669828 | 0.668258 | 0.664442 |
| 0.66346 | 0.662369 | 0.662544 | 0.673561 | 0.673409 | 0.666667 | 0.659912 | 0.663822 | 0.660841 |
| 0.664135 | 0.660351 | 0.661938 | 0.668453 | 0.66708 | 0.666448 | 0.661923 | 0.662133 | 0.658109 |
| 0.667774 | 0.658192 | 0.664904 | 0.668787 | 0.669447 | 0.670894 | 0.664515 | 0.663685 | 0.657066 |
| 0.671467 | 0.670281 | 0.671544 | 0.675832 | 0.673927 | 0.678439 | 0.669809 | 0.664266 | 0.670579 |
| 0.668149 | 0.669612 | 0.666751 | 0.678634 | 0.671625 | 0.668641 | 0.664999 | 0.66439 | 0.666144 |
| 0.663723 | 0.666581 | 0.663061 | 0.668577 | 0.666667 | 0.669918 | 0.669236 | 0.65949 | 0.658404 |
| 0.66082 | 0.661105 | 0.668838 | 0.672905 | 0.668312 | 0.672737 | 0.671294 | 0.66254 | 0.663652 |
| 0.667685 | 0.663939 | 0.666416 | 0.671597 | 0.671873 | 0.672763 | 0.666667 | 0.660435 | 0.660351 |
| 0.66585 | 0.667353 | 0.663529 | 0.673894 | 0.666377 | 0.670182 | 0.666801 | 0.662633 | 0.661497 |
| 0.656759 | 0.657331 | 0.662942 | 0.663484 | 0.666412 | 0.669982 | 0.661631 | 0.658557 | 0.657487 |
| 0.663016 | 0.670161 | 0.668368 | 0.671412 | 0.669852 | 0.674045 | 0.667643 | 0.655895 | 0.666041 |
| 0.671273 | 0.671224 | 0.67184 | 0.675103 | 0.676522 | 0.677513 | 0.671313 | 0.666267 | 0.666926 |
| 0.675976 | 0.671672 | 0.677286 | 0.686907 | 0.677487 | 0.68114 | 0.675498 | 0.674853 | 0.675606 |
| 0.672084 | 0.666038 | 0.667244 | 0.675374 | 0.671856 | 0.672767 | 0.669561 | 0.664275 | 0.669252 |
| 0.672107 | 0.671547 | 0.674343 | 0.681965 | 0.67691 | 0.675506 | 0.67503 | 0.667464 | 0.667611 |
| 0.668523 | 0.667445 | 0.664981 | 0.673108 | 0.675927 | 0.674199 | 0.668017 | 0.665229 | 0.665711 |
| 0.668391 | 0.672295 | 0.670055 | 0.675028 | 0.670797 | 0.672443 | 0.673845 | 0.662966 | 0.669547 |
| 0.671544 | 0.665634 | 0.670162 | 0.677041 | 0.669753 | 0.673651 | 0.672769 | 0.663485 | 0.66645 |
| 0.67027 | 0.668688 | 0.670599 | 0.675936 | 0.669615 | 0.674125 | 0.674063 | 0.665862 | 0.664203 |
| 0.675439 | 0.67245 | 0.670547 | 0.679859 | 0.671618 | 0.680016 | 0.673458 | 0.66698 | 0.669751 |
| 0.672946 | 0.676436 | 0.672129 | 0.673849 | 0.670976 | 0.675992 | 0.666667 | 0.667576 | 0.666404 |
| 0.673062 | 0.672715 | 0.673766 | 0.683012 | 0.673707 | 0.67762 | 0.676707 | 0.672995 | 0.667377 |
| 0.761454 | 0.752897 | 0.771075 | 0.7757 | 0.751514 | 0.769896 | 0.744086 | 0.768521 | 0.750912 |
| 0.663025 | 0.666624 | 0.666625 | 0.674648 | 0.674743 | 0.66977 | 0.672494 | 0.654329 | 0.666966 |
| 0.666828 | 0.666878 | 0.668706 | 0.675272 | 0.677407 | 0.669554 | 0.673013 | 0.660029 | 0.666367 |
| 0.668333 | 0.67206 | 0.671317 | 0.677664 | 0.669468 | 0.67336 | 0.675037 | 0.668584 | 0.670432 |
| 0.668203 | 0.670725 | 0.672297 | 0.674433 | 0.673967 | 0.671443 | 0.670409 | 0.659544 | 0.662734 |
| 0.670424 | 0.667185 | 0.671948 | 0.676141 | 0.672595 | 0.677269 | 0.674058 | 0.662468 | 0.669894 |
| 0.669501 | 0.668824 | 0.670426 | 0.673592 | 0.676367 | 0.67408 | 0.675152 | 0.660317 | 0.666104 |
| 0.665851 | 0.668896 | 0.668549 | 0.673675 | 0.68013 | 0.668325 | 0.672052 | 0.659274 | 0.664818 |
| 0.665596 | 0.669828 | 0.668482 | 0.67242 | 0.67361 | 0.666092 | 0.673886 | 0.663751 | 0.664056 |
| 0.661953 | 0.664973 | 0.665732 | 0.67037 | 0.672216 | 0.666402 | 0.67328 | 0.662287 | 0.665619 |
| 0.660826 | 0.663957 | 0.66491 | 0.673766 | 0.672584 | 0.666272 | 0.666712 | 0.658032 | 0.661644 |
| 0.670493 | 0.677894 | 0.67056 | 0.67504 | 0.673781 | 0.678226 | 0.673667 | 0.664894 | 0.667012 |
| 0.672435 | 0.669051 | 0.670877 | 0.679597 | 0.674916 | 0.670049 | 0.674254 | 0.663361 | 0.672095 |
| 0.669382 | 0.670644 | 0.667042 | 0.677359 | 0.675582 | 0.677635 | 0.668231 | 0.66538 | 0.666839 |
| 0.670794 | 0.666363 | 0.667722 | 0.675403 | 0.673888 | 0.672592 | 0.67146 | 0.667654 | 0.671803 |
| 0.664044 | 0.659468 | 0.667255 | 0.669666 | 0.667953 | 0.66579 | 0.66987 | 0.654958 | 0.667318 |
| 0.665681 | 0.668485 | 0.671029 | 0.677346 | 0.673779 | 0.674391 | 0.677468 | 0.66389 | 0.669912 |
| 0.667823 | 0.664817 | 0.666751 | 0.675567 | 0.673695 | 0.671654 | 0.672915 | 0.665192 | 0.669517 |
| 0.666869 | 0.667134 | 0.665213 | 0.671616 | 0.673419 | 0.668141 | 0.67327 | 0.663707 | 0.663326 |
| 0.671296 | 0.672129 | 0.671173 | 0.682817 | 0.677431 | 0.680979 | 0.67362 | 0.662792 | 0.66821 |
| 0.671057 | 0.667221 | 0.667251 | 0.674072 | 0.679911 | 0.674918 | 0.670383 | 0.662706 | 0.66508 |

| | | | | | | | | |
|---|---|---|---|---|---|---|---|---|
| 0.66353 | 0.665394 | 0.663807 | 0.674332 | 0.667932 | 0.672486 | 0.668086 | 0.656693 | 0.659291 |
| 0.66932 | 0.667257 | 0.668684 | 0.677321 | 0.672319 | 0.674855 | 0.669808 | 0.662217 | 0.66773 |
| 0.664757 | 0.665047 | 0.66809 | 0.674477 | 0.671683 | 0.672996 | 0.671573 | 0.663525 | 0.661111 |
| 0.655451 | 0.661559 | 0.6658 | 0.66358 | 0.663167 | 0.659173 | 0.664414 | 0.643172 | 0.651203 |
| 0.66723 | 0.667935 | 0.665592 | 0.675212 | 0.672574 | 0.672534 | 0.667947 | 0.655204 | 0.659694 |
| 0.667801 | 0.66722 | 0.668712 | 0.66877 | 0.666749 | 0.66989 | 0.666399 | 0.65874 | 0.664821 |
| 0.670637 | 0.674713 | 0.671783 | 0.674833 | 0.674238 | 0.674616 | 0.676178 | 0.666667 | 0.664534 |
| 0.663858 | 0.670685 | 0.665883 | 0.676101 | 0.671112 | 0.675016 | 0.66981 | 0.659139 | 0.664752 |
| 0.671998 | 0.670831 | 0.673452 | 0.683943 | 0.679301 | 0.677436 | 0.672241 | 0.663463 | 0.668459 |
| 0.665297 | 0.671474 | 0.668121 | 0.676999 | 0.676997 | 0.674264 | 0.671982 | 0.66363 | 0.661842 |
| 0.660792 | 0.671947 | 0.670376 | 0.673429 | 0.670262 | 0.676639 | 0.669376 | 0.663892 | 0.669712 |
| 0.665855 | 0.672555 | 0.673703 | 0.67973 | 0.675478 | 0.67722 | 0.671804 | 0.664502 | 0.666839 |
| 0.671598 | 0.672491 | 0.674731 | 0.681587 | 0.678899 | 0.673419 | 0.674297 | 0.666359 | 0.668424 |
| 0.670174 | 0.666667 | 0.668407 | 0.671994 | 0.670094 | 0.673851 | 0.665961 | 0.657502 | 0.661725 |
| 0.669698 | 0.669671 | 0.672784 | 0.676194 | 0.673635 | 0.678255 | 0.670701 | 0.664208 | 0.666581 |
| 0.661945 | 0.666751 | 0.664673 | 0.672476 | 0.668345 | 0.670659 | 0.661784 | 0.660474 | 0.65631 |
| 0.668757 | 0.668096 | 0.671498 | 0.673726 | 0.674923 | 0.680837 | 0.672395 | 0.663344 | 0.674445 |
| 0.672428 | 0.666624 | 0.668077 | 0.67364 | 0.671978 | 0.669814 | 0.671267 | 0.655894 | 0.66782 |
| 0.664616 | 0.663494 | 0.66783 | 0.671671 | 0.669128 | 0.669997 | 0.671088 | 0.656283 | 0.657908 |
| 0.666425 | 0.667175 | 0.666418 | 0.671761 | 0.668992 | 0.673766 | 0.665335 | 0.654248 | 0.663973 |
| 0.672078 | 0.667813 | 0.674913 | 0.679351 | 0.674264 | 0.680574 | 0.66952 | 0.663069 | 0.664875 |
| 0.674308 | 0.672363 | 0.675344 | 0.677607 | 0.675285 | 0.680628 | 0.677108 | 0.66552 | 0.668812 |
| 0.690848 | 0.695164 | 0.693511 | 0.697304 | 0.694185 | 0.697333 | 0.69876 | 0.695991 | 0.68916 |
| 0.727347 | 0.711949 | 0.723422 | 0.732113 | 0.696568 | 0.733199 | 0.712984 | 0.732107 | 0.734796 |
| 0.742912 | 0.751522 | 0.747786 | 0.746 | 0.73377 | 0.748685 | 0.742815 | 0.736308 | 0.763547 |
| 0.730951 | 0.738578 | 0.747204 | 0.744908 | 0.734839 | 0.738 | 0.727983 | 0.988932 | 0.73778 |
| 0.654395 | 0.662966 | 0.663366 | 0.661259 | 0.660305 | 0.666114 | 0.664254 | 0.655141 | 0.657419 |
| 0.660941 | 0.659439 | 0.668475 | 0.668037 | 0.654168 | 0.666365 | 0.671227 | 0.653048 | 0.666593 |
| 0.657679 | 0.652749 | 0.662121 | 0.660711 | 0.658132 | 0.66312 | 0.652944 | 0.650898 | 0.660507 |
| 0.663073 | 0.671271 | 0.664235 | 0.667687 | 0.655138 | 0.666753 | 0.668138 | 0.652741 | 0.669483 |
| 0.654919 | 0.653685 | 0.651856 | 0.674015 | 0.652141 | 0.660222 | 0.65672 | 0.64905 | 0.655674 |
| 0.63926 | 0.650016 | 0.652456 | 0.651797 | 0.63951 | 0.642338 | 0.651965 | 0.633864 | 0.646231 |
| 0.652202 | 0.640385 | 0.650753 | 0.648448 | 0.635105 | 0.646245 | 0.648185 | 0.643093 | 0.637859 |
| 0.736754 | 0.738656 | 0.74502 | 0.734476 | 0.752056 | 0.731627 | 0.744831 | 0.728722 | 0.744636 |
| 0.669059 | 0.665225 | 0.663192 | 0.669335 | 0.671469 | 0.667703 | 0.666312 | 0.659348 | 0.667481 |
| 0.662354 | 0.658145 | 0.666907 | 0.669427 | 0.66486 | 0.669998 | 0.664583 | 0.660146 | 0.662393 |
| 0.660552 | 0.659146 | 0.662575 | 0.673518 | 0.658901 | 0.670052 | 0.663497 | 0.662415 | 0.665267 |
| 0.649482 | 0.656501 | 0.648638 | 0.656899 | 0.655101 | 0.653589 | 0.662315 | 0.64488 | 0.652862 |
| 0.658934 | 0.662574 | 0.662106 | 0.671371 | 0.667476 | 0.668995 | 0.67035 | 0.661027 | 0.658145 |
| 0.655838 | 0.659258 | 0.661817 | 0.668931 | 0.665348 | 0.667362 | 0.663252 | 0.660196 | 0.653608 |
| 0.651078 | 0.657273 | 0.651329 | 0.659351 | 0.661775 | 0.662954 | 0.662191 | 0.647979 | 0.652796 |
| 0.650757 | 0.654806 | 0.66123 | 0.662274 | 0.658758 | 0.665469 | 0.663361 | 0.654871 | 0.653275 |
| 0.647932 | 0.657377 | 0.648945 | 0.657842 | 0.654294 | 0.658372 | 0.656666 | 0.650523 | 0.65235 |
| 0.663232 | 0.65857 | 0.659398 | 0.664139 | 0.657679 | 0.662952 | 0.662526 | 0.650033 | 0.659271 |
| 0.646512 | 0.668936 | 0.662878 | 0.659676 | 0.651233 | 0.668903 | 0.662665 | 0.655868 | 0.658659 |
| 0.642208 | 0.646685 | 0.649635 | 0.655755 | 0.636301 | 0.658098 | 0.645138 | 0.64623 | 0.641875 |

| | | | | | | | | |
|---|---|---|---|---|---|---|---|---|
| 0.672923 | 0.669228 | 0.673024 | 0.674721 | 0.670522 | 0.678072 | 0.66948 | 0.670222 | 0.670032 |
| 0.646013 | 0.655509 | 0.653377 | 0.658799 | 0.653846 | 0.655966 | 0.64715 | 0.645721 | 0.651635 |
| 0.655918 | 0.653894 | 0.658934 | 0.655471 | 0.653528 | 0.663164 | 0.661329 | 0.654604 | 0.656041 |
| 0.6539 | 0.659711 | 0.658875 | 0.657343 | 0.654797 | 0.665861 | 0.666855 | 0.65656 | 0.663077 |
| 0.655632 | 0.662156 | 0.662742 | 0.662629 | 0.653292 | 0.662818 | 0.656821 | 0.660041 | 0.66493 |
| 0.662816 | 0.665638 | 0.671081 | 0.667939 | 0.660317 | 0.671578 | 0.661936 | 0.664638 | 0.660197 |
| 0.654431 | 0.664151 | 0.654654 | 0.66112 | 0.659802 | 0.655247 | 0.65625 | 0.658367 | 0.654983 |
| 0.648215 | 0.649985 | 0.654639 | 0.655947 | 0.646578 | 0.660731 | 0.656003 | 0.648988 | 0.651354 |
| 0.64018 | 0.65599 | 0.645604 | 0.646356 | 0.645347 | 0.651814 | 0.653375 | 0.646178 | 0.642879 |
| 0.67051 | 0.670506 | 0.667664 | 0.671325 | 0.673921 | 0.673221 | 0.668297 | 0.669185 | 0.666132 |
| 0.668622 | 0.667437 | 0.668078 | 0.672991 | 0.673772 | 0.675676 | 0.673379 | 0.665599 | 0.671246 |
| 0.664704 | 0.669581 | 0.670362 | 0.672459 | 0.673594 | 0.672786 | 0.67187 | 0.664419 | 0.670103 |
| 0.667397 | 0.666211 | 0.667896 | 0.672439 | 0.669833 | 0.672116 | 0.66719 | 0.665744 | 0.668612 |
| 0.666865 | 0.66615 | 0.664849 | 0.676689 | 0.670835 | 0.67506 | 0.668651 | 0.662658 | 0.663459 |
| 0.661336 | 0.670381 | 0.664822 | 0.668504 | 0.667556 | 0.663746 | 0.665977 | 0.66204 | 0.659254 |

**INERA Coffee Collection**

| Collection_ | Collection_ | Collection_ | Collection_ | Collection_ | Collection_ | Collection_ | Collection_ | Collection_ |
|---|---|---|---|---|---|---|---|---|
| 0.74043 | 0.743088 | 0.762999 | 0.718359 | 0.766893 | 0.744848 | 0.788517 | 0.772775 | 0.75 |
| 0.745941 | 0.758024 | 0.765324 | 0.726138 | 0.784185 | 0.742201 | 0.765131 | 0.769444 | 0.758744 |
| 0.760478 | 0.748555 | 0.762135 | 0.719867 | 0.761231 | 0.749033 | 0.765587 | 0.766161 | 0.750763 |
| 0.73818 | 0.747157 | 0.754584 | 0.723063 | 0.758975 | 0.743163 | 0.767893 | 0.778257 | 0.763133 |
| 0.735008 | 0.751296 | 0.743177 | 0.727438 | 0.738218 | 0.740451 | 0.778719 | 0.768032 | 0.744273 |
| 0.760074 | 0.738565 | 0.74275 | 0.7307 | 0.740517 | 0.750913 | 0.761136 | 0.77167 | 0.759268 |
| 0.666621 | 0.664413 | 0.662765 | 0.660992 | 0.671437 | 0.655098 | 0.697263 | 0.70856 | 0.687309 |
| 0.663384 | 0.661302 | 0.663777 | 0.659877 | 0.657987 | 0.650907 | 0.695221 | 0.706831 | 0.679416 |
| 0.772767 | 0.746036 | 0.748615 | 0.733597 | 0.753092 | 0.768687 | 0.773299 | 0.77836 | 0.767558 |
| 0.735428 | 0.75159 | 0.752565 | 0.721607 | 0.764986 | 0.736392 | 0.79044 | 0.769731 | 0.748912 |
| 0.741163 | 0.742711 | 0.754324 | 0.723779 | 0.745513 | 0.745772 | 0.786624 | 0.770361 | 0.748566 |
| 0.73252 | 0.76337 | 0.782131 | 0.748965 | 0.757171 | 0.743206 | 0.781572 | 0.77039 | 0.747325 |
| 0.752736 | 0.752443 | 0.746824 | 0.737647 | 0.785403 | 0.74154 | 0.758131 | 0.767472 | 0.753664 |
| 0.742906 | 0.74585 | 0.75911 | 0.720042 | 0.745573 | 0.744315 | 0.773267 | 0.765064 | 0.747567 |
| 0.725661 | 0.738293 | 0.738861 | 0.713069 | 0.73786 | 0.720619 | 0.75724 | 0.757184 | 0.740728 |
| 0.757689 | 0.735182 | 0.769469 | 0.736633 | 0.745959 | 0.752639 | 0.784022 | 0.771259 | 0.744643 |
| 0.732869 | 0.734205 | 0.758631 | 0.719227 | 0.739243 | 0.736695 | 0.793183 | 0.76328 | 0.743113 |
| 0.71486 | 0.734343 | 0.721939 | 0.698689 | 0.736999 | 0.710535 | 0.748353 | 0.747778 | 0.721577 |
| 0.741829 | 0.752334 | 0.786226 | 0.752092 | 0.787247 | 0.737469 | 0.764485 | 0.77056 | 0.749823 |
| 0.744346 | 0.761096 | 0.758357 | 0.730863 | 0.776725 | 0.742777 | 0.770881 | 0.77886 | 0.758861 |
| 0.735541 | 0.754123 | 0.764226 | 0.726333 | 0.75498 | 0.741025 | 0.772557 | 0.781171 | 0.760303 |
| 0.744766 | 0.760776 | 0.773112 | 0.726077 | 0.76092 | 0.743742 | 0.791211 | 0.774867 | 0.747883 |
| 0.751036 | 0.757791 | 0.763081 | 0.730886 | 0.778103 | 0.757708 | 0.784219 | 0.781756 | 0.769231 |
| 0.753277 | 0.760802 | 0.769549 | 0.748 | 0.765944 | 0.747337 | 0.783927 | 0.775557 | 0.761967 |
| 0.742935 | 0.756137 | 0.761743 | 0.7443 | 0.759877 | 0.746895 | 0.768277 | 0.779383 | 0.755656 |
| 0.742715 | 0.749411 | 0.769129 | 0.729705 | 0.757191 | 0.749913 | 0.765849 | 0.775363 | 0.757829 |
| 0.73417 | 0.746531 | 0.761602 | 0.7304 | 0.769883 | 0.737263 | 0.75774 | 0.771536 | 0.742158 |
| 0.747336 | 0.755457 | 0.753068 | 0.728261 | 0.756344 | 0.744448 | 0.769679 | 0.774578 | 0.769077 |
| 0.748614 | 0.759805 | 0.757953 | 0.727437 | 0.745933 | 0.745742 | 0.769181 | 0.785144 | 0.765656 |
| 0.734609 | 0.755721 | 0.756533 | 0.732606 | 0.769862 | 0.736962 | 0.78659 | 0.76668 | 0.744304 |
| 0.744129 | 0.741973 | 0.747445 | 0.727562 | 0.745714 | 0.746862 | 0.767286 | 0.775248 | 0.761826 |
| 0.741945 | 0.739924 | 0.74894 | 0.725393 | 0.744119 | 0.74 | 0.767961 | 0.775006 | 0.756976 |
| 0.741035 | 0.740616 | 0.758116 | 0.730166 | 0.758084 | 0.735787 | 0.76197 | 0.776475 | 0.754772 |
| 0.740709 | 0.735165 | 0.742587 | 0.720174 | 0.737384 | 0.728913 | 0.755146 | 0.772864 | 0.749036 |
| 0.736759 | 0.731764 | 0.744332 | 0.73373 | 0.740701 | 0.734892 | 0.753933 | 0.766105 | 0.749498 |
| 0.748777 | 0.744028 | 0.768056 | 0.738259 | 0.767923 | 0.746704 | 0.766147 | 0.773823 | 0.742891 |
| 0.741611 | 0.745115 | 0.761266 | 0.722757 | 0.751292 | 0.738062 | 0.765951 | 0.768628 | 0.748013 |
| 0.734379 | 0.732467 | 0.743382 | 0.73054 | 0.739257 | 0.732713 | 0.759013 | 0.771889 | 0.754634 |
| 0.736349 | 0.738608 | 0.741514 | 0.727561 | 0.747946 | 0.737864 | 0.765869 | 0.775984 | 0.753756 |

| | | | | | | | | |
|---|---|---|---|---|---|---|---|---|
| 0.7125 | 0.713183 | 0.73253 | 0.706897 | 0.70853 | 0.709546 | 0.757552 | 0.748308 | 0.724874 |
| 0.740788 | 0.753065 | 0.750255 | 0.713233 | 0.779616 | 0.744186 | 0.795587 | 0.76382 | 0.74672 |
| 0.748745 | 0.756549 | 0.776948 | 0.73989 | 0.754106 | 0.749035 | 0.798464 | 0.771233 | 0.749088 |
| 0.742772 | 0.75337 | 0.768825 | 0.742653 | 0.776839 | 0.747696 | 0.790527 | 0.774899 | 0.752079 |
| 0.747535 | 0.752656 | 0.768289 | 0.737378 | 0.758953 | 0.748209 | 0.787452 | 0.775683 | 0.75285 |
| 0.747227 | 0.751733 | 0.757393 | 0.734339 | 0.755576 | 0.742747 | 0.781827 | 0.773443 | 0.761341 |
| 0.742523 | 0.745992 | 0.736636 | 0.724336 | 0.742715 | 0.741964 | 0.763592 | 0.772687 | 0.757972 |
| 0.745126 | 0.75637 | 0.760018 | 0.729533 | 0.790265 | 0.751535 | 0.767389 | 0.782734 | 0.759479 |
| 0.743925 | 0.744877 | 0.753953 | 0.727579 | 0.743637 | 0.746429 | 0.767695 | 0.772328 | 0.760779 |
| 0.739495 | 0.742487 | 0.741344 | 0.71923 | 0.737456 | 0.739667 | 0.765758 | 0.777959 | 0.776366 |
| 0.748814 | 0.754025 | 0.776866 | 0.735299 | 0.76714 | 0.742687 | 0.793146 | 0.765695 | 0.745437 |
| 0.749387 | 0.760785 | 0.769345 | 0.732508 | 0.774545 | 0.746305 | 0.773225 | 0.77101 | 0.743802 |
| 0.746856 | 0.752662 | 0.771967 | 0.731399 | 0.771625 | 0.746501 | 0.786589 | 0.771915 | 0.743917 |
| 0.7667 | 0.764205 | 0.762173 | 0.72801 | 0.753965 | 0.758316 | 0.78292 | 0.765605 | 0.746959 |
| 0.734368 | 0.758543 | 0.764294 | 0.726158 | 0.75884 | 0.731464 | 0.769401 | 0.770629 | 0.748186 |
| 0.735629 | 0.75304 | 0.772495 | 0.726937 | 0.753418 | 0.735445 | 0.770414 | 0.766854 | 0.738662 |
| 0.73757 | 0.737978 | 0.773718 | 0.720566 | 0.74666 | 0.732959 | 0.776758 | 0.763559 | 0.736637 |
| 0.74903 | 0.75799 | 0.783167 | 0.728074 | 0.781581 | 0.747272 | 0.788015 | 0.773831 | 0.745653 |
| 0.744902 | 0.768967 | 0.758793 | 0.727508 | 0.766815 | 0.740527 | 0.779802 | 0.773276 | 0.751149 |
| 0.745968 | 0.75716 | 0.769516 | 0.73202 | 0.782949 | 0.749691 | 0.784467 | 0.76954 | 0.750266 |
| 0.757725 | 0.743151 | 0.79598 | 0.726715 | 0.736834 | 0.745772 | 0.797641 | 0.770052 | 0.740627 |
| 0.743062 | 0.748329 | 0.757 | 0.725118 | 0.774595 | 0.742562 | 0.785667 | 0.766776 | 0.742076 |
| 0.758245 | 0.763612 | 0.756268 | 0.728492 | 0.76682 | 0.751225 | 0.766583 | 0.777579 | 0.743114 |
| 0.740256 | 0.740229 | 0.743381 | 0.723388 | 0.743104 | 0.743201 | 0.756938 | 0.78126 | 0.772198 |
| 0.725587 | 0.730529 | 0.728719 | 0.71618 | 0.733634 | 0.723801 | 0.760694 | 0.764283 | 0.756026 |
| 0.6663 | 0.664719 | 0.664605 | 0.66015 | 0.65755 | 0.65232 | 0.696781 | 0.706153 | 0.680446 |
| 0.671858 | 0.670775 | 0.66388 | 0.666374 | 0.668221 | 0.660227 | 0.699273 | 0.710073 | 0.687131 |
| 0.666897 | 0.671906 | 0.666667 | 0.66568 | 0.669072 | 0.656008 | 0.699835 | 0.710144 | 0.690209 |
| 0.732785 | 0.753982 | 0.74164 | 0.728711 | 0.78703 | 0.741017 | 0.757093 | 0.76785 | 0.747749 |
| 0.736487 | 0.762902 | 0.74683 | 0.724212 | 0.782718 | 0.739291 | 0.756035 | 0.771046 | 0.75 |
| 0.728018 | 0.741893 | 0.729091 | 0.711829 | 0.758647 | 0.731661 | 0.786114 | 0.761085 | 0.738237 |
| 0.728606 | 0.758417 | 0.747281 | 0.727027 | 0.789661 | 0.740356 | 0.760314 | 0.764246 | 0.753906 |
| 0.743652 | 0.784615 | 0.800452 | 0.728793 | 0.844245 | 0.746543 | 0.776659 | 0.77625 | 0.760862 |
| 0.749892 | 0.750815 | 0.817217 | 0.766739 | 0.757688 | 0.743127 | 0.771822 | 0.777349 | 0.745912 |
| 0.74291 | 0.777026 | 0.812102 | 0.736794 | 0.841676 | 0.741039 | 0.770763 | 0.775876 | 0.75 |
| 0.743732 | 0.741232 | 0.837735 | 0.736834 | 0.75459 | 0.738478 | 0.77194 | 0.775755 | 0.745395 |
| 0.84566 | 0.795061 | 0.756037 | 0.733617 | 0.843581 | 0.793583 | 0.772016 | 0.776253 | 0.76497 |
| 0.781853 | 0.797615 | 0.802132 | 0.754778 | 0.84478 | 0.778355 | 0.814232 | 0.795678 | 0.777261 |
| 0.735665 | 0.764168 | 0.793213 | 0.725179 | 0.819422 | 0.74336 | 0.762976 | 0.774606 | 0.754445 |
| 0.740998 | 0.772383 | 0.756895 | 0.734907 | 0.752355 | 0.74856 | 0.772393 | 0.78696 | 0.751432 |
| 0.746551 | 0.746684 | 0.799225 | 0.72822 | 0.749051 | 0.736659 | 0.770032 | 0.77655 | 0.74177 |
| 0.777827 | 0.788666 | 0.802083 | 0.752323 | 0.83499 | 0.782232 | 0.816916 | 0.793814 | 0.776793 |
| 0.742418 | 0.755211 | 0.754758 | 0.732424 | 0.753009 | 0.748519 | 0.817863 | 0.776813 | 0.752433 |
| 0.741455 | 0.758174 | 0.794468 | 0.745967 | 0.748937 | 0.746051 | 0.807045 | 0.771862 | 0.751609 |
| 0.75126 | 0.759507 | 0.755222 | 0.739044 | 0.763966 | 0.747106 | 0.773107 | 0.779298 | 0.750323 |
| 0.740662 | 0.746387 | 0.789918 | 0.730839 | 0.757887 | 0.747776 | 0.765665 | 0.776704 | 0.758381 |

| | | | | | | | | |
|---|---|---|---|---|---|---|---|---|
| 0.761699 | 0.767023 | 0.752193 | 0.733829 | 0.75038 | 0.755552 | 0.82378 | 0.778033 | 0.757601 |
| 0.785071 | 0.757223 | 0.746243 | 0.726545 | 0.74787 | 0.838522 | 0.768502 | 0.772117 | 0.789133 |
| 0.84402 | 0.773134 | 0.757833 | 0.736938 | 0.749427 | 0.803812 | 0.784286 | 0.779264 | 0.764775 |
| 0.775451 | 0.777478 | 0.774889 | 0.747056 | 0.812352 | 0.765028 | 0.797562 | 0.78559 | 0.768856 |
| 0.819522 | 0.772491 | 0.7591 | 0.772454 | 0.759384 | 0.778719 | 0.776146 | 0.77729 | 0.769403 |
| 0.741069 | 0.75579 | 0.76262 | 0.730043 | 0.751298 | 0.750571 | 0.7722 | 0.772041 | 0.752773 |
| 0.782155 | 0.780182 | 0.750581 | 0.726435 | 0.838357 | 0.753484 | 0.770224 | 0.777886 | 0.768858 |
| 0.841282 | 0.756439 | 0.744466 | 0.727314 | 0.75269 | 0.803765 | 0.769904 | 0.775729 | 0.765154 |
| 0.750827 | 0.747428 | 0.752019 | 0.729833 | 0.757464 | 0.748372 | 0.765539 | 0.775107 | 0.746313 |
| 0.732312 | 0.732096 | 0.751981 | 0.757207 | 0.738121 | 0.727115 | 0.762962 | 0.771149 | 0.741761 |
| 0.747944 | 0.744566 | 0.755717 | 0.741677 | 0.749585 | 0.74541 | 0.773643 | 0.779094 | 0.747351 |
| 0.741133 | 0.753015 | 0.765496 | 0.747769 | 0.750496 | 0.746177 | 0.766393 | 0.773509 | 0.757619 |
| 0.748847 | 0.756313 | 0.753322 | 0.731984 | 0.749385 | 0.753047 | 0.768215 | 0.774585 | 0.753177 |
| 0.748275 | 0.749741 | 0.767235 | 0.766972 | 0.741553 | 0.745873 | 0.769282 | 0.772821 | 0.749144 |
| 0.744166 | 0.741826 | 0.743091 | 0.732137 | 0.749542 | 0.739138 | 0.767793 | 0.779075 | 0.74331 |
| 0.7612 | 0.750663 | 0.750876 | 0.729976 | 0.742023 | 0.755245 | 0.764253 | 0.770344 | 0.750706 |
| 0.750613 | 0.751894 | 0.747 | 0.735523 | 0.759242 | 0.749558 | 0.766934 | 0.771227 | 0.755063 |
| 0.740961 | 0.740638 | 0.790206 | 0.729881 | 0.745003 | 0.742322 | 0.763683 | 0.774817 | 0.754483 |
| 0.748895 | 0.752952 | 0.765754 | 0.750191 | 0.760441 | 0.752401 | 0.774476 | 0.777113 | 0.760061 |
| 0.742546 | 0.749074 | 0.743951 | 0.726154 | 0.751073 | 0.745573 | 0.76483 | 0.781106 | 0.752723 |
| 0.760773 | 0.754979 | 0.756717 | 0.73201 | 0.751713 | 0.759888 | 0.764068 | 0.771517 | 0.750879 |
| 0.745729 | 0.743913 | 0.75276 | 0.730132 | 0.752432 | 0.750786 | 0.766694 | 0.777059 | 0.762604 |
| 0.775247 | 0.750627 | 0.759082 | 0.736649 | 0.74741 | 0.780267 | 0.774122 | 0.773841 | 0.777592 |
| 0.743447 | 0.747245 | 0.763162 | 0.842474 | 0.749013 | 0.744077 | 0.768134 | 0.771873 | 0.744259 |
| 0.769623 | 0.776019 | 0.749963 | 0.72629 | 0.811241 | 0.755521 | 0.764021 | 0.774231 | 0.749237 |
| 0.770092 | 0.791842 | 0.752924 | 0.725275 | 0.851686 | 0.75729 | 0.775055 | 0.770307 | 0.786105 |
| 0.774338 | 0.767794 | 0.756883 | 0.733162 | 0.818752 | 0.758303 | 0.772201 | 0.773838 | 0.75632 |
| 0.74052 | 0.73775 | 0.746802 | 0.71898 | 0.746434 | 0.743126 | 0.767934 | 0.772406 | 0.752237 |
| 0.737402 | 0.746321 | 0.754132 | 0.731533 | 0.749733 | 0.742372 | 0.768919 | 0.774309 | 0.755911 |
| 0.795074 | 0.817767 | 0.747152 | 0.733273 | 0.809262 | 0.759972 | 0.76878 | 0.772916 | 0.744208 |
| 0.759376 | 0.73934 | 0.757886 | 0.739763 | 0.747579 | 0.744951 | 0.763795 | 0.772183 | 0.752292 |
| 0.762299 | 0.747587 | 0.745217 | 0.726673 | 0.749093 | 0.763176 | 0.761183 | 0.775702 | 0.757778 |
| 0.755572 | 0.776529 | 0.754739 | 0.727148 | 0.847712 | 0.743729 | 0.773821 | 0.781629 | 0.758175 |
| 0.747352 | 0.756681 | 0.770932 | 0.744378 | 0.780216 | 0.75092 | 0.790206 | 0.774334 | 0.756568 |
| 0.752929 | 0.756529 | 0.75495 | 0.740049 | 0.754749 | 0.75856 | 0.770124 | 0.786033 | 0.755073 |
| 0.745006 | 0.748008 | 0.775063 | 0.725182 | 0.754206 | 0.748209 | 0.778064 | 0.776949 | 0.754266 |
| 0.746398 | 0.749831 | 0.759607 | 0.744609 | 0.739802 | 0.753591 | 0.765684 | 0.777457 | 0.743065 |
| 0.754596 | 0.740153 | 0.743077 | 0.774035 | 0.735256 | 0.782841 | 0.7435 | 0.76079 | 0.766314 |
| 0.743527 | 0.76057 | 0.753011 | 0.731042 | 0.754894 | 0.747182 | 0.754705 | 0.752773 | 0.742081 |
| 0.749883 | 0.751724 | 0.737431 | 0.718374 | 0.753314 | 0.745434 | 0.738761 | 0.75908 | 0.750588 |
| 0.743243 | 0.747819 | 0.790741 | 0.734286 | 0.748473 | 0.747631 | 0.747816 | 0.751114 | 0.738854 |
| 0.735657 | 0.727861 | 0.775622 | 0.777698 | 0.726641 | 0.734839 | 0.736515 | 0.733822 | 0.734241 |
| 0.751102 | 0.751084 | 0.765935 | 0.747559 | 0.750589 | 0.750654 | 0.789883 | 0.742286 | 0.740768 |
| 0.769284 | 0.753134 | 0.789483 | 0.729504 | 0.756182 | 0.796144 | 0.743785 | 0.738642 | 0.756574 |
| 0.752463 | 0.749323 | 0.751023 | 0.727857 | 0.76 | 0.74359 | 0.737287 | 0.749536 | 0.743336 |
| 0.784073 | 0.774754 | 0.77001 | 0.731351 | 0.805075 | 0.84122 | 0.74201 | 0.744974 | 0.793819 |
| 0.742013 | 0.740331 | 0.799433 | 0.725147 | 0.749817 | 0.743751 | 0.738351 | 0.747446 | 0.736293 |

| | | | | | | | | |
|---|---|---|---|---|---|---|---|---|
| 0.740868 | 0.73144 | 0.727942 | 0.728193 | 0.733491 | 0.736008 | 0.725235 | 0.767733 | 0.736633 |
| 0.740269 | 0.771209 | 0.810684 | 0.733044 | 0.840877 | 0.742399 | 0.743891 | 0.748151 | 0.745003 |
| 0.747003 | 0.73511 | 0.740938 | 0.719667 | 0.738979 | 0.745086 | 0.72484 | 0.740006 | 0.749406 |
| 0.7473 | 0.756373 | 0.740814 | 0.722781 | 0.737724 | 0.742064 | 0.781813 | 0.739915 | 0.742844 |
| 0.745194 | 0.77363 | 0.774756 | 0.765616 | 0.842744 | 0.744611 | 0.74005 | 0.743293 | 0.750114 |
| 1 | 0.782516 | 0.739951 | 0.728286 | 0.741369 | 0.836335 | 0.742232 | 0.743692 | 0.761397 |
| 0.782516 | 1 | 0.745897 | 0.725844 | 0.809051 | 0.775062 | 0.749927 | 0.741473 | 0.757576 |
| 0.739951 | 0.745897 | 1 | 0.749559 | 0.753361 | 0.743318 | 0.738486 | 0.739879 | 0.741103 |
| 0.728286 | 0.725844 | 0.749559 | 1 | 0.724237 | 0.72709 | 0.724112 | 0.729305 | 0.723813 |
| 0.741369 | 0.809051 | 0.753361 | 0.724237 | 1 | 0.744847 | 0.745893 | 0.743718 | 0.747617 |
| 0.836335 | 0.775062 | 0.743318 | 0.72709 | 0.744847 | 1 | 0.738231 | 0.74056 | 0.838382 |
| 0.742232 | 0.749927 | 0.738486 | 0.724112 | 0.745893 | 0.738231 | 1 | 0.75992 | 0.742078 |
| 0.743692 | 0.741473 | 0.739879 | 0.729305 | 0.743718 | 0.74056 | 0.75992 | 1 | 0.752283 |
| 0.761397 | 0.757576 | 0.741103 | 0.723813 | 0.747617 | 0.838382 | 0.742078 | 0.752283 | 1 |
| 0.74414 | 0.747196 | 0.764654 | 0.737003 | 0.744707 | 0.737402 | 0.772199 | 0.769967 | 0.749207 |
| 0.734876 | 0.743412 | 0.758256 | 0.749693 | 0.741636 | 0.738187 | 0.758604 | 0.769551 | 0.741321 |
| 0.74414 | 0.745431 | 0.74456 | 0.732426 | 0.754213 | 0.741311 | 0.770328 | 0.769411 | 0.747816 |
| 0.737703 | 0.738477 | 0.741997 | 0.72372 | 0.74592 | 0.747861 | 0.763184 | 0.772664 | 0.75566 |
| 0.77198 | 0.730603 | 0.723932 | 0.702858 | 0.723651 | 0.750516 | 0.758275 | 0.756904 | 0.734537 |
| 0.740585 | 0.735009 | 0.7632 | 0.748149 | 0.741877 | 0.742506 | 0.762796 | 0.77713 | 0.803943 |
| 0.828164 | 0.748653 | 0.737061 | 0.718529 | 0.734064 | 0.784341 | 0.74271 | 0.750277 | 0.794061 |
| 0.735265 | 0.743144 | 0.737819 | 0.723986 | 0.747442 | 0.741385 | 0.764127 | 0.769979 | 0.755692 |
| 0.73322 | 0.738144 | 0.736475 | 0.717549 | 0.741822 | 0.742718 | 0.760506 | 0.771193 | 0.760208 |
| 0.72583 | 0.7511 | 0.743748 | 0.713871 | 0.737129 | 0.739189 | 0.734641 | 0.740402 | 0.761198 |
| 0.707947 | 0.726618 | 0.733586 | 0.749961 | 0.76622 | 0.706742 | 0.734007 | 0.737023 | 0.721434 |
| 0.75648 | 0.756258 | 0.775725 | 0.737714 | 0.745524 | 0.769503 | 0.748298 | 0.748837 | 0.767065 |
| 0.781343 | 0.751546 | 0.727286 | 0.711998 | 0.724223 | 0.791371 | 0.754945 | 0.760123 | 0.790732 |
| 0.736611 | 0.735739 | 0.74 | 0.717307 | 0.74059 | 0.74751 | 0.762197 | 0.768021 | 0.757629 |
| 0.717775 | 0.721237 | 0.713964 | 0.694011 | 0.725314 | 0.710955 | 0.739323 | 0.753585 | 0.730919 |
| 0.733764 | 0.754511 | 0.733543 | 0.715941 | 0.825106 | 0.727693 | 0.757966 | 0.771147 | 0.760578 |
| 0.728731 | 0.735634 | 0.727376 | 0.724869 | 0.746877 | 0.733345 | 0.752715 | 0.76699 | 0.750782 |
| 0.72749 | 0.738661 | 0.7397 | 0.70778 | 0.75599 | 0.724435 | 0.769338 | 0.756971 | 0.732609 |
| 0.722074 | 0.741251 | 0.741679 | 0.71208 | 0.756584 | 0.723364 | 0.768264 | 0.757236 | 0.736247 |
| 0.727618 | 0.736113 | 0.73707 | 0.70755 | 0.74097 | 0.731328 | 0.760894 | 0.776403 | 0.839005 |
| 0.734923 | 0.718942 | 0.722112 | 0.723886 | 0.725406 | 0.731586 | 0.748143 | 0.777697 | 0.75162 |
| 0.73032 | 0.737442 | 0.742043 | 0.716829 | 0.738271 | 0.734542 | 0.758668 | 0.76581 | 0.749124 |
| 0.717344 | 0.698961 | 0.701684 | 0.681549 | 0.691651 | 0.705774 | 0.73247 | 0.730721 | 0.710042 |
| 0.722246 | 0.718196 | 0.725718 | 0.704148 | 0.725816 | 0.716939 | 0.74565 | 0.754311 | 0.729287 |
| 0.716326 | 0.696406 | 0.69218 | 0.682414 | 0.694191 | 0.735064 | 0.728252 | 0.727124 | 0.732522 |
| 0.744109 | 0.697063 | 0.671835 | 0.663767 | 0.678287 | 0.743538 | 0.728966 | 0.717603 | 0.716994 |
| 0.740071 | 0.746565 | 0.745489 | 0.734145 | 0.750417 | 0.733586 | 0.756402 | 0.766219 | 0.745064 |
| 0.72501 | 0.705752 | 0.677847 | 0.668224 | 0.677665 | 0.716642 | 0.727976 | 0.717144 | 0.709373 |
| 0.684029 | 0.692318 | 0.699012 | 0.664926 | 0.720203 | 0.675291 | 0.722013 | 0.712995 | 0.6827 |
| 0.6731 | 0.700834 | 0.67934 | 0.658433 | 0.731679 | 0.663946 | 0.707758 | 0.708868 | 0.682291 |
| 0.733034 | 0.745089 | 0.729521 | 0.710765 | 0.752226 | 0.73275 | 0.771994 | 0.761786 | 0.736714 |
| 0.663778 | 0.675303 | 0.687441 | 0.654785 | 0.718475 | 0.658974 | 0.707838 | 0.706013 | 0.673334 |
| 0.733887 | 0.738569 | 0.735366 | 0.719319 | 0.751619 | 0.734483 | 0.759092 | 0.76421 | 0.743358 |

| | | | | | | | | |
|---|---|---|---|---|---|---|---|---|
| 0.724725 | 0.731158 | 0.726561 | 0.714137 | 0.748565 | 0.721359 | 0.755055 | 0.760936 | 0.736757 |
| 0.737925 | 0.742521 | 0.738058 | 0.722888 | 0.753582 | 0.73667 | 0.762008 | 0.766734 | 0.742519 |
| 0.717785 | 0.719816 | 0.71146 | 0.705386 | 0.718833 | 0.711519 | 0.746205 | 0.748888 | 0.729503 |
| 0.664058 | 0.668467 | 0.679159 | 0.644819 | 0.699292 | 0.65045 | 0.700723 | 0.700647 | 0.658578 |
| 0.687669 | 0.690978 | 0.67156 | 0.649371 | 0.719618 | 0.684802 | 0.705921 | 0.694919 | 0.684546 |
| 0.742164 | 0.740047 | 0.744836 | 0.721665 | 0.747244 | 0.734149 | 0.7606 | 0.759742 | 0.740367 |
| 0.653332 | 0.655541 | 0.662184 | 0.644343 | 0.657372 | 0.65774 | 0.705053 | 0.699093 | 0.676249 |
| 0.688963 | 0.689205 | 0.694413 | 0.695978 | 0.694581 | 0.690492 | 0.723809 | 0.736812 | 0.711882 |
| 0.727898 | 0.727866 | 0.739853 | 0.714071 | 0.734154 | 0.730802 | 0.764443 | 0.756224 | 0.732993 |
| 0.676381 | 0.673308 | 0.680399 | 0.679707 | 0.677817 | 0.671086 | 0.711216 | 0.725307 | 0.695029 |
| 0.708551 | 0.721012 | 0.717722 | 0.700961 | 0.735339 | 0.706959 | 0.744942 | 0.744447 | 0.724644 |
| 0.666667 | 0.641892 | 0.676692 | 0.655629 | 0.676923 | 0.664286 | 0.628049 | 0.631579 | 0.651613 |
| 0.660024 | 0.664402 | 0.650648 | 0.651352 | 0.647619 | 0.65369 | 0.65205 | 0.655166 | 0.665413 |
| 0.656874 | 0.658713 | 0.649034 | 0.652339 | 0.649084 | 0.643796 | 0.650493 | 0.650276 | 0.666892 |
| 0.66952 | 0.658061 | 0.659203 | 0.657292 | 0.653619 | 0.655515 | 0.651371 | 0.659003 | 0.680528 |
| 0.655284 | 0.660808 | 0.650069 | 0.657417 | 0.660474 | 0.649416 | 0.651771 | 0.66246 | 0.669189 |
| 0.602476 | 0.607843 | 0.606335 | 0.593072 | 0.606574 | 0.60499 | 0.633315 | 0.622171 | 0.619911 |
| 0.654406 | 0.65593 | 0.659417 | 0.654412 | 0.655234 | 0.654495 | 0.651928 | 0.662144 | 0.672725 |
| 0.703823 | 0.716171 | 0.715457 | 0.692228 | 0.727847 | 0.699095 | 0.732244 | 0.739387 | 0.71772 |
| 0.698949 | 0.705162 | 0.71348 | 0.693988 | 0.711232 | 0.70289 | 0.734671 | 0.739864 | 0.716891 |
| 0.707829 | 0.720701 | 0.712531 | 0.690359 | 0.731429 | 0.702315 | 0.735621 | 0.7452 | 0.721989 |
| 0.58836 | 0.578174 | 0.564037 | 0.559859 | 0.564221 | 0.568832 | 0.635228 | 0.617672 | 0.585416 |
| 0.557865 | 0.555771 | 0.546245 | 0.548077 | 0.551282 | 0.547921 | 0.614787 | 0.6003 | 0.567468 |
| 0.663997 | 0.664526 | 0.660228 | 0.65953 | 0.65838 | 0.656303 | 0.694849 | 0.705972 | 0.684548 |
| 0.666208 | 0.663216 | 0.661519 | 0.663619 | 0.660438 | 0.657966 | 0.69535 | 0.70249 | 0.684145 |
| 0.655514 | 0.660778 | 0.648872 | 0.658043 | 0.650461 | 0.645474 | 0.688276 | 0.698604 | 0.676122 |
| 0.659243 | 0.658819 | 0.656263 | 0.65752 | 0.654103 | 0.647742 | 0.692043 | 0.701572 | 0.685103 |
| 0.665981 | 0.664642 | 0.660053 | 0.66043 | 0.656977 | 0.660131 | 0.694843 | 0.706054 | 0.687179 |
| 0.663925 | 0.66421 | 0.662703 | 0.664211 | 0.66027 | 0.659903 | 0.698396 | 0.70913 | 0.685633 |
| 0.663308 | 0.665446 | 0.665508 | 0.657359 | 0.664738 | 0.649761 | 0.698352 | 0.70416 | 0.683913 |
| 0.665265 | 0.668219 | 0.669103 | 0.661858 | 0.664227 | 0.656485 | 0.698814 | 0.705523 | 0.682216 |
| 0.673866 | 0.674721 | 0.674069 | 0.671008 | 0.671814 | 0.665224 | 0.704563 | 0.712829 | 0.693208 |
| 0.668812 | 0.667328 | 0.66447 | 0.661104 | 0.665984 | 0.66007 | 0.700707 | 0.710189 | 0.684893 |
| 0.66135 | 0.662316 | 0.669223 | 0.666228 | 0.664063 | 0.658403 | 0.697799 | 0.710654 | 0.6869 |
| 0.668272 | 0.665673 | 0.665407 | 0.668586 | 0.671265 | 0.66386 | 0.704479 | 0.711984 | 0.690444 |
| 0.65896 | 0.658122 | 0.659919 | 0.660845 | 0.659562 | 0.652312 | 0.697533 | 0.707139 | 0.684337 |
| 0.664614 | 0.667329 | 0.664565 | 0.664954 | 0.66463 | 0.66192 | 0.6989 | 0.70838 | 0.6877 |
| 0.666027 | 0.665585 | 0.666013 | 0.669164 | 0.662115 | 0.658154 | 0.698945 | 0.708986 | 0.685282 |
| 0.663819 | 0.663357 | 0.659116 | 0.660389 | 0.66544 | 0.657806 | 0.69759 | 0.705539 | 0.681547 |
| 0.663399 | 0.666103 | 0.665127 | 0.666422 | 0.667592 | 0.658598 | 0.701852 | 0.710704 | 0.685254 |
| 0.660892 | 0.65672 | 0.649823 | 0.658589 | 0.648222 | 0.654537 | 0.650075 | 0.656093 | 0.670465 |
| 0.666592 | 0.661546 | 0.6601 | 0.661758 | 0.657863 | 0.65981 | 0.659899 | 0.661591 | 0.669519 |
| 0.67145 | 0.669711 | 0.664128 | 0.66473 | 0.669384 | 0.663944 | 0.702123 | 0.712083 | 0.686597 |
| 0.673961 | 0.671252 | 0.666061 | 0.670495 | 0.674177 | 0.669253 | 0.701974 | 0.71362 | 0.694158 |
| 0.635105 | 0.648649 | 0.639585 | 0.647387 | 0.641671 | 0.642563 | 0.637355 | 0.646005 | 0.654764 |
| 0.66548 | 0.670017 | 0.661734 | 0.666325 | 0.66798 | 0.661401 | 0.702762 | 0.710868 | 0.685023 |
| 0.655513 | 0.652418 | 0.652411 | 0.654516 | 0.658433 | 0.65165 | 0.687728 | 0.698921 | 0.678349 |

| | | | | | | | | |
|---|---|---|---|---|---|---|---|---|
| 0.666074 | 0.668924 | 0.661968 | 0.65977 | 0.667927 | 0.660667 | 0.698759 | 0.707919 | 0.682579 |
| 0.663722 | 0.670316 | 0.660603 | 0.664826 | 0.663915 | 0.660854 | 0.700039 | 0.70579 | 0.681557 |
| 0.668512 | 0.667613 | 0.666714 | 0.667158 | 0.666177 | 0.657267 | 0.701581 | 0.710409 | 0.689358 |
| 0.666393 | 0.676207 | 0.667649 | 0.662349 | 0.673063 | 0.660848 | 0.69881 | 0.708952 | 0.684246 |
| 0.672819 | 0.676038 | 0.671425 | 0.66989 | 0.67577 | 0.667667 | 0.708219 | 0.71802 | 0.694217 |
| 0.667626 | 0.667793 | 0.661315 | 0.66603 | 0.66894 | 0.658321 | 0.695796 | 0.7057 | 0.684211 |
| 0.664761 | 0.66977 | 0.659875 | 0.659078 | 0.667539 | 0.657734 | 0.698673 | 0.705896 | 0.683456 |
| 0.656834 | 0.660242 | 0.654227 | 0.657934 | 0.656001 | 0.652867 | 0.693465 | 0.704531 | 0.674732 |
| 0.662966 | 0.660619 | 0.661742 | 0.658533 | 0.663654 | 0.649734 | 0.697587 | 0.706333 | 0.680045 |
| 0.660599 | 0.659629 | 0.664076 | 0.66218 | 0.659141 | 0.655746 | 0.696103 | 0.704921 | 0.677489 |
| 0.665936 | 0.665538 | 0.664341 | 0.665493 | 0.666035 | 0.655236 | 0.701 | 0.710936 | 0.684515 |
| 0.667925 | 0.66824 | 0.662676 | 0.658461 | 0.662274 | 0.6614 | 0.693591 | 0.703878 | 0.682719 |
| 0.65531 | 0.653644 | 0.652571 | 0.657269 | 0.656886 | 0.655974 | 0.690078 | 0.701116 | 0.675157 |
| 0.658428 | 0.659045 | 0.655408 | 0.653778 | 0.659037 | 0.654555 | 0.689368 | 0.698612 | 0.679138 |
| 0.67513 | 0.672589 | 0.672653 | 0.667545 | 0.667732 | 0.664385 | 0.704145 | 0.71176 | 0.687352 |
| 0.666303 | 0.668175 | 0.667878 | 0.664622 | 0.668227 | 0.6573 | 0.697714 | 0.710139 | 0.684407 |
| 0.665745 | 0.668389 | 0.659749 | 0.663712 | 0.660527 | 0.65508 | 0.693384 | 0.699405 | 0.689783 |
| 0.659504 | 0.66296 | 0.661727 | 0.66068 | 0.660097 | 0.652827 | 0.695307 | 0.703194 | 0.680055 |
| 0.661368 | 0.663037 | 0.659333 | 0.657474 | 0.655941 | 0.653936 | 0.691774 | 0.703838 | 0.683791 |
| 0.661138 | 0.665435 | 0.660459 | 0.658937 | 0.655769 | 0.651089 | 0.692428 | 0.700252 | 0.686174 |
| 0.656483 | 0.657538 | 0.652998 | 0.655126 | 0.655229 | 0.649355 | 0.691345 | 0.697514 | 0.676254 |
| 0.661362 | 0.667433 | 0.656409 | 0.665123 | 0.652397 | 0.653331 | 0.691508 | 0.70016 | 0.682289 |
| 0.66535 | 0.669397 | 0.661944 | 0.662615 | 0.657564 | 0.654491 | 0.694188 | 0.702523 | 0.689683 |
| 0.661158 | 0.664164 | 0.662558 | 0.663042 | 0.664918 | 0.650942 | 0.698652 | 0.706453 | 0.682369 |
| 0.658925 | 0.664816 | 0.660499 | 0.658353 | 0.661487 | 0.654958 | 0.694167 | 0.703594 | 0.683469 |
| 0.660305 | 0.661178 | 0.657973 | 0.658515 | 0.657005 | 0.650226 | 0.688842 | 0.700063 | 0.6792 |
| 0.654234 | 0.658662 | 0.663257 | 0.656829 | 0.66097 | 0.648267 | 0.692655 | 0.704427 | 0.679848 |
| 0.656518 | 0.655556 | 0.655378 | 0.663878 | 0.654352 | 0.652416 | 0.691241 | 0.699487 | 0.681259 |
| 0.662038 | 0.660281 | 0.657924 | 0.663127 | 0.660604 | 0.653827 | 0.693144 | 0.697601 | 0.681393 |
| 0.661135 | 0.662283 | 0.661597 | 0.665292 | 0.660196 | 0.652864 | 0.694976 | 0.703096 | 0.684607 |
| 0.668542 | 0.669361 | 0.663765 | 0.661421 | 0.664473 | 0.661171 | 0.699976 | 0.709005 | 0.687893 |
| 0.664959 | 0.6729 | 0.665769 | 0.66622 | 0.67018 | 0.658438 | 0.702479 | 0.706342 | 0.687695 |
| 0.668345 | 0.671221 | 0.668155 | 0.668662 | 0.667005 | 0.663845 | 0.701213 | 0.709586 | 0.690895 |
| 0.664791 | 0.671824 | 0.668499 | 0.670265 | 0.666423 | 0.658102 | 0.703349 | 0.712941 | 0.688712 |
| 0.660377 | 0.657727 | 0.655839 | 0.659088 | 0.656054 | 0.652437 | 0.690499 | 0.700673 | 0.678576 |
| 0.663222 | 0.668611 | 0.666338 | 0.664944 | 0.664805 | 0.657353 | 0.694519 | 0.706529 | 0.685672 |
| 0.663518 | 0.670356 | 0.663333 | 0.66115 | 0.66334 | 0.654685 | 0.698189 | 0.702389 | 0.684577 |
| 0.661224 | 0.661338 | 0.659431 | 0.661032 | 0.657272 | 0.648496 | 0.688742 | 0.698816 | 0.685027 |
| 0.655225 | 0.659651 | 0.657094 | 0.659792 | 0.654305 | 0.64527 | 0.688505 | 0.696695 | 0.677698 |
| 0.660088 | 0.666476 | 0.656994 | 0.661163 | 0.656044 | 0.649008 | 0.688539 | 0.691801 | 0.681578 |
| 0.65947 | 0.661049 | 0.658299 | 0.654072 | 0.652943 | 0.649357 | 0.684079 | 0.693947 | 0.679281 |
| 0.658284 | 0.664876 | 0.654811 | 0.660564 | 0.655826 | 0.653381 | 0.685013 | 0.695667 | 0.682324 |
| 0.65595 | 0.660957 | 0.657982 | 0.652694 | 0.658753 | 0.649717 | 0.688976 | 0.697425 | 0.678563 |
| 0.654715 | 0.658135 | 0.655343 | 0.655072 | 0.646864 | 0.643339 | 0.678632 | 0.688342 | 0.677827 |
| 0.662486 | 0.663924 | 0.660431 | 0.662732 | 0.655876 | 0.655846 | 0.691537 | 0.699188 | 0.684443 |
| 0.653022 | 0.661503 | 0.659279 | 0.654378 | 0.66352 | 0.646644 | 0.684626 | 0.693698 | 0.673946 |

| | | | | | | | | |
|---|---|---|---|---|---|---|---|---|
| 0.675231 | 0.672382 | 0.666526 | 0.665976 | 0.664766 | 0.662899 | 0.696677 | 0.706316 | 0.687186 |
| 0.668415 | 0.671327 | 0.661567 | 0.663524 | 0.664221 | 0.660624 | 0.699547 | 0.707943 | 0.6877 |
| 0.657502 | 0.660337 | 0.658877 | 0.660786 | 0.659354 | 0.65011 | 0.691747 | 0.701294 | 0.674951 |
| 0.668121 | 0.668554 | 0.656791 | 0.658796 | 0.663673 | 0.660783 | 0.701901 | 0.711003 | 0.682663 |
| 0.671775 | 0.673768 | 0.67318 | 0.66975 | 0.666229 | 0.667169 | 0.706418 | 0.714908 | 0.691019 |
| 0.665078 | 0.668744 | 0.668948 | 0.662056 | 0.669288 | 0.660359 | 0.700031 | 0.709182 | 0.684525 |
| 0.663989 | 0.661558 | 0.663648 | 0.664783 | 0.663554 | 0.655196 | 0.696223 | 0.701434 | 0.681702 |
| 0.662214 | 0.664733 | 0.66012 | 0.659453 | 0.66555 | 0.654582 | 0.70025 | 0.706086 | 0.682623 |
| 0.669124 | 0.669831 | 0.668987 | 0.665013 | 0.672268 | 0.662141 | 0.693609 | 0.700181 | 0.686414 |
| 0.662834 | 0.666808 | 0.659838 | 0.659194 | 0.654648 | 0.65457 | 0.697819 | 0.705279 | 0.681385 |
| 0.664899 | 0.666854 | 0.662187 | 0.663417 | 0.673205 | 0.660913 | 0.699775 | 0.706657 | 0.686272 |
| 0.664898 | 0.667707 | 0.665407 | 0.665008 | 0.666957 | 0.663656 | 0.699992 | 0.708323 | 0.686769 |
| 0.666804 | 0.666479 | 0.662483 | 0.657991 | 0.666327 | 0.655132 | 0.696311 | 0.70366 | 0.681854 |
| 0.664308 | 0.67119 | 0.660064 | 0.665203 | 0.661022 | 0.659557 | 0.699616 | 0.706698 | 0.685202 |
| 0.668255 | 0.671667 | 0.6649 | 0.659329 | 0.670301 | 0.657646 | 0.69909 | 0.709392 | 0.689628 |
| 0.665659 | 0.663699 | 0.664748 | 0.664018 | 0.664963 | 0.657691 | 0.700477 | 0.705383 | 0.686176 |
| 0.66703 | 0.66409 | 0.666667 | 0.661923 | 0.672822 | 0.656816 | 0.698736 | 0.709708 | 0.688716 |
| 0.664585 | 0.67087 | 0.662889 | 0.656467 | 0.670976 | 0.658759 | 0.70069 | 0.710711 | 0.683909 |
| 0.663838 | 0.670495 | 0.672115 | 0.661439 | 0.664681 | 0.663089 | 0.70194 | 0.714139 | 0.68802 |
| 0.660678 | 0.665683 | 0.66634 | 0.662273 | 0.66866 | 0.66214 | 0.696544 | 0.707405 | 0.684801 |
| 0.662582 | 0.66723 | 0.663589 | 0.663985 | 0.65836 | 0.649877 | 0.698029 | 0.705784 | 0.685536 |
| 0.668126 | 0.666714 | 0.668778 | 0.662465 | 0.666667 | 0.654134 | 0.698058 | 0.704291 | 0.683977 |
| 0.665533 | 0.669717 | 0.662341 | 0.66428 | 0.670943 | 0.657711 | 0.701919 | 0.712218 | 0.68864 |
| 0.670267 | 0.666292 | 0.664539 | 0.662407 | 0.66851 | 0.66137 | 0.699674 | 0.713223 | 0.690387 |
| 0.665308 | 0.662852 | 0.659298 | 0.661258 | 0.66196 | 0.655354 | 0.698926 | 0.706248 | 0.68114 |
| 0.666713 | 0.665241 | 0.661137 | 0.662454 | 0.666521 | 0.657828 | 0.698022 | 0.703779 | 0.679468 |
| 0.659864 | 0.663048 | 0.655106 | 0.656377 | 0.659843 | 0.65182 | 0.699517 | 0.705299 | 0.680221 |
| 0.666349 | 0.667042 | 0.6632 | 0.660769 | 0.664689 | 0.65624 | 0.701225 | 0.702503 | 0.679807 |
| 0.667666 | 0.669057 | 0.66444 | 0.664289 | 0.665746 | 0.661025 | 0.699953 | 0.709817 | 0.687599 |
| 0.672992 | 0.673434 | 0.668512 | 0.672378 | 0.671138 | 0.672153 | 0.698718 | 0.711623 | 0.691136 |
| 0.66538 | 0.66605 | 0.664042 | 0.665089 | 0.661786 | 0.661683 | 0.697066 | 0.704942 | 0.686744 |
| 0.671519 | 0.673672 | 0.670217 | 0.666422 | 0.672682 | 0.668458 | 0.702703 | 0.71201 | 0.691799 |
| 0.669583 | 0.667419 | 0.662844 | 0.666227 | 0.664626 | 0.668689 | 0.700008 | 0.708794 | 0.688508 |
| 0.664239 | 0.665528 | 0.663014 | 0.664307 | 0.665238 | 0.656799 | 0.694144 | 0.705983 | 0.687155 |
| 0.666068 | 0.670146 | 0.669029 | 0.663974 | 0.66302 | 0.656166 | 0.696713 | 0.707852 | 0.68579 |
| 0.669496 | 0.675781 | 0.667332 | 0.663747 | 0.666518 | 0.662171 | 0.696967 | 0.703792 | 0.681509 |
| 0.672762 | 0.666949 | 0.668729 | 0.668437 | 0.672714 | 0.662653 | 0.706792 | 0.715279 | 0.689812 |
| 0.664231 | 0.671685 | 0.664794 | 0.664298 | 0.668087 | 0.660269 | 0.697932 | 0.709462 | 0.686821 |
| 0.667259 | 0.667841 | 0.663932 | 0.669689 | 0.666958 | 0.659098 | 0.700455 | 0.710774 | 0.683758 |
| 0.669611 | 0.668421 | 0.663448 | 0.663085 | 0.666667 | 0.659041 | 0.698539 | 0.711559 | 0.686871 |
| 0.669895 | 0.66793 | 0.671536 | 0.665356 | 0.67112 | 0.665395 | 0.704725 | 0.715928 | 0.691123 |
| 0.669002 | 0.669399 | 0.667878 | 0.664567 | 0.67254 | 0.660969 | 0.698343 | 0.712649 | 0.689459 |
| 0.670307 | 0.671519 | 0.666435 | 0.665159 | 0.672942 | 0.65977 | 0.702068 | 0.714297 | 0.685505 |
| 0.66393 | 0.67378 | 0.665214 | 0.658851 | 0.66531 | 0.66007 | 0.697689 | 0.707398 | 0.683728 |
| 0.671866 | 0.672443 | 0.660274 | 0.665734 | 0.667251 | 0.66487 | 0.701377 | 0.708066 | 0.690014 |
| 0.661089 | 0.663125 | 0.659324 | 0.650824 | 0.661267 | 0.653147 | 0.693579 | 0.704653 | 0.684482 |

| | | | | | | | | |
|---|---|---|---|---|---|---|---|---|
| 0.659612 | 0.661902 | 0.661896 | 0.657209 | 0.658087 | 0.649726 | 0.6832 | 0.699408 | 0.677325 |
| 0.710095 | 0.710621 | 0.717078 | 0.697747 | 0.712511 | 0.699357 | 0.738718 | 0.749306 | 0.743158 |
| 0.664737 | 0.670047 | 0.66421 | 0.663862 | 0.667157 | 0.65549 | 0.695027 | 0.707058 | 0.685051 |
| 0.670133 | 0.671917 | 0.66792 | 0.66327 | 0.667294 | 0.664274 | 0.699588 | 0.712706 | 0.684727 |
| 0.665598 | 0.664272 | 0.662961 | 0.662083 | 0.661386 | 0.660099 | 0.692764 | 0.705506 | 0.678348 |
| 0.665976 | 0.669528 | 0.659203 | 0.657887 | 0.667699 | 0.65857 | 0.698291 | 0.708699 | 0.680348 |
| 0.669889 | 0.668542 | 0.665953 | 0.658707 | 0.663939 | 0.660583 | 0.690854 | 0.70655 | 0.687354 |
| 0.66437 | 0.666327 | 0.662878 | 0.658801 | 0.664666 | 0.660072 | 0.689048 | 0.701925 | 0.683257 |
| 0.662652 | 0.66264 | 0.652529 | 0.659028 | 0.661792 | 0.654633 | 0.69972 | 0.707447 | 0.684103 |
| 0.66855 | 0.67682 | 0.674091 | 0.668286 | 0.675187 | 0.665318 | 0.703972 | 0.712585 | 0.691631 |
| 0.663208 | 0.668269 | 0.668965 | 0.66466 | 0.668614 | 0.663371 | 0.702204 | 0.708928 | 0.688773 |
| 0.669835 | 0.664879 | 0.671979 | 0.664951 | 0.668129 | 0.664647 | 0.701836 | 0.708076 | 0.690096 |
| 0.67015 | 0.663524 | 0.673028 | 0.670199 | 0.672469 | 0.661209 | 0.703529 | 0.711927 | 0.688068 |
| 0.664347 | 0.670663 | 0.667923 | 0.66477 | 0.671911 | 0.664539 | 0.702292 | 0.709958 | 0.693282 |
| 0.668182 | 0.667802 | 0.664933 | 0.668344 | 0.67069 | 0.663241 | 0.69906 | 0.706354 | 0.686806 |
| 0.661301 | 0.663093 | 0.668375 | 0.660975 | 0.662677 | 0.660262 | 0.700162 | 0.71043 | 0.689367 |
| 0.668322 | 0.670927 | 0.664753 | 0.666912 | 0.671393 | 0.664138 | 0.700094 | 0.710991 | 0.689722 |
| 0.667861 | 0.666619 | 0.668917 | 0.664753 | 0.671495 | 0.667172 | 0.70204 | 0.704187 | 0.687072 |
| 0.673375 | 0.666379 | 0.66762 | 0.665769 | 0.672065 | 0.662362 | 0.692943 | 0.701781 | 0.686242 |
| 0.662495 | 0.668208 | 0.661832 | 0.658678 | 0.66577 | 0.656896 | 0.69427 | 0.701545 | 0.681945 |
| 0.668085 | 0.67026 | 0.668023 | 0.661922 | 0.669643 | 0.662802 | 0.699232 | 0.709258 | 0.687364 |
| 0.650787 | 0.658487 | 0.655801 | 0.652611 | 0.656768 | 0.646959 | 0.688298 | 0.701861 | 0.673014 |
| 0.661376 | 0.662652 | 0.663537 | 0.659041 | 0.661168 | 0.662237 | 0.694742 | 0.706073 | 0.680967 |
| 0.663154 | 0.662893 | 0.663173 | 0.65838 | 0.663163 | 0.66045 | 0.698061 | 0.709417 | 0.685428 |
| 0.667863 | 0.670612 | 0.668931 | 0.66446 | 0.671429 | 0.661678 | 0.70008 | 0.712091 | 0.689893 |
| 0.663413 | 0.665287 | 0.664552 | 0.66218 | 0.66452 | 0.655271 | 0.696813 | 0.710361 | 0.681121 |
| 0.659527 | 0.664455 | 0.660496 | 0.658914 | 0.663194 | 0.645353 | 0.692893 | 0.702398 | 0.680502 |
| 0.662237 | 0.662706 | 0.662457 | 0.663347 | 0.664254 | 0.65232 | 0.692077 | 0.702758 | 0.680359 |
| 0.660989 | 0.665895 | 0.664322 | 0.659692 | 0.671546 | 0.65444 | 0.690807 | 0.704085 | 0.67903 |
| 0.732026 | 0.74552 | 0.741951 | 0.721296 | 0.743058 | 0.730443 | 0.748383 | 0.774301 | 0.740448 |
| 0.750649 | 0.763893 | 0.752791 | 0.725726 | 0.836457 | 0.74515 | 0.769142 | 0.773114 | 0.747403 |
| 0.75909 | 0.75861 | 0.741506 | 0.733099 | 0.749271 | 0.839897 | 0.769471 | 0.775888 | 0.79373 |
| 0.812327 | 0.767352 | 0.743936 | 0.73136 | 0.748125 | 0.833175 | 0.791347 | 0.762836 | 0.77951 |
| 0.814073 | 0.792608 | 0.746799 | 0.72916 | 0.834737 | 0.837867 | 0.772764 | 0.775734 | 0.79057 |
| 0.656505 | 0.653721 | 0.65772 | 0.654017 | 0.658476 | 0.653178 | 0.691692 | 0.7052 | 0.677067 |
| 0.6591 | 0.661872 | 0.65989 | 0.660701 | 0.662835 | 0.654576 | 0.696124 | 0.707069 | 0.679795 |
| 0.665015 | 0.662867 | 0.660794 | 0.664949 | 0.665152 | 0.654704 | 0.697612 | 0.707811 | 0.683394 |
| 0.665656 | 0.662367 | 0.65918 | 0.657625 | 0.661992 | 0.659825 | 0.700331 | 0.708668 | 0.683402 |
| 0.661279 | 0.661634 | 0.661071 | 0.658343 | 0.659788 | 0.655941 | 0.696666 | 0.706636 | 0.682882 |
| 0.666118 | 0.665066 | 0.663863 | 0.660568 | 0.667644 | 0.66005 | 0.700442 | 0.712965 | 0.687251 |
| 0.664144 | 0.662212 | 0.664082 | 0.661309 | 0.658316 | 0.658442 | 0.697242 | 0.707342 | 0.681188 |
| 0.657671 | 0.660474 | 0.6591 | 0.658515 | 0.660003 | 0.652326 | 0.695059 | 0.708471 | 0.678584 |
| 0.658822 | 0.659417 | 0.656804 | 0.65445 | 0.657083 | 0.655368 | 0.687784 | 0.700217 | 0.679223 |
| 0.657705 | 0.657472 | 0.659593 | 0.659067 | 0.654043 | 0.653773 | 0.689911 | 0.699116 | 0.680547 |
| 0.664016 | 0.664119 | 0.662012 | 0.663423 | 0.664042 | 0.651653 | 0.692166 | 0.705391 | 0.680171 |
| 0.665602 | 0.664328 | 0.657231 | 0.662076 | 0.663029 | 0.653653 | 0.696777 | 0.70999 | 0.682877 |

| | | | | | | | | |
|---|---|---|---|---|---|---|---|---|
| 0.665421 | 0.665857 | 0.666902 | 0.661939 | 0.666716 | 0.659524 | 0.699021 | 0.710215 | 0.686037 |
| 0.662957 | 0.65953 | 0.662367 | 0.659559 | 0.663549 | 0.652986 | 0.696492 | 0.707558 | 0.68329 |
| 0.665709 | 0.666241 | 0.666714 | 0.667259 | 0.670556 | 0.660137 | 0.700087 | 0.711855 | 0.686467 |
| 0.657209 | 0.662872 | 0.657732 | 0.657569 | 0.663196 | 0.655941 | 0.696686 | 0.706204 | 0.680902 |
| 0.660273 | 0.661631 | 0.656929 | 0.652457 | 0.661359 | 0.6543 | 0.694567 | 0.705186 | 0.679256 |
| 0.664822 | 0.664479 | 0.660948 | 0.65248 | 0.662986 | 0.656008 | 0.694922 | 0.702548 | 0.681771 |
| 0.666298 | 0.667097 | 0.668175 | 0.66469 | 0.67001 | 0.659456 | 0.700113 | 0.710526 | 0.689663 |
| 0.670171 | 0.670583 | 0.664769 | 0.66274 | 0.66785 | 0.661664 | 0.697922 | 0.708929 | 0.686018 |
| 0.66311 | 0.669053 | 0.655991 | 0.661283 | 0.659481 | 0.65457 | 0.692536 | 0.707072 | 0.68198 |
| 0.665796 | 0.664912 | 0.661752 | 0.660656 | 0.664908 | 0.658524 | 0.695023 | 0.7084 | 0.683838 |
| 0.663769 | 0.665475 | 0.661281 | 0.658062 | 0.666079 | 0.656591 | 0.697574 | 0.705037 | 0.68488 |
| 0.663253 | 0.668383 | 0.659661 | 0.658638 | 0.664947 | 0.658664 | 0.697137 | 0.704397 | 0.686485 |
| 0.655973 | 0.661066 | 0.657337 | 0.654388 | 0.66241 | 0.652823 | 0.686298 | 0.696902 | 0.681089 |
| 0.665709 | 0.667552 | 0.664181 | 0.662332 | 0.66692 | 0.658635 | 0.68606 | 0.699763 | 0.687477 |
| 0.663071 | 0.667095 | 0.664639 | 0.667312 | 0.665926 | 0.66206 | 0.699295 | 0.706291 | 0.686938 |
| 0.672557 | 0.672836 | 0.674422 | 0.672881 | 0.67289 | 0.668071 | 0.702664 | 0.710213 | 0.694336 |
| 0.669661 | 0.664105 | 0.663483 | 0.669438 | 0.664451 | 0.663505 | 0.697885 | 0.706774 | 0.690705 |
| 0.667492 | 0.667849 | 0.665263 | 0.670899 | 0.66394 | 0.665715 | 0.701205 | 0.708409 | 0.690699 |
| 0.667272 | 0.667913 | 0.661569 | 0.658969 | 0.668005 | 0.659549 | 0.696176 | 0.701587 | 0.685447 |
| 0.665643 | 0.666475 | 0.667474 | 0.664578 | 0.665387 | 0.654625 | 0.693066 | 0.703078 | 0.686747 |
| 0.666341 | 0.668055 | 0.663539 | 0.664179 | 0.667213 | 0.662107 | 0.693593 | 0.699598 | 0.687956 |
| 0.66625 | 0.670563 | 0.663316 | 0.662722 | 0.66943 | 0.659895 | 0.698448 | 0.706916 | 0.688999 |
| 0.670817 | 0.669349 | 0.66728 | 0.667014 | 0.672337 | 0.66502 | 0.700219 | 0.706757 | 0.689555 |
| 0.670255 | 0.671381 | 0.666619 | 0.662998 | 0.672075 | 0.66375 | 0.696952 | 0.703334 | 0.683814 |
| 0.672421 | 0.673526 | 0.666474 | 0.663784 | 0.671958 | 0.668974 | 0.696737 | 0.706916 | 0.692445 |
| 0.748595 | 0.766916 | 0.776324 | 0.731793 | 0.752762 | 0.761468 | 0.792056 | 0.777898 | 0.759129 |
| 0.665349 | 0.666432 | 0.663833 | 0.664086 | 0.666909 | 0.658683 | 0.697397 | 0.70814 | 0.681946 |
| 0.667948 | 0.665774 | 0.665032 | 0.662614 | 0.663037 | 0.662054 | 0.699157 | 0.711472 | 0.683417 |
| 0.669155 | 0.673008 | 0.66916 | 0.665827 | 0.664324 | 0.658125 | 0.707513 | 0.709079 | 0.688123 |
| 0.6707 | 0.666714 | 0.660364 | 0.66583 | 0.663517 | 0.665382 | 0.698826 | 0.705436 | 0.685994 |
| 0.672346 | 0.66839 | 0.672269 | 0.665824 | 0.665725 | 0.663289 | 0.693708 | 0.706516 | 0.684606 |
| 0.665051 | 0.669535 | 0.663083 | 0.661195 | 0.669574 | 0.662146 | 0.694175 | 0.706369 | 0.683669 |
| 0.664686 | 0.663596 | 0.660599 | 0.653666 | 0.662762 | 0.659015 | 0.695527 | 0.70789 | 0.682182 |
| 0.667505 | 0.664551 | 0.663775 | 0.664184 | 0.659897 | 0.662704 | 0.697498 | 0.702705 | 0.682757 |
| 0.667273 | 0.666618 | 0.66547 | 0.663012 | 0.662772 | 0.657872 | 0.692467 | 0.702507 | 0.684291 |
| 0.667832 | 0.663481 | 0.658599 | 0.664181 | 0.657402 | 0.658055 | 0.693639 | 0.707841 | 0.681305 |
| 0.673098 | 0.666667 | 0.669445 | 0.668197 | 0.670304 | 0.668417 | 0.701499 | 0.710632 | 0.690607 |
| 0.66685 | 0.671083 | 0.663261 | 0.666765 | 0.672709 | 0.663812 | 0.701902 | 0.708808 | 0.687721 |
| 0.665704 | 0.662283 | 0.668217 | 0.666225 | 0.669062 | 0.653436 | 0.699556 | 0.706245 | 0.680291 |
| 0.669244 | 0.671681 | 0.665332 | 0.666467 | 0.668448 | 0.666268 | 0.698926 | 0.70581 | 0.691976 |
| 0.663993 | 0.664855 | 0.666336 | 0.66031 | 0.662277 | 0.663561 | 0.693237 | 0.703517 | 0.685467 |
| 0.668345 | 0.669987 | 0.661825 | 0.662277 | 0.66503 | 0.66336 | 0.701822 | 0.70837 | 0.691236 |
| 0.665229 | 0.669258 | 0.664305 | 0.664237 | 0.669647 | 0.663455 | 0.693652 | 0.70632 | 0.686806 |
| 0.664098 | 0.66828 | 0.663902 | 0.663718 | 0.668135 | 0.65746 | 0.698178 | 0.707384 | 0.68115 |
| 0.668314 | 0.669506 | 0.672612 | 0.663339 | 0.672022 | 0.662272 | 0.698354 | 0.711858 | 0.688551 |
| 0.669333 | 0.669787 | 0.663622 | 0.665829 | 0.662227 | 0.660786 | 0.699521 | 0.713707 | 0.686021 |

| | | | | | | | | |
|---|---|---|---|---|---|---|---|---|
| 0.664562 | 0.662525 | 0.662147 | 0.665883 | 0.660213 | 0.658843 | 0.699601 | 0.706538 | 0.685643 |
| 0.662235 | 0.664698 | 0.665417 | 0.665743 | 0.665988 | 0.660196 | 0.700719 | 0.709799 | 0.687712 |
| 0.66662 | 0.668234 | 0.662523 | 0.661217 | 0.664165 | 0.658063 | 0.698404 | 0.704965 | 0.683019 |
| 0.653321 | 0.660085 | 0.654834 | 0.656386 | 0.64313 | 0.664163 | 0.663668 | 0.66841 | 0.673527 |
| 0.665214 | 0.666901 | 0.665226 | 0.664665 | 0.666473 | 0.653419 | 0.697446 | 0.70851 | 0.687847 |
| 0.663042 | 0.662982 | 0.65956 | 0.660297 | 0.65816 | 0.651826 | 0.6971 | 0.707421 | 0.680323 |
| 0.666301 | 0.666619 | 0.665683 | 0.668932 | 0.659822 | 0.65797 | 0.698234 | 0.710778 | 0.690294 |
| 0.666347 | 0.661446 | 0.659145 | 0.659822 | 0.659957 | 0.653251 | 0.701795 | 0.708274 | 0.684831 |
| 0.672602 | 0.667374 | 0.671646 | 0.66515 | 0.669215 | 0.668917 | 0.703449 | 0.715604 | 0.69222 |
| 0.671172 | 0.669739 | 0.665036 | 0.663243 | 0.667008 | 0.663642 | 0.701812 | 0.709038 | 0.685744 |
| 0.66621 | 0.669888 | 0.662734 | 0.656825 | 0.666423 | 0.656443 | 0.696711 | 0.710528 | 0.685938 |
| 0.671229 | 0.67222 | 0.668642 | 0.666617 | 0.665886 | 0.657983 | 0.702698 | 0.714717 | 0.692202 |
| 0.674066 | 0.674706 | 0.668676 | 0.663832 | 0.670771 | 0.665267 | 0.701262 | 0.712005 | 0.691667 |
| 0.664577 | 0.663291 | 0.66569 | 0.660176 | 0.659861 | 0.657186 | 0.700062 | 0.707446 | 0.687018 |
| 0.668133 | 0.670339 | 0.66876 | 0.667352 | 0.665262 | 0.657697 | 0.704355 | 0.710153 | 0.691525 |
| 0.66078 | 0.661723 | 0.66048 | 0.658974 | 0.653483 | 0.648056 | 0.696656 | 0.704554 | 0.683189 |
| 0.669672 | 0.669758 | 0.671429 | 0.667105 | 0.669534 | 0.660013 | 0.703384 | 0.71157 | 0.685108 |
| 0.663385 | 0.664413 | 0.667786 | 0.663596 | 0.660533 | 0.653597 | 0.699517 | 0.710074 | 0.688239 |
| 0.663379 | 0.664688 | 0.660246 | 0.661715 | 0.664767 | 0.6583 | 0.692859 | 0.70346 | 0.682871 |
| 0.660524 | 0.667043 | 0.666946 | 0.659743 | 0.66264 | 0.654718 | 0.698532 | 0.70726 | 0.689423 |
| 0.666575 | 0.667325 | 0.666713 | 0.660474 | 0.664103 | 0.661477 | 0.702614 | 0.71013 | 0.692036 |
| 0.673661 | 0.674247 | 0.670703 | 0.668582 | 0.669892 | 0.664145 | 0.701732 | 0.711831 | 0.693379 |
| 0.692588 | 0.692467 | 0.696783 | 0.684501 | 0.688509 | 0.6875 | 0.680094 | 0.694698 | 0.69743 |
| 0.718777 | 0.728483 | 0.729336 | 0.717333 | 0.71729 | 0.710927 | 0.724903 | 0.704386 | 0.726579 |
| 0.738197 | 0.748737 | 0.77663 | 0.738931 | 0.753162 | 0.737175 | 0.744279 | 0.753067 | 0.743685 |
| 0.741784 | 0.75613 | 0.735376 | 0.718227 | 0.733762 | 0.741247 | 0.78288 | 0.737209 | 0.732106 |
| 0.659053 | 0.659661 | 0.656691 | 0.657483 | 0.65425 | 0.659719 | 0.656165 | 0.665419 | 0.669177 |
| 0.654699 | 0.660224 | 0.655592 | 0.655197 | 0.664904 | 0.653529 | 0.650454 | 0.661943 | 0.679163 |
| 0.656469 | 0.662326 | 0.647881 | 0.660386 | 0.650142 | 0.646731 | 0.652256 | 0.660736 | 0.671772 |
| 0.660096 | 0.667783 | 0.655711 | 0.659983 | 0.666265 | 0.65375 | 0.66899 | 0.673482 | 0.679294 |
| 0.653315 | 0.655778 | 0.648575 | 0.641758 | 0.658451 | 0.651678 | 0.653459 | 0.649606 | 0.656379 |
| 0.642116 | 0.656427 | 0.637345 | 0.643359 | 0.646127 | 0.645232 | 0.633912 | 0.640519 | 0.657957 |
| 0.636274 | 0.64467 | 0.644768 | 0.644948 | 0.632719 | 0.641553 | 0.646992 | 0.648656 | 0.665235 |
| 0.767189 | 0.737697 | 0.743246 | 0.729335 | 0.731185 | 0.753927 | 0.714857 | 0.726537 | 0.723666 |
| 0.66484 | 0.664595 | 0.659474 | 0.667402 | 0.66285 | 0.655585 | 0.693116 | 0.703823 | 0.685219 |
| 0.6688 | 0.668429 | 0.65628 | 0.661724 | 0.655822 | 0.661155 | 0.663721 | 0.67957 | 0.674944 |
| 0.661612 | 0.666372 | 0.662985 | 0.656807 | 0.655535 | 0.658005 | 0.664395 | 0.669778 | 0.674847 |
| 0.650929 | 0.657246 | 0.648649 | 0.658326 | 0.652132 | 0.649334 | 0.645778 | 0.655068 | 0.660259 |
| 0.670132 | 0.666774 | 0.658796 | 0.661629 | 0.664432 | 0.654808 | 0.663284 | 0.672316 | 0.682119 |
| 0.66451 | 0.666382 | 0.657435 | 0.658164 | 0.65339 | 0.656524 | 0.659572 | 0.669793 | 0.671431 |
| 0.653461 | 0.662124 | 0.64836 | 0.650458 | 0.655727 | 0.649875 | 0.652436 | 0.660665 | 0.666149 |
| 0.660179 | 0.655112 | 0.659167 | 0.661248 | 0.649189 | 0.650442 | 0.65546 | 0.664911 | 0.667854 |
| 0.647625 | 0.653435 | 0.647605 | 0.658839 | 0.651519 | 0.650348 | 0.650652 | 0.658068 | 0.664768 |
| 0.658788 | 0.665484 | 0.65132 | 0.661337 | 0.657308 | 0.653948 | 0.650038 | 0.658486 | 0.676127 |
| 0.665003 | 0.661169 | 0.661189 | 0.665664 | 0.658331 | 0.665455 | 0.653285 | 0.666584 | 0.673138 |
| 0.649142 | 0.651481 | 0.639823 | 0.649888 | 0.647736 | 0.6423 | 0.635076 | 0.636221 | 0.656448 |

| | | | | | | | | |
|---|---|---|---|---|---|---|---|---|
| 0.666283 | 0.666914 | 0.667104 | 0.66758 | 0.662463 | 0.662594 | 0.68003 | 0.692814 | 0.682734 |
| 0.655439 | 0.653127 | 0.6525 | 0.655299 | 0.648225 | 0.652597 | 0.646184 | 0.659508 | 0.665742 |
| 0.665349 | 0.657845 | 0.658868 | 0.661378 | 0.650192 | 0.652541 | 0.654049 | 0.661397 | 0.670588 |
| 0.659904 | 0.660552 | 0.656774 | 0.656969 | 0.657764 | 0.651002 | 0.651594 | 0.658901 | 0.670359 |
| 0.664433 | 0.654906 | 0.656295 | 0.657638 | 0.648813 | 0.660862 | 0.656699 | 0.652503 | 0.668158 |
| 0.665708 | 0.674766 | 0.664366 | 0.658508 | 0.671633 | 0.67287 | 0.653381 | 0.663165 | 0.671518 |
| 0.65935 | 0.651768 | 0.661952 | 0.654416 | 0.65604 | 0.66399 | 0.644806 | 0.661744 | 0.672654 |
| 0.658902 | 0.661679 | 0.654896 | 0.641753 | 0.644166 | 0.65 | 0.631607 | 0.648612 | 0.656747 |
| 0.647905 | 0.653608 | 0.649481 | 0.647527 | 0.647018 | 0.641962 | 0.647233 | 0.657909 | 0.655723 |
| 0.668006 | 0.666075 | 0.66111 | 0.668086 | 0.662381 | 0.659817 | 0.684151 | 0.690767 | 0.6829 |
| 0.66773 | 0.669257 | 0.665008 | 0.672096 | 0.665105 | 0.656979 | 0.67913 | 0.688249 | 0.685512 |
| 0.67089 | 0.66906 | 0.668213 | 0.669416 | 0.664273 | 0.668004 | 0.67514 | 0.681176 | 0.684429 |
| 0.668761 | 0.669906 | 0.667098 | 0.665161 | 0.663042 | 0.658657 | 0.669013 | 0.673946 | 0.676919 |
| 0.667048 | 0.667621 | 0.667278 | 0.666082 | 0.661871 | 0.665889 | 0.673322 | 0.681262 | 0.685888 |
| 0.657588 | 0.664382 | 0.656348 | 0.664159 | 0.661636 | 0.660974 | 0.666979 | 0.675918 | 0.677667 |

| Collection_ | Collection_ | Collection_ | Collection_ | Collection_ | Collection_ | Collection_ | Collection_ | Collection_ |
|---|---|---|---|---|---|---|---|---|
| 0.778472 | 0.767888 | 0.772192 | 0.769358 | 0.749577 | 0.774262 | 0.747209 | 0.771377 | 0.774379 |
| 0.772304 | 0.768668 | 0.779765 | 0.772287 | 0.750564 | 0.770647 | 0.754274 | 0.772462 | 0.769041 |
| 0.768848 | 0.757991 | 0.77737 | 0.774036 | 0.756877 | 0.764078 | 0.750279 | 0.764143 | 0.767585 |
| 0.78001 | 0.768428 | 0.772809 | 0.785868 | 0.754926 | 0.783832 | 0.759174 | 0.773058 | 0.775006 |
| 0.775675 | 0.770622 | 0.77844 | 0.772218 | 0.757058 | 0.769162 | 0.736752 | 0.775443 | 0.772761 |
| 0.774948 | 0.761808 | 0.769866 | 0.77929 | 0.7625 | 0.780091 | 0.766016 | 0.778124 | 0.777038 |
| 0.716624 | 0.703422 | 0.714511 | 0.721923 | 0.695428 | 0.71832 | 0.693482 | 0.717334 | 0.720757 |
| 0.714412 | 0.705831 | 0.714184 | 0.716808 | 0.692752 | 0.714793 | 0.687988 | 0.718287 | 0.719151 |
| 0.776301 | 0.769212 | 0.777523 | 0.782316 | 0.763955 | 0.774183 | 0.776269 | 0.774066 | 0.776108 |
| 0.778723 | 0.754991 | 0.775006 | 0.775271 | 0.753607 | 0.769381 | 0.741428 | 0.774055 | 0.775807 |
| 0.783661 | 0.758643 | 0.771288 | 0.774338 | 0.757731 | 0.783555 | 0.752944 | 0.76819 | 0.768351 |
| 0.773662 | 0.774691 | 0.781863 | 0.776509 | 0.755145 | 0.790231 | 0.743013 | 0.774672 | 0.767904 |
| 0.792987 | 0.774948 | 0.781147 | 0.782674 | 0.751673 | 0.795969 | 0.752395 | 0.771003 | 0.767408 |
| 0.779548 | 0.758166 | 0.769519 | 0.776216 | 0.749577 | 0.776437 | 0.75204 | 0.768051 | 0.77235 |
| 0.764459 | 0.753072 | 0.770727 | 0.773976 | 0.74686 | 0.766895 | 0.743978 | 0.764299 | 0.767598 |
| 0.781427 | 0.765957 | 0.778886 | 0.779166 | 0.761878 | 0.771802 | 0.754902 | 0.772272 | 0.780405 |
| 0.777387 | 0.756216 | 0.772828 | 0.775289 | 0.745882 | 0.770608 | 0.743485 | 0.76555 | 0.772284 |
| 0.752966 | 0.740862 | 0.758521 | 0.754142 | 0.734884 | 0.751807 | 0.728742 | 0.757593 | 0.748296 |
| 0.782525 | 0.772269 | 0.776718 | 0.775424 | 0.753867 | 0.791268 | 0.738225 | 0.770394 | 0.777925 |
| 0.774 | 0.770679 | 0.782677 | 0.781076 | 0.757157 | 0.768718 | 0.743091 | 0.775949 | 0.773864 |
| 0.771034 | 0.76775 | 0.776232 | 0.779988 | 0.757946 | 0.769824 | 0.747085 | 0.775146 | 0.776469 |
| 0.784583 | 0.763834 | 0.775041 | 0.780271 | 0.761739 | 0.772876 | 0.748265 | 0.773826 | 0.767826 |
| 0.77824 | 0.775436 | 0.781483 | 0.787672 | 0.759648 | 0.790493 | 0.760065 | 0.780637 | 0.775839 |
| 0.778946 | 0.769414 | 0.788594 | 0.777529 | 0.758139 | 0.799273 | 0.744831 | 0.778934 | 0.776047 |
| 0.786676 | 0.771846 | 0.783509 | 0.785881 | 0.760198 | 0.789899 | 0.749127 | 0.783782 | 0.778708 |
| 0.765486 | 0.769884 | 0.775335 | 0.779098 | 0.762024 | 0.777553 | 0.755641 | 0.770551 | 0.773356 |
| 0.760945 | 0.761803 | 0.772976 | 0.771183 | 0.748165 | 0.776589 | 0.735439 | 0.76937 | 0.766579 |
| 0.776892 | 0.761579 | 0.779173 | 0.786514 | 0.76273 | 0.772797 | 0.755427 | 0.773871 | 0.774865 |
| 0.776187 | 0.776395 | 0.785896 | 0.781071 | 0.759156 | 0.795827 | 0.759011 | 0.779116 | 0.772862 |
| 0.779325 | 0.762427 | 0.780647 | 0.772289 | 0.746466 | 0.778498 | 0.741783 | 0.773408 | 0.770813 |
| 0.770884 | 0.760937 | 0.774408 | 0.781586 | 0.756048 | 0.778717 | 0.754996 | 0.775644 | 0.77082 |
| 0.768618 | 0.765446 | 0.7739 | 0.770078 | 0.759964 | 0.77313 | 0.746388 | 0.765086 | 0.767902 |
| 0.774515 | 0.771818 | 0.77609 | 0.780851 | 0.751888 | 0.78702 | 0.745091 | 0.779461 | 0.770908 |
| 0.76995 | 0.768966 | 0.77756 | 0.779368 | 0.756879 | 0.774846 | 0.750802 | 0.776079 | 0.770647 |
| 0.764023 | 0.767728 | 0.766949 | 0.770866 | 0.75111 | 0.775614 | 0.743796 | 0.76446 | 0.770098 |
| 0.774483 | 0.767651 | 0.777038 | 0.775245 | 0.758929 | 0.785462 | 0.744319 | 0.76601 | 0.773204 |
| 0.776795 | 0.761716 | 0.777353 | 0.782777 | 0.758266 | 0.773551 | 0.743398 | 0.777214 | 0.77101 |
| 0.765169 | 0.759993 | 0.766271 | 0.775435 | 0.748475 | 0.787352 | 0.753227 | 0.769244 | 0.772425 |
| 0.771991 | 0.760601 | 0.7693 | 0.780833 | 0.754919 | 0.7792 | 0.755517 | 0.775604 | 0.77334 |

| | | | | | | | | |
|---|---|---|---|---|---|---|---|---|
| 0.756113 | 0.743074 | 0.759682 | 0.757605 | 0.73326 | 0.761821 | 0.73076 | 0.752515 | 0.756107 |
| 0.769435 | 0.755094 | 0.77855 | 0.771157 | 0.754537 | 0.767953 | 0.745767 | 0.76434 | 0.765872 |
| 0.775188 | 0.773981 | 0.786092 | 0.784553 | 0.752421 | 0.78751 | 0.751028 | 0.773275 | 0.769737 |
| 0.784231 | 0.775253 | 0.779232 | 0.778875 | 0.753961 | 0.787382 | 0.745513 | 0.771007 | 0.775143 |
| 0.778388 | 0.76831 | 0.778295 | 0.779712 | 0.75884 | 0.785289 | 0.753111 | 0.775719 | 0.769477 |
| 0.780998 | 0.771495 | 0.779799 | 0.775699 | 0.755907 | 0.788631 | 0.753237 | 0.779699 | 0.777676 |
| 0.769607 | 0.767276 | 0.770764 | 0.776896 | 0.755529 | 0.767361 | 0.753999 | 0.779826 | 0.77439 |
| 0.778263 | 0.770223 | 0.780405 | 0.783318 | 0.756375 | 0.782476 | 0.746765 | 0.777023 | 0.778312 |
| 0.771974 | 0.76522 | 0.775665 | 0.778474 | 0.7561 | 0.77218 | 0.747847 | 0.776683 | 0.782457 |
| 0.769791 | 0.757522 | 0.77355 | 0.771252 | 0.752128 | 0.780518 | 0.766766 | 0.77215 | 0.769867 |
| 0.78319 | 0.764623 | 0.778794 | 0.770003 | 0.75866 | 0.776193 | 0.747574 | 0.772423 | 0.771946 |
| 0.775751 | 0.76792 | 0.781881 | 0.779947 | 0.75998 | 0.771573 | 0.747794 | 0.778561 | 0.771785 |
| 0.780621 | 0.761815 | 0.787627 | 0.78411 | 0.757966 | 0.772686 | 0.750063 | 0.776431 | 0.773361 |
| 0.789909 | 0.766796 | 0.776789 | 0.779262 | 0.763577 | 0.775517 | 0.754406 | 0.772212 | 0.769638 |
| 0.77524 | 0.767236 | 0.776345 | 0.771835 | 0.762341 | 0.764971 | 0.736587 | 0.767286 | 0.765942 |
| 0.772704 | 0.765444 | 0.779943 | 0.773302 | 0.760873 | 0.760471 | 0.73735 | 0.767156 | 0.767708 |
| 0.776615 | 0.755971 | 0.775993 | 0.769969 | 0.75394 | 0.768548 | 0.739114 | 0.759264 | 0.762696 |
| 0.788856 | 0.768522 | 0.781176 | 0.780899 | 0.765364 | 0.774236 | 0.746802 | 0.779024 | 0.776305 |
| 0.777833 | 0.766533 | 0.781596 | 0.774263 | 0.754443 | 0.784805 | 0.750309 | 0.772311 | 0.770528 |
| 0.776173 | 0.766223 | 0.780669 | 0.77573 | 0.756609 | 0.777211 | 0.741763 | 0.770428 | 0.772071 |
| 0.778925 | 0.761167 | 0.779825 | 0.770651 | 0.753897 | 0.764994 | 0.754793 | 0.771218 | 0.766606 |
| 0.773043 | 0.762295 | 0.772899 | 0.770628 | 0.751152 | 0.764318 | 0.746577 | 0.762585 | 0.765634 |
| 0.772065 | 0.763303 | 0.775691 | 0.774798 | 0.770513 | 0.76482 | 0.748336 | 0.773848 | 0.770085 |
| 0.77192 | 0.758094 | 0.770366 | 0.788038 | 0.763443 | 0.789365 | 0.765498 | 0.771862 | 0.772908 |
| 0.769046 | 0.752536 | 0.76554 | 0.790682 | 0.742676 | 0.785586 | 0.757328 | 0.790085 | 0.791252 |
| 0.716216 | 0.706457 | 0.715005 | 0.71839 | 0.692653 | 0.716781 | 0.691497 | 0.718679 | 0.719376 |
| 0.716415 | 0.706284 | 0.715452 | 0.721892 | 0.699454 | 0.719291 | 0.689655 | 0.719189 | 0.722627 |
| 0.717298 | 0.707673 | 0.716683 | 0.722308 | 0.698421 | 0.72335 | 0.695018 | 0.719236 | 0.724199 |
| 0.776029 | 0.769983 | 0.766786 | 0.77985 | 0.75514 | 0.773659 | 0.735017 | 0.770746 | 0.773132 |
| 0.779966 | 0.765799 | 0.771179 | 0.783986 | 0.749294 | 0.770314 | 0.740232 | 0.774043 | 0.774705 |
| 0.781122 | 0.761542 | 0.772579 | 0.785316 | 0.743408 | 0.766895 | 0.737753 | 0.772937 | 0.779584 |
| 0.76956 | 0.761507 | 0.771998 | 0.773653 | 0.757059 | 0.770962 | 0.747588 | 0.772277 | 0.769122 |
| 0.783235 | 0.767018 | 0.779671 | 0.786094 | 0.756359 | 0.774676 | 0.752128 | 0.774818 | 0.773505 |
| 0.788767 | 0.788442 | 0.782238 | 0.781617 | 0.75599 | 0.820858 | 0.745052 | 0.772185 | 0.774191 |
| 0.785241 | 0.776811 | 0.77404 | 0.774514 | 0.754117 | 0.773306 | 0.744438 | 0.768667 | 0.773298 |
| 0.793287 | 0.767482 | 0.778181 | 0.778214 | 0.760442 | 0.771111 | 0.748569 | 0.767143 | 0.7749 |
| 0.773669 | 0.771025 | 0.779803 | 0.777384 | 0.772757 | 0.778192 | 0.77892 | 0.770084 | 0.771781 |
| 0.80736 | 0.785495 | 0.804487 | 0.797054 | 0.780636 | 0.796907 | 0.779477 | 0.7871 | 0.789351 |
| 0.777181 | 0.762858 | 0.765934 | 0.776771 | 0.752732 | 0.767902 | 0.740759 | 0.766584 | 0.7712 |
| 0.782064 | 0.780562 | 0.783951 | 0.78017 | 0.781703 | 0.772005 | 0.744358 | 0.77683 | 0.773416 |
| 0.784961 | 0.764902 | 0.773596 | 0.771104 | 0.752764 | 0.766489 | 0.741641 | 0.768904 | 0.764149 |
| 0.805402 | 0.784742 | 0.811691 | 0.801958 | 0.776156 | 0.796238 | 0.771883 | 0.794341 | 0.795133 |
| 0.785397 | 0.771125 | 0.782348 | 0.780065 | 0.763284 | 0.776615 | 0.749691 | 0.776144 | 0.770899 |
| 0.778321 | 0.784884 | 0.787398 | 0.777494 | 0.750801 | 0.787421 | 0.747684 | 0.776748 | 0.772604 |
| 0.779524 | 0.771221 | 0.782655 | 0.77787 | 0.784309 | 0.773073 | 0.750652 | 0.775392 | 0.774681 |
| 0.777321 | 0.767053 | 0.771759 | 0.777508 | 0.760592 | 0.772009 | 0.743358 | 0.766526 | 0.776432 |

| | | | | | | | | |
|---|---|---|---|---|---|---|---|---|
| 0.780996 | 0.771633 | 0.780289 | 0.772166 | 0.760631 | 0.772245 | 0.753699 | 0.77287 | 0.766103 |
| 0.778564 | 0.768941 | 0.778086 | 0.776749 | 0.773544 | 0.768931 | 0.759027 | 0.778647 | 0.775954 |
| 0.776187 | 0.774494 | 0.776093 | 0.778814 | 0.782393 | 0.778824 | 0.783845 | 0.771952 | 0.77093 |
| 0.797943 | 0.778881 | 0.79166 | 0.788561 | 0.770789 | 0.787086 | 0.774528 | 0.783213 | 0.781326 |
| 0.782913 | 0.779528 | 0.782022 | 0.776731 | 0.779696 | 0.791994 | 0.792569 | 0.783115 | 0.771996 |
| 0.781479 | 0.778156 | 0.780599 | 0.770154 | 0.761351 | 0.771337 | 0.735974 | 0.771433 | 0.764458 |
| 0.774148 | 0.772186 | 0.775781 | 0.780693 | 0.765587 | 0.790753 | 0.799116 | 0.769601 | 0.767611 |
| 0.776097 | 0.76848 | 0.774568 | 0.779236 | 0.839102 | 0.774135 | 0.802738 | 0.776283 | 0.775909 |
| 0.779862 | 0.770492 | 0.777906 | 0.784259 | 0.76382 | 0.770644 | 0.744818 | 0.78375 | 0.784415 |
| 0.772481 | 0.779695 | 0.774739 | 0.777354 | 0.750262 | 0.779032 | 0.728919 | 0.773277 | 0.774574 |
| 0.778155 | 0.775188 | 0.775621 | 0.776635 | 0.76359 | 0.769753 | 0.747881 | 0.776676 | 0.772701 |
| 0.777104 | 0.789615 | 0.771183 | 0.772541 | 0.756564 | 0.786047 | 0.745502 | 0.774062 | 0.769519 |
| 0.774191 | 0.778765 | 0.781433 | 0.772742 | 0.761616 | 0.782336 | 0.74344 | 0.778092 | 0.767559 |
| 0.777797 | 0.794764 | 0.777473 | 0.770013 | 0.75377 | 0.795643 | 0.738543 | 0.765534 | 0.769901 |
| 0.767116 | 0.768648 | 0.780472 | 0.778314 | 0.755273 | 0.763325 | 0.746063 | 0.772535 | 0.771728 |
| 0.777667 | 0.762517 | 0.7821 | 0.776666 | 0.755525 | 0.778086 | 0.746515 | 0.771989 | 0.775896 |
| 0.769327 | 0.771819 | 0.790202 | 0.781425 | 0.767257 | 0.7766 | 0.75285 | 0.776643 | 0.769803 |
| 0.784898 | 0.771276 | 0.770656 | 0.785589 | 0.755003 | 0.771459 | 0.746489 | 0.773375 | 0.779892 |
| 0.77966 | 0.786621 | 0.779114 | 0.787998 | 0.765195 | 0.790978 | 0.749105 | 0.775843 | 0.775534 |
| 0.77375 | 0.764706 | 0.77601 | 0.7872 | 0.761258 | 0.772019 | 0.740089 | 0.780031 | 0.782334 |
| 0.774552 | 0.772056 | 0.779198 | 0.772172 | 0.760544 | 0.773869 | 0.756137 | 0.774661 | 0.768945 |
| 0.768014 | 0.768997 | 0.772779 | 0.787884 | 0.75785 | 0.773427 | 0.741212 | 0.76834 | 0.78009 |
| 0.774899 | 0.767363 | 0.77673 | 0.771367 | 0.76192 | 0.779609 | 0.753728 | 0.768779 | 0.775196 |
| 0.778258 | 0.789561 | 0.776942 | 0.771242 | 0.764806 | 0.781944 | 0.738986 | 0.773015 | 0.76576 |
| 0.770283 | 0.768355 | 0.771965 | 0.771098 | 0.764962 | 0.771925 | 0.755511 | 0.772382 | 0.770057 |
| 0.77153 | 0.770832 | 0.775364 | 0.79092 | 0.767925 | 0.788688 | 0.78732 | 0.782612 | 0.782805 |
| 0.770364 | 0.765656 | 0.775083 | 0.773675 | 0.765561 | 0.771965 | 0.757258 | 0.768841 | 0.773033 |
| 0.772746 | 0.772161 | 0.769306 | 0.781899 | 0.757632 | 0.773525 | 0.745383 | 0.779959 | 0.778958 |
| 0.771483 | 0.769969 | 0.779679 | 0.777161 | 0.754967 | 0.775061 | 0.742043 | 0.776898 | 0.771112 |
| 0.768933 | 0.76452 | 0.773071 | 0.772421 | 0.76626 | 0.769396 | 0.772875 | 0.768698 | 0.768628 |
| 0.776594 | 0.769243 | 0.772226 | 0.776682 | 0.762308 | 0.783601 | 0.757066 | 0.776576 | 0.768072 |
| 0.76968 | 0.77345 | 0.773677 | 0.774713 | 0.767421 | 0.770687 | 0.754894 | 0.769062 | 0.77363 |
| 0.779642 | 0.770594 | 0.780652 | 0.777675 | 0.760745 | 0.773614 | 0.755617 | 0.773396 | 0.774404 |
| 0.786315 | 0.774804 | 0.774981 | 0.773548 | 0.753841 | 0.789336 | 0.746269 | 0.767118 | 0.770941 |
| 0.78163 | 0.77489 | 0.775144 | 0.776925 | 0.760535 | 0.783254 | 0.748552 | 0.774573 | 0.772471 |
| 0.78204 | 0.769006 | 0.771732 | 0.776703 | 0.758972 | 0.787816 | 0.758556 | 0.772085 | 0.775132 |
| 0.777863 | 0.774118 | 0.776647 | 0.780504 | 0.757002 | 0.778554 | 0.737568 | 0.775672 | 0.774145 |
| 0.767908 | 0.748309 | 0.750052 | 0.752498 | 0.749741 | 0.765524 | 0.756632 | 0.750996 | 0.748872 |
| 0.76559 | 0.755227 | 0.758004 | 0.755719 | 0.767261 | 0.750181 | 0.739795 | 0.757221 | 0.749817 |
| 0.750133 | 0.739801 | 0.747304 | 0.754818 | 0.732098 | 0.750053 | 0.751589 | 0.753792 | 0.751834 |
| 0.764692 | 0.748022 | 0.748257 | 0.751791 | 0.738254 | 0.744967 | 0.736326 | 0.744093 | 0.74287 |
| 0.756704 | 0.774784 | 0.742785 | 0.741186 | 0.717003 | 0.839879 | 0.720363 | 0.731531 | 0.736804 |
| 0.759318 | 0.76408 | 0.754388 | 0.743835 | 0.731689 | 0.761899 | 0.743961 | 0.743303 | 0.734873 |
| 0.754399 | 0.734985 | 0.744396 | 0.747917 | 0.732501 | 0.739591 | 0.745763 | 0.738531 | 0.738966 |
| 0.748509 | 0.744886 | 0.744348 | 0.743535 | 0.732918 | 0.743489 | 0.738118 | 0.743254 | 0.73453 |
| 0.750195 | 0.738503 | 0.742101 | 0.754322 | 0.740094 | 0.745123 | 0.75941 | 0.741564 | 0.745558 |
| 0.760724 | 0.741837 | 0.739878 | 0.746473 | 0.734642 | 0.738801 | 0.74081 | 0.737077 | 0.741642 |

| | | | | | | | | |
|---|---|---|---|---|---|---|---|---|
| 0.739607 | 0.736702 | 0.739919 | 0.742923 | 0.735228 | 0.736571 | 0.727175 | 0.753707 | 0.733817 |
| 0.753393 | 0.746923 | 0.740484 | 0.751873 | 0.728451 | 0.751238 | 0.742871 | 0.742745 | 0.744312 |
| 0.739401 | 0.738122 | 0.7339 | 0.749025 | 0.721803 | 0.743425 | 0.746013 | 0.738968 | 0.74355 |
| 0.752589 | 0.730533 | 0.753007 | 0.738092 | 0.721158 | 0.735589 | 0.741492 | 0.741798 | 0.730595 |
| 0.754464 | 0.764571 | 0.752812 | 0.743891 | 0.723598 | 0.804312 | 0.731836 | 0.743321 | 0.74433 |
| 0.74414 | 0.734876 | 0.74414 | 0.737703 | 0.77198 | 0.740585 | 0.828164 | 0.735265 | 0.73322 |
| 0.747196 | 0.743412 | 0.745431 | 0.738477 | 0.730603 | 0.735009 | 0.748653 | 0.743144 | 0.738144 |
| 0.764654 | 0.758256 | 0.74456 | 0.741997 | 0.723932 | 0.7632 | 0.737061 | 0.737819 | 0.736475 |
| 0.737003 | 0.749693 | 0.732426 | 0.72372 | 0.702858 | 0.748149 | 0.718529 | 0.723986 | 0.717549 |
| 0.744707 | 0.741636 | 0.754213 | 0.74592 | 0.723651 | 0.741877 | 0.734064 | 0.747442 | 0.741822 |
| 0.737402 | 0.738187 | 0.741311 | 0.747861 | 0.750516 | 0.742506 | 0.784341 | 0.741385 | 0.742718 |
| 0.772199 | 0.758604 | 0.770328 | 0.763184 | 0.758275 | 0.762796 | 0.74271 | 0.764127 | 0.760506 |
| 0.769967 | 0.769551 | 0.769411 | 0.772664 | 0.756904 | 0.77713 | 0.750277 | 0.769979 | 0.771193 |
| 0.749207 | 0.741321 | 0.747816 | 0.75566 | 0.734537 | 0.803943 | 0.794061 | 0.755692 | 0.760208 |
| 1 | 0.77242 | 0.775761 | 0.783412 | 0.752902 | 0.790305 | 0.756498 | 0.769973 | 0.77209 |
| 0.77242 | 1 | 0.767538 | 0.761516 | 0.752383 | 0.783387 | 0.733128 | 0.761787 | 0.761948 |
| 0.775761 | 0.767538 | 1 | 0.78767 | 0.757907 | 0.774586 | 0.745026 | 0.783013 | 0.773775 |
| 0.783412 | 0.761516 | 0.78767 | 1 | 0.756444 | 0.780825 | 0.759275 | 0.797549 | 0.837419 |
| 0.752902 | 0.752383 | 0.757907 | 0.756444 | 1 | 0.752017 | 0.749939 | 0.753399 | 0.748503 |
| 0.790305 | 0.783387 | 0.774586 | 0.780825 | 0.752017 | 1 | 0.799119 | 0.775002 | 0.775382 |
| 0.756498 | 0.733128 | 0.745026 | 0.759275 | 0.749939 | 0.799119 | 1 | 0.743602 | 0.746308 |
| 0.769973 | 0.761787 | 0.783013 | 0.797549 | 0.753399 | 0.775002 | 0.743602 | 1 | 0.808246 |
| 0.77209 | 0.761948 | 0.773775 | 0.837419 | 0.748503 | 0.775382 | 0.746308 | 0.808246 | 1 |
| 0.752889 | 0.726112 | 0.745177 | 0.751895 | 0.721931 | 0.76061 | 0.776881 | 0.739343 | 0.740036 |
| 0.754863 | 0.753216 | 0.747488 | 0.755916 | 0.726417 | 0.74334 | 0.718638 | 0.736044 | 0.735441 |
| 0.762792 | 0.748375 | 0.755202 | 0.757326 | 0.734741 | 0.75279 | 0.753204 | 0.74475 | 0.746466 |
| 0.760903 | 0.749416 | 0.754105 | 0.764637 | 0.749711 | 0.803459 | 0.828447 | 0.756332 | 0.76073 |
| 0.770084 | 0.759312 | 0.78044 | 0.779902 | 0.753079 | 0.779133 | 0.764253 | 0.773706 | 0.770031 |
| 0.747166 | 0.756328 | 0.741052 | 0.748508 | 0.739474 | 0.752027 | 0.715889 | 0.739728 | 0.744132 |
| 0.786893 | 0.765598 | 0.768989 | 0.778563 | 0.745821 | 0.785008 | 0.753552 | 0.771664 | 0.783709 |
| 0.768816 | 0.753542 | 0.774225 | 0.774333 | 0.753338 | 0.772089 | 0.740151 | 0.773415 | 0.776705 |
| 0.765317 | 0.753247 | 0.759547 | 0.758872 | 0.756111 | 0.754105 | 0.73237 | 0.753011 | 0.754527 |
| 0.762239 | 0.765606 | 0.75787 | 0.75397 | 0.758632 | 0.757498 | 0.72754 | 0.750669 | 0.754028 |
| 0.776088 | 0.756834 | 0.761051 | 0.778634 | 0.747556 | 0.85885 | 0.841888 | 0.771992 | 0.773934 |
| 0.78332 | 0.758563 | 0.760463 | 0.768969 | 0.750119 | 0.776416 | 0.748661 | 0.774196 | 0.770783 |
| 0.774623 | 0.753852 | 0.759238 | 0.767581 | 0.752549 | 0.769262 | 0.742969 | 0.771726 | 0.767735 |
| 0.72344 | 0.733253 | 0.721052 | 0.720388 | 0.733725 | 0.721999 | 0.705186 | 0.728751 | 0.723087 |
| 0.752521 | 0.753036 | 0.752531 | 0.761581 | 0.740826 | 0.754029 | 0.734524 | 0.755579 | 0.754928 |
| 0.723084 | 0.73346 | 0.72627 | 0.731216 | 0.745152 | 0.72568 | 0.710087 | 0.723989 | 0.724704 |
| 0.717088 | 0.720756 | 0.716881 | 0.721817 | 0.734082 | 0.716747 | 0.711093 | 0.717921 | 0.717927 |
| 0.774924 | 0.763656 | 0.767603 | 0.768286 | 0.755274 | 0.772379 | 0.742066 | 0.765631 | 0.760402 |
| 0.717591 | 0.716817 | 0.713426 | 0.714029 | 0.721831 | 0.712719 | 0.703367 | 0.714319 | 0.715075 |
| 0.710241 | 0.716109 | 0.713759 | 0.713152 | 0.731496 | 0.712428 | 0.67943 | 0.709927 | 0.707524 |
| 0.706394 | 0.713522 | 0.704415 | 0.705502 | 0.709838 | 0.702889 | 0.674536 | 0.706585 | 0.704686 |
| 0.763007 | 0.752037 | 0.764606 | 0.764016 | 0.748739 | 0.766355 | 0.747831 | 0.755593 | 0.75468 |
| 0.703268 | 0.712942 | 0.707858 | 0.706292 | 0.715885 | 0.705559 | 0.66808 | 0.701128 | 0.705111 |
| 0.768639 | 0.756247 | 0.764792 | 0.760952 | 0.754529 | 0.766783 | 0.738829 | 0.762366 | 0.758446 |

| | | | | | | | | |
|---|---|---|---|---|---|---|---|---|
| 0.761195 | 0.74994 | 0.766629 | 0.758076 | 0.738399 | 0.76007 | 0.735241 | 0.756027 | 0.756912 |
| 0.763151 | 0.75668 | 0.764102 | 0.761263 | 0.751483 | 0.761195 | 0.734257 | 0.762768 | 0.760602 |
| 0.753878 | 0.740979 | 0.74783 | 0.755746 | 0.739445 | 0.755934 | 0.729539 | 0.750143 | 0.756242 |
| 0.696142 | 0.697541 | 0.694732 | 0.70023 | 0.711283 | 0.690396 | 0.657117 | 0.698358 | 0.695567 |
| 0.69457 | 0.705458 | 0.696928 | 0.693596 | 0.716834 | 0.698485 | 0.678694 | 0.690655 | 0.694088 |
| 0.767507 | 0.758083 | 0.7675 | 0.766836 | 0.74626 | 0.774089 | 0.750952 | 0.758003 | 0.758138 |
| 0.691167 | 0.705123 | 0.692714 | 0.696452 | 0.719682 | 0.693099 | 0.65592 | 0.696713 | 0.693749 |
| 0.737059 | 0.728476 | 0.73398 | 0.742462 | 0.7144 | 0.737722 | 0.709666 | 0.739568 | 0.740871 |
| 0.764035 | 0.746775 | 0.760721 | 0.758519 | 0.749516 | 0.759935 | 0.734211 | 0.762397 | 0.758646 |
| 0.720043 | 0.712761 | 0.722227 | 0.729821 | 0.706653 | 0.725222 | 0.702452 | 0.734266 | 0.728074 |
| 0.747753 | 0.740271 | 0.752438 | 0.746086 | 0.72924 | 0.751961 | 0.72342 | 0.745312 | 0.743565 |
| 0.627451 | 0.588235 | 0.636905 | 0.651899 | 0.623457 | 0.66879 | 0.653846 | 0.576687 | 0.619355 |
| 0.661589 | 0.655418 | 0.659436 | 0.664286 | 0.645747 | 0.666667 | 0.676777 | 0.658184 | 0.660611 |
| 0.668339 | 0.65601 | 0.653168 | 0.665508 | 0.642425 | 0.666219 | 0.666197 | 0.66459 | 0.663882 |
| 0.674806 | 0.662368 | 0.671683 | 0.668945 | 0.656206 | 0.67927 | 0.680082 | 0.670169 | 0.677108 |
| 0.669349 | 0.661793 | 0.666839 | 0.671607 | 0.652774 | 0.671981 | 0.666509 | 0.66671 | 0.670651 |
| 0.62558 | 0.634165 | 0.61716 | 0.619607 | 0.656371 | 0.62091 | 0.609263 | 0.620519 | 0.615789 |
| 0.672999 | 0.662778 | 0.670634 | 0.676808 | 0.652516 | 0.67996 | 0.679573 | 0.671295 | 0.673616 |
| 0.745483 | 0.737395 | 0.739014 | 0.738735 | 0.725412 | 0.750729 | 0.721203 | 0.739655 | 0.738072 |
| 0.74225 | 0.728271 | 0.740674 | 0.738904 | 0.723368 | 0.74328 | 0.719043 | 0.735938 | 0.730239 |
| 0.748777 | 0.738606 | 0.751122 | 0.750103 | 0.731005 | 0.749858 | 0.723577 | 0.744327 | 0.747109 |
| 0.611072 | 0.628641 | 0.614499 | 0.616563 | 0.643175 | 0.619725 | 0.585006 | 0.613992 | 0.619029 |
| 0.595257 | 0.613311 | 0.594565 | 0.598308 | 0.625539 | 0.601132 | 0.56168 | 0.599411 | 0.600559 |
| 0.715093 | 0.704958 | 0.707206 | 0.71382 | 0.693863 | 0.717749 | 0.688706 | 0.71417 | 0.718424 |
| 0.716769 | 0.703189 | 0.708271 | 0.715873 | 0.696051 | 0.718547 | 0.688262 | 0.715671 | 0.71642 |
| 0.70914 | 0.697706 | 0.699905 | 0.708722 | 0.687048 | 0.707255 | 0.678632 | 0.708757 | 0.710473 |
| 0.711521 | 0.701516 | 0.701187 | 0.712852 | 0.691851 | 0.710039 | 0.681578 | 0.712527 | 0.71408 |
| 0.718712 | 0.703144 | 0.709046 | 0.717992 | 0.698113 | 0.719194 | 0.69356 | 0.716799 | 0.721064 |
| 0.71754 | 0.704328 | 0.70751 | 0.714274 | 0.691499 | 0.720462 | 0.68756 | 0.714574 | 0.716986 |
| 0.718537 | 0.704096 | 0.709749 | 0.715908 | 0.695693 | 0.722753 | 0.687947 | 0.71345 | 0.717761 |
| 0.715022 | 0.706372 | 0.708829 | 0.71808 | 0.694645 | 0.720212 | 0.691475 | 0.71521 | 0.719741 |
| 0.721789 | 0.709529 | 0.719196 | 0.720858 | 0.702926 | 0.722931 | 0.697032 | 0.722869 | 0.723325 |
| 0.715683 | 0.705996 | 0.71271 | 0.721337 | 0.698098 | 0.718367 | 0.697514 | 0.716831 | 0.721456 |
| 0.715497 | 0.70547 | 0.715262 | 0.723972 | 0.698376 | 0.720446 | 0.689713 | 0.716875 | 0.725293 |
| 0.717667 | 0.709788 | 0.71789 | 0.726259 | 0.704447 | 0.723347 | 0.696129 | 0.718435 | 0.725273 |
| 0.715133 | 0.702153 | 0.707788 | 0.720311 | 0.693203 | 0.714773 | 0.689971 | 0.717383 | 0.720778 |
| 0.716024 | 0.703881 | 0.710834 | 0.718543 | 0.694429 | 0.721099 | 0.691566 | 0.716984 | 0.721104 |
| 0.717758 | 0.70684 | 0.713181 | 0.720325 | 0.696863 | 0.722222 | 0.692326 | 0.715804 | 0.722841 |
| 0.710668 | 0.699788 | 0.707063 | 0.716243 | 0.692097 | 0.715892 | 0.693086 | 0.714803 | 0.718033 |
| 0.718092 | 0.706764 | 0.713348 | 0.722173 | 0.695405 | 0.720996 | 0.695453 | 0.717764 | 0.72212 |
| 0.660158 | 0.655956 | 0.665993 | 0.661211 | 0.646583 | 0.673473 | 0.670973 | 0.664065 | 0.662276 |
| 0.671963 | 0.668072 | 0.668481 | 0.670016 | 0.649778 | 0.673745 | 0.673986 | 0.665693 | 0.6688 |
| 0.717411 | 0.707911 | 0.717884 | 0.721335 | 0.696698 | 0.720205 | 0.691977 | 0.718563 | 0.722729 |
| 0.721209 | 0.708884 | 0.716851 | 0.721568 | 0.699403 | 0.722081 | 0.695825 | 0.720682 | 0.72265 |
| 0.653099 | 0.647364 | 0.653981 | 0.659345 | 0.626543 | 0.656142 | 0.663105 | 0.650307 | 0.656122 |
| 0.718522 | 0.707878 | 0.714411 | 0.723442 | 0.696171 | 0.72289 | 0.691762 | 0.71808 | 0.723242 |
| 0.708487 | 0.694714 | 0.704587 | 0.713175 | 0.686641 | 0.712384 | 0.681895 | 0.706931 | 0.71194 |

| | | | | | | | | |
|---|---|---|---|---|---|---|---|---|
| 0.714008 | 0.701207 | 0.709739 | 0.71771 | 0.695181 | 0.718514 | 0.69192 | 0.713406 | 0.719958 |
| 0.717142 | 0.707562 | 0.713014 | 0.722805 | 0.696149 | 0.720839 | 0.688439 | 0.719328 | 0.72539 |
| 0.722512 | 0.705882 | 0.712342 | 0.720419 | 0.699135 | 0.723973 | 0.697762 | 0.715057 | 0.720634 |
| 0.718366 | 0.705578 | 0.713121 | 0.721931 | 0.697904 | 0.719708 | 0.692843 | 0.715656 | 0.723922 |
| 0.726934 | 0.71731 | 0.718983 | 0.725952 | 0.704585 | 0.729747 | 0.704718 | 0.71915 | 0.722107 |
| 0.714878 | 0.70191 | 0.711422 | 0.720042 | 0.691059 | 0.715183 | 0.693028 | 0.714907 | 0.718533 |
| 0.713723 | 0.705286 | 0.711658 | 0.719705 | 0.695024 | 0.719836 | 0.688847 | 0.713432 | 0.71808 |
| 0.711479 | 0.699591 | 0.705134 | 0.713587 | 0.688933 | 0.71146 | 0.684063 | 0.709195 | 0.713754 |
| 0.714586 | 0.705114 | 0.711973 | 0.716829 | 0.693876 | 0.7173 | 0.687523 | 0.714778 | 0.720787 |
| 0.713718 | 0.702068 | 0.711113 | 0.718214 | 0.694811 | 0.715933 | 0.686421 | 0.714263 | 0.716296 |
| 0.71844 | 0.704236 | 0.715425 | 0.723035 | 0.700477 | 0.720047 | 0.689676 | 0.718745 | 0.719636 |
| 0.709947 | 0.701283 | 0.70798 | 0.713837 | 0.689161 | 0.712609 | 0.689773 | 0.709855 | 0.7174 |
| 0.708153 | 0.695969 | 0.702094 | 0.708509 | 0.686773 | 0.70884 | 0.681741 | 0.705583 | 0.712305 |
| 0.710294 | 0.696247 | 0.702047 | 0.713097 | 0.684146 | 0.709353 | 0.682442 | 0.70738 | 0.710796 |
| 0.721396 | 0.711537 | 0.715061 | 0.724653 | 0.702148 | 0.72733 | 0.701188 | 0.720615 | 0.724842 |
| 0.717971 | 0.710698 | 0.713383 | 0.719043 | 0.698016 | 0.722117 | 0.690619 | 0.714058 | 0.72256 |
| 0.712114 | 0.70055 | 0.706854 | 0.713811 | 0.692252 | 0.715058 | 0.688399 | 0.707752 | 0.713813 |
| 0.712746 | 0.703011 | 0.708459 | 0.716039 | 0.69197 | 0.713537 | 0.68454 | 0.712031 | 0.720019 |
| 0.715532 | 0.69985 | 0.709649 | 0.716889 | 0.698257 | 0.71574 | 0.687432 | 0.713807 | 0.719218 |
| 0.711244 | 0.700612 | 0.708294 | 0.71687 | 0.690424 | 0.715842 | 0.689464 | 0.714217 | 0.71905 |
| 0.71091 | 0.700907 | 0.706922 | 0.711715 | 0.69216 | 0.713018 | 0.684796 | 0.71052 | 0.717548 |
| 0.710972 | 0.698883 | 0.705395 | 0.715411 | 0.693584 | 0.711803 | 0.68309 | 0.712294 | 0.717823 |
| 0.714802 | 0.70243 | 0.71108 | 0.719696 | 0.693984 | 0.716711 | 0.69277 | 0.717402 | 0.721402 |
| 0.718196 | 0.706402 | 0.712471 | 0.717693 | 0.696123 | 0.717074 | 0.687958 | 0.713886 | 0.722733 |
| 0.714075 | 0.701288 | 0.707577 | 0.715763 | 0.692137 | 0.717046 | 0.687568 | 0.710907 | 0.717874 |
| 0.71231 | 0.698824 | 0.706383 | 0.712227 | 0.692192 | 0.711161 | 0.683567 | 0.713686 | 0.712628 |
| 0.713712 | 0.702829 | 0.705972 | 0.715077 | 0.691244 | 0.713872 | 0.684473 | 0.710861 | 0.720463 |
| 0.707058 | 0.697827 | 0.708037 | 0.711721 | 0.690996 | 0.709587 | 0.683877 | 0.708323 | 0.71598 |
| 0.714413 | 0.704167 | 0.707961 | 0.713757 | 0.693151 | 0.718884 | 0.686699 | 0.715305 | 0.718223 |
| 0.714632 | 0.701745 | 0.709657 | 0.718652 | 0.69309 | 0.716333 | 0.689069 | 0.717454 | 0.721522 |
| 0.715512 | 0.707689 | 0.710255 | 0.719919 | 0.694512 | 0.717957 | 0.686702 | 0.717471 | 0.723786 |
| 0.719226 | 0.707307 | 0.71377 | 0.720782 | 0.693744 | 0.716721 | 0.692804 | 0.719521 | 0.72356 |
| 0.722665 | 0.709239 | 0.712863 | 0.721516 | 0.701381 | 0.722476 | 0.693164 | 0.719723 | 0.723869 |
| 0.723194 | 0.710759 | 0.713076 | 0.720348 | 0.701862 | 0.723651 | 0.692818 | 0.717409 | 0.72159 |
| 0.712394 | 0.697284 | 0.706883 | 0.714679 | 0.687788 | 0.712787 | 0.683752 | 0.716069 | 0.716426 |
| 0.715599 | 0.704467 | 0.708086 | 0.716292 | 0.692631 | 0.716079 | 0.690349 | 0.714193 | 0.7189 |
| 0.71939 | 0.70387 | 0.712617 | 0.717231 | 0.691937 | 0.719635 | 0.687817 | 0.716419 | 0.718717 |
| 0.712463 | 0.697947 | 0.706278 | 0.714007 | 0.689186 | 0.714446 | 0.685201 | 0.714888 | 0.714216 |
| 0.709022 | 0.696686 | 0.701668 | 0.707925 | 0.688073 | 0.707816 | 0.681918 | 0.706691 | 0.712108 |
| 0.706528 | 0.694234 | 0.698706 | 0.706452 | 0.685698 | 0.70766 | 0.68215 | 0.706164 | 0.709944 |
| 0.703472 | 0.697309 | 0.701416 | 0.708583 | 0.684279 | 0.70786 | 0.680748 | 0.704825 | 0.709451 |
| 0.704339 | 0.692768 | 0.700421 | 0.708394 | 0.683405 | 0.70813 | 0.6835 | 0.705722 | 0.710367 |
| 0.706962 | 0.697443 | 0.704222 | 0.71113 | 0.685434 | 0.712005 | 0.682251 | 0.709407 | 0.715162 |
| 0.700855 | 0.686288 | 0.694893 | 0.702287 | 0.68011 | 0.700216 | 0.676504 | 0.699187 | 0.704168 |
| 0.711137 | 0.698564 | 0.702379 | 0.715167 | 0.688044 | 0.710028 | 0.68709 | 0.71384 | 0.716332 |
| 0.704537 | 0.692868 | 0.700946 | 0.705346 | 0.680975 | 0.70259 | 0.679587 | 0.705574 | 0.708579 |

| | | | | | | | | |
|---|---|---|---|---|---|---|---|---|
| 0.718089 | 0.704642 | 0.713616 | 0.72141 | 0.69641 | 0.721342 | 0.695088 | 0.717671 | 0.720325 |
| 0.714694 | 0.705484 | 0.712443 | 0.716772 | 0.698754 | 0.720547 | 0.692019 | 0.716452 | 0.718463 |
| 0.711466 | 0.70045 | 0.713792 | 0.715149 | 0.691612 | 0.714892 | 0.685889 | 0.713208 | 0.713904 |
| 0.715215 | 0.708679 | 0.716362 | 0.720794 | 0.698769 | 0.717753 | 0.689556 | 0.718128 | 0.723992 |
| 0.721916 | 0.710399 | 0.722037 | 0.725119 | 0.70663 | 0.726309 | 0.696981 | 0.722485 | 0.72515 |
| 0.716211 | 0.706489 | 0.717522 | 0.719852 | 0.696944 | 0.722794 | 0.689729 | 0.719002 | 0.721138 |
| 0.71377 | 0.702989 | 0.709932 | 0.71357 | 0.694949 | 0.710479 | 0.684159 | 0.712311 | 0.71654 |
| 0.71333 | 0.704271 | 0.713743 | 0.71954 | 0.693137 | 0.719312 | 0.688463 | 0.717136 | 0.72016 |
| 0.712902 | 0.697614 | 0.709288 | 0.714122 | 0.686614 | 0.715067 | 0.688401 | 0.713495 | 0.714373 |
| 0.713164 | 0.70584 | 0.713184 | 0.718243 | 0.696965 | 0.717902 | 0.686601 | 0.715905 | 0.718446 |
| 0.717058 | 0.707021 | 0.715905 | 0.71705 | 0.696381 | 0.722397 | 0.682055 | 0.718708 | 0.722169 |
| 0.716828 | 0.706802 | 0.712726 | 0.71871 | 0.699945 | 0.721975 | 0.687021 | 0.715303 | 0.721937 |
| 0.711367 | 0.704136 | 0.710855 | 0.721712 | 0.696619 | 0.716331 | 0.684846 | 0.71602 | 0.720535 |
| 0.714528 | 0.703504 | 0.714308 | 0.718316 | 0.69673 | 0.71842 | 0.691966 | 0.718199 | 0.718415 |
| 0.717961 | 0.705789 | 0.717099 | 0.721146 | 0.69598 | 0.721234 | 0.687388 | 0.720573 | 0.722811 |
| 0.715946 | 0.705449 | 0.713269 | 0.722258 | 0.696209 | 0.720105 | 0.688803 | 0.718488 | 0.726622 |
| 0.718339 | 0.704667 | 0.716584 | 0.722608 | 0.697212 | 0.720141 | 0.694019 | 0.718571 | 0.723291 |
| 0.713625 | 0.704822 | 0.716911 | 0.719096 | 0.698823 | 0.719474 | 0.688675 | 0.7153 | 0.721318 |
| 0.722752 | 0.707729 | 0.717908 | 0.723663 | 0.698645 | 0.720702 | 0.691355 | 0.718985 | 0.723343 |
| 0.71648 | 0.704833 | 0.713173 | 0.720887 | 0.695398 | 0.716294 | 0.689288 | 0.717115 | 0.722719 |
| 0.716747 | 0.705638 | 0.713137 | 0.722484 | 0.698205 | 0.719974 | 0.68771 | 0.719249 | 0.722159 |
| 0.714007 | 0.702673 | 0.712406 | 0.719682 | 0.693415 | 0.718558 | 0.690171 | 0.714424 | 0.721682 |
| 0.715179 | 0.705813 | 0.716619 | 0.725191 | 0.697104 | 0.720723 | 0.69584 | 0.719593 | 0.72563 |
| 0.716108 | 0.707523 | 0.719621 | 0.719914 | 0.701771 | 0.718981 | 0.691995 | 0.722023 | 0.722576 |
| 0.711295 | 0.708697 | 0.714677 | 0.720057 | 0.695525 | 0.716737 | 0.686773 | 0.717681 | 0.722512 |
| 0.71257 | 0.705056 | 0.713754 | 0.719011 | 0.698237 | 0.72098 | 0.691129 | 0.716056 | 0.723734 |
| 0.712248 | 0.703427 | 0.712414 | 0.716587 | 0.695409 | 0.713983 | 0.685462 | 0.714195 | 0.721448 |
| 0.709711 | 0.704723 | 0.707265 | 0.716853 | 0.694656 | 0.71587 | 0.684017 | 0.711412 | 0.714909 |
| 0.714423 | 0.704843 | 0.717628 | 0.72094 | 0.701724 | 0.72117 | 0.686537 | 0.716369 | 0.721847 |
| 0.715589 | 0.707129 | 0.719748 | 0.721858 | 0.703219 | 0.724172 | 0.695461 | 0.716938 | 0.724839 |
| 0.711696 | 0.703408 | 0.715678 | 0.717413 | 0.694129 | 0.716029 | 0.686645 | 0.715282 | 0.719304 |
| 0.717899 | 0.708267 | 0.721189 | 0.723129 | 0.703011 | 0.720562 | 0.69431 | 0.721894 | 0.725411 |
| 0.717486 | 0.708383 | 0.718339 | 0.7222 | 0.699258 | 0.721753 | 0.689556 | 0.719594 | 0.721431 |
| 0.712105 | 0.703051 | 0.70851 | 0.71399 | 0.691092 | 0.71815 | 0.687143 | 0.711277 | 0.711876 |
| 0.714131 | 0.703474 | 0.712535 | 0.718343 | 0.697953 | 0.715704 | 0.692204 | 0.716332 | 0.719792 |
| 0.706982 | 0.700149 | 0.709413 | 0.715944 | 0.692226 | 0.713528 | 0.687615 | 0.709858 | 0.715173 |
| 0.719741 | 0.708717 | 0.723585 | 0.726533 | 0.704291 | 0.72463 | 0.696196 | 0.722053 | 0.726544 |
| 0.713288 | 0.70458 | 0.712582 | 0.71999 | 0.697567 | 0.718079 | 0.690337 | 0.715367 | 0.717815 |
| 0.718991 | 0.702102 | 0.714263 | 0.71798 | 0.698526 | 0.718123 | 0.688924 | 0.716344 | 0.716528 |
| 0.716324 | 0.70654 | 0.711875 | 0.720781 | 0.696541 | 0.720023 | 0.690755 | 0.716467 | 0.718546 |
| 0.720683 | 0.711315 | 0.719434 | 0.72216 | 0.706421 | 0.723993 | 0.693423 | 0.719586 | 0.725421 |
| 0.71826 | 0.708343 | 0.71636 | 0.723149 | 0.700926 | 0.723367 | 0.693242 | 0.717842 | 0.721386 |
| 0.720986 | 0.706457 | 0.7174 | 0.720912 | 0.700436 | 0.72272 | 0.697733 | 0.717865 | 0.718958 |
| 0.713686 | 0.701092 | 0.713997 | 0.720608 | 0.69643 | 0.716407 | 0.692503 | 0.714809 | 0.71842 |
| 0.716003 | 0.706592 | 0.715265 | 0.724012 | 0.699069 | 0.720622 | 0.693685 | 0.721452 | 0.724525 |
| 0.710643 | 0.700395 | 0.712003 | 0.721089 | 0.69551 | 0.717813 | 0.691738 | 0.716739 | 0.716344 |

| | | | | | | | | |
|---|---|---|---|---|---|---|---|---|
| 0.700703 | 0.688956 | 0.702317 | 0.709062 | 0.685419 | 0.701662 | 0.685417 | 0.702855 | 0.710077 |
| 0.750561 | 0.734499 | 0.747222 | 0.75901 | 0.729771 | 0.759892 | 0.740736 | 0.746916 | 0.747083 |
| 0.711341 | 0.705239 | 0.710223 | 0.712396 | 0.695185 | 0.719045 | 0.692083 | 0.712797 | 0.716857 |
| 0.718429 | 0.709025 | 0.718671 | 0.719415 | 0.698544 | 0.722485 | 0.696744 | 0.719115 | 0.72159 |
| 0.709518 | 0.702214 | 0.706847 | 0.712176 | 0.694696 | 0.718219 | 0.688213 | 0.70995 | 0.716439 |
| 0.713108 | 0.703689 | 0.713165 | 0.719791 | 0.694708 | 0.718314 | 0.690349 | 0.716356 | 0.720662 |
| 0.708344 | 0.702154 | 0.710738 | 0.711001 | 0.693501 | 0.717218 | 0.69112 | 0.71168 | 0.710491 |
| 0.703874 | 0.697782 | 0.704644 | 0.710057 | 0.687912 | 0.714958 | 0.685868 | 0.706707 | 0.71215 |
| 0.712828 | 0.704646 | 0.714432 | 0.720394 | 0.695246 | 0.720793 | 0.689878 | 0.71062 | 0.720472 |
| 0.721675 | 0.707076 | 0.719336 | 0.726534 | 0.699086 | 0.724165 | 0.692846 | 0.720377 | 0.723574 |
| 0.71826 | 0.705189 | 0.713622 | 0.722227 | 0.698712 | 0.724536 | 0.691145 | 0.714762 | 0.71971 |
| 0.717347 | 0.709585 | 0.71506 | 0.726896 | 0.69837 | 0.725156 | 0.692866 | 0.71877 | 0.722016 |
| 0.721934 | 0.710001 | 0.720317 | 0.723707 | 0.699757 | 0.724431 | 0.690593 | 0.721946 | 0.7266 |
| 0.719526 | 0.707928 | 0.716157 | 0.724578 | 0.698394 | 0.726115 | 0.690573 | 0.72248 | 0.724108 |
| 0.712397 | 0.703895 | 0.711744 | 0.721379 | 0.69465 | 0.722074 | 0.692634 | 0.712174 | 0.720463 |
| 0.716289 | 0.705823 | 0.710697 | 0.718882 | 0.700535 | 0.721862 | 0.690564 | 0.715252 | 0.718246 |
| 0.717538 | 0.705601 | 0.716437 | 0.722132 | 0.698401 | 0.722559 | 0.689805 | 0.717852 | 0.722321 |
| 0.715527 | 0.704373 | 0.715187 | 0.721132 | 0.69873 | 0.719904 | 0.689822 | 0.714435 | 0.719055 |
| 0.711105 | 0.70037 | 0.704893 | 0.716235 | 0.690751 | 0.715637 | 0.692576 | 0.708825 | 0.713913 |
| 0.708461 | 0.695234 | 0.704644 | 0.711972 | 0.689669 | 0.714884 | 0.688148 | 0.711081 | 0.713633 |
| 0.714812 | 0.705617 | 0.712135 | 0.719331 | 0.696554 | 0.724374 | 0.69477 | 0.714656 | 0.717976 |
| 0.704802 | 0.696159 | 0.697865 | 0.708344 | 0.687782 | 0.710882 | 0.678281 | 0.70977 | 0.714628 |
| 0.707021 | 0.700531 | 0.712172 | 0.71385 | 0.692206 | 0.715358 | 0.686666 | 0.715564 | 0.715358 |
| 0.713417 | 0.703766 | 0.716131 | 0.71585 | 0.694854 | 0.717932 | 0.692958 | 0.718933 | 0.71819 |
| 0.715608 | 0.704029 | 0.716413 | 0.719467 | 0.699413 | 0.720065 | 0.692495 | 0.718812 | 0.721009 |
| 0.709493 | 0.701466 | 0.713398 | 0.714961 | 0.693439 | 0.715146 | 0.685725 | 0.718134 | 0.717671 |
| 0.709398 | 0.696376 | 0.708251 | 0.715411 | 0.695754 | 0.717037 | 0.688349 | 0.70988 | 0.71294 |
| 0.711742 | 0.698789 | 0.708741 | 0.715775 | 0.692783 | 0.712888 | 0.687661 | 0.709332 | 0.71361 |
| 0.708546 | 0.698673 | 0.703574 | 0.712537 | 0.690052 | 0.710983 | 0.678598 | 0.706802 | 0.715582 |
| 0.766959 | 0.756081 | 0.764628 | 0.759355 | 0.745687 | 0.756164 | 0.728281 | 0.765852 | 0.75382 |
| 0.773174 | 0.76723 | 0.783009 | 0.777992 | 0.772825 | 0.768687 | 0.743357 | 0.779252 | 0.769673 |
| 0.773825 | 0.770818 | 0.777851 | 0.779241 | 0.75639 | 0.770648 | 0.742889 | 0.778381 | 0.776944 |
| 0.772249 | 0.755187 | 0.772503 | 0.768262 | 0.762094 | 0.762171 | 0.773427 | 0.763206 | 0.768948 |
| 0.777085 | 0.768804 | 0.782335 | 0.787432 | 0.77183 | 0.778628 | 0.774585 | 0.774487 | 0.776317 |
| 0.707654 | 0.697467 | 0.704432 | 0.712719 | 0.688514 | 0.712967 | 0.681191 | 0.711874 | 0.713317 |
| 0.709509 | 0.702073 | 0.70826 | 0.717331 | 0.695597 | 0.714148 | 0.682032 | 0.711032 | 0.715685 |
| 0.713506 | 0.705068 | 0.711662 | 0.717982 | 0.692149 | 0.72017 | 0.688623 | 0.714344 | 0.719485 |
| 0.712687 | 0.703772 | 0.711309 | 0.721314 | 0.695467 | 0.717832 | 0.690942 | 0.717901 | 0.720923 |
| 0.712962 | 0.703653 | 0.706468 | 0.720225 | 0.692689 | 0.715037 | 0.688276 | 0.714933 | 0.716839 |
| 0.716618 | 0.707374 | 0.70939 | 0.717565 | 0.697233 | 0.723499 | 0.687755 | 0.718964 | 0.718259 |
| 0.71273 | 0.704457 | 0.710617 | 0.716406 | 0.690288 | 0.716857 | 0.683994 | 0.712391 | 0.717672 |
| 0.70973 | 0.701318 | 0.710881 | 0.717286 | 0.694044 | 0.714758 | 0.681906 | 0.712936 | 0.718684 |
| 0.706178 | 0.695594 | 0.705611 | 0.711496 | 0.686636 | 0.710847 | 0.684971 | 0.713643 | 0.710974 |
| 0.706347 | 0.692682 | 0.702113 | 0.708708 | 0.688036 | 0.709584 | 0.683565 | 0.708315 | 0.714458 |
| 0.713005 | 0.701968 | 0.709057 | 0.715736 | 0.695677 | 0.711465 | 0.683999 | 0.714227 | 0.715968 |
| 0.709456 | 0.703318 | 0.712105 | 0.720396 | 0.697056 | 0.717383 | 0.6888 | 0.713349 | 0.719802 |

| | | | | | | | | |
|---|---|---|---|---|---|---|---|---|
| 0.715562 | 0.702841 | 0.710622 | 0.718601 | 0.695778 | 0.718113 | 0.695764 | 0.714672 | 0.717606 |
| 0.712695 | 0.700831 | 0.711522 | 0.717486 | 0.697516 | 0.718194 | 0.68905 | 0.714619 | 0.719961 |
| 0.714929 | 0.704542 | 0.716873 | 0.719285 | 0.701376 | 0.715487 | 0.69672 | 0.717374 | 0.721178 |
| 0.713856 | 0.700862 | 0.712002 | 0.718175 | 0.692139 | 0.715762 | 0.687122 | 0.714803 | 0.715606 |
| 0.710705 | 0.698308 | 0.70765 | 0.714487 | 0.690341 | 0.714622 | 0.68731 | 0.708623 | 0.71283 |
| 0.712052 | 0.701544 | 0.708402 | 0.717208 | 0.693776 | 0.712172 | 0.68648 | 0.710387 | 0.715639 |
| 0.712971 | 0.706296 | 0.710678 | 0.719596 | 0.696296 | 0.71846 | 0.694275 | 0.720781 | 0.719229 |
| 0.716643 | 0.704854 | 0.713192 | 0.721067 | 0.695715 | 0.718692 | 0.689799 | 0.715694 | 0.716939 |
| 0.711511 | 0.702152 | 0.711082 | 0.717023 | 0.690407 | 0.712886 | 0.688485 | 0.717513 | 0.717425 |
| 0.713621 | 0.704222 | 0.713048 | 0.718005 | 0.693685 | 0.718567 | 0.687606 | 0.715188 | 0.721042 |
| 0.715499 | 0.699685 | 0.712001 | 0.714858 | 0.693773 | 0.715217 | 0.691754 | 0.713576 | 0.715777 |
| 0.715558 | 0.701021 | 0.711834 | 0.714215 | 0.695242 | 0.717518 | 0.691474 | 0.714987 | 0.714589 |
| 0.702743 | 0.692031 | 0.702195 | 0.704626 | 0.687516 | 0.707056 | 0.687798 | 0.702626 | 0.706719 |
| 0.708995 | 0.69255 | 0.703233 | 0.711607 | 0.68769 | 0.710837 | 0.691404 | 0.707906 | 0.715226 |
| 0.718464 | 0.705388 | 0.714437 | 0.719198 | 0.69938 | 0.723248 | 0.695833 | 0.718127 | 0.718712 |
| 0.719632 | 0.710176 | 0.717701 | 0.724158 | 0.700016 | 0.723827 | 0.697878 | 0.719182 | 0.723219 |
| 0.715052 | 0.705929 | 0.712111 | 0.718386 | 0.700343 | 0.72203 | 0.691693 | 0.717383 | 0.717764 |
| 0.720068 | 0.710062 | 0.717291 | 0.724241 | 0.701175 | 0.72232 | 0.695647 | 0.719412 | 0.724694 |
| 0.717824 | 0.701384 | 0.709254 | 0.714751 | 0.691602 | 0.717296 | 0.693463 | 0.712011 | 0.715388 |
| 0.714274 | 0.703138 | 0.706964 | 0.715403 | 0.692158 | 0.720112 | 0.688442 | 0.710657 | 0.714605 |
| 0.710679 | 0.69882 | 0.706173 | 0.714525 | 0.688298 | 0.718463 | 0.689613 | 0.713607 | 0.712726 |
| 0.715901 | 0.703094 | 0.713793 | 0.722136 | 0.695235 | 0.722173 | 0.69553 | 0.718631 | 0.722687 |
| 0.714967 | 0.707568 | 0.712161 | 0.719937 | 0.69817 | 0.720747 | 0.697428 | 0.714416 | 0.720159 |
| 0.713593 | 0.703379 | 0.706781 | 0.717446 | 0.694066 | 0.713959 | 0.688623 | 0.715601 | 0.713967 |
| 0.716439 | 0.704024 | 0.712085 | 0.715362 | 0.693417 | 0.717942 | 0.696974 | 0.711107 | 0.719476 |
| 0.780252 | 0.772992 | 0.786365 | 0.781695 | 0.757789 | 0.771766 | 0.744021 | 0.780042 | 0.783175 |
| 0.713436 | 0.704801 | 0.713508 | 0.714879 | 0.694208 | 0.719545 | 0.688918 | 0.713751 | 0.71714 |
| 0.714874 | 0.709211 | 0.712749 | 0.721689 | 0.700023 | 0.720323 | 0.68646 | 0.717992 | 0.723992 |
| 0.71754 | 0.709677 | 0.720176 | 0.719899 | 0.699549 | 0.721481 | 0.695969 | 0.721288 | 0.722897 |
| 0.715621 | 0.705023 | 0.710537 | 0.720181 | 0.697151 | 0.71869 | 0.691583 | 0.716032 | 0.720413 |
| 0.710533 | 0.703215 | 0.708857 | 0.711559 | 0.689446 | 0.716031 | 0.691513 | 0.712896 | 0.717358 |
| 0.711425 | 0.701248 | 0.709863 | 0.715428 | 0.695391 | 0.7176 | 0.688353 | 0.714807 | 0.714806 |
| 0.711417 | 0.702988 | 0.711696 | 0.720511 | 0.698494 | 0.719825 | 0.6888 | 0.716553 | 0.722441 |
| 0.714056 | 0.700208 | 0.711006 | 0.715189 | 0.693651 | 0.714962 | 0.687907 | 0.716582 | 0.716368 |
| 0.710049 | 0.701721 | 0.709821 | 0.712954 | 0.692181 | 0.717195 | 0.688925 | 0.709677 | 0.715544 |
| 0.710093 | 0.701603 | 0.709389 | 0.714097 | 0.695719 | 0.716747 | 0.682847 | 0.713279 | 0.718634 |
| 0.713393 | 0.707533 | 0.716244 | 0.719153 | 0.699174 | 0.719314 | 0.695467 | 0.718035 | 0.721603 |
| 0.716194 | 0.705604 | 0.716871 | 0.717003 | 0.699564 | 0.723842 | 0.695857 | 0.718011 | 0.721768 |
| 0.718165 | 0.705418 | 0.713324 | 0.719354 | 0.699801 | 0.719267 | 0.691283 | 0.716309 | 0.719717 |
| 0.715491 | 0.704655 | 0.712902 | 0.715751 | 0.695086 | 0.717768 | 0.694527 | 0.718928 | 0.716713 |
| 0.709946 | 0.701807 | 0.708292 | 0.709823 | 0.692913 | 0.712375 | 0.686585 | 0.714214 | 0.715434 |
| 0.721202 | 0.709172 | 0.716775 | 0.722651 | 0.699183 | 0.722879 | 0.695935 | 0.717878 | 0.723366 |
| 0.7113 | 0.700571 | 0.710854 | 0.713127 | 0.690521 | 0.716973 | 0.692091 | 0.712561 | 0.720837 |
| 0.714217 | 0.704643 | 0.709632 | 0.716463 | 0.697164 | 0.716643 | 0.685172 | 0.714286 | 0.716568 |
| 0.715702 | 0.70553 | 0.712772 | 0.716018 | 0.695909 | 0.720661 | 0.693724 | 0.720048 | 0.717881 |
| 0.718917 | 0.70678 | 0.712935 | 0.716323 | 0.69898 | 0.722566 | 0.690824 | 0.719589 | 0.721727 |

| | | | | | | | | |
|---|---|---|---|---|---|---|---|---|
| 0.710854 | 0.704704 | 0.710572 | 0.716522 | 0.695639 | 0.719098 | 0.687005 | 0.715196 | 0.717838 |
| 0.717263 | 0.705347 | 0.710385 | 0.713897 | 0.697556 | 0.722205 | 0.689417 | 0.714206 | 0.71895 |
| 0.712631 | 0.703004 | 0.711191 | 0.71476 | 0.699992 | 0.714138 | 0.683855 | 0.712154 | 0.718188 |
| 0.675211 | 0.666783 | 0.676573 | 0.678068 | 0.653557 | 0.68487 | 0.668506 | 0.676594 | 0.67546 |
| 0.71479 | 0.699708 | 0.709863 | 0.716148 | 0.694494 | 0.718001 | 0.68623 | 0.713421 | 0.71771 |
| 0.713939 | 0.703036 | 0.70832 | 0.714596 | 0.699576 | 0.71844 | 0.686638 | 0.713053 | 0.712845 |
| 0.712087 | 0.705812 | 0.711645 | 0.718632 | 0.698596 | 0.718067 | 0.689481 | 0.716414 | 0.716215 |
| 0.71609 | 0.705322 | 0.710151 | 0.718479 | 0.698053 | 0.721774 | 0.687103 | 0.711796 | 0.718633 |
| 0.719331 | 0.707192 | 0.717827 | 0.720399 | 0.701739 | 0.724012 | 0.698624 | 0.720099 | 0.721111 |
| 0.716884 | 0.707243 | 0.710716 | 0.715295 | 0.69938 | 0.71728 | 0.691203 | 0.717971 | 0.717224 |
| 0.719744 | 0.707482 | 0.715421 | 0.716 | 0.697941 | 0.717825 | 0.68735 | 0.711803 | 0.718301 |
| 0.721284 | 0.706131 | 0.718628 | 0.725423 | 0.703044 | 0.723592 | 0.698858 | 0.715683 | 0.719663 |
| 0.718138 | 0.709162 | 0.715102 | 0.719352 | 0.699507 | 0.721588 | 0.689602 | 0.715202 | 0.719805 |
| 0.711316 | 0.701856 | 0.709797 | 0.712029 | 0.698178 | 0.719991 | 0.691561 | 0.713445 | 0.719776 |
| 0.721051 | 0.709344 | 0.716077 | 0.719665 | 0.701872 | 0.724215 | 0.69278 | 0.720064 | 0.724027 |
| 0.712843 | 0.70119 | 0.71106 | 0.716276 | 0.696759 | 0.720178 | 0.688027 | 0.71311 | 0.718753 |
| 0.717517 | 0.706369 | 0.713579 | 0.720201 | 0.700897 | 0.723379 | 0.693432 | 0.716067 | 0.722898 |
| 0.716828 | 0.709406 | 0.71767 | 0.716004 | 0.698122 | 0.725305 | 0.69156 | 0.720994 | 0.719272 |
| 0.712259 | 0.700616 | 0.707535 | 0.715559 | 0.698368 | 0.718659 | 0.687349 | 0.713761 | 0.717715 |
| 0.714067 | 0.7057 | 0.713416 | 0.712705 | 0.697553 | 0.725085 | 0.688972 | 0.715405 | 0.719035 |
| 0.717657 | 0.705021 | 0.712175 | 0.720087 | 0.697935 | 0.72362 | 0.691983 | 0.711778 | 0.723974 |
| 0.718034 | 0.708456 | 0.715609 | 0.721873 | 0.702238 | 0.724157 | 0.692021 | 0.715883 | 0.727733 |
| 0.698921 | 0.694536 | 0.698161 | 0.703016 | 0.672682 | 0.695479 | 0.696309 | 0.698713 | 0.703574 |
| 0.765883 | 0.71488 | 0.712797 | 0.725307 | 0.697006 | 0.723014 | 0.720644 | 0.70669 | 0.724002 |
| 0.757555 | 0.774725 | 0.748557 | 0.751568 | 0.73266 | 0.758897 | 0.737127 | 0.744045 | 0.743521 |
| 0.747081 | 0.722845 | 0.747657 | 0.727195 | 0.711724 | 0.725572 | 0.737679 | 0.736446 | 0.729707 |
| 0.670881 | 0.665079 | 0.6719 | 0.676954 | 0.655021 | 0.674282 | 0.672705 | 0.674527 | 0.673796 |
| 0.667285 | 0.660566 | 0.661716 | 0.667533 | 0.639812 | 0.67033 | 0.67425 | 0.66741 | 0.665989 |
| 0.672562 | 0.66064 | 0.665346 | 0.670157 | 0.651463 | 0.682102 | 0.670461 | 0.666901 | 0.663127 |
| 0.688041 | 0.676139 | 0.685997 | 0.689774 | 0.66627 | 0.688725 | 0.691705 | 0.684317 | 0.683046 |
| 0.666001 | 0.655952 | 0.646892 | 0.666368 | 0.641591 | 0.666012 | 0.668603 | 0.657455 | 0.666593 |
| 0.658033 | 0.645497 | 0.655961 | 0.654162 | 0.638326 | 0.653165 | 0.667677 | 0.652198 | 0.64873 |
| 0.652374 | 0.649052 | 0.652581 | 0.657274 | 0.63583 | 0.6566 | 0.662128 | 0.649886 | 0.656963 |
| 0.735936 | 0.732285 | 0.727572 | 0.724386 | 0.732811 | 0.728699 | 0.744637 | 0.727021 | 0.723231 |
| 0.711098 | 0.700706 | 0.702776 | 0.709028 | 0.691634 | 0.7165 | 0.688294 | 0.707792 | 0.708684 |
| 0.685402 | 0.672882 | 0.68282 | 0.686319 | 0.667734 | 0.685014 | 0.682885 | 0.683919 | 0.682685 |
| 0.6783 | 0.665668 | 0.672425 | 0.675304 | 0.655045 | 0.679049 | 0.683121 | 0.674879 | 0.673195 |
| 0.662938 | 0.66094 | 0.656 | 0.663667 | 0.643973 | 0.660151 | 0.668581 | 0.658862 | 0.665108 |
| 0.690948 | 0.674385 | 0.678599 | 0.688039 | 0.664898 | 0.687164 | 0.685141 | 0.683228 | 0.686777 |
| 0.67639 | 0.667657 | 0.670436 | 0.677311 | 0.65453 | 0.676608 | 0.679795 | 0.670504 | 0.675679 |
| 0.669723 | 0.65414 | 0.664988 | 0.67275 | 0.654184 | 0.669148 | 0.673282 | 0.666 | 0.668825 |
| 0.670089 | 0.666013 | 0.66587 | 0.678185 | 0.65332 | 0.671542 | 0.675054 | 0.665811 | 0.671308 |
| 0.663582 | 0.657366 | 0.66435 | 0.666085 | 0.641074 | 0.663979 | 0.670277 | 0.660259 | 0.670056 |
| 0.660259 | 0.660252 | 0.664649 | 0.666349 | 0.648613 | 0.665663 | 0.671327 | 0.668259 | 0.664826 |
| 0.664688 | 0.663087 | 0.657521 | 0.67156 | 0.641231 | 0.673035 | 0.67694 | 0.660429 | 0.663843 |
| 0.645767 | 0.641141 | 0.653055 | 0.652146 | 0.626431 | 0.657008 | 0.66328 | 0.645933 | 0.655084 |

| | | | | | | | | |
|---|---|---|---|---|---|---|---|---|
| 0.698769 | 0.685701 | 0.691391 | 0.701995 | 0.680324 | 0.7002 | 0.69066 | 0.696654 | 0.700968 |
| 0.668027 | 0.658269 | 0.66104 | 0.667977 | 0.647495 | 0.671858 | 0.672769 | 0.671392 | 0.665807 |
| 0.670741 | 0.660712 | 0.668158 | 0.677065 | 0.65022 | 0.674322 | 0.676851 | 0.672719 | 0.674829 |
| 0.674985 | 0.654032 | 0.667694 | 0.667355 | 0.653256 | 0.675573 | 0.679737 | 0.670028 | 0.670658 |
| 0.66575 | 0.654542 | 0.662257 | 0.673977 | 0.642625 | 0.668414 | 0.673201 | 0.663327 | 0.667785 |
| 0.667475 | 0.663046 | 0.668588 | 0.678832 | 0.648833 | 0.6673 | 0.679827 | 0.664402 | 0.672714 |
| 0.657371 | 0.65717 | 0.658554 | 0.663925 | 0.645095 | 0.668988 | 0.675439 | 0.658973 | 0.658365 |
| 0.652304 | 0.651182 | 0.651081 | 0.66201 | 0.635977 | 0.663784 | 0.67337 | 0.659757 | 0.662188 |
| 0.660695 | 0.648671 | 0.652906 | 0.659592 | 0.638808 | 0.659085 | 0.667269 | 0.655476 | 0.65202 |
| 0.700594 | 0.686032 | 0.691721 | 0.702099 | 0.683458 | 0.701598 | 0.697152 | 0.694334 | 0.701028 |
| 0.699348 | 0.684794 | 0.689268 | 0.694995 | 0.679579 | 0.703119 | 0.692328 | 0.694077 | 0.695512 |
| 0.694657 | 0.681299 | 0.685532 | 0.69064 | 0.671952 | 0.69478 | 0.687329 | 0.689729 | 0.691497 |
| 0.68299 | 0.676284 | 0.685329 | 0.684849 | 0.665328 | 0.686359 | 0.681024 | 0.682616 | 0.686254 |
| 0.686636 | 0.680646 | 0.68929 | 0.688638 | 0.668365 | 0.696713 | 0.689732 | 0.684173 | 0.691138 |
| 0.681758 | 0.670018 | 0.675963 | 0.682165 | 0.657907 | 0.683214 | 0.682306 | 0.680579 | 0.686842 |

| Collection_ | Collection_ | Collection_ | Collection_ | Collection_ | Collection_ | Collection_ | Collection_ | Collection_ |
|---|---|---|---|---|---|---|---|---|
| 0.743994 | 0.753383 | 0.761899 | 0.761654 | 0.774423 | 0.743575 | 0.787844 | 0.765532 | 0.773952 |
| 0.759067 | 0.752511 | 0.768445 | 0.76147 | 0.778937 | 0.743753 | 0.779455 | 0.769392 | 0.766193 |
| 0.7358 | 0.756391 | 0.762209 | 0.76713 | 0.770908 | 0.740829 | 0.766853 | 0.769324 | 0.765947 |
| 0.757088 | 0.75214 | 0.760767 | 0.777694 | 0.78237 | 0.749741 | 0.780706 | 0.764241 | 0.76448 |
| 0.734624 | 0.742802 | 0.748514 | 0.760113 | 0.770725 | 0.749261 | 0.764141 | 0.763858 | 0.755335 |
| 0.742646 | 0.742908 | 0.759168 | 0.775755 | 0.777932 | 0.742494 | 0.770112 | 0.76513 | 0.74812 |
| 0.688029 | 0.68574 | 0.69287 | 0.70834 | 0.723552 | 0.685381 | 0.723445 | 0.727416 | 0.694945 |
| 0.68492 | 0.685886 | 0.688543 | 0.708255 | 0.715102 | 0.683396 | 0.720917 | 0.725603 | 0.693865 |
| 0.762118 | 0.745669 | 0.763913 | 0.779506 | 0.791801 | 0.750921 | 0.770782 | 0.771666 | 0.75388 |
| 0.734869 | 0.749342 | 0.744984 | 0.755391 | 0.764797 | 0.74401 | 0.790816 | 0.763801 | 0.771727 |
| 0.75087 | 0.748173 | 0.752106 | 0.762796 | 0.772824 | 0.737807 | 0.770297 | 0.76215 | 0.763563 |
| 0.757993 | 0.74934 | 0.760324 | 0.760955 | 0.777705 | 0.740785 | 0.763973 | 0.765534 | 0.763527 |
| 0.761098 | 0.757389 | 0.752666 | 0.762491 | 0.772079 | 0.741683 | 0.797576 | 0.765672 | 0.758414 |
| 0.73465 | 0.744477 | 0.759859 | 0.763055 | 0.766212 | 0.746232 | 0.777914 | 0.771943 | 0.761149 |
| 0.73668 | 0.736094 | 0.746971 | 0.755038 | 0.761289 | 0.732362 | 0.769014 | 0.764894 | 0.754316 |
| 0.741839 | 0.754818 | 0.773498 | 0.764804 | 0.775454 | 0.75118 | 0.775114 | 0.764654 | 0.767686 |
| 0.732122 | 0.748656 | 0.758986 | 0.758563 | 0.765487 | 0.738524 | 0.768492 | 0.764093 | 0.76932 |
| 0.745998 | 0.729355 | 0.733698 | 0.73946 | 0.764739 | 0.724149 | 0.763257 | 0.75712 | 0.742642 |
| 0.746721 | 0.755788 | 0.758897 | 0.749311 | 0.77667 | 0.74803 | 0.774117 | 0.765947 | 0.761477 |
| 0.765674 | 0.756193 | 0.757903 | 0.763972 | 0.78424 | 0.74914 | 0.774729 | 0.776732 | 0.76272 |
| 0.777453 | 0.74983 | 0.768483 | 0.766697 | 0.788506 | 0.749319 | 0.770075 | 0.779445 | 0.758037 |
| 0.761295 | 0.753172 | 0.767174 | 0.761789 | 0.777213 | 0.750581 | 0.77221 | 0.76956 | 0.770644 |
| 0.750808 | 0.755574 | 0.768171 | 0.77146 | 0.783717 | 0.753993 | 0.773122 | 0.770301 | 0.769205 |
| 0.765818 | 0.758217 | 0.769423 | 0.762988 | 0.787659 | 0.750962 | 0.775469 | 0.772166 | 0.766932 |
| 0.75778 | 0.750619 | 0.762328 | 0.764745 | 0.785974 | 0.749417 | 0.772297 | 0.772458 | 0.757753 |
| 0.771095 | 0.755474 | 0.76206 | 0.765934 | 0.782094 | 0.742212 | 0.768685 | 0.765463 | 0.764495 |
| 0.738585 | 0.754014 | 0.752248 | 0.751728 | 0.7684 | 0.744783 | 0.765607 | 0.760289 | 0.761193 |
| 0.754609 | 0.751443 | 0.760867 | 0.769609 | 0.781079 | 0.751716 | 0.768739 | 0.773374 | 0.763128 |
| 0.754803 | 0.744874 | 0.764821 | 0.766195 | 0.778625 | 0.749359 | 0.769343 | 0.774939 | 0.764092 |
| 0.742706 | 0.749018 | 0.759011 | 0.75496 | 0.771215 | 0.746201 | 0.77714 | 0.764353 | 0.759657 |
| 0.766997 | 0.743119 | 0.750974 | 0.773092 | 0.782281 | 0.743182 | 0.764677 | 0.767205 | 0.757325 |
| 0.752698 | 0.742303 | 0.751521 | 0.759427 | 0.771715 | 0.744116 | 0.767087 | 0.765766 | 0.751808 |
| 0.738903 | 0.751992 | 0.754345 | 0.765157 | 0.778723 | 0.749621 | 0.769231 | 0.769905 | 0.76395 |
| 0.740732 | 0.744467 | 0.75582 | 0.75377 | 0.77138 | 0.745185 | 0.763135 | 0.767526 | 0.760683 |
| 0.735147 | 0.744591 | 0.749675 | 0.758445 | 0.76764 | 0.742848 | 0.760434 | 0.762798 | 0.75239 |
| 0.73839 | 0.753484 | 0.756289 | 0.756804 | 0.777604 | 0.745425 | 0.781956 | 0.762685 | 0.770868 |
| 0.765619 | 0.751882 | 0.761595 | 0.758824 | 0.78337 | 0.744899 | 0.766597 | 0.773081 | 0.760345 |
| 0.741134 | 0.734258 | 0.747872 | 0.765632 | 0.769752 | 0.737797 | 0.759357 | 0.76494 | 0.746858 |
| 0.755489 | 0.74585 | 0.758706 | 0.771237 | 0.776262 | 0.74828 | 0.774456 | 0.772958 | 0.755904 |

| | | | | | | | | |
|---|---|---|---|---|---|---|---|---|
| 0.725387 | 0.733023 | 0.7374 | 0.743588 | 0.7551 | 0.726298 | 0.766453 | 0.756697 | 0.747005 |
| 0.74192 | 0.754592 | 0.756454 | 0.760084 | 0.7723 | 0.739638 | 0.779941 | 0.764971 | 0.777137 |
| 0.750481 | 0.750298 | 0.77002 | 0.763251 | 0.774569 | 0.744842 | 0.76461 | 0.765308 | 0.766368 |
| 0.741299 | 0.751901 | 0.7574 | 0.760664 | 0.775816 | 0.747659 | 0.787057 | 0.771426 | 0.776648 |
| 0.75635 | 0.74818 | 0.762869 | 0.762096 | 0.791347 | 0.744423 | 0.773059 | 0.768495 | 0.764566 |
| 0.752739 | 0.748655 | 0.761661 | 0.772109 | 0.782967 | 0.752192 | 0.772412 | 0.779778 | 0.761113 |
| 0.741805 | 0.737661 | 0.747387 | 0.758113 | 0.771799 | 0.744455 | 0.76885 | 0.763628 | 0.751915 |
| 0.756917 | 0.749317 | 0.760912 | 0.758875 | 0.782767 | 0.7543 | 0.786889 | 0.77488 | 0.767922 |
| 0.741169 | 0.746461 | 0.758784 | 0.764765 | 0.77499 | 0.751534 | 0.767051 | 0.772642 | 0.752849 |
| 0.773063 | 0.73372 | 0.762016 | 0.784536 | 0.781009 | 0.740514 | 0.771847 | 0.770868 | 0.750427 |
| 0.752751 | 0.749829 | 0.769231 | 0.764311 | 0.779776 | 0.744154 | 0.770687 | 0.77119 | 0.767661 |
| 0.753627 | 0.755914 | 0.763348 | 0.766027 | 0.785514 | 0.74677 | 0.774244 | 0.76661 | 0.764022 |
| 0.746915 | 0.760566 | 0.762213 | 0.758192 | 0.777566 | 0.746472 | 0.784865 | 0.769068 | 0.768327 |
| 0.745216 | 0.749487 | 0.76643 | 0.769672 | 0.775021 | 0.749738 | 0.760299 | 0.765602 | 0.75615 |
| 0.751971 | 0.759026 | 0.755985 | 0.754741 | 0.774311 | 0.741213 | 0.768014 | 0.763139 | 0.765705 |
| 0.758096 | 0.75458 | 0.7544 | 0.756155 | 0.773264 | 0.743571 | 0.758621 | 0.761249 | 0.764789 |
| 0.735958 | 0.75125 | 0.76367 | 0.75543 | 0.766066 | 0.737432 | 0.769035 | 0.761507 | 0.761873 |
| 0.748497 | 0.770367 | 0.762854 | 0.767695 | 0.776221 | 0.752265 | 0.788241 | 0.774507 | 0.783005 |
| 0.754363 | 0.742591 | 0.755483 | 0.760793 | 0.773826 | 0.748378 | 0.776729 | 0.767899 | 0.763711 |
| 0.754591 | 0.754197 | 0.762212 | 0.755338 | 0.777952 | 0.746554 | 0.776465 | 0.765959 | 0.76491 |
| 0.742634 | 0.750963 | 0.775645 | 0.763293 | 0.774839 | 0.744525 | 0.761266 | 0.761689 | 0.771912 |
| 0.744479 | 0.752346 | 0.755099 | 0.763491 | 0.773429 | 0.739141 | 0.782138 | 0.766963 | 0.767394 |
| 0.759793 | 0.757661 | 0.761322 | 0.757241 | 0.778195 | 0.744075 | 0.781258 | 0.765262 | 0.759697 |
| 0.764309 | 0.743387 | 0.755799 | 0.774335 | 0.774721 | 0.740187 | 0.774172 | 0.776002 | 0.753396 |
| 0.758277 | 0.73292 | 0.750647 | 0.761948 | 0.780224 | 0.735062 | 0.770611 | 0.772438 | 0.744098 |
| 0.685995 | 0.688086 | 0.688173 | 0.708084 | 0.717252 | 0.684009 | 0.721552 | 0.725362 | 0.693406 |
| 0.686086 | 0.690765 | 0.686543 | 0.709383 | 0.720193 | 0.689061 | 0.724996 | 0.728078 | 0.700665 |
| 0.688484 | 0.693716 | 0.687516 | 0.709219 | 0.718076 | 0.690872 | 0.724755 | 0.72809 | 0.701466 |
| 0.751346 | 0.750787 | 0.747485 | 0.759876 | 0.772977 | 0.745051 | 0.786616 | 0.764048 | 0.762331 |
| 0.761138 | 0.743322 | 0.752529 | 0.755365 | 0.769367 | 0.739248 | 0.777927 | 0.767185 | 0.755833 |
| 0.766024 | 0.743029 | 0.749903 | 0.753043 | 0.772201 | 0.743419 | 0.787869 | 0.778654 | 0.764522 |
| 0.766719 | 0.74556 | 0.754845 | 0.757924 | 0.775653 | 0.737524 | 0.780762 | 0.769577 | 0.757318 |
| 0.756819 | 0.790218 | 0.781518 | 0.7656 | 0.78009 | 0.754124 | 0.804087 | 0.769658 | 0.786567 |
| 0.743279 | 0.766392 | 0.777484 | 0.758885 | 0.782475 | 0.753805 | 0.771457 | 0.770319 | 0.774967 |
| 0.748675 | 0.777206 | 0.790797 | 0.755168 | 0.774778 | 0.755012 | 0.80521 | 0.768834 | 0.77898 |
| 0.745339 | 0.779143 | 0.801199 | 0.758094 | 0.775443 | 0.753063 | 0.772163 | 0.762197 | 0.788776 |
| 0.746965 | 0.761313 | 0.75863 | 0.783913 | 0.773333 | 0.753681 | 0.811677 | 0.768603 | 0.767948 |
| 0.771491 | 0.794472 | 0.790094 | 0.784029 | 0.795349 | 0.774327 | 0.82202 | 0.788005 | 0.857069 |
| 0.744504 | 0.759884 | 0.769844 | 0.752492 | 0.763366 | 0.778653 | 0.79526 | 0.763665 | 0.771208 |
| 0.772382 | 0.746275 | 0.764683 | 0.756728 | 0.779099 | 0.750457 | 0.773542 | 0.769995 | 0.760345 |
| 0.737531 | 0.752621 | 0.776407 | 0.756564 | 0.773357 | 0.750658 | 0.762984 | 0.756735 | 0.775869 |
| 0.770484 | 0.791674 | 0.792584 | 0.785047 | 0.798751 | 0.772986 | 0.814846 | 0.78904 | 0.835302 |
| 0.740957 | 0.754153 | 0.756877 | 0.762904 | 0.77343 | 0.74836 | 0.771738 | 0.766021 | 0.779111 |
| 0.747131 | 0.748541 | 0.756161 | 0.760302 | 0.773997 | 0.749043 | 0.763446 | 0.770454 | 0.76374 |
| 0.74715 | 0.751345 | 0.759843 | 0.765819 | 0.783113 | 0.753405 | 0.776259 | 0.780524 | 0.758326 |
| 0.745119 | 0.757245 | 0.791422 | 0.759089 | 0.769823 | 0.80163 | 0.771607 | 0.766879 | 0.771041 |

| | | | | | | | | |
|---|---|---|---|---|---|---|---|---|
| 0.747486 | 0.749893 | 0.762284 | 0.768155 | 0.776935 | 0.753482 | 0.766805 | 0.764255 | 0.77549 |
| 0.784278 | 0.743978 | 0.764813 | 0.781558 | 0.792198 | 0.745555 | 0.765286 | 0.769775 | 0.752972 |
| 0.737979 | 0.749209 | 0.767463 | 0.792382 | 0.768252 | 0.752797 | 0.77091 | 0.765189 | 0.76003 |
| 0.762112 | 0.775886 | 0.779663 | 0.777522 | 0.787135 | 0.765726 | 0.805453 | 0.781019 | 0.825738 |
| 0.764275 | 0.753968 | 0.770866 | 0.799796 | 0.786852 | 0.754955 | 0.782025 | 0.770745 | 0.761815 |
| 0.770907 | 0.743982 | 0.765362 | 0.754999 | 0.786999 | 0.74996 | 0.766865 | 0.765351 | 0.753019 |
| 0.762759 | 0.771004 | 0.759439 | 0.801967 | 0.779072 | 0.762614 | 0.817169 | 0.774411 | 0.771064 |
| 0.743633 | 0.744645 | 0.761917 | 0.783604 | 0.779964 | 0.751151 | 0.769212 | 0.7697 | 0.759795 |
| 0.748735 | 0.748273 | 0.760881 | 0.765977 | 0.77426 | 0.755186 | 0.773521 | 0.767081 | 0.759726 |
| 0.731556 | 0.745686 | 0.752249 | 0.751289 | 0.768854 | 0.74408 | 0.76383 | 0.770318 | 0.74949 |
| 0.742134 | 0.748487 | 0.763777 | 0.760561 | 0.771033 | 0.758703 | 0.772052 | 0.769052 | 0.767923 |
| 0.735554 | 0.746462 | 0.756905 | 0.755728 | 0.767862 | 0.74755 | 0.768023 | 0.765436 | 0.754739 |
| 0.744578 | 0.750342 | 0.752455 | 0.759161 | 0.77662 | 0.752205 | 0.762763 | 0.773643 | 0.763173 |
| 0.74199 | 0.752093 | 0.758834 | 0.755349 | 0.767004 | 0.743369 | 0.765035 | 0.759419 | 0.750145 |
| 0.737942 | 0.750299 | 0.763224 | 0.757662 | 0.778199 | 0.752823 | 0.764687 | 0.772403 | 0.754616 |
| 0.73767 | 0.744625 | 0.758167 | 0.767007 | 0.77199 | 0.747368 | 0.765281 | 0.770513 | 0.761012 |
| 0.742981 | 0.74953 | 0.757954 | 0.767827 | 0.776428 | 0.74843 | 0.770355 | 0.769873 | 0.753883 |
| 0.744729 | 0.76325 | 0.771481 | 0.758264 | 0.768784 | 0.748883 | 0.773826 | 0.762169 | 0.774606 |
| 0.738722 | 0.74919 | 0.759719 | 0.764498 | 0.772649 | 0.757334 | 0.774794 | 0.770066 | 0.755572 |
| 0.741611 | 0.745419 | 0.758951 | 0.756542 | 0.776415 | 0.753478 | 0.778697 | 0.771119 | 0.763835 |
| 0.742374 | 0.740063 | 0.756192 | 0.768887 | 0.773667 | 0.744473 | 0.77048 | 0.77279 | 0.755162 |
| 0.743237 | 0.746971 | 0.775713 | 0.760988 | 0.771029 | 0.79236 | 0.767804 | 0.764105 | 0.768085 |
| 0.745785 | 0.745458 | 0.772178 | 0.773522 | 0.772025 | 0.753962 | 0.769874 | 0.763405 | 0.762681 |
| 0.762482 | 0.77474 | 0.757466 | 0.75481 | 0.782099 | 0.745608 | 0.768738 | 0.761432 | 0.755857 |
| 0.736509 | 0.764604 | 0.75 | 0.765394 | 0.765075 | 0.744225 | 0.797269 | 0.769174 | 0.7594 |
| 0.762419 | 0.768398 | 0.755708 | 0.802137 | 0.787077 | 0.756035 | 0.81932 | 0.776255 | 0.769685 |
| 0.737123 | 0.760088 | 0.751613 | 0.765874 | 0.769983 | 0.753999 | 0.795367 | 0.765526 | 0.764595 |
| 0.738599 | 0.74001 | 0.753437 | 0.762275 | 0.772482 | 0.752025 | 0.768653 | 0.76928 | 0.759791 |
| 0.742385 | 0.741535 | 0.756746 | 0.763292 | 0.774025 | 0.751981 | 0.768238 | 0.771493 | 0.759676 |
| 0.739615 | 0.762567 | 0.756835 | 0.766905 | 0.769691 | 0.743532 | 0.785815 | 0.76617 | 0.76232 |
| 0.742314 | 0.741092 | 0.756856 | 0.75962 | 0.793995 | 0.745595 | 0.769426 | 0.772194 | 0.756755 |
| 0.739711 | 0.74666 | 0.758237 | 0.766994 | 0.786942 | 0.752321 | 0.769607 | 0.772054 | 0.759057 |
| 0.749849 | 0.77829 | 0.762123 | 0.762433 | 0.776596 | 0.759814 | 0.815951 | 0.772511 | 0.775616 |
| 0.741363 | 0.751974 | 0.760477 | 0.757863 | 0.771513 | 0.748279 | 0.789181 | 0.768008 | 0.778548 |
| 0.761669 | 0.751173 | 0.762152 | 0.760981 | 0.782576 | 0.75372 | 0.768045 | 0.770225 | 0.766904 |
| 0.768262 | 0.745637 | 0.768968 | 0.763466 | 0.773654 | 0.745093 | 0.767551 | 0.764068 | 0.761905 |
| 0.735489 | 0.750941 | 0.756234 | 0.753834 | 0.769202 | 0.754052 | 0.767012 | 0.77156 | 0.760904 |
| 0.737461 | 0.732244 | 0.758326 | 0.751181 | 0.749948 | 0.725051 | 0.762514 | 0.744744 | 0.731652 |
| 0.78599 | 0.731805 | 0.760547 | 0.734651 | 0.767206 | 0.726494 | 0.753916 | 0.751894 | 0.743924 |
| 0.734058 | 0.713419 | 0.744776 | 0.748506 | 0.745659 | 0.727451 | 0.746067 | 0.745894 | 0.734428 |
| 0.738703 | 0.751022 | 0.780379 | 0.737472 | 0.75539 | 0.725075 | 0.744943 | 0.742469 | 0.751292 |
| 0.710368 | 0.717159 | 0.738522 | 0.719244 | 0.728277 | 0.711158 | 0.724342 | 0.726584 | 0.717123 |
| 0.73568 | 0.738291 | 0.75309 | 0.730136 | 0.740453 | 0.718796 | 0.736628 | 0.732834 | 0.744463 |
| 0.729529 | 0.727124 | 0.799869 | 0.745239 | 0.740569 | 0.72328 | 0.74212 | 0.73147 | 0.735915 |
| 0.729042 | 0.712733 | 0.75412 | 0.728188 | 0.738242 | 0.717572 | 0.742827 | 0.750588 | 0.724138 |
| 0.739846 | 0.738154 | 0.772436 | 0.749768 | 0.741495 | 0.721009 | 0.760984 | 0.738672 | 0.745103 |
| 0.737547 | 0.738745 | 0.771933 | 0.73046 | 0.74696 | 0.720688 | 0.742623 | 0.73597 | 0.749009 |

| | | | | | | | | |
|---|---|---|---|---|---|---|---|---|
| 0.722793 | 0.714594 | 0.739784 | 0.711908 | 0.740074 | 0.713599 | 0.736598 | 0.73554 | 0.724665 |
| 0.744406 | 0.771405 | 0.784378 | 0.732373 | 0.744174 | 0.722986 | 0.782335 | 0.740225 | 0.759538 |
| 0.735656 | 0.712439 | 0.751456 | 0.721911 | 0.73552 | 0.716044 | 0.742998 | 0.74137 | 0.727709 |
| 0.730431 | 0.720044 | 0.74038 | 0.730545 | 0.736354 | 0.711419 | 0.727029 | 0.731694 | 0.747263 |
| 0.733965 | 0.748151 | 0.753856 | 0.727557 | 0.732266 | 0.723355 | 0.782971 | 0.739136 | 0.741123 |
| 0.72583 | 0.707947 | 0.75648 | 0.781343 | 0.736611 | 0.717775 | 0.733764 | 0.728731 | 0.72749 |
| 0.7511 | 0.726618 | 0.756258 | 0.751546 | 0.735739 | 0.721237 | 0.754511 | 0.735634 | 0.738661 |
| 0.743748 | 0.733586 | 0.775725 | 0.727286 | 0.74 | 0.713964 | 0.733543 | 0.727376 | 0.7397 |
| 0.713871 | 0.749961 | 0.737714 | 0.711998 | 0.717307 | 0.694011 | 0.715941 | 0.724869 | 0.70778 |
| 0.737129 | 0.76622 | 0.745524 | 0.724223 | 0.74059 | 0.725314 | 0.825106 | 0.746877 | 0.75599 |
| 0.739189 | 0.706742 | 0.769503 | 0.791371 | 0.74751 | 0.710955 | 0.727693 | 0.733345 | 0.724435 |
| 0.734641 | 0.734007 | 0.748298 | 0.754945 | 0.762197 | 0.739323 | 0.757966 | 0.752715 | 0.769338 |
| 0.740402 | 0.737023 | 0.748837 | 0.760123 | 0.768021 | 0.753585 | 0.771147 | 0.76699 | 0.756971 |
| 0.761198 | 0.721434 | 0.767065 | 0.790732 | 0.757629 | 0.730919 | 0.760578 | 0.750782 | 0.732609 |
| 0.752889 | 0.754863 | 0.762792 | 0.760903 | 0.770084 | 0.747166 | 0.786893 | 0.768816 | 0.765317 |
| 0.726112 | 0.753216 | 0.748375 | 0.749416 | 0.759312 | 0.756328 | 0.765598 | 0.753542 | 0.753247 |
| 0.745177 | 0.747488 | 0.755202 | 0.754105 | 0.78044 | 0.741052 | 0.768989 | 0.774225 | 0.759547 |
| 0.751895 | 0.755916 | 0.757326 | 0.764637 | 0.779902 | 0.748508 | 0.778563 | 0.774333 | 0.758872 |
| 0.721931 | 0.726417 | 0.734741 | 0.749711 | 0.753079 | 0.739474 | 0.745821 | 0.753338 | 0.756111 |
| 0.76061 | 0.74334 | 0.75279 | 0.803459 | 0.779133 | 0.752027 | 0.785008 | 0.772089 | 0.754105 |
| 0.776881 | 0.718638 | 0.753204 | 0.828447 | 0.764253 | 0.715889 | 0.753552 | 0.740151 | 0.73237 |
| 0.739343 | 0.736044 | 0.74475 | 0.756332 | 0.773706 | 0.739728 | 0.771664 | 0.773415 | 0.753011 |
| 0.740036 | 0.735441 | 0.746466 | 0.76073 | 0.770031 | 0.744132 | 0.783709 | 0.776705 | 0.754527 |
| 1 | 0.716832 | 0.756067 | 0.749296 | 0.831491 | 0.703551 | 0.754209 | 0.732243 | 0.726264 |
| 0.716832 | 1 | 0.732438 | 0.724179 | 0.74349 | 0.742216 | 0.767527 | 0.733129 | 0.768264 |
| 0.756067 | 0.732438 | 1 | 0.744417 | 0.758974 | 0.736143 | 0.743649 | 0.743119 | 0.748894 |
| 0.749296 | 0.724179 | 0.744417 | 1 | 0.768522 | 0.733491 | 0.763706 | 0.757775 | 0.740691 |
| 0.831491 | 0.74349 | 0.758974 | 0.768522 | 1 | 0.740635 | 0.771271 | 0.762902 | 0.752609 |
| 0.703551 | 0.742216 | 0.736143 | 0.733491 | 0.740635 | 1 | 0.748 | 0.739291 | 0.763162 |
| 0.754209 | 0.767527 | 0.743649 | 0.763706 | 0.771271 | 0.748 | 1 | 0.777367 | 0.764543 |
| 0.732243 | 0.733129 | 0.743119 | 0.757775 | 0.762902 | 0.739291 | 0.777367 | 1 | 0.746121 |
| 0.726264 | 0.768264 | 0.748894 | 0.740691 | 0.752609 | 0.763162 | 0.764543 | 0.746121 | 1 |
| 0.726133 | 0.762981 | 0.748255 | 0.74185 | 0.755038 | 0.763809 | 0.766298 | 0.75012 | 0.888285 |
| 0.781443 | 0.740124 | 0.744856 | 0.844396 | 0.789565 | 0.750439 | 0.786139 | 0.775517 | 0.749311 |
| 0.73767 | 0.73565 | 0.744052 | 0.750774 | 0.764838 | 0.745466 | 0.805179 | 0.77405 | 0.74928 |
| 0.751579 | 0.736664 | 0.749579 | 0.752529 | 0.77007 | 0.739912 | 0.778958 | 0.775254 | 0.749612 |
| 0.690569 | 0.72392 | 0.709689 | 0.72478 | 0.724001 | 0.73225 | 0.726124 | 0.721691 | 0.741928 |
| 0.723314 | 0.732452 | 0.740103 | 0.737812 | 0.758551 | 0.739406 | 0.756963 | 0.756244 | 0.746625 |
| 0.693975 | 0.715911 | 0.715266 | 0.733879 | 0.720122 | 0.731628 | 0.723762 | 0.721162 | 0.758972 |
| 0.682697 | 0.705183 | 0.696404 | 0.727113 | 0.717583 | 0.718868 | 0.707407 | 0.712476 | 0.741333 |
| 0.752741 | 0.742687 | 0.755852 | 0.749382 | 0.769187 | 0.738811 | 0.768928 | 0.755935 | 0.758054 |
| 0.693681 | 0.707622 | 0.697861 | 0.721413 | 0.722615 | 0.713444 | 0.713333 | 0.707455 | 0.738695 |
| 0.675475 | 0.720668 | 0.699037 | 0.691614 | 0.707308 | 0.718418 | 0.73189 | 0.709435 | 0.745858 |
| 0.668368 | 0.735865 | 0.68924 | 0.68756 | 0.69979 | 0.722378 | 0.728028 | 0.703441 | 0.747646 |
| 0.7481 | 0.739549 | 0.740945 | 0.752377 | 0.768237 | 0.736061 | 0.77552 | 0.758788 | 0.75587 |
| 0.671315 | 0.719258 | 0.687197 | 0.686079 | 0.706193 | 0.709717 | 0.725999 | 0.70496 | 0.736404 |
| 0.745381 | 0.735768 | 0.748268 | 0.745732 | 0.764067 | 0.737913 | 0.777561 | 0.759183 | 0.760108 |

| | | | | | | | | |
|---|---|---|---|---|---|---|---|---|
| 0.739171 | 0.733472 | 0.741091 | 0.745706 | 0.762259 | 0.7337 | 0.774594 | 0.755246 | 0.753075 |
| 0.739807 | 0.738011 | 0.74955 | 0.74767 | 0.76098 | 0.73538 | 0.774196 | 0.762859 | 0.758542 |
| 0.722754 | 0.729058 | 0.73893 | 0.74119 | 0.758288 | 0.725742 | 0.751655 | 0.758434 | 0.740617 |
| 0.65554 | 0.715134 | 0.678201 | 0.679899 | 0.694746 | 0.71025 | 0.709522 | 0.698931 | 0.741688 |
| 0.662659 | 0.727725 | 0.689509 | 0.692417 | 0.690834 | 0.71955 | 0.731993 | 0.694198 | 0.739058 |
| 0.738961 | 0.747114 | 0.755886 | 0.751948 | 0.766139 | 0.735373 | 0.770177 | 0.761131 | 0.753254 |
| 0.654147 | 0.691942 | 0.674566 | 0.676036 | 0.68816 | 0.71907 | 0.691781 | 0.689653 | 0.816759 |
| 0.702278 | 0.707561 | 0.71346 | 0.727876 | 0.74379 | 0.709726 | 0.73878 | 0.733925 | 0.716809 |
| 0.731152 | 0.740928 | 0.758339 | 0.74408 | 0.752815 | 0.732453 | 0.763147 | 0.756937 | 0.751344 |
| 0.691981 | 0.695459 | 0.704118 | 0.715491 | 0.735313 | 0.698758 | 0.726651 | 0.726044 | 0.706898 |
| 0.719098 | 0.723641 | 0.738532 | 0.739622 | 0.741951 | 0.723538 | 0.765133 | 0.74795 | 0.739091 |
| 0.631579 | 0.608974 | 0.638298 | 0.628378 | 0.611111 | 0.610063 | 0.664516 | 0.672956 | 0.666667 |
| 0.678284 | 0.637692 | 0.687238 | 0.659172 | 0.66937 | 0.635349 | 0.674271 | 0.676251 | 0.648092 |
| 0.665634 | 0.637401 | 0.674401 | 0.66029 | 0.667523 | 0.641462 | 0.669756 | 0.676992 | 0.651009 |
| 0.675924 | 0.6494 | 0.679249 | 0.665036 | 0.672717 | 0.64658 | 0.681936 | 0.682881 | 0.658246 |
| 0.668602 | 0.650584 | 0.696059 | 0.662926 | 0.67302 | 0.640698 | 0.681842 | 0.677647 | 0.655773 |
| 0.606214 | 0.63109 | 0.615329 | 0.617693 | 0.614983 | 0.638731 | 0.620349 | 0.616429 | 0.710284 |
| 0.672528 | 0.650559 | 0.682489 | 0.665382 | 0.677464 | 0.645185 | 0.684539 | 0.681699 | 0.658967 |
| 0.718034 | 0.721321 | 0.733831 | 0.731596 | 0.740524 | 0.718671 | 0.750061 | 0.740376 | 0.726323 |
| 0.706015 | 0.716457 | 0.726116 | 0.731367 | 0.736303 | 0.718258 | 0.751721 | 0.731601 | 0.732971 |
| 0.714964 | 0.727959 | 0.733013 | 0.732235 | 0.742701 | 0.722789 | 0.768198 | 0.750323 | 0.731167 |
| 0.570063 | 0.635434 | 0.58554 | 0.610792 | 0.61619 | 0.648466 | 0.615791 | 0.616759 | 0.663397 |
| 0.553301 | 0.620119 | 0.567657 | 0.592161 | 0.597814 | 0.631357 | 0.60126 | 0.599609 | 0.639925 |
| 0.686341 | 0.683654 | 0.692701 | 0.703521 | 0.715024 | 0.684524 | 0.723915 | 0.722635 | 0.692209 |
| 0.67758 | 0.686294 | 0.696012 | 0.704501 | 0.712233 | 0.687243 | 0.724245 | 0.724878 | 0.696499 |
| 0.676828 | 0.681543 | 0.683575 | 0.697046 | 0.70729 | 0.680309 | 0.71901 | 0.713685 | 0.688367 |
| 0.677386 | 0.681124 | 0.67867 | 0.701436 | 0.710558 | 0.681481 | 0.722293 | 0.717937 | 0.692442 |
| 0.685625 | 0.688487 | 0.692629 | 0.706817 | 0.71584 | 0.688347 | 0.724897 | 0.72363 | 0.697368 |
| 0.687814 | 0.68493 | 0.691204 | 0.702805 | 0.712692 | 0.685893 | 0.725845 | 0.718713 | 0.699485 |
| 0.685582 | 0.686462 | 0.692879 | 0.703942 | 0.712717 | 0.685588 | 0.725631 | 0.722881 | 0.698494 |
| 0.685861 | 0.689317 | 0.691312 | 0.705847 | 0.714977 | 0.685719 | 0.721723 | 0.725786 | 0.695188 |
| 0.68719 | 0.694563 | 0.69465 | 0.714923 | 0.724239 | 0.68886 | 0.727775 | 0.732489 | 0.701111 |
| 0.686968 | 0.68832 | 0.69223 | 0.706839 | 0.722191 | 0.686119 | 0.721347 | 0.726023 | 0.698742 |
| 0.684843 | 0.691715 | 0.690829 | 0.706655 | 0.721123 | 0.68961 | 0.72442 | 0.73151 | 0.697165 |
| 0.68785 | 0.693 | 0.693813 | 0.714321 | 0.722923 | 0.688358 | 0.730977 | 0.73308 | 0.698615 |
| 0.681337 | 0.684882 | 0.692866 | 0.706657 | 0.717202 | 0.68513 | 0.721527 | 0.725445 | 0.693654 |
| 0.68312 | 0.687981 | 0.693386 | 0.706651 | 0.719324 | 0.686143 | 0.724009 | 0.728451 | 0.696258 |
| 0.686178 | 0.69178 | 0.693593 | 0.71283 | 0.718042 | 0.6893 | 0.723534 | 0.727065 | 0.696563 |
| 0.679888 | 0.683354 | 0.686821 | 0.705961 | 0.713386 | 0.682657 | 0.722363 | 0.721573 | 0.68956 |
| 0.686524 | 0.689638 | 0.687163 | 0.707357 | 0.720745 | 0.687133 | 0.724614 | 0.729879 | 0.695954 |
| 0.66133 | 0.63803 | 0.685131 | 0.654019 | 0.660104 | 0.640078 | 0.67349 | 0.672721 | 0.648188 |
| 0.668654 | 0.647016 | 0.682046 | 0.655956 | 0.676234 | 0.640345 | 0.683953 | 0.673344 | 0.656239 |
| 0.684289 | 0.691883 | 0.693949 | 0.709714 | 0.720282 | 0.688359 | 0.72561 | 0.732638 | 0.70134 |
| 0.693368 | 0.691584 | 0.69881 | 0.711902 | 0.723868 | 0.694576 | 0.728776 | 0.731177 | 0.703999 |
| 0.650765 | 0.634083 | 0.662829 | 0.641055 | 0.651596 | 0.625803 | 0.660128 | 0.661029 | 0.640256 |
| 0.682918 | 0.687645 | 0.690368 | 0.710209 | 0.71867 | 0.687685 | 0.723463 | 0.726539 | 0.700781 |
| 0.672008 | 0.681606 | 0.677947 | 0.700985 | 0.708284 | 0.67462 | 0.714579 | 0.715996 | 0.689742 |

| | | | | | | | | |
|---|---|---|---|---|---|---|---|---|
| 0.683905 | 0.685594 | 0.69187 | 0.708255 | 0.714786 | 0.685595 | 0.723529 | 0.727071 | 0.696339 |
| 0.679756 | 0.689469 | 0.691623 | 0.705849 | 0.715043 | 0.686995 | 0.722559 | 0.726688 | 0.69951 |
| 0.687265 | 0.688355 | 0.692895 | 0.712025 | 0.717512 | 0.686723 | 0.726719 | 0.726164 | 0.699278 |
| 0.680221 | 0.688186 | 0.689607 | 0.709788 | 0.717637 | 0.687864 | 0.724853 | 0.727007 | 0.698032 |
| 0.692997 | 0.697823 | 0.697713 | 0.712381 | 0.723097 | 0.694594 | 0.732956 | 0.731399 | 0.704835 |
| 0.684253 | 0.688147 | 0.687311 | 0.707002 | 0.717908 | 0.685168 | 0.721903 | 0.725048 | 0.695236 |
| 0.68719 | 0.686413 | 0.689339 | 0.705074 | 0.715565 | 0.686748 | 0.725589 | 0.726614 | 0.697006 |
| 0.673171 | 0.67875 | 0.686908 | 0.703228 | 0.708156 | 0.678594 | 0.716221 | 0.716389 | 0.688257 |
| 0.678295 | 0.685611 | 0.685185 | 0.704468 | 0.712797 | 0.684868 | 0.721242 | 0.723972 | 0.695184 |
| 0.678517 | 0.684228 | 0.688056 | 0.706372 | 0.717709 | 0.682769 | 0.716238 | 0.722279 | 0.693446 |
| 0.682085 | 0.68772 | 0.690176 | 0.707453 | 0.719787 | 0.687336 | 0.723166 | 0.723072 | 0.697864 |
| 0.678676 | 0.685154 | 0.688167 | 0.705651 | 0.713243 | 0.682192 | 0.719383 | 0.722565 | 0.693349 |
| 0.670627 | 0.680276 | 0.677223 | 0.701084 | 0.707597 | 0.677672 | 0.711996 | 0.715914 | 0.688145 |
| 0.676509 | 0.680014 | 0.681357 | 0.700593 | 0.707081 | 0.679177 | 0.714085 | 0.71486 | 0.688992 |
| 0.69271 | 0.695348 | 0.691622 | 0.712191 | 0.728766 | 0.691525 | 0.730176 | 0.730839 | 0.70344 |
| 0.680273 | 0.691826 | 0.694458 | 0.709432 | 0.716391 | 0.689519 | 0.729854 | 0.728018 | 0.701767 |
| 0.680961 | 0.681664 | 0.690772 | 0.704062 | 0.711347 | 0.679177 | 0.717249 | 0.722213 | 0.693323 |
| 0.677092 | 0.680942 | 0.682337 | 0.70668 | 0.714092 | 0.685033 | 0.719498 | 0.724075 | 0.694436 |
| 0.680619 | 0.68543 | 0.687967 | 0.701373 | 0.714921 | 0.685338 | 0.719668 | 0.724793 | 0.694189 |
| 0.6852 | 0.685015 | 0.687737 | 0.706382 | 0.714217 | 0.684935 | 0.725246 | 0.723778 | 0.690548 |
| 0.674 | 0.683156 | 0.681975 | 0.703249 | 0.709458 | 0.683761 | 0.717976 | 0.716727 | 0.692171 |
| 0.6774 | 0.685525 | 0.687316 | 0.70175 | 0.712493 | 0.681962 | 0.719179 | 0.721766 | 0.694155 |
| 0.689559 | 0.689389 | 0.690829 | 0.706799 | 0.71803 | 0.686326 | 0.725841 | 0.726852 | 0.699154 |
| 0.681839 | 0.686101 | 0.688844 | 0.711165 | 0.717478 | 0.686608 | 0.726281 | 0.725047 | 0.696087 |
| 0.685142 | 0.684338 | 0.685627 | 0.7029 | 0.714988 | 0.68553 | 0.720984 | 0.722076 | 0.694496 |
| 0.681378 | 0.684784 | 0.684237 | 0.704432 | 0.710122 | 0.683331 | 0.717275 | 0.7213 | 0.694369 |
| 0.676744 | 0.681337 | 0.686325 | 0.704699 | 0.712395 | 0.682714 | 0.721963 | 0.718553 | 0.688287 |
| 0.677757 | 0.681594 | 0.685366 | 0.701902 | 0.708472 | 0.678312 | 0.719359 | 0.720348 | 0.68926 |
| 0.676836 | 0.683898 | 0.686066 | 0.708652 | 0.710046 | 0.682286 | 0.721592 | 0.72302 | 0.691089 |
| 0.68304 | 0.687515 | 0.689148 | 0.702291 | 0.714751 | 0.68625 | 0.721236 | 0.725209 | 0.694482 |
| 0.677599 | 0.685907 | 0.690865 | 0.708011 | 0.716448 | 0.687321 | 0.723057 | 0.723814 | 0.698023 |
| 0.683265 | 0.687594 | 0.691118 | 0.709954 | 0.717635 | 0.689346 | 0.723396 | 0.724 | 0.700926 |
| 0.691101 | 0.691291 | 0.700126 | 0.710288 | 0.721017 | 0.694056 | 0.728419 | 0.730014 | 0.705594 |
| 0.688083 | 0.688619 | 0.69449 | 0.712168 | 0.720417 | 0.693589 | 0.730616 | 0.727065 | 0.703904 |
| 0.675148 | 0.681551 | 0.692818 | 0.700586 | 0.708979 | 0.681991 | 0.718091 | 0.721401 | 0.689998 |
| 0.682417 | 0.687531 | 0.687524 | 0.706962 | 0.713583 | 0.685061 | 0.722365 | 0.722009 | 0.696086 |
| 0.684609 | 0.686032 | 0.690537 | 0.708942 | 0.718041 | 0.683587 | 0.721582 | 0.72466 | 0.695336 |
| 0.68256 | 0.685418 | 0.687627 | 0.703004 | 0.713286 | 0.68345 | 0.719808 | 0.72284 | 0.693745 |
| 0.672977 | 0.680582 | 0.679709 | 0.699438 | 0.705235 | 0.679956 | 0.714009 | 0.713782 | 0.68814 |
| 0.683108 | 0.678742 | 0.687308 | 0.695785 | 0.705858 | 0.677608 | 0.715119 | 0.714298 | 0.685778 |
| 0.674358 | 0.678067 | 0.678041 | 0.69594 | 0.703461 | 0.677246 | 0.713454 | 0.712971 | 0.684279 |
| 0.681485 | 0.680576 | 0.682161 | 0.698302 | 0.706758 | 0.675469 | 0.714105 | 0.713425 | 0.686286 |
| 0.673577 | 0.679317 | 0.676707 | 0.702108 | 0.707847 | 0.677903 | 0.716488 | 0.715168 | 0.686258 |
| 0.670314 | 0.673639 | 0.67963 | 0.692157 | 0.70155 | 0.668076 | 0.707667 | 0.709215 | 0.67677 |
| 0.684561 | 0.682002 | 0.688418 | 0.705923 | 0.710383 | 0.68278 | 0.71863 | 0.718851 | 0.692096 |
| 0.67667 | 0.674366 | 0.67945 | 0.697264 | 0.704465 | 0.67341 | 0.712679 | 0.714441 | 0.684568 |

| | | | | | | | | |
|---|---|---|---|---|---|---|---|---|
| 0.687025 | 0.690265 | 0.694621 | 0.711488 | 0.713126 | 0.690787 | 0.724099 | 0.72772 | 0.700622 |
| 0.682591 | 0.686778 | 0.695122 | 0.707723 | 0.718995 | 0.687299 | 0.722151 | 0.723386 | 0.696975 |
| 0.676156 | 0.681231 | 0.683418 | 0.706789 | 0.713883 | 0.680824 | 0.719891 | 0.721734 | 0.690083 |
| 0.678493 | 0.68321 | 0.689369 | 0.703743 | 0.719997 | 0.690012 | 0.724396 | 0.728322 | 0.700055 |
| 0.69309 | 0.693167 | 0.697392 | 0.712646 | 0.725638 | 0.691895 | 0.727359 | 0.734239 | 0.704187 |
| 0.681067 | 0.684875 | 0.692802 | 0.708235 | 0.717955 | 0.689394 | 0.720378 | 0.731249 | 0.698216 |
| 0.676887 | 0.684282 | 0.687941 | 0.70333 | 0.715328 | 0.683223 | 0.719476 | 0.725982 | 0.689459 |
| 0.67858 | 0.685015 | 0.68801 | 0.704417 | 0.71957 | 0.68678 | 0.721787 | 0.724752 | 0.700906 |
| 0.685822 | 0.683308 | 0.690892 | 0.69906 | 0.711844 | 0.681429 | 0.715611 | 0.723515 | 0.694252 |
| 0.68 | 0.68266 | 0.688361 | 0.710075 | 0.717709 | 0.686826 | 0.723144 | 0.727836 | 0.695504 |
| 0.679793 | 0.684013 | 0.693872 | 0.702121 | 0.719527 | 0.688874 | 0.724802 | 0.732611 | 0.69721 |
| 0.688182 | 0.683019 | 0.693241 | 0.705267 | 0.719895 | 0.688648 | 0.721825 | 0.728439 | 0.697757 |
| 0.680157 | 0.682699 | 0.689372 | 0.702939 | 0.7179 | 0.684903 | 0.723155 | 0.723465 | 0.693536 |
| 0.683411 | 0.687366 | 0.687973 | 0.702591 | 0.718454 | 0.682099 | 0.723629 | 0.725678 | 0.695738 |
| 0.680988 | 0.690316 | 0.68851 | 0.707701 | 0.719098 | 0.687322 | 0.724268 | 0.72916 | 0.696967 |
| 0.685116 | 0.686694 | 0.689083 | 0.704139 | 0.719178 | 0.686044 | 0.72378 | 0.726054 | 0.698616 |
| 0.688343 | 0.686715 | 0.696177 | 0.710624 | 0.725018 | 0.687947 | 0.724972 | 0.729715 | 0.696927 |
| 0.683746 | 0.687627 | 0.687804 | 0.709338 | 0.717745 | 0.688541 | 0.725384 | 0.726522 | 0.697782 |
| 0.685303 | 0.693069 | 0.693524 | 0.705916 | 0.721523 | 0.68824 | 0.727222 | 0.731301 | 0.69932 |
| 0.681718 | 0.688845 | 0.690602 | 0.70454 | 0.718453 | 0.686413 | 0.724132 | 0.729949 | 0.694486 |
| 0.683832 | 0.690674 | 0.688199 | 0.709202 | 0.721141 | 0.686746 | 0.727926 | 0.724986 | 0.698372 |
| 0.679769 | 0.688061 | 0.689742 | 0.70297 | 0.718509 | 0.681723 | 0.723939 | 0.723294 | 0.699352 |
| 0.689332 | 0.694646 | 0.690856 | 0.714993 | 0.72413 | 0.690621 | 0.730322 | 0.72929 | 0.700213 |
| 0.684641 | 0.691166 | 0.693783 | 0.70828 | 0.724635 | 0.690559 | 0.725418 | 0.73044 | 0.7 |
| 0.683257 | 0.686764 | 0.687617 | 0.705991 | 0.720963 | 0.686993 | 0.723511 | 0.725579 | 0.70094 |
| 0.682822 | 0.685343 | 0.687106 | 0.707717 | 0.720099 | 0.686146 | 0.724283 | 0.724606 | 0.699318 |
| 0.675011 | 0.685102 | 0.687367 | 0.703945 | 0.715968 | 0.684486 | 0.722353 | 0.724381 | 0.696032 |
| 0.679797 | 0.683088 | 0.688917 | 0.705048 | 0.71622 | 0.685659 | 0.718642 | 0.722888 | 0.693777 |
| 0.685609 | 0.686198 | 0.693293 | 0.710388 | 0.722306 | 0.687289 | 0.723143 | 0.729706 | 0.702076 |
| 0.69606 | 0.692171 | 0.697184 | 0.71165 | 0.728097 | 0.689749 | 0.724459 | 0.728656 | 0.699447 |
| 0.6838 | 0.687615 | 0.689638 | 0.703185 | 0.71975 | 0.686009 | 0.719274 | 0.724235 | 0.695108 |
| 0.695169 | 0.692148 | 0.699149 | 0.707176 | 0.724867 | 0.690686 | 0.725357 | 0.729226 | 0.702048 |
| 0.690759 | 0.689009 | 0.696195 | 0.706582 | 0.723063 | 0.688195 | 0.724045 | 0.732448 | 0.699296 |
| 0.680169 | 0.685193 | 0.691041 | 0.706381 | 0.712486 | 0.680484 | 0.721618 | 0.722854 | 0.694199 |
| 0.684543 | 0.692752 | 0.694094 | 0.704847 | 0.721077 | 0.686961 | 0.725839 | 0.727428 | 0.699396 |
| 0.68787 | 0.68422 | 0.697283 | 0.700255 | 0.715083 | 0.68317 | 0.719518 | 0.720129 | 0.693942 |
| 0.692084 | 0.696069 | 0.694648 | 0.712385 | 0.7272 | 0.694773 | 0.731579 | 0.737509 | 0.702227 |
| 0.685952 | 0.684114 | 0.692532 | 0.706534 | 0.720143 | 0.683842 | 0.721076 | 0.72665 | 0.696167 |
| 0.684423 | 0.689977 | 0.690336 | 0.70616 | 0.718909 | 0.687442 | 0.72254 | 0.728044 | 0.702981 |
| 0.685874 | 0.68783 | 0.690775 | 0.706628 | 0.71824 | 0.68836 | 0.724986 | 0.724243 | 0.70021 |
| 0.693003 | 0.695144 | 0.698325 | 0.712403 | 0.727425 | 0.692415 | 0.73022 | 0.729617 | 0.705277 |
| 0.688198 | 0.691065 | 0.69435 | 0.710693 | 0.722554 | 0.69096 | 0.725939 | 0.732446 | 0.701421 |
| 0.692451 | 0.690858 | 0.69315 | 0.707773 | 0.723088 | 0.690093 | 0.724713 | 0.73391 | 0.700567 |
| 0.684358 | 0.687006 | 0.687246 | 0.704166 | 0.715373 | 0.686157 | 0.722222 | 0.728139 | 0.697058 |
| 0.688982 | 0.692583 | 0.695294 | 0.709139 | 0.72001 | 0.690759 | 0.723901 | 0.729213 | 0.700735 |
| 0.687419 | 0.684431 | 0.684959 | 0.704013 | 0.71661 | 0.687687 | 0.723152 | 0.727514 | 0.693794 |

| | | | | | | | | |
|---|---|---|---|---|---|---|---|---|
| 0.682029 | 0.677504 | 0.687936 | 0.696409 | 0.705478 | 0.674795 | 0.714103 | 0.714312 | 0.683647 |
| 0.737886 | 0.723139 | 0.736537 | 0.755083 | 0.759872 | 0.717625 | 0.755292 | 0.758923 | 0.732565 |
| 0.68112 | 0.684503 | 0.691336 | 0.707254 | 0.715546 | 0.681898 | 0.724792 | 0.727539 | 0.69658 |
| 0.685642 | 0.689734 | 0.696206 | 0.711663 | 0.719991 | 0.685962 | 0.726872 | 0.728029 | 0.69722 |
| 0.682476 | 0.683249 | 0.685071 | 0.704817 | 0.713884 | 0.681087 | 0.717459 | 0.72371 | 0.690574 |
| 0.680908 | 0.685992 | 0.691432 | 0.708553 | 0.71682 | 0.683758 | 0.722196 | 0.723408 | 0.697419 |
| 0.685993 | 0.684749 | 0.691669 | 0.705242 | 0.715202 | 0.682489 | 0.721317 | 0.724341 | 0.694617 |
| 0.683159 | 0.678958 | 0.68764 | 0.701069 | 0.710806 | 0.676072 | 0.718528 | 0.722364 | 0.689201 |
| 0.683625 | 0.689304 | 0.685366 | 0.707641 | 0.718958 | 0.688185 | 0.72605 | 0.723151 | 0.695841 |
| 0.689325 | 0.694057 | 0.703453 | 0.71521 | 0.725712 | 0.691618 | 0.730246 | 0.731252 | 0.702481 |
| 0.686042 | 0.687515 | 0.690638 | 0.71046 | 0.723803 | 0.690473 | 0.726634 | 0.72377 | 0.699353 |
| 0.687223 | 0.692206 | 0.696211 | 0.710057 | 0.723784 | 0.689488 | 0.729007 | 0.729195 | 0.700853 |
| 0.682676 | 0.694495 | 0.696004 | 0.708651 | 0.722945 | 0.694395 | 0.729646 | 0.729764 | 0.703607 |
| 0.688056 | 0.68995 | 0.696638 | 0.709589 | 0.72618 | 0.691517 | 0.729552 | 0.732418 | 0.700545 |
| 0.683255 | 0.68838 | 0.69413 | 0.706641 | 0.717492 | 0.688596 | 0.720851 | 0.725209 | 0.699871 |
| 0.683636 | 0.689857 | 0.692337 | 0.709354 | 0.720586 | 0.690289 | 0.727109 | 0.725399 | 0.699922 |
| 0.688108 | 0.693889 | 0.693927 | 0.71038 | 0.721919 | 0.688177 | 0.728189 | 0.727532 | 0.698424 |
| 0.684033 | 0.692052 | 0.692571 | 0.705628 | 0.721766 | 0.686929 | 0.722949 | 0.727367 | 0.696478 |
| 0.681038 | 0.685923 | 0.694665 | 0.695231 | 0.708884 | 0.683803 | 0.721 | 0.715715 | 0.693602 |
| 0.680143 | 0.683024 | 0.689432 | 0.701078 | 0.713204 | 0.68034 | 0.718024 | 0.719379 | 0.692739 |
| 0.684493 | 0.689169 | 0.695023 | 0.70857 | 0.720019 | 0.688245 | 0.724221 | 0.724848 | 0.698506 |
| 0.674543 | 0.678858 | 0.681292 | 0.696894 | 0.707993 | 0.677422 | 0.712741 | 0.712528 | 0.685209 |
| 0.685634 | 0.688371 | 0.688297 | 0.705901 | 0.717677 | 0.685076 | 0.718515 | 0.720608 | 0.695311 |
| 0.690559 | 0.690161 | 0.69223 | 0.709999 | 0.720726 | 0.688031 | 0.726983 | 0.722705 | 0.700853 |
| 0.687669 | 0.690611 | 0.699295 | 0.707139 | 0.721746 | 0.688806 | 0.72629 | 0.723935 | 0.700252 |
| 0.689846 | 0.692075 | 0.693789 | 0.705014 | 0.718179 | 0.688993 | 0.72121 | 0.721462 | 0.69677 |
| 0.684286 | 0.683028 | 0.68304 | 0.702183 | 0.716594 | 0.682485 | 0.723154 | 0.716944 | 0.695071 |
| 0.68247 | 0.683694 | 0.687708 | 0.70085 | 0.713009 | 0.68447 | 0.719105 | 0.715864 | 0.693815 |
| 0.685121 | 0.683313 | 0.692509 | 0.698354 | 0.710936 | 0.67947 | 0.719209 | 0.719593 | 0.693286 |
| 0.77819 | 0.732789 | 0.744899 | 0.737796 | 0.782717 | 0.738627 | 0.759457 | 0.752094 | 0.743421 |
| 0.769325 | 0.762994 | 0.753828 | 0.758105 | 0.784015 | 0.744169 | 0.813273 | 0.76888 | 0.770481 |
| 0.789721 | 0.742864 | 0.765695 | 0.778421 | 0.797322 | 0.747454 | 0.765727 | 0.764845 | 0.754425 |
| 0.737722 | 0.742403 | 0.759221 | 0.787762 | 0.765646 | 0.739594 | 0.75766 | 0.758738 | 0.767614 |
| 0.742242 | 0.763115 | 0.771907 | 0.799949 | 0.775194 | 0.753576 | 0.795185 | 0.769681 | 0.774401 |
| 0.675698 | 0.678994 | 0.685159 | 0.70198 | 0.713957 | 0.679312 | 0.715966 | 0.718264 | 0.690873 |
| 0.678688 | 0.683948 | 0.688182 | 0.69977 | 0.715727 | 0.68573 | 0.720035 | 0.718563 | 0.693661 |
| 0.685082 | 0.688489 | 0.686047 | 0.709073 | 0.715975 | 0.687623 | 0.723304 | 0.72389 | 0.695586 |
| 0.681214 | 0.684946 | 0.689546 | 0.705781 | 0.71888 | 0.689419 | 0.723426 | 0.724556 | 0.698562 |
| 0.675735 | 0.687225 | 0.687484 | 0.69919 | 0.713312 | 0.683807 | 0.722316 | 0.720419 | 0.697314 |
| 0.687318 | 0.691562 | 0.688187 | 0.706007 | 0.720813 | 0.68986 | 0.723486 | 0.725632 | 0.700232 |
| 0.681234 | 0.68701 | 0.687143 | 0.70564 | 0.716015 | 0.686213 | 0.718617 | 0.72153 | 0.695785 |
| 0.682146 | 0.68604 | 0.688014 | 0.703759 | 0.714529 | 0.687723 | 0.718251 | 0.721861 | 0.695561 |
| 0.677098 | 0.677715 | 0.685624 | 0.700198 | 0.706416 | 0.676127 | 0.713999 | 0.715691 | 0.687724 |
| 0.672659 | 0.676992 | 0.68905 | 0.69808 | 0.70739 | 0.678123 | 0.714359 | 0.715146 | 0.688585 |
| 0.682125 | 0.686481 | 0.689438 | 0.701366 | 0.717484 | 0.68289 | 0.717867 | 0.717312 | 0.696351 |
| 0.680515 | 0.687244 | 0.685743 | 0.705514 | 0.71904 | 0.688352 | 0.721087 | 0.721738 | 0.69811 |

| | | | | | | | | |
|---|---|---|---|---|---|---|---|---|
| 0.688697 | 0.686364 | 0.696449 | 0.708625 | 0.718016 | 0.685185 | 0.720575 | 0.7258 | 0.695974 |
| 0.685151 | 0.685598 | 0.688531 | 0.705748 | 0.716667 | 0.688398 | 0.724061 | 0.724769 | 0.695624 |
| 0.687507 | 0.686116 | 0.698358 | 0.709326 | 0.723077 | 0.688345 | 0.724074 | 0.725212 | 0.701968 |
| 0.68097 | 0.683409 | 0.688279 | 0.706363 | 0.715167 | 0.68695 | 0.720061 | 0.725915 | 0.697324 |
| 0.683068 | 0.685095 | 0.683788 | 0.701789 | 0.714531 | 0.683439 | 0.720693 | 0.722668 | 0.693588 |
| 0.680556 | 0.682383 | 0.684392 | 0.704857 | 0.713744 | 0.68168 | 0.716628 | 0.720523 | 0.694305 |
| 0.687219 | 0.689393 | 0.691943 | 0.708135 | 0.718873 | 0.690356 | 0.727555 | 0.728823 | 0.699641 |
| 0.687774 | 0.686179 | 0.694523 | 0.709998 | 0.718323 | 0.686339 | 0.724163 | 0.723908 | 0.695132 |
| 0.686476 | 0.686708 | 0.689836 | 0.705352 | 0.718169 | 0.683574 | 0.72058 | 0.726653 | 0.693413 |
| 0.683466 | 0.682747 | 0.69081 | 0.706807 | 0.71878 | 0.688265 | 0.722249 | 0.727171 | 0.695227 |
| 0.683033 | 0.686483 | 0.690898 | 0.709503 | 0.718732 | 0.68548 | 0.723363 | 0.726716 | 0.696607 |
| 0.68031 | 0.683512 | 0.690774 | 0.707187 | 0.7149 | 0.683895 | 0.723194 | 0.724653 | 0.695677 |
| 0.67607 | 0.677493 | 0.690269 | 0.698007 | 0.707111 | 0.678138 | 0.714727 | 0.716387 | 0.690073 |
| 0.684861 | 0.676738 | 0.694865 | 0.701604 | 0.710815 | 0.678391 | 0.714612 | 0.718725 | 0.688134 |
| 0.688333 | 0.688059 | 0.693885 | 0.707195 | 0.716895 | 0.68932 | 0.72395 | 0.725692 | 0.700196 |
| 0.692209 | 0.690626 | 0.695873 | 0.71267 | 0.721256 | 0.690445 | 0.729834 | 0.729885 | 0.702467 |
| 0.684636 | 0.689122 | 0.689418 | 0.70906 | 0.717793 | 0.688696 | 0.725833 | 0.728605 | 0.699803 |
| 0.687011 | 0.69211 | 0.695569 | 0.712312 | 0.721946 | 0.689513 | 0.728934 | 0.733736 | 0.702487 |
| 0.683975 | 0.688005 | 0.69208 | 0.706339 | 0.71451 | 0.682289 | 0.719565 | 0.721767 | 0.694734 |
| 0.685491 | 0.685802 | 0.690657 | 0.703952 | 0.715769 | 0.684704 | 0.722921 | 0.723224 | 0.691321 |
| 0.684361 | 0.684508 | 0.687685 | 0.702085 | 0.711686 | 0.682009 | 0.718943 | 0.721068 | 0.691454 |
| 0.687964 | 0.691729 | 0.692033 | 0.709133 | 0.720744 | 0.687451 | 0.726592 | 0.72539 | 0.696785 |
| 0.689831 | 0.691076 | 0.696005 | 0.710828 | 0.720046 | 0.689019 | 0.727205 | 0.72766 | 0.700041 |
| 0.685707 | 0.686448 | 0.691836 | 0.707032 | 0.717083 | 0.682761 | 0.719776 | 0.721273 | 0.69583 |
| 0.688935 | 0.685903 | 0.694821 | 0.707086 | 0.714629 | 0.685652 | 0.723217 | 0.723401 | 0.695185 |
| 0.76755 | 0.745433 | 0.766377 | 0.769028 | 0.788391 | 0.742718 | 0.765548 | 0.761947 | 0.768155 |
| 0.68342 | 0.688874 | 0.693577 | 0.704992 | 0.717579 | 0.685261 | 0.723144 | 0.726336 | 0.696064 |
| 0.686439 | 0.690665 | 0.695119 | 0.70635 | 0.722461 | 0.690913 | 0.725775 | 0.727639 | 0.700738 |
| 0.685597 | 0.691866 | 0.696722 | 0.707011 | 0.722908 | 0.689337 | 0.727499 | 0.7296 | 0.702646 |
| 0.690858 | 0.687583 | 0.692639 | 0.707167 | 0.719541 | 0.686994 | 0.723951 | 0.723463 | 0.700032 |
| 0.683637 | 0.685996 | 0.695253 | 0.707079 | 0.715863 | 0.682982 | 0.719243 | 0.720923 | 0.695309 |
| 0.688068 | 0.686868 | 0.692021 | 0.702292 | 0.717079 | 0.685788 | 0.722541 | 0.722754 | 0.69358 |
| 0.688494 | 0.685278 | 0.687786 | 0.70843 | 0.717278 | 0.687856 | 0.720919 | 0.728073 | 0.695388 |
| 0.689532 | 0.684784 | 0.690683 | 0.702155 | 0.714417 | 0.679212 | 0.718631 | 0.723892 | 0.698526 |
| 0.683708 | 0.683541 | 0.691869 | 0.70649 | 0.714755 | 0.680973 | 0.71823 | 0.718693 | 0.696601 |
| 0.681042 | 0.686915 | 0.691339 | 0.70005 | 0.716913 | 0.682647 | 0.721259 | 0.723981 | 0.69678 |
| 0.687368 | 0.686988 | 0.695435 | 0.71008 | 0.722558 | 0.689202 | 0.726985 | 0.725688 | 0.698549 |
| 0.689687 | 0.688953 | 0.691369 | 0.709754 | 0.724207 | 0.690077 | 0.72497 | 0.726613 | 0.698928 |
| 0.685213 | 0.689313 | 0.690066 | 0.706942 | 0.716602 | 0.691262 | 0.72365 | 0.72606 | 0.69938 |
| 0.689363 | 0.685383 | 0.691909 | 0.707218 | 0.718589 | 0.688104 | 0.719609 | 0.723116 | 0.697258 |
| 0.680572 | 0.683599 | 0.689447 | 0.704657 | 0.71292 | 0.680741 | 0.719459 | 0.724614 | 0.691457 |
| 0.687937 | 0.687018 | 0.688173 | 0.710893 | 0.725167 | 0.690788 | 0.725882 | 0.726922 | 0.699063 |
| 0.686228 | 0.686651 | 0.694409 | 0.704062 | 0.716531 | 0.683687 | 0.721508 | 0.723289 | 0.697765 |
| 0.679717 | 0.689385 | 0.685837 | 0.701529 | 0.714263 | 0.687766 | 0.724204 | 0.722615 | 0.698465 |
| 0.685174 | 0.691804 | 0.693436 | 0.708708 | 0.716836 | 0.689137 | 0.723806 | 0.724414 | 0.697573 |
| 0.679986 | 0.691113 | 0.689276 | 0.710707 | 0.714707 | 0.690023 | 0.72639 | 0.724372 | 0.700915 |

| | | | | | | | | |
|---|---|---|---|---|---|---|---|---|
| 0.682876 | 0.688506 | 0.689113 | 0.703479 | 0.714918 | 0.686184 | 0.722683 | 0.722131 | 0.698366 |
| 0.689635 | 0.690018 | 0.688825 | 0.707436 | 0.715384 | 0.687679 | 0.729917 | 0.72695 | 0.698845 |
| 0.682196 | 0.68764 | 0.688662 | 0.701275 | 0.714148 | 0.686571 | 0.721365 | 0.720956 | 0.69887 |
| 0.667224 | 0.647524 | 0.678058 | 0.666192 | 0.679047 | 0.655079 | 0.684156 | 0.682692 | 0.65846 |
| 0.686529 | 0.688549 | 0.692751 | 0.701299 | 0.712457 | 0.682844 | 0.721246 | 0.720069 | 0.694574 |
| 0.684437 | 0.686618 | 0.690189 | 0.706953 | 0.714843 | 0.684822 | 0.723984 | 0.723525 | 0.696559 |
| 0.688077 | 0.692518 | 0.698235 | 0.704226 | 0.716457 | 0.687113 | 0.722015 | 0.724081 | 0.697406 |
| 0.681119 | 0.687654 | 0.692037 | 0.705992 | 0.718979 | 0.689581 | 0.72746 | 0.724211 | 0.69966 |
| 0.690507 | 0.69182 | 0.698963 | 0.711707 | 0.720643 | 0.693868 | 0.729119 | 0.727724 | 0.703404 |
| 0.683972 | 0.688307 | 0.691636 | 0.706451 | 0.720811 | 0.692846 | 0.724356 | 0.723418 | 0.700763 |
| 0.686803 | 0.687613 | 0.688758 | 0.703361 | 0.71718 | 0.687185 | 0.725557 | 0.723657 | 0.702094 |
| 0.687215 | 0.694246 | 0.699008 | 0.70783 | 0.720505 | 0.692523 | 0.728354 | 0.725461 | 0.701729 |
| 0.685754 | 0.689851 | 0.6941 | 0.707216 | 0.717253 | 0.692516 | 0.724487 | 0.72227 | 0.701922 |
| 0.679655 | 0.685698 | 0.694048 | 0.709387 | 0.715259 | 0.68721 | 0.719167 | 0.722109 | 0.697116 |
| 0.683378 | 0.692536 | 0.697373 | 0.709732 | 0.717181 | 0.693575 | 0.723515 | 0.72543 | 0.703586 |
| 0.676402 | 0.686789 | 0.687587 | 0.705524 | 0.709408 | 0.686585 | 0.721645 | 0.719556 | 0.697484 |
| 0.688606 | 0.693834 | 0.696128 | 0.709196 | 0.720035 | 0.69052 | 0.726722 | 0.723556 | 0.700118 |
| 0.682848 | 0.690417 | 0.693731 | 0.709866 | 0.719415 | 0.690292 | 0.726912 | 0.725367 | 0.701156 |
| 0.677211 | 0.684609 | 0.686445 | 0.702782 | 0.712066 | 0.685354 | 0.719345 | 0.719588 | 0.695752 |
| 0.679219 | 0.687141 | 0.687981 | 0.708513 | 0.713119 | 0.686138 | 0.721458 | 0.721645 | 0.698599 |
| 0.684094 | 0.68961 | 0.697819 | 0.70853 | 0.716094 | 0.689867 | 0.727489 | 0.723948 | 0.70092 |
| 0.685362 | 0.69338 | 0.698178 | 0.707948 | 0.718028 | 0.691384 | 0.724942 | 0.728083 | 0.702825 |
| 0.695545 | 0.669397 | 0.699345 | 0.686548 | 0.705395 | 0.671378 | 0.70645 | 0.698175 | 0.688686 |
| 0.728487 | 0.705495 | 0.734379 | 0.695191 | 0.716203 | 0.689176 | 0.736432 | 0.705812 | 0.718012 |
| 0.730626 | 0.729577 | 0.756388 | 0.729656 | 0.742502 | 0.728606 | 0.741693 | 0.744699 | 0.73983 |
| 0.729584 | 0.715641 | 0.733917 | 0.721059 | 0.730161 | 0.708272 | 0.722972 | 0.725998 | 0.741473 |
| 0.665833 | 0.651947 | 0.684364 | 0.667454 | 0.668484 | 0.640724 | 0.68649 | 0.683756 | 0.658302 |
| 0.667291 | 0.639027 | 0.687193 | 0.661905 | 0.673864 | 0.640519 | 0.674291 | 0.673387 | 0.652918 |
| 0.664093 | 0.646905 | 0.675183 | 0.65928 | 0.666204 | 0.638502 | 0.681024 | 0.682553 | 0.656837 |
| 0.679451 | 0.663943 | 0.691942 | 0.678365 | 0.687538 | 0.653892 | 0.699372 | 0.697347 | 0.66976 |
| 0.655762 | 0.640407 | 0.674614 | 0.649234 | 0.660698 | 0.629803 | 0.675099 | 0.671435 | 0.650498 |
| 0.655546 | 0.626047 | 0.668137 | 0.644686 | 0.646788 | 0.623692 | 0.669487 | 0.664855 | 0.646866 |
| 0.64921 | 0.626158 | 0.672385 | 0.641618 | 0.652161 | 0.622802 | 0.649059 | 0.653187 | 0.633408 |
| 0.720996 | 0.704541 | 0.738847 | 0.713856 | 0.723708 | 0.706208 | 0.732127 | 0.730414 | 0.710709 |
| 0.681099 | 0.677094 | 0.696743 | 0.703331 | 0.711876 | 0.685501 | 0.71632 | 0.717033 | 0.694288 |
| 0.676623 | 0.660275 | 0.688387 | 0.671289 | 0.681828 | 0.65251 | 0.69211 | 0.690439 | 0.671014 |
| 0.674616 | 0.650221 | 0.685813 | 0.66463 | 0.675521 | 0.644641 | 0.68323 | 0.684756 | 0.660336 |
| 0.670705 | 0.63575 | 0.675064 | 0.6501 | 0.661651 | 0.636018 | 0.667732 | 0.670165 | 0.649378 |
| 0.676035 | 0.658086 | 0.687126 | 0.670921 | 0.68235 | 0.656322 | 0.692316 | 0.692153 | 0.667375 |
| 0.669832 | 0.655764 | 0.68977 | 0.663729 | 0.671147 | 0.648792 | 0.684295 | 0.683149 | 0.662591 |
| 0.669449 | 0.639717 | 0.682685 | 0.656495 | 0.664018 | 0.638556 | 0.683677 | 0.676632 | 0.651336 |
| 0.66447 | 0.647796 | 0.681988 | 0.661536 | 0.667068 | 0.641636 | 0.679581 | 0.679575 | 0.662451 |
| 0.666471 | 0.640714 | 0.673427 | 0.64841 | 0.667304 | 0.635024 | 0.67728 | 0.67583 | 0.651963 |
| 0.660941 | 0.641921 | 0.681808 | 0.656971 | 0.662811 | 0.635311 | 0.675671 | 0.676705 | 0.653046 |
| 0.665037 | 0.645327 | 0.684456 | 0.655998 | 0.662142 | 0.643062 | 0.671337 | 0.668302 | 0.651611 |
| 0.654985 | 0.630449 | 0.671005 | 0.643768 | 0.643084 | 0.624769 | 0.659292 | 0.6541 | 0.644916 |

| | | | | | | | | |
|---|---|---|---|---|---|---|---|---|
| 0.685483 | 0.670093 | 0.691963 | 0.688414 | 0.698819 | 0.66586 | 0.706939 | 0.70902 | 0.68 |
| 0.663309 | 0.643098 | 0.668243 | 0.662099 | 0.665584 | 0.644903 | 0.678125 | 0.6779 | 0.653392 |
| 0.660726 | 0.648947 | 0.678655 | 0.661114 | 0.671488 | 0.646695 | 0.673484 | 0.67549 | 0.655212 |
| 0.66501 | 0.647094 | 0.678414 | 0.664041 | 0.668675 | 0.642816 | 0.677282 | 0.680558 | 0.658476 |
| 0.669711 | 0.634627 | 0.677674 | 0.659009 | 0.669557 | 0.637911 | 0.669835 | 0.670739 | 0.651862 |
| 0.672961 | 0.645246 | 0.689861 | 0.654954 | 0.672539 | 0.638683 | 0.675077 | 0.676197 | 0.659118 |
| 0.660852 | 0.641294 | 0.677675 | 0.653435 | 0.658576 | 0.636797 | 0.669844 | 0.666589 | 0.646596 |
| 0.655255 | 0.635113 | 0.674065 | 0.637721 | 0.65648 | 0.627925 | 0.662261 | 0.667208 | 0.639547 |
| 0.652644 | 0.62961 | 0.664815 | 0.641651 | 0.653867 | 0.633669 | 0.661334 | 0.662524 | 0.6503 |
| 0.681382 | 0.668788 | 0.690152 | 0.692233 | 0.698748 | 0.668501 | 0.708518 | 0.706065 | 0.678944 |
| 0.682159 | 0.667558 | 0.691836 | 0.687824 | 0.696152 | 0.665725 | 0.706316 | 0.704532 | 0.683613 |
| 0.679614 | 0.66587 | 0.688489 | 0.678914 | 0.693609 | 0.662858 | 0.701494 | 0.698376 | 0.679316 |
| 0.679256 | 0.660257 | 0.68656 | 0.674814 | 0.683742 | 0.653246 | 0.689583 | 0.694419 | 0.670781 |
| 0.684142 | 0.66361 | 0.689061 | 0.676921 | 0.69427 | 0.660835 | 0.694575 | 0.696487 | 0.669477 |
| 0.673368 | 0.655742 | 0.680406 | 0.669045 | 0.680944 | 0.651661 | 0.690619 | 0.689647 | 0.667178 |

| Collection_ | Collection_ | Collection_ | Collection_ | Collection_ | Collection_ | Collection_ | Collection_ | Collection_ |
|---|---|---|---|---|---|---|---|---|
| 0.774461 | 0.769861 | 0.769807 | 0.766434 | 0.721715 | 0.753372 | 0.726586 | 0.721017 | 0.769193 |
| 0.767759 | 0.769533 | 0.761748 | 0.769693 | 0.721542 | 0.758498 | 0.723951 | 0.7215 | 0.768999 |
| 0.765425 | 0.758872 | 0.756651 | 0.759464 | 0.723224 | 0.751892 | 0.733449 | 0.727152 | 0.763385 |
| 0.7615 | 0.78603 | 0.763982 | 0.770516 | 0.722442 | 0.760333 | 0.731903 | 0.717344 | 0.776307 |
| 0.765419 | 0.765985 | 0.763814 | 0.765062 | 0.719604 | 0.751391 | 0.72977 | 0.726319 | 0.763296 |
| 0.755457 | 0.778122 | 0.766208 | 0.769803 | 0.724097 | 0.763982 | 0.733863 | 0.724475 | 0.765552 |
| 0.696583 | 0.725736 | 0.723125 | 0.724528 | 0.677499 | 0.718824 | 0.673611 | 0.671175 | 0.72173 |
| 0.693666 | 0.721311 | 0.717274 | 0.720205 | 0.670559 | 0.719197 | 0.673134 | 0.669037 | 0.717467 |
| 0.755233 | 0.784555 | 0.765192 | 0.769675 | 0.736155 | 0.757134 | 0.732175 | 0.72996 | 0.770521 |
| 0.773565 | 0.766501 | 0.769669 | 0.767545 | 0.716491 | 0.752747 | 0.722109 | 0.711593 | 0.766089 |
| 0.759259 | 0.775224 | 0.772239 | 0.765868 | 0.725216 | 0.749555 | 0.728954 | 0.71962 | 0.766018 |
| 0.766217 | 0.764639 | 0.756337 | 0.773245 | 0.721176 | 0.755365 | 0.721843 | 0.714186 | 0.78979 |
| 0.757428 | 0.767468 | 0.771297 | 0.76762 | 0.714716 | 0.745709 | 0.726407 | 0.715116 | 0.783618 |
| 0.760429 | 0.76908 | 0.760964 | 0.769756 | 0.718987 | 0.76013 | 0.725433 | 0.721705 | 0.776265 |
| 0.750221 | 0.768182 | 0.757884 | 0.760822 | 0.716265 | 0.755099 | 0.720097 | 0.711354 | 0.766895 |
| 0.765304 | 0.770908 | 0.766718 | 0.764123 | 0.730266 | 0.760975 | 0.732274 | 0.727422 | 0.769992 |
| 0.767092 | 0.767556 | 0.765552 | 0.75772 | 0.719122 | 0.753949 | 0.725004 | 0.709133 | 0.765919 |
| 0.746037 | 0.754372 | 0.751279 | 0.764768 | 0.706184 | 0.751931 | 0.710427 | 0.714973 | 0.789499 |
| 0.766486 | 0.759138 | 0.759398 | 0.76252 | 0.720579 | 0.75684 | 0.725472 | 0.714018 | 0.782469 |
| 0.767195 | 0.771157 | 0.76316 | 0.767042 | 0.725692 | 0.759172 | 0.734699 | 0.724464 | 0.771242 |
| 0.759964 | 0.77113 | 0.766693 | 0.772948 | 0.730484 | 0.755064 | 0.728582 | 0.724243 | 0.774249 |
| 0.769867 | 0.766858 | 0.759423 | 0.770232 | 0.730926 | 0.763094 | 0.727809 | 0.727949 | 0.779475 |
| 0.765684 | 0.784022 | 0.763636 | 0.76882 | 0.73034 | 0.763633 | 0.732714 | 0.729105 | 0.77766 |
| 0.770238 | 0.768217 | 0.771979 | 0.775943 | 0.732129 | 0.756847 | 0.734355 | 0.727237 | 0.777868 |
| 0.757114 | 0.775694 | 0.764929 | 0.769523 | 0.728101 | 0.759521 | 0.730398 | 0.726261 | 0.776985 |
| 0.76442 | 0.773865 | 0.7706 | 0.767083 | 0.720585 | 0.755393 | 0.724267 | 0.720452 | 0.780198 |
| 0.767136 | 0.760854 | 0.752539 | 0.766889 | 0.71475 | 0.751964 | 0.727222 | 0.718692 | 0.77103 |
| 0.762657 | 0.779104 | 0.762414 | 0.766808 | 0.724113 | 0.755899 | 0.731814 | 0.729555 | 0.765043 |
| 0.759031 | 0.783935 | 0.764512 | 0.766746 | 0.730052 | 0.759659 | 0.7288 | 0.723939 | 0.770601 |
| 0.759951 | 0.762841 | 0.777236 | 0.766956 | 0.720272 | 0.748745 | 0.719402 | 0.710663 | 0.76766 |
| 0.75608 | 0.781743 | 0.777435 | 0.769796 | 0.726436 | 0.764076 | 0.723054 | 0.718726 | 0.76708 |
| 0.751008 | 0.775557 | 0.772702 | 0.768216 | 0.720491 | 0.756072 | 0.725229 | 0.718211 | 0.764872 |
| 0.76844 | 0.78155 | 0.765036 | 0.76486 | 0.721738 | 0.757573 | 0.724443 | 0.71217 | 0.766648 |
| 0.75778 | 0.776171 | 0.775209 | 0.764658 | 0.720978 | 0.755217 | 0.72426 | 0.721697 | 0.764844 |
| 0.755572 | 0.76647 | 0.754114 | 0.759029 | 0.715293 | 0.749091 | 0.726915 | 0.715816 | 0.766111 |
| 0.769952 | 0.765562 | 0.770656 | 0.768165 | 0.720006 | 0.75529 | 0.728231 | 0.721965 | 0.772139 |
| 0.759935 | 0.770112 | 0.759987 | 0.774597 | 0.725617 | 0.756709 | 0.72516 | 0.715982 | 0.773739 |
| 0.746938 | 0.784074 | 0.759165 | 0.761795 | 0.721385 | 0.757437 | 0.715683 | 0.711111 | 0.759199 |
| 0.752794 | 0.786943 | 0.764376 | 0.767175 | 0.724163 | 0.765322 | 0.722401 | 0.714308 | 0.766294 |

| | | | | | | | | |
|---|---|---|---|---|---|---|---|---|
| 0.74874 | 0.757168 | 0.759753 | 0.752594 | 0.711507 | 0.74994 | 0.708766 | 0.704753 | 0.791023 |
| 0.781712 | 0.767045 | 0.758433 | 0.758722 | 0.716153 | 0.75006 | 0.731846 | 0.715669 | 0.766175 |
| 0.763946 | 0.759951 | 0.763474 | 0.76628 | 0.72866 | 0.754198 | 0.727731 | 0.724612 | 0.784518 |
| 0.775532 | 0.762578 | 0.778564 | 0.77197 | 0.724536 | 0.758266 | 0.727124 | 0.723047 | 0.783619 |
| 0.76211 | 0.769162 | 0.761056 | 0.767459 | 0.726673 | 0.753589 | 0.724284 | 0.722865 | 0.785369 |
| 0.762704 | 0.786413 | 0.763578 | 0.770547 | 0.730309 | 0.759421 | 0.724629 | 0.720979 | 0.774571 |
| 0.758055 | 0.779739 | 0.76805 | 0.761691 | 0.731021 | 0.755412 | 0.728404 | 0.719929 | 0.763423 |
| 0.773886 | 0.769199 | 0.76566 | 0.768996 | 0.723346 | 0.762867 | 0.732461 | 0.720491 | 0.773895 |
| 0.75715 | 0.778191 | 0.765468 | 0.766595 | 0.732116 | 0.75924 | 0.730833 | 0.712124 | 0.767701 |
| 0.751859 | 0.795372 | 0.76151 | 0.762733 | 0.724474 | 0.75171 | 0.723906 | 0.714407 | 0.760235 |
| 0.772095 | 0.764412 | 0.759034 | 0.76789 | 0.720208 | 0.753876 | 0.720186 | 0.716079 | 0.776575 |
| 0.768612 | 0.762968 | 0.760793 | 0.774717 | 0.723647 | 0.754591 | 0.727539 | 0.719338 | 0.783933 |
| 0.770063 | 0.765117 | 0.758866 | 0.767348 | 0.723229 | 0.757957 | 0.730458 | 0.718985 | 0.768815 |
| 0.765398 | 0.766319 | 0.760194 | 0.770894 | 0.724707 | 0.752014 | 0.729308 | 0.721209 | 0.774588 |
| 0.765604 | 0.760424 | 0.76307 | 0.756444 | 0.71837 | 0.750897 | 0.728797 | 0.718588 | 0.771229 |
| 0.764149 | 0.763025 | 0.760091 | 0.759721 | 0.719106 | 0.750964 | 0.727862 | 0.717717 | 0.76758 |
| 0.765778 | 0.761042 | 0.761754 | 0.764823 | 0.715857 | 0.741676 | 0.723957 | 0.712797 | 0.764045 |
| 0.781466 | 0.771522 | 0.777509 | 0.770874 | 0.727019 | 0.76163 | 0.728561 | 0.726506 | 0.782214 |
| 0.767187 | 0.770338 | 0.764115 | 0.773473 | 0.722853 | 0.75887 | 0.72892 | 0.724407 | 0.772472 |
| 0.765821 | 0.760348 | 0.758079 | 0.767039 | 0.719785 | 0.755054 | 0.725016 | 0.720352 | 0.772429 |
| 0.770416 | 0.760574 | 0.757075 | 0.758524 | 0.722663 | 0.748883 | 0.727929 | 0.720022 | 0.767026 |
| 0.763404 | 0.761948 | 0.764 | 0.761193 | 0.718772 | 0.748787 | 0.721541 | 0.714663 | 0.767713 |
| 0.758794 | 0.764319 | 0.766486 | 0.778271 | 0.729329 | 0.755911 | 0.729147 | 0.730685 | 0.774348 |
| 0.753827 | 0.790844 | 0.776102 | 0.773501 | 0.726349 | 0.7552 | 0.727711 | 0.714464 | 0.770367 |
| 0.745798 | 0.796917 | 0.762152 | 0.768966 | 0.717916 | 0.757972 | 0.72217 | 0.71264 | 0.758479 |
| 0.693471 | 0.721901 | 0.718686 | 0.720763 | 0.670146 | 0.720205 | 0.672748 | 0.669055 | 0.717229 |
| 0.700335 | 0.727625 | 0.724135 | 0.723313 | 0.683354 | 0.723214 | 0.682857 | 0.676161 | 0.723696 |
| 0.701597 | 0.730142 | 0.722121 | 0.725449 | 0.682639 | 0.723684 | 0.681442 | 0.673142 | 0.725578 |
| 0.758138 | 0.762731 | 0.75965 | 0.765625 | 0.722737 | 0.751459 | 0.735948 | 0.7277 | 0.770747 |
| 0.754745 | 0.762718 | 0.761369 | 0.765984 | 0.71863 | 0.749958 | 0.723984 | 0.719319 | 0.768508 |
| 0.766607 | 0.7606 | 0.76575 | 0.770619 | 0.71618 | 0.746834 | 0.722432 | 0.714119 | 0.765012 |
| 0.759427 | 0.76862 | 0.760815 | 0.768034 | 0.721676 | 0.75645 | 0.726896 | 0.719438 | 0.772825 |
| 0.784257 | 0.777049 | 0.769626 | 0.775642 | 0.732669 | 0.766711 | 0.731963 | 0.717118 | 0.781705 |
| 0.77649 | 0.762669 | 0.765178 | 0.767565 | 0.726675 | 0.76228 | 0.725852 | 0.726264 | 0.791574 |
| 0.783397 | 0.771497 | 0.760553 | 0.7712 | 0.730523 | 0.764779 | 0.731848 | 0.71142 | 0.791663 |
| 0.786152 | 0.770688 | 0.766109 | 0.771336 | 0.728379 | 0.759863 | 0.730425 | 0.713829 | 0.779601 |
| 0.76835 | 0.766448 | 0.764532 | 0.76994 | 0.731202 | 0.76854 | 0.741254 | 0.736879 | 0.775809 |
| 0.850745 | 0.794672 | 0.7837 | 0.789342 | 0.744375 | 0.781808 | 0.754738 | 0.736855 | 0.801306 |
| 0.773033 | 0.769408 | 0.759672 | 0.766194 | 0.725176 | 0.755411 | 0.736764 | 0.709188 | 0.77086 |
| 0.763982 | 0.763403 | 0.7657 | 0.777933 | 0.726432 | 0.760528 | 0.733841 | 0.73103 | 0.778021 |
| 0.773902 | 0.758587 | 0.761814 | 0.765215 | 0.727193 | 0.758495 | 0.723726 | 0.712084 | 0.775722 |
| 0.830782 | 0.791107 | 0.784986 | 0.785637 | 0.747567 | 0.778106 | 0.753765 | 0.736759 | 0.794669 |
| 0.778162 | 0.772164 | 0.765995 | 0.77752 | 0.727338 | 0.759131 | 0.736205 | 0.71865 | 0.772843 |
| 0.770063 | 0.767985 | 0.756129 | 0.77177 | 0.725174 | 0.75235 | 0.726533 | 0.720666 | 0.773726 |
| 0.762123 | 0.765453 | 0.765172 | 0.775556 | 0.726208 | 0.763459 | 0.72649 | 0.720846 | 0.765839 |
| 0.777152 | 0.776222 | 0.7669 | 0.772428 | 0.728715 | 0.759613 | 0.753796 | 0.714879 | 0.775801 |

| | | | | | | | | |
|---|---|---|---|---|---|---|---|---|
| 0.772608 | 0.771384 | 0.762776 | 0.767735 | 0.729312 | 0.754869 | 0.733897 | 0.723391 | 0.769619 |
| 0.755613 | 0.760065 | 0.764009 | 0.771264 | 0.733983 | 0.757975 | 0.744096 | 0.746596 | 0.778466 |
| 0.757225 | 0.766645 | 0.767194 | 0.767446 | 0.737723 | 0.75502 | 0.740933 | 0.7499 | 0.769035 |
| 0.817407 | 0.784707 | 0.779024 | 0.784384 | 0.734219 | 0.77233 | 0.744975 | 0.73354 | 0.790369 |
| 0.766848 | 0.791321 | 0.76973 | 0.774035 | 0.74085 | 0.766073 | 0.728592 | 0.736171 | 0.782881 |
| 0.760505 | 0.765789 | 0.763545 | 0.770845 | 0.729728 | 0.763173 | 0.727699 | 0.728148 | 0.774093 |
| 0.770049 | 0.819242 | 0.764004 | 0.767921 | 0.737782 | 0.766074 | 0.733632 | 0.72362 | 0.774978 |
| 0.757675 | 0.767371 | 0.778782 | 0.765359 | 0.740498 | 0.760825 | 0.738004 | 0.745877 | 0.77514 |
| 0.758226 | 0.766417 | 0.765417 | 0.767791 | 0.727675 | 0.763229 | 0.728044 | 0.717945 | 0.764289 |
| 0.755409 | 0.760365 | 0.76437 | 0.762957 | 0.717507 | 0.751049 | 0.725145 | 0.717809 | 0.768437 |
| 0.763226 | 0.766602 | 0.771193 | 0.768051 | 0.734463 | 0.768348 | 0.732524 | 0.728894 | 0.771659 |
| 0.754635 | 0.768759 | 0.76598 | 0.769968 | 0.725604 | 0.758975 | 0.729004 | 0.720699 | 0.764315 |
| 0.766922 | 0.765466 | 0.763804 | 0.762491 | 0.723793 | 0.761737 | 0.731764 | 0.722057 | 0.761307 |
| 0.757806 | 0.755216 | 0.763861 | 0.760615 | 0.727457 | 0.753015 | 0.723921 | 0.720218 | 0.767842 |
| 0.755802 | 0.761951 | 0.763411 | 0.764997 | 0.730464 | 0.761055 | 0.727129 | 0.718428 | 0.76755 |
| 0.756113 | 0.76831 | 0.761676 | 0.764282 | 0.729283 | 0.758079 | 0.72848 | 0.727656 | 0.763345 |
| 0.754706 | 0.768725 | 0.766567 | 0.769958 | 0.727595 | 0.756876 | 0.731353 | 0.723494 | 0.766422 |
| 0.768401 | 0.770733 | 0.773722 | 0.76887 | 0.725999 | 0.759877 | 0.728999 | 0.712547 | 0.771043 |
| 0.759113 | 0.772578 | 0.761786 | 0.769199 | 0.725902 | 0.757241 | 0.731707 | 0.722417 | 0.774465 |
| 0.757731 | 0.766006 | 0.775568 | 0.760187 | 0.720952 | 0.757952 | 0.724752 | 0.720143 | 0.765395 |
| 0.754712 | 0.76377 | 0.768857 | 0.768831 | 0.73519 | 0.757221 | 0.727215 | 0.719276 | 0.766126 |
| 0.773559 | 0.776702 | 0.772724 | 0.767068 | 0.726417 | 0.754368 | 0.751257 | 0.717063 | 0.767587 |
| 0.767101 | 0.773516 | 0.76807 | 0.769231 | 0.726223 | 0.758261 | 0.74291 | 0.731297 | 0.771112 |
| 0.763915 | 0.758151 | 0.765688 | 0.768411 | 0.727541 | 0.757752 | 0.725738 | 0.721186 | 0.7845 |
| 0.757738 | 0.759297 | 0.765659 | 0.76387 | 0.728483 | 0.764658 | 0.722519 | 0.72069 | 0.769104 |
| 0.771059 | 0.820342 | 0.769434 | 0.772273 | 0.730644 | 0.761329 | 0.734298 | 0.732136 | 0.777283 |
| 0.766615 | 0.766253 | 0.767766 | 0.764178 | 0.732628 | 0.765253 | 0.725743 | 0.720084 | 0.770015 |
| 0.754475 | 0.766482 | 0.761764 | 0.763149 | 0.72359 | 0.758228 | 0.722987 | 0.72303 | 0.768346 |
| 0.760778 | 0.767514 | 0.763881 | 0.771097 | 0.729868 | 0.764496 | 0.731936 | 0.722145 | 0.768537 |
| 0.76299 | 0.761401 | 0.759417 | 0.762827 | 0.730953 | 0.760382 | 0.724504 | 0.724487 | 0.774706 |
| 0.760538 | 0.766232 | 0.763711 | 0.773082 | 0.731629 | 0.764932 | 0.725537 | 0.719279 | 0.7724 |
| 0.757096 | 0.769486 | 0.767007 | 0.765862 | 0.722468 | 0.759843 | 0.738711 | 0.720758 | 0.769714 |
| 0.773613 | 0.772449 | 0.770957 | 0.768585 | 0.730477 | 0.769682 | 0.729471 | 0.71626 | 0.772934 |
| 0.77803 | 0.765409 | 0.780504 | 0.772624 | 0.723555 | 0.756992 | 0.727324 | 0.719039 | 0.786051 |
| 0.765795 | 0.76346 | 0.763377 | 0.770293 | 0.728001 | 0.756909 | 0.733307 | 0.72907 | 0.773077 |
| 0.763003 | 0.779657 | 0.758559 | 0.775836 | 0.721389 | 0.760837 | 0.727862 | 0.723765 | 0.769561 |
| 0.763691 | 0.762333 | 0.761579 | 0.762395 | 0.727083 | 0.754083 | 0.728409 | 0.72104 | 0.76988 |
| 0.735233 | 0.746835 | 0.796508 | 0.749817 | 0.706715 | 0.731313 | 0.721904 | 0.701598 | 0.74552 |
| 0.747362 | 0.740607 | 0.743265 | 0.763051 | 0.706076 | 0.739184 | 0.707856 | 0.700443 | 0.764645 |
| 0.734687 | 0.74992 | 0.740985 | 0.74261 | 0.702651 | 0.731295 | 0.712117 | 0.706991 | 0.744688 |
| 0.754019 | 0.740471 | 0.741108 | 0.750143 | 0.712068 | 0.73549 | 0.705045 | 0.687493 | 0.757185 |
| 0.726563 | 0.717179 | 0.728386 | 0.72639 | 0.684459 | 0.717839 | 0.686131 | 0.685809 | 0.754758 |
| 0.748555 | 0.734414 | 0.735628 | 0.736849 | 0.703306 | 0.723452 | 0.698375 | 0.686701 | 0.744151 |
| 0.736203 | 0.732389 | 0.730727 | 0.741448 | 0.701327 | 0.725267 | 0.713819 | 0.692088 | 0.747407 |
| 0.724574 | 0.735125 | 0.725467 | 0.732752 | 0.690685 | 0.730729 | 0.693034 | 0.685889 | 0.738596 |
| 0.748344 | 0.742259 | 0.743531 | 0.741202 | 0.704914 | 0.726705 | 0.714197 | 0.719068 | 0.745448 |
| 0.751631 | 0.739247 | 0.742396 | 0.743398 | 0.699195 | 0.735551 | 0.700268 | 0.676748 | 0.74714 |

| | | | | | | | | |
|---|---|---|---|---|---|---|---|---|
| 0.727284 | 0.723466 | 0.835205 | 0.733451 | 0.687228 | 0.721301 | 0.690301 | 0.682807 | 0.729883 |
| 0.761464 | 0.75223 | 0.728919 | 0.7524 | 0.699277 | 0.731347 | 0.69847 | 0.684197 | 0.752382 |
| 0.723893 | 0.746516 | 0.743919 | 0.73137 | 0.692105 | 0.726963 | 0.688522 | 0.67852 | 0.734711 |
| 0.746991 | 0.730441 | 0.725956 | 0.727493 | 0.690327 | 0.717189 | 0.689079 | 0.678622 | 0.736577 |
| 0.743029 | 0.733254 | 0.731101 | 0.737995 | 0.696893 | 0.72652 | 0.68983 | 0.680176 | 0.758487 |
| 0.722074 | 0.727618 | 0.734923 | 0.73032 | 0.717344 | 0.722246 | 0.716326 | 0.744109 | 0.740071 |
| 0.741251 | 0.736113 | 0.718942 | 0.737442 | 0.698961 | 0.718196 | 0.696406 | 0.697063 | 0.746565 |
| 0.741679 | 0.73707 | 0.722112 | 0.742043 | 0.701684 | 0.725718 | 0.69218 | 0.671835 | 0.745489 |
| 0.71208 | 0.70755 | 0.723886 | 0.716829 | 0.681549 | 0.704148 | 0.682414 | 0.663767 | 0.734145 |
| 0.756584 | 0.74097 | 0.725406 | 0.738271 | 0.691651 | 0.725816 | 0.694191 | 0.678287 | 0.750417 |
| 0.723364 | 0.731328 | 0.731586 | 0.734542 | 0.705774 | 0.716939 | 0.735064 | 0.743538 | 0.733586 |
| 0.768264 | 0.760894 | 0.748143 | 0.758668 | 0.73247 | 0.74565 | 0.728252 | 0.728966 | 0.756402 |
| 0.757236 | 0.776403 | 0.777697 | 0.76581 | 0.730721 | 0.754311 | 0.727124 | 0.717603 | 0.766219 |
| 0.736247 | 0.839005 | 0.75162 | 0.749124 | 0.710042 | 0.729287 | 0.732522 | 0.716994 | 0.745064 |
| 0.762239 | 0.776088 | 0.78332 | 0.774623 | 0.72344 | 0.752521 | 0.723084 | 0.717088 | 0.774924 |
| 0.765606 | 0.756834 | 0.758563 | 0.753852 | 0.733253 | 0.753036 | 0.73346 | 0.720756 | 0.763656 |
| 0.75787 | 0.761051 | 0.760463 | 0.759238 | 0.721052 | 0.752531 | 0.72627 | 0.716881 | 0.767603 |
| 0.75397 | 0.778634 | 0.768969 | 0.767581 | 0.720388 | 0.761581 | 0.731216 | 0.721817 | 0.768286 |
| 0.758632 | 0.747556 | 0.750119 | 0.752549 | 0.733725 | 0.740826 | 0.745152 | 0.734082 | 0.755274 |
| 0.757498 | 0.85885 | 0.776416 | 0.769262 | 0.721999 | 0.754029 | 0.72568 | 0.716747 | 0.772379 |
| 0.72754 | 0.841888 | 0.748661 | 0.742969 | 0.705186 | 0.734524 | 0.710087 | 0.711093 | 0.742066 |
| 0.750669 | 0.771992 | 0.774196 | 0.771726 | 0.728751 | 0.755579 | 0.723989 | 0.717921 | 0.765631 |
| 0.754028 | 0.773934 | 0.770783 | 0.767735 | 0.723087 | 0.754928 | 0.724704 | 0.717927 | 0.760402 |
| 0.726133 | 0.781443 | 0.73767 | 0.751579 | 0.690569 | 0.723314 | 0.693975 | 0.682697 | 0.752741 |
| 0.762981 | 0.740124 | 0.73565 | 0.736664 | 0.72392 | 0.732452 | 0.715911 | 0.705183 | 0.742687 |
| 0.748255 | 0.744856 | 0.744052 | 0.749579 | 0.709689 | 0.740103 | 0.715266 | 0.696404 | 0.755852 |
| 0.74185 | 0.844396 | 0.750774 | 0.752529 | 0.72478 | 0.737812 | 0.733879 | 0.727113 | 0.749382 |
| 0.755038 | 0.789565 | 0.764838 | 0.77007 | 0.724001 | 0.758551 | 0.720122 | 0.717583 | 0.769187 |
| 0.763809 | 0.750439 | 0.745466 | 0.739912 | 0.73225 | 0.739406 | 0.731628 | 0.718868 | 0.738811 |
| 0.766298 | 0.786139 | 0.805179 | 0.778958 | 0.726124 | 0.756963 | 0.723762 | 0.707407 | 0.768928 |
| 0.75012 | 0.775517 | 0.77405 | 0.775254 | 0.721691 | 0.756244 | 0.721162 | 0.712476 | 0.755935 |
| 0.888285 | 0.749311 | 0.74928 | 0.749612 | 0.741928 | 0.746625 | 0.758972 | 0.741333 | 0.758054 |
| 1 | 0.752889 | 0.749801 | 0.75267 | 0.740257 | 0.741827 | 0.7703 | 0.734515 | 0.7536 |
| 0.752889 | 1 | 0.776392 | 0.768506 | 0.722213 | 0.751086 | 0.727987 | 0.710109 | 0.761229 |
| 0.749801 | 0.776392 | 1 | 0.773198 | 0.72025 | 0.753156 | 0.727111 | 0.712776 | 0.761662 |
| 0.75267 | 0.768506 | 0.773198 | 1 | 0.718511 | 0.758629 | 0.724067 | 0.714052 | 0.766696 |
| 0.740257 | 0.722213 | 0.72025 | 0.718511 | 1 | 0.711186 | 0.720555 | 0.734155 | 0.720157 |
| 0.741827 | 0.751086 | 0.753156 | 0.758629 | 0.711186 | 1 | 0.714264 | 0.703093 | 0.751279 |
| 0.7703 | 0.727987 | 0.727111 | 0.724067 | 0.720555 | 0.714264 | 1 | 0.736927 | 0.720724 |
| 0.734515 | 0.710109 | 0.712776 | 0.714052 | 0.734155 | 0.703093 | 0.736927 | 1 | 0.716157 |
| 0.7536 | 0.761229 | 0.761662 | 0.766696 | 0.720157 | 0.751279 | 0.720724 | 0.716157 | 1 |
| 0.735548 | 0.710175 | 0.712174 | 0.720456 | 0.745735 | 0.706397 | 0.738657 | 0.843823 | 0.718417 |
| 0.744607 | 0.705338 | 0.70606 | 0.707697 | 0.71605 | 0.700496 | 0.705363 | 0.754811 | 0.71611 |
| 0.752826 | 0.701711 | 0.69391 | 0.704663 | 0.723398 | 0.691048 | 0.722231 | 0.768497 | 0.705267 |
| 0.75727 | 0.765434 | 0.7593 | 0.768597 | 0.716379 | 0.753316 | 0.717391 | 0.709002 | 0.806017 |
| 0.7378 | 0.700496 | 0.699368 | 0.702154 | 0.719719 | 0.691217 | 0.710146 | 0.732281 | 0.708902 |
| 0.755713 | 0.755099 | 0.759132 | 0.76884 | 0.71284 | 0.750485 | 0.718286 | 0.712313 | 0.847997 |

| | | | | | | | | |
|---|---|---|---|---|---|---|---|---|
| 0.74986 | 0.75838 | 0.759081 | 0.762241 | 0.714286 | 0.746745 | 0.715504 | 0.707567 | 0.826335 |
| 0.753587 | 0.756576 | 0.758662 | 0.763454 | 0.725085 | 0.751425 | 0.716316 | 0.714043 | 0.854088 |
| 0.740744 | 0.757974 | 0.74998 | 0.760312 | 0.709501 | 0.749008 | 0.720543 | 0.70554 | 0.754809 |
| 0.737754 | 0.688518 | 0.689681 | 0.690456 | 0.718409 | 0.683593 | 0.717062 | 0.748617 | 0.700256 |
| 0.743431 | 0.69134 | 0.690699 | 0.6934 | 0.723192 | 0.689655 | 0.718656 | 0.762716 | 0.69176 |
| 0.754036 | 0.761781 | 0.761988 | 0.764557 | 0.723726 | 0.755851 | 0.719293 | 0.712944 | 0.813234 |
| 0.821504 | 0.68907 | 0.692709 | 0.687851 | 0.718125 | 0.685751 | 0.763345 | 0.719388 | 0.690891 |
| 0.718895 | 0.740614 | 0.731566 | 0.739645 | 0.693 | 0.76113 | 0.695567 | 0.689663 | 0.730936 |
| 0.748283 | 0.755518 | 0.758635 | 0.762903 | 0.7164 | 0.755206 | 0.719041 | 0.708391 | 0.784404 |
| 0.702592 | 0.72962 | 0.723811 | 0.731423 | 0.686907 | 0.760105 | 0.68532 | 0.679898 | 0.72312 |
| 0.742251 | 0.751302 | 0.750241 | 0.74659 | 0.710407 | 0.739158 | 0.70327 | 0.694971 | 0.804606 |
| 0.638158 | 0.679487 | 0.662338 | 0.675159 | 0.666667 | 0.664516 | 0.677852 | 0.565476 | 0.636943 |
| 0.644013 | 0.676522 | 0.669667 | 0.670842 | 0.634005 | 0.676059 | 0.626953 | 0.615906 | 0.664294 |
| 0.649351 | 0.674872 | 0.666965 | 0.672492 | 0.630211 | 0.669532 | 0.62275 | 0.612696 | 0.667419 |
| 0.655557 | 0.686972 | 0.678012 | 0.685664 | 0.634835 | 0.678184 | 0.631208 | 0.619264 | 0.679109 |
| 0.660087 | 0.67827 | 0.679907 | 0.680983 | 0.633843 | 0.678655 | 0.630335 | 0.623833 | 0.675738 |
| 0.71036 | 0.623399 | 0.618708 | 0.623217 | 0.660176 | 0.613641 | 0.783777 | 0.661893 | 0.61988 |
| 0.658116 | 0.686764 | 0.676193 | 0.685804 | 0.638257 | 0.681545 | 0.63509 | 0.620387 | 0.675986 |
| 0.729398 | 0.744526 | 0.741611 | 0.748812 | 0.698798 | 0.737058 | 0.701015 | 0.695048 | 0.805261 |
| 0.730691 | 0.747039 | 0.738422 | 0.741935 | 0.700392 | 0.738732 | 0.699889 | 0.693301 | 0.793077 |
| 0.73239 | 0.753616 | 0.745612 | 0.748237 | 0.707315 | 0.738266 | 0.700601 | 0.693444 | 0.799817 |
| 0.66664 | 0.613025 | 0.614492 | 0.616309 | 0.670046 | 0.613961 | 0.657917 | 0.722993 | 0.614972 |
| 0.646293 | 0.599794 | 0.598673 | 0.600637 | 0.653351 | 0.597712 | 0.642072 | 0.687358 | 0.600268 |
| 0.694544 | 0.723554 | 0.719669 | 0.723118 | 0.67515 | 0.722061 | 0.674388 | 0.670054 | 0.730324 |
| 0.696662 | 0.720188 | 0.719846 | 0.724577 | 0.677161 | 0.724446 | 0.674549 | 0.667373 | 0.73477 |
| 0.688826 | 0.716902 | 0.717114 | 0.719919 | 0.669741 | 0.711339 | 0.668825 | 0.657839 | 0.720651 |
| 0.695198 | 0.723012 | 0.720031 | 0.722343 | 0.675917 | 0.717452 | 0.674726 | 0.664877 | 0.724009 |
| 0.697259 | 0.727745 | 0.725415 | 0.726645 | 0.676861 | 0.720834 | 0.677015 | 0.666893 | 0.728985 |
| 0.698781 | 0.725642 | 0.721259 | 0.727495 | 0.675355 | 0.719016 | 0.675462 | 0.667076 | 0.72897 |
| 0.697506 | 0.726952 | 0.723265 | 0.725675 | 0.67507 | 0.722728 | 0.674568 | 0.665455 | 0.730822 |
| 0.697975 | 0.724588 | 0.718231 | 0.725338 | 0.67534 | 0.716339 | 0.674904 | 0.667853 | 0.717618 |
| 0.702728 | 0.730111 | 0.727643 | 0.731172 | 0.682187 | 0.72347 | 0.682852 | 0.680218 | 0.724551 |
| 0.699276 | 0.72474 | 0.723697 | 0.724799 | 0.678339 | 0.719487 | 0.67928 | 0.671876 | 0.721033 |
| 0.698128 | 0.726437 | 0.720292 | 0.726411 | 0.67662 | 0.721597 | 0.677392 | 0.668415 | 0.721427 |
| 0.702545 | 0.731176 | 0.723745 | 0.728493 | 0.678006 | 0.72127 | 0.681675 | 0.671385 | 0.723673 |
| 0.696454 | 0.724626 | 0.716853 | 0.726823 | 0.673845 | 0.719843 | 0.677696 | 0.667223 | 0.718952 |
| 0.69724 | 0.726542 | 0.722938 | 0.722724 | 0.678359 | 0.721729 | 0.678059 | 0.671212 | 0.715748 |
| 0.698075 | 0.726002 | 0.721452 | 0.725449 | 0.678519 | 0.722231 | 0.675653 | 0.6708 | 0.720634 |
| 0.692183 | 0.721775 | 0.718163 | 0.723097 | 0.673922 | 0.71551 | 0.672236 | 0.664983 | 0.716124 |
| 0.697609 | 0.725223 | 0.725373 | 0.726522 | 0.675826 | 0.722681 | 0.674213 | 0.668375 | 0.717189 |
| 0.647548 | 0.676933 | 0.679522 | 0.678782 | 0.631434 | 0.675447 | 0.622176 | 0.612939 | 0.67182 |
| 0.65269 | 0.680754 | 0.676338 | 0.681587 | 0.631063 | 0.677648 | 0.627806 | 0.615582 | 0.682407 |
| 0.702134 | 0.724914 | 0.726112 | 0.729912 | 0.678409 | 0.723182 | 0.677789 | 0.674877 | 0.72705 |
| 0.700314 | 0.730791 | 0.726865 | 0.732193 | 0.68271 | 0.725243 | 0.682112 | 0.672836 | 0.725669 |
| 0.638294 | 0.66568 | 0.659731 | 0.667839 | 0.624821 | 0.661675 | 0.616923 | 0.598604 | 0.663216 |
| 0.697948 | 0.727105 | 0.723054 | 0.727888 | 0.676576 | 0.717957 | 0.676294 | 0.670589 | 0.722311 |
| 0.68999 | 0.718549 | 0.713061 | 0.716257 | 0.668492 | 0.711453 | 0.667553 | 0.66303 | 0.709715 |

| | | | | | | | | |
|---|---|---|---|---|---|---|---|---|
| 0.695371 | 0.724331 | 0.718802 | 0.726328 | 0.675554 | 0.718628 | 0.673675 | 0.667374 | 0.719604 |
| 0.700902 | 0.724423 | 0.725252 | 0.727862 | 0.676407 | 0.72195 | 0.677483 | 0.670656 | 0.724113 |
| 0.701436 | 0.732786 | 0.723582 | 0.727564 | 0.677691 | 0.721057 | 0.67538 | 0.66695 | 0.724701 |
| 0.698398 | 0.729929 | 0.723131 | 0.729256 | 0.678969 | 0.720897 | 0.678741 | 0.671494 | 0.72318 |
| 0.705394 | 0.734954 | 0.728728 | 0.734832 | 0.680404 | 0.729861 | 0.680877 | 0.67362 | 0.734583 |
| 0.694867 | 0.72504 | 0.716642 | 0.723645 | 0.672879 | 0.716785 | 0.673968 | 0.666895 | 0.715776 |
| 0.696276 | 0.726837 | 0.722889 | 0.72728 | 0.676691 | 0.719671 | 0.676421 | 0.668147 | 0.718587 |
| 0.68977 | 0.719101 | 0.714626 | 0.721811 | 0.668496 | 0.71198 | 0.670516 | 0.66325 | 0.716287 |
| 0.694475 | 0.723588 | 0.721099 | 0.723415 | 0.673159 | 0.716977 | 0.673946 | 0.669373 | 0.716503 |
| 0.693073 | 0.718665 | 0.715411 | 0.721327 | 0.674473 | 0.715094 | 0.671689 | 0.667072 | 0.717089 |
| 0.697412 | 0.724174 | 0.722509 | 0.727791 | 0.677049 | 0.722024 | 0.676006 | 0.669632 | 0.72267 |
| 0.694546 | 0.72399 | 0.717154 | 0.723716 | 0.672858 | 0.716592 | 0.671813 | 0.666641 | 0.715543 |
| 0.690122 | 0.715496 | 0.709968 | 0.714795 | 0.669073 | 0.709494 | 0.668011 | 0.662251 | 0.711223 |
| 0.687239 | 0.716939 | 0.713374 | 0.718475 | 0.668117 | 0.709389 | 0.669673 | 0.659286 | 0.709443 |
| 0.703091 | 0.731157 | 0.728254 | 0.731422 | 0.684643 | 0.726761 | 0.680454 | 0.67592 | 0.726921 |
| 0.702696 | 0.72741 | 0.722366 | 0.723774 | 0.680505 | 0.726241 | 0.679942 | 0.67072 | 0.72829 |
| 0.694755 | 0.72274 | 0.713718 | 0.722851 | 0.671259 | 0.715649 | 0.667823 | 0.663384 | 0.719813 |
| 0.696023 | 0.721608 | 0.715612 | 0.722101 | 0.677707 | 0.719958 | 0.672043 | 0.667377 | 0.723335 |
| 0.693752 | 0.720951 | 0.716928 | 0.721678 | 0.677046 | 0.720566 | 0.672265 | 0.664126 | 0.725736 |
| 0.69446 | 0.721844 | 0.721083 | 0.720331 | 0.67576 | 0.718468 | 0.672389 | 0.664417 | 0.723491 |
| 0.690715 | 0.719201 | 0.714549 | 0.715276 | 0.672478 | 0.714702 | 0.670725 | 0.66489 | 0.721381 |
| 0.694164 | 0.718928 | 0.716298 | 0.718298 | 0.673702 | 0.716057 | 0.672512 | 0.662969 | 0.729055 |
| 0.698489 | 0.725942 | 0.725121 | 0.724243 | 0.676939 | 0.718224 | 0.679224 | 0.668424 | 0.727964 |
| 0.698517 | 0.72763 | 0.717836 | 0.724586 | 0.6787 | 0.720847 | 0.675184 | 0.66831 | 0.723587 |
| 0.693536 | 0.724793 | 0.719547 | 0.722326 | 0.673378 | 0.71811 | 0.67396 | 0.665799 | 0.7267 |
| 0.690735 | 0.720135 | 0.715608 | 0.717841 | 0.672622 | 0.713657 | 0.670932 | 0.663642 | 0.732228 |
| 0.689922 | 0.722569 | 0.715708 | 0.714471 | 0.673721 | 0.713539 | 0.673029 | 0.665668 | 0.717125 |
| 0.689976 | 0.719948 | 0.716469 | 0.717798 | 0.67191 | 0.712135 | 0.671149 | 0.664916 | 0.7192 |
| 0.693381 | 0.718476 | 0.71498 | 0.716625 | 0.674737 | 0.718596 | 0.675159 | 0.670926 | 0.731101 |
| 0.696001 | 0.722968 | 0.721413 | 0.718818 | 0.67752 | 0.716694 | 0.673526 | 0.666918 | 0.726051 |
| 0.69651 | 0.726828 | 0.718799 | 0.723242 | 0.679931 | 0.718392 | 0.675383 | 0.670993 | 0.725738 |
| 0.699202 | 0.723776 | 0.720222 | 0.723692 | 0.679905 | 0.721413 | 0.678336 | 0.669168 | 0.724357 |
| 0.703579 | 0.728571 | 0.724895 | 0.73124 | 0.681195 | 0.729822 | 0.679766 | 0.673872 | 0.738757 |
| 0.705088 | 0.734018 | 0.725529 | 0.730878 | 0.681645 | 0.726838 | 0.680457 | 0.671945 | 0.730089 |
| 0.692253 | 0.715667 | 0.717047 | 0.718267 | 0.671581 | 0.716794 | 0.670428 | 0.664665 | 0.729095 |
| 0.696744 | 0.722808 | 0.716862 | 0.721515 | 0.67602 | 0.720003 | 0.676276 | 0.667154 | 0.730588 |
| 0.696156 | 0.722352 | 0.718725 | 0.724945 | 0.676431 | 0.719544 | 0.669667 | 0.665853 | 0.729029 |
| 0.695898 | 0.720639 | 0.720832 | 0.722619 | 0.672867 | 0.714389 | 0.672134 | 0.665773 | 0.723209 |
| 0.687215 | 0.713091 | 0.707447 | 0.71233 | 0.671524 | 0.713025 | 0.667992 | 0.662834 | 0.718474 |
| 0.687309 | 0.716664 | 0.71291 | 0.718721 | 0.66999 | 0.707441 | 0.664665 | 0.655611 | 0.721717 |
| 0.684185 | 0.715549 | 0.709706 | 0.712461 | 0.670205 | 0.707687 | 0.664169 | 0.660574 | 0.723885 |
| 0.685626 | 0.713013 | 0.711083 | 0.709085 | 0.670002 | 0.704762 | 0.665879 | 0.657168 | 0.722783 |
| 0.68651 | 0.715384 | 0.707724 | 0.711664 | 0.670845 | 0.71178 | 0.66632 | 0.660433 | 0.719707 |
| 0.678399 | 0.71102 | 0.705482 | 0.703915 | 0.66227 | 0.702376 | 0.660131 | 0.649662 | 0.713686 |
| 0.691303 | 0.719678 | 0.718724 | 0.719666 | 0.67017 | 0.712236 | 0.671441 | 0.661746 | 0.72086 |
| 0.682617 | 0.711422 | 0.702537 | 0.712325 | 0.662081 | 0.706959 | 0.661761 | 0.652815 | 0.712679 |

| | | | | | | | | |
|---|---|---|---|---|---|---|---|---|
| 0.699334 | 0.725712 | 0.720793 | 0.723296 | 0.677276 | 0.720259 | 0.673258 | 0.671181 | 0.74709 |
| 0.695957 | 0.728071 | 0.72148 | 0.721779 | 0.67955 | 0.71972 | 0.67627 | 0.667765 | 0.716725 |
| 0.694194 | 0.721908 | 0.716297 | 0.720427 | 0.673178 | 0.716026 | 0.669971 | 0.663223 | 0.716771 |
| 0.700564 | 0.722928 | 0.721607 | 0.722116 | 0.677912 | 0.71929 | 0.675495 | 0.671259 | 0.722231 |
| 0.704263 | 0.73167 | 0.728162 | 0.727543 | 0.684399 | 0.727135 | 0.682206 | 0.672199 | 0.729417 |
| 0.696729 | 0.728442 | 0.723267 | 0.723455 | 0.677847 | 0.720146 | 0.676129 | 0.669962 | 0.722764 |
| 0.688854 | 0.719615 | 0.718556 | 0.722594 | 0.674548 | 0.71652 | 0.671606 | 0.665015 | 0.716073 |
| 0.698498 | 0.72421 | 0.719931 | 0.723269 | 0.677063 | 0.719209 | 0.677553 | 0.667246 | 0.71988 |
| 0.691446 | 0.717001 | 0.713659 | 0.716128 | 0.670525 | 0.713839 | 0.670792 | 0.661508 | 0.712974 |
| 0.694563 | 0.723031 | 0.722424 | 0.723142 | 0.6752 | 0.717324 | 0.674 | 0.670488 | 0.717243 |
| 0.698492 | 0.72574 | 0.7274 | 0.72615 | 0.678068 | 0.719921 | 0.675461 | 0.67152 | 0.719868 |
| 0.697817 | 0.72558 | 0.722641 | 0.725805 | 0.675867 | 0.717231 | 0.675705 | 0.669212 | 0.722482 |
| 0.697251 | 0.719319 | 0.718686 | 0.725476 | 0.675831 | 0.71663 | 0.674001 | 0.668611 | 0.718493 |
| 0.695488 | 0.721668 | 0.723741 | 0.722919 | 0.676541 | 0.71432 | 0.677219 | 0.668937 | 0.718879 |
| 0.696189 | 0.725785 | 0.722517 | 0.724928 | 0.67717 | 0.721959 | 0.681787 | 0.672842 | 0.720683 |
| 0.695727 | 0.723608 | 0.720788 | 0.724643 | 0.678452 | 0.718193 | 0.673686 | 0.671913 | 0.719889 |
| 0.698876 | 0.727986 | 0.725513 | 0.725275 | 0.678544 | 0.720227 | 0.67623 | 0.672647 | 0.72637 |
| 0.69947 | 0.724582 | 0.722606 | 0.723468 | 0.673622 | 0.722043 | 0.673744 | 0.671807 | 0.724095 |
| 0.70181 | 0.727505 | 0.724178 | 0.72599 | 0.682105 | 0.722817 | 0.679433 | 0.672536 | 0.723649 |
| 0.697747 | 0.722328 | 0.719853 | 0.727716 | 0.677355 | 0.721989 | 0.672544 | 0.671937 | 0.720363 |
| 0.70004 | 0.729808 | 0.723608 | 0.723923 | 0.677891 | 0.718683 | 0.678238 | 0.667502 | 0.718613 |
| 0.696429 | 0.723529 | 0.721553 | 0.723912 | 0.676565 | 0.716776 | 0.676797 | 0.670966 | 0.717922 |
| 0.701366 | 0.728185 | 0.726931 | 0.73306 | 0.677713 | 0.721234 | 0.678236 | 0.67378 | 0.725529 |
| 0.700368 | 0.727287 | 0.727906 | 0.725231 | 0.678798 | 0.725396 | 0.675171 | 0.672986 | 0.722986 |
| 0.7 | 0.722997 | 0.720784 | 0.725345 | 0.678637 | 0.717463 | 0.6762 | 0.671204 | 0.723754 |
| 0.698584 | 0.722834 | 0.721398 | 0.720429 | 0.676788 | 0.72034 | 0.677367 | 0.670328 | 0.71938 |
| 0.696462 | 0.720193 | 0.717087 | 0.722683 | 0.675301 | 0.720697 | 0.67269 | 0.667518 | 0.719202 |
| 0.695265 | 0.72169 | 0.721845 | 0.719863 | 0.676012 | 0.720864 | 0.676305 | 0.671106 | 0.719584 |
| 0.70277 | 0.725228 | 0.721889 | 0.724815 | 0.676877 | 0.723394 | 0.678838 | 0.671754 | 0.72234 |
| 0.698642 | 0.729558 | 0.725827 | 0.72859 | 0.681665 | 0.723791 | 0.679053 | 0.674179 | 0.723571 |
| 0.693497 | 0.723109 | 0.720667 | 0.720365 | 0.678698 | 0.717395 | 0.674633 | 0.669152 | 0.718526 |
| 0.702632 | 0.727258 | 0.723633 | 0.725457 | 0.682942 | 0.724733 | 0.678403 | 0.676174 | 0.721952 |
| 0.697497 | 0.729205 | 0.72475 | 0.729371 | 0.678995 | 0.72034 | 0.677816 | 0.673842 | 0.723048 |
| 0.693355 | 0.720223 | 0.715877 | 0.722126 | 0.677138 | 0.719639 | 0.674191 | 0.666667 | 0.718301 |
| 0.700317 | 0.723883 | 0.723216 | 0.72444 | 0.678917 | 0.723261 | 0.674433 | 0.671037 | 0.719738 |
| 0.691096 | 0.718413 | 0.713441 | 0.719545 | 0.671908 | 0.720487 | 0.670347 | 0.665504 | 0.717568 |
| 0.701897 | 0.728302 | 0.727416 | 0.730165 | 0.682469 | 0.72559 | 0.680742 | 0.676935 | 0.728928 |
| 0.700056 | 0.725014 | 0.72426 | 0.721078 | 0.674791 | 0.723998 | 0.676182 | 0.666436 | 0.719574 |
| 0.70316 | 0.722697 | 0.720333 | 0.726475 | 0.680128 | 0.719008 | 0.677745 | 0.667196 | 0.723856 |
| 0.698512 | 0.724031 | 0.721331 | 0.720796 | 0.679411 | 0.721816 | 0.676289 | 0.672233 | 0.718929 |
| 0.705632 | 0.730152 | 0.72406 | 0.72752 | 0.684174 | 0.724042 | 0.683162 | 0.67687 | 0.724201 |
| 0.705246 | 0.727994 | 0.726035 | 0.72283 | 0.679046 | 0.723394 | 0.680043 | 0.672448 | 0.724267 |
| 0.703666 | 0.727525 | 0.72701 | 0.72744 | 0.679123 | 0.723742 | 0.678743 | 0.671496 | 0.725968 |
| 0.696583 | 0.719629 | 0.720211 | 0.723906 | 0.672618 | 0.721771 | 0.675469 | 0.668621 | 0.720071 |
| 0.700445 | 0.727874 | 0.725498 | 0.727957 | 0.677409 | 0.726434 | 0.681572 | 0.676856 | 0.723481 |
| 0.696105 | 0.722832 | 0.7224 | 0.72562 | 0.675885 | 0.718148 | 0.676535 | 0.668593 | 0.714745 |

| | | | | | | | | |
|---|---|---|---|---|---|---|---|---|
| 0.683517 | 0.712079 | 0.708816 | 0.709952 | 0.663148 | 0.707092 | 0.660996 | 0.653295 | 0.705288 |
| 0.736915 | 0.776078 | 0.748732 | 0.752496 | 0.709724 | 0.741781 | 0.706829 | 0.695817 | 0.755679 |
| 0.697473 | 0.721458 | 0.721019 | 0.71954 | 0.679432 | 0.718223 | 0.674596 | 0.670007 | 0.720904 |
| 0.696461 | 0.725693 | 0.722244 | 0.725866 | 0.680657 | 0.721204 | 0.675936 | 0.671129 | 0.724669 |
| 0.691975 | 0.71889 | 0.717963 | 0.716434 | 0.676243 | 0.717531 | 0.671356 | 0.664963 | 0.714403 |
| 0.696109 | 0.722307 | 0.721551 | 0.722312 | 0.674068 | 0.724196 | 0.674841 | 0.672116 | 0.720843 |
| 0.692282 | 0.722176 | 0.715013 | 0.718295 | 0.673496 | 0.716463 | 0.676816 | 0.667488 | 0.716718 |
| 0.690508 | 0.713929 | 0.713004 | 0.716121 | 0.670925 | 0.710585 | 0.669332 | 0.663102 | 0.711031 |
| 0.697742 | 0.72799 | 0.72099 | 0.72328 | 0.677726 | 0.72176 | 0.676226 | 0.666742 | 0.715466 |
| 0.702857 | 0.73006 | 0.723515 | 0.72789 | 0.680735 | 0.725542 | 0.677814 | 0.672066 | 0.724242 |
| 0.698208 | 0.727417 | 0.722717 | 0.724152 | 0.680452 | 0.722284 | 0.678754 | 0.671121 | 0.721658 |
| 0.702857 | 0.729643 | 0.725771 | 0.724917 | 0.682801 | 0.721165 | 0.678384 | 0.674874 | 0.725022 |
| 0.706385 | 0.727033 | 0.724929 | 0.727595 | 0.682247 | 0.724647 | 0.6812 | 0.673814 | 0.72585 |
| 0.701028 | 0.73007 | 0.726917 | 0.72525 | 0.684137 | 0.722768 | 0.681599 | 0.673263 | 0.724483 |
| 0.697994 | 0.724158 | 0.71878 | 0.724285 | 0.675755 | 0.721258 | 0.673871 | 0.668488 | 0.716756 |
| 0.702325 | 0.729188 | 0.721265 | 0.723854 | 0.680939 | 0.721783 | 0.678771 | 0.670052 | 0.722931 |
| 0.700198 | 0.727822 | 0.726287 | 0.726143 | 0.678359 | 0.719962 | 0.675946 | 0.670397 | 0.723725 |
| 0.697302 | 0.723439 | 0.721584 | 0.724806 | 0.676275 | 0.721216 | 0.674973 | 0.66781 | 0.721987 |
| 0.693445 | 0.715902 | 0.718998 | 0.722062 | 0.674743 | 0.712219 | 0.671897 | 0.658589 | 0.713708 |
| 0.689961 | 0.717455 | 0.714333 | 0.719012 | 0.66964 | 0.713806 | 0.669168 | 0.661108 | 0.716128 |
| 0.697982 | 0.729292 | 0.720445 | 0.719475 | 0.677353 | 0.718139 | 0.675087 | 0.668428 | 0.71872 |
| 0.686199 | 0.711928 | 0.71281 | 0.714215 | 0.672216 | 0.710271 | 0.666929 | 0.661165 | 0.71261 |
| 0.696617 | 0.717218 | 0.721829 | 0.719276 | 0.677111 | 0.71424 | 0.674757 | 0.669192 | 0.716512 |
| 0.699114 | 0.726696 | 0.724484 | 0.729133 | 0.676044 | 0.719728 | 0.676342 | 0.671764 | 0.720274 |
| 0.698051 | 0.726454 | 0.724174 | 0.72559 | 0.677673 | 0.724678 | 0.675887 | 0.673795 | 0.722096 |
| 0.697849 | 0.720794 | 0.720383 | 0.722938 | 0.679229 | 0.717783 | 0.676274 | 0.670135 | 0.718974 |
| 0.693114 | 0.721644 | 0.713036 | 0.7181 | 0.67377 | 0.713547 | 0.671621 | 0.663205 | 0.71541 |
| 0.691059 | 0.715151 | 0.714332 | 0.71998 | 0.672925 | 0.712121 | 0.66956 | 0.663397 | 0.712567 |
| 0.690093 | 0.715286 | 0.710287 | 0.720461 | 0.666803 | 0.715689 | 0.670389 | 0.663309 | 0.713939 |
| 0.744785 | 0.746183 | 0.793989 | 0.762439 | 0.707201 | 0.741191 | 0.714299 | 0.708693 | 0.759114 |
| 0.770689 | 0.761294 | 0.760088 | 0.771191 | 0.721009 | 0.756724 | 0.725501 | 0.723798 | 0.780302 |
| 0.758476 | 0.760732 | 0.764482 | 0.766321 | 0.726379 | 0.749918 | 0.739315 | 0.741659 | 0.774271 |
| 0.766134 | 0.75919 | 0.751965 | 0.759413 | 0.719266 | 0.739485 | 0.739552 | 0.742576 | 0.76117 |
| 0.76873 | 0.764596 | 0.766413 | 0.768726 | 0.731245 | 0.762885 | 0.750974 | 0.755031 | 0.776685 |
| 0.691237 | 0.721708 | 0.71454 | 0.717879 | 0.667679 | 0.714789 | 0.67096 | 0.662588 | 0.713574 |
| 0.693068 | 0.719761 | 0.718501 | 0.721608 | 0.673068 | 0.717397 | 0.671759 | 0.667689 | 0.717528 |
| 0.692565 | 0.723901 | 0.720423 | 0.723409 | 0.675873 | 0.714446 | 0.67452 | 0.669529 | 0.716744 |
| 0.696946 | 0.72367 | 0.722042 | 0.724725 | 0.673584 | 0.721888 | 0.674644 | 0.669128 | 0.71948 |
| 0.697601 | 0.718402 | 0.720571 | 0.721101 | 0.674288 | 0.719971 | 0.674293 | 0.667712 | 0.717836 |
| 0.701159 | 0.727258 | 0.723778 | 0.72898 | 0.679887 | 0.720897 | 0.680118 | 0.668903 | 0.725708 |
| 0.694396 | 0.721655 | 0.720105 | 0.722091 | 0.676027 | 0.720919 | 0.673705 | 0.667946 | 0.719997 |
| 0.695126 | 0.72108 | 0.719408 | 0.724889 | 0.674931 | 0.717638 | 0.672818 | 0.666744 | 0.717688 |
| 0.684712 | 0.713901 | 0.711822 | 0.715266 | 0.669287 | 0.710851 | 0.665377 | 0.662136 | 0.713339 |
| 0.688279 | 0.71474 | 0.715484 | 0.718478 | 0.664007 | 0.714359 | 0.666996 | 0.661855 | 0.712391 |
| 0.695896 | 0.718783 | 0.714481 | 0.723288 | 0.673301 | 0.718362 | 0.671223 | 0.667419 | 0.718867 |
| 0.695878 | 0.725039 | 0.718255 | 0.722222 | 0.673785 | 0.719787 | 0.674442 | 0.669328 | 0.721331 |

| | | | | | | | | |
|---|---|---|---|---|---|---|---|---|
| 0.698497 | 0.725222 | 0.717856 | 0.719721 | 0.676756 | 0.718961 | 0.674269 | 0.670159 | 0.717441 |
| 0.695973 | 0.723165 | 0.718298 | 0.720259 | 0.676567 | 0.718381 | 0.67275 | 0.667098 | 0.719297 |
| 0.698042 | 0.721823 | 0.719599 | 0.724093 | 0.680038 | 0.72418 | 0.67572 | 0.671844 | 0.723735 |
| 0.694069 | 0.720131 | 0.718681 | 0.719368 | 0.674988 | 0.7195 | 0.671133 | 0.66809 | 0.717196 |
| 0.690134 | 0.718707 | 0.719106 | 0.718101 | 0.668991 | 0.716744 | 0.671378 | 0.663361 | 0.713393 |
| 0.693899 | 0.719547 | 0.715214 | 0.718579 | 0.673761 | 0.717775 | 0.67207 | 0.662105 | 0.71779 |
| 0.698339 | 0.726099 | 0.722586 | 0.723845 | 0.67968 | 0.719432 | 0.675006 | 0.670041 | 0.719644 |
| 0.693983 | 0.722997 | 0.719852 | 0.724459 | 0.677499 | 0.71784 | 0.673457 | 0.668585 | 0.722181 |
| 0.693451 | 0.720439 | 0.717938 | 0.720505 | 0.675542 | 0.717729 | 0.67248 | 0.667884 | 0.718625 |
| 0.694343 | 0.722424 | 0.71836 | 0.719409 | 0.678501 | 0.718192 | 0.673371 | 0.66945 | 0.720569 |
| 0.6941 | 0.723171 | 0.719901 | 0.723173 | 0.675584 | 0.717733 | 0.6716 | 0.669579 | 0.715 |
| 0.69568 | 0.722824 | 0.719442 | 0.721426 | 0.673651 | 0.718398 | 0.671532 | 0.668806 | 0.714482 |
| 0.687027 | 0.712631 | 0.710643 | 0.710558 | 0.667448 | 0.712843 | 0.666391 | 0.65882 | 0.708512 |
| 0.686166 | 0.716859 | 0.712719 | 0.716909 | 0.668097 | 0.709774 | 0.667569 | 0.661077 | 0.71411 |
| 0.699223 | 0.724076 | 0.721886 | 0.727145 | 0.68107 | 0.720085 | 0.676789 | 0.67304 | 0.722674 |
| 0.706495 | 0.728619 | 0.728087 | 0.726949 | 0.681743 | 0.724832 | 0.681176 | 0.674344 | 0.726641 |
| 0.702026 | 0.726804 | 0.723896 | 0.727374 | 0.682925 | 0.719994 | 0.680375 | 0.675177 | 0.721916 |
| 0.701448 | 0.729091 | 0.729633 | 0.731062 | 0.684272 | 0.722342 | 0.678091 | 0.677929 | 0.723106 |
| 0.693991 | 0.721702 | 0.717708 | 0.719521 | 0.674774 | 0.715404 | 0.672006 | 0.666614 | 0.716347 |
| 0.693316 | 0.724218 | 0.720065 | 0.720053 | 0.673845 | 0.720951 | 0.673356 | 0.668886 | 0.715517 |
| 0.692446 | 0.721351 | 0.716342 | 0.716796 | 0.672474 | 0.715479 | 0.671444 | 0.668035 | 0.714509 |
| 0.698658 | 0.728571 | 0.724705 | 0.724894 | 0.677938 | 0.721882 | 0.678577 | 0.670195 | 0.718128 |
| 0.701289 | 0.728781 | 0.721693 | 0.725004 | 0.678428 | 0.724928 | 0.678509 | 0.674734 | 0.722907 |
| 0.694526 | 0.721119 | 0.716286 | 0.72091 | 0.675287 | 0.71828 | 0.674723 | 0.663772 | 0.717783 |
| 0.697749 | 0.723074 | 0.719481 | 0.72329 | 0.675123 | 0.724933 | 0.675533 | 0.667864 | 0.720384 |
| 0.77619 | 0.765559 | 0.765916 | 0.765288 | 0.723732 | 0.751044 | 0.732864 | 0.728456 | 0.781058 |
| 0.699984 | 0.722448 | 0.720685 | 0.723898 | 0.679282 | 0.720314 | 0.677646 | 0.669354 | 0.719043 |
| 0.70052 | 0.719514 | 0.726344 | 0.729053 | 0.681949 | 0.723485 | 0.67829 | 0.671107 | 0.721822 |
| 0.703509 | 0.727478 | 0.72231 | 0.727509 | 0.680289 | 0.726415 | 0.676846 | 0.673117 | 0.725764 |
| 0.700555 | 0.719349 | 0.721405 | 0.729167 | 0.680172 | 0.723754 | 0.67527 | 0.674332 | 0.722324 |
| 0.695534 | 0.721789 | 0.719763 | 0.719936 | 0.672605 | 0.719338 | 0.672771 | 0.667461 | 0.715021 |
| 0.69132 | 0.71941 | 0.71829 | 0.721742 | 0.674465 | 0.720095 | 0.673371 | 0.668027 | 0.716309 |
| 0.693248 | 0.723429 | 0.722597 | 0.726623 | 0.677348 | 0.721761 | 0.674963 | 0.669325 | 0.719497 |
| 0.695897 | 0.717987 | 0.717269 | 0.717026 | 0.675259 | 0.71456 | 0.67165 | 0.666587 | 0.715855 |
| 0.695012 | 0.721072 | 0.721143 | 0.713139 | 0.675385 | 0.717979 | 0.67057 | 0.665309 | 0.714393 |
| 0.694308 | 0.716867 | 0.722509 | 0.719115 | 0.678386 | 0.71902 | 0.672318 | 0.668425 | 0.716209 |
| 0.700588 | 0.728581 | 0.724443 | 0.723713 | 0.675172 | 0.718988 | 0.67576 | 0.671366 | 0.719156 |
| 0.698989 | 0.731134 | 0.72276 | 0.724099 | 0.681124 | 0.722427 | 0.67285 | 0.670691 | 0.722351 |
| 0.699832 | 0.722029 | 0.723435 | 0.724059 | 0.676318 | 0.7225 | 0.677645 | 0.668686 | 0.720051 |
| 0.696773 | 0.723504 | 0.7181 | 0.72404 | 0.678071 | 0.719262 | 0.676322 | 0.6673 | 0.722508 |
| 0.693861 | 0.718252 | 0.715422 | 0.7174 | 0.674443 | 0.718979 | 0.669223 | 0.667353 | 0.715725 |
| 0.702385 | 0.726075 | 0.723684 | 0.72279 | 0.674729 | 0.724648 | 0.67724 | 0.671432 | 0.724001 |
| 0.695248 | 0.723564 | 0.719215 | 0.722297 | 0.676704 | 0.719064 | 0.673533 | 0.665615 | 0.716155 |
| 0.694984 | 0.722147 | 0.718992 | 0.722066 | 0.6747 | 0.717536 | 0.675397 | 0.662951 | 0.720383 |
| 0.701413 | 0.729899 | 0.725051 | 0.724495 | 0.678962 | 0.722338 | 0.679092 | 0.668317 | 0.721191 |
| 0.701035 | 0.726371 | 0.728227 | 0.72469 | 0.678861 | 0.720815 | 0.679606 | 0.671888 | 0.724045 |

| | | | | | | | | |
|---|---|---|---|---|---|---|---|---|
| 0.69743 | 0.723582 | 0.718999 | 0.724038 | 0.679957 | 0.722509 | 0.677579 | 0.668377 | 0.720057 |
| 0.697494 | 0.728332 | 0.724479 | 0.729207 | 0.678253 | 0.720101 | 0.679154 | 0.668404 | 0.723192 |
| 0.697052 | 0.720908 | 0.718264 | 0.720781 | 0.678549 | 0.717612 | 0.675278 | 0.669167 | 0.717298 |
| 0.651075 | 0.691549 | 0.678228 | 0.692281 | 0.642304 | 0.674532 | 0.634146 | 0.623631 | 0.680194 |
| 0.697392 | 0.72288 | 0.720278 | 0.721504 | 0.677792 | 0.717039 | 0.675997 | 0.662613 | 0.71912 |
| 0.698076 | 0.722573 | 0.721146 | 0.724944 | 0.677497 | 0.718208 | 0.677395 | 0.666869 | 0.720448 |
| 0.699945 | 0.724199 | 0.725092 | 0.72336 | 0.683808 | 0.716957 | 0.681054 | 0.669873 | 0.721914 |
| 0.699112 | 0.724701 | 0.724382 | 0.727069 | 0.677632 | 0.723862 | 0.680009 | 0.669163 | 0.718725 |
| 0.705141 | 0.733328 | 0.727836 | 0.727739 | 0.687041 | 0.724587 | 0.683477 | 0.673888 | 0.725113 |
| 0.700016 | 0.724413 | 0.723688 | 0.725155 | 0.681747 | 0.724713 | 0.676495 | 0.670027 | 0.722827 |
| 0.701487 | 0.721268 | 0.72182 | 0.720376 | 0.67801 | 0.721855 | 0.678423 | 0.671054 | 0.720779 |
| 0.701897 | 0.729771 | 0.723267 | 0.731137 | 0.683285 | 0.725066 | 0.679041 | 0.672756 | 0.724278 |
| 0.702856 | 0.723632 | 0.723832 | 0.722561 | 0.680679 | 0.724893 | 0.680771 | 0.671317 | 0.72652 |
| 0.69743 | 0.725518 | 0.720213 | 0.722897 | 0.67933 | 0.721681 | 0.676666 | 0.671808 | 0.722723 |
| 0.703765 | 0.728705 | 0.721939 | 0.728552 | 0.682062 | 0.720832 | 0.677615 | 0.671237 | 0.725881 |
| 0.696788 | 0.721929 | 0.717792 | 0.722826 | 0.675396 | 0.721223 | 0.674989 | 0.665256 | 0.722041 |
| 0.705205 | 0.728514 | 0.721406 | 0.728224 | 0.681558 | 0.722178 | 0.677819 | 0.66739 | 0.72574 |
| 0.701746 | 0.728745 | 0.723857 | 0.725471 | 0.681309 | 0.723056 | 0.677055 | 0.670493 | 0.720502 |
| 0.696653 | 0.721027 | 0.716288 | 0.72302 | 0.676935 | 0.71805 | 0.674847 | 0.663898 | 0.718012 |
| 0.697674 | 0.725805 | 0.718322 | 0.725347 | 0.67676 | 0.717939 | 0.676642 | 0.668883 | 0.719621 |
| 0.702134 | 0.726286 | 0.725271 | 0.731458 | 0.680778 | 0.722311 | 0.677826 | 0.671311 | 0.724478 |
| 0.703882 | 0.728665 | 0.720956 | 0.730515 | 0.68597 | 0.71734 | 0.681477 | 0.671944 | 0.728262 |
| 0.68535 | 0.702229 | 0.696538 | 0.698638 | 0.657715 | 0.722664 | 0.657513 | 0.640628 | 0.696159 |
| 0.713275 | 0.724003 | 0.716699 | 0.719137 | 0.669785 | 0.714182 | 0.679021 | 0.651884 | 0.734081 |
| 0.74182 | 0.741179 | 0.736405 | 0.746039 | 0.700324 | 0.729665 | 0.710441 | 0.696577 | 0.757895 |
| 0.746133 | 0.719322 | 0.722176 | 0.726905 | 0.683657 | 0.710999 | 0.68234 | 0.670228 | 0.730492 |
| 0.651163 | 0.681365 | 0.679725 | 0.68914 | 0.638405 | 0.678206 | 0.640284 | 0.628819 | 0.6809 |
| 0.649306 | 0.683405 | 0.678951 | 0.682292 | 0.633911 | 0.675671 | 0.627673 | 0.61486 | 0.67481 |
| 0.655743 | 0.676768 | 0.680532 | 0.675095 | 0.635448 | 0.67568 | 0.632759 | 0.615258 | 0.671039 |
| 0.669952 | 0.703673 | 0.690481 | 0.696723 | 0.643361 | 0.67845 | 0.638926 | 0.634841 | 0.685155 |
| 0.646296 | 0.667996 | 0.669298 | 0.674651 | 0.620901 | 0.668584 | 0.62093 | 0.607955 | 0.668652 |
| 0.639063 | 0.660675 | 0.659207 | 0.659245 | 0.623085 | 0.653731 | 0.610532 | 0.60232 | 0.652632 |
| 0.630006 | 0.668003 | 0.662048 | 0.668768 | 0.620093 | 0.649662 | 0.615301 | 0.603845 | 0.659387 |
| 0.714024 | 0.713582 | 0.747696 | 0.719832 | 0.68935 | 0.709019 | 0.672225 | 0.671011 | 0.723696 |
| 0.690915 | 0.71859 | 0.717191 | 0.720298 | 0.669849 | 0.71538 | 0.673789 | 0.664104 | 0.71592 |
| 0.667858 | 0.695696 | 0.689889 | 0.69604 | 0.647921 | 0.690766 | 0.640134 | 0.636121 | 0.693733 |
| 0.658399 | 0.6883 | 0.68176 | 0.685737 | 0.638586 | 0.681599 | 0.63591 | 0.624106 | 0.681205 |
| 0.64519 | 0.670839 | 0.662963 | 0.673788 | 0.622531 | 0.675729 | 0.621354 | 0.61438 | 0.66839 |
| 0.670282 | 0.695598 | 0.69442 | 0.696613 | 0.648905 | 0.686125 | 0.646486 | 0.634201 | 0.68611 |
| 0.658725 | 0.683372 | 0.688046 | 0.682824 | 0.639223 | 0.683063 | 0.635265 | 0.62547 | 0.685484 |
| 0.644518 | 0.678089 | 0.673479 | 0.681356 | 0.638266 | 0.674354 | 0.62637 | 0.618222 | 0.674977 |
| 0.654951 | 0.681241 | 0.676996 | 0.678073 | 0.634253 | 0.680539 | 0.631192 | 0.617024 | 0.68052 |
| 0.643237 | 0.671878 | 0.669374 | 0.672683 | 0.629717 | 0.667359 | 0.619574 | 0.611558 | 0.675178 |
| 0.652182 | 0.680598 | 0.671881 | 0.671267 | 0.627261 | 0.670465 | 0.626987 | 0.613735 | 0.675409 |
| 0.646026 | 0.677524 | 0.674063 | 0.683114 | 0.634597 | 0.669537 | 0.628154 | 0.614162 | 0.672597 |
| 0.639406 | 0.660618 | 0.65869 | 0.663593 | 0.623805 | 0.652201 | 0.611922 | 0.593928 | 0.661563 |

| | | | | | | | | |
|---|---|---|---|---|---|---|---|---|
| 0.680137 | 0.7051 | 0.702087 | 0.706844 | 0.659758 | 0.703636 | 0.657293 | 0.648012 | 0.702876 |
| 0.648353 | 0.678892 | 0.677051 | 0.682404 | 0.632541 | 0.672516 | 0.630311 | 0.62135 | 0.674027 |
| 0.651894 | 0.679174 | 0.673571 | 0.68243 | 0.636211 | 0.68074 | 0.628228 | 0.615984 | 0.677601 |
| 0.655611 | 0.684601 | 0.675537 | 0.684857 | 0.631333 | 0.676835 | 0.631171 | 0.617385 | 0.677253 |
| 0.647127 | 0.67679 | 0.676328 | 0.675855 | 0.62603 | 0.67287 | 0.624069 | 0.617631 | 0.674696 |
| 0.653541 | 0.669863 | 0.665013 | 0.676323 | 0.636555 | 0.680774 | 0.629536 | 0.621005 | 0.679182 |
| 0.644869 | 0.676595 | 0.670613 | 0.674637 | 0.629087 | 0.673791 | 0.628016 | 0.605572 | 0.671352 |
| 0.638739 | 0.665393 | 0.654877 | 0.666398 | 0.622537 | 0.66821 | 0.617854 | 0.606452 | 0.669473 |
| 0.638949 | 0.668949 | 0.657174 | 0.670492 | 0.619147 | 0.664313 | 0.615204 | 0.610925 | 0.661321 |
| 0.679813 | 0.708262 | 0.703274 | 0.709607 | 0.660284 | 0.704623 | 0.657275 | 0.648294 | 0.701424 |
| 0.682541 | 0.706418 | 0.702677 | 0.705635 | 0.656476 | 0.701618 | 0.656495 | 0.643969 | 0.702682 |
| 0.67815 | 0.702407 | 0.70243 | 0.701211 | 0.653656 | 0.694852 | 0.653811 | 0.64603 | 0.697793 |
| 0.670542 | 0.693921 | 0.687442 | 0.691687 | 0.646482 | 0.689308 | 0.644611 | 0.631136 | 0.690426 |
| 0.673515 | 0.700154 | 0.693162 | 0.695878 | 0.650279 | 0.690684 | 0.65 | 0.634111 | 0.688735 |
| 0.665179 | 0.69273 | 0.686606 | 0.693723 | 0.648605 | 0.685284 | 0.64366 | 0.629106 | 0.686228 |

| Collection_ | Collection_ | Collection_ | Collection_ | Collection_ | Collection_ | Collection_ | Collection_ | Collection_ |
|---|---|---|---|---|---|---|---|---|
| 0.716822 | 0.715575 | 0.707707 | 0.766545 | 0.716247 | 0.764927 | 0.761635 | 0.767854 | 0.746113 |
| 0.719526 | 0.725375 | 0.721651 | 0.765791 | 0.724117 | 0.770724 | 0.765688 | 0.762373 | 0.747757 |
| 0.713346 | 0.727153 | 0.709529 | 0.767334 | 0.715177 | 0.76147 | 0.763994 | 0.763962 | 0.74588 |
| 0.714927 | 0.719285 | 0.709087 | 0.770731 | 0.709455 | 0.770575 | 0.772536 | 0.765715 | 0.749838 |
| 0.720692 | 0.716209 | 0.709554 | 0.765196 | 0.708359 | 0.759288 | 0.756655 | 0.757791 | 0.750081 |
| 0.718547 | 0.705671 | 0.702949 | 0.75595 | 0.702727 | 0.761029 | 0.763251 | 0.758957 | 0.755592 |
| 0.672853 | 0.662794 | 0.659275 | 0.729125 | 0.664143 | 0.720418 | 0.731329 | 0.724494 | 0.748577 |
| 0.667322 | 0.658052 | 0.655456 | 0.724877 | 0.659099 | 0.718301 | 0.727121 | 0.723309 | 0.742069 |
| 0.722456 | 0.715576 | 0.706463 | 0.771291 | 0.710075 | 0.769667 | 0.76422 | 0.768534 | 0.747291 |
| 0.709735 | 0.727118 | 0.715251 | 0.793123 | 0.714286 | 0.777778 | 0.769618 | 0.776571 | 0.751061 |
| 0.710496 | 0.716397 | 0.703956 | 0.791488 | 0.710549 | 0.762763 | 0.756512 | 0.768841 | 0.748367 |
| 0.719305 | 0.723849 | 0.708765 | 0.770473 | 0.713789 | 0.778116 | 0.774729 | 0.773388 | 0.748226 |
| 0.713155 | 0.720077 | 0.715399 | 0.771868 | 0.713802 | 0.777347 | 0.773046 | 0.767499 | 0.745763 |
| 0.713678 | 0.713351 | 0.713086 | 0.780043 | 0.711762 | 0.776013 | 0.783686 | 0.777732 | 0.756026 |
| 0.70824 | 0.715897 | 0.705219 | 0.782765 | 0.70805 | 0.779077 | 0.779363 | 0.793016 | 0.760075 |
| 0.718667 | 0.726463 | 0.709229 | 0.774951 | 0.719639 | 0.772348 | 0.772133 | 0.776135 | 0.754346 |
| 0.711395 | 0.718908 | 0.70856 | 0.781309 | 0.710157 | 0.770404 | 0.767907 | 0.769635 | 0.751335 |
| 0.710327 | 0.708137 | 0.701302 | 0.797365 | 0.702474 | 0.788933 | 0.811145 | 0.804846 | 0.762167 |
| 0.709034 | 0.720782 | 0.719588 | 0.76798 | 0.727315 | 0.782551 | 0.762435 | 0.771492 | 0.742974 |
| 0.721595 | 0.726411 | 0.715838 | 0.770874 | 0.709767 | 0.771823 | 0.762783 | 0.767594 | 0.754321 |
| 0.726697 | 0.719737 | 0.710052 | 0.765769 | 0.709095 | 0.770868 | 0.761374 | 0.771538 | 0.749307 |
| 0.724151 | 0.727138 | 0.71113 | 0.774154 | 0.716186 | 0.778365 | 0.772076 | 0.783023 | 0.751452 |
| 0.725481 | 0.723022 | 0.718066 | 0.781577 | 0.720813 | 0.777458 | 0.771459 | 0.780566 | 0.75737 |
| 0.729618 | 0.726286 | 0.714555 | 0.773783 | 0.727824 | 0.778409 | 0.777126 | 0.781319 | 0.755503 |
| 0.727244 | 0.722564 | 0.716011 | 0.768407 | 0.715692 | 0.772589 | 0.762067 | 0.769785 | 0.760423 |
| 0.714308 | 0.714516 | 0.711798 | 0.766107 | 0.71271 | 0.775297 | 0.766591 | 0.774149 | 0.748237 |
| 0.711979 | 0.722089 | 0.714612 | 0.755815 | 0.709178 | 0.765631 | 0.756283 | 0.767659 | 0.743964 |
| 0.725689 | 0.716106 | 0.712743 | 0.767711 | 0.715942 | 0.757826 | 0.760752 | 0.765649 | 0.758891 |
| 0.723769 | 0.716692 | 0.705136 | 0.766496 | 0.710132 | 0.767768 | 0.774015 | 0.774209 | 0.756856 |
| 0.706093 | 0.721802 | 0.719987 | 0.767105 | 0.721409 | 0.769421 | 0.760455 | 0.774936 | 0.746059 |
| 0.716613 | 0.711589 | 0.705688 | 0.76472 | 0.703456 | 0.76001 | 0.759708 | 0.758789 | 0.750777 |
| 0.713821 | 0.715122 | 0.701431 | 0.758265 | 0.708151 | 0.756801 | 0.760364 | 0.755974 | 0.749491 |
| 0.712194 | 0.714593 | 0.712288 | 0.764844 | 0.712553 | 0.761932 | 0.763404 | 0.765459 | 0.750386 |
| 0.719026 | 0.714637 | 0.708789 | 0.755452 | 0.706617 | 0.75541 | 0.758394 | 0.756145 | 0.75596 |
| 0.714852 | 0.717794 | 0.708523 | 0.754751 | 0.709183 | 0.75928 | 0.754939 | 0.75452 | 0.739615 |
| 0.717251 | 0.720675 | 0.717204 | 0.76737 | 0.719216 | 0.771843 | 0.765421 | 0.769231 | 0.747218 |
| 0.71541 | 0.719975 | 0.70834 | 0.766527 | 0.718702 | 0.766352 | 0.759244 | 0.766624 | 0.747653 |
| 0.710467 | 0.708048 | 0.697303 | 0.760157 | 0.705417 | 0.755406 | 0.7581 | 0.761455 | 0.749526 |
| 0.712718 | 0.715342 | 0.706563 | 0.771901 | 0.709886 | 0.770985 | 0.760817 | 0.767084 | 0.751711 |

| | | | | | | | | |
|---|---|---|---|---|---|---|---|---|
| 0.705043 | 0.706052 | 0.701523 | 0.799804 | 0.70548 | 0.782609 | 0.791606 | 0.797285 | 0.756905 |
| 0.715514 | 0.715916 | 0.715609 | 0.772168 | 0.710828 | 0.767282 | 0.762213 | 0.768717 | 0.754148 |
| 0.723384 | 0.718828 | 0.707797 | 0.768334 | 0.713877 | 0.770703 | 0.766903 | 0.787627 | 0.75418 |
| 0.720887 | 0.734294 | 0.723534 | 0.79493 | 0.720769 | 0.788026 | 0.777463 | 0.782233 | 0.755832 |
| 0.717412 | 0.722897 | 0.711168 | 0.766277 | 0.714737 | 0.774478 | 0.766033 | 0.778096 | 0.753728 |
| 0.719019 | 0.718451 | 0.710632 | 0.764663 | 0.712871 | 0.770832 | 0.770367 | 0.77285 | 0.753598 |
| 0.718898 | 0.715543 | 0.705238 | 0.755235 | 0.706401 | 0.764663 | 0.756959 | 0.761654 | 0.747112 |
| 0.719236 | 0.725513 | 0.713553 | 0.763158 | 0.721751 | 0.76918 | 0.763272 | 0.766598 | 0.744966 |
| 0.712582 | 0.712213 | 0.704696 | 0.760928 | 0.712612 | 0.759054 | 0.759271 | 0.765067 | 0.756899 |
| 0.712785 | 0.706959 | 0.695848 | 0.757524 | 0.69808 | 0.761034 | 0.755015 | 0.760664 | 0.755031 |
| 0.717042 | 0.72051 | 0.71017 | 0.774573 | 0.71097 | 0.772153 | 0.768772 | 0.767491 | 0.747482 |
| 0.721201 | 0.725629 | 0.713982 | 0.771515 | 0.712309 | 0.775488 | 0.763438 | 0.77418 | 0.746809 |
| 0.712243 | 0.727016 | 0.719914 | 0.776049 | 0.724842 | 0.773752 | 0.763918 | 0.764292 | 0.749466 |
| 0.719743 | 0.717963 | 0.710618 | 0.769578 | 0.708362 | 0.771837 | 0.753139 | 0.772448 | 0.748728 |
| 0.716701 | 0.723414 | 0.713763 | 0.76868 | 0.715099 | 0.777722 | 0.76781 | 0.771203 | 0.749423 |
| 0.716259 | 0.713593 | 0.706932 | 0.764473 | 0.707191 | 0.763503 | 0.76268 | 0.762365 | 0.751327 |
| 0.710242 | 0.714542 | 0.706139 | 0.7678 | 0.707321 | 0.766854 | 0.759314 | 0.762433 | 0.745303 |
| 0.721764 | 0.73701 | 0.724514 | 0.780634 | 0.72471 | 0.781469 | 0.765954 | 0.774483 | 0.756732 |
| 0.722395 | 0.726885 | 0.714916 | 0.777869 | 0.718694 | 0.768108 | 0.774312 | 0.769086 | 0.751122 |
| 0.716903 | 0.728548 | 0.718869 | 0.770599 | 0.727358 | 0.782417 | 0.769375 | 0.774257 | 0.750288 |
| 0.716656 | 0.729542 | 0.714263 | 0.774011 | 0.714896 | 0.765635 | 0.768621 | 0.768052 | 0.743665 |
| 0.712066 | 0.718392 | 0.712014 | 0.787334 | 0.717692 | 0.763674 | 0.767552 | 0.768218 | 0.749242 |
| 0.723424 | 0.72896 | 0.723117 | 0.772388 | 0.724865 | 0.774867 | 0.765556 | 0.775951 | 0.748033 |
| 0.710083 | 0.715072 | 0.703599 | 0.763094 | 0.699908 | 0.768902 | 0.756173 | 0.767412 | 0.753337 |
| 0.707691 | 0.702201 | 0.701238 | 0.757231 | 0.69853 | 0.754417 | 0.755774 | 0.756001 | 0.765978 |
| 0.66743 | 0.658432 | 0.65397 | 0.725913 | 0.658715 | 0.718878 | 0.7279 | 0.723792 | 0.743296 |
| 0.67541 | 0.663939 | 0.663454 | 0.726411 | 0.665808 | 0.72081 | 0.731264 | 0.723825 | 0.743869 |
| 0.675163 | 0.663395 | 0.659705 | 0.72906 | 0.664779 | 0.720801 | 0.736182 | 0.727039 | 0.744529 |
| 0.721793 | 0.72439 | 0.715365 | 0.763917 | 0.71611 | 0.762151 | 0.763923 | 0.762037 | 0.751062 |
| 0.717226 | 0.715991 | 0.71491 | 0.759395 | 0.717745 | 0.763204 | 0.75877 | 0.761279 | 0.746536 |
| 0.719152 | 0.712619 | 0.707692 | 0.764931 | 0.710242 | 0.76849 | 0.760535 | 0.770735 | 0.753258 |
| 0.717162 | 0.717051 | 0.713877 | 0.770183 | 0.716414 | 0.763123 | 0.769736 | 0.764386 | 0.755453 |
| 0.718566 | 0.761539 | 0.741613 | 0.780567 | 0.748905 | 0.776981 | 0.781165 | 0.787622 | 0.756243 |
| 0.723486 | 0.735276 | 0.715383 | 0.767611 | 0.729098 | 0.788081 | 0.765547 | 0.777338 | 0.748825 |
| 0.716492 | 0.754417 | 0.752843 | 0.780622 | 0.745929 | 0.784918 | 0.780391 | 0.786196 | 0.750767 |
| 0.719647 | 0.751006 | 0.728365 | 0.770497 | 0.735814 | 0.779494 | 0.777132 | 0.775907 | 0.752688 |
| 0.736789 | 0.733499 | 0.731009 | 0.776515 | 0.72583 | 0.772093 | 0.774632 | 0.776722 | 0.752617 |
| 0.736714 | 0.750473 | 0.743286 | 0.808226 | 0.739431 | 0.8014 | 0.792462 | 0.799204 | 0.773895 |
| 0.712835 | 0.743924 | 0.724625 | 0.771031 | 0.730719 | 0.772289 | 0.761151 | 0.775234 | 0.744117 |
| 0.732898 | 0.721424 | 0.711699 | 0.769231 | 0.708782 | 0.770927 | 0.766892 | 0.773063 | 0.754058 |
| 0.717628 | 0.732364 | 0.715827 | 0.774476 | 0.717087 | 0.772301 | 0.771087 | 0.77182 | 0.747752 |
| 0.73477 | 0.752846 | 0.738936 | 0.805861 | 0.743192 | 0.795668 | 0.792799 | 0.798604 | 0.776864 |
| 0.719418 | 0.720323 | 0.709688 | 0.782827 | 0.708929 | 0.775445 | 0.770221 | 0.772411 | 0.762285 |
| 0.71671 | 0.71377 | 0.70746 | 0.777651 | 0.709972 | 0.776084 | 0.763177 | 0.777299 | 0.753127 |
| 0.718542 | 0.717873 | 0.710177 | 0.766743 | 0.712931 | 0.768977 | 0.765257 | 0.759739 | 0.753407 |
| 0.717219 | 0.746199 | 0.725989 | 0.769465 | 0.725327 | 0.776052 | 0.766525 | 0.769717 | 0.749117 |

| | | | | | | | | |
|---|---|---|---|---|---|---|---|---|
| 0.725041 | 0.719643 | 0.711867 | 0.78338 | 0.710752 | 0.774462 | 0.76858 | 0.773397 | 0.754528 |
| 0.741584 | 0.714417 | 0.706507 | 0.766469 | 0.706576 | 0.773257 | 0.758284 | 0.775755 | 0.745965 |
| 0.737426 | 0.717744 | 0.71126 | 0.766814 | 0.702615 | 0.766239 | 0.765601 | 0.768453 | 0.751666 |
| 0.731291 | 0.738766 | 0.728628 | 0.791871 | 0.724656 | 0.788857 | 0.781232 | 0.784278 | 0.769616 |
| 0.737175 | 0.72185 | 0.712508 | 0.77069 | 0.708758 | 0.776392 | 0.769479 | 0.772052 | 0.753176 |
| 0.726379 | 0.716799 | 0.708561 | 0.768341 | 0.704979 | 0.771377 | 0.772418 | 0.772503 | 0.750822 |
| 0.721638 | 0.727609 | 0.731978 | 0.778968 | 0.722812 | 0.774636 | 0.770109 | 0.772536 | 0.752617 |
| 0.729493 | 0.725203 | 0.707369 | 0.766847 | 0.710209 | 0.772517 | 0.756956 | 0.770027 | 0.751355 |
| 0.715582 | 0.716474 | 0.704021 | 0.768878 | 0.706655 | 0.764537 | 0.760714 | 0.76732 | 0.748687 |
| 0.713066 | 0.714286 | 0.707413 | 0.761341 | 0.709777 | 0.771342 | 0.754344 | 0.762995 | 0.748407 |
| 0.723983 | 0.720884 | 0.709881 | 0.769823 | 0.71282 | 0.768223 | 0.759494 | 0.768007 | 0.753584 |
| 0.716999 | 0.715817 | 0.70977 | 0.763831 | 0.707992 | 0.76357 | 0.756285 | 0.762128 | 0.761655 |
| 0.717123 | 0.715395 | 0.710874 | 0.763115 | 0.717119 | 0.762588 | 0.759632 | 0.760838 | 0.750555 |
| 0.718202 | 0.71108 | 0.706635 | 0.758961 | 0.697877 | 0.760641 | 0.755796 | 0.761336 | 0.746591 |
| 0.721497 | 0.716011 | 0.708573 | 0.766454 | 0.713512 | 0.762288 | 0.762679 | 0.768374 | 0.74957 |
| 0.720411 | 0.71478 | 0.707906 | 0.759237 | 0.707857 | 0.763792 | 0.755895 | 0.760353 | 0.752938 |
| 0.721359 | 0.713055 | 0.705378 | 0.770406 | 0.711222 | 0.764342 | 0.761622 | 0.767522 | 0.752655 |
| 0.713866 | 0.726786 | 0.71173 | 0.758827 | 0.720031 | 0.769413 | 0.767012 | 0.768537 | 0.748837 |
| 0.718791 | 0.720824 | 0.71022 | 0.763601 | 0.708869 | 0.768465 | 0.760211 | 0.766074 | 0.755395 |
| 0.718297 | 0.716387 | 0.71351 | 0.761994 | 0.712045 | 0.765628 | 0.762233 | 0.757381 | 0.750551 |
| 0.717608 | 0.71484 | 0.70391 | 0.767679 | 0.700578 | 0.763378 | 0.756479 | 0.765283 | 0.750041 |
| 0.715096 | 0.716009 | 0.707873 | 0.763328 | 0.70464 | 0.762473 | 0.756094 | 0.765479 | 0.752511 |
| 0.727573 | 0.718602 | 0.712458 | 0.767959 | 0.70834 | 0.768939 | 0.766217 | 0.770597 | 0.753902 |
| 0.724122 | 0.719653 | 0.710522 | 0.762371 | 0.707413 | 0.770813 | 0.764977 | 0.773817 | 0.745639 |
| 0.716688 | 0.728503 | 0.724986 | 0.769751 | 0.714787 | 0.767511 | 0.766305 | 0.76998 | 0.749403 |
| 0.72248 | 0.737701 | 0.74283 | 0.778491 | 0.736736 | 0.776105 | 0.771313 | 0.777136 | 0.754656 |
| 0.714903 | 0.729531 | 0.720757 | 0.772267 | 0.709119 | 0.769469 | 0.762853 | 0.773525 | 0.753314 |
| 0.719584 | 0.714165 | 0.70647 | 0.761693 | 0.712924 | 0.766778 | 0.765396 | 0.763448 | 0.751753 |
| 0.716947 | 0.714902 | 0.707181 | 0.761766 | 0.707051 | 0.764408 | 0.76031 | 0.766201 | 0.75244 |
| 0.724751 | 0.724369 | 0.725896 | 0.774437 | 0.715306 | 0.770082 | 0.76443 | 0.770747 | 0.752922 |
| 0.722853 | 0.717388 | 0.709472 | 0.765745 | 0.70769 | 0.767346 | 0.759538 | 0.767017 | 0.754753 |
| 0.718204 | 0.717534 | 0.709156 | 0.76393 | 0.701958 | 0.768008 | 0.755351 | 0.764399 | 0.751981 |
| 0.718082 | 0.734983 | 0.73344 | 0.776253 | 0.729992 | 0.77403 | 0.768958 | 0.771452 | 0.751137 |
| 0.720233 | 0.734052 | 0.721904 | 0.795401 | 0.717597 | 0.788333 | 0.777044 | 0.782092 | 0.756576 |
| 0.726809 | 0.721211 | 0.709204 | 0.763927 | 0.711289 | 0.765809 | 0.763029 | 0.766013 | 0.754031 |
| 0.719273 | 0.717605 | 0.707189 | 0.763546 | 0.710223 | 0.7656 | 0.762559 | 0.771109 | 0.752776 |
| 0.716693 | 0.719048 | 0.708561 | 0.757914 | 0.706578 | 0.764497 | 0.757423 | 0.761989 | 0.753391 |
| 0.697491 | 0.688741 | 0.679898 | 0.745112 | 0.678939 | 0.74302 | 0.737767 | 0.736913 | 0.724177 |
| 0.706529 | 0.696916 | 0.689176 | 0.751365 | 0.687281 | 0.755971 | 0.744076 | 0.748407 | 0.725843 |
| 0.695274 | 0.690978 | 0.682815 | 0.738621 | 0.686195 | 0.742214 | 0.735654 | 0.738105 | 0.732684 |
| 0.69711 | 0.705482 | 0.689443 | 0.744659 | 0.694334 | 0.753192 | 0.743963 | 0.747789 | 0.728246 |
| 0.678818 | 0.676791 | 0.664417 | 0.717827 | 0.665061 | 0.741178 | 0.718631 | 0.736685 | 0.710828 |
| 0.687882 | 0.683144 | 0.676905 | 0.750467 | 0.672742 | 0.743398 | 0.738522 | 0.742697 | 0.727127 |
| 0.691003 | 0.698013 | 0.675466 | 0.73181 | 0.680318 | 0.735167 | 0.73269 | 0.737129 | 0.712537 |
| 0.682071 | 0.690576 | 0.676984 | 0.732614 | 0.681404 | 0.733697 | 0.72449 | 0.727627 | 0.726178 |
| 0.701022 | 0.711013 | 0.696911 | 0.743167 | 0.696407 | 0.741882 | 0.732505 | 0.744882 | 0.72331 |
| 0.682956 | 0.698263 | 0.68127 | 0.736162 | 0.688809 | 0.749936 | 0.743073 | 0.744773 | 0.725919 |

| | | | | | | | | |
|---|---|---|---|---|---|---|---|---|
| 0.678954 | 0.680158 | 0.669902 | 0.724047 | 0.665014 | 0.725262 | 0.725714 | 0.726428 | 0.717356 |
| 0.686426 | 0.720715 | 0.71885 | 0.752013 | 0.710031 | 0.752887 | 0.742595 | 0.757149 | 0.730376 |
| 0.673818 | 0.667988 | 0.662795 | 0.726197 | 0.663221 | 0.724382 | 0.723315 | 0.728224 | 0.719092 |
| 0.678841 | 0.674448 | 0.665737 | 0.761709 | 0.656002 | 0.742369 | 0.735468 | 0.744186 | 0.723296 |
| 0.681116 | 0.704393 | 0.7054 | 0.741829 | 0.693662 | 0.747315 | 0.73364 | 0.752351 | 0.712933 |
| 0.72501 | 0.684029 | 0.6731 | 0.733034 | 0.663778 | 0.733887 | 0.724725 | 0.737925 | 0.717785 |
| 0.705752 | 0.692318 | 0.700834 | 0.745089 | 0.675303 | 0.738569 | 0.731158 | 0.742521 | 0.719816 |
| 0.677847 | 0.699012 | 0.67934 | 0.729521 | 0.687441 | 0.735366 | 0.726561 | 0.738058 | 0.71146 |
| 0.668224 | 0.664926 | 0.658433 | 0.710765 | 0.654785 | 0.719319 | 0.714137 | 0.722888 | 0.705386 |
| 0.677665 | 0.720203 | 0.731679 | 0.752226 | 0.718475 | 0.751619 | 0.748565 | 0.753582 | 0.718833 |
| 0.716642 | 0.675291 | 0.663946 | 0.73275 | 0.658974 | 0.734483 | 0.721359 | 0.73667 | 0.711519 |
| 0.727976 | 0.722013 | 0.707758 | 0.771994 | 0.707838 | 0.759092 | 0.755055 | 0.762008 | 0.746205 |
| 0.717144 | 0.712995 | 0.708868 | 0.761786 | 0.706013 | 0.76421 | 0.760936 | 0.766734 | 0.748888 |
| 0.709373 | 0.6827 | 0.682291 | 0.736714 | 0.673334 | 0.743358 | 0.736757 | 0.742519 | 0.729503 |
| 0.717591 | 0.710241 | 0.706394 | 0.763007 | 0.703268 | 0.768639 | 0.761195 | 0.763151 | 0.753878 |
| 0.716817 | 0.716109 | 0.713522 | 0.752037 | 0.712942 | 0.756247 | 0.74994 | 0.75668 | 0.740979 |
| 0.713426 | 0.713759 | 0.704415 | 0.764606 | 0.707858 | 0.764792 | 0.766629 | 0.764102 | 0.74783 |
| 0.714029 | 0.713152 | 0.705502 | 0.764016 | 0.706292 | 0.760952 | 0.758076 | 0.761263 | 0.755746 |
| 0.721831 | 0.731496 | 0.709838 | 0.748739 | 0.715885 | 0.754529 | 0.738399 | 0.751483 | 0.739445 |
| 0.712719 | 0.712428 | 0.702889 | 0.766355 | 0.705559 | 0.766783 | 0.76007 | 0.761195 | 0.755934 |
| 0.703367 | 0.67943 | 0.674536 | 0.747831 | 0.66808 | 0.738829 | 0.735241 | 0.734257 | 0.729539 |
| 0.714319 | 0.709927 | 0.706585 | 0.755593 | 0.701128 | 0.762366 | 0.756027 | 0.762768 | 0.750143 |
| 0.715075 | 0.707524 | 0.704686 | 0.75468 | 0.705111 | 0.758446 | 0.756912 | 0.760602 | 0.756242 |
| 0.693681 | 0.675475 | 0.668368 | 0.7481 | 0.671315 | 0.745381 | 0.739171 | 0.739807 | 0.722754 |
| 0.707622 | 0.720668 | 0.735865 | 0.739549 | 0.719258 | 0.735768 | 0.733472 | 0.738011 | 0.729058 |
| 0.697861 | 0.699037 | 0.68924 | 0.740945 | 0.687197 | 0.748268 | 0.741091 | 0.74955 | 0.73893 |
| 0.721413 | 0.691614 | 0.68756 | 0.752377 | 0.686079 | 0.745732 | 0.745706 | 0.74767 | 0.74119 |
| 0.722615 | 0.707308 | 0.69979 | 0.768237 | 0.706193 | 0.764067 | 0.762259 | 0.76098 | 0.758288 |
| 0.713444 | 0.718418 | 0.722378 | 0.736061 | 0.709717 | 0.737913 | 0.7337 | 0.73538 | 0.725742 |
| 0.713333 | 0.73189 | 0.728028 | 0.77552 | 0.725999 | 0.777561 | 0.774594 | 0.774196 | 0.751655 |
| 0.707455 | 0.709435 | 0.703441 | 0.758788 | 0.70496 | 0.759183 | 0.755246 | 0.762859 | 0.758434 |
| 0.738695 | 0.745858 | 0.747646 | 0.75587 | 0.736404 | 0.760108 | 0.753075 | 0.758542 | 0.740617 |
| 0.735548 | 0.744607 | 0.752826 | 0.75727 | 0.7378 | 0.755713 | 0.74986 | 0.753587 | 0.740744 |
| 0.710175 | 0.705338 | 0.701711 | 0.765434 | 0.700496 | 0.755099 | 0.75838 | 0.756576 | 0.757974 |
| 0.712174 | 0.70606 | 0.69391 | 0.7593 | 0.699368 | 0.759132 | 0.759081 | 0.758662 | 0.74998 |
| 0.720456 | 0.707697 | 0.704663 | 0.768597 | 0.702154 | 0.76884 | 0.762241 | 0.763454 | 0.760312 |
| 0.745735 | 0.71605 | 0.723398 | 0.716379 | 0.719719 | 0.71284 | 0.714286 | 0.725085 | 0.709501 |
| 0.706397 | 0.700496 | 0.691048 | 0.753316 | 0.691217 | 0.750485 | 0.746745 | 0.751425 | 0.749008 |
| 0.738657 | 0.705363 | 0.722231 | 0.717391 | 0.710146 | 0.718286 | 0.715504 | 0.716316 | 0.720543 |
| 0.843823 | 0.754811 | 0.768497 | 0.709002 | 0.732281 | 0.712313 | 0.707567 | 0.714043 | 0.70554 |
| 0.718417 | 0.71611 | 0.705267 | 0.806017 | 0.708902 | 0.847997 | 0.826335 | 0.854088 | 0.754809 |
| 1 | 0.750192 | 0.763528 | 0.711278 | 0.740495 | 0.715949 | 0.710765 | 0.717543 | 0.707546 |
| 0.750192 | 1 | 0.80359 | 0.714659 | 0.848932 | 0.718661 | 0.711486 | 0.723044 | 0.695114 |
| 0.763528 | 0.80359 | 1 | 0.706893 | 0.798009 | 0.707727 | 0.70399 | 0.711128 | 0.696124 |
| 0.711278 | 0.714659 | 0.706893 | 1 | 0.710331 | 0.834424 | 0.810964 | 0.809602 | 0.757711 |
| 0.740495 | 0.848932 | 0.798009 | 0.710331 | 1 | 0.709389 | 0.703231 | 0.714242 | 0.689359 |
| 0.715949 | 0.718661 | 0.707727 | 0.834424 | 0.709389 | 1 | 0.802872 | 0.831149 | 0.753352 |

| | | | | | | | | |
|---|---|---|---|---|---|---|---|---|
| 0.710765 | 0.711486 | 0.70399 | 0.810964 | 0.703231 | 0.802872 | 1 | 0.821248 | 0.7595 |
| 0.717543 | 0.723044 | 0.711128 | 0.809602 | 0.714242 | 0.831149 | 0.821248 | 1 | 0.759684 |
| 0.707546 | 0.695114 | 0.696124 | 0.757711 | 0.689359 | 0.753352 | 0.7595 | 0.759684 | 1 |
| 0.739571 | 0.847616 | 0.806441 | 0.698146 | 0.834935 | 0.701003 | 0.701181 | 0.706671 | 0.683185 |
| 0.765048 | 0.795526 | 0.789262 | 0.69616 | 0.787889 | 0.694663 | 0.692184 | 0.69446 | 0.684407 |
| 0.718289 | 0.713825 | 0.705128 | 0.805599 | 0.707931 | 0.802191 | 0.807172 | 0.815912 | 0.756983 |
| 0.721792 | 0.720607 | 0.726524 | 0.683638 | 0.718366 | 0.685587 | 0.686566 | 0.687697 | 0.684993 |
| 0.687336 | 0.679484 | 0.678006 | 0.733677 | 0.684873 | 0.728709 | 0.738138 | 0.729523 | 0.736911 |
| 0.709237 | 0.717895 | 0.705491 | 0.814309 | 0.709036 | 0.815706 | 0.797298 | 0.813473 | 0.758346 |
| 0.681387 | 0.671426 | 0.664343 | 0.726679 | 0.670788 | 0.727043 | 0.729597 | 0.728082 | 0.736942 |
| 0.694815 | 0.705261 | 0.701593 | 0.793315 | 0.702436 | 0.804639 | 0.801873 | 0.818692 | 0.745279 |
| 0.617284 | 0.635762 | 0.598802 | 0.649351 | 0.562874 | 0.704698 | 0.642857 | 0.658065 | 0.674699 |
| 0.619682 | 0.605244 | 0.605121 | 0.675437 | 0.603192 | 0.674343 | 0.673708 | 0.66808 | 0.69971 |
| 0.621433 | 0.60568 | 0.604622 | 0.677094 | 0.603049 | 0.674191 | 0.676696 | 0.66978 | 0.702348 |
| 0.623876 | 0.614789 | 0.612732 | 0.691266 | 0.611969 | 0.686828 | 0.690704 | 0.68315 | 0.707877 |
| 0.625502 | 0.620736 | 0.620799 | 0.68907 | 0.615963 | 0.68139 | 0.684408 | 0.684156 | 0.704347 |
| 0.671063 | 0.646882 | 0.670056 | 0.614952 | 0.661392 | 0.616777 | 0.609914 | 0.61417 | 0.625957 |
| 0.622699 | 0.618722 | 0.614249 | 0.691882 | 0.61487 | 0.686871 | 0.688263 | 0.68392 | 0.712541 |
| 0.693815 | 0.698398 | 0.692115 | 0.809588 | 0.693663 | 0.804905 | 0.813221 | 0.802869 | 0.747791 |
| 0.696477 | 0.700148 | 0.691925 | 0.807666 | 0.691584 | 0.807656 | 0.807417 | 0.799158 | 0.743547 |
| 0.692551 | 0.704662 | 0.698069 | 0.810933 | 0.702987 | 0.809914 | 0.796993 | 0.814871 | 0.749384 |
| 0.731672 | 0.751231 | 0.735249 | 0.616342 | 0.740973 | 0.610978 | 0.612573 | 0.614706 | 0.615331 |
| 0.690297 | 0.746277 | 0.71076 | 0.598945 | 0.762685 | 0.596359 | 0.59672 | 0.601261 | 0.599618 |
| 0.671061 | 0.661687 | 0.659797 | 0.735409 | 0.657052 | 0.733393 | 0.747471 | 0.736145 | 0.746387 |
| 0.668763 | 0.662508 | 0.656894 | 0.73951 | 0.657972 | 0.731966 | 0.748512 | 0.736686 | 0.742594 |
| 0.661317 | 0.65468 | 0.654846 | 0.729998 | 0.651086 | 0.722937 | 0.733441 | 0.725304 | 0.741433 |
| 0.668258 | 0.661169 | 0.659901 | 0.730868 | 0.657704 | 0.724868 | 0.731995 | 0.724813 | 0.743043 |
| 0.668981 | 0.66059 | 0.659667 | 0.737132 | 0.659605 | 0.726705 | 0.738683 | 0.732408 | 0.746103 |
| 0.668451 | 0.66067 | 0.659372 | 0.735909 | 0.658322 | 0.725937 | 0.745469 | 0.730695 | 0.751031 |
| 0.670183 | 0.664955 | 0.659633 | 0.738763 | 0.659675 | 0.730831 | 0.746146 | 0.733317 | 0.746675 |
| 0.66635 | 0.660213 | 0.656225 | 0.726773 | 0.658531 | 0.719113 | 0.730186 | 0.722721 | 0.749083 |
| 0.680052 | 0.667413 | 0.662947 | 0.734191 | 0.667789 | 0.724176 | 0.731009 | 0.726337 | 0.758873 |
| 0.672369 | 0.663432 | 0.655424 | 0.727009 | 0.663302 | 0.719403 | 0.728194 | 0.721994 | 0.75279 |
| 0.668945 | 0.664555 | 0.6599 | 0.72611 | 0.664531 | 0.719804 | 0.731578 | 0.724239 | 0.755828 |
| 0.672086 | 0.665239 | 0.663325 | 0.732264 | 0.664561 | 0.722485 | 0.73481 | 0.726533 | 0.752816 |
| 0.666946 | 0.661027 | 0.656056 | 0.729454 | 0.663094 | 0.71936 | 0.73042 | 0.718795 | 0.746354 |
| 0.671812 | 0.661694 | 0.657278 | 0.7268 | 0.659991 | 0.721104 | 0.728982 | 0.721792 | 0.752505 |
| 0.671948 | 0.663766 | 0.658222 | 0.730368 | 0.662969 | 0.721649 | 0.731713 | 0.723411 | 0.753997 |
| 0.665939 | 0.65707 | 0.65143 | 0.723577 | 0.658891 | 0.716337 | 0.726662 | 0.724217 | 0.747906 |
| 0.669998 | 0.662852 | 0.658364 | 0.724406 | 0.662127 | 0.718964 | 0.727698 | 0.721429 | 0.751019 |
| 0.610603 | 0.608607 | 0.602524 | 0.683519 | 0.605565 | 0.676471 | 0.685395 | 0.679763 | 0.704929 |
| 0.619178 | 0.616123 | 0.613331 | 0.692774 | 0.609785 | 0.689971 | 0.693292 | 0.682008 | 0.705958 |
| 0.674494 | 0.667091 | 0.66082 | 0.731991 | 0.666567 | 0.725638 | 0.734108 | 0.729502 | 0.752096 |
| 0.674325 | 0.665263 | 0.66465 | 0.743064 | 0.669179 | 0.729046 | 0.740973 | 0.734037 | 0.753186 |
| 0.609408 | 0.595961 | 0.600869 | 0.671927 | 0.590628 | 0.664458 | 0.678704 | 0.664449 | 0.70756 |
| 0.671636 | 0.664834 | 0.660858 | 0.729818 | 0.66506 | 0.722209 | 0.732516 | 0.725349 | 0.753132 |
| 0.661674 | 0.654848 | 0.649724 | 0.723136 | 0.655928 | 0.711162 | 0.720361 | 0.715281 | 0.738852 |

| | | | | | | | | |
|---|---|---|---|---|---|---|---|---|
| 0.668946 | 0.660678 | 0.657112 | 0.730948 | 0.659788 | 0.721808 | 0.729928 | 0.726227 | 0.748898 |
| 0.670813 | 0.66185 | 0.660667 | 0.728562 | 0.665321 | 0.721043 | 0.733254 | 0.727186 | 0.754352 |
| 0.667311 | 0.663239 | 0.659369 | 0.731474 | 0.661141 | 0.724664 | 0.731882 | 0.728603 | 0.754703 |
| 0.673508 | 0.665839 | 0.661292 | 0.731354 | 0.664884 | 0.722975 | 0.736648 | 0.728537 | 0.751757 |
| 0.676556 | 0.669983 | 0.666896 | 0.738322 | 0.668073 | 0.732877 | 0.742008 | 0.738562 | 0.760327 |
| 0.666489 | 0.659875 | 0.657204 | 0.728809 | 0.66042 | 0.722182 | 0.729347 | 0.721311 | 0.750718 |
| 0.669362 | 0.664022 | 0.65839 | 0.730631 | 0.663493 | 0.72096 | 0.73276 | 0.722306 | 0.749594 |
| 0.662784 | 0.657162 | 0.651351 | 0.720515 | 0.656688 | 0.713535 | 0.722767 | 0.717713 | 0.737766 |
| 0.67116 | 0.661922 | 0.6577 | 0.725917 | 0.661409 | 0.71984 | 0.729356 | 0.725355 | 0.74904 |
| 0.664741 | 0.657169 | 0.652244 | 0.725526 | 0.658285 | 0.715969 | 0.727666 | 0.720811 | 0.740285 |
| 0.669438 | 0.663121 | 0.657643 | 0.727143 | 0.662182 | 0.720914 | 0.73389 | 0.727042 | 0.752749 |
| 0.668308 | 0.65904 | 0.657656 | 0.727101 | 0.660322 | 0.716604 | 0.72787 | 0.721333 | 0.74725 |
| 0.659524 | 0.652451 | 0.647376 | 0.720229 | 0.653248 | 0.712832 | 0.71699 | 0.713358 | 0.738905 |
| 0.660931 | 0.653163 | 0.651319 | 0.719688 | 0.654684 | 0.712894 | 0.717793 | 0.714416 | 0.737181 |
| 0.677029 | 0.668063 | 0.664713 | 0.733784 | 0.665967 | 0.727005 | 0.734671 | 0.730165 | 0.756193 |
| 0.672186 | 0.665144 | 0.661767 | 0.74747 | 0.664771 | 0.737452 | 0.74726 | 0.739712 | 0.750407 |
| 0.668943 | 0.6595 | 0.654064 | 0.728637 | 0.657087 | 0.718727 | 0.733328 | 0.725941 | 0.745091 |
| 0.667377 | 0.662294 | 0.658088 | 0.735289 | 0.662091 | 0.728521 | 0.741506 | 0.731721 | 0.742761 |
| 0.667145 | 0.658101 | 0.657696 | 0.735845 | 0.658214 | 0.724994 | 0.742318 | 0.730972 | 0.743362 |
| 0.666667 | 0.658821 | 0.658411 | 0.737729 | 0.659291 | 0.727794 | 0.73961 | 0.734418 | 0.750389 |
| 0.663918 | 0.655928 | 0.654893 | 0.740313 | 0.657436 | 0.730636 | 0.735249 | 0.731046 | 0.737108 |
| 0.666255 | 0.657774 | 0.655387 | 0.731681 | 0.657531 | 0.723225 | 0.73891 | 0.730851 | 0.743175 |
| 0.669984 | 0.66336 | 0.661743 | 0.733317 | 0.663055 | 0.725634 | 0.737794 | 0.733615 | 0.749899 |
| 0.670745 | 0.665962 | 0.659703 | 0.73406 | 0.664231 | 0.728621 | 0.737028 | 0.727993 | 0.750471 |
| 0.667024 | 0.659186 | 0.658652 | 0.734972 | 0.660743 | 0.727836 | 0.73964 | 0.734667 | 0.746626 |
| 0.664656 | 0.65754 | 0.656067 | 0.738905 | 0.657257 | 0.724587 | 0.745586 | 0.737848 | 0.73933 |
| 0.663216 | 0.658207 | 0.657085 | 0.730479 | 0.658173 | 0.723652 | 0.735208 | 0.723063 | 0.742317 |
| 0.666178 | 0.656092 | 0.652353 | 0.724287 | 0.656802 | 0.72113 | 0.729974 | 0.720819 | 0.737134 |
| 0.669075 | 0.65728 | 0.657611 | 0.740529 | 0.657076 | 0.731656 | 0.748853 | 0.739683 | 0.737817 |
| 0.670961 | 0.659707 | 0.657871 | 0.733456 | 0.66294 | 0.725056 | 0.740364 | 0.729094 | 0.747042 |
| 0.671474 | 0.66288 | 0.661135 | 0.738309 | 0.662431 | 0.733479 | 0.740497 | 0.736728 | 0.743686 |
| 0.670579 | 0.662849 | 0.661872 | 0.73854 | 0.6634 | 0.726019 | 0.736578 | 0.733755 | 0.74812 |
| 0.675825 | 0.668212 | 0.663273 | 0.74239 | 0.664429 | 0.738688 | 0.749321 | 0.739401 | 0.751806 |
| 0.677331 | 0.668725 | 0.666156 | 0.738372 | 0.667897 | 0.732974 | 0.74064 | 0.736383 | 0.752874 |
| 0.665753 | 0.654908 | 0.654392 | 0.73991 | 0.656436 | 0.72602 | 0.748504 | 0.735963 | 0.737864 |
| 0.670051 | 0.662289 | 0.658929 | 0.741403 | 0.659714 | 0.731456 | 0.746664 | 0.737075 | 0.744563 |
| 0.669137 | 0.662041 | 0.656784 | 0.732456 | 0.65976 | 0.725932 | 0.740477 | 0.734061 | 0.745888 |
| 0.668022 | 0.658592 | 0.656057 | 0.726943 | 0.657721 | 0.721713 | 0.734583 | 0.726421 | 0.74068 |
| 0.663015 | 0.657296 | 0.653687 | 0.728858 | 0.656214 | 0.72532 | 0.736412 | 0.729092 | 0.737249 |
| 0.660735 | 0.654031 | 0.651296 | 0.726595 | 0.651272 | 0.713357 | 0.734992 | 0.725428 | 0.7349 |
| 0.660988 | 0.651067 | 0.652504 | 0.73299 | 0.651648 | 0.726916 | 0.745467 | 0.734613 | 0.731799 |
| 0.662718 | 0.651035 | 0.650692 | 0.725234 | 0.65147 | 0.721387 | 0.735616 | 0.726173 | 0.734022 |
| 0.660103 | 0.654723 | 0.650379 | 0.725676 | 0.65196 | 0.723248 | 0.734965 | 0.725393 | 0.739691 |
| 0.65364 | 0.645992 | 0.645277 | 0.719383 | 0.643741 | 0.711778 | 0.727297 | 0.718523 | 0.733482 |
| 0.664253 | 0.657008 | 0.650324 | 0.723962 | 0.654313 | 0.721207 | 0.728579 | 0.724271 | 0.738539 |
| 0.657125 | 0.64969 | 0.646771 | 0.72067 | 0.650992 | 0.712684 | 0.719751 | 0.715018 | 0.738062 |

| | | | | | | | | |
|---|---|---|---|---|---|---|---|---|
| 0.672004 | 0.663022 | 0.659893 | 0.753329 | 0.662392 | 0.743331 | 0.765984 | 0.755142 | 0.743058 |
| 0.672806 | 0.660986 | 0.659981 | 0.731371 | 0.664608 | 0.718917 | 0.731344 | 0.724055 | 0.747746 |
| 0.666334 | 0.658928 | 0.656533 | 0.725471 | 0.658563 | 0.716082 | 0.728331 | 0.720006 | 0.741645 |
| 0.670963 | 0.66259 | 0.657813 | 0.727194 | 0.663831 | 0.720875 | 0.730507 | 0.725428 | 0.746985 |
| 0.676845 | 0.667847 | 0.663248 | 0.732641 | 0.667745 | 0.723713 | 0.735271 | 0.730305 | 0.753336 |
| 0.670526 | 0.662931 | 0.657751 | 0.729059 | 0.662182 | 0.721459 | 0.731911 | 0.725799 | 0.753513 |
| 0.66475 | 0.65817 | 0.654492 | 0.729484 | 0.659362 | 0.717673 | 0.725593 | 0.721378 | 0.753747 |
| 0.668131 | 0.664317 | 0.658388 | 0.729308 | 0.665715 | 0.719057 | 0.731884 | 0.725181 | 0.749368 |
| 0.664125 | 0.655385 | 0.652794 | 0.720534 | 0.657394 | 0.712189 | 0.727177 | 0.718439 | 0.747656 |
| 0.669253 | 0.661423 | 0.657152 | 0.727047 | 0.661998 | 0.719187 | 0.727127 | 0.724141 | 0.741967 |
| 0.667643 | 0.662154 | 0.659368 | 0.727754 | 0.664529 | 0.722147 | 0.728672 | 0.727294 | 0.754019 |
| 0.669572 | 0.663634 | 0.659058 | 0.7256 | 0.66377 | 0.720163 | 0.729492 | 0.724508 | 0.752534 |
| 0.669371 | 0.663884 | 0.659096 | 0.723653 | 0.663763 | 0.717544 | 0.726629 | 0.722716 | 0.74165 |
| 0.668943 | 0.666767 | 0.660124 | 0.725606 | 0.66594 | 0.718405 | 0.728816 | 0.720257 | 0.743915 |
| 0.673383 | 0.663008 | 0.657996 | 0.727488 | 0.663461 | 0.722868 | 0.730891 | 0.725631 | 0.747592 |
| 0.671971 | 0.667193 | 0.661675 | 0.726955 | 0.664095 | 0.721588 | 0.730007 | 0.72506 | 0.744377 |
| 0.674913 | 0.664186 | 0.66134 | 0.732346 | 0.665922 | 0.723818 | 0.734182 | 0.728271 | 0.748567 |
| 0.672917 | 0.666171 | 0.660152 | 0.731392 | 0.667088 | 0.720164 | 0.733323 | 0.727566 | 0.742839 |
| 0.672904 | 0.66212 | 0.662184 | 0.728712 | 0.663745 | 0.722817 | 0.734407 | 0.726154 | 0.74669 |
| 0.674248 | 0.663758 | 0.661903 | 0.726166 | 0.663284 | 0.721353 | 0.728035 | 0.724897 | 0.746725 |
| 0.669055 | 0.664698 | 0.659271 | 0.726974 | 0.664549 | 0.718019 | 0.728963 | 0.727849 | 0.743533 |
| 0.66822 | 0.662484 | 0.656317 | 0.724534 | 0.663592 | 0.718937 | 0.724204 | 0.721068 | 0.749116 |
| 0.673138 | 0.665145 | 0.659629 | 0.733217 | 0.666991 | 0.725778 | 0.738107 | 0.729071 | 0.744314 |
| 0.67156 | 0.664036 | 0.662569 | 0.730544 | 0.664083 | 0.72497 | 0.732524 | 0.727524 | 0.744846 |
| 0.672218 | 0.664711 | 0.660463 | 0.728524 | 0.663043 | 0.726733 | 0.731734 | 0.728735 | 0.745016 |
| 0.672313 | 0.66514 | 0.660891 | 0.726158 | 0.662991 | 0.722382 | 0.728421 | 0.722588 | 0.74373 |
| 0.669049 | 0.659414 | 0.656186 | 0.723958 | 0.660781 | 0.719418 | 0.726326 | 0.719502 | 0.74024 |
| 0.671876 | 0.660317 | 0.658786 | 0.724577 | 0.662338 | 0.716222 | 0.729871 | 0.720919 | 0.742804 |
| 0.671646 | 0.663513 | 0.658967 | 0.733058 | 0.664258 | 0.72356 | 0.731104 | 0.727093 | 0.745676 |
| 0.672441 | 0.667493 | 0.664442 | 0.733877 | 0.664993 | 0.724771 | 0.733119 | 0.728819 | 0.747241 |
| 0.670285 | 0.660567 | 0.657542 | 0.726373 | 0.661891 | 0.720533 | 0.726296 | 0.719181 | 0.742662 |
| 0.678637 | 0.666541 | 0.665 | 0.73067 | 0.667271 | 0.724432 | 0.73176 | 0.728596 | 0.74257 |
| 0.673676 | 0.666892 | 0.659516 | 0.730344 | 0.667492 | 0.728677 | 0.73327 | 0.726471 | 0.748048 |
| 0.664458 | 0.657257 | 0.655099 | 0.725867 | 0.660119 | 0.719136 | 0.728999 | 0.722058 | 0.739963 |
| 0.672645 | 0.663855 | 0.664463 | 0.729299 | 0.663708 | 0.72053 | 0.728967 | 0.725808 | 0.741736 |
| 0.666322 | 0.658749 | 0.655654 | 0.726306 | 0.659489 | 0.719273 | 0.728781 | 0.721708 | 0.740596 |
| 0.677243 | 0.668997 | 0.665544 | 0.735733 | 0.672318 | 0.72782 | 0.73896 | 0.730202 | 0.751896 |
| 0.666949 | 0.661367 | 0.654227 | 0.726293 | 0.661499 | 0.720661 | 0.731644 | 0.721854 | 0.745152 |
| 0.670695 | 0.665939 | 0.660318 | 0.729771 | 0.666416 | 0.722587 | 0.731939 | 0.72625 | 0.74351 |
| 0.672467 | 0.662228 | 0.658359 | 0.727805 | 0.664547 | 0.72044 | 0.72894 | 0.722922 | 0.744815 |
| 0.676794 | 0.667541 | 0.663877 | 0.730513 | 0.669161 | 0.728065 | 0.733979 | 0.728621 | 0.747607 |
| 0.674426 | 0.667143 | 0.663382 | 0.728467 | 0.670962 | 0.724074 | 0.734041 | 0.728709 | 0.744861 |
| 0.673196 | 0.666866 | 0.660789 | 0.728834 | 0.668411 | 0.726324 | 0.737331 | 0.729509 | 0.747509 |
| 0.666718 | 0.660608 | 0.657938 | 0.729231 | 0.662085 | 0.721388 | 0.73111 | 0.724894 | 0.743477 |
| 0.675748 | 0.664902 | 0.661314 | 0.730911 | 0.665209 | 0.726149 | 0.73268 | 0.727535 | 0.745684 |
| 0.667952 | 0.659273 | 0.657037 | 0.727516 | 0.660476 | 0.722562 | 0.729894 | 0.724765 | 0.743549 |

| | | | | | | | | |
|---|---|---|---|---|---|---|---|---|
| 0.654923 | 0.649586 | 0.64932 | 0.712446 | 0.648371 | 0.706657 | 0.718188 | 0.71077 | 0.736361 |
| 0.697866 | 0.693994 | 0.688132 | 0.75545 | 0.691678 | 0.749386 | 0.754674 | 0.756497 | 0.760527 |
| 0.670455 | 0.663424 | 0.659468 | 0.725814 | 0.663096 | 0.720512 | 0.725345 | 0.723484 | 0.744099 |
| 0.670655 | 0.66329 | 0.662111 | 0.731392 | 0.665052 | 0.725384 | 0.73591 | 0.729406 | 0.746926 |
| 0.665225 | 0.657354 | 0.651407 | 0.722768 | 0.65793 | 0.716428 | 0.72219 | 0.718634 | 0.73666 |
| 0.67089 | 0.66118 | 0.656358 | 0.723221 | 0.660153 | 0.720598 | 0.727178 | 0.723741 | 0.742238 |
| 0.666984 | 0.658943 | 0.654955 | 0.724049 | 0.661788 | 0.715635 | 0.726749 | 0.71989 | 0.744123 |
| 0.663084 | 0.654826 | 0.65138 | 0.714286 | 0.65525 | 0.712958 | 0.716752 | 0.714176 | 0.736603 |
| 0.668654 | 0.659374 | 0.656162 | 0.728832 | 0.662133 | 0.72015 | 0.731506 | 0.725016 | 0.746431 |
| 0.6732 | 0.664206 | 0.663394 | 0.732916 | 0.664856 | 0.725451 | 0.730265 | 0.727935 | 0.748039 |
| 0.671044 | 0.665993 | 0.661016 | 0.732829 | 0.663732 | 0.724311 | 0.735051 | 0.726919 | 0.747139 |
| 0.67252 | 0.665132 | 0.660725 | 0.726242 | 0.666188 | 0.725909 | 0.732723 | 0.727833 | 0.748281 |
| 0.676213 | 0.668099 | 0.664113 | 0.732635 | 0.667269 | 0.729247 | 0.736543 | 0.728208 | 0.750184 |
| 0.675119 | 0.664923 | 0.660889 | 0.732713 | 0.665572 | 0.726458 | 0.732793 | 0.727778 | 0.749939 |
| 0.668569 | 0.662994 | 0.658246 | 0.728654 | 0.663021 | 0.725647 | 0.728717 | 0.722078 | 0.744558 |
| 0.672183 | 0.66289 | 0.660012 | 0.72708 | 0.6622 | 0.721337 | 0.729912 | 0.729349 | 0.743647 |
| 0.67126 | 0.662651 | 0.663002 | 0.727774 | 0.664324 | 0.723916 | 0.7288 | 0.72552 | 0.746264 |
| 0.667736 | 0.661406 | 0.658482 | 0.73198 | 0.661304 | 0.723885 | 0.726821 | 0.726965 | 0.745047 |
| 0.667271 | 0.656036 | 0.655758 | 0.72643 | 0.654942 | 0.72016 | 0.727635 | 0.721123 | 0.743136 |
| 0.661669 | 0.652721 | 0.65432 | 0.722718 | 0.658587 | 0.720261 | 0.725261 | 0.720246 | 0.740163 |
| 0.66879 | 0.660991 | 0.659683 | 0.731323 | 0.664992 | 0.723206 | 0.730184 | 0.726055 | 0.746623 |
| 0.66143 | 0.65617 | 0.648513 | 0.717437 | 0.65484 | 0.713969 | 0.716281 | 0.714835 | 0.731833 |
| 0.670586 | 0.663636 | 0.656302 | 0.72508 | 0.662191 | 0.717576 | 0.722849 | 0.723558 | 0.73808 |
| 0.671135 | 0.663475 | 0.659761 | 0.727949 | 0.664982 | 0.721003 | 0.729425 | 0.724788 | 0.746498 |
| 0.673698 | 0.663743 | 0.661319 | 0.730344 | 0.666641 | 0.721926 | 0.731578 | 0.727273 | 0.740467 |
| 0.668566 | 0.66353 | 0.660143 | 0.726588 | 0.662725 | 0.719265 | 0.729662 | 0.722056 | 0.740799 |
| 0.660857 | 0.659718 | 0.653271 | 0.722926 | 0.661221 | 0.712942 | 0.724823 | 0.717888 | 0.737691 |
| 0.662478 | 0.654346 | 0.65082 | 0.719498 | 0.65698 | 0.717507 | 0.728027 | 0.717339 | 0.741028 |
| 0.664962 | 0.659286 | 0.655012 | 0.721261 | 0.660836 | 0.715155 | 0.720948 | 0.718976 | 0.735488 |
| 0.717479 | 0.702426 | 0.69893 | 0.752699 | 0.693723 | 0.756642 | 0.751425 | 0.757245 | 0.733439 |
| 0.715237 | 0.729273 | 0.723597 | 0.771586 | 0.730138 | 0.776326 | 0.769319 | 0.776483 | 0.7479 |
| 0.733397 | 0.711883 | 0.701956 | 0.76472 | 0.702571 | 0.773132 | 0.763121 | 0.76989 | 0.744817 |
| 0.725594 | 0.70839 | 0.698121 | 0.775867 | 0.699072 | 0.764222 | 0.761093 | 0.769545 | 0.749141 |
| 0.738338 | 0.728264 | 0.727481 | 0.779424 | 0.718671 | 0.774885 | 0.768453 | 0.775726 | 0.75218 |
| 0.664063 | 0.654495 | 0.651727 | 0.720816 | 0.656829 | 0.712921 | 0.719682 | 0.718044 | 0.741737 |
| 0.665593 | 0.657835 | 0.653621 | 0.723442 | 0.660198 | 0.717194 | 0.730069 | 0.72189 | 0.742138 |
| 0.669099 | 0.660243 | 0.655598 | 0.726781 | 0.66311 | 0.715437 | 0.727727 | 0.724411 | 0.742939 |
| 0.669643 | 0.659515 | 0.658073 | 0.729213 | 0.664353 | 0.720984 | 0.727215 | 0.724755 | 0.743951 |
| 0.670365 | 0.659204 | 0.659961 | 0.724713 | 0.661334 | 0.716091 | 0.725821 | 0.726615 | 0.745952 |
| 0.669946 | 0.66109 | 0.659772 | 0.73336 | 0.663817 | 0.725068 | 0.730026 | 0.729891 | 0.747681 |
| 0.665388 | 0.657849 | 0.655612 | 0.728546 | 0.660287 | 0.717368 | 0.728301 | 0.723076 | 0.743069 |
| 0.668369 | 0.658204 | 0.655156 | 0.727961 | 0.659561 | 0.715621 | 0.728826 | 0.718963 | 0.742304 |
| 0.665223 | 0.654303 | 0.65201 | 0.719164 | 0.656738 | 0.713067 | 0.720249 | 0.718784 | 0.740957 |
| 0.663344 | 0.653806 | 0.650835 | 0.718788 | 0.65443 | 0.709838 | 0.719609 | 0.716042 | 0.737602 |
| 0.667498 | 0.662363 | 0.657217 | 0.722145 | 0.662795 | 0.714966 | 0.728502 | 0.723237 | 0.736882 |
| 0.67144 | 0.661012 | 0.655396 | 0.729254 | 0.663089 | 0.7195 | 0.733982 | 0.725478 | 0.747018 |

| | | | | | | | | |
|---|---|---|---|---|---|---|---|---|
| 0.667054 | 0.660488 | 0.661311 | 0.730505 | 0.665919 | 0.720525 | 0.727169 | 0.726417 | 0.746661 |
| 0.670638 | 0.664544 | 0.659506 | 0.725223 | 0.66042 | 0.720767 | 0.726055 | 0.723351 | 0.743384 |
| 0.672314 | 0.663192 | 0.660138 | 0.730118 | 0.664225 | 0.724177 | 0.730104 | 0.728523 | 0.745376 |
| 0.667886 | 0.65905 | 0.654655 | 0.727616 | 0.661483 | 0.720906 | 0.725278 | 0.721342 | 0.743695 |
| 0.666098 | 0.655744 | 0.654895 | 0.723535 | 0.658638 | 0.719803 | 0.722821 | 0.715983 | 0.740262 |
| 0.664915 | 0.659102 | 0.654867 | 0.720392 | 0.659692 | 0.715385 | 0.721614 | 0.723323 | 0.738674 |
| 0.669262 | 0.662503 | 0.661898 | 0.726471 | 0.664659 | 0.720634 | 0.729012 | 0.724546 | 0.743041 |
| 0.672584 | 0.662191 | 0.657905 | 0.730633 | 0.661678 | 0.723469 | 0.732791 | 0.723162 | 0.738892 |
| 0.668717 | 0.658875 | 0.653513 | 0.726312 | 0.662886 | 0.718227 | 0.723621 | 0.722812 | 0.740241 |
| 0.668815 | 0.661139 | 0.655635 | 0.726539 | 0.658307 | 0.717361 | 0.724598 | 0.724013 | 0.741301 |
| 0.668703 | 0.661591 | 0.657551 | 0.726075 | 0.659612 | 0.721114 | 0.72456 | 0.720023 | 0.743724 |
| 0.667748 | 0.659796 | 0.653742 | 0.726199 | 0.657822 | 0.720248 | 0.728132 | 0.720523 | 0.743185 |
| 0.658946 | 0.652993 | 0.651027 | 0.716513 | 0.651649 | 0.711585 | 0.716289 | 0.713303 | 0.734094 |
| 0.662021 | 0.650472 | 0.647251 | 0.717998 | 0.653014 | 0.71718 | 0.719198 | 0.718017 | 0.735979 |
| 0.670426 | 0.662591 | 0.65787 | 0.733262 | 0.661022 | 0.726226 | 0.729569 | 0.727176 | 0.740572 |
| 0.673401 | 0.666641 | 0.661551 | 0.736384 | 0.668811 | 0.728985 | 0.737731 | 0.734199 | 0.748891 |
| 0.672166 | 0.663444 | 0.656106 | 0.730152 | 0.664036 | 0.722403 | 0.732031 | 0.727589 | 0.745168 |
| 0.675525 | 0.666138 | 0.660613 | 0.731828 | 0.667347 | 0.723286 | 0.737155 | 0.728606 | 0.749877 |
| 0.666929 | 0.660209 | 0.653315 | 0.727898 | 0.659578 | 0.721426 | 0.728237 | 0.721032 | 0.744512 |
| 0.668235 | 0.661666 | 0.654795 | 0.72596 | 0.66249 | 0.722994 | 0.733471 | 0.722012 | 0.740632 |
| 0.665088 | 0.661094 | 0.655781 | 0.723211 | 0.660433 | 0.719205 | 0.726428 | 0.719099 | 0.736744 |
| 0.671969 | 0.659692 | 0.657736 | 0.728241 | 0.660296 | 0.722727 | 0.731217 | 0.725577 | 0.745362 |
| 0.670467 | 0.664252 | 0.660537 | 0.731927 | 0.663634 | 0.72068 | 0.733973 | 0.731467 | 0.746189 |
| 0.666315 | 0.65795 | 0.654128 | 0.723654 | 0.658009 | 0.718485 | 0.723873 | 0.722969 | 0.737984 |
| 0.667157 | 0.65661 | 0.6526 | 0.728406 | 0.659245 | 0.721528 | 0.730495 | 0.725116 | 0.745969 |
| 0.725841 | 0.717511 | 0.707274 | 0.772254 | 0.713124 | 0.779515 | 0.769542 | 0.782507 | 0.750228 |
| 0.671684 | 0.662276 | 0.660874 | 0.72728 | 0.663353 | 0.720092 | 0.724522 | 0.72317 | 0.742521 |
| 0.673316 | 0.66569 | 0.661324 | 0.732278 | 0.665617 | 0.722965 | 0.731054 | 0.726846 | 0.741718 |
| 0.675328 | 0.667501 | 0.665894 | 0.736455 | 0.668719 | 0.727015 | 0.735908 | 0.729227 | 0.750021 |
| 0.674805 | 0.66397 | 0.661053 | 0.726124 | 0.665986 | 0.722948 | 0.732354 | 0.724497 | 0.740016 |
| 0.666773 | 0.65851 | 0.654925 | 0.725046 | 0.658583 | 0.721122 | 0.72339 | 0.720324 | 0.742205 |
| 0.667372 | 0.659632 | 0.656442 | 0.726258 | 0.662117 | 0.721614 | 0.727452 | 0.721087 | 0.74221 |
| 0.668346 | 0.659522 | 0.657303 | 0.728162 | 0.664335 | 0.726385 | 0.726785 | 0.724186 | 0.744741 |
| 0.667622 | 0.659155 | 0.657886 | 0.724897 | 0.659197 | 0.714896 | 0.723573 | 0.720221 | 0.741275 |
| 0.666427 | 0.65743 | 0.655244 | 0.724391 | 0.658284 | 0.716647 | 0.725819 | 0.720101 | 0.735734 |
| 0.66765 | 0.661941 | 0.654059 | 0.7244 | 0.660175 | 0.716065 | 0.725698 | 0.723338 | 0.738837 |
| 0.670771 | 0.661331 | 0.660055 | 0.727813 | 0.662147 | 0.721484 | 0.729706 | 0.726747 | 0.745424 |
| 0.669649 | 0.66137 | 0.659886 | 0.732623 | 0.666161 | 0.726191 | 0.736129 | 0.72865 | 0.746363 |
| 0.66682 | 0.660396 | 0.657317 | 0.728396 | 0.662996 | 0.720893 | 0.732019 | 0.726335 | 0.751802 |
| 0.669046 | 0.660878 | 0.657407 | 0.726992 | 0.662611 | 0.721902 | 0.728443 | 0.727182 | 0.741498 |
| 0.665981 | 0.660597 | 0.653546 | 0.720619 | 0.658771 | 0.714405 | 0.725912 | 0.720522 | 0.744113 |
| 0.670893 | 0.663327 | 0.657258 | 0.730134 | 0.662502 | 0.720915 | 0.733648 | 0.72731 | 0.748588 |
| 0.664695 | 0.656683 | 0.656607 | 0.72703 | 0.658026 | 0.720164 | 0.731728 | 0.724253 | 0.741127 |
| 0.665988 | 0.660774 | 0.657903 | 0.729371 | 0.66038 | 0.716551 | 0.727518 | 0.722403 | 0.740042 |
| 0.670251 | 0.667096 | 0.662843 | 0.732487 | 0.666414 | 0.723068 | 0.731447 | 0.727988 | 0.747909 |
| 0.672273 | 0.666491 | 0.661248 | 0.73127 | 0.664124 | 0.721434 | 0.735351 | 0.727433 | 0.746868 |

| | | | | | | | | |
|---|---|---|---|---|---|---|---|---|
| 0.671176 | 0.663848 | 0.664451 | 0.725171 | 0.663075 | 0.721534 | 0.727006 | 0.723131 | 0.744501 |
| 0.671588 | 0.663144 | 0.658996 | 0.733434 | 0.663621 | 0.724474 | 0.732316 | 0.728741 | 0.745174 |
| 0.666819 | 0.665277 | 0.659187 | 0.729062 | 0.661896 | 0.719322 | 0.723817 | 0.722213 | 0.743012 |
| 0.612305 | 0.609732 | 0.613659 | 0.693408 | 0.611872 | 0.675779 | 0.680903 | 0.675545 | 0.718839 |
| 0.668276 | 0.663278 | 0.658379 | 0.727735 | 0.660104 | 0.719975 | 0.729719 | 0.723452 | 0.735546 |
| 0.670265 | 0.659304 | 0.657132 | 0.73047 | 0.659168 | 0.721673 | 0.728766 | 0.726619 | 0.738808 |
| 0.671 | 0.662137 | 0.658628 | 0.729592 | 0.662382 | 0.722049 | 0.731701 | 0.724859 | 0.744935 |
| 0.67176 | 0.6624 | 0.658301 | 0.726781 | 0.663022 | 0.721478 | 0.732055 | 0.722399 | 0.744524 |
| 0.673916 | 0.668599 | 0.664198 | 0.731758 | 0.668571 | 0.728577 | 0.7355 | 0.728849 | 0.750183 |
| 0.673961 | 0.665613 | 0.661415 | 0.726759 | 0.662236 | 0.721111 | 0.726832 | 0.726961 | 0.74432 |
| 0.67202 | 0.667171 | 0.663191 | 0.726895 | 0.664027 | 0.722365 | 0.727657 | 0.724053 | 0.741752 |
| 0.675244 | 0.667625 | 0.662139 | 0.732468 | 0.663929 | 0.725435 | 0.730942 | 0.727738 | 0.745358 |
| 0.673252 | 0.668801 | 0.662005 | 0.732277 | 0.666867 | 0.725008 | 0.730622 | 0.727839 | 0.741551 |
| 0.673667 | 0.663449 | 0.65916 | 0.725535 | 0.662398 | 0.724826 | 0.731688 | 0.725787 | 0.743982 |
| 0.673634 | 0.667318 | 0.663022 | 0.728909 | 0.667971 | 0.727681 | 0.732609 | 0.728309 | 0.745515 |
| 0.667147 | 0.662382 | 0.656147 | 0.728763 | 0.657988 | 0.719155 | 0.728853 | 0.722382 | 0.740717 |
| 0.668994 | 0.666542 | 0.665172 | 0.734637 | 0.665497 | 0.722227 | 0.733238 | 0.727589 | 0.751033 |
| 0.672292 | 0.666817 | 0.661387 | 0.728884 | 0.66392 | 0.723733 | 0.72988 | 0.723068 | 0.745384 |
| 0.668256 | 0.658851 | 0.655519 | 0.722982 | 0.659739 | 0.718413 | 0.728815 | 0.722152 | 0.7375 |
| 0.670271 | 0.665742 | 0.656252 | 0.730264 | 0.663368 | 0.721298 | 0.728031 | 0.723678 | 0.741563 |
| 0.673647 | 0.667118 | 0.663328 | 0.7341 | 0.664639 | 0.723986 | 0.734248 | 0.728542 | 0.748208 |
| 0.67531 | 0.669548 | 0.667254 | 0.733371 | 0.668673 | 0.724428 | 0.733678 | 0.730401 | 0.74492 |
| 0.646674 | 0.640644 | 0.639263 | 0.700219 | 0.637528 | 0.698802 | 0.703165 | 0.686183 | 0.694771 |
| 0.659361 | 0.65301 | 0.655063 | 0.708444 | 0.644898 | 0.723368 | 0.713837 | 0.718637 | 0.703544 |
| 0.704897 | 0.69854 | 0.688099 | 0.737217 | 0.688287 | 0.741184 | 0.734802 | 0.747571 | 0.726309 |
| 0.671168 | 0.66879 | 0.656225 | 0.754762 | 0.650801 | 0.735646 | 0.736142 | 0.737328 | 0.717565 |
| 0.631867 | 0.614489 | 0.613995 | 0.691873 | 0.614822 | 0.687002 | 0.696749 | 0.682514 | 0.707363 |
| 0.61913 | 0.616615 | 0.613155 | 0.686278 | 0.608801 | 0.682329 | 0.683099 | 0.677592 | 0.703517 |
| 0.621106 | 0.607166 | 0.606496 | 0.683273 | 0.605796 | 0.678769 | 0.683553 | 0.677855 | 0.700331 |
| 0.636173 | 0.6283 | 0.62987 | 0.698561 | 0.625544 | 0.695688 | 0.6935 | 0.690548 | 0.705751 |
| 0.619347 | 0.61295 | 0.605741 | 0.681788 | 0.6065 | 0.675454 | 0.67085 | 0.672936 | 0.703312 |
| 0.60082 | 0.590018 | 0.588728 | 0.667933 | 0.586129 | 0.663152 | 0.664677 | 0.655331 | 0.689523 |
| 0.607075 | 0.596727 | 0.590291 | 0.670094 | 0.59194 | 0.662316 | 0.674706 | 0.656901 | 0.695942 |
| 0.678274 | 0.66379 | 0.656271 | 0.719285 | 0.651492 | 0.724012 | 0.719455 | 0.715371 | 0.707185 |
| 0.666508 | 0.656925 | 0.654048 | 0.72725 | 0.656297 | 0.719863 | 0.724731 | 0.720217 | 0.738063 |
| 0.635194 | 0.631077 | 0.627185 | 0.699842 | 0.627196 | 0.69299 | 0.697612 | 0.692969 | 0.714707 |
| 0.62761 | 0.618838 | 0.614597 | 0.694658 | 0.616302 | 0.688232 | 0.693614 | 0.685688 | 0.710371 |
| 0.618687 | 0.605091 | 0.60446 | 0.679296 | 0.604146 | 0.674236 | 0.67439 | 0.670777 | 0.699134 |
| 0.636335 | 0.624736 | 0.622107 | 0.700146 | 0.625645 | 0.694692 | 0.699052 | 0.694763 | 0.719134 |
| 0.633283 | 0.62388 | 0.619661 | 0.695847 | 0.623906 | 0.689271 | 0.690132 | 0.68302 | 0.710855 |
| 0.623969 | 0.612107 | 0.605672 | 0.690326 | 0.611167 | 0.684194 | 0.686034 | 0.679048 | 0.701624 |
| 0.626314 | 0.616464 | 0.613234 | 0.687614 | 0.618285 | 0.684848 | 0.685495 | 0.68445 | 0.706758 |
| 0.615862 | 0.604082 | 0.60058 | 0.676328 | 0.60342 | 0.676757 | 0.688241 | 0.673856 | 0.704774 |
| 0.624865 | 0.60952 | 0.610953 | 0.682452 | 0.60813 | 0.685115 | 0.682085 | 0.673913 | 0.694439 |
| 0.62254 | 0.605611 | 0.601145 | 0.68789 | 0.60383 | 0.67911 | 0.675037 | 0.683042 | 0.700673 |
| 0.60524 | 0.598442 | 0.599402 | 0.675048 | 0.590329 | 0.665609 | 0.661071 | 0.667199 | 0.678208 |

| | | | | | | | | |
|---|---|---|---|---|---|---|---|---|
| 0.649022 | 0.642683 | 0.639068 | 0.711113 | 0.641561 | 0.706182 | 0.710292 | 0.707957 | 0.725446 |
| 0.624393 | 0.613345 | 0.609599 | 0.68456 | 0.608426 | 0.676753 | 0.684503 | 0.676784 | 0.706108 |
| 0.618491 | 0.610346 | 0.612899 | 0.683511 | 0.609895 | 0.680626 | 0.68559 | 0.682705 | 0.703022 |
| 0.622691 | 0.616918 | 0.609558 | 0.684086 | 0.613313 | 0.683673 | 0.687039 | 0.68253 | 0.705725 |
| 0.625023 | 0.610194 | 0.603661 | 0.683448 | 0.603932 | 0.676373 | 0.69337 | 0.680709 | 0.705615 |
| 0.629689 | 0.609569 | 0.608861 | 0.685569 | 0.606719 | 0.68149 | 0.685728 | 0.673661 | 0.69935 |
| 0.611465 | 0.601184 | 0.602003 | 0.677049 | 0.602716 | 0.669618 | 0.678755 | 0.664104 | 0.695519 |
| 0.604204 | 0.598985 | 0.605216 | 0.668287 | 0.598267 | 0.668838 | 0.677508 | 0.670688 | 0.689101 |
| 0.609107 | 0.608953 | 0.607106 | 0.675683 | 0.596227 | 0.669471 | 0.675734 | 0.667485 | 0.696895 |
| 0.65281 | 0.639886 | 0.640524 | 0.716996 | 0.641459 | 0.706268 | 0.712105 | 0.707275 | 0.729973 |
| 0.649866 | 0.640235 | 0.636055 | 0.709997 | 0.637575 | 0.705423 | 0.713143 | 0.708459 | 0.727721 |
| 0.647834 | 0.63459 | 0.63298 | 0.706625 | 0.635587 | 0.702173 | 0.707257 | 0.706444 | 0.723491 |
| 0.632797 | 0.623904 | 0.627276 | 0.699251 | 0.623195 | 0.692533 | 0.700247 | 0.694591 | 0.717209 |
| 0.638586 | 0.629605 | 0.626968 | 0.701651 | 0.627072 | 0.695069 | 0.699176 | 0.69433 | 0.71809 |
| 0.632233 | 0.624304 | 0.622351 | 0.697286 | 0.625264 | 0.689178 | 0.691508 | 0.692153 | 0.711302 |

| Collection_ | Collection_ | Collection_ | Collection_ | Collection_ | Collection_ | Collection_ | Collection_ | Collection_ |
|---|---|---|---|---|---|---|---|---|
| 0.704312 | 0.703276 | 0.77745 | 0.687514 | 0.72996 | 0.763575 | 0.718164 | 0.749654 | 0.611111 |
| 0.711549 | 0.714853 | 0.777268 | 0.693687 | 0.734498 | 0.75935 | 0.72293 | 0.750285 | 0.659722 |
| 0.714918 | 0.704195 | 0.76833 | 0.689092 | 0.728298 | 0.761368 | 0.71843 | 0.750611 | 0.643836 |
| 0.702909 | 0.696282 | 0.773356 | 0.693175 | 0.745996 | 0.762738 | 0.729266 | 0.753826 | 0.655844 |
| 0.697548 | 0.692008 | 0.762767 | 0.694524 | 0.734046 | 0.760877 | 0.722308 | 0.745359 | 0.623529 |
| 0.697257 | 0.696289 | 0.766669 | 0.688937 | 0.74108 | 0.755662 | 0.734047 | 0.75136 | 0.625 |
| 0.654261 | 0.655393 | 0.735021 | 0.651955 | 0.740229 | 0.728046 | 0.738352 | 0.733603 | 0.716216 |
| 0.64979 | 0.651428 | 0.726953 | 0.652232 | 0.732624 | 0.730927 | 0.732285 | 0.730999 | 0.671053 |
| 0.702584 | 0.702424 | 0.771667 | 0.688781 | 0.737775 | 0.765058 | 0.725382 | 0.756659 | 0.6125 |
| 0.708029 | 0.698092 | 0.768849 | 0.69119 | 0.729773 | 0.763611 | 0.720635 | 0.763571 | 0.610778 |
| 0.702034 | 0.69795 | 0.768739 | 0.692495 | 0.730994 | 0.760592 | 0.71573 | 0.753196 | 0.621118 |
| 0.70966 | 0.695656 | 0.77113 | 0.686583 | 0.728023 | 0.763078 | 0.717182 | 0.750326 | 0.597561 |
| 0.706365 | 0.713613 | 0.771246 | 0.685902 | 0.73183 | 0.762817 | 0.720981 | 0.757509 | 0.595092 |
| 0.700455 | 0.699869 | 0.785802 | 0.690345 | 0.74248 | 0.772269 | 0.725032 | 0.773596 | 0.649007 |
| 0.703975 | 0.694913 | 0.790479 | 0.683174 | 0.741509 | 0.793943 | 0.734841 | 0.788943 | 0.689189 |
| 0.715255 | 0.699193 | 0.799983 | 0.692115 | 0.739865 | 0.82552 | 0.724679 | 0.757318 | 0.624204 |
| 0.711084 | 0.688419 | 0.783905 | 0.688887 | 0.730474 | 0.784677 | 0.722816 | 0.762538 | 0.684211 |
| 0.695834 | 0.687831 | 0.806951 | 0.683147 | 0.741142 | 0.802549 | 0.73792 | 0.805148 | 0.643312 |
| 0.718158 | 0.70792 | 0.770837 | 0.691489 | 0.731453 | 0.77254 | 0.713591 | 0.756657 | 0.652482 |
| 0.705528 | 0.701103 | 0.771141 | 0.694726 | 0.737581 | 0.759001 | 0.719925 | 0.75096 | 0.602484 |
| 0.709253 | 0.695098 | 0.769702 | 0.694265 | 0.732134 | 0.755368 | 0.720622 | 0.748494 | 0.586826 |
| 0.71243 | 0.694608 | 0.771135 | 0.69302 | 0.736324 | 0.767602 | 0.729314 | 0.758235 | 0.606061 |
| 0.715171 | 0.701055 | 0.775353 | 0.697264 | 0.741284 | 0.764691 | 0.733165 | 0.762841 | 0.633987 |
| 0.716383 | 0.711554 | 0.788296 | 0.695418 | 0.74015 | 0.766759 | 0.726741 | 0.762276 | 0.568862 |
| 0.707028 | 0.701632 | 0.773 | 0.694529 | 0.738804 | 0.765117 | 0.729483 | 0.748755 | 0.594771 |
| 0.701327 | 0.704638 | 0.76753 | 0.688834 | 0.732839 | 0.768014 | 0.721488 | 0.747591 | 0.624204 |
| 0.707163 | 0.705752 | 0.757894 | 0.688196 | 0.72787 | 0.754831 | 0.716216 | 0.744846 | 0.570552 |
| 0.705037 | 0.703834 | 0.766537 | 0.696811 | 0.738286 | 0.754285 | 0.724865 | 0.745085 | 0.641892 |
| 0.702409 | 0.699746 | 0.772114 | 0.695649 | 0.733652 | 0.758241 | 0.726306 | 0.752265 | 0.605263 |
| 0.711643 | 0.70934 | 0.771177 | 0.683356 | 0.729363 | 0.774673 | 0.718938 | 0.760662 | 0.653061 |
| 0.697306 | 0.694943 | 0.758041 | 0.687842 | 0.734795 | 0.757789 | 0.719606 | 0.740313 | 0.6375 |
| 0.700423 | 0.693416 | 0.75902 | 0.687458 | 0.736701 | 0.75641 | 0.723219 | 0.740965 | 0.632911 |
| 0.70932 | 0.699707 | 0.767344 | 0.695004 | 0.73445 | 0.760451 | 0.721506 | 0.746168 | 0.628931 |
| 0.702893 | 0.696256 | 0.75395 | 0.692092 | 0.737881 | 0.753069 | 0.726022 | 0.739215 | 0.608696 |
| 0.711272 | 0.695827 | 0.762847 | 0.691899 | 0.72999 | 0.760226 | 0.717652 | 0.742624 | 0.616438 |
| 0.711252 | 0.704907 | 0.764513 | 0.696476 | 0.733377 | 0.765676 | 0.721463 | 0.762593 | 0.646259 |
| 0.701032 | 0.697671 | 0.768127 | 0.689655 | 0.741025 | 0.754196 | 0.724216 | 0.750486 | 0.590909 |
| 0.694453 | 0.692674 | 0.761944 | 0.685147 | 0.73621 | 0.747829 | 0.727825 | 0.745586 | 0.601266 |
| 0.702834 | 0.69691 | 0.768836 | 0.68845 | 0.740635 | 0.762503 | 0.727295 | 0.750571 | 0.6125 |

| | | | | | | | | |
|---|---|---|---|---|---|---|---|---|
| 0.688653 | 0.694563 | 0.813252 | 0.684678 | 0.742724 | 0.805748 | 0.737774 | 0.858201 | 0.660256 |
| 0.704962 | 0.714074 | 0.771569 | 0.691871 | 0.72717 | 0.756902 | 0.714954 | 0.752428 | 0.690141 |
| 0.708614 | 0.701602 | 0.794174 | 0.692963 | 0.731828 | 0.765281 | 0.717876 | 0.751988 | 0.569697 |
| 0.717795 | 0.701913 | 0.799951 | 0.690541 | 0.738255 | 0.773172 | 0.723242 | 0.767818 | 0.6 |
| 0.707789 | 0.699199 | 0.769281 | 0.690145 | 0.734247 | 0.766475 | 0.721844 | 0.747741 | 0.625 |
| 0.707696 | 0.694971 | 0.766134 | 0.689527 | 0.735922 | 0.768876 | 0.725742 | 0.754067 | 0.642384 |
| 0.701425 | 0.695716 | 0.754674 | 0.686298 | 0.730064 | 0.759365 | 0.721469 | 0.742727 | 0.623377 |
| 0.714823 | 0.71237 | 0.771584 | 0.700069 | 0.737032 | 0.765393 | 0.723058 | 0.758183 | 0.616766 |
| 0.698664 | 0.699333 | 0.759355 | 0.692049 | 0.735962 | 0.756936 | 0.725883 | 0.745684 | 0.640523 |
| 0.693557 | 0.690784 | 0.757559 | 0.6905 | 0.73397 | 0.756005 | 0.723286 | 0.745603 | 0.613924 |
| 0.7079 | 0.697352 | 0.773452 | 0.685326 | 0.729662 | 0.771773 | 0.716995 | 0.75115 | 0.651007 |
| 0.719324 | 0.703889 | 0.781899 | 0.690511 | 0.732681 | 0.764233 | 0.722344 | 0.756242 | 0.602564 |
| 0.71378 | 0.71217 | 0.775804 | 0.691033 | 0.735106 | 0.769193 | 0.720862 | 0.757491 | 0.631206 |
| 0.704301 | 0.698688 | 0.780369 | 0.688252 | 0.731061 | 0.76459 | 0.72058 | 0.755351 | 0.627329 |
| 0.711541 | 0.698561 | 0.774535 | 0.688615 | 0.730325 | 0.756997 | 0.717588 | 0.747753 | 0.587097 |
| 0.706418 | 0.690837 | 0.772527 | 0.685626 | 0.728365 | 0.755859 | 0.719279 | 0.743844 | 0.633987 |
| 0.700307 | 0.692136 | 0.771433 | 0.68508 | 0.728346 | 0.765799 | 0.71474 | 0.749381 | 0.591195 |
| 0.724651 | 0.71132 | 0.78032 | 0.694506 | 0.735918 | 0.776636 | 0.72077 | 0.758663 | 0.612903 |
| 0.705583 | 0.704777 | 0.779757 | 0.692577 | 0.738398 | 0.767011 | 0.72175 | 0.762054 | 0.607595 |
| 0.712221 | 0.712684 | 0.76868 | 0.690376 | 0.73329 | 0.777123 | 0.722231 | 0.754015 | 0.607595 |
| 0.709129 | 0.695174 | 0.779412 | 0.6899 | 0.72594 | 0.776517 | 0.714089 | 0.750821 | 0.613924 |
| 0.70577 | 0.70597 | 0.777137 | 0.685201 | 0.731352 | 0.766716 | 0.715627 | 0.756675 | 0.580247 |
| 0.709589 | 0.703864 | 0.767764 | 0.690782 | 0.731126 | 0.764802 | 0.720039 | 0.75555 | 0.634731 |
| 0.700213 | 0.693478 | 0.762527 | 0.688506 | 0.738697 | 0.763566 | 0.725985 | 0.753014 | 0.608696 |
| 0.691221 | 0.688691 | 0.762664 | 0.682456 | 0.740211 | 0.755987 | 0.733274 | 0.741002 | 0.647799 |
| 0.649242 | 0.65173 | 0.728678 | 0.652541 | 0.732549 | 0.73217 | 0.732156 | 0.731126 | 0.675497 |
| 0.656336 | 0.65469 | 0.730806 | 0.658639 | 0.733186 | 0.727295 | 0.731373 | 0.730245 | 0.709459 |
| 0.654719 | 0.654582 | 0.732726 | 0.658325 | 0.733989 | 0.733485 | 0.732238 | 0.734033 | 0.702703 |
| 0.708054 | 0.709287 | 0.760371 | 0.69958 | 0.730186 | 0.752891 | 0.717617 | 0.744104 | 0.634615 |
| 0.705237 | 0.698708 | 0.762706 | 0.68676 | 0.729714 | 0.747519 | 0.718374 | 0.74295 | 0.626667 |
| 0.694337 | 0.693521 | 0.77659 | 0.685063 | 0.735512 | 0.760052 | 0.718639 | 0.75217 | 0.679245 |
| 0.70692 | 0.703578 | 0.764405 | 0.689352 | 0.749834 | 0.753847 | 0.735304 | 0.747486 | 0.637584 |
| 0.744468 | 0.72927 | 0.785964 | 0.694686 | 0.736059 | 0.781878 | 0.720758 | 0.769006 | 0.635762 |
| 0.720096 | 0.704761 | 0.781597 | 0.694131 | 0.733765 | 0.781091 | 0.719043 | 0.760711 | 0.601307 |
| 0.740939 | 0.72093 | 0.791317 | 0.695609 | 0.732552 | 0.784103 | 0.716373 | 0.779953 | 0.632258 |
| 0.737067 | 0.711785 | 0.78885 | 0.696248 | 0.732833 | 0.791118 | 0.718024 | 0.769365 | 0.629139 |
| 0.716773 | 0.73921 | 0.779062 | 0.690577 | 0.732108 | 0.763258 | 0.721355 | 0.753161 | 0.670968 |
| 0.733514 | 0.740512 | 0.797479 | 0.710363 | 0.749365 | 0.790731 | 0.734079 | 0.782464 | 0.651007 |
| 0.726615 | 0.717565 | 0.772503 | 0.697444 | 0.729963 | 0.770691 | 0.709824 | 0.764907 | 0.675676 |
| 0.700053 | 0.703239 | 0.772473 | 0.695846 | 0.73285 | 0.763524 | 0.723899 | 0.749273 | 0.607595 |
| 0.713516 | 0.704625 | 0.785913 | 0.692267 | 0.72919 | 0.772107 | 0.723069 | 0.758076 | 0.636364 |
| 0.732878 | 0.732847 | 0.792937 | 0.713955 | 0.755953 | 0.796473 | 0.737498 | 0.781666 | 0.675676 |
| 0.698935 | 0.699428 | 0.782856 | 0.692518 | 0.739542 | 0.772552 | 0.725917 | 0.760279 | 0.645161 |
| 0.695537 | 0.693376 | 0.777134 | 0.690097 | 0.734773 | 0.764452 | 0.722459 | 0.755067 | 0.611111 |
| 0.699397 | 0.705451 | 0.770532 | 0.69181 | 0.736347 | 0.765455 | 0.726262 | 0.744954 | 0.647436 |
| 0.726331 | 0.707706 | 0.769104 | 0.715939 | 0.731665 | 0.777473 | 0.717739 | 0.761716 | 0.615385 |

| | | | | | | | | |
|---|---|---|---|---|---|---|---|---|
| 0.700684 | 0.700698 | 0.776928 | 0.691096 | 0.73143 | 0.769092 | 0.720826 | 0.751185 | 0.639241 |
| 0.698371 | 0.705461 | 0.767529 | 0.696327 | 0.739431 | 0.759684 | 0.727126 | 0.749023 | 0.589744 |
| 0.699648 | 0.714155 | 0.771947 | 0.6913 | 0.730492 | 0.764322 | 0.718549 | 0.75138 | 0.625767 |
| 0.722382 | 0.72563 | 0.783862 | 0.713264 | 0.739116 | 0.776676 | 0.730195 | 0.768103 | 0.68 |
| 0.703878 | 0.708705 | 0.77968 | 0.693361 | 0.738669 | 0.768151 | 0.723617 | 0.757363 | 0.652695 |
| 0.698884 | 0.696539 | 0.768724 | 0.694521 | 0.738123 | 0.759681 | 0.72737 | 0.749918 | 0.627451 |
| 0.713192 | 0.728082 | 0.779932 | 0.690659 | 0.734043 | 0.762452 | 0.721757 | 0.759002 | 0.625 |
| 0.705166 | 0.710017 | 0.769985 | 0.695233 | 0.733524 | 0.764526 | 0.729026 | 0.747084 | 0.62963 |
| 0.703505 | 0.700272 | 0.768782 | 0.68749 | 0.73642 | 0.763286 | 0.724382 | 0.751027 | 0.596026 |
| 0.702588 | 0.697242 | 0.764165 | 0.693476 | 0.733138 | 0.759302 | 0.718798 | 0.745199 | 0.636986 |
| 0.702958 | 0.704838 | 0.767017 | 0.69972 | 0.730618 | 0.766483 | 0.724471 | 0.752937 | 0.666667 |
| 0.697726 | 0.701091 | 0.763542 | 0.694182 | 0.734201 | 0.760803 | 0.721994 | 0.745736 | 0.664516 |
| 0.702994 | 0.701882 | 0.762283 | 0.694258 | 0.73607 | 0.760921 | 0.720881 | 0.747823 | 0.605096 |
| 0.69044 | 0.697401 | 0.763482 | 0.691555 | 0.729122 | 0.758469 | 0.715691 | 0.743372 | 0.62 |
| 0.701204 | 0.700098 | 0.764423 | 0.69116 | 0.730488 | 0.759511 | 0.723409 | 0.751391 | 0.623377 |
| 0.701429 | 0.699584 | 0.760078 | 0.693828 | 0.732561 | 0.760896 | 0.722919 | 0.745173 | 0.655629 |
| 0.696214 | 0.706393 | 0.768996 | 0.692607 | 0.734717 | 0.756963 | 0.722272 | 0.750267 | 0.61039 |
| 0.7218 | 0.70067 | 0.778124 | 0.694045 | 0.732117 | 0.777339 | 0.719356 | 0.755196 | 0.633333 |
| 0.701203 | 0.702288 | 0.766119 | 0.692905 | 0.735445 | 0.759797 | 0.720055 | 0.749039 | 0.625 |
| 0.702452 | 0.699441 | 0.763506 | 0.697414 | 0.734765 | 0.759374 | 0.7221 | 0.750409 | 0.607843 |
| 0.694273 | 0.697072 | 0.760349 | 0.690787 | 0.730669 | 0.759561 | 0.724163 | 0.747306 | 0.675862 |
| 0.697955 | 0.694019 | 0.764816 | 0.72452 | 0.729042 | 0.757662 | 0.716903 | 0.746482 | 0.660377 |
| 0.700634 | 0.700807 | 0.765398 | 0.697657 | 0.729668 | 0.759221 | 0.717808 | 0.751728 | 0.655629 |
| 0.701908 | 0.699175 | 0.77187 | 0.694118 | 0.738173 | 0.762165 | 0.720281 | 0.747015 | 0.62963 |
| 0.708948 | 0.71781 | 0.771682 | 0.688274 | 0.728973 | 0.757728 | 0.718801 | 0.753518 | 0.616352 |
| 0.720866 | 0.728669 | 0.7801 | 0.694166 | 0.742103 | 0.766849 | 0.722012 | 0.764643 | 0.625806 |
| 0.706114 | 0.717037 | 0.772047 | 0.690589 | 0.727273 | 0.76027 | 0.718511 | 0.755725 | 0.664384 |
| 0.701241 | 0.699135 | 0.76873 | 0.695942 | 0.731567 | 0.758432 | 0.723008 | 0.75059 | 0.623457 |
| 0.698925 | 0.693028 | 0.766553 | 0.696484 | 0.745044 | 0.761218 | 0.731819 | 0.753257 | 0.613497 |
| 0.704693 | 0.710932 | 0.771058 | 0.691353 | 0.731279 | 0.759152 | 0.724351 | 0.752424 | 0.620482 |
| 0.699763 | 0.697663 | 0.768725 | 0.695423 | 0.737896 | 0.758793 | 0.729023 | 0.74438 | 0.641975 |
| 0.701443 | 0.699716 | 0.76327 | 0.695983 | 0.730185 | 0.759977 | 0.728897 | 0.747463 | 0.613924 |
| 0.720398 | 0.729247 | 0.777916 | 0.694442 | 0.734936 | 0.767316 | 0.724572 | 0.759213 | 0.627451 |
| 0.711725 | 0.701737 | 0.800357 | 0.690239 | 0.735994 | 0.773267 | 0.719082 | 0.769049 | 0.612121 |
| 0.69955 | 0.699327 | 0.76556 | 0.694767 | 0.737947 | 0.754892 | 0.716528 | 0.751561 | 0.635762 |
| 0.697815 | 0.69328 | 0.767291 | 0.689719 | 0.735549 | 0.760046 | 0.723632 | 0.748113 | 0.617284 |
| 0.70026 | 0.696114 | 0.762652 | 0.692175 | 0.727928 | 0.754261 | 0.718439 | 0.748304 | 0.605634 |
| 0.66689 | 0.67482 | 0.74587 | 0.668524 | 0.710271 | 0.736395 | 0.693871 | 0.723444 | 0.61875 |
| 0.678174 | 0.680832 | 0.752297 | 0.676414 | 0.716009 | 0.742138 | 0.70273 | 0.724508 | 0.62 |
| 0.677697 | 0.675297 | 0.736842 | 0.672123 | 0.703894 | 0.735861 | 0.693339 | 0.722158 | 0.625 |
| 0.69158 | 0.678415 | 0.753127 | 0.666282 | 0.707736 | 0.751601 | 0.69095 | 0.739876 | 0.630137 |
| 0.651468 | 0.661437 | 0.73586 | 0.656357 | 0.691164 | 0.71878 | 0.67616 | 0.712138 | 0.59375 |
| 0.660002 | 0.666112 | 0.75395 | 0.6581 | 0.692157 | 0.734635 | 0.68027 | 0.72456 | 0.655629 |
| 0.680181 | 0.679035 | 0.745145 | 0.666667 | 0.697435 | 0.741964 | 0.682638 | 0.727056 | 0.61875 |
| 0.666061 | 0.673411 | 0.731404 | 0.652758 | 0.700651 | 0.73431 | 0.68519 | 0.71451 | 0.673469 |
| 0.677776 | 0.701818 | 0.75226 | 0.666351 | 0.695928 | 0.739532 | 0.682989 | 0.72575 | 0.715328 |
| 0.687426 | 0.673608 | 0.749936 | 0.656621 | 0.698895 | 0.745315 | 0.683589 | 0.727145 | 0.654135 |

| | | | | | | | | |
|---|---|---|---|---|---|---|---|---|
| 0.655992 | 0.66497 | 0.727857 | 0.657502 | 0.693298 | 0.725545 | 0.682034 | 0.711614 | 0.646667 |
| 0.704835 | 0.716083 | 0.756968 | 0.66044 | 0.699205 | 0.752465 | 0.680354 | 0.743805 | 0.673913 |
| 0.651546 | 0.659862 | 0.734358 | 0.65329 | 0.697813 | 0.719112 | 0.692088 | 0.716644 | 0.597315 |
| 0.655125 | 0.658411 | 0.745318 | 0.648615 | 0.687819 | 0.742517 | 0.684218 | 0.715156 | 0.643357 |
| 0.674081 | 0.696524 | 0.752602 | 0.657738 | 0.692982 | 0.726126 | 0.672914 | 0.729089 | 0.648649 |
| 0.664058 | 0.687669 | 0.742164 | 0.653332 | 0.688963 | 0.727898 | 0.676381 | 0.708551 | 0.666667 |
| 0.668467 | 0.690978 | 0.740047 | 0.655541 | 0.689205 | 0.727866 | 0.673308 | 0.721012 | 0.641892 |
| 0.679159 | 0.67156 | 0.744836 | 0.662184 | 0.694413 | 0.739853 | 0.680399 | 0.717722 | 0.676692 |
| 0.644819 | 0.649371 | 0.721665 | 0.644343 | 0.695978 | 0.714071 | 0.679707 | 0.700961 | 0.655629 |
| 0.699292 | 0.719618 | 0.747244 | 0.657372 | 0.694581 | 0.734154 | 0.677817 | 0.735339 | 0.676923 |
| 0.65045 | 0.684802 | 0.734149 | 0.65774 | 0.690492 | 0.730802 | 0.671086 | 0.706959 | 0.664286 |
| 0.700723 | 0.705921 | 0.7606 | 0.705053 | 0.723809 | 0.764443 | 0.711216 | 0.744942 | 0.628049 |
| 0.700647 | 0.694919 | 0.759742 | 0.699093 | 0.736812 | 0.756224 | 0.725307 | 0.744447 | 0.631579 |
| 0.658578 | 0.684546 | 0.740367 | 0.676249 | 0.711882 | 0.732993 | 0.695029 | 0.724644 | 0.651613 |
| 0.696142 | 0.69457 | 0.767507 | 0.691167 | 0.737059 | 0.764035 | 0.720043 | 0.747753 | 0.627451 |
| 0.697541 | 0.705458 | 0.758083 | 0.705123 | 0.728476 | 0.746775 | 0.712761 | 0.740271 | 0.588235 |
| 0.694732 | 0.696928 | 0.7675 | 0.692714 | 0.73398 | 0.760721 | 0.722227 | 0.752438 | 0.636905 |
| 0.70023 | 0.693596 | 0.766836 | 0.696452 | 0.742462 | 0.758519 | 0.729821 | 0.746086 | 0.651899 |
| 0.711283 | 0.716834 | 0.74626 | 0.719682 | 0.7144 | 0.749516 | 0.706653 | 0.72924 | 0.623457 |
| 0.690396 | 0.698485 | 0.774089 | 0.693099 | 0.737722 | 0.759935 | 0.725222 | 0.751961 | 0.66879 |
| 0.657117 | 0.678694 | 0.750952 | 0.65592 | 0.709666 | 0.734211 | 0.702452 | 0.72342 | 0.653846 |
| 0.698358 | 0.690655 | 0.758003 | 0.696713 | 0.739568 | 0.762397 | 0.734266 | 0.745312 | 0.576687 |
| 0.695567 | 0.694088 | 0.758138 | 0.693749 | 0.740871 | 0.758646 | 0.728074 | 0.743565 | 0.619355 |
| 0.65554 | 0.662659 | 0.738961 | 0.654147 | 0.702278 | 0.731152 | 0.691981 | 0.719098 | 0.631579 |
| 0.715134 | 0.727725 | 0.747114 | 0.691942 | 0.707561 | 0.740928 | 0.695459 | 0.723641 | 0.608974 |
| 0.678201 | 0.689509 | 0.755886 | 0.674566 | 0.71346 | 0.758339 | 0.704118 | 0.738532 | 0.638298 |
| 0.679899 | 0.692417 | 0.751948 | 0.676036 | 0.727876 | 0.74408 | 0.715491 | 0.739622 | 0.628378 |
| 0.694746 | 0.690834 | 0.766139 | 0.68816 | 0.74379 | 0.752815 | 0.735313 | 0.741951 | 0.611111 |
| 0.71025 | 0.71955 | 0.735373 | 0.71907 | 0.709726 | 0.732453 | 0.698758 | 0.723538 | 0.610063 |
| 0.709522 | 0.731993 | 0.770177 | 0.691781 | 0.73878 | 0.763147 | 0.726651 | 0.765133 | 0.664516 |
| 0.698931 | 0.694198 | 0.761131 | 0.689653 | 0.733925 | 0.756937 | 0.726044 | 0.74795 | 0.672956 |
| 0.741688 | 0.739058 | 0.753254 | 0.816759 | 0.716809 | 0.751344 | 0.706898 | 0.739091 | 0.666667 |
| 0.737754 | 0.743431 | 0.754036 | 0.821504 | 0.718895 | 0.748283 | 0.702592 | 0.742251 | 0.638158 |
| 0.688518 | 0.69134 | 0.761781 | 0.68907 | 0.740614 | 0.755518 | 0.72962 | 0.751302 | 0.679487 |
| 0.689681 | 0.690699 | 0.761988 | 0.692709 | 0.731566 | 0.758635 | 0.723811 | 0.750241 | 0.662338 |
| 0.690456 | 0.6934 | 0.764557 | 0.687851 | 0.739645 | 0.762903 | 0.731423 | 0.74659 | 0.675159 |
| 0.718409 | 0.723192 | 0.723726 | 0.718125 | 0.693 | 0.7164 | 0.686907 | 0.710407 | 0.666667 |
| 0.683593 | 0.689655 | 0.755851 | 0.685751 | 0.76113 | 0.755206 | 0.760105 | 0.739158 | 0.664516 |
| 0.717062 | 0.718656 | 0.719293 | 0.763345 | 0.695567 | 0.719041 | 0.68532 | 0.70327 | 0.677852 |
| 0.748617 | 0.762716 | 0.712944 | 0.719388 | 0.689663 | 0.708391 | 0.679898 | 0.694971 | 0.565476 |
| 0.700256 | 0.69176 | 0.813234 | 0.690891 | 0.730936 | 0.784404 | 0.72312 | 0.804606 | 0.636943 |
| 0.739571 | 0.765048 | 0.718289 | 0.721792 | 0.687336 | 0.709237 | 0.681387 | 0.694815 | 0.617284 |
| 0.847616 | 0.795526 | 0.713825 | 0.720607 | 0.679484 | 0.717895 | 0.671426 | 0.705261 | 0.635762 |
| 0.806441 | 0.789262 | 0.705128 | 0.726524 | 0.678006 | 0.705491 | 0.664343 | 0.701593 | 0.598802 |
| 0.698146 | 0.69616 | 0.805599 | 0.683638 | 0.733677 | 0.814309 | 0.726679 | 0.793315 | 0.649351 |
| 0.834935 | 0.787889 | 0.707931 | 0.718366 | 0.684873 | 0.709036 | 0.670788 | 0.702436 | 0.562874 |
| 0.701003 | 0.694663 | 0.802191 | 0.685587 | 0.728709 | 0.815706 | 0.727043 | 0.804639 | 0.704698 |

| | | | | | | | | |
|---|---|---|---|---|---|---|---|---|
| 0.701181 | 0.692184 | 0.807172 | 0.686566 | 0.738138 | 0.797298 | 0.729597 | 0.801873 | 0.642857 |
| 0.706671 | 0.69446 | 0.815912 | 0.687697 | 0.729523 | 0.813473 | 0.728082 | 0.818692 | 0.658065 |
| 0.683185 | 0.684407 | 0.756983 | 0.684993 | 0.736911 | 0.758346 | 0.736942 | 0.745279 | 0.674699 |
| 1 | 0.786414 | 0.706328 | 0.721403 | 0.671614 | 0.702838 | 0.662032 | 0.692005 | 0.579618 |
| 0.786414 | 1 | 0.699667 | 0.719412 | 0.670359 | 0.690108 | 0.657258 | 0.685845 | 0.625806 |
| 0.706328 | 0.699667 | 1 | 0.685721 | 0.738616 | 0.810106 | 0.724845 | 0.814504 | 0.673469 |
| 0.721403 | 0.719412 | 0.685721 | 1 | 0.669943 | 0.686083 | 0.661334 | 0.679767 | 0.624204 |
| 0.671614 | 0.670359 | 0.738616 | 0.669943 | 1 | 0.738066 | 0.796177 | 0.724315 | 0.707483 |
| 0.702838 | 0.690108 | 0.810106 | 0.686083 | 0.738066 | 1 | 0.735985 | 0.815027 | 0.697183 |
| 0.662032 | 0.657258 | 0.724845 | 0.661334 | 0.796177 | 0.735985 | 1 | 0.724698 | 0.702128 |
| 0.692005 | 0.685845 | 0.814504 | 0.679767 | 0.724315 | 0.815027 | 0.724698 | 1 | 0.716312 |
| 0.579618 | 0.625806 | 0.673469 | 0.624204 | 0.707483 | 0.697183 | 0.702128 | 0.716312 | 1 |
| 0.59235 | 0.604834 | 0.682604 | 0.599646 | 0.6848 | 0.685603 | 0.691604 | 0.685996 | 0.821705 |
| 0.59368 | 0.608752 | 0.682294 | 0.601196 | 0.679387 | 0.683146 | 0.681452 | 0.678101 | 0.895161 |
| 0.599787 | 0.613887 | 0.692836 | 0.611111 | 0.69107 | 0.698855 | 0.690636 | 0.692251 | 0.804348 |
| 0.605701 | 0.614869 | 0.688281 | 0.608384 | 0.688977 | 0.693072 | 0.695914 | 0.696754 | 0.738255 |
| 0.667923 | 0.664114 | 0.615101 | 0.756364 | 0.607053 | 0.617413 | 0.599978 | 0.608967 | 0.566265 |
| 0.604904 | 0.612019 | 0.691461 | 0.611773 | 0.694363 | 0.696127 | 0.69453 | 0.695574 | 0.792857 |
| 0.684656 | 0.678551 | 0.81461 | 0.673567 | 0.731614 | 0.815391 | 0.72104 | 0.82884 | 0.696552 |
| 0.684352 | 0.683051 | 0.818267 | 0.674101 | 0.727175 | 0.815587 | 0.726391 | 0.841652 | 0.654088 |
| 0.690529 | 0.684649 | 0.815687 | 0.677839 | 0.727868 | 0.81938 | 0.731585 | 0.884683 | 0.675497 |
| 0.751976 | 0.788035 | 0.61591 | 0.681259 | 0.605887 | 0.612951 | 0.598808 | 0.608128 | 0.580838 |
| 0.757772 | 0.724612 | 0.598846 | 0.670454 | 0.59366 | 0.596377 | 0.588926 | 0.591595 | 0.512048 |
| 0.649981 | 0.64956 | 0.744961 | 0.652305 | 0.731579 | 0.74699 | 0.73371 | 0.75519 | 0.698113 |
| 0.649627 | 0.651378 | 0.748646 | 0.652755 | 0.728247 | 0.74572 | 0.735299 | 0.758884 | 0.7 |
| 0.642252 | 0.644498 | 0.729988 | 0.646185 | 0.7215 | 0.729365 | 0.726615 | 0.738467 | 0.662338 |
| 0.649712 | 0.646412 | 0.72992 | 0.651254 | 0.729158 | 0.734146 | 0.73347 | 0.741108 | 0.735099 |
| 0.647621 | 0.649188 | 0.735254 | 0.654049 | 0.728858 | 0.740226 | 0.733464 | 0.743427 | 0.739437 |
| 0.650239 | 0.650665 | 0.740768 | 0.656947 | 0.730086 | 0.740565 | 0.736215 | 0.750456 | 0.746575 |
| 0.651335 | 0.650267 | 0.743179 | 0.657716 | 0.72871 | 0.746089 | 0.731623 | 0.746872 | 0.729032 |
| 0.648132 | 0.653683 | 0.728739 | 0.652416 | 0.727435 | 0.729109 | 0.726197 | 0.730209 | 0.703448 |
| 0.657418 | 0.658111 | 0.735798 | 0.657881 | 0.740444 | 0.736053 | 0.739484 | 0.734416 | 0.695364 |
| 0.655572 | 0.654079 | 0.731084 | 0.65221 | 0.732814 | 0.731977 | 0.737598 | 0.733164 | 0.660377 |
| 0.654306 | 0.655446 | 0.732559 | 0.651956 | 0.738505 | 0.727186 | 0.733889 | 0.728086 | 0.709459 |
| 0.653437 | 0.657097 | 0.734002 | 0.658417 | 0.739746 | 0.736655 | 0.739269 | 0.736137 | 0.721088 |
| 0.650108 | 0.651978 | 0.729521 | 0.653343 | 0.731382 | 0.727875 | 0.735282 | 0.731737 | 0.718954 |
| 0.650608 | 0.65267 | 0.733972 | 0.652086 | 0.737217 | 0.731424 | 0.735002 | 0.731658 | 0.731034 |
| 0.653656 | 0.657262 | 0.730431 | 0.65414 | 0.733685 | 0.731933 | 0.737311 | 0.733204 | 0.711538 |
| 0.647019 | 0.648761 | 0.731328 | 0.65048 | 0.73473 | 0.726467 | 0.733777 | 0.727613 | 0.693878 |
| 0.651423 | 0.652451 | 0.728891 | 0.653097 | 0.733901 | 0.73127 | 0.73207 | 0.728047 | 0.746377 |
| 0.591562 | 0.606012 | 0.688541 | 0.60438 | 0.688448 | 0.688305 | 0.687273 | 0.690258 | 0.796748 |
| 0.597665 | 0.613919 | 0.686106 | 0.609257 | 0.684298 | 0.697679 | 0.689862 | 0.685271 | 0.764286 |
| 0.657588 | 0.656312 | 0.737234 | 0.655724 | 0.737987 | 0.73418 | 0.739908 | 0.731364 | 0.723684 |
| 0.660255 | 0.659009 | 0.735631 | 0.656313 | 0.735872 | 0.738709 | 0.740844 | 0.744539 | 0.72973 |
| 0.583432 | 0.59786 | 0.680127 | 0.593633 | 0.680136 | 0.67382 | 0.676499 | 0.681698 | 0.740157 |
| 0.654659 | 0.653397 | 0.732369 | 0.657612 | 0.734553 | 0.730896 | 0.736777 | 0.73179 | 0.694805 |
| 0.645389 | 0.64502 | 0.724138 | 0.646091 | 0.72428 | 0.723375 | 0.729925 | 0.725131 | 0.72549 |

| | | | | | | | | |
|---|---|---|---|---|---|---|---|---|
| 0.653889 | 0.648987 | 0.732488 | 0.652015 | 0.734463 | 0.731014 | 0.734293 | 0.736684 | 0.726667 |
| 0.654805 | 0.65506 | 0.731602 | 0.654816 | 0.735394 | 0.731918 | 0.736633 | 0.736206 | 0.736486 |
| 0.65201 | 0.654332 | 0.732497 | 0.657188 | 0.733433 | 0.732717 | 0.735555 | 0.732535 | 0.816176 |
| 0.657383 | 0.656728 | 0.735311 | 0.654504 | 0.732125 | 0.736153 | 0.729611 | 0.734656 | 0.75 |
| 0.65786 | 0.660202 | 0.742162 | 0.658622 | 0.740744 | 0.740364 | 0.739433 | 0.741729 | 0.758389 |
| 0.650818 | 0.64903 | 0.727595 | 0.648822 | 0.72986 | 0.730632 | 0.732165 | 0.731624 | 0.710526 |
| 0.655798 | 0.653221 | 0.729114 | 0.651881 | 0.733657 | 0.72986 | 0.73085 | 0.733823 | 0.768212 |
| 0.646962 | 0.648079 | 0.724737 | 0.648158 | 0.728445 | 0.723775 | 0.731399 | 0.725031 | 0.723684 |
| 0.653238 | 0.651133 | 0.730096 | 0.654329 | 0.732629 | 0.727705 | 0.729584 | 0.729543 | 0.699346 |
| 0.645867 | 0.64838 | 0.731748 | 0.651375 | 0.72862 | 0.7308 | 0.731447 | 0.727295 | 0.700637 |
| 0.65343 | 0.653545 | 0.73456 | 0.656252 | 0.735344 | 0.731995 | 0.735493 | 0.73269 | 0.741497 |
| 0.648801 | 0.651829 | 0.729687 | 0.65078 | 0.733104 | 0.728646 | 0.733772 | 0.730183 | 0.742857 |
| 0.641138 | 0.644149 | 0.722033 | 0.645733 | 0.723198 | 0.723273 | 0.728062 | 0.724961 | 0.700637 |
| 0.645174 | 0.644699 | 0.718633 | 0.644812 | 0.723392 | 0.721747 | 0.724258 | 0.72583 | 0.706667 |
| 0.654702 | 0.657951 | 0.738141 | 0.658161 | 0.738687 | 0.737079 | 0.73815 | 0.735715 | 0.77931 |
| 0.652511 | 0.655582 | 0.742311 | 0.658649 | 0.734581 | 0.748419 | 0.736679 | 0.757858 | 0.736486 |
| 0.649686 | 0.648547 | 0.732951 | 0.649904 | 0.731461 | 0.732655 | 0.730474 | 0.734403 | 0.755245 |
| 0.6525 | 0.650164 | 0.736987 | 0.651154 | 0.734625 | 0.741295 | 0.733832 | 0.742486 | 0.714286 |
| 0.64816 | 0.649421 | 0.739179 | 0.651356 | 0.733035 | 0.740923 | 0.733883 | 0.745586 | 0.714286 |
| 0.649844 | 0.648821 | 0.737772 | 0.651114 | 0.732006 | 0.738259 | 0.733459 | 0.747643 | 0.727891 |
| 0.649781 | 0.646109 | 0.735826 | 0.648448 | 0.726574 | 0.749543 | 0.729087 | 0.744901 | 0.71831 |
| 0.647904 | 0.648962 | 0.736579 | 0.651216 | 0.728769 | 0.738976 | 0.731431 | 0.746944 | 0.704698 |
| 0.654517 | 0.652647 | 0.736691 | 0.654915 | 0.73196 | 0.73724 | 0.735089 | 0.744819 | 0.685535 |
| 0.655516 | 0.653009 | 0.73605 | 0.654367 | 0.737568 | 0.738577 | 0.736583 | 0.740138 | 0.701987 |
| 0.650983 | 0.648478 | 0.736297 | 0.652027 | 0.731903 | 0.743704 | 0.733825 | 0.745309 | 0.727273 |
| 0.648606 | 0.649518 | 0.740693 | 0.649843 | 0.72763 | 0.746435 | 0.729728 | 0.756017 | 0.697368 |
| 0.650105 | 0.648959 | 0.730126 | 0.651938 | 0.72996 | 0.736444 | 0.728633 | 0.739285 | 0.712329 |
| 0.647792 | 0.647226 | 0.727154 | 0.64826 | 0.725388 | 0.729956 | 0.72964 | 0.738399 | 0.713333 |
| 0.648726 | 0.650236 | 0.743658 | 0.649473 | 0.726763 | 0.749288 | 0.731851 | 0.75842 | 0.707006 |
| 0.651786 | 0.650667 | 0.741093 | 0.654967 | 0.732916 | 0.737289 | 0.734902 | 0.745727 | 0.712418 |
| 0.652904 | 0.65006 | 0.736846 | 0.654117 | 0.733611 | 0.746402 | 0.737982 | 0.74743 | 0.707006 |
| 0.656798 | 0.65431 | 0.736081 | 0.655141 | 0.732933 | 0.745338 | 0.736929 | 0.741474 | 0.701299 |
| 0.654848 | 0.659676 | 0.75014 | 0.658209 | 0.73955 | 0.752807 | 0.741326 | 0.752735 | 0.685897 |
| 0.657156 | 0.658214 | 0.739462 | 0.661396 | 0.739662 | 0.74169 | 0.735823 | 0.747987 | 0.758621 |
| 0.64784 | 0.645913 | 0.742769 | 0.649274 | 0.724534 | 0.742025 | 0.732632 | 0.752511 | 0.708861 |
| 0.652014 | 0.653456 | 0.738232 | 0.650745 | 0.730367 | 0.743791 | 0.734065 | 0.745079 | 0.710526 |
| 0.651704 | 0.652335 | 0.738051 | 0.653843 | 0.729393 | 0.734315 | 0.733141 | 0.746093 | 0.666667 |
| 0.651389 | 0.647504 | 0.734329 | 0.649553 | 0.727833 | 0.733922 | 0.731412 | 0.739756 | 0.7 |
| 0.64725 | 0.645701 | 0.733826 | 0.644495 | 0.723685 | 0.738542 | 0.729181 | 0.744709 | 0.72549 |
| 0.640883 | 0.641622 | 0.735421 | 0.643527 | 0.71968 | 0.7281 | 0.722265 | 0.737646 | 0.712418 |
| 0.646341 | 0.643363 | 0.743238 | 0.643981 | 0.723184 | 0.748395 | 0.72344 | 0.755361 | 0.684211 |
| 0.641024 | 0.640782 | 0.733436 | 0.645743 | 0.724343 | 0.733775 | 0.72634 | 0.744342 | 0.705882 |
| 0.644618 | 0.6442 | 0.733726 | 0.644675 | 0.724513 | 0.733767 | 0.727179 | 0.747468 | 0.647059 |
| 0.635922 | 0.635704 | 0.725445 | 0.637013 | 0.718703 | 0.724248 | 0.720072 | 0.735952 | 0.717241 |
| 0.644746 | 0.647127 | 0.732797 | 0.647238 | 0.728018 | 0.731678 | 0.728303 | 0.738494 | 0.677215 |
| 0.637934 | 0.641212 | 0.72105 | 0.641335 | 0.720086 | 0.720373 | 0.724377 | 0.72662 | 0.668831 |

| | | | | | | | | |
|---|---|---|---|---|---|---|---|---|
| 0.654911 | 0.655591 | 0.760281 | 0.654457 | 0.730199 | 0.76584 | 0.7314 | 0.782612 | 0.698113 |
| 0.653494 | 0.654229 | 0.733279 | 0.652589 | 0.730969 | 0.730979 | 0.734966 | 0.733306 | 0.75 |
| 0.645896 | 0.648671 | 0.726417 | 0.649962 | 0.729179 | 0.729048 | 0.727535 | 0.727571 | 0.756944 |
| 0.651826 | 0.65215 | 0.731057 | 0.654284 | 0.737643 | 0.729866 | 0.738037 | 0.727513 | 0.717105 |
| 0.656919 | 0.655272 | 0.740279 | 0.658537 | 0.743546 | 0.738384 | 0.741917 | 0.736625 | 0.782609 |
| 0.653804 | 0.651318 | 0.733264 | 0.65172 | 0.735633 | 0.734894 | 0.735126 | 0.732897 | 0.716129 |
| 0.64881 | 0.648497 | 0.729445 | 0.648776 | 0.732584 | 0.728609 | 0.732172 | 0.725128 | 0.690141 |
| 0.65309 | 0.652503 | 0.730201 | 0.655695 | 0.736073 | 0.730649 | 0.736683 | 0.731372 | 0.726667 |
| 0.644444 | 0.647242 | 0.727505 | 0.648811 | 0.726097 | 0.725266 | 0.726649 | 0.723952 | 0.724138 |
| 0.652981 | 0.650766 | 0.727922 | 0.653159 | 0.731392 | 0.732045 | 0.731611 | 0.727501 | 0.7 |
| 0.656882 | 0.652956 | 0.734033 | 0.655718 | 0.735021 | 0.733232 | 0.732541 | 0.730126 | 0.748299 |
| 0.65237 | 0.651858 | 0.726924 | 0.652825 | 0.73703 | 0.731305 | 0.735205 | 0.73108 | 0.732877 |
| 0.653633 | 0.653198 | 0.73112 | 0.650337 | 0.731781 | 0.728082 | 0.735344 | 0.728026 | 0.72 |
| 0.656838 | 0.654643 | 0.731909 | 0.656371 | 0.728908 | 0.728604 | 0.732632 | 0.727786 | 0.705128 |
| 0.654866 | 0.653917 | 0.735879 | 0.656463 | 0.737281 | 0.736614 | 0.737517 | 0.734932 | 0.688742 |
| 0.654778 | 0.653466 | 0.732682 | 0.6539 | 0.734893 | 0.732465 | 0.737422 | 0.731129 | 0.763514 |
| 0.65471 | 0.656506 | 0.736963 | 0.654188 | 0.742165 | 0.730245 | 0.740566 | 0.736166 | 0.726027 |
| 0.654714 | 0.654557 | 0.735352 | 0.656009 | 0.733119 | 0.731599 | 0.735706 | 0.732147 | 0.705479 |
| 0.654966 | 0.656252 | 0.734434 | 0.654638 | 0.736863 | 0.73206 | 0.737899 | 0.72891 | 0.746479 |
| 0.654384 | 0.655442 | 0.730928 | 0.656847 | 0.734178 | 0.73229 | 0.740231 | 0.735161 | 0.75 |
| 0.652449 | 0.653843 | 0.73027 | 0.658796 | 0.732348 | 0.728937 | 0.73209 | 0.733453 | 0.713376 |
| 0.653617 | 0.652796 | 0.728328 | 0.655467 | 0.732016 | 0.727523 | 0.733607 | 0.726136 | 0.688312 |
| 0.65564 | 0.655966 | 0.73494 | 0.65888 | 0.734679 | 0.736377 | 0.735913 | 0.733425 | 0.751724 |
| 0.656358 | 0.659749 | 0.731579 | 0.655424 | 0.737131 | 0.734939 | 0.735027 | 0.734538 | 0.757962 |
| 0.655914 | 0.653548 | 0.73121 | 0.654351 | 0.734586 | 0.731261 | 0.732124 | 0.733605 | 0.774648 |
| 0.656107 | 0.655382 | 0.731709 | 0.655134 | 0.73263 | 0.729867 | 0.732297 | 0.734141 | 0.728477 |
| 0.650355 | 0.648163 | 0.724602 | 0.652806 | 0.724284 | 0.729669 | 0.733939 | 0.726735 | 0.717241 |
| 0.651393 | 0.653171 | 0.726378 | 0.653334 | 0.729791 | 0.730431 | 0.73234 | 0.72704 | 0.746753 |
| 0.653288 | 0.655774 | 0.733938 | 0.657516 | 0.738972 | 0.731992 | 0.740566 | 0.734951 | 0.757353 |
| 0.65606 | 0.657996 | 0.735746 | 0.65582 | 0.735459 | 0.734079 | 0.737302 | 0.733638 | 0.74 |
| 0.650386 | 0.654463 | 0.728456 | 0.652413 | 0.731131 | 0.729663 | 0.730027 | 0.729874 | 0.740506 |
| 0.657946 | 0.659122 | 0.729452 | 0.655862 | 0.734495 | 0.735667 | 0.736508 | 0.732189 | 0.72973 |
| 0.655939 | 0.657506 | 0.733509 | 0.657605 | 0.732529 | 0.73724 | 0.737345 | 0.737544 | 0.708075 |
| 0.650225 | 0.649554 | 0.730125 | 0.649794 | 0.732787 | 0.72962 | 0.728265 | 0.731369 | 0.72973 |
| 0.653518 | 0.65818 | 0.731766 | 0.655526 | 0.732392 | 0.732929 | 0.734694 | 0.732644 | 0.76129 |
| 0.648225 | 0.650042 | 0.728389 | 0.648042 | 0.729454 | 0.729682 | 0.728608 | 0.731102 | 0.685185 |
| 0.661897 | 0.661001 | 0.73846 | 0.660441 | 0.742043 | 0.742277 | 0.739201 | 0.73906 | 0.755102 |
| 0.649876 | 0.653918 | 0.728434 | 0.654561 | 0.735899 | 0.730228 | 0.742051 | 0.730716 | 0.779412 |
| 0.654407 | 0.653216 | 0.733977 | 0.654696 | 0.733827 | 0.733894 | 0.731476 | 0.731167 | 0.75 |
| 0.652807 | 0.655094 | 0.73156 | 0.657035 | 0.733069 | 0.731397 | 0.735375 | 0.734309 | 0.726027 |
| 0.658643 | 0.661238 | 0.734915 | 0.661997 | 0.736345 | 0.737151 | 0.735982 | 0.739879 | 0.707483 |
| 0.661056 | 0.65717 | 0.737346 | 0.660383 | 0.74017 | 0.73084 | 0.738128 | 0.733215 | 0.724138 |
| 0.659651 | 0.657997 | 0.73576 | 0.658603 | 0.737856 | 0.733525 | 0.735837 | 0.736584 | 0.75 |
| 0.652482 | 0.652354 | 0.730103 | 0.654639 | 0.727663 | 0.732659 | 0.729575 | 0.733499 | 0.733813 |
| 0.658976 | 0.653017 | 0.735522 | 0.658398 | 0.735143 | 0.735526 | 0.736851 | 0.737108 | 0.712418 |
| 0.651456 | 0.651379 | 0.729462 | 0.654108 | 0.730218 | 0.729194 | 0.732736 | 0.732675 | 0.707792 |

| | | | | | | | | |
|---|---|---|---|---|---|---|---|---|
| 0.640588 | 0.640249 | 0.715123 | 0.642388 | 0.722321 | 0.71923 | 0.725141 | 0.719728 | 0.688742 |
| 0.684531 | 0.680101 | 0.758762 | 0.676927 | 0.737652 | 0.754097 | 0.732844 | 0.748185 | 0.701389 |
| 0.651193 | 0.654028 | 0.728251 | 0.655478 | 0.729281 | 0.728308 | 0.731235 | 0.728343 | 0.634615 |
| 0.652976 | 0.653116 | 0.735881 | 0.655055 | 0.737795 | 0.737476 | 0.734812 | 0.733847 | 0.673333 |
| 0.644392 | 0.646782 | 0.723406 | 0.65148 | 0.723958 | 0.726269 | 0.726889 | 0.721595 | 0.682759 |
| 0.651272 | 0.652079 | 0.727589 | 0.653177 | 0.732972 | 0.730049 | 0.734055 | 0.728676 | 0.687075 |
| 0.650077 | 0.651462 | 0.730029 | 0.650982 | 0.725698 | 0.727878 | 0.726575 | 0.728265 | 0.739437 |
| 0.643595 | 0.645887 | 0.726769 | 0.648543 | 0.721922 | 0.72378 | 0.722685 | 0.725229 | 0.72 |
| 0.651512 | 0.651835 | 0.734784 | 0.657067 | 0.737073 | 0.731698 | 0.735356 | 0.732737 | 0.662651 |
| 0.657251 | 0.659535 | 0.737877 | 0.654741 | 0.736911 | 0.735385 | 0.737486 | 0.73448 | 0.675676 |
| 0.655763 | 0.655722 | 0.731807 | 0.659152 | 0.736518 | 0.734159 | 0.738613 | 0.733484 | 0.72956 |
| 0.656085 | 0.657354 | 0.734375 | 0.656677 | 0.740095 | 0.731279 | 0.738666 | 0.732678 | 0.721429 |
| 0.657811 | 0.66045 | 0.738575 | 0.661172 | 0.740686 | 0.736315 | 0.739009 | 0.737551 | 0.700637 |
| 0.655279 | 0.655405 | 0.737478 | 0.658136 | 0.737578 | 0.732479 | 0.735297 | 0.735629 | 0.704698 |
| 0.652131 | 0.65047 | 0.730255 | 0.653931 | 0.734353 | 0.731261 | 0.731493 | 0.728946 | 0.716216 |
| 0.65469 | 0.654058 | 0.731249 | 0.657365 | 0.73623 | 0.731841 | 0.733194 | 0.731711 | 0.710345 |
| 0.654029 | 0.655159 | 0.734347 | 0.654966 | 0.736239 | 0.73363 | 0.734592 | 0.73186 | 0.692308 |
| 0.652197 | 0.651535 | 0.732563 | 0.654543 | 0.732106 | 0.73641 | 0.733514 | 0.730707 | 0.695946 |
| 0.644914 | 0.652748 | 0.729468 | 0.649457 | 0.727057 | 0.731571 | 0.72556 | 0.73103 | 0.774648 |
| 0.647687 | 0.648285 | 0.727996 | 0.645804 | 0.725081 | 0.728216 | 0.730201 | 0.726946 | 0.682119 |
| 0.652857 | 0.653716 | 0.732008 | 0.654565 | 0.735413 | 0.7315 | 0.733537 | 0.729051 | 0.77551 |
| 0.644789 | 0.641975 | 0.719252 | 0.646655 | 0.727569 | 0.720185 | 0.727632 | 0.723176 | 0.727891 |
| 0.651037 | 0.651066 | 0.728079 | 0.653046 | 0.733366 | 0.727959 | 0.731139 | 0.725127 | 0.748344 |
| 0.652571 | 0.653313 | 0.732791 | 0.65653 | 0.73642 | 0.732982 | 0.735627 | 0.727803 | 0.72549 |
| 0.65396 | 0.655127 | 0.733377 | 0.654897 | 0.736715 | 0.732937 | 0.736763 | 0.729766 | 0.699346 |
| 0.655087 | 0.65517 | 0.728929 | 0.654639 | 0.733807 | 0.729869 | 0.731075 | 0.727907 | 0.794326 |
| 0.647054 | 0.646686 | 0.727408 | 0.653075 | 0.727479 | 0.722772 | 0.723151 | 0.726506 | 0.722973 |
| 0.646018 | 0.646512 | 0.72284 | 0.650513 | 0.72973 | 0.726755 | 0.728136 | 0.726921 | 0.758865 |
| 0.646595 | 0.651492 | 0.722622 | 0.646884 | 0.726306 | 0.725297 | 0.725355 | 0.721753 | 0.711268 |
| 0.689257 | 0.687808 | 0.754265 | 0.679692 | 0.719573 | 0.747201 | 0.708329 | 0.734282 | 0.627586 |
| 0.711082 | 0.716418 | 0.773381 | 0.692006 | 0.733339 | 0.760076 | 0.718558 | 0.75322 | 0.662162 |
| 0.696051 | 0.69901 | 0.760332 | 0.696237 | 0.729796 | 0.754951 | 0.718074 | 0.748659 | 0.578313 |
| 0.685738 | 0.698286 | 0.778063 | 0.683816 | 0.720169 | 0.757213 | 0.708311 | 0.741915 | 0.607595 |
| 0.71384 | 0.733582 | 0.779593 | 0.687792 | 0.731238 | 0.763055 | 0.717784 | 0.760775 | 0.630573 |
| 0.64665 | 0.644483 | 0.721952 | 0.649019 | 0.731654 | 0.724355 | 0.7292 | 0.721157 | 0.666667 |
| 0.649585 | 0.649028 | 0.728926 | 0.654232 | 0.734119 | 0.728444 | 0.736816 | 0.728868 | 0.701299 |
| 0.650814 | 0.647981 | 0.729108 | 0.652367 | 0.734041 | 0.727539 | 0.736694 | 0.729495 | 0.732026 |
| 0.649341 | 0.648322 | 0.72911 | 0.65534 | 0.735128 | 0.731492 | 0.737725 | 0.730092 | 0.660131 |
| 0.649955 | 0.653369 | 0.726789 | 0.65292 | 0.73369 | 0.729437 | 0.734348 | 0.728279 | 0.736842 |
| 0.651935 | 0.652711 | 0.734496 | 0.654752 | 0.733628 | 0.735512 | 0.735807 | 0.734622 | 0.705882 |
| 0.650372 | 0.651396 | 0.730838 | 0.653483 | 0.734445 | 0.731307 | 0.732505 | 0.731711 | 0.692308 |
| 0.649323 | 0.648638 | 0.727384 | 0.650351 | 0.731853 | 0.727258 | 0.73148 | 0.726221 | 0.769784 |
| 0.644947 | 0.646793 | 0.725357 | 0.643958 | 0.728007 | 0.72507 | 0.728859 | 0.724872 | 0.731544 |
| 0.640717 | 0.641979 | 0.720766 | 0.646381 | 0.725452 | 0.718361 | 0.729892 | 0.718294 | 0.72 |
| 0.652313 | 0.651509 | 0.730046 | 0.651631 | 0.73257 | 0.726266 | 0.732483 | 0.727567 | 0.66443 |
| 0.653257 | 0.647961 | 0.732345 | 0.654928 | 0.732645 | 0.729175 | 0.734096 | 0.729389 | 0.707006 |

| | | | | | | | | |
|---|---|---|---|---|---|---|---|---|
| 0.653557 | 0.652195 | 0.729366 | 0.656041 | 0.73395 | 0.732937 | 0.738265 | 0.731456 | 0.695035 |
| 0.655314 | 0.651975 | 0.726782 | 0.653308 | 0.733077 | 0.73091 | 0.730826 | 0.729822 | 0.732877 |
| 0.654305 | 0.65332 | 0.728209 | 0.654799 | 0.7333 | 0.735164 | 0.733229 | 0.736674 | 0.679739 |
| 0.65168 | 0.647597 | 0.725407 | 0.653826 | 0.730961 | 0.731203 | 0.731255 | 0.730704 | 0.736486 |
| 0.647522 | 0.648779 | 0.725088 | 0.64908 | 0.730705 | 0.725908 | 0.730369 | 0.728504 | 0.74026 |
| 0.649812 | 0.647455 | 0.724075 | 0.652144 | 0.728303 | 0.724035 | 0.727088 | 0.725705 | 0.696429 |
| 0.656279 | 0.654066 | 0.729311 | 0.657228 | 0.732716 | 0.728636 | 0.733043 | 0.727258 | 0.751773 |
| 0.649315 | 0.651673 | 0.729456 | 0.652906 | 0.729613 | 0.732119 | 0.730888 | 0.733272 | 0.724638 |
| 0.649397 | 0.649653 | 0.726945 | 0.651605 | 0.725106 | 0.72519 | 0.729877 | 0.728553 | 0.726115 |
| 0.649187 | 0.650621 | 0.728934 | 0.652167 | 0.731663 | 0.72762 | 0.732755 | 0.730952 | 0.755396 |
| 0.651454 | 0.652538 | 0.726991 | 0.652548 | 0.731091 | 0.730972 | 0.731134 | 0.731225 | 0.707483 |
| 0.648557 | 0.651877 | 0.72728 | 0.652851 | 0.733777 | 0.732341 | 0.730151 | 0.729975 | 0.71519 |
| 0.641974 | 0.64212 | 0.718506 | 0.645568 | 0.7219 | 0.720316 | 0.725442 | 0.722887 | 0.771429 |
| 0.642472 | 0.646803 | 0.725522 | 0.646134 | 0.723029 | 0.727964 | 0.720395 | 0.723016 | 0.736486 |
| 0.649411 | 0.653717 | 0.731685 | 0.657407 | 0.733433 | 0.735776 | 0.730025 | 0.732212 | 0.757143 |
| 0.658653 | 0.659243 | 0.739141 | 0.65921 | 0.74068 | 0.742499 | 0.738883 | 0.740535 | 0.719178 |
| 0.653182 | 0.654935 | 0.738509 | 0.657743 | 0.736953 | 0.735979 | 0.731553 | 0.734956 | 0.727848 |
| 0.656385 | 0.65788 | 0.732942 | 0.659788 | 0.733708 | 0.733684 | 0.736631 | 0.738344 | 0.709459 |
| 0.649084 | 0.652019 | 0.725868 | 0.648044 | 0.727556 | 0.728608 | 0.727512 | 0.730136 | 0.722222 |
| 0.650696 | 0.651866 | 0.726828 | 0.651598 | 0.736838 | 0.731058 | 0.734594 | 0.732936 | 0.707792 |
| 0.650046 | 0.651803 | 0.723383 | 0.648876 | 0.732733 | 0.727379 | 0.732248 | 0.727739 | 0.726667 |
| 0.65258 | 0.654005 | 0.730106 | 0.652851 | 0.731892 | 0.730581 | 0.73506 | 0.728016 | 0.743243 |
| 0.655275 | 0.659749 | 0.736267 | 0.658243 | 0.738037 | 0.736961 | 0.737593 | 0.738204 | 0.756757 |
| 0.649996 | 0.65273 | 0.727085 | 0.652018 | 0.729067 | 0.728119 | 0.733257 | 0.728852 | 0.733766 |
| 0.648773 | 0.653564 | 0.730361 | 0.651354 | 0.73296 | 0.73168 | 0.732019 | 0.733077 | 0.715278 |
| 0.704538 | 0.695372 | 0.783617 | 0.690615 | 0.727743 | 0.762608 | 0.716391 | 0.747521 | 0.655172 |
| 0.653702 | 0.654027 | 0.728878 | 0.657493 | 0.7324 | 0.733933 | 0.737177 | 0.730293 | 0.700637 |
| 0.653733 | 0.653097 | 0.730754 | 0.657179 | 0.734845 | 0.736838 | 0.735546 | 0.734918 | 0.743243 |
| 0.660379 | 0.658438 | 0.736795 | 0.658266 | 0.736535 | 0.740816 | 0.739249 | 0.740115 | 0.711111 |
| 0.654338 | 0.653581 | 0.733011 | 0.654598 | 0.737079 | 0.737625 | 0.735398 | 0.733653 | 0.74 |
| 0.64833 | 0.649634 | 0.725907 | 0.652374 | 0.729861 | 0.727058 | 0.731132 | 0.729326 | 0.693333 |
| 0.648591 | 0.6534 | 0.728691 | 0.650622 | 0.732026 | 0.728614 | 0.731633 | 0.728581 | 0.724832 |
| 0.650173 | 0.653312 | 0.730335 | 0.651715 | 0.730906 | 0.731513 | 0.731994 | 0.734305 | 0.697987 |
| 0.64849 | 0.652576 | 0.725229 | 0.652846 | 0.729106 | 0.73087 | 0.731624 | 0.729831 | 0.75817 |
| 0.646339 | 0.648015 | 0.723797 | 0.650125 | 0.725677 | 0.728479 | 0.728745 | 0.727785 | 0.729167 |
| 0.650167 | 0.649629 | 0.726521 | 0.654559 | 0.72897 | 0.729019 | 0.732234 | 0.727528 | 0.736111 |
| 0.651754 | 0.65258 | 0.732869 | 0.657564 | 0.738471 | 0.731999 | 0.736328 | 0.731953 | 0.726667 |
| 0.652245 | 0.653386 | 0.734013 | 0.654526 | 0.74024 | 0.732775 | 0.737746 | 0.733937 | 0.701299 |
| 0.652034 | 0.652437 | 0.728883 | 0.655916 | 0.735801 | 0.734342 | 0.735005 | 0.727681 | 0.724832 |
| 0.650802 | 0.652625 | 0.735237 | 0.652481 | 0.733226 | 0.729614 | 0.729546 | 0.731779 | 0.731034 |
| 0.648457 | 0.652412 | 0.726422 | 0.651897 | 0.729386 | 0.723765 | 0.728842 | 0.727649 | 0.753425 |
| 0.652259 | 0.654084 | 0.737213 | 0.65499 | 0.735594 | 0.731331 | 0.736201 | 0.733333 | 0.729167 |
| 0.648613 | 0.647879 | 0.726946 | 0.651561 | 0.732575 | 0.7311 | 0.731655 | 0.730104 | 0.677852 |
| 0.646661 | 0.653726 | 0.728319 | 0.657306 | 0.731844 | 0.73101 | 0.728527 | 0.729353 | 0.716129 |
| 0.652442 | 0.657969 | 0.736267 | 0.656196 | 0.737691 | 0.735475 | 0.734177 | 0.735772 | 0.77551 |
| 0.652287 | 0.656546 | 0.736287 | 0.656919 | 0.738193 | 0.734603 | 0.732766 | 0.736081 | 0.690789 |

| | | | | | | | | |
|---|---|---|---|---|---|---|---|---|
| 0.652421 | 0.656442 | 0.729659 | 0.657694 | 0.732286 | 0.729459 | 0.729774 | 0.731487 | 0.714286 |
| 0.649416 | 0.655494 | 0.736074 | 0.656042 | 0.736062 | 0.73736 | 0.73121 | 0.738035 | 0.773973 |
| 0.650227 | 0.655301 | 0.730321 | 0.65702 | 0.730798 | 0.734511 | 0.72946 | 0.730216 | 0.708075 |
| 0.595914 | 0.613825 | 0.681834 | 0.610365 | 0.692088 | 0.689222 | 0.678471 | 0.68386 | 0.753425 |
| 0.64989 | 0.651925 | 0.729426 | 0.655499 | 0.727397 | 0.734873 | 0.726375 | 0.729301 | 0.746667 |
| 0.647862 | 0.651457 | 0.733628 | 0.657196 | 0.730418 | 0.733226 | 0.728521 | 0.732446 | 0.703226 |
| 0.652601 | 0.654425 | 0.734777 | 0.65744 | 0.732057 | 0.735555 | 0.729382 | 0.734973 | 0.79562 |
| 0.650942 | 0.654506 | 0.732488 | 0.653775 | 0.732506 | 0.731356 | 0.732133 | 0.729344 | 0.731544 |
| 0.656617 | 0.65898 | 0.737028 | 0.6603 | 0.73501 | 0.737771 | 0.731925 | 0.737489 | 0.722973 |
| 0.652241 | 0.654704 | 0.731781 | 0.658387 | 0.731079 | 0.732748 | 0.7328 | 0.729518 | 0.748299 |
| 0.656071 | 0.657905 | 0.732253 | 0.655203 | 0.729422 | 0.729562 | 0.728578 | 0.725554 | 0.713287 |
| 0.653058 | 0.655026 | 0.733387 | 0.655545 | 0.73373 | 0.735361 | 0.730263 | 0.733355 | 0.781022 |
| 0.655769 | 0.657268 | 0.732986 | 0.657942 | 0.730065 | 0.731745 | 0.734677 | 0.732352 | 0.718121 |
| 0.649636 | 0.654134 | 0.732768 | 0.657746 | 0.729944 | 0.737198 | 0.732227 | 0.73586 | 0.772414 |
| 0.655523 | 0.656942 | 0.734827 | 0.658314 | 0.73038 | 0.73306 | 0.729382 | 0.739215 | 0.687117 |
| 0.649612 | 0.65372 | 0.729736 | 0.656753 | 0.729897 | 0.731686 | 0.731719 | 0.733431 | 0.733766 |
| 0.654648 | 0.657483 | 0.732682 | 0.659384 | 0.733527 | 0.737103 | 0.733015 | 0.737299 | 0.732026 |
| 0.653508 | 0.65652 | 0.732444 | 0.655229 | 0.736675 | 0.732316 | 0.732617 | 0.735275 | 0.75 |
| 0.646453 | 0.650582 | 0.731905 | 0.653316 | 0.732889 | 0.731648 | 0.731888 | 0.735113 | 0.771429 |
| 0.650962 | 0.654785 | 0.734643 | 0.656839 | 0.73066 | 0.734811 | 0.72969 | 0.73522 | 0.76259 |
| 0.655804 | 0.659798 | 0.737278 | 0.657685 | 0.738825 | 0.738516 | 0.734892 | 0.737793 | 0.743243 |
| 0.656615 | 0.660175 | 0.735101 | 0.66045 | 0.732803 | 0.738013 | 0.729148 | 0.743494 | 0.762238 |
| 0.626413 | 0.633727 | 0.695668 | 0.623009 | 0.762122 | 0.695398 | 0.743294 | 0.688811 | 0.681159 |
| 0.638174 | 0.645939 | 0.72326 | 0.639428 | 0.698072 | 0.714528 | 0.668595 | 0.698757 | 0.655738 |
| 0.680513 | 0.682644 | 0.751287 | 0.671555 | 0.705402 | 0.739305 | 0.686626 | 0.713969 | 0.660194 |
| 0.643742 | 0.650745 | 0.741979 | 0.642071 | 0.683481 | 0.731844 | 0.676825 | 0.700737 | 0.623288 |
| 0.600505 | 0.615608 | 0.692386 | 0.617021 | 0.69645 | 0.694226 | 0.701804 | 0.692376 | 0.832 |
| 0.599283 | 0.613604 | 0.686319 | 0.604894 | 0.686699 | 0.686882 | 0.687618 | 0.688172 | 0.759398 |
| 0.591681 | 0.612292 | 0.684929 | 0.606742 | 0.692676 | 0.686199 | 0.690896 | 0.688893 | 0.80916 |
| 0.615442 | 0.624951 | 0.694929 | 0.623496 | 0.701379 | 0.704587 | 0.696421 | 0.69575 | 0.752212 |
| 0.591549 | 0.604307 | 0.680988 | 0.600369 | 0.678395 | 0.678041 | 0.679233 | 0.676702 | 0.727273 |
| 0.575012 | 0.594792 | 0.670219 | 0.590486 | 0.674819 | 0.671658 | 0.679299 | 0.673769 | 0.784314 |
| 0.579518 | 0.592512 | 0.678758 | 0.589939 | 0.682466 | 0.679919 | 0.683667 | 0.671689 | 0.801887 |
| 0.642353 | 0.654372 | 0.724132 | 0.639658 | 0.68256 | 0.712157 | 0.670539 | 0.698494 | 0.705426 |
| 0.644907 | 0.649126 | 0.729873 | 0.64976 | 0.72553 | 0.728325 | 0.726962 | 0.727465 | 0.732484 |
| 0.614057 | 0.624877 | 0.703329 | 0.623564 | 0.704965 | 0.702638 | 0.704169 | 0.704958 | 0.706667 |
| 0.60622 | 0.617125 | 0.694693 | 0.61444 | 0.700533 | 0.695292 | 0.700416 | 0.694105 | 0.756757 |
| 0.591508 | 0.605679 | 0.675184 | 0.598257 | 0.687923 | 0.680755 | 0.683892 | 0.682141 | 0.79661 |
| 0.614914 | 0.622881 | 0.703973 | 0.623356 | 0.700711 | 0.705006 | 0.704443 | 0.702591 | 0.755245 |
| 0.608015 | 0.619359 | 0.697878 | 0.613722 | 0.695316 | 0.700966 | 0.692796 | 0.695465 | 0.774648 |
| 0.601075 | 0.608466 | 0.690602 | 0.600509 | 0.691995 | 0.692224 | 0.693986 | 0.688314 | 0.766423 |
| 0.601581 | 0.617191 | 0.695407 | 0.612333 | 0.694735 | 0.695798 | 0.69435 | 0.695833 | 0.761905 |
| 0.590641 | 0.604578 | 0.684137 | 0.598756 | 0.686229 | 0.675882 | 0.682701 | 0.683931 | 0.75969 |
| 0.595719 | 0.606548 | 0.692634 | 0.607778 | 0.689202 | 0.692441 | 0.685312 | 0.692864 | 0.768657 |
| 0.592288 | 0.615312 | 0.687924 | 0.598353 | 0.691061 | 0.691884 | 0.687643 | 0.689655 | 0.760331 |
| 0.580334 | 0.597332 | 0.670294 | 0.596028 | 0.667313 | 0.678846 | 0.665688 | 0.673821 | 0.747664 |

| | | | | | | | | |
|---|---|---|---|---|---|---|---|---|
| 0.628744 | 0.632546 | 0.712137 | 0.63742 | 0.713868 | 0.716873 | 0.715551 | 0.717387 | 0.777778 |
| 0.597984 | 0.608729 | 0.68814 | 0.609726 | 0.687845 | 0.689095 | 0.68743 | 0.692957 | 0.783784 |
| 0.597817 | 0.606711 | 0.691299 | 0.61019 | 0.693989 | 0.692466 | 0.68737 | 0.693763 | 0.81746 |
| 0.600924 | 0.615221 | 0.689423 | 0.60972 | 0.694063 | 0.68789 | 0.694313 | 0.691043 | 0.736486 |
| 0.591804 | 0.610647 | 0.68446 | 0.603451 | 0.685555 | 0.687177 | 0.686594 | 0.685954 | 0.795455 |
| 0.597035 | 0.612053 | 0.688867 | 0.60887 | 0.690406 | 0.695175 | 0.68784 | 0.695299 | 0.821705 |
| 0.592239 | 0.604181 | 0.675441 | 0.601471 | 0.678275 | 0.678098 | 0.677296 | 0.676204 | 0.785047 |
| 0.583105 | 0.596469 | 0.679537 | 0.59109 | 0.675235 | 0.679484 | 0.67652 | 0.680548 | 0.806452 |
| 0.590152 | 0.597269 | 0.672527 | 0.592768 | 0.67438 | 0.676334 | 0.675676 | 0.671107 | 0.783019 |
| 0.626829 | 0.637859 | 0.711551 | 0.637835 | 0.716244 | 0.717706 | 0.71991 | 0.719392 | 0.727891 |
| 0.62937 | 0.633219 | 0.712042 | 0.636446 | 0.713875 | 0.715215 | 0.715599 | 0.71532 | 0.728477 |
| 0.620312 | 0.630117 | 0.708975 | 0.635592 | 0.706491 | 0.713189 | 0.709277 | 0.716923 | 0.794521 |
| 0.613158 | 0.622548 | 0.697792 | 0.624959 | 0.701675 | 0.702604 | 0.696817 | 0.702945 | 0.751773 |
| 0.615893 | 0.622436 | 0.699384 | 0.630068 | 0.701809 | 0.71073 | 0.707959 | 0.702836 | 0.776119 |
| 0.613937 | 0.620962 | 0.694732 | 0.619837 | 0.694703 | 0.704409 | 0.70264 | 0.702187 | 0.792857 |

| Collection_ | Collection_ | Collection_ | Collection_ | Collection_ | Collection_ | Collection_ | Collection_ | Collection_ |
|---|---|---|---|---|---|---|---|---|
| 0.644146 | 0.654254 | 0.664536 | 0.658055 | 0.619723 | 0.657767 | 0.737777 | 0.732755 | 0.742359 |
| 0.65848 | 0.665391 | 0.674231 | 0.666449 | 0.62072 | 0.676982 | 0.744483 | 0.740667 | 0.744366 |
| 0.64387 | 0.650784 | 0.657629 | 0.657117 | 0.624986 | 0.659935 | 0.732903 | 0.735684 | 0.741497 |
| 0.661449 | 0.656694 | 0.667828 | 0.668922 | 0.623583 | 0.668672 | 0.743031 | 0.745706 | 0.749206 |
| 0.657945 | 0.652265 | 0.662912 | 0.658094 | 0.626071 | 0.663661 | 0.734432 | 0.734701 | 0.741377 |
| 0.663793 | 0.661919 | 0.668243 | 0.669416 | 0.618902 | 0.670411 | 0.745924 | 0.741579 | 0.746586 |
| 0.724378 | 0.725582 | 0.729242 | 0.728614 | 0.590337 | 0.728607 | 0.737657 | 0.736501 | 0.74249 |
| 0.722543 | 0.7199 | 0.722897 | 0.733683 | 0.587097 | 0.728597 | 0.732988 | 0.735998 | 0.738364 |
| 0.661836 | 0.659146 | 0.663048 | 0.665845 | 0.617818 | 0.672927 | 0.739187 | 0.740152 | 0.75382 |
| 0.656455 | 0.647572 | 0.663945 | 0.660336 | 0.616643 | 0.667002 | 0.749206 | 0.738816 | 0.747528 |
| 0.656851 | 0.654746 | 0.667344 | 0.668479 | 0.626661 | 0.668378 | 0.740883 | 0.744933 | 0.749365 |
| 0.64637 | 0.644391 | 0.657673 | 0.65486 | 0.616124 | 0.657522 | 0.747691 | 0.740475 | 0.739765 |
| 0.659459 | 0.65895 | 0.673843 | 0.665282 | 0.621839 | 0.66847 | 0.748815 | 0.745352 | 0.756071 |
| 0.672565 | 0.666616 | 0.675417 | 0.678669 | 0.624973 | 0.677383 | 0.759039 | 0.764275 | 0.768456 |
| 0.689305 | 0.687882 | 0.69472 | 0.696863 | 0.617782 | 0.698635 | 0.785785 | 0.784347 | 0.789692 |
| 0.668093 | 0.666869 | 0.670438 | 0.672349 | 0.620493 | 0.673261 | 0.746301 | 0.749773 | 0.756334 |
| 0.658937 | 0.656155 | 0.666085 | 0.66472 | 0.616197 | 0.669322 | 0.741988 | 0.748422 | 0.753837 |
| 0.691196 | 0.682831 | 0.694513 | 0.704002 | 0.612855 | 0.706653 | 0.814324 | 0.810113 | 0.810003 |
| 0.650658 | 0.646785 | 0.660026 | 0.658075 | 0.614912 | 0.663061 | 0.74246 | 0.740195 | 0.748479 |
| 0.661014 | 0.649481 | 0.672595 | 0.661965 | 0.628051 | 0.667085 | 0.742636 | 0.738725 | 0.750205 |
| 0.653824 | 0.654677 | 0.662276 | 0.659948 | 0.621851 | 0.662316 | 0.739636 | 0.731965 | 0.745919 |
| 0.6543 | 0.654855 | 0.660887 | 0.660931 | 0.620606 | 0.664371 | 0.74336 | 0.743069 | 0.750348 |
| 0.656238 | 0.661183 | 0.672627 | 0.665894 | 0.623558 | 0.671377 | 0.752939 | 0.746641 | 0.758445 |
| 0.651913 | 0.656524 | 0.660683 | 0.663293 | 0.620773 | 0.66629 | 0.7527 | 0.746908 | 0.75244 |
| 0.652805 | 0.661695 | 0.670455 | 0.668835 | 0.623347 | 0.671966 | 0.748135 | 0.745024 | 0.753269 |
| 0.654849 | 0.655922 | 0.663792 | 0.662632 | 0.61793 | 0.668169 | 0.740471 | 0.734208 | 0.747116 |
| 0.648628 | 0.649376 | 0.660255 | 0.660143 | 0.621315 | 0.659812 | 0.739368 | 0.731747 | 0.74049 |
| 0.654948 | 0.652911 | 0.671637 | 0.66028 | 0.622596 | 0.661854 | 0.741894 | 0.735311 | 0.746123 |
| 0.661343 | 0.660466 | 0.671016 | 0.669048 | 0.619982 | 0.669553 | 0.74462 | 0.742664 | 0.747486 |
| 0.659144 | 0.662755 | 0.667543 | 0.665319 | 0.615885 | 0.667425 | 0.753244 | 0.740243 | 0.754081 |
| 0.660151 | 0.656588 | 0.66768 | 0.661761 | 0.618142 | 0.666792 | 0.741678 | 0.733416 | 0.735932 |
| 0.652614 | 0.650022 | 0.65498 | 0.654206 | 0.617786 | 0.660067 | 0.735928 | 0.733895 | 0.738668 |
| 0.655939 | 0.658727 | 0.660196 | 0.660385 | 0.618106 | 0.663757 | 0.738007 | 0.736575 | 0.741071 |
| 0.649571 | 0.656306 | 0.665803 | 0.661446 | 0.623263 | 0.6665 | 0.735099 | 0.735128 | 0.737243 |
| 0.646463 | 0.649655 | 0.660026 | 0.657276 | 0.623709 | 0.66365 | 0.733668 | 0.728912 | 0.744256 |
| 0.651896 | 0.652357 | 0.662495 | 0.667801 | 0.619815 | 0.665365 | 0.745592 | 0.741395 | 0.75446 |
| 0.659812 | 0.661356 | 0.671154 | 0.669897 | 0.618037 | 0.669082 | 0.745343 | 0.743668 | 0.743831 |
| 0.656384 | 0.654568 | 0.67097 | 0.666405 | 0.610457 | 0.670391 | 0.742079 | 0.741267 | 0.743839 |
| 0.666925 | 0.65795 | 0.670571 | 0.663952 | 0.614838 | 0.670133 | 0.74512 | 0.747108 | 0.745143 |

| | | | | | | | | |
|---|---|---|---|---|---|---|---|---|
| 0.707647 | 0.695135 | 0.7079 | 0.707755 | 0.614826 | 0.711477 | 0.809631 | 0.861809 | 0.85801 |
| 0.646946 | 0.649647 | 0.660944 | 0.653708 | 0.621898 | 0.660928 | 0.739092 | 0.741058 | 0.747301 |
| 0.651989 | 0.655073 | 0.660484 | 0.659494 | 0.619873 | 0.660236 | 0.745321 | 0.74131 | 0.745818 |
| 0.659778 | 0.661208 | 0.671334 | 0.667655 | 0.617474 | 0.675016 | 0.750932 | 0.757318 | 0.759072 |
| 0.653742 | 0.660118 | 0.663915 | 0.662063 | 0.620584 | 0.667797 | 0.745825 | 0.7415 | 0.740046 |
| 0.663295 | 0.659778 | 0.668949 | 0.666321 | 0.616274 | 0.665829 | 0.741416 | 0.738739 | 0.745708 |
| 0.650019 | 0.648786 | 0.66225 | 0.654672 | 0.616968 | 0.655001 | 0.737729 | 0.731869 | 0.739766 |
| 0.650732 | 0.65972 | 0.666181 | 0.669437 | 0.627468 | 0.671376 | 0.749712 | 0.742497 | 0.749116 |
| 0.659401 | 0.658544 | 0.677267 | 0.66436 | 0.625277 | 0.669876 | 0.739961 | 0.736698 | 0.744266 |
| 0.671442 | 0.665404 | 0.673135 | 0.666581 | 0.618985 | 0.675091 | 0.736591 | 0.734876 | 0.7454 |
| 0.652772 | 0.653018 | 0.660977 | 0.656891 | 0.612101 | 0.659411 | 0.746391 | 0.7357 | 0.745195 |
| 0.644351 | 0.649758 | 0.663268 | 0.658546 | 0.617559 | 0.656906 | 0.741176 | 0.737111 | 0.748849 |
| 0.653013 | 0.653343 | 0.662121 | 0.658314 | 0.621481 | 0.66358 | 0.739801 | 0.746978 | 0.748885 |
| 0.654362 | 0.65342 | 0.662438 | 0.659212 | 0.619766 | 0.6644 | 0.744463 | 0.743792 | 0.751795 |
| 0.65 | 0.65279 | 0.66155 | 0.65995 | 0.618262 | 0.665309 | 0.743795 | 0.73387 | 0.746075 |
| 0.650828 | 0.650307 | 0.665341 | 0.660919 | 0.618788 | 0.664286 | 0.740349 | 0.7337 | 0.743771 |
| 0.645575 | 0.649595 | 0.655001 | 0.657038 | 0.615444 | 0.6588 | 0.737223 | 0.740954 | 0.740292 |
| 0.657885 | 0.652365 | 0.666084 | 0.663204 | 0.622401 | 0.664484 | 0.745666 | 0.755342 | 0.753586 |
| 0.653234 | 0.659263 | 0.668212 | 0.66395 | 0.621642 | 0.665746 | 0.749673 | 0.74515 | 0.752032 |
| 0.644573 | 0.654121 | 0.662782 | 0.655704 | 0.617254 | 0.660692 | 0.746326 | 0.737652 | 0.74868 |
| 0.64508 | 0.651771 | 0.660309 | 0.658355 | 0.613697 | 0.659987 | 0.739706 | 0.74281 | 0.743737 |
| 0.653876 | 0.652517 | 0.661574 | 0.663082 | 0.614254 | 0.661761 | 0.742988 | 0.743899 | 0.75247 |
| 0.661272 | 0.652655 | 0.660632 | 0.658577 | 0.618605 | 0.665297 | 0.743505 | 0.739013 | 0.754611 |
| 0.662217 | 0.662306 | 0.673348 | 0.670899 | 0.615545 | 0.673341 | 0.744981 | 0.74124 | 0.752986 |
| 0.67485 | 0.675713 | 0.682346 | 0.675978 | 0.618892 | 0.686198 | 0.740015 | 0.737649 | 0.744927 |
| 0.722078 | 0.71968 | 0.724102 | 0.733252 | 0.589844 | 0.72974 | 0.734701 | 0.73655 | 0.739527 |
| 0.728766 | 0.731772 | 0.738874 | 0.736792 | 0.597788 | 0.733412 | 0.736192 | 0.734062 | 0.737326 |
| 0.724664 | 0.725134 | 0.736394 | 0.73056 | 0.599891 | 0.736038 | 0.738739 | 0.739011 | 0.744707 |
| 0.653735 | 0.656077 | 0.663724 | 0.658301 | 0.631249 | 0.664454 | 0.738483 | 0.732972 | 0.737353 |
| 0.651208 | 0.654308 | 0.662131 | 0.660333 | 0.620867 | 0.662109 | 0.738207 | 0.733654 | 0.736749 |
| 0.664414 | 0.665452 | 0.667438 | 0.663776 | 0.616395 | 0.669848 | 0.747665 | 0.740454 | 0.7457 |
| 0.670402 | 0.66697 | 0.675774 | 0.675091 | 0.623983 | 0.673431 | 0.7446 | 0.739848 | 0.7403 |
| 0.653586 | 0.655919 | 0.665455 | 0.666797 | 0.621293 | 0.66873 | 0.760534 | 0.755294 | 0.77146 |
| 0.650482 | 0.654933 | 0.664287 | 0.665365 | 0.620185 | 0.665577 | 0.754614 | 0.753302 | 0.759161 |
| 0.653081 | 0.653957 | 0.666909 | 0.666103 | 0.623329 | 0.667254 | 0.76225 | 0.764264 | 0.771258 |
| 0.652575 | 0.654895 | 0.665989 | 0.666753 | 0.623693 | 0.671432 | 0.75259 | 0.763292 | 0.764696 |
| 0.658162 | 0.658496 | 0.667293 | 0.666882 | 0.617946 | 0.672067 | 0.74767 | 0.742782 | 0.75379 |
| 0.678858 | 0.675507 | 0.687739 | 0.684211 | 0.641505 | 0.684104 | 0.762363 | 0.766036 | 0.769581 |
| 0.651627 | 0.653465 | 0.663988 | 0.659982 | 0.617938 | 0.663412 | 0.752769 | 0.750366 | 0.757013 |
| 0.655113 | 0.652239 | 0.665802 | 0.656774 | 0.626176 | 0.66107 | 0.742608 | 0.741101 | 0.746323 |
| 0.647708 | 0.649723 | 0.66044 | 0.661334 | 0.617689 | 0.664586 | 0.747772 | 0.752283 | 0.753526 |
| 0.673883 | 0.675651 | 0.684932 | 0.681222 | 0.641212 | 0.68362 | 0.76207 | 0.764902 | 0.7705 |
| 0.654053 | 0.662477 | 0.669432 | 0.667401 | 0.624738 | 0.662592 | 0.743548 | 0.746527 | 0.749815 |
| 0.648432 | 0.649492 | 0.663004 | 0.658678 | 0.621171 | 0.660192 | 0.742811 | 0.744697 | 0.747475 |
| 0.65613 | 0.65345 | 0.665654 | 0.662143 | 0.621526 | 0.662649 | 0.743143 | 0.738768 | 0.744257 |
| 0.656731 | 0.656962 | 0.665314 | 0.663868 | 0.620868 | 0.668848 | 0.749155 | 0.747583 | 0.759004 |

| | | | | | | | | |
|---|---|---|---|---|---|---|---|---|
| 0.653573 | 0.658312 | 0.662609 | 0.660044 | 0.621087 | 0.664828 | 0.74117 | 0.743391 | 0.746374 |
| 0.653323 | 0.656409 | 0.667053 | 0.663524 | 0.628166 | 0.67114 | 0.743567 | 0.736747 | 0.745515 |
| 0.650517 | 0.656499 | 0.667921 | 0.662758 | 0.618531 | 0.665749 | 0.741443 | 0.741898 | 0.747813 |
| 0.671653 | 0.66885 | 0.676505 | 0.677005 | 0.638451 | 0.678196 | 0.752499 | 0.754646 | 0.75869 |
| 0.667568 | 0.664706 | 0.676275 | 0.67348 | 0.620389 | 0.670429 | 0.748904 | 0.745926 | 0.751018 |
| 0.655026 | 0.655967 | 0.66861 | 0.6589 | 0.617805 | 0.666248 | 0.744884 | 0.737186 | 0.747838 |
| 0.661907 | 0.662898 | 0.672899 | 0.668561 | 0.618316 | 0.673897 | 0.75441 | 0.745206 | 0.75542 |
| 0.662388 | 0.658481 | 0.67383 | 0.666884 | 0.62021 | 0.673038 | 0.739565 | 0.741369 | 0.745618 |
| 0.6555 | 0.6586 | 0.660621 | 0.660519 | 0.61541 | 0.670759 | 0.741199 | 0.741446 | 0.747365 |
| 0.650693 | 0.648571 | 0.658878 | 0.657513 | 0.622658 | 0.65999 | 0.737261 | 0.734234 | 0.738423 |
| 0.65433 | 0.656236 | 0.661182 | 0.663076 | 0.623212 | 0.661757 | 0.741007 | 0.74049 | 0.74591 |
| 0.657436 | 0.659545 | 0.669886 | 0.66671 | 0.625733 | 0.667592 | 0.74476 | 0.738468 | 0.744704 |
| 0.655977 | 0.654353 | 0.665152 | 0.664621 | 0.621666 | 0.6684 | 0.740401 | 0.739152 | 0.74613 |
| 0.651486 | 0.654895 | 0.66735 | 0.665103 | 0.61743 | 0.671877 | 0.745426 | 0.732158 | 0.740885 |
| 0.655485 | 0.656651 | 0.664535 | 0.667622 | 0.620238 | 0.666625 | 0.740904 | 0.740401 | 0.745265 |
| 0.652266 | 0.649236 | 0.662279 | 0.655062 | 0.619292 | 0.658291 | 0.734956 | 0.736652 | 0.740614 |
| 0.654436 | 0.654775 | 0.665886 | 0.665661 | 0.619389 | 0.669286 | 0.746978 | 0.738295 | 0.746481 |
| 0.653987 | 0.656241 | 0.662873 | 0.663968 | 0.622563 | 0.667084 | 0.744924 | 0.74493 | 0.751518 |
| 0.663123 | 0.661562 | 0.671462 | 0.668221 | 0.62511 | 0.671389 | 0.744365 | 0.739163 | 0.747704 |
| 0.658154 | 0.664285 | 0.668807 | 0.661303 | 0.620807 | 0.668392 | 0.740705 | 0.739658 | 0.744287 |
| 0.660294 | 0.659607 | 0.669678 | 0.668829 | 0.619053 | 0.667501 | 0.743143 | 0.738843 | 0.743569 |
| 0.653186 | 0.656133 | 0.659349 | 0.654696 | 0.617188 | 0.659762 | 0.740181 | 0.736146 | 0.745103 |
| 0.655564 | 0.662775 | 0.66856 | 0.665674 | 0.62683 | 0.666124 | 0.744602 | 0.739264 | 0.744665 |
| 0.654134 | 0.659587 | 0.672131 | 0.665032 | 0.618813 | 0.665958 | 0.744057 | 0.737831 | 0.743993 |
| 0.655352 | 0.65885 | 0.665453 | 0.669886 | 0.619861 | 0.668887 | 0.751633 | 0.739098 | 0.750166 |
| 0.663071 | 0.662414 | 0.666619 | 0.671049 | 0.621857 | 0.670948 | 0.75986 | 0.749199 | 0.756155 |
| 0.656363 | 0.660513 | 0.664507 | 0.665722 | 0.618778 | 0.670659 | 0.752665 | 0.746161 | 0.750326 |
| 0.649378 | 0.650675 | 0.668655 | 0.66071 | 0.620391 | 0.663458 | 0.740548 | 0.738248 | 0.742259 |
| 0.65504 | 0.655898 | 0.670435 | 0.669214 | 0.618694 | 0.665291 | 0.743533 | 0.741951 | 0.748754 |
| 0.662408 | 0.662822 | 0.667343 | 0.671044 | 0.618353 | 0.671114 | 0.752468 | 0.742329 | 0.753018 |
| 0.654089 | 0.663362 | 0.671998 | 0.669776 | 0.615561 | 0.667293 | 0.742672 | 0.738374 | 0.743326 |
| 0.655534 | 0.654328 | 0.665712 | 0.663247 | 0.616556 | 0.664513 | 0.743342 | 0.738812 | 0.746947 |
| 0.660341 | 0.662616 | 0.674466 | 0.669837 | 0.624018 | 0.675621 | 0.755812 | 0.745913 | 0.760837 |
| 0.659423 | 0.659834 | 0.674385 | 0.669397 | 0.620181 | 0.674304 | 0.751423 | 0.757447 | 0.758929 |
| 0.652589 | 0.660512 | 0.662156 | 0.662321 | 0.617824 | 0.663668 | 0.744932 | 0.740064 | 0.746875 |
| 0.661568 | 0.661513 | 0.669334 | 0.661481 | 0.619121 | 0.67116 | 0.746795 | 0.742568 | 0.745613 |
| 0.641133 | 0.650026 | 0.665448 | 0.655561 | 0.619982 | 0.657351 | 0.742211 | 0.735602 | 0.743666 |
| 0.661517 | 0.658935 | 0.668107 | 0.663282 | 0.613532 | 0.661022 | 0.713873 | 0.714123 | 0.718015 |
| 0.663366 | 0.658216 | 0.670896 | 0.661005 | 0.619094 | 0.66789 | 0.723968 | 0.719464 | 0.722573 |
| 0.653145 | 0.651934 | 0.66331 | 0.657273 | 0.617716 | 0.657515 | 0.719245 | 0.713923 | 0.719476 |
| 0.657485 | 0.654764 | 0.662537 | 0.659699 | 0.615607 | 0.660968 | 0.723009 | 0.726316 | 0.737 |
| 0.644763 | 0.644239 | 0.656826 | 0.644029 | 0.596312 | 0.651862 | 0.711828 | 0.693807 | 0.704055 |
| 0.651228 | 0.649249 | 0.659348 | 0.651555 | 0.600028 | 0.653724 | 0.708544 | 0.709327 | 0.713984 |
| 0.653956 | 0.64354 | 0.65596 | 0.656329 | 0.607396 | 0.657534 | 0.715261 | 0.721404 | 0.717734 |
| 0.657781 | 0.649139 | 0.659411 | 0.655105 | 0.603717 | 0.65177 | 0.708252 | 0.704498 | 0.71453 |
| 0.659663 | 0.655939 | 0.660156 | 0.657377 | 0.608274 | 0.663072 | 0.718638 | 0.714801 | 0.723404 |
| 0.659321 | 0.65494 | 0.662052 | 0.658174 | 0.611282 | 0.664515 | 0.716864 | 0.717544 | 0.722796 |

| | | | | | | | | |
|---|---|---|---|---|---|---|---|---|
| 0.647239 | 0.649318 | 0.661977 | 0.658164 | 0.602473 | 0.653039 | 0.701346 | 0.702522 | 0.70826 |
| 0.663433 | 0.655904 | 0.665757 | 0.660825 | 0.609999 | 0.662971 | 0.736566 | 0.733821 | 0.744909 |
| 0.654892 | 0.648131 | 0.657569 | 0.658533 | 0.599404 | 0.658921 | 0.704082 | 0.702236 | 0.711673 |
| 0.65799 | 0.650994 | 0.656256 | 0.653184 | 0.597132 | 0.655806 | 0.706547 | 0.704363 | 0.706248 |
| 0.653915 | 0.65171 | 0.659315 | 0.653285 | 0.601733 | 0.657066 | 0.725056 | 0.7061 | 0.719457 |
| 0.660024 | 0.656874 | 0.66952 | 0.655284 | 0.602476 | 0.654406 | 0.703823 | 0.698949 | 0.707829 |
| 0.664402 | 0.658713 | 0.658061 | 0.660808 | 0.607843 | 0.65593 | 0.716171 | 0.705162 | 0.720701 |
| 0.650648 | 0.649034 | 0.659203 | 0.650069 | 0.606335 | 0.659417 | 0.715457 | 0.71348 | 0.712531 |
| 0.651352 | 0.652339 | 0.657292 | 0.657417 | 0.593072 | 0.654412 | 0.692228 | 0.693988 | 0.690359 |
| 0.647619 | 0.649084 | 0.653619 | 0.660474 | 0.606574 | 0.655234 | 0.727847 | 0.711232 | 0.731429 |
| 0.65369 | 0.643796 | 0.655515 | 0.649416 | 0.60499 | 0.654495 | 0.699095 | 0.70289 | 0.702315 |
| 0.65205 | 0.650493 | 0.651371 | 0.651771 | 0.633315 | 0.651928 | 0.732244 | 0.734671 | 0.735621 |
| 0.655166 | 0.650276 | 0.659003 | 0.66246 | 0.622171 | 0.662144 | 0.739387 | 0.739864 | 0.7452 |
| 0.665413 | 0.666892 | 0.680528 | 0.669189 | 0.619911 | 0.672725 | 0.71772 | 0.716891 | 0.721989 |
| 0.661589 | 0.668339 | 0.674806 | 0.669349 | 0.62558 | 0.672999 | 0.745483 | 0.74225 | 0.748777 |
| 0.655418 | 0.65601 | 0.662368 | 0.661793 | 0.634165 | 0.662778 | 0.737395 | 0.728271 | 0.738606 |
| 0.659436 | 0.653168 | 0.671683 | 0.666839 | 0.61716 | 0.670634 | 0.739014 | 0.740674 | 0.751122 |
| 0.664286 | 0.665508 | 0.668945 | 0.671607 | 0.619607 | 0.676808 | 0.738735 | 0.738904 | 0.750103 |
| 0.645747 | 0.642425 | 0.656206 | 0.652774 | 0.656371 | 0.652516 | 0.725412 | 0.723368 | 0.731005 |
| 0.666667 | 0.666219 | 0.67927 | 0.671981 | 0.62091 | 0.67996 | 0.750729 | 0.74328 | 0.749858 |
| 0.676777 | 0.666197 | 0.680082 | 0.666509 | 0.609263 | 0.679573 | 0.721203 | 0.719043 | 0.723577 |
| 0.658184 | 0.66459 | 0.670169 | 0.66671 | 0.620519 | 0.671295 | 0.739655 | 0.735938 | 0.744327 |
| 0.660611 | 0.663882 | 0.677108 | 0.670651 | 0.615789 | 0.673616 | 0.738072 | 0.730239 | 0.747109 |
| 0.678284 | 0.665634 | 0.675924 | 0.668602 | 0.606214 | 0.672528 | 0.718034 | 0.706015 | 0.714964 |
| 0.637692 | 0.637401 | 0.6494 | 0.650584 | 0.63109 | 0.650559 | 0.721321 | 0.716457 | 0.727959 |
| 0.687238 | 0.674401 | 0.679249 | 0.696059 | 0.615329 | 0.682489 | 0.733831 | 0.726116 | 0.733013 |
| 0.659172 | 0.66029 | 0.665036 | 0.662926 | 0.617693 | 0.665382 | 0.731596 | 0.731367 | 0.732235 |
| 0.66937 | 0.667523 | 0.672717 | 0.67302 | 0.614983 | 0.677464 | 0.740524 | 0.736303 | 0.742701 |
| 0.635349 | 0.641462 | 0.64658 | 0.640698 | 0.638731 | 0.645185 | 0.718671 | 0.718258 | 0.722789 |
| 0.674271 | 0.669756 | 0.681936 | 0.681842 | 0.620349 | 0.684539 | 0.750061 | 0.751721 | 0.768198 |
| 0.676251 | 0.676992 | 0.682881 | 0.677647 | 0.616429 | 0.681699 | 0.740376 | 0.731601 | 0.750323 |
| 0.648092 | 0.651009 | 0.658246 | 0.655773 | 0.710284 | 0.658967 | 0.726323 | 0.732971 | 0.731167 |
| 0.644013 | 0.649351 | 0.655557 | 0.660087 | 0.71036 | 0.658116 | 0.729398 | 0.730691 | 0.73239 |
| 0.676522 | 0.674872 | 0.686972 | 0.67827 | 0.623399 | 0.686764 | 0.744526 | 0.747039 | 0.753616 |
| 0.669667 | 0.666965 | 0.678012 | 0.679907 | 0.618708 | 0.676193 | 0.741611 | 0.738422 | 0.745612 |
| 0.670842 | 0.672492 | 0.685664 | 0.680983 | 0.623217 | 0.685804 | 0.748812 | 0.741935 | 0.748237 |
| 0.634005 | 0.630211 | 0.634835 | 0.633843 | 0.660176 | 0.638257 | 0.698798 | 0.700392 | 0.707315 |
| 0.676059 | 0.669532 | 0.678184 | 0.678655 | 0.613641 | 0.681545 | 0.737058 | 0.738732 | 0.738266 |
| 0.626953 | 0.62275 | 0.631208 | 0.630335 | 0.783777 | 0.63509 | 0.701015 | 0.699889 | 0.700601 |
| 0.615906 | 0.612696 | 0.619264 | 0.623833 | 0.661893 | 0.620387 | 0.695048 | 0.693301 | 0.693444 |
| 0.664294 | 0.667419 | 0.679109 | 0.675738 | 0.61988 | 0.675986 | 0.805261 | 0.793077 | 0.799817 |
| 0.619682 | 0.621433 | 0.623876 | 0.625502 | 0.671063 | 0.622699 | 0.693815 | 0.696477 | 0.692551 |
| 0.605244 | 0.60568 | 0.614789 | 0.620736 | 0.646882 | 0.618722 | 0.698398 | 0.700148 | 0.704662 |
| 0.605121 | 0.604622 | 0.612732 | 0.620799 | 0.670056 | 0.614249 | 0.692115 | 0.691925 | 0.698069 |
| 0.675437 | 0.677094 | 0.691266 | 0.68907 | 0.614952 | 0.691882 | 0.809588 | 0.807666 | 0.810933 |
| 0.603192 | 0.603049 | 0.611969 | 0.615963 | 0.661392 | 0.61487 | 0.693663 | 0.691584 | 0.702987 |
| 0.674343 | 0.674191 | 0.686828 | 0.68139 | 0.616777 | 0.686871 | 0.804905 | 0.807656 | 0.809914 |

| | | | | | | | | |
|---|---|---|---|---|---|---|---|---|
| 0.673708 | 0.676696 | 0.690704 | 0.684408 | 0.609914 | 0.688263 | 0.813221 | 0.807417 | 0.796993 |
| 0.66808 | 0.66978 | 0.68315 | 0.684156 | 0.61417 | 0.68392 | 0.802869 | 0.799158 | 0.814871 |
| 0.69971 | 0.702348 | 0.707877 | 0.704347 | 0.625957 | 0.712541 | 0.747791 | 0.743547 | 0.749384 |
| 0.59235 | 0.59368 | 0.599787 | 0.605701 | 0.667923 | 0.604904 | 0.684656 | 0.684352 | 0.690529 |
| 0.604834 | 0.608752 | 0.613887 | 0.614869 | 0.664114 | 0.612019 | 0.678551 | 0.683051 | 0.684649 |
| 0.682604 | 0.682294 | 0.692836 | 0.688281 | 0.615101 | 0.691461 | 0.81461 | 0.818267 | 0.815687 |
| 0.599646 | 0.601196 | 0.611111 | 0.608384 | 0.756364 | 0.611773 | 0.673567 | 0.674101 | 0.677839 |
| 0.6848 | 0.679387 | 0.69107 | 0.688977 | 0.607053 | 0.694363 | 0.731614 | 0.727175 | 0.727868 |
| 0.685603 | 0.683146 | 0.698855 | 0.693072 | 0.617413 | 0.696127 | 0.815391 | 0.815587 | 0.81938 |
| 0.691604 | 0.681452 | 0.690636 | 0.695914 | 0.599978 | 0.69453 | 0.72104 | 0.726391 | 0.731585 |
| 0.685996 | 0.678101 | 0.692251 | 0.696754 | 0.608967 | 0.695574 | 0.82884 | 0.841652 | 0.884683 |
| 0.821705 | 0.895161 | 0.804348 | 0.738255 | 0.566265 | 0.792857 | 0.696552 | 0.654088 | 0.675497 |
| 1 | 0.789942 | 0.756709 | 0.751231 | 0.569971 | 0.807738 | 0.694592 | 0.69535 | 0.697772 |
| 0.789942 | 1 | 0.745533 | 0.743658 | 0.56794 | 0.801921 | 0.691739 | 0.691818 | 0.692081 |
| 0.756709 | 0.745533 | 1 | 0.746721 | 0.573812 | 0.749909 | 0.697926 | 0.699077 | 0.697413 |
| 0.751231 | 0.743658 | 0.746721 | 1 | 0.570959 | 0.751955 | 0.701233 | 0.698601 | 0.700532 |
| 0.569971 | 0.56794 | 0.573812 | 0.570959 | 1 | 0.571927 | 0.609711 | 0.606462 | 0.609882 |
| 0.807738 | 0.801921 | 0.749909 | 0.751955 | 0.571927 | 1 | 0.707323 | 0.704593 | 0.706489 |
| 0.694592 | 0.691739 | 0.697926 | 0.701233 | 0.609711 | 0.707323 | 1 | 0.846657 | 0.832261 |
| 0.69535 | 0.691818 | 0.699077 | 0.698601 | 0.606462 | 0.704593 | 0.846657 | 1 | 0.841655 |
| 0.697772 | 0.692081 | 0.697413 | 0.700532 | 0.609882 | 0.706489 | 0.832261 | 0.841655 | 1 |
| 0.528247 | 0.536221 | 0.540596 | 0.546085 | 0.646509 | 0.541722 | 0.60507 | 0.608876 | 0.609141 |
| 0.518081 | 0.521021 | 0.523896 | 0.5333 | 0.647842 | 0.529405 | 0.591624 | 0.592539 | 0.593151 |
| 0.720491 | 0.719623 | 0.725434 | 0.723817 | 0.590716 | 0.73292 | 0.768618 | 0.763312 | 0.767092 |
| 0.722178 | 0.722178 | 0.721578 | 0.725466 | 0.593373 | 0.735391 | 0.771097 | 0.763041 | 0.764113 |
| 0.71863 | 0.707493 | 0.714911 | 0.718628 | 0.581773 | 0.723755 | 0.744893 | 0.743248 | 0.747079 |
| 0.72067 | 0.712733 | 0.722982 | 0.719423 | 0.588166 | 0.732159 | 0.741893 | 0.743372 | 0.750164 |
| 0.717668 | 0.718794 | 0.721781 | 0.722937 | 0.588337 | 0.726883 | 0.753205 | 0.752153 | 0.754751 |
| 0.727218 | 0.720433 | 0.731689 | 0.728412 | 0.590075 | 0.740231 | 0.75222 | 0.754457 | 0.754411 |
| 0.723645 | 0.724631 | 0.732239 | 0.728245 | 0.592597 | 0.7298 | 0.751931 | 0.758051 | 0.754709 |
| 0.721624 | 0.718665 | 0.729004 | 0.7251 | 0.590919 | 0.727503 | 0.728924 | 0.72887 | 0.735417 |
| 0.721836 | 0.721823 | 0.738652 | 0.734541 | 0.599718 | 0.735256 | 0.739614 | 0.73568 | 0.739085 |
| 0.721544 | 0.718085 | 0.728956 | 0.723007 | 0.591504 | 0.723067 | 0.734443 | 0.734585 | 0.738301 |
| 0.709544 | 0.712931 | 0.717918 | 0.720806 | 0.58956 | 0.724552 | 0.730871 | 0.732921 | 0.736927 |
| 0.719872 | 0.71404 | 0.734434 | 0.732948 | 0.597028 | 0.728811 | 0.740385 | 0.737788 | 0.73979 |
| 0.717964 | 0.715692 | 0.731774 | 0.725752 | 0.596846 | 0.728791 | 0.735155 | 0.734632 | 0.737062 |
| 0.718775 | 0.712218 | 0.726917 | 0.723168 | 0.590973 | 0.72805 | 0.734276 | 0.731944 | 0.736102 |
| 0.716278 | 0.724638 | 0.735201 | 0.729214 | 0.593407 | 0.729818 | 0.734397 | 0.735724 | 0.736068 |
| 0.718881 | 0.722794 | 0.725853 | 0.721243 | 0.590351 | 0.725361 | 0.730542 | 0.733745 | 0.7363 |
| 0.712614 | 0.718706 | 0.726167 | 0.715426 | 0.590111 | 0.724089 | 0.731892 | 0.730907 | 0.735938 |
| 0.740217 | 0.731446 | 0.731473 | 0.731143 | 0.570296 | 0.741302 | 0.694612 | 0.692922 | 0.694877 |
| 0.735465 | 0.727098 | 0.739506 | 0.744155 | 0.571931 | 0.72867 | 0.694218 | 0.700977 | 0.701031 |
| 0.708383 | 0.723939 | 0.73131 | 0.72674 | 0.595481 | 0.725255 | 0.737716 | 0.737696 | 0.737034 |
| 0.722794 | 0.717881 | 0.734657 | 0.733073 | 0.597897 | 0.733463 | 0.747935 | 0.741027 | 0.748546 |
| 0.71412 | 0.702087 | 0.715248 | 0.709131 | 0.566568 | 0.714755 | 0.683704 | 0.67968 | 0.68522 |
| 0.722666 | 0.72264 | 0.726835 | 0.729435 | 0.592761 | 0.728855 | 0.736791 | 0.734946 | 0.736567 |
| 0.709851 | 0.709094 | 0.723158 | 0.719581 | 0.586021 | 0.722136 | 0.731794 | 0.728899 | 0.730937 |

| | | | | | | | | |
|---|---|---|---|---|---|---|---|---|
| 0.714257 | 0.716843 | 0.728319 | 0.724671 | 0.591668 | 0.730994 | 0.737575 | 0.736104 | 0.740018 |
| 0.715923 | 0.720613 | 0.725987 | 0.730759 | 0.591153 | 0.730234 | 0.735425 | 0.734727 | 0.738525 |
| 0.719576 | 0.717094 | 0.725378 | 0.726302 | 0.593869 | 0.730393 | 0.736659 | 0.738691 | 0.740911 |
| 0.721803 | 0.723548 | 0.726756 | 0.735691 | 0.59625 | 0.728135 | 0.740385 | 0.736088 | 0.741537 |
| 0.728571 | 0.724699 | 0.736226 | 0.735943 | 0.59599 | 0.740052 | 0.743719 | 0.745789 | 0.747774 |
| 0.722532 | 0.72185 | 0.723947 | 0.728392 | 0.589274 | 0.734363 | 0.733431 | 0.730693 | 0.736385 |
| 0.719881 | 0.714308 | 0.723385 | 0.729621 | 0.590071 | 0.72461 | 0.736778 | 0.731508 | 0.734777 |
| 0.716989 | 0.70799 | 0.716231 | 0.719325 | 0.58492 | 0.71547 | 0.726544 | 0.730824 | 0.730393 |
| 0.720656 | 0.719499 | 0.726291 | 0.726481 | 0.588824 | 0.727793 | 0.730096 | 0.731631 | 0.734627 |
| 0.710853 | 0.708891 | 0.720121 | 0.718001 | 0.58467 | 0.72463 | 0.733208 | 0.737068 | 0.737501 |
| 0.717551 | 0.717577 | 0.726374 | 0.729119 | 0.592445 | 0.72934 | 0.738147 | 0.736338 | 0.738598 |
| 0.714802 | 0.722028 | 0.727912 | 0.726096 | 0.58888 | 0.728673 | 0.735644 | 0.737029 | 0.736895 |
| 0.706094 | 0.708669 | 0.716528 | 0.71334 | 0.585302 | 0.721727 | 0.72793 | 0.727491 | 0.731575 |
| 0.711484 | 0.707797 | 0.71854 | 0.721902 | 0.587268 | 0.717745 | 0.728005 | 0.724285 | 0.730237 |
| 0.722588 | 0.723949 | 0.726524 | 0.725247 | 0.593198 | 0.730118 | 0.738387 | 0.739198 | 0.742572 |
| 0.72542 | 0.719385 | 0.733413 | 0.734209 | 0.595048 | 0.734659 | 0.759673 | 0.761054 | 0.765247 |
| 0.729724 | 0.718367 | 0.731409 | 0.726644 | 0.588325 | 0.73165 | 0.741044 | 0.741412 | 0.739726 |
| 0.721292 | 0.712764 | 0.724913 | 0.718809 | 0.586595 | 0.730603 | 0.750762 | 0.757198 | 0.746261 |
| 0.718619 | 0.713751 | 0.719167 | 0.72462 | 0.589264 | 0.730925 | 0.755935 | 0.755982 | 0.761284 |
| 0.720512 | 0.718186 | 0.719538 | 0.726526 | 0.58569 | 0.732637 | 0.751091 | 0.753464 | 0.745607 |
| 0.712725 | 0.706235 | 0.714221 | 0.719744 | 0.587657 | 0.722391 | 0.755291 | 0.760238 | 0.755076 |
| 0.718068 | 0.708831 | 0.717041 | 0.722169 | 0.589041 | 0.729043 | 0.751377 | 0.750782 | 0.755563 |
| 0.730362 | 0.719178 | 0.728524 | 0.731883 | 0.590709 | 0.733593 | 0.747746 | 0.749052 | 0.752122 |
| 0.722334 | 0.720238 | 0.726414 | 0.720037 | 0.590753 | 0.730334 | 0.748299 | 0.748774 | 0.747209 |
| 0.723396 | 0.726211 | 0.724728 | 0.721331 | 0.586285 | 0.731193 | 0.74799 | 0.75735 | 0.753562 |
| 0.714856 | 0.711021 | 0.720723 | 0.726631 | 0.587398 | 0.728918 | 0.765417 | 0.758876 | 0.764524 |
| 0.710707 | 0.709243 | 0.717626 | 0.710055 | 0.587585 | 0.721227 | 0.750811 | 0.749642 | 0.746011 |
| 0.710378 | 0.703982 | 0.71749 | 0.716815 | 0.587107 | 0.724347 | 0.741774 | 0.744785 | 0.747221 |
| 0.718335 | 0.71323 | 0.719245 | 0.715933 | 0.588815 | 0.731063 | 0.773334 | 0.768928 | 0.769154 |
| 0.714058 | 0.712768 | 0.72285 | 0.723764 | 0.588369 | 0.731787 | 0.759242 | 0.759406 | 0.758985 |
| 0.717687 | 0.718343 | 0.724185 | 0.725037 | 0.590535 | 0.732122 | 0.75621 | 0.761738 | 0.752343 |
| 0.71697 | 0.718834 | 0.723918 | 0.722509 | 0.5954 | 0.728026 | 0.747742 | 0.750649 | 0.747125 |
| 0.72915 | 0.728981 | 0.728755 | 0.729404 | 0.597087 | 0.73594 | 0.761652 | 0.760627 | 0.759726 |
| 0.722568 | 0.722275 | 0.730478 | 0.730142 | 0.600696 | 0.732064 | 0.752559 | 0.751991 | 0.754822 |
| 0.707487 | 0.710794 | 0.717378 | 0.718259 | 0.588363 | 0.728455 | 0.769506 | 0.759699 | 0.763488 |
| 0.722991 | 0.71776 | 0.7322 | 0.724036 | 0.590212 | 0.727785 | 0.753944 | 0.763484 | 0.752609 |
| 0.717008 | 0.713391 | 0.719598 | 0.716555 | 0.587955 | 0.723225 | 0.750165 | 0.749352 | 0.751801 |
| 0.720847 | 0.715313 | 0.726397 | 0.724642 | 0.589406 | 0.723584 | 0.745129 | 0.739947 | 0.744317 |
| 0.709003 | 0.703132 | 0.71034 | 0.716823 | 0.584899 | 0.720406 | 0.74912 | 0.754349 | 0.747113 |
| 0.714457 | 0.707378 | 0.723156 | 0.727297 | 0.586592 | 0.725056 | 0.744154 | 0.74123 | 0.743385 |
| 0.710124 | 0.706042 | 0.709199 | 0.715912 | 0.582014 | 0.720679 | 0.767141 | 0.767371 | 0.761999 |
| 0.719209 | 0.712375 | 0.717149 | 0.721275 | 0.584347 | 0.724428 | 0.751464 | 0.745283 | 0.750306 |
| 0.71232 | 0.704374 | 0.709911 | 0.710432 | 0.580918 | 0.719051 | 0.751098 | 0.754542 | 0.746796 |
| 0.714988 | 0.708522 | 0.718642 | 0.71912 | 0.582349 | 0.722569 | 0.739091 | 0.742067 | 0.745371 |
| 0.717474 | 0.714986 | 0.728626 | 0.726962 | 0.588704 | 0.725524 | 0.736865 | 0.740317 | 0.741922 |
| 0.707286 | 0.710856 | 0.715006 | 0.705737 | 0.582152 | 0.718887 | 0.73137 | 0.729935 | 0.733114 |

| | | | | | | | | |
|---|---|---|---|---|---|---|---|---|
| 0.718505 | 0.714599 | 0.724868 | 0.727616 | 0.588025 | 0.730447 | 0.788692 | 0.790199 | 0.787879 |
| 0.738231 | 0.736142 | 0.732919 | 0.728057 | 0.595212 | 0.737683 | 0.733881 | 0.737851 | 0.735537 |
| 0.717793 | 0.71922 | 0.722604 | 0.722207 | 0.584209 | 0.735259 | 0.736642 | 0.733455 | 0.73591 |
| 0.71718 | 0.728867 | 0.723398 | 0.722318 | 0.588657 | 0.727801 | 0.736626 | 0.73317 | 0.738184 |
| 0.73332 | 0.736842 | 0.739933 | 0.738204 | 0.597985 | 0.741316 | 0.743734 | 0.740903 | 0.744543 |
| 0.731864 | 0.723923 | 0.735764 | 0.727613 | 0.587038 | 0.733828 | 0.739742 | 0.739291 | 0.738573 |
| 0.727971 | 0.726379 | 0.726429 | 0.721687 | 0.587588 | 0.734551 | 0.735608 | 0.729039 | 0.735663 |
| 0.720697 | 0.719831 | 0.730427 | 0.718079 | 0.589892 | 0.726989 | 0.734727 | 0.730864 | 0.73802 |
| 0.724342 | 0.724803 | 0.732702 | 0.72721 | 0.591166 | 0.730018 | 0.727604 | 0.726939 | 0.728737 |
| 0.718 | 0.714397 | 0.714796 | 0.720768 | 0.58833 | 0.72609 | 0.734541 | 0.731643 | 0.738302 |
| 0.714821 | 0.716476 | 0.722718 | 0.722407 | 0.588916 | 0.728117 | 0.733076 | 0.730507 | 0.737542 |
| 0.720644 | 0.721902 | 0.722364 | 0.728064 | 0.589589 | 0.731935 | 0.736001 | 0.733536 | 0.737163 |
| 0.726655 | 0.722031 | 0.731036 | 0.728993 | 0.589917 | 0.738636 | 0.73907 | 0.736189 | 0.738546 |
| 0.725119 | 0.726304 | 0.739961 | 0.72989 | 0.594674 | 0.730949 | 0.736722 | 0.737167 | 0.737804 |
| 0.731198 | 0.729802 | 0.737685 | 0.731391 | 0.594039 | 0.733066 | 0.74 | 0.735992 | 0.740108 |
| 0.722077 | 0.718843 | 0.735165 | 0.726523 | 0.591874 | 0.731211 | 0.739626 | 0.736379 | 0.741109 |
| 0.726193 | 0.728224 | 0.731126 | 0.731511 | 0.591916 | 0.72882 | 0.739449 | 0.740029 | 0.744246 |
| 0.728016 | 0.722524 | 0.729319 | 0.733485 | 0.589341 | 0.734593 | 0.741045 | 0.740096 | 0.742413 |
| 0.724138 | 0.729236 | 0.736644 | 0.73463 | 0.594922 | 0.739575 | 0.739014 | 0.736993 | 0.738014 |
| 0.731668 | 0.727843 | 0.73249 | 0.734447 | 0.593125 | 0.736657 | 0.735513 | 0.737984 | 0.739885 |
| 0.720294 | 0.720118 | 0.728449 | 0.717863 | 0.597575 | 0.732684 | 0.735292 | 0.736264 | 0.742808 |
| 0.721158 | 0.730418 | 0.728518 | 0.728688 | 0.59482 | 0.739564 | 0.733699 | 0.732732 | 0.73514 |
| 0.721106 | 0.724181 | 0.725549 | 0.728306 | 0.593504 | 0.733169 | 0.743419 | 0.741515 | 0.742664 |
| 0.720699 | 0.720652 | 0.730585 | 0.732532 | 0.592648 | 0.730963 | 0.735843 | 0.731747 | 0.737837 |
| 0.734026 | 0.726891 | 0.738618 | 0.735857 | 0.590565 | 0.738228 | 0.7368 | 0.73773 | 0.74401 |
| 0.726267 | 0.731049 | 0.737901 | 0.731799 | 0.59291 | 0.739153 | 0.737141 | 0.73772 | 0.739919 |
| 0.727327 | 0.723896 | 0.729997 | 0.731439 | 0.585009 | 0.733342 | 0.731444 | 0.732508 | 0.734092 |
| 0.722047 | 0.72192 | 0.732036 | 0.730579 | 0.591441 | 0.732842 | 0.730389 | 0.734831 | 0.736961 |
| 0.726673 | 0.730504 | 0.73826 | 0.734147 | 0.591518 | 0.743014 | 0.737513 | 0.736739 | 0.743376 |
| 0.743153 | 0.729213 | 0.742019 | 0.735606 | 0.596441 | 0.737319 | 0.73889 | 0.738623 | 0.742161 |
| 0.724699 | 0.718838 | 0.730152 | 0.726326 | 0.59315 | 0.728296 | 0.731155 | 0.733796 | 0.736189 |
| 0.732247 | 0.726445 | 0.735604 | 0.735119 | 0.59689 | 0.737816 | 0.737939 | 0.736381 | 0.743337 |
| 0.734322 | 0.729251 | 0.734065 | 0.736757 | 0.594131 | 0.738301 | 0.740431 | 0.736605 | 0.745616 |
| 0.726485 | 0.730806 | 0.7358 | 0.734913 | 0.590939 | 0.738272 | 0.73364 | 0.735573 | 0.739203 |
| 0.727491 | 0.721402 | 0.734112 | 0.734724 | 0.596316 | 0.733631 | 0.733406 | 0.732449 | 0.738631 |
| 0.730987 | 0.725242 | 0.739443 | 0.733443 | 0.590338 | 0.738019 | 0.733747 | 0.735018 | 0.737135 |
| 0.729746 | 0.730328 | 0.741155 | 0.738503 | 0.59335 | 0.742302 | 0.742769 | 0.74333 | 0.746002 |
| 0.730022 | 0.730279 | 0.733887 | 0.735612 | 0.594956 | 0.743886 | 0.734446 | 0.736714 | 0.736002 |
| 0.735773 | 0.719495 | 0.738322 | 0.731074 | 0.593801 | 0.73493 | 0.735697 | 0.737678 | 0.740656 |
| 0.731446 | 0.724295 | 0.734678 | 0.734472 | 0.595163 | 0.737597 | 0.735707 | 0.733506 | 0.739355 |
| 0.730111 | 0.727032 | 0.73636 | 0.732788 | 0.599396 | 0.738204 | 0.739141 | 0.743554 | 0.745988 |
| 0.72864 | 0.72553 | 0.735352 | 0.739325 | 0.595551 | 0.743243 | 0.741108 | 0.735654 | 0.740122 |
| 0.728163 | 0.731252 | 0.742736 | 0.739588 | 0.595938 | 0.741334 | 0.739608 | 0.738932 | 0.742178 |
| 0.724565 | 0.720138 | 0.729811 | 0.730671 | 0.590909 | 0.734862 | 0.737021 | 0.73761 | 0.739452 |
| 0.731909 | 0.72626 | 0.736492 | 0.73772 | 0.600522 | 0.734649 | 0.739483 | 0.739188 | 0.744877 |
| 0.720832 | 0.729477 | 0.738026 | 0.733651 | 0.587409 | 0.731711 | 0.736232 | 0.736732 | 0.741834 |

| | | | | | | | | |
|---|---|---|---|---|---|---|---|---|
| 0.727138 | 0.726937 | 0.733258 | 0.729179 | 0.58223 | 0.73106 | 0.72308 | 0.724713 | 0.726728 |
| 0.707856 | 0.711197 | 0.714455 | 0.716543 | 0.611475 | 0.716072 | 0.75261 | 0.747201 | 0.74971 |
| 0.727932 | 0.720732 | 0.730074 | 0.728591 | 0.59391 | 0.731881 | 0.737861 | 0.732536 | 0.735326 |
| 0.726639 | 0.73039 | 0.745835 | 0.731652 | 0.590831 | 0.736561 | 0.739029 | 0.738546 | 0.741001 |
| 0.724306 | 0.723513 | 0.731595 | 0.725573 | 0.590634 | 0.733475 | 0.729198 | 0.729307 | 0.732528 |
| 0.718429 | 0.72195 | 0.73919 | 0.732599 | 0.590494 | 0.733817 | 0.735577 | 0.73345 | 0.733394 |
| 0.733715 | 0.732939 | 0.742338 | 0.734341 | 0.592465 | 0.740706 | 0.73249 | 0.731536 | 0.736398 |
| 0.72644 | 0.724852 | 0.736998 | 0.729569 | 0.588585 | 0.735089 | 0.726011 | 0.726768 | 0.729108 |
| 0.730461 | 0.726006 | 0.739964 | 0.734691 | 0.592469 | 0.735979 | 0.736748 | 0.737933 | 0.738004 |
| 0.731776 | 0.732501 | 0.739639 | 0.730022 | 0.594041 | 0.738897 | 0.74068 | 0.739278 | 0.740059 |
| 0.729317 | 0.730436 | 0.735695 | 0.734078 | 0.59521 | 0.737945 | 0.739173 | 0.739697 | 0.74134 |
| 0.72499 | 0.727444 | 0.733858 | 0.733162 | 0.595065 | 0.741303 | 0.737452 | 0.739037 | 0.739433 |
| 0.735431 | 0.7303 | 0.73848 | 0.74435 | 0.59745 | 0.743506 | 0.744914 | 0.742497 | 0.74299 |
| 0.727146 | 0.728623 | 0.744884 | 0.737303 | 0.599611 | 0.745001 | 0.74105 | 0.736877 | 0.740596 |
| 0.727872 | 0.727731 | 0.735089 | 0.731434 | 0.594828 | 0.736614 | 0.735182 | 0.73374 | 0.736376 |
| 0.727615 | 0.723715 | 0.730781 | 0.735274 | 0.592399 | 0.735264 | 0.733749 | 0.735904 | 0.737036 |
| 0.726434 | 0.72604 | 0.732567 | 0.730899 | 0.590658 | 0.737511 | 0.735337 | 0.739285 | 0.739644 |
| 0.7268 | 0.729738 | 0.736164 | 0.728191 | 0.587017 | 0.736539 | 0.733932 | 0.736777 | 0.736277 |
| 0.739592 | 0.735115 | 0.742804 | 0.739035 | 0.592465 | 0.740498 | 0.733902 | 0.732055 | 0.736847 |
| 0.725308 | 0.724408 | 0.735544 | 0.725571 | 0.588745 | 0.732644 | 0.731548 | 0.72954 | 0.733535 |
| 0.723188 | 0.728993 | 0.733938 | 0.731349 | 0.590047 | 0.734338 | 0.73796 | 0.732957 | 0.738502 |
| 0.724933 | 0.721802 | 0.727441 | 0.730552 | 0.5847 | 0.727841 | 0.728077 | 0.725403 | 0.728314 |
| 0.727767 | 0.733607 | 0.734366 | 0.73525 | 0.590481 | 0.73397 | 0.730987 | 0.732667 | 0.730053 |
| 0.728422 | 0.739234 | 0.731788 | 0.7355 | 0.591528 | 0.744326 | 0.737117 | 0.734843 | 0.733643 |
| 0.745354 | 0.738586 | 0.733515 | 0.738474 | 0.591697 | 0.747289 | 0.738071 | 0.736842 | 0.736622 |
| 0.733912 | 0.731139 | 0.730868 | 0.736785 | 0.59173 | 0.739388 | 0.735432 | 0.733445 | 0.732492 |
| 0.72411 | 0.730324 | 0.73612 | 0.725855 | 0.588027 | 0.727103 | 0.73025 | 0.732184 | 0.732113 |
| 0.735133 | 0.734965 | 0.739051 | 0.740279 | 0.588351 | 0.735474 | 0.729794 | 0.73668 | 0.732297 |
| 0.732296 | 0.727754 | 0.73818 | 0.732113 | 0.591559 | 0.733564 | 0.725942 | 0.725392 | 0.72914 |
| 0.649827 | 0.652299 | 0.659594 | 0.657186 | 0.613956 | 0.661207 | 0.728405 | 0.722159 | 0.730237 |
| 0.656682 | 0.653303 | 0.664436 | 0.662371 | 0.619011 | 0.670296 | 0.742643 | 0.739333 | 0.751138 |
| 0.65188 | 0.655115 | 0.661902 | 0.656743 | 0.624086 | 0.664898 | 0.737274 | 0.734063 | 0.742132 |
| 0.651265 | 0.654629 | 0.662176 | 0.656604 | 0.616413 | 0.659435 | 0.727937 | 0.732731 | 0.737249 |
| 0.656118 | 0.650926 | 0.664781 | 0.65838 | 0.618271 | 0.665626 | 0.751323 | 0.743592 | 0.753309 |
| 0.727199 | 0.736003 | 0.752504 | 0.737897 | 0.587076 | 0.737421 | 0.73018 | 0.727219 | 0.726631 |
| 0.732617 | 0.737412 | 0.746073 | 0.743352 | 0.588913 | 0.741141 | 0.734119 | 0.734662 | 0.735165 |
| 0.747479 | 0.740227 | 0.748671 | 0.748742 | 0.592283 | 0.748054 | 0.735152 | 0.733112 | 0.734776 |
| 0.741296 | 0.745348 | 0.753734 | 0.747167 | 0.592171 | 0.748352 | 0.739113 | 0.734187 | 0.734757 |
| 0.738719 | 0.737449 | 0.747526 | 0.748775 | 0.588818 | 0.745235 | 0.732155 | 0.732131 | 0.73451 |
| 0.731957 | 0.744681 | 0.749164 | 0.746454 | 0.594156 | 0.748752 | 0.738846 | 0.737857 | 0.742857 |
| 0.734343 | 0.736909 | 0.74577 | 0.738418 | 0.590706 | 0.741799 | 0.734697 | 0.738985 | 0.73743 |
| 0.734435 | 0.73825 | 0.745011 | 0.737442 | 0.589066 | 0.738282 | 0.730366 | 0.730782 | 0.73155 |
| 0.735923 | 0.741925 | 0.748907 | 0.744134 | 0.586957 | 0.736927 | 0.731066 | 0.727935 | 0.731525 |
| 0.728995 | 0.735729 | 0.748043 | 0.743389 | 0.587409 | 0.732913 | 0.724889 | 0.723616 | 0.726096 |
| 0.730738 | 0.7392 | 0.741189 | 0.737335 | 0.592922 | 0.738396 | 0.732544 | 0.731961 | 0.730666 |
| 0.736607 | 0.742029 | 0.74655 | 0.747976 | 0.591997 | 0.749168 | 0.737594 | 0.733827 | 0.734593 |

| | | | | | | | | |
|---|---|---|---|---|---|---|---|---|
| 0.732928 | 0.742229 | 0.753163 | 0.750513 | 0.589822 | 0.750497 | 0.737756 | 0.73575 | 0.736644 |
| 0.740418 | 0.734484 | 0.75674 | 0.752838 | 0.588982 | 0.75593 | 0.734914 | 0.737871 | 0.735732 |
| 0.74406 | 0.746155 | 0.76288 | 0.751088 | 0.59577 | 0.757269 | 0.735117 | 0.734082 | 0.741013 |
| 0.733172 | 0.738281 | 0.740295 | 0.748629 | 0.591082 | 0.746859 | 0.732089 | 0.734136 | 0.73733 |
| 0.742329 | 0.738486 | 0.744264 | 0.752653 | 0.589102 | 0.746758 | 0.728942 | 0.731164 | 0.732431 |
| 0.740197 | 0.743974 | 0.749045 | 0.755237 | 0.593362 | 0.752192 | 0.729714 | 0.729365 | 0.732416 |
| 0.739695 | 0.737287 | 0.757422 | 0.753336 | 0.59335 | 0.751029 | 0.73525 | 0.734536 | 0.733923 |
| 0.742845 | 0.744033 | 0.757225 | 0.75097 | 0.590128 | 0.7456 | 0.737735 | 0.737406 | 0.73551 |
| 0.744962 | 0.740214 | 0.757538 | 0.746125 | 0.589791 | 0.753928 | 0.732319 | 0.734428 | 0.738247 |
| 0.734545 | 0.741915 | 0.7479 | 0.752053 | 0.58899 | 0.755175 | 0.734921 | 0.736934 | 0.736175 |
| 0.737705 | 0.734264 | 0.743114 | 0.750137 | 0.595422 | 0.75179 | 0.732733 | 0.73183 | 0.73498 |
| 0.733793 | 0.732665 | 0.744388 | 0.745846 | 0.593654 | 0.751128 | 0.736842 | 0.735664 | 0.735902 |
| 0.743484 | 0.735547 | 0.747398 | 0.744878 | 0.589471 | 0.748207 | 0.725277 | 0.721687 | 0.726833 |
| 0.747401 | 0.737221 | 0.755808 | 0.74469 | 0.590289 | 0.752576 | 0.73011 | 0.729237 | 0.730558 |
| 0.724492 | 0.72253 | 0.738685 | 0.723886 | 0.593729 | 0.731251 | 0.739902 | 0.73986 | 0.739337 |
| 0.733845 | 0.732839 | 0.738189 | 0.737647 | 0.597778 | 0.743596 | 0.745522 | 0.743894 | 0.747777 |
| 0.725595 | 0.722136 | 0.736282 | 0.728735 | 0.595848 | 0.731827 | 0.741297 | 0.738637 | 0.741628 |
| 0.726415 | 0.721926 | 0.736397 | 0.732795 | 0.596942 | 0.737484 | 0.741471 | 0.742822 | 0.739244 |
| 0.728175 | 0.725347 | 0.737997 | 0.734117 | 0.591029 | 0.740234 | 0.734565 | 0.736252 | 0.737792 |
| 0.730675 | 0.727374 | 0.734069 | 0.727746 | 0.59108 | 0.739102 | 0.736633 | 0.733935 | 0.74232 |
| 0.732589 | 0.724029 | 0.739217 | 0.740135 | 0.590214 | 0.735714 | 0.733761 | 0.729681 | 0.735109 |
| 0.722943 | 0.724127 | 0.736391 | 0.73166 | 0.590451 | 0.730856 | 0.737993 | 0.733555 | 0.735781 |
| 0.728927 | 0.73102 | 0.739338 | 0.738862 | 0.596701 | 0.742086 | 0.743187 | 0.740481 | 0.742765 |
| 0.747174 | 0.74818 | 0.757292 | 0.757576 | 0.5935 | 0.752669 | 0.734385 | 0.733116 | 0.736373 |
| 0.736647 | 0.736156 | 0.739762 | 0.736585 | 0.595047 | 0.75164 | 0.741059 | 0.737453 | 0.736805 |
| 0.648534 | 0.651685 | 0.652444 | 0.656161 | 0.61996 | 0.652934 | 0.736859 | 0.736482 | 0.739785 |
| 0.731615 | 0.726588 | 0.737977 | 0.734274 | 0.595089 | 0.738801 | 0.734076 | 0.733983 | 0.738103 |
| 0.732745 | 0.729347 | 0.739433 | 0.736607 | 0.592585 | 0.739807 | 0.738682 | 0.737141 | 0.742783 |
| 0.727768 | 0.725995 | 0.73303 | 0.734866 | 0.593196 | 0.742951 | 0.741167 | 0.742572 | 0.747464 |
| 0.735412 | 0.730733 | 0.743749 | 0.731454 | 0.591504 | 0.741413 | 0.735846 | 0.738979 | 0.739587 |
| 0.733198 | 0.733365 | 0.735371 | 0.736929 | 0.59104 | 0.737038 | 0.731956 | 0.7281 | 0.734303 |
| 0.741215 | 0.731893 | 0.739682 | 0.738619 | 0.59138 | 0.74361 | 0.737242 | 0.735141 | 0.734265 |
| 0.725506 | 0.724654 | 0.73677 | 0.732314 | 0.591812 | 0.73853 | 0.740476 | 0.736369 | 0.742485 |
| 0.732563 | 0.7282 | 0.73681 | 0.733897 | 0.586469 | 0.737389 | 0.728799 | 0.730845 | 0.735868 |
| 0.724138 | 0.727521 | 0.739265 | 0.728558 | 0.588744 | 0.738431 | 0.732036 | 0.731775 | 0.733833 |
| 0.730285 | 0.725528 | 0.732012 | 0.734288 | 0.58928 | 0.73531 | 0.731719 | 0.730132 | 0.735053 |
| 0.745625 | 0.731108 | 0.740301 | 0.732503 | 0.590978 | 0.744012 | 0.738228 | 0.734422 | 0.741148 |
| 0.727656 | 0.727001 | 0.73592 | 0.735938 | 0.587975 | 0.736215 | 0.73846 | 0.741469 | 0.741182 |
| 0.729801 | 0.727832 | 0.734539 | 0.736864 | 0.595002 | 0.736233 | 0.735903 | 0.736001 | 0.736161 |
| 0.730049 | 0.728489 | 0.736286 | 0.732199 | 0.590643 | 0.739241 | 0.735072 | 0.738304 | 0.738582 |
| 0.72581 | 0.724361 | 0.725809 | 0.725156 | 0.590849 | 0.736592 | 0.728104 | 0.728159 | 0.731941 |
| 0.730463 | 0.72794 | 0.726814 | 0.728958 | 0.587694 | 0.736 | 0.738745 | 0.741343 | 0.741027 |
| 0.729268 | 0.725282 | 0.735982 | 0.736345 | 0.589419 | 0.737149 | 0.736167 | 0.733173 | 0.737505 |
| 0.717 | 0.720743 | 0.728094 | 0.731994 | 0.589572 | 0.731439 | 0.731766 | 0.734737 | 0.735997 |
| 0.739095 | 0.735113 | 0.747679 | 0.741082 | 0.595719 | 0.738351 | 0.741447 | 0.743634 | 0.741666 |
| 0.724228 | 0.730787 | 0.735357 | 0.734891 | 0.593808 | 0.734862 | 0.742196 | 0.739686 | 0.741386 |

| | | | | | | | | |
|---|---|---|---|---|---|---|---|---|
| 0.732892 | 0.722214 | 0.737145 | 0.738978 | 0.598028 | 0.743367 | 0.728958 | 0.734086 | 0.734408 |
| 0.742604 | 0.739962 | 0.750339 | 0.749696 | 0.595282 | 0.742271 | 0.743154 | 0.742394 | 0.741762 |
| 0.732685 | 0.721376 | 0.729403 | 0.737036 | 0.593747 | 0.737119 | 0.734456 | 0.734372 | 0.739267 |
| 0.705882 | 0.706922 | 0.717008 | 0.713778 | 0.565846 | 0.717926 | 0.695178 | 0.689594 | 0.687127 |
| 0.725826 | 0.729793 | 0.737135 | 0.735634 | 0.594615 | 0.737273 | 0.733963 | 0.737493 | 0.735858 |
| 0.727236 | 0.727545 | 0.735419 | 0.727162 | 0.596382 | 0.734242 | 0.737451 | 0.740071 | 0.736975 |
| 0.733213 | 0.732379 | 0.742979 | 0.740173 | 0.596242 | 0.742565 | 0.740955 | 0.742407 | 0.73877 |
| 0.73182 | 0.725466 | 0.741255 | 0.745551 | 0.588254 | 0.746926 | 0.735613 | 0.740832 | 0.738364 |
| 0.738545 | 0.732829 | 0.742694 | 0.74368 | 0.596427 | 0.74604 | 0.739561 | 0.739752 | 0.741144 |
| 0.738611 | 0.73269 | 0.739631 | 0.741194 | 0.593682 | 0.74593 | 0.736462 | 0.734872 | 0.736295 |
| 0.738554 | 0.728475 | 0.74329 | 0.735818 | 0.591589 | 0.74282 | 0.733496 | 0.72961 | 0.732813 |
| 0.73301 | 0.733481 | 0.744788 | 0.739035 | 0.589763 | 0.739474 | 0.743978 | 0.741765 | 0.738697 |
| 0.736058 | 0.724095 | 0.742195 | 0.741143 | 0.5985 | 0.739746 | 0.737714 | 0.735111 | 0.738198 |
| 0.731435 | 0.732123 | 0.742507 | 0.740517 | 0.594802 | 0.744558 | 0.739653 | 0.736593 | 0.74046 |
| 0.740145 | 0.743671 | 0.745292 | 0.746347 | 0.592681 | 0.745371 | 0.743909 | 0.741077 | 0.742561 |
| 0.723684 | 0.728477 | 0.731397 | 0.738913 | 0.59116 | 0.736464 | 0.735621 | 0.738425 | 0.737189 |
| 0.734106 | 0.737646 | 0.73978 | 0.745381 | 0.597049 | 0.739362 | 0.743731 | 0.744381 | 0.744386 |
| 0.736303 | 0.73724 | 0.743004 | 0.744859 | 0.593575 | 0.747473 | 0.739236 | 0.740881 | 0.740912 |
| 0.731064 | 0.725662 | 0.740393 | 0.737876 | 0.590458 | 0.737131 | 0.735402 | 0.736232 | 0.737325 |
| 0.737626 | 0.734267 | 0.733967 | 0.736892 | 0.593514 | 0.736231 | 0.737605 | 0.741821 | 0.740539 |
| 0.733012 | 0.738959 | 0.741032 | 0.74473 | 0.593314 | 0.744482 | 0.744275 | 0.744799 | 0.745645 |
| 0.742713 | 0.73785 | 0.744257 | 0.750851 | 0.599241 | 0.743728 | 0.743225 | 0.743577 | 0.744676 |
| 0.681151 | 0.671716 | 0.676135 | 0.676781 | 0.589226 | 0.678693 | 0.682274 | 0.691385 | 0.687684 |
| 0.67099 | 0.643486 | 0.656313 | 0.660832 | 0.585275 | 0.648764 | 0.69876 | 0.688445 | 0.691448 |
| 0.644409 | 0.639572 | 0.649431 | 0.640538 | 0.61279 | 0.645215 | 0.711958 | 0.709171 | 0.720294 |
| 0.657068 | 0.639426 | 0.643737 | 0.647736 | 0.586544 | 0.64495 | 0.694932 | 0.70027 | 0.693405 |
| 0.753741 | 0.74625 | 0.759149 | 0.766673 | 0.575377 | 0.746635 | 0.700797 | 0.702429 | 0.702438 |
| 0.76579 | 0.742474 | 0.752656 | 0.751959 | 0.576102 | 0.747037 | 0.693939 | 0.693901 | 0.691769 |
| 0.74348 | 0.740591 | 0.744496 | 0.746174 | 0.564778 | 0.742623 | 0.699129 | 0.693186 | 0.696539 |
| 0.745474 | 0.740387 | 0.753254 | 0.758029 | 0.576871 | 0.751501 | 0.698329 | 0.697496 | 0.698931 |
| 0.754605 | 0.731714 | 0.740644 | 0.752418 | 0.560582 | 0.754489 | 0.683206 | 0.682788 | 0.693822 |
| 0.742996 | 0.737102 | 0.729877 | 0.757375 | 0.554299 | 0.739695 | 0.674599 | 0.67713 | 0.676267 |
| 0.753978 | 0.73185 | 0.732579 | 0.750256 | 0.562128 | 0.746488 | 0.677065 | 0.679543 | 0.681871 |
| 0.648774 | 0.647234 | 0.646109 | 0.654339 | 0.583743 | 0.647348 | 0.691891 | 0.686163 | 0.693375 |
| 0.745429 | 0.743777 | 0.754118 | 0.757931 | 0.59322 | 0.752169 | 0.730443 | 0.734826 | 0.733965 |
| 0.752346 | 0.738478 | 0.75234 | 0.755398 | 0.581774 | 0.753739 | 0.709256 | 0.706406 | 0.711761 |
| 0.751465 | 0.745154 | 0.758756 | 0.745892 | 0.57351 | 0.756053 | 0.703876 | 0.705645 | 0.711466 |
| 0.745963 | 0.749213 | 0.745986 | 0.749777 | 0.563216 | 0.743319 | 0.68699 | 0.683537 | 0.687968 |
| 0.741146 | 0.740308 | 0.748758 | 0.747294 | 0.584015 | 0.749811 | 0.711658 | 0.707495 | 0.708852 |
| 0.736854 | 0.730649 | 0.737673 | 0.743337 | 0.577008 | 0.737825 | 0.701236 | 0.706883 | 0.703313 |
| 0.737533 | 0.725789 | 0.731952 | 0.734372 | 0.567972 | 0.73309 | 0.700141 | 0.702498 | 0.697319 |
| 0.747276 | 0.72872 | 0.743024 | 0.736418 | 0.574612 | 0.739778 | 0.694811 | 0.703472 | 0.702657 |
| 0.743191 | 0.73252 | 0.741053 | 0.734498 | 0.567263 | 0.734005 | 0.693369 | 0.694075 | 0.690434 |
| 0.748422 | 0.737134 | 0.747012 | 0.741399 | 0.568945 | 0.738381 | 0.699162 | 0.695005 | 0.695151 |
| 0.744358 | 0.743722 | 0.741505 | 0.74933 | 0.561525 | 0.745396 | 0.68994 | 0.696738 | 0.698875 |
| 0.730337 | 0.726559 | 0.738516 | 0.724314 | 0.552137 | 0.736383 | 0.678053 | 0.680274 | 0.680208 |

| | | | | | | | | |
|---|---|---|---|---|---|---|---|---|
| 0.759732 | 0.750041 | 0.760914 | 0.757304 | 0.590395 | 0.767555 | 0.716624 | 0.720292 | 0.72161 |
| 0.744325 | 0.744396 | 0.750808 | 0.746514 | 0.570049 | 0.750861 | 0.694494 | 0.697067 | 0.697833 |
| 0.741654 | 0.735367 | 0.751869 | 0.742958 | 0.565682 | 0.750228 | 0.69672 | 0.698082 | 0.703158 |
| 0.744551 | 0.741686 | 0.760959 | 0.748404 | 0.570029 | 0.75657 | 0.695423 | 0.698665 | 0.700258 |
| 0.754354 | 0.743946 | 0.745237 | 0.745779 | 0.561235 | 0.751992 | 0.695251 | 0.694652 | 0.684672 |
| 0.743382 | 0.747669 | 0.74977 | 0.75205 | 0.572433 | 0.751332 | 0.691819 | 0.696582 | 0.698499 |
| 0.757895 | 0.74488 | 0.749102 | 0.744935 | 0.57039 | 0.758936 | 0.679962 | 0.687817 | 0.686303 |
| 0.749104 | 0.753363 | 0.752206 | 0.739879 | 0.56351 | 0.750871 | 0.684867 | 0.686512 | 0.68605 |
| 0.753545 | 0.730115 | 0.742282 | 0.749181 | 0.560102 | 0.744031 | 0.680407 | 0.683049 | 0.682601 |
| 0.742857 | 0.740548 | 0.752766 | 0.74692 | 0.590532 | 0.749658 | 0.723182 | 0.722937 | 0.727414 |
| 0.73706 | 0.737387 | 0.750079 | 0.74278 | 0.591352 | 0.740997 | 0.721764 | 0.716702 | 0.720434 |
| 0.748009 | 0.743643 | 0.75323 | 0.73889 | 0.589063 | 0.741973 | 0.71903 | 0.718302 | 0.717699 |
| 0.745871 | 0.733417 | 0.743911 | 0.737731 | 0.584845 | 0.74434 | 0.711651 | 0.705485 | 0.70896 |
| 0.745309 | 0.738249 | 0.754178 | 0.744144 | 0.582531 | 0.742983 | 0.707953 | 0.707966 | 0.711487 |
| 0.745939 | 0.742095 | 0.743935 | 0.742686 | 0.581201 | 0.748065 | 0.704329 | 0.702519 | 0.70814 |

| Collection_ | Collection_ | Collection_ | Collection_ | Collection_ | Collection_ | Collection_ | Collection_ | Collection_ |
|---|---|---|---|---|---|---|---|---|
| 0.618364 | 0.601559 | 0.705206 | 0.708304 | 0.698548 | 0.701006 | 0.706302 | 0.704051 | 0.706927 |
| 0.617729 | 0.601051 | 0.711419 | 0.711851 | 0.706275 | 0.707441 | 0.710779 | 0.711244 | 0.709896 |
| 0.619287 | 0.599508 | 0.69817 | 0.700039 | 0.693772 | 0.702008 | 0.703543 | 0.701582 | 0.70019 |
| 0.618461 | 0.600364 | 0.709654 | 0.708551 | 0.702574 | 0.706946 | 0.712379 | 0.710623 | 0.707868 |
| 0.624053 | 0.605117 | 0.705533 | 0.703777 | 0.697726 | 0.701968 | 0.707681 | 0.70592 | 0.704322 |
| 0.620868 | 0.59891 | 0.713333 | 0.717407 | 0.707109 | 0.714478 | 0.718142 | 0.715134 | 0.714661 |
| 0.591931 | 0.580863 | 0.763402 | 0.765694 | 0.760254 | 0.761088 | 0.762452 | 0.765334 | 0.766382 |
| 0.594366 | 0.580954 | 0.759407 | 0.762551 | 0.752527 | 0.756651 | 0.758879 | 0.767054 | 0.763644 |
| 0.625262 | 0.601224 | 0.707813 | 0.70831 | 0.701704 | 0.706732 | 0.708142 | 0.707644 | 0.711912 |
| 0.618412 | 0.599603 | 0.704606 | 0.705445 | 0.700128 | 0.704281 | 0.704535 | 0.704022 | 0.70434 |
| 0.6201 | 0.599555 | 0.706604 | 0.708327 | 0.702022 | 0.704281 | 0.709036 | 0.708185 | 0.707389 |
| 0.614625 | 0.598746 | 0.699149 | 0.70087 | 0.692791 | 0.696132 | 0.699921 | 0.700465 | 0.698107 |
| 0.619169 | 0.597538 | 0.709456 | 0.710239 | 0.703062 | 0.705583 | 0.711468 | 0.712571 | 0.715476 |
| 0.62196 | 0.602526 | 0.731497 | 0.728644 | 0.720427 | 0.721395 | 0.7296 | 0.726073 | 0.726184 |
| 0.61439 | 0.596848 | 0.743726 | 0.748971 | 0.732589 | 0.735734 | 0.744263 | 0.741616 | 0.745935 |
| 0.619753 | 0.602846 | 0.71454 | 0.71366 | 0.702437 | 0.711348 | 0.712687 | 0.713871 | 0.713375 |
| 0.61676 | 0.598499 | 0.713702 | 0.712942 | 0.703647 | 0.709344 | 0.710958 | 0.713915 | 0.713057 |
| 0.616673 | 0.595125 | 0.761905 | 0.762012 | 0.741385 | 0.74634 | 0.75843 | 0.7527 | 0.756574 |
| 0.615928 | 0.597145 | 0.699661 | 0.702498 | 0.6942 | 0.700032 | 0.702151 | 0.703128 | 0.703323 |
| 0.621928 | 0.602802 | 0.707205 | 0.708151 | 0.701347 | 0.707375 | 0.707099 | 0.709135 | 0.707589 |
| 0.618011 | 0.596806 | 0.704878 | 0.705845 | 0.699522 | 0.703434 | 0.707058 | 0.704828 | 0.701974 |
| 0.617734 | 0.599889 | 0.708121 | 0.709356 | 0.70108 | 0.705317 | 0.706587 | 0.710312 | 0.709045 |
| 0.619084 | 0.602921 | 0.711545 | 0.714115 | 0.703275 | 0.709222 | 0.711636 | 0.714562 | 0.714834 |
| 0.623196 | 0.603999 | 0.708837 | 0.707917 | 0.699129 | 0.704006 | 0.709823 | 0.711473 | 0.707441 |
| 0.622493 | 0.603389 | 0.710772 | 0.714675 | 0.703185 | 0.707735 | 0.708589 | 0.711752 | 0.710444 |
| 0.619264 | 0.596482 | 0.711769 | 0.706598 | 0.701916 | 0.706499 | 0.710024 | 0.710933 | 0.709288 |
| 0.618896 | 0.600541 | 0.702381 | 0.701772 | 0.694176 | 0.69871 | 0.704787 | 0.703278 | 0.702378 |
| 0.618502 | 0.600965 | 0.707405 | 0.707675 | 0.700978 | 0.705771 | 0.709527 | 0.709938 | 0.708931 |
| 0.620738 | 0.600953 | 0.708145 | 0.708429 | 0.704491 | 0.708784 | 0.712103 | 0.712269 | 0.71064 |
| 0.619189 | 0.599264 | 0.706239 | 0.707792 | 0.69826 | 0.703506 | 0.704414 | 0.706668 | 0.705225 |
| 0.615963 | 0.598949 | 0.709132 | 0.707888 | 0.70209 | 0.705166 | 0.70914 | 0.708805 | 0.705248 |
| 0.617334 | 0.596934 | 0.707223 | 0.705501 | 0.697939 | 0.701633 | 0.705743 | 0.708159 | 0.706045 |
| 0.616841 | 0.596919 | 0.706468 | 0.704405 | 0.696338 | 0.700631 | 0.70521 | 0.705506 | 0.704191 |
| 0.620153 | 0.599462 | 0.712052 | 0.70956 | 0.704467 | 0.707555 | 0.711975 | 0.71097 | 0.710832 |
| 0.627375 | 0.603745 | 0.70316 | 0.702761 | 0.695132 | 0.699945 | 0.704556 | 0.703537 | 0.701146 |
| 0.620716 | 0.602587 | 0.708837 | 0.706796 | 0.697684 | 0.70502 | 0.708217 | 0.704577 | 0.71022 |
| 0.620568 | 0.6 | 0.712702 | 0.714286 | 0.708094 | 0.710909 | 0.715924 | 0.709592 | 0.714979 |
| 0.614155 | 0.59923 | 0.708999 | 0.707896 | 0.700379 | 0.709623 | 0.711764 | 0.713052 | 0.709309 |
| 0.614205 | 0.599555 | 0.713628 | 0.712353 | 0.706796 | 0.710796 | 0.715807 | 0.714977 | 0.713284 |

| | | | | | | | | |
|---|---|---|---|---|---|---|---|---|
| 0.616095 | 0.59785 | 0.769301 | 0.764115 | 0.746517 | 0.750483 | 0.760151 | 0.757801 | 0.761399 |
| 0.618269 | 0.599714 | 0.698493 | 0.696358 | 0.691032 | 0.697425 | 0.701887 | 0.697615 | 0.697474 |
| 0.620214 | 0.597901 | 0.700692 | 0.703098 | 0.693886 | 0.697544 | 0.702193 | 0.704196 | 0.70275 |
| 0.61863 | 0.598959 | 0.711992 | 0.712012 | 0.704281 | 0.709143 | 0.710035 | 0.713001 | 0.711366 |
| 0.616131 | 0.60046 | 0.704125 | 0.708 | 0.698247 | 0.704473 | 0.705859 | 0.704116 | 0.703962 |
| 0.619185 | 0.601727 | 0.709821 | 0.712026 | 0.699438 | 0.705915 | 0.708473 | 0.711994 | 0.708643 |
| 0.619816 | 0.598416 | 0.705363 | 0.703036 | 0.695815 | 0.70109 | 0.70578 | 0.703349 | 0.704394 |
| 0.620988 | 0.603703 | 0.71033 | 0.712453 | 0.701312 | 0.703973 | 0.71156 | 0.71092 | 0.708534 |
| 0.61957 | 0.602465 | 0.707052 | 0.70771 | 0.699301 | 0.705474 | 0.70815 | 0.710697 | 0.706693 |
| 0.616006 | 0.594416 | 0.710776 | 0.708188 | 0.699976 | 0.703748 | 0.705794 | 0.709734 | 0.7108 |
| 0.611515 | 0.594726 | 0.69895 | 0.697265 | 0.692652 | 0.697661 | 0.699682 | 0.700992 | 0.700279 |
| 0.616731 | 0.596925 | 0.702358 | 0.700833 | 0.696411 | 0.700397 | 0.702506 | 0.704762 | 0.703373 |
| 0.619734 | 0.601136 | 0.703551 | 0.707563 | 0.697426 | 0.702075 | 0.705193 | 0.705701 | 0.704779 |
| 0.618302 | 0.594674 | 0.705706 | 0.704651 | 0.696979 | 0.700955 | 0.702776 | 0.702777 | 0.702346 |
| 0.615612 | 0.598267 | 0.702872 | 0.702404 | 0.698618 | 0.704518 | 0.706137 | 0.701524 | 0.70208 |
| 0.614692 | 0.595165 | 0.703667 | 0.701867 | 0.696388 | 0.700056 | 0.709906 | 0.701344 | 0.701089 |
| 0.609747 | 0.589072 | 0.698265 | 0.697621 | 0.691558 | 0.698002 | 0.701935 | 0.700495 | 0.698916 |
| 0.623914 | 0.604142 | 0.708491 | 0.709554 | 0.704751 | 0.705724 | 0.70948 | 0.706345 | 0.708496 |
| 0.618827 | 0.601064 | 0.714297 | 0.712785 | 0.705562 | 0.70949 | 0.713376 | 0.710927 | 0.707123 |
| 0.618632 | 0.59984 | 0.706417 | 0.706974 | 0.698367 | 0.704032 | 0.704573 | 0.705578 | 0.704873 |
| 0.615905 | 0.596455 | 0.699984 | 0.700119 | 0.696549 | 0.69865 | 0.702617 | 0.699068 | 0.698547 |
| 0.618193 | 0.597769 | 0.702849 | 0.70358 | 0.698709 | 0.701303 | 0.703833 | 0.703208 | 0.705487 |
| 0.619888 | 0.600348 | 0.704707 | 0.704533 | 0.696027 | 0.703996 | 0.708422 | 0.705953 | 0.707435 |
| 0.614386 | 0.59497 | 0.712423 | 0.712233 | 0.707777 | 0.712219 | 0.714331 | 0.716544 | 0.713279 |
| 0.611883 | 0.596304 | 0.721912 | 0.719551 | 0.711653 | 0.715818 | 0.721112 | 0.724806 | 0.722432 |
| 0.593591 | 0.579581 | 0.76007 | 0.76324 | 0.753191 | 0.756497 | 0.759717 | 0.767942 | 0.765154 |
| 0.600201 | 0.58465 | 0.758399 | 0.755267 | 0.75301 | 0.7635 | 0.763851 | 0.766139 | 0.763193 |
| 0.601388 | 0.585923 | 0.766232 | 0.764148 | 0.75871 | 0.769364 | 0.770045 | 0.765745 | 0.764361 |
| 0.621351 | 0.604256 | 0.702979 | 0.702716 | 0.696277 | 0.702385 | 0.70669 | 0.706019 | 0.705315 |
| 0.61563 | 0.596735 | 0.702738 | 0.702612 | 0.697521 | 0.701635 | 0.70805 | 0.70494 | 0.700164 |
| 0.617576 | 0.594687 | 0.711634 | 0.71351 | 0.702136 | 0.707067 | 0.712567 | 0.709059 | 0.710036 |
| 0.613205 | 0.599984 | 0.712079 | 0.71371 | 0.705224 | 0.711613 | 0.712055 | 0.713411 | 0.716058 |
| 0.618118 | 0.602881 | 0.706746 | 0.705448 | 0.700332 | 0.704682 | 0.706912 | 0.706605 | 0.707641 |
| 0.621413 | 0.600352 | 0.704577 | 0.705379 | 0.695148 | 0.699538 | 0.701074 | 0.704882 | 0.706038 |
| 0.616254 | 0.600581 | 0.703474 | 0.705289 | 0.697686 | 0.700698 | 0.702842 | 0.705503 | 0.707613 |
| 0.618629 | 0.603085 | 0.708171 | 0.707309 | 0.701381 | 0.704969 | 0.709559 | 0.708364 | 0.708283 |
| 0.631118 | 0.605888 | 0.706164 | 0.709316 | 0.701199 | 0.702654 | 0.708868 | 0.708011 | 0.709187 |
| 0.630348 | 0.611652 | 0.718872 | 0.72329 | 0.713378 | 0.71716 | 0.721513 | 0.723634 | 0.723498 |
| 0.614237 | 0.60109 | 0.699439 | 0.700553 | 0.691541 | 0.698574 | 0.700033 | 0.699226 | 0.69905 |
| 0.615237 | 0.602145 | 0.70934 | 0.710238 | 0.701809 | 0.705517 | 0.707457 | 0.708719 | 0.709394 |
| 0.618085 | 0.60098 | 0.70831 | 0.705901 | 0.701328 | 0.703099 | 0.706963 | 0.702079 | 0.70544 |
| 0.634213 | 0.614457 | 0.718194 | 0.719395 | 0.711938 | 0.71967 | 0.722778 | 0.721442 | 0.722985 |
| 0.620127 | 0.60183 | 0.703169 | 0.709356 | 0.701122 | 0.704737 | 0.710135 | 0.707303 | 0.710318 |
| 0.61818 | 0.598283 | 0.701504 | 0.704651 | 0.695635 | 0.699757 | 0.701611 | 0.704756 | 0.703476 |
| 0.621167 | 0.603172 | 0.709486 | 0.707261 | 0.70082 | 0.702912 | 0.706715 | 0.708855 | 0.705939 |
| 0.618739 | 0.604319 | 0.705318 | 0.706524 | 0.7005 | 0.7056 | 0.709442 | 0.706207 | 0.704254 |

| | | | | | | | | |
|---|---|---|---|---|---|---|---|---|
| 0.620341 | 0.59992 | 0.70678 | 0.702862 | 0.699872 | 0.704569 | 0.70626 | 0.70422 | 0.702847 |
| 0.623027 | 0.601823 | 0.709532 | 0.711737 | 0.703296 | 0.70767 | 0.709708 | 0.70823 | 0.705691 |
| 0.623698 | 0.600048 | 0.709249 | 0.712596 | 0.701137 | 0.705948 | 0.707036 | 0.709377 | 0.707507 |
| 0.630989 | 0.612407 | 0.712124 | 0.714424 | 0.707404 | 0.710846 | 0.716153 | 0.714554 | 0.715812 |
| 0.625087 | 0.605906 | 0.710695 | 0.716331 | 0.705986 | 0.712392 | 0.714896 | 0.713861 | 0.713627 |
| 0.614475 | 0.601298 | 0.707765 | 0.710699 | 0.70133 | 0.706598 | 0.706835 | 0.707446 | 0.706961 |
| 0.619652 | 0.600473 | 0.711387 | 0.711733 | 0.706652 | 0.707965 | 0.711855 | 0.712212 | 0.710622 |
| 0.632996 | 0.60478 | 0.706354 | 0.710537 | 0.703345 | 0.705506 | 0.708566 | 0.7068 | 0.705934 |
| 0.617043 | 0.600288 | 0.706838 | 0.706695 | 0.697038 | 0.701841 | 0.709543 | 0.706924 | 0.705035 |
| 0.619446 | 0.603035 | 0.704926 | 0.703378 | 0.694535 | 0.697748 | 0.703071 | 0.704622 | 0.702172 |
| 0.623477 | 0.605929 | 0.705911 | 0.708409 | 0.700681 | 0.705508 | 0.709573 | 0.706754 | 0.704658 |
| 0.620576 | 0.608225 | 0.706651 | 0.709493 | 0.701402 | 0.704001 | 0.708002 | 0.708045 | 0.709644 |
| 0.622659 | 0.60573 | 0.708408 | 0.708253 | 0.701039 | 0.704607 | 0.713552 | 0.706665 | 0.705953 |
| 0.61937 | 0.596361 | 0.704968 | 0.705109 | 0.700716 | 0.704367 | 0.70619 | 0.706286 | 0.703148 |
| 0.622935 | 0.603322 | 0.704987 | 0.70846 | 0.6998 | 0.703857 | 0.708698 | 0.706029 | 0.705991 |
| 0.620588 | 0.602428 | 0.705268 | 0.703204 | 0.697956 | 0.701255 | 0.706718 | 0.699545 | 0.702429 |
| 0.623168 | 0.603639 | 0.712516 | 0.712852 | 0.703426 | 0.704138 | 0.711463 | 0.712058 | 0.708895 |
| 0.616402 | 0.600238 | 0.706865 | 0.704191 | 0.698437 | 0.702786 | 0.711549 | 0.709511 | 0.705873 |
| 0.618744 | 0.602952 | 0.706449 | 0.70741 | 0.698889 | 0.703942 | 0.705948 | 0.709261 | 0.706286 |
| 0.622759 | 0.601387 | 0.706044 | 0.70841 | 0.701402 | 0.707793 | 0.707966 | 0.706446 | 0.704578 |
| 0.613121 | 0.597735 | 0.707511 | 0.709984 | 0.701277 | 0.706174 | 0.710094 | 0.70753 | 0.707009 |
| 0.616794 | 0.600318 | 0.702184 | 0.702921 | 0.698404 | 0.700427 | 0.708978 | 0.703947 | 0.704239 |
| 0.624619 | 0.60325 | 0.709329 | 0.710115 | 0.703772 | 0.703597 | 0.713931 | 0.707723 | 0.712354 |
| 0.621689 | 0.604296 | 0.707509 | 0.710777 | 0.704829 | 0.708426 | 0.709833 | 0.713779 | 0.707777 |
| 0.616613 | 0.598815 | 0.707546 | 0.71122 | 0.700598 | 0.704429 | 0.712096 | 0.707781 | 0.70969 |
| 0.624701 | 0.599779 | 0.710817 | 0.709792 | 0.705246 | 0.708995 | 0.71458 | 0.709886 | 0.714002 |
| 0.616564 | 0.59951 | 0.707599 | 0.711097 | 0.704414 | 0.704553 | 0.711393 | 0.708883 | 0.709516 |
| 0.622755 | 0.605299 | 0.703597 | 0.70572 | 0.702578 | 0.706479 | 0.709575 | 0.704442 | 0.708546 |
| 0.617443 | 0.599826 | 0.708552 | 0.709913 | 0.702284 | 0.705397 | 0.709634 | 0.707888 | 0.706986 |
| 0.620248 | 0.599324 | 0.711723 | 0.715202 | 0.703276 | 0.705726 | 0.714263 | 0.712754 | 0.71248 |
| 0.616149 | 0.60024 | 0.710586 | 0.71307 | 0.702788 | 0.707451 | 0.71072 | 0.710242 | 0.710729 |
| 0.618409 | 0.599843 | 0.707977 | 0.706382 | 0.702392 | 0.704963 | 0.709538 | 0.705826 | 0.709155 |
| 0.620391 | 0.601591 | 0.708214 | 0.711299 | 0.702841 | 0.704108 | 0.710372 | 0.707299 | 0.711413 |
| 0.613113 | 0.596145 | 0.710104 | 0.71108 | 0.703743 | 0.709116 | 0.709704 | 0.711089 | 0.711189 |
| 0.618973 | 0.601435 | 0.711371 | 0.70794 | 0.701571 | 0.701633 | 0.709994 | 0.707903 | 0.707926 |
| 0.609483 | 0.595297 | 0.709371 | 0.708054 | 0.701194 | 0.70962 | 0.709745 | 0.712086 | 0.712168 |
| 0.615066 | 0.600798 | 0.706618 | 0.706839 | 0.699133 | 0.701128 | 0.70862 | 0.708406 | 0.707448 |
| 0.584548 | 0.569967 | 0.680432 | 0.680896 | 0.67302 | 0.677276 | 0.682971 | 0.682382 | 0.679723 |
| 0.590191 | 0.573782 | 0.68907 | 0.690187 | 0.680524 | 0.684402 | 0.686719 | 0.691994 | 0.688053 |
| 0.587836 | 0.573957 | 0.680664 | 0.679716 | 0.671337 | 0.678027 | 0.681073 | 0.676382 | 0.680033 |
| 0.584278 | 0.567423 | 0.676208 | 0.677849 | 0.668741 | 0.672641 | 0.677987 | 0.680652 | 0.680164 |
| 0.576099 | 0.560822 | 0.661794 | 0.663468 | 0.652804 | 0.656152 | 0.66007 | 0.664787 | 0.65629 |
| 0.571678 | 0.553841 | 0.666301 | 0.666828 | 0.661893 | 0.663621 | 0.66973 | 0.666344 | 0.666027 |
| 0.568831 | 0.555727 | 0.666218 | 0.665207 | 0.658269 | 0.660589 | 0.664128 | 0.665408 | 0.663383 |
| 0.573477 | 0.558012 | 0.666283 | 0.667857 | 0.656713 | 0.661161 | 0.666624 | 0.669766 | 0.669541 |
| 0.575798 | 0.557549 | 0.661171 | 0.663086 | 0.656551 | 0.66108 | 0.669865 | 0.668082 | 0.666749 |
| 0.572784 | 0.554313 | 0.670484 | 0.670301 | 0.659175 | 0.665517 | 0.673916 | 0.675904 | 0.675104 |

| | | | | | | | | |
|---|---|---|---|---|---|---|---|---|
| 0.568292 | 0.551204 | 0.67047 | 0.673497 | 0.662828 | 0.666093 | 0.671682 | 0.67302 | 0.671795 |
| 0.574906 | 0.560423 | 0.671474 | 0.670493 | 0.663438 | 0.667186 | 0.669267 | 0.674642 | 0.673702 |
| 0.562556 | 0.550389 | 0.665595 | 0.668098 | 0.666487 | 0.666355 | 0.669955 | 0.673239 | 0.667107 |
| 0.565383 | 0.548039 | 0.6618 | 0.660377 | 0.65676 | 0.657162 | 0.663807 | 0.659316 | 0.66322 |
| 0.565755 | 0.55352 | 0.660411 | 0.667096 | 0.653772 | 0.656027 | 0.662434 | 0.660586 | 0.664416 |
| 0.58836 | 0.557865 | 0.663997 | 0.666208 | 0.655514 | 0.659243 | 0.665981 | 0.663925 | 0.663308 |
| 0.578174 | 0.555771 | 0.664526 | 0.663216 | 0.660778 | 0.658819 | 0.664642 | 0.66421 | 0.665446 |
| 0.564037 | 0.546245 | 0.660228 | 0.661519 | 0.648872 | 0.656263 | 0.660053 | 0.662703 | 0.665508 |
| 0.559859 | 0.548077 | 0.65953 | 0.663619 | 0.658043 | 0.65752 | 0.66043 | 0.664211 | 0.657359 |
| 0.564221 | 0.551282 | 0.65838 | 0.660438 | 0.650461 | 0.654103 | 0.656977 | 0.66027 | 0.664738 |
| 0.568832 | 0.547921 | 0.656303 | 0.657966 | 0.645474 | 0.647742 | 0.660131 | 0.659903 | 0.649761 |
| 0.635228 | 0.614787 | 0.694849 | 0.69535 | 0.688276 | 0.692043 | 0.694843 | 0.698396 | 0.698352 |
| 0.617672 | 0.6003 | 0.705972 | 0.70249 | 0.698604 | 0.701572 | 0.706054 | 0.70913 | 0.70416 |
| 0.585416 | 0.567468 | 0.684548 | 0.684145 | 0.676122 | 0.685103 | 0.687179 | 0.685633 | 0.683913 |
| 0.611072 | 0.595257 | 0.715093 | 0.716769 | 0.70914 | 0.711521 | 0.718712 | 0.71754 | 0.718537 |
| 0.628641 | 0.613311 | 0.704958 | 0.703189 | 0.697706 | 0.701516 | 0.703144 | 0.704328 | 0.704096 |
| 0.614499 | 0.594565 | 0.707206 | 0.708271 | 0.699905 | 0.701187 | 0.709046 | 0.70751 | 0.709749 |
| 0.616563 | 0.598308 | 0.71382 | 0.715873 | 0.708722 | 0.712852 | 0.717992 | 0.714274 | 0.715908 |
| 0.643175 | 0.625539 | 0.693863 | 0.696051 | 0.687048 | 0.691851 | 0.698113 | 0.691499 | 0.695693 |
| 0.619725 | 0.601132 | 0.717749 | 0.718547 | 0.707255 | 0.710039 | 0.719194 | 0.720462 | 0.722753 |
| 0.585006 | 0.56168 | 0.688706 | 0.688262 | 0.678632 | 0.681578 | 0.69356 | 0.68756 | 0.687947 |
| 0.613992 | 0.599411 | 0.71417 | 0.715671 | 0.708757 | 0.712527 | 0.716799 | 0.714574 | 0.71345 |
| 0.619029 | 0.600559 | 0.718424 | 0.71642 | 0.710473 | 0.71408 | 0.721064 | 0.716986 | 0.717761 |
| 0.570063 | 0.553301 | 0.686341 | 0.67758 | 0.676828 | 0.677386 | 0.685625 | 0.687814 | 0.685582 |
| 0.635434 | 0.620119 | 0.683654 | 0.686294 | 0.681543 | 0.681124 | 0.688487 | 0.68493 | 0.686462 |
| 0.58554 | 0.567657 | 0.692701 | 0.696012 | 0.683575 | 0.67867 | 0.692629 | 0.691204 | 0.692879 |
| 0.610792 | 0.592161 | 0.703521 | 0.704501 | 0.697046 | 0.701436 | 0.706817 | 0.702805 | 0.703942 |
| 0.61619 | 0.597814 | 0.715024 | 0.712233 | 0.70729 | 0.710558 | 0.71584 | 0.712692 | 0.712717 |
| 0.648466 | 0.631357 | 0.684524 | 0.687243 | 0.680309 | 0.681481 | 0.688347 | 0.685893 | 0.685588 |
| 0.615791 | 0.60126 | 0.723915 | 0.724245 | 0.71901 | 0.722293 | 0.724897 | 0.725845 | 0.725631 |
| 0.616759 | 0.599609 | 0.722635 | 0.724878 | 0.713685 | 0.717937 | 0.72363 | 0.718713 | 0.722881 |
| 0.663397 | 0.639925 | 0.692209 | 0.696499 | 0.688367 | 0.692442 | 0.697368 | 0.699485 | 0.698494 |
| 0.66664 | 0.646293 | 0.694544 | 0.696662 | 0.688826 | 0.695198 | 0.697259 | 0.698781 | 0.697506 |
| 0.613025 | 0.599794 | 0.723554 | 0.720188 | 0.716902 | 0.723012 | 0.727745 | 0.725642 | 0.726952 |
| 0.614492 | 0.598673 | 0.719669 | 0.719846 | 0.717114 | 0.720031 | 0.725415 | 0.721259 | 0.723265 |
| 0.616309 | 0.600637 | 0.723118 | 0.724577 | 0.719919 | 0.722343 | 0.726645 | 0.727495 | 0.725675 |
| 0.670046 | 0.653351 | 0.67515 | 0.677161 | 0.669741 | 0.675917 | 0.676861 | 0.675355 | 0.67507 |
| 0.613961 | 0.597712 | 0.722061 | 0.724446 | 0.711339 | 0.717452 | 0.720834 | 0.719016 | 0.722728 |
| 0.657917 | 0.642072 | 0.674388 | 0.674549 | 0.668825 | 0.674726 | 0.677015 | 0.675462 | 0.674568 |
| 0.722993 | 0.687358 | 0.670054 | 0.667373 | 0.657839 | 0.664877 | 0.666893 | 0.667076 | 0.665455 |
| 0.614972 | 0.600268 | 0.730324 | 0.73477 | 0.720651 | 0.724009 | 0.728985 | 0.72897 | 0.730822 |
| 0.731672 | 0.690297 | 0.671061 | 0.668763 | 0.661317 | 0.668258 | 0.668981 | 0.668451 | 0.670183 |
| 0.751231 | 0.746277 | 0.661687 | 0.662508 | 0.65468 | 0.661169 | 0.66059 | 0.66067 | 0.664955 |
| 0.735249 | 0.71076 | 0.659797 | 0.656894 | 0.654846 | 0.659901 | 0.659667 | 0.659372 | 0.659633 |
| 0.616342 | 0.598945 | 0.735409 | 0.73951 | 0.729998 | 0.730868 | 0.737132 | 0.735909 | 0.738763 |
| 0.740973 | 0.762685 | 0.657052 | 0.657972 | 0.651086 | 0.657704 | 0.659605 | 0.658322 | 0.659675 |
| 0.610978 | 0.596359 | 0.733393 | 0.731966 | 0.722937 | 0.724868 | 0.726705 | 0.725937 | 0.730831 |

| | | | | | | | | |
|---|---|---|---|---|---|---|---|---|
| 0.612573 | 0.59672 | 0.747471 | 0.748512 | 0.733441 | 0.731995 | 0.738683 | 0.745469 | 0.746146 |
| 0.614706 | 0.601261 | 0.736145 | 0.736686 | 0.725304 | 0.724813 | 0.732408 | 0.730695 | 0.733317 |
| 0.615331 | 0.599618 | 0.746387 | 0.742594 | 0.741433 | 0.743043 | 0.746103 | 0.751031 | 0.746675 |
| 0.751976 | 0.757772 | 0.649981 | 0.649627 | 0.642252 | 0.649712 | 0.647621 | 0.650239 | 0.651335 |
| 0.788035 | 0.724612 | 0.64956 | 0.651378 | 0.644498 | 0.646412 | 0.649188 | 0.650665 | 0.650267 |
| 0.61591 | 0.598846 | 0.744961 | 0.748646 | 0.729988 | 0.72992 | 0.735254 | 0.740768 | 0.743179 |
| 0.681259 | 0.670454 | 0.652305 | 0.652755 | 0.646185 | 0.651254 | 0.654049 | 0.656947 | 0.657716 |
| 0.605887 | 0.59366 | 0.731579 | 0.728247 | 0.7215 | 0.729158 | 0.728858 | 0.730086 | 0.72871 |
| 0.612951 | 0.596377 | 0.74699 | 0.74572 | 0.729365 | 0.734146 | 0.740226 | 0.740565 | 0.746089 |
| 0.598808 | 0.588926 | 0.73371 | 0.735299 | 0.726615 | 0.73347 | 0.733464 | 0.736215 | 0.731623 |
| 0.608128 | 0.591595 | 0.75519 | 0.758884 | 0.738467 | 0.741108 | 0.743427 | 0.750456 | 0.746872 |
| 0.580838 | 0.512048 | 0.698113 | 0.7 | 0.662338 | 0.735099 | 0.739437 | 0.746575 | 0.729032 |
| 0.528247 | 0.518081 | 0.720491 | 0.722178 | 0.71863 | 0.72067 | 0.717668 | 0.727218 | 0.723645 |
| 0.536221 | 0.521021 | 0.719623 | 0.722178 | 0.707493 | 0.712733 | 0.718794 | 0.720433 | 0.724631 |
| 0.540596 | 0.523896 | 0.725434 | 0.721578 | 0.714911 | 0.722982 | 0.721781 | 0.731689 | 0.732239 |
| 0.546085 | 0.5333 | 0.723817 | 0.725466 | 0.718628 | 0.719423 | 0.722937 | 0.728412 | 0.728245 |
| 0.646509 | 0.647842 | 0.590716 | 0.593373 | 0.581773 | 0.588166 | 0.588337 | 0.590075 | 0.592597 |
| 0.541722 | 0.529405 | 0.73292 | 0.735391 | 0.723755 | 0.732159 | 0.726883 | 0.740231 | 0.7298 |
| 0.60507 | 0.591624 | 0.768618 | 0.771097 | 0.744893 | 0.741893 | 0.753205 | 0.75222 | 0.751931 |
| 0.608876 | 0.592539 | 0.763312 | 0.763041 | 0.743248 | 0.743372 | 0.752153 | 0.754457 | 0.758051 |
| 0.609141 | 0.593151 | 0.767092 | 0.764113 | 0.747079 | 0.750164 | 0.754751 | 0.754411 | 0.754709 |
| 1 | 0.798965 | 0.592377 | 0.594163 | 0.587567 | 0.592703 | 0.594555 | 0.591695 | 0.592638 |
| 0.798965 | 1 | 0.579702 | 0.581725 | 0.575408 | 0.581598 | 0.581377 | 0.578697 | 0.581787 |
| 0.592377 | 0.579702 | 1 | 0.861407 | 0.813783 | 0.811356 | 0.821051 | 0.823646 | 0.818439 |
| 0.594163 | 0.581725 | 0.861407 | 1 | 0.800692 | 0.801386 | 0.815982 | 0.815395 | 0.821374 |
| 0.587567 | 0.575408 | 0.813783 | 0.800692 | 1 | 0.862252 | 0.849027 | 0.797551 | 0.795736 |
| 0.592703 | 0.581598 | 0.811356 | 0.801386 | 0.862252 | 1 | 0.842159 | 0.803514 | 0.799983 |
| 0.594555 | 0.581377 | 0.821051 | 0.815982 | 0.849027 | 0.842159 | 1 | 0.802042 | 0.811226 |
| 0.591695 | 0.578697 | 0.823646 | 0.815395 | 0.797551 | 0.803514 | 0.802042 | 1 | 0.819067 |
| 0.592638 | 0.581787 | 0.818439 | 0.821374 | 0.795736 | 0.799983 | 0.811226 | 0.819067 | 1 |
| 0.593543 | 0.580906 | 0.758749 | 0.759555 | 0.749441 | 0.755766 | 0.759123 | 0.767149 | 0.767992 |
| 0.601536 | 0.585156 | 0.763677 | 0.764414 | 0.761924 | 0.766952 | 0.773317 | 0.773208 | 0.77167 |
| 0.597677 | 0.585032 | 0.757904 | 0.758811 | 0.752537 | 0.758635 | 0.761547 | 0.759145 | 0.761628 |
| 0.598725 | 0.584304 | 0.763277 | 0.762577 | 0.753757 | 0.75748 | 0.759851 | 0.763989 | 0.761668 |
| 0.594226 | 0.582724 | 0.765763 | 0.765 | 0.758905 | 0.763867 | 0.765072 | 0.769077 | 0.768882 |
| 0.596049 | 0.582 | 0.758244 | 0.76112 | 0.756725 | 0.761281 | 0.760571 | 0.761746 | 0.761454 |
| 0.598414 | 0.58117 | 0.764662 | 0.764391 | 0.760976 | 0.766447 | 0.766297 | 0.769571 | 0.773701 |
| 0.603173 | 0.58733 | 0.765136 | 0.766762 | 0.761779 | 0.764701 | 0.76627 | 0.767411 | 0.770911 |
| 0.592928 | 0.580716 | 0.756241 | 0.762367 | 0.750976 | 0.757127 | 0.757841 | 0.762173 | 0.762227 |
| 0.596152 | 0.581998 | 0.761277 | 0.762437 | 0.755462 | 0.759798 | 0.763939 | 0.765047 | 0.765807 |
| 0.533099 | 0.519546 | 0.722535 | 0.72308 | 0.710873 | 0.714634 | 0.717922 | 0.73411 | 0.722569 |
| 0.537043 | 0.522513 | 0.724308 | 0.72389 | 0.711759 | 0.726164 | 0.723763 | 0.736484 | 0.732569 |
| 0.598932 | 0.586958 | 0.761267 | 0.763708 | 0.752602 | 0.758854 | 0.762841 | 0.768664 | 0.768027 |
| 0.598807 | 0.584962 | 0.782729 | 0.785702 | 0.776528 | 0.784059 | 0.785756 | 0.793497 | 0.786844 |
| 0.525113 | 0.50764 | 0.718362 | 0.725707 | 0.711137 | 0.718424 | 0.723092 | 0.721932 | 0.722945 |
| 0.598496 | 0.584636 | 0.762774 | 0.764942 | 0.76054 | 0.765924 | 0.766207 | 0.764848 | 0.767411 |
| 0.588984 | 0.579397 | 0.754602 | 0.76425 | 0.747547 | 0.757988 | 0.761629 | 0.760043 | 0.764885 |

| | | | | | | | | |
|---|---|---|---|---|---|---|---|---|
| 0.595515 | 0.58148 | 0.760608 | 0.766355 | 0.757553 | 0.763413 | 0.765868 | 0.764955 | 0.764769 |
| 0.598142 | 0.584464 | 0.765912 | 0.7685 | 0.759816 | 0.764835 | 0.768748 | 0.767246 | 0.766058 |
| 0.594086 | 0.582771 | 0.761872 | 0.764706 | 0.756657 | 0.763323 | 0.766814 | 0.766338 | 0.76753 |
| 0.598028 | 0.584577 | 0.76665 | 0.76722 | 0.757753 | 0.764349 | 0.769983 | 0.771605 | 0.766872 |
| 0.60072 | 0.586172 | 0.768412 | 0.770409 | 0.764886 | 0.772332 | 0.77195 | 0.772199 | 0.774382 |
| 0.592157 | 0.577142 | 0.760394 | 0.766758 | 0.753796 | 0.762648 | 0.760743 | 0.764086 | 0.767777 |
| 0.596407 | 0.583051 | 0.764345 | 0.766361 | 0.75633 | 0.763218 | 0.765719 | 0.762643 | 0.769623 |
| 0.593781 | 0.57992 | 0.750335 | 0.757167 | 0.749605 | 0.752373 | 0.757829 | 0.75469 | 0.758216 |
| 0.597089 | 0.58141 | 0.760499 | 0.764377 | 0.755306 | 0.760125 | 0.760631 | 0.76268 | 0.765482 |
| 0.592187 | 0.577169 | 0.753431 | 0.760968 | 0.747938 | 0.753933 | 0.757871 | 0.756833 | 0.76455 |
| 0.596831 | 0.584555 | 0.763356 | 0.766509 | 0.757611 | 0.759912 | 0.762807 | 0.765746 | 0.775216 |
| 0.594671 | 0.580219 | 0.763167 | 0.766432 | 0.76014 | 0.764626 | 0.767579 | 0.769514 | 0.768041 |
| 0.589126 | 0.579346 | 0.749853 | 0.751095 | 0.745134 | 0.755443 | 0.753483 | 0.757601 | 0.757314 |
| 0.588577 | 0.576024 | 0.750865 | 0.756304 | 0.748904 | 0.755425 | 0.752831 | 0.760445 | 0.758789 |
| 0.599558 | 0.585328 | 0.76511 | 0.769878 | 0.760482 | 0.770398 | 0.773428 | 0.764323 | 0.778199 |
| 0.594985 | 0.585917 | 0.799527 | 0.807615 | 0.796523 | 0.802898 | 0.824771 | 0.795451 | 0.815438 |
| 0.592072 | 0.579656 | 0.795747 | 0.803272 | 0.797055 | 0.789026 | 0.794446 | 0.807167 | 0.806741 |
| 0.592035 | 0.586453 | 0.809862 | 0.805595 | 0.810002 | 0.809784 | 0.802961 | 0.795197 | 0.799983 |
| 0.594284 | 0.580554 | 0.832373 | 0.822368 | 0.821438 | 0.815995 | 0.891431 | 0.800354 | 0.802896 |
| 0.593166 | 0.580757 | 0.801534 | 0.804983 | 0.833635 | 0.836761 | 0.842762 | 0.789695 | 0.805195 |
| 0.590895 | 0.584265 | 0.795839 | 0.79942 | 0.782279 | 0.798688 | 0.812975 | 0.780577 | 0.78955 |
| 0.593638 | 0.577546 | 0.817151 | 0.805484 | 0.827178 | 0.839653 | 0.864111 | 0.801481 | 0.801196 |
| 0.596084 | 0.582961 | 0.804427 | 0.794598 | 0.875141 | 0.877648 | 0.885857 | 0.797922 | 0.808416 |
| 0.595986 | 0.584512 | 0.808196 | 0.803144 | 0.812277 | 0.80422 | 0.801034 | 0.793833 | 0.806917 |
| 0.593963 | 0.583407 | 0.817867 | 0.806468 | 0.793118 | 0.799594 | 0.807239 | 0.804648 | 0.801753 |
| 0.592518 | 0.578152 | 0.820148 | 0.801671 | 0.830929 | 0.836437 | 0.847688 | 0.800494 | 0.808877 |
| 0.59013 | 0.579228 | 0.804809 | 0.808565 | 0.794457 | 0.800068 | 0.810193 | 0.785921 | 0.802504 |
| 0.588592 | 0.578577 | 0.815295 | 0.797921 | 0.839162 | 0.832278 | 0.815758 | 0.812587 | 0.790563 |
| 0.594665 | 0.58128 | 0.831164 | 0.852992 | 0.799797 | 0.799716 | 0.806325 | 0.798478 | 0.791656 |
| 0.594198 | 0.582704 | 0.819139 | 0.810107 | 0.812903 | 0.815703 | 0.82627 | 0.805151 | 0.789925 |
| 0.594194 | 0.585631 | 0.802413 | 0.809424 | 0.80707 | 0.816067 | 0.81199 | 0.799306 | 0.797202 |
| 0.597986 | 0.585802 | 0.804538 | 0.806903 | 0.806488 | 0.800321 | 0.794805 | 0.784001 | 0.801134 |
| 0.597436 | 0.586712 | 0.85671 | 0.885384 | 0.80129 | 0.800695 | 0.827375 | 0.815752 | 0.841749 |
| 0.597819 | 0.590417 | 0.806263 | 0.807535 | 0.832192 | 0.822187 | 0.814232 | 0.804304 | 0.814042 |
| 0.590707 | 0.577995 | 0.854796 | 0.889016 | 0.802939 | 0.801935 | 0.815438 | 0.816454 | 0.80044 |
| 0.5931 | 0.581137 | 0.809862 | 0.816047 | 0.785909 | 0.794516 | 0.822752 | 0.799038 | 0.812013 |
| 0.59341 | 0.580942 | 0.822163 | 0.799444 | 0.812217 | 0.799562 | 0.801787 | 0.801056 | 0.794878 |
| 0.59086 | 0.579688 | 0.797187 | 0.790839 | 0.826162 | 0.832933 | 0.834492 | 0.806697 | 0.803326 |
| 0.591514 | 0.582007 | 0.801556 | 0.802222 | 0.794105 | 0.800871 | 0.810646 | 0.777948 | 0.790619 |
| 0.584238 | 0.573701 | 0.795485 | 0.794503 | 0.828941 | 0.850686 | 0.829179 | 0.799965 | 0.794892 |
| 0.588254 | 0.577468 | 0.804091 | 0.8029 | 0.783624 | 0.797404 | 0.807886 | 0.78038 | 0.783211 |
| 0.58332 | 0.568004 | 0.821311 | 0.807055 | 0.832464 | 0.835492 | 0.837201 | 0.805897 | 0.796927 |
| 0.586094 | 0.574719 | 0.798668 | 0.794341 | 0.8031 | 0.79428 | 0.806046 | 0.778421 | 0.786829 |
| 0.579918 | 0.56556 | 0.801751 | 0.791139 | 0.821026 | 0.83048 | 0.854295 | 0.785234 | 0.790442 |
| 0.58856 | 0.577717 | 0.790943 | 0.787611 | 0.831396 | 0.825222 | 0.832547 | 0.790692 | 0.800369 |
| 0.581081 | 0.56765 | 0.78757 | 0.783864 | 0.790988 | 0.783287 | 0.778201 | 0.777927 | 0.788747 |

| | | | | | | | | |
|---|---|---|---|---|---|---|---|---|
| 0.595461 | 0.583142 | 0.827937 | 0.814105 | 0.786989 | 0.787689 | 0.80488 | 0.801783 | 0.800942 |
| 0.60288 | 0.585192 | 0.764825 | 0.763552 | 0.757075 | 0.761522 | 0.763404 | 0.764765 | 0.767034 |
| 0.591846 | 0.581297 | 0.760576 | 0.763125 | 0.752465 | 0.757588 | 0.756269 | 0.765426 | 0.762688 |
| 0.599238 | 0.584444 | 0.755951 | 0.760634 | 0.751184 | 0.755129 | 0.75804 | 0.766121 | 0.760932 |
| 0.604418 | 0.58732 | 0.768115 | 0.771323 | 0.760738 | 0.76568 | 0.770328 | 0.76941 | 0.769807 |
| 0.596947 | 0.582751 | 0.761036 | 0.76448 | 0.757673 | 0.759912 | 0.761238 | 0.764433 | 0.768314 |
| 0.59296 | 0.577005 | 0.763119 | 0.765719 | 0.757503 | 0.759901 | 0.765219 | 0.762769 | 0.764827 |
| 0.594797 | 0.584559 | 0.759837 | 0.764855 | 0.760819 | 0.769104 | 0.761552 | 0.76675 | 0.767051 |
| 0.586118 | 0.573754 | 0.752808 | 0.756915 | 0.747386 | 0.749844 | 0.758859 | 0.761096 | 0.763127 |
| 0.594379 | 0.582075 | 0.755587 | 0.763244 | 0.752503 | 0.760525 | 0.761451 | 0.765659 | 0.760435 |
| 0.598692 | 0.582787 | 0.763793 | 0.765457 | 0.756642 | 0.761017 | 0.764806 | 0.769669 | 0.768545 |
| 0.593671 | 0.580597 | 0.762205 | 0.763035 | 0.751393 | 0.765818 | 0.764538 | 0.767533 | 0.769034 |
| 0.599647 | 0.582023 | 0.765841 | 0.763414 | 0.756881 | 0.758719 | 0.759197 | 0.763345 | 0.762225 |
| 0.600046 | 0.586939 | 0.759305 | 0.763592 | 0.753209 | 0.759987 | 0.762279 | 0.765415 | 0.762881 |
| 0.597205 | 0.585598 | 0.760552 | 0.759273 | 0.751654 | 0.762598 | 0.760845 | 0.767113 | 0.764135 |
| 0.600107 | 0.584802 | 0.76715 | 0.765178 | 0.755804 | 0.760861 | 0.762317 | 0.767743 | 0.763884 |
| 0.594959 | 0.580851 | 0.76506 | 0.766523 | 0.7616 | 0.764285 | 0.765416 | 0.765707 | 0.767945 |
| 0.598022 | 0.581691 | 0.761721 | 0.761601 | 0.753799 | 0.76092 | 0.761053 | 0.766388 | 0.761897 |
| 0.594757 | 0.581395 | 0.768216 | 0.765721 | 0.759502 | 0.765741 | 0.76908 | 0.768956 | 0.771087 |
| 0.601251 | 0.585766 | 0.766752 | 0.765728 | 0.758952 | 0.764485 | 0.764873 | 0.7667 | 0.772156 |
| 0.597111 | 0.582398 | 0.757717 | 0.761611 | 0.749567 | 0.757352 | 0.76142 | 0.761492 | 0.75811 |
| 0.59721 | 0.582798 | 0.759702 | 0.764001 | 0.756658 | 0.764147 | 0.764231 | 0.7708 | 0.764409 |
| 0.599453 | 0.585107 | 0.767484 | 0.771927 | 0.756803 | 0.762296 | 0.767783 | 0.764444 | 0.764095 |
| 0.600944 | 0.584175 | 0.760961 | 0.763434 | 0.752835 | 0.761078 | 0.76269 | 0.769009 | 0.762575 |
| 0.597323 | 0.584711 | 0.763885 | 0.759303 | 0.756244 | 0.761597 | 0.76519 | 0.764383 | 0.76771 |
| 0.598242 | 0.583958 | 0.762371 | 0.76254 | 0.753621 | 0.760804 | 0.762581 | 0.767307 | 0.763711 |
| 0.595767 | 0.581938 | 0.762042 | 0.759786 | 0.752103 | 0.75864 | 0.76335 | 0.762697 | 0.7623 |
| 0.59757 | 0.58394 | 0.760185 | 0.758915 | 0.752922 | 0.758915 | 0.759611 | 0.764992 | 0.761509 |
| 0.598661 | 0.580932 | 0.764516 | 0.763286 | 0.756635 | 0.761295 | 0.76658 | 0.767104 | 0.767337 |
| 0.599764 | 0.583293 | 0.763228 | 0.76357 | 0.759517 | 0.765336 | 0.7696 | 0.769612 | 0.76685 |
| 0.594574 | 0.581696 | 0.75867 | 0.752274 | 0.750898 | 0.765514 | 0.756209 | 0.762113 | 0.752758 |
| 0.602855 | 0.589287 | 0.766717 | 0.764429 | 0.757556 | 0.768277 | 0.769842 | 0.768999 | 0.767059 |
| 0.597807 | 0.587837 | 0.766924 | 0.764682 | 0.753463 | 0.766122 | 0.763083 | 0.767721 | 0.764167 |
| 0.592371 | 0.576693 | 0.761032 | 0.763704 | 0.754499 | 0.759083 | 0.759045 | 0.766267 | 0.760041 |
| 0.597984 | 0.582557 | 0.762634 | 0.762142 | 0.755988 | 0.760793 | 0.758887 | 0.765243 | 0.762536 |
| 0.592999 | 0.578705 | 0.761263 | 0.760718 | 0.75151 | 0.760625 | 0.757013 | 0.766847 | 0.762417 |
| 0.603944 | 0.588976 | 0.766348 | 0.767876 | 0.760794 | 0.767818 | 0.767923 | 0.768516 | 0.768959 |
| 0.593577 | 0.58005 | 0.76511 | 0.767432 | 0.753243 | 0.757236 | 0.759353 | 0.767416 | 0.767508 |
| 0.599939 | 0.583576 | 0.760748 | 0.761928 | 0.757243 | 0.764218 | 0.767126 | 0.769327 | 0.76598 |
| 0.597008 | 0.581807 | 0.761425 | 0.761921 | 0.756182 | 0.759613 | 0.759315 | 0.769981 | 0.765076 |
| 0.601037 | 0.584849 | 0.763282 | 0.76204 | 0.75689 | 0.759756 | 0.766373 | 0.766741 | 0.764624 |
| 0.600352 | 0.588231 | 0.76428 | 0.768744 | 0.757446 | 0.766637 | 0.766389 | 0.77091 | 0.76695 |
| 0.602715 | 0.589639 | 0.766209 | 0.769639 | 0.759403 | 0.768134 | 0.768393 | 0.774559 | 0.769999 |
| 0.595896 | 0.579878 | 0.761921 | 0.762028 | 0.752877 | 0.759213 | 0.762825 | 0.760576 | 0.764291 |
| 0.596955 | 0.583934 | 0.767395 | 0.763693 | 0.758283 | 0.761971 | 0.766194 | 0.768357 | 0.767526 |
| 0.592868 | 0.579468 | 0.762074 | 0.762561 | 0.752867 | 0.76045 | 0.759265 | 0.766557 | 0.76447 |

| | | | | | | | | |
|---|---|---|---|---|---|---|---|---|
| 0.582383 | 0.56667 | 0.750167 | 0.75229 | 0.743541 | 0.745701 | 0.753106 | 0.750403 | 0.751206 |
| 0.607087 | 0.590144 | 0.743844 | 0.742862 | 0.735539 | 0.741212 | 0.743846 | 0.744462 | 0.74604 |
| 0.596341 | 0.582202 | 0.754303 | 0.759169 | 0.749937 | 0.754772 | 0.760278 | 0.759704 | 0.759018 |
| 0.595895 | 0.584461 | 0.760667 | 0.759565 | 0.750651 | 0.759469 | 0.758197 | 0.769072 | 0.760283 |
| 0.587463 | 0.575563 | 0.750858 | 0.753713 | 0.742753 | 0.753371 | 0.755054 | 0.75691 | 0.754974 |
| 0.59528 | 0.580227 | 0.756263 | 0.761822 | 0.748111 | 0.755304 | 0.75891 | 0.76342 | 0.759086 |
| 0.59368 | 0.579354 | 0.759725 | 0.759928 | 0.752547 | 0.76022 | 0.757413 | 0.761339 | 0.763826 |
| 0.589986 | 0.576985 | 0.753362 | 0.750948 | 0.74473 | 0.752644 | 0.751407 | 0.757422 | 0.754392 |
| 0.597104 | 0.585467 | 0.765447 | 0.762648 | 0.758433 | 0.765109 | 0.765765 | 0.766476 | 0.766967 |
| 0.59464 | 0.581261 | 0.761445 | 0.760465 | 0.753506 | 0.761846 | 0.767831 | 0.766342 | 0.765648 |
| 0.597456 | 0.584073 | 0.761074 | 0.765234 | 0.759425 | 0.770547 | 0.768735 | 0.771918 | 0.769029 |
| 0.598029 | 0.585385 | 0.759444 | 0.760788 | 0.753803 | 0.761517 | 0.761194 | 0.76663 | 0.768237 |
| 0.600629 | 0.586969 | 0.76768 | 0.766429 | 0.755546 | 0.76279 | 0.767491 | 0.766681 | 0.76642 |
| 0.599832 | 0.585601 | 0.762314 | 0.76163 | 0.758128 | 0.762626 | 0.764826 | 0.768893 | 0.762379 |
| 0.595631 | 0.582492 | 0.760927 | 0.7624 | 0.75495 | 0.763307 | 0.761374 | 0.769315 | 0.766792 |
| 0.595315 | 0.583975 | 0.762279 | 0.760567 | 0.759294 | 0.766888 | 0.767033 | 0.76743 | 0.765364 |
| 0.594688 | 0.579783 | 0.761742 | 0.761631 | 0.758564 | 0.758923 | 0.763016 | 0.76179 | 0.76837 |
| 0.589598 | 0.576227 | 0.760084 | 0.754246 | 0.753297 | 0.755364 | 0.758734 | 0.757543 | 0.761152 |
| 0.586478 | 0.574804 | 0.758297 | 0.760738 | 0.754683 | 0.756496 | 0.759389 | 0.768189 | 0.763838 |
| 0.588373 | 0.572748 | 0.759245 | 0.760792 | 0.753009 | 0.7584 | 0.762559 | 0.762234 | 0.763124 |
| 0.594699 | 0.580668 | 0.759728 | 0.760702 | 0.754624 | 0.762 | 0.760007 | 0.763783 | 0.765087 |
| 0.592406 | 0.578657 | 0.749936 | 0.750924 | 0.739086 | 0.748092 | 0.74631 | 0.752062 | 0.750126 |
| 0.598024 | 0.582481 | 0.756772 | 0.757241 | 0.746764 | 0.753381 | 0.756845 | 0.758424 | 0.755852 |
| 0.597469 | 0.584789 | 0.762203 | 0.759629 | 0.7513 | 0.757591 | 0.759895 | 0.762963 | 0.763618 |
| 0.595318 | 0.585317 | 0.764721 | 0.766415 | 0.752632 | 0.760127 | 0.756982 | 0.764246 | 0.768077 |
| 0.599315 | 0.584442 | 0.760906 | 0.759524 | 0.743259 | 0.753104 | 0.75312 | 0.764126 | 0.758598 |
| 0.590295 | 0.582118 | 0.757664 | 0.757596 | 0.74849 | 0.755195 | 0.752819 | 0.76175 | 0.763885 |
| 0.586556 | 0.574196 | 0.760712 | 0.758826 | 0.74876 | 0.761601 | 0.762745 | 0.760469 | 0.7612 |
| 0.589194 | 0.580173 | 0.756066 | 0.757096 | 0.746183 | 0.75649 | 0.753387 | 0.762147 | 0.762311 |
| 0.604278 | 0.585135 | 0.695146 | 0.694177 | 0.682938 | 0.691243 | 0.693352 | 0.694264 | 0.693584 |
| 0.614677 | 0.596351 | 0.7006 | 0.704556 | 0.697209 | 0.700016 | 0.705991 | 0.703836 | 0.707091 |
| 0.617013 | 0.599567 | 0.704606 | 0.706052 | 0.696516 | 0.698843 | 0.704776 | 0.699855 | 0.700545 |
| 0.612532 | 0.5873 | 0.695687 | 0.695299 | 0.686136 | 0.690184 | 0.696823 | 0.694092 | 0.69199 |
| 0.632139 | 0.600079 | 0.701398 | 0.708129 | 0.697203 | 0.699992 | 0.70604 | 0.702806 | 0.705887 |
| 0.586641 | 0.576948 | 0.753658 | 0.758463 | 0.747896 | 0.756713 | 0.756934 | 0.762198 | 0.759082 |
| 0.59383 | 0.57867 | 0.761905 | 0.760143 | 0.750187 | 0.758692 | 0.759814 | 0.76476 | 0.761469 |
| 0.594004 | 0.581512 | 0.760319 | 0.762438 | 0.753921 | 0.767993 | 0.761609 | 0.762301 | 0.763031 |
| 0.592788 | 0.57854 | 0.760067 | 0.762662 | 0.753359 | 0.75859 | 0.759325 | 0.768261 | 0.765567 |
| 0.592615 | 0.583227 | 0.759732 | 0.762905 | 0.753362 | 0.760472 | 0.763978 | 0.761988 | 0.763742 |
| 0.595697 | 0.581963 | 0.765106 | 0.767859 | 0.755567 | 0.766323 | 0.765592 | 0.767721 | 0.767376 |
| 0.593825 | 0.579609 | 0.759027 | 0.762237 | 0.75547 | 0.763759 | 0.764149 | 0.766101 | 0.766341 |
| 0.592515 | 0.577599 | 0.760787 | 0.762191 | 0.74954 | 0.758058 | 0.760253 | 0.759646 | 0.760786 |
| 0.586778 | 0.579263 | 0.754032 | 0.757584 | 0.741193 | 0.75333 | 0.755965 | 0.753161 | 0.756834 |
| 0.585864 | 0.572237 | 0.75456 | 0.757678 | 0.748668 | 0.756363 | 0.755937 | 0.758685 | 0.754965 |
| 0.592052 | 0.578743 | 0.76073 | 0.756857 | 0.749218 | 0.756732 | 0.75475 | 0.761365 | 0.760741 |
| 0.593396 | 0.582033 | 0.762109 | 0.763193 | 0.754643 | 0.761379 | 0.762043 | 0.766442 | 0.765727 |

| | | | | | | | | |
|---|---|---|---|---|---|---|---|---|
| 0.59552 | 0.583233 | 0.767446 | 0.76103 | 0.758183 | 0.763516 | 0.765742 | 0.766959 | 0.765361 |
| 0.59793 | 0.582336 | 0.760291 | 0.758184 | 0.750021 | 0.758678 | 0.757262 | 0.763026 | 0.762263 |
| 0.595806 | 0.58395 | 0.765023 | 0.767236 | 0.752645 | 0.757571 | 0.761286 | 0.764256 | 0.763408 |
| 0.591285 | 0.580936 | 0.75818 | 0.759579 | 0.7563 | 0.75657 | 0.757183 | 0.762934 | 0.764068 |
| 0.589424 | 0.575064 | 0.758906 | 0.754247 | 0.748904 | 0.756259 | 0.755121 | 0.763644 | 0.763025 |
| 0.589936 | 0.576464 | 0.756638 | 0.758943 | 0.751795 | 0.758193 | 0.759015 | 0.76389 | 0.763268 |
| 0.594383 | 0.580929 | 0.764037 | 0.763475 | 0.753429 | 0.75621 | 0.759507 | 0.765814 | 0.766759 |
| 0.592798 | 0.580029 | 0.763079 | 0.760299 | 0.752622 | 0.756748 | 0.761881 | 0.768738 | 0.769289 |
| 0.593195 | 0.582902 | 0.754759 | 0.757515 | 0.750611 | 0.755331 | 0.757117 | 0.757795 | 0.759183 |
| 0.592478 | 0.578977 | 0.757497 | 0.760663 | 0.75125 | 0.755559 | 0.758669 | 0.760382 | 0.762296 |
| 0.592448 | 0.580226 | 0.762371 | 0.762422 | 0.755956 | 0.760955 | 0.757866 | 0.763603 | 0.766605 |
| 0.593879 | 0.581688 | 0.766485 | 0.763469 | 0.758755 | 0.761606 | 0.765395 | 0.765853 | 0.765675 |
| 0.581173 | 0.571187 | 0.758683 | 0.75687 | 0.748571 | 0.751414 | 0.754364 | 0.760244 | 0.757495 |
| 0.585072 | 0.572217 | 0.759128 | 0.754696 | 0.746362 | 0.753008 | 0.753479 | 0.761502 | 0.758082 |
| 0.594955 | 0.580438 | 0.76014 | 0.757781 | 0.753682 | 0.761613 | 0.761519 | 0.761848 | 0.761094 |
| 0.602702 | 0.588315 | 0.765134 | 0.767096 | 0.760043 | 0.766284 | 0.765611 | 0.77163 | 0.768268 |
| 0.597409 | 0.582565 | 0.764444 | 0.766943 | 0.75598 | 0.760211 | 0.76078 | 0.766022 | 0.76349 |
| 0.599555 | 0.586411 | 0.763274 | 0.764686 | 0.755419 | 0.760973 | 0.76155 | 0.768302 | 0.763953 |
| 0.5912 | 0.58317 | 0.759518 | 0.762505 | 0.749893 | 0.758638 | 0.752851 | 0.767725 | 0.76015 |
| 0.593913 | 0.580086 | 0.759797 | 0.760829 | 0.757242 | 0.763292 | 0.764412 | 0.766076 | 0.766201 |
| 0.591619 | 0.578836 | 0.759343 | 0.759692 | 0.753766 | 0.762507 | 0.764465 | 0.760069 | 0.764771 |
| 0.594975 | 0.582949 | 0.760181 | 0.755777 | 0.75204 | 0.757834 | 0.758316 | 0.762972 | 0.759585 |
| 0.599809 | 0.583306 | 0.76809 | 0.770246 | 0.761815 | 0.764496 | 0.76648 | 0.775588 | 0.769054 |
| 0.589637 | 0.576515 | 0.762236 | 0.756859 | 0.752609 | 0.761539 | 0.756847 | 0.764648 | 0.762106 |
| 0.593866 | 0.578961 | 0.763931 | 0.763972 | 0.755109 | 0.764768 | 0.762716 | 0.770044 | 0.764017 |
| 0.613656 | 0.5977 | 0.696255 | 0.69714 | 0.691655 | 0.697509 | 0.702631 | 0.700642 | 0.700685 |
| 0.595567 | 0.584334 | 0.764833 | 0.764461 | 0.757508 | 0.76028 | 0.762746 | 0.766515 | 0.766508 |
| 0.600614 | 0.584534 | 0.760496 | 0.761924 | 0.756603 | 0.763281 | 0.760608 | 0.763746 | 0.761628 |
| 0.601221 | 0.58931 | 0.765763 | 0.76403 | 0.752901 | 0.759555 | 0.76668 | 0.77238 | 0.767274 |
| 0.600678 | 0.5846 | 0.759947 | 0.762461 | 0.753849 | 0.761018 | 0.762578 | 0.765668 | 0.762424 |
| 0.590333 | 0.580256 | 0.760367 | 0.760806 | 0.750906 | 0.760151 | 0.756805 | 0.764619 | 0.765579 |
| 0.593167 | 0.579738 | 0.763713 | 0.766032 | 0.75096 | 0.756738 | 0.75879 | 0.762584 | 0.768392 |
| 0.595216 | 0.579904 | 0.765246 | 0.764054 | 0.750441 | 0.754523 | 0.757495 | 0.764492 | 0.763989 |
| 0.592961 | 0.574859 | 0.754909 | 0.766428 | 0.756553 | 0.760757 | 0.76371 | 0.763568 | 0.762615 |
| 0.591769 | 0.575076 | 0.751832 | 0.755077 | 0.749042 | 0.755358 | 0.757102 | 0.759519 | 0.759549 |
| 0.594421 | 0.580193 | 0.753195 | 0.755082 | 0.747476 | 0.755524 | 0.755454 | 0.762775 | 0.759484 |
| 0.595121 | 0.581136 | 0.764153 | 0.76298 | 0.756352 | 0.761543 | 0.761881 | 0.769087 | 0.764898 |
| 0.596333 | 0.58458 | 0.764869 | 0.762999 | 0.756469 | 0.764054 | 0.767581 | 0.767654 | 0.770377 |
| 0.596237 | 0.582982 | 0.765038 | 0.764345 | 0.758575 | 0.76824 | 0.768797 | 0.769574 | 0.764838 |
| 0.594659 | 0.581966 | 0.761752 | 0.763576 | 0.756537 | 0.760685 | 0.762455 | 0.765475 | 0.766096 |
| 0.591812 | 0.575916 | 0.757794 | 0.759223 | 0.750945 | 0.760459 | 0.757431 | 0.763902 | 0.759788 |
| 0.596106 | 0.580306 | 0.766409 | 0.768342 | 0.757186 | 0.764019 | 0.766614 | 0.770667 | 0.768702 |
| 0.590139 | 0.575 | 0.756845 | 0.758694 | 0.752386 | 0.759829 | 0.761823 | 0.762498 | 0.761004 |
| 0.592897 | 0.580402 | 0.759157 | 0.760547 | 0.755132 | 0.755919 | 0.761339 | 0.760819 | 0.765092 |
| 0.597108 | 0.584037 | 0.765956 | 0.767929 | 0.757659 | 0.763822 | 0.768869 | 0.769263 | 0.771042 |
| 0.596772 | 0.584361 | 0.765899 | 0.766197 | 0.753481 | 0.764392 | 0.76703 | 0.763232 | 0.77072 |

| | | | | | | | | |
|---|---|---|---|---|---|---|---|---|
| 0.59723 | 0.588082 | 0.764384 | 0.764591 | 0.759389 | 0.762161 | 0.764677 | 0.767001 | 0.767159 |
| 0.596535 | 0.586069 | 0.761964 | 0.766243 | 0.758638 | 0.760815 | 0.765754 | 0.770564 | 0.774034 |
| 0.596863 | 0.583763 | 0.765234 | 0.761219 | 0.75341 | 0.757686 | 0.765392 | 0.76337 | 0.767709 |
| 0.539251 | 0.525378 | 0.728249 | 0.733285 | 0.717251 | 0.726949 | 0.729331 | 0.730879 | 0.722874 |
| 0.59414 | 0.581654 | 0.758848 | 0.760434 | 0.752151 | 0.759301 | 0.762693 | 0.763857 | 0.766896 |
| 0.592777 | 0.580017 | 0.763259 | 0.76529 | 0.75326 | 0.758072 | 0.76488 | 0.76792 | 0.765118 |
| 0.596082 | 0.585443 | 0.769212 | 0.771732 | 0.761475 | 0.767335 | 0.767387 | 0.773976 | 0.770835 |
| 0.593664 | 0.582868 | 0.760235 | 0.766476 | 0.759713 | 0.759694 | 0.76497 | 0.765266 | 0.771457 |
| 0.599755 | 0.587258 | 0.765619 | 0.765851 | 0.760207 | 0.764735 | 0.768226 | 0.768995 | 0.767822 |
| 0.597931 | 0.587161 | 0.760709 | 0.766003 | 0.764404 | 0.765356 | 0.762965 | 0.769563 | 0.76906 |
| 0.595357 | 0.584861 | 0.764226 | 0.762987 | 0.756518 | 0.760551 | 0.762069 | 0.767368 | 0.770923 |
| 0.595707 | 0.583202 | 0.767127 | 0.767806 | 0.754667 | 0.765112 | 0.765594 | 0.766048 | 0.767164 |
| 0.597185 | 0.587111 | 0.761655 | 0.766092 | 0.757423 | 0.764407 | 0.764311 | 0.765998 | 0.769705 |
| 0.596233 | 0.583811 | 0.761795 | 0.765326 | 0.757259 | 0.762002 | 0.764533 | 0.765002 | 0.766437 |
| 0.597686 | 0.587023 | 0.759784 | 0.767709 | 0.759614 | 0.766721 | 0.765455 | 0.764745 | 0.768927 |
| 0.59452 | 0.581317 | 0.763835 | 0.767291 | 0.757152 | 0.764036 | 0.760085 | 0.766972 | 0.77 |
| 0.598252 | 0.586748 | 0.763143 | 0.766082 | 0.754611 | 0.763184 | 0.763424 | 0.766329 | 0.77441 |
| 0.597692 | 0.586297 | 0.769974 | 0.77352 | 0.759918 | 0.766459 | 0.768715 | 0.770207 | 0.772671 |
| 0.593919 | 0.580691 | 0.761277 | 0.760726 | 0.752136 | 0.757502 | 0.760809 | 0.765825 | 0.765213 |
| 0.594667 | 0.584909 | 0.765352 | 0.767583 | 0.755433 | 0.763547 | 0.76435 | 0.771933 | 0.769547 |
| 0.597812 | 0.584529 | 0.766897 | 0.768417 | 0.758598 | 0.765221 | 0.769798 | 0.772364 | 0.773691 |
| 0.604405 | 0.594415 | 0.767008 | 0.767667 | 0.758731 | 0.767939 | 0.76828 | 0.766299 | 0.769795 |
| 0.548084 | 0.534754 | 0.68121 | 0.675228 | 0.673342 | 0.677302 | 0.680634 | 0.685546 | 0.684029 |
| 0.54527 | 0.529667 | 0.665146 | 0.667947 | 0.652725 | 0.653637 | 0.659833 | 0.662291 | 0.66024 |
| 0.593737 | 0.57621 | 0.670445 | 0.672794 | 0.657691 | 0.659329 | 0.663952 | 0.674206 | 0.674688 |
| 0.555067 | 0.533512 | 0.651803 | 0.649902 | 0.644849 | 0.64772 | 0.651485 | 0.64753 | 0.650446 |
| 0.545192 | 0.527217 | 0.729148 | 0.727956 | 0.72122 | 0.727939 | 0.72855 | 0.730451 | 0.72971 |
| 0.539677 | 0.523818 | 0.723101 | 0.720904 | 0.707536 | 0.722594 | 0.72117 | 0.725415 | 0.72539 |
| 0.527651 | 0.514718 | 0.72145 | 0.711377 | 0.712307 | 0.723649 | 0.725662 | 0.730296 | 0.725901 |
| 0.550903 | 0.534335 | 0.722534 | 0.729461 | 0.723227 | 0.727817 | 0.731587 | 0.73738 | 0.73669 |
| 0.52881 | 0.513836 | 0.705909 | 0.712652 | 0.702746 | 0.713733 | 0.717222 | 0.720276 | 0.719936 |
| 0.515981 | 0.498062 | 0.709204 | 0.709122 | 0.70697 | 0.704316 | 0.714407 | 0.71762 | 0.716653 |
| 0.513616 | 0.505362 | 0.705675 | 0.699532 | 0.697496 | 0.711118 | 0.70583 | 0.720379 | 0.716089 |
| 0.548029 | 0.526757 | 0.654067 | 0.654506 | 0.641236 | 0.645298 | 0.658969 | 0.659372 | 0.658946 |
| 0.591094 | 0.57809 | 0.755411 | 0.754531 | 0.753393 | 0.761146 | 0.758683 | 0.759575 | 0.758128 |
| 0.553585 | 0.542567 | 0.737064 | 0.735548 | 0.731101 | 0.735695 | 0.73331 | 0.746601 | 0.739785 |
| 0.541543 | 0.52798 | 0.735053 | 0.722823 | 0.723148 | 0.735207 | 0.729475 | 0.737298 | 0.734131 |
| 0.529511 | 0.515307 | 0.717728 | 0.708994 | 0.717662 | 0.718738 | 0.717138 | 0.73469 | 0.71809 |
| 0.554186 | 0.545256 | 0.727676 | 0.729472 | 0.728516 | 0.735373 | 0.73863 | 0.738723 | 0.739965 |
| 0.547453 | 0.533477 | 0.72125 | 0.725565 | 0.717341 | 0.72544 | 0.723347 | 0.732214 | 0.73313 |
| 0.53863 | 0.525281 | 0.721725 | 0.7251 | 0.720595 | 0.716754 | 0.723316 | 0.732417 | 0.729572 |
| 0.539671 | 0.531638 | 0.722335 | 0.720563 | 0.710825 | 0.718556 | 0.725109 | 0.731868 | 0.729341 |
| 0.531573 | 0.517477 | 0.724582 | 0.729159 | 0.711007 | 0.71951 | 0.721161 | 0.734924 | 0.722737 |
| 0.533898 | 0.521875 | 0.707279 | 0.720388 | 0.704375 | 0.717015 | 0.713386 | 0.719402 | 0.724037 |
| 0.526983 | 0.507501 | 0.719938 | 0.716261 | 0.720942 | 0.715057 | 0.720235 | 0.73537 | 0.729481 |
| 0.520531 | 0.504698 | 0.711303 | 0.706155 | 0.701894 | 0.700654 | 0.714804 | 0.727545 | 0.706942 |

| | | | | | | | | |
|---|---|---|---|---|---|---|---|---|
| 0.563602 | 0.554257 | 0.743239 | 0.745145 | 0.740143 | 0.750025 | 0.747032 | 0.75035 | 0.751185 |
| 0.535851 | 0.525108 | 0.727157 | 0.728984 | 0.717606 | 0.727963 | 0.72596 | 0.731019 | 0.731075 |
| 0.536535 | 0.524707 | 0.725265 | 0.727506 | 0.717473 | 0.721799 | 0.728237 | 0.731571 | 0.727747 |
| 0.543796 | 0.525546 | 0.72834 | 0.727217 | 0.714439 | 0.722138 | 0.722146 | 0.728874 | 0.730594 |
| 0.530926 | 0.518567 | 0.725349 | 0.727254 | 0.714402 | 0.720439 | 0.729741 | 0.723174 | 0.725 |
| 0.541619 | 0.519244 | 0.725697 | 0.730507 | 0.710912 | 0.715318 | 0.715633 | 0.73573 | 0.719057 |
| 0.521903 | 0.511325 | 0.712524 | 0.706464 | 0.710134 | 0.718826 | 0.712938 | 0.72038 | 0.71408 |
| 0.516425 | 0.502845 | 0.721036 | 0.714043 | 0.705882 | 0.709442 | 0.714847 | 0.726292 | 0.719978 |
| 0.51718 | 0.509723 | 0.721096 | 0.712657 | 0.705269 | 0.713761 | 0.716098 | 0.720215 | 0.711926 |
| 0.568434 | 0.557712 | 0.754704 | 0.755481 | 0.74279 | 0.755963 | 0.75493 | 0.755657 | 0.754745 |
| 0.567793 | 0.554252 | 0.74972 | 0.752579 | 0.741374 | 0.751664 | 0.751612 | 0.752236 | 0.753096 |
| 0.564672 | 0.550904 | 0.740258 | 0.744127 | 0.738362 | 0.740737 | 0.744235 | 0.746831 | 0.748401 |
| 0.550632 | 0.538592 | 0.734788 | 0.737277 | 0.725488 | 0.732697 | 0.733325 | 0.741071 | 0.739838 |
| 0.555132 | 0.542445 | 0.735045 | 0.738177 | 0.730826 | 0.743747 | 0.737861 | 0.74963 | 0.741681 |
| 0.554838 | 0.540587 | 0.730945 | 0.735138 | 0.725046 | 0.73852 | 0.738778 | 0.738636 | 0.737561 |

| | | | | | | | | |
|---|---|---|---|---|---|---|---|---|
| | plot 01 | | | | | | | |
| Collection_ | Wild_plot_ | Wild_plot_ | Wild_plot_ | Wild_plot_ | Wild_plot_ | Wild_plot_ | Wild_plot_ | Wild_plot_ |
| 0.706181 | 0.711827 | 0.709002 | 0.709644 | 0.716041 | 0.706549 | 0.712652 | 0.709455 | 0.704872 |
| 0.710665 | 0.717772 | 0.717605 | 0.715268 | 0.719437 | 0.713679 | 0.719434 | 0.716587 | 0.710339 |
| 0.702524 | 0.708258 | 0.703349 | 0.705062 | 0.706393 | 0.703192 | 0.704861 | 0.70806 | 0.701137 |
| 0.710487 | 0.72017 | 0.714766 | 0.715117 | 0.717357 | 0.711806 | 0.714889 | 0.717471 | 0.709438 |
| 0.703927 | 0.711887 | 0.711984 | 0.707246 | 0.709999 | 0.706149 | 0.710145 | 0.710042 | 0.703887 |
| 0.713622 | 0.724943 | 0.716644 | 0.717953 | 0.721776 | 0.716564 | 0.716105 | 0.720631 | 0.713576 |
| 0.76053 | 0.771559 | 0.766755 | 0.765742 | 0.768003 | 0.759649 | 0.768356 | 0.766531 | 0.759071 |
| 0.752258 | 0.767917 | 0.756761 | 0.7644 | 0.769212 | 0.760612 | 0.760272 | 0.762883 | 0.763551 |
| 0.709239 | 0.721084 | 0.713295 | 0.717764 | 0.719225 | 0.715032 | 0.712823 | 0.713877 | 0.708412 |
| 0.704103 | 0.714894 | 0.713242 | 0.711595 | 0.71466 | 0.705488 | 0.708719 | 0.711423 | 0.704555 |
| 0.706714 | 0.714637 | 0.711475 | 0.710674 | 0.713795 | 0.709245 | 0.711063 | 0.711932 | 0.706499 |
| 0.704866 | 0.709817 | 0.708039 | 0.703004 | 0.709624 | 0.701879 | 0.706998 | 0.71001 | 0.700147 |
| 0.711193 | 0.718222 | 0.712769 | 0.715912 | 0.714843 | 0.709564 | 0.711924 | 0.714036 | 0.710231 |
| 0.718528 | 0.724359 | 0.719751 | 0.724431 | 0.725606 | 0.72145 | 0.724833 | 0.72294 | 0.721066 |
| 0.731239 | 0.736508 | 0.737856 | 0.73421 | 0.738783 | 0.732974 | 0.733264 | 0.737818 | 0.733366 |
| 0.71519 | 0.723033 | 0.715743 | 0.71819 | 0.719407 | 0.712836 | 0.716535 | 0.719459 | 0.712286 |
| 0.714958 | 0.721499 | 0.713628 | 0.720437 | 0.719274 | 0.711583 | 0.718427 | 0.715098 | 0.712401 |
| 0.736097 | 0.743111 | 0.740786 | 0.7391 | 0.746441 | 0.736872 | 0.744532 | 0.741425 | 0.737932 |
| 0.703321 | 0.710038 | 0.707697 | 0.708475 | 0.709191 | 0.70643 | 0.705553 | 0.707548 | 0.703328 |
| 0.710035 | 0.717309 | 0.710273 | 0.712548 | 0.716823 | 0.709326 | 0.711452 | 0.713206 | 0.708163 |
| 0.703103 | 0.709876 | 0.710446 | 0.710175 | 0.713103 | 0.704314 | 0.711262 | 0.71094 | 0.702524 |
| 0.706928 | 0.713552 | 0.712791 | 0.710613 | 0.712647 | 0.709353 | 0.707387 | 0.710643 | 0.706243 |
| 0.713191 | 0.719744 | 0.714861 | 0.719061 | 0.721038 | 0.715237 | 0.71798 | 0.717398 | 0.712568 |
| 0.710255 | 0.719099 | 0.712355 | 0.712519 | 0.71792 | 0.709649 | 0.715189 | 0.719081 | 0.706594 |
| 0.710454 | 0.717377 | 0.717073 | 0.714456 | 0.718632 | 0.714274 | 0.716641 | 0.715564 | 0.709058 |
| 0.707351 | 0.716717 | 0.714366 | 0.713155 | 0.716755 | 0.709131 | 0.71473 | 0.711387 | 0.704844 |
| 0.700573 | 0.710955 | 0.706585 | 0.706756 | 0.707334 | 0.704413 | 0.707687 | 0.705412 | 0.701772 |
| 0.710008 | 0.715837 | 0.711902 | 0.715598 | 0.714694 | 0.714115 | 0.713786 | 0.714173 | 0.710962 |
| 0.712775 | 0.719882 | 0.712518 | 0.716729 | 0.718436 | 0.712957 | 0.713384 | 0.711719 | 0.710653 |
| 0.708709 | 0.714796 | 0.712542 | 0.713176 | 0.715578 | 0.712031 | 0.712209 | 0.711253 | 0.708234 |
| 0.703658 | 0.715558 | 0.711888 | 0.711044 | 0.711764 | 0.705849 | 0.708967 | 0.710019 | 0.709013 |
| 0.702808 | 0.714342 | 0.707545 | 0.708997 | 0.712654 | 0.70915 | 0.71016 | 0.706731 | 0.700405 |
| 0.703348 | 0.71316 | 0.709804 | 0.710031 | 0.709981 | 0.704677 | 0.711798 | 0.709234 | 0.704594 |
| 0.708752 | 0.718454 | 0.709827 | 0.713298 | 0.712871 | 0.707554 | 0.714217 | 0.714694 | 0.707967 |
| 0.700033 | 0.711519 | 0.705953 | 0.708363 | 0.711095 | 0.7077 | 0.706299 | 0.706722 | 0.703997 |
| 0.708825 | 0.715181 | 0.708333 | 0.711468 | 0.716663 | 0.706363 | 0.709677 | 0.709523 | 0.706545 |
| 0.714203 | 0.720483 | 0.71152 | 0.717067 | 0.718847 | 0.715355 | 0.716977 | 0.715369 | 0.709698 |
| 0.709884 | 0.71836 | 0.710287 | 0.714081 | 0.715699 | 0.710139 | 0.713385 | 0.712574 | 0.709883 |
| 0.713737 | 0.720612 | 0.719027 | 0.718992 | 0.721708 | 0.714822 | 0.720054 | 0.718238 | 0.714251 |

| | | | | | | | | |
|---|---|---|---|---|---|---|---|---|
| 0.742208 | 0.748667 | 0.750976 | 0.746054 | 0.749639 | 0.744061 | 0.746881 | 0.746627 | 0.744535 |
| 0.69764 | 0.706368 | 0.704145 | 0.705527 | 0.708621 | 0.701535 | 0.70473 | 0.70359 | 0.699537 |
| 0.703522 | 0.712502 | 0.706043 | 0.706857 | 0.707922 | 0.705596 | 0.705535 | 0.70783 | 0.703822 |
| 0.714321 | 0.719965 | 0.714686 | 0.717979 | 0.715021 | 0.714502 | 0.714024 | 0.714669 | 0.709141 |
| 0.702473 | 0.717359 | 0.711991 | 0.711902 | 0.712881 | 0.705564 | 0.711324 | 0.713549 | 0.706516 |
| 0.711232 | 0.716157 | 0.714851 | 0.715713 | 0.717348 | 0.712666 | 0.713183 | 0.714868 | 0.706519 |
| 0.706756 | 0.714714 | 0.710708 | 0.708579 | 0.712798 | 0.705466 | 0.706657 | 0.712758 | 0.704732 |
| 0.707378 | 0.714774 | 0.712527 | 0.712994 | 0.714663 | 0.708729 | 0.70947 | 0.711659 | 0.70796 |
| 0.706483 | 0.716154 | 0.714205 | 0.715921 | 0.717918 | 0.713559 | 0.716731 | 0.715407 | 0.707067 |
| 0.708237 | 0.71772 | 0.713016 | 0.716008 | 0.719309 | 0.711063 | 0.711462 | 0.715736 | 0.710133 |
| 0.701627 | 0.710353 | 0.706333 | 0.709156 | 0.707379 | 0.700625 | 0.706524 | 0.705545 | 0.700884 |
| 0.700857 | 0.708251 | 0.70654 | 0.708337 | 0.708765 | 0.700678 | 0.704455 | 0.708131 | 0.704202 |
| 0.706104 | 0.714263 | 0.712389 | 0.715242 | 0.715926 | 0.707474 | 0.709265 | 0.709056 | 0.703251 |
| 0.70366 | 0.712014 | 0.708152 | 0.707651 | 0.712239 | 0.704679 | 0.707835 | 0.707177 | 0.702323 |
| 0.702306 | 0.711798 | 0.702764 | 0.707936 | 0.709341 | 0.70274 | 0.708693 | 0.707423 | 0.699441 |
| 0.705306 | 0.713368 | 0.70442 | 0.711292 | 0.708653 | 0.704961 | 0.710479 | 0.707754 | 0.698751 |
| 0.703109 | 0.706126 | 0.706499 | 0.709281 | 0.71132 | 0.702694 | 0.707337 | 0.704222 | 0.700832 |
| 0.709224 | 0.716242 | 0.711358 | 0.71486 | 0.716935 | 0.708283 | 0.712166 | 0.713605 | 0.704621 |
| 0.712117 | 0.720659 | 0.71582 | 0.716501 | 0.719329 | 0.714079 | 0.714583 | 0.712664 | 0.713757 |
| 0.705129 | 0.712501 | 0.709672 | 0.710261 | 0.712385 | 0.70418 | 0.708115 | 0.707122 | 0.706018 |
| 0.702126 | 0.709375 | 0.703722 | 0.70831 | 0.705671 | 0.705009 | 0.707006 | 0.705812 | 0.698249 |
| 0.705294 | 0.710861 | 0.706611 | 0.709193 | 0.708519 | 0.703736 | 0.704762 | 0.711549 | 0.701325 |
| 0.705693 | 0.715804 | 0.710806 | 0.714128 | 0.713268 | 0.709752 | 0.708654 | 0.71178 | 0.703935 |
| 0.709877 | 0.724673 | 0.718558 | 0.720603 | 0.723078 | 0.71486 | 0.716236 | 0.718995 | 0.712783 |
| 0.719746 | 0.72684 | 0.722854 | 0.726675 | 0.72662 | 0.722672 | 0.723713 | 0.720481 | 0.721658 |
| 0.751457 | 0.769794 | 0.757959 | 0.764586 | 0.76932 | 0.760828 | 0.760046 | 0.762571 | 0.763698 |
| 0.762937 | 0.77533 | 0.770087 | 0.765357 | 0.776051 | 0.766179 | 0.76948 | 0.770443 | 0.767396 |
| 0.76396 | 0.772399 | 0.770296 | 0.767474 | 0.775554 | 0.768975 | 0.76863 | 0.767545 | 0.765595 |
| 0.701612 | 0.714274 | 0.707154 | 0.70695 | 0.708211 | 0.705069 | 0.706982 | 0.707516 | 0.703531 |
| 0.705958 | 0.71376 | 0.709376 | 0.70815 | 0.711846 | 0.706313 | 0.708089 | 0.7108 | 0.704438 |
| 0.711783 | 0.721001 | 0.716096 | 0.719485 | 0.719562 | 0.715286 | 0.718401 | 0.71652 | 0.714603 |
| 0.713371 | 0.722817 | 0.717012 | 0.717193 | 0.717066 | 0.716981 | 0.715804 | 0.717299 | 0.715829 |
| 0.707073 | 0.7115 | 0.708587 | 0.710539 | 0.714021 | 0.707335 | 0.707183 | 0.710723 | 0.702787 |
| 0.706292 | 0.712383 | 0.710155 | 0.71187 | 0.71252 | 0.705216 | 0.708635 | 0.711697 | 0.702794 |
| 0.703738 | 0.71165 | 0.705902 | 0.710018 | 0.709597 | 0.708584 | 0.706439 | 0.708817 | 0.70521 |
| 0.707403 | 0.712073 | 0.708838 | 0.711715 | 0.711429 | 0.708491 | 0.709778 | 0.710291 | 0.701095 |
| 0.706649 | 0.711813 | 0.707735 | 0.710148 | 0.713147 | 0.705728 | 0.70912 | 0.707511 | 0.706887 |
| 0.722464 | 0.725935 | 0.722685 | 0.723418 | 0.724401 | 0.719182 | 0.719453 | 0.721021 | 0.717177 |
| 0.702458 | 0.704969 | 0.702085 | 0.706037 | 0.704773 | 0.705574 | 0.699495 | 0.700948 | 0.698083 |
| 0.708271 | 0.714611 | 0.71092 | 0.708668 | 0.715066 | 0.706884 | 0.708961 | 0.71058 | 0.7016 |
| 0.702692 | 0.709473 | 0.70336 | 0.705864 | 0.707877 | 0.705579 | 0.706498 | 0.708932 | 0.701733 |
| 0.721979 | 0.729218 | 0.726564 | 0.729114 | 0.726893 | 0.722814 | 0.724683 | 0.722794 | 0.718421 |
| 0.709182 | 0.71651 | 0.71222 | 0.710574 | 0.713261 | 0.708922 | 0.710339 | 0.71158 | 0.708135 |
| 0.706388 | 0.712288 | 0.709404 | 0.710708 | 0.709084 | 0.709231 | 0.706237 | 0.706692 | 0.703445 |
| 0.708066 | 0.716378 | 0.712 | 0.714411 | 0.716969 | 0.70828 | 0.714263 | 0.716024 | 0.704687 |
| 0.704926 | 0.715006 | 0.706025 | 0.710598 | 0.708828 | 0.70829 | 0.705953 | 0.709952 | 0.699606 |

| | | | | | | | | |
|---|---|---|---|---|---|---|---|---|
| 0.710013 | 0.712768 | 0.70999 | 0.709529 | 0.71464 | 0.708006 | 0.708697 | 0.709154 | 0.707049 |
| 0.707165 | 0.719368 | 0.713072 | 0.712702 | 0.716115 | 0.706388 | 0.709508 | 0.712078 | 0.708082 |
| 0.706177 | 0.716012 | 0.711515 | 0.713192 | 0.713998 | 0.704628 | 0.707801 | 0.712302 | 0.703402 |
| 0.718864 | 0.719445 | 0.719217 | 0.716881 | 0.720711 | 0.715073 | 0.713094 | 0.717176 | 0.712367 |
| 0.71652 | 0.72066 | 0.713897 | 0.718385 | 0.718929 | 0.712133 | 0.713101 | 0.715607 | 0.712733 |
| 0.707887 | 0.715359 | 0.710629 | 0.708978 | 0.710632 | 0.70936 | 0.708166 | 0.709344 | 0.70431 |
| 0.711139 | 0.716057 | 0.710572 | 0.712666 | 0.717815 | 0.709035 | 0.7099 | 0.709533 | 0.707765 |
| 0.708916 | 0.722658 | 0.714748 | 0.715778 | 0.720406 | 0.707192 | 0.711263 | 0.711348 | 0.708721 |
| 0.71074 | 0.712856 | 0.714714 | 0.711557 | 0.716693 | 0.710088 | 0.710695 | 0.713908 | 0.705989 |
| 0.702864 | 0.712186 | 0.705717 | 0.705961 | 0.711069 | 0.706591 | 0.707716 | 0.707947 | 0.70333 |
| 0.711122 | 0.714834 | 0.711439 | 0.70835 | 0.71221 | 0.709064 | 0.711792 | 0.707482 | 0.702353 |
| 0.705833 | 0.713071 | 0.711772 | 0.71036 | 0.714586 | 0.707491 | 0.70668 | 0.709085 | 0.706297 |
| 0.70958 | 0.718313 | 0.713013 | 0.713728 | 0.716834 | 0.707974 | 0.713203 | 0.714939 | 0.704704 |
| 0.707766 | 0.711307 | 0.708153 | 0.705554 | 0.711845 | 0.703212 | 0.707234 | 0.709698 | 0.703292 |
| 0.708821 | 0.716487 | 0.71055 | 0.710447 | 0.715842 | 0.710137 | 0.710438 | 0.714673 | 0.706591 |
| 0.705063 | 0.712892 | 0.709264 | 0.710547 | 0.71287 | 0.706135 | 0.705943 | 0.710241 | 0.703122 |
| 0.709289 | 0.717079 | 0.715085 | 0.717142 | 0.719034 | 0.713363 | 0.712227 | 0.718507 | 0.712081 |
| 0.705536 | 0.712999 | 0.70989 | 0.710327 | 0.711866 | 0.707598 | 0.707626 | 0.710777 | 0.702478 |
| 0.708066 | 0.71261 | 0.709849 | 0.709922 | 0.715966 | 0.707649 | 0.708477 | 0.710656 | 0.705285 |
| 0.707532 | 0.716019 | 0.715001 | 0.711157 | 0.71522 | 0.706588 | 0.711904 | 0.713034 | 0.70608 |
| 0.706943 | 0.715502 | 0.711276 | 0.712643 | 0.716821 | 0.704642 | 0.71129 | 0.709729 | 0.707695 |
| 0.703811 | 0.708913 | 0.708132 | 0.708107 | 0.710001 | 0.703633 | 0.704939 | 0.706666 | 0.699592 |
| 0.710641 | 0.717023 | 0.70891 | 0.708799 | 0.715652 | 0.709409 | 0.710467 | 0.712076 | 0.703408 |
| 0.711294 | 0.717395 | 0.707686 | 0.714207 | 0.712752 | 0.709443 | 0.708928 | 0.711659 | 0.705733 |
| 0.709577 | 0.714069 | 0.709593 | 0.713623 | 0.7147 | 0.708644 | 0.710718 | 0.710499 | 0.707525 |
| 0.710345 | 0.720328 | 0.716521 | 0.716216 | 0.722969 | 0.713798 | 0.714797 | 0.717364 | 0.710728 |
| 0.710051 | 0.71586 | 0.708662 | 0.711569 | 0.716839 | 0.706821 | 0.711774 | 0.710159 | 0.706465 |
| 0.707248 | 0.714871 | 0.711574 | 0.713544 | 0.717355 | 0.70912 | 0.710029 | 0.715966 | 0.705354 |
| 0.708579 | 0.717939 | 0.71362 | 0.710949 | 0.716413 | 0.710165 | 0.712142 | 0.715377 | 0.708742 |
| 0.712678 | 0.715445 | 0.710164 | 0.711628 | 0.716389 | 0.707931 | 0.712786 | 0.714182 | 0.707404 |
| 0.711687 | 0.71856 | 0.715978 | 0.716816 | 0.721142 | 0.708675 | 0.714228 | 0.713323 | 0.709295 |
| 0.7075 | 0.715827 | 0.708746 | 0.711738 | 0.716857 | 0.709989 | 0.710364 | 0.710376 | 0.705174 |
| 0.711902 | 0.714915 | 0.710704 | 0.713315 | 0.716107 | 0.709876 | 0.710398 | 0.711848 | 0.707553 |
| 0.712735 | 0.716551 | 0.711313 | 0.713209 | 0.712368 | 0.711407 | 0.710692 | 0.710584 | 0.705868 |
| 0.704779 | 0.71316 | 0.708835 | 0.708106 | 0.711686 | 0.703043 | 0.708993 | 0.710423 | 0.703414 |
| 0.709588 | 0.716513 | 0.710821 | 0.709484 | 0.715653 | 0.709834 | 0.712126 | 0.708827 | 0.705005 |
| 0.7058 | 0.710443 | 0.713223 | 0.707956 | 0.713792 | 0.702325 | 0.706636 | 0.709023 | 0.705712 |
| 0.683039 | 0.689673 | 0.686312 | 0.681805 | 0.69161 | 0.681148 | 0.687557 | 0.684599 | 0.679922 |
| 0.694331 | 0.697248 | 0.691534 | 0.691466 | 0.693167 | 0.68818 | 0.68757 | 0.688685 | 0.682538 |
| 0.68136 | 0.687083 | 0.682369 | 0.676542 | 0.683603 | 0.67465 | 0.680987 | 0.685531 | 0.680092 |
| 0.678231 | 0.682814 | 0.678211 | 0.678862 | 0.678758 | 0.677186 | 0.680196 | 0.682938 | 0.674062 |
| 0.668663 | 0.67027 | 0.664639 | 0.662302 | 0.66729 | 0.658027 | 0.664599 | 0.662739 | 0.661484 |
| 0.671631 | 0.676065 | 0.667952 | 0.670357 | 0.672253 | 0.66566 | 0.670109 | 0.669657 | 0.664555 |
| 0.669404 | 0.672751 | 0.667923 | 0.664545 | 0.668041 | 0.666101 | 0.669583 | 0.671759 | 0.664101 |
| 0.672225 | 0.675772 | 0.670739 | 0.670076 | 0.668577 | 0.665433 | 0.671642 | 0.672862 | 0.668122 |
| 0.670507 | 0.677947 | 0.673149 | 0.671796 | 0.673405 | 0.666916 | 0.668374 | 0.675238 | 0.668009 |
| 0.674489 | 0.679822 | 0.674021 | 0.67543 | 0.674597 | 0.67248 | 0.675856 | 0.679553 | 0.668738 |

| | | | | | | | | |
|---|---|---|---|---|---|---|---|---|
| 0.674981 | 0.675898 | 0.677387 | 0.673357 | 0.677172 | 0.667648 | 0.674935 | 0.669524 | 0.672684 |
| 0.675398 | 0.679382 | 0.67558 | 0.677155 | 0.680136 | 0.675768 | 0.677436 | 0.673814 | 0.673999 |
| 0.6697 | 0.678238 | 0.675295 | 0.670834 | 0.672827 | 0.664798 | 0.672375 | 0.674624 | 0.670441 |
| 0.673895 | 0.673556 | 0.663283 | 0.665791 | 0.671815 | 0.661235 | 0.670209 | 0.670876 | 0.669741 |
| 0.66469 | 0.666497 | 0.661785 | 0.663567 | 0.664778 | 0.656374 | 0.665426 | 0.661757 | 0.659827 |
| 0.665265 | 0.673866 | 0.668812 | 0.66135 | 0.668272 | 0.65896 | 0.664614 | 0.666027 | 0.663819 |
| 0.668219 | 0.674721 | 0.667328 | 0.662316 | 0.665673 | 0.658122 | 0.667329 | 0.665585 | 0.663357 |
| 0.669103 | 0.674069 | 0.66447 | 0.669223 | 0.665407 | 0.659919 | 0.664565 | 0.666013 | 0.659116 |
| 0.661858 | 0.671008 | 0.661104 | 0.666228 | 0.668586 | 0.660845 | 0.664954 | 0.669164 | 0.660389 |
| 0.664227 | 0.671814 | 0.665984 | 0.664063 | 0.671265 | 0.659562 | 0.66463 | 0.662115 | 0.66544 |
| 0.656485 | 0.665224 | 0.66007 | 0.658403 | 0.66386 | 0.652312 | 0.66192 | 0.658154 | 0.657806 |
| 0.698814 | 0.704563 | 0.700707 | 0.697799 | 0.704479 | 0.697533 | 0.6989 | 0.698945 | 0.69759 |
| 0.705523 | 0.712829 | 0.710189 | 0.710654 | 0.711984 | 0.707139 | 0.70838 | 0.708986 | 0.705539 |
| 0.682216 | 0.693208 | 0.684893 | 0.6869 | 0.690444 | 0.684337 | 0.6877 | 0.685282 | 0.681547 |
| 0.715022 | 0.721789 | 0.715683 | 0.715497 | 0.717667 | 0.715133 | 0.716024 | 0.717758 | 0.710668 |
| 0.706372 | 0.709529 | 0.705996 | 0.70547 | 0.709788 | 0.702153 | 0.703881 | 0.70684 | 0.699788 |
| 0.708829 | 0.719196 | 0.71271 | 0.715262 | 0.71789 | 0.707788 | 0.710834 | 0.713181 | 0.707063 |
| 0.71808 | 0.720858 | 0.721337 | 0.723972 | 0.726259 | 0.720311 | 0.718543 | 0.720325 | 0.716243 |
| 0.694645 | 0.702926 | 0.698098 | 0.698376 | 0.704447 | 0.693203 | 0.694429 | 0.696863 | 0.692097 |
| 0.720212 | 0.722931 | 0.718367 | 0.720446 | 0.723347 | 0.714773 | 0.721099 | 0.722222 | 0.715892 |
| 0.691475 | 0.697032 | 0.697514 | 0.689713 | 0.696129 | 0.689971 | 0.691566 | 0.692326 | 0.693086 |
| 0.71521 | 0.722869 | 0.716831 | 0.716875 | 0.718435 | 0.717383 | 0.716984 | 0.715804 | 0.714803 |
| 0.719741 | 0.723325 | 0.721456 | 0.725293 | 0.725273 | 0.720778 | 0.721104 | 0.722841 | 0.718033 |
| 0.685861 | 0.68719 | 0.686968 | 0.684843 | 0.68785 | 0.681337 | 0.68312 | 0.686178 | 0.679888 |
| 0.689317 | 0.694563 | 0.68832 | 0.691715 | 0.693 | 0.684882 | 0.687981 | 0.69178 | 0.683354 |
| 0.691312 | 0.69465 | 0.69223 | 0.690829 | 0.693813 | 0.692866 | 0.693386 | 0.693593 | 0.686821 |
| 0.705847 | 0.714923 | 0.706839 | 0.706655 | 0.714321 | 0.706657 | 0.706651 | 0.71283 | 0.705961 |
| 0.714977 | 0.724239 | 0.722191 | 0.721123 | 0.722923 | 0.717202 | 0.719324 | 0.718042 | 0.713386 |
| 0.685719 | 0.68886 | 0.686119 | 0.68961 | 0.688358 | 0.68513 | 0.686143 | 0.6893 | 0.682657 |
| 0.721723 | 0.727775 | 0.721347 | 0.72442 | 0.730977 | 0.721527 | 0.724009 | 0.723534 | 0.722363 |
| 0.725786 | 0.732489 | 0.726023 | 0.73151 | 0.73308 | 0.725445 | 0.728451 | 0.727065 | 0.721573 |
| 0.695188 | 0.701111 | 0.698742 | 0.697165 | 0.698615 | 0.693654 | 0.696258 | 0.696563 | 0.68956 |
| 0.697975 | 0.702728 | 0.699276 | 0.698128 | 0.702545 | 0.696454 | 0.69724 | 0.698075 | 0.692183 |
| 0.724588 | 0.730111 | 0.72474 | 0.726437 | 0.731176 | 0.724626 | 0.726542 | 0.726002 | 0.721775 |
| 0.718231 | 0.727643 | 0.723697 | 0.720292 | 0.723745 | 0.716853 | 0.722938 | 0.721452 | 0.718163 |
| 0.725338 | 0.731172 | 0.724799 | 0.726411 | 0.728493 | 0.726823 | 0.722724 | 0.725449 | 0.723097 |
| 0.67534 | 0.682187 | 0.678339 | 0.67662 | 0.678006 | 0.673845 | 0.678359 | 0.678519 | 0.673922 |
| 0.716339 | 0.72347 | 0.719487 | 0.721597 | 0.72127 | 0.719843 | 0.721729 | 0.722231 | 0.71551 |
| 0.674904 | 0.682852 | 0.67928 | 0.677392 | 0.681675 | 0.677696 | 0.678059 | 0.675653 | 0.672236 |
| 0.667853 | 0.680218 | 0.671876 | 0.668415 | 0.671385 | 0.667223 | 0.671212 | 0.6708 | 0.664983 |
| 0.717618 | 0.724551 | 0.721033 | 0.721427 | 0.723673 | 0.718952 | 0.715748 | 0.720634 | 0.716124 |
| 0.66635 | 0.680052 | 0.672369 | 0.668945 | 0.672086 | 0.666946 | 0.671812 | 0.671948 | 0.665939 |
| 0.660213 | 0.667413 | 0.663432 | 0.664555 | 0.665239 | 0.661027 | 0.661694 | 0.663766 | 0.65707 |
| 0.656225 | 0.662947 | 0.655424 | 0.6599 | 0.663325 | 0.656056 | 0.657278 | 0.658222 | 0.65143 |
| 0.726773 | 0.734191 | 0.727009 | 0.72611 | 0.732264 | 0.729454 | 0.7268 | 0.730368 | 0.723577 |
| 0.658531 | 0.667789 | 0.663302 | 0.664531 | 0.664561 | 0.663094 | 0.659991 | 0.662969 | 0.658891 |
| 0.719113 | 0.724176 | 0.719403 | 0.719804 | 0.722485 | 0.71936 | 0.721104 | 0.721649 | 0.716337 |

| | | | | | | | | |
|---|---|---|---|---|---|---|---|---|
| 0.730186 | 0.731009 | 0.728194 | 0.731578 | 0.73481 | 0.73042 | 0.728982 | 0.731713 | 0.726662 |
| 0.722721 | 0.726337 | 0.721994 | 0.724239 | 0.726533 | 0.718795 | 0.721792 | 0.723411 | 0.724217 |
| 0.749083 | 0.758873 | 0.75279 | 0.755828 | 0.752816 | 0.746354 | 0.752505 | 0.753997 | 0.747906 |
| 0.648132 | 0.657418 | 0.655572 | 0.654306 | 0.653437 | 0.650108 | 0.650608 | 0.653656 | 0.647019 |
| 0.653683 | 0.658111 | 0.654079 | 0.655446 | 0.657097 | 0.651978 | 0.65267 | 0.657262 | 0.648761 |
| 0.728739 | 0.735798 | 0.731084 | 0.732559 | 0.734002 | 0.729521 | 0.733972 | 0.730431 | 0.731328 |
| 0.652416 | 0.657881 | 0.65221 | 0.651956 | 0.658417 | 0.653343 | 0.652086 | 0.65414 | 0.65048 |
| 0.727435 | 0.740444 | 0.732814 | 0.738505 | 0.739746 | 0.731382 | 0.737217 | 0.733685 | 0.73473 |
| 0.729109 | 0.736053 | 0.731977 | 0.727186 | 0.736655 | 0.727875 | 0.731424 | 0.731933 | 0.726467 |
| 0.726197 | 0.739484 | 0.737598 | 0.733889 | 0.739269 | 0.735282 | 0.735002 | 0.737311 | 0.733777 |
| 0.730209 | 0.734416 | 0.733164 | 0.728086 | 0.736137 | 0.731737 | 0.731658 | 0.733204 | 0.727613 |
| 0.703448 | 0.695364 | 0.660377 | 0.709459 | 0.721088 | 0.718954 | 0.731034 | 0.711538 | 0.693878 |
| 0.721624 | 0.721836 | 0.721544 | 0.709544 | 0.719872 | 0.717964 | 0.718775 | 0.716278 | 0.718881 |
| 0.718665 | 0.721823 | 0.718085 | 0.712931 | 0.71404 | 0.715692 | 0.712218 | 0.724638 | 0.722794 |
| 0.729004 | 0.738652 | 0.728956 | 0.717918 | 0.734434 | 0.731774 | 0.726917 | 0.735201 | 0.725853 |
| 0.7251 | 0.734541 | 0.723007 | 0.720806 | 0.732948 | 0.725752 | 0.723168 | 0.729214 | 0.721243 |
| 0.590919 | 0.599718 | 0.591504 | 0.58956 | 0.597028 | 0.596846 | 0.590973 | 0.593407 | 0.590351 |
| 0.727503 | 0.735256 | 0.723067 | 0.724552 | 0.728811 | 0.728791 | 0.72805 | 0.729818 | 0.725361 |
| 0.728924 | 0.739614 | 0.734443 | 0.730871 | 0.740385 | 0.735155 | 0.734276 | 0.734397 | 0.730542 |
| 0.72887 | 0.73568 | 0.734585 | 0.732921 | 0.737788 | 0.734632 | 0.731944 | 0.735724 | 0.733745 |
| 0.735417 | 0.739085 | 0.738301 | 0.736927 | 0.73979 | 0.737062 | 0.736102 | 0.736068 | 0.7363 |
| 0.593543 | 0.601536 | 0.597677 | 0.598725 | 0.594226 | 0.596049 | 0.598414 | 0.603173 | 0.592928 |
| 0.580906 | 0.585156 | 0.585032 | 0.584304 | 0.582724 | 0.582 | 0.58117 | 0.58733 | 0.580716 |
| 0.758749 | 0.763677 | 0.757904 | 0.763277 | 0.765763 | 0.758244 | 0.764662 | 0.765136 | 0.756241 |
| 0.759555 | 0.764414 | 0.758811 | 0.762577 | 0.765 | 0.76112 | 0.764391 | 0.766762 | 0.762367 |
| 0.749441 | 0.761924 | 0.752537 | 0.753757 | 0.758905 | 0.756725 | 0.760976 | 0.761779 | 0.750976 |
| 0.755766 | 0.766952 | 0.758635 | 0.75748 | 0.763867 | 0.761281 | 0.766447 | 0.764701 | 0.757127 |
| 0.759123 | 0.773317 | 0.761547 | 0.759851 | 0.765072 | 0.760571 | 0.766297 | 0.76627 | 0.757841 |
| 0.767149 | 0.773208 | 0.759145 | 0.763989 | 0.769077 | 0.761746 | 0.769571 | 0.767411 | 0.762173 |
| 0.767992 | 0.77167 | 0.761628 | 0.761668 | 0.768882 | 0.761454 | 0.773701 | 0.770911 | 0.762227 |
| 1 | 0.785892 | 0.786546 | 0.782351 | 0.786544 | 0.790395 | 0.781974 | 0.808339 | 0.788063 |
| 0.785892 | 1 | 0.803741 | 0.791582 | 0.79793 | 0.789221 | 0.805299 | 0.790698 | 0.79376 |
| 0.786546 | 0.803741 | 1 | 0.789701 | 0.791027 | 0.796536 | 0.784202 | 0.795901 | 0.798546 |
| 0.782351 | 0.791582 | 0.789701 | 1 | 0.800694 | 0.817953 | 0.788141 | 0.786189 | 0.795203 |
| 0.786544 | 0.79793 | 0.791027 | 0.800694 | 1 | 0.79777 | 0.806097 | 0.792001 | 0.789271 |
| 0.790395 | 0.789221 | 0.796536 | 0.817953 | 0.79777 | 1 | 0.792115 | 0.793823 | 0.789786 |
| 0.781974 | 0.805299 | 0.784202 | 0.788141 | 0.806097 | 0.792115 | 1 | 0.789491 | 0.793744 |
| 0.808339 | 0.790698 | 0.795901 | 0.786189 | 0.792001 | 0.793823 | 0.789491 | 1 | 0.787392 |
| 0.788063 | 0.79376 | 0.798546 | 0.795203 | 0.789271 | 0.789786 | 0.793744 | 0.787392 | 1 |
| 0.790521 | 0.791318 | 0.804093 | 0.797204 | 0.795893 | 0.788492 | 0.79725 | 0.800599 | 0.826147 |
| 0.720072 | 0.72955 | 0.71008 | 0.721294 | 0.718168 | 0.712148 | 0.716049 | 0.716925 | 0.716287 |
| 0.722053 | 0.727684 | 0.72604 | 0.719174 | 0.731481 | 0.726655 | 0.725428 | 0.731329 | 0.728852 |
| 0.77385 | 0.784399 | 0.778138 | 0.774115 | 0.781278 | 0.7726 | 0.780909 | 0.774323 | 0.77631 |
| 0.774002 | 0.789743 | 0.775719 | 0.775492 | 0.784259 | 0.777483 | 0.779083 | 0.781307 | 0.778316 |
| 0.747475 | 0.75146 | 0.749462 | 0.739232 | 0.748331 | 0.732425 | 0.748588 | 0.748383 | 0.745583 |
| 0.786968 | 0.791646 | 0.787477 | 0.787236 | 0.792868 | 0.782514 | 0.783473 | 0.789233 | 0.79331 |
| 0.777214 | 0.788613 | 0.78407 | 0.776519 | 0.788282 | 0.778303 | 0.784861 | 0.782219 | 0.782002 |

| | | | | | | | | |
|---|---|---|---|---|---|---|---|---|
| 0.786211 | 0.794334 | 0.784378 | 0.788758 | 0.793863 | 0.784137 | 0.783948 | 0.792356 | 0.790375 |
| 0.785394 | 0.791592 | 0.787783 | 0.791152 | 0.793736 | 0.78488 | 0.788318 | 0.789075 | 0.788784 |
| 0.781656 | 0.794103 | 0.782226 | 0.78647 | 0.796329 | 0.786734 | 0.791037 | 0.781806 | 0.78627 |
| 0.790362 | 0.801508 | 0.784268 | 0.795183 | 0.794394 | 0.782111 | 0.781728 | 0.798401 | 0.791388 |
| 0.788759 | 0.798449 | 0.779496 | 0.789763 | 0.797642 | 0.784859 | 0.796375 | 0.791129 | 0.787031 |
| 0.784456 | 0.793292 | 0.778607 | 0.782989 | 0.795133 | 0.777287 | 0.78917 | 0.791249 | 0.786904 |
| 0.777491 | 0.794783 | 0.775613 | 0.785936 | 0.793568 | 0.77896 | 0.785228 | 0.78943 | 0.784767 |
| 0.777564 | 0.777732 | 0.785757 | 0.768628 | 0.782208 | 0.775393 | 0.781229 | 0.779587 | 0.778141 |
| 0.779576 | 0.790195 | 0.780013 | 0.785879 | 0.788998 | 0.779209 | 0.783152 | 0.78867 | 0.782501 |
| 0.776216 | 0.784188 | 0.775594 | 0.775929 | 0.783266 | 0.774328 | 0.779434 | 0.775386 | 0.778376 |
| 0.778027 | 0.795981 | 0.782069 | 0.789155 | 0.79113 | 0.781842 | 0.788845 | 0.79039 | 0.78848 |
| 0.788025 | 0.789281 | 0.785899 | 0.786682 | 0.792584 | 0.780939 | 0.788859 | 0.791945 | 0.787895 |
| 0.773804 | 0.783478 | 0.775378 | 0.774963 | 0.786709 | 0.773353 | 0.778881 | 0.778354 | 0.778743 |
| 0.775993 | 0.782848 | 0.776829 | 0.773205 | 0.78634 | 0.773212 | 0.778038 | 0.777079 | 0.778617 |
| 0.789093 | 0.792832 | 0.782912 | 0.790295 | 0.791564 | 0.782731 | 0.800583 | 0.79525 | 0.790775 |
| 0.76929 | 0.777315 | 0.768135 | 0.761327 | 0.772306 | 0.764232 | 0.769674 | 0.774313 | 0.768308 |
| 0.762602 | 0.774589 | 0.76528 | 0.763012 | 0.77357 | 0.764661 | 0.775135 | 0.771939 | 0.76528 |
| 0.760529 | 0.76993 | 0.760924 | 0.763072 | 0.767584 | 0.765292 | 0.767789 | 0.769218 | 0.759458 |
| 0.76065 | 0.774083 | 0.764419 | 0.762878 | 0.767581 | 0.760305 | 0.768311 | 0.772208 | 0.760354 |
| 0.760754 | 0.767923 | 0.765968 | 0.769571 | 0.77024 | 0.764745 | 0.773195 | 0.770523 | 0.764167 |
| 0.758609 | 0.763278 | 0.759434 | 0.755436 | 0.763673 | 0.757113 | 0.766001 | 0.765541 | 0.757563 |
| 0.756997 | 0.76822 | 0.757795 | 0.760754 | 0.759191 | 0.760333 | 0.766113 | 0.766146 | 0.759667 |
| 0.761726 | 0.774702 | 0.769339 | 0.767938 | 0.766734 | 0.768807 | 0.772045 | 0.774186 | 0.765239 |
| 0.763282 | 0.777067 | 0.762402 | 0.762733 | 0.772383 | 0.764153 | 0.771166 | 0.773784 | 0.763133 |
| 0.763658 | 0.768219 | 0.760831 | 0.765668 | 0.764911 | 0.762705 | 0.768463 | 0.767005 | 0.761852 |
| 0.757367 | 0.763659 | 0.757863 | 0.75598 | 0.76001 | 0.751693 | 0.760486 | 0.763857 | 0.75631 |
| 0.758186 | 0.767956 | 0.757045 | 0.758641 | 0.764515 | 0.757256 | 0.766815 | 0.765361 | 0.757677 |
| 0.750283 | 0.764994 | 0.757596 | 0.76132 | 0.766091 | 0.755239 | 0.766008 | 0.763063 | 0.757911 |
| 0.755516 | 0.765392 | 0.760017 | 0.760407 | 0.765644 | 0.756088 | 0.767794 | 0.76612 | 0.754858 |
| 0.753861 | 0.767461 | 0.764126 | 0.762303 | 0.766921 | 0.756963 | 0.769891 | 0.763212 | 0.760013 |
| 0.761127 | 0.768438 | 0.761692 | 0.764586 | 0.771267 | 0.762101 | 0.767901 | 0.769484 | 0.762178 |
| 0.76231 | 0.772069 | 0.762885 | 0.76595 | 0.772629 | 0.764559 | 0.77026 | 0.77013 | 0.761113 |
| 0.774362 | 0.775809 | 0.768554 | 0.770816 | 0.77145 | 0.771954 | 0.776362 | 0.778505 | 0.77207 |
| 0.769304 | 0.778241 | 0.769717 | 0.767705 | 0.776169 | 0.771506 | 0.773722 | 0.774154 | 0.770102 |
| 0.751865 | 0.764081 | 0.752442 | 0.757154 | 0.763364 | 0.756075 | 0.763991 | 0.761846 | 0.755721 |
| 0.76193 | 0.77027 | 0.760514 | 0.760683 | 0.766481 | 0.76637 | 0.765363 | 0.771145 | 0.764004 |
| 0.753693 | 0.765767 | 0.759357 | 0.7593 | 0.766866 | 0.759648 | 0.766575 | 0.765231 | 0.75484 |
| 0.757875 | 0.773054 | 0.764253 | 0.763346 | 0.765779 | 0.763705 | 0.768352 | 0.763343 | 0.757599 |
| 0.755715 | 0.762749 | 0.754711 | 0.758158 | 0.759408 | 0.752701 | 0.763615 | 0.760754 | 0.750771 |
| 0.748332 | 0.761138 | 0.748447 | 0.754299 | 0.761248 | 0.752417 | 0.760768 | 0.755493 | 0.750878 |
| 0.750773 | 0.752959 | 0.75299 | 0.750775 | 0.756899 | 0.742833 | 0.752416 | 0.758271 | 0.746517 |
| 0.74726 | 0.759951 | 0.752041 | 0.753568 | 0.754972 | 0.750371 | 0.759786 | 0.759606 | 0.752679 |
| 0.755875 | 0.761462 | 0.757319 | 0.755873 | 0.763149 | 0.751012 | 0.760963 | 0.760014 | 0.75077 |
| 0.748576 | 0.758936 | 0.752757 | 0.751047 | 0.755546 | 0.75455 | 0.752904 | 0.758559 | 0.748048 |
| 0.754545 | 0.765753 | 0.758973 | 0.758935 | 0.76022 | 0.754424 | 0.759941 | 0.761148 | 0.756852 |
| 0.751638 | 0.762597 | 0.749125 | 0.752005 | 0.760712 | 0.757766 | 0.761238 | 0.759928 | 0.749783 |

| | | | | | | | | |
|---|---|---|---|---|---|---|---|---|
| 0.758363 | 0.766972 | 0.759927 | 0.76246 | 0.76167 | 0.756527 | 0.76615 | 0.764773 | 0.758562 |
| 0.764834 | 0.774774 | 0.763892 | 0.768406 | 0.777156 | 0.765698 | 0.773966 | 0.767783 | 0.767368 |
| 0.759156 | 0.768696 | 0.760222 | 0.767038 | 0.772875 | 0.760746 | 0.766203 | 0.759474 | 0.762697 |
| 0.763686 | 0.772296 | 0.766403 | 0.766303 | 0.775614 | 0.76586 | 0.763706 | 0.765539 | 0.760077 |
| 0.773276 | 0.780644 | 0.772308 | 0.774853 | 0.776332 | 0.769015 | 0.768398 | 0.773131 | 0.774451 |
| 0.770764 | 0.783611 | 0.76874 | 0.769562 | 0.772041 | 0.768153 | 0.776308 | 0.772366 | 0.7714 |
| 0.765484 | 0.779744 | 0.768003 | 0.773998 | 0.775812 | 0.771315 | 0.76612 | 0.770954 | 0.765186 |
| 0.764272 | 0.771366 | 0.763084 | 0.76678 | 0.774869 | 0.767791 | 0.772727 | 0.770206 | 0.767281 |
| 0.771585 | 0.77469 | 0.767567 | 0.773299 | 0.774022 | 0.765259 | 0.771234 | 0.774578 | 0.763717 |
| 0.762843 | 0.773955 | 0.764383 | 0.765876 | 0.769453 | 0.766446 | 0.770507 | 0.766376 | 0.764566 |
| 0.771961 | 0.784805 | 0.771426 | 0.771809 | 0.774959 | 0.771412 | 0.778325 | 0.78184 | 0.776468 |
| 0.772735 | 0.779769 | 0.773417 | 0.774643 | 0.779309 | 0.768939 | 0.779592 | 0.771623 | 0.775259 |
| 0.759226 | 0.769664 | 0.765875 | 0.761195 | 0.765916 | 0.757919 | 0.762436 | 0.768166 | 0.760504 |
| 0.759288 | 0.769218 | 0.760862 | 0.763184 | 0.76524 | 0.760941 | 0.76368 | 0.766773 | 0.76078 |
| 0.765252 | 0.775729 | 0.766766 | 0.765265 | 0.768987 | 0.76613 | 0.767707 | 0.770616 | 0.766482 |
| 0.763307 | 0.769149 | 0.765285 | 0.761939 | 0.768821 | 0.761125 | 0.765147 | 0.76229 | 0.763808 |
| 0.762042 | 0.772676 | 0.767587 | 0.767562 | 0.769337 | 0.761277 | 0.769627 | 0.768082 | 0.760987 |
| 0.762405 | 0.774764 | 0.763154 | 0.762027 | 0.769542 | 0.763278 | 0.764151 | 0.768402 | 0.768691 |
| 0.761156 | 0.77543 | 0.771365 | 0.76413 | 0.772121 | 0.763593 | 0.769743 | 0.773692 | 0.764167 |
| 0.766197 | 0.77644 | 0.768546 | 0.765408 | 0.776168 | 0.765607 | 0.769465 | 0.774647 | 0.768557 |
| 0.76084 | 0.771187 | 0.765269 | 0.766086 | 0.76795 | 0.760778 | 0.763654 | 0.76722 | 0.762384 |
| 0.760825 | 0.765462 | 0.765908 | 0.761457 | 0.766931 | 0.764189 | 0.765515 | 0.76781 | 0.766896 |
| 0.760402 | 0.770469 | 0.76458 | 0.765932 | 0.772104 | 0.767509 | 0.764196 | 0.766119 | 0.763028 |
| 0.765223 | 0.771818 | 0.762968 | 0.766188 | 0.771937 | 0.770247 | 0.770864 | 0.767638 | 0.763623 |
| 0.761901 | 0.771341 | 0.764095 | 0.769818 | 0.773732 | 0.768917 | 0.768252 | 0.768857 | 0.766158 |
| 0.760509 | 0.768999 | 0.759308 | 0.764129 | 0.765831 | 0.758607 | 0.763087 | 0.768009 | 0.757644 |
| 0.758134 | 0.768614 | 0.761655 | 0.763758 | 0.766531 | 0.760545 | 0.762173 | 0.764677 | 0.764301 |
| 0.760892 | 0.772395 | 0.757209 | 0.759058 | 0.767609 | 0.759987 | 0.762014 | 0.762639 | 0.757982 |
| 0.76563 | 0.771826 | 0.767691 | 0.769721 | 0.774652 | 0.770931 | 0.766432 | 0.773368 | 0.76388 |
| 0.761821 | 0.772991 | 0.764651 | 0.765782 | 0.764282 | 0.766161 | 0.767921 | 0.767139 | 0.764752 |
| 0.757336 | 0.762168 | 0.756373 | 0.763875 | 0.764388 | 0.757318 | 0.758353 | 0.766016 | 0.754712 |
| 0.762858 | 0.776258 | 0.766055 | 0.767472 | 0.777824 | 0.765087 | 0.769659 | 0.769478 | 0.768719 |
| 0.765281 | 0.770005 | 0.76254 | 0.766897 | 0.767429 | 0.762517 | 0.768011 | 0.769835 | 0.763543 |
| 0.758621 | 0.771327 | 0.759079 | 0.759163 | 0.765954 | 0.757281 | 0.758691 | 0.763822 | 0.758889 |
| 0.763044 | 0.76934 | 0.759966 | 0.761017 | 0.768302 | 0.76516 | 0.764492 | 0.76341 | 0.759638 |
| 0.761008 | 0.768816 | 0.756344 | 0.762107 | 0.770032 | 0.76314 | 0.763534 | 0.765453 | 0.76014 |
| 0.76549 | 0.774312 | 0.763754 | 0.769962 | 0.776318 | 0.769599 | 0.767373 | 0.770835 | 0.765792 |
| 0.759336 | 0.772153 | 0.763012 | 0.764852 | 0.771682 | 0.764262 | 0.768721 | 0.770471 | 0.764469 |
| 0.763533 | 0.772831 | 0.762657 | 0.763299 | 0.770792 | 0.763621 | 0.768521 | 0.769193 | 0.761151 |
| 0.762984 | 0.768521 | 0.764949 | 0.768322 | 0.767222 | 0.767161 | 0.764207 | 0.76788 | 0.765784 |
| 0.7668 | 0.772344 | 0.769529 | 0.765048 | 0.772742 | 0.767064 | 0.765187 | 0.770711 | 0.765234 |
| 0.759202 | 0.776579 | 0.767037 | 0.7642 | 0.775337 | 0.766315 | 0.768799 | 0.76716 | 0.764378 |
| 0.76082 | 0.775514 | 0.769573 | 0.766204 | 0.775657 | 0.767745 | 0.771576 | 0.77066 | 0.767649 |
| 0.759777 | 0.771868 | 0.762702 | 0.762479 | 0.767951 | 0.767644 | 0.759645 | 0.767908 | 0.762447 |
| 0.763837 | 0.773479 | 0.76562 | 0.768442 | 0.768071 | 0.767832 | 0.768171 | 0.769726 | 0.764396 |
| 0.758972 | 0.771056 | 0.761188 | 0.762748 | 0.76756 | 0.763725 | 0.766129 | 0.768203 | 0.763717 |

| | | | | | | | | |
|---|---|---|---|---|---|---|---|---|
| 0.748458 | 0.763051 | 0.750213 | 0.757678 | 0.760526 | 0.750142 | 0.750825 | 0.76118 | 0.74976 |
| 0.742146 | 0.756247 | 0.746059 | 0.745242 | 0.751091 | 0.744481 | 0.741435 | 0.750555 | 0.742759 |
| 0.756828 | 0.772666 | 0.759892 | 0.761107 | 0.765678 | 0.762955 | 0.762885 | 0.767054 | 0.756599 |
| 0.761084 | 0.773904 | 0.769345 | 0.762307 | 0.769738 | 0.762691 | 0.766509 | 0.774054 | 0.765001 |
| 0.754069 | 0.763398 | 0.762048 | 0.757851 | 0.760692 | 0.758747 | 0.755764 | 0.762293 | 0.758266 |
| 0.756773 | 0.769028 | 0.758598 | 0.759075 | 0.770017 | 0.762373 | 0.758672 | 0.764181 | 0.756337 |
| 0.762371 | 0.769928 | 0.761913 | 0.760163 | 0.770808 | 0.760514 | 0.769397 | 0.765412 | 0.758645 |
| 0.753195 | 0.766761 | 0.756048 | 0.753936 | 0.765003 | 0.756805 | 0.758367 | 0.75987 | 0.750539 |
| 0.767649 | 0.771849 | 0.765157 | 0.767387 | 0.772218 | 0.762033 | 0.766486 | 0.776349 | 0.765879 |
| 0.761022 | 0.768841 | 0.768379 | 0.769268 | 0.770871 | 0.761524 | 0.765455 | 0.771248 | 0.762966 |
| 0.76714 | 0.774657 | 0.769053 | 0.763338 | 0.772371 | 0.764865 | 0.767451 | 0.773485 | 0.767536 |
| 0.767877 | 0.770876 | 0.768718 | 0.768137 | 0.765257 | 0.763931 | 0.76051 | 0.771539 | 0.762731 |
| 0.769211 | 0.771844 | 0.769205 | 0.769005 | 0.777238 | 0.773122 | 0.768546 | 0.771648 | 0.766145 |
| 0.766305 | 0.775191 | 0.767223 | 0.771334 | 0.76977 | 0.769231 | 0.763522 | 0.770857 | 0.765121 |
| 0.765508 | 0.766587 | 0.761037 | 0.760228 | 0.76477 | 0.758477 | 0.76073 | 0.771159 | 0.763728 |
| 0.760239 | 0.765012 | 0.760949 | 0.761396 | 0.766575 | 0.762523 | 0.758841 | 0.767266 | 0.758824 |
| 0.759067 | 0.77382 | 0.765352 | 0.765879 | 0.77254 | 0.76146 | 0.766449 | 0.767056 | 0.758615 |
| 0.75529 | 0.767535 | 0.758678 | 0.763846 | 0.768901 | 0.758701 | 0.762157 | 0.764754 | 0.757044 |
| 0.759063 | 0.767347 | 0.761862 | 0.755318 | 0.7652 | 0.759688 | 0.763485 | 0.764758 | 0.759534 |
| 0.752542 | 0.768341 | 0.760805 | 0.75962 | 0.76514 | 0.755888 | 0.766087 | 0.765578 | 0.760291 |
| 0.762247 | 0.768768 | 0.762986 | 0.76431 | 0.76744 | 0.76221 | 0.766322 | 0.77029 | 0.761633 |
| 0.74395 | 0.760968 | 0.752759 | 0.748858 | 0.758432 | 0.754387 | 0.753383 | 0.755981 | 0.746145 |
| 0.754527 | 0.769413 | 0.758291 | 0.760434 | 0.762076 | 0.759316 | 0.754819 | 0.763433 | 0.754097 |
| 0.761043 | 0.772626 | 0.762664 | 0.765215 | 0.770875 | 0.763938 | 0.761209 | 0.770397 | 0.761796 |
| 0.75811 | 0.771636 | 0.760906 | 0.765191 | 0.768312 | 0.765529 | 0.763439 | 0.765228 | 0.763939 |
| 0.758294 | 0.768618 | 0.759292 | 0.757374 | 0.765815 | 0.762635 | 0.755537 | 0.76475 | 0.754843 |
| 0.751301 | 0.764825 | 0.754473 | 0.75739 | 0.764956 | 0.758071 | 0.762322 | 0.759936 | 0.755175 |
| 0.752174 | 0.765439 | 0.762052 | 0.758437 | 0.765449 | 0.757169 | 0.759884 | 0.762391 | 0.763013 |
| 0.751603 | 0.765722 | 0.7546 | 0.756708 | 0.759629 | 0.758333 | 0.75709 | 0.76422 | 0.758584 |
| 0.691276 | 0.701381 | 0.699574 | 0.698239 | 0.698228 | 0.693462 | 0.69655 | 0.694984 | 0.693321 |
| 0.705231 | 0.713692 | 0.708959 | 0.711497 | 0.711336 | 0.707731 | 0.70838 | 0.70703 | 0.702843 |
| 0.702875 | 0.711835 | 0.706725 | 0.706916 | 0.709739 | 0.701271 | 0.704399 | 0.706979 | 0.702467 |
| 0.695644 | 0.704432 | 0.700178 | 0.698854 | 0.703236 | 0.694165 | 0.700372 | 0.700704 | 0.693797 |
| 0.702498 | 0.71405 | 0.706323 | 0.709639 | 0.715362 | 0.707441 | 0.70987 | 0.709428 | 0.70515 |
| 0.756281 | 0.767583 | 0.75627 | 0.760524 | 0.768726 | 0.758451 | 0.760722 | 0.76272 | 0.760477 |
| 0.760911 | 0.767103 | 0.761689 | 0.758771 | 0.7665 | 0.759753 | 0.7609 | 0.760709 | 0.760592 |
| 0.764782 | 0.773779 | 0.764302 | 0.759554 | 0.766667 | 0.764215 | 0.766625 | 0.76973 | 0.760825 |
| 0.761408 | 0.772768 | 0.763044 | 0.76533 | 0.775861 | 0.767614 | 0.766669 | 0.768418 | 0.765455 |
| 0.765795 | 0.7646 | 0.761145 | 0.7596 | 0.767879 | 0.759179 | 0.763123 | 0.763744 | 0.761824 |
| 0.766101 | 0.769609 | 0.762055 | 0.757295 | 0.768304 | 0.764258 | 0.762393 | 0.765796 | 0.764101 |
| 0.763479 | 0.774489 | 0.76395 | 0.765216 | 0.772299 | 0.762259 | 0.766669 | 0.767552 | 0.76289 |
| 0.757244 | 0.767746 | 0.758739 | 0.75845 | 0.766759 | 0.760202 | 0.761813 | 0.75877 | 0.766879 |
| 0.754475 | 0.76637 | 0.755672 | 0.754906 | 0.764281 | 0.753535 | 0.75905 | 0.758391 | 0.75555 |
| 0.755785 | 0.765644 | 0.758382 | 0.756166 | 0.759594 | 0.754692 | 0.764429 | 0.761762 | 0.757856 |
| 0.756217 | 0.765975 | 0.756182 | 0.75652 | 0.760742 | 0.758408 | 0.76 | 0.76051 | 0.75756 |
| 0.767012 | 0.767836 | 0.762773 | 0.756326 | 0.771343 | 0.763252 | 0.766683 | 0.764183 | 0.764938 |

| | | | | | | | | |
|---|---|---|---|---|---|---|---|---|
| 0.760403 | 0.7705 | 0.76322 | 0.763589 | 0.771383 | 0.761969 | 0.768561 | 0.772651 | 0.764146 |
| 0.758335 | 0.764633 | 0.756768 | 0.758736 | 0.769174 | 0.759407 | 0.762098 | 0.764711 | 0.758772 |
| 0.759955 | 0.767621 | 0.765398 | 0.760394 | 0.769448 | 0.761722 | 0.763476 | 0.763608 | 0.761378 |
| 0.754139 | 0.76351 | 0.75903 | 0.754553 | 0.760105 | 0.757077 | 0.762043 | 0.764275 | 0.758221 |
| 0.756518 | 0.763011 | 0.759454 | 0.757977 | 0.763151 | 0.756652 | 0.761081 | 0.75701 | 0.758657 |
| 0.757883 | 0.763071 | 0.756748 | 0.755116 | 0.764451 | 0.756982 | 0.763095 | 0.760047 | 0.7609 |
| 0.759121 | 0.767175 | 0.763122 | 0.758727 | 0.76718 | 0.760851 | 0.766373 | 0.763332 | 0.759707 |
| 0.759367 | 0.765317 | 0.759346 | 0.753175 | 0.765596 | 0.756912 | 0.76 | 0.764449 | 0.757534 |
| 0.758394 | 0.765973 | 0.754813 | 0.75745 | 0.765017 | 0.759036 | 0.759283 | 0.754777 | 0.757824 |
| 0.757043 | 0.766837 | 0.755151 | 0.754333 | 0.769461 | 0.761269 | 0.759349 | 0.761913 | 0.757094 |
| 0.755966 | 0.765788 | 0.761427 | 0.75548 | 0.765271 | 0.75699 | 0.763626 | 0.761147 | 0.75617 |
| 0.757134 | 0.766853 | 0.759683 | 0.756077 | 0.766446 | 0.759046 | 0.764909 | 0.763737 | 0.757123 |
| 0.748647 | 0.754074 | 0.755404 | 0.751975 | 0.757764 | 0.752511 | 0.752818 | 0.751833 | 0.747496 |
| 0.754884 | 0.758948 | 0.758877 | 0.754474 | 0.76345 | 0.755056 | 0.754082 | 0.75493 | 0.755306 |
| 0.762235 | 0.770023 | 0.763695 | 0.761316 | 0.768241 | 0.762652 | 0.759116 | 0.768656 | 0.756564 |
| 0.764559 | 0.775525 | 0.764437 | 0.769023 | 0.771093 | 0.769103 | 0.768459 | 0.772386 | 0.76676 |
| 0.76218 | 0.770351 | 0.762178 | 0.766454 | 0.766523 | 0.76436 | 0.762514 | 0.765899 | 0.761288 |
| 0.761806 | 0.775791 | 0.765957 | 0.767322 | 0.773513 | 0.76997 | 0.763136 | 0.769351 | 0.764771 |
| 0.761888 | 0.773234 | 0.764181 | 0.766111 | 0.770229 | 0.767889 | 0.765533 | 0.769722 | 0.761534 |
| 0.759361 | 0.768801 | 0.759175 | 0.765187 | 0.76537 | 0.763827 | 0.763567 | 0.767083 | 0.75848 |
| 0.756709 | 0.765056 | 0.765968 | 0.759597 | 0.767892 | 0.760924 | 0.766583 | 0.761836 | 0.761942 |
| 0.758665 | 0.767038 | 0.758074 | 0.760604 | 0.764896 | 0.759364 | 0.757825 | 0.768176 | 0.755601 |
| 0.764059 | 0.771908 | 0.762593 | 0.769562 | 0.771101 | 0.765584 | 0.766525 | 0.768127 | 0.763777 |
| 0.755229 | 0.764201 | 0.760447 | 0.750954 | 0.763635 | 0.758864 | 0.76111 | 0.756894 | 0.760946 |
| 0.764749 | 0.772775 | 0.759961 | 0.770522 | 0.770361 | 0.765009 | 0.768999 | 0.769933 | 0.764281 |
| 0.703504 | 0.710235 | 0.702744 | 0.709252 | 0.70682 | 0.705391 | 0.705256 | 0.703404 | 0.700867 |
| 0.760984 | 0.773256 | 0.760441 | 0.762447 | 0.773207 | 0.765359 | 0.76487 | 0.765793 | 0.760436 |
| 0.768518 | 0.774302 | 0.761889 | 0.764229 | 0.77538 | 0.763128 | 0.765339 | 0.764399 | 0.763382 |
| 0.765873 | 0.773244 | 0.764927 | 0.768769 | 0.774863 | 0.767711 | 0.771901 | 0.764735 | 0.760787 |
| 0.763759 | 0.772866 | 0.767395 | 0.762188 | 0.773596 | 0.765554 | 0.769595 | 0.763363 | 0.764571 |
| 0.763623 | 0.771267 | 0.760301 | 0.757581 | 0.76699 | 0.760017 | 0.765133 | 0.763366 | 0.758693 |
| 0.76018 | 0.771475 | 0.763318 | 0.762425 | 0.769722 | 0.767112 | 0.761503 | 0.766607 | 0.762963 |
| 0.760993 | 0.767446 | 0.760169 | 0.763846 | 0.767724 | 0.761905 | 0.76626 | 0.766427 | 0.760832 |
| 0.760043 | 0.768604 | 0.760461 | 0.757694 | 0.772763 | 0.760898 | 0.764996 | 0.764579 | 0.760018 |
| 0.758031 | 0.762719 | 0.756604 | 0.758031 | 0.772174 | 0.753621 | 0.762269 | 0.759664 | 0.753133 |
| 0.757573 | 0.768405 | 0.755918 | 0.7626 | 0.77397 | 0.760335 | 0.762753 | 0.759692 | 0.755854 |
| 0.761705 | 0.771562 | 0.76308 | 0.762702 | 0.774907 | 0.76903 | 0.768538 | 0.768709 | 0.762101 |
| 0.759421 | 0.773257 | 0.763955 | 0.766699 | 0.771246 | 0.763953 | 0.768482 | 0.772923 | 0.766323 |
| 0.760349 | 0.77201 | 0.76477 | 0.767388 | 0.774925 | 0.767917 | 0.769701 | 0.770482 | 0.771132 |
| 0.757983 | 0.767499 | 0.76316 | 0.763534 | 0.769237 | 0.760316 | 0.766287 | 0.765164 | 0.760446 |
| 0.756925 | 0.771353 | 0.758466 | 0.76387 | 0.770957 | 0.760823 | 0.76422 | 0.766039 | 0.759339 |
| 0.764316 | 0.771609 | 0.768073 | 0.767839 | 0.776631 | 0.763906 | 0.769224 | 0.7673 | 0.768461 |
| 0.76594 | 0.76973 | 0.757675 | 0.763348 | 0.766624 | 0.760789 | 0.765895 | 0.761295 | 0.761457 |
| 0.758412 | 0.7627 | 0.755857 | 0.757361 | 0.766425 | 0.756622 | 0.757935 | 0.763201 | 0.757586 |
| 0.766641 | 0.774544 | 0.76265 | 0.764235 | 0.774952 | 0.76474 | 0.771829 | 0.766958 | 0.764462 |
| 0.763627 | 0.769074 | 0.759726 | 0.762493 | 0.768858 | 0.761736 | 0.764793 | 0.76404 | 0.762831 |

| | | | | | | | | |
|---|---|---|---|---|---|---|---|---|
| 0.759588 | 0.76955 | 0.765382 | 0.758654 | 0.767867 | 0.762065 | 0.764248 | 0.767962 | 0.759586 |
| 0.763734 | 0.770407 | 0.759875 | 0.758412 | 0.767358 | 0.762007 | 0.763805 | 0.76732 | 0.760464 |
| 0.757563 | 0.765436 | 0.76026 | 0.756969 | 0.765276 | 0.754558 | 0.763053 | 0.760681 | 0.754632 |
| 0.721098 | 0.722104 | 0.726686 | 0.723915 | 0.73191 | 0.714286 | 0.715404 | 0.733429 | 0.713525 |
| 0.762805 | 0.767701 | 0.755257 | 0.756539 | 0.762741 | 0.757405 | 0.761975 | 0.763626 | 0.758401 |
| 0.760718 | 0.770332 | 0.757139 | 0.757825 | 0.760626 | 0.760559 | 0.760198 | 0.760633 | 0.757192 |
| 0.765044 | 0.77064 | 0.756474 | 0.7562 | 0.766519 | 0.760628 | 0.75907 | 0.760933 | 0.756998 |
| 0.766218 | 0.767342 | 0.760555 | 0.759603 | 0.765628 | 0.7652 | 0.764022 | 0.762495 | 0.760279 |
| 0.764171 | 0.772853 | 0.763491 | 0.769712 | 0.769489 | 0.766828 | 0.766234 | 0.766051 | 0.762699 |
| 0.759178 | 0.771741 | 0.761464 | 0.763109 | 0.767347 | 0.76158 | 0.765994 | 0.763321 | 0.755675 |
| 0.758683 | 0.770137 | 0.758957 | 0.755548 | 0.76037 | 0.761378 | 0.762211 | 0.763736 | 0.753637 |
| 0.762585 | 0.76615 | 0.762142 | 0.760767 | 0.766283 | 0.768854 | 0.762117 | 0.767476 | 0.754047 |
| 0.758627 | 0.773076 | 0.763561 | 0.761334 | 0.763483 | 0.764366 | 0.759661 | 0.76372 | 0.75782 |
| 0.764098 | 0.768668 | 0.763621 | 0.761261 | 0.762695 | 0.76693 | 0.763378 | 0.765359 | 0.756852 |
| 0.762993 | 0.768993 | 0.764171 | 0.764362 | 0.766956 | 0.763364 | 0.76303 | 0.766563 | 0.763391 |
| 0.759678 | 0.761947 | 0.758024 | 0.757824 | 0.764012 | 0.762483 | 0.76344 | 0.764928 | 0.756814 |
| 0.762576 | 0.768901 | 0.763745 | 0.766618 | 0.770063 | 0.762515 | 0.763463 | 0.767747 | 0.763246 |
| 0.763117 | 0.770385 | 0.763429 | 0.7617 | 0.768589 | 0.766068 | 0.763035 | 0.766697 | 0.75913 |
| 0.757068 | 0.763536 | 0.759464 | 0.757306 | 0.764171 | 0.758306 | 0.758756 | 0.76408 | 0.755184 |
| 0.761115 | 0.761773 | 0.761873 | 0.757346 | 0.765269 | 0.76347 | 0.766442 | 0.766874 | 0.763327 |
| 0.767226 | 0.772776 | 0.76399 | 0.765963 | 0.769533 | 0.768609 | 0.762972 | 0.766233 | 0.766376 |
| 0.764097 | 0.773109 | 0.763489 | 0.767434 | 0.772291 | 0.761286 | 0.762687 | 0.763457 | 0.761833 |
| 0.681512 | 0.682089 | 0.68372 | 0.677373 | 0.687776 | 0.671561 | 0.674359 | 0.678806 | 0.678532 |
| 0.656573 | 0.660642 | 0.649587 | 0.660266 | 0.661955 | 0.647492 | 0.648222 | 0.658171 | 0.653992 |
| 0.673165 | 0.678649 | 0.675904 | 0.672871 | 0.67675 | 0.675906 | 0.674837 | 0.681393 | 0.673804 |
| 0.663085 | 0.661791 | 0.654161 | 0.656423 | 0.660817 | 0.651718 | 0.657925 | 0.661859 | 0.658565 |
| 0.724804 | 0.738501 | 0.724869 | 0.72019 | 0.735014 | 0.724375 | 0.723537 | 0.728102 | 0.722606 |
| 0.720125 | 0.728862 | 0.718135 | 0.7119 | 0.722137 | 0.720721 | 0.726577 | 0.725014 | 0.714478 |
| 0.729319 | 0.722462 | 0.717531 | 0.720601 | 0.728094 | 0.72026 | 0.718542 | 0.724659 | 0.715646 |
| 0.737572 | 0.73765 | 0.728438 | 0.721993 | 0.742677 | 0.729188 | 0.732959 | 0.734238 | 0.727924 |
| 0.719188 | 0.722718 | 0.706162 | 0.703562 | 0.717849 | 0.70277 | 0.711091 | 0.714416 | 0.705733 |
| 0.70523 | 0.710898 | 0.704309 | 0.704941 | 0.714245 | 0.70048 | 0.703631 | 0.719238 | 0.697243 |
| 0.71791 | 0.720627 | 0.715296 | 0.692441 | 0.713823 | 0.709535 | 0.715543 | 0.709123 | 0.708482 |
| 0.659112 | 0.665299 | 0.661881 | 0.660876 | 0.662491 | 0.650802 | 0.658266 | 0.65611 | 0.653116 |
| 0.758392 | 0.76532 | 0.757628 | 0.754899 | 0.762118 | 0.756016 | 0.753627 | 0.758668 | 0.758393 |
| 0.740541 | 0.746146 | 0.736511 | 0.729298 | 0.7415 | 0.72915 | 0.734279 | 0.740922 | 0.738264 |
| 0.735951 | 0.742965 | 0.728094 | 0.732071 | 0.733008 | 0.733523 | 0.733072 | 0.730541 | 0.724759 |
| 0.723364 | 0.721657 | 0.714233 | 0.709254 | 0.719657 | 0.709487 | 0.716261 | 0.725093 | 0.717549 |
| 0.736312 | 0.740732 | 0.733746 | 0.728575 | 0.741066 | 0.738584 | 0.738571 | 0.738471 | 0.731044 |
| 0.729963 | 0.735031 | 0.726748 | 0.729212 | 0.735839 | 0.727168 | 0.729726 | 0.727996 | 0.726485 |
| 0.720588 | 0.726595 | 0.721082 | 0.72 | 0.725693 | 0.722054 | 0.726925 | 0.719549 | 0.72255 |
| 0.734375 | 0.731206 | 0.724359 | 0.727216 | 0.733103 | 0.724653 | 0.729831 | 0.730589 | 0.731067 |
| 0.724488 | 0.727338 | 0.723324 | 0.725148 | 0.730205 | 0.717246 | 0.722842 | 0.723715 | 0.716209 |
| 0.71865 | 0.724618 | 0.720542 | 0.706534 | 0.715907 | 0.708698 | 0.716974 | 0.720134 | 0.718349 |
| 0.718419 | 0.727881 | 0.714395 | 0.712652 | 0.728208 | 0.710211 | 0.716463 | 0.720082 | 0.720978 |
| 0.697429 | 0.727303 | 0.694652 | 0.70026 | 0.710958 | 0.708706 | 0.715702 | 0.708017 | 0.700197 |

| | | | | | | | | |
|---|---|---|---|---|---|---|---|---|
| 0.745662 | 0.749459 | 0.744528 | 0.742331 | 0.750124 | 0.743138 | 0.744455 | 0.749205 | 0.747453 |
| 0.72285 | 0.719833 | 0.713966 | 0.716989 | 0.722682 | 0.72507 | 0.718829 | 0.726699 | 0.715899 |
| 0.723225 | 0.729528 | 0.721493 | 0.714899 | 0.728347 | 0.726766 | 0.722057 | 0.72715 | 0.728386 |
| 0.729509 | 0.735249 | 0.723776 | 0.711936 | 0.728289 | 0.727273 | 0.726464 | 0.734154 | 0.717674 |
| 0.726213 | 0.726478 | 0.7164 | 0.708543 | 0.725932 | 0.715096 | 0.721311 | 0.720798 | 0.722845 |
| 0.72658 | 0.715483 | 0.706863 | 0.718076 | 0.723821 | 0.714712 | 0.715171 | 0.715451 | 0.720371 |
| 0.721444 | 0.721077 | 0.701607 | 0.703178 | 0.717756 | 0.706689 | 0.712052 | 0.713834 | 0.718086 |
| 0.711642 | 0.706588 | 0.705193 | 0.696523 | 0.710092 | 0.707399 | 0.710681 | 0.71043 | 0.70729 |
| 0.722936 | 0.719085 | 0.700334 | 0.703003 | 0.709218 | 0.710497 | 0.71076 | 0.71614 | 0.717324 |
| 0.748534 | 0.756823 | 0.750025 | 0.744413 | 0.757864 | 0.749803 | 0.748161 | 0.75539 | 0.747464 |
| 0.744416 | 0.755018 | 0.741845 | 0.74129 | 0.752697 | 0.742918 | 0.747154 | 0.749523 | 0.748824 |
| 0.735832 | 0.747573 | 0.734873 | 0.738429 | 0.743905 | 0.735803 | 0.739529 | 0.742178 | 0.737606 |
| 0.730836 | 0.742556 | 0.729101 | 0.728921 | 0.736727 | 0.731492 | 0.732415 | 0.732664 | 0.731075 |
| 0.73797 | 0.747652 | 0.736816 | 0.737059 | 0.739104 | 0.740327 | 0.733512 | 0.740763 | 0.739302 |
| 0.730173 | 0.742331 | 0.735833 | 0.731562 | 0.738723 | 0.731455 | 0.734003 | 0.734151 | 0.732978 |

| Wild_plot_ | Wild_plot_ | Wild_plot_ | Wild_plot_ | Wild_plot_ | Wild_plot_ | Wild_plot_ | Wild_plot_ | Wild_plot_ |
|---|---|---|---|---|---|---|---|---|
| 0.709534 | 0.65758 | 0.653972 | 0.710722 | 0.712104 | 0.642305 | 0.7088 | 0.700284 | 0.705238 |
| 0.715172 | 0.663723 | 0.66489 | 0.719656 | 0.718223 | 0.649805 | 0.715795 | 0.706582 | 0.712051 |
| 0.704748 | 0.650548 | 0.657914 | 0.708639 | 0.708686 | 0.636081 | 0.703698 | 0.696668 | 0.702658 |
| 0.713271 | 0.659317 | 0.666059 | 0.716121 | 0.720353 | 0.654903 | 0.716433 | 0.705234 | 0.710462 |
| 0.710963 | 0.657128 | 0.666849 | 0.712308 | 0.717881 | 0.645351 | 0.712785 | 0.699064 | 0.707375 |
| 0.716144 | 0.667507 | 0.666667 | 0.722147 | 0.724703 | 0.653988 | 0.721275 | 0.709403 | 0.714172 |
| 0.761544 | 0.72538 | 0.727136 | 0.766686 | 0.772623 | 0.713438 | 0.763569 | 0.761532 | 0.765544 |
| 0.759329 | 0.716467 | 0.724864 | 0.760673 | 0.768594 | 0.715881 | 0.7633 | 0.756965 | 0.760408 |
| 0.710266 | 0.664467 | 0.667031 | 0.717694 | 0.714457 | 0.651257 | 0.715578 | 0.705719 | 0.712803 |
| 0.71172 | 0.652489 | 0.659675 | 0.709504 | 0.712609 | 0.645914 | 0.711023 | 0.703119 | 0.709908 |
| 0.709636 | 0.658849 | 0.669408 | 0.710938 | 0.716687 | 0.647331 | 0.714377 | 0.702468 | 0.710358 |
| 0.703821 | 0.650708 | 0.655574 | 0.708109 | 0.708598 | 0.63789 | 0.707995 | 0.696528 | 0.702892 |
| 0.715785 | 0.661877 | 0.665147 | 0.717215 | 0.716769 | 0.654204 | 0.716055 | 0.706591 | 0.714058 |
| 0.722368 | 0.675199 | 0.67441 | 0.7251 | 0.728782 | 0.66263 | 0.721005 | 0.715988 | 0.718436 |
| 0.732639 | 0.686118 | 0.687192 | 0.741956 | 0.743066 | 0.681852 | 0.735535 | 0.72854 | 0.734664 |
| 0.717965 | 0.664014 | 0.670088 | 0.720267 | 0.718524 | 0.65994 | 0.718358 | 0.7086 | 0.712763 |
| 0.716448 | 0.659963 | 0.665814 | 0.719123 | 0.719585 | 0.657207 | 0.720436 | 0.710453 | 0.717437 |
| 0.738872 | 0.69552 | 0.696533 | 0.748087 | 0.754237 | 0.687952 | 0.741048 | 0.736909 | 0.74311 |
| 0.708768 | 0.649874 | 0.658862 | 0.712025 | 0.707858 | 0.63851 | 0.70834 | 0.700175 | 0.706093 |
| 0.713593 | 0.6549 | 0.665754 | 0.714592 | 0.714035 | 0.64379 | 0.711164 | 0.704769 | 0.708628 |
| 0.711844 | 0.65175 | 0.658847 | 0.710593 | 0.711572 | 0.6432 | 0.709898 | 0.698811 | 0.704867 |
| 0.711392 | 0.657278 | 0.662406 | 0.713651 | 0.715949 | 0.643425 | 0.711159 | 0.70049 | 0.707239 |
| 0.716915 | 0.664763 | 0.668852 | 0.723241 | 0.726567 | 0.649853 | 0.720013 | 0.707599 | 0.715459 |
| 0.715215 | 0.661479 | 0.667155 | 0.715907 | 0.719185 | 0.648694 | 0.716181 | 0.703258 | 0.713181 |
| 0.71555 | 0.664974 | 0.669844 | 0.719559 | 0.719158 | 0.650298 | 0.714962 | 0.707498 | 0.715111 |
| 0.71217 | 0.657555 | 0.658651 | 0.716848 | 0.714274 | 0.648514 | 0.714115 | 0.705559 | 0.710505 |
| 0.705534 | 0.64636 | 0.659455 | 0.709274 | 0.706606 | 0.64381 | 0.709464 | 0.696102 | 0.703521 |
| 0.717448 | 0.664027 | 0.659378 | 0.716329 | 0.71768 | 0.650902 | 0.715716 | 0.704596 | 0.711672 |
| 0.714865 | 0.666441 | 0.664552 | 0.71626 | 0.719866 | 0.648838 | 0.713875 | 0.703179 | 0.709942 |
| 0.711681 | 0.667294 | 0.667402 | 0.713512 | 0.712082 | 0.645845 | 0.715905 | 0.703242 | 0.712477 |
| 0.710144 | 0.658693 | 0.665577 | 0.716236 | 0.716125 | 0.656297 | 0.716121 | 0.704551 | 0.710425 |
| 0.708462 | 0.65287 | 0.658138 | 0.712793 | 0.71086 | 0.647417 | 0.713627 | 0.700196 | 0.706809 |
| 0.713054 | 0.655743 | 0.666062 | 0.711529 | 0.712588 | 0.650267 | 0.709326 | 0.701517 | 0.708333 |
| 0.712501 | 0.655276 | 0.664078 | 0.71502 | 0.718208 | 0.648962 | 0.716996 | 0.704585 | 0.71315 |
| 0.705845 | 0.650479 | 0.657995 | 0.710789 | 0.708806 | 0.647681 | 0.708778 | 0.695741 | 0.701397 |
| 0.71424 | 0.664289 | 0.663312 | 0.716259 | 0.713097 | 0.647092 | 0.712932 | 0.702293 | 0.709159 |
| 0.713746 | 0.662855 | 0.662157 | 0.717945 | 0.719291 | 0.654294 | 0.714874 | 0.704529 | 0.711599 |
| 0.712178 | 0.663712 | 0.668452 | 0.716877 | 0.719981 | 0.648432 | 0.716297 | 0.705892 | 0.711299 |
| 0.719936 | 0.660969 | 0.667882 | 0.72091 | 0.721872 | 0.658129 | 0.722712 | 0.714558 | 0.716986 |

| | | | | | | | | |
|---|---|---|---|---|---|---|---|---|
| 0.746365 | 0.697892 | 0.700714 | 0.747405 | 0.756198 | 0.696362 | 0.74593 | 0.743097 | 0.750886 |
| 0.704897 | 0.641707 | 0.648795 | 0.711569 | 0.707362 | 0.634897 | 0.706835 | 0.695974 | 0.701282 |
| 0.707039 | 0.656165 | 0.659074 | 0.708673 | 0.710315 | 0.641764 | 0.709226 | 0.694215 | 0.703018 |
| 0.71432 | 0.664764 | 0.670663 | 0.721427 | 0.719711 | 0.65223 | 0.713878 | 0.705457 | 0.711941 |
| 0.712813 | 0.654072 | 0.662668 | 0.714229 | 0.712709 | 0.644531 | 0.712531 | 0.700228 | 0.708085 |
| 0.715678 | 0.663669 | 0.670398 | 0.716281 | 0.718785 | 0.64755 | 0.716236 | 0.7029 | 0.708227 |
| 0.709624 | 0.647784 | 0.654076 | 0.713743 | 0.712517 | 0.643699 | 0.708919 | 0.698334 | 0.708948 |
| 0.713498 | 0.659333 | 0.667642 | 0.715049 | 0.715385 | 0.650847 | 0.713987 | 0.701697 | 0.708144 |
| 0.714343 | 0.663655 | 0.665564 | 0.716565 | 0.718356 | 0.650037 | 0.715644 | 0.703654 | 0.712302 |
| 0.718222 | 0.664315 | 0.664305 | 0.712334 | 0.714965 | 0.6565 | 0.715492 | 0.70055 | 0.711676 |
| 0.700761 | 0.652005 | 0.659216 | 0.707876 | 0.708027 | 0.633717 | 0.705634 | 0.695319 | 0.701696 |
| 0.704012 | 0.645253 | 0.657504 | 0.709357 | 0.711177 | 0.635041 | 0.707635 | 0.695216 | 0.70263 |
| 0.712994 | 0.651746 | 0.659908 | 0.714686 | 0.713804 | 0.639993 | 0.711454 | 0.701224 | 0.70418 |
| 0.70641 | 0.652299 | 0.655066 | 0.710617 | 0.708902 | 0.636953 | 0.707916 | 0.697875 | 0.706616 |
| 0.708347 | 0.655785 | 0.658505 | 0.708004 | 0.708206 | 0.64064 | 0.708656 | 0.698025 | 0.704259 |
| 0.702575 | 0.655308 | 0.653655 | 0.710378 | 0.709113 | 0.63905 | 0.703393 | 0.697566 | 0.699236 |
| 0.70608 | 0.650179 | 0.654995 | 0.710123 | 0.70893 | 0.636584 | 0.708921 | 0.69786 | 0.701754 |
| 0.712645 | 0.655045 | 0.659629 | 0.713766 | 0.714809 | 0.64238 | 0.714741 | 0.702839 | 0.710748 |
| 0.718571 | 0.663055 | 0.66169 | 0.71432 | 0.715946 | 0.648124 | 0.718292 | 0.705664 | 0.7095 |
| 0.715865 | 0.650397 | 0.656619 | 0.712587 | 0.709396 | 0.644118 | 0.711463 | 0.698149 | 0.70616 |
| 0.702662 | 0.654809 | 0.651677 | 0.707872 | 0.708506 | 0.632105 | 0.706216 | 0.694273 | 0.700717 |
| 0.707864 | 0.654146 | 0.657842 | 0.710114 | 0.708606 | 0.636464 | 0.708536 | 0.698307 | 0.703801 |
| 0.710244 | 0.649269 | 0.658545 | 0.712888 | 0.712625 | 0.639678 | 0.710361 | 0.699623 | 0.708412 |
| 0.71908 | 0.6652 | 0.669641 | 0.721779 | 0.723693 | 0.653667 | 0.722634 | 0.710011 | 0.717936 |
| 0.721931 | 0.673281 | 0.680743 | 0.727352 | 0.72832 | 0.66333 | 0.725859 | 0.713178 | 0.724358 |
| 0.758745 | 0.716693 | 0.725649 | 0.761698 | 0.76951 | 0.717949 | 0.763783 | 0.756175 | 0.761885 |
| 0.764851 | 0.728206 | 0.727632 | 0.76817 | 0.768855 | 0.730313 | 0.775042 | 0.767017 | 0.76644 |
| 0.764566 | 0.730578 | 0.731758 | 0.770447 | 0.772897 | 0.725352 | 0.77218 | 0.766875 | 0.771578 |
| 0.70837 | 0.658405 | 0.66134 | 0.712544 | 0.710344 | 0.644751 | 0.70757 | 0.69947 | 0.703051 |
| 0.709965 | 0.65529 | 0.657856 | 0.712428 | 0.710708 | 0.642964 | 0.708104 | 0.699821 | 0.706471 |
| 0.716003 | 0.662504 | 0.662921 | 0.722409 | 0.719686 | 0.653065 | 0.720794 | 0.708069 | 0.716872 |
| 0.717563 | 0.666106 | 0.669575 | 0.721157 | 0.720742 | 0.650656 | 0.71572 | 0.706118 | 0.712741 |
| 0.711418 | 0.658073 | 0.659683 | 0.709964 | 0.711258 | 0.646504 | 0.70619 | 0.70048 | 0.706826 |
| 0.711491 | 0.659603 | 0.662392 | 0.711558 | 0.711767 | 0.645383 | 0.706377 | 0.701244 | 0.704407 |
| 0.706956 | 0.656888 | 0.663495 | 0.709256 | 0.709758 | 0.645518 | 0.70805 | 0.695683 | 0.705394 |
| 0.707806 | 0.658941 | 0.663131 | 0.70938 | 0.709366 | 0.642897 | 0.708938 | 0.699857 | 0.705552 |
| 0.710238 | 0.657868 | 0.661483 | 0.710681 | 0.714297 | 0.644936 | 0.709473 | 0.697008 | 0.706978 |
| 0.720581 | 0.678828 | 0.683977 | 0.726016 | 0.728646 | 0.667354 | 0.723744 | 0.715744 | 0.721718 |
| 0.703695 | 0.655172 | 0.654177 | 0.703816 | 0.703685 | 0.643269 | 0.700455 | 0.693299 | 0.702475 |
| 0.707754 | 0.661582 | 0.658955 | 0.711132 | 0.71211 | 0.641565 | 0.709851 | 0.7004 | 0.709996 |
| 0.705411 | 0.65696 | 0.659942 | 0.70812 | 0.709996 | 0.636996 | 0.70538 | 0.693766 | 0.700923 |
| 0.725786 | 0.677995 | 0.679919 | 0.732118 | 0.73197 | 0.662655 | 0.728343 | 0.71944 | 0.727647 |
| 0.710029 | 0.663502 | 0.663275 | 0.715469 | 0.716102 | 0.645929 | 0.716592 | 0.701663 | 0.710831 |
| 0.708258 | 0.654946 | 0.662543 | 0.711646 | 0.711057 | 0.642792 | 0.709998 | 0.697869 | 0.703718 |
| 0.712128 | 0.660512 | 0.656363 | 0.715851 | 0.715879 | 0.649528 | 0.714562 | 0.703109 | 0.711663 |
| 0.707266 | 0.662561 | 0.663937 | 0.710408 | 0.709558 | 0.647937 | 0.708793 | 0.700128 | 0.706861 |

| | | | | | | | | |
|---|---|---|---|---|---|---|---|---|
| 0.7094 | 0.658865 | 0.662844 | 0.712286 | 0.712492 | 0.645209 | 0.711978 | 0.69834 | 0.706393 |
| 0.714037 | 0.661471 | 0.657757 | 0.713812 | 0.715306 | 0.641447 | 0.713079 | 0.701786 | 0.712072 |
| 0.708363 | 0.660789 | 0.663404 | 0.71502 | 0.71479 | 0.640659 | 0.71318 | 0.699308 | 0.705373 |
| 0.716954 | 0.672393 | 0.674355 | 0.719052 | 0.722582 | 0.659777 | 0.721361 | 0.709998 | 0.718007 |
| 0.718718 | 0.66953 | 0.673544 | 0.717539 | 0.722943 | 0.650137 | 0.718355 | 0.706638 | 0.714445 |
| 0.710324 | 0.65875 | 0.667767 | 0.711535 | 0.715272 | 0.647233 | 0.708444 | 0.696977 | 0.706005 |
| 0.714489 | 0.664076 | 0.668005 | 0.712601 | 0.714602 | 0.655364 | 0.713538 | 0.70345 | 0.710412 |
| 0.711625 | 0.663277 | 0.668748 | 0.716769 | 0.713382 | 0.642278 | 0.712163 | 0.705028 | 0.70999 |
| 0.711122 | 0.655574 | 0.659536 | 0.710719 | 0.713806 | 0.643612 | 0.712805 | 0.703766 | 0.706988 |
| 0.708762 | 0.652824 | 0.659625 | 0.713504 | 0.707702 | 0.639396 | 0.708798 | 0.697656 | 0.703519 |
| 0.706515 | 0.653122 | 0.655364 | 0.714128 | 0.713136 | 0.649936 | 0.709588 | 0.702637 | 0.704948 |
| 0.711297 | 0.661695 | 0.666605 | 0.712964 | 0.713778 | 0.649394 | 0.710922 | 0.697962 | 0.702993 |
| 0.712745 | 0.658028 | 0.662011 | 0.714446 | 0.715169 | 0.651623 | 0.713008 | 0.703716 | 0.710683 |
| 0.708005 | 0.654724 | 0.663183 | 0.707915 | 0.711185 | 0.644486 | 0.709586 | 0.698451 | 0.705977 |
| 0.711044 | 0.653342 | 0.661936 | 0.71299 | 0.711993 | 0.648035 | 0.713586 | 0.70222 | 0.707356 |
| 0.712498 | 0.652422 | 0.658311 | 0.712161 | 0.712722 | 0.642018 | 0.710908 | 0.701267 | 0.704298 |
| 0.71432 | 0.657483 | 0.661151 | 0.71615 | 0.71914 | 0.650233 | 0.715136 | 0.7045 | 0.712222 |
| 0.708729 | 0.65523 | 0.664609 | 0.711352 | 0.712331 | 0.644616 | 0.709531 | 0.698211 | 0.70436 |
| 0.710365 | 0.658094 | 0.662194 | 0.713139 | 0.712735 | 0.645876 | 0.709313 | 0.698845 | 0.700667 |
| 0.710779 | 0.656076 | 0.666178 | 0.717201 | 0.713511 | 0.65346 | 0.713452 | 0.705402 | 0.708105 |
| 0.71144 | 0.659542 | 0.668185 | 0.713761 | 0.714343 | 0.6509 | 0.71013 | 0.699374 | 0.708153 |
| 0.704803 | 0.652224 | 0.655682 | 0.710301 | 0.713205 | 0.644167 | 0.707507 | 0.696364 | 0.703674 |
| 0.711042 | 0.664192 | 0.663024 | 0.712485 | 0.71377 | 0.645209 | 0.710097 | 0.699595 | 0.708814 |
| 0.712668 | 0.663071 | 0.664367 | 0.715668 | 0.716493 | 0.642779 | 0.714525 | 0.701743 | 0.708386 |
| 0.709812 | 0.653419 | 0.665209 | 0.715734 | 0.71705 | 0.648613 | 0.714621 | 0.70326 | 0.71199 |
| 0.715638 | 0.657431 | 0.667756 | 0.718871 | 0.718299 | 0.652038 | 0.715276 | 0.705026 | 0.713934 |
| 0.708955 | 0.658516 | 0.661436 | 0.714906 | 0.718036 | 0.648926 | 0.713378 | 0.702953 | 0.712595 |
| 0.712262 | 0.659596 | 0.661757 | 0.713495 | 0.718061 | 0.645451 | 0.714012 | 0.702901 | 0.711189 |
| 0.710749 | 0.657582 | 0.663152 | 0.717372 | 0.718247 | 0.645791 | 0.711885 | 0.703873 | 0.708773 |
| 0.710934 | 0.656377 | 0.668129 | 0.715853 | 0.720562 | 0.653001 | 0.715259 | 0.704019 | 0.715189 |
| 0.715006 | 0.658754 | 0.663575 | 0.72009 | 0.721272 | 0.652254 | 0.714332 | 0.705939 | 0.712551 |
| 0.710196 | 0.658675 | 0.659414 | 0.714634 | 0.712303 | 0.640531 | 0.711648 | 0.700094 | 0.708448 |
| 0.712939 | 0.658277 | 0.673462 | 0.715712 | 0.712949 | 0.650491 | 0.709373 | 0.700857 | 0.708196 |
| 0.709962 | 0.668191 | 0.670255 | 0.717421 | 0.714801 | 0.655172 | 0.712016 | 0.701764 | 0.707476 |
| 0.708473 | 0.661733 | 0.657576 | 0.710423 | 0.711129 | 0.64695 | 0.708901 | 0.698387 | 0.707591 |
| 0.711556 | 0.662887 | 0.665452 | 0.711138 | 0.712457 | 0.648059 | 0.71147 | 0.70056 | 0.707165 |
| 0.70848 | 0.649714 | 0.65827 | 0.710748 | 0.710827 | 0.63944 | 0.708326 | 0.700764 | 0.702378 |
| 0.688163 | 0.662725 | 0.665141 | 0.692206 | 0.689932 | 0.647906 | 0.685897 | 0.672903 | 0.682984 |
| 0.69216 | 0.66181 | 0.662874 | 0.693767 | 0.693935 | 0.641367 | 0.689221 | 0.67622 | 0.687343 |
| 0.680796 | 0.656854 | 0.666211 | 0.681139 | 0.686959 | 0.642395 | 0.683518 | 0.669497 | 0.681183 |
| 0.680093 | 0.652886 | 0.654447 | 0.682725 | 0.685426 | 0.642726 | 0.678686 | 0.673451 | 0.676451 |
| 0.669778 | 0.648119 | 0.656327 | 0.668448 | 0.66593 | 0.637082 | 0.66866 | 0.656388 | 0.661172 |
| 0.668399 | 0.651346 | 0.655087 | 0.674598 | 0.675705 | 0.637657 | 0.675911 | 0.658678 | 0.665825 |
| 0.668845 | 0.658766 | 0.660158 | 0.672323 | 0.670961 | 0.639351 | 0.668242 | 0.656657 | 0.66881 |
| 0.668742 | 0.658972 | 0.661054 | 0.675683 | 0.674116 | 0.647488 | 0.673308 | 0.659631 | 0.672519 |
| 0.671318 | 0.659914 | 0.663855 | 0.675286 | 0.671822 | 0.647491 | 0.672037 | 0.660743 | 0.667577 |
| 0.673985 | 0.660871 | 0.666101 | 0.679245 | 0.674763 | 0.647911 | 0.673991 | 0.660071 | 0.670919 |

| | | | | | | | | |
|---|---|---|---|---|---|---|---|---|
| 0.676959 | 0.661089 | 0.664553 | 0.679396 | 0.677133 | 0.643552 | 0.674754 | 0.666051 | 0.670747 |
| 0.674422 | 0.658833 | 0.66933 | 0.676627 | 0.677344 | 0.652001 | 0.673811 | 0.667056 | 0.67636 |
| 0.669405 | 0.65684 | 0.66077 | 0.678961 | 0.677678 | 0.651634 | 0.671457 | 0.654509 | 0.67164 |
| 0.668561 | 0.650813 | 0.666161 | 0.675354 | 0.671695 | 0.640898 | 0.671612 | 0.658148 | 0.662066 |
| 0.667739 | 0.656675 | 0.663671 | 0.668455 | 0.670434 | 0.64245 | 0.665471 | 0.654285 | 0.663988 |
| 0.663399 | 0.660892 | 0.666592 | 0.67145 | 0.673961 | 0.635105 | 0.66548 | 0.655513 | 0.666074 |
| 0.666103 | 0.65672 | 0.661546 | 0.669711 | 0.671252 | 0.648649 | 0.670017 | 0.652418 | 0.668924 |
| 0.665127 | 0.649823 | 0.6601 | 0.664128 | 0.666061 | 0.639585 | 0.661734 | 0.652411 | 0.661968 |
| 0.666422 | 0.658589 | 0.661758 | 0.66473 | 0.670495 | 0.647387 | 0.666325 | 0.654516 | 0.65977 |
| 0.667592 | 0.648222 | 0.657863 | 0.669384 | 0.674177 | 0.641671 | 0.66798 | 0.658433 | 0.667927 |
| 0.658598 | 0.654537 | 0.65981 | 0.663944 | 0.669253 | 0.642563 | 0.661401 | 0.65165 | 0.660667 |
| 0.701852 | 0.650075 | 0.659899 | 0.702123 | 0.701974 | 0.637355 | 0.702762 | 0.687728 | 0.698759 |
| 0.710704 | 0.656093 | 0.661591 | 0.712083 | 0.71362 | 0.646005 | 0.710868 | 0.698921 | 0.707919 |
| 0.685254 | 0.670465 | 0.669519 | 0.686597 | 0.694158 | 0.654764 | 0.685023 | 0.678349 | 0.682579 |
| 0.718092 | 0.660158 | 0.671963 | 0.717411 | 0.721209 | 0.653099 | 0.718522 | 0.708487 | 0.714008 |
| 0.706764 | 0.655956 | 0.668072 | 0.707911 | 0.708884 | 0.647364 | 0.707878 | 0.694714 | 0.701207 |
| 0.713348 | 0.665993 | 0.668481 | 0.717884 | 0.716851 | 0.653981 | 0.714411 | 0.704587 | 0.709739 |
| 0.722173 | 0.661211 | 0.670016 | 0.721335 | 0.721568 | 0.659345 | 0.723442 | 0.713175 | 0.71771 |
| 0.695405 | 0.646583 | 0.649778 | 0.696698 | 0.699403 | 0.626543 | 0.696171 | 0.686641 | 0.695181 |
| 0.720996 | 0.673473 | 0.673745 | 0.720205 | 0.722081 | 0.656142 | 0.72289 | 0.712384 | 0.718514 |
| 0.695453 | 0.670973 | 0.673986 | 0.691977 | 0.695825 | 0.663105 | 0.691762 | 0.681895 | 0.69192 |
| 0.717764 | 0.664065 | 0.665693 | 0.718563 | 0.720682 | 0.650307 | 0.71808 | 0.706931 | 0.713406 |
| 0.72212 | 0.662276 | 0.6688 | 0.722729 | 0.72265 | 0.656122 | 0.723242 | 0.71194 | 0.719958 |
| 0.686524 | 0.66133 | 0.668654 | 0.684289 | 0.693368 | 0.650765 | 0.682918 | 0.672008 | 0.683905 |
| 0.689638 | 0.63803 | 0.647016 | 0.691883 | 0.691584 | 0.634083 | 0.687645 | 0.681606 | 0.685594 |
| 0.687163 | 0.685131 | 0.682046 | 0.693949 | 0.69881 | 0.662829 | 0.690368 | 0.677947 | 0.69187 |
| 0.707357 | 0.654019 | 0.655956 | 0.709714 | 0.711902 | 0.641055 | 0.710209 | 0.700985 | 0.708255 |
| 0.720745 | 0.660104 | 0.676234 | 0.720282 | 0.723868 | 0.651596 | 0.71867 | 0.708284 | 0.714786 |
| 0.687133 | 0.640078 | 0.640345 | 0.688359 | 0.694576 | 0.625803 | 0.687685 | 0.67462 | 0.685595 |
| 0.724614 | 0.67349 | 0.683953 | 0.72561 | 0.728776 | 0.660128 | 0.723463 | 0.714579 | 0.723529 |
| 0.729879 | 0.672721 | 0.673344 | 0.732638 | 0.731177 | 0.661029 | 0.726539 | 0.715996 | 0.727071 |
| 0.695954 | 0.648188 | 0.656239 | 0.70134 | 0.703999 | 0.640256 | 0.700781 | 0.689742 | 0.696339 |
| 0.697609 | 0.647548 | 0.65269 | 0.702134 | 0.700314 | 0.638294 | 0.697948 | 0.68999 | 0.695371 |
| 0.725223 | 0.676933 | 0.680754 | 0.724914 | 0.730791 | 0.66568 | 0.727105 | 0.718549 | 0.724331 |
| 0.725373 | 0.679522 | 0.676338 | 0.726112 | 0.726865 | 0.659731 | 0.723054 | 0.713061 | 0.718802 |
| 0.726522 | 0.678782 | 0.681587 | 0.729912 | 0.732193 | 0.667839 | 0.727888 | 0.716257 | 0.726328 |
| 0.675826 | 0.631434 | 0.631063 | 0.678409 | 0.68271 | 0.624821 | 0.676576 | 0.668492 | 0.675554 |
| 0.722681 | 0.675447 | 0.677648 | 0.723182 | 0.725243 | 0.661675 | 0.717957 | 0.711453 | 0.718628 |
| 0.674213 | 0.622176 | 0.627806 | 0.677789 | 0.682112 | 0.616923 | 0.676294 | 0.667553 | 0.673675 |
| 0.668375 | 0.612939 | 0.615582 | 0.674877 | 0.672836 | 0.598604 | 0.670589 | 0.66303 | 0.667374 |
| 0.717189 | 0.67182 | 0.682407 | 0.72705 | 0.725669 | 0.663216 | 0.722311 | 0.709715 | 0.719604 |
| 0.669998 | 0.610603 | 0.619178 | 0.674494 | 0.674325 | 0.609408 | 0.671636 | 0.661674 | 0.668946 |
| 0.662852 | 0.608607 | 0.616123 | 0.667091 | 0.665263 | 0.595961 | 0.664834 | 0.654848 | 0.660678 |
| 0.658364 | 0.602524 | 0.613331 | 0.66082 | 0.66465 | 0.600869 | 0.660858 | 0.649724 | 0.657112 |
| 0.724406 | 0.683519 | 0.692774 | 0.731991 | 0.743064 | 0.671927 | 0.729818 | 0.723136 | 0.730948 |
| 0.662127 | 0.605565 | 0.609785 | 0.666567 | 0.669179 | 0.590628 | 0.66506 | 0.655928 | 0.659788 |
| 0.718964 | 0.676471 | 0.689971 | 0.725638 | 0.729046 | 0.664458 | 0.722209 | 0.711162 | 0.721808 |

| | | | | | | | | |
|---|---|---|---|---|---|---|---|---|
| 0.727698 | 0.685395 | 0.693292 | 0.734108 | 0.740973 | 0.678704 | 0.732516 | 0.720361 | 0.729928 |
| 0.721429 | 0.679763 | 0.682008 | 0.729502 | 0.734037 | 0.664449 | 0.725349 | 0.715281 | 0.726227 |
| 0.751019 | 0.704929 | 0.705958 | 0.752096 | 0.753186 | 0.70756 | 0.753132 | 0.738852 | 0.748898 |
| 0.651423 | 0.591562 | 0.597665 | 0.657588 | 0.660255 | 0.583432 | 0.654659 | 0.645389 | 0.653889 |
| 0.652451 | 0.606012 | 0.613919 | 0.656312 | 0.659009 | 0.59786 | 0.653397 | 0.64502 | 0.648987 |
| 0.728891 | 0.688541 | 0.686106 | 0.737234 | 0.735631 | 0.680127 | 0.732369 | 0.724138 | 0.732488 |
| 0.653097 | 0.60438 | 0.609257 | 0.655724 | 0.656313 | 0.593633 | 0.657612 | 0.646091 | 0.652015 |
| 0.733901 | 0.688448 | 0.684298 | 0.737987 | 0.735872 | 0.680136 | 0.734553 | 0.72428 | 0.734463 |
| 0.73127 | 0.688305 | 0.697679 | 0.73418 | 0.738709 | 0.67382 | 0.730896 | 0.723375 | 0.731014 |
| 0.73207 | 0.687273 | 0.689862 | 0.739908 | 0.740844 | 0.676499 | 0.736777 | 0.729925 | 0.734293 |
| 0.728047 | 0.690258 | 0.685271 | 0.731364 | 0.744539 | 0.681698 | 0.73179 | 0.725131 | 0.736684 |
| 0.746377 | 0.796748 | 0.764286 | 0.723684 | 0.72973 | 0.740157 | 0.694805 | 0.72549 | 0.726667 |
| 0.712614 | 0.740217 | 0.735465 | 0.708383 | 0.722794 | 0.71412 | 0.722666 | 0.709851 | 0.714257 |
| 0.718706 | 0.731446 | 0.727098 | 0.723939 | 0.717881 | 0.702087 | 0.72264 | 0.709094 | 0.716843 |
| 0.726167 | 0.731473 | 0.739506 | 0.73131 | 0.734657 | 0.715248 | 0.726835 | 0.723158 | 0.728319 |
| 0.715426 | 0.731143 | 0.744155 | 0.72674 | 0.733073 | 0.709131 | 0.729435 | 0.719581 | 0.724671 |
| 0.590111 | 0.570296 | 0.571931 | 0.595481 | 0.597897 | 0.566568 | 0.592761 | 0.586021 | 0.591668 |
| 0.724089 | 0.741302 | 0.72867 | 0.725255 | 0.733463 | 0.714755 | 0.728855 | 0.722136 | 0.730994 |
| 0.731892 | 0.694612 | 0.694218 | 0.737716 | 0.747935 | 0.683704 | 0.736791 | 0.731794 | 0.737575 |
| 0.730907 | 0.692922 | 0.700977 | 0.737696 | 0.741027 | 0.67968 | 0.734946 | 0.728899 | 0.736104 |
| 0.735938 | 0.694877 | 0.701031 | 0.737034 | 0.748546 | 0.68522 | 0.736567 | 0.730937 | 0.740018 |
| 0.596152 | 0.533099 | 0.537043 | 0.598932 | 0.598807 | 0.525113 | 0.598496 | 0.588984 | 0.595515 |
| 0.581998 | 0.519546 | 0.522513 | 0.586958 | 0.584962 | 0.50764 | 0.584636 | 0.579397 | 0.58148 |
| 0.761277 | 0.722535 | 0.724308 | 0.761267 | 0.782729 | 0.718362 | 0.762774 | 0.754602 | 0.760608 |
| 0.762437 | 0.72308 | 0.72389 | 0.763708 | 0.785702 | 0.725707 | 0.764942 | 0.76425 | 0.766355 |
| 0.755462 | 0.710873 | 0.711759 | 0.752602 | 0.776528 | 0.711137 | 0.76054 | 0.747547 | 0.757553 |
| 0.759798 | 0.714634 | 0.726164 | 0.758854 | 0.784059 | 0.718424 | 0.765924 | 0.757988 | 0.763413 |
| 0.763939 | 0.717922 | 0.723763 | 0.762841 | 0.785756 | 0.723092 | 0.766207 | 0.761629 | 0.765868 |
| 0.765047 | 0.73411 | 0.736484 | 0.768664 | 0.793497 | 0.721932 | 0.764848 | 0.760043 | 0.764955 |
| 0.765807 | 0.722569 | 0.732569 | 0.768027 | 0.786844 | 0.722945 | 0.767411 | 0.764885 | 0.764769 |
| 0.790521 | 0.720072 | 0.722053 | 0.77385 | 0.774002 | 0.747475 | 0.786968 | 0.777214 | 0.786211 |
| 0.791318 | 0.72955 | 0.727684 | 0.784399 | 0.789743 | 0.75146 | 0.791646 | 0.788613 | 0.794334 |
| 0.804093 | 0.71008 | 0.72604 | 0.778138 | 0.775719 | 0.749462 | 0.787477 | 0.78407 | 0.784378 |
| 0.797204 | 0.721294 | 0.719174 | 0.774115 | 0.775492 | 0.739232 | 0.787236 | 0.776519 | 0.788758 |
| 0.795893 | 0.718168 | 0.731481 | 0.781278 | 0.784259 | 0.748331 | 0.792868 | 0.788282 | 0.793863 |
| 0.788492 | 0.712148 | 0.726655 | 0.7726 | 0.777483 | 0.732425 | 0.782514 | 0.778303 | 0.784137 |
| 0.79725 | 0.716049 | 0.725428 | 0.780909 | 0.779083 | 0.748588 | 0.783473 | 0.784861 | 0.783948 |
| 0.800599 | 0.716925 | 0.731329 | 0.774323 | 0.781307 | 0.748383 | 0.789233 | 0.782219 | 0.792356 |
| 0.826147 | 0.716287 | 0.728852 | 0.77631 | 0.778316 | 0.745583 | 0.79331 | 0.782002 | 0.790375 |
| 1 | 0.711894 | 0.72215 | 0.778278 | 0.772757 | 0.74228 | 0.788543 | 0.779244 | 0.785389 |
| 0.711894 | 1 | 0.740799 | 0.713542 | 0.719659 | 0.708518 | 0.71906 | 0.71187 | 0.717369 |
| 0.72215 | 0.740799 | 1 | 0.730928 | 0.740741 | 0.733091 | 0.735682 | 0.721438 | 0.73395 |
| 0.778278 | 0.713542 | 0.730928 | 1 | 0.782186 | 0.756751 | 0.798736 | 0.809425 | 0.800984 |
| 0.772757 | 0.719659 | 0.740741 | 0.782186 | 1 | 0.75117 | 0.789224 | 0.796679 | 0.800117 |
| 0.74228 | 0.708518 | 0.733091 | 0.756751 | 0.75117 | 1 | 0.80012 | 0.835459 | 0.806768 |
| 0.788543 | 0.71906 | 0.735682 | 0.798736 | 0.789224 | 0.80012 | 1 | 0.815342 | 0.850317 |
| 0.779244 | 0.71187 | 0.721438 | 0.809425 | 0.796679 | 0.835459 | 0.815342 | 1 | 0.837459 |

| | | | | | | | | |
|---|---|---|---|---|---|---|---|---|
| 0.785389 | 0.717369 | 0.73395 | 0.800984 | 0.800117 | 0.806768 | 0.850317 | 0.837459 | 1 |
| 0.789422 | 0.721084 | 0.726882 | 0.807533 | 0.785973 | 0.812826 | 0.857021 | 0.83641 | 0.858377 |
| 0.787718 | 0.72489 | 0.721062 | 0.802709 | 0.790806 | 0.777489 | 0.817829 | 0.835747 | 0.817884 |
| 0.785667 | 0.718231 | 0.732318 | 0.789609 | 0.798847 | 0.787637 | 0.837905 | 0.800383 | 0.8536 |
| 0.788026 | 0.725147 | 0.738482 | 0.787122 | 0.787999 | 0.819369 | 0.831416 | 0.808007 | 0.835981 |
| 0.78215 | 0.719965 | 0.729831 | 0.78118 | 0.818736 | 0.796925 | 0.816484 | 0.812395 | 0.831648 |
| 0.779658 | 0.719134 | 0.734325 | 0.78088 | 0.824876 | 0.778129 | 0.827461 | 0.808603 | 0.857302 |
| 0.776242 | 0.696335 | 0.715838 | 0.796871 | 0.78954 | 0.766834 | 0.838623 | 0.824102 | 0.815163 |
| 0.778583 | 0.712116 | 0.728561 | 0.776781 | 0.788035 | 0.783741 | 0.838043 | 0.796788 | 0.83955 |
| 0.775935 | 0.709056 | 0.725349 | 0.820736 | 0.785014 | 0.77119 | 0.811112 | 0.846732 | 0.823204 |
| 0.787138 | 0.717024 | 0.732244 | 0.794183 | 0.78542 | 0.786537 | 0.838389 | 0.822706 | 0.839387 |
| 0.785446 | 0.711996 | 0.730195 | 0.802232 | 0.795281 | 0.817718 | 0.836944 | 0.83606 | 0.87692 |
| 0.780137 | 0.70739 | 0.713292 | 0.802535 | 0.793976 | 0.784749 | 0.814707 | 0.889124 | 0.829646 |
| 0.7716 | 0.717503 | 0.721602 | 0.781476 | 0.856127 | 0.757521 | 0.798253 | 0.832332 | 0.827751 |
| 0.794418 | 0.709905 | 0.728775 | 0.782933 | 0.779407 | 0.794953 | 0.811408 | 0.789335 | 0.803362 |
| 0.767672 | 0.725231 | 0.729436 | 0.76563 | 0.784907 | 0.728347 | 0.776067 | 0.764845 | 0.76814 |
| 0.760523 | 0.727851 | 0.730514 | 0.766845 | 0.78281 | 0.725578 | 0.77234 | 0.766593 | 0.770239 |
| 0.76308 | 0.72141 | 0.719404 | 0.76036 | 0.778423 | 0.718774 | 0.769841 | 0.758835 | 0.77237 |
| 0.761416 | 0.722378 | 0.723224 | 0.76879 | 0.785567 | 0.721802 | 0.768725 | 0.763364 | 0.769553 |
| 0.769212 | 0.717627 | 0.726826 | 0.763997 | 0.781258 | 0.729025 | 0.769478 | 0.761312 | 0.767922 |
| 0.758767 | 0.712343 | 0.721446 | 0.762377 | 0.770218 | 0.718927 | 0.764218 | 0.757898 | 0.764676 |
| 0.760539 | 0.716326 | 0.722757 | 0.763103 | 0.78095 | 0.722635 | 0.766253 | 0.758486 | 0.76024 |
| 0.771412 | 0.720236 | 0.729806 | 0.766756 | 0.786636 | 0.730658 | 0.772861 | 0.762596 | 0.76721 |
| 0.768247 | 0.71893 | 0.723276 | 0.764836 | 0.783782 | 0.72146 | 0.774324 | 0.767095 | 0.774547 |
| 0.765217 | 0.723055 | 0.729309 | 0.763879 | 0.781014 | 0.718896 | 0.767042 | 0.759385 | 0.766196 |
| 0.760481 | 0.71311 | 0.721739 | 0.761615 | 0.78384 | 0.719876 | 0.763567 | 0.756104 | 0.76032 |
| 0.760862 | 0.715574 | 0.719916 | 0.758857 | 0.775344 | 0.725324 | 0.766964 | 0.760743 | 0.766519 |
| 0.757853 | 0.71235 | 0.71601 | 0.760412 | 0.78323 | 0.707671 | 0.760063 | 0.753759 | 0.760398 |
| 0.760546 | 0.718936 | 0.72402 | 0.760522 | 0.78636 | 0.718853 | 0.761936 | 0.758683 | 0.76096 |
| 0.762421 | 0.721764 | 0.727788 | 0.766762 | 0.786982 | 0.722763 | 0.769998 | 0.76206 | 0.768687 |
| 0.764691 | 0.723993 | 0.725372 | 0.765197 | 0.784179 | 0.723235 | 0.768927 | 0.764754 | 0.770917 |
| 0.766951 | 0.722665 | 0.729119 | 0.764356 | 0.782187 | 0.725671 | 0.77092 | 0.765519 | 0.773615 |
| 0.774091 | 0.732286 | 0.734349 | 0.773852 | 0.789994 | 0.732972 | 0.773701 | 0.772416 | 0.778133 |
| 0.768475 | 0.727 | 0.733422 | 0.770511 | 0.790524 | 0.730986 | 0.778301 | 0.77 | 0.77753 |
| 0.755746 | 0.718772 | 0.717056 | 0.760312 | 0.788947 | 0.708947 | 0.75601 | 0.752635 | 0.759086 |
| 0.767105 | 0.722495 | 0.732525 | 0.763136 | 0.782217 | 0.729751 | 0.767273 | 0.760325 | 0.767387 |
| 0.757314 | 0.715132 | 0.718339 | 0.761329 | 0.779843 | 0.717152 | 0.767492 | 0.758706 | 0.764102 |
| 0.759397 | 0.712742 | 0.728625 | 0.763423 | 0.784519 | 0.724705 | 0.763322 | 0.753311 | 0.761814 |
| 0.757164 | 0.711945 | 0.717176 | 0.75485 | 0.77189 | 0.712947 | 0.758894 | 0.754322 | 0.757662 |
| 0.752194 | 0.716912 | 0.71885 | 0.749681 | 0.768199 | 0.716934 | 0.75513 | 0.746695 | 0.755072 |
| 0.755071 | 0.707408 | 0.725358 | 0.755786 | 0.770604 | 0.711625 | 0.754422 | 0.747188 | 0.756718 |
| 0.757792 | 0.713753 | 0.721086 | 0.757078 | 0.778052 | 0.724655 | 0.758669 | 0.749848 | 0.759306 |
| 0.755531 | 0.711259 | 0.713266 | 0.75356 | 0.769277 | 0.716397 | 0.766362 | 0.752109 | 0.762578 |
| 0.746455 | 0.711941 | 0.714588 | 0.753247 | 0.772517 | 0.715228 | 0.754928 | 0.751279 | 0.754975 |
| 0.755064 | 0.722242 | 0.726035 | 0.759208 | 0.778695 | 0.724104 | 0.765249 | 0.755067 | 0.761601 |
| 0.753046 | 0.709019 | 0.712811 | 0.751004 | 0.774184 | 0.705797 | 0.755907 | 0.756299 | 0.762409 |

| | | | | | | | | |
|---|---|---|---|---|---|---|---|---|
| 0.761971 | 0.715013 | 0.731517 | 0.764257 | 0.784156 | 0.7157 | 0.763785 | 0.757864 | 0.761592 |
| 0.769161 | 0.733475 | 0.733067 | 0.767455 | 0.772595 | 0.731478 | 0.768897 | 0.762577 | 0.768153 |
| 0.758729 | 0.719821 | 0.717787 | 0.761826 | 0.769148 | 0.711685 | 0.761347 | 0.758318 | 0.761157 |
| 0.765256 | 0.714459 | 0.723034 | 0.768999 | 0.769206 | 0.71464 | 0.771953 | 0.756881 | 0.768942 |
| 0.77525 | 0.736165 | 0.738754 | 0.774268 | 0.773957 | 0.734111 | 0.782058 | 0.764975 | 0.774831 |
| 0.767556 | 0.72913 | 0.730603 | 0.771994 | 0.767645 | 0.73166 | 0.777383 | 0.765721 | 0.771548 |
| 0.770988 | 0.725115 | 0.72748 | 0.763652 | 0.772501 | 0.720405 | 0.770365 | 0.759466 | 0.767689 |
| 0.770619 | 0.721355 | 0.724651 | 0.765876 | 0.773387 | 0.719461 | 0.771335 | 0.762153 | 0.767093 |
| 0.768016 | 0.731571 | 0.728571 | 0.763864 | 0.76334 | 0.725595 | 0.772641 | 0.75769 | 0.763648 |
| 0.763082 | 0.709593 | 0.723493 | 0.763214 | 0.768559 | 0.720378 | 0.770537 | 0.759032 | 0.766411 |
| 0.771793 | 0.721456 | 0.727885 | 0.772321 | 0.776216 | 0.725273 | 0.779877 | 0.765978 | 0.776832 |
| 0.775373 | 0.724908 | 0.728555 | 0.773328 | 0.770241 | 0.725596 | 0.7808 | 0.767835 | 0.775534 |
| 0.759977 | 0.717063 | 0.724228 | 0.768973 | 0.767654 | 0.719217 | 0.767998 | 0.762216 | 0.762162 |
| 0.761017 | 0.724969 | 0.727443 | 0.764922 | 0.767259 | 0.725993 | 0.766136 | 0.762359 | 0.765189 |
| 0.763868 | 0.721497 | 0.741662 | 0.767889 | 0.774722 | 0.716584 | 0.768671 | 0.763905 | 0.771828 |
| 0.759798 | 0.715185 | 0.731925 | 0.769533 | 0.767543 | 0.718901 | 0.767444 | 0.763467 | 0.762006 |
| 0.762124 | 0.72533 | 0.729209 | 0.769044 | 0.773223 | 0.712687 | 0.765174 | 0.763126 | 0.767552 |
| 0.761587 | 0.724748 | 0.73241 | 0.767462 | 0.771359 | 0.716492 | 0.764754 | 0.758002 | 0.765475 |
| 0.76553 | 0.726124 | 0.727221 | 0.766044 | 0.776108 | 0.725464 | 0.768562 | 0.766286 | 0.769768 |
| 0.76339 | 0.727591 | 0.736145 | 0.768955 | 0.767069 | 0.716887 | 0.768659 | 0.764921 | 0.766936 |
| 0.762229 | 0.717207 | 0.729547 | 0.761316 | 0.767809 | 0.720207 | 0.767058 | 0.760905 | 0.760718 |
| 0.76257 | 0.718614 | 0.732642 | 0.764991 | 0.767776 | 0.725101 | 0.768518 | 0.759806 | 0.765711 |
| 0.765279 | 0.723874 | 0.725182 | 0.770164 | 0.772334 | 0.720752 | 0.76628 | 0.759955 | 0.768929 |
| 0.763343 | 0.724612 | 0.731589 | 0.767267 | 0.774739 | 0.714613 | 0.771545 | 0.761712 | 0.768754 |
| 0.760714 | 0.721111 | 0.733596 | 0.764 | 0.767106 | 0.71705 | 0.770238 | 0.757732 | 0.762601 |
| 0.760215 | 0.720238 | 0.727736 | 0.764956 | 0.767438 | 0.721169 | 0.765082 | 0.758388 | 0.762665 |
| 0.759583 | 0.722513 | 0.727803 | 0.762663 | 0.769494 | 0.714203 | 0.767901 | 0.762514 | 0.767372 |
| 0.760431 | 0.724126 | 0.730582 | 0.765589 | 0.768279 | 0.712441 | 0.762443 | 0.755985 | 0.766088 |
| 0.764863 | 0.739314 | 0.736002 | 0.774333 | 0.770315 | 0.720391 | 0.772508 | 0.765705 | 0.768315 |
| 0.761533 | 0.738434 | 0.73894 | 0.768753 | 0.77097 | 0.724312 | 0.771545 | 0.76112 | 0.766737 |
| 0.755695 | 0.71749 | 0.72729 | 0.756922 | 0.760302 | 0.715061 | 0.757538 | 0.752392 | 0.758621 |
| 0.766979 | 0.727337 | 0.730683 | 0.768148 | 0.76939 | 0.716237 | 0.763154 | 0.760256 | 0.767157 |
| 0.763201 | 0.729445 | 0.737181 | 0.765687 | 0.769055 | 0.725223 | 0.766315 | 0.759825 | 0.770423 |
| 0.758195 | 0.732814 | 0.730147 | 0.760265 | 0.766839 | 0.72084 | 0.766692 | 0.760199 | 0.764676 |
| 0.758716 | 0.736065 | 0.737182 | 0.766198 | 0.768438 | 0.721931 | 0.7624 | 0.75944 | 0.760958 |
| 0.755991 | 0.729259 | 0.736692 | 0.76261 | 0.766635 | 0.718513 | 0.761643 | 0.754413 | 0.760432 |
| 0.765607 | 0.72882 | 0.737031 | 0.771711 | 0.776736 | 0.718954 | 0.771797 | 0.761835 | 0.775896 |
| 0.762415 | 0.733216 | 0.730834 | 0.772186 | 0.771815 | 0.72856 | 0.767672 | 0.766069 | 0.768195 |
| 0.761055 | 0.728944 | 0.73459 | 0.768212 | 0.77444 | 0.716695 | 0.764986 | 0.761391 | 0.772315 |
| 0.767434 | 0.726281 | 0.728374 | 0.763184 | 0.771311 | 0.720917 | 0.769708 | 0.757916 | 0.766686 |
| 0.761037 | 0.726512 | 0.727765 | 0.763728 | 0.773699 | 0.719067 | 0.76837 | 0.759093 | 0.76808 |
| 0.765599 | 0.732699 | 0.734775 | 0.767423 | 0.773205 | 0.722868 | 0.773448 | 0.759889 | 0.771927 |
| 0.767248 | 0.737027 | 0.737804 | 0.771266 | 0.77193 | 0.722104 | 0.773602 | 0.760359 | 0.770682 |
| 0.762372 | 0.723659 | 0.732424 | 0.761059 | 0.767354 | 0.723532 | 0.767524 | 0.757994 | 0.765089 |
| 0.764745 | 0.727 | 0.730558 | 0.764194 | 0.771026 | 0.725131 | 0.769167 | 0.762507 | 0.772483 |
| 0.759486 | 0.72163 | 0.727512 | 0.766332 | 0.771354 | 0.719258 | 0.765909 | 0.758114 | 0.766929 |

| | | | | | | | | |
|---|---|---|---|---|---|---|---|---|
| 0.7519 | 0.719949 | 0.729258 | 0.754551 | 0.761202 | 0.721586 | 0.755193 | 0.750118 | 0.755175 |
| 0.742624 | 0.706784 | 0.713835 | 0.748077 | 0.749343 | 0.698013 | 0.748392 | 0.738049 | 0.743082 |
| 0.757783 | 0.727979 | 0.728376 | 0.762662 | 0.762588 | 0.718334 | 0.764061 | 0.758713 | 0.760289 |
| 0.763829 | 0.735459 | 0.737637 | 0.767527 | 0.764792 | 0.71954 | 0.768056 | 0.759981 | 0.770959 |
| 0.757922 | 0.716944 | 0.719589 | 0.758772 | 0.759929 | 0.706579 | 0.759786 | 0.751772 | 0.757591 |
| 0.758094 | 0.718476 | 0.726721 | 0.764585 | 0.763647 | 0.716412 | 0.768591 | 0.757157 | 0.760374 |
| 0.758574 | 0.731319 | 0.737397 | 0.762528 | 0.765954 | 0.726593 | 0.762023 | 0.759288 | 0.762171 |
| 0.75046 | 0.723492 | 0.719177 | 0.755864 | 0.760979 | 0.721091 | 0.761968 | 0.753792 | 0.7593 |
| 0.769789 | 0.726395 | 0.736535 | 0.768392 | 0.775801 | 0.717534 | 0.76806 | 0.760718 | 0.766691 |
| 0.766132 | 0.730912 | 0.735456 | 0.767265 | 0.770789 | 0.728299 | 0.769517 | 0.760978 | 0.764924 |
| 0.766542 | 0.731485 | 0.737991 | 0.768103 | 0.771685 | 0.728688 | 0.77341 | 0.763431 | 0.770031 |
| 0.766036 | 0.728311 | 0.732349 | 0.765579 | 0.774671 | 0.71916 | 0.767384 | 0.758869 | 0.766697 |
| 0.765959 | 0.732982 | 0.738068 | 0.77062 | 0.769123 | 0.718504 | 0.771309 | 0.760666 | 0.770991 |
| 0.768643 | 0.734408 | 0.742944 | 0.772253 | 0.773121 | 0.727798 | 0.773387 | 0.765762 | 0.771716 |
| 0.764691 | 0.730403 | 0.7411 | 0.764447 | 0.769218 | 0.727167 | 0.768721 | 0.757525 | 0.765099 |
| 0.761156 | 0.721232 | 0.739872 | 0.764597 | 0.775181 | 0.718267 | 0.762451 | 0.755465 | 0.760335 |
| 0.76513 | 0.725735 | 0.724695 | 0.767339 | 0.767231 | 0.720466 | 0.768341 | 0.75894 | 0.768843 |
| 0.760843 | 0.724658 | 0.722108 | 0.767383 | 0.765065 | 0.71784 | 0.76258 | 0.754712 | 0.76347 |
| 0.761304 | 0.737002 | 0.747543 | 0.76248 | 0.765399 | 0.72381 | 0.761233 | 0.751283 | 0.761684 |
| 0.757159 | 0.729701 | 0.732037 | 0.76463 | 0.767537 | 0.719127 | 0.763204 | 0.754673 | 0.76087 |
| 0.760755 | 0.729364 | 0.73244 | 0.766174 | 0.767777 | 0.723664 | 0.768776 | 0.761822 | 0.764077 |
| 0.746206 | 0.720797 | 0.728359 | 0.754212 | 0.758772 | 0.695754 | 0.751888 | 0.749267 | 0.749811 |
| 0.756006 | 0.730058 | 0.723123 | 0.766266 | 0.763321 | 0.706245 | 0.762197 | 0.75798 | 0.757064 |
| 0.763775 | 0.728715 | 0.733006 | 0.765937 | 0.770181 | 0.714203 | 0.76673 | 0.761374 | 0.765336 |
| 0.762128 | 0.733734 | 0.74177 | 0.764814 | 0.767035 | 0.721663 | 0.764666 | 0.756323 | 0.764938 |
| 0.757874 | 0.727657 | 0.728407 | 0.763722 | 0.761932 | 0.707645 | 0.765569 | 0.757224 | 0.760502 |
| 0.751684 | 0.719693 | 0.723042 | 0.762066 | 0.765489 | 0.710335 | 0.760844 | 0.753753 | 0.763635 |
| 0.757298 | 0.728829 | 0.738009 | 0.762561 | 0.767317 | 0.719526 | 0.761142 | 0.760184 | 0.759758 |
| 0.757421 | 0.730255 | 0.738251 | 0.757257 | 0.760633 | 0.708277 | 0.760662 | 0.749204 | 0.754348 |
| 0.699991 | 0.66254 | 0.660275 | 0.698227 | 0.702464 | 0.638002 | 0.695948 | 0.686158 | 0.695704 |
| 0.70997 | 0.653437 | 0.659153 | 0.711909 | 0.712163 | 0.646094 | 0.710961 | 0.697903 | 0.707362 |
| 0.708153 | 0.649379 | 0.653559 | 0.706957 | 0.708767 | 0.640244 | 0.704755 | 0.697787 | 0.705218 |
| 0.694067 | 0.658246 | 0.656104 | 0.703283 | 0.702813 | 0.635912 | 0.69985 | 0.690145 | 0.695641 |
| 0.708915 | 0.653659 | 0.655141 | 0.716875 | 0.711915 | 0.640951 | 0.709652 | 0.697961 | 0.708422 |
| 0.759002 | 0.73293 | 0.74079 | 0.758847 | 0.763764 | 0.714729 | 0.764891 | 0.759191 | 0.759195 |
| 0.759366 | 0.72801 | 0.750285 | 0.765734 | 0.763357 | 0.713623 | 0.764045 | 0.758951 | 0.762222 |
| 0.760591 | 0.734153 | 0.74249 | 0.765931 | 0.771293 | 0.712011 | 0.768802 | 0.761838 | 0.765411 |
| 0.765175 | 0.736479 | 0.739221 | 0.767723 | 0.768989 | 0.719244 | 0.768354 | 0.765103 | 0.766614 |
| 0.759648 | 0.734518 | 0.740503 | 0.763383 | 0.761972 | 0.721441 | 0.762563 | 0.756408 | 0.759518 |
| 0.758178 | 0.734397 | 0.752137 | 0.764247 | 0.775493 | 0.71519 | 0.765801 | 0.759498 | 0.7644 |
| 0.764184 | 0.736018 | 0.743097 | 0.765804 | 0.769327 | 0.718533 | 0.764828 | 0.763086 | 0.760667 |
| 0.753851 | 0.726347 | 0.739698 | 0.763648 | 0.769552 | 0.711253 | 0.764824 | 0.760859 | 0.760906 |
| 0.75506 | 0.726471 | 0.73951 | 0.759117 | 0.761888 | 0.713215 | 0.761715 | 0.760132 | 0.758154 |
| 0.757392 | 0.72438 | 0.740059 | 0.757203 | 0.759944 | 0.720135 | 0.76241 | 0.754289 | 0.759379 |
| 0.755706 | 0.734375 | 0.735835 | 0.75934 | 0.765241 | 0.700835 | 0.75998 | 0.757596 | 0.760034 |
| 0.763971 | 0.734126 | 0.736752 | 0.76548 | 0.763677 | 0.717277 | 0.768433 | 0.760596 | 0.763138 |

| | | | | | | | | |
|---|---|---|---|---|---|---|---|---|
| 0.761714 | 0.746617 | 0.739635 | 0.763407 | 0.777253 | 0.721388 | 0.768038 | 0.763764 | 0.766838 |
| 0.756402 | 0.736553 | 0.73351 | 0.760413 | 0.768995 | 0.716557 | 0.763375 | 0.753357 | 0.762164 |
| 0.759696 | 0.732971 | 0.738297 | 0.762688 | 0.770176 | 0.720631 | 0.76585 | 0.758626 | 0.764265 |
| 0.7541 | 0.738066 | 0.736822 | 0.756143 | 0.763959 | 0.700115 | 0.756817 | 0.75364 | 0.756886 |
| 0.751435 | 0.735152 | 0.740126 | 0.75645 | 0.760094 | 0.713069 | 0.764036 | 0.750792 | 0.756159 |
| 0.755235 | 0.731531 | 0.744665 | 0.758615 | 0.761767 | 0.716886 | 0.758942 | 0.754201 | 0.754675 |
| 0.757265 | 0.749732 | 0.746414 | 0.759045 | 0.767012 | 0.713897 | 0.764825 | 0.756488 | 0.762269 |
| 0.758344 | 0.739721 | 0.739992 | 0.762483 | 0.768926 | 0.716009 | 0.767183 | 0.760632 | 0.76086 |
| 0.760488 | 0.734033 | 0.733169 | 0.761439 | 0.767248 | 0.713458 | 0.759778 | 0.755392 | 0.758502 |
| 0.753965 | 0.733757 | 0.731343 | 0.761025 | 0.771201 | 0.712451 | 0.76068 | 0.752944 | 0.756714 |
| 0.753832 | 0.733392 | 0.733396 | 0.757573 | 0.766249 | 0.714838 | 0.762459 | 0.75467 | 0.757896 |
| 0.758641 | 0.742056 | 0.745027 | 0.758767 | 0.767304 | 0.713869 | 0.762248 | 0.754469 | 0.760201 |
| 0.747009 | 0.733477 | 0.735351 | 0.75291 | 0.761038 | 0.71603 | 0.754234 | 0.744613 | 0.752412 |
| 0.755518 | 0.743201 | 0.741631 | 0.75036 | 0.761053 | 0.705093 | 0.760418 | 0.74219 | 0.755431 |
| 0.758609 | 0.723352 | 0.729385 | 0.76994 | 0.763676 | 0.713235 | 0.76362 | 0.756764 | 0.758618 |
| 0.763335 | 0.733929 | 0.739426 | 0.774802 | 0.77551 | 0.717622 | 0.772313 | 0.764836 | 0.767453 |
| 0.760996 | 0.719903 | 0.725205 | 0.771807 | 0.76885 | 0.716221 | 0.767357 | 0.76473 | 0.76496 |
| 0.763992 | 0.72689 | 0.729827 | 0.773147 | 0.771516 | 0.714671 | 0.771381 | 0.761667 | 0.766694 |
| 0.75914 | 0.723589 | 0.738123 | 0.766749 | 0.767158 | 0.719538 | 0.766909 | 0.760561 | 0.762225 |
| 0.7592 | 0.729585 | 0.734557 | 0.769433 | 0.770383 | 0.719084 | 0.77006 | 0.761256 | 0.761209 |
| 0.75738 | 0.731777 | 0.737235 | 0.764235 | 0.768421 | 0.717124 | 0.765824 | 0.756697 | 0.75903 |
| 0.753471 | 0.719363 | 0.729843 | 0.767134 | 0.766026 | 0.716013 | 0.765821 | 0.755707 | 0.762236 |
| 0.764512 | 0.719845 | 0.733168 | 0.775919 | 0.772111 | 0.726437 | 0.77426 | 0.765634 | 0.772359 |
| 0.75505 | 0.738826 | 0.754067 | 0.758554 | 0.769034 | 0.720362 | 0.762791 | 0.753305 | 0.756529 |
| 0.766728 | 0.734164 | 0.742818 | 0.766344 | 0.775109 | 0.727147 | 0.769231 | 0.760074 | 0.768782 |
| 0.706034 | 0.650986 | 0.661237 | 0.708167 | 0.707156 | 0.643572 | 0.707033 | 0.696725 | 0.704527 |
| 0.762087 | 0.730037 | 0.726401 | 0.766634 | 0.771125 | 0.71511 | 0.768248 | 0.758956 | 0.766222 |
| 0.76146 | 0.73139 | 0.735183 | 0.769018 | 0.769206 | 0.721033 | 0.765895 | 0.762976 | 0.766118 |
| 0.765424 | 0.729141 | 0.730529 | 0.767285 | 0.770071 | 0.7206 | 0.769684 | 0.76234 | 0.771452 |
| 0.76748 | 0.732663 | 0.744394 | 0.766228 | 0.769161 | 0.716232 | 0.772321 | 0.761452 | 0.766702 |
| 0.758795 | 0.728625 | 0.730286 | 0.764661 | 0.768316 | 0.711621 | 0.762474 | 0.75395 | 0.759873 |
| 0.762348 | 0.721987 | 0.741316 | 0.763283 | 0.765066 | 0.724225 | 0.769914 | 0.755214 | 0.763605 |
| 0.760337 | 0.726919 | 0.73356 | 0.765168 | 0.769244 | 0.721074 | 0.768034 | 0.757144 | 0.767628 |
| 0.761231 | 0.732127 | 0.737922 | 0.766406 | 0.767934 | 0.727094 | 0.765313 | 0.75736 | 0.76348 |
| 0.755144 | 0.725735 | 0.733629 | 0.760193 | 0.757387 | 0.715602 | 0.757335 | 0.752023 | 0.755234 |
| 0.758364 | 0.726228 | 0.724816 | 0.764711 | 0.76123 | 0.717709 | 0.761969 | 0.757166 | 0.758846 |
| 0.763579 | 0.727867 | 0.74153 | 0.769559 | 0.770477 | 0.728708 | 0.771222 | 0.767859 | 0.770551 |
| 0.763106 | 0.729426 | 0.741098 | 0.765868 | 0.773546 | 0.721942 | 0.7715 | 0.761783 | 0.767122 |
| 0.767505 | 0.729141 | 0.734485 | 0.767232 | 0.770471 | 0.723561 | 0.768584 | 0.762469 | 0.765107 |
| 0.763872 | 0.71636 | 0.734049 | 0.767649 | 0.767669 | 0.71993 | 0.76739 | 0.758723 | 0.764766 |
| 0.758221 | 0.722437 | 0.719579 | 0.762563 | 0.76797 | 0.718939 | 0.761746 | 0.75817 | 0.762023 |
| 0.761826 | 0.72899 | 0.728637 | 0.765895 | 0.773526 | 0.721654 | 0.772563 | 0.763827 | 0.767043 |
| 0.759908 | 0.726146 | 0.739022 | 0.76141 | 0.765114 | 0.720545 | 0.75999 | 0.753973 | 0.76199 |
| 0.758498 | 0.729229 | 0.736164 | 0.760817 | 0.763913 | 0.719512 | 0.758279 | 0.756382 | 0.755859 |
| 0.762584 | 0.737868 | 0.7432 | 0.769566 | 0.766417 | 0.723125 | 0.768881 | 0.763763 | 0.76657 |
| 0.761986 | 0.735802 | 0.735545 | 0.76656 | 0.765349 | 0.718424 | 0.767626 | 0.761138 | 0.765849 |

| | | | | | | | | |
|---|---|---|---|---|---|---|---|---|
| 0.76325 | 0.735393 | 0.734405 | 0.762308 | 0.765359 | 0.712754 | 0.765313 | 0.757868 | 0.766344 |
| 0.7623 | 0.741352 | 0.748627 | 0.763796 | 0.767594 | 0.720146 | 0.764937 | 0.760691 | 0.762434 |
| 0.75632 | 0.733511 | 0.73913 | 0.760228 | 0.763928 | 0.721649 | 0.75918 | 0.756399 | 0.758569 |
| 0.716767 | 0.723599 | 0.714604 | 0.726331 | 0.734118 | 0.705233 | 0.731214 | 0.709872 | 0.70985 |
| 0.757458 | 0.732377 | 0.729405 | 0.759063 | 0.764351 | 0.714258 | 0.762083 | 0.751434 | 0.760825 |
| 0.756014 | 0.730762 | 0.731471 | 0.760167 | 0.769503 | 0.71423 | 0.76366 | 0.753479 | 0.759803 |
| 0.756147 | 0.733087 | 0.738634 | 0.760537 | 0.769667 | 0.714918 | 0.761678 | 0.757401 | 0.761371 |
| 0.756814 | 0.734059 | 0.732754 | 0.757998 | 0.76665 | 0.718047 | 0.763037 | 0.758172 | 0.767535 |
| 0.762083 | 0.740354 | 0.738949 | 0.765266 | 0.770327 | 0.718768 | 0.766577 | 0.75958 | 0.76751 |
| 0.758284 | 0.739262 | 0.744146 | 0.763162 | 0.7679 | 0.724245 | 0.763587 | 0.757287 | 0.762162 |
| 0.758292 | 0.729174 | 0.738176 | 0.756882 | 0.767239 | 0.721451 | 0.763817 | 0.7595 | 0.765336 |
| 0.757198 | 0.731664 | 0.735658 | 0.764595 | 0.769839 | 0.717615 | 0.764526 | 0.757717 | 0.764203 |
| 0.756438 | 0.729185 | 0.737628 | 0.759142 | 0.771053 | 0.722233 | 0.763451 | 0.758276 | 0.764817 |
| 0.758657 | 0.731789 | 0.735621 | 0.763747 | 0.76845 | 0.72282 | 0.76761 | 0.758921 | 0.762629 |
| 0.761617 | 0.733556 | 0.736377 | 0.763856 | 0.770599 | 0.727994 | 0.769547 | 0.761547 | 0.766508 |
| 0.756374 | 0.724544 | 0.73167 | 0.757837 | 0.770832 | 0.717959 | 0.7625 | 0.756832 | 0.758059 |
| 0.766375 | 0.739856 | 0.744089 | 0.762571 | 0.773577 | 0.723819 | 0.770419 | 0.759835 | 0.767055 |
| 0.758769 | 0.73637 | 0.737001 | 0.765266 | 0.773751 | 0.724797 | 0.765575 | 0.760632 | 0.766404 |
| 0.754426 | 0.733017 | 0.733813 | 0.761082 | 0.764176 | 0.723585 | 0.768307 | 0.759726 | 0.764214 |
| 0.760875 | 0.734834 | 0.734125 | 0.761823 | 0.765408 | 0.718672 | 0.762888 | 0.758702 | 0.765778 |
| 0.760382 | 0.735604 | 0.740825 | 0.764792 | 0.77331 | 0.719707 | 0.769136 | 0.761784 | 0.765917 |
| 0.759071 | 0.731094 | 0.739213 | 0.760924 | 0.774986 | 0.725966 | 0.766475 | 0.763374 | 0.769585 |
| 0.673503 | 0.67838 | 0.688418 | 0.688946 | 0.692336 | 0.669338 | 0.683118 | 0.675419 | 0.678105 |
| 0.658773 | 0.65133 | 0.66857 | 0.663901 | 0.665 | 0.64084 | 0.656375 | 0.645131 | 0.654353 |
| 0.676969 | 0.638481 | 0.652943 | 0.681378 | 0.68492 | 0.644485 | 0.673509 | 0.665451 | 0.66997 |
| 0.656863 | 0.649311 | 0.657701 | 0.663134 | 0.663002 | 0.637406 | 0.661089 | 0.648968 | 0.654655 |
| 0.716066 | 0.743357 | 0.746089 | 0.724342 | 0.736724 | 0.71571 | 0.731126 | 0.718099 | 0.728476 |
| 0.718934 | 0.733819 | 0.740267 | 0.720535 | 0.731034 | 0.717298 | 0.723664 | 0.720653 | 0.719232 |
| 0.717578 | 0.75 | 0.742679 | 0.724156 | 0.733803 | 0.716761 | 0.730922 | 0.723473 | 0.726114 |
| 0.729562 | 0.720381 | 0.747297 | 0.741086 | 0.745155 | 0.704221 | 0.737739 | 0.728333 | 0.733472 |
| 0.708296 | 0.724883 | 0.730348 | 0.717431 | 0.723995 | 0.694642 | 0.717202 | 0.708485 | 0.706507 |
| 0.692997 | 0.724509 | 0.727031 | 0.706425 | 0.721274 | 0.701248 | 0.709459 | 0.707684 | 0.700893 |
| 0.704366 | 0.740295 | 0.720976 | 0.705061 | 0.719407 | 0.702534 | 0.706967 | 0.699649 | 0.706208 |
| 0.660283 | 0.655504 | 0.660126 | 0.662706 | 0.665712 | 0.650769 | 0.657843 | 0.647667 | 0.662698 |
| 0.75309 | 0.745541 | 0.736166 | 0.760003 | 0.765979 | 0.714231 | 0.762659 | 0.752944 | 0.758452 |
| 0.728425 | 0.72684 | 0.734439 | 0.737536 | 0.744897 | 0.715648 | 0.743226 | 0.731606 | 0.739439 |
| 0.726051 | 0.742698 | 0.745132 | 0.732553 | 0.742202 | 0.72015 | 0.733181 | 0.726595 | 0.733171 |
| 0.713653 | 0.732194 | 0.751238 | 0.711802 | 0.719298 | 0.7122 | 0.71837 | 0.705524 | 0.714496 |
| 0.73235 | 0.737024 | 0.745335 | 0.733943 | 0.74443 | 0.715356 | 0.73911 | 0.731177 | 0.728874 |
| 0.728898 | 0.724056 | 0.734342 | 0.730917 | 0.735765 | 0.723409 | 0.73217 | 0.731836 | 0.731177 |
| 0.722933 | 0.724942 | 0.731384 | 0.719163 | 0.727693 | 0.711754 | 0.72358 | 0.719226 | 0.727157 |
| 0.724676 | 0.734769 | 0.737869 | 0.722257 | 0.732084 | 0.721756 | 0.722811 | 0.723218 | 0.724887 |
| 0.72043 | 0.731388 | 0.739084 | 0.727126 | 0.738247 | 0.714324 | 0.725995 | 0.718727 | 0.722582 |
| 0.711332 | 0.732047 | 0.743335 | 0.727344 | 0.731187 | 0.714737 | 0.725186 | 0.712968 | 0.715259 |
| 0.72553 | 0.736509 | 0.743553 | 0.727179 | 0.737196 | 0.72244 | 0.716871 | 0.720072 | 0.720285 |
| 0.705999 | 0.73102 | 0.73182 | 0.715374 | 0.719391 | 0.699914 | 0.716694 | 0.702488 | 0.703594 |

| | | | | | | | | |
|---|---|---|---|---|---|---|---|---|
| 0.74428 | 0.739687 | 0.744403 | 0.742056 | 0.755909 | 0.721949 | 0.750544 | 0.741875 | 0.747748 |
| 0.715753 | 0.730655 | 0.731149 | 0.724605 | 0.729263 | 0.712783 | 0.726674 | 0.71769 | 0.720324 |
| 0.720608 | 0.727594 | 0.739995 | 0.730327 | 0.734356 | 0.714012 | 0.729932 | 0.725019 | 0.729369 |
| 0.720463 | 0.732203 | 0.739287 | 0.728624 | 0.728302 | 0.710255 | 0.725209 | 0.71997 | 0.72439 |
| 0.712249 | 0.729927 | 0.737881 | 0.734506 | 0.731249 | 0.719344 | 0.725872 | 0.717489 | 0.725244 |
| 0.710882 | 0.741967 | 0.737491 | 0.716007 | 0.727861 | 0.714627 | 0.721703 | 0.71238 | 0.713792 |
| 0.710089 | 0.732154 | 0.742657 | 0.715665 | 0.715876 | 0.712187 | 0.710173 | 0.711333 | 0.711971 |
| 0.70845 | 0.72779 | 0.735878 | 0.708567 | 0.71364 | 0.713313 | 0.711317 | 0.699637 | 0.70392 |
| 0.708602 | 0.733976 | 0.741119 | 0.712528 | 0.726604 | 0.713789 | 0.710003 | 0.70906 | 0.715695 |
| 0.746179 | 0.776586 | 0.760747 | 0.752244 | 0.759366 | 0.719493 | 0.75498 | 0.748144 | 0.748952 |
| 0.747216 | 0.781417 | 0.752489 | 0.743439 | 0.756819 | 0.71319 | 0.753779 | 0.741224 | 0.747616 |
| 0.740435 | 0.782137 | 0.759894 | 0.73622 | 0.74906 | 0.715886 | 0.743194 | 0.730533 | 0.740037 |
| 0.731593 | 0.795135 | 0.74979 | 0.732716 | 0.741104 | 0.719585 | 0.731122 | 0.727284 | 0.7383 |
| 0.736356 | 0.800207 | 0.752321 | 0.734432 | 0.746378 | 0.718249 | 0.7432 | 0.73672 | 0.742815 |
| 0.72969 | 0.781444 | 0.753536 | 0.734041 | 0.739743 | 0.72135 | 0.740285 | 0.732932 | 0.737789 |

| | | plot 02 | | | | | | |
|---|---|---|---|---|---|---|---|---|
| Wild_plot_ | Wild_plot_ | Wild_plot_ | Wild_plot_ | Wild_plot_ | Wild_plot_ | Wild_plot_ | Wild_plot_ | Wild_plot_ |
| 0.707361 | 0.710586 | 0.706615 | 0.715263 | 0.704402 | 0.705339 | 0.699992 | 0.700514 | 0.705658 |
| 0.716439 | 0.713202 | 0.716042 | 0.716882 | 0.710915 | 0.714774 | 0.709993 | 0.711362 | 0.711715 |
| 0.70391 | 0.705378 | 0.701703 | 0.708103 | 0.699548 | 0.701256 | 0.69838 | 0.698014 | 0.702092 |
| 0.713029 | 0.712546 | 0.714297 | 0.719122 | 0.712268 | 0.711519 | 0.706663 | 0.710334 | 0.709183 |
| 0.712507 | 0.708671 | 0.708756 | 0.713754 | 0.707055 | 0.706151 | 0.70307 | 0.706409 | 0.70585 |
| 0.718457 | 0.718717 | 0.722784 | 0.721591 | 0.716126 | 0.716138 | 0.709763 | 0.714955 | 0.715265 |
| 0.766713 | 0.762 | 0.76689 | 0.76918 | 0.766551 | 0.768271 | 0.756636 | 0.765173 | 0.75879 |
| 0.766134 | 0.758681 | 0.764441 | 0.764773 | 0.763292 | 0.7604 | 0.752514 | 0.757132 | 0.756615 |
| 0.714455 | 0.715079 | 0.716568 | 0.722031 | 0.715767 | 0.711644 | 0.707954 | 0.70895 | 0.713702 |
| 0.713586 | 0.710824 | 0.712195 | 0.715876 | 0.708656 | 0.711519 | 0.701528 | 0.709811 | 0.703998 |
| 0.713526 | 0.710094 | 0.714069 | 0.714628 | 0.706546 | 0.714604 | 0.703742 | 0.709004 | 0.707963 |
| 0.70891 | 0.703962 | 0.705178 | 0.710299 | 0.70253 | 0.705283 | 0.700569 | 0.701951 | 0.704182 |
| 0.715487 | 0.711772 | 0.715574 | 0.716789 | 0.713002 | 0.712198 | 0.709553 | 0.711646 | 0.713954 |
| 0.723617 | 0.725047 | 0.724182 | 0.729334 | 0.716427 | 0.722487 | 0.715382 | 0.718972 | 0.718181 |
| 0.738251 | 0.736091 | 0.736088 | 0.738006 | 0.734983 | 0.732299 | 0.72948 | 0.732826 | 0.734743 |
| 0.716385 | 0.716927 | 0.716706 | 0.721434 | 0.712563 | 0.712851 | 0.711534 | 0.712094 | 0.716777 |
| 0.718558 | 0.719865 | 0.717204 | 0.725316 | 0.714008 | 0.717405 | 0.705758 | 0.71424 | 0.713192 |
| 0.747693 | 0.739481 | 0.744448 | 0.747266 | 0.742758 | 0.742943 | 0.731094 | 0.739068 | 0.739472 |
| 0.709944 | 0.70944 | 0.709049 | 0.7115 | 0.705693 | 0.703921 | 0.699139 | 0.702647 | 0.705736 |
| 0.711619 | 0.712038 | 0.712057 | 0.716761 | 0.708484 | 0.70768 | 0.704023 | 0.70969 | 0.710451 |
| 0.707924 | 0.708915 | 0.706977 | 0.714661 | 0.704902 | 0.705628 | 0.701219 | 0.700822 | 0.704142 |
| 0.709769 | 0.711006 | 0.710505 | 0.715868 | 0.708701 | 0.708997 | 0.704251 | 0.704071 | 0.704814 |
| 0.718421 | 0.715403 | 0.717787 | 0.719857 | 0.714297 | 0.717464 | 0.710319 | 0.711652 | 0.717682 |
| 0.71501 | 0.713424 | 0.715675 | 0.721548 | 0.711939 | 0.715659 | 0.70609 | 0.712202 | 0.711807 |
| 0.715523 | 0.713293 | 0.715384 | 0.71837 | 0.712309 | 0.714685 | 0.708975 | 0.712007 | 0.717274 |
| 0.714241 | 0.710199 | 0.714207 | 0.718311 | 0.710762 | 0.708002 | 0.706639 | 0.711446 | 0.712018 |
| 0.707261 | 0.701099 | 0.707102 | 0.709865 | 0.699411 | 0.70344 | 0.697661 | 0.703713 | 0.703101 |
| 0.71459 | 0.711809 | 0.714602 | 0.717117 | 0.71078 | 0.710941 | 0.704527 | 0.710364 | 0.710424 |
| 0.713608 | 0.714659 | 0.713557 | 0.716492 | 0.71014 | 0.713457 | 0.707537 | 0.710598 | 0.712798 |
| 0.714479 | 0.713043 | 0.711469 | 0.715759 | 0.707884 | 0.71253 | 0.707362 | 0.709639 | 0.7082 |
| 0.714047 | 0.711122 | 0.713371 | 0.716869 | 0.710325 | 0.710133 | 0.704905 | 0.705568 | 0.711738 |
| 0.71184 | 0.709116 | 0.71203 | 0.716065 | 0.703966 | 0.70999 | 0.702021 | 0.70686 | 0.707975 |
| 0.710877 | 0.709029 | 0.709549 | 0.715544 | 0.708725 | 0.709766 | 0.702761 | 0.707894 | 0.70878 |
| 0.715209 | 0.715004 | 0.715877 | 0.720712 | 0.711028 | 0.712975 | 0.706184 | 0.710128 | 0.710235 |
| 0.706429 | 0.70572 | 0.706274 | 0.70928 | 0.701619 | 0.702592 | 0.699526 | 0.702548 | 0.706601 |
| 0.711279 | 0.711131 | 0.712527 | 0.715419 | 0.705255 | 0.709438 | 0.70463 | 0.710349 | 0.706782 |
| 0.717343 | 0.71246 | 0.716414 | 0.718243 | 0.712063 | 0.713925 | 0.703078 | 0.709774 | 0.710785 |
| 0.71456 | 0.71155 | 0.713422 | 0.715559 | 0.709826 | 0.713885 | 0.709838 | 0.709402 | 0.713812 |
| 0.720307 | 0.716669 | 0.7182 | 0.721355 | 0.716439 | 0.716333 | 0.713076 | 0.715504 | 0.715725 |

| | | | | | | | | |
|---|---|---|---|---|---|---|---|---|
| 0.751815 | 0.746944 | 0.748195 | 0.751825 | 0.746646 | 0.749259 | 0.741365 | 0.742075 | 0.743559 |
| 0.706624 | 0.704959 | 0.705487 | 0.708604 | 0.70233 | 0.700944 | 0.699064 | 0.700531 | 0.702554 |
| 0.707462 | 0.705935 | 0.708085 | 0.712829 | 0.704493 | 0.704883 | 0.699534 | 0.703977 | 0.705939 |
| 0.714623 | 0.715526 | 0.715136 | 0.723145 | 0.7131 | 0.71459 | 0.709264 | 0.711831 | 0.714206 |
| 0.710323 | 0.711907 | 0.713207 | 0.715295 | 0.707821 | 0.708893 | 0.702508 | 0.704419 | 0.708041 |
| 0.714786 | 0.711504 | 0.713829 | 0.71416 | 0.711469 | 0.71283 | 0.706566 | 0.710707 | 0.706942 |
| 0.70986 | 0.707742 | 0.708548 | 0.710947 | 0.704397 | 0.70956 | 0.703327 | 0.706675 | 0.704172 |
| 0.713911 | 0.711268 | 0.715134 | 0.717139 | 0.708739 | 0.708429 | 0.705312 | 0.708517 | 0.709157 |
| 0.71521 | 0.713864 | 0.715501 | 0.72145 | 0.71319 | 0.711932 | 0.70573 | 0.710151 | 0.713192 |
| 0.711992 | 0.71032 | 0.713448 | 0.718765 | 0.709605 | 0.709807 | 0.706057 | 0.710543 | 0.710706 |
| 0.702096 | 0.703655 | 0.703813 | 0.711692 | 0.702757 | 0.702448 | 0.698046 | 0.699921 | 0.700886 |
| 0.705836 | 0.704075 | 0.709796 | 0.709683 | 0.703772 | 0.702242 | 0.700548 | 0.703503 | 0.702129 |
| 0.709952 | 0.707333 | 0.707315 | 0.710463 | 0.706541 | 0.70636 | 0.705096 | 0.703225 | 0.707649 |
| 0.708 | 0.705882 | 0.70652 | 0.710946 | 0.705291 | 0.703604 | 0.697588 | 0.703429 | 0.70748 |
| 0.707722 | 0.706734 | 0.709151 | 0.712388 | 0.703357 | 0.705093 | 0.69698 | 0.705462 | 0.704214 |
| 0.706189 | 0.7078 | 0.707421 | 0.710881 | 0.701826 | 0.701952 | 0.695995 | 0.701756 | 0.70284 |
| 0.70812 | 0.70346 | 0.705377 | 0.708939 | 0.701348 | 0.703893 | 0.699489 | 0.701314 | 0.705783 |
| 0.712627 | 0.708435 | 0.713194 | 0.716184 | 0.708103 | 0.710493 | 0.705193 | 0.710794 | 0.709194 |
| 0.716107 | 0.716294 | 0.714183 | 0.720645 | 0.710463 | 0.715023 | 0.706441 | 0.711955 | 0.711259 |
| 0.709665 | 0.705099 | 0.708253 | 0.710341 | 0.706689 | 0.703203 | 0.706107 | 0.703269 | 0.706005 |
| 0.703413 | 0.70439 | 0.702731 | 0.707507 | 0.701202 | 0.700728 | 0.696065 | 0.700851 | 0.703051 |
| 0.707174 | 0.707391 | 0.705031 | 0.713408 | 0.703555 | 0.705188 | 0.701241 | 0.703125 | 0.708263 |
| 0.711349 | 0.710258 | 0.711842 | 0.716371 | 0.707067 | 0.70676 | 0.702905 | 0.708779 | 0.707013 |
| 0.719605 | 0.718012 | 0.722294 | 0.72515 | 0.715526 | 0.715514 | 0.712607 | 0.718747 | 0.714618 |
| 0.725167 | 0.720143 | 0.727897 | 0.727527 | 0.723245 | 0.723367 | 0.71668 | 0.721355 | 0.719158 |
| 0.765561 | 0.758862 | 0.76565 | 0.765229 | 0.764321 | 0.761039 | 0.751038 | 0.758425 | 0.756218 |
| 0.772218 | 0.768046 | 0.771047 | 0.770625 | 0.770783 | 0.768792 | 0.760782 | 0.76809 | 0.765464 |
| 0.772447 | 0.769108 | 0.774867 | 0.774319 | 0.768879 | 0.773339 | 0.76334 | 0.769613 | 0.766769 |
| 0.708686 | 0.705265 | 0.707519 | 0.712019 | 0.706666 | 0.706377 | 0.699205 | 0.704187 | 0.705338 |
| 0.709706 | 0.708571 | 0.70835 | 0.715108 | 0.707893 | 0.708177 | 0.700207 | 0.704949 | 0.704944 |
| 0.722581 | 0.718245 | 0.716238 | 0.725317 | 0.715551 | 0.717312 | 0.713256 | 0.715121 | 0.717031 |
| 0.716383 | 0.717504 | 0.716986 | 0.719535 | 0.712431 | 0.712484 | 0.709593 | 0.709553 | 0.714019 |
| 0.707438 | 0.710994 | 0.708484 | 0.717642 | 0.706539 | 0.707743 | 0.699594 | 0.703585 | 0.702648 |
| 0.707561 | 0.708101 | 0.706823 | 0.715145 | 0.704996 | 0.706965 | 0.701945 | 0.70501 | 0.709508 |
| 0.706076 | 0.707846 | 0.705906 | 0.716192 | 0.705372 | 0.707464 | 0.699902 | 0.704208 | 0.704954 |
| 0.706517 | 0.709348 | 0.707097 | 0.715704 | 0.702536 | 0.705434 | 0.700105 | 0.705906 | 0.707393 |
| 0.707661 | 0.708162 | 0.709448 | 0.715221 | 0.707358 | 0.707872 | 0.698911 | 0.707142 | 0.70194 |
| 0.723364 | 0.725833 | 0.722441 | 0.731947 | 0.721635 | 0.723685 | 0.716157 | 0.718094 | 0.719537 |
| 0.7011 | 0.706519 | 0.699637 | 0.709811 | 0.698425 | 0.700559 | 0.695761 | 0.69658 | 0.699171 |
| 0.71143 | 0.711183 | 0.711127 | 0.71591 | 0.704787 | 0.707954 | 0.701154 | 0.707893 | 0.708419 |
| 0.699571 | 0.705128 | 0.705567 | 0.712391 | 0.69977 | 0.701655 | 0.698925 | 0.698946 | 0.702998 |
| 0.724176 | 0.726703 | 0.725476 | 0.732499 | 0.724169 | 0.723476 | 0.71728 | 0.7211 | 0.724734 |
| 0.711217 | 0.710776 | 0.713316 | 0.718268 | 0.709945 | 0.713273 | 0.702248 | 0.708636 | 0.708731 |
| 0.705601 | 0.706877 | 0.707421 | 0.712493 | 0.706126 | 0.707145 | 0.700455 | 0.702309 | 0.704 |
| 0.714366 | 0.712475 | 0.713908 | 0.718727 | 0.709506 | 0.710017 | 0.704279 | 0.712361 | 0.70735 |
| 0.707325 | 0.709265 | 0.707895 | 0.714814 | 0.703038 | 0.702823 | 0.700908 | 0.706506 | 0.707107 |

| | | | | | | | | |
|---|---|---|---|---|---|---|---|---|
| 0.707793 | 0.713035 | 0.709376 | 0.715744 | 0.708558 | 0.706745 | 0.702966 | 0.704974 | 0.706631 |
| 0.712157 | 0.711122 | 0.712382 | 0.716189 | 0.707772 | 0.709596 | 0.702763 | 0.706544 | 0.708588 |
| 0.711195 | 0.709234 | 0.712514 | 0.716354 | 0.706702 | 0.709361 | 0.701791 | 0.707415 | 0.705783 |
| 0.718231 | 0.720794 | 0.718596 | 0.724707 | 0.71793 | 0.71463 | 0.708903 | 0.710929 | 0.716071 |
| 0.715479 | 0.716279 | 0.716123 | 0.725058 | 0.713783 | 0.716282 | 0.706462 | 0.71382 | 0.710173 |
| 0.709004 | 0.707866 | 0.708491 | 0.715512 | 0.704196 | 0.708353 | 0.69874 | 0.706005 | 0.705097 |
| 0.711787 | 0.712529 | 0.71424 | 0.718963 | 0.709257 | 0.712193 | 0.704342 | 0.708963 | 0.70829 |
| 0.715232 | 0.71206 | 0.712646 | 0.718552 | 0.710136 | 0.711163 | 0.704213 | 0.710081 | 0.71288 |
| 0.710088 | 0.710716 | 0.710067 | 0.716338 | 0.706516 | 0.708287 | 0.702585 | 0.706833 | 0.707372 |
| 0.708438 | 0.706208 | 0.707207 | 0.71179 | 0.702679 | 0.703739 | 0.702837 | 0.704536 | 0.701174 |
| 0.710376 | 0.710955 | 0.708287 | 0.714139 | 0.707462 | 0.705482 | 0.701925 | 0.704141 | 0.709187 |
| 0.709564 | 0.710236 | 0.708246 | 0.715715 | 0.705606 | 0.70665 | 0.705954 | 0.702746 | 0.704667 |
| 0.713271 | 0.712914 | 0.714389 | 0.721131 | 0.709586 | 0.712745 | 0.705696 | 0.71074 | 0.708478 |
| 0.708631 | 0.707791 | 0.70804 | 0.717758 | 0.706713 | 0.707474 | 0.700245 | 0.703417 | 0.705283 |
| 0.711867 | 0.710619 | 0.710354 | 0.71601 | 0.707155 | 0.706689 | 0.705873 | 0.705253 | 0.709419 |
| 0.707691 | 0.709704 | 0.708446 | 0.710655 | 0.701284 | 0.705966 | 0.703558 | 0.706554 | 0.708914 |
| 0.715129 | 0.715575 | 0.717362 | 0.720174 | 0.708999 | 0.712085 | 0.704827 | 0.713274 | 0.710859 |
| 0.708873 | 0.709559 | 0.710088 | 0.71424 | 0.701044 | 0.705079 | 0.700819 | 0.703274 | 0.706135 |
| 0.709001 | 0.711443 | 0.706997 | 0.717436 | 0.704808 | 0.705417 | 0.699582 | 0.703078 | 0.705666 |
| 0.710974 | 0.709478 | 0.712475 | 0.715711 | 0.710084 | 0.70898 | 0.701593 | 0.708469 | 0.705807 |
| 0.708032 | 0.711964 | 0.708134 | 0.71865 | 0.709296 | 0.706275 | 0.703722 | 0.70592 | 0.707577 |
| 0.70553 | 0.70881 | 0.706405 | 0.710754 | 0.70384 | 0.700961 | 0.698341 | 0.701927 | 0.703651 |
| 0.711974 | 0.712423 | 0.709719 | 0.714938 | 0.70654 | 0.708688 | 0.701601 | 0.706567 | 0.707102 |
| 0.712214 | 0.710637 | 0.711991 | 0.719516 | 0.711529 | 0.71027 | 0.701331 | 0.711054 | 0.7084 |
| 0.714194 | 0.716685 | 0.711799 | 0.718306 | 0.707919 | 0.711879 | 0.703081 | 0.709334 | 0.711109 |
| 0.716843 | 0.712469 | 0.718388 | 0.717788 | 0.716772 | 0.715075 | 0.708033 | 0.714887 | 0.710071 |
| 0.71157 | 0.712599 | 0.708402 | 0.717334 | 0.708702 | 0.709492 | 0.705821 | 0.707773 | 0.709654 |
| 0.715314 | 0.711176 | 0.711349 | 0.716527 | 0.711029 | 0.711643 | 0.702277 | 0.70874 | 0.709757 |
| 0.714399 | 0.711811 | 0.711653 | 0.717845 | 0.710299 | 0.709529 | 0.703641 | 0.709629 | 0.708978 |
| 0.712655 | 0.713107 | 0.717296 | 0.721511 | 0.712891 | 0.714366 | 0.705887 | 0.710973 | 0.707681 |
| 0.712055 | 0.713595 | 0.713139 | 0.719453 | 0.7152 | 0.714229 | 0.706178 | 0.708792 | 0.71222 |
| 0.710083 | 0.712259 | 0.712002 | 0.717937 | 0.706585 | 0.708226 | 0.702104 | 0.709426 | 0.709002 |
| 0.70889 | 0.712011 | 0.712958 | 0.72179 | 0.707295 | 0.709328 | 0.699366 | 0.707953 | 0.705958 |
| 0.711349 | 0.715494 | 0.711224 | 0.723346 | 0.708617 | 0.710134 | 0.705805 | 0.707578 | 0.710898 |
| 0.708764 | 0.70818 | 0.710307 | 0.715272 | 0.703951 | 0.705069 | 0.699814 | 0.70501 | 0.70346 |
| 0.707317 | 0.710568 | 0.710452 | 0.715151 | 0.706011 | 0.70776 | 0.703036 | 0.703826 | 0.705925 |
| 0.706692 | 0.708936 | 0.704759 | 0.712136 | 0.70177 | 0.701895 | 0.700475 | 0.702348 | 0.704306 |
| 0.684057 | 0.688095 | 0.687399 | 0.692105 | 0.683158 | 0.686273 | 0.67173 | 0.685578 | 0.680823 |
| 0.689448 | 0.692268 | 0.68877 | 0.697098 | 0.689512 | 0.687669 | 0.673374 | 0.685116 | 0.688072 |
| 0.682809 | 0.681861 | 0.68345 | 0.686011 | 0.680541 | 0.682269 | 0.671522 | 0.678406 | 0.675793 |
| 0.677979 | 0.68043 | 0.680451 | 0.688687 | 0.67648 | 0.676138 | 0.671144 | 0.673447 | 0.674839 |
| 0.663546 | 0.667884 | 0.666279 | 0.678083 | 0.664302 | 0.664307 | 0.657416 | 0.662551 | 0.662096 |
| 0.668902 | 0.668856 | 0.669888 | 0.674651 | 0.667149 | 0.668022 | 0.660988 | 0.664376 | 0.664037 |
| 0.667712 | 0.666097 | 0.670303 | 0.674824 | 0.666384 | 0.667267 | 0.659593 | 0.66594 | 0.664798 |
| 0.674095 | 0.676312 | 0.675229 | 0.684036 | 0.671193 | 0.670077 | 0.663817 | 0.665145 | 0.667434 |
| 0.66716 | 0.672312 | 0.671269 | 0.678286 | 0.667581 | 0.669315 | 0.655099 | 0.660994 | 0.663953 |
| 0.672129 | 0.673642 | 0.673037 | 0.684002 | 0.672448 | 0.674262 | 0.661184 | 0.669585 | 0.671931 |

| | | | | | | | | |
|---|---|---|---|---|---|---|---|---|
| 0.676489 | 0.675468 | 0.676055 | 0.680597 | 0.671129 | 0.675679 | 0.658873 | 0.668911 | 0.673421 |
| 0.675655 | 0.675697 | 0.674023 | 0.688024 | 0.673735 | 0.673443 | 0.665306 | 0.665933 | 0.67401 |
| 0.668738 | 0.664928 | 0.673317 | 0.675568 | 0.668263 | 0.67245 | 0.660416 | 0.666223 | 0.661058 |
| 0.666667 | 0.672621 | 0.668122 | 0.677189 | 0.664515 | 0.665048 | 0.66059 | 0.664998 | 0.659317 |
| 0.663952 | 0.667183 | 0.666667 | 0.673938 | 0.66607 | 0.666709 | 0.655706 | 0.664912 | 0.655581 |
| 0.663722 | 0.668512 | 0.666393 | 0.672819 | 0.667626 | 0.664761 | 0.656834 | 0.662966 | 0.660599 |
| 0.670316 | 0.667613 | 0.676207 | 0.676038 | 0.667793 | 0.66977 | 0.660242 | 0.660619 | 0.659629 |
| 0.660603 | 0.666714 | 0.667649 | 0.671425 | 0.661315 | 0.659875 | 0.654227 | 0.661742 | 0.664076 |
| 0.664826 | 0.667158 | 0.662349 | 0.66989 | 0.66603 | 0.659078 | 0.657934 | 0.658533 | 0.66218 |
| 0.663915 | 0.666177 | 0.673063 | 0.67577 | 0.66894 | 0.667539 | 0.656001 | 0.663654 | 0.659141 |
| 0.660854 | 0.657267 | 0.660848 | 0.667667 | 0.658321 | 0.657734 | 0.652867 | 0.649734 | 0.655746 |
| 0.700039 | 0.701581 | 0.69881 | 0.708219 | 0.695796 | 0.698673 | 0.693465 | 0.697587 | 0.696103 |
| 0.70579 | 0.710409 | 0.708952 | 0.71802 | 0.7057 | 0.705896 | 0.704531 | 0.706333 | 0.704921 |
| 0.681557 | 0.689358 | 0.684246 | 0.694217 | 0.684211 | 0.683456 | 0.674732 | 0.680045 | 0.677489 |
| 0.717142 | 0.722512 | 0.718366 | 0.726934 | 0.714878 | 0.713723 | 0.711479 | 0.714586 | 0.713718 |
| 0.707562 | 0.705882 | 0.705578 | 0.71731 | 0.70191 | 0.705286 | 0.699591 | 0.705114 | 0.702068 |
| 0.713014 | 0.712342 | 0.713121 | 0.718983 | 0.711422 | 0.711658 | 0.705134 | 0.711973 | 0.711113 |
| 0.722805 | 0.720419 | 0.721931 | 0.725952 | 0.720042 | 0.719705 | 0.713587 | 0.716829 | 0.718214 |
| 0.696149 | 0.699135 | 0.697904 | 0.704585 | 0.691059 | 0.695024 | 0.688933 | 0.693876 | 0.694811 |
| 0.720839 | 0.723973 | 0.719708 | 0.729747 | 0.715183 | 0.719836 | 0.71146 | 0.7173 | 0.715933 |
| 0.688439 | 0.697762 | 0.692843 | 0.704718 | 0.693028 | 0.688847 | 0.684063 | 0.687523 | 0.686421 |
| 0.719328 | 0.715057 | 0.715656 | 0.71915 | 0.714907 | 0.713432 | 0.709195 | 0.714778 | 0.714263 |
| 0.72539 | 0.720634 | 0.723922 | 0.722107 | 0.718533 | 0.71808 | 0.713754 | 0.720787 | 0.716296 |
| 0.679756 | 0.687265 | 0.680221 | 0.692997 | 0.684253 | 0.68719 | 0.673171 | 0.678295 | 0.678517 |
| 0.689469 | 0.688355 | 0.688186 | 0.697823 | 0.688147 | 0.686413 | 0.67875 | 0.685611 | 0.684228 |
| 0.691623 | 0.692895 | 0.689607 | 0.697713 | 0.687311 | 0.689339 | 0.686908 | 0.685185 | 0.688056 |
| 0.705849 | 0.712025 | 0.709788 | 0.712381 | 0.707002 | 0.705074 | 0.703228 | 0.704468 | 0.706372 |
| 0.715043 | 0.717512 | 0.717637 | 0.723097 | 0.717908 | 0.715565 | 0.708156 | 0.712797 | 0.717709 |
| 0.686995 | 0.686723 | 0.687864 | 0.694594 | 0.685168 | 0.686748 | 0.678594 | 0.684868 | 0.682769 |
| 0.722559 | 0.726719 | 0.724853 | 0.732956 | 0.721903 | 0.725589 | 0.716221 | 0.721242 | 0.716238 |
| 0.726688 | 0.726164 | 0.727007 | 0.731399 | 0.725048 | 0.726614 | 0.716389 | 0.723972 | 0.722279 |
| 0.69951 | 0.699278 | 0.698032 | 0.704835 | 0.695236 | 0.697006 | 0.688257 | 0.695184 | 0.693446 |
| 0.700902 | 0.701436 | 0.698398 | 0.705394 | 0.694867 | 0.696276 | 0.68977 | 0.694475 | 0.693073 |
| 0.724423 | 0.732786 | 0.729929 | 0.734954 | 0.72504 | 0.726837 | 0.719101 | 0.723588 | 0.718665 |
| 0.725252 | 0.723582 | 0.723131 | 0.728728 | 0.716642 | 0.722889 | 0.714626 | 0.721099 | 0.715411 |
| 0.727862 | 0.727564 | 0.729256 | 0.734832 | 0.723645 | 0.72728 | 0.721811 | 0.723415 | 0.721327 |
| 0.676407 | 0.677691 | 0.678969 | 0.680404 | 0.672879 | 0.676691 | 0.668496 | 0.673159 | 0.674473 |
| 0.72195 | 0.721057 | 0.720897 | 0.729861 | 0.716785 | 0.719671 | 0.71198 | 0.716977 | 0.715094 |
| 0.677483 | 0.67538 | 0.678741 | 0.680877 | 0.673968 | 0.676421 | 0.670516 | 0.673946 | 0.671689 |
| 0.670656 | 0.66695 | 0.671494 | 0.67362 | 0.666895 | 0.668147 | 0.66325 | 0.669373 | 0.667072 |
| 0.724113 | 0.724701 | 0.72318 | 0.734583 | 0.715776 | 0.718587 | 0.716287 | 0.716503 | 0.717089 |
| 0.670813 | 0.667311 | 0.673508 | 0.676556 | 0.666489 | 0.669362 | 0.662784 | 0.67116 | 0.664741 |
| 0.66185 | 0.663239 | 0.665839 | 0.669983 | 0.659875 | 0.664022 | 0.657162 | 0.661922 | 0.657169 |
| 0.660667 | 0.659369 | 0.661292 | 0.666896 | 0.657204 | 0.65839 | 0.651351 | 0.6577 | 0.652244 |
| 0.728562 | 0.731474 | 0.731354 | 0.738322 | 0.728809 | 0.730631 | 0.720515 | 0.725917 | 0.725526 |
| 0.665321 | 0.661141 | 0.664884 | 0.668073 | 0.66042 | 0.663493 | 0.656688 | 0.661409 | 0.658285 |
| 0.721043 | 0.724664 | 0.722975 | 0.732877 | 0.722182 | 0.72096 | 0.713535 | 0.71984 | 0.715969 |

| | | | | | | | | |
|---|---|---|---|---|---|---|---|---|
| 0.733254 | 0.731882 | 0.736648 | 0.742008 | 0.729347 | 0.73276 | 0.722767 | 0.729356 | 0.727666 |
| 0.727186 | 0.728603 | 0.728537 | 0.738562 | 0.721311 | 0.722306 | 0.717713 | 0.725355 | 0.720811 |
| 0.754352 | 0.754703 | 0.751757 | 0.760327 | 0.750718 | 0.749594 | 0.737766 | 0.74904 | 0.740285 |
| 0.654805 | 0.65201 | 0.657383 | 0.65786 | 0.650818 | 0.655798 | 0.646962 | 0.653238 | 0.645867 |
| 0.65506 | 0.654332 | 0.656728 | 0.660202 | 0.64903 | 0.653221 | 0.648079 | 0.651133 | 0.64838 |
| 0.731602 | 0.732497 | 0.735311 | 0.742162 | 0.727595 | 0.729114 | 0.724737 | 0.730096 | 0.731748 |
| 0.654816 | 0.657188 | 0.654504 | 0.658622 | 0.648822 | 0.651881 | 0.648158 | 0.654329 | 0.651375 |
| 0.735394 | 0.733433 | 0.732125 | 0.740744 | 0.72986 | 0.733657 | 0.728445 | 0.732629 | 0.72862 |
| 0.731918 | 0.732717 | 0.736153 | 0.740364 | 0.730632 | 0.72986 | 0.723775 | 0.727705 | 0.7308 |
| 0.736633 | 0.735555 | 0.729611 | 0.739433 | 0.732165 | 0.73085 | 0.731399 | 0.729584 | 0.731447 |
| 0.736206 | 0.732535 | 0.734656 | 0.741729 | 0.731624 | 0.733823 | 0.725031 | 0.729543 | 0.727295 |
| 0.736486 | 0.816176 | 0.75 | 0.758389 | 0.710526 | 0.768212 | 0.723684 | 0.699346 | 0.700637 |
| 0.715923 | 0.719576 | 0.721803 | 0.728571 | 0.722532 | 0.719881 | 0.716989 | 0.720656 | 0.710853 |
| 0.720613 | 0.717094 | 0.723548 | 0.724699 | 0.72185 | 0.714308 | 0.70799 | 0.719499 | 0.708891 |
| 0.725987 | 0.725378 | 0.726756 | 0.736226 | 0.723947 | 0.723385 | 0.716231 | 0.726291 | 0.720121 |
| 0.730759 | 0.726302 | 0.735691 | 0.735943 | 0.728392 | 0.729621 | 0.719325 | 0.726481 | 0.718001 |
| 0.591153 | 0.593869 | 0.59625 | 0.59599 | 0.589274 | 0.590071 | 0.58492 | 0.588824 | 0.58467 |
| 0.730234 | 0.730393 | 0.728135 | 0.740052 | 0.734363 | 0.72461 | 0.71547 | 0.727793 | 0.72463 |
| 0.735425 | 0.736659 | 0.740385 | 0.743719 | 0.733431 | 0.736778 | 0.726544 | 0.730096 | 0.733208 |
| 0.734727 | 0.738691 | 0.736088 | 0.745789 | 0.730693 | 0.731508 | 0.730824 | 0.731631 | 0.737068 |
| 0.738525 | 0.740911 | 0.741537 | 0.747774 | 0.736385 | 0.734777 | 0.730393 | 0.734627 | 0.737501 |
| 0.598142 | 0.594086 | 0.598028 | 0.60072 | 0.592157 | 0.596407 | 0.593781 | 0.597089 | 0.592187 |
| 0.584464 | 0.582771 | 0.584577 | 0.586172 | 0.577142 | 0.583051 | 0.57992 | 0.58141 | 0.577169 |
| 0.765912 | 0.761872 | 0.76665 | 0.768412 | 0.760394 | 0.764345 | 0.750335 | 0.760499 | 0.753431 |
| 0.7685 | 0.764706 | 0.76722 | 0.770409 | 0.766758 | 0.766361 | 0.757167 | 0.764377 | 0.760968 |
| 0.759816 | 0.756657 | 0.757753 | 0.764886 | 0.753796 | 0.75633 | 0.749605 | 0.755306 | 0.747938 |
| 0.764835 | 0.763323 | 0.764349 | 0.772332 | 0.762648 | 0.763218 | 0.752373 | 0.760125 | 0.753933 |
| 0.768748 | 0.766814 | 0.769983 | 0.77195 | 0.760743 | 0.765719 | 0.757829 | 0.760631 | 0.757871 |
| 0.767246 | 0.766338 | 0.771605 | 0.772199 | 0.764086 | 0.762643 | 0.75469 | 0.76268 | 0.756833 |
| 0.766058 | 0.76753 | 0.766872 | 0.774382 | 0.767777 | 0.769623 | 0.758216 | 0.765482 | 0.76455 |
| 0.785394 | 0.781656 | 0.790362 | 0.788759 | 0.784456 | 0.777491 | 0.777564 | 0.779576 | 0.776216 |
| 0.791592 | 0.794103 | 0.801508 | 0.798449 | 0.793292 | 0.794783 | 0.777732 | 0.790195 | 0.784188 |
| 0.787783 | 0.782226 | 0.784268 | 0.779496 | 0.778607 | 0.775613 | 0.785757 | 0.780013 | 0.775594 |
| 0.791152 | 0.78647 | 0.795183 | 0.789763 | 0.782989 | 0.785936 | 0.768628 | 0.785879 | 0.775929 |
| 0.793736 | 0.796329 | 0.794394 | 0.797642 | 0.795133 | 0.793568 | 0.782208 | 0.788998 | 0.783266 |
| 0.78488 | 0.786734 | 0.782111 | 0.784859 | 0.777287 | 0.77896 | 0.775393 | 0.779209 | 0.774328 |
| 0.788318 | 0.791037 | 0.781728 | 0.796375 | 0.78917 | 0.785228 | 0.781229 | 0.783152 | 0.779434 |
| 0.789075 | 0.781806 | 0.798401 | 0.791129 | 0.791249 | 0.78943 | 0.779587 | 0.78867 | 0.775386 |
| 0.788784 | 0.78627 | 0.791388 | 0.787031 | 0.786904 | 0.784767 | 0.778141 | 0.782501 | 0.778376 |
| 0.789422 | 0.787718 | 0.785667 | 0.788026 | 0.78215 | 0.779658 | 0.776242 | 0.778583 | 0.775935 |
| 0.721084 | 0.72489 | 0.718231 | 0.725147 | 0.719965 | 0.719134 | 0.696335 | 0.712116 | 0.709056 |
| 0.726882 | 0.721062 | 0.732318 | 0.738482 | 0.729831 | 0.734325 | 0.715838 | 0.728561 | 0.725349 |
| 0.807533 | 0.802709 | 0.789609 | 0.787122 | 0.78118 | 0.78088 | 0.796871 | 0.776781 | 0.820736 |
| 0.785973 | 0.790806 | 0.798847 | 0.787999 | 0.818736 | 0.824876 | 0.78954 | 0.788035 | 0.785014 |
| 0.812826 | 0.777489 | 0.787637 | 0.819369 | 0.796925 | 0.778129 | 0.766834 | 0.783741 | 0.77119 |
| 0.857021 | 0.817829 | 0.837905 | 0.831416 | 0.816484 | 0.827461 | 0.838623 | 0.838043 | 0.811112 |
| 0.83641 | 0.835747 | 0.800383 | 0.808007 | 0.812395 | 0.808603 | 0.824102 | 0.796788 | 0.846732 |

| | | | | | | | | |
|---|---|---|---|---|---|---|---|---|
| 0.858377 | 0.817884 | 0.8536 | 0.835981 | 0.831648 | 0.857302 | 0.815163 | 0.83955 | 0.823204 |
| 1 | 0.82175 | 0.834529 | 0.821193 | 0.813551 | 0.829058 | 0.806558 | 0.840905 | 0.829178 |
| 0.82175 | 1 | 0.802536 | 0.870305 | 0.810375 | 0.805961 | 0.825847 | 0.816171 | 0.825962 |
| 0.834529 | 0.802536 | 1 | 0.818036 | 0.820674 | 0.853588 | 0.791951 | 0.835247 | 0.792536 |
| 0.821193 | 0.870305 | 0.818036 | 1 | 0.835478 | 0.824962 | 0.803484 | 0.827813 | 0.799682 |
| 0.813551 | 0.810375 | 0.820674 | 0.835478 | 1 | 0.842753 | 0.797719 | 0.803204 | 0.794792 |
| 0.829058 | 0.805961 | 0.853588 | 0.824962 | 0.842753 | 1 | 0.787211 | 0.839154 | 0.793675 |
| 0.806558 | 0.825847 | 0.791951 | 0.803484 | 0.797719 | 0.787211 | 1 | 0.78258 | 0.828934 |
| 0.840905 | 0.816171 | 0.835247 | 0.827813 | 0.803204 | 0.839154 | 0.78258 | 1 | 0.780224 |
| 0.829178 | 0.825962 | 0.792536 | 0.799682 | 0.794792 | 0.793675 | 0.828934 | 0.780224 | 1 |
| 0.855947 | 0.811313 | 0.826211 | 0.817292 | 0.796193 | 0.836432 | 0.798704 | 0.823217 | 0.837148 |
| 0.864805 | 0.814802 | 0.835155 | 0.834398 | 0.83708 | 0.846928 | 0.813451 | 0.842045 | 0.812663 |
| 0.820201 | 0.834215 | 0.798966 | 0.808745 | 0.804491 | 0.805518 | 0.827137 | 0.777004 | 0.850065 |
| 0.804388 | 0.817577 | 0.81438 | 0.799076 | 0.858201 | 0.858714 | 0.809134 | 0.796319 | 0.805384 |
| 0.802984 | 0.797615 | 0.801761 | 0.842816 | 0.834407 | 0.797408 | 0.792606 | 0.802701 | 0.792202 |
| 0.770641 | 0.767757 | 0.77554 | 0.777285 | 0.773529 | 0.769727 | 0.758174 | 0.767923 | 0.761409 |
| 0.774139 | 0.76996 | 0.774051 | 0.77021 | 0.773473 | 0.771581 | 0.76323 | 0.773932 | 0.763936 |
| 0.769963 | 0.761118 | 0.772686 | 0.76528 | 0.770357 | 0.765585 | 0.753726 | 0.765803 | 0.756832 |
| 0.771274 | 0.767932 | 0.768524 | 0.771799 | 0.766529 | 0.767598 | 0.758161 | 0.763119 | 0.762978 |
| 0.772308 | 0.762797 | 0.772491 | 0.768795 | 0.767361 | 0.769629 | 0.753313 | 0.767638 | 0.758572 |
| 0.764725 | 0.75835 | 0.770006 | 0.764529 | 0.758702 | 0.762875 | 0.750505 | 0.760904 | 0.758624 |
| 0.766262 | 0.764298 | 0.765897 | 0.767434 | 0.760846 | 0.761003 | 0.757883 | 0.758379 | 0.758658 |
| 0.773557 | 0.766328 | 0.770298 | 0.776479 | 0.769022 | 0.771724 | 0.75709 | 0.764002 | 0.761523 |
| 0.775232 | 0.770169 | 0.773814 | 0.772432 | 0.774062 | 0.774621 | 0.75635 | 0.774122 | 0.761893 |
| 0.765431 | 0.762831 | 0.768929 | 0.76745 | 0.760536 | 0.759074 | 0.753753 | 0.767166 | 0.761037 |
| 0.764474 | 0.759777 | 0.764287 | 0.769402 | 0.763807 | 0.76525 | 0.753667 | 0.756886 | 0.756267 |
| 0.765594 | 0.757576 | 0.768404 | 0.763934 | 0.764454 | 0.765537 | 0.749101 | 0.764721 | 0.758845 |
| 0.76521 | 0.757153 | 0.762649 | 0.765084 | 0.759328 | 0.76175 | 0.747794 | 0.757016 | 0.754766 |
| 0.76194 | 0.758557 | 0.765102 | 0.766492 | 0.762003 | 0.760578 | 0.749629 | 0.762116 | 0.756386 |
| 0.768722 | 0.76374 | 0.767022 | 0.768235 | 0.765108 | 0.76634 | 0.756309 | 0.764097 | 0.759714 |
| 0.769848 | 0.76614 | 0.7706 | 0.766817 | 0.766153 | 0.766102 | 0.757784 | 0.770881 | 0.76862 |
| 0.773258 | 0.765486 | 0.774331 | 0.766476 | 0.767672 | 0.769186 | 0.760124 | 0.772435 | 0.763858 |
| 0.776339 | 0.776002 | 0.779307 | 0.777216 | 0.773251 | 0.775308 | 0.763603 | 0.774312 | 0.771501 |
| 0.778252 | 0.773428 | 0.776048 | 0.776534 | 0.772517 | 0.773868 | 0.764569 | 0.774295 | 0.767016 |
| 0.763311 | 0.754796 | 0.76474 | 0.763238 | 0.759275 | 0.759662 | 0.751119 | 0.760149 | 0.755331 |
| 0.767085 | 0.761448 | 0.771283 | 0.768557 | 0.764805 | 0.767038 | 0.751111 | 0.768105 | 0.75982 |
| 0.766371 | 0.758522 | 0.768231 | 0.7678 | 0.76386 | 0.767733 | 0.752663 | 0.764531 | 0.758903 |
| 0.76295 | 0.758126 | 0.766736 | 0.768797 | 0.764167 | 0.763988 | 0.748352 | 0.760597 | 0.758586 |
| 0.760273 | 0.756289 | 0.761574 | 0.762868 | 0.756804 | 0.758581 | 0.747035 | 0.760251 | 0.757754 |
| 0.757612 | 0.754345 | 0.754735 | 0.768031 | 0.757238 | 0.757576 | 0.743406 | 0.756259 | 0.747143 |
| 0.756841 | 0.752402 | 0.761593 | 0.761251 | 0.75731 | 0.75606 | 0.746546 | 0.754555 | 0.753632 |
| 0.75825 | 0.75427 | 0.758368 | 0.758275 | 0.754334 | 0.759089 | 0.747539 | 0.75617 | 0.750262 |
| 0.762455 | 0.753326 | 0.765809 | 0.761639 | 0.762946 | 0.765075 | 0.747368 | 0.761536 | 0.751162 |
| 0.754719 | 0.749908 | 0.755738 | 0.757889 | 0.751306 | 0.7536 | 0.742852 | 0.748564 | 0.750813 |
| 0.765331 | 0.757159 | 0.760279 | 0.7642 | 0.760343 | 0.762313 | 0.753495 | 0.759642 | 0.757665 |
| 0.759831 | 0.75649 | 0.760652 | 0.759017 | 0.757851 | 0.764062 | 0.745189 | 0.759625 | 0.750095 |

| | | | | | | | | |
|---|---|---|---|---|---|---|---|---|
| 0.763623 | 0.762005 | 0.76809 | 0.770549 | 0.763498 | 0.763972 | 0.75753 | 0.762533 | 0.757581 |
| 0.769014 | 0.766035 | 0.764672 | 0.772996 | 0.765696 | 0.769945 | 0.760239 | 0.769122 | 0.760417 |
| 0.761587 | 0.758957 | 0.760535 | 0.766287 | 0.760639 | 0.761519 | 0.753325 | 0.76196 | 0.756538 |
| 0.766025 | 0.762527 | 0.768086 | 0.766227 | 0.759797 | 0.766838 | 0.760574 | 0.770484 | 0.758846 |
| 0.776822 | 0.770486 | 0.776346 | 0.775132 | 0.77141 | 0.774403 | 0.765077 | 0.774572 | 0.766491 |
| 0.773688 | 0.771837 | 0.772153 | 0.77619 | 0.769834 | 0.768823 | 0.760447 | 0.774066 | 0.765316 |
| 0.769807 | 0.76487 | 0.771794 | 0.770258 | 0.765722 | 0.767291 | 0.75568 | 0.772141 | 0.755726 |
| 0.77367 | 0.76545 | 0.768581 | 0.770113 | 0.768202 | 0.766928 | 0.756772 | 0.770135 | 0.761064 |
| 0.768852 | 0.764053 | 0.767373 | 0.769481 | 0.762094 | 0.766121 | 0.756106 | 0.764865 | 0.754663 |
| 0.767103 | 0.760727 | 0.767248 | 0.769262 | 0.765655 | 0.764605 | 0.761817 | 0.765215 | 0.758683 |
| 0.778436 | 0.770609 | 0.779569 | 0.780038 | 0.774042 | 0.773724 | 0.761333 | 0.77285 | 0.765105 |
| 0.778315 | 0.769903 | 0.77743 | 0.778061 | 0.77387 | 0.769206 | 0.763697 | 0.769161 | 0.766645 |
| 0.769782 | 0.765895 | 0.768222 | 0.767815 | 0.765322 | 0.762626 | 0.757769 | 0.759641 | 0.762872 |
| 0.7663 | 0.763681 | 0.765612 | 0.768342 | 0.766158 | 0.764006 | 0.755183 | 0.762467 | 0.760923 |
| 0.770166 | 0.764407 | 0.773961 | 0.770051 | 0.768416 | 0.767234 | 0.76054 | 0.765527 | 0.762399 |
| 0.772456 | 0.764499 | 0.765933 | 0.766866 | 0.766007 | 0.762894 | 0.755431 | 0.765053 | 0.760823 |
| 0.767634 | 0.763138 | 0.767381 | 0.771326 | 0.766541 | 0.769867 | 0.759092 | 0.767082 | 0.759948 |
| 0.763139 | 0.76124 | 0.764611 | 0.771271 | 0.766004 | 0.762866 | 0.758883 | 0.763333 | 0.758383 |
| 0.770755 | 0.764765 | 0.774035 | 0.775191 | 0.768408 | 0.774471 | 0.758284 | 0.762742 | 0.761526 |
| 0.772081 | 0.762987 | 0.769155 | 0.76834 | 0.766398 | 0.76651 | 0.760706 | 0.763895 | 0.765111 |
| 0.769559 | 0.761553 | 0.763675 | 0.768781 | 0.766824 | 0.76208 | 0.756667 | 0.762375 | 0.756007 |
| 0.767239 | 0.75998 | 0.766499 | 0.768242 | 0.763487 | 0.762409 | 0.757495 | 0.763642 | 0.756255 |
| 0.771995 | 0.760183 | 0.768189 | 0.767559 | 0.767039 | 0.764873 | 0.755119 | 0.763512 | 0.761326 |
| 0.773643 | 0.766628 | 0.771871 | 0.770633 | 0.766868 | 0.768857 | 0.758813 | 0.769137 | 0.760222 |
| 0.766024 | 0.763223 | 0.76983 | 0.767664 | 0.761487 | 0.764687 | 0.754405 | 0.765578 | 0.759474 |
| 0.765592 | 0.760351 | 0.765952 | 0.764994 | 0.762567 | 0.764701 | 0.758898 | 0.762205 | 0.76167 |
| 0.766909 | 0.759495 | 0.768811 | 0.767415 | 0.759595 | 0.763286 | 0.758083 | 0.766185 | 0.761559 |
| 0.766233 | 0.765378 | 0.762001 | 0.767533 | 0.762634 | 0.764587 | 0.750143 | 0.761951 | 0.759063 |
| 0.772993 | 0.767554 | 0.770384 | 0.775525 | 0.769711 | 0.769087 | 0.761042 | 0.766142 | 0.766951 |
| 0.767152 | 0.762342 | 0.771016 | 0.769679 | 0.764323 | 0.765307 | 0.758048 | 0.762867 | 0.760506 |
| 0.762251 | 0.755845 | 0.760731 | 0.759685 | 0.760381 | 0.760215 | 0.747301 | 0.755197 | 0.75029 |
| 0.770446 | 0.765257 | 0.767959 | 0.766285 | 0.768214 | 0.768162 | 0.755925 | 0.761093 | 0.760794 |
| 0.769937 | 0.764854 | 0.770497 | 0.772129 | 0.767894 | 0.766157 | 0.755711 | 0.761155 | 0.760337 |
| 0.766967 | 0.760507 | 0.766802 | 0.769205 | 0.762333 | 0.764795 | 0.757423 | 0.76022 | 0.757517 |
| 0.765427 | 0.765315 | 0.765346 | 0.769947 | 0.763644 | 0.760191 | 0.75828 | 0.760448 | 0.762413 |
| 0.765068 | 0.759342 | 0.766096 | 0.768246 | 0.765087 | 0.761042 | 0.751602 | 0.760182 | 0.757581 |
| 0.773912 | 0.768432 | 0.776463 | 0.776447 | 0.770876 | 0.774654 | 0.76043 | 0.774988 | 0.763862 |
| 0.772225 | 0.762687 | 0.769977 | 0.77037 | 0.76863 | 0.767939 | 0.763323 | 0.766279 | 0.758814 |
| 0.768873 | 0.764365 | 0.773083 | 0.77052 | 0.769313 | 0.767059 | 0.755499 | 0.763363 | 0.762973 |
| 0.764511 | 0.761657 | 0.769314 | 0.76846 | 0.768557 | 0.767186 | 0.755947 | 0.763008 | 0.75822 |
| 0.768682 | 0.766062 | 0.769155 | 0.769921 | 0.767385 | 0.766791 | 0.759921 | 0.760351 | 0.762844 |
| 0.771015 | 0.767333 | 0.770362 | 0.772944 | 0.768589 | 0.771726 | 0.763058 | 0.766155 | 0.76362 |
| 0.772119 | 0.764374 | 0.773162 | 0.772412 | 0.768569 | 0.769987 | 0.761842 | 0.77074 | 0.765216 |
| 0.764174 | 0.76013 | 0.769012 | 0.765922 | 0.768342 | 0.761395 | 0.757254 | 0.764465 | 0.759742 |
| 0.773161 | 0.765737 | 0.771478 | 0.772205 | 0.770582 | 0.770681 | 0.762784 | 0.766951 | 0.764364 |
| 0.771177 | 0.759248 | 0.771146 | 0.767 | 0.765012 | 0.767674 | 0.756714 | 0.764765 | 0.759281 |

| | | | | | | | | |
|---|---|---|---|---|---|---|---|---|
| 0.757838 | 0.75782 | 0.75773 | 0.758881 | 0.756857 | 0.753926 | 0.749093 | 0.75224 | 0.752083 |
| 0.74812 | 0.744168 | 0.746634 | 0.750021 | 0.743801 | 0.741509 | 0.73737 | 0.741351 | 0.740145 |
| 0.763718 | 0.760323 | 0.763412 | 0.768001 | 0.760175 | 0.759753 | 0.755137 | 0.76386 | 0.756609 |
| 0.766343 | 0.767211 | 0.769626 | 0.772946 | 0.764682 | 0.765006 | 0.758001 | 0.770765 | 0.761761 |
| 0.758417 | 0.756843 | 0.760105 | 0.762673 | 0.755885 | 0.758916 | 0.751071 | 0.760085 | 0.754364 |
| 0.762687 | 0.755991 | 0.763428 | 0.76493 | 0.760768 | 0.758575 | 0.756849 | 0.763356 | 0.756336 |
| 0.763113 | 0.761604 | 0.764438 | 0.769383 | 0.763099 | 0.763372 | 0.753826 | 0.766649 | 0.758199 |
| 0.754527 | 0.754395 | 0.762886 | 0.764113 | 0.758177 | 0.756513 | 0.751831 | 0.757306 | 0.749801 |
| 0.77211 | 0.764637 | 0.773686 | 0.774381 | 0.768851 | 0.765766 | 0.757073 | 0.769874 | 0.760245 |
| 0.768695 | 0.763403 | 0.770643 | 0.774478 | 0.768638 | 0.766757 | 0.754434 | 0.767474 | 0.759944 |
| 0.769823 | 0.763724 | 0.771646 | 0.771457 | 0.770294 | 0.768874 | 0.764755 | 0.763958 | 0.766568 |
| 0.768601 | 0.76474 | 0.77084 | 0.772829 | 0.762936 | 0.765928 | 0.757367 | 0.76772 | 0.757613 |
| 0.771748 | 0.760606 | 0.777659 | 0.770367 | 0.769632 | 0.772683 | 0.759066 | 0.76887 | 0.760663 |
| 0.772128 | 0.768464 | 0.772564 | 0.772226 | 0.762459 | 0.768442 | 0.76049 | 0.767914 | 0.768264 |
| 0.769544 | 0.758893 | 0.770028 | 0.769822 | 0.766804 | 0.761802 | 0.753849 | 0.76474 | 0.758065 |
| 0.761821 | 0.765352 | 0.766599 | 0.774434 | 0.760149 | 0.759176 | 0.752885 | 0.76483 | 0.759165 |
| 0.769778 | 0.763975 | 0.771129 | 0.772076 | 0.761826 | 0.767063 | 0.753644 | 0.767013 | 0.763399 |
| 0.765419 | 0.757558 | 0.767063 | 0.766324 | 0.759503 | 0.765557 | 0.751059 | 0.762926 | 0.758441 |
| 0.761174 | 0.76203 | 0.762096 | 0.767384 | 0.75967 | 0.757149 | 0.749215 | 0.756976 | 0.754985 |
| 0.765959 | 0.758438 | 0.761554 | 0.767297 | 0.761362 | 0.761769 | 0.749258 | 0.760738 | 0.756468 |
| 0.76937 | 0.763127 | 0.773967 | 0.769141 | 0.76347 | 0.765604 | 0.756147 | 0.764149 | 0.756708 |
| 0.755485 | 0.752708 | 0.758868 | 0.756618 | 0.75237 | 0.754112 | 0.743405 | 0.751896 | 0.750021 |
| 0.764912 | 0.757143 | 0.761337 | 0.761567 | 0.757332 | 0.760834 | 0.753493 | 0.757846 | 0.755485 |
| 0.76706 | 0.765256 | 0.767183 | 0.769174 | 0.765082 | 0.766798 | 0.7602 | 0.765425 | 0.761028 |
| 0.766021 | 0.762712 | 0.771482 | 0.769773 | 0.765599 | 0.766313 | 0.755725 | 0.764839 | 0.755548 |
| 0.762247 | 0.756461 | 0.765504 | 0.764784 | 0.760884 | 0.759854 | 0.757005 | 0.758543 | 0.758132 |
| 0.764736 | 0.762694 | 0.764576 | 0.766907 | 0.759335 | 0.763182 | 0.750768 | 0.760897 | 0.759656 |
| 0.761284 | 0.761382 | 0.763216 | 0.767289 | 0.766489 | 0.762175 | 0.750515 | 0.760764 | 0.757138 |
| 0.757578 | 0.7539 | 0.760947 | 0.763089 | 0.760818 | 0.757557 | 0.750459 | 0.753849 | 0.749589 |
| 0.700201 | 0.699888 | 0.701388 | 0.69901 | 0.694242 | 0.700733 | 0.683341 | 0.695744 | 0.692464 |
| 0.709974 | 0.709152 | 0.708935 | 0.71652 | 0.709424 | 0.707637 | 0.698657 | 0.708779 | 0.705206 |
| 0.70674 | 0.705892 | 0.705779 | 0.710412 | 0.702579 | 0.699754 | 0.697601 | 0.699369 | 0.702948 |
| 0.69764 | 0.701348 | 0.696405 | 0.704642 | 0.696568 | 0.695896 | 0.694311 | 0.694856 | 0.695683 |
| 0.710481 | 0.707667 | 0.708825 | 0.715214 | 0.705335 | 0.706595 | 0.701958 | 0.703762 | 0.705901 |
| 0.76089 | 0.757218 | 0.765065 | 0.761237 | 0.760133 | 0.76215 | 0.753356 | 0.759211 | 0.757501 |
| 0.762109 | 0.760831 | 0.763655 | 0.765859 | 0.765376 | 0.759874 | 0.754931 | 0.760486 | 0.76177 |
| 0.767476 | 0.766037 | 0.766595 | 0.76941 | 0.768304 | 0.764541 | 0.757817 | 0.765596 | 0.761192 |
| 0.767425 | 0.763829 | 0.772803 | 0.770799 | 0.769685 | 0.76668 | 0.760543 | 0.764007 | 0.763654 |
| 0.76587 | 0.760182 | 0.761499 | 0.767814 | 0.758686 | 0.756695 | 0.755934 | 0.752101 | 0.755078 |
| 0.766948 | 0.766546 | 0.765813 | 0.768452 | 0.765116 | 0.763716 | 0.758333 | 0.762505 | 0.762741 |
| 0.764886 | 0.762878 | 0.765225 | 0.769026 | 0.76691 | 0.763996 | 0.757581 | 0.763951 | 0.75931 |
| 0.761723 | 0.761495 | 0.765829 | 0.767868 | 0.765088 | 0.761287 | 0.760643 | 0.763913 | 0.761362 |
| 0.764843 | 0.75794 | 0.761971 | 0.761167 | 0.754843 | 0.757539 | 0.754952 | 0.757202 | 0.75583 |
| 0.762901 | 0.760378 | 0.761791 | 0.76802 | 0.762074 | 0.758308 | 0.753275 | 0.759099 | 0.751612 |
| 0.759933 | 0.760065 | 0.762209 | 0.762484 | 0.760517 | 0.758753 | 0.751845 | 0.760263 | 0.756422 |
| 0.766432 | 0.765618 | 0.769533 | 0.767799 | 0.767698 | 0.761983 | 0.755539 | 0.764133 | 0.765701 |

| | | | | | | | | |
|---|---|---|---|---|---|---|---|---|
| 0.77091 | 0.760676 | 0.769956 | 0.769677 | 0.768585 | 0.769838 | 0.754725 | 0.769796 | 0.758699 |
| 0.764107 | 0.762915 | 0.762701 | 0.77 | 0.762474 | 0.764342 | 0.754971 | 0.765926 | 0.757536 |
| 0.765887 | 0.759313 | 0.762398 | 0.769557 | 0.760437 | 0.76353 | 0.753184 | 0.758366 | 0.756817 |
| 0.758199 | 0.755401 | 0.757907 | 0.764319 | 0.761581 | 0.763318 | 0.748075 | 0.758396 | 0.754659 |
| 0.759757 | 0.758428 | 0.760507 | 0.763964 | 0.758915 | 0.758373 | 0.749429 | 0.75682 | 0.756443 |
| 0.756893 | 0.758908 | 0.755875 | 0.76236 | 0.760773 | 0.759081 | 0.748783 | 0.759878 | 0.753738 |
| 0.763083 | 0.761757 | 0.762296 | 0.769224 | 0.762669 | 0.765367 | 0.752414 | 0.762919 | 0.760352 |
| 0.764967 | 0.761798 | 0.767353 | 0.766077 | 0.762537 | 0.759893 | 0.754562 | 0.764706 | 0.756891 |
| 0.762339 | 0.75907 | 0.761605 | 0.762424 | 0.760732 | 0.760486 | 0.750655 | 0.757176 | 0.753641 |
| 0.762392 | 0.759679 | 0.758812 | 0.764409 | 0.758669 | 0.760823 | 0.750167 | 0.758501 | 0.754145 |
| 0.760186 | 0.756585 | 0.758692 | 0.767749 | 0.761421 | 0.760645 | 0.749622 | 0.756148 | 0.754834 |
| 0.761291 | 0.759952 | 0.762786 | 0.767635 | 0.758082 | 0.761417 | 0.750506 | 0.757333 | 0.756757 |
| 0.753297 | 0.746227 | 0.754714 | 0.757178 | 0.754008 | 0.755344 | 0.74078 | 0.75102 | 0.745408 |
| 0.7596 | 0.754307 | 0.75745 | 0.760631 | 0.756966 | 0.756303 | 0.748147 | 0.756471 | 0.745869 |
| 0.766672 | 0.76233 | 0.768178 | 0.767075 | 0.760242 | 0.759333 | 0.751945 | 0.763513 | 0.75675 |
| 0.771773 | 0.76798 | 0.773807 | 0.772667 | 0.768605 | 0.768073 | 0.76104 | 0.766845 | 0.767507 |
| 0.769094 | 0.767636 | 0.769934 | 0.768399 | 0.767651 | 0.762817 | 0.755829 | 0.766411 | 0.761446 |
| 0.76947 | 0.767369 | 0.774636 | 0.773046 | 0.764166 | 0.767057 | 0.757246 | 0.768736 | 0.761689 |
| 0.766012 | 0.765626 | 0.770448 | 0.770738 | 0.766752 | 0.762289 | 0.757005 | 0.763695 | 0.759606 |
| 0.767361 | 0.765976 | 0.767157 | 0.771126 | 0.763194 | 0.762742 | 0.754478 | 0.766383 | 0.758236 |
| 0.761553 | 0.761357 | 0.764993 | 0.769382 | 0.763391 | 0.764496 | 0.755634 | 0.759115 | 0.759043 |
| 0.767127 | 0.758419 | 0.767534 | 0.76551 | 0.758718 | 0.761818 | 0.752104 | 0.75956 | 0.759088 |
| 0.776355 | 0.769693 | 0.773976 | 0.773514 | 0.767935 | 0.770418 | 0.760278 | 0.769657 | 0.76615 |
| 0.760161 | 0.756728 | 0.758675 | 0.761733 | 0.755129 | 0.758853 | 0.751972 | 0.757142 | 0.75361 |
| 0.772002 | 0.769782 | 0.773243 | 0.775773 | 0.766158 | 0.766948 | 0.759836 | 0.765758 | 0.762366 |
| 0.707184 | 0.703859 | 0.707272 | 0.710849 | 0.704192 | 0.705873 | 0.697272 | 0.703954 | 0.702742 |
| 0.766742 | 0.761925 | 0.769187 | 0.768069 | 0.766505 | 0.765423 | 0.758248 | 0.761932 | 0.756219 |
| 0.767349 | 0.76212 | 0.769679 | 0.766427 | 0.765684 | 0.765079 | 0.752575 | 0.766155 | 0.763128 |
| 0.774066 | 0.765522 | 0.769014 | 0.771265 | 0.76725 | 0.768641 | 0.757923 | 0.769531 | 0.762791 |
| 0.769805 | 0.7671 | 0.771152 | 0.77481 | 0.76792 | 0.768477 | 0.764122 | 0.769123 | 0.765633 |
| 0.760936 | 0.759958 | 0.76179 | 0.763388 | 0.75865 | 0.759504 | 0.749957 | 0.760292 | 0.751801 |
| 0.763639 | 0.756586 | 0.765275 | 0.767252 | 0.763951 | 0.766562 | 0.750919 | 0.760326 | 0.754954 |
| 0.767072 | 0.758926 | 0.766893 | 0.76655 | 0.764121 | 0.767486 | 0.752999 | 0.763912 | 0.757698 |
| 0.76589 | 0.762878 | 0.762412 | 0.765866 | 0.76441 | 0.76564 | 0.757386 | 0.757844 | 0.758677 |
| 0.760306 | 0.755131 | 0.761917 | 0.759226 | 0.756106 | 0.755997 | 0.743935 | 0.757507 | 0.754016 |
| 0.761302 | 0.756162 | 0.767638 | 0.76119 | 0.758757 | 0.758552 | 0.749303 | 0.759852 | 0.756223 |
| 0.771338 | 0.768465 | 0.766689 | 0.773855 | 0.771134 | 0.76448 | 0.761925 | 0.765639 | 0.758774 |
| 0.771688 | 0.765368 | 0.77304 | 0.775928 | 0.769327 | 0.774081 | 0.761315 | 0.770572 | 0.763797 |
| 0.76817 | 0.765576 | 0.768796 | 0.770984 | 0.764883 | 0.764643 | 0.763081 | 0.763259 | 0.760526 |
| 0.768351 | 0.762416 | 0.764731 | 0.766378 | 0.758917 | 0.762716 | 0.757436 | 0.76014 | 0.757183 |
| 0.763308 | 0.755811 | 0.764464 | 0.765048 | 0.756984 | 0.765874 | 0.75147 | 0.761084 | 0.75543 |
| 0.769263 | 0.763655 | 0.767667 | 0.77102 | 0.766445 | 0.770868 | 0.761397 | 0.76474 | 0.761642 |
| 0.766644 | 0.759826 | 0.764706 | 0.767055 | 0.765857 | 0.763903 | 0.753556 | 0.759791 | 0.7552 |
| 0.759679 | 0.764627 | 0.76194 | 0.769224 | 0.760023 | 0.759237 | 0.749096 | 0.759108 | 0.754779 |
| 0.771177 | 0.770544 | 0.768659 | 0.773091 | 0.76517 | 0.766317 | 0.76314 | 0.769899 | 0.762007 |
| 0.765782 | 0.766883 | 0.762944 | 0.774116 | 0.762003 | 0.761259 | 0.757785 | 0.763277 | 0.755981 |

| | | | | | | | | |
|---|---|---|---|---|---|---|---|---|
| 0.765577 | 0.764293 | 0.766429 | 0.770578 | 0.758347 | 0.75975 | 0.753435 | 0.764937 | 0.754967 |
| 0.758802 | 0.765806 | 0.763816 | 0.772422 | 0.761803 | 0.762807 | 0.755191 | 0.759687 | 0.75509 |
| 0.756945 | 0.764103 | 0.759067 | 0.770621 | 0.757583 | 0.758014 | 0.75407 | 0.758458 | 0.754165 |
| 0.72316 | 0.71014 | 0.724262 | 0.727758 | 0.71317 | 0.722401 | 0.714491 | 0.708199 | 0.709861 |
| 0.761804 | 0.763116 | 0.763418 | 0.770505 | 0.757457 | 0.757564 | 0.751312 | 0.758963 | 0.751121 |
| 0.758724 | 0.761063 | 0.759633 | 0.767348 | 0.75617 | 0.756479 | 0.751862 | 0.759662 | 0.755057 |
| 0.763505 | 0.762869 | 0.764581 | 0.772489 | 0.760725 | 0.762149 | 0.751173 | 0.762483 | 0.755663 |
| 0.762933 | 0.769186 | 0.761971 | 0.771686 | 0.760889 | 0.764936 | 0.761629 | 0.762684 | 0.760738 |
| 0.768905 | 0.768825 | 0.767054 | 0.777048 | 0.767723 | 0.768345 | 0.755166 | 0.763662 | 0.763928 |
| 0.765822 | 0.76734 | 0.766787 | 0.772936 | 0.760569 | 0.762401 | 0.753798 | 0.764431 | 0.762187 |
| 0.762359 | 0.76635 | 0.766231 | 0.771037 | 0.761842 | 0.762532 | 0.752482 | 0.764377 | 0.758683 |
| 0.760646 | 0.766818 | 0.764035 | 0.773334 | 0.760324 | 0.763823 | 0.750972 | 0.761509 | 0.759479 |
| 0.765616 | 0.768112 | 0.765498 | 0.771616 | 0.76298 | 0.764019 | 0.753847 | 0.756388 | 0.757404 |
| 0.760513 | 0.767987 | 0.766568 | 0.771801 | 0.764817 | 0.764811 | 0.758826 | 0.762956 | 0.754785 |
| 0.767063 | 0.770802 | 0.7674 | 0.772242 | 0.770247 | 0.767271 | 0.758883 | 0.760544 | 0.760043 |
| 0.762285 | 0.766199 | 0.762499 | 0.769938 | 0.762117 | 0.764149 | 0.754815 | 0.758313 | 0.754055 |
| 0.76849 | 0.769385 | 0.769676 | 0.775854 | 0.768134 | 0.768801 | 0.760372 | 0.761278 | 0.762421 |
| 0.769043 | 0.765706 | 0.766642 | 0.771562 | 0.766159 | 0.768879 | 0.754371 | 0.762826 | 0.757678 |
| 0.76871 | 0.763735 | 0.762135 | 0.770279 | 0.762019 | 0.760602 | 0.754878 | 0.76056 | 0.758233 |
| 0.765327 | 0.766153 | 0.766232 | 0.771459 | 0.7654 | 0.765018 | 0.756422 | 0.763924 | 0.758776 |
| 0.767947 | 0.767352 | 0.768504 | 0.774562 | 0.76572 | 0.765652 | 0.758895 | 0.764696 | 0.761787 |
| 0.769406 | 0.768263 | 0.769307 | 0.773608 | 0.767574 | 0.766245 | 0.755163 | 0.762916 | 0.760082 |
| 0.676264 | 0.686019 | 0.679949 | 0.690146 | 0.682719 | 0.678196 | 0.671465 | 0.676753 | 0.676043 |
| 0.658192 | 0.662997 | 0.660109 | 0.670069 | 0.649494 | 0.654181 | 0.645679 | 0.65642 | 0.651207 |
| 0.676244 | 0.678329 | 0.669953 | 0.686996 | 0.668677 | 0.676123 | 0.667392 | 0.664315 | 0.668821 |
| 0.654704 | 0.659143 | 0.657566 | 0.665303 | 0.659833 | 0.65789 | 0.652261 | 0.655291 | 0.652988 |
| 0.729375 | 0.722081 | 0.728115 | 0.734553 | 0.729895 | 0.734501 | 0.718769 | 0.7242 | 0.715356 |
| 0.721283 | 0.72511 | 0.720008 | 0.733627 | 0.727533 | 0.729533 | 0.71459 | 0.717031 | 0.713023 |
| 0.73179 | 0.729099 | 0.730424 | 0.735696 | 0.72509 | 0.723503 | 0.707739 | 0.717935 | 0.719279 |
| 0.730928 | 0.738265 | 0.731612 | 0.744792 | 0.735221 | 0.730244 | 0.719265 | 0.727726 | 0.722464 |
| 0.711352 | 0.715499 | 0.714026 | 0.719234 | 0.712563 | 0.710865 | 0.709947 | 0.706576 | 0.710448 |
| 0.707378 | 0.705549 | 0.707042 | 0.713122 | 0.704016 | 0.701931 | 0.708189 | 0.699944 | 0.695946 |
| 0.71299 | 0.710464 | 0.710434 | 0.708986 | 0.71091 | 0.705539 | 0.696296 | 0.704472 | 0.695869 |
| 0.653995 | 0.657777 | 0.655075 | 0.668267 | 0.661328 | 0.665305 | 0.648038 | 0.650213 | 0.654046 |
| 0.76127 | 0.76188 | 0.761942 | 0.763034 | 0.75503 | 0.757449 | 0.749848 | 0.754396 | 0.75435 |
| 0.73997 | 0.737813 | 0.740444 | 0.738949 | 0.737619 | 0.736927 | 0.729532 | 0.735748 | 0.733677 |
| 0.731498 | 0.738436 | 0.733649 | 0.744557 | 0.734884 | 0.732741 | 0.723605 | 0.730692 | 0.725645 |
| 0.720832 | 0.718954 | 0.718281 | 0.724755 | 0.717491 | 0.720044 | 0.709098 | 0.708241 | 0.703348 |
| 0.73405 | 0.740902 | 0.732946 | 0.7444 | 0.735746 | 0.732463 | 0.729002 | 0.736536 | 0.728326 |
| 0.733503 | 0.73779 | 0.729255 | 0.739003 | 0.729681 | 0.727041 | 0.724474 | 0.730962 | 0.726284 |
| 0.725249 | 0.726996 | 0.72908 | 0.733704 | 0.72359 | 0.730654 | 0.706454 | 0.723624 | 0.713898 |
| 0.726313 | 0.73327 | 0.724063 | 0.734691 | 0.719017 | 0.720415 | 0.716793 | 0.722144 | 0.714286 |
| 0.725641 | 0.724232 | 0.724802 | 0.731461 | 0.729842 | 0.723866 | 0.723264 | 0.713873 | 0.71535 |
| 0.714119 | 0.725377 | 0.71967 | 0.731717 | 0.72549 | 0.718939 | 0.715936 | 0.716833 | 0.709754 |
| 0.720783 | 0.727131 | 0.723758 | 0.727089 | 0.716578 | 0.717057 | 0.715325 | 0.710761 | 0.71355 |
| 0.711118 | 0.71438 | 0.716239 | 0.715824 | 0.706981 | 0.706326 | 0.702496 | 0.700751 | 0.70363 |

| | | | | | | | | |
|---|---|---|---|---|---|---|---|---|
| 0.746211 | 0.747618 | 0.748591 | 0.755885 | 0.74826 | 0.749778 | 0.736763 | 0.748688 | 0.738001 |
| 0.724321 | 0.731069 | 0.720859 | 0.728518 | 0.719085 | 0.722624 | 0.71265 | 0.723461 | 0.713906 |
| 0.725409 | 0.72268 | 0.727668 | 0.73492 | 0.726335 | 0.725534 | 0.718451 | 0.726551 | 0.722054 |
| 0.72154 | 0.726205 | 0.722603 | 0.729833 | 0.729326 | 0.718096 | 0.717758 | 0.716366 | 0.721023 |
| 0.72731 | 0.728239 | 0.726072 | 0.735857 | 0.729035 | 0.730612 | 0.715907 | 0.72264 | 0.713672 |
| 0.715929 | 0.712732 | 0.715307 | 0.72562 | 0.716123 | 0.711515 | 0.701238 | 0.705103 | 0.713076 |
| 0.716112 | 0.717211 | 0.71321 | 0.718245 | 0.710875 | 0.703721 | 0.699682 | 0.707892 | 0.706227 |
| 0.713367 | 0.708555 | 0.708928 | 0.719089 | 0.698124 | 0.70302 | 0.696721 | 0.705669 | 0.701606 |
| 0.713408 | 0.710259 | 0.719989 | 0.722844 | 0.705422 | 0.711297 | 0.697055 | 0.701725 | 0.709375 |
| 0.750947 | 0.752145 | 0.753053 | 0.758819 | 0.756232 | 0.7564 | 0.749181 | 0.75122 | 0.748011 |
| 0.747376 | 0.746285 | 0.75098 | 0.752757 | 0.747462 | 0.753542 | 0.742172 | 0.745238 | 0.738302 |
| 0.738845 | 0.74205 | 0.743771 | 0.7544 | 0.743434 | 0.745126 | 0.730748 | 0.736076 | 0.733754 |
| 0.738204 | 0.733893 | 0.736836 | 0.742351 | 0.738326 | 0.739749 | 0.715276 | 0.730128 | 0.727273 |
| 0.738084 | 0.74192 | 0.741111 | 0.749878 | 0.740475 | 0.737417 | 0.725403 | 0.735957 | 0.730445 |
| 0.734592 | 0.737924 | 0.739756 | 0.741731 | 0.739838 | 0.737837 | 0.724343 | 0.732005 | 0.733515 |

| Wild_plot_ | Wild_plot_ | Wild_plot_ | Wild_plot_ | Wild_plot_ | Wild_plot_ | Wild_plot_ | Wild_plot_ | Wild_plot_ |
|---|---|---|---|---|---|---|---|---|
| 0.706602 | 0.700767 | 0.700538 | 0.697828 | 0.713099 | 0.706152 | 0.703034 | 0.705231 | 0.704281 |
| 0.715454 | 0.712182 | 0.708902 | 0.706866 | 0.717711 | 0.711455 | 0.708121 | 0.710783 | 0.709913 |
| 0.706731 | 0.701911 | 0.697292 | 0.69547 | 0.70622 | 0.707248 | 0.700204 | 0.701341 | 0.703739 |
| 0.713441 | 0.709727 | 0.70087 | 0.707275 | 0.718782 | 0.711963 | 0.706311 | 0.712455 | 0.710406 |
| 0.70855 | 0.706776 | 0.696402 | 0.700322 | 0.71298 | 0.70943 | 0.704411 | 0.707972 | 0.707767 |
| 0.718192 | 0.711548 | 0.709362 | 0.70932 | 0.721389 | 0.716726 | 0.713407 | 0.719464 | 0.717072 |
| 0.763786 | 0.766672 | 0.756835 | 0.758267 | 0.764438 | 0.768729 | 0.764786 | 0.761678 | 0.764639 |
| 0.764202 | 0.762826 | 0.752974 | 0.75234 | 0.76132 | 0.763349 | 0.763136 | 0.760272 | 0.758983 |
| 0.713299 | 0.71014 | 0.707183 | 0.705951 | 0.720714 | 0.714013 | 0.707849 | 0.711475 | 0.709772 |
| 0.711745 | 0.70631 | 0.700698 | 0.70192 | 0.707377 | 0.709887 | 0.702548 | 0.708187 | 0.70484 |
| 0.712922 | 0.70783 | 0.698567 | 0.699743 | 0.714638 | 0.710646 | 0.70499 | 0.706805 | 0.706466 |
| 0.704739 | 0.702049 | 0.697473 | 0.696091 | 0.709466 | 0.704395 | 0.70233 | 0.702004 | 0.701455 |
| 0.714558 | 0.707934 | 0.705484 | 0.707412 | 0.716314 | 0.714513 | 0.709988 | 0.709076 | 0.709867 |
| 0.725745 | 0.718229 | 0.71282 | 0.714536 | 0.725813 | 0.7238 | 0.720486 | 0.720122 | 0.728256 |
| 0.738596 | 0.733086 | 0.725992 | 0.725327 | 0.737553 | 0.748332 | 0.738174 | 0.746421 | 0.747304 |
| 0.718363 | 0.713302 | 0.706275 | 0.702399 | 0.720035 | 0.715623 | 0.710882 | 0.712701 | 0.714331 |
| 0.718107 | 0.715043 | 0.706612 | 0.707148 | 0.718475 | 0.713761 | 0.711959 | 0.712917 | 0.710144 |
| 0.746345 | 0.744347 | 0.734011 | 0.730683 | 0.750219 | 0.763794 | 0.745283 | 0.752398 | 0.765171 |
| 0.708234 | 0.702636 | 0.697397 | 0.69774 | 0.709192 | 0.707552 | 0.700915 | 0.702124 | 0.699944 |
| 0.714342 | 0.706949 | 0.703635 | 0.702025 | 0.713209 | 0.710817 | 0.704361 | 0.705915 | 0.709154 |
| 0.707304 | 0.703622 | 0.699039 | 0.698026 | 0.710225 | 0.706552 | 0.697889 | 0.702079 | 0.704949 |
| 0.710799 | 0.70541 | 0.700129 | 0.702586 | 0.711722 | 0.709391 | 0.704938 | 0.707239 | 0.7098 |
| 0.71972 | 0.712366 | 0.709729 | 0.71012 | 0.718054 | 0.714172 | 0.7105 | 0.710518 | 0.711042 |
| 0.713889 | 0.713003 | 0.70053 | 0.702806 | 0.718678 | 0.713411 | 0.710952 | 0.710996 | 0.706674 |
| 0.716462 | 0.709752 | 0.708232 | 0.706726 | 0.718177 | 0.715667 | 0.707339 | 0.709639 | 0.707498 |
| 0.715233 | 0.707434 | 0.704106 | 0.7007 | 0.717307 | 0.708494 | 0.706908 | 0.708479 | 0.707972 |
| 0.707097 | 0.702831 | 0.695914 | 0.693763 | 0.706128 | 0.707021 | 0.70276 | 0.702397 | 0.70221 |
| 0.714985 | 0.707373 | 0.701822 | 0.703658 | 0.716392 | 0.714625 | 0.707764 | 0.711846 | 0.710443 |
| 0.71568 | 0.708798 | 0.706195 | 0.707536 | 0.715792 | 0.715105 | 0.708165 | 0.708526 | 0.710485 |
| 0.712502 | 0.707257 | 0.703212 | 0.701933 | 0.71246 | 0.709657 | 0.706466 | 0.708006 | 0.704042 |
| 0.714695 | 0.704868 | 0.702855 | 0.70249 | 0.716034 | 0.707565 | 0.706334 | 0.711509 | 0.707587 |
| 0.71284 | 0.704481 | 0.696929 | 0.696124 | 0.71162 | 0.706383 | 0.701169 | 0.703325 | 0.706505 |
| 0.712017 | 0.705815 | 0.699704 | 0.702493 | 0.712167 | 0.707567 | 0.705318 | 0.704085 | 0.703975 |
| 0.715551 | 0.709519 | 0.704593 | 0.704812 | 0.712281 | 0.709504 | 0.706003 | 0.707288 | 0.710631 |
| 0.708941 | 0.701232 | 0.696943 | 0.698319 | 0.706997 | 0.704465 | 0.696408 | 0.703166 | 0.704183 |
| 0.710376 | 0.705354 | 0.701621 | 0.701556 | 0.713616 | 0.714594 | 0.707214 | 0.706047 | 0.708383 |
| 0.716717 | 0.710226 | 0.70474 | 0.703576 | 0.718698 | 0.714951 | 0.705474 | 0.712674 | 0.711646 |
| 0.713669 | 0.709883 | 0.703559 | 0.703741 | 0.715028 | 0.711339 | 0.707202 | 0.70961 | 0.71144 |
| 0.720295 | 0.713806 | 0.710259 | 0.709026 | 0.720543 | 0.716194 | 0.712285 | 0.715829 | 0.713934 |

| | | | | | | | | |
|---|---|---|---|---|---|---|---|---|
| 0.7503 | 0.750849 | 0.738866 | 0.741765 | 0.750299 | 0.768055 | 0.749626 | 0.761387 | 0.770402 |
| 0.70533 | 0.699788 | 0.696351 | 0.693233 | 0.706068 | 0.703242 | 0.697996 | 0.699126 | 0.701534 |
| 0.707746 | 0.704019 | 0.695635 | 0.695069 | 0.710947 | 0.704132 | 0.701306 | 0.703166 | 0.701574 |
| 0.713914 | 0.710973 | 0.70663 | 0.705887 | 0.71987 | 0.715121 | 0.711135 | 0.713014 | 0.710056 |
| 0.711213 | 0.706233 | 0.699656 | 0.698588 | 0.716614 | 0.709029 | 0.705287 | 0.707789 | 0.707964 |
| 0.71391 | 0.708299 | 0.701364 | 0.703463 | 0.71894 | 0.710309 | 0.708295 | 0.709295 | 0.709436 |
| 0.709158 | 0.70799 | 0.700537 | 0.697381 | 0.711763 | 0.707001 | 0.701759 | 0.704018 | 0.701602 |
| 0.710296 | 0.708069 | 0.701932 | 0.701223 | 0.714639 | 0.710713 | 0.703667 | 0.709325 | 0.712447 |
| 0.714617 | 0.707935 | 0.707172 | 0.70955 | 0.715667 | 0.711969 | 0.707095 | 0.709019 | 0.709368 |
| 0.712064 | 0.709042 | 0.701248 | 0.703909 | 0.715908 | 0.711071 | 0.706885 | 0.708575 | 0.709392 |
| 0.704428 | 0.700016 | 0.695127 | 0.692977 | 0.70746 | 0.705135 | 0.698494 | 0.699736 | 0.700302 |
| 0.706223 | 0.699951 | 0.695747 | 0.695778 | 0.709294 | 0.707191 | 0.699786 | 0.704183 | 0.700293 |
| 0.710045 | 0.705344 | 0.700874 | 0.698743 | 0.70987 | 0.710789 | 0.70128 | 0.706005 | 0.70508 |
| 0.707564 | 0.70325 | 0.696994 | 0.697835 | 0.710775 | 0.707117 | 0.70066 | 0.705339 | 0.703282 |
| 0.706606 | 0.702324 | 0.69867 | 0.697117 | 0.710082 | 0.705078 | 0.701898 | 0.703725 | 0.703351 |
| 0.705722 | 0.698526 | 0.697229 | 0.695087 | 0.710543 | 0.706531 | 0.701592 | 0.702071 | 0.705395 |
| 0.706451 | 0.703152 | 0.69785 | 0.698482 | 0.705478 | 0.705704 | 0.696023 | 0.700587 | 0.698905 |
| 0.713549 | 0.707163 | 0.699664 | 0.700828 | 0.713755 | 0.71245 | 0.703868 | 0.707269 | 0.708581 |
| 0.715829 | 0.709273 | 0.705674 | 0.701763 | 0.719758 | 0.714706 | 0.707847 | 0.709842 | 0.711528 |
| 0.710007 | 0.702262 | 0.699199 | 0.697529 | 0.711845 | 0.706692 | 0.703056 | 0.705807 | 0.702341 |
| 0.703724 | 0.701371 | 0.693506 | 0.694523 | 0.706386 | 0.704272 | 0.698648 | 0.700224 | 0.698508 |
| 0.705477 | 0.70214 | 0.70046 | 0.696457 | 0.710714 | 0.70407 | 0.700412 | 0.702724 | 0.703945 |
| 0.71086 | 0.704185 | 0.698755 | 0.699597 | 0.716696 | 0.708593 | 0.703235 | 0.704342 | 0.705957 |
| 0.721769 | 0.713463 | 0.70799 | 0.710092 | 0.718521 | 0.715391 | 0.712964 | 0.714744 | 0.71367 |
| 0.723374 | 0.720108 | 0.712757 | 0.716631 | 0.728161 | 0.722505 | 0.71779 | 0.720118 | 0.719831 |
| 0.763203 | 0.760563 | 0.751997 | 0.753907 | 0.760275 | 0.764295 | 0.765148 | 0.761727 | 0.760988 |
| 0.772356 | 0.771707 | 0.764193 | 0.764025 | 0.768997 | 0.766488 | 0.762962 | 0.760507 | 0.762247 |
| 0.771499 | 0.770306 | 0.768366 | 0.763619 | 0.768738 | 0.766782 | 0.769356 | 0.763367 | 0.768774 |
| 0.707796 | 0.702362 | 0.700434 | 0.699329 | 0.713832 | 0.705868 | 0.702119 | 0.70514 | 0.705752 |
| 0.708361 | 0.705253 | 0.699121 | 0.698748 | 0.71173 | 0.706257 | 0.701078 | 0.706691 | 0.705902 |
| 0.718795 | 0.715923 | 0.708269 | 0.706858 | 0.719866 | 0.715836 | 0.710498 | 0.710876 | 0.711672 |
| 0.715674 | 0.709284 | 0.709293 | 0.706653 | 0.718348 | 0.716083 | 0.715536 | 0.713962 | 0.714597 |
| 0.707282 | 0.705736 | 0.69935 | 0.700855 | 0.714091 | 0.709478 | 0.703661 | 0.707325 | 0.704179 |
| 0.707473 | 0.700927 | 0.699838 | 0.700483 | 0.716011 | 0.70829 | 0.703259 | 0.705339 | 0.702079 |
| 0.707374 | 0.703253 | 0.699389 | 0.697754 | 0.71355 | 0.709459 | 0.700848 | 0.702183 | 0.701973 |
| 0.709893 | 0.705306 | 0.698284 | 0.697431 | 0.711879 | 0.70976 | 0.704185 | 0.706713 | 0.705887 |
| 0.706963 | 0.704243 | 0.699045 | 0.695688 | 0.71623 | 0.714387 | 0.703997 | 0.705333 | 0.70516 |
| 0.723074 | 0.718804 | 0.715756 | 0.713403 | 0.728491 | 0.726597 | 0.718592 | 0.719544 | 0.719782 |
| 0.700214 | 0.696083 | 0.693347 | 0.693 | 0.706483 | 0.703215 | 0.694791 | 0.698796 | 0.697826 |
| 0.711413 | 0.705062 | 0.700518 | 0.698907 | 0.711531 | 0.709234 | 0.701603 | 0.704879 | 0.704724 |
| 0.703506 | 0.699131 | 0.693899 | 0.696546 | 0.709602 | 0.704038 | 0.699421 | 0.70351 | 0.704821 |
| 0.726559 | 0.721446 | 0.719678 | 0.717923 | 0.729587 | 0.726342 | 0.719463 | 0.72435 | 0.721729 |
| 0.712262 | 0.710821 | 0.702918 | 0.702867 | 0.713765 | 0.71089 | 0.705176 | 0.707219 | 0.703992 |
| 0.707733 | 0.703671 | 0.697011 | 0.697929 | 0.711123 | 0.707231 | 0.70293 | 0.706965 | 0.70228 |
| 0.71513 | 0.710353 | 0.699346 | 0.700983 | 0.716417 | 0.708904 | 0.706815 | 0.709683 | 0.710192 |
| 0.707707 | 0.701317 | 0.69893 | 0.69697 | 0.712549 | 0.709105 | 0.700935 | 0.705138 | 0.705392 |

| | | | | | | | | |
|---|---|---|---|---|---|---|---|---|
| 0.708555 | 0.705998 | 0.700966 | 0.702586 | 0.711717 | 0.707559 | 0.704354 | 0.705097 | 0.702559 |
| 0.713562 | 0.706835 | 0.702276 | 0.700935 | 0.713464 | 0.71475 | 0.70634 | 0.708893 | 0.707776 |
| 0.711462 | 0.706538 | 0.699636 | 0.6996 | 0.716486 | 0.713016 | 0.704968 | 0.706603 | 0.704027 |
| 0.717951 | 0.712449 | 0.710552 | 0.708071 | 0.722361 | 0.721594 | 0.713797 | 0.713938 | 0.713548 |
| 0.715425 | 0.712057 | 0.706297 | 0.707753 | 0.721239 | 0.715155 | 0.712453 | 0.712273 | 0.711982 |
| 0.708018 | 0.704986 | 0.697109 | 0.697477 | 0.713978 | 0.709104 | 0.702196 | 0.705212 | 0.704839 |
| 0.710597 | 0.70703 | 0.703739 | 0.701515 | 0.719113 | 0.715559 | 0.706546 | 0.708621 | 0.707271 |
| 0.715029 | 0.707793 | 0.705262 | 0.701601 | 0.718297 | 0.710969 | 0.705464 | 0.709054 | 0.709309 |
| 0.709531 | 0.705572 | 0.700388 | 0.699451 | 0.71257 | 0.711157 | 0.704321 | 0.707652 | 0.707457 |
| 0.709124 | 0.701215 | 0.698821 | 0.695986 | 0.710139 | 0.706846 | 0.703147 | 0.702888 | 0.701907 |
| 0.709677 | 0.704125 | 0.703916 | 0.701547 | 0.712269 | 0.711919 | 0.705008 | 0.705585 | 0.706068 |
| 0.710936 | 0.702841 | 0.702575 | 0.698191 | 0.715031 | 0.713042 | 0.703306 | 0.70728 | 0.705556 |
| 0.712995 | 0.70884 | 0.704223 | 0.701227 | 0.717811 | 0.711514 | 0.706375 | 0.71007 | 0.708786 |
| 0.707241 | 0.704949 | 0.696716 | 0.697799 | 0.714022 | 0.706592 | 0.700108 | 0.699282 | 0.700232 |
| 0.712124 | 0.705641 | 0.702539 | 0.699344 | 0.713197 | 0.710996 | 0.706871 | 0.708974 | 0.707929 |
| 0.709853 | 0.704885 | 0.701981 | 0.699184 | 0.709528 | 0.70922 | 0.703634 | 0.710021 | 0.705394 |
| 0.714799 | 0.707917 | 0.703776 | 0.702199 | 0.719385 | 0.715696 | 0.708008 | 0.707979 | 0.711471 |
| 0.709547 | 0.702392 | 0.698325 | 0.697122 | 0.713024 | 0.708749 | 0.700508 | 0.706089 | 0.707273 |
| 0.707222 | 0.699738 | 0.700662 | 0.697362 | 0.710743 | 0.708741 | 0.703121 | 0.704607 | 0.702709 |
| 0.71323 | 0.70635 | 0.700563 | 0.700464 | 0.713041 | 0.70924 | 0.704261 | 0.7067 | 0.708028 |
| 0.709937 | 0.703296 | 0.69904 | 0.698317 | 0.712432 | 0.710126 | 0.707398 | 0.708858 | 0.705082 |
| 0.70532 | 0.700196 | 0.696334 | 0.696355 | 0.713471 | 0.706287 | 0.699492 | 0.700842 | 0.702525 |
| 0.710531 | 0.708106 | 0.702091 | 0.701459 | 0.713227 | 0.71368 | 0.701463 | 0.707666 | 0.707073 |
| 0.712759 | 0.707435 | 0.703117 | 0.702779 | 0.716713 | 0.712122 | 0.704437 | 0.705279 | 0.705817 |
| 0.713 | 0.709932 | 0.706108 | 0.704954 | 0.714103 | 0.715101 | 0.701068 | 0.706087 | 0.707284 |
| 0.714951 | 0.713948 | 0.704154 | 0.70547 | 0.720518 | 0.713801 | 0.710569 | 0.712801 | 0.71151 |
| 0.708733 | 0.71069 | 0.706718 | 0.704974 | 0.715184 | 0.713778 | 0.705589 | 0.707854 | 0.706921 |
| 0.712477 | 0.70532 | 0.703942 | 0.701737 | 0.714726 | 0.712148 | 0.703355 | 0.70748 | 0.710875 |
| 0.71047 | 0.709463 | 0.701594 | 0.703109 | 0.714851 | 0.71122 | 0.703828 | 0.711482 | 0.712085 |
| 0.711018 | 0.71039 | 0.70555 | 0.704511 | 0.718521 | 0.716835 | 0.704443 | 0.711005 | 0.709389 |
| 0.715086 | 0.708684 | 0.705214 | 0.707309 | 0.719744 | 0.71271 | 0.708592 | 0.711193 | 0.707802 |
| 0.711139 | 0.708205 | 0.701487 | 0.700057 | 0.716984 | 0.710469 | 0.702965 | 0.706361 | 0.705011 |
| 0.70851 | 0.705775 | 0.699341 | 0.700629 | 0.716019 | 0.712791 | 0.702797 | 0.705569 | 0.707519 |
| 0.710398 | 0.70643 | 0.704018 | 0.703414 | 0.718421 | 0.713087 | 0.707515 | 0.710678 | 0.707506 |
| 0.706478 | 0.705373 | 0.698829 | 0.696718 | 0.713671 | 0.710818 | 0.703449 | 0.707343 | 0.706906 |
| 0.711754 | 0.703648 | 0.700277 | 0.701253 | 0.71165 | 0.714643 | 0.705218 | 0.706044 | 0.706636 |
| 0.70895 | 0.70323 | 0.697772 | 0.693895 | 0.715047 | 0.707942 | 0.698343 | 0.704267 | 0.705701 |
| 0.684625 | 0.680322 | 0.67195 | 0.677453 | 0.6915 | 0.682757 | 0.678046 | 0.680122 | 0.677299 |
| 0.689294 | 0.682837 | 0.679355 | 0.681785 | 0.69624 | 0.688104 | 0.684671 | 0.682274 | 0.685027 |
| 0.675662 | 0.676301 | 0.67043 | 0.673444 | 0.68734 | 0.683251 | 0.675342 | 0.679212 | 0.677184 |
| 0.678935 | 0.675171 | 0.672875 | 0.669148 | 0.684942 | 0.683721 | 0.677769 | 0.675778 | 0.676343 |
| 0.662024 | 0.659451 | 0.657176 | 0.657405 | 0.675196 | 0.665271 | 0.659662 | 0.659535 | 0.659748 |
| 0.666506 | 0.669575 | 0.661172 | 0.662102 | 0.67221 | 0.673065 | 0.665529 | 0.666423 | 0.6653 |
| 0.666747 | 0.666955 | 0.65701 | 0.661785 | 0.673425 | 0.672881 | 0.666174 | 0.663943 | 0.664446 |
| 0.672558 | 0.669809 | 0.662075 | 0.659814 | 0.681911 | 0.669418 | 0.664999 | 0.671825 | 0.665989 |
| 0.670229 | 0.67018 | 0.663186 | 0.659623 | 0.673543 | 0.671388 | 0.665243 | 0.665329 | 0.662443 |
| 0.675925 | 0.669204 | 0.662606 | 0.663567 | 0.679719 | 0.671341 | 0.673587 | 0.666501 | 0.672982 |

| | | | | | | | | |
|---|---|---|---|---|---|---|---|---|
| 0.678171 | 0.671122 | 0.66291 | 0.662997 | 0.678413 | 0.676438 | 0.666749 | 0.667079 | 0.666093 |
| 0.673953 | 0.674625 | 0.667498 | 0.667508 | 0.684313 | 0.675247 | 0.67222 | 0.669421 | 0.668055 |
| 0.672492 | 0.665631 | 0.655705 | 0.662819 | 0.679728 | 0.673711 | 0.674331 | 0.673954 | 0.667779 |
| 0.665482 | 0.665326 | 0.656923 | 0.661399 | 0.671379 | 0.662727 | 0.662006 | 0.658795 | 0.661209 |
| 0.661656 | 0.662771 | 0.65408 | 0.65994 | 0.670075 | 0.668679 | 0.66307 | 0.657939 | 0.658218 |
| 0.665936 | 0.667925 | 0.65531 | 0.658428 | 0.67513 | 0.666303 | 0.665745 | 0.659504 | 0.661368 |
| 0.665538 | 0.66824 | 0.653644 | 0.659045 | 0.672589 | 0.668175 | 0.668389 | 0.66296 | 0.663037 |
| 0.664341 | 0.662676 | 0.652571 | 0.655408 | 0.672653 | 0.667878 | 0.659749 | 0.661727 | 0.659333 |
| 0.665493 | 0.658461 | 0.657269 | 0.653778 | 0.667545 | 0.664622 | 0.663712 | 0.66068 | 0.657474 |
| 0.666035 | 0.662274 | 0.656886 | 0.659037 | 0.667732 | 0.668227 | 0.660527 | 0.660097 | 0.655941 |
| 0.655236 | 0.6614 | 0.655974 | 0.654555 | 0.664385 | 0.6573 | 0.65508 | 0.652827 | 0.653936 |
| 0.701 | 0.693591 | 0.690078 | 0.689368 | 0.704145 | 0.697714 | 0.693384 | 0.695307 | 0.691774 |
| 0.710936 | 0.703878 | 0.701116 | 0.698612 | 0.71176 | 0.710139 | 0.699405 | 0.703194 | 0.703838 |
| 0.684515 | 0.682719 | 0.675157 | 0.679138 | 0.687352 | 0.684407 | 0.689783 | 0.680055 | 0.683791 |
| 0.71844 | 0.709947 | 0.708153 | 0.710294 | 0.721396 | 0.717971 | 0.712114 | 0.712746 | 0.715532 |
| 0.704236 | 0.701283 | 0.695969 | 0.696247 | 0.711537 | 0.710698 | 0.70055 | 0.703011 | 0.69985 |
| 0.715425 | 0.70798 | 0.702094 | 0.702047 | 0.715061 | 0.713383 | 0.706854 | 0.708459 | 0.709649 |
| 0.723035 | 0.713837 | 0.708509 | 0.713097 | 0.724653 | 0.719043 | 0.713811 | 0.716039 | 0.716889 |
| 0.700477 | 0.689161 | 0.686773 | 0.684146 | 0.702148 | 0.698016 | 0.692252 | 0.69197 | 0.698257 |
| 0.720047 | 0.712609 | 0.70884 | 0.709353 | 0.72733 | 0.722117 | 0.715058 | 0.713537 | 0.71574 |
| 0.689676 | 0.689773 | 0.681741 | 0.682442 | 0.701188 | 0.690619 | 0.688399 | 0.68454 | 0.687432 |
| 0.718745 | 0.709855 | 0.705583 | 0.70738 | 0.720615 | 0.714058 | 0.707752 | 0.712031 | 0.713807 |
| 0.719636 | 0.7174 | 0.712305 | 0.710796 | 0.724842 | 0.72256 | 0.713813 | 0.720019 | 0.719218 |
| 0.682085 | 0.678676 | 0.670627 | 0.676509 | 0.69271 | 0.680273 | 0.680961 | 0.677092 | 0.680619 |
| 0.68772 | 0.685154 | 0.680276 | 0.680014 | 0.695348 | 0.691826 | 0.681664 | 0.680942 | 0.68543 |
| 0.690176 | 0.688167 | 0.677223 | 0.681357 | 0.691622 | 0.694458 | 0.690772 | 0.682337 | 0.687967 |
| 0.707453 | 0.705651 | 0.701084 | 0.700593 | 0.712191 | 0.709432 | 0.704062 | 0.70668 | 0.701373 |
| 0.719787 | 0.713243 | 0.707597 | 0.707081 | 0.728766 | 0.716391 | 0.711347 | 0.714092 | 0.714921 |
| 0.687336 | 0.682192 | 0.677672 | 0.679177 | 0.691525 | 0.689519 | 0.679177 | 0.685033 | 0.685338 |
| 0.723166 | 0.719383 | 0.711996 | 0.714085 | 0.730176 | 0.729854 | 0.717249 | 0.719498 | 0.719668 |
| 0.723072 | 0.722565 | 0.715914 | 0.71486 | 0.730839 | 0.728018 | 0.722213 | 0.724075 | 0.724793 |
| 0.697864 | 0.693349 | 0.688145 | 0.688992 | 0.70344 | 0.701767 | 0.693323 | 0.694436 | 0.694189 |
| 0.697412 | 0.694546 | 0.690122 | 0.687239 | 0.703091 | 0.702696 | 0.694755 | 0.696023 | 0.693752 |
| 0.724174 | 0.72399 | 0.715496 | 0.716939 | 0.731157 | 0.72741 | 0.72274 | 0.721608 | 0.720951 |
| 0.722509 | 0.717154 | 0.709968 | 0.713374 | 0.728254 | 0.722366 | 0.713718 | 0.715612 | 0.716928 |
| 0.727791 | 0.723716 | 0.714795 | 0.718475 | 0.731422 | 0.723774 | 0.722851 | 0.722101 | 0.721678 |
| 0.677049 | 0.672858 | 0.669073 | 0.668117 | 0.684643 | 0.680505 | 0.671259 | 0.677707 | 0.677046 |
| 0.722024 | 0.716592 | 0.709494 | 0.709389 | 0.726761 | 0.726241 | 0.715649 | 0.719958 | 0.720566 |
| 0.676006 | 0.671813 | 0.668011 | 0.669673 | 0.680454 | 0.679942 | 0.667823 | 0.672043 | 0.672265 |
| 0.669632 | 0.666641 | 0.662251 | 0.659286 | 0.67592 | 0.67072 | 0.663384 | 0.667377 | 0.664126 |
| 0.72267 | 0.715543 | 0.711223 | 0.709443 | 0.726921 | 0.72829 | 0.719813 | 0.723335 | 0.725736 |
| 0.669438 | 0.668308 | 0.659524 | 0.660931 | 0.677029 | 0.672186 | 0.668943 | 0.667377 | 0.667145 |
| 0.663121 | 0.65904 | 0.652451 | 0.653163 | 0.668063 | 0.665144 | 0.6595 | 0.662294 | 0.658101 |
| 0.657643 | 0.657656 | 0.647376 | 0.651319 | 0.664713 | 0.661767 | 0.654064 | 0.658088 | 0.657696 |
| 0.727143 | 0.727101 | 0.720229 | 0.719688 | 0.733784 | 0.74747 | 0.728637 | 0.735289 | 0.735845 |
| 0.662182 | 0.660322 | 0.653248 | 0.654684 | 0.665967 | 0.664771 | 0.657087 | 0.662091 | 0.658214 |
| 0.720914 | 0.716604 | 0.712832 | 0.712894 | 0.727005 | 0.737452 | 0.718727 | 0.728521 | 0.724994 |

| | | | | | | | | |
|---|---|---|---|---|---|---|---|---|
| 0.73389 | 0.72787 | 0.71699 | 0.717793 | 0.734671 | 0.74726 | 0.733328 | 0.741506 | 0.742318 |
| 0.727042 | 0.721333 | 0.713358 | 0.714416 | 0.730165 | 0.739712 | 0.725941 | 0.731721 | 0.730972 |
| 0.752749 | 0.74725 | 0.738905 | 0.737181 | 0.756193 | 0.750407 | 0.745091 | 0.742761 | 0.743362 |
| 0.65343 | 0.648801 | 0.641138 | 0.645174 | 0.654702 | 0.652511 | 0.649686 | 0.6525 | 0.64816 |
| 0.653545 | 0.651829 | 0.644149 | 0.644699 | 0.657951 | 0.655582 | 0.648547 | 0.650164 | 0.649421 |
| 0.73456 | 0.729687 | 0.722033 | 0.718633 | 0.738141 | 0.742311 | 0.732951 | 0.736987 | 0.739179 |
| 0.656252 | 0.65078 | 0.645733 | 0.644812 | 0.658161 | 0.658649 | 0.649904 | 0.651154 | 0.651356 |
| 0.735344 | 0.733104 | 0.723198 | 0.723392 | 0.738687 | 0.734581 | 0.731461 | 0.734625 | 0.733035 |
| 0.731995 | 0.728646 | 0.723273 | 0.721747 | 0.737079 | 0.748419 | 0.732655 | 0.741295 | 0.740923 |
| 0.735493 | 0.733772 | 0.728062 | 0.724258 | 0.73815 | 0.736679 | 0.730474 | 0.733832 | 0.733883 |
| 0.73269 | 0.730183 | 0.724961 | 0.72583 | 0.735715 | 0.757858 | 0.734403 | 0.742486 | 0.745586 |
| 0.741497 | 0.742857 | 0.700637 | 0.706667 | 0.77931 | 0.736486 | 0.755245 | 0.714286 | 0.714286 |
| 0.717551 | 0.714802 | 0.706094 | 0.711484 | 0.722588 | 0.72542 | 0.729724 | 0.721292 | 0.718619 |
| 0.717577 | 0.722028 | 0.708669 | 0.707797 | 0.723949 | 0.719385 | 0.718367 | 0.712764 | 0.713751 |
| 0.726374 | 0.727912 | 0.716528 | 0.71854 | 0.726524 | 0.733413 | 0.731409 | 0.724913 | 0.719167 |
| 0.729119 | 0.726096 | 0.71334 | 0.721902 | 0.725247 | 0.734209 | 0.726644 | 0.718809 | 0.72462 |
| 0.592445 | 0.58888 | 0.585302 | 0.587268 | 0.593198 | 0.595048 | 0.588325 | 0.586595 | 0.589264 |
| 0.72934 | 0.728673 | 0.721727 | 0.717745 | 0.730118 | 0.734659 | 0.73165 | 0.730603 | 0.730925 |
| 0.738147 | 0.735644 | 0.72793 | 0.728005 | 0.738387 | 0.759673 | 0.741044 | 0.750762 | 0.755935 |
| 0.736338 | 0.737029 | 0.727491 | 0.724285 | 0.739198 | 0.761054 | 0.741412 | 0.757198 | 0.755982 |
| 0.738598 | 0.736895 | 0.731575 | 0.730237 | 0.742572 | 0.765247 | 0.739726 | 0.746261 | 0.761284 |
| 0.596831 | 0.594671 | 0.589126 | 0.588577 | 0.599558 | 0.594985 | 0.592072 | 0.592035 | 0.594284 |
| 0.584555 | 0.580219 | 0.579346 | 0.576024 | 0.585328 | 0.585917 | 0.579656 | 0.586453 | 0.580554 |
| 0.763356 | 0.763167 | 0.749853 | 0.750865 | 0.76511 | 0.799527 | 0.795747 | 0.809862 | 0.832373 |
| 0.766509 | 0.766432 | 0.751095 | 0.756304 | 0.769878 | 0.807615 | 0.803272 | 0.805595 | 0.822368 |
| 0.757611 | 0.76014 | 0.745134 | 0.748904 | 0.760482 | 0.796523 | 0.797055 | 0.810002 | 0.821438 |
| 0.759912 | 0.764626 | 0.755443 | 0.755425 | 0.770398 | 0.802898 | 0.789026 | 0.809784 | 0.815995 |
| 0.762807 | 0.767579 | 0.753483 | 0.752831 | 0.773428 | 0.824771 | 0.794446 | 0.802961 | 0.891431 |
| 0.765746 | 0.769514 | 0.757601 | 0.760445 | 0.764323 | 0.795451 | 0.807167 | 0.795197 | 0.800354 |
| 0.775216 | 0.768041 | 0.757314 | 0.758789 | 0.778199 | 0.815438 | 0.806741 | 0.799983 | 0.802896 |
| 0.778027 | 0.788025 | 0.773804 | 0.775993 | 0.789093 | 0.76929 | 0.762602 | 0.760529 | 0.76065 |
| 0.795981 | 0.789281 | 0.783478 | 0.782848 | 0.792832 | 0.777315 | 0.774589 | 0.76993 | 0.774083 |
| 0.782069 | 0.785899 | 0.775378 | 0.776829 | 0.782912 | 0.768135 | 0.76528 | 0.760924 | 0.764419 |
| 0.789155 | 0.786682 | 0.774963 | 0.773205 | 0.790295 | 0.761327 | 0.763012 | 0.763072 | 0.762878 |
| 0.79113 | 0.792584 | 0.786709 | 0.78634 | 0.791564 | 0.772306 | 0.77357 | 0.767584 | 0.767581 |
| 0.781842 | 0.780939 | 0.773353 | 0.773212 | 0.782731 | 0.764232 | 0.764661 | 0.765292 | 0.760305 |
| 0.788845 | 0.788859 | 0.778881 | 0.778038 | 0.800583 | 0.769674 | 0.775135 | 0.767789 | 0.768311 |
| 0.79039 | 0.791945 | 0.778354 | 0.777079 | 0.79525 | 0.774313 | 0.771939 | 0.769218 | 0.772208 |
| 0.78848 | 0.787895 | 0.778743 | 0.778617 | 0.790775 | 0.768308 | 0.76528 | 0.759458 | 0.760354 |
| 0.787138 | 0.785446 | 0.780137 | 0.7716 | 0.794418 | 0.767672 | 0.760523 | 0.76308 | 0.761416 |
| 0.717024 | 0.711996 | 0.70739 | 0.717503 | 0.709905 | 0.725231 | 0.727851 | 0.72141 | 0.722378 |
| 0.732244 | 0.730195 | 0.713292 | 0.721602 | 0.728775 | 0.729436 | 0.730514 | 0.719404 | 0.723224 |
| 0.794183 | 0.802232 | 0.802535 | 0.781476 | 0.782933 | 0.76563 | 0.766845 | 0.76036 | 0.76879 |
| 0.78542 | 0.795281 | 0.793976 | 0.856127 | 0.779407 | 0.784907 | 0.78281 | 0.778423 | 0.785567 |
| 0.786537 | 0.817718 | 0.784749 | 0.757521 | 0.794953 | 0.728347 | 0.725578 | 0.718774 | 0.721802 |
| 0.838389 | 0.836944 | 0.814707 | 0.798253 | 0.811408 | 0.776067 | 0.77234 | 0.769841 | 0.768725 |
| 0.822706 | 0.83606 | 0.889124 | 0.832332 | 0.789335 | 0.764845 | 0.766593 | 0.758835 | 0.763364 |

| | | | | | | | | |
|---|---|---|---|---|---|---|---|---|
| 0.839387 | 0.87692 | 0.829646 | 0.827751 | 0.803362 | 0.76814 | 0.770239 | 0.77237 | 0.769553 |
| 0.855947 | 0.864805 | 0.820201 | 0.804388 | 0.802984 | 0.770641 | 0.774139 | 0.769963 | 0.771274 |
| 0.811313 | 0.814802 | 0.834215 | 0.817577 | 0.797615 | 0.767757 | 0.76996 | 0.761118 | 0.767932 |
| 0.826211 | 0.835155 | 0.798966 | 0.81438 | 0.801761 | 0.77554 | 0.774051 | 0.772686 | 0.768524 |
| 0.817292 | 0.834398 | 0.808745 | 0.799076 | 0.842816 | 0.777285 | 0.77021 | 0.76528 | 0.771799 |
| 0.796193 | 0.83708 | 0.804491 | 0.858201 | 0.834407 | 0.773529 | 0.773473 | 0.770357 | 0.766529 |
| 0.836432 | 0.846928 | 0.805518 | 0.858714 | 0.797408 | 0.769727 | 0.771581 | 0.765585 | 0.767598 |
| 0.798704 | 0.813451 | 0.827137 | 0.809134 | 0.792606 | 0.758174 | 0.76323 | 0.753726 | 0.758161 |
| 0.823217 | 0.842045 | 0.777004 | 0.796319 | 0.802701 | 0.767923 | 0.773932 | 0.765803 | 0.763119 |
| 0.837148 | 0.812663 | 0.850065 | 0.805384 | 0.792202 | 0.761409 | 0.763936 | 0.756832 | 0.762978 |
| 1 | 0.823427 | 0.814263 | 0.787368 | 0.79596 | 0.768403 | 0.76996 | 0.766642 | 0.768012 |
| 0.823427 | 1 | 0.818918 | 0.818955 | 0.809454 | 0.770597 | 0.772942 | 0.766712 | 0.771184 |
| 0.814263 | 0.818918 | 1 | 0.830981 | 0.780145 | 0.758279 | 0.760372 | 0.754068 | 0.7566 |
| 0.787368 | 0.818955 | 0.830981 | 1 | 0.77732 | 0.763187 | 0.764289 | 0.754389 | 0.759634 |
| 0.79596 | 0.809454 | 0.780145 | 0.77732 | 1 | 0.779133 | 0.774692 | 0.77082 | 0.77308 |
| 0.768403 | 0.770597 | 0.758279 | 0.763187 | 0.779133 | 1 | 0.794065 | 0.814637 | 0.820916 |
| 0.76996 | 0.772942 | 0.760372 | 0.764289 | 0.774692 | 0.794065 | 1 | 0.82894 | 0.800173 |
| 0.766642 | 0.766712 | 0.754068 | 0.754389 | 0.77082 | 0.814637 | 0.82894 | 1 | 0.815745 |
| 0.768012 | 0.771184 | 0.7566 | 0.759634 | 0.77308 | 0.820916 | 0.800173 | 0.815745 | 1 |
| 0.769497 | 0.772173 | 0.754564 | 0.754943 | 0.774459 | 0.821124 | 0.800763 | 0.831351 | 0.821719 |
| 0.764407 | 0.761176 | 0.753633 | 0.751259 | 0.764261 | 0.827277 | 0.801852 | 0.836804 | 0.819902 |
| 0.762186 | 0.764153 | 0.755235 | 0.757067 | 0.767817 | 0.807955 | 0.786713 | 0.797188 | 0.895813 |
| 0.769806 | 0.773488 | 0.758521 | 0.758847 | 0.774935 | 0.815568 | 0.796638 | 0.809245 | 0.837541 |
| 0.774347 | 0.771625 | 0.761379 | 0.760662 | 0.771305 | 0.7996 | 0.841599 | 0.887414 | 0.806746 |
| 0.765652 | 0.761803 | 0.757466 | 0.753841 | 0.766791 | 0.807343 | 0.810627 | 0.832744 | 0.823157 |
| 0.765938 | 0.764681 | 0.749127 | 0.753524 | 0.763653 | 0.799614 | 0.783625 | 0.794522 | 0.83856 |
| 0.765189 | 0.764296 | 0.753173 | 0.752381 | 0.768294 | 0.822677 | 0.810885 | 0.865062 | 0.814672 |
| 0.762472 | 0.761925 | 0.752004 | 0.752611 | 0.764393 | 0.794332 | 0.802857 | 0.804918 | 0.817557 |
| 0.765298 | 0.760748 | 0.752606 | 0.753354 | 0.761521 | 0.813801 | 0.794174 | 0.814583 | 0.825591 |
| 0.766533 | 0.772322 | 0.756551 | 0.757076 | 0.765195 | 0.81077 | 0.802635 | 0.819214 | 0.845271 |
| 0.771555 | 0.767696 | 0.75923 | 0.762129 | 0.771394 | 0.82095 | 0.822813 | 0.852276 | 0.821191 |
| 0.7715 | 0.770202 | 0.758933 | 0.762381 | 0.77158 | 0.815192 | 0.822743 | 0.861467 | 0.810008 |
| 0.77717 | 0.773138 | 0.765692 | 0.766948 | 0.77917 | 0.817431 | 0.826181 | 0.837257 | 0.831083 |
| 0.773454 | 0.774962 | 0.75993 | 0.764994 | 0.778878 | 0.812249 | 0.856292 | 0.853673 | 0.805818 |
| 0.761156 | 0.761302 | 0.747639 | 0.751502 | 0.758967 | 0.800754 | 0.797758 | 0.797976 | 0.826065 |
| 0.766963 | 0.764095 | 0.754231 | 0.756176 | 0.768068 | 0.818555 | 0.804313 | 0.82399 | 0.827539 |
| 0.768362 | 0.762168 | 0.753563 | 0.753327 | 0.764856 | 0.795519 | 0.828211 | 0.813989 | 0.81714 |
| 0.764394 | 0.763221 | 0.749395 | 0.755588 | 0.76709 | 0.803561 | 0.79986 | 0.801326 | 0.812075 |
| 0.75991 | 0.756473 | 0.749051 | 0.749393 | 0.76288 | 0.824159 | 0.798241 | 0.86622 | 0.826083 |
| 0.756338 | 0.756623 | 0.744613 | 0.746187 | 0.762664 | 0.793043 | 0.802008 | 0.800831 | 0.803818 |
| 0.752689 | 0.755695 | 0.742901 | 0.751697 | 0.761013 | 0.809976 | 0.775325 | 0.814659 | 0.822644 |
| 0.756311 | 0.758744 | 0.745871 | 0.748396 | 0.754865 | 0.801424 | 0.792267 | 0.796736 | 0.826899 |
| 0.762052 | 0.759706 | 0.74948 | 0.75185 | 0.764676 | 0.82913 | 0.807556 | 0.844815 | 0.821009 |
| 0.748482 | 0.754129 | 0.745984 | 0.749604 | 0.757396 | 0.810846 | 0.78265 | 0.797561 | 0.883474 |
| 0.761803 | 0.76081 | 0.751718 | 0.756745 | 0.763038 | 0.793644 | 0.816308 | 0.809089 | 0.808199 |
| 0.756676 | 0.758188 | 0.747461 | 0.754802 | 0.756422 | 0.781689 | 0.818925 | 0.83126 | 0.789003 |

| | | | | | | | | |
|---|---|---|---|---|---|---|---|---|
| 0.765289 | 0.763861 | 0.751371 | 0.755987 | 0.769199 | 0.802242 | 0.77978 | 0.797292 | 0.811959 |
| 0.765748 | 0.769937 | 0.759061 | 0.762136 | 0.767455 | 0.767579 | 0.768787 | 0.763583 | 0.765867 |
| 0.758521 | 0.763304 | 0.751805 | 0.753981 | 0.768988 | 0.760109 | 0.764386 | 0.763153 | 0.760718 |
| 0.766196 | 0.765152 | 0.75002 | 0.755331 | 0.766183 | 0.762404 | 0.763486 | 0.758744 | 0.762338 |
| 0.772291 | 0.773044 | 0.762751 | 0.761709 | 0.77502 | 0.772645 | 0.76959 | 0.768768 | 0.773801 |
| 0.769809 | 0.768984 | 0.762699 | 0.758773 | 0.771698 | 0.769608 | 0.768636 | 0.764276 | 0.764792 |
| 0.761282 | 0.768052 | 0.753769 | 0.756195 | 0.771312 | 0.767801 | 0.769803 | 0.766093 | 0.768492 |
| 0.769551 | 0.766861 | 0.756779 | 0.760821 | 0.771216 | 0.76637 | 0.770727 | 0.767323 | 0.764326 |
| 0.762489 | 0.766999 | 0.756781 | 0.758476 | 0.764 | 0.764648 | 0.764472 | 0.75632 | 0.761135 |
| 0.768568 | 0.764132 | 0.757063 | 0.754723 | 0.770246 | 0.763963 | 0.765554 | 0.76438 | 0.763618 |
| 0.775447 | 0.773957 | 0.762188 | 0.761804 | 0.778148 | 0.772712 | 0.764953 | 0.76619 | 0.769043 |
| 0.776399 | 0.773974 | 0.765 | 0.76217 | 0.775629 | 0.775493 | 0.768799 | 0.769028 | 0.767587 |
| 0.765761 | 0.763768 | 0.756197 | 0.755725 | 0.762505 | 0.763787 | 0.766053 | 0.760722 | 0.766154 |
| 0.766727 | 0.763405 | 0.756542 | 0.756861 | 0.766863 | 0.764614 | 0.763149 | 0.760616 | 0.764277 |
| 0.767672 | 0.766485 | 0.760775 | 0.76053 | 0.7689 | 0.768911 | 0.763883 | 0.763828 | 0.764926 |
| 0.764687 | 0.768775 | 0.756401 | 0.757482 | 0.76562 | 0.766225 | 0.761159 | 0.763257 | 0.766503 |
| 0.765353 | 0.768075 | 0.759198 | 0.759572 | 0.765583 | 0.771053 | 0.766142 | 0.763293 | 0.766556 |
| 0.760058 | 0.763991 | 0.755705 | 0.756248 | 0.764342 | 0.76538 | 0.764647 | 0.759961 | 0.766718 |
| 0.767768 | 0.768073 | 0.759997 | 0.760961 | 0.767861 | 0.768728 | 0.771414 | 0.768566 | 0.770069 |
| 0.768448 | 0.768077 | 0.758922 | 0.75922 | 0.770154 | 0.771361 | 0.771387 | 0.767949 | 0.76771 |
| 0.767337 | 0.763766 | 0.753244 | 0.754412 | 0.760532 | 0.763007 | 0.759314 | 0.759482 | 0.764062 |
| 0.765844 | 0.764821 | 0.753581 | 0.752943 | 0.767665 | 0.764159 | 0.766428 | 0.761692 | 0.769205 |
| 0.766509 | 0.765299 | 0.757771 | 0.755926 | 0.764763 | 0.764859 | 0.764309 | 0.76384 | 0.769621 |
| 0.769643 | 0.766261 | 0.757511 | 0.758509 | 0.766844 | 0.769556 | 0.764646 | 0.760016 | 0.765304 |
| 0.764015 | 0.764209 | 0.752997 | 0.755081 | 0.766141 | 0.767113 | 0.766345 | 0.766645 | 0.765971 |
| 0.763638 | 0.764706 | 0.75566 | 0.756957 | 0.766865 | 0.766359 | 0.763962 | 0.76176 | 0.765514 |
| 0.76522 | 0.768196 | 0.758584 | 0.756031 | 0.76383 | 0.760936 | 0.761447 | 0.757687 | 0.761712 |
| 0.765355 | 0.76121 | 0.756666 | 0.756392 | 0.762882 | 0.765231 | 0.762102 | 0.760242 | 0.763104 |
| 0.772842 | 0.767528 | 0.758042 | 0.761249 | 0.771618 | 0.771256 | 0.768238 | 0.767129 | 0.767669 |
| 0.766664 | 0.764675 | 0.759082 | 0.758182 | 0.771364 | 0.771373 | 0.768266 | 0.761696 | 0.765291 |
| 0.756373 | 0.757179 | 0.74662 | 0.751038 | 0.760615 | 0.757876 | 0.756738 | 0.754851 | 0.758812 |
| 0.765044 | 0.764096 | 0.758339 | 0.759315 | 0.764788 | 0.769801 | 0.762538 | 0.761121 | 0.770064 |
| 0.764118 | 0.767238 | 0.755939 | 0.757744 | 0.768162 | 0.767683 | 0.764189 | 0.761569 | 0.762963 |
| 0.762041 | 0.761756 | 0.756387 | 0.760318 | 0.762935 | 0.76235 | 0.761668 | 0.759437 | 0.761973 |
| 0.765264 | 0.762687 | 0.758212 | 0.756771 | 0.768519 | 0.764973 | 0.766387 | 0.762735 | 0.760643 |
| 0.763061 | 0.763267 | 0.752747 | 0.755579 | 0.764344 | 0.764242 | 0.764571 | 0.765165 | 0.761061 |
| 0.771765 | 0.769829 | 0.758472 | 0.7652 | 0.775357 | 0.771045 | 0.772221 | 0.7688 | 0.768069 |
| 0.770492 | 0.768152 | 0.761301 | 0.760059 | 0.76824 | 0.76452 | 0.759487 | 0.761574 | 0.76272 |
| 0.767773 | 0.767395 | 0.761255 | 0.76175 | 0.762342 | 0.768827 | 0.764002 | 0.761117 | 0.769439 |
| 0.763594 | 0.7646 | 0.755533 | 0.760292 | 0.766207 | 0.77033 | 0.760993 | 0.762133 | 0.76423 |
| 0.768717 | 0.765151 | 0.760215 | 0.759918 | 0.766707 | 0.773846 | 0.762648 | 0.76472 | 0.769168 |
| 0.769401 | 0.766056 | 0.755668 | 0.762235 | 0.77091 | 0.768998 | 0.766463 | 0.763825 | 0.766939 |
| 0.773264 | 0.765095 | 0.758246 | 0.762651 | 0.769287 | 0.774396 | 0.765753 | 0.76425 | 0.771573 |
| 0.761944 | 0.765708 | 0.755402 | 0.753407 | 0.764476 | 0.764657 | 0.759143 | 0.759458 | 0.760603 |
| 0.770085 | 0.768909 | 0.762729 | 0.761211 | 0.773259 | 0.772251 | 0.766385 | 0.763388 | 0.767348 |
| 0.769122 | 0.766928 | 0.759133 | 0.75725 | 0.764823 | 0.764208 | 0.763131 | 0.759736 | 0.763066 |

| | | | | | | | | |
|---|---|---|---|---|---|---|---|---|
| 0.756024 | 0.749928 | 0.746744 | 0.747102 | 0.755436 | 0.753399 | 0.752275 | 0.75026 | 0.753441 |
| 0.747771 | 0.743193 | 0.734837 | 0.730675 | 0.748303 | 0.748381 | 0.748131 | 0.741237 | 0.744905 |
| 0.765617 | 0.762003 | 0.753714 | 0.752534 | 0.765612 | 0.759441 | 0.760722 | 0.755772 | 0.762809 |
| 0.769443 | 0.766463 | 0.757103 | 0.755157 | 0.770022 | 0.764763 | 0.76293 | 0.764653 | 0.764706 |
| 0.762244 | 0.755614 | 0.750043 | 0.750816 | 0.756745 | 0.758133 | 0.758733 | 0.754143 | 0.759189 |
| 0.76331 | 0.757578 | 0.754613 | 0.754097 | 0.760226 | 0.763225 | 0.759537 | 0.756741 | 0.761123 |
| 0.768709 | 0.760623 | 0.755588 | 0.755678 | 0.762473 | 0.761542 | 0.763282 | 0.758312 | 0.762955 |
| 0.761544 | 0.754879 | 0.747237 | 0.748401 | 0.760165 | 0.75645 | 0.761492 | 0.753558 | 0.756343 |
| 0.765184 | 0.766302 | 0.754428 | 0.758779 | 0.77292 | 0.767538 | 0.767825 | 0.762586 | 0.765492 |
| 0.770589 | 0.766203 | 0.757971 | 0.761605 | 0.765763 | 0.768871 | 0.76772 | 0.764128 | 0.76885 |
| 0.768818 | 0.768188 | 0.758772 | 0.760831 | 0.770742 | 0.767217 | 0.770399 | 0.764331 | 0.767229 |
| 0.765434 | 0.764945 | 0.756755 | 0.758264 | 0.770151 | 0.765755 | 0.770006 | 0.763004 | 0.762727 |
| 0.769639 | 0.774572 | 0.759236 | 0.760855 | 0.768083 | 0.766686 | 0.764342 | 0.764232 | 0.767617 |
| 0.771917 | 0.769841 | 0.761225 | 0.759336 | 0.770881 | 0.766946 | 0.767351 | 0.763578 | 0.767848 |
| 0.766235 | 0.764706 | 0.755147 | 0.755208 | 0.763864 | 0.764466 | 0.770592 | 0.761877 | 0.763279 |
| 0.763331 | 0.758145 | 0.75222 | 0.754695 | 0.767246 | 0.763878 | 0.764647 | 0.758359 | 0.760822 |
| 0.769161 | 0.767244 | 0.757465 | 0.756114 | 0.763924 | 0.765502 | 0.765432 | 0.765956 | 0.767166 |
| 0.762475 | 0.760722 | 0.75578 | 0.756433 | 0.762528 | 0.762601 | 0.75928 | 0.761058 | 0.761036 |
| 0.762362 | 0.759584 | 0.752493 | 0.755236 | 0.760781 | 0.766978 | 0.762201 | 0.75404 | 0.758384 |
| 0.764462 | 0.760796 | 0.754206 | 0.755429 | 0.763523 | 0.761217 | 0.762389 | 0.761884 | 0.767161 |
| 0.764667 | 0.765653 | 0.757964 | 0.757312 | 0.765471 | 0.766995 | 0.767733 | 0.760163 | 0.7614 |
| 0.756895 | 0.747903 | 0.745858 | 0.747698 | 0.748522 | 0.751509 | 0.7586 | 0.750822 | 0.749497 |
| 0.758076 | 0.755296 | 0.751032 | 0.753731 | 0.762242 | 0.759188 | 0.758694 | 0.756903 | 0.758629 |
| 0.763559 | 0.761965 | 0.7564 | 0.757482 | 0.76653 | 0.764122 | 0.761593 | 0.760979 | 0.760923 |
| 0.765685 | 0.764406 | 0.75309 | 0.757039 | 0.761921 | 0.76446 | 0.766791 | 0.765296 | 0.762092 |
| 0.762393 | 0.758261 | 0.751555 | 0.754202 | 0.762149 | 0.764178 | 0.762126 | 0.762993 | 0.756025 |
| 0.762927 | 0.76079 | 0.75206 | 0.756203 | 0.758812 | 0.759275 | 0.757976 | 0.755343 | 0.756322 |
| 0.761658 | 0.759797 | 0.75599 | 0.756206 | 0.762397 | 0.762986 | 0.761817 | 0.759774 | 0.764289 |
| 0.757291 | 0.753709 | 0.750393 | 0.752382 | 0.758543 | 0.758779 | 0.759271 | 0.760428 | 0.759524 |
| 0.699606 | 0.69658 | 0.68615 | 0.689518 | 0.700688 | 0.695907 | 0.690706 | 0.690774 | 0.691554 |
| 0.711665 | 0.703258 | 0.697739 | 0.697688 | 0.712183 | 0.709216 | 0.700372 | 0.701068 | 0.704385 |
| 0.705385 | 0.700858 | 0.698406 | 0.696524 | 0.70769 | 0.706262 | 0.697292 | 0.701918 | 0.701173 |
| 0.699253 | 0.698517 | 0.690056 | 0.688657 | 0.701255 | 0.698962 | 0.695664 | 0.695852 | 0.69463 |
| 0.708484 | 0.704777 | 0.700272 | 0.695848 | 0.712543 | 0.710464 | 0.700757 | 0.701579 | 0.70396 |
| 0.759852 | 0.755164 | 0.754514 | 0.755794 | 0.762423 | 0.75922 | 0.764917 | 0.757779 | 0.757314 |
| 0.761716 | 0.756466 | 0.757553 | 0.751533 | 0.767236 | 0.765873 | 0.767951 | 0.762616 | 0.761972 |
| 0.760818 | 0.76339 | 0.759426 | 0.757932 | 0.767413 | 0.768466 | 0.769867 | 0.765667 | 0.763969 |
| 0.763013 | 0.764076 | 0.761093 | 0.761765 | 0.767551 | 0.769599 | 0.76563 | 0.762677 | 0.762128 |
| 0.760978 | 0.753012 | 0.75514 | 0.752363 | 0.766248 | 0.7639 | 0.75845 | 0.756797 | 0.760496 |
| 0.759755 | 0.760702 | 0.757953 | 0.760041 | 0.770501 | 0.76968 | 0.767523 | 0.763149 | 0.76782 |
| 0.763966 | 0.76066 | 0.760172 | 0.758074 | 0.769742 | 0.767658 | 0.767302 | 0.765089 | 0.767931 |
| 0.759316 | 0.762583 | 0.757284 | 0.756251 | 0.769622 | 0.762599 | 0.764981 | 0.759317 | 0.7632 |
| 0.76225 | 0.753374 | 0.754275 | 0.750315 | 0.754858 | 0.760311 | 0.759196 | 0.755278 | 0.754846 |
| 0.760166 | 0.756984 | 0.749733 | 0.749338 | 0.761641 | 0.756389 | 0.757639 | 0.754251 | 0.754526 |
| 0.757077 | 0.755434 | 0.752697 | 0.755018 | 0.761119 | 0.758205 | 0.765234 | 0.75787 | 0.760926 |
| 0.760651 | 0.761188 | 0.756793 | 0.755068 | 0.768147 | 0.763182 | 0.766595 | 0.760967 | 0.765782 |

| | | | | | | | | |
|---|---|---|---|---|---|---|---|---|
| 0.766711 | 0.765317 | 0.759358 | 0.761437 | 0.766834 | 0.766025 | 0.772375 | 0.766816 | 0.766558 |
| 0.763459 | 0.762428 | 0.748588 | 0.754232 | 0.763084 | 0.760365 | 0.762213 | 0.760692 | 0.756422 |
| 0.765027 | 0.757537 | 0.755869 | 0.755398 | 0.762545 | 0.766401 | 0.760955 | 0.761029 | 0.76678 |
| 0.763557 | 0.75819 | 0.74836 | 0.7532 | 0.762217 | 0.761248 | 0.760825 | 0.756258 | 0.758096 |
| 0.760084 | 0.757683 | 0.749725 | 0.755233 | 0.757454 | 0.761268 | 0.760766 | 0.757155 | 0.758667 |
| 0.759662 | 0.75593 | 0.749615 | 0.752161 | 0.761025 | 0.764301 | 0.762486 | 0.759032 | 0.761561 |
| 0.765066 | 0.759972 | 0.755644 | 0.756622 | 0.764021 | 0.764134 | 0.763018 | 0.756832 | 0.763238 |
| 0.764587 | 0.760623 | 0.754824 | 0.755131 | 0.762458 | 0.76576 | 0.765707 | 0.762851 | 0.763255 |
| 0.761638 | 0.757811 | 0.752654 | 0.752401 | 0.757396 | 0.762995 | 0.758227 | 0.755868 | 0.757243 |
| 0.759714 | 0.757413 | 0.752698 | 0.752909 | 0.76078 | 0.762565 | 0.758074 | 0.755302 | 0.759402 |
| 0.7612 | 0.755034 | 0.750421 | 0.754517 | 0.761683 | 0.765383 | 0.763003 | 0.761622 | 0.760083 |
| 0.760861 | 0.758609 | 0.751499 | 0.751999 | 0.760423 | 0.765237 | 0.763502 | 0.761597 | 0.765124 |
| 0.753419 | 0.749161 | 0.745266 | 0.747174 | 0.752248 | 0.759901 | 0.755813 | 0.752379 | 0.755682 |
| 0.75377 | 0.752362 | 0.742985 | 0.749812 | 0.751575 | 0.758664 | 0.759044 | 0.752678 | 0.753938 |
| 0.761746 | 0.762557 | 0.754872 | 0.751656 | 0.763745 | 0.761777 | 0.757712 | 0.762588 | 0.760355 |
| 0.772634 | 0.768137 | 0.766047 | 0.763539 | 0.76665 | 0.766867 | 0.763839 | 0.763404 | 0.768071 |
| 0.763811 | 0.763639 | 0.761626 | 0.758638 | 0.768054 | 0.76371 | 0.761107 | 0.759993 | 0.76182 |
| 0.767579 | 0.767622 | 0.761297 | 0.75914 | 0.768759 | 0.7682 | 0.765126 | 0.761552 | 0.764658 |
| 0.76496 | 0.765808 | 0.75849 | 0.755882 | 0.769192 | 0.762115 | 0.761859 | 0.761302 | 0.760807 |
| 0.765272 | 0.76279 | 0.758947 | 0.758117 | 0.763797 | 0.765115 | 0.766385 | 0.761344 | 0.769598 |
| 0.765463 | 0.761537 | 0.755644 | 0.754729 | 0.764116 | 0.760283 | 0.766899 | 0.760418 | 0.764958 |
| 0.765221 | 0.760857 | 0.755546 | 0.755469 | 0.760273 | 0.764128 | 0.759321 | 0.761553 | 0.758791 |
| 0.770614 | 0.769339 | 0.766952 | 0.761428 | 0.765893 | 0.76488 | 0.765087 | 0.763598 | 0.767142 |
| 0.759615 | 0.753868 | 0.747105 | 0.754248 | 0.758244 | 0.760441 | 0.759537 | 0.756147 | 0.76045 |
| 0.770778 | 0.768116 | 0.762229 | 0.757806 | 0.765001 | 0.764737 | 0.76657 | 0.760598 | 0.76031 |
| 0.703413 | 0.701157 | 0.69525 | 0.695838 | 0.707692 | 0.70183 | 0.699792 | 0.70231 | 0.698919 |
| 0.763817 | 0.763238 | 0.756788 | 0.755085 | 0.768331 | 0.766697 | 0.766896 | 0.761154 | 0.764461 |
| 0.764428 | 0.760407 | 0.754726 | 0.760791 | 0.768974 | 0.767539 | 0.765555 | 0.759027 | 0.763141 |
| 0.770139 | 0.770446 | 0.760725 | 0.760069 | 0.767624 | 0.765187 | 0.769383 | 0.766173 | 0.769754 |
| 0.772922 | 0.76389 | 0.758376 | 0.756775 | 0.774273 | 0.768281 | 0.765881 | 0.762982 | 0.762465 |
| 0.761161 | 0.755702 | 0.750583 | 0.749622 | 0.76387 | 0.763752 | 0.766282 | 0.76047 | 0.760458 |
| 0.767809 | 0.759839 | 0.750834 | 0.756087 | 0.765671 | 0.763229 | 0.763043 | 0.761075 | 0.76117 |
| 0.767772 | 0.764797 | 0.756983 | 0.759822 | 0.763948 | 0.764657 | 0.764074 | 0.766133 | 0.762795 |
| 0.767304 | 0.765613 | 0.754374 | 0.759355 | 0.766741 | 0.766031 | 0.762453 | 0.762451 | 0.767037 |
| 0.756882 | 0.75397 | 0.74769 | 0.745612 | 0.760089 | 0.76 | 0.754995 | 0.753338 | 0.756897 |
| 0.762863 | 0.755862 | 0.75174 | 0.748933 | 0.767309 | 0.762136 | 0.759522 | 0.753186 | 0.754167 |
| 0.769752 | 0.768042 | 0.763147 | 0.759459 | 0.772727 | 0.767779 | 0.765098 | 0.763169 | 0.765696 |
| 0.769415 | 0.76554 | 0.75659 | 0.757821 | 0.77224 | 0.77097 | 0.770247 | 0.7643 | 0.767563 |
| 0.768104 | 0.762222 | 0.757474 | 0.75762 | 0.766691 | 0.765855 | 0.764408 | 0.761881 | 0.771048 |
| 0.76808 | 0.765142 | 0.75634 | 0.755469 | 0.765092 | 0.765008 | 0.766814 | 0.766658 | 0.765693 |
| 0.767366 | 0.761452 | 0.752856 | 0.753001 | 0.762853 | 0.760498 | 0.757961 | 0.761156 | 0.759624 |
| 0.767107 | 0.766615 | 0.760064 | 0.763539 | 0.76918 | 0.769976 | 0.76675 | 0.763138 | 0.769083 |
| 0.763082 | 0.761115 | 0.75332 | 0.753889 | 0.763178 | 0.765861 | 0.762551 | 0.756706 | 0.760347 |
| 0.758299 | 0.756015 | 0.750989 | 0.753964 | 0.764754 | 0.760903 | 0.757965 | 0.751305 | 0.756983 |
| 0.770688 | 0.765011 | 0.761627 | 0.762196 | 0.771738 | 0.765818 | 0.76902 | 0.760803 | 0.765656 |
| 0.766771 | 0.761744 | 0.754225 | 0.753338 | 0.770324 | 0.76877 | 0.762459 | 0.759395 | 0.76375 |

| | | | | | | | | |
|---|---|---|---|---|---|---|---|---|
| 0.759627 | 0.760276 | 0.754497 | 0.754598 | 0.767304 | 0.763945 | 0.763566 | 0.75981 | 0.762227 |
| 0.764998 | 0.754863 | 0.757731 | 0.754142 | 0.766342 | 0.769883 | 0.766441 | 0.761885 | 0.766334 |
| 0.758995 | 0.754661 | 0.756159 | 0.7563 | 0.76212 | 0.765625 | 0.757061 | 0.754969 | 0.762962 |
| 0.718952 | 0.714389 | 0.706884 | 0.710061 | 0.731171 | 0.733046 | 0.728035 | 0.722502 | 0.731059 |
| 0.759006 | 0.760471 | 0.749877 | 0.750382 | 0.760052 | 0.763188 | 0.760449 | 0.753504 | 0.760664 |
| 0.759228 | 0.756141 | 0.752575 | 0.750812 | 0.763585 | 0.761582 | 0.758571 | 0.752395 | 0.761126 |
| 0.76425 | 0.759133 | 0.754134 | 0.754498 | 0.761364 | 0.763914 | 0.768216 | 0.763089 | 0.765531 |
| 0.763992 | 0.762262 | 0.753903 | 0.758038 | 0.767181 | 0.763719 | 0.762659 | 0.758745 | 0.761653 |
| 0.769971 | 0.764542 | 0.759983 | 0.760726 | 0.769906 | 0.769212 | 0.763308 | 0.763869 | 0.767482 |
| 0.765789 | 0.758975 | 0.755911 | 0.752088 | 0.767251 | 0.766634 | 0.763609 | 0.757191 | 0.762962 |
| 0.765337 | 0.759993 | 0.755845 | 0.755366 | 0.766294 | 0.764186 | 0.763144 | 0.758723 | 0.762417 |
| 0.764629 | 0.756567 | 0.751405 | 0.754344 | 0.764432 | 0.765057 | 0.764906 | 0.761034 | 0.763402 |
| 0.767077 | 0.760677 | 0.756799 | 0.753025 | 0.767675 | 0.76496 | 0.761061 | 0.755996 | 0.765753 |
| 0.762751 | 0.764751 | 0.753326 | 0.75764 | 0.766455 | 0.76947 | 0.762942 | 0.754642 | 0.761064 |
| 0.763781 | 0.763438 | 0.761802 | 0.760633 | 0.771961 | 0.771806 | 0.763488 | 0.761306 | 0.764027 |
| 0.760284 | 0.75641 | 0.753545 | 0.758671 | 0.761707 | 0.763068 | 0.76163 | 0.758041 | 0.755461 |
| 0.761874 | 0.764331 | 0.757715 | 0.760632 | 0.771061 | 0.769737 | 0.766863 | 0.75891 | 0.761407 |
| 0.762999 | 0.761579 | 0.755078 | 0.756642 | 0.766859 | 0.771559 | 0.767359 | 0.76162 | 0.767398 |
| 0.765478 | 0.759534 | 0.753702 | 0.755945 | 0.762767 | 0.763819 | 0.760335 | 0.750795 | 0.758956 |
| 0.759512 | 0.762816 | 0.754528 | 0.757304 | 0.764931 | 0.764078 | 0.760455 | 0.759964 | 0.760424 |
| 0.769275 | 0.764159 | 0.760555 | 0.760907 | 0.766726 | 0.767759 | 0.767323 | 0.764401 | 0.765457 |
| 0.764394 | 0.763616 | 0.757032 | 0.76252 | 0.767385 | 0.76852 | 0.766483 | 0.761515 | 0.76486 |
| 0.687283 | 0.67761 | 0.673353 | 0.677395 | 0.682993 | 0.687717 | 0.678571 | 0.681649 | 0.68008 |
| 0.662233 | 0.654874 | 0.647594 | 0.652059 | 0.658554 | 0.662857 | 0.654342 | 0.65153 | 0.6596 |
| 0.670114 | 0.664474 | 0.670437 | 0.666519 | 0.678794 | 0.665314 | 0.664262 | 0.660856 | 0.662945 |
| 0.655442 | 0.654875 | 0.649951 | 0.653048 | 0.660075 | 0.653138 | 0.652639 | 0.649802 | 0.649134 |
| 0.727777 | 0.727687 | 0.717752 | 0.717507 | 0.725394 | 0.731247 | 0.736195 | 0.721795 | 0.725191 |
| 0.719232 | 0.722868 | 0.713075 | 0.716287 | 0.718066 | 0.720855 | 0.725494 | 0.719522 | 0.717387 |
| 0.727534 | 0.721121 | 0.716121 | 0.713557 | 0.724853 | 0.729507 | 0.727686 | 0.717297 | 0.728721 |
| 0.725713 | 0.723898 | 0.719841 | 0.731509 | 0.733597 | 0.730524 | 0.736509 | 0.728109 | 0.736842 |
| 0.702727 | 0.713888 | 0.708534 | 0.708012 | 0.708832 | 0.72031 | 0.714121 | 0.70607 | 0.709008 |
| 0.699663 | 0.699829 | 0.696789 | 0.705525 | 0.713762 | 0.715896 | 0.709004 | 0.696388 | 0.706347 |
| 0.705019 | 0.703979 | 0.701387 | 0.707288 | 0.703487 | 0.725082 | 0.724148 | 0.705447 | 0.705015 |
| 0.66096 | 0.653846 | 0.647597 | 0.65245 | 0.662405 | 0.655822 | 0.660724 | 0.651507 | 0.653481 |
| 0.763232 | 0.756231 | 0.750869 | 0.753008 | 0.763765 | 0.761349 | 0.755028 | 0.748948 | 0.753472 |
| 0.737661 | 0.736352 | 0.723064 | 0.729865 | 0.734218 | 0.74177 | 0.741436 | 0.73345 | 0.734566 |
| 0.728166 | 0.732619 | 0.720071 | 0.728821 | 0.732984 | 0.738005 | 0.733006 | 0.726654 | 0.729369 |
| 0.718067 | 0.721154 | 0.707976 | 0.709471 | 0.713104 | 0.726651 | 0.718523 | 0.711517 | 0.716809 |
| 0.736114 | 0.726887 | 0.725228 | 0.731315 | 0.738893 | 0.737999 | 0.739862 | 0.727124 | 0.73481 |
| 0.726532 | 0.727273 | 0.721426 | 0.718363 | 0.727701 | 0.731368 | 0.732284 | 0.714213 | 0.722364 |
| 0.72603 | 0.718464 | 0.714172 | 0.720495 | 0.72398 | 0.727591 | 0.728264 | 0.715083 | 0.723581 |
| 0.72313 | 0.724346 | 0.716418 | 0.713698 | 0.724047 | 0.73004 | 0.728838 | 0.721586 | 0.719187 |
| 0.715746 | 0.723474 | 0.712471 | 0.725667 | 0.726765 | 0.727584 | 0.73161 | 0.719486 | 0.724493 |
| 0.719253 | 0.719305 | 0.706185 | 0.7204 | 0.717849 | 0.72904 | 0.723211 | 0.720939 | 0.71623 |
| 0.722265 | 0.718361 | 0.70976 | 0.72403 | 0.719294 | 0.722591 | 0.722107 | 0.713623 | 0.72175 |
| 0.710692 | 0.703125 | 0.691196 | 0.704447 | 0.703486 | 0.716123 | 0.716617 | 0.700925 | 0.704463 |

| | | | | | | | | |
|---|---|---|---|---|---|---|---|---|
| 0.754199 | 0.743077 | 0.736154 | 0.741853 | 0.751381 | 0.752277 | 0.745055 | 0.742817 | 0.744101 |
| 0.727681 | 0.720176 | 0.71207 | 0.710736 | 0.729479 | 0.730748 | 0.727916 | 0.725429 | 0.724463 |
| 0.727749 | 0.723133 | 0.721776 | 0.720519 | 0.72257 | 0.728103 | 0.730485 | 0.718773 | 0.71985 |
| 0.724117 | 0.722171 | 0.714858 | 0.718132 | 0.727134 | 0.731752 | 0.730745 | 0.72194 | 0.723557 |
| 0.722718 | 0.730491 | 0.709902 | 0.722384 | 0.726152 | 0.733823 | 0.727851 | 0.718858 | 0.71847 |
| 0.717154 | 0.718977 | 0.705664 | 0.710725 | 0.708354 | 0.727295 | 0.726861 | 0.716328 | 0.713117 |
| 0.713107 | 0.704074 | 0.704236 | 0.710278 | 0.709515 | 0.729406 | 0.718591 | 0.70804 | 0.713664 |
| 0.709024 | 0.705615 | 0.698225 | 0.699517 | 0.707774 | 0.725634 | 0.723217 | 0.707096 | 0.711917 |
| 0.707017 | 0.712034 | 0.69797 | 0.708417 | 0.71581 | 0.716923 | 0.715909 | 0.707145 | 0.719457 |
| 0.751658 | 0.753831 | 0.744034 | 0.750354 | 0.75175 | 0.76022 | 0.756453 | 0.749336 | 0.752448 |
| 0.750503 | 0.747768 | 0.738821 | 0.742366 | 0.746704 | 0.751257 | 0.753475 | 0.743178 | 0.751665 |
| 0.742212 | 0.739057 | 0.728855 | 0.735576 | 0.741791 | 0.746007 | 0.742458 | 0.734642 | 0.743089 |
| 0.737128 | 0.733989 | 0.723633 | 0.728667 | 0.734328 | 0.740126 | 0.73615 | 0.731425 | 0.73343 |
| 0.739703 | 0.740129 | 0.727929 | 0.730726 | 0.7395 | 0.749267 | 0.7423 | 0.740481 | 0.743489 |
| 0.739771 | 0.734901 | 0.728105 | 0.731401 | 0.739689 | 0.737787 | 0.739527 | 0.727778 | 0.737808 |

| Wild_plot_ | Wild_plot_ | Wild_plot_ | Wild_plot_ | Wild_plot_ | Wild_plot_ | Wild_plot_ | Wild_plot_ | Wild_plot_ |
|---|---|---|---|---|---|---|---|---|
| 0.705676 | 0.699782 | 0.703832 | 0.707174 | 0.70736 | 0.705523 | 0.701726 | 0.702709 | 0.700476 |
| 0.713244 | 0.705635 | 0.709162 | 0.715158 | 0.712646 | 0.710713 | 0.706619 | 0.708243 | 0.708554 |
| 0.702274 | 0.694957 | 0.703354 | 0.707235 | 0.703595 | 0.700809 | 0.700375 | 0.697479 | 0.698776 |
| 0.710098 | 0.707055 | 0.707467 | 0.714511 | 0.713232 | 0.712013 | 0.708759 | 0.709755 | 0.706994 |
| 0.705093 | 0.700395 | 0.704863 | 0.706003 | 0.710116 | 0.70711 | 0.702845 | 0.707272 | 0.700742 |
| 0.715866 | 0.713131 | 0.714713 | 0.718052 | 0.720653 | 0.715526 | 0.712333 | 0.716668 | 0.711963 |
| 0.76319 | 0.756705 | 0.761476 | 0.768442 | 0.764908 | 0.764153 | 0.763794 | 0.757034 | 0.755336 |
| 0.760216 | 0.757487 | 0.756122 | 0.763291 | 0.760964 | 0.762256 | 0.76161 | 0.752791 | 0.755131 |
| 0.706758 | 0.704566 | 0.70706 | 0.709626 | 0.71205 | 0.710107 | 0.707892 | 0.706614 | 0.704082 |
| 0.704554 | 0.703584 | 0.701951 | 0.706753 | 0.708933 | 0.706374 | 0.701876 | 0.704279 | 0.699927 |
| 0.706715 | 0.702953 | 0.704921 | 0.709808 | 0.711565 | 0.703994 | 0.706751 | 0.704269 | 0.704558 |
| 0.699905 | 0.6984 | 0.699339 | 0.703296 | 0.701427 | 0.702522 | 0.696883 | 0.700121 | 0.695954 |
| 0.708074 | 0.70743 | 0.708019 | 0.710626 | 0.714229 | 0.710367 | 0.707708 | 0.709029 | 0.707335 |
| 0.722811 | 0.72033 | 0.728217 | 0.727962 | 0.725096 | 0.725325 | 0.733295 | 0.720689 | 0.719053 |
| 0.745195 | 0.752331 | 0.742346 | 0.743336 | 0.73955 | 0.745309 | 0.744141 | 0.740635 | 0.733529 |
| 0.710621 | 0.711062 | 0.713242 | 0.714716 | 0.715242 | 0.715778 | 0.709597 | 0.711829 | 0.706725 |
| 0.710141 | 0.707275 | 0.711231 | 0.714217 | 0.714688 | 0.714111 | 0.706196 | 0.711263 | 0.710986 |
| 0.758966 | 0.762592 | 0.758822 | 0.75974 | 0.750281 | 0.764537 | 0.766283 | 0.74817 | 0.748835 |
| 0.70216 | 0.70087 | 0.699716 | 0.70599 | 0.705099 | 0.702138 | 0.702614 | 0.702676 | 0.696501 |
| 0.706337 | 0.703803 | 0.707691 | 0.711363 | 0.710205 | 0.706499 | 0.705528 | 0.704077 | 0.705615 |
| 0.704994 | 0.698789 | 0.703455 | 0.705248 | 0.703962 | 0.70393 | 0.703384 | 0.700972 | 0.701179 |
| 0.705502 | 0.702821 | 0.707904 | 0.709438 | 0.708111 | 0.709706 | 0.703766 | 0.702707 | 0.704286 |
| 0.713395 | 0.709999 | 0.710286 | 0.715891 | 0.715004 | 0.713974 | 0.708935 | 0.711632 | 0.710294 |
| 0.710507 | 0.708912 | 0.707248 | 0.712626 | 0.713441 | 0.711134 | 0.706819 | 0.708676 | 0.703887 |
| 0.712691 | 0.707706 | 0.70625 | 0.710838 | 0.712201 | 0.710752 | 0.704547 | 0.708982 | 0.70753 |
| 0.707024 | 0.703552 | 0.705221 | 0.712948 | 0.709123 | 0.709889 | 0.706215 | 0.707565 | 0.702816 |
| 0.701233 | 0.70097 | 0.700663 | 0.705265 | 0.703021 | 0.705541 | 0.701928 | 0.699839 | 0.696367 |
| 0.70736 | 0.704789 | 0.70513 | 0.712934 | 0.711817 | 0.709714 | 0.706028 | 0.708102 | 0.703313 |
| 0.709544 | 0.707971 | 0.710848 | 0.714218 | 0.714823 | 0.710673 | 0.709732 | 0.710237 | 0.706092 |
| 0.706551 | 0.704605 | 0.706016 | 0.707795 | 0.711287 | 0.705407 | 0.702158 | 0.707679 | 0.701186 |
| 0.707159 | 0.703008 | 0.705023 | 0.709163 | 0.711703 | 0.707242 | 0.704177 | 0.708495 | 0.704782 |
| 0.705667 | 0.698931 | 0.703552 | 0.707085 | 0.706237 | 0.706585 | 0.701936 | 0.702599 | 0.70497 |
| 0.706186 | 0.699236 | 0.701076 | 0.708865 | 0.709082 | 0.700008 | 0.704582 | 0.70317 | 0.696527 |
| 0.709623 | 0.705322 | 0.707421 | 0.713124 | 0.710635 | 0.711607 | 0.708403 | 0.70893 | 0.706601 |
| 0.702353 | 0.700517 | 0.701441 | 0.70632 | 0.702853 | 0.703355 | 0.701057 | 0.699019 | 0.69841 |
| 0.706382 | 0.703966 | 0.706313 | 0.709075 | 0.710375 | 0.706353 | 0.704926 | 0.70915 | 0.697782 |
| 0.713172 | 0.708286 | 0.709166 | 0.716242 | 0.71338 | 0.710396 | 0.711767 | 0.710133 | 0.705916 |
| 0.708858 | 0.704095 | 0.708719 | 0.712333 | 0.713698 | 0.707991 | 0.706524 | 0.70914 | 0.705416 |
| 0.713989 | 0.706977 | 0.711428 | 0.713913 | 0.716876 | 0.712341 | 0.710727 | 0.71207 | 0.707608 |

| | | | | | | | | |
|---|---|---|---|---|---|---|---|---|
| 0.758903 | 0.764322 | 0.762022 | 0.761691 | 0.758002 | 0.766103 | 0.771159 | 0.754748 | 0.755594 |
| 0.700389 | 0.694806 | 0.699612 | 0.70296 | 0.702332 | 0.699254 | 0.697366 | 0.696933 | 0.696339 |
| 0.700621 | 0.699733 | 0.699968 | 0.705158 | 0.706463 | 0.701883 | 0.698579 | 0.701596 | 0.698682 |
| 0.70913 | 0.705949 | 0.707784 | 0.713319 | 0.714263 | 0.709605 | 0.708107 | 0.709414 | 0.706617 |
| 0.707708 | 0.702035 | 0.7045 | 0.710601 | 0.709962 | 0.706498 | 0.700956 | 0.704417 | 0.699847 |
| 0.710858 | 0.708286 | 0.707536 | 0.711159 | 0.711897 | 0.71213 | 0.707967 | 0.70935 | 0.705442 |
| 0.701672 | 0.70191 | 0.701239 | 0.707 | 0.705934 | 0.703386 | 0.701577 | 0.702874 | 0.698792 |
| 0.707581 | 0.703948 | 0.709598 | 0.712818 | 0.712769 | 0.707226 | 0.707746 | 0.707085 | 0.703187 |
| 0.705379 | 0.703354 | 0.709788 | 0.709978 | 0.7116 | 0.708182 | 0.706352 | 0.705348 | 0.705844 |
| 0.707106 | 0.703808 | 0.704778 | 0.708589 | 0.709708 | 0.704862 | 0.706056 | 0.704206 | 0.706244 |
| 0.697626 | 0.697493 | 0.698346 | 0.701106 | 0.702571 | 0.702417 | 0.696394 | 0.699055 | 0.697459 |
| 0.697804 | 0.697639 | 0.699814 | 0.703791 | 0.706643 | 0.704226 | 0.697706 | 0.701041 | 0.699635 |
| 0.705821 | 0.707221 | 0.701413 | 0.708025 | 0.707728 | 0.706939 | 0.702328 | 0.707859 | 0.69978 |
| 0.699753 | 0.698871 | 0.699627 | 0.701541 | 0.705943 | 0.705773 | 0.698742 | 0.701869 | 0.701049 |
| 0.704906 | 0.703725 | 0.704737 | 0.708526 | 0.707401 | 0.704557 | 0.70121 | 0.70214 | 0.703002 |
| 0.703312 | 0.699082 | 0.704601 | 0.706906 | 0.704191 | 0.705459 | 0.700957 | 0.702149 | 0.697486 |
| 0.701636 | 0.698281 | 0.69746 | 0.702752 | 0.70479 | 0.702813 | 0.696508 | 0.700691 | 0.697401 |
| 0.706466 | 0.706712 | 0.708816 | 0.712885 | 0.71006 | 0.709831 | 0.703556 | 0.704567 | 0.705582 |
| 0.710865 | 0.710173 | 0.70935 | 0.717315 | 0.713258 | 0.714598 | 0.708919 | 0.709651 | 0.706516 |
| 0.70152 | 0.702317 | 0.703079 | 0.708018 | 0.708153 | 0.704949 | 0.700684 | 0.70304 | 0.703392 |
| 0.700631 | 0.699017 | 0.699895 | 0.702414 | 0.700686 | 0.701623 | 0.698941 | 0.700494 | 0.697327 |
| 0.702043 | 0.697999 | 0.705806 | 0.705091 | 0.707385 | 0.70083 | 0.700875 | 0.700654 | 0.697999 |
| 0.704346 | 0.701675 | 0.70237 | 0.708301 | 0.705985 | 0.706429 | 0.704187 | 0.700048 | 0.703003 |
| 0.715032 | 0.710597 | 0.709394 | 0.716684 | 0.717536 | 0.715514 | 0.713572 | 0.712298 | 0.712595 |
| 0.720772 | 0.716365 | 0.720625 | 0.723257 | 0.722453 | 0.72394 | 0.717965 | 0.720058 | 0.716909 |
| 0.762165 | 0.758656 | 0.758543 | 0.763803 | 0.762313 | 0.762407 | 0.762523 | 0.755163 | 0.755755 |
| 0.761621 | 0.755085 | 0.764899 | 0.7697 | 0.766299 | 0.761905 | 0.762531 | 0.757182 | 0.759445 |
| 0.76699 | 0.756807 | 0.76475 | 0.773962 | 0.766284 | 0.765423 | 0.768358 | 0.756883 | 0.762461 |
| 0.701393 | 0.699386 | 0.702362 | 0.707481 | 0.708094 | 0.707264 | 0.70208 | 0.704328 | 0.698237 |
| 0.701869 | 0.69811 | 0.704111 | 0.706371 | 0.708227 | 0.70281 | 0.702406 | 0.700356 | 0.697564 |
| 0.710613 | 0.711491 | 0.709835 | 0.71212 | 0.715833 | 0.711615 | 0.708961 | 0.711506 | 0.707081 |
| 0.712379 | 0.705068 | 0.715087 | 0.716388 | 0.716783 | 0.716004 | 0.711287 | 0.707436 | 0.709709 |
| 0.706646 | 0.703 | 0.704281 | 0.708985 | 0.709047 | 0.708521 | 0.703353 | 0.703758 | 0.701071 |
| 0.704697 | 0.705435 | 0.699642 | 0.707662 | 0.707758 | 0.706011 | 0.700886 | 0.702698 | 0.698918 |
| 0.701275 | 0.700607 | 0.702998 | 0.705887 | 0.705236 | 0.703456 | 0.700374 | 0.700367 | 0.69697 |
| 0.704077 | 0.703764 | 0.704839 | 0.710952 | 0.70844 | 0.706996 | 0.702218 | 0.704016 | 0.702221 |
| 0.704655 | 0.702449 | 0.701918 | 0.708147 | 0.70859 | 0.70783 | 0.70419 | 0.704338 | 0.702104 |
| 0.719119 | 0.715042 | 0.717377 | 0.723184 | 0.7206 | 0.720925 | 0.716756 | 0.714598 | 0.713215 |
| 0.697341 | 0.695214 | 0.696484 | 0.701813 | 0.701448 | 0.697688 | 0.694027 | 0.694927 | 0.692804 |
| 0.706138 | 0.700056 | 0.703834 | 0.70785 | 0.708228 | 0.704469 | 0.700673 | 0.705562 | 0.701099 |
| 0.703768 | 0.697885 | 0.702757 | 0.707658 | 0.705794 | 0.704555 | 0.701728 | 0.704442 | 0.699133 |
| 0.721819 | 0.718997 | 0.720611 | 0.726214 | 0.725867 | 0.723989 | 0.716876 | 0.719721 | 0.715619 |
| 0.70646 | 0.704212 | 0.701011 | 0.709759 | 0.707961 | 0.705119 | 0.70092 | 0.705003 | 0.700762 |
| 0.703821 | 0.699695 | 0.70221 | 0.703762 | 0.708567 | 0.703455 | 0.70105 | 0.703784 | 0.698339 |
| 0.710203 | 0.703803 | 0.705202 | 0.707685 | 0.71028 | 0.711088 | 0.70422 | 0.704273 | 0.703432 |
| 0.707511 | 0.702246 | 0.704088 | 0.707914 | 0.706373 | 0.705591 | 0.701162 | 0.704011 | 0.700948 |

| | | | | | | | | |
|---|---|---|---|---|---|---|---|---|
| 0.701826 | 0.6972 | 0.701668 | 0.705822 | 0.706754 | 0.702249 | 0.700745 | 0.700435 | 0.699109 |
| 0.710124 | 0.704946 | 0.706794 | 0.711852 | 0.712159 | 0.710023 | 0.706399 | 0.708296 | 0.705498 |
| 0.704042 | 0.701143 | 0.703121 | 0.70467 | 0.708478 | 0.703491 | 0.701079 | 0.70334 | 0.707193 |
| 0.712012 | 0.709322 | 0.710809 | 0.717625 | 0.712693 | 0.712591 | 0.709882 | 0.708255 | 0.706703 |
| 0.710656 | 0.707191 | 0.709831 | 0.712653 | 0.715729 | 0.710799 | 0.70886 | 0.707443 | 0.709962 |
| 0.705433 | 0.698844 | 0.705 | 0.708782 | 0.71063 | 0.706608 | 0.704137 | 0.702657 | 0.702877 |
| 0.708535 | 0.705173 | 0.706213 | 0.712087 | 0.710114 | 0.709133 | 0.706177 | 0.705802 | 0.706319 |
| 0.707768 | 0.705065 | 0.707154 | 0.708641 | 0.710006 | 0.70906 | 0.704796 | 0.70833 | 0.703927 |
| 0.70913 | 0.70333 | 0.702947 | 0.7073 | 0.709468 | 0.706582 | 0.704375 | 0.705143 | 0.701764 |
| 0.703707 | 0.698934 | 0.700709 | 0.70437 | 0.704729 | 0.705764 | 0.697083 | 0.70503 | 0.702948 |
| 0.707836 | 0.70032 | 0.703603 | 0.709131 | 0.70862 | 0.707512 | 0.705191 | 0.70458 | 0.702818 |
| 0.70603 | 0.702685 | 0.703716 | 0.704088 | 0.707067 | 0.704033 | 0.702552 | 0.704188 | 0.70159 |
| 0.708782 | 0.707377 | 0.70831 | 0.709089 | 0.711941 | 0.711332 | 0.705617 | 0.706786 | 0.700204 |
| 0.701803 | 0.696791 | 0.700827 | 0.705001 | 0.706034 | 0.701166 | 0.697571 | 0.700065 | 0.702996 |
| 0.708153 | 0.706912 | 0.704588 | 0.708079 | 0.710031 | 0.708587 | 0.705664 | 0.708398 | 0.704605 |
| 0.704899 | 0.702781 | 0.701159 | 0.707331 | 0.709847 | 0.707266 | 0.700876 | 0.705641 | 0.700807 |
| 0.710162 | 0.70787 | 0.707854 | 0.714388 | 0.711358 | 0.713216 | 0.70752 | 0.70853 | 0.707999 |
| 0.702858 | 0.703428 | 0.70465 | 0.705873 | 0.706894 | 0.705508 | 0.705354 | 0.702701 | 0.698934 |
| 0.705967 | 0.702727 | 0.701554 | 0.705934 | 0.704549 | 0.708306 | 0.701807 | 0.704979 | 0.702615 |
| 0.706925 | 0.704444 | 0.704031 | 0.710158 | 0.707793 | 0.707475 | 0.706265 | 0.705583 | 0.704455 |
| 0.70721 | 0.702259 | 0.703445 | 0.708655 | 0.708273 | 0.704248 | 0.701895 | 0.706383 | 0.700008 |
| 0.703521 | 0.700048 | 0.697787 | 0.704495 | 0.702308 | 0.70295 | 0.699537 | 0.698949 | 0.700381 |
| 0.707031 | 0.703326 | 0.706281 | 0.70969 | 0.711059 | 0.706793 | 0.70584 | 0.706739 | 0.705252 |
| 0.706262 | 0.701814 | 0.707206 | 0.70941 | 0.710434 | 0.706057 | 0.703716 | 0.703846 | 0.704503 |
| 0.708418 | 0.707067 | 0.705993 | 0.70815 | 0.708219 | 0.707078 | 0.706073 | 0.705301 | 0.703112 |
| 0.713207 | 0.709163 | 0.709173 | 0.714691 | 0.715294 | 0.709954 | 0.70822 | 0.709957 | 0.709133 |
| 0.708796 | 0.704353 | 0.703308 | 0.709966 | 0.70898 | 0.706807 | 0.70708 | 0.70436 | 0.700899 |
| 0.70901 | 0.705982 | 0.707909 | 0.712921 | 0.709491 | 0.709091 | 0.704591 | 0.703769 | 0.702598 |
| 0.711129 | 0.707258 | 0.708104 | 0.7114 | 0.713183 | 0.711396 | 0.708554 | 0.706398 | 0.707319 |
| 0.710146 | 0.705602 | 0.707518 | 0.709646 | 0.710624 | 0.708862 | 0.706624 | 0.704289 | 0.700049 |
| 0.708812 | 0.708085 | 0.705202 | 0.712484 | 0.712977 | 0.707796 | 0.707852 | 0.708476 | 0.704976 |
| 0.703108 | 0.702068 | 0.702908 | 0.70682 | 0.709178 | 0.705544 | 0.706279 | 0.705901 | 0.702859 |
| 0.70602 | 0.703022 | 0.704305 | 0.710189 | 0.708794 | 0.703942 | 0.704262 | 0.702072 | 0.702105 |
| 0.705065 | 0.702225 | 0.705887 | 0.7094 | 0.708834 | 0.706028 | 0.704383 | 0.705407 | 0.704257 |
| 0.705181 | 0.702166 | 0.703484 | 0.706485 | 0.708598 | 0.704066 | 0.70511 | 0.704319 | 0.699691 |
| 0.705058 | 0.702958 | 0.70536 | 0.709475 | 0.709338 | 0.712936 | 0.704472 | 0.703664 | 0.703039 |
| 0.703053 | 0.696087 | 0.7022 | 0.706179 | 0.703996 | 0.697439 | 0.703169 | 0.702177 | 0.697692 |
| 0.677364 | 0.673132 | 0.679437 | 0.685359 | 0.681612 | 0.67871 | 0.67469 | 0.677175 | 0.674409 |
| 0.681291 | 0.678334 | 0.684803 | 0.68706 | 0.688473 | 0.682243 | 0.680529 | 0.681711 | 0.684788 |
| 0.678568 | 0.679486 | 0.677251 | 0.679886 | 0.678593 | 0.677359 | 0.675062 | 0.67486 | 0.67158 |
| 0.676233 | 0.668386 | 0.675293 | 0.680835 | 0.679386 | 0.672887 | 0.671953 | 0.673412 | 0.672333 |
| 0.662338 | 0.661332 | 0.660206 | 0.66291 | 0.662738 | 0.66176 | 0.659517 | 0.655992 | 0.653629 |
| 0.666586 | 0.65903 | 0.664059 | 0.669545 | 0.668599 | 0.661224 | 0.662658 | 0.665604 | 0.664996 |
| 0.661878 | 0.659211 | 0.663189 | 0.667068 | 0.666343 | 0.659177 | 0.660407 | 0.658126 | 0.659988 |
| 0.666837 | 0.667446 | 0.66512 | 0.663215 | 0.668792 | 0.667094 | 0.661768 | 0.665506 | 0.659126 |
| 0.663267 | 0.656664 | 0.664524 | 0.668068 | 0.666252 | 0.667548 | 0.663041 | 0.66168 | 0.660505 |
| 0.670696 | 0.666416 | 0.668417 | 0.675819 | 0.6688 | 0.67153 | 0.664977 | 0.665168 | 0.665462 |

| | | | | | | | | |
|---|---|---|---|---|---|---|---|---|
| 0.668721 | 0.662889 | 0.666459 | 0.670711 | 0.665359 | 0.673589 | 0.66576 | 0.666086 | 0.667125 |
| 0.668447 | 0.667855 | 0.668825 | 0.674413 | 0.675683 | 0.673598 | 0.666317 | 0.666228 | 0.667811 |
| 0.66885 | 0.663151 | 0.664817 | 0.675938 | 0.676779 | 0.667788 | 0.666264 | 0.667208 | 0.670722 |
| 0.664595 | 0.651861 | 0.663765 | 0.663342 | 0.664596 | 0.659546 | 0.658248 | 0.659458 | 0.654313 |
| 0.664948 | 0.657816 | 0.661167 | 0.665216 | 0.664009 | 0.660636 | 0.660333 | 0.660984 | 0.651865 |
| 0.661138 | 0.656483 | 0.661362 | 0.66535 | 0.661158 | 0.658925 | 0.660305 | 0.654234 | 0.656518 |
| 0.665435 | 0.657538 | 0.667433 | 0.669397 | 0.664164 | 0.664816 | 0.661178 | 0.658662 | 0.655556 |
| 0.660459 | 0.652998 | 0.656409 | 0.661944 | 0.662558 | 0.660499 | 0.657973 | 0.663257 | 0.655378 |
| 0.658937 | 0.655126 | 0.665123 | 0.662615 | 0.663042 | 0.658353 | 0.658515 | 0.656829 | 0.663878 |
| 0.655769 | 0.655229 | 0.652397 | 0.657564 | 0.664918 | 0.661487 | 0.657005 | 0.66097 | 0.654352 |
| 0.651089 | 0.649355 | 0.653331 | 0.654491 | 0.650942 | 0.654958 | 0.650226 | 0.648267 | 0.652416 |
| 0.692428 | 0.691345 | 0.691508 | 0.694188 | 0.698652 | 0.694167 | 0.688842 | 0.692655 | 0.691241 |
| 0.700252 | 0.697514 | 0.70016 | 0.702523 | 0.706453 | 0.703594 | 0.700063 | 0.704427 | 0.699487 |
| 0.686174 | 0.676254 | 0.682289 | 0.689683 | 0.682369 | 0.683469 | 0.6792 | 0.679848 | 0.681259 |
| 0.711244 | 0.71091 | 0.710972 | 0.714802 | 0.718196 | 0.714075 | 0.71231 | 0.713712 | 0.707058 |
| 0.700612 | 0.700907 | 0.698883 | 0.70243 | 0.706402 | 0.701288 | 0.698824 | 0.702829 | 0.697827 |
| 0.708294 | 0.706922 | 0.705395 | 0.71108 | 0.712471 | 0.707577 | 0.706383 | 0.705972 | 0.708037 |
| 0.71687 | 0.711715 | 0.715411 | 0.719696 | 0.717693 | 0.715763 | 0.712227 | 0.715077 | 0.711721 |
| 0.690424 | 0.69216 | 0.693584 | 0.693984 | 0.696123 | 0.692137 | 0.692192 | 0.691244 | 0.690996 |
| 0.715842 | 0.713018 | 0.711803 | 0.716711 | 0.717074 | 0.717046 | 0.711161 | 0.713872 | 0.709587 |
| 0.689464 | 0.684796 | 0.68309 | 0.69277 | 0.687958 | 0.687568 | 0.683567 | 0.684473 | 0.683877 |
| 0.714217 | 0.71052 | 0.712294 | 0.717402 | 0.713886 | 0.710907 | 0.713686 | 0.710861 | 0.708323 |
| 0.71905 | 0.717548 | 0.717823 | 0.721402 | 0.722733 | 0.717874 | 0.712628 | 0.720463 | 0.71598 |
| 0.6852 | 0.674 | 0.6774 | 0.689559 | 0.681839 | 0.685142 | 0.681378 | 0.676744 | 0.677757 |
| 0.685015 | 0.683156 | 0.685525 | 0.689389 | 0.686101 | 0.684338 | 0.684784 | 0.681337 | 0.681594 |
| 0.687737 | 0.681975 | 0.687316 | 0.690829 | 0.688844 | 0.685627 | 0.684237 | 0.686325 | 0.685366 |
| 0.706382 | 0.703249 | 0.70175 | 0.706799 | 0.711165 | 0.7029 | 0.704432 | 0.704699 | 0.701902 |
| 0.714217 | 0.709458 | 0.712493 | 0.71803 | 0.717478 | 0.714988 | 0.710122 | 0.712395 | 0.708472 |
| 0.684935 | 0.683761 | 0.681962 | 0.686326 | 0.686608 | 0.68553 | 0.683331 | 0.682714 | 0.678312 |
| 0.725246 | 0.717976 | 0.719179 | 0.725841 | 0.726281 | 0.720984 | 0.717275 | 0.721963 | 0.719359 |
| 0.723778 | 0.716727 | 0.721766 | 0.726852 | 0.725047 | 0.722076 | 0.7213 | 0.718553 | 0.720348 |
| 0.690548 | 0.692171 | 0.694155 | 0.699154 | 0.696087 | 0.694496 | 0.694369 | 0.688287 | 0.68926 |
| 0.69446 | 0.690715 | 0.694164 | 0.698489 | 0.698517 | 0.693536 | 0.690735 | 0.689922 | 0.689976 |
| 0.721844 | 0.719201 | 0.718928 | 0.725942 | 0.72763 | 0.724793 | 0.720135 | 0.722569 | 0.719948 |
| 0.721083 | 0.714549 | 0.716298 | 0.725121 | 0.717836 | 0.719547 | 0.715608 | 0.715708 | 0.716469 |
| 0.720331 | 0.715276 | 0.718298 | 0.724243 | 0.724586 | 0.722326 | 0.717841 | 0.714471 | 0.717798 |
| 0.67576 | 0.672478 | 0.673702 | 0.676939 | 0.6787 | 0.673378 | 0.672622 | 0.673721 | 0.67191 |
| 0.718468 | 0.714702 | 0.716057 | 0.718224 | 0.720847 | 0.71811 | 0.713657 | 0.713539 | 0.712135 |
| 0.672389 | 0.670725 | 0.672512 | 0.679224 | 0.675184 | 0.67396 | 0.670932 | 0.673029 | 0.671149 |
| 0.664417 | 0.66489 | 0.662969 | 0.668424 | 0.66831 | 0.665799 | 0.663642 | 0.665668 | 0.664916 |
| 0.723491 | 0.721381 | 0.729055 | 0.727964 | 0.723587 | 0.7267 | 0.732228 | 0.717125 | 0.7192 |
| 0.666667 | 0.663918 | 0.666255 | 0.669984 | 0.670745 | 0.667024 | 0.664656 | 0.663216 | 0.666178 |
| 0.658821 | 0.655928 | 0.657774 | 0.66336 | 0.665962 | 0.659186 | 0.65754 | 0.658207 | 0.656092 |
| 0.658411 | 0.654893 | 0.655387 | 0.661743 | 0.659703 | 0.658652 | 0.656067 | 0.657085 | 0.652353 |
| 0.737729 | 0.740313 | 0.731681 | 0.733317 | 0.73406 | 0.734972 | 0.738905 | 0.730479 | 0.724287 |
| 0.659291 | 0.657436 | 0.657531 | 0.663055 | 0.664231 | 0.660743 | 0.657257 | 0.658173 | 0.656802 |
| 0.727794 | 0.730636 | 0.723225 | 0.725634 | 0.728621 | 0.727836 | 0.724587 | 0.723652 | 0.72113 |

| | | | | | | | | |
|---|---|---|---|---|---|---|---|---|
| 0.73961 | 0.735249 | 0.73891 | 0.737794 | 0.737028 | 0.73964 | 0.745586 | 0.735208 | 0.729974 |
| 0.734418 | 0.731046 | 0.730851 | 0.733615 | 0.727993 | 0.734667 | 0.737848 | 0.723063 | 0.720819 |
| 0.750389 | 0.737108 | 0.743175 | 0.749899 | 0.750471 | 0.746626 | 0.73933 | 0.742317 | 0.737134 |
| 0.649844 | 0.649781 | 0.647904 | 0.654517 | 0.655516 | 0.650983 | 0.648606 | 0.650105 | 0.647792 |
| 0.648821 | 0.646109 | 0.648962 | 0.652647 | 0.653009 | 0.648478 | 0.649518 | 0.648959 | 0.647226 |
| 0.737772 | 0.735826 | 0.736579 | 0.736691 | 0.73605 | 0.736297 | 0.740693 | 0.730126 | 0.727154 |
| 0.651114 | 0.648448 | 0.651216 | 0.654915 | 0.654367 | 0.652027 | 0.649843 | 0.651938 | 0.64826 |
| 0.732006 | 0.726574 | 0.728769 | 0.73196 | 0.737568 | 0.731903 | 0.72763 | 0.72996 | 0.725388 |
| 0.738259 | 0.749543 | 0.738976 | 0.73724 | 0.738577 | 0.743704 | 0.746435 | 0.736444 | 0.729956 |
| 0.733459 | 0.729087 | 0.731431 | 0.735089 | 0.736583 | 0.733825 | 0.729728 | 0.728633 | 0.72964 |
| 0.747643 | 0.744901 | 0.746944 | 0.744819 | 0.740138 | 0.745309 | 0.756017 | 0.739285 | 0.738399 |
| 0.727891 | 0.71831 | 0.704698 | 0.685535 | 0.701987 | 0.727273 | 0.697368 | 0.712329 | 0.713333 |
| 0.720512 | 0.712725 | 0.718068 | 0.730362 | 0.722334 | 0.723396 | 0.714856 | 0.710707 | 0.710378 |
| 0.718186 | 0.706235 | 0.708831 | 0.719178 | 0.720238 | 0.726211 | 0.711021 | 0.709243 | 0.703982 |
| 0.719538 | 0.714221 | 0.717041 | 0.728524 | 0.726414 | 0.724728 | 0.720723 | 0.717626 | 0.71749 |
| 0.726526 | 0.719744 | 0.722169 | 0.731883 | 0.720037 | 0.721331 | 0.726631 | 0.710055 | 0.716815 |
| 0.58569 | 0.587657 | 0.589041 | 0.590709 | 0.590753 | 0.586285 | 0.587398 | 0.587585 | 0.587107 |
| 0.732637 | 0.722391 | 0.729043 | 0.733593 | 0.730334 | 0.731193 | 0.728918 | 0.721227 | 0.724347 |
| 0.751091 | 0.755291 | 0.751377 | 0.747746 | 0.748299 | 0.74799 | 0.765417 | 0.750811 | 0.741774 |
| 0.753464 | 0.760238 | 0.750782 | 0.749052 | 0.748774 | 0.75735 | 0.758876 | 0.749642 | 0.744785 |
| 0.745607 | 0.755076 | 0.755563 | 0.752122 | 0.747209 | 0.753562 | 0.764524 | 0.746011 | 0.747221 |
| 0.593166 | 0.590895 | 0.593638 | 0.596084 | 0.595986 | 0.593963 | 0.592518 | 0.59013 | 0.588592 |
| 0.580757 | 0.584265 | 0.577546 | 0.582961 | 0.584512 | 0.583407 | 0.578152 | 0.579228 | 0.578577 |
| 0.801534 | 0.795839 | 0.817151 | 0.804427 | 0.808196 | 0.817867 | 0.820148 | 0.804809 | 0.815295 |
| 0.804983 | 0.79942 | 0.805484 | 0.794598 | 0.803144 | 0.806468 | 0.801671 | 0.808565 | 0.797921 |
| 0.833635 | 0.782279 | 0.827178 | 0.875141 | 0.812277 | 0.793118 | 0.830929 | 0.794457 | 0.839162 |
| 0.836761 | 0.798688 | 0.839653 | 0.877648 | 0.80422 | 0.799594 | 0.836437 | 0.800068 | 0.832278 |
| 0.842762 | 0.812975 | 0.864111 | 0.885857 | 0.801034 | 0.807239 | 0.847688 | 0.810193 | 0.815758 |
| 0.789695 | 0.780577 | 0.801481 | 0.797922 | 0.793833 | 0.804648 | 0.800494 | 0.785921 | 0.812587 |
| 0.805195 | 0.78955 | 0.801196 | 0.808416 | 0.806917 | 0.801753 | 0.808877 | 0.802504 | 0.790563 |
| 0.760754 | 0.758609 | 0.756997 | 0.761726 | 0.763282 | 0.763658 | 0.757367 | 0.758186 | 0.750283 |
| 0.767923 | 0.763278 | 0.76822 | 0.774702 | 0.777067 | 0.768219 | 0.763659 | 0.767956 | 0.764994 |
| 0.765968 | 0.759434 | 0.757795 | 0.769339 | 0.762402 | 0.760831 | 0.757863 | 0.757045 | 0.757596 |
| 0.769571 | 0.755436 | 0.760754 | 0.767938 | 0.762733 | 0.765668 | 0.75598 | 0.758641 | 0.76132 |
| 0.77024 | 0.763673 | 0.759191 | 0.766734 | 0.772383 | 0.764911 | 0.76001 | 0.764515 | 0.766091 |
| 0.764745 | 0.757113 | 0.760333 | 0.768807 | 0.764153 | 0.762705 | 0.751693 | 0.757256 | 0.755239 |
| 0.773195 | 0.766001 | 0.766113 | 0.772045 | 0.771166 | 0.768463 | 0.760486 | 0.766815 | 0.766008 |
| 0.770523 | 0.765541 | 0.766146 | 0.774186 | 0.773784 | 0.767005 | 0.763857 | 0.765361 | 0.763063 |
| 0.764167 | 0.757563 | 0.759667 | 0.765239 | 0.763133 | 0.761852 | 0.75631 | 0.757677 | 0.757911 |
| 0.769212 | 0.758767 | 0.760539 | 0.771412 | 0.768247 | 0.765217 | 0.760481 | 0.760862 | 0.757853 |
| 0.717627 | 0.712343 | 0.716326 | 0.720236 | 0.71893 | 0.723055 | 0.71311 | 0.715574 | 0.71235 |
| 0.726826 | 0.721446 | 0.722757 | 0.729806 | 0.723276 | 0.729309 | 0.721739 | 0.719916 | 0.71601 |
| 0.763997 | 0.762377 | 0.763103 | 0.766756 | 0.764836 | 0.763879 | 0.761615 | 0.758857 | 0.760412 |
| 0.781258 | 0.770218 | 0.78095 | 0.786636 | 0.783782 | 0.781014 | 0.78384 | 0.775344 | 0.78323 |
| 0.729025 | 0.718927 | 0.722635 | 0.730658 | 0.72146 | 0.718896 | 0.719876 | 0.725324 | 0.707671 |
| 0.769478 | 0.764218 | 0.766253 | 0.772861 | 0.774324 | 0.767042 | 0.763567 | 0.766964 | 0.760063 |
| 0.761312 | 0.757898 | 0.758486 | 0.762596 | 0.767095 | 0.759385 | 0.756104 | 0.760743 | 0.753759 |

| | | | | | | | | |
|---|---|---|---|---|---|---|---|---|
| 0.767922 | 0.764676 | 0.76024 | 0.76721 | 0.774547 | 0.766196 | 0.76032 | 0.766519 | 0.760398 |
| 0.772308 | 0.764725 | 0.766262 | 0.773557 | 0.775232 | 0.765431 | 0.764474 | 0.765594 | 0.76521 |
| 0.762797 | 0.75835 | 0.764298 | 0.766328 | 0.770169 | 0.762831 | 0.759777 | 0.757576 | 0.757153 |
| 0.772491 | 0.770006 | 0.765897 | 0.770298 | 0.773814 | 0.768929 | 0.764287 | 0.768404 | 0.762649 |
| 0.768795 | 0.764529 | 0.767434 | 0.776479 | 0.772432 | 0.76745 | 0.769402 | 0.763934 | 0.765084 |
| 0.767361 | 0.758702 | 0.760846 | 0.769022 | 0.774062 | 0.760536 | 0.763807 | 0.764454 | 0.759328 |
| 0.769629 | 0.762875 | 0.761003 | 0.771724 | 0.774621 | 0.759074 | 0.76525 | 0.765537 | 0.76175 |
| 0.753313 | 0.750505 | 0.757883 | 0.75709 | 0.75635 | 0.753753 | 0.753667 | 0.749101 | 0.747794 |
| 0.767638 | 0.760904 | 0.758379 | 0.764002 | 0.774122 | 0.767166 | 0.756886 | 0.764721 | 0.757016 |
| 0.758572 | 0.758624 | 0.758658 | 0.761523 | 0.761893 | 0.761037 | 0.756267 | 0.758845 | 0.754766 |
| 0.769497 | 0.764407 | 0.762186 | 0.769806 | 0.774347 | 0.765652 | 0.765938 | 0.765189 | 0.762472 |
| 0.772173 | 0.761176 | 0.764153 | 0.773488 | 0.771625 | 0.761803 | 0.764681 | 0.764296 | 0.761925 |
| 0.754564 | 0.753633 | 0.755235 | 0.758521 | 0.761379 | 0.757466 | 0.749127 | 0.753173 | 0.752004 |
| 0.754943 | 0.751259 | 0.757067 | 0.758847 | 0.760662 | 0.753841 | 0.753524 | 0.752381 | 0.752611 |
| 0.774459 | 0.764261 | 0.767817 | 0.774935 | 0.771305 | 0.766791 | 0.763653 | 0.768294 | 0.764393 |
| 0.821124 | 0.827277 | 0.807955 | 0.815568 | 0.7996 | 0.807343 | 0.799614 | 0.822677 | 0.794332 |
| 0.800763 | 0.801852 | 0.786713 | 0.796638 | 0.841599 | 0.810627 | 0.783625 | 0.810885 | 0.802857 |
| 0.831351 | 0.836804 | 0.797188 | 0.809245 | 0.887414 | 0.832744 | 0.794522 | 0.865062 | 0.804918 |
| 0.821719 | 0.819902 | 0.895813 | 0.837541 | 0.806746 | 0.823157 | 0.83856 | 0.814672 | 0.817557 |
| 1 | 0.830868 | 0.833434 | 0.879692 | 0.80536 | 0.818696 | 0.823454 | 0.831027 | 0.80366 |
| 0.830868 | 1 | 0.804587 | 0.807728 | 0.795847 | 0.839008 | 0.801478 | 0.834652 | 0.807958 |
| 0.833434 | 0.804587 | 1 | 0.872372 | 0.797567 | 0.813091 | 0.844427 | 0.798757 | 0.812303 |
| 0.879692 | 0.807728 | 0.872372 | 1 | 0.808006 | 0.810683 | 0.866356 | 0.8058 | 0.833233 |
| 0.80536 | 0.795847 | 0.797567 | 0.808006 | 1 | 0.81012 | 0.793656 | 0.864645 | 0.815803 |
| 0.818696 | 0.839008 | 0.813091 | 0.810683 | 0.81012 | 1 | 0.800751 | 0.827169 | 0.790188 |
| 0.823454 | 0.801478 | 0.844427 | 0.866356 | 0.793656 | 0.800751 | 1 | 0.79616 | 0.807351 |
| 0.831027 | 0.834652 | 0.798757 | 0.8058 | 0.864645 | 0.827169 | 0.79616 | 1 | 0.801145 |
| 0.80366 | 0.807958 | 0.812303 | 0.833233 | 0.815803 | 0.790188 | 0.807351 | 0.801145 | 1 |
| 0.812897 | 0.835417 | 0.807376 | 0.801698 | 0.800335 | 0.817372 | 0.815593 | 0.811308 | 0.810112 |
| 0.807624 | 0.809503 | 0.823347 | 0.827296 | 0.826043 | 0.808248 | 0.8251 | 0.818384 | 0.848532 |
| 0.828491 | 0.847316 | 0.800324 | 0.8065 | 0.820933 | 0.83108 | 0.804381 | 0.826974 | 0.823758 |
| 0.828958 | 0.835932 | 0.79084 | 0.797806 | 0.835458 | 0.831521 | 0.794245 | 0.873103 | 0.797709 |
| 0.823708 | 0.821151 | 0.816868 | 0.806784 | 0.830404 | 0.835413 | 0.801973 | 0.841418 | 0.788822 |
| 0.817468 | 0.821241 | 0.799813 | 0.81863 | 0.881021 | 0.813145 | 0.790051 | 0.83506 | 0.836125 |
| 0.799409 | 0.795534 | 0.807312 | 0.803 | 0.793366 | 0.807952 | 0.823035 | 0.7947 | 0.808708 |
| 0.836356 | 0.836303 | 0.819087 | 0.809879 | 0.805069 | 0.850339 | 0.803329 | 0.832575 | 0.792868 |
| 0.797784 | 0.800636 | 0.80441 | 0.800084 | 0.831425 | 0.804151 | 0.80252 | 0.814445 | 0.816535 |
| 0.821945 | 0.800828 | 0.820353 | 0.868049 | 0.797481 | 0.798903 | 0.832554 | 0.797029 | 0.826804 |
| 0.849745 | 0.87491 | 0.806193 | 0.808404 | 0.813876 | 0.847017 | 0.80338 | 0.843338 | 0.79293 |
| 0.815949 | 0.786674 | 0.816614 | 0.859338 | 0.798155 | 0.788851 | 0.822682 | 0.790493 | 0.82007 |
| 0.814059 | 0.847203 | 0.803942 | 0.799604 | 0.778808 | 0.811805 | 0.833511 | 0.808154 | 0.789566 |
| 0.836367 | 0.79455 | 0.845571 | 0.864653 | 0.796298 | 0.802526 | 0.84715 | 0.797704 | 0.824089 |
| 0.846908 | 0.836954 | 0.804939 | 0.808442 | 0.812511 | 0.8066 | 0.795549 | 0.852335 | 0.796415 |
| 0.826669 | 0.7982 | 0.881576 | 0.870607 | 0.788938 | 0.796429 | 0.827339 | 0.794257 | 0.801497 |
| 0.825707 | 0.791318 | 0.807934 | 0.865661 | 0.810787 | 0.802823 | 0.818466 | 0.800893 | 0.812641 |
| 0.795834 | 0.780562 | 0.776853 | 0.78469 | 0.882953 | 0.793067 | 0.777115 | 0.839757 | 0.796907 |

| | | | | | | | | |
|---|---|---|---|---|---|---|---|---|
| 0.797663 | 0.813469 | 0.806529 | 0.799833 | 0.7831 | 0.809263 | 0.865106 | 0.787725 | 0.791901 |
| 0.764466 | 0.758743 | 0.762812 | 0.769759 | 0.764745 | 0.768647 | 0.766544 | 0.761752 | 0.76576 |
| 0.760154 | 0.755307 | 0.756899 | 0.759571 | 0.768222 | 0.762536 | 0.761633 | 0.757752 | 0.75599 |
| 0.757146 | 0.753193 | 0.755888 | 0.760423 | 0.76712 | 0.762743 | 0.755057 | 0.757016 | 0.757591 |
| 0.772101 | 0.764711 | 0.769385 | 0.774945 | 0.774742 | 0.769212 | 0.770994 | 0.764084 | 0.766409 |
| 0.768831 | 0.760513 | 0.757649 | 0.767992 | 0.768549 | 0.769973 | 0.764322 | 0.761968 | 0.76011 |
| 0.765104 | 0.760464 | 0.760233 | 0.767472 | 0.7696 | 0.768383 | 0.765117 | 0.768058 | 0.765763 |
| 0.769668 | 0.759319 | 0.761675 | 0.770813 | 0.769805 | 0.760915 | 0.76059 | 0.760642 | 0.761849 |
| 0.761042 | 0.749431 | 0.754875 | 0.761083 | 0.761712 | 0.760291 | 0.756197 | 0.751477 | 0.748069 |
| 0.761194 | 0.755379 | 0.758946 | 0.763696 | 0.76777 | 0.758804 | 0.757978 | 0.759133 | 0.759407 |
| 0.768935 | 0.760733 | 0.760287 | 0.771371 | 0.769488 | 0.767869 | 0.763283 | 0.762806 | 0.761597 |
| 0.770091 | 0.758661 | 0.762886 | 0.770786 | 0.770544 | 0.766445 | 0.764065 | 0.76409 | 0.760203 |
| 0.768038 | 0.758178 | 0.762504 | 0.766072 | 0.768327 | 0.762256 | 0.760821 | 0.761838 | 0.761698 |
| 0.767206 | 0.753779 | 0.762318 | 0.766898 | 0.764199 | 0.763298 | 0.760073 | 0.753831 | 0.759064 |
| 0.766153 | 0.75796 | 0.760814 | 0.765044 | 0.767618 | 0.764322 | 0.76162 | 0.762803 | 0.760994 |
| 0.76472 | 0.756299 | 0.762358 | 0.767042 | 0.768776 | 0.762059 | 0.762834 | 0.760477 | 0.76171 |
| 0.763848 | 0.75938 | 0.763557 | 0.769225 | 0.766824 | 0.764483 | 0.765842 | 0.760309 | 0.758039 |
| 0.760207 | 0.757022 | 0.760365 | 0.767292 | 0.764496 | 0.765604 | 0.761804 | 0.757593 | 0.75549 |
| 0.767813 | 0.76032 | 0.763966 | 0.770337 | 0.774225 | 0.766562 | 0.769841 | 0.76589 | 0.766195 |
| 0.771006 | 0.762095 | 0.762774 | 0.769443 | 0.770528 | 0.766678 | 0.766743 | 0.765843 | 0.760346 |
| 0.760218 | 0.759469 | 0.759578 | 0.7625 | 0.763829 | 0.76169 | 0.757767 | 0.756494 | 0.757614 |
| 0.769071 | 0.761503 | 0.765335 | 0.770365 | 0.76773 | 0.761881 | 0.764324 | 0.757434 | 0.76281 |
| 0.765741 | 0.758446 | 0.764167 | 0.770311 | 0.76736 | 0.766841 | 0.766949 | 0.758891 | 0.763261 |
| 0.762575 | 0.758103 | 0.760632 | 0.765272 | 0.763002 | 0.763557 | 0.762848 | 0.75788 | 0.754775 |
| 0.769852 | 0.761873 | 0.762372 | 0.768004 | 0.768519 | 0.765561 | 0.762716 | 0.757922 | 0.758865 |
| 0.762827 | 0.756361 | 0.761568 | 0.765777 | 0.764103 | 0.766249 | 0.764881 | 0.755119 | 0.755996 |
| 0.763106 | 0.75157 | 0.759371 | 0.76573 | 0.76156 | 0.758061 | 0.760381 | 0.755832 | 0.757305 |
| 0.76048 | 0.754815 | 0.759646 | 0.762539 | 0.763799 | 0.760326 | 0.759921 | 0.755762 | 0.758589 |
| 0.767611 | 0.759486 | 0.763979 | 0.76788 | 0.765349 | 0.765458 | 0.764035 | 0.761869 | 0.758635 |
| 0.765248 | 0.760528 | 0.759904 | 0.771285 | 0.768178 | 0.761511 | 0.7653 | 0.757465 | 0.763176 |
| 0.762422 | 0.75484 | 0.756085 | 0.763286 | 0.754762 | 0.754477 | 0.757323 | 0.751007 | 0.75499 |
| 0.765528 | 0.759063 | 0.764988 | 0.770957 | 0.768371 | 0.76691 | 0.765082 | 0.762654 | 0.763473 |
| 0.770485 | 0.760296 | 0.760106 | 0.768674 | 0.76185 | 0.763558 | 0.765462 | 0.758202 | 0.760432 |
| 0.761111 | 0.754431 | 0.759035 | 0.764394 | 0.76592 | 0.758102 | 0.75611 | 0.75337 | 0.759169 |
| 0.757711 | 0.754076 | 0.757755 | 0.765446 | 0.764785 | 0.762949 | 0.758916 | 0.754468 | 0.757202 |
| 0.760244 | 0.757908 | 0.755488 | 0.761653 | 0.760948 | 0.760524 | 0.758963 | 0.755501 | 0.759196 |
| 0.769509 | 0.75991 | 0.767135 | 0.771605 | 0.771529 | 0.770709 | 0.769212 | 0.768249 | 0.763377 |
| 0.761106 | 0.753615 | 0.75818 | 0.766385 | 0.764313 | 0.763455 | 0.761773 | 0.755107 | 0.759871 |
| 0.764386 | 0.758788 | 0.762744 | 0.770117 | 0.763445 | 0.762609 | 0.767684 | 0.756466 | 0.761469 |
| 0.765618 | 0.759949 | 0.76018 | 0.766104 | 0.759907 | 0.76213 | 0.765351 | 0.754032 | 0.76216 |
| 0.767033 | 0.761014 | 0.764276 | 0.769825 | 0.76497 | 0.763428 | 0.767995 | 0.757073 | 0.766186 |
| 0.76629 | 0.764037 | 0.762965 | 0.772073 | 0.768352 | 0.76966 | 0.76781 | 0.760156 | 0.760047 |
| 0.769168 | 0.764056 | 0.767135 | 0.775116 | 0.768108 | 0.771888 | 0.768763 | 0.762632 | 0.763285 |
| 0.766127 | 0.754309 | 0.756341 | 0.763374 | 0.761921 | 0.757698 | 0.756306 | 0.755257 | 0.75623 |
| 0.764989 | 0.762859 | 0.762919 | 0.767793 | 0.764202 | 0.766447 | 0.763636 | 0.756926 | 0.765261 |
| 0.762012 | 0.755622 | 0.756688 | 0.765813 | 0.762973 | 0.762512 | 0.758274 | 0.755501 | 0.758246 |

| | | | | | | | | |
|---|---|---|---|---|---|---|---|---|
| 0.752741 | 0.743821 | 0.749833 | 0.756139 | 0.755226 | 0.756132 | 0.748602 | 0.747129 | 0.743953 |
| 0.745243 | 0.737066 | 0.74569 | 0.751988 | 0.745991 | 0.743047 | 0.743713 | 0.737975 | 0.737014 |
| 0.756001 | 0.753092 | 0.757558 | 0.759863 | 0.7621 | 0.759061 | 0.754337 | 0.755015 | 0.758058 |
| 0.76147 | 0.758284 | 0.760139 | 0.764331 | 0.76527 | 0.767644 | 0.758167 | 0.757636 | 0.76087 |
| 0.755332 | 0.748428 | 0.751185 | 0.756174 | 0.75875 | 0.755106 | 0.751704 | 0.748863 | 0.750992 |
| 0.759373 | 0.751575 | 0.756761 | 0.762065 | 0.760816 | 0.75717 | 0.757996 | 0.752065 | 0.751448 |
| 0.758469 | 0.751479 | 0.760957 | 0.767157 | 0.761942 | 0.761087 | 0.759302 | 0.753275 | 0.757798 |
| 0.753872 | 0.746576 | 0.747094 | 0.755433 | 0.75771 | 0.752764 | 0.750265 | 0.745033 | 0.752716 |
| 0.766771 | 0.757327 | 0.763224 | 0.771224 | 0.768069 | 0.767326 | 0.768252 | 0.760209 | 0.760978 |
| 0.762954 | 0.760175 | 0.762919 | 0.767537 | 0.770381 | 0.766171 | 0.76612 | 0.765418 | 0.76327 |
| 0.767347 | 0.760192 | 0.761842 | 0.770393 | 0.772154 | 0.762137 | 0.764227 | 0.76043 | 0.76837 |
| 0.760696 | 0.753383 | 0.757826 | 0.766983 | 0.769625 | 0.763281 | 0.758669 | 0.758015 | 0.757581 |
| 0.769878 | 0.764716 | 0.765045 | 0.771125 | 0.769952 | 0.772568 | 0.765441 | 0.761925 | 0.764519 |
| 0.762017 | 0.755318 | 0.763544 | 0.765401 | 0.767408 | 0.762349 | 0.759237 | 0.756813 | 0.75905 |
| 0.762004 | 0.754237 | 0.761141 | 0.768605 | 0.76972 | 0.762831 | 0.760544 | 0.759107 | 0.757084 |
| 0.762063 | 0.751842 | 0.758822 | 0.765905 | 0.759948 | 0.763294 | 0.76055 | 0.754081 | 0.75659 |
| 0.765896 | 0.756018 | 0.758621 | 0.765959 | 0.764769 | 0.766352 | 0.761818 | 0.762569 | 0.762639 |
| 0.760886 | 0.751739 | 0.753635 | 0.760818 | 0.761273 | 0.759706 | 0.757272 | 0.754542 | 0.758548 |
| 0.763363 | 0.752625 | 0.751487 | 0.761354 | 0.75518 | 0.758745 | 0.756614 | 0.748473 | 0.753771 |
| 0.763876 | 0.755389 | 0.763351 | 0.76631 | 0.763733 | 0.761674 | 0.765191 | 0.756235 | 0.7587 |
| 0.759502 | 0.754766 | 0.758775 | 0.766625 | 0.765694 | 0.760369 | 0.758885 | 0.75576 | 0.758577 |
| 0.746493 | 0.749358 | 0.744991 | 0.752172 | 0.755623 | 0.753879 | 0.750318 | 0.746167 | 0.74364 |
| 0.758259 | 0.752284 | 0.751535 | 0.758229 | 0.758869 | 0.756949 | 0.754759 | 0.750454 | 0.749004 |
| 0.759252 | 0.758739 | 0.75999 | 0.764629 | 0.765642 | 0.762661 | 0.759075 | 0.754704 | 0.75502 |
| 0.760285 | 0.756333 | 0.758136 | 0.764569 | 0.765625 | 0.762595 | 0.756973 | 0.759935 | 0.757989 |
| 0.756158 | 0.755187 | 0.752342 | 0.758774 | 0.762095 | 0.761215 | 0.756514 | 0.753435 | 0.750359 |
| 0.755916 | 0.75263 | 0.752755 | 0.760091 | 0.756876 | 0.754638 | 0.757374 | 0.749658 | 0.75796 |
| 0.761852 | 0.758902 | 0.758953 | 0.76521 | 0.760747 | 0.762662 | 0.759248 | 0.753341 | 0.756806 |
| 0.757439 | 0.752468 | 0.754662 | 0.759245 | 0.759511 | 0.759206 | 0.752392 | 0.75 | 0.752421 |
| 0.694375 | 0.687995 | 0.6921 | 0.694437 | 0.694702 | 0.695419 | 0.688875 | 0.688558 | 0.6902 |
| 0.703179 | 0.699844 | 0.701387 | 0.706159 | 0.704602 | 0.70393 | 0.699911 | 0.702815 | 0.702665 |
| 0.702317 | 0.699853 | 0.701018 | 0.704137 | 0.706057 | 0.704921 | 0.69884 | 0.702711 | 0.696999 |
| 0.694021 | 0.687701 | 0.695157 | 0.697428 | 0.694696 | 0.695115 | 0.692832 | 0.694534 | 0.689248 |
| 0.704191 | 0.700563 | 0.703564 | 0.70663 | 0.705589 | 0.704276 | 0.700938 | 0.70141 | 0.697519 |
| 0.758226 | 0.752539 | 0.754022 | 0.758377 | 0.766077 | 0.758508 | 0.753441 | 0.759016 | 0.754277 |
| 0.761063 | 0.753949 | 0.756465 | 0.761339 | 0.766136 | 0.758841 | 0.761319 | 0.755966 | 0.757371 |
| 0.768427 | 0.757008 | 0.761053 | 0.769307 | 0.769564 | 0.765829 | 0.765708 | 0.76074 | 0.756926 |
| 0.760683 | 0.759711 | 0.75873 | 0.76217 | 0.767025 | 0.760647 | 0.757167 | 0.762726 | 0.760494 |
| 0.757302 | 0.755786 | 0.757685 | 0.763391 | 0.761206 | 0.757451 | 0.757621 | 0.758415 | 0.749853 |
| 0.763169 | 0.756012 | 0.761153 | 0.763839 | 0.764929 | 0.763255 | 0.758915 | 0.755685 | 0.758511 |
| 0.766312 | 0.761824 | 0.765279 | 0.766118 | 0.768806 | 0.764265 | 0.759759 | 0.759808 | 0.761275 |
| 0.75943 | 0.752632 | 0.760227 | 0.763573 | 0.766159 | 0.761341 | 0.756571 | 0.755283 | 0.754523 |
| 0.755778 | 0.750264 | 0.751253 | 0.755546 | 0.758151 | 0.752573 | 0.751066 | 0.752506 | 0.748136 |
| 0.757557 | 0.752776 | 0.752307 | 0.760144 | 0.759328 | 0.755137 | 0.753801 | 0.75324 | 0.755426 |
| 0.757142 | 0.753239 | 0.758706 | 0.760321 | 0.762591 | 0.758674 | 0.753971 | 0.753787 | 0.75411 |
| 0.766304 | 0.755768 | 0.763216 | 0.768014 | 0.765881 | 0.764096 | 0.760141 | 0.760088 | 0.75963 |

| | | | | | | | | |
|---|---|---|---|---|---|---|---|---|
| 0.765599 | 0.760698 | 0.76343 | 0.769759 | 0.767289 | 0.764536 | 0.758757 | 0.761119 | 0.755607 |
| 0.762505 | 0.756349 | 0.751107 | 0.761858 | 0.763108 | 0.762474 | 0.756211 | 0.755041 | 0.752514 |
| 0.761742 | 0.758621 | 0.761647 | 0.762948 | 0.766026 | 0.762996 | 0.760868 | 0.754008 | 0.75979 |
| 0.75999 | 0.752684 | 0.753344 | 0.759526 | 0.763162 | 0.761233 | 0.757849 | 0.755635 | 0.757449 |
| 0.756545 | 0.751621 | 0.757815 | 0.762355 | 0.758783 | 0.760899 | 0.757399 | 0.750318 | 0.759023 |
| 0.760095 | 0.751706 | 0.755684 | 0.766173 | 0.765272 | 0.761769 | 0.757025 | 0.75589 | 0.754097 |
| 0.75815 | 0.7534 | 0.756949 | 0.761988 | 0.76429 | 0.761893 | 0.756675 | 0.753382 | 0.76037 |
| 0.758499 | 0.757357 | 0.758859 | 0.762801 | 0.763842 | 0.763943 | 0.757568 | 0.756023 | 0.755268 |
| 0.754153 | 0.753631 | 0.753151 | 0.759897 | 0.759275 | 0.758218 | 0.753548 | 0.753824 | 0.755121 |
| 0.761199 | 0.75318 | 0.753699 | 0.763672 | 0.759847 | 0.761641 | 0.759957 | 0.753729 | 0.755563 |
| 0.76091 | 0.755637 | 0.75629 | 0.763662 | 0.765276 | 0.758094 | 0.759287 | 0.755111 | 0.761519 |
| 0.763244 | 0.756888 | 0.761941 | 0.768012 | 0.767295 | 0.76498 | 0.761897 | 0.757098 | 0.761194 |
| 0.757184 | 0.751036 | 0.751169 | 0.755305 | 0.754685 | 0.757239 | 0.750623 | 0.752407 | 0.754038 |
| 0.756299 | 0.754811 | 0.748763 | 0.756786 | 0.757842 | 0.758836 | 0.751917 | 0.748877 | 0.745741 |
| 0.761096 | 0.757506 | 0.755335 | 0.76405 | 0.760624 | 0.757383 | 0.760855 | 0.754112 | 0.7561 |
| 0.768784 | 0.757852 | 0.764647 | 0.770994 | 0.767094 | 0.764794 | 0.767595 | 0.758155 | 0.762433 |
| 0.764188 | 0.752885 | 0.760794 | 0.767457 | 0.762675 | 0.760996 | 0.760141 | 0.757119 | 0.758045 |
| 0.766763 | 0.759615 | 0.761849 | 0.767373 | 0.764416 | 0.771381 | 0.756451 | 0.75971 | 0.759483 |
| 0.766141 | 0.760545 | 0.755891 | 0.762597 | 0.760122 | 0.765026 | 0.755375 | 0.755753 | 0.761079 |
| 0.765616 | 0.754652 | 0.766745 | 0.767552 | 0.764091 | 0.764017 | 0.762729 | 0.762569 | 0.758414 |
| 0.763232 | 0.753359 | 0.760953 | 0.76948 | 0.765411 | 0.760685 | 0.758721 | 0.753992 | 0.758627 |
| 0.760642 | 0.754736 | 0.755613 | 0.763891 | 0.762793 | 0.759604 | 0.757424 | 0.755699 | 0.75667 |
| 0.766621 | 0.755715 | 0.763892 | 0.770247 | 0.768916 | 0.76634 | 0.768379 | 0.760825 | 0.762222 |
| 0.762266 | 0.752474 | 0.759196 | 0.764258 | 0.760081 | 0.759776 | 0.760731 | 0.753685 | 0.753817 |
| 0.763793 | 0.75195 | 0.762598 | 0.766977 | 0.76752 | 0.761335 | 0.762884 | 0.758303 | 0.760007 |
| 0.700282 | 0.69643 | 0.699762 | 0.702947 | 0.704958 | 0.701671 | 0.695421 | 0.700448 | 0.69689 |
| 0.765082 | 0.755798 | 0.760884 | 0.764734 | 0.76447 | 0.762244 | 0.760977 | 0.757243 | 0.757924 |
| 0.76594 | 0.755718 | 0.758641 | 0.76851 | 0.762477 | 0.762525 | 0.764157 | 0.755767 | 0.757735 |
| 0.768827 | 0.759976 | 0.7648 | 0.76944 | 0.771571 | 0.76275 | 0.763903 | 0.759605 | 0.759125 |
| 0.767552 | 0.759352 | 0.758245 | 0.769079 | 0.765172 | 0.765822 | 0.76164 | 0.756748 | 0.75951 |
| 0.762953 | 0.755447 | 0.759615 | 0.763719 | 0.764462 | 0.763828 | 0.755134 | 0.757956 | 0.755257 |
| 0.764646 | 0.757395 | 0.759312 | 0.762258 | 0.766001 | 0.76419 | 0.759009 | 0.753983 | 0.754761 |
| 0.762453 | 0.753917 | 0.758368 | 0.763027 | 0.768423 | 0.766965 | 0.757354 | 0.758993 | 0.758747 |
| 0.764915 | 0.756686 | 0.759923 | 0.768948 | 0.764959 | 0.760923 | 0.759251 | 0.75869 | 0.757449 |
| 0.76101 | 0.748637 | 0.75415 | 0.761113 | 0.754189 | 0.757882 | 0.756663 | 0.747686 | 0.752153 |
| 0.762013 | 0.750106 | 0.75102 | 0.761512 | 0.757289 | 0.756553 | 0.757106 | 0.75155 | 0.751401 |
| 0.766958 | 0.758709 | 0.760997 | 0.768495 | 0.766935 | 0.761337 | 0.761817 | 0.759338 | 0.762214 |
| 0.76378 | 0.763098 | 0.761267 | 0.766283 | 0.767338 | 0.765423 | 0.762891 | 0.759578 | 0.763327 |
| 0.770434 | 0.762183 | 0.766026 | 0.771452 | 0.769308 | 0.769549 | 0.769889 | 0.763235 | 0.763418 |
| 0.766954 | 0.76116 | 0.761353 | 0.76427 | 0.767989 | 0.768613 | 0.758841 | 0.761905 | 0.762697 |
| 0.75881 | 0.753128 | 0.753534 | 0.763557 | 0.760144 | 0.760821 | 0.757224 | 0.753959 | 0.758482 |
| 0.768545 | 0.762689 | 0.764741 | 0.769308 | 0.767249 | 0.770486 | 0.765555 | 0.756542 | 0.764049 |
| 0.762199 | 0.754117 | 0.758091 | 0.76397 | 0.760522 | 0.7616 | 0.76084 | 0.756937 | 0.759777 |
| 0.755877 | 0.748925 | 0.758279 | 0.761723 | 0.759282 | 0.758193 | 0.756114 | 0.74895 | 0.75359 |
| 0.761432 | 0.756786 | 0.760963 | 0.763763 | 0.764876 | 0.762541 | 0.756999 | 0.760214 | 0.757512 |
| 0.761999 | 0.757244 | 0.762016 | 0.763134 | 0.765409 | 0.763559 | 0.759848 | 0.758552 | 0.756782 |

| | | | | | | | | |
|---|---|---|---|---|---|---|---|---|
| 0.761667 | 0.755051 | 0.760506 | 0.764085 | 0.763937 | 0.756569 | 0.75299 | 0.757021 | 0.758919 |
| 0.762772 | 0.758962 | 0.760882 | 0.765432 | 0.766214 | 0.762945 | 0.761285 | 0.759695 | 0.756867 |
| 0.758561 | 0.753129 | 0.761311 | 0.764236 | 0.760315 | 0.76062 | 0.759474 | 0.753224 | 0.750672 |
| 0.72688 | 0.721755 | 0.7213 | 0.730756 | 0.716732 | 0.716524 | 0.722562 | 0.71657 | 0.719109 |
| 0.757806 | 0.748405 | 0.76009 | 0.763505 | 0.758587 | 0.754969 | 0.756537 | 0.753987 | 0.752752 |
| 0.758448 | 0.751293 | 0.757599 | 0.75917 | 0.75617 | 0.759694 | 0.761486 | 0.751906 | 0.751477 |
| 0.761506 | 0.759162 | 0.763413 | 0.768327 | 0.767665 | 0.764745 | 0.765866 | 0.764936 | 0.76112 |
| 0.763943 | 0.756418 | 0.76005 | 0.764801 | 0.760706 | 0.758738 | 0.7565 | 0.757425 | 0.758678 |
| 0.763871 | 0.758692 | 0.765133 | 0.769062 | 0.764634 | 0.763313 | 0.761979 | 0.75492 | 0.760853 |
| 0.763026 | 0.755248 | 0.761285 | 0.766477 | 0.763428 | 0.763948 | 0.761917 | 0.755934 | 0.759252 |
| 0.757245 | 0.755327 | 0.759061 | 0.764053 | 0.76577 | 0.760047 | 0.759785 | 0.755104 | 0.756707 |
| 0.759137 | 0.758248 | 0.759946 | 0.764884 | 0.766324 | 0.763885 | 0.758199 | 0.76011 | 0.758395 |
| 0.759306 | 0.753837 | 0.75879 | 0.762571 | 0.763609 | 0.758326 | 0.759145 | 0.752192 | 0.759392 |
| 0.761249 | 0.752114 | 0.75991 | 0.764272 | 0.761442 | 0.758419 | 0.761321 | 0.755987 | 0.757105 |
| 0.767473 | 0.755102 | 0.765016 | 0.767315 | 0.764036 | 0.762585 | 0.76238 | 0.755764 | 0.760708 |
| 0.760478 | 0.753165 | 0.754607 | 0.759164 | 0.759888 | 0.758102 | 0.757601 | 0.757114 | 0.754417 |
| 0.762538 | 0.756346 | 0.758965 | 0.765105 | 0.764338 | 0.763359 | 0.762544 | 0.760336 | 0.758923 |
| 0.765054 | 0.758474 | 0.76262 | 0.770008 | 0.769521 | 0.763147 | 0.765318 | 0.759233 | 0.762076 |
| 0.754689 | 0.751703 | 0.75715 | 0.759265 | 0.754969 | 0.757721 | 0.758228 | 0.748845 | 0.754196 |
| 0.766447 | 0.754029 | 0.760944 | 0.765644 | 0.762473 | 0.760232 | 0.759203 | 0.756757 | 0.757434 |
| 0.763824 | 0.762164 | 0.764445 | 0.769049 | 0.770268 | 0.76907 | 0.764462 | 0.763642 | 0.761622 |
| 0.765823 | 0.76022 | 0.764201 | 0.769363 | 0.7655 | 0.765483 | 0.766286 | 0.761976 | 0.760844 |
| 0.679583 | 0.677294 | 0.674359 | 0.680174 | 0.67792 | 0.692039 | 0.677254 | 0.674728 | 0.677969 |
| 0.650225 | 0.655131 | 0.653013 | 0.655587 | 0.652297 | 0.65523 | 0.654472 | 0.654707 | 0.651588 |
| 0.657985 | 0.657564 | 0.66565 | 0.666024 | 0.670043 | 0.66537 | 0.65914 | 0.654518 | 0.669211 |
| 0.651889 | 0.646815 | 0.652462 | 0.65524 | 0.65097 | 0.652495 | 0.646557 | 0.648658 | 0.643564 |
| 0.731238 | 0.71368 | 0.719632 | 0.72757 | 0.728882 | 0.731368 | 0.726945 | 0.717247 | 0.71435 |
| 0.718882 | 0.713814 | 0.715085 | 0.72641 | 0.726783 | 0.71617 | 0.712344 | 0.716198 | 0.71346 |
| 0.731265 | 0.713712 | 0.725455 | 0.733129 | 0.727387 | 0.721689 | 0.726465 | 0.716388 | 0.717879 |
| 0.727818 | 0.72005 | 0.725753 | 0.73595 | 0.733695 | 0.725993 | 0.730858 | 0.722803 | 0.728895 |
| 0.713926 | 0.705692 | 0.711445 | 0.71332 | 0.711517 | 0.708333 | 0.702308 | 0.704404 | 0.707339 |
| 0.70271 | 0.692068 | 0.704134 | 0.713246 | 0.704443 | 0.698166 | 0.705483 | 0.697357 | 0.700876 |
| 0.704787 | 0.699376 | 0.700442 | 0.704952 | 0.710192 | 0.704378 | 0.707296 | 0.712085 | 0.702222 |
| 0.654229 | 0.646315 | 0.651908 | 0.655491 | 0.646787 | 0.650135 | 0.651055 | 0.646667 | 0.650822 |
| 0.755557 | 0.752118 | 0.752948 | 0.762381 | 0.757602 | 0.758235 | 0.754501 | 0.747542 | 0.754771 |
| 0.735671 | 0.727101 | 0.730616 | 0.736392 | 0.737111 | 0.733614 | 0.735129 | 0.722756 | 0.728776 |
| 0.730161 | 0.720985 | 0.728721 | 0.737288 | 0.73346 | 0.723627 | 0.729495 | 0.721538 | 0.729527 |
| 0.723042 | 0.711078 | 0.714847 | 0.723954 | 0.716803 | 0.717002 | 0.715642 | 0.706257 | 0.714179 |
| 0.736541 | 0.730255 | 0.733255 | 0.738949 | 0.730291 | 0.733193 | 0.734452 | 0.724862 | 0.729365 |
| 0.725819 | 0.714785 | 0.723696 | 0.728404 | 0.719974 | 0.716357 | 0.727706 | 0.714267 | 0.714249 |
| 0.71875 | 0.715207 | 0.717374 | 0.725512 | 0.722682 | 0.718929 | 0.726356 | 0.713324 | 0.724077 |
| 0.726209 | 0.713205 | 0.719767 | 0.728342 | 0.718998 | 0.717676 | 0.721145 | 0.718021 | 0.71201 |
| 0.720872 | 0.716396 | 0.722638 | 0.724335 | 0.719305 | 0.720783 | 0.719899 | 0.711664 | 0.715998 |
| 0.717828 | 0.709207 | 0.709091 | 0.719922 | 0.723107 | 0.712451 | 0.711804 | 0.711477 | 0.711283 |
| 0.719528 | 0.713619 | 0.714763 | 0.722322 | 0.713555 | 0.72 | 0.715865 | 0.708301 | 0.718799 |
| 0.700426 | 0.702513 | 0.703704 | 0.713399 | 0.713481 | 0.710943 | 0.714666 | 0.705803 | 0.713194 |

| | | | | | | | | |
|---|---|---|---|---|---|---|---|---|
| 0.744983 | 0.737598 | 0.743703 | 0.747225 | 0.748221 | 0.741085 | 0.740042 | 0.736974 | 0.74011 |
| 0.73241 | 0.723319 | 0.723302 | 0.731995 | 0.728771 | 0.72523 | 0.720721 | 0.716847 | 0.71594 |
| 0.722582 | 0.716157 | 0.716012 | 0.729948 | 0.722189 | 0.721004 | 0.728346 | 0.716781 | 0.716139 |
| 0.717381 | 0.714086 | 0.715473 | 0.723217 | 0.72567 | 0.721811 | 0.718923 | 0.717706 | 0.71768 |
| 0.724356 | 0.716124 | 0.720066 | 0.728545 | 0.723089 | 0.715514 | 0.724343 | 0.707272 | 0.719869 |
| 0.713143 | 0.71134 | 0.713102 | 0.720701 | 0.720808 | 0.718384 | 0.713858 | 0.709903 | 0.711649 |
| 0.718735 | 0.698872 | 0.712299 | 0.722397 | 0.71584 | 0.709935 | 0.709215 | 0.705031 | 0.704838 |
| 0.713962 | 0.698833 | 0.713838 | 0.717697 | 0.710393 | 0.713154 | 0.714854 | 0.70151 | 0.701467 |
| 0.717905 | 0.708536 | 0.723465 | 0.724118 | 0.72068 | 0.710741 | 0.724335 | 0.704881 | 0.714123 |
| 0.753716 | 0.744431 | 0.755465 | 0.759026 | 0.748289 | 0.752455 | 0.752689 | 0.746989 | 0.746455 |
| 0.750759 | 0.738563 | 0.74954 | 0.754734 | 0.745458 | 0.747966 | 0.746003 | 0.740268 | 0.744533 |
| 0.743145 | 0.73683 | 0.744416 | 0.748787 | 0.735926 | 0.742633 | 0.738971 | 0.733452 | 0.737635 |
| 0.731974 | 0.728612 | 0.731213 | 0.741456 | 0.731341 | 0.735161 | 0.7315 | 0.725101 | 0.726884 |
| 0.738774 | 0.733845 | 0.740461 | 0.745585 | 0.741392 | 0.739978 | 0.74136 | 0.734941 | 0.735567 |
| 0.736978 | 0.729406 | 0.733913 | 0.735269 | 0.727962 | 0.735462 | 0.7339 | 0.721135 | 0.726406 |

| | | | plot 03 | | | | | |
|---|---|---|---|---|---|---|---|---|
| Wild_plot_ | Wild_plot_ | Wild_plot_ | Wild_plot_ | Wild_plot_ | Wild_plot_ | Wild_plot_ | Wild_plot_ | Wild_plot_ |
| 0.701639 | 0.708615 | 0.707561 | 0.708209 | 0.710833 | 0.710608 | 0.705101 | 0.704646 | 0.70843 |
| 0.708877 | 0.71489 | 0.713351 | 0.714931 | 0.717197 | 0.718346 | 0.708407 | 0.707027 | 0.712542 |
| 0.699953 | 0.700426 | 0.703598 | 0.705351 | 0.707509 | 0.707184 | 0.698043 | 0.700714 | 0.7046 |
| 0.707872 | 0.712366 | 0.712575 | 0.714721 | 0.715602 | 0.715158 | 0.704884 | 0.710147 | 0.712298 |
| 0.70412 | 0.708314 | 0.709381 | 0.710752 | 0.709594 | 0.711046 | 0.702889 | 0.707246 | 0.707454 |
| 0.717001 | 0.716433 | 0.720013 | 0.720502 | 0.723377 | 0.722452 | 0.713273 | 0.715137 | 0.716264 |
| 0.761573 | 0.762029 | 0.762598 | 0.763321 | 0.767444 | 0.768538 | 0.762222 | 0.761308 | 0.7661 |
| 0.761211 | 0.762497 | 0.760203 | 0.758925 | 0.767784 | 0.766855 | 0.760691 | 0.760375 | 0.759364 |
| 0.706753 | 0.710819 | 0.713989 | 0.712133 | 0.715474 | 0.715357 | 0.707605 | 0.706713 | 0.710799 |
| 0.701825 | 0.704542 | 0.705929 | 0.708771 | 0.708875 | 0.708839 | 0.705097 | 0.705441 | 0.707611 |
| 0.706585 | 0.707955 | 0.71051 | 0.709292 | 0.710835 | 0.712698 | 0.706877 | 0.7065 | 0.7099 |
| 0.698735 | 0.703146 | 0.704592 | 0.707025 | 0.706109 | 0.706579 | 0.69903 | 0.703107 | 0.704642 |
| 0.71211 | 0.708039 | 0.710642 | 0.714655 | 0.716938 | 0.716327 | 0.710297 | 0.709044 | 0.715796 |
| 0.725922 | 0.726894 | 0.725136 | 0.723739 | 0.729799 | 0.72833 | 0.729728 | 0.725931 | 0.726258 |
| 0.752754 | 0.745333 | 0.750464 | 0.747233 | 0.751584 | 0.74518 | 0.74482 | 0.74809 | 0.739137 |
| 0.711629 | 0.713445 | 0.717822 | 0.716173 | 0.719642 | 0.717402 | 0.711969 | 0.714818 | 0.712509 |
| 0.710358 | 0.713255 | 0.714791 | 0.715688 | 0.717957 | 0.719099 | 0.710314 | 0.710039 | 0.712138 |
| 0.773289 | 0.761878 | 0.762574 | 0.752653 | 0.76087 | 0.752885 | 0.764639 | 0.764064 | 0.755148 |
| 0.702316 | 0.702885 | 0.707378 | 0.707063 | 0.710314 | 0.708151 | 0.699984 | 0.704516 | 0.704325 |
| 0.706013 | 0.71083 | 0.709834 | 0.711684 | 0.713731 | 0.712567 | 0.704206 | 0.70829 | 0.711251 |
| 0.703011 | 0.70516 | 0.707247 | 0.710446 | 0.709494 | 0.707466 | 0.703155 | 0.705665 | 0.704103 |
| 0.704172 | 0.710514 | 0.710327 | 0.711825 | 0.714795 | 0.713603 | 0.706555 | 0.707451 | 0.707382 |
| 0.713263 | 0.716003 | 0.71619 | 0.715668 | 0.71792 | 0.7182 | 0.710766 | 0.711827 | 0.71479 |
| 0.708668 | 0.712059 | 0.713568 | 0.714251 | 0.716539 | 0.716292 | 0.710057 | 0.708555 | 0.713288 |
| 0.711702 | 0.711889 | 0.715523 | 0.71513 | 0.714172 | 0.716461 | 0.709952 | 0.71325 | 0.710439 |
| 0.707086 | 0.707675 | 0.712849 | 0.712466 | 0.713668 | 0.713539 | 0.704913 | 0.709849 | 0.709261 |
| 0.700571 | 0.701995 | 0.705293 | 0.705703 | 0.710116 | 0.708363 | 0.698173 | 0.70399 | 0.705194 |
| 0.707641 | 0.709367 | 0.715536 | 0.71274 | 0.71358 | 0.714874 | 0.706405 | 0.710414 | 0.710535 |
| 0.709521 | 0.712464 | 0.712748 | 0.71371 | 0.714681 | 0.715905 | 0.70789 | 0.712078 | 0.711118 |
| 0.705756 | 0.708959 | 0.70875 | 0.713071 | 0.712429 | 0.715305 | 0.705279 | 0.707236 | 0.705104 |
| 0.706521 | 0.710833 | 0.710966 | 0.711766 | 0.714308 | 0.713075 | 0.706259 | 0.70613 | 0.711819 |
| 0.703991 | 0.707119 | 0.706585 | 0.708547 | 0.706224 | 0.709897 | 0.702799 | 0.706469 | 0.705864 |
| 0.70256 | 0.705957 | 0.706334 | 0.707011 | 0.710985 | 0.709279 | 0.703821 | 0.707211 | 0.706135 |
| 0.70584 | 0.709213 | 0.710903 | 0.711969 | 0.713353 | 0.714467 | 0.703789 | 0.708454 | 0.710745 |
| 0.702003 | 0.701642 | 0.70434 | 0.70646 | 0.707525 | 0.7057 | 0.699411 | 0.702565 | 0.70176 |
| 0.707943 | 0.705653 | 0.7122 | 0.71439 | 0.715253 | 0.711844 | 0.706161 | 0.709898 | 0.710265 |
| 0.711006 | 0.712781 | 0.713199 | 0.715604 | 0.718285 | 0.715564 | 0.707798 | 0.711108 | 0.713032 |
| 0.706822 | 0.71184 | 0.712515 | 0.711012 | 0.71344 | 0.716811 | 0.707008 | 0.710262 | 0.708973 |
| 0.714422 | 0.713593 | 0.715469 | 0.718553 | 0.717813 | 0.718368 | 0.711875 | 0.71257 | 0.716834 |

| | | | | | | | | |
|---|---|---|---|---|---|---|---|---|
| 0.774798 | 0.772772 | 0.763982 | 0.759768 | 0.762709 | 0.757768 | 0.771765 | 0.767392 | 0.760815 |
| 0.696212 | 0.70148 | 0.700673 | 0.701544 | 0.703861 | 0.705322 | 0.694669 | 0.7 | 0.701519 |
| 0.700254 | 0.704869 | 0.704695 | 0.707056 | 0.708007 | 0.708755 | 0.700734 | 0.705963 | 0.706496 |
| 0.708805 | 0.71301 | 0.713493 | 0.716585 | 0.717234 | 0.718941 | 0.710813 | 0.709714 | 0.712483 |
| 0.706302 | 0.709396 | 0.709293 | 0.709341 | 0.714274 | 0.712617 | 0.706337 | 0.707439 | 0.707204 |
| 0.713565 | 0.710264 | 0.715168 | 0.71395 | 0.716162 | 0.712951 | 0.708394 | 0.713821 | 0.712705 |
| 0.703466 | 0.703523 | 0.706315 | 0.709843 | 0.709581 | 0.708934 | 0.701326 | 0.705196 | 0.703063 |
| 0.708667 | 0.71352 | 0.710461 | 0.710384 | 0.715707 | 0.715476 | 0.706973 | 0.705615 | 0.711837 |
| 0.707676 | 0.709206 | 0.713152 | 0.713371 | 0.713432 | 0.714309 | 0.706964 | 0.70814 | 0.711258 |
| 0.709337 | 0.709075 | 0.711099 | 0.711881 | 0.713634 | 0.71491 | 0.709068 | 0.706577 | 0.710785 |
| 0.699769 | 0.702232 | 0.705422 | 0.705925 | 0.703551 | 0.707165 | 0.698345 | 0.703464 | 0.704472 |
| 0.699309 | 0.703548 | 0.706009 | 0.706527 | 0.706594 | 0.710102 | 0.69913 | 0.702989 | 0.706176 |
| 0.705195 | 0.706432 | 0.711795 | 0.713973 | 0.713535 | 0.710758 | 0.703055 | 0.706577 | 0.704283 |
| 0.703323 | 0.708243 | 0.707943 | 0.708856 | 0.708327 | 0.709275 | 0.700953 | 0.705204 | 0.70739 |
| 0.704825 | 0.70593 | 0.708008 | 0.710255 | 0.707709 | 0.711295 | 0.701569 | 0.705187 | 0.706249 |
| 0.702071 | 0.701697 | 0.70471 | 0.709063 | 0.708223 | 0.709602 | 0.700461 | 0.703341 | 0.705489 |
| 0.699047 | 0.703084 | 0.704206 | 0.704953 | 0.704823 | 0.706208 | 0.697316 | 0.702178 | 0.701825 |
| 0.70606 | 0.710432 | 0.712535 | 0.712174 | 0.714906 | 0.714388 | 0.706727 | 0.708068 | 0.71238 |
| 0.711839 | 0.711638 | 0.714914 | 0.713607 | 0.716288 | 0.717398 | 0.710169 | 0.711777 | 0.711049 |
| 0.70504 | 0.705831 | 0.708878 | 0.712881 | 0.711433 | 0.714205 | 0.705363 | 0.704049 | 0.707599 |
| 0.701958 | 0.702569 | 0.701761 | 0.707569 | 0.704006 | 0.70599 | 0.698378 | 0.701514 | 0.701904 |
| 0.699849 | 0.703598 | 0.704625 | 0.707054 | 0.709317 | 0.710717 | 0.701596 | 0.70176 | 0.704642 |
| 0.705245 | 0.706903 | 0.708627 | 0.707226 | 0.70867 | 0.711109 | 0.704852 | 0.705552 | 0.708104 |
| 0.713077 | 0.716547 | 0.716595 | 0.716572 | 0.718891 | 0.720766 | 0.711449 | 0.71417 | 0.713331 |
| 0.722728 | 0.721319 | 0.722817 | 0.72298 | 0.724943 | 0.72408 | 0.716959 | 0.720829 | 0.722312 |
| 0.761629 | 0.763293 | 0.760769 | 0.760734 | 0.768156 | 0.767096 | 0.761001 | 0.762688 | 0.76162 |
| 0.755589 | 0.764433 | 0.761995 | 0.762592 | 0.764754 | 0.765799 | 0.751982 | 0.763979 | 0.761294 |
| 0.763171 | 0.769256 | 0.763773 | 0.764995 | 0.769042 | 0.770031 | 0.760704 | 0.761949 | 0.765024 |
| 0.701635 | 0.706191 | 0.706018 | 0.707021 | 0.710791 | 0.708466 | 0.701514 | 0.707343 | 0.703463 |
| 0.6992 | 0.704328 | 0.70555 | 0.706145 | 0.709683 | 0.7089 | 0.699885 | 0.703891 | 0.705455 |
| 0.711831 | 0.713955 | 0.713475 | 0.717741 | 0.71966 | 0.716548 | 0.711236 | 0.710686 | 0.71605 |
| 0.709516 | 0.717812 | 0.716006 | 0.714881 | 0.718825 | 0.719311 | 0.711203 | 0.713431 | 0.715659 |
| 0.701849 | 0.705467 | 0.707237 | 0.713469 | 0.713278 | 0.714666 | 0.701662 | 0.705517 | 0.704389 |
| 0.706647 | 0.704115 | 0.709219 | 0.70951 | 0.712157 | 0.709752 | 0.703933 | 0.70697 | 0.708606 |
| 0.698247 | 0.703287 | 0.706738 | 0.707377 | 0.71183 | 0.712358 | 0.699217 | 0.704616 | 0.704791 |
| 0.70433 | 0.707678 | 0.708993 | 0.709425 | 0.713038 | 0.714297 | 0.702672 | 0.706646 | 0.707237 |
| 0.706314 | 0.707461 | 0.709994 | 0.708882 | 0.712662 | 0.713553 | 0.703525 | 0.704795 | 0.707406 |
| 0.717026 | 0.72001 | 0.719904 | 0.720932 | 0.730751 | 0.728141 | 0.715236 | 0.719548 | 0.720669 |
| 0.695111 | 0.697307 | 0.701159 | 0.702601 | 0.706728 | 0.706457 | 0.694555 | 0.696742 | 0.696139 |
| 0.704683 | 0.705595 | 0.706258 | 0.70978 | 0.714745 | 0.713609 | 0.703315 | 0.706717 | 0.7092 |
| 0.702 | 0.705813 | 0.70494 | 0.709639 | 0.713668 | 0.711759 | 0.700443 | 0.704835 | 0.705976 |
| 0.718795 | 0.723558 | 0.724066 | 0.724826 | 0.728881 | 0.728434 | 0.717708 | 0.723845 | 0.722375 |
| 0.703165 | 0.70686 | 0.71125 | 0.713039 | 0.715477 | 0.717614 | 0.701645 | 0.708252 | 0.705618 |
| 0.702311 | 0.706105 | 0.706747 | 0.710369 | 0.709308 | 0.711171 | 0.702504 | 0.706307 | 0.704974 |
| 0.706854 | 0.706853 | 0.713586 | 0.713129 | 0.714718 | 0.71297 | 0.706099 | 0.706392 | 0.710458 |
| 0.70409 | 0.705835 | 0.708022 | 0.709585 | 0.711063 | 0.712356 | 0.700836 | 0.703384 | 0.706453 |

| | | | | | | | | |
|---|---|---|---|---|---|---|---|---|
| 0.700739 | 0.704044 | 0.707864 | 0.709508 | 0.710828 | 0.712588 | 0.700048 | 0.704282 | 0.705594 |
| 0.711753 | 0.710407 | 0.713308 | 0.712509 | 0.716483 | 0.714898 | 0.709166 | 0.712813 | 0.714343 |
| 0.710535 | 0.708033 | 0.711941 | 0.710498 | 0.712849 | 0.712679 | 0.707641 | 0.709793 | 0.711196 |
| 0.709763 | 0.712065 | 0.71521 | 0.715265 | 0.723577 | 0.721701 | 0.707591 | 0.717719 | 0.712203 |
| 0.711566 | 0.711586 | 0.715545 | 0.714767 | 0.720945 | 0.7219 | 0.708383 | 0.715024 | 0.714766 |
| 0.707097 | 0.708063 | 0.708668 | 0.710759 | 0.714013 | 0.714584 | 0.708317 | 0.706864 | 0.710127 |
| 0.706988 | 0.708888 | 0.711705 | 0.713027 | 0.716925 | 0.720463 | 0.704091 | 0.711261 | 0.71044 |
| 0.709111 | 0.711424 | 0.712096 | 0.713207 | 0.71506 | 0.713588 | 0.705655 | 0.71037 | 0.709318 |
| 0.706377 | 0.707354 | 0.709773 | 0.711023 | 0.713033 | 0.713622 | 0.703196 | 0.709211 | 0.707187 |
| 0.705733 | 0.70385 | 0.705649 | 0.70816 | 0.706914 | 0.70997 | 0.702675 | 0.703081 | 0.708567 |
| 0.707246 | 0.704924 | 0.710248 | 0.709556 | 0.71414 | 0.713756 | 0.703145 | 0.706043 | 0.708108 |
| 0.704032 | 0.70592 | 0.709094 | 0.710196 | 0.714664 | 0.712627 | 0.703542 | 0.706036 | 0.706126 |
| 0.707093 | 0.708377 | 0.709574 | 0.713532 | 0.714991 | 0.711961 | 0.705254 | 0.70979 | 0.708162 |
| 0.704832 | 0.70371 | 0.702402 | 0.706094 | 0.709623 | 0.711314 | 0.701434 | 0.702444 | 0.706328 |
| 0.705488 | 0.70958 | 0.710606 | 0.713731 | 0.715799 | 0.713645 | 0.706988 | 0.710018 | 0.71062 |
| 0.704751 | 0.706907 | 0.70835 | 0.709685 | 0.70966 | 0.711406 | 0.702874 | 0.706095 | 0.706201 |
| 0.711807 | 0.711241 | 0.713746 | 0.712685 | 0.716392 | 0.716135 | 0.710953 | 0.711661 | 0.71306 |
| 0.702866 | 0.708599 | 0.709701 | 0.709836 | 0.712231 | 0.711173 | 0.703677 | 0.707665 | 0.706566 |
| 0.709136 | 0.705289 | 0.70978 | 0.709852 | 0.715399 | 0.712441 | 0.701446 | 0.705559 | 0.706019 |
| 0.707858 | 0.711242 | 0.711108 | 0.71018 | 0.709871 | 0.712999 | 0.705469 | 0.706948 | 0.70827 |
| 0.703337 | 0.707144 | 0.709499 | 0.710795 | 0.716103 | 0.714606 | 0.700648 | 0.706813 | 0.707729 |
| 0.700451 | 0.701821 | 0.70619 | 0.705802 | 0.709708 | 0.711 | 0.69717 | 0.701239 | 0.701934 |
| 0.708012 | 0.708587 | 0.713251 | 0.711381 | 0.715163 | 0.715468 | 0.707345 | 0.707535 | 0.709855 |
| 0.706568 | 0.708932 | 0.707846 | 0.70846 | 0.716464 | 0.716216 | 0.706557 | 0.709581 | 0.712943 |
| 0.708491 | 0.709929 | 0.711264 | 0.709843 | 0.713554 | 0.714205 | 0.705654 | 0.709764 | 0.705302 |
| 0.71071 | 0.71241 | 0.715487 | 0.715935 | 0.718545 | 0.717904 | 0.708115 | 0.714573 | 0.710258 |
| 0.706068 | 0.705952 | 0.711106 | 0.710179 | 0.715038 | 0.71605 | 0.704385 | 0.708548 | 0.706079 |
| 0.707276 | 0.710257 | 0.710928 | 0.71154 | 0.712607 | 0.711377 | 0.702782 | 0.710645 | 0.708243 |
| 0.708879 | 0.711798 | 0.714685 | 0.712267 | 0.715538 | 0.713752 | 0.706899 | 0.710499 | 0.711824 |
| 0.708151 | 0.708599 | 0.712003 | 0.71126 | 0.714997 | 0.717247 | 0.707058 | 0.71082 | 0.707605 |
| 0.710811 | 0.711549 | 0.71401 | 0.717945 | 0.717779 | 0.720354 | 0.706544 | 0.713388 | 0.712194 |
| 0.703085 | 0.707143 | 0.707409 | 0.709475 | 0.712645 | 0.712811 | 0.704102 | 0.706269 | 0.707818 |
| 0.707272 | 0.70575 | 0.706727 | 0.710374 | 0.715868 | 0.714459 | 0.703731 | 0.706508 | 0.707967 |
| 0.705033 | 0.709245 | 0.711605 | 0.713623 | 0.715879 | 0.717634 | 0.707766 | 0.706958 | 0.709247 |
| 0.704442 | 0.705831 | 0.70838 | 0.71257 | 0.713228 | 0.712648 | 0.701908 | 0.705037 | 0.708098 |
| 0.704248 | 0.706541 | 0.709195 | 0.712976 | 0.715171 | 0.716474 | 0.705115 | 0.707483 | 0.708537 |
| 0.700777 | 0.703614 | 0.706731 | 0.70578 | 0.713771 | 0.708378 | 0.700426 | 0.702564 | 0.70493 |
| 0.678387 | 0.68185 | 0.681224 | 0.682762 | 0.688824 | 0.686478 | 0.673447 | 0.677839 | 0.677852 |
| 0.684487 | 0.685902 | 0.685882 | 0.689316 | 0.693995 | 0.694464 | 0.685332 | 0.687786 | 0.691728 |
| 0.675146 | 0.672463 | 0.681299 | 0.68684 | 0.686379 | 0.686398 | 0.676206 | 0.678974 | 0.675542 |
| 0.671513 | 0.678627 | 0.678919 | 0.682247 | 0.683554 | 0.684515 | 0.67066 | 0.675887 | 0.676903 |
| 0.663539 | 0.660593 | 0.66123 | 0.664307 | 0.668101 | 0.666667 | 0.660207 | 0.664316 | 0.667805 |
| 0.66376 | 0.666344 | 0.66856 | 0.67378 | 0.670834 | 0.674815 | 0.661204 | 0.666991 | 0.667113 |
| 0.661572 | 0.664082 | 0.669622 | 0.666218 | 0.670573 | 0.668972 | 0.660139 | 0.662248 | 0.668861 |
| 0.662212 | 0.665733 | 0.669442 | 0.67214 | 0.676684 | 0.675163 | 0.661907 | 0.66833 | 0.664035 |
| 0.66061 | 0.663888 | 0.669834 | 0.66755 | 0.669605 | 0.671296 | 0.659489 | 0.664418 | 0.661533 |
| 0.664157 | 0.669595 | 0.669416 | 0.671002 | 0.676953 | 0.67114 | 0.665384 | 0.673113 | 0.670759 |

| | | | | | | | | |
|---|---|---|---|---|---|---|---|---|
| 0.668352 | 0.670243 | 0.673972 | 0.673495 | 0.67759 | 0.675716 | 0.669308 | 0.671258 | 0.664946 |
| 0.66532 | 0.673384 | 0.674294 | 0.673416 | 0.679886 | 0.677782 | 0.667453 | 0.670548 | 0.671936 |
| 0.663863 | 0.666089 | 0.672773 | 0.67474 | 0.677493 | 0.675781 | 0.663663 | 0.670467 | 0.67497 |
| 0.655513 | 0.661295 | 0.660037 | 0.666488 | 0.665835 | 0.667197 | 0.6554 | 0.664363 | 0.663926 |
| 0.660409 | 0.658026 | 0.663578 | 0.66502 | 0.669744 | 0.664865 | 0.663001 | 0.662021 | 0.666969 |
| 0.662038 | 0.661135 | 0.668542 | 0.664959 | 0.668345 | 0.664791 | 0.660377 | 0.663222 | 0.663518 |
| 0.660281 | 0.662283 | 0.669361 | 0.6729 | 0.671221 | 0.671824 | 0.657727 | 0.668611 | 0.670356 |
| 0.657924 | 0.661597 | 0.663765 | 0.665769 | 0.668155 | 0.668499 | 0.655839 | 0.666338 | 0.663333 |
| 0.663127 | 0.665292 | 0.661421 | 0.66622 | 0.668662 | 0.670265 | 0.659088 | 0.664944 | 0.66115 |
| 0.660604 | 0.660196 | 0.664473 | 0.67018 | 0.667005 | 0.666423 | 0.656054 | 0.664805 | 0.66334 |
| 0.653827 | 0.652864 | 0.661171 | 0.658438 | 0.663845 | 0.658102 | 0.652437 | 0.657353 | 0.654685 |
| 0.693144 | 0.694976 | 0.699976 | 0.702479 | 0.701213 | 0.703349 | 0.690499 | 0.694519 | 0.698189 |
| 0.697601 | 0.703096 | 0.709005 | 0.706342 | 0.709586 | 0.712941 | 0.700673 | 0.706529 | 0.702389 |
| 0.681393 | 0.684607 | 0.687893 | 0.687695 | 0.690895 | 0.688712 | 0.678576 | 0.685672 | 0.684577 |
| 0.714413 | 0.714632 | 0.715512 | 0.719226 | 0.722665 | 0.723194 | 0.712394 | 0.715599 | 0.71939 |
| 0.704167 | 0.701745 | 0.707689 | 0.707307 | 0.709239 | 0.710759 | 0.697284 | 0.704467 | 0.70387 |
| 0.707961 | 0.709657 | 0.710255 | 0.71377 | 0.712863 | 0.713076 | 0.706883 | 0.708086 | 0.712617 |
| 0.713757 | 0.718652 | 0.719919 | 0.720782 | 0.721516 | 0.720348 | 0.714679 | 0.716292 | 0.717231 |
| 0.693151 | 0.69309 | 0.694512 | 0.693744 | 0.701381 | 0.701862 | 0.687788 | 0.692631 | 0.691937 |
| 0.718884 | 0.716333 | 0.717957 | 0.716721 | 0.722476 | 0.723651 | 0.712787 | 0.716079 | 0.719635 |
| 0.686699 | 0.689069 | 0.686702 | 0.692804 | 0.693164 | 0.692818 | 0.683752 | 0.690349 | 0.687817 |
| 0.715305 | 0.717454 | 0.717471 | 0.719521 | 0.719723 | 0.717409 | 0.716069 | 0.714193 | 0.716419 |
| 0.718223 | 0.721522 | 0.723786 | 0.72356 | 0.723869 | 0.72159 | 0.716426 | 0.7189 | 0.718717 |
| 0.676836 | 0.68304 | 0.677599 | 0.683265 | 0.691101 | 0.688083 | 0.675148 | 0.682417 | 0.684609 |
| 0.683898 | 0.687515 | 0.685907 | 0.687594 | 0.691291 | 0.688619 | 0.681551 | 0.687531 | 0.686032 |
| 0.686066 | 0.689148 | 0.690865 | 0.691118 | 0.700126 | 0.69449 | 0.692818 | 0.687524 | 0.690537 |
| 0.708652 | 0.702291 | 0.708011 | 0.709954 | 0.710288 | 0.712168 | 0.700586 | 0.706962 | 0.708942 |
| 0.710046 | 0.714751 | 0.716448 | 0.717635 | 0.721017 | 0.720417 | 0.708979 | 0.713583 | 0.718041 |
| 0.682286 | 0.68625 | 0.687321 | 0.689346 | 0.694056 | 0.693589 | 0.681991 | 0.685061 | 0.683587 |
| 0.721592 | 0.721236 | 0.723057 | 0.723396 | 0.728419 | 0.730616 | 0.718091 | 0.722365 | 0.721582 |
| 0.72302 | 0.725209 | 0.723814 | 0.724 | 0.730014 | 0.727065 | 0.721401 | 0.722009 | 0.72466 |
| 0.691089 | 0.694482 | 0.698023 | 0.700926 | 0.705594 | 0.703904 | 0.689998 | 0.696086 | 0.695336 |
| 0.693381 | 0.696001 | 0.69651 | 0.699202 | 0.703579 | 0.705088 | 0.692253 | 0.696744 | 0.696156 |
| 0.718476 | 0.722968 | 0.726828 | 0.723776 | 0.728571 | 0.734018 | 0.715667 | 0.722808 | 0.722352 |
| 0.71498 | 0.721413 | 0.718799 | 0.720222 | 0.724895 | 0.725529 | 0.717047 | 0.716862 | 0.718725 |
| 0.716625 | 0.718818 | 0.723242 | 0.723692 | 0.73124 | 0.730878 | 0.718267 | 0.721515 | 0.724945 |
| 0.674737 | 0.67752 | 0.679931 | 0.679905 | 0.681195 | 0.681645 | 0.671581 | 0.67602 | 0.676431 |
| 0.718596 | 0.716694 | 0.718392 | 0.721413 | 0.729822 | 0.726838 | 0.716794 | 0.720003 | 0.719544 |
| 0.675159 | 0.673526 | 0.675383 | 0.678336 | 0.679766 | 0.680457 | 0.670428 | 0.676276 | 0.669667 |
| 0.670926 | 0.666918 | 0.670993 | 0.669168 | 0.673872 | 0.671945 | 0.664665 | 0.667154 | 0.665853 |
| 0.731101 | 0.726051 | 0.725738 | 0.724357 | 0.738757 | 0.730089 | 0.729095 | 0.730588 | 0.729029 |
| 0.669075 | 0.670961 | 0.671474 | 0.670579 | 0.675825 | 0.677331 | 0.665753 | 0.670051 | 0.669137 |
| 0.65728 | 0.659707 | 0.66288 | 0.662849 | 0.668212 | 0.668725 | 0.654908 | 0.662289 | 0.662041 |
| 0.657611 | 0.657871 | 0.661135 | 0.661872 | 0.663273 | 0.666156 | 0.654392 | 0.658929 | 0.656784 |
| 0.740529 | 0.733456 | 0.738309 | 0.73854 | 0.74239 | 0.738372 | 0.73991 | 0.741403 | 0.732456 |
| 0.657076 | 0.66294 | 0.662431 | 0.6634 | 0.664429 | 0.667897 | 0.656436 | 0.659714 | 0.65976 |
| 0.731656 | 0.725056 | 0.733479 | 0.726019 | 0.738688 | 0.732974 | 0.72602 | 0.731456 | 0.725932 |

| | | | | | | | | |
|---|---|---|---|---|---|---|---|---|
| 0.748853 | 0.740364 | 0.740497 | 0.736578 | 0.749321 | 0.74064 | 0.748504 | 0.746664 | 0.740477 |
| 0.739683 | 0.729094 | 0.736728 | 0.733755 | 0.739401 | 0.736383 | 0.735963 | 0.737075 | 0.734061 |
| 0.737817 | 0.747042 | 0.743686 | 0.74812 | 0.751806 | 0.752874 | 0.737864 | 0.744563 | 0.745888 |
| 0.648726 | 0.651786 | 0.652904 | 0.656798 | 0.654848 | 0.657156 | 0.64784 | 0.652014 | 0.651704 |
| 0.650236 | 0.650667 | 0.65006 | 0.65431 | 0.659676 | 0.658214 | 0.645913 | 0.653456 | 0.652335 |
| 0.743658 | 0.741093 | 0.736846 | 0.736081 | 0.75014 | 0.739462 | 0.742769 | 0.738232 | 0.738051 |
| 0.649473 | 0.654967 | 0.654117 | 0.655141 | 0.658209 | 0.661396 | 0.649274 | 0.650745 | 0.653843 |
| 0.726763 | 0.732916 | 0.733611 | 0.732933 | 0.73955 | 0.739662 | 0.724534 | 0.730367 | 0.729393 |
| 0.749288 | 0.737289 | 0.746402 | 0.745338 | 0.752807 | 0.74169 | 0.742025 | 0.743791 | 0.734315 |
| 0.731851 | 0.734902 | 0.737982 | 0.736929 | 0.741326 | 0.735823 | 0.732632 | 0.734065 | 0.733141 |
| 0.75842 | 0.745727 | 0.74743 | 0.741474 | 0.752735 | 0.747987 | 0.752511 | 0.745079 | 0.746093 |
| 0.707006 | 0.712418 | 0.707006 | 0.701299 | 0.685897 | 0.758621 | 0.708861 | 0.710526 | 0.666667 |
| 0.718335 | 0.714058 | 0.717687 | 0.71697 | 0.72915 | 0.722568 | 0.707487 | 0.722991 | 0.717008 |
| 0.71323 | 0.712768 | 0.718343 | 0.718834 | 0.728981 | 0.722275 | 0.710794 | 0.71776 | 0.713391 |
| 0.719245 | 0.72285 | 0.724185 | 0.723918 | 0.728755 | 0.730478 | 0.717378 | 0.7322 | 0.719598 |
| 0.715933 | 0.723764 | 0.725037 | 0.722509 | 0.729404 | 0.730142 | 0.718259 | 0.724036 | 0.716555 |
| 0.588815 | 0.588369 | 0.590535 | 0.5954 | 0.597087 | 0.600696 | 0.588363 | 0.590212 | 0.587955 |
| 0.731063 | 0.731787 | 0.732122 | 0.728026 | 0.73594 | 0.732064 | 0.728455 | 0.727785 | 0.723225 |
| 0.773334 | 0.759242 | 0.75621 | 0.747742 | 0.761652 | 0.752559 | 0.769506 | 0.753944 | 0.750165 |
| 0.768928 | 0.759406 | 0.761738 | 0.750649 | 0.760627 | 0.751991 | 0.759699 | 0.763484 | 0.749352 |
| 0.769154 | 0.758985 | 0.752343 | 0.747125 | 0.759726 | 0.754822 | 0.763488 | 0.752609 | 0.751801 |
| 0.594665 | 0.594198 | 0.594194 | 0.597986 | 0.597436 | 0.597819 | 0.590707 | 0.5931 | 0.59341 |
| 0.58128 | 0.582704 | 0.585631 | 0.585802 | 0.586712 | 0.590417 | 0.577995 | 0.581137 | 0.580942 |
| 0.831164 | 0.819139 | 0.802413 | 0.804538 | 0.85671 | 0.806263 | 0.854796 | 0.809862 | 0.822163 |
| 0.852992 | 0.810107 | 0.809424 | 0.806903 | 0.885384 | 0.807535 | 0.889016 | 0.816047 | 0.799444 |
| 0.799797 | 0.812903 | 0.80707 | 0.806488 | 0.80129 | 0.832192 | 0.802939 | 0.785909 | 0.812217 |
| 0.799716 | 0.815703 | 0.816067 | 0.800321 | 0.800695 | 0.822187 | 0.801935 | 0.794516 | 0.799562 |
| 0.806325 | 0.82627 | 0.81199 | 0.794805 | 0.827375 | 0.814232 | 0.815438 | 0.822752 | 0.801787 |
| 0.798478 | 0.805151 | 0.799306 | 0.784001 | 0.815752 | 0.804304 | 0.816454 | 0.799038 | 0.801056 |
| 0.791656 | 0.789925 | 0.797202 | 0.801134 | 0.841749 | 0.814042 | 0.80044 | 0.812013 | 0.794878 |
| 0.755516 | 0.753861 | 0.761127 | 0.76231 | 0.774362 | 0.769304 | 0.751865 | 0.76193 | 0.753693 |
| 0.765392 | 0.767461 | 0.768438 | 0.772069 | 0.775809 | 0.778241 | 0.764081 | 0.77027 | 0.765767 |
| 0.760017 | 0.764126 | 0.761692 | 0.762885 | 0.768554 | 0.769717 | 0.752442 | 0.760514 | 0.759357 |
| 0.760407 | 0.762303 | 0.764586 | 0.76595 | 0.770816 | 0.767705 | 0.757154 | 0.760683 | 0.7593 |
| 0.765644 | 0.766921 | 0.771267 | 0.772629 | 0.77145 | 0.776169 | 0.763364 | 0.766481 | 0.766866 |
| 0.756088 | 0.756963 | 0.762101 | 0.764559 | 0.771954 | 0.771506 | 0.756075 | 0.76637 | 0.759648 |
| 0.767794 | 0.769891 | 0.767901 | 0.77026 | 0.776362 | 0.773722 | 0.763991 | 0.765363 | 0.766575 |
| 0.76612 | 0.763212 | 0.769484 | 0.77013 | 0.778505 | 0.774154 | 0.761846 | 0.771145 | 0.765231 |
| 0.754858 | 0.760013 | 0.762178 | 0.761113 | 0.77207 | 0.770102 | 0.755721 | 0.764004 | 0.75484 |
| 0.760546 | 0.762421 | 0.764691 | 0.766951 | 0.774091 | 0.768475 | 0.755746 | 0.767105 | 0.757314 |
| 0.718936 | 0.721764 | 0.723993 | 0.722665 | 0.732286 | 0.727 | 0.718772 | 0.722495 | 0.715132 |
| 0.72402 | 0.727788 | 0.725372 | 0.729119 | 0.734349 | 0.733422 | 0.717056 | 0.732525 | 0.718339 |
| 0.760522 | 0.766762 | 0.765197 | 0.764356 | 0.773852 | 0.770511 | 0.760312 | 0.763136 | 0.761329 |
| 0.78636 | 0.786982 | 0.784179 | 0.782187 | 0.789994 | 0.790524 | 0.788947 | 0.782217 | 0.779843 |
| 0.718853 | 0.722763 | 0.723235 | 0.725671 | 0.732972 | 0.730986 | 0.708947 | 0.729751 | 0.717152 |
| 0.761936 | 0.769998 | 0.768927 | 0.77092 | 0.773701 | 0.778301 | 0.75601 | 0.767273 | 0.767492 |
| 0.758683 | 0.76206 | 0.764754 | 0.765519 | 0.772416 | 0.77 | 0.752635 | 0.760325 | 0.758706 |

| | | | | | | | | |
|---|---|---|---|---|---|---|---|---|
| 0.76096 | 0.768687 | 0.770917 | 0.773615 | 0.778133 | 0.77753 | 0.759086 | 0.767387 | 0.764102 |
| 0.76194 | 0.768722 | 0.769848 | 0.773258 | 0.776339 | 0.778252 | 0.763311 | 0.767085 | 0.766371 |
| 0.758557 | 0.76374 | 0.76614 | 0.765486 | 0.776002 | 0.773428 | 0.754796 | 0.761448 | 0.758522 |
| 0.765102 | 0.767022 | 0.7706 | 0.774331 | 0.779307 | 0.776048 | 0.76474 | 0.771283 | 0.768231 |
| 0.766492 | 0.768235 | 0.766817 | 0.766476 | 0.777216 | 0.776534 | 0.763238 | 0.768557 | 0.7678 |
| 0.762003 | 0.765108 | 0.766153 | 0.767672 | 0.773251 | 0.772517 | 0.759275 | 0.764805 | 0.76386 |
| 0.760578 | 0.76634 | 0.766102 | 0.769186 | 0.775308 | 0.773868 | 0.759662 | 0.767038 | 0.767733 |
| 0.749629 | 0.756309 | 0.757784 | 0.760124 | 0.763603 | 0.764569 | 0.751119 | 0.751111 | 0.752663 |
| 0.762116 | 0.764097 | 0.770881 | 0.772435 | 0.774312 | 0.774295 | 0.760149 | 0.768105 | 0.764531 |
| 0.756386 | 0.759714 | 0.76862 | 0.763858 | 0.771501 | 0.767016 | 0.755331 | 0.75982 | 0.758903 |
| 0.765298 | 0.766533 | 0.771555 | 0.7715 | 0.77717 | 0.773454 | 0.761156 | 0.766963 | 0.768362 |
| 0.760748 | 0.772322 | 0.767696 | 0.770202 | 0.773138 | 0.774962 | 0.761302 | 0.764095 | 0.762168 |
| 0.752606 | 0.756551 | 0.75923 | 0.758933 | 0.765692 | 0.75993 | 0.747639 | 0.754231 | 0.753563 |
| 0.753354 | 0.757076 | 0.762129 | 0.762381 | 0.766948 | 0.764994 | 0.751502 | 0.756176 | 0.753327 |
| 0.761521 | 0.765195 | 0.771394 | 0.77158 | 0.77917 | 0.778878 | 0.758967 | 0.768068 | 0.764856 |
| 0.813801 | 0.81077 | 0.82095 | 0.815192 | 0.817431 | 0.812249 | 0.800754 | 0.818555 | 0.795519 |
| 0.794174 | 0.802635 | 0.822813 | 0.822743 | 0.826181 | 0.856292 | 0.797758 | 0.804313 | 0.828211 |
| 0.814583 | 0.819214 | 0.852276 | 0.861467 | 0.837257 | 0.853673 | 0.797976 | 0.82399 | 0.813989 |
| 0.825591 | 0.845271 | 0.821191 | 0.810008 | 0.831083 | 0.805818 | 0.826065 | 0.827539 | 0.81714 |
| 0.812897 | 0.807624 | 0.828491 | 0.828958 | 0.823708 | 0.817468 | 0.799409 | 0.836356 | 0.797784 |
| 0.835417 | 0.809503 | 0.847316 | 0.835932 | 0.821151 | 0.821241 | 0.795534 | 0.836303 | 0.800636 |
| 0.807376 | 0.823347 | 0.800324 | 0.79084 | 0.816868 | 0.799813 | 0.807312 | 0.819087 | 0.80441 |
| 0.801698 | 0.827296 | 0.8065 | 0.797806 | 0.806784 | 0.81863 | 0.803 | 0.809879 | 0.800084 |
| 0.800335 | 0.826043 | 0.820933 | 0.835458 | 0.830404 | 0.881021 | 0.793366 | 0.805069 | 0.831425 |
| 0.817372 | 0.808248 | 0.83108 | 0.831521 | 0.835413 | 0.813145 | 0.807952 | 0.850339 | 0.804151 |
| 0.815593 | 0.8251 | 0.804381 | 0.794245 | 0.801973 | 0.790051 | 0.823035 | 0.803329 | 0.80252 |
| 0.811308 | 0.818384 | 0.826974 | 0.873103 | 0.841418 | 0.83506 | 0.7947 | 0.832575 | 0.814445 |
| 0.810112 | 0.848532 | 0.823758 | 0.797709 | 0.788822 | 0.836125 | 0.808708 | 0.792868 | 0.816535 |
| 1 | 0.822337 | 0.830094 | 0.819609 | 0.815346 | 0.797361 | 0.878888 | 0.825157 | 0.817587 |
| 0.822337 | 1 | 0.826146 | 0.804329 | 0.807449 | 0.817524 | 0.82297 | 0.798204 | 0.817271 |
| 0.830094 | 0.826146 | 1 | 0.86215 | 0.827839 | 0.824475 | 0.805218 | 0.835581 | 0.813965 |
| 0.819609 | 0.804329 | 0.86215 | 1 | 0.83822 | 0.833519 | 0.80017 | 0.832593 | 0.818268 |
| 0.815346 | 0.807449 | 0.827839 | 0.83822 | 1 | 0.838539 | 0.822717 | 0.846855 | 0.817033 |
| 0.797361 | 0.817524 | 0.824475 | 0.833519 | 0.838539 | 1 | 0.792409 | 0.811934 | 0.825976 |
| 0.878888 | 0.82297 | 0.805218 | 0.80017 | 0.822717 | 0.792409 | 1 | 0.808943 | 0.816963 |
| 0.825157 | 0.798204 | 0.835581 | 0.832593 | 0.846855 | 0.811934 | 0.808943 | 1 | 0.804587 |
| 0.817587 | 0.817271 | 0.813965 | 0.818268 | 0.817033 | 0.825976 | 0.816963 | 0.804587 | 1 |
| 0.795925 | 0.819956 | 0.809851 | 0.795106 | 0.805979 | 0.799848 | 0.795972 | 0.796158 | 0.803997 |
| 0.842978 | 0.81146 | 0.869615 | 0.858041 | 0.827899 | 0.811198 | 0.79803 | 0.844205 | 0.80574 |
| 0.795305 | 0.813758 | 0.800176 | 0.793972 | 0.794253 | 0.808415 | 0.803079 | 0.785879 | 0.831221 |
| 0.840049 | 0.809014 | 0.828411 | 0.811427 | 0.795862 | 0.778254 | 0.811848 | 0.803496 | 0.790275 |
| 0.805127 | 0.820797 | 0.808734 | 0.787914 | 0.799699 | 0.79655 | 0.81684 | 0.805722 | 0.793827 |
| 0.815224 | 0.803256 | 0.840611 | 0.840884 | 0.830368 | 0.822116 | 0.792534 | 0.82683 | 0.832901 |
| 0.79071 | 0.824028 | 0.800221 | 0.790049 | 0.809029 | 0.797019 | 0.792318 | 0.805535 | 0.799313 |
| 0.787597 | 0.80891 | 0.814211 | 0.821422 | 0.812736 | 0.827276 | 0.792627 | 0.797387 | 0.804744 |
| 0.779478 | 0.788811 | 0.809103 | 0.841917 | 0.827831 | 0.848094 | 0.774258 | 0.798286 | 0.814435 |

| | | | | | | | | |
|---|---|---|---|---|---|---|---|---|
| 0.852645 | 0.803571 | 0.812089 | 0.796495 | 0.804321 | 0.783418 | 0.834304 | 0.813971 | 0.802439 |
| 0.763154 | 0.765511 | 0.769982 | 0.769341 | 0.771568 | 0.770462 | 0.760533 | 0.767473 | 0.76632 |
| 0.765359 | 0.761849 | 0.764848 | 0.764067 | 0.766755 | 0.766536 | 0.76173 | 0.761555 | 0.762272 |
| 0.757369 | 0.764179 | 0.764188 | 0.764541 | 0.770315 | 0.76903 | 0.757993 | 0.766512 | 0.764425 |
| 0.766595 | 0.772354 | 0.77421 | 0.771312 | 0.778904 | 0.780892 | 0.767259 | 0.770034 | 0.767841 |
| 0.76683 | 0.764341 | 0.771508 | 0.767912 | 0.776463 | 0.772029 | 0.759731 | 0.76943 | 0.761709 |
| 0.762605 | 0.767243 | 0.76803 | 0.767327 | 0.769048 | 0.773254 | 0.760302 | 0.76174 | 0.762414 |
| 0.761188 | 0.768493 | 0.766932 | 0.770068 | 0.770541 | 0.773756 | 0.758666 | 0.762654 | 0.761881 |
| 0.755617 | 0.756713 | 0.763799 | 0.761632 | 0.764369 | 0.764674 | 0.750944 | 0.759062 | 0.753811 |
| 0.756076 | 0.759859 | 0.759635 | 0.767519 | 0.772332 | 0.776693 | 0.753353 | 0.760037 | 0.757498 |
| 0.76382 | 0.76398 | 0.768202 | 0.771268 | 0.776966 | 0.774894 | 0.761154 | 0.768708 | 0.763727 |
| 0.75859 | 0.76838 | 0.769123 | 0.770954 | 0.773946 | 0.776187 | 0.759037 | 0.769712 | 0.764016 |
| 0.762927 | 0.767904 | 0.765222 | 0.765709 | 0.769275 | 0.770254 | 0.762995 | 0.760136 | 0.765484 |
| 0.761452 | 0.763055 | 0.762953 | 0.763682 | 0.766447 | 0.767173 | 0.759352 | 0.763799 | 0.761182 |
| 0.763025 | 0.76861 | 0.768264 | 0.769148 | 0.769978 | 0.769093 | 0.760222 | 0.763465 | 0.765049 |
| 0.763968 | 0.767907 | 0.764093 | 0.765331 | 0.768261 | 0.768695 | 0.763364 | 0.757247 | 0.764929 |
| 0.76197 | 0.763656 | 0.765549 | 0.764001 | 0.769336 | 0.769349 | 0.762165 | 0.763821 | 0.767522 |
| 0.758267 | 0.763507 | 0.76263 | 0.762407 | 0.769825 | 0.768049 | 0.761248 | 0.763795 | 0.765061 |
| 0.768536 | 0.773602 | 0.770272 | 0.773861 | 0.7694 | 0.770194 | 0.765218 | 0.768476 | 0.772011 |
| 0.767594 | 0.765573 | 0.76847 | 0.777085 | 0.770279 | 0.773574 | 0.762204 | 0.769973 | 0.762272 |
| 0.759694 | 0.762579 | 0.763353 | 0.764642 | 0.768934 | 0.764812 | 0.756224 | 0.76545 | 0.761798 |
| 0.760488 | 0.766744 | 0.765769 | 0.766441 | 0.768921 | 0.768163 | 0.762287 | 0.764617 | 0.766714 |
| 0.768526 | 0.766376 | 0.766732 | 0.765317 | 0.770847 | 0.769992 | 0.76906 | 0.768617 | 0.769358 |
| 0.758803 | 0.760866 | 0.764132 | 0.767166 | 0.767647 | 0.765734 | 0.76281 | 0.764901 | 0.765358 |
| 0.76021 | 0.767695 | 0.764687 | 0.763903 | 0.767853 | 0.771171 | 0.75821 | 0.763753 | 0.768556 |
| 0.761831 | 0.764215 | 0.760822 | 0.761103 | 0.767274 | 0.760933 | 0.75807 | 0.764293 | 0.762585 |
| 0.756208 | 0.758436 | 0.760557 | 0.757571 | 0.763315 | 0.761264 | 0.758652 | 0.760819 | 0.758462 |
| 0.759246 | 0.763447 | 0.763767 | 0.766126 | 0.768123 | 0.765696 | 0.758216 | 0.760321 | 0.76059 |
| 0.758968 | 0.765772 | 0.771808 | 0.767895 | 0.774538 | 0.770333 | 0.761022 | 0.769844 | 0.765207 |
| 0.762118 | 0.763362 | 0.767493 | 0.762972 | 0.770237 | 0.77126 | 0.759632 | 0.763911 | 0.766091 |
| 0.75666 | 0.758192 | 0.75993 | 0.757543 | 0.75999 | 0.756241 | 0.753575 | 0.754768 | 0.758355 |
| 0.764269 | 0.766408 | 0.767242 | 0.768324 | 0.772503 | 0.768577 | 0.761365 | 0.768946 | 0.765686 |
| 0.762531 | 0.765186 | 0.768864 | 0.766361 | 0.771726 | 0.765589 | 0.759544 | 0.766251 | 0.762758 |
| 0.761836 | 0.757972 | 0.763269 | 0.758279 | 0.765954 | 0.764333 | 0.757506 | 0.758627 | 0.759626 |
| 0.75937 | 0.756636 | 0.762294 | 0.760801 | 0.766237 | 0.765538 | 0.760836 | 0.760082 | 0.76366 |
| 0.757833 | 0.764964 | 0.762068 | 0.763903 | 0.768276 | 0.764276 | 0.759603 | 0.758227 | 0.759299 |
| 0.767085 | 0.768897 | 0.771626 | 0.772094 | 0.776334 | 0.772366 | 0.767944 | 0.771684 | 0.767783 |
| 0.763351 | 0.760754 | 0.760233 | 0.761268 | 0.771134 | 0.768068 | 0.763434 | 0.760014 | 0.761149 |
| 0.757281 | 0.766522 | 0.765697 | 0.766412 | 0.768195 | 0.766174 | 0.758799 | 0.762987 | 0.761315 |
| 0.764356 | 0.765345 | 0.768902 | 0.763498 | 0.768222 | 0.762413 | 0.759135 | 0.764766 | 0.759936 |
| 0.764691 | 0.768439 | 0.769092 | 0.762653 | 0.76963 | 0.764739 | 0.760458 | 0.763225 | 0.764334 |
| 0.762697 | 0.768316 | 0.768177 | 0.765772 | 0.77259 | 0.772216 | 0.764847 | 0.763971 | 0.762522 |
| 0.763175 | 0.7712 | 0.771431 | 0.767413 | 0.770877 | 0.771941 | 0.767246 | 0.768899 | 0.768346 |
| 0.758436 | 0.764553 | 0.762685 | 0.761514 | 0.766772 | 0.764642 | 0.760275 | 0.762079 | 0.758184 |
| 0.765566 | 0.768132 | 0.764346 | 0.76476 | 0.768154 | 0.76781 | 0.761195 | 0.765834 | 0.767277 |
| 0.759077 | 0.768047 | 0.76108 | 0.759768 | 0.765086 | 0.768227 | 0.757647 | 0.761748 | 0.762261 |

| | | | | | | | | |
|---|---|---|---|---|---|---|---|---|
| 0.751275 | 0.750354 | 0.753906 | 0.754436 | 0.757986 | 0.754658 | 0.74721 | 0.752027 | 0.749149 |
| 0.741051 | 0.74273 | 0.745592 | 0.739674 | 0.747125 | 0.749404 | 0.742593 | 0.746702 | 0.744209 |
| 0.755375 | 0.762939 | 0.760087 | 0.758425 | 0.766708 | 0.76649 | 0.755539 | 0.757444 | 0.761816 |
| 0.759789 | 0.763693 | 0.766044 | 0.759237 | 0.76887 | 0.76713 | 0.758188 | 0.767122 | 0.762166 |
| 0.752969 | 0.757529 | 0.757329 | 0.754045 | 0.761132 | 0.760109 | 0.749257 | 0.753436 | 0.757106 |
| 0.759464 | 0.759593 | 0.762438 | 0.755937 | 0.765996 | 0.762699 | 0.758421 | 0.762101 | 0.761615 |
| 0.760107 | 0.760498 | 0.757589 | 0.760505 | 0.765339 | 0.762964 | 0.757062 | 0.758118 | 0.758954 |
| 0.751745 | 0.757819 | 0.756289 | 0.754942 | 0.758727 | 0.761017 | 0.750686 | 0.751001 | 0.757587 |
| 0.766304 | 0.768073 | 0.765585 | 0.76614 | 0.769156 | 0.769086 | 0.762704 | 0.766819 | 0.764372 |
| 0.763674 | 0.769326 | 0.764178 | 0.765358 | 0.765379 | 0.769662 | 0.760733 | 0.766781 | 0.768604 |
| 0.763407 | 0.77052 | 0.768289 | 0.763091 | 0.767406 | 0.775637 | 0.766585 | 0.762702 | 0.763879 |
| 0.761466 | 0.767423 | 0.764498 | 0.763147 | 0.77181 | 0.769998 | 0.757102 | 0.761869 | 0.759252 |
| 0.767436 | 0.769982 | 0.767741 | 0.766802 | 0.772215 | 0.772735 | 0.767079 | 0.768768 | 0.768697 |
| 0.757029 | 0.767102 | 0.76935 | 0.765668 | 0.770117 | 0.771067 | 0.755889 | 0.765023 | 0.766052 |
| 0.760677 | 0.765806 | 0.763564 | 0.761539 | 0.765538 | 0.770453 | 0.759742 | 0.759814 | 0.762481 |
| 0.757429 | 0.763937 | 0.762158 | 0.762142 | 0.768552 | 0.771796 | 0.756088 | 0.762903 | 0.759292 |
| 0.764487 | 0.768552 | 0.767216 | 0.767039 | 0.771115 | 0.763889 | 0.756741 | 0.765913 | 0.761348 |
| 0.756274 | 0.763648 | 0.765678 | 0.762525 | 0.765741 | 0.764289 | 0.754123 | 0.761338 | 0.757171 |
| 0.757414 | 0.760442 | 0.761007 | 0.760458 | 0.769046 | 0.766628 | 0.755416 | 0.763391 | 0.753502 |
| 0.76176 | 0.766467 | 0.758746 | 0.760418 | 0.763165 | 0.769318 | 0.760017 | 0.762257 | 0.758254 |
| 0.759479 | 0.763784 | 0.763952 | 0.760033 | 0.764076 | 0.765843 | 0.759874 | 0.757047 | 0.759159 |
| 0.749916 | 0.749601 | 0.753765 | 0.749745 | 0.756104 | 0.756297 | 0.744333 | 0.752965 | 0.751122 |
| 0.757274 | 0.75652 | 0.75948 | 0.75592 | 0.761094 | 0.759935 | 0.752131 | 0.758099 | 0.75812 |
| 0.756188 | 0.762344 | 0.762676 | 0.760944 | 0.766015 | 0.767538 | 0.757053 | 0.761218 | 0.758969 |
| 0.762481 | 0.764223 | 0.769483 | 0.766766 | 0.768685 | 0.76691 | 0.763311 | 0.761189 | 0.761961 |
| 0.759434 | 0.757259 | 0.76395 | 0.75956 | 0.76509 | 0.762269 | 0.756134 | 0.760892 | 0.757867 |
| 0.756933 | 0.761937 | 0.761468 | 0.75855 | 0.766203 | 0.760998 | 0.756095 | 0.755517 | 0.756895 |
| 0.760072 | 0.763576 | 0.761264 | 0.761252 | 0.768826 | 0.762721 | 0.757316 | 0.760599 | 0.759377 |
| 0.75559 | 0.756666 | 0.764031 | 0.758569 | 0.766644 | 0.759761 | 0.753952 | 0.756106 | 0.752908 |
| 0.692824 | 0.695333 | 0.69705 | 0.699357 | 0.699753 | 0.697039 | 0.692811 | 0.696741 | 0.695584 |
| 0.704768 | 0.704799 | 0.704073 | 0.707301 | 0.709325 | 0.708865 | 0.701615 | 0.704362 | 0.706873 |
| 0.704282 | 0.702257 | 0.704812 | 0.706989 | 0.708821 | 0.708166 | 0.702761 | 0.704682 | 0.705706 |
| 0.694978 | 0.693944 | 0.700186 | 0.703372 | 0.703073 | 0.698669 | 0.693758 | 0.696573 | 0.697007 |
| 0.704283 | 0.703756 | 0.710079 | 0.709615 | 0.711783 | 0.709262 | 0.702313 | 0.707352 | 0.705826 |
| 0.750717 | 0.757612 | 0.762584 | 0.76135 | 0.765816 | 0.767244 | 0.752349 | 0.75887 | 0.753188 |
| 0.755647 | 0.761758 | 0.763907 | 0.761853 | 0.765973 | 0.767378 | 0.758632 | 0.759518 | 0.75992 |
| 0.760189 | 0.76368 | 0.767678 | 0.76585 | 0.769898 | 0.766999 | 0.761547 | 0.765471 | 0.764928 |
| 0.756854 | 0.763978 | 0.763371 | 0.764285 | 0.767678 | 0.770476 | 0.758495 | 0.761378 | 0.75992 |
| 0.75325 | 0.75847 | 0.758156 | 0.759106 | 0.766048 | 0.767715 | 0.754798 | 0.76006 | 0.758179 |
| 0.761379 | 0.770275 | 0.762442 | 0.763392 | 0.77322 | 0.772528 | 0.760864 | 0.76355 | 0.762696 |
| 0.759777 | 0.768513 | 0.767769 | 0.764661 | 0.769791 | 0.768404 | 0.759097 | 0.765009 | 0.76216 |
| 0.75858 | 0.762931 | 0.765169 | 0.75838 | 0.766642 | 0.764234 | 0.758904 | 0.7598 | 0.760553 |
| 0.753632 | 0.755199 | 0.757271 | 0.758077 | 0.760796 | 0.7609 | 0.753066 | 0.752856 | 0.755877 |
| 0.754918 | 0.758328 | 0.759389 | 0.758695 | 0.760652 | 0.762082 | 0.752765 | 0.753058 | 0.753156 |
| 0.754286 | 0.756582 | 0.762247 | 0.759718 | 0.765218 | 0.765421 | 0.75722 | 0.756611 | 0.755631 |
| 0.757151 | 0.763874 | 0.764681 | 0.76378 | 0.770678 | 0.769661 | 0.759633 | 0.766342 | 0.760388 |

| | | | | | | | | |
|---|---|---|---|---|---|---|---|---|
| 0.763903 | 0.763795 | 0.767779 | 0.76725 | 0.769946 | 0.769982 | 0.760893 | 0.766921 | 0.766004 |
| 0.76137 | 0.758995 | 0.76475 | 0.760737 | 0.765197 | 0.765085 | 0.758418 | 0.761673 | 0.759337 |
| 0.762965 | 0.760992 | 0.765911 | 0.764736 | 0.767857 | 0.765899 | 0.763468 | 0.765736 | 0.764052 |
| 0.756368 | 0.758482 | 0.761237 | 0.761282 | 0.766814 | 0.767269 | 0.758136 | 0.761052 | 0.761984 |
| 0.758163 | 0.758018 | 0.760323 | 0.757948 | 0.76142 | 0.762782 | 0.756085 | 0.758025 | 0.759159 |
| 0.756486 | 0.754864 | 0.761138 | 0.758544 | 0.762929 | 0.765357 | 0.755357 | 0.758606 | 0.76015 |
| 0.758948 | 0.755727 | 0.759195 | 0.75903 | 0.76954 | 0.768686 | 0.762199 | 0.760903 | 0.764671 |
| 0.757476 | 0.764533 | 0.763946 | 0.763799 | 0.768399 | 0.76542 | 0.761393 | 0.763794 | 0.761194 |
| 0.754758 | 0.764255 | 0.758878 | 0.757956 | 0.75834 | 0.758525 | 0.754517 | 0.755444 | 0.75513 |
| 0.760229 | 0.759215 | 0.763899 | 0.758759 | 0.76518 | 0.760542 | 0.759359 | 0.761051 | 0.754701 |
| 0.761829 | 0.761535 | 0.762207 | 0.760421 | 0.766686 | 0.76809 | 0.75918 | 0.760037 | 0.761949 |
| 0.762307 | 0.762635 | 0.764039 | 0.766582 | 0.766561 | 0.767746 | 0.760624 | 0.767079 | 0.762593 |
| 0.753864 | 0.755194 | 0.755593 | 0.755599 | 0.760279 | 0.757587 | 0.752648 | 0.758323 | 0.755175 |
| 0.755078 | 0.754213 | 0.75837 | 0.756104 | 0.761325 | 0.761464 | 0.75093 | 0.754862 | 0.752411 |
| 0.762005 | 0.760436 | 0.76358 | 0.762054 | 0.767033 | 0.767012 | 0.757576 | 0.762817 | 0.760604 |
| 0.767063 | 0.769791 | 0.765697 | 0.763217 | 0.770372 | 0.770026 | 0.763539 | 0.7632 | 0.76245 |
| 0.762794 | 0.761414 | 0.763053 | 0.760987 | 0.772316 | 0.765657 | 0.760997 | 0.763473 | 0.760356 |
| 0.76311 | 0.766406 | 0.768622 | 0.764721 | 0.769734 | 0.76487 | 0.760771 | 0.767417 | 0.766141 |
| 0.758255 | 0.761876 | 0.767444 | 0.765558 | 0.768207 | 0.766778 | 0.755775 | 0.764741 | 0.75857 |
| 0.761217 | 0.768068 | 0.76248 | 0.766391 | 0.768469 | 0.765726 | 0.758956 | 0.763503 | 0.762663 |
| 0.757338 | 0.765245 | 0.763856 | 0.760059 | 0.766641 | 0.763142 | 0.75323 | 0.762809 | 0.759011 |
| 0.761387 | 0.762232 | 0.758686 | 0.760757 | 0.766057 | 0.765101 | 0.757279 | 0.761135 | 0.760548 |
| 0.768306 | 0.770332 | 0.764226 | 0.763268 | 0.773858 | 0.772071 | 0.765665 | 0.762565 | 0.762854 |
| 0.757013 | 0.758199 | 0.763251 | 0.761243 | 0.762262 | 0.763462 | 0.754451 | 0.760848 | 0.761799 |
| 0.765128 | 0.767116 | 0.760987 | 0.762325 | 0.767227 | 0.76573 | 0.764064 | 0.759603 | 0.762307 |
| 0.699107 | 0.697829 | 0.703883 | 0.707532 | 0.705309 | 0.709376 | 0.696884 | 0.703161 | 0.706112 |
| 0.758295 | 0.76269 | 0.760359 | 0.763145 | 0.767359 | 0.771113 | 0.762466 | 0.763191 | 0.760884 |
| 0.763031 | 0.760571 | 0.766041 | 0.762077 | 0.768993 | 0.766083 | 0.758208 | 0.762075 | 0.756116 |
| 0.76417 | 0.766678 | 0.768784 | 0.771568 | 0.771892 | 0.768874 | 0.759977 | 0.765087 | 0.764308 |
| 0.764498 | 0.762262 | 0.765021 | 0.763406 | 0.76884 | 0.76595 | 0.762054 | 0.764164 | 0.75948 |
| 0.756592 | 0.759842 | 0.760107 | 0.763195 | 0.767324 | 0.767809 | 0.756245 | 0.759613 | 0.759226 |
| 0.760449 | 0.760739 | 0.762216 | 0.762223 | 0.771073 | 0.76676 | 0.758999 | 0.761593 | 0.762011 |
| 0.759042 | 0.762898 | 0.761921 | 0.761675 | 0.767097 | 0.765351 | 0.759317 | 0.758008 | 0.759259 |
| 0.7604 | 0.758413 | 0.762656 | 0.763321 | 0.768492 | 0.765511 | 0.759579 | 0.75955 | 0.761756 |
| 0.751771 | 0.750496 | 0.759078 | 0.753986 | 0.759794 | 0.757516 | 0.750975 | 0.758439 | 0.748742 |
| 0.751797 | 0.754405 | 0.758734 | 0.756775 | 0.762408 | 0.760111 | 0.753314 | 0.758465 | 0.751179 |
| 0.76208 | 0.763233 | 0.766457 | 0.764054 | 0.770976 | 0.769501 | 0.758277 | 0.761004 | 0.759417 |
| 0.76454 | 0.767026 | 0.770399 | 0.765696 | 0.76992 | 0.772042 | 0.763817 | 0.768006 | 0.767827 |
| 0.769872 | 0.770064 | 0.767277 | 0.765549 | 0.773596 | 0.772788 | 0.764083 | 0.765411 | 0.76574 |
| 0.761215 | 0.76618 | 0.766649 | 0.769197 | 0.7727 | 0.771537 | 0.76213 | 0.765813 | 0.766027 |
| 0.758103 | 0.758029 | 0.762543 | 0.759952 | 0.764611 | 0.763809 | 0.758816 | 0.761024 | 0.760954 |
| 0.76543 | 0.764593 | 0.770844 | 0.76652 | 0.771048 | 0.768896 | 0.764528 | 0.768407 | 0.763187 |
| 0.761819 | 0.762584 | 0.762908 | 0.763187 | 0.763633 | 0.768664 | 0.759701 | 0.763516 | 0.762608 |
| 0.757586 | 0.758227 | 0.759788 | 0.756643 | 0.764267 | 0.767727 | 0.753572 | 0.755129 | 0.758638 |
| 0.762537 | 0.761732 | 0.765906 | 0.76837 | 0.770937 | 0.773337 | 0.763326 | 0.761287 | 0.763164 |
| 0.765982 | 0.762253 | 0.762666 | 0.761806 | 0.77151 | 0.770192 | 0.760934 | 0.762835 | 0.763402 |

| | | | | | | | | |
|---|---|---|---|---|---|---|---|---|
| 0.757057 | 0.762956 | 0.764821 | 0.763731 | 0.770006 | 0.77127 | 0.756838 | 0.758409 | 0.761285 |
| 0.763281 | 0.763037 | 0.763556 | 0.765717 | 0.770664 | 0.773387 | 0.759898 | 0.764657 | 0.761145 |
| 0.760477 | 0.757164 | 0.758187 | 0.758923 | 0.766538 | 0.767382 | 0.753267 | 0.759476 | 0.761524 |
| 0.716284 | 0.731556 | 0.712251 | 0.717022 | 0.735357 | 0.738529 | 0.728587 | 0.722023 | 0.718188 |
| 0.754473 | 0.756697 | 0.756792 | 0.755763 | 0.763782 | 0.764922 | 0.750956 | 0.754558 | 0.756622 |
| 0.754683 | 0.760321 | 0.759109 | 0.757558 | 0.770738 | 0.763895 | 0.754216 | 0.757362 | 0.754947 |
| 0.759476 | 0.763016 | 0.767031 | 0.767973 | 0.778142 | 0.772742 | 0.76048 | 0.762918 | 0.760959 |
| 0.761289 | 0.761345 | 0.763458 | 0.761346 | 0.774998 | 0.772124 | 0.758527 | 0.764613 | 0.757725 |
| 0.762688 | 0.764547 | 0.762806 | 0.76222 | 0.770838 | 0.77025 | 0.762229 | 0.759259 | 0.760421 |
| 0.759594 | 0.759954 | 0.763792 | 0.762879 | 0.771954 | 0.772241 | 0.758697 | 0.760793 | 0.762585 |
| 0.760448 | 0.764691 | 0.764788 | 0.759599 | 0.774434 | 0.769168 | 0.75853 | 0.762498 | 0.760056 |
| 0.762046 | 0.765728 | 0.766777 | 0.762718 | 0.774183 | 0.772255 | 0.759293 | 0.760452 | 0.757957 |
| 0.760903 | 0.76502 | 0.762384 | 0.757713 | 0.769344 | 0.769155 | 0.759974 | 0.760616 | 0.757766 |
| 0.759397 | 0.758076 | 0.761839 | 0.760844 | 0.773135 | 0.7661 | 0.754759 | 0.758469 | 0.761568 |
| 0.763089 | 0.765034 | 0.766031 | 0.764359 | 0.773173 | 0.7699 | 0.758898 | 0.76152 | 0.763342 |
| 0.757591 | 0.760187 | 0.758367 | 0.762134 | 0.770828 | 0.768314 | 0.759191 | 0.75892 | 0.762847 |
| 0.758637 | 0.762013 | 0.767795 | 0.76474 | 0.771396 | 0.773342 | 0.755617 | 0.764239 | 0.763309 |
| 0.76747 | 0.764499 | 0.762938 | 0.762215 | 0.777134 | 0.773297 | 0.762069 | 0.764006 | 0.76557 |
| 0.755179 | 0.759271 | 0.76057 | 0.757205 | 0.765556 | 0.764817 | 0.752948 | 0.757066 | 0.756061 |
| 0.757927 | 0.762251 | 0.763709 | 0.762741 | 0.772987 | 0.766353 | 0.760105 | 0.762952 | 0.762428 |
| 0.764508 | 0.763911 | 0.766429 | 0.768292 | 0.77422 | 0.7743 | 0.760546 | 0.766533 | 0.764201 |
| 0.763741 | 0.765656 | 0.766233 | 0.768771 | 0.771831 | 0.77445 | 0.761662 | 0.763272 | 0.765162 |
| 0.671884 | 0.683786 | 0.682998 | 0.680354 | 0.693024 | 0.690989 | 0.667092 | 0.687904 | 0.680952 |
| 0.664118 | 0.659341 | 0.660932 | 0.658325 | 0.671574 | 0.667065 | 0.658164 | 0.66054 | 0.65782 |
| 0.660349 | 0.664166 | 0.666882 | 0.661548 | 0.681025 | 0.674378 | 0.663147 | 0.659218 | 0.667379 |
| 0.645884 | 0.653701 | 0.652687 | 0.66006 | 0.657171 | 0.656327 | 0.647987 | 0.650575 | 0.655121 |
| 0.724651 | 0.726809 | 0.729114 | 0.731633 | 0.731675 | 0.735498 | 0.72541 | 0.733364 | 0.724655 |
| 0.718816 | 0.718469 | 0.721346 | 0.725029 | 0.725434 | 0.734148 | 0.714972 | 0.720177 | 0.718888 |
| 0.713542 | 0.721653 | 0.722952 | 0.723486 | 0.725134 | 0.734627 | 0.712129 | 0.729951 | 0.724206 |
| 0.730761 | 0.734151 | 0.735068 | 0.732427 | 0.735203 | 0.745366 | 0.727634 | 0.732914 | 0.731813 |
| 0.70564 | 0.711182 | 0.713501 | 0.708256 | 0.730108 | 0.7213 | 0.703254 | 0.719991 | 0.701798 |
| 0.702307 | 0.699441 | 0.706911 | 0.703863 | 0.711669 | 0.719213 | 0.696398 | 0.706754 | 0.708661 |
| 0.705079 | 0.710822 | 0.712722 | 0.720881 | 0.711006 | 0.720707 | 0.700824 | 0.710847 | 0.714793 |
| 0.646975 | 0.654337 | 0.656484 | 0.661112 | 0.667321 | 0.663002 | 0.647048 | 0.661558 | 0.6485 |
| 0.753548 | 0.756267 | 0.756408 | 0.755723 | 0.761018 | 0.76461 | 0.750043 | 0.752042 | 0.752921 |
| 0.730079 | 0.737033 | 0.736114 | 0.733536 | 0.742025 | 0.745878 | 0.72996 | 0.733934 | 0.734233 |
| 0.724133 | 0.735698 | 0.729792 | 0.723372 | 0.730749 | 0.741544 | 0.724902 | 0.727581 | 0.723947 |
| 0.711528 | 0.713092 | 0.717896 | 0.720917 | 0.72211 | 0.724581 | 0.711078 | 0.720112 | 0.718901 |
| 0.731753 | 0.731616 | 0.735915 | 0.733373 | 0.734948 | 0.746656 | 0.728175 | 0.741278 | 0.732547 |
| 0.719173 | 0.725042 | 0.728273 | 0.725198 | 0.730979 | 0.73461 | 0.719954 | 0.728248 | 0.724872 |
| 0.721576 | 0.719217 | 0.724769 | 0.72175 | 0.729157 | 0.729084 | 0.724972 | 0.724552 | 0.721418 |
| 0.714397 | 0.719285 | 0.726021 | 0.724209 | 0.732423 | 0.733877 | 0.716698 | 0.730606 | 0.721937 |
| 0.717093 | 0.719409 | 0.723746 | 0.723307 | 0.727289 | 0.734597 | 0.717972 | 0.719978 | 0.721288 |
| 0.717736 | 0.725153 | 0.723216 | 0.716081 | 0.721994 | 0.727578 | 0.709449 | 0.719181 | 0.719589 |
| 0.715203 | 0.714322 | 0.726432 | 0.720229 | 0.73074 | 0.731308 | 0.708611 | 0.72066 | 0.709801 |
| 0.703569 | 0.707631 | 0.713485 | 0.712601 | 0.719987 | 0.714761 | 0.698008 | 0.713386 | 0.711836 |

| | | | | | | | | |
|---|---|---|---|---|---|---|---|---|
| 0.739814 | 0.744746 | 0.749157 | 0.742517 | 0.752985 | 0.755273 | 0.740328 | 0.743495 | 0.739587 |
| 0.721542 | 0.721948 | 0.727427 | 0.727376 | 0.73522 | 0.738082 | 0.719177 | 0.725697 | 0.717169 |
| 0.721754 | 0.721798 | 0.727136 | 0.726885 | 0.734221 | 0.724071 | 0.717964 | 0.725889 | 0.721199 |
| 0.71573 | 0.72269 | 0.725358 | 0.724786 | 0.729322 | 0.737352 | 0.720376 | 0.71989 | 0.724238 |
| 0.726563 | 0.720277 | 0.722392 | 0.72637 | 0.732549 | 0.728461 | 0.726019 | 0.720016 | 0.720342 |
| 0.727889 | 0.714891 | 0.72002 | 0.722573 | 0.731518 | 0.723058 | 0.727617 | 0.727432 | 0.719721 |
| 0.70493 | 0.711959 | 0.715433 | 0.714462 | 0.714286 | 0.720127 | 0.701424 | 0.706453 | 0.702872 |
| 0.714205 | 0.711788 | 0.712475 | 0.712238 | 0.727067 | 0.713881 | 0.706633 | 0.720397 | 0.721591 |
| 0.716719 | 0.7176 | 0.712598 | 0.71502 | 0.721519 | 0.726781 | 0.708558 | 0.714042 | 0.712398 |
| 0.753275 | 0.751475 | 0.753138 | 0.749454 | 0.759187 | 0.757271 | 0.750911 | 0.756349 | 0.749852 |
| 0.746159 | 0.751185 | 0.749344 | 0.747092 | 0.753152 | 0.752273 | 0.746627 | 0.754177 | 0.742326 |
| 0.742623 | 0.737984 | 0.741339 | 0.741885 | 0.744973 | 0.746374 | 0.73845 | 0.747597 | 0.738066 |
| 0.733966 | 0.736554 | 0.738512 | 0.736268 | 0.739188 | 0.737757 | 0.730432 | 0.735065 | 0.727482 |
| 0.739488 | 0.739024 | 0.744488 | 0.74274 | 0.747079 | 0.746775 | 0.734411 | 0.745342 | 0.736887 |
| 0.732292 | 0.733729 | 0.736447 | 0.734641 | 0.737311 | 0.73838 | 0.733103 | 0.740745 | 0.72715 |

| Wild_plot_ | Wild_plot_ | Wild_plot_ | Wild_plot_ | Wild_plot_ | Wild_plot_ | Wild_plot_ | Wild_plot_ | Wild_plot_ |
|---|---|---|---|---|---|---|---|---|
| 0.703656 | 0.697155 | 0.697502 | 0.69244 | 0.696967 | 0.69748 | 0.690807 | 0.701435 | 0.698077 |
| 0.70831 | 0.703758 | 0.698892 | 0.701688 | 0.704383 | 0.708393 | 0.698889 | 0.706469 | 0.704933 |
| 0.700665 | 0.696016 | 0.693734 | 0.689276 | 0.696875 | 0.69722 | 0.690838 | 0.698769 | 0.692162 |
| 0.710942 | 0.704421 | 0.700707 | 0.702102 | 0.69979 | 0.707538 | 0.697561 | 0.710234 | 0.698945 |
| 0.704567 | 0.700121 | 0.695472 | 0.695503 | 0.697084 | 0.699545 | 0.693162 | 0.701463 | 0.6967 |
| 0.713013 | 0.711061 | 0.70292 | 0.705897 | 0.705134 | 0.712017 | 0.703292 | 0.714799 | 0.705947 |
| 0.760882 | 0.7548 | 0.75999 | 0.755821 | 0.753753 | 0.752971 | 0.750202 | 0.76087 | 0.751754 |
| 0.756927 | 0.755284 | 0.7524 | 0.748483 | 0.750977 | 0.75316 | 0.748722 | 0.753882 | 0.74476 |
| 0.708159 | 0.702819 | 0.699434 | 0.700025 | 0.700692 | 0.70334 | 0.696978 | 0.707211 | 0.700119 |
| 0.702586 | 0.698631 | 0.696371 | 0.695969 | 0.696681 | 0.698099 | 0.691091 | 0.701357 | 0.694687 |
| 0.702507 | 0.698148 | 0.700367 | 0.699153 | 0.700763 | 0.702167 | 0.692716 | 0.705414 | 0.700092 |
| 0.701291 | 0.698669 | 0.691196 | 0.690336 | 0.692612 | 0.70119 | 0.687283 | 0.699317 | 0.694609 |
| 0.708343 | 0.701811 | 0.700584 | 0.701069 | 0.702337 | 0.701963 | 0.695645 | 0.703728 | 0.703439 |
| 0.722452 | 0.715462 | 0.713221 | 0.721403 | 0.720918 | 0.716079 | 0.709955 | 0.722213 | 0.711111 |
| 0.739212 | 0.743452 | 0.728104 | 0.743916 | 0.736766 | 0.74147 | 0.729069 | 0.733861 | 0.724883 |
| 0.711849 | 0.706853 | 0.703645 | 0.705268 | 0.705604 | 0.707614 | 0.700551 | 0.70908 | 0.704764 |
| 0.709769 | 0.706569 | 0.702914 | 0.702112 | 0.703811 | 0.709181 | 0.697781 | 0.708826 | 0.702847 |
| 0.748926 | 0.751025 | 0.744356 | 0.762276 | 0.753455 | 0.74958 | 0.739031 | 0.743165 | 0.73015 |
| 0.699638 | 0.697139 | 0.691942 | 0.694911 | 0.694762 | 0.698475 | 0.688683 | 0.6972 | 0.696131 |
| 0.704445 | 0.700712 | 0.698932 | 0.696895 | 0.699619 | 0.701617 | 0.695478 | 0.706164 | 0.701017 |
| 0.69881 | 0.696706 | 0.692881 | 0.69727 | 0.696656 | 0.698581 | 0.691699 | 0.701268 | 0.69355 |
| 0.704724 | 0.701938 | 0.700477 | 0.696097 | 0.69741 | 0.702 | 0.696438 | 0.704492 | 0.700366 |
| 0.710877 | 0.708144 | 0.705637 | 0.703761 | 0.703801 | 0.706892 | 0.698981 | 0.711987 | 0.703029 |
| 0.708876 | 0.702827 | 0.702312 | 0.701355 | 0.700651 | 0.705319 | 0.695133 | 0.705247 | 0.700137 |
| 0.705119 | 0.7042 | 0.698413 | 0.70364 | 0.701095 | 0.703376 | 0.695804 | 0.707311 | 0.702954 |
| 0.709017 | 0.701617 | 0.701476 | 0.700311 | 0.701368 | 0.70157 | 0.694118 | 0.702479 | 0.699463 |
| 0.700721 | 0.696017 | 0.6928 | 0.694907 | 0.692592 | 0.698452 | 0.690329 | 0.699026 | 0.696381 |
| 0.706414 | 0.704489 | 0.702465 | 0.700196 | 0.702138 | 0.703774 | 0.697043 | 0.705159 | 0.700282 |
| 0.704662 | 0.705104 | 0.703608 | 0.701323 | 0.70397 | 0.70309 | 0.699564 | 0.705116 | 0.703164 |
| 0.70408 | 0.698581 | 0.698777 | 0.696636 | 0.699339 | 0.700375 | 0.694158 | 0.705793 | 0.699917 |
| 0.70547 | 0.701991 | 0.700527 | 0.698502 | 0.696567 | 0.702106 | 0.693146 | 0.705912 | 0.699478 |
| 0.703105 | 0.696585 | 0.695749 | 0.695482 | 0.693527 | 0.699693 | 0.691526 | 0.696834 | 0.69448 |
| 0.703177 | 0.696145 | 0.692829 | 0.695261 | 0.69995 | 0.7015 | 0.691921 | 0.701512 | 0.701194 |
| 0.706967 | 0.699122 | 0.69956 | 0.698308 | 0.69968 | 0.700207 | 0.697412 | 0.706103 | 0.700456 |
| 0.69891 | 0.697738 | 0.694239 | 0.695656 | 0.696371 | 0.697476 | 0.691905 | 0.702362 | 0.692546 |
| 0.701819 | 0.70126 | 0.696929 | 0.699389 | 0.697822 | 0.705389 | 0.694955 | 0.702748 | 0.698156 |
| 0.709014 | 0.703989 | 0.704672 | 0.701273 | 0.704433 | 0.705025 | 0.698404 | 0.709156 | 0.701574 |
| 0.704976 | 0.702952 | 0.701247 | 0.702424 | 0.701512 | 0.7025 | 0.697421 | 0.708094 | 0.701719 |
| 0.712742 | 0.707556 | 0.703122 | 0.707305 | 0.701734 | 0.707532 | 0.698382 | 0.710498 | 0.706702 |

| | | | | | | | | |
|---|---|---|---|---|---|---|---|---|
| 0.752421 | 0.757631 | 0.752293 | 0.764374 | 0.756687 | 0.756967 | 0.750887 | 0.755114 | 0.739587 |
| 0.696556 | 0.692023 | 0.688913 | 0.689224 | 0.692003 | 0.693382 | 0.685615 | 0.693978 | 0.69195 |
| 0.701326 | 0.696358 | 0.693575 | 0.693646 | 0.695612 | 0.699509 | 0.688891 | 0.699599 | 0.696826 |
| 0.710533 | 0.702831 | 0.702582 | 0.702518 | 0.702491 | 0.703716 | 0.697574 | 0.707341 | 0.70337 |
| 0.704758 | 0.700915 | 0.696519 | 0.696893 | 0.699451 | 0.700806 | 0.692936 | 0.701851 | 0.697304 |
| 0.705935 | 0.704449 | 0.701224 | 0.704356 | 0.701248 | 0.704481 | 0.694391 | 0.706175 | 0.696836 |
| 0.702473 | 0.697714 | 0.696756 | 0.697619 | 0.691491 | 0.698485 | 0.690487 | 0.700465 | 0.692119 |
| 0.705455 | 0.701033 | 0.70047 | 0.699752 | 0.698708 | 0.702883 | 0.69753 | 0.709072 | 0.700166 |
| 0.709866 | 0.703758 | 0.700931 | 0.701266 | 0.700136 | 0.703266 | 0.698456 | 0.705591 | 0.70023 |
| 0.704744 | 0.698238 | 0.698869 | 0.701073 | 0.700455 | 0.702723 | 0.693441 | 0.703682 | 0.69585 |
| 0.697893 | 0.693846 | 0.691962 | 0.692985 | 0.690818 | 0.69558 | 0.686835 | 0.696845 | 0.69247 |
| 0.697337 | 0.695459 | 0.689787 | 0.692528 | 0.692139 | 0.698972 | 0.686175 | 0.699547 | 0.695026 |
| 0.702602 | 0.700603 | 0.697379 | 0.697653 | 0.697101 | 0.703943 | 0.690472 | 0.701657 | 0.697235 |
| 0.700895 | 0.696188 | 0.695103 | 0.694424 | 0.691748 | 0.697935 | 0.689923 | 0.698072 | 0.693006 |
| 0.703911 | 0.700033 | 0.695835 | 0.69758 | 0.695745 | 0.6992 | 0.693845 | 0.700447 | 0.693363 |
| 0.699846 | 0.698016 | 0.693306 | 0.693196 | 0.69351 | 0.696389 | 0.690895 | 0.700229 | 0.694712 |
| 0.696634 | 0.691687 | 0.687385 | 0.69211 | 0.693895 | 0.695088 | 0.687445 | 0.698018 | 0.692599 |
| 0.707125 | 0.701113 | 0.700667 | 0.699877 | 0.699189 | 0.703497 | 0.695379 | 0.707362 | 0.700238 |
| 0.711048 | 0.703575 | 0.703911 | 0.700279 | 0.702149 | 0.706767 | 0.699764 | 0.708445 | 0.701289 |
| 0.701008 | 0.699439 | 0.696484 | 0.693997 | 0.698248 | 0.699322 | 0.688353 | 0.702054 | 0.698426 |
| 0.698728 | 0.695522 | 0.691307 | 0.693037 | 0.690185 | 0.696008 | 0.688237 | 0.697704 | 0.691674 |
| 0.700177 | 0.694133 | 0.694741 | 0.691652 | 0.696171 | 0.697607 | 0.69294 | 0.700588 | 0.695449 |
| 0.704103 | 0.698106 | 0.697807 | 0.697925 | 0.695539 | 0.699992 | 0.692141 | 0.701045 | 0.693861 |
| 0.712207 | 0.708299 | 0.705011 | 0.706679 | 0.708231 | 0.707303 | 0.700218 | 0.709942 | 0.704908 |
| 0.720926 | 0.71645 | 0.712302 | 0.71218 | 0.709571 | 0.715252 | 0.711468 | 0.718412 | 0.712466 |
| 0.757947 | 0.756516 | 0.752634 | 0.749596 | 0.750852 | 0.754826 | 0.750904 | 0.754321 | 0.746204 |
| 0.763767 | 0.753422 | 0.756841 | 0.750873 | 0.75272 | 0.750689 | 0.752458 | 0.760277 | 0.751912 |
| 0.76844 | 0.756222 | 0.760939 | 0.755869 | 0.757562 | 0.752601 | 0.758013 | 0.766423 | 0.756264 |
| 0.700353 | 0.696017 | 0.690561 | 0.693919 | 0.694912 | 0.696731 | 0.688877 | 0.699816 | 0.694912 |
| 0.700198 | 0.697083 | 0.694456 | 0.692094 | 0.69633 | 0.694601 | 0.692294 | 0.701116 | 0.696856 |
| 0.70943 | 0.704151 | 0.703524 | 0.701457 | 0.703148 | 0.704687 | 0.697658 | 0.708603 | 0.703944 |
| 0.713072 | 0.705199 | 0.704768 | 0.699884 | 0.705711 | 0.704224 | 0.703111 | 0.712808 | 0.702698 |
| 0.704901 | 0.701483 | 0.697782 | 0.698886 | 0.696819 | 0.699253 | 0.693575 | 0.702271 | 0.697436 |
| 0.701196 | 0.700033 | 0.693389 | 0.698463 | 0.694553 | 0.698645 | 0.689266 | 0.702251 | 0.700212 |
| 0.703495 | 0.698819 | 0.695587 | 0.693353 | 0.695563 | 0.695904 | 0.689482 | 0.699366 | 0.697096 |
| 0.707147 | 0.699667 | 0.697797 | 0.698599 | 0.697631 | 0.697407 | 0.692979 | 0.705254 | 0.696814 |
| 0.7036 | 0.700225 | 0.698519 | 0.698793 | 0.69727 | 0.702173 | 0.689096 | 0.700058 | 0.697594 |
| 0.719229 | 0.712879 | 0.71256 | 0.708014 | 0.710171 | 0.712195 | 0.701234 | 0.71473 | 0.708173 |
| 0.695967 | 0.692475 | 0.691126 | 0.692839 | 0.688632 | 0.691647 | 0.686571 | 0.697483 | 0.689804 |
| 0.703305 | 0.697637 | 0.69749 | 0.695543 | 0.696698 | 0.699281 | 0.691897 | 0.701531 | 0.694517 |
| 0.702413 | 0.697723 | 0.696223 | 0.696747 | 0.696101 | 0.69862 | 0.691296 | 0.698102 | 0.694314 |
| 0.720789 | 0.714588 | 0.711462 | 0.710424 | 0.714921 | 0.714453 | 0.705919 | 0.719743 | 0.712889 |
| 0.703611 | 0.703131 | 0.698823 | 0.694935 | 0.693859 | 0.704509 | 0.690547 | 0.701991 | 0.696548 |
| 0.702511 | 0.697777 | 0.696217 | 0.694107 | 0.695087 | 0.699092 | 0.687667 | 0.698096 | 0.697968 |
| 0.702755 | 0.704105 | 0.697081 | 0.702084 | 0.697696 | 0.704355 | 0.693974 | 0.704145 | 0.699559 |
| 0.704555 | 0.700188 | 0.697501 | 0.696543 | 0.696477 | 0.69734 | 0.692477 | 0.700742 | 0.695812 |

| | | | | | | | | |
|---|---|---|---|---|---|---|---|---|
| 0.702186 | 0.698574 | 0.699967 | 0.690921 | 0.695523 | 0.698768 | 0.691624 | 0.698867 | 0.694982 |
| 0.708581 | 0.703988 | 0.703883 | 0.701543 | 0.700379 | 0.705188 | 0.696354 | 0.706456 | 0.699927 |
| 0.702076 | 0.699886 | 0.696293 | 0.698098 | 0.694816 | 0.703019 | 0.689782 | 0.702748 | 0.696752 |
| 0.710997 | 0.709149 | 0.703978 | 0.700398 | 0.702904 | 0.707134 | 0.702417 | 0.710825 | 0.702795 |
| 0.709974 | 0.703268 | 0.705589 | 0.702269 | 0.703804 | 0.70375 | 0.6978 | 0.710017 | 0.702597 |
| 0.703332 | 0.697502 | 0.700075 | 0.696011 | 0.698656 | 0.69923 | 0.691209 | 0.703881 | 0.694646 |
| 0.707272 | 0.702822 | 0.705457 | 0.699021 | 0.702907 | 0.702886 | 0.692946 | 0.706861 | 0.700791 |
| 0.70463 | 0.702306 | 0.70036 | 0.699678 | 0.698703 | 0.704232 | 0.694982 | 0.704965 | 0.699761 |
| 0.706091 | 0.70228 | 0.695244 | 0.698976 | 0.697526 | 0.701554 | 0.694427 | 0.70376 | 0.698148 |
| 0.700328 | 0.697581 | 0.697576 | 0.695684 | 0.694393 | 0.698631 | 0.690669 | 0.700725 | 0.694855 |
| 0.703159 | 0.701124 | 0.693112 | 0.695875 | 0.694096 | 0.701709 | 0.693012 | 0.703085 | 0.697725 |
| 0.699141 | 0.697823 | 0.696438 | 0.697531 | 0.694596 | 0.70118 | 0.692906 | 0.69989 | 0.695043 |
| 0.702508 | 0.707177 | 0.697097 | 0.701542 | 0.697821 | 0.707955 | 0.696267 | 0.700008 | 0.697463 |
| 0.699325 | 0.692113 | 0.699046 | 0.69596 | 0.693661 | 0.695245 | 0.689707 | 0.699568 | 0.692436 |
| 0.703316 | 0.703092 | 0.696593 | 0.701026 | 0.699873 | 0.703409 | 0.694335 | 0.704772 | 0.699034 |
| 0.701766 | 0.701718 | 0.695526 | 0.696198 | 0.697945 | 0.700969 | 0.689428 | 0.702113 | 0.698478 |
| 0.709384 | 0.70368 | 0.699303 | 0.701532 | 0.699141 | 0.707741 | 0.694766 | 0.707523 | 0.700359 |
| 0.702909 | 0.702849 | 0.698561 | 0.699458 | 0.695199 | 0.697194 | 0.691205 | 0.701429 | 0.69408 |
| 0.701076 | 0.698164 | 0.696808 | 0.699185 | 0.698342 | 0.70005 | 0.689076 | 0.701547 | 0.696057 |
| 0.705072 | 0.701998 | 0.700008 | 0.698054 | 0.699246 | 0.698178 | 0.699342 | 0.702624 | 0.697391 |
| 0.704379 | 0.700439 | 0.698923 | 0.693924 | 0.694858 | 0.703577 | 0.691678 | 0.703626 | 0.696928 |
| 0.700441 | 0.694773 | 0.692939 | 0.693342 | 0.691358 | 0.694246 | 0.69003 | 0.700067 | 0.69186 |
| 0.703278 | 0.701312 | 0.701281 | 0.697549 | 0.703786 | 0.703458 | 0.690896 | 0.703878 | 0.69879 |
| 0.70759 | 0.698492 | 0.700615 | 0.697844 | 0.699705 | 0.69917 | 0.692154 | 0.704322 | 0.696931 |
| 0.70561 | 0.702435 | 0.698797 | 0.696769 | 0.699581 | 0.706648 | 0.694701 | 0.703047 | 0.699796 |
| 0.708966 | 0.706959 | 0.702436 | 0.702751 | 0.704798 | 0.707366 | 0.697164 | 0.708861 | 0.702033 |
| 0.705281 | 0.700628 | 0.699436 | 0.699584 | 0.701001 | 0.704775 | 0.69595 | 0.702602 | 0.696453 |
| 0.706679 | 0.704033 | 0.698377 | 0.699681 | 0.697856 | 0.703082 | 0.696034 | 0.70761 | 0.697636 |
| 0.708199 | 0.702941 | 0.698361 | 0.699811 | 0.700034 | 0.703107 | 0.696685 | 0.707007 | 0.699245 |
| 0.705335 | 0.701723 | 0.698259 | 0.700959 | 0.701517 | 0.705251 | 0.697457 | 0.704001 | 0.698771 |
| 0.704617 | 0.706767 | 0.700025 | 0.701284 | 0.701163 | 0.70793 | 0.695801 | 0.704683 | 0.701472 |
| 0.703207 | 0.698438 | 0.700182 | 0.6986 | 0.695445 | 0.700551 | 0.692778 | 0.699134 | 0.697853 |
| 0.703399 | 0.699828 | 0.701027 | 0.699118 | 0.696482 | 0.700622 | 0.694518 | 0.703309 | 0.697031 |
| 0.707959 | 0.698963 | 0.70185 | 0.700463 | 0.699888 | 0.698327 | 0.695629 | 0.704608 | 0.698284 |
| 0.702834 | 0.70166 | 0.700192 | 0.695993 | 0.698826 | 0.698982 | 0.689342 | 0.700109 | 0.695253 |
| 0.709905 | 0.699918 | 0.701418 | 0.700234 | 0.699057 | 0.699141 | 0.695502 | 0.707537 | 0.699501 |
| 0.700055 | 0.694447 | 0.69501 | 0.691826 | 0.697535 | 0.699754 | 0.688884 | 0.696187 | 0.693201 |
| 0.679261 | 0.668911 | 0.676546 | 0.670605 | 0.677812 | 0.672956 | 0.671084 | 0.67823 | 0.670933 |
| 0.68583 | 0.676378 | 0.683359 | 0.676428 | 0.67682 | 0.675005 | 0.674077 | 0.685759 | 0.680158 |
| 0.674704 | 0.674035 | 0.670874 | 0.670069 | 0.673202 | 0.675699 | 0.669596 | 0.676807 | 0.671609 |
| 0.675598 | 0.671044 | 0.670333 | 0.668675 | 0.671669 | 0.670509 | 0.668032 | 0.677386 | 0.671218 |
| 0.661355 | 0.656239 | 0.657 | 0.660588 | 0.658598 | 0.658704 | 0.654233 | 0.659974 | 0.657607 |
| 0.663089 | 0.659515 | 0.663813 | 0.656223 | 0.660045 | 0.661879 | 0.663973 | 0.667119 | 0.661425 |
| 0.665529 | 0.657824 | 0.663796 | 0.66114 | 0.661558 | 0.659652 | 0.658249 | 0.663639 | 0.66036 |
| 0.660234 | 0.663526 | 0.659362 | 0.664365 | 0.657481 | 0.65965 | 0.65525 | 0.664072 | 0.659075 |
| 0.665082 | 0.659879 | 0.656836 | 0.661095 | 0.660894 | 0.65999 | 0.655829 | 0.665477 | 0.657225 |
| 0.669556 | 0.663465 | 0.663968 | 0.663681 | 0.662714 | 0.658524 | 0.66465 | 0.67129 | 0.65782 |

| | | | | | | | | |
|---|---|---|---|---|---|---|---|---|
| 0.670334 | 0.663645 | 0.664174 | 0.659515 | 0.661252 | 0.658015 | 0.660377 | 0.669481 | 0.661826 |
| 0.669405 | 0.667108 | 0.669399 | 0.666042 | 0.667516 | 0.664475 | 0.661436 | 0.670587 | 0.665662 |
| 0.670989 | 0.665853 | 0.665319 | 0.660263 | 0.664791 | 0.665522 | 0.663372 | 0.674199 | 0.666473 |
| 0.659929 | 0.654441 | 0.658653 | 0.651922 | 0.656718 | 0.658537 | 0.656947 | 0.658092 | 0.657332 |
| 0.660309 | 0.657243 | 0.659886 | 0.655448 | 0.658296 | 0.657396 | 0.650558 | 0.661208 | 0.658437 |
| 0.661224 | 0.655225 | 0.660088 | 0.65947 | 0.658284 | 0.65595 | 0.654715 | 0.662486 | 0.653022 |
| 0.661338 | 0.659651 | 0.666476 | 0.661049 | 0.664876 | 0.660957 | 0.658135 | 0.663924 | 0.661503 |
| 0.659431 | 0.657094 | 0.656994 | 0.658299 | 0.654811 | 0.657982 | 0.655343 | 0.660431 | 0.659279 |
| 0.661032 | 0.659792 | 0.661163 | 0.654072 | 0.660564 | 0.652694 | 0.655072 | 0.662732 | 0.654378 |
| 0.657272 | 0.654305 | 0.656044 | 0.652943 | 0.655826 | 0.658753 | 0.646864 | 0.655876 | 0.66352 |
| 0.648496 | 0.64527 | 0.649008 | 0.649357 | 0.653381 | 0.649717 | 0.643339 | 0.655846 | 0.646644 |
| 0.688742 | 0.688505 | 0.688539 | 0.684079 | 0.685013 | 0.688976 | 0.678632 | 0.691537 | 0.684626 |
| 0.698816 | 0.696695 | 0.691801 | 0.693947 | 0.695667 | 0.697425 | 0.688342 | 0.699188 | 0.693698 |
| 0.685027 | 0.677698 | 0.681578 | 0.679281 | 0.682324 | 0.678563 | 0.677827 | 0.684443 | 0.673946 |
| 0.712463 | 0.709022 | 0.706528 | 0.703472 | 0.704339 | 0.706962 | 0.700855 | 0.711137 | 0.704537 |
| 0.697947 | 0.696686 | 0.694234 | 0.697309 | 0.692768 | 0.697443 | 0.686288 | 0.698564 | 0.692868 |
| 0.706278 | 0.701668 | 0.698706 | 0.701416 | 0.700421 | 0.704222 | 0.694893 | 0.702379 | 0.700946 |
| 0.714007 | 0.707925 | 0.706452 | 0.708583 | 0.708394 | 0.71113 | 0.702287 | 0.715167 | 0.705346 |
| 0.689186 | 0.688073 | 0.685698 | 0.684279 | 0.683405 | 0.685434 | 0.68011 | 0.688044 | 0.680975 |
| 0.714446 | 0.707816 | 0.70766 | 0.70786 | 0.70813 | 0.712005 | 0.700216 | 0.710028 | 0.70259 |
| 0.685201 | 0.681918 | 0.68215 | 0.680748 | 0.6835 | 0.682251 | 0.676504 | 0.68709 | 0.679587 |
| 0.714888 | 0.706691 | 0.706164 | 0.704825 | 0.705722 | 0.709407 | 0.699187 | 0.71384 | 0.705574 |
| 0.714216 | 0.712108 | 0.709944 | 0.709451 | 0.710367 | 0.715162 | 0.704168 | 0.716332 | 0.708579 |
| 0.68256 | 0.672977 | 0.683108 | 0.674358 | 0.681485 | 0.673577 | 0.670314 | 0.684561 | 0.67667 |
| 0.685418 | 0.680582 | 0.678742 | 0.678067 | 0.680576 | 0.679317 | 0.673639 | 0.682002 | 0.674366 |
| 0.687627 | 0.679709 | 0.687308 | 0.678041 | 0.682161 | 0.676707 | 0.67963 | 0.688418 | 0.67945 |
| 0.703004 | 0.699438 | 0.695785 | 0.69594 | 0.698302 | 0.702108 | 0.692157 | 0.705923 | 0.697264 |
| 0.713286 | 0.705235 | 0.705858 | 0.703461 | 0.706758 | 0.707847 | 0.70155 | 0.710383 | 0.704465 |
| 0.68345 | 0.679956 | 0.677608 | 0.677246 | 0.675469 | 0.677903 | 0.668076 | 0.68278 | 0.67341 |
| 0.719808 | 0.714009 | 0.715119 | 0.713454 | 0.714105 | 0.716488 | 0.707667 | 0.71863 | 0.712679 |
| 0.72284 | 0.713782 | 0.714298 | 0.712971 | 0.713425 | 0.715168 | 0.709215 | 0.718851 | 0.714441 |
| 0.693745 | 0.68814 | 0.685778 | 0.684279 | 0.686286 | 0.686258 | 0.67677 | 0.692096 | 0.684568 |
| 0.695898 | 0.687215 | 0.687309 | 0.684185 | 0.685626 | 0.68651 | 0.678399 | 0.691303 | 0.682617 |
| 0.720639 | 0.713091 | 0.716664 | 0.715549 | 0.713013 | 0.715384 | 0.71102 | 0.719678 | 0.711422 |
| 0.720832 | 0.707447 | 0.71291 | 0.709706 | 0.711083 | 0.707724 | 0.705482 | 0.718724 | 0.702537 |
| 0.722619 | 0.71233 | 0.718721 | 0.712461 | 0.709085 | 0.711664 | 0.703915 | 0.719666 | 0.712325 |
| 0.672867 | 0.671524 | 0.66999 | 0.670205 | 0.670002 | 0.670845 | 0.66227 | 0.67017 | 0.662081 |
| 0.714389 | 0.713025 | 0.707441 | 0.707687 | 0.704762 | 0.71178 | 0.702376 | 0.712236 | 0.706959 |
| 0.672134 | 0.667992 | 0.664665 | 0.664169 | 0.665879 | 0.66632 | 0.660131 | 0.671441 | 0.661761 |
| 0.665773 | 0.662834 | 0.655611 | 0.660574 | 0.657168 | 0.660433 | 0.649662 | 0.661746 | 0.652815 |
| 0.723209 | 0.718474 | 0.721717 | 0.723885 | 0.722783 | 0.719707 | 0.713686 | 0.72086 | 0.712679 |
| 0.668022 | 0.663015 | 0.660735 | 0.660988 | 0.662718 | 0.660103 | 0.65364 | 0.664253 | 0.657125 |
| 0.658592 | 0.657296 | 0.654031 | 0.651067 | 0.651035 | 0.654723 | 0.645992 | 0.657008 | 0.64969 |
| 0.656057 | 0.653687 | 0.651296 | 0.652504 | 0.650692 | 0.650379 | 0.645277 | 0.650324 | 0.646771 |
| 0.726943 | 0.728858 | 0.726595 | 0.73299 | 0.725234 | 0.725676 | 0.719383 | 0.723962 | 0.72067 |
| 0.657721 | 0.656214 | 0.651272 | 0.651648 | 0.65147 | 0.65196 | 0.643741 | 0.654313 | 0.650992 |
| 0.721713 | 0.72532 | 0.713357 | 0.726916 | 0.721387 | 0.723248 | 0.711778 | 0.721207 | 0.712684 |

| | | | | | | | | |
|---|---|---|---|---|---|---|---|---|
| 0.734583 | 0.736412 | 0.734992 | 0.745467 | 0.735616 | 0.734965 | 0.727297 | 0.728579 | 0.719751 |
| 0.726421 | 0.729092 | 0.725428 | 0.734613 | 0.726173 | 0.725393 | 0.718523 | 0.724271 | 0.715018 |
| 0.74068 | 0.737249 | 0.7349 | 0.731799 | 0.734022 | 0.739691 | 0.733482 | 0.738539 | 0.738062 |
| 0.651389 | 0.64725 | 0.640883 | 0.646341 | 0.641024 | 0.644618 | 0.635922 | 0.644746 | 0.637934 |
| 0.647504 | 0.645701 | 0.641622 | 0.643363 | 0.640782 | 0.6442 | 0.635704 | 0.647127 | 0.641212 |
| 0.734329 | 0.733826 | 0.735421 | 0.743238 | 0.733436 | 0.733726 | 0.725445 | 0.732797 | 0.72105 |
| 0.649553 | 0.644495 | 0.643527 | 0.643981 | 0.645743 | 0.644675 | 0.637013 | 0.647238 | 0.641335 |
| 0.727833 | 0.723685 | 0.71968 | 0.723184 | 0.724343 | 0.724513 | 0.718703 | 0.728018 | 0.720086 |
| 0.733922 | 0.738542 | 0.7281 | 0.748395 | 0.733775 | 0.733767 | 0.724248 | 0.731678 | 0.720373 |
| 0.731412 | 0.729181 | 0.722265 | 0.72344 | 0.72634 | 0.727179 | 0.720072 | 0.728303 | 0.724377 |
| 0.739756 | 0.744709 | 0.737646 | 0.755361 | 0.744342 | 0.747468 | 0.735952 | 0.738494 | 0.72662 |
| 0.7 | 0.72549 | 0.712418 | 0.684211 | 0.705882 | 0.647059 | 0.717241 | 0.677215 | 0.668831 |
| 0.720847 | 0.709003 | 0.714457 | 0.710124 | 0.719209 | 0.71232 | 0.714988 | 0.717474 | 0.707286 |
| 0.715313 | 0.703132 | 0.707378 | 0.706042 | 0.712375 | 0.704374 | 0.708522 | 0.714986 | 0.710856 |
| 0.726397 | 0.71034 | 0.723156 | 0.709199 | 0.717149 | 0.709911 | 0.718642 | 0.728626 | 0.715006 |
| 0.724642 | 0.716823 | 0.727297 | 0.715912 | 0.721275 | 0.710432 | 0.71912 | 0.726962 | 0.705737 |
| 0.589406 | 0.584899 | 0.586592 | 0.582014 | 0.584347 | 0.580918 | 0.582349 | 0.588704 | 0.582152 |
| 0.723584 | 0.720406 | 0.725056 | 0.720679 | 0.724428 | 0.719051 | 0.722569 | 0.725524 | 0.718887 |
| 0.745129 | 0.74912 | 0.744154 | 0.767141 | 0.751464 | 0.751098 | 0.739091 | 0.736865 | 0.73137 |
| 0.739947 | 0.754349 | 0.74123 | 0.767371 | 0.745283 | 0.754542 | 0.742067 | 0.740317 | 0.729935 |
| 0.744317 | 0.747113 | 0.743385 | 0.761999 | 0.750306 | 0.746796 | 0.745371 | 0.741922 | 0.733114 |
| 0.59086 | 0.591514 | 0.584238 | 0.588254 | 0.58332 | 0.586094 | 0.579918 | 0.58856 | 0.581081 |
| 0.579688 | 0.582007 | 0.573701 | 0.577468 | 0.568004 | 0.574719 | 0.56556 | 0.577717 | 0.56765 |
| 0.797187 | 0.801556 | 0.795485 | 0.804091 | 0.821311 | 0.798668 | 0.801751 | 0.790943 | 0.78757 |
| 0.790839 | 0.802222 | 0.794503 | 0.8029 | 0.807055 | 0.794341 | 0.791139 | 0.787611 | 0.783864 |
| 0.826162 | 0.794105 | 0.828941 | 0.783624 | 0.832464 | 0.8031 | 0.821026 | 0.831396 | 0.790988 |
| 0.832933 | 0.800871 | 0.850686 | 0.797404 | 0.835492 | 0.79428 | 0.83048 | 0.825222 | 0.783287 |
| 0.834492 | 0.810646 | 0.829179 | 0.807886 | 0.837201 | 0.806046 | 0.854295 | 0.832547 | 0.778201 |
| 0.806697 | 0.777948 | 0.799965 | 0.78038 | 0.805897 | 0.778421 | 0.785234 | 0.790692 | 0.777927 |
| 0.803326 | 0.790619 | 0.794892 | 0.783211 | 0.796927 | 0.786829 | 0.790442 | 0.800369 | 0.788747 |
| 0.757875 | 0.755715 | 0.748332 | 0.750773 | 0.74726 | 0.755875 | 0.748576 | 0.754545 | 0.751638 |
| 0.773054 | 0.762749 | 0.761138 | 0.752959 | 0.759951 | 0.761462 | 0.758936 | 0.765753 | 0.762597 |
| 0.764253 | 0.754711 | 0.748447 | 0.75299 | 0.752041 | 0.757319 | 0.752757 | 0.758973 | 0.749125 |
| 0.763346 | 0.758158 | 0.754299 | 0.750775 | 0.753568 | 0.755873 | 0.751047 | 0.758935 | 0.752005 |
| 0.765779 | 0.759408 | 0.761248 | 0.756899 | 0.754972 | 0.763149 | 0.755546 | 0.76022 | 0.760712 |
| 0.763705 | 0.752701 | 0.752417 | 0.742833 | 0.750371 | 0.751012 | 0.75455 | 0.754424 | 0.757766 |
| 0.768352 | 0.763615 | 0.760768 | 0.752416 | 0.759786 | 0.760963 | 0.752904 | 0.759941 | 0.761238 |
| 0.763343 | 0.760754 | 0.755493 | 0.758271 | 0.759606 | 0.760014 | 0.758559 | 0.761148 | 0.759928 |
| 0.757599 | 0.750771 | 0.750878 | 0.746517 | 0.752679 | 0.75077 | 0.748048 | 0.756852 | 0.749783 |
| 0.759397 | 0.757164 | 0.752194 | 0.755071 | 0.757792 | 0.755531 | 0.746455 | 0.755064 | 0.753046 |
| 0.712742 | 0.711945 | 0.716912 | 0.707408 | 0.713753 | 0.711259 | 0.711941 | 0.722242 | 0.709019 |
| 0.728625 | 0.717176 | 0.71885 | 0.725358 | 0.721086 | 0.713266 | 0.714588 | 0.726035 | 0.712811 |
| 0.763423 | 0.75485 | 0.749681 | 0.755786 | 0.757078 | 0.75356 | 0.753247 | 0.759208 | 0.751004 |
| 0.784519 | 0.77189 | 0.768199 | 0.770604 | 0.778052 | 0.769277 | 0.772517 | 0.778695 | 0.774184 |
| 0.724705 | 0.712947 | 0.716934 | 0.711625 | 0.724655 | 0.716397 | 0.715228 | 0.724104 | 0.705797 |
| 0.763322 | 0.758894 | 0.75513 | 0.754422 | 0.758669 | 0.766362 | 0.754928 | 0.765249 | 0.755907 |
| 0.753311 | 0.754322 | 0.746695 | 0.747188 | 0.749848 | 0.752109 | 0.751279 | 0.755067 | 0.756299 |

| | | | | | | | | |
|---|---|---|---|---|---|---|---|---|
| 0.761814 | 0.757662 | 0.755072 | 0.756718 | 0.759306 | 0.762578 | 0.754975 | 0.761601 | 0.762409 |
| 0.76295 | 0.760273 | 0.757612 | 0.756841 | 0.75825 | 0.762455 | 0.754719 | 0.765331 | 0.759831 |
| 0.758126 | 0.756289 | 0.754345 | 0.752402 | 0.75427 | 0.753326 | 0.749908 | 0.757159 | 0.75649 |
| 0.766736 | 0.761574 | 0.754735 | 0.761593 | 0.758368 | 0.765809 | 0.755738 | 0.760279 | 0.760652 |
| 0.768797 | 0.762868 | 0.768031 | 0.761251 | 0.758275 | 0.761639 | 0.757889 | 0.7642 | 0.759017 |
| 0.764167 | 0.756804 | 0.757238 | 0.75731 | 0.754334 | 0.762946 | 0.751306 | 0.760343 | 0.757851 |
| 0.763988 | 0.758581 | 0.757576 | 0.75606 | 0.759089 | 0.765075 | 0.7536 | 0.762313 | 0.764062 |
| 0.748352 | 0.747035 | 0.743406 | 0.746546 | 0.747539 | 0.747368 | 0.742852 | 0.753495 | 0.745189 |
| 0.760597 | 0.760251 | 0.756259 | 0.754555 | 0.75617 | 0.761536 | 0.748564 | 0.759642 | 0.759625 |
| 0.758586 | 0.757754 | 0.747143 | 0.753632 | 0.750262 | 0.751162 | 0.750813 | 0.757665 | 0.750095 |
| 0.764394 | 0.75991 | 0.756338 | 0.752689 | 0.756311 | 0.762052 | 0.748482 | 0.761803 | 0.756676 |
| 0.763221 | 0.756473 | 0.756623 | 0.755695 | 0.758744 | 0.759706 | 0.754129 | 0.76081 | 0.758188 |
| 0.749395 | 0.749051 | 0.744613 | 0.742901 | 0.745871 | 0.74948 | 0.745984 | 0.751718 | 0.747461 |
| 0.755588 | 0.749393 | 0.746187 | 0.751697 | 0.748396 | 0.75185 | 0.749604 | 0.756745 | 0.754802 |
| 0.76709 | 0.76288 | 0.762664 | 0.761013 | 0.754865 | 0.764676 | 0.757396 | 0.763038 | 0.756422 |
| 0.803561 | 0.824159 | 0.793043 | 0.809976 | 0.801424 | 0.82913 | 0.810846 | 0.793644 | 0.781689 |
| 0.79986 | 0.798241 | 0.802008 | 0.775325 | 0.792267 | 0.807556 | 0.78265 | 0.816308 | 0.818925 |
| 0.801326 | 0.86622 | 0.800831 | 0.814659 | 0.796736 | 0.844815 | 0.797561 | 0.809089 | 0.83126 |
| 0.812075 | 0.826083 | 0.803818 | 0.822644 | 0.826899 | 0.821009 | 0.883474 | 0.808199 | 0.789003 |
| 0.821945 | 0.849745 | 0.815949 | 0.814059 | 0.836367 | 0.846908 | 0.826669 | 0.825707 | 0.795834 |
| 0.800828 | 0.87491 | 0.786674 | 0.847203 | 0.79455 | 0.836954 | 0.7982 | 0.791318 | 0.780562 |
| 0.820353 | 0.806193 | 0.816614 | 0.803942 | 0.845571 | 0.804939 | 0.881576 | 0.807934 | 0.776853 |
| 0.868049 | 0.808404 | 0.859338 | 0.799604 | 0.864653 | 0.808442 | 0.870607 | 0.865661 | 0.78469 |
| 0.797481 | 0.813876 | 0.798155 | 0.778808 | 0.796298 | 0.812511 | 0.788938 | 0.810787 | 0.882953 |
| 0.798903 | 0.847017 | 0.788851 | 0.811805 | 0.802526 | 0.8066 | 0.796429 | 0.802823 | 0.793067 |
| 0.832554 | 0.80338 | 0.822682 | 0.833511 | 0.84715 | 0.795549 | 0.827339 | 0.818466 | 0.777115 |
| 0.797029 | 0.843338 | 0.790493 | 0.808154 | 0.797704 | 0.852335 | 0.794257 | 0.800893 | 0.839757 |
| 0.826804 | 0.79293 | 0.82007 | 0.789566 | 0.824089 | 0.796415 | 0.801497 | 0.812641 | 0.796907 |
| 0.795925 | 0.842978 | 0.795305 | 0.840049 | 0.805127 | 0.815224 | 0.79071 | 0.787597 | 0.779478 |
| 0.819956 | 0.81146 | 0.813758 | 0.809014 | 0.820797 | 0.803256 | 0.824028 | 0.80891 | 0.788811 |
| 0.809851 | 0.869615 | 0.800176 | 0.828411 | 0.808734 | 0.840611 | 0.800221 | 0.814211 | 0.809103 |
| 0.795106 | 0.858041 | 0.793972 | 0.811427 | 0.787914 | 0.840884 | 0.790049 | 0.821422 | 0.841917 |
| 0.805979 | 0.827899 | 0.794253 | 0.795862 | 0.799699 | 0.830368 | 0.809029 | 0.812736 | 0.827831 |
| 0.799848 | 0.811198 | 0.808415 | 0.778254 | 0.79655 | 0.822116 | 0.797019 | 0.827276 | 0.848094 |
| 0.795972 | 0.79803 | 0.803079 | 0.811848 | 0.81684 | 0.792534 | 0.792318 | 0.792627 | 0.774258 |
| 0.796158 | 0.844205 | 0.785879 | 0.803496 | 0.805722 | 0.82683 | 0.805535 | 0.797387 | 0.798286 |
| 0.803997 | 0.80574 | 0.831221 | 0.790275 | 0.793827 | 0.832901 | 0.799313 | 0.804744 | 0.814435 |
| 1 | 0.796552 | 0.834673 | 0.785975 | 0.820543 | 0.790527 | 0.815836 | 0.837696 | 0.785645 |
| 0.796552 | 1 | 0.795085 | 0.843437 | 0.794203 | 0.853993 | 0.80637 | 0.807273 | 0.804991 |
| 0.834673 | 0.795085 | 1 | 0.784722 | 0.814693 | 0.785786 | 0.821827 | 0.831643 | 0.781827 |
| 0.785975 | 0.843437 | 0.784722 | 1 | 0.806544 | 0.819927 | 0.795015 | 0.78496 | 0.755096 |
| 0.820543 | 0.794203 | 0.814693 | 0.806544 | 1 | 0.792551 | 0.822941 | 0.794919 | 0.774874 |
| 0.790527 | 0.853993 | 0.785786 | 0.819927 | 0.792551 | 1 | 0.805949 | 0.804079 | 0.805441 |
| 0.815836 | 0.80637 | 0.821827 | 0.795015 | 0.822941 | 0.805949 | 1 | 0.817725 | 0.770762 |
| 0.837696 | 0.807273 | 0.831643 | 0.78496 | 0.794919 | 0.804079 | 0.817725 | 1 | 0.810065 |
| 0.785645 | 0.804991 | 0.781827 | 0.755096 | 0.774874 | 0.805441 | 0.770762 | 0.810065 | 1 |

| | | | | | | | | |
|---|---|---|---|---|---|---|---|---|
| 0.792633 | 0.811752 | 0.788203 | 0.870429 | 0.808971 | 0.796254 | 0.785024 | 0.781731 | 0.760849 |
| 0.766437 | 0.754322 | 0.757071 | 0.75685 | 0.75845 | 0.757513 | 0.756654 | 0.763685 | 0.758015 |
| 0.758067 | 0.755035 | 0.752937 | 0.758716 | 0.752871 | 0.751564 | 0.746127 | 0.753487 | 0.751956 |
| 0.75357 | 0.750145 | 0.753626 | 0.750883 | 0.752336 | 0.748364 | 0.746789 | 0.758394 | 0.755958 |
| 0.768061 | 0.76 | 0.761371 | 0.760779 | 0.763914 | 0.760034 | 0.762931 | 0.772076 | 0.758683 |
| 0.76296 | 0.758454 | 0.757849 | 0.754966 | 0.752837 | 0.758388 | 0.75339 | 0.763611 | 0.758702 |
| 0.763408 | 0.756297 | 0.756696 | 0.755558 | 0.761801 | 0.756998 | 0.754791 | 0.763599 | 0.759332 |
| 0.765569 | 0.756365 | 0.755567 | 0.753318 | 0.756856 | 0.757399 | 0.753347 | 0.759656 | 0.754467 |
| 0.754724 | 0.751319 | 0.750697 | 0.748456 | 0.746475 | 0.752052 | 0.753155 | 0.758466 | 0.754157 |
| 0.756466 | 0.752463 | 0.750773 | 0.750763 | 0.749782 | 0.74865 | 0.749798 | 0.759089 | 0.752807 |
| 0.764376 | 0.76153 | 0.75924 | 0.754538 | 0.756815 | 0.759452 | 0.753904 | 0.76671 | 0.760026 |
| 0.764569 | 0.758207 | 0.760333 | 0.753384 | 0.75761 | 0.758939 | 0.754106 | 0.768168 | 0.760634 |
| 0.759155 | 0.759332 | 0.756525 | 0.755652 | 0.751782 | 0.756597 | 0.753079 | 0.760724 | 0.751945 |
| 0.758478 | 0.756727 | 0.753371 | 0.75184 | 0.756646 | 0.752932 | 0.753896 | 0.756056 | 0.748452 |
| 0.760599 | 0.759714 | 0.755879 | 0.755827 | 0.752851 | 0.755272 | 0.757019 | 0.75888 | 0.752779 |
| 0.761593 | 0.755041 | 0.758621 | 0.755527 | 0.756294 | 0.754578 | 0.752498 | 0.762348 | 0.754609 |
| 0.76185 | 0.754996 | 0.760693 | 0.758319 | 0.757175 | 0.75465 | 0.752546 | 0.763102 | 0.752958 |
| 0.758463 | 0.754983 | 0.754469 | 0.752083 | 0.753247 | 0.75355 | 0.754455 | 0.757123 | 0.749699 |
| 0.766141 | 0.765153 | 0.757127 | 0.760444 | 0.758314 | 0.760275 | 0.75541 | 0.762649 | 0.753256 |
| 0.76313 | 0.763515 | 0.758945 | 0.760247 | 0.758363 | 0.760489 | 0.754631 | 0.760918 | 0.760913 |
| 0.760589 | 0.758317 | 0.751076 | 0.752251 | 0.752182 | 0.752734 | 0.751939 | 0.755146 | 0.748497 |
| 0.76179 | 0.755186 | 0.758158 | 0.754495 | 0.759365 | 0.753947 | 0.754893 | 0.759674 | 0.751934 |
| 0.760651 | 0.759033 | 0.756305 | 0.754922 | 0.756529 | 0.7546 | 0.754303 | 0.759383 | 0.754781 |
| 0.759678 | 0.755234 | 0.751616 | 0.752868 | 0.753954 | 0.750741 | 0.749086 | 0.75898 | 0.750186 |
| 0.765579 | 0.757995 | 0.759609 | 0.751063 | 0.758017 | 0.755815 | 0.753608 | 0.761401 | 0.755614 |
| 0.761055 | 0.756827 | 0.754894 | 0.752807 | 0.752409 | 0.751536 | 0.752634 | 0.758968 | 0.751198 |
| 0.757558 | 0.750021 | 0.75324 | 0.74715 | 0.750065 | 0.752017 | 0.749978 | 0.756902 | 0.75098 |
| 0.760671 | 0.751531 | 0.752944 | 0.750776 | 0.748512 | 0.751233 | 0.752395 | 0.759334 | 0.753719 |
| 0.762567 | 0.755888 | 0.761259 | 0.755274 | 0.758839 | 0.75643 | 0.755292 | 0.761037 | 0.757981 |
| 0.766687 | 0.752736 | 0.760803 | 0.75625 | 0.759971 | 0.752942 | 0.753683 | 0.76359 | 0.750072 |
| 0.753826 | 0.752047 | 0.752805 | 0.74938 | 0.749496 | 0.747381 | 0.748204 | 0.755436 | 0.749359 |
| 0.764686 | 0.758088 | 0.75761 | 0.759024 | 0.755839 | 0.752709 | 0.755429 | 0.761517 | 0.755131 |
| 0.764823 | 0.758796 | 0.756935 | 0.756163 | 0.753864 | 0.750809 | 0.755005 | 0.763568 | 0.754887 |
| 0.756895 | 0.750792 | 0.750878 | 0.749066 | 0.751943 | 0.744102 | 0.746716 | 0.752844 | 0.747177 |
| 0.762144 | 0.751582 | 0.751753 | 0.748291 | 0.751691 | 0.746931 | 0.747407 | 0.755084 | 0.749569 |
| 0.75955 | 0.751546 | 0.756798 | 0.750265 | 0.749344 | 0.751007 | 0.749185 | 0.757396 | 0.752301 |
| 0.767656 | 0.762366 | 0.760395 | 0.755966 | 0.761345 | 0.761733 | 0.755322 | 0.76465 | 0.760068 |
| 0.759248 | 0.749979 | 0.751413 | 0.751653 | 0.75541 | 0.751457 | 0.751322 | 0.758582 | 0.747835 |
| 0.763825 | 0.756761 | 0.754788 | 0.753508 | 0.756311 | 0.752205 | 0.755245 | 0.762949 | 0.748945 |
| 0.763735 | 0.757395 | 0.755 | 0.756398 | 0.753409 | 0.751937 | 0.750251 | 0.760641 | 0.750452 |
| 0.768098 | 0.75522 | 0.755313 | 0.757494 | 0.756766 | 0.753636 | 0.752348 | 0.758692 | 0.74807 |
| 0.765972 | 0.757429 | 0.759456 | 0.760007 | 0.754067 | 0.753766 | 0.752562 | 0.764579 | 0.75096 |
| 0.768787 | 0.760826 | 0.763309 | 0.761053 | 0.756848 | 0.757746 | 0.760702 | 0.76687 | 0.752938 |
| 0.75742 | 0.750761 | 0.756485 | 0.750988 | 0.752431 | 0.750392 | 0.745918 | 0.756223 | 0.74765 |
| 0.761563 | 0.755833 | 0.761075 | 0.754529 | 0.752973 | 0.751589 | 0.759677 | 0.761785 | 0.750942 |
| 0.759738 | 0.752644 | 0.757955 | 0.753096 | 0.750749 | 0.750978 | 0.754243 | 0.760723 | 0.749192 |

| | | | | | | | | |
|---|---|---|---|---|---|---|---|---|
| 0.748839 | 0.744522 | 0.743149 | 0.741444 | 0.746543 | 0.739325 | 0.739406 | 0.750293 | 0.741054 |
| 0.745096 | 0.735129 | 0.73631 | 0.734259 | 0.738201 | 0.734069 | 0.736411 | 0.742194 | 0.731949 |
| 0.757635 | 0.748001 | 0.75237 | 0.744442 | 0.746081 | 0.748056 | 0.750941 | 0.749212 | 0.75399 |
| 0.764517 | 0.756001 | 0.752957 | 0.751236 | 0.750818 | 0.75138 | 0.750089 | 0.758954 | 0.752847 |
| 0.75523 | 0.744432 | 0.748672 | 0.740738 | 0.744675 | 0.742464 | 0.746136 | 0.754494 | 0.745537 |
| 0.757791 | 0.748574 | 0.752292 | 0.745407 | 0.747812 | 0.749159 | 0.745432 | 0.753885 | 0.748981 |
| 0.761834 | 0.748499 | 0.752989 | 0.748104 | 0.75102 | 0.747434 | 0.747761 | 0.76036 | 0.747223 |
| 0.755231 | 0.743684 | 0.746387 | 0.740051 | 0.746382 | 0.74048 | 0.742863 | 0.748397 | 0.746799 |
| 0.767271 | 0.755146 | 0.76033 | 0.756789 | 0.758345 | 0.752603 | 0.753427 | 0.760443 | 0.752576 |
| 0.764412 | 0.755273 | 0.752886 | 0.758392 | 0.757645 | 0.752106 | 0.751307 | 0.760194 | 0.755587 |
| 0.765901 | 0.756992 | 0.759997 | 0.75925 | 0.756057 | 0.755182 | 0.754939 | 0.765895 | 0.755587 |
| 0.762713 | 0.752025 | 0.754239 | 0.748604 | 0.75299 | 0.751374 | 0.749527 | 0.760764 | 0.753608 |
| 0.767289 | 0.759513 | 0.759023 | 0.758808 | 0.760339 | 0.758547 | 0.756214 | 0.762709 | 0.762094 |
| 0.762583 | 0.753048 | 0.754573 | 0.752501 | 0.753574 | 0.751574 | 0.749531 | 0.762031 | 0.751613 |
| 0.7649 | 0.754215 | 0.754522 | 0.75478 | 0.758663 | 0.749349 | 0.751275 | 0.761842 | 0.753629 |
| 0.756415 | 0.751169 | 0.754208 | 0.750708 | 0.751933 | 0.748321 | 0.748159 | 0.755928 | 0.743592 |
| 0.763777 | 0.757769 | 0.753289 | 0.753776 | 0.753955 | 0.754213 | 0.750991 | 0.759654 | 0.748706 |
| 0.753368 | 0.751088 | 0.751708 | 0.74983 | 0.749497 | 0.749548 | 0.747815 | 0.755257 | 0.748512 |
| 0.75493 | 0.751062 | 0.75172 | 0.748742 | 0.751326 | 0.745118 | 0.749234 | 0.754748 | 0.74296 |
| 0.761117 | 0.753818 | 0.753094 | 0.748951 | 0.756994 | 0.75036 | 0.755333 | 0.75828 | 0.753158 |
| 0.755882 | 0.749727 | 0.754877 | 0.752205 | 0.754905 | 0.748082 | 0.748735 | 0.756345 | 0.74858 |
| 0.750742 | 0.745482 | 0.739279 | 0.742422 | 0.737843 | 0.741576 | 0.737421 | 0.746284 | 0.740616 |
| 0.754121 | 0.749107 | 0.747049 | 0.74701 | 0.744863 | 0.745676 | 0.743955 | 0.749786 | 0.7426 |
| 0.757257 | 0.750836 | 0.750795 | 0.751463 | 0.751132 | 0.747944 | 0.747306 | 0.756897 | 0.746595 |
| 0.756215 | 0.756824 | 0.750044 | 0.749827 | 0.751639 | 0.754185 | 0.750618 | 0.760183 | 0.750622 |
| 0.754459 | 0.752724 | 0.747054 | 0.753004 | 0.743421 | 0.746904 | 0.741821 | 0.752951 | 0.746109 |
| 0.756118 | 0.748198 | 0.749558 | 0.742266 | 0.750604 | 0.747288 | 0.745609 | 0.755624 | 0.75075 |
| 0.758023 | 0.754427 | 0.748958 | 0.747371 | 0.755016 | 0.75071 | 0.749375 | 0.756605 | 0.748229 |
| 0.757279 | 0.755673 | 0.749414 | 0.749197 | 0.750114 | 0.745162 | 0.74861 | 0.754654 | 0.745185 |
| 0.694393 | 0.686577 | 0.68626 | 0.681951 | 0.684996 | 0.685725 | 0.681751 | 0.688768 | 0.686362 |
| 0.703529 | 0.696691 | 0.692527 | 0.696849 | 0.693437 | 0.696177 | 0.690871 | 0.697761 | 0.694672 |
| 0.699603 | 0.695964 | 0.694069 | 0.695274 | 0.694289 | 0.698382 | 0.689417 | 0.69759 | 0.691779 |
| 0.692512 | 0.689089 | 0.689289 | 0.686448 | 0.687696 | 0.690527 | 0.681194 | 0.693569 | 0.68826 |
| 0.701858 | 0.696907 | 0.694093 | 0.694852 | 0.697336 | 0.699909 | 0.689085 | 0.697854 | 0.694875 |
| 0.755867 | 0.747771 | 0.748109 | 0.741686 | 0.75222 | 0.747776 | 0.745763 | 0.757275 | 0.756529 |
| 0.761121 | 0.751203 | 0.750759 | 0.749101 | 0.75101 | 0.747729 | 0.745853 | 0.756787 | 0.751858 |
| 0.76256 | 0.756734 | 0.754064 | 0.752585 | 0.75849 | 0.756963 | 0.752243 | 0.760483 | 0.757042 |
| 0.760296 | 0.754651 | 0.75397 | 0.750234 | 0.754702 | 0.75433 | 0.750203 | 0.756188 | 0.754424 |
| 0.756536 | 0.749623 | 0.754777 | 0.747762 | 0.747923 | 0.749655 | 0.746929 | 0.755421 | 0.748201 |
| 0.762715 | 0.752572 | 0.755525 | 0.750127 | 0.757581 | 0.752625 | 0.753207 | 0.759381 | 0.753852 |
| 0.76378 | 0.75662 | 0.75815 | 0.751197 | 0.756268 | 0.751905 | 0.753779 | 0.763112 | 0.757509 |
| 0.760892 | 0.750382 | 0.752001 | 0.748712 | 0.752538 | 0.750065 | 0.750342 | 0.754594 | 0.752968 |
| 0.753931 | 0.747326 | 0.746876 | 0.746901 | 0.750367 | 0.74594 | 0.742108 | 0.749549 | 0.746721 |
| 0.756504 | 0.747941 | 0.749427 | 0.740503 | 0.749118 | 0.74571 | 0.740702 | 0.752941 | 0.743979 |
| 0.756323 | 0.749488 | 0.748241 | 0.745868 | 0.749041 | 0.745751 | 0.748391 | 0.75619 | 0.748446 |
| 0.760221 | 0.753423 | 0.75349 | 0.747733 | 0.757504 | 0.752738 | 0.75109 | 0.755414 | 0.75702 |

| | | | | | | | | |
|---|---|---|---|---|---|---|---|---|
| 0.764061 | 0.758644 | 0.753416 | 0.754029 | 0.753286 | 0.756027 | 0.751144 | 0.765382 | 0.756151 |
| 0.759409 | 0.756981 | 0.747656 | 0.751149 | 0.749651 | 0.753547 | 0.744339 | 0.760534 | 0.747687 |
| 0.759624 | 0.753811 | 0.75225 | 0.753259 | 0.756567 | 0.75403 | 0.751519 | 0.754336 | 0.753799 |
| 0.757566 | 0.748372 | 0.747392 | 0.746437 | 0.748864 | 0.749614 | 0.743414 | 0.756155 | 0.75276 |
| 0.755601 | 0.751216 | 0.751274 | 0.749892 | 0.75307 | 0.744214 | 0.744691 | 0.750351 | 0.747726 |
| 0.758221 | 0.750991 | 0.750728 | 0.747986 | 0.749127 | 0.750222 | 0.748152 | 0.754918 | 0.748187 |
| 0.75981 | 0.748742 | 0.752501 | 0.745294 | 0.7514 | 0.749453 | 0.74547 | 0.757344 | 0.75531 |
| 0.757282 | 0.750768 | 0.75581 | 0.749113 | 0.757117 | 0.750591 | 0.748345 | 0.758834 | 0.752502 |
| 0.761706 | 0.749425 | 0.750526 | 0.74896 | 0.749401 | 0.744605 | 0.741535 | 0.752916 | 0.752575 |
| 0.757363 | 0.753451 | 0.747209 | 0.750512 | 0.752859 | 0.747732 | 0.744054 | 0.753714 | 0.749192 |
| 0.758822 | 0.751165 | 0.750567 | 0.75043 | 0.753334 | 0.749544 | 0.747878 | 0.755999 | 0.752535 |
| 0.760207 | 0.753187 | 0.75192 | 0.751574 | 0.759299 | 0.752096 | 0.749863 | 0.758491 | 0.757945 |
| 0.750693 | 0.743776 | 0.745404 | 0.744962 | 0.750117 | 0.745746 | 0.744086 | 0.746688 | 0.742528 |
| 0.75267 | 0.750944 | 0.744911 | 0.742844 | 0.747265 | 0.746107 | 0.742002 | 0.752564 | 0.746401 |
| 0.758198 | 0.75643 | 0.751639 | 0.752095 | 0.754062 | 0.751424 | 0.749635 | 0.755118 | 0.75024 |
| 0.765193 | 0.757906 | 0.756652 | 0.756568 | 0.758385 | 0.754585 | 0.75951 | 0.764869 | 0.752507 |
| 0.763385 | 0.756913 | 0.754556 | 0.754578 | 0.753689 | 0.754821 | 0.752947 | 0.756046 | 0.74793 |
| 0.763639 | 0.760131 | 0.756295 | 0.75542 | 0.755769 | 0.758027 | 0.75477 | 0.756871 | 0.75132 |
| 0.760492 | 0.758049 | 0.749735 | 0.750241 | 0.751571 | 0.751772 | 0.750162 | 0.751662 | 0.753785 |
| 0.761002 | 0.754147 | 0.752433 | 0.758478 | 0.756942 | 0.756292 | 0.754578 | 0.757719 | 0.75029 |
| 0.763939 | 0.753341 | 0.758174 | 0.748749 | 0.754427 | 0.751044 | 0.754056 | 0.760301 | 0.752264 |
| 0.757581 | 0.753263 | 0.751062 | 0.749403 | 0.751536 | 0.752706 | 0.749319 | 0.757004 | 0.74884 |
| 0.764079 | 0.755018 | 0.756936 | 0.758699 | 0.760184 | 0.756663 | 0.756754 | 0.763146 | 0.753437 |
| 0.757638 | 0.753348 | 0.751915 | 0.752495 | 0.74866 | 0.748788 | 0.749189 | 0.757034 | 0.746395 |
| 0.762474 | 0.755084 | 0.75732 | 0.749206 | 0.756593 | 0.750528 | 0.752991 | 0.760442 | 0.749975 |
| 0.699392 | 0.693932 | 0.692049 | 0.691944 | 0.691981 | 0.695842 | 0.685072 | 0.698085 | 0.693078 |
| 0.759921 | 0.749565 | 0.753425 | 0.749053 | 0.753778 | 0.750659 | 0.75175 | 0.759681 | 0.749392 |
| 0.758441 | 0.753698 | 0.754822 | 0.75103 | 0.755482 | 0.749062 | 0.747386 | 0.757451 | 0.747918 |
| 0.762068 | 0.758507 | 0.756897 | 0.754054 | 0.75562 | 0.754516 | 0.756828 | 0.76566 | 0.756318 |
| 0.76008 | 0.755295 | 0.755 | 0.754717 | 0.754725 | 0.751693 | 0.750293 | 0.759277 | 0.750447 |
| 0.760128 | 0.748582 | 0.750335 | 0.745571 | 0.751568 | 0.74886 | 0.749158 | 0.754604 | 0.750414 |
| 0.757532 | 0.75086 | 0.749802 | 0.751201 | 0.750447 | 0.751831 | 0.751464 | 0.757587 | 0.751769 |
| 0.759142 | 0.757181 | 0.753471 | 0.755323 | 0.753393 | 0.753584 | 0.7516 | 0.75785 | 0.751986 |
| 0.765203 | 0.751394 | 0.753872 | 0.747855 | 0.755612 | 0.751856 | 0.755974 | 0.762721 | 0.752079 |
| 0.751763 | 0.747522 | 0.748 | 0.747657 | 0.754703 | 0.743859 | 0.743443 | 0.751577 | 0.742467 |
| 0.755829 | 0.750766 | 0.748101 | 0.744887 | 0.749135 | 0.742934 | 0.740231 | 0.753835 | 0.744535 |
| 0.764358 | 0.755523 | 0.75489 | 0.751786 | 0.756776 | 0.754412 | 0.752748 | 0.756464 | 0.751147 |
| 0.761022 | 0.759251 | 0.758522 | 0.752749 | 0.755986 | 0.757233 | 0.75641 | 0.760014 | 0.756269 |
| 0.763778 | 0.760679 | 0.756593 | 0.755548 | 0.760011 | 0.756383 | 0.755499 | 0.758624 | 0.751989 |
| 0.758668 | 0.759745 | 0.753789 | 0.752228 | 0.75589 | 0.757056 | 0.75304 | 0.760054 | 0.756188 |
| 0.758258 | 0.752821 | 0.753564 | 0.746311 | 0.748512 | 0.7476 | 0.748209 | 0.755038 | 0.752761 |
| 0.764865 | 0.756883 | 0.760685 | 0.757123 | 0.759676 | 0.755431 | 0.75695 | 0.761354 | 0.752248 |
| 0.761588 | 0.751539 | 0.755875 | 0.749076 | 0.752958 | 0.752153 | 0.753677 | 0.756723 | 0.748276 |
| 0.756381 | 0.746937 | 0.748804 | 0.74286 | 0.749264 | 0.743764 | 0.747272 | 0.749487 | 0.750635 |
| 0.764362 | 0.752655 | 0.758052 | 0.750999 | 0.753912 | 0.751158 | 0.75061 | 0.758811 | 0.751748 |
| 0.760814 | 0.756605 | 0.756406 | 0.753607 | 0.753666 | 0.752592 | 0.747238 | 0.759023 | 0.752643 |

| | | | | | | | | |
|---|---|---|---|---|---|---|---|---|
| 0.760999 | 0.754502 | 0.754514 | 0.747939 | 0.754039 | 0.753624 | 0.749149 | 0.754174 | 0.753332 |
| 0.761968 | 0.753698 | 0.755797 | 0.751012 | 0.753721 | 0.754691 | 0.748616 | 0.757713 | 0.749789 |
| 0.758244 | 0.752247 | 0.75123 | 0.747402 | 0.749715 | 0.748641 | 0.750749 | 0.754465 | 0.747037 |
| 0.728629 | 0.709457 | 0.726944 | 0.712464 | 0.722509 | 0.70923 | 0.71476 | 0.719738 | 0.711864 |
| 0.756608 | 0.746696 | 0.750681 | 0.747186 | 0.751949 | 0.745006 | 0.746054 | 0.755472 | 0.742055 |
| 0.755443 | 0.749709 | 0.749249 | 0.75036 | 0.750414 | 0.74479 | 0.747214 | 0.75213 | 0.740953 |
| 0.764589 | 0.757578 | 0.760635 | 0.751838 | 0.757787 | 0.753124 | 0.752518 | 0.756676 | 0.756132 |
| 0.759346 | 0.75585 | 0.751859 | 0.750548 | 0.753801 | 0.752312 | 0.745974 | 0.756817 | 0.749272 |
| 0.76159 | 0.75205 | 0.758304 | 0.751643 | 0.753221 | 0.751645 | 0.752847 | 0.760554 | 0.748478 |
| 0.76284 | 0.752958 | 0.759325 | 0.752515 | 0.756206 | 0.747843 | 0.751011 | 0.760151 | 0.748289 |
| 0.760864 | 0.75326 | 0.752815 | 0.751528 | 0.756941 | 0.752832 | 0.745611 | 0.755663 | 0.749835 |
| 0.76179 | 0.752406 | 0.757277 | 0.750572 | 0.756941 | 0.749957 | 0.754073 | 0.754375 | 0.751062 |
| 0.761214 | 0.748878 | 0.752831 | 0.746079 | 0.7561 | 0.7506 | 0.750135 | 0.75466 | 0.746693 |
| 0.762642 | 0.748449 | 0.754507 | 0.745635 | 0.754063 | 0.749489 | 0.749173 | 0.758127 | 0.751032 |
| 0.765082 | 0.754752 | 0.758597 | 0.748926 | 0.755809 | 0.753535 | 0.748856 | 0.761179 | 0.752047 |
| 0.758446 | 0.750812 | 0.755009 | 0.744263 | 0.750044 | 0.753348 | 0.743049 | 0.753352 | 0.750613 |
| 0.761752 | 0.752288 | 0.756203 | 0.753188 | 0.755273 | 0.752592 | 0.751451 | 0.761628 | 0.753971 |
| 0.765276 | 0.753298 | 0.757819 | 0.749726 | 0.757897 | 0.751664 | 0.749081 | 0.760576 | 0.755223 |
| 0.756837 | 0.748051 | 0.750021 | 0.745958 | 0.749174 | 0.746679 | 0.747779 | 0.755232 | 0.743486 |
| 0.761372 | 0.754328 | 0.7555 | 0.750506 | 0.757214 | 0.753591 | 0.751463 | 0.759041 | 0.752167 |
| 0.765972 | 0.758063 | 0.76072 | 0.753816 | 0.756976 | 0.753469 | 0.755405 | 0.760217 | 0.754872 |
| 0.765487 | 0.756203 | 0.761434 | 0.754373 | 0.758834 | 0.756525 | 0.751391 | 0.762302 | 0.750514 |
| 0.68256 | 0.675222 | 0.675482 | 0.666728 | 0.677755 | 0.678989 | 0.675849 | 0.684554 | 0.676279 |
| 0.647951 | 0.650973 | 0.65687 | 0.64545 | 0.655647 | 0.65409 | 0.648008 | 0.650916 | 0.649331 |
| 0.66394 | 0.655531 | 0.663243 | 0.647949 | 0.660942 | 0.65172 | 0.652567 | 0.658167 | 0.658171 |
| 0.649223 | 0.644026 | 0.647243 | 0.644374 | 0.648033 | 0.647864 | 0.645342 | 0.651055 | 0.646314 |
| 0.722928 | 0.711895 | 0.724535 | 0.711521 | 0.718346 | 0.708532 | 0.71344 | 0.729693 | 0.72573 |
| 0.717753 | 0.708986 | 0.720602 | 0.699825 | 0.71121 | 0.70771 | 0.708465 | 0.722428 | 0.715544 |
| 0.720129 | 0.716236 | 0.723362 | 0.709748 | 0.724037 | 0.718067 | 0.718867 | 0.725375 | 0.721193 |
| 0.732667 | 0.72264 | 0.728821 | 0.723757 | 0.723152 | 0.72227 | 0.720381 | 0.731937 | 0.728038 |
| 0.708087 | 0.701613 | 0.698974 | 0.696503 | 0.708497 | 0.702373 | 0.704918 | 0.707917 | 0.708598 |
| 0.716879 | 0.692505 | 0.709142 | 0.6849 | 0.703067 | 0.689557 | 0.696812 | 0.719656 | 0.704456 |
| 0.704899 | 0.696566 | 0.711969 | 0.695158 | 0.708031 | 0.7059 | 0.696612 | 0.714115 | 0.704959 |
| 0.653764 | 0.649186 | 0.650313 | 0.644065 | 0.651589 | 0.647999 | 0.644071 | 0.656545 | 0.650588 |
| 0.752528 | 0.748471 | 0.749911 | 0.743281 | 0.749007 | 0.741889 | 0.742642 | 0.753523 | 0.743379 |
| 0.735006 | 0.724748 | 0.728073 | 0.72145 | 0.732391 | 0.7233 | 0.724729 | 0.733905 | 0.725297 |
| 0.730421 | 0.71754 | 0.727521 | 0.715466 | 0.731761 | 0.713832 | 0.724768 | 0.727849 | 0.714494 |
| 0.714867 | 0.706832 | 0.716462 | 0.699397 | 0.712786 | 0.709358 | 0.711391 | 0.726703 | 0.710327 |
| 0.734734 | 0.726317 | 0.733136 | 0.721845 | 0.72736 | 0.719342 | 0.730128 | 0.735035 | 0.725067 |
| 0.725297 | 0.710968 | 0.727743 | 0.715109 | 0.721521 | 0.715699 | 0.717137 | 0.728069 | 0.710211 |
| 0.72356 | 0.707567 | 0.721027 | 0.714704 | 0.720273 | 0.706446 | 0.714451 | 0.724384 | 0.713624 |
| 0.724509 | 0.713835 | 0.721183 | 0.719528 | 0.726143 | 0.716793 | 0.717687 | 0.723299 | 0.719709 |
| 0.714642 | 0.70856 | 0.72199 | 0.709017 | 0.720206 | 0.711631 | 0.721796 | 0.718927 | 0.710486 |
| 0.713306 | 0.708135 | 0.712404 | 0.71013 | 0.71875 | 0.69982 | 0.714836 | 0.71026 | 0.710521 |
| 0.715162 | 0.712536 | 0.718061 | 0.704995 | 0.716476 | 0.702022 | 0.709763 | 0.720864 | 0.699755 |
| 0.707309 | 0.700134 | 0.707575 | 0.697962 | 0.715292 | 0.697235 | 0.699088 | 0.707415 | 0.709587 |

| | | | | | | | | |
|---|---|---|---|---|---|---|---|---|
| 0.742533 | 0.736879 | 0.746054 | 0.733077 | 0.738061 | 0.728497 | 0.738639 | 0.744374 | 0.733555 |
| 0.719714 | 0.72049 | 0.724294 | 0.715984 | 0.72129 | 0.715442 | 0.719321 | 0.729486 | 0.720374 |
| 0.723121 | 0.716545 | 0.723175 | 0.714329 | 0.723571 | 0.713293 | 0.713952 | 0.723339 | 0.711377 |
| 0.718023 | 0.711289 | 0.719339 | 0.706534 | 0.713841 | 0.713658 | 0.716358 | 0.726833 | 0.718429 |
| 0.720978 | 0.709598 | 0.724759 | 0.718002 | 0.719305 | 0.711194 | 0.710758 | 0.727725 | 0.711986 |
| 0.713547 | 0.714898 | 0.720248 | 0.710392 | 0.714214 | 0.71275 | 0.712055 | 0.715795 | 0.707555 |
| 0.713214 | 0.704545 | 0.707973 | 0.702483 | 0.712844 | 0.704138 | 0.700197 | 0.716356 | 0.70793 |
| 0.71296 | 0.699035 | 0.718274 | 0.697395 | 0.714902 | 0.702054 | 0.711914 | 0.716867 | 0.703804 |
| 0.713124 | 0.705158 | 0.716714 | 0.699138 | 0.721868 | 0.706223 | 0.717003 | 0.718885 | 0.708333 |
| 0.744934 | 0.743084 | 0.746215 | 0.741147 | 0.750637 | 0.739736 | 0.743919 | 0.746962 | 0.740137 |
| 0.747264 | 0.73882 | 0.744872 | 0.733724 | 0.744854 | 0.732819 | 0.739855 | 0.749896 | 0.733131 |
| 0.743896 | 0.730184 | 0.740328 | 0.730308 | 0.734745 | 0.727314 | 0.734794 | 0.739333 | 0.7256 |
| 0.731403 | 0.724016 | 0.726816 | 0.721198 | 0.73109 | 0.721162 | 0.727112 | 0.7334 | 0.721656 |
| 0.737713 | 0.732726 | 0.732781 | 0.726661 | 0.732449 | 0.73187 | 0.735548 | 0.736604 | 0.735473 |
| 0.734155 | 0.72356 | 0.729286 | 0.72422 | 0.727571 | 0.720672 | 0.723259 | 0.733872 | 0.720726 |

| | | | | | | | | |
|---|---|---|---|---|---|---|---|---|
| | | | | | | plot 04 | | |
| Wild_plot_ | Wild_plot_ | Wild_plot_ | Wild_plot_ | Wild_plot_ | Wild_plot_ | Wild_plot_ | Wild_plot_ | Wild_plot_ |
| 0.708131 | 0.704108 | 0.703246 | 0.709227 | 0.71341 | 0.710311 | 0.704957 | 0.707338 | 0.700948 |
| 0.7114 | 0.712498 | 0.711138 | 0.716372 | 0.719862 | 0.714434 | 0.709664 | 0.712007 | 0.707566 |
| 0.705026 | 0.703244 | 0.70315 | 0.704792 | 0.708541 | 0.702275 | 0.700214 | 0.705139 | 0.695521 |
| 0.712866 | 0.715378 | 0.708427 | 0.715242 | 0.722059 | 0.714737 | 0.711093 | 0.7151 | 0.707942 |
| 0.709549 | 0.710709 | 0.704893 | 0.710852 | 0.714683 | 0.711271 | 0.705276 | 0.711099 | 0.699359 |
| 0.718104 | 0.716999 | 0.712223 | 0.718577 | 0.724765 | 0.717837 | 0.712776 | 0.72149 | 0.710487 |
| 0.765107 | 0.775609 | 0.772595 | 0.765887 | 0.775232 | 0.770318 | 0.767825 | 0.763312 | 0.768389 |
| 0.761212 | 0.771626 | 0.772193 | 0.766408 | 0.77972 | 0.764658 | 0.761317 | 0.76171 | 0.761014 |
| 0.715947 | 0.710427 | 0.710202 | 0.711945 | 0.71867 | 0.715161 | 0.713313 | 0.710503 | 0.705944 |
| 0.706475 | 0.706623 | 0.705727 | 0.709281 | 0.714593 | 0.707972 | 0.707618 | 0.710014 | 0.702362 |
| 0.709757 | 0.708918 | 0.702764 | 0.707958 | 0.715769 | 0.710392 | 0.707548 | 0.711275 | 0.704213 |
| 0.704099 | 0.706128 | 0.698798 | 0.703147 | 0.712013 | 0.710245 | 0.701049 | 0.702564 | 0.698654 |
| 0.715958 | 0.710486 | 0.709482 | 0.71206 | 0.718264 | 0.711543 | 0.708053 | 0.711166 | 0.708745 |
| 0.741614 | 0.723127 | 0.72106 | 0.723839 | 0.728278 | 0.722603 | 0.722741 | 0.723176 | 0.717416 |
| 0.761571 | 0.734106 | 0.733268 | 0.73423 | 0.743264 | 0.734237 | 0.730074 | 0.735174 | 0.726947 |
| 0.717365 | 0.715356 | 0.714217 | 0.718597 | 0.719426 | 0.718952 | 0.711966 | 0.714104 | 0.70771 |
| 0.716695 | 0.717181 | 0.712041 | 0.714115 | 0.719475 | 0.715543 | 0.710594 | 0.713624 | 0.708187 |
| 0.789452 | 0.743389 | 0.741113 | 0.746836 | 0.751563 | 0.744545 | 0.742633 | 0.741248 | 0.738245 |
| 0.706226 | 0.704173 | 0.702655 | 0.705612 | 0.709698 | 0.706322 | 0.705589 | 0.705396 | 0.702078 |
| 0.709381 | 0.710167 | 0.70611 | 0.710745 | 0.717249 | 0.71172 | 0.704497 | 0.711529 | 0.705017 |
| 0.709577 | 0.708139 | 0.70566 | 0.709955 | 0.717109 | 0.712993 | 0.70367 | 0.708587 | 0.702771 |
| 0.709443 | 0.708096 | 0.70752 | 0.711705 | 0.719262 | 0.711065 | 0.705318 | 0.706498 | 0.702191 |
| 0.71537 | 0.716226 | 0.712902 | 0.717051 | 0.724016 | 0.720134 | 0.710677 | 0.717124 | 0.709088 |
| 0.716194 | 0.715509 | 0.709125 | 0.714987 | 0.720964 | 0.716524 | 0.709426 | 0.712583 | 0.706128 |
| 0.713301 | 0.713868 | 0.709931 | 0.713945 | 0.718572 | 0.716969 | 0.70815 | 0.714845 | 0.70974 |
| 0.712698 | 0.712109 | 0.706432 | 0.710723 | 0.718134 | 0.711717 | 0.706976 | 0.709462 | 0.702911 |
| 0.709819 | 0.706081 | 0.701117 | 0.706482 | 0.712668 | 0.709288 | 0.701495 | 0.706274 | 0.701325 |
| 0.710338 | 0.709321 | 0.70867 | 0.7136 | 0.717469 | 0.714229 | 0.708842 | 0.714681 | 0.704628 |
| 0.714331 | 0.714343 | 0.711481 | 0.71329 | 0.719981 | 0.716846 | 0.709302 | 0.713706 | 0.708261 |
| 0.705586 | 0.713926 | 0.707828 | 0.715878 | 0.717152 | 0.715335 | 0.711065 | 0.713384 | 0.709867 |
| 0.707861 | 0.708609 | 0.705977 | 0.710765 | 0.716528 | 0.712635 | 0.705375 | 0.710734 | 0.705096 |
| 0.710042 | 0.707088 | 0.700855 | 0.708494 | 0.712312 | 0.709978 | 0.70142 | 0.706216 | 0.699792 |
| 0.708635 | 0.709584 | 0.705892 | 0.709998 | 0.715886 | 0.715304 | 0.706188 | 0.710391 | 0.70412 |
| 0.709685 | 0.714493 | 0.707781 | 0.712878 | 0.720769 | 0.714105 | 0.708388 | 0.712591 | 0.70673 |
| 0.705826 | 0.706787 | 0.703224 | 0.707392 | 0.711612 | 0.707052 | 0.701743 | 0.704932 | 0.700244 |
| 0.712885 | 0.714297 | 0.709424 | 0.713022 | 0.716256 | 0.709927 | 0.705026 | 0.711374 | 0.706011 |
| 0.713378 | 0.71589 | 0.71219 | 0.717121 | 0.722213 | 0.720472 | 0.710509 | 0.711934 | 0.710156 |
| 0.709958 | 0.71073 | 0.710269 | 0.712233 | 0.719179 | 0.714584 | 0.707668 | 0.71647 | 0.708363 |
| 0.717017 | 0.712504 | 0.712296 | 0.715675 | 0.721131 | 0.715578 | 0.710837 | 0.718312 | 0.709388 |

| | | | | | | | | |
|---|---|---|---|---|---|---|---|---|
| 0.796154 | 0.748882 | 0.743867 | 0.748068 | 0.754767 | 0.75004 | 0.748199 | 0.74616 | 0.739237 |
| 0.701502 | 0.703338 | 0.6981 | 0.704281 | 0.710215 | 0.704078 | 0.698287 | 0.706148 | 0.694435 |
| 0.707 | 0.708515 | 0.702218 | 0.705971 | 0.713909 | 0.707913 | 0.702783 | 0.707608 | 0.695509 |
| 0.714683 | 0.713988 | 0.707557 | 0.714218 | 0.723039 | 0.718001 | 0.710364 | 0.714003 | 0.711528 |
| 0.708469 | 0.709031 | 0.708122 | 0.711884 | 0.717548 | 0.714433 | 0.704741 | 0.713072 | 0.704334 |
| 0.718974 | 0.713081 | 0.710603 | 0.716029 | 0.717757 | 0.714365 | 0.710581 | 0.711436 | 0.706242 |
| 0.709754 | 0.706631 | 0.701344 | 0.709446 | 0.714763 | 0.711834 | 0.704425 | 0.708686 | 0.702665 |
| 0.712161 | 0.708276 | 0.708654 | 0.711697 | 0.72019 | 0.712859 | 0.705203 | 0.711264 | 0.705003 |
| 0.710947 | 0.713612 | 0.710783 | 0.70967 | 0.721301 | 0.715235 | 0.710173 | 0.715784 | 0.709578 |
| 0.711392 | 0.711084 | 0.706297 | 0.712505 | 0.718004 | 0.714014 | 0.708498 | 0.713122 | 0.705698 |
| 0.704753 | 0.705122 | 0.700194 | 0.70689 | 0.707858 | 0.707164 | 0.702651 | 0.70038 | 0.700655 |
| 0.702425 | 0.70405 | 0.702047 | 0.705757 | 0.712229 | 0.707092 | 0.700265 | 0.705612 | 0.699321 |
| 0.710433 | 0.710479 | 0.706064 | 0.710761 | 0.718058 | 0.70959 | 0.704988 | 0.707282 | 0.701256 |
| 0.706354 | 0.70833 | 0.702281 | 0.705216 | 0.713301 | 0.70629 | 0.70077 | 0.706037 | 0.700166 |
| 0.708007 | 0.705255 | 0.702451 | 0.702651 | 0.714193 | 0.709908 | 0.701067 | 0.705995 | 0.698126 |
| 0.708023 | 0.705911 | 0.704009 | 0.705326 | 0.712214 | 0.705267 | 0.702998 | 0.703448 | 0.698201 |
| 0.704958 | 0.703845 | 0.702747 | 0.704129 | 0.710755 | 0.707022 | 0.702386 | 0.705024 | 0.693772 |
| 0.711017 | 0.711365 | 0.707613 | 0.711 | 0.718349 | 0.712698 | 0.707509 | 0.706538 | 0.701677 |
| 0.716334 | 0.715199 | 0.709994 | 0.712606 | 0.720125 | 0.716709 | 0.711642 | 0.715002 | 0.710377 |
| 0.706038 | 0.707709 | 0.706417 | 0.70921 | 0.715879 | 0.713999 | 0.704814 | 0.711203 | 0.701109 |
| 0.704837 | 0.703259 | 0.698445 | 0.703344 | 0.714377 | 0.70255 | 0.702105 | 0.702593 | 0.696744 |
| 0.70553 | 0.703984 | 0.703226 | 0.705351 | 0.712883 | 0.706797 | 0.703172 | 0.704672 | 0.699564 |
| 0.709629 | 0.704576 | 0.705572 | 0.708546 | 0.713278 | 0.709515 | 0.706337 | 0.709968 | 0.69756 |
| 0.720463 | 0.718057 | 0.71344 | 0.717761 | 0.724614 | 0.717687 | 0.715797 | 0.717388 | 0.708253 |
| 0.72504 | 0.725154 | 0.720071 | 0.724647 | 0.731748 | 0.729481 | 0.720123 | 0.725449 | 0.720084 |
| 0.762379 | 0.772422 | 0.770482 | 0.766615 | 0.780514 | 0.766186 | 0.762268 | 0.763114 | 0.761085 |
| 0.760781 | 0.77512 | 0.769794 | 0.768566 | 0.774896 | 0.771509 | 0.771343 | 0.770651 | 0.764238 |
| 0.766722 | 0.778021 | 0.771683 | 0.768171 | 0.775863 | 0.772874 | 0.771843 | 0.773159 | 0.767366 |
| 0.707002 | 0.704677 | 0.703266 | 0.708422 | 0.713887 | 0.707798 | 0.701659 | 0.70672 | 0.700991 |
| 0.705873 | 0.706462 | 0.702662 | 0.706821 | 0.711371 | 0.704922 | 0.698206 | 0.705484 | 0.701373 |
| 0.713977 | 0.715107 | 0.711836 | 0.719095 | 0.721915 | 0.716974 | 0.709629 | 0.71539 | 0.707602 |
| 0.714679 | 0.715453 | 0.71251 | 0.718243 | 0.724144 | 0.715356 | 0.713421 | 0.71424 | 0.710644 |
| 0.711345 | 0.705074 | 0.70499 | 0.705699 | 0.711453 | 0.708286 | 0.70415 | 0.705897 | 0.698604 |
| 0.710543 | 0.709586 | 0.704448 | 0.709575 | 0.716699 | 0.710928 | 0.706054 | 0.70848 | 0.705201 |
| 0.706711 | 0.703448 | 0.703943 | 0.705991 | 0.710624 | 0.707398 | 0.704095 | 0.704138 | 0.701798 |
| 0.710814 | 0.709035 | 0.705416 | 0.709539 | 0.717685 | 0.708804 | 0.705418 | 0.705256 | 0.699747 |
| 0.711058 | 0.705845 | 0.709273 | 0.709921 | 0.71248 | 0.708075 | 0.704988 | 0.705596 | 0.703425 |
| 0.724227 | 0.720798 | 0.71609 | 0.724014 | 0.729312 | 0.72137 | 0.717573 | 0.721555 | 0.717008 |
| 0.700083 | 0.69844 | 0.697883 | 0.697602 | 0.705912 | 0.701697 | 0.696558 | 0.699316 | 0.69425 |
| 0.707622 | 0.707571 | 0.706493 | 0.71012 | 0.719546 | 0.710444 | 0.704396 | 0.708861 | 0.701261 |
| 0.707844 | 0.702459 | 0.700614 | 0.70413 | 0.713618 | 0.707582 | 0.701682 | 0.703008 | 0.697972 |
| 0.725793 | 0.723731 | 0.718798 | 0.724683 | 0.730145 | 0.724713 | 0.718088 | 0.721281 | 0.719922 |
| 0.706357 | 0.714239 | 0.704799 | 0.712889 | 0.717821 | 0.710719 | 0.707635 | 0.710673 | 0.70225 |
| 0.704495 | 0.706597 | 0.703325 | 0.709195 | 0.712701 | 0.707325 | 0.701916 | 0.710416 | 0.698328 |
| 0.714068 | 0.711517 | 0.70612 | 0.711003 | 0.719713 | 0.715727 | 0.709885 | 0.712058 | 0.705517 |
| 0.70749 | 0.705983 | 0.707003 | 0.705939 | 0.715046 | 0.70817 | 0.702321 | 0.706844 | 0.698478 |

| | | | | | | | | |
|---|---|---|---|---|---|---|---|---|
| 0.704858 | 0.710484 | 0.704164 | 0.706898 | 0.716173 | 0.712306 | 0.703473 | 0.708303 | 0.702315 |
| 0.714696 | 0.710861 | 0.706071 | 0.715133 | 0.717626 | 0.712165 | 0.705287 | 0.711347 | 0.70751 |
| 0.713824 | 0.710226 | 0.704767 | 0.710113 | 0.713769 | 0.708746 | 0.70679 | 0.706561 | 0.700473 |
| 0.716966 | 0.713436 | 0.710073 | 0.717161 | 0.722636 | 0.71806 | 0.712265 | 0.712701 | 0.710721 |
| 0.71895 | 0.714791 | 0.714816 | 0.716622 | 0.72482 | 0.719017 | 0.709704 | 0.714794 | 0.709773 |
| 0.710306 | 0.709966 | 0.708788 | 0.708062 | 0.719336 | 0.710873 | 0.707433 | 0.709869 | 0.702367 |
| 0.71106 | 0.710946 | 0.71094 | 0.713904 | 0.717886 | 0.714252 | 0.708165 | 0.708979 | 0.705327 |
| 0.712813 | 0.713821 | 0.709363 | 0.71926 | 0.716705 | 0.715212 | 0.711195 | 0.713007 | 0.709136 |
| 0.710979 | 0.710028 | 0.711493 | 0.709403 | 0.715593 | 0.712357 | 0.709677 | 0.709284 | 0.704018 |
| 0.707327 | 0.706762 | 0.704704 | 0.708515 | 0.713774 | 0.710174 | 0.704545 | 0.706325 | 0.701442 |
| 0.709826 | 0.70834 | 0.704785 | 0.710322 | 0.715258 | 0.71136 | 0.704287 | 0.706354 | 0.70326 |
| 0.708897 | 0.710004 | 0.705786 | 0.707259 | 0.716343 | 0.711057 | 0.704218 | 0.707486 | 0.705025 |
| 0.711892 | 0.713286 | 0.714274 | 0.712487 | 0.713748 | 0.713018 | 0.710038 | 0.712099 | 0.706043 |
| 0.704444 | 0.709247 | 0.70049 | 0.708637 | 0.713742 | 0.709007 | 0.706249 | 0.708024 | 0.704867 |
| 0.71094 | 0.712374 | 0.707528 | 0.708294 | 0.718206 | 0.714012 | 0.706709 | 0.71081 | 0.70082 |
| 0.705985 | 0.709457 | 0.704294 | 0.710821 | 0.715364 | 0.711718 | 0.705024 | 0.710606 | 0.700892 |
| 0.715902 | 0.712637 | 0.70781 | 0.713112 | 0.718017 | 0.712776 | 0.709338 | 0.713725 | 0.70748 |
| 0.711882 | 0.706829 | 0.706204 | 0.708041 | 0.7161 | 0.708656 | 0.701121 | 0.706116 | 0.701642 |
| 0.709197 | 0.712216 | 0.707643 | 0.709741 | 0.717778 | 0.714149 | 0.706302 | 0.71259 | 0.70413 |
| 0.709922 | 0.711364 | 0.707376 | 0.717262 | 0.717914 | 0.714434 | 0.710032 | 0.712682 | 0.708819 |
| 0.707012 | 0.709604 | 0.702509 | 0.709754 | 0.715305 | 0.709549 | 0.704214 | 0.709088 | 0.702549 |
| 0.704222 | 0.70656 | 0.703742 | 0.706454 | 0.708847 | 0.704198 | 0.700619 | 0.705962 | 0.699254 |
| 0.710484 | 0.71021 | 0.707194 | 0.709885 | 0.714068 | 0.71123 | 0.702842 | 0.707867 | 0.698942 |
| 0.712646 | 0.713548 | 0.705443 | 0.711581 | 0.721592 | 0.713038 | 0.705625 | 0.712526 | 0.705065 |
| 0.711469 | 0.709283 | 0.708867 | 0.711323 | 0.716373 | 0.712015 | 0.708679 | 0.708755 | 0.705598 |
| 0.713583 | 0.713702 | 0.713116 | 0.716178 | 0.72164 | 0.718379 | 0.710273 | 0.717045 | 0.709092 |
| 0.712742 | 0.706508 | 0.708096 | 0.710189 | 0.716749 | 0.712434 | 0.707253 | 0.707972 | 0.70439 |
| 0.710088 | 0.710839 | 0.705161 | 0.713642 | 0.716319 | 0.712138 | 0.71011 | 0.711285 | 0.708668 |
| 0.71187 | 0.715508 | 0.709422 | 0.712045 | 0.718743 | 0.714558 | 0.710755 | 0.713776 | 0.709222 |
| 0.714077 | 0.709701 | 0.708047 | 0.711636 | 0.715686 | 0.714286 | 0.706365 | 0.707578 | 0.706808 |
| 0.712457 | 0.713588 | 0.710119 | 0.714799 | 0.721509 | 0.713131 | 0.707037 | 0.711475 | 0.700044 |
| 0.710543 | 0.707199 | 0.705324 | 0.711508 | 0.717292 | 0.709662 | 0.706152 | 0.707692 | 0.701428 |
| 0.708192 | 0.708847 | 0.708442 | 0.70787 | 0.715151 | 0.711968 | 0.705572 | 0.71141 | 0.70432 |
| 0.711222 | 0.710827 | 0.704841 | 0.711163 | 0.720355 | 0.714718 | 0.706231 | 0.709712 | 0.708508 |
| 0.710605 | 0.707362 | 0.705949 | 0.709963 | 0.715031 | 0.708134 | 0.704532 | 0.705174 | 0.702797 |
| 0.708743 | 0.709372 | 0.707678 | 0.709457 | 0.717749 | 0.712061 | 0.705561 | 0.705688 | 0.702415 |
| 0.70361 | 0.700292 | 0.703619 | 0.704384 | 0.711951 | 0.709625 | 0.701205 | 0.702262 | 0.700884 |
| 0.679688 | 0.687615 | 0.677635 | 0.68633 | 0.690817 | 0.691108 | 0.68225 | 0.680795 | 0.687727 |
| 0.689403 | 0.689112 | 0.684351 | 0.686811 | 0.697264 | 0.688533 | 0.687403 | 0.686051 | 0.690002 |
| 0.680302 | 0.678948 | 0.676124 | 0.678013 | 0.683517 | 0.686659 | 0.680547 | 0.680799 | 0.683041 |
| 0.67951 | 0.681117 | 0.67754 | 0.681003 | 0.691244 | 0.680945 | 0.674521 | 0.680009 | 0.679463 |
| 0.665102 | 0.66698 | 0.656084 | 0.665003 | 0.67115 | 0.670742 | 0.664315 | 0.665732 | 0.671611 |
| 0.665536 | 0.673288 | 0.660465 | 0.670768 | 0.676104 | 0.663317 | 0.666012 | 0.668033 | 0.673186 |
| 0.668661 | 0.667967 | 0.659431 | 0.668111 | 0.67511 | 0.668438 | 0.668484 | 0.662165 | 0.667928 |
| 0.674502 | 0.671113 | 0.663163 | 0.662845 | 0.678754 | 0.672374 | 0.662921 | 0.670062 | 0.670369 |
| 0.667622 | 0.667419 | 0.664256 | 0.663782 | 0.672058 | 0.668034 | 0.662217 | 0.664175 | 0.669373 |
| 0.675323 | 0.671675 | 0.664502 | 0.671986 | 0.681762 | 0.676681 | 0.669407 | 0.670358 | 0.678232 |

| | | | | | | | | |
|---|---|---|---|---|---|---|---|---|
| 0.66774 | 0.674349 | 0.664717 | 0.669721 | 0.679457 | 0.674932 | 0.667912 | 0.666993 | 0.676721 |
| 0.674087 | 0.672193 | 0.665299 | 0.671119 | 0.674772 | 0.676765 | 0.670011 | 0.670173 | 0.673098 |
| 0.670253 | 0.67155 | 0.663634 | 0.670432 | 0.67783 | 0.670785 | 0.669973 | 0.671923 | 0.675125 |
| 0.664989 | 0.667465 | 0.656794 | 0.662157 | 0.669402 | 0.668377 | 0.661239 | 0.668741 | 0.661262 |
| 0.667011 | 0.665721 | 0.660544 | 0.662258 | 0.667052 | 0.670004 | 0.662449 | 0.665215 | 0.668742 |
| 0.675231 | 0.668415 | 0.657502 | 0.668121 | 0.671775 | 0.665078 | 0.663989 | 0.662214 | 0.669124 |
| 0.672382 | 0.671327 | 0.660337 | 0.668554 | 0.673768 | 0.668744 | 0.661558 | 0.664733 | 0.669831 |
| 0.666526 | 0.661567 | 0.658877 | 0.656791 | 0.67318 | 0.668948 | 0.663648 | 0.66012 | 0.668987 |
| 0.665976 | 0.663524 | 0.660786 | 0.658796 | 0.66975 | 0.662056 | 0.664783 | 0.659453 | 0.665013 |
| 0.664766 | 0.664221 | 0.659354 | 0.663673 | 0.666229 | 0.669288 | 0.663554 | 0.66555 | 0.672268 |
| 0.662899 | 0.660624 | 0.65011 | 0.660783 | 0.667169 | 0.660359 | 0.655196 | 0.654582 | 0.662141 |
| 0.696677 | 0.699547 | 0.691747 | 0.701901 | 0.706418 | 0.700031 | 0.696223 | 0.70025 | 0.693609 |
| 0.706316 | 0.707943 | 0.701294 | 0.711003 | 0.714908 | 0.709182 | 0.701434 | 0.706086 | 0.700181 |
| 0.687186 | 0.6877 | 0.674951 | 0.682663 | 0.691019 | 0.684525 | 0.681702 | 0.682623 | 0.686414 |
| 0.718089 | 0.714694 | 0.711466 | 0.715215 | 0.721916 | 0.716211 | 0.71377 | 0.71333 | 0.712902 |
| 0.704642 | 0.705484 | 0.70045 | 0.708679 | 0.710399 | 0.706489 | 0.702989 | 0.704271 | 0.697614 |
| 0.713616 | 0.712443 | 0.713792 | 0.716362 | 0.722037 | 0.717522 | 0.709932 | 0.713743 | 0.709288 |
| 0.72141 | 0.716772 | 0.715149 | 0.720794 | 0.725119 | 0.719852 | 0.71357 | 0.71954 | 0.714122 |
| 0.69641 | 0.698754 | 0.691612 | 0.698769 | 0.70663 | 0.696944 | 0.694949 | 0.693137 | 0.686614 |
| 0.721342 | 0.720547 | 0.714892 | 0.717753 | 0.726309 | 0.722794 | 0.710479 | 0.719312 | 0.715067 |
| 0.695088 | 0.692019 | 0.685889 | 0.689556 | 0.696981 | 0.689729 | 0.684159 | 0.688463 | 0.688401 |
| 0.717671 | 0.716452 | 0.713208 | 0.718128 | 0.722485 | 0.719002 | 0.712311 | 0.717136 | 0.713495 |
| 0.720325 | 0.718463 | 0.713904 | 0.723992 | 0.72515 | 0.721138 | 0.71654 | 0.72016 | 0.714373 |
| 0.687025 | 0.682591 | 0.676156 | 0.678493 | 0.69309 | 0.681067 | 0.676887 | 0.67858 | 0.685822 |
| 0.690265 | 0.686778 | 0.681231 | 0.68321 | 0.693167 | 0.684875 | 0.684282 | 0.685015 | 0.683308 |
| 0.694621 | 0.695122 | 0.683418 | 0.689369 | 0.697392 | 0.692802 | 0.687941 | 0.68801 | 0.690892 |
| 0.711488 | 0.707723 | 0.706789 | 0.703743 | 0.712646 | 0.708235 | 0.70333 | 0.704417 | 0.69906 |
| 0.713126 | 0.718995 | 0.713883 | 0.719997 | 0.725638 | 0.717955 | 0.715328 | 0.71957 | 0.711844 |
| 0.690787 | 0.687299 | 0.680824 | 0.690012 | 0.691895 | 0.689394 | 0.683223 | 0.68678 | 0.681429 |
| 0.724099 | 0.722151 | 0.719891 | 0.724396 | 0.727359 | 0.720378 | 0.719476 | 0.721787 | 0.715611 |
| 0.72772 | 0.723386 | 0.721734 | 0.728322 | 0.734239 | 0.731249 | 0.725982 | 0.724752 | 0.723515 |
| 0.700622 | 0.696975 | 0.690083 | 0.700055 | 0.704187 | 0.698216 | 0.689459 | 0.700906 | 0.694252 |
| 0.699334 | 0.695957 | 0.694194 | 0.700564 | 0.704263 | 0.696729 | 0.688854 | 0.698498 | 0.691446 |
| 0.725712 | 0.728071 | 0.721908 | 0.722928 | 0.73167 | 0.728442 | 0.719615 | 0.72421 | 0.717001 |
| 0.720793 | 0.72148 | 0.716297 | 0.721607 | 0.728162 | 0.723267 | 0.718556 | 0.719931 | 0.713659 |
| 0.723296 | 0.721779 | 0.720427 | 0.722116 | 0.727543 | 0.723455 | 0.722594 | 0.723269 | 0.716128 |
| 0.677276 | 0.67955 | 0.673178 | 0.677912 | 0.684399 | 0.677847 | 0.674548 | 0.677063 | 0.670525 |
| 0.720259 | 0.71972 | 0.716026 | 0.71929 | 0.727135 | 0.720146 | 0.71652 | 0.719209 | 0.713839 |
| 0.673258 | 0.67627 | 0.669971 | 0.675495 | 0.682206 | 0.676129 | 0.671606 | 0.677553 | 0.670792 |
| 0.671181 | 0.667765 | 0.663223 | 0.671259 | 0.672199 | 0.669962 | 0.665015 | 0.667246 | 0.661508 |
| 0.74709 | 0.716725 | 0.716771 | 0.722231 | 0.729417 | 0.722764 | 0.716073 | 0.71988 | 0.712974 |
| 0.672004 | 0.672806 | 0.666334 | 0.670963 | 0.676845 | 0.670526 | 0.66475 | 0.668131 | 0.664125 |
| 0.663022 | 0.660986 | 0.658928 | 0.66259 | 0.667847 | 0.662931 | 0.65817 | 0.664317 | 0.655385 |
| 0.659893 | 0.659981 | 0.656533 | 0.657813 | 0.663248 | 0.657751 | 0.654492 | 0.658388 | 0.652794 |
| 0.753329 | 0.731371 | 0.725471 | 0.727194 | 0.732641 | 0.729059 | 0.729484 | 0.729308 | 0.720534 |
| 0.662392 | 0.664608 | 0.658563 | 0.663831 | 0.667745 | 0.662182 | 0.659362 | 0.665715 | 0.657394 |
| 0.743331 | 0.718917 | 0.716082 | 0.720875 | 0.723713 | 0.721459 | 0.717673 | 0.719057 | 0.712189 |

| | | | | | | | | |
|---|---|---|---|---|---|---|---|---|
| 0.765984 | 0.731344 | 0.728331 | 0.730507 | 0.735271 | 0.731911 | 0.725593 | 0.731884 | 0.727177 |
| 0.755142 | 0.724055 | 0.720006 | 0.725428 | 0.730305 | 0.725799 | 0.721378 | 0.725181 | 0.718439 |
| 0.743058 | 0.747746 | 0.741645 | 0.746985 | 0.753336 | 0.753513 | 0.753747 | 0.749368 | 0.747656 |
| 0.654911 | 0.653494 | 0.645896 | 0.651826 | 0.656919 | 0.653804 | 0.64881 | 0.65309 | 0.644444 |
| 0.655591 | 0.654229 | 0.648671 | 0.65215 | 0.655272 | 0.651318 | 0.648497 | 0.652503 | 0.647242 |
| 0.760281 | 0.733279 | 0.726417 | 0.731057 | 0.740279 | 0.733264 | 0.729445 | 0.730201 | 0.727505 |
| 0.654457 | 0.652589 | 0.649962 | 0.654284 | 0.658537 | 0.65172 | 0.648776 | 0.655695 | 0.648811 |
| 0.730199 | 0.730969 | 0.729179 | 0.737643 | 0.743546 | 0.735633 | 0.732584 | 0.736073 | 0.726097 |
| 0.76584 | 0.730979 | 0.729048 | 0.729866 | 0.738384 | 0.734894 | 0.728609 | 0.730649 | 0.725266 |
| 0.7314 | 0.734966 | 0.727535 | 0.738037 | 0.741917 | 0.735126 | 0.732172 | 0.736683 | 0.726649 |
| 0.782612 | 0.733306 | 0.727571 | 0.727513 | 0.736625 | 0.732897 | 0.725128 | 0.731372 | 0.723952 |
| 0.698113 | 0.75 | 0.756944 | 0.717105 | 0.782609 | 0.716129 | 0.690141 | 0.726667 | 0.724138 |
| 0.718505 | 0.738231 | 0.717793 | 0.71718 | 0.73332 | 0.731864 | 0.727971 | 0.720697 | 0.724342 |
| 0.714599 | 0.736142 | 0.71922 | 0.728867 | 0.736842 | 0.723923 | 0.726379 | 0.719831 | 0.724803 |
| 0.724868 | 0.732919 | 0.722604 | 0.723398 | 0.739933 | 0.735764 | 0.726429 | 0.730427 | 0.732702 |
| 0.727616 | 0.728057 | 0.722207 | 0.722318 | 0.738204 | 0.727613 | 0.721687 | 0.718079 | 0.72721 |
| 0.588025 | 0.595212 | 0.584209 | 0.588657 | 0.597985 | 0.587038 | 0.587588 | 0.589892 | 0.591166 |
| 0.730447 | 0.737683 | 0.735259 | 0.727801 | 0.741316 | 0.733828 | 0.734551 | 0.726989 | 0.730018 |
| 0.788692 | 0.733881 | 0.736642 | 0.736626 | 0.743734 | 0.739742 | 0.735608 | 0.734727 | 0.727604 |
| 0.790199 | 0.737851 | 0.733455 | 0.73317 | 0.740903 | 0.739291 | 0.729039 | 0.730864 | 0.726939 |
| 0.787879 | 0.735537 | 0.73591 | 0.738184 | 0.744543 | 0.738573 | 0.735663 | 0.73802 | 0.728737 |
| 0.595461 | 0.60288 | 0.591846 | 0.599238 | 0.604418 | 0.596947 | 0.59296 | 0.594797 | 0.586118 |
| 0.583142 | 0.585192 | 0.581297 | 0.584444 | 0.58732 | 0.582751 | 0.577005 | 0.584559 | 0.573754 |
| 0.827937 | 0.764825 | 0.760576 | 0.755951 | 0.768115 | 0.761036 | 0.763119 | 0.759837 | 0.752808 |
| 0.814105 | 0.763552 | 0.763125 | 0.760634 | 0.771323 | 0.76448 | 0.765719 | 0.764855 | 0.756915 |
| 0.786989 | 0.757075 | 0.752465 | 0.751184 | 0.760738 | 0.757673 | 0.757503 | 0.760819 | 0.747386 |
| 0.787689 | 0.761522 | 0.757588 | 0.755129 | 0.76568 | 0.759912 | 0.759901 | 0.769104 | 0.749844 |
| 0.80488 | 0.763404 | 0.756269 | 0.75804 | 0.770328 | 0.761238 | 0.765219 | 0.761552 | 0.758859 |
| 0.801783 | 0.764765 | 0.765426 | 0.766121 | 0.76941 | 0.764433 | 0.762769 | 0.76675 | 0.761096 |
| 0.800942 | 0.767034 | 0.762688 | 0.760932 | 0.769807 | 0.768314 | 0.764827 | 0.767051 | 0.763127 |
| 0.758363 | 0.764834 | 0.759156 | 0.763686 | 0.773276 | 0.770764 | 0.765484 | 0.764272 | 0.771585 |
| 0.766972 | 0.774774 | 0.768696 | 0.772296 | 0.780644 | 0.783611 | 0.779744 | 0.771366 | 0.77469 |
| 0.759927 | 0.763892 | 0.760222 | 0.766403 | 0.772308 | 0.76874 | 0.768003 | 0.763084 | 0.767567 |
| 0.76246 | 0.768406 | 0.767038 | 0.766303 | 0.774853 | 0.769562 | 0.773998 | 0.76678 | 0.773299 |
| 0.76167 | 0.777156 | 0.772875 | 0.775614 | 0.776332 | 0.772041 | 0.775812 | 0.774869 | 0.774022 |
| 0.756527 | 0.765698 | 0.760746 | 0.76586 | 0.769015 | 0.768153 | 0.771315 | 0.767791 | 0.765259 |
| 0.76615 | 0.773966 | 0.766203 | 0.763706 | 0.768398 | 0.776308 | 0.76612 | 0.772727 | 0.771234 |
| 0.764773 | 0.767783 | 0.759474 | 0.765539 | 0.773131 | 0.772366 | 0.770954 | 0.770206 | 0.774578 |
| 0.758562 | 0.767368 | 0.762697 | 0.760077 | 0.774451 | 0.7714 | 0.765186 | 0.767281 | 0.763717 |
| 0.761971 | 0.769161 | 0.758729 | 0.765256 | 0.77525 | 0.767556 | 0.770988 | 0.770619 | 0.768016 |
| 0.715013 | 0.733475 | 0.719821 | 0.714459 | 0.736165 | 0.72913 | 0.725115 | 0.721355 | 0.731571 |
| 0.731517 | 0.733067 | 0.717787 | 0.723034 | 0.738754 | 0.730603 | 0.72748 | 0.724651 | 0.728571 |
| 0.764257 | 0.767455 | 0.761826 | 0.768999 | 0.774268 | 0.771994 | 0.763652 | 0.765876 | 0.763864 |
| 0.784156 | 0.772595 | 0.769148 | 0.769206 | 0.773957 | 0.767645 | 0.772501 | 0.773387 | 0.76334 |
| 0.7157 | 0.731478 | 0.711685 | 0.71464 | 0.734111 | 0.73166 | 0.720405 | 0.719461 | 0.725595 |
| 0.763785 | 0.768897 | 0.761347 | 0.771953 | 0.782058 | 0.777383 | 0.770365 | 0.771335 | 0.772641 |
| 0.757864 | 0.762577 | 0.758318 | 0.756881 | 0.764975 | 0.765721 | 0.759466 | 0.762153 | 0.75769 |

| | | | | | | | | |
|---|---|---|---|---|---|---|---|---|
| 0.761592 | 0.768153 | 0.761157 | 0.768942 | 0.774831 | 0.771548 | 0.767689 | 0.767093 | 0.763648 |
| 0.763623 | 0.769014 | 0.761587 | 0.766025 | 0.776822 | 0.773688 | 0.769807 | 0.77367 | 0.768852 |
| 0.762005 | 0.766035 | 0.758957 | 0.762527 | 0.770486 | 0.771837 | 0.76487 | 0.76545 | 0.764053 |
| 0.76809 | 0.764672 | 0.760535 | 0.768086 | 0.776346 | 0.772153 | 0.771794 | 0.768581 | 0.767373 |
| 0.770549 | 0.772996 | 0.766287 | 0.766227 | 0.775132 | 0.77619 | 0.770258 | 0.770113 | 0.769481 |
| 0.763498 | 0.765696 | 0.760639 | 0.759797 | 0.77141 | 0.769834 | 0.765722 | 0.768202 | 0.762094 |
| 0.763972 | 0.769945 | 0.761519 | 0.766838 | 0.774403 | 0.768823 | 0.767291 | 0.766928 | 0.766121 |
| 0.75753 | 0.760239 | 0.753325 | 0.760574 | 0.765077 | 0.760447 | 0.75568 | 0.756772 | 0.756106 |
| 0.762533 | 0.769122 | 0.76196 | 0.770484 | 0.774572 | 0.774066 | 0.772141 | 0.770135 | 0.764865 |
| 0.757581 | 0.760417 | 0.756538 | 0.758846 | 0.766491 | 0.765316 | 0.755726 | 0.761064 | 0.754663 |
| 0.765289 | 0.765748 | 0.758521 | 0.766196 | 0.772291 | 0.769809 | 0.761282 | 0.769551 | 0.762489 |
| 0.763861 | 0.769937 | 0.763304 | 0.765152 | 0.773044 | 0.768984 | 0.768052 | 0.766861 | 0.766999 |
| 0.751371 | 0.759061 | 0.751805 | 0.75002 | 0.762751 | 0.762699 | 0.753769 | 0.756779 | 0.756781 |
| 0.755987 | 0.762136 | 0.753981 | 0.755331 | 0.761709 | 0.758773 | 0.756195 | 0.760821 | 0.758476 |
| 0.769199 | 0.767455 | 0.768988 | 0.766183 | 0.77502 | 0.771698 | 0.771312 | 0.771216 | 0.764 |
| 0.802242 | 0.767579 | 0.760109 | 0.762404 | 0.772645 | 0.769608 | 0.767801 | 0.76637 | 0.764648 |
| 0.77978 | 0.768787 | 0.764386 | 0.763486 | 0.76959 | 0.768636 | 0.769803 | 0.770727 | 0.764472 |
| 0.797292 | 0.763583 | 0.763153 | 0.758744 | 0.768768 | 0.764276 | 0.766093 | 0.767323 | 0.75632 |
| 0.811959 | 0.765867 | 0.760718 | 0.762338 | 0.773801 | 0.764792 | 0.768492 | 0.764326 | 0.761135 |
| 0.797663 | 0.764466 | 0.760154 | 0.757146 | 0.772101 | 0.768831 | 0.765104 | 0.769668 | 0.761042 |
| 0.813469 | 0.758743 | 0.755307 | 0.753193 | 0.764711 | 0.760513 | 0.760464 | 0.759319 | 0.749431 |
| 0.806529 | 0.762812 | 0.756899 | 0.755888 | 0.769385 | 0.757649 | 0.760233 | 0.761675 | 0.754875 |
| 0.799833 | 0.769759 | 0.759571 | 0.760423 | 0.774945 | 0.767992 | 0.767472 | 0.770813 | 0.761083 |
| 0.7831 | 0.764745 | 0.768222 | 0.76712 | 0.774742 | 0.768549 | 0.7696 | 0.769805 | 0.761712 |
| 0.809263 | 0.768647 | 0.762536 | 0.762743 | 0.769212 | 0.769973 | 0.768383 | 0.760915 | 0.760291 |
| 0.865106 | 0.766544 | 0.761633 | 0.755057 | 0.770994 | 0.764322 | 0.765117 | 0.76059 | 0.756197 |
| 0.787725 | 0.761752 | 0.757752 | 0.757016 | 0.764084 | 0.761968 | 0.768058 | 0.760642 | 0.751477 |
| 0.791901 | 0.76576 | 0.75599 | 0.757591 | 0.766409 | 0.76011 | 0.765763 | 0.761849 | 0.748069 |
| 0.852645 | 0.763154 | 0.765359 | 0.757369 | 0.766595 | 0.76683 | 0.762605 | 0.761188 | 0.755617 |
| 0.803571 | 0.765511 | 0.761849 | 0.764179 | 0.772354 | 0.764341 | 0.767243 | 0.768493 | 0.756713 |
| 0.812089 | 0.769982 | 0.764848 | 0.764188 | 0.77421 | 0.771508 | 0.76803 | 0.766932 | 0.763799 |
| 0.796495 | 0.769341 | 0.764067 | 0.764541 | 0.771312 | 0.767912 | 0.767327 | 0.770068 | 0.761632 |
| 0.804321 | 0.771568 | 0.766755 | 0.770315 | 0.778904 | 0.776463 | 0.769048 | 0.770541 | 0.764369 |
| 0.783418 | 0.770462 | 0.766536 | 0.76903 | 0.780892 | 0.772029 | 0.773254 | 0.773756 | 0.764674 |
| 0.834304 | 0.760533 | 0.76173 | 0.757993 | 0.767259 | 0.759731 | 0.760302 | 0.758666 | 0.750944 |
| 0.813971 | 0.767473 | 0.761555 | 0.766512 | 0.770034 | 0.76943 | 0.76174 | 0.762654 | 0.759062 |
| 0.802439 | 0.76632 | 0.762272 | 0.764425 | 0.767841 | 0.761709 | 0.762414 | 0.761881 | 0.753811 |
| 0.792633 | 0.766437 | 0.758067 | 0.75357 | 0.768061 | 0.76296 | 0.763408 | 0.765569 | 0.754724 |
| 0.811752 | 0.754322 | 0.755035 | 0.750145 | 0.76 | 0.758454 | 0.756297 | 0.756365 | 0.751319 |
| 0.788203 | 0.757071 | 0.752937 | 0.753626 | 0.761371 | 0.757849 | 0.756696 | 0.755567 | 0.750697 |
| 0.870429 | 0.75685 | 0.758716 | 0.750883 | 0.760779 | 0.754966 | 0.755558 | 0.753318 | 0.748456 |
| 0.808971 | 0.75845 | 0.752871 | 0.752336 | 0.763914 | 0.752837 | 0.761801 | 0.756856 | 0.746475 |
| 0.796254 | 0.757513 | 0.751564 | 0.748364 | 0.760034 | 0.758388 | 0.756998 | 0.757399 | 0.752052 |
| 0.785024 | 0.756654 | 0.746127 | 0.746789 | 0.762931 | 0.75339 | 0.754791 | 0.753347 | 0.753155 |
| 0.781731 | 0.763685 | 0.753487 | 0.758394 | 0.772076 | 0.763611 | 0.763599 | 0.759656 | 0.758466 |
| 0.760849 | 0.758015 | 0.751956 | 0.755958 | 0.758683 | 0.758702 | 0.759332 | 0.754467 | 0.754157 |

| | | | | | | | | |
|---|---|---|---|---|---|---|---|---|
| 1 | 0.766483 | 0.764967 | 0.760742 | 0.767707 | 0.762566 | 0.763287 | 0.762018 | 0.753537 |
| 0.766483 | 1 | 0.781657 | 0.812072 | 0.813805 | 0.802946 | 0.797206 | 0.791688 | 0.790927 |
| 0.764967 | 0.781657 | 1 | 0.781542 | 0.78593 | 0.777834 | 0.7803 | 0.778771 | 0.768681 |
| 0.760742 | 0.812072 | 0.781542 | 1 | 0.823184 | 0.799387 | 0.792144 | 0.788267 | 0.786814 |
| 0.767707 | 0.813805 | 0.78593 | 0.823184 | 1 | 0.80913 | 0.792107 | 0.792206 | 0.799616 |
| 0.762566 | 0.802946 | 0.777834 | 0.799387 | 0.80913 | 1 | 0.791861 | 0.781584 | 0.809498 |
| 0.763287 | 0.797206 | 0.7803 | 0.792144 | 0.792107 | 0.791861 | 1 | 0.780318 | 0.786962 |
| 0.762018 | 0.791688 | 0.778771 | 0.788267 | 0.792206 | 0.781584 | 0.780318 | 1 | 0.776273 |
| 0.753537 | 0.790927 | 0.768681 | 0.786814 | 0.799616 | 0.809498 | 0.786962 | 0.776273 | 1 |
| 0.75466 | 0.783 | 0.773512 | 0.801817 | 0.812553 | 0.776887 | 0.775718 | 0.8 | 0.764436 |
| 0.758786 | 0.78144 | 0.770588 | 0.785738 | 0.783254 | 0.814676 | 0.791853 | 0.784556 | 0.782429 |
| 0.761667 | 0.783802 | 0.773266 | 0.802056 | 0.802434 | 0.828526 | 0.780356 | 0.798056 | 0.775322 |
| 0.762578 | 0.772678 | 0.770799 | 0.771307 | 0.779832 | 0.768431 | 0.760712 | 0.765264 | 0.762718 |
| 0.762199 | 0.773399 | 0.769749 | 0.772582 | 0.776716 | 0.76978 | 0.768319 | 0.767354 | 0.762648 |
| 0.764206 | 0.773827 | 0.76925 | 0.775507 | 0.777705 | 0.773904 | 0.768526 | 0.77522 | 0.762784 |
| 0.76266 | 0.773623 | 0.768624 | 0.770606 | 0.779316 | 0.774355 | 0.76569 | 0.768154 | 0.762762 |
| 0.766912 | 0.777262 | 0.773629 | 0.766376 | 0.775514 | 0.773418 | 0.769179 | 0.764773 | 0.770923 |
| 0.76035 | 0.767665 | 0.765996 | 0.768734 | 0.77691 | 0.769162 | 0.761785 | 0.767011 | 0.756558 |
| 0.771473 | 0.776304 | 0.770259 | 0.77388 | 0.778892 | 0.77099 | 0.766823 | 0.775344 | 0.764447 |
| 0.768651 | 0.777593 | 0.774566 | 0.773968 | 0.778453 | 0.776796 | 0.771924 | 0.768661 | 0.767561 |
| 0.7615 | 0.772777 | 0.766126 | 0.773389 | 0.774153 | 0.76752 | 0.767692 | 0.762838 | 0.761625 |
| 0.765422 | 0.770715 | 0.769734 | 0.77236 | 0.775769 | 0.775492 | 0.766288 | 0.767551 | 0.765031 |
| 0.767128 | 0.779239 | 0.769822 | 0.774149 | 0.780787 | 0.775866 | 0.770083 | 0.773183 | 0.769632 |
| 0.763555 | 0.776238 | 0.771381 | 0.773924 | 0.785168 | 0.776143 | 0.772603 | 0.772664 | 0.764979 |
| 0.765695 | 0.773499 | 0.765126 | 0.769342 | 0.777914 | 0.770319 | 0.77272 | 0.768727 | 0.765224 |
| 0.760766 | 0.772754 | 0.770807 | 0.772969 | 0.778887 | 0.769086 | 0.77126 | 0.765249 | 0.764305 |
| 0.75834 | 0.77211 | 0.762516 | 0.768825 | 0.777578 | 0.775769 | 0.772173 | 0.768317 | 0.762647 |
| 0.756493 | 0.767922 | 0.768123 | 0.766659 | 0.771956 | 0.76821 | 0.767648 | 0.767493 | 0.760947 |
| 0.767375 | 0.778147 | 0.767011 | 0.775177 | 0.778661 | 0.775359 | 0.777474 | 0.773553 | 0.774057 |
| 0.765183 | 0.772014 | 0.769706 | 0.777374 | 0.784187 | 0.777618 | 0.776836 | 0.769341 | 0.770264 |
| 0.757873 | 0.769777 | 0.760061 | 0.76072 | 0.769064 | 0.768431 | 0.764527 | 0.763291 | 0.763271 |
| 0.766841 | 0.773882 | 0.769838 | 0.769186 | 0.777307 | 0.770532 | 0.768595 | 0.765665 | 0.764972 |
| 0.76567 | 0.77652 | 0.767469 | 0.768216 | 0.780355 | 0.773637 | 0.773202 | 0.77339 | 0.764236 |
| 0.761472 | 0.764796 | 0.764242 | 0.769393 | 0.774509 | 0.769042 | 0.769638 | 0.764508 | 0.758732 |
| 0.761328 | 0.771927 | 0.762888 | 0.76381 | 0.776882 | 0.768732 | 0.766981 | 0.766434 | 0.766673 |
| 0.757933 | 0.769969 | 0.768017 | 0.765385 | 0.774838 | 0.772166 | 0.761761 | 0.766392 | 0.76356 |
| 0.76839 | 0.780874 | 0.771112 | 0.777851 | 0.782236 | 0.779969 | 0.777425 | 0.778298 | 0.768064 |
| 0.767424 | 0.774875 | 0.765671 | 0.769812 | 0.777554 | 0.77083 | 0.769479 | 0.770623 | 0.768486 |
| 0.767557 | 0.775864 | 0.7713 | 0.769438 | 0.77676 | 0.772058 | 0.769974 | 0.771764 | 0.765472 |
| 0.768972 | 0.777654 | 0.771266 | 0.766372 | 0.780278 | 0.771419 | 0.766252 | 0.765265 | 0.767816 |
| 0.771131 | 0.774824 | 0.770588 | 0.770794 | 0.781486 | 0.770254 | 0.766953 | 0.770108 | 0.763747 |
| 0.767862 | 0.777564 | 0.771674 | 0.775176 | 0.778622 | 0.773947 | 0.772396 | 0.774048 | 0.772024 |
| 0.768947 | 0.777335 | 0.773905 | 0.776611 | 0.782334 | 0.777623 | 0.773386 | 0.774831 | 0.775422 |
| 0.762855 | 0.771184 | 0.764383 | 0.76733 | 0.777403 | 0.771429 | 0.767517 | 0.765064 | 0.75698 |
| 0.76565 | 0.773982 | 0.770455 | 0.771459 | 0.780702 | 0.774575 | 0.76576 | 0.768774 | 0.768178 |
| 0.762307 | 0.774166 | 0.762849 | 0.767112 | 0.777796 | 0.771715 | 0.764359 | 0.767225 | 0.763757 |

| | | | | | | | | |
|---|---|---|---|---|---|---|---|---|
| 0.751279 | 0.75915 | 0.756487 | 0.76009 | 0.768149 | 0.762813 | 0.759281 | 0.751435 | 0.756488 |
| 0.746574 | 0.75393 | 0.745835 | 0.752007 | 0.756667 | 0.751417 | 0.747628 | 0.749549 | 0.746841 |
| 0.754747 | 0.771954 | 0.764751 | 0.766318 | 0.778551 | 0.771779 | 0.76745 | 0.764242 | 0.762298 |
| 0.763446 | 0.775147 | 0.770174 | 0.771832 | 0.778442 | 0.774841 | 0.76944 | 0.765518 | 0.76278 |
| 0.751466 | 0.761774 | 0.764833 | 0.759469 | 0.766357 | 0.767438 | 0.758861 | 0.757365 | 0.757556 |
| 0.757075 | 0.770345 | 0.763985 | 0.770191 | 0.778864 | 0.770249 | 0.763459 | 0.761426 | 0.763411 |
| 0.759677 | 0.775805 | 0.768271 | 0.766812 | 0.779291 | 0.775734 | 0.772205 | 0.767588 | 0.768337 |
| 0.751633 | 0.766179 | 0.760254 | 0.761775 | 0.771434 | 0.764706 | 0.76048 | 0.757266 | 0.759606 |
| 0.768335 | 0.779396 | 0.771433 | 0.767468 | 0.778197 | 0.775033 | 0.773932 | 0.772049 | 0.768878 |
| 0.76525 | 0.778067 | 0.77157 | 0.765307 | 0.780673 | 0.768624 | 0.773611 | 0.765498 | 0.767501 |
| 0.765923 | 0.779155 | 0.767649 | 0.770236 | 0.779365 | 0.775687 | 0.772304 | 0.771938 | 0.766457 |
| 0.75998 | 0.774502 | 0.767995 | 0.773396 | 0.7774 | 0.775095 | 0.769489 | 0.76679 | 0.764791 |
| 0.768477 | 0.77762 | 0.769794 | 0.775738 | 0.782939 | 0.778509 | 0.772914 | 0.769099 | 0.770219 |
| 0.762663 | 0.777343 | 0.768918 | 0.776157 | 0.78143 | 0.773035 | 0.772015 | 0.769638 | 0.771911 |
| 0.759106 | 0.776993 | 0.769244 | 0.766058 | 0.778896 | 0.773936 | 0.769565 | 0.765585 | 0.767557 |
| 0.762746 | 0.773962 | 0.769528 | 0.770576 | 0.775169 | 0.770021 | 0.770622 | 0.761836 | 0.765714 |
| 0.761995 | 0.7697 | 0.765831 | 0.769533 | 0.778727 | 0.76744 | 0.771267 | 0.761776 | 0.766495 |
| 0.76087 | 0.771865 | 0.76478 | 0.768283 | 0.779237 | 0.767436 | 0.769145 | 0.763984 | 0.763835 |
| 0.757692 | 0.768543 | 0.760529 | 0.770523 | 0.778327 | 0.772723 | 0.763416 | 0.763593 | 0.767243 |
| 0.761315 | 0.772372 | 0.766025 | 0.760731 | 0.775161 | 0.767849 | 0.767234 | 0.762218 | 0.76071 |
| 0.763239 | 0.774812 | 0.767589 | 0.769676 | 0.777861 | 0.770251 | 0.77194 | 0.770017 | 0.766419 |
| 0.749095 | 0.758261 | 0.753699 | 0.757553 | 0.759074 | 0.755864 | 0.75188 | 0.754076 | 0.757036 |
| 0.756257 | 0.76868 | 0.760269 | 0.76292 | 0.766399 | 0.764846 | 0.760539 | 0.762971 | 0.758645 |
| 0.7603 | 0.773408 | 0.767405 | 0.766134 | 0.772772 | 0.768387 | 0.764468 | 0.763481 | 0.762681 |
| 0.764018 | 0.775695 | 0.764506 | 0.764627 | 0.775471 | 0.771757 | 0.767816 | 0.764893 | 0.76933 |
| 0.763004 | 0.767658 | 0.766901 | 0.765779 | 0.771261 | 0.765181 | 0.76157 | 0.76075 | 0.759536 |
| 0.75689 | 0.767646 | 0.755569 | 0.757989 | 0.769572 | 0.765484 | 0.752966 | 0.760436 | 0.75849 |
| 0.758379 | 0.771114 | 0.759543 | 0.76335 | 0.771964 | 0.765542 | 0.758672 | 0.764019 | 0.757971 |
| 0.756129 | 0.761938 | 0.761695 | 0.760345 | 0.766481 | 0.765211 | 0.761049 | 0.760128 | 0.757598 |
| 0.695043 | 0.695094 | 0.69371 | 0.695581 | 0.701777 | 0.700938 | 0.69065 | 0.695927 | 0.695092 |
| 0.709633 | 0.704457 | 0.70659 | 0.70833 | 0.714102 | 0.708998 | 0.706305 | 0.707194 | 0.703674 |
| 0.708119 | 0.703695 | 0.702122 | 0.709181 | 0.711931 | 0.708463 | 0.698345 | 0.704446 | 0.699098 |
| 0.696394 | 0.697515 | 0.688466 | 0.698636 | 0.701518 | 0.697636 | 0.690526 | 0.697959 | 0.689342 |
| 0.710326 | 0.707087 | 0.706278 | 0.711099 | 0.71338 | 0.71116 | 0.703664 | 0.706017 | 0.702345 |
| 0.752116 | 0.7656 | 0.759763 | 0.762037 | 0.770507 | 0.768086 | 0.759568 | 0.765589 | 0.762358 |
| 0.758403 | 0.768952 | 0.765927 | 0.766269 | 0.772993 | 0.771612 | 0.762728 | 0.765555 | 0.761433 |
| 0.764922 | 0.771749 | 0.767021 | 0.765109 | 0.778836 | 0.776602 | 0.769911 | 0.766457 | 0.763977 |
| 0.756401 | 0.773112 | 0.76907 | 0.769313 | 0.77472 | 0.773655 | 0.765302 | 0.768692 | 0.763962 |
| 0.757874 | 0.766407 | 0.765972 | 0.765693 | 0.772327 | 0.767273 | 0.762779 | 0.765003 | 0.764253 |
| 0.761535 | 0.774099 | 0.762805 | 0.769041 | 0.777178 | 0.77017 | 0.764035 | 0.769218 | 0.763791 |
| 0.76034 | 0.77197 | 0.765697 | 0.768262 | 0.774718 | 0.772178 | 0.766366 | 0.767107 | 0.765354 |
| 0.760261 | 0.767138 | 0.759466 | 0.768776 | 0.769726 | 0.772149 | 0.760958 | 0.765725 | 0.768998 |
| 0.754895 | 0.760715 | 0.758457 | 0.759914 | 0.768253 | 0.76468 | 0.760104 | 0.762487 | 0.756325 |
| 0.752867 | 0.761361 | 0.759032 | 0.76014 | 0.765621 | 0.759284 | 0.757385 | 0.759989 | 0.758571 |
| 0.75634 | 0.767543 | 0.764162 | 0.761622 | 0.7709 | 0.7682 | 0.762794 | 0.764211 | 0.759728 |
| 0.763061 | 0.766866 | 0.765138 | 0.7698 | 0.780547 | 0.77333 | 0.762871 | 0.768828 | 0.764192 |

| | | | | | | | | |
|---|---|---|---|---|---|---|---|---|
| 0.763662 | 0.769564 | 0.765291 | 0.765705 | 0.776135 | 0.770563 | 0.765961 | 0.768921 | 0.759368 |
| 0.763715 | 0.7661 | 0.762642 | 0.768965 | 0.773964 | 0.767572 | 0.758056 | 0.760263 | 0.760407 |
| 0.760879 | 0.768569 | 0.765429 | 0.764429 | 0.773819 | 0.768552 | 0.762822 | 0.767158 | 0.760572 |
| 0.754568 | 0.770075 | 0.75998 | 0.763589 | 0.768843 | 0.765089 | 0.756949 | 0.764696 | 0.759953 |
| 0.760695 | 0.763205 | 0.763402 | 0.762471 | 0.769853 | 0.768933 | 0.759045 | 0.76533 | 0.76018 |
| 0.758744 | 0.764361 | 0.762656 | 0.761909 | 0.771282 | 0.764551 | 0.759796 | 0.766468 | 0.758009 |
| 0.759515 | 0.768279 | 0.763908 | 0.762954 | 0.76962 | 0.767454 | 0.765875 | 0.763721 | 0.758179 |
| 0.764303 | 0.770778 | 0.7634 | 0.767456 | 0.778268 | 0.761283 | 0.761967 | 0.761861 | 0.763216 |
| 0.75824 | 0.767349 | 0.76094 | 0.760645 | 0.770387 | 0.767542 | 0.762484 | 0.763423 | 0.759806 |
| 0.762178 | 0.769802 | 0.762272 | 0.767184 | 0.771743 | 0.766412 | 0.761235 | 0.763679 | 0.758543 |
| 0.759387 | 0.769969 | 0.761197 | 0.763265 | 0.771881 | 0.766275 | 0.757124 | 0.764067 | 0.758975 |
| 0.760702 | 0.765813 | 0.761731 | 0.764392 | 0.770707 | 0.7667 | 0.759951 | 0.763933 | 0.757493 |
| 0.751644 | 0.758364 | 0.756898 | 0.752931 | 0.763303 | 0.762095 | 0.754016 | 0.755465 | 0.75478 |
| 0.755388 | 0.758488 | 0.757439 | 0.762953 | 0.763504 | 0.76109 | 0.757891 | 0.74991 | 0.750848 |
| 0.764037 | 0.77158 | 0.763367 | 0.767372 | 0.771123 | 0.773631 | 0.761229 | 0.76605 | 0.760293 |
| 0.7676 | 0.777796 | 0.771778 | 0.773258 | 0.782804 | 0.774679 | 0.774614 | 0.772098 | 0.771343 |
| 0.763977 | 0.773868 | 0.76508 | 0.763766 | 0.773253 | 0.769375 | 0.765757 | 0.76758 | 0.764423 |
| 0.765504 | 0.774347 | 0.764873 | 0.772097 | 0.780664 | 0.774701 | 0.769489 | 0.770325 | 0.767328 |
| 0.759141 | 0.770243 | 0.764268 | 0.76761 | 0.773152 | 0.768805 | 0.763635 | 0.766267 | 0.764957 |
| 0.765223 | 0.770015 | 0.76743 | 0.762806 | 0.774246 | 0.768159 | 0.764576 | 0.766522 | 0.760823 |
| 0.757713 | 0.76747 | 0.759131 | 0.758767 | 0.769152 | 0.769099 | 0.765244 | 0.760419 | 0.761869 |
| 0.758618 | 0.767166 | 0.762813 | 0.76491 | 0.773163 | 0.767047 | 0.768204 | 0.763441 | 0.759651 |
| 0.768836 | 0.777451 | 0.771137 | 0.771412 | 0.77947 | 0.772321 | 0.770839 | 0.770304 | 0.770168 |
| 0.761623 | 0.763714 | 0.76196 | 0.766187 | 0.773851 | 0.766975 | 0.759669 | 0.758832 | 0.762647 |
| 0.76174 | 0.770284 | 0.765774 | 0.770534 | 0.775664 | 0.774262 | 0.770556 | 0.767085 | 0.764386 |
| 0.703264 | 0.705224 | 0.701525 | 0.708533 | 0.713073 | 0.708189 | 0.701272 | 0.705438 | 0.697989 |
| 0.759402 | 0.769129 | 0.772523 | 0.767574 | 0.776792 | 0.769608 | 0.769856 | 0.770086 | 0.764174 |
| 0.763863 | 0.774075 | 0.765929 | 0.770794 | 0.780552 | 0.775611 | 0.766274 | 0.767832 | 0.767486 |
| 0.76518 | 0.775586 | 0.772941 | 0.77172 | 0.784532 | 0.775476 | 0.770473 | 0.773635 | 0.768587 |
| 0.766893 | 0.770328 | 0.768799 | 0.770053 | 0.781497 | 0.77305 | 0.767372 | 0.768863 | 0.765509 |
| 0.757116 | 0.766923 | 0.764146 | 0.767872 | 0.771767 | 0.768534 | 0.769576 | 0.767025 | 0.760512 |
| 0.761125 | 0.771825 | 0.763657 | 0.767621 | 0.777683 | 0.76974 | 0.764108 | 0.767473 | 0.770402 |
| 0.763395 | 0.771849 | 0.768975 | 0.766731 | 0.771858 | 0.772312 | 0.76294 | 0.766831 | 0.761104 |
| 0.758009 | 0.768902 | 0.765318 | 0.767144 | 0.776526 | 0.771303 | 0.763893 | 0.761099 | 0.766442 |
| 0.759041 | 0.768278 | 0.756057 | 0.763085 | 0.769204 | 0.769004 | 0.76047 | 0.764008 | 0.760767 |
| 0.756645 | 0.76887 | 0.755781 | 0.768837 | 0.774218 | 0.767812 | 0.762113 | 0.766577 | 0.758112 |
| 0.766737 | 0.772457 | 0.763006 | 0.766756 | 0.78044 | 0.772693 | 0.766397 | 0.773593 | 0.762395 |
| 0.764716 | 0.77609 | 0.766655 | 0.769611 | 0.779675 | 0.772276 | 0.761303 | 0.766747 | 0.76371 |
| 0.766039 | 0.770134 | 0.767522 | 0.76774 | 0.783852 | 0.773037 | 0.768437 | 0.768286 | 0.768454 |
| 0.758993 | 0.776702 | 0.767203 | 0.770027 | 0.773569 | 0.769917 | 0.764727 | 0.768025 | 0.76344 |
| 0.757389 | 0.766843 | 0.763526 | 0.768632 | 0.773843 | 0.765103 | 0.766442 | 0.763926 | 0.7651 |
| 0.765292 | 0.770014 | 0.765109 | 0.767957 | 0.776324 | 0.768705 | 0.767982 | 0.767504 | 0.765106 |
| 0.761267 | 0.773127 | 0.762087 | 0.768429 | 0.778363 | 0.770683 | 0.760557 | 0.767788 | 0.761945 |
| 0.759038 | 0.76081 | 0.762469 | 0.761607 | 0.768493 | 0.76607 | 0.763582 | 0.761975 | 0.756089 |
| 0.760447 | 0.771017 | 0.769192 | 0.76673 | 0.771259 | 0.773039 | 0.766583 | 0.767632 | 0.76372 |
| 0.764177 | 0.769333 | 0.766268 | 0.76374 | 0.77173 | 0.771403 | 0.765872 | 0.768125 | 0.759486 |

| | | | | | | | | |
|---|---|---|---|---|---|---|---|---|
| 0.754652 | 0.769787 | 0.761215 | 0.76429 | 0.77041 | 0.767165 | 0.759725 | 0.764956 | 0.75888 |
| 0.762222 | 0.769671 | 0.76704 | 0.766572 | 0.771218 | 0.773924 | 0.761557 | 0.769281 | 0.760061 |
| 0.756795 | 0.764987 | 0.7631 | 0.759338 | 0.767151 | 0.763793 | 0.762439 | 0.76064 | 0.759851 |
| 0.719971 | 0.724617 | 0.714912 | 0.724777 | 0.74293 | 0.725847 | 0.715217 | 0.725413 | 0.731959 |
| 0.756209 | 0.7649 | 0.766266 | 0.762614 | 0.768406 | 0.763599 | 0.75891 | 0.765755 | 0.757792 |
| 0.759743 | 0.770664 | 0.766605 | 0.764143 | 0.768667 | 0.76445 | 0.761697 | 0.768591 | 0.757589 |
| 0.761296 | 0.766495 | 0.763855 | 0.7631 | 0.768056 | 0.76901 | 0.75928 | 0.764936 | 0.758286 |
| 0.758288 | 0.764623 | 0.763335 | 0.757068 | 0.77431 | 0.770281 | 0.764374 | 0.761676 | 0.759595 |
| 0.76244 | 0.774102 | 0.76902 | 0.767415 | 0.777046 | 0.772679 | 0.766786 | 0.765931 | 0.763014 |
| 0.763744 | 0.770262 | 0.771488 | 0.765181 | 0.774018 | 0.772567 | 0.764444 | 0.76485 | 0.763189 |
| 0.760875 | 0.765186 | 0.765201 | 0.767668 | 0.772862 | 0.769858 | 0.765295 | 0.764122 | 0.763151 |
| 0.758882 | 0.767193 | 0.765851 | 0.767497 | 0.774924 | 0.769186 | 0.764519 | 0.763041 | 0.758822 |
| 0.756584 | 0.768539 | 0.766536 | 0.764346 | 0.770198 | 0.767147 | 0.76232 | 0.76208 | 0.759734 |
| 0.756229 | 0.768714 | 0.765159 | 0.763961 | 0.77417 | 0.770129 | 0.760879 | 0.765917 | 0.758859 |
| 0.75951 | 0.768263 | 0.767308 | 0.76751 | 0.771457 | 0.769306 | 0.763344 | 0.767341 | 0.760975 |
| 0.757294 | 0.768144 | 0.765212 | 0.759268 | 0.767922 | 0.765395 | 0.760646 | 0.763434 | 0.75987 |
| 0.763186 | 0.770771 | 0.772566 | 0.76938 | 0.775872 | 0.77134 | 0.765189 | 0.769543 | 0.766148 |
| 0.761674 | 0.771167 | 0.765668 | 0.768484 | 0.777759 | 0.770111 | 0.760914 | 0.770088 | 0.767712 |
| 0.757414 | 0.767388 | 0.760282 | 0.76532 | 0.766448 | 0.768487 | 0.759058 | 0.760813 | 0.760506 |
| 0.762755 | 0.768406 | 0.769397 | 0.764792 | 0.772563 | 0.765422 | 0.761096 | 0.765192 | 0.763343 |
| 0.763337 | 0.769391 | 0.768512 | 0.766503 | 0.768586 | 0.769862 | 0.766644 | 0.769975 | 0.763726 |
| 0.766038 | 0.770165 | 0.770799 | 0.76972 | 0.778171 | 0.770515 | 0.767331 | 0.769338 | 0.767369 |
| 0.676401 | 0.691896 | 0.675696 | 0.682701 | 0.692857 | 0.691647 | 0.681674 | 0.675504 | 0.684161 |
| 0.664341 | 0.664202 | 0.652805 | 0.661524 | 0.657366 | 0.656975 | 0.654463 | 0.651887 | 0.665856 |
| 0.667821 | 0.676041 | 0.672959 | 0.675536 | 0.683326 | 0.677447 | 0.673565 | 0.678007 | 0.675964 |
| 0.655695 | 0.660116 | 0.648415 | 0.652803 | 0.664689 | 0.66065 | 0.653923 | 0.65783 | 0.654957 |
| 0.729998 | 0.737836 | 0.731544 | 0.733283 | 0.745128 | 0.738896 | 0.732347 | 0.723741 | 0.73716 |
| 0.71445 | 0.731306 | 0.718556 | 0.726163 | 0.732769 | 0.723509 | 0.719798 | 0.723742 | 0.724328 |
| 0.719206 | 0.734705 | 0.712543 | 0.73241 | 0.734938 | 0.7283 | 0.72681 | 0.717544 | 0.739726 |
| 0.731281 | 0.745251 | 0.736343 | 0.742009 | 0.736123 | 0.740718 | 0.734282 | 0.741019 | 0.733866 |
| 0.710912 | 0.720968 | 0.713254 | 0.715974 | 0.72872 | 0.724697 | 0.717225 | 0.708979 | 0.724777 |
| 0.70386 | 0.719467 | 0.704623 | 0.711658 | 0.724472 | 0.710157 | 0.702526 | 0.711023 | 0.718277 |
| 0.705621 | 0.72377 | 0.721936 | 0.724342 | 0.732617 | 0.726145 | 0.712202 | 0.712766 | 0.708831 |
| 0.659079 | 0.659494 | 0.65219 | 0.656395 | 0.667563 | 0.658359 | 0.651657 | 0.653442 | 0.662818 |
| 0.755999 | 0.760135 | 0.756025 | 0.759561 | 0.762839 | 0.757159 | 0.75124 | 0.757508 | 0.757816 |
| 0.733909 | 0.74285 | 0.733396 | 0.736313 | 0.752671 | 0.744154 | 0.739177 | 0.735342 | 0.737512 |
| 0.72891 | 0.733569 | 0.732563 | 0.729111 | 0.741062 | 0.738719 | 0.728834 | 0.733433 | 0.729117 |
| 0.704815 | 0.723034 | 0.710877 | 0.7186 | 0.724907 | 0.721214 | 0.71857 | 0.7228 | 0.721442 |
| 0.735425 | 0.740628 | 0.735131 | 0.73454 | 0.740322 | 0.745236 | 0.735936 | 0.732303 | 0.73886 |
| 0.728278 | 0.735788 | 0.725884 | 0.7216 | 0.735759 | 0.735042 | 0.731291 | 0.728298 | 0.728672 |
| 0.725832 | 0.738191 | 0.728097 | 0.719174 | 0.731419 | 0.728497 | 0.726411 | 0.723996 | 0.72484 |
| 0.726062 | 0.739453 | 0.725681 | 0.72346 | 0.735221 | 0.73723 | 0.726292 | 0.725246 | 0.730274 |
| 0.720448 | 0.734586 | 0.719758 | 0.728749 | 0.739336 | 0.737604 | 0.732107 | 0.722553 | 0.734587 |
| 0.716315 | 0.727417 | 0.722991 | 0.725598 | 0.739156 | 0.729187 | 0.726145 | 0.725277 | 0.73188 |
| 0.726067 | 0.733663 | 0.718303 | 0.725354 | 0.733126 | 0.731105 | 0.720197 | 0.735036 | 0.716649 |
| 0.708223 | 0.727303 | 0.713901 | 0.714617 | 0.7229 | 0.723566 | 0.716522 | 0.707591 | 0.709656 |

| | | | | | | | | |
|---|---|---|---|---|---|---|---|---|
| 0.741299 | 0.750425 | 0.748114 | 0.74958 | 0.75895 | 0.755283 | 0.746576 | 0.747573 | 0.751399 |
| 0.724771 | 0.73136 | 0.725846 | 0.726369 | 0.738129 | 0.731186 | 0.72739 | 0.723581 | 0.730819 |
| 0.726436 | 0.729876 | 0.720884 | 0.723657 | 0.737705 | 0.735263 | 0.720054 | 0.724891 | 0.732941 |
| 0.719932 | 0.73281 | 0.721266 | 0.727813 | 0.738903 | 0.734609 | 0.723893 | 0.73225 | 0.730181 |
| 0.728471 | 0.73024 | 0.724683 | 0.722177 | 0.733647 | 0.72934 | 0.723954 | 0.719772 | 0.732861 |
| 0.71738 | 0.721637 | 0.718797 | 0.718189 | 0.733584 | 0.72462 | 0.709718 | 0.714712 | 0.724682 |
| 0.709356 | 0.722398 | 0.715027 | 0.709554 | 0.725547 | 0.722601 | 0.717088 | 0.721816 | 0.714286 |
| 0.709586 | 0.7246 | 0.714616 | 0.716167 | 0.721102 | 0.718254 | 0.708381 | 0.717477 | 0.721644 |
| 0.714693 | 0.72273 | 0.715234 | 0.717861 | 0.733238 | 0.726016 | 0.712984 | 0.718065 | 0.727588 |
| 0.757429 | 0.752912 | 0.751043 | 0.746383 | 0.761679 | 0.755503 | 0.75267 | 0.752029 | 0.74922 |
| 0.749596 | 0.749036 | 0.746409 | 0.743536 | 0.761306 | 0.753947 | 0.752953 | 0.748368 | 0.75 |
| 0.740918 | 0.74986 | 0.741617 | 0.740044 | 0.75478 | 0.747894 | 0.739581 | 0.741277 | 0.742363 |
| 0.733228 | 0.741067 | 0.732694 | 0.7255 | 0.743892 | 0.742538 | 0.734165 | 0.741658 | 0.743638 |
| 0.738057 | 0.742706 | 0.734578 | 0.737862 | 0.750704 | 0.74388 | 0.736554 | 0.741541 | 0.740471 |
| 0.737532 | 0.742925 | 0.736064 | 0.736115 | 0.744825 | 0.738162 | 0.732708 | 0.732648 | 0.738779 |

| | | | | | | | | plot 05 |
|---|---|---|---|---|---|---|---|---|
| Wild_plot_ | Wild_plot_ | Wild_plot_ | Wild_plot_ | Wild_plot_ | Wild_plot_ | Wild_plot_ | Wild_plot_ | Wild_plot_ |
| 0.710615 | 0.709578 | 0.708849 | 0.707136 | 0.708108 | 0.707723 | 0.708201 | 0.711563 | 0.709 |
| 0.715314 | 0.720291 | 0.716176 | 0.712174 | 0.715129 | 0.717945 | 0.713003 | 0.720687 | 0.717484 |
| 0.706541 | 0.704179 | 0.70658 | 0.699717 | 0.705654 | 0.706662 | 0.703607 | 0.708291 | 0.70474 |
| 0.715606 | 0.714106 | 0.718333 | 0.710899 | 0.711324 | 0.715928 | 0.714546 | 0.717666 | 0.711375 |
| 0.711508 | 0.713297 | 0.714695 | 0.708742 | 0.710189 | 0.710978 | 0.709848 | 0.711277 | 0.708708 |
| 0.716695 | 0.717937 | 0.721239 | 0.718138 | 0.718172 | 0.720875 | 0.717412 | 0.721192 | 0.717328 |
| 0.767037 | 0.772513 | 0.77234 | 0.787113 | 0.779964 | 0.777841 | 0.77796 | 0.986099 | 0.781456 |
| 0.766839 | 0.767967 | 0.766585 | 0.779727 | 0.778061 | 0.784484 | 0.779196 | 0.787737 | 0.780697 |
| 0.711532 | 0.716174 | 0.71252 | 0.711077 | 0.712055 | 0.712579 | 0.712103 | 0.717197 | 0.714387 |
| 0.708588 | 0.712497 | 0.710409 | 0.708099 | 0.708551 | 0.709423 | 0.713242 | 0.713082 | 0.710906 |
| 0.707074 | 0.713382 | 0.710748 | 0.70833 | 0.710952 | 0.711918 | 0.712682 | 0.71657 | 0.712442 |
| 0.703548 | 0.705392 | 0.705239 | 0.701753 | 0.702375 | 0.705855 | 0.707099 | 0.705818 | 0.705574 |
| 0.713051 | 0.717589 | 0.716371 | 0.713411 | 0.71342 | 0.716097 | 0.715628 | 0.71657 | 0.71709 |
| 0.720368 | 0.722152 | 0.720713 | 0.720051 | 0.720688 | 0.722985 | 0.721845 | 0.726782 | 0.720454 |
| 0.7339 | 0.736746 | 0.733381 | 0.734244 | 0.733732 | 0.73797 | 0.735177 | 0.737553 | 0.736492 |
| 0.717763 | 0.71691 | 0.719547 | 0.714843 | 0.71432 | 0.720301 | 0.715873 | 0.717926 | 0.714768 |
| 0.714263 | 0.717339 | 0.717821 | 0.713632 | 0.716387 | 0.71506 | 0.717782 | 0.717402 | 0.710587 |
| 0.740978 | 0.743343 | 0.741493 | 0.742342 | 0.741382 | 0.744919 | 0.744091 | 0.744149 | 0.742625 |
| 0.703911 | 0.709912 | 0.712524 | 0.704891 | 0.702505 | 0.708845 | 0.7065 | 0.709537 | 0.706123 |
| 0.707593 | 0.715279 | 0.713694 | 0.709785 | 0.709536 | 0.715304 | 0.713104 | 0.716104 | 0.713858 |
| 0.709672 | 0.711381 | 0.710579 | 0.706376 | 0.709398 | 0.711143 | 0.706152 | 0.714815 | 0.707189 |
| 0.709307 | 0.711974 | 0.711525 | 0.709795 | 0.708977 | 0.710339 | 0.714411 | 0.714557 | 0.712176 |
| 0.71497 | 0.719343 | 0.720726 | 0.715217 | 0.714059 | 0.721284 | 0.718096 | 0.721228 | 0.720404 |
| 0.713238 | 0.716045 | 0.716451 | 0.709417 | 0.713477 | 0.714873 | 0.709634 | 0.720978 | 0.714398 |
| 0.710227 | 0.71475 | 0.715601 | 0.710327 | 0.713829 | 0.712871 | 0.715565 | 0.718263 | 0.71477 |
| 0.710151 | 0.712147 | 0.710243 | 0.705929 | 0.708574 | 0.71105 | 0.707392 | 0.715149 | 0.706029 |
| 0.705396 | 0.705642 | 0.70628 | 0.703 | 0.704813 | 0.708511 | 0.706798 | 0.707025 | 0.705601 |
| 0.713355 | 0.712223 | 0.716099 | 0.709721 | 0.708893 | 0.713522 | 0.711997 | 0.713681 | 0.715002 |
| 0.712755 | 0.718131 | 0.716918 | 0.712376 | 0.712475 | 0.716995 | 0.714468 | 0.718611 | 0.716834 |
| 0.71275 | 0.715634 | 0.716577 | 0.708493 | 0.710863 | 0.715693 | 0.71139 | 0.716908 | 0.711181 |
| 0.708673 | 0.709308 | 0.70914 | 0.707257 | 0.709458 | 0.711227 | 0.712518 | 0.713799 | 0.710043 |
| 0.708366 | 0.711151 | 0.712416 | 0.703797 | 0.707257 | 0.708938 | 0.709953 | 0.708379 | 0.70742 |
| 0.710744 | 0.713769 | 0.713707 | 0.706779 | 0.70506 | 0.711992 | 0.707452 | 0.716364 | 0.709488 |
| 0.711502 | 0.71745 | 0.715227 | 0.709005 | 0.712768 | 0.713915 | 0.712961 | 0.712569 | 0.713156 |
| 0.705329 | 0.707559 | 0.70619 | 0.698484 | 0.705962 | 0.705748 | 0.705509 | 0.708694 | 0.704312 |
| 0.712452 | 0.710858 | 0.710974 | 0.710022 | 0.708337 | 0.713514 | 0.714892 | 0.714501 | 0.710657 |
| 0.711872 | 0.714857 | 0.715358 | 0.713078 | 0.71746 | 0.715587 | 0.716092 | 0.7174 | 0.715245 |
| 0.710905 | 0.711416 | 0.714045 | 0.710393 | 0.710076 | 0.715678 | 0.713942 | 0.718322 | 0.714036 |
| 0.714149 | 0.716878 | 0.717433 | 0.71441 | 0.714479 | 0.715141 | 0.718616 | 0.721427 | 0.718263 |

| | | | | | | | | |
|---|---|---|---|---|---|---|---|---|
| 0.74497 | 0.745769 | 0.744056 | 0.745723 | 0.744516 | 0.746082 | 0.748439 | 0.748947 | 0.747901 |
| 0.702741 | 0.70446 | 0.699653 | 0.70165 | 0.700981 | 0.703152 | 0.70279 | 0.705435 | 0.704165 |
| 0.703904 | 0.709204 | 0.70829 | 0.702568 | 0.707344 | 0.708258 | 0.705468 | 0.71223 | 0.706937 |
| 0.71558 | 0.719635 | 0.717709 | 0.712774 | 0.717063 | 0.7146 | 0.717689 | 0.7221 | 0.717584 |
| 0.710619 | 0.712162 | 0.713364 | 0.709206 | 0.713774 | 0.711003 | 0.712439 | 0.714274 | 0.711814 |
| 0.713896 | 0.717122 | 0.717486 | 0.717906 | 0.712446 | 0.715274 | 0.717327 | 0.717255 | 0.715741 |
| 0.707181 | 0.710802 | 0.707236 | 0.703029 | 0.705366 | 0.708373 | 0.709435 | 0.711806 | 0.709609 |
| 0.711967 | 0.716923 | 0.714434 | 0.714343 | 0.71002 | 0.713184 | 0.715796 | 0.718355 | 0.715291 |
| 0.715795 | 0.713049 | 0.713576 | 0.710535 | 0.712547 | 0.71782 | 0.715842 | 0.716058 | 0.715859 |
| 0.713785 | 0.712569 | 0.712213 | 0.708073 | 0.711012 | 0.716291 | 0.709915 | 0.712513 | 0.712554 |
| 0.706019 | 0.704131 | 0.706375 | 0.701447 | 0.70438 | 0.706778 | 0.707364 | 0.706142 | 0.705947 |
| 0.704344 | 0.705553 | 0.705513 | 0.707739 | 0.705011 | 0.70605 | 0.710182 | 0.710593 | 0.708242 |
| 0.707948 | 0.710556 | 0.709566 | 0.708937 | 0.711796 | 0.710405 | 0.709422 | 0.714047 | 0.7094 |
| 0.706278 | 0.708732 | 0.70802 | 0.707617 | 0.705859 | 0.709659 | 0.7084 | 0.70864 | 0.705933 |
| 0.706511 | 0.707509 | 0.707272 | 0.705048 | 0.706732 | 0.706973 | 0.707982 | 0.708333 | 0.708069 |
| 0.703557 | 0.706094 | 0.704339 | 0.705678 | 0.70874 | 0.7082 | 0.70968 | 0.708573 | 0.707175 |
| 0.70803 | 0.708327 | 0.707047 | 0.703487 | 0.703805 | 0.705836 | 0.704398 | 0.711891 | 0.70598 |
| 0.71235 | 0.712564 | 0.713706 | 0.711383 | 0.714331 | 0.715088 | 0.712686 | 0.716679 | 0.713882 |
| 0.71295 | 0.716772 | 0.716258 | 0.714888 | 0.71316 | 0.716573 | 0.716015 | 0.715606 | 0.71331 |
| 0.707006 | 0.712842 | 0.713208 | 0.703695 | 0.708517 | 0.714889 | 0.709087 | 0.710489 | 0.714422 |
| 0.704649 | 0.706422 | 0.704562 | 0.702916 | 0.704466 | 0.705133 | 0.708111 | 0.712088 | 0.707552 |
| 0.706536 | 0.710958 | 0.70934 | 0.704629 | 0.705967 | 0.708729 | 0.709536 | 0.712409 | 0.709888 |
| 0.70713 | 0.711118 | 0.709833 | 0.705526 | 0.709877 | 0.711127 | 0.707622 | 0.714767 | 0.710934 |
| 0.718612 | 0.719219 | 0.717988 | 0.7169 | 0.715084 | 0.7168 | 0.718293 | 0.720149 | 0.716585 |
| 0.724729 | 0.725499 | 0.727811 | 0.722718 | 0.724493 | 0.725022 | 0.723399 | 0.729091 | 0.724429 |
| 0.768745 | 0.770127 | 0.76796 | 0.780443 | 0.778257 | 0.785549 | 0.78017 | 0.788907 | 0.782348 |
| 0.769711 | 0.769206 | 0.765847 | 0.775941 | 0.777611 | 0.778825 | 0.771947 | 0.778474 | 0.773163 |
| 0.768931 | 0.76973 | 0.769747 | 0.777732 | 0.776017 | 0.777164 | 0.772359 | 0.778024 | 0.777769 |
| 0.705957 | 0.709285 | 0.709159 | 0.706135 | 0.706051 | 0.709522 | 0.709065 | 0.709565 | 0.709665 |
| 0.707686 | 0.711368 | 0.709292 | 0.706743 | 0.70648 | 0.709167 | 0.70545 | 0.708225 | 0.706834 |
| 0.714331 | 0.717059 | 0.718094 | 0.713306 | 0.717969 | 0.717053 | 0.717055 | 0.7164 | 0.719294 |
| 0.714159 | 0.719512 | 0.713332 | 0.716602 | 0.716457 | 0.720122 | 0.717525 | 0.722931 | 0.715922 |
| 0.705745 | 0.708895 | 0.709316 | 0.703051 | 0.70669 | 0.70815 | 0.705242 | 0.711904 | 0.710152 |
| 0.710902 | 0.713808 | 0.711626 | 0.708196 | 0.71167 | 0.710715 | 0.711662 | 0.714775 | 0.71089 |
| 0.703793 | 0.711997 | 0.709173 | 0.704845 | 0.706771 | 0.7086 | 0.70435 | 0.708836 | 0.709623 |
| 0.709244 | 0.708689 | 0.71036 | 0.705774 | 0.710106 | 0.709998 | 0.708513 | 0.714897 | 0.708146 |
| 0.702253 | 0.708884 | 0.7071 | 0.706369 | 0.70734 | 0.708486 | 0.706511 | 0.712752 | 0.708998 |
| 0.719756 | 0.723965 | 0.722903 | 0.721133 | 0.72212 | 0.723629 | 0.721566 | 0.725727 | 0.720451 |
| 0.698999 | 0.702913 | 0.701246 | 0.698639 | 0.700693 | 0.703716 | 0.700833 | 0.70507 | 0.703979 |
| 0.71053 | 0.712028 | 0.711143 | 0.709137 | 0.710391 | 0.70969 | 0.710769 | 0.712528 | 0.711368 |
| 0.704091 | 0.708617 | 0.706907 | 0.702051 | 0.704124 | 0.704722 | 0.705679 | 0.710225 | 0.701125 |
| 0.723596 | 0.726911 | 0.724996 | 0.721622 | 0.722703 | 0.726441 | 0.721663 | 0.731042 | 0.724739 |
| 0.709855 | 0.71337 | 0.710936 | 0.708625 | 0.70841 | 0.71226 | 0.712657 | 0.71268 | 0.708088 |
| 0.709238 | 0.706472 | 0.705627 | 0.708231 | 0.708656 | 0.709915 | 0.712774 | 0.712313 | 0.708639 |
| 0.711641 | 0.71507 | 0.716522 | 0.71118 | 0.711209 | 0.712266 | 0.711385 | 0.715035 | 0.712964 |
| 0.706715 | 0.710405 | 0.709548 | 0.70454 | 0.710222 | 0.709174 | 0.70961 | 0.71202 | 0.709406 |

| | | | | | | | | |
|---|---|---|---|---|---|---|---|---|
| 0.708043 | 0.708596 | 0.709886 | 0.705854 | 0.705578 | 0.708297 | 0.710152 | 0.711848 | 0.703837 |
| 0.710694 | 0.712399 | 0.716875 | 0.710107 | 0.708241 | 0.711844 | 0.710903 | 0.713388 | 0.710967 |
| 0.708286 | 0.707573 | 0.710324 | 0.709478 | 0.710258 | 0.710195 | 0.710862 | 0.711885 | 0.709657 |
| 0.713904 | 0.716966 | 0.719083 | 0.713066 | 0.717636 | 0.718517 | 0.714907 | 0.722089 | 0.717638 |
| 0.716618 | 0.714421 | 0.71875 | 0.714297 | 0.712539 | 0.717173 | 0.716179 | 0.717444 | 0.715762 |
| 0.709151 | 0.709939 | 0.711236 | 0.704005 | 0.709266 | 0.707437 | 0.708543 | 0.710121 | 0.710351 |
| 0.710399 | 0.713368 | 0.711872 | 0.708422 | 0.709897 | 0.712975 | 0.709517 | 0.712979 | 0.713651 |
| 0.713113 | 0.711384 | 0.714824 | 0.713288 | 0.714148 | 0.71341 | 0.714137 | 0.715093 | 0.715192 |
| 0.712221 | 0.714229 | 0.710791 | 0.712203 | 0.714458 | 0.708687 | 0.714491 | 0.71407 | 0.711734 |
| 0.707459 | 0.70573 | 0.710062 | 0.705715 | 0.7066 | 0.711724 | 0.710861 | 0.708546 | 0.709134 |
| 0.706449 | 0.710288 | 0.710261 | 0.706882 | 0.708524 | 0.70749 | 0.709571 | 0.710724 | 0.707618 |
| 0.707872 | 0.713232 | 0.709192 | 0.709615 | 0.706491 | 0.711965 | 0.711903 | 0.7096 | 0.711828 |
| 0.712788 | 0.715366 | 0.714503 | 0.711984 | 0.712796 | 0.712544 | 0.714938 | 0.718743 | 0.713833 |
| 0.708769 | 0.708343 | 0.705631 | 0.704553 | 0.706628 | 0.708056 | 0.708051 | 0.70894 | 0.70494 |
| 0.713415 | 0.712213 | 0.71104 | 0.709462 | 0.709791 | 0.710357 | 0.710035 | 0.714625 | 0.713349 |
| 0.706965 | 0.710638 | 0.709864 | 0.704996 | 0.705264 | 0.710061 | 0.707517 | 0.70948 | 0.708738 |
| 0.710162 | 0.712538 | 0.714722 | 0.71369 | 0.712979 | 0.71358 | 0.713829 | 0.713425 | 0.712375 |
| 0.708755 | 0.708415 | 0.70881 | 0.708254 | 0.709649 | 0.709011 | 0.710179 | 0.711484 | 0.708582 |
| 0.712112 | 0.71137 | 0.711554 | 0.712573 | 0.711329 | 0.712305 | 0.713738 | 0.71424 | 0.710878 |
| 0.71 | 0.714263 | 0.713282 | 0.710407 | 0.71432 | 0.713719 | 0.710901 | 0.712683 | 0.709437 |
| 0.710157 | 0.710109 | 0.711543 | 0.707513 | 0.707247 | 0.709961 | 0.712643 | 0.709484 | 0.710296 |
| 0.704473 | 0.705799 | 0.705663 | 0.703334 | 0.702773 | 0.707761 | 0.705915 | 0.706979 | 0.706952 |
| 0.705402 | 0.709736 | 0.709445 | 0.707284 | 0.711177 | 0.710009 | 0.708566 | 0.712194 | 0.709609 |
| 0.70898 | 0.712902 | 0.71399 | 0.709064 | 0.709798 | 0.712457 | 0.712155 | 0.714184 | 0.708562 |
| 0.707233 | 0.712335 | 0.712205 | 0.707601 | 0.708293 | 0.713897 | 0.710277 | 0.715348 | 0.712085 |
| 0.714206 | 0.718229 | 0.71554 | 0.713101 | 0.714705 | 0.715993 | 0.715957 | 0.720274 | 0.718161 |
| 0.706944 | 0.710157 | 0.710075 | 0.705363 | 0.707286 | 0.71127 | 0.707486 | 0.712752 | 0.711016 |
| 0.709762 | 0.711041 | 0.712177 | 0.711076 | 0.711746 | 0.709865 | 0.712316 | 0.717757 | 0.71217 |
| 0.711058 | 0.714893 | 0.712732 | 0.709844 | 0.710668 | 0.71353 | 0.713018 | 0.712321 | 0.712783 |
| 0.708624 | 0.711312 | 0.712151 | 0.710414 | 0.710741 | 0.712396 | 0.711357 | 0.715077 | 0.716879 |
| 0.710706 | 0.712197 | 0.713827 | 0.708383 | 0.712507 | 0.716076 | 0.712088 | 0.715059 | 0.717136 |
| 0.708738 | 0.709203 | 0.710835 | 0.709258 | 0.708435 | 0.710058 | 0.708677 | 0.714118 | 0.708453 |
| 0.707614 | 0.713372 | 0.712728 | 0.710043 | 0.711317 | 0.713302 | 0.711893 | 0.715737 | 0.717562 |
| 0.712273 | 0.716318 | 0.715199 | 0.709312 | 0.712882 | 0.711535 | 0.713596 | 0.717711 | 0.713859 |
| 0.709398 | 0.706306 | 0.706848 | 0.704555 | 0.70711 | 0.708081 | 0.706239 | 0.712062 | 0.710612 |
| 0.711691 | 0.710042 | 0.712624 | 0.708024 | 0.709912 | 0.708567 | 0.713974 | 0.714503 | 0.709526 |
| 0.702147 | 0.707892 | 0.704404 | 0.703513 | 0.703276 | 0.705044 | 0.708805 | 0.709643 | 0.704598 |
| 0.682131 | 0.685795 | 0.689121 | 0.680351 | 0.684423 | 0.68365 | 0.684707 | 0.688544 | 0.683788 |
| 0.687802 | 0.690964 | 0.691618 | 0.685441 | 0.689003 | 0.692416 | 0.68954 | 0.693012 | 0.689714 |
| 0.677002 | 0.682737 | 0.682689 | 0.675983 | 0.682197 | 0.682069 | 0.678447 | 0.682787 | 0.677573 |
| 0.681869 | 0.68283 | 0.680964 | 0.677676 | 0.680448 | 0.67959 | 0.679876 | 0.68355 | 0.683667 |
| 0.665539 | 0.668485 | 0.670876 | 0.664687 | 0.662599 | 0.66628 | 0.671088 | 0.666705 | 0.666821 |
| 0.666506 | 0.663528 | 0.667148 | 0.665501 | 0.666747 | 0.666707 | 0.669048 | 0.668743 | 0.667861 |
| 0.667675 | 0.669634 | 0.668721 | 0.666343 | 0.668887 | 0.668716 | 0.666019 | 0.670277 | 0.665988 |
| 0.666328 | 0.676392 | 0.673269 | 0.665696 | 0.666371 | 0.671268 | 0.67023 | 0.669871 | 0.668638 |
| 0.66505 | 0.671095 | 0.667492 | 0.660639 | 0.669025 | 0.664027 | 0.66596 | 0.66831 | 0.667283 |
| 0.671763 | 0.676084 | 0.674631 | 0.666054 | 0.670266 | 0.66977 | 0.669179 | 0.677688 | 0.673253 |

| | | | | | | | | |
|---|---|---|---|---|---|---|---|---|
| 0.668966 | 0.67704 | 0.675018 | 0.669982 | 0.675919 | 0.673387 | 0.667775 | 0.674908 | 0.664404 |
| 0.670166 | 0.676764 | 0.674639 | 0.669863 | 0.671302 | 0.677792 | 0.67083 | 0.675742 | 0.672376 |
| 0.668266 | 0.67312 | 0.67427 | 0.667997 | 0.665341 | 0.669006 | 0.671453 | 0.677826 | 0.667325 |
| 0.660266 | 0.663792 | 0.667458 | 0.660827 | 0.665259 | 0.661872 | 0.666534 | 0.665357 | 0.66037 |
| 0.660298 | 0.668877 | 0.666326 | 0.661693 | 0.665043 | 0.669017 | 0.667482 | 0.668541 | 0.669135 |
| 0.662834 | 0.664899 | 0.664898 | 0.666804 | 0.664308 | 0.668255 | 0.665659 | 0.66703 | 0.664585 |
| 0.666808 | 0.666854 | 0.667707 | 0.666479 | 0.67119 | 0.671667 | 0.663699 | 0.66409 | 0.67087 |
| 0.659838 | 0.662187 | 0.665407 | 0.662483 | 0.660064 | 0.6649 | 0.664748 | 0.666667 | 0.662889 |
| 0.659194 | 0.663417 | 0.665008 | 0.657991 | 0.665203 | 0.659329 | 0.664018 | 0.661923 | 0.656467 |
| 0.654648 | 0.673205 | 0.666957 | 0.666327 | 0.661022 | 0.670301 | 0.664963 | 0.672822 | 0.670976 |
| 0.65457 | 0.660913 | 0.663656 | 0.655132 | 0.659557 | 0.657646 | 0.657691 | 0.656816 | 0.658759 |
| 0.697819 | 0.699775 | 0.699992 | 0.696311 | 0.699616 | 0.69909 | 0.700477 | 0.698736 | 0.70069 |
| 0.705279 | 0.706657 | 0.708323 | 0.70366 | 0.706698 | 0.709392 | 0.705383 | 0.709708 | 0.710711 |
| 0.681385 | 0.686272 | 0.686769 | 0.681854 | 0.685202 | 0.689628 | 0.686176 | 0.688716 | 0.683909 |
| 0.713164 | 0.717058 | 0.716828 | 0.711367 | 0.714528 | 0.717961 | 0.715946 | 0.718339 | 0.713625 |
| 0.70584 | 0.707021 | 0.706802 | 0.704136 | 0.703504 | 0.705789 | 0.705449 | 0.704667 | 0.704822 |
| 0.713184 | 0.715905 | 0.712726 | 0.710855 | 0.714308 | 0.717099 | 0.713269 | 0.716584 | 0.716911 |
| 0.718243 | 0.71705 | 0.71871 | 0.721712 | 0.718316 | 0.721146 | 0.722258 | 0.722608 | 0.719096 |
| 0.696965 | 0.696381 | 0.699945 | 0.696619 | 0.69673 | 0.69598 | 0.696209 | 0.697212 | 0.698823 |
| 0.717902 | 0.722397 | 0.721975 | 0.716331 | 0.71842 | 0.721234 | 0.720105 | 0.720141 | 0.719474 |
| 0.686601 | 0.682055 | 0.687021 | 0.684846 | 0.691966 | 0.687388 | 0.688803 | 0.694019 | 0.688675 |
| 0.715905 | 0.718708 | 0.715303 | 0.71602 | 0.718199 | 0.720573 | 0.718488 | 0.718571 | 0.7153 |
| 0.718446 | 0.722169 | 0.721937 | 0.720535 | 0.718415 | 0.722811 | 0.726622 | 0.723291 | 0.721318 |
| 0.68 | 0.679793 | 0.688182 | 0.680157 | 0.683411 | 0.680988 | 0.685116 | 0.688343 | 0.683746 |
| 0.68266 | 0.684013 | 0.683019 | 0.682699 | 0.687366 | 0.690316 | 0.686694 | 0.686715 | 0.687627 |
| 0.688361 | 0.693872 | 0.693241 | 0.689372 | 0.687973 | 0.68851 | 0.689083 | 0.696177 | 0.687804 |
| 0.710075 | 0.702121 | 0.705267 | 0.702939 | 0.702591 | 0.707701 | 0.704139 | 0.710624 | 0.709338 |
| 0.717709 | 0.719527 | 0.719895 | 0.7179 | 0.718454 | 0.719098 | 0.719178 | 0.725018 | 0.717745 |
| 0.686826 | 0.688874 | 0.688648 | 0.684903 | 0.682099 | 0.687322 | 0.686044 | 0.687947 | 0.688541 |
| 0.723144 | 0.724802 | 0.721825 | 0.723155 | 0.723629 | 0.724268 | 0.72378 | 0.724972 | 0.725384 |
| 0.727836 | 0.732611 | 0.728439 | 0.723465 | 0.725678 | 0.72916 | 0.726054 | 0.729715 | 0.726522 |
| 0.695504 | 0.69721 | 0.697757 | 0.693536 | 0.695738 | 0.696967 | 0.698616 | 0.696927 | 0.697782 |
| 0.694563 | 0.698492 | 0.697817 | 0.697251 | 0.695488 | 0.696189 | 0.695727 | 0.698876 | 0.69947 |
| 0.723031 | 0.72574 | 0.72558 | 0.719319 | 0.721668 | 0.725785 | 0.723608 | 0.727986 | 0.724582 |
| 0.722424 | 0.7274 | 0.722641 | 0.718686 | 0.723741 | 0.722517 | 0.720788 | 0.725513 | 0.722606 |
| 0.723142 | 0.72615 | 0.725805 | 0.725476 | 0.722919 | 0.724928 | 0.724643 | 0.725275 | 0.723468 |
| 0.6752 | 0.678068 | 0.675867 | 0.675831 | 0.676541 | 0.67717 | 0.678452 | 0.678544 | 0.673622 |
| 0.717324 | 0.719921 | 0.717231 | 0.71663 | 0.71432 | 0.721959 | 0.718193 | 0.720227 | 0.722043 |
| 0.674 | 0.675461 | 0.675705 | 0.674001 | 0.677219 | 0.681787 | 0.673686 | 0.67623 | 0.673744 |
| 0.670488 | 0.67152 | 0.669212 | 0.668611 | 0.668937 | 0.672842 | 0.671913 | 0.672647 | 0.671807 |
| 0.717243 | 0.719868 | 0.722482 | 0.718493 | 0.718879 | 0.720683 | 0.719889 | 0.72637 | 0.724095 |
| 0.669253 | 0.667643 | 0.669572 | 0.669371 | 0.668943 | 0.673383 | 0.671971 | 0.674913 | 0.672917 |
| 0.661423 | 0.662154 | 0.663634 | 0.663884 | 0.666767 | 0.663008 | 0.667193 | 0.664186 | 0.666171 |
| 0.657152 | 0.659368 | 0.659058 | 0.659096 | 0.660124 | 0.657996 | 0.661675 | 0.66134 | 0.660152 |
| 0.727047 | 0.727754 | 0.7256 | 0.723653 | 0.725606 | 0.727488 | 0.726955 | 0.732346 | 0.731392 |
| 0.661998 | 0.664529 | 0.66377 | 0.663763 | 0.66594 | 0.663461 | 0.664095 | 0.665922 | 0.667088 |
| 0.719187 | 0.722147 | 0.720163 | 0.717544 | 0.718405 | 0.722868 | 0.721588 | 0.723818 | 0.720164 |

| | | | | | | | | |
|---|---|---|---|---|---|---|---|---|
| 0.727127 | 0.728672 | 0.729492 | 0.726629 | 0.728816 | 0.730891 | 0.730007 | 0.734182 | 0.733323 |
| 0.724141 | 0.727294 | 0.724508 | 0.722716 | 0.720257 | 0.725631 | 0.72506 | 0.728271 | 0.727566 |
| 0.741967 | 0.754019 | 0.752534 | 0.74165 | 0.743915 | 0.747592 | 0.744377 | 0.748567 | 0.742839 |
| 0.652981 | 0.656882 | 0.65237 | 0.653633 | 0.656838 | 0.654866 | 0.654778 | 0.65471 | 0.654714 |
| 0.650766 | 0.652956 | 0.651858 | 0.653198 | 0.654643 | 0.653917 | 0.653466 | 0.656506 | 0.654557 |
| 0.727922 | 0.734033 | 0.726924 | 0.73112 | 0.731909 | 0.735879 | 0.732682 | 0.736963 | 0.735352 |
| 0.653159 | 0.655718 | 0.652825 | 0.650337 | 0.656371 | 0.656463 | 0.6539 | 0.654188 | 0.656009 |
| 0.731392 | 0.735021 | 0.73703 | 0.731781 | 0.728908 | 0.737281 | 0.734893 | 0.742165 | 0.733119 |
| 0.732045 | 0.733232 | 0.731305 | 0.728082 | 0.728604 | 0.736614 | 0.732465 | 0.730245 | 0.731599 |
| 0.731611 | 0.732541 | 0.735205 | 0.735344 | 0.732632 | 0.737517 | 0.737422 | 0.740566 | 0.735706 |
| 0.727501 | 0.730126 | 0.73108 | 0.728026 | 0.727786 | 0.734932 | 0.731129 | 0.736166 | 0.732147 |
| 0.7 | 0.748299 | 0.732877 | 0.72 | 0.705128 | 0.688742 | 0.763514 | 0.726027 | 0.705479 |
| 0.718 | 0.714821 | 0.720644 | 0.726655 | 0.725119 | 0.731198 | 0.722077 | 0.726193 | 0.728016 |
| 0.714397 | 0.716476 | 0.721902 | 0.722031 | 0.726304 | 0.729802 | 0.718843 | 0.728224 | 0.722524 |
| 0.714796 | 0.722718 | 0.722364 | 0.731036 | 0.739961 | 0.737685 | 0.735165 | 0.731126 | 0.729319 |
| 0.720768 | 0.722407 | 0.728064 | 0.728993 | 0.72989 | 0.731391 | 0.726523 | 0.731511 | 0.733485 |
| 0.58833 | 0.588916 | 0.589589 | 0.589917 | 0.594674 | 0.594039 | 0.591874 | 0.591916 | 0.589341 |
| 0.72609 | 0.728117 | 0.731935 | 0.738636 | 0.730949 | 0.733066 | 0.731211 | 0.72882 | 0.734593 |
| 0.734541 | 0.733076 | 0.736001 | 0.73907 | 0.736722 | 0.74 | 0.739626 | 0.739449 | 0.741045 |
| 0.731643 | 0.730507 | 0.733536 | 0.736189 | 0.737167 | 0.735992 | 0.736379 | 0.740029 | 0.740096 |
| 0.738302 | 0.737542 | 0.737163 | 0.738546 | 0.737804 | 0.740108 | 0.741109 | 0.744246 | 0.742413 |
| 0.594379 | 0.598692 | 0.593671 | 0.599647 | 0.600046 | 0.597205 | 0.600107 | 0.594959 | 0.598022 |
| 0.582075 | 0.582787 | 0.580597 | 0.582023 | 0.586939 | 0.585598 | 0.584802 | 0.580851 | 0.581691 |
| 0.755587 | 0.763793 | 0.762205 | 0.765841 | 0.759305 | 0.760552 | 0.76715 | 0.76506 | 0.761721 |
| 0.763244 | 0.765457 | 0.763035 | 0.763414 | 0.763592 | 0.759273 | 0.765178 | 0.766523 | 0.761601 |
| 0.752503 | 0.756642 | 0.751393 | 0.756881 | 0.753209 | 0.751654 | 0.755804 | 0.7616 | 0.753799 |
| 0.760525 | 0.761017 | 0.765818 | 0.758719 | 0.759987 | 0.762598 | 0.760861 | 0.764285 | 0.76092 |
| 0.761451 | 0.764806 | 0.764538 | 0.759197 | 0.762279 | 0.760845 | 0.762317 | 0.765416 | 0.761053 |
| 0.765659 | 0.769669 | 0.767533 | 0.763345 | 0.765415 | 0.767113 | 0.767743 | 0.765707 | 0.766388 |
| 0.760435 | 0.768545 | 0.769034 | 0.762225 | 0.762881 | 0.764135 | 0.763884 | 0.767945 | 0.761897 |
| 0.762843 | 0.771961 | 0.772735 | 0.759226 | 0.759288 | 0.765252 | 0.763307 | 0.762042 | 0.762405 |
| 0.773955 | 0.784805 | 0.779769 | 0.769664 | 0.769218 | 0.775729 | 0.769149 | 0.772676 | 0.774764 |
| 0.764383 | 0.771426 | 0.773417 | 0.765875 | 0.760862 | 0.766766 | 0.765285 | 0.767587 | 0.763154 |
| 0.765876 | 0.771809 | 0.774643 | 0.761195 | 0.763184 | 0.765265 | 0.761939 | 0.767562 | 0.762027 |
| 0.769453 | 0.774959 | 0.779309 | 0.765916 | 0.76524 | 0.768987 | 0.768821 | 0.769337 | 0.769542 |
| 0.766446 | 0.771412 | 0.768939 | 0.757919 | 0.760941 | 0.76613 | 0.761125 | 0.761277 | 0.763278 |
| 0.770507 | 0.778325 | 0.779592 | 0.762436 | 0.76368 | 0.767707 | 0.765147 | 0.769627 | 0.764151 |
| 0.766376 | 0.78184 | 0.771623 | 0.768166 | 0.766773 | 0.770616 | 0.76229 | 0.768082 | 0.768402 |
| 0.764566 | 0.776468 | 0.775259 | 0.760504 | 0.76078 | 0.766482 | 0.763808 | 0.760987 | 0.768691 |
| 0.763082 | 0.771793 | 0.775373 | 0.759977 | 0.761017 | 0.763868 | 0.759798 | 0.762124 | 0.761587 |
| 0.709593 | 0.721456 | 0.724908 | 0.717063 | 0.724969 | 0.721497 | 0.715185 | 0.72533 | 0.724748 |
| 0.723493 | 0.727885 | 0.728555 | 0.724228 | 0.727443 | 0.741662 | 0.731925 | 0.729209 | 0.73241 |
| 0.763214 | 0.772321 | 0.773328 | 0.768973 | 0.764922 | 0.767889 | 0.769533 | 0.769044 | 0.767462 |
| 0.768559 | 0.776216 | 0.770241 | 0.767654 | 0.767259 | 0.774722 | 0.767543 | 0.773223 | 0.771359 |
| 0.720378 | 0.725273 | 0.725596 | 0.719217 | 0.725993 | 0.716584 | 0.718901 | 0.712687 | 0.716492 |
| 0.770537 | 0.779877 | 0.7808 | 0.767998 | 0.766136 | 0.768671 | 0.767444 | 0.765174 | 0.764754 |
| 0.759032 | 0.765978 | 0.767835 | 0.762216 | 0.762359 | 0.763905 | 0.763467 | 0.763126 | 0.758002 |

| | | | | | | | | |
|---|---|---|---|---|---|---|---|---|
| 0.766411 | 0.776832 | 0.775534 | 0.762162 | 0.765189 | 0.771828 | 0.762006 | 0.767552 | 0.765475 |
| 0.767103 | 0.778436 | 0.778315 | 0.769782 | 0.7663 | 0.770166 | 0.772456 | 0.767634 | 0.763139 |
| 0.760727 | 0.770609 | 0.769903 | 0.765895 | 0.763681 | 0.764407 | 0.764499 | 0.763138 | 0.76124 |
| 0.767248 | 0.779569 | 0.77743 | 0.768222 | 0.765612 | 0.773961 | 0.765933 | 0.767381 | 0.764611 |
| 0.769262 | 0.780038 | 0.778061 | 0.767815 | 0.768342 | 0.770051 | 0.766866 | 0.771326 | 0.771271 |
| 0.765655 | 0.774042 | 0.77387 | 0.765322 | 0.766158 | 0.768416 | 0.766007 | 0.766541 | 0.766004 |
| 0.764605 | 0.773724 | 0.769206 | 0.762626 | 0.764006 | 0.767234 | 0.762894 | 0.769867 | 0.762866 |
| 0.761817 | 0.761333 | 0.763697 | 0.757769 | 0.755183 | 0.76054 | 0.755431 | 0.759092 | 0.758883 |
| 0.765215 | 0.77285 | 0.769161 | 0.759641 | 0.762467 | 0.765527 | 0.765053 | 0.767082 | 0.763333 |
| 0.758683 | 0.765105 | 0.766645 | 0.762872 | 0.760923 | 0.762399 | 0.760823 | 0.759948 | 0.758383 |
| 0.768568 | 0.775447 | 0.776399 | 0.765761 | 0.766727 | 0.767672 | 0.764687 | 0.765353 | 0.760058 |
| 0.764132 | 0.773957 | 0.773974 | 0.763768 | 0.763405 | 0.766485 | 0.768775 | 0.768075 | 0.763991 |
| 0.757063 | 0.762188 | 0.765 | 0.756197 | 0.756542 | 0.760775 | 0.756401 | 0.759198 | 0.755705 |
| 0.754723 | 0.761804 | 0.76217 | 0.755725 | 0.756861 | 0.76053 | 0.757482 | 0.759572 | 0.756248 |
| 0.770246 | 0.778148 | 0.775629 | 0.762505 | 0.766863 | 0.7689 | 0.76562 | 0.765583 | 0.764342 |
| 0.763963 | 0.772712 | 0.775493 | 0.763787 | 0.764614 | 0.768911 | 0.766225 | 0.771053 | 0.76538 |
| 0.765554 | 0.764953 | 0.768799 | 0.766053 | 0.763149 | 0.763883 | 0.761159 | 0.766142 | 0.764647 |
| 0.76438 | 0.76619 | 0.769028 | 0.760722 | 0.760616 | 0.763828 | 0.763257 | 0.763293 | 0.759961 |
| 0.763618 | 0.769043 | 0.767587 | 0.766154 | 0.764277 | 0.764926 | 0.766503 | 0.766556 | 0.766718 |
| 0.761194 | 0.768935 | 0.770091 | 0.768038 | 0.767206 | 0.766153 | 0.76472 | 0.763848 | 0.760207 |
| 0.755379 | 0.760733 | 0.758661 | 0.758178 | 0.753779 | 0.75796 | 0.756299 | 0.75938 | 0.757022 |
| 0.758946 | 0.760287 | 0.762886 | 0.762504 | 0.762318 | 0.760814 | 0.762358 | 0.763557 | 0.760365 |
| 0.763696 | 0.771371 | 0.770786 | 0.766072 | 0.766898 | 0.765044 | 0.767042 | 0.769225 | 0.767292 |
| 0.76777 | 0.769488 | 0.770544 | 0.768327 | 0.764199 | 0.767618 | 0.768776 | 0.766824 | 0.764496 |
| 0.758804 | 0.767869 | 0.766445 | 0.762256 | 0.763298 | 0.764322 | 0.762059 | 0.764483 | 0.765604 |
| 0.757978 | 0.763283 | 0.764065 | 0.760821 | 0.760073 | 0.76162 | 0.762834 | 0.765842 | 0.761804 |
| 0.759133 | 0.762806 | 0.76409 | 0.761838 | 0.753831 | 0.762803 | 0.760477 | 0.760309 | 0.757593 |
| 0.759407 | 0.761597 | 0.760203 | 0.761698 | 0.759064 | 0.760994 | 0.76171 | 0.758039 | 0.75549 |
| 0.756076 | 0.76382 | 0.75859 | 0.762927 | 0.761452 | 0.763025 | 0.763968 | 0.76197 | 0.758267 |
| 0.759859 | 0.76398 | 0.76838 | 0.767904 | 0.763055 | 0.76861 | 0.767907 | 0.763656 | 0.763507 |
| 0.759635 | 0.768202 | 0.769123 | 0.765222 | 0.762953 | 0.768264 | 0.764093 | 0.765549 | 0.76263 |
| 0.767519 | 0.771268 | 0.770954 | 0.765709 | 0.763682 | 0.769148 | 0.765331 | 0.764001 | 0.762407 |
| 0.772332 | 0.776966 | 0.773946 | 0.769275 | 0.766447 | 0.769978 | 0.768261 | 0.769336 | 0.769825 |
| 0.776693 | 0.774894 | 0.776187 | 0.770254 | 0.767173 | 0.769093 | 0.768695 | 0.769349 | 0.768049 |
| 0.753353 | 0.761154 | 0.759037 | 0.762995 | 0.759352 | 0.760222 | 0.763364 | 0.762165 | 0.761248 |
| 0.760037 | 0.768708 | 0.769712 | 0.760136 | 0.763799 | 0.763465 | 0.757247 | 0.763821 | 0.763795 |
| 0.757498 | 0.763727 | 0.764016 | 0.765484 | 0.761182 | 0.765049 | 0.764929 | 0.767522 | 0.765061 |
| 0.756466 | 0.764376 | 0.764569 | 0.759155 | 0.758478 | 0.760599 | 0.761593 | 0.76185 | 0.758463 |
| 0.752463 | 0.76153 | 0.758207 | 0.759332 | 0.756727 | 0.759714 | 0.755041 | 0.754996 | 0.754983 |
| 0.750773 | 0.75924 | 0.760333 | 0.756525 | 0.753371 | 0.755879 | 0.758621 | 0.760693 | 0.754469 |
| 0.750763 | 0.754538 | 0.753384 | 0.755652 | 0.75184 | 0.755827 | 0.755527 | 0.758319 | 0.752083 |
| 0.749782 | 0.756815 | 0.75761 | 0.751782 | 0.756646 | 0.752851 | 0.756294 | 0.757175 | 0.753247 |
| 0.74865 | 0.759452 | 0.758939 | 0.756597 | 0.752932 | 0.755272 | 0.754578 | 0.75465 | 0.75355 |
| 0.749798 | 0.753904 | 0.754106 | 0.753079 | 0.753896 | 0.757019 | 0.752498 | 0.752546 | 0.754455 |
| 0.759089 | 0.76671 | 0.768168 | 0.760724 | 0.756056 | 0.75888 | 0.762348 | 0.763102 | 0.757123 |
| 0.752807 | 0.760026 | 0.760634 | 0.751945 | 0.748452 | 0.752779 | 0.754609 | 0.752958 | 0.749699 |

| | | | | | | | | |
|---|---|---|---|---|---|---|---|---|
| 0.75466 | 0.758786 | 0.761667 | 0.762578 | 0.762199 | 0.764206 | 0.76266 | 0.766912 | 0.76035 |
| 0.783 | 0.78144 | 0.783802 | 0.772678 | 0.773399 | 0.773827 | 0.773623 | 0.777262 | 0.767665 |
| 0.773512 | 0.770588 | 0.773266 | 0.770799 | 0.769749 | 0.76925 | 0.768624 | 0.773629 | 0.765996 |
| 0.801817 | 0.785738 | 0.802056 | 0.771307 | 0.772582 | 0.775507 | 0.770606 | 0.766376 | 0.768734 |
| 0.812553 | 0.783254 | 0.802434 | 0.779832 | 0.776716 | 0.777705 | 0.779316 | 0.775514 | 0.77691 |
| 0.776887 | 0.814676 | 0.828526 | 0.768431 | 0.76978 | 0.773904 | 0.774355 | 0.773418 | 0.769162 |
| 0.775718 | 0.791853 | 0.780356 | 0.760712 | 0.768319 | 0.768526 | 0.76569 | 0.769179 | 0.761785 |
| 0.8 | 0.784556 | 0.798056 | 0.765264 | 0.767354 | 0.77522 | 0.768154 | 0.764773 | 0.767011 |
| 0.764436 | 0.782429 | 0.775322 | 0.762718 | 0.762648 | 0.762784 | 0.762762 | 0.770923 | 0.756558 |
| 1 | 0.783392 | 0.819233 | 0.764377 | 0.768363 | 0.765134 | 0.762073 | 0.768652 | 0.76848 |
| 0.783392 | 1 | 0.834622 | 0.771424 | 0.764425 | 0.772606 | 0.777188 | 0.776051 | 0.77058 |
| 0.819233 | 0.834622 | 1 | 0.766601 | 0.76891 | 0.770743 | 0.771513 | 0.773511 | 0.767812 |
| 0.764377 | 0.771424 | 0.766601 | 1 | 0.783339 | 0.778223 | 0.876306 | 0.788105 | 0.785223 |
| 0.768363 | 0.764425 | 0.76891 | 0.783339 | 1 | 0.784539 | 0.785064 | 0.781462 | 0.782488 |
| 0.765134 | 0.772606 | 0.770743 | 0.778223 | 0.784539 | 1 | 0.784159 | 0.778871 | 0.781068 |
| 0.762073 | 0.777188 | 0.771513 | 0.876306 | 0.785064 | 0.784159 | 1 | 0.778711 | 0.781454 |
| 0.768652 | 0.776051 | 0.773511 | 0.788105 | 0.781462 | 0.778871 | 0.778711 | 1 | 0.784322 |
| 0.76848 | 0.77058 | 0.767812 | 0.785223 | 0.782488 | 0.781068 | 0.781454 | 0.784322 | 1 |
| 0.769636 | 0.77485 | 0.774201 | 0.790257 | 0.783025 | 0.790274 | 0.784585 | 0.784761 | 0.778761 |
| 0.765442 | 0.774199 | 0.773161 | 0.785773 | 0.783757 | 0.787646 | 0.789049 | 0.78292 | 0.780464 |
| 0.767924 | 0.765105 | 0.765533 | 0.783613 | 0.816502 | 0.786134 | 0.78736 | 0.771714 | 0.778852 |
| 0.770553 | 0.767788 | 0.771566 | 0.78801 | 0.818443 | 0.790516 | 0.800672 | 0.776173 | 0.775798 |
| 0.763381 | 0.769187 | 0.771706 | 0.782991 | 0.803351 | 0.78838 | 0.787211 | 0.781054 | 0.779669 |
| 0.775456 | 0.773501 | 0.774555 | 0.771881 | 0.777405 | 0.773427 | 0.776118 | 0.779922 | 0.771311 |
| 0.765799 | 0.773664 | 0.772613 | 0.78074 | 0.776203 | 0.782616 | 0.780784 | 0.776157 | 0.779868 |
| 0.768388 | 0.776345 | 0.768948 | 0.778353 | 0.77181 | 0.77707 | 0.778866 | 0.775301 | 0.773203 |
| 0.763233 | 0.770047 | 0.769519 | 0.768436 | 0.770867 | 0.779258 | 0.769137 | 0.771794 | 0.769949 |
| 0.769319 | 0.769505 | 0.76629 | 0.778351 | 0.773343 | 0.774141 | 0.777669 | 0.775725 | 0.779272 |
| 0.768413 | 0.772459 | 0.770723 | 0.776471 | 0.775699 | 0.783667 | 0.77836 | 0.782422 | 0.775109 |
| 0.771463 | 0.775591 | 0.769653 | 0.774125 | 0.774922 | 0.778358 | 0.773812 | 0.779894 | 0.776129 |
| 0.760023 | 0.769301 | 0.767688 | 0.767897 | 0.76756 | 0.773554 | 0.770317 | 0.773036 | 0.766776 |
| 0.771937 | 0.767648 | 0.775245 | 0.774536 | 0.771247 | 0.776751 | 0.775236 | 0.780486 | 0.780878 |
| 0.764239 | 0.773947 | 0.772895 | 0.776009 | 0.77253 | 0.78148 | 0.773289 | 0.777308 | 0.773293 |
| 0.765804 | 0.766692 | 0.763817 | 0.773654 | 0.770098 | 0.774996 | 0.771848 | 0.779099 | 0.773345 |
| 0.766613 | 0.766034 | 0.765261 | 0.773962 | 0.774196 | 0.774577 | 0.776619 | 0.771984 | 0.774183 |
| 0.762328 | 0.765892 | 0.767492 | 0.775831 | 0.77185 | 0.769507 | 0.773574 | 0.774713 | 0.770539 |
| 0.767192 | 0.777932 | 0.775592 | 0.782995 | 0.780778 | 0.782174 | 0.783479 | 0.786574 | 0.777308 |
| 0.762975 | 0.769914 | 0.76882 | 0.7706 | 0.769186 | 0.777509 | 0.771996 | 0.78063 | 0.776048 |
| 0.765359 | 0.770911 | 0.770525 | 0.779532 | 0.771595 | 0.772545 | 0.776715 | 0.772031 | 0.773307 |
| 0.768843 | 0.773706 | 0.770157 | 0.773037 | 0.775269 | 0.777722 | 0.77129 | 0.779233 | 0.772543 |
| 0.768481 | 0.771243 | 0.770789 | 0.774826 | 0.77765 | 0.780434 | 0.776459 | 0.781873 | 0.777031 |
| 0.769548 | 0.774244 | 0.772242 | 0.782526 | 0.775401 | 0.781488 | 0.776452 | 0.778804 | 0.773565 |
| 0.770055 | 0.776095 | 0.770656 | 0.781127 | 0.776552 | 0.778923 | 0.77707 | 0.779834 | 0.771157 |
| 0.76719 | 0.764735 | 0.763751 | 0.77478 | 0.767809 | 0.772659 | 0.770409 | 0.771246 | 0.770785 |
| 0.768014 | 0.768783 | 0.771745 | 0.778192 | 0.776894 | 0.780198 | 0.778901 | 0.778498 | 0.784892 |
| 0.760523 | 0.768102 | 0.763855 | 0.777414 | 0.770111 | 0.773499 | 0.774104 | 0.774552 | 0.777335 |

| | | | | | | | | |
|---|---|---|---|---|---|---|---|---|
| 0.758614 | 0.753416 | 0.75783 | 0.767091 | 0.762352 | 0.765322 | 0.7678 | 0.765304 | 0.768992 |
| 0.745516 | 0.749694 | 0.750988 | 0.75074 | 0.749979 | 0.75533 | 0.751091 | 0.75826 | 0.759215 |
| 0.762743 | 0.765192 | 0.763801 | 0.769574 | 0.771624 | 0.772544 | 0.77158 | 0.770124 | 0.77289 |
| 0.767181 | 0.775805 | 0.773292 | 0.77315 | 0.775552 | 0.777751 | 0.778215 | 0.778499 | 0.77454 |
| 0.75404 | 0.761591 | 0.762613 | 0.764961 | 0.771257 | 0.76817 | 0.764224 | 0.767937 | 0.765628 |
| 0.757292 | 0.770035 | 0.763343 | 0.774051 | 0.772807 | 0.776372 | 0.778204 | 0.771593 | 0.772627 |
| 0.764026 | 0.765644 | 0.769237 | 0.768516 | 0.769317 | 0.779127 | 0.769734 | 0.778283 | 0.768989 |
| 0.756393 | 0.760025 | 0.760021 | 0.765179 | 0.771939 | 0.770489 | 0.764722 | 0.76775 | 0.76348 |
| 0.763556 | 0.771591 | 0.765353 | 0.777586 | 0.774575 | 0.777208 | 0.778133 | 0.778184 | 0.775107 |
| 0.762752 | 0.770743 | 0.769433 | 0.772228 | 0.776003 | 0.777211 | 0.7702 | 0.778224 | 0.770638 |
| 0.767665 | 0.773979 | 0.774812 | 0.769068 | 0.77091 | 0.779343 | 0.769293 | 0.773843 | 0.770956 |
| 0.767378 | 0.767009 | 0.772344 | 0.771299 | 0.771584 | 0.772877 | 0.774862 | 0.775059 | 0.777841 |
| 0.768832 | 0.76955 | 0.774618 | 0.777934 | 0.775169 | 0.782452 | 0.777467 | 0.778495 | 0.780488 |
| 0.767995 | 0.772978 | 0.770801 | 0.770549 | 0.775632 | 0.777334 | 0.771303 | 0.776624 | 0.773891 |
| 0.76509 | 0.768403 | 0.768891 | 0.766403 | 0.769179 | 0.774118 | 0.767763 | 0.776496 | 0.775375 |
| 0.763169 | 0.767703 | 0.767453 | 0.768964 | 0.766494 | 0.775843 | 0.768405 | 0.773986 | 0.774165 |
| 0.763969 | 0.769841 | 0.768032 | 0.769408 | 0.768685 | 0.774844 | 0.769186 | 0.773278 | 0.775437 |
| 0.760697 | 0.767935 | 0.766031 | 0.77083 | 0.766529 | 0.772873 | 0.771785 | 0.768627 | 0.771742 |
| 0.758999 | 0.766516 | 0.76369 | 0.766102 | 0.768103 | 0.775212 | 0.766932 | 0.765377 | 0.770819 |
| 0.758534 | 0.76654 | 0.763491 | 0.767076 | 0.770478 | 0.772322 | 0.770276 | 0.772145 | 0.769985 |
| 0.767008 | 0.770043 | 0.767467 | 0.770944 | 0.76782 | 0.778506 | 0.773144 | 0.774913 | 0.770602 |
| 0.754438 | 0.752375 | 0.75271 | 0.756206 | 0.759165 | 0.760496 | 0.757078 | 0.765233 | 0.753925 |
| 0.7604 | 0.757838 | 0.756215 | 0.766238 | 0.765676 | 0.765602 | 0.768531 | 0.77222 | 0.766076 |
| 0.763193 | 0.764595 | 0.76297 | 0.769725 | 0.769662 | 0.768477 | 0.772251 | 0.773189 | 0.77127 |
| 0.761253 | 0.762237 | 0.765228 | 0.76905 | 0.769929 | 0.772324 | 0.767836 | 0.770619 | 0.764384 |
| 0.761984 | 0.763416 | 0.759844 | 0.767477 | 0.765461 | 0.774921 | 0.768513 | 0.769333 | 0.766194 |
| 0.756139 | 0.762338 | 0.758501 | 0.76522 | 0.761412 | 0.767742 | 0.760108 | 0.76476 | 0.763607 |
| 0.760749 | 0.760226 | 0.764681 | 0.766345 | 0.767752 | 0.769335 | 0.765344 | 0.768449 | 0.765143 |
| 0.75435 | 0.762585 | 0.763532 | 0.763121 | 0.764629 | 0.767534 | 0.76417 | 0.762621 | 0.766179 |
| 0.697389 | 0.69856 | 0.69854 | 0.696394 | 0.697687 | 0.699042 | 0.695253 | 0.702678 | 0.695206 |
| 0.709818 | 0.711756 | 0.710452 | 0.70931 | 0.709234 | 0.710686 | 0.708654 | 0.713146 | 0.710668 |
| 0.703683 | 0.708578 | 0.708233 | 0.705355 | 0.702588 | 0.707199 | 0.706099 | 0.706857 | 0.705109 |
| 0.697746 | 0.697313 | 0.696874 | 0.696463 | 0.697365 | 0.697544 | 0.697191 | 0.698673 | 0.698093 |
| 0.707406 | 0.710638 | 0.709769 | 0.706616 | 0.705386 | 0.707755 | 0.707965 | 0.709824 | 0.709984 |
| 0.759176 | 0.763841 | 0.764741 | 0.762647 | 0.763269 | 0.770525 | 0.758982 | 0.766099 | 0.764972 |
| 0.758377 | 0.76687 | 0.767902 | 0.767157 | 0.770199 | 0.770455 | 0.766012 | 0.772517 | 0.768707 |
| 0.761592 | 0.76776 | 0.768323 | 0.767568 | 0.770347 | 0.775596 | 0.773376 | 0.773425 | 0.76896 |
| 0.765241 | 0.769294 | 0.771492 | 0.768013 | 0.771026 | 0.770429 | 0.773652 | 0.774024 | 0.774093 |
| 0.757849 | 0.765617 | 0.764628 | 0.767715 | 0.761716 | 0.769828 | 0.76174 | 0.77196 | 0.768424 |
| 0.764039 | 0.768028 | 0.767726 | 0.765915 | 0.769765 | 0.774045 | 0.76805 | 0.771349 | 0.771695 |
| 0.76042 | 0.770402 | 0.769404 | 0.771017 | 0.767692 | 0.774028 | 0.76959 | 0.768964 | 0.773936 |
| 0.758996 | 0.768228 | 0.767646 | 0.764819 | 0.76621 | 0.771785 | 0.770558 | 0.765704 | 0.766785 |
| 0.756409 | 0.766534 | 0.75991 | 0.762177 | 0.760607 | 0.762155 | 0.76169 | 0.766104 | 0.759391 |
| 0.753161 | 0.759853 | 0.758323 | 0.762595 | 0.767884 | 0.763339 | 0.760057 | 0.766444 | 0.763414 |
| 0.754693 | 0.765758 | 0.762579 | 0.766318 | 0.765932 | 0.767884 | 0.7654 | 0.769392 | 0.770162 |
| 0.762842 | 0.766303 | 0.771016 | 0.768326 | 0.767661 | 0.773134 | 0.772872 | 0.772031 | 0.77446 |

| | | | | | | | | |
|---|---|---|---|---|---|---|---|---|
| 0.761604 | 0.767172 | 0.766007 | 0.765911 | 0.76936 | 0.771905 | 0.769289 | 0.770689 | 0.768911 |
| 0.7591 | 0.762753 | 0.763032 | 0.766647 | 0.765274 | 0.770272 | 0.769199 | 0.766037 | 0.771195 |
| 0.758678 | 0.764469 | 0.765628 | 0.771303 | 0.771517 | 0.77243 | 0.77314 | 0.772761 | 0.772178 |
| 0.760373 | 0.762196 | 0.76196 | 0.768989 | 0.76429 | 0.77414 | 0.763121 | 0.764792 | 0.7663 |
| 0.758595 | 0.763653 | 0.762665 | 0.769497 | 0.768672 | 0.772177 | 0.767338 | 0.765673 | 0.761328 |
| 0.763385 | 0.761641 | 0.763663 | 0.768437 | 0.766718 | 0.767145 | 0.76625 | 0.76713 | 0.772386 |
| 0.761844 | 0.768168 | 0.765805 | 0.766431 | 0.765225 | 0.770463 | 0.768382 | 0.767512 | 0.769906 |
| 0.757903 | 0.761619 | 0.759729 | 0.769718 | 0.766698 | 0.771438 | 0.770912 | 0.770675 | 0.769724 |
| 0.757619 | 0.760393 | 0.761857 | 0.764137 | 0.763375 | 0.76936 | 0.766765 | 0.767108 | 0.765556 |
| 0.760305 | 0.760834 | 0.763873 | 0.765862 | 0.76691 | 0.770508 | 0.766044 | 0.762201 | 0.769742 |
| 0.757229 | 0.764993 | 0.765721 | 0.769759 | 0.766201 | 0.767187 | 0.768169 | 0.765635 | 0.768677 |
| 0.757048 | 0.767595 | 0.766262 | 0.765697 | 0.763878 | 0.771171 | 0.76905 | 0.764935 | 0.766981 |
| 0.751021 | 0.757065 | 0.757527 | 0.763275 | 0.760036 | 0.763572 | 0.758182 | 0.764342 | 0.756695 |
| 0.749345 | 0.7564 | 0.756855 | 0.761283 | 0.757507 | 0.76264 | 0.761065 | 0.76077 | 0.762503 |
| 0.760081 | 0.775073 | 0.76721 | 0.766038 | 0.765412 | 0.773689 | 0.772341 | 0.772044 | 0.771533 |
| 0.767536 | 0.773736 | 0.77354 | 0.775431 | 0.773822 | 0.779602 | 0.777778 | 0.781535 | 0.781181 |
| 0.764403 | 0.773744 | 0.770151 | 0.772423 | 0.767111 | 0.775487 | 0.775362 | 0.782111 | 0.773952 |
| 0.772513 | 0.772534 | 0.772615 | 0.771158 | 0.768059 | 0.779823 | 0.771944 | 0.776187 | 0.773261 |
| 0.764575 | 0.77 | 0.767291 | 0.768338 | 0.769342 | 0.771509 | 0.768357 | 0.768789 | 0.766911 |
| 0.758899 | 0.76864 | 0.764341 | 0.774317 | 0.767357 | 0.774262 | 0.771665 | 0.773888 | 0.76892 |
| 0.758182 | 0.767787 | 0.763792 | 0.766801 | 0.7646 | 0.767594 | 0.768697 | 0.767527 | 0.769361 |
| 0.75979 | 0.767731 | 0.761685 | 0.76777 | 0.766697 | 0.7774 | 0.770811 | 0.775406 | 0.771316 |
| 0.763454 | 0.773342 | 0.771803 | 0.775606 | 0.76927 | 0.778968 | 0.774331 | 0.779493 | 0.776354 |
| 0.758885 | 0.75983 | 0.759912 | 0.768402 | 0.763939 | 0.76942 | 0.766878 | 0.763254 | 0.764814 |
| 0.762957 | 0.768034 | 0.770569 | 0.772788 | 0.767174 | 0.776912 | 0.770595 | 0.77208 | 0.773995 |
| 0.703931 | 0.706484 | 0.707331 | 0.703352 | 0.705328 | 0.707596 | 0.70346 | 0.707862 | 0.702692 |
| 0.76392 | 0.771211 | 0.770396 | 0.7745 | 0.765824 | 0.782178 | 0.768585 | 0.77851 | 0.776697 |
| 0.766795 | 0.773097 | 0.771487 | 0.773427 | 0.774552 | 0.77246 | 0.773794 | 0.776121 | 0.770425 |
| 0.766302 | 0.773258 | 0.770423 | 0.776837 | 0.770773 | 0.777952 | 0.779802 | 0.775401 | 0.774077 |
| 0.76545 | 0.772552 | 0.771551 | 0.77248 | 0.769937 | 0.778666 | 0.774636 | 0.776528 | 0.774814 |
| 0.76232 | 0.764591 | 0.770213 | 0.770326 | 0.76347 | 0.775372 | 0.766058 | 0.764395 | 0.771114 |
| 0.764961 | 0.768913 | 0.768851 | 0.771421 | 0.768602 | 0.772727 | 0.765194 | 0.775464 | 0.767725 |
| 0.767234 | 0.770745 | 0.771908 | 0.767068 | 0.770238 | 0.774191 | 0.768341 | 0.77319 | 0.776724 |
| 0.762813 | 0.765712 | 0.766715 | 0.775893 | 0.764216 | 0.772794 | 0.774617 | 0.776689 | 0.767674 |
| 0.754694 | 0.763638 | 0.764196 | 0.766387 | 0.764297 | 0.766782 | 0.767642 | 0.772425 | 0.761289 |
| 0.760815 | 0.768447 | 0.764079 | 0.765997 | 0.760715 | 0.766244 | 0.768797 | 0.769828 | 0.766534 |
| 0.765002 | 0.771557 | 0.76743 | 0.776396 | 0.770605 | 0.780792 | 0.777834 | 0.781627 | 0.778648 |
| 0.767504 | 0.773997 | 0.774301 | 0.779935 | 0.776484 | 0.782277 | 0.773406 | 0.790137 | 0.784359 |
| 0.766811 | 0.773325 | 0.773246 | 0.781305 | 0.777267 | 0.780686 | 0.777001 | 0.785105 | 0.78078 |
| 0.762466 | 0.769736 | 0.768651 | 0.783074 | 0.775159 | 0.781143 | 0.780294 | 0.782907 | 0.779066 |
| 0.761741 | 0.765366 | 0.765756 | 0.774604 | 0.770977 | 0.775594 | 0.781304 | 0.775737 | 0.773261 |
| 0.769257 | 0.771571 | 0.770338 | 0.782766 | 0.783079 | 0.782587 | 0.780422 | 0.787859 | 0.781933 |
| 0.766117 | 0.769637 | 0.767672 | 0.776975 | 0.76811 | 0.775043 | 0.773274 | 0.782992 | 0.781634 |
| 0.751119 | 0.765468 | 0.761959 | 0.76274 | 0.763403 | 0.769806 | 0.763703 | 0.767691 | 0.766565 |
| 0.759944 | 0.768929 | 0.767317 | 0.769085 | 0.76953 | 0.773501 | 0.76676 | 0.771555 | 0.766561 |
| 0.760371 | 0.766013 | 0.763119 | 0.768541 | 0.766738 | 0.770591 | 0.770325 | 0.769814 | 0.767582 |

| | | | | | | | | |
|---|---|---|---|---|---|---|---|---|
| 0.757722 | 0.762451 | 0.762094 | 0.768689 | 0.76632 | 0.767351 | 0.768632 | 0.768981 | 0.766419 |
| 0.760672 | 0.764749 | 0.765589 | 0.769054 | 0.770227 | 0.770996 | 0.766825 | 0.771918 | 0.76956 |
| 0.756233 | 0.766272 | 0.758817 | 0.761728 | 0.762997 | 0.768646 | 0.76321 | 0.768055 | 0.766367 |
| 0.728336 | 0.71956 | 0.72973 | 0.724508 | 0.737586 | 0.723548 | 0.731034 | 0.725259 | 0.719633 |
| 0.754639 | 0.760558 | 0.764337 | 0.768998 | 0.762751 | 0.768582 | 0.762866 | 0.76872 | 0.766957 |
| 0.754719 | 0.756746 | 0.760478 | 0.76487 | 0.767243 | 0.770883 | 0.76613 | 0.766735 | 0.767644 |
| 0.756914 | 0.765284 | 0.766615 | 0.767539 | 0.768156 | 0.771324 | 0.768568 | 0.769075 | 0.765367 |
| 0.760475 | 0.763632 | 0.765173 | 0.764576 | 0.762235 | 0.772787 | 0.762013 | 0.767779 | 0.762358 |
| 0.764392 | 0.763339 | 0.76714 | 0.770852 | 0.768776 | 0.776887 | 0.766084 | 0.769623 | 0.770557 |
| 0.760377 | 0.768717 | 0.764624 | 0.768593 | 0.76719 | 0.773482 | 0.767202 | 0.773441 | 0.770067 |
| 0.758561 | 0.766018 | 0.763056 | 0.765377 | 0.764701 | 0.770263 | 0.763662 | 0.768392 | 0.765666 |
| 0.762509 | 0.760605 | 0.762635 | 0.769884 | 0.77035 | 0.769691 | 0.767769 | 0.769946 | 0.769945 |
| 0.762816 | 0.768117 | 0.767816 | 0.76609 | 0.766424 | 0.770743 | 0.762306 | 0.773476 | 0.764977 |
| 0.759173 | 0.761485 | 0.766272 | 0.765642 | 0.767682 | 0.775179 | 0.768929 | 0.772642 | 0.769771 |
| 0.764324 | 0.762075 | 0.766577 | 0.766189 | 0.77016 | 0.770813 | 0.767982 | 0.770188 | 0.77074 |
| 0.751226 | 0.76026 | 0.761148 | 0.761834 | 0.769624 | 0.765032 | 0.764344 | 0.766726 | 0.76386 |
| 0.764538 | 0.768821 | 0.770954 | 0.767043 | 0.768805 | 0.770557 | 0.764935 | 0.771062 | 0.771916 |
| 0.76054 | 0.765412 | 0.766568 | 0.771938 | 0.769408 | 0.775842 | 0.771168 | 0.773763 | 0.773413 |
| 0.754541 | 0.760223 | 0.759395 | 0.769161 | 0.765849 | 0.766694 | 0.767404 | 0.767103 | 0.760084 |
| 0.756748 | 0.765776 | 0.763711 | 0.768336 | 0.767041 | 0.76841 | 0.769079 | 0.768549 | 0.771341 |
| 0.75978 | 0.768066 | 0.76685 | 0.773091 | 0.775431 | 0.773616 | 0.77163 | 0.770557 | 0.771222 |
| 0.763151 | 0.76878 | 0.770442 | 0.769799 | 0.769944 | 0.776008 | 0.766097 | 0.774222 | 0.771544 |
| 0.676 | 0.680678 | 0.681984 | 0.676867 | 0.683282 | 0.687238 | 0.678997 | 0.682016 | 0.669705 |
| 0.646795 | 0.655734 | 0.666348 | 0.649645 | 0.655645 | 0.656955 | 0.654472 | 0.658267 | 0.659934 |
| 0.67054 | 0.676635 | 0.670971 | 0.676065 | 0.678564 | 0.674379 | 0.67533 | 0.67508 | 0.676144 |
| 0.650481 | 0.6551 | 0.65669 | 0.650294 | 0.653877 | 0.655376 | 0.655428 | 0.654774 | 0.650177 |
| 0.727778 | 0.726719 | 0.729572 | 0.733985 | 0.737626 | 0.735872 | 0.73239 | 0.735768 | 0.737101 |
| 0.717978 | 0.714096 | 0.718347 | 0.72839 | 0.722878 | 0.731173 | 0.728843 | 0.725714 | 0.716846 |
| 0.722771 | 0.723014 | 0.724888 | 0.726376 | 0.739617 | 0.731874 | 0.735482 | 0.720369 | 0.726837 |
| 0.724252 | 0.740079 | 0.731596 | 0.733707 | 0.743729 | 0.739112 | 0.733874 | 0.742946 | 0.731172 |
| 0.709891 | 0.718864 | 0.713831 | 0.706709 | 0.717421 | 0.725006 | 0.711366 | 0.709329 | 0.713704 |
| 0.703767 | 0.709858 | 0.708968 | 0.699637 | 0.708228 | 0.714885 | 0.71222 | 0.719234 | 0.6977 |
| 0.71378 | 0.70566 | 0.718833 | 0.715755 | 0.713992 | 0.723541 | 0.72254 | 0.716965 | 0.719319 |
| 0.652716 | 0.655111 | 0.658865 | 0.655356 | 0.657041 | 0.657666 | 0.655401 | 0.661522 | 0.64936 |
| 0.754613 | 0.756699 | 0.757817 | 0.76497 | 0.761076 | 0.763624 | 0.763142 | 0.763255 | 0.758007 |
| 0.732543 | 0.734576 | 0.737232 | 0.744387 | 0.738353 | 0.743348 | 0.740922 | 0.74111 | 0.742738 |
| 0.724231 | 0.732661 | 0.734699 | 0.730028 | 0.732676 | 0.74207 | 0.735262 | 0.734057 | 0.739838 |
| 0.712435 | 0.719582 | 0.723735 | 0.717806 | 0.715257 | 0.718785 | 0.724125 | 0.725263 | 0.724157 |
| 0.725029 | 0.73187 | 0.740361 | 0.734511 | 0.740806 | 0.739418 | 0.739817 | 0.737413 | 0.73907 |
| 0.726876 | 0.727319 | 0.737579 | 0.728041 | 0.731056 | 0.734949 | 0.730008 | 0.724408 | 0.725572 |
| 0.722178 | 0.719931 | 0.732188 | 0.729618 | 0.731322 | 0.733037 | 0.72624 | 0.727344 | 0.725157 |
| 0.722526 | 0.717468 | 0.729785 | 0.722845 | 0.727287 | 0.731886 | 0.726603 | 0.726233 | 0.722171 |
| 0.71637 | 0.727534 | 0.733829 | 0.72765 | 0.729208 | 0.734088 | 0.723627 | 0.724273 | 0.721096 |
| 0.727002 | 0.720749 | 0.7262 | 0.730018 | 0.732716 | 0.729825 | 0.73177 | 0.726651 | 0.72777 |
| 0.713079 | 0.72002 | 0.721441 | 0.722493 | 0.729521 | 0.72848 | 0.726732 | 0.720978 | 0.715949 |
| 0.707171 | 0.700913 | 0.716189 | 0.706211 | 0.720992 | 0.713485 | 0.714618 | 0.716975 | 0.706037 |

| | | | | | | | | |
|---|---|---|---|---|---|---|---|---|
| 0.745016 | 0.742574 | 0.74758 | 0.749677 | 0.751865 | 0.755086 | 0.747545 | 0.753279 | 0.755468 |
| 0.723177 | 0.72782 | 0.728614 | 0.726774 | 0.726085 | 0.733847 | 0.722543 | 0.730769 | 0.730834 |
| 0.716843 | 0.720781 | 0.722803 | 0.720719 | 0.729774 | 0.73112 | 0.719615 | 0.730671 | 0.72703 |
| 0.72281 | 0.72364 | 0.72791 | 0.72698 | 0.727717 | 0.735353 | 0.731244 | 0.736666 | 0.730939 |
| 0.717338 | 0.727328 | 0.726384 | 0.728886 | 0.729774 | 0.734531 | 0.732923 | 0.736574 | 0.731495 |
| 0.706743 | 0.711898 | 0.718031 | 0.720278 | 0.72371 | 0.717702 | 0.714572 | 0.72418 | 0.726185 |
| 0.714911 | 0.715868 | 0.715494 | 0.713698 | 0.716872 | 0.724079 | 0.718162 | 0.718915 | 0.7212 |
| 0.704757 | 0.71341 | 0.710386 | 0.723621 | 0.714005 | 0.715527 | 0.720884 | 0.719989 | 0.719136 |
| 0.717324 | 0.718093 | 0.721781 | 0.721907 | 0.724548 | 0.717483 | 0.721888 | 0.717863 | 0.723374 |
| 0.743838 | 0.752231 | 0.752831 | 0.752344 | 0.74656 | 0.756786 | 0.746199 | 0.756449 | 0.753877 |
| 0.736837 | 0.750499 | 0.755591 | 0.74902 | 0.743585 | 0.751203 | 0.745548 | 0.752254 | 0.751794 |
| 0.732703 | 0.742321 | 0.744109 | 0.741505 | 0.74185 | 0.745904 | 0.739943 | 0.746952 | 0.751016 |
| 0.729658 | 0.733782 | 0.733967 | 0.73368 | 0.741826 | 0.73871 | 0.736855 | 0.737803 | 0.744781 |
| 0.733957 | 0.741001 | 0.742777 | 0.740429 | 0.736682 | 0.752256 | 0.7408 | 0.747476 | 0.737857 |
| 0.729683 | 0.734371 | 0.739237 | 0.74003 | 0.74467 | 0.743078 | 0.735856 | 0.739152 | 0.736253 |

| Wild_plot_ | Wild_plot_ | Wild_plot_ | Wild_plot_ | Wild_plot_ | Wild_plot_ | Wild_plot_ | Wild_plot_ | Wild_plot_ |
|---|---|---|---|---|---|---|---|---|
| 0.712676 | 0.708478 | 0.708866 | 0.706541 | 0.708906 | 0.71146 | 0.708164 | 0.709156 | 0.706827 |
| 0.718713 | 0.716742 | 0.712147 | 0.713953 | 0.715871 | 0.717989 | 0.713679 | 0.715878 | 0.713369 |
| 0.70942 | 0.708736 | 0.702748 | 0.702258 | 0.704285 | 0.706963 | 0.704946 | 0.705279 | 0.70198 |
| 0.719451 | 0.717137 | 0.713273 | 0.709654 | 0.718074 | 0.716232 | 0.714934 | 0.713924 | 0.710027 |
| 0.715553 | 0.710002 | 0.709621 | 0.708752 | 0.711113 | 0.711711 | 0.709839 | 0.706403 | 0.707704 |
| 0.721452 | 0.717739 | 0.717525 | 0.718043 | 0.720363 | 0.718199 | 0.716981 | 0.716392 | 0.715734 |
| 0.782641 | 0.781273 | 0.76899 | 0.774401 | 0.778925 | 0.777062 | 0.77457 | 0.774241 | 0.769493 |
| 0.780877 | 0.785843 | 0.774622 | 0.773329 | 0.779053 | 0.7756 | 0.77433 | 0.769256 | 0.766992 |
| 0.718429 | 0.712553 | 0.713062 | 0.710159 | 0.715182 | 0.718658 | 0.712378 | 0.710024 | 0.711088 |
| 0.714819 | 0.710759 | 0.710818 | 0.709242 | 0.712932 | 0.71315 | 0.707468 | 0.71046 | 0.709223 |
| 0.713944 | 0.709197 | 0.709452 | 0.706716 | 0.713763 | 0.714082 | 0.708769 | 0.710759 | 0.708442 |
| 0.709485 | 0.704115 | 0.700659 | 0.699165 | 0.70787 | 0.704572 | 0.703226 | 0.706241 | 0.703564 |
| 0.716563 | 0.714342 | 0.714938 | 0.710533 | 0.715021 | 0.712705 | 0.711245 | 0.711624 | 0.711476 |
| 0.727615 | 0.722496 | 0.723103 | 0.719524 | 0.726193 | 0.722447 | 0.721789 | 0.723799 | 0.720941 |
| 0.739089 | 0.735804 | 0.733955 | 0.736028 | 0.739813 | 0.737876 | 0.734175 | 0.739426 | 0.731779 |
| 0.720013 | 0.717257 | 0.71408 | 0.714297 | 0.716611 | 0.717888 | 0.713715 | 0.714569 | 0.712542 |
| 0.721847 | 0.716598 | 0.715588 | 0.710932 | 0.71677 | 0.715093 | 0.714116 | 0.712444 | 0.708975 |
| 0.744323 | 0.744444 | 0.739909 | 0.740247 | 0.752196 | 0.744153 | 0.740764 | 0.742384 | 0.738626 |
| 0.711038 | 0.707818 | 0.703618 | 0.703829 | 0.708747 | 0.713696 | 0.706779 | 0.704287 | 0.702626 |
| 0.715253 | 0.713459 | 0.71023 | 0.70738 | 0.71424 | 0.712974 | 0.712993 | 0.710591 | 0.710558 |
| 0.711108 | 0.70444 | 0.707802 | 0.703742 | 0.711781 | 0.711783 | 0.707143 | 0.704228 | 0.710769 |
| 0.717685 | 0.709532 | 0.709476 | 0.709111 | 0.714444 | 0.716945 | 0.707026 | 0.708502 | 0.711101 |
| 0.721051 | 0.718728 | 0.71859 | 0.71504 | 0.722372 | 0.719731 | 0.716627 | 0.713593 | 0.713981 |
| 0.717784 | 0.712874 | 0.711674 | 0.712316 | 0.715315 | 0.712843 | 0.71132 | 0.712879 | 0.709168 |
| 0.71783 | 0.714331 | 0.71501 | 0.710975 | 0.717435 | 0.717839 | 0.712653 | 0.710459 | 0.710798 |
| 0.715824 | 0.711602 | 0.713398 | 0.709978 | 0.713466 | 0.7135 | 0.711201 | 0.709652 | 0.710528 |
| 0.707454 | 0.705762 | 0.7063 | 0.703855 | 0.707668 | 0.708396 | 0.7087 | 0.703041 | 0.706022 |
| 0.719446 | 0.713016 | 0.713044 | 0.708867 | 0.713319 | 0.714949 | 0.710742 | 0.708646 | 0.708565 |
| 0.717838 | 0.714785 | 0.714069 | 0.712393 | 0.715656 | 0.715598 | 0.712631 | 0.710236 | 0.712501 |
| 0.71318 | 0.71042 | 0.712555 | 0.713364 | 0.715014 | 0.713231 | 0.712978 | 0.707965 | 0.709025 |
| 0.715221 | 0.708373 | 0.712507 | 0.70817 | 0.714683 | 0.712939 | 0.708183 | 0.709094 | 0.70631 |
| 0.716257 | 0.707846 | 0.707519 | 0.705995 | 0.708929 | 0.712328 | 0.705557 | 0.707728 | 0.706298 |
| 0.719142 | 0.706627 | 0.710967 | 0.708257 | 0.709614 | 0.714027 | 0.707569 | 0.70996 | 0.708036 |
| 0.71731 | 0.713148 | 0.716601 | 0.712836 | 0.717321 | 0.713859 | 0.712341 | 0.711377 | 0.713026 |
| 0.705226 | 0.709417 | 0.705028 | 0.703256 | 0.708406 | 0.710215 | 0.708422 | 0.707185 | 0.704855 |
| 0.712848 | 0.712059 | 0.710907 | 0.711082 | 0.714559 | 0.71716 | 0.711637 | 0.709178 | 0.70897 |
| 0.717523 | 0.713848 | 0.716847 | 0.714422 | 0.715376 | 0.714745 | 0.710661 | 0.712842 | 0.712393 |
| 0.716238 | 0.713453 | 0.713963 | 0.712976 | 0.718895 | 0.714616 | 0.710923 | 0.710419 | 0.710281 |
| 0.722355 | 0.718678 | 0.717906 | 0.711933 | 0.721613 | 0.718427 | 0.717675 | 0.71289 | 0.717324 |

| | | | | | | | | |
|---|---|---|---|---|---|---|---|---|
| 0.74716 | 0.750399 | 0.750525 | 0.746744 | 0.750999 | 0.743689 | 0.752817 | 0.748299 | 0.744225 |
| 0.707238 | 0.701618 | 0.700215 | 0.700295 | 0.703075 | 0.708179 | 0.704219 | 0.703686 | 0.700581 |
| 0.708998 | 0.7081 | 0.703677 | 0.705355 | 0.709297 | 0.706963 | 0.705177 | 0.708005 | 0.704282 |
| 0.718673 | 0.711763 | 0.712686 | 0.71359 | 0.718604 | 0.71845 | 0.716376 | 0.717286 | 0.712703 |
| 0.716987 | 0.709728 | 0.708967 | 0.711134 | 0.716618 | 0.710976 | 0.708871 | 0.711735 | 0.706155 |
| 0.718587 | 0.713717 | 0.713998 | 0.710664 | 0.713047 | 0.715895 | 0.716464 | 0.718432 | 0.71262 |
| 0.71381 | 0.709167 | 0.708824 | 0.707958 | 0.710458 | 0.712013 | 0.707176 | 0.707244 | 0.707006 |
| 0.712074 | 0.713057 | 0.70911 | 0.709138 | 0.712991 | 0.712303 | 0.709609 | 0.709999 | 0.708955 |
| 0.716412 | 0.712534 | 0.71394 | 0.712282 | 0.720925 | 0.717379 | 0.714749 | 0.712605 | 0.711143 |
| 0.716597 | 0.715652 | 0.714628 | 0.712082 | 0.718377 | 0.716776 | 0.714902 | 0.709575 | 0.710907 |
| 0.711388 | 0.703223 | 0.702675 | 0.700953 | 0.708363 | 0.706297 | 0.703634 | 0.706126 | 0.701745 |
| 0.711384 | 0.705785 | 0.703991 | 0.703203 | 0.70945 | 0.711426 | 0.704175 | 0.70905 | 0.705375 |
| 0.715592 | 0.706265 | 0.709404 | 0.707641 | 0.713738 | 0.711603 | 0.711438 | 0.711369 | 0.70876 |
| 0.711129 | 0.704074 | 0.703505 | 0.702742 | 0.710158 | 0.70734 | 0.708094 | 0.705544 | 0.704932 |
| 0.710961 | 0.70835 | 0.708596 | 0.704476 | 0.707897 | 0.709253 | 0.702359 | 0.703301 | 0.701591 |
| 0.711821 | 0.707732 | 0.707647 | 0.707097 | 0.708541 | 0.705403 | 0.702863 | 0.703704 | 0.700917 |
| 0.710607 | 0.701818 | 0.702733 | 0.702026 | 0.707321 | 0.705188 | 0.706498 | 0.700998 | 0.702265 |
| 0.716456 | 0.712671 | 0.710547 | 0.712386 | 0.711942 | 0.713812 | 0.711988 | 0.711053 | 0.713551 |
| 0.715525 | 0.714807 | 0.713954 | 0.713369 | 0.718926 | 0.714173 | 0.713309 | 0.714002 | 0.711552 |
| 0.712137 | 0.707674 | 0.703612 | 0.704067 | 0.711258 | 0.711684 | 0.710514 | 0.708627 | 0.705966 |
| 0.710957 | 0.704536 | 0.705387 | 0.702196 | 0.705262 | 0.705505 | 0.701052 | 0.706371 | 0.701589 |
| 0.709958 | 0.705789 | 0.706028 | 0.707141 | 0.707891 | 0.709121 | 0.706854 | 0.705639 | 0.706133 |
| 0.716636 | 0.708193 | 0.711805 | 0.706579 | 0.711434 | 0.71469 | 0.710719 | 0.706763 | 0.704691 |
| 0.721862 | 0.716537 | 0.715904 | 0.714274 | 0.718969 | 0.722408 | 0.716327 | 0.715495 | 0.717431 |
| 0.727403 | 0.724833 | 0.725885 | 0.724799 | 0.72754 | 0.726912 | 0.725107 | 0.725493 | 0.723156 |
| 0.782461 | 0.786241 | 0.775843 | 0.774301 | 0.778999 | 0.776422 | 0.776066 | 0.771095 | 0.768573 |
| 0.779746 | 0.777227 | 0.774417 | 0.775004 | 0.777916 | 0.782059 | 0.780648 | 0.780685 | 0.781623 |
| 0.781418 | 0.77938 | 0.773965 | 0.774877 | 0.780766 | 0.777915 | 0.776479 | 0.778027 | 0.782003 |
| 0.708396 | 0.708277 | 0.706876 | 0.70569 | 0.711828 | 0.709331 | 0.707267 | 0.706153 | 0.701046 |
| 0.709144 | 0.705734 | 0.705921 | 0.705722 | 0.709152 | 0.711249 | 0.703987 | 0.703214 | 0.70605 |
| 0.71737 | 0.713333 | 0.718184 | 0.715968 | 0.722067 | 0.715974 | 0.719662 | 0.715369 | 0.711528 |
| 0.71941 | 0.715985 | 0.713603 | 0.714634 | 0.720312 | 0.718858 | 0.714706 | 0.716774 | 0.712353 |
| 0.713354 | 0.708483 | 0.705445 | 0.705801 | 0.709404 | 0.709852 | 0.706732 | 0.705344 | 0.701926 |
| 0.713999 | 0.707822 | 0.707321 | 0.707297 | 0.712322 | 0.710871 | 0.708488 | 0.70811 | 0.70606 |
| 0.71066 | 0.707949 | 0.706699 | 0.704073 | 0.70762 | 0.712992 | 0.707694 | 0.707733 | 0.70325 |
| 0.715931 | 0.710809 | 0.708822 | 0.707537 | 0.711468 | 0.709667 | 0.708287 | 0.707951 | 0.704936 |
| 0.713442 | 0.707537 | 0.704445 | 0.705299 | 0.711326 | 0.710594 | 0.708013 | 0.707792 | 0.703629 |
| 0.725002 | 0.719695 | 0.722581 | 0.720113 | 0.72433 | 0.727251 | 0.725631 | 0.723787 | 0.721559 |
| 0.701679 | 0.700912 | 0.702202 | 0.701015 | 0.70032 | 0.705694 | 0.702013 | 0.70014 | 0.695638 |
| 0.709477 | 0.70454 | 0.710166 | 0.708264 | 0.710904 | 0.708852 | 0.706234 | 0.705341 | 0.706789 |
| 0.711604 | 0.704339 | 0.704106 | 0.704704 | 0.708238 | 0.706546 | 0.702518 | 0.704208 | 0.700586 |
| 0.728324 | 0.723709 | 0.725541 | 0.724227 | 0.727425 | 0.729006 | 0.725316 | 0.723169 | 0.722744 |
| 0.71216 | 0.707485 | 0.709737 | 0.708253 | 0.713855 | 0.711675 | 0.709119 | 0.709939 | 0.708828 |
| 0.714557 | 0.707486 | 0.709731 | 0.70469 | 0.711969 | 0.708634 | 0.708457 | 0.705002 | 0.706928 |
| 0.71408 | 0.708314 | 0.709797 | 0.711509 | 0.712524 | 0.715957 | 0.710957 | 0.711897 | 0.711258 |
| 0.712349 | 0.709188 | 0.709547 | 0.706306 | 0.708972 | 0.708967 | 0.709258 | 0.706052 | 0.705208 |

| | | | | | | | | |
|---|---|---|---|---|---|---|---|---|
| 0.709464 | 0.701504 | 0.709662 | 0.70844 | 0.711969 | 0.709277 | 0.707587 | 0.707554 | 0.704729 |
| 0.71389 | 0.713248 | 0.710143 | 0.707795 | 0.716125 | 0.714759 | 0.713158 | 0.711407 | 0.708713 |
| 0.713473 | 0.708752 | 0.710295 | 0.706751 | 0.710665 | 0.713226 | 0.712633 | 0.710081 | 0.708396 |
| 0.716694 | 0.712172 | 0.715387 | 0.715065 | 0.718241 | 0.72108 | 0.716619 | 0.7163 | 0.714927 |
| 0.718705 | 0.711644 | 0.713955 | 0.713555 | 0.716293 | 0.71665 | 0.717856 | 0.714093 | 0.715187 |
| 0.713026 | 0.70932 | 0.711009 | 0.706585 | 0.710734 | 0.710212 | 0.709058 | 0.708784 | 0.70762 |
| 0.713621 | 0.707143 | 0.712102 | 0.707383 | 0.716977 | 0.714387 | 0.710945 | 0.709266 | 0.712549 |
| 0.714823 | 0.713921 | 0.713421 | 0.712475 | 0.720737 | 0.718041 | 0.713257 | 0.714962 | 0.709215 |
| 0.714583 | 0.710286 | 0.71289 | 0.711591 | 0.716089 | 0.714161 | 0.713857 | 0.710121 | 0.709496 |
| 0.712297 | 0.710229 | 0.709412 | 0.70848 | 0.713913 | 0.709408 | 0.708242 | 0.708657 | 0.703564 |
| 0.712125 | 0.711392 | 0.710105 | 0.706531 | 0.711148 | 0.712236 | 0.707289 | 0.706373 | 0.705689 |
| 0.713652 | 0.706923 | 0.709443 | 0.705711 | 0.71323 | 0.712606 | 0.708446 | 0.707129 | 0.706051 |
| 0.712309 | 0.710623 | 0.712317 | 0.709235 | 0.714309 | 0.714161 | 0.712555 | 0.708867 | 0.709523 |
| 0.710943 | 0.705326 | 0.710422 | 0.706494 | 0.705039 | 0.710214 | 0.705611 | 0.703531 | 0.707196 |
| 0.71367 | 0.711002 | 0.708206 | 0.707491 | 0.7144 | 0.713697 | 0.710308 | 0.711436 | 0.707377 |
| 0.716054 | 0.711629 | 0.707461 | 0.702709 | 0.711742 | 0.71151 | 0.711733 | 0.70806 | 0.707127 |
| 0.715385 | 0.712511 | 0.714562 | 0.712622 | 0.715632 | 0.7149 | 0.712489 | 0.711757 | 0.709465 |
| 0.711584 | 0.709502 | 0.708902 | 0.708463 | 0.711642 | 0.709045 | 0.708947 | 0.70689 | 0.705341 |
| 0.713921 | 0.711102 | 0.709892 | 0.71063 | 0.713968 | 0.711391 | 0.710576 | 0.708752 | 0.707051 |
| 0.714308 | 0.710946 | 0.711001 | 0.709507 | 0.714637 | 0.709971 | 0.708723 | 0.70829 | 0.708604 |
| 0.709919 | 0.707725 | 0.708658 | 0.706989 | 0.709821 | 0.713229 | 0.710327 | 0.710114 | 0.709329 |
| 0.708901 | 0.706741 | 0.708267 | 0.703793 | 0.707814 | 0.705623 | 0.706113 | 0.705617 | 0.703584 |
| 0.712252 | 0.710063 | 0.709703 | 0.708411 | 0.717294 | 0.709972 | 0.707369 | 0.709592 | 0.704686 |
| 0.716286 | 0.707801 | 0.710863 | 0.708497 | 0.71204 | 0.714195 | 0.711226 | 0.708069 | 0.704504 |
| 0.712839 | 0.708027 | 0.711129 | 0.710571 | 0.715591 | 0.713302 | 0.710995 | 0.709354 | 0.708217 |
| 0.716307 | 0.714004 | 0.716542 | 0.711681 | 0.71952 | 0.721093 | 0.715112 | 0.716912 | 0.714645 |
| 0.713879 | 0.708758 | 0.709464 | 0.709675 | 0.713847 | 0.711635 | 0.710473 | 0.71023 | 0.705633 |
| 0.717396 | 0.709573 | 0.711902 | 0.710331 | 0.718411 | 0.713057 | 0.711709 | 0.708205 | 0.708684 |
| 0.715761 | 0.713281 | 0.713098 | 0.709616 | 0.713011 | 0.712796 | 0.710965 | 0.713934 | 0.714704 |
| 0.713617 | 0.70926 | 0.709437 | 0.710458 | 0.713746 | 0.711585 | 0.711621 | 0.710955 | 0.710353 |
| 0.717811 | 0.711019 | 0.713676 | 0.711756 | 0.715677 | 0.716566 | 0.711878 | 0.715615 | 0.707993 |
| 0.71213 | 0.705139 | 0.71227 | 0.708971 | 0.712049 | 0.713436 | 0.708353 | 0.708026 | 0.706327 |
| 0.714735 | 0.710054 | 0.710665 | 0.714518 | 0.713519 | 0.713805 | 0.710918 | 0.710124 | 0.71141 |
| 0.71508 | 0.709869 | 0.70973 | 0.710613 | 0.712613 | 0.713392 | 0.713269 | 0.713701 | 0.7081 |
| 0.706459 | 0.704315 | 0.705325 | 0.70611 | 0.712008 | 0.71106 | 0.707023 | 0.70842 | 0.704655 |
| 0.714147 | 0.710945 | 0.712786 | 0.707095 | 0.714297 | 0.713023 | 0.707941 | 0.705404 | 0.704964 |
| 0.708085 | 0.702771 | 0.704958 | 0.707957 | 0.705322 | 0.707895 | 0.706576 | 0.707813 | 0.701534 |
| 0.687273 | 0.684163 | 0.688425 | 0.684733 | 0.688857 | 0.686271 | 0.682563 | 0.68307 | 0.678654 |
| 0.689481 | 0.686477 | 0.687424 | 0.687759 | 0.692261 | 0.690694 | 0.687319 | 0.68583 | 0.685867 |
| 0.680221 | 0.678659 | 0.680394 | 0.678157 | 0.684848 | 0.684243 | 0.679437 | 0.680814 | 0.680057 |
| 0.686418 | 0.68094 | 0.677039 | 0.679548 | 0.683256 | 0.679141 | 0.676641 | 0.68136 | 0.674664 |
| 0.666043 | 0.662602 | 0.663727 | 0.663517 | 0.665504 | 0.669144 | 0.662522 | 0.667174 | 0.660371 |
| 0.671905 | 0.66056 | 0.667829 | 0.666024 | 0.664672 | 0.670602 | 0.66834 | 0.672408 | 0.666023 |
| 0.67588 | 0.66526 | 0.668038 | 0.667513 | 0.669566 | 0.667749 | 0.664468 | 0.665859 | 0.664182 |
| 0.670881 | 0.667514 | 0.666033 | 0.668234 | 0.665614 | 0.676042 | 0.665368 | 0.66743 | 0.659515 |
| 0.67053 | 0.66675 | 0.666832 | 0.668241 | 0.664199 | 0.667695 | 0.662744 | 0.665461 | 0.658618 |
| 0.680079 | 0.6706 | 0.672224 | 0.675592 | 0.671117 | 0.678087 | 0.669431 | 0.676656 | 0.6663 |

| | | | | | | | | |
|---|---|---|---|---|---|---|---|---|
| 0.674212 | 0.668582 | 0.672023 | 0.672042 | 0.674116 | 0.678493 | 0.668977 | 0.672224 | 0.66923 |
| 0.680267 | 0.676164 | 0.675707 | 0.670909 | 0.674732 | 0.674919 | 0.671331 | 0.673873 | 0.664174 |
| 0.670921 | 0.666002 | 0.667952 | 0.673311 | 0.672831 | 0.671296 | 0.667896 | 0.669209 | 0.666091 |
| 0.670863 | 0.65994 | 0.664342 | 0.66434 | 0.664346 | 0.667983 | 0.662108 | 0.66543 | 0.66128 |
| 0.667568 | 0.662998 | 0.668768 | 0.66564 | 0.667947 | 0.665605 | 0.662145 | 0.664998 | 0.660628 |
| 0.663838 | 0.660678 | 0.662582 | 0.668126 | 0.665533 | 0.670267 | 0.665308 | 0.666713 | 0.659864 |
| 0.670495 | 0.665683 | 0.66723 | 0.666714 | 0.669717 | 0.666292 | 0.662852 | 0.665241 | 0.663048 |
| 0.672115 | 0.66634 | 0.663589 | 0.668778 | 0.662341 | 0.664539 | 0.659298 | 0.661137 | 0.655106 |
| 0.661439 | 0.662273 | 0.663985 | 0.662465 | 0.66428 | 0.662407 | 0.661258 | 0.662454 | 0.656377 |
| 0.664681 | 0.66866 | 0.65836 | 0.666667 | 0.670943 | 0.66851 | 0.66196 | 0.666521 | 0.659843 |
| 0.663089 | 0.66214 | 0.649877 | 0.654134 | 0.657711 | 0.66137 | 0.655354 | 0.657828 | 0.65182 |
| 0.70194 | 0.696544 | 0.698029 | 0.698058 | 0.701919 | 0.699674 | 0.698926 | 0.698022 | 0.699517 |
| 0.714139 | 0.707405 | 0.705784 | 0.704291 | 0.712218 | 0.713223 | 0.706248 | 0.703779 | 0.705299 |
| 0.68802 | 0.684801 | 0.685536 | 0.683977 | 0.68864 | 0.690387 | 0.68114 | 0.679468 | 0.680221 |
| 0.722752 | 0.71648 | 0.716747 | 0.714007 | 0.715179 | 0.716108 | 0.711295 | 0.71257 | 0.712248 |
| 0.707729 | 0.704833 | 0.705638 | 0.702673 | 0.705813 | 0.707523 | 0.708697 | 0.705056 | 0.703427 |
| 0.717908 | 0.713173 | 0.713137 | 0.712406 | 0.716619 | 0.719621 | 0.714677 | 0.713754 | 0.712414 |
| 0.723663 | 0.720887 | 0.722484 | 0.719682 | 0.725191 | 0.719914 | 0.720057 | 0.719011 | 0.716587 |
| 0.698645 | 0.695398 | 0.698205 | 0.693415 | 0.697104 | 0.701771 | 0.695525 | 0.698237 | 0.695409 |
| 0.720702 | 0.716294 | 0.719974 | 0.718558 | 0.720723 | 0.718981 | 0.716737 | 0.72098 | 0.713983 |
| 0.691355 | 0.689288 | 0.68771 | 0.690171 | 0.69584 | 0.691995 | 0.686773 | 0.691129 | 0.685462 |
| 0.718985 | 0.717115 | 0.719249 | 0.714424 | 0.719593 | 0.722023 | 0.717681 | 0.716056 | 0.714195 |
| 0.723343 | 0.722719 | 0.722159 | 0.721682 | 0.72563 | 0.722576 | 0.722512 | 0.723734 | 0.721448 |
| 0.685303 | 0.681718 | 0.683832 | 0.679769 | 0.689332 | 0.684641 | 0.683257 | 0.682822 | 0.675011 |
| 0.693069 | 0.688845 | 0.690674 | 0.688061 | 0.694646 | 0.691166 | 0.686764 | 0.685343 | 0.685102 |
| 0.693524 | 0.690602 | 0.688199 | 0.689742 | 0.690856 | 0.693783 | 0.687617 | 0.687106 | 0.687367 |
| 0.705916 | 0.70454 | 0.709202 | 0.70297 | 0.714993 | 0.70828 | 0.705991 | 0.707717 | 0.703945 |
| 0.721523 | 0.718453 | 0.721141 | 0.718509 | 0.72413 | 0.724635 | 0.720963 | 0.720099 | 0.715968 |
| 0.68824 | 0.686413 | 0.686746 | 0.681723 | 0.690621 | 0.690559 | 0.686993 | 0.686146 | 0.684486 |
| 0.727222 | 0.724132 | 0.727926 | 0.723939 | 0.730322 | 0.725418 | 0.723511 | 0.724283 | 0.722353 |
| 0.731301 | 0.729949 | 0.724986 | 0.723294 | 0.72929 | 0.73044 | 0.725579 | 0.724606 | 0.724381 |
| 0.69932 | 0.694486 | 0.698372 | 0.699352 | 0.700213 | 0.7 | 0.70094 | 0.699318 | 0.696032 |
| 0.70181 | 0.697747 | 0.70004 | 0.696429 | 0.701366 | 0.700368 | 0.7 | 0.698584 | 0.696462 |
| 0.727505 | 0.722328 | 0.729808 | 0.723529 | 0.728185 | 0.727287 | 0.722997 | 0.722834 | 0.720193 |
| 0.724178 | 0.719853 | 0.723608 | 0.721553 | 0.726931 | 0.727906 | 0.720784 | 0.721398 | 0.717087 |
| 0.72599 | 0.727716 | 0.723923 | 0.723912 | 0.73306 | 0.725231 | 0.725345 | 0.720429 | 0.722683 |
| 0.682105 | 0.677355 | 0.677891 | 0.676565 | 0.677713 | 0.678798 | 0.678637 | 0.676788 | 0.675301 |
| 0.722817 | 0.721989 | 0.718683 | 0.716776 | 0.721234 | 0.725396 | 0.717463 | 0.72034 | 0.720697 |
| 0.679433 | 0.672544 | 0.678238 | 0.676797 | 0.678236 | 0.675171 | 0.6762 | 0.677367 | 0.67269 |
| 0.672536 | 0.671937 | 0.667502 | 0.670966 | 0.67378 | 0.672986 | 0.671204 | 0.670328 | 0.667518 |
| 0.723649 | 0.720363 | 0.718613 | 0.717922 | 0.725529 | 0.722986 | 0.723754 | 0.71938 | 0.719202 |
| 0.672904 | 0.674248 | 0.669055 | 0.66822 | 0.673138 | 0.67156 | 0.672218 | 0.672313 | 0.669049 |
| 0.66212 | 0.663758 | 0.664698 | 0.662484 | 0.665145 | 0.664036 | 0.664711 | 0.66514 | 0.659414 |
| 0.662184 | 0.661903 | 0.659271 | 0.656317 | 0.659629 | 0.662569 | 0.660463 | 0.660891 | 0.656186 |
| 0.728712 | 0.726166 | 0.726974 | 0.724534 | 0.733217 | 0.730544 | 0.728524 | 0.726158 | 0.723958 |
| 0.663745 | 0.663284 | 0.664549 | 0.663592 | 0.666991 | 0.664083 | 0.663043 | 0.662991 | 0.660781 |
| 0.722817 | 0.721353 | 0.718019 | 0.718937 | 0.725778 | 0.72497 | 0.726733 | 0.722382 | 0.719418 |

| | | | | | | | | |
|---|---|---|---|---|---|---|---|---|
| 0.734407 | 0.728035 | 0.728963 | 0.724204 | 0.738107 | 0.732524 | 0.731734 | 0.728421 | 0.726326 |
| 0.726154 | 0.724897 | 0.727849 | 0.721068 | 0.729071 | 0.727524 | 0.728735 | 0.722588 | 0.719502 |
| 0.74669 | 0.746725 | 0.743533 | 0.749116 | 0.744314 | 0.744846 | 0.745016 | 0.74373 | 0.74024 |
| 0.654966 | 0.654384 | 0.652449 | 0.653617 | 0.65564 | 0.656358 | 0.655914 | 0.656107 | 0.650355 |
| 0.656252 | 0.655442 | 0.653843 | 0.652796 | 0.655966 | 0.659749 | 0.653548 | 0.655382 | 0.648163 |
| 0.734434 | 0.730928 | 0.73027 | 0.728328 | 0.73494 | 0.731579 | 0.73121 | 0.731709 | 0.724602 |
| 0.654638 | 0.656847 | 0.658796 | 0.655467 | 0.65888 | 0.655424 | 0.654351 | 0.655134 | 0.652806 |
| 0.736863 | 0.734178 | 0.732348 | 0.732016 | 0.734679 | 0.737131 | 0.734586 | 0.73263 | 0.724284 |
| 0.73206 | 0.73229 | 0.728937 | 0.727523 | 0.736377 | 0.734939 | 0.731261 | 0.729867 | 0.729669 |
| 0.737899 | 0.740231 | 0.73209 | 0.733607 | 0.735913 | 0.735027 | 0.732124 | 0.732297 | 0.733939 |
| 0.72891 | 0.735161 | 0.733453 | 0.726136 | 0.733425 | 0.734538 | 0.733605 | 0.734141 | 0.726735 |
| 0.746479 | 0.75 | 0.713376 | 0.688312 | 0.751724 | 0.757962 | 0.774648 | 0.728477 | 0.717241 |
| 0.724138 | 0.731668 | 0.720294 | 0.721158 | 0.721106 | 0.720699 | 0.734026 | 0.726267 | 0.727327 |
| 0.729236 | 0.727843 | 0.720118 | 0.730418 | 0.724181 | 0.720652 | 0.726891 | 0.731049 | 0.723896 |
| 0.736644 | 0.73249 | 0.728449 | 0.728518 | 0.725549 | 0.730585 | 0.738618 | 0.737901 | 0.729997 |
| 0.73463 | 0.734447 | 0.717863 | 0.728688 | 0.728306 | 0.732532 | 0.735857 | 0.731799 | 0.731439 |
| 0.594922 | 0.593125 | 0.597575 | 0.59482 | 0.593504 | 0.592648 | 0.590565 | 0.59291 | 0.585009 |
| 0.739575 | 0.736657 | 0.732684 | 0.739564 | 0.733169 | 0.730963 | 0.738228 | 0.739153 | 0.733342 |
| 0.739014 | 0.735513 | 0.735292 | 0.733699 | 0.743419 | 0.735843 | 0.7368 | 0.737141 | 0.731444 |
| 0.736993 | 0.737984 | 0.736264 | 0.732732 | 0.741515 | 0.731747 | 0.73773 | 0.73772 | 0.732508 |
| 0.738014 | 0.739885 | 0.742808 | 0.73514 | 0.742664 | 0.737837 | 0.74401 | 0.739919 | 0.734092 |
| 0.594757 | 0.601251 | 0.597111 | 0.59721 | 0.599453 | 0.600944 | 0.597323 | 0.598242 | 0.595767 |
| 0.581395 | 0.585766 | 0.582398 | 0.582798 | 0.585107 | 0.584175 | 0.584711 | 0.583958 | 0.581938 |
| 0.768216 | 0.766752 | 0.757717 | 0.759702 | 0.767484 | 0.760961 | 0.763885 | 0.762371 | 0.762042 |
| 0.765721 | 0.765728 | 0.761611 | 0.764001 | 0.771927 | 0.763434 | 0.759303 | 0.76254 | 0.759786 |
| 0.759502 | 0.758952 | 0.749567 | 0.756658 | 0.756803 | 0.752835 | 0.756244 | 0.753621 | 0.752103 |
| 0.765741 | 0.764485 | 0.757352 | 0.764147 | 0.762296 | 0.761078 | 0.761597 | 0.760804 | 0.75864 |
| 0.76908 | 0.764873 | 0.76142 | 0.764231 | 0.767783 | 0.76269 | 0.76519 | 0.762581 | 0.76335 |
| 0.768956 | 0.7667 | 0.761492 | 0.7708 | 0.764444 | 0.769009 | 0.764383 | 0.767307 | 0.762697 |
| 0.771087 | 0.772156 | 0.75811 | 0.764409 | 0.764095 | 0.762575 | 0.76771 | 0.763711 | 0.7623 |
| 0.761156 | 0.766197 | 0.76084 | 0.760825 | 0.760402 | 0.765223 | 0.761901 | 0.760509 | 0.758134 |
| 0.77543 | 0.77644 | 0.771187 | 0.765462 | 0.770469 | 0.771818 | 0.771341 | 0.768999 | 0.768614 |
| 0.771365 | 0.768546 | 0.765269 | 0.765908 | 0.76458 | 0.762968 | 0.764095 | 0.759308 | 0.761655 |
| 0.76413 | 0.765408 | 0.766086 | 0.761457 | 0.765932 | 0.766188 | 0.769818 | 0.764129 | 0.763758 |
| 0.772121 | 0.776168 | 0.76795 | 0.766931 | 0.772104 | 0.771937 | 0.773732 | 0.765831 | 0.766531 |
| 0.763593 | 0.765607 | 0.760778 | 0.764189 | 0.767509 | 0.770247 | 0.768917 | 0.758607 | 0.760545 |
| 0.769743 | 0.769465 | 0.763654 | 0.765515 | 0.764196 | 0.770864 | 0.768252 | 0.763087 | 0.762173 |
| 0.773692 | 0.774647 | 0.76722 | 0.76781 | 0.766119 | 0.767638 | 0.768857 | 0.768009 | 0.764677 |
| 0.764167 | 0.768557 | 0.762384 | 0.766896 | 0.763028 | 0.763623 | 0.766158 | 0.757644 | 0.764301 |
| 0.76553 | 0.76339 | 0.762229 | 0.76257 | 0.765279 | 0.763343 | 0.760714 | 0.760215 | 0.759583 |
| 0.726124 | 0.727591 | 0.717207 | 0.718614 | 0.723874 | 0.724612 | 0.721111 | 0.720238 | 0.722513 |
| 0.727221 | 0.736145 | 0.729547 | 0.732642 | 0.725182 | 0.731589 | 0.733596 | 0.727736 | 0.727803 |
| 0.766044 | 0.768955 | 0.761316 | 0.764991 | 0.770164 | 0.767267 | 0.764 | 0.764956 | 0.762663 |
| 0.776108 | 0.767069 | 0.767809 | 0.767776 | 0.772334 | 0.774739 | 0.767106 | 0.767438 | 0.769494 |
| 0.725464 | 0.716887 | 0.720207 | 0.725101 | 0.720752 | 0.714613 | 0.71705 | 0.721169 | 0.714203 |
| 0.768562 | 0.768659 | 0.767058 | 0.768518 | 0.76628 | 0.771545 | 0.770238 | 0.765082 | 0.767901 |
| 0.766286 | 0.764921 | 0.760905 | 0.759806 | 0.759955 | 0.761712 | 0.757732 | 0.758388 | 0.762514 |

| | | | | | | | | |
|---|---|---|---|---|---|---|---|---|
| 0.769768 | 0.766936 | 0.760718 | 0.765711 | 0.768929 | 0.768754 | 0.762601 | 0.762665 | 0.767372 |
| 0.770755 | 0.772081 | 0.769559 | 0.767239 | 0.771995 | 0.773643 | 0.766024 | 0.765592 | 0.766909 |
| 0.764765 | 0.762987 | 0.761553 | 0.75998 | 0.760183 | 0.766628 | 0.763223 | 0.760351 | 0.759495 |
| 0.774035 | 0.769155 | 0.763675 | 0.766499 | 0.768189 | 0.771871 | 0.76983 | 0.765952 | 0.768811 |
| 0.775191 | 0.76834 | 0.768781 | 0.768242 | 0.767559 | 0.770633 | 0.767664 | 0.764994 | 0.767415 |
| 0.768408 | 0.766398 | 0.766824 | 0.763487 | 0.767039 | 0.766868 | 0.761487 | 0.762567 | 0.759595 |
| 0.774471 | 0.76651 | 0.76208 | 0.762409 | 0.764873 | 0.768857 | 0.764687 | 0.764701 | 0.763286 |
| 0.758284 | 0.760706 | 0.756667 | 0.757495 | 0.755119 | 0.758813 | 0.754405 | 0.758898 | 0.758083 |
| 0.762742 | 0.763895 | 0.762375 | 0.763642 | 0.763512 | 0.769137 | 0.765578 | 0.762205 | 0.766185 |
| 0.761526 | 0.765111 | 0.756007 | 0.756255 | 0.761326 | 0.760222 | 0.759474 | 0.76167 | 0.761559 |
| 0.767768 | 0.768448 | 0.767337 | 0.765844 | 0.766509 | 0.769643 | 0.764015 | 0.763638 | 0.76522 |
| 0.768073 | 0.768077 | 0.763766 | 0.764821 | 0.765299 | 0.766261 | 0.764209 | 0.764706 | 0.768196 |
| 0.759997 | 0.758922 | 0.753244 | 0.753581 | 0.757771 | 0.757511 | 0.752997 | 0.75566 | 0.758584 |
| 0.760961 | 0.75922 | 0.754412 | 0.752943 | 0.755926 | 0.758509 | 0.755081 | 0.756957 | 0.756031 |
| 0.767861 | 0.770154 | 0.760532 | 0.767665 | 0.764763 | 0.766844 | 0.766141 | 0.766865 | 0.76383 |
| 0.768728 | 0.771361 | 0.763007 | 0.764159 | 0.764859 | 0.769556 | 0.767113 | 0.766359 | 0.760936 |
| 0.771414 | 0.771387 | 0.759314 | 0.766428 | 0.764309 | 0.764646 | 0.766345 | 0.763962 | 0.761447 |
| 0.768566 | 0.767949 | 0.759482 | 0.761692 | 0.76384 | 0.760016 | 0.766645 | 0.76176 | 0.757687 |
| 0.770069 | 0.76771 | 0.764062 | 0.769205 | 0.769621 | 0.765304 | 0.765971 | 0.765514 | 0.761712 |
| 0.767813 | 0.771006 | 0.760218 | 0.769071 | 0.765741 | 0.762575 | 0.769852 | 0.762827 | 0.763106 |
| 0.76032 | 0.762095 | 0.759469 | 0.761503 | 0.758446 | 0.758103 | 0.761873 | 0.756361 | 0.75157 |
| 0.763966 | 0.762774 | 0.759578 | 0.765335 | 0.764167 | 0.760632 | 0.762372 | 0.761568 | 0.759371 |
| 0.770337 | 0.769443 | 0.7625 | 0.770365 | 0.770311 | 0.765272 | 0.768004 | 0.765777 | 0.76573 |
| 0.774225 | 0.770528 | 0.763829 | 0.76773 | 0.76736 | 0.763002 | 0.768519 | 0.764103 | 0.76156 |
| 0.766562 | 0.766678 | 0.76169 | 0.761881 | 0.766841 | 0.763557 | 0.765561 | 0.766249 | 0.758061 |
| 0.769841 | 0.766743 | 0.757767 | 0.764324 | 0.766949 | 0.762848 | 0.762716 | 0.764881 | 0.760381 |
| 0.76589 | 0.765843 | 0.756494 | 0.757434 | 0.758891 | 0.75788 | 0.757922 | 0.755119 | 0.755832 |
| 0.766195 | 0.760346 | 0.757614 | 0.76281 | 0.763261 | 0.754775 | 0.758865 | 0.755996 | 0.757305 |
| 0.768536 | 0.767594 | 0.759694 | 0.760488 | 0.768526 | 0.758803 | 0.76021 | 0.761831 | 0.756208 |
| 0.773602 | 0.765573 | 0.762579 | 0.766744 | 0.766376 | 0.760866 | 0.767695 | 0.764215 | 0.758436 |
| 0.770272 | 0.76847 | 0.763353 | 0.765769 | 0.766732 | 0.764132 | 0.764687 | 0.760822 | 0.760557 |
| 0.773861 | 0.777085 | 0.764642 | 0.766441 | 0.765317 | 0.767166 | 0.763903 | 0.761103 | 0.757571 |
| 0.7694 | 0.770279 | 0.768934 | 0.768921 | 0.770847 | 0.767647 | 0.767853 | 0.767274 | 0.763315 |
| 0.770194 | 0.773574 | 0.764812 | 0.768163 | 0.769992 | 0.765734 | 0.771171 | 0.760933 | 0.761264 |
| 0.765218 | 0.762204 | 0.756224 | 0.762287 | 0.76906 | 0.76281 | 0.75821 | 0.75807 | 0.758652 |
| 0.768476 | 0.769973 | 0.76545 | 0.764617 | 0.768617 | 0.764901 | 0.763753 | 0.764293 | 0.760819 |
| 0.772011 | 0.762272 | 0.761798 | 0.766714 | 0.769358 | 0.765358 | 0.768556 | 0.762585 | 0.758462 |
| 0.766141 | 0.76313 | 0.760589 | 0.76179 | 0.760651 | 0.759678 | 0.765579 | 0.761055 | 0.757558 |
| 0.765153 | 0.763515 | 0.758317 | 0.755186 | 0.759033 | 0.755234 | 0.757995 | 0.756827 | 0.750021 |
| 0.757127 | 0.758945 | 0.751076 | 0.758158 | 0.756305 | 0.751616 | 0.759609 | 0.754894 | 0.75324 |
| 0.760444 | 0.760247 | 0.752251 | 0.754495 | 0.754922 | 0.752868 | 0.751063 | 0.752807 | 0.74715 |
| 0.758314 | 0.758363 | 0.752182 | 0.759365 | 0.756529 | 0.753954 | 0.758017 | 0.752409 | 0.750065 |
| 0.760275 | 0.760489 | 0.752734 | 0.753947 | 0.7546 | 0.750741 | 0.755815 | 0.751536 | 0.752017 |
| 0.75541 | 0.754631 | 0.751939 | 0.754893 | 0.754303 | 0.749086 | 0.753608 | 0.752634 | 0.749978 |
| 0.762649 | 0.760918 | 0.755146 | 0.759674 | 0.759383 | 0.75898 | 0.761401 | 0.758968 | 0.756902 |
| 0.753256 | 0.760913 | 0.748497 | 0.751934 | 0.754781 | 0.750186 | 0.755614 | 0.751198 | 0.75098 |

| | | | | | | | | |
|---|---|---|---|---|---|---|---|---|
| 0.771473 | 0.768651 | 0.7615 | 0.765422 | 0.767128 | 0.763555 | 0.765695 | 0.760766 | 0.75834 |
| 0.776304 | 0.777593 | 0.772777 | 0.770715 | 0.779239 | 0.776238 | 0.773499 | 0.772754 | 0.77211 |
| 0.770259 | 0.774566 | 0.766126 | 0.769734 | 0.769822 | 0.771381 | 0.765126 | 0.770807 | 0.762516 |
| 0.77388 | 0.773968 | 0.773389 | 0.77236 | 0.774149 | 0.773924 | 0.769342 | 0.772969 | 0.768825 |
| 0.778892 | 0.778453 | 0.774153 | 0.775769 | 0.780787 | 0.785168 | 0.777914 | 0.778887 | 0.777578 |
| 0.77099 | 0.776796 | 0.76752 | 0.775492 | 0.775866 | 0.776143 | 0.770319 | 0.769086 | 0.775769 |
| 0.766823 | 0.771924 | 0.767692 | 0.766288 | 0.770083 | 0.772603 | 0.77272 | 0.77126 | 0.772173 |
| 0.775344 | 0.768661 | 0.762838 | 0.767551 | 0.773183 | 0.772664 | 0.768727 | 0.765249 | 0.768317 |
| 0.764447 | 0.767561 | 0.761625 | 0.765031 | 0.769632 | 0.764979 | 0.765224 | 0.764305 | 0.762647 |
| 0.769636 | 0.765442 | 0.767924 | 0.770553 | 0.763381 | 0.775456 | 0.765799 | 0.768388 | 0.763233 |
| 0.77485 | 0.774199 | 0.765105 | 0.767788 | 0.769187 | 0.773501 | 0.773664 | 0.776345 | 0.770047 |
| 0.774201 | 0.773161 | 0.765533 | 0.771566 | 0.771706 | 0.774555 | 0.772613 | 0.768948 | 0.769519 |
| 0.790257 | 0.785773 | 0.783613 | 0.78801 | 0.782991 | 0.771881 | 0.78074 | 0.778353 | 0.768436 |
| 0.783025 | 0.783757 | 0.816502 | 0.818443 | 0.803351 | 0.777405 | 0.776203 | 0.77181 | 0.770867 |
| 0.790274 | 0.787646 | 0.786134 | 0.790516 | 0.78838 | 0.773427 | 0.782616 | 0.77707 | 0.779258 |
| 0.784585 | 0.789049 | 0.78736 | 0.800672 | 0.787211 | 0.776118 | 0.780784 | 0.778866 | 0.769137 |
| 0.784761 | 0.78292 | 0.771714 | 0.776173 | 0.781054 | 0.779922 | 0.776157 | 0.775301 | 0.771794 |
| 0.778761 | 0.780464 | 0.778852 | 0.775798 | 0.779669 | 0.771311 | 0.779868 | 0.773203 | 0.769949 |
| 1 | 0.792771 | 0.780185 | 0.779896 | 0.784254 | 0.775796 | 0.776679 | 0.775854 | 0.771217 |
| 0.792771 | 1 | 0.786593 | 0.780074 | 0.790656 | 0.775159 | 0.772936 | 0.775102 | 0.773159 |
| 0.780185 | 0.786593 | 1 | 0.825564 | 0.802202 | 0.773817 | 0.774273 | 0.770823 | 0.76779 |
| 0.779896 | 0.780074 | 0.825564 | 1 | 0.8001 | 0.775551 | 0.776874 | 0.775929 | 0.771931 |
| 0.784254 | 0.790656 | 0.802202 | 0.8001 | 1 | 0.778139 | 0.776743 | 0.772865 | 0.774514 |
| 0.775796 | 0.775159 | 0.773817 | 0.775551 | 0.778139 | 1 | 0.786735 | 0.788881 | 0.787402 |
| 0.776679 | 0.772936 | 0.774273 | 0.776874 | 0.776743 | 0.786735 | 1 | 0.803327 | 0.803357 |
| 0.775854 | 0.775102 | 0.770823 | 0.775929 | 0.772865 | 0.788881 | 0.803327 | 1 | 0.782267 |
| 0.771217 | 0.773159 | 0.76779 | 0.771931 | 0.774514 | 0.787402 | 0.803357 | 0.782267 | 1 |
| 0.771064 | 0.771823 | 0.769174 | 0.775436 | 0.770793 | 0.782712 | 0.780624 | 0.786037 | 0.783191 |
| 0.782541 | 0.783995 | 0.778455 | 0.779052 | 0.774896 | 0.790635 | 0.788957 | 0.783607 | 0.783261 |
| 0.775411 | 0.775335 | 0.771712 | 0.770087 | 0.778405 | 0.798284 | 0.786677 | 0.780956 | 0.784752 |
| 0.771153 | 0.769633 | 0.764627 | 0.768785 | 0.766197 | 0.812886 | 0.789702 | 0.809625 | 0.779111 |
| 0.777861 | 0.777393 | 0.772194 | 0.770153 | 0.771405 | 0.798668 | 0.790283 | 0.785101 | 0.790698 |
| 0.776795 | 0.777433 | 0.772937 | 0.7741 | 0.77433 | 0.796557 | 0.818875 | 0.791488 | 0.785562 |
| 0.776032 | 0.775617 | 0.765835 | 0.769913 | 0.770897 | 0.797377 | 0.77583 | 0.775843 | 0.78347 |
| 0.777024 | 0.775517 | 0.770579 | 0.769774 | 0.770668 | 0.825194 | 0.781969 | 0.788591 | 0.78763 |
| 0.771149 | 0.770765 | 0.767922 | 0.771292 | 0.773149 | 0.779849 | 0.786488 | 0.781626 | 0.781317 |
| 0.78448 | 0.783821 | 0.779421 | 0.78098 | 0.786519 | 0.78582 | 0.787045 | 0.785992 | 0.785937 |
| 0.779017 | 0.779522 | 0.771229 | 0.766984 | 0.772016 | 0.776629 | 0.780101 | 0.781577 | 0.779575 |
| 0.772597 | 0.770707 | 0.77246 | 0.769053 | 0.772634 | 0.779069 | 0.78084 | 0.778556 | 0.778633 |
| 0.776687 | 0.777852 | 0.773682 | 0.771436 | 0.77761 | 0.781454 | 0.777519 | 0.775522 | 0.778084 |
| 0.783589 | 0.7797 | 0.775811 | 0.774146 | 0.776936 | 0.777941 | 0.782218 | 0.784745 | 0.780186 |
| 0.780038 | 0.775362 | 0.772637 | 0.774766 | 0.775984 | 0.789088 | 0.785109 | 0.786954 | 0.785538 |
| 0.782321 | 0.778917 | 0.771934 | 0.774589 | 0.778699 | 0.787498 | 0.783885 | 0.785182 | 0.787119 |
| 0.774937 | 0.780359 | 0.769846 | 0.772099 | 0.771586 | 0.775689 | 0.777676 | 0.775394 | 0.785385 |
| 0.781704 | 0.777686 | 0.773751 | 0.771208 | 0.779842 | 0.783463 | 0.785027 | 0.783013 | 0.788872 |
| 0.782335 | 0.777926 | 0.768366 | 0.763312 | 0.770484 | 0.778592 | 0.775691 | 0.778644 | 0.782663 |

| | | | | | | | | |
|---|---|---|---|---|---|---|---|---|
| 0.771176 | 0.766963 | 0.767446 | 0.765222 | 0.771309 | 0.760399 | 0.762788 | 0.766803 | 0.764002 |
| 0.754506 | 0.746994 | 0.748905 | 0.744758 | 0.755437 | 0.756571 | 0.752396 | 0.752185 | 0.752657 |
| 0.77443 | 0.771488 | 0.76647 | 0.771988 | 0.771712 | 0.768084 | 0.774525 | 0.774093 | 0.771369 |
| 0.774585 | 0.776196 | 0.775058 | 0.775983 | 0.776531 | 0.771193 | 0.770523 | 0.77588 | 0.775408 |
| 0.770049 | 0.767243 | 0.767306 | 0.766652 | 0.773199 | 0.765124 | 0.770911 | 0.770576 | 0.763515 |
| 0.77572 | 0.767825 | 0.770064 | 0.771708 | 0.777695 | 0.773061 | 0.774273 | 0.770864 | 0.775008 |
| 0.773137 | 0.7754 | 0.772381 | 0.768864 | 0.767114 | 0.771705 | 0.770316 | 0.771584 | 0.769329 |
| 0.768904 | 0.768755 | 0.769669 | 0.765177 | 0.769434 | 0.761637 | 0.770724 | 0.769231 | 0.767923 |
| 0.777123 | 0.779892 | 0.771163 | 0.773579 | 0.773897 | 0.771568 | 0.773455 | 0.775505 | 0.7689 |
| 0.778053 | 0.775021 | 0.772132 | 0.770911 | 0.776096 | 0.773391 | 0.772709 | 0.7678 | 0.765166 |
| 0.780452 | 0.775915 | 0.771567 | 0.772536 | 0.773511 | 0.772122 | 0.777868 | 0.772345 | 0.777335 |
| 0.775764 | 0.780574 | 0.771161 | 0.773826 | 0.771227 | 0.773226 | 0.774784 | 0.775141 | 0.768085 |
| 0.778015 | 0.780993 | 0.777584 | 0.780376 | 0.775205 | 0.775408 | 0.781528 | 0.777595 | 0.773198 |
| 0.778789 | 0.777443 | 0.775662 | 0.773678 | 0.774774 | 0.774322 | 0.777517 | 0.77605 | 0.774361 |
| 0.772735 | 0.778037 | 0.767728 | 0.768016 | 0.770842 | 0.767758 | 0.767235 | 0.768852 | 0.764716 |
| 0.771934 | 0.771546 | 0.771605 | 0.765724 | 0.77351 | 0.773485 | 0.775175 | 0.770283 | 0.766132 |
| 0.773638 | 0.774209 | 0.768555 | 0.766189 | 0.770943 | 0.771044 | 0.772924 | 0.768654 | 0.762569 |
| 0.769485 | 0.774798 | 0.765173 | 0.766274 | 0.772488 | 0.772653 | 0.769457 | 0.768634 | 0.766626 |
| 0.771681 | 0.773201 | 0.761977 | 0.767227 | 0.770314 | 0.768711 | 0.767244 | 0.762614 | 0.764487 |
| 0.772502 | 0.773284 | 0.764947 | 0.768253 | 0.77096 | 0.768421 | 0.766272 | 0.77061 | 0.76959 |
| 0.773746 | 0.775539 | 0.768994 | 0.770292 | 0.769243 | 0.774122 | 0.769843 | 0.769352 | 0.767994 |
| 0.762394 | 0.758219 | 0.756745 | 0.760592 | 0.760849 | 0.762595 | 0.766143 | 0.756777 | 0.761817 |
| 0.769313 | 0.767637 | 0.763626 | 0.767336 | 0.76438 | 0.769513 | 0.770798 | 0.765019 | 0.766054 |
| 0.771992 | 0.768563 | 0.766863 | 0.770793 | 0.770031 | 0.774469 | 0.771654 | 0.766384 | 0.766887 |
| 0.771601 | 0.773504 | 0.773953 | 0.770328 | 0.772301 | 0.77523 | 0.771908 | 0.768243 | 0.769006 |
| 0.773243 | 0.770253 | 0.767181 | 0.767839 | 0.769314 | 0.77411 | 0.771045 | 0.764544 | 0.767758 |
| 0.766938 | 0.766791 | 0.763398 | 0.763169 | 0.766247 | 0.765549 | 0.76255 | 0.762712 | 0.763054 |
| 0.770395 | 0.770995 | 0.765144 | 0.769204 | 0.767206 | 0.767992 | 0.76602 | 0.76424 | 0.766994 |
| 0.765486 | 0.765751 | 0.76229 | 0.763438 | 0.760148 | 0.766857 | 0.763548 | 0.760464 | 0.757495 |
| 0.69742 | 0.693591 | 0.698394 | 0.693998 | 0.700284 | 0.700675 | 0.692996 | 0.694732 | 0.690747 |
| 0.711213 | 0.705789 | 0.707715 | 0.707912 | 0.70713 | 0.708835 | 0.707555 | 0.707073 | 0.705297 |
| 0.707516 | 0.705035 | 0.705126 | 0.699863 | 0.709214 | 0.70671 | 0.707628 | 0.706052 | 0.704318 |
| 0.701134 | 0.695401 | 0.696887 | 0.693386 | 0.704862 | 0.696567 | 0.695781 | 0.697478 | 0.694892 |
| 0.709316 | 0.708222 | 0.705672 | 0.707053 | 0.71244 | 0.714634 | 0.709428 | 0.706253 | 0.706205 |
| 0.770733 | 0.768177 | 0.762816 | 0.766829 | 0.762221 | 0.768796 | 0.768953 | 0.770419 | 0.765127 |
| 0.769563 | 0.772456 | 0.764412 | 0.768482 | 0.764482 | 0.770664 | 0.767184 | 0.76772 | 0.764735 |
| 0.771863 | 0.769396 | 0.768903 | 0.77012 | 0.771014 | 0.775735 | 0.777502 | 0.773105 | 0.776912 |
| 0.775431 | 0.773652 | 0.769001 | 0.77289 | 0.767963 | 0.771694 | 0.774531 | 0.769142 | 0.769218 |
| 0.766791 | 0.772031 | 0.765353 | 0.766461 | 0.765928 | 0.767713 | 0.771918 | 0.769728 | 0.766181 |
| 0.772844 | 0.772858 | 0.769595 | 0.770507 | 0.770002 | 0.772451 | 0.773125 | 0.771965 | 0.769692 |
| 0.772481 | 0.771735 | 0.768921 | 0.768456 | 0.767662 | 0.769682 | 0.774209 | 0.770965 | 0.769294 |
| 0.771722 | 0.76868 | 0.7652 | 0.768972 | 0.766975 | 0.773254 | 0.768082 | 0.769025 | 0.765155 |
| 0.761686 | 0.768085 | 0.761921 | 0.761088 | 0.760062 | 0.766031 | 0.767448 | 0.765645 | 0.767482 |
| 0.764321 | 0.765713 | 0.765428 | 0.761452 | 0.762952 | 0.765301 | 0.766407 | 0.765592 | 0.762114 |
| 0.769769 | 0.771325 | 0.765333 | 0.76486 | 0.761885 | 0.769341 | 0.766448 | 0.766097 | 0.765219 |
| 0.770346 | 0.773756 | 0.770721 | 0.771306 | 0.770565 | 0.775987 | 0.770897 | 0.767448 | 0.76839 |

| | | | | | | | | |
|---|---|---|---|---|---|---|---|---|
| 0.772586 | 0.773211 | 0.767924 | 0.774951 | 0.771378 | 0.770208 | 0.77111 | 0.768817 | 0.775746 |
| 0.771466 | 0.765237 | 0.760123 | 0.763243 | 0.766987 | 0.771758 | 0.777969 | 0.770591 | 0.767406 |
| 0.773343 | 0.772855 | 0.764407 | 0.771532 | 0.775089 | 0.772562 | 0.77656 | 0.776192 | 0.769161 |
| 0.769085 | 0.767377 | 0.760013 | 0.768706 | 0.766809 | 0.764119 | 0.769946 | 0.768317 | 0.763362 |
| 0.770399 | 0.769528 | 0.761812 | 0.766562 | 0.769101 | 0.768941 | 0.773487 | 0.766305 | 0.765463 |
| 0.767142 | 0.766576 | 0.759028 | 0.768193 | 0.763807 | 0.767948 | 0.769944 | 0.765106 | 0.766204 |
| 0.772471 | 0.771565 | 0.765625 | 0.771028 | 0.766759 | 0.77203 | 0.768392 | 0.764607 | 0.765261 |
| 0.772137 | 0.766982 | 0.759939 | 0.769407 | 0.767385 | 0.770294 | 0.768751 | 0.764815 | 0.768457 |
| 0.769334 | 0.764809 | 0.761611 | 0.763686 | 0.765795 | 0.770242 | 0.770038 | 0.766095 | 0.763197 |
| 0.764385 | 0.762837 | 0.762159 | 0.769791 | 0.764158 | 0.768347 | 0.768486 | 0.765771 | 0.766702 |
| 0.769237 | 0.7695 | 0.762847 | 0.768303 | 0.768916 | 0.769832 | 0.774036 | 0.772365 | 0.766531 |
| 0.76842 | 0.772077 | 0.762308 | 0.771255 | 0.769276 | 0.771753 | 0.771616 | 0.766837 | 0.770162 |
| 0.762479 | 0.761112 | 0.754635 | 0.760969 | 0.76249 | 0.764086 | 0.764546 | 0.762659 | 0.761374 |
| 0.761767 | 0.763505 | 0.755878 | 0.761848 | 0.759159 | 0.760267 | 0.764701 | 0.758175 | 0.758717 |
| 0.770457 | 0.77353 | 0.766339 | 0.770012 | 0.774777 | 0.770774 | 0.767496 | 0.764376 | 0.764613 |
| 0.778586 | 0.782648 | 0.771751 | 0.77635 | 0.779407 | 0.782025 | 0.776659 | 0.779535 | 0.773082 |
| 0.769811 | 0.77611 | 0.763708 | 0.771849 | 0.774516 | 0.779212 | 0.777321 | 0.778858 | 0.772172 |
| 0.774738 | 0.774627 | 0.769282 | 0.770201 | 0.77027 | 0.77436 | 0.776161 | 0.774657 | 0.771337 |
| 0.76948 | 0.772221 | 0.766966 | 0.768716 | 0.767126 | 0.772044 | 0.770105 | 0.77013 | 0.769577 |
| 0.774298 | 0.773881 | 0.773136 | 0.769686 | 0.770385 | 0.772088 | 0.772743 | 0.767473 | 0.765929 |
| 0.770988 | 0.768273 | 0.765778 | 0.763841 | 0.765056 | 0.767297 | 0.771753 | 0.767209 | 0.765927 |
| 0.772014 | 0.773435 | 0.768544 | 0.766683 | 0.77158 | 0.770497 | 0.766093 | 0.76999 | 0.763647 |
| 0.776387 | 0.779269 | 0.767833 | 0.777443 | 0.777816 | 0.778004 | 0.774827 | 0.777844 | 0.777929 |
| 0.765368 | 0.76681 | 0.756308 | 0.762637 | 0.765041 | 0.766065 | 0.769191 | 0.764401 | 0.767501 |
| 0.779129 | 0.779068 | 0.768998 | 0.770937 | 0.774299 | 0.770938 | 0.770522 | 0.775157 | 0.772836 |
| 0.710868 | 0.702904 | 0.704229 | 0.701715 | 0.708066 | 0.705948 | 0.706578 | 0.699752 | 0.701333 |
| 0.774336 | 0.773319 | 0.764658 | 0.773317 | 0.773112 | 0.7822 | 0.782584 | 0.781504 | 0.782733 |
| 0.778837 | 0.774906 | 0.767738 | 0.769205 | 0.772534 | 0.786745 | 0.783885 | 0.780074 | 0.783246 |
| 0.776536 | 0.775863 | 0.772243 | 0.775033 | 0.776436 | 0.780415 | 0.785644 | 0.786519 | 0.788599 |
| 0.774143 | 0.776149 | 0.773381 | 0.772549 | 0.775177 | 0.776462 | 0.781812 | 0.778597 | 0.778757 |
| 0.770801 | 0.77175 | 0.762859 | 0.765766 | 0.766193 | 0.781798 | 0.77852 | 0.778807 | 0.7739 |
| 0.772279 | 0.770292 | 0.767586 | 0.769824 | 0.766889 | 0.779963 | 0.774754 | 0.78334 | 0.774481 |
| 0.772865 | 0.768846 | 0.767561 | 0.769789 | 0.774655 | 0.779779 | 0.778962 | 0.779555 | 0.778048 |
| 0.771059 | 0.772438 | 0.770063 | 0.771622 | 0.772645 | 0.777414 | 0.782553 | 0.774475 | 0.782489 |
| 0.762658 | 0.765935 | 0.760313 | 0.761818 | 0.763394 | 0.776805 | 0.772485 | 0.773454 | 0.777267 |
| 0.775447 | 0.769076 | 0.760923 | 0.761499 | 0.765173 | 0.782213 | 0.776693 | 0.773233 | 0.7753 |
| 0.780725 | 0.775331 | 0.773195 | 0.769569 | 0.778969 | 0.782332 | 0.783211 | 0.782473 | 0.780788 |
| 0.778728 | 0.778394 | 0.780081 | 0.78088 | 0.779553 | 0.779292 | 0.78575 | 0.777491 | 0.778154 |
| 0.779517 | 0.781393 | 0.773191 | 0.781313 | 0.783525 | 0.779409 | 0.781245 | 0.779619 | 0.782627 |
| 0.778638 | 0.773015 | 0.773557 | 0.776487 | 0.777911 | 0.774014 | 0.778052 | 0.779551 | 0.771658 |
| 0.779119 | 0.786059 | 0.769178 | 0.77544 | 0.778674 | 0.776664 | 0.774372 | 0.768377 | 0.769973 |
| 0.780807 | 0.783303 | 0.77774 | 0.778022 | 0.780206 | 0.778251 | 0.778829 | 0.780793 | 0.774301 |
| 0.772712 | 0.77244 | 0.767588 | 0.770689 | 0.780769 | 0.77237 | 0.776027 | 0.773692 | 0.772413 |
| 0.767321 | 0.768223 | 0.760829 | 0.764144 | 0.766941 | 0.766759 | 0.767799 | 0.764278 | 0.765971 |
| 0.769155 | 0.770338 | 0.768937 | 0.774547 | 0.771857 | 0.773048 | 0.770304 | 0.763995 | 0.768695 |
| 0.770305 | 0.764792 | 0.768535 | 0.770105 | 0.77272 | 0.771335 | 0.771245 | 0.766302 | 0.766405 |

| | | | | | | | | |
|---|---|---|---|---|---|---|---|---|
| 0.770855 | 0.773466 | 0.767951 | 0.771098 | 0.768386 | 0.76617 | 0.771232 | 0.768828 | 0.766247 |
| 0.773996 | 0.774826 | 0.769009 | 0.774247 | 0.767548 | 0.770332 | 0.772912 | 0.771494 | 0.76655 |
| 0.765987 | 0.771417 | 0.762605 | 0.762024 | 0.766549 | 0.765626 | 0.766306 | 0.765947 | 0.764749 |
| 0.723792 | 0.727143 | 0.739701 | 0.741039 | 0.734291 | 0.714386 | 0.728307 | 0.71836 | 0.722302 |
| 0.766786 | 0.766092 | 0.763339 | 0.768933 | 0.772135 | 0.770173 | 0.766932 | 0.766531 | 0.766434 |
| 0.766702 | 0.768859 | 0.763911 | 0.764755 | 0.768626 | 0.769061 | 0.77118 | 0.768041 | 0.770057 |
| 0.770517 | 0.769218 | 0.766422 | 0.766326 | 0.768096 | 0.766621 | 0.767449 | 0.770379 | 0.762408 |
| 0.770689 | 0.766287 | 0.763672 | 0.769345 | 0.765859 | 0.770884 | 0.773738 | 0.768005 | 0.765876 |
| 0.776185 | 0.771871 | 0.763928 | 0.772445 | 0.772467 | 0.77211 | 0.771819 | 0.772511 | 0.771508 |
| 0.773134 | 0.771358 | 0.767096 | 0.772761 | 0.771099 | 0.771375 | 0.769455 | 0.769527 | 0.763305 |
| 0.767736 | 0.769048 | 0.763386 | 0.768285 | 0.762033 | 0.772306 | 0.770652 | 0.769256 | 0.768621 |
| 0.772944 | 0.773187 | 0.765714 | 0.77159 | 0.769667 | 0.778728 | 0.769794 | 0.76688 | 0.76721 |
| 0.770533 | 0.771615 | 0.767409 | 0.76725 | 0.769947 | 0.770372 | 0.77326 | 0.767724 | 0.76756 |
| 0.768904 | 0.772006 | 0.765987 | 0.769669 | 0.772053 | 0.772561 | 0.772897 | 0.770319 | 0.770815 |
| 0.774684 | 0.7723 | 0.76365 | 0.770078 | 0.773934 | 0.769675 | 0.769517 | 0.770708 | 0.765857 |
| 0.77036 | 0.767444 | 0.761748 | 0.772001 | 0.769313 | 0.767251 | 0.767653 | 0.766193 | 0.764725 |
| 0.774756 | 0.776666 | 0.768607 | 0.772205 | 0.773557 | 0.77151 | 0.771265 | 0.766545 | 0.771752 |
| 0.775064 | 0.770713 | 0.763922 | 0.768525 | 0.771317 | 0.772086 | 0.772392 | 0.765984 | 0.771484 |
| 0.764826 | 0.76652 | 0.765491 | 0.765312 | 0.770513 | 0.765059 | 0.766319 | 0.7685 | 0.766157 |
| 0.768765 | 0.773392 | 0.762195 | 0.768499 | 0.769193 | 0.769988 | 0.768296 | 0.769294 | 0.765041 |
| 0.773958 | 0.772158 | 0.766897 | 0.773658 | 0.773871 | 0.770302 | 0.772018 | 0.775891 | 0.771822 |
| 0.775094 | 0.776396 | 0.767488 | 0.772467 | 0.770891 | 0.770382 | 0.775698 | 0.770656 | 0.771557 |
| 0.680972 | 0.685215 | 0.679753 | 0.681777 | 0.67569 | 0.680036 | 0.68546 | 0.687443 | 0.674732 |
| 0.662127 | 0.658438 | 0.657957 | 0.655532 | 0.66508 | 0.665404 | 0.661252 | 0.655311 | 0.649707 |
| 0.678951 | 0.669675 | 0.666524 | 0.674989 | 0.678225 | 0.676237 | 0.669283 | 0.670344 | 0.670452 |
| 0.661944 | 0.65283 | 0.65 | 0.654706 | 0.654317 | 0.656133 | 0.653933 | 0.653423 | 0.651424 |
| 0.746099 | 0.73592 | 0.732638 | 0.736519 | 0.73991 | 0.743597 | 0.7397 | 0.738513 | 0.74112 |
| 0.740333 | 0.724971 | 0.72019 | 0.725742 | 0.727048 | 0.724797 | 0.725565 | 0.717287 | 0.718406 |
| 0.734239 | 0.728948 | 0.72938 | 0.732007 | 0.728762 | 0.732361 | 0.729846 | 0.733224 | 0.723953 |
| 0.732751 | 0.750889 | 0.724882 | 0.749737 | 0.740879 | 0.741269 | 0.738825 | 0.746526 | 0.736154 |
| 0.719431 | 0.724943 | 0.70914 | 0.712107 | 0.716727 | 0.720428 | 0.716265 | 0.716899 | 0.71458 |
| 0.709522 | 0.715537 | 0.707215 | 0.718785 | 0.712079 | 0.719547 | 0.721298 | 0.713639 | 0.715333 |
| 0.725268 | 0.716184 | 0.713821 | 0.73226 | 0.719673 | 0.720571 | 0.719118 | 0.719574 | 0.706816 |
| 0.659605 | 0.650709 | 0.658506 | 0.65615 | 0.655063 | 0.654855 | 0.657259 | 0.659975 | 0.657123 |
| 0.765616 | 0.763521 | 0.757693 | 0.760903 | 0.759365 | 0.763399 | 0.762018 | 0.758544 | 0.763473 |
| 0.745533 | 0.744281 | 0.731036 | 0.738092 | 0.733402 | 0.744224 | 0.737024 | 0.746078 | 0.734852 |
| 0.735005 | 0.73481 | 0.731967 | 0.732467 | 0.7351 | 0.742009 | 0.734503 | 0.733965 | 0.730372 |
| 0.715683 | 0.72246 | 0.720643 | 0.723377 | 0.727374 | 0.727222 | 0.728653 | 0.726698 | 0.72181 |
| 0.741838 | 0.740517 | 0.738001 | 0.737106 | 0.735737 | 0.745822 | 0.742405 | 0.744306 | 0.740288 |
| 0.734972 | 0.738434 | 0.723647 | 0.738951 | 0.728811 | 0.734866 | 0.732646 | 0.728731 | 0.731536 |
| 0.735322 | 0.733078 | 0.726479 | 0.736255 | 0.732593 | 0.735219 | 0.732103 | 0.730506 | 0.727504 |
| 0.730069 | 0.735837 | 0.72818 | 0.732988 | 0.721009 | 0.729373 | 0.729839 | 0.724828 | 0.728732 |
| 0.732476 | 0.728375 | 0.723549 | 0.729837 | 0.728202 | 0.72742 | 0.732377 | 0.733321 | 0.726945 |
| 0.73869 | 0.727255 | 0.729767 | 0.729474 | 0.723367 | 0.72875 | 0.730596 | 0.727903 | 0.722691 |
| 0.737471 | 0.737442 | 0.714249 | 0.729786 | 0.723812 | 0.726553 | 0.733025 | 0.731289 | 0.722821 |
| 0.713858 | 0.719008 | 0.709287 | 0.717898 | 0.720938 | 0.715364 | 0.714052 | 0.720588 | 0.708815 |

| | | | | | | | | |
|---|---|---|---|---|---|---|---|---|
| 0.753057 | 0.749061 | 0.745053 | 0.750868 | 0.750295 | 0.752494 | 0.753982 | 0.755264 | 0.753635 |
| 0.732506 | 0.739364 | 0.723238 | 0.729939 | 0.72529 | 0.737777 | 0.735459 | 0.734552 | 0.736233 |
| 0.729429 | 0.729336 | 0.719168 | 0.722164 | 0.724683 | 0.726082 | 0.730518 | 0.724283 | 0.734194 |
| 0.730551 | 0.736543 | 0.718278 | 0.729071 | 0.726511 | 0.73775 | 0.732837 | 0.727538 | 0.734907 |
| 0.73749 | 0.738414 | 0.727808 | 0.729675 | 0.724791 | 0.723099 | 0.730512 | 0.739489 | 0.723469 |
| 0.730063 | 0.730567 | 0.717182 | 0.720268 | 0.72112 | 0.720873 | 0.727655 | 0.728432 | 0.718835 |
| 0.720885 | 0.724539 | 0.711803 | 0.720292 | 0.718171 | 0.727273 | 0.724105 | 0.720107 | 0.71619 |
| 0.719184 | 0.731241 | 0.717876 | 0.71213 | 0.715941 | 0.719358 | 0.725169 | 0.726439 | 0.715254 |
| 0.721953 | 0.722504 | 0.715491 | 0.717495 | 0.728454 | 0.722767 | 0.727579 | 0.721056 | 0.722144 |
| 0.753882 | 0.748713 | 0.748218 | 0.752402 | 0.749781 | 0.755363 | 0.757994 | 0.757171 | 0.757211 |
| 0.752486 | 0.752742 | 0.749095 | 0.744751 | 0.747376 | 0.75358 | 0.753191 | 0.750781 | 0.753665 |
| 0.747673 | 0.746662 | 0.738082 | 0.745746 | 0.745351 | 0.748433 | 0.746 | 0.746227 | 0.743491 |
| 0.741272 | 0.741081 | 0.73533 | 0.736919 | 0.735283 | 0.73833 | 0.734091 | 0.733196 | 0.735052 |
| 0.743219 | 0.741444 | 0.738877 | 0.738347 | 0.744067 | 0.745024 | 0.742258 | 0.744328 | 0.742861 |
| 0.74217 | 0.74 | 0.739519 | 0.738309 | 0.740854 | 0.744767 | 0.741492 | 0.738601 | 0.74311 |

| | | | | | | | | |
|---|---|---|---|---|---|---|---|---|
| | | plot 06 | | | | | | |
| Wild_plot_ | Wild_plot_ | Wild_plot_ | Wild_plot_ | Wild_plot_ | Wild_plot_ | Wild_plot_ | Wild_plot_ | Wild_plot_ |
| 0.705535 | 0.711749 | 0.709931 | 0.705295 | 0.711202 | 0.710047 | 0.704508 | 0.706897 | 0.702761 |
| 0.709693 | 0.718095 | 0.717559 | 0.714714 | 0.717479 | 0.719421 | 0.709349 | 0.715225 | 0.710866 |
| 0.699329 | 0.704209 | 0.707444 | 0.701885 | 0.708529 | 0.706796 | 0.699172 | 0.707265 | 0.697563 |
| 0.713971 | 0.717176 | 0.714553 | 0.712365 | 0.719284 | 0.717142 | 0.710475 | 0.715535 | 0.709135 |
| 0.706611 | 0.712048 | 0.708272 | 0.708857 | 0.71384 | 0.712918 | 0.706853 | 0.70918 | 0.706502 |
| 0.71542 | 0.717444 | 0.720955 | 0.714929 | 0.721211 | 0.718866 | 0.713388 | 0.717194 | 0.710025 |
| 0.773699 | 0.780773 | 0.778662 | 0.771398 | 0.776226 | 0.775933 | 0.776611 | 0.769711 | 0.772857 |
| 0.773985 | 0.775053 | 0.766933 | 0.764681 | 0.768768 | 0.770728 | 0.771409 | 0.768964 | 0.770598 |
| 0.710458 | 0.719574 | 0.716373 | 0.71276 | 0.713588 | 0.715788 | 0.708937 | 0.714896 | 0.708139 |
| 0.707379 | 0.709908 | 0.712152 | 0.70974 | 0.711786 | 0.712272 | 0.706748 | 0.709522 | 0.704121 |
| 0.711805 | 0.711523 | 0.715837 | 0.708333 | 0.711382 | 0.713934 | 0.705458 | 0.708525 | 0.707264 |
| 0.703132 | 0.704922 | 0.708871 | 0.703894 | 0.705971 | 0.705989 | 0.705117 | 0.702593 | 0.700422 |
| 0.710184 | 0.713011 | 0.717436 | 0.711964 | 0.716374 | 0.716218 | 0.70936 | 0.713922 | 0.7109 |
| 0.720915 | 0.724102 | 0.724416 | 0.722466 | 0.725926 | 0.723835 | 0.720161 | 0.72499 | 0.717718 |
| 0.734708 | 0.738553 | 0.734965 | 0.736307 | 0.740624 | 0.739694 | 0.729937 | 0.737691 | 0.734646 |
| 0.713745 | 0.714567 | 0.717254 | 0.71499 | 0.71654 | 0.71544 | 0.710397 | 0.714895 | 0.713106 |
| 0.710234 | 0.716855 | 0.715064 | 0.711742 | 0.715704 | 0.71464 | 0.714061 | 0.714675 | 0.70928 |
| 0.73668 | 0.746349 | 0.751072 | 0.739296 | 0.746739 | 0.74526 | 0.738709 | 0.742341 | 0.739875 |
| 0.702408 | 0.706867 | 0.709675 | 0.707173 | 0.710757 | 0.707459 | 0.703928 | 0.708748 | 0.705794 |
| 0.711099 | 0.712954 | 0.714896 | 0.710805 | 0.713781 | 0.715673 | 0.70847 | 0.714297 | 0.708903 |
| 0.706192 | 0.709028 | 0.711298 | 0.706417 | 0.712377 | 0.710096 | 0.707269 | 0.707439 | 0.705589 |
| 0.707963 | 0.712222 | 0.713268 | 0.70923 | 0.714206 | 0.712572 | 0.710932 | 0.712222 | 0.709325 |
| 0.713767 | 0.719858 | 0.720365 | 0.711744 | 0.721267 | 0.718107 | 0.713723 | 0.714075 | 0.713892 |
| 0.709352 | 0.713464 | 0.717432 | 0.711278 | 0.715916 | 0.71363 | 0.710768 | 0.7106 | 0.71 |
| 0.711963 | 0.716634 | 0.721791 | 0.712489 | 0.717791 | 0.718825 | 0.711475 | 0.714615 | 0.709008 |
| 0.707922 | 0.711685 | 0.713983 | 0.70828 | 0.714468 | 0.709891 | 0.705433 | 0.708401 | 0.705464 |
| 0.701464 | 0.70831 | 0.709034 | 0.701776 | 0.70663 | 0.706478 | 0.701153 | 0.701648 | 0.700498 |
| 0.711132 | 0.713287 | 0.714495 | 0.708263 | 0.716856 | 0.71415 | 0.708507 | 0.715359 | 0.709942 |
| 0.712217 | 0.719038 | 0.717522 | 0.711233 | 0.71541 | 0.718072 | 0.711894 | 0.713617 | 0.713008 |
| 0.711132 | 0.713697 | 0.718071 | 0.710567 | 0.715853 | 0.713216 | 0.708059 | 0.709877 | 0.710026 |
| 0.708271 | 0.711872 | 0.711179 | 0.707075 | 0.714791 | 0.711569 | 0.706813 | 0.710343 | 0.704411 |
| 0.707045 | 0.709931 | 0.707953 | 0.707768 | 0.712565 | 0.710853 | 0.702641 | 0.708632 | 0.704427 |
| 0.706247 | 0.708487 | 0.71205 | 0.710253 | 0.713624 | 0.712501 | 0.703887 | 0.712302 | 0.702238 |
| 0.713284 | 0.714241 | 0.712955 | 0.712157 | 0.715963 | 0.716232 | 0.705213 | 0.712105 | 0.711089 |
| 0.705089 | 0.708123 | 0.710652 | 0.704683 | 0.707375 | 0.707908 | 0.703952 | 0.702326 | 0.701603 |
| 0.709552 | 0.713866 | 0.710662 | 0.711048 | 0.716154 | 0.71408 | 0.706677 | 0.713826 | 0.707182 |
| 0.707727 | 0.713345 | 0.719271 | 0.712051 | 0.718777 | 0.717596 | 0.70942 | 0.713484 | 0.709976 |
| 0.709098 | 0.714514 | 0.717774 | 0.711323 | 0.71542 | 0.713919 | 0.708371 | 0.713065 | 0.708505 |
| 0.712731 | 0.719427 | 0.719139 | 0.716562 | 0.721512 | 0.720035 | 0.711783 | 0.713981 | 0.714975 |

| | | | | | | | | |
|---|---|---|---|---|---|---|---|---|
| 0.744422 | 0.752033 | 0.751466 | 0.747149 | 0.749737 | 0.752886 | 0.745697 | 0.745315 | 0.749264 |
| 0.701608 | 0.706743 | 0.707773 | 0.702247 | 0.707247 | 0.705316 | 0.699153 | 0.703015 | 0.700224 |
| 0.704447 | 0.70831 | 0.705806 | 0.706795 | 0.710463 | 0.708525 | 0.703073 | 0.705791 | 0.70521 |
| 0.713759 | 0.720032 | 0.719957 | 0.716091 | 0.718513 | 0.718036 | 0.714064 | 0.715722 | 0.707359 |
| 0.708674 | 0.711484 | 0.710604 | 0.707529 | 0.71287 | 0.712949 | 0.708286 | 0.707307 | 0.708427 |
| 0.712911 | 0.716917 | 0.719775 | 0.71518 | 0.719345 | 0.719162 | 0.710728 | 0.712535 | 0.710312 |
| 0.710147 | 0.710228 | 0.710939 | 0.70593 | 0.712516 | 0.709525 | 0.706871 | 0.710234 | 0.707473 |
| 0.712956 | 0.711465 | 0.712286 | 0.708084 | 0.717173 | 0.713579 | 0.70846 | 0.711675 | 0.711917 |
| 0.711393 | 0.716787 | 0.720315 | 0.715265 | 0.721347 | 0.720308 | 0.710324 | 0.716593 | 0.711948 |
| 0.712677 | 0.714848 | 0.714892 | 0.713078 | 0.717748 | 0.716926 | 0.713841 | 0.714368 | 0.713823 |
| 0.701753 | 0.706431 | 0.704191 | 0.703105 | 0.706798 | 0.707296 | 0.698673 | 0.70249 | 0.699658 |
| 0.70359 | 0.709565 | 0.710686 | 0.706573 | 0.711281 | 0.710526 | 0.702393 | 0.707964 | 0.70161 |
| 0.707253 | 0.709875 | 0.712964 | 0.71143 | 0.714343 | 0.712617 | 0.709608 | 0.710696 | 0.707106 |
| 0.707375 | 0.708241 | 0.707692 | 0.703922 | 0.707664 | 0.709939 | 0.701802 | 0.705839 | 0.701624 |
| 0.702892 | 0.706757 | 0.708237 | 0.704766 | 0.709774 | 0.708547 | 0.703587 | 0.708816 | 0.699774 |
| 0.703763 | 0.7087 | 0.708824 | 0.702888 | 0.708073 | 0.708904 | 0.700514 | 0.704845 | 0.703161 |
| 0.703612 | 0.707729 | 0.709122 | 0.706094 | 0.711563 | 0.710788 | 0.700711 | 0.703337 | 0.705506 |
| 0.710394 | 0.713677 | 0.716769 | 0.708646 | 0.717827 | 0.714989 | 0.707958 | 0.711721 | 0.709197 |
| 0.710869 | 0.716989 | 0.719569 | 0.713535 | 0.717638 | 0.712894 | 0.711466 | 0.712271 | 0.710891 |
| 0.707013 | 0.708595 | 0.711223 | 0.70721 | 0.71143 | 0.713167 | 0.702881 | 0.709675 | 0.707603 |
| 0.701161 | 0.705153 | 0.707442 | 0.703395 | 0.707437 | 0.704704 | 0.697619 | 0.70457 | 0.698286 |
| 0.703946 | 0.705645 | 0.70995 | 0.703644 | 0.71193 | 0.708569 | 0.70045 | 0.705477 | 0.703392 |
| 0.70533 | 0.713926 | 0.714193 | 0.705977 | 0.712442 | 0.711006 | 0.70517 | 0.711893 | 0.703538 |
| 0.717175 | 0.720827 | 0.718451 | 0.71439 | 0.721657 | 0.72068 | 0.717084 | 0.718484 | 0.712529 |
| 0.723484 | 0.72217 | 0.725883 | 0.722915 | 0.726431 | 0.727345 | 0.720869 | 0.724634 | 0.725831 |
| 0.773728 | 0.776559 | 0.769852 | 0.764474 | 0.769928 | 0.773434 | 0.772588 | 0.768701 | 0.771297 |
| 0.778946 | 0.782137 | 0.781199 | 0.773451 | 0.774695 | 0.773423 | 0.77286 | 0.780267 | 0.77804 |
| 0.779207 | 0.78244 | 0.78102 | 0.768721 | 0.775764 | 0.776264 | 0.776445 | 0.779573 | 0.778 |
| 0.706537 | 0.708724 | 0.710267 | 0.704943 | 0.709187 | 0.70911 | 0.703846 | 0.706744 | 0.704252 |
| 0.703192 | 0.710205 | 0.709333 | 0.704959 | 0.711124 | 0.711066 | 0.70313 | 0.704841 | 0.702926 |
| 0.710903 | 0.719234 | 0.719141 | 0.710294 | 0.720607 | 0.71605 | 0.710831 | 0.714603 | 0.71141 |
| 0.717304 | 0.718785 | 0.720581 | 0.713084 | 0.717575 | 0.719445 | 0.71277 | 0.715768 | 0.712818 |
| 0.705892 | 0.707682 | 0.708695 | 0.705983 | 0.712897 | 0.709437 | 0.700709 | 0.708999 | 0.704314 |
| 0.707635 | 0.71028 | 0.709146 | 0.70852 | 0.712935 | 0.713587 | 0.705654 | 0.711748 | 0.708183 |
| 0.706985 | 0.706136 | 0.7088 | 0.705156 | 0.713308 | 0.705991 | 0.701936 | 0.709466 | 0.703526 |
| 0.706139 | 0.709557 | 0.710585 | 0.708059 | 0.714113 | 0.713601 | 0.700008 | 0.711027 | 0.702885 |
| 0.706665 | 0.713793 | 0.712928 | 0.70536 | 0.713683 | 0.710355 | 0.704837 | 0.709032 | 0.707283 |
| 0.719765 | 0.724201 | 0.726411 | 0.719197 | 0.72606 | 0.723216 | 0.71925 | 0.725805 | 0.718952 |
| 0.697711 | 0.701073 | 0.705927 | 0.701823 | 0.705579 | 0.702663 | 0.69638 | 0.70254 | 0.696637 |
| 0.703964 | 0.710897 | 0.713646 | 0.706354 | 0.710629 | 0.714094 | 0.706862 | 0.704737 | 0.705746 |
| 0.70318 | 0.706085 | 0.705052 | 0.701271 | 0.707128 | 0.709134 | 0.697985 | 0.704927 | 0.700652 |
| 0.721344 | 0.72605 | 0.72659 | 0.721969 | 0.730414 | 0.724662 | 0.721015 | 0.727145 | 0.721466 |
| 0.709634 | 0.713414 | 0.713476 | 0.708539 | 0.709269 | 0.710489 | 0.706259 | 0.708973 | 0.707617 |
| 0.7055 | 0.711964 | 0.710001 | 0.705219 | 0.711453 | 0.711132 | 0.703677 | 0.705887 | 0.704267 |
| 0.712706 | 0.710724 | 0.712231 | 0.707837 | 0.716016 | 0.714469 | 0.709035 | 0.708875 | 0.707327 |
| 0.70681 | 0.710551 | 0.711628 | 0.709241 | 0.712308 | 0.711093 | 0.702077 | 0.707933 | 0.705157 |

| | | | | | | | | |
|---|---|---|---|---|---|---|---|---|
| 0.705957 | 0.709088 | 0.70907 | 0.706196 | 0.711571 | 0.710721 | 0.702302 | 0.70662 | 0.703929 |
| 0.707237 | 0.713904 | 0.715919 | 0.71318 | 0.716665 | 0.716902 | 0.707174 | 0.711069 | 0.707779 |
| 0.710315 | 0.712975 | 0.714274 | 0.708126 | 0.713639 | 0.714974 | 0.708227 | 0.70674 | 0.708481 |
| 0.714629 | 0.719646 | 0.721127 | 0.713747 | 0.718806 | 0.718929 | 0.710884 | 0.717983 | 0.711806 |
| 0.71315 | 0.715892 | 0.71851 | 0.714689 | 0.719872 | 0.719025 | 0.711697 | 0.715385 | 0.713538 |
| 0.70535 | 0.711955 | 0.714853 | 0.708995 | 0.711425 | 0.717386 | 0.705005 | 0.708454 | 0.707186 |
| 0.71032 | 0.712028 | 0.71637 | 0.7093 | 0.715071 | 0.713115 | 0.707012 | 0.710599 | 0.705338 |
| 0.714343 | 0.718096 | 0.715948 | 0.711671 | 0.720156 | 0.715496 | 0.709118 | 0.713283 | 0.709141 |
| 0.708996 | 0.715344 | 0.71318 | 0.711167 | 0.714332 | 0.716554 | 0.706663 | 0.710615 | 0.709144 |
| 0.705368 | 0.708425 | 0.712471 | 0.702392 | 0.709634 | 0.708699 | 0.704737 | 0.710692 | 0.704381 |
| 0.706094 | 0.711233 | 0.708945 | 0.70807 | 0.716025 | 0.711046 | 0.702939 | 0.709154 | 0.706198 |
| 0.706861 | 0.710674 | 0.711443 | 0.710978 | 0.71112 | 0.713813 | 0.70541 | 0.708834 | 0.70562 |
| 0.711636 | 0.71563 | 0.716202 | 0.708896 | 0.715916 | 0.713817 | 0.708474 | 0.708911 | 0.708519 |
| 0.703677 | 0.708884 | 0.709648 | 0.706167 | 0.710939 | 0.711231 | 0.706459 | 0.706107 | 0.702047 |
| 0.710086 | 0.714229 | 0.715841 | 0.710052 | 0.71664 | 0.715635 | 0.703619 | 0.713333 | 0.706355 |
| 0.705859 | 0.713802 | 0.712398 | 0.71119 | 0.713486 | 0.712229 | 0.704455 | 0.711553 | 0.705487 |
| 0.706679 | 0.714422 | 0.713519 | 0.713344 | 0.719031 | 0.716909 | 0.707367 | 0.712447 | 0.707293 |
| 0.707839 | 0.709052 | 0.708561 | 0.707012 | 0.715135 | 0.712255 | 0.704042 | 0.709993 | 0.705549 |
| 0.708208 | 0.711236 | 0.715812 | 0.709547 | 0.714998 | 0.714047 | 0.707265 | 0.710239 | 0.707234 |
| 0.705437 | 0.711511 | 0.714485 | 0.71072 | 0.716414 | 0.712937 | 0.706573 | 0.710377 | 0.706687 |
| 0.706797 | 0.715184 | 0.714992 | 0.709011 | 0.712343 | 0.713932 | 0.706222 | 0.712462 | 0.708093 |
| 0.703147 | 0.708064 | 0.707826 | 0.704853 | 0.708486 | 0.706834 | 0.698981 | 0.705638 | 0.703039 |
| 0.709385 | 0.714274 | 0.712907 | 0.711347 | 0.712404 | 0.710188 | 0.70436 | 0.708856 | 0.706674 |
| 0.706442 | 0.709581 | 0.716772 | 0.710654 | 0.711395 | 0.713527 | 0.709173 | 0.712343 | 0.704802 |
| 0.710148 | 0.712905 | 0.713814 | 0.708221 | 0.713602 | 0.711109 | 0.706287 | 0.712391 | 0.707407 |
| 0.713612 | 0.718961 | 0.719869 | 0.715781 | 0.721339 | 0.717845 | 0.71023 | 0.715568 | 0.71329 |
| 0.707877 | 0.710994 | 0.713123 | 0.707601 | 0.7135 | 0.709456 | 0.706567 | 0.712014 | 0.706515 |
| 0.707898 | 0.71213 | 0.714473 | 0.709353 | 0.71432 | 0.714773 | 0.70848 | 0.709448 | 0.707226 |
| 0.710804 | 0.717113 | 0.716639 | 0.713755 | 0.717486 | 0.714897 | 0.711042 | 0.711304 | 0.709471 |
| 0.713057 | 0.710602 | 0.713836 | 0.705979 | 0.715077 | 0.711229 | 0.707161 | 0.714869 | 0.709026 |
| 0.710742 | 0.71538 | 0.715348 | 0.71046 | 0.715558 | 0.712975 | 0.710474 | 0.713672 | 0.706656 |
| 0.705428 | 0.709982 | 0.708953 | 0.709484 | 0.71296 | 0.715253 | 0.703177 | 0.711638 | 0.705679 |
| 0.708733 | 0.71248 | 0.714877 | 0.706968 | 0.714867 | 0.711034 | 0.707331 | 0.71228 | 0.70925 |
| 0.710929 | 0.715575 | 0.715293 | 0.711023 | 0.713344 | 0.714788 | 0.709721 | 0.712651 | 0.703817 |
| 0.705304 | 0.70869 | 0.710214 | 0.704542 | 0.709836 | 0.70967 | 0.706143 | 0.704699 | 0.706888 |
| 0.705022 | 0.7094 | 0.71287 | 0.707604 | 0.71382 | 0.715568 | 0.707599 | 0.706414 | 0.706469 |
| 0.703126 | 0.706284 | 0.702163 | 0.705584 | 0.710142 | 0.710729 | 0.701202 | 0.703214 | 0.700683 |
| 0.683064 | 0.681199 | 0.690896 | 0.683748 | 0.686841 | 0.688821 | 0.683282 | 0.685269 | 0.682886 |
| 0.68298 | 0.689399 | 0.691358 | 0.687468 | 0.692988 | 0.694051 | 0.683388 | 0.689659 | 0.691429 |
| 0.679191 | 0.68274 | 0.686367 | 0.683418 | 0.689326 | 0.687526 | 0.676299 | 0.681363 | 0.684177 |
| 0.676374 | 0.681083 | 0.684051 | 0.679686 | 0.683936 | 0.687054 | 0.674455 | 0.685528 | 0.679215 |
| 0.664234 | 0.668252 | 0.664894 | 0.66757 | 0.669478 | 0.665968 | 0.67006 | 0.664424 | 0.669063 |
| 0.668825 | 0.663791 | 0.672989 | 0.66752 | 0.672569 | 0.668919 | 0.670662 | 0.668293 | 0.672275 |
| 0.666347 | 0.668873 | 0.674387 | 0.668623 | 0.673792 | 0.670664 | 0.662759 | 0.671082 | 0.671126 |
| 0.667931 | 0.670249 | 0.672294 | 0.661609 | 0.671345 | 0.670822 | 0.66799 | 0.671379 | 0.674458 |
| 0.663997 | 0.669472 | 0.675508 | 0.667503 | 0.672188 | 0.669324 | 0.668678 | 0.668508 | 0.667634 |
| 0.669728 | 0.673886 | 0.68031 | 0.672705 | 0.684289 | 0.67957 | 0.672691 | 0.681279 | 0.673282 |

| | | | | | | | | |
|---|---|---|---|---|---|---|---|---|
| 0.672269 | 0.675506 | 0.674404 | 0.67577 | 0.678479 | 0.675009 | 0.668983 | 0.674089 | 0.670547 |
| 0.669459 | 0.671173 | 0.679046 | 0.671254 | 0.679132 | 0.672463 | 0.668896 | 0.678977 | 0.673062 |
| 0.666711 | 0.677346 | 0.673036 | 0.676077 | 0.676822 | 0.678638 | 0.669001 | 0.674431 | 0.670187 |
| 0.660358 | 0.664354 | 0.671083 | 0.661365 | 0.666931 | 0.665172 | 0.659735 | 0.66751 | 0.664839 |
| 0.660109 | 0.66374 | 0.671776 | 0.662115 | 0.670785 | 0.662701 | 0.66887 | 0.667785 | 0.667841 |
| 0.666349 | 0.667666 | 0.672992 | 0.66538 | 0.671519 | 0.669583 | 0.664239 | 0.666068 | 0.669496 |
| 0.667042 | 0.669057 | 0.673434 | 0.66605 | 0.673672 | 0.667419 | 0.665528 | 0.670146 | 0.675781 |
| 0.6632 | 0.66444 | 0.668512 | 0.664042 | 0.670217 | 0.662844 | 0.663014 | 0.669029 | 0.667332 |
| 0.660769 | 0.664289 | 0.672378 | 0.665089 | 0.666422 | 0.666227 | 0.664307 | 0.663974 | 0.663747 |
| 0.664689 | 0.665746 | 0.671138 | 0.661786 | 0.672682 | 0.664626 | 0.665238 | 0.66302 | 0.666518 |
| 0.65624 | 0.661025 | 0.672153 | 0.661683 | 0.668458 | 0.668689 | 0.656799 | 0.656166 | 0.662171 |
| 0.701225 | 0.699953 | 0.698718 | 0.697066 | 0.702703 | 0.700008 | 0.694144 | 0.696713 | 0.696967 |
| 0.702503 | 0.709817 | 0.711623 | 0.704942 | 0.71201 | 0.708794 | 0.705983 | 0.707852 | 0.703792 |
| 0.679807 | 0.687599 | 0.691136 | 0.686744 | 0.691799 | 0.688508 | 0.687155 | 0.68579 | 0.681509 |
| 0.709711 | 0.714423 | 0.715589 | 0.711696 | 0.717899 | 0.717486 | 0.712105 | 0.714131 | 0.706982 |
| 0.704723 | 0.704843 | 0.707129 | 0.703408 | 0.708267 | 0.708383 | 0.703051 | 0.703474 | 0.700149 |
| 0.707265 | 0.717628 | 0.719748 | 0.715678 | 0.721189 | 0.718339 | 0.70851 | 0.712535 | 0.709413 |
| 0.716853 | 0.72094 | 0.721858 | 0.717413 | 0.723129 | 0.7222 | 0.71399 | 0.718343 | 0.715944 |
| 0.694656 | 0.701724 | 0.703219 | 0.694129 | 0.703011 | 0.699258 | 0.691092 | 0.697953 | 0.692226 |
| 0.71587 | 0.72117 | 0.724172 | 0.716029 | 0.720562 | 0.721753 | 0.71815 | 0.715704 | 0.713528 |
| 0.684017 | 0.686537 | 0.695461 | 0.686645 | 0.69431 | 0.689556 | 0.687143 | 0.692204 | 0.687615 |
| 0.711412 | 0.716369 | 0.716938 | 0.715282 | 0.721894 | 0.719594 | 0.711277 | 0.716332 | 0.709858 |
| 0.714909 | 0.721847 | 0.724839 | 0.719304 | 0.725411 | 0.721431 | 0.711876 | 0.719792 | 0.715173 |
| 0.679797 | 0.685609 | 0.69606 | 0.6838 | 0.695169 | 0.690759 | 0.680169 | 0.684543 | 0.68787 |
| 0.683088 | 0.686198 | 0.692171 | 0.687615 | 0.692148 | 0.689009 | 0.685193 | 0.692752 | 0.68422 |
| 0.688917 | 0.693293 | 0.697184 | 0.689638 | 0.699149 | 0.696195 | 0.691041 | 0.694094 | 0.697283 |
| 0.705048 | 0.710388 | 0.71165 | 0.703185 | 0.707176 | 0.706582 | 0.706381 | 0.704847 | 0.700255 |
| 0.71622 | 0.722306 | 0.728097 | 0.71975 | 0.724867 | 0.723063 | 0.712486 | 0.721077 | 0.715083 |
| 0.685659 | 0.687289 | 0.689749 | 0.686009 | 0.690686 | 0.688195 | 0.680484 | 0.686961 | 0.68317 |
| 0.718642 | 0.723143 | 0.724459 | 0.719274 | 0.725357 | 0.724045 | 0.721618 | 0.725839 | 0.719518 |
| 0.722888 | 0.729706 | 0.728656 | 0.724235 | 0.729226 | 0.732448 | 0.722854 | 0.727428 | 0.720129 |
| 0.693777 | 0.702076 | 0.699447 | 0.695108 | 0.702048 | 0.699296 | 0.694199 | 0.699396 | 0.693942 |
| 0.695265 | 0.70277 | 0.698642 | 0.693497 | 0.702632 | 0.697497 | 0.693355 | 0.700317 | 0.691096 |
| 0.72169 | 0.725228 | 0.729558 | 0.723109 | 0.727258 | 0.729205 | 0.720223 | 0.723883 | 0.718413 |
| 0.721845 | 0.721889 | 0.725827 | 0.720667 | 0.723633 | 0.72475 | 0.715877 | 0.723216 | 0.713441 |
| 0.719863 | 0.724815 | 0.72859 | 0.720365 | 0.725457 | 0.729371 | 0.722126 | 0.72444 | 0.719545 |
| 0.676012 | 0.676877 | 0.681665 | 0.678698 | 0.682942 | 0.678995 | 0.677138 | 0.678917 | 0.671908 |
| 0.720864 | 0.723394 | 0.723791 | 0.717395 | 0.724733 | 0.72034 | 0.719639 | 0.723261 | 0.720487 |
| 0.676305 | 0.678838 | 0.679053 | 0.674633 | 0.678403 | 0.677816 | 0.674191 | 0.674433 | 0.670347 |
| 0.671106 | 0.671754 | 0.674179 | 0.669152 | 0.676174 | 0.673842 | 0.666667 | 0.671037 | 0.665504 |
| 0.719584 | 0.72234 | 0.723571 | 0.718526 | 0.721952 | 0.723048 | 0.718301 | 0.719738 | 0.717568 |
| 0.671876 | 0.671646 | 0.672441 | 0.670285 | 0.678637 | 0.673676 | 0.664458 | 0.672645 | 0.666322 |
| 0.660317 | 0.663513 | 0.667493 | 0.660567 | 0.666541 | 0.666892 | 0.657257 | 0.663855 | 0.658749 |
| 0.658786 | 0.658967 | 0.664442 | 0.657542 | 0.665 | 0.659516 | 0.655099 | 0.664463 | 0.655654 |
| 0.724577 | 0.733058 | 0.733877 | 0.726373 | 0.73067 | 0.730344 | 0.725867 | 0.729299 | 0.726306 |
| 0.662338 | 0.664258 | 0.664993 | 0.661891 | 0.667271 | 0.667492 | 0.660119 | 0.663708 | 0.659489 |
| 0.716222 | 0.72356 | 0.724771 | 0.720533 | 0.724432 | 0.728677 | 0.719136 | 0.72053 | 0.719273 |

| | | | | | | | | |
|---|---|---|---|---|---|---|---|---|
| 0.729871 | 0.731104 | 0.733119 | 0.726296 | 0.73176 | 0.73327 | 0.728999 | 0.728967 | 0.728781 |
| 0.720919 | 0.727093 | 0.728819 | 0.719181 | 0.728596 | 0.726471 | 0.722058 | 0.725808 | 0.721708 |
| 0.742804 | 0.745676 | 0.747241 | 0.742662 | 0.74257 | 0.748048 | 0.739963 | 0.741736 | 0.740596 |
| 0.651393 | 0.653288 | 0.65606 | 0.650386 | 0.657946 | 0.655939 | 0.650225 | 0.653518 | 0.648225 |
| 0.653171 | 0.655774 | 0.657996 | 0.654463 | 0.659122 | 0.657506 | 0.649554 | 0.65818 | 0.650042 |
| 0.726378 | 0.733938 | 0.735746 | 0.728456 | 0.729452 | 0.733509 | 0.730125 | 0.731766 | 0.728389 |
| 0.653334 | 0.657516 | 0.65582 | 0.652413 | 0.655862 | 0.657605 | 0.649794 | 0.655526 | 0.648042 |
| 0.729791 | 0.738972 | 0.735459 | 0.731131 | 0.734495 | 0.732529 | 0.732787 | 0.732392 | 0.729454 |
| 0.730431 | 0.731992 | 0.734079 | 0.729663 | 0.735667 | 0.73724 | 0.72962 | 0.732929 | 0.729682 |
| 0.73234 | 0.740566 | 0.737302 | 0.730027 | 0.736508 | 0.737345 | 0.728265 | 0.734694 | 0.728608 |
| 0.72704 | 0.734951 | 0.733638 | 0.729874 | 0.732189 | 0.737544 | 0.731369 | 0.732644 | 0.731102 |
| 0.746753 | 0.757353 | 0.74 | 0.740506 | 0.72973 | 0.708075 | 0.72973 | 0.76129 | 0.685185 |
| 0.722047 | 0.726673 | 0.743153 | 0.724699 | 0.732247 | 0.734322 | 0.726485 | 0.727491 | 0.730987 |
| 0.72192 | 0.730504 | 0.729213 | 0.718838 | 0.726445 | 0.729251 | 0.730806 | 0.721402 | 0.725242 |
| 0.732036 | 0.73826 | 0.742019 | 0.730152 | 0.735604 | 0.734065 | 0.7358 | 0.734112 | 0.739443 |
| 0.730579 | 0.734147 | 0.735606 | 0.726326 | 0.735119 | 0.736757 | 0.734913 | 0.734724 | 0.733443 |
| 0.591441 | 0.591518 | 0.596441 | 0.59315 | 0.59689 | 0.594131 | 0.590939 | 0.596316 | 0.590338 |
| 0.732842 | 0.743014 | 0.737319 | 0.728296 | 0.737816 | 0.738301 | 0.738272 | 0.733631 | 0.738019 |
| 0.730389 | 0.737513 | 0.73889 | 0.731155 | 0.737939 | 0.740431 | 0.73364 | 0.733406 | 0.733747 |
| 0.734831 | 0.736739 | 0.738623 | 0.733796 | 0.736381 | 0.736605 | 0.735573 | 0.732449 | 0.735018 |
| 0.736961 | 0.743376 | 0.742161 | 0.736189 | 0.743337 | 0.745616 | 0.739203 | 0.738631 | 0.737135 |
| 0.59757 | 0.598661 | 0.599764 | 0.594574 | 0.602855 | 0.597807 | 0.592371 | 0.597984 | 0.592999 |
| 0.58394 | 0.580932 | 0.583293 | 0.581696 | 0.589287 | 0.587837 | 0.576693 | 0.582557 | 0.578705 |
| 0.760185 | 0.764516 | 0.763228 | 0.75867 | 0.766717 | 0.766924 | 0.761032 | 0.762634 | 0.761263 |
| 0.758915 | 0.763286 | 0.76357 | 0.752274 | 0.764429 | 0.764682 | 0.763704 | 0.762142 | 0.760718 |
| 0.752922 | 0.756635 | 0.759517 | 0.750898 | 0.757556 | 0.753463 | 0.754499 | 0.755988 | 0.75151 |
| 0.758915 | 0.761295 | 0.765336 | 0.765514 | 0.768277 | 0.766122 | 0.759083 | 0.760793 | 0.760625 |
| 0.759611 | 0.76658 | 0.7696 | 0.756209 | 0.769842 | 0.763083 | 0.759045 | 0.758887 | 0.757013 |
| 0.764992 | 0.767104 | 0.769612 | 0.762113 | 0.768999 | 0.767721 | 0.766267 | 0.765243 | 0.766847 |
| 0.761509 | 0.767337 | 0.76685 | 0.752758 | 0.767059 | 0.764167 | 0.760041 | 0.762536 | 0.762417 |
| 0.760892 | 0.76563 | 0.761821 | 0.757336 | 0.762858 | 0.765281 | 0.758621 | 0.763044 | 0.761008 |
| 0.772395 | 0.771826 | 0.772991 | 0.762168 | 0.776258 | 0.770005 | 0.771327 | 0.76934 | 0.768816 |
| 0.757209 | 0.767691 | 0.764651 | 0.756373 | 0.766055 | 0.76254 | 0.759079 | 0.759966 | 0.756344 |
| 0.759058 | 0.769721 | 0.765782 | 0.763875 | 0.767472 | 0.766897 | 0.759163 | 0.761017 | 0.762107 |
| 0.767609 | 0.774652 | 0.764282 | 0.764388 | 0.777824 | 0.767429 | 0.765954 | 0.768302 | 0.770032 |
| 0.759987 | 0.770931 | 0.766161 | 0.757318 | 0.765087 | 0.762517 | 0.757281 | 0.76516 | 0.76314 |
| 0.762014 | 0.766432 | 0.767921 | 0.758353 | 0.769659 | 0.768011 | 0.758691 | 0.764492 | 0.763534 |
| 0.762639 | 0.773368 | 0.767139 | 0.766016 | 0.769478 | 0.769835 | 0.763822 | 0.76341 | 0.765453 |
| 0.757982 | 0.76388 | 0.764752 | 0.754712 | 0.768719 | 0.763543 | 0.758889 | 0.759638 | 0.76014 |
| 0.760431 | 0.764863 | 0.761533 | 0.755695 | 0.766979 | 0.763201 | 0.758195 | 0.758716 | 0.755991 |
| 0.724126 | 0.739314 | 0.738434 | 0.71749 | 0.727337 | 0.729445 | 0.732814 | 0.736065 | 0.729259 |
| 0.730582 | 0.736002 | 0.73894 | 0.72729 | 0.730683 | 0.737181 | 0.730147 | 0.737182 | 0.736692 |
| 0.765589 | 0.774333 | 0.768753 | 0.756922 | 0.768148 | 0.765687 | 0.760265 | 0.766198 | 0.76261 |
| 0.768279 | 0.770315 | 0.77097 | 0.760302 | 0.76939 | 0.769055 | 0.766839 | 0.768438 | 0.766635 |
| 0.712441 | 0.720391 | 0.724312 | 0.715061 | 0.716237 | 0.725223 | 0.72084 | 0.721931 | 0.718513 |
| 0.762443 | 0.772508 | 0.771545 | 0.757538 | 0.763154 | 0.766315 | 0.766692 | 0.7624 | 0.761643 |
| 0.755985 | 0.765705 | 0.76112 | 0.752392 | 0.760256 | 0.759825 | 0.760199 | 0.75944 | 0.754413 |

| | | | | | | | | |
|---|---|---|---|---|---|---|---|---|
| 0.766088 | 0.768315 | 0.766737 | 0.758621 | 0.767157 | 0.770423 | 0.764676 | 0.760958 | 0.760432 |
| 0.766233 | 0.772993 | 0.767152 | 0.762251 | 0.770446 | 0.769937 | 0.766967 | 0.765427 | 0.765068 |
| 0.765378 | 0.767554 | 0.762342 | 0.755845 | 0.765257 | 0.764854 | 0.760507 | 0.765315 | 0.759342 |
| 0.762001 | 0.770384 | 0.771016 | 0.760731 | 0.767959 | 0.770497 | 0.766802 | 0.765346 | 0.766096 |
| 0.767533 | 0.775525 | 0.769679 | 0.759685 | 0.766285 | 0.772129 | 0.769205 | 0.769947 | 0.768246 |
| 0.762634 | 0.769711 | 0.764323 | 0.760381 | 0.768214 | 0.767894 | 0.762333 | 0.763644 | 0.765087 |
| 0.764587 | 0.769087 | 0.765307 | 0.760215 | 0.768162 | 0.766157 | 0.764795 | 0.760191 | 0.761042 |
| 0.750143 | 0.761042 | 0.758048 | 0.747301 | 0.755925 | 0.755711 | 0.757423 | 0.75828 | 0.751602 |
| 0.761951 | 0.766142 | 0.762867 | 0.755197 | 0.761093 | 0.761155 | 0.76022 | 0.760448 | 0.760182 |
| 0.759063 | 0.766951 | 0.760506 | 0.75029 | 0.760794 | 0.760337 | 0.757517 | 0.762413 | 0.757581 |
| 0.765355 | 0.772842 | 0.766664 | 0.756373 | 0.765044 | 0.764118 | 0.762041 | 0.765264 | 0.763061 |
| 0.76121 | 0.767528 | 0.764675 | 0.757179 | 0.764096 | 0.767238 | 0.761756 | 0.762687 | 0.763267 |
| 0.756666 | 0.758042 | 0.759082 | 0.74662 | 0.758339 | 0.755939 | 0.756387 | 0.758212 | 0.752747 |
| 0.756392 | 0.761249 | 0.758182 | 0.751038 | 0.759315 | 0.757744 | 0.760318 | 0.756771 | 0.755579 |
| 0.762882 | 0.771618 | 0.771364 | 0.760615 | 0.764788 | 0.768162 | 0.762935 | 0.768519 | 0.764344 |
| 0.765231 | 0.771256 | 0.771373 | 0.757876 | 0.769801 | 0.767683 | 0.76235 | 0.764973 | 0.764242 |
| 0.762102 | 0.768238 | 0.768266 | 0.756738 | 0.762538 | 0.764189 | 0.761668 | 0.766387 | 0.764571 |
| 0.760242 | 0.767129 | 0.761696 | 0.754851 | 0.761121 | 0.761569 | 0.759437 | 0.762735 | 0.765165 |
| 0.763104 | 0.767669 | 0.765291 | 0.758812 | 0.770064 | 0.762963 | 0.761973 | 0.760643 | 0.761061 |
| 0.76048 | 0.767611 | 0.765248 | 0.762422 | 0.765528 | 0.770485 | 0.761111 | 0.757711 | 0.760244 |
| 0.754815 | 0.759486 | 0.760528 | 0.75484 | 0.759063 | 0.760296 | 0.754431 | 0.754076 | 0.757908 |
| 0.759646 | 0.763979 | 0.759904 | 0.756085 | 0.764988 | 0.760106 | 0.759035 | 0.757755 | 0.755488 |
| 0.762539 | 0.76788 | 0.771285 | 0.763286 | 0.770957 | 0.768674 | 0.764394 | 0.765446 | 0.761653 |
| 0.763799 | 0.765349 | 0.768178 | 0.754762 | 0.768371 | 0.76185 | 0.76592 | 0.764785 | 0.760948 |
| 0.760326 | 0.765458 | 0.761511 | 0.754477 | 0.76691 | 0.763558 | 0.758102 | 0.762949 | 0.760524 |
| 0.759921 | 0.764035 | 0.7653 | 0.757323 | 0.765082 | 0.765462 | 0.75611 | 0.758916 | 0.758963 |
| 0.755762 | 0.761869 | 0.757465 | 0.751007 | 0.762654 | 0.758202 | 0.75337 | 0.754468 | 0.755501 |
| 0.758589 | 0.758635 | 0.763176 | 0.75499 | 0.763473 | 0.760432 | 0.759169 | 0.757202 | 0.759196 |
| 0.759246 | 0.758968 | 0.762118 | 0.75666 | 0.764269 | 0.762531 | 0.761836 | 0.75937 | 0.757833 |
| 0.763447 | 0.765772 | 0.763362 | 0.758192 | 0.766408 | 0.765186 | 0.757972 | 0.756636 | 0.764964 |
| 0.763767 | 0.771808 | 0.767493 | 0.75993 | 0.767242 | 0.768864 | 0.763269 | 0.762294 | 0.762068 |
| 0.766126 | 0.767895 | 0.762972 | 0.757543 | 0.768324 | 0.766361 | 0.758279 | 0.760801 | 0.763903 |
| 0.768123 | 0.774538 | 0.770237 | 0.75999 | 0.772503 | 0.771726 | 0.765954 | 0.766237 | 0.768276 |
| 0.765696 | 0.770333 | 0.77126 | 0.756241 | 0.768577 | 0.765589 | 0.764333 | 0.765538 | 0.764276 |
| 0.758216 | 0.761022 | 0.759632 | 0.753575 | 0.761365 | 0.759544 | 0.757506 | 0.760836 | 0.759603 |
| 0.760321 | 0.769844 | 0.763911 | 0.754768 | 0.768946 | 0.766251 | 0.758627 | 0.760082 | 0.758227 |
| 0.76059 | 0.765207 | 0.766091 | 0.758355 | 0.765686 | 0.762758 | 0.759626 | 0.76366 | 0.759299 |
| 0.760671 | 0.762567 | 0.766687 | 0.753826 | 0.764686 | 0.764823 | 0.756895 | 0.762144 | 0.75955 |
| 0.751531 | 0.755888 | 0.752736 | 0.752047 | 0.758088 | 0.758796 | 0.750792 | 0.751582 | 0.751546 |
| 0.752944 | 0.761259 | 0.760803 | 0.752805 | 0.75761 | 0.756935 | 0.750878 | 0.751753 | 0.756798 |
| 0.750776 | 0.755274 | 0.75625 | 0.74938 | 0.759024 | 0.756163 | 0.749066 | 0.748291 | 0.750265 |
| 0.748512 | 0.758839 | 0.759971 | 0.749496 | 0.755839 | 0.753864 | 0.751943 | 0.751691 | 0.749344 |
| 0.751233 | 0.75643 | 0.752942 | 0.747381 | 0.752709 | 0.750809 | 0.744102 | 0.746931 | 0.751007 |
| 0.752395 | 0.755292 | 0.753683 | 0.748204 | 0.755429 | 0.755005 | 0.746716 | 0.747407 | 0.749185 |
| 0.759334 | 0.761037 | 0.76359 | 0.755436 | 0.761517 | 0.763568 | 0.752844 | 0.755084 | 0.757396 |
| 0.753719 | 0.757981 | 0.750072 | 0.749359 | 0.755131 | 0.754887 | 0.747177 | 0.749569 | 0.752301 |

| | | | | | | | | |
|---|---|---|---|---|---|---|---|---|
| 0.756493 | 0.767375 | 0.765183 | 0.757873 | 0.766841 | 0.76567 | 0.761472 | 0.761328 | 0.757933 |
| 0.767922 | 0.778147 | 0.772014 | 0.769777 | 0.773882 | 0.77652 | 0.764796 | 0.771927 | 0.769969 |
| 0.768123 | 0.767011 | 0.769706 | 0.760061 | 0.769838 | 0.767469 | 0.764242 | 0.762888 | 0.768017 |
| 0.766659 | 0.775177 | 0.777374 | 0.76072 | 0.769186 | 0.768216 | 0.769393 | 0.76381 | 0.765385 |
| 0.771956 | 0.778661 | 0.784187 | 0.769064 | 0.777307 | 0.780355 | 0.774509 | 0.776882 | 0.774838 |
| 0.76821 | 0.775359 | 0.777618 | 0.768431 | 0.770532 | 0.773637 | 0.769042 | 0.768732 | 0.772166 |
| 0.767648 | 0.777474 | 0.776836 | 0.764527 | 0.768595 | 0.773202 | 0.769638 | 0.766981 | 0.761761 |
| 0.767493 | 0.773553 | 0.769341 | 0.763291 | 0.765665 | 0.77339 | 0.764508 | 0.766434 | 0.766392 |
| 0.760947 | 0.774057 | 0.770264 | 0.763271 | 0.764972 | 0.764236 | 0.758732 | 0.766673 | 0.76356 |
| 0.769319 | 0.768413 | 0.771463 | 0.760023 | 0.771937 | 0.764239 | 0.765804 | 0.766613 | 0.762328 |
| 0.769505 | 0.772459 | 0.775591 | 0.769301 | 0.767648 | 0.773947 | 0.766692 | 0.766034 | 0.765892 |
| 0.76629 | 0.770723 | 0.769653 | 0.767688 | 0.775245 | 0.772895 | 0.763817 | 0.765261 | 0.767492 |
| 0.778351 | 0.776471 | 0.774125 | 0.767897 | 0.774536 | 0.776009 | 0.773654 | 0.773962 | 0.775831 |
| 0.773343 | 0.775699 | 0.774922 | 0.76756 | 0.771247 | 0.77253 | 0.770098 | 0.774196 | 0.77185 |
| 0.774141 | 0.783667 | 0.778358 | 0.773554 | 0.776751 | 0.78148 | 0.774996 | 0.774577 | 0.769507 |
| 0.777669 | 0.77836 | 0.773812 | 0.770317 | 0.775236 | 0.773289 | 0.771848 | 0.776619 | 0.773574 |
| 0.775725 | 0.782422 | 0.779894 | 0.773036 | 0.780486 | 0.777308 | 0.779099 | 0.771984 | 0.774713 |
| 0.779272 | 0.775109 | 0.776129 | 0.766776 | 0.780878 | 0.773293 | 0.773345 | 0.774183 | 0.770539 |
| 0.771064 | 0.782541 | 0.775411 | 0.771153 | 0.777861 | 0.776795 | 0.776032 | 0.777024 | 0.771149 |
| 0.771823 | 0.783995 | 0.775335 | 0.769633 | 0.777393 | 0.777433 | 0.775617 | 0.775517 | 0.770765 |
| 0.769174 | 0.778455 | 0.771712 | 0.764627 | 0.772194 | 0.772937 | 0.765835 | 0.770579 | 0.767922 |
| 0.775436 | 0.779052 | 0.770087 | 0.768785 | 0.770153 | 0.7741 | 0.769913 | 0.769774 | 0.771292 |
| 0.770793 | 0.774896 | 0.778405 | 0.766197 | 0.771405 | 0.77433 | 0.770897 | 0.770668 | 0.773149 |
| 0.782712 | 0.790635 | 0.798284 | 0.812886 | 0.798668 | 0.796557 | 0.797377 | 0.825194 | 0.779849 |
| 0.780624 | 0.788957 | 0.786677 | 0.789702 | 0.790283 | 0.818875 | 0.77583 | 0.781969 | 0.786488 |
| 0.786037 | 0.783607 | 0.780956 | 0.809625 | 0.785101 | 0.791488 | 0.775843 | 0.788591 | 0.781626 |
| 0.783191 | 0.783261 | 0.784752 | 0.779111 | 0.790698 | 0.785562 | 0.78347 | 0.78763 | 0.781317 |
| 1 | 0.775037 | 0.778958 | 0.781856 | 0.783855 | 0.782944 | 0.778698 | 0.780352 | 0.81657 |
| 0.775037 | 1 | 0.787984 | 0.792806 | 0.802253 | 0.797722 | 0.782255 | 0.790932 | 0.783791 |
| 0.778958 | 0.787984 | 1 | 0.785229 | 0.792078 | 0.788927 | 0.789723 | 0.79294 | 0.782083 |
| 0.781856 | 0.792806 | 0.785229 | 1 | 0.800494 | 0.862651 | 0.779522 | 0.816859 | 0.781044 |
| 0.783855 | 0.802253 | 0.792078 | 0.800494 | 1 | 0.801979 | 0.777074 | 0.800933 | 0.782974 |
| 0.782944 | 0.797722 | 0.788927 | 0.862651 | 0.801979 | 1 | 0.779173 | 0.789411 | 0.792087 |
| 0.778698 | 0.782255 | 0.789723 | 0.779522 | 0.777074 | 0.779173 | 1 | 0.793711 | 0.783149 |
| 0.780352 | 0.790932 | 0.79294 | 0.816859 | 0.800933 | 0.789411 | 0.793711 | 1 | 0.777778 |
| 0.81657 | 0.783791 | 0.782083 | 0.781044 | 0.782974 | 0.792087 | 0.783149 | 0.777778 | 1 |
| 0.779488 | 0.787824 | 0.7805 | 0.785403 | 0.782363 | 0.788476 | 0.781841 | 0.78863 | 0.780939 |
| 0.772822 | 0.780645 | 0.781724 | 0.776037 | 0.781777 | 0.779388 | 0.776509 | 0.778295 | 0.782544 |
| 0.780176 | 0.779618 | 0.779 | 0.774956 | 0.781273 | 0.783039 | 0.776234 | 0.775604 | 0.774773 |
| 0.769634 | 0.784989 | 0.779549 | 0.768956 | 0.778484 | 0.778944 | 0.776652 | 0.780847 | 0.77427 |
| 0.775455 | 0.788211 | 0.781673 | 0.773238 | 0.780504 | 0.779859 | 0.775455 | 0.783452 | 0.774944 |
| 0.779707 | 0.784854 | 0.785052 | 0.780035 | 0.784186 | 0.78552 | 0.780124 | 0.780751 | 0.779927 |
| 0.783702 | 0.785551 | 0.790103 | 0.778611 | 0.785466 | 0.787054 | 0.781747 | 0.77818 | 0.784866 |
| 0.777128 | 0.780217 | 0.779662 | 0.772627 | 0.778323 | 0.78016 | 0.78046 | 0.776293 | 0.776286 |
| 0.779756 | 0.789805 | 0.783622 | 0.78204 | 0.782921 | 0.783095 | 0.779949 | 0.783018 | 0.783243 |
| 0.774075 | 0.78108 | 0.781315 | 0.778383 | 0.778125 | 0.78208 | 0.777691 | 0.778045 | 0.779667 |

| | | | | | | | | |
|---|---|---|---|---|---|---|---|---|
| 0.761479 | 0.766525 | 0.766337 | 0.758012 | 0.76024 | 0.76625 | 0.762788 | 0.760008 | 0.758833 |
| 0.75356 | 0.754314 | 0.757391 | 0.748126 | 0.754534 | 0.754871 | 0.748623 | 0.752598 | 0.749096 |
| 0.764087 | 0.77479 | 0.77513 | 0.770145 | 0.770589 | 0.77171 | 0.771085 | 0.772755 | 0.774954 |
| 0.769611 | 0.775697 | 0.77385 | 0.767592 | 0.77507 | 0.774152 | 0.772835 | 0.771695 | 0.771318 |
| 0.762078 | 0.768461 | 0.770921 | 0.763433 | 0.76463 | 0.768054 | 0.764102 | 0.766807 | 0.767164 |
| 0.770837 | 0.77467 | 0.775491 | 0.766104 | 0.774816 | 0.771466 | 0.769474 | 0.771443 | 0.76749 |
| 0.77088 | 0.775507 | 0.771613 | 0.770238 | 0.772026 | 0.775939 | 0.771638 | 0.773592 | 0.766368 |
| 0.762871 | 0.777335 | 0.771434 | 0.765786 | 0.769053 | 0.772582 | 0.769399 | 0.774618 | 0.763306 |
| 0.773215 | 0.775788 | 0.774101 | 0.768886 | 0.77272 | 0.773799 | 0.768938 | 0.768507 | 0.772547 |
| 0.7678 | 0.774252 | 0.778864 | 0.766034 | 0.7738 | 0.776095 | 0.766367 | 0.773628 | 0.770315 |
| 0.770997 | 0.77738 | 0.777862 | 0.767469 | 0.776411 | 0.778413 | 0.772209 | 0.775696 | 0.767906 |
| 0.770246 | 0.77957 | 0.77511 | 0.765536 | 0.769716 | 0.770647 | 0.767205 | 0.77243 | 0.767424 |
| 0.775622 | 0.778277 | 0.779161 | 0.769948 | 0.776562 | 0.775045 | 0.775115 | 0.775583 | 0.775648 |
| 0.772934 | 0.777669 | 0.779365 | 0.767342 | 0.772761 | 0.773205 | 0.768343 | 0.773844 | 0.765325 |
| 0.766175 | 0.77306 | 0.774335 | 0.763033 | 0.766641 | 0.76968 | 0.769502 | 0.769746 | 0.767174 |
| 0.770145 | 0.772837 | 0.772276 | 0.763711 | 0.771193 | 0.773982 | 0.766956 | 0.768033 | 0.764501 |
| 0.767029 | 0.775787 | 0.770798 | 0.765705 | 0.765982 | 0.775871 | 0.769683 | 0.76957 | 0.766066 |
| 0.766781 | 0.771126 | 0.770945 | 0.764814 | 0.764009 | 0.772948 | 0.764055 | 0.770816 | 0.763955 |
| 0.761172 | 0.767287 | 0.773922 | 0.761785 | 0.765284 | 0.771362 | 0.765605 | 0.764523 | 0.763769 |
| 0.766531 | 0.773112 | 0.776528 | 0.765038 | 0.767133 | 0.77191 | 0.772153 | 0.768716 | 0.772038 |
| 0.767172 | 0.775728 | 0.773566 | 0.764612 | 0.767617 | 0.768505 | 0.772217 | 0.771433 | 0.766368 |
| 0.764142 | 0.760154 | 0.762161 | 0.755873 | 0.763343 | 0.767812 | 0.751388 | 0.761162 | 0.757008 |
| 0.763344 | 0.768904 | 0.768438 | 0.761211 | 0.768087 | 0.767236 | 0.760912 | 0.767747 | 0.7637 |
| 0.773146 | 0.771405 | 0.770789 | 0.760637 | 0.771704 | 0.767067 | 0.76689 | 0.769961 | 0.769497 |
| 0.770692 | 0.77343 | 0.773639 | 0.766474 | 0.771547 | 0.779634 | 0.768471 | 0.768375 | 0.772263 |
| 0.769671 | 0.77083 | 0.769594 | 0.76382 | 0.77134 | 0.773588 | 0.766466 | 0.771173 | 0.768928 |
| 0.764174 | 0.766622 | 0.766286 | 0.761307 | 0.763695 | 0.769185 | 0.757304 | 0.76369 | 0.763351 |
| 0.76835 | 0.767665 | 0.772002 | 0.768431 | 0.768742 | 0.772875 | 0.757507 | 0.762589 | 0.770984 |
| 0.761929 | 0.770368 | 0.766776 | 0.765624 | 0.769258 | 0.769846 | 0.763097 | 0.767485 | 0.769607 |
| 0.693686 | 0.702772 | 0.6964 | 0.697735 | 0.702313 | 0.700688 | 0.693855 | 0.696811 | 0.696118 |
| 0.706636 | 0.708705 | 0.708959 | 0.70469 | 0.710722 | 0.70837 | 0.704814 | 0.707734 | 0.705467 |
| 0.701896 | 0.708492 | 0.708646 | 0.709012 | 0.709914 | 0.712761 | 0.70385 | 0.704837 | 0.699891 |
| 0.694202 | 0.698812 | 0.703172 | 0.697148 | 0.70147 | 0.698895 | 0.693176 | 0.696981 | 0.692329 |
| 0.708791 | 0.711336 | 0.712389 | 0.706628 | 0.714799 | 0.710087 | 0.706884 | 0.708147 | 0.705756 |
| 0.765063 | 0.77204 | 0.768137 | 0.765394 | 0.765263 | 0.76867 | 0.760254 | 0.767826 | 0.762494 |
| 0.76825 | 0.772249 | 0.771841 | 0.769433 | 0.772632 | 0.7717 | 0.761334 | 0.774432 | 0.768848 |
| 0.772799 | 0.775168 | 0.77405 | 0.769244 | 0.772739 | 0.776752 | 0.77209 | 0.776164 | 0.769271 |
| 0.772306 | 0.775396 | 0.771346 | 0.767607 | 0.773928 | 0.775968 | 0.764991 | 0.769329 | 0.767868 |
| 0.764725 | 0.772667 | 0.76674 | 0.763441 | 0.768788 | 0.767319 | 0.762972 | 0.768273 | 0.766302 |
| 0.768272 | 0.773732 | 0.772293 | 0.768701 | 0.772329 | 0.776886 | 0.764561 | 0.772588 | 0.765102 |
| 0.770925 | 0.772264 | 0.773839 | 0.766551 | 0.774747 | 0.775684 | 0.764499 | 0.769297 | 0.770003 |
| 0.76792 | 0.770522 | 0.769099 | 0.766154 | 0.768628 | 0.772727 | 0.761799 | 0.766303 | 0.764783 |
| 0.763367 | 0.766897 | 0.767757 | 0.764085 | 0.760313 | 0.767928 | 0.76061 | 0.763542 | 0.76348 |
| 0.762733 | 0.768322 | 0.765799 | 0.762533 | 0.763902 | 0.768161 | 0.755535 | 0.764061 | 0.762567 |
| 0.765387 | 0.77076 | 0.770379 | 0.762858 | 0.767473 | 0.769989 | 0.763064 | 0.763276 | 0.765895 |
| 0.771061 | 0.774392 | 0.773991 | 0.767422 | 0.774129 | 0.77185 | 0.763986 | 0.770174 | 0.767383 |

| | | | | | | | | |
|---|---|---|---|---|---|---|---|---|
| 0.772309 | 0.772868 | 0.777749 | 0.764279 | 0.772414 | 0.769366 | 0.765956 | 0.77311 | 0.772623 |
| 0.767009 | 0.771189 | 0.770769 | 0.768985 | 0.771927 | 0.770538 | 0.763725 | 0.769453 | 0.764091 |
| 0.769275 | 0.772269 | 0.776263 | 0.762729 | 0.770558 | 0.773433 | 0.76557 | 0.769862 | 0.771153 |
| 0.765154 | 0.771727 | 0.765806 | 0.762466 | 0.762604 | 0.768306 | 0.761294 | 0.766194 | 0.764309 |
| 0.766725 | 0.766228 | 0.768885 | 0.759976 | 0.767535 | 0.769948 | 0.761698 | 0.766996 | 0.761804 |
| 0.766994 | 0.763943 | 0.769881 | 0.762065 | 0.767667 | 0.767675 | 0.760276 | 0.765651 | 0.767833 |
| 0.766146 | 0.775017 | 0.768218 | 0.759656 | 0.776868 | 0.767522 | 0.764251 | 0.768831 | 0.769142 |
| 0.768818 | 0.774526 | 0.76969 | 0.761036 | 0.766971 | 0.770267 | 0.766713 | 0.766574 | 0.766717 |
| 0.771369 | 0.770926 | 0.768812 | 0.765098 | 0.766892 | 0.769826 | 0.754546 | 0.765493 | 0.767302 |
| 0.768869 | 0.765452 | 0.768173 | 0.763473 | 0.764947 | 0.768715 | 0.764203 | 0.768185 | 0.764977 |
| 0.763651 | 0.76633 | 0.771721 | 0.761042 | 0.76651 | 0.771333 | 0.764691 | 0.76695 | 0.764737 |
| 0.764559 | 0.764349 | 0.768496 | 0.761123 | 0.766689 | 0.769534 | 0.761227 | 0.766069 | 0.767244 |
| 0.761976 | 0.762157 | 0.765926 | 0.753809 | 0.764158 | 0.765632 | 0.760649 | 0.761463 | 0.763688 |
| 0.756876 | 0.755868 | 0.764249 | 0.754928 | 0.765377 | 0.762931 | 0.760048 | 0.755019 | 0.761167 |
| 0.766633 | 0.769077 | 0.775128 | 0.763511 | 0.771784 | 0.772601 | 0.763394 | 0.769982 | 0.768574 |
| 0.777074 | 0.776536 | 0.780952 | 0.774744 | 0.780278 | 0.779228 | 0.775895 | 0.781416 | 0.77594 |
| 0.776659 | 0.773092 | 0.778888 | 0.77211 | 0.774292 | 0.776322 | 0.773949 | 0.778461 | 0.774443 |
| 0.770867 | 0.776239 | 0.779241 | 0.771971 | 0.774537 | 0.78173 | 0.771779 | 0.777759 | 0.77378 |
| 0.766774 | 0.772391 | 0.768465 | 0.767116 | 0.774961 | 0.773588 | 0.768886 | 0.774233 | 0.771551 |
| 0.76592 | 0.772658 | 0.77293 | 0.761974 | 0.773621 | 0.771606 | 0.768712 | 0.768315 | 0.763767 |
| 0.763522 | 0.769492 | 0.770001 | 0.758922 | 0.769847 | 0.767577 | 0.767694 | 0.764355 | 0.765394 |
| 0.769891 | 0.774808 | 0.769481 | 0.769231 | 0.770544 | 0.7735 | 0.766687 | 0.771504 | 0.769956 |
| 0.775562 | 0.776777 | 0.778585 | 0.771722 | 0.778712 | 0.776662 | 0.776163 | 0.780197 | 0.775304 |
| 0.764155 | 0.765292 | 0.770106 | 0.758293 | 0.767576 | 0.768404 | 0.758941 | 0.768279 | 0.764771 |
| 0.765776 | 0.769935 | 0.779319 | 0.766971 | 0.777211 | 0.774211 | 0.774199 | 0.777626 | 0.770375 |
| 0.700295 | 0.706122 | 0.708716 | 0.704202 | 0.709882 | 0.707076 | 0.699884 | 0.704398 | 0.702337 |
| 0.777361 | 0.784622 | 0.784109 | 0.775459 | 0.779402 | 0.77982 | 0.779868 | 0.782022 | 0.780572 |
| 0.777406 | 0.78366 | 0.786606 | 0.774094 | 0.782421 | 0.780868 | 0.77843 | 0.780151 | 0.781773 |
| 0.783297 | 0.783073 | 0.78688 | 0.780467 | 0.781229 | 0.784947 | 0.781341 | 0.785451 | 0.781499 |
| 0.779977 | 0.780411 | 0.777136 | 0.772888 | 0.783248 | 0.781939 | 0.773081 | 0.786326 | 0.786869 |
| 0.777024 | 0.778897 | 0.777581 | 0.776385 | 0.780073 | 0.781077 | 0.771984 | 0.780076 | 0.778732 |
| 0.771474 | 0.782057 | 0.776854 | 0.778106 | 0.77395 | 0.775441 | 0.77522 | 0.779181 | 0.780318 |
| 0.775318 | 0.779509 | 0.780184 | 0.773483 | 0.776347 | 0.778189 | 0.775856 | 0.784233 | 0.776809 |
| 0.77665 | 0.781733 | 0.779209 | 0.777661 | 0.783137 | 0.784605 | 0.774521 | 0.777659 | 0.784041 |
| 0.765882 | 0.775125 | 0.777559 | 0.768579 | 0.777143 | 0.774808 | 0.768084 | 0.771426 | 0.772901 |
| 0.770992 | 0.777451 | 0.781769 | 0.771421 | 0.77956 | 0.774183 | 0.774401 | 0.775411 | 0.770182 |
| 0.779609 | 0.789571 | 0.786659 | 0.77867 | 0.785186 | 0.787062 | 0.780437 | 0.784121 | 0.781785 |
| 0.777383 | 0.785958 | 0.77906 | 0.775044 | 0.772451 | 0.779206 | 0.775831 | 0.776466 | 0.779017 |
| 0.784928 | 0.782182 | 0.77896 | 0.773913 | 0.782426 | 0.780157 | 0.782549 | 0.772603 | 0.777933 |
| 0.770823 | 0.776391 | 0.772315 | 0.770585 | 0.774448 | 0.775642 | 0.771618 | 0.772234 | 0.772226 |
| 0.77071 | 0.776512 | 0.77089 | 0.765104 | 0.773242 | 0.77426 | 0.77319 | 0.772499 | 0.767224 |
| 0.777026 | 0.783969 | 0.778354 | 0.771822 | 0.780945 | 0.784648 | 0.776238 | 0.773622 | 0.774903 |
| 0.774812 | 0.776446 | 0.776926 | 0.767234 | 0.776853 | 0.774618 | 0.771998 | 0.769637 | 0.777548 |
| 0.765308 | 0.769394 | 0.768519 | 0.764031 | 0.769814 | 0.770136 | 0.760415 | 0.760363 | 0.762729 |
| 0.767299 | 0.76988 | 0.775644 | 0.766496 | 0.765707 | 0.773054 | 0.767739 | 0.769878 | 0.771627 |
| 0.772079 | 0.768534 | 0.7727 | 0.765941 | 0.764745 | 0.772253 | 0.763957 | 0.76674 | 0.773377 |

| | | | | | | | | |
|---|---|---|---|---|---|---|---|---|
| 0.766829 | 0.771605 | 0.770053 | 0.761842 | 0.767744 | 0.771354 | 0.762154 | 0.763867 | 0.761019 |
| 0.769412 | 0.771043 | 0.767387 | 0.765998 | 0.769884 | 0.776084 | 0.768607 | 0.770809 | 0.77495 |
| 0.765891 | 0.768222 | 0.768857 | 0.760769 | 0.77037 | 0.766334 | 0.763284 | 0.76858 | 0.766206 |
| 0.723132 | 0.731439 | 0.729866 | 0.724014 | 0.722202 | 0.727796 | 0.734392 | 0.744355 | 0.736096 |
| 0.765006 | 0.769162 | 0.770274 | 0.76515 | 0.765044 | 0.769908 | 0.764137 | 0.768764 | 0.762243 |
| 0.766138 | 0.772321 | 0.771662 | 0.761778 | 0.766109 | 0.771058 | 0.763649 | 0.76687 | 0.770608 |
| 0.762799 | 0.765987 | 0.7699 | 0.767403 | 0.767394 | 0.769017 | 0.76316 | 0.767601 | 0.76805 |
| 0.770545 | 0.771828 | 0.766852 | 0.766237 | 0.767235 | 0.771405 | 0.765715 | 0.769231 | 0.76721 |
| 0.768318 | 0.774115 | 0.774897 | 0.770186 | 0.770532 | 0.774133 | 0.768445 | 0.773564 | 0.771966 |
| 0.770815 | 0.774489 | 0.774366 | 0.770087 | 0.773193 | 0.771871 | 0.765099 | 0.772083 | 0.769311 |
| 0.764365 | 0.770531 | 0.778648 | 0.765892 | 0.76763 | 0.770666 | 0.765556 | 0.768951 | 0.765285 |
| 0.762375 | 0.773364 | 0.778308 | 0.769089 | 0.769467 | 0.771626 | 0.770386 | 0.771657 | 0.77177 |
| 0.765418 | 0.771617 | 0.77675 | 0.766547 | 0.770987 | 0.774262 | 0.763361 | 0.767829 | 0.76998 |
| 0.773733 | 0.771947 | 0.774032 | 0.76475 | 0.771207 | 0.772419 | 0.767874 | 0.773158 | 0.770297 |
| 0.767701 | 0.774431 | 0.767597 | 0.760859 | 0.772994 | 0.771096 | 0.7657 | 0.769601 | 0.770352 |
| 0.764648 | 0.771312 | 0.766523 | 0.76372 | 0.764643 | 0.77084 | 0.761131 | 0.766173 | 0.762769 |
| 0.768808 | 0.773018 | 0.774573 | 0.759822 | 0.771403 | 0.773611 | 0.765322 | 0.768798 | 0.772268 |
| 0.769018 | 0.774656 | 0.774106 | 0.764079 | 0.770495 | 0.770959 | 0.767239 | 0.769634 | 0.771268 |
| 0.762519 | 0.764859 | 0.767738 | 0.759854 | 0.763582 | 0.766781 | 0.763102 | 0.763497 | 0.763815 |
| 0.764601 | 0.776041 | 0.768504 | 0.763848 | 0.76925 | 0.773356 | 0.763853 | 0.764721 | 0.767224 |
| 0.771485 | 0.775906 | 0.774286 | 0.764549 | 0.772648 | 0.769187 | 0.764686 | 0.770943 | 0.769476 |
| 0.772445 | 0.774299 | 0.772773 | 0.766739 | 0.774041 | 0.775825 | 0.768501 | 0.77272 | 0.76841 |
| 0.684048 | 0.683972 | 0.694347 | 0.685908 | 0.681645 | 0.683692 | 0.682487 | 0.682291 | 0.684939 |
| 0.652205 | 0.6547 | 0.66867 | 0.656677 | 0.659458 | 0.653018 | 0.66283 | 0.662849 | 0.653491 |
| 0.67231 | 0.671811 | 0.674622 | 0.673198 | 0.671997 | 0.675535 | 0.677899 | 0.67376 | 0.670187 |
| 0.652343 | 0.653633 | 0.659402 | 0.657677 | 0.65811 | 0.656858 | 0.649427 | 0.660265 | 0.656385 |
| 0.741762 | 0.740275 | 0.740033 | 0.734725 | 0.741304 | 0.745902 | 0.740684 | 0.744742 | 0.738117 |
| 0.726478 | 0.728642 | 0.727835 | 0.718624 | 0.725163 | 0.733244 | 0.727518 | 0.733359 | 0.725897 |
| 0.731223 | 0.732793 | 0.743087 | 0.728545 | 0.735914 | 0.733611 | 0.731305 | 0.733985 | 0.728618 |
| 0.738503 | 0.749062 | 0.747889 | 0.732997 | 0.742212 | 0.742307 | 0.740435 | 0.745073 | 0.74296 |
| 0.721014 | 0.71969 | 0.730636 | 0.718226 | 0.714806 | 0.724494 | 0.720968 | 0.73304 | 0.72778 |
| 0.708626 | 0.716573 | 0.724525 | 0.709586 | 0.721269 | 0.715789 | 0.720584 | 0.718937 | 0.711604 |
| 0.722451 | 0.710896 | 0.729204 | 0.710029 | 0.713949 | 0.719158 | 0.719104 | 0.715896 | 0.713989 |
| 0.654953 | 0.660515 | 0.664332 | 0.655958 | 0.659936 | 0.660613 | 0.659548 | 0.664336 | 0.662403 |
| 0.763512 | 0.767147 | 0.765933 | 0.755847 | 0.764471 | 0.766345 | 0.763545 | 0.762831 | 0.764669 |
| 0.74457 | 0.745467 | 0.751808 | 0.736626 | 0.74122 | 0.746093 | 0.744117 | 0.74435 | 0.746036 |
| 0.723798 | 0.737835 | 0.741173 | 0.736936 | 0.740766 | 0.742194 | 0.739386 | 0.744416 | 0.734452 |
| 0.731784 | 0.719993 | 0.72306 | 0.724883 | 0.730381 | 0.724465 | 0.724028 | 0.727765 | 0.728249 |
| 0.740248 | 0.748306 | 0.746626 | 0.742171 | 0.75227 | 0.747009 | 0.742844 | 0.749466 | 0.752161 |
| 0.737178 | 0.735313 | 0.735565 | 0.72938 | 0.738368 | 0.732203 | 0.728133 | 0.729674 | 0.736849 |
| 0.734442 | 0.732242 | 0.738633 | 0.729639 | 0.734335 | 0.73408 | 0.73199 | 0.736944 | 0.735366 |
| 0.729684 | 0.734424 | 0.7317 | 0.722561 | 0.727798 | 0.731493 | 0.725971 | 0.731108 | 0.729219 |
| 0.724181 | 0.734844 | 0.731925 | 0.730692 | 0.730602 | 0.732211 | 0.724932 | 0.728048 | 0.737199 |
| 0.723375 | 0.733557 | 0.736327 | 0.727038 | 0.733728 | 0.737773 | 0.731809 | 0.736257 | 0.726748 |
| 0.721532 | 0.726224 | 0.732453 | 0.724023 | 0.726197 | 0.730233 | 0.722752 | 0.7307 | 0.730381 |
| 0.702826 | 0.707597 | 0.725249 | 0.709464 | 0.719973 | 0.717219 | 0.721909 | 0.720544 | 0.718469 |

| | | | | | | | | |
|---|---|---|---|---|---|---|---|---|
| 0.748107 | 0.762193 | 0.762127 | 0.750701 | 0.753487 | 0.757612 | 0.755663 | 0.755351 | 0.751834 |
| 0.735348 | 0.739707 | 0.740455 | 0.733646 | 0.733841 | 0.740457 | 0.731902 | 0.740071 | 0.738054 |
| 0.726719 | 0.728019 | 0.731343 | 0.725585 | 0.726591 | 0.73085 | 0.734091 | 0.730101 | 0.734351 |
| 0.726624 | 0.737575 | 0.748199 | 0.726448 | 0.726161 | 0.732853 | 0.732623 | 0.737021 | 0.731164 |
| 0.724235 | 0.737983 | 0.740037 | 0.730422 | 0.725228 | 0.737316 | 0.7331 | 0.74023 | 0.730469 |
| 0.735228 | 0.731354 | 0.740329 | 0.725101 | 0.725223 | 0.725378 | 0.724888 | 0.733683 | 0.744221 |
| 0.72301 | 0.725724 | 0.737102 | 0.719581 | 0.719912 | 0.728867 | 0.725514 | 0.732222 | 0.726675 |
| 0.722222 | 0.734112 | 0.731693 | 0.714813 | 0.717637 | 0.722854 | 0.721541 | 0.728199 | 0.727376 |
| 0.716505 | 0.723529 | 0.736345 | 0.726059 | 0.723332 | 0.730769 | 0.727456 | 0.72843 | 0.729923 |
| 0.756062 | 0.759871 | 0.763456 | 0.750074 | 0.755377 | 0.758117 | 0.755773 | 0.756565 | 0.757849 |
| 0.747279 | 0.754389 | 0.757134 | 0.746175 | 0.754629 | 0.757228 | 0.749718 | 0.748775 | 0.751575 |
| 0.744007 | 0.751656 | 0.752678 | 0.741447 | 0.749917 | 0.752689 | 0.748959 | 0.752727 | 0.745216 |
| 0.743376 | 0.739487 | 0.74766 | 0.736617 | 0.739896 | 0.744254 | 0.738971 | 0.747505 | 0.742315 |
| 0.741638 | 0.747297 | 0.751772 | 0.739467 | 0.746874 | 0.745579 | 0.737063 | 0.744212 | 0.745191 |
| 0.741438 | 0.742133 | 0.752355 | 0.741919 | 0.742744 | 0.748205 | 0.737944 | 0.745088 | 0.746566 |

| | | | | | plot 07 | | | |
|---|---|---|---|---|---|---|---|---|
| Wild_plot_ | Wild_plot_ | Wild_plot_ | Wild_plot_ | Wild_plot_ | Wild_plot_ | Wild_plot_ | Wild_plot_ | Wild_plot_ |
| 0.714728 | 0.706216 | 0.709175 | 0.709453 | 0.713336 | 0.712435 | 0.711059 | 0.710028 | 0.709994 |
| 0.72137 | 0.716178 | 0.715665 | 0.714867 | 0.719276 | 0.721987 | 0.722081 | 0.713137 | 0.714492 |
| 0.709976 | 0.701807 | 0.701982 | 0.706205 | 0.710096 | 0.704829 | 0.707905 | 0.707667 | 0.708824 |
| 0.722165 | 0.713444 | 0.718261 | 0.713743 | 0.720534 | 0.716469 | 0.716813 | 0.711602 | 0.714297 |
| 0.712432 | 0.706816 | 0.706221 | 0.710063 | 0.713948 | 0.709688 | 0.711394 | 0.70787 | 0.711871 |
| 0.725314 | 0.718364 | 0.720149 | 0.714931 | 0.719779 | 0.718785 | 0.71863 | 0.712867 | 0.717922 |
| 0.78585 | 0.777183 | 0.769704 | 0.778383 | 0.779223 | 0.777183 | 0.776806 | 0.769745 | 0.776303 |
| 0.77958 | 0.770195 | 0.768529 | 0.768562 | 0.76815 | 0.771074 | 0.774523 | 0.773482 | 0.776666 |
| 0.722435 | 0.710906 | 0.714743 | 0.712906 | 0.716043 | 0.717081 | 0.71909 | 0.711848 | 0.715825 |
| 0.716542 | 0.708179 | 0.708489 | 0.708431 | 0.713755 | 0.709922 | 0.711151 | 0.709458 | 0.707531 |
| 0.718573 | 0.709222 | 0.710808 | 0.711506 | 0.71618 | 0.714503 | 0.714988 | 0.709926 | 0.710239 |
| 0.709608 | 0.704522 | 0.707092 | 0.705141 | 0.710067 | 0.709196 | 0.708054 | 0.704599 | 0.708004 |
| 0.721318 | 0.713301 | 0.715699 | 0.712998 | 0.719657 | 0.713072 | 0.717503 | 0.70962 | 0.714882 |
| 0.729814 | 0.720013 | 0.724912 | 0.720785 | 0.725264 | 0.725884 | 0.723875 | 0.720316 | 0.723129 |
| 0.739294 | 0.736975 | 0.736369 | 0.73572 | 0.739748 | 0.739047 | 0.740223 | 0.73778 | 0.737318 |
| 0.721476 | 0.711495 | 0.714728 | 0.710582 | 0.717316 | 0.71669 | 0.717182 | 0.715456 | 0.719349 |
| 0.721229 | 0.710736 | 0.712173 | 0.714075 | 0.717154 | 0.712459 | 0.714354 | 0.71298 | 0.714055 |
| 0.751642 | 0.739874 | 0.740425 | 0.743269 | 0.749005 | 0.745333 | 0.748372 | 0.741836 | 0.746039 |
| 0.715439 | 0.704516 | 0.707971 | 0.707079 | 0.709829 | 0.714377 | 0.714252 | 0.709309 | 0.708856 |
| 0.718882 | 0.708975 | 0.714765 | 0.713484 | 0.715647 | 0.71473 | 0.716062 | 0.710046 | 0.71519 |
| 0.71373 | 0.705764 | 0.711819 | 0.708084 | 0.709161 | 0.713411 | 0.709733 | 0.705312 | 0.71184 |
| 0.716912 | 0.709649 | 0.711612 | 0.707699 | 0.715658 | 0.713614 | 0.714173 | 0.709455 | 0.710709 |
| 0.722617 | 0.716153 | 0.717227 | 0.719541 | 0.723626 | 0.719505 | 0.719234 | 0.715505 | 0.718249 |
| 0.72043 | 0.711318 | 0.713468 | 0.713032 | 0.71657 | 0.716129 | 0.715561 | 0.712884 | 0.715705 |
| 0.721929 | 0.713495 | 0.714137 | 0.715172 | 0.720373 | 0.715634 | 0.720271 | 0.712459 | 0.715654 |
| 0.716774 | 0.711095 | 0.712118 | 0.709329 | 0.714398 | 0.713451 | 0.716897 | 0.70838 | 0.711639 |
| 0.708776 | 0.704788 | 0.708476 | 0.706278 | 0.708534 | 0.710121 | 0.70923 | 0.70324 | 0.710406 |
| 0.718949 | 0.710617 | 0.712296 | 0.71275 | 0.716336 | 0.714037 | 0.714241 | 0.712246 | 0.712239 |
| 0.72091 | 0.716909 | 0.714491 | 0.71475 | 0.718499 | 0.718623 | 0.720638 | 0.711521 | 0.715477 |
| 0.718345 | 0.713599 | 0.71052 | 0.712362 | 0.713502 | 0.713295 | 0.717839 | 0.711156 | 0.711316 |
| 0.717494 | 0.70859 | 0.71106 | 0.709044 | 0.715364 | 0.71456 | 0.716701 | 0.71076 | 0.710864 |
| 0.711776 | 0.708145 | 0.705868 | 0.707134 | 0.710628 | 0.709183 | 0.710764 | 0.705721 | 0.708227 |
| 0.715633 | 0.711154 | 0.711406 | 0.710507 | 0.713395 | 0.715238 | 0.714555 | 0.709321 | 0.713498 |
| 0.718864 | 0.713156 | 0.713674 | 0.712923 | 0.715923 | 0.71704 | 0.720372 | 0.708925 | 0.712212 |
| 0.711855 | 0.708772 | 0.707416 | 0.70485 | 0.711134 | 0.710796 | 0.711545 | 0.706426 | 0.70645 |
| 0.718119 | 0.712304 | 0.714618 | 0.713982 | 0.715413 | 0.71629 | 0.714819 | 0.706083 | 0.71107 |
| 0.722323 | 0.715322 | 0.715066 | 0.713134 | 0.720183 | 0.71626 | 0.717651 | 0.711215 | 0.720457 |
| 0.719727 | 0.713105 | 0.711791 | 0.713254 | 0.719759 | 0.717746 | 0.717848 | 0.71019 | 0.714563 |
| 0.723343 | 0.718009 | 0.716628 | 0.718166 | 0.720441 | 0.720395 | 0.725014 | 0.714529 | 0.717296 |

| | | | | | | | | |
|---|---|---|---|---|---|---|---|---|
| 0.756005 | 0.747558 | 0.747774 | 0.749448 | 0.756889 | 0.7494 | 0.750279 | 0.749241 | 0.750869 |
| 0.710649 | 0.701021 | 0.704223 | 0.705682 | 0.710458 | 0.702188 | 0.703663 | 0.702447 | 0.706239 |
| 0.713876 | 0.704404 | 0.707637 | 0.704273 | 0.707853 | 0.70833 | 0.710778 | 0.706278 | 0.709087 |
| 0.722161 | 0.712801 | 0.716256 | 0.714147 | 0.718863 | 0.72047 | 0.71861 | 0.712461 | 0.714479 |
| 0.716282 | 0.7087 | 0.710439 | 0.709137 | 0.714071 | 0.712553 | 0.712745 | 0.711626 | 0.712 |
| 0.720067 | 0.713496 | 0.713817 | 0.709788 | 0.718077 | 0.715919 | 0.717181 | 0.711867 | 0.712199 |
| 0.714818 | 0.706171 | 0.710338 | 0.706267 | 0.709546 | 0.710053 | 0.711844 | 0.706224 | 0.71187 |
| 0.719414 | 0.713252 | 0.714205 | 0.711656 | 0.715847 | 0.716297 | 0.716412 | 0.71073 | 0.714988 |
| 0.721673 | 0.71333 | 0.713826 | 0.715068 | 0.720768 | 0.717173 | 0.719116 | 0.712328 | 0.713502 |
| 0.721335 | 0.715257 | 0.714184 | 0.71637 | 0.720127 | 0.717079 | 0.72112 | 0.713696 | 0.711316 |
| 0.710004 | 0.703538 | 0.703819 | 0.707567 | 0.708905 | 0.708889 | 0.706516 | 0.704898 | 0.70832 |
| 0.711298 | 0.703293 | 0.705915 | 0.706998 | 0.712945 | 0.707658 | 0.708712 | 0.707192 | 0.711566 |
| 0.715053 | 0.708938 | 0.70952 | 0.710862 | 0.714548 | 0.714859 | 0.714877 | 0.7123 | 0.711222 |
| 0.711106 | 0.705959 | 0.709525 | 0.702538 | 0.710211 | 0.711317 | 0.711113 | 0.70693 | 0.711728 |
| 0.712007 | 0.706718 | 0.706562 | 0.707385 | 0.711466 | 0.707606 | 0.705195 | 0.705469 | 0.708956 |
| 0.710812 | 0.705501 | 0.705868 | 0.705592 | 0.709374 | 0.708676 | 0.709011 | 0.703746 | 0.706975 |
| 0.70822 | 0.704451 | 0.70653 | 0.704158 | 0.70846 | 0.70729 | 0.708058 | 0.704067 | 0.705773 |
| 0.718601 | 0.710364 | 0.715948 | 0.711946 | 0.718069 | 0.716936 | 0.718202 | 0.710247 | 0.715121 |
| 0.719873 | 0.716281 | 0.713693 | 0.711968 | 0.714184 | 0.718501 | 0.71709 | 0.712802 | 0.711623 |
| 0.717384 | 0.708866 | 0.706343 | 0.707558 | 0.712384 | 0.713438 | 0.711389 | 0.710425 | 0.71134 |
| 0.710002 | 0.699733 | 0.705117 | 0.705495 | 0.708456 | 0.705457 | 0.706124 | 0.701632 | 0.705099 |
| 0.71424 | 0.704372 | 0.708659 | 0.708615 | 0.7129 | 0.709731 | 0.709963 | 0.708188 | 0.707038 |
| 0.716295 | 0.708343 | 0.708551 | 0.709542 | 0.713197 | 0.711925 | 0.712583 | 0.707833 | 0.714252 |
| 0.722382 | 0.717738 | 0.722883 | 0.719902 | 0.720854 | 0.722155 | 0.724259 | 0.715498 | 0.719452 |
| 0.732507 | 0.72487 | 0.725681 | 0.724902 | 0.73061 | 0.725503 | 0.729685 | 0.721667 | 0.724777 |
| 0.779041 | 0.770546 | 0.768652 | 0.769633 | 0.77043 | 0.771689 | 0.775699 | 0.773877 | 0.776666 |
| 0.783042 | 0.778549 | 0.777002 | 0.780231 | 0.783092 | 0.778074 | 0.779657 | 0.780339 | 0.787945 |
| 0.782926 | 0.776132 | 0.783368 | 0.785411 | 0.784981 | 0.782396 | 0.785555 | 0.77882 | 0.793815 |
| 0.715685 | 0.710266 | 0.706597 | 0.709869 | 0.71286 | 0.709251 | 0.707906 | 0.706035 | 0.709453 |
| 0.710473 | 0.705599 | 0.709431 | 0.706815 | 0.7139 | 0.709825 | 0.708628 | 0.705799 | 0.710911 |
| 0.725208 | 0.712073 | 0.715543 | 0.716706 | 0.719898 | 0.717457 | 0.717212 | 0.717613 | 0.716375 |
| 0.726101 | 0.715036 | 0.718998 | 0.717166 | 0.723701 | 0.718228 | 0.71862 | 0.714052 | 0.716862 |
| 0.715519 | 0.705541 | 0.707433 | 0.709174 | 0.714057 | 0.709561 | 0.707871 | 0.708511 | 0.711356 |
| 0.714549 | 0.708843 | 0.708958 | 0.712367 | 0.714137 | 0.715341 | 0.711751 | 0.711315 | 0.711197 |
| 0.713536 | 0.702477 | 0.710773 | 0.707455 | 0.712538 | 0.711425 | 0.710919 | 0.707709 | 0.709816 |
| 0.713442 | 0.706246 | 0.71117 | 0.709226 | 0.716975 | 0.714503 | 0.714308 | 0.708374 | 0.70918 |
| 0.717663 | 0.708447 | 0.706911 | 0.709174 | 0.715095 | 0.711176 | 0.710833 | 0.706818 | 0.7083 |
| 0.728395 | 0.721797 | 0.725062 | 0.72229 | 0.729423 | 0.72576 | 0.724847 | 0.721433 | 0.724589 |
| 0.708975 | 0.70005 | 0.701982 | 0.701676 | 0.707287 | 0.705029 | 0.703203 | 0.701328 | 0.703633 |
| 0.716022 | 0.70845 | 0.7121 | 0.710518 | 0.712067 | 0.711787 | 0.7128 | 0.711466 | 0.71115 |
| 0.712076 | 0.704212 | 0.70604 | 0.704617 | 0.709475 | 0.707736 | 0.708897 | 0.702327 | 0.702833 |
| 0.731335 | 0.726967 | 0.727024 | 0.726609 | 0.733488 | 0.727834 | 0.729704 | 0.724451 | 0.727597 |
| 0.718311 | 0.707098 | 0.711431 | 0.707619 | 0.714047 | 0.709256 | 0.7098 | 0.7076 | 0.711512 |
| 0.713936 | 0.705108 | 0.706664 | 0.706332 | 0.711655 | 0.706921 | 0.708275 | 0.708883 | 0.710964 |
| 0.717212 | 0.714938 | 0.71556 | 0.711287 | 0.718238 | 0.716529 | 0.715557 | 0.710819 | 0.708746 |
| 0.716955 | 0.707608 | 0.711093 | 0.7095 | 0.715533 | 0.713057 | 0.710363 | 0.708279 | 0.712413 |

| | | | | | | | | |
|---|---|---|---|---|---|---|---|---|
| 0.711964 | 0.706951 | 0.707841 | 0.706503 | 0.712958 | 0.705719 | 0.707793 | 0.705682 | 0.70791 |
| 0.717737 | 0.712786 | 0.713006 | 0.713845 | 0.71459 | 0.715818 | 0.71222 | 0.710903 | 0.716728 |
| 0.71625 | 0.70868 | 0.706932 | 0.709954 | 0.714069 | 0.709274 | 0.711191 | 0.709039 | 0.711091 |
| 0.723576 | 0.717553 | 0.719369 | 0.716177 | 0.72152 | 0.718939 | 0.720087 | 0.714333 | 0.719415 |
| 0.722845 | 0.712685 | 0.716545 | 0.718334 | 0.71987 | 0.717588 | 0.717698 | 0.713858 | 0.714343 |
| 0.714171 | 0.708235 | 0.707352 | 0.708996 | 0.711299 | 0.710168 | 0.710899 | 0.708773 | 0.712074 |
| 0.717469 | 0.708009 | 0.710395 | 0.712973 | 0.717189 | 0.711595 | 0.713155 | 0.711145 | 0.712659 |
| 0.72105 | 0.712907 | 0.712772 | 0.713923 | 0.718246 | 0.717299 | 0.717372 | 0.71221 | 0.716016 |
| 0.717238 | 0.708659 | 0.71258 | 0.709135 | 0.714627 | 0.715626 | 0.713196 | 0.708364 | 0.711545 |
| 0.711737 | 0.70556 | 0.706209 | 0.707773 | 0.71303 | 0.709693 | 0.712449 | 0.706505 | 0.709879 |
| 0.714714 | 0.70617 | 0.708952 | 0.70697 | 0.714589 | 0.711982 | 0.712785 | 0.706119 | 0.710412 |
| 0.717593 | 0.706926 | 0.710399 | 0.708217 | 0.715078 | 0.710698 | 0.712683 | 0.713144 | 0.710548 |
| 0.718326 | 0.711305 | 0.71087 | 0.709078 | 0.713329 | 0.719897 | 0.717367 | 0.712497 | 0.71189 |
| 0.715395 | 0.70327 | 0.706148 | 0.70716 | 0.712161 | 0.712059 | 0.714056 | 0.705689 | 0.708645 |
| 0.718867 | 0.711321 | 0.709376 | 0.711559 | 0.71382 | 0.712811 | 0.71582 | 0.71128 | 0.715115 |
| 0.713099 | 0.708122 | 0.70545 | 0.709625 | 0.714274 | 0.707879 | 0.711425 | 0.708195 | 0.710631 |
| 0.722401 | 0.714988 | 0.715341 | 0.713911 | 0.717348 | 0.717519 | 0.720317 | 0.715421 | 0.716133 |
| 0.715648 | 0.706053 | 0.709416 | 0.708842 | 0.711274 | 0.712547 | 0.710495 | 0.710015 | 0.707492 |
| 0.719082 | 0.711071 | 0.710203 | 0.711333 | 0.715309 | 0.712816 | 0.716316 | 0.705816 | 0.712196 |
| 0.717294 | 0.70945 | 0.707346 | 0.714251 | 0.718999 | 0.712998 | 0.713403 | 0.710336 | 0.715458 |
| 0.717339 | 0.712135 | 0.708533 | 0.710812 | 0.716568 | 0.713268 | 0.715002 | 0.707761 | 0.71455 |
| 0.712256 | 0.707589 | 0.705537 | 0.709193 | 0.712938 | 0.708419 | 0.710016 | 0.704869 | 0.707851 |
| 0.718818 | 0.711022 | 0.708504 | 0.710468 | 0.716376 | 0.712936 | 0.712409 | 0.707908 | 0.711002 |
| 0.718308 | 0.710356 | 0.713341 | 0.71114 | 0.712502 | 0.711659 | 0.711249 | 0.707925 | 0.710866 |
| 0.716159 | 0.710045 | 0.708186 | 0.711778 | 0.71557 | 0.711645 | 0.711072 | 0.710095 | 0.71306 |
| 0.72246 | 0.7169 | 0.712083 | 0.713558 | 0.722165 | 0.717902 | 0.715889 | 0.71816 | 0.717159 |
| 0.714998 | 0.710158 | 0.708156 | 0.71039 | 0.714241 | 0.711101 | 0.710347 | 0.713628 | 0.709818 |
| 0.721502 | 0.709711 | 0.712357 | 0.712481 | 0.717199 | 0.713664 | 0.714432 | 0.710488 | 0.7144 |
| 0.717352 | 0.710573 | 0.711019 | 0.712954 | 0.715404 | 0.715853 | 0.716452 | 0.714611 | 0.714092 |
| 0.720817 | 0.710788 | 0.710662 | 0.709975 | 0.716092 | 0.716906 | 0.716578 | 0.712144 | 0.714599 |
| 0.722249 | 0.71515 | 0.713048 | 0.716837 | 0.71829 | 0.71674 | 0.718862 | 0.709159 | 0.713536 |
| 0.715457 | 0.712132 | 0.707551 | 0.708935 | 0.7147 | 0.714049 | 0.712488 | 0.705477 | 0.71068 |
| 0.7178 | 0.709651 | 0.712417 | 0.713838 | 0.717291 | 0.714955 | 0.712859 | 0.71033 | 0.714797 |
| 0.717661 | 0.710188 | 0.71493 | 0.708486 | 0.715308 | 0.717953 | 0.715932 | 0.710051 | 0.710038 |
| 0.715052 | 0.709312 | 0.708924 | 0.70718 | 0.710145 | 0.713358 | 0.712026 | 0.707249 | 0.710448 |
| 0.717994 | 0.711356 | 0.70916 | 0.710976 | 0.714539 | 0.71432 | 0.71562 | 0.710324 | 0.710293 |
| 0.715752 | 0.705428 | 0.706581 | 0.707531 | 0.718948 | 0.709371 | 0.707113 | 0.701131 | 0.708193 |
| 0.692777 | 0.685685 | 0.683998 | 0.682737 | 0.686275 | 0.682361 | 0.686391 | 0.681027 | 0.688665 |
| 0.69105 | 0.690029 | 0.691733 | 0.691475 | 0.693717 | 0.692688 | 0.694232 | 0.689068 | 0.6918 |
| 0.689591 | 0.678997 | 0.681267 | 0.680931 | 0.68289 | 0.682112 | 0.682041 | 0.677503 | 0.685602 |
| 0.681698 | 0.682711 | 0.680929 | 0.679292 | 0.68584 | 0.685756 | 0.685496 | 0.679915 | 0.684445 |
| 0.667367 | 0.666627 | 0.667134 | 0.667058 | 0.66934 | 0.670994 | 0.66806 | 0.666275 | 0.668661 |
| 0.672631 | 0.667555 | 0.671933 | 0.669299 | 0.672756 | 0.670138 | 0.67133 | 0.664564 | 0.666425 |
| 0.672476 | 0.668979 | 0.667954 | 0.666991 | 0.674475 | 0.666425 | 0.666065 | 0.666424 | 0.673643 |
| 0.672018 | 0.672239 | 0.663894 | 0.668545 | 0.67363 | 0.668406 | 0.668944 | 0.667347 | 0.669309 |
| 0.671313 | 0.669248 | 0.666625 | 0.670839 | 0.677043 | 0.669979 | 0.665431 | 0.669758 | 0.670957 |
| 0.677304 | 0.675723 | 0.675596 | 0.674975 | 0.684431 | 0.674387 | 0.674598 | 0.677095 | 0.67607 |

| | | | | | | | | |
|---|---|---|---|---|---|---|---|---|
| 0.680841 | 0.675662 | 0.6723 | 0.671409 | 0.671188 | 0.674141 | 0.675851 | 0.667283 | 0.674071 |
| 0.675363 | 0.673132 | 0.674159 | 0.674437 | 0.683418 | 0.676941 | 0.674395 | 0.680084 | 0.676925 |
| 0.675928 | 0.675141 | 0.66844 | 0.674879 | 0.676964 | 0.672846 | 0.675611 | 0.663866 | 0.673095 |
| 0.670363 | 0.66649 | 0.663861 | 0.661811 | 0.668641 | 0.668346 | 0.669486 | 0.667197 | 0.663761 |
| 0.66769 | 0.663238 | 0.665475 | 0.66942 | 0.668364 | 0.664107 | 0.667392 | 0.664735 | 0.66916 |
| 0.672762 | 0.664231 | 0.667259 | 0.669611 | 0.669895 | 0.669002 | 0.670307 | 0.66393 | 0.671866 |
| 0.666949 | 0.671685 | 0.667841 | 0.668421 | 0.66793 | 0.669399 | 0.671519 | 0.67378 | 0.672443 |
| 0.668729 | 0.664794 | 0.663932 | 0.663448 | 0.671536 | 0.667878 | 0.666435 | 0.665214 | 0.660274 |
| 0.668437 | 0.664298 | 0.669689 | 0.663085 | 0.665356 | 0.664567 | 0.665159 | 0.658851 | 0.665734 |
| 0.672714 | 0.668087 | 0.666958 | 0.666667 | 0.67112 | 0.67254 | 0.672942 | 0.66531 | 0.667251 |
| 0.662653 | 0.660269 | 0.659098 | 0.659041 | 0.665395 | 0.660969 | 0.65977 | 0.66007 | 0.66487 |
| 0.706792 | 0.697932 | 0.700455 | 0.698539 | 0.704725 | 0.698343 | 0.702068 | 0.697689 | 0.701377 |
| 0.715279 | 0.709462 | 0.710774 | 0.711559 | 0.715928 | 0.712649 | 0.714297 | 0.707398 | 0.708066 |
| 0.689812 | 0.686821 | 0.683758 | 0.686871 | 0.691123 | 0.689459 | 0.685505 | 0.683728 | 0.690014 |
| 0.719741 | 0.713288 | 0.718991 | 0.716324 | 0.720683 | 0.71826 | 0.720986 | 0.713686 | 0.716003 |
| 0.708717 | 0.70458 | 0.702102 | 0.70654 | 0.711315 | 0.708343 | 0.706457 | 0.701092 | 0.706592 |
| 0.723585 | 0.712582 | 0.714263 | 0.711875 | 0.719434 | 0.71636 | 0.7174 | 0.713997 | 0.715265 |
| 0.726533 | 0.71999 | 0.71798 | 0.720781 | 0.72216 | 0.723149 | 0.720912 | 0.720608 | 0.724012 |
| 0.704291 | 0.697567 | 0.698526 | 0.696541 | 0.706421 | 0.700926 | 0.700436 | 0.69643 | 0.699069 |
| 0.72463 | 0.718079 | 0.718123 | 0.720023 | 0.723993 | 0.723367 | 0.72272 | 0.716407 | 0.720622 |
| 0.696196 | 0.690337 | 0.688924 | 0.690755 | 0.693423 | 0.693242 | 0.697733 | 0.692503 | 0.693685 |
| 0.722053 | 0.715367 | 0.716344 | 0.716467 | 0.719586 | 0.717842 | 0.717865 | 0.714809 | 0.721452 |
| 0.726544 | 0.717815 | 0.716528 | 0.718546 | 0.725421 | 0.721386 | 0.718958 | 0.71842 | 0.724525 |
| 0.692084 | 0.685952 | 0.684423 | 0.685874 | 0.693003 | 0.688198 | 0.692451 | 0.684358 | 0.688982 |
| 0.696069 | 0.684114 | 0.689977 | 0.68783 | 0.695144 | 0.691065 | 0.690858 | 0.687006 | 0.692583 |
| 0.694648 | 0.692532 | 0.690336 | 0.690775 | 0.698325 | 0.69435 | 0.69315 | 0.687246 | 0.695294 |
| 0.712385 | 0.706534 | 0.70616 | 0.706628 | 0.712403 | 0.710693 | 0.707773 | 0.704166 | 0.709139 |
| 0.7272 | 0.720143 | 0.718909 | 0.71824 | 0.727425 | 0.722554 | 0.723088 | 0.715373 | 0.72001 |
| 0.694773 | 0.683842 | 0.687442 | 0.68836 | 0.692415 | 0.69096 | 0.690093 | 0.686157 | 0.690759 |
| 0.731579 | 0.721076 | 0.72254 | 0.724986 | 0.73022 | 0.725939 | 0.724713 | 0.722222 | 0.723901 |
| 0.737509 | 0.72665 | 0.728044 | 0.724243 | 0.729617 | 0.732446 | 0.73391 | 0.728139 | 0.729213 |
| 0.702227 | 0.696167 | 0.702981 | 0.70021 | 0.705277 | 0.701421 | 0.700567 | 0.697058 | 0.700735 |
| 0.701897 | 0.700056 | 0.70316 | 0.698512 | 0.705632 | 0.705246 | 0.703666 | 0.696583 | 0.700445 |
| 0.728302 | 0.725014 | 0.722697 | 0.724031 | 0.730152 | 0.727994 | 0.727525 | 0.719629 | 0.727874 |
| 0.727416 | 0.72426 | 0.720333 | 0.721331 | 0.72406 | 0.726035 | 0.72701 | 0.720211 | 0.725498 |
| 0.730165 | 0.721078 | 0.726475 | 0.720796 | 0.72752 | 0.72283 | 0.72744 | 0.723906 | 0.727957 |
| 0.682469 | 0.674791 | 0.680128 | 0.679411 | 0.684174 | 0.679046 | 0.679123 | 0.672618 | 0.677409 |
| 0.72559 | 0.723998 | 0.719008 | 0.721816 | 0.724042 | 0.723394 | 0.723742 | 0.721771 | 0.726434 |
| 0.680742 | 0.676182 | 0.677745 | 0.676289 | 0.683162 | 0.680043 | 0.678743 | 0.675469 | 0.681572 |
| 0.676935 | 0.666436 | 0.667196 | 0.672233 | 0.67687 | 0.672448 | 0.671496 | 0.668621 | 0.676856 |
| 0.728928 | 0.719574 | 0.723856 | 0.718929 | 0.724201 | 0.724267 | 0.725968 | 0.720071 | 0.723481 |
| 0.677243 | 0.666949 | 0.670695 | 0.672467 | 0.676794 | 0.674426 | 0.673196 | 0.666718 | 0.675748 |
| 0.668997 | 0.661367 | 0.665939 | 0.662228 | 0.667541 | 0.667143 | 0.666866 | 0.660608 | 0.664902 |
| 0.665544 | 0.654227 | 0.660318 | 0.658359 | 0.663877 | 0.663382 | 0.660789 | 0.657938 | 0.661314 |
| 0.735733 | 0.726293 | 0.729771 | 0.727805 | 0.730513 | 0.728467 | 0.728834 | 0.729231 | 0.730911 |
| 0.672318 | 0.661499 | 0.666416 | 0.664547 | 0.669161 | 0.670962 | 0.668411 | 0.662085 | 0.665209 |
| 0.72782 | 0.720661 | 0.722587 | 0.72044 | 0.728065 | 0.724074 | 0.726324 | 0.721388 | 0.726149 |

| | | | | | | | | |
|---|---|---|---|---|---|---|---|---|
| 0.73896 | 0.731644 | 0.731939 | 0.72894 | 0.733979 | 0.734041 | 0.737331 | 0.73111 | 0.73268 |
| 0.730202 | 0.721854 | 0.72625 | 0.722922 | 0.728621 | 0.728709 | 0.729509 | 0.724894 | 0.727535 |
| 0.751896 | 0.745152 | 0.74351 | 0.744815 | 0.747607 | 0.744861 | 0.747509 | 0.743477 | 0.745684 |
| 0.661897 | 0.649876 | 0.654407 | 0.652807 | 0.658643 | 0.661056 | 0.659651 | 0.652482 | 0.658976 |
| 0.661001 | 0.653918 | 0.653216 | 0.655094 | 0.661238 | 0.65717 | 0.657997 | 0.652354 | 0.653017 |
| 0.73846 | 0.728434 | 0.733977 | 0.73156 | 0.734915 | 0.737346 | 0.73576 | 0.730103 | 0.735522 |
| 0.660441 | 0.654561 | 0.654696 | 0.657035 | 0.661997 | 0.660383 | 0.658603 | 0.654639 | 0.658398 |
| 0.742043 | 0.735899 | 0.733827 | 0.733069 | 0.736345 | 0.74017 | 0.737856 | 0.727663 | 0.735143 |
| 0.742277 | 0.730228 | 0.733894 | 0.731397 | 0.737151 | 0.73084 | 0.733525 | 0.732659 | 0.735526 |
| 0.739201 | 0.742051 | 0.731476 | 0.735375 | 0.735982 | 0.738128 | 0.735837 | 0.729575 | 0.736851 |
| 0.73906 | 0.730716 | 0.731167 | 0.734309 | 0.739879 | 0.733215 | 0.736584 | 0.733499 | 0.737108 |
| 0.755102 | 0.779412 | 0.75 | 0.726027 | 0.707483 | 0.724138 | 0.75 | 0.733813 | 0.712418 |
| 0.729746 | 0.730022 | 0.735773 | 0.731446 | 0.730111 | 0.72864 | 0.728163 | 0.724565 | 0.731909 |
| 0.730328 | 0.730279 | 0.719495 | 0.724295 | 0.727032 | 0.72553 | 0.731252 | 0.720138 | 0.72626 |
| 0.741155 | 0.733887 | 0.738322 | 0.734678 | 0.73636 | 0.735352 | 0.742736 | 0.729811 | 0.736492 |
| 0.738503 | 0.735612 | 0.731074 | 0.734472 | 0.732788 | 0.739325 | 0.739588 | 0.730671 | 0.73772 |
| 0.59335 | 0.594956 | 0.593801 | 0.595163 | 0.599396 | 0.595551 | 0.595938 | 0.590909 | 0.600522 |
| 0.742302 | 0.743886 | 0.73493 | 0.737597 | 0.738204 | 0.743243 | 0.741334 | 0.734862 | 0.734649 |
| 0.742769 | 0.734446 | 0.735697 | 0.735707 | 0.739141 | 0.741108 | 0.739608 | 0.737021 | 0.739483 |
| 0.74333 | 0.736714 | 0.737678 | 0.733506 | 0.743554 | 0.735654 | 0.738932 | 0.73761 | 0.739188 |
| 0.746002 | 0.736002 | 0.740656 | 0.739355 | 0.745988 | 0.740122 | 0.742178 | 0.739452 | 0.744877 |
| 0.603944 | 0.593577 | 0.599939 | 0.597008 | 0.601037 | 0.600352 | 0.602715 | 0.595896 | 0.596955 |
| 0.588976 | 0.58005 | 0.583576 | 0.581807 | 0.584849 | 0.588231 | 0.589639 | 0.579878 | 0.583934 |
| 0.766348 | 0.76511 | 0.760748 | 0.761425 | 0.763282 | 0.76428 | 0.766209 | 0.761921 | 0.767395 |
| 0.767876 | 0.767432 | 0.761928 | 0.761921 | 0.76204 | 0.768744 | 0.769639 | 0.762028 | 0.763693 |
| 0.760794 | 0.753243 | 0.757243 | 0.756182 | 0.75689 | 0.757446 | 0.759403 | 0.752877 | 0.758283 |
| 0.767818 | 0.757236 | 0.764218 | 0.759613 | 0.759756 | 0.766637 | 0.768134 | 0.759213 | 0.761971 |
| 0.767923 | 0.759353 | 0.767126 | 0.759315 | 0.766373 | 0.766389 | 0.768393 | 0.762825 | 0.766194 |
| 0.768516 | 0.767416 | 0.769327 | 0.769981 | 0.766741 | 0.77091 | 0.774559 | 0.760576 | 0.768357 |
| 0.768959 | 0.767508 | 0.76598 | 0.765076 | 0.764624 | 0.76695 | 0.769999 | 0.764291 | 0.767526 |
| 0.76549 | 0.759336 | 0.763533 | 0.762984 | 0.7668 | 0.759202 | 0.76082 | 0.759777 | 0.763837 |
| 0.774312 | 0.772153 | 0.772831 | 0.768521 | 0.772344 | 0.776579 | 0.775514 | 0.771868 | 0.773479 |
| 0.763754 | 0.763012 | 0.762657 | 0.764949 | 0.769529 | 0.767037 | 0.769573 | 0.762702 | 0.76562 |
| 0.769962 | 0.764852 | 0.763299 | 0.768322 | 0.765048 | 0.7642 | 0.766204 | 0.762479 | 0.768442 |
| 0.776318 | 0.771682 | 0.770792 | 0.767222 | 0.772742 | 0.775337 | 0.775657 | 0.767951 | 0.768071 |
| 0.769599 | 0.764262 | 0.763621 | 0.767161 | 0.767064 | 0.766315 | 0.767745 | 0.767644 | 0.767832 |
| 0.767373 | 0.768721 | 0.768521 | 0.764207 | 0.765187 | 0.768799 | 0.771576 | 0.759645 | 0.768171 |
| 0.770835 | 0.770471 | 0.769193 | 0.76788 | 0.770711 | 0.76716 | 0.77066 | 0.767908 | 0.769726 |
| 0.765792 | 0.764469 | 0.761151 | 0.765784 | 0.765234 | 0.764378 | 0.767649 | 0.762447 | 0.764396 |
| 0.765607 | 0.762415 | 0.761055 | 0.767434 | 0.761037 | 0.765599 | 0.767248 | 0.762372 | 0.764745 |
| 0.72882 | 0.733216 | 0.728944 | 0.726281 | 0.726512 | 0.732699 | 0.737027 | 0.723659 | 0.727 |
| 0.737031 | 0.730834 | 0.73459 | 0.728374 | 0.727765 | 0.734775 | 0.737804 | 0.732424 | 0.730558 |
| 0.771711 | 0.772186 | 0.768212 | 0.763184 | 0.763728 | 0.767423 | 0.771266 | 0.761059 | 0.764194 |
| 0.776736 | 0.771815 | 0.77444 | 0.771311 | 0.773699 | 0.773205 | 0.77193 | 0.767354 | 0.771026 |
| 0.718954 | 0.72856 | 0.716695 | 0.720917 | 0.719067 | 0.722868 | 0.722104 | 0.723532 | 0.725131 |
| 0.771797 | 0.767672 | 0.764986 | 0.769708 | 0.76837 | 0.773448 | 0.773602 | 0.767524 | 0.769167 |
| 0.761835 | 0.766069 | 0.761391 | 0.757916 | 0.759093 | 0.759889 | 0.760359 | 0.757994 | 0.762507 |

| | | | | | | | | |
|---|---|---|---|---|---|---|---|---|
| 0.775896 | 0.768195 | 0.772315 | 0.766686 | 0.76808 | 0.771927 | 0.770682 | 0.765089 | 0.772483 |
| 0.773912 | 0.772225 | 0.768873 | 0.764511 | 0.768682 | 0.771015 | 0.772119 | 0.764174 | 0.773161 |
| 0.768432 | 0.762687 | 0.764365 | 0.761657 | 0.766062 | 0.767333 | 0.764374 | 0.76013 | 0.765737 |
| 0.776463 | 0.769977 | 0.773083 | 0.769314 | 0.769155 | 0.770362 | 0.773162 | 0.769012 | 0.771478 |
| 0.776447 | 0.77037 | 0.77052 | 0.76846 | 0.769921 | 0.772944 | 0.772412 | 0.765922 | 0.772205 |
| 0.770876 | 0.76863 | 0.769313 | 0.768557 | 0.767385 | 0.768589 | 0.768569 | 0.768342 | 0.770582 |
| 0.774654 | 0.767939 | 0.767059 | 0.767186 | 0.766791 | 0.771726 | 0.769987 | 0.761395 | 0.770681 |
| 0.76043 | 0.763323 | 0.755499 | 0.755947 | 0.759921 | 0.763058 | 0.761842 | 0.757254 | 0.762784 |
| 0.774988 | 0.766279 | 0.763363 | 0.763008 | 0.760351 | 0.766155 | 0.77074 | 0.764465 | 0.766951 |
| 0.763862 | 0.758814 | 0.762973 | 0.75822 | 0.762844 | 0.76362 | 0.765216 | 0.759742 | 0.764364 |
| 0.771765 | 0.770492 | 0.767773 | 0.763594 | 0.768717 | 0.769401 | 0.773264 | 0.761944 | 0.770085 |
| 0.769829 | 0.768152 | 0.767395 | 0.7646 | 0.765151 | 0.766056 | 0.765095 | 0.765708 | 0.768909 |
| 0.758472 | 0.761301 | 0.761255 | 0.755533 | 0.760215 | 0.755668 | 0.758246 | 0.755402 | 0.762729 |
| 0.7652 | 0.760059 | 0.76175 | 0.760292 | 0.759918 | 0.762235 | 0.762651 | 0.753407 | 0.761211 |
| 0.775357 | 0.76824 | 0.762342 | 0.766207 | 0.766707 | 0.77091 | 0.769287 | 0.764476 | 0.773259 |
| 0.771045 | 0.76452 | 0.768827 | 0.77033 | 0.773846 | 0.768998 | 0.774396 | 0.764657 | 0.772251 |
| 0.772221 | 0.759487 | 0.764002 | 0.760993 | 0.762648 | 0.766463 | 0.765753 | 0.759143 | 0.766385 |
| 0.7688 | 0.761574 | 0.761117 | 0.762133 | 0.76472 | 0.763825 | 0.76425 | 0.759458 | 0.763388 |
| 0.768069 | 0.76272 | 0.769439 | 0.76423 | 0.769168 | 0.766939 | 0.771573 | 0.760603 | 0.767348 |
| 0.769509 | 0.761106 | 0.764386 | 0.765618 | 0.767033 | 0.76629 | 0.769168 | 0.766127 | 0.764989 |
| 0.75991 | 0.753615 | 0.758788 | 0.759949 | 0.761014 | 0.764037 | 0.764056 | 0.754309 | 0.762859 |
| 0.767135 | 0.75818 | 0.762744 | 0.76018 | 0.764276 | 0.762965 | 0.767135 | 0.756341 | 0.762919 |
| 0.771605 | 0.766385 | 0.770117 | 0.766104 | 0.769825 | 0.772073 | 0.775116 | 0.763374 | 0.767793 |
| 0.771529 | 0.764313 | 0.763445 | 0.759907 | 0.76497 | 0.768352 | 0.768108 | 0.761921 | 0.764202 |
| 0.770709 | 0.763455 | 0.762609 | 0.76213 | 0.763428 | 0.76966 | 0.771888 | 0.757698 | 0.766447 |
| 0.769212 | 0.761773 | 0.767684 | 0.765351 | 0.767995 | 0.76781 | 0.768763 | 0.756306 | 0.763636 |
| 0.768249 | 0.755107 | 0.756466 | 0.754032 | 0.757073 | 0.760156 | 0.762632 | 0.755257 | 0.756926 |
| 0.763377 | 0.759871 | 0.761469 | 0.76216 | 0.766186 | 0.760047 | 0.763285 | 0.75623 | 0.765261 |
| 0.767085 | 0.763351 | 0.757281 | 0.764356 | 0.764691 | 0.762697 | 0.763175 | 0.758436 | 0.765566 |
| 0.768897 | 0.760754 | 0.766522 | 0.765345 | 0.768439 | 0.768316 | 0.7712 | 0.764553 | 0.768132 |
| 0.771626 | 0.760233 | 0.765697 | 0.768902 | 0.769092 | 0.768177 | 0.771431 | 0.762685 | 0.764346 |
| 0.772094 | 0.761268 | 0.766412 | 0.763498 | 0.762653 | 0.765772 | 0.767413 | 0.761514 | 0.76476 |
| 0.776334 | 0.771134 | 0.768195 | 0.768222 | 0.76963 | 0.77259 | 0.770877 | 0.766772 | 0.768154 |
| 0.772366 | 0.768068 | 0.766174 | 0.762413 | 0.764739 | 0.772216 | 0.771941 | 0.764642 | 0.76781 |
| 0.767944 | 0.763434 | 0.758799 | 0.759135 | 0.760458 | 0.764847 | 0.767246 | 0.760275 | 0.761195 |
| 0.771684 | 0.760014 | 0.762987 | 0.764766 | 0.763225 | 0.763971 | 0.768899 | 0.762079 | 0.765834 |
| 0.767783 | 0.761149 | 0.761315 | 0.759936 | 0.764334 | 0.762522 | 0.768346 | 0.758184 | 0.767277 |
| 0.767656 | 0.759248 | 0.763825 | 0.763735 | 0.768098 | 0.765972 | 0.768787 | 0.75742 | 0.761563 |
| 0.762366 | 0.749979 | 0.756761 | 0.757395 | 0.75522 | 0.757429 | 0.760826 | 0.750761 | 0.755833 |
| 0.760395 | 0.751413 | 0.754788 | 0.755 | 0.755313 | 0.759456 | 0.763309 | 0.756485 | 0.761075 |
| 0.755966 | 0.751653 | 0.753508 | 0.756398 | 0.757494 | 0.760007 | 0.761053 | 0.750988 | 0.754529 |
| 0.761345 | 0.75541 | 0.756311 | 0.753409 | 0.756766 | 0.754067 | 0.756848 | 0.752431 | 0.752973 |
| 0.761733 | 0.751457 | 0.752205 | 0.751937 | 0.753636 | 0.753766 | 0.757746 | 0.750392 | 0.751589 |
| 0.755322 | 0.751322 | 0.755245 | 0.750251 | 0.752348 | 0.752562 | 0.760702 | 0.745918 | 0.759677 |
| 0.76465 | 0.758582 | 0.762949 | 0.760641 | 0.758692 | 0.764579 | 0.76687 | 0.756223 | 0.761785 |
| 0.760068 | 0.747835 | 0.748945 | 0.750452 | 0.74807 | 0.75096 | 0.752938 | 0.74765 | 0.750942 |

| | | | | | | | | |
|---|---|---|---|---|---|---|---|---|
| 0.76839 | 0.767424 | 0.767557 | 0.768972 | 0.771131 | 0.767862 | 0.768947 | 0.762855 | 0.76565 |
| 0.780874 | 0.774875 | 0.775864 | 0.777654 | 0.774824 | 0.777564 | 0.777335 | 0.771184 | 0.773982 |
| 0.771112 | 0.765671 | 0.7713 | 0.771266 | 0.770588 | 0.771674 | 0.773905 | 0.764383 | 0.770455 |
| 0.777851 | 0.769812 | 0.769438 | 0.766372 | 0.770794 | 0.775176 | 0.776611 | 0.76733 | 0.771459 |
| 0.782236 | 0.777554 | 0.77676 | 0.780278 | 0.781486 | 0.778622 | 0.782334 | 0.777403 | 0.780702 |
| 0.779969 | 0.77083 | 0.772058 | 0.771419 | 0.770254 | 0.773947 | 0.777623 | 0.771429 | 0.774575 |
| 0.777425 | 0.769479 | 0.769974 | 0.766252 | 0.766953 | 0.772396 | 0.773386 | 0.767517 | 0.76576 |
| 0.778298 | 0.770623 | 0.771764 | 0.765265 | 0.770108 | 0.774048 | 0.774831 | 0.765064 | 0.768774 |
| 0.768064 | 0.768486 | 0.765472 | 0.767816 | 0.763747 | 0.772024 | 0.775422 | 0.75698 | 0.768178 |
| 0.767192 | 0.762975 | 0.765359 | 0.768843 | 0.768481 | 0.769548 | 0.770055 | 0.76719 | 0.768014 |
| 0.777932 | 0.769914 | 0.770911 | 0.773706 | 0.771243 | 0.774244 | 0.776095 | 0.764735 | 0.768783 |
| 0.775592 | 0.76882 | 0.770525 | 0.770157 | 0.770789 | 0.772242 | 0.770656 | 0.763751 | 0.771745 |
| 0.782995 | 0.7706 | 0.779532 | 0.773037 | 0.774826 | 0.782526 | 0.781127 | 0.77478 | 0.778192 |
| 0.780778 | 0.769186 | 0.771595 | 0.775269 | 0.77765 | 0.775401 | 0.776552 | 0.767809 | 0.776894 |
| 0.782174 | 0.777509 | 0.772545 | 0.777722 | 0.780434 | 0.781488 | 0.778923 | 0.772659 | 0.780198 |
| 0.783479 | 0.771996 | 0.776715 | 0.77129 | 0.776459 | 0.776452 | 0.77707 | 0.770409 | 0.778901 |
| 0.786574 | 0.78063 | 0.772031 | 0.779233 | 0.781873 | 0.778804 | 0.779834 | 0.771246 | 0.778498 |
| 0.777308 | 0.776048 | 0.773307 | 0.772543 | 0.777031 | 0.773565 | 0.771157 | 0.770785 | 0.784892 |
| 0.78448 | 0.779017 | 0.772597 | 0.776687 | 0.783589 | 0.780038 | 0.782321 | 0.774937 | 0.781704 |
| 0.783821 | 0.779522 | 0.770707 | 0.777852 | 0.7797 | 0.775362 | 0.778917 | 0.780359 | 0.777686 |
| 0.779421 | 0.771229 | 0.77246 | 0.773682 | 0.775811 | 0.772637 | 0.771934 | 0.769846 | 0.773751 |
| 0.78098 | 0.766984 | 0.769053 | 0.771436 | 0.774146 | 0.774766 | 0.774589 | 0.772099 | 0.771208 |
| 0.786519 | 0.772016 | 0.772634 | 0.77761 | 0.776936 | 0.775984 | 0.778699 | 0.771586 | 0.779842 |
| 0.78582 | 0.776629 | 0.779069 | 0.781454 | 0.777941 | 0.789088 | 0.787498 | 0.775689 | 0.783463 |
| 0.787045 | 0.780101 | 0.78084 | 0.777519 | 0.782218 | 0.785109 | 0.783885 | 0.777676 | 0.785027 |
| 0.785992 | 0.781577 | 0.778556 | 0.775522 | 0.784745 | 0.786954 | 0.785182 | 0.775394 | 0.783013 |
| 0.785937 | 0.779575 | 0.778633 | 0.778084 | 0.780186 | 0.785538 | 0.787119 | 0.785385 | 0.788872 |
| 0.779488 | 0.772822 | 0.780176 | 0.769634 | 0.775455 | 0.779707 | 0.783702 | 0.777128 | 0.779756 |
| 0.787824 | 0.780645 | 0.779618 | 0.784989 | 0.788211 | 0.784854 | 0.785551 | 0.780217 | 0.789805 |
| 0.7805 | 0.781724 | 0.779 | 0.779549 | 0.781673 | 0.785052 | 0.790103 | 0.779662 | 0.783622 |
| 0.785403 | 0.776037 | 0.774956 | 0.768956 | 0.773238 | 0.780035 | 0.778611 | 0.772627 | 0.78204 |
| 0.782363 | 0.781777 | 0.781273 | 0.778484 | 0.780504 | 0.784186 | 0.785466 | 0.778323 | 0.782921 |
| 0.788476 | 0.779388 | 0.783039 | 0.778944 | 0.779859 | 0.78552 | 0.787054 | 0.78016 | 0.783095 |
| 0.781841 | 0.776509 | 0.776234 | 0.776652 | 0.775455 | 0.780124 | 0.781747 | 0.78046 | 0.779949 |
| 0.78863 | 0.778295 | 0.775604 | 0.780847 | 0.783452 | 0.780751 | 0.77818 | 0.776293 | 0.783018 |
| 0.780939 | 0.782544 | 0.774773 | 0.77427 | 0.774944 | 0.779927 | 0.784866 | 0.776286 | 0.783243 |
| 1 | 0.781934 | 0.780211 | 0.784866 | 0.785386 | 0.788352 | 0.790131 | 0.782675 | 0.789276 |
| 0.781934 | 1 | 0.777135 | 0.782013 | 0.803565 | 0.823772 | 0.816987 | 0.782467 | 0.783869 |
| 0.780211 | 0.777135 | 1 | 0.780903 | 0.787929 | 0.82477 | 0.818705 | 0.782184 | 0.784162 |
| 0.784866 | 0.782013 | 0.780903 | 1 | 0.863613 | 0.786124 | 0.787423 | 0.7876 | 0.813603 |
| 0.785386 | 0.803565 | 0.787929 | 0.863613 | 1 | 0.785861 | 0.784786 | 0.792629 | 0.797268 |
| 0.788352 | 0.823772 | 0.82477 | 0.786124 | 0.785861 | 1 | 0.883146 | 0.783033 | 0.786005 |
| 0.790131 | 0.816987 | 0.818705 | 0.787423 | 0.784786 | 0.883146 | 1 | 0.785469 | 0.790441 |
| 0.782675 | 0.782467 | 0.782184 | 0.7876 | 0.792629 | 0.783033 | 0.785469 | 1 | 0.782141 |
| 0.789276 | 0.783869 | 0.784162 | 0.813603 | 0.797268 | 0.786005 | 0.790441 | 0.782141 | 1 |
| 0.788256 | 0.800743 | 0.782974 | 0.79693 | 0.791443 | 0.805414 | 0.810891 | 0.793971 | 0.860911 |

| | | | | | | | | |
|---|---|---|---|---|---|---|---|---|
| 0.771825 | 0.768233 | 0.764183 | 0.766932 | 0.770104 | 0.775998 | 0.782778 | 0.769364 | 0.78001 |
| 0.759964 | 0.757163 | 0.760929 | 0.755925 | 0.759819 | 0.765399 | 0.761963 | 0.756194 | 0.756195 |
| 0.781607 | 0.777424 | 0.772071 | 0.775415 | 0.780862 | 0.780703 | 0.777226 | 0.768286 | 0.776822 |
| 0.786194 | 0.774438 | 0.771888 | 0.78103 | 0.78112 | 0.778296 | 0.779006 | 0.776962 | 0.781668 |
| 0.776451 | 0.770144 | 0.766019 | 0.76998 | 0.773129 | 0.772731 | 0.774259 | 0.762639 | 0.772825 |
| 0.781328 | 0.776904 | 0.774027 | 0.778686 | 0.779412 | 0.77862 | 0.778283 | 0.770619 | 0.775291 |
| 0.780793 | 0.774109 | 0.774324 | 0.770417 | 0.773397 | 0.778392 | 0.778493 | 0.773243 | 0.778502 |
| 0.777373 | 0.772473 | 0.772275 | 0.773432 | 0.774134 | 0.772808 | 0.773448 | 0.772744 | 0.775802 |
| 0.781117 | 0.774161 | 0.776446 | 0.771864 | 0.774336 | 0.778351 | 0.780675 | 0.771753 | 0.77667 |
| 0.779118 | 0.771996 | 0.77153 | 0.76696 | 0.774289 | 0.772499 | 0.779678 | 0.772371 | 0.774983 |
| 0.780822 | 0.774392 | 0.77203 | 0.775878 | 0.777769 | 0.777823 | 0.777769 | 0.774757 | 0.777494 |
| 0.776725 | 0.778947 | 0.770582 | 0.772666 | 0.7769 | 0.774851 | 0.776976 | 0.771657 | 0.775531 |
| 0.787267 | 0.78259 | 0.777769 | 0.776777 | 0.777696 | 0.780281 | 0.779841 | 0.774362 | 0.780459 |
| 0.780768 | 0.771778 | 0.774117 | 0.770103 | 0.774417 | 0.77796 | 0.780593 | 0.769384 | 0.775717 |
| 0.777713 | 0.772031 | 0.767065 | 0.77062 | 0.770492 | 0.772263 | 0.7752 | 0.768577 | 0.774042 |
| 0.779222 | 0.774058 | 0.769661 | 0.771223 | 0.770825 | 0.775083 | 0.776318 | 0.764575 | 0.775235 |
| 0.775561 | 0.768946 | 0.769288 | 0.765727 | 0.768695 | 0.769826 | 0.770746 | 0.765559 | 0.768409 |
| 0.775876 | 0.769864 | 0.768729 | 0.764944 | 0.770244 | 0.77178 | 0.773847 | 0.76732 | 0.769954 |
| 0.77041 | 0.766208 | 0.765262 | 0.771028 | 0.769751 | 0.767998 | 0.770865 | 0.764418 | 0.771045 |
| 0.775253 | 0.767417 | 0.767351 | 0.768816 | 0.765787 | 0.769866 | 0.774071 | 0.764349 | 0.771065 |
| 0.778499 | 0.774434 | 0.771871 | 0.770793 | 0.774234 | 0.776748 | 0.777052 | 0.769543 | 0.775856 |
| 0.76124 | 0.765396 | 0.755976 | 0.760316 | 0.767128 | 0.764074 | 0.762983 | 0.757821 | 0.759363 |
| 0.768878 | 0.768942 | 0.761733 | 0.766954 | 0.771155 | 0.767087 | 0.769375 | 0.768702 | 0.763597 |
| 0.774866 | 0.770683 | 0.767179 | 0.772352 | 0.776523 | 0.774204 | 0.775421 | 0.7696 | 0.769014 |
| 0.776636 | 0.775423 | 0.768234 | 0.7724 | 0.775055 | 0.778013 | 0.776463 | 0.770297 | 0.771919 |
| 0.77122 | 0.770906 | 0.764672 | 0.769694 | 0.77364 | 0.772815 | 0.773264 | 0.764181 | 0.769509 |
| 0.769146 | 0.768559 | 0.767717 | 0.764284 | 0.765297 | 0.766585 | 0.764874 | 0.764726 | 0.765198 |
| 0.771777 | 0.77122 | 0.770844 | 0.766467 | 0.773379 | 0.771455 | 0.772375 | 0.769625 | 0.766835 |
| 0.76841 | 0.767126 | 0.765442 | 0.768264 | 0.769558 | 0.769505 | 0.767612 | 0.763353 | 0.768759 |
| 0.703316 | 0.702108 | 0.696872 | 0.697969 | 0.700923 | 0.699257 | 0.70158 | 0.698606 | 0.699963 |
| 0.715756 | 0.706074 | 0.709527 | 0.709408 | 0.712243 | 0.709874 | 0.711419 | 0.705335 | 0.708686 |
| 0.711826 | 0.704656 | 0.708223 | 0.706444 | 0.711856 | 0.70961 | 0.709911 | 0.706982 | 0.710749 |
| 0.703551 | 0.695037 | 0.695893 | 0.69592 | 0.702593 | 0.695874 | 0.693038 | 0.697348 | 0.699061 |
| 0.715846 | 0.706729 | 0.708011 | 0.70975 | 0.715738 | 0.714014 | 0.712818 | 0.707447 | 0.709939 |
| 0.769335 | 0.763962 | 0.767046 | 0.765765 | 0.769549 | 0.766327 | 0.765841 | 0.760062 | 0.769573 |
| 0.775702 | 0.769413 | 0.770482 | 0.772066 | 0.774547 | 0.769212 | 0.768589 | 0.768193 | 0.77746 |
| 0.775874 | 0.773871 | 0.773745 | 0.774013 | 0.774635 | 0.775133 | 0.777879 | 0.77083 | 0.777312 |
| 0.77545 | 0.774234 | 0.773062 | 0.769716 | 0.773644 | 0.776653 | 0.776234 | 0.768693 | 0.776298 |
| 0.771975 | 0.766352 | 0.76694 | 0.762653 | 0.768399 | 0.7676 | 0.771063 | 0.762194 | 0.773385 |
| 0.77263 | 0.773908 | 0.774348 | 0.771395 | 0.776943 | 0.773702 | 0.773124 | 0.767459 | 0.777824 |
| 0.77361 | 0.774128 | 0.77487 | 0.768754 | 0.772637 | 0.774848 | 0.775819 | 0.77122 | 0.778049 |
| 0.771379 | 0.768178 | 0.771555 | 0.76576 | 0.768355 | 0.768903 | 0.768923 | 0.765891 | 0.769535 |
| 0.769565 | 0.76242 | 0.7666 | 0.764306 | 0.769038 | 0.762408 | 0.763986 | 0.75788 | 0.764213 |
| 0.767894 | 0.764965 | 0.765039 | 0.762872 | 0.766535 | 0.763416 | 0.766182 | 0.760813 | 0.766409 |
| 0.774526 | 0.767236 | 0.76731 | 0.76457 | 0.769535 | 0.768614 | 0.765799 | 0.764378 | 0.770073 |
| 0.775123 | 0.767751 | 0.774016 | 0.772127 | 0.771213 | 0.771726 | 0.772938 | 0.767031 | 0.773639 |

| | | | | | | | | |
|---|---|---|---|---|---|---|---|---|
| 0.780502 | 0.772992 | 0.772785 | 0.770762 | 0.775334 | 0.773604 | 0.776855 | 0.768874 | 0.770488 |
| 0.775329 | 0.769923 | 0.772302 | 0.766425 | 0.76851 | 0.772637 | 0.774586 | 0.766714 | 0.76773 |
| 0.77575 | 0.767672 | 0.770988 | 0.76755 | 0.771779 | 0.770278 | 0.77535 | 0.76479 | 0.773329 |
| 0.768996 | 0.768114 | 0.766403 | 0.762715 | 0.768619 | 0.76958 | 0.77359 | 0.760762 | 0.767076 |
| 0.770403 | 0.768182 | 0.771436 | 0.767099 | 0.768431 | 0.769607 | 0.769244 | 0.763406 | 0.763767 |
| 0.769165 | 0.769271 | 0.771275 | 0.766283 | 0.76551 | 0.769277 | 0.77363 | 0.765336 | 0.769198 |
| 0.770819 | 0.77041 | 0.772056 | 0.769707 | 0.774034 | 0.767956 | 0.774664 | 0.766372 | 0.76942 |
| 0.775343 | 0.764701 | 0.770927 | 0.7676 | 0.767024 | 0.767299 | 0.767847 | 0.768426 | 0.769315 |
| 0.773237 | 0.768599 | 0.769426 | 0.763819 | 0.766809 | 0.768667 | 0.770197 | 0.762703 | 0.766867 |
| 0.774314 | 0.768658 | 0.768994 | 0.763957 | 0.768977 | 0.776989 | 0.774655 | 0.765248 | 0.764966 |
| 0.770633 | 0.764771 | 0.76758 | 0.764536 | 0.770013 | 0.769649 | 0.77155 | 0.758755 | 0.76947 |
| 0.773154 | 0.767501 | 0.769909 | 0.76424 | 0.773278 | 0.769411 | 0.77355 | 0.768106 | 0.768674 |
| 0.768984 | 0.764249 | 0.760957 | 0.759513 | 0.762027 | 0.764266 | 0.768241 | 0.759553 | 0.76364 |
| 0.762855 | 0.761511 | 0.763506 | 0.763573 | 0.76419 | 0.766946 | 0.769342 | 0.762242 | 0.760553 |
| 0.776814 | 0.767119 | 0.769127 | 0.767767 | 0.770563 | 0.771337 | 0.772864 | 0.762974 | 0.772473 |
| 0.778513 | 0.777198 | 0.779615 | 0.785538 | 0.783701 | 0.781042 | 0.785744 | 0.777627 | 0.784156 |
| 0.784878 | 0.776061 | 0.770855 | 0.778028 | 0.780756 | 0.774678 | 0.778382 | 0.771502 | 0.782061 |
| 0.779972 | 0.775659 | 0.772124 | 0.774433 | 0.77413 | 0.776017 | 0.779073 | 0.771371 | 0.776974 |
| 0.775937 | 0.772913 | 0.777987 | 0.780019 | 0.775623 | 0.77392 | 0.774951 | 0.77498 | 0.778928 |
| 0.777702 | 0.772053 | 0.770031 | 0.774449 | 0.76901 | 0.771782 | 0.77579 | 0.763212 | 0.772493 |
| 0.772813 | 0.767967 | 0.76538 | 0.766527 | 0.769277 | 0.770558 | 0.770855 | 0.761599 | 0.769323 |
| 0.771822 | 0.770447 | 0.772352 | 0.772136 | 0.770901 | 0.771946 | 0.774034 | 0.766515 | 0.770835 |
| 0.783066 | 0.779458 | 0.774001 | 0.776599 | 0.778553 | 0.780267 | 0.780801 | 0.778039 | 0.782067 |
| 0.768317 | 0.768762 | 0.768264 | 0.761773 | 0.766477 | 0.770909 | 0.774415 | 0.762544 | 0.765758 |
| 0.781634 | 0.778008 | 0.775396 | 0.775806 | 0.776158 | 0.777482 | 0.781516 | 0.775462 | 0.781923 |
| 0.710863 | 0.706997 | 0.70548 | 0.702891 | 0.708752 | 0.711245 | 0.707921 | 0.706785 | 0.707554 |
| 0.782648 | 0.774172 | 0.776363 | 0.779914 | 0.780753 | 0.780913 | 0.780956 | 0.778504 | 0.782375 |
| 0.78515 | 0.781646 | 0.779151 | 0.788206 | 0.784776 | 0.786729 | 0.789893 | 0.78193 | 0.782209 |
| 0.793464 | 0.784292 | 0.787273 | 0.784947 | 0.785061 | 0.785756 | 0.787814 | 0.781459 | 0.785756 |
| 0.78543 | 0.782623 | 0.780897 | 0.785751 | 0.786082 | 0.786672 | 0.787959 | 0.77775 | 0.77991 |
| 0.773952 | 0.772098 | 0.775471 | 0.770329 | 0.777232 | 0.777384 | 0.777138 | 0.771956 | 0.777981 |
| 0.776161 | 0.782201 | 0.773258 | 0.779897 | 0.78137 | 0.785476 | 0.791442 | 0.77627 | 0.774927 |
| 0.783565 | 0.780389 | 0.776881 | 0.775409 | 0.780754 | 0.785436 | 0.787784 | 0.775201 | 0.776638 |
| 0.78141 | 0.777113 | 0.778712 | 0.769649 | 0.777615 | 0.778866 | 0.783634 | 0.770917 | 0.77867 |
| 0.775746 | 0.776339 | 0.773775 | 0.7729 | 0.773637 | 0.77863 | 0.783697 | 0.77025 | 0.769486 |
| 0.781438 | 0.774371 | 0.775205 | 0.77953 | 0.775055 | 0.780473 | 0.781608 | 0.775933 | 0.778147 |
| 0.782506 | 0.783329 | 0.782009 | 0.789011 | 0.790647 | 0.78627 | 0.787805 | 0.779253 | 0.785308 |
| 0.783174 | 0.778803 | 0.782259 | 0.778277 | 0.784568 | 0.780376 | 0.784559 | 0.777928 | 0.784149 |
| 0.781612 | 0.782091 | 0.779005 | 0.785305 | 0.783854 | 0.784176 | 0.787248 | 0.778287 | 0.782638 |
| 0.781261 | 0.774295 | 0.776015 | 0.777914 | 0.779993 | 0.774711 | 0.774138 | 0.771941 | 0.771438 |
| 0.779798 | 0.772858 | 0.772782 | 0.770579 | 0.775157 | 0.775852 | 0.772374 | 0.768996 | 0.775556 |
| 0.783388 | 0.775675 | 0.774096 | 0.780074 | 0.77907 | 0.779859 | 0.783219 | 0.774991 | 0.780288 |
| 0.779376 | 0.774825 | 0.773408 | 0.768923 | 0.772355 | 0.778266 | 0.783232 | 0.775028 | 0.775434 |
| 0.766289 | 0.768588 | 0.76671 | 0.762214 | 0.766829 | 0.769376 | 0.770369 | 0.761188 | 0.769288 |
| 0.777465 | 0.770318 | 0.769936 | 0.76575 | 0.770761 | 0.774746 | 0.773944 | 0.767642 | 0.769953 |
| 0.77595 | 0.767292 | 0.768117 | 0.769703 | 0.77097 | 0.774994 | 0.774793 | 0.766664 | 0.771509 |

| | | | | | | | | |
|---|---|---|---|---|---|---|---|---|
| 0.771473 | 0.766794 | 0.767853 | 0.769565 | 0.773533 | 0.77037 | 0.769312 | 0.768974 | 0.769788 |
| 0.772489 | 0.770174 | 0.771107 | 0.766133 | 0.773139 | 0.772455 | 0.773973 | 0.770041 | 0.770289 |
| 0.769581 | 0.772685 | 0.772508 | 0.764096 | 0.768665 | 0.770093 | 0.772701 | 0.762985 | 0.770837 |
| 0.731375 | 0.72358 | 0.730394 | 0.730366 | 0.727305 | 0.727632 | 0.728086 | 0.718761 | 0.730187 |
| 0.770166 | 0.770571 | 0.772116 | 0.766097 | 0.770738 | 0.772095 | 0.772687 | 0.765073 | 0.771017 |
| 0.771107 | 0.770566 | 0.76816 | 0.76623 | 0.771129 | 0.770099 | 0.767956 | 0.761506 | 0.768099 |
| 0.771553 | 0.766403 | 0.76736 | 0.766105 | 0.770794 | 0.768871 | 0.769275 | 0.764141 | 0.769123 |
| 0.774777 | 0.776587 | 0.767388 | 0.768633 | 0.771471 | 0.77576 | 0.771927 | 0.763929 | 0.769826 |
| 0.775217 | 0.773648 | 0.772306 | 0.774837 | 0.777442 | 0.780345 | 0.780304 | 0.770325 | 0.772757 |
| 0.773915 | 0.769784 | 0.771574 | 0.772609 | 0.774685 | 0.777108 | 0.77318 | 0.766293 | 0.771282 |
| 0.772742 | 0.769998 | 0.76925 | 0.766742 | 0.768802 | 0.774074 | 0.773043 | 0.767422 | 0.767788 |
| 0.773338 | 0.767005 | 0.766656 | 0.769419 | 0.774928 | 0.775014 | 0.774479 | 0.766781 | 0.767521 |
| 0.77278 | 0.769713 | 0.767161 | 0.769991 | 0.764419 | 0.770775 | 0.771792 | 0.766366 | 0.767994 |
| 0.775954 | 0.771004 | 0.771353 | 0.764593 | 0.768167 | 0.772817 | 0.772882 | 0.768012 | 0.7683 |
| 0.771193 | 0.773931 | 0.769705 | 0.767508 | 0.773935 | 0.772387 | 0.775635 | 0.767158 | 0.768514 |
| 0.769086 | 0.765384 | 0.765093 | 0.761499 | 0.765277 | 0.763848 | 0.768109 | 0.765101 | 0.766584 |
| 0.772302 | 0.772652 | 0.768629 | 0.772746 | 0.775199 | 0.774693 | 0.77604 | 0.766495 | 0.772586 |
| 0.772902 | 0.773943 | 0.77534 | 0.765663 | 0.775053 | 0.775259 | 0.777978 | 0.766155 | 0.767333 |
| 0.769477 | 0.767452 | 0.766516 | 0.765605 | 0.769155 | 0.770722 | 0.771678 | 0.763178 | 0.771062 |
| 0.772265 | 0.765652 | 0.768007 | 0.766135 | 0.770803 | 0.771661 | 0.773168 | 0.768709 | 0.768445 |
| 0.77986 | 0.77307 | 0.769573 | 0.769063 | 0.772166 | 0.774992 | 0.777096 | 0.765817 | 0.769675 |
| 0.770363 | 0.774307 | 0.774646 | 0.77224 | 0.775047 | 0.777009 | 0.77885 | 0.771797 | 0.77437 |
| 0.687865 | 0.693976 | 0.68107 | 0.682476 | 0.685367 | 0.685192 | 0.687102 | 0.675582 | 0.687454 |
| 0.669214 | 0.662868 | 0.654443 | 0.663348 | 0.66517 | 0.665553 | 0.66857 | 0.655968 | 0.664921 |
| 0.675412 | 0.671814 | 0.668809 | 0.671885 | 0.679031 | 0.678953 | 0.681906 | 0.674707 | 0.674111 |
| 0.658041 | 0.658272 | 0.653465 | 0.657683 | 0.660953 | 0.65563 | 0.659269 | 0.656127 | 0.658292 |
| 0.742401 | 0.741066 | 0.739496 | 0.744564 | 0.741926 | 0.737794 | 0.74068 | 0.730428 | 0.741132 |
| 0.727203 | 0.728849 | 0.72677 | 0.729131 | 0.730828 | 0.733359 | 0.731168 | 0.722553 | 0.730311 |
| 0.730061 | 0.732653 | 0.733285 | 0.732982 | 0.727859 | 0.730081 | 0.732696 | 0.728654 | 0.731787 |
| 0.760194 | 0.751958 | 0.748268 | 0.742647 | 0.741379 | 0.747267 | 0.754927 | 0.739709 | 0.745798 |
| 0.727607 | 0.730309 | 0.72619 | 0.726119 | 0.723696 | 0.727691 | 0.733042 | 0.714383 | 0.726335 |
| 0.723108 | 0.723374 | 0.719114 | 0.72 | 0.714166 | 0.713229 | 0.719898 | 0.717985 | 0.715695 |
| 0.72563 | 0.727164 | 0.714957 | 0.722754 | 0.719588 | 0.722535 | 0.72517 | 0.722354 | 0.72118 |
| 0.664456 | 0.65996 | 0.66087 | 0.662312 | 0.663186 | 0.662778 | 0.658228 | 0.662845 | 0.658506 |
| 0.765712 | 0.765943 | 0.766066 | 0.762485 | 0.76641 | 0.7705 | 0.772458 | 0.760325 | 0.764563 |
| 0.745812 | 0.745621 | 0.747113 | 0.746642 | 0.750787 | 0.744132 | 0.748052 | 0.744076 | 0.743238 |
| 0.734318 | 0.742217 | 0.733154 | 0.736162 | 0.732519 | 0.738395 | 0.740911 | 0.741255 | 0.739644 |
| 0.727138 | 0.728499 | 0.7175 | 0.732022 | 0.728471 | 0.725831 | 0.718762 | 0.721129 | 0.723906 |
| 0.749737 | 0.750148 | 0.741337 | 0.746131 | 0.743846 | 0.745741 | 0.742638 | 0.748434 | 0.747499 |
| 0.740812 | 0.730507 | 0.733095 | 0.728962 | 0.731122 | 0.735808 | 0.737279 | 0.734385 | 0.732212 |
| 0.737794 | 0.73597 | 0.73363 | 0.729673 | 0.733567 | 0.730831 | 0.732973 | 0.729466 | 0.729459 |
| 0.73569 | 0.732067 | 0.732394 | 0.733449 | 0.731278 | 0.737024 | 0.740286 | 0.729925 | 0.728742 |
| 0.733515 | 0.734426 | 0.737607 | 0.730972 | 0.726376 | 0.73887 | 0.740338 | 0.729412 | 0.725777 |
| 0.730301 | 0.734059 | 0.731467 | 0.725946 | 0.726614 | 0.732291 | 0.728487 | 0.725559 | 0.726233 |
| 0.73581 | 0.727626 | 0.721156 | 0.729366 | 0.734747 | 0.728686 | 0.733093 | 0.720183 | 0.732759 |
| 0.715471 | 0.725025 | 0.70918 | 0.715367 | 0.719697 | 0.725876 | 0.729215 | 0.723369 | 0.714004 |

| | | | | | | | | |
|---|---|---|---|---|---|---|---|---|
| 0.759182 | 0.751378 | 0.75144 | 0.7498 | 0.757006 | 0.75358 | 0.754767 | 0.752483 | 0.751843 |
| 0.736768 | 0.739397 | 0.740589 | 0.733475 | 0.735253 | 0.739229 | 0.74117 | 0.73269 | 0.7382 |
| 0.738074 | 0.730932 | 0.72821 | 0.733364 | 0.732047 | 0.730042 | 0.735484 | 0.733434 | 0.732528 |
| 0.733927 | 0.741345 | 0.728868 | 0.736899 | 0.734227 | 0.736404 | 0.737773 | 0.734955 | 0.736793 |
| 0.725538 | 0.74113 | 0.73527 | 0.73614 | 0.736469 | 0.739847 | 0.740132 | 0.727698 | 0.734167 |
| 0.724095 | 0.727543 | 0.724112 | 0.736261 | 0.733201 | 0.723372 | 0.734184 | 0.725726 | 0.724431 |
| 0.720603 | 0.731863 | 0.725176 | 0.717291 | 0.719383 | 0.728096 | 0.72871 | 0.704442 | 0.716164 |
| 0.727143 | 0.728437 | 0.717883 | 0.719943 | 0.724544 | 0.724148 | 0.728103 | 0.725406 | 0.723242 |
| 0.727169 | 0.720812 | 0.720123 | 0.728673 | 0.720695 | 0.728208 | 0.735277 | 0.724737 | 0.735849 |
| 0.760897 | 0.761687 | 0.759339 | 0.75384 | 0.756433 | 0.765164 | 0.761942 | 0.759431 | 0.756413 |
| 0.753117 | 0.757752 | 0.753648 | 0.753768 | 0.753036 | 0.759179 | 0.756654 | 0.751018 | 0.751848 |
| 0.747562 | 0.749104 | 0.746049 | 0.750391 | 0.753314 | 0.752763 | 0.754481 | 0.750729 | 0.749361 |
| 0.744048 | 0.746541 | 0.74349 | 0.735812 | 0.741877 | 0.740615 | 0.753899 | 0.744326 | 0.744348 |
| 0.747862 | 0.748611 | 0.74223 | 0.744243 | 0.742455 | 0.749172 | 0.752869 | 0.745627 | 0.749908 |
| 0.743187 | 0.747261 | 0.744956 | 0.743466 | 0.735573 | 0.746969 | 0.748029 | 0.742424 | 0.743119 |

| | | | plot 08 | | | | | |
|---|---|---|---|---|---|---|---|---|
| Wild_plot_ | Wild_plot_ | Wild_plot_ | Wild_plot_ | Wild_plot_ | Wild_plot_ | Wild_plot_ | Wild_plot_ | Wild_plot_ |
| 0.703331 | 0.693725 | 0.750041 | 0.706934 | 0.711975 | 0.704796 | 0.708394 | 0.704187 | 0.701897 |
| 0.712272 | 0.698322 | 0.753497 | 0.709496 | 0.712268 | 0.708233 | 0.711458 | 0.710104 | 0.710189 |
| 0.7039 | 0.690294 | 0.745167 | 0.701661 | 0.706976 | 0.699109 | 0.701498 | 0.701375 | 0.697949 |
| 0.711869 | 0.700191 | 0.760332 | 0.710849 | 0.714263 | 0.705091 | 0.710484 | 0.711499 | 0.704606 |
| 0.7074 | 0.695057 | 0.743996 | 0.708676 | 0.71252 | 0.706057 | 0.709978 | 0.706543 | 0.704786 |
| 0.715218 | 0.70647 | 0.763728 | 0.716582 | 0.722292 | 0.713862 | 0.718278 | 0.71608 | 0.710367 |
| 0.772177 | 0.764684 | 0.755422 | 0.768423 | 0.775434 | 0.76678 | 0.770338 | 0.776294 | 0.765635 |
| 0.771281 | 0.767822 | 0.749014 | 0.761845 | 0.772764 | 0.762663 | 0.767061 | 0.767086 | 0.762503 |
| 0.711822 | 0.700623 | 0.756911 | 0.709159 | 0.717965 | 0.705167 | 0.709234 | 0.710862 | 0.706475 |
| 0.702117 | 0.697396 | 0.742787 | 0.705543 | 0.706552 | 0.70454 | 0.706953 | 0.704306 | 0.700239 |
| 0.707972 | 0.699377 | 0.750577 | 0.709293 | 0.709642 | 0.705702 | 0.7091 | 0.704326 | 0.698692 |
| 0.704435 | 0.689463 | 0.742927 | 0.700162 | 0.704308 | 0.7 | 0.703903 | 0.69779 | 0.694888 |
| 0.711819 | 0.700679 | 0.748521 | 0.715411 | 0.717739 | 0.706888 | 0.712226 | 0.707632 | 0.703945 |
| 0.71911 | 0.705148 | 0.758666 | 0.71998 | 0.723571 | 0.718804 | 0.720815 | 0.720971 | 0.712214 |
| 0.736967 | 0.721823 | 0.764009 | 0.733542 | 0.738764 | 0.730138 | 0.731381 | 0.728999 | 0.726405 |
| 0.710662 | 0.706309 | 0.753532 | 0.714951 | 0.718253 | 0.710444 | 0.713391 | 0.710662 | 0.707208 |
| 0.709866 | 0.701274 | 0.74423 | 0.712875 | 0.714445 | 0.710629 | 0.711277 | 0.710842 | 0.707887 |
| 0.742198 | 0.727256 | 0.765865 | 0.739778 | 0.744429 | 0.732756 | 0.736084 | 0.738528 | 0.731674 |
| 0.704684 | 0.693021 | 0.749277 | 0.706774 | 0.708756 | 0.69944 | 0.704765 | 0.701779 | 0.69697 |
| 0.708769 | 0.696903 | 0.752182 | 0.710068 | 0.715497 | 0.70783 | 0.71149 | 0.708775 | 0.704514 |
| 0.705042 | 0.695898 | 0.755055 | 0.703851 | 0.709256 | 0.702122 | 0.708266 | 0.702449 | 0.699932 |
| 0.706842 | 0.695947 | 0.747887 | 0.71064 | 0.712921 | 0.706082 | 0.710249 | 0.702407 | 0.701549 |
| 0.714829 | 0.701855 | 0.760531 | 0.71785 | 0.721056 | 0.713043 | 0.71478 | 0.713273 | 0.710766 |
| 0.712192 | 0.698251 | 0.755416 | 0.711069 | 0.716975 | 0.710722 | 0.71455 | 0.708764 | 0.706266 |
| 0.711001 | 0.699057 | 0.760335 | 0.712762 | 0.718315 | 0.711154 | 0.71402 | 0.711651 | 0.705017 |
| 0.705548 | 0.697986 | 0.752289 | 0.710443 | 0.713166 | 0.706274 | 0.708912 | 0.706165 | 0.701216 |
| 0.704219 | 0.695086 | 0.749588 | 0.702775 | 0.708517 | 0.699374 | 0.705764 | 0.701807 | 0.699171 |
| 0.707791 | 0.698845 | 0.760118 | 0.708455 | 0.712951 | 0.705578 | 0.709482 | 0.706685 | 0.702364 |
| 0.711694 | 0.698326 | 0.754976 | 0.712791 | 0.717806 | 0.712556 | 0.712822 | 0.713018 | 0.708616 |
| 0.710871 | 0.6983 | 0.746422 | 0.710895 | 0.713923 | 0.706541 | 0.711253 | 0.70513 | 0.702635 |
| 0.704781 | 0.699395 | 0.758487 | 0.706219 | 0.712756 | 0.707026 | 0.708119 | 0.703602 | 0.703218 |
| 0.703608 | 0.696183 | 0.752255 | 0.705768 | 0.711478 | 0.70579 | 0.707171 | 0.706141 | 0.696638 |
| 0.710151 | 0.697772 | 0.757795 | 0.707373 | 0.710522 | 0.703244 | 0.705699 | 0.707113 | 0.700469 |
| 0.710756 | 0.699563 | 0.750921 | 0.710629 | 0.712937 | 0.708165 | 0.709881 | 0.704572 | 0.701639 |
| 0.701032 | 0.69783 | 0.749712 | 0.706472 | 0.711339 | 0.705221 | 0.7076 | 0.703821 | 0.702483 |
| 0.709112 | 0.697546 | 0.755692 | 0.708866 | 0.71295 | 0.702647 | 0.70961 | 0.709028 | 0.702989 |
| 0.715045 | 0.70134 | 0.753761 | 0.712941 | 0.71786 | 0.708724 | 0.713085 | 0.710596 | 0.707344 |
| 0.711796 | 0.697423 | 0.756467 | 0.711869 | 0.718712 | 0.711931 | 0.713425 | 0.711209 | 0.707895 |
| 0.7141 | 0.706222 | 0.759799 | 0.713097 | 0.720114 | 0.708671 | 0.713103 | 0.711598 | 0.705359 |

| | | | | | | | | |
|---|---|---|---|---|---|---|---|---|
| 0.745716 | 0.735503 | 0.761496 | 0.744559 | 0.749801 | 0.740449 | 0.742462 | 0.744547 | 0.740114 |
| 0.700386 | 0.688075 | 0.742088 | 0.701537 | 0.703011 | 0.694857 | 0.700912 | 0.699229 | 0.690767 |
| 0.702241 | 0.688348 | 0.747885 | 0.704401 | 0.71077 | 0.698839 | 0.702926 | 0.699252 | 0.695723 |
| 0.710243 | 0.696843 | 0.747881 | 0.710853 | 0.716466 | 0.707435 | 0.715778 | 0.710884 | 0.705421 |
| 0.707073 | 0.695814 | 0.754619 | 0.705452 | 0.713982 | 0.70745 | 0.710159 | 0.706621 | 0.702481 |
| 0.710952 | 0.698513 | 0.763757 | 0.711647 | 0.715979 | 0.708898 | 0.712635 | 0.711691 | 0.705953 |
| 0.706181 | 0.693909 | 0.748181 | 0.707014 | 0.711517 | 0.70207 | 0.705901 | 0.702242 | 0.697305 |
| 0.710492 | 0.699411 | 0.751159 | 0.710466 | 0.71508 | 0.706569 | 0.710237 | 0.707593 | 0.705462 |
| 0.710646 | 0.697988 | 0.744972 | 0.713568 | 0.716027 | 0.707208 | 0.71381 | 0.709813 | 0.704893 |
| 0.707742 | 0.699918 | 0.763434 | 0.712282 | 0.71478 | 0.707016 | 0.712033 | 0.710064 | 0.702405 |
| 0.703863 | 0.689113 | 0.742458 | 0.702658 | 0.706899 | 0.699711 | 0.701862 | 0.698757 | 0.697331 |
| 0.703665 | 0.693473 | 0.746869 | 0.702588 | 0.707342 | 0.700182 | 0.705925 | 0.702399 | 0.696367 |
| 0.706361 | 0.692555 | 0.749132 | 0.706872 | 0.714081 | 0.705159 | 0.712244 | 0.70604 | 0.704059 |
| 0.706948 | 0.693262 | 0.74666 | 0.702838 | 0.708168 | 0.699561 | 0.705085 | 0.703118 | 0.697988 |
| 0.703824 | 0.689585 | 0.741154 | 0.704493 | 0.706637 | 0.70231 | 0.705539 | 0.702555 | 0.69753 |
| 0.702906 | 0.690793 | 0.741153 | 0.703888 | 0.704763 | 0.700619 | 0.702514 | 0.702418 | 0.698197 |
| 0.701068 | 0.690542 | 0.742379 | 0.701431 | 0.70546 | 0.698496 | 0.706657 | 0.700151 | 0.6973 |
| 0.710737 | 0.692644 | 0.749836 | 0.714204 | 0.714511 | 0.707333 | 0.712701 | 0.709747 | 0.705524 |
| 0.710168 | 0.700046 | 0.74881 | 0.712196 | 0.713315 | 0.70768 | 0.709184 | 0.708354 | 0.706963 |
| 0.70815 | 0.691764 | 0.74604 | 0.707522 | 0.711366 | 0.699561 | 0.709361 | 0.702913 | 0.69818 |
| 0.699587 | 0.688678 | 0.743393 | 0.699257 | 0.704439 | 0.69803 | 0.701351 | 0.698458 | 0.694168 |
| 0.70323 | 0.692082 | 0.744927 | 0.704696 | 0.710099 | 0.699637 | 0.703781 | 0.702092 | 0.697868 |
| 0.705656 | 0.695921 | 0.747302 | 0.705944 | 0.709688 | 0.704193 | 0.706235 | 0.703743 | 0.699474 |
| 0.718465 | 0.706185 | 0.76537 | 0.715342 | 0.719376 | 0.712777 | 0.717419 | 0.715916 | 0.705508 |
| 0.722173 | 0.715047 | 0.766562 | 0.721731 | 0.721679 | 0.717649 | 0.724915 | 0.720204 | 0.715329 |
| 0.770388 | 0.769165 | 0.749959 | 0.764063 | 0.773702 | 0.763666 | 0.769334 | 0.767815 | 0.763306 |
| 0.785216 | 0.768569 | 0.752943 | 0.785048 | 0.776377 | 0.778775 | 0.783683 | 0.778425 | 0.777489 |
| 0.785502 | 0.770616 | 0.754505 | 0.785452 | 0.784595 | 0.781982 | 0.786339 | 0.780839 | 0.779133 |
| 0.704857 | 0.689087 | 0.747288 | 0.708701 | 0.711109 | 0.70421 | 0.707299 | 0.703239 | 0.698102 |
| 0.704816 | 0.693632 | 0.74458 | 0.707022 | 0.712015 | 0.703224 | 0.710087 | 0.705113 | 0.699974 |
| 0.712419 | 0.701027 | 0.747941 | 0.712426 | 0.717778 | 0.709076 | 0.713581 | 0.710577 | 0.704101 |
| 0.709812 | 0.704189 | 0.757447 | 0.719744 | 0.722679 | 0.71207 | 0.712473 | 0.715494 | 0.704506 |
| 0.704167 | 0.695829 | 0.756849 | 0.706237 | 0.711944 | 0.704699 | 0.705261 | 0.703139 | 0.697171 |
| 0.706794 | 0.694623 | 0.74971 | 0.707501 | 0.713853 | 0.706879 | 0.709806 | 0.705724 | 0.701265 |
| 0.7058 | 0.694255 | 0.752299 | 0.707186 | 0.710455 | 0.701009 | 0.707259 | 0.706022 | 0.698892 |
| 0.706322 | 0.698978 | 0.752807 | 0.702824 | 0.709457 | 0.70319 | 0.706874 | 0.703868 | 0.699803 |
| 0.703896 | 0.696526 | 0.744605 | 0.707637 | 0.71236 | 0.706143 | 0.708792 | 0.703481 | 0.703142 |
| 0.72054 | 0.707019 | 0.767241 | 0.718157 | 0.721261 | 0.714084 | 0.721124 | 0.715588 | 0.711197 |
| 0.698199 | 0.687195 | 0.741675 | 0.697565 | 0.701466 | 0.696119 | 0.698518 | 0.695499 | 0.6943 |
| 0.704858 | 0.695242 | 0.746383 | 0.706653 | 0.712214 | 0.706307 | 0.705069 | 0.704934 | 0.699406 |
| 0.699712 | 0.690141 | 0.747571 | 0.702281 | 0.705206 | 0.698511 | 0.700835 | 0.698961 | 0.692529 |
| 0.722644 | 0.710176 | 0.770604 | 0.721074 | 0.727802 | 0.717761 | 0.721745 | 0.719614 | 0.715668 |
| 0.70523 | 0.691429 | 0.747262 | 0.703619 | 0.708409 | 0.702788 | 0.707352 | 0.707527 | 0.6977 |
| 0.704214 | 0.691748 | 0.751247 | 0.703902 | 0.709326 | 0.699256 | 0.703565 | 0.703271 | 0.698088 |
| 0.70646 | 0.699339 | 0.748493 | 0.708149 | 0.714104 | 0.70833 | 0.711383 | 0.706729 | 0.705051 |
| 0.705767 | 0.695792 | 0.751183 | 0.70698 | 0.706009 | 0.704824 | 0.705267 | 0.705291 | 0.70163 |

| | | | | | | | | |
|---|---|---|---|---|---|---|---|---|
| 0.701894 | 0.692237 | 0.745943 | 0.705421 | 0.707517 | 0.703888 | 0.704463 | 0.703722 | 0.697234 |
| 0.709192 | 0.696523 | 0.743025 | 0.709423 | 0.714782 | 0.706973 | 0.710674 | 0.706606 | 0.703738 |
| 0.70716 | 0.693245 | 0.7432 | 0.705713 | 0.713759 | 0.705284 | 0.70947 | 0.700886 | 0.701753 |
| 0.714885 | 0.702693 | 0.757811 | 0.711356 | 0.716392 | 0.709318 | 0.715415 | 0.712528 | 0.706153 |
| 0.713951 | 0.70296 | 0.757646 | 0.712935 | 0.717807 | 0.709651 | 0.713851 | 0.708596 | 0.707617 |
| 0.707749 | 0.694365 | 0.747081 | 0.705607 | 0.71152 | 0.704461 | 0.708114 | 0.704919 | 0.700465 |
| 0.706302 | 0.700757 | 0.757874 | 0.710054 | 0.712192 | 0.702479 | 0.711508 | 0.705688 | 0.705913 |
| 0.712583 | 0.700508 | 0.747048 | 0.710736 | 0.718433 | 0.709747 | 0.714507 | 0.712832 | 0.706512 |
| 0.705285 | 0.694911 | 0.74767 | 0.710241 | 0.715355 | 0.710861 | 0.712711 | 0.704696 | 0.70301 |
| 0.702997 | 0.695465 | 0.742624 | 0.709306 | 0.713394 | 0.701744 | 0.708692 | 0.707679 | 0.701011 |
| 0.704202 | 0.695353 | 0.743861 | 0.708151 | 0.710217 | 0.703446 | 0.707412 | 0.702019 | 0.699788 |
| 0.707777 | 0.695957 | 0.745269 | 0.708457 | 0.711001 | 0.706241 | 0.708483 | 0.706046 | 0.702656 |
| 0.710029 | 0.697868 | 0.744928 | 0.709324 | 0.712807 | 0.70825 | 0.710177 | 0.709254 | 0.701974 |
| 0.70554 | 0.692308 | 0.741399 | 0.706155 | 0.711716 | 0.703601 | 0.70706 | 0.702205 | 0.699618 |
| 0.70484 | 0.701482 | 0.746992 | 0.706836 | 0.713447 | 0.706072 | 0.708691 | 0.708368 | 0.702954 |
| 0.706579 | 0.6938 | 0.750533 | 0.706771 | 0.711598 | 0.704246 | 0.707485 | 0.703777 | 0.701006 |
| 0.712079 | 0.701182 | 0.752671 | 0.709831 | 0.717327 | 0.7089 | 0.710364 | 0.710201 | 0.706259 |
| 0.706515 | 0.695867 | 0.748567 | 0.70781 | 0.712267 | 0.704845 | 0.706765 | 0.70428 | 0.699795 |
| 0.706516 | 0.699432 | 0.751587 | 0.709477 | 0.714524 | 0.70469 | 0.708501 | 0.706 | 0.702389 |
| 0.71006 | 0.697992 | 0.746422 | 0.707668 | 0.715306 | 0.706286 | 0.707458 | 0.707602 | 0.705426 |
| 0.707636 | 0.696709 | 0.744349 | 0.709865 | 0.713888 | 0.707412 | 0.708051 | 0.70716 | 0.705066 |
| 0.704026 | 0.692055 | 0.739028 | 0.701693 | 0.706875 | 0.701118 | 0.703416 | 0.701588 | 0.69543 |
| 0.705863 | 0.693647 | 0.747581 | 0.707213 | 0.706447 | 0.707581 | 0.711234 | 0.705029 | 0.700974 |
| 0.708525 | 0.693809 | 0.744951 | 0.707536 | 0.715844 | 0.704936 | 0.709088 | 0.705677 | 0.699991 |
| 0.706595 | 0.701091 | 0.746332 | 0.709915 | 0.713167 | 0.708595 | 0.711028 | 0.704414 | 0.702242 |
| 0.712755 | 0.705593 | 0.762394 | 0.716017 | 0.721714 | 0.710277 | 0.714206 | 0.710289 | 0.707242 |
| 0.708333 | 0.697182 | 0.744728 | 0.709379 | 0.709296 | 0.703971 | 0.707851 | 0.701662 | 0.702991 |
| 0.705353 | 0.69842 | 0.747443 | 0.711365 | 0.716281 | 0.709086 | 0.711058 | 0.706429 | 0.706724 |
| 0.71039 | 0.697872 | 0.75304 | 0.709283 | 0.713723 | 0.706595 | 0.713619 | 0.709519 | 0.707426 |
| 0.708511 | 0.702074 | 0.745734 | 0.708409 | 0.713988 | 0.707811 | 0.713117 | 0.703763 | 0.704887 |
| 0.708136 | 0.699265 | 0.750021 | 0.710444 | 0.714491 | 0.710163 | 0.712742 | 0.708783 | 0.705127 |
| 0.71112 | 0.693966 | 0.749796 | 0.706714 | 0.710364 | 0.703341 | 0.708771 | 0.704143 | 0.698723 |
| 0.705372 | 0.697855 | 0.750519 | 0.709587 | 0.712893 | 0.706345 | 0.70926 | 0.707175 | 0.703607 |
| 0.707032 | 0.693923 | 0.74464 | 0.706859 | 0.712211 | 0.704731 | 0.710237 | 0.707911 | 0.702239 |
| 0.708567 | 0.695254 | 0.74383 | 0.707788 | 0.713422 | 0.705624 | 0.704771 | 0.703028 | 0.699837 |
| 0.705661 | 0.696732 | 0.74896 | 0.707079 | 0.712909 | 0.704413 | 0.706635 | 0.70622 | 0.706594 |
| 0.70415 | 0.690122 | 0.743877 | 0.702155 | 0.712263 | 0.70647 | 0.708891 | 0.703429 | 0.696209 |
| 0.679336 | 0.675759 | 0.722463 | 0.683012 | 0.689395 | 0.690578 | 0.681869 | 0.686338 | 0.683732 |
| 0.686824 | 0.680568 | 0.727732 | 0.688546 | 0.687682 | 0.683807 | 0.683461 | 0.684046 | 0.681958 |
| 0.677666 | 0.67518 | 0.728202 | 0.678166 | 0.685574 | 0.67478 | 0.681515 | 0.680122 | 0.675597 |
| 0.675358 | 0.670828 | 0.727696 | 0.682381 | 0.678199 | 0.674151 | 0.678379 | 0.680438 | 0.676692 |
| 0.660647 | 0.65846 | 0.703933 | 0.666272 | 0.672101 | 0.669698 | 0.666627 | 0.664948 | 0.664684 |
| 0.658744 | 0.658018 | 0.712683 | 0.667929 | 0.670398 | 0.668315 | 0.665974 | 0.673389 | 0.666417 |
| 0.663711 | 0.661402 | 0.714639 | 0.662087 | 0.668394 | 0.662842 | 0.667074 | 0.666584 | 0.661396 |
| 0.666667 | 0.663116 | 0.719745 | 0.671151 | 0.668519 | 0.663556 | 0.65989 | 0.664249 | 0.666316 |
| 0.661333 | 0.660707 | 0.723152 | 0.67012 | 0.67167 | 0.670268 | 0.665913 | 0.666709 | 0.665939 |
| 0.674308 | 0.670328 | 0.71743 | 0.674404 | 0.674911 | 0.673208 | 0.674349 | 0.674154 | 0.666582 |

| | | | | | | | | |
|---|---|---|---|---|---|---|---|---|
| 0.662849 | 0.664329 | 0.705824 | 0.667286 | 0.676215 | 0.668467 | 0.667533 | 0.667335 | 0.666582 |
| 0.66885 | 0.670925 | 0.721758 | 0.671222 | 0.678052 | 0.668253 | 0.672935 | 0.670133 | 0.667432 |
| 0.668275 | 0.668484 | 0.714518 | 0.668102 | 0.672055 | 0.667711 | 0.670085 | 0.669616 | 0.660447 |
| 0.659501 | 0.658098 | 0.707981 | 0.66041 | 0.663056 | 0.6596 | 0.656491 | 0.65917 | 0.656313 |
| 0.660749 | 0.661631 | 0.709703 | 0.669 | 0.670206 | 0.665227 | 0.666494 | 0.668858 | 0.661809 |
| 0.661089 | 0.659612 | 0.710095 | 0.664737 | 0.670133 | 0.665598 | 0.665976 | 0.669889 | 0.66437 |
| 0.663125 | 0.661902 | 0.710621 | 0.670047 | 0.671917 | 0.664272 | 0.669528 | 0.668542 | 0.666327 |
| 0.659324 | 0.661896 | 0.717078 | 0.66421 | 0.66792 | 0.662961 | 0.659203 | 0.665953 | 0.662878 |
| 0.650824 | 0.657209 | 0.697747 | 0.663862 | 0.66327 | 0.662083 | 0.657887 | 0.658707 | 0.658801 |
| 0.661267 | 0.658087 | 0.712511 | 0.667157 | 0.667294 | 0.661386 | 0.667699 | 0.663939 | 0.664666 |
| 0.653147 | 0.649726 | 0.699357 | 0.65549 | 0.664274 | 0.660099 | 0.65857 | 0.660583 | 0.660072 |
| 0.693579 | 0.6832 | 0.738718 | 0.695027 | 0.699588 | 0.692764 | 0.698291 | 0.690854 | 0.689048 |
| 0.704653 | 0.699408 | 0.749306 | 0.707058 | 0.712706 | 0.705506 | 0.708699 | 0.70655 | 0.701925 |
| 0.684482 | 0.677325 | 0.743158 | 0.685051 | 0.684727 | 0.678348 | 0.680348 | 0.687354 | 0.683257 |
| 0.710643 | 0.700703 | 0.750561 | 0.711341 | 0.718429 | 0.709518 | 0.713108 | 0.708344 | 0.703874 |
| 0.700395 | 0.688956 | 0.734499 | 0.705239 | 0.709025 | 0.702214 | 0.703689 | 0.702154 | 0.697782 |
| 0.712003 | 0.702317 | 0.747222 | 0.710223 | 0.718671 | 0.706847 | 0.713165 | 0.710738 | 0.704644 |
| 0.721089 | 0.709062 | 0.75901 | 0.712396 | 0.719415 | 0.712176 | 0.719791 | 0.711001 | 0.710057 |
| 0.69551 | 0.685419 | 0.729771 | 0.695185 | 0.698544 | 0.694696 | 0.694708 | 0.693501 | 0.687912 |
| 0.717813 | 0.701662 | 0.759892 | 0.719045 | 0.722485 | 0.718219 | 0.718314 | 0.717218 | 0.714958 |
| 0.691738 | 0.685417 | 0.740736 | 0.692083 | 0.696744 | 0.688213 | 0.690349 | 0.69112 | 0.685868 |
| 0.716739 | 0.702855 | 0.746916 | 0.712797 | 0.719115 | 0.70995 | 0.716356 | 0.71168 | 0.706707 |
| 0.716344 | 0.710077 | 0.747083 | 0.716857 | 0.72159 | 0.716439 | 0.720662 | 0.710491 | 0.71215 |
| 0.687419 | 0.682029 | 0.737886 | 0.68112 | 0.685642 | 0.682476 | 0.680908 | 0.685993 | 0.683159 |
| 0.684431 | 0.677504 | 0.723139 | 0.684503 | 0.689734 | 0.683249 | 0.685992 | 0.684749 | 0.678958 |
| 0.684959 | 0.687936 | 0.736537 | 0.691336 | 0.696206 | 0.685071 | 0.691432 | 0.691669 | 0.68764 |
| 0.704013 | 0.696409 | 0.755083 | 0.707254 | 0.711663 | 0.704817 | 0.708553 | 0.705242 | 0.701069 |
| 0.71661 | 0.705478 | 0.759872 | 0.715546 | 0.719991 | 0.713884 | 0.71682 | 0.715202 | 0.710806 |
| 0.687687 | 0.674795 | 0.717625 | 0.681898 | 0.685962 | 0.681087 | 0.683758 | 0.682489 | 0.676072 |
| 0.723152 | 0.714103 | 0.755292 | 0.724792 | 0.726872 | 0.717459 | 0.722196 | 0.721317 | 0.718528 |
| 0.727514 | 0.714312 | 0.758923 | 0.727539 | 0.728029 | 0.72371 | 0.723408 | 0.724341 | 0.722364 |
| 0.693794 | 0.683647 | 0.732565 | 0.69658 | 0.69722 | 0.690574 | 0.697419 | 0.694617 | 0.689201 |
| 0.696105 | 0.683517 | 0.736915 | 0.697473 | 0.696461 | 0.691975 | 0.696109 | 0.692282 | 0.690508 |
| 0.722832 | 0.712079 | 0.776078 | 0.721458 | 0.725693 | 0.71889 | 0.722307 | 0.722176 | 0.713929 |
| 0.7224 | 0.708816 | 0.748732 | 0.721019 | 0.722244 | 0.717963 | 0.721551 | 0.715013 | 0.713004 |
| 0.72562 | 0.709952 | 0.752496 | 0.71954 | 0.725866 | 0.716434 | 0.722312 | 0.718295 | 0.716121 |
| 0.675885 | 0.663148 | 0.709724 | 0.679432 | 0.680657 | 0.676243 | 0.674068 | 0.673496 | 0.670925 |
| 0.718148 | 0.707092 | 0.741781 | 0.718223 | 0.721204 | 0.717531 | 0.724196 | 0.716463 | 0.710585 |
| 0.676535 | 0.660996 | 0.706829 | 0.674596 | 0.675936 | 0.671356 | 0.674841 | 0.676816 | 0.669332 |
| 0.668593 | 0.653295 | 0.695817 | 0.670007 | 0.671129 | 0.664963 | 0.672116 | 0.667488 | 0.663102 |
| 0.714745 | 0.705288 | 0.755679 | 0.720904 | 0.724669 | 0.714403 | 0.720843 | 0.716718 | 0.711031 |
| 0.667952 | 0.654923 | 0.697866 | 0.670455 | 0.670655 | 0.665225 | 0.67089 | 0.666984 | 0.663084 |
| 0.659273 | 0.649586 | 0.693994 | 0.663424 | 0.66329 | 0.657354 | 0.66118 | 0.658943 | 0.654826 |
| 0.657037 | 0.64932 | 0.688132 | 0.659468 | 0.662111 | 0.651407 | 0.656358 | 0.654955 | 0.65138 |
| 0.727516 | 0.712446 | 0.75545 | 0.725814 | 0.731392 | 0.722768 | 0.723221 | 0.724049 | 0.714286 |
| 0.660476 | 0.648371 | 0.691678 | 0.663096 | 0.665052 | 0.65793 | 0.660153 | 0.661788 | 0.65525 |
| 0.722562 | 0.706657 | 0.749386 | 0.720512 | 0.725384 | 0.716428 | 0.720598 | 0.715635 | 0.712958 |

| | | | | | | | | |
|---|---|---|---|---|---|---|---|---|
| 0.729894 | 0.718188 | 0.754674 | 0.725345 | 0.73591 | 0.72219 | 0.727178 | 0.726749 | 0.716752 |
| 0.724765 | 0.71077 | 0.756497 | 0.723484 | 0.729406 | 0.718634 | 0.723741 | 0.71989 | 0.714176 |
| 0.743549 | 0.736361 | 0.760527 | 0.744099 | 0.746926 | 0.73666 | 0.742238 | 0.744123 | 0.736603 |
| 0.651456 | 0.640588 | 0.684531 | 0.651193 | 0.652976 | 0.644392 | 0.651272 | 0.650077 | 0.643595 |
| 0.651379 | 0.640249 | 0.680101 | 0.654028 | 0.653116 | 0.646782 | 0.652079 | 0.651462 | 0.645887 |
| 0.729462 | 0.715123 | 0.758762 | 0.728251 | 0.735881 | 0.723406 | 0.727589 | 0.730029 | 0.726769 |
| 0.654108 | 0.642388 | 0.676927 | 0.655478 | 0.655055 | 0.65148 | 0.653177 | 0.650982 | 0.648543 |
| 0.730218 | 0.722321 | 0.737652 | 0.729281 | 0.737795 | 0.723958 | 0.732972 | 0.725698 | 0.721922 |
| 0.729194 | 0.71923 | 0.754097 | 0.728308 | 0.737476 | 0.726269 | 0.730049 | 0.727878 | 0.72378 |
| 0.732736 | 0.725141 | 0.732844 | 0.731235 | 0.734812 | 0.726889 | 0.734055 | 0.726575 | 0.722685 |
| 0.732675 | 0.719728 | 0.748185 | 0.728343 | 0.733847 | 0.721595 | 0.728676 | 0.728265 | 0.725229 |
| 0.707792 | 0.688742 | 0.701389 | 0.634615 | 0.673333 | 0.682759 | 0.687075 | 0.739437 | 0.72 |
| 0.720832 | 0.727138 | 0.707856 | 0.727932 | 0.726639 | 0.724306 | 0.718429 | 0.733715 | 0.72644 |
| 0.729477 | 0.726937 | 0.711197 | 0.720732 | 0.73039 | 0.723513 | 0.72195 | 0.732939 | 0.724852 |
| 0.738026 | 0.733258 | 0.714455 | 0.730074 | 0.745835 | 0.731595 | 0.73919 | 0.742338 | 0.736998 |
| 0.733651 | 0.729179 | 0.716543 | 0.728591 | 0.731652 | 0.725573 | 0.732599 | 0.734341 | 0.729569 |
| 0.587409 | 0.58223 | 0.611475 | 0.59391 | 0.590831 | 0.590634 | 0.590494 | 0.592465 | 0.588585 |
| 0.731711 | 0.73106 | 0.716072 | 0.731881 | 0.736561 | 0.733475 | 0.733817 | 0.740706 | 0.735089 |
| 0.736232 | 0.72308 | 0.75261 | 0.737861 | 0.739029 | 0.729198 | 0.735577 | 0.73249 | 0.726011 |
| 0.736732 | 0.724713 | 0.747201 | 0.732536 | 0.738546 | 0.729307 | 0.73345 | 0.731536 | 0.726768 |
| 0.741834 | 0.726728 | 0.74971 | 0.735326 | 0.741001 | 0.732528 | 0.733394 | 0.736398 | 0.729108 |
| 0.592868 | 0.582383 | 0.607087 | 0.596341 | 0.595895 | 0.587463 | 0.59528 | 0.59368 | 0.589986 |
| 0.579468 | 0.56667 | 0.590144 | 0.582202 | 0.584461 | 0.575563 | 0.580227 | 0.579354 | 0.576985 |
| 0.762074 | 0.750167 | 0.743844 | 0.754303 | 0.760667 | 0.750858 | 0.756263 | 0.759725 | 0.753362 |
| 0.762561 | 0.75229 | 0.742862 | 0.759169 | 0.759565 | 0.753713 | 0.761822 | 0.759928 | 0.750948 |
| 0.752867 | 0.743541 | 0.735539 | 0.749937 | 0.750651 | 0.742753 | 0.748111 | 0.752547 | 0.74473 |
| 0.76045 | 0.745701 | 0.741212 | 0.754772 | 0.759469 | 0.753371 | 0.755304 | 0.76022 | 0.752644 |
| 0.759265 | 0.753106 | 0.743846 | 0.760278 | 0.758197 | 0.755054 | 0.75891 | 0.757413 | 0.751407 |
| 0.766557 | 0.750403 | 0.744462 | 0.759704 | 0.769072 | 0.75691 | 0.76342 | 0.761339 | 0.757422 |
| 0.76447 | 0.751206 | 0.74604 | 0.759018 | 0.760283 | 0.754974 | 0.759086 | 0.763826 | 0.754392 |
| 0.758972 | 0.748458 | 0.742146 | 0.756828 | 0.761084 | 0.754069 | 0.756773 | 0.762371 | 0.753195 |
| 0.771056 | 0.763051 | 0.756247 | 0.772666 | 0.773904 | 0.763398 | 0.769028 | 0.769928 | 0.766761 |
| 0.761188 | 0.750213 | 0.746059 | 0.759892 | 0.769345 | 0.762048 | 0.758598 | 0.761913 | 0.756048 |
| 0.762748 | 0.757678 | 0.745242 | 0.761107 | 0.762307 | 0.757851 | 0.759075 | 0.760163 | 0.753936 |
| 0.76756 | 0.760526 | 0.751091 | 0.765678 | 0.769738 | 0.760692 | 0.770017 | 0.770808 | 0.765003 |
| 0.763725 | 0.750142 | 0.744481 | 0.762955 | 0.762691 | 0.758747 | 0.762373 | 0.760514 | 0.756805 |
| 0.766129 | 0.750825 | 0.741435 | 0.762885 | 0.766509 | 0.755764 | 0.758672 | 0.769397 | 0.758367 |
| 0.768203 | 0.76118 | 0.750555 | 0.767054 | 0.774054 | 0.762293 | 0.764181 | 0.765412 | 0.75987 |
| 0.763717 | 0.74976 | 0.742759 | 0.756599 | 0.765001 | 0.758266 | 0.756337 | 0.758645 | 0.750539 |
| 0.759486 | 0.7519 | 0.742624 | 0.757783 | 0.763829 | 0.757922 | 0.758094 | 0.758574 | 0.75046 |
| 0.72163 | 0.719949 | 0.706784 | 0.727979 | 0.735459 | 0.716944 | 0.718476 | 0.731319 | 0.723492 |
| 0.727512 | 0.729258 | 0.713835 | 0.728376 | 0.737637 | 0.719589 | 0.726721 | 0.737397 | 0.719177 |
| 0.766332 | 0.754551 | 0.748077 | 0.762662 | 0.767527 | 0.758772 | 0.764585 | 0.762528 | 0.755864 |
| 0.771354 | 0.761202 | 0.749343 | 0.762588 | 0.764792 | 0.759929 | 0.763647 | 0.765954 | 0.760979 |
| 0.719258 | 0.721586 | 0.698013 | 0.718334 | 0.71954 | 0.706579 | 0.716412 | 0.726593 | 0.721091 |
| 0.765909 | 0.755193 | 0.748392 | 0.764061 | 0.768056 | 0.759786 | 0.768591 | 0.762023 | 0.761968 |
| 0.758114 | 0.750118 | 0.738049 | 0.758713 | 0.759981 | 0.751772 | 0.757157 | 0.759288 | 0.753792 |

| | | | | | | | | |
|---|---|---|---|---|---|---|---|---|
| 0.766929 | 0.755175 | 0.743082 | 0.760289 | 0.770959 | 0.757591 | 0.760374 | 0.762171 | 0.7593 |
| 0.771177 | 0.757838 | 0.74812 | 0.763718 | 0.766343 | 0.758417 | 0.762687 | 0.763113 | 0.754527 |
| 0.759248 | 0.75782 | 0.744168 | 0.760323 | 0.767211 | 0.756843 | 0.755991 | 0.761604 | 0.754395 |
| 0.771146 | 0.75773 | 0.746634 | 0.763412 | 0.769626 | 0.760105 | 0.763428 | 0.764438 | 0.762886 |
| 0.767 | 0.758881 | 0.750021 | 0.768001 | 0.772946 | 0.762673 | 0.76493 | 0.769383 | 0.764113 |
| 0.765012 | 0.756857 | 0.743801 | 0.760175 | 0.764682 | 0.755885 | 0.760768 | 0.763099 | 0.758177 |
| 0.767674 | 0.753926 | 0.741509 | 0.759753 | 0.765006 | 0.758916 | 0.758575 | 0.763372 | 0.756513 |
| 0.756714 | 0.749093 | 0.73737 | 0.755137 | 0.758001 | 0.751071 | 0.756849 | 0.753826 | 0.751831 |
| 0.764765 | 0.75224 | 0.741351 | 0.76386 | 0.770765 | 0.760085 | 0.763356 | 0.766649 | 0.757306 |
| 0.759281 | 0.752083 | 0.740145 | 0.756609 | 0.761761 | 0.754364 | 0.756336 | 0.758199 | 0.749801 |
| 0.769122 | 0.756024 | 0.747771 | 0.765617 | 0.769443 | 0.762244 | 0.76331 | 0.768709 | 0.761544 |
| 0.766928 | 0.749928 | 0.743193 | 0.762003 | 0.766463 | 0.755614 | 0.757578 | 0.760623 | 0.754879 |
| 0.759133 | 0.746744 | 0.734837 | 0.753714 | 0.757103 | 0.750043 | 0.754613 | 0.755588 | 0.747237 |
| 0.75725 | 0.747102 | 0.730675 | 0.752534 | 0.755157 | 0.750816 | 0.754097 | 0.755678 | 0.748401 |
| 0.764823 | 0.755436 | 0.748303 | 0.765612 | 0.770022 | 0.756745 | 0.760226 | 0.762473 | 0.760165 |
| 0.764208 | 0.753399 | 0.748381 | 0.759441 | 0.764763 | 0.758133 | 0.763225 | 0.761542 | 0.75645 |
| 0.763131 | 0.752275 | 0.748131 | 0.760722 | 0.76293 | 0.758733 | 0.759537 | 0.763282 | 0.761492 |
| 0.759736 | 0.75026 | 0.741237 | 0.755772 | 0.764653 | 0.754143 | 0.756741 | 0.758312 | 0.753558 |
| 0.763066 | 0.753441 | 0.744905 | 0.762809 | 0.764706 | 0.759189 | 0.761123 | 0.762955 | 0.756343 |
| 0.762012 | 0.752741 | 0.745243 | 0.756001 | 0.76147 | 0.755332 | 0.759373 | 0.758469 | 0.753872 |
| 0.755622 | 0.743821 | 0.737066 | 0.753092 | 0.758284 | 0.748428 | 0.751575 | 0.751479 | 0.746576 |
| 0.756688 | 0.749833 | 0.74569 | 0.757558 | 0.760139 | 0.751185 | 0.756761 | 0.760957 | 0.747094 |
| 0.765813 | 0.756139 | 0.751988 | 0.759863 | 0.764331 | 0.756174 | 0.762065 | 0.767157 | 0.755433 |
| 0.762973 | 0.755226 | 0.745991 | 0.7621 | 0.76527 | 0.75875 | 0.760816 | 0.761942 | 0.75771 |
| 0.762512 | 0.756132 | 0.743047 | 0.759061 | 0.767644 | 0.755106 | 0.75717 | 0.761087 | 0.752764 |
| 0.758274 | 0.748602 | 0.743713 | 0.754337 | 0.758167 | 0.751704 | 0.757996 | 0.759302 | 0.750265 |
| 0.755501 | 0.747129 | 0.737975 | 0.755015 | 0.757636 | 0.748863 | 0.752065 | 0.753275 | 0.745033 |
| 0.758246 | 0.743953 | 0.737014 | 0.758058 | 0.76087 | 0.750992 | 0.751448 | 0.757798 | 0.752716 |
| 0.759077 | 0.751275 | 0.741051 | 0.755375 | 0.759789 | 0.752969 | 0.759464 | 0.760107 | 0.751745 |
| 0.768047 | 0.750354 | 0.74273 | 0.762939 | 0.763693 | 0.757529 | 0.759593 | 0.760498 | 0.757819 |
| 0.76108 | 0.753906 | 0.745592 | 0.760087 | 0.766044 | 0.757329 | 0.762438 | 0.757589 | 0.756289 |
| 0.759768 | 0.754436 | 0.739674 | 0.758425 | 0.759237 | 0.754045 | 0.755937 | 0.760505 | 0.754942 |
| 0.765086 | 0.757986 | 0.747125 | 0.766708 | 0.76887 | 0.761132 | 0.765996 | 0.765339 | 0.758727 |
| 0.768227 | 0.754658 | 0.749404 | 0.76649 | 0.76713 | 0.760109 | 0.762699 | 0.762964 | 0.761017 |
| 0.757647 | 0.74721 | 0.742593 | 0.755539 | 0.758188 | 0.749257 | 0.758421 | 0.757062 | 0.750686 |
| 0.761748 | 0.752027 | 0.746702 | 0.757444 | 0.767122 | 0.753436 | 0.762101 | 0.758118 | 0.751001 |
| 0.762261 | 0.749149 | 0.744209 | 0.761816 | 0.762166 | 0.757106 | 0.761615 | 0.758954 | 0.757587 |
| 0.759738 | 0.748839 | 0.745096 | 0.757635 | 0.764517 | 0.75523 | 0.757791 | 0.761834 | 0.755231 |
| 0.752644 | 0.744522 | 0.735129 | 0.748001 | 0.756001 | 0.744432 | 0.748574 | 0.748499 | 0.743684 |
| 0.757955 | 0.743149 | 0.73631 | 0.75237 | 0.752957 | 0.748672 | 0.752292 | 0.752989 | 0.746387 |
| 0.753096 | 0.741444 | 0.734259 | 0.744442 | 0.751236 | 0.740738 | 0.745407 | 0.748104 | 0.740051 |
| 0.750749 | 0.746543 | 0.738201 | 0.746081 | 0.750818 | 0.744675 | 0.747812 | 0.75102 | 0.746382 |
| 0.750978 | 0.739325 | 0.734069 | 0.748056 | 0.75138 | 0.742464 | 0.749159 | 0.747434 | 0.74048 |
| 0.754243 | 0.739406 | 0.736411 | 0.750941 | 0.750089 | 0.746136 | 0.745432 | 0.747761 | 0.742863 |
| 0.760723 | 0.750293 | 0.742194 | 0.749212 | 0.758954 | 0.754494 | 0.753885 | 0.76036 | 0.748397 |
| 0.749192 | 0.741054 | 0.731949 | 0.75399 | 0.752847 | 0.745537 | 0.748981 | 0.747223 | 0.746799 |

| | | | | | | | | |
|---|---|---|---|---|---|---|---|---|
| 0.762307 | 0.751279 | 0.746574 | 0.754747 | 0.763446 | 0.751466 | 0.757075 | 0.759677 | 0.751633 |
| 0.774166 | 0.75915 | 0.75393 | 0.771954 | 0.775147 | 0.761774 | 0.770345 | 0.775805 | 0.766179 |
| 0.762849 | 0.756487 | 0.745835 | 0.764751 | 0.770174 | 0.764833 | 0.763985 | 0.768271 | 0.760254 |
| 0.767112 | 0.76009 | 0.752007 | 0.766318 | 0.771832 | 0.759469 | 0.770191 | 0.766812 | 0.761775 |
| 0.777796 | 0.768149 | 0.756667 | 0.778551 | 0.778442 | 0.766357 | 0.778864 | 0.779291 | 0.771434 |
| 0.771715 | 0.762813 | 0.751417 | 0.771779 | 0.774841 | 0.767438 | 0.770249 | 0.775734 | 0.764706 |
| 0.764359 | 0.759281 | 0.747628 | 0.76745 | 0.76944 | 0.758861 | 0.763459 | 0.772205 | 0.76048 |
| 0.767225 | 0.751435 | 0.749549 | 0.764242 | 0.765518 | 0.757365 | 0.761426 | 0.767588 | 0.757266 |
| 0.763757 | 0.756488 | 0.746841 | 0.762298 | 0.76278 | 0.757556 | 0.763411 | 0.768337 | 0.759606 |
| 0.760523 | 0.758614 | 0.745516 | 0.762743 | 0.767181 | 0.75404 | 0.757292 | 0.764026 | 0.756393 |
| 0.768102 | 0.753416 | 0.749694 | 0.765192 | 0.775805 | 0.761591 | 0.770035 | 0.765644 | 0.760025 |
| 0.763855 | 0.75783 | 0.750988 | 0.763801 | 0.773292 | 0.762613 | 0.763343 | 0.769237 | 0.760021 |
| 0.777414 | 0.767091 | 0.75074 | 0.769574 | 0.77315 | 0.764961 | 0.774051 | 0.768516 | 0.765179 |
| 0.770111 | 0.762352 | 0.749979 | 0.771624 | 0.775552 | 0.771257 | 0.772807 | 0.769317 | 0.771939 |
| 0.773499 | 0.765322 | 0.75533 | 0.772544 | 0.777751 | 0.76817 | 0.776372 | 0.779127 | 0.770489 |
| 0.774104 | 0.7678 | 0.751091 | 0.77158 | 0.778215 | 0.764224 | 0.778204 | 0.769734 | 0.764722 |
| 0.774552 | 0.765304 | 0.75826 | 0.770124 | 0.778499 | 0.767937 | 0.771593 | 0.778283 | 0.76775 |
| 0.777335 | 0.768992 | 0.759215 | 0.77289 | 0.77454 | 0.765628 | 0.772627 | 0.768989 | 0.76348 |
| 0.782335 | 0.771176 | 0.754506 | 0.77443 | 0.774585 | 0.770049 | 0.77572 | 0.773137 | 0.768904 |
| 0.777926 | 0.766963 | 0.746994 | 0.771488 | 0.776196 | 0.767243 | 0.767825 | 0.7754 | 0.768755 |
| 0.768366 | 0.767446 | 0.748905 | 0.76647 | 0.775058 | 0.767306 | 0.770064 | 0.772381 | 0.769669 |
| 0.763312 | 0.765222 | 0.744758 | 0.771988 | 0.775983 | 0.766652 | 0.771708 | 0.768864 | 0.765177 |
| 0.770484 | 0.771309 | 0.755437 | 0.771712 | 0.776531 | 0.773199 | 0.777695 | 0.767114 | 0.769434 |
| 0.778592 | 0.760399 | 0.756571 | 0.768084 | 0.771193 | 0.765124 | 0.773061 | 0.771705 | 0.761637 |
| 0.775691 | 0.762788 | 0.752396 | 0.774525 | 0.770523 | 0.770911 | 0.774273 | 0.770316 | 0.770724 |
| 0.778644 | 0.766803 | 0.752185 | 0.774093 | 0.77588 | 0.770576 | 0.770864 | 0.771584 | 0.769231 |
| 0.782663 | 0.764002 | 0.752657 | 0.771369 | 0.775408 | 0.763515 | 0.775008 | 0.769329 | 0.767923 |
| 0.774075 | 0.761479 | 0.75356 | 0.764087 | 0.769611 | 0.762078 | 0.770837 | 0.77088 | 0.762871 |
| 0.78108 | 0.766525 | 0.754314 | 0.77479 | 0.775697 | 0.768461 | 0.77467 | 0.775507 | 0.777335 |
| 0.781315 | 0.766337 | 0.757391 | 0.77513 | 0.77385 | 0.770921 | 0.775491 | 0.771613 | 0.771434 |
| 0.778383 | 0.758012 | 0.748126 | 0.770145 | 0.767592 | 0.763433 | 0.766104 | 0.770238 | 0.765786 |
| 0.778125 | 0.76024 | 0.754534 | 0.770589 | 0.77507 | 0.76463 | 0.774816 | 0.772026 | 0.769053 |
| 0.78208 | 0.76625 | 0.754871 | 0.77171 | 0.774152 | 0.768054 | 0.771466 | 0.775939 | 0.772582 |
| 0.777691 | 0.762788 | 0.748623 | 0.771085 | 0.772835 | 0.764102 | 0.769474 | 0.771638 | 0.769399 |
| 0.778045 | 0.760008 | 0.752598 | 0.772755 | 0.771695 | 0.766807 | 0.771443 | 0.773592 | 0.774618 |
| 0.779667 | 0.758833 | 0.749096 | 0.774954 | 0.771318 | 0.767164 | 0.76749 | 0.766368 | 0.763306 |
| 0.788256 | 0.771825 | 0.759964 | 0.781607 | 0.786194 | 0.776451 | 0.781328 | 0.780793 | 0.777373 |
| 0.800743 | 0.768233 | 0.757163 | 0.777424 | 0.774438 | 0.770144 | 0.776904 | 0.774109 | 0.772473 |
| 0.782974 | 0.764183 | 0.760929 | 0.772071 | 0.771888 | 0.766019 | 0.774027 | 0.774324 | 0.772275 |
| 0.79693 | 0.766932 | 0.755925 | 0.775415 | 0.78103 | 0.76998 | 0.778686 | 0.770417 | 0.773432 |
| 0.791443 | 0.770104 | 0.759819 | 0.780862 | 0.78112 | 0.773129 | 0.779412 | 0.773397 | 0.774134 |
| 0.805414 | 0.775998 | 0.765399 | 0.780703 | 0.778296 | 0.772731 | 0.77862 | 0.778392 | 0.772808 |
| 0.810891 | 0.782778 | 0.761963 | 0.777226 | 0.779006 | 0.774259 | 0.778283 | 0.778493 | 0.773448 |
| 0.793971 | 0.769364 | 0.756194 | 0.768286 | 0.776962 | 0.762639 | 0.770619 | 0.773243 | 0.772744 |
| 0.860911 | 0.78001 | 0.756195 | 0.776822 | 0.781668 | 0.772825 | 0.775291 | 0.778502 | 0.775802 |
| 1 | 0.768328 | 0.75834 | 0.778256 | 0.779287 | 0.768209 | 0.774706 | 0.772552 | 0.775226 |

| | | | | | | | | |
|---|---|---|---|---|---|---|---|---|
| 0.768328 | 1 | 0.74609 | 0.761025 | 0.76217 | 0.756914 | 0.76039 | 0.758897 | 0.761991 |
| 0.75834 | 0.74609 | 1 | 0.750148 | 0.754584 | 0.744958 | 0.748259 | 0.749419 | 0.741944 |
| 0.778256 | 0.761025 | 0.750148 | 1 | 0.800679 | 0.850882 | 0.820403 | 0.795875 | 0.806106 |
| 0.779287 | 0.76217 | 0.754584 | 0.800679 | 1 | 0.811781 | 0.802744 | 0.803918 | 0.792977 |
| 0.768209 | 0.756914 | 0.744958 | 0.850882 | 0.811781 | 1 | 0.839868 | 0.788968 | 0.809185 |
| 0.774706 | 0.76039 | 0.748259 | 0.820403 | 0.802744 | 0.839868 | 1 | 0.781212 | 0.790394 |
| 0.772552 | 0.758897 | 0.749419 | 0.795875 | 0.803918 | 0.788968 | 0.781212 | 1 | 0.804745 |
| 0.775226 | 0.761991 | 0.741944 | 0.806106 | 0.792977 | 0.809185 | 0.790394 | 0.804745 | 1 |
| 0.776322 | 0.765139 | 0.752358 | 0.774952 | 0.779309 | 0.774213 | 0.776737 | 0.775584 | 0.772059 |
| 0.770438 | 0.765125 | 0.751113 | 0.776778 | 0.78219 | 0.771772 | 0.772009 | 0.774954 | 0.769437 |
| 0.77144 | 0.766225 | 0.74737 | 0.770861 | 0.78081 | 0.77332 | 0.776859 | 0.773551 | 0.767752 |
| 0.773147 | 0.762371 | 0.751675 | 0.768906 | 0.777695 | 0.767556 | 0.770147 | 0.774171 | 0.769422 |
| 0.780353 | 0.766046 | 0.754677 | 0.777317 | 0.785509 | 0.770065 | 0.780596 | 0.778278 | 0.778995 |
| 0.769666 | 0.770026 | 0.752558 | 0.770024 | 0.779553 | 0.76893 | 0.774017 | 0.777539 | 0.770572 |
| 0.770394 | 0.759536 | 0.750356 | 0.771039 | 0.777648 | 0.764431 | 0.770752 | 0.77525 | 0.763966 |
| 0.768153 | 0.768384 | 0.74925 | 0.771424 | 0.77724 | 0.764053 | 0.769306 | 0.773556 | 0.76711 |
| 0.762909 | 0.761611 | 0.749278 | 0.77125 | 0.774286 | 0.763523 | 0.76893 | 0.771438 | 0.763389 |
| 0.763043 | 0.763015 | 0.750372 | 0.771611 | 0.771157 | 0.767791 | 0.771101 | 0.768117 | 0.764848 |
| 0.768254 | 0.764914 | 0.749465 | 0.762664 | 0.776707 | 0.76046 | 0.764928 | 0.767517 | 0.756347 |
| 0.768052 | 0.757328 | 0.744823 | 0.771581 | 0.773716 | 0.764331 | 0.768449 | 0.769824 | 0.763733 |
| 0.77373 | 0.767529 | 0.753582 | 0.772688 | 0.776036 | 0.766851 | 0.769179 | 0.772332 | 0.763807 |
| 0.759297 | 0.748548 | 0.739904 | 0.757479 | 0.756142 | 0.750022 | 0.753122 | 0.755814 | 0.749299 |
| 0.762631 | 0.757335 | 0.746916 | 0.761322 | 0.762249 | 0.75855 | 0.760159 | 0.760912 | 0.754536 |
| 0.767298 | 0.758869 | 0.752401 | 0.766914 | 0.767326 | 0.761372 | 0.764324 | 0.767876 | 0.757104 |
| 0.772957 | 0.759012 | 0.754299 | 0.765318 | 0.771455 | 0.763137 | 0.766933 | 0.767816 | 0.761081 |
| 0.770833 | 0.754433 | 0.75069 | 0.765972 | 0.767598 | 0.760528 | 0.766588 | 0.767105 | 0.759771 |
| 0.763727 | 0.75693 | 0.742741 | 0.760909 | 0.764503 | 0.759455 | 0.764938 | 0.759158 | 0.756941 |
| 0.765801 | 0.762935 | 0.743803 | 0.765846 | 0.771443 | 0.759534 | 0.764681 | 0.765959 | 0.76265 |
| 0.764183 | 0.753259 | 0.747056 | 0.764508 | 0.768545 | 0.759606 | 0.760683 | 0.761549 | 0.757142 |
| 0.694029 | 0.68551 | 0.730715 | 0.693866 | 0.696559 | 0.690138 | 0.690104 | 0.693228 | 0.687307 |
| 0.703661 | 0.69484 | 0.748094 | 0.706975 | 0.709953 | 0.704197 | 0.706574 | 0.702465 | 0.70057 |
| 0.703189 | 0.690282 | 0.7406 | 0.706056 | 0.709469 | 0.704902 | 0.705492 | 0.703122 | 0.698268 |
| 0.689903 | 0.681993 | 0.734685 | 0.692852 | 0.696959 | 0.691848 | 0.694222 | 0.692595 | 0.689222 |
| 0.704985 | 0.69228 | 0.748118 | 0.706446 | 0.712319 | 0.704443 | 0.707685 | 0.705006 | 0.702106 |
| 0.764858 | 0.755193 | 0.743776 | 0.759938 | 0.76642 | 0.756236 | 0.761129 | 0.761602 | 0.752484 |
| 0.76694 | 0.757553 | 0.743083 | 0.765584 | 0.768734 | 0.761995 | 0.766403 | 0.76297 | 0.761845 |
| 0.772272 | 0.755905 | 0.746141 | 0.763091 | 0.76951 | 0.761409 | 0.767842 | 0.761517 | 0.764237 |
| 0.77074 | 0.75882 | 0.749731 | 0.766134 | 0.770753 | 0.764812 | 0.766797 | 0.768154 | 0.757678 |
| 0.764641 | 0.754276 | 0.744446 | 0.758257 | 0.766773 | 0.755687 | 0.759537 | 0.765409 | 0.757144 |
| 0.771977 | 0.759673 | 0.746897 | 0.765975 | 0.769798 | 0.762937 | 0.768588 | 0.764417 | 0.762977 |
| 0.771623 | 0.762618 | 0.748521 | 0.76839 | 0.77217 | 0.762776 | 0.767334 | 0.765341 | 0.761688 |
| 0.766831 | 0.752669 | 0.746742 | 0.760424 | 0.769199 | 0.757918 | 0.765312 | 0.763873 | 0.757573 |
| 0.760889 | 0.751863 | 0.740357 | 0.762091 | 0.766485 | 0.756992 | 0.76251 | 0.760065 | 0.757593 |
| 0.761037 | 0.749627 | 0.739226 | 0.7603 | 0.763915 | 0.75387 | 0.759226 | 0.759169 | 0.752843 |
| 0.763971 | 0.758897 | 0.74722 | 0.761367 | 0.76737 | 0.754671 | 0.761488 | 0.760871 | 0.753541 |
| 0.770093 | 0.759215 | 0.748584 | 0.765193 | 0.77249 | 0.760403 | 0.765986 | 0.766428 | 0.764701 |

| | | | | | | | | |
|---|---|---|---|---|---|---|---|---|
| 0.771078 | 0.759275 | 0.74792 | 0.767899 | 0.773203 | 0.762991 | 0.767069 | 0.770392 | 0.762381 |
| 0.768196 | 0.755977 | 0.751704 | 0.765744 | 0.769022 | 0.758515 | 0.768935 | 0.767755 | 0.757244 |
| 0.768426 | 0.759143 | 0.751913 | 0.769388 | 0.77391 | 0.765466 | 0.766521 | 0.770746 | 0.764065 |
| 0.767694 | 0.756359 | 0.742478 | 0.764587 | 0.765596 | 0.758541 | 0.760196 | 0.765571 | 0.754046 |
| 0.765345 | 0.749015 | 0.745354 | 0.760941 | 0.766134 | 0.756501 | 0.764831 | 0.764939 | 0.75757 |
| 0.762459 | 0.755508 | 0.746958 | 0.762398 | 0.768299 | 0.760412 | 0.766285 | 0.771612 | 0.755723 |
| 0.764776 | 0.758614 | 0.750105 | 0.761085 | 0.768967 | 0.761146 | 0.763786 | 0.766057 | 0.758991 |
| 0.765487 | 0.757722 | 0.749326 | 0.760965 | 0.771314 | 0.76001 | 0.765999 | 0.771835 | 0.759931 |
| 0.765491 | 0.756227 | 0.750444 | 0.766707 | 0.771997 | 0.757647 | 0.763129 | 0.771739 | 0.760542 |
| 0.764755 | 0.75589 | 0.746513 | 0.767687 | 0.770722 | 0.76188 | 0.76805 | 0.770728 | 0.762253 |
| 0.767462 | 0.752938 | 0.749707 | 0.762879 | 0.765686 | 0.76056 | 0.765475 | 0.766996 | 0.759035 |
| 0.763185 | 0.757945 | 0.74916 | 0.760045 | 0.770545 | 0.758971 | 0.767934 | 0.765195 | 0.75775 |
| 0.763557 | 0.746379 | 0.741684 | 0.760617 | 0.759065 | 0.754053 | 0.756909 | 0.765622 | 0.755805 |
| 0.760363 | 0.749237 | 0.737126 | 0.756219 | 0.762925 | 0.758389 | 0.756135 | 0.762999 | 0.756994 |
| 0.769547 | 0.757188 | 0.746012 | 0.772205 | 0.772689 | 0.765818 | 0.771091 | 0.765197 | 0.762984 |
| 0.7823 | 0.768043 | 0.757135 | 0.77791 | 0.777384 | 0.77147 | 0.77748 | 0.775657 | 0.769395 |
| 0.780522 | 0.761932 | 0.752105 | 0.775813 | 0.770835 | 0.766342 | 0.777668 | 0.767571 | 0.766169 |
| 0.771539 | 0.758156 | 0.752889 | 0.77247 | 0.774299 | 0.763178 | 0.770362 | 0.768082 | 0.767719 |
| 0.7736 | 0.757101 | 0.750768 | 0.768902 | 0.772535 | 0.76468 | 0.772582 | 0.769402 | 0.765573 |
| 0.768132 | 0.758124 | 0.745879 | 0.76911 | 0.770425 | 0.763888 | 0.772876 | 0.769579 | 0.763545 |
| 0.769513 | 0.755092 | 0.739042 | 0.764588 | 0.76505 | 0.762052 | 0.766244 | 0.770492 | 0.762136 |
| 0.773678 | 0.754663 | 0.74898 | 0.771643 | 0.776388 | 0.761095 | 0.769921 | 0.769988 | 0.760722 |
| 0.778057 | 0.764425 | 0.752187 | 0.773458 | 0.775235 | 0.765326 | 0.775026 | 0.772986 | 0.764352 |
| 0.762372 | 0.754647 | 0.747473 | 0.7628 | 0.766296 | 0.757276 | 0.763092 | 0.770183 | 0.759519 |
| 0.77822 | 0.763731 | 0.74958 | 0.773509 | 0.77231 | 0.7625 | 0.770094 | 0.770831 | 0.761428 |
| 0.701633 | 0.690296 | 0.745559 | 0.703829 | 0.710147 | 0.699733 | 0.703195 | 0.701353 | 0.695521 |
| 0.779543 | 0.76585 | 0.75374 | 0.77452 | 0.777098 | 0.767367 | 0.77098 | 0.769231 | 0.770226 |
| 0.776847 | 0.76267 | 0.757556 | 0.776554 | 0.768874 | 0.77051 | 0.773572 | 0.767969 | 0.771026 |
| 0.783377 | 0.767097 | 0.756278 | 0.775268 | 0.777668 | 0.769026 | 0.776036 | 0.769846 | 0.771475 |
| 0.774636 | 0.761194 | 0.756023 | 0.775871 | 0.773295 | 0.769422 | 0.773328 | 0.772195 | 0.767991 |
| 0.776188 | 0.763388 | 0.750366 | 0.768525 | 0.767394 | 0.766257 | 0.762688 | 0.768016 | 0.76489 |
| 0.776948 | 0.765887 | 0.75017 | 0.766402 | 0.767049 | 0.766382 | 0.76962 | 0.775051 | 0.765381 |
| 0.777291 | 0.762629 | 0.750358 | 0.766437 | 0.769314 | 0.76656 | 0.769296 | 0.766063 | 0.763381 |
| 0.774216 | 0.769933 | 0.751271 | 0.773708 | 0.767854 | 0.76468 | 0.768574 | 0.767229 | 0.767097 |
| 0.768048 | 0.755108 | 0.749435 | 0.762955 | 0.763428 | 0.762794 | 0.764001 | 0.766507 | 0.76698 |
| 0.774474 | 0.754032 | 0.753613 | 0.769337 | 0.763827 | 0.764726 | 0.769047 | 0.762357 | 0.762799 |
| 0.784056 | 0.766056 | 0.759619 | 0.77398 | 0.777685 | 0.763728 | 0.768397 | 0.773374 | 0.771931 |
| 0.784505 | 0.763517 | 0.757946 | 0.77385 | 0.77953 | 0.773034 | 0.779443 | 0.767637 | 0.772898 |
| 0.775773 | 0.76734 | 0.750686 | 0.775334 | 0.778542 | 0.774702 | 0.781878 | 0.773271 | 0.768317 |
| 0.772282 | 0.762288 | 0.749829 | 0.768297 | 0.777825 | 0.769826 | 0.776955 | 0.773315 | 0.763486 |
| 0.770737 | 0.764181 | 0.74701 | 0.767578 | 0.772529 | 0.761812 | 0.767579 | 0.767718 | 0.766489 |
| 0.771996 | 0.768673 | 0.752199 | 0.773754 | 0.774623 | 0.76675 | 0.772593 | 0.771914 | 0.76706 |
| 0.776308 | 0.756438 | 0.753375 | 0.769522 | 0.77174 | 0.767897 | 0.776724 | 0.768055 | 0.767256 |
| 0.763939 | 0.75621 | 0.743405 | 0.75954 | 0.760569 | 0.756677 | 0.762361 | 0.763104 | 0.76095 |
| 0.770849 | 0.76083 | 0.746781 | 0.766399 | 0.77099 | 0.764196 | 0.767936 | 0.770559 | 0.760444 |
| 0.768019 | 0.754353 | 0.746992 | 0.763273 | 0.772225 | 0.764524 | 0.765526 | 0.768201 | 0.761778 |

| | | | | | | | | |
|---|---|---|---|---|---|---|---|---|
| 0.764955 | 0.760571 | 0.744526 | 0.764607 | 0.769981 | 0.760547 | 0.762615 | 0.768993 | 0.76227 |
| 0.771813 | 0.761945 | 0.747829 | 0.769774 | 0.775198 | 0.770058 | 0.770021 | 0.774976 | 0.765777 |
| 0.770759 | 0.759672 | 0.742687 | 0.760316 | 0.765761 | 0.758157 | 0.765254 | 0.769914 | 0.75969 |
| 0.721786 | 0.735936 | 0.709666 | 0.731261 | 0.730906 | 0.726147 | 0.729956 | 0.732841 | 0.740421 |
| 0.764999 | 0.758803 | 0.746647 | 0.763006 | 0.769568 | 0.761393 | 0.765362 | 0.768413 | 0.759673 |
| 0.767678 | 0.751537 | 0.746226 | 0.764557 | 0.769363 | 0.760506 | 0.763791 | 0.771819 | 0.759126 |
| 0.762055 | 0.754892 | 0.751505 | 0.764745 | 0.771557 | 0.762986 | 0.763347 | 0.772018 | 0.758124 |
| 0.766794 | 0.756323 | 0.744418 | 0.768259 | 0.766259 | 0.761561 | 0.766918 | 0.767923 | 0.761644 |
| 0.770475 | 0.758722 | 0.752321 | 0.772033 | 0.772131 | 0.766088 | 0.766156 | 0.773256 | 0.764883 |
| 0.768826 | 0.758363 | 0.748396 | 0.769517 | 0.770713 | 0.764389 | 0.766012 | 0.767258 | 0.760556 |
| 0.768567 | 0.755551 | 0.748353 | 0.76981 | 0.769451 | 0.762096 | 0.768533 | 0.767414 | 0.763265 |
| 0.767781 | 0.75658 | 0.747591 | 0.768273 | 0.769885 | 0.760581 | 0.767047 | 0.768805 | 0.763999 |
| 0.767638 | 0.751983 | 0.752928 | 0.765538 | 0.768104 | 0.7645 | 0.761987 | 0.769131 | 0.763048 |
| 0.771109 | 0.75923 | 0.744629 | 0.770294 | 0.770245 | 0.760882 | 0.770143 | 0.775181 | 0.769605 |
| 0.76559 | 0.759966 | 0.745617 | 0.76583 | 0.769362 | 0.763692 | 0.765949 | 0.771254 | 0.762532 |
| 0.762894 | 0.758333 | 0.742362 | 0.764746 | 0.764175 | 0.761138 | 0.76473 | 0.764868 | 0.758508 |
| 0.771704 | 0.763039 | 0.74717 | 0.772024 | 0.768356 | 0.765703 | 0.763098 | 0.770297 | 0.764494 |
| 0.767979 | 0.758383 | 0.749979 | 0.766998 | 0.768162 | 0.765793 | 0.766863 | 0.777586 | 0.766965 |
| 0.766075 | 0.757768 | 0.739625 | 0.764033 | 0.76866 | 0.760677 | 0.765456 | 0.76464 | 0.758971 |
| 0.767217 | 0.754174 | 0.747432 | 0.761909 | 0.769275 | 0.757948 | 0.764345 | 0.770245 | 0.76481 |
| 0.763603 | 0.758043 | 0.750802 | 0.770928 | 0.775029 | 0.767267 | 0.76847 | 0.769496 | 0.767019 |
| 0.769449 | 0.760398 | 0.750123 | 0.768946 | 0.766933 | 0.766863 | 0.767836 | 0.771451 | 0.762006 |
| 0.682068 | 0.676471 | 0.690975 | 0.681284 | 0.696272 | 0.6811 | 0.68064 | 0.680682 | 0.678832 |
| 0.656546 | 0.658573 | 0.703106 | 0.659732 | 0.661122 | 0.652477 | 0.65613 | 0.654589 | 0.651096 |
| 0.673123 | 0.662114 | 0.719101 | 0.677922 | 0.678104 | 0.673094 | 0.669495 | 0.676464 | 0.665456 |
| 0.654498 | 0.648883 | 0.703388 | 0.65143 | 0.654492 | 0.654746 | 0.650247 | 0.649768 | 0.647864 |
| 0.7326 | 0.726433 | 0.715783 | 0.726065 | 0.74695 | 0.731039 | 0.736147 | 0.747033 | 0.738277 |
| 0.72936 | 0.73591 | 0.715123 | 0.722768 | 0.725404 | 0.721673 | 0.724725 | 0.725223 | 0.723728 |
| 0.727355 | 0.722509 | 0.719482 | 0.717029 | 0.730549 | 0.717626 | 0.723018 | 0.72516 | 0.72256 |
| 0.743331 | 0.728403 | 0.721857 | 0.735431 | 0.740988 | 0.733662 | 0.744093 | 0.742845 | 0.735435 |
| 0.722872 | 0.716611 | 0.703272 | 0.71966 | 0.721005 | 0.721216 | 0.727632 | 0.721652 | 0.727681 |
| 0.711691 | 0.721925 | 0.697405 | 0.705034 | 0.716042 | 0.707499 | 0.716495 | 0.721581 | 0.710924 |
| 0.71155 | 0.716847 | 0.708418 | 0.706213 | 0.725648 | 0.712112 | 0.714712 | 0.722524 | 0.710494 |
| 0.64992 | 0.662802 | 0.691042 | 0.653317 | 0.656438 | 0.65708 | 0.660633 | 0.655228 | 0.659124 |
| 0.763261 | 0.750024 | 0.743625 | 0.765501 | 0.763243 | 0.761968 | 0.768312 | 0.7628 | 0.756234 |
| 0.743744 | 0.740671 | 0.723409 | 0.731619 | 0.740201 | 0.730928 | 0.738748 | 0.748504 | 0.737135 |
| 0.733623 | 0.726671 | 0.714608 | 0.733897 | 0.74452 | 0.735927 | 0.737481 | 0.746304 | 0.73543 |
| 0.720075 | 0.725195 | 0.6992 | 0.709767 | 0.730534 | 0.708738 | 0.722473 | 0.724484 | 0.71163 |
| 0.744633 | 0.740989 | 0.726696 | 0.739485 | 0.740792 | 0.738975 | 0.745721 | 0.742457 | 0.734037 |
| 0.728516 | 0.729204 | 0.717358 | 0.730309 | 0.734856 | 0.729232 | 0.732866 | 0.740971 | 0.72673 |
| 0.72473 | 0.728822 | 0.711027 | 0.727097 | 0.731773 | 0.718659 | 0.726498 | 0.739074 | 0.72772 |
| 0.7269 | 0.727092 | 0.71356 | 0.725959 | 0.732504 | 0.721835 | 0.721029 | 0.731058 | 0.720378 |
| 0.725394 | 0.722382 | 0.71089 | 0.733859 | 0.73013 | 0.725854 | 0.723763 | 0.733198 | 0.731205 |
| 0.727488 | 0.731907 | 0.708922 | 0.732072 | 0.731534 | 0.728763 | 0.720428 | 0.739596 | 0.735401 |
| 0.729982 | 0.725173 | 0.701312 | 0.719104 | 0.728325 | 0.723735 | 0.721164 | 0.732002 | 0.730373 |
| 0.709964 | 0.723865 | 0.703211 | 0.709529 | 0.727394 | 0.713289 | 0.725385 | 0.714716 | 0.722846 |

| | | | | | | | | |
|---|---|---|---|---|---|---|---|---|
| 0.749124 | 0.750083 | 0.734503 | 0.756122 | 0.754947 | 0.747326 | 0.750526 | 0.759024 | 0.755267 |
| 0.737404 | 0.725288 | 0.711916 | 0.728099 | 0.732723 | 0.724996 | 0.72628 | 0.740656 | 0.730775 |
| 0.72844 | 0.722576 | 0.707989 | 0.723893 | 0.733562 | 0.72256 | 0.72916 | 0.73976 | 0.726625 |
| 0.737127 | 0.729637 | 0.714847 | 0.721456 | 0.73602 | 0.721324 | 0.726108 | 0.735621 | 0.721786 |
| 0.730605 | 0.729318 | 0.705425 | 0.731974 | 0.737249 | 0.724803 | 0.726019 | 0.737435 | 0.727102 |
| 0.720988 | 0.719162 | 0.709229 | 0.716915 | 0.732232 | 0.713534 | 0.719166 | 0.732466 | 0.718916 |
| 0.716618 | 0.732326 | 0.704518 | 0.727004 | 0.734949 | 0.718404 | 0.714951 | 0.739021 | 0.720676 |
| 0.717137 | 0.719065 | 0.702583 | 0.713839 | 0.724711 | 0.713474 | 0.715466 | 0.736305 | 0.71531 |
| 0.728522 | 0.722895 | 0.70253 | 0.715503 | 0.729538 | 0.719523 | 0.721009 | 0.723521 | 0.71601 |
| 0.754887 | 0.745934 | 0.735735 | 0.752979 | 0.763553 | 0.753146 | 0.752202 | 0.753594 | 0.750619 |
| 0.750994 | 0.744946 | 0.733407 | 0.747528 | 0.757506 | 0.748939 | 0.748781 | 0.754835 | 0.749022 |
| 0.747762 | 0.740285 | 0.728651 | 0.740749 | 0.754557 | 0.742421 | 0.744539 | 0.745726 | 0.743002 |
| 0.739301 | 0.738154 | 0.72103 | 0.737429 | 0.742325 | 0.736614 | 0.736977 | 0.741292 | 0.73539 |
| 0.745086 | 0.735306 | 0.724363 | 0.741747 | 0.750061 | 0.74014 | 0.74013 | 0.748397 | 0.742642 |
| 0.739233 | 0.729988 | 0.721507 | 0.734412 | 0.746098 | 0.736478 | 0.735636 | 0.743402 | 0.742758 |

| | | | | | | plot 09 | | |
|---|---|---|---|---|---|---|---|---|
| Wild_plot_ | Wild_plot_ | Wild_plot_ | Wild_plot_ | Wild_plot_ | Wild_plot_ | Wild_plot_ | Wild_plot_ | Wild_plot_ |
| 0.705404 | 0.714058 | 0.710774 | 0.710133 | 0.712274 | 0.709379 | 0.706689 | 0.708768 | 0.710838 |
| 0.712025 | 0.72116 | 0.717545 | 0.718123 | 0.720566 | 0.719733 | 0.71468 | 0.713489 | 0.716542 |
| 0.704225 | 0.708049 | 0.704824 | 0.707216 | 0.708729 | 0.705836 | 0.704677 | 0.704188 | 0.706949 |
| 0.711267 | 0.717148 | 0.716372 | 0.714343 | 0.718983 | 0.717432 | 0.713215 | 0.714163 | 0.715512 |
| 0.712489 | 0.716269 | 0.711824 | 0.711941 | 0.714399 | 0.711884 | 0.708374 | 0.70766 | 0.714183 |
| 0.718881 | 0.725573 | 0.716652 | 0.721664 | 0.721772 | 0.721708 | 0.715724 | 0.714174 | 0.719465 |
| 0.776986 | 0.776386 | 0.771382 | 0.772705 | 0.776126 | 0.775464 | 0.77379 | 0.773945 | 0.770899 |
| 0.769795 | 0.772634 | 0.774088 | 0.769738 | 0.772143 | 0.771795 | 0.770816 | 0.76999 | 0.77003 |
| 0.709315 | 0.716038 | 0.714422 | 0.71867 | 0.718785 | 0.71669 | 0.710998 | 0.710912 | 0.714663 |
| 0.704245 | 0.713865 | 0.70873 | 0.713614 | 0.709501 | 0.713665 | 0.708384 | 0.707985 | 0.712133 |
| 0.706801 | 0.710421 | 0.710958 | 0.712599 | 0.7103 | 0.713605 | 0.70863 | 0.710634 | 0.713289 |
| 0.703689 | 0.709641 | 0.706138 | 0.709907 | 0.710671 | 0.706793 | 0.70468 | 0.70557 | 0.706248 |
| 0.717428 | 0.720173 | 0.719132 | 0.719393 | 0.721609 | 0.715713 | 0.713927 | 0.712044 | 0.720116 |
| 0.721117 | 0.729257 | 0.726535 | 0.724096 | 0.729519 | 0.727316 | 0.721983 | 0.721225 | 0.723087 |
| 0.736649 | 0.740851 | 0.735807 | 0.737681 | 0.740319 | 0.736917 | 0.73284 | 0.73519 | 0.740272 |
| 0.713642 | 0.723826 | 0.716013 | 0.718212 | 0.718631 | 0.720626 | 0.715963 | 0.716437 | 0.720878 |
| 0.711042 | 0.720624 | 0.714821 | 0.718056 | 0.719646 | 0.71882 | 0.718456 | 0.71509 | 0.71943 |
| 0.74578 | 0.74421 | 0.745149 | 0.743505 | 0.74762 | 0.74326 | 0.737611 | 0.740508 | 0.740975 |
| 0.707946 | 0.715241 | 0.710162 | 0.714113 | 0.711306 | 0.711747 | 0.709468 | 0.709983 | 0.712343 |
| 0.70833 | 0.719224 | 0.717112 | 0.712684 | 0.714627 | 0.718745 | 0.710622 | 0.711726 | 0.7162 |
| 0.70389 | 0.715141 | 0.710578 | 0.709657 | 0.714365 | 0.713008 | 0.709401 | 0.70878 | 0.710971 |
| 0.707876 | 0.715979 | 0.716326 | 0.713965 | 0.714718 | 0.71337 | 0.711109 | 0.71032 | 0.71093 |
| 0.715881 | 0.722584 | 0.719965 | 0.719564 | 0.721468 | 0.718814 | 0.718003 | 0.715318 | 0.718302 |
| 0.712061 | 0.718029 | 0.717493 | 0.717881 | 0.721188 | 0.717672 | 0.715062 | 0.712361 | 0.715691 |
| 0.715049 | 0.719955 | 0.715208 | 0.719862 | 0.720308 | 0.720979 | 0.71909 | 0.71459 | 0.720606 |
| 0.712114 | 0.718026 | 0.715073 | 0.716038 | 0.71754 | 0.714601 | 0.71302 | 0.710789 | 0.714252 |
| 0.704883 | 0.711816 | 0.707807 | 0.71248 | 0.708972 | 0.71106 | 0.709461 | 0.706246 | 0.710436 |
| 0.709388 | 0.716195 | 0.711354 | 0.712888 | 0.716395 | 0.71351 | 0.710384 | 0.711507 | 0.715082 |
| 0.71161 | 0.718169 | 0.714274 | 0.71684 | 0.717893 | 0.716566 | 0.713949 | 0.713163 | 0.715845 |
| 0.712864 | 0.717728 | 0.714502 | 0.718112 | 0.721364 | 0.714922 | 0.715322 | 0.708461 | 0.713184 |
| 0.706339 | 0.714492 | 0.71136 | 0.713873 | 0.714331 | 0.713866 | 0.71092 | 0.709827 | 0.714837 |
| 0.705365 | 0.713205 | 0.709738 | 0.712759 | 0.713198 | 0.709944 | 0.707907 | 0.709733 | 0.709824 |
| 0.708415 | 0.713081 | 0.709594 | 0.715435 | 0.712408 | 0.71397 | 0.708562 | 0.707889 | 0.708883 |
| 0.709881 | 0.718847 | 0.716535 | 0.71539 | 0.716512 | 0.715855 | 0.712935 | 0.713069 | 0.715072 |
| 0.705534 | 0.710579 | 0.707995 | 0.70765 | 0.71299 | 0.708244 | 0.706101 | 0.707978 | 0.710381 |
| 0.707478 | 0.715962 | 0.712297 | 0.713918 | 0.717484 | 0.71588 | 0.713738 | 0.708835 | 0.713368 |
| 0.714668 | 0.721539 | 0.714905 | 0.718675 | 0.717512 | 0.715645 | 0.717256 | 0.714063 | 0.720656 |
| 0.710713 | 0.716791 | 0.714686 | 0.719008 | 0.719022 | 0.715925 | 0.71541 | 0.710316 | 0.716065 |
| 0.713856 | 0.722967 | 0.719316 | 0.720071 | 0.717941 | 0.72167 | 0.716854 | 0.7159 | 0.718312 |

| | | | | | | | | |
|---|---|---|---|---|---|---|---|---|
| 0.747567 | 0.7523 | 0.749641 | 0.750605 | 0.751446 | 0.748563 | 0.746648 | 0.745417 | 0.752019 |
| 0.700016 | 0.708615 | 0.704549 | 0.70284 | 0.709875 | 0.705451 | 0.700443 | 0.702823 | 0.705985 |
| 0.704432 | 0.713451 | 0.706166 | 0.709899 | 0.708887 | 0.710337 | 0.70885 | 0.704732 | 0.707857 |
| 0.711892 | 0.722763 | 0.716111 | 0.719263 | 0.722611 | 0.71838 | 0.714217 | 0.714509 | 0.72162 |
| 0.711834 | 0.718088 | 0.711616 | 0.716664 | 0.715104 | 0.71511 | 0.710637 | 0.71616 | 0.711848 |
| 0.711978 | 0.719144 | 0.714695 | 0.716196 | 0.720292 | 0.718765 | 0.711265 | 0.714692 | 0.716369 |
| 0.708084 | 0.713546 | 0.710999 | 0.70939 | 0.714002 | 0.712006 | 0.705343 | 0.709114 | 0.710598 |
| 0.711332 | 0.720532 | 0.716152 | 0.718114 | 0.718936 | 0.716566 | 0.712865 | 0.714173 | 0.716045 |
| 0.713283 | 0.723452 | 0.71424 | 0.717246 | 0.719283 | 0.716545 | 0.713784 | 0.713099 | 0.712966 |
| 0.714139 | 0.719221 | 0.716959 | 0.716966 | 0.718934 | 0.717823 | 0.712339 | 0.712933 | 0.713772 |
| 0.704262 | 0.711056 | 0.704961 | 0.709747 | 0.709958 | 0.707864 | 0.705361 | 0.702486 | 0.705496 |
| 0.707429 | 0.713358 | 0.709179 | 0.712303 | 0.709842 | 0.709979 | 0.709481 | 0.705471 | 0.711213 |
| 0.708652 | 0.716131 | 0.708967 | 0.711213 | 0.714091 | 0.71187 | 0.708557 | 0.712082 | 0.713618 |
| 0.705163 | 0.712002 | 0.708056 | 0.710087 | 0.711099 | 0.707956 | 0.706508 | 0.704149 | 0.708347 |
| 0.704096 | 0.713039 | 0.70828 | 0.710762 | 0.71202 | 0.71069 | 0.707774 | 0.706557 | 0.710424 |
| 0.706365 | 0.71287 | 0.705755 | 0.712002 | 0.710097 | 0.709119 | 0.705179 | 0.707849 | 0.709884 |
| 0.702077 | 0.711388 | 0.705606 | 0.70574 | 0.709535 | 0.705252 | 0.703202 | 0.704623 | 0.709266 |
| 0.710682 | 0.720477 | 0.716561 | 0.716263 | 0.718082 | 0.716857 | 0.712258 | 0.713031 | 0.719331 |
| 0.713618 | 0.717754 | 0.713008 | 0.716909 | 0.717091 | 0.713969 | 0.711612 | 0.711743 | 0.713418 |
| 0.70768 | 0.715102 | 0.712249 | 0.7106 | 0.712922 | 0.711293 | 0.708785 | 0.707683 | 0.711111 |
| 0.699019 | 0.712886 | 0.704502 | 0.704757 | 0.71051 | 0.706004 | 0.703325 | 0.703002 | 0.704696 |
| 0.707095 | 0.71518 | 0.709434 | 0.709375 | 0.713166 | 0.70794 | 0.706336 | 0.705249 | 0.71036 |
| 0.706903 | 0.715342 | 0.71195 | 0.712209 | 0.71187 | 0.716067 | 0.705963 | 0.710688 | 0.714081 |
| 0.716677 | 0.721972 | 0.716864 | 0.719955 | 0.721937 | 0.720808 | 0.716355 | 0.716324 | 0.721475 |
| 0.725177 | 0.729634 | 0.72385 | 0.727967 | 0.731974 | 0.729444 | 0.723505 | 0.722388 | 0.727755 |
| 0.772173 | 0.772689 | 0.77595 | 0.770762 | 0.774515 | 0.771986 | 0.77303 | 0.769795 | 0.769702 |
| 0.776089 | 0.776508 | 0.778081 | 0.775573 | 0.778064 | 0.780419 | 0.772585 | 0.773441 | 0.773189 |
| 0.774151 | 0.780212 | 0.782702 | 0.77264 | 0.779042 | 0.779959 | 0.771114 | 0.77529 | 0.774177 |
| 0.705761 | 0.717757 | 0.711002 | 0.7084 | 0.714422 | 0.710216 | 0.708995 | 0.708825 | 0.711562 |
| 0.707351 | 0.713402 | 0.710832 | 0.707728 | 0.712853 | 0.711488 | 0.705462 | 0.707972 | 0.710411 |
| 0.713208 | 0.723615 | 0.715749 | 0.717938 | 0.722494 | 0.718937 | 0.713438 | 0.712712 | 0.71766 |
| 0.714882 | 0.724427 | 0.720537 | 0.719422 | 0.721395 | 0.720373 | 0.71904 | 0.715857 | 0.720757 |
| 0.705023 | 0.714448 | 0.711255 | 0.712505 | 0.712869 | 0.711206 | 0.706627 | 0.710556 | 0.710473 |
| 0.706888 | 0.715968 | 0.71253 | 0.713766 | 0.713758 | 0.714103 | 0.708018 | 0.711674 | 0.713905 |
| 0.702945 | 0.714855 | 0.710207 | 0.709499 | 0.712851 | 0.711582 | 0.704052 | 0.710904 | 0.711848 |
| 0.705527 | 0.716473 | 0.710396 | 0.70997 | 0.7144 | 0.712126 | 0.706595 | 0.711692 | 0.711402 |
| 0.708232 | 0.715512 | 0.708076 | 0.712789 | 0.708769 | 0.710584 | 0.710579 | 0.709116 | 0.711034 |
| 0.72066 | 0.730168 | 0.723014 | 0.725587 | 0.728477 | 0.726373 | 0.723614 | 0.723871 | 0.724432 |
| 0.695471 | 0.707465 | 0.703372 | 0.701354 | 0.704876 | 0.704364 | 0.699288 | 0.702549 | 0.703369 |
| 0.707327 | 0.714614 | 0.709124 | 0.71296 | 0.712157 | 0.710919 | 0.710499 | 0.709735 | 0.712023 |
| 0.701441 | 0.710702 | 0.705044 | 0.707155 | 0.713008 | 0.708949 | 0.70473 | 0.706969 | 0.708462 |
| 0.722027 | 0.731944 | 0.7267 | 0.729894 | 0.728573 | 0.729198 | 0.725405 | 0.726344 | 0.727824 |
| 0.707924 | 0.713634 | 0.712789 | 0.711068 | 0.713373 | 0.711692 | 0.707307 | 0.712127 | 0.711061 |
| 0.705799 | 0.712266 | 0.706523 | 0.712073 | 0.714897 | 0.711328 | 0.709789 | 0.707414 | 0.708796 |
| 0.709588 | 0.715896 | 0.714001 | 0.715426 | 0.715523 | 0.714707 | 0.711055 | 0.711185 | 0.714033 |
| 0.705817 | 0.715048 | 0.712031 | 0.707994 | 0.712205 | 0.712308 | 0.707299 | 0.711203 | 0.71011 |

| | | | | | | | | |
|---|---|---|---|---|---|---|---|---|
| 0.705179 | 0.714343 | 0.709846 | 0.709066 | 0.713007 | 0.708086 | 0.706827 | 0.709794 | 0.709962 |
| 0.707678 | 0.713556 | 0.711919 | 0.715711 | 0.715068 | 0.715394 | 0.715269 | 0.709148 | 0.713398 |
| 0.70844 | 0.713065 | 0.709464 | 0.713132 | 0.71485 | 0.713726 | 0.71035 | 0.711136 | 0.711627 |
| 0.713667 | 0.722479 | 0.718559 | 0.720728 | 0.720582 | 0.721191 | 0.716411 | 0.719291 | 0.718773 |
| 0.713496 | 0.720105 | 0.71742 | 0.719347 | 0.720487 | 0.71968 | 0.714794 | 0.712769 | 0.716578 |
| 0.708626 | 0.714401 | 0.711858 | 0.714609 | 0.714228 | 0.71221 | 0.710264 | 0.709502 | 0.708588 |
| 0.712016 | 0.717626 | 0.715017 | 0.717504 | 0.716673 | 0.715253 | 0.713894 | 0.714408 | 0.716663 |
| 0.712245 | 0.718962 | 0.715244 | 0.720424 | 0.716671 | 0.717038 | 0.710552 | 0.71584 | 0.718614 |
| 0.708443 | 0.717237 | 0.713841 | 0.712765 | 0.713473 | 0.71432 | 0.706776 | 0.709181 | 0.712265 |
| 0.707984 | 0.710405 | 0.709257 | 0.712165 | 0.715117 | 0.710892 | 0.708869 | 0.707065 | 0.708752 |
| 0.70568 | 0.713302 | 0.709175 | 0.709315 | 0.71068 | 0.711106 | 0.705355 | 0.708608 | 0.71263 |
| 0.708995 | 0.712087 | 0.711097 | 0.711517 | 0.714736 | 0.713586 | 0.707423 | 0.711346 | 0.711063 |
| 0.711584 | 0.715991 | 0.711322 | 0.715357 | 0.717767 | 0.712919 | 0.709851 | 0.714331 | 0.714989 |
| 0.70755 | 0.711293 | 0.707618 | 0.712499 | 0.712495 | 0.709836 | 0.707858 | 0.708977 | 0.710084 |
| 0.70801 | 0.713357 | 0.711007 | 0.711579 | 0.714092 | 0.715448 | 0.711249 | 0.711867 | 0.715655 |
| 0.71067 | 0.714411 | 0.711535 | 0.714024 | 0.713571 | 0.715912 | 0.708014 | 0.710118 | 0.712078 |
| 0.712942 | 0.720757 | 0.715084 | 0.718177 | 0.716439 | 0.720324 | 0.714684 | 0.720267 | 0.717584 |
| 0.70519 | 0.714537 | 0.70755 | 0.711216 | 0.715139 | 0.710558 | 0.705963 | 0.709655 | 0.714149 |
| 0.711115 | 0.714469 | 0.710733 | 0.713071 | 0.717388 | 0.713286 | 0.709565 | 0.711365 | 0.712854 |
| 0.711069 | 0.716029 | 0.714195 | 0.713186 | 0.713943 | 0.715522 | 0.710612 | 0.709078 | 0.713552 |
| 0.707149 | 0.716791 | 0.713137 | 0.715483 | 0.711794 | 0.715662 | 0.709221 | 0.712686 | 0.712174 |
| 0.703721 | 0.711752 | 0.706793 | 0.709732 | 0.71257 | 0.708458 | 0.704872 | 0.706633 | 0.706629 |
| 0.708044 | 0.713998 | 0.708602 | 0.711283 | 0.716113 | 0.710682 | 0.708245 | 0.708353 | 0.710605 |
| 0.711145 | 0.71408 | 0.710385 | 0.714229 | 0.717904 | 0.714206 | 0.709432 | 0.712058 | 0.713227 |
| 0.706479 | 0.716218 | 0.710444 | 0.711131 | 0.714286 | 0.711892 | 0.711288 | 0.708591 | 0.713626 |
| 0.714038 | 0.723465 | 0.718688 | 0.719528 | 0.721371 | 0.719946 | 0.716637 | 0.715 | 0.722009 |
| 0.706972 | 0.715798 | 0.709741 | 0.710876 | 0.714739 | 0.710566 | 0.709125 | 0.706605 | 0.710297 |
| 0.71101 | 0.718473 | 0.713462 | 0.716342 | 0.714331 | 0.71468 | 0.712575 | 0.710708 | 0.716948 |
| 0.713935 | 0.715084 | 0.715124 | 0.714708 | 0.720473 | 0.717446 | 0.710971 | 0.709935 | 0.71195 |
| 0.711848 | 0.718038 | 0.712586 | 0.713438 | 0.717419 | 0.711761 | 0.711316 | 0.710378 | 0.714669 |
| 0.71086 | 0.719013 | 0.716078 | 0.717274 | 0.715855 | 0.716664 | 0.712405 | 0.71589 | 0.715798 |
| 0.708709 | 0.7138 | 0.709907 | 0.711507 | 0.714398 | 0.71151 | 0.702853 | 0.70919 | 0.712296 |
| 0.710547 | 0.716189 | 0.714849 | 0.715993 | 0.715371 | 0.715938 | 0.710827 | 0.712765 | 0.715212 |
| 0.706494 | 0.717561 | 0.711853 | 0.714286 | 0.718235 | 0.716269 | 0.711122 | 0.713203 | 0.718195 |
| 0.707375 | 0.713643 | 0.708148 | 0.71234 | 0.713759 | 0.710323 | 0.705897 | 0.714467 | 0.710734 |
| 0.706095 | 0.713472 | 0.710172 | 0.710191 | 0.71483 | 0.709799 | 0.708217 | 0.712566 | 0.710641 |
| 0.708773 | 0.716426 | 0.707166 | 0.713776 | 0.711333 | 0.711656 | 0.706212 | 0.707298 | 0.709483 |
| 0.682615 | 0.692457 | 0.684706 | 0.691374 | 0.686453 | 0.68961 | 0.686901 | 0.686255 | 0.685503 |
| 0.68669 | 0.694004 | 0.691087 | 0.695714 | 0.692401 | 0.692254 | 0.693426 | 0.690217 | 0.689945 |
| 0.680442 | 0.688582 | 0.685667 | 0.68088 | 0.685673 | 0.68233 | 0.68163 | 0.679984 | 0.685886 |
| 0.677473 | 0.686748 | 0.685342 | 0.683027 | 0.685673 | 0.687708 | 0.680491 | 0.684673 | 0.681843 |
| 0.662255 | 0.668268 | 0.670675 | 0.671487 | 0.673178 | 0.668139 | 0.673327 | 0.668707 | 0.671259 |
| 0.660395 | 0.675617 | 0.667832 | 0.672588 | 0.676738 | 0.670077 | 0.670717 | 0.665591 | 0.673028 |
| 0.659906 | 0.67529 | 0.667474 | 0.673823 | 0.673855 | 0.670705 | 0.671603 | 0.668262 | 0.671046 |
| 0.659585 | 0.675596 | 0.67378 | 0.669694 | 0.673305 | 0.673783 | 0.6688 | 0.669018 | 0.675475 |
| 0.661792 | 0.677176 | 0.667702 | 0.671761 | 0.670777 | 0.673352 | 0.669122 | 0.665806 | 0.673728 |
| 0.668262 | 0.678222 | 0.679009 | 0.674456 | 0.679411 | 0.676781 | 0.674213 | 0.677218 | 0.676555 |

| | | | | | | | | |
|---|---|---|---|---|---|---|---|---|
| 0.667687 | 0.678876 | 0.670189 | 0.67812 | 0.675825 | 0.677281 | 0.673275 | 0.671206 | 0.673628 |
| 0.667703 | 0.683187 | 0.677142 | 0.678557 | 0.676574 | 0.675704 | 0.672024 | 0.672239 | 0.678361 |
| 0.664896 | 0.678221 | 0.673221 | 0.673374 | 0.67128 | 0.678367 | 0.672886 | 0.667938 | 0.679062 |
| 0.659661 | 0.677015 | 0.669316 | 0.668929 | 0.668032 | 0.672316 | 0.666446 | 0.658562 | 0.670477 |
| 0.659466 | 0.671268 | 0.666923 | 0.666924 | 0.66829 | 0.666667 | 0.666237 | 0.664559 | 0.668605 |
| 0.662652 | 0.66855 | 0.663208 | 0.669835 | 0.67015 | 0.664347 | 0.668182 | 0.661301 | 0.668322 |
| 0.66264 | 0.67682 | 0.668269 | 0.664879 | 0.663524 | 0.670663 | 0.667802 | 0.663093 | 0.670927 |
| 0.652529 | 0.674091 | 0.668965 | 0.671979 | 0.673028 | 0.667923 | 0.664933 | 0.668375 | 0.664753 |
| 0.659028 | 0.668286 | 0.66466 | 0.664951 | 0.670199 | 0.66477 | 0.668344 | 0.660975 | 0.666912 |
| 0.661792 | 0.675187 | 0.668614 | 0.668129 | 0.672469 | 0.671911 | 0.67069 | 0.662677 | 0.671393 |
| 0.654633 | 0.665318 | 0.663371 | 0.664647 | 0.661209 | 0.664539 | 0.663241 | 0.660262 | 0.664138 |
| 0.69972 | 0.703972 | 0.702204 | 0.701836 | 0.703529 | 0.702292 | 0.69906 | 0.700162 | 0.700094 |
| 0.707447 | 0.712585 | 0.708928 | 0.708076 | 0.711927 | 0.709958 | 0.706354 | 0.71043 | 0.710991 |
| 0.684103 | 0.691631 | 0.688773 | 0.690096 | 0.688068 | 0.693282 | 0.686806 | 0.689367 | 0.689722 |
| 0.712828 | 0.721675 | 0.71826 | 0.717347 | 0.721934 | 0.719526 | 0.712397 | 0.716289 | 0.717538 |
| 0.704646 | 0.707076 | 0.705189 | 0.709585 | 0.710001 | 0.707928 | 0.703895 | 0.705823 | 0.705601 |
| 0.714432 | 0.719336 | 0.713622 | 0.71506 | 0.720317 | 0.716157 | 0.711744 | 0.710697 | 0.716437 |
| 0.720394 | 0.726534 | 0.722227 | 0.726896 | 0.723707 | 0.724578 | 0.721379 | 0.718882 | 0.722132 |
| 0.695246 | 0.699086 | 0.698712 | 0.69837 | 0.699757 | 0.698394 | 0.69465 | 0.700535 | 0.698401 |
| 0.720793 | 0.724165 | 0.724536 | 0.725156 | 0.724431 | 0.726115 | 0.722074 | 0.721862 | 0.722559 |
| 0.689878 | 0.692846 | 0.691145 | 0.692866 | 0.690593 | 0.690573 | 0.692634 | 0.690564 | 0.689805 |
| 0.71062 | 0.720377 | 0.714762 | 0.71877 | 0.721946 | 0.72248 | 0.712174 | 0.715252 | 0.717852 |
| 0.720472 | 0.723574 | 0.71971 | 0.722016 | 0.7266 | 0.724108 | 0.720463 | 0.718246 | 0.722321 |
| 0.683625 | 0.689325 | 0.686042 | 0.687223 | 0.682676 | 0.688056 | 0.683255 | 0.683636 | 0.688108 |
| 0.689304 | 0.694057 | 0.687515 | 0.692206 | 0.694495 | 0.68995 | 0.68838 | 0.689857 | 0.693889 |
| 0.685366 | 0.703453 | 0.690638 | 0.696211 | 0.696004 | 0.696638 | 0.69413 | 0.692337 | 0.693927 |
| 0.707641 | 0.71521 | 0.71046 | 0.710057 | 0.708651 | 0.709589 | 0.706641 | 0.709354 | 0.71038 |
| 0.718958 | 0.725712 | 0.723803 | 0.723784 | 0.722945 | 0.72618 | 0.717492 | 0.720586 | 0.721919 |
| 0.688185 | 0.691618 | 0.690473 | 0.689488 | 0.694395 | 0.691517 | 0.688596 | 0.690289 | 0.688177 |
| 0.72605 | 0.730246 | 0.726634 | 0.729007 | 0.729646 | 0.729552 | 0.720851 | 0.727109 | 0.728189 |
| 0.723151 | 0.731252 | 0.72377 | 0.729195 | 0.729764 | 0.732418 | 0.725209 | 0.725399 | 0.727532 |
| 0.695841 | 0.702481 | 0.699353 | 0.700853 | 0.703607 | 0.700545 | 0.699871 | 0.699922 | 0.698424 |
| 0.697742 | 0.702857 | 0.698208 | 0.702857 | 0.706385 | 0.701028 | 0.697994 | 0.702325 | 0.700198 |
| 0.72799 | 0.73006 | 0.727417 | 0.729643 | 0.727033 | 0.73007 | 0.724158 | 0.729188 | 0.727822 |
| 0.72099 | 0.723515 | 0.722717 | 0.725771 | 0.724929 | 0.726917 | 0.71878 | 0.721265 | 0.726287 |
| 0.72328 | 0.72789 | 0.724152 | 0.724917 | 0.727595 | 0.72525 | 0.724285 | 0.723854 | 0.726143 |
| 0.677726 | 0.680735 | 0.680452 | 0.682801 | 0.682247 | 0.684137 | 0.675755 | 0.680939 | 0.678359 |
| 0.72176 | 0.725542 | 0.722284 | 0.721165 | 0.724647 | 0.722768 | 0.721258 | 0.721783 | 0.719962 |
| 0.676226 | 0.677814 | 0.678754 | 0.678384 | 0.6812 | 0.681599 | 0.673871 | 0.678771 | 0.675946 |
| 0.666742 | 0.672066 | 0.671121 | 0.674874 | 0.673814 | 0.673263 | 0.668488 | 0.670052 | 0.670397 |
| 0.715466 | 0.724242 | 0.721658 | 0.725022 | 0.72585 | 0.724483 | 0.716756 | 0.722931 | 0.723725 |
| 0.668654 | 0.6732 | 0.671044 | 0.67252 | 0.676213 | 0.675119 | 0.668569 | 0.672183 | 0.67126 |
| 0.659374 | 0.664206 | 0.665993 | 0.665132 | 0.668099 | 0.664923 | 0.662994 | 0.66289 | 0.662651 |
| 0.656162 | 0.663394 | 0.661016 | 0.660725 | 0.664113 | 0.660889 | 0.658246 | 0.660012 | 0.663002 |
| 0.728832 | 0.732916 | 0.732829 | 0.726242 | 0.732635 | 0.732713 | 0.728654 | 0.72708 | 0.727774 |
| 0.662133 | 0.664856 | 0.663732 | 0.666188 | 0.667269 | 0.665572 | 0.663021 | 0.6622 | 0.664324 |
| 0.72015 | 0.725451 | 0.724311 | 0.725909 | 0.729247 | 0.726458 | 0.725647 | 0.721337 | 0.723916 |

| | | | | | | | | |
|---|---|---|---|---|---|---|---|---|
| 0.731506 | 0.730265 | 0.735051 | 0.732723 | 0.736543 | 0.732793 | 0.728717 | 0.729912 | 0.7288 |
| 0.725016 | 0.727935 | 0.726919 | 0.727833 | 0.728208 | 0.727778 | 0.722078 | 0.729349 | 0.72552 |
| 0.746431 | 0.748039 | 0.747139 | 0.748281 | 0.750184 | 0.749939 | 0.744558 | 0.743647 | 0.746264 |
| 0.651512 | 0.657251 | 0.655763 | 0.656085 | 0.657811 | 0.655279 | 0.652131 | 0.65469 | 0.654029 |
| 0.651835 | 0.659535 | 0.655722 | 0.657354 | 0.66045 | 0.655405 | 0.65047 | 0.654058 | 0.655159 |
| 0.734784 | 0.737877 | 0.731807 | 0.734375 | 0.738575 | 0.737478 | 0.730255 | 0.731249 | 0.734347 |
| 0.657067 | 0.654741 | 0.659152 | 0.656677 | 0.661172 | 0.658136 | 0.653931 | 0.657365 | 0.654966 |
| 0.737073 | 0.736911 | 0.736518 | 0.740095 | 0.740686 | 0.737578 | 0.734353 | 0.73623 | 0.736239 |
| 0.731698 | 0.735385 | 0.734159 | 0.731279 | 0.736315 | 0.732479 | 0.731261 | 0.731841 | 0.73363 |
| 0.735356 | 0.737486 | 0.738613 | 0.738666 | 0.739009 | 0.735297 | 0.731493 | 0.733194 | 0.734592 |
| 0.732737 | 0.73448 | 0.733484 | 0.732678 | 0.737551 | 0.735629 | 0.728946 | 0.731711 | 0.73186 |
| 0.662651 | 0.675676 | 0.72956 | 0.721429 | 0.700637 | 0.704698 | 0.716216 | 0.710345 | 0.692308 |
| 0.730461 | 0.731776 | 0.729317 | 0.72499 | 0.735431 | 0.727146 | 0.727872 | 0.727615 | 0.726434 |
| 0.726006 | 0.732501 | 0.730436 | 0.727444 | 0.7303 | 0.728623 | 0.727731 | 0.723715 | 0.72604 |
| 0.739964 | 0.739639 | 0.735695 | 0.733858 | 0.73848 | 0.744884 | 0.735089 | 0.730781 | 0.732567 |
| 0.734691 | 0.730022 | 0.734078 | 0.733162 | 0.74435 | 0.737303 | 0.731434 | 0.735274 | 0.730899 |
| 0.592469 | 0.594041 | 0.59521 | 0.595065 | 0.59745 | 0.599611 | 0.594828 | 0.592399 | 0.590658 |
| 0.735979 | 0.738897 | 0.737945 | 0.741303 | 0.743506 | 0.745001 | 0.736614 | 0.735264 | 0.737511 |
| 0.736748 | 0.74068 | 0.739173 | 0.737452 | 0.744914 | 0.74105 | 0.735182 | 0.733749 | 0.735337 |
| 0.737933 | 0.739278 | 0.739697 | 0.739037 | 0.742497 | 0.736877 | 0.73374 | 0.735904 | 0.739285 |
| 0.738004 | 0.740059 | 0.74134 | 0.739433 | 0.74299 | 0.740596 | 0.736376 | 0.737036 | 0.739644 |
| 0.597104 | 0.59464 | 0.597456 | 0.598029 | 0.600629 | 0.599832 | 0.595631 | 0.595315 | 0.594688 |
| 0.585467 | 0.581261 | 0.584073 | 0.585385 | 0.586969 | 0.585601 | 0.582492 | 0.583975 | 0.579783 |
| 0.765447 | 0.761445 | 0.761074 | 0.759444 | 0.76768 | 0.762314 | 0.760927 | 0.762279 | 0.761742 |
| 0.762648 | 0.760465 | 0.765234 | 0.760788 | 0.766429 | 0.76163 | 0.7624 | 0.760567 | 0.761631 |
| 0.758433 | 0.753506 | 0.759425 | 0.753803 | 0.755546 | 0.758128 | 0.75495 | 0.759294 | 0.758564 |
| 0.765109 | 0.761846 | 0.770547 | 0.761517 | 0.76279 | 0.762626 | 0.763307 | 0.766888 | 0.758923 |
| 0.765765 | 0.767831 | 0.768735 | 0.761194 | 0.767491 | 0.764826 | 0.761374 | 0.767033 | 0.763016 |
| 0.766476 | 0.766342 | 0.771918 | 0.76663 | 0.766681 | 0.768893 | 0.769315 | 0.76743 | 0.76179 |
| 0.766967 | 0.765648 | 0.769029 | 0.768237 | 0.76642 | 0.762379 | 0.766792 | 0.765364 | 0.76837 |
| 0.767649 | 0.761022 | 0.76714 | 0.767877 | 0.769211 | 0.766305 | 0.765508 | 0.760239 | 0.759067 |
| 0.771849 | 0.768841 | 0.774657 | 0.770876 | 0.771844 | 0.775191 | 0.766587 | 0.765012 | 0.77382 |
| 0.765157 | 0.768379 | 0.769053 | 0.768718 | 0.769205 | 0.767223 | 0.761037 | 0.760949 | 0.765352 |
| 0.767387 | 0.769268 | 0.763338 | 0.768137 | 0.769005 | 0.771334 | 0.760228 | 0.761396 | 0.765879 |
| 0.772218 | 0.770871 | 0.772371 | 0.765257 | 0.777238 | 0.76977 | 0.76477 | 0.766575 | 0.77254 |
| 0.762033 | 0.761524 | 0.764865 | 0.763931 | 0.773122 | 0.769231 | 0.758477 | 0.762523 | 0.76146 |
| 0.766486 | 0.765455 | 0.767451 | 0.76051 | 0.768546 | 0.763522 | 0.76073 | 0.758841 | 0.766449 |
| 0.776349 | 0.771248 | 0.773485 | 0.771539 | 0.771648 | 0.770857 | 0.771159 | 0.767266 | 0.767056 |
| 0.765879 | 0.762966 | 0.767536 | 0.762731 | 0.766145 | 0.765121 | 0.763728 | 0.758824 | 0.758615 |
| 0.769789 | 0.766132 | 0.766542 | 0.766036 | 0.765959 | 0.768643 | 0.764691 | 0.761156 | 0.76513 |
| 0.726395 | 0.730912 | 0.731485 | 0.728311 | 0.732982 | 0.734408 | 0.730403 | 0.721232 | 0.725735 |
| 0.736535 | 0.735456 | 0.737991 | 0.732349 | 0.738068 | 0.742944 | 0.7411 | 0.739872 | 0.724695 |
| 0.768392 | 0.767265 | 0.768103 | 0.765579 | 0.77062 | 0.772253 | 0.764447 | 0.764597 | 0.767339 |
| 0.775801 | 0.770789 | 0.771685 | 0.774671 | 0.769123 | 0.773121 | 0.769218 | 0.775181 | 0.767231 |
| 0.717534 | 0.728299 | 0.728688 | 0.71916 | 0.718504 | 0.727798 | 0.727167 | 0.718267 | 0.720466 |
| 0.76806 | 0.769517 | 0.77341 | 0.767384 | 0.771309 | 0.773387 | 0.768721 | 0.762451 | 0.768341 |
| 0.760718 | 0.760978 | 0.763431 | 0.758869 | 0.760666 | 0.765762 | 0.757525 | 0.755465 | 0.75894 |

| | | | | | | | | |
|---|---|---|---|---|---|---|---|---|
| 0.766691 | 0.764924 | 0.770031 | 0.766697 | 0.770991 | 0.771716 | 0.765099 | 0.760335 | 0.768843 |
| 0.77211 | 0.768695 | 0.769823 | 0.768601 | 0.771748 | 0.772128 | 0.769544 | 0.761821 | 0.769778 |
| 0.764637 | 0.763403 | 0.763724 | 0.76474 | 0.760606 | 0.768464 | 0.758893 | 0.765352 | 0.763975 |
| 0.773686 | 0.770643 | 0.771646 | 0.77084 | 0.777659 | 0.772564 | 0.770028 | 0.766599 | 0.771129 |
| 0.774381 | 0.774478 | 0.771457 | 0.772829 | 0.770367 | 0.772226 | 0.769822 | 0.774434 | 0.772076 |
| 0.768851 | 0.768638 | 0.770294 | 0.762936 | 0.769632 | 0.762459 | 0.766804 | 0.760149 | 0.761826 |
| 0.765766 | 0.766757 | 0.768874 | 0.765928 | 0.772683 | 0.768442 | 0.761802 | 0.759176 | 0.767063 |
| 0.757073 | 0.754434 | 0.764755 | 0.757367 | 0.759066 | 0.76049 | 0.753849 | 0.752885 | 0.753644 |
| 0.769874 | 0.767474 | 0.763958 | 0.76772 | 0.76887 | 0.767914 | 0.76474 | 0.76483 | 0.767013 |
| 0.760245 | 0.759944 | 0.766568 | 0.757613 | 0.760663 | 0.768264 | 0.758065 | 0.759165 | 0.763399 |
| 0.765184 | 0.770589 | 0.768818 | 0.765434 | 0.769639 | 0.771917 | 0.766235 | 0.763331 | 0.769161 |
| 0.766302 | 0.766203 | 0.768188 | 0.764945 | 0.774572 | 0.769841 | 0.764706 | 0.758145 | 0.767244 |
| 0.754428 | 0.757971 | 0.758772 | 0.756755 | 0.759236 | 0.761225 | 0.755147 | 0.75222 | 0.757465 |
| 0.758779 | 0.761605 | 0.760831 | 0.758264 | 0.760855 | 0.759336 | 0.755208 | 0.754695 | 0.756114 |
| 0.77292 | 0.765763 | 0.770742 | 0.770151 | 0.768083 | 0.770881 | 0.763864 | 0.767246 | 0.763924 |
| 0.767538 | 0.768871 | 0.767217 | 0.765755 | 0.766686 | 0.766946 | 0.764466 | 0.763878 | 0.765502 |
| 0.767825 | 0.76772 | 0.770399 | 0.770006 | 0.764342 | 0.767351 | 0.770592 | 0.764647 | 0.765432 |
| 0.762586 | 0.764128 | 0.764331 | 0.763004 | 0.764232 | 0.763578 | 0.761877 | 0.758359 | 0.765956 |
| 0.765492 | 0.76885 | 0.767229 | 0.762727 | 0.767617 | 0.767848 | 0.763279 | 0.760822 | 0.767166 |
| 0.766771 | 0.762954 | 0.767347 | 0.760696 | 0.769878 | 0.762017 | 0.762004 | 0.762063 | 0.765896 |
| 0.757327 | 0.760175 | 0.760192 | 0.753383 | 0.764716 | 0.755318 | 0.754237 | 0.751842 | 0.756018 |
| 0.763224 | 0.762919 | 0.761842 | 0.757826 | 0.765045 | 0.763544 | 0.761141 | 0.758822 | 0.758621 |
| 0.771224 | 0.767537 | 0.770393 | 0.766983 | 0.771125 | 0.765401 | 0.768605 | 0.765905 | 0.765959 |
| 0.768069 | 0.770381 | 0.772154 | 0.769625 | 0.769952 | 0.767408 | 0.76972 | 0.759948 | 0.764769 |
| 0.767326 | 0.766171 | 0.762137 | 0.763281 | 0.772568 | 0.762349 | 0.762831 | 0.763294 | 0.766352 |
| 0.768252 | 0.76612 | 0.764227 | 0.758669 | 0.765441 | 0.759237 | 0.760544 | 0.76055 | 0.761818 |
| 0.760209 | 0.765418 | 0.76043 | 0.758015 | 0.761925 | 0.756813 | 0.759107 | 0.754081 | 0.762569 |
| 0.760978 | 0.76327 | 0.76837 | 0.757581 | 0.764519 | 0.75905 | 0.757084 | 0.75659 | 0.762639 |
| 0.766304 | 0.763674 | 0.763407 | 0.761466 | 0.767436 | 0.757029 | 0.760677 | 0.757429 | 0.764487 |
| 0.768073 | 0.769326 | 0.77052 | 0.767423 | 0.769982 | 0.767102 | 0.765806 | 0.763937 | 0.768552 |
| 0.765585 | 0.764178 | 0.768289 | 0.764498 | 0.767741 | 0.76935 | 0.763564 | 0.762158 | 0.767216 |
| 0.76614 | 0.765358 | 0.763091 | 0.763147 | 0.766802 | 0.765668 | 0.761539 | 0.762142 | 0.767039 |
| 0.769156 | 0.765379 | 0.767406 | 0.77181 | 0.772215 | 0.770117 | 0.765538 | 0.768552 | 0.771115 |
| 0.769086 | 0.769662 | 0.775637 | 0.769998 | 0.772735 | 0.771067 | 0.770453 | 0.771796 | 0.763889 |
| 0.762704 | 0.760733 | 0.766585 | 0.757102 | 0.767079 | 0.755889 | 0.759742 | 0.756088 | 0.756741 |
| 0.766819 | 0.766781 | 0.762702 | 0.761869 | 0.768768 | 0.765023 | 0.759814 | 0.762903 | 0.765913 |
| 0.764372 | 0.768604 | 0.763879 | 0.759252 | 0.768697 | 0.766052 | 0.762481 | 0.759292 | 0.761348 |
| 0.767271 | 0.764412 | 0.765901 | 0.762713 | 0.767289 | 0.762583 | 0.7649 | 0.756415 | 0.763777 |
| 0.755146 | 0.755273 | 0.756992 | 0.752025 | 0.759513 | 0.753048 | 0.754215 | 0.751169 | 0.757769 |
| 0.76033 | 0.752886 | 0.759997 | 0.754239 | 0.759023 | 0.754573 | 0.754522 | 0.754208 | 0.753289 |
| 0.756789 | 0.758392 | 0.75925 | 0.748604 | 0.758808 | 0.752501 | 0.75478 | 0.750708 | 0.753776 |
| 0.758345 | 0.757645 | 0.756057 | 0.75299 | 0.760339 | 0.753574 | 0.758663 | 0.751933 | 0.753955 |
| 0.752603 | 0.752106 | 0.755182 | 0.751374 | 0.758547 | 0.751574 | 0.749349 | 0.748321 | 0.754213 |
| 0.753427 | 0.751307 | 0.754939 | 0.749527 | 0.756214 | 0.749531 | 0.751275 | 0.748159 | 0.750991 |
| 0.760443 | 0.760194 | 0.765895 | 0.760764 | 0.762709 | 0.762031 | 0.761842 | 0.755928 | 0.759654 |
| 0.752576 | 0.755587 | 0.755587 | 0.753608 | 0.762094 | 0.751613 | 0.753629 | 0.743592 | 0.748706 |

| | | | | | | | | |
|---|---|---|---|---|---|---|---|---|
| 0.768335 | 0.76525 | 0.765923 | 0.75998 | 0.768477 | 0.762663 | 0.759106 | 0.762746 | 0.761995 |
| 0.779396 | 0.778067 | 0.779155 | 0.774502 | 0.77762 | 0.777343 | 0.776993 | 0.773962 | 0.7697 |
| 0.771433 | 0.77157 | 0.767649 | 0.767995 | 0.769794 | 0.768918 | 0.769244 | 0.769528 | 0.765831 |
| 0.767468 | 0.765307 | 0.770236 | 0.773396 | 0.775738 | 0.776157 | 0.766058 | 0.770576 | 0.769533 |
| 0.778197 | 0.780673 | 0.779365 | 0.7774 | 0.782939 | 0.78143 | 0.778896 | 0.775169 | 0.778727 |
| 0.775033 | 0.768624 | 0.775687 | 0.775095 | 0.778509 | 0.773035 | 0.773936 | 0.770021 | 0.76744 |
| 0.773932 | 0.773611 | 0.772304 | 0.769489 | 0.772914 | 0.772015 | 0.769565 | 0.770622 | 0.771267 |
| 0.772049 | 0.765498 | 0.771938 | 0.76679 | 0.769099 | 0.769638 | 0.765585 | 0.761836 | 0.761776 |
| 0.768878 | 0.767501 | 0.766457 | 0.764791 | 0.770219 | 0.771911 | 0.767557 | 0.765714 | 0.766495 |
| 0.763556 | 0.762752 | 0.767665 | 0.767378 | 0.768832 | 0.767995 | 0.76509 | 0.763169 | 0.763969 |
| 0.771591 | 0.770743 | 0.773979 | 0.767009 | 0.76955 | 0.772978 | 0.768403 | 0.767703 | 0.769841 |
| 0.765353 | 0.769433 | 0.774812 | 0.772344 | 0.774618 | 0.770801 | 0.768891 | 0.767453 | 0.768032 |
| 0.777586 | 0.772228 | 0.769068 | 0.771299 | 0.777934 | 0.770549 | 0.766403 | 0.768964 | 0.769408 |
| 0.774575 | 0.776003 | 0.77091 | 0.771584 | 0.775169 | 0.775632 | 0.769179 | 0.766494 | 0.768685 |
| 0.777208 | 0.777211 | 0.779343 | 0.772877 | 0.782452 | 0.777334 | 0.774118 | 0.775843 | 0.774844 |
| 0.778133 | 0.7702 | 0.769293 | 0.774862 | 0.777467 | 0.771303 | 0.767763 | 0.768405 | 0.769186 |
| 0.778184 | 0.778224 | 0.773843 | 0.775059 | 0.778495 | 0.776624 | 0.776496 | 0.773986 | 0.773278 |
| 0.775107 | 0.770638 | 0.770956 | 0.777841 | 0.780488 | 0.773891 | 0.775375 | 0.774165 | 0.775437 |
| 0.777123 | 0.778053 | 0.780452 | 0.775764 | 0.778015 | 0.778789 | 0.772735 | 0.771934 | 0.773638 |
| 0.779892 | 0.775021 | 0.775915 | 0.780574 | 0.780993 | 0.777443 | 0.778037 | 0.771546 | 0.774209 |
| 0.771163 | 0.772132 | 0.771567 | 0.771161 | 0.777584 | 0.775662 | 0.767728 | 0.771605 | 0.768555 |
| 0.773579 | 0.770911 | 0.772536 | 0.773826 | 0.780376 | 0.773678 | 0.768016 | 0.765724 | 0.766189 |
| 0.773897 | 0.776096 | 0.773511 | 0.771227 | 0.775205 | 0.774774 | 0.770842 | 0.77351 | 0.770943 |
| 0.771568 | 0.773391 | 0.772122 | 0.773226 | 0.775408 | 0.774322 | 0.767758 | 0.773485 | 0.771044 |
| 0.773455 | 0.772709 | 0.777868 | 0.774784 | 0.781528 | 0.777517 | 0.767235 | 0.775175 | 0.772924 |
| 0.775505 | 0.7678 | 0.772345 | 0.775141 | 0.777595 | 0.77605 | 0.768852 | 0.770283 | 0.768654 |
| 0.7689 | 0.765166 | 0.777335 | 0.768085 | 0.773198 | 0.774361 | 0.764716 | 0.766132 | 0.762569 |
| 0.773215 | 0.7678 | 0.770997 | 0.770246 | 0.775622 | 0.772934 | 0.766175 | 0.770145 | 0.767029 |
| 0.775788 | 0.774252 | 0.77738 | 0.77957 | 0.778277 | 0.777669 | 0.77306 | 0.772837 | 0.775787 |
| 0.774101 | 0.778864 | 0.777862 | 0.77511 | 0.779161 | 0.779365 | 0.774335 | 0.772276 | 0.770798 |
| 0.768886 | 0.766034 | 0.767469 | 0.765536 | 0.769948 | 0.767342 | 0.763033 | 0.763711 | 0.765705 |
| 0.77272 | 0.7738 | 0.776411 | 0.769716 | 0.776562 | 0.772761 | 0.766641 | 0.771193 | 0.765982 |
| 0.773799 | 0.776095 | 0.778413 | 0.770647 | 0.775045 | 0.773205 | 0.76968 | 0.773982 | 0.775871 |
| 0.768938 | 0.766367 | 0.772209 | 0.767205 | 0.775115 | 0.768343 | 0.769502 | 0.766956 | 0.769683 |
| 0.768507 | 0.773628 | 0.775696 | 0.77243 | 0.775583 | 0.773844 | 0.769746 | 0.768033 | 0.76957 |
| 0.772547 | 0.770315 | 0.767906 | 0.767424 | 0.775648 | 0.765325 | 0.767174 | 0.764501 | 0.766066 |
| 0.781117 | 0.779118 | 0.780822 | 0.776725 | 0.787267 | 0.780768 | 0.777713 | 0.779222 | 0.775561 |
| 0.774161 | 0.771996 | 0.774392 | 0.778947 | 0.78259 | 0.771778 | 0.772031 | 0.774058 | 0.768946 |
| 0.776446 | 0.77153 | 0.77203 | 0.770582 | 0.777769 | 0.774117 | 0.767065 | 0.769661 | 0.769288 |
| 0.771864 | 0.76696 | 0.775878 | 0.772666 | 0.776777 | 0.770103 | 0.77062 | 0.771223 | 0.765727 |
| 0.774336 | 0.774289 | 0.777769 | 0.7769 | 0.777696 | 0.774417 | 0.770492 | 0.770825 | 0.768695 |
| 0.778351 | 0.772499 | 0.777823 | 0.774851 | 0.780281 | 0.77796 | 0.772263 | 0.775083 | 0.769826 |
| 0.780675 | 0.779678 | 0.777769 | 0.776976 | 0.779841 | 0.780593 | 0.7752 | 0.776318 | 0.770746 |
| 0.771753 | 0.772371 | 0.774757 | 0.771657 | 0.774362 | 0.769384 | 0.768577 | 0.764575 | 0.765559 |
| 0.77667 | 0.774983 | 0.777494 | 0.775531 | 0.780459 | 0.775717 | 0.774042 | 0.775235 | 0.768409 |
| 0.776322 | 0.770438 | 0.77144 | 0.773147 | 0.780353 | 0.769666 | 0.770394 | 0.768153 | 0.762909 |

| | | | | | | | | |
|---|---|---|---|---|---|---|---|---|
| 0.765139 | 0.765125 | 0.766225 | 0.762371 | 0.766046 | 0.770026 | 0.759536 | 0.768384 | 0.761611 |
| 0.752358 | 0.751113 | 0.74737 | 0.751675 | 0.754677 | 0.752558 | 0.750356 | 0.74925 | 0.749278 |
| 0.774952 | 0.776778 | 0.770861 | 0.768906 | 0.777317 | 0.770024 | 0.771039 | 0.771424 | 0.77125 |
| 0.779309 | 0.78219 | 0.78081 | 0.777695 | 0.785509 | 0.779553 | 0.777648 | 0.77724 | 0.774286 |
| 0.774213 | 0.771772 | 0.77332 | 0.767556 | 0.770065 | 0.76893 | 0.764431 | 0.764053 | 0.763523 |
| 0.776737 | 0.772009 | 0.776859 | 0.770147 | 0.780596 | 0.774017 | 0.770752 | 0.769306 | 0.76893 |
| 0.775584 | 0.774954 | 0.773551 | 0.774171 | 0.778278 | 0.777539 | 0.77525 | 0.773556 | 0.771438 |
| 0.772059 | 0.769437 | 0.767752 | 0.769422 | 0.778995 | 0.770572 | 0.763966 | 0.76711 | 0.763389 |
| 1 | 0.806357 | 0.797246 | 0.818075 | 0.784927 | 0.801916 | 0.824027 | 0.808437 | 0.788032 |
| 0.806357 | 1 | 0.803424 | 0.805009 | 0.780121 | 0.825834 | 0.817448 | 0.808181 | 0.836379 |
| 0.797246 | 0.803424 | 1 | 0.793888 | 0.782512 | 0.826632 | 0.794496 | 0.784648 | 0.783952 |
| 0.818075 | 0.805009 | 0.793888 | 1 | 0.778164 | 0.84924 | 0.83546 | 0.813196 | 0.821295 |
| 0.784927 | 0.780121 | 0.782512 | 0.778164 | 1 | 0.782355 | 0.781132 | 0.780329 | 0.780814 |
| 0.801916 | 0.825834 | 0.826632 | 0.84924 | 0.782355 | 1 | 0.799882 | 0.826123 | 0.815452 |
| 0.824027 | 0.817448 | 0.794496 | 0.83546 | 0.781132 | 0.799882 | 1 | 0.780569 | 0.800256 |
| 0.808437 | 0.808181 | 0.784648 | 0.813196 | 0.780329 | 0.826123 | 0.780569 | 1 | 0.825206 |
| 0.788032 | 0.836379 | 0.783952 | 0.821295 | 0.780814 | 0.815452 | 0.800256 | 0.825206 | 1 |
| 0.792851 | 0.831964 | 0.777392 | 0.818483 | 0.778166 | 0.823514 | 0.792719 | 0.826171 | 0.867165 |
| 0.776665 | 0.777048 | 0.776616 | 0.775209 | 0.775358 | 0.776071 | 0.778933 | 0.777847 | 0.781456 |
| 0.789114 | 0.829356 | 0.77654 | 0.774196 | 0.779322 | 0.77579 | 0.79871 | 0.779043 | 0.81002 |
| 0.809291 | 0.792876 | 0.794179 | 0.798442 | 0.779409 | 0.790168 | 0.803265 | 0.783793 | 0.785582 |
| 0.760641 | 0.763887 | 0.757517 | 0.761662 | 0.762782 | 0.758386 | 0.760371 | 0.753972 | 0.757168 |
| 0.769596 | 0.76716 | 0.769501 | 0.767812 | 0.769193 | 0.763744 | 0.766642 | 0.762101 | 0.76445 |
| 0.772477 | 0.772467 | 0.771201 | 0.773478 | 0.77055 | 0.772395 | 0.771316 | 0.768256 | 0.769333 |
| 0.777021 | 0.773759 | 0.773818 | 0.774218 | 0.775476 | 0.775365 | 0.772817 | 0.769581 | 0.772566 |
| 0.768668 | 0.770296 | 0.767436 | 0.772029 | 0.768222 | 0.768484 | 0.767511 | 0.766076 | 0.768188 |
| 0.767395 | 0.764681 | 0.764239 | 0.766092 | 0.761563 | 0.766486 | 0.758659 | 0.76239 | 0.768319 |
| 0.771304 | 0.770254 | 0.76948 | 0.769792 | 0.772427 | 0.76682 | 0.766811 | 0.761057 | 0.76669 |
| 0.772174 | 0.767962 | 0.770583 | 0.76444 | 0.770968 | 0.769177 | 0.766293 | 0.764671 | 0.764074 |
| 0.692861 | 0.700969 | 0.696072 | 0.702175 | 0.699724 | 0.700411 | 0.697445 | 0.692669 | 0.696088 |
| 0.706591 | 0.712524 | 0.708197 | 0.712958 | 0.71432 | 0.709649 | 0.707793 | 0.707726 | 0.714759 |
| 0.703186 | 0.7098 | 0.70523 | 0.710276 | 0.709771 | 0.707911 | 0.704632 | 0.70328 | 0.707519 |
| 0.691416 | 0.702523 | 0.699114 | 0.701461 | 0.705315 | 0.699413 | 0.699143 | 0.693148 | 0.701551 |
| 0.707005 | 0.714991 | 0.710611 | 0.712876 | 0.713222 | 0.71136 | 0.710046 | 0.70662 | 0.712468 |
| 0.772442 | 0.763534 | 0.773155 | 0.772377 | 0.771514 | 0.771769 | 0.766529 | 0.761432 | 0.764496 |
| 0.773029 | 0.769848 | 0.77619 | 0.7683 | 0.775096 | 0.772022 | 0.769752 | 0.765183 | 0.768858 |
| 0.778573 | 0.773126 | 0.778987 | 0.774663 | 0.780563 | 0.776633 | 0.77558 | 0.767303 | 0.770823 |
| 0.777924 | 0.774577 | 0.775761 | 0.77069 | 0.776176 | 0.771375 | 0.77471 | 0.764472 | 0.77377 |
| 0.77517 | 0.767204 | 0.768857 | 0.769686 | 0.774633 | 0.769339 | 0.76695 | 0.770867 | 0.764472 |
| 0.776024 | 0.769965 | 0.772205 | 0.771661 | 0.771923 | 0.770457 | 0.768746 | 0.772368 | 0.765231 |
| 0.776638 | 0.774169 | 0.780616 | 0.772136 | 0.779042 | 0.775293 | 0.772249 | 0.769136 | 0.769662 |
| 0.771171 | 0.767811 | 0.771514 | 0.768726 | 0.773667 | 0.768654 | 0.768713 | 0.766455 | 0.764898 |
| 0.769611 | 0.763036 | 0.765346 | 0.763839 | 0.765073 | 0.767013 | 0.762005 | 0.764288 | 0.761143 |
| 0.770531 | 0.765574 | 0.769312 | 0.765193 | 0.770351 | 0.764659 | 0.767722 | 0.761589 | 0.76364 |
| 0.769645 | 0.768163 | 0.770793 | 0.7726 | 0.769497 | 0.767418 | 0.768596 | 0.76476 | 0.765501 |
| 0.776595 | 0.765798 | 0.775359 | 0.774275 | 0.775081 | 0.771659 | 0.773388 | 0.768731 | 0.769526 |

| | | | | | | | | |
|---|---|---|---|---|---|---|---|---|
| 0.774074 | 0.775642 | 0.77398 | 0.774727 | 0.778265 | 0.775594 | 0.773204 | 0.77097 | 0.775752 |
| 0.768787 | 0.772531 | 0.777363 | 0.772026 | 0.772264 | 0.774345 | 0.770302 | 0.764778 | 0.768975 |
| 0.774022 | 0.771909 | 0.772212 | 0.769591 | 0.776997 | 0.770652 | 0.766352 | 0.76598 | 0.766917 |
| 0.77245 | 0.768618 | 0.771583 | 0.766054 | 0.770465 | 0.769237 | 0.76361 | 0.764505 | 0.764857 |
| 0.770739 | 0.768736 | 0.770587 | 0.767479 | 0.769516 | 0.768194 | 0.766903 | 0.764501 | 0.767706 |
| 0.767552 | 0.768606 | 0.769153 | 0.766564 | 0.769126 | 0.770417 | 0.762291 | 0.763482 | 0.764937 |
| 0.772051 | 0.772812 | 0.770576 | 0.77398 | 0.770538 | 0.772601 | 0.770178 | 0.767891 | 0.768148 |
| 0.770342 | 0.770091 | 0.775916 | 0.769752 | 0.773855 | 0.770523 | 0.766019 | 0.76394 | 0.770574 |
| 0.767208 | 0.767044 | 0.767418 | 0.768821 | 0.76475 | 0.76805 | 0.765686 | 0.762813 | 0.770021 |
| 0.767198 | 0.769981 | 0.771798 | 0.769751 | 0.77501 | 0.769963 | 0.76572 | 0.764264 | 0.765244 |
| 0.770658 | 0.770285 | 0.76774 | 0.768481 | 0.772212 | 0.768565 | 0.76567 | 0.763708 | 0.769179 |
| 0.769571 | 0.770677 | 0.768935 | 0.768551 | 0.775437 | 0.766286 | 0.764926 | 0.762723 | 0.769354 |
| 0.766622 | 0.763196 | 0.762014 | 0.762933 | 0.767844 | 0.757448 | 0.761213 | 0.7586 | 0.757703 |
| 0.766878 | 0.76064 | 0.765089 | 0.759386 | 0.766294 | 0.762666 | 0.759119 | 0.755405 | 0.757179 |
| 0.772103 | 0.766563 | 0.770015 | 0.771943 | 0.772001 | 0.771799 | 0.770038 | 0.7653 | 0.768115 |
| 0.780452 | 0.779836 | 0.773146 | 0.778296 | 0.777851 | 0.778641 | 0.778218 | 0.776271 | 0.774881 |
| 0.776656 | 0.769394 | 0.768919 | 0.775211 | 0.771927 | 0.774586 | 0.774025 | 0.771759 | 0.766678 |
| 0.775068 | 0.77211 | 0.771517 | 0.774712 | 0.777539 | 0.776855 | 0.776057 | 0.773385 | 0.773966 |
| 0.770462 | 0.771625 | 0.772511 | 0.769 | 0.775205 | 0.769407 | 0.768651 | 0.768336 | 0.770472 |
| 0.773878 | 0.773917 | 0.771674 | 0.775722 | 0.77812 | 0.777541 | 0.773413 | 0.7722 | 0.770577 |
| 0.77447 | 0.774091 | 0.775533 | 0.768272 | 0.773328 | 0.773154 | 0.772815 | 0.768113 | 0.766744 |
| 0.777224 | 0.775817 | 0.772284 | 0.773714 | 0.77763 | 0.778147 | 0.775023 | 0.770324 | 0.770819 |
| 0.776994 | 0.775997 | 0.772519 | 0.777217 | 0.776706 | 0.772785 | 0.773421 | 0.775196 | 0.771297 |
| 0.769427 | 0.772885 | 0.766956 | 0.766823 | 0.770011 | 0.771178 | 0.764674 | 0.761537 | 0.769319 |
| 0.776344 | 0.777748 | 0.772735 | 0.774773 | 0.777265 | 0.775862 | 0.771513 | 0.774001 | 0.775109 |
| 0.70159 | 0.709216 | 0.707986 | 0.705069 | 0.711814 | 0.707489 | 0.705464 | 0.704136 | 0.705911 |
| 0.775069 | 0.775233 | 0.769981 | 0.768618 | 0.774069 | 0.771235 | 0.767494 | 0.767303 | 0.766087 |
| 0.770033 | 0.770403 | 0.771853 | 0.773928 | 0.769935 | 0.777269 | 0.768559 | 0.767145 | 0.768212 |
| 0.777759 | 0.777425 | 0.775023 | 0.777537 | 0.779119 | 0.776145 | 0.777956 | 0.772156 | 0.771669 |
| 0.771584 | 0.769065 | 0.773549 | 0.773679 | 0.775635 | 0.773577 | 0.765959 | 0.776463 | 0.768013 |
| 0.767284 | 0.770395 | 0.764746 | 0.767617 | 0.769729 | 0.766123 | 0.764897 | 0.763868 | 0.766042 |
| 0.771484 | 0.768123 | 0.772438 | 0.768987 | 0.772231 | 0.771349 | 0.766776 | 0.766038 | 0.762537 |
| 0.772884 | 0.769134 | 0.768858 | 0.7708 | 0.77345 | 0.772065 | 0.77136 | 0.766647 | 0.771057 |
| 0.770473 | 0.769171 | 0.769753 | 0.76703 | 0.772468 | 0.774091 | 0.765255 | 0.766633 | 0.765398 |
| 0.765199 | 0.763614 | 0.76436 | 0.765557 | 0.768629 | 0.766491 | 0.758242 | 0.76578 | 0.761847 |
| 0.768527 | 0.762081 | 0.767124 | 0.765625 | 0.766739 | 0.768011 | 0.764184 | 0.758618 | 0.75934 |
| 0.775483 | 0.774905 | 0.769905 | 0.775733 | 0.776255 | 0.771581 | 0.773859 | 0.774159 | 0.770015 |
| 0.775561 | 0.775583 | 0.77707 | 0.76785 | 0.783428 | 0.77299 | 0.772409 | 0.773919 | 0.771486 |
| 0.774073 | 0.776948 | 0.773235 | 0.773476 | 0.779711 | 0.769938 | 0.774511 | 0.769641 | 0.775257 |
| 0.772392 | 0.773492 | 0.767289 | 0.768892 | 0.777481 | 0.769737 | 0.771878 | 0.770169 | 0.772747 |
| 0.770559 | 0.767929 | 0.772416 | 0.766087 | 0.772263 | 0.770474 | 0.765504 | 0.771038 | 0.766876 |
| 0.775468 | 0.775684 | 0.774942 | 0.775706 | 0.782462 | 0.774385 | 0.769924 | 0.775744 | 0.772693 |
| 0.765939 | 0.769131 | 0.769205 | 0.767132 | 0.778603 | 0.765057 | 0.766321 | 0.763695 | 0.763705 |
| 0.768072 | 0.769256 | 0.76606 | 0.763904 | 0.770272 | 0.767954 | 0.762194 | 0.769969 | 0.76474 |
| 0.770667 | 0.772799 | 0.774353 | 0.771799 | 0.776242 | 0.772622 | 0.770246 | 0.773202 | 0.770636 |
| 0.769243 | 0.772242 | 0.773554 | 0.770842 | 0.774431 | 0.770623 | 0.767588 | 0.77324 | 0.767334 |

| | | | | | | | | |
|---|---|---|---|---|---|---|---|---|
| 0.770052 | 0.770768 | 0.768448 | 0.771612 | 0.773614 | 0.771175 | 0.76452 | 0.772834 | 0.768605 |
| 0.775176 | 0.771995 | 0.775524 | 0.767728 | 0.777979 | 0.771529 | 0.76562 | 0.772481 | 0.76743 |
| 0.771128 | 0.771178 | 0.767974 | 0.767119 | 0.775083 | 0.768419 | 0.766186 | 0.769628 | 0.766188 |
| 0.735199 | 0.731234 | 0.730727 | 0.713112 | 0.732188 | 0.732136 | 0.722883 | 0.73129 | 0.723727 |
| 0.769607 | 0.771053 | 0.769787 | 0.767812 | 0.77365 | 0.76935 | 0.766539 | 0.770125 | 0.767988 |
| 0.773026 | 0.768309 | 0.769962 | 0.77144 | 0.76996 | 0.772738 | 0.762709 | 0.7692 | 0.767246 |
| 0.768158 | 0.76823 | 0.769074 | 0.769047 | 0.772753 | 0.772082 | 0.758204 | 0.769386 | 0.762268 |
| 0.76712 | 0.769705 | 0.769921 | 0.769079 | 0.772339 | 0.770367 | 0.759698 | 0.765947 | 0.767889 |
| 0.772571 | 0.776713 | 0.776143 | 0.77211 | 0.775138 | 0.77427 | 0.76986 | 0.768407 | 0.770294 |
| 0.771183 | 0.774041 | 0.777933 | 0.768755 | 0.778531 | 0.773367 | 0.768087 | 0.770044 | 0.769936 |
| 0.768979 | 0.767959 | 0.770442 | 0.774772 | 0.772409 | 0.774305 | 0.766879 | 0.7716 | 0.767062 |
| 0.772641 | 0.772037 | 0.768411 | 0.769428 | 0.774236 | 0.772806 | 0.767159 | 0.767895 | 0.77132 |
| 0.773155 | 0.77071 | 0.775306 | 0.774983 | 0.771661 | 0.779281 | 0.762186 | 0.773914 | 0.768222 |
| 0.769996 | 0.767799 | 0.767516 | 0.769933 | 0.772828 | 0.771326 | 0.763348 | 0.770211 | 0.765929 |
| 0.775002 | 0.768624 | 0.767611 | 0.771179 | 0.773371 | 0.770104 | 0.767028 | 0.771405 | 0.764812 |
| 0.770019 | 0.766461 | 0.763577 | 0.76425 | 0.7722 | 0.763478 | 0.75787 | 0.768121 | 0.76331 |
| 0.775528 | 0.773267 | 0.773573 | 0.772065 | 0.775972 | 0.773299 | 0.768625 | 0.772621 | 0.765924 |
| 0.77321 | 0.768725 | 0.767597 | 0.770073 | 0.774485 | 0.772342 | 0.768942 | 0.77512 | 0.762631 |
| 0.766237 | 0.766477 | 0.76221 | 0.762987 | 0.76942 | 0.767153 | 0.760718 | 0.768845 | 0.765693 |
| 0.774323 | 0.765377 | 0.769024 | 0.766215 | 0.773064 | 0.767677 | 0.766895 | 0.770685 | 0.76458 |
| 0.772098 | 0.771497 | 0.772634 | 0.774151 | 0.773064 | 0.773613 | 0.769359 | 0.769904 | 0.765626 |
| 0.773631 | 0.773935 | 0.771072 | 0.772885 | 0.776241 | 0.774975 | 0.768447 | 0.770753 | 0.769763 |
| 0.676727 | 0.68541 | 0.682508 | 0.697584 | 0.683172 | 0.685138 | 0.687777 | 0.690731 | 0.690415 |
| 0.650483 | 0.663113 | 0.655784 | 0.659843 | 0.659938 | 0.666031 | 0.656599 | 0.672209 | 0.672143 |
| 0.672982 | 0.683843 | 0.678486 | 0.678859 | 0.672994 | 0.675804 | 0.679504 | 0.66504 | 0.679523 |
| 0.646639 | 0.66446 | 0.660638 | 0.657852 | 0.659273 | 0.656872 | 0.656898 | 0.65033 | 0.655223 |
| 0.736763 | 0.733755 | 0.740393 | 0.736159 | 0.750417 | 0.733453 | 0.735188 | 0.734452 | 0.733645 |
| 0.725138 | 0.729766 | 0.732525 | 0.73094 | 0.728042 | 0.733384 | 0.730688 | 0.727601 | 0.727536 |
| 0.736756 | 0.737253 | 0.730838 | 0.732127 | 0.732033 | 0.735665 | 0.731359 | 0.730563 | 0.732065 |
| 0.737769 | 0.739987 | 0.747595 | 0.735943 | 0.742352 | 0.743041 | 0.73788 | 0.744321 | 0.736328 |
| 0.722184 | 0.719274 | 0.730883 | 0.723963 | 0.725504 | 0.725023 | 0.731054 | 0.719791 | 0.721484 |
| 0.709795 | 0.722492 | 0.712155 | 0.711331 | 0.712997 | 0.727453 | 0.712155 | 0.709732 | 0.715925 |
| 0.716264 | 0.72204 | 0.727381 | 0.71598 | 0.722766 | 0.722436 | 0.723284 | 0.716431 | 0.720812 |
| 0.644951 | 0.665286 | 0.658212 | 0.664393 | 0.66405 | 0.660106 | 0.658537 | 0.654846 | 0.66074 |
| 0.767328 | 0.761917 | 0.768023 | 0.760732 | 0.768499 | 0.764394 | 0.763888 | 0.765847 | 0.764335 |
| 0.74598 | 0.74368 | 0.743748 | 0.738309 | 0.746706 | 0.743215 | 0.739845 | 0.741018 | 0.739201 |
| 0.745813 | 0.738238 | 0.739566 | 0.735698 | 0.739824 | 0.734247 | 0.73984 | 0.735077 | 0.732926 |
| 0.728842 | 0.722295 | 0.725683 | 0.724022 | 0.733271 | 0.729775 | 0.728678 | 0.720666 | 0.720692 |
| 0.743868 | 0.743536 | 0.744621 | 0.743272 | 0.751345 | 0.74497 | 0.749793 | 0.743729 | 0.737659 |
| 0.733103 | 0.738651 | 0.729899 | 0.73729 | 0.734529 | 0.734785 | 0.731538 | 0.733147 | 0.740238 |
| 0.731327 | 0.734828 | 0.733504 | 0.7368 | 0.7317 | 0.727646 | 0.733355 | 0.725173 | 0.731453 |
| 0.72757 | 0.73066 | 0.724836 | 0.734034 | 0.735952 | 0.736505 | 0.731746 | 0.731228 | 0.736892 |
| 0.734262 | 0.727339 | 0.734046 | 0.730294 | 0.733478 | 0.724539 | 0.736535 | 0.734977 | 0.724475 |
| 0.732844 | 0.728653 | 0.73033 | 0.72609 | 0.734868 | 0.730187 | 0.733704 | 0.73358 | 0.730335 |
| 0.735499 | 0.72884 | 0.723481 | 0.722882 | 0.727577 | 0.735104 | 0.729424 | 0.728965 | 0.726848 |
| 0.715838 | 0.72393 | 0.717168 | 0.718315 | 0.712541 | 0.716127 | 0.723496 | 0.719057 | 0.723645 |

| | | | | | | | | |
|---|---|---|---|---|---|---|---|---|
| 0.755507 | 0.756188 | 0.749333 | 0.754919 | 0.754879 | 0.750271 | 0.752187 | 0.754309 | 0.752537 |
| 0.730634 | 0.729432 | 0.731972 | 0.732693 | 0.741296 | 0.729412 | 0.731069 | 0.7285 | 0.730197 |
| 0.727953 | 0.733424 | 0.732951 | 0.730966 | 0.734305 | 0.729185 | 0.728289 | 0.729111 | 0.726848 |
| 0.729738 | 0.729846 | 0.733129 | 0.729925 | 0.73893 | 0.728532 | 0.731978 | 0.729697 | 0.72751 |
| 0.728752 | 0.736463 | 0.74081 | 0.722517 | 0.73889 | 0.722662 | 0.725331 | 0.72632 | 0.731396 |
| 0.731781 | 0.730721 | 0.733034 | 0.730846 | 0.721522 | 0.727386 | 0.72614 | 0.718497 | 0.72842 |
| 0.721189 | 0.720488 | 0.727096 | 0.722101 | 0.723643 | 0.722894 | 0.720448 | 0.72261 | 0.726626 |
| 0.718936 | 0.719613 | 0.724246 | 0.727195 | 0.715011 | 0.716288 | 0.721088 | 0.725049 | 0.714692 |
| 0.72114 | 0.732133 | 0.729714 | 0.733466 | 0.729606 | 0.732614 | 0.730923 | 0.724341 | 0.726545 |
| 0.756667 | 0.761919 | 0.760776 | 0.754795 | 0.767307 | 0.762021 | 0.755108 | 0.754444 | 0.749388 |
| 0.753907 | 0.755011 | 0.757686 | 0.752551 | 0.756918 | 0.756558 | 0.752626 | 0.753806 | 0.749495 |
| 0.748317 | 0.748138 | 0.752543 | 0.749334 | 0.755612 | 0.74725 | 0.746187 | 0.747317 | 0.745539 |
| 0.741289 | 0.75083 | 0.743867 | 0.743149 | 0.742542 | 0.746336 | 0.743527 | 0.741468 | 0.740426 |
| 0.746846 | 0.74917 | 0.750581 | 0.743455 | 0.747646 | 0.749511 | 0.746512 | 0.743964 | 0.747692 |
| 0.74376 | 0.747148 | 0.744752 | 0.735301 | 0.74701 | 0.74243 | 0.738705 | 0.739262 | 0.745633 |

| Yangambi rainforest | | | | | | | | |
|---|---|---|---|---|---|---|---|---|
| Wild_plot_ | Wild_plot_ | Wild_plot_ | Wild_plot_ | Wild_plot_ | Wild_plot_ | Wild_plot_ | Wild_plot_ | Wild_plot_ |
| 0.70921 | 0.701576 | 0.705086 | 0.709044 | 0.694501 | 0.706615 | 0.708981 | 0.710528 | 0.708034 |
| 0.714 | 0.70956 | 0.710298 | 0.714367 | 0.703646 | 0.710265 | 0.71456 | 0.719476 | 0.714912 |
| 0.704918 | 0.698667 | 0.698821 | 0.705628 | 0.696642 | 0.702609 | 0.707797 | 0.710238 | 0.703885 |
| 0.714286 | 0.704637 | 0.708844 | 0.715651 | 0.703195 | 0.714184 | 0.715175 | 0.714448 | 0.716377 |
| 0.712415 | 0.701653 | 0.706567 | 0.710084 | 0.699366 | 0.706678 | 0.708704 | 0.71146 | 0.708851 |
| 0.71718 | 0.706818 | 0.712144 | 0.713621 | 0.706055 | 0.714704 | 0.71915 | 0.721083 | 0.715104 |
| 0.76804 | 0.764455 | 0.771488 | 0.771886 | 0.764017 | 0.77002 | 0.77058 | 0.769882 | 0.767399 |
| 0.764029 | 0.764669 | 0.769846 | 0.767322 | 0.758548 | 0.758972 | 0.761149 | 0.770149 | 0.760864 |
| 0.714792 | 0.703777 | 0.710382 | 0.7108 | 0.701013 | 0.714948 | 0.715593 | 0.715582 | 0.71337 |
| 0.711744 | 0.698934 | 0.701488 | 0.710245 | 0.704212 | 0.706902 | 0.711272 | 0.712868 | 0.711219 |
| 0.709763 | 0.701329 | 0.703305 | 0.71175 | 0.701934 | 0.708197 | 0.709806 | 0.711051 | 0.710183 |
| 0.704839 | 0.696235 | 0.700444 | 0.705107 | 0.695564 | 0.699297 | 0.706248 | 0.710128 | 0.704749 |
| 0.71794 | 0.707153 | 0.708606 | 0.716516 | 0.70513 | 0.708714 | 0.711448 | 0.713819 | 0.712213 |
| 0.719184 | 0.714038 | 0.716929 | 0.724269 | 0.710116 | 0.718354 | 0.727214 | 0.723441 | 0.72409 |
| 0.736918 | 0.7322 | 0.730558 | 0.734294 | 0.72698 | 0.731411 | 0.73545 | 0.736855 | 0.736063 |
| 0.716554 | 0.706055 | 0.715396 | 0.717121 | 0.705561 | 0.71253 | 0.714388 | 0.715066 | 0.715938 |
| 0.717928 | 0.709632 | 0.711211 | 0.714844 | 0.701373 | 0.709512 | 0.713538 | 0.714133 | 0.714776 |
| 0.739695 | 0.736058 | 0.736598 | 0.742209 | 0.733229 | 0.740322 | 0.74524 | 0.745315 | 0.742296 |
| 0.7098 | 0.701463 | 0.699882 | 0.708481 | 0.700968 | 0.707247 | 0.706386 | 0.709532 | 0.707925 |
| 0.715193 | 0.705923 | 0.706402 | 0.707193 | 0.701372 | 0.707474 | 0.711729 | 0.714998 | 0.713486 |
| 0.711415 | 0.702467 | 0.70159 | 0.705472 | 0.697234 | 0.706167 | 0.710071 | 0.712496 | 0.707987 |
| 0.712509 | 0.701863 | 0.70604 | 0.714447 | 0.704944 | 0.709324 | 0.714023 | 0.714344 | 0.713407 |
| 0.717428 | 0.711826 | 0.71121 | 0.722533 | 0.706041 | 0.714161 | 0.718878 | 0.721301 | 0.717599 |
| 0.715328 | 0.702607 | 0.707473 | 0.716051 | 0.70493 | 0.711831 | 0.714651 | 0.715346 | 0.713997 |
| 0.718077 | 0.708866 | 0.7109 | 0.71779 | 0.709401 | 0.711718 | 0.717404 | 0.719546 | 0.715927 |
| 0.711591 | 0.703244 | 0.704621 | 0.712832 | 0.698962 | 0.711036 | 0.714706 | 0.713106 | 0.712193 |
| 0.708487 | 0.697988 | 0.699539 | 0.707005 | 0.695652 | 0.704545 | 0.707146 | 0.707795 | 0.706158 |
| 0.711923 | 0.704076 | 0.709864 | 0.71556 | 0.702058 | 0.709358 | 0.710067 | 0.714738 | 0.711113 |
| 0.712041 | 0.705872 | 0.706432 | 0.712776 | 0.704982 | 0.713261 | 0.718002 | 0.717712 | 0.715468 |
| 0.715196 | 0.70353 | 0.70692 | 0.712456 | 0.70437 | 0.707759 | 0.714217 | 0.713957 | 0.713879 |
| 0.715012 | 0.703572 | 0.702632 | 0.71029 | 0.700008 | 0.708633 | 0.714883 | 0.715067 | 0.71214 |
| 0.709034 | 0.697079 | 0.703335 | 0.708504 | 0.695642 | 0.705403 | 0.708519 | 0.708482 | 0.709088 |
| 0.711913 | 0.700523 | 0.70164 | 0.712116 | 0.697614 | 0.707516 | 0.711644 | 0.713173 | 0.7088 |
| 0.712666 | 0.703748 | 0.708253 | 0.715424 | 0.701675 | 0.706826 | 0.711841 | 0.714367 | 0.708956 |
| 0.709062 | 0.702521 | 0.700729 | 0.706824 | 0.69938 | 0.705602 | 0.709603 | 0.709833 | 0.709322 |
| 0.711732 | 0.704161 | 0.700833 | 0.71171 | 0.699484 | 0.707835 | 0.711134 | 0.710225 | 0.70726 |
| 0.715152 | 0.705368 | 0.70864 | 0.716363 | 0.706691 | 0.71554 | 0.719305 | 0.717934 | 0.713998 |
| 0.712399 | 0.708945 | 0.707826 | 0.714774 | 0.705483 | 0.711252 | 0.71402 | 0.717201 | 0.714613 |
| 0.71361 | 0.704857 | 0.709037 | 0.71977 | 0.709847 | 0.713015 | 0.714845 | 0.716355 | 0.715324 |

| | | | | | | | | |
|---|---|---|---|---|---|---|---|---|
| 0.747275 | 0.743592 | 0.741796 | 0.746409 | 0.732316 | 0.740859 | 0.744154 | 0.748256 | 0.742973 |
| 0.704113 | 0.69267 | 0.69845 | 0.705542 | 0.693475 | 0.706148 | 0.705953 | 0.701183 | 0.705697 |
| 0.707472 | 0.697592 | 0.700395 | 0.707769 | 0.69578 | 0.706182 | 0.705868 | 0.707069 | 0.706487 |
| 0.72101 | 0.708567 | 0.712452 | 0.714698 | 0.706978 | 0.713427 | 0.714569 | 0.716745 | 0.717491 |
| 0.714755 | 0.700207 | 0.70564 | 0.712756 | 0.701839 | 0.710234 | 0.712312 | 0.712602 | 0.711725 |
| 0.711147 | 0.70505 | 0.709798 | 0.714529 | 0.708931 | 0.714641 | 0.7151 | 0.716713 | 0.715205 |
| 0.709088 | 0.700851 | 0.704342 | 0.707912 | 0.697735 | 0.707317 | 0.711669 | 0.710652 | 0.711031 |
| 0.715001 | 0.707533 | 0.709778 | 0.711719 | 0.70343 | 0.713268 | 0.71494 | 0.71293 | 0.713159 |
| 0.715151 | 0.709059 | 0.710917 | 0.71568 | 0.702222 | 0.706831 | 0.712634 | 0.714262 | 0.712308 |
| 0.712554 | 0.707139 | 0.709954 | 0.716179 | 0.701767 | 0.710428 | 0.715345 | 0.712211 | 0.711168 |
| 0.705523 | 0.694334 | 0.700438 | 0.705536 | 0.694213 | 0.702617 | 0.706165 | 0.70606 | 0.702517 |
| 0.710653 | 0.700329 | 0.702974 | 0.707738 | 0.696762 | 0.70447 | 0.702849 | 0.706559 | 0.704327 |
| 0.711917 | 0.698351 | 0.704407 | 0.711715 | 0.698488 | 0.706423 | 0.70898 | 0.711122 | 0.707561 |
| 0.709101 | 0.697904 | 0.703722 | 0.70835 | 0.696404 | 0.704073 | 0.705505 | 0.709152 | 0.707115 |
| 0.709163 | 0.702893 | 0.703316 | 0.706002 | 0.702725 | 0.7055 | 0.707851 | 0.709871 | 0.709124 |
| 0.709384 | 0.697469 | 0.700145 | 0.703333 | 0.698517 | 0.705133 | 0.707143 | 0.706848 | 0.709895 |
| 0.710234 | 0.698084 | 0.696546 | 0.704849 | 0.694678 | 0.704286 | 0.707772 | 0.706899 | 0.705806 |
| 0.716995 | 0.703138 | 0.706303 | 0.711308 | 0.701232 | 0.710729 | 0.712607 | 0.71368 | 0.710788 |
| 0.713872 | 0.705171 | 0.705739 | 0.712066 | 0.707543 | 0.712557 | 0.716794 | 0.719392 | 0.714518 |
| 0.708886 | 0.698838 | 0.704278 | 0.708728 | 0.69638 | 0.706709 | 0.708701 | 0.711142 | 0.708493 |
| 0.705604 | 0.694379 | 0.69984 | 0.704775 | 0.694124 | 0.702821 | 0.707251 | 0.703991 | 0.704957 |
| 0.710952 | 0.697904 | 0.703092 | 0.707862 | 0.694347 | 0.703356 | 0.708377 | 0.706563 | 0.703276 |
| 0.710124 | 0.698107 | 0.702671 | 0.70911 | 0.700244 | 0.708248 | 0.710061 | 0.71316 | 0.709012 |
| 0.716625 | 0.710972 | 0.710582 | 0.716019 | 0.710035 | 0.714913 | 0.719116 | 0.722894 | 0.721258 |
| 0.725501 | 0.716733 | 0.721659 | 0.723907 | 0.714145 | 0.720348 | 0.725056 | 0.727801 | 0.723942 |
| 0.766537 | 0.76578 | 0.771631 | 0.768349 | 0.760367 | 0.761187 | 0.764501 | 0.770877 | 0.763187 |
| 0.77129 | 0.774182 | 0.772121 | 0.776974 | 0.761167 | 0.767373 | 0.768713 | 0.769002 | 0.766709 |
| 0.773289 | 0.774147 | 0.774866 | 0.774788 | 0.761953 | 0.768538 | 0.77112 | 0.772623 | 0.76903 |
| 0.708901 | 0.700517 | 0.704595 | 0.707641 | 0.699461 | 0.704177 | 0.710459 | 0.713527 | 0.708263 |
| 0.71083 | 0.701483 | 0.702861 | 0.710559 | 0.696499 | 0.703023 | 0.70797 | 0.708469 | 0.709731 |
| 0.717033 | 0.709719 | 0.708965 | 0.715709 | 0.705655 | 0.71152 | 0.716417 | 0.719081 | 0.716141 |
| 0.719844 | 0.709832 | 0.714895 | 0.718666 | 0.708302 | 0.7176 | 0.720306 | 0.721928 | 0.721064 |
| 0.709604 | 0.70451 | 0.702932 | 0.71202 | 0.697801 | 0.705443 | 0.709956 | 0.714734 | 0.707279 |
| 0.714274 | 0.703282 | 0.70299 | 0.709063 | 0.702098 | 0.70854 | 0.708812 | 0.713652 | 0.706786 |
| 0.708814 | 0.705004 | 0.703814 | 0.706462 | 0.696318 | 0.702983 | 0.706891 | 0.710548 | 0.706436 |
| 0.711442 | 0.704599 | 0.703435 | 0.708458 | 0.698558 | 0.705554 | 0.706372 | 0.711217 | 0.708117 |
| 0.710145 | 0.704616 | 0.705494 | 0.710886 | 0.698503 | 0.7053 | 0.708934 | 0.711915 | 0.708253 |
| 0.722871 | 0.718027 | 0.718607 | 0.722475 | 0.708873 | 0.718902 | 0.723199 | 0.723465 | 0.720088 |
| 0.699809 | 0.699087 | 0.693632 | 0.703055 | 0.692301 | 0.697617 | 0.701642 | 0.706111 | 0.70156 |
| 0.707861 | 0.704314 | 0.700841 | 0.706883 | 0.697398 | 0.706077 | 0.709887 | 0.712507 | 0.710178 |
| 0.710076 | 0.699152 | 0.699085 | 0.703311 | 0.69232 | 0.701022 | 0.70297 | 0.707353 | 0.703203 |
| 0.726883 | 0.716874 | 0.72017 | 0.724691 | 0.712782 | 0.721474 | 0.72819 | 0.728253 | 0.722856 |
| 0.713354 | 0.700673 | 0.702083 | 0.708364 | 0.702663 | 0.706861 | 0.710871 | 0.713421 | 0.710326 |
| 0.709878 | 0.696198 | 0.7002 | 0.706739 | 0.700268 | 0.706086 | 0.708528 | 0.709931 | 0.709371 |
| 0.712476 | 0.702426 | 0.702128 | 0.714077 | 0.698777 | 0.708927 | 0.710533 | 0.71452 | 0.708726 |
| 0.707866 | 0.704737 | 0.700618 | 0.708756 | 0.701676 | 0.705897 | 0.708518 | 0.711616 | 0.706693 |

| | | | | | | | | |
|---|---|---|---|---|---|---|---|---|
| 0.711631 | 0.702289 | 0.703046 | 0.708189 | 0.69638 | 0.703739 | 0.709771 | 0.710567 | 0.707921 |
| 0.710043 | 0.701367 | 0.702314 | 0.70813 | 0.704027 | 0.713028 | 0.713819 | 0.714414 | 0.711681 |
| 0.712133 | 0.706826 | 0.703501 | 0.708097 | 0.699325 | 0.708817 | 0.708964 | 0.711235 | 0.707487 |
| 0.714993 | 0.710896 | 0.710605 | 0.71537 | 0.702743 | 0.712721 | 0.715569 | 0.716998 | 0.714075 |
| 0.718129 | 0.709939 | 0.707974 | 0.714722 | 0.704114 | 0.711866 | 0.716701 | 0.71539 | 0.712742 |
| 0.708005 | 0.704942 | 0.703219 | 0.703794 | 0.702275 | 0.709022 | 0.711362 | 0.715273 | 0.712523 |
| 0.713103 | 0.705377 | 0.704667 | 0.715327 | 0.702536 | 0.707098 | 0.710608 | 0.712239 | 0.70825 |
| 0.71622 | 0.708065 | 0.707842 | 0.710064 | 0.703463 | 0.709845 | 0.713424 | 0.715425 | 0.712958 |
| 0.712386 | 0.701269 | 0.704703 | 0.710155 | 0.701949 | 0.709765 | 0.711894 | 0.712692 | 0.712213 |
| 0.707105 | 0.697564 | 0.700402 | 0.710342 | 0.698401 | 0.705761 | 0.706567 | 0.710803 | 0.707654 |
| 0.709677 | 0.702137 | 0.700416 | 0.707811 | 0.697266 | 0.703418 | 0.709016 | 0.709399 | 0.707524 |
| 0.711684 | 0.706519 | 0.703234 | 0.706793 | 0.697613 | 0.706242 | 0.70799 | 0.708364 | 0.707433 |
| 0.713374 | 0.702813 | 0.704718 | 0.713219 | 0.70136 | 0.710176 | 0.712918 | 0.711095 | 0.708405 |
| 0.709811 | 0.701984 | 0.702102 | 0.705183 | 0.699461 | 0.705127 | 0.707103 | 0.708199 | 0.706881 |
| 0.711236 | 0.703813 | 0.706528 | 0.71359 | 0.700565 | 0.7076 | 0.71101 | 0.715783 | 0.711596 |
| 0.707862 | 0.700705 | 0.703514 | 0.711333 | 0.701783 | 0.709134 | 0.711278 | 0.708607 | 0.709797 |
| 0.712623 | 0.703488 | 0.709094 | 0.716729 | 0.699318 | 0.711893 | 0.713906 | 0.713546 | 0.713212 |
| 0.71083 | 0.704823 | 0.703273 | 0.707733 | 0.700734 | 0.708383 | 0.708669 | 0.71213 | 0.706595 |
| 0.71279 | 0.706025 | 0.702825 | 0.711893 | 0.701889 | 0.706769 | 0.70821 | 0.711612 | 0.710864 |
| 0.711599 | 0.701876 | 0.705793 | 0.706545 | 0.702431 | 0.709089 | 0.712924 | 0.7134 | 0.709278 |
| 0.710594 | 0.707046 | 0.702666 | 0.708306 | 0.701799 | 0.708629 | 0.710229 | 0.708643 | 0.709725 |
| 0.706136 | 0.699218 | 0.696311 | 0.705901 | 0.691481 | 0.697485 | 0.704455 | 0.713095 | 0.704105 |
| 0.709735 | 0.704593 | 0.702253 | 0.70503 | 0.69748 | 0.705722 | 0.709488 | 0.709463 | 0.704663 |
| 0.712923 | 0.709624 | 0.70394 | 0.709633 | 0.699292 | 0.709074 | 0.713315 | 0.713075 | 0.710698 |
| 0.709926 | 0.708999 | 0.705528 | 0.710102 | 0.705131 | 0.712479 | 0.713222 | 0.710262 | 0.709767 |
| 0.717002 | 0.706155 | 0.710456 | 0.719874 | 0.703499 | 0.712874 | 0.718392 | 0.721639 | 0.714366 |
| 0.709621 | 0.70795 | 0.703763 | 0.708383 | 0.701362 | 0.708511 | 0.709708 | 0.706789 | 0.708353 |
| 0.714709 | 0.704585 | 0.706291 | 0.713103 | 0.702943 | 0.710819 | 0.713068 | 0.709599 | 0.710752 |
| 0.711943 | 0.7081 | 0.707327 | 0.711907 | 0.704672 | 0.710248 | 0.713784 | 0.718523 | 0.717961 |
| 0.713273 | 0.708876 | 0.708245 | 0.712343 | 0.701674 | 0.71018 | 0.714425 | 0.711702 | 0.713276 |
| 0.71342 | 0.7097 | 0.708132 | 0.710671 | 0.704482 | 0.71179 | 0.712276 | 0.714815 | 0.712366 |
| 0.712039 | 0.702938 | 0.701445 | 0.708463 | 0.699992 | 0.705855 | 0.71159 | 0.712382 | 0.709922 |
| 0.710601 | 0.708789 | 0.705687 | 0.714601 | 0.70133 | 0.709906 | 0.710724 | 0.715905 | 0.710995 |
| 0.716285 | 0.708988 | 0.70658 | 0.710788 | 0.702872 | 0.709496 | 0.711602 | 0.713009 | 0.714012 |
| 0.708925 | 0.700156 | 0.704194 | 0.709223 | 0.697007 | 0.705023 | 0.708859 | 0.709564 | 0.70665 |
| 0.70832 | 0.705846 | 0.702073 | 0.709941 | 0.702629 | 0.70797 | 0.711503 | 0.714463 | 0.713087 |
| 0.710333 | 0.703142 | 0.70311 | 0.709253 | 0.698606 | 0.703519 | 0.700415 | 0.705588 | 0.703292 |
| 0.685153 | 0.684521 | 0.680698 | 0.685302 | 0.673583 | 0.681583 | 0.684952 | 0.687628 | 0.683499 |
| 0.691752 | 0.690748 | 0.685518 | 0.688033 | 0.676085 | 0.687399 | 0.689543 | 0.696203 | 0.693664 |
| 0.685179 | 0.683478 | 0.680089 | 0.680033 | 0.671122 | 0.676093 | 0.680561 | 0.685866 | 0.680771 |
| 0.683988 | 0.680959 | 0.679194 | 0.683077 | 0.669945 | 0.678195 | 0.67835 | 0.684944 | 0.678432 |
| 0.67078 | 0.668597 | 0.660592 | 0.667446 | 0.658876 | 0.666159 | 0.661537 | 0.667414 | 0.664938 |
| 0.672301 | 0.667286 | 0.667284 | 0.667715 | 0.656755 | 0.6645 | 0.665579 | 0.670604 | 0.667234 |
| 0.674138 | 0.670957 | 0.666048 | 0.669747 | 0.658552 | 0.667193 | 0.665779 | 0.672631 | 0.669889 |
| 0.67117 | 0.66928 | 0.663775 | 0.669943 | 0.657628 | 0.667391 | 0.669439 | 0.670727 | 0.670276 |
| 0.674746 | 0.669556 | 0.663913 | 0.671207 | 0.664028 | 0.664504 | 0.671059 | 0.674808 | 0.668505 |
| 0.673487 | 0.67232 | 0.671337 | 0.674755 | 0.663355 | 0.668558 | 0.669137 | 0.676291 | 0.674744 |

| | | | | | | | | |
|---|---|---|---|---|---|---|---|---|
| 0.672367 | 0.676609 | 0.67085 | 0.672646 | 0.660183 | 0.671758 | 0.667647 | 0.676245 | 0.671832 |
| 0.679775 | 0.677691 | 0.670083 | 0.676371 | 0.665467 | 0.671796 | 0.671867 | 0.67835 | 0.674734 |
| 0.679606 | 0.671353 | 0.671866 | 0.673286 | 0.66248 | 0.669168 | 0.670631 | 0.674074 | 0.667249 |
| 0.667728 | 0.662665 | 0.660142 | 0.662171 | 0.650821 | 0.659892 | 0.663672 | 0.665514 | 0.660695 |
| 0.668039 | 0.668943 | 0.666054 | 0.668426 | 0.65506 | 0.660912 | 0.66671 | 0.671982 | 0.661681 |
| 0.667861 | 0.673375 | 0.662495 | 0.668085 | 0.650787 | 0.661376 | 0.663154 | 0.667863 | 0.663413 |
| 0.666619 | 0.666379 | 0.668208 | 0.67026 | 0.658487 | 0.662652 | 0.662893 | 0.670612 | 0.665287 |
| 0.668917 | 0.66762 | 0.661832 | 0.668023 | 0.655801 | 0.663537 | 0.663173 | 0.668931 | 0.664552 |
| 0.664753 | 0.665769 | 0.658678 | 0.661922 | 0.652611 | 0.659041 | 0.65838 | 0.66446 | 0.66218 |
| 0.671495 | 0.672065 | 0.66577 | 0.669643 | 0.656768 | 0.661168 | 0.663163 | 0.671429 | 0.66452 |
| 0.667172 | 0.662362 | 0.656896 | 0.662802 | 0.646959 | 0.662237 | 0.66045 | 0.661678 | 0.655271 |
| 0.70204 | 0.692943 | 0.69427 | 0.699232 | 0.688298 | 0.694742 | 0.698061 | 0.70008 | 0.696813 |
| 0.704187 | 0.701781 | 0.701545 | 0.709258 | 0.701861 | 0.706073 | 0.709417 | 0.712091 | 0.710361 |
| 0.687072 | 0.686242 | 0.681945 | 0.687364 | 0.673014 | 0.680967 | 0.685428 | 0.689893 | 0.681121 |
| 0.715527 | 0.711105 | 0.708461 | 0.714812 | 0.704802 | 0.707021 | 0.713417 | 0.715608 | 0.709493 |
| 0.704373 | 0.70037 | 0.695234 | 0.705617 | 0.696159 | 0.700531 | 0.703766 | 0.704029 | 0.701466 |
| 0.715187 | 0.704893 | 0.704644 | 0.712135 | 0.697865 | 0.712172 | 0.716131 | 0.716413 | 0.713398 |
| 0.721132 | 0.716235 | 0.711972 | 0.719331 | 0.708344 | 0.71385 | 0.71585 | 0.719467 | 0.714961 |
| 0.69873 | 0.690751 | 0.689669 | 0.696554 | 0.687782 | 0.692206 | 0.694854 | 0.699413 | 0.693439 |
| 0.719904 | 0.715637 | 0.714884 | 0.724374 | 0.710882 | 0.715358 | 0.717932 | 0.720065 | 0.715146 |
| 0.689822 | 0.692576 | 0.688148 | 0.69477 | 0.678281 | 0.686666 | 0.692958 | 0.692495 | 0.685725 |
| 0.714435 | 0.708825 | 0.711081 | 0.714656 | 0.70977 | 0.715564 | 0.718933 | 0.718812 | 0.718134 |
| 0.719055 | 0.713913 | 0.713633 | 0.717976 | 0.714628 | 0.715358 | 0.71819 | 0.721009 | 0.717671 |
| 0.684033 | 0.681038 | 0.680143 | 0.684493 | 0.674543 | 0.685634 | 0.690559 | 0.687669 | 0.689846 |
| 0.692052 | 0.685923 | 0.683024 | 0.689169 | 0.678858 | 0.688371 | 0.690161 | 0.690611 | 0.692075 |
| 0.692571 | 0.694665 | 0.689432 | 0.695023 | 0.681292 | 0.688297 | 0.69223 | 0.699295 | 0.693789 |
| 0.705628 | 0.695231 | 0.701078 | 0.70857 | 0.696894 | 0.705901 | 0.709999 | 0.707139 | 0.705014 |
| 0.721766 | 0.708884 | 0.713204 | 0.720019 | 0.707993 | 0.717677 | 0.720726 | 0.721746 | 0.718179 |
| 0.686929 | 0.683803 | 0.68034 | 0.688245 | 0.677422 | 0.685076 | 0.688031 | 0.688806 | 0.688993 |
| 0.722949 | 0.721 | 0.718024 | 0.724221 | 0.712741 | 0.718515 | 0.726983 | 0.72629 | 0.72121 |
| 0.727367 | 0.715715 | 0.719379 | 0.724848 | 0.712528 | 0.720608 | 0.722705 | 0.723935 | 0.721462 |
| 0.696478 | 0.693602 | 0.692739 | 0.698506 | 0.685209 | 0.695311 | 0.700853 | 0.700252 | 0.69677 |
| 0.697302 | 0.693445 | 0.689961 | 0.697982 | 0.686199 | 0.696617 | 0.699114 | 0.698051 | 0.697849 |
| 0.723439 | 0.715902 | 0.717455 | 0.729292 | 0.711928 | 0.717218 | 0.726696 | 0.726454 | 0.720794 |
| 0.721584 | 0.718998 | 0.714333 | 0.720445 | 0.71281 | 0.721829 | 0.724484 | 0.724174 | 0.720383 |
| 0.724806 | 0.722062 | 0.719012 | 0.719475 | 0.714215 | 0.719276 | 0.729133 | 0.72559 | 0.722938 |
| 0.676275 | 0.674743 | 0.66964 | 0.677353 | 0.672216 | 0.677111 | 0.676044 | 0.677673 | 0.679229 |
| 0.721216 | 0.712219 | 0.713806 | 0.718139 | 0.710271 | 0.71424 | 0.719728 | 0.724678 | 0.717783 |
| 0.674973 | 0.671897 | 0.669168 | 0.675087 | 0.666929 | 0.674757 | 0.676342 | 0.675887 | 0.676274 |
| 0.66781 | 0.658589 | 0.661108 | 0.668428 | 0.661165 | 0.669192 | 0.671764 | 0.673795 | 0.670135 |
| 0.721987 | 0.713708 | 0.716128 | 0.71872 | 0.71261 | 0.716512 | 0.720274 | 0.722096 | 0.718974 |
| 0.667736 | 0.667271 | 0.661669 | 0.66879 | 0.66143 | 0.670586 | 0.671135 | 0.673698 | 0.668566 |
| 0.661406 | 0.656036 | 0.652721 | 0.660991 | 0.65617 | 0.663636 | 0.663475 | 0.663743 | 0.66353 |
| 0.658482 | 0.655758 | 0.65432 | 0.659683 | 0.648513 | 0.656302 | 0.659761 | 0.661319 | 0.660143 |
| 0.73198 | 0.72643 | 0.722718 | 0.731323 | 0.717437 | 0.72508 | 0.727949 | 0.730344 | 0.726588 |
| 0.661304 | 0.654942 | 0.658587 | 0.664992 | 0.65484 | 0.662191 | 0.664982 | 0.666641 | 0.662725 |
| 0.723885 | 0.72016 | 0.720261 | 0.723206 | 0.713969 | 0.717576 | 0.721003 | 0.721926 | 0.719265 |

| | | | | | | | | |
|---|---|---|---|---|---|---|---|---|
| 0.726821 | 0.727635 | 0.725261 | 0.730184 | 0.716281 | 0.722849 | 0.729425 | 0.731578 | 0.729662 |
| 0.726965 | 0.721123 | 0.720246 | 0.726055 | 0.714835 | 0.723558 | 0.724788 | 0.727273 | 0.722056 |
| 0.745047 | 0.743136 | 0.740163 | 0.746623 | 0.731833 | 0.73808 | 0.746498 | 0.740467 | 0.740799 |
| 0.652197 | 0.644914 | 0.647687 | 0.652857 | 0.644789 | 0.651037 | 0.652571 | 0.65396 | 0.655087 |
| 0.651535 | 0.652748 | 0.648285 | 0.653716 | 0.641975 | 0.651066 | 0.653313 | 0.655127 | 0.65517 |
| 0.732563 | 0.729468 | 0.727996 | 0.732008 | 0.719252 | 0.728079 | 0.732791 | 0.733377 | 0.728929 |
| 0.654543 | 0.649457 | 0.645804 | 0.654565 | 0.646655 | 0.653046 | 0.65653 | 0.654897 | 0.654639 |
| 0.732106 | 0.727057 | 0.725081 | 0.735413 | 0.727569 | 0.733366 | 0.73642 | 0.736715 | 0.733807 |
| 0.73641 | 0.731571 | 0.728216 | 0.7315 | 0.720185 | 0.727959 | 0.732982 | 0.732937 | 0.729869 |
| 0.733514 | 0.72556 | 0.730201 | 0.733537 | 0.727632 | 0.731139 | 0.735627 | 0.736763 | 0.731075 |
| 0.730707 | 0.73103 | 0.726946 | 0.729051 | 0.723176 | 0.725127 | 0.727803 | 0.729766 | 0.727907 |
| 0.695946 | 0.774648 | 0.682119 | 0.77551 | 0.727891 | 0.748344 | 0.72549 | 0.699346 | 0.794326 |
| 0.7268 | 0.739592 | 0.725308 | 0.723188 | 0.724933 | 0.727767 | 0.728422 | 0.745354 | 0.733912 |
| 0.729738 | 0.735115 | 0.724408 | 0.728993 | 0.721802 | 0.733607 | 0.739234 | 0.738586 | 0.731139 |
| 0.736164 | 0.742804 | 0.735544 | 0.733938 | 0.727441 | 0.734366 | 0.731788 | 0.733515 | 0.730868 |
| 0.728191 | 0.739035 | 0.725571 | 0.731349 | 0.730552 | 0.73525 | 0.7355 | 0.738474 | 0.736785 |
| 0.587017 | 0.592465 | 0.588745 | 0.590047 | 0.5847 | 0.590481 | 0.591528 | 0.591697 | 0.59173 |
| 0.736539 | 0.740498 | 0.732644 | 0.734338 | 0.727841 | 0.73397 | 0.744326 | 0.747289 | 0.739388 |
| 0.733932 | 0.733902 | 0.731548 | 0.73796 | 0.728077 | 0.730987 | 0.737117 | 0.738071 | 0.735432 |
| 0.736777 | 0.732055 | 0.72954 | 0.732957 | 0.725403 | 0.732667 | 0.734843 | 0.736842 | 0.733445 |
| 0.736277 | 0.736847 | 0.733535 | 0.738502 | 0.728314 | 0.730053 | 0.733643 | 0.736622 | 0.732492 |
| 0.589598 | 0.586478 | 0.588373 | 0.594699 | 0.592406 | 0.598024 | 0.597469 | 0.595318 | 0.599315 |
| 0.576227 | 0.574804 | 0.572748 | 0.580668 | 0.578657 | 0.582481 | 0.584789 | 0.585317 | 0.584442 |
| 0.760084 | 0.758297 | 0.759245 | 0.759728 | 0.749936 | 0.756772 | 0.762203 | 0.764721 | 0.760906 |
| 0.754246 | 0.760738 | 0.760792 | 0.760702 | 0.750924 | 0.757241 | 0.759629 | 0.766415 | 0.759524 |
| 0.753297 | 0.754683 | 0.753009 | 0.754624 | 0.739086 | 0.746764 | 0.7513 | 0.752632 | 0.743259 |
| 0.755364 | 0.756496 | 0.7584 | 0.762 | 0.748092 | 0.753381 | 0.757591 | 0.760127 | 0.753104 |
| 0.758734 | 0.759389 | 0.762559 | 0.760007 | 0.74631 | 0.756845 | 0.759895 | 0.756982 | 0.75312 |
| 0.757543 | 0.768189 | 0.762234 | 0.763783 | 0.752062 | 0.758424 | 0.762963 | 0.764246 | 0.764126 |
| 0.761152 | 0.763838 | 0.763124 | 0.765087 | 0.750126 | 0.755852 | 0.763618 | 0.768077 | 0.758598 |
| 0.75529 | 0.759063 | 0.752542 | 0.762247 | 0.74395 | 0.754527 | 0.761043 | 0.75811 | 0.758294 |
| 0.767535 | 0.767347 | 0.768341 | 0.768768 | 0.760968 | 0.769413 | 0.772626 | 0.771636 | 0.768618 |
| 0.758678 | 0.761862 | 0.760805 | 0.762986 | 0.752759 | 0.758291 | 0.762664 | 0.760906 | 0.759292 |
| 0.763846 | 0.755318 | 0.75962 | 0.76431 | 0.748858 | 0.760434 | 0.765215 | 0.765191 | 0.757374 |
| 0.768901 | 0.7652 | 0.76514 | 0.76744 | 0.758432 | 0.762076 | 0.770875 | 0.768312 | 0.765815 |
| 0.758701 | 0.759688 | 0.755888 | 0.76221 | 0.754387 | 0.759316 | 0.763938 | 0.765529 | 0.762635 |
| 0.762157 | 0.763485 | 0.766087 | 0.766322 | 0.753383 | 0.754819 | 0.761209 | 0.763439 | 0.755537 |
| 0.764754 | 0.764758 | 0.765578 | 0.77029 | 0.755981 | 0.763433 | 0.770397 | 0.765228 | 0.76475 |
| 0.757044 | 0.759534 | 0.760291 | 0.761633 | 0.746145 | 0.754097 | 0.761796 | 0.763939 | 0.754843 |
| 0.760843 | 0.761304 | 0.757159 | 0.760755 | 0.746206 | 0.756006 | 0.763775 | 0.762128 | 0.757874 |
| 0.724658 | 0.737002 | 0.729701 | 0.729364 | 0.720797 | 0.730058 | 0.728715 | 0.733734 | 0.727657 |
| 0.722108 | 0.747543 | 0.732037 | 0.73244 | 0.728359 | 0.723123 | 0.733006 | 0.74177 | 0.728407 |
| 0.767383 | 0.76248 | 0.76463 | 0.766174 | 0.754212 | 0.766266 | 0.765937 | 0.764814 | 0.763722 |
| 0.765065 | 0.765399 | 0.767537 | 0.767777 | 0.758772 | 0.763321 | 0.770181 | 0.767035 | 0.761932 |
| 0.71784 | 0.72381 | 0.719127 | 0.723664 | 0.695754 | 0.706245 | 0.714203 | 0.721663 | 0.707645 |
| 0.76258 | 0.761233 | 0.763204 | 0.768776 | 0.751888 | 0.762197 | 0.76673 | 0.764666 | 0.765569 |
| 0.754712 | 0.751283 | 0.754673 | 0.761822 | 0.749267 | 0.75798 | 0.761374 | 0.756323 | 0.757224 |

| | | | | | | | | |
|---|---|---|---|---|---|---|---|---|
| 0.76347 | 0.761684 | 0.76087 | 0.764077 | 0.749811 | 0.757064 | 0.765336 | 0.764938 | 0.760502 |
| 0.765419 | 0.761174 | 0.765959 | 0.76937 | 0.755485 | 0.764912 | 0.76706 | 0.766021 | 0.762247 |
| 0.757558 | 0.76203 | 0.758438 | 0.763127 | 0.752708 | 0.757143 | 0.765256 | 0.762712 | 0.756461 |
| 0.767063 | 0.762096 | 0.761554 | 0.773967 | 0.758868 | 0.761337 | 0.767183 | 0.771482 | 0.765504 |
| 0.766324 | 0.767384 | 0.767297 | 0.769141 | 0.756618 | 0.761567 | 0.769174 | 0.769773 | 0.764784 |
| 0.759503 | 0.75967 | 0.761362 | 0.76347 | 0.75237 | 0.757332 | 0.765082 | 0.765599 | 0.760884 |
| 0.765557 | 0.757149 | 0.761769 | 0.765604 | 0.754112 | 0.760834 | 0.766798 | 0.766313 | 0.759854 |
| 0.751059 | 0.749215 | 0.749258 | 0.756147 | 0.743405 | 0.753493 | 0.7602 | 0.755725 | 0.757005 |
| 0.762926 | 0.756976 | 0.760738 | 0.764149 | 0.751896 | 0.757846 | 0.765425 | 0.764839 | 0.758543 |
| 0.758441 | 0.754985 | 0.756468 | 0.756708 | 0.750021 | 0.755485 | 0.761028 | 0.755548 | 0.758132 |
| 0.762475 | 0.762362 | 0.764462 | 0.764667 | 0.756895 | 0.758076 | 0.763559 | 0.765685 | 0.762393 |
| 0.760722 | 0.759584 | 0.760796 | 0.765653 | 0.747903 | 0.755296 | 0.761965 | 0.764406 | 0.758261 |
| 0.75578 | 0.752493 | 0.754206 | 0.757964 | 0.745858 | 0.751032 | 0.7564 | 0.75309 | 0.751555 |
| 0.756433 | 0.755236 | 0.755429 | 0.757312 | 0.747698 | 0.753731 | 0.757482 | 0.757039 | 0.754202 |
| 0.762528 | 0.760781 | 0.763523 | 0.765471 | 0.748522 | 0.762242 | 0.76653 | 0.761921 | 0.762149 |
| 0.762601 | 0.766978 | 0.761217 | 0.766995 | 0.751509 | 0.759188 | 0.764122 | 0.76446 | 0.764178 |
| 0.75928 | 0.762201 | 0.762389 | 0.767733 | 0.7586 | 0.758694 | 0.761593 | 0.766791 | 0.762126 |
| 0.761058 | 0.75404 | 0.761884 | 0.760163 | 0.750822 | 0.756903 | 0.760979 | 0.765296 | 0.762993 |
| 0.761036 | 0.758384 | 0.767161 | 0.7614 | 0.749497 | 0.758629 | 0.760923 | 0.762092 | 0.756025 |
| 0.760886 | 0.763363 | 0.763876 | 0.759502 | 0.746493 | 0.758259 | 0.759252 | 0.760285 | 0.756158 |
| 0.751739 | 0.752625 | 0.755389 | 0.754766 | 0.749358 | 0.752284 | 0.758739 | 0.756333 | 0.755187 |
| 0.753635 | 0.751487 | 0.763351 | 0.758775 | 0.744991 | 0.751535 | 0.75999 | 0.758136 | 0.752342 |
| 0.760818 | 0.761354 | 0.76631 | 0.766625 | 0.752172 | 0.758229 | 0.764629 | 0.764569 | 0.758774 |
| 0.761273 | 0.75518 | 0.763733 | 0.765694 | 0.755623 | 0.758869 | 0.765642 | 0.765625 | 0.762095 |
| 0.759706 | 0.758745 | 0.761674 | 0.760369 | 0.753879 | 0.756949 | 0.762661 | 0.762595 | 0.761215 |
| 0.757272 | 0.756614 | 0.765191 | 0.758885 | 0.750318 | 0.754759 | 0.759075 | 0.756973 | 0.756514 |
| 0.754542 | 0.748473 | 0.756235 | 0.75576 | 0.746167 | 0.750454 | 0.754704 | 0.759935 | 0.753435 |
| 0.758548 | 0.753771 | 0.7587 | 0.758577 | 0.74364 | 0.749004 | 0.75502 | 0.757989 | 0.750359 |
| 0.756274 | 0.757414 | 0.76176 | 0.759479 | 0.749916 | 0.757274 | 0.756188 | 0.762481 | 0.759434 |
| 0.763648 | 0.760442 | 0.766467 | 0.763784 | 0.749601 | 0.75652 | 0.762344 | 0.764223 | 0.757259 |
| 0.765678 | 0.761007 | 0.758746 | 0.763952 | 0.753765 | 0.75948 | 0.762676 | 0.769483 | 0.76395 |
| 0.762525 | 0.760458 | 0.760418 | 0.760033 | 0.749745 | 0.75592 | 0.760944 | 0.766766 | 0.75956 |
| 0.765741 | 0.769046 | 0.763165 | 0.764076 | 0.756104 | 0.761094 | 0.766015 | 0.768685 | 0.76509 |
| 0.764289 | 0.766628 | 0.769318 | 0.765843 | 0.756297 | 0.759935 | 0.767538 | 0.76691 | 0.762269 |
| 0.754123 | 0.755416 | 0.760017 | 0.759874 | 0.744333 | 0.752131 | 0.757053 | 0.763311 | 0.756134 |
| 0.761338 | 0.763391 | 0.762257 | 0.757047 | 0.752965 | 0.758099 | 0.761218 | 0.761189 | 0.760892 |
| 0.757171 | 0.753502 | 0.758254 | 0.759159 | 0.751122 | 0.75812 | 0.758969 | 0.761961 | 0.757867 |
| 0.753368 | 0.75493 | 0.761117 | 0.755882 | 0.750742 | 0.754121 | 0.757257 | 0.756215 | 0.754459 |
| 0.751088 | 0.751062 | 0.753818 | 0.749727 | 0.745482 | 0.749107 | 0.750836 | 0.756824 | 0.752724 |
| 0.751708 | 0.75172 | 0.753094 | 0.754877 | 0.739279 | 0.747049 | 0.750795 | 0.750044 | 0.747054 |
| 0.74983 | 0.748742 | 0.748951 | 0.752205 | 0.742422 | 0.74701 | 0.751463 | 0.749827 | 0.753004 |
| 0.749497 | 0.751326 | 0.756994 | 0.754905 | 0.737843 | 0.744863 | 0.751132 | 0.751639 | 0.743421 |
| 0.749548 | 0.745118 | 0.75036 | 0.748082 | 0.741576 | 0.745676 | 0.747944 | 0.754185 | 0.746904 |
| 0.747815 | 0.749234 | 0.755333 | 0.748735 | 0.737421 | 0.743955 | 0.747306 | 0.750618 | 0.741821 |
| 0.755257 | 0.754748 | 0.75828 | 0.756345 | 0.746284 | 0.749786 | 0.756897 | 0.760183 | 0.752951 |
| 0.748512 | 0.74296 | 0.753158 | 0.74858 | 0.740616 | 0.7426 | 0.746595 | 0.750622 | 0.746109 |

| | | | | | | | | |
|---|---|---|---|---|---|---|---|---|
| 0.76087 | 0.757692 | 0.761315 | 0.763239 | 0.749095 | 0.756257 | 0.7603 | 0.764018 | 0.763004 |
| 0.771865 | 0.768543 | 0.772372 | 0.774812 | 0.758261 | 0.76868 | 0.773408 | 0.775695 | 0.767658 |
| 0.76478 | 0.760529 | 0.766025 | 0.767589 | 0.753699 | 0.760269 | 0.767405 | 0.764506 | 0.766901 |
| 0.768283 | 0.770523 | 0.760731 | 0.769676 | 0.757553 | 0.76292 | 0.766134 | 0.764627 | 0.765779 |
| 0.779237 | 0.778327 | 0.775161 | 0.777861 | 0.759074 | 0.766399 | 0.772772 | 0.775471 | 0.771261 |
| 0.767436 | 0.772723 | 0.767849 | 0.770251 | 0.755864 | 0.764846 | 0.768387 | 0.771757 | 0.765181 |
| 0.769145 | 0.763416 | 0.767234 | 0.77194 | 0.75188 | 0.760539 | 0.764468 | 0.767816 | 0.76157 |
| 0.763984 | 0.763593 | 0.762218 | 0.770017 | 0.754076 | 0.762971 | 0.763481 | 0.764893 | 0.76075 |
| 0.763835 | 0.767243 | 0.76071 | 0.766419 | 0.757036 | 0.758645 | 0.762681 | 0.76933 | 0.759536 |
| 0.760697 | 0.758999 | 0.758534 | 0.767008 | 0.754438 | 0.7604 | 0.763193 | 0.761253 | 0.761984 |
| 0.767935 | 0.766516 | 0.76654 | 0.770043 | 0.752375 | 0.757838 | 0.764595 | 0.762237 | 0.763416 |
| 0.766031 | 0.76369 | 0.763491 | 0.767467 | 0.75271 | 0.756215 | 0.76297 | 0.765228 | 0.759844 |
| 0.77083 | 0.766102 | 0.767076 | 0.770944 | 0.756206 | 0.766238 | 0.769725 | 0.76905 | 0.767477 |
| 0.766529 | 0.768103 | 0.770478 | 0.76782 | 0.759165 | 0.765676 | 0.769662 | 0.769929 | 0.765461 |
| 0.772873 | 0.775212 | 0.772322 | 0.778506 | 0.760496 | 0.765602 | 0.768477 | 0.772324 | 0.774921 |
| 0.771785 | 0.766932 | 0.770276 | 0.773144 | 0.757078 | 0.768531 | 0.772251 | 0.767836 | 0.768513 |
| 0.768627 | 0.765377 | 0.772145 | 0.774913 | 0.765233 | 0.77222 | 0.773189 | 0.770619 | 0.769333 |
| 0.771742 | 0.770819 | 0.769985 | 0.770602 | 0.753925 | 0.766076 | 0.77127 | 0.764384 | 0.766194 |
| 0.769485 | 0.771681 | 0.772502 | 0.773746 | 0.762394 | 0.769313 | 0.771992 | 0.771601 | 0.773243 |
| 0.774798 | 0.773201 | 0.773284 | 0.775539 | 0.758219 | 0.767637 | 0.768563 | 0.773504 | 0.770253 |
| 0.765173 | 0.761977 | 0.764947 | 0.768994 | 0.756745 | 0.763626 | 0.766863 | 0.773953 | 0.767181 |
| 0.766274 | 0.767227 | 0.768253 | 0.770292 | 0.760592 | 0.767336 | 0.770793 | 0.770328 | 0.767839 |
| 0.772488 | 0.770314 | 0.77096 | 0.769243 | 0.760849 | 0.76438 | 0.770031 | 0.772301 | 0.769314 |
| 0.772653 | 0.768711 | 0.768421 | 0.774122 | 0.762595 | 0.769513 | 0.774469 | 0.77523 | 0.77411 |
| 0.769457 | 0.767244 | 0.766272 | 0.769843 | 0.766143 | 0.770798 | 0.771654 | 0.771908 | 0.771045 |
| 0.768634 | 0.762614 | 0.77061 | 0.769352 | 0.756777 | 0.765019 | 0.766384 | 0.768243 | 0.764544 |
| 0.766626 | 0.764487 | 0.76959 | 0.767994 | 0.761817 | 0.766054 | 0.766887 | 0.769006 | 0.767758 |
| 0.766781 | 0.761172 | 0.766531 | 0.767172 | 0.764142 | 0.763344 | 0.773146 | 0.770692 | 0.769671 |
| 0.771126 | 0.767287 | 0.773112 | 0.775728 | 0.760154 | 0.768904 | 0.771405 | 0.77343 | 0.77083 |
| 0.770945 | 0.773922 | 0.776528 | 0.773566 | 0.762161 | 0.768438 | 0.770789 | 0.773639 | 0.769594 |
| 0.764814 | 0.761785 | 0.765038 | 0.764612 | 0.755873 | 0.761211 | 0.760637 | 0.766474 | 0.76382 |
| 0.764009 | 0.765284 | 0.767133 | 0.767617 | 0.763343 | 0.768087 | 0.771704 | 0.771547 | 0.77134 |
| 0.772948 | 0.771362 | 0.77191 | 0.768505 | 0.767812 | 0.767236 | 0.767067 | 0.779634 | 0.773588 |
| 0.764055 | 0.765605 | 0.772153 | 0.772217 | 0.751388 | 0.760912 | 0.76689 | 0.768471 | 0.766466 |
| 0.770816 | 0.764523 | 0.768716 | 0.771433 | 0.761162 | 0.767747 | 0.769961 | 0.768375 | 0.771173 |
| 0.763955 | 0.763769 | 0.772038 | 0.766368 | 0.757008 | 0.7637 | 0.769497 | 0.772263 | 0.768928 |
| 0.775876 | 0.77041 | 0.775253 | 0.778499 | 0.76124 | 0.768878 | 0.774866 | 0.776636 | 0.77122 |
| 0.769864 | 0.766208 | 0.767417 | 0.774434 | 0.765396 | 0.768942 | 0.770683 | 0.775423 | 0.770906 |
| 0.768729 | 0.765262 | 0.767351 | 0.771871 | 0.755976 | 0.761733 | 0.767179 | 0.768234 | 0.764672 |
| 0.764944 | 0.771028 | 0.768816 | 0.770793 | 0.760316 | 0.766954 | 0.772352 | 0.7724 | 0.769694 |
| 0.770244 | 0.769751 | 0.765787 | 0.774234 | 0.767128 | 0.771155 | 0.776523 | 0.775055 | 0.77364 |
| 0.77178 | 0.767998 | 0.769866 | 0.776748 | 0.764074 | 0.767087 | 0.774204 | 0.778013 | 0.772815 |
| 0.773847 | 0.770865 | 0.774071 | 0.777052 | 0.762983 | 0.769375 | 0.775421 | 0.776463 | 0.773264 |
| 0.76732 | 0.764418 | 0.764349 | 0.769543 | 0.757821 | 0.768702 | 0.7696 | 0.770297 | 0.764181 |
| 0.769954 | 0.771045 | 0.771065 | 0.775856 | 0.759363 | 0.763597 | 0.769014 | 0.771919 | 0.769509 |
| 0.763043 | 0.768254 | 0.768052 | 0.77373 | 0.759297 | 0.762631 | 0.767298 | 0.772957 | 0.770833 |

| | | | | | | | | |
|---|---|---|---|---|---|---|---|---|
| 0.763015 | 0.764914 | 0.757328 | 0.767529 | 0.748548 | 0.757335 | 0.758869 | 0.759012 | 0.754433 |
| 0.750372 | 0.749465 | 0.744823 | 0.753582 | 0.739904 | 0.746916 | 0.752401 | 0.754299 | 0.75069 |
| 0.771611 | 0.762664 | 0.771581 | 0.772688 | 0.757479 | 0.761322 | 0.766914 | 0.765318 | 0.765972 |
| 0.771157 | 0.776707 | 0.773716 | 0.776036 | 0.756142 | 0.762249 | 0.767326 | 0.771455 | 0.767598 |
| 0.767791 | 0.76046 | 0.764331 | 0.766851 | 0.750022 | 0.75855 | 0.761372 | 0.763137 | 0.760528 |
| 0.771101 | 0.764928 | 0.768449 | 0.769179 | 0.753122 | 0.760159 | 0.764324 | 0.766933 | 0.766588 |
| 0.768117 | 0.767517 | 0.769824 | 0.772332 | 0.755814 | 0.760912 | 0.767876 | 0.767816 | 0.767105 |
| 0.764848 | 0.756347 | 0.763733 | 0.763807 | 0.749299 | 0.754536 | 0.757104 | 0.761081 | 0.759771 |
| 0.792851 | 0.776665 | 0.789114 | 0.809291 | 0.760641 | 0.769596 | 0.772477 | 0.777021 | 0.768668 |
| 0.831964 | 0.777048 | 0.829356 | 0.792876 | 0.763887 | 0.76716 | 0.772467 | 0.773759 | 0.770296 |
| 0.777392 | 0.776616 | 0.77654 | 0.794179 | 0.757517 | 0.769501 | 0.771201 | 0.773818 | 0.767436 |
| 0.818483 | 0.775209 | 0.774196 | 0.798442 | 0.761662 | 0.767812 | 0.773478 | 0.774218 | 0.772029 |
| 0.778166 | 0.775358 | 0.779322 | 0.779409 | 0.762782 | 0.769193 | 0.77055 | 0.775476 | 0.768222 |
| 0.823514 | 0.776071 | 0.77579 | 0.790168 | 0.758386 | 0.763744 | 0.772395 | 0.775365 | 0.768484 |
| 0.792719 | 0.778933 | 0.79871 | 0.803265 | 0.760371 | 0.766642 | 0.771316 | 0.772817 | 0.767511 |
| 0.826171 | 0.777847 | 0.779043 | 0.783793 | 0.753972 | 0.762101 | 0.768256 | 0.769581 | 0.766076 |
| 0.867165 | 0.781456 | 0.81002 | 0.785582 | 0.757168 | 0.76445 | 0.769333 | 0.772566 | 0.768188 |
| 1 | 0.776606 | 0.80949 | 0.776088 | 0.752832 | 0.761269 | 0.768246 | 0.767367 | 0.766524 |
| 0.776606 | 1 | 0.775878 | 0.780289 | 0.754519 | 0.7601 | 0.762411 | 0.770358 | 0.766431 |
| 0.80949 | 0.775878 | 1 | 0.777748 | 0.752916 | 0.763379 | 0.765198 | 0.767178 | 0.764198 |
| 0.776088 | 0.780289 | 0.777748 | 1 | 0.760208 | 0.766072 | 0.772211 | 0.767833 | 0.766506 |
| 0.752832 | 0.754519 | 0.752916 | 0.760208 | 1 | 0.848653 | 0.805899 | 0.789681 | 0.825456 |
| 0.761269 | 0.7601 | 0.763379 | 0.766072 | 0.848653 | 1 | 0.869776 | 0.786424 | 0.865879 |
| 0.768246 | 0.762411 | 0.765198 | 0.772211 | 0.805899 | 0.869776 | 1 | 0.815962 | 0.874118 |
| 0.767367 | 0.770358 | 0.767178 | 0.767833 | 0.789681 | 0.786424 | 0.815962 | 1 | 0.831074 |
| 0.766524 | 0.766431 | 0.764198 | 0.766506 | 0.825456 | 0.865879 | 0.874118 | 0.831074 | 1 |
| 0.763171 | 0.760055 | 0.759671 | 0.764121 | 0.778551 | 0.786118 | 0.79744 | 0.799032 | 0.789506 |
| 0.762529 | 0.761962 | 0.762007 | 0.767612 | 0.789151 | 0.782066 | 0.792358 | 0.798529 | 0.792943 |
| 0.759548 | 0.763721 | 0.763403 | 0.762518 | 0.787269 | 0.78344 | 0.795806 | 0.83283 | 0.806408 |
| 0.696866 | 0.691192 | 0.690238 | 0.695684 | 0.68602 | 0.695764 | 0.697157 | 0.7018 | 0.698368 |
| 0.712286 | 0.701626 | 0.703349 | 0.708201 | 0.696025 | 0.706666 | 0.711066 | 0.712635 | 0.708762 |
| 0.704743 | 0.691955 | 0.697576 | 0.704282 | 0.69916 | 0.707149 | 0.707769 | 0.709558 | 0.705077 |
| 0.700896 | 0.687816 | 0.692131 | 0.695001 | 0.688769 | 0.698105 | 0.700831 | 0.700396 | 0.698806 |
| 0.710862 | 0.702104 | 0.704137 | 0.710215 | 0.698556 | 0.706705 | 0.708518 | 0.709502 | 0.705934 |
| 0.761495 | 0.761063 | 0.762411 | 0.768438 | 0.774366 | 0.775222 | 0.776894 | 0.782821 | 0.774954 |
| 0.765405 | 0.761746 | 0.76423 | 0.766479 | 0.775688 | 0.779351 | 0.782199 | 0.78203 | 0.775739 |
| 0.766853 | 0.767352 | 0.763192 | 0.770882 | 0.784258 | 0.781891 | 0.786664 | 0.790869 | 0.785769 |
| 0.766631 | 0.763144 | 0.771149 | 0.768069 | 0.782542 | 0.783317 | 0.786527 | 0.788629 | 0.784777 |
| 0.760193 | 0.76631 | 0.757051 | 0.763978 | 0.769776 | 0.77358 | 0.780106 | 0.781186 | 0.775384 |
| 0.76773 | 0.7687 | 0.760313 | 0.767027 | 0.786359 | 0.793562 | 0.788732 | 0.790932 | 0.788641 |
| 0.764371 | 0.763829 | 0.763927 | 0.769172 | 0.767989 | 0.773936 | 0.781987 | 0.785156 | 0.773066 |
| 0.767127 | 0.763279 | 0.760441 | 0.764621 | 0.784179 | 0.788427 | 0.791277 | 0.789907 | 0.790262 |
| 0.759565 | 0.760345 | 0.753127 | 0.766699 | 0.77178 | 0.777068 | 0.774295 | 0.777708 | 0.772715 |
| 0.757882 | 0.759483 | 0.754878 | 0.764281 | 0.771731 | 0.775919 | 0.781186 | 0.782043 | 0.772801 |
| 0.760774 | 0.75981 | 0.758012 | 0.765002 | 0.777729 | 0.774057 | 0.779113 | 0.787125 | 0.776269 |
| 0.765297 | 0.765115 | 0.762627 | 0.767949 | 0.776896 | 0.778783 | 0.78622 | 0.789428 | 0.776917 |

| | | | | | | | | |
|---|---|---|---|---|---|---|---|---|
| 0.775824 | 0.761957 | 0.771287 | 0.77621 | 0.763438 | 0.768448 | 0.774232 | 0.776789 | 0.772622 |
| 0.766675 | 0.760928 | 0.76322 | 0.767259 | 0.764761 | 0.764691 | 0.768899 | 0.770606 | 0.769742 |
| 0.765286 | 0.761755 | 0.765524 | 0.767991 | 0.768801 | 0.772545 | 0.776771 | 0.773733 | 0.775833 |
| 0.762713 | 0.762272 | 0.762249 | 0.765771 | 0.761277 | 0.759023 | 0.765364 | 0.769544 | 0.768959 |
| 0.761973 | 0.764813 | 0.766014 | 0.76593 | 0.761504 | 0.761513 | 0.767047 | 0.777161 | 0.769877 |
| 0.762744 | 0.760947 | 0.760943 | 0.76094 | 0.76367 | 0.764071 | 0.768744 | 0.771336 | 0.768672 |
| 0.766338 | 0.764152 | 0.764309 | 0.767961 | 0.760804 | 0.770042 | 0.773275 | 0.770884 | 0.770655 |
| 0.766289 | 0.765773 | 0.766043 | 0.769933 | 0.764022 | 0.76901 | 0.77444 | 0.77211 | 0.774144 |
| 0.758624 | 0.757165 | 0.760914 | 0.765766 | 0.763977 | 0.766793 | 0.771954 | 0.771256 | 0.771049 |
| 0.766572 | 0.762375 | 0.758004 | 0.7663 | 0.762365 | 0.767965 | 0.771625 | 0.767357 | 0.769432 |
| 0.764128 | 0.761594 | 0.766057 | 0.765943 | 0.760712 | 0.764509 | 0.768869 | 0.773552 | 0.769707 |
| 0.765212 | 0.763199 | 0.765756 | 0.767765 | 0.762896 | 0.765922 | 0.769762 | 0.772477 | 0.770983 |
| 0.75473 | 0.756489 | 0.757246 | 0.755663 | 0.758283 | 0.761247 | 0.767721 | 0.768367 | 0.766567 |
| 0.755582 | 0.759327 | 0.759954 | 0.760681 | 0.759944 | 0.763165 | 0.767834 | 0.76797 | 0.764749 |
| 0.767591 | 0.767518 | 0.760587 | 0.769335 | 0.758606 | 0.76254 | 0.766537 | 0.772077 | 0.765657 |
| 0.778288 | 0.772986 | 0.775099 | 0.777405 | 0.768122 | 0.771426 | 0.775214 | 0.77225 | 0.776114 |
| 0.772511 | 0.76591 | 0.760099 | 0.771223 | 0.758854 | 0.766824 | 0.770755 | 0.766708 | 0.768104 |
| 0.769639 | 0.769251 | 0.763494 | 0.779017 | 0.761437 | 0.767857 | 0.773964 | 0.776164 | 0.769906 |
| 0.766581 | 0.772375 | 0.769799 | 0.771217 | 0.75974 | 0.764781 | 0.772462 | 0.776571 | 0.769832 |
| 0.768862 | 0.77168 | 0.767096 | 0.775194 | 0.755212 | 0.76282 | 0.765959 | 0.773732 | 0.764807 |
| 0.766328 | 0.767457 | 0.762836 | 0.774507 | 0.757797 | 0.762677 | 0.773159 | 0.769797 | 0.765073 |
| 0.769353 | 0.7667 | 0.773164 | 0.772743 | 0.757826 | 0.760927 | 0.771006 | 0.76961 | 0.767673 |
| 0.772256 | 0.777316 | 0.774127 | 0.77456 | 0.761538 | 0.769264 | 0.775507 | 0.774523 | 0.772079 |
| 0.762258 | 0.763835 | 0.762834 | 0.7631 | 0.756686 | 0.765267 | 0.769156 | 0.7747 | 0.770537 |
| 0.772077 | 0.768931 | 0.772232 | 0.772662 | 0.761213 | 0.762178 | 0.771238 | 0.774129 | 0.768672 |
| 0.703419 | 0.695026 | 0.699941 | 0.705418 | 0.694639 | 0.703713 | 0.706853 | 0.710035 | 0.705027 |
| 0.765552 | 0.76834 | 0.765163 | 0.770257 | 0.761845 | 0.768326 | 0.770933 | 0.77365 | 0.772587 |
| 0.768939 | 0.769039 | 0.765567 | 0.77108 | 0.761628 | 0.767059 | 0.772176 | 0.771726 | 0.76866 |
| 0.775439 | 0.767915 | 0.77171 | 0.77508 | 0.7586 | 0.767153 | 0.773712 | 0.776523 | 0.773975 |
| 0.768478 | 0.765238 | 0.762765 | 0.769385 | 0.759838 | 0.76932 | 0.777852 | 0.771092 | 0.770991 |
| 0.759561 | 0.763389 | 0.761952 | 0.76558 | 0.75507 | 0.76391 | 0.768107 | 0.770388 | 0.766474 |
| 0.765582 | 0.770136 | 0.769473 | 0.766302 | 0.755383 | 0.764346 | 0.768345 | 0.776296 | 0.767808 |
| 0.768555 | 0.767505 | 0.7684 | 0.771747 | 0.758439 | 0.766032 | 0.768206 | 0.770168 | 0.768515 |
| 0.765 | 0.769933 | 0.766712 | 0.76887 | 0.758328 | 0.766991 | 0.766753 | 0.772439 | 0.767931 |
| 0.762752 | 0.760896 | 0.759276 | 0.763248 | 0.75675 | 0.759533 | 0.764747 | 0.766664 | 0.763795 |
| 0.75688 | 0.764426 | 0.759497 | 0.764731 | 0.757329 | 0.764112 | 0.766588 | 0.76708 | 0.762282 |
| 0.767389 | 0.767634 | 0.766737 | 0.77113 | 0.75762 | 0.769779 | 0.774597 | 0.775133 | 0.769237 |
| 0.767136 | 0.769155 | 0.77686 | 0.775599 | 0.759439 | 0.768249 | 0.770485 | 0.774883 | 0.768441 |
| 0.768264 | 0.766883 | 0.772932 | 0.773056 | 0.75669 | 0.765719 | 0.769224 | 0.771156 | 0.765703 |
| 0.771364 | 0.763172 | 0.770563 | 0.770432 | 0.755107 | 0.765307 | 0.769583 | 0.770454 | 0.764127 |
| 0.767175 | 0.765814 | 0.768583 | 0.765639 | 0.753354 | 0.762299 | 0.765324 | 0.763043 | 0.760662 |
| 0.773567 | 0.765872 | 0.770118 | 0.770155 | 0.760481 | 0.769733 | 0.771253 | 0.772135 | 0.766412 |
| 0.763373 | 0.759147 | 0.771318 | 0.765628 | 0.753912 | 0.760864 | 0.767594 | 0.771905 | 0.765315 |
| 0.767916 | 0.764994 | 0.758466 | 0.762215 | 0.756384 | 0.7629 | 0.766261 | 0.765554 | 0.768269 |
| 0.766268 | 0.770798 | 0.766678 | 0.774442 | 0.76237 | 0.774076 | 0.775287 | 0.774194 | 0.771222 |
| 0.764794 | 0.772364 | 0.769674 | 0.767161 | 0.762199 | 0.773091 | 0.769822 | 0.770749 | 0.769707 |

| | | | | | | | | |
|---|---|---|---|---|---|---|---|---|
| 0.765205 | 0.76967 | 0.764726 | 0.766722 | 0.761124 | 0.76104 | 0.765549 | 0.770784 | 0.765038 |
| 0.761478 | 0.770796 | 0.767567 | 0.768129 | 0.766931 | 0.770146 | 0.773345 | 0.777572 | 0.776329 |
| 0.766293 | 0.767591 | 0.763428 | 0.76479 | 0.759338 | 0.761188 | 0.765426 | 0.770277 | 0.767297 |
| 0.722787 | 0.736288 | 0.731278 | 0.733696 | 0.720888 | 0.725108 | 0.728832 | 0.72425 | 0.727141 |
| 0.761788 | 0.7622 | 0.762854 | 0.765118 | 0.758276 | 0.760932 | 0.766099 | 0.768819 | 0.763915 |
| 0.766217 | 0.7601 | 0.76111 | 0.764207 | 0.760695 | 0.764657 | 0.767 | 0.765072 | 0.767974 |
| 0.766015 | 0.764617 | 0.759854 | 0.763097 | 0.763647 | 0.765612 | 0.769307 | 0.768043 | 0.770939 |
| 0.765642 | 0.764979 | 0.763545 | 0.765587 | 0.758147 | 0.761625 | 0.767007 | 0.769619 | 0.763405 |
| 0.766713 | 0.767712 | 0.771294 | 0.768217 | 0.761599 | 0.768594 | 0.775389 | 0.772861 | 0.769673 |
| 0.76857 | 0.762447 | 0.768165 | 0.765811 | 0.763392 | 0.772406 | 0.775298 | 0.771077 | 0.773236 |
| 0.766606 | 0.764806 | 0.757741 | 0.7652 | 0.759192 | 0.765809 | 0.768226 | 0.765695 | 0.766976 |
| 0.772716 | 0.769763 | 0.765924 | 0.764236 | 0.761076 | 0.771344 | 0.772912 | 0.774741 | 0.770779 |
| 0.768341 | 0.766809 | 0.76388 | 0.763005 | 0.760139 | 0.766443 | 0.768596 | 0.770409 | 0.770285 |
| 0.763188 | 0.764151 | 0.766101 | 0.765986 | 0.764041 | 0.768165 | 0.769699 | 0.769909 | 0.76941 |
| 0.763674 | 0.769004 | 0.764925 | 0.767799 | 0.763902 | 0.767495 | 0.77314 | 0.770781 | 0.769475 |
| 0.75947 | 0.764133 | 0.761971 | 0.763525 | 0.757698 | 0.761631 | 0.765856 | 0.766047 | 0.765335 |
| 0.767194 | 0.774377 | 0.768176 | 0.772303 | 0.764864 | 0.765985 | 0.772846 | 0.770291 | 0.769593 |
| 0.764031 | 0.772353 | 0.764578 | 0.77079 | 0.765397 | 0.770868 | 0.774207 | 0.773553 | 0.772333 |
| 0.762655 | 0.764758 | 0.761162 | 0.76338 | 0.759103 | 0.767108 | 0.76574 | 0.766524 | 0.767966 |
| 0.764172 | 0.765755 | 0.761418 | 0.767621 | 0.759612 | 0.764499 | 0.767459 | 0.769825 | 0.766586 |
| 0.767686 | 0.76941 | 0.766461 | 0.765982 | 0.769276 | 0.771027 | 0.775782 | 0.774302 | 0.773975 |
| 0.768271 | 0.77673 | 0.76537 | 0.774506 | 0.760917 | 0.764874 | 0.769686 | 0.773814 | 0.770373 |
| 0.686551 | 0.681516 | 0.681911 | 0.689261 | 0.67809 | 0.676856 | 0.674672 | 0.683367 | 0.676159 |
| 0.666906 | 0.661103 | 0.659879 | 0.663462 | 0.65455 | 0.65401 | 0.661874 | 0.663874 | 0.658065 |
| 0.675342 | 0.673543 | 0.670149 | 0.675699 | 0.66221 | 0.668539 | 0.663358 | 0.672955 | 0.669787 |
| 0.655907 | 0.656823 | 0.653245 | 0.653493 | 0.645107 | 0.649672 | 0.656347 | 0.662689 | 0.652654 |
| 0.723698 | 0.745198 | 0.738146 | 0.735032 | 0.744443 | 0.745316 | 0.742559 | 0.749392 | 0.749695 |
| 0.718027 | 0.736114 | 0.726547 | 0.730903 | 0.734409 | 0.731081 | 0.729497 | 0.743228 | 0.731494 |
| 0.72453 | 0.736183 | 0.733554 | 0.72972 | 0.735556 | 0.7375 | 0.741217 | 0.734557 | 0.744331 |
| 0.735116 | 0.752597 | 0.746784 | 0.745443 | 0.734432 | 0.740338 | 0.73738 | 0.742539 | 0.735865 |
| 0.717037 | 0.73363 | 0.717543 | 0.719274 | 0.716946 | 0.707556 | 0.714873 | 0.719161 | 0.704457 |
| 0.708688 | 0.724296 | 0.710451 | 0.721888 | 0.711869 | 0.720824 | 0.715093 | 0.720744 | 0.718037 |
| 0.710868 | 0.731047 | 0.718797 | 0.723983 | 0.726335 | 0.722784 | 0.726272 | 0.732227 | 0.732772 |
| 0.657782 | 0.660119 | 0.65471 | 0.662121 | 0.642082 | 0.653402 | 0.654885 | 0.659173 | 0.653564 |
| 0.764711 | 0.761544 | 0.760978 | 0.766 | 0.752802 | 0.754482 | 0.764966 | 0.767591 | 0.762667 |
| 0.739574 | 0.754316 | 0.742032 | 0.746106 | 0.738669 | 0.742637 | 0.746494 | 0.748734 | 0.743659 |
| 0.737563 | 0.742786 | 0.738909 | 0.744345 | 0.729561 | 0.737006 | 0.739655 | 0.747509 | 0.732798 |
| 0.721373 | 0.731703 | 0.726828 | 0.726664 | 0.725785 | 0.725219 | 0.728186 | 0.735355 | 0.723396 |
| 0.735075 | 0.752485 | 0.743946 | 0.744459 | 0.74306 | 0.743875 | 0.750471 | 0.748283 | 0.742016 |
| 0.734407 | 0.736711 | 0.739966 | 0.733651 | 0.726675 | 0.733798 | 0.734941 | 0.734974 | 0.731571 |
| 0.731398 | 0.736102 | 0.732221 | 0.735915 | 0.726381 | 0.727666 | 0.727258 | 0.734586 | 0.726836 |
| 0.734402 | 0.735158 | 0.73295 | 0.731566 | 0.721531 | 0.729353 | 0.730129 | 0.735971 | 0.734007 |
| 0.720318 | 0.742455 | 0.735776 | 0.732668 | 0.729132 | 0.728644 | 0.72798 | 0.738377 | 0.730629 |
| 0.729072 | 0.737001 | 0.728603 | 0.730067 | 0.728548 | 0.723661 | 0.725281 | 0.735883 | 0.729237 |
| 0.727884 | 0.739472 | 0.741765 | 0.735614 | 0.721709 | 0.729129 | 0.724994 | 0.729102 | 0.726102 |
| 0.71755 | 0.72551 | 0.720993 | 0.727426 | 0.714286 | 0.713573 | 0.71556 | 0.719855 | 0.716617 |

| | | | | | | | | |
|---|---|---|---|---|---|---|---|---|
| 0.753223 | 0.753236 | 0.75144 | 0.751472 | 0.752089 | 0.753901 | 0.756835 | 0.761052 | 0.754174 |
| 0.728412 | 0.739118 | 0.729803 | 0.7304 | 0.726252 | 0.726189 | 0.726966 | 0.740202 | 0.729237 |
| 0.722929 | 0.73964 | 0.728733 | 0.732562 | 0.721459 | 0.729352 | 0.735582 | 0.73574 | 0.730223 |
| 0.727511 | 0.743727 | 0.725829 | 0.737459 | 0.728968 | 0.728668 | 0.730587 | 0.737095 | 0.727259 |
| 0.728615 | 0.752599 | 0.732354 | 0.728163 | 0.726741 | 0.726382 | 0.732409 | 0.731722 | 0.731533 |
| 0.723104 | 0.737069 | 0.722152 | 0.718415 | 0.715441 | 0.724009 | 0.725915 | 0.736802 | 0.719564 |
| 0.715639 | 0.734472 | 0.720491 | 0.724933 | 0.717656 | 0.7238 | 0.728669 | 0.735417 | 0.723581 |
| 0.712472 | 0.73047 | 0.717795 | 0.712913 | 0.709164 | 0.71696 | 0.715459 | 0.72996 | 0.718652 |
| 0.723471 | 0.733007 | 0.736295 | 0.729668 | 0.723157 | 0.722994 | 0.725529 | 0.736317 | 0.726627 |
| 0.748309 | 0.758134 | 0.751645 | 0.753298 | 0.748092 | 0.757799 | 0.756451 | 0.761696 | 0.753967 |
| 0.749495 | 0.756665 | 0.747763 | 0.751945 | 0.747589 | 0.755819 | 0.754283 | 0.761088 | 0.753931 |
| 0.743846 | 0.754617 | 0.750171 | 0.748664 | 0.746153 | 0.740303 | 0.744622 | 0.755851 | 0.744593 |
| 0.741833 | 0.751006 | 0.747817 | 0.741979 | 0.733226 | 0.740709 | 0.743934 | 0.747483 | 0.74328 |
| 0.741647 | 0.754259 | 0.745886 | 0.746531 | 0.742341 | 0.745595 | 0.751354 | 0.751706 | 0.749351 |
| 0.739672 | 0.745791 | 0.743262 | 0.743178 | 0.739653 | 0.741422 | 0.743881 | 0.7495 | 0.742871 |

| | plot 10 | | | | | | | |
|---|---|---|---|---|---|---|---|---|
| Wild_plot_ | Wild_plot_ | Wild_plot_ | Wild_plot_ | Wild_plot_ | Wild_plot_ | Wild_plot_ | Wild_plot_ | Wild_plot_ |
| 0.702705 | 0.702983 | 0.70041 | 0.76263 | 0.788616 | 0.776487 | 0.793909 | 0.791423 | 0.702667 |
| 0.715018 | 0.709172 | 0.711677 | 0.769481 | 0.797597 | 0.779137 | 0.780721 | 0.792473 | 0.707608 |
| 0.703231 | 0.700443 | 0.699357 | 0.758043 | 0.785994 | 0.772276 | 0.790293 | 0.789863 | 0.696091 |
| 0.709966 | 0.707726 | 0.708082 | 0.760242 | 0.782425 | 0.776143 | 0.780702 | 0.789707 | 0.706027 |
| 0.703667 | 0.701363 | 0.70316 | 0.769868 | 0.78262 | 0.778385 | 0.787628 | 0.781194 | 0.702291 |
| 0.709253 | 0.711978 | 0.710707 | 0.759079 | 0.779947 | 0.774239 | 0.773341 | 0.788164 | 0.709733 |
| 0.763229 | 0.766851 | 0.763144 | 0.699807 | 0.709135 | 0.706291 | 0.697521 | 0.708991 | 0.765866 |
| 0.761252 | 0.759864 | 0.759969 | 0.693145 | 0.706565 | 0.703106 | 0.693677 | 0.703336 | 0.763987 |
| 0.707997 | 0.70297 | 0.70617 | 0.766396 | 0.792041 | 0.791077 | 0.783137 | 0.804143 | 0.706672 |
| 0.703334 | 0.705829 | 0.703729 | 0.75702 | 0.785804 | 0.775971 | 0.802225 | 0.785433 | 0.698386 |
| 0.706494 | 0.706661 | 0.705783 | 0.7596 | 0.776248 | 0.775833 | 0.786518 | 0.77682 | 0.701633 |
| 0.699206 | 0.699745 | 0.702149 | 0.774539 | 0.789535 | 0.793121 | 0.790276 | 0.792713 | 0.694712 |
| 0.707181 | 0.706617 | 0.708928 | 0.763982 | 0.801684 | 0.781349 | 0.762204 | 0.799834 | 0.705152 |
| 0.715499 | 0.715979 | 0.717736 | 0.760762 | 0.778543 | 0.771223 | 0.786462 | 0.783839 | 0.714886 |
| 0.731877 | 0.733139 | 0.727265 | 0.751759 | 0.771134 | 0.760779 | 0.757407 | 0.76905 | 0.72656 |
| 0.707975 | 0.709424 | 0.70726 | 0.761589 | 0.778938 | 0.773764 | 0.799741 | 0.78704 | 0.710247 |
| 0.708579 | 0.70908 | 0.705055 | 0.753378 | 0.773088 | 0.764848 | 0.793305 | 0.773318 | 0.708464 |
| 0.737873 | 0.740679 | 0.737764 | 0.754953 | 0.77003 | 0.764069 | 0.756631 | 0.767434 | 0.732686 |
| 0.701221 | 0.700224 | 0.702395 | 0.768929 | 0.792897 | 0.775454 | 0.765585 | 0.792187 | 0.69949 |
| 0.705553 | 0.704953 | 0.709108 | 0.784049 | 0.795254 | 0.795129 | 0.776312 | 0.793588 | 0.70628 |
| 0.70479 | 0.704003 | 0.70251 | 0.777203 | 0.792243 | 0.789703 | 0.7779 | 0.785121 | 0.703124 |
| 0.710672 | 0.707166 | 0.707385 | 0.776041 | 0.793141 | 0.785118 | 0.789537 | 0.796619 | 0.703479 |
| 0.710676 | 0.709001 | 0.711448 | 0.780663 | 0.79921 | 0.791872 | 0.784079 | 0.801159 | 0.706882 |
| 0.710217 | 0.70916 | 0.706526 | 0.780953 | 0.798065 | 0.789588 | 0.791963 | 0.792065 | 0.707223 |
| 0.713113 | 0.711259 | 0.713149 | 0.781042 | 0.802491 | 0.794576 | 0.769323 | 0.787333 | 0.708547 |
| 0.706412 | 0.705385 | 0.705582 | 0.776742 | 0.795049 | 0.783463 | 0.775758 | 0.786885 | 0.702128 |
| 0.704078 | 0.700767 | 0.701563 | 0.765515 | 0.784053 | 0.773875 | 0.7619 | 0.781917 | 0.700197 |
| 0.707415 | 0.705969 | 0.706001 | 0.768931 | 0.780366 | 0.78332 | 0.782657 | 0.789331 | 0.702857 |
| 0.711799 | 0.707903 | 0.711057 | 0.772831 | 0.781573 | 0.783315 | 0.774927 | 0.781245 | 0.705467 |
| 0.705907 | 0.706517 | 0.705351 | 0.765279 | 0.788054 | 0.773307 | 0.781689 | 0.782267 | 0.704574 |
| 0.711683 | 0.70289 | 0.706775 | 0.775038 | 0.781786 | 0.787115 | 0.771439 | 0.781044 | 0.700049 |
| 0.702756 | 0.701603 | 0.703638 | 0.767784 | 0.777483 | 0.778625 | 0.765112 | 0.774772 | 0.698573 |
| 0.706935 | 0.706086 | 0.700852 | 0.76381 | 0.783307 | 0.778091 | 0.770389 | 0.788544 | 0.705337 |
| 0.705926 | 0.706294 | 0.707063 | 0.771453 | 0.774263 | 0.775782 | 0.766312 | 0.778485 | 0.704052 |
| 0.705094 | 0.704512 | 0.700134 | 0.756259 | 0.76839 | 0.766892 | 0.760889 | 0.770801 | 0.697545 |
| 0.703853 | 0.705843 | 0.701669 | 0.762437 | 0.787107 | 0.775733 | 0.771909 | 0.792835 | 0.702366 |
| 0.709854 | 0.708128 | 0.709715 | 0.777334 | 0.788778 | 0.778575 | 0.77193 | 0.781449 | 0.70818 |
| 0.704655 | 0.705107 | 0.708362 | 0.751959 | 0.774166 | 0.765114 | 0.759022 | 0.768546 | 0.70555 |
| 0.709025 | 0.706712 | 0.707694 | 0.763476 | 0.777096 | 0.773257 | 0.767416 | 0.779297 | 0.710515 |

| | | | | | | | | |
|---|---|---|---|---|---|---|---|---|
| 0.743466 | 0.743588 | 0.738851 | 0.743046 | 0.751561 | 0.751236 | 0.755544 | 0.751299 | 0.736038 |
| 0.699787 | 0.696713 | 0.696476 | 0.761109 | 0.793514 | 0.774944 | 0.795945 | 0.790884 | 0.696504 |
| 0.701593 | 0.701569 | 0.702341 | 0.781767 | 0.791313 | 0.79763 | 0.795706 | 0.79275 | 0.700213 |
| 0.710705 | 0.711635 | 0.708887 | 0.765692 | 0.793101 | 0.786388 | 0.815493 | 0.805281 | 0.707391 |
| 0.707315 | 0.706605 | 0.705098 | 0.775941 | 0.79705 | 0.788251 | 0.778562 | 0.788906 | 0.707648 |
| 0.708216 | 0.708247 | 0.712217 | 0.768751 | 0.784685 | 0.785822 | 0.777102 | 0.782752 | 0.706309 |
| 0.703115 | 0.702213 | 0.704212 | 0.764082 | 0.77237 | 0.783608 | 0.766906 | 0.775996 | 0.702053 |
| 0.706106 | 0.70335 | 0.708147 | 0.777949 | 0.809026 | 0.785882 | 0.771497 | 0.805862 | 0.703511 |
| 0.708813 | 0.70817 | 0.704271 | 0.764289 | 0.776254 | 0.778782 | 0.77219 | 0.781178 | 0.700882 |
| 0.711463 | 0.706845 | 0.705384 | 0.759258 | 0.778423 | 0.775294 | 0.766112 | 0.779474 | 0.706167 |
| 0.703026 | 0.696942 | 0.697283 | 0.772043 | 0.794125 | 0.791786 | 0.803136 | 0.788973 | 0.698388 |
| 0.701137 | 0.697055 | 0.701187 | 0.783259 | 0.80891 | 0.796398 | 0.795332 | 0.796333 | 0.699268 |
| 0.704369 | 0.704493 | 0.702522 | 0.775613 | 0.797898 | 0.789159 | 0.795896 | 0.785815 | 0.703642 |
| 0.701154 | 0.701036 | 0.698569 | 0.777178 | 0.783918 | 0.791575 | 0.804987 | 0.787821 | 0.699424 |
| 0.706323 | 0.701633 | 0.700466 | 0.784765 | 0.797791 | 0.78691 | 0.781683 | 0.779861 | 0.700613 |
| 0.703722 | 0.70112 | 0.699157 | 0.774712 | 0.793691 | 0.77878 | 0.793213 | 0.774086 | 0.699369 |
| 0.698706 | 0.698859 | 0.696736 | 0.755664 | 0.779294 | 0.772033 | 0.788314 | 0.773615 | 0.699203 |
| 0.708998 | 0.706334 | 0.708488 | 0.772675 | 0.803048 | 0.781001 | 0.797813 | 0.799967 | 0.704378 |
| 0.706844 | 0.7067 | 0.704064 | 0.77863 | 0.800504 | 0.787062 | 0.78721 | 0.79614 | 0.706037 |
| 0.701702 | 0.701861 | 0.70236 | 0.783025 | 0.792243 | 0.793639 | 0.787865 | 0.793 | 0.701048 |
| 0.7007 | 0.699917 | 0.697767 | 0.761577 | 0.771862 | 0.769801 | 0.801235 | 0.777064 | 0.697169 |
| 0.700675 | 0.700488 | 0.698293 | 0.763414 | 0.790791 | 0.777546 | 0.793333 | 0.793272 | 0.700806 |
| 0.705147 | 0.69776 | 0.7034 | 0.770064 | 0.792601 | 0.788857 | 0.779835 | 0.792495 | 0.701214 |
| 0.716012 | 0.713977 | 0.711873 | 0.758565 | 0.780455 | 0.773604 | 0.76024 | 0.78062 | 0.712152 |
| 0.718351 | 0.717348 | 0.717248 | 0.749365 | 0.767246 | 0.760481 | 0.748558 | 0.764944 | 0.719528 |
| 0.762185 | 0.761815 | 0.761087 | 0.693263 | 0.706774 | 0.703364 | 0.694813 | 0.705107 | 0.765098 |
| 0.764283 | 0.770054 | 0.765859 | 0.700361 | 0.712234 | 0.704791 | 0.695725 | 0.709713 | 0.769112 |
| 0.764887 | 0.769045 | 0.767189 | 0.70038 | 0.711041 | 0.708953 | 0.697267 | 0.713021 | 0.76858 |
| 0.703451 | 0.704457 | 0.70795 | 0.764745 | 0.798132 | 0.778065 | 0.767338 | 0.796815 | 0.702035 |
| 0.701595 | 0.700195 | 0.702306 | 0.76727 | 0.786764 | 0.776227 | 0.764952 | 0.792487 | 0.699571 |
| 0.710079 | 0.70934 | 0.712096 | 0.779265 | 0.785314 | 0.796179 | 0.796514 | 0.780145 | 0.708045 |
| 0.71273 | 0.710722 | 0.717592 | 0.77049 | 0.797893 | 0.77889 | 0.768594 | 0.790705 | 0.712146 |
| 0.706034 | 0.702209 | 0.707993 | 0.766939 | 0.817971 | 0.777264 | 0.781413 | 0.807349 | 0.702835 |
| 0.706765 | 0.705305 | 0.708853 | 0.768088 | 0.784517 | 0.782052 | 0.771053 | 0.787859 | 0.703621 |
| 0.703034 | 0.700242 | 0.705531 | 0.759613 | 0.804532 | 0.77316 | 0.774921 | 0.808235 | 0.700939 |
| 0.705425 | 0.703314 | 0.701744 | 0.764082 | 0.780535 | 0.767235 | 0.772598 | 0.778374 | 0.700658 |
| 0.702379 | 0.699681 | 0.703212 | 0.766819 | 0.821483 | 0.789478 | 0.801163 | 0.859101 | 0.698603 |
| 0.716802 | 0.716293 | 0.719245 | 0.782943 | 0.829824 | 0.791614 | 0.825191 | 0.830616 | 0.715301 |
| 0.697599 | 0.694551 | 0.69762 | 0.762646 | 0.791012 | 0.770607 | 0.761932 | 0.797132 | 0.693351 |
| 0.704068 | 0.704488 | 0.703148 | 0.804245 | 0.821842 | 0.821386 | 0.771085 | 0.785819 | 0.701821 |
| 0.702568 | 0.700328 | 0.700584 | 0.76509 | 0.779598 | 0.769155 | 0.78192 | 0.780839 | 0.695585 |
| 0.720861 | 0.720287 | 0.720626 | 0.783677 | 0.836517 | 0.797539 | 0.823346 | 0.83442 | 0.717851 |
| 0.70364 | 0.704179 | 0.704741 | 0.768996 | 0.783259 | 0.778963 | 0.821751 | 0.783304 | 0.703206 |
| 0.701078 | 0.698414 | 0.705065 | 0.764912 | 0.779932 | 0.781848 | 0.823804 | 0.78113 | 0.701597 |
| 0.70803 | 0.706186 | 0.705843 | 0.779137 | 0.803658 | 0.783119 | 0.776489 | 0.791722 | 0.702486 |
| 0.705257 | 0.705083 | 0.705043 | 0.767941 | 0.774954 | 0.775399 | 0.766079 | 0.77996 | 0.70359 |

| | | | | | | | | |
|---|---|---|---|---|---|---|---|---|
| 0.702252 | 0.703223 | 0.700345 | 0.7671 | 0.784845 | 0.780323 | 0.823143 | 0.78987 | 0.700658 |
| 0.706812 | 0.705669 | 0.708935 | 0.824007 | 0.805806 | 0.889375 | 0.816808 | 0.808753 | 0.706022 |
| 0.703701 | 0.702053 | 0.701673 | 0.768034 | 0.784955 | 0.790859 | 0.812746 | 0.812258 | 0.69874 |
| 0.70938 | 0.710009 | 0.710733 | 0.776083 | 0.81294 | 0.786862 | 0.81113 | 0.813462 | 0.706282 |
| 0.713263 | 0.711111 | 0.709236 | 0.779708 | 0.808842 | 0.788396 | 0.792055 | 0.798995 | 0.710358 |
| 0.707655 | 0.701144 | 0.708185 | 0.80197 | 0.800119 | 0.820166 | 0.766262 | 0.777015 | 0.70201 |
| 0.705729 | 0.703334 | 0.705813 | 0.75636 | 0.815293 | 0.777366 | 0.783588 | 0.82008 | 0.700602 |
| 0.710884 | 0.708961 | 0.706181 | 0.768619 | 0.784909 | 0.786026 | 0.805761 | 0.820212 | 0.705464 |
| 0.703633 | 0.702492 | 0.705668 | 0.770363 | 0.78581 | 0.780077 | 0.772061 | 0.789465 | 0.699383 |
| 0.70308 | 0.702406 | 0.702462 | 0.764521 | 0.775977 | 0.776855 | 0.761414 | 0.77198 | 0.702603 |
| 0.705192 | 0.702997 | 0.702298 | 0.762448 | 0.777429 | 0.777282 | 0.768499 | 0.779808 | 0.700993 |
| 0.702885 | 0.7057 | 0.705897 | 0.759774 | 0.776411 | 0.776889 | 0.766399 | 0.773746 | 0.703098 |
| 0.709101 | 0.705124 | 0.706706 | 0.769423 | 0.789718 | 0.776597 | 0.772367 | 0.784263 | 0.701453 |
| 0.7003 | 0.70103 | 0.699983 | 0.768052 | 0.781787 | 0.783951 | 0.762809 | 0.781767 | 0.698274 |
| 0.706412 | 0.70457 | 0.709175 | 0.766971 | 0.78168 | 0.77838 | 0.772673 | 0.779847 | 0.705421 |
| 0.703798 | 0.702767 | 0.702648 | 0.764025 | 0.777437 | 0.782108 | 0.775073 | 0.784413 | 0.700834 |
| 0.706992 | 0.705687 | 0.704742 | 0.764176 | 0.784145 | 0.779127 | 0.76967 | 0.791563 | 0.701546 |
| 0.707778 | 0.704156 | 0.706209 | 0.762474 | 0.774421 | 0.769454 | 0.768183 | 0.774386 | 0.701113 |
| 0.704231 | 0.704854 | 0.708756 | 0.761025 | 0.785349 | 0.784663 | 0.771216 | 0.783549 | 0.703238 |
| 0.703445 | 0.705941 | 0.706483 | 0.767939 | 0.785332 | 0.781472 | 0.76802 | 0.784689 | 0.702469 |
| 0.707571 | 0.705022 | 0.703863 | 0.760286 | 0.771338 | 0.780151 | 0.774661 | 0.782048 | 0.704168 |
| 0.701059 | 0.698831 | 0.699648 | 0.764913 | 0.773228 | 0.778886 | 0.767782 | 0.781805 | 0.696483 |
| 0.707422 | 0.703405 | 0.704172 | 0.773645 | 0.777927 | 0.798272 | 0.785998 | 0.797294 | 0.701512 |
| 0.708813 | 0.705664 | 0.704563 | 0.79204 | 0.816477 | 0.79467 | 0.766292 | 0.777614 | 0.704749 |
| 0.706753 | 0.703571 | 0.70663 | 0.765362 | 0.798381 | 0.782106 | 0.77888 | 0.812083 | 0.702075 |
| 0.710375 | 0.709717 | 0.710467 | 0.766803 | 0.819941 | 0.783185 | 0.785531 | 0.828727 | 0.708555 |
| 0.704503 | 0.701638 | 0.702209 | 0.761193 | 0.802384 | 0.782681 | 0.782692 | 0.810009 | 0.701183 |
| 0.703461 | 0.706511 | 0.705793 | 0.764092 | 0.781234 | 0.778027 | 0.76774 | 0.779595 | 0.708652 |
| 0.710366 | 0.705049 | 0.709713 | 0.75975 | 0.783149 | 0.777815 | 0.770289 | 0.782951 | 0.705844 |
| 0.707871 | 0.705592 | 0.707363 | 0.758142 | 0.789729 | 0.776079 | 0.779446 | 0.814705 | 0.703177 |
| 0.708714 | 0.708392 | 0.703629 | 0.758929 | 0.776819 | 0.774407 | 0.771415 | 0.78002 | 0.702956 |
| 0.702114 | 0.704512 | 0.703633 | 0.755926 | 0.777192 | 0.773157 | 0.779278 | 0.787716 | 0.701559 |
| 0.706157 | 0.704373 | 0.708769 | 0.770473 | 0.822915 | 0.776561 | 0.775321 | 0.819754 | 0.702792 |
| 0.708608 | 0.708901 | 0.705614 | 0.762024 | 0.790238 | 0.782082 | 0.812073 | 0.80109 | 0.7029 |
| 0.70378 | 0.703421 | 0.699402 | 0.768551 | 0.7884 | 0.782492 | 0.77077 | 0.786325 | 0.698814 |
| 0.706167 | 0.704969 | 0.704609 | 0.769216 | 0.782031 | 0.774435 | 0.777623 | 0.78569 | 0.701607 |
| 0.702936 | 0.701819 | 0.698819 | 0.763484 | 0.778105 | 0.780247 | 0.763342 | 0.778566 | 0.694156 |
| 0.682143 | 0.681931 | 0.680693 | 0.778349 | 0.753835 | 0.786817 | 0.766355 | 0.772435 | 0.679125 |
| 0.687864 | 0.688257 | 0.689601 | 0.798162 | 0.853269 | 0.780202 | 0.755002 | 0.755872 | 0.68259 |
| 0.679096 | 0.679451 | 0.681124 | 0.74542 | 0.752793 | 0.756111 | 0.7498 | 0.769504 | 0.671733 |
| 0.675497 | 0.677846 | 0.679853 | 0.747219 | 0.762846 | 0.757344 | 0.760413 | 0.75983 | 0.67585 |
| 0.663149 | 0.664721 | 0.661689 | 0.730592 | 0.738458 | 0.745862 | 0.743273 | 0.749181 | 0.659728 |
| 0.660877 | 0.666135 | 0.665838 | 0.749693 | 0.763599 | 0.756163 | 0.801164 | 0.755987 | 0.660108 |
| 0.660679 | 0.660521 | 0.666876 | 0.735482 | 0.768431 | 0.767445 | 0.773793 | 0.775233 | 0.663351 |
| 0.665105 | 0.665331 | 0.668365 | 0.73777 | 0.749867 | 0.738284 | 0.74616 | 0.755181 | 0.6646 |
| 0.66314 | 0.660755 | 0.668112 | 0.740813 | 0.77956 | 0.800053 | 0.803656 | 0.815354 | 0.661302 |
| 0.666204 | 0.671466 | 0.672658 | 0.743965 | 0.760021 | 0.745078 | 0.756038 | 0.747954 | 0.667957 |

| | | | | | | | | |
|---|---|---|---|---|---|---|---|---|
| 0.665541 | 0.672422 | 0.668605 | 0.832447 | 0.745224 | 0.743125 | 0.737997 | 0.747202 | 0.664142 |
| 0.673619 | 0.675672 | 0.678798 | 0.735976 | 0.777349 | 0.745343 | 0.753609 | 0.794354 | 0.667982 |
| 0.664991 | 0.664597 | 0.670333 | 0.73649 | 0.748388 | 0.739922 | 0.737992 | 0.74469 | 0.666936 |
| 0.658664 | 0.664262 | 0.657919 | 0.735614 | 0.743877 | 0.743657 | 0.802788 | 0.752271 | 0.659776 |
| 0.66213 | 0.661959 | 0.667585 | 0.742051 | 0.786215 | 0.751691 | 0.741375 | 0.796759 | 0.660304 |
| 0.659527 | 0.662237 | 0.660989 | 0.732026 | 0.750649 | 0.75909 | 0.812327 | 0.814073 | 0.656505 |
| 0.664455 | 0.662706 | 0.665895 | 0.74552 | 0.763893 | 0.75861 | 0.767352 | 0.792608 | 0.653721 |
| 0.660496 | 0.662457 | 0.664322 | 0.741951 | 0.752791 | 0.741506 | 0.743936 | 0.746799 | 0.65772 |
| 0.658914 | 0.663347 | 0.659692 | 0.721296 | 0.725726 | 0.733099 | 0.73136 | 0.72916 | 0.654017 |
| 0.663194 | 0.664254 | 0.671546 | 0.743058 | 0.836457 | 0.749271 | 0.748125 | 0.834737 | 0.658476 |
| 0.645353 | 0.65232 | 0.65444 | 0.730443 | 0.74515 | 0.839897 | 0.833175 | 0.837867 | 0.653178 |
| 0.692893 | 0.692077 | 0.690807 | 0.748383 | 0.769142 | 0.769471 | 0.791347 | 0.772764 | 0.691692 |
| 0.702398 | 0.702758 | 0.704085 | 0.774301 | 0.773114 | 0.775888 | 0.762836 | 0.775734 | 0.7052 |
| 0.680502 | 0.680359 | 0.67903 | 0.740448 | 0.747403 | 0.79373 | 0.77951 | 0.79057 | 0.677067 |
| 0.709398 | 0.711742 | 0.708546 | 0.766959 | 0.773174 | 0.773825 | 0.772249 | 0.777085 | 0.707654 |
| 0.696376 | 0.698789 | 0.698673 | 0.756081 | 0.76723 | 0.770818 | 0.755187 | 0.768804 | 0.697467 |
| 0.708251 | 0.708741 | 0.703574 | 0.764628 | 0.783009 | 0.777851 | 0.772503 | 0.782335 | 0.704432 |
| 0.715411 | 0.715775 | 0.712537 | 0.759355 | 0.777992 | 0.779241 | 0.768262 | 0.787432 | 0.712719 |
| 0.695754 | 0.692783 | 0.690052 | 0.745687 | 0.772825 | 0.75639 | 0.762094 | 0.77183 | 0.688514 |
| 0.717037 | 0.712888 | 0.710983 | 0.756164 | 0.768687 | 0.770648 | 0.762171 | 0.778628 | 0.712967 |
| 0.688349 | 0.687661 | 0.678598 | 0.728281 | 0.743357 | 0.742889 | 0.773427 | 0.774585 | 0.681191 |
| 0.70988 | 0.709332 | 0.706802 | 0.765852 | 0.779252 | 0.778381 | 0.763206 | 0.774487 | 0.711874 |
| 0.71294 | 0.71361 | 0.715582 | 0.75382 | 0.769673 | 0.776944 | 0.768948 | 0.776317 | 0.713317 |
| 0.684286 | 0.68247 | 0.685121 | 0.77819 | 0.769325 | 0.789721 | 0.737722 | 0.742242 | 0.675698 |
| 0.683028 | 0.683694 | 0.683313 | 0.732789 | 0.762994 | 0.742864 | 0.742403 | 0.763115 | 0.678994 |
| 0.68304 | 0.687708 | 0.692509 | 0.744899 | 0.753828 | 0.765695 | 0.759221 | 0.771907 | 0.685159 |
| 0.702183 | 0.70085 | 0.698354 | 0.737796 | 0.758105 | 0.778421 | 0.787762 | 0.799949 | 0.70198 |
| 0.716594 | 0.713009 | 0.710936 | 0.782717 | 0.784015 | 0.797322 | 0.765646 | 0.775194 | 0.713957 |
| 0.682485 | 0.68447 | 0.67947 | 0.738627 | 0.744169 | 0.747454 | 0.739594 | 0.753576 | 0.679312 |
| 0.723154 | 0.719105 | 0.719209 | 0.759457 | 0.813273 | 0.765727 | 0.75766 | 0.795185 | 0.715966 |
| 0.716944 | 0.715864 | 0.719593 | 0.752094 | 0.76888 | 0.764845 | 0.758738 | 0.769681 | 0.718264 |
| 0.695071 | 0.693815 | 0.693286 | 0.743421 | 0.770481 | 0.754425 | 0.767614 | 0.774401 | 0.690873 |
| 0.693114 | 0.691059 | 0.690093 | 0.744785 | 0.770689 | 0.758476 | 0.766134 | 0.76873 | 0.691237 |
| 0.721644 | 0.715151 | 0.715286 | 0.746183 | 0.761294 | 0.760732 | 0.75919 | 0.764596 | 0.721708 |
| 0.713036 | 0.714332 | 0.710287 | 0.793989 | 0.760088 | 0.764482 | 0.751965 | 0.766413 | 0.71454 |
| 0.7181 | 0.71998 | 0.720461 | 0.762439 | 0.771191 | 0.766321 | 0.759413 | 0.768726 | 0.717879 |
| 0.67377 | 0.672925 | 0.666803 | 0.707201 | 0.721009 | 0.726379 | 0.719266 | 0.731245 | 0.667679 |
| 0.713547 | 0.712121 | 0.715689 | 0.741191 | 0.756724 | 0.749918 | 0.739485 | 0.762885 | 0.714789 |
| 0.671621 | 0.66956 | 0.670389 | 0.714299 | 0.725501 | 0.739315 | 0.739552 | 0.750974 | 0.67096 |
| 0.663205 | 0.663397 | 0.663309 | 0.708693 | 0.723798 | 0.741659 | 0.742576 | 0.755031 | 0.662588 |
| 0.71541 | 0.712567 | 0.713939 | 0.759114 | 0.780302 | 0.774271 | 0.76117 | 0.776685 | 0.713574 |
| 0.660857 | 0.662478 | 0.664962 | 0.717479 | 0.715237 | 0.733397 | 0.725594 | 0.738338 | 0.664063 |
| 0.659718 | 0.654346 | 0.659286 | 0.702426 | 0.729273 | 0.711883 | 0.70839 | 0.728264 | 0.654495 |
| 0.653271 | 0.65082 | 0.655012 | 0.69893 | 0.723597 | 0.701956 | 0.698121 | 0.727481 | 0.651727 |
| 0.722926 | 0.719498 | 0.721261 | 0.752699 | 0.771586 | 0.76472 | 0.775867 | 0.779424 | 0.720816 |
| 0.661221 | 0.65698 | 0.660836 | 0.693723 | 0.730138 | 0.702571 | 0.699072 | 0.718671 | 0.656829 |
| 0.712942 | 0.717507 | 0.715155 | 0.756642 | 0.776326 | 0.773132 | 0.764222 | 0.774885 | 0.712921 |

| | | | | | | | | |
|---|---|---|---|---|---|---|---|---|
| 0.724823 | 0.728027 | 0.720948 | 0.751425 | 0.769319 | 0.763121 | 0.761093 | 0.768453 | 0.719682 |
| 0.717888 | 0.717339 | 0.718976 | 0.757245 | 0.776483 | 0.76989 | 0.769545 | 0.775726 | 0.718044 |
| 0.737691 | 0.741028 | 0.735488 | 0.733439 | 0.7479 | 0.744817 | 0.749141 | 0.75218 | 0.741737 |
| 0.647054 | 0.646018 | 0.646595 | 0.689257 | 0.711082 | 0.696051 | 0.685738 | 0.71384 | 0.64665 |
| 0.646686 | 0.646512 | 0.651492 | 0.687808 | 0.716418 | 0.69901 | 0.698286 | 0.733582 | 0.644483 |
| 0.727408 | 0.72284 | 0.722622 | 0.754265 | 0.773381 | 0.760332 | 0.778063 | 0.779593 | 0.721952 |
| 0.653075 | 0.650513 | 0.646884 | 0.679692 | 0.692006 | 0.696237 | 0.683816 | 0.687792 | 0.649019 |
| 0.727479 | 0.72973 | 0.726306 | 0.719573 | 0.733339 | 0.729796 | 0.720169 | 0.731238 | 0.731654 |
| 0.722772 | 0.726755 | 0.725297 | 0.747201 | 0.760076 | 0.754951 | 0.757213 | 0.763055 | 0.724355 |
| 0.723151 | 0.728136 | 0.725355 | 0.708329 | 0.718558 | 0.718074 | 0.708311 | 0.717784 | 0.7292 |
| 0.726506 | 0.726921 | 0.721753 | 0.734282 | 0.75322 | 0.748659 | 0.741915 | 0.760775 | 0.721157 |
| 0.722973 | 0.758865 | 0.711268 | 0.627586 | 0.662162 | 0.578313 | 0.607595 | 0.630573 | 0.666667 |
| 0.72411 | 0.735133 | 0.732296 | 0.649827 | 0.656682 | 0.65188 | 0.651265 | 0.656118 | 0.727199 |
| 0.730324 | 0.734965 | 0.727754 | 0.652299 | 0.653303 | 0.655115 | 0.654629 | 0.650926 | 0.736003 |
| 0.73612 | 0.739051 | 0.73818 | 0.659594 | 0.664436 | 0.661902 | 0.662176 | 0.664781 | 0.752504 |
| 0.725855 | 0.740279 | 0.732113 | 0.657186 | 0.662371 | 0.656743 | 0.656604 | 0.65838 | 0.737897 |
| 0.588027 | 0.588351 | 0.591559 | 0.613956 | 0.619011 | 0.624086 | 0.616413 | 0.618271 | 0.587076 |
| 0.727103 | 0.735474 | 0.733564 | 0.661207 | 0.670296 | 0.664898 | 0.659435 | 0.665626 | 0.737421 |
| 0.73025 | 0.729794 | 0.725942 | 0.728405 | 0.742643 | 0.737274 | 0.727937 | 0.751323 | 0.73018 |
| 0.732184 | 0.73668 | 0.725392 | 0.722159 | 0.739333 | 0.734063 | 0.732731 | 0.743592 | 0.727219 |
| 0.732113 | 0.732297 | 0.72914 | 0.730237 | 0.751138 | 0.742132 | 0.737249 | 0.753309 | 0.726631 |
| 0.590295 | 0.586556 | 0.589194 | 0.604278 | 0.614677 | 0.617013 | 0.612532 | 0.632139 | 0.586641 |
| 0.582118 | 0.574196 | 0.580173 | 0.585135 | 0.596351 | 0.599567 | 0.5873 | 0.600079 | 0.576948 |
| 0.757664 | 0.760712 | 0.756066 | 0.695146 | 0.7006 | 0.704606 | 0.695687 | 0.701398 | 0.753658 |
| 0.757596 | 0.758826 | 0.757096 | 0.694177 | 0.704556 | 0.706052 | 0.695299 | 0.708129 | 0.758463 |
| 0.74849 | 0.74876 | 0.746183 | 0.682938 | 0.697209 | 0.696516 | 0.686136 | 0.697203 | 0.747896 |
| 0.755195 | 0.761601 | 0.75649 | 0.691243 | 0.700016 | 0.698843 | 0.690184 | 0.699992 | 0.756713 |
| 0.752819 | 0.762745 | 0.753387 | 0.693352 | 0.705991 | 0.704776 | 0.696823 | 0.70604 | 0.756934 |
| 0.76175 | 0.760469 | 0.762147 | 0.694264 | 0.703836 | 0.699855 | 0.694092 | 0.702806 | 0.762198 |
| 0.763885 | 0.7612 | 0.762311 | 0.693584 | 0.707091 | 0.700545 | 0.69199 | 0.705887 | 0.759082 |
| 0.751301 | 0.752174 | 0.751603 | 0.691276 | 0.705231 | 0.702875 | 0.695644 | 0.702498 | 0.756281 |
| 0.764825 | 0.765439 | 0.765722 | 0.701381 | 0.713692 | 0.711835 | 0.704432 | 0.71405 | 0.767583 |
| 0.754473 | 0.762052 | 0.7546 | 0.699574 | 0.708959 | 0.706725 | 0.700178 | 0.706323 | 0.75627 |
| 0.75739 | 0.758437 | 0.756708 | 0.698239 | 0.711497 | 0.706916 | 0.698854 | 0.709639 | 0.760524 |
| 0.764956 | 0.765449 | 0.759629 | 0.698228 | 0.711336 | 0.709739 | 0.703236 | 0.715362 | 0.768726 |
| 0.758071 | 0.757169 | 0.758333 | 0.693462 | 0.707731 | 0.701271 | 0.694165 | 0.707441 | 0.758451 |
| 0.762322 | 0.759884 | 0.75709 | 0.69655 | 0.70838 | 0.704399 | 0.700372 | 0.70987 | 0.760722 |
| 0.759936 | 0.762391 | 0.76422 | 0.694984 | 0.70703 | 0.706979 | 0.700704 | 0.709428 | 0.76272 |
| 0.755175 | 0.763013 | 0.758584 | 0.693321 | 0.702843 | 0.702467 | 0.693797 | 0.70515 | 0.760477 |
| 0.751684 | 0.757298 | 0.757421 | 0.699991 | 0.70997 | 0.708153 | 0.694067 | 0.708915 | 0.759002 |
| 0.719693 | 0.728829 | 0.730255 | 0.66254 | 0.653437 | 0.649379 | 0.658246 | 0.653659 | 0.73293 |
| 0.723042 | 0.738009 | 0.738251 | 0.660275 | 0.659153 | 0.653559 | 0.656104 | 0.655141 | 0.74079 |
| 0.762066 | 0.762561 | 0.757257 | 0.698227 | 0.711909 | 0.706957 | 0.703283 | 0.716875 | 0.758847 |
| 0.765489 | 0.767317 | 0.760633 | 0.702464 | 0.712163 | 0.708767 | 0.702813 | 0.711915 | 0.763764 |
| 0.710335 | 0.719526 | 0.708277 | 0.638002 | 0.646094 | 0.640244 | 0.635912 | 0.640951 | 0.714729 |
| 0.760844 | 0.761142 | 0.760662 | 0.695948 | 0.710961 | 0.704755 | 0.69985 | 0.709652 | 0.764891 |
| 0.753753 | 0.760184 | 0.749204 | 0.686158 | 0.697903 | 0.697787 | 0.690145 | 0.697961 | 0.759191 |

| | | | | | | | | |
|---|---|---|---|---|---|---|---|---|
| 0.763635 | 0.759758 | 0.754348 | 0.695704 | 0.707362 | 0.705218 | 0.695641 | 0.708422 | 0.759195 |
| 0.764736 | 0.761284 | 0.757578 | 0.700201 | 0.709974 | 0.70674 | 0.69764 | 0.710481 | 0.76089 |
| 0.762694 | 0.761382 | 0.7539 | 0.699888 | 0.709152 | 0.705892 | 0.701348 | 0.707667 | 0.757218 |
| 0.764576 | 0.763216 | 0.760947 | 0.701388 | 0.708935 | 0.705779 | 0.696405 | 0.708825 | 0.765065 |
| 0.766907 | 0.767289 | 0.763089 | 0.69901 | 0.71652 | 0.710412 | 0.704642 | 0.715214 | 0.761237 |
| 0.759335 | 0.766489 | 0.760818 | 0.694242 | 0.709424 | 0.702579 | 0.696568 | 0.705335 | 0.760133 |
| 0.763182 | 0.762175 | 0.757557 | 0.700733 | 0.707637 | 0.699754 | 0.695896 | 0.706595 | 0.76215 |
| 0.750768 | 0.750515 | 0.750459 | 0.683341 | 0.698657 | 0.697601 | 0.694311 | 0.701958 | 0.753356 |
| 0.760897 | 0.760764 | 0.753849 | 0.695744 | 0.708779 | 0.699369 | 0.694856 | 0.703762 | 0.759211 |
| 0.759656 | 0.757138 | 0.749589 | 0.692464 | 0.705206 | 0.702948 | 0.695683 | 0.705901 | 0.757501 |
| 0.762927 | 0.761658 | 0.757291 | 0.699606 | 0.711665 | 0.705385 | 0.699253 | 0.708484 | 0.759852 |
| 0.76079 | 0.759797 | 0.753709 | 0.69658 | 0.703258 | 0.700858 | 0.698517 | 0.704777 | 0.755164 |
| 0.75206 | 0.75599 | 0.750393 | 0.68615 | 0.697739 | 0.698406 | 0.690056 | 0.700272 | 0.754514 |
| 0.756203 | 0.756206 | 0.752382 | 0.689518 | 0.697688 | 0.696524 | 0.688657 | 0.695848 | 0.755794 |
| 0.758812 | 0.762397 | 0.758543 | 0.700688 | 0.712183 | 0.70769 | 0.701255 | 0.712543 | 0.762423 |
| 0.759275 | 0.762986 | 0.758779 | 0.695907 | 0.709216 | 0.706262 | 0.698962 | 0.710464 | 0.75922 |
| 0.757976 | 0.761817 | 0.759271 | 0.690706 | 0.700372 | 0.697292 | 0.695664 | 0.700757 | 0.764917 |
| 0.755343 | 0.759774 | 0.760428 | 0.690774 | 0.701068 | 0.701918 | 0.695852 | 0.701579 | 0.757779 |
| 0.756322 | 0.764289 | 0.759524 | 0.691554 | 0.704385 | 0.701173 | 0.69463 | 0.70396 | 0.757314 |
| 0.755916 | 0.761852 | 0.757439 | 0.694375 | 0.703179 | 0.702317 | 0.694021 | 0.704191 | 0.758226 |
| 0.75263 | 0.758902 | 0.752468 | 0.687995 | 0.699844 | 0.699853 | 0.687701 | 0.700563 | 0.752539 |
| 0.752755 | 0.758953 | 0.754662 | 0.6921 | 0.701387 | 0.701018 | 0.695157 | 0.703564 | 0.754022 |
| 0.760091 | 0.76521 | 0.759245 | 0.694437 | 0.706159 | 0.704137 | 0.697428 | 0.70663 | 0.758377 |
| 0.756876 | 0.760747 | 0.759511 | 0.694702 | 0.704602 | 0.706057 | 0.694696 | 0.705589 | 0.766077 |
| 0.754638 | 0.762662 | 0.759206 | 0.695419 | 0.70393 | 0.704921 | 0.695115 | 0.704276 | 0.758508 |
| 0.757374 | 0.759248 | 0.752392 | 0.688875 | 0.699911 | 0.69884 | 0.692832 | 0.700938 | 0.753441 |
| 0.749658 | 0.753341 | 0.75 | 0.688558 | 0.702815 | 0.702711 | 0.694534 | 0.70141 | 0.759016 |
| 0.75796 | 0.756806 | 0.752421 | 0.6902 | 0.702665 | 0.696999 | 0.689248 | 0.697519 | 0.754277 |
| 0.756933 | 0.760072 | 0.75559 | 0.692824 | 0.704768 | 0.704282 | 0.694978 | 0.704283 | 0.750717 |
| 0.761937 | 0.763576 | 0.756666 | 0.695333 | 0.704799 | 0.702257 | 0.693944 | 0.703756 | 0.757612 |
| 0.761468 | 0.761264 | 0.764031 | 0.69705 | 0.704073 | 0.704812 | 0.700186 | 0.710079 | 0.762584 |
| 0.75855 | 0.761252 | 0.758569 | 0.699357 | 0.707301 | 0.706989 | 0.703372 | 0.709615 | 0.76135 |
| 0.766203 | 0.768826 | 0.766644 | 0.699753 | 0.709325 | 0.708821 | 0.703073 | 0.711783 | 0.765816 |
| 0.760998 | 0.762721 | 0.759761 | 0.697039 | 0.708865 | 0.708166 | 0.698669 | 0.709262 | 0.767244 |
| 0.756095 | 0.757316 | 0.753952 | 0.692811 | 0.701615 | 0.702761 | 0.693758 | 0.702313 | 0.752349 |
| 0.755517 | 0.760599 | 0.756106 | 0.696741 | 0.704362 | 0.704682 | 0.696573 | 0.707352 | 0.75887 |
| 0.756895 | 0.759377 | 0.752908 | 0.695584 | 0.706873 | 0.705706 | 0.697007 | 0.705826 | 0.753188 |
| 0.756118 | 0.758023 | 0.757279 | 0.694393 | 0.703529 | 0.699603 | 0.692512 | 0.701858 | 0.755867 |
| 0.748198 | 0.754427 | 0.755673 | 0.686577 | 0.696691 | 0.695964 | 0.689089 | 0.696907 | 0.747771 |
| 0.749558 | 0.748958 | 0.749414 | 0.68626 | 0.692527 | 0.694069 | 0.689289 | 0.694093 | 0.748109 |
| 0.742266 | 0.747371 | 0.749197 | 0.681951 | 0.696849 | 0.695274 | 0.686448 | 0.694852 | 0.741686 |
| 0.750604 | 0.755016 | 0.750114 | 0.684996 | 0.693437 | 0.694289 | 0.687696 | 0.697336 | 0.75222 |
| 0.747288 | 0.75071 | 0.745162 | 0.685725 | 0.696177 | 0.698382 | 0.690527 | 0.699909 | 0.747776 |
| 0.745609 | 0.749375 | 0.74861 | 0.681751 | 0.690871 | 0.689417 | 0.681194 | 0.689085 | 0.745763 |
| 0.755624 | 0.756605 | 0.754654 | 0.688768 | 0.697761 | 0.69759 | 0.693569 | 0.697854 | 0.757275 |
| 0.75075 | 0.748229 | 0.745185 | 0.686362 | 0.694672 | 0.691779 | 0.68826 | 0.694875 | 0.756529 |

| | | | | | | | | |
|---|---|---|---|---|---|---|---|---|
| 0.75689 | 0.758379 | 0.756129 | 0.695043 | 0.709633 | 0.708119 | 0.696394 | 0.710326 | 0.752116 |
| 0.767646 | 0.771114 | 0.761938 | 0.695094 | 0.704457 | 0.703695 | 0.697515 | 0.707087 | 0.7656 |
| 0.755569 | 0.759543 | 0.761695 | 0.69371 | 0.70659 | 0.702122 | 0.688466 | 0.706278 | 0.759763 |
| 0.757989 | 0.76335 | 0.760345 | 0.695581 | 0.70833 | 0.709181 | 0.698636 | 0.711099 | 0.762037 |
| 0.769572 | 0.771964 | 0.766481 | 0.701777 | 0.714102 | 0.711931 | 0.701518 | 0.71338 | 0.770507 |
| 0.765484 | 0.765542 | 0.765211 | 0.700938 | 0.708998 | 0.708463 | 0.697636 | 0.71116 | 0.768086 |
| 0.752966 | 0.758672 | 0.761049 | 0.69065 | 0.706305 | 0.698345 | 0.690526 | 0.703664 | 0.759568 |
| 0.760436 | 0.764019 | 0.760128 | 0.695927 | 0.707194 | 0.704446 | 0.697959 | 0.706017 | 0.765589 |
| 0.75849 | 0.757971 | 0.757598 | 0.695092 | 0.703674 | 0.699098 | 0.689342 | 0.702345 | 0.762358 |
| 0.756139 | 0.760749 | 0.75435 | 0.697389 | 0.709818 | 0.703683 | 0.697746 | 0.707406 | 0.759176 |
| 0.762338 | 0.760226 | 0.762585 | 0.69856 | 0.711756 | 0.708578 | 0.697313 | 0.710638 | 0.763841 |
| 0.758501 | 0.764681 | 0.763532 | 0.69854 | 0.710452 | 0.708233 | 0.696874 | 0.709769 | 0.764741 |
| 0.76522 | 0.766345 | 0.763121 | 0.696394 | 0.70931 | 0.705355 | 0.696463 | 0.706616 | 0.762647 |
| 0.761412 | 0.767752 | 0.764629 | 0.697687 | 0.709234 | 0.702588 | 0.697365 | 0.705386 | 0.763269 |
| 0.767742 | 0.769335 | 0.767534 | 0.699042 | 0.710686 | 0.707199 | 0.697544 | 0.707755 | 0.770525 |
| 0.760108 | 0.765344 | 0.76417 | 0.695253 | 0.708654 | 0.706099 | 0.697191 | 0.707965 | 0.758982 |
| 0.76476 | 0.768449 | 0.762621 | 0.702678 | 0.713146 | 0.706857 | 0.698673 | 0.709824 | 0.766099 |
| 0.763607 | 0.765143 | 0.766179 | 0.695206 | 0.710668 | 0.705109 | 0.698093 | 0.709984 | 0.764972 |
| 0.766938 | 0.770395 | 0.765486 | 0.69742 | 0.711213 | 0.707516 | 0.701134 | 0.709316 | 0.770733 |
| 0.766791 | 0.770995 | 0.765751 | 0.693591 | 0.705789 | 0.705035 | 0.695401 | 0.708222 | 0.768177 |
| 0.763398 | 0.765144 | 0.76229 | 0.698394 | 0.707715 | 0.705126 | 0.696887 | 0.705672 | 0.762816 |
| 0.763169 | 0.769204 | 0.763438 | 0.693998 | 0.707912 | 0.699863 | 0.693386 | 0.707053 | 0.766829 |
| 0.766247 | 0.767206 | 0.760148 | 0.700284 | 0.70713 | 0.709214 | 0.704862 | 0.71244 | 0.762221 |
| 0.765549 | 0.767992 | 0.766857 | 0.700675 | 0.708835 | 0.70671 | 0.696567 | 0.714634 | 0.768796 |
| 0.76255 | 0.76602 | 0.763548 | 0.692996 | 0.707555 | 0.707628 | 0.695781 | 0.709428 | 0.768953 |
| 0.762712 | 0.76424 | 0.760464 | 0.694732 | 0.707073 | 0.706052 | 0.697478 | 0.706253 | 0.770419 |
| 0.763054 | 0.766994 | 0.757495 | 0.690747 | 0.705297 | 0.704318 | 0.694892 | 0.706205 | 0.765127 |
| 0.764174 | 0.76835 | 0.761929 | 0.693686 | 0.706636 | 0.701896 | 0.694202 | 0.708791 | 0.765063 |
| 0.766622 | 0.767665 | 0.770368 | 0.702772 | 0.708705 | 0.708492 | 0.698812 | 0.711336 | 0.77204 |
| 0.766286 | 0.772002 | 0.766776 | 0.6964 | 0.708959 | 0.708646 | 0.703172 | 0.712389 | 0.768137 |
| 0.761307 | 0.768431 | 0.765624 | 0.697735 | 0.70469 | 0.709012 | 0.697148 | 0.706628 | 0.765394 |
| 0.763695 | 0.768742 | 0.769258 | 0.702313 | 0.710722 | 0.709914 | 0.70147 | 0.714799 | 0.765263 |
| 0.769185 | 0.772875 | 0.769846 | 0.700688 | 0.70837 | 0.712761 | 0.698895 | 0.710087 | 0.76867 |
| 0.757304 | 0.757507 | 0.763097 | 0.693855 | 0.704814 | 0.70385 | 0.693176 | 0.706884 | 0.760254 |
| 0.76369 | 0.762589 | 0.767485 | 0.696811 | 0.707734 | 0.704837 | 0.696981 | 0.708147 | 0.767826 |
| 0.763351 | 0.770984 | 0.769607 | 0.696118 | 0.705467 | 0.699891 | 0.692329 | 0.705756 | 0.762494 |
| 0.769146 | 0.771777 | 0.76841 | 0.703316 | 0.715756 | 0.711826 | 0.703551 | 0.715846 | 0.769335 |
| 0.768559 | 0.77122 | 0.767126 | 0.702108 | 0.706074 | 0.704656 | 0.695037 | 0.706729 | 0.763962 |
| 0.767717 | 0.770844 | 0.765442 | 0.696872 | 0.709527 | 0.708223 | 0.695893 | 0.708011 | 0.767046 |
| 0.764284 | 0.766467 | 0.768264 | 0.697969 | 0.709408 | 0.706444 | 0.69592 | 0.70975 | 0.765765 |
| 0.765297 | 0.773379 | 0.769558 | 0.700923 | 0.712243 | 0.711856 | 0.702593 | 0.715738 | 0.769549 |
| 0.766585 | 0.771455 | 0.769505 | 0.699257 | 0.709874 | 0.70961 | 0.695874 | 0.714014 | 0.766327 |
| 0.764874 | 0.772375 | 0.767612 | 0.70158 | 0.711419 | 0.709911 | 0.693038 | 0.712818 | 0.765841 |
| 0.764726 | 0.769625 | 0.763353 | 0.698606 | 0.705335 | 0.706982 | 0.697348 | 0.707447 | 0.760062 |
| 0.765198 | 0.766835 | 0.768759 | 0.699963 | 0.708686 | 0.710749 | 0.699061 | 0.709939 | 0.769573 |
| 0.763727 | 0.765801 | 0.764183 | 0.694029 | 0.703661 | 0.703189 | 0.689903 | 0.704985 | 0.764858 |

| | | | | | | | | |
|---|---|---|---|---|---|---|---|---|
| 0.75693 | 0.762935 | 0.753259 | 0.68551 | 0.69484 | 0.690282 | 0.681993 | 0.69228 | 0.755193 |
| 0.742741 | 0.743803 | 0.747056 | 0.730715 | 0.748094 | 0.7406 | 0.734685 | 0.748118 | 0.743776 |
| 0.760909 | 0.765846 | 0.764508 | 0.693866 | 0.706975 | 0.706056 | 0.692852 | 0.706446 | 0.759938 |
| 0.764503 | 0.771443 | 0.768545 | 0.696559 | 0.709953 | 0.709469 | 0.696959 | 0.712319 | 0.76642 |
| 0.759455 | 0.759534 | 0.759606 | 0.690138 | 0.704197 | 0.704902 | 0.691848 | 0.704443 | 0.756236 |
| 0.764938 | 0.764681 | 0.760683 | 0.690104 | 0.706574 | 0.705492 | 0.694222 | 0.707685 | 0.761129 |
| 0.759158 | 0.765959 | 0.761549 | 0.693228 | 0.702465 | 0.703122 | 0.692595 | 0.705006 | 0.761602 |
| 0.756941 | 0.76265 | 0.757142 | 0.687307 | 0.70057 | 0.698268 | 0.689222 | 0.702106 | 0.752484 |
| 0.767395 | 0.771304 | 0.772174 | 0.692861 | 0.706591 | 0.703186 | 0.691416 | 0.707005 | 0.772442 |
| 0.764681 | 0.770254 | 0.767962 | 0.700969 | 0.712524 | 0.7098 | 0.702523 | 0.714991 | 0.763534 |
| 0.764239 | 0.76948 | 0.770583 | 0.696072 | 0.708197 | 0.70523 | 0.699114 | 0.710611 | 0.773155 |
| 0.766092 | 0.769792 | 0.76444 | 0.702175 | 0.712958 | 0.710276 | 0.701461 | 0.712876 | 0.772377 |
| 0.761563 | 0.772427 | 0.770968 | 0.699724 | 0.71432 | 0.709771 | 0.705315 | 0.713222 | 0.771514 |
| 0.766486 | 0.76682 | 0.769177 | 0.700411 | 0.709649 | 0.707911 | 0.699413 | 0.71136 | 0.771769 |
| 0.758659 | 0.766811 | 0.766293 | 0.697445 | 0.707793 | 0.704632 | 0.699143 | 0.710046 | 0.766529 |
| 0.76239 | 0.761057 | 0.764671 | 0.692669 | 0.707726 | 0.70328 | 0.693148 | 0.70662 | 0.761432 |
| 0.768319 | 0.76669 | 0.764074 | 0.696088 | 0.714759 | 0.707519 | 0.701551 | 0.712468 | 0.764496 |
| 0.763171 | 0.762529 | 0.759548 | 0.696866 | 0.712286 | 0.704743 | 0.700896 | 0.710862 | 0.761495 |
| 0.760055 | 0.761962 | 0.763721 | 0.691192 | 0.701626 | 0.691955 | 0.687816 | 0.702104 | 0.761063 |
| 0.759671 | 0.762007 | 0.763403 | 0.690238 | 0.703349 | 0.697576 | 0.692131 | 0.704137 | 0.762411 |
| 0.764121 | 0.767612 | 0.762518 | 0.695684 | 0.708201 | 0.704282 | 0.695001 | 0.710215 | 0.768438 |
| 0.778551 | 0.789151 | 0.787269 | 0.68602 | 0.696025 | 0.69916 | 0.688769 | 0.698556 | 0.774366 |
| 0.786118 | 0.782066 | 0.78344 | 0.695764 | 0.706666 | 0.707149 | 0.698105 | 0.706705 | 0.775222 |
| 0.79744 | 0.792358 | 0.795806 | 0.697157 | 0.711066 | 0.707769 | 0.700831 | 0.708518 | 0.776894 |
| 0.799032 | 0.798529 | 0.83283 | 0.7018 | 0.712635 | 0.709558 | 0.700396 | 0.709502 | 0.782821 |
| 0.789506 | 0.792943 | 0.806408 | 0.698368 | 0.708762 | 0.705077 | 0.698806 | 0.705934 | 0.774954 |
| 1 | 0.846189 | 0.78231 | 0.694195 | 0.705711 | 0.701176 | 0.692064 | 0.702703 | 0.77668 |
| 0.846189 | 1 | 0.796572 | 0.696893 | 0.70346 | 0.700532 | 0.691848 | 0.70032 | 0.772779 |
| 0.78231 | 0.796572 | 1 | 0.693643 | 0.704785 | 0.70279 | 0.696961 | 0.707239 | 0.780319 |
| 0.694195 | 0.696893 | 0.693643 | 1 | 0.785777 | 0.816588 | 0.75551 | 0.766733 | 0.691621 |
| 0.705711 | 0.70346 | 0.704785 | 0.785777 | 1 | 0.794055 | 0.77466 | 0.814605 | 0.699926 |
| 0.701176 | 0.700532 | 0.70279 | 0.816588 | 0.794055 | 1 | 0.79761 | 0.811329 | 0.69693 |
| 0.692064 | 0.691848 | 0.696961 | 0.75551 | 0.77466 | 0.79761 | 1 | 0.819774 | 0.693093 |
| 0.702703 | 0.70032 | 0.707239 | 0.766733 | 0.814605 | 0.811329 | 0.819774 | 1 | 0.696169 |
| 0.77668 | 0.772779 | 0.780319 | 0.691621 | 0.699926 | 0.69693 | 0.693093 | 0.696169 | 1 |
| 0.775932 | 0.781367 | 0.779207 | 0.691433 | 0.705777 | 0.699207 | 0.695324 | 0.702701 | 0.828076 |
| 0.782726 | 0.787805 | 0.780559 | 0.693161 | 0.707353 | 0.703797 | 0.699572 | 0.707571 | 0.812034 |
| 0.777959 | 0.778373 | 0.782888 | 0.694421 | 0.709711 | 0.703793 | 0.700879 | 0.705742 | 0.867227 |
| 0.768862 | 0.7702 | 0.77363 | 0.687793 | 0.702371 | 0.699984 | 0.69207 | 0.703361 | 0.832273 |
| 0.781495 | 0.785999 | 0.786984 | 0.698995 | 0.711025 | 0.707705 | 0.699194 | 0.707437 | 0.820077 |
| 0.774132 | 0.780057 | 0.780229 | 0.694049 | 0.706863 | 0.702597 | 0.696678 | 0.706861 | 0.824223 |
| 0.783652 | 0.785 | 0.781821 | 0.697488 | 0.706926 | 0.702496 | 0.697027 | 0.703686 | 0.80774 |
| 0.768906 | 0.770764 | 0.776115 | 0.686748 | 0.698501 | 0.695773 | 0.691952 | 0.69888 | 0.834273 |
| 0.775497 | 0.771612 | 0.775999 | 0.689541 | 0.698235 | 0.699233 | 0.691464 | 0.69604 | 0.8408 |
| 0.778158 | 0.778316 | 0.780081 | 0.694145 | 0.708695 | 0.704652 | 0.693655 | 0.702635 | 0.812807 |
| 0.782462 | 0.782775 | 0.783301 | 0.697339 | 0.706524 | 0.705208 | 0.700832 | 0.705802 | 0.821388 |

| | | | | | | | | |
|---|---|---|---|---|---|---|---|---|
| 0.77069 | 0.772099 | 0.766003 | 0.695774 | 0.706646 | 0.703336 | 0.69682 | 0.703342 | 0.774778 |
| 0.765047 | 0.764015 | 0.767768 | 0.692258 | 0.704044 | 0.704474 | 0.697064 | 0.705284 | 0.773994 |
| 0.772657 | 0.77255 | 0.767152 | 0.696995 | 0.709565 | 0.706569 | 0.698044 | 0.7073 | 0.775822 |
| 0.761423 | 0.76888 | 0.762478 | 0.695933 | 0.707731 | 0.702601 | 0.699849 | 0.705314 | 0.769277 |
| 0.762954 | 0.764876 | 0.767957 | 0.691919 | 0.699573 | 0.700049 | 0.697262 | 0.703224 | 0.7677 |
| 0.764095 | 0.76855 | 0.7693 | 0.692257 | 0.705153 | 0.701501 | 0.696427 | 0.703001 | 0.771237 |
| 0.764409 | 0.765628 | 0.767576 | 0.699101 | 0.710424 | 0.702321 | 0.698066 | 0.70902 | 0.770886 |
| 0.767066 | 0.769695 | 0.76683 | 0.690032 | 0.704304 | 0.705452 | 0.699521 | 0.707615 | 0.774768 |
| 0.767867 | 0.769278 | 0.76849 | 0.691561 | 0.70459 | 0.703071 | 0.693713 | 0.704315 | 0.772739 |
| 0.762623 | 0.765834 | 0.761331 | 0.695086 | 0.709389 | 0.705368 | 0.699902 | 0.709375 | 0.767099 |
| 0.763003 | 0.768253 | 0.76738 | 0.693876 | 0.704312 | 0.702103 | 0.694732 | 0.70538 | 0.76749 |
| 0.767502 | 0.772329 | 0.767675 | 0.695012 | 0.704255 | 0.705334 | 0.693368 | 0.705802 | 0.768098 |
| 0.759703 | 0.76631 | 0.760246 | 0.687019 | 0.698497 | 0.694644 | 0.687588 | 0.696737 | 0.762625 |
| 0.761579 | 0.765738 | 0.764493 | 0.694212 | 0.699345 | 0.698681 | 0.690822 | 0.700831 | 0.76638 |
| 0.763663 | 0.765029 | 0.76372 | 0.697026 | 0.709502 | 0.705062 | 0.698774 | 0.711421 | 0.761317 |
| 0.76831 | 0.772547 | 0.766894 | 0.700175 | 0.716451 | 0.710237 | 0.701372 | 0.715152 | 0.774865 |
| 0.758499 | 0.761704 | 0.759091 | 0.695506 | 0.709039 | 0.706681 | 0.696547 | 0.71342 | 0.766366 |
| 0.763646 | 0.769845 | 0.764578 | 0.696036 | 0.711684 | 0.708622 | 0.700468 | 0.712263 | 0.770883 |
| 0.765946 | 0.77104 | 0.76764 | 0.696432 | 0.70716 | 0.702721 | 0.695759 | 0.706216 | 0.766728 |
| 0.767593 | 0.764478 | 0.765675 | 0.694237 | 0.705936 | 0.701347 | 0.691967 | 0.70521 | 0.761971 |
| 0.762031 | 0.762314 | 0.762282 | 0.692913 | 0.702541 | 0.699335 | 0.693482 | 0.702343 | 0.770496 |
| 0.760324 | 0.768148 | 0.767579 | 0.697674 | 0.706193 | 0.707998 | 0.694015 | 0.709396 | 0.761787 |
| 0.763574 | 0.763769 | 0.766843 | 0.697095 | 0.709251 | 0.705546 | 0.700182 | 0.713453 | 0.769582 |
| 0.762129 | 0.767751 | 0.769444 | 0.695737 | 0.707378 | 0.703659 | 0.698823 | 0.705079 | 0.769447 |
| 0.764057 | 0.767942 | 0.768374 | 0.697303 | 0.70667 | 0.700344 | 0.696022 | 0.708098 | 0.766473 |
| 0.699062 | 0.698702 | 0.698957 | 0.783891 | 0.788724 | 0.803756 | 0.803771 | 0.792892 | 0.697514 |
| 0.76482 | 0.772448 | 0.767295 | 0.698525 | 0.707917 | 0.701495 | 0.69407 | 0.708192 | 0.766703 |
| 0.763975 | 0.767079 | 0.766889 | 0.698156 | 0.710229 | 0.711939 | 0.701092 | 0.711002 | 0.771625 |
| 0.773988 | 0.770492 | 0.768139 | 0.697737 | 0.710751 | 0.705645 | 0.699164 | 0.712367 | 0.770503 |
| 0.767235 | 0.769317 | 0.766516 | 0.697477 | 0.71082 | 0.708649 | 0.695437 | 0.711159 | 0.770152 |
| 0.75842 | 0.764083 | 0.761728 | 0.695079 | 0.703722 | 0.698303 | 0.69476 | 0.707467 | 0.765405 |
| 0.763622 | 0.764877 | 0.763024 | 0.698386 | 0.706728 | 0.702922 | 0.693505 | 0.704872 | 0.766965 |
| 0.766424 | 0.766339 | 0.766044 | 0.703025 | 0.708111 | 0.70427 | 0.694952 | 0.708249 | 0.762053 |
| 0.766409 | 0.770063 | 0.76301 | 0.693953 | 0.704779 | 0.703921 | 0.69465 | 0.703553 | 0.766581 |
| 0.756913 | 0.759767 | 0.758552 | 0.691955 | 0.70313 | 0.702888 | 0.693633 | 0.7064 | 0.764506 |
| 0.757736 | 0.760708 | 0.758358 | 0.694898 | 0.702315 | 0.703224 | 0.693949 | 0.705963 | 0.762697 |
| 0.769558 | 0.772521 | 0.76979 | 0.700955 | 0.709122 | 0.710861 | 0.699095 | 0.712471 | 0.768179 |
| 0.767394 | 0.766148 | 0.763983 | 0.6982 | 0.712988 | 0.708185 | 0.698979 | 0.710912 | 0.76952 |
| 0.764847 | 0.771087 | 0.765005 | 0.695717 | 0.709515 | 0.705229 | 0.695849 | 0.70539 | 0.763995 |
| 0.764789 | 0.766056 | 0.761347 | 0.700685 | 0.705035 | 0.707489 | 0.698453 | 0.709669 | 0.76519 |
| 0.759679 | 0.761062 | 0.75366 | 0.690128 | 0.701471 | 0.702457 | 0.690688 | 0.702761 | 0.757276 |
| 0.769497 | 0.766374 | 0.765211 | 0.697375 | 0.710164 | 0.710375 | 0.699051 | 0.71206 | 0.768166 |
| 0.7587 | 0.765312 | 0.763533 | 0.697917 | 0.705853 | 0.704658 | 0.696663 | 0.705202 | 0.759831 |
| 0.759611 | 0.761614 | 0.763208 | 0.691653 | 0.708904 | 0.700637 | 0.69365 | 0.704624 | 0.764766 |
| 0.774487 | 0.771741 | 0.77189 | 0.695941 | 0.710303 | 0.705001 | 0.69637 | 0.709739 | 0.776985 |
| 0.766508 | 0.76825 | 0.768007 | 0.695192 | 0.709831 | 0.703336 | 0.697935 | 0.711035 | 0.776105 |

| | | | | | | | | |
|---|---|---|---|---|---|---|---|---|
| 0.763187 | 0.763142 | 0.766366 | 0.693947 | 0.708223 | 0.706522 | 0.693595 | 0.704531 | 0.766252 |
| 0.772627 | 0.772665 | 0.76721 | 0.696499 | 0.706847 | 0.704469 | 0.695283 | 0.707565 | 0.780655 |
| 0.764964 | 0.767534 | 0.762235 | 0.69287 | 0.708523 | 0.704911 | 0.694971 | 0.706033 | 0.764676 |
| 0.710612 | 0.72583 | 0.730147 | 0.657885 | 0.658714 | 0.6642 | 0.656318 | 0.665742 | 0.731681 |
| 0.763883 | 0.761796 | 0.760701 | 0.696656 | 0.706498 | 0.700794 | 0.692079 | 0.703309 | 0.761344 |
| 0.764566 | 0.765763 | 0.763819 | 0.690896 | 0.703501 | 0.706533 | 0.692396 | 0.704971 | 0.767099 |
| 0.764801 | 0.767039 | 0.761047 | 0.695812 | 0.703188 | 0.704278 | 0.696372 | 0.707234 | 0.77061 |
| 0.769814 | 0.76595 | 0.761765 | 0.692934 | 0.704703 | 0.699459 | 0.693653 | 0.70699 | 0.765783 |
| 0.769355 | 0.771118 | 0.766652 | 0.698359 | 0.711935 | 0.708093 | 0.696339 | 0.7109 | 0.773493 |
| 0.766438 | 0.76585 | 0.766571 | 0.696069 | 0.706217 | 0.704751 | 0.698613 | 0.71047 | 0.772448 |
| 0.767672 | 0.765999 | 0.764849 | 0.694907 | 0.705428 | 0.703119 | 0.694315 | 0.707848 | 0.769408 |
| 0.767949 | 0.771168 | 0.767529 | 0.700398 | 0.708801 | 0.706903 | 0.697053 | 0.711868 | 0.770881 |
| 0.768301 | 0.767244 | 0.767492 | 0.698868 | 0.711662 | 0.708487 | 0.698326 | 0.711407 | 0.766998 |
| 0.761776 | 0.76971 | 0.765512 | 0.693157 | 0.706781 | 0.706582 | 0.69876 | 0.707615 | 0.772098 |
| 0.771545 | 0.772766 | 0.767067 | 0.699844 | 0.711631 | 0.709975 | 0.698065 | 0.710111 | 0.77029 |
| 0.762552 | 0.765602 | 0.763803 | 0.691919 | 0.705873 | 0.702327 | 0.688762 | 0.704136 | 0.763771 |
| 0.770842 | 0.769042 | 0.76864 | 0.697415 | 0.709562 | 0.706999 | 0.694408 | 0.711068 | 0.769198 |
| 0.768161 | 0.773907 | 0.766175 | 0.697996 | 0.707233 | 0.708964 | 0.6964 | 0.705531 | 0.769753 |
| 0.76627 | 0.765945 | 0.760779 | 0.686359 | 0.702162 | 0.701285 | 0.691768 | 0.704051 | 0.767175 |
| 0.765703 | 0.765959 | 0.765991 | 0.694895 | 0.705267 | 0.703902 | 0.694493 | 0.708714 | 0.768544 |
| 0.765715 | 0.769783 | 0.76742 | 0.694681 | 0.711328 | 0.706104 | 0.698775 | 0.70787 | 0.772153 |
| 0.772921 | 0.77555 | 0.768882 | 0.699789 | 0.70807 | 0.706456 | 0.695313 | 0.710072 | 0.768101 |
| 0.674921 | 0.679055 | 0.678135 | 0.68767 | 0.692774 | 0.692209 | 0.685084 | 0.6875 | 0.68098 |
| 0.656 | 0.658058 | 0.650565 | 0.713771 | 0.713548 | 0.720228 | 0.725051 | 0.705669 | 0.64626 |
| 0.673094 | 0.668712 | 0.674685 | 0.742166 | 0.759591 | 0.755927 | 0.747962 | 0.761137 | 0.675078 |
| 0.653538 | 0.659384 | 0.649725 | 0.733674 | 0.742975 | 0.741002 | 0.80233 | 0.748116 | 0.651901 |
| 0.741573 | 0.7449 | 0.744747 | 0.656032 | 0.659959 | 0.659574 | 0.661596 | 0.663751 | 0.747554 |
| 0.727166 | 0.73121 | 0.728697 | 0.655369 | 0.65836 | 0.659432 | 0.657331 | 0.659707 | 0.740972 |
| 0.735658 | 0.745891 | 0.737936 | 0.660926 | 0.660328 | 0.656157 | 0.652925 | 0.652988 | 0.732854 |
| 0.737324 | 0.729377 | 0.737465 | 0.65728 | 0.680065 | 0.668548 | 0.669452 | 0.676459 | 0.749894 |
| 0.715888 | 0.722635 | 0.72337 | 0.654045 | 0.656091 | 0.645584 | 0.642417 | 0.655507 | 0.728631 |
| 0.721658 | 0.718394 | 0.715069 | 0.634621 | 0.643265 | 0.639967 | 0.639841 | 0.638986 | 0.728843 |
| 0.72374 | 0.720923 | 0.721208 | 0.64307 | 0.635641 | 0.64494 | 0.639528 | 0.63593 | 0.731141 |
| 0.653498 | 0.660384 | 0.654691 | 0.736204 | 0.740011 | 0.725134 | 0.743105 | 0.733616 | 0.648915 |
| 0.762697 | 0.765887 | 0.766454 | 0.68944 | 0.700572 | 0.699043 | 0.692343 | 0.697981 | 0.767326 |
| 0.747246 | 0.754753 | 0.75012 | 0.672268 | 0.674427 | 0.66881 | 0.668642 | 0.67117 | 0.75 |
| 0.739586 | 0.748029 | 0.736326 | 0.667748 | 0.665446 | 0.665709 | 0.664802 | 0.663392 | 0.743969 |
| 0.728032 | 0.728778 | 0.734771 | 0.650066 | 0.650458 | 0.649409 | 0.651141 | 0.645881 | 0.73872 |
| 0.744844 | 0.748775 | 0.745793 | 0.669239 | 0.670993 | 0.668741 | 0.670735 | 0.671894 | 0.751824 |
| 0.7282 | 0.733229 | 0.729865 | 0.664826 | 0.665627 | 0.661296 | 0.66497 | 0.668227 | 0.733169 |
| 0.734482 | 0.739832 | 0.730826 | 0.660458 | 0.655803 | 0.649801 | 0.651029 | 0.658816 | 0.730312 |
| 0.725087 | 0.732775 | 0.735922 | 0.652508 | 0.666768 | 0.655193 | 0.658323 | 0.661119 | 0.729956 |
| 0.731203 | 0.736081 | 0.739703 | 0.649483 | 0.652947 | 0.653338 | 0.649211 | 0.646584 | 0.72898 |
| 0.731678 | 0.733693 | 0.732114 | 0.658586 | 0.657834 | 0.651752 | 0.657459 | 0.659328 | 0.730769 |
| 0.720615 | 0.729568 | 0.731433 | 0.652731 | 0.653357 | 0.651616 | 0.65736 | 0.653429 | 0.731612 |
| 0.72585 | 0.723411 | 0.718521 | 0.64191 | 0.640764 | 0.635407 | 0.646445 | 0.640227 | 0.727364 |

| | | | | | | | | |
|---|---|---|---|---|---|---|---|---|
| 0.750433 | 0.755104 | 0.758763 | 0.680482 | 0.687784 | 0.685012 | 0.683055 | 0.683965 | 0.761987 |
| 0.727534 | 0.733438 | 0.731373 | 0.653907 | 0.653763 | 0.652156 | 0.64827 | 0.658058 | 0.739734 |
| 0.734775 | 0.738706 | 0.732821 | 0.654568 | 0.660125 | 0.655641 | 0.657414 | 0.6616 | 0.733596 |
| 0.730926 | 0.734992 | 0.729942 | 0.656077 | 0.655249 | 0.654283 | 0.66017 | 0.660756 | 0.74193 |
| 0.732614 | 0.740367 | 0.733762 | 0.652401 | 0.651789 | 0.646325 | 0.662942 | 0.650301 | 0.733636 |
| 0.735347 | 0.735968 | 0.727067 | 0.667688 | 0.661825 | 0.658478 | 0.662594 | 0.658152 | 0.730187 |
| 0.721728 | 0.724663 | 0.723441 | 0.649472 | 0.648585 | 0.648105 | 0.645764 | 0.651244 | 0.740292 |
| 0.724667 | 0.720534 | 0.721526 | 0.63803 | 0.645908 | 0.637524 | 0.646266 | 0.641067 | 0.726961 |
| 0.732501 | 0.732117 | 0.731326 | 0.643367 | 0.651374 | 0.640893 | 0.641504 | 0.641525 | 0.743162 |
| 0.753574 | 0.755504 | 0.754073 | 0.683667 | 0.687231 | 0.682425 | 0.686343 | 0.685663 | 0.761603 |
| 0.749432 | 0.755271 | 0.756397 | 0.682132 | 0.685378 | 0.681849 | 0.681414 | 0.686894 | 0.75482 |
| 0.748413 | 0.750311 | 0.748573 | 0.675692 | 0.678875 | 0.677354 | 0.677011 | 0.680387 | 0.751384 |
| 0.74086 | 0.744923 | 0.742775 | 0.673488 | 0.676551 | 0.675106 | 0.674991 | 0.672114 | 0.737508 |
| 0.746278 | 0.748251 | 0.745093 | 0.67914 | 0.673802 | 0.677886 | 0.676463 | 0.675717 | 0.745113 |
| 0.743392 | 0.745544 | 0.742647 | 0.673698 | 0.668054 | 0.669083 | 0.671873 | 0.666354 | 0.743721 |

| plot 11 | | | | | | | | |
|---|---|---|---|---|---|---|---|---|
| Wild_plot_ | Wild_plot_ | Wild_plot_ | Wild_plot_ | Wild_plot_ | Wild_plot_ | Wild_plot_ | Wild_plot_ | Wild_plot_ |
| 0.703135 | 0.705156 | 0.710352 | 0.703124 | 0.709546 | 0.704918 | 0.70653 | 0.698374 | 0.700615 |
| 0.71179 | 0.712481 | 0.711301 | 0.709693 | 0.715078 | 0.711874 | 0.71101 | 0.705803 | 0.704047 |
| 0.699784 | 0.703345 | 0.701989 | 0.700231 | 0.70143 | 0.699369 | 0.700289 | 0.695235 | 0.696368 |
| 0.709062 | 0.711444 | 0.71249 | 0.706121 | 0.711816 | 0.711001 | 0.70921 | 0.706213 | 0.706891 |
| 0.704871 | 0.706622 | 0.708675 | 0.703916 | 0.706773 | 0.706553 | 0.705086 | 0.700939 | 0.701238 |
| 0.712851 | 0.714962 | 0.719821 | 0.715671 | 0.716438 | 0.713689 | 0.715233 | 0.707758 | 0.706088 |
| 0.770993 | 0.771341 | 0.77165 | 0.770211 | 0.769903 | 0.768796 | 0.763353 | 0.764038 | 0.76569 |
| 0.770592 | 0.769942 | 0.770081 | 0.766334 | 0.767987 | 0.767628 | 0.759563 | 0.760138 | 0.760664 |
| 0.709863 | 0.712012 | 0.714446 | 0.707822 | 0.713711 | 0.710675 | 0.708384 | 0.705247 | 0.702916 |
| 0.705193 | 0.703189 | 0.708244 | 0.702263 | 0.707492 | 0.704535 | 0.705308 | 0.69736 | 0.701597 |
| 0.704152 | 0.707183 | 0.707799 | 0.70324 | 0.708996 | 0.707081 | 0.707206 | 0.703238 | 0.701723 |
| 0.698877 | 0.70201 | 0.701605 | 0.69639 | 0.70302 | 0.698032 | 0.69946 | 0.695361 | 0.694544 |
| 0.708307 | 0.711937 | 0.714045 | 0.70954 | 0.712303 | 0.71387 | 0.709764 | 0.702716 | 0.706054 |
| 0.720697 | 0.720915 | 0.721504 | 0.719752 | 0.720251 | 0.720642 | 0.720667 | 0.713852 | 0.715381 |
| 0.726862 | 0.731903 | 0.734617 | 0.731963 | 0.732496 | 0.732397 | 0.733388 | 0.726127 | 0.723604 |
| 0.708096 | 0.712655 | 0.714994 | 0.70983 | 0.713623 | 0.710075 | 0.713846 | 0.706853 | 0.705197 |
| 0.710067 | 0.710908 | 0.713445 | 0.708905 | 0.714182 | 0.71121 | 0.713772 | 0.705504 | 0.70395 |
| 0.740178 | 0.742068 | 0.740753 | 0.738868 | 0.746172 | 0.741316 | 0.740407 | 0.736567 | 0.730614 |
| 0.703234 | 0.703047 | 0.703591 | 0.702119 | 0.705102 | 0.70091 | 0.70141 | 0.699078 | 0.696618 |
| 0.705915 | 0.708625 | 0.713577 | 0.70729 | 0.713106 | 0.709493 | 0.707121 | 0.700008 | 0.701616 |
| 0.703481 | 0.707715 | 0.710997 | 0.70339 | 0.705709 | 0.704195 | 0.705095 | 0.69872 | 0.697502 |
| 0.708565 | 0.712719 | 0.713919 | 0.705717 | 0.712582 | 0.708384 | 0.707274 | 0.701824 | 0.705266 |
| 0.708846 | 0.714816 | 0.715051 | 0.71083 | 0.715499 | 0.711971 | 0.709427 | 0.706625 | 0.704124 |
| 0.712521 | 0.712614 | 0.713473 | 0.707962 | 0.712031 | 0.710369 | 0.712711 | 0.706756 | 0.704224 |
| 0.709125 | 0.710261 | 0.712693 | 0.709488 | 0.715173 | 0.710732 | 0.709007 | 0.70337 | 0.708441 |
| 0.708233 | 0.709206 | 0.708588 | 0.70358 | 0.712056 | 0.708687 | 0.709363 | 0.701009 | 0.699041 |
| 0.703554 | 0.704998 | 0.706821 | 0.702354 | 0.705262 | 0.703784 | 0.703227 | 0.6977 | 0.698512 |
| 0.707221 | 0.708243 | 0.709264 | 0.707977 | 0.708864 | 0.70838 | 0.70886 | 0.702757 | 0.698572 |
| 0.707315 | 0.710249 | 0.713714 | 0.708654 | 0.715996 | 0.714286 | 0.711145 | 0.705724 | 0.703431 |
| 0.707453 | 0.706961 | 0.710414 | 0.706763 | 0.70924 | 0.707471 | 0.710473 | 0.704896 | 0.705209 |
| 0.703611 | 0.704889 | 0.706056 | 0.70387 | 0.707206 | 0.705233 | 0.708597 | 0.699294 | 0.700086 |
| 0.703582 | 0.703612 | 0.706611 | 0.70247 | 0.706285 | 0.705854 | 0.705427 | 0.696825 | 0.698563 |
| 0.705223 | 0.705685 | 0.711382 | 0.70624 | 0.709654 | 0.707621 | 0.707715 | 0.699407 | 0.701434 |
| 0.706325 | 0.707916 | 0.709252 | 0.706 | 0.710596 | 0.707698 | 0.708642 | 0.703601 | 0.701632 |
| 0.699968 | 0.704198 | 0.704593 | 0.700496 | 0.705868 | 0.701437 | 0.699088 | 0.695331 | 0.697078 |
| 0.704123 | 0.708192 | 0.708136 | 0.705338 | 0.7091 | 0.708087 | 0.70972 | 0.703018 | 0.699159 |
| 0.71086 | 0.713178 | 0.717977 | 0.713123 | 0.716268 | 0.713241 | 0.715579 | 0.707235 | 0.705852 |
| 0.710528 | 0.709586 | 0.712233 | 0.711337 | 0.713429 | 0.710803 | 0.709839 | 0.705952 | 0.70466 |
| 0.709698 | 0.708965 | 0.713909 | 0.71028 | 0.713553 | 0.71031 | 0.708468 | 0.706586 | 0.705571 |

| | | | | | | | | |
|---|---|---|---|---|---|---|---|---|
| 0.744475 | 0.745138 | 0.743013 | 0.742391 | 0.747175 | 0.75053 | 0.74306 | 0.741338 | 0.733602 |
| 0.70196 | 0.701986 | 0.702598 | 0.69559 | 0.703034 | 0.701829 | 0.700701 | 0.694827 | 0.692563 |
| 0.703486 | 0.706138 | 0.708247 | 0.70352 | 0.70824 | 0.704157 | 0.702222 | 0.699782 | 0.699082 |
| 0.711434 | 0.714023 | 0.716471 | 0.710572 | 0.715276 | 0.711387 | 0.713917 | 0.704019 | 0.705071 |
| 0.706356 | 0.708658 | 0.71035 | 0.707623 | 0.712087 | 0.70754 | 0.707623 | 0.704099 | 0.700513 |
| 0.708681 | 0.710714 | 0.713332 | 0.706972 | 0.714366 | 0.711318 | 0.711316 | 0.705634 | 0.705659 |
| 0.702932 | 0.706203 | 0.708052 | 0.703562 | 0.707676 | 0.706062 | 0.705001 | 0.699448 | 0.699464 |
| 0.710161 | 0.706959 | 0.709887 | 0.7068 | 0.712075 | 0.708908 | 0.707256 | 0.704933 | 0.703091 |
| 0.705759 | 0.712325 | 0.70844 | 0.705428 | 0.711785 | 0.710371 | 0.711557 | 0.704011 | 0.700611 |
| 0.710913 | 0.713014 | 0.711357 | 0.704061 | 0.712929 | 0.71013 | 0.710182 | 0.705631 | 0.702829 |
| 0.701227 | 0.701352 | 0.704751 | 0.698362 | 0.704135 | 0.701556 | 0.701321 | 0.694634 | 0.696913 |
| 0.700346 | 0.702414 | 0.706915 | 0.7002 | 0.703401 | 0.701303 | 0.704289 | 0.69749 | 0.697431 |
| 0.70646 | 0.704902 | 0.707898 | 0.705579 | 0.705097 | 0.706397 | 0.703471 | 0.700745 | 0.701545 |
| 0.70075 | 0.702858 | 0.70828 | 0.701245 | 0.704859 | 0.705492 | 0.700917 | 0.696676 | 0.696447 |
| 0.702959 | 0.70739 | 0.705338 | 0.703758 | 0.70916 | 0.703912 | 0.705061 | 0.70079 | 0.698188 |
| 0.704312 | 0.704918 | 0.70603 | 0.702389 | 0.706843 | 0.705815 | 0.70199 | 0.699123 | 0.696112 |
| 0.70026 | 0.702426 | 0.705778 | 0.701471 | 0.700483 | 0.698435 | 0.700662 | 0.697317 | 0.697282 |
| 0.708199 | 0.706563 | 0.711483 | 0.707243 | 0.707438 | 0.707182 | 0.709544 | 0.703128 | 0.702765 |
| 0.709154 | 0.709532 | 0.713703 | 0.71026 | 0.71235 | 0.711845 | 0.712973 | 0.705392 | 0.701754 |
| 0.704872 | 0.704604 | 0.708943 | 0.701305 | 0.705703 | 0.707559 | 0.705164 | 0.699467 | 0.698392 |
| 0.701477 | 0.704074 | 0.703051 | 0.700313 | 0.703727 | 0.700185 | 0.701135 | 0.696903 | 0.694114 |
| 0.703047 | 0.706186 | 0.707802 | 0.702158 | 0.705543 | 0.70342 | 0.704118 | 0.697168 | 0.700575 |
| 0.70434 | 0.706164 | 0.708462 | 0.704096 | 0.70779 | 0.705892 | 0.706873 | 0.699232 | 0.700196 |
| 0.71147 | 0.716133 | 0.7201 | 0.713814 | 0.719368 | 0.712473 | 0.713936 | 0.707173 | 0.710116 |
| 0.722523 | 0.721133 | 0.725493 | 0.719589 | 0.723065 | 0.721533 | 0.722516 | 0.715527 | 0.716346 |
| 0.7729 | 0.770276 | 0.771902 | 0.764417 | 0.76868 | 0.768011 | 0.762245 | 0.761726 | 0.76278 |
| 0.772161 | 0.771591 | 0.772478 | 0.773135 | 0.769686 | 0.774637 | 0.766133 | 0.770181 | 0.763847 |
| 0.773442 | 0.773764 | 0.775726 | 0.77027 | 0.772981 | 0.774088 | 0.768123 | 0.770503 | 0.765335 |
| 0.704827 | 0.70689 | 0.706723 | 0.703944 | 0.707979 | 0.706898 | 0.70495 | 0.695151 | 0.699402 |
| 0.703217 | 0.70404 | 0.705776 | 0.700986 | 0.707341 | 0.703682 | 0.701536 | 0.697696 | 0.699055 |
| 0.712088 | 0.718186 | 0.718793 | 0.71208 | 0.715893 | 0.713869 | 0.713112 | 0.704404 | 0.706348 |
| 0.714344 | 0.717196 | 0.715631 | 0.712764 | 0.715097 | 0.717987 | 0.715767 | 0.707417 | 0.709166 |
| 0.703912 | 0.70352 | 0.708559 | 0.707353 | 0.709779 | 0.704233 | 0.705112 | 0.70251 | 0.701677 |
| 0.705587 | 0.707432 | 0.705689 | 0.705764 | 0.709677 | 0.706707 | 0.705359 | 0.701642 | 0.70031 |
| 0.70235 | 0.701203 | 0.70365 | 0.702471 | 0.709407 | 0.70352 | 0.703106 | 0.700077 | 0.696435 |
| 0.704884 | 0.705175 | 0.706562 | 0.707103 | 0.708842 | 0.704278 | 0.7054 | 0.701443 | 0.699458 |
| 0.701698 | 0.705498 | 0.705938 | 0.704621 | 0.71249 | 0.708067 | 0.701714 | 0.699042 | 0.698421 |
| 0.71739 | 0.720376 | 0.721426 | 0.720213 | 0.724408 | 0.719435 | 0.71968 | 0.71338 | 0.71351 |
| 0.698357 | 0.696289 | 0.699038 | 0.699701 | 0.702947 | 0.695739 | 0.698365 | 0.694889 | 0.690659 |
| 0.705709 | 0.706684 | 0.708881 | 0.706817 | 0.713264 | 0.704892 | 0.708711 | 0.698013 | 0.699839 |
| 0.699537 | 0.700804 | 0.705262 | 0.701458 | 0.70788 | 0.703033 | 0.70349 | 0.696083 | 0.694987 |
| 0.72056 | 0.723283 | 0.724524 | 0.721989 | 0.72579 | 0.723094 | 0.723291 | 0.715689 | 0.713978 |
| 0.705497 | 0.707016 | 0.709639 | 0.704146 | 0.711889 | 0.705555 | 0.707181 | 0.701553 | 0.700523 |
| 0.703639 | 0.706056 | 0.707443 | 0.702795 | 0.707701 | 0.704842 | 0.707162 | 0.696558 | 0.698394 |
| 0.705539 | 0.709059 | 0.708183 | 0.703665 | 0.712293 | 0.708134 | 0.71234 | 0.699713 | 0.700112 |
| 0.705118 | 0.707569 | 0.706797 | 0.706172 | 0.707692 | 0.704466 | 0.705167 | 0.700874 | 0.703512 |

| | | | | | | | | |
|---|---|---|---|---|---|---|---|---|
| 0.704445 | 0.703898 | 0.708878 | 0.707068 | 0.707323 | 0.704663 | 0.705692 | 0.695422 | 0.701613 |
| 0.707292 | 0.711694 | 0.714537 | 0.706705 | 0.71408 | 0.708976 | 0.708884 | 0.704984 | 0.703672 |
| 0.704573 | 0.709698 | 0.709956 | 0.708138 | 0.710605 | 0.710925 | 0.705777 | 0.699272 | 0.698908 |
| 0.711556 | 0.714507 | 0.715697 | 0.713786 | 0.718851 | 0.713608 | 0.712572 | 0.70693 | 0.703761 |
| 0.710659 | 0.712049 | 0.716236 | 0.711437 | 0.71743 | 0.715254 | 0.712222 | 0.70834 | 0.706516 |
| 0.703461 | 0.706216 | 0.711152 | 0.705669 | 0.715094 | 0.706947 | 0.7078 | 0.69866 | 0.701915 |
| 0.705003 | 0.706431 | 0.707864 | 0.705442 | 0.711788 | 0.706005 | 0.704266 | 0.700827 | 0.698649 |
| 0.710139 | 0.712687 | 0.713065 | 0.70869 | 0.71074 | 0.711986 | 0.707142 | 0.705618 | 0.706167 |
| 0.705468 | 0.70861 | 0.70825 | 0.704848 | 0.706237 | 0.71001 | 0.70884 | 0.700743 | 0.69969 |
| 0.704228 | 0.7068 | 0.706998 | 0.703662 | 0.704252 | 0.703344 | 0.705935 | 0.701824 | 0.702687 |
| 0.703565 | 0.702976 | 0.707768 | 0.703721 | 0.709601 | 0.703313 | 0.705038 | 0.699142 | 0.700093 |
| 0.703317 | 0.707295 | 0.712376 | 0.707382 | 0.710096 | 0.706866 | 0.704823 | 0.700449 | 0.700623 |
| 0.704568 | 0.705025 | 0.708869 | 0.704914 | 0.706176 | 0.705406 | 0.707277 | 0.698024 | 0.697722 |
| 0.701591 | 0.704547 | 0.706955 | 0.703595 | 0.708249 | 0.703837 | 0.70282 | 0.699267 | 0.697727 |
| 0.706343 | 0.707098 | 0.711717 | 0.707024 | 0.708641 | 0.706802 | 0.708631 | 0.703419 | 0.698755 |
| 0.705651 | 0.705566 | 0.706641 | 0.70228 | 0.706965 | 0.707731 | 0.703674 | 0.699288 | 0.696381 |
| 0.709174 | 0.706653 | 0.707755 | 0.707362 | 0.710953 | 0.709095 | 0.702345 | 0.701398 | 0.698993 |
| 0.705944 | 0.705166 | 0.708536 | 0.708661 | 0.71249 | 0.704761 | 0.707132 | 0.699446 | 0.699906 |
| 0.705759 | 0.708283 | 0.712953 | 0.708772 | 0.710885 | 0.708961 | 0.704862 | 0.701496 | 0.698144 |
| 0.705664 | 0.706048 | 0.710385 | 0.704381 | 0.709864 | 0.709029 | 0.707181 | 0.701955 | 0.702582 |
| 0.707112 | 0.710064 | 0.706279 | 0.708827 | 0.712103 | 0.710059 | 0.706939 | 0.698779 | 0.703309 |
| 0.701302 | 0.702425 | 0.703255 | 0.703271 | 0.704614 | 0.699188 | 0.702569 | 0.694454 | 0.69618 |
| 0.702305 | 0.705645 | 0.706181 | 0.705617 | 0.709579 | 0.707301 | 0.706616 | 0.700711 | 0.697531 |
| 0.707258 | 0.709812 | 0.711833 | 0.709623 | 0.712617 | 0.709449 | 0.708455 | 0.699239 | 0.703196 |
| 0.706711 | 0.711448 | 0.710705 | 0.710507 | 0.711774 | 0.705705 | 0.706757 | 0.700449 | 0.698915 |
| 0.712132 | 0.711598 | 0.716498 | 0.709001 | 0.714559 | 0.714768 | 0.711097 | 0.706581 | 0.705837 |
| 0.702698 | 0.707161 | 0.709772 | 0.708663 | 0.712429 | 0.70355 | 0.705155 | 0.700217 | 0.698766 |
| 0.70729 | 0.710037 | 0.713098 | 0.70789 | 0.709886 | 0.71063 | 0.708168 | 0.704154 | 0.701119 |
| 0.707229 | 0.705661 | 0.715923 | 0.708 | 0.710699 | 0.709579 | 0.707785 | 0.703284 | 0.706442 |
| 0.708313 | 0.709293 | 0.711881 | 0.711795 | 0.714669 | 0.709002 | 0.706412 | 0.703606 | 0.701961 |
| 0.712924 | 0.709319 | 0.710857 | 0.707984 | 0.715797 | 0.710616 | 0.70948 | 0.701701 | 0.702395 |
| 0.703018 | 0.704407 | 0.70918 | 0.707877 | 0.710655 | 0.706568 | 0.704609 | 0.695706 | 0.698418 |
| 0.708306 | 0.709341 | 0.708984 | 0.710667 | 0.71446 | 0.708272 | 0.709929 | 0.705185 | 0.702745 |
| 0.707394 | 0.710485 | 0.712586 | 0.708731 | 0.71391 | 0.708306 | 0.71076 | 0.700627 | 0.702445 |
| 0.70458 | 0.706206 | 0.704164 | 0.70474 | 0.707807 | 0.705073 | 0.707347 | 0.695004 | 0.696527 |
| 0.706663 | 0.708858 | 0.709265 | 0.706643 | 0.713785 | 0.704748 | 0.706482 | 0.699369 | 0.700859 |
| 0.698764 | 0.698852 | 0.702506 | 0.698213 | 0.701724 | 0.699086 | 0.702907 | 0.689511 | 0.690783 |
| 0.68265 | 0.680847 | 0.68174 | 0.680069 | 0.686554 | 0.680982 | 0.678299 | 0.678112 | 0.681389 |
| 0.686019 | 0.6915 | 0.688824 | 0.68533 | 0.695295 | 0.685912 | 0.689888 | 0.680976 | 0.682034 |
| 0.678256 | 0.680099 | 0.682179 | 0.67722 | 0.682005 | 0.680485 | 0.680357 | 0.674126 | 0.676005 |
| 0.676251 | 0.679786 | 0.678096 | 0.672458 | 0.680827 | 0.679786 | 0.679431 | 0.672642 | 0.676912 |
| 0.664623 | 0.663384 | 0.662304 | 0.663354 | 0.664915 | 0.665568 | 0.662284 | 0.66152 | 0.662888 |
| 0.662505 | 0.660839 | 0.665378 | 0.662687 | 0.668354 | 0.66772 | 0.661505 | 0.657104 | 0.665374 |
| 0.660792 | 0.662671 | 0.669013 | 0.661076 | 0.665738 | 0.663586 | 0.665002 | 0.660047 | 0.662506 |
| 0.665511 | 0.667306 | 0.662548 | 0.663155 | 0.668445 | 0.66641 | 0.670255 | 0.665219 | 0.662622 |
| 0.6629 | 0.662928 | 0.66621 | 0.66252 | 0.668534 | 0.66579 | 0.660307 | 0.663159 | 0.664517 |
| 0.669396 | 0.669342 | 0.667609 | 0.663349 | 0.674174 | 0.669145 | 0.669646 | 0.667092 | 0.665558 |

| | | | | | | | | |
|---|---|---|---|---|---|---|---|---|
| 0.671336 | 0.669588 | 0.668716 | 0.663888 | 0.673442 | 0.669845 | 0.66972 | 0.661552 | 0.668371 |
| 0.669679 | 0.668968 | 0.66671 | 0.666494 | 0.675796 | 0.66541 | 0.672996 | 0.667964 | 0.66888 |
| 0.663137 | 0.665017 | 0.670371 | 0.670645 | 0.670136 | 0.666087 | 0.667068 | 0.660643 | 0.663859 |
| 0.665649 | 0.65853 | 0.663672 | 0.658241 | 0.667282 | 0.663218 | 0.66272 | 0.655691 | 0.658619 |
| 0.663177 | 0.662344 | 0.663801 | 0.658636 | 0.66581 | 0.665289 | 0.6601 | 0.660521 | 0.657989 |
| 0.6591 | 0.665015 | 0.665656 | 0.661279 | 0.666118 | 0.664144 | 0.657671 | 0.658822 | 0.657705 |
| 0.661872 | 0.662867 | 0.662367 | 0.661634 | 0.665066 | 0.662212 | 0.660474 | 0.659417 | 0.657472 |
| 0.65989 | 0.660794 | 0.65918 | 0.661071 | 0.663863 | 0.664082 | 0.6591 | 0.656804 | 0.659593 |
| 0.660701 | 0.664949 | 0.657625 | 0.658343 | 0.660568 | 0.661309 | 0.658515 | 0.65445 | 0.659067 |
| 0.662835 | 0.665152 | 0.661992 | 0.659788 | 0.667644 | 0.658316 | 0.660003 | 0.657083 | 0.654043 |
| 0.654576 | 0.654704 | 0.659825 | 0.655941 | 0.66005 | 0.658442 | 0.652326 | 0.655368 | 0.653773 |
| 0.696124 | 0.697612 | 0.700331 | 0.696666 | 0.700442 | 0.697242 | 0.695059 | 0.687784 | 0.689911 |
| 0.707069 | 0.707811 | 0.708668 | 0.706636 | 0.712965 | 0.707342 | 0.708471 | 0.700217 | 0.699116 |
| 0.679795 | 0.683394 | 0.683402 | 0.682882 | 0.687251 | 0.681188 | 0.678584 | 0.679223 | 0.680547 |
| 0.709509 | 0.713506 | 0.712687 | 0.712962 | 0.716618 | 0.71273 | 0.70973 | 0.706178 | 0.706347 |
| 0.702073 | 0.705068 | 0.703772 | 0.703653 | 0.707374 | 0.704457 | 0.701318 | 0.695594 | 0.692682 |
| 0.70826 | 0.711662 | 0.711309 | 0.706468 | 0.70939 | 0.710617 | 0.710881 | 0.705611 | 0.702113 |
| 0.717331 | 0.717982 | 0.721314 | 0.720225 | 0.717565 | 0.716406 | 0.717286 | 0.711496 | 0.708708 |
| 0.695597 | 0.692149 | 0.695467 | 0.692689 | 0.697233 | 0.690288 | 0.694044 | 0.686636 | 0.688036 |
| 0.714148 | 0.72017 | 0.717832 | 0.715037 | 0.723499 | 0.716857 | 0.714758 | 0.710847 | 0.709584 |
| 0.682032 | 0.688623 | 0.690942 | 0.688276 | 0.687755 | 0.683994 | 0.681906 | 0.684971 | 0.683565 |
| 0.711032 | 0.714344 | 0.717901 | 0.714933 | 0.718964 | 0.712391 | 0.712936 | 0.713643 | 0.708315 |
| 0.715685 | 0.719485 | 0.720923 | 0.716839 | 0.718259 | 0.717672 | 0.718684 | 0.710974 | 0.714458 |
| 0.678688 | 0.685082 | 0.681214 | 0.675735 | 0.687318 | 0.681234 | 0.682146 | 0.677098 | 0.672659 |
| 0.683948 | 0.688489 | 0.684946 | 0.687225 | 0.691562 | 0.68701 | 0.68604 | 0.677715 | 0.676992 |
| 0.688182 | 0.686047 | 0.689546 | 0.687484 | 0.688187 | 0.687143 | 0.688014 | 0.685624 | 0.68905 |
| 0.69977 | 0.709073 | 0.705781 | 0.69919 | 0.706007 | 0.70564 | 0.703759 | 0.700198 | 0.69808 |
| 0.715727 | 0.715975 | 0.71888 | 0.713312 | 0.720813 | 0.716015 | 0.714529 | 0.706416 | 0.70739 |
| 0.68573 | 0.687623 | 0.689419 | 0.683807 | 0.68986 | 0.686213 | 0.687723 | 0.676127 | 0.678123 |
| 0.720035 | 0.723304 | 0.723426 | 0.722316 | 0.723486 | 0.718617 | 0.718251 | 0.713999 | 0.714359 |
| 0.718563 | 0.72389 | 0.724556 | 0.720419 | 0.725632 | 0.72153 | 0.721861 | 0.715691 | 0.715146 |
| 0.693661 | 0.695586 | 0.698562 | 0.697314 | 0.700232 | 0.695785 | 0.695561 | 0.687724 | 0.688585 |
| 0.693068 | 0.692565 | 0.696946 | 0.697601 | 0.701159 | 0.694396 | 0.695126 | 0.684712 | 0.688279 |
| 0.719761 | 0.723901 | 0.72367 | 0.718402 | 0.727258 | 0.721655 | 0.72108 | 0.713901 | 0.71474 |
| 0.718501 | 0.720423 | 0.722042 | 0.720571 | 0.723778 | 0.720105 | 0.719408 | 0.711822 | 0.715484 |
| 0.721608 | 0.723409 | 0.724725 | 0.721101 | 0.72898 | 0.722091 | 0.724889 | 0.715266 | 0.718478 |
| 0.673068 | 0.675873 | 0.673584 | 0.674288 | 0.679887 | 0.676027 | 0.674931 | 0.669287 | 0.664007 |
| 0.717397 | 0.714446 | 0.721888 | 0.719971 | 0.720897 | 0.720919 | 0.717638 | 0.710851 | 0.714359 |
| 0.671759 | 0.67452 | 0.674644 | 0.674293 | 0.680118 | 0.673705 | 0.672818 | 0.665377 | 0.666996 |
| 0.667689 | 0.669529 | 0.669128 | 0.667712 | 0.668903 | 0.667946 | 0.666744 | 0.662136 | 0.661855 |
| 0.717528 | 0.716744 | 0.71948 | 0.717836 | 0.725708 | 0.719997 | 0.717688 | 0.713339 | 0.712391 |
| 0.665593 | 0.669099 | 0.669643 | 0.670365 | 0.669946 | 0.665388 | 0.668369 | 0.665223 | 0.663344 |
| 0.657835 | 0.660243 | 0.659515 | 0.659204 | 0.66109 | 0.657849 | 0.658204 | 0.654303 | 0.653806 |
| 0.653621 | 0.655598 | 0.658073 | 0.659961 | 0.659772 | 0.655612 | 0.655156 | 0.65201 | 0.650835 |
| 0.723442 | 0.726781 | 0.729213 | 0.724713 | 0.73336 | 0.728546 | 0.727961 | 0.719164 | 0.718788 |
| 0.660198 | 0.66311 | 0.664353 | 0.661334 | 0.663817 | 0.660287 | 0.659561 | 0.656738 | 0.65443 |
| 0.717194 | 0.715437 | 0.720984 | 0.716091 | 0.725068 | 0.717368 | 0.715621 | 0.713067 | 0.709838 |

| | | | | | | | | |
|---|---|---|---|---|---|---|---|---|
| 0.730069 | 0.727727 | 0.727215 | 0.725821 | 0.730026 | 0.728301 | 0.728826 | 0.720249 | 0.719609 |
| 0.72189 | 0.724411 | 0.724755 | 0.726615 | 0.729891 | 0.723076 | 0.718963 | 0.718784 | 0.716042 |
| 0.742138 | 0.742939 | 0.743951 | 0.745952 | 0.747681 | 0.743069 | 0.742304 | 0.740957 | 0.737602 |
| 0.649585 | 0.650814 | 0.649341 | 0.649955 | 0.651935 | 0.650372 | 0.649323 | 0.644947 | 0.640717 |
| 0.649028 | 0.647981 | 0.648322 | 0.653369 | 0.652711 | 0.651396 | 0.648638 | 0.646793 | 0.641979 |
| 0.728926 | 0.729108 | 0.72911 | 0.726789 | 0.734496 | 0.730838 | 0.727384 | 0.725357 | 0.720766 |
| 0.654232 | 0.652367 | 0.65534 | 0.65292 | 0.654752 | 0.653483 | 0.650351 | 0.643958 | 0.646381 |
| 0.734119 | 0.734041 | 0.735128 | 0.73369 | 0.733628 | 0.734445 | 0.731853 | 0.728007 | 0.725452 |
| 0.728444 | 0.727539 | 0.731492 | 0.729437 | 0.735512 | 0.731307 | 0.727258 | 0.72507 | 0.718361 |
| 0.736816 | 0.736694 | 0.737725 | 0.734348 | 0.735807 | 0.732505 | 0.73148 | 0.728859 | 0.729892 |
| 0.728868 | 0.729495 | 0.730092 | 0.728279 | 0.734622 | 0.731711 | 0.726221 | 0.724872 | 0.718294 |
| 0.701299 | 0.732026 | 0.660131 | 0.736842 | 0.705882 | 0.692308 | 0.769784 | 0.731544 | 0.72 |
| 0.732617 | 0.747479 | 0.741296 | 0.738719 | 0.731957 | 0.734343 | 0.734435 | 0.735923 | 0.728995 |
| 0.737412 | 0.740227 | 0.745348 | 0.737449 | 0.744681 | 0.736909 | 0.73825 | 0.741925 | 0.735729 |
| 0.746073 | 0.748671 | 0.753734 | 0.747526 | 0.749164 | 0.74577 | 0.745011 | 0.748907 | 0.748043 |
| 0.743352 | 0.748742 | 0.747167 | 0.748775 | 0.746454 | 0.738418 | 0.737442 | 0.744134 | 0.743389 |
| 0.588913 | 0.592283 | 0.592171 | 0.588818 | 0.594156 | 0.590706 | 0.589066 | 0.586957 | 0.587409 |
| 0.741141 | 0.748054 | 0.748352 | 0.745235 | 0.748752 | 0.741799 | 0.738282 | 0.736927 | 0.732913 |
| 0.734119 | 0.735152 | 0.739113 | 0.732155 | 0.738846 | 0.734697 | 0.730366 | 0.731066 | 0.724889 |
| 0.734662 | 0.733112 | 0.734187 | 0.732131 | 0.737857 | 0.738985 | 0.730782 | 0.727935 | 0.723616 |
| 0.735165 | 0.734776 | 0.734757 | 0.73451 | 0.742857 | 0.73743 | 0.73155 | 0.731525 | 0.726096 |
| 0.59383 | 0.594004 | 0.592788 | 0.592615 | 0.595697 | 0.593825 | 0.592515 | 0.586778 | 0.585864 |
| 0.57867 | 0.581512 | 0.57854 | 0.583227 | 0.581963 | 0.579609 | 0.577599 | 0.579263 | 0.572237 |
| 0.761905 | 0.760319 | 0.760067 | 0.759732 | 0.765106 | 0.759027 | 0.760787 | 0.754032 | 0.75456 |
| 0.760143 | 0.762438 | 0.762662 | 0.762905 | 0.767859 | 0.762237 | 0.762191 | 0.757584 | 0.757678 |
| 0.750187 | 0.753921 | 0.753359 | 0.753362 | 0.755567 | 0.75547 | 0.74954 | 0.741193 | 0.748668 |
| 0.758692 | 0.767993 | 0.75859 | 0.760472 | 0.766323 | 0.763759 | 0.758058 | 0.75333 | 0.756363 |
| 0.759814 | 0.761609 | 0.759325 | 0.763978 | 0.765592 | 0.764149 | 0.760253 | 0.755965 | 0.755937 |
| 0.76476 | 0.762301 | 0.768261 | 0.761988 | 0.767721 | 0.766101 | 0.759646 | 0.753161 | 0.758685 |
| 0.761469 | 0.763031 | 0.765567 | 0.763742 | 0.767376 | 0.766341 | 0.760786 | 0.756834 | 0.754965 |
| 0.760911 | 0.764782 | 0.761408 | 0.765795 | 0.766101 | 0.763479 | 0.757244 | 0.754475 | 0.755785 |
| 0.767103 | 0.773779 | 0.772768 | 0.7646 | 0.769609 | 0.774489 | 0.767746 | 0.76637 | 0.765644 |
| 0.761689 | 0.764302 | 0.763044 | 0.761145 | 0.762055 | 0.76395 | 0.758739 | 0.755672 | 0.758382 |
| 0.758771 | 0.759554 | 0.76533 | 0.7596 | 0.757295 | 0.765216 | 0.75845 | 0.754906 | 0.756166 |
| 0.7665 | 0.766667 | 0.775861 | 0.767879 | 0.768304 | 0.772299 | 0.766759 | 0.764281 | 0.759594 |
| 0.759753 | 0.764215 | 0.767614 | 0.759179 | 0.764258 | 0.762259 | 0.760202 | 0.753535 | 0.754692 |
| 0.7609 | 0.766625 | 0.766669 | 0.763123 | 0.762393 | 0.766669 | 0.761813 | 0.75905 | 0.764429 |
| 0.760709 | 0.76973 | 0.768418 | 0.763744 | 0.765796 | 0.767552 | 0.75877 | 0.758391 | 0.761762 |
| 0.760592 | 0.760825 | 0.765455 | 0.761824 | 0.764101 | 0.76289 | 0.766879 | 0.75555 | 0.757856 |
| 0.759366 | 0.760591 | 0.765175 | 0.759648 | 0.758178 | 0.764184 | 0.753851 | 0.75506 | 0.757392 |
| 0.72801 | 0.734153 | 0.736479 | 0.734518 | 0.734397 | 0.736018 | 0.726347 | 0.726471 | 0.72438 |
| 0.750285 | 0.74249 | 0.739221 | 0.740503 | 0.752137 | 0.743097 | 0.739698 | 0.73951 | 0.740059 |
| 0.765734 | 0.765931 | 0.767723 | 0.763383 | 0.764247 | 0.765804 | 0.763648 | 0.759117 | 0.757203 |
| 0.763357 | 0.771293 | 0.768989 | 0.761972 | 0.775493 | 0.769327 | 0.769552 | 0.761888 | 0.759944 |
| 0.713623 | 0.712011 | 0.719244 | 0.721441 | 0.71519 | 0.718533 | 0.711253 | 0.713215 | 0.720135 |
| 0.764045 | 0.768802 | 0.768354 | 0.762563 | 0.765801 | 0.764828 | 0.764824 | 0.761715 | 0.76241 |
| 0.758951 | 0.761838 | 0.765103 | 0.756408 | 0.759498 | 0.763086 | 0.760859 | 0.760132 | 0.754289 |

| | | | | | | | | |
|---|---|---|---|---|---|---|---|---|
| 0.762222 | 0.765411 | 0.766614 | 0.759518 | 0.7644 | 0.760667 | 0.760906 | 0.758154 | 0.759379 |
| 0.762109 | 0.767476 | 0.767425 | 0.76587 | 0.766948 | 0.764886 | 0.761723 | 0.764843 | 0.762901 |
| 0.760831 | 0.766037 | 0.763829 | 0.760182 | 0.766546 | 0.762878 | 0.761495 | 0.75794 | 0.760378 |
| 0.763655 | 0.766595 | 0.772803 | 0.761499 | 0.765813 | 0.765225 | 0.765829 | 0.761971 | 0.761791 |
| 0.765859 | 0.76941 | 0.770799 | 0.767814 | 0.768452 | 0.769026 | 0.767868 | 0.761167 | 0.76802 |
| 0.765376 | 0.768304 | 0.769685 | 0.758686 | 0.765116 | 0.76691 | 0.765088 | 0.754843 | 0.762074 |
| 0.759874 | 0.764541 | 0.76668 | 0.756695 | 0.763716 | 0.763996 | 0.761287 | 0.757539 | 0.758308 |
| 0.754931 | 0.757817 | 0.760543 | 0.755934 | 0.758333 | 0.757581 | 0.760643 | 0.754952 | 0.753275 |
| 0.760486 | 0.765596 | 0.764007 | 0.752101 | 0.762505 | 0.763951 | 0.763913 | 0.757202 | 0.759099 |
| 0.76177 | 0.761192 | 0.763654 | 0.755078 | 0.762741 | 0.75931 | 0.761362 | 0.75583 | 0.751612 |
| 0.761716 | 0.760818 | 0.763013 | 0.760978 | 0.759755 | 0.763966 | 0.759316 | 0.76225 | 0.760166 |
| 0.756466 | 0.76339 | 0.764076 | 0.753012 | 0.760702 | 0.76066 | 0.762583 | 0.753374 | 0.756984 |
| 0.757553 | 0.759426 | 0.761093 | 0.75514 | 0.757953 | 0.760172 | 0.757284 | 0.754275 | 0.749733 |
| 0.751533 | 0.757932 | 0.761765 | 0.752363 | 0.760041 | 0.758074 | 0.756251 | 0.750315 | 0.749338 |
| 0.767236 | 0.767413 | 0.767551 | 0.766248 | 0.770501 | 0.769742 | 0.769622 | 0.754858 | 0.761641 |
| 0.765873 | 0.768466 | 0.769599 | 0.7639 | 0.76968 | 0.767658 | 0.762599 | 0.760311 | 0.756389 |
| 0.767951 | 0.769867 | 0.76563 | 0.75845 | 0.767523 | 0.767302 | 0.764981 | 0.759196 | 0.757639 |
| 0.762616 | 0.765667 | 0.762677 | 0.756797 | 0.763149 | 0.765089 | 0.759317 | 0.755278 | 0.754251 |
| 0.761972 | 0.763969 | 0.762128 | 0.760496 | 0.76782 | 0.767931 | 0.7632 | 0.754846 | 0.754526 |
| 0.761063 | 0.768427 | 0.760683 | 0.757302 | 0.763169 | 0.766312 | 0.75943 | 0.755778 | 0.757557 |
| 0.753949 | 0.757008 | 0.759711 | 0.755786 | 0.756012 | 0.761824 | 0.752632 | 0.750264 | 0.752776 |
| 0.756465 | 0.761053 | 0.75873 | 0.757685 | 0.761153 | 0.765279 | 0.760227 | 0.751253 | 0.752307 |
| 0.761339 | 0.769307 | 0.76217 | 0.763391 | 0.763839 | 0.766118 | 0.763573 | 0.755546 | 0.760144 |
| 0.766136 | 0.769564 | 0.767025 | 0.761206 | 0.764929 | 0.768806 | 0.766159 | 0.758151 | 0.759328 |
| 0.758841 | 0.765829 | 0.760647 | 0.757451 | 0.763255 | 0.764265 | 0.761341 | 0.752573 | 0.755137 |
| 0.761319 | 0.765708 | 0.757167 | 0.757621 | 0.758915 | 0.759759 | 0.756571 | 0.751066 | 0.753801 |
| 0.755966 | 0.76074 | 0.762726 | 0.758415 | 0.755685 | 0.759808 | 0.755283 | 0.752506 | 0.75324 |
| 0.757371 | 0.756926 | 0.760494 | 0.749853 | 0.758511 | 0.761275 | 0.754523 | 0.748136 | 0.755426 |
| 0.755647 | 0.760189 | 0.756854 | 0.75325 | 0.761379 | 0.759777 | 0.75858 | 0.753632 | 0.754918 |
| 0.761758 | 0.76368 | 0.763978 | 0.75847 | 0.770275 | 0.768513 | 0.762931 | 0.755199 | 0.758328 |
| 0.763907 | 0.767678 | 0.763371 | 0.758156 | 0.762442 | 0.767769 | 0.765169 | 0.757271 | 0.759389 |
| 0.761853 | 0.76585 | 0.764285 | 0.759106 | 0.763392 | 0.764661 | 0.75838 | 0.758077 | 0.758695 |
| 0.765973 | 0.769898 | 0.767678 | 0.766048 | 0.77322 | 0.769791 | 0.766642 | 0.760796 | 0.760652 |
| 0.767378 | 0.766999 | 0.770476 | 0.767715 | 0.772528 | 0.768404 | 0.764234 | 0.7609 | 0.762082 |
| 0.758632 | 0.761547 | 0.758495 | 0.754798 | 0.760864 | 0.759097 | 0.758904 | 0.753066 | 0.752765 |
| 0.759518 | 0.765471 | 0.761378 | 0.76006 | 0.76355 | 0.765009 | 0.7598 | 0.752856 | 0.753058 |
| 0.75992 | 0.764928 | 0.75992 | 0.758179 | 0.762696 | 0.76216 | 0.760553 | 0.755877 | 0.753156 |
| 0.761121 | 0.76256 | 0.760296 | 0.756536 | 0.762715 | 0.76378 | 0.760892 | 0.753931 | 0.756504 |
| 0.751203 | 0.756734 | 0.754651 | 0.749623 | 0.752572 | 0.75662 | 0.750382 | 0.747326 | 0.747941 |
| 0.750759 | 0.754064 | 0.75397 | 0.754777 | 0.755525 | 0.75815 | 0.752001 | 0.746876 | 0.749427 |
| 0.749101 | 0.752585 | 0.750234 | 0.747762 | 0.750127 | 0.751197 | 0.748712 | 0.746901 | 0.740503 |
| 0.75101 | 0.75849 | 0.754702 | 0.747923 | 0.757581 | 0.756268 | 0.752538 | 0.750367 | 0.749118 |
| 0.747729 | 0.756963 | 0.75433 | 0.749655 | 0.752625 | 0.751905 | 0.750065 | 0.74594 | 0.74571 |
| 0.745853 | 0.752243 | 0.750203 | 0.746929 | 0.753207 | 0.753779 | 0.750342 | 0.742108 | 0.740702 |
| 0.756787 | 0.760483 | 0.756188 | 0.755421 | 0.759381 | 0.763112 | 0.754594 | 0.749549 | 0.752941 |
| 0.751858 | 0.757042 | 0.754424 | 0.748201 | 0.753852 | 0.757509 | 0.752968 | 0.746721 | 0.743979 |

| | | | | | | | | |
|---|---|---|---|---|---|---|---|---|
| 0.758403 | 0.764922 | 0.756401 | 0.757874 | 0.761535 | 0.76034 | 0.760261 | 0.754895 | 0.752867 |
| 0.768952 | 0.771749 | 0.773112 | 0.766407 | 0.774099 | 0.77197 | 0.767138 | 0.760715 | 0.761361 |
| 0.765927 | 0.767021 | 0.76907 | 0.765972 | 0.762805 | 0.765697 | 0.759466 | 0.758457 | 0.759032 |
| 0.766269 | 0.765109 | 0.769313 | 0.765693 | 0.769041 | 0.768262 | 0.768776 | 0.759914 | 0.76014 |
| 0.772993 | 0.778836 | 0.77472 | 0.772327 | 0.777178 | 0.774718 | 0.769726 | 0.768253 | 0.765621 |
| 0.771612 | 0.776602 | 0.773655 | 0.767273 | 0.77017 | 0.772178 | 0.772149 | 0.76468 | 0.759284 |
| 0.762728 | 0.769911 | 0.765302 | 0.762779 | 0.764035 | 0.766366 | 0.760958 | 0.760104 | 0.757385 |
| 0.765555 | 0.766457 | 0.768692 | 0.765003 | 0.769218 | 0.767107 | 0.765725 | 0.762487 | 0.759989 |
| 0.761433 | 0.763977 | 0.763962 | 0.764253 | 0.763791 | 0.765354 | 0.768998 | 0.756325 | 0.758571 |
| 0.758377 | 0.761592 | 0.765241 | 0.757849 | 0.764039 | 0.76042 | 0.758996 | 0.756409 | 0.753161 |
| 0.76687 | 0.76776 | 0.769294 | 0.765617 | 0.768028 | 0.770402 | 0.768228 | 0.766534 | 0.759853 |
| 0.767902 | 0.768323 | 0.771492 | 0.764628 | 0.767726 | 0.769404 | 0.767646 | 0.75991 | 0.758323 |
| 0.767157 | 0.767568 | 0.768013 | 0.767715 | 0.765915 | 0.771017 | 0.764819 | 0.762177 | 0.762595 |
| 0.770199 | 0.770347 | 0.771026 | 0.761716 | 0.769765 | 0.767692 | 0.76621 | 0.760607 | 0.767884 |
| 0.770455 | 0.775596 | 0.770429 | 0.769828 | 0.774045 | 0.774028 | 0.771785 | 0.762155 | 0.763339 |
| 0.766012 | 0.773376 | 0.773652 | 0.76174 | 0.76805 | 0.76959 | 0.770558 | 0.76169 | 0.760057 |
| 0.772517 | 0.773425 | 0.774024 | 0.77196 | 0.771349 | 0.768964 | 0.765704 | 0.766104 | 0.766444 |
| 0.768707 | 0.76896 | 0.774093 | 0.768424 | 0.771695 | 0.773936 | 0.766785 | 0.759391 | 0.763414 |
| 0.769563 | 0.771863 | 0.775431 | 0.766791 | 0.772844 | 0.772481 | 0.771722 | 0.761686 | 0.764321 |
| 0.772456 | 0.769396 | 0.773652 | 0.772031 | 0.772858 | 0.771735 | 0.76868 | 0.768085 | 0.765713 |
| 0.764412 | 0.768903 | 0.769001 | 0.765353 | 0.769595 | 0.768921 | 0.7652 | 0.761921 | 0.765428 |
| 0.768482 | 0.77012 | 0.77289 | 0.766461 | 0.770507 | 0.768456 | 0.768972 | 0.761088 | 0.761452 |
| 0.764482 | 0.771014 | 0.767963 | 0.765928 | 0.770002 | 0.767662 | 0.766975 | 0.760062 | 0.762952 |
| 0.770664 | 0.775735 | 0.771694 | 0.767713 | 0.772451 | 0.769682 | 0.773254 | 0.766031 | 0.765301 |
| 0.767184 | 0.777502 | 0.774531 | 0.771918 | 0.773125 | 0.774209 | 0.768082 | 0.767448 | 0.766407 |
| 0.76772 | 0.773105 | 0.769142 | 0.769728 | 0.771965 | 0.770965 | 0.769025 | 0.765645 | 0.765592 |
| 0.764735 | 0.776912 | 0.769218 | 0.766181 | 0.769692 | 0.769294 | 0.765155 | 0.767482 | 0.762114 |
| 0.76825 | 0.772799 | 0.772306 | 0.764725 | 0.768272 | 0.770925 | 0.76792 | 0.763367 | 0.762733 |
| 0.772249 | 0.775168 | 0.775396 | 0.772667 | 0.773732 | 0.772264 | 0.770522 | 0.766897 | 0.768322 |
| 0.771841 | 0.77405 | 0.771346 | 0.76674 | 0.772293 | 0.773839 | 0.769099 | 0.767757 | 0.765799 |
| 0.769433 | 0.769244 | 0.767607 | 0.763441 | 0.768701 | 0.766551 | 0.766154 | 0.764085 | 0.762533 |
| 0.772632 | 0.772739 | 0.773928 | 0.768788 | 0.772329 | 0.774747 | 0.768628 | 0.760313 | 0.763902 |
| 0.7717 | 0.776752 | 0.775968 | 0.767319 | 0.776886 | 0.775684 | 0.772727 | 0.767928 | 0.768161 |
| 0.761334 | 0.77209 | 0.764991 | 0.762972 | 0.764561 | 0.764499 | 0.761799 | 0.76061 | 0.755535 |
| 0.774432 | 0.776164 | 0.769329 | 0.768273 | 0.772588 | 0.769297 | 0.766303 | 0.763542 | 0.764061 |
| 0.768848 | 0.769271 | 0.767868 | 0.766302 | 0.765102 | 0.770003 | 0.764783 | 0.76348 | 0.762567 |
| 0.775702 | 0.775874 | 0.77545 | 0.771975 | 0.77263 | 0.77361 | 0.771379 | 0.769565 | 0.767894 |
| 0.769413 | 0.773871 | 0.774234 | 0.766352 | 0.773908 | 0.774128 | 0.768178 | 0.76242 | 0.764965 |
| 0.770482 | 0.773745 | 0.773062 | 0.76694 | 0.774348 | 0.77487 | 0.771555 | 0.7666 | 0.765039 |
| 0.772066 | 0.774013 | 0.769716 | 0.762653 | 0.771395 | 0.768754 | 0.76576 | 0.764306 | 0.762872 |
| 0.774547 | 0.774635 | 0.773644 | 0.768399 | 0.776943 | 0.772637 | 0.768355 | 0.769038 | 0.766535 |
| 0.769212 | 0.775133 | 0.776653 | 0.7676 | 0.773702 | 0.774848 | 0.768903 | 0.762408 | 0.763416 |
| 0.768589 | 0.777879 | 0.776234 | 0.771063 | 0.773124 | 0.775819 | 0.768923 | 0.763986 | 0.766182 |
| 0.768193 | 0.77083 | 0.768693 | 0.762194 | 0.767459 | 0.77122 | 0.765891 | 0.75788 | 0.760813 |
| 0.77746 | 0.777312 | 0.776298 | 0.773385 | 0.777824 | 0.778049 | 0.769535 | 0.764213 | 0.766409 |
| 0.76694 | 0.772272 | 0.77074 | 0.764641 | 0.771977 | 0.771623 | 0.766831 | 0.760889 | 0.761037 |

| | | | | | | | | |
|---|---|---|---|---|---|---|---|---|
| 0.757553 | 0.755905 | 0.75882 | 0.754276 | 0.759673 | 0.762618 | 0.752669 | 0.751863 | 0.749627 |
| 0.743083 | 0.746141 | 0.749731 | 0.744446 | 0.746897 | 0.748521 | 0.746742 | 0.740357 | 0.739226 |
| 0.765584 | 0.763091 | 0.766134 | 0.758257 | 0.765975 | 0.76839 | 0.760424 | 0.762091 | 0.7603 |
| 0.768734 | 0.76951 | 0.770753 | 0.766773 | 0.769798 | 0.77217 | 0.769199 | 0.766485 | 0.763915 |
| 0.761995 | 0.761409 | 0.764812 | 0.755687 | 0.762937 | 0.762776 | 0.757918 | 0.756992 | 0.75387 |
| 0.766403 | 0.767842 | 0.766797 | 0.759537 | 0.768588 | 0.767334 | 0.765312 | 0.76251 | 0.759226 |
| 0.76297 | 0.761517 | 0.768154 | 0.765409 | 0.764417 | 0.765341 | 0.763873 | 0.760065 | 0.759169 |
| 0.761845 | 0.764237 | 0.757678 | 0.757144 | 0.762977 | 0.761688 | 0.757573 | 0.757593 | 0.752843 |
| 0.773029 | 0.778573 | 0.777924 | 0.77517 | 0.776024 | 0.776638 | 0.771171 | 0.769611 | 0.770531 |
| 0.769848 | 0.773126 | 0.774577 | 0.767204 | 0.769965 | 0.774169 | 0.767811 | 0.763036 | 0.765574 |
| 0.77619 | 0.778987 | 0.775761 | 0.768857 | 0.772205 | 0.780616 | 0.771514 | 0.765346 | 0.769312 |
| 0.7683 | 0.774663 | 0.77069 | 0.769686 | 0.771661 | 0.772136 | 0.768726 | 0.763839 | 0.765193 |
| 0.775096 | 0.780563 | 0.776176 | 0.774633 | 0.771923 | 0.779042 | 0.773667 | 0.765073 | 0.770351 |
| 0.772022 | 0.776633 | 0.771375 | 0.769339 | 0.770457 | 0.775293 | 0.768654 | 0.767013 | 0.764659 |
| 0.769752 | 0.77558 | 0.77471 | 0.76695 | 0.768746 | 0.772249 | 0.768713 | 0.762005 | 0.767722 |
| 0.765183 | 0.767303 | 0.764472 | 0.770867 | 0.772368 | 0.769136 | 0.766455 | 0.764288 | 0.761589 |
| 0.768858 | 0.770823 | 0.77377 | 0.764472 | 0.765231 | 0.769662 | 0.764898 | 0.761143 | 0.76364 |
| 0.765405 | 0.766853 | 0.766631 | 0.760193 | 0.76773 | 0.764371 | 0.767127 | 0.759565 | 0.757882 |
| 0.761746 | 0.767352 | 0.763144 | 0.76631 | 0.7687 | 0.763829 | 0.763279 | 0.760345 | 0.759483 |
| 0.76423 | 0.763192 | 0.771149 | 0.757051 | 0.760313 | 0.763927 | 0.760441 | 0.753127 | 0.754878 |
| 0.766479 | 0.770882 | 0.768069 | 0.763978 | 0.767027 | 0.769172 | 0.764621 | 0.766699 | 0.764281 |
| 0.775688 | 0.784258 | 0.782542 | 0.769776 | 0.786359 | 0.767989 | 0.784179 | 0.77178 | 0.771731 |
| 0.779351 | 0.781891 | 0.783317 | 0.77358 | 0.793562 | 0.773936 | 0.788427 | 0.777068 | 0.775919 |
| 0.782199 | 0.786664 | 0.786527 | 0.780106 | 0.788732 | 0.781987 | 0.791277 | 0.774295 | 0.781186 |
| 0.78203 | 0.790869 | 0.788629 | 0.781186 | 0.790932 | 0.785156 | 0.789907 | 0.777708 | 0.782043 |
| 0.775739 | 0.785769 | 0.784777 | 0.775384 | 0.788641 | 0.773066 | 0.790262 | 0.772715 | 0.772801 |
| 0.775932 | 0.782726 | 0.777959 | 0.768862 | 0.781495 | 0.774132 | 0.783652 | 0.768906 | 0.775497 |
| 0.781367 | 0.787805 | 0.778373 | 0.7702 | 0.785999 | 0.780057 | 0.785 | 0.770764 | 0.771612 |
| 0.779207 | 0.780559 | 0.782888 | 0.77363 | 0.786984 | 0.780229 | 0.781821 | 0.776115 | 0.775999 |
| 0.691433 | 0.693161 | 0.694421 | 0.687793 | 0.698995 | 0.694049 | 0.697488 | 0.686748 | 0.689541 |
| 0.705777 | 0.707353 | 0.709711 | 0.702371 | 0.711025 | 0.706863 | 0.706926 | 0.698501 | 0.698235 |
| 0.699207 | 0.703797 | 0.703793 | 0.699984 | 0.707705 | 0.702597 | 0.702496 | 0.695773 | 0.699233 |
| 0.695324 | 0.699572 | 0.700879 | 0.69207 | 0.699194 | 0.696678 | 0.697027 | 0.691952 | 0.691464 |
| 0.702701 | 0.707571 | 0.705742 | 0.703361 | 0.707437 | 0.706861 | 0.703686 | 0.69888 | 0.69604 |
| 0.828076 | 0.812034 | 0.867227 | 0.832273 | 0.820077 | 0.824223 | 0.80774 | 0.834273 | 0.8408 |
| 1 | 0.834553 | 0.826049 | 0.795831 | 0.833607 | 0.839331 | 0.817669 | 0.798906 | 0.800911 |
| 0.834553 | 1 | 0.820363 | 0.811884 | 0.836622 | 0.836957 | 0.825781 | 0.809921 | 0.797033 |
| 0.826049 | 0.820363 | 1 | 0.843434 | 0.824812 | 0.820634 | 0.810911 | 0.82298 | 0.835249 |
| 0.795831 | 0.811884 | 0.843434 | 1 | 0.80278 | 0.804216 | 0.793449 | 0.832007 | 0.831609 |
| 0.833607 | 0.836622 | 0.824812 | 0.80278 | 1 | 0.841909 | 0.841402 | 0.804185 | 0.799258 |
| 0.839331 | 0.836957 | 0.820634 | 0.804216 | 0.841909 | 1 | 0.823458 | 0.802692 | 0.798396 |
| 0.817669 | 0.825781 | 0.810911 | 0.793449 | 0.841402 | 0.823458 | 1 | 0.788771 | 0.791872 |
| 0.798906 | 0.809921 | 0.82298 | 0.832007 | 0.804185 | 0.802692 | 0.788771 | 1 | 0.830487 |
| 0.800911 | 0.797033 | 0.835249 | 0.831609 | 0.799258 | 0.798396 | 0.791872 | 0.830487 | 1 |
| 0.828078 | 0.835446 | 0.817053 | 0.799342 | 0.831712 | 0.815955 | 0.827882 | 0.800345 | 0.797783 |
| 0.862837 | 0.83842 | 0.834588 | 0.814288 | 0.822037 | 0.829504 | 0.83147 | 0.803581 | 0.806003 |

| | | | | | | | | |
|---|---|---|---|---|---|---|---|---|
| 0.775259 | 0.780982 | 0.780693 | 0.777305 | 0.779364 | 0.778633 | 0.776796 | 0.77338 | 0.775111 |
| 0.77327 | 0.778592 | 0.777222 | 0.776351 | 0.778512 | 0.778068 | 0.77418 | 0.776163 | 0.768731 |
| 0.773057 | 0.778644 | 0.784334 | 0.774655 | 0.774169 | 0.777618 | 0.773417 | 0.77517 | 0.774542 |
| 0.7714 | 0.774912 | 0.777314 | 0.772435 | 0.771645 | 0.769211 | 0.770495 | 0.768921 | 0.762854 |
| 0.768146 | 0.774848 | 0.774805 | 0.768255 | 0.771145 | 0.769205 | 0.76867 | 0.768887 | 0.765062 |
| 0.777181 | 0.777095 | 0.781307 | 0.772426 | 0.773192 | 0.775256 | 0.770654 | 0.769286 | 0.767922 |
| 0.771649 | 0.775186 | 0.782382 | 0.774378 | 0.774862 | 0.779747 | 0.768623 | 0.769745 | 0.768012 |
| 0.775107 | 0.779177 | 0.786411 | 0.778225 | 0.775649 | 0.777826 | 0.768034 | 0.779432 | 0.771598 |
| 0.772591 | 0.775465 | 0.775846 | 0.774367 | 0.771928 | 0.776822 | 0.769898 | 0.779901 | 0.769133 |
| 0.771294 | 0.777609 | 0.771722 | 0.768572 | 0.773032 | 0.776554 | 0.769608 | 0.767395 | 0.762335 |
| 0.769891 | 0.775151 | 0.776381 | 0.770402 | 0.770645 | 0.772673 | 0.768752 | 0.769401 | 0.767352 |
| 0.769538 | 0.774183 | 0.777307 | 0.770281 | 0.772869 | 0.773583 | 0.770615 | 0.765585 | 0.765363 |
| 0.76724 | 0.773056 | 0.767606 | 0.765597 | 0.764853 | 0.765863 | 0.768534 | 0.760474 | 0.763673 |
| 0.765128 | 0.772153 | 0.770588 | 0.768673 | 0.77174 | 0.772505 | 0.766476 | 0.764706 | 0.755748 |
| 0.770201 | 0.770534 | 0.770979 | 0.763116 | 0.770572 | 0.770478 | 0.768329 | 0.763893 | 0.76009 |
| 0.778329 | 0.776605 | 0.781788 | 0.769731 | 0.781801 | 0.776815 | 0.770111 | 0.769069 | 0.768854 |
| 0.77028 | 0.770432 | 0.774453 | 0.763453 | 0.77055 | 0.771911 | 0.766484 | 0.763668 | 0.756066 |
| 0.772104 | 0.772986 | 0.777343 | 0.77504 | 0.771924 | 0.773321 | 0.770252 | 0.771023 | 0.769621 |
| 0.771577 | 0.773216 | 0.772135 | 0.76874 | 0.767173 | 0.772582 | 0.767123 | 0.762732 | 0.768188 |
| 0.766362 | 0.766443 | 0.765928 | 0.767434 | 0.764434 | 0.768613 | 0.759529 | 0.764299 | 0.756629 |
| 0.768603 | 0.767924 | 0.771076 | 0.767842 | 0.767468 | 0.772125 | 0.763078 | 0.768079 | 0.765467 |
| 0.763173 | 0.766241 | 0.771297 | 0.760879 | 0.763984 | 0.761088 | 0.761254 | 0.758615 | 0.764216 |
| 0.774571 | 0.771956 | 0.776471 | 0.769892 | 0.774639 | 0.772877 | 0.767763 | 0.763863 | 0.764872 |
| 0.773237 | 0.77739 | 0.776769 | 0.772128 | 0.777304 | 0.775516 | 0.771695 | 0.769595 | 0.762811 |
| 0.769563 | 0.771357 | 0.772821 | 0.763623 | 0.775552 | 0.770846 | 0.766685 | 0.762029 | 0.763407 |
| 0.701253 | 0.703466 | 0.705306 | 0.702323 | 0.705739 | 0.703508 | 0.703798 | 0.696648 | 0.696271 |
| 0.766142 | 0.776781 | 0.772686 | 0.768142 | 0.766648 | 0.77049 | 0.763606 | 0.76704 | 0.759986 |
| 0.774662 | 0.778176 | 0.777319 | 0.768436 | 0.774407 | 0.77841 | 0.771909 | 0.761999 | 0.767286 |
| 0.775696 | 0.774544 | 0.77421 | 0.77057 | 0.77404 | 0.775198 | 0.768017 | 0.765396 | 0.762283 |
| 0.771999 | 0.77595 | 0.775835 | 0.772348 | 0.775908 | 0.770985 | 0.769458 | 0.760424 | 0.767694 |
| 0.761826 | 0.773881 | 0.773019 | 0.768436 | 0.767418 | 0.767828 | 0.76362 | 0.765299 | 0.760836 |
| 0.76754 | 0.77408 | 0.772026 | 0.772688 | 0.771192 | 0.766348 | 0.767442 | 0.761186 | 0.763676 |
| 0.766542 | 0.772379 | 0.766838 | 0.767613 | 0.77118 | 0.769755 | 0.764525 | 0.762107 | 0.75687 |
| 0.76342 | 0.773689 | 0.768082 | 0.766849 | 0.770327 | 0.773603 | 0.768778 | 0.765432 | 0.763408 |
| 0.764312 | 0.769549 | 0.770609 | 0.761581 | 0.764151 | 0.768082 | 0.7637 | 0.753952 | 0.760738 |
| 0.765333 | 0.769966 | 0.769758 | 0.760785 | 0.764487 | 0.766678 | 0.76208 | 0.756251 | 0.757748 |
| 0.769133 | 0.774642 | 0.77549 | 0.772388 | 0.770923 | 0.770372 | 0.768343 | 0.765632 | 0.762942 |
| 0.772391 | 0.770894 | 0.773634 | 0.770351 | 0.773273 | 0.779064 | 0.770212 | 0.764685 | 0.768155 |
| 0.769296 | 0.773606 | 0.77347 | 0.769489 | 0.768145 | 0.770709 | 0.763951 | 0.765916 | 0.769397 |
| 0.766041 | 0.763224 | 0.771315 | 0.764584 | 0.764209 | 0.773297 | 0.763036 | 0.762288 | 0.759691 |
| 0.759882 | 0.765547 | 0.766635 | 0.760286 | 0.76382 | 0.766296 | 0.759639 | 0.757031 | 0.761213 |
| 0.77272 | 0.774396 | 0.773698 | 0.77005 | 0.775398 | 0.772569 | 0.767507 | 0.763685 | 0.764791 |
| 0.762515 | 0.76493 | 0.764523 | 0.763445 | 0.763445 | 0.770535 | 0.758349 | 0.754291 | 0.756495 |
| 0.771015 | 0.770818 | 0.773764 | 0.767708 | 0.774039 | 0.772413 | 0.769192 | 0.758483 | 0.760064 |
| 0.773731 | 0.775002 | 0.777935 | 0.774919 | 0.783384 | 0.774161 | 0.776164 | 0.773007 | 0.770919 |
| 0.770696 | 0.776523 | 0.778925 | 0.775716 | 0.777408 | 0.773424 | 0.773558 | 0.771964 | 0.770952 |

| | | | | | | | | |
|---|---|---|---|---|---|---|---|---|
| 0.762385 | 0.771583 | 0.771198 | 0.769071 | 0.76944 | 0.765772 | 0.7673 | 0.76695 | 0.760215 |
| 0.769858 | 0.774271 | 0.778587 | 0.77538 | 0.779998 | 0.777973 | 0.772492 | 0.775761 | 0.771525 |
| 0.770723 | 0.769418 | 0.770136 | 0.767769 | 0.771854 | 0.767763 | 0.768957 | 0.75971 | 0.763221 |
| 0.736726 | 0.726131 | 0.737319 | 0.732156 | 0.752735 | 0.726686 | 0.743337 | 0.724854 | 0.730176 |
| 0.762098 | 0.770759 | 0.766459 | 0.765561 | 0.771833 | 0.76314 | 0.762419 | 0.762316 | 0.761737 |
| 0.765293 | 0.771782 | 0.772701 | 0.773266 | 0.776298 | 0.772193 | 0.765086 | 0.764034 | 0.764685 |
| 0.772644 | 0.770394 | 0.773758 | 0.771741 | 0.774648 | 0.767981 | 0.768702 | 0.768172 | 0.766729 |
| 0.769802 | 0.775277 | 0.769212 | 0.770509 | 0.774356 | 0.763798 | 0.770079 | 0.764236 | 0.766791 |
| 0.771721 | 0.774948 | 0.778201 | 0.772678 | 0.777355 | 0.771531 | 0.771433 | 0.770891 | 0.775365 |
| 0.774072 | 0.778131 | 0.775862 | 0.774333 | 0.776933 | 0.773563 | 0.771887 | 0.765697 | 0.768193 |
| 0.772279 | 0.77487 | 0.771841 | 0.773118 | 0.775017 | 0.773074 | 0.767013 | 0.764824 | 0.764601 |
| 0.772331 | 0.772264 | 0.774869 | 0.774745 | 0.780372 | 0.769244 | 0.769787 | 0.765364 | 0.765434 |
| 0.767426 | 0.770927 | 0.77272 | 0.772875 | 0.777437 | 0.770739 | 0.766388 | 0.760135 | 0.764529 |
| 0.770596 | 0.773944 | 0.776746 | 0.770889 | 0.776518 | 0.773599 | 0.773142 | 0.764956 | 0.765231 |
| 0.770851 | 0.777945 | 0.774642 | 0.775458 | 0.778294 | 0.773911 | 0.770488 | 0.767432 | 0.767469 |
| 0.763344 | 0.772758 | 0.769014 | 0.77051 | 0.773583 | 0.768717 | 0.763233 | 0.765393 | 0.763543 |
| 0.773932 | 0.780211 | 0.774241 | 0.772847 | 0.780849 | 0.775169 | 0.768066 | 0.766632 | 0.760365 |
| 0.76955 | 0.77689 | 0.779676 | 0.775087 | 0.777125 | 0.772871 | 0.768382 | 0.769887 | 0.772053 |
| 0.768367 | 0.768668 | 0.774135 | 0.770906 | 0.771551 | 0.768429 | 0.763644 | 0.763534 | 0.763585 |
| 0.764379 | 0.772644 | 0.77016 | 0.772826 | 0.775836 | 0.770641 | 0.767071 | 0.763995 | 0.766227 |
| 0.769866 | 0.774681 | 0.777207 | 0.774073 | 0.775365 | 0.774979 | 0.773029 | 0.766313 | 0.767697 |
| 0.768265 | 0.773676 | 0.774615 | 0.77668 | 0.775461 | 0.769 | 0.769886 | 0.768989 | 0.766619 |
| 0.678454 | 0.683834 | 0.682293 | 0.677078 | 0.685121 | 0.676503 | 0.683269 | 0.683346 | 0.68283 |
| 0.654576 | 0.659021 | 0.647765 | 0.654148 | 0.654309 | 0.651958 | 0.65239 | 0.654872 | 0.653317 |
| 0.664996 | 0.670998 | 0.672459 | 0.670771 | 0.672395 | 0.675016 | 0.667682 | 0.663835 | 0.667354 |
| 0.658115 | 0.652046 | 0.657514 | 0.648494 | 0.65783 | 0.656065 | 0.652138 | 0.652996 | 0.653853 |
| 0.746203 | 0.74947 | 0.75072 | 0.749962 | 0.749131 | 0.743438 | 0.747902 | 0.74415 | 0.740654 |
| 0.737473 | 0.739768 | 0.741313 | 0.738963 | 0.742272 | 0.737515 | 0.736984 | 0.731017 | 0.735939 |
| 0.730162 | 0.738436 | 0.744944 | 0.734905 | 0.741075 | 0.738226 | 0.736088 | 0.740286 | 0.738211 |
| 0.747475 | 0.751049 | 0.759115 | 0.753953 | 0.749738 | 0.743767 | 0.740927 | 0.749623 | 0.742732 |
| 0.724936 | 0.732486 | 0.741898 | 0.737907 | 0.734444 | 0.719835 | 0.732345 | 0.73376 | 0.738715 |
| 0.722143 | 0.721845 | 0.734174 | 0.73553 | 0.721515 | 0.715179 | 0.722761 | 0.72969 | 0.738018 |
| 0.721874 | 0.726893 | 0.735321 | 0.730631 | 0.735329 | 0.723943 | 0.735719 | 0.728927 | 0.744064 |
| 0.651621 | 0.652151 | 0.653221 | 0.645726 | 0.660246 | 0.65602 | 0.649822 | 0.646201 | 0.652841 |
| 0.769318 | 0.771361 | 0.770707 | 0.769284 | 0.770438 | 0.76832 | 0.769366 | 0.76653 | 0.762365 |
| 0.750235 | 0.751934 | 0.754029 | 0.745274 | 0.754717 | 0.753225 | 0.751123 | 0.750121 | 0.747417 |
| 0.747912 | 0.746598 | 0.751538 | 0.743024 | 0.752452 | 0.744164 | 0.739692 | 0.743371 | 0.747786 |
| 0.731657 | 0.73316 | 0.741911 | 0.737647 | 0.743912 | 0.733271 | 0.734694 | 0.736058 | 0.732318 |
| 0.748907 | 0.752466 | 0.755254 | 0.749148 | 0.758495 | 0.750532 | 0.754032 | 0.746441 | 0.743408 |
| 0.736406 | 0.733752 | 0.735665 | 0.732719 | 0.739192 | 0.733867 | 0.733497 | 0.728782 | 0.731041 |
| 0.728059 | 0.731591 | 0.733643 | 0.734272 | 0.735819 | 0.733002 | 0.73175 | 0.727065 | 0.728934 |
| 0.731988 | 0.738488 | 0.737157 | 0.734953 | 0.739943 | 0.729899 | 0.737413 | 0.729764 | 0.725932 |
| 0.729254 | 0.732095 | 0.73469 | 0.731933 | 0.746589 | 0.73523 | 0.732847 | 0.725501 | 0.726515 |
| 0.734625 | 0.746448 | 0.735973 | 0.729142 | 0.73826 | 0.734017 | 0.742412 | 0.726942 | 0.734623 |
| 0.734476 | 0.73189 | 0.742976 | 0.73702 | 0.738581 | 0.741708 | 0.732935 | 0.72948 | 0.728733 |
| 0.72076 | 0.718472 | 0.726818 | 0.725705 | 0.740566 | 0.724826 | 0.709911 | 0.718443 | 0.726004 |

| | | | | | | | | |
|---|---|---|---|---|---|---|---|---|
| 0.764468 | 0.763673 | 0.76666 | 0.766898 | 0.764929 | 0.760528 | 0.760351 | 0.756564 | 0.760833 |
| 0.736015 | 0.742514 | 0.743044 | 0.743448 | 0.744657 | 0.740308 | 0.737493 | 0.742511 | 0.74404 |
| 0.734858 | 0.747991 | 0.73747 | 0.735117 | 0.746226 | 0.738648 | 0.734922 | 0.740873 | 0.740527 |
| 0.738088 | 0.747677 | 0.747946 | 0.74209 | 0.745595 | 0.736801 | 0.732628 | 0.735632 | 0.738501 |
| 0.736008 | 0.737544 | 0.737712 | 0.740412 | 0.744301 | 0.745391 | 0.735018 | 0.738958 | 0.744924 |
| 0.733082 | 0.731169 | 0.734985 | 0.731871 | 0.7335 | 0.725461 | 0.731059 | 0.729957 | 0.728619 |
| 0.730882 | 0.7295 | 0.731078 | 0.732535 | 0.736881 | 0.737217 | 0.721614 | 0.733881 | 0.736341 |
| 0.723902 | 0.729676 | 0.738129 | 0.735908 | 0.731437 | 0.727662 | 0.725457 | 0.727273 | 0.727246 |
| 0.7402 | 0.741046 | 0.735973 | 0.737532 | 0.736423 | 0.734607 | 0.731812 | 0.732009 | 0.738925 |
| 0.760008 | 0.757354 | 0.7653 | 0.762854 | 0.763168 | 0.761393 | 0.754372 | 0.754584 | 0.757045 |
| 0.752624 | 0.755971 | 0.756281 | 0.758292 | 0.757981 | 0.758829 | 0.754994 | 0.750498 | 0.750583 |
| 0.75462 | 0.752481 | 0.759655 | 0.751753 | 0.755709 | 0.755807 | 0.748465 | 0.746472 | 0.749653 |
| 0.741447 | 0.742669 | 0.742707 | 0.741065 | 0.748512 | 0.748346 | 0.744121 | 0.741094 | 0.740101 |
| 0.740309 | 0.743401 | 0.751815 | 0.74785 | 0.753234 | 0.754804 | 0.744704 | 0.747113 | 0.749648 |
| 0.748786 | 0.745398 | 0.754338 | 0.747587 | 0.752044 | 0.745258 | 0.74273 | 0.738954 | 0.753719 |

| | | | | | | | | plot 12 |
|---|---|---|---|---|---|---|---|---|
| Wild_plot_ | Wild_plot_ | Wild_plot_ | Wild_plot_ | Wild_plot_ | Wild_plot_ | Wild_plot_ | Wild_plot_ | Wild_plot_ |
| 0.704072 | 0.706588 | 0.705514 | 0.705699 | 0.707476 | 0.705831 | 0.705695 | 0.702894 | 0.705892 |
| 0.71364 | 0.713835 | 0.71542 | 0.713368 | 0.720191 | 0.714332 | 0.711491 | 0.711758 | 0.716794 |
| 0.698994 | 0.702222 | 0.702085 | 0.703209 | 0.706822 | 0.697956 | 0.699052 | 0.703807 | 0.706318 |
| 0.709672 | 0.71158 | 0.714506 | 0.712858 | 0.71547 | 0.710392 | 0.708455 | 0.710989 | 0.71544 |
| 0.704174 | 0.708608 | 0.706274 | 0.702984 | 0.710395 | 0.708307 | 0.704861 | 0.705222 | 0.708818 |
| 0.715348 | 0.717589 | 0.719456 | 0.715494 | 0.716816 | 0.714936 | 0.71225 | 0.710799 | 0.717581 |
| 0.768733 | 0.768975 | 0.767667 | 0.763011 | 0.770681 | 0.763283 | 0.764232 | 0.766154 | 0.764316 |
| 0.765945 | 0.767246 | 0.768106 | 0.765041 | 0.772447 | 0.767194 | 0.762121 | 0.764156 | 0.765052 |
| 0.707832 | 0.711167 | 0.711556 | 0.708223 | 0.713188 | 0.707793 | 0.703773 | 0.704439 | 0.710895 |
| 0.705351 | 0.705944 | 0.707119 | 0.706253 | 0.709924 | 0.707391 | 0.702841 | 0.706467 | 0.710312 |
| 0.705805 | 0.707349 | 0.70676 | 0.712449 | 0.712158 | 0.710068 | 0.707146 | 0.705955 | 0.709736 |
| 0.69678 | 0.701189 | 0.701548 | 0.702472 | 0.704881 | 0.701654 | 0.697922 | 0.697291 | 0.703975 |
| 0.707781 | 0.711677 | 0.711733 | 0.714113 | 0.715429 | 0.712694 | 0.70684 | 0.707624 | 0.714344 |
| 0.716451 | 0.721394 | 0.721218 | 0.719312 | 0.723699 | 0.719836 | 0.71632 | 0.718509 | 0.722838 |
| 0.728414 | 0.730685 | 0.737603 | 0.733317 | 0.737124 | 0.735169 | 0.728823 | 0.729911 | 0.733153 |
| 0.709251 | 0.712443 | 0.720163 | 0.714057 | 0.717458 | 0.717296 | 0.709935 | 0.710487 | 0.717061 |
| 0.713318 | 0.712798 | 0.718735 | 0.712579 | 0.715833 | 0.711844 | 0.708601 | 0.710352 | 0.714592 |
| 0.736786 | 0.740687 | 0.742434 | 0.739941 | 0.742573 | 0.740435 | 0.738041 | 0.735524 | 0.740738 |
| 0.702729 | 0.704027 | 0.704436 | 0.705278 | 0.706206 | 0.703393 | 0.700065 | 0.703125 | 0.706253 |
| 0.707577 | 0.70937 | 0.711815 | 0.709938 | 0.71491 | 0.711779 | 0.706654 | 0.706178 | 0.712403 |
| 0.70924 | 0.709073 | 0.707287 | 0.704743 | 0.710334 | 0.706365 | 0.702522 | 0.705088 | 0.705217 |
| 0.70864 | 0.710094 | 0.712398 | 0.710758 | 0.716587 | 0.708882 | 0.707462 | 0.70876 | 0.712168 |
| 0.71314 | 0.71371 | 0.717631 | 0.715788 | 0.717707 | 0.717229 | 0.711999 | 0.714676 | 0.717831 |
| 0.713337 | 0.713743 | 0.713411 | 0.713828 | 0.715349 | 0.708879 | 0.7057 | 0.709637 | 0.714695 |
| 0.709762 | 0.71381 | 0.712943 | 0.711791 | 0.715596 | 0.714895 | 0.709461 | 0.712981 | 0.71691 |
| 0.706083 | 0.709776 | 0.712292 | 0.708221 | 0.713039 | 0.706279 | 0.706945 | 0.706104 | 0.710475 |
| 0.704018 | 0.707859 | 0.704327 | 0.704774 | 0.707725 | 0.701068 | 0.698529 | 0.698823 | 0.705028 |
| 0.709814 | 0.711441 | 0.707278 | 0.711131 | 0.713368 | 0.709421 | 0.706945 | 0.706128 | 0.71325 |
| 0.710345 | 0.717539 | 0.713352 | 0.711035 | 0.720705 | 0.711711 | 0.709762 | 0.711394 | 0.71856 |
| 0.705285 | 0.708448 | 0.712144 | 0.708598 | 0.713845 | 0.711599 | 0.706987 | 0.707826 | 0.713499 |
| 0.704306 | 0.709206 | 0.70763 | 0.707874 | 0.712455 | 0.708594 | 0.706094 | 0.705605 | 0.708921 |
| 0.701305 | 0.703458 | 0.708465 | 0.704603 | 0.710493 | 0.706299 | 0.701743 | 0.70292 | 0.707454 |
| 0.707506 | 0.710988 | 0.710131 | 0.708323 | 0.712607 | 0.708121 | 0.704923 | 0.701899 | 0.711465 |
| 0.705667 | 0.711436 | 0.710088 | 0.712141 | 0.717525 | 0.71269 | 0.706527 | 0.709563 | 0.712113 |
| 0.701753 | 0.705963 | 0.701604 | 0.703147 | 0.707594 | 0.700773 | 0.700203 | 0.703008 | 0.707499 |
| 0.708059 | 0.710516 | 0.710369 | 0.71172 | 0.713403 | 0.713354 | 0.706147 | 0.706778 | 0.712721 |
| 0.712495 | 0.714274 | 0.714089 | 0.714468 | 0.716537 | 0.712244 | 0.708208 | 0.709907 | 0.715772 |
| 0.707546 | 0.714146 | 0.712942 | 0.712099 | 0.716458 | 0.713026 | 0.70964 | 0.709121 | 0.713576 |
| 0.710473 | 0.714793 | 0.711743 | 0.71573 | 0.716878 | 0.713908 | 0.708442 | 0.712647 | 0.719341 |

| | | | | | | | | |
|---|---|---|---|---|---|---|---|---|
| 0.743269 | 0.74379 | 0.748274 | 0.743416 | 0.749206 | 0.741684 | 0.739163 | 0.740897 | 0.744167 |
| 0.698725 | 0.701196 | 0.703091 | 0.702421 | 0.704013 | 0.701042 | 0.700154 | 0.699245 | 0.703828 |
| 0.702864 | 0.707116 | 0.705049 | 0.702077 | 0.705792 | 0.706108 | 0.700423 | 0.701039 | 0.706012 |
| 0.708841 | 0.714894 | 0.714796 | 0.712533 | 0.718287 | 0.716518 | 0.709476 | 0.709696 | 0.714646 |
| 0.703436 | 0.707894 | 0.708797 | 0.708107 | 0.712121 | 0.709024 | 0.701199 | 0.70229 | 0.711779 |
| 0.713684 | 0.712986 | 0.713537 | 0.710677 | 0.713475 | 0.711954 | 0.707699 | 0.709712 | 0.716764 |
| 0.702962 | 0.705222 | 0.705549 | 0.704738 | 0.708089 | 0.705142 | 0.699515 | 0.701146 | 0.704794 |
| 0.707689 | 0.708938 | 0.710058 | 0.710658 | 0.714971 | 0.70845 | 0.707556 | 0.706004 | 0.711396 |
| 0.707309 | 0.714413 | 0.714684 | 0.711423 | 0.715726 | 0.710715 | 0.711534 | 0.709696 | 0.714392 |
| 0.711631 | 0.714816 | 0.71273 | 0.712674 | 0.717744 | 0.710922 | 0.709423 | 0.709508 | 0.713192 |
| 0.700147 | 0.701234 | 0.703383 | 0.703549 | 0.705821 | 0.706133 | 0.698549 | 0.699876 | 0.704232 |
| 0.704769 | 0.703838 | 0.70705 | 0.706657 | 0.70864 | 0.704837 | 0.698606 | 0.701673 | 0.705911 |
| 0.700805 | 0.706958 | 0.708611 | 0.709032 | 0.709048 | 0.707158 | 0.706332 | 0.70596 | 0.708131 |
| 0.705482 | 0.703854 | 0.704266 | 0.704752 | 0.7083 | 0.706303 | 0.70482 | 0.704504 | 0.704605 |
| 0.706857 | 0.705772 | 0.707431 | 0.703695 | 0.709851 | 0.705773 | 0.700484 | 0.701702 | 0.707976 |
| 0.705428 | 0.70515 | 0.70511 | 0.700534 | 0.707656 | 0.707028 | 0.702024 | 0.703846 | 0.707567 |
| 0.698632 | 0.702885 | 0.705553 | 0.702325 | 0.705926 | 0.70124 | 0.70084 | 0.700607 | 0.703795 |
| 0.706973 | 0.711335 | 0.709381 | 0.711154 | 0.713282 | 0.707924 | 0.706596 | 0.70489 | 0.708901 |
| 0.710989 | 0.714077 | 0.712421 | 0.710967 | 0.71417 | 0.711711 | 0.70617 | 0.708804 | 0.714742 |
| 0.701747 | 0.707281 | 0.712099 | 0.707058 | 0.709478 | 0.708139 | 0.704793 | 0.701828 | 0.710182 |
| 0.700647 | 0.704051 | 0.705729 | 0.703299 | 0.706462 | 0.703855 | 0.70242 | 0.701363 | 0.705098 |
| 0.705165 | 0.70836 | 0.707162 | 0.704092 | 0.709028 | 0.705566 | 0.703988 | 0.699769 | 0.710485 |
| 0.707093 | 0.705045 | 0.707901 | 0.706466 | 0.711981 | 0.706845 | 0.698799 | 0.704068 | 0.70782 |
| 0.715954 | 0.716465 | 0.715925 | 0.713849 | 0.715926 | 0.713713 | 0.714333 | 0.712689 | 0.714239 |
| 0.721674 | 0.723146 | 0.725268 | 0.719311 | 0.723435 | 0.716598 | 0.71797 | 0.716465 | 0.724728 |
| 0.767689 | 0.770123 | 0.768267 | 0.766575 | 0.77251 | 0.767393 | 0.762937 | 0.764907 | 0.767292 |
| 0.762817 | 0.768837 | 0.773697 | 0.768954 | 0.766616 | 0.76675 | 0.765131 | 0.76933 | 0.767416 |
| 0.766757 | 0.773251 | 0.771579 | 0.773038 | 0.770373 | 0.764769 | 0.764861 | 0.771996 | 0.768937 |
| 0.703523 | 0.706837 | 0.708894 | 0.706523 | 0.705088 | 0.707664 | 0.703477 | 0.706421 | 0.708616 |
| 0.703844 | 0.703468 | 0.709748 | 0.703999 | 0.709726 | 0.704849 | 0.702857 | 0.702364 | 0.705942 |
| 0.711115 | 0.713151 | 0.71202 | 0.715148 | 0.716297 | 0.716132 | 0.712417 | 0.709733 | 0.712914 |
| 0.716564 | 0.717086 | 0.716269 | 0.712847 | 0.720354 | 0.715107 | 0.711948 | 0.712233 | 0.714109 |
| 0.705333 | 0.705112 | 0.706245 | 0.707588 | 0.710799 | 0.704621 | 0.703143 | 0.703065 | 0.710598 |
| 0.705519 | 0.709683 | 0.710658 | 0.709125 | 0.709785 | 0.70783 | 0.70292 | 0.707358 | 0.712507 |
| 0.704014 | 0.702848 | 0.705034 | 0.707212 | 0.710625 | 0.704809 | 0.702136 | 0.70219 | 0.711719 |
| 0.706463 | 0.708013 | 0.708787 | 0.708048 | 0.711655 | 0.703772 | 0.702625 | 0.705839 | 0.710567 |
| 0.700785 | 0.703222 | 0.70666 | 0.70741 | 0.709584 | 0.707418 | 0.705454 | 0.705387 | 0.709557 |
| 0.718475 | 0.720169 | 0.723114 | 0.718572 | 0.725369 | 0.720818 | 0.717957 | 0.716262 | 0.723569 |
| 0.695999 | 0.6977 | 0.700742 | 0.705569 | 0.703911 | 0.696026 | 0.697753 | 0.699102 | 0.703236 |
| 0.707993 | 0.709073 | 0.706475 | 0.705635 | 0.708722 | 0.708651 | 0.702487 | 0.70111 | 0.706225 |
| 0.70315 | 0.70478 | 0.703274 | 0.701331 | 0.706592 | 0.702442 | 0.698509 | 0.701134 | 0.704902 |
| 0.720727 | 0.7253 | 0.72662 | 0.724118 | 0.727829 | 0.725222 | 0.72186 | 0.721741 | 0.72368 |
| 0.707039 | 0.709117 | 0.708799 | 0.70816 | 0.712968 | 0.709189 | 0.70569 | 0.70783 | 0.708982 |
| 0.704269 | 0.708587 | 0.706449 | 0.703818 | 0.7083 | 0.707579 | 0.701666 | 0.70421 | 0.706164 |
| 0.705105 | 0.707594 | 0.710953 | 0.709467 | 0.711873 | 0.708611 | 0.706713 | 0.704497 | 0.709977 |
| 0.705979 | 0.706787 | 0.70868 | 0.709917 | 0.70999 | 0.704457 | 0.701419 | 0.706662 | 0.710972 |

| | | | | | | | | |
|---|---|---|---|---|---|---|---|---|
| 0.705237 | 0.70583 | 0.706953 | 0.703203 | 0.713754 | 0.708106 | 0.703971 | 0.70374 | 0.704902 |
| 0.710348 | 0.710463 | 0.710355 | 0.709976 | 0.711852 | 0.710572 | 0.705414 | 0.708156 | 0.713399 |
| 0.701429 | 0.707204 | 0.708916 | 0.708502 | 0.712098 | 0.708747 | 0.706414 | 0.704391 | 0.709106 |
| 0.713409 | 0.713003 | 0.715785 | 0.710753 | 0.718553 | 0.713881 | 0.710944 | 0.710915 | 0.714534 |
| 0.713095 | 0.713883 | 0.713844 | 0.712269 | 0.717028 | 0.713204 | 0.70738 | 0.71133 | 0.716247 |
| 0.706638 | 0.707815 | 0.709603 | 0.706723 | 0.710454 | 0.703739 | 0.702614 | 0.703071 | 0.704849 |
| 0.705177 | 0.707472 | 0.711284 | 0.709613 | 0.712968 | 0.710146 | 0.70664 | 0.704191 | 0.712347 |
| 0.708906 | 0.715345 | 0.710565 | 0.712739 | 0.717731 | 0.709948 | 0.707742 | 0.710176 | 0.713685 |
| 0.707179 | 0.705868 | 0.711812 | 0.70691 | 0.713311 | 0.708374 | 0.707826 | 0.708219 | 0.712307 |
| 0.706531 | 0.708754 | 0.703752 | 0.70836 | 0.708732 | 0.705329 | 0.702771 | 0.706197 | 0.705782 |
| 0.703074 | 0.70639 | 0.707782 | 0.705859 | 0.711791 | 0.70871 | 0.702605 | 0.703117 | 0.706763 |
| 0.706068 | 0.70849 | 0.711534 | 0.706598 | 0.712625 | 0.712754 | 0.706257 | 0.705214 | 0.710635 |
| 0.707115 | 0.709848 | 0.708104 | 0.708162 | 0.714647 | 0.706847 | 0.703953 | 0.706614 | 0.711931 |
| 0.702676 | 0.705502 | 0.70771 | 0.701604 | 0.70594 | 0.703538 | 0.701412 | 0.699967 | 0.704692 |
| 0.707844 | 0.70967 | 0.710132 | 0.706902 | 0.715411 | 0.709996 | 0.705892 | 0.704566 | 0.711729 |
| 0.704723 | 0.705362 | 0.709877 | 0.704423 | 0.710459 | 0.705494 | 0.703149 | 0.706332 | 0.707722 |
| 0.707235 | 0.712148 | 0.710431 | 0.708776 | 0.715276 | 0.708521 | 0.704903 | 0.705502 | 0.712498 |
| 0.709714 | 0.708276 | 0.709627 | 0.706985 | 0.709667 | 0.705033 | 0.700357 | 0.704792 | 0.710189 |
| 0.706422 | 0.708926 | 0.711414 | 0.709654 | 0.716107 | 0.708771 | 0.705767 | 0.707204 | 0.710918 |
| 0.706041 | 0.709907 | 0.708806 | 0.709652 | 0.712272 | 0.712451 | 0.706594 | 0.706675 | 0.711321 |
| 0.706442 | 0.708992 | 0.710836 | 0.708839 | 0.712361 | 0.706558 | 0.703764 | 0.70507 | 0.711387 |
| 0.701694 | 0.705176 | 0.706759 | 0.705652 | 0.707783 | 0.70316 | 0.698133 | 0.701324 | 0.703526 |
| 0.706807 | 0.70679 | 0.707879 | 0.705282 | 0.711462 | 0.709005 | 0.70415 | 0.707374 | 0.70758 |
| 0.707888 | 0.710944 | 0.710483 | 0.710954 | 0.711616 | 0.709389 | 0.701746 | 0.70642 | 0.710834 |
| 0.705378 | 0.707692 | 0.710526 | 0.706834 | 0.713492 | 0.70659 | 0.705611 | 0.709064 | 0.713185 |
| 0.70935 | 0.714688 | 0.719234 | 0.713909 | 0.717907 | 0.714103 | 0.708316 | 0.713298 | 0.717606 |
| 0.702582 | 0.705145 | 0.706788 | 0.707538 | 0.710029 | 0.707145 | 0.70239 | 0.704177 | 0.708222 |
| 0.70744 | 0.710391 | 0.713715 | 0.706576 | 0.714044 | 0.70828 | 0.705705 | 0.702982 | 0.714111 |
| 0.707909 | 0.71042 | 0.710782 | 0.709835 | 0.715266 | 0.708769 | 0.704414 | 0.708175 | 0.709162 |
| 0.706609 | 0.709006 | 0.710931 | 0.707424 | 0.711935 | 0.709523 | 0.705882 | 0.706378 | 0.71285 |
| 0.710878 | 0.713995 | 0.713229 | 0.712324 | 0.714646 | 0.712717 | 0.706128 | 0.711392 | 0.714498 |
| 0.703425 | 0.705896 | 0.705986 | 0.705478 | 0.707888 | 0.70572 | 0.702368 | 0.702458 | 0.708994 |
| 0.707228 | 0.709836 | 0.711464 | 0.707972 | 0.712967 | 0.712025 | 0.709269 | 0.705157 | 0.71393 |
| 0.706642 | 0.711656 | 0.710022 | 0.708306 | 0.713813 | 0.71074 | 0.706064 | 0.705063 | 0.711663 |
| 0.704221 | 0.70482 | 0.705565 | 0.705181 | 0.71108 | 0.703867 | 0.703361 | 0.702108 | 0.707385 |
| 0.707197 | 0.708832 | 0.714132 | 0.708208 | 0.712088 | 0.707236 | 0.70397 | 0.703018 | 0.708879 |
| 0.701688 | 0.701144 | 0.704775 | 0.701339 | 0.710748 | 0.704589 | 0.701738 | 0.702969 | 0.707185 |
| 0.680174 | 0.684065 | 0.681963 | 0.682082 | 0.68794 | 0.679772 | 0.678825 | 0.682062 | 0.68624 |
| 0.687906 | 0.688239 | 0.688439 | 0.684504 | 0.693713 | 0.688174 | 0.681753 | 0.684314 | 0.690969 |
| 0.675546 | 0.678616 | 0.678553 | 0.674387 | 0.678542 | 0.676258 | 0.67169 | 0.673766 | 0.679125 |
| 0.681412 | 0.679423 | 0.680811 | 0.676769 | 0.683859 | 0.676143 | 0.674096 | 0.67754 | 0.683273 |
| 0.660772 | 0.664029 | 0.663152 | 0.664941 | 0.669254 | 0.668632 | 0.658473 | 0.661562 | 0.668449 |
| 0.661637 | 0.664152 | 0.668215 | 0.667071 | 0.669778 | 0.668253 | 0.667319 | 0.664261 | 0.66402 |
| 0.664039 | 0.659342 | 0.67027 | 0.663474 | 0.670729 | 0.66346 | 0.664135 | 0.667774 | 0.671467 |
| 0.666753 | 0.666966 | 0.666495 | 0.667219 | 0.671805 | 0.662369 | 0.660351 | 0.658192 | 0.670281 |
| 0.664257 | 0.665954 | 0.667002 | 0.668252 | 0.667748 | 0.662544 | 0.661938 | 0.664904 | 0.671544 |
| 0.672373 | 0.67236 | 0.673 | 0.671871 | 0.677147 | 0.673561 | 0.668453 | 0.668787 | 0.675832 |

| | | | | | | | | |
|---|---|---|---|---|---|---|---|---|
| 0.667499 | 0.669102 | 0.669151 | 0.672965 | 0.679224 | 0.673409 | 0.66708 | 0.669447 | 0.673927 |
| 0.666182 | 0.665097 | 0.675069 | 0.670898 | 0.675386 | 0.666667 | 0.666448 | 0.670894 | 0.678439 |
| 0.657547 | 0.664025 | 0.666265 | 0.666444 | 0.669828 | 0.659912 | 0.661923 | 0.664515 | 0.669809 |
| 0.662766 | 0.664891 | 0.663659 | 0.660977 | 0.668258 | 0.663822 | 0.662133 | 0.663685 | 0.664266 |
| 0.660086 | 0.663476 | 0.664033 | 0.665291 | 0.664442 | 0.660841 | 0.658109 | 0.657066 | 0.670579 |
| 0.664016 | 0.665602 | 0.665421 | 0.662957 | 0.665709 | 0.657209 | 0.660273 | 0.664822 | 0.666298 |
| 0.664119 | 0.664328 | 0.665857 | 0.65953 | 0.666241 | 0.662872 | 0.661631 | 0.664479 | 0.667097 |
| 0.662012 | 0.657231 | 0.666902 | 0.662367 | 0.666714 | 0.657732 | 0.656929 | 0.660948 | 0.668175 |
| 0.663423 | 0.662076 | 0.661939 | 0.659559 | 0.667259 | 0.657569 | 0.652457 | 0.65248 | 0.66469 |
| 0.664042 | 0.663029 | 0.666716 | 0.663549 | 0.670556 | 0.663196 | 0.661359 | 0.662986 | 0.67001 |
| 0.651653 | 0.653653 | 0.659524 | 0.652986 | 0.660137 | 0.655941 | 0.6543 | 0.656008 | 0.659456 |
| 0.692166 | 0.696777 | 0.699021 | 0.696492 | 0.700087 | 0.696686 | 0.694567 | 0.694922 | 0.700113 |
| 0.705391 | 0.70999 | 0.710215 | 0.707558 | 0.711855 | 0.706204 | 0.705186 | 0.702548 | 0.710526 |
| 0.680171 | 0.682877 | 0.686037 | 0.68329 | 0.686467 | 0.680902 | 0.679256 | 0.681771 | 0.689663 |
| 0.713005 | 0.709456 | 0.715562 | 0.712695 | 0.714929 | 0.713856 | 0.710705 | 0.712052 | 0.712971 |
| 0.701968 | 0.703318 | 0.702841 | 0.700831 | 0.704542 | 0.700862 | 0.698308 | 0.701544 | 0.706296 |
| 0.709057 | 0.712105 | 0.710622 | 0.711522 | 0.716873 | 0.712002 | 0.70765 | 0.708402 | 0.710678 |
| 0.715736 | 0.720396 | 0.718601 | 0.717486 | 0.719285 | 0.718175 | 0.714487 | 0.717208 | 0.719596 |
| 0.695677 | 0.697056 | 0.695778 | 0.697516 | 0.701376 | 0.692139 | 0.690341 | 0.693776 | 0.696296 |
| 0.711465 | 0.717383 | 0.718113 | 0.718194 | 0.715487 | 0.715762 | 0.714622 | 0.712172 | 0.71846 |
| 0.683999 | 0.6888 | 0.695764 | 0.68905 | 0.69672 | 0.687122 | 0.68731 | 0.68648 | 0.694275 |
| 0.714227 | 0.713349 | 0.714672 | 0.714619 | 0.717374 | 0.714803 | 0.708623 | 0.710387 | 0.720781 |
| 0.715968 | 0.719802 | 0.717606 | 0.719961 | 0.721178 | 0.715606 | 0.71283 | 0.715639 | 0.719229 |
| 0.682125 | 0.680515 | 0.688697 | 0.685151 | 0.687507 | 0.68097 | 0.683068 | 0.680556 | 0.687219 |
| 0.686481 | 0.687244 | 0.686364 | 0.685598 | 0.686116 | 0.683409 | 0.685095 | 0.682383 | 0.689393 |
| 0.689438 | 0.685743 | 0.696449 | 0.688531 | 0.698358 | 0.688279 | 0.683788 | 0.684392 | 0.691943 |
| 0.701366 | 0.705514 | 0.708625 | 0.705748 | 0.709326 | 0.706363 | 0.701789 | 0.704857 | 0.708135 |
| 0.717484 | 0.71904 | 0.718016 | 0.716667 | 0.723077 | 0.715167 | 0.714531 | 0.713744 | 0.718873 |
| 0.68289 | 0.688352 | 0.685185 | 0.688398 | 0.688345 | 0.68695 | 0.683439 | 0.68168 | 0.690356 |
| 0.717867 | 0.721087 | 0.720575 | 0.724061 | 0.724074 | 0.720061 | 0.720693 | 0.716628 | 0.727555 |
| 0.717312 | 0.721738 | 0.7258 | 0.724769 | 0.725212 | 0.725915 | 0.722668 | 0.720523 | 0.728823 |
| 0.696351 | 0.69811 | 0.695974 | 0.695624 | 0.701968 | 0.697324 | 0.693588 | 0.694305 | 0.699641 |
| 0.695896 | 0.695878 | 0.698497 | 0.695973 | 0.698042 | 0.694069 | 0.690134 | 0.693899 | 0.698339 |
| 0.718783 | 0.725039 | 0.725222 | 0.723165 | 0.721823 | 0.720131 | 0.718707 | 0.719547 | 0.726099 |
| 0.714481 | 0.718255 | 0.717856 | 0.718298 | 0.719599 | 0.718681 | 0.719106 | 0.715214 | 0.722586 |
| 0.723288 | 0.722222 | 0.719721 | 0.720259 | 0.724093 | 0.719368 | 0.718101 | 0.718579 | 0.723845 |
| 0.673301 | 0.673785 | 0.676756 | 0.676567 | 0.680038 | 0.674988 | 0.668991 | 0.673761 | 0.67968 |
| 0.718362 | 0.719787 | 0.718961 | 0.718381 | 0.72418 | 0.7195 | 0.716744 | 0.717775 | 0.719432 |
| 0.671223 | 0.674442 | 0.674269 | 0.67275 | 0.67572 | 0.671133 | 0.671378 | 0.67207 | 0.675006 |
| 0.667419 | 0.669328 | 0.670159 | 0.667098 | 0.671844 | 0.66809 | 0.663361 | 0.662105 | 0.670041 |
| 0.718867 | 0.721331 | 0.717441 | 0.719297 | 0.723735 | 0.717196 | 0.713393 | 0.71779 | 0.719644 |
| 0.667498 | 0.67144 | 0.667054 | 0.670638 | 0.672314 | 0.667886 | 0.666098 | 0.664915 | 0.669262 |
| 0.662363 | 0.661012 | 0.660488 | 0.664544 | 0.663192 | 0.65905 | 0.655744 | 0.659102 | 0.662503 |
| 0.657217 | 0.655396 | 0.661311 | 0.659506 | 0.660138 | 0.654655 | 0.654895 | 0.654867 | 0.661898 |
| 0.722145 | 0.729254 | 0.730505 | 0.725223 | 0.730118 | 0.727616 | 0.723535 | 0.720392 | 0.726471 |
| 0.662795 | 0.663089 | 0.665919 | 0.66042 | 0.664225 | 0.661483 | 0.658638 | 0.659692 | 0.664659 |
| 0.714966 | 0.7195 | 0.720525 | 0.720767 | 0.724177 | 0.720906 | 0.719803 | 0.715385 | 0.720634 |

| | | | | | | | | |
|---|---|---|---|---|---|---|---|---|
| 0.728502 | 0.733982 | 0.727169 | 0.726055 | 0.730104 | 0.725278 | 0.722821 | 0.721614 | 0.729012 |
| 0.723237 | 0.725478 | 0.726417 | 0.723351 | 0.728523 | 0.721342 | 0.715983 | 0.723323 | 0.724546 |
| 0.736882 | 0.747018 | 0.746661 | 0.743384 | 0.745376 | 0.743695 | 0.740262 | 0.738674 | 0.743041 |
| 0.652313 | 0.653257 | 0.653557 | 0.655314 | 0.654305 | 0.65168 | 0.647522 | 0.649812 | 0.656279 |
| 0.651509 | 0.647961 | 0.652195 | 0.651975 | 0.65332 | 0.647597 | 0.648779 | 0.647455 | 0.654066 |
| 0.730046 | 0.732345 | 0.729366 | 0.726782 | 0.728209 | 0.725407 | 0.725088 | 0.724075 | 0.729311 |
| 0.651631 | 0.654928 | 0.656041 | 0.653308 | 0.654799 | 0.653826 | 0.64908 | 0.652144 | 0.657228 |
| 0.73257 | 0.732645 | 0.73395 | 0.733077 | 0.7333 | 0.730961 | 0.730705 | 0.728303 | 0.732716 |
| 0.726266 | 0.729175 | 0.732937 | 0.73091 | 0.735164 | 0.731203 | 0.725908 | 0.724035 | 0.728636 |
| 0.732483 | 0.734096 | 0.738265 | 0.730826 | 0.733229 | 0.731255 | 0.730369 | 0.727088 | 0.733043 |
| 0.727567 | 0.729389 | 0.731456 | 0.729822 | 0.736674 | 0.730704 | 0.728504 | 0.725705 | 0.727258 |
| 0.66443 | 0.707006 | 0.695035 | 0.732877 | 0.679739 | 0.736486 | 0.74026 | 0.696429 | 0.751773 |
| 0.730738 | 0.736607 | 0.732928 | 0.740418 | 0.74406 | 0.733172 | 0.742329 | 0.740197 | 0.739695 |
| 0.7392 | 0.742029 | 0.742229 | 0.734484 | 0.746155 | 0.738281 | 0.738486 | 0.743974 | 0.737287 |
| 0.741189 | 0.74655 | 0.753163 | 0.75674 | 0.76288 | 0.740295 | 0.744264 | 0.749045 | 0.757422 |
| 0.737335 | 0.747976 | 0.750513 | 0.752838 | 0.751088 | 0.748629 | 0.752653 | 0.755237 | 0.753336 |
| 0.592922 | 0.591997 | 0.589822 | 0.588982 | 0.59577 | 0.591082 | 0.589102 | 0.593362 | 0.59335 |
| 0.738396 | 0.749168 | 0.750497 | 0.75593 | 0.757269 | 0.746859 | 0.746758 | 0.752192 | 0.751029 |
| 0.732544 | 0.737594 | 0.737756 | 0.734914 | 0.735117 | 0.732089 | 0.728942 | 0.729714 | 0.73525 |
| 0.731961 | 0.733827 | 0.73575 | 0.737871 | 0.734082 | 0.734136 | 0.731164 | 0.729365 | 0.734536 |
| 0.730666 | 0.734593 | 0.736644 | 0.735732 | 0.741013 | 0.73733 | 0.732431 | 0.732416 | 0.733923 |
| 0.592052 | 0.593396 | 0.59552 | 0.59793 | 0.595806 | 0.591285 | 0.589424 | 0.589936 | 0.594383 |
| 0.578743 | 0.582033 | 0.583233 | 0.582336 | 0.58395 | 0.580936 | 0.575064 | 0.576464 | 0.580929 |
| 0.76073 | 0.762109 | 0.767446 | 0.760291 | 0.765023 | 0.75818 | 0.758906 | 0.756638 | 0.764037 |
| 0.756857 | 0.763193 | 0.76103 | 0.758184 | 0.767236 | 0.759579 | 0.754247 | 0.758943 | 0.763475 |
| 0.749218 | 0.754643 | 0.758183 | 0.750021 | 0.752645 | 0.7563 | 0.748904 | 0.751795 | 0.753429 |
| 0.756732 | 0.761379 | 0.763516 | 0.758678 | 0.757571 | 0.75657 | 0.756259 | 0.758193 | 0.75621 |
| 0.75475 | 0.762043 | 0.765742 | 0.757262 | 0.761286 | 0.757183 | 0.755121 | 0.759015 | 0.759507 |
| 0.761365 | 0.766442 | 0.766959 | 0.763026 | 0.764256 | 0.762934 | 0.763644 | 0.76389 | 0.765814 |
| 0.760741 | 0.765727 | 0.765361 | 0.762263 | 0.763408 | 0.764068 | 0.763025 | 0.763268 | 0.766759 |
| 0.756217 | 0.767012 | 0.760403 | 0.758335 | 0.759955 | 0.754139 | 0.756518 | 0.757883 | 0.759121 |
| 0.765975 | 0.767836 | 0.7705 | 0.764633 | 0.767621 | 0.76351 | 0.763011 | 0.763071 | 0.767175 |
| 0.756182 | 0.762773 | 0.76322 | 0.756768 | 0.765398 | 0.75903 | 0.759454 | 0.756748 | 0.763122 |
| 0.75652 | 0.756326 | 0.763589 | 0.758736 | 0.760394 | 0.754553 | 0.757977 | 0.755116 | 0.758727 |
| 0.760742 | 0.771343 | 0.771383 | 0.769174 | 0.769448 | 0.760105 | 0.763151 | 0.764451 | 0.76718 |
| 0.758408 | 0.763252 | 0.761969 | 0.759407 | 0.761722 | 0.757077 | 0.756652 | 0.756982 | 0.760851 |
| 0.76 | 0.766683 | 0.768561 | 0.762098 | 0.763476 | 0.762043 | 0.761081 | 0.763095 | 0.766373 |
| 0.76051 | 0.764183 | 0.772651 | 0.764711 | 0.763608 | 0.764275 | 0.75701 | 0.760047 | 0.763332 |
| 0.75756 | 0.764938 | 0.764146 | 0.758772 | 0.761378 | 0.758221 | 0.758657 | 0.7609 | 0.759707 |
| 0.755706 | 0.763971 | 0.761714 | 0.756402 | 0.759696 | 0.7541 | 0.751435 | 0.755235 | 0.757265 |
| 0.734375 | 0.734126 | 0.746617 | 0.736553 | 0.732971 | 0.738066 | 0.735152 | 0.731531 | 0.749732 |
| 0.735835 | 0.736752 | 0.739635 | 0.73351 | 0.738297 | 0.736822 | 0.740126 | 0.744665 | 0.746414 |
| 0.75934 | 0.76548 | 0.763407 | 0.760413 | 0.762688 | 0.756143 | 0.75645 | 0.758615 | 0.759045 |
| 0.765241 | 0.763677 | 0.777253 | 0.768995 | 0.770176 | 0.763959 | 0.760094 | 0.761767 | 0.767012 |
| 0.700835 | 0.717277 | 0.721388 | 0.716557 | 0.720631 | 0.700115 | 0.713069 | 0.716886 | 0.713897 |
| 0.75998 | 0.768433 | 0.768038 | 0.763375 | 0.76585 | 0.756817 | 0.764036 | 0.758942 | 0.764825 |
| 0.757596 | 0.760596 | 0.763764 | 0.753357 | 0.758626 | 0.75364 | 0.750792 | 0.754201 | 0.756488 |

| | | | | | | | | |
|---|---|---|---|---|---|---|---|---|
| 0.760034 | 0.763138 | 0.766838 | 0.762164 | 0.764265 | 0.756886 | 0.756159 | 0.754675 | 0.762269 |
| 0.759933 | 0.766432 | 0.77091 | 0.764107 | 0.765887 | 0.758199 | 0.759757 | 0.756893 | 0.763083 |
| 0.760065 | 0.765618 | 0.760676 | 0.762915 | 0.759313 | 0.755401 | 0.758428 | 0.758908 | 0.761757 |
| 0.762209 | 0.769533 | 0.769956 | 0.762701 | 0.762398 | 0.757907 | 0.760507 | 0.755875 | 0.762296 |
| 0.762484 | 0.767799 | 0.769677 | 0.77 | 0.769557 | 0.764319 | 0.763964 | 0.76236 | 0.769224 |
| 0.760517 | 0.767698 | 0.768585 | 0.762474 | 0.760437 | 0.761581 | 0.758915 | 0.760773 | 0.762669 |
| 0.758753 | 0.761983 | 0.769838 | 0.764342 | 0.76353 | 0.763318 | 0.758373 | 0.759081 | 0.765367 |
| 0.751845 | 0.755539 | 0.754725 | 0.754971 | 0.753184 | 0.748075 | 0.749429 | 0.748783 | 0.752414 |
| 0.760263 | 0.764133 | 0.769796 | 0.765926 | 0.758366 | 0.758396 | 0.75682 | 0.759878 | 0.762919 |
| 0.756422 | 0.765701 | 0.758699 | 0.757536 | 0.756817 | 0.754659 | 0.756443 | 0.753738 | 0.760352 |
| 0.757077 | 0.760651 | 0.766711 | 0.763459 | 0.765027 | 0.763557 | 0.760084 | 0.759662 | 0.765066 |
| 0.755434 | 0.761188 | 0.765317 | 0.762428 | 0.757537 | 0.75819 | 0.757683 | 0.75593 | 0.759972 |
| 0.752697 | 0.756793 | 0.759358 | 0.748588 | 0.755869 | 0.74836 | 0.749725 | 0.749615 | 0.755644 |
| 0.755018 | 0.755068 | 0.761437 | 0.754232 | 0.755398 | 0.7532 | 0.755233 | 0.752161 | 0.756622 |
| 0.761119 | 0.768147 | 0.766834 | 0.763084 | 0.762545 | 0.762217 | 0.757454 | 0.761025 | 0.764021 |
| 0.758205 | 0.763182 | 0.766025 | 0.760365 | 0.766401 | 0.761248 | 0.761268 | 0.764301 | 0.764134 |
| 0.765234 | 0.766595 | 0.772375 | 0.762213 | 0.760955 | 0.760825 | 0.760766 | 0.762486 | 0.763018 |
| 0.75787 | 0.760967 | 0.766816 | 0.760692 | 0.761029 | 0.756258 | 0.757155 | 0.759032 | 0.756832 |
| 0.760926 | 0.765782 | 0.766558 | 0.756422 | 0.76678 | 0.758096 | 0.758667 | 0.761561 | 0.763238 |
| 0.757142 | 0.766304 | 0.765599 | 0.762505 | 0.761742 | 0.75999 | 0.756545 | 0.760095 | 0.75815 |
| 0.753239 | 0.755768 | 0.760698 | 0.756349 | 0.758621 | 0.752684 | 0.751621 | 0.751706 | 0.7534 |
| 0.758706 | 0.763216 | 0.76343 | 0.751107 | 0.761647 | 0.753344 | 0.757815 | 0.755684 | 0.756949 |
| 0.760321 | 0.768014 | 0.769759 | 0.761858 | 0.762948 | 0.759526 | 0.762355 | 0.766173 | 0.761988 |
| 0.762591 | 0.765881 | 0.767289 | 0.763108 | 0.766026 | 0.763162 | 0.758783 | 0.765272 | 0.76429 |
| 0.758674 | 0.764096 | 0.764536 | 0.762474 | 0.762996 | 0.761233 | 0.760899 | 0.761769 | 0.761893 |
| 0.753971 | 0.760141 | 0.758757 | 0.756211 | 0.760868 | 0.757849 | 0.757399 | 0.757025 | 0.756675 |
| 0.753787 | 0.760088 | 0.761119 | 0.755041 | 0.754008 | 0.755635 | 0.750318 | 0.75589 | 0.753382 |
| 0.75411 | 0.75963 | 0.755607 | 0.752514 | 0.75979 | 0.757449 | 0.759023 | 0.754097 | 0.76037 |
| 0.754286 | 0.757151 | 0.763903 | 0.76137 | 0.762965 | 0.756368 | 0.758163 | 0.756486 | 0.758948 |
| 0.756582 | 0.763874 | 0.763795 | 0.758995 | 0.760992 | 0.758482 | 0.758018 | 0.754864 | 0.755727 |
| 0.762247 | 0.764681 | 0.767779 | 0.76475 | 0.765911 | 0.761237 | 0.760323 | 0.761138 | 0.759195 |
| 0.759718 | 0.76378 | 0.76725 | 0.760737 | 0.764736 | 0.761282 | 0.757948 | 0.758544 | 0.75903 |
| 0.765218 | 0.770678 | 0.769946 | 0.765197 | 0.767857 | 0.766814 | 0.76142 | 0.762929 | 0.76954 |
| 0.765421 | 0.769661 | 0.769982 | 0.765085 | 0.765899 | 0.767269 | 0.762782 | 0.765357 | 0.768686 |
| 0.75722 | 0.759633 | 0.760893 | 0.758418 | 0.763468 | 0.758136 | 0.756085 | 0.755357 | 0.762199 |
| 0.756611 | 0.766342 | 0.766921 | 0.761673 | 0.765736 | 0.761052 | 0.758025 | 0.758606 | 0.760903 |
| 0.755631 | 0.760388 | 0.766004 | 0.759337 | 0.764052 | 0.761984 | 0.759159 | 0.76015 | 0.764671 |
| 0.756323 | 0.760221 | 0.764061 | 0.759409 | 0.759624 | 0.757566 | 0.755601 | 0.758221 | 0.75981 |
| 0.749488 | 0.753423 | 0.758644 | 0.756981 | 0.753811 | 0.748372 | 0.751216 | 0.750991 | 0.748742 |
| 0.748241 | 0.75349 | 0.753416 | 0.747656 | 0.75225 | 0.747392 | 0.751274 | 0.750728 | 0.752501 |
| 0.745868 | 0.747733 | 0.754029 | 0.751149 | 0.753259 | 0.746437 | 0.749892 | 0.747986 | 0.745294 |
| 0.749041 | 0.757504 | 0.753286 | 0.749651 | 0.756567 | 0.748864 | 0.75307 | 0.749127 | 0.7514 |
| 0.745751 | 0.752738 | 0.756027 | 0.753547 | 0.75403 | 0.749614 | 0.744214 | 0.750222 | 0.749453 |
| 0.748391 | 0.75109 | 0.751144 | 0.744339 | 0.751519 | 0.743414 | 0.744691 | 0.748152 | 0.74547 |
| 0.75619 | 0.755414 | 0.765382 | 0.760534 | 0.754336 | 0.756155 | 0.750351 | 0.754918 | 0.757344 |
| 0.748446 | 0.75702 | 0.756151 | 0.747687 | 0.753799 | 0.75276 | 0.747726 | 0.748187 | 0.75531 |

| | | | | | | | | |
|---|---|---|---|---|---|---|---|---|
| 0.75634 | 0.763061 | 0.763662 | 0.763715 | 0.760879 | 0.754568 | 0.760695 | 0.758744 | 0.759515 |
| 0.767543 | 0.766866 | 0.769564 | 0.7661 | 0.768569 | 0.770075 | 0.763205 | 0.764361 | 0.768279 |
| 0.764162 | 0.765138 | 0.765291 | 0.762642 | 0.765429 | 0.75998 | 0.763402 | 0.762656 | 0.763908 |
| 0.761622 | 0.7698 | 0.765705 | 0.768965 | 0.764429 | 0.763589 | 0.762471 | 0.761909 | 0.762954 |
| 0.7709 | 0.780547 | 0.776135 | 0.773964 | 0.773819 | 0.768843 | 0.769853 | 0.771282 | 0.76962 |
| 0.7682 | 0.77333 | 0.770563 | 0.767572 | 0.768552 | 0.765089 | 0.768933 | 0.764551 | 0.767454 |
| 0.762794 | 0.762871 | 0.765961 | 0.758056 | 0.762822 | 0.756949 | 0.759045 | 0.759796 | 0.765875 |
| 0.764211 | 0.768828 | 0.768921 | 0.760263 | 0.767158 | 0.764696 | 0.76533 | 0.766468 | 0.763721 |
| 0.759728 | 0.764192 | 0.759368 | 0.760407 | 0.760572 | 0.759953 | 0.76018 | 0.758009 | 0.758179 |
| 0.754693 | 0.762842 | 0.761604 | 0.7591 | 0.758678 | 0.760373 | 0.758595 | 0.763385 | 0.761844 |
| 0.765758 | 0.766303 | 0.767172 | 0.762753 | 0.764469 | 0.762196 | 0.763653 | 0.761641 | 0.768168 |
| 0.762579 | 0.771016 | 0.766007 | 0.763032 | 0.765628 | 0.76196 | 0.762665 | 0.763663 | 0.765805 |
| 0.766318 | 0.768326 | 0.765911 | 0.766647 | 0.771303 | 0.768989 | 0.769497 | 0.768437 | 0.766431 |
| 0.765932 | 0.767661 | 0.76936 | 0.765274 | 0.771517 | 0.76429 | 0.768672 | 0.766718 | 0.765225 |
| 0.767884 | 0.773134 | 0.771905 | 0.770272 | 0.77243 | 0.77414 | 0.772177 | 0.767145 | 0.770463 |
| 0.7654 | 0.772872 | 0.769289 | 0.769199 | 0.77314 | 0.763121 | 0.767338 | 0.76625 | 0.768382 |
| 0.769392 | 0.772031 | 0.770689 | 0.766037 | 0.772761 | 0.764792 | 0.765673 | 0.76713 | 0.767512 |
| 0.770162 | 0.77446 | 0.768911 | 0.771195 | 0.772178 | 0.7663 | 0.761328 | 0.772386 | 0.769906 |
| 0.769769 | 0.770346 | 0.772586 | 0.771466 | 0.773343 | 0.769085 | 0.770399 | 0.767142 | 0.772471 |
| 0.771325 | 0.773756 | 0.773211 | 0.765237 | 0.772855 | 0.767377 | 0.769528 | 0.766576 | 0.771565 |
| 0.765333 | 0.770721 | 0.767924 | 0.760123 | 0.764407 | 0.760013 | 0.761812 | 0.759028 | 0.765625 |
| 0.76486 | 0.771306 | 0.774951 | 0.763243 | 0.771532 | 0.768706 | 0.766562 | 0.768193 | 0.771028 |
| 0.761885 | 0.770565 | 0.771378 | 0.766987 | 0.775089 | 0.766809 | 0.769101 | 0.763807 | 0.766759 |
| 0.769341 | 0.775987 | 0.770208 | 0.771758 | 0.772562 | 0.764119 | 0.768941 | 0.767948 | 0.77203 |
| 0.766448 | 0.770897 | 0.77111 | 0.777969 | 0.77656 | 0.769946 | 0.773487 | 0.769944 | 0.768392 |
| 0.766097 | 0.767448 | 0.768817 | 0.770591 | 0.776192 | 0.768317 | 0.766305 | 0.765106 | 0.764607 |
| 0.765219 | 0.76839 | 0.775746 | 0.767406 | 0.769161 | 0.763362 | 0.765463 | 0.766204 | 0.765261 |
| 0.765387 | 0.771061 | 0.772309 | 0.767009 | 0.769275 | 0.765154 | 0.766725 | 0.766994 | 0.766146 |
| 0.77076 | 0.774392 | 0.772868 | 0.771189 | 0.772269 | 0.771727 | 0.766228 | 0.763943 | 0.775017 |
| 0.770379 | 0.773991 | 0.777749 | 0.770769 | 0.776263 | 0.765806 | 0.768885 | 0.769881 | 0.768218 |
| 0.762858 | 0.767422 | 0.764279 | 0.768985 | 0.762729 | 0.762466 | 0.759976 | 0.762065 | 0.759656 |
| 0.767473 | 0.774129 | 0.772414 | 0.771927 | 0.770558 | 0.762604 | 0.767535 | 0.767667 | 0.776868 |
| 0.769989 | 0.77185 | 0.769366 | 0.770538 | 0.773433 | 0.768306 | 0.769948 | 0.767675 | 0.767522 |
| 0.763064 | 0.763986 | 0.765956 | 0.763725 | 0.76557 | 0.761294 | 0.761698 | 0.760276 | 0.764251 |
| 0.763276 | 0.770174 | 0.77311 | 0.769453 | 0.769862 | 0.766194 | 0.766996 | 0.765651 | 0.768831 |
| 0.765895 | 0.767383 | 0.772623 | 0.764091 | 0.771153 | 0.764309 | 0.761804 | 0.767833 | 0.769142 |
| 0.774526 | 0.775123 | 0.780502 | 0.775329 | 0.77575 | 0.768996 | 0.770403 | 0.769165 | 0.770819 |
| 0.767236 | 0.767751 | 0.772992 | 0.769923 | 0.767672 | 0.768114 | 0.768182 | 0.769271 | 0.77041 |
| 0.76731 | 0.774016 | 0.772785 | 0.772302 | 0.770988 | 0.766403 | 0.771436 | 0.771275 | 0.772056 |
| 0.76457 | 0.772127 | 0.770762 | 0.766425 | 0.76755 | 0.762715 | 0.767099 | 0.766283 | 0.769707 |
| 0.769535 | 0.771213 | 0.775334 | 0.76851 | 0.771779 | 0.768619 | 0.768431 | 0.76551 | 0.774034 |
| 0.768614 | 0.771726 | 0.773604 | 0.772637 | 0.770278 | 0.76958 | 0.769607 | 0.769277 | 0.767956 |
| 0.765799 | 0.772938 | 0.776855 | 0.774586 | 0.77535 | 0.77359 | 0.769244 | 0.77363 | 0.774664 |
| 0.764378 | 0.767031 | 0.768874 | 0.766714 | 0.76479 | 0.760762 | 0.763406 | 0.765336 | 0.766372 |
| 0.770073 | 0.773639 | 0.770488 | 0.76773 | 0.773329 | 0.767076 | 0.763767 | 0.769198 | 0.76942 |
| 0.763971 | 0.770093 | 0.771078 | 0.768196 | 0.768426 | 0.767694 | 0.765345 | 0.762459 | 0.764776 |

| | | | | | | | | |
|---|---|---|---|---|---|---|---|---|
| 0.758897 | 0.759215 | 0.759275 | 0.755977 | 0.759143 | 0.756359 | 0.749015 | 0.755508 | 0.758614 |
| 0.74722 | 0.748584 | 0.74792 | 0.751704 | 0.751913 | 0.742478 | 0.745354 | 0.746958 | 0.750105 |
| 0.761367 | 0.765193 | 0.767899 | 0.765744 | 0.769388 | 0.764587 | 0.760941 | 0.762398 | 0.761085 |
| 0.76737 | 0.77249 | 0.773203 | 0.769022 | 0.77391 | 0.765596 | 0.766134 | 0.768299 | 0.768967 |
| 0.754671 | 0.760403 | 0.762991 | 0.758515 | 0.765466 | 0.758541 | 0.756501 | 0.760412 | 0.761146 |
| 0.761488 | 0.765986 | 0.767069 | 0.768935 | 0.766521 | 0.760196 | 0.764831 | 0.766285 | 0.763786 |
| 0.760871 | 0.766428 | 0.770392 | 0.767755 | 0.770746 | 0.765571 | 0.764939 | 0.771612 | 0.766057 |
| 0.753541 | 0.764701 | 0.762381 | 0.757244 | 0.764065 | 0.754046 | 0.75757 | 0.755723 | 0.758991 |
| 0.769645 | 0.776595 | 0.774074 | 0.768787 | 0.774022 | 0.77245 | 0.770739 | 0.767552 | 0.772051 |
| 0.768163 | 0.765798 | 0.775642 | 0.772531 | 0.771909 | 0.768618 | 0.768736 | 0.768606 | 0.772812 |
| 0.770793 | 0.775359 | 0.77398 | 0.777363 | 0.772212 | 0.771583 | 0.770587 | 0.769153 | 0.770576 |
| 0.7726 | 0.774275 | 0.774727 | 0.772026 | 0.769591 | 0.766054 | 0.767479 | 0.766564 | 0.77398 |
| 0.769497 | 0.775081 | 0.778265 | 0.772264 | 0.776997 | 0.770465 | 0.769516 | 0.769126 | 0.770538 |
| 0.767418 | 0.771659 | 0.775594 | 0.774345 | 0.770652 | 0.769237 | 0.768194 | 0.770417 | 0.772601 |
| 0.768596 | 0.773388 | 0.773204 | 0.770302 | 0.766352 | 0.76361 | 0.766903 | 0.762291 | 0.770178 |
| 0.76476 | 0.768731 | 0.77097 | 0.764778 | 0.76598 | 0.764505 | 0.764501 | 0.763482 | 0.767891 |
| 0.765501 | 0.769526 | 0.775752 | 0.768975 | 0.766917 | 0.764857 | 0.767706 | 0.764937 | 0.768148 |
| 0.760774 | 0.765297 | 0.775824 | 0.766675 | 0.765286 | 0.762713 | 0.761973 | 0.762744 | 0.766338 |
| 0.75981 | 0.765115 | 0.761957 | 0.760928 | 0.761755 | 0.762272 | 0.764813 | 0.760947 | 0.764152 |
| 0.758012 | 0.762627 | 0.771287 | 0.76322 | 0.765524 | 0.762249 | 0.766014 | 0.760943 | 0.764309 |
| 0.765002 | 0.767949 | 0.77621 | 0.767259 | 0.767991 | 0.765771 | 0.76593 | 0.76094 | 0.767961 |
| 0.777729 | 0.776896 | 0.763438 | 0.764761 | 0.768801 | 0.761277 | 0.761504 | 0.76367 | 0.760804 |
| 0.774057 | 0.778783 | 0.768448 | 0.764691 | 0.772545 | 0.759023 | 0.761513 | 0.764071 | 0.770042 |
| 0.779113 | 0.78622 | 0.774232 | 0.768899 | 0.776771 | 0.765364 | 0.767047 | 0.768744 | 0.773275 |
| 0.787125 | 0.789428 | 0.776789 | 0.770606 | 0.773733 | 0.769544 | 0.777161 | 0.771336 | 0.770884 |
| 0.776269 | 0.776917 | 0.772622 | 0.769742 | 0.775833 | 0.768959 | 0.769877 | 0.768672 | 0.770655 |
| 0.778158 | 0.782462 | 0.77069 | 0.765047 | 0.772657 | 0.761423 | 0.762954 | 0.764095 | 0.764409 |
| 0.778316 | 0.782775 | 0.772099 | 0.764015 | 0.77255 | 0.76888 | 0.764876 | 0.76855 | 0.765628 |
| 0.780081 | 0.783301 | 0.766003 | 0.767768 | 0.767152 | 0.762478 | 0.767957 | 0.7693 | 0.767576 |
| 0.694145 | 0.697339 | 0.695774 | 0.692258 | 0.696995 | 0.695933 | 0.691919 | 0.692257 | 0.699101 |
| 0.708695 | 0.706524 | 0.706646 | 0.704044 | 0.709565 | 0.707731 | 0.699573 | 0.705153 | 0.710424 |
| 0.704652 | 0.705208 | 0.703336 | 0.704474 | 0.706569 | 0.702601 | 0.700049 | 0.701501 | 0.702321 |
| 0.693655 | 0.700832 | 0.69682 | 0.697064 | 0.698044 | 0.699849 | 0.697262 | 0.696427 | 0.698066 |
| 0.702635 | 0.705802 | 0.703342 | 0.705284 | 0.7073 | 0.705314 | 0.703224 | 0.703001 | 0.70902 |
| 0.812807 | 0.821388 | 0.774778 | 0.773994 | 0.775822 | 0.769277 | 0.7677 | 0.771237 | 0.770886 |
| 0.828078 | 0.862837 | 0.775259 | 0.77327 | 0.773057 | 0.7714 | 0.768146 | 0.777181 | 0.771649 |
| 0.835446 | 0.83842 | 0.780982 | 0.778592 | 0.778644 | 0.774912 | 0.774848 | 0.777095 | 0.775186 |
| 0.817053 | 0.834588 | 0.780693 | 0.777222 | 0.784334 | 0.777314 | 0.774805 | 0.781307 | 0.782382 |
| 0.799342 | 0.814288 | 0.777305 | 0.776351 | 0.774655 | 0.772435 | 0.768255 | 0.772426 | 0.774378 |
| 0.831712 | 0.822037 | 0.779364 | 0.778512 | 0.774169 | 0.771645 | 0.771145 | 0.773192 | 0.774862 |
| 0.815955 | 0.829504 | 0.778633 | 0.778068 | 0.777618 | 0.769211 | 0.769205 | 0.775256 | 0.779747 |
| 0.827882 | 0.83147 | 0.776796 | 0.77418 | 0.773417 | 0.770495 | 0.76867 | 0.770654 | 0.768623 |
| 0.800345 | 0.803581 | 0.77338 | 0.776163 | 0.77517 | 0.768921 | 0.768887 | 0.769286 | 0.769745 |
| 0.797783 | 0.806003 | 0.775111 | 0.768731 | 0.774542 | 0.762854 | 0.765062 | 0.767922 | 0.768012 |
| 1 | 0.846318 | 0.774395 | 0.767793 | 0.775622 | 0.764825 | 0.760673 | 0.773839 | 0.770927 |
| 0.846318 | 1 | 0.779456 | 0.776689 | 0.780032 | 0.77789 | 0.773928 | 0.779754 | 0.777472 |

| | | | | | | | | |
|---|---|---|---|---|---|---|---|---|
| 0.774395 | 0.779456 | 1 | 0.79058 | 0.791581 | 0.789352 | 0.789382 | 0.788533 | 0.792962 |
| 0.767793 | 0.776689 | 0.79058 | 1 | 0.790836 | 0.78638 | 0.794662 | 0.800506 | 0.789194 |
| 0.775622 | 0.780032 | 0.791581 | 0.790836 | 1 | 0.819974 | 0.804853 | 0.799334 | 0.803466 |
| 0.764825 | 0.77789 | 0.789352 | 0.78638 | 0.819974 | 1 | 0.820379 | 0.798612 | 0.794108 |
| 0.760673 | 0.773928 | 0.789382 | 0.794662 | 0.804853 | 0.820379 | 1 | 0.803402 | 0.795029 |
| 0.773839 | 0.779754 | 0.788533 | 0.800506 | 0.799334 | 0.798612 | 0.803402 | 1 | 0.82169 |
| 0.770927 | 0.777472 | 0.792962 | 0.789194 | 0.803466 | 0.794108 | 0.795029 | 0.82169 | 1 |
| 0.771496 | 0.784012 | 0.78661 | 0.796013 | 0.791803 | 0.781551 | 0.788596 | 0.782318 | 0.787477 |
| 0.767468 | 0.776701 | 0.782329 | 0.783292 | 0.792141 | 0.785556 | 0.781247 | 0.784224 | 0.792932 |
| 0.767653 | 0.773172 | 0.788529 | 0.839304 | 0.785908 | 0.778627 | 0.784713 | 0.782982 | 0.781874 |
| 0.767558 | 0.773164 | 0.789866 | 0.793743 | 0.864656 | 0.862949 | 0.858015 | 0.822442 | 0.814685 |
| 0.773211 | 0.774232 | 0.787876 | 0.784076 | 0.828457 | 0.811812 | 0.81044 | 0.794673 | 0.806446 |
| 0.762117 | 0.767543 | 0.790119 | 0.782896 | 0.864418 | 0.8125 | 0.823276 | 0.799177 | 0.801722 |
| 0.759055 | 0.766621 | 0.779666 | 0.79894 | 0.779315 | 0.775664 | 0.778797 | 0.778449 | 0.79263 |
| 0.763522 | 0.767985 | 0.771847 | 0.764348 | 0.768081 | 0.759822 | 0.760785 | 0.760139 | 0.764549 |
| 0.771808 | 0.780093 | 0.781317 | 0.769224 | 0.775728 | 0.770443 | 0.76858 | 0.772953 | 0.773854 |
| 0.767163 | 0.772558 | 0.771295 | 0.767347 | 0.770096 | 0.764039 | 0.762718 | 0.765896 | 0.763573 |
| 0.770227 | 0.777796 | 0.777025 | 0.771229 | 0.774383 | 0.768893 | 0.770526 | 0.765941 | 0.769537 |
| 0.761776 | 0.773922 | 0.771144 | 0.768636 | 0.771048 | 0.765255 | 0.768645 | 0.768065 | 0.767211 |
| 0.757599 | 0.768026 | 0.766531 | 0.765974 | 0.765499 | 0.762092 | 0.764762 | 0.76482 | 0.767649 |
| 0.763052 | 0.77157 | 0.767452 | 0.765227 | 0.765772 | 0.762875 | 0.76603 | 0.759757 | 0.77 |
| 0.764836 | 0.766639 | 0.770326 | 0.76399 | 0.767188 | 0.760436 | 0.760962 | 0.766119 | 0.765742 |
| 0.770456 | 0.771251 | 0.775867 | 0.768604 | 0.771232 | 0.767517 | 0.768587 | 0.767733 | 0.768479 |
| 0.768299 | 0.778276 | 0.782593 | 0.794872 | 0.790285 | 0.787936 | 0.796949 | 0.811046 | 0.787961 |
| 0.767613 | 0.768789 | 0.775326 | 0.765895 | 0.771355 | 0.764936 | 0.76907 | 0.771822 | 0.769175 |
| 0.700835 | 0.704492 | 0.704301 | 0.704656 | 0.705001 | 0.702294 | 0.701258 | 0.704472 | 0.703088 |
| 0.768907 | 0.767139 | 0.773452 | 0.769662 | 0.769562 | 0.762828 | 0.766216 | 0.770087 | 0.765243 |
| 0.771847 | 0.774654 | 0.769845 | 0.772177 | 0.771868 | 0.762709 | 0.76679 | 0.766908 | 0.767579 |
| 0.770399 | 0.774251 | 0.782074 | 0.772173 | 0.773537 | 0.768214 | 0.766774 | 0.769297 | 0.773537 |
| 0.771528 | 0.773866 | 0.771591 | 0.774662 | 0.767424 | 0.76738 | 0.765119 | 0.773391 | 0.766509 |
| 0.770176 | 0.764757 | 0.770023 | 0.764578 | 0.762996 | 0.759959 | 0.761809 | 0.766809 | 0.763306 |
| 0.764649 | 0.76576 | 0.77184 | 0.770764 | 0.768735 | 0.76639 | 0.766568 | 0.766042 | 0.766844 |
| 0.762857 | 0.769028 | 0.770374 | 0.771318 | 0.770888 | 0.769393 | 0.762268 | 0.773028 | 0.767928 |
| 0.767141 | 0.76917 | 0.767624 | 0.769726 | 0.770425 | 0.767075 | 0.758814 | 0.763807 | 0.766725 |
| 0.762353 | 0.766808 | 0.764014 | 0.761983 | 0.764582 | 0.753806 | 0.760039 | 0.758283 | 0.76057 |
| 0.763044 | 0.768646 | 0.764923 | 0.761905 | 0.764881 | 0.75453 | 0.756775 | 0.76159 | 0.762591 |
| 0.767526 | 0.770534 | 0.77443 | 0.769133 | 0.773484 | 0.765688 | 0.773132 | 0.770748 | 0.774104 |
| 0.764976 | 0.770201 | 0.774558 | 0.770411 | 0.771995 | 0.770579 | 0.76529 | 0.764342 | 0.76523 |
| 0.768475 | 0.770478 | 0.775971 | 0.769829 | 0.771017 | 0.766327 | 0.76792 | 0.768309 | 0.766735 |
| 0.762859 | 0.769808 | 0.767826 | 0.768494 | 0.767286 | 0.762958 | 0.766816 | 0.765865 | 0.76437 |
| 0.759586 | 0.766211 | 0.766892 | 0.762586 | 0.769197 | 0.765504 | 0.761129 | 0.76119 | 0.764432 |
| 0.769363 | 0.772654 | 0.771924 | 0.76752 | 0.774734 | 0.769548 | 0.768001 | 0.767664 | 0.768748 |
| 0.761022 | 0.763386 | 0.764721 | 0.760226 | 0.764497 | 0.766126 | 0.756172 | 0.760929 | 0.761938 |
| 0.76763 | 0.772208 | 0.773152 | 0.764037 | 0.766007 | 0.768299 | 0.76778 | 0.764516 | 0.765773 |
| 0.775198 | 0.77158 | 0.771982 | 0.770625 | 0.773066 | 0.772474 | 0.774554 | 0.766473 | 0.770357 |
| 0.773357 | 0.771079 | 0.775699 | 0.76909 | 0.76782 | 0.763666 | 0.768879 | 0.765564 | 0.769902 |

| | | | | | | | | |
|---|---|---|---|---|---|---|---|---|
| 0.769263 | 0.767964 | 0.772609 | 0.767388 | 0.770132 | 0.76302 | 0.767949 | 0.769951 | 0.771809 |
| 0.771527 | 0.774398 | 0.77629 | 0.77445 | 0.775492 | 0.773554 | 0.776538 | 0.772445 | 0.780168 |
| 0.767618 | 0.771776 | 0.775192 | 0.767182 | 0.77015 | 0.768396 | 0.768072 | 0.768059 | 0.770924 |
| 0.725712 | 0.741246 | 0.729009 | 0.730296 | 0.72688 | 0.717508 | 0.717829 | 0.71894 | 0.725235 |
| 0.766034 | 0.764672 | 0.770068 | 0.769034 | 0.765859 | 0.76217 | 0.764661 | 0.75721 | 0.764519 |
| 0.765677 | 0.765771 | 0.771373 | 0.764779 | 0.769346 | 0.766468 | 0.762778 | 0.75798 | 0.762887 |
| 0.770572 | 0.774234 | 0.770312 | 0.769091 | 0.772118 | 0.766848 | 0.76479 | 0.766539 | 0.768562 |
| 0.76753 | 0.769911 | 0.767257 | 0.774735 | 0.770348 | 0.767683 | 0.769808 | 0.766189 | 0.772689 |
| 0.771933 | 0.77316 | 0.779939 | 0.773871 | 0.77736 | 0.770519 | 0.770893 | 0.771592 | 0.774586 |
| 0.770791 | 0.776448 | 0.77349 | 0.771582 | 0.77457 | 0.772085 | 0.77277 | 0.775465 | 0.769127 |
| 0.769146 | 0.772624 | 0.771758 | 0.768287 | 0.773323 | 0.764517 | 0.766962 | 0.765576 | 0.769244 |
| 0.768554 | 0.771254 | 0.771163 | 0.7756 | 0.773786 | 0.769377 | 0.77372 | 0.769713 | 0.772716 |
| 0.770752 | 0.771619 | 0.769619 | 0.769186 | 0.768373 | 0.766554 | 0.769647 | 0.768916 | 0.771378 |
| 0.770127 | 0.77316 | 0.774752 | 0.771488 | 0.77392 | 0.768964 | 0.76912 | 0.768348 | 0.775342 |
| 0.766773 | 0.774546 | 0.769877 | 0.77287 | 0.775044 | 0.772637 | 0.772162 | 0.775182 | 0.776875 |
| 0.762927 | 0.766441 | 0.767507 | 0.769122 | 0.768109 | 0.766557 | 0.76676 | 0.767766 | 0.777005 |
| 0.770365 | 0.77322 | 0.774571 | 0.769464 | 0.773081 | 0.76609 | 0.770605 | 0.766994 | 0.771199 |
| 0.765993 | 0.775015 | 0.772559 | 0.775605 | 0.77636 | 0.77212 | 0.777269 | 0.772821 | 0.77535 |
| 0.765625 | 0.769788 | 0.766896 | 0.763064 | 0.766437 | 0.768493 | 0.764855 | 0.764806 | 0.764622 |
| 0.768451 | 0.773516 | 0.770078 | 0.77082 | 0.769237 | 0.768729 | 0.768907 | 0.766048 | 0.773134 |
| 0.76914 | 0.771906 | 0.776847 | 0.776121 | 0.776624 | 0.772897 | 0.777325 | 0.769565 | 0.776658 |
| 0.766054 | 0.77211 | 0.775345 | 0.77068 | 0.776851 | 0.76892 | 0.76995 | 0.771207 | 0.770709 |
| 0.677312 | 0.678159 | 0.682136 | 0.679494 | 0.69045 | 0.679919 | 0.681215 | 0.677684 | 0.682119 |
| 0.65253 | 0.647881 | 0.654321 | 0.65625 | 0.668735 | 0.65719 | 0.64843 | 0.661627 | 0.659952 |
| 0.666812 | 0.66855 | 0.676302 | 0.657186 | 0.67342 | 0.669262 | 0.669519 | 0.671878 | 0.673703 |
| 0.652152 | 0.653426 | 0.657275 | 0.653095 | 0.657312 | 0.653606 | 0.648017 | 0.65226 | 0.655532 |
| 0.744369 | 0.747243 | 0.757202 | 0.747764 | 0.753452 | 0.750153 | 0.756923 | 0.747599 | 0.755757 |
| 0.743494 | 0.739798 | 0.745964 | 0.740604 | 0.741482 | 0.732525 | 0.745647 | 0.741373 | 0.741527 |
| 0.732446 | 0.744229 | 0.738915 | 0.736058 | 0.745957 | 0.74093 | 0.739043 | 0.742951 | 0.747628 |
| 0.752425 | 0.750317 | 0.743337 | 0.741672 | 0.766025 | 0.744475 | 0.752233 | 0.75746 | 0.754203 |
| 0.727443 | 0.733178 | 0.722068 | 0.73125 | 0.73666 | 0.731178 | 0.725576 | 0.739272 | 0.727315 |
| 0.719723 | 0.721203 | 0.732582 | 0.723921 | 0.729469 | 0.714651 | 0.721699 | 0.729683 | 0.722957 |
| 0.724572 | 0.726999 | 0.743389 | 0.731032 | 0.738046 | 0.734639 | 0.731824 | 0.732244 | 0.734295 |
| 0.655091 | 0.653535 | 0.664309 | 0.65508 | 0.658884 | 0.650116 | 0.651529 | 0.65375 | 0.659624 |
| 0.767919 | 0.774559 | 0.777168 | 0.777467 | 0.770478 | 0.772131 | 0.769869 | 0.76934 | 0.776506 |
| 0.748091 | 0.751419 | 0.759671 | 0.757526 | 0.766438 | 0.746795 | 0.754116 | 0.754071 | 0.750917 |
| 0.741891 | 0.74425 | 0.746533 | 0.748631 | 0.751637 | 0.744561 | 0.752215 | 0.747634 | 0.748596 |
| 0.732536 | 0.738109 | 0.737318 | 0.740057 | 0.740299 | 0.727662 | 0.734648 | 0.741102 | 0.739621 |
| 0.757601 | 0.757953 | 0.753713 | 0.758261 | 0.751708 | 0.751449 | 0.753765 | 0.75656 | 0.757129 |
| 0.735256 | 0.738058 | 0.745343 | 0.739192 | 0.741644 | 0.729298 | 0.741952 | 0.737497 | 0.742422 |
| 0.728731 | 0.739208 | 0.735992 | 0.735422 | 0.740948 | 0.738011 | 0.74227 | 0.738877 | 0.740962 |
| 0.737271 | 0.742161 | 0.741864 | 0.743715 | 0.737893 | 0.725991 | 0.737337 | 0.73831 | 0.74517 |
| 0.731519 | 0.739849 | 0.736437 | 0.743091 | 0.737584 | 0.723868 | 0.732883 | 0.736525 | 0.739967 |
| 0.734223 | 0.744873 | 0.732667 | 0.736097 | 0.741942 | 0.722906 | 0.72884 | 0.734238 | 0.729633 |
| 0.737856 | 0.734391 | 0.747042 | 0.737984 | 0.734552 | 0.728029 | 0.738164 | 0.734921 | 0.738238 |
| 0.714858 | 0.725008 | 0.721317 | 0.724184 | 0.730196 | 0.716413 | 0.713671 | 0.719934 | 0.727909 |

| | | | | | | | | |
|---|---|---|---|---|---|---|---|---|
| 0.759498 | 0.763264 | 0.769998 | 0.766294 | 0.766216 | 0.764937 | 0.762528 | 0.771937 | 0.767837 |
| 0.741141 | 0.744831 | 0.746626 | 0.756115 | 0.753313 | 0.743155 | 0.744146 | 0.745232 | 0.747965 |
| 0.731532 | 0.742438 | 0.748467 | 0.747907 | 0.758074 | 0.747903 | 0.743895 | 0.754486 | 0.743543 |
| 0.737312 | 0.745409 | 0.751252 | 0.749186 | 0.74454 | 0.742964 | 0.754019 | 0.756693 | 0.752619 |
| 0.738259 | 0.731918 | 0.745956 | 0.748292 | 0.743742 | 0.735409 | 0.748705 | 0.747729 | 0.744268 |
| 0.726334 | 0.731837 | 0.748473 | 0.741159 | 0.746737 | 0.74625 | 0.744456 | 0.74584 | 0.749809 |
| 0.735009 | 0.739763 | 0.7438 | 0.744094 | 0.748217 | 0.731994 | 0.737749 | 0.753652 | 0.737596 |
| 0.723178 | 0.726184 | 0.736071 | 0.734326 | 0.742498 | 0.738484 | 0.739966 | 0.750573 | 0.7409 |
| 0.732818 | 0.740006 | 0.738395 | 0.733485 | 0.754501 | 0.741245 | 0.743014 | 0.749281 | 0.748489 |
| 0.751906 | 0.757922 | 0.767153 | 0.766581 | 0.766325 | 0.760773 | 0.765799 | 0.765713 | 0.770639 |
| 0.749227 | 0.753598 | 0.770118 | 0.760867 | 0.760645 | 0.755235 | 0.765339 | 0.761791 | 0.768008 |
| 0.754 | 0.751399 | 0.764094 | 0.760789 | 0.759536 | 0.758949 | 0.757032 | 0.754877 | 0.764939 |
| 0.736855 | 0.747966 | 0.751999 | 0.749816 | 0.750917 | 0.750092 | 0.755949 | 0.745721 | 0.756556 |
| 0.740695 | 0.74758 | 0.760588 | 0.75314 | 0.760261 | 0.750737 | 0.753598 | 0.753471 | 0.762196 |
| 0.73675 | 0.749281 | 0.757397 | 0.742649 | 0.753399 | 0.74643 | 0.750562 | 0.748343 | 0.765529 |

| Wild_plot_ | Wild_plot_ | Wild_plot_ | Wild_plot_ | Wild_plot_ | Wild_plot_ | Wild_plot_ | Wild_plot_ | Wild_plot_ |
|---|---|---|---|---|---|---|---|---|
| 0.70594 | 0.704597 | 0.707331 | 0.70267 | 0.701655 | 0.698264 | 0.699085 | 0.7077 | 0.71256 |
| 0.715658 | 0.712627 | 0.713537 | 0.714007 | 0.713424 | 0.709831 | 0.7054 | 0.714484 | 0.722884 |
| 0.703338 | 0.696916 | 0.703624 | 0.699863 | 0.700572 | 0.693472 | 0.693075 | 0.705963 | 0.711085 |
| 0.712796 | 0.708354 | 0.717493 | 0.711375 | 0.711097 | 0.704739 | 0.704807 | 0.717763 | 0.719811 |
| 0.709063 | 0.705787 | 0.707221 | 0.705092 | 0.706591 | 0.699846 | 0.700351 | 0.705749 | 0.713326 |
| 0.713774 | 0.712893 | 0.716613 | 0.71296 | 0.712496 | 0.703545 | 0.707825 | 0.719381 | 0.722093 |
| 0.769929 | 0.763922 | 0.76 | 0.765015 | 0.76252 | 0.762234 | 0.759299 | 0.768708 | 0.778467 |
| 0.766978 | 0.768764 | 0.766788 | 0.76421 | 0.76419 | 0.757013 | 0.756608 | 0.76646 | 0.780749 |
| 0.711978 | 0.705743 | 0.710904 | 0.708621 | 0.709961 | 0.7005 | 0.703994 | 0.709888 | 0.718149 |
| 0.707722 | 0.705456 | 0.708434 | 0.70391 | 0.704345 | 0.699031 | 0.700754 | 0.707268 | 0.712052 |
| 0.710916 | 0.704547 | 0.710984 | 0.710708 | 0.705261 | 0.697032 | 0.701136 | 0.711718 | 0.713919 |
| 0.70152 | 0.701277 | 0.704334 | 0.700823 | 0.701457 | 0.690391 | 0.692955 | 0.7038 | 0.711047 |
| 0.715622 | 0.709989 | 0.709955 | 0.711028 | 0.712436 | 0.702691 | 0.704241 | 0.715442 | 0.721489 |
| 0.723327 | 0.718671 | 0.720384 | 0.718816 | 0.717788 | 0.710495 | 0.712164 | 0.722927 | 0.731083 |
| 0.732846 | 0.733344 | 0.733646 | 0.732149 | 0.730509 | 0.724684 | 0.728665 | 0.734186 | 0.742827 |
| 0.716532 | 0.712245 | 0.712316 | 0.715666 | 0.711759 | 0.70673 | 0.706307 | 0.717166 | 0.719263 |
| 0.713426 | 0.706808 | 0.714123 | 0.710391 | 0.711073 | 0.704301 | 0.708485 | 0.716436 | 0.715865 |
| 0.742351 | 0.739696 | 0.740514 | 0.742255 | 0.739254 | 0.729776 | 0.733944 | 0.746234 | 0.75139 |
| 0.701987 | 0.69991 | 0.705203 | 0.702633 | 0.701706 | 0.694145 | 0.696342 | 0.709949 | 0.714194 |
| 0.710255 | 0.70618 | 0.710304 | 0.711256 | 0.710173 | 0.705369 | 0.699956 | 0.711906 | 0.715214 |
| 0.708823 | 0.702654 | 0.705557 | 0.703698 | 0.706559 | 0.697441 | 0.698054 | 0.707004 | 0.711259 |
| 0.713793 | 0.707208 | 0.708853 | 0.709215 | 0.708387 | 0.701406 | 0.701882 | 0.708642 | 0.717815 |
| 0.714531 | 0.714087 | 0.715321 | 0.715215 | 0.714599 | 0.707813 | 0.706644 | 0.717695 | 0.722222 |
| 0.71126 | 0.70849 | 0.71233 | 0.710242 | 0.707904 | 0.702542 | 0.702453 | 0.715044 | 0.717414 |
| 0.714733 | 0.713558 | 0.71216 | 0.713156 | 0.713223 | 0.705358 | 0.70594 | 0.714438 | 0.724185 |
| 0.711183 | 0.705625 | 0.708007 | 0.711172 | 0.709789 | 0.701463 | 0.701014 | 0.707167 | 0.716231 |
| 0.704878 | 0.702982 | 0.704802 | 0.702078 | 0.702703 | 0.693482 | 0.699216 | 0.703267 | 0.710419 |
| 0.71361 | 0.711545 | 0.712554 | 0.7095 | 0.707392 | 0.701759 | 0.703512 | 0.711833 | 0.718495 |
| 0.712964 | 0.714403 | 0.710677 | 0.713368 | 0.713505 | 0.704394 | 0.707306 | 0.720726 | 0.723046 |
| 0.710611 | 0.709006 | 0.709521 | 0.70988 | 0.709738 | 0.703145 | 0.702662 | 0.711899 | 0.718443 |
| 0.708566 | 0.70787 | 0.707453 | 0.710715 | 0.708011 | 0.699285 | 0.701579 | 0.705297 | 0.714446 |
| 0.706016 | 0.705327 | 0.705948 | 0.707467 | 0.707639 | 0.698158 | 0.697221 | 0.704959 | 0.709942 |
| 0.708323 | 0.708486 | 0.708343 | 0.7083 | 0.706457 | 0.697818 | 0.69894 | 0.707719 | 0.715347 |
| 0.711719 | 0.708255 | 0.710138 | 0.712978 | 0.711311 | 0.700103 | 0.702558 | 0.710161 | 0.715503 |
| 0.703948 | 0.703441 | 0.703234 | 0.701464 | 0.703599 | 0.695562 | 0.701444 | 0.705581 | 0.712444 |
| 0.709987 | 0.708039 | 0.709101 | 0.712258 | 0.708959 | 0.700606 | 0.701387 | 0.713242 | 0.715634 |
| 0.714274 | 0.711053 | 0.715407 | 0.712465 | 0.712522 | 0.705282 | 0.702552 | 0.713568 | 0.720984 |
| 0.710446 | 0.712691 | 0.711936 | 0.711007 | 0.71237 | 0.700557 | 0.700027 | 0.711923 | 0.718958 |
| 0.718079 | 0.712409 | 0.715783 | 0.709904 | 0.713194 | 0.702998 | 0.705566 | 0.713902 | 0.719865 |

| | | | | | | | | |
|---|---|---|---|---|---|---|---|---|
| 0.744198 | 0.743164 | 0.746109 | 0.741999 | 0.743133 | 0.733345 | 0.736642 | 0.748295 | 0.756827 |
| 0.700853 | 0.697124 | 0.702878 | 0.698776 | 0.697858 | 0.692499 | 0.691125 | 0.704116 | 0.707809 |
| 0.704198 | 0.699748 | 0.701228 | 0.704389 | 0.702083 | 0.694444 | 0.695427 | 0.708119 | 0.712921 |
| 0.713601 | 0.70633 | 0.715432 | 0.711245 | 0.711176 | 0.703561 | 0.70758 | 0.716013 | 0.721378 |
| 0.706892 | 0.706371 | 0.706881 | 0.704164 | 0.706606 | 0.697529 | 0.702216 | 0.715021 | 0.717025 |
| 0.713486 | 0.712527 | 0.712121 | 0.711584 | 0.712929 | 0.703688 | 0.704592 | 0.716563 | 0.719761 |
| 0.703791 | 0.705386 | 0.705171 | 0.703567 | 0.705198 | 0.694998 | 0.697589 | 0.705406 | 0.71416 |
| 0.710027 | 0.708412 | 0.710922 | 0.708174 | 0.706567 | 0.70083 | 0.701503 | 0.713253 | 0.718838 |
| 0.711453 | 0.709489 | 0.712847 | 0.712662 | 0.710769 | 0.703974 | 0.706822 | 0.712489 | 0.720492 |
| 0.711942 | 0.712221 | 0.711464 | 0.71131 | 0.712362 | 0.703898 | 0.705453 | 0.711342 | 0.716154 |
| 0.70223 | 0.699584 | 0.705375 | 0.702422 | 0.699545 | 0.692321 | 0.698726 | 0.702039 | 0.710628 |
| 0.703365 | 0.704088 | 0.705745 | 0.7021 | 0.701844 | 0.69364 | 0.699154 | 0.70546 | 0.71277 |
| 0.709058 | 0.705684 | 0.706941 | 0.705239 | 0.704378 | 0.698586 | 0.698725 | 0.709097 | 0.715984 |
| 0.704229 | 0.702795 | 0.704403 | 0.705523 | 0.70535 | 0.696894 | 0.696128 | 0.707764 | 0.708618 |
| 0.702133 | 0.703557 | 0.704085 | 0.705402 | 0.702667 | 0.697438 | 0.695156 | 0.705433 | 0.716872 |
| 0.703183 | 0.702618 | 0.704736 | 0.702067 | 0.701116 | 0.692214 | 0.697965 | 0.705494 | 0.714997 |
| 0.702436 | 0.698644 | 0.701272 | 0.703154 | 0.700402 | 0.694338 | 0.691599 | 0.707373 | 0.710656 |
| 0.711834 | 0.708388 | 0.712093 | 0.709612 | 0.706044 | 0.701567 | 0.700114 | 0.710839 | 0.713464 |
| 0.709622 | 0.709303 | 0.709787 | 0.708769 | 0.710652 | 0.702854 | 0.703143 | 0.71326 | 0.718793 |
| 0.708546 | 0.70241 | 0.709123 | 0.707341 | 0.705926 | 0.698881 | 0.699239 | 0.707731 | 0.714101 |
| 0.706929 | 0.700172 | 0.701328 | 0.700796 | 0.700162 | 0.693291 | 0.695553 | 0.706696 | 0.711356 |
| 0.707347 | 0.701412 | 0.704461 | 0.704049 | 0.706026 | 0.694653 | 0.695128 | 0.708129 | 0.715299 |
| 0.708732 | 0.708252 | 0.705723 | 0.705612 | 0.704617 | 0.698333 | 0.698313 | 0.708256 | 0.714877 |
| 0.719772 | 0.717035 | 0.714274 | 0.712925 | 0.712727 | 0.704927 | 0.708592 | 0.716803 | 0.726655 |
| 0.723985 | 0.720894 | 0.721441 | 0.718196 | 0.720504 | 0.712637 | 0.72056 | 0.720985 | 0.726388 |
| 0.769486 | 0.770225 | 0.767802 | 0.765057 | 0.766388 | 0.758627 | 0.759898 | 0.767452 | 0.780371 |
| 0.767083 | 0.764123 | 0.769315 | 0.767228 | 0.760793 | 0.761619 | 0.759291 | 0.777258 | 0.785091 |
| 0.771069 | 0.767458 | 0.772167 | 0.769185 | 0.766818 | 0.7635 | 0.766817 | 0.779507 | 0.786685 |
| 0.707114 | 0.704659 | 0.706294 | 0.704558 | 0.706121 | 0.69607 | 0.697142 | 0.709998 | 0.717594 |
| 0.708817 | 0.703626 | 0.707532 | 0.705661 | 0.706049 | 0.69737 | 0.699531 | 0.710352 | 0.710578 |
| 0.71492 | 0.712398 | 0.716054 | 0.713948 | 0.712702 | 0.705512 | 0.707479 | 0.717441 | 0.721153 |
| 0.71618 | 0.712951 | 0.715952 | 0.71533 | 0.715685 | 0.707271 | 0.708866 | 0.716507 | 0.721847 |
| 0.706004 | 0.702534 | 0.705152 | 0.704906 | 0.705766 | 0.699521 | 0.699006 | 0.709491 | 0.713718 |
| 0.709023 | 0.706661 | 0.705297 | 0.707677 | 0.706787 | 0.695972 | 0.701897 | 0.709192 | 0.716798 |
| 0.70744 | 0.700831 | 0.70762 | 0.704975 | 0.7058 | 0.697869 | 0.700695 | 0.707841 | 0.714437 |
| 0.708357 | 0.702039 | 0.70602 | 0.706303 | 0.706961 | 0.697694 | 0.698747 | 0.709693 | 0.71447 |
| 0.707533 | 0.707335 | 0.709654 | 0.70619 | 0.70412 | 0.697933 | 0.700858 | 0.706599 | 0.714854 |
| 0.718781 | 0.718496 | 0.718978 | 0.721763 | 0.717342 | 0.71379 | 0.71249 | 0.721834 | 0.729662 |
| 0.701289 | 0.697885 | 0.702903 | 0.699111 | 0.69821 | 0.695278 | 0.693373 | 0.700371 | 0.708129 |
| 0.705315 | 0.703647 | 0.704592 | 0.705179 | 0.707033 | 0.69405 | 0.702215 | 0.711324 | 0.714116 |
| 0.705406 | 0.699387 | 0.7035 | 0.70122 | 0.70267 | 0.693757 | 0.697597 | 0.702419 | 0.710436 |
| 0.723594 | 0.721445 | 0.722339 | 0.722845 | 0.720982 | 0.717081 | 0.714526 | 0.722008 | 0.732771 |
| 0.707944 | 0.704673 | 0.707009 | 0.707734 | 0.705558 | 0.697548 | 0.698775 | 0.712078 | 0.713231 |
| 0.706269 | 0.70374 | 0.705497 | 0.705518 | 0.701969 | 0.69452 | 0.699372 | 0.70443 | 0.710786 |
| 0.71049 | 0.708542 | 0.70697 | 0.706102 | 0.708194 | 0.699628 | 0.701757 | 0.712878 | 0.715126 |
| 0.709712 | 0.705863 | 0.707572 | 0.706345 | 0.703089 | 0.699064 | 0.697755 | 0.708385 | 0.714101 |

| | | | | | | | | |
|---|---|---|---|---|---|---|---|---|
| 0.705267 | 0.701654 | 0.702555 | 0.704077 | 0.704549 | 0.697238 | 0.696383 | 0.708476 | 0.714539 |
| 0.710533 | 0.705696 | 0.711715 | 0.70728 | 0.711147 | 0.70129 | 0.702216 | 0.714996 | 0.716467 |
| 0.711421 | 0.709116 | 0.708455 | 0.706232 | 0.705868 | 0.698936 | 0.70328 | 0.709391 | 0.71617 |
| 0.71327 | 0.712986 | 0.711706 | 0.713065 | 0.713488 | 0.70483 | 0.707289 | 0.717319 | 0.721906 |
| 0.713808 | 0.712067 | 0.711367 | 0.710484 | 0.712076 | 0.702775 | 0.707326 | 0.714425 | 0.719328 |
| 0.704544 | 0.704232 | 0.70811 | 0.703209 | 0.704097 | 0.696084 | 0.699389 | 0.712746 | 0.719148 |
| 0.707368 | 0.708178 | 0.708808 | 0.711408 | 0.708555 | 0.703173 | 0.701469 | 0.70894 | 0.717983 |
| 0.713861 | 0.708244 | 0.711875 | 0.710073 | 0.710103 | 0.702426 | 0.704509 | 0.712301 | 0.718247 |
| 0.710959 | 0.70782 | 0.707993 | 0.708686 | 0.706776 | 0.701575 | 0.701485 | 0.709585 | 0.713929 |
| 0.70862 | 0.704223 | 0.705689 | 0.703023 | 0.704966 | 0.695853 | 0.695751 | 0.707107 | 0.713049 |
| 0.710738 | 0.707457 | 0.705939 | 0.707863 | 0.706219 | 0.697926 | 0.697918 | 0.711197 | 0.711456 |
| 0.707537 | 0.709029 | 0.709409 | 0.709547 | 0.708649 | 0.69946 | 0.704738 | 0.708457 | 0.713832 |
| 0.708429 | 0.712635 | 0.708036 | 0.706832 | 0.707036 | 0.698061 | 0.702607 | 0.708425 | 0.716466 |
| 0.702696 | 0.70321 | 0.70619 | 0.701871 | 0.705649 | 0.691559 | 0.698563 | 0.70805 | 0.712457 |
| 0.710492 | 0.711179 | 0.708859 | 0.707257 | 0.705546 | 0.699861 | 0.700898 | 0.710647 | 0.71539 |
| 0.706404 | 0.703217 | 0.709292 | 0.703068 | 0.704797 | 0.698519 | 0.69887 | 0.706377 | 0.71552 |
| 0.710126 | 0.707485 | 0.71088 | 0.70796 | 0.708082 | 0.698522 | 0.702152 | 0.712689 | 0.719644 |
| 0.70619 | 0.704294 | 0.705514 | 0.705777 | 0.708878 | 0.69823 | 0.702188 | 0.706299 | 0.714733 |
| 0.708565 | 0.706479 | 0.710779 | 0.70926 | 0.706769 | 0.701493 | 0.70164 | 0.711691 | 0.714228 |
| 0.71139 | 0.705092 | 0.712127 | 0.708249 | 0.708971 | 0.698354 | 0.701965 | 0.709299 | 0.712454 |
| 0.707617 | 0.704911 | 0.709912 | 0.705744 | 0.705815 | 0.699422 | 0.701058 | 0.710468 | 0.712617 |
| 0.704588 | 0.701032 | 0.70635 | 0.701929 | 0.701359 | 0.695552 | 0.695047 | 0.704038 | 0.705366 |
| 0.707054 | 0.703755 | 0.707042 | 0.707066 | 0.708047 | 0.702215 | 0.698526 | 0.711583 | 0.717307 |
| 0.709403 | 0.707893 | 0.709103 | 0.708644 | 0.7076 | 0.699191 | 0.700272 | 0.710481 | 0.717897 |
| 0.7066 | 0.708881 | 0.710252 | 0.709934 | 0.706771 | 0.702252 | 0.703034 | 0.712587 | 0.716257 |
| 0.7123 | 0.712323 | 0.713575 | 0.712164 | 0.710344 | 0.702619 | 0.704889 | 0.716283 | 0.723782 |
| 0.70645 | 0.705068 | 0.710269 | 0.705026 | 0.703948 | 0.701019 | 0.70042 | 0.711117 | 0.712534 |
| 0.711729 | 0.709349 | 0.711678 | 0.708788 | 0.709411 | 0.700688 | 0.705396 | 0.71211 | 0.718961 |
| 0.709567 | 0.710591 | 0.71276 | 0.708414 | 0.708523 | 0.701521 | 0.699553 | 0.711717 | 0.71801 |
| 0.708558 | 0.71015 | 0.710036 | 0.710095 | 0.707939 | 0.700427 | 0.704133 | 0.712075 | 0.71582 |
| 0.714686 | 0.712172 | 0.712119 | 0.712196 | 0.711167 | 0.705698 | 0.70463 | 0.712256 | 0.716943 |
| 0.705056 | 0.705678 | 0.706838 | 0.706392 | 0.703968 | 0.697581 | 0.700523 | 0.708216 | 0.711007 |
| 0.707769 | 0.710067 | 0.707498 | 0.709109 | 0.707451 | 0.700915 | 0.70415 | 0.709187 | 0.715923 |
| 0.711225 | 0.70205 | 0.711651 | 0.708784 | 0.707311 | 0.701683 | 0.702577 | 0.712733 | 0.717885 |
| 0.705593 | 0.706632 | 0.707657 | 0.707257 | 0.710831 | 0.6967 | 0.697436 | 0.709225 | 0.711942 |
| 0.711622 | 0.70649 | 0.707949 | 0.707167 | 0.706972 | 0.700105 | 0.70511 | 0.707077 | 0.715101 |
| 0.703659 | 0.702826 | 0.707695 | 0.705497 | 0.705485 | 0.692705 | 0.696014 | 0.707919 | 0.712707 |
| 0.685319 | 0.681015 | 0.681684 | 0.683693 | 0.67927 | 0.676234 | 0.681603 | 0.686408 | 0.692221 |
| 0.690094 | 0.684599 | 0.686997 | 0.687117 | 0.689836 | 0.6797 | 0.686516 | 0.695769 | 0.699021 |
| 0.679051 | 0.674711 | 0.677166 | 0.675098 | 0.67533 | 0.671187 | 0.677652 | 0.681135 | 0.688586 |
| 0.680618 | 0.675745 | 0.678051 | 0.67914 | 0.678507 | 0.673701 | 0.676554 | 0.687444 | 0.689221 |
| 0.669181 | 0.660522 | 0.66205 | 0.662506 | 0.665049 | 0.659717 | 0.664106 | 0.667771 | 0.674832 |
| 0.673467 | 0.661137 | 0.66833 | 0.667685 | 0.664106 | 0.661898 | 0.661688 | 0.67168 | 0.67582 |
| 0.668149 | 0.663723 | 0.66082 | 0.667685 | 0.66585 | 0.656759 | 0.663016 | 0.671273 | 0.675976 |
| 0.669612 | 0.666581 | 0.661105 | 0.663939 | 0.667353 | 0.657331 | 0.670161 | 0.671224 | 0.671672 |
| 0.666751 | 0.663061 | 0.668838 | 0.666416 | 0.663529 | 0.662942 | 0.668368 | 0.67184 | 0.677286 |
| 0.678634 | 0.668577 | 0.672905 | 0.671597 | 0.673894 | 0.663484 | 0.671412 | 0.675103 | 0.686907 |

| | | | | | | | | |
|---|---|---|---|---|---|---|---|---|
| 0.671625 | 0.666667 | 0.668312 | 0.671873 | 0.666377 | 0.666412 | 0.669852 | 0.676522 | 0.677487 |
| 0.668641 | 0.669918 | 0.672737 | 0.672763 | 0.670182 | 0.669982 | 0.674045 | 0.677513 | 0.68114 |
| 0.664999 | 0.669236 | 0.671294 | 0.666667 | 0.666801 | 0.661631 | 0.667643 | 0.671313 | 0.675498 |
| 0.66439 | 0.65949 | 0.66254 | 0.660435 | 0.662633 | 0.658557 | 0.655895 | 0.666267 | 0.674853 |
| 0.666144 | 0.658404 | 0.663652 | 0.660351 | 0.661497 | 0.657487 | 0.666041 | 0.666926 | 0.675606 |
| 0.670171 | 0.66311 | 0.665796 | 0.663769 | 0.663253 | 0.655973 | 0.665709 | 0.663071 | 0.672557 |
| 0.670583 | 0.669053 | 0.664912 | 0.665475 | 0.668383 | 0.661066 | 0.667552 | 0.667095 | 0.672836 |
| 0.664769 | 0.655991 | 0.661752 | 0.661281 | 0.659661 | 0.657337 | 0.664181 | 0.664639 | 0.674422 |
| 0.66274 | 0.661283 | 0.660656 | 0.658062 | 0.658638 | 0.654388 | 0.662332 | 0.667312 | 0.672881 |
| 0.66785 | 0.659481 | 0.664908 | 0.666079 | 0.664947 | 0.66241 | 0.66692 | 0.665926 | 0.67289 |
| 0.661664 | 0.65457 | 0.658524 | 0.656591 | 0.658664 | 0.652823 | 0.658635 | 0.66206 | 0.668071 |
| 0.697922 | 0.692536 | 0.695023 | 0.697574 | 0.697137 | 0.686298 | 0.68606 | 0.699295 | 0.702664 |
| 0.708929 | 0.707072 | 0.7084 | 0.705037 | 0.704397 | 0.696902 | 0.699763 | 0.706291 | 0.710213 |
| 0.686018 | 0.68198 | 0.683838 | 0.68488 | 0.686485 | 0.681089 | 0.687477 | 0.686938 | 0.694336 |
| 0.716643 | 0.711511 | 0.713621 | 0.715499 | 0.715558 | 0.702743 | 0.708995 | 0.718464 | 0.719632 |
| 0.704854 | 0.702152 | 0.704222 | 0.699685 | 0.701021 | 0.692031 | 0.69255 | 0.705388 | 0.710176 |
| 0.713192 | 0.711082 | 0.713048 | 0.712001 | 0.711834 | 0.702195 | 0.703233 | 0.714437 | 0.717701 |
| 0.721067 | 0.717023 | 0.718005 | 0.714858 | 0.714215 | 0.704626 | 0.711607 | 0.719198 | 0.724158 |
| 0.695715 | 0.690407 | 0.693685 | 0.693773 | 0.695242 | 0.687516 | 0.68769 | 0.69938 | 0.700016 |
| 0.718692 | 0.712886 | 0.718567 | 0.715217 | 0.717518 | 0.707056 | 0.710837 | 0.723248 | 0.723827 |
| 0.689799 | 0.688485 | 0.687606 | 0.691754 | 0.691474 | 0.687798 | 0.691404 | 0.695833 | 0.697878 |
| 0.715694 | 0.717513 | 0.715188 | 0.713576 | 0.714987 | 0.702626 | 0.707906 | 0.718127 | 0.719182 |
| 0.716939 | 0.717425 | 0.721042 | 0.715777 | 0.714589 | 0.706719 | 0.715226 | 0.718712 | 0.723219 |
| 0.687774 | 0.686476 | 0.683466 | 0.683033 | 0.68031 | 0.67607 | 0.684861 | 0.688333 | 0.692209 |
| 0.686179 | 0.686708 | 0.682747 | 0.686483 | 0.683512 | 0.677493 | 0.676738 | 0.688059 | 0.690626 |
| 0.694523 | 0.689836 | 0.69081 | 0.690898 | 0.690774 | 0.690269 | 0.694865 | 0.693885 | 0.695873 |
| 0.709998 | 0.705352 | 0.706807 | 0.709503 | 0.707187 | 0.698007 | 0.701604 | 0.707195 | 0.71267 |
| 0.718323 | 0.718169 | 0.71878 | 0.718732 | 0.7149 | 0.707111 | 0.710815 | 0.716895 | 0.721256 |
| 0.686339 | 0.683574 | 0.688265 | 0.68548 | 0.683895 | 0.678138 | 0.678391 | 0.68932 | 0.690445 |
| 0.724163 | 0.72058 | 0.722249 | 0.723363 | 0.723194 | 0.714727 | 0.714612 | 0.72395 | 0.729834 |
| 0.723908 | 0.726653 | 0.727171 | 0.726716 | 0.724653 | 0.716387 | 0.718725 | 0.725692 | 0.729885 |
| 0.695132 | 0.693413 | 0.695227 | 0.696607 | 0.695677 | 0.690073 | 0.688134 | 0.700196 | 0.702467 |
| 0.693983 | 0.693451 | 0.694343 | 0.6941 | 0.69568 | 0.687027 | 0.686166 | 0.699223 | 0.706495 |
| 0.722997 | 0.720439 | 0.722424 | 0.723171 | 0.722824 | 0.712631 | 0.716859 | 0.724076 | 0.728619 |
| 0.719852 | 0.717938 | 0.71836 | 0.719901 | 0.719442 | 0.710643 | 0.712719 | 0.721886 | 0.728087 |
| 0.724459 | 0.720505 | 0.719409 | 0.723173 | 0.721426 | 0.710558 | 0.716909 | 0.727145 | 0.726949 |
| 0.677499 | 0.675542 | 0.678501 | 0.675584 | 0.673651 | 0.667448 | 0.668097 | 0.68107 | 0.681743 |
| 0.71784 | 0.717729 | 0.718192 | 0.717733 | 0.718398 | 0.712843 | 0.709774 | 0.720085 | 0.724832 |
| 0.673457 | 0.67248 | 0.673371 | 0.6716 | 0.671532 | 0.666391 | 0.667569 | 0.676789 | 0.681176 |
| 0.668585 | 0.667884 | 0.66945 | 0.669579 | 0.668806 | 0.65882 | 0.661077 | 0.67304 | 0.674344 |
| 0.722181 | 0.718625 | 0.720569 | 0.715 | 0.714482 | 0.708512 | 0.71411 | 0.722674 | 0.726641 |
| 0.672584 | 0.668717 | 0.668815 | 0.668703 | 0.667748 | 0.658946 | 0.662021 | 0.670426 | 0.673401 |
| 0.662191 | 0.658875 | 0.661139 | 0.661591 | 0.659796 | 0.652993 | 0.650472 | 0.662591 | 0.666641 |
| 0.657905 | 0.653513 | 0.655635 | 0.657551 | 0.653742 | 0.651027 | 0.647251 | 0.65787 | 0.661551 |
| 0.730633 | 0.726312 | 0.726539 | 0.726075 | 0.726199 | 0.716513 | 0.717998 | 0.733262 | 0.736384 |
| 0.661678 | 0.662886 | 0.658307 | 0.659612 | 0.657822 | 0.651649 | 0.653014 | 0.661022 | 0.668811 |
| 0.723469 | 0.718227 | 0.717361 | 0.721114 | 0.720248 | 0.711585 | 0.71718 | 0.726226 | 0.728985 |

| | | | | | | | | |
|---|---|---|---|---|---|---|---|---|
| 0.732791 | 0.723621 | 0.724598 | 0.72456 | 0.728132 | 0.716289 | 0.719198 | 0.729569 | 0.737731 |
| 0.723162 | 0.722812 | 0.724013 | 0.720023 | 0.720523 | 0.713303 | 0.718017 | 0.727176 | 0.734199 |
| 0.738892 | 0.740241 | 0.741301 | 0.743724 | 0.743185 | 0.734094 | 0.735979 | 0.740572 | 0.748891 |
| 0.649315 | 0.649397 | 0.649187 | 0.651454 | 0.648557 | 0.641974 | 0.642472 | 0.649411 | 0.658653 |
| 0.651673 | 0.649653 | 0.650621 | 0.652538 | 0.651877 | 0.64212 | 0.646803 | 0.653717 | 0.659243 |
| 0.729456 | 0.726945 | 0.728934 | 0.726991 | 0.72728 | 0.718506 | 0.725522 | 0.731685 | 0.739141 |
| 0.652906 | 0.651605 | 0.652167 | 0.652548 | 0.652851 | 0.645568 | 0.646134 | 0.657407 | 0.65921 |
| 0.729613 | 0.725106 | 0.731663 | 0.731091 | 0.733777 | 0.7219 | 0.723029 | 0.733433 | 0.74068 |
| 0.732119 | 0.72519 | 0.72762 | 0.730972 | 0.732341 | 0.720316 | 0.727964 | 0.735776 | 0.742499 |
| 0.730888 | 0.729877 | 0.732755 | 0.731134 | 0.730151 | 0.725442 | 0.720395 | 0.730025 | 0.738883 |
| 0.733272 | 0.728553 | 0.730952 | 0.731225 | 0.729975 | 0.722887 | 0.723016 | 0.732212 | 0.740535 |
| 0.724638 | 0.726115 | 0.755396 | 0.707483 | 0.71519 | 0.771429 | 0.736486 | 0.757143 | 0.719178 |
| 0.742845 | 0.744962 | 0.734545 | 0.737705 | 0.733793 | 0.743484 | 0.747401 | 0.724492 | 0.733845 |
| 0.744033 | 0.740214 | 0.741915 | 0.734264 | 0.732665 | 0.735547 | 0.737221 | 0.72253 | 0.732839 |
| 0.757225 | 0.757538 | 0.7479 | 0.743114 | 0.744388 | 0.747398 | 0.755808 | 0.738685 | 0.738189 |
| 0.75097 | 0.746125 | 0.752053 | 0.750137 | 0.745846 | 0.744878 | 0.74469 | 0.723886 | 0.737647 |
| 0.590128 | 0.589791 | 0.58899 | 0.595422 | 0.593654 | 0.589471 | 0.590289 | 0.593729 | 0.597778 |
| 0.7456 | 0.753928 | 0.755175 | 0.75179 | 0.751128 | 0.748207 | 0.752576 | 0.731251 | 0.743596 |
| 0.737735 | 0.732319 | 0.734921 | 0.732733 | 0.736842 | 0.725277 | 0.73011 | 0.739902 | 0.745522 |
| 0.737406 | 0.734428 | 0.736934 | 0.73183 | 0.735664 | 0.721687 | 0.729237 | 0.73986 | 0.743894 |
| 0.73551 | 0.738247 | 0.736175 | 0.73498 | 0.735902 | 0.726833 | 0.730558 | 0.739337 | 0.747777 |
| 0.592798 | 0.593195 | 0.592478 | 0.592448 | 0.593879 | 0.581173 | 0.585072 | 0.594955 | 0.602702 |
| 0.580029 | 0.582902 | 0.578977 | 0.580226 | 0.581688 | 0.571187 | 0.572217 | 0.580438 | 0.588315 |
| 0.763079 | 0.754759 | 0.757497 | 0.762371 | 0.766485 | 0.758683 | 0.759128 | 0.76014 | 0.765134 |
| 0.760299 | 0.757515 | 0.760663 | 0.762422 | 0.763469 | 0.75687 | 0.754696 | 0.757781 | 0.767096 |
| 0.752622 | 0.750611 | 0.75125 | 0.755956 | 0.758755 | 0.748571 | 0.746362 | 0.753682 | 0.760043 |
| 0.756748 | 0.755331 | 0.755559 | 0.760955 | 0.761606 | 0.751414 | 0.753008 | 0.761613 | 0.766284 |
| 0.761881 | 0.757117 | 0.758669 | 0.757866 | 0.765395 | 0.754364 | 0.753479 | 0.761519 | 0.765611 |
| 0.768738 | 0.757795 | 0.760382 | 0.763603 | 0.765853 | 0.760244 | 0.761502 | 0.761848 | 0.77163 |
| 0.769289 | 0.759183 | 0.762296 | 0.766605 | 0.765675 | 0.757495 | 0.758082 | 0.761094 | 0.768268 |
| 0.759367 | 0.758394 | 0.757043 | 0.755966 | 0.757134 | 0.748647 | 0.754884 | 0.762235 | 0.764559 |
| 0.765317 | 0.765973 | 0.766837 | 0.765788 | 0.766853 | 0.754074 | 0.758948 | 0.770023 | 0.775525 |
| 0.759346 | 0.754813 | 0.755151 | 0.761427 | 0.759683 | 0.755404 | 0.758877 | 0.763695 | 0.764437 |
| 0.753175 | 0.75745 | 0.754333 | 0.75548 | 0.756077 | 0.751975 | 0.754474 | 0.761316 | 0.769023 |
| 0.765596 | 0.765017 | 0.769461 | 0.765271 | 0.766446 | 0.757764 | 0.76345 | 0.768241 | 0.771093 |
| 0.756912 | 0.759036 | 0.761269 | 0.75699 | 0.759046 | 0.752511 | 0.755056 | 0.762652 | 0.769103 |
| 0.76 | 0.759283 | 0.759349 | 0.763626 | 0.764909 | 0.752818 | 0.754082 | 0.759116 | 0.768459 |
| 0.764449 | 0.754777 | 0.761913 | 0.761147 | 0.763737 | 0.751833 | 0.75493 | 0.768656 | 0.772386 |
| 0.757534 | 0.757824 | 0.757094 | 0.75617 | 0.757123 | 0.747496 | 0.755306 | 0.756564 | 0.76676 |
| 0.758344 | 0.760488 | 0.753965 | 0.753832 | 0.758641 | 0.747009 | 0.755518 | 0.758609 | 0.763335 |
| 0.739721 | 0.734033 | 0.733757 | 0.733392 | 0.742056 | 0.733477 | 0.743201 | 0.723352 | 0.733929 |
| 0.739992 | 0.733169 | 0.731343 | 0.733396 | 0.745027 | 0.735351 | 0.741631 | 0.729385 | 0.739426 |
| 0.762483 | 0.761439 | 0.761025 | 0.757573 | 0.758767 | 0.75291 | 0.75036 | 0.76994 | 0.774802 |
| 0.768926 | 0.767248 | 0.771201 | 0.766249 | 0.767304 | 0.761038 | 0.761053 | 0.763676 | 0.77551 |
| 0.716009 | 0.713458 | 0.712451 | 0.714838 | 0.713869 | 0.71603 | 0.705093 | 0.713235 | 0.717622 |
| 0.767183 | 0.759778 | 0.76068 | 0.762459 | 0.762248 | 0.754234 | 0.760418 | 0.76362 | 0.772313 |
| 0.760632 | 0.755392 | 0.752944 | 0.75467 | 0.754469 | 0.744613 | 0.74219 | 0.756764 | 0.764836 |

| | | | | | | | | |
|---|---|---|---|---|---|---|---|---|
| 0.76086 | 0.758502 | 0.756714 | 0.757896 | 0.760201 | 0.752412 | 0.755431 | 0.758618 | 0.767453 |
| 0.764967 | 0.762339 | 0.762392 | 0.760186 | 0.761291 | 0.753297 | 0.7596 | 0.766672 | 0.771773 |
| 0.761798 | 0.75907 | 0.759679 | 0.756585 | 0.759952 | 0.746227 | 0.754307 | 0.76233 | 0.76798 |
| 0.767353 | 0.761605 | 0.758812 | 0.758692 | 0.762786 | 0.754714 | 0.75745 | 0.768178 | 0.773807 |
| 0.766077 | 0.762424 | 0.764409 | 0.767749 | 0.767635 | 0.757178 | 0.760631 | 0.767075 | 0.772667 |
| 0.762537 | 0.760732 | 0.758669 | 0.761421 | 0.758082 | 0.754008 | 0.756966 | 0.760242 | 0.768605 |
| 0.759893 | 0.760486 | 0.760823 | 0.760645 | 0.761417 | 0.755344 | 0.756303 | 0.759333 | 0.768073 |
| 0.754562 | 0.750655 | 0.750167 | 0.749622 | 0.750506 | 0.74078 | 0.748147 | 0.751945 | 0.76104 |
| 0.764706 | 0.757176 | 0.758501 | 0.756148 | 0.757333 | 0.75102 | 0.756471 | 0.763513 | 0.766845 |
| 0.756891 | 0.753641 | 0.754145 | 0.754834 | 0.756757 | 0.745408 | 0.745869 | 0.75675 | 0.767507 |
| 0.764587 | 0.761638 | 0.759714 | 0.7612 | 0.760861 | 0.753419 | 0.75377 | 0.761746 | 0.772634 |
| 0.760623 | 0.757811 | 0.757413 | 0.755034 | 0.758609 | 0.749161 | 0.752362 | 0.762557 | 0.768137 |
| 0.754824 | 0.752654 | 0.752698 | 0.750421 | 0.751499 | 0.745266 | 0.742985 | 0.754872 | 0.766047 |
| 0.755131 | 0.752401 | 0.752909 | 0.754517 | 0.751999 | 0.747174 | 0.749812 | 0.751656 | 0.763539 |
| 0.762458 | 0.757396 | 0.76078 | 0.761683 | 0.760423 | 0.752248 | 0.751575 | 0.763745 | 0.76665 |
| 0.76576 | 0.762995 | 0.762565 | 0.765383 | 0.765237 | 0.759901 | 0.758664 | 0.761777 | 0.766867 |
| 0.765707 | 0.758227 | 0.758074 | 0.763003 | 0.763502 | 0.755813 | 0.759044 | 0.757712 | 0.763839 |
| 0.762851 | 0.755868 | 0.755302 | 0.761622 | 0.761597 | 0.752379 | 0.752678 | 0.762588 | 0.763404 |
| 0.763255 | 0.757243 | 0.759402 | 0.760083 | 0.765124 | 0.755682 | 0.753938 | 0.760355 | 0.768071 |
| 0.758499 | 0.754153 | 0.761199 | 0.76091 | 0.763244 | 0.757184 | 0.756299 | 0.761096 | 0.768784 |
| 0.757357 | 0.753631 | 0.75318 | 0.755637 | 0.756888 | 0.751036 | 0.754811 | 0.757506 | 0.757852 |
| 0.758859 | 0.753151 | 0.753699 | 0.75629 | 0.761941 | 0.751169 | 0.748763 | 0.755335 | 0.764647 |
| 0.762801 | 0.759897 | 0.763672 | 0.763662 | 0.768012 | 0.755305 | 0.756786 | 0.76405 | 0.770994 |
| 0.763842 | 0.759275 | 0.759847 | 0.765276 | 0.767295 | 0.754685 | 0.757842 | 0.760624 | 0.767094 |
| 0.763943 | 0.758218 | 0.761641 | 0.758094 | 0.76498 | 0.757239 | 0.758836 | 0.757383 | 0.764794 |
| 0.757568 | 0.753548 | 0.759957 | 0.759287 | 0.761897 | 0.750623 | 0.751917 | 0.760855 | 0.767595 |
| 0.756023 | 0.753824 | 0.753729 | 0.755111 | 0.757098 | 0.752407 | 0.748877 | 0.754112 | 0.758155 |
| 0.755268 | 0.755121 | 0.755563 | 0.761519 | 0.761194 | 0.754038 | 0.745741 | 0.7561 | 0.762433 |
| 0.757476 | 0.754758 | 0.760229 | 0.761829 | 0.762307 | 0.753864 | 0.755078 | 0.762005 | 0.767063 |
| 0.764533 | 0.764255 | 0.759215 | 0.761535 | 0.762635 | 0.755194 | 0.754213 | 0.760436 | 0.769791 |
| 0.763946 | 0.758878 | 0.763899 | 0.762207 | 0.764039 | 0.755593 | 0.75837 | 0.76358 | 0.765697 |
| 0.763799 | 0.757956 | 0.758759 | 0.760421 | 0.766582 | 0.755599 | 0.756104 | 0.762054 | 0.763217 |
| 0.768399 | 0.75834 | 0.76518 | 0.766686 | 0.766561 | 0.760279 | 0.761325 | 0.767033 | 0.770372 |
| 0.76542 | 0.758525 | 0.760542 | 0.76809 | 0.767746 | 0.757587 | 0.761464 | 0.767012 | 0.770026 |
| 0.761393 | 0.754517 | 0.759359 | 0.75918 | 0.760624 | 0.752648 | 0.75093 | 0.757576 | 0.763539 |
| 0.763794 | 0.755444 | 0.761051 | 0.760037 | 0.767079 | 0.758323 | 0.754862 | 0.762817 | 0.7632 |
| 0.761194 | 0.75513 | 0.754701 | 0.761949 | 0.762593 | 0.755175 | 0.752411 | 0.760604 | 0.76245 |
| 0.757282 | 0.761706 | 0.757363 | 0.758822 | 0.760207 | 0.750693 | 0.75267 | 0.758198 | 0.765193 |
| 0.750768 | 0.749425 | 0.753451 | 0.751165 | 0.753187 | 0.743776 | 0.750944 | 0.75643 | 0.757906 |
| 0.75581 | 0.750526 | 0.747209 | 0.750567 | 0.75192 | 0.745404 | 0.744911 | 0.751639 | 0.756652 |
| 0.749113 | 0.74896 | 0.750512 | 0.75043 | 0.751574 | 0.744962 | 0.742844 | 0.752095 | 0.756568 |
| 0.757117 | 0.749401 | 0.752859 | 0.753334 | 0.759299 | 0.750117 | 0.747265 | 0.754062 | 0.758385 |
| 0.750591 | 0.744605 | 0.747732 | 0.749544 | 0.752096 | 0.745746 | 0.746107 | 0.751424 | 0.754585 |
| 0.748345 | 0.741535 | 0.744054 | 0.747878 | 0.749863 | 0.744086 | 0.742002 | 0.749635 | 0.75951 |
| 0.758834 | 0.752916 | 0.753714 | 0.755999 | 0.758491 | 0.746688 | 0.752564 | 0.755118 | 0.764869 |
| 0.752502 | 0.752575 | 0.749192 | 0.752535 | 0.757945 | 0.742528 | 0.746401 | 0.75024 | 0.752507 |

| | | | | | | | | |
|---|---|---|---|---|---|---|---|---|
| 0.764303 | 0.75824 | 0.762178 | 0.759387 | 0.760702 | 0.751644 | 0.755388 | 0.764037 | 0.7676 |
| 0.770778 | 0.767349 | 0.769802 | 0.769969 | 0.765813 | 0.758364 | 0.758488 | 0.77158 | 0.777796 |
| 0.7634 | 0.76094 | 0.762272 | 0.761197 | 0.761731 | 0.756898 | 0.757439 | 0.763367 | 0.771778 |
| 0.767456 | 0.760645 | 0.767184 | 0.763265 | 0.764392 | 0.752931 | 0.762953 | 0.767372 | 0.773258 |
| 0.778268 | 0.770387 | 0.771743 | 0.771881 | 0.770707 | 0.763303 | 0.763504 | 0.771123 | 0.782804 |
| 0.761283 | 0.767542 | 0.766412 | 0.766275 | 0.7667 | 0.762095 | 0.76109 | 0.773631 | 0.774679 |
| 0.761967 | 0.762484 | 0.761235 | 0.757124 | 0.759951 | 0.754016 | 0.757891 | 0.761229 | 0.774614 |
| 0.761861 | 0.763423 | 0.763679 | 0.764067 | 0.763933 | 0.755465 | 0.74991 | 0.76605 | 0.772098 |
| 0.763216 | 0.759806 | 0.758543 | 0.758975 | 0.757493 | 0.75478 | 0.750848 | 0.760293 | 0.771343 |
| 0.757903 | 0.757619 | 0.760305 | 0.757229 | 0.757048 | 0.751021 | 0.749345 | 0.760081 | 0.767536 |
| 0.761619 | 0.760393 | 0.760834 | 0.764993 | 0.767595 | 0.757065 | 0.7564 | 0.775073 | 0.773736 |
| 0.759729 | 0.761857 | 0.763873 | 0.765721 | 0.766262 | 0.757527 | 0.756855 | 0.76721 | 0.77354 |
| 0.769718 | 0.764137 | 0.765862 | 0.769759 | 0.765697 | 0.763275 | 0.761283 | 0.766038 | 0.775431 |
| 0.766698 | 0.763375 | 0.76691 | 0.766201 | 0.763878 | 0.760036 | 0.757507 | 0.765412 | 0.773822 |
| 0.771438 | 0.76936 | 0.770508 | 0.767187 | 0.771171 | 0.763572 | 0.76264 | 0.773689 | 0.779602 |
| 0.770912 | 0.766765 | 0.766044 | 0.768169 | 0.76905 | 0.758182 | 0.761065 | 0.772341 | 0.777778 |
| 0.770675 | 0.767108 | 0.762201 | 0.765635 | 0.764935 | 0.764342 | 0.76077 | 0.772044 | 0.781535 |
| 0.769724 | 0.765556 | 0.769742 | 0.768677 | 0.766981 | 0.756695 | 0.762503 | 0.771533 | 0.781181 |
| 0.772137 | 0.769334 | 0.764385 | 0.769237 | 0.76842 | 0.762479 | 0.761767 | 0.770457 | 0.778586 |
| 0.766982 | 0.764809 | 0.762837 | 0.7695 | 0.772077 | 0.761112 | 0.763505 | 0.77353 | 0.782648 |
| 0.759939 | 0.761611 | 0.762159 | 0.762847 | 0.762308 | 0.754635 | 0.755878 | 0.766339 | 0.771751 |
| 0.769407 | 0.763686 | 0.769791 | 0.768303 | 0.771255 | 0.760969 | 0.761848 | 0.770012 | 0.77635 |
| 0.767385 | 0.765795 | 0.764158 | 0.768916 | 0.769276 | 0.76249 | 0.759159 | 0.774777 | 0.779407 |
| 0.770294 | 0.770242 | 0.768347 | 0.769832 | 0.771753 | 0.764086 | 0.760267 | 0.770774 | 0.782025 |
| 0.768751 | 0.770038 | 0.768486 | 0.774036 | 0.771616 | 0.764546 | 0.764701 | 0.767496 | 0.776659 |
| 0.764815 | 0.766095 | 0.765771 | 0.772365 | 0.766837 | 0.762659 | 0.758175 | 0.764376 | 0.779535 |
| 0.768457 | 0.763197 | 0.766702 | 0.766531 | 0.770162 | 0.761374 | 0.758717 | 0.764613 | 0.773082 |
| 0.768818 | 0.771369 | 0.768869 | 0.763651 | 0.764559 | 0.761976 | 0.756876 | 0.766633 | 0.777074 |
| 0.774526 | 0.770926 | 0.765452 | 0.76633 | 0.764349 | 0.762157 | 0.755868 | 0.769077 | 0.776536 |
| 0.76969 | 0.768812 | 0.768173 | 0.771721 | 0.768496 | 0.765926 | 0.764249 | 0.775128 | 0.780952 |
| 0.761036 | 0.765098 | 0.763473 | 0.761042 | 0.761123 | 0.753809 | 0.754928 | 0.763511 | 0.774744 |
| 0.766971 | 0.766892 | 0.764947 | 0.76651 | 0.766689 | 0.764158 | 0.765377 | 0.771784 | 0.780278 |
| 0.770267 | 0.769826 | 0.768715 | 0.771333 | 0.769534 | 0.765632 | 0.762931 | 0.772601 | 0.779228 |
| 0.766713 | 0.754546 | 0.764203 | 0.764691 | 0.761227 | 0.760649 | 0.760048 | 0.763394 | 0.775895 |
| 0.766574 | 0.765493 | 0.768185 | 0.76695 | 0.766069 | 0.761463 | 0.755019 | 0.769982 | 0.781416 |
| 0.766717 | 0.767302 | 0.764977 | 0.764737 | 0.767244 | 0.763688 | 0.761167 | 0.768574 | 0.77594 |
| 0.775343 | 0.773237 | 0.774314 | 0.770633 | 0.773154 | 0.768984 | 0.762855 | 0.776814 | 0.778513 |
| 0.764701 | 0.768599 | 0.768658 | 0.764771 | 0.767501 | 0.764249 | 0.761511 | 0.767119 | 0.777198 |
| 0.770927 | 0.769426 | 0.768994 | 0.76758 | 0.769909 | 0.760957 | 0.763506 | 0.769127 | 0.779615 |
| 0.7676 | 0.763819 | 0.763957 | 0.764536 | 0.76424 | 0.759513 | 0.763573 | 0.767767 | 0.785538 |
| 0.767024 | 0.766809 | 0.768977 | 0.770013 | 0.773278 | 0.762027 | 0.76419 | 0.770563 | 0.783701 |
| 0.767299 | 0.768667 | 0.776989 | 0.769649 | 0.769411 | 0.764266 | 0.766946 | 0.771337 | 0.781042 |
| 0.767847 | 0.770197 | 0.774655 | 0.77155 | 0.77355 | 0.768241 | 0.769342 | 0.772864 | 0.785744 |
| 0.768426 | 0.762703 | 0.765248 | 0.758755 | 0.768106 | 0.759553 | 0.762242 | 0.762974 | 0.777627 |
| 0.769315 | 0.766867 | 0.764966 | 0.76947 | 0.768674 | 0.76364 | 0.760553 | 0.772473 | 0.784156 |
| 0.765487 | 0.765491 | 0.764755 | 0.767462 | 0.763185 | 0.763557 | 0.760363 | 0.769547 | 0.7823 |

| | | | | | | | | |
|---|---|---|---|---|---|---|---|---|
| 0.757722 | 0.756227 | 0.75589 | 0.752938 | 0.757945 | 0.746379 | 0.749237 | 0.757188 | 0.768043 |
| 0.749326 | 0.750444 | 0.746513 | 0.749707 | 0.74916 | 0.741684 | 0.737126 | 0.746012 | 0.757135 |
| 0.760965 | 0.766707 | 0.767687 | 0.762879 | 0.760045 | 0.760617 | 0.756219 | 0.772205 | 0.77791 |
| 0.771314 | 0.771997 | 0.770722 | 0.765686 | 0.770545 | 0.759065 | 0.762925 | 0.772689 | 0.777384 |
| 0.76001 | 0.757647 | 0.76188 | 0.76056 | 0.758971 | 0.754053 | 0.758389 | 0.765818 | 0.77147 |
| 0.765999 | 0.763129 | 0.76805 | 0.765475 | 0.767934 | 0.756909 | 0.756135 | 0.771091 | 0.77748 |
| 0.771835 | 0.771739 | 0.770728 | 0.766996 | 0.765195 | 0.765622 | 0.762999 | 0.765197 | 0.775657 |
| 0.759931 | 0.760542 | 0.762253 | 0.759035 | 0.75775 | 0.755805 | 0.756994 | 0.762984 | 0.769395 |
| 0.770342 | 0.767208 | 0.767198 | 0.770658 | 0.769571 | 0.766622 | 0.766878 | 0.772103 | 0.780452 |
| 0.770091 | 0.767044 | 0.769981 | 0.770285 | 0.770677 | 0.763196 | 0.76064 | 0.766563 | 0.779836 |
| 0.775916 | 0.767418 | 0.771798 | 0.76774 | 0.768935 | 0.762014 | 0.765089 | 0.770015 | 0.773146 |
| 0.769752 | 0.768821 | 0.769751 | 0.768481 | 0.768551 | 0.762933 | 0.759386 | 0.771943 | 0.778296 |
| 0.773855 | 0.76475 | 0.77501 | 0.772212 | 0.775437 | 0.767844 | 0.766294 | 0.772001 | 0.777851 |
| 0.770523 | 0.76805 | 0.769963 | 0.768565 | 0.766286 | 0.757448 | 0.762666 | 0.771799 | 0.778641 |
| 0.766019 | 0.765686 | 0.76572 | 0.76567 | 0.764926 | 0.761213 | 0.759119 | 0.770038 | 0.778218 |
| 0.76394 | 0.762813 | 0.764264 | 0.763708 | 0.762723 | 0.7586 | 0.755405 | 0.7653 | 0.776271 |
| 0.770574 | 0.770021 | 0.765244 | 0.769179 | 0.769354 | 0.757703 | 0.757179 | 0.768115 | 0.774881 |
| 0.766289 | 0.758624 | 0.766572 | 0.764128 | 0.765212 | 0.75473 | 0.755582 | 0.767591 | 0.778288 |
| 0.765773 | 0.757165 | 0.762375 | 0.761594 | 0.763199 | 0.756489 | 0.759327 | 0.767518 | 0.772986 |
| 0.766043 | 0.760914 | 0.758004 | 0.766057 | 0.765756 | 0.757246 | 0.759954 | 0.760587 | 0.775099 |
| 0.769933 | 0.765766 | 0.7663 | 0.765943 | 0.767765 | 0.755663 | 0.760681 | 0.769335 | 0.777405 |
| 0.764022 | 0.763977 | 0.762365 | 0.760712 | 0.762896 | 0.758283 | 0.759944 | 0.758606 | 0.768122 |
| 0.76901 | 0.766793 | 0.767965 | 0.764509 | 0.765922 | 0.761247 | 0.763165 | 0.76254 | 0.771426 |
| 0.77444 | 0.771954 | 0.771625 | 0.768869 | 0.769762 | 0.767721 | 0.767834 | 0.766537 | 0.775214 |
| 0.77211 | 0.771256 | 0.767357 | 0.773552 | 0.772477 | 0.768367 | 0.76797 | 0.772077 | 0.77225 |
| 0.774144 | 0.771049 | 0.769432 | 0.769707 | 0.770983 | 0.766567 | 0.764749 | 0.765657 | 0.776114 |
| 0.767066 | 0.767867 | 0.762623 | 0.763003 | 0.767502 | 0.759703 | 0.761579 | 0.763663 | 0.76831 |
| 0.769695 | 0.769278 | 0.765834 | 0.768253 | 0.772329 | 0.76631 | 0.765738 | 0.765029 | 0.772547 |
| 0.76683 | 0.76849 | 0.761331 | 0.76738 | 0.767675 | 0.760246 | 0.764493 | 0.76372 | 0.766894 |
| 0.690032 | 0.691561 | 0.695086 | 0.693876 | 0.695012 | 0.687019 | 0.694212 | 0.697026 | 0.700175 |
| 0.704304 | 0.70459 | 0.709389 | 0.704312 | 0.704255 | 0.698497 | 0.699345 | 0.709502 | 0.716451 |
| 0.705452 | 0.703071 | 0.705368 | 0.702103 | 0.705334 | 0.694644 | 0.698681 | 0.705062 | 0.710237 |
| 0.699521 | 0.693713 | 0.699902 | 0.694732 | 0.693368 | 0.687588 | 0.690822 | 0.698774 | 0.701372 |
| 0.707615 | 0.704315 | 0.709375 | 0.70538 | 0.705802 | 0.696737 | 0.700831 | 0.711421 | 0.715152 |
| 0.774768 | 0.772739 | 0.767099 | 0.76749 | 0.768098 | 0.762625 | 0.76638 | 0.761317 | 0.774865 |
| 0.775107 | 0.772591 | 0.771294 | 0.769891 | 0.769538 | 0.76724 | 0.765128 | 0.770201 | 0.778329 |
| 0.779177 | 0.775465 | 0.777609 | 0.775151 | 0.774183 | 0.773056 | 0.772153 | 0.770534 | 0.776605 |
| 0.786411 | 0.775846 | 0.771722 | 0.776381 | 0.777307 | 0.767606 | 0.770588 | 0.770979 | 0.781788 |
| 0.778225 | 0.774367 | 0.768572 | 0.770402 | 0.770281 | 0.765597 | 0.768673 | 0.763116 | 0.769731 |
| 0.775649 | 0.771928 | 0.773032 | 0.770645 | 0.772869 | 0.764853 | 0.77174 | 0.770572 | 0.781801 |
| 0.777826 | 0.776822 | 0.776554 | 0.772673 | 0.773583 | 0.765863 | 0.772505 | 0.770478 | 0.776815 |
| 0.768034 | 0.769898 | 0.769608 | 0.768752 | 0.770615 | 0.768534 | 0.766476 | 0.768329 | 0.770111 |
| 0.779432 | 0.779901 | 0.767395 | 0.769401 | 0.765585 | 0.760474 | 0.764706 | 0.763893 | 0.769069 |
| 0.771598 | 0.769133 | 0.762335 | 0.767352 | 0.765363 | 0.763673 | 0.755748 | 0.76009 | 0.768854 |
| 0.771496 | 0.767468 | 0.767653 | 0.767558 | 0.773211 | 0.762117 | 0.759055 | 0.763522 | 0.771808 |
| 0.784012 | 0.776701 | 0.773172 | 0.773164 | 0.774232 | 0.767543 | 0.766621 | 0.767985 | 0.780093 |

| | | | | | | | | |
|---|---|---|---|---|---|---|---|---|
| 0.78661 | 0.782329 | 0.788529 | 0.789866 | 0.787876 | 0.790119 | 0.779666 | 0.771847 | 0.781317 |
| 0.796013 | 0.783292 | 0.839304 | 0.793743 | 0.784076 | 0.782896 | 0.79894 | 0.764348 | 0.769224 |
| 0.791803 | 0.792141 | 0.785908 | 0.864656 | 0.828457 | 0.864418 | 0.779315 | 0.768081 | 0.775728 |
| 0.781551 | 0.785556 | 0.778627 | 0.862949 | 0.811812 | 0.8125 | 0.775664 | 0.759822 | 0.770443 |
| 0.788596 | 0.781247 | 0.784713 | 0.858015 | 0.81044 | 0.823276 | 0.778797 | 0.760785 | 0.76858 |
| 0.782318 | 0.784224 | 0.782982 | 0.822442 | 0.794673 | 0.799177 | 0.778449 | 0.760139 | 0.772953 |
| 0.787477 | 0.792932 | 0.781874 | 0.814685 | 0.806446 | 0.801722 | 0.79263 | 0.764549 | 0.773854 |
| 1 | 0.797107 | 0.783665 | 0.782342 | 0.780619 | 0.778919 | 0.784517 | 0.76464 | 0.77353 |
| 0.797107 | 1 | 0.778415 | 0.786369 | 0.787766 | 0.782125 | 0.774939 | 0.760357 | 0.769218 |
| 0.783665 | 0.778415 | 1 | 0.779462 | 0.777731 | 0.78011 | 0.793286 | 0.764222 | 0.768601 |
| 0.782342 | 0.786369 | 0.779462 | 1 | 0.853767 | 0.859886 | 0.777335 | 0.763706 | 0.772425 |
| 0.780619 | 0.787766 | 0.777731 | 0.853767 | 1 | 0.812784 | 0.787664 | 0.766007 | 0.774134 |
| 0.778919 | 0.782125 | 0.78011 | 0.859886 | 0.812784 | 1 | 0.774263 | 0.758877 | 0.766081 |
| 0.784517 | 0.774939 | 0.793286 | 0.777335 | 0.787664 | 0.774263 | 1 | 0.755752 | 0.770047 |
| 0.76464 | 0.760357 | 0.764222 | 0.763706 | 0.766007 | 0.758877 | 0.755752 | 1 | 0.800343 |
| 0.77353 | 0.769218 | 0.768601 | 0.772425 | 0.774134 | 0.766081 | 0.770047 | 0.800343 | 1 |
| 0.761297 | 0.763764 | 0.771226 | 0.761861 | 0.765389 | 0.760709 | 0.758627 | 0.817226 | 0.866075 |
| 0.771089 | 0.765959 | 0.770526 | 0.766837 | 0.772137 | 0.762848 | 0.762621 | 0.822228 | 0.816819 |
| 0.769163 | 0.763378 | 0.764124 | 0.767034 | 0.771777 | 0.762354 | 0.762342 | 0.796442 | 0.786976 |
| 0.768064 | 0.764193 | 0.762334 | 0.767547 | 0.764874 | 0.760366 | 0.75474 | 0.797547 | 0.798332 |
| 0.769271 | 0.762209 | 0.765452 | 0.764051 | 0.763142 | 0.756039 | 0.758879 | 0.784763 | 0.789067 |
| 0.769126 | 0.761727 | 0.764346 | 0.763776 | 0.768364 | 0.761409 | 0.757606 | 0.784401 | 0.794306 |
| 0.768271 | 0.76628 | 0.769654 | 0.766817 | 0.766459 | 0.763613 | 0.765349 | 0.798661 | 0.872224 |
| 0.788221 | 0.778945 | 0.786716 | 0.78915 | 0.782699 | 0.779103 | 0.791914 | 0.762178 | 0.771386 |
| 0.769314 | 0.766579 | 0.762713 | 0.771338 | 0.771707 | 0.765198 | 0.76124 | 0.78589 | 0.858918 |
| 0.705566 | 0.702823 | 0.707472 | 0.699731 | 0.69959 | 0.691741 | 0.699029 | 0.706722 | 0.709126 |
| 0.767874 | 0.76445 | 0.768798 | 0.765656 | 0.771306 | 0.765098 | 0.760797 | 0.77107 | 0.783419 |
| 0.768474 | 0.765932 | 0.772984 | 0.767529 | 0.76806 | 0.763275 | 0.758511 | 0.772471 | 0.781499 |
| 0.774315 | 0.771375 | 0.772132 | 0.765667 | 0.768047 | 0.762706 | 0.767833 | 0.771869 | 0.783298 |
| 0.769309 | 0.765647 | 0.767153 | 0.767079 | 0.771576 | 0.761215 | 0.762598 | 0.767964 | 0.778222 |
| 0.763077 | 0.764032 | 0.764685 | 0.761688 | 0.769048 | 0.761535 | 0.764734 | 0.764644 | 0.780869 |
| 0.767724 | 0.761764 | 0.768429 | 0.765311 | 0.763244 | 0.7637 | 0.762759 | 0.764823 | 0.781563 |
| 0.765202 | 0.765176 | 0.770676 | 0.766041 | 0.767563 | 0.76106 | 0.762398 | 0.771122 | 0.781758 |
| 0.765359 | 0.763195 | 0.766548 | 0.765439 | 0.76628 | 0.757341 | 0.759376 | 0.768497 | 0.779312 |
| 0.761444 | 0.756971 | 0.763165 | 0.760797 | 0.76371 | 0.754685 | 0.749175 | 0.765358 | 0.771704 |
| 0.764289 | 0.758742 | 0.762656 | 0.759833 | 0.764429 | 0.755427 | 0.755887 | 0.765034 | 0.77874 |
| 0.769899 | 0.768833 | 0.768662 | 0.771043 | 0.772003 | 0.763108 | 0.763891 | 0.773098 | 0.783873 |
| 0.768412 | 0.764721 | 0.762708 | 0.767089 | 0.767729 | 0.757479 | 0.754681 | 0.771111 | 0.781956 |
| 0.770906 | 0.764301 | 0.769489 | 0.771678 | 0.773315 | 0.763335 | 0.762515 | 0.770423 | 0.783686 |
| 0.764964 | 0.764375 | 0.767408 | 0.763907 | 0.766827 | 0.764424 | 0.759707 | 0.768711 | 0.78039 |
| 0.762255 | 0.766225 | 0.763386 | 0.763936 | 0.767729 | 0.761356 | 0.756501 | 0.767903 | 0.775538 |
| 0.765371 | 0.762113 | 0.767202 | 0.768386 | 0.771531 | 0.76541 | 0.76052 | 0.774994 | 0.782642 |
| 0.765543 | 0.762704 | 0.76077 | 0.763866 | 0.767489 | 0.757401 | 0.751987 | 0.765896 | 0.77641 |
| 0.768427 | 0.762747 | 0.767272 | 0.766784 | 0.759973 | 0.758783 | 0.75771 | 0.764288 | 0.768616 |
| 0.772924 | 0.773052 | 0.769186 | 0.768293 | 0.770427 | 0.764996 | 0.770366 | 0.766526 | 0.776553 |
| 0.77329 | 0.766515 | 0.768057 | 0.765159 | 0.766277 | 0.760068 | 0.765351 | 0.771296 | 0.774346 |

| | | | | | | | | |
|---|---|---|---|---|---|---|---|---|
| 0.768752 | 0.767497 | 0.76733 | 0.765559 | 0.765064 | 0.763137 | 0.76285 | 0.768992 | 0.769642 |
| 0.77476 | 0.772076 | 0.772501 | 0.772898 | 0.775366 | 0.767659 | 0.774132 | 0.766736 | 0.77329 |
| 0.762676 | 0.766567 | 0.768448 | 0.769231 | 0.765362 | 0.760384 | 0.762171 | 0.762801 | 0.76925 |
| 0.737518 | 0.730893 | 0.730769 | 0.729964 | 0.721252 | 0.725533 | 0.720434 | 0.732312 | 0.733574 |
| 0.765954 | 0.767934 | 0.769511 | 0.764222 | 0.763624 | 0.754809 | 0.760524 | 0.764175 | 0.772118 |
| 0.761194 | 0.76235 | 0.764716 | 0.766728 | 0.767096 | 0.763812 | 0.76237 | 0.765267 | 0.770442 |
| 0.765909 | 0.767262 | 0.767455 | 0.768124 | 0.766896 | 0.758229 | 0.766157 | 0.762372 | 0.76836 |
| 0.768492 | 0.765087 | 0.77249 | 0.767932 | 0.771652 | 0.764256 | 0.766773 | 0.757961 | 0.773746 |
| 0.771527 | 0.772624 | 0.771652 | 0.771978 | 0.771088 | 0.768635 | 0.765364 | 0.768083 | 0.779196 |
| 0.771349 | 0.76754 | 0.768668 | 0.772605 | 0.772922 | 0.76636 | 0.763309 | 0.764656 | 0.775579 |
| 0.765495 | 0.76318 | 0.76941 | 0.770704 | 0.767286 | 0.761798 | 0.765371 | 0.762976 | 0.772705 |
| 0.774641 | 0.769061 | 0.769488 | 0.77418 | 0.77058 | 0.768519 | 0.769188 | 0.763348 | 0.767602 |
| 0.768056 | 0.765592 | 0.768319 | 0.7656 | 0.759158 | 0.76196 | 0.764508 | 0.765695 | 0.773419 |
| 0.772819 | 0.769625 | 0.767482 | 0.765383 | 0.76965 | 0.767641 | 0.767115 | 0.768946 | 0.771973 |
| 0.774183 | 0.769458 | 0.773981 | 0.771165 | 0.77184 | 0.767384 | 0.770121 | 0.767442 | 0.772332 |
| 0.770331 | 0.764359 | 0.767937 | 0.764499 | 0.765827 | 0.761468 | 0.764062 | 0.763213 | 0.767648 |
| 0.773254 | 0.768761 | 0.769845 | 0.769939 | 0.766562 | 0.762761 | 0.769865 | 0.767219 | 0.774551 |
| 0.775565 | 0.765041 | 0.775541 | 0.773721 | 0.771008 | 0.766996 | 0.773461 | 0.764602 | 0.772727 |
| 0.763733 | 0.761238 | 0.763783 | 0.765892 | 0.764134 | 0.757568 | 0.760363 | 0.760301 | 0.768399 |
| 0.767818 | 0.764874 | 0.770375 | 0.766199 | 0.766988 | 0.759269 | 0.763883 | 0.766362 | 0.768399 |
| 0.770341 | 0.764983 | 0.774089 | 0.775183 | 0.772391 | 0.768537 | 0.770514 | 0.763595 | 0.778304 |
| 0.774886 | 0.770035 | 0.770885 | 0.771151 | 0.771359 | 0.765067 | 0.768587 | 0.7707 | 0.773272 |
| 0.687942 | 0.680206 | 0.68463 | 0.678096 | 0.676535 | 0.680493 | 0.674007 | 0.688089 | 0.690655 |
| 0.666989 | 0.653168 | 0.656205 | 0.655718 | 0.649156 | 0.657664 | 0.651456 | 0.657958 | 0.668422 |
| 0.669236 | 0.673272 | 0.669251 | 0.670063 | 0.677106 | 0.665401 | 0.67423 | 0.671609 | 0.678517 |
| 0.658586 | 0.652418 | 0.654093 | 0.65014 | 0.649186 | 0.648803 | 0.65049 | 0.654195 | 0.665347 |
| 0.753035 | 0.757156 | 0.756369 | 0.751569 | 0.747863 | 0.755524 | 0.759777 | 0.735325 | 0.736984 |
| 0.744222 | 0.745898 | 0.731317 | 0.744801 | 0.740806 | 0.735306 | 0.74834 | 0.727608 | 0.731533 |
| 0.751384 | 0.740314 | 0.73102 | 0.736544 | 0.734025 | 0.733235 | 0.73713 | 0.724401 | 0.740029 |
| 0.756665 | 0.746234 | 0.739929 | 0.757846 | 0.756201 | 0.758146 | 0.744896 | 0.736831 | 0.744181 |
| 0.735845 | 0.721721 | 0.730501 | 0.729398 | 0.73134 | 0.730543 | 0.731497 | 0.715826 | 0.720393 |
| 0.727325 | 0.709065 | 0.726749 | 0.727821 | 0.73395 | 0.717691 | 0.72644 | 0.711299 | 0.723301 |
| 0.742355 | 0.735653 | 0.729454 | 0.726426 | 0.727273 | 0.723835 | 0.740707 | 0.708086 | 0.71382 |
| 0.658726 | 0.658651 | 0.651046 | 0.653522 | 0.654011 | 0.654245 | 0.657841 | 0.658335 | 0.659274 |
| 0.776149 | 0.77134 | 0.77297 | 0.769995 | 0.767964 | 0.765813 | 0.769442 | 0.76065 | 0.76467 |
| 0.753936 | 0.749615 | 0.752565 | 0.752961 | 0.753927 | 0.754223 | 0.761988 | 0.738587 | 0.749765 |
| 0.752286 | 0.744672 | 0.749658 | 0.746097 | 0.745342 | 0.745775 | 0.75025 | 0.733879 | 0.746246 |
| 0.752055 | 0.727835 | 0.735835 | 0.728611 | 0.729377 | 0.728814 | 0.741085 | 0.726828 | 0.729078 |
| 0.748984 | 0.751455 | 0.753519 | 0.751424 | 0.752469 | 0.74768 | 0.754178 | 0.73952 | 0.7505 |
| 0.73595 | 0.742786 | 0.737555 | 0.735522 | 0.734876 | 0.741508 | 0.735433 | 0.727026 | 0.74053 |
| 0.740656 | 0.742645 | 0.737383 | 0.739501 | 0.738739 | 0.7403 | 0.73255 | 0.720923 | 0.732545 |
| 0.737647 | 0.7407 | 0.736701 | 0.734019 | 0.734326 | 0.734347 | 0.735973 | 0.730216 | 0.733271 |
| 0.745996 | 0.735558 | 0.741282 | 0.733503 | 0.739442 | 0.732169 | 0.745676 | 0.721728 | 0.727734 |
| 0.745955 | 0.734942 | 0.73577 | 0.732178 | 0.729708 | 0.731559 | 0.735143 | 0.731887 | 0.741242 |
| 0.744764 | 0.732709 | 0.735957 | 0.724929 | 0.729183 | 0.730515 | 0.737503 | 0.728725 | 0.740078 |
| 0.724681 | 0.720294 | 0.721279 | 0.722018 | 0.725312 | 0.721047 | 0.728043 | 0.726875 | 0.720715 |

| | | | | | | | | |
|---|---|---|---|---|---|---|---|---|
| 0.769663 | 0.766024 | 0.762153 | 0.762638 | 0.757799 | 0.76084 | 0.765707 | 0.749471 | 0.761991 |
| 0.748629 | 0.748069 | 0.747143 | 0.748429 | 0.745816 | 0.743534 | 0.745269 | 0.7315 | 0.736479 |
| 0.752776 | 0.746104 | 0.753015 | 0.750229 | 0.744271 | 0.745517 | 0.743252 | 0.728388 | 0.731301 |
| 0.756271 | 0.750117 | 0.748092 | 0.747318 | 0.745318 | 0.747607 | 0.745627 | 0.726909 | 0.734308 |
| 0.752194 | 0.743117 | 0.749845 | 0.739905 | 0.738996 | 0.73809 | 0.744324 | 0.726843 | 0.734423 |
| 0.753582 | 0.732915 | 0.741522 | 0.743927 | 0.735481 | 0.742666 | 0.750065 | 0.723565 | 0.73243 |
| 0.745568 | 0.742454 | 0.744958 | 0.737568 | 0.736558 | 0.737767 | 0.741724 | 0.721408 | 0.727096 |
| 0.741344 | 0.728988 | 0.74568 | 0.734116 | 0.728543 | 0.739194 | 0.743787 | 0.714853 | 0.721497 |
| 0.735547 | 0.736662 | 0.742407 | 0.746169 | 0.745172 | 0.746073 | 0.731083 | 0.721573 | 0.725834 |
| 0.760112 | 0.760648 | 0.763837 | 0.758138 | 0.760707 | 0.756057 | 0.766184 | 0.751592 | 0.759283 |
| 0.758244 | 0.759206 | 0.761129 | 0.762691 | 0.758638 | 0.756585 | 0.766844 | 0.744768 | 0.759421 |
| 0.759973 | 0.754609 | 0.753244 | 0.757831 | 0.759649 | 0.750232 | 0.76729 | 0.745287 | 0.755494 |
| 0.754275 | 0.745926 | 0.744331 | 0.752797 | 0.753548 | 0.745379 | 0.751543 | 0.736784 | 0.749817 |
| 0.758893 | 0.755769 | 0.751539 | 0.752627 | 0.752795 | 0.749616 | 0.759199 | 0.743822 | 0.745802 |
| 0.75428 | 0.750847 | 0.739899 | 0.746178 | 0.749439 | 0.742941 | 0.744906 | 0.73017 | 0.746723 |

| plot 13 | | | | | | | | |
|---|---|---|---|---|---|---|---|---|
| Wild_plot_ | Wild_plot_ | Wild_plot_ | Wild_plot_ | Wild_plot_ | Wild_plot_ | Wild_plot_ | Wild_plot_ | Wild_plot_ |
| 0.710091 | 0.71082 | 0.702367 | 0.705351 | 0.701718 | 0.705387 | 0.708559 | 0.705562 | 0.70553 |
| 0.715822 | 0.71807 | 0.712855 | 0.715965 | 0.711264 | 0.717818 | 0.719506 | 0.714004 | 0.715419 |
| 0.706709 | 0.711713 | 0.70207 | 0.701242 | 0.701657 | 0.704809 | 0.711288 | 0.703169 | 0.702224 |
| 0.716081 | 0.718872 | 0.714509 | 0.709622 | 0.708519 | 0.713885 | 0.714391 | 0.709834 | 0.710421 |
| 0.709655 | 0.714559 | 0.703646 | 0.706945 | 0.704782 | 0.7077 | 0.70849 | 0.705723 | 0.703461 |
| 0.720602 | 0.724416 | 0.712782 | 0.713631 | 0.709895 | 0.719345 | 0.719651 | 0.71352 | 0.719283 |
| 0.778004 | 0.772241 | 0.768786 | 0.771927 | 0.766986 | 0.773127 | 0.777683 | 0.76239 | 0.772239 |
| 0.770509 | 0.775694 | 0.768784 | 0.76834 | 0.76649 | 0.769563 | 0.774199 | 0.759068 | 0.771188 |
| 0.712289 | 0.711674 | 0.705654 | 0.711032 | 0.706292 | 0.713604 | 0.715338 | 0.708822 | 0.711646 |
| 0.711155 | 0.714536 | 0.702063 | 0.707067 | 0.706433 | 0.708243 | 0.712065 | 0.703707 | 0.706696 |
| 0.713257 | 0.714618 | 0.705202 | 0.707317 | 0.704636 | 0.710168 | 0.712845 | 0.70723 | 0.70665 |
| 0.708238 | 0.708781 | 0.700247 | 0.698576 | 0.697732 | 0.704064 | 0.704312 | 0.701201 | 0.70231 |
| 0.713496 | 0.716321 | 0.709952 | 0.712371 | 0.708451 | 0.715544 | 0.717595 | 0.711043 | 0.711212 |
| 0.722561 | 0.725327 | 0.718183 | 0.722034 | 0.718149 | 0.720808 | 0.728494 | 0.72313 | 0.720762 |
| 0.737645 | 0.740818 | 0.732658 | 0.734266 | 0.727551 | 0.736411 | 0.741739 | 0.731605 | 0.734298 |
| 0.712418 | 0.717457 | 0.713483 | 0.71331 | 0.710654 | 0.714309 | 0.716371 | 0.712506 | 0.713305 |
| 0.717044 | 0.718159 | 0.712668 | 0.711089 | 0.710945 | 0.714541 | 0.717522 | 0.710332 | 0.712235 |
| 0.747936 | 0.748855 | 0.741935 | 0.740814 | 0.73591 | 0.739877 | 0.75079 | 0.741362 | 0.743992 |
| 0.708347 | 0.705051 | 0.704675 | 0.705839 | 0.698686 | 0.707574 | 0.70905 | 0.703871 | 0.705746 |
| 0.712335 | 0.716183 | 0.710957 | 0.711007 | 0.708427 | 0.711627 | 0.713353 | 0.708159 | 0.708205 |
| 0.706196 | 0.711209 | 0.70562 | 0.70553 | 0.702187 | 0.70835 | 0.708886 | 0.70353 | 0.708351 |
| 0.712683 | 0.71326 | 0.708837 | 0.708443 | 0.708105 | 0.709145 | 0.71366 | 0.711273 | 0.708398 |
| 0.716837 | 0.718445 | 0.714179 | 0.714498 | 0.710147 | 0.717698 | 0.717781 | 0.713919 | 0.713403 |
| 0.712155 | 0.71878 | 0.711159 | 0.712637 | 0.706936 | 0.714032 | 0.714085 | 0.707067 | 0.713057 |
| 0.713583 | 0.716227 | 0.714393 | 0.714499 | 0.710678 | 0.71527 | 0.714842 | 0.711628 | 0.713459 |
| 0.710953 | 0.716217 | 0.708686 | 0.710887 | 0.703704 | 0.712616 | 0.713383 | 0.708773 | 0.707587 |
| 0.706364 | 0.708602 | 0.70238 | 0.702581 | 0.698822 | 0.705613 | 0.710277 | 0.699498 | 0.705999 |
| 0.715072 | 0.716742 | 0.707143 | 0.709799 | 0.707687 | 0.714538 | 0.716024 | 0.707773 | 0.71133 |
| 0.715688 | 0.720986 | 0.712001 | 0.715149 | 0.712217 | 0.716676 | 0.718377 | 0.709575 | 0.714052 |
| 0.71209 | 0.715594 | 0.709121 | 0.711966 | 0.708212 | 0.71151 | 0.715235 | 0.7103 | 0.710908 |
| 0.708044 | 0.713645 | 0.706889 | 0.708175 | 0.704026 | 0.709696 | 0.706963 | 0.707956 | 0.707206 |
| 0.708742 | 0.711607 | 0.703261 | 0.705391 | 0.701547 | 0.707456 | 0.707637 | 0.702187 | 0.707067 |
| 0.710269 | 0.712603 | 0.705264 | 0.708576 | 0.703914 | 0.709191 | 0.71263 | 0.704626 | 0.70843 |
| 0.711628 | 0.713119 | 0.707494 | 0.707941 | 0.707824 | 0.711775 | 0.71351 | 0.709834 | 0.707219 |
| 0.706081 | 0.708234 | 0.701935 | 0.703473 | 0.703456 | 0.705367 | 0.707751 | 0.70193 | 0.704466 |
| 0.710483 | 0.718496 | 0.709375 | 0.711542 | 0.709976 | 0.713069 | 0.712177 | 0.708144 | 0.70815 |
| 0.716496 | 0.719787 | 0.710664 | 0.712936 | 0.711978 | 0.71257 | 0.719243 | 0.713229 | 0.714237 |
| 0.711963 | 0.71678 | 0.711195 | 0.713081 | 0.710202 | 0.714623 | 0.715737 | 0.709246 | 0.711645 |
| 0.717039 | 0.722774 | 0.710691 | 0.711288 | 0.712533 | 0.714366 | 0.721362 | 0.711396 | 0.714765 |

| | | | | | | | | |
|---|---|---|---|---|---|---|---|---|
| 0.749363 | 0.75262 | 0.746437 | 0.745554 | 0.742672 | 0.74165 | 0.755238 | 0.745101 | 0.750325 |
| 0.705356 | 0.704984 | 0.701767 | 0.699264 | 0.698609 | 0.701944 | 0.707726 | 0.700959 | 0.703475 |
| 0.708962 | 0.709925 | 0.701532 | 0.702994 | 0.698157 | 0.707695 | 0.710031 | 0.703833 | 0.703113 |
| 0.718249 | 0.71879 | 0.710981 | 0.710632 | 0.708125 | 0.714812 | 0.719223 | 0.711205 | 0.711345 |
| 0.709969 | 0.713967 | 0.706705 | 0.70654 | 0.705892 | 0.707208 | 0.710617 | 0.705021 | 0.706824 |
| 0.716636 | 0.717582 | 0.711032 | 0.71342 | 0.710274 | 0.712931 | 0.715469 | 0.709824 | 0.712724 |
| 0.711814 | 0.712537 | 0.702829 | 0.703209 | 0.701061 | 0.710869 | 0.708922 | 0.701002 | 0.708255 |
| 0.714501 | 0.714435 | 0.711942 | 0.711583 | 0.705267 | 0.713288 | 0.71457 | 0.711924 | 0.70953 |
| 0.713524 | 0.71493 | 0.709932 | 0.711709 | 0.707077 | 0.715492 | 0.716783 | 0.710676 | 0.715004 |
| 0.714218 | 0.717914 | 0.70916 | 0.709683 | 0.705689 | 0.714366 | 0.713992 | 0.709931 | 0.710475 |
| 0.705664 | 0.70746 | 0.702172 | 0.699447 | 0.69695 | 0.704663 | 0.707517 | 0.701351 | 0.701476 |
| 0.706801 | 0.707947 | 0.701917 | 0.702345 | 0.697448 | 0.707897 | 0.706602 | 0.702304 | 0.702533 |
| 0.712501 | 0.713171 | 0.706567 | 0.707248 | 0.703494 | 0.709594 | 0.715085 | 0.706044 | 0.709781 |
| 0.708916 | 0.709409 | 0.703376 | 0.706198 | 0.702599 | 0.706934 | 0.708133 | 0.7041 | 0.702047 |
| 0.710189 | 0.708642 | 0.706093 | 0.702862 | 0.704069 | 0.709767 | 0.709852 | 0.703054 | 0.705887 |
| 0.708377 | 0.708313 | 0.704107 | 0.703515 | 0.70156 | 0.705219 | 0.710434 | 0.70477 | 0.705488 |
| 0.707218 | 0.710015 | 0.699775 | 0.70214 | 0.699567 | 0.703986 | 0.708462 | 0.701514 | 0.702567 |
| 0.712438 | 0.715243 | 0.706139 | 0.707977 | 0.706267 | 0.712985 | 0.713435 | 0.708914 | 0.709293 |
| 0.717077 | 0.716214 | 0.708443 | 0.713341 | 0.708561 | 0.711836 | 0.717193 | 0.712158 | 0.711654 |
| 0.70936 | 0.710979 | 0.70693 | 0.709375 | 0.70602 | 0.707564 | 0.709977 | 0.705781 | 0.706217 |
| 0.706469 | 0.709251 | 0.703115 | 0.703391 | 0.701801 | 0.704454 | 0.705337 | 0.70233 | 0.701695 |
| 0.709484 | 0.713073 | 0.704822 | 0.704055 | 0.702853 | 0.704584 | 0.712593 | 0.706573 | 0.706632 |
| 0.705717 | 0.7123 | 0.707447 | 0.707706 | 0.702643 | 0.709193 | 0.706522 | 0.706781 | 0.70667 |
| 0.721115 | 0.720308 | 0.71348 | 0.715339 | 0.713573 | 0.71891 | 0.72231 | 0.711214 | 0.717307 |
| 0.722961 | 0.725809 | 0.720698 | 0.72066 | 0.720288 | 0.723397 | 0.721938 | 0.724704 | 0.720155 |
| 0.773763 | 0.777963 | 0.768173 | 0.770882 | 0.767322 | 0.76954 | 0.775307 | 0.762671 | 0.771112 |
| 0.784744 | 0.783639 | 0.785374 | 0.780509 | 0.768083 | 0.773483 | 0.77884 | 0.766325 | 0.775098 |
| 0.785744 | 0.779715 | 0.780071 | 0.777481 | 0.766638 | 0.772525 | 0.780938 | 0.76166 | 0.777599 |
| 0.710223 | 0.712502 | 0.70628 | 0.703256 | 0.703593 | 0.710182 | 0.714985 | 0.704019 | 0.710235 |
| 0.707493 | 0.710833 | 0.704418 | 0.70672 | 0.701943 | 0.708416 | 0.710721 | 0.706382 | 0.705846 |
| 0.71999 | 0.718522 | 0.711421 | 0.712101 | 0.708153 | 0.71567 | 0.718108 | 0.715637 | 0.714397 |
| 0.713167 | 0.720494 | 0.712804 | 0.714823 | 0.70965 | 0.721693 | 0.718551 | 0.710812 | 0.71539 |
| 0.708094 | 0.710169 | 0.706349 | 0.707287 | 0.706246 | 0.709659 | 0.712711 | 0.706531 | 0.707675 |
| 0.709495 | 0.711689 | 0.707384 | 0.708943 | 0.706604 | 0.7103 | 0.713012 | 0.706842 | 0.707819 |
| 0.70577 | 0.710294 | 0.704295 | 0.705053 | 0.70367 | 0.707945 | 0.709613 | 0.708456 | 0.708196 |
| 0.708337 | 0.715216 | 0.706576 | 0.706189 | 0.705398 | 0.708114 | 0.711087 | 0.707135 | 0.706156 |
| 0.708672 | 0.708389 | 0.706827 | 0.705176 | 0.702652 | 0.709353 | 0.713478 | 0.704205 | 0.71001 |
| 0.725517 | 0.726644 | 0.720173 | 0.718222 | 0.715735 | 0.726393 | 0.723471 | 0.722667 | 0.720747 |
| 0.703946 | 0.705658 | 0.696298 | 0.699634 | 0.697573 | 0.70067 | 0.705491 | 0.703628 | 0.699664 |
| 0.710664 | 0.713324 | 0.708022 | 0.709202 | 0.701016 | 0.71122 | 0.711723 | 0.703169 | 0.707502 |
| 0.705754 | 0.710309 | 0.701515 | 0.701059 | 0.702367 | 0.704921 | 0.704323 | 0.701829 | 0.701044 |
| 0.725386 | 0.730107 | 0.723585 | 0.721548 | 0.718028 | 0.728013 | 0.725087 | 0.723685 | 0.724631 |
| 0.712139 | 0.71217 | 0.704904 | 0.702005 | 0.703228 | 0.7107 | 0.710013 | 0.708686 | 0.705312 |
| 0.708836 | 0.711723 | 0.702858 | 0.70465 | 0.697638 | 0.7086 | 0.70817 | 0.705538 | 0.704766 |
| 0.714206 | 0.712267 | 0.706536 | 0.707951 | 0.704551 | 0.710759 | 0.71122 | 0.708012 | 0.709224 |
| 0.711618 | 0.712087 | 0.704076 | 0.705541 | 0.704041 | 0.706766 | 0.71189 | 0.708481 | 0.705201 |

| | | | | | | | | |
|---|---|---|---|---|---|---|---|---|
| 0.708034 | 0.713209 | 0.70743 | 0.704082 | 0.705219 | 0.707225 | 0.709576 | 0.704281 | 0.707957 |
| 0.71459 | 0.713876 | 0.710604 | 0.710193 | 0.709296 | 0.711063 | 0.712805 | 0.708834 | 0.707999 |
| 0.712602 | 0.714205 | 0.708121 | 0.707495 | 0.703349 | 0.70855 | 0.711862 | 0.708606 | 0.708852 |
| 0.716421 | 0.721015 | 0.71431 | 0.712489 | 0.7113 | 0.718433 | 0.717213 | 0.713435 | 0.714323 |
| 0.718972 | 0.719592 | 0.710732 | 0.711357 | 0.710381 | 0.715182 | 0.714156 | 0.71065 | 0.711937 |
| 0.711989 | 0.712606 | 0.705936 | 0.708084 | 0.704049 | 0.714635 | 0.711144 | 0.705716 | 0.710096 |
| 0.7132 | 0.712655 | 0.705993 | 0.707724 | 0.706936 | 0.712434 | 0.717985 | 0.709011 | 0.710922 |
| 0.716141 | 0.718977 | 0.710693 | 0.709002 | 0.708243 | 0.716041 | 0.716115 | 0.712427 | 0.712381 |
| 0.71019 | 0.712921 | 0.7078 | 0.708275 | 0.706485 | 0.712097 | 0.712828 | 0.706559 | 0.707629 |
| 0.707595 | 0.711882 | 0.70382 | 0.702971 | 0.699325 | 0.710878 | 0.708416 | 0.70491 | 0.704376 |
| 0.710807 | 0.712283 | 0.706619 | 0.706531 | 0.703849 | 0.707668 | 0.709333 | 0.704672 | 0.704613 |
| 0.709175 | 0.712701 | 0.706325 | 0.708007 | 0.70494 | 0.708701 | 0.709314 | 0.706543 | 0.708993 |
| 0.711657 | 0.716577 | 0.708216 | 0.712112 | 0.709284 | 0.713392 | 0.712184 | 0.706308 | 0.705908 |
| 0.708033 | 0.71169 | 0.706326 | 0.70383 | 0.700377 | 0.705421 | 0.707915 | 0.703976 | 0.707081 |
| 0.712214 | 0.716685 | 0.709476 | 0.70825 | 0.70694 | 0.708556 | 0.712673 | 0.705923 | 0.705654 |
| 0.706988 | 0.713071 | 0.703603 | 0.706383 | 0.703789 | 0.712006 | 0.70661 | 0.70454 | 0.707246 |
| 0.716313 | 0.717442 | 0.708825 | 0.710823 | 0.70499 | 0.713139 | 0.71412 | 0.708229 | 0.711073 |
| 0.709639 | 0.714206 | 0.704716 | 0.705293 | 0.703884 | 0.705826 | 0.712515 | 0.704391 | 0.707847 |
| 0.710306 | 0.715764 | 0.705382 | 0.708533 | 0.706053 | 0.711733 | 0.711933 | 0.707104 | 0.708247 |
| 0.709398 | 0.715693 | 0.704395 | 0.709222 | 0.70579 | 0.711871 | 0.710796 | 0.708383 | 0.708426 |
| 0.70966 | 0.715494 | 0.707119 | 0.70475 | 0.706152 | 0.708735 | 0.710539 | 0.707781 | 0.711245 |
| 0.705517 | 0.708915 | 0.698146 | 0.701541 | 0.70123 | 0.703258 | 0.703139 | 0.702719 | 0.701102 |
| 0.712512 | 0.713284 | 0.706294 | 0.709055 | 0.702018 | 0.710757 | 0.713799 | 0.703196 | 0.708542 |
| 0.714974 | 0.713636 | 0.70611 | 0.712436 | 0.707854 | 0.708698 | 0.711236 | 0.707542 | 0.705295 |
| 0.711502 | 0.710772 | 0.707906 | 0.70618 | 0.703627 | 0.713597 | 0.71671 | 0.704711 | 0.711161 |
| 0.717912 | 0.716168 | 0.709826 | 0.712518 | 0.710401 | 0.717786 | 0.722937 | 0.714554 | 0.716913 |
| 0.709192 | 0.710616 | 0.707055 | 0.705757 | 0.70274 | 0.714148 | 0.714075 | 0.705658 | 0.706183 |
| 0.714083 | 0.714456 | 0.706623 | 0.708751 | 0.705897 | 0.711344 | 0.713803 | 0.706952 | 0.71143 |
| 0.714162 | 0.714024 | 0.706442 | 0.708354 | 0.707394 | 0.713745 | 0.714427 | 0.708213 | 0.706144 |
| 0.713015 | 0.714205 | 0.710537 | 0.710632 | 0.706914 | 0.71404 | 0.716454 | 0.707397 | 0.712842 |
| 0.714934 | 0.715321 | 0.71174 | 0.711452 | 0.709001 | 0.713624 | 0.712839 | 0.71004 | 0.710125 |
| 0.710861 | 0.714263 | 0.708221 | 0.706045 | 0.700628 | 0.708931 | 0.710786 | 0.703904 | 0.708457 |
| 0.710921 | 0.71171 | 0.709737 | 0.707796 | 0.703118 | 0.713982 | 0.715278 | 0.711542 | 0.710978 |
| 0.714657 | 0.715119 | 0.706716 | 0.706679 | 0.706012 | 0.711047 | 0.714867 | 0.70907 | 0.708202 |
| 0.709329 | 0.712631 | 0.705228 | 0.704566 | 0.702301 | 0.709134 | 0.708164 | 0.707219 | 0.704223 |
| 0.713323 | 0.713973 | 0.708882 | 0.710244 | 0.705038 | 0.711235 | 0.714011 | 0.707631 | 0.707493 |
| 0.709608 | 0.709812 | 0.702264 | 0.705246 | 0.697821 | 0.708821 | 0.708228 | 0.703474 | 0.704944 |
| 0.686428 | 0.694853 | 0.686549 | 0.682577 | 0.683628 | 0.685577 | 0.684254 | 0.684776 | 0.68756 |
| 0.69146 | 0.691558 | 0.690632 | 0.69202 | 0.689172 | 0.69159 | 0.693398 | 0.68979 | 0.69285 |
| 0.679399 | 0.684162 | 0.680739 | 0.676973 | 0.679962 | 0.683176 | 0.683879 | 0.680295 | 0.678367 |
| 0.683308 | 0.689482 | 0.680041 | 0.678648 | 0.680335 | 0.67904 | 0.685313 | 0.680317 | 0.684058 |
| 0.664779 | 0.671355 | 0.667262 | 0.670418 | 0.671533 | 0.668955 | 0.669448 | 0.665821 | 0.667318 |
| 0.67162 | 0.671752 | 0.669861 | 0.671941 | 0.671636 | 0.671697 | 0.67516 | 0.675144 | 0.673333 |
| 0.672084 | 0.672107 | 0.668523 | 0.668391 | 0.671544 | 0.67027 | 0.675439 | 0.672946 | 0.673062 |
| 0.666038 | 0.671547 | 0.667445 | 0.672295 | 0.665634 | 0.668688 | 0.67245 | 0.676436 | 0.672715 |
| 0.667244 | 0.674343 | 0.664981 | 0.670055 | 0.670162 | 0.670599 | 0.670547 | 0.672129 | 0.673766 |
| 0.675374 | 0.681965 | 0.673108 | 0.675028 | 0.677041 | 0.675936 | 0.679859 | 0.673849 | 0.683012 |

| | | | | | | | | |
|---|---|---|---|---|---|---|---|---|
| 0.671856 | 0.67691 | 0.675927 | 0.670797 | 0.669753 | 0.669615 | 0.671618 | 0.670976 | 0.673707 |
| 0.672767 | 0.675506 | 0.674199 | 0.672443 | 0.673651 | 0.674125 | 0.680016 | 0.675992 | 0.67762 |
| 0.669561 | 0.67503 | 0.668017 | 0.673845 | 0.672769 | 0.674063 | 0.673458 | 0.666667 | 0.676707 |
| 0.664275 | 0.667464 | 0.665229 | 0.662966 | 0.663485 | 0.665862 | 0.66698 | 0.667576 | 0.672995 |
| 0.669252 | 0.667611 | 0.665711 | 0.669547 | 0.66645 | 0.664203 | 0.669751 | 0.666404 | 0.667377 |
| 0.669661 | 0.667492 | 0.667272 | 0.665643 | 0.666341 | 0.66625 | 0.670817 | 0.670255 | 0.672421 |
| 0.664105 | 0.667849 | 0.667913 | 0.666475 | 0.668055 | 0.670563 | 0.669349 | 0.671381 | 0.673526 |
| 0.663483 | 0.665263 | 0.661569 | 0.667474 | 0.663539 | 0.663316 | 0.66728 | 0.666619 | 0.666474 |
| 0.669438 | 0.670899 | 0.658969 | 0.664578 | 0.664179 | 0.662722 | 0.667014 | 0.662998 | 0.663784 |
| 0.664451 | 0.66394 | 0.668005 | 0.665387 | 0.667213 | 0.66943 | 0.672337 | 0.672075 | 0.671958 |
| 0.663505 | 0.665715 | 0.659549 | 0.654625 | 0.662107 | 0.659895 | 0.66502 | 0.66375 | 0.668974 |
| 0.697885 | 0.701205 | 0.696176 | 0.693066 | 0.693593 | 0.698448 | 0.700219 | 0.696952 | 0.696737 |
| 0.706774 | 0.708409 | 0.701587 | 0.703078 | 0.699598 | 0.706916 | 0.706757 | 0.703334 | 0.706916 |
| 0.690705 | 0.690699 | 0.685447 | 0.686747 | 0.687956 | 0.688999 | 0.689555 | 0.683814 | 0.692445 |
| 0.715052 | 0.720068 | 0.717824 | 0.714274 | 0.710679 | 0.715901 | 0.714967 | 0.713593 | 0.716439 |
| 0.705929 | 0.710062 | 0.701384 | 0.703138 | 0.69882 | 0.703094 | 0.707568 | 0.703379 | 0.704024 |
| 0.712111 | 0.717291 | 0.709254 | 0.706964 | 0.706173 | 0.713793 | 0.712161 | 0.706781 | 0.712085 |
| 0.718386 | 0.724241 | 0.714751 | 0.715403 | 0.714525 | 0.722136 | 0.719937 | 0.717446 | 0.715362 |
| 0.700343 | 0.701175 | 0.691602 | 0.692158 | 0.688298 | 0.695235 | 0.69817 | 0.694066 | 0.693417 |
| 0.72203 | 0.72232 | 0.717296 | 0.720112 | 0.718463 | 0.722173 | 0.720747 | 0.713959 | 0.717942 |
| 0.691693 | 0.695647 | 0.693463 | 0.688442 | 0.689613 | 0.69553 | 0.697428 | 0.688623 | 0.696974 |
| 0.717383 | 0.719412 | 0.712011 | 0.710657 | 0.713607 | 0.718631 | 0.714416 | 0.715601 | 0.711107 |
| 0.717764 | 0.724694 | 0.715388 | 0.714605 | 0.712726 | 0.722687 | 0.720159 | 0.713967 | 0.719476 |
| 0.684636 | 0.687011 | 0.683975 | 0.685491 | 0.684361 | 0.687964 | 0.689831 | 0.685707 | 0.688935 |
| 0.689122 | 0.69211 | 0.688005 | 0.685802 | 0.684508 | 0.691729 | 0.691076 | 0.686448 | 0.685903 |
| 0.689418 | 0.695569 | 0.69208 | 0.690657 | 0.687685 | 0.692033 | 0.696005 | 0.691836 | 0.694821 |
| 0.70906 | 0.712312 | 0.706339 | 0.703952 | 0.702085 | 0.709133 | 0.710828 | 0.707032 | 0.707086 |
| 0.717793 | 0.721946 | 0.71451 | 0.715769 | 0.711686 | 0.720744 | 0.720046 | 0.717083 | 0.714629 |
| 0.688696 | 0.689513 | 0.682289 | 0.684704 | 0.682009 | 0.687451 | 0.689019 | 0.682761 | 0.685652 |
| 0.725833 | 0.728934 | 0.719565 | 0.722921 | 0.718943 | 0.726592 | 0.727205 | 0.719776 | 0.723217 |
| 0.728605 | 0.733736 | 0.721767 | 0.723224 | 0.721068 | 0.72539 | 0.72766 | 0.721273 | 0.723401 |
| 0.699803 | 0.702487 | 0.694734 | 0.691321 | 0.691454 | 0.696785 | 0.700041 | 0.69583 | 0.695185 |
| 0.702026 | 0.701448 | 0.693991 | 0.693316 | 0.692446 | 0.698658 | 0.701289 | 0.694526 | 0.697749 |
| 0.726804 | 0.729091 | 0.721702 | 0.724218 | 0.721351 | 0.728571 | 0.728781 | 0.721119 | 0.723074 |
| 0.723896 | 0.729633 | 0.717708 | 0.720065 | 0.716342 | 0.724705 | 0.721693 | 0.716286 | 0.719481 |
| 0.727374 | 0.731062 | 0.719521 | 0.720053 | 0.716796 | 0.724894 | 0.725004 | 0.72091 | 0.72329 |
| 0.682925 | 0.684272 | 0.674774 | 0.673845 | 0.672474 | 0.677938 | 0.678428 | 0.675287 | 0.675123 |
| 0.719994 | 0.722342 | 0.715404 | 0.720951 | 0.715479 | 0.721882 | 0.724928 | 0.71828 | 0.724933 |
| 0.680375 | 0.678091 | 0.672006 | 0.673356 | 0.671444 | 0.678577 | 0.678509 | 0.674723 | 0.675533 |
| 0.675177 | 0.677929 | 0.666614 | 0.668886 | 0.668035 | 0.670195 | 0.674734 | 0.663772 | 0.667864 |
| 0.721916 | 0.723106 | 0.716347 | 0.715517 | 0.714509 | 0.718128 | 0.722907 | 0.717783 | 0.720384 |
| 0.672166 | 0.675525 | 0.666929 | 0.668235 | 0.665088 | 0.671969 | 0.670467 | 0.666315 | 0.667157 |
| 0.663444 | 0.666138 | 0.660209 | 0.661666 | 0.661094 | 0.659692 | 0.664252 | 0.65795 | 0.65661 |
| 0.656106 | 0.660613 | 0.653315 | 0.654795 | 0.655781 | 0.657736 | 0.660537 | 0.654128 | 0.6526 |
| 0.730152 | 0.731828 | 0.727898 | 0.72596 | 0.723211 | 0.728241 | 0.731927 | 0.723654 | 0.728406 |
| 0.664036 | 0.667347 | 0.659578 | 0.66249 | 0.660433 | 0.660296 | 0.663634 | 0.658009 | 0.659245 |
| 0.722403 | 0.723286 | 0.721426 | 0.722994 | 0.719205 | 0.722727 | 0.72068 | 0.718485 | 0.721528 |

| | | | | | | | | |
|---|---|---|---|---|---|---|---|---|
| 0.732031 | 0.737155 | 0.728237 | 0.733471 | 0.726428 | 0.731217 | 0.733973 | 0.723873 | 0.730495 |
| 0.727589 | 0.728606 | 0.721032 | 0.722012 | 0.719099 | 0.725577 | 0.731467 | 0.722969 | 0.725116 |
| 0.745168 | 0.749877 | 0.744512 | 0.740632 | 0.736744 | 0.745362 | 0.746189 | 0.737984 | 0.745969 |
| 0.653182 | 0.656385 | 0.649084 | 0.650696 | 0.650046 | 0.65258 | 0.655275 | 0.649996 | 0.648773 |
| 0.654935 | 0.65788 | 0.652019 | 0.651866 | 0.651803 | 0.654005 | 0.659749 | 0.65273 | 0.653564 |
| 0.738509 | 0.732942 | 0.725868 | 0.726828 | 0.723383 | 0.730106 | 0.736267 | 0.727085 | 0.730361 |
| 0.657743 | 0.659788 | 0.648044 | 0.651598 | 0.648876 | 0.652851 | 0.658243 | 0.652018 | 0.651354 |
| 0.736953 | 0.733708 | 0.727556 | 0.736838 | 0.732733 | 0.731892 | 0.738037 | 0.729067 | 0.73296 |
| 0.735979 | 0.733684 | 0.728608 | 0.731058 | 0.727379 | 0.730581 | 0.736961 | 0.728119 | 0.73168 |
| 0.731553 | 0.736631 | 0.727512 | 0.734594 | 0.732248 | 0.73506 | 0.737593 | 0.733257 | 0.732019 |
| 0.734956 | 0.738344 | 0.730136 | 0.732936 | 0.727739 | 0.728016 | 0.738204 | 0.728852 | 0.733077 |
| 0.727848 | 0.709459 | 0.722222 | 0.707792 | 0.726667 | 0.743243 | 0.756757 | 0.733766 | 0.715278 |
| 0.725595 | 0.726415 | 0.728175 | 0.730675 | 0.732589 | 0.722943 | 0.728927 | 0.747174 | 0.736647 |
| 0.722136 | 0.721926 | 0.725347 | 0.727374 | 0.724029 | 0.724127 | 0.73102 | 0.74818 | 0.736156 |
| 0.736282 | 0.736397 | 0.737997 | 0.734069 | 0.739217 | 0.736391 | 0.739338 | 0.757292 | 0.739762 |
| 0.728735 | 0.732795 | 0.734117 | 0.727746 | 0.740135 | 0.73166 | 0.738862 | 0.757576 | 0.736585 |
| 0.595848 | 0.596942 | 0.591029 | 0.59108 | 0.590214 | 0.590451 | 0.596701 | 0.5935 | 0.595047 |
| 0.731827 | 0.737484 | 0.740234 | 0.739102 | 0.735714 | 0.730856 | 0.742086 | 0.752669 | 0.75164 |
| 0.741297 | 0.741471 | 0.734565 | 0.736633 | 0.733761 | 0.737993 | 0.743187 | 0.734385 | 0.741059 |
| 0.738637 | 0.742822 | 0.736252 | 0.733935 | 0.729681 | 0.733555 | 0.740481 | 0.733116 | 0.737453 |
| 0.741628 | 0.739244 | 0.737792 | 0.74232 | 0.735109 | 0.735781 | 0.742765 | 0.736373 | 0.736805 |
| 0.597409 | 0.599555 | 0.5912 | 0.593913 | 0.591619 | 0.594975 | 0.599809 | 0.589637 | 0.593866 |
| 0.582565 | 0.586411 | 0.58317 | 0.580086 | 0.578836 | 0.582949 | 0.583306 | 0.576515 | 0.578961 |
| 0.764444 | 0.763274 | 0.759518 | 0.759797 | 0.759343 | 0.760181 | 0.76809 | 0.762236 | 0.763931 |
| 0.766943 | 0.764686 | 0.762505 | 0.760829 | 0.759692 | 0.755777 | 0.770246 | 0.756859 | 0.763972 |
| 0.75598 | 0.755419 | 0.749893 | 0.757242 | 0.753766 | 0.75204 | 0.761815 | 0.752609 | 0.755109 |
| 0.760211 | 0.760973 | 0.758638 | 0.763292 | 0.762507 | 0.757834 | 0.764496 | 0.761539 | 0.764768 |
| 0.76078 | 0.76155 | 0.752851 | 0.764412 | 0.764465 | 0.758316 | 0.76648 | 0.756847 | 0.762716 |
| 0.766022 | 0.768302 | 0.767725 | 0.766076 | 0.760069 | 0.762972 | 0.775588 | 0.764648 | 0.770044 |
| 0.76349 | 0.763953 | 0.76015 | 0.766201 | 0.764771 | 0.759585 | 0.769054 | 0.762106 | 0.764017 |
| 0.76218 | 0.761806 | 0.761888 | 0.759361 | 0.756709 | 0.758665 | 0.764059 | 0.755229 | 0.764749 |
| 0.770351 | 0.775791 | 0.773234 | 0.768801 | 0.765056 | 0.767038 | 0.771908 | 0.764201 | 0.772775 |
| 0.762178 | 0.765957 | 0.764181 | 0.759175 | 0.765968 | 0.758074 | 0.762593 | 0.760447 | 0.759961 |
| 0.766454 | 0.767322 | 0.766111 | 0.765187 | 0.759597 | 0.760604 | 0.769562 | 0.750954 | 0.770522 |
| 0.766523 | 0.773513 | 0.770229 | 0.76537 | 0.767892 | 0.764896 | 0.771101 | 0.763635 | 0.770361 |
| 0.76436 | 0.76997 | 0.767889 | 0.763827 | 0.760924 | 0.759364 | 0.765584 | 0.758864 | 0.765009 |
| 0.762514 | 0.763136 | 0.765533 | 0.763567 | 0.766583 | 0.757825 | 0.766525 | 0.76111 | 0.768999 |
| 0.765899 | 0.769351 | 0.769722 | 0.767083 | 0.761836 | 0.768176 | 0.768127 | 0.756894 | 0.769933 |
| 0.761288 | 0.764771 | 0.761534 | 0.75848 | 0.761942 | 0.755601 | 0.763777 | 0.760946 | 0.764281 |
| 0.760996 | 0.763992 | 0.75914 | 0.7592 | 0.75738 | 0.753471 | 0.764512 | 0.75505 | 0.766728 |
| 0.719903 | 0.72689 | 0.723589 | 0.729585 | 0.731777 | 0.719363 | 0.719845 | 0.738826 | 0.734164 |
| 0.725205 | 0.729827 | 0.738123 | 0.734557 | 0.737235 | 0.729843 | 0.733168 | 0.754067 | 0.742818 |
| 0.771807 | 0.773147 | 0.766749 | 0.769433 | 0.764235 | 0.767134 | 0.775919 | 0.758554 | 0.766344 |
| 0.76885 | 0.771516 | 0.767158 | 0.770383 | 0.768421 | 0.766026 | 0.772111 | 0.769034 | 0.775109 |
| 0.716221 | 0.714671 | 0.719538 | 0.719084 | 0.717124 | 0.716013 | 0.726437 | 0.720362 | 0.727147 |
| 0.767357 | 0.771381 | 0.766909 | 0.77006 | 0.765824 | 0.765821 | 0.77426 | 0.762791 | 0.769231 |
| 0.76473 | 0.761667 | 0.760561 | 0.761256 | 0.756697 | 0.755707 | 0.765634 | 0.753305 | 0.760074 |

| | | | | | | | | |
|---|---|---|---|---|---|---|---|---|
| 0.76496 | 0.766694 | 0.762225 | 0.761209 | 0.75903 | 0.762236 | 0.772359 | 0.756529 | 0.768782 |
| 0.769094 | 0.76947 | 0.766012 | 0.767361 | 0.761553 | 0.767127 | 0.776355 | 0.760161 | 0.772002 |
| 0.767636 | 0.767369 | 0.765626 | 0.765976 | 0.761357 | 0.758419 | 0.769693 | 0.756728 | 0.769782 |
| 0.769934 | 0.774636 | 0.770448 | 0.767157 | 0.764993 | 0.767534 | 0.773976 | 0.758675 | 0.773243 |
| 0.768399 | 0.773046 | 0.770738 | 0.771126 | 0.769382 | 0.76551 | 0.773514 | 0.761733 | 0.775773 |
| 0.767651 | 0.764166 | 0.766752 | 0.763194 | 0.763391 | 0.758718 | 0.767935 | 0.755129 | 0.766158 |
| 0.762817 | 0.767057 | 0.762289 | 0.762742 | 0.764496 | 0.761818 | 0.770418 | 0.758853 | 0.766948 |
| 0.755829 | 0.757246 | 0.757005 | 0.754478 | 0.755634 | 0.752104 | 0.760278 | 0.751972 | 0.759836 |
| 0.766411 | 0.768736 | 0.763695 | 0.766383 | 0.759115 | 0.75956 | 0.769657 | 0.757142 | 0.765758 |
| 0.761446 | 0.761689 | 0.759606 | 0.758236 | 0.759043 | 0.759088 | 0.76615 | 0.75361 | 0.762366 |
| 0.763811 | 0.767579 | 0.76496 | 0.765272 | 0.765463 | 0.765221 | 0.770614 | 0.759615 | 0.770778 |
| 0.763639 | 0.767622 | 0.765808 | 0.76279 | 0.761537 | 0.760857 | 0.769339 | 0.753868 | 0.768116 |
| 0.761626 | 0.761297 | 0.75849 | 0.758947 | 0.755644 | 0.755546 | 0.766952 | 0.747105 | 0.762229 |
| 0.758638 | 0.75914 | 0.755882 | 0.758117 | 0.754729 | 0.755469 | 0.761428 | 0.754248 | 0.757806 |
| 0.768054 | 0.768759 | 0.769192 | 0.763797 | 0.764116 | 0.760273 | 0.765893 | 0.758244 | 0.765001 |
| 0.76371 | 0.7682 | 0.762115 | 0.765115 | 0.760283 | 0.764128 | 0.76488 | 0.760441 | 0.764737 |
| 0.761107 | 0.765126 | 0.761859 | 0.766385 | 0.766899 | 0.759321 | 0.765087 | 0.759537 | 0.76657 |
| 0.759993 | 0.761552 | 0.761302 | 0.761344 | 0.760418 | 0.761553 | 0.763598 | 0.756147 | 0.760598 |
| 0.76182 | 0.764658 | 0.760807 | 0.769598 | 0.764958 | 0.758791 | 0.767142 | 0.76045 | 0.76031 |
| 0.764188 | 0.766763 | 0.766141 | 0.765616 | 0.763232 | 0.760642 | 0.766621 | 0.762266 | 0.763793 |
| 0.752885 | 0.759615 | 0.760545 | 0.754652 | 0.753359 | 0.754736 | 0.755715 | 0.752474 | 0.75195 |
| 0.760794 | 0.761849 | 0.755891 | 0.766745 | 0.760953 | 0.755613 | 0.763892 | 0.759196 | 0.762598 |
| 0.767457 | 0.767373 | 0.762597 | 0.767552 | 0.76948 | 0.763891 | 0.770247 | 0.764258 | 0.766977 |
| 0.762675 | 0.764416 | 0.760122 | 0.764091 | 0.765411 | 0.762793 | 0.768916 | 0.760081 | 0.76752 |
| 0.760996 | 0.771381 | 0.765026 | 0.764017 | 0.760685 | 0.759604 | 0.76634 | 0.759776 | 0.761335 |
| 0.760141 | 0.756451 | 0.755375 | 0.762729 | 0.758721 | 0.757424 | 0.768379 | 0.760731 | 0.762884 |
| 0.757119 | 0.75971 | 0.755753 | 0.762569 | 0.753992 | 0.755699 | 0.760825 | 0.753685 | 0.758303 |
| 0.758045 | 0.759483 | 0.761079 | 0.758414 | 0.758627 | 0.75667 | 0.762222 | 0.753817 | 0.760007 |
| 0.762794 | 0.76311 | 0.758255 | 0.761217 | 0.757338 | 0.761387 | 0.768306 | 0.757013 | 0.765128 |
| 0.761414 | 0.766406 | 0.761876 | 0.768068 | 0.765245 | 0.762232 | 0.770332 | 0.758199 | 0.767116 |
| 0.763053 | 0.768622 | 0.767444 | 0.76248 | 0.763856 | 0.758686 | 0.764226 | 0.763251 | 0.760987 |
| 0.760987 | 0.764721 | 0.765558 | 0.766391 | 0.760059 | 0.760757 | 0.763268 | 0.761243 | 0.762325 |
| 0.772316 | 0.769734 | 0.768207 | 0.768469 | 0.766641 | 0.766057 | 0.773858 | 0.762262 | 0.767227 |
| 0.765657 | 0.76487 | 0.766778 | 0.765726 | 0.763142 | 0.765101 | 0.772071 | 0.763462 | 0.76573 |
| 0.760997 | 0.760771 | 0.755775 | 0.758956 | 0.75323 | 0.757279 | 0.765665 | 0.754451 | 0.764064 |
| 0.763473 | 0.767417 | 0.764741 | 0.763503 | 0.762809 | 0.761135 | 0.762565 | 0.760848 | 0.759603 |
| 0.760356 | 0.766141 | 0.75857 | 0.762663 | 0.759011 | 0.760548 | 0.762854 | 0.761799 | 0.762307 |
| 0.763385 | 0.763639 | 0.760492 | 0.761002 | 0.763939 | 0.757581 | 0.764079 | 0.757638 | 0.762474 |
| 0.756913 | 0.760131 | 0.758049 | 0.754147 | 0.753341 | 0.753263 | 0.755018 | 0.753348 | 0.755084 |
| 0.754556 | 0.756295 | 0.749735 | 0.752433 | 0.758174 | 0.751062 | 0.756936 | 0.751915 | 0.75732 |
| 0.754578 | 0.75542 | 0.750241 | 0.758478 | 0.748749 | 0.749403 | 0.758699 | 0.752495 | 0.749206 |
| 0.753689 | 0.755769 | 0.751571 | 0.756942 | 0.754427 | 0.751536 | 0.760184 | 0.74866 | 0.756593 |
| 0.754821 | 0.758027 | 0.751772 | 0.756292 | 0.751044 | 0.752706 | 0.756663 | 0.748788 | 0.750528 |
| 0.752947 | 0.75477 | 0.750162 | 0.754578 | 0.754056 | 0.749319 | 0.756754 | 0.749189 | 0.752991 |
| 0.756046 | 0.756871 | 0.751662 | 0.757719 | 0.760301 | 0.757004 | 0.763146 | 0.757034 | 0.760442 |
| 0.74793 | 0.75132 | 0.753785 | 0.75029 | 0.752264 | 0.74884 | 0.753437 | 0.746395 | 0.749975 |

| | | | | | | | | |
|---|---|---|---|---|---|---|---|---|
| 0.763977 | 0.765504 | 0.759141 | 0.765223 | 0.757713 | 0.758618 | 0.768836 | 0.761623 | 0.76174 |
| 0.773868 | 0.774347 | 0.770243 | 0.770015 | 0.76747 | 0.767166 | 0.777451 | 0.763714 | 0.770284 |
| 0.76508 | 0.764873 | 0.764268 | 0.76743 | 0.759131 | 0.762813 | 0.771137 | 0.76196 | 0.765774 |
| 0.763766 | 0.772097 | 0.76761 | 0.762806 | 0.758767 | 0.76491 | 0.771412 | 0.766187 | 0.770534 |
| 0.773253 | 0.780664 | 0.773152 | 0.774246 | 0.769152 | 0.773163 | 0.77947 | 0.773851 | 0.775664 |
| 0.769375 | 0.774701 | 0.768805 | 0.768159 | 0.769099 | 0.767047 | 0.772321 | 0.766975 | 0.774262 |
| 0.765757 | 0.769489 | 0.763635 | 0.764576 | 0.765244 | 0.768204 | 0.770839 | 0.759669 | 0.770556 |
| 0.76758 | 0.770325 | 0.766267 | 0.766522 | 0.760419 | 0.763441 | 0.770304 | 0.758832 | 0.767085 |
| 0.764423 | 0.767328 | 0.764957 | 0.760823 | 0.761869 | 0.759651 | 0.770168 | 0.762647 | 0.764386 |
| 0.764403 | 0.772513 | 0.764575 | 0.758899 | 0.758182 | 0.75979 | 0.763454 | 0.758885 | 0.762957 |
| 0.773744 | 0.772534 | 0.77 | 0.76864 | 0.767787 | 0.767731 | 0.773342 | 0.75983 | 0.768034 |
| 0.770151 | 0.772615 | 0.767291 | 0.764341 | 0.763792 | 0.761685 | 0.771803 | 0.759912 | 0.770569 |
| 0.772423 | 0.771158 | 0.768338 | 0.774317 | 0.766801 | 0.76777 | 0.775606 | 0.768402 | 0.772788 |
| 0.767111 | 0.768059 | 0.769342 | 0.767357 | 0.7646 | 0.766697 | 0.76927 | 0.763939 | 0.767174 |
| 0.775487 | 0.779823 | 0.771509 | 0.774262 | 0.767594 | 0.7774 | 0.778968 | 0.76942 | 0.776912 |
| 0.775362 | 0.771944 | 0.768357 | 0.771665 | 0.768697 | 0.770811 | 0.774331 | 0.766878 | 0.770595 |
| 0.782111 | 0.776187 | 0.768789 | 0.773888 | 0.767527 | 0.775406 | 0.779493 | 0.763254 | 0.77208 |
| 0.773952 | 0.773261 | 0.766911 | 0.76892 | 0.769361 | 0.771316 | 0.776354 | 0.764814 | 0.773995 |
| 0.769811 | 0.774738 | 0.76948 | 0.774298 | 0.770988 | 0.772014 | 0.776387 | 0.765368 | 0.779129 |
| 0.77611 | 0.774627 | 0.772221 | 0.773881 | 0.768273 | 0.773435 | 0.779269 | 0.76681 | 0.779068 |
| 0.763708 | 0.769282 | 0.766966 | 0.773136 | 0.765778 | 0.768544 | 0.767833 | 0.756308 | 0.768998 |
| 0.771849 | 0.770201 | 0.768716 | 0.769686 | 0.763841 | 0.766683 | 0.777443 | 0.762637 | 0.770937 |
| 0.774516 | 0.77027 | 0.767126 | 0.770385 | 0.765056 | 0.77158 | 0.777816 | 0.765041 | 0.774299 |
| 0.779212 | 0.77436 | 0.772044 | 0.772088 | 0.767297 | 0.770497 | 0.778004 | 0.766065 | 0.770938 |
| 0.777321 | 0.776161 | 0.770105 | 0.772743 | 0.771753 | 0.766093 | 0.774827 | 0.769191 | 0.770522 |
| 0.778858 | 0.774657 | 0.77013 | 0.767473 | 0.767209 | 0.76999 | 0.777844 | 0.764401 | 0.775157 |
| 0.772172 | 0.771337 | 0.769577 | 0.765929 | 0.765927 | 0.763647 | 0.777929 | 0.767501 | 0.772836 |
| 0.776659 | 0.770867 | 0.766774 | 0.76592 | 0.763522 | 0.769891 | 0.775562 | 0.764155 | 0.765776 |
| 0.773092 | 0.776239 | 0.772391 | 0.772658 | 0.769492 | 0.774808 | 0.776777 | 0.765292 | 0.769935 |
| 0.778888 | 0.779241 | 0.768465 | 0.77293 | 0.770001 | 0.769481 | 0.778585 | 0.770106 | 0.779319 |
| 0.77211 | 0.771971 | 0.767116 | 0.761974 | 0.758922 | 0.769231 | 0.771722 | 0.758293 | 0.766971 |
| 0.774292 | 0.774537 | 0.774961 | 0.773621 | 0.769847 | 0.770544 | 0.778712 | 0.767576 | 0.777211 |
| 0.776322 | 0.78173 | 0.773588 | 0.771606 | 0.767577 | 0.7735 | 0.776662 | 0.768404 | 0.774211 |
| 0.773949 | 0.771779 | 0.768886 | 0.768712 | 0.767694 | 0.766687 | 0.776163 | 0.758941 | 0.774199 |
| 0.778461 | 0.777759 | 0.774233 | 0.768315 | 0.764355 | 0.771504 | 0.780197 | 0.768279 | 0.777626 |
| 0.774443 | 0.77378 | 0.771551 | 0.763767 | 0.765394 | 0.769956 | 0.775304 | 0.764771 | 0.770375 |
| 0.784878 | 0.779972 | 0.775937 | 0.777702 | 0.772813 | 0.771822 | 0.783066 | 0.768317 | 0.781634 |
| 0.776061 | 0.775659 | 0.772913 | 0.772053 | 0.767967 | 0.770447 | 0.779458 | 0.768762 | 0.778008 |
| 0.770855 | 0.772124 | 0.777987 | 0.770031 | 0.76538 | 0.772352 | 0.774001 | 0.768264 | 0.775396 |
| 0.778028 | 0.774433 | 0.780019 | 0.774449 | 0.766527 | 0.772136 | 0.776599 | 0.761773 | 0.775806 |
| 0.780756 | 0.77413 | 0.775623 | 0.76901 | 0.769277 | 0.770901 | 0.778553 | 0.766477 | 0.776158 |
| 0.774678 | 0.776017 | 0.77392 | 0.771782 | 0.770558 | 0.771946 | 0.780267 | 0.770909 | 0.777482 |
| 0.778382 | 0.779073 | 0.774951 | 0.77579 | 0.770855 | 0.774034 | 0.780801 | 0.774415 | 0.781516 |
| 0.771502 | 0.771371 | 0.77498 | 0.763212 | 0.761599 | 0.766515 | 0.778039 | 0.762544 | 0.775462 |
| 0.782061 | 0.776974 | 0.778928 | 0.772493 | 0.769323 | 0.770835 | 0.782067 | 0.765758 | 0.781923 |
| 0.780522 | 0.771539 | 0.7736 | 0.768132 | 0.769513 | 0.773678 | 0.778057 | 0.762372 | 0.77822 |

| | | | | | | | | |
|---|---|---|---|---|---|---|---|---|
| 0.761932 | 0.758156 | 0.757101 | 0.758124 | 0.755092 | 0.754663 | 0.764425 | 0.754647 | 0.763731 |
| 0.752105 | 0.752889 | 0.750768 | 0.745879 | 0.739042 | 0.74898 | 0.752187 | 0.747473 | 0.74958 |
| 0.775813 | 0.77247 | 0.768902 | 0.76911 | 0.764588 | 0.771643 | 0.773458 | 0.7628 | 0.773509 |
| 0.770835 | 0.774299 | 0.772535 | 0.770425 | 0.76505 | 0.776388 | 0.775235 | 0.766296 | 0.77231 |
| 0.766342 | 0.763178 | 0.76468 | 0.763888 | 0.762052 | 0.761095 | 0.765326 | 0.757276 | 0.7625 |
| 0.777668 | 0.770362 | 0.772582 | 0.772876 | 0.766244 | 0.769921 | 0.775026 | 0.763092 | 0.770094 |
| 0.767571 | 0.768082 | 0.769402 | 0.769579 | 0.770492 | 0.769988 | 0.772986 | 0.770183 | 0.770831 |
| 0.766169 | 0.767719 | 0.765573 | 0.763545 | 0.762136 | 0.760722 | 0.764352 | 0.759519 | 0.761428 |
| 0.776656 | 0.775068 | 0.770462 | 0.773878 | 0.77447 | 0.777224 | 0.776994 | 0.769427 | 0.776344 |
| 0.769394 | 0.77211 | 0.771625 | 0.773917 | 0.774091 | 0.775817 | 0.775997 | 0.772885 | 0.777748 |
| 0.768919 | 0.771517 | 0.772511 | 0.771674 | 0.775533 | 0.772284 | 0.772519 | 0.766956 | 0.772735 |
| 0.775211 | 0.774712 | 0.769 | 0.775722 | 0.768272 | 0.773714 | 0.777217 | 0.766823 | 0.774773 |
| 0.771927 | 0.777539 | 0.775205 | 0.77812 | 0.773328 | 0.77763 | 0.776706 | 0.770011 | 0.777265 |
| 0.774586 | 0.776855 | 0.769407 | 0.777541 | 0.773154 | 0.778147 | 0.772785 | 0.771178 | 0.775862 |
| 0.774025 | 0.776057 | 0.768651 | 0.773413 | 0.772815 | 0.775023 | 0.773421 | 0.764674 | 0.771513 |
| 0.771759 | 0.773385 | 0.768336 | 0.7722 | 0.768113 | 0.770324 | 0.775196 | 0.761537 | 0.774001 |
| 0.766678 | 0.773966 | 0.770472 | 0.770577 | 0.766744 | 0.770819 | 0.771297 | 0.769319 | 0.775109 |
| 0.772511 | 0.769639 | 0.766581 | 0.768862 | 0.766328 | 0.769353 | 0.772256 | 0.762258 | 0.772077 |
| 0.76591 | 0.769251 | 0.772375 | 0.77168 | 0.767457 | 0.7667 | 0.777316 | 0.763835 | 0.768931 |
| 0.760099 | 0.763494 | 0.769799 | 0.767096 | 0.762836 | 0.773164 | 0.774127 | 0.762834 | 0.772232 |
| 0.771223 | 0.779017 | 0.771217 | 0.775194 | 0.774507 | 0.772743 | 0.77456 | 0.7631 | 0.772662 |
| 0.758854 | 0.761437 | 0.75974 | 0.755212 | 0.757797 | 0.757826 | 0.761538 | 0.756686 | 0.761213 |
| 0.766824 | 0.767857 | 0.764781 | 0.76282 | 0.762677 | 0.760927 | 0.769264 | 0.765267 | 0.762178 |
| 0.770755 | 0.773964 | 0.772462 | 0.765959 | 0.773159 | 0.771006 | 0.775507 | 0.769156 | 0.771238 |
| 0.766708 | 0.776164 | 0.776571 | 0.773732 | 0.769797 | 0.76961 | 0.774523 | 0.7747 | 0.774129 |
| 0.768104 | 0.769906 | 0.769832 | 0.764807 | 0.765073 | 0.767673 | 0.772079 | 0.770537 | 0.768672 |
| 0.758499 | 0.763646 | 0.765946 | 0.767593 | 0.762031 | 0.760324 | 0.763574 | 0.762129 | 0.764057 |
| 0.761704 | 0.769845 | 0.77104 | 0.764478 | 0.762314 | 0.768148 | 0.763769 | 0.767751 | 0.767942 |
| 0.759091 | 0.764578 | 0.76764 | 0.765675 | 0.762282 | 0.767579 | 0.766843 | 0.769444 | 0.768374 |
| 0.695506 | 0.696036 | 0.696432 | 0.694237 | 0.692913 | 0.697674 | 0.697095 | 0.695737 | 0.697303 |
| 0.709039 | 0.711684 | 0.70716 | 0.705936 | 0.702541 | 0.706193 | 0.709251 | 0.707378 | 0.70667 |
| 0.706681 | 0.708622 | 0.702721 | 0.701347 | 0.699335 | 0.707998 | 0.705546 | 0.703659 | 0.700344 |
| 0.696547 | 0.700468 | 0.695759 | 0.691967 | 0.693482 | 0.694015 | 0.700182 | 0.698823 | 0.696022 |
| 0.71342 | 0.712263 | 0.706216 | 0.70521 | 0.702343 | 0.709396 | 0.713453 | 0.705079 | 0.708098 |
| 0.766366 | 0.770883 | 0.766728 | 0.761971 | 0.770496 | 0.761787 | 0.769582 | 0.769447 | 0.766473 |
| 0.77028 | 0.772104 | 0.771577 | 0.766362 | 0.768603 | 0.763173 | 0.774571 | 0.773237 | 0.769563 |
| 0.770432 | 0.772986 | 0.773216 | 0.766443 | 0.767924 | 0.766241 | 0.771956 | 0.77739 | 0.771357 |
| 0.774453 | 0.777343 | 0.772135 | 0.765928 | 0.771076 | 0.771297 | 0.776471 | 0.776769 | 0.772821 |
| 0.763453 | 0.77504 | 0.76874 | 0.767434 | 0.767842 | 0.760879 | 0.769892 | 0.772128 | 0.763623 |
| 0.77055 | 0.771924 | 0.767173 | 0.764434 | 0.767468 | 0.763984 | 0.774639 | 0.777304 | 0.775552 |
| 0.771911 | 0.773321 | 0.772582 | 0.768613 | 0.772125 | 0.761088 | 0.772877 | 0.775516 | 0.770846 |
| 0.766484 | 0.770252 | 0.767123 | 0.759529 | 0.763078 | 0.761254 | 0.767763 | 0.771695 | 0.766685 |
| 0.763668 | 0.771023 | 0.762732 | 0.764299 | 0.768079 | 0.758615 | 0.763863 | 0.769595 | 0.762029 |
| 0.756066 | 0.769621 | 0.768188 | 0.756629 | 0.765467 | 0.764216 | 0.764872 | 0.762811 | 0.763407 |
| 0.767163 | 0.770227 | 0.761776 | 0.757599 | 0.763052 | 0.764836 | 0.770456 | 0.768299 | 0.767613 |
| 0.772558 | 0.777796 | 0.773922 | 0.768026 | 0.77157 | 0.766639 | 0.771251 | 0.778276 | 0.768789 |

| | | | | | | | | |
|---|---|---|---|---|---|---|---|---|
| 0.771295 | 0.777025 | 0.771144 | 0.766531 | 0.767452 | 0.770326 | 0.775867 | 0.782593 | 0.775326 |
| 0.767347 | 0.771229 | 0.768636 | 0.765974 | 0.765227 | 0.76399 | 0.768604 | 0.794872 | 0.765895 |
| 0.770096 | 0.774383 | 0.771048 | 0.765499 | 0.765772 | 0.767188 | 0.771232 | 0.790285 | 0.771355 |
| 0.764039 | 0.768893 | 0.765255 | 0.762092 | 0.762875 | 0.760436 | 0.767517 | 0.787936 | 0.764936 |
| 0.762718 | 0.770526 | 0.768645 | 0.764762 | 0.76603 | 0.760962 | 0.768587 | 0.796949 | 0.76907 |
| 0.765896 | 0.765941 | 0.768065 | 0.76482 | 0.759757 | 0.766119 | 0.767733 | 0.811046 | 0.771822 |
| 0.763573 | 0.769537 | 0.767211 | 0.767649 | 0.77 | 0.765742 | 0.768479 | 0.787961 | 0.769175 |
| 0.761297 | 0.771089 | 0.769163 | 0.768064 | 0.769271 | 0.769126 | 0.768271 | 0.788221 | 0.769314 |
| 0.763764 | 0.765959 | 0.763378 | 0.764193 | 0.762209 | 0.761727 | 0.76628 | 0.778945 | 0.766579 |
| 0.771226 | 0.770526 | 0.764124 | 0.762334 | 0.765452 | 0.764346 | 0.769654 | 0.786716 | 0.762713 |
| 0.761861 | 0.766837 | 0.767034 | 0.767547 | 0.764051 | 0.763776 | 0.766817 | 0.78915 | 0.771338 |
| 0.765389 | 0.772137 | 0.771777 | 0.764874 | 0.763142 | 0.768364 | 0.766459 | 0.782699 | 0.771707 |
| 0.760709 | 0.762848 | 0.762354 | 0.760366 | 0.756039 | 0.761409 | 0.763613 | 0.779103 | 0.765198 |
| 0.758627 | 0.762621 | 0.762342 | 0.75474 | 0.758879 | 0.757606 | 0.765349 | 0.791914 | 0.76124 |
| 0.817226 | 0.822228 | 0.796442 | 0.797547 | 0.784763 | 0.784401 | 0.798661 | 0.762178 | 0.78589 |
| 0.866075 | 0.816819 | 0.786976 | 0.798332 | 0.789067 | 0.794306 | 0.872224 | 0.771386 | 0.858918 |
| 1 | 0.837726 | 0.78834 | 0.78924 | 0.778383 | 0.77708 | 0.866732 | 0.760132 | 0.779567 |
| 0.837726 | 1 | 0.81268 | 0.802746 | 0.793928 | 0.794199 | 0.811175 | 0.768229 | 0.791741 |
| 0.78834 | 0.81268 | 1 | 0.795853 | 0.785785 | 0.787581 | 0.782802 | 0.767577 | 0.787234 |
| 0.78924 | 0.802746 | 0.795853 | 1 | 0.845095 | 0.801575 | 0.799089 | 0.764327 | 0.799001 |
| 0.778383 | 0.793928 | 0.785785 | 0.845095 | 1 | 0.782879 | 0.783142 | 0.762931 | 0.780357 |
| 0.77708 | 0.794199 | 0.787581 | 0.801575 | 0.782879 | 1 | 0.788136 | 0.761935 | 0.790525 |
| 0.866732 | 0.811175 | 0.782802 | 0.799089 | 0.783142 | 0.788136 | 1 | 0.764733 | 0.859032 |
| 0.760132 | 0.768229 | 0.767577 | 0.764327 | 0.762931 | 0.761935 | 0.764733 | 1 | 0.769332 |
| 0.779567 | 0.791741 | 0.787234 | 0.799001 | 0.780357 | 0.790525 | 0.859032 | 0.769332 | 1 |
| 0.704676 | 0.70866 | 0.704301 | 0.7045 | 0.699666 | 0.700268 | 0.707035 | 0.703305 | 0.704441 |
| 0.7823 | 0.772664 | 0.769899 | 0.770208 | 0.763053 | 0.769479 | 0.782026 | 0.763818 | 0.77705 |
| 0.777814 | 0.775023 | 0.770227 | 0.769165 | 0.766083 | 0.772595 | 0.77704 | 0.766094 | 0.772414 |
| 0.779282 | 0.777537 | 0.775426 | 0.774338 | 0.772012 | 0.774996 | 0.783056 | 0.769251 | 0.782218 |
| 0.772228 | 0.77264 | 0.771149 | 0.772977 | 0.76712 | 0.773743 | 0.776055 | 0.770335 | 0.77563 |
| 0.775631 | 0.774768 | 0.766687 | 0.766711 | 0.762644 | 0.766199 | 0.780818 | 0.762354 | 0.773144 |
| 0.772997 | 0.769598 | 0.770376 | 0.771713 | 0.766183 | 0.768112 | 0.777944 | 0.768923 | 0.770615 |
| 0.774266 | 0.775386 | 0.77168 | 0.766545 | 0.76023 | 0.769865 | 0.778146 | 0.762842 | 0.773149 |
| 0.773392 | 0.769638 | 0.767214 | 0.767365 | 0.76337 | 0.766329 | 0.778744 | 0.766526 | 0.773762 |
| 0.767241 | 0.771578 | 0.765858 | 0.762185 | 0.759524 | 0.764283 | 0.771202 | 0.763774 | 0.768811 |
| 0.775764 | 0.766031 | 0.763348 | 0.762344 | 0.759902 | 0.770353 | 0.774179 | 0.760866 | 0.768773 |
| 0.781632 | 0.773485 | 0.770822 | 0.77939 | 0.767006 | 0.774323 | 0.782453 | 0.768098 | 0.774855 |
| 0.774055 | 0.78005 | 0.773888 | 0.769752 | 0.769569 | 0.771225 | 0.778782 | 0.767663 | 0.776574 |
| 0.783719 | 0.77775 | 0.775971 | 0.773049 | 0.768745 | 0.776118 | 0.784953 | 0.764784 | 0.777348 |
| 0.771533 | 0.77535 | 0.769867 | 0.768288 | 0.767137 | 0.766834 | 0.775847 | 0.764453 | 0.774188 |
| 0.773446 | 0.770618 | 0.767135 | 0.765174 | 0.762112 | 0.764854 | 0.770049 | 0.757603 | 0.768713 |
| 0.778677 | 0.77936 | 0.773873 | 0.777312 | 0.767982 | 0.773344 | 0.779307 | 0.76557 | 0.775951 |
| 0.773193 | 0.77162 | 0.770413 | 0.764503 | 0.757887 | 0.762871 | 0.773462 | 0.75965 | 0.770413 |
| 0.764534 | 0.770097 | 0.765492 | 0.769774 | 0.763025 | 0.763364 | 0.767622 | 0.761315 | 0.76214 |
| 0.770611 | 0.771227 | 0.771057 | 0.768888 | 0.763872 | 0.768961 | 0.774171 | 0.771226 | 0.771092 |
| 0.769797 | 0.77272 | 0.769534 | 0.769507 | 0.764847 | 0.766115 | 0.769948 | 0.766434 | 0.767223 |

| | | | | | | | | |
|---|---|---|---|---|---|---|---|---|
| 0.767346 | 0.773618 | 0.767869 | 0.766313 | 0.763503 | 0.767715 | 0.770902 | 0.765669 | 0.766349 |
| 0.770325 | 0.771541 | 0.77222 | 0.767546 | 0.766084 | 0.765013 | 0.775354 | 0.770767 | 0.769659 |
| 0.764619 | 0.770727 | 0.766816 | 0.767992 | 0.760631 | 0.761064 | 0.765216 | 0.765653 | 0.762733 |
| 0.722597 | 0.730645 | 0.718356 | 0.723054 | 0.722364 | 0.727273 | 0.72576 | 0.720058 | 0.736223 |
| 0.766672 | 0.770746 | 0.763484 | 0.764616 | 0.762075 | 0.763125 | 0.76886 | 0.762877 | 0.766077 |
| 0.770578 | 0.770947 | 0.76788 | 0.768094 | 0.75974 | 0.76179 | 0.770979 | 0.768481 | 0.764669 |
| 0.76678 | 0.767341 | 0.765344 | 0.764821 | 0.759127 | 0.765163 | 0.771901 | 0.768911 | 0.762224 |
| 0.771843 | 0.767721 | 0.764046 | 0.76629 | 0.765445 | 0.767808 | 0.7712 | 0.768688 | 0.768898 |
| 0.77245 | 0.770643 | 0.771052 | 0.766327 | 0.767464 | 0.769926 | 0.773779 | 0.770195 | 0.770608 |
| 0.770127 | 0.771384 | 0.76979 | 0.764746 | 0.76465 | 0.768642 | 0.772999 | 0.769319 | 0.770051 |
| 0.764701 | 0.767904 | 0.764264 | 0.763055 | 0.760393 | 0.763451 | 0.768221 | 0.766493 | 0.769792 |
| 0.769728 | 0.771774 | 0.765377 | 0.769211 | 0.767172 | 0.769744 | 0.765787 | 0.770817 | 0.764612 |
| 0.767546 | 0.771839 | 0.768784 | 0.769807 | 0.766558 | 0.768035 | 0.7731 | 0.765391 | 0.77065 |
| 0.768749 | 0.768474 | 0.770826 | 0.767734 | 0.765324 | 0.77312 | 0.771426 | 0.769928 | 0.770421 |
| 0.76555 | 0.769756 | 0.772646 | 0.771173 | 0.767537 | 0.766736 | 0.769663 | 0.773523 | 0.77118 |
| 0.764495 | 0.770177 | 0.769553 | 0.767484 | 0.763449 | 0.765979 | 0.768451 | 0.762182 | 0.765729 |
| 0.764644 | 0.771431 | 0.766396 | 0.771076 | 0.767355 | 0.768166 | 0.773008 | 0.773186 | 0.769034 |
| 0.767674 | 0.772041 | 0.773775 | 0.76925 | 0.76552 | 0.770392 | 0.77039 | 0.774513 | 0.766649 |
| 0.763675 | 0.768635 | 0.764345 | 0.765907 | 0.757289 | 0.765306 | 0.763811 | 0.767528 | 0.763121 |
| 0.765484 | 0.771885 | 0.768422 | 0.77102 | 0.762734 | 0.765617 | 0.769969 | 0.766796 | 0.767448 |
| 0.770013 | 0.777695 | 0.772335 | 0.773767 | 0.765249 | 0.771752 | 0.773809 | 0.771593 | 0.773795 |
| 0.766515 | 0.775603 | 0.770584 | 0.770961 | 0.765027 | 0.772251 | 0.773332 | 0.770765 | 0.768931 |
| 0.684324 | 0.685412 | 0.680428 | 0.683439 | 0.685852 | 0.689162 | 0.689009 | 0.682106 | 0.686215 |
| 0.657183 | 0.662607 | 0.662476 | 0.657876 | 0.656992 | 0.659266 | 0.662497 | 0.666424 | 0.67275 |
| 0.672421 | 0.675687 | 0.670726 | 0.676484 | 0.678619 | 0.676406 | 0.671731 | 0.666296 | 0.672933 |
| 0.657993 | 0.659627 | 0.658716 | 0.65543 | 0.658557 | 0.658561 | 0.660257 | 0.659099 | 0.661667 |
| 0.727219 | 0.742452 | 0.733109 | 0.73489 | 0.750038 | 0.732471 | 0.740673 | 0.761534 | 0.739422 |
| 0.720948 | 0.725456 | 0.72759 | 0.723011 | 0.734982 | 0.725384 | 0.726181 | 0.746609 | 0.73035 |
| 0.73105 | 0.735932 | 0.735919 | 0.732611 | 0.741846 | 0.733871 | 0.732737 | 0.750884 | 0.731011 |
| 0.741882 | 0.743547 | 0.744319 | 0.740756 | 0.743557 | 0.733947 | 0.741963 | 0.763735 | 0.736466 |
| 0.717024 | 0.727315 | 0.724468 | 0.719035 | 0.732833 | 0.716964 | 0.725349 | 0.746141 | 0.722549 |
| 0.72241 | 0.711783 | 0.70906 | 0.704616 | 0.717772 | 0.710594 | 0.721523 | 0.742438 | 0.71686 |
| 0.709191 | 0.712732 | 0.717312 | 0.717411 | 0.716212 | 0.722172 | 0.721739 | 0.738513 | 0.719878 |
| 0.656559 | 0.665169 | 0.661578 | 0.657443 | 0.657364 | 0.661591 | 0.659398 | 0.658426 | 0.66642 |
| 0.759251 | 0.764188 | 0.761926 | 0.764118 | 0.762953 | 0.761764 | 0.76048 | 0.777778 | 0.759842 |
| 0.739699 | 0.743406 | 0.747937 | 0.744602 | 0.744455 | 0.741175 | 0.745949 | 0.759927 | 0.749848 |
| 0.731825 | 0.738823 | 0.74 | 0.742636 | 0.741678 | 0.739089 | 0.739323 | 0.762753 | 0.743139 |
| 0.724766 | 0.729252 | 0.733763 | 0.728657 | 0.720674 | 0.716939 | 0.725155 | 0.747848 | 0.730217 |
| 0.737818 | 0.741637 | 0.745905 | 0.740931 | 0.747036 | 0.741512 | 0.742433 | 0.757009 | 0.747314 |
| 0.728709 | 0.729949 | 0.729733 | 0.732767 | 0.730473 | 0.73131 | 0.736986 | 0.739569 | 0.741432 |
| 0.724084 | 0.723913 | 0.728009 | 0.727757 | 0.730352 | 0.728341 | 0.733624 | 0.740844 | 0.727958 |
| 0.723562 | 0.736111 | 0.733684 | 0.72904 | 0.734191 | 0.723296 | 0.731178 | 0.744715 | 0.734928 |
| 0.721406 | 0.731938 | 0.73168 | 0.738307 | 0.726974 | 0.725253 | 0.735912 | 0.746038 | 0.739106 |
| 0.732066 | 0.72645 | 0.738177 | 0.733387 | 0.737286 | 0.729384 | 0.728417 | 0.74578 | 0.730675 |
| 0.733008 | 0.729716 | 0.733753 | 0.724895 | 0.729737 | 0.725552 | 0.737559 | 0.746959 | 0.737975 |
| 0.713303 | 0.72123 | 0.72027 | 0.721489 | 0.713856 | 0.719639 | 0.718112 | 0.731121 | 0.727459 |

| | | | | | | | | |
|---|---|---|---|---|---|---|---|---|
| 0.751846 | 0.752041 | 0.752986 | 0.749516 | 0.756247 | 0.746963 | 0.759763 | 0.771486 | 0.761134 |
| 0.730475 | 0.735793 | 0.738448 | 0.734688 | 0.737378 | 0.728478 | 0.734991 | 0.758939 | 0.734004 |
| 0.722972 | 0.732477 | 0.72954 | 0.728229 | 0.739377 | 0.718154 | 0.73247 | 0.756719 | 0.733023 |
| 0.729693 | 0.734722 | 0.737312 | 0.72186 | 0.730481 | 0.726298 | 0.738293 | 0.761512 | 0.739234 |
| 0.726522 | 0.733894 | 0.736853 | 0.734369 | 0.730205 | 0.725669 | 0.743302 | 0.757001 | 0.74338 |
| 0.721352 | 0.733833 | 0.718719 | 0.721746 | 0.729892 | 0.722827 | 0.73473 | 0.767028 | 0.740072 |
| 0.721555 | 0.724062 | 0.718164 | 0.719882 | 0.724602 | 0.717509 | 0.725649 | 0.75944 | 0.722608 |
| 0.718662 | 0.717617 | 0.707289 | 0.712493 | 0.720216 | 0.707499 | 0.722397 | 0.756339 | 0.726824 |
| 0.718976 | 0.723235 | 0.723307 | 0.714326 | 0.729576 | 0.722397 | 0.726204 | 0.751092 | 0.735028 |
| 0.743899 | 0.760556 | 0.756245 | 0.749249 | 0.76164 | 0.754529 | 0.753287 | 0.76644 | 0.757996 |
| 0.744554 | 0.760329 | 0.748865 | 0.752769 | 0.755465 | 0.751913 | 0.751598 | 0.764944 | 0.756728 |
| 0.745467 | 0.753844 | 0.74697 | 0.739391 | 0.749662 | 0.737571 | 0.752601 | 0.760598 | 0.758313 |
| 0.73744 | 0.750214 | 0.74371 | 0.734867 | 0.750839 | 0.737502 | 0.745509 | 0.756944 | 0.752756 |
| 0.735645 | 0.745026 | 0.740213 | 0.736178 | 0.746438 | 0.743292 | 0.743574 | 0.758529 | 0.751082 |
| 0.732365 | 0.742653 | 0.740018 | 0.74271 | 0.750567 | 0.729797 | 0.74369 | 0.755459 | 0.747981 |

| | plot 14 | | | | | | | |
|---|---|---|---|---|---|---|---|---|
| Wild_plot_ | Wild_plot_ | Wild_plot_ | Wild_plot_ | Wild_plot_ | Wild_plot_ | Wild_plot_ | Wild_plot_ | Wild_plot_ |
| 0.80424 | 0.709797 | 0.711925 | 0.711913 | 0.710077 | 0.705828 | 0.704661 | 0.707313 | 0.704337 |
| 0.785882 | 0.718026 | 0.715149 | 0.715389 | 0.714046 | 0.714549 | 0.712471 | 0.717015 | 0.7131 |
| 0.788381 | 0.704506 | 0.703222 | 0.707455 | 0.703127 | 0.701842 | 0.699517 | 0.703345 | 0.700648 |
| 0.794108 | 0.711134 | 0.713857 | 0.716159 | 0.714138 | 0.712083 | 0.710447 | 0.713615 | 0.711915 |
| 0.790731 | 0.707909 | 0.709446 | 0.710522 | 0.709469 | 0.70515 | 0.70425 | 0.710825 | 0.706861 |
| 0.777224 | 0.716023 | 0.715131 | 0.722548 | 0.721587 | 0.712031 | 0.714415 | 0.717076 | 0.714262 |
| 0.706491 | 0.775517 | 0.773781 | 0.774095 | 0.77373 | 0.76348 | 0.773141 | 0.77204 | 0.774414 |
| 0.703591 | 0.766438 | 0.766982 | 0.773349 | 0.766716 | 0.760319 | 0.771436 | 0.765553 | 0.762812 |
| 0.782857 | 0.710453 | 0.715482 | 0.714182 | 0.712671 | 0.709942 | 0.709268 | 0.708394 | 0.712095 |
| 0.784686 | 0.710684 | 0.709249 | 0.714366 | 0.707305 | 0.707539 | 0.705926 | 0.707205 | 0.705089 |
| 0.788192 | 0.710194 | 0.70918 | 0.716481 | 0.712794 | 0.710575 | 0.710852 | 0.71146 | 0.71096 |
| 0.796139 | 0.705696 | 0.701943 | 0.706216 | 0.702207 | 0.701991 | 0.699459 | 0.700315 | 0.701282 |
| 0.774723 | 0.711907 | 0.714775 | 0.714998 | 0.71416 | 0.707046 | 0.710635 | 0.714833 | 0.712383 |
| 0.789738 | 0.721687 | 0.720226 | 0.726405 | 0.720906 | 0.717725 | 0.718211 | 0.723451 | 0.714644 |
| 0.759031 | 0.731444 | 0.73788 | 0.740959 | 0.736787 | 0.731883 | 0.731158 | 0.738099 | 0.733003 |
| 0.797846 | 0.714783 | 0.715898 | 0.719939 | 0.716883 | 0.709906 | 0.709606 | 0.712605 | 0.71264 |
| 0.796862 | 0.715085 | 0.71495 | 0.717837 | 0.713606 | 0.710013 | 0.712341 | 0.711682 | 0.715046 |
| 0.771234 | 0.742292 | 0.743404 | 0.74659 | 0.740281 | 0.741517 | 0.738736 | 0.742521 | 0.738854 |
| 0.797647 | 0.705453 | 0.70556 | 0.712889 | 0.704383 | 0.701911 | 0.702285 | 0.707383 | 0.70235 |
| 0.787739 | 0.712904 | 0.713207 | 0.717566 | 0.709092 | 0.70982 | 0.708388 | 0.711659 | 0.707382 |
| 0.798127 | 0.708725 | 0.710626 | 0.714205 | 0.709724 | 0.703316 | 0.705482 | 0.706587 | 0.706422 |
| 0.8 | 0.711718 | 0.714661 | 0.716716 | 0.712958 | 0.707804 | 0.711162 | 0.708858 | 0.707268 |
| 0.78969 | 0.714161 | 0.72044 | 0.722827 | 0.71764 | 0.713868 | 0.714486 | 0.718156 | 0.714095 |
| 0.804353 | 0.712803 | 0.715081 | 0.714079 | 0.712927 | 0.708278 | 0.708577 | 0.712093 | 0.709753 |
| 0.782105 | 0.715402 | 0.716641 | 0.715888 | 0.714711 | 0.707842 | 0.712441 | 0.716675 | 0.711897 |
| 0.793034 | 0.70812 | 0.714829 | 0.714604 | 0.714479 | 0.706175 | 0.708967 | 0.712862 | 0.70825 |
| 0.778588 | 0.707957 | 0.705705 | 0.704584 | 0.706221 | 0.702347 | 0.702166 | 0.701624 | 0.700067 |
| 0.791001 | 0.710673 | 0.715656 | 0.714297 | 0.716159 | 0.711742 | 0.71131 | 0.715063 | 0.710776 |
| 0.791912 | 0.713175 | 0.717397 | 0.717792 | 0.712376 | 0.71006 | 0.709378 | 0.714344 | 0.712753 |
| 0.78601 | 0.710016 | 0.712917 | 0.717395 | 0.71162 | 0.707166 | 0.706669 | 0.712265 | 0.710323 |
| 0.778141 | 0.708942 | 0.7096 | 0.714389 | 0.711239 | 0.703973 | 0.706943 | 0.710689 | 0.70461 |
| 0.777704 | 0.707223 | 0.704118 | 0.708047 | 0.708562 | 0.703731 | 0.70361 | 0.704263 | 0.702451 |
| 0.791555 | 0.709708 | 0.71067 | 0.712378 | 0.707313 | 0.70712 | 0.702524 | 0.708084 | 0.704827 |
| 0.777075 | 0.712293 | 0.711224 | 0.71319 | 0.709243 | 0.706224 | 0.708624 | 0.713845 | 0.710307 |
| 0.774831 | 0.708356 | 0.707884 | 0.70848 | 0.706866 | 0.704466 | 0.700796 | 0.707911 | 0.701192 |
| 0.788591 | 0.710681 | 0.71189 | 0.713397 | 0.710292 | 0.706616 | 0.706106 | 0.709559 | 0.706331 |
| 0.78669 | 0.711747 | 0.715539 | 0.717029 | 0.715696 | 0.709886 | 0.714789 | 0.717252 | 0.713157 |
| 0.771736 | 0.710827 | 0.714171 | 0.7169 | 0.711447 | 0.707001 | 0.709249 | 0.715887 | 0.710266 |
| 0.77377 | 0.712805 | 0.716387 | 0.719537 | 0.714766 | 0.711376 | 0.715569 | 0.71726 | 0.71413 |

| | | | | | | | | |
|---|---|---|---|---|---|---|---|---|
| 0.762362 | 0.746177 | 0.748957 | 0.754035 | 0.748105 | 0.742155 | 0.748336 | 0.746769 | 0.747406 |
| 0.798651 | 0.702093 | 0.704748 | 0.70848 | 0.701174 | 0.700374 | 0.697999 | 0.705173 | 0.699444 |
| 0.807493 | 0.705719 | 0.706923 | 0.70997 | 0.708807 | 0.701289 | 0.702871 | 0.704496 | 0.703663 |
| 0.797894 | 0.712545 | 0.715842 | 0.72158 | 0.712244 | 0.709677 | 0.712294 | 0.712566 | 0.71318 |
| 0.792439 | 0.708804 | 0.70863 | 0.711348 | 0.710711 | 0.705275 | 0.707604 | 0.711804 | 0.703173 |
| 0.790321 | 0.710604 | 0.710629 | 0.719258 | 0.711896 | 0.707064 | 0.709077 | 0.712931 | 0.708239 |
| 0.782348 | 0.706879 | 0.709448 | 0.712592 | 0.710329 | 0.703194 | 0.707069 | 0.708394 | 0.707215 |
| 0.788457 | 0.709154 | 0.713326 | 0.717934 | 0.712216 | 0.708347 | 0.709288 | 0.713268 | 0.707485 |
| 0.777414 | 0.713032 | 0.714514 | 0.718621 | 0.714758 | 0.709808 | 0.7122 | 0.71384 | 0.708064 |
| 0.776282 | 0.710771 | 0.714716 | 0.717001 | 0.712269 | 0.708257 | 0.709553 | 0.714228 | 0.712496 |
| 0.806988 | 0.702872 | 0.703816 | 0.710388 | 0.702577 | 0.700451 | 0.699481 | 0.700146 | 0.701089 |
| 0.793112 | 0.705766 | 0.705775 | 0.710624 | 0.706184 | 0.702099 | 0.701987 | 0.708146 | 0.700753 |
| 0.796177 | 0.707958 | 0.711403 | 0.71077 | 0.711726 | 0.704098 | 0.705643 | 0.708678 | 0.706333 |
| 0.799252 | 0.705747 | 0.705003 | 0.707392 | 0.70601 | 0.702094 | 0.701361 | 0.703854 | 0.702477 |
| 0.80434 | 0.703949 | 0.707999 | 0.708849 | 0.706372 | 0.70084 | 0.704636 | 0.707467 | 0.702737 |
| 0.795662 | 0.703894 | 0.705129 | 0.707547 | 0.704505 | 0.703738 | 0.703682 | 0.70232 | 0.700778 |
| 0.792398 | 0.700605 | 0.706704 | 0.710031 | 0.702296 | 0.698774 | 0.698584 | 0.704601 | 0.702061 |
| 0.804944 | 0.709644 | 0.710814 | 0.715973 | 0.712346 | 0.707817 | 0.711082 | 0.710099 | 0.706702 |
| 0.792392 | 0.708847 | 0.714683 | 0.719619 | 0.712618 | 0.710192 | 0.708292 | 0.71225 | 0.713339 |
| 0.80467 | 0.704677 | 0.709435 | 0.712818 | 0.706412 | 0.703276 | 0.705118 | 0.708841 | 0.705205 |
| 0.80471 | 0.702555 | 0.707971 | 0.70831 | 0.702796 | 0.701089 | 0.700157 | 0.702588 | 0.702154 |
| 0.789815 | 0.705482 | 0.70641 | 0.707784 | 0.705661 | 0.701428 | 0.702819 | 0.704008 | 0.704413 |
| 0.786442 | 0.709591 | 0.711444 | 0.709798 | 0.712363 | 0.706131 | 0.705155 | 0.709471 | 0.704832 |
| 0.771583 | 0.716574 | 0.719 | 0.721378 | 0.71708 | 0.714513 | 0.716093 | 0.71716 | 0.715848 |
| 0.760244 | 0.72179 | 0.726859 | 0.727419 | 0.726285 | 0.718411 | 0.720727 | 0.725192 | 0.724337 |
| 0.703914 | 0.767934 | 0.767419 | 0.77272 | 0.76767 | 0.760999 | 0.772583 | 0.766633 | 0.764284 |
| 0.708381 | 0.776709 | 0.778119 | 0.782664 | 0.777276 | 0.778514 | 0.777787 | 0.777542 | 0.776682 |
| 0.706364 | 0.77686 | 0.779584 | 0.781981 | 0.77534 | 0.780991 | 0.774243 | 0.774625 | 0.775668 |
| 0.77762 | 0.706776 | 0.710562 | 0.710452 | 0.710564 | 0.703124 | 0.704946 | 0.707543 | 0.701449 |
| 0.777692 | 0.70742 | 0.707072 | 0.706135 | 0.705708 | 0.702151 | 0.703203 | 0.706476 | 0.702398 |
| 0.790285 | 0.713957 | 0.722462 | 0.717959 | 0.717269 | 0.709982 | 0.714109 | 0.714858 | 0.71455 |
| 0.778265 | 0.715157 | 0.716776 | 0.715366 | 0.715744 | 0.714213 | 0.714763 | 0.716639 | 0.714117 |
| 0.796285 | 0.704803 | 0.707975 | 0.710847 | 0.70601 | 0.705016 | 0.703654 | 0.70637 | 0.701988 |
| 0.798929 | 0.70829 | 0.711268 | 0.710763 | 0.707576 | 0.707947 | 0.703829 | 0.708996 | 0.705261 |
| 0.80362 | 0.706085 | 0.705683 | 0.71214 | 0.705111 | 0.70425 | 0.701281 | 0.706033 | 0.699933 |
| 0.814451 | 0.706823 | 0.710404 | 0.712301 | 0.70898 | 0.704135 | 0.701919 | 0.706608 | 0.705119 |
| 0.78384 | 0.708134 | 0.709472 | 0.713943 | 0.708033 | 0.708551 | 0.704703 | 0.706678 | 0.704743 |
| 0.822571 | 0.723827 | 0.725122 | 0.728599 | 0.722889 | 0.720104 | 0.717359 | 0.72089 | 0.720313 |
| 0.78756 | 0.700427 | 0.701905 | 0.707115 | 0.700737 | 0.699699 | 0.700374 | 0.701287 | 0.693955 |
| 0.798105 | 0.708081 | 0.706834 | 0.711557 | 0.704849 | 0.703701 | 0.705993 | 0.71 | 0.706019 |
| 0.805858 | 0.701087 | 0.706197 | 0.707942 | 0.703121 | 0.7013 | 0.69895 | 0.703477 | 0.701011 |
| 0.828004 | 0.726779 | 0.728486 | 0.730157 | 0.729653 | 0.722255 | 0.720023 | 0.723101 | 0.721427 |
| 0.802254 | 0.708152 | 0.710088 | 0.714862 | 0.705911 | 0.704914 | 0.705819 | 0.708245 | 0.705547 |
| 0.814068 | 0.704619 | 0.708606 | 0.711932 | 0.704655 | 0.705028 | 0.702897 | 0.706015 | 0.703108 |
| 0.788007 | 0.711154 | 0.713509 | 0.713721 | 0.707343 | 0.706794 | 0.709672 | 0.712557 | 0.705498 |
| 0.786864 | 0.708617 | 0.712297 | 0.712051 | 0.706461 | 0.705625 | 0.704152 | 0.709064 | 0.70131 |

| | | | | | | | | |
|---|---|---|---|---|---|---|---|---|
| 0.806803 | 0.70414 | 0.709564 | 0.709716 | 0.70808 | 0.701864 | 0.702392 | 0.705347 | 0.703351 |
| 0.798128 | 0.71159 | 0.715815 | 0.709763 | 0.712484 | 0.707021 | 0.711668 | 0.710121 | 0.70673 |
| 0.782399 | 0.711355 | 0.70997 | 0.713868 | 0.708306 | 0.709514 | 0.706185 | 0.708976 | 0.703666 |
| 0.805029 | 0.71625 | 0.72054 | 0.719267 | 0.716977 | 0.713936 | 0.712228 | 0.71586 | 0.714552 |
| 0.790281 | 0.713914 | 0.717393 | 0.718545 | 0.713307 | 0.709782 | 0.711586 | 0.715839 | 0.710011 |
| 0.794996 | 0.709262 | 0.708612 | 0.71045 | 0.7087 | 0.704906 | 0.706218 | 0.709918 | 0.70371 |
| 0.779439 | 0.711634 | 0.713733 | 0.714685 | 0.708092 | 0.708371 | 0.709783 | 0.712011 | 0.705785 |
| 0.78557 | 0.712323 | 0.716109 | 0.716153 | 0.716582 | 0.712068 | 0.712219 | 0.718056 | 0.709378 |
| 0.784653 | 0.712242 | 0.714974 | 0.714551 | 0.711261 | 0.707976 | 0.7088 | 0.711556 | 0.704374 |
| 0.777973 | 0.707406 | 0.706254 | 0.70876 | 0.708157 | 0.704341 | 0.70286 | 0.705577 | 0.701997 |
| 0.782259 | 0.708752 | 0.712945 | 0.710329 | 0.711187 | 0.705999 | 0.705254 | 0.705367 | 0.703928 |
| 0.775201 | 0.706727 | 0.712131 | 0.712021 | 0.705749 | 0.707406 | 0.703704 | 0.708608 | 0.703595 |
| 0.779001 | 0.711916 | 0.711003 | 0.712447 | 0.709896 | 0.708249 | 0.707649 | 0.713442 | 0.705476 |
| 0.776404 | 0.706502 | 0.711852 | 0.71055 | 0.707458 | 0.704981 | 0.704195 | 0.707269 | 0.705837 |
| 0.779995 | 0.710535 | 0.712701 | 0.712996 | 0.709525 | 0.707446 | 0.709016 | 0.710655 | 0.707374 |
| 0.780168 | 0.708034 | 0.710693 | 0.711841 | 0.711801 | 0.705355 | 0.704711 | 0.707035 | 0.703137 |
| 0.783012 | 0.711881 | 0.712321 | 0.715593 | 0.714067 | 0.710599 | 0.709126 | 0.713677 | 0.709729 |
| 0.793719 | 0.705022 | 0.70948 | 0.713368 | 0.708407 | 0.701769 | 0.703543 | 0.707539 | 0.704154 |
| 0.777461 | 0.708677 | 0.71109 | 0.715197 | 0.708651 | 0.705317 | 0.70833 | 0.710003 | 0.704291 |
| 0.779837 | 0.706856 | 0.710166 | 0.710275 | 0.710493 | 0.702588 | 0.707285 | 0.709858 | 0.705318 |
| 0.779671 | 0.708519 | 0.712836 | 0.712637 | 0.709623 | 0.707528 | 0.7058 | 0.708357 | 0.706582 |
| 0.779468 | 0.703809 | 0.70667 | 0.707853 | 0.705319 | 0.704589 | 0.700819 | 0.703812 | 0.698986 |
| 0.788917 | 0.709551 | 0.708639 | 0.710014 | 0.708692 | 0.705069 | 0.707165 | 0.710488 | 0.707622 |
| 0.788392 | 0.711146 | 0.711407 | 0.714469 | 0.710552 | 0.705056 | 0.703858 | 0.706645 | 0.705677 |
| 0.778927 | 0.71276 | 0.712929 | 0.717326 | 0.710027 | 0.708358 | 0.710113 | 0.712958 | 0.705436 |
| 0.782366 | 0.714466 | 0.713564 | 0.71975 | 0.715629 | 0.710778 | 0.712214 | 0.715292 | 0.710112 |
| 0.77971 | 0.710168 | 0.709853 | 0.712323 | 0.707424 | 0.707374 | 0.706613 | 0.709784 | 0.70227 |
| 0.783698 | 0.71213 | 0.712632 | 0.714057 | 0.710989 | 0.707599 | 0.706022 | 0.710475 | 0.704962 |
| 0.783687 | 0.713099 | 0.714445 | 0.718423 | 0.710953 | 0.708854 | 0.709643 | 0.712903 | 0.705926 |
| 0.774869 | 0.711676 | 0.711955 | 0.718334 | 0.709654 | 0.705654 | 0.70954 | 0.710657 | 0.704599 |
| 0.77145 | 0.7115 | 0.712959 | 0.717344 | 0.711841 | 0.709317 | 0.709873 | 0.711362 | 0.705848 |
| 0.779747 | 0.707099 | 0.710326 | 0.708839 | 0.708577 | 0.70597 | 0.706265 | 0.708849 | 0.705834 |
| 0.780731 | 0.710558 | 0.714067 | 0.713995 | 0.708546 | 0.707626 | 0.707688 | 0.709797 | 0.70623 |
| 0.79338 | 0.7081 | 0.711167 | 0.714427 | 0.707563 | 0.706564 | 0.710031 | 0.709138 | 0.708412 |
| 0.78193 | 0.708609 | 0.706629 | 0.712298 | 0.705022 | 0.707577 | 0.701837 | 0.707779 | 0.70041 |
| 0.788693 | 0.707724 | 0.711473 | 0.712211 | 0.706368 | 0.706914 | 0.709845 | 0.708014 | 0.707061 |
| 0.779304 | 0.709209 | 0.705479 | 0.712284 | 0.703281 | 0.702356 | 0.704005 | 0.708603 | 0.703482 |
| 0.755753 | 0.68695 | 0.68514 | 0.685703 | 0.687214 | 0.682228 | 0.684118 | 0.687087 | 0.683622 |
| 0.776432 | 0.689248 | 0.688976 | 0.691391 | 0.688437 | 0.687935 | 0.692101 | 0.690239 | 0.687826 |
| 0.756171 | 0.680916 | 0.683333 | 0.684205 | 0.683499 | 0.681948 | 0.683172 | 0.68074 | 0.681032 |
| 0.766787 | 0.681044 | 0.681833 | 0.680156 | 0.681003 | 0.683547 | 0.677869 | 0.682842 | 0.678007 |
| 0.743084 | 0.664336 | 0.666939 | 0.667531 | 0.663164 | 0.672139 | 0.664885 | 0.665136 | 0.664632 |
| 0.773551 | 0.666145 | 0.669728 | 0.672557 | 0.667071 | 0.6705 | 0.667319 | 0.667194 | 0.671728 |
| 0.761454 | 0.663025 | 0.666828 | 0.668333 | 0.668203 | 0.670424 | 0.669501 | 0.665851 | 0.665596 |
| 0.752897 | 0.666624 | 0.666878 | 0.67206 | 0.670725 | 0.667185 | 0.668824 | 0.668896 | 0.669828 |
| 0.771075 | 0.666625 | 0.668706 | 0.671317 | 0.672297 | 0.671948 | 0.670426 | 0.668549 | 0.668482 |
| 0.7757 | 0.674648 | 0.675272 | 0.677664 | 0.674433 | 0.676141 | 0.673592 | 0.673675 | 0.67242 |

| | | | | | | | | |
|---|---|---|---|---|---|---|---|---|
| 0.751514 | 0.674743 | 0.677407 | 0.669468 | 0.673967 | 0.672595 | 0.676367 | 0.68013 | 0.67361 |
| 0.769896 | 0.66977 | 0.669554 | 0.67336 | 0.671443 | 0.677269 | 0.67408 | 0.668325 | 0.666092 |
| 0.744086 | 0.672494 | 0.673013 | 0.675037 | 0.670409 | 0.674058 | 0.675152 | 0.672052 | 0.673886 |
| 0.768521 | 0.654329 | 0.660029 | 0.668584 | 0.659544 | 0.662468 | 0.660317 | 0.659274 | 0.663751 |
| 0.750912 | 0.666966 | 0.666367 | 0.670432 | 0.662734 | 0.669894 | 0.666104 | 0.664818 | 0.664056 |
| 0.748595 | 0.665349 | 0.667948 | 0.669155 | 0.6707 | 0.672346 | 0.665051 | 0.664686 | 0.667505 |
| 0.766916 | 0.666432 | 0.665774 | 0.673008 | 0.666714 | 0.66839 | 0.669535 | 0.663596 | 0.664551 |
| 0.776324 | 0.663833 | 0.665032 | 0.66916 | 0.660364 | 0.672269 | 0.663083 | 0.660599 | 0.663775 |
| 0.731793 | 0.664086 | 0.662614 | 0.665827 | 0.66583 | 0.665824 | 0.661195 | 0.653666 | 0.664184 |
| 0.752762 | 0.666909 | 0.663037 | 0.664324 | 0.663517 | 0.665725 | 0.669574 | 0.662762 | 0.659897 |
| 0.761468 | 0.658683 | 0.662054 | 0.658125 | 0.665382 | 0.663289 | 0.662146 | 0.659015 | 0.662704 |
| 0.792056 | 0.697397 | 0.699157 | 0.707513 | 0.698826 | 0.693708 | 0.694175 | 0.695527 | 0.697498 |
| 0.777898 | 0.70814 | 0.711472 | 0.709079 | 0.705436 | 0.706516 | 0.706369 | 0.70789 | 0.702705 |
| 0.759129 | 0.681946 | 0.683417 | 0.688123 | 0.685994 | 0.684606 | 0.683669 | 0.682182 | 0.682757 |
| 0.780252 | 0.713436 | 0.714874 | 0.71754 | 0.715621 | 0.710533 | 0.711425 | 0.711417 | 0.714056 |
| 0.772992 | 0.704801 | 0.709211 | 0.709677 | 0.705023 | 0.703215 | 0.701248 | 0.702988 | 0.700208 |
| 0.786365 | 0.713508 | 0.712749 | 0.720176 | 0.710537 | 0.708857 | 0.709863 | 0.711696 | 0.711006 |
| 0.781695 | 0.714879 | 0.721689 | 0.719899 | 0.720181 | 0.711559 | 0.715428 | 0.720511 | 0.715189 |
| 0.757789 | 0.694208 | 0.700023 | 0.699549 | 0.697151 | 0.689446 | 0.695391 | 0.698494 | 0.693651 |
| 0.771766 | 0.719545 | 0.720323 | 0.721481 | 0.71869 | 0.716031 | 0.7176 | 0.719825 | 0.714962 |
| 0.744021 | 0.688918 | 0.68646 | 0.695969 | 0.691583 | 0.691513 | 0.688353 | 0.6888 | 0.687907 |
| 0.780042 | 0.713751 | 0.717992 | 0.721288 | 0.716032 | 0.712896 | 0.714807 | 0.716553 | 0.716582 |
| 0.783175 | 0.71714 | 0.723992 | 0.722897 | 0.720413 | 0.717358 | 0.714806 | 0.722441 | 0.716368 |
| 0.76755 | 0.68342 | 0.686439 | 0.685597 | 0.690858 | 0.683637 | 0.688068 | 0.688494 | 0.689532 |
| 0.745433 | 0.688874 | 0.690665 | 0.691866 | 0.687583 | 0.685996 | 0.686868 | 0.685278 | 0.684784 |
| 0.766377 | 0.693577 | 0.695119 | 0.696722 | 0.692639 | 0.695253 | 0.692021 | 0.687786 | 0.690683 |
| 0.769028 | 0.704992 | 0.70635 | 0.707011 | 0.707167 | 0.707079 | 0.702292 | 0.70843 | 0.702155 |
| 0.788391 | 0.717579 | 0.722461 | 0.722908 | 0.719541 | 0.715863 | 0.717079 | 0.717278 | 0.714417 |
| 0.742718 | 0.685261 | 0.690913 | 0.689337 | 0.686994 | 0.682982 | 0.685788 | 0.687856 | 0.679212 |
| 0.765548 | 0.723144 | 0.725775 | 0.727499 | 0.723951 | 0.719243 | 0.722541 | 0.720919 | 0.718631 |
| 0.761947 | 0.726336 | 0.727639 | 0.7296 | 0.723463 | 0.720923 | 0.722754 | 0.728073 | 0.723892 |
| 0.768155 | 0.696064 | 0.700738 | 0.702646 | 0.700032 | 0.695309 | 0.69358 | 0.695388 | 0.698526 |
| 0.77619 | 0.699984 | 0.70052 | 0.703509 | 0.700555 | 0.695534 | 0.69132 | 0.693248 | 0.695897 |
| 0.765559 | 0.722448 | 0.719514 | 0.727478 | 0.719349 | 0.721789 | 0.71941 | 0.723429 | 0.717987 |
| 0.765916 | 0.720685 | 0.726344 | 0.72231 | 0.721405 | 0.719763 | 0.71829 | 0.722597 | 0.717269 |
| 0.765288 | 0.723898 | 0.729053 | 0.727509 | 0.729167 | 0.719936 | 0.721742 | 0.726623 | 0.717026 |
| 0.723732 | 0.679282 | 0.681949 | 0.680289 | 0.680172 | 0.672605 | 0.674465 | 0.677348 | 0.675259 |
| 0.751044 | 0.720314 | 0.723485 | 0.726415 | 0.723754 | 0.719338 | 0.720095 | 0.721761 | 0.71456 |
| 0.732864 | 0.677646 | 0.67829 | 0.676846 | 0.67527 | 0.672771 | 0.673371 | 0.674963 | 0.67165 |
| 0.728456 | 0.669354 | 0.671107 | 0.673117 | 0.674332 | 0.667461 | 0.668027 | 0.669325 | 0.666587 |
| 0.781058 | 0.719043 | 0.721822 | 0.725764 | 0.722324 | 0.715021 | 0.716309 | 0.719497 | 0.715855 |
| 0.725841 | 0.671684 | 0.673316 | 0.675328 | 0.674805 | 0.666773 | 0.667372 | 0.668346 | 0.667622 |
| 0.717511 | 0.662276 | 0.66569 | 0.667501 | 0.66397 | 0.65851 | 0.659632 | 0.659522 | 0.659155 |
| 0.707274 | 0.660874 | 0.661324 | 0.665894 | 0.661053 | 0.654925 | 0.656442 | 0.657303 | 0.657886 |
| 0.772254 | 0.72728 | 0.732278 | 0.736455 | 0.726124 | 0.725046 | 0.726258 | 0.728162 | 0.724897 |
| 0.713124 | 0.663353 | 0.665617 | 0.668719 | 0.665986 | 0.658583 | 0.662117 | 0.664335 | 0.659197 |
| 0.779515 | 0.720092 | 0.722965 | 0.727015 | 0.722948 | 0.721122 | 0.721614 | 0.726385 | 0.714896 |

| | | | | | | | | |
|---|---|---|---|---|---|---|---|---|
| 0.769542 | 0.724522 | 0.731054 | 0.735908 | 0.732354 | 0.72339 | 0.727452 | 0.726785 | 0.723573 |
| 0.782507 | 0.72317 | 0.726846 | 0.729227 | 0.724497 | 0.720324 | 0.721087 | 0.724186 | 0.720221 |
| 0.750228 | 0.742521 | 0.741718 | 0.750021 | 0.740016 | 0.742205 | 0.74221 | 0.744741 | 0.741275 |
| 0.704538 | 0.653702 | 0.653733 | 0.660379 | 0.654338 | 0.64833 | 0.648591 | 0.650173 | 0.64849 |
| 0.695372 | 0.654027 | 0.653097 | 0.658438 | 0.653581 | 0.649634 | 0.6534 | 0.653312 | 0.652576 |
| 0.783617 | 0.728878 | 0.730754 | 0.736795 | 0.733011 | 0.725907 | 0.728691 | 0.730335 | 0.725229 |
| 0.690615 | 0.657493 | 0.657179 | 0.658266 | 0.654598 | 0.652374 | 0.650622 | 0.651715 | 0.652846 |
| 0.727743 | 0.7324 | 0.734845 | 0.736535 | 0.737079 | 0.729861 | 0.732026 | 0.730906 | 0.729106 |
| 0.762608 | 0.733933 | 0.736838 | 0.740816 | 0.737625 | 0.727058 | 0.728614 | 0.731513 | 0.73087 |
| 0.716391 | 0.737177 | 0.735546 | 0.739249 | 0.735398 | 0.731132 | 0.731633 | 0.731994 | 0.731624 |
| 0.747521 | 0.730293 | 0.734918 | 0.740115 | 0.733653 | 0.729326 | 0.728581 | 0.734305 | 0.729831 |
| 0.655172 | 0.700637 | 0.743243 | 0.711111 | 0.74 | 0.693333 | 0.724832 | 0.697987 | 0.75817 |
| 0.648534 | 0.731615 | 0.732745 | 0.727768 | 0.735412 | 0.733198 | 0.741215 | 0.725506 | 0.732563 |
| 0.651685 | 0.726588 | 0.729347 | 0.725995 | 0.730733 | 0.733365 | 0.731893 | 0.724654 | 0.7282 |
| 0.652444 | 0.737977 | 0.739433 | 0.73303 | 0.743749 | 0.735371 | 0.739682 | 0.73677 | 0.73681 |
| 0.656161 | 0.734274 | 0.736607 | 0.734866 | 0.731454 | 0.736929 | 0.738619 | 0.732314 | 0.733897 |
| 0.61996 | 0.595089 | 0.592585 | 0.593196 | 0.591504 | 0.59104 | 0.59138 | 0.591812 | 0.586469 |
| 0.652934 | 0.738801 | 0.739807 | 0.742951 | 0.741413 | 0.737038 | 0.74361 | 0.73853 | 0.737389 |
| 0.736859 | 0.734076 | 0.738682 | 0.741167 | 0.735846 | 0.731956 | 0.737242 | 0.740476 | 0.728799 |
| 0.736482 | 0.733983 | 0.737141 | 0.742572 | 0.738979 | 0.7281 | 0.735141 | 0.736369 | 0.730845 |
| 0.739785 | 0.738103 | 0.742783 | 0.747464 | 0.739587 | 0.734303 | 0.734265 | 0.742485 | 0.735868 |
| 0.613656 | 0.595567 | 0.600614 | 0.601221 | 0.600678 | 0.590333 | 0.593167 | 0.595216 | 0.592961 |
| 0.5977 | 0.584334 | 0.584534 | 0.58931 | 0.5846 | 0.580256 | 0.579738 | 0.579904 | 0.574859 |
| 0.696255 | 0.764833 | 0.760496 | 0.765763 | 0.759947 | 0.760367 | 0.763713 | 0.765246 | 0.754909 |
| 0.69714 | 0.764461 | 0.761924 | 0.76403 | 0.762461 | 0.760806 | 0.766032 | 0.764054 | 0.766428 |
| 0.691655 | 0.757508 | 0.756603 | 0.752901 | 0.753849 | 0.750906 | 0.75096 | 0.750441 | 0.756553 |
| 0.697509 | 0.76028 | 0.763281 | 0.759555 | 0.761018 | 0.760151 | 0.756738 | 0.754523 | 0.760757 |
| 0.702631 | 0.762746 | 0.760608 | 0.76668 | 0.762578 | 0.756805 | 0.75879 | 0.757495 | 0.76371 |
| 0.700642 | 0.766515 | 0.763746 | 0.77238 | 0.765668 | 0.764619 | 0.762584 | 0.764492 | 0.763568 |
| 0.700685 | 0.766508 | 0.761628 | 0.767274 | 0.762424 | 0.765579 | 0.768392 | 0.763989 | 0.762615 |
| 0.703504 | 0.760984 | 0.768518 | 0.765873 | 0.763759 | 0.763623 | 0.76018 | 0.760993 | 0.760043 |
| 0.710235 | 0.773256 | 0.774302 | 0.773244 | 0.772866 | 0.771267 | 0.771475 | 0.767446 | 0.768604 |
| 0.702744 | 0.760441 | 0.761889 | 0.764927 | 0.767395 | 0.760301 | 0.763318 | 0.760169 | 0.760461 |
| 0.709252 | 0.762447 | 0.764229 | 0.768769 | 0.762188 | 0.757581 | 0.762425 | 0.763846 | 0.757694 |
| 0.70682 | 0.773207 | 0.77538 | 0.774863 | 0.773596 | 0.76699 | 0.769722 | 0.767724 | 0.772763 |
| 0.705391 | 0.765359 | 0.763128 | 0.767711 | 0.765554 | 0.760017 | 0.767112 | 0.761905 | 0.760898 |
| 0.705256 | 0.76487 | 0.765339 | 0.771901 | 0.769595 | 0.765133 | 0.761503 | 0.76626 | 0.764996 |
| 0.703404 | 0.765793 | 0.764399 | 0.764735 | 0.763363 | 0.763366 | 0.766607 | 0.766427 | 0.764579 |
| 0.700867 | 0.760436 | 0.763382 | 0.760787 | 0.764571 | 0.758693 | 0.762963 | 0.760832 | 0.760018 |
| 0.706034 | 0.762087 | 0.76146 | 0.765424 | 0.76748 | 0.758795 | 0.762348 | 0.760337 | 0.761231 |
| 0.650986 | 0.730037 | 0.73139 | 0.729141 | 0.732663 | 0.728625 | 0.721987 | 0.726919 | 0.732127 |
| 0.661237 | 0.726401 | 0.735183 | 0.730529 | 0.744394 | 0.730286 | 0.741316 | 0.73356 | 0.737922 |
| 0.708167 | 0.766634 | 0.769018 | 0.767285 | 0.766228 | 0.764661 | 0.763283 | 0.765168 | 0.766406 |
| 0.707156 | 0.771125 | 0.769206 | 0.770071 | 0.769161 | 0.768316 | 0.765066 | 0.769244 | 0.767934 |
| 0.643572 | 0.71511 | 0.721033 | 0.7206 | 0.716232 | 0.711621 | 0.724225 | 0.721074 | 0.727094 |
| 0.707033 | 0.768248 | 0.765895 | 0.769684 | 0.772321 | 0.762474 | 0.769914 | 0.768034 | 0.765313 |
| 0.696725 | 0.758956 | 0.762976 | 0.76234 | 0.761452 | 0.75395 | 0.755214 | 0.757144 | 0.75736 |

| | | | | | | | | |
|---|---|---|---|---|---|---|---|---|
| 0.704527 | 0.766222 | 0.766118 | 0.771452 | 0.766702 | 0.759873 | 0.763605 | 0.767628 | 0.76348 |
| 0.707184 | 0.766742 | 0.767349 | 0.774066 | 0.769805 | 0.760936 | 0.763639 | 0.767072 | 0.76589 |
| 0.703859 | 0.761925 | 0.76212 | 0.765522 | 0.7671 | 0.759958 | 0.756586 | 0.758926 | 0.762878 |
| 0.707272 | 0.769187 | 0.769679 | 0.769014 | 0.771152 | 0.76179 | 0.765275 | 0.766893 | 0.762412 |
| 0.710849 | 0.768069 | 0.766427 | 0.771265 | 0.77481 | 0.763388 | 0.767252 | 0.76655 | 0.765866 |
| 0.704192 | 0.766505 | 0.765684 | 0.76725 | 0.76792 | 0.75865 | 0.763951 | 0.764121 | 0.76441 |
| 0.705873 | 0.765423 | 0.765079 | 0.768641 | 0.768477 | 0.759504 | 0.766562 | 0.767486 | 0.76564 |
| 0.697272 | 0.758248 | 0.752575 | 0.757923 | 0.764122 | 0.749957 | 0.750919 | 0.752999 | 0.757386 |
| 0.703954 | 0.761932 | 0.766155 | 0.769531 | 0.769123 | 0.760292 | 0.760326 | 0.763912 | 0.757844 |
| 0.702742 | 0.756219 | 0.763128 | 0.762791 | 0.765633 | 0.751801 | 0.754954 | 0.757698 | 0.758677 |
| 0.703413 | 0.763817 | 0.764428 | 0.770139 | 0.772922 | 0.761161 | 0.767809 | 0.767772 | 0.767304 |
| 0.701157 | 0.763238 | 0.760407 | 0.770446 | 0.76389 | 0.755702 | 0.759839 | 0.764797 | 0.765613 |
| 0.69525 | 0.756788 | 0.754726 | 0.760725 | 0.758376 | 0.750583 | 0.750834 | 0.756983 | 0.754374 |
| 0.695838 | 0.755085 | 0.760791 | 0.760069 | 0.756775 | 0.749622 | 0.756087 | 0.759822 | 0.759355 |
| 0.707692 | 0.768331 | 0.768974 | 0.767624 | 0.774273 | 0.76387 | 0.765671 | 0.763948 | 0.766741 |
| 0.70183 | 0.766697 | 0.767539 | 0.765187 | 0.768281 | 0.763752 | 0.763229 | 0.764657 | 0.766031 |
| 0.699792 | 0.766896 | 0.765555 | 0.769383 | 0.765881 | 0.766282 | 0.763043 | 0.764074 | 0.762453 |
| 0.70231 | 0.761154 | 0.759027 | 0.766173 | 0.762982 | 0.76047 | 0.761075 | 0.766133 | 0.762451 |
| 0.698919 | 0.764461 | 0.763141 | 0.769754 | 0.762465 | 0.760458 | 0.76117 | 0.762795 | 0.767037 |
| 0.700282 | 0.765082 | 0.76594 | 0.768827 | 0.767552 | 0.762953 | 0.764646 | 0.762453 | 0.764915 |
| 0.69643 | 0.755798 | 0.755718 | 0.759976 | 0.759352 | 0.755447 | 0.757395 | 0.753917 | 0.756686 |
| 0.699762 | 0.760884 | 0.758641 | 0.7648 | 0.758245 | 0.759615 | 0.759312 | 0.758368 | 0.759923 |
| 0.702947 | 0.764734 | 0.76851 | 0.76944 | 0.769079 | 0.763719 | 0.762258 | 0.763027 | 0.768948 |
| 0.704958 | 0.76447 | 0.762477 | 0.771571 | 0.765172 | 0.764462 | 0.766001 | 0.768423 | 0.764959 |
| 0.701671 | 0.762244 | 0.762525 | 0.76275 | 0.765822 | 0.763828 | 0.76419 | 0.766965 | 0.760923 |
| 0.695421 | 0.760977 | 0.764157 | 0.763903 | 0.76164 | 0.755134 | 0.759009 | 0.757354 | 0.759251 |
| 0.700448 | 0.757243 | 0.755767 | 0.759605 | 0.756748 | 0.757956 | 0.753983 | 0.758993 | 0.75869 |
| 0.69689 | 0.757924 | 0.757735 | 0.759125 | 0.75951 | 0.755257 | 0.754761 | 0.758747 | 0.757449 |
| 0.699107 | 0.758295 | 0.763031 | 0.76417 | 0.764498 | 0.756592 | 0.760449 | 0.759042 | 0.7604 |
| 0.697829 | 0.76269 | 0.760571 | 0.766678 | 0.762262 | 0.759842 | 0.760739 | 0.762898 | 0.758413 |
| 0.703883 | 0.760359 | 0.766041 | 0.768784 | 0.765021 | 0.760107 | 0.762216 | 0.761921 | 0.762656 |
| 0.707532 | 0.763145 | 0.762077 | 0.771568 | 0.763406 | 0.763195 | 0.762223 | 0.761675 | 0.763321 |
| 0.705309 | 0.767359 | 0.768993 | 0.771892 | 0.76884 | 0.767324 | 0.771073 | 0.767097 | 0.768492 |
| 0.709376 | 0.771113 | 0.766083 | 0.768874 | 0.76595 | 0.767809 | 0.76676 | 0.765351 | 0.765511 |
| 0.696884 | 0.762466 | 0.758208 | 0.759977 | 0.762054 | 0.756245 | 0.758999 | 0.759317 | 0.759579 |
| 0.703161 | 0.763191 | 0.762075 | 0.765087 | 0.764164 | 0.759613 | 0.761593 | 0.758008 | 0.75955 |
| 0.706112 | 0.760884 | 0.756116 | 0.764308 | 0.75948 | 0.759226 | 0.762011 | 0.759259 | 0.761756 |
| 0.699392 | 0.759921 | 0.758441 | 0.762068 | 0.76008 | 0.760128 | 0.757532 | 0.759142 | 0.765203 |
| 0.693932 | 0.749565 | 0.753698 | 0.758507 | 0.755295 | 0.748582 | 0.75086 | 0.757181 | 0.751394 |
| 0.692049 | 0.753425 | 0.754822 | 0.756897 | 0.755 | 0.750335 | 0.749802 | 0.753471 | 0.753872 |
| 0.691944 | 0.749053 | 0.75103 | 0.754054 | 0.754717 | 0.745571 | 0.751201 | 0.755323 | 0.747855 |
| 0.691981 | 0.753778 | 0.755482 | 0.75562 | 0.754725 | 0.751568 | 0.750447 | 0.753393 | 0.755612 |
| 0.695842 | 0.750659 | 0.749062 | 0.754516 | 0.751693 | 0.74886 | 0.751831 | 0.753584 | 0.751856 |
| 0.685072 | 0.75175 | 0.747386 | 0.756828 | 0.750293 | 0.749158 | 0.751464 | 0.7516 | 0.755974 |
| 0.698085 | 0.759681 | 0.757451 | 0.76566 | 0.759277 | 0.754604 | 0.757587 | 0.75785 | 0.762721 |
| 0.693078 | 0.749392 | 0.747918 | 0.756318 | 0.750447 | 0.750414 | 0.751769 | 0.751986 | 0.752079 |

| | | | | | | | | |
|---|---|---|---|---|---|---|---|---|
| 0.703264 | 0.759402 | 0.763863 | 0.76518 | 0.766893 | 0.757116 | 0.761125 | 0.763395 | 0.758009 |
| 0.705224 | 0.769129 | 0.774075 | 0.775586 | 0.770328 | 0.766923 | 0.771825 | 0.771849 | 0.768902 |
| 0.701525 | 0.772523 | 0.765929 | 0.772941 | 0.768799 | 0.764146 | 0.763657 | 0.768975 | 0.765318 |
| 0.708533 | 0.767574 | 0.770794 | 0.77172 | 0.770053 | 0.767872 | 0.767621 | 0.766731 | 0.767144 |
| 0.713073 | 0.776792 | 0.780552 | 0.784532 | 0.781497 | 0.771767 | 0.777683 | 0.771858 | 0.776526 |
| 0.708189 | 0.769608 | 0.775611 | 0.775476 | 0.77305 | 0.768534 | 0.76974 | 0.772312 | 0.771303 |
| 0.701272 | 0.769856 | 0.766274 | 0.770473 | 0.767372 | 0.769576 | 0.764108 | 0.76294 | 0.763893 |
| 0.705438 | 0.770086 | 0.767832 | 0.773635 | 0.768863 | 0.767025 | 0.767473 | 0.766831 | 0.761099 |
| 0.697989 | 0.764174 | 0.767486 | 0.768587 | 0.765509 | 0.760512 | 0.770402 | 0.761104 | 0.766442 |
| 0.703931 | 0.76392 | 0.766795 | 0.766302 | 0.76545 | 0.76232 | 0.764961 | 0.767234 | 0.762813 |
| 0.706484 | 0.771211 | 0.773097 | 0.773258 | 0.772552 | 0.764591 | 0.768913 | 0.770745 | 0.765712 |
| 0.707331 | 0.770396 | 0.771487 | 0.770423 | 0.771551 | 0.770213 | 0.768851 | 0.771908 | 0.766715 |
| 0.703352 | 0.7745 | 0.773427 | 0.776837 | 0.77248 | 0.770326 | 0.771421 | 0.767068 | 0.775893 |
| 0.705328 | 0.765824 | 0.774552 | 0.770773 | 0.769937 | 0.76347 | 0.768602 | 0.770238 | 0.764216 |
| 0.707596 | 0.782178 | 0.77246 | 0.777952 | 0.778666 | 0.775372 | 0.772727 | 0.774191 | 0.772794 |
| 0.70346 | 0.768585 | 0.773794 | 0.779802 | 0.774636 | 0.766058 | 0.765194 | 0.768341 | 0.774617 |
| 0.707862 | 0.77851 | 0.776121 | 0.775401 | 0.776528 | 0.764395 | 0.775464 | 0.77319 | 0.776689 |
| 0.702692 | 0.776697 | 0.770425 | 0.774077 | 0.774814 | 0.771114 | 0.767725 | 0.776724 | 0.767674 |
| 0.710868 | 0.774336 | 0.778837 | 0.776536 | 0.774143 | 0.770801 | 0.772279 | 0.772865 | 0.771059 |
| 0.702904 | 0.773319 | 0.774906 | 0.775863 | 0.776149 | 0.77175 | 0.770292 | 0.768846 | 0.772438 |
| 0.704229 | 0.764658 | 0.767738 | 0.772243 | 0.773381 | 0.762859 | 0.767586 | 0.767561 | 0.770063 |
| 0.701715 | 0.773317 | 0.769205 | 0.775033 | 0.772549 | 0.765766 | 0.769824 | 0.769789 | 0.771622 |
| 0.708066 | 0.773112 | 0.772534 | 0.776436 | 0.775177 | 0.766193 | 0.766889 | 0.774655 | 0.772645 |
| 0.705948 | 0.7822 | 0.786745 | 0.780415 | 0.776462 | 0.781798 | 0.779963 | 0.779779 | 0.777414 |
| 0.706578 | 0.782584 | 0.783885 | 0.785644 | 0.781812 | 0.77852 | 0.774754 | 0.778962 | 0.782553 |
| 0.699752 | 0.781504 | 0.780074 | 0.786519 | 0.778597 | 0.778807 | 0.78334 | 0.779555 | 0.774475 |
| 0.701333 | 0.782733 | 0.783246 | 0.788599 | 0.778757 | 0.7739 | 0.774481 | 0.778048 | 0.782489 |
| 0.700295 | 0.777361 | 0.777406 | 0.783297 | 0.779977 | 0.777024 | 0.771474 | 0.775318 | 0.77665 |
| 0.706122 | 0.784622 | 0.78366 | 0.783073 | 0.780411 | 0.778897 | 0.782057 | 0.779509 | 0.781733 |
| 0.708716 | 0.784109 | 0.786606 | 0.78688 | 0.777136 | 0.777581 | 0.776854 | 0.780184 | 0.779209 |
| 0.704202 | 0.775459 | 0.774094 | 0.780467 | 0.772888 | 0.776385 | 0.778106 | 0.773483 | 0.777661 |
| 0.709882 | 0.779402 | 0.782421 | 0.781229 | 0.783248 | 0.780073 | 0.77395 | 0.776347 | 0.783137 |
| 0.707076 | 0.77982 | 0.780868 | 0.784947 | 0.781939 | 0.781077 | 0.775441 | 0.778189 | 0.784605 |
| 0.699884 | 0.779868 | 0.77843 | 0.781341 | 0.773081 | 0.771984 | 0.77522 | 0.775856 | 0.774521 |
| 0.704398 | 0.782022 | 0.780151 | 0.785451 | 0.786326 | 0.780076 | 0.779181 | 0.784233 | 0.777659 |
| 0.702337 | 0.780572 | 0.781773 | 0.781499 | 0.786869 | 0.778732 | 0.780318 | 0.776809 | 0.784041 |
| 0.710863 | 0.782648 | 0.78515 | 0.793464 | 0.78543 | 0.773952 | 0.776161 | 0.783565 | 0.78141 |
| 0.706997 | 0.774172 | 0.781646 | 0.784292 | 0.782623 | 0.772098 | 0.782201 | 0.780389 | 0.777113 |
| 0.70548 | 0.776363 | 0.779151 | 0.787273 | 0.780897 | 0.775471 | 0.773258 | 0.776881 | 0.778712 |
| 0.702891 | 0.779914 | 0.788206 | 0.784947 | 0.785751 | 0.770329 | 0.779897 | 0.775409 | 0.769649 |
| 0.708752 | 0.780753 | 0.784776 | 0.785061 | 0.786082 | 0.777232 | 0.78137 | 0.780754 | 0.777615 |
| 0.711245 | 0.780913 | 0.786729 | 0.785756 | 0.786672 | 0.777384 | 0.785476 | 0.785436 | 0.778866 |
| 0.707921 | 0.780956 | 0.789893 | 0.787814 | 0.787959 | 0.777138 | 0.791442 | 0.787784 | 0.783634 |
| 0.706785 | 0.778504 | 0.78193 | 0.781459 | 0.77775 | 0.771956 | 0.77627 | 0.775201 | 0.770917 |
| 0.707554 | 0.782375 | 0.782209 | 0.785756 | 0.77991 | 0.777981 | 0.774927 | 0.776638 | 0.77867 |
| 0.701633 | 0.779543 | 0.776847 | 0.783377 | 0.774636 | 0.776188 | 0.776948 | 0.777291 | 0.774216 |

| | | | | | | | | |
|---|---|---|---|---|---|---|---|---|
| 0.690296 | 0.76585 | 0.76267 | 0.767097 | 0.761194 | 0.763388 | 0.765887 | 0.762629 | 0.769933 |
| 0.745559 | 0.75374 | 0.757556 | 0.756278 | 0.756023 | 0.750366 | 0.75017 | 0.750358 | 0.751271 |
| 0.703829 | 0.77452 | 0.776554 | 0.775268 | 0.775871 | 0.768525 | 0.766402 | 0.766437 | 0.773708 |
| 0.710147 | 0.777098 | 0.768874 | 0.777668 | 0.773295 | 0.767394 | 0.767049 | 0.769314 | 0.767854 |
| 0.699733 | 0.767367 | 0.77051 | 0.769026 | 0.769422 | 0.766257 | 0.766382 | 0.76656 | 0.76468 |
| 0.703195 | 0.77098 | 0.773572 | 0.776036 | 0.773328 | 0.762688 | 0.76962 | 0.769296 | 0.768574 |
| 0.701353 | 0.769231 | 0.767969 | 0.769846 | 0.772195 | 0.768016 | 0.775051 | 0.766063 | 0.767229 |
| 0.695521 | 0.770226 | 0.771026 | 0.771475 | 0.767991 | 0.76489 | 0.765381 | 0.763381 | 0.767097 |
| 0.70159 | 0.775069 | 0.770033 | 0.777759 | 0.771584 | 0.767284 | 0.771484 | 0.772884 | 0.770473 |
| 0.709216 | 0.775233 | 0.770403 | 0.777425 | 0.769065 | 0.770395 | 0.768123 | 0.769134 | 0.769171 |
| 0.707986 | 0.769981 | 0.771853 | 0.775023 | 0.773549 | 0.764746 | 0.772438 | 0.768858 | 0.769753 |
| 0.705069 | 0.768618 | 0.773928 | 0.777537 | 0.773679 | 0.767617 | 0.768987 | 0.7708 | 0.76703 |
| 0.711814 | 0.774069 | 0.769935 | 0.779119 | 0.775635 | 0.769729 | 0.772231 | 0.77345 | 0.772468 |
| 0.707489 | 0.771235 | 0.777269 | 0.776145 | 0.773577 | 0.766123 | 0.771349 | 0.772065 | 0.774091 |
| 0.705464 | 0.767494 | 0.768559 | 0.777956 | 0.765959 | 0.764897 | 0.766776 | 0.77136 | 0.765255 |
| 0.704136 | 0.767303 | 0.767145 | 0.772156 | 0.776463 | 0.763868 | 0.766038 | 0.766647 | 0.766633 |
| 0.705911 | 0.766087 | 0.768212 | 0.771669 | 0.768013 | 0.766042 | 0.762537 | 0.771057 | 0.765398 |
| 0.703419 | 0.765552 | 0.768939 | 0.775439 | 0.768478 | 0.759561 | 0.765582 | 0.768555 | 0.765 |
| 0.695026 | 0.76834 | 0.769039 | 0.767915 | 0.765238 | 0.763389 | 0.770136 | 0.767505 | 0.769933 |
| 0.699941 | 0.765163 | 0.765567 | 0.77171 | 0.762765 | 0.761952 | 0.769473 | 0.7684 | 0.766712 |
| 0.705418 | 0.770257 | 0.77108 | 0.77508 | 0.769385 | 0.76558 | 0.766302 | 0.771747 | 0.76887 |
| 0.694639 | 0.761845 | 0.761628 | 0.7586 | 0.759838 | 0.75507 | 0.755383 | 0.758439 | 0.758328 |
| 0.703713 | 0.768326 | 0.767059 | 0.767153 | 0.76932 | 0.76391 | 0.764346 | 0.766032 | 0.766991 |
| 0.706853 | 0.770933 | 0.772176 | 0.773712 | 0.777852 | 0.768107 | 0.768345 | 0.768206 | 0.766753 |
| 0.710035 | 0.77365 | 0.771726 | 0.776523 | 0.771092 | 0.770388 | 0.776296 | 0.770168 | 0.772439 |
| 0.705027 | 0.772587 | 0.76866 | 0.773975 | 0.770991 | 0.766474 | 0.767808 | 0.768515 | 0.767931 |
| 0.699062 | 0.76482 | 0.763975 | 0.773988 | 0.767235 | 0.75842 | 0.763622 | 0.766424 | 0.766409 |
| 0.698702 | 0.772448 | 0.767079 | 0.770492 | 0.769317 | 0.764083 | 0.764877 | 0.766339 | 0.770063 |
| 0.698957 | 0.767295 | 0.766889 | 0.768139 | 0.766516 | 0.761728 | 0.763024 | 0.766044 | 0.76301 |
| 0.783891 | 0.698525 | 0.698156 | 0.697737 | 0.697477 | 0.695079 | 0.698386 | 0.703025 | 0.693953 |
| 0.788724 | 0.707917 | 0.710229 | 0.710751 | 0.71082 | 0.703722 | 0.706728 | 0.708111 | 0.704779 |
| 0.803756 | 0.701495 | 0.711939 | 0.705645 | 0.708649 | 0.698303 | 0.702922 | 0.70427 | 0.703921 |
| 0.803771 | 0.69407 | 0.701092 | 0.699164 | 0.695437 | 0.69476 | 0.693505 | 0.694952 | 0.69465 |
| 0.792892 | 0.708192 | 0.711002 | 0.712367 | 0.711159 | 0.707467 | 0.704872 | 0.708249 | 0.703553 |
| 0.697514 | 0.766703 | 0.771625 | 0.770503 | 0.770152 | 0.765405 | 0.766965 | 0.762053 | 0.766581 |
| 0.701253 | 0.766142 | 0.774662 | 0.775696 | 0.771999 | 0.761826 | 0.76754 | 0.766542 | 0.76342 |
| 0.703466 | 0.776781 | 0.778176 | 0.774544 | 0.77595 | 0.773881 | 0.77408 | 0.772379 | 0.773689 |
| 0.705306 | 0.772686 | 0.777319 | 0.77421 | 0.775835 | 0.773019 | 0.772026 | 0.766838 | 0.768082 |
| 0.702323 | 0.768142 | 0.768436 | 0.77057 | 0.772348 | 0.768436 | 0.772688 | 0.767613 | 0.766849 |
| 0.705739 | 0.766648 | 0.774407 | 0.77404 | 0.775908 | 0.767418 | 0.771192 | 0.77118 | 0.770327 |
| 0.703508 | 0.77049 | 0.77841 | 0.775198 | 0.770985 | 0.767828 | 0.766348 | 0.769755 | 0.773603 |
| 0.703798 | 0.763606 | 0.771909 | 0.768017 | 0.769458 | 0.76362 | 0.767442 | 0.764525 | 0.768778 |
| 0.696648 | 0.76704 | 0.761999 | 0.765396 | 0.760424 | 0.765299 | 0.761186 | 0.762107 | 0.765432 |
| 0.696271 | 0.759986 | 0.767286 | 0.762283 | 0.767694 | 0.760836 | 0.763676 | 0.75687 | 0.763408 |
| 0.700835 | 0.768907 | 0.771847 | 0.770399 | 0.771528 | 0.770176 | 0.764649 | 0.762857 | 0.767141 |
| 0.704492 | 0.767139 | 0.774654 | 0.774251 | 0.773866 | 0.764757 | 0.76576 | 0.769028 | 0.76917 |

| | | | | | | | | |
|---|---|---|---|---|---|---|---|---|
| 0.704301 | 0.773452 | 0.769845 | 0.782074 | 0.771591 | 0.770023 | 0.77184 | 0.770374 | 0.767624 |
| 0.704656 | 0.769662 | 0.772177 | 0.772173 | 0.774662 | 0.764578 | 0.770764 | 0.771318 | 0.769726 |
| 0.705001 | 0.769562 | 0.771868 | 0.773537 | 0.767424 | 0.762996 | 0.768735 | 0.770888 | 0.770425 |
| 0.702294 | 0.762828 | 0.762709 | 0.768214 | 0.76738 | 0.759959 | 0.76639 | 0.769393 | 0.767075 |
| 0.701258 | 0.766216 | 0.76679 | 0.766774 | 0.765119 | 0.761809 | 0.766568 | 0.762268 | 0.758814 |
| 0.704472 | 0.770087 | 0.766908 | 0.769297 | 0.773391 | 0.766809 | 0.766042 | 0.773028 | 0.763807 |
| 0.703088 | 0.765243 | 0.767579 | 0.773537 | 0.766509 | 0.763306 | 0.766844 | 0.767928 | 0.766725 |
| 0.705566 | 0.767874 | 0.768474 | 0.774315 | 0.769309 | 0.763077 | 0.767724 | 0.765202 | 0.765359 |
| 0.702823 | 0.76445 | 0.765932 | 0.771375 | 0.765647 | 0.764032 | 0.761764 | 0.765176 | 0.763195 |
| 0.707472 | 0.768798 | 0.772984 | 0.772132 | 0.767153 | 0.764685 | 0.768429 | 0.770676 | 0.766548 |
| 0.699731 | 0.765656 | 0.767529 | 0.765667 | 0.767079 | 0.761688 | 0.765311 | 0.766041 | 0.765439 |
| 0.69959 | 0.771306 | 0.76806 | 0.768047 | 0.771576 | 0.769048 | 0.763244 | 0.767563 | 0.76628 |
| 0.691741 | 0.765098 | 0.763275 | 0.762706 | 0.761215 | 0.761535 | 0.7637 | 0.76106 | 0.757341 |
| 0.699029 | 0.760797 | 0.758511 | 0.767833 | 0.762598 | 0.764734 | 0.762759 | 0.762398 | 0.759376 |
| 0.706722 | 0.77107 | 0.772471 | 0.771869 | 0.767964 | 0.764644 | 0.764823 | 0.771122 | 0.768497 |
| 0.709126 | 0.783419 | 0.781499 | 0.783298 | 0.778222 | 0.780869 | 0.781563 | 0.781758 | 0.779312 |
| 0.704676 | 0.7823 | 0.777814 | 0.779282 | 0.772228 | 0.775631 | 0.772997 | 0.774266 | 0.773392 |
| 0.70866 | 0.772664 | 0.775023 | 0.777537 | 0.77264 | 0.774768 | 0.769598 | 0.775386 | 0.769638 |
| 0.704301 | 0.769899 | 0.770227 | 0.775426 | 0.771149 | 0.766687 | 0.770376 | 0.77168 | 0.767214 |
| 0.7045 | 0.770208 | 0.769165 | 0.774338 | 0.772977 | 0.766711 | 0.771713 | 0.766545 | 0.767365 |
| 0.699666 | 0.763053 | 0.766083 | 0.772012 | 0.76712 | 0.762644 | 0.766183 | 0.76023 | 0.76337 |
| 0.700268 | 0.769479 | 0.772595 | 0.774996 | 0.773743 | 0.766199 | 0.768112 | 0.769865 | 0.766329 |
| 0.707035 | 0.782026 | 0.77704 | 0.783056 | 0.776055 | 0.780818 | 0.777944 | 0.778146 | 0.778744 |
| 0.703305 | 0.763818 | 0.766094 | 0.769251 | 0.770335 | 0.762354 | 0.768923 | 0.762842 | 0.766526 |
| 0.704441 | 0.77705 | 0.772414 | 0.782218 | 0.77563 | 0.773144 | 0.770615 | 0.773149 | 0.773762 |
| 1 | 0.703988 | 0.70433 | 0.71031 | 0.703964 | 0.702151 | 0.69849 | 0.702323 | 0.701747 |
| 0.703988 | 1 | 0.785591 | 0.787468 | 0.782389 | 0.874288 | 0.789329 | 0.780223 | 0.784675 |
| 0.70433 | 0.785591 | 1 | 0.787667 | 0.837405 | 0.779164 | 0.786705 | 0.777983 | 0.799195 |
| 0.71031 | 0.787468 | 0.787667 | 1 | 0.784935 | 0.776114 | 0.787343 | 0.78951 | 0.781825 |
| 0.703964 | 0.782389 | 0.837405 | 0.784935 | 1 | 0.776967 | 0.785936 | 0.778625 | 0.801199 |
| 0.702151 | 0.874288 | 0.779164 | 0.776114 | 0.776967 | 1 | 0.787392 | 0.778287 | 0.776743 |
| 0.69849 | 0.789329 | 0.786705 | 0.787343 | 0.785936 | 0.787392 | 1 | 0.831351 | 0.780573 |
| 0.702323 | 0.780223 | 0.777983 | 0.78951 | 0.778625 | 0.778287 | 0.831351 | 1 | 0.776279 |
| 0.701747 | 0.784675 | 0.799195 | 0.781825 | 0.801199 | 0.776743 | 0.780573 | 0.776279 | 1 |
| 0.697217 | 0.773299 | 0.875331 | 0.779098 | 0.818842 | 0.769189 | 0.781431 | 0.772562 | 0.813335 |
| 0.700462 | 0.780728 | 0.886724 | 0.78355 | 0.813587 | 0.771781 | 0.779463 | 0.775813 | 0.78864 |
| 0.706008 | 0.794781 | 0.789665 | 0.787579 | 0.786887 | 0.788209 | 0.78607 | 0.78058 | 0.782985 |
| 0.704965 | 0.777262 | 0.776136 | 0.775835 | 0.77788 | 0.773017 | 0.778583 | 0.774913 | 0.776092 |
| 0.702617 | 0.780561 | 0.782983 | 0.78207 | 0.78023 | 0.773811 | 0.77699 | 0.77446 | 0.781094 |
| 0.702923 | 0.773034 | 0.773173 | 0.775798 | 0.773452 | 0.770102 | 0.774575 | 0.774044 | 0.770979 |
| 0.699397 | 0.769356 | 0.772575 | 0.779237 | 0.774029 | 0.76594 | 0.76936 | 0.774041 | 0.774431 |
| 0.70794 | 0.777871 | 0.780437 | 0.781197 | 0.785297 | 0.772927 | 0.776464 | 0.771541 | 0.78112 |
| 0.700754 | 0.771602 | 0.7775 | 0.771798 | 0.774684 | 0.77116 | 0.773228 | 0.770024 | 0.772911 |
| 0.701572 | 0.769513 | 0.768339 | 0.771388 | 0.76349 | 0.764085 | 0.762395 | 0.761162 | 0.768833 |
| 0.703728 | 0.771417 | 0.773129 | 0.775611 | 0.77201 | 0.769751 | 0.773366 | 0.773369 | 0.767244 |
| 0.704558 | 0.772522 | 0.77196 | 0.773324 | 0.768868 | 0.766457 | 0.766272 | 0.766955 | 0.758873 |

| | | | | | | | | |
|---|---|---|---|---|---|---|---|---|
| 0.701992 | 0.772601 | 0.769703 | 0.772194 | 0.768118 | 0.768807 | 0.766678 | 0.763762 | 0.763877 |
| 0.701889 | 0.770694 | 0.769017 | 0.777234 | 0.772772 | 0.769595 | 0.769407 | 0.771843 | 0.7672 |
| 0.702299 | 0.771417 | 0.766744 | 0.776367 | 0.767564 | 0.764987 | 0.768081 | 0.766873 | 0.768112 |
| 0.667373 | 0.723121 | 0.720285 | 0.725824 | 0.73169 | 0.719048 | 0.724982 | 0.719786 | 0.728131 |
| 0.702526 | 0.768317 | 0.762427 | 0.774915 | 0.766957 | 0.767346 | 0.763036 | 0.765463 | 0.762773 |
| 0.701754 | 0.769243 | 0.767028 | 0.769161 | 0.769594 | 0.762475 | 0.768843 | 0.770726 | 0.768376 |
| 0.706754 | 0.771349 | 0.767086 | 0.771827 | 0.768761 | 0.774398 | 0.767678 | 0.766323 | 0.76353 |
| 0.703864 | 0.766428 | 0.771106 | 0.775598 | 0.772465 | 0.767418 | 0.764497 | 0.767974 | 0.773644 |
| 0.711202 | 0.776546 | 0.776244 | 0.782786 | 0.771563 | 0.771915 | 0.776658 | 0.776459 | 0.770848 |
| 0.7062 | 0.77326 | 0.771743 | 0.774089 | 0.773793 | 0.766859 | 0.772144 | 0.772498 | 0.767361 |
| 0.705403 | 0.773048 | 0.770523 | 0.779938 | 0.769729 | 0.771965 | 0.766251 | 0.770438 | 0.762458 |
| 0.70762 | 0.769547 | 0.774146 | 0.777564 | 0.772055 | 0.770908 | 0.768113 | 0.772132 | 0.767561 |
| 0.708044 | 0.770823 | 0.767455 | 0.776594 | 0.770718 | 0.769843 | 0.768144 | 0.769057 | 0.767845 |
| 0.702538 | 0.771781 | 0.772259 | 0.776039 | 0.771544 | 0.76757 | 0.769329 | 0.765965 | 0.771665 |
| 0.708165 | 0.775299 | 0.77269 | 0.772203 | 0.774076 | 0.772028 | 0.770603 | 0.765272 | 0.769816 |
| 0.698611 | 0.76784 | 0.761648 | 0.771923 | 0.763623 | 0.765274 | 0.76574 | 0.761889 | 0.763162 |
| 0.707874 | 0.778077 | 0.774562 | 0.776945 | 0.777083 | 0.774532 | 0.773997 | 0.769519 | 0.773805 |
| 0.706015 | 0.775967 | 0.772526 | 0.775329 | 0.772416 | 0.769549 | 0.770715 | 0.767889 | 0.771224 |
| 0.698511 | 0.769489 | 0.768198 | 0.770136 | 0.76828 | 0.7617 | 0.763772 | 0.761315 | 0.76466 |
| 0.701698 | 0.76903 | 0.76605 | 0.771774 | 0.768247 | 0.767402 | 0.766805 | 0.766731 | 0.764073 |
| 0.705707 | 0.777805 | 0.773679 | 0.774961 | 0.773521 | 0.775669 | 0.774712 | 0.773079 | 0.768693 |
| 0.708122 | 0.776494 | 0.774042 | 0.777436 | 0.777245 | 0.767143 | 0.770089 | 0.769515 | 0.768316 |
| 0.692873 | 0.679709 | 0.679476 | 0.685867 | 0.678748 | 0.679165 | 0.682457 | 0.676164 | 0.680163 |
| 0.727792 | 0.652461 | 0.657976 | 0.653251 | 0.652994 | 0.651594 | 0.660124 | 0.653609 | 0.659518 |
| 0.766811 | 0.669819 | 0.671268 | 0.676066 | 0.675918 | 0.67634 | 0.681698 | 0.67066 | 0.66674 |
| 0.766661 | 0.650943 | 0.65626 | 0.662238 | 0.653051 | 0.654491 | 0.653872 | 0.653916 | 0.656125 |
| 0.663289 | 0.742788 | 0.740707 | 0.73486 | 0.741834 | 0.745661 | 0.742 | 0.735325 | 0.744282 |
| 0.655704 | 0.730902 | 0.72458 | 0.72501 | 0.727866 | 0.73157 | 0.732353 | 0.730925 | 0.720534 |
| 0.655503 | 0.733874 | 0.728762 | 0.733565 | 0.735145 | 0.731944 | 0.730243 | 0.733927 | 0.730397 |
| 0.664377 | 0.743611 | 0.74144 | 0.74203 | 0.753211 | 0.738298 | 0.74469 | 0.743465 | 0.733546 |
| 0.642337 | 0.731409 | 0.724611 | 0.724698 | 0.729748 | 0.726193 | 0.729065 | 0.719487 | 0.715443 |
| 0.636862 | 0.718952 | 0.713314 | 0.709024 | 0.714407 | 0.728377 | 0.72055 | 0.711874 | 0.711758 |
| 0.640273 | 0.726381 | 0.710179 | 0.72322 | 0.719081 | 0.727465 | 0.717038 | 0.711929 | 0.720443 |
| 0.743729 | 0.66129 | 0.656677 | 0.662225 | 0.654877 | 0.657985 | 0.657205 | 0.653715 | 0.655729 |
| 0.695736 | 0.761386 | 0.767296 | 0.769378 | 0.767488 | 0.762178 | 0.762378 | 0.763139 | 0.762552 |
| 0.670276 | 0.742562 | 0.743294 | 0.751684 | 0.744561 | 0.742624 | 0.749702 | 0.740924 | 0.737373 |
| 0.664268 | 0.738115 | 0.740409 | 0.739627 | 0.739378 | 0.744228 | 0.738738 | 0.737143 | 0.734806 |
| 0.648102 | 0.727611 | 0.727442 | 0.728379 | 0.731163 | 0.725446 | 0.731096 | 0.72809 | 0.726262 |
| 0.669688 | 0.745518 | 0.743011 | 0.745603 | 0.751146 | 0.747707 | 0.744382 | 0.744389 | 0.747989 |
| 0.660354 | 0.740303 | 0.733701 | 0.733428 | 0.732846 | 0.738391 | 0.734491 | 0.732928 | 0.732472 |
| 0.658635 | 0.736355 | 0.722002 | 0.729063 | 0.736909 | 0.734228 | 0.729639 | 0.733301 | 0.738374 |
| 0.656335 | 0.732584 | 0.727117 | 0.733827 | 0.725074 | 0.73852 | 0.727344 | 0.729101 | 0.730739 |
| 0.657904 | 0.732091 | 0.732594 | 0.732302 | 0.727322 | 0.732944 | 0.730896 | 0.726828 | 0.731023 |
| 0.656691 | 0.736582 | 0.72738 | 0.727976 | 0.731881 | 0.733174 | 0.734213 | 0.730174 | 0.734269 |
| 0.65568 | 0.728979 | 0.729161 | 0.727249 | 0.727273 | 0.734076 | 0.729919 | 0.726522 | 0.732157 |
| 0.638995 | 0.720676 | 0.720266 | 0.724545 | 0.724024 | 0.723504 | 0.72035 | 0.712261 | 0.724126 |

| | | | | | | | | |
|---|---|---|---|---|---|---|---|---|
| 0.68263 | 0.754843 | 0.756294 | 0.75564 | 0.755529 | 0.758055 | 0.755852 | 0.750552 | 0.754001 |
| 0.649939 | 0.73181 | 0.735257 | 0.738288 | 0.737407 | 0.741165 | 0.737973 | 0.733087 | 0.737863 |
| 0.657172 | 0.73003 | 0.731891 | 0.734252 | 0.732831 | 0.732999 | 0.735901 | 0.731461 | 0.732388 |
| 0.658569 | 0.740889 | 0.738326 | 0.731836 | 0.739747 | 0.740347 | 0.74292 | 0.727608 | 0.736569 |
| 0.650235 | 0.728703 | 0.731712 | 0.739992 | 0.73276 | 0.732606 | 0.734339 | 0.729963 | 0.726731 |
| 0.660382 | 0.731726 | 0.740276 | 0.728608 | 0.738346 | 0.737417 | 0.737468 | 0.722042 | 0.728737 |
| 0.642191 | 0.727162 | 0.728134 | 0.725533 | 0.721792 | 0.721243 | 0.721937 | 0.718849 | 0.725553 |
| 0.641334 | 0.717311 | 0.724109 | 0.730991 | 0.728702 | 0.727794 | 0.727743 | 0.717804 | 0.723598 |
| 0.642896 | 0.720943 | 0.737158 | 0.729529 | 0.726602 | 0.725949 | 0.736311 | 0.724711 | 0.728392 |
| 0.687243 | 0.755168 | 0.75814 | 0.754998 | 0.759002 | 0.753803 | 0.759237 | 0.753096 | 0.755046 |
| 0.685591 | 0.748897 | 0.752542 | 0.755946 | 0.757551 | 0.748579 | 0.754359 | 0.750025 | 0.750494 |
| 0.680336 | 0.751162 | 0.751219 | 0.753836 | 0.755082 | 0.744874 | 0.751669 | 0.745609 | 0.74776 |
| 0.67351 | 0.741826 | 0.740082 | 0.739734 | 0.742624 | 0.742458 | 0.745782 | 0.737549 | 0.741892 |
| 0.677213 | 0.745241 | 0.741581 | 0.748401 | 0.742046 | 0.745527 | 0.744881 | 0.738122 | 0.742814 |
| 0.668172 | 0.737745 | 0.745098 | 0.741375 | 0.746104 | 0.740416 | 0.748143 | 0.739794 | 0.743279 |

| | | | plot 15 | | | | | |
|---|---|---|---|---|---|---|---|---|
| Wild_plot_ | Wild_plot_ | Wild_plot_ | Wild_plot_ | Wild_plot_ | Wild_plot_ | Wild_plot_ | Wild_plot_ | Wild_plot_ |
| 0.704744 | 0.7064 | 0.708006 | 0.712114 | 0.709139 | 0.708773 | 0.70335 | 0.711246 | 0.703912 |
| 0.708797 | 0.710089 | 0.717052 | 0.717903 | 0.712904 | 0.712071 | 0.708091 | 0.715156 | 0.711942 |
| 0.700008 | 0.70017 | 0.707234 | 0.703621 | 0.706118 | 0.703364 | 0.698098 | 0.706924 | 0.699324 |
| 0.709391 | 0.708594 | 0.715085 | 0.717456 | 0.714618 | 0.714357 | 0.708654 | 0.715623 | 0.712544 |
| 0.703673 | 0.703163 | 0.709913 | 0.712786 | 0.710649 | 0.708838 | 0.706951 | 0.713443 | 0.704344 |
| 0.711795 | 0.712568 | 0.71728 | 0.722605 | 0.715054 | 0.716631 | 0.712781 | 0.720229 | 0.717761 |
| 0.771722 | 0.768067 | 0.780005 | 0.788306 | 0.782482 | 0.781871 | 0.774482 | 0.786495 | 0.781255 |
| 0.756561 | 0.75877 | 0.770346 | 0.779478 | 0.773277 | 0.777471 | 0.778448 | 0.779207 | 0.772794 |
| 0.708555 | 0.710935 | 0.715406 | 0.715956 | 0.714782 | 0.711272 | 0.710065 | 0.716264 | 0.709123 |
| 0.701391 | 0.705701 | 0.711028 | 0.71037 | 0.70543 | 0.709889 | 0.702073 | 0.707127 | 0.708032 |
| 0.710244 | 0.706719 | 0.711646 | 0.714746 | 0.711459 | 0.707613 | 0.703556 | 0.71332 | 0.710147 |
| 0.696227 | 0.700545 | 0.705597 | 0.70824 | 0.701746 | 0.702716 | 0.695943 | 0.704162 | 0.70348 |
| 0.708411 | 0.709198 | 0.711021 | 0.71563 | 0.712476 | 0.710088 | 0.71021 | 0.718475 | 0.707281 |
| 0.717743 | 0.713983 | 0.722277 | 0.723696 | 0.720155 | 0.719807 | 0.719155 | 0.725078 | 0.717002 |
| 0.733905 | 0.731709 | 0.738117 | 0.738522 | 0.739166 | 0.734542 | 0.730442 | 0.741244 | 0.734168 |
| 0.708872 | 0.712619 | 0.713694 | 0.715488 | 0.716946 | 0.715 | 0.711356 | 0.717739 | 0.714107 |
| 0.710176 | 0.710784 | 0.717641 | 0.715003 | 0.711746 | 0.714741 | 0.711923 | 0.714239 | 0.710934 |
| 0.735401 | 0.73553 | 0.74254 | 0.743393 | 0.743218 | 0.742057 | 0.739562 | 0.747374 | 0.73987 |
| 0.700825 | 0.703613 | 0.708347 | 0.710586 | 0.707933 | 0.708629 | 0.698663 | 0.709649 | 0.702914 |
| 0.707519 | 0.707383 | 0.712726 | 0.715748 | 0.710554 | 0.711733 | 0.704073 | 0.709343 | 0.712009 |
| 0.699674 | 0.70717 | 0.706996 | 0.710999 | 0.708712 | 0.707315 | 0.702286 | 0.709154 | 0.708949 |
| 0.707436 | 0.708868 | 0.711093 | 0.712757 | 0.712162 | 0.710174 | 0.704276 | 0.715644 | 0.708191 |
| 0.709265 | 0.714286 | 0.715956 | 0.721272 | 0.716382 | 0.71669 | 0.708506 | 0.718205 | 0.716261 |
| 0.707688 | 0.710209 | 0.715378 | 0.715801 | 0.711494 | 0.711222 | 0.70597 | 0.714668 | 0.712154 |
| 0.71048 | 0.712351 | 0.715315 | 0.717479 | 0.712979 | 0.716046 | 0.704682 | 0.716152 | 0.712329 |
| 0.707023 | 0.707357 | 0.715431 | 0.715429 | 0.709786 | 0.706862 | 0.70469 | 0.714182 | 0.706382 |
| 0.699085 | 0.703568 | 0.707062 | 0.705812 | 0.704919 | 0.702274 | 0.700732 | 0.707902 | 0.69995 |
| 0.708207 | 0.709743 | 0.712643 | 0.714045 | 0.710697 | 0.711554 | 0.7045 | 0.713985 | 0.711257 |
| 0.709835 | 0.713046 | 0.715276 | 0.717405 | 0.713963 | 0.716123 | 0.708368 | 0.715969 | 0.711595 |
| 0.706646 | 0.708751 | 0.711617 | 0.713709 | 0.708347 | 0.708104 | 0.706187 | 0.712634 | 0.708897 |
| 0.704306 | 0.704523 | 0.70974 | 0.714228 | 0.709925 | 0.707841 | 0.704598 | 0.710595 | 0.710603 |
| 0.701119 | 0.700956 | 0.709633 | 0.708989 | 0.707022 | 0.706881 | 0.700594 | 0.710618 | 0.704346 |
| 0.703784 | 0.708841 | 0.711738 | 0.710129 | 0.710963 | 0.708337 | 0.703169 | 0.707101 | 0.707992 |
| 0.705873 | 0.70603 | 0.709385 | 0.710539 | 0.712346 | 0.710805 | 0.707557 | 0.713695 | 0.708796 |
| 0.697721 | 0.703825 | 0.710458 | 0.706836 | 0.703971 | 0.705482 | 0.702555 | 0.709772 | 0.703106 |
| 0.706991 | 0.709823 | 0.712699 | 0.713974 | 0.713927 | 0.705745 | 0.706402 | 0.715009 | 0.70891 |
| 0.707101 | 0.710339 | 0.714898 | 0.718187 | 0.714629 | 0.716562 | 0.710448 | 0.717676 | 0.713931 |
| 0.705822 | 0.710537 | 0.712762 | 0.71845 | 0.713004 | 0.712566 | 0.709924 | 0.715765 | 0.712506 |
| 0.712133 | 0.708771 | 0.717053 | 0.719278 | 0.714654 | 0.717532 | 0.711845 | 0.716373 | 0.715792 |

| | | | | | | | | |
|---|---|---|---|---|---|---|---|---|
| 0.741299 | 0.740154 | 0.750369 | 0.751174 | 0.749004 | 0.745529 | 0.741239 | 0.747676 | 0.74424 |
| 0.698005 | 0.700869 | 0.706898 | 0.706743 | 0.704711 | 0.70425 | 0.69676 | 0.703877 | 0.702084 |
| 0.701843 | 0.703359 | 0.70978 | 0.709745 | 0.70684 | 0.707398 | 0.702398 | 0.708168 | 0.705516 |
| 0.707999 | 0.709787 | 0.715418 | 0.71898 | 0.712305 | 0.713931 | 0.709565 | 0.714654 | 0.712785 |
| 0.705344 | 0.705532 | 0.714018 | 0.711051 | 0.709439 | 0.7096 | 0.707141 | 0.714935 | 0.708029 |
| 0.710274 | 0.707705 | 0.71383 | 0.716839 | 0.71366 | 0.711921 | 0.706815 | 0.718272 | 0.714095 |
| 0.705147 | 0.704154 | 0.708671 | 0.710263 | 0.70627 | 0.705235 | 0.703453 | 0.70834 | 0.707984 |
| 0.704473 | 0.705767 | 0.712979 | 0.712269 | 0.712121 | 0.712228 | 0.706118 | 0.713737 | 0.707236 |
| 0.707867 | 0.71129 | 0.716053 | 0.718217 | 0.715797 | 0.712204 | 0.709996 | 0.717259 | 0.71063 |
| 0.708442 | 0.711098 | 0.718862 | 0.717107 | 0.710299 | 0.710758 | 0.705625 | 0.715246 | 0.712687 |
| 0.698112 | 0.699281 | 0.707247 | 0.708988 | 0.705916 | 0.702763 | 0.699086 | 0.706479 | 0.704201 |
| 0.700968 | 0.700497 | 0.705973 | 0.70637 | 0.704778 | 0.705691 | 0.699138 | 0.709069 | 0.702527 |
| 0.702829 | 0.707439 | 0.712857 | 0.714043 | 0.711431 | 0.710458 | 0.702915 | 0.712394 | 0.710487 |
| 0.699217 | 0.701323 | 0.705039 | 0.707923 | 0.704078 | 0.70778 | 0.702479 | 0.708771 | 0.703241 |
| 0.702216 | 0.701094 | 0.708634 | 0.710353 | 0.705629 | 0.708715 | 0.69808 | 0.709767 | 0.704863 |
| 0.700085 | 0.697249 | 0.709576 | 0.706772 | 0.704338 | 0.704494 | 0.699471 | 0.708525 | 0.702002 |
| 0.698195 | 0.702224 | 0.705617 | 0.707711 | 0.705457 | 0.69969 | 0.700931 | 0.707337 | 0.702103 |
| 0.704801 | 0.706246 | 0.713368 | 0.712488 | 0.712816 | 0.710275 | 0.705619 | 0.715086 | 0.70636 |
| 0.708295 | 0.707144 | 0.715332 | 0.715839 | 0.712847 | 0.718442 | 0.709675 | 0.715772 | 0.714763 |
| 0.700338 | 0.704512 | 0.711009 | 0.713568 | 0.711151 | 0.705273 | 0.704722 | 0.710359 | 0.70786 |
| 0.697204 | 0.699869 | 0.707303 | 0.704005 | 0.702369 | 0.704686 | 0.699325 | 0.70748 | 0.698678 |
| 0.702241 | 0.702147 | 0.705303 | 0.709542 | 0.706257 | 0.704095 | 0.705318 | 0.707776 | 0.70672 |
| 0.705311 | 0.704801 | 0.709873 | 0.711822 | 0.705995 | 0.705417 | 0.704944 | 0.709548 | 0.707525 |
| 0.710738 | 0.715714 | 0.720023 | 0.717683 | 0.714713 | 0.713414 | 0.709395 | 0.718604 | 0.714047 |
| 0.720932 | 0.721227 | 0.726433 | 0.725151 | 0.727052 | 0.720073 | 0.719773 | 0.725778 | 0.718885 |
| 0.757494 | 0.760051 | 0.772446 | 0.779997 | 0.774775 | 0.778347 | 0.778289 | 0.780759 | 0.775549 |
| 0.771476 | 0.772642 | 0.783286 | 0.777209 | 0.780691 | 0.776696 | 0.777007 | 0.781563 | 0.771725 |
| 0.773233 | 0.772299 | 0.781261 | 0.778364 | 0.780995 | 0.777884 | 0.779661 | 0.784194 | 0.777181 |
| 0.703084 | 0.705081 | 0.708286 | 0.708481 | 0.708652 | 0.707425 | 0.702226 | 0.712907 | 0.706872 |
| 0.701621 | 0.701038 | 0.706402 | 0.706105 | 0.705464 | 0.705006 | 0.703057 | 0.708692 | 0.703912 |
| 0.71477 | 0.714567 | 0.717871 | 0.715922 | 0.714794 | 0.71561 | 0.711941 | 0.718074 | 0.715659 |
| 0.712218 | 0.710943 | 0.715486 | 0.716844 | 0.718678 | 0.713554 | 0.713949 | 0.718503 | 0.715273 |
| 0.699687 | 0.703088 | 0.710734 | 0.712612 | 0.708795 | 0.705498 | 0.7029 | 0.711658 | 0.704384 |
| 0.702538 | 0.707078 | 0.70977 | 0.712607 | 0.711364 | 0.709356 | 0.70418 | 0.711927 | 0.704642 |
| 0.700374 | 0.701441 | 0.710838 | 0.711191 | 0.706084 | 0.705917 | 0.701377 | 0.708005 | 0.70338 |
| 0.703211 | 0.706365 | 0.70977 | 0.711433 | 0.707842 | 0.706128 | 0.70366 | 0.710917 | 0.704984 |
| 0.704669 | 0.701257 | 0.711383 | 0.709775 | 0.707257 | 0.707567 | 0.704362 | 0.713274 | 0.704097 |
| 0.717171 | 0.720142 | 0.723881 | 0.724893 | 0.722402 | 0.720318 | 0.718133 | 0.726143 | 0.722679 |
| 0.692381 | 0.695528 | 0.705188 | 0.703568 | 0.703614 | 0.70161 | 0.69531 | 0.704365 | 0.700567 |
| 0.702021 | 0.701869 | 0.713582 | 0.708911 | 0.709526 | 0.704603 | 0.699836 | 0.707708 | 0.704691 |
| 0.697433 | 0.70085 | 0.706775 | 0.703972 | 0.703718 | 0.704422 | 0.699803 | 0.706719 | 0.699294 |
| 0.719997 | 0.724411 | 0.727153 | 0.727465 | 0.728615 | 0.723099 | 0.72079 | 0.72705 | 0.723984 |
| 0.701886 | 0.702117 | 0.708882 | 0.710399 | 0.707584 | 0.709141 | 0.705441 | 0.712495 | 0.709285 |
| 0.699799 | 0.704707 | 0.708667 | 0.708463 | 0.708894 | 0.707228 | 0.703334 | 0.712279 | 0.705329 |
| 0.705196 | 0.708066 | 0.71561 | 0.713848 | 0.714483 | 0.707681 | 0.705406 | 0.715349 | 0.708901 |
| 0.700448 | 0.705389 | 0.711647 | 0.712395 | 0.707856 | 0.706163 | 0.70268 | 0.709015 | 0.705863 |

| | | | | | | | | |
|---|---|---|---|---|---|---|---|---|
| 0.702853 | 0.702674 | 0.708581 | 0.708551 | 0.706375 | 0.70645 | 0.699636 | 0.711703 | 0.707656 |
| 0.707217 | 0.709567 | 0.71668 | 0.712079 | 0.70866 | 0.712805 | 0.70673 | 0.714378 | 0.704202 |
| 0.70509 | 0.706672 | 0.709155 | 0.713787 | 0.710977 | 0.708072 | 0.704285 | 0.712971 | 0.705238 |
| 0.711323 | 0.714534 | 0.717894 | 0.717455 | 0.717553 | 0.714346 | 0.708824 | 0.718316 | 0.713529 |
| 0.710617 | 0.712018 | 0.719086 | 0.717095 | 0.713758 | 0.714844 | 0.71009 | 0.717546 | 0.712204 |
| 0.703381 | 0.706588 | 0.713384 | 0.710015 | 0.708039 | 0.70439 | 0.703948 | 0.708868 | 0.705302 |
| 0.707356 | 0.705335 | 0.716115 | 0.713647 | 0.71267 | 0.711118 | 0.707438 | 0.71218 | 0.708313 |
| 0.709705 | 0.712261 | 0.716916 | 0.715247 | 0.714588 | 0.711845 | 0.710588 | 0.718235 | 0.71194 |
| 0.70407 | 0.707834 | 0.711004 | 0.715686 | 0.70896 | 0.712827 | 0.706378 | 0.711281 | 0.713362 |
| 0.700921 | 0.702014 | 0.707359 | 0.709804 | 0.708625 | 0.709511 | 0.699545 | 0.710827 | 0.705897 |
| 0.70359 | 0.706544 | 0.709917 | 0.709441 | 0.710077 | 0.710146 | 0.703457 | 0.712409 | 0.705486 |
| 0.704671 | 0.707652 | 0.71257 | 0.711446 | 0.709777 | 0.708421 | 0.70494 | 0.711926 | 0.707661 |
| 0.708673 | 0.706752 | 0.712003 | 0.712156 | 0.710136 | 0.709458 | 0.706065 | 0.71306 | 0.709826 |
| 0.702666 | 0.707024 | 0.709803 | 0.707051 | 0.704799 | 0.707136 | 0.700982 | 0.71045 | 0.702453 |
| 0.704954 | 0.707961 | 0.709672 | 0.712959 | 0.710095 | 0.710566 | 0.704511 | 0.715864 | 0.706643 |
| 0.70198 | 0.706226 | 0.709889 | 0.711343 | 0.710292 | 0.709506 | 0.705522 | 0.711374 | 0.706609 |
| 0.709382 | 0.708303 | 0.71848 | 0.712205 | 0.712491 | 0.709937 | 0.709796 | 0.716228 | 0.712295 |
| 0.703297 | 0.705518 | 0.708709 | 0.708852 | 0.708511 | 0.703249 | 0.703781 | 0.712917 | 0.705917 |
| 0.701345 | 0.709213 | 0.710494 | 0.713167 | 0.710741 | 0.711703 | 0.705725 | 0.716465 | 0.705056 |
| 0.705049 | 0.705024 | 0.714997 | 0.714182 | 0.712738 | 0.711903 | 0.70652 | 0.715681 | 0.705677 |
| 0.704572 | 0.707025 | 0.708686 | 0.709082 | 0.70957 | 0.705843 | 0.704273 | 0.70927 | 0.705657 |
| 0.700033 | 0.702206 | 0.706513 | 0.704812 | 0.704197 | 0.704585 | 0.699313 | 0.709343 | 0.705692 |
| 0.702776 | 0.706153 | 0.712797 | 0.710605 | 0.708117 | 0.707321 | 0.704394 | 0.71015 | 0.707289 |
| 0.703598 | 0.706791 | 0.711682 | 0.713381 | 0.709395 | 0.712017 | 0.701718 | 0.710277 | 0.707012 |
| 0.704407 | 0.704713 | 0.712607 | 0.715542 | 0.712163 | 0.708071 | 0.703202 | 0.712587 | 0.707683 |
| 0.710449 | 0.708023 | 0.718717 | 0.720147 | 0.714813 | 0.713117 | 0.712936 | 0.718052 | 0.714854 |
| 0.702466 | 0.702779 | 0.712945 | 0.71125 | 0.70843 | 0.706889 | 0.704308 | 0.711702 | 0.704326 |
| 0.705094 | 0.708781 | 0.714019 | 0.716516 | 0.709152 | 0.710039 | 0.707625 | 0.715466 | 0.708942 |
| 0.704761 | 0.709175 | 0.714553 | 0.713758 | 0.713031 | 0.708471 | 0.709806 | 0.713811 | 0.708316 |
| 0.705542 | 0.70494 | 0.711904 | 0.712515 | 0.712986 | 0.710449 | 0.705221 | 0.715123 | 0.708337 |
| 0.702534 | 0.709948 | 0.713432 | 0.712217 | 0.713601 | 0.711251 | 0.705735 | 0.717648 | 0.708232 |
| 0.704052 | 0.705427 | 0.711892 | 0.708459 | 0.706983 | 0.706681 | 0.701264 | 0.710193 | 0.703661 |
| 0.706628 | 0.704804 | 0.71232 | 0.717063 | 0.711006 | 0.70999 | 0.706904 | 0.714592 | 0.712068 |
| 0.70396 | 0.706767 | 0.711276 | 0.71543 | 0.708003 | 0.711373 | 0.704772 | 0.712344 | 0.709213 |
| 0.702691 | 0.700229 | 0.708962 | 0.708787 | 0.708988 | 0.703833 | 0.701239 | 0.712344 | 0.707692 |
| 0.703351 | 0.705085 | 0.71194 | 0.711625 | 0.709962 | 0.707801 | 0.702453 | 0.711302 | 0.710189 |
| 0.695528 | 0.701738 | 0.707572 | 0.708489 | 0.706764 | 0.704331 | 0.698879 | 0.709144 | 0.706238 |
| 0.684089 | 0.681599 | 0.687948 | 0.687157 | 0.686582 | 0.685249 | 0.679245 | 0.68894 | 0.688658 |
| 0.684645 | 0.683518 | 0.689458 | 0.691979 | 0.688042 | 0.688135 | 0.6847 | 0.693116 | 0.688255 |
| 0.676708 | 0.676825 | 0.678186 | 0.683345 | 0.685459 | 0.682476 | 0.680286 | 0.685177 | 0.679277 |
| 0.673965 | 0.677256 | 0.681491 | 0.680701 | 0.682006 | 0.679314 | 0.675682 | 0.681066 | 0.679814 |
| 0.665025 | 0.666587 | 0.666863 | 0.668704 | 0.67102 | 0.670389 | 0.664098 | 0.669766 | 0.666227 |
| 0.668495 | 0.667646 | 0.673133 | 0.669259 | 0.666747 | 0.674635 | 0.667077 | 0.667775 | 0.672463 |
| 0.661953 | 0.660826 | 0.670493 | 0.672435 | 0.669382 | 0.670794 | 0.664044 | 0.665681 | 0.667823 |
| 0.664973 | 0.663957 | 0.677894 | 0.669051 | 0.670644 | 0.666363 | 0.659468 | 0.668485 | 0.664817 |
| 0.665732 | 0.66491 | 0.67056 | 0.670877 | 0.667042 | 0.667722 | 0.667255 | 0.671029 | 0.666751 |
| 0.67037 | 0.673766 | 0.67504 | 0.679597 | 0.677359 | 0.675403 | 0.669666 | 0.677346 | 0.675567 |

| | | | | | | | | |
|---|---|---|---|---|---|---|---|---|
| 0.672216 | 0.672584 | 0.673781 | 0.674916 | 0.675582 | 0.673888 | 0.667953 | 0.673779 | 0.673695 |
| 0.666402 | 0.666272 | 0.678226 | 0.670049 | 0.677635 | 0.672592 | 0.66579 | 0.674391 | 0.671654 |
| 0.67328 | 0.666712 | 0.673667 | 0.674254 | 0.668231 | 0.67146 | 0.66987 | 0.677468 | 0.672915 |
| 0.662287 | 0.658032 | 0.664894 | 0.663361 | 0.66538 | 0.667654 | 0.654958 | 0.66389 | 0.665192 |
| 0.665619 | 0.661644 | 0.667012 | 0.672095 | 0.666839 | 0.671803 | 0.667318 | 0.669912 | 0.669517 |
| 0.667273 | 0.667832 | 0.673098 | 0.66685 | 0.665704 | 0.669244 | 0.663993 | 0.668345 | 0.665229 |
| 0.666618 | 0.663481 | 0.666667 | 0.671083 | 0.662283 | 0.671681 | 0.664855 | 0.669987 | 0.669258 |
| 0.66547 | 0.658599 | 0.669445 | 0.663261 | 0.668217 | 0.665332 | 0.666336 | 0.661825 | 0.664305 |
| 0.663012 | 0.664181 | 0.668197 | 0.666765 | 0.666225 | 0.666467 | 0.66031 | 0.662277 | 0.664237 |
| 0.662772 | 0.657402 | 0.670304 | 0.672709 | 0.669062 | 0.668448 | 0.662277 | 0.66503 | 0.669647 |
| 0.657872 | 0.658055 | 0.668417 | 0.663812 | 0.653436 | 0.666268 | 0.663561 | 0.66336 | 0.663455 |
| 0.692467 | 0.693639 | 0.701499 | 0.701902 | 0.699556 | 0.698926 | 0.693237 | 0.701822 | 0.693652 |
| 0.702507 | 0.707841 | 0.710632 | 0.708808 | 0.706245 | 0.70581 | 0.703517 | 0.70837 | 0.70632 |
| 0.684291 | 0.681305 | 0.690607 | 0.687721 | 0.680291 | 0.691976 | 0.685467 | 0.691236 | 0.686806 |
| 0.710049 | 0.710093 | 0.713393 | 0.716194 | 0.718165 | 0.715491 | 0.709946 | 0.721202 | 0.7113 |
| 0.701721 | 0.701603 | 0.707533 | 0.705604 | 0.705418 | 0.704655 | 0.701807 | 0.709172 | 0.700571 |
| 0.709821 | 0.709389 | 0.716244 | 0.716871 | 0.713324 | 0.712902 | 0.708292 | 0.716775 | 0.710854 |
| 0.712954 | 0.714097 | 0.719153 | 0.717003 | 0.719354 | 0.715751 | 0.709823 | 0.722651 | 0.713127 |
| 0.692181 | 0.695719 | 0.699174 | 0.699564 | 0.699801 | 0.695086 | 0.692913 | 0.699183 | 0.690521 |
| 0.717195 | 0.716747 | 0.719314 | 0.723842 | 0.719267 | 0.717768 | 0.712375 | 0.722879 | 0.716973 |
| 0.688925 | 0.682847 | 0.695467 | 0.695857 | 0.691283 | 0.694527 | 0.686585 | 0.695935 | 0.692091 |
| 0.709677 | 0.713279 | 0.718035 | 0.718011 | 0.716309 | 0.718928 | 0.714214 | 0.717878 | 0.712561 |
| 0.715544 | 0.718634 | 0.721603 | 0.721768 | 0.719717 | 0.716713 | 0.715434 | 0.723366 | 0.720837 |
| 0.683708 | 0.681042 | 0.687368 | 0.689687 | 0.685213 | 0.689363 | 0.680572 | 0.687937 | 0.686228 |
| 0.683541 | 0.686915 | 0.686988 | 0.688953 | 0.689313 | 0.685383 | 0.683599 | 0.687018 | 0.686651 |
| 0.691869 | 0.691339 | 0.695435 | 0.691369 | 0.690066 | 0.691909 | 0.689447 | 0.688173 | 0.694409 |
| 0.70649 | 0.70005 | 0.71008 | 0.709754 | 0.706942 | 0.707218 | 0.704657 | 0.710893 | 0.704062 |
| 0.714755 | 0.716913 | 0.722558 | 0.724207 | 0.716602 | 0.718589 | 0.71292 | 0.725167 | 0.716531 |
| 0.680973 | 0.682647 | 0.689202 | 0.690077 | 0.691262 | 0.688104 | 0.680741 | 0.690788 | 0.683687 |
| 0.71823 | 0.721259 | 0.726985 | 0.72497 | 0.72365 | 0.719609 | 0.719459 | 0.725882 | 0.721508 |
| 0.718693 | 0.723981 | 0.725688 | 0.726613 | 0.72606 | 0.723116 | 0.724614 | 0.726922 | 0.723289 |
| 0.696601 | 0.69678 | 0.698549 | 0.698928 | 0.69938 | 0.697258 | 0.691457 | 0.699063 | 0.697765 |
| 0.695012 | 0.694308 | 0.700588 | 0.698989 | 0.699832 | 0.696773 | 0.693861 | 0.702385 | 0.695248 |
| 0.721072 | 0.716867 | 0.728581 | 0.731134 | 0.722029 | 0.723504 | 0.718252 | 0.726075 | 0.723564 |
| 0.721143 | 0.722509 | 0.724443 | 0.72276 | 0.723435 | 0.7181 | 0.715422 | 0.723684 | 0.719215 |
| 0.713139 | 0.719115 | 0.723713 | 0.724099 | 0.724059 | 0.72404 | 0.7174 | 0.72279 | 0.722297 |
| 0.675385 | 0.678386 | 0.675172 | 0.681124 | 0.676318 | 0.678071 | 0.674443 | 0.674729 | 0.676704 |
| 0.717979 | 0.71902 | 0.718988 | 0.722427 | 0.7225 | 0.719262 | 0.718979 | 0.724648 | 0.719064 |
| 0.67057 | 0.672318 | 0.67576 | 0.67285 | 0.677645 | 0.676322 | 0.669223 | 0.67724 | 0.673533 |
| 0.665309 | 0.668425 | 0.671366 | 0.670691 | 0.668686 | 0.6673 | 0.667353 | 0.671432 | 0.665615 |
| 0.714393 | 0.716209 | 0.719156 | 0.722351 | 0.720051 | 0.722508 | 0.715725 | 0.724001 | 0.716155 |
| 0.666427 | 0.66765 | 0.670771 | 0.669649 | 0.66682 | 0.669046 | 0.665981 | 0.670893 | 0.664695 |
| 0.65743 | 0.661941 | 0.661331 | 0.66137 | 0.660396 | 0.660878 | 0.660597 | 0.663327 | 0.656683 |
| 0.655244 | 0.654059 | 0.660055 | 0.659886 | 0.657317 | 0.657407 | 0.653546 | 0.657258 | 0.656607 |
| 0.724391 | 0.7244 | 0.727813 | 0.732623 | 0.728396 | 0.726992 | 0.720619 | 0.730134 | 0.72703 |
| 0.658284 | 0.660175 | 0.662147 | 0.666161 | 0.662996 | 0.662611 | 0.658771 | 0.662502 | 0.658026 |
| 0.716647 | 0.716065 | 0.721484 | 0.726191 | 0.720893 | 0.721902 | 0.714405 | 0.720915 | 0.720164 |

| | | | | | | | | |
|---|---|---|---|---|---|---|---|---|
| 0.725819 | 0.725698 | 0.729706 | 0.736129 | 0.732019 | 0.728443 | 0.725912 | 0.733648 | 0.731728 |
| 0.720101 | 0.723338 | 0.726747 | 0.72865 | 0.726335 | 0.727182 | 0.720522 | 0.72731 | 0.724253 |
| 0.735734 | 0.738837 | 0.745424 | 0.746363 | 0.751802 | 0.741498 | 0.744113 | 0.748588 | 0.741127 |
| 0.646339 | 0.650167 | 0.651754 | 0.652245 | 0.652034 | 0.650802 | 0.648457 | 0.652259 | 0.648613 |
| 0.648015 | 0.649629 | 0.65258 | 0.653386 | 0.652437 | 0.652625 | 0.652412 | 0.654084 | 0.647879 |
| 0.723797 | 0.726521 | 0.732869 | 0.734013 | 0.728883 | 0.735237 | 0.726422 | 0.737213 | 0.726946 |
| 0.650125 | 0.654559 | 0.657564 | 0.654526 | 0.655916 | 0.652481 | 0.651897 | 0.65499 | 0.651561 |
| 0.725677 | 0.72897 | 0.738471 | 0.74024 | 0.735801 | 0.733226 | 0.729386 | 0.735594 | 0.732575 |
| 0.728479 | 0.729019 | 0.731999 | 0.732775 | 0.734342 | 0.729614 | 0.723765 | 0.731331 | 0.7311 |
| 0.728745 | 0.732234 | 0.736328 | 0.737746 | 0.735005 | 0.729546 | 0.728842 | 0.736201 | 0.731655 |
| 0.727785 | 0.727528 | 0.731953 | 0.733937 | 0.727681 | 0.731779 | 0.727649 | 0.733333 | 0.730104 |
| 0.729167 | 0.736111 | 0.726667 | 0.701299 | 0.724832 | 0.731034 | 0.753425 | 0.729167 | 0.677852 |
| 0.724138 | 0.730285 | 0.745625 | 0.727656 | 0.729801 | 0.730049 | 0.72581 | 0.730463 | 0.729268 |
| 0.727521 | 0.725528 | 0.731108 | 0.727001 | 0.727832 | 0.728489 | 0.724361 | 0.72794 | 0.725282 |
| 0.739265 | 0.732012 | 0.740301 | 0.73592 | 0.734539 | 0.736286 | 0.725809 | 0.726814 | 0.735982 |
| 0.728558 | 0.734288 | 0.732503 | 0.735938 | 0.736864 | 0.732199 | 0.725156 | 0.728958 | 0.736345 |
| 0.588744 | 0.58928 | 0.590978 | 0.587975 | 0.595002 | 0.590643 | 0.590849 | 0.587694 | 0.589419 |
| 0.738431 | 0.73531 | 0.744012 | 0.736215 | 0.736233 | 0.739241 | 0.736592 | 0.736 | 0.737149 |
| 0.732036 | 0.731719 | 0.738228 | 0.73846 | 0.735903 | 0.735072 | 0.728104 | 0.738745 | 0.736167 |
| 0.731775 | 0.730132 | 0.734422 | 0.741469 | 0.736001 | 0.738304 | 0.728159 | 0.741343 | 0.733173 |
| 0.733833 | 0.735053 | 0.741148 | 0.741182 | 0.736161 | 0.738582 | 0.731941 | 0.741027 | 0.737505 |
| 0.591769 | 0.594421 | 0.595121 | 0.596333 | 0.596237 | 0.594659 | 0.591812 | 0.596106 | 0.590139 |
| 0.575076 | 0.580193 | 0.581136 | 0.58458 | 0.582982 | 0.581966 | 0.575916 | 0.580306 | 0.575 |
| 0.751832 | 0.753195 | 0.764153 | 0.764869 | 0.765038 | 0.761752 | 0.757794 | 0.766409 | 0.756845 |
| 0.755077 | 0.755082 | 0.76298 | 0.762999 | 0.764345 | 0.763576 | 0.759223 | 0.768342 | 0.758694 |
| 0.749042 | 0.747476 | 0.756352 | 0.756469 | 0.758575 | 0.756537 | 0.750945 | 0.757186 | 0.752386 |
| 0.755358 | 0.755524 | 0.761543 | 0.764054 | 0.76824 | 0.760685 | 0.760459 | 0.764019 | 0.759829 |
| 0.757102 | 0.755454 | 0.761881 | 0.767581 | 0.768797 | 0.762455 | 0.757431 | 0.766614 | 0.761823 |
| 0.759519 | 0.762775 | 0.769087 | 0.767654 | 0.769574 | 0.765475 | 0.763902 | 0.770667 | 0.762498 |
| 0.759549 | 0.759484 | 0.764898 | 0.770377 | 0.764838 | 0.766096 | 0.759788 | 0.768702 | 0.761004 |
| 0.758031 | 0.757573 | 0.761705 | 0.759421 | 0.760349 | 0.757983 | 0.756925 | 0.764316 | 0.76594 |
| 0.762719 | 0.768405 | 0.771562 | 0.773257 | 0.77201 | 0.767499 | 0.771353 | 0.771609 | 0.76973 |
| 0.756604 | 0.755918 | 0.76308 | 0.763955 | 0.76477 | 0.76316 | 0.758466 | 0.768073 | 0.757675 |
| 0.758031 | 0.7626 | 0.762702 | 0.766699 | 0.767388 | 0.763534 | 0.76387 | 0.767839 | 0.763348 |
| 0.772174 | 0.77397 | 0.774907 | 0.771246 | 0.774925 | 0.769237 | 0.770957 | 0.776631 | 0.766624 |
| 0.753621 | 0.760335 | 0.76903 | 0.763953 | 0.767917 | 0.760316 | 0.760823 | 0.763906 | 0.760789 |
| 0.762269 | 0.762753 | 0.768538 | 0.768482 | 0.769701 | 0.766287 | 0.76422 | 0.769224 | 0.765895 |
| 0.759664 | 0.759692 | 0.768709 | 0.772923 | 0.770482 | 0.765164 | 0.766039 | 0.7673 | 0.761295 |
| 0.753133 | 0.755854 | 0.762101 | 0.766323 | 0.771132 | 0.760446 | 0.759339 | 0.768461 | 0.761457 |
| 0.755144 | 0.758364 | 0.763579 | 0.763106 | 0.767505 | 0.763872 | 0.758221 | 0.761826 | 0.759908 |
| 0.725735 | 0.726228 | 0.727867 | 0.729426 | 0.729141 | 0.71636 | 0.722437 | 0.72899 | 0.726146 |
| 0.733629 | 0.724816 | 0.74153 | 0.741098 | 0.734485 | 0.734049 | 0.719579 | 0.728637 | 0.739022 |
| 0.760193 | 0.764711 | 0.769559 | 0.765868 | 0.767232 | 0.767649 | 0.762563 | 0.765895 | 0.76141 |
| 0.757387 | 0.76123 | 0.770477 | 0.773546 | 0.770471 | 0.767669 | 0.76797 | 0.773526 | 0.765114 |
| 0.715602 | 0.717709 | 0.728708 | 0.721942 | 0.723561 | 0.71993 | 0.718939 | 0.721654 | 0.720545 |
| 0.757335 | 0.761969 | 0.771222 | 0.7715 | 0.768584 | 0.76739 | 0.761746 | 0.772563 | 0.75999 |
| 0.752023 | 0.757166 | 0.767859 | 0.761783 | 0.762469 | 0.758723 | 0.75817 | 0.763827 | 0.753973 |

| | | | | | | | | |
|---|---|---|---|---|---|---|---|---|
| 0.755234 | 0.758846 | 0.770551 | 0.767122 | 0.765107 | 0.764766 | 0.762023 | 0.767043 | 0.76199 |
| 0.760306 | 0.761302 | 0.771338 | 0.771688 | 0.76817 | 0.768351 | 0.763308 | 0.769263 | 0.766644 |
| 0.755131 | 0.756162 | 0.768465 | 0.765368 | 0.765576 | 0.762416 | 0.755811 | 0.763655 | 0.759826 |
| 0.761917 | 0.767638 | 0.766689 | 0.77304 | 0.768796 | 0.764731 | 0.764464 | 0.767667 | 0.764706 |
| 0.759226 | 0.76119 | 0.773855 | 0.775928 | 0.770984 | 0.766378 | 0.765048 | 0.77102 | 0.767055 |
| 0.756106 | 0.758757 | 0.771134 | 0.769327 | 0.764883 | 0.758917 | 0.756984 | 0.766445 | 0.765857 |
| 0.755997 | 0.758552 | 0.76448 | 0.774081 | 0.764643 | 0.762716 | 0.765874 | 0.770868 | 0.763903 |
| 0.743935 | 0.749303 | 0.761925 | 0.761315 | 0.763081 | 0.757436 | 0.75147 | 0.761397 | 0.753556 |
| 0.757507 | 0.759852 | 0.765639 | 0.770572 | 0.763259 | 0.76014 | 0.761084 | 0.76474 | 0.759791 |
| 0.754016 | 0.756223 | 0.758774 | 0.763797 | 0.760526 | 0.757183 | 0.75543 | 0.761642 | 0.7552 |
| 0.756882 | 0.762863 | 0.769752 | 0.769415 | 0.768104 | 0.76808 | 0.767366 | 0.767107 | 0.763082 |
| 0.75397 | 0.755862 | 0.768042 | 0.76554 | 0.762222 | 0.765142 | 0.761452 | 0.766615 | 0.761115 |
| 0.74769 | 0.75174 | 0.763147 | 0.75659 | 0.757474 | 0.75634 | 0.752856 | 0.760064 | 0.75332 |
| 0.745612 | 0.748933 | 0.759459 | 0.757821 | 0.75762 | 0.755469 | 0.753001 | 0.763539 | 0.753889 |
| 0.760089 | 0.767309 | 0.772727 | 0.77224 | 0.766691 | 0.765092 | 0.762853 | 0.76918 | 0.763178 |
| 0.76 | 0.762136 | 0.767779 | 0.77097 | 0.765855 | 0.765008 | 0.760498 | 0.769976 | 0.765861 |
| 0.754995 | 0.759522 | 0.765098 | 0.770247 | 0.764408 | 0.766814 | 0.757961 | 0.76675 | 0.762551 |
| 0.753338 | 0.753186 | 0.763169 | 0.7643 | 0.761881 | 0.766658 | 0.761156 | 0.763138 | 0.756706 |
| 0.756897 | 0.754167 | 0.765696 | 0.767563 | 0.771048 | 0.765693 | 0.759624 | 0.769083 | 0.760347 |
| 0.76101 | 0.762013 | 0.766958 | 0.76378 | 0.770434 | 0.766954 | 0.75881 | 0.768545 | 0.762199 |
| 0.748637 | 0.750106 | 0.758709 | 0.763098 | 0.762183 | 0.76116 | 0.753128 | 0.762689 | 0.754117 |
| 0.75415 | 0.75102 | 0.760997 | 0.761267 | 0.766026 | 0.761353 | 0.753534 | 0.764741 | 0.758091 |
| 0.761113 | 0.761512 | 0.768495 | 0.766283 | 0.771452 | 0.76427 | 0.763557 | 0.769308 | 0.76397 |
| 0.754189 | 0.757289 | 0.766935 | 0.767338 | 0.769308 | 0.767989 | 0.760144 | 0.767249 | 0.760522 |
| 0.757882 | 0.756553 | 0.761337 | 0.765423 | 0.769549 | 0.768613 | 0.760821 | 0.770486 | 0.7616 |
| 0.756663 | 0.757106 | 0.761817 | 0.762891 | 0.769889 | 0.758841 | 0.757224 | 0.765555 | 0.76084 |
| 0.747686 | 0.75155 | 0.759338 | 0.759578 | 0.763235 | 0.761905 | 0.753959 | 0.756542 | 0.756937 |
| 0.752153 | 0.751401 | 0.762214 | 0.763327 | 0.763418 | 0.762697 | 0.758482 | 0.764049 | 0.759777 |
| 0.751771 | 0.751797 | 0.76208 | 0.76454 | 0.769872 | 0.761215 | 0.758103 | 0.76543 | 0.761819 |
| 0.750496 | 0.754405 | 0.763233 | 0.767026 | 0.770064 | 0.76618 | 0.758029 | 0.764593 | 0.762584 |
| 0.759078 | 0.758734 | 0.766457 | 0.770399 | 0.767277 | 0.766649 | 0.762543 | 0.770844 | 0.762908 |
| 0.753986 | 0.756775 | 0.764054 | 0.765696 | 0.765549 | 0.769197 | 0.759952 | 0.76652 | 0.763187 |
| 0.759794 | 0.762408 | 0.770976 | 0.76992 | 0.773596 | 0.7727 | 0.764611 | 0.771048 | 0.763633 |
| 0.757516 | 0.760111 | 0.769501 | 0.772042 | 0.772788 | 0.771537 | 0.763809 | 0.768896 | 0.768664 |
| 0.750975 | 0.753314 | 0.758277 | 0.763817 | 0.764083 | 0.76213 | 0.758816 | 0.764528 | 0.759701 |
| 0.758439 | 0.758465 | 0.761004 | 0.768006 | 0.765411 | 0.765813 | 0.761024 | 0.768407 | 0.763516 |
| 0.748742 | 0.751179 | 0.759417 | 0.767827 | 0.76574 | 0.766027 | 0.760954 | 0.763187 | 0.762608 |
| 0.751763 | 0.755829 | 0.764358 | 0.761022 | 0.763778 | 0.758668 | 0.758258 | 0.764865 | 0.761588 |
| 0.747522 | 0.750766 | 0.755523 | 0.759251 | 0.760679 | 0.759745 | 0.752821 | 0.756883 | 0.751539 |
| 0.748 | 0.748101 | 0.75489 | 0.758522 | 0.756593 | 0.753789 | 0.753564 | 0.760685 | 0.755875 |
| 0.747657 | 0.744887 | 0.751786 | 0.752749 | 0.755548 | 0.752228 | 0.746311 | 0.757123 | 0.749076 |
| 0.754703 | 0.749135 | 0.756776 | 0.755986 | 0.760011 | 0.75589 | 0.748512 | 0.759676 | 0.752958 |
| 0.743859 | 0.742934 | 0.754412 | 0.757233 | 0.756383 | 0.757056 | 0.7476 | 0.755431 | 0.752153 |
| 0.743443 | 0.740231 | 0.752748 | 0.75641 | 0.755499 | 0.75304 | 0.748209 | 0.75695 | 0.753677 |
| 0.751577 | 0.753835 | 0.756464 | 0.760014 | 0.758624 | 0.760054 | 0.755038 | 0.761354 | 0.756723 |
| 0.742467 | 0.744535 | 0.751147 | 0.756269 | 0.751989 | 0.756188 | 0.752761 | 0.752248 | 0.748276 |

| | | | | | | | | |
|---|---|---|---|---|---|---|---|---|
| 0.759041 | 0.756645 | 0.766737 | 0.764716 | 0.766039 | 0.758993 | 0.757389 | 0.765292 | 0.761267 |
| 0.768278 | 0.76887 | 0.772457 | 0.77609 | 0.770134 | 0.776702 | 0.766843 | 0.770014 | 0.773127 |
| 0.756057 | 0.755781 | 0.763006 | 0.766655 | 0.767522 | 0.767203 | 0.763526 | 0.765109 | 0.762087 |
| 0.763085 | 0.768837 | 0.766756 | 0.769611 | 0.76774 | 0.770027 | 0.768632 | 0.767957 | 0.768429 |
| 0.769204 | 0.774218 | 0.78044 | 0.779675 | 0.783852 | 0.773569 | 0.773843 | 0.776324 | 0.778363 |
| 0.769004 | 0.767812 | 0.772693 | 0.772276 | 0.773037 | 0.769917 | 0.765103 | 0.768705 | 0.770683 |
| 0.76047 | 0.762113 | 0.766397 | 0.761303 | 0.768437 | 0.764727 | 0.766442 | 0.767982 | 0.760557 |
| 0.764008 | 0.766577 | 0.773593 | 0.766747 | 0.768286 | 0.768025 | 0.763926 | 0.767504 | 0.767788 |
| 0.760767 | 0.758112 | 0.762395 | 0.76371 | 0.768454 | 0.76344 | 0.7651 | 0.765106 | 0.761945 |
| 0.754694 | 0.760815 | 0.765002 | 0.767504 | 0.766811 | 0.762466 | 0.761741 | 0.769257 | 0.766117 |
| 0.763638 | 0.768447 | 0.771557 | 0.773997 | 0.773325 | 0.769736 | 0.765366 | 0.771571 | 0.769637 |
| 0.764196 | 0.764079 | 0.76743 | 0.774301 | 0.773246 | 0.768651 | 0.765756 | 0.770338 | 0.767672 |
| 0.766387 | 0.765997 | 0.776396 | 0.779935 | 0.781305 | 0.783074 | 0.774604 | 0.782766 | 0.776975 |
| 0.764297 | 0.760715 | 0.770605 | 0.776484 | 0.777267 | 0.775159 | 0.770977 | 0.783079 | 0.76811 |
| 0.766782 | 0.766244 | 0.780792 | 0.782277 | 0.780686 | 0.781143 | 0.775594 | 0.782587 | 0.775043 |
| 0.767642 | 0.768797 | 0.777834 | 0.773406 | 0.777001 | 0.780294 | 0.781304 | 0.780422 | 0.773274 |
| 0.772425 | 0.769828 | 0.781627 | 0.790137 | 0.785105 | 0.782907 | 0.775737 | 0.787859 | 0.782992 |
| 0.761289 | 0.766534 | 0.778648 | 0.784359 | 0.78078 | 0.779066 | 0.773261 | 0.781933 | 0.781634 |
| 0.762658 | 0.775447 | 0.780725 | 0.778728 | 0.779517 | 0.778638 | 0.779119 | 0.780807 | 0.772712 |
| 0.765935 | 0.769076 | 0.775331 | 0.778394 | 0.781393 | 0.773015 | 0.786059 | 0.783303 | 0.77244 |
| 0.760313 | 0.760923 | 0.773195 | 0.780081 | 0.773191 | 0.773557 | 0.769178 | 0.77774 | 0.767588 |
| 0.761818 | 0.761499 | 0.769569 | 0.78088 | 0.781313 | 0.776487 | 0.77544 | 0.778022 | 0.770689 |
| 0.763394 | 0.765173 | 0.778969 | 0.779553 | 0.783525 | 0.777911 | 0.778674 | 0.780206 | 0.780769 |
| 0.776805 | 0.782213 | 0.782332 | 0.779292 | 0.779409 | 0.774014 | 0.776664 | 0.778251 | 0.77237 |
| 0.772485 | 0.776693 | 0.783211 | 0.78575 | 0.781245 | 0.778052 | 0.774372 | 0.778829 | 0.776027 |
| 0.773454 | 0.773233 | 0.782473 | 0.777491 | 0.779619 | 0.779551 | 0.768377 | 0.780793 | 0.773692 |
| 0.777267 | 0.7753 | 0.780788 | 0.778154 | 0.782627 | 0.771658 | 0.769973 | 0.774301 | 0.772413 |
| 0.765882 | 0.770992 | 0.779609 | 0.777383 | 0.784928 | 0.770823 | 0.77071 | 0.777026 | 0.774812 |
| 0.775125 | 0.777451 | 0.789571 | 0.785958 | 0.782182 | 0.776391 | 0.776512 | 0.783969 | 0.776446 |
| 0.777559 | 0.781769 | 0.786659 | 0.77906 | 0.77896 | 0.772315 | 0.77089 | 0.778354 | 0.776926 |
| 0.768579 | 0.771421 | 0.77867 | 0.775044 | 0.773913 | 0.770585 | 0.765104 | 0.771822 | 0.767234 |
| 0.777143 | 0.77956 | 0.785186 | 0.772451 | 0.782426 | 0.774448 | 0.773242 | 0.780945 | 0.776853 |
| 0.774808 | 0.774183 | 0.787062 | 0.779206 | 0.780157 | 0.775642 | 0.77426 | 0.784648 | 0.774618 |
| 0.768084 | 0.774401 | 0.780437 | 0.775831 | 0.782549 | 0.771618 | 0.77319 | 0.776238 | 0.771998 |
| 0.771426 | 0.775411 | 0.784121 | 0.776466 | 0.772603 | 0.772234 | 0.772499 | 0.773622 | 0.769637 |
| 0.772901 | 0.770182 | 0.781785 | 0.779017 | 0.777933 | 0.772226 | 0.767224 | 0.774903 | 0.777548 |
| 0.775746 | 0.781438 | 0.782506 | 0.783174 | 0.781612 | 0.781261 | 0.779798 | 0.783388 | 0.779376 |
| 0.776339 | 0.774371 | 0.783329 | 0.778803 | 0.782091 | 0.774295 | 0.772858 | 0.775675 | 0.774825 |
| 0.773775 | 0.775205 | 0.782009 | 0.782259 | 0.779005 | 0.776015 | 0.772782 | 0.774096 | 0.773408 |
| 0.7729 | 0.77953 | 0.789011 | 0.778277 | 0.785305 | 0.777914 | 0.770579 | 0.780074 | 0.768923 |
| 0.773637 | 0.775055 | 0.790647 | 0.784568 | 0.783854 | 0.779993 | 0.775157 | 0.77907 | 0.772355 |
| 0.77863 | 0.780473 | 0.78627 | 0.780376 | 0.784176 | 0.774711 | 0.775852 | 0.779859 | 0.778266 |
| 0.783697 | 0.781608 | 0.787805 | 0.784559 | 0.787248 | 0.774138 | 0.772374 | 0.783219 | 0.783232 |
| 0.77025 | 0.775933 | 0.779253 | 0.777928 | 0.778287 | 0.771941 | 0.768996 | 0.774991 | 0.775028 |
| 0.769486 | 0.778147 | 0.785308 | 0.784149 | 0.782638 | 0.771438 | 0.775556 | 0.780288 | 0.775434 |
| 0.768048 | 0.774474 | 0.784056 | 0.784505 | 0.775773 | 0.772282 | 0.770737 | 0.771996 | 0.776308 |

| | | | | | | | | |
|---|---|---|---|---|---|---|---|---|
| 0.755108 | 0.754032 | 0.766056 | 0.763517 | 0.76734 | 0.762288 | 0.764181 | 0.768673 | 0.756438 |
| 0.749435 | 0.753613 | 0.759619 | 0.757946 | 0.750686 | 0.749829 | 0.74701 | 0.752199 | 0.753375 |
| 0.762955 | 0.769337 | 0.77398 | 0.77385 | 0.775334 | 0.768297 | 0.767578 | 0.773754 | 0.769522 |
| 0.763428 | 0.763827 | 0.777685 | 0.77953 | 0.778542 | 0.777825 | 0.772529 | 0.774623 | 0.77174 |
| 0.762794 | 0.764726 | 0.763728 | 0.773034 | 0.774702 | 0.769826 | 0.761812 | 0.76675 | 0.767897 |
| 0.764001 | 0.769047 | 0.768397 | 0.779443 | 0.781878 | 0.776955 | 0.767579 | 0.772593 | 0.776724 |
| 0.766507 | 0.762357 | 0.773374 | 0.767637 | 0.773271 | 0.773315 | 0.767718 | 0.771914 | 0.768055 |
| 0.76698 | 0.762799 | 0.771931 | 0.772898 | 0.768317 | 0.763486 | 0.766489 | 0.76706 | 0.767256 |
| 0.765199 | 0.768527 | 0.775483 | 0.775561 | 0.774073 | 0.772392 | 0.770559 | 0.775468 | 0.765939 |
| 0.763614 | 0.762081 | 0.774905 | 0.775583 | 0.776948 | 0.773492 | 0.767929 | 0.775684 | 0.769131 |
| 0.76436 | 0.767124 | 0.769905 | 0.77707 | 0.773235 | 0.767289 | 0.772416 | 0.774942 | 0.769205 |
| 0.765557 | 0.765625 | 0.775733 | 0.76785 | 0.773476 | 0.768892 | 0.766087 | 0.775706 | 0.767132 |
| 0.768629 | 0.766739 | 0.776255 | 0.783428 | 0.779711 | 0.777481 | 0.772263 | 0.782462 | 0.778603 |
| 0.766491 | 0.768011 | 0.771581 | 0.77299 | 0.769938 | 0.769737 | 0.770474 | 0.774385 | 0.765057 |
| 0.758242 | 0.764184 | 0.773859 | 0.772409 | 0.774511 | 0.771878 | 0.765504 | 0.769924 | 0.766321 |
| 0.76578 | 0.758618 | 0.774159 | 0.773919 | 0.769641 | 0.770169 | 0.771038 | 0.775744 | 0.763695 |
| 0.761847 | 0.75934 | 0.770015 | 0.771486 | 0.775257 | 0.772747 | 0.766876 | 0.772693 | 0.763705 |
| 0.762752 | 0.75688 | 0.767389 | 0.767136 | 0.768264 | 0.771364 | 0.767175 | 0.773567 | 0.763373 |
| 0.760896 | 0.764426 | 0.767634 | 0.769155 | 0.766883 | 0.763172 | 0.765814 | 0.765872 | 0.759147 |
| 0.759276 | 0.759497 | 0.766737 | 0.77686 | 0.772932 | 0.770563 | 0.768583 | 0.770118 | 0.771318 |
| 0.763248 | 0.764731 | 0.77113 | 0.775599 | 0.773056 | 0.770432 | 0.765639 | 0.770155 | 0.765628 |
| 0.75675 | 0.757329 | 0.75762 | 0.759439 | 0.75669 | 0.755107 | 0.753354 | 0.760481 | 0.753912 |
| 0.759533 | 0.764112 | 0.769779 | 0.768249 | 0.765719 | 0.765307 | 0.762299 | 0.769733 | 0.760864 |
| 0.764747 | 0.766588 | 0.774597 | 0.770485 | 0.769224 | 0.769583 | 0.765324 | 0.771253 | 0.767594 |
| 0.766664 | 0.76708 | 0.775133 | 0.774883 | 0.771156 | 0.770454 | 0.763043 | 0.772135 | 0.771905 |
| 0.763795 | 0.762282 | 0.769237 | 0.768441 | 0.765703 | 0.764127 | 0.760662 | 0.766412 | 0.765315 |
| 0.756913 | 0.757736 | 0.769558 | 0.767394 | 0.764847 | 0.764789 | 0.759679 | 0.769497 | 0.7587 |
| 0.759767 | 0.760708 | 0.772521 | 0.766148 | 0.771087 | 0.766056 | 0.761062 | 0.766374 | 0.765312 |
| 0.758552 | 0.758358 | 0.76979 | 0.763983 | 0.765005 | 0.761347 | 0.75366 | 0.765211 | 0.763533 |
| 0.691955 | 0.694898 | 0.700955 | 0.6982 | 0.695717 | 0.700685 | 0.690128 | 0.697375 | 0.697917 |
| 0.70313 | 0.702315 | 0.709122 | 0.712988 | 0.709515 | 0.705035 | 0.701471 | 0.710164 | 0.705853 |
| 0.702888 | 0.703224 | 0.710861 | 0.708185 | 0.705229 | 0.707489 | 0.702457 | 0.710375 | 0.704658 |
| 0.693633 | 0.693949 | 0.699095 | 0.698979 | 0.695849 | 0.698453 | 0.690688 | 0.699051 | 0.696663 |
| 0.7064 | 0.705963 | 0.712471 | 0.710912 | 0.70539 | 0.709669 | 0.702761 | 0.71206 | 0.705202 |
| 0.764506 | 0.762697 | 0.768179 | 0.76952 | 0.763995 | 0.76519 | 0.757276 | 0.768166 | 0.759831 |
| 0.764312 | 0.765333 | 0.769133 | 0.772391 | 0.769296 | 0.766041 | 0.759882 | 0.77272 | 0.762515 |
| 0.769549 | 0.769966 | 0.774642 | 0.770894 | 0.773606 | 0.763224 | 0.765547 | 0.774396 | 0.76493 |
| 0.770609 | 0.769758 | 0.77549 | 0.773634 | 0.77347 | 0.771315 | 0.766635 | 0.773698 | 0.764523 |
| 0.761581 | 0.760785 | 0.772388 | 0.770351 | 0.769489 | 0.764584 | 0.760286 | 0.77005 | 0.763445 |
| 0.764151 | 0.764487 | 0.770923 | 0.773273 | 0.768145 | 0.764209 | 0.76382 | 0.775398 | 0.763445 |
| 0.768082 | 0.766678 | 0.770372 | 0.779064 | 0.770709 | 0.773297 | 0.766296 | 0.772569 | 0.770535 |
| 0.7637 | 0.76208 | 0.768343 | 0.770212 | 0.763951 | 0.763036 | 0.759639 | 0.767507 | 0.758349 |
| 0.753952 | 0.756251 | 0.765632 | 0.764685 | 0.765916 | 0.762288 | 0.757031 | 0.763685 | 0.754291 |
| 0.760738 | 0.757748 | 0.762942 | 0.768155 | 0.769397 | 0.759691 | 0.761213 | 0.764791 | 0.756495 |
| 0.762353 | 0.763044 | 0.767526 | 0.764976 | 0.768475 | 0.762859 | 0.759586 | 0.769363 | 0.761022 |
| 0.766808 | 0.768646 | 0.770534 | 0.770201 | 0.770478 | 0.769808 | 0.766211 | 0.772654 | 0.763386 |

| | | | | | | | | |
|---|---|---|---|---|---|---|---|---|
| 0.764014 | 0.764923 | 0.77443 | 0.774558 | 0.775971 | 0.767826 | 0.766892 | 0.771924 | 0.764721 |
| 0.761983 | 0.761905 | 0.769133 | 0.770411 | 0.769829 | 0.768494 | 0.762586 | 0.76752 | 0.760226 |
| 0.764582 | 0.764881 | 0.773484 | 0.771995 | 0.771017 | 0.767286 | 0.769197 | 0.774734 | 0.764497 |
| 0.753806 | 0.75453 | 0.765688 | 0.770579 | 0.766327 | 0.762958 | 0.765504 | 0.769548 | 0.766126 |
| 0.760039 | 0.756775 | 0.773132 | 0.76529 | 0.76792 | 0.766816 | 0.761129 | 0.768001 | 0.756172 |
| 0.758283 | 0.76159 | 0.770748 | 0.764342 | 0.768309 | 0.765865 | 0.76119 | 0.767664 | 0.760929 |
| 0.76057 | 0.762591 | 0.774104 | 0.76523 | 0.766735 | 0.76437 | 0.764432 | 0.768748 | 0.761938 |
| 0.761444 | 0.764289 | 0.769899 | 0.768412 | 0.770906 | 0.764964 | 0.762255 | 0.765371 | 0.765543 |
| 0.756971 | 0.758742 | 0.768833 | 0.764721 | 0.764301 | 0.764375 | 0.766225 | 0.762113 | 0.762704 |
| 0.763165 | 0.762656 | 0.768662 | 0.762708 | 0.769489 | 0.767408 | 0.763386 | 0.767202 | 0.76077 |
| 0.760797 | 0.759833 | 0.771043 | 0.767089 | 0.771678 | 0.763907 | 0.763936 | 0.768386 | 0.763866 |
| 0.76371 | 0.764429 | 0.772003 | 0.767729 | 0.773315 | 0.766827 | 0.767729 | 0.771531 | 0.767489 |
| 0.754685 | 0.755427 | 0.763108 | 0.757479 | 0.763335 | 0.764424 | 0.761356 | 0.76541 | 0.757401 |
| 0.749175 | 0.755887 | 0.763891 | 0.754681 | 0.762515 | 0.759707 | 0.756501 | 0.76052 | 0.751987 |
| 0.765358 | 0.765034 | 0.773098 | 0.771111 | 0.770423 | 0.768711 | 0.767903 | 0.774994 | 0.765896 |
| 0.771704 | 0.77874 | 0.783873 | 0.781956 | 0.783686 | 0.78039 | 0.775538 | 0.782642 | 0.77641 |
| 0.767241 | 0.775764 | 0.781632 | 0.774055 | 0.783719 | 0.771533 | 0.773446 | 0.778677 | 0.773193 |
| 0.771578 | 0.766031 | 0.773485 | 0.78005 | 0.77775 | 0.77535 | 0.770618 | 0.77936 | 0.77162 |
| 0.765858 | 0.763348 | 0.770822 | 0.773888 | 0.775971 | 0.769867 | 0.767135 | 0.773873 | 0.770413 |
| 0.762185 | 0.762344 | 0.77939 | 0.769752 | 0.773049 | 0.768288 | 0.765174 | 0.777312 | 0.764503 |
| 0.759524 | 0.759902 | 0.767006 | 0.769569 | 0.768745 | 0.767137 | 0.762112 | 0.767982 | 0.757887 |
| 0.764283 | 0.770353 | 0.774323 | 0.771225 | 0.776118 | 0.766834 | 0.764854 | 0.773344 | 0.762871 |
| 0.771202 | 0.774179 | 0.782453 | 0.778782 | 0.784953 | 0.775847 | 0.770049 | 0.779307 | 0.773462 |
| 0.763774 | 0.760866 | 0.768098 | 0.767663 | 0.764784 | 0.764453 | 0.757603 | 0.76557 | 0.75965 |
| 0.768811 | 0.768773 | 0.774855 | 0.776574 | 0.777348 | 0.774188 | 0.768713 | 0.775951 | 0.770413 |
| 0.697217 | 0.700462 | 0.706008 | 0.704965 | 0.702617 | 0.702923 | 0.699397 | 0.70794 | 0.700754 |
| 0.773299 | 0.780728 | 0.794781 | 0.777262 | 0.780561 | 0.773034 | 0.769356 | 0.777871 | 0.771602 |
| 0.875331 | 0.886724 | 0.789665 | 0.776136 | 0.782983 | 0.773173 | 0.772575 | 0.780437 | 0.7775 |
| 0.779098 | 0.78355 | 0.787579 | 0.775835 | 0.78207 | 0.775798 | 0.779237 | 0.781197 | 0.771798 |
| 0.818842 | 0.813587 | 0.786887 | 0.77788 | 0.78023 | 0.773452 | 0.774029 | 0.785297 | 0.774684 |
| 0.769189 | 0.771781 | 0.788209 | 0.773017 | 0.773811 | 0.770102 | 0.76594 | 0.772927 | 0.77116 |
| 0.781431 | 0.779463 | 0.78607 | 0.778583 | 0.77699 | 0.774575 | 0.76936 | 0.776464 | 0.773228 |
| 0.772562 | 0.775813 | 0.78058 | 0.774913 | 0.77446 | 0.774044 | 0.774041 | 0.771541 | 0.770024 |
| 0.813335 | 0.78864 | 0.782985 | 0.776092 | 0.781094 | 0.770979 | 0.774431 | 0.78112 | 0.772911 |
| 1 | 0.83341 | 0.780323 | 0.770927 | 0.770489 | 0.769523 | 0.76605 | 0.777946 | 0.771361 |
| 0.83341 | 1 | 0.783683 | 0.773219 | 0.776764 | 0.769603 | 0.76717 | 0.77286 | 0.771637 |
| 0.780323 | 0.783683 | 1 | 0.778542 | 0.78314 | 0.774858 | 0.767219 | 0.778073 | 0.775175 |
| 0.770927 | 0.773219 | 0.778542 | 1 | 0.786131 | 0.790197 | 0.77376 | 0.790916 | 0.830009 |
| 0.770489 | 0.776764 | 0.78314 | 0.786131 | 1 | 0.7947 | 0.780012 | 0.78777 | 0.783022 |
| 0.769523 | 0.769603 | 0.774858 | 0.790197 | 0.7947 | 1 | 0.771025 | 0.780342 | 0.789635 |
| 0.76605 | 0.76717 | 0.767219 | 0.77376 | 0.780012 | 0.771025 | 1 | 0.81713 | 0.775454 |
| 0.777946 | 0.77286 | 0.778073 | 0.790916 | 0.78777 | 0.780342 | 0.81713 | 1 | 0.78702 |
| 0.771361 | 0.771637 | 0.775175 | 0.830009 | 0.783022 | 0.789635 | 0.775454 | 0.78702 | 1 |
| 0.758981 | 0.757328 | 0.764716 | 0.768099 | 0.76712 | 0.762926 | 0.759481 | 0.768115 | 0.76214 |
| 0.762237 | 0.765304 | 0.768881 | 0.776634 | 0.773146 | 0.767476 | 0.765268 | 0.778013 | 0.767446 |
| 0.762964 | 0.763587 | 0.770364 | 0.772077 | 0.77513 | 0.764352 | 0.766661 | 0.77236 | 0.768477 |

| | | | | | | | | |
|---|---|---|---|---|---|---|---|---|
| 0.763411 | 0.761486 | 0.76762 | 0.77246 | 0.767928 | 0.764686 | 0.760041 | 0.769558 | 0.763851 |
| 0.759307 | 0.762615 | 0.770023 | 0.774419 | 0.774424 | 0.769053 | 0.769574 | 0.774556 | 0.767068 |
| 0.761792 | 0.761041 | 0.769022 | 0.770252 | 0.764376 | 0.764747 | 0.755083 | 0.771876 | 0.761046 |
| 0.722751 | 0.726647 | 0.724299 | 0.73644 | 0.728222 | 0.735598 | 0.722861 | 0.726545 | 0.73353 |
| 0.756383 | 0.757171 | 0.771096 | 0.770651 | 0.766485 | 0.766484 | 0.765133 | 0.765279 | 0.765694 |
| 0.759979 | 0.760286 | 0.771691 | 0.766279 | 0.767277 | 0.762907 | 0.763907 | 0.767358 | 0.758278 |
| 0.756635 | 0.760289 | 0.772785 | 0.766617 | 0.766284 | 0.76346 | 0.762895 | 0.768115 | 0.765174 |
| 0.764395 | 0.761961 | 0.77099 | 0.763071 | 0.771201 | 0.76155 | 0.762372 | 0.768723 | 0.761641 |
| 0.768215 | 0.771303 | 0.773128 | 0.76654 | 0.776049 | 0.769647 | 0.764716 | 0.770999 | 0.765149 |
| 0.767088 | 0.766103 | 0.774068 | 0.76702 | 0.767712 | 0.769901 | 0.76466 | 0.776142 | 0.759887 |
| 0.76195 | 0.762883 | 0.773325 | 0.766791 | 0.76496 | 0.764923 | 0.765479 | 0.772108 | 0.758968 |
| 0.763901 | 0.766393 | 0.772616 | 0.770719 | 0.764907 | 0.765304 | 0.760856 | 0.769166 | 0.761835 |
| 0.760455 | 0.759174 | 0.770807 | 0.771902 | 0.767869 | 0.763903 | 0.765771 | 0.776834 | 0.765755 |
| 0.765027 | 0.767061 | 0.773365 | 0.771741 | 0.772874 | 0.769586 | 0.767886 | 0.769622 | 0.762351 |
| 0.76571 | 0.765741 | 0.771306 | 0.766474 | 0.770466 | 0.766484 | 0.767925 | 0.770884 | 0.767095 |
| 0.755598 | 0.754412 | 0.766905 | 0.768285 | 0.768558 | 0.76823 | 0.765445 | 0.772338 | 0.759846 |
| 0.767823 | 0.766917 | 0.775699 | 0.769508 | 0.772464 | 0.768333 | 0.769553 | 0.773668 | 0.765762 |
| 0.764854 | 0.765772 | 0.775113 | 0.773119 | 0.774325 | 0.769832 | 0.769626 | 0.774083 | 0.768559 |
| 0.764614 | 0.762709 | 0.765523 | 0.765635 | 0.767395 | 0.766421 | 0.768511 | 0.770463 | 0.758462 |
| 0.756979 | 0.757474 | 0.76735 | 0.771248 | 0.770088 | 0.766116 | 0.769052 | 0.770146 | 0.760892 |
| 0.771391 | 0.765745 | 0.772933 | 0.774476 | 0.767876 | 0.771084 | 0.769899 | 0.779574 | 0.769191 |
| 0.768147 | 0.763041 | 0.770983 | 0.772769 | 0.771659 | 0.766202 | 0.770507 | 0.773882 | 0.770055 |
| 0.674643 | 0.676764 | 0.678879 | 0.682655 | 0.682316 | 0.692006 | 0.673634 | 0.681462 | 0.685045 |
| 0.652988 | 0.648951 | 0.668261 | 0.660166 | 0.659107 | 0.658566 | 0.660269 | 0.667558 | 0.66186 |
| 0.677485 | 0.669305 | 0.674786 | 0.66977 | 0.6717 | 0.672707 | 0.669166 | 0.679034 | 0.670936 |
| 0.652588 | 0.649085 | 0.657721 | 0.654941 | 0.66363 | 0.662659 | 0.647039 | 0.657372 | 0.654145 |
| 0.734725 | 0.737395 | 0.74197 | 0.743182 | 0.736 | 0.733803 | 0.73766 | 0.74148 | 0.736705 |
| 0.720117 | 0.721315 | 0.736517 | 0.722594 | 0.728476 | 0.7257 | 0.726765 | 0.729978 | 0.718326 |
| 0.734011 | 0.725812 | 0.736679 | 0.731103 | 0.736823 | 0.7246 | 0.734002 | 0.727572 | 0.73664 |
| 0.738879 | 0.729979 | 0.743781 | 0.738931 | 0.75216 | 0.741302 | 0.733642 | 0.746548 | 0.744568 |
| 0.719606 | 0.713039 | 0.731656 | 0.724319 | 0.723288 | 0.724452 | 0.715763 | 0.724468 | 0.719339 |
| 0.714492 | 0.714245 | 0.716634 | 0.720193 | 0.719637 | 0.720266 | 0.711544 | 0.720023 | 0.716367 |
| 0.706477 | 0.70731 | 0.727382 | 0.718252 | 0.719094 | 0.722521 | 0.721448 | 0.719479 | 0.717385 |
| 0.656108 | 0.651829 | 0.660252 | 0.660205 | 0.653324 | 0.659066 | 0.650116 | 0.658594 | 0.663291 |
| 0.75721 | 0.758976 | 0.7622 | 0.75974 | 0.766347 | 0.766667 | 0.760733 | 0.767951 | 0.7564 |
| 0.738408 | 0.737195 | 0.746868 | 0.74953 | 0.740042 | 0.743376 | 0.746196 | 0.742453 | 0.74718 |
| 0.730859 | 0.735153 | 0.74274 | 0.74374 | 0.742904 | 0.735326 | 0.735101 | 0.734287 | 0.745001 |
| 0.722254 | 0.72454 | 0.731739 | 0.726126 | 0.727103 | 0.726651 | 0.721785 | 0.729436 | 0.731233 |
| 0.743043 | 0.739993 | 0.745352 | 0.733628 | 0.748411 | 0.744038 | 0.741989 | 0.744092 | 0.738194 |
| 0.732612 | 0.728864 | 0.738565 | 0.736414 | 0.735446 | 0.736268 | 0.729186 | 0.736365 | 0.736393 |
| 0.725255 | 0.721023 | 0.736317 | 0.73001 | 0.736131 | 0.734558 | 0.726192 | 0.730308 | 0.732007 |
| 0.731121 | 0.725738 | 0.740694 | 0.72953 | 0.737899 | 0.734173 | 0.731433 | 0.738941 | 0.725269 |
| 0.726201 | 0.727339 | 0.736861 | 0.732175 | 0.729514 | 0.72619 | 0.727918 | 0.734175 | 0.730713 |
| 0.728978 | 0.727345 | 0.733797 | 0.72645 | 0.729458 | 0.737427 | 0.726157 | 0.728407 | 0.727998 |
| 0.730021 | 0.722165 | 0.734161 | 0.730077 | 0.730819 | 0.728725 | 0.735522 | 0.73015 | 0.725019 |
| 0.721373 | 0.721643 | 0.718553 | 0.71438 | 0.72849 | 0.720514 | 0.71178 | 0.724786 | 0.723397 |

| | | | | | | | | |
|---|---|---|---|---|---|---|---|---|
| 0.751084 | 0.753824 | 0.756154 | 0.754625 | 0.755792 | 0.757342 | 0.749822 | 0.756261 | 0.750994 |
| 0.725809 | 0.73235 | 0.739328 | 0.732458 | 0.739285 | 0.735231 | 0.726511 | 0.734044 | 0.731397 |
| 0.72698 | 0.724408 | 0.735201 | 0.732124 | 0.726916 | 0.732565 | 0.728703 | 0.730093 | 0.727314 |
| 0.733386 | 0.732653 | 0.741561 | 0.730722 | 0.735248 | 0.730572 | 0.729726 | 0.728565 | 0.732804 |
| 0.731057 | 0.730864 | 0.738611 | 0.733238 | 0.734391 | 0.727666 | 0.736418 | 0.738822 | 0.73658 |
| 0.739086 | 0.725872 | 0.731677 | 0.72673 | 0.726841 | 0.71997 | 0.731242 | 0.732517 | 0.729382 |
| 0.724752 | 0.722562 | 0.723238 | 0.710488 | 0.729498 | 0.715898 | 0.722615 | 0.720247 | 0.719482 |
| 0.717241 | 0.723979 | 0.725719 | 0.721781 | 0.720467 | 0.716486 | 0.717854 | 0.714988 | 0.717137 |
| 0.726173 | 0.718803 | 0.729553 | 0.722946 | 0.726136 | 0.71754 | 0.716266 | 0.726984 | 0.729071 |
| 0.748204 | 0.754458 | 0.759844 | 0.757773 | 0.757956 | 0.757242 | 0.75891 | 0.761065 | 0.753572 |
| 0.741442 | 0.750564 | 0.753039 | 0.755686 | 0.752942 | 0.758876 | 0.759913 | 0.756135 | 0.748508 |
| 0.744438 | 0.745852 | 0.74578 | 0.746088 | 0.748245 | 0.745467 | 0.744558 | 0.750565 | 0.748555 |
| 0.737803 | 0.734214 | 0.74164 | 0.745663 | 0.741211 | 0.740888 | 0.735594 | 0.743984 | 0.742475 |
| 0.734109 | 0.740332 | 0.744649 | 0.748558 | 0.745209 | 0.744931 | 0.743871 | 0.748754 | 0.744596 |
| 0.736663 | 0.737713 | 0.744609 | 0.746591 | 0.739614 | 0.73803 | 0.743258 | 0.747074 | 0.743055 |

| plot 16 | | | | | | | plot 17 | |
|---|---|---|---|---|---|---|---|---|
| Wild_plot_ | Wild_plot_ | Wild_plot_ | Wild_plot_ | Wild_plot_ | Wild_plot_ | Wild_plot_ | Wild_plot_ | Wild_plot_ |
| 0.704183 | 0.707469 | 0.710992 | 0.70406 | 0.705538 | 0.699912 | 0.65652 | 0.701393 | 0.704867 |
| 0.711758 | 0.71495 | 0.711782 | 0.712744 | 0.712062 | 0.712271 | 0.67428 | 0.711277 | 0.709935 |
| 0.702094 | 0.705261 | 0.708221 | 0.703456 | 0.705061 | 0.704293 | 0.658511 | 0.702363 | 0.700815 |
| 0.7113 | 0.716074 | 0.713063 | 0.713351 | 0.71255 | 0.710265 | 0.669705 | 0.711304 | 0.710535 |
| 0.705326 | 0.707707 | 0.706992 | 0.706095 | 0.71145 | 0.703461 | 0.663212 | 0.704633 | 0.705327 |
| 0.714342 | 0.721574 | 0.720181 | 0.714365 | 0.717579 | 0.712699 | 0.673043 | 0.715334 | 0.715555 |
| 0.766176 | 0.768433 | 0.767834 | 0.766912 | 0.768679 | 0.766146 | 0.724534 | 0.766243 | 0.76472 |
| 0.761245 | 0.769713 | 0.766267 | 0.766585 | 0.765786 | 0.76185 | 0.718706 | 0.76671 | 0.767518 |
| 0.708492 | 0.711124 | 0.712701 | 0.709823 | 0.711238 | 0.706841 | 0.664348 | 0.708993 | 0.708483 |
| 0.704131 | 0.708124 | 0.710541 | 0.706291 | 0.707304 | 0.704053 | 0.656423 | 0.706229 | 0.706158 |
| 0.706918 | 0.708764 | 0.709474 | 0.707536 | 0.708191 | 0.704904 | 0.668635 | 0.707564 | 0.706847 |
| 0.699126 | 0.702914 | 0.701603 | 0.703014 | 0.701444 | 0.703158 | 0.659052 | 0.698587 | 0.699446 |
| 0.711813 | 0.712705 | 0.712543 | 0.711226 | 0.714683 | 0.710982 | 0.676419 | 0.707747 | 0.704962 |
| 0.716968 | 0.721767 | 0.720157 | 0.718126 | 0.723827 | 0.720673 | 0.686473 | 0.719346 | 0.716966 |
| 0.72921 | 0.734161 | 0.731378 | 0.729138 | 0.737301 | 0.731219 | 0.696422 | 0.730409 | 0.730609 |
| 0.71058 | 0.716482 | 0.716473 | 0.714444 | 0.717554 | 0.715724 | 0.680042 | 0.709286 | 0.712453 |
| 0.71001 | 0.716411 | 0.714206 | 0.713691 | 0.715316 | 0.713405 | 0.664921 | 0.710468 | 0.711708 |
| 0.732837 | 0.738549 | 0.741111 | 0.73665 | 0.743612 | 0.737915 | 0.696641 | 0.738254 | 0.737924 |
| 0.700856 | 0.70398 | 0.70428 | 0.703106 | 0.705593 | 0.702544 | 0.662583 | 0.70187 | 0.702912 |
| 0.709657 | 0.71393 | 0.714286 | 0.713185 | 0.712967 | 0.712973 | 0.667482 | 0.709997 | 0.710405 |
| 0.703304 | 0.706613 | 0.7078 | 0.708508 | 0.703683 | 0.704738 | 0.658766 | 0.705209 | 0.706326 |
| 0.709017 | 0.710044 | 0.709495 | 0.70551 | 0.70935 | 0.706175 | 0.660384 | 0.705984 | 0.710701 |
| 0.711799 | 0.716508 | 0.715712 | 0.711595 | 0.713788 | 0.709398 | 0.672963 | 0.711991 | 0.711251 |
| 0.708904 | 0.712468 | 0.712885 | 0.711465 | 0.712665 | 0.711823 | 0.673043 | 0.709458 | 0.709464 |
| 0.711528 | 0.714642 | 0.715158 | 0.711927 | 0.711906 | 0.711428 | 0.67507 | 0.711037 | 0.712867 |
| 0.713687 | 0.709427 | 0.710629 | 0.710636 | 0.707704 | 0.711134 | 0.66204 | 0.707469 | 0.708762 |
| 0.702281 | 0.703547 | 0.706782 | 0.704214 | 0.703967 | 0.7006 | 0.656153 | 0.702148 | 0.70406 |
| 0.711343 | 0.712009 | 0.711577 | 0.711025 | 0.71108 | 0.712865 | 0.672504 | 0.708396 | 0.709856 |
| 0.71256 | 0.71597 | 0.716218 | 0.711097 | 0.713479 | 0.709307 | 0.670744 | 0.713276 | 0.710415 |
| 0.70782 | 0.70836 | 0.712727 | 0.709128 | 0.710495 | 0.708578 | 0.678282 | 0.706723 | 0.704406 |
| 0.709888 | 0.709644 | 0.70884 | 0.706246 | 0.712352 | 0.708625 | 0.675157 | 0.704844 | 0.705409 |
| 0.707646 | 0.704612 | 0.705887 | 0.704039 | 0.707837 | 0.704061 | 0.65884 | 0.702395 | 0.703013 |
| 0.703783 | 0.710688 | 0.71084 | 0.706865 | 0.710111 | 0.706888 | 0.66368 | 0.705425 | 0.707894 |
| 0.709685 | 0.709444 | 0.712957 | 0.705401 | 0.710368 | 0.706211 | 0.654898 | 0.708389 | 0.708327 |
| 0.703258 | 0.704061 | 0.70366 | 0.702598 | 0.700868 | 0.704502 | 0.660417 | 0.701478 | 0.70613 |
| 0.708828 | 0.711874 | 0.710978 | 0.708499 | 0.71285 | 0.706409 | 0.664806 | 0.707969 | 0.709696 |
| 0.711673 | 0.717016 | 0.714988 | 0.711735 | 0.718594 | 0.710717 | 0.671384 | 0.711224 | 0.711642 |
| 0.71165 | 0.718968 | 0.717505 | 0.70936 | 0.71106 | 0.708962 | 0.669346 | 0.70837 | 0.709722 |
| 0.716099 | 0.714251 | 0.715606 | 0.714048 | 0.717141 | 0.715607 | 0.66852 | 0.713573 | 0.711931 |

| | | | | | | | | |
|---|---|---|---|---|---|---|---|---|
| 0.742463 | 0.746895 | 0.747371 | 0.744984 | 0.750862 | 0.745586 | 0.69972 | 0.745501 | 0.744464 |
| 0.699677 | 0.703624 | 0.700864 | 0.699119 | 0.701645 | 0.699107 | 0.655817 | 0.698443 | 0.700443 |
| 0.70098 | 0.705652 | 0.704349 | 0.703648 | 0.706717 | 0.703371 | 0.66655 | 0.703677 | 0.702492 |
| 0.712217 | 0.714992 | 0.716839 | 0.713509 | 0.714229 | 0.714788 | 0.664932 | 0.708768 | 0.711414 |
| 0.706287 | 0.709227 | 0.710018 | 0.70635 | 0.709813 | 0.704575 | 0.668772 | 0.705743 | 0.707251 |
| 0.708098 | 0.708494 | 0.710514 | 0.71109 | 0.712678 | 0.709828 | 0.66713 | 0.709079 | 0.7108 |
| 0.704411 | 0.704396 | 0.708015 | 0.703298 | 0.704549 | 0.702923 | 0.667821 | 0.704476 | 0.708085 |
| 0.70952 | 0.712825 | 0.716169 | 0.70979 | 0.710757 | 0.70979 | 0.671016 | 0.706154 | 0.709194 |
| 0.712723 | 0.712918 | 0.712237 | 0.713111 | 0.708489 | 0.70955 | 0.667489 | 0.709829 | 0.706089 |
| 0.711236 | 0.712558 | 0.713888 | 0.708764 | 0.712265 | 0.710922 | 0.666435 | 0.710294 | 0.710627 |
| 0.701263 | 0.704256 | 0.705365 | 0.702377 | 0.700412 | 0.699743 | 0.649686 | 0.700768 | 0.697348 |
| 0.698249 | 0.705171 | 0.703426 | 0.703888 | 0.704109 | 0.700577 | 0.66156 | 0.699258 | 0.703063 |
| 0.70893 | 0.70875 | 0.709796 | 0.706262 | 0.708018 | 0.708249 | 0.671353 | 0.705541 | 0.708243 |
| 0.700643 | 0.708086 | 0.707696 | 0.703201 | 0.707358 | 0.703083 | 0.660695 | 0.70152 | 0.706255 |
| 0.700869 | 0.705526 | 0.706696 | 0.700668 | 0.705727 | 0.703245 | 0.65979 | 0.702927 | 0.703793 |
| 0.703052 | 0.705348 | 0.70425 | 0.701256 | 0.703267 | 0.704847 | 0.652189 | 0.698906 | 0.705061 |
| 0.700503 | 0.703151 | 0.706412 | 0.699268 | 0.704186 | 0.70226 | 0.659067 | 0.697451 | 0.703859 |
| 0.707991 | 0.712092 | 0.711534 | 0.704554 | 0.711241 | 0.709652 | 0.672498 | 0.706012 | 0.709162 |
| 0.713186 | 0.711769 | 0.714388 | 0.711051 | 0.712847 | 0.713021 | 0.668171 | 0.71037 | 0.712224 |
| 0.705292 | 0.7093 | 0.7107 | 0.702924 | 0.709888 | 0.705811 | 0.66016 | 0.704089 | 0.702567 |
| 0.699905 | 0.704137 | 0.704571 | 0.700736 | 0.701754 | 0.703119 | 0.661982 | 0.697003 | 0.700487 |
| 0.705901 | 0.70844 | 0.70827 | 0.703071 | 0.706958 | 0.702012 | 0.661437 | 0.70205 | 0.705464 |
| 0.703088 | 0.708655 | 0.709376 | 0.704918 | 0.707884 | 0.705148 | 0.655483 | 0.70364 | 0.706175 |
| 0.712004 | 0.71836 | 0.716918 | 0.714172 | 0.719287 | 0.715611 | 0.67993 | 0.714104 | 0.715236 |
| 0.72108 | 0.724243 | 0.722698 | 0.718237 | 0.723257 | 0.720848 | 0.685806 | 0.720559 | 0.718401 |
| 0.761924 | 0.769474 | 0.767897 | 0.766623 | 0.76699 | 0.762504 | 0.719929 | 0.768554 | 0.768823 |
| 0.760414 | 0.771697 | 0.770222 | 0.765922 | 0.772351 | 0.766817 | 0.71832 | 0.766615 | 0.765578 |
| 0.761576 | 0.771158 | 0.768116 | 0.770482 | 0.773602 | 0.766805 | 0.7307 | 0.768238 | 0.764944 |
| 0.703633 | 0.707241 | 0.707441 | 0.703168 | 0.706853 | 0.703273 | 0.659508 | 0.703426 | 0.702969 |
| 0.704954 | 0.70457 | 0.704948 | 0.701832 | 0.705128 | 0.704179 | 0.660174 | 0.70721 | 0.70446 |
| 0.711037 | 0.714355 | 0.7144 | 0.713421 | 0.715117 | 0.712259 | 0.66307 | 0.712584 | 0.712026 |
| 0.712929 | 0.720381 | 0.717552 | 0.711864 | 0.714953 | 0.713925 | 0.671384 | 0.712132 | 0.711787 |
| 0.709592 | 0.715479 | 0.711545 | 0.709315 | 0.709566 | 0.709504 | 0.667597 | 0.708074 | 0.706944 |
| 0.704681 | 0.70899 | 0.709768 | 0.708337 | 0.708786 | 0.707287 | 0.666899 | 0.703374 | 0.706827 |
| 0.705395 | 0.713182 | 0.711046 | 0.709787 | 0.709631 | 0.708415 | 0.665977 | 0.705246 | 0.707288 |
| 0.70685 | 0.713917 | 0.711232 | 0.708784 | 0.712097 | 0.707873 | 0.676896 | 0.707807 | 0.709737 |
| 0.707983 | 0.712979 | 0.713007 | 0.709138 | 0.70853 | 0.709011 | 0.660384 | 0.70818 | 0.706239 |
| 0.724132 | 0.727832 | 0.728224 | 0.72513 | 0.724944 | 0.725235 | 0.694159 | 0.722156 | 0.722195 |
| 0.701806 | 0.706366 | 0.707147 | 0.703279 | 0.704946 | 0.703895 | 0.669831 | 0.702732 | 0.700943 |
| 0.707319 | 0.71016 | 0.709408 | 0.71014 | 0.710514 | 0.711074 | 0.671652 | 0.709933 | 0.710238 |
| 0.703164 | 0.709114 | 0.705664 | 0.702862 | 0.701879 | 0.704665 | 0.65804 | 0.70186 | 0.702784 |
| 0.724966 | 0.725216 | 0.727675 | 0.726523 | 0.727032 | 0.723917 | 0.689864 | 0.723493 | 0.72207 |
| 0.706694 | 0.708193 | 0.710774 | 0.709798 | 0.711433 | 0.710029 | 0.671223 | 0.70716 | 0.707352 |
| 0.702361 | 0.706831 | 0.705282 | 0.70591 | 0.706387 | 0.704493 | 0.666667 | 0.703448 | 0.7022 |
| 0.709791 | 0.711147 | 0.710211 | 0.705041 | 0.706574 | 0.709317 | 0.663205 | 0.707174 | 0.706169 |
| 0.709328 | 0.711377 | 0.714851 | 0.709396 | 0.711003 | 0.707173 | 0.678534 | 0.706014 | 0.708407 |

| | | | | | | | | |
|---|---|---|---|---|---|---|---|---|
| 0.706457 | 0.709905 | 0.710754 | 0.706652 | 0.708095 | 0.702843 | 0.665162 | 0.704515 | 0.704983 |
| 0.708113 | 0.711738 | 0.709852 | 0.710647 | 0.710481 | 0.70927 | 0.661485 | 0.708589 | 0.711662 |
| 0.707793 | 0.712728 | 0.710539 | 0.70751 | 0.708247 | 0.709855 | 0.666783 | 0.708846 | 0.704977 |
| 0.715894 | 0.717936 | 0.719064 | 0.715065 | 0.717901 | 0.717026 | 0.692775 | 0.715375 | 0.717451 |
| 0.716877 | 0.715732 | 0.718349 | 0.715822 | 0.714873 | 0.714183 | 0.677793 | 0.713113 | 0.716026 |
| 0.708223 | 0.710181 | 0.709015 | 0.707008 | 0.71134 | 0.709086 | 0.669346 | 0.707434 | 0.707214 |
| 0.712586 | 0.712391 | 0.712659 | 0.708144 | 0.714984 | 0.710017 | 0.673754 | 0.710978 | 0.711317 |
| 0.7102 | 0.711899 | 0.712965 | 0.710873 | 0.710991 | 0.709503 | 0.665502 | 0.710294 | 0.711177 |
| 0.70858 | 0.71138 | 0.714217 | 0.709716 | 0.711219 | 0.709166 | 0.667831 | 0.70773 | 0.707642 |
| 0.703739 | 0.706407 | 0.710738 | 0.709212 | 0.704011 | 0.703609 | 0.658377 | 0.702839 | 0.705307 |
| 0.708271 | 0.71066 | 0.712907 | 0.709455 | 0.707348 | 0.707929 | 0.662795 | 0.705979 | 0.706314 |
| 0.707117 | 0.709578 | 0.711683 | 0.708815 | 0.707555 | 0.710619 | 0.667487 | 0.706641 | 0.708649 |
| 0.710387 | 0.713191 | 0.711749 | 0.711466 | 0.708529 | 0.711381 | 0.673303 | 0.706462 | 0.710613 |
| 0.704095 | 0.706635 | 0.711523 | 0.711714 | 0.707533 | 0.705681 | 0.664916 | 0.704863 | 0.703719 |
| 0.705251 | 0.707966 | 0.708007 | 0.708446 | 0.709126 | 0.706115 | 0.660515 | 0.707121 | 0.707294 |
| 0.706736 | 0.70978 | 0.708495 | 0.706829 | 0.708317 | 0.705995 | 0.667129 | 0.706082 | 0.707762 |
| 0.710265 | 0.711607 | 0.714769 | 0.710948 | 0.710002 | 0.710514 | 0.669741 | 0.710188 | 0.710389 |
| 0.706613 | 0.713189 | 0.712201 | 0.709396 | 0.711726 | 0.707724 | 0.667017 | 0.710152 | 0.708327 |
| 0.711791 | 0.713804 | 0.71448 | 0.711288 | 0.713569 | 0.710828 | 0.677351 | 0.709429 | 0.707928 |
| 0.708794 | 0.709065 | 0.710394 | 0.707296 | 0.710597 | 0.705953 | 0.668763 | 0.705976 | 0.711408 |
| 0.709947 | 0.71279 | 0.711973 | 0.707846 | 0.708977 | 0.706806 | 0.668531 | 0.706694 | 0.711697 |
| 0.706805 | 0.706182 | 0.710539 | 0.70745 | 0.707759 | 0.705246 | 0.665059 | 0.703914 | 0.704491 |
| 0.705114 | 0.712293 | 0.713137 | 0.711372 | 0.711532 | 0.708643 | 0.673305 | 0.710636 | 0.707919 |
| 0.714569 | 0.710636 | 0.715494 | 0.711911 | 0.710766 | 0.711477 | 0.669583 | 0.711231 | 0.712466 |
| 0.70779 | 0.715258 | 0.715025 | 0.707796 | 0.711909 | 0.708215 | 0.651826 | 0.709422 | 0.710635 |
| 0.712742 | 0.716862 | 0.715409 | 0.711843 | 0.713451 | 0.712676 | 0.660281 | 0.71182 | 0.710101 |
| 0.707402 | 0.712717 | 0.710108 | 0.708915 | 0.712146 | 0.706862 | 0.656541 | 0.709317 | 0.712493 |
| 0.709968 | 0.711649 | 0.714513 | 0.713515 | 0.709323 | 0.711052 | 0.6597 | 0.708071 | 0.707702 |
| 0.709289 | 0.711797 | 0.710959 | 0.711812 | 0.712624 | 0.710094 | 0.665977 | 0.708639 | 0.707296 |
| 0.712201 | 0.711216 | 0.711248 | 0.705458 | 0.71146 | 0.712151 | 0.664011 | 0.707434 | 0.709223 |
| 0.710234 | 0.71618 | 0.7119 | 0.710768 | 0.712288 | 0.711063 | 0.675061 | 0.709069 | 0.71292 |
| 0.708716 | 0.708759 | 0.712139 | 0.709435 | 0.709857 | 0.705943 | 0.669077 | 0.706857 | 0.710522 |
| 0.71394 | 0.712976 | 0.711599 | 0.712584 | 0.713297 | 0.714565 | 0.666667 | 0.709282 | 0.708253 |
| 0.712254 | 0.714133 | 0.715551 | 0.712502 | 0.713517 | 0.713589 | 0.665503 | 0.708462 | 0.711965 |
| 0.709809 | 0.709644 | 0.71229 | 0.706658 | 0.70906 | 0.710518 | 0.669212 | 0.705508 | 0.710674 |
| 0.709802 | 0.711884 | 0.713196 | 0.70901 | 0.712869 | 0.706985 | 0.660912 | 0.708404 | 0.707855 |
| 0.705162 | 0.705476 | 0.709938 | 0.705623 | 0.709071 | 0.702883 | 0.669361 | 0.704114 | 0.703035 |
| 0.683933 | 0.687201 | 0.687475 | 0.684088 | 0.688156 | 0.683845 | 0.666033 | 0.681178 | 0.684766 |
| 0.691255 | 0.690292 | 0.68911 | 0.686164 | 0.685511 | 0.687946 | 0.664618 | 0.691348 | 0.687588 |
| 0.68145 | 0.683277 | 0.685048 | 0.677396 | 0.684427 | 0.680397 | 0.653488 | 0.676606 | 0.679274 |
| 0.678378 | 0.68148 | 0.678825 | 0.677878 | 0.682612 | 0.678898 | 0.659367 | 0.678646 | 0.678277 |
| 0.662815 | 0.667135 | 0.669328 | 0.665965 | 0.665778 | 0.662078 | 0.64472 | 0.662642 | 0.665377 |
| 0.665662 | 0.67205 | 0.666505 | 0.667149 | 0.665392 | 0.667356 | 0.650987 | 0.662795 | 0.664889 |
| 0.666869 | 0.671296 | 0.671057 | 0.66353 | 0.66932 | 0.664757 | 0.655451 | 0.66723 | 0.667801 |
| 0.667134 | 0.672129 | 0.667221 | 0.665394 | 0.667257 | 0.665047 | 0.661559 | 0.667935 | 0.66722 |
| 0.665213 | 0.671173 | 0.667251 | 0.663807 | 0.668684 | 0.66809 | 0.6658 | 0.665592 | 0.668712 |
| 0.671616 | 0.682817 | 0.674072 | 0.674332 | 0.677321 | 0.674477 | 0.66358 | 0.675212 | 0.66877 |

| | | | | | | | | |
|---|---|---|---|---|---|---|---|---|
| 0.673419 | 0.677431 | 0.679911 | 0.667932 | 0.672319 | 0.671683 | 0.663167 | 0.672574 | 0.666749 |
| 0.668141 | 0.680979 | 0.674918 | 0.672486 | 0.674855 | 0.672996 | 0.659173 | 0.672534 | 0.66989 |
| 0.67327 | 0.67362 | 0.670383 | 0.668086 | 0.669808 | 0.671573 | 0.664414 | 0.667947 | 0.666399 |
| 0.663707 | 0.662792 | 0.662706 | 0.656693 | 0.662217 | 0.663525 | 0.643172 | 0.655204 | 0.65874 |
| 0.663326 | 0.66821 | 0.66508 | 0.659291 | 0.66773 | 0.661111 | 0.651203 | 0.659694 | 0.664821 |
| 0.664098 | 0.668314 | 0.669333 | 0.664562 | 0.662235 | 0.66662 | 0.653321 | 0.665214 | 0.663042 |
| 0.66828 | 0.669506 | 0.669787 | 0.662525 | 0.664698 | 0.668234 | 0.660085 | 0.666901 | 0.662982 |
| 0.663902 | 0.672612 | 0.663622 | 0.662147 | 0.665417 | 0.662523 | 0.654834 | 0.665226 | 0.65956 |
| 0.663718 | 0.663339 | 0.665829 | 0.665883 | 0.665743 | 0.661217 | 0.656386 | 0.664665 | 0.660297 |
| 0.668135 | 0.672022 | 0.662227 | 0.660213 | 0.665988 | 0.664165 | 0.64313 | 0.666473 | 0.65816 |
| 0.65746 | 0.662272 | 0.660786 | 0.658843 | 0.660196 | 0.658063 | 0.664163 | 0.653419 | 0.651826 |
| 0.698178 | 0.698354 | 0.699521 | 0.699601 | 0.700719 | 0.698404 | 0.663668 | 0.697446 | 0.6971 |
| 0.707384 | 0.711858 | 0.713707 | 0.706538 | 0.709799 | 0.704965 | 0.66841 | 0.70851 | 0.707421 |
| 0.68115 | 0.688551 | 0.686021 | 0.685643 | 0.687712 | 0.683019 | 0.673527 | 0.687847 | 0.680323 |
| 0.714217 | 0.715702 | 0.718917 | 0.710854 | 0.717263 | 0.712631 | 0.675211 | 0.71479 | 0.713939 |
| 0.704643 | 0.70553 | 0.70678 | 0.704704 | 0.705347 | 0.703004 | 0.666783 | 0.699708 | 0.703036 |
| 0.709632 | 0.712772 | 0.712935 | 0.710572 | 0.710385 | 0.711191 | 0.676573 | 0.709863 | 0.70832 |
| 0.716463 | 0.716018 | 0.716323 | 0.716522 | 0.713897 | 0.71476 | 0.678068 | 0.716148 | 0.714596 |
| 0.697164 | 0.695909 | 0.69898 | 0.695639 | 0.697556 | 0.699992 | 0.653557 | 0.694494 | 0.699576 |
| 0.716643 | 0.720661 | 0.722566 | 0.719098 | 0.722205 | 0.714138 | 0.68487 | 0.718001 | 0.71844 |
| 0.685172 | 0.693724 | 0.690824 | 0.687005 | 0.689417 | 0.683855 | 0.668506 | 0.68623 | 0.686638 |
| 0.714286 | 0.720048 | 0.719589 | 0.715196 | 0.714206 | 0.712154 | 0.676594 | 0.713421 | 0.713053 |
| 0.716568 | 0.717881 | 0.721727 | 0.717838 | 0.71895 | 0.718188 | 0.67546 | 0.71771 | 0.712845 |
| 0.679717 | 0.685174 | 0.679986 | 0.682876 | 0.689635 | 0.682196 | 0.667224 | 0.686529 | 0.684437 |
| 0.689385 | 0.691804 | 0.691113 | 0.688506 | 0.690018 | 0.68764 | 0.647524 | 0.688549 | 0.686618 |
| 0.685837 | 0.693436 | 0.689276 | 0.689113 | 0.688825 | 0.688662 | 0.678058 | 0.692751 | 0.690189 |
| 0.701529 | 0.708708 | 0.710707 | 0.703479 | 0.707436 | 0.701275 | 0.666192 | 0.701299 | 0.706953 |
| 0.714263 | 0.716836 | 0.714707 | 0.714918 | 0.715384 | 0.714148 | 0.679047 | 0.712457 | 0.714843 |
| 0.687766 | 0.689137 | 0.690023 | 0.686184 | 0.687679 | 0.686571 | 0.655079 | 0.682844 | 0.684822 |
| 0.724204 | 0.723806 | 0.72639 | 0.722683 | 0.729917 | 0.721365 | 0.684156 | 0.721246 | 0.723984 |
| 0.722615 | 0.724414 | 0.724372 | 0.722131 | 0.72695 | 0.720956 | 0.682692 | 0.720069 | 0.723525 |
| 0.698465 | 0.697573 | 0.700915 | 0.698366 | 0.698845 | 0.69887 | 0.65846 | 0.694574 | 0.696559 |
| 0.694984 | 0.701413 | 0.701035 | 0.69743 | 0.697494 | 0.697052 | 0.651075 | 0.697392 | 0.698076 |
| 0.722147 | 0.729899 | 0.726371 | 0.723582 | 0.728332 | 0.720908 | 0.691549 | 0.72288 | 0.722573 |
| 0.718992 | 0.725051 | 0.728227 | 0.718999 | 0.724479 | 0.718264 | 0.678228 | 0.720278 | 0.721146 |
| 0.722066 | 0.724495 | 0.72469 | 0.724038 | 0.729207 | 0.720781 | 0.692281 | 0.721504 | 0.724944 |
| 0.6747 | 0.678962 | 0.678861 | 0.679957 | 0.678253 | 0.678549 | 0.642304 | 0.677792 | 0.677497 |
| 0.717536 | 0.722338 | 0.720815 | 0.722509 | 0.720101 | 0.717612 | 0.674532 | 0.717039 | 0.718208 |
| 0.675397 | 0.679092 | 0.679606 | 0.677579 | 0.679154 | 0.675278 | 0.634146 | 0.675997 | 0.677395 |
| 0.662951 | 0.668317 | 0.671888 | 0.668377 | 0.668404 | 0.669167 | 0.623631 | 0.662613 | 0.666869 |
| 0.720383 | 0.721191 | 0.724045 | 0.720057 | 0.723192 | 0.717298 | 0.680194 | 0.71912 | 0.720448 |
| 0.665988 | 0.670251 | 0.672273 | 0.671176 | 0.671588 | 0.666819 | 0.612305 | 0.668276 | 0.670265 |
| 0.660774 | 0.667096 | 0.666491 | 0.663848 | 0.663144 | 0.665277 | 0.609732 | 0.663278 | 0.659304 |
| 0.657903 | 0.662843 | 0.661248 | 0.664451 | 0.658996 | 0.659187 | 0.613659 | 0.658379 | 0.657132 |
| 0.729371 | 0.732487 | 0.73127 | 0.725171 | 0.733434 | 0.729062 | 0.693408 | 0.727735 | 0.73047 |
| 0.66038 | 0.666414 | 0.664124 | 0.663075 | 0.663621 | 0.661896 | 0.611872 | 0.660104 | 0.659168 |
| 0.716551 | 0.723068 | 0.721434 | 0.721534 | 0.724474 | 0.719322 | 0.675779 | 0.719975 | 0.721673 |

| | | | | | | | | |
|---|---|---|---|---|---|---|---|---|
| 0.727518 | 0.731447 | 0.735351 | 0.727006 | 0.732316 | 0.723817 | 0.680903 | 0.729719 | 0.728766 |
| 0.722403 | 0.727988 | 0.727433 | 0.723131 | 0.728741 | 0.722213 | 0.675545 | 0.723452 | 0.726619 |
| 0.740042 | 0.747909 | 0.746868 | 0.744501 | 0.745174 | 0.743012 | 0.718839 | 0.735546 | 0.738808 |
| 0.646661 | 0.652442 | 0.652287 | 0.652421 | 0.649416 | 0.650227 | 0.595914 | 0.64989 | 0.647862 |
| 0.653726 | 0.657969 | 0.656546 | 0.656442 | 0.655494 | 0.655301 | 0.613825 | 0.651925 | 0.651457 |
| 0.728319 | 0.736267 | 0.736287 | 0.729659 | 0.736074 | 0.730321 | 0.681834 | 0.729426 | 0.733628 |
| 0.657306 | 0.656196 | 0.656919 | 0.657694 | 0.656042 | 0.65702 | 0.610365 | 0.655499 | 0.657196 |
| 0.731844 | 0.737691 | 0.738193 | 0.732286 | 0.736062 | 0.730798 | 0.692088 | 0.727397 | 0.730418 |
| 0.73101 | 0.735475 | 0.734603 | 0.729459 | 0.73736 | 0.734511 | 0.689222 | 0.734873 | 0.733226 |
| 0.728527 | 0.734177 | 0.732766 | 0.729774 | 0.73121 | 0.72946 | 0.678471 | 0.726375 | 0.728521 |
| 0.729353 | 0.735772 | 0.736081 | 0.731487 | 0.738035 | 0.730216 | 0.68386 | 0.729301 | 0.732446 |
| 0.716129 | 0.77551 | 0.690789 | 0.714286 | 0.773973 | 0.708075 | 0.753425 | 0.746667 | 0.703226 |
| 0.717 | 0.739095 | 0.724228 | 0.732892 | 0.742604 | 0.732685 | 0.705882 | 0.725826 | 0.727236 |
| 0.720743 | 0.735113 | 0.730787 | 0.722214 | 0.739962 | 0.721376 | 0.706922 | 0.729793 | 0.727545 |
| 0.728094 | 0.747679 | 0.735357 | 0.737145 | 0.750339 | 0.729403 | 0.717008 | 0.737135 | 0.735419 |
| 0.731994 | 0.741082 | 0.734891 | 0.738978 | 0.749696 | 0.737036 | 0.713778 | 0.735634 | 0.727162 |
| 0.589572 | 0.595719 | 0.593808 | 0.598028 | 0.595282 | 0.593747 | 0.565846 | 0.594615 | 0.596382 |
| 0.731439 | 0.738351 | 0.734862 | 0.743367 | 0.742271 | 0.737119 | 0.717926 | 0.737273 | 0.734242 |
| 0.731766 | 0.741447 | 0.742196 | 0.728958 | 0.743154 | 0.734456 | 0.695178 | 0.733963 | 0.737451 |
| 0.734737 | 0.743634 | 0.739686 | 0.734086 | 0.742394 | 0.734372 | 0.689594 | 0.737493 | 0.740071 |
| 0.735997 | 0.741666 | 0.741386 | 0.734408 | 0.741762 | 0.739267 | 0.687127 | 0.735858 | 0.736975 |
| 0.592897 | 0.597108 | 0.596772 | 0.59723 | 0.596535 | 0.596863 | 0.539251 | 0.59414 | 0.592777 |
| 0.580402 | 0.584037 | 0.584361 | 0.588082 | 0.586069 | 0.583763 | 0.525378 | 0.581654 | 0.580017 |
| 0.759157 | 0.765956 | 0.765899 | 0.764384 | 0.761964 | 0.765234 | 0.728249 | 0.758848 | 0.763259 |
| 0.760547 | 0.767929 | 0.766197 | 0.764591 | 0.766243 | 0.761219 | 0.733285 | 0.760434 | 0.76529 |
| 0.755132 | 0.757659 | 0.753481 | 0.759389 | 0.758638 | 0.75341 | 0.717251 | 0.752151 | 0.75326 |
| 0.755919 | 0.763822 | 0.764392 | 0.762161 | 0.760815 | 0.757686 | 0.726949 | 0.759301 | 0.758072 |
| 0.761339 | 0.768869 | 0.76703 | 0.764677 | 0.765754 | 0.765392 | 0.729331 | 0.762693 | 0.76488 |
| 0.760819 | 0.769263 | 0.763232 | 0.767001 | 0.770564 | 0.76337 | 0.730879 | 0.763857 | 0.76792 |
| 0.765092 | 0.771042 | 0.77072 | 0.767159 | 0.774034 | 0.767709 | 0.722874 | 0.766896 | 0.765118 |
| 0.758412 | 0.766641 | 0.763627 | 0.759588 | 0.763734 | 0.757563 | 0.721098 | 0.762805 | 0.760718 |
| 0.7627 | 0.774544 | 0.769074 | 0.76955 | 0.770407 | 0.765436 | 0.722104 | 0.767701 | 0.770332 |
| 0.755857 | 0.76265 | 0.759726 | 0.765382 | 0.759875 | 0.76026 | 0.726686 | 0.755257 | 0.757139 |
| 0.757361 | 0.764235 | 0.762493 | 0.758654 | 0.758412 | 0.756969 | 0.723915 | 0.756539 | 0.757825 |
| 0.766425 | 0.774952 | 0.768858 | 0.767867 | 0.767358 | 0.765276 | 0.73191 | 0.762741 | 0.760626 |
| 0.756622 | 0.76474 | 0.761736 | 0.762065 | 0.762007 | 0.754558 | 0.714286 | 0.757405 | 0.760559 |
| 0.757935 | 0.771829 | 0.764793 | 0.764248 | 0.763805 | 0.763053 | 0.715404 | 0.761975 | 0.760198 |
| 0.763201 | 0.766958 | 0.76404 | 0.767962 | 0.76732 | 0.760681 | 0.733429 | 0.763626 | 0.760633 |
| 0.757586 | 0.764462 | 0.762831 | 0.759586 | 0.760464 | 0.754632 | 0.713525 | 0.758401 | 0.757192 |
| 0.758498 | 0.762584 | 0.761986 | 0.76325 | 0.7623 | 0.75632 | 0.716767 | 0.757458 | 0.756014 |
| 0.729229 | 0.737868 | 0.735802 | 0.735393 | 0.741352 | 0.733511 | 0.723599 | 0.732377 | 0.730762 |
| 0.736164 | 0.7432 | 0.735545 | 0.734405 | 0.748627 | 0.73913 | 0.714604 | 0.729405 | 0.731471 |
| 0.760817 | 0.769566 | 0.76656 | 0.762308 | 0.763796 | 0.760228 | 0.726331 | 0.759063 | 0.760167 |
| 0.763913 | 0.766417 | 0.765349 | 0.765359 | 0.767594 | 0.763928 | 0.734118 | 0.764351 | 0.769503 |
| 0.719512 | 0.723125 | 0.718424 | 0.712754 | 0.720146 | 0.721649 | 0.705233 | 0.714258 | 0.71423 |
| 0.758279 | 0.768881 | 0.767626 | 0.765313 | 0.764937 | 0.75918 | 0.731214 | 0.762083 | 0.76366 |
| 0.756382 | 0.763763 | 0.761138 | 0.757868 | 0.760691 | 0.756399 | 0.709872 | 0.751434 | 0.753479 |

| | | | | | | | | |
|---|---|---|---|---|---|---|---|---|
| 0.755859 | 0.76657 | 0.765849 | 0.766344 | 0.762434 | 0.758569 | 0.70985 | 0.760825 | 0.759803 |
| 0.759679 | 0.771177 | 0.765782 | 0.765577 | 0.758802 | 0.756945 | 0.72316 | 0.761804 | 0.758724 |
| 0.764627 | 0.770544 | 0.766883 | 0.764293 | 0.765806 | 0.764103 | 0.71014 | 0.763116 | 0.761063 |
| 0.76194 | 0.768659 | 0.762944 | 0.766429 | 0.763816 | 0.759067 | 0.724262 | 0.763418 | 0.759633 |
| 0.769224 | 0.773091 | 0.774116 | 0.770578 | 0.772422 | 0.770621 | 0.727758 | 0.770505 | 0.767348 |
| 0.760023 | 0.76517 | 0.762003 | 0.758347 | 0.761803 | 0.757583 | 0.71317 | 0.757457 | 0.75617 |
| 0.759237 | 0.766317 | 0.761259 | 0.75975 | 0.762807 | 0.758014 | 0.722401 | 0.757564 | 0.756479 |
| 0.749096 | 0.76314 | 0.757785 | 0.753435 | 0.755191 | 0.75407 | 0.714491 | 0.751312 | 0.751862 |
| 0.759108 | 0.769899 | 0.763277 | 0.764937 | 0.759687 | 0.758458 | 0.708199 | 0.758963 | 0.759662 |
| 0.754779 | 0.762007 | 0.755981 | 0.754967 | 0.75509 | 0.754165 | 0.709861 | 0.751121 | 0.755057 |
| 0.758299 | 0.770688 | 0.766771 | 0.759627 | 0.764998 | 0.758995 | 0.718952 | 0.759006 | 0.759228 |
| 0.756015 | 0.765011 | 0.761744 | 0.760276 | 0.754863 | 0.754661 | 0.714389 | 0.760471 | 0.756141 |
| 0.750989 | 0.761627 | 0.754225 | 0.754497 | 0.757731 | 0.756159 | 0.706884 | 0.749877 | 0.752575 |
| 0.753964 | 0.762196 | 0.753338 | 0.754598 | 0.754142 | 0.7563 | 0.710061 | 0.750382 | 0.750812 |
| 0.764754 | 0.771738 | 0.770324 | 0.767304 | 0.766342 | 0.76212 | 0.731171 | 0.760052 | 0.763585 |
| 0.760903 | 0.765818 | 0.76877 | 0.763945 | 0.769883 | 0.765625 | 0.733046 | 0.763188 | 0.761582 |
| 0.757965 | 0.76902 | 0.762459 | 0.763566 | 0.766441 | 0.757061 | 0.728035 | 0.760449 | 0.758571 |
| 0.751305 | 0.760803 | 0.759395 | 0.75981 | 0.761885 | 0.754969 | 0.722502 | 0.753504 | 0.752395 |
| 0.756983 | 0.765656 | 0.76375 | 0.762227 | 0.766334 | 0.762962 | 0.731059 | 0.760664 | 0.761126 |
| 0.755877 | 0.761432 | 0.761999 | 0.761667 | 0.762772 | 0.758561 | 0.72688 | 0.757806 | 0.758448 |
| 0.748925 | 0.756786 | 0.757244 | 0.755051 | 0.758962 | 0.753129 | 0.721755 | 0.748405 | 0.751293 |
| 0.758279 | 0.760963 | 0.762016 | 0.760506 | 0.760882 | 0.761311 | 0.7213 | 0.76009 | 0.757599 |
| 0.761723 | 0.763763 | 0.763134 | 0.764085 | 0.765432 | 0.764236 | 0.730756 | 0.763505 | 0.75917 |
| 0.759282 | 0.764876 | 0.765409 | 0.763937 | 0.766214 | 0.760315 | 0.716732 | 0.758587 | 0.75617 |
| 0.758193 | 0.762541 | 0.763559 | 0.756569 | 0.762945 | 0.76062 | 0.716524 | 0.754969 | 0.759694 |
| 0.756114 | 0.756999 | 0.759848 | 0.75299 | 0.761285 | 0.759474 | 0.722562 | 0.756537 | 0.761486 |
| 0.74895 | 0.760214 | 0.758552 | 0.757021 | 0.759695 | 0.753224 | 0.71657 | 0.753987 | 0.751906 |
| 0.75359 | 0.757512 | 0.756782 | 0.758919 | 0.756867 | 0.750672 | 0.719109 | 0.752752 | 0.751477 |
| 0.757586 | 0.762537 | 0.765982 | 0.757057 | 0.763281 | 0.760477 | 0.716284 | 0.754473 | 0.754683 |
| 0.758227 | 0.761732 | 0.762253 | 0.762956 | 0.763037 | 0.757164 | 0.731556 | 0.756697 | 0.760321 |
| 0.759788 | 0.765906 | 0.762666 | 0.764821 | 0.763556 | 0.758187 | 0.712251 | 0.756792 | 0.759109 |
| 0.756643 | 0.76837 | 0.761806 | 0.763731 | 0.765717 | 0.758923 | 0.717022 | 0.755763 | 0.757558 |
| 0.764267 | 0.770937 | 0.77151 | 0.770006 | 0.770664 | 0.766538 | 0.735357 | 0.763782 | 0.770738 |
| 0.767727 | 0.773337 | 0.770192 | 0.77127 | 0.773387 | 0.767382 | 0.738529 | 0.764922 | 0.763895 |
| 0.753572 | 0.763326 | 0.760934 | 0.756838 | 0.759898 | 0.753267 | 0.728587 | 0.750956 | 0.754216 |
| 0.755129 | 0.761287 | 0.762835 | 0.758409 | 0.764657 | 0.759476 | 0.722023 | 0.754558 | 0.757362 |
| 0.758638 | 0.763164 | 0.763402 | 0.761285 | 0.761145 | 0.761524 | 0.718188 | 0.756622 | 0.754947 |
| 0.756381 | 0.764362 | 0.760814 | 0.760999 | 0.761968 | 0.758244 | 0.728629 | 0.756608 | 0.755443 |
| 0.746937 | 0.752655 | 0.756605 | 0.754502 | 0.753698 | 0.752247 | 0.709457 | 0.746696 | 0.749709 |
| 0.748804 | 0.758052 | 0.756406 | 0.754514 | 0.755797 | 0.75123 | 0.726944 | 0.750681 | 0.749249 |
| 0.74286 | 0.750999 | 0.753607 | 0.747939 | 0.751012 | 0.747402 | 0.712464 | 0.747186 | 0.75036 |
| 0.749264 | 0.753912 | 0.753666 | 0.754039 | 0.753721 | 0.749715 | 0.722509 | 0.751949 | 0.750414 |
| 0.743764 | 0.751158 | 0.752592 | 0.753624 | 0.754691 | 0.748641 | 0.70923 | 0.745006 | 0.74479 |
| 0.747272 | 0.75061 | 0.747238 | 0.749149 | 0.748616 | 0.750749 | 0.71476 | 0.746054 | 0.747214 |
| 0.749487 | 0.758811 | 0.759023 | 0.754174 | 0.757713 | 0.754465 | 0.719738 | 0.755472 | 0.75213 |
| 0.750635 | 0.751748 | 0.752643 | 0.753332 | 0.749789 | 0.747037 | 0.711864 | 0.742055 | 0.740953 |

| | | | | | | | | |
|---|---|---|---|---|---|---|---|---|
| 0.759038 | 0.760447 | 0.764177 | 0.754652 | 0.762222 | 0.756795 | 0.719971 | 0.756209 | 0.759743 |
| 0.76081 | 0.771017 | 0.769333 | 0.769787 | 0.769671 | 0.764987 | 0.724617 | 0.7649 | 0.770664 |
| 0.762469 | 0.769192 | 0.766268 | 0.761215 | 0.76704 | 0.7631 | 0.714912 | 0.766266 | 0.766605 |
| 0.761607 | 0.76673 | 0.76374 | 0.76429 | 0.766572 | 0.759338 | 0.724777 | 0.762614 | 0.764143 |
| 0.768493 | 0.771259 | 0.77173 | 0.77041 | 0.771218 | 0.767151 | 0.74293 | 0.768406 | 0.768667 |
| 0.76607 | 0.773039 | 0.771403 | 0.767165 | 0.773924 | 0.763793 | 0.725847 | 0.763599 | 0.76445 |
| 0.763582 | 0.766583 | 0.765872 | 0.759725 | 0.761557 | 0.762439 | 0.715217 | 0.75891 | 0.761697 |
| 0.761975 | 0.767632 | 0.768125 | 0.764956 | 0.769281 | 0.76064 | 0.725413 | 0.765755 | 0.768591 |
| 0.756089 | 0.76372 | 0.759486 | 0.75888 | 0.760061 | 0.759851 | 0.731959 | 0.757792 | 0.757589 |
| 0.751119 | 0.759944 | 0.760371 | 0.757722 | 0.760672 | 0.756233 | 0.728336 | 0.754639 | 0.754719 |
| 0.765468 | 0.768929 | 0.766013 | 0.762451 | 0.764749 | 0.766272 | 0.71956 | 0.760558 | 0.756746 |
| 0.761959 | 0.767317 | 0.763119 | 0.762094 | 0.765589 | 0.758817 | 0.72973 | 0.764337 | 0.760478 |
| 0.76274 | 0.769085 | 0.768541 | 0.768689 | 0.769054 | 0.761728 | 0.724508 | 0.768998 | 0.76487 |
| 0.763403 | 0.76953 | 0.766738 | 0.76632 | 0.770227 | 0.762997 | 0.737586 | 0.762751 | 0.767243 |
| 0.769806 | 0.773501 | 0.770591 | 0.767351 | 0.770996 | 0.768646 | 0.723548 | 0.768582 | 0.770883 |
| 0.763703 | 0.76676 | 0.770325 | 0.768632 | 0.766825 | 0.76321 | 0.731034 | 0.762866 | 0.76613 |
| 0.767691 | 0.771555 | 0.769814 | 0.768981 | 0.771918 | 0.768055 | 0.725259 | 0.76872 | 0.766735 |
| 0.766565 | 0.766561 | 0.767582 | 0.766419 | 0.76956 | 0.766367 | 0.719633 | 0.766957 | 0.767644 |
| 0.767321 | 0.769155 | 0.770305 | 0.770855 | 0.773996 | 0.765987 | 0.723792 | 0.766786 | 0.766702 |
| 0.768223 | 0.770338 | 0.764792 | 0.773466 | 0.774826 | 0.771417 | 0.727143 | 0.766092 | 0.768859 |
| 0.760829 | 0.768937 | 0.768535 | 0.767951 | 0.769009 | 0.762605 | 0.739701 | 0.763339 | 0.763911 |
| 0.764144 | 0.774547 | 0.770105 | 0.771098 | 0.774247 | 0.762024 | 0.741039 | 0.768933 | 0.764755 |
| 0.766941 | 0.771857 | 0.77272 | 0.768386 | 0.767548 | 0.766549 | 0.734291 | 0.772135 | 0.768626 |
| 0.766759 | 0.773048 | 0.771335 | 0.76617 | 0.770332 | 0.765626 | 0.714386 | 0.770173 | 0.769061 |
| 0.767799 | 0.770304 | 0.771245 | 0.771232 | 0.772912 | 0.766306 | 0.728307 | 0.766932 | 0.77118 |
| 0.764278 | 0.763995 | 0.766302 | 0.768828 | 0.771494 | 0.765947 | 0.71836 | 0.766531 | 0.768041 |
| 0.765971 | 0.768695 | 0.766405 | 0.766247 | 0.76655 | 0.764749 | 0.722302 | 0.766434 | 0.770057 |
| 0.765308 | 0.767299 | 0.772079 | 0.766829 | 0.769412 | 0.765891 | 0.723132 | 0.765006 | 0.766138 |
| 0.769394 | 0.76988 | 0.768534 | 0.771605 | 0.771043 | 0.768222 | 0.731439 | 0.769162 | 0.772321 |
| 0.768519 | 0.775644 | 0.7727 | 0.770053 | 0.767387 | 0.768857 | 0.729866 | 0.770274 | 0.771662 |
| 0.764031 | 0.766496 | 0.765941 | 0.761842 | 0.765998 | 0.760769 | 0.724014 | 0.76515 | 0.761778 |
| 0.769814 | 0.765707 | 0.764745 | 0.767744 | 0.769884 | 0.77037 | 0.722202 | 0.765044 | 0.766109 |
| 0.770136 | 0.773054 | 0.772253 | 0.771354 | 0.776084 | 0.766334 | 0.727796 | 0.769908 | 0.771058 |
| 0.760415 | 0.767739 | 0.763957 | 0.762154 | 0.768607 | 0.763284 | 0.734392 | 0.764137 | 0.763649 |
| 0.760363 | 0.769878 | 0.76674 | 0.763867 | 0.770809 | 0.76858 | 0.744355 | 0.768764 | 0.76687 |
| 0.762729 | 0.771627 | 0.773377 | 0.761019 | 0.77495 | 0.766206 | 0.736096 | 0.762243 | 0.770608 |
| 0.766289 | 0.777465 | 0.77595 | 0.771473 | 0.772489 | 0.769581 | 0.731375 | 0.770166 | 0.771107 |
| 0.768588 | 0.770318 | 0.767292 | 0.766794 | 0.770174 | 0.772685 | 0.72358 | 0.770571 | 0.770566 |
| 0.76671 | 0.769936 | 0.768117 | 0.767853 | 0.771107 | 0.772508 | 0.730394 | 0.772116 | 0.76816 |
| 0.762214 | 0.76575 | 0.769703 | 0.769565 | 0.766133 | 0.764096 | 0.730366 | 0.766097 | 0.76623 |
| 0.766829 | 0.770761 | 0.77097 | 0.773533 | 0.773139 | 0.768665 | 0.727305 | 0.770738 | 0.771129 |
| 0.769376 | 0.774746 | 0.774994 | 0.77037 | 0.772455 | 0.770093 | 0.727632 | 0.772095 | 0.770099 |
| 0.770369 | 0.773944 | 0.774793 | 0.769312 | 0.773973 | 0.772701 | 0.728086 | 0.772687 | 0.767956 |
| 0.761188 | 0.767642 | 0.766664 | 0.768974 | 0.770041 | 0.762985 | 0.718761 | 0.765073 | 0.761506 |
| 0.769288 | 0.769953 | 0.771509 | 0.769788 | 0.770289 | 0.770837 | 0.730187 | 0.771017 | 0.768099 |
| 0.763939 | 0.770849 | 0.768019 | 0.764955 | 0.771813 | 0.770759 | 0.721786 | 0.764999 | 0.767678 |

| | | | | | | | | |
|---|---|---|---|---|---|---|---|---|
| 0.75621 | 0.76083 | 0.754353 | 0.760571 | 0.761945 | 0.759672 | 0.735936 | 0.758803 | 0.751537 |
| 0.743405 | 0.746781 | 0.746992 | 0.744526 | 0.747829 | 0.742687 | 0.709666 | 0.746647 | 0.746226 |
| 0.75954 | 0.766399 | 0.763273 | 0.764607 | 0.769774 | 0.760316 | 0.731261 | 0.763006 | 0.764557 |
| 0.760569 | 0.77099 | 0.772225 | 0.769981 | 0.775198 | 0.765761 | 0.730906 | 0.769568 | 0.769363 |
| 0.756677 | 0.764196 | 0.764524 | 0.760547 | 0.770058 | 0.758157 | 0.726147 | 0.761393 | 0.760506 |
| 0.762361 | 0.767936 | 0.765526 | 0.762615 | 0.770021 | 0.765254 | 0.729956 | 0.765362 | 0.763791 |
| 0.763104 | 0.770559 | 0.768201 | 0.768993 | 0.774976 | 0.769914 | 0.732841 | 0.768413 | 0.771819 |
| 0.76095 | 0.760444 | 0.761778 | 0.76227 | 0.765777 | 0.75969 | 0.740421 | 0.759673 | 0.759126 |
| 0.768072 | 0.770667 | 0.769243 | 0.770052 | 0.775176 | 0.771128 | 0.735199 | 0.769607 | 0.773026 |
| 0.769256 | 0.772799 | 0.772242 | 0.770768 | 0.771995 | 0.771178 | 0.731234 | 0.771053 | 0.768309 |
| 0.76606 | 0.774353 | 0.773554 | 0.768448 | 0.775524 | 0.767974 | 0.730727 | 0.769787 | 0.769962 |
| 0.763904 | 0.771799 | 0.770842 | 0.771612 | 0.767728 | 0.767119 | 0.713112 | 0.767812 | 0.77144 |
| 0.770272 | 0.776242 | 0.774431 | 0.773614 | 0.777979 | 0.775083 | 0.732188 | 0.77365 | 0.76996 |
| 0.767954 | 0.772622 | 0.770623 | 0.771175 | 0.771529 | 0.768419 | 0.732136 | 0.76935 | 0.772738 |
| 0.762194 | 0.770246 | 0.767588 | 0.76452 | 0.76562 | 0.766186 | 0.722883 | 0.766539 | 0.762709 |
| 0.769969 | 0.773202 | 0.77324 | 0.772834 | 0.772481 | 0.769628 | 0.73129 | 0.770125 | 0.7692 |
| 0.76474 | 0.770636 | 0.767334 | 0.768605 | 0.76743 | 0.766188 | 0.723727 | 0.767988 | 0.767246 |
| 0.767916 | 0.766268 | 0.764794 | 0.765205 | 0.761478 | 0.766293 | 0.722787 | 0.761788 | 0.766217 |
| 0.764994 | 0.770798 | 0.772364 | 0.76967 | 0.770796 | 0.767591 | 0.736288 | 0.7622 | 0.7601 |
| 0.758466 | 0.766678 | 0.769674 | 0.764726 | 0.767567 | 0.763428 | 0.731278 | 0.762854 | 0.76111 |
| 0.762215 | 0.774442 | 0.767161 | 0.766722 | 0.768129 | 0.76479 | 0.733696 | 0.765118 | 0.764207 |
| 0.756384 | 0.76237 | 0.762199 | 0.761124 | 0.766931 | 0.759338 | 0.720888 | 0.758276 | 0.760695 |
| 0.7629 | 0.774076 | 0.773091 | 0.76104 | 0.770146 | 0.761188 | 0.725108 | 0.760932 | 0.764657 |
| 0.766261 | 0.775287 | 0.769822 | 0.765549 | 0.773345 | 0.765426 | 0.728832 | 0.766099 | 0.767 |
| 0.765554 | 0.774194 | 0.770749 | 0.770784 | 0.777572 | 0.770277 | 0.72425 | 0.768819 | 0.765072 |
| 0.768269 | 0.771222 | 0.769707 | 0.765038 | 0.776329 | 0.767297 | 0.727141 | 0.763915 | 0.767974 |
| 0.759611 | 0.774487 | 0.766508 | 0.763187 | 0.772627 | 0.764964 | 0.710612 | 0.763883 | 0.764566 |
| 0.761614 | 0.771741 | 0.76825 | 0.763142 | 0.772665 | 0.767534 | 0.72583 | 0.761796 | 0.765763 |
| 0.763208 | 0.77189 | 0.768007 | 0.766366 | 0.76721 | 0.762235 | 0.730147 | 0.760701 | 0.763819 |
| 0.691653 | 0.695941 | 0.695192 | 0.693947 | 0.696499 | 0.69287 | 0.657885 | 0.696656 | 0.690896 |
| 0.708904 | 0.710303 | 0.709831 | 0.708223 | 0.706847 | 0.708523 | 0.658714 | 0.706498 | 0.703501 |
| 0.700637 | 0.705001 | 0.703336 | 0.706522 | 0.704469 | 0.704911 | 0.6642 | 0.700794 | 0.706533 |
| 0.69365 | 0.69637 | 0.697935 | 0.693595 | 0.695283 | 0.694971 | 0.656318 | 0.692079 | 0.692396 |
| 0.704624 | 0.709739 | 0.711035 | 0.704531 | 0.707565 | 0.706033 | 0.665742 | 0.703309 | 0.704971 |
| 0.764766 | 0.776985 | 0.776105 | 0.766252 | 0.780655 | 0.764676 | 0.731681 | 0.761344 | 0.767099 |
| 0.771015 | 0.773731 | 0.770696 | 0.762385 | 0.769858 | 0.770723 | 0.736726 | 0.762098 | 0.765293 |
| 0.770818 | 0.775002 | 0.776523 | 0.771583 | 0.774271 | 0.769418 | 0.726131 | 0.770759 | 0.771782 |
| 0.773764 | 0.777935 | 0.778925 | 0.771198 | 0.778587 | 0.770136 | 0.737319 | 0.766459 | 0.772701 |
| 0.767708 | 0.774919 | 0.775716 | 0.769071 | 0.77538 | 0.767769 | 0.732156 | 0.765561 | 0.773266 |
| 0.774039 | 0.783384 | 0.777408 | 0.76944 | 0.779998 | 0.771854 | 0.752735 | 0.771833 | 0.776298 |
| 0.772413 | 0.774161 | 0.773424 | 0.765772 | 0.777973 | 0.767763 | 0.726686 | 0.76314 | 0.772193 |
| 0.769192 | 0.776164 | 0.773558 | 0.7673 | 0.772492 | 0.768957 | 0.743337 | 0.762419 | 0.765086 |
| 0.758483 | 0.773007 | 0.771964 | 0.76695 | 0.775761 | 0.75971 | 0.724854 | 0.762316 | 0.764034 |
| 0.760064 | 0.770919 | 0.770952 | 0.760215 | 0.771525 | 0.763221 | 0.730176 | 0.761737 | 0.764685 |
| 0.76763 | 0.775198 | 0.773357 | 0.769263 | 0.771527 | 0.767618 | 0.725712 | 0.766034 | 0.765677 |
| 0.772208 | 0.77158 | 0.771079 | 0.767964 | 0.774398 | 0.771776 | 0.741246 | 0.764672 | 0.765771 |

| | | | | | | | | |
|---|---|---|---|---|---|---|---|---|
| 0.773152 | 0.771982 | 0.775699 | 0.772609 | 0.77629 | 0.775192 | 0.729009 | 0.770068 | 0.771373 |
| 0.764037 | 0.770625 | 0.76909 | 0.767388 | 0.77445 | 0.767182 | 0.730296 | 0.769034 | 0.764779 |
| 0.766007 | 0.773066 | 0.76782 | 0.770132 | 0.775492 | 0.77015 | 0.72688 | 0.765859 | 0.769346 |
| 0.768299 | 0.772474 | 0.763666 | 0.76302 | 0.773554 | 0.768396 | 0.717508 | 0.76217 | 0.766468 |
| 0.76778 | 0.774554 | 0.768879 | 0.767949 | 0.776538 | 0.768072 | 0.717829 | 0.764661 | 0.762778 |
| 0.764516 | 0.766473 | 0.765564 | 0.769951 | 0.772445 | 0.768059 | 0.71894 | 0.75721 | 0.75798 |
| 0.765773 | 0.770357 | 0.769902 | 0.771809 | 0.780168 | 0.770924 | 0.725235 | 0.764519 | 0.762887 |
| 0.768427 | 0.772924 | 0.77329 | 0.768752 | 0.77476 | 0.762676 | 0.737518 | 0.765954 | 0.761194 |
| 0.762747 | 0.773052 | 0.766515 | 0.767497 | 0.772076 | 0.766567 | 0.730893 | 0.767934 | 0.76235 |
| 0.767272 | 0.769186 | 0.768057 | 0.76733 | 0.772501 | 0.768448 | 0.730769 | 0.769511 | 0.764716 |
| 0.766784 | 0.768293 | 0.765159 | 0.765559 | 0.772898 | 0.769231 | 0.729964 | 0.764222 | 0.766728 |
| 0.759973 | 0.770427 | 0.766277 | 0.765064 | 0.775366 | 0.765362 | 0.721252 | 0.763624 | 0.767096 |
| 0.758783 | 0.764996 | 0.760068 | 0.763137 | 0.767659 | 0.760384 | 0.725533 | 0.754809 | 0.763812 |
| 0.75771 | 0.770366 | 0.765351 | 0.76285 | 0.774132 | 0.762171 | 0.720434 | 0.760524 | 0.76237 |
| 0.764288 | 0.766526 | 0.771296 | 0.768992 | 0.766736 | 0.762801 | 0.732312 | 0.764175 | 0.765267 |
| 0.768616 | 0.776553 | 0.774346 | 0.769642 | 0.77329 | 0.76925 | 0.733574 | 0.772118 | 0.770442 |
| 0.764534 | 0.770611 | 0.769797 | 0.767346 | 0.770325 | 0.764619 | 0.722597 | 0.766672 | 0.770578 |
| 0.770097 | 0.771227 | 0.77272 | 0.773618 | 0.771541 | 0.770727 | 0.730645 | 0.770746 | 0.770947 |
| 0.765492 | 0.771057 | 0.769534 | 0.767869 | 0.77222 | 0.766816 | 0.718356 | 0.763484 | 0.76788 |
| 0.769774 | 0.768888 | 0.769507 | 0.766313 | 0.767546 | 0.767992 | 0.723054 | 0.764616 | 0.768094 |
| 0.763025 | 0.763872 | 0.764847 | 0.763503 | 0.766084 | 0.760631 | 0.722364 | 0.762075 | 0.75974 |
| 0.763364 | 0.768961 | 0.766115 | 0.767715 | 0.765013 | 0.761064 | 0.727273 | 0.763125 | 0.76179 |
| 0.767622 | 0.774171 | 0.769948 | 0.770902 | 0.775354 | 0.765216 | 0.72576 | 0.76886 | 0.770979 |
| 0.761315 | 0.771226 | 0.766434 | 0.765669 | 0.770767 | 0.765653 | 0.720058 | 0.762877 | 0.768481 |
| 0.76214 | 0.771092 | 0.767223 | 0.766349 | 0.769659 | 0.762733 | 0.736223 | 0.766077 | 0.764669 |
| 0.701572 | 0.703728 | 0.704558 | 0.701992 | 0.701889 | 0.702299 | 0.667373 | 0.702526 | 0.701754 |
| 0.769513 | 0.771417 | 0.772522 | 0.772601 | 0.770694 | 0.771417 | 0.723121 | 0.768317 | 0.769243 |
| 0.768339 | 0.773129 | 0.77196 | 0.769703 | 0.769017 | 0.766744 | 0.720285 | 0.762427 | 0.767028 |
| 0.771388 | 0.775611 | 0.773324 | 0.772194 | 0.777234 | 0.776367 | 0.725824 | 0.774915 | 0.769161 |
| 0.76349 | 0.77201 | 0.768868 | 0.768118 | 0.772772 | 0.767564 | 0.73169 | 0.766957 | 0.769594 |
| 0.764085 | 0.769751 | 0.766457 | 0.768807 | 0.769595 | 0.764987 | 0.719048 | 0.767346 | 0.762475 |
| 0.762395 | 0.773366 | 0.766272 | 0.766678 | 0.769407 | 0.768081 | 0.724982 | 0.763036 | 0.768843 |
| 0.761162 | 0.773369 | 0.766955 | 0.763762 | 0.771843 | 0.766873 | 0.719786 | 0.765463 | 0.770726 |
| 0.768833 | 0.767244 | 0.758873 | 0.763877 | 0.7672 | 0.768112 | 0.728131 | 0.762773 | 0.768376 |
| 0.758981 | 0.762237 | 0.762964 | 0.763411 | 0.759307 | 0.761792 | 0.722751 | 0.756383 | 0.759979 |
| 0.757328 | 0.765304 | 0.763587 | 0.761486 | 0.762615 | 0.761041 | 0.726647 | 0.757171 | 0.760286 |
| 0.764716 | 0.768881 | 0.770364 | 0.76762 | 0.770023 | 0.769022 | 0.724299 | 0.771096 | 0.771691 |
| 0.768099 | 0.776634 | 0.772077 | 0.77246 | 0.774419 | 0.770252 | 0.73644 | 0.770651 | 0.766279 |
| 0.76712 | 0.773146 | 0.77513 | 0.767928 | 0.774424 | 0.764376 | 0.728222 | 0.766485 | 0.767277 |
| 0.762926 | 0.767476 | 0.764352 | 0.764686 | 0.769053 | 0.764747 | 0.735598 | 0.766484 | 0.762907 |
| 0.759481 | 0.765268 | 0.766661 | 0.760041 | 0.769574 | 0.755083 | 0.722861 | 0.765133 | 0.763907 |
| 0.768115 | 0.778013 | 0.77236 | 0.769558 | 0.774556 | 0.771876 | 0.726545 | 0.765279 | 0.767358 |
| 0.76214 | 0.767446 | 0.768477 | 0.763851 | 0.767068 | 0.761046 | 0.73353 | 0.765694 | 0.758278 |
| 1 | 0.803299 | 0.803439 | 0.80544 | 0.803953 | 0.859891 | 0.760264 | 0.788386 | 0.777283 |
| 0.803299 | 1 | 0.846908 | 0.806657 | 0.823201 | 0.809228 | 0.77104 | 0.798498 | 0.794564 |
| 0.803439 | 0.846908 | 1 | 0.804206 | 0.813649 | 0.802745 | 0.77557 | 0.785266 | 0.792424 |

| | | | | | | | | |
|---|---|---|---|---|---|---|---|---|
| 0.80544 | 0.806657 | 0.804206 | 1 | 0.808989 | 0.811751 | 0.768279 | 0.791241 | 0.783583 |
| 0.803953 | 0.823201 | 0.813649 | 0.808989 | 1 | 0.800017 | 0.774873 | 0.800479 | 0.791269 |
| 0.859891 | 0.809228 | 0.802745 | 0.811751 | 0.800017 | 1 | 0.771018 | 0.794425 | 0.784578 |
| 0.760264 | 0.77104 | 0.77557 | 0.768279 | 0.774873 | 0.771018 | 1 | 0.742694 | 0.73949 |
| 0.788386 | 0.798498 | 0.785266 | 0.791241 | 0.800479 | 0.794425 | 0.742694 | 1 | 0.80085 |
| 0.777283 | 0.794564 | 0.792424 | 0.783583 | 0.791269 | 0.784578 | 0.73949 | 0.80085 | 1 |
| 0.784889 | 0.795015 | 0.790562 | 0.788586 | 0.79784 | 0.786061 | 0.738758 | 0.80596 | 0.815161 |
| 0.770837 | 0.782364 | 0.780106 | 0.774975 | 0.779619 | 0.777474 | 0.730797 | 0.779462 | 0.783764 |
| 0.777951 | 0.786195 | 0.789657 | 0.783635 | 0.789365 | 0.781718 | 0.743423 | 0.788989 | 0.788323 |
| 0.77691 | 0.783214 | 0.789915 | 0.776719 | 0.788725 | 0.77578 | 0.741679 | 0.784627 | 0.785963 |
| 0.779178 | 0.782225 | 0.780665 | 0.779298 | 0.787271 | 0.780512 | 0.728089 | 0.780839 | 0.783425 |
| 0.781198 | 0.783142 | 0.784544 | 0.781041 | 0.785483 | 0.779417 | 0.737261 | 0.778373 | 0.78972 |
| 0.781775 | 0.783795 | 0.784857 | 0.781779 | 0.780964 | 0.778925 | 0.735178 | 0.781342 | 0.782389 |
| 0.783138 | 0.79616 | 0.790181 | 0.793349 | 0.789822 | 0.788251 | 0.747308 | 0.789513 | 0.786393 |
| 0.785791 | 0.79413 | 0.789238 | 0.791677 | 0.800532 | 0.79145 | 0.74452 | 0.79249 | 0.788302 |
| 0.78882 | 0.790752 | 0.788402 | 0.79534 | 0.798184 | 0.790328 | 0.74659 | 0.788621 | 0.783077 |
| 0.788878 | 0.798804 | 0.788156 | 0.786509 | 0.79784 | 0.790613 | 0.753498 | 0.785802 | 0.78114 |
| 0.789717 | 0.791139 | 0.794665 | 0.791333 | 0.794157 | 0.789808 | 0.7557 | 0.787286 | 0.790116 |
| 0.788641 | 0.794765 | 0.794882 | 0.797196 | 0.791815 | 0.796936 | 0.738534 | 0.783174 | 0.778475 |
| 0.788301 | 0.788209 | 0.791687 | 0.792796 | 0.79402 | 0.79025 | 0.747754 | 0.78715 | 0.788047 |
| 0.790013 | 0.79669 | 0.793853 | 0.79912 | 0.800897 | 0.796317 | 0.75662 | 0.795577 | 0.790916 |
| 0.790482 | 0.797698 | 0.798018 | 0.797619 | 0.801862 | 0.794236 | 0.748108 | 0.790246 | 0.786721 |
| 0.678087 | 0.682152 | 0.684635 | 0.680322 | 0.685902 | 0.683795 | 0.666482 | 0.672196 | 0.682429 |
| 0.662509 | 0.663667 | 0.664666 | 0.648924 | 0.662796 | 0.655132 | 0.655795 | 0.66018 | 0.664428 |
| 0.670987 | 0.675816 | 0.676977 | 0.673791 | 0.679063 | 0.670726 | 0.653727 | 0.669241 | 0.672582 |
| 0.654554 | 0.65625 | 0.653132 | 0.648891 | 0.655352 | 0.65325 | 0.644327 | 0.649226 | 0.642276 |
| 0.740382 | 0.746417 | 0.737923 | 0.741648 | 0.750301 | 0.743157 | 0.715125 | 0.742076 | 0.737879 |
| 0.725729 | 0.74012 | 0.737679 | 0.737847 | 0.744611 | 0.728569 | 0.727603 | 0.73425 | 0.722678 |
| 0.731424 | 0.738345 | 0.733152 | 0.727716 | 0.737894 | 0.740334 | 0.72924 | 0.728823 | 0.727587 |
| 0.741051 | 0.74728 | 0.747784 | 0.747494 | 0.754156 | 0.750423 | 0.722883 | 0.74215 | 0.747894 |
| 0.731034 | 0.736818 | 0.73218 | 0.734548 | 0.742712 | 0.731685 | 0.732872 | 0.733287 | 0.724138 |
| 0.715909 | 0.722317 | 0.719303 | 0.731078 | 0.73587 | 0.733564 | 0.699462 | 0.723217 | 0.721905 |
| 0.728274 | 0.738576 | 0.728062 | 0.730324 | 0.744283 | 0.729722 | 0.709302 | 0.725987 | 0.715135 |
| 0.654376 | 0.660057 | 0.661439 | 0.651887 | 0.653907 | 0.651631 | 0.651515 | 0.656616 | 0.657005 |
| 0.759457 | 0.771263 | 0.76753 | 0.76718 | 0.773111 | 0.764588 | 0.735049 | 0.769065 | 0.764711 |
| 0.743841 | 0.753954 | 0.746659 | 0.749796 | 0.758621 | 0.747559 | 0.723608 | 0.74582 | 0.74783 |
| 0.740816 | 0.746053 | 0.740821 | 0.740144 | 0.752101 | 0.740336 | 0.723338 | 0.74712 | 0.738254 |
| 0.720684 | 0.73948 | 0.731173 | 0.727425 | 0.73691 | 0.730588 | 0.721339 | 0.730406 | 0.727751 |
| 0.747239 | 0.757165 | 0.753893 | 0.751407 | 0.760563 | 0.75341 | 0.734581 | 0.750817 | 0.748999 |
| 0.735142 | 0.74569 | 0.741548 | 0.740484 | 0.744201 | 0.739762 | 0.722682 | 0.739142 | 0.745144 |
| 0.736234 | 0.734824 | 0.730197 | 0.733917 | 0.746034 | 0.735701 | 0.720326 | 0.735731 | 0.736435 |
| 0.732547 | 0.747951 | 0.74029 | 0.741768 | 0.742172 | 0.73893 | 0.720098 | 0.738229 | 0.737739 |
| 0.73739 | 0.7423 | 0.734023 | 0.741788 | 0.743571 | 0.745231 | 0.739989 | 0.736405 | 0.725515 |
| 0.737229 | 0.742527 | 0.739633 | 0.734019 | 0.746195 | 0.73778 | 0.72515 | 0.739925 | 0.737525 |
| 0.738269 | 0.737115 | 0.736883 | 0.745511 | 0.744252 | 0.740412 | 0.732075 | 0.725852 | 0.732152 |
| 0.715474 | 0.722001 | 0.715143 | 0.722259 | 0.72392 | 0.72849 | 0.716312 | 0.719351 | 0.726138 |

| | | | | | | | | |
|---|---|---|---|---|---|---|---|---|
| 0.752432 | 0.760416 | 0.758376 | 0.753425 | 0.765491 | 0.754159 | 0.743699 | 0.759322 | 0.757057 |
| 0.73842 | 0.743251 | 0.74025 | 0.734821 | 0.748513 | 0.74348 | 0.727525 | 0.737308 | 0.737509 |
| 0.734978 | 0.743971 | 0.736866 | 0.729925 | 0.747582 | 0.733283 | 0.71898 | 0.730109 | 0.73913 |
| 0.732286 | 0.745697 | 0.743848 | 0.734466 | 0.75 | 0.732223 | 0.727549 | 0.728795 | 0.735735 |
| 0.73432 | 0.748031 | 0.737914 | 0.732665 | 0.746807 | 0.737005 | 0.711895 | 0.727899 | 0.726095 |
| 0.726276 | 0.730836 | 0.731126 | 0.732546 | 0.737092 | 0.727318 | 0.721118 | 0.734958 | 0.725044 |
| 0.723617 | 0.741197 | 0.72587 | 0.729432 | 0.737098 | 0.724054 | 0.720028 | 0.725448 | 0.727965 |
| 0.723392 | 0.737625 | 0.731476 | 0.728465 | 0.737397 | 0.721414 | 0.720285 | 0.719461 | 0.73142 |
| 0.726129 | 0.733447 | 0.730539 | 0.732026 | 0.740311 | 0.727868 | 0.708562 | 0.724662 | 0.721841 |
| 0.753925 | 0.766363 | 0.76153 | 0.755682 | 0.766903 | 0.75597 | 0.734435 | 0.760297 | 0.75732 |
| 0.752067 | 0.760903 | 0.757919 | 0.753476 | 0.763812 | 0.750507 | 0.724565 | 0.756426 | 0.752144 |
| 0.746509 | 0.755284 | 0.753195 | 0.747035 | 0.7596 | 0.752182 | 0.725506 | 0.746308 | 0.754388 |
| 0.733543 | 0.748746 | 0.748773 | 0.738385 | 0.75417 | 0.735161 | 0.707795 | 0.740597 | 0.742644 |
| 0.74856 | 0.751976 | 0.749142 | 0.755082 | 0.758591 | 0.752659 | 0.723719 | 0.745722 | 0.743835 |
| 0.741028 | 0.750186 | 0.745828 | 0.745963 | 0.753076 | 0.744876 | 0.7247 | 0.74271 | 0.741907 |

| | plot 18 | | | | | | | |
|---|---|---|---|---|---|---|---|---|
| Wild_plot_ | Wild_plot_ | Wild_plot_ | Wild_plot_ | Wild_plot_ | Wild_plot_ | Wild_plot_ | Wild_plot_ | Wild_plot_ |
| 0.70425 | 0.704687 | 0.709808 | 0.705035 | 0.708217 | 0.708294 | 0.710037 | 0.704912 | 0.711297 |
| 0.712354 | 0.709882 | 0.715764 | 0.714252 | 0.713098 | 0.71519 | 0.716157 | 0.712936 | 0.719133 |
| 0.706952 | 0.702263 | 0.707623 | 0.702 | 0.704328 | 0.705257 | 0.706115 | 0.70447 | 0.708076 |
| 0.713449 | 0.71398 | 0.714974 | 0.712656 | 0.712505 | 0.714843 | 0.714422 | 0.712886 | 0.716575 |
| 0.705093 | 0.71042 | 0.707785 | 0.708238 | 0.708926 | 0.711844 | 0.713957 | 0.708067 | 0.709529 |
| 0.718081 | 0.718852 | 0.719594 | 0.717083 | 0.713707 | 0.720613 | 0.716411 | 0.715346 | 0.718844 |
| 0.767037 | 0.764514 | 0.76898 | 0.772887 | 0.767577 | 0.768643 | 0.771743 | 0.768697 | 0.768859 |
| 0.765602 | 0.761653 | 0.773642 | 0.766475 | 0.765054 | 0.766804 | 0.767137 | 0.772422 | 0.768839 |
| 0.713716 | 0.710503 | 0.718748 | 0.715676 | 0.71207 | 0.715178 | 0.714195 | 0.709782 | 0.715368 |
| 0.707278 | 0.704619 | 0.711863 | 0.706381 | 0.707429 | 0.710338 | 0.712128 | 0.704844 | 0.710522 |
| 0.709079 | 0.70828 | 0.711198 | 0.708078 | 0.70817 | 0.709198 | 0.712757 | 0.706394 | 0.714343 |
| 0.700815 | 0.70093 | 0.702718 | 0.699109 | 0.703186 | 0.706098 | 0.707218 | 0.702833 | 0.705472 |
| 0.711629 | 0.711688 | 0.714615 | 0.708049 | 0.710831 | 0.712458 | 0.71083 | 0.710835 | 0.713353 |
| 0.717786 | 0.717041 | 0.725812 | 0.723802 | 0.71884 | 0.722138 | 0.724036 | 0.718395 | 0.72472 |
| 0.732224 | 0.733062 | 0.732446 | 0.732251 | 0.730766 | 0.738134 | 0.734842 | 0.729302 | 0.734736 |
| 0.716424 | 0.714455 | 0.719241 | 0.713775 | 0.71749 | 0.714947 | 0.716492 | 0.716681 | 0.7173 |
| 0.715652 | 0.711581 | 0.718956 | 0.713128 | 0.713768 | 0.712172 | 0.71566 | 0.717579 | 0.715293 |
| 0.737385 | 0.737055 | 0.744784 | 0.735229 | 0.73904 | 0.740847 | 0.740054 | 0.737406 | 0.742791 |
| 0.706032 | 0.704345 | 0.70839 | 0.706792 | 0.707059 | 0.706978 | 0.709448 | 0.705663 | 0.709922 |
| 0.713773 | 0.709143 | 0.715378 | 0.708608 | 0.710724 | 0.713576 | 0.7127 | 0.710898 | 0.714707 |
| 0.707508 | 0.706561 | 0.71165 | 0.708856 | 0.706051 | 0.708287 | 0.710167 | 0.704066 | 0.711836 |
| 0.710853 | 0.708396 | 0.712923 | 0.712191 | 0.712324 | 0.711671 | 0.714001 | 0.711173 | 0.713489 |
| 0.712085 | 0.713538 | 0.720354 | 0.717528 | 0.714787 | 0.718068 | 0.7189 | 0.714399 | 0.718683 |
| 0.712628 | 0.710067 | 0.716947 | 0.707433 | 0.714594 | 0.715602 | 0.714957 | 0.712307 | 0.715266 |
| 0.710772 | 0.712579 | 0.717744 | 0.713064 | 0.71338 | 0.715741 | 0.717595 | 0.714388 | 0.719554 |
| 0.710508 | 0.708993 | 0.715653 | 0.711754 | 0.710222 | 0.710822 | 0.714546 | 0.708481 | 0.712726 |
| 0.70344 | 0.70592 | 0.708148 | 0.706724 | 0.706028 | 0.705671 | 0.706603 | 0.703698 | 0.708184 |
| 0.710821 | 0.710829 | 0.718708 | 0.712082 | 0.713832 | 0.714694 | 0.715964 | 0.711232 | 0.716382 |
| 0.713443 | 0.71205 | 0.717395 | 0.712886 | 0.716653 | 0.716552 | 0.716326 | 0.710925 | 0.717353 |
| 0.708633 | 0.710049 | 0.716377 | 0.709167 | 0.708603 | 0.716719 | 0.711618 | 0.70918 | 0.716059 |
| 0.709081 | 0.70728 | 0.712054 | 0.709268 | 0.71016 | 0.715723 | 0.714172 | 0.711134 | 0.712609 |
| 0.706789 | 0.703266 | 0.712841 | 0.706045 | 0.705524 | 0.711265 | 0.709723 | 0.701984 | 0.708241 |
| 0.711317 | 0.707521 | 0.71187 | 0.709902 | 0.708905 | 0.708641 | 0.7109 | 0.701562 | 0.710954 |
| 0.706505 | 0.711013 | 0.714252 | 0.710814 | 0.712502 | 0.713251 | 0.716099 | 0.70831 | 0.715488 |
| 0.70349 | 0.704301 | 0.710873 | 0.705826 | 0.705251 | 0.706014 | 0.707816 | 0.703023 | 0.706274 |
| 0.709659 | 0.710665 | 0.712253 | 0.709275 | 0.709215 | 0.711605 | 0.715317 | 0.70906 | 0.713897 |
| 0.714624 | 0.711237 | 0.717345 | 0.715632 | 0.71174 | 0.715657 | 0.715279 | 0.715606 | 0.719851 |
| 0.711489 | 0.710185 | 0.717334 | 0.712138 | 0.71095 | 0.714689 | 0.714999 | 0.708833 | 0.716208 |
| 0.712947 | 0.715556 | 0.717149 | 0.714933 | 0.712192 | 0.716008 | 0.716722 | 0.711774 | 0.716685 |

| | | | | | | | | |
|---|---|---|---|---|---|---|---|---|
| 0.744798 | 0.744957 | 0.74996 | 0.740081 | 0.742158 | 0.747464 | 0.746736 | 0.74551 | 0.749236 |
| 0.699203 | 0.699172 | 0.703982 | 0.702003 | 0.704295 | 0.704324 | 0.709275 | 0.700677 | 0.70488 |
| 0.705952 | 0.703554 | 0.707402 | 0.707369 | 0.707195 | 0.710667 | 0.712442 | 0.707093 | 0.708884 |
| 0.714715 | 0.712519 | 0.713655 | 0.713313 | 0.71115 | 0.714569 | 0.716425 | 0.710148 | 0.714104 |
| 0.707811 | 0.706918 | 0.712486 | 0.71231 | 0.708956 | 0.715884 | 0.716029 | 0.709764 | 0.713661 |
| 0.71148 | 0.707338 | 0.717084 | 0.711237 | 0.710758 | 0.714171 | 0.716218 | 0.712418 | 0.715292 |
| 0.707838 | 0.705252 | 0.709598 | 0.706175 | 0.707532 | 0.711716 | 0.710189 | 0.703421 | 0.709172 |
| 0.710295 | 0.709554 | 0.714605 | 0.7132 | 0.710008 | 0.712289 | 0.712777 | 0.707298 | 0.711292 |
| 0.711069 | 0.712528 | 0.715567 | 0.713598 | 0.712219 | 0.716691 | 0.714079 | 0.71335 | 0.718499 |
| 0.713492 | 0.71257 | 0.718695 | 0.712475 | 0.711755 | 0.71358 | 0.715308 | 0.710913 | 0.713515 |
| 0.701676 | 0.700515 | 0.704578 | 0.703314 | 0.702985 | 0.701276 | 0.705104 | 0.700641 | 0.705249 |
| 0.703866 | 0.69801 | 0.706965 | 0.703222 | 0.701194 | 0.704129 | 0.705667 | 0.702816 | 0.70754 |
| 0.710082 | 0.705643 | 0.711326 | 0.707337 | 0.70819 | 0.713527 | 0.71269 | 0.707965 | 0.710938 |
| 0.703978 | 0.702763 | 0.709107 | 0.705006 | 0.705234 | 0.710872 | 0.711289 | 0.704331 | 0.706488 |
| 0.70837 | 0.70569 | 0.70866 | 0.706345 | 0.706261 | 0.708176 | 0.706571 | 0.703997 | 0.704747 |
| 0.705485 | 0.703356 | 0.710444 | 0.70631 | 0.707619 | 0.709008 | 0.70998 | 0.705401 | 0.703141 |
| 0.705349 | 0.702126 | 0.705529 | 0.702722 | 0.701052 | 0.708112 | 0.7076 | 0.70277 | 0.703849 |
| 0.709718 | 0.710257 | 0.714682 | 0.713604 | 0.711283 | 0.715587 | 0.713945 | 0.710343 | 0.713865 |
| 0.712287 | 0.708858 | 0.715797 | 0.711702 | 0.711471 | 0.712779 | 0.717204 | 0.713006 | 0.71506 |
| 0.707557 | 0.707229 | 0.711595 | 0.709205 | 0.709677 | 0.710235 | 0.710404 | 0.705162 | 0.709858 |
| 0.703356 | 0.701272 | 0.70591 | 0.705203 | 0.707676 | 0.711885 | 0.71193 | 0.702949 | 0.707565 |
| 0.705592 | 0.704257 | 0.708131 | 0.707938 | 0.705882 | 0.707216 | 0.709742 | 0.703625 | 0.708975 |
| 0.70446 | 0.706267 | 0.712108 | 0.70914 | 0.711881 | 0.709844 | 0.713448 | 0.704774 | 0.7094 |
| 0.719763 | 0.714968 | 0.720746 | 0.71521 | 0.715248 | 0.71881 | 0.717327 | 0.713376 | 0.720365 |
| 0.725803 | 0.723452 | 0.726561 | 0.724646 | 0.722586 | 0.724584 | 0.725464 | 0.724896 | 0.729725 |
| 0.766314 | 0.761858 | 0.773017 | 0.767842 | 0.766098 | 0.767695 | 0.767773 | 0.772494 | 0.7689 |
| 0.76777 | 0.763968 | 0.7731 | 0.770492 | 0.770655 | 0.768559 | 0.770747 | 0.769822 | 0.769912 |
| 0.766969 | 0.76297 | 0.774431 | 0.770373 | 0.76748 | 0.770861 | 0.771686 | 0.774207 | 0.770802 |
| 0.708138 | 0.703944 | 0.712388 | 0.708615 | 0.706821 | 0.708825 | 0.707426 | 0.705798 | 0.708638 |
| 0.705623 | 0.705406 | 0.709009 | 0.707469 | 0.707092 | 0.707726 | 0.707303 | 0.70581 | 0.709461 |
| 0.713863 | 0.712626 | 0.715118 | 0.71472 | 0.713326 | 0.717625 | 0.714868 | 0.712943 | 0.713966 |
| 0.715115 | 0.715264 | 0.717657 | 0.715415 | 0.71678 | 0.72253 | 0.719686 | 0.714596 | 0.717942 |
| 0.711249 | 0.709424 | 0.714343 | 0.710522 | 0.708615 | 0.712255 | 0.711365 | 0.71119 | 0.71515 |
| 0.70797 | 0.708896 | 0.712456 | 0.7113 | 0.706426 | 0.710412 | 0.713333 | 0.710699 | 0.711343 |
| 0.710596 | 0.708072 | 0.712775 | 0.709636 | 0.70926 | 0.711237 | 0.712621 | 0.70961 | 0.711904 |
| 0.711509 | 0.709436 | 0.715417 | 0.713085 | 0.710375 | 0.71339 | 0.714457 | 0.711551 | 0.713954 |
| 0.706975 | 0.710319 | 0.712384 | 0.711274 | 0.712068 | 0.715294 | 0.713529 | 0.706892 | 0.708287 |
| 0.72476 | 0.727906 | 0.729424 | 0.726485 | 0.726696 | 0.730496 | 0.729736 | 0.723443 | 0.730744 |
| 0.704435 | 0.704701 | 0.710182 | 0.704895 | 0.706467 | 0.70876 | 0.707071 | 0.7058 | 0.705702 |
| 0.71101 | 0.707608 | 0.714196 | 0.709888 | 0.709267 | 0.712758 | 0.714331 | 0.708598 | 0.713159 |
| 0.707952 | 0.707426 | 0.710137 | 0.708442 | 0.70401 | 0.709363 | 0.71169 | 0.702569 | 0.707064 |
| 0.725459 | 0.724688 | 0.728381 | 0.724813 | 0.724766 | 0.728985 | 0.727836 | 0.723465 | 0.728647 |
| 0.711904 | 0.708495 | 0.712067 | 0.708961 | 0.712061 | 0.716002 | 0.714297 | 0.71291 | 0.715062 |
| 0.705492 | 0.706169 | 0.707801 | 0.706667 | 0.708482 | 0.710166 | 0.709935 | 0.705171 | 0.708785 |
| 0.705099 | 0.707467 | 0.712084 | 0.710537 | 0.707059 | 0.710596 | 0.711431 | 0.708124 | 0.710558 |
| 0.708137 | 0.70852 | 0.714894 | 0.708932 | 0.708679 | 0.715431 | 0.714746 | 0.70929 | 0.712275 |

| | | | | | | | | |
|---|---|---|---|---|---|---|---|---|
| 0.70932 | 0.708118 | 0.709537 | 0.707624 | 0.707738 | 0.710072 | 0.709055 | 0.706573 | 0.709307 |
| 0.708383 | 0.704874 | 0.714908 | 0.712413 | 0.710489 | 0.711731 | 0.71365 | 0.712592 | 0.717421 |
| 0.711822 | 0.707138 | 0.711997 | 0.710778 | 0.710868 | 0.712857 | 0.715618 | 0.711874 | 0.713437 |
| 0.715391 | 0.718521 | 0.719894 | 0.717591 | 0.718871 | 0.720097 | 0.721696 | 0.714653 | 0.721764 |
| 0.714966 | 0.717496 | 0.720228 | 0.719085 | 0.71499 | 0.722529 | 0.719344 | 0.717178 | 0.718934 |
| 0.709136 | 0.709639 | 0.714755 | 0.711719 | 0.709116 | 0.712756 | 0.713598 | 0.708842 | 0.714297 |
| 0.709766 | 0.711923 | 0.719304 | 0.710501 | 0.71252 | 0.714887 | 0.710955 | 0.710435 | 0.71327 |
| 0.715204 | 0.713782 | 0.718274 | 0.71594 | 0.715771 | 0.71904 | 0.716663 | 0.7115 | 0.716341 |
| 0.709353 | 0.708506 | 0.713749 | 0.71039 | 0.713275 | 0.717363 | 0.715606 | 0.709353 | 0.711858 |
| 0.705771 | 0.706711 | 0.710773 | 0.706897 | 0.70434 | 0.713188 | 0.710436 | 0.704379 | 0.709042 |
| 0.71041 | 0.709037 | 0.710303 | 0.712177 | 0.706936 | 0.715023 | 0.71041 | 0.708775 | 0.710744 |
| 0.709161 | 0.711025 | 0.715426 | 0.712198 | 0.71196 | 0.711971 | 0.713824 | 0.707395 | 0.714401 |
| 0.709597 | 0.708313 | 0.714743 | 0.711381 | 0.711515 | 0.71355 | 0.713427 | 0.708078 | 0.713129 |
| 0.707258 | 0.708273 | 0.711261 | 0.708236 | 0.707625 | 0.709281 | 0.71352 | 0.705174 | 0.71121 |
| 0.70763 | 0.708661 | 0.712882 | 0.707821 | 0.707483 | 0.712922 | 0.710665 | 0.707707 | 0.713783 |
| 0.709146 | 0.706393 | 0.71217 | 0.707408 | 0.708135 | 0.71234 | 0.711856 | 0.710006 | 0.710232 |
| 0.712912 | 0.710722 | 0.718013 | 0.716696 | 0.713516 | 0.714965 | 0.715308 | 0.710295 | 0.715905 |
| 0.712045 | 0.712801 | 0.715569 | 0.711279 | 0.711056 | 0.713521 | 0.714661 | 0.710155 | 0.714035 |
| 0.7106 | 0.710489 | 0.715642 | 0.71368 | 0.709935 | 0.715317 | 0.716482 | 0.709775 | 0.713043 |
| 0.708516 | 0.709228 | 0.714263 | 0.711106 | 0.7112 | 0.714114 | 0.715601 | 0.708095 | 0.712415 |
| 0.711784 | 0.713351 | 0.714445 | 0.710967 | 0.71211 | 0.716817 | 0.71496 | 0.713066 | 0.715157 |
| 0.706511 | 0.703955 | 0.711836 | 0.707369 | 0.707736 | 0.71194 | 0.70914 | 0.705632 | 0.70939 |
| 0.709367 | 0.712628 | 0.714469 | 0.711028 | 0.708357 | 0.71643 | 0.714538 | 0.708756 | 0.710668 |
| 0.712742 | 0.710331 | 0.716369 | 0.710579 | 0.711352 | 0.717977 | 0.717691 | 0.709349 | 0.715116 |
| 0.709097 | 0.713231 | 0.714792 | 0.712824 | 0.713191 | 0.715997 | 0.714401 | 0.712007 | 0.711665 |
| 0.711181 | 0.714365 | 0.718634 | 0.71327 | 0.712066 | 0.718053 | 0.715587 | 0.711824 | 0.71432 |
| 0.709425 | 0.71127 | 0.71512 | 0.711185 | 0.710884 | 0.716208 | 0.715384 | 0.710054 | 0.712118 |
| 0.710372 | 0.70929 | 0.712468 | 0.709997 | 0.711242 | 0.716442 | 0.713083 | 0.707177 | 0.714422 |
| 0.711134 | 0.711984 | 0.718255 | 0.71148 | 0.709311 | 0.714936 | 0.712787 | 0.710124 | 0.713026 |
| 0.709469 | 0.711066 | 0.713835 | 0.711882 | 0.710752 | 0.714274 | 0.713501 | 0.71264 | 0.712831 |
| 0.713713 | 0.717137 | 0.718056 | 0.712751 | 0.711263 | 0.716866 | 0.717671 | 0.711363 | 0.713931 |
| 0.712758 | 0.714117 | 0.71276 | 0.712521 | 0.711027 | 0.717524 | 0.711787 | 0.706928 | 0.711786 |
| 0.713121 | 0.710115 | 0.716675 | 0.714978 | 0.71267 | 0.718271 | 0.713225 | 0.710934 | 0.71379 |
| 0.714753 | 0.711831 | 0.713411 | 0.712744 | 0.710755 | 0.714122 | 0.717051 | 0.708245 | 0.711543 |
| 0.711858 | 0.710373 | 0.715268 | 0.710602 | 0.709149 | 0.714056 | 0.712044 | 0.707071 | 0.711669 |
| 0.711655 | 0.712949 | 0.717642 | 0.713615 | 0.713534 | 0.714007 | 0.714274 | 0.709524 | 0.713904 |
| 0.712234 | 0.707194 | 0.711429 | 0.707609 | 0.707932 | 0.713037 | 0.712962 | 0.704337 | 0.710359 |
| 0.687063 | 0.681512 | 0.689914 | 0.687145 | 0.686916 | 0.688934 | 0.69501 | 0.683833 | 0.689313 |
| 0.68921 | 0.689964 | 0.693532 | 0.689732 | 0.689694 | 0.690813 | 0.693789 | 0.686609 | 0.695349 |
| 0.685263 | 0.678637 | 0.685427 | 0.678505 | 0.677823 | 0.681935 | 0.682163 | 0.677538 | 0.678648 |
| 0.681591 | 0.68015 | 0.687417 | 0.68066 | 0.680585 | 0.683847 | 0.688827 | 0.679872 | 0.682902 |
| 0.66608 | 0.664808 | 0.666279 | 0.666823 | 0.666199 | 0.670813 | 0.668337 | 0.663098 | 0.669155 |
| 0.670097 | 0.666426 | 0.675355 | 0.670661 | 0.669727 | 0.673703 | 0.673037 | 0.662509 | 0.672714 |
| 0.670637 | 0.663858 | 0.671998 | 0.665297 | 0.660792 | 0.665855 | 0.671598 | 0.670174 | 0.669698 |
| 0.674713 | 0.670685 | 0.670831 | 0.671474 | 0.671947 | 0.672555 | 0.672491 | 0.666667 | 0.669671 |
| 0.671783 | 0.665883 | 0.673452 | 0.668121 | 0.670376 | 0.673703 | 0.674731 | 0.668407 | 0.672784 |
| 0.674833 | 0.676101 | 0.683943 | 0.676999 | 0.673429 | 0.67973 | 0.681587 | 0.671994 | 0.676194 |

| | | | | | | | | |
|---|---|---|---|---|---|---|---|---|
| 0.674238 | 0.671112 | 0.679301 | 0.676997 | 0.670262 | 0.675478 | 0.678899 | 0.670094 | 0.673635 |
| 0.674616 | 0.675016 | 0.677436 | 0.674264 | 0.676639 | 0.67722 | 0.673419 | 0.673851 | 0.678255 |
| 0.676178 | 0.66981 | 0.672241 | 0.671982 | 0.669376 | 0.671804 | 0.674297 | 0.665961 | 0.670701 |
| 0.666667 | 0.659139 | 0.663463 | 0.66363 | 0.663892 | 0.664502 | 0.666359 | 0.657502 | 0.664208 |
| 0.664534 | 0.664752 | 0.668459 | 0.661842 | 0.669712 | 0.666839 | 0.668424 | 0.661725 | 0.666581 |
| 0.666301 | 0.666347 | 0.672602 | 0.671172 | 0.66621 | 0.671229 | 0.674066 | 0.664577 | 0.668133 |
| 0.666619 | 0.661446 | 0.667374 | 0.669739 | 0.669888 | 0.67222 | 0.674706 | 0.663291 | 0.670339 |
| 0.665683 | 0.659145 | 0.671646 | 0.665036 | 0.662734 | 0.668642 | 0.668676 | 0.66569 | 0.66876 |
| 0.668932 | 0.659822 | 0.66515 | 0.663243 | 0.656825 | 0.666617 | 0.663832 | 0.660176 | 0.667352 |
| 0.659822 | 0.659957 | 0.669215 | 0.667008 | 0.666423 | 0.665886 | 0.670771 | 0.659861 | 0.665262 |
| 0.65797 | 0.653251 | 0.668917 | 0.663642 | 0.656443 | 0.657983 | 0.665267 | 0.657186 | 0.657697 |
| 0.698234 | 0.701795 | 0.703449 | 0.701812 | 0.696711 | 0.702698 | 0.701262 | 0.700062 | 0.704355 |
| 0.710778 | 0.708274 | 0.715604 | 0.709038 | 0.710528 | 0.714717 | 0.712005 | 0.707446 | 0.710153 |
| 0.690294 | 0.684831 | 0.69222 | 0.685744 | 0.685938 | 0.692202 | 0.691667 | 0.687018 | 0.691525 |
| 0.712087 | 0.71609 | 0.719331 | 0.716884 | 0.719744 | 0.721284 | 0.718138 | 0.711316 | 0.721051 |
| 0.705812 | 0.705322 | 0.707192 | 0.707243 | 0.707482 | 0.706131 | 0.709162 | 0.701856 | 0.709344 |
| 0.711645 | 0.710151 | 0.717827 | 0.710716 | 0.715421 | 0.718628 | 0.715102 | 0.709797 | 0.716077 |
| 0.718632 | 0.718479 | 0.720399 | 0.715295 | 0.716 | 0.725423 | 0.719352 | 0.712029 | 0.719665 |
| 0.698596 | 0.698053 | 0.701739 | 0.69938 | 0.697941 | 0.703044 | 0.699507 | 0.698178 | 0.701872 |
| 0.718067 | 0.721774 | 0.724012 | 0.71728 | 0.717825 | 0.723592 | 0.721588 | 0.719991 | 0.724215 |
| 0.689481 | 0.687103 | 0.698624 | 0.691203 | 0.68735 | 0.698858 | 0.689602 | 0.691561 | 0.69278 |
| 0.716414 | 0.711796 | 0.720099 | 0.717971 | 0.711803 | 0.715683 | 0.715202 | 0.713445 | 0.720064 |
| 0.716215 | 0.718633 | 0.721111 | 0.717224 | 0.718301 | 0.719663 | 0.719805 | 0.719776 | 0.724027 |
| 0.688077 | 0.681119 | 0.690507 | 0.683972 | 0.686803 | 0.687215 | 0.685754 | 0.679655 | 0.683378 |
| 0.692518 | 0.687654 | 0.69182 | 0.688307 | 0.687613 | 0.694246 | 0.689851 | 0.685698 | 0.692536 |
| 0.698235 | 0.692037 | 0.698963 | 0.691636 | 0.688758 | 0.699008 | 0.6941 | 0.694048 | 0.697373 |
| 0.704226 | 0.705992 | 0.711707 | 0.706451 | 0.703361 | 0.70783 | 0.707216 | 0.709387 | 0.709732 |
| 0.716457 | 0.718979 | 0.720643 | 0.720811 | 0.71718 | 0.720505 | 0.717253 | 0.715259 | 0.717181 |
| 0.687113 | 0.689581 | 0.693868 | 0.692846 | 0.687185 | 0.692523 | 0.692516 | 0.68721 | 0.693575 |
| 0.722015 | 0.72746 | 0.729119 | 0.724356 | 0.725557 | 0.728354 | 0.724487 | 0.719167 | 0.723515 |
| 0.724081 | 0.724211 | 0.727724 | 0.723418 | 0.723657 | 0.725461 | 0.72227 | 0.722109 | 0.72543 |
| 0.697406 | 0.69966 | 0.703404 | 0.700763 | 0.702094 | 0.701729 | 0.701922 | 0.697116 | 0.703586 |
| 0.699945 | 0.699112 | 0.705141 | 0.700016 | 0.701487 | 0.701897 | 0.702856 | 0.69743 | 0.703765 |
| 0.724199 | 0.724701 | 0.733328 | 0.724413 | 0.721268 | 0.729771 | 0.723632 | 0.725518 | 0.728705 |
| 0.725092 | 0.724382 | 0.727836 | 0.723688 | 0.72182 | 0.723267 | 0.723832 | 0.720213 | 0.721939 |
| 0.72336 | 0.727069 | 0.727739 | 0.725155 | 0.720376 | 0.731137 | 0.722561 | 0.722897 | 0.728552 |
| 0.683808 | 0.677632 | 0.687041 | 0.681747 | 0.67801 | 0.683285 | 0.680679 | 0.67933 | 0.682062 |
| 0.716957 | 0.723862 | 0.724587 | 0.724713 | 0.721855 | 0.725066 | 0.724893 | 0.721681 | 0.720832 |
| 0.681054 | 0.680009 | 0.683477 | 0.676495 | 0.678423 | 0.679041 | 0.680771 | 0.676666 | 0.677615 |
| 0.669873 | 0.669163 | 0.673888 | 0.670027 | 0.671054 | 0.672756 | 0.671317 | 0.671808 | 0.671237 |
| 0.721914 | 0.718725 | 0.725113 | 0.722827 | 0.720779 | 0.724278 | 0.72652 | 0.722723 | 0.725881 |
| 0.671 | 0.67176 | 0.673916 | 0.673961 | 0.67202 | 0.675244 | 0.673252 | 0.673667 | 0.673634 |
| 0.662137 | 0.6624 | 0.668599 | 0.665613 | 0.667171 | 0.667625 | 0.668801 | 0.663449 | 0.667318 |
| 0.658628 | 0.658301 | 0.664198 | 0.661415 | 0.663191 | 0.662139 | 0.662005 | 0.65916 | 0.663022 |
| 0.729592 | 0.726781 | 0.731758 | 0.726759 | 0.726895 | 0.732468 | 0.732277 | 0.725535 | 0.728909 |
| 0.662382 | 0.663022 | 0.668571 | 0.662236 | 0.664027 | 0.663929 | 0.666867 | 0.662398 | 0.667971 |
| 0.722049 | 0.721478 | 0.728577 | 0.721111 | 0.722365 | 0.725435 | 0.725008 | 0.724826 | 0.727681 |

| | | | | | | | | |
|---|---|---|---|---|---|---|---|---|
| 0.731701 | 0.732055 | 0.7355 | 0.726832 | 0.727657 | 0.730942 | 0.730622 | 0.731688 | 0.732609 |
| 0.724859 | 0.722399 | 0.728849 | 0.726961 | 0.724053 | 0.727738 | 0.727839 | 0.725787 | 0.728309 |
| 0.744935 | 0.744524 | 0.750183 | 0.74432 | 0.741752 | 0.745358 | 0.741551 | 0.743982 | 0.745515 |
| 0.652601 | 0.650942 | 0.656617 | 0.652241 | 0.656071 | 0.653058 | 0.655769 | 0.649636 | 0.655523 |
| 0.654425 | 0.654506 | 0.65898 | 0.654704 | 0.657905 | 0.655026 | 0.657268 | 0.654134 | 0.656942 |
| 0.734777 | 0.732488 | 0.737028 | 0.731781 | 0.732253 | 0.733387 | 0.732986 | 0.732768 | 0.734827 |
| 0.65744 | 0.653775 | 0.6603 | 0.658387 | 0.655203 | 0.655545 | 0.657942 | 0.657746 | 0.658314 |
| 0.732057 | 0.732506 | 0.73501 | 0.731079 | 0.729422 | 0.73373 | 0.730065 | 0.729944 | 0.73038 |
| 0.735555 | 0.731356 | 0.737771 | 0.732748 | 0.729562 | 0.735361 | 0.731745 | 0.737198 | 0.73306 |
| 0.729382 | 0.732133 | 0.731925 | 0.7328 | 0.728578 | 0.730263 | 0.734677 | 0.732227 | 0.729382 |
| 0.734973 | 0.729344 | 0.737489 | 0.729518 | 0.725554 | 0.733355 | 0.732352 | 0.73586 | 0.739215 |
| 0.79562 | 0.731544 | 0.722973 | 0.748299 | 0.713287 | 0.781022 | 0.718121 | 0.772414 | 0.687117 |
| 0.733213 | 0.73182 | 0.738545 | 0.738611 | 0.738554 | 0.73301 | 0.736058 | 0.731435 | 0.740145 |
| 0.732379 | 0.725466 | 0.732829 | 0.73269 | 0.728475 | 0.733481 | 0.724095 | 0.732123 | 0.743671 |
| 0.742979 | 0.741255 | 0.742694 | 0.739631 | 0.74329 | 0.744788 | 0.742195 | 0.742507 | 0.745292 |
| 0.740173 | 0.745551 | 0.74368 | 0.741194 | 0.735818 | 0.739035 | 0.741143 | 0.740517 | 0.746347 |
| 0.596242 | 0.588254 | 0.596427 | 0.593682 | 0.591589 | 0.589763 | 0.5985 | 0.594802 | 0.592681 |
| 0.742565 | 0.746926 | 0.74604 | 0.74593 | 0.74282 | 0.739474 | 0.739746 | 0.744558 | 0.745371 |
| 0.740955 | 0.735613 | 0.739561 | 0.736462 | 0.733496 | 0.743978 | 0.737714 | 0.739653 | 0.743909 |
| 0.742407 | 0.740832 | 0.739752 | 0.734872 | 0.72961 | 0.741765 | 0.735111 | 0.736593 | 0.741077 |
| 0.73877 | 0.738364 | 0.741144 | 0.736295 | 0.732813 | 0.738697 | 0.738198 | 0.74046 | 0.742561 |
| 0.596082 | 0.593664 | 0.599755 | 0.597931 | 0.595357 | 0.595707 | 0.597185 | 0.596233 | 0.597686 |
| 0.585443 | 0.582868 | 0.587258 | 0.587161 | 0.584861 | 0.583202 | 0.587111 | 0.583811 | 0.587023 |
| 0.769212 | 0.760235 | 0.765619 | 0.760709 | 0.764226 | 0.767127 | 0.761655 | 0.761795 | 0.759784 |
| 0.771732 | 0.766476 | 0.765851 | 0.766003 | 0.762987 | 0.767806 | 0.766092 | 0.765326 | 0.767709 |
| 0.761475 | 0.759713 | 0.760207 | 0.764404 | 0.756518 | 0.754667 | 0.757423 | 0.757259 | 0.759614 |
| 0.767335 | 0.759694 | 0.764735 | 0.765356 | 0.760551 | 0.765112 | 0.764407 | 0.762002 | 0.766721 |
| 0.767387 | 0.76497 | 0.768226 | 0.762965 | 0.762069 | 0.765594 | 0.764311 | 0.764533 | 0.765455 |
| 0.773976 | 0.765266 | 0.768995 | 0.769563 | 0.767368 | 0.766048 | 0.765998 | 0.765002 | 0.764745 |
| 0.770835 | 0.771457 | 0.767822 | 0.76906 | 0.770923 | 0.767164 | 0.769705 | 0.766437 | 0.768927 |
| 0.765044 | 0.766218 | 0.764171 | 0.759178 | 0.758683 | 0.762585 | 0.758627 | 0.764098 | 0.762993 |
| 0.77064 | 0.767342 | 0.772853 | 0.771741 | 0.770137 | 0.76615 | 0.773076 | 0.768668 | 0.768993 |
| 0.756474 | 0.760555 | 0.763491 | 0.761464 | 0.758957 | 0.762142 | 0.763561 | 0.763621 | 0.764171 |
| 0.7562 | 0.759603 | 0.769712 | 0.763109 | 0.755548 | 0.760767 | 0.761334 | 0.761261 | 0.764362 |
| 0.766519 | 0.765628 | 0.769489 | 0.767347 | 0.76037 | 0.766283 | 0.763483 | 0.762695 | 0.766956 |
| 0.760628 | 0.7652 | 0.766828 | 0.76158 | 0.761378 | 0.768854 | 0.764366 | 0.76693 | 0.763364 |
| 0.75907 | 0.764022 | 0.766234 | 0.765994 | 0.762211 | 0.762117 | 0.759661 | 0.763378 | 0.76303 |
| 0.760933 | 0.762495 | 0.766051 | 0.763321 | 0.763736 | 0.767476 | 0.76372 | 0.765359 | 0.766563 |
| 0.756998 | 0.760279 | 0.762699 | 0.755675 | 0.753637 | 0.754047 | 0.75782 | 0.756852 | 0.763391 |
| 0.756147 | 0.756814 | 0.762083 | 0.758284 | 0.758292 | 0.757198 | 0.756438 | 0.758657 | 0.761617 |
| 0.733087 | 0.734059 | 0.740354 | 0.739262 | 0.729174 | 0.731664 | 0.729185 | 0.731789 | 0.733556 |
| 0.738634 | 0.732754 | 0.738949 | 0.744146 | 0.738176 | 0.735658 | 0.737628 | 0.735621 | 0.736377 |
| 0.760537 | 0.757998 | 0.765266 | 0.763162 | 0.756882 | 0.764595 | 0.759142 | 0.763747 | 0.763856 |
| 0.769667 | 0.76665 | 0.770327 | 0.7679 | 0.767239 | 0.769839 | 0.771053 | 0.76845 | 0.770599 |
| 0.714918 | 0.718047 | 0.718768 | 0.724245 | 0.721451 | 0.717615 | 0.722233 | 0.72282 | 0.727994 |
| 0.761678 | 0.763037 | 0.766577 | 0.763587 | 0.763817 | 0.764526 | 0.763451 | 0.76761 | 0.769547 |
| 0.757401 | 0.758172 | 0.75958 | 0.757287 | 0.7595 | 0.757717 | 0.758276 | 0.758921 | 0.761547 |

| | | | | | | | | |
|---|---|---|---|---|---|---|---|---|
| 0.761371 | 0.767535 | 0.76751 | 0.762162 | 0.765336 | 0.764203 | 0.764817 | 0.762629 | 0.766508 |
| 0.763505 | 0.762933 | 0.768905 | 0.765822 | 0.762359 | 0.760646 | 0.765616 | 0.760513 | 0.767063 |
| 0.762869 | 0.769186 | 0.768825 | 0.76734 | 0.76635 | 0.766818 | 0.768112 | 0.767987 | 0.770802 |
| 0.764581 | 0.761971 | 0.767054 | 0.766787 | 0.766231 | 0.764035 | 0.765498 | 0.766568 | 0.7674 |
| 0.772489 | 0.771686 | 0.777048 | 0.772936 | 0.771037 | 0.773334 | 0.771616 | 0.771801 | 0.772242 |
| 0.760725 | 0.760889 | 0.767723 | 0.760569 | 0.761842 | 0.760324 | 0.76298 | 0.764817 | 0.770247 |
| 0.762149 | 0.764936 | 0.768345 | 0.762401 | 0.762532 | 0.763823 | 0.764019 | 0.764811 | 0.767271 |
| 0.751173 | 0.761629 | 0.755166 | 0.753798 | 0.752482 | 0.750972 | 0.753847 | 0.758826 | 0.758883 |
| 0.762483 | 0.762684 | 0.763662 | 0.764431 | 0.764377 | 0.761509 | 0.756388 | 0.762956 | 0.760544 |
| 0.755663 | 0.760738 | 0.763928 | 0.762187 | 0.758683 | 0.759479 | 0.757404 | 0.754785 | 0.760043 |
| 0.76425 | 0.763992 | 0.769971 | 0.765789 | 0.765337 | 0.764629 | 0.767077 | 0.762751 | 0.763781 |
| 0.759133 | 0.762262 | 0.764542 | 0.758975 | 0.759993 | 0.756567 | 0.760677 | 0.764751 | 0.763438 |
| 0.754134 | 0.753903 | 0.759983 | 0.755911 | 0.755845 | 0.751405 | 0.756799 | 0.753326 | 0.761802 |
| 0.754498 | 0.758038 | 0.760726 | 0.752088 | 0.755366 | 0.754344 | 0.753025 | 0.75764 | 0.760633 |
| 0.761364 | 0.767181 | 0.769906 | 0.767251 | 0.766294 | 0.764432 | 0.767675 | 0.766455 | 0.771961 |
| 0.763914 | 0.763719 | 0.769212 | 0.766634 | 0.764186 | 0.765057 | 0.76496 | 0.76947 | 0.771806 |
| 0.768216 | 0.762659 | 0.763308 | 0.763609 | 0.763144 | 0.764906 | 0.761061 | 0.762942 | 0.763488 |
| 0.763089 | 0.758745 | 0.763869 | 0.757191 | 0.758723 | 0.761034 | 0.755996 | 0.754642 | 0.761306 |
| 0.765531 | 0.761653 | 0.767482 | 0.762962 | 0.762417 | 0.763402 | 0.765753 | 0.761064 | 0.764027 |
| 0.761506 | 0.763943 | 0.763871 | 0.763026 | 0.757245 | 0.759137 | 0.759306 | 0.761249 | 0.767473 |
| 0.759162 | 0.756418 | 0.758692 | 0.755248 | 0.755327 | 0.758248 | 0.753837 | 0.752114 | 0.755102 |
| 0.763413 | 0.76005 | 0.765133 | 0.761285 | 0.759061 | 0.759946 | 0.75879 | 0.75991 | 0.765016 |
| 0.768327 | 0.764801 | 0.769062 | 0.766477 | 0.764053 | 0.764884 | 0.762571 | 0.764272 | 0.767315 |
| 0.767665 | 0.760706 | 0.764634 | 0.763428 | 0.76577 | 0.766324 | 0.763609 | 0.761442 | 0.764036 |
| 0.764745 | 0.758738 | 0.763313 | 0.763948 | 0.760047 | 0.763885 | 0.758326 | 0.758419 | 0.762585 |
| 0.765866 | 0.7565 | 0.761979 | 0.761917 | 0.759785 | 0.758199 | 0.759145 | 0.761321 | 0.76238 |
| 0.764936 | 0.757425 | 0.75492 | 0.755934 | 0.755104 | 0.76011 | 0.752192 | 0.755987 | 0.755764 |
| 0.76112 | 0.758678 | 0.760853 | 0.759252 | 0.756707 | 0.758395 | 0.759392 | 0.757105 | 0.760708 |
| 0.759476 | 0.761289 | 0.762688 | 0.759594 | 0.760448 | 0.762046 | 0.760903 | 0.759397 | 0.763089 |
| 0.763016 | 0.761345 | 0.764547 | 0.759954 | 0.764691 | 0.765728 | 0.76502 | 0.758076 | 0.765034 |
| 0.767031 | 0.763458 | 0.762806 | 0.763792 | 0.764788 | 0.766777 | 0.762384 | 0.761839 | 0.766031 |
| 0.767973 | 0.761346 | 0.76222 | 0.762879 | 0.759599 | 0.762718 | 0.757713 | 0.760844 | 0.764359 |
| 0.778142 | 0.774998 | 0.770838 | 0.771954 | 0.774434 | 0.774183 | 0.769344 | 0.773135 | 0.773173 |
| 0.772742 | 0.772124 | 0.77025 | 0.772241 | 0.769168 | 0.772255 | 0.769155 | 0.7661 | 0.7699 |
| 0.76048 | 0.758527 | 0.762229 | 0.758697 | 0.75853 | 0.759293 | 0.759974 | 0.754759 | 0.758898 |
| 0.762918 | 0.764613 | 0.759259 | 0.760793 | 0.762498 | 0.760452 | 0.760616 | 0.758469 | 0.76152 |
| 0.760959 | 0.757725 | 0.760421 | 0.762585 | 0.760056 | 0.757957 | 0.757766 | 0.761568 | 0.763342 |
| 0.764589 | 0.759346 | 0.76159 | 0.76284 | 0.760864 | 0.76179 | 0.761214 | 0.762642 | 0.765082 |
| 0.757578 | 0.75585 | 0.75205 | 0.752958 | 0.75326 | 0.752406 | 0.748878 | 0.748449 | 0.754752 |
| 0.760635 | 0.751859 | 0.758304 | 0.759325 | 0.752815 | 0.757277 | 0.752831 | 0.754507 | 0.758597 |
| 0.751838 | 0.750548 | 0.751643 | 0.752515 | 0.751528 | 0.750572 | 0.746079 | 0.745635 | 0.748926 |
| 0.757787 | 0.753801 | 0.753221 | 0.756206 | 0.756941 | 0.756941 | 0.7561 | 0.754063 | 0.755809 |
| 0.753124 | 0.752312 | 0.751645 | 0.747843 | 0.752832 | 0.749957 | 0.7506 | 0.749489 | 0.753535 |
| 0.752518 | 0.745974 | 0.752847 | 0.751011 | 0.745611 | 0.754073 | 0.750135 | 0.749173 | 0.748856 |
| 0.756676 | 0.756817 | 0.760554 | 0.760151 | 0.755663 | 0.754375 | 0.75466 | 0.758127 | 0.761179 |
| 0.756132 | 0.749272 | 0.748478 | 0.748289 | 0.749835 | 0.751062 | 0.746693 | 0.751032 | 0.752047 |

| | | | | | | | | |
|---|---|---|---|---|---|---|---|---|
| 0.761296 | 0.758288 | 0.76244 | 0.763744 | 0.760875 | 0.758882 | 0.756584 | 0.756229 | 0.75951 |
| 0.766495 | 0.764623 | 0.774102 | 0.770262 | 0.765186 | 0.767193 | 0.768539 | 0.768714 | 0.768263 |
| 0.763855 | 0.763335 | 0.76902 | 0.771488 | 0.765201 | 0.765851 | 0.766536 | 0.765159 | 0.767308 |
| 0.7631 | 0.757068 | 0.767415 | 0.765181 | 0.767668 | 0.767497 | 0.764346 | 0.763961 | 0.76751 |
| 0.768056 | 0.77431 | 0.777046 | 0.774018 | 0.772862 | 0.774924 | 0.770198 | 0.77417 | 0.771457 |
| 0.76901 | 0.770281 | 0.772679 | 0.772567 | 0.769858 | 0.769186 | 0.767147 | 0.770129 | 0.769306 |
| 0.75928 | 0.764374 | 0.766786 | 0.764444 | 0.765295 | 0.764519 | 0.76232 | 0.760879 | 0.763344 |
| 0.764936 | 0.761676 | 0.765931 | 0.76485 | 0.764122 | 0.763041 | 0.76208 | 0.765917 | 0.767341 |
| 0.758286 | 0.759595 | 0.763014 | 0.763189 | 0.763151 | 0.758822 | 0.759734 | 0.758859 | 0.760975 |
| 0.756914 | 0.760475 | 0.764392 | 0.760377 | 0.758561 | 0.762509 | 0.762816 | 0.759173 | 0.764324 |
| 0.765284 | 0.763632 | 0.763339 | 0.768717 | 0.766018 | 0.760605 | 0.768117 | 0.761485 | 0.762075 |
| 0.766615 | 0.765173 | 0.76714 | 0.764624 | 0.763056 | 0.762635 | 0.767816 | 0.766272 | 0.766577 |
| 0.767539 | 0.764576 | 0.770852 | 0.768593 | 0.765377 | 0.769884 | 0.76609 | 0.765642 | 0.766189 |
| 0.768156 | 0.762235 | 0.768776 | 0.76719 | 0.764701 | 0.77035 | 0.766424 | 0.767682 | 0.77016 |
| 0.771324 | 0.772787 | 0.776887 | 0.773482 | 0.770263 | 0.769691 | 0.770743 | 0.775179 | 0.770813 |
| 0.768568 | 0.762013 | 0.766084 | 0.767202 | 0.763662 | 0.767769 | 0.762306 | 0.768929 | 0.767982 |
| 0.769075 | 0.767779 | 0.769623 | 0.773441 | 0.768392 | 0.769946 | 0.773476 | 0.772642 | 0.770188 |
| 0.765367 | 0.762358 | 0.770557 | 0.770067 | 0.765666 | 0.769945 | 0.764977 | 0.769771 | 0.77074 |
| 0.770517 | 0.770689 | 0.776185 | 0.773134 | 0.767736 | 0.772944 | 0.770533 | 0.768904 | 0.774684 |
| 0.769218 | 0.766287 | 0.771871 | 0.771358 | 0.769048 | 0.773187 | 0.771615 | 0.772006 | 0.7723 |
| 0.766422 | 0.763672 | 0.763928 | 0.767096 | 0.763386 | 0.765714 | 0.767409 | 0.765987 | 0.76365 |
| 0.766326 | 0.769345 | 0.772445 | 0.772761 | 0.768285 | 0.77159 | 0.76725 | 0.769669 | 0.770078 |
| 0.768096 | 0.765859 | 0.772467 | 0.771099 | 0.762033 | 0.769667 | 0.769947 | 0.772053 | 0.773934 |
| 0.766621 | 0.770884 | 0.77211 | 0.771375 | 0.772306 | 0.778728 | 0.770372 | 0.772561 | 0.769675 |
| 0.767449 | 0.773738 | 0.771819 | 0.769455 | 0.770652 | 0.769794 | 0.77326 | 0.772897 | 0.769517 |
| 0.770379 | 0.768005 | 0.772511 | 0.769527 | 0.769256 | 0.76688 | 0.767724 | 0.770319 | 0.770708 |
| 0.762408 | 0.765876 | 0.771508 | 0.763305 | 0.768621 | 0.76721 | 0.76756 | 0.770815 | 0.765857 |
| 0.762799 | 0.770545 | 0.768318 | 0.770815 | 0.764365 | 0.762375 | 0.765418 | 0.773733 | 0.767701 |
| 0.765987 | 0.771828 | 0.774115 | 0.774489 | 0.770531 | 0.773364 | 0.771617 | 0.771947 | 0.774431 |
| 0.7699 | 0.766852 | 0.774897 | 0.774366 | 0.778648 | 0.778308 | 0.77675 | 0.774032 | 0.767597 |
| 0.767403 | 0.766237 | 0.770186 | 0.770087 | 0.765892 | 0.769089 | 0.766547 | 0.76475 | 0.760859 |
| 0.767394 | 0.767235 | 0.770532 | 0.773193 | 0.76763 | 0.769467 | 0.770987 | 0.771207 | 0.772994 |
| 0.769017 | 0.771405 | 0.774133 | 0.771871 | 0.770666 | 0.771626 | 0.774262 | 0.772419 | 0.771096 |
| 0.76316 | 0.765715 | 0.768445 | 0.765099 | 0.765556 | 0.770386 | 0.763361 | 0.767874 | 0.7657 |
| 0.767601 | 0.769231 | 0.773564 | 0.772083 | 0.768951 | 0.771657 | 0.767829 | 0.773158 | 0.769601 |
| 0.76805 | 0.76721 | 0.771966 | 0.769311 | 0.765285 | 0.77177 | 0.76998 | 0.770297 | 0.770352 |
| 0.771553 | 0.774777 | 0.775217 | 0.773915 | 0.772742 | 0.773338 | 0.77278 | 0.775954 | 0.771193 |
| 0.766403 | 0.776587 | 0.773648 | 0.769784 | 0.769998 | 0.767005 | 0.769713 | 0.771004 | 0.773931 |
| 0.76736 | 0.767388 | 0.772306 | 0.771574 | 0.76925 | 0.766656 | 0.767161 | 0.771353 | 0.769705 |
| 0.766105 | 0.768633 | 0.774837 | 0.772609 | 0.766742 | 0.769419 | 0.769991 | 0.764593 | 0.767508 |
| 0.770794 | 0.771471 | 0.777442 | 0.774685 | 0.768802 | 0.774928 | 0.764419 | 0.768167 | 0.773935 |
| 0.768871 | 0.77576 | 0.780345 | 0.777108 | 0.774074 | 0.775014 | 0.770775 | 0.772817 | 0.772387 |
| 0.769275 | 0.771927 | 0.780304 | 0.77318 | 0.773043 | 0.774479 | 0.771792 | 0.772882 | 0.775635 |
| 0.764141 | 0.763929 | 0.770325 | 0.766293 | 0.767422 | 0.766781 | 0.766366 | 0.768012 | 0.767158 |
| 0.769123 | 0.769826 | 0.772757 | 0.771282 | 0.767788 | 0.767521 | 0.767994 | 0.7683 | 0.768514 |
| 0.762055 | 0.766794 | 0.770475 | 0.768826 | 0.768567 | 0.767781 | 0.767638 | 0.771109 | 0.76559 |

| | | | | | | | | |
|---|---|---|---|---|---|---|---|---|
| 0.754892 | 0.756323 | 0.758722 | 0.758363 | 0.755551 | 0.75658 | 0.751983 | 0.75923 | 0.759966 |
| 0.751505 | 0.744418 | 0.752321 | 0.748396 | 0.748353 | 0.747591 | 0.752928 | 0.744629 | 0.745617 |
| 0.764745 | 0.768259 | 0.772033 | 0.769517 | 0.76981 | 0.768273 | 0.765538 | 0.770294 | 0.76583 |
| 0.771557 | 0.766259 | 0.772131 | 0.770713 | 0.769451 | 0.769885 | 0.768104 | 0.770245 | 0.769362 |
| 0.762986 | 0.761561 | 0.766088 | 0.764389 | 0.762096 | 0.760581 | 0.7645 | 0.760882 | 0.763692 |
| 0.763347 | 0.766918 | 0.766156 | 0.766012 | 0.768533 | 0.767047 | 0.761987 | 0.770143 | 0.765949 |
| 0.772018 | 0.767923 | 0.773256 | 0.767258 | 0.767414 | 0.768805 | 0.769131 | 0.775181 | 0.771254 |
| 0.758124 | 0.761644 | 0.764883 | 0.760556 | 0.763265 | 0.763999 | 0.763048 | 0.769605 | 0.762532 |
| 0.768158 | 0.76712 | 0.772571 | 0.771183 | 0.768979 | 0.772641 | 0.773155 | 0.769996 | 0.775002 |
| 0.76823 | 0.769705 | 0.776713 | 0.774041 | 0.767959 | 0.772037 | 0.77071 | 0.767799 | 0.768624 |
| 0.769074 | 0.769921 | 0.776143 | 0.777933 | 0.770442 | 0.768411 | 0.775306 | 0.767516 | 0.767611 |
| 0.769047 | 0.769079 | 0.77211 | 0.768755 | 0.774772 | 0.769428 | 0.774983 | 0.769933 | 0.771179 |
| 0.772753 | 0.772339 | 0.775138 | 0.778531 | 0.772409 | 0.774236 | 0.771661 | 0.772828 | 0.773371 |
| 0.772082 | 0.770367 | 0.77427 | 0.773367 | 0.774305 | 0.772806 | 0.779281 | 0.771326 | 0.770104 |
| 0.758204 | 0.759698 | 0.76986 | 0.768087 | 0.766879 | 0.767159 | 0.762186 | 0.763348 | 0.767028 |
| 0.769386 | 0.765947 | 0.768407 | 0.770044 | 0.7716 | 0.767895 | 0.773914 | 0.770211 | 0.771405 |
| 0.762268 | 0.767889 | 0.770294 | 0.769936 | 0.767062 | 0.77132 | 0.768222 | 0.765929 | 0.764812 |
| 0.766015 | 0.765642 | 0.766713 | 0.76857 | 0.766606 | 0.772716 | 0.768341 | 0.763188 | 0.763674 |
| 0.764617 | 0.764979 | 0.767712 | 0.762447 | 0.764806 | 0.769763 | 0.766809 | 0.764151 | 0.769004 |
| 0.759854 | 0.763545 | 0.771294 | 0.768165 | 0.757741 | 0.765924 | 0.76388 | 0.766101 | 0.764925 |
| 0.763097 | 0.765587 | 0.768217 | 0.765811 | 0.7652 | 0.764236 | 0.763005 | 0.765986 | 0.767799 |
| 0.763647 | 0.758147 | 0.761599 | 0.763392 | 0.759192 | 0.761076 | 0.760139 | 0.764041 | 0.763902 |
| 0.765612 | 0.761625 | 0.768594 | 0.772406 | 0.765809 | 0.771344 | 0.766443 | 0.768165 | 0.767495 |
| 0.769307 | 0.767007 | 0.775389 | 0.775298 | 0.768226 | 0.772912 | 0.768596 | 0.769699 | 0.77314 |
| 0.768043 | 0.769619 | 0.772861 | 0.771077 | 0.765695 | 0.774741 | 0.770409 | 0.769909 | 0.770781 |
| 0.770939 | 0.763405 | 0.769673 | 0.773236 | 0.766976 | 0.770779 | 0.770285 | 0.76941 | 0.769475 |
| 0.764801 | 0.769814 | 0.769355 | 0.766438 | 0.767672 | 0.767949 | 0.768301 | 0.761776 | 0.771545 |
| 0.767039 | 0.76595 | 0.771118 | 0.76585 | 0.765999 | 0.771168 | 0.767244 | 0.76971 | 0.772766 |
| 0.761047 | 0.761765 | 0.766652 | 0.766571 | 0.764849 | 0.767529 | 0.767492 | 0.765512 | 0.767067 |
| 0.695812 | 0.692934 | 0.698359 | 0.696069 | 0.694907 | 0.700398 | 0.698868 | 0.693157 | 0.699844 |
| 0.703188 | 0.704703 | 0.711935 | 0.706217 | 0.705428 | 0.708801 | 0.711662 | 0.706781 | 0.711631 |
| 0.704278 | 0.699459 | 0.708093 | 0.704751 | 0.703119 | 0.706903 | 0.708487 | 0.706582 | 0.709975 |
| 0.696372 | 0.693653 | 0.696339 | 0.698613 | 0.694315 | 0.697053 | 0.698326 | 0.69876 | 0.698065 |
| 0.707234 | 0.70699 | 0.7109 | 0.71047 | 0.707848 | 0.711868 | 0.711407 | 0.707615 | 0.710111 |
| 0.77061 | 0.765783 | 0.773493 | 0.772448 | 0.769408 | 0.770881 | 0.766998 | 0.772098 | 0.77029 |
| 0.772644 | 0.769802 | 0.771721 | 0.774072 | 0.772279 | 0.772331 | 0.767426 | 0.770596 | 0.770851 |
| 0.770394 | 0.775277 | 0.774948 | 0.778131 | 0.77487 | 0.772264 | 0.770927 | 0.773944 | 0.777945 |
| 0.773758 | 0.769212 | 0.778201 | 0.775862 | 0.771841 | 0.774869 | 0.77272 | 0.776746 | 0.774642 |
| 0.771741 | 0.770509 | 0.772678 | 0.774333 | 0.773118 | 0.774745 | 0.772875 | 0.770889 | 0.775458 |
| 0.774648 | 0.774356 | 0.777355 | 0.776933 | 0.775017 | 0.780372 | 0.777437 | 0.776518 | 0.778294 |
| 0.767981 | 0.763798 | 0.771531 | 0.773563 | 0.773074 | 0.769244 | 0.770739 | 0.773599 | 0.773911 |
| 0.768702 | 0.770079 | 0.771433 | 0.771887 | 0.767013 | 0.769787 | 0.766388 | 0.773142 | 0.770488 |
| 0.768172 | 0.764236 | 0.770891 | 0.765697 | 0.764824 | 0.765364 | 0.760135 | 0.764956 | 0.767432 |
| 0.766729 | 0.766791 | 0.775365 | 0.768193 | 0.764601 | 0.765434 | 0.764529 | 0.765231 | 0.767469 |
| 0.770572 | 0.76753 | 0.771933 | 0.770791 | 0.769146 | 0.768554 | 0.770752 | 0.770127 | 0.766773 |
| 0.774234 | 0.769911 | 0.77316 | 0.776448 | 0.772624 | 0.771254 | 0.771619 | 0.77316 | 0.774546 |

| | | | | | | | | |
|---|---|---|---|---|---|---|---|---|
| 0.770312 | 0.767257 | 0.779939 | 0.77349 | 0.771758 | 0.771163 | 0.769619 | 0.774752 | 0.769877 |
| 0.769091 | 0.774735 | 0.773871 | 0.771582 | 0.768287 | 0.7756 | 0.769186 | 0.771488 | 0.77287 |
| 0.772118 | 0.770348 | 0.77736 | 0.77457 | 0.773323 | 0.773786 | 0.768373 | 0.77392 | 0.775044 |
| 0.766848 | 0.767683 | 0.770519 | 0.772085 | 0.764517 | 0.769377 | 0.766554 | 0.768964 | 0.772637 |
| 0.76479 | 0.769808 | 0.770893 | 0.77277 | 0.766962 | 0.77372 | 0.769647 | 0.76912 | 0.772162 |
| 0.766539 | 0.766189 | 0.771592 | 0.775465 | 0.765576 | 0.769713 | 0.768916 | 0.768348 | 0.775182 |
| 0.768562 | 0.772689 | 0.774586 | 0.769127 | 0.769244 | 0.772716 | 0.771378 | 0.775342 | 0.776875 |
| 0.765909 | 0.768492 | 0.771527 | 0.771349 | 0.765495 | 0.774641 | 0.768056 | 0.772819 | 0.774183 |
| 0.767262 | 0.765087 | 0.772624 | 0.76754 | 0.76318 | 0.769061 | 0.765592 | 0.769625 | 0.769458 |
| 0.767455 | 0.77249 | 0.771652 | 0.768668 | 0.76941 | 0.769488 | 0.768319 | 0.767482 | 0.773981 |
| 0.768124 | 0.767932 | 0.771978 | 0.772605 | 0.770704 | 0.77418 | 0.7656 | 0.765383 | 0.771165 |
| 0.766896 | 0.771652 | 0.771088 | 0.772922 | 0.767286 | 0.77058 | 0.759158 | 0.76965 | 0.77184 |
| 0.758229 | 0.764256 | 0.768635 | 0.76636 | 0.761798 | 0.768519 | 0.76196 | 0.767641 | 0.767384 |
| 0.766157 | 0.766773 | 0.765364 | 0.763309 | 0.765371 | 0.769188 | 0.764508 | 0.767115 | 0.770121 |
| 0.762372 | 0.757961 | 0.768083 | 0.764656 | 0.762976 | 0.763348 | 0.765695 | 0.768946 | 0.767442 |
| 0.76836 | 0.773746 | 0.779196 | 0.775579 | 0.772705 | 0.767602 | 0.773419 | 0.771973 | 0.772332 |
| 0.76678 | 0.771843 | 0.77245 | 0.770127 | 0.764701 | 0.769728 | 0.767546 | 0.768749 | 0.76555 |
| 0.767341 | 0.767721 | 0.770643 | 0.771384 | 0.767904 | 0.771774 | 0.771839 | 0.768474 | 0.769756 |
| 0.765344 | 0.764046 | 0.771052 | 0.76979 | 0.764264 | 0.765377 | 0.768784 | 0.770826 | 0.772646 |
| 0.764821 | 0.76629 | 0.766327 | 0.764746 | 0.763055 | 0.769211 | 0.769807 | 0.767734 | 0.771173 |
| 0.759127 | 0.765445 | 0.767464 | 0.76465 | 0.760393 | 0.767172 | 0.766558 | 0.765324 | 0.767537 |
| 0.765163 | 0.767808 | 0.769926 | 0.768642 | 0.763451 | 0.769744 | 0.768035 | 0.77312 | 0.766736 |
| 0.771901 | 0.7712 | 0.773779 | 0.772999 | 0.768221 | 0.765787 | 0.7731 | 0.771426 | 0.769663 |
| 0.768911 | 0.768688 | 0.770195 | 0.769319 | 0.766493 | 0.770817 | 0.765391 | 0.769928 | 0.773523 |
| 0.762224 | 0.768898 | 0.770608 | 0.770051 | 0.769792 | 0.764612 | 0.77065 | 0.770421 | 0.77118 |
| 0.706754 | 0.703864 | 0.711202 | 0.7062 | 0.705403 | 0.70762 | 0.708044 | 0.702538 | 0.708165 |
| 0.771349 | 0.766428 | 0.776546 | 0.77326 | 0.773048 | 0.769547 | 0.770823 | 0.771781 | 0.775299 |
| 0.767086 | 0.771106 | 0.776244 | 0.771743 | 0.770523 | 0.774146 | 0.767455 | 0.772259 | 0.77269 |
| 0.771827 | 0.775598 | 0.782786 | 0.774089 | 0.779938 | 0.777564 | 0.776594 | 0.776039 | 0.772203 |
| 0.768761 | 0.772465 | 0.771563 | 0.773793 | 0.769729 | 0.772055 | 0.770718 | 0.771544 | 0.774076 |
| 0.774398 | 0.767418 | 0.771915 | 0.766859 | 0.771965 | 0.770908 | 0.769843 | 0.76757 | 0.772028 |
| 0.767678 | 0.764497 | 0.776658 | 0.772144 | 0.766251 | 0.768113 | 0.768144 | 0.769329 | 0.770603 |
| 0.766323 | 0.767974 | 0.776459 | 0.772498 | 0.770438 | 0.772132 | 0.769057 | 0.765965 | 0.765272 |
| 0.76353 | 0.773644 | 0.770848 | 0.767361 | 0.762458 | 0.767561 | 0.767845 | 0.771665 | 0.769816 |
| 0.756635 | 0.764395 | 0.768215 | 0.767088 | 0.76195 | 0.763901 | 0.760455 | 0.765027 | 0.76571 |
| 0.760289 | 0.761961 | 0.771303 | 0.766103 | 0.762883 | 0.766393 | 0.759174 | 0.767061 | 0.765741 |
| 0.772785 | 0.77099 | 0.773128 | 0.774068 | 0.773325 | 0.772616 | 0.770807 | 0.773365 | 0.771306 |
| 0.766617 | 0.763071 | 0.76654 | 0.76702 | 0.766791 | 0.770719 | 0.771902 | 0.771741 | 0.766474 |
| 0.766284 | 0.771201 | 0.776049 | 0.767712 | 0.76496 | 0.764907 | 0.767869 | 0.772874 | 0.770466 |
| 0.76346 | 0.76155 | 0.769647 | 0.769901 | 0.764923 | 0.765304 | 0.763903 | 0.769586 | 0.766484 |
| 0.762895 | 0.762372 | 0.764716 | 0.76466 | 0.765479 | 0.760856 | 0.765771 | 0.767886 | 0.767925 |
| 0.768115 | 0.768723 | 0.770999 | 0.776142 | 0.772108 | 0.769166 | 0.776834 | 0.769622 | 0.770884 |
| 0.765174 | 0.761641 | 0.765149 | 0.759887 | 0.758968 | 0.761835 | 0.765755 | 0.762351 | 0.767095 |
| 0.784889 | 0.770837 | 0.777951 | 0.77691 | 0.779178 | 0.781198 | 0.781775 | 0.783138 | 0.785791 |
| 0.795015 | 0.782364 | 0.786195 | 0.783214 | 0.782225 | 0.783142 | 0.783795 | 0.79616 | 0.79413 |
| 0.790562 | 0.780106 | 0.789657 | 0.789915 | 0.780665 | 0.784544 | 0.784857 | 0.790181 | 0.789238 |

| | | | | | | | | |
|---|---|---|---|---|---|---|---|---|
| 0.788586 | 0.774975 | 0.783635 | 0.776719 | 0.779298 | 0.781041 | 0.781779 | 0.793349 | 0.791677 |
| 0.79784 | 0.779619 | 0.789365 | 0.788725 | 0.787271 | 0.785483 | 0.780964 | 0.789822 | 0.800532 |
| 0.786061 | 0.777474 | 0.781718 | 0.77578 | 0.780512 | 0.779417 | 0.778925 | 0.788251 | 0.79145 |
| 0.738758 | 0.730797 | 0.743423 | 0.741679 | 0.728089 | 0.737261 | 0.735178 | 0.747308 | 0.74452 |
| 0.80596 | 0.779462 | 0.788989 | 0.784627 | 0.780839 | 0.778373 | 0.781342 | 0.789513 | 0.79249 |
| 0.815161 | 0.783764 | 0.788323 | 0.785963 | 0.783425 | 0.78972 | 0.782389 | 0.786393 | 0.788302 |
| 1 | 0.78674 | 0.795199 | 0.790958 | 0.790703 | 0.794147 | 0.783326 | 0.78708 | 0.790246 |
| 0.78674 | 1 | 0.803804 | 0.806215 | 0.792837 | 0.800918 | 0.799218 | 0.778982 | 0.782193 |
| 0.795199 | 0.803804 | 1 | 0.817444 | 0.811098 | 0.809971 | 0.804852 | 0.787692 | 0.785031 |
| 0.790958 | 0.806215 | 0.817444 | 1 | 0.83175 | 0.811163 | 0.814228 | 0.780542 | 0.781286 |
| 0.790703 | 0.792837 | 0.811098 | 0.83175 | 1 | 0.805878 | 0.850867 | 0.777092 | 0.783343 |
| 0.794147 | 0.800918 | 0.809971 | 0.811163 | 0.805878 | 1 | 0.80563 | 0.779661 | 0.77901 |
| 0.783326 | 0.799218 | 0.804852 | 0.814228 | 0.850867 | 0.80563 | 1 | 0.781371 | 0.784121 |
| 0.78708 | 0.778982 | 0.787692 | 0.780542 | 0.777092 | 0.779661 | 0.781371 | 1 | 0.844989 |
| 0.790246 | 0.782193 | 0.785031 | 0.781286 | 0.783343 | 0.77901 | 0.784121 | 0.844989 | 1 |
| 0.785756 | 0.773769 | 0.781389 | 0.779465 | 0.780621 | 0.776545 | 0.779863 | 0.839963 | 0.844651 |
| 0.786476 | 0.781549 | 0.785158 | 0.783866 | 0.781091 | 0.778954 | 0.784262 | 0.79855 | 0.834286 |
| 0.788222 | 0.783135 | 0.785638 | 0.784089 | 0.781492 | 0.78093 | 0.780329 | 0.837349 | 0.859918 |
| 0.7867 | 0.773272 | 0.781314 | 0.77841 | 0.773775 | 0.777217 | 0.772888 | 0.816422 | 0.805049 |
| 0.787869 | 0.779605 | 0.781068 | 0.778464 | 0.776687 | 0.780684 | 0.78058 | 0.82948 | 0.848593 |
| 0.791055 | 0.789692 | 0.788363 | 0.78683 | 0.785033 | 0.788136 | 0.784385 | 0.819471 | 0.816083 |
| 0.791429 | 0.785373 | 0.784341 | 0.783503 | 0.783645 | 0.78171 | 0.787899 | 0.80259 | 0.853242 |
| 0.685861 | 0.681369 | 0.683776 | 0.682476 | 0.674541 | 0.687581 | 0.683864 | 0.681202 | 0.683625 |
| 0.658566 | 0.649836 | 0.664833 | 0.658126 | 0.65564 | 0.658425 | 0.660766 | 0.659328 | 0.664614 |
| 0.669462 | 0.668382 | 0.681454 | 0.670677 | 0.674464 | 0.674555 | 0.672 | 0.664539 | 0.675384 |
| 0.655716 | 0.64922 | 0.658101 | 0.659431 | 0.654653 | 0.653498 | 0.660526 | 0.650597 | 0.653852 |
| 0.740097 | 0.739802 | 0.748639 | 0.742136 | 0.74159 | 0.744034 | 0.739569 | 0.748112 | 0.747765 |
| 0.729719 | 0.729183 | 0.731478 | 0.733525 | 0.72825 | 0.738142 | 0.73054 | 0.737868 | 0.738237 |
| 0.729012 | 0.731672 | 0.730457 | 0.733454 | 0.723828 | 0.730993 | 0.728059 | 0.73373 | 0.734141 |
| 0.75079 | 0.744868 | 0.746605 | 0.74692 | 0.746928 | 0.745752 | 0.741336 | 0.745992 | 0.753733 |
| 0.734168 | 0.735057 | 0.729355 | 0.732932 | 0.729648 | 0.729686 | 0.726897 | 0.732118 | 0.737781 |
| 0.732092 | 0.725206 | 0.724928 | 0.725843 | 0.719422 | 0.72393 | 0.720057 | 0.721298 | 0.730168 |
| 0.736985 | 0.723505 | 0.726703 | 0.725677 | 0.724933 | 0.718638 | 0.719905 | 0.722977 | 0.729923 |
| 0.65949 | 0.65334 | 0.663558 | 0.66009 | 0.653832 | 0.658069 | 0.660698 | 0.654971 | 0.655368 |
| 0.769562 | 0.767558 | 0.77626 | 0.774732 | 0.769184 | 0.771089 | 0.768057 | 0.773854 | 0.770212 |
| 0.754202 | 0.745482 | 0.756766 | 0.750556 | 0.753916 | 0.754677 | 0.746626 | 0.746451 | 0.755683 |
| 0.751165 | 0.745215 | 0.74908 | 0.745427 | 0.742764 | 0.748419 | 0.748029 | 0.743086 | 0.748944 |
| 0.729957 | 0.723125 | 0.738911 | 0.734194 | 0.732133 | 0.734498 | 0.739783 | 0.735701 | 0.739869 |
| 0.755208 | 0.746439 | 0.757084 | 0.755071 | 0.745499 | 0.755676 | 0.746985 | 0.749883 | 0.750439 |
| 0.737187 | 0.739208 | 0.740273 | 0.733974 | 0.734145 | 0.745992 | 0.731748 | 0.737629 | 0.73913 |
| 0.735831 | 0.737629 | 0.73872 | 0.736302 | 0.731408 | 0.739419 | 0.732918 | 0.728862 | 0.738894 |
| 0.740519 | 0.741141 | 0.74185 | 0.74342 | 0.736233 | 0.742161 | 0.73224 | 0.744369 | 0.739638 |
| 0.737663 | 0.733008 | 0.73947 | 0.737396 | 0.731324 | 0.729538 | 0.73429 | 0.739217 | 0.741912 |
| 0.740637 | 0.736423 | 0.733834 | 0.731326 | 0.728636 | 0.737577 | 0.734443 | 0.735782 | 0.741993 |
| 0.739402 | 0.743849 | 0.73869 | 0.737115 | 0.73071 | 0.741952 | 0.729479 | 0.737899 | 0.736856 |
| 0.739336 | 0.720452 | 0.723482 | 0.733378 | 0.722792 | 0.729711 | 0.716324 | 0.720159 | 0.725588 |

| | | | | | | | | |
|---|---|---|---|---|---|---|---|---|
| 0.767037 | 0.756286 | 0.760255 | 0.763431 | 0.758024 | 0.762926 | 0.755611 | 0.763425 | 0.761297 |
| 0.743812 | 0.739492 | 0.744212 | 0.737587 | 0.735079 | 0.737953 | 0.74146 | 0.737525 | 0.74083 |
| 0.742948 | 0.73769 | 0.745948 | 0.741429 | 0.732913 | 0.739988 | 0.734949 | 0.73826 | 0.748333 |
| 0.735692 | 0.736389 | 0.74587 | 0.73745 | 0.743614 | 0.739528 | 0.73646 | 0.740638 | 0.741414 |
| 0.741536 | 0.739522 | 0.744056 | 0.74002 | 0.741497 | 0.741016 | 0.736336 | 0.731837 | 0.736648 |
| 0.737001 | 0.732036 | 0.732899 | 0.731512 | 0.728979 | 0.739219 | 0.732317 | 0.73755 | 0.739391 |
| 0.731563 | 0.729106 | 0.737555 | 0.719815 | 0.721105 | 0.724087 | 0.719055 | 0.735101 | 0.737717 |
| 0.7374 | 0.722977 | 0.727713 | 0.725825 | 0.723103 | 0.726231 | 0.732877 | 0.728919 | 0.731659 |
| 0.738506 | 0.733504 | 0.741434 | 0.734 | 0.722473 | 0.739193 | 0.729131 | 0.740582 | 0.733069 |
| 0.760393 | 0.764217 | 0.76146 | 0.763971 | 0.760094 | 0.761341 | 0.766104 | 0.768868 | 0.76226 |
| 0.755997 | 0.755652 | 0.759184 | 0.758254 | 0.755296 | 0.753753 | 0.759445 | 0.762922 | 0.759289 |
| 0.752949 | 0.749807 | 0.755365 | 0.754082 | 0.752499 | 0.75265 | 0.75294 | 0.752772 | 0.75058 |
| 0.74458 | 0.742421 | 0.749299 | 0.743163 | 0.738614 | 0.74641 | 0.744523 | 0.745241 | 0.740893 |
| 0.747266 | 0.749237 | 0.750092 | 0.751628 | 0.746698 | 0.749106 | 0.751623 | 0.749695 | 0.746883 |
| 0.741804 | 0.744999 | 0.747778 | 0.739371 | 0.736778 | 0.746217 | 0.74708 | 0.748244 | 0.744283 |

| plot 19 | | | | | | | plot | |
|---|---|---|---|---|---|---|---|---|
| Wild_plot_ | Wild_plot_ | Wild_plot_ | Wild_plot_ | Wild_plot_ | Wild_plot_ | Wild_plot_ | Wild_plot_ | Wild_plot_ |
| 0.700832 | 0.706354 | 0.708248 | 0.698535 | 0.707271 | 0.705579 | 0.706511 | 0.690922 | 0.72468 |
| 0.710719 | 0.715009 | 0.713638 | 0.707966 | 0.712443 | 0.714856 | 0.716084 | 0.701016 | 0.715479 |
| 0.702372 | 0.704809 | 0.704667 | 0.70089 | 0.705349 | 0.708347 | 0.707259 | 0.684652 | 0.713237 |
| 0.711196 | 0.715284 | 0.716242 | 0.710024 | 0.712027 | 0.713811 | 0.716854 | 0.709071 | 0.715753 |
| 0.702799 | 0.706971 | 0.709412 | 0.701535 | 0.705961 | 0.707267 | 0.708732 | 0.680014 | 0.715505 |
| 0.709726 | 0.715632 | 0.719499 | 0.715803 | 0.713088 | 0.717924 | 0.717908 | 0.694263 | 0.715741 |
| 0.765452 | 0.770168 | 0.770462 | 0.764365 | 0.765869 | 0.767872 | 0.770847 | 0.681802 | 0.65962 |
| 0.768883 | 0.767712 | 0.774759 | 0.763803 | 0.768197 | 0.772073 | 0.769546 | 0.67417 | 0.658583 |
| 0.705047 | 0.713822 | 0.712331 | 0.707083 | 0.712061 | 0.713568 | 0.714513 | 0.697619 | 0.71537 |
| 0.704421 | 0.706541 | 0.708491 | 0.706805 | 0.706813 | 0.710189 | 0.712789 | 0.69543 | 0.723749 |
| 0.708337 | 0.712162 | 0.711681 | 0.702107 | 0.709754 | 0.709981 | 0.711689 | 0.681039 | 0.720624 |
| 0.700079 | 0.700894 | 0.703958 | 0.697445 | 0.703519 | 0.701543 | 0.702692 | 0.69356 | 0.724951 |
| 0.712475 | 0.714083 | 0.71094 | 0.709777 | 0.715833 | 0.715938 | 0.714354 | 0.691489 | 0.725185 |
| 0.716758 | 0.724195 | 0.721386 | 0.716127 | 0.71933 | 0.722787 | 0.727033 | 0.710628 | 0.731875 |
| 0.732815 | 0.734338 | 0.734062 | 0.730754 | 0.733669 | 0.738796 | 0.737941 | 0.704313 | 0.715677 |
| 0.710721 | 0.715086 | 0.714694 | 0.710426 | 0.712929 | 0.718503 | 0.714399 | 0.699502 | 0.734467 |
| 0.710946 | 0.711916 | 0.714686 | 0.710348 | 0.714366 | 0.717353 | 0.714984 | 0.69514 | 0.726737 |
| 0.73502 | 0.745632 | 0.741802 | 0.736139 | 0.740558 | 0.743033 | 0.743144 | 0.698933 | 0.705798 |
| 0.701767 | 0.707773 | 0.705864 | 0.702486 | 0.702349 | 0.708317 | 0.707577 | 0.692789 | 0.723232 |
| 0.709644 | 0.710283 | 0.712957 | 0.707373 | 0.709483 | 0.709436 | 0.712228 | 0.694103 | 0.718551 |
| 0.703777 | 0.704899 | 0.710516 | 0.704169 | 0.704768 | 0.708842 | 0.7093 | 0.693614 | 0.721963 |
| 0.707094 | 0.708809 | 0.709189 | 0.70515 | 0.710374 | 0.71199 | 0.711418 | 0.695064 | 0.722685 |
| 0.713716 | 0.716204 | 0.719402 | 0.711554 | 0.713221 | 0.715843 | 0.718188 | 0.695636 | 0.724079 |
| 0.708168 | 0.711698 | 0.712185 | 0.707422 | 0.711742 | 0.711844 | 0.711416 | 0.694031 | 0.714495 |
| 0.714183 | 0.714841 | 0.718068 | 0.710696 | 0.710296 | 0.715361 | 0.715518 | 0.693309 | 0.72001 |
| 0.70744 | 0.707737 | 0.708333 | 0.703599 | 0.706262 | 0.709461 | 0.711478 | 0.69855 | 0.723655 |
| 0.701841 | 0.704102 | 0.707292 | 0.700918 | 0.70508 | 0.708559 | 0.70668 | 0.699262 | 0.7139 |
| 0.706426 | 0.714062 | 0.714816 | 0.707907 | 0.710342 | 0.71398 | 0.714829 | 0.691233 | 0.714563 |
| 0.71129 | 0.714908 | 0.716389 | 0.710708 | 0.712716 | 0.71595 | 0.719288 | 0.693307 | 0.722562 |
| 0.707387 | 0.712091 | 0.71309 | 0.702604 | 0.710163 | 0.710556 | 0.712089 | 0.689699 | 0.719961 |
| 0.705967 | 0.710101 | 0.711123 | 0.706973 | 0.704805 | 0.712289 | 0.711951 | 0.690707 | 0.716064 |
| 0.703136 | 0.706081 | 0.705492 | 0.700819 | 0.703386 | 0.705395 | 0.70702 | 0.690701 | 0.710061 |
| 0.702905 | 0.709272 | 0.709731 | 0.703859 | 0.703718 | 0.707622 | 0.710318 | 0.700604 | 0.71737 |
| 0.708849 | 0.710752 | 0.713019 | 0.704649 | 0.709874 | 0.712054 | 0.713439 | 0.693248 | 0.702102 |
| 0.700807 | 0.703669 | 0.70354 | 0.701677 | 0.70285 | 0.70892 | 0.704871 | 0.691039 | 0.705596 |
| 0.706897 | 0.711432 | 0.713395 | 0.705934 | 0.711572 | 0.709876 | 0.712486 | 0.692364 | 0.7195 |
| 0.713822 | 0.713714 | 0.719477 | 0.711622 | 0.715469 | 0.714444 | 0.717928 | 0.697318 | 0.724096 |
| 0.709312 | 0.713567 | 0.716594 | 0.709947 | 0.712049 | 0.715737 | 0.714905 | 0.686652 | 0.71516 |
| 0.710917 | 0.716132 | 0.717765 | 0.710775 | 0.713708 | 0.715125 | 0.718385 | 0.697258 | 0.711431 |

| | | | | | | | | |
|---|---|---|---|---|---|---|---|---|
| 0.741894 | 0.749502 | 0.749379 | 0.742039 | 0.747738 | 0.752071 | 0.750742 | 0.704512 | 0.699041 |
| 0.697978 | 0.703183 | 0.702393 | 0.696929 | 0.699007 | 0.702506 | 0.699109 | 0.686953 | 0.724691 |
| 0.703031 | 0.705841 | 0.705766 | 0.700609 | 0.703806 | 0.708211 | 0.707758 | 0.686428 | 0.714042 |
| 0.71006 | 0.713098 | 0.716827 | 0.709677 | 0.710122 | 0.714173 | 0.715562 | 0.700329 | 0.735872 |
| 0.704988 | 0.712427 | 0.713991 | 0.705985 | 0.70982 | 0.712109 | 0.713651 | 0.685041 | 0.729776 |
| 0.710964 | 0.71207 | 0.712601 | 0.70776 | 0.710927 | 0.711901 | 0.717774 | 0.70203 | 0.72396 |
| 0.705836 | 0.705767 | 0.711266 | 0.701395 | 0.702905 | 0.705752 | 0.708838 | 0.686221 | 0.716411 |
| 0.705681 | 0.712585 | 0.711251 | 0.704146 | 0.709505 | 0.710823 | 0.710648 | 0.696702 | 0.717841 |
| 0.70661 | 0.716149 | 0.714343 | 0.705573 | 0.710621 | 0.713703 | 0.717079 | 0.690353 | 0.707197 |
| 0.70686 | 0.713511 | 0.714387 | 0.708442 | 0.710002 | 0.710751 | 0.714535 | 0.697882 | 0.705385 |
| 0.699101 | 0.701207 | 0.700928 | 0.697595 | 0.70085 | 0.701805 | 0.702538 | 0.696077 | 0.727878 |
| 0.700755 | 0.705381 | 0.704206 | 0.70046 | 0.703812 | 0.705102 | 0.704621 | 0.691558 | 0.727048 |
| 0.704509 | 0.709818 | 0.709224 | 0.70584 | 0.707249 | 0.709411 | 0.708686 | 0.6897 | 0.727681 |
| 0.701602 | 0.701918 | 0.708184 | 0.700199 | 0.701869 | 0.703288 | 0.705456 | 0.692195 | 0.733449 |
| 0.701483 | 0.708323 | 0.705403 | 0.701522 | 0.700614 | 0.70728 | 0.705534 | 0.689292 | 0.721685 |
| 0.703486 | 0.704482 | 0.705996 | 0.697268 | 0.703186 | 0.70678 | 0.705428 | 0.693653 | 0.728907 |
| 0.700048 | 0.706472 | 0.705119 | 0.699177 | 0.701245 | 0.705585 | 0.705544 | 0.689841 | 0.722775 |
| 0.706706 | 0.711186 | 0.709956 | 0.707358 | 0.709578 | 0.713389 | 0.712032 | 0.698089 | 0.732986 |
| 0.712746 | 0.709771 | 0.715924 | 0.708356 | 0.714399 | 0.713603 | 0.714036 | 0.699872 | 0.719242 |
| 0.702796 | 0.710516 | 0.707705 | 0.702649 | 0.704683 | 0.704854 | 0.707605 | 0.694922 | 0.719303 |
| 0.703125 | 0.703894 | 0.704354 | 0.69973 | 0.701525 | 0.706154 | 0.705037 | 0.690006 | 0.72004 |
| 0.704719 | 0.708936 | 0.705854 | 0.699612 | 0.70703 | 0.707527 | 0.70721 | 0.68294 | 0.721159 |
| 0.703295 | 0.710423 | 0.706688 | 0.703497 | 0.705873 | 0.708041 | 0.709532 | 0.695779 | 0.72175 |
| 0.713429 | 0.720768 | 0.71867 | 0.715246 | 0.714696 | 0.720571 | 0.721431 | 0.694561 | 0.713902 |
| 0.721064 | 0.725676 | 0.725056 | 0.721304 | 0.721748 | 0.725421 | 0.729438 | 0.695676 | 0.728589 |
| 0.768969 | 0.767668 | 0.775498 | 0.763325 | 0.767651 | 0.772905 | 0.770198 | 0.674791 | 0.657476 |
| 0.766213 | 0.768852 | 0.768124 | 0.761826 | 0.766303 | 0.771367 | 0.766513 | 0.68022 | 0.664147 |
| 0.767346 | 0.768299 | 0.774007 | 0.765002 | 0.76777 | 0.772848 | 0.768602 | 0.680973 | 0.659752 |
| 0.703023 | 0.70713 | 0.707251 | 0.701322 | 0.706247 | 0.708562 | 0.708734 | 0.689567 | 0.726054 |
| 0.704611 | 0.705725 | 0.70581 | 0.702329 | 0.702291 | 0.707451 | 0.709979 | 0.688398 | 0.727544 |
| 0.708855 | 0.714727 | 0.715869 | 0.7144 | 0.709403 | 0.715782 | 0.715197 | 0.686684 | 0.724822 |
| 0.710423 | 0.716046 | 0.717815 | 0.710383 | 0.715633 | 0.719835 | 0.71538 | 0.704771 | 0.722874 |
| 0.707647 | 0.7111 | 0.711819 | 0.704717 | 0.710571 | 0.714136 | 0.71175 | 0.698211 | 0.727273 |
| 0.70704 | 0.711384 | 0.710141 | 0.705854 | 0.709543 | 0.710343 | 0.712273 | 0.699336 | 0.724316 |
| 0.708639 | 0.712681 | 0.710798 | 0.70628 | 0.707804 | 0.710609 | 0.7123 | 0.703307 | 0.740208 |
| 0.706211 | 0.711465 | 0.712406 | 0.705391 | 0.708373 | 0.710398 | 0.711813 | 0.695364 | 0.73483 |
| 0.704869 | 0.708327 | 0.710155 | 0.704335 | 0.707938 | 0.709553 | 0.708045 | 0.694171 | 0.72105 |
| 0.722267 | 0.725085 | 0.728218 | 0.721556 | 0.725136 | 0.728499 | 0.727367 | 0.718715 | 0.760151 |
| 0.701853 | 0.703643 | 0.703414 | 0.696631 | 0.700354 | 0.707843 | 0.707627 | 0.69136 | 0.724138 |
| 0.707943 | 0.71395 | 0.714049 | 0.705758 | 0.70871 | 0.71201 | 0.711834 | 0.691818 | 0.71803 |
| 0.702526 | 0.706461 | 0.707841 | 0.699835 | 0.704062 | 0.706322 | 0.707039 | 0.695811 | 0.723707 |
| 0.720677 | 0.725107 | 0.726274 | 0.718743 | 0.724838 | 0.728223 | 0.727477 | 0.718494 | 0.754506 |
| 0.70837 | 0.71053 | 0.714821 | 0.704896 | 0.711523 | 0.712859 | 0.713819 | 0.699872 | 0.749191 |
| 0.703061 | 0.705652 | 0.70699 | 0.701359 | 0.704921 | 0.705215 | 0.708685 | 0.691217 | 0.71917 |
| 0.70537 | 0.709396 | 0.70906 | 0.706174 | 0.707282 | 0.707927 | 0.710861 | 0.698186 | 0.721187 |
| 0.707064 | 0.711148 | 0.711143 | 0.704396 | 0.706263 | 0.712127 | 0.713643 | 0.694556 | 0.718591 |

| | | | | | | | | |
|---|---|---|---|---|---|---|---|---|
| 0.703822 | 0.706854 | 0.706587 | 0.705159 | 0.70398 | 0.712142 | 0.70978 | 0.693672 | 0.725846 |
| 0.710472 | 0.710504 | 0.715295 | 0.706115 | 0.709744 | 0.714411 | 0.712329 | 0.697888 | 0.723994 |
| 0.708615 | 0.711272 | 0.711776 | 0.711259 | 0.709881 | 0.710911 | 0.710295 | 0.694038 | 0.719238 |
| 0.711927 | 0.719436 | 0.719253 | 0.714758 | 0.714596 | 0.721701 | 0.720229 | 0.709522 | 0.747439 |
| 0.71549 | 0.715892 | 0.71927 | 0.712613 | 0.715666 | 0.716824 | 0.718482 | 0.706713 | 0.736132 |
| 0.709657 | 0.714297 | 0.712878 | 0.704442 | 0.712161 | 0.710303 | 0.711973 | 0.70711 | 0.730104 |
| 0.707886 | 0.714521 | 0.712665 | 0.70941 | 0.711239 | 0.711538 | 0.71406 | 0.699196 | 0.725567 |
| 0.708571 | 0.715378 | 0.712343 | 0.711494 | 0.710629 | 0.710451 | 0.712797 | 0.692237 | 0.722456 |
| 0.70847 | 0.710579 | 0.71051 | 0.707734 | 0.709335 | 0.712957 | 0.71384 | 0.693923 | 0.71275 |
| 0.702033 | 0.707081 | 0.708703 | 0.700963 | 0.705603 | 0.707606 | 0.708023 | 0.690198 | 0.724028 |
| 0.705919 | 0.709148 | 0.710914 | 0.706086 | 0.706585 | 0.710673 | 0.712013 | 0.69778 | 0.718704 |
| 0.712082 | 0.712264 | 0.712851 | 0.708666 | 0.707607 | 0.711478 | 0.712929 | 0.695957 | 0.722031 |
| 0.709202 | 0.710984 | 0.7124 | 0.709463 | 0.711337 | 0.711335 | 0.715443 | 0.691682 | 0.708864 |
| 0.705574 | 0.711475 | 0.711192 | 0.70439 | 0.706573 | 0.708505 | 0.7114 | 0.697153 | 0.711716 |
| 0.707103 | 0.710666 | 0.712368 | 0.707473 | 0.709289 | 0.713337 | 0.715381 | 0.70397 | 0.709184 |
| 0.708749 | 0.708565 | 0.708607 | 0.703756 | 0.707651 | 0.710176 | 0.708515 | 0.695144 | 0.714216 |
| 0.708718 | 0.712292 | 0.712573 | 0.707019 | 0.709592 | 0.713335 | 0.716049 | 0.692606 | 0.713866 |
| 0.710152 | 0.713103 | 0.711183 | 0.709362 | 0.707582 | 0.711143 | 0.710829 | 0.689285 | 0.732315 |
| 0.708589 | 0.713358 | 0.714548 | 0.709204 | 0.710619 | 0.713134 | 0.715038 | 0.702593 | 0.719363 |
| 0.707513 | 0.71122 | 0.713523 | 0.703827 | 0.708264 | 0.712255 | 0.71199 | 0.70549 | 0.719665 |
| 0.710079 | 0.710193 | 0.714457 | 0.709325 | 0.710831 | 0.708712 | 0.714252 | 0.693747 | 0.70952 |
| 0.706527 | 0.711227 | 0.705757 | 0.703317 | 0.702669 | 0.707944 | 0.711413 | 0.696341 | 0.721629 |
| 0.706165 | 0.710583 | 0.708739 | 0.701704 | 0.706854 | 0.712201 | 0.713667 | 0.690524 | 0.730523 |
| 0.710984 | 0.712643 | 0.715979 | 0.70631 | 0.711116 | 0.714286 | 0.713413 | 0.696246 | 0.729478 |
| 0.709446 | 0.71112 | 0.710782 | 0.708621 | 0.709193 | 0.713377 | 0.712484 | 0.692026 | 0.721497 |
| 0.707709 | 0.714118 | 0.714026 | 0.708415 | 0.712899 | 0.714851 | 0.715663 | 0.702287 | 0.723227 |
| 0.70836 | 0.711242 | 0.712076 | 0.705984 | 0.708616 | 0.712795 | 0.711269 | 0.691944 | 0.718948 |
| 0.707781 | 0.710334 | 0.710354 | 0.708558 | 0.709935 | 0.713605 | 0.713775 | 0.68926 | 0.708313 |
| 0.709042 | 0.714477 | 0.714547 | 0.712821 | 0.713911 | 0.713831 | 0.713152 | 0.703303 | 0.710194 |
| 0.707531 | 0.714 | 0.713894 | 0.709992 | 0.70988 | 0.714899 | 0.7111 | 0.695429 | 0.714843 |
| 0.708196 | 0.715673 | 0.714834 | 0.708973 | 0.707788 | 0.712819 | 0.714366 | 0.705364 | 0.725893 |
| 0.705411 | 0.710438 | 0.710543 | 0.705638 | 0.708735 | 0.709842 | 0.713892 | 0.711634 | 0.717181 |
| 0.710584 | 0.71416 | 0.714712 | 0.706941 | 0.710484 | 0.717521 | 0.712608 | 0.704181 | 0.725711 |
| 0.70765 | 0.711495 | 0.714309 | 0.708265 | 0.708676 | 0.712734 | 0.715197 | 0.701964 | 0.739506 |
| 0.707102 | 0.712708 | 0.712056 | 0.705929 | 0.707356 | 0.709879 | 0.713041 | 0.697315 | 0.723133 |
| 0.704745 | 0.71533 | 0.710541 | 0.705982 | 0.708551 | 0.711425 | 0.717696 | 0.696307 | 0.728801 |
| 0.701715 | 0.706553 | 0.711801 | 0.701988 | 0.704801 | 0.70724 | 0.709339 | 0.699495 | 0.706543 |
| 0.684724 | 0.686387 | 0.688133 | 0.685292 | 0.684788 | 0.688272 | 0.688589 | 0.69256 | 0.72122 |
| 0.689254 | 0.692879 | 0.688889 | 0.684988 | 0.686928 | 0.689613 | 0.693501 | 0.695105 | 0.730195 |
| 0.675192 | 0.681619 | 0.682232 | 0.679257 | 0.677085 | 0.683496 | 0.682631 | 0.690448 | 0.717705 |
| 0.67677 | 0.680598 | 0.678867 | 0.676909 | 0.675851 | 0.684695 | 0.68328 | 0.693388 | 0.723797 |
| 0.663202 | 0.667597 | 0.667365 | 0.664172 | 0.666939 | 0.666082 | 0.670187 | 0.675373 | 0.715024 |
| 0.660014 | 0.674144 | 0.670574 | 0.665781 | 0.666185 | 0.668354 | 0.674232 | 0.688083 | 0.73531 |
| 0.661945 | 0.668757 | 0.672428 | 0.664616 | 0.666425 | 0.672078 | 0.674308 | 0.690848 | 0.727347 |
| 0.666751 | 0.668096 | 0.666624 | 0.663494 | 0.667175 | 0.667813 | 0.672363 | 0.695164 | 0.711949 |
| 0.664673 | 0.671498 | 0.668077 | 0.66783 | 0.666418 | 0.674913 | 0.675344 | 0.693511 | 0.723422 |
| 0.672476 | 0.673726 | 0.67364 | 0.671671 | 0.671761 | 0.679351 | 0.677607 | 0.697304 | 0.732113 |

| | | | | | | | | |
|---|---|---|---|---|---|---|---|---|
| 0.668345 | 0.674923 | 0.671978 | 0.669128 | 0.668992 | 0.674264 | 0.675285 | 0.694185 | 0.696568 |
| 0.670659 | 0.680837 | 0.669814 | 0.669997 | 0.673766 | 0.680574 | 0.680628 | 0.697333 | 0.733199 |
| 0.661784 | 0.672395 | 0.671267 | 0.671088 | 0.665335 | 0.66952 | 0.677108 | 0.69876 | 0.712984 |
| 0.660474 | 0.663344 | 0.655894 | 0.656283 | 0.654248 | 0.663069 | 0.66552 | 0.695991 | 0.732107 |
| 0.65631 | 0.674445 | 0.66782 | 0.657908 | 0.663973 | 0.664875 | 0.668812 | 0.68916 | 0.734796 |
| 0.66078 | 0.669672 | 0.663385 | 0.663379 | 0.660524 | 0.666575 | 0.673661 | 0.692588 | 0.718777 |
| 0.661723 | 0.669758 | 0.664413 | 0.664688 | 0.667043 | 0.667325 | 0.674247 | 0.692467 | 0.728483 |
| 0.66048 | 0.671429 | 0.667786 | 0.660246 | 0.666946 | 0.666713 | 0.670703 | 0.696783 | 0.729336 |
| 0.658974 | 0.667105 | 0.663596 | 0.661715 | 0.659743 | 0.660474 | 0.668582 | 0.684501 | 0.717333 |
| 0.653483 | 0.669534 | 0.660533 | 0.664767 | 0.66264 | 0.664103 | 0.669892 | 0.688509 | 0.71729 |
| 0.648056 | 0.660013 | 0.653597 | 0.6583 | 0.654718 | 0.661477 | 0.664145 | 0.6875 | 0.710927 |
| 0.696656 | 0.703384 | 0.699517 | 0.692859 | 0.698532 | 0.702614 | 0.701732 | 0.680094 | 0.724903 |
| 0.704554 | 0.71157 | 0.710074 | 0.70346 | 0.70726 | 0.71013 | 0.711831 | 0.694698 | 0.704386 |
| 0.683189 | 0.685108 | 0.688239 | 0.682871 | 0.689423 | 0.692036 | 0.693379 | 0.69743 | 0.726579 |
| 0.712843 | 0.717517 | 0.716828 | 0.712259 | 0.714067 | 0.717657 | 0.718034 | 0.698921 | 0.765883 |
| 0.70119 | 0.706369 | 0.709406 | 0.700616 | 0.7057 | 0.705021 | 0.708456 | 0.694536 | 0.71488 |
| 0.71106 | 0.713579 | 0.71767 | 0.707535 | 0.713416 | 0.712175 | 0.715609 | 0.698161 | 0.712797 |
| 0.716276 | 0.720201 | 0.716004 | 0.715559 | 0.712705 | 0.720087 | 0.721873 | 0.703016 | 0.725307 |
| 0.696759 | 0.700897 | 0.698122 | 0.698368 | 0.697553 | 0.697935 | 0.702238 | 0.672682 | 0.697006 |
| 0.720178 | 0.723379 | 0.725305 | 0.718659 | 0.725085 | 0.72362 | 0.724157 | 0.695479 | 0.723014 |
| 0.688027 | 0.693432 | 0.69156 | 0.687349 | 0.688972 | 0.691983 | 0.692021 | 0.696309 | 0.720644 |
| 0.71311 | 0.716067 | 0.720994 | 0.713761 | 0.715405 | 0.711778 | 0.715883 | 0.698713 | 0.70669 |
| 0.718753 | 0.722898 | 0.719272 | 0.717715 | 0.719035 | 0.723974 | 0.727733 | 0.703574 | 0.724002 |
| 0.676402 | 0.688606 | 0.682848 | 0.677211 | 0.679219 | 0.684094 | 0.685362 | 0.695545 | 0.728487 |
| 0.686789 | 0.693834 | 0.690417 | 0.684609 | 0.687141 | 0.68961 | 0.69338 | 0.669397 | 0.705495 |
| 0.687587 | 0.696128 | 0.693731 | 0.686445 | 0.687981 | 0.697819 | 0.698178 | 0.699345 | 0.734379 |
| 0.705524 | 0.709196 | 0.709866 | 0.702782 | 0.708513 | 0.70853 | 0.707948 | 0.686548 | 0.695191 |
| 0.709408 | 0.720035 | 0.719415 | 0.712066 | 0.713119 | 0.716094 | 0.718028 | 0.705395 | 0.716203 |
| 0.686585 | 0.69052 | 0.690292 | 0.685354 | 0.686138 | 0.689867 | 0.691384 | 0.671378 | 0.689176 |
| 0.721645 | 0.726722 | 0.726912 | 0.719345 | 0.721458 | 0.727489 | 0.724942 | 0.70645 | 0.736432 |
| 0.719556 | 0.723556 | 0.725367 | 0.719588 | 0.721645 | 0.723948 | 0.728083 | 0.698175 | 0.705812 |
| 0.697484 | 0.700118 | 0.701156 | 0.695752 | 0.698599 | 0.70092 | 0.702825 | 0.688686 | 0.718012 |
| 0.696788 | 0.705205 | 0.701746 | 0.696653 | 0.697674 | 0.702134 | 0.703882 | 0.68535 | 0.713275 |
| 0.721929 | 0.728514 | 0.728745 | 0.721027 | 0.725805 | 0.726286 | 0.728665 | 0.702229 | 0.724003 |
| 0.717792 | 0.721406 | 0.723857 | 0.716288 | 0.718322 | 0.725271 | 0.720956 | 0.696538 | 0.716699 |
| 0.722826 | 0.728224 | 0.725471 | 0.72302 | 0.725347 | 0.731458 | 0.730515 | 0.698638 | 0.719137 |
| 0.675396 | 0.681558 | 0.681309 | 0.676935 | 0.67676 | 0.680778 | 0.68597 | 0.657715 | 0.669785 |
| 0.721223 | 0.722178 | 0.723056 | 0.71805 | 0.717939 | 0.722311 | 0.71734 | 0.722664 | 0.714182 |
| 0.674989 | 0.677819 | 0.677055 | 0.674847 | 0.676642 | 0.677826 | 0.681477 | 0.657513 | 0.679021 |
| 0.665256 | 0.66739 | 0.670493 | 0.663898 | 0.668883 | 0.671311 | 0.671944 | 0.640628 | 0.651884 |
| 0.722041 | 0.72574 | 0.720502 | 0.718012 | 0.719621 | 0.724478 | 0.728262 | 0.696159 | 0.734081 |
| 0.667147 | 0.668994 | 0.672292 | 0.668256 | 0.670271 | 0.673647 | 0.67531 | 0.646674 | 0.659361 |
| 0.662382 | 0.666542 | 0.666817 | 0.658851 | 0.665742 | 0.667118 | 0.669548 | 0.640644 | 0.65301 |
| 0.656147 | 0.665172 | 0.661387 | 0.655519 | 0.656252 | 0.663328 | 0.667254 | 0.639263 | 0.655063 |
| 0.728763 | 0.734637 | 0.728884 | 0.722982 | 0.730264 | 0.7341 | 0.733371 | 0.700219 | 0.708444 |
| 0.657988 | 0.665497 | 0.66392 | 0.659739 | 0.663368 | 0.664639 | 0.668673 | 0.637528 | 0.644898 |
| 0.719155 | 0.722227 | 0.723733 | 0.718413 | 0.721298 | 0.723986 | 0.724428 | 0.698802 | 0.723368 |

| | | | | | | | | |
|---|---|---|---|---|---|---|---|---|
| 0.728853 | 0.733238 | 0.72988 | 0.728815 | 0.728031 | 0.734248 | 0.733678 | 0.703165 | 0.713837 |
| 0.722382 | 0.727589 | 0.723068 | 0.722152 | 0.723678 | 0.728542 | 0.730401 | 0.686183 | 0.718637 |
| 0.740717 | 0.751033 | 0.745384 | 0.7375 | 0.741563 | 0.748208 | 0.74492 | 0.694771 | 0.703544 |
| 0.649612 | 0.654648 | 0.653508 | 0.646453 | 0.650962 | 0.655804 | 0.656615 | 0.626413 | 0.638174 |
| 0.65372 | 0.657483 | 0.65652 | 0.650582 | 0.654785 | 0.659798 | 0.660175 | 0.633727 | 0.645939 |
| 0.729736 | 0.732682 | 0.732444 | 0.731905 | 0.734643 | 0.737278 | 0.735101 | 0.695668 | 0.72326 |
| 0.656753 | 0.659384 | 0.655229 | 0.653316 | 0.656839 | 0.657685 | 0.66045 | 0.623009 | 0.639428 |
| 0.729897 | 0.733527 | 0.736675 | 0.732889 | 0.73066 | 0.738825 | 0.732803 | 0.762122 | 0.698072 |
| 0.731686 | 0.737103 | 0.732316 | 0.731648 | 0.734811 | 0.738516 | 0.738013 | 0.695398 | 0.714528 |
| 0.731719 | 0.733015 | 0.732617 | 0.731888 | 0.72969 | 0.734892 | 0.729148 | 0.743294 | 0.668595 |
| 0.733431 | 0.737299 | 0.735275 | 0.735113 | 0.73522 | 0.737793 | 0.743494 | 0.688811 | 0.698757 |
| 0.733766 | 0.732026 | 0.75 | 0.771429 | 0.76259 | 0.743243 | 0.762238 | 0.681159 | 0.655738 |
| 0.723684 | 0.734106 | 0.736303 | 0.731064 | 0.737626 | 0.733012 | 0.742713 | 0.681151 | 0.67099 |
| 0.728477 | 0.737646 | 0.73724 | 0.725662 | 0.734267 | 0.738959 | 0.73785 | 0.671716 | 0.643486 |
| 0.731397 | 0.73978 | 0.743004 | 0.740393 | 0.733967 | 0.741032 | 0.744257 | 0.676135 | 0.656313 |
| 0.738913 | 0.745381 | 0.744859 | 0.737876 | 0.736892 | 0.74473 | 0.750851 | 0.676781 | 0.660832 |
| 0.59116 | 0.597049 | 0.593575 | 0.590458 | 0.593514 | 0.593314 | 0.599241 | 0.589226 | 0.585275 |
| 0.736464 | 0.739362 | 0.747473 | 0.737131 | 0.736231 | 0.744482 | 0.743728 | 0.678693 | 0.648764 |
| 0.735621 | 0.743731 | 0.739236 | 0.735402 | 0.737605 | 0.744275 | 0.743225 | 0.682274 | 0.69876 |
| 0.738425 | 0.744381 | 0.740881 | 0.736232 | 0.741821 | 0.744799 | 0.743577 | 0.691385 | 0.688445 |
| 0.737189 | 0.744386 | 0.740912 | 0.737325 | 0.740539 | 0.745645 | 0.744676 | 0.687684 | 0.691448 |
| 0.59452 | 0.598252 | 0.597692 | 0.593919 | 0.594667 | 0.597812 | 0.604405 | 0.548084 | 0.54527 |
| 0.581317 | 0.586748 | 0.586297 | 0.580691 | 0.584909 | 0.584529 | 0.594415 | 0.534754 | 0.529667 |
| 0.763835 | 0.763143 | 0.769974 | 0.761277 | 0.765352 | 0.766897 | 0.767008 | 0.68121 | 0.665146 |
| 0.767291 | 0.766082 | 0.77352 | 0.760726 | 0.767583 | 0.768417 | 0.767667 | 0.675228 | 0.667947 |
| 0.757152 | 0.754611 | 0.759918 | 0.752136 | 0.755433 | 0.758598 | 0.758731 | 0.673342 | 0.652725 |
| 0.764036 | 0.763184 | 0.766459 | 0.757502 | 0.763547 | 0.765221 | 0.767939 | 0.677302 | 0.653637 |
| 0.760085 | 0.763424 | 0.768715 | 0.760809 | 0.76435 | 0.769798 | 0.76828 | 0.680634 | 0.659833 |
| 0.766972 | 0.766329 | 0.770207 | 0.765825 | 0.771933 | 0.772364 | 0.766299 | 0.685546 | 0.662291 |
| 0.77 | 0.77441 | 0.772671 | 0.765213 | 0.769547 | 0.773691 | 0.769795 | 0.684029 | 0.66024 |
| 0.759678 | 0.762576 | 0.763117 | 0.757068 | 0.761115 | 0.767226 | 0.764097 | 0.681512 | 0.656573 |
| 0.761947 | 0.768901 | 0.770385 | 0.763536 | 0.761773 | 0.772776 | 0.773109 | 0.682089 | 0.660642 |
| 0.758024 | 0.763745 | 0.763429 | 0.759464 | 0.761873 | 0.76399 | 0.763489 | 0.68372 | 0.649587 |
| 0.757824 | 0.766618 | 0.7617 | 0.757306 | 0.757346 | 0.765963 | 0.767434 | 0.677373 | 0.660266 |
| 0.764012 | 0.770063 | 0.768589 | 0.764171 | 0.765269 | 0.769533 | 0.772291 | 0.687776 | 0.661955 |
| 0.762483 | 0.762515 | 0.766068 | 0.758306 | 0.76347 | 0.768609 | 0.761286 | 0.671561 | 0.647492 |
| 0.76344 | 0.763463 | 0.763035 | 0.758756 | 0.766442 | 0.762972 | 0.762687 | 0.674359 | 0.648222 |
| 0.764928 | 0.767747 | 0.766697 | 0.76408 | 0.766874 | 0.766233 | 0.763457 | 0.678806 | 0.658171 |
| 0.756814 | 0.763246 | 0.75913 | 0.755184 | 0.763327 | 0.766376 | 0.761833 | 0.678532 | 0.653992 |
| 0.756374 | 0.766375 | 0.758769 | 0.754426 | 0.760875 | 0.760382 | 0.759071 | 0.673503 | 0.658773 |
| 0.724544 | 0.739856 | 0.73637 | 0.733017 | 0.734834 | 0.735604 | 0.731094 | 0.67838 | 0.65133 |
| 0.73167 | 0.744089 | 0.737001 | 0.733813 | 0.734125 | 0.740825 | 0.739213 | 0.688418 | 0.66857 |
| 0.757837 | 0.762571 | 0.765266 | 0.761082 | 0.761823 | 0.764792 | 0.760924 | 0.688946 | 0.663901 |
| 0.770832 | 0.773577 | 0.773751 | 0.764176 | 0.765408 | 0.77331 | 0.774986 | 0.692336 | 0.665 |
| 0.717959 | 0.723819 | 0.724797 | 0.723585 | 0.718672 | 0.719707 | 0.725966 | 0.669338 | 0.64084 |
| 0.7625 | 0.770419 | 0.765575 | 0.768307 | 0.762888 | 0.769136 | 0.766475 | 0.683118 | 0.656375 |
| 0.756832 | 0.759835 | 0.760632 | 0.759726 | 0.758702 | 0.761784 | 0.763374 | 0.675419 | 0.645131 |

| | | | | | | | | |
|---|---|---|---|---|---|---|---|---|
| 0.758059 | 0.767055 | 0.766404 | 0.764214 | 0.765778 | 0.765917 | 0.769585 | 0.678105 | 0.654353 |
| 0.762285 | 0.76849 | 0.769043 | 0.76871 | 0.765327 | 0.767947 | 0.769406 | 0.676264 | 0.658192 |
| 0.766199 | 0.769385 | 0.765706 | 0.763735 | 0.766153 | 0.767352 | 0.768263 | 0.686019 | 0.662997 |
| 0.762499 | 0.769676 | 0.766642 | 0.762135 | 0.766232 | 0.768504 | 0.769307 | 0.679949 | 0.660109 |
| 0.769938 | 0.775854 | 0.771562 | 0.770279 | 0.771459 | 0.774562 | 0.773608 | 0.690146 | 0.670069 |
| 0.762117 | 0.768134 | 0.766159 | 0.762019 | 0.7654 | 0.76572 | 0.767574 | 0.682719 | 0.649494 |
| 0.764149 | 0.768801 | 0.768879 | 0.760602 | 0.765018 | 0.765652 | 0.766245 | 0.678196 | 0.654181 |
| 0.754815 | 0.760372 | 0.754371 | 0.754878 | 0.756422 | 0.758895 | 0.755163 | 0.671465 | 0.645679 |
| 0.758313 | 0.761278 | 0.762826 | 0.76056 | 0.763924 | 0.764696 | 0.762916 | 0.676753 | 0.65642 |
| 0.754055 | 0.762421 | 0.757678 | 0.758233 | 0.758776 | 0.761787 | 0.760082 | 0.676043 | 0.651207 |
| 0.760284 | 0.761874 | 0.762999 | 0.765478 | 0.759512 | 0.769275 | 0.764394 | 0.687283 | 0.662233 |
| 0.75641 | 0.764331 | 0.761579 | 0.759534 | 0.762816 | 0.764159 | 0.763616 | 0.67761 | 0.654874 |
| 0.753545 | 0.757715 | 0.755078 | 0.753702 | 0.754528 | 0.760555 | 0.757032 | 0.673353 | 0.647594 |
| 0.758671 | 0.760632 | 0.756642 | 0.755945 | 0.757304 | 0.760907 | 0.76252 | 0.677395 | 0.652059 |
| 0.761707 | 0.771061 | 0.766859 | 0.762767 | 0.764931 | 0.766726 | 0.767385 | 0.682993 | 0.658554 |
| 0.763068 | 0.769737 | 0.771559 | 0.763819 | 0.764078 | 0.767759 | 0.76852 | 0.687717 | 0.662857 |
| 0.76163 | 0.766863 | 0.767359 | 0.760335 | 0.760455 | 0.767323 | 0.766483 | 0.678571 | 0.654342 |
| 0.758041 | 0.75891 | 0.76162 | 0.750795 | 0.759964 | 0.764401 | 0.761515 | 0.681649 | 0.65153 |
| 0.755461 | 0.761407 | 0.767398 | 0.758956 | 0.760424 | 0.765457 | 0.76486 | 0.68008 | 0.6596 |
| 0.760478 | 0.762538 | 0.765054 | 0.754689 | 0.766447 | 0.763824 | 0.765823 | 0.679583 | 0.650225 |
| 0.753165 | 0.756346 | 0.758474 | 0.751703 | 0.754029 | 0.762164 | 0.76022 | 0.677294 | 0.655131 |
| 0.754607 | 0.758965 | 0.76262 | 0.75715 | 0.760944 | 0.764445 | 0.764201 | 0.674359 | 0.653013 |
| 0.759164 | 0.765105 | 0.770008 | 0.759265 | 0.765644 | 0.769049 | 0.769363 | 0.680174 | 0.655587 |
| 0.759888 | 0.764338 | 0.769521 | 0.754969 | 0.762473 | 0.770268 | 0.7655 | 0.67792 | 0.652297 |
| 0.758102 | 0.763359 | 0.763147 | 0.757721 | 0.760232 | 0.76907 | 0.765483 | 0.692039 | 0.65523 |
| 0.757601 | 0.762544 | 0.765318 | 0.758228 | 0.759203 | 0.764462 | 0.766286 | 0.677254 | 0.654472 |
| 0.757114 | 0.760336 | 0.759233 | 0.748845 | 0.756757 | 0.763642 | 0.761976 | 0.674728 | 0.654707 |
| 0.754417 | 0.758923 | 0.762076 | 0.754196 | 0.757434 | 0.761622 | 0.760844 | 0.677969 | 0.651588 |
| 0.757591 | 0.758637 | 0.76747 | 0.755179 | 0.757927 | 0.764508 | 0.763741 | 0.671884 | 0.664118 |
| 0.760187 | 0.762013 | 0.764499 | 0.759271 | 0.762251 | 0.763911 | 0.765656 | 0.683786 | 0.659341 |
| 0.758367 | 0.767795 | 0.762938 | 0.76057 | 0.763709 | 0.766429 | 0.766233 | 0.682998 | 0.660932 |
| 0.762134 | 0.76474 | 0.762215 | 0.757205 | 0.762741 | 0.768292 | 0.768771 | 0.680354 | 0.658325 |
| 0.770828 | 0.771396 | 0.777134 | 0.765556 | 0.772987 | 0.77422 | 0.771831 | 0.693024 | 0.671574 |
| 0.768314 | 0.773342 | 0.773297 | 0.764817 | 0.766353 | 0.7743 | 0.77445 | 0.690989 | 0.667065 |
| 0.759191 | 0.755617 | 0.762069 | 0.752948 | 0.760105 | 0.760546 | 0.761662 | 0.667092 | 0.658164 |
| 0.75892 | 0.764239 | 0.764006 | 0.757066 | 0.762952 | 0.766533 | 0.763272 | 0.687904 | 0.66054 |
| 0.762847 | 0.763309 | 0.76557 | 0.756061 | 0.762428 | 0.764201 | 0.765162 | 0.680952 | 0.65782 |
| 0.758446 | 0.761752 | 0.765276 | 0.756837 | 0.761372 | 0.765972 | 0.765487 | 0.68256 | 0.647951 |
| 0.750812 | 0.752288 | 0.753298 | 0.748051 | 0.754328 | 0.758063 | 0.756203 | 0.675222 | 0.650973 |
| 0.755009 | 0.756203 | 0.757819 | 0.750021 | 0.7555 | 0.76072 | 0.761434 | 0.675482 | 0.65687 |
| 0.744263 | 0.753188 | 0.749726 | 0.745958 | 0.750506 | 0.753816 | 0.754373 | 0.666728 | 0.64545 |
| 0.750044 | 0.755273 | 0.757897 | 0.749174 | 0.757214 | 0.756976 | 0.758834 | 0.677755 | 0.655647 |
| 0.753348 | 0.752592 | 0.751664 | 0.746679 | 0.753591 | 0.753469 | 0.756525 | 0.678989 | 0.65409 |
| 0.743049 | 0.751451 | 0.749081 | 0.747779 | 0.751463 | 0.755405 | 0.751391 | 0.675849 | 0.648008 |
| 0.753352 | 0.761628 | 0.760576 | 0.755232 | 0.759041 | 0.760217 | 0.762302 | 0.684554 | 0.650916 |
| 0.750613 | 0.753971 | 0.755223 | 0.743486 | 0.752167 | 0.754872 | 0.750514 | 0.676279 | 0.649331 |

| | | | | | | | | |
|---|---|---|---|---|---|---|---|---|
| 0.757294 | 0.763186 | 0.761674 | 0.757414 | 0.762755 | 0.763337 | 0.766038 | 0.676401 | 0.664341 |
| 0.768144 | 0.770771 | 0.771167 | 0.767388 | 0.768406 | 0.769391 | 0.770165 | 0.691896 | 0.664202 |
| 0.765212 | 0.772566 | 0.765668 | 0.760282 | 0.769397 | 0.768512 | 0.770799 | 0.675696 | 0.652805 |
| 0.759268 | 0.76938 | 0.768484 | 0.76532 | 0.764792 | 0.766503 | 0.76972 | 0.682701 | 0.661524 |
| 0.767922 | 0.775872 | 0.777759 | 0.766448 | 0.772563 | 0.768586 | 0.778171 | 0.692857 | 0.657366 |
| 0.765395 | 0.77134 | 0.770111 | 0.768487 | 0.765422 | 0.769862 | 0.770515 | 0.691647 | 0.656975 |
| 0.760646 | 0.765189 | 0.760914 | 0.759058 | 0.761096 | 0.766644 | 0.767331 | 0.681674 | 0.654463 |
| 0.763434 | 0.769543 | 0.770088 | 0.760813 | 0.765192 | 0.769975 | 0.769338 | 0.675504 | 0.651887 |
| 0.75987 | 0.766148 | 0.767712 | 0.760506 | 0.763343 | 0.763726 | 0.767369 | 0.684161 | 0.665856 |
| 0.751226 | 0.764538 | 0.76054 | 0.754541 | 0.756748 | 0.75978 | 0.763151 | 0.676 | 0.646795 |
| 0.76026 | 0.768821 | 0.765412 | 0.760223 | 0.765776 | 0.768066 | 0.76878 | 0.680678 | 0.655734 |
| 0.761148 | 0.770954 | 0.766568 | 0.759395 | 0.763711 | 0.76685 | 0.770442 | 0.681984 | 0.666348 |
| 0.761834 | 0.767043 | 0.771938 | 0.769161 | 0.768336 | 0.773091 | 0.769799 | 0.676867 | 0.649645 |
| 0.769624 | 0.768805 | 0.769408 | 0.765849 | 0.767041 | 0.775431 | 0.769944 | 0.683282 | 0.655645 |
| 0.765032 | 0.770557 | 0.775842 | 0.766694 | 0.76841 | 0.773616 | 0.776008 | 0.687238 | 0.656955 |
| 0.764344 | 0.764935 | 0.771168 | 0.767404 | 0.769079 | 0.77163 | 0.766097 | 0.678997 | 0.654472 |
| 0.766726 | 0.771062 | 0.773763 | 0.767103 | 0.768549 | 0.770557 | 0.774222 | 0.682016 | 0.658267 |
| 0.76386 | 0.771916 | 0.773413 | 0.760084 | 0.771341 | 0.771222 | 0.771544 | 0.669705 | 0.659934 |
| 0.77036 | 0.774756 | 0.775064 | 0.764826 | 0.768765 | 0.773958 | 0.775094 | 0.680972 | 0.662127 |
| 0.767444 | 0.776666 | 0.770713 | 0.76652 | 0.773392 | 0.772158 | 0.776396 | 0.685215 | 0.658438 |
| 0.761748 | 0.768607 | 0.763922 | 0.765491 | 0.762195 | 0.766897 | 0.767488 | 0.679753 | 0.657957 |
| 0.772001 | 0.772205 | 0.768525 | 0.765312 | 0.768499 | 0.773658 | 0.772467 | 0.681777 | 0.655532 |
| 0.769313 | 0.773557 | 0.771317 | 0.770513 | 0.769193 | 0.773871 | 0.770891 | 0.67569 | 0.66508 |
| 0.767251 | 0.77151 | 0.772086 | 0.765059 | 0.769988 | 0.770302 | 0.770382 | 0.680036 | 0.665404 |
| 0.767653 | 0.771265 | 0.772392 | 0.766319 | 0.768296 | 0.772018 | 0.775698 | 0.68546 | 0.661252 |
| 0.766193 | 0.766545 | 0.765984 | 0.7685 | 0.769294 | 0.775891 | 0.770656 | 0.687443 | 0.655311 |
| 0.764725 | 0.771752 | 0.771484 | 0.766157 | 0.765041 | 0.771822 | 0.771557 | 0.674732 | 0.649707 |
| 0.764648 | 0.768808 | 0.769018 | 0.762519 | 0.764601 | 0.771485 | 0.772445 | 0.684048 | 0.652205 |
| 0.771312 | 0.773018 | 0.774656 | 0.764859 | 0.776041 | 0.775906 | 0.774299 | 0.683972 | 0.6547 |
| 0.766523 | 0.774573 | 0.774106 | 0.767738 | 0.768504 | 0.774286 | 0.772773 | 0.694347 | 0.66867 |
| 0.76372 | 0.759822 | 0.764079 | 0.759854 | 0.763848 | 0.764549 | 0.766739 | 0.685908 | 0.656677 |
| 0.764643 | 0.771403 | 0.770495 | 0.763582 | 0.76925 | 0.772648 | 0.774041 | 0.681645 | 0.659458 |
| 0.77084 | 0.773611 | 0.770959 | 0.766781 | 0.773356 | 0.769187 | 0.775825 | 0.683692 | 0.653018 |
| 0.761131 | 0.765322 | 0.767239 | 0.763102 | 0.763853 | 0.764686 | 0.768501 | 0.682487 | 0.66283 |
| 0.766173 | 0.768798 | 0.769634 | 0.763497 | 0.764721 | 0.770943 | 0.77272 | 0.682291 | 0.662849 |
| 0.762769 | 0.772268 | 0.771268 | 0.763815 | 0.767224 | 0.769476 | 0.76841 | 0.684939 | 0.653491 |
| 0.769086 | 0.772302 | 0.772902 | 0.769477 | 0.772265 | 0.77986 | 0.770363 | 0.687865 | 0.669214 |
| 0.765384 | 0.772652 | 0.773943 | 0.767452 | 0.765652 | 0.77307 | 0.774307 | 0.693976 | 0.662868 |
| 0.765093 | 0.768629 | 0.77534 | 0.766516 | 0.768007 | 0.769573 | 0.774646 | 0.68107 | 0.654443 |
| 0.761499 | 0.772746 | 0.765663 | 0.765605 | 0.766135 | 0.769063 | 0.77224 | 0.682476 | 0.663348 |
| 0.765277 | 0.775199 | 0.775053 | 0.769155 | 0.770803 | 0.772166 | 0.775047 | 0.685367 | 0.66517 |
| 0.763848 | 0.774693 | 0.775259 | 0.770722 | 0.771661 | 0.774992 | 0.777009 | 0.685192 | 0.665553 |
| 0.768109 | 0.77604 | 0.777978 | 0.771678 | 0.773168 | 0.777096 | 0.77885 | 0.687102 | 0.66857 |
| 0.765101 | 0.766495 | 0.766155 | 0.763178 | 0.768709 | 0.765817 | 0.771797 | 0.675582 | 0.655968 |
| 0.766584 | 0.772586 | 0.767333 | 0.771062 | 0.768445 | 0.769675 | 0.77437 | 0.687454 | 0.664921 |
| 0.762894 | 0.771704 | 0.767979 | 0.766075 | 0.767217 | 0.763603 | 0.769449 | 0.682068 | 0.656546 |

| | | | | | | | | |
|---|---|---|---|---|---|---|---|---|
| 0.758333 | 0.763039 | 0.758383 | 0.757768 | 0.754174 | 0.758043 | 0.760398 | 0.676471 | 0.658573 |
| 0.742362 | 0.74717 | 0.749979 | 0.739625 | 0.747432 | 0.750802 | 0.750123 | 0.690975 | 0.703106 |
| 0.764746 | 0.772024 | 0.766998 | 0.764033 | 0.761909 | 0.770928 | 0.768946 | 0.681284 | 0.659732 |
| 0.764175 | 0.768356 | 0.768162 | 0.76866 | 0.769275 | 0.775029 | 0.766933 | 0.696272 | 0.661122 |
| 0.761138 | 0.765703 | 0.765793 | 0.760677 | 0.757948 | 0.767267 | 0.766863 | 0.6811 | 0.652477 |
| 0.76473 | 0.763098 | 0.766863 | 0.765456 | 0.764345 | 0.76847 | 0.767836 | 0.68064 | 0.65613 |
| 0.764868 | 0.770297 | 0.777586 | 0.76464 | 0.770245 | 0.769496 | 0.771451 | 0.680682 | 0.654589 |
| 0.758508 | 0.764494 | 0.766965 | 0.758971 | 0.76481 | 0.767019 | 0.762006 | 0.678832 | 0.651096 |
| 0.770019 | 0.775528 | 0.77321 | 0.766237 | 0.774323 | 0.772098 | 0.773631 | 0.676727 | 0.650483 |
| 0.766461 | 0.773267 | 0.768725 | 0.766477 | 0.765377 | 0.771497 | 0.773935 | 0.68541 | 0.663113 |
| 0.763577 | 0.773573 | 0.767597 | 0.76221 | 0.769024 | 0.772634 | 0.771072 | 0.682508 | 0.655784 |
| 0.76425 | 0.772065 | 0.770073 | 0.762987 | 0.766215 | 0.774151 | 0.772885 | 0.697584 | 0.659843 |
| 0.7722 | 0.775972 | 0.774485 | 0.76942 | 0.773064 | 0.773064 | 0.776241 | 0.683172 | 0.659938 |
| 0.763478 | 0.773299 | 0.772342 | 0.767153 | 0.767677 | 0.773613 | 0.774975 | 0.685138 | 0.666031 |
| 0.75787 | 0.768625 | 0.768942 | 0.760718 | 0.766895 | 0.769359 | 0.768447 | 0.687777 | 0.656599 |
| 0.768121 | 0.772621 | 0.77512 | 0.768845 | 0.770685 | 0.769904 | 0.770753 | 0.690731 | 0.672209 |
| 0.76331 | 0.765924 | 0.762631 | 0.765693 | 0.76458 | 0.765626 | 0.769763 | 0.690415 | 0.672143 |
| 0.75947 | 0.767194 | 0.764031 | 0.762655 | 0.764172 | 0.767686 | 0.768271 | 0.686551 | 0.666906 |
| 0.764133 | 0.774377 | 0.772353 | 0.764758 | 0.765755 | 0.76941 | 0.77673 | 0.681516 | 0.661103 |
| 0.761971 | 0.768176 | 0.764578 | 0.761162 | 0.761418 | 0.766461 | 0.76537 | 0.681911 | 0.659879 |
| 0.763525 | 0.772303 | 0.77079 | 0.76338 | 0.767621 | 0.765982 | 0.774506 | 0.689261 | 0.663462 |
| 0.757698 | 0.764864 | 0.765397 | 0.759103 | 0.759612 | 0.769276 | 0.760917 | 0.67809 | 0.65455 |
| 0.761631 | 0.765985 | 0.770868 | 0.767108 | 0.764499 | 0.771027 | 0.764874 | 0.676856 | 0.65401 |
| 0.765856 | 0.772846 | 0.774207 | 0.76574 | 0.767459 | 0.775782 | 0.769686 | 0.674672 | 0.661874 |
| 0.766047 | 0.770291 | 0.773553 | 0.766524 | 0.769825 | 0.774302 | 0.773814 | 0.683367 | 0.663874 |
| 0.765335 | 0.769593 | 0.772333 | 0.767966 | 0.766586 | 0.773975 | 0.770373 | 0.676159 | 0.658065 |
| 0.762552 | 0.770842 | 0.768161 | 0.76627 | 0.765703 | 0.765715 | 0.772921 | 0.674921 | 0.656 |
| 0.765602 | 0.769042 | 0.773907 | 0.765945 | 0.765959 | 0.769783 | 0.77555 | 0.679055 | 0.658058 |
| 0.763803 | 0.76864 | 0.766175 | 0.760779 | 0.765991 | 0.76742 | 0.768882 | 0.678135 | 0.650565 |
| 0.691919 | 0.697415 | 0.697996 | 0.686359 | 0.694895 | 0.694681 | 0.699789 | 0.68767 | 0.713771 |
| 0.705873 | 0.709562 | 0.707233 | 0.702162 | 0.705267 | 0.711328 | 0.70807 | 0.692774 | 0.713548 |
| 0.702327 | 0.706999 | 0.708964 | 0.701285 | 0.703902 | 0.706104 | 0.706456 | 0.692209 | 0.720228 |
| 0.688762 | 0.694408 | 0.6964 | 0.691768 | 0.694493 | 0.698775 | 0.695313 | 0.685084 | 0.725051 |
| 0.704136 | 0.711068 | 0.705531 | 0.704051 | 0.708714 | 0.70787 | 0.710072 | 0.6875 | 0.705669 |
| 0.763771 | 0.769198 | 0.769753 | 0.767175 | 0.768544 | 0.772153 | 0.768101 | 0.68098 | 0.64626 |
| 0.763344 | 0.773932 | 0.76955 | 0.768367 | 0.764379 | 0.769866 | 0.768265 | 0.678454 | 0.654576 |
| 0.772758 | 0.780211 | 0.77689 | 0.768668 | 0.772644 | 0.774681 | 0.773676 | 0.683834 | 0.659021 |
| 0.769014 | 0.774241 | 0.779676 | 0.774135 | 0.77016 | 0.777207 | 0.774615 | 0.682293 | 0.647765 |
| 0.77051 | 0.772847 | 0.775087 | 0.770906 | 0.772826 | 0.774073 | 0.77668 | 0.677078 | 0.654148 |
| 0.773583 | 0.780849 | 0.777125 | 0.771551 | 0.775836 | 0.775365 | 0.775461 | 0.685121 | 0.654309 |
| 0.768717 | 0.775169 | 0.772871 | 0.768429 | 0.770641 | 0.774979 | 0.769 | 0.676503 | 0.651958 |
| 0.763233 | 0.768066 | 0.768382 | 0.763644 | 0.767071 | 0.773029 | 0.769886 | 0.683269 | 0.65239 |
| 0.765393 | 0.766632 | 0.769887 | 0.763534 | 0.763995 | 0.766313 | 0.768989 | 0.683346 | 0.654872 |
| 0.763543 | 0.760365 | 0.772053 | 0.763585 | 0.766227 | 0.767697 | 0.766619 | 0.68283 | 0.653317 |
| 0.762927 | 0.770365 | 0.765993 | 0.765625 | 0.768451 | 0.76914 | 0.766054 | 0.677312 | 0.65253 |
| 0.766441 | 0.77322 | 0.775015 | 0.769788 | 0.773516 | 0.771906 | 0.77211 | 0.678159 | 0.647881 |

| | | | | | | | | |
|---|---|---|---|---|---|---|---|---|
| 0.767507 | 0.774571 | 0.772559 | 0.766896 | 0.770078 | 0.776847 | 0.775345 | 0.682136 | 0.654321 |
| 0.769122 | 0.769464 | 0.775605 | 0.763064 | 0.77082 | 0.776121 | 0.77068 | 0.679494 | 0.65625 |
| 0.768109 | 0.773081 | 0.77636 | 0.766437 | 0.769237 | 0.776624 | 0.776851 | 0.69045 | 0.668735 |
| 0.766557 | 0.76609 | 0.77212 | 0.768493 | 0.768729 | 0.772897 | 0.76892 | 0.679919 | 0.65719 |
| 0.76676 | 0.770605 | 0.777269 | 0.764855 | 0.768907 | 0.777325 | 0.76995 | 0.681215 | 0.64843 |
| 0.767766 | 0.766994 | 0.772821 | 0.764806 | 0.766048 | 0.769565 | 0.771207 | 0.677684 | 0.661627 |
| 0.777005 | 0.771199 | 0.77535 | 0.764622 | 0.773134 | 0.776658 | 0.770709 | 0.682119 | 0.659952 |
| 0.770331 | 0.773254 | 0.775565 | 0.763733 | 0.767818 | 0.770341 | 0.774886 | 0.687942 | 0.666989 |
| 0.764359 | 0.768761 | 0.765041 | 0.761238 | 0.764874 | 0.764983 | 0.770035 | 0.680206 | 0.653168 |
| 0.767937 | 0.769845 | 0.775541 | 0.763783 | 0.770375 | 0.774089 | 0.770885 | 0.68463 | 0.656205 |
| 0.764499 | 0.769939 | 0.773721 | 0.765892 | 0.766199 | 0.775183 | 0.771151 | 0.678096 | 0.655718 |
| 0.765827 | 0.766562 | 0.771008 | 0.764134 | 0.766988 | 0.772391 | 0.771359 | 0.676535 | 0.649156 |
| 0.761468 | 0.762761 | 0.766996 | 0.757568 | 0.759269 | 0.768537 | 0.765067 | 0.680493 | 0.657664 |
| 0.764062 | 0.769865 | 0.773461 | 0.760363 | 0.763883 | 0.770514 | 0.768587 | 0.674007 | 0.651456 |
| 0.763213 | 0.767219 | 0.764602 | 0.760301 | 0.766362 | 0.763595 | 0.7707 | 0.688089 | 0.657958 |
| 0.767648 | 0.774551 | 0.772727 | 0.768399 | 0.768399 | 0.778304 | 0.773272 | 0.690655 | 0.668422 |
| 0.764495 | 0.764644 | 0.767674 | 0.763675 | 0.765484 | 0.770013 | 0.766515 | 0.684324 | 0.657183 |
| 0.770177 | 0.771431 | 0.772041 | 0.768635 | 0.771885 | 0.777695 | 0.775603 | 0.685412 | 0.662607 |
| 0.769553 | 0.766396 | 0.773775 | 0.764345 | 0.768422 | 0.772335 | 0.770584 | 0.680428 | 0.662476 |
| 0.767484 | 0.771076 | 0.76925 | 0.765907 | 0.77102 | 0.773767 | 0.770961 | 0.683439 | 0.657876 |
| 0.763449 | 0.767355 | 0.76552 | 0.757289 | 0.762734 | 0.765249 | 0.765027 | 0.685852 | 0.656992 |
| 0.765979 | 0.768166 | 0.770392 | 0.765306 | 0.765617 | 0.771752 | 0.772251 | 0.689162 | 0.659266 |
| 0.768451 | 0.773008 | 0.77039 | 0.763811 | 0.769969 | 0.773809 | 0.773332 | 0.689009 | 0.662497 |
| 0.762182 | 0.773186 | 0.774513 | 0.767528 | 0.766796 | 0.771593 | 0.770765 | 0.682106 | 0.666424 |
| 0.765729 | 0.769034 | 0.766649 | 0.763121 | 0.767448 | 0.773795 | 0.768931 | 0.686215 | 0.67275 |
| 0.698611 | 0.707874 | 0.706015 | 0.698511 | 0.701698 | 0.705707 | 0.708122 | 0.692873 | 0.727792 |
| 0.76784 | 0.778077 | 0.775967 | 0.769489 | 0.76903 | 0.777805 | 0.776494 | 0.679709 | 0.652461 |
| 0.761648 | 0.774562 | 0.772526 | 0.768198 | 0.76605 | 0.773679 | 0.774042 | 0.679476 | 0.657976 |
| 0.771923 | 0.776945 | 0.775329 | 0.770136 | 0.771774 | 0.774961 | 0.777436 | 0.685867 | 0.653251 |
| 0.763623 | 0.777083 | 0.772416 | 0.76828 | 0.768247 | 0.773521 | 0.777245 | 0.678748 | 0.652994 |
| 0.765274 | 0.774532 | 0.769549 | 0.7617 | 0.767402 | 0.775669 | 0.767143 | 0.679165 | 0.651594 |
| 0.76574 | 0.773997 | 0.770715 | 0.763772 | 0.766805 | 0.774712 | 0.770089 | 0.682457 | 0.660124 |
| 0.761889 | 0.769519 | 0.767889 | 0.761315 | 0.766731 | 0.773079 | 0.769515 | 0.676164 | 0.653609 |
| 0.763162 | 0.773805 | 0.771224 | 0.76466 | 0.764073 | 0.768693 | 0.768316 | 0.680163 | 0.659518 |
| 0.755598 | 0.767823 | 0.764854 | 0.764614 | 0.756979 | 0.771391 | 0.768147 | 0.674643 | 0.652988 |
| 0.754412 | 0.766917 | 0.765772 | 0.762709 | 0.757474 | 0.765745 | 0.763041 | 0.676764 | 0.648951 |
| 0.766905 | 0.775699 | 0.775113 | 0.765523 | 0.76735 | 0.772933 | 0.770983 | 0.678879 | 0.668261 |
| 0.768285 | 0.769508 | 0.773119 | 0.765635 | 0.771248 | 0.774476 | 0.772769 | 0.682655 | 0.660166 |
| 0.768558 | 0.772464 | 0.774325 | 0.767395 | 0.770088 | 0.767876 | 0.771659 | 0.682316 | 0.659107 |
| 0.76823 | 0.768333 | 0.769832 | 0.766421 | 0.766116 | 0.771084 | 0.766202 | 0.692006 | 0.658566 |
| 0.765445 | 0.769553 | 0.769626 | 0.768511 | 0.769052 | 0.769899 | 0.770507 | 0.673634 | 0.660269 |
| 0.772338 | 0.773668 | 0.774083 | 0.770463 | 0.770146 | 0.779574 | 0.773882 | 0.681462 | 0.667558 |
| 0.759846 | 0.765762 | 0.768559 | 0.758462 | 0.760892 | 0.769191 | 0.770055 | 0.685045 | 0.66186 |
| 0.78882 | 0.788878 | 0.789717 | 0.788641 | 0.788301 | 0.790013 | 0.790482 | 0.678087 | 0.662509 |
| 0.790752 | 0.798804 | 0.791139 | 0.794765 | 0.788209 | 0.79669 | 0.797698 | 0.682152 | 0.663667 |
| 0.788402 | 0.788156 | 0.794665 | 0.794882 | 0.791687 | 0.793853 | 0.798018 | 0.684635 | 0.664666 |

| | | | | | | | | |
|---|---|---|---|---|---|---|---|---|
| 0.79534 | 0.786509 | 0.791333 | 0.797196 | 0.792796 | 0.79912 | 0.797619 | 0.680322 | 0.648924 |
| 0.798184 | 0.79784 | 0.794157 | 0.791815 | 0.79402 | 0.800897 | 0.801862 | 0.685902 | 0.662796 |
| 0.790328 | 0.790613 | 0.789808 | 0.796936 | 0.79025 | 0.796317 | 0.794236 | 0.683795 | 0.655132 |
| 0.74659 | 0.753498 | 0.7557 | 0.738534 | 0.747754 | 0.75662 | 0.748108 | 0.666482 | 0.655795 |
| 0.788621 | 0.785802 | 0.787286 | 0.783174 | 0.78715 | 0.795577 | 0.790246 | 0.672196 | 0.66018 |
| 0.783077 | 0.78114 | 0.790116 | 0.778475 | 0.788047 | 0.790916 | 0.786721 | 0.682429 | 0.664428 |
| 0.785756 | 0.786476 | 0.788222 | 0.7867 | 0.787869 | 0.791055 | 0.791429 | 0.685861 | 0.658566 |
| 0.773769 | 0.781549 | 0.783135 | 0.773272 | 0.779605 | 0.789692 | 0.785373 | 0.681369 | 0.649836 |
| 0.781389 | 0.785158 | 0.785638 | 0.781314 | 0.781068 | 0.788363 | 0.784341 | 0.683776 | 0.664833 |
| 0.779465 | 0.783866 | 0.784089 | 0.77841 | 0.778464 | 0.78683 | 0.783503 | 0.682476 | 0.658126 |
| 0.780621 | 0.781091 | 0.781492 | 0.773775 | 0.776687 | 0.785033 | 0.783645 | 0.674541 | 0.65564 |
| 0.776545 | 0.778954 | 0.78093 | 0.777217 | 0.780684 | 0.788136 | 0.78171 | 0.687581 | 0.658425 |
| 0.779863 | 0.784262 | 0.780329 | 0.772888 | 0.78058 | 0.784385 | 0.787899 | 0.683864 | 0.660766 |
| 0.839963 | 0.79855 | 0.837349 | 0.816422 | 0.82948 | 0.819471 | 0.80259 | 0.681202 | 0.659328 |
| 0.844651 | 0.834286 | 0.859918 | 0.805049 | 0.848593 | 0.816083 | 0.853242 | 0.683625 | 0.664614 |
| 1 | 0.799238 | 0.827811 | 0.812967 | 0.853062 | 0.818274 | 0.806829 | 0.682467 | 0.661183 |
| 0.799238 | 1 | 0.808631 | 0.796105 | 0.80504 | 0.795546 | 0.835656 | 0.682257 | 0.660231 |
| 0.827811 | 0.808631 | 1 | 0.805588 | 0.83939 | 0.809544 | 0.81544 | 0.684979 | 0.667141 |
| 0.812967 | 0.796105 | 0.805588 | 1 | 0.805277 | 0.809835 | 0.803649 | 0.684249 | 0.661106 |
| 0.853062 | 0.80504 | 0.83939 | 0.805277 | 1 | 0.813271 | 0.806581 | 0.683294 | 0.661363 |
| 0.818274 | 0.795546 | 0.809544 | 0.809835 | 0.813271 | 1 | 0.812814 | 0.688558 | 0.657658 |
| 0.806829 | 0.835656 | 0.81544 | 0.803649 | 0.806581 | 0.812814 | 1 | 0.689553 | 0.663736 |
| 0.682467 | 0.682257 | 0.684979 | 0.684249 | 0.683294 | 0.688558 | 0.689553 | 1 | 0.709931 |
| 0.661183 | 0.660231 | 0.667141 | 0.661106 | 0.661363 | 0.657658 | 0.663736 | 0.709931 | 1 |
| 0.674092 | 0.676244 | 0.670188 | 0.675594 | 0.673272 | 0.677043 | 0.675359 | 0.696667 | 0.708702 |
| 0.653298 | 0.653978 | 0.650708 | 0.65152 | 0.645979 | 0.651019 | 0.653808 | 0.695423 | 0.72522 |
| 0.739786 | 0.750414 | 0.745187 | 0.739176 | 0.738987 | 0.748181 | 0.745208 | 0.684025 | 0.662591 |
| 0.731881 | 0.730189 | 0.735645 | 0.725081 | 0.735943 | 0.733897 | 0.737819 | 0.685786 | 0.672332 |
| 0.731175 | 0.744565 | 0.736909 | 0.730394 | 0.734175 | 0.734491 | 0.737473 | 0.674357 | 0.660191 |
| 0.747158 | 0.747869 | 0.747961 | 0.745127 | 0.745713 | 0.745287 | 0.75972 | 0.680277 | 0.648849 |
| 0.733241 | 0.742554 | 0.733441 | 0.724494 | 0.730743 | 0.740861 | 0.735709 | 0.672219 | 0.660543 |
| 0.720659 | 0.73195 | 0.726989 | 0.716515 | 0.729054 | 0.718218 | 0.732605 | 0.671004 | 0.653796 |
| 0.722781 | 0.735303 | 0.730126 | 0.726593 | 0.72989 | 0.72214 | 0.734976 | 0.668213 | 0.652154 |
| 0.651345 | 0.657927 | 0.655313 | 0.656016 | 0.648601 | 0.64947 | 0.663254 | 0.687713 | 0.71135 |
| 0.765106 | 0.771659 | 0.772114 | 0.764205 | 0.767886 | 0.773895 | 0.768363 | 0.683884 | 0.66643 |
| 0.742052 | 0.757389 | 0.753792 | 0.742555 | 0.749971 | 0.75475 | 0.751839 | 0.678593 | 0.659596 |
| 0.738355 | 0.748175 | 0.746744 | 0.733008 | 0.747747 | 0.741244 | 0.74635 | 0.678395 | 0.663859 |
| 0.726884 | 0.737729 | 0.736576 | 0.728172 | 0.735694 | 0.738821 | 0.731207 | 0.683903 | 0.655924 |
| 0.750587 | 0.757679 | 0.750993 | 0.74918 | 0.750645 | 0.752374 | 0.754156 | 0.687462 | 0.662062 |
| 0.736136 | 0.747619 | 0.736815 | 0.7371 | 0.738515 | 0.744042 | 0.743702 | 0.685023 | 0.654484 |
| 0.732545 | 0.740764 | 0.738444 | 0.724984 | 0.735506 | 0.743812 | 0.743169 | 0.680923 | 0.657674 |
| 0.739329 | 0.750942 | 0.738144 | 0.737057 | 0.736619 | 0.742947 | 0.745771 | 0.688583 | 0.657372 |
| 0.734395 | 0.742485 | 0.750046 | 0.727207 | 0.736041 | 0.74 | 0.738773 | 0.682523 | 0.675949 |
| 0.729687 | 0.751344 | 0.745234 | 0.735137 | 0.73512 | 0.743396 | 0.748443 | 0.675862 | 0.653007 |
| 0.732903 | 0.738163 | 0.73885 | 0.735034 | 0.73885 | 0.740337 | 0.752806 | 0.689381 | 0.672485 |
| 0.721946 | 0.736194 | 0.718263 | 0.718677 | 0.724801 | 0.725242 | 0.735166 | 0.691645 | 0.660099 |

| | | | | | | | | |
|---|---|---|---|---|---|---|---|---|
| 0.756209 | 0.762907 | 0.766296 | 0.750273 | 0.755891 | 0.760655 | 0.758923 | 0.684072 | 0.665377 |
| 0.735718 | 0.743434 | 0.741876 | 0.73551 | 0.737191 | 0.744239 | 0.741665 | 0.674754 | 0.655828 |
| 0.731546 | 0.740602 | 0.742379 | 0.726619 | 0.736621 | 0.739483 | 0.744034 | 0.688234 | 0.657013 |
| 0.734691 | 0.747038 | 0.739877 | 0.741906 | 0.737335 | 0.734591 | 0.745477 | 0.680484 | 0.662311 |
| 0.72255 | 0.73232 | 0.740809 | 0.729945 | 0.730675 | 0.739765 | 0.736356 | 0.684994 | 0.676305 |
| 0.725935 | 0.735411 | 0.736038 | 0.724807 | 0.734679 | 0.740787 | 0.743635 | 0.685042 | 0.679662 |
| 0.726074 | 0.739152 | 0.73878 | 0.729412 | 0.727893 | 0.735807 | 0.732515 | 0.676355 | 0.654369 |
| 0.722285 | 0.730845 | 0.731874 | 0.721939 | 0.730051 | 0.737939 | 0.732744 | 0.692564 | 0.666089 |
| 0.719977 | 0.731326 | 0.737489 | 0.72771 | 0.731458 | 0.739565 | 0.734127 | 0.664766 | 0.655496 |
| 0.761184 | 0.762189 | 0.764862 | 0.760778 | 0.761198 | 0.768852 | 0.763047 | 0.685875 | 0.664562 |
| 0.754873 | 0.759068 | 0.760924 | 0.761639 | 0.757405 | 0.761765 | 0.760417 | 0.68559 | 0.664312 |
| 0.743399 | 0.756017 | 0.746334 | 0.749668 | 0.748502 | 0.755032 | 0.75425 | 0.685558 | 0.666833 |
| 0.742282 | 0.751758 | 0.748755 | 0.745771 | 0.746379 | 0.751004 | 0.747867 | 0.678968 | 0.664275 |
| 0.74767 | 0.748298 | 0.749573 | 0.748045 | 0.753025 | 0.754775 | 0.751751 | 0.688268 | 0.657596 |
| 0.74219 | 0.747696 | 0.749168 | 0.745392 | 0.745625 | 0.749414 | 0.749506 | 0.682145 | 0.659326 |

| t 20 | | plot 21 | | | | | | |
|---|---|---|---|---|---|---|---|---|
| Wild_plot_ | Wild_plot_ | Wild_plot_ | Wild_plot_ | Wild_plot_ | Wild_plot_ | Wild_plot_ | Wild_plot_ | Wild_plot_ |
| 0.742266 | 0.760727 | 0.663535 | 0.65636 | 0.661223 | 0.669754 | 0.648356 | 0.645135 | 0.637809 |
| 0.756092 | 0.734634 | 0.669143 | 0.66979 | 0.664452 | 0.684971 | 0.661477 | 0.649671 | 0.650115 |
| 0.748885 | 0.746498 | 0.659317 | 0.653938 | 0.654238 | 0.669747 | 0.643956 | 0.635259 | 0.64212 |
| 0.759883 | 0.742789 | 0.669358 | 0.670009 | 0.662843 | 0.680886 | 0.657604 | 0.647814 | 0.649343 |
| 0.755923 | 0.757373 | 0.659822 | 0.662635 | 0.662277 | 0.6736 | 0.646724 | 0.641422 | 0.64043 |
| 0.746673 | 0.726325 | 0.671864 | 0.666113 | 0.67036 | 0.677562 | 0.655346 | 0.64958 | 0.647875 |
| 0.672879 | 0.653827 | 0.733992 | 0.725791 | 0.71767 | 0.73988 | 0.711753 | 0.718043 | 0.714707 |
| 0.670334 | 0.647997 | 0.733263 | 0.721819 | 0.728553 | 0.734592 | 0.71751 | 0.7136 | 0.715205 |
| 0.755331 | 0.741913 | 0.670052 | 0.669015 | 0.660829 | 0.679032 | 0.655821 | 0.652233 | 0.644684 |
| 0.743304 | 0.77182 | 0.664399 | 0.665057 | 0.653398 | 0.681468 | 0.655494 | 0.647768 | 0.643444 |
| 0.751741 | 0.74471 | 0.665703 | 0.66537 | 0.665974 | 0.669954 | 0.658256 | 0.650946 | 0.643182 |
| 0.758119 | 0.756547 | 0.654839 | 0.653141 | 0.65465 | 0.66485 | 0.638639 | 0.632421 | 0.638214 |
| 0.75524 | 0.736609 | 0.671395 | 0.664189 | 0.66829 | 0.686803 | 0.661611 | 0.654641 | 0.649332 |
| 0.750728 | 0.744884 | 0.683236 | 0.678292 | 0.672208 | 0.681882 | 0.660607 | 0.664103 | 0.660679 |
| 0.741528 | 0.72553 | 0.693708 | 0.687 | 0.693942 | 0.700851 | 0.678253 | 0.675979 | 0.678706 |
| 0.764825 | 0.765954 | 0.670231 | 0.668026 | 0.665102 | 0.686358 | 0.659207 | 0.650082 | 0.656849 |
| 0.745243 | 0.748714 | 0.66982 | 0.667723 | 0.666959 | 0.676979 | 0.655196 | 0.648819 | 0.648997 |
| 0.729034 | 0.715211 | 0.705573 | 0.697635 | 0.697975 | 0.698231 | 0.683882 | 0.681532 | 0.678348 |
| 0.766735 | 0.735274 | 0.656232 | 0.65808 | 0.653292 | 0.669825 | 0.647215 | 0.632973 | 0.640011 |
| 0.758155 | 0.74351 | 0.66831 | 0.671279 | 0.66423 | 0.680243 | 0.648678 | 0.649089 | 0.650771 |
| 0.755735 | 0.741536 | 0.661614 | 0.661977 | 0.656017 | 0.667536 | 0.651569 | 0.639228 | 0.636236 |
| 0.756537 | 0.755176 | 0.670001 | 0.663648 | 0.658082 | 0.680984 | 0.65754 | 0.647811 | 0.65261 |
| 0.76578 | 0.749316 | 0.672682 | 0.671808 | 0.668634 | 0.680718 | 0.658035 | 0.645388 | 0.653451 |
| 0.765158 | 0.757158 | 0.668605 | 0.6603 | 0.661603 | 0.674848 | 0.641834 | 0.646738 | 0.643718 |
| 0.758108 | 0.73768 | 0.674236 | 0.67075 | 0.663174 | 0.679211 | 0.659537 | 0.647802 | 0.644647 |
| 0.761104 | 0.742057 | 0.664418 | 0.660081 | 0.657414 | 0.680648 | 0.654228 | 0.640389 | 0.645721 |
| 0.75134 | 0.727133 | 0.66044 | 0.660185 | 0.661327 | 0.67163 | 0.650376 | 0.640391 | 0.636003 |
| 0.760054 | 0.741156 | 0.66638 | 0.66193 | 0.657532 | 0.676065 | 0.651239 | 0.635827 | 0.63758 |
| 0.763371 | 0.740224 | 0.675923 | 0.668837 | 0.672381 | 0.679976 | 0.655051 | 0.652884 | 0.64798 |
| 0.751807 | 0.756794 | 0.667835 | 0.667659 | 0.668308 | 0.680456 | 0.647797 | 0.647991 | 0.646604 |
| 0.753597 | 0.739443 | 0.664691 | 0.67103 | 0.661153 | 0.673569 | 0.647952 | 0.641026 | 0.64818 |
| 0.759131 | 0.733696 | 0.662635 | 0.661036 | 0.655953 | 0.673363 | 0.648964 | 0.626298 | 0.638944 |
| 0.75663 | 0.734942 | 0.660924 | 0.661876 | 0.658617 | 0.673188 | 0.647136 | 0.641046 | 0.642336 |
| 0.749059 | 0.730554 | 0.663522 | 0.665133 | 0.657433 | 0.6796 | 0.64458 | 0.641835 | 0.644897 |
| 0.741503 | 0.731282 | 0.663669 | 0.656932 | 0.660745 | 0.67179 | 0.646838 | 0.635428 | 0.637916 |
| 0.760417 | 0.74208 | 0.663821 | 0.660037 | 0.664041 | 0.680782 | 0.652184 | 0.643806 | 0.639561 |
| 0.745446 | 0.732871 | 0.668926 | 0.667465 | 0.672585 | 0.678119 | 0.652499 | 0.647297 | 0.651865 |
| 0.757465 | 0.72592 | 0.671197 | 0.668035 | 0.664269 | 0.677723 | 0.650475 | 0.643289 | 0.653468 |
| 0.739516 | 0.732339 | 0.670423 | 0.671013 | 0.661867 | 0.678078 | 0.652775 | 0.64831 | 0.649093 |

| | | | | | | | | |
|---|---|---|---|---|---|---|---|---|
| 0.718285 | 0.712787 | 0.71175 | 0.699608 | 0.706973 | 0.70813 | 0.689033 | 0.688619 | 0.683832 |
| 0.749106 | 0.765315 | 0.659599 | 0.65211 | 0.66196 | 0.670076 | 0.639308 | 0.630691 | 0.63285 |
| 0.772129 | 0.765598 | 0.660317 | 0.655853 | 0.662026 | 0.671705 | 0.650232 | 0.638258 | 0.639228 |
| 0.760179 | 0.776316 | 0.67534 | 0.669309 | 0.670091 | 0.67541 | 0.667108 | 0.652529 | 0.647943 |
| 0.759118 | 0.747304 | 0.661707 | 0.661237 | 0.665279 | 0.674157 | 0.651685 | 0.641622 | 0.63796 |
| 0.759568 | 0.742416 | 0.663384 | 0.666729 | 0.662581 | 0.676715 | 0.650993 | 0.638453 | 0.64622 |
| 0.751738 | 0.740918 | 0.657534 | 0.659496 | 0.661299 | 0.671282 | 0.643731 | 0.637389 | 0.634222 |
| 0.753829 | 0.737446 | 0.662522 | 0.661964 | 0.663929 | 0.681198 | 0.660667 | 0.653668 | 0.649624 |
| 0.749101 | 0.741355 | 0.667251 | 0.664314 | 0.66353 | 0.68118 | 0.649402 | 0.643033 | 0.649288 |
| 0.750112 | 0.731546 | 0.671203 | 0.665868 | 0.670028 | 0.683109 | 0.656986 | 0.656011 | 0.647743 |
| 0.763624 | 0.780839 | 0.658636 | 0.652399 | 0.654776 | 0.666937 | 0.644612 | 0.634847 | 0.640104 |
| 0.759074 | 0.751293 | 0.656436 | 0.656459 | 0.653772 | 0.66998 | 0.644337 | 0.634279 | 0.636467 |
| 0.748933 | 0.767948 | 0.655187 | 0.656513 | 0.657328 | 0.677005 | 0.64032 | 0.63287 | 0.642426 |
| 0.755134 | 0.757723 | 0.656941 | 0.656899 | 0.655635 | 0.676125 | 0.650323 | 0.640851 | 0.63934 |
| 0.750396 | 0.756959 | 0.657853 | 0.654833 | 0.655046 | 0.670672 | 0.652603 | 0.636314 | 0.641217 |
| 0.747171 | 0.765304 | 0.661871 | 0.653471 | 0.659593 | 0.671292 | 0.64439 | 0.637653 | 0.638139 |
| 0.743192 | 0.760603 | 0.654842 | 0.649869 | 0.649694 | 0.666127 | 0.638501 | 0.624763 | 0.632513 |
| 0.762742 | 0.775201 | 0.665896 | 0.659041 | 0.659012 | 0.6739 | 0.656174 | 0.641979 | 0.64306 |
| 0.751344 | 0.76087 | 0.66287 | 0.662528 | 0.662385 | 0.674203 | 0.65483 | 0.640845 | 0.649631 |
| 0.760722 | 0.766829 | 0.655854 | 0.658405 | 0.657307 | 0.667201 | 0.648971 | 0.635501 | 0.631817 |
| 0.748083 | 0.770888 | 0.66079 | 0.651366 | 0.655106 | 0.671675 | 0.643 | 0.635767 | 0.632486 |
| 0.749269 | 0.767985 | 0.663665 | 0.653363 | 0.660456 | 0.665462 | 0.647862 | 0.635208 | 0.633644 |
| 0.76153 | 0.743301 | 0.665995 | 0.65431 | 0.658726 | 0.67482 | 0.642142 | 0.642395 | 0.639201 |
| 0.744857 | 0.727319 | 0.675632 | 0.669941 | 0.671312 | 0.689366 | 0.658365 | 0.654098 | 0.657012 |
| 0.738301 | 0.716675 | 0.684614 | 0.676247 | 0.675182 | 0.695095 | 0.672456 | 0.665027 | 0.655309 |
| 0.669913 | 0.650484 | 0.734601 | 0.721594 | 0.729019 | 0.735539 | 0.7168 | 0.714326 | 0.718137 |
| 0.67588 | 0.660703 | 0.734213 | 0.728733 | 0.735236 | 0.736897 | 0.714123 | 0.716615 | 0.715416 |
| 0.684833 | 0.65491 | 0.73685 | 0.731314 | 0.723864 | 0.73852 | 0.717541 | 0.708829 | 0.708198 |
| 0.750339 | 0.719772 | 0.665032 | 0.66463 | 0.662574 | 0.676465 | 0.648286 | 0.642993 | 0.640497 |
| 0.757169 | 0.731078 | 0.6622 | 0.6555 | 0.657188 | 0.672846 | 0.650524 | 0.643618 | 0.641048 |
| 0.749777 | 0.747599 | 0.670971 | 0.666667 | 0.671252 | 0.684807 | 0.66049 | 0.653428 | 0.654023 |
| 0.755097 | 0.731567 | 0.674395 | 0.670495 | 0.666435 | 0.683074 | 0.661009 | 0.655823 | 0.653454 |
| 0.762966 | 0.746034 | 0.667832 | 0.662069 | 0.661762 | 0.67378 | 0.654332 | 0.649329 | 0.637138 |
| 0.786685 | 0.747506 | 0.663182 | 0.656129 | 0.662945 | 0.670423 | 0.65072 | 0.647187 | 0.632174 |
| 0.764198 | 0.749225 | 0.665942 | 0.666729 | 0.661132 | 0.67538 | 0.659858 | 0.638791 | 0.641185 |
| 0.756087 | 0.744612 | 0.662755 | 0.660923 | 0.657817 | 0.675396 | 0.648034 | 0.644105 | 0.641033 |
| 0.758343 | 0.741523 | 0.669551 | 0.670189 | 0.662892 | 0.678888 | 0.655907 | 0.641888 | 0.638143 |
| 0.780265 | 0.805546 | 0.681487 | 0.685882 | 0.683451 | 0.700329 | 0.684891 | 0.665469 | 0.664439 |
| 0.745214 | 0.731236 | 0.659962 | 0.663801 | 0.652773 | 0.666463 | 0.653564 | 0.644621 | 0.63706 |
| 0.768608 | 0.737889 | 0.660202 | 0.657803 | 0.665568 | 0.671291 | 0.650558 | 0.648554 | 0.646015 |
| 0.756201 | 0.835208 | 0.661332 | 0.65379 | 0.654125 | 0.668202 | 0.645669 | 0.635996 | 0.639298 |
| 0.783329 | 0.805052 | 0.678884 | 0.681329 | 0.681514 | 0.695456 | 0.677055 | 0.666481 | 0.661346 |
| 0.764226 | 0.785166 | 0.669507 | 0.670268 | 0.661859 | 0.676387 | 0.652318 | 0.643167 | 0.647344 |
| 0.789007 | 0.776526 | 0.661249 | 0.654552 | 0.656266 | 0.669214 | 0.647188 | 0.634527 | 0.645061 |
| 0.756763 | 0.743647 | 0.668749 | 0.665369 | 0.662552 | 0.680572 | 0.65 | 0.6469 | 0.645426 |
| 0.753666 | 0.734227 | 0.664355 | 0.662297 | 0.657271 | 0.681559 | 0.651495 | 0.642954 | 0.646421 |

| | | | | | | | | |
|---|---|---|---|---|---|---|---|---|
| 0.756318 | 0.850693 | 0.666619 | 0.659125 | 0.659615 | 0.670372 | 0.652935 | 0.638005 | 0.645537 |
| 0.757996 | 0.738401 | 0.666427 | 0.660291 | 0.665854 | 0.671747 | 0.65735 | 0.643535 | 0.636109 |
| 0.753604 | 0.758045 | 0.670322 | 0.657909 | 0.659079 | 0.678478 | 0.653557 | 0.64884 | 0.646259 |
| 0.777396 | 0.790694 | 0.677971 | 0.676745 | 0.682442 | 0.695019 | 0.682679 | 0.663251 | 0.657028 |
| 0.760494 | 0.756181 | 0.672006 | 0.667713 | 0.667535 | 0.682671 | 0.662786 | 0.651404 | 0.648725 |
| 0.762896 | 0.744018 | 0.667875 | 0.661281 | 0.663636 | 0.673147 | 0.652088 | 0.639858 | 0.648308 |
| 0.762205 | 0.74093 | 0.669356 | 0.669813 | 0.664646 | 0.683471 | 0.66209 | 0.651585 | 0.645189 |
| 0.76147 | 0.738787 | 0.670753 | 0.661891 | 0.672414 | 0.683428 | 0.662564 | 0.645969 | 0.650243 |
| 0.756026 | 0.740499 | 0.664299 | 0.665482 | 0.651759 | 0.67226 | 0.652145 | 0.633117 | 0.649928 |
| 0.766742 | 0.739693 | 0.663399 | 0.663311 | 0.657372 | 0.67567 | 0.652953 | 0.638292 | 0.64 |
| 0.760412 | 0.742061 | 0.66123 | 0.668689 | 0.662707 | 0.677542 | 0.656736 | 0.64485 | 0.651669 |
| 0.781739 | 0.738788 | 0.666618 | 0.663324 | 0.66064 | 0.675444 | 0.654 | 0.643368 | 0.636932 |
| 0.756787 | 0.745546 | 0.665309 | 0.665418 | 0.66415 | 0.675501 | 0.649666 | 0.64171 | 0.646007 |
| 0.779743 | 0.739361 | 0.659682 | 0.663005 | 0.662531 | 0.676644 | 0.648763 | 0.643013 | 0.637941 |
| 0.758824 | 0.752632 | 0.666909 | 0.662715 | 0.659549 | 0.679161 | 0.654117 | 0.646305 | 0.647259 |
| 0.748936 | 0.741803 | 0.659019 | 0.660155 | 0.660597 | 0.669263 | 0.647598 | 0.639024 | 0.64375 |
| 0.753646 | 0.741958 | 0.666618 | 0.663931 | 0.659962 | 0.670043 | 0.653265 | 0.645223 | 0.644355 |
| 0.757242 | 0.736288 | 0.664937 | 0.657316 | 0.65884 | 0.67408 | 0.651619 | 0.644637 | 0.646623 |
| 0.7808 | 0.73502 | 0.670325 | 0.662362 | 0.668583 | 0.68612 | 0.659593 | 0.651581 | 0.65037 |
| 0.755006 | 0.736292 | 0.669474 | 0.663803 | 0.663226 | 0.68308 | 0.657684 | 0.642469 | 0.651263 |
| 0.758753 | 0.738805 | 0.669132 | 0.67026 | 0.658718 | 0.678968 | 0.651819 | 0.652316 | 0.644915 |
| 0.759015 | 0.735543 | 0.660753 | 0.658214 | 0.65376 | 0.673809 | 0.648725 | 0.640675 | 0.63672 |
| 0.754518 | 0.747889 | 0.666137 | 0.660447 | 0.668757 | 0.677491 | 0.656534 | 0.646883 | 0.650825 |
| 0.764851 | 0.746113 | 0.666185 | 0.66138 | 0.660446 | 0.670861 | 0.649072 | 0.642644 | 0.642028 |
| 0.749265 | 0.732119 | 0.672303 | 0.662797 | 0.668244 | 0.685499 | 0.661333 | 0.645504 | 0.639943 |
| 0.760582 | 0.731716 | 0.669824 | 0.667158 | 0.665747 | 0.685967 | 0.661173 | 0.646348 | 0.641632 |
| 0.754797 | 0.74591 | 0.667528 | 0.664404 | 0.668985 | 0.679101 | 0.666373 | 0.643145 | 0.636466 |
| 0.751339 | 0.727472 | 0.664398 | 0.666543 | 0.665159 | 0.682617 | 0.652499 | 0.647876 | 0.651083 |
| 0.75156 | 0.735955 | 0.669603 | 0.657798 | 0.664187 | 0.674522 | 0.652949 | 0.637416 | 0.651163 |
| 0.751817 | 0.744484 | 0.671402 | 0.667596 | 0.668589 | 0.684264 | 0.663938 | 0.654843 | 0.64539 |
| 0.763027 | 0.736555 | 0.668399 | 0.668695 | 0.663939 | 0.686239 | 0.65837 | 0.645022 | 0.645734 |
| 0.751839 | 0.744182 | 0.666905 | 0.659909 | 0.65439 | 0.678336 | 0.65245 | 0.646649 | 0.649167 |
| 0.758204 | 0.746182 | 0.670556 | 0.664564 | 0.668941 | 0.686974 | 0.658233 | 0.655675 | 0.646638 |
| 0.758769 | 0.778048 | 0.675026 | 0.669266 | 0.668825 | 0.675228 | 0.669715 | 0.654755 | 0.648417 |
| 0.76009 | 0.743485 | 0.665942 | 0.668405 | 0.659862 | 0.676204 | 0.651168 | 0.645126 | 0.651839 |
| 0.754547 | 0.74532 | 0.66739 | 0.665677 | 0.655633 | 0.678687 | 0.653838 | 0.648832 | 0.655457 |
| 0.770869 | 0.735497 | 0.658141 | 0.651148 | 0.65998 | 0.678263 | 0.645802 | 0.641832 | 0.64224 |
| 0.736921 | 0.732327 | 0.658807 | 0.654894 | 0.668207 | 0.667546 | 0.655268 | 0.644733 | 0.648889 |
| 0.752988 | 0.744735 | 0.665417 | 0.663116 | 0.66468 | 0.669598 | 0.658968 | 0.644274 | 0.645759 |
| 0.747692 | 0.735273 | 0.660826 | 0.661017 | 0.65431 | 0.672529 | 0.646165 | 0.645431 | 0.639737 |
| 0.754377 | 0.73649 | 0.666556 | 0.659261 | 0.653425 | 0.670035 | 0.6543 | 0.647253 | 0.642614 |
| 0.77371 | 0.728002 | 0.651147 | 0.646984 | 0.652434 | 0.655205 | 0.639871 | 0.628201 | 0.62313 |
| 0.758776 | 0.786771 | 0.661063 | 0.654279 | 0.657603 | 0.663391 | 0.655849 | 0.641469 | 0.64443 |
| 0.742912 | 0.730951 | 0.654395 | 0.660941 | 0.657679 | 0.663073 | 0.654919 | 0.63926 | 0.652202 |
| 0.751522 | 0.738578 | 0.662966 | 0.659439 | 0.652749 | 0.671271 | 0.653685 | 0.650016 | 0.640385 |
| 0.747786 | 0.747204 | 0.663366 | 0.668475 | 0.662121 | 0.664235 | 0.651856 | 0.652456 | 0.650753 |
| 0.746 | 0.744908 | 0.661259 | 0.668037 | 0.660711 | 0.667687 | 0.674015 | 0.651797 | 0.648448 |

| | | | | | | | | |
|---|---|---|---|---|---|---|---|---|
| 0.73377 | 0.734839 | 0.660305 | 0.654168 | 0.658132 | 0.655138 | 0.652141 | 0.63951 | 0.635105 |
| 0.748685 | 0.738 | 0.666114 | 0.666365 | 0.66312 | 0.666753 | 0.660222 | 0.642338 | 0.646245 |
| 0.742815 | 0.727983 | 0.664254 | 0.671227 | 0.652944 | 0.668138 | 0.65672 | 0.651965 | 0.648185 |
| 0.736308 | 0.988932 | 0.655141 | 0.653048 | 0.650898 | 0.652741 | 0.64905 | 0.633864 | 0.643093 |
| 0.763547 | 0.73778 | 0.657419 | 0.666593 | 0.660507 | 0.669483 | 0.655674 | 0.646231 | 0.637859 |
| 0.738197 | 0.741784 | 0.659053 | 0.654699 | 0.656469 | 0.660096 | 0.653315 | 0.642116 | 0.636274 |
| 0.748737 | 0.75613 | 0.659661 | 0.660224 | 0.662326 | 0.667783 | 0.655778 | 0.656427 | 0.64467 |
| 0.77663 | 0.735376 | 0.656691 | 0.655592 | 0.647881 | 0.655711 | 0.648575 | 0.637345 | 0.644768 |
| 0.738931 | 0.718227 | 0.657483 | 0.655197 | 0.660386 | 0.659983 | 0.641758 | 0.643359 | 0.644948 |
| 0.753162 | 0.733762 | 0.65425 | 0.664904 | 0.650142 | 0.666265 | 0.658451 | 0.646127 | 0.632719 |
| 0.737175 | 0.741247 | 0.659719 | 0.653529 | 0.646731 | 0.65375 | 0.651678 | 0.645232 | 0.641553 |
| 0.744279 | 0.78288 | 0.656165 | 0.650454 | 0.652256 | 0.66899 | 0.653459 | 0.633912 | 0.646992 |
| 0.753067 | 0.737209 | 0.665419 | 0.661943 | 0.660736 | 0.673482 | 0.649606 | 0.640519 | 0.648656 |
| 0.743685 | 0.732106 | 0.669177 | 0.679163 | 0.671772 | 0.679294 | 0.656379 | 0.657957 | 0.665235 |
| 0.757555 | 0.747081 | 0.670881 | 0.667285 | 0.672562 | 0.688041 | 0.666001 | 0.658033 | 0.652374 |
| 0.774725 | 0.722845 | 0.665079 | 0.660566 | 0.66064 | 0.676139 | 0.655952 | 0.645497 | 0.649052 |
| 0.748557 | 0.747657 | 0.6719 | 0.661716 | 0.665346 | 0.685997 | 0.646892 | 0.655961 | 0.652581 |
| 0.751568 | 0.727195 | 0.676954 | 0.667533 | 0.670157 | 0.689774 | 0.666368 | 0.654162 | 0.657274 |
| 0.73266 | 0.711724 | 0.655021 | 0.639812 | 0.651463 | 0.66627 | 0.641591 | 0.638326 | 0.63583 |
| 0.758897 | 0.725572 | 0.674282 | 0.67033 | 0.682102 | 0.688725 | 0.666012 | 0.653165 | 0.6566 |
| 0.737127 | 0.737679 | 0.672705 | 0.67425 | 0.670461 | 0.691705 | 0.668603 | 0.667677 | 0.662128 |
| 0.744045 | 0.736446 | 0.674527 | 0.66741 | 0.666901 | 0.684317 | 0.657455 | 0.652198 | 0.649886 |
| 0.743521 | 0.729707 | 0.673796 | 0.665989 | 0.663127 | 0.683046 | 0.666593 | 0.64873 | 0.656963 |
| 0.730626 | 0.729584 | 0.665833 | 0.667291 | 0.664093 | 0.679451 | 0.655762 | 0.655546 | 0.64921 |
| 0.729577 | 0.715641 | 0.651947 | 0.639027 | 0.646905 | 0.663943 | 0.640407 | 0.626047 | 0.626158 |
| 0.756388 | 0.733917 | 0.684364 | 0.687193 | 0.675183 | 0.691942 | 0.674614 | 0.668137 | 0.672385 |
| 0.729656 | 0.721059 | 0.667454 | 0.661905 | 0.65928 | 0.678365 | 0.649234 | 0.644686 | 0.641618 |
| 0.742502 | 0.730161 | 0.668484 | 0.673864 | 0.666204 | 0.687538 | 0.660698 | 0.646788 | 0.652161 |
| 0.728606 | 0.708272 | 0.640724 | 0.640519 | 0.638502 | 0.653892 | 0.629803 | 0.623692 | 0.622802 |
| 0.741693 | 0.722972 | 0.68649 | 0.674291 | 0.681024 | 0.699372 | 0.675099 | 0.669487 | 0.649059 |
| 0.744699 | 0.725998 | 0.683756 | 0.673387 | 0.682553 | 0.697347 | 0.671435 | 0.664855 | 0.653187 |
| 0.73983 | 0.741473 | 0.658302 | 0.652918 | 0.656837 | 0.66976 | 0.650498 | 0.646866 | 0.633408 |
| 0.74182 | 0.746133 | 0.651163 | 0.649306 | 0.655743 | 0.669952 | 0.646296 | 0.639063 | 0.630006 |
| 0.741179 | 0.719322 | 0.681365 | 0.683405 | 0.676768 | 0.703673 | 0.667996 | 0.660675 | 0.668003 |
| 0.736405 | 0.722176 | 0.679725 | 0.678951 | 0.680532 | 0.690481 | 0.669298 | 0.659207 | 0.662048 |
| 0.746039 | 0.726905 | 0.68914 | 0.682292 | 0.675095 | 0.696723 | 0.674651 | 0.659245 | 0.668768 |
| 0.700324 | 0.683657 | 0.638405 | 0.633911 | 0.635448 | 0.643361 | 0.620901 | 0.623085 | 0.620093 |
| 0.729665 | 0.710999 | 0.678206 | 0.675671 | 0.67568 | 0.67845 | 0.668584 | 0.653731 | 0.649662 |
| 0.710441 | 0.68234 | 0.640284 | 0.627673 | 0.632759 | 0.638926 | 0.62093 | 0.610532 | 0.615301 |
| 0.696577 | 0.670228 | 0.628819 | 0.61486 | 0.615258 | 0.634841 | 0.607955 | 0.60232 | 0.603845 |
| 0.757895 | 0.730492 | 0.6809 | 0.67481 | 0.671039 | 0.685155 | 0.668652 | 0.652632 | 0.659387 |
| 0.704897 | 0.671168 | 0.631867 | 0.61913 | 0.621106 | 0.636173 | 0.619347 | 0.60082 | 0.607075 |
| 0.69854 | 0.66879 | 0.614489 | 0.616615 | 0.607166 | 0.6283 | 0.61295 | 0.590018 | 0.596727 |
| 0.688099 | 0.656225 | 0.613995 | 0.613155 | 0.606496 | 0.62987 | 0.605741 | 0.588728 | 0.590291 |
| 0.737217 | 0.754762 | 0.691873 | 0.686278 | 0.683273 | 0.698561 | 0.681788 | 0.667933 | 0.670094 |
| 0.688287 | 0.650801 | 0.614822 | 0.608801 | 0.605796 | 0.625544 | 0.6065 | 0.586129 | 0.59194 |
| 0.741184 | 0.735646 | 0.687002 | 0.682329 | 0.678769 | 0.695688 | 0.675454 | 0.663152 | 0.662316 |

| | | | | | | | | |
|---|---|---|---|---|---|---|---|---|
| 0.734802 | 0.736142 | 0.696749 | 0.683099 | 0.683553 | 0.6935 | 0.67085 | 0.664677 | 0.674706 |
| 0.747571 | 0.737328 | 0.682514 | 0.677592 | 0.677855 | 0.690548 | 0.672936 | 0.655331 | 0.656901 |
| 0.726309 | 0.717565 | 0.707363 | 0.703517 | 0.700331 | 0.705751 | 0.703312 | 0.689523 | 0.695942 |
| 0.680513 | 0.643742 | 0.600505 | 0.599283 | 0.591681 | 0.615442 | 0.591549 | 0.575012 | 0.579518 |
| 0.682644 | 0.650745 | 0.615608 | 0.613604 | 0.612292 | 0.624951 | 0.604307 | 0.594792 | 0.592512 |
| 0.751287 | 0.741979 | 0.692386 | 0.686319 | 0.684929 | 0.694929 | 0.680988 | 0.670219 | 0.678758 |
| 0.671555 | 0.642071 | 0.617021 | 0.604894 | 0.606742 | 0.623496 | 0.600369 | 0.590486 | 0.589939 |
| 0.705402 | 0.683481 | 0.69645 | 0.686699 | 0.692676 | 0.701379 | 0.678395 | 0.674819 | 0.682466 |
| 0.739305 | 0.731844 | 0.694226 | 0.686882 | 0.686199 | 0.704587 | 0.678041 | 0.671658 | 0.679919 |
| 0.686626 | 0.676825 | 0.701804 | 0.687618 | 0.690896 | 0.696421 | 0.679233 | 0.679299 | 0.683667 |
| 0.713969 | 0.700737 | 0.692376 | 0.688172 | 0.688893 | 0.69575 | 0.676702 | 0.673769 | 0.671689 |
| 0.660194 | 0.623288 | 0.832 | 0.759398 | 0.80916 | 0.752212 | 0.727273 | 0.784314 | 0.801887 |
| 0.644409 | 0.657068 | 0.753741 | 0.76579 | 0.74348 | 0.745474 | 0.754605 | 0.742996 | 0.753978 |
| 0.639572 | 0.639426 | 0.74625 | 0.742474 | 0.740591 | 0.740387 | 0.731714 | 0.737102 | 0.73185 |
| 0.649431 | 0.643737 | 0.759149 | 0.752656 | 0.744496 | 0.753254 | 0.740644 | 0.729877 | 0.732579 |
| 0.640538 | 0.647736 | 0.766673 | 0.751959 | 0.746174 | 0.758029 | 0.752418 | 0.757375 | 0.750256 |
| 0.61279 | 0.586544 | 0.575377 | 0.576102 | 0.564778 | 0.576871 | 0.560582 | 0.554299 | 0.562128 |
| 0.645215 | 0.64495 | 0.746635 | 0.747037 | 0.742623 | 0.751501 | 0.754489 | 0.739695 | 0.746488 |
| 0.711958 | 0.694932 | 0.700797 | 0.693939 | 0.699129 | 0.698329 | 0.683206 | 0.674599 | 0.677065 |
| 0.709171 | 0.70027 | 0.702429 | 0.693901 | 0.693186 | 0.697496 | 0.682788 | 0.67713 | 0.679543 |
| 0.720294 | 0.693405 | 0.702438 | 0.691769 | 0.696539 | 0.698931 | 0.693822 | 0.676267 | 0.681871 |
| 0.593737 | 0.555067 | 0.545192 | 0.539677 | 0.527651 | 0.550903 | 0.52881 | 0.515981 | 0.513616 |
| 0.57621 | 0.533512 | 0.527217 | 0.523818 | 0.514718 | 0.534335 | 0.513836 | 0.498062 | 0.505362 |
| 0.670445 | 0.651803 | 0.729148 | 0.723101 | 0.72145 | 0.722534 | 0.705909 | 0.709204 | 0.705675 |
| 0.672794 | 0.649902 | 0.727956 | 0.720904 | 0.711377 | 0.729461 | 0.712652 | 0.709122 | 0.699532 |
| 0.657691 | 0.644849 | 0.72122 | 0.707536 | 0.712307 | 0.723227 | 0.702746 | 0.70697 | 0.697496 |
| 0.659329 | 0.64772 | 0.727939 | 0.722594 | 0.723649 | 0.727817 | 0.713733 | 0.704316 | 0.711118 |
| 0.663952 | 0.651485 | 0.72855 | 0.72117 | 0.725662 | 0.731587 | 0.717222 | 0.714407 | 0.70583 |
| 0.674206 | 0.64753 | 0.730451 | 0.725415 | 0.730296 | 0.73738 | 0.720276 | 0.71762 | 0.720379 |
| 0.674688 | 0.650446 | 0.72971 | 0.72539 | 0.725901 | 0.73669 | 0.719936 | 0.716653 | 0.716089 |
| 0.673165 | 0.663085 | 0.724804 | 0.720125 | 0.729319 | 0.737572 | 0.719188 | 0.70523 | 0.71791 |
| 0.678649 | 0.661791 | 0.738501 | 0.728862 | 0.722462 | 0.73765 | 0.722718 | 0.710898 | 0.720627 |
| 0.675904 | 0.654161 | 0.724869 | 0.718135 | 0.717531 | 0.728438 | 0.706162 | 0.704309 | 0.715296 |
| 0.672871 | 0.656423 | 0.72019 | 0.7119 | 0.720601 | 0.721993 | 0.703562 | 0.704941 | 0.692441 |
| 0.67675 | 0.660817 | 0.735014 | 0.722137 | 0.728094 | 0.742677 | 0.717849 | 0.714245 | 0.713823 |
| 0.675906 | 0.651718 | 0.724375 | 0.720721 | 0.72026 | 0.729188 | 0.70277 | 0.70048 | 0.709535 |
| 0.674837 | 0.657925 | 0.723537 | 0.726577 | 0.718542 | 0.732959 | 0.711091 | 0.703631 | 0.715543 |
| 0.681393 | 0.661859 | 0.728102 | 0.725014 | 0.724659 | 0.734238 | 0.714416 | 0.719238 | 0.709123 |
| 0.673804 | 0.658565 | 0.722606 | 0.714478 | 0.715646 | 0.727924 | 0.705733 | 0.697243 | 0.708482 |
| 0.676969 | 0.656863 | 0.716066 | 0.718934 | 0.717578 | 0.729562 | 0.708296 | 0.692997 | 0.704366 |
| 0.638481 | 0.649311 | 0.743357 | 0.733819 | 0.75 | 0.720381 | 0.724883 | 0.724509 | 0.740295 |
| 0.652943 | 0.657701 | 0.746089 | 0.740267 | 0.742679 | 0.747297 | 0.730348 | 0.727031 | 0.720976 |
| 0.681378 | 0.663134 | 0.724342 | 0.720535 | 0.724156 | 0.741086 | 0.717431 | 0.706425 | 0.705061 |
| 0.68492 | 0.663002 | 0.736724 | 0.731034 | 0.733803 | 0.745155 | 0.723995 | 0.721274 | 0.719407 |
| 0.644485 | 0.637406 | 0.71571 | 0.717298 | 0.716761 | 0.704221 | 0.694642 | 0.701248 | 0.702534 |
| 0.673509 | 0.661089 | 0.731126 | 0.723664 | 0.730922 | 0.737739 | 0.717202 | 0.709459 | 0.706967 |
| 0.665451 | 0.648968 | 0.718099 | 0.720653 | 0.723473 | 0.728333 | 0.708485 | 0.707684 | 0.699649 |

| | | | | | | | | |
|---|---|---|---|---|---|---|---|---|
| 0.66997 | 0.654655 | 0.728476 | 0.719232 | 0.726114 | 0.733472 | 0.706507 | 0.700893 | 0.706208 |
| 0.676244 | 0.654704 | 0.729375 | 0.721283 | 0.73179 | 0.730928 | 0.711352 | 0.707378 | 0.71299 |
| 0.678329 | 0.659143 | 0.722081 | 0.72511 | 0.729099 | 0.738265 | 0.715499 | 0.705549 | 0.710464 |
| 0.669953 | 0.657566 | 0.728115 | 0.720008 | 0.730424 | 0.731612 | 0.714026 | 0.707042 | 0.710434 |
| 0.686996 | 0.665303 | 0.734553 | 0.733627 | 0.735696 | 0.744792 | 0.719234 | 0.713122 | 0.708986 |
| 0.668677 | 0.659833 | 0.729895 | 0.727533 | 0.72509 | 0.735221 | 0.712563 | 0.704016 | 0.71091 |
| 0.676123 | 0.65789 | 0.734501 | 0.729533 | 0.723503 | 0.730244 | 0.710865 | 0.701931 | 0.705539 |
| 0.667392 | 0.652261 | 0.718769 | 0.71459 | 0.707739 | 0.719265 | 0.709947 | 0.708189 | 0.696296 |
| 0.664315 | 0.655291 | 0.7242 | 0.717031 | 0.717935 | 0.727726 | 0.706576 | 0.699944 | 0.704472 |
| 0.668821 | 0.652988 | 0.715356 | 0.713023 | 0.719279 | 0.722464 | 0.710448 | 0.695946 | 0.695869 |
| 0.670114 | 0.655442 | 0.727777 | 0.719232 | 0.727534 | 0.725713 | 0.702727 | 0.699663 | 0.705019 |
| 0.664474 | 0.654875 | 0.727687 | 0.722868 | 0.721121 | 0.723898 | 0.713888 | 0.699829 | 0.703979 |
| 0.670437 | 0.649951 | 0.717752 | 0.713075 | 0.716121 | 0.719841 | 0.708534 | 0.696789 | 0.701387 |
| 0.666519 | 0.653048 | 0.717507 | 0.716287 | 0.713557 | 0.731509 | 0.708012 | 0.705525 | 0.707288 |
| 0.678794 | 0.660075 | 0.725394 | 0.718066 | 0.724853 | 0.733597 | 0.708832 | 0.713762 | 0.703487 |
| 0.665314 | 0.653138 | 0.731247 | 0.720855 | 0.729507 | 0.730524 | 0.72031 | 0.715896 | 0.725082 |
| 0.664262 | 0.652639 | 0.736195 | 0.725494 | 0.727686 | 0.736509 | 0.714121 | 0.709004 | 0.724148 |
| 0.660856 | 0.649802 | 0.721795 | 0.719522 | 0.717297 | 0.728109 | 0.70607 | 0.696388 | 0.705447 |
| 0.662945 | 0.649134 | 0.725191 | 0.717387 | 0.728721 | 0.736842 | 0.709008 | 0.706347 | 0.705015 |
| 0.657985 | 0.651889 | 0.731238 | 0.718882 | 0.731265 | 0.727818 | 0.713926 | 0.70271 | 0.704787 |
| 0.657564 | 0.646815 | 0.71368 | 0.713814 | 0.713712 | 0.72005 | 0.705692 | 0.692068 | 0.699376 |
| 0.66565 | 0.652462 | 0.719632 | 0.715085 | 0.725455 | 0.725753 | 0.711445 | 0.704134 | 0.700442 |
| 0.666024 | 0.65524 | 0.72757 | 0.72641 | 0.733129 | 0.73595 | 0.71332 | 0.713246 | 0.704952 |
| 0.670043 | 0.65097 | 0.728882 | 0.726783 | 0.727387 | 0.733695 | 0.711517 | 0.704443 | 0.710192 |
| 0.66537 | 0.652495 | 0.731368 | 0.71617 | 0.721689 | 0.725993 | 0.708333 | 0.698166 | 0.704378 |
| 0.65914 | 0.646557 | 0.726945 | 0.712344 | 0.726465 | 0.730858 | 0.702308 | 0.705483 | 0.707296 |
| 0.654518 | 0.648658 | 0.717247 | 0.716198 | 0.716388 | 0.722803 | 0.704404 | 0.697357 | 0.712085 |
| 0.669211 | 0.643564 | 0.71435 | 0.71346 | 0.717879 | 0.728895 | 0.707339 | 0.700876 | 0.702222 |
| 0.660349 | 0.645884 | 0.724651 | 0.718816 | 0.713542 | 0.730761 | 0.70564 | 0.702307 | 0.705079 |
| 0.664166 | 0.653701 | 0.726809 | 0.718469 | 0.721653 | 0.734151 | 0.711182 | 0.699441 | 0.710822 |
| 0.666882 | 0.652687 | 0.729114 | 0.721346 | 0.722952 | 0.735068 | 0.713501 | 0.706911 | 0.712722 |
| 0.661548 | 0.66006 | 0.731633 | 0.725029 | 0.723486 | 0.732427 | 0.708256 | 0.703863 | 0.720881 |
| 0.681025 | 0.657171 | 0.731675 | 0.725434 | 0.725134 | 0.735203 | 0.730108 | 0.711669 | 0.711006 |
| 0.674378 | 0.656327 | 0.735498 | 0.734148 | 0.734627 | 0.745366 | 0.7213 | 0.719213 | 0.720707 |
| 0.663147 | 0.647987 | 0.72541 | 0.714972 | 0.712129 | 0.727634 | 0.703254 | 0.696398 | 0.700824 |
| 0.659218 | 0.650575 | 0.733364 | 0.720177 | 0.729951 | 0.732914 | 0.719991 | 0.706754 | 0.710847 |
| 0.667379 | 0.655121 | 0.724655 | 0.718888 | 0.724206 | 0.731813 | 0.701798 | 0.708661 | 0.714793 |
| 0.66394 | 0.649223 | 0.722928 | 0.717753 | 0.720129 | 0.732667 | 0.708087 | 0.716879 | 0.704899 |
| 0.655531 | 0.644026 | 0.711895 | 0.708986 | 0.716236 | 0.72264 | 0.701613 | 0.692505 | 0.696566 |
| 0.663243 | 0.647243 | 0.724535 | 0.720602 | 0.723362 | 0.728821 | 0.698974 | 0.709142 | 0.711969 |
| 0.647949 | 0.644374 | 0.711521 | 0.699825 | 0.709748 | 0.723757 | 0.696503 | 0.6849 | 0.695158 |
| 0.660942 | 0.648033 | 0.718346 | 0.71121 | 0.724037 | 0.723152 | 0.708497 | 0.703067 | 0.708031 |
| 0.65172 | 0.647864 | 0.708532 | 0.70771 | 0.718067 | 0.72227 | 0.702373 | 0.689557 | 0.7059 |
| 0.652567 | 0.645342 | 0.71344 | 0.708465 | 0.718867 | 0.720381 | 0.704918 | 0.696812 | 0.696612 |
| 0.658167 | 0.651055 | 0.729693 | 0.722428 | 0.725375 | 0.731937 | 0.707917 | 0.719656 | 0.714115 |
| 0.658171 | 0.646314 | 0.72573 | 0.715544 | 0.721193 | 0.728038 | 0.708598 | 0.704456 | 0.704959 |

| | | | | | | | | |
|---|---|---|---|---|---|---|---|---|
| 0.667821 | 0.655695 | 0.729998 | 0.71445 | 0.719206 | 0.731281 | 0.710912 | 0.70386 | 0.705621 |
| 0.676041 | 0.660116 | 0.737836 | 0.731306 | 0.734705 | 0.745251 | 0.720968 | 0.719467 | 0.72377 |
| 0.672959 | 0.648415 | 0.731544 | 0.718556 | 0.712543 | 0.736343 | 0.713254 | 0.704623 | 0.721936 |
| 0.675536 | 0.652803 | 0.733283 | 0.726163 | 0.73241 | 0.742009 | 0.715974 | 0.711658 | 0.724342 |
| 0.683326 | 0.664689 | 0.745128 | 0.732769 | 0.734938 | 0.736123 | 0.72872 | 0.724472 | 0.732617 |
| 0.677447 | 0.66065 | 0.738896 | 0.723509 | 0.7283 | 0.740718 | 0.724697 | 0.710157 | 0.726145 |
| 0.673565 | 0.653923 | 0.732347 | 0.719798 | 0.72681 | 0.734282 | 0.717225 | 0.702526 | 0.712202 |
| 0.678007 | 0.65783 | 0.723741 | 0.723742 | 0.717544 | 0.741019 | 0.708979 | 0.711023 | 0.712766 |
| 0.675964 | 0.654957 | 0.73716 | 0.724328 | 0.739726 | 0.733866 | 0.724777 | 0.718277 | 0.708831 |
| 0.67054 | 0.650481 | 0.727778 | 0.717978 | 0.722771 | 0.724252 | 0.709891 | 0.703767 | 0.71378 |
| 0.676635 | 0.6551 | 0.726719 | 0.714096 | 0.723014 | 0.740079 | 0.718864 | 0.709858 | 0.70566 |
| 0.670971 | 0.65669 | 0.729572 | 0.718347 | 0.724888 | 0.731596 | 0.713831 | 0.708968 | 0.718833 |
| 0.676065 | 0.650294 | 0.733985 | 0.72839 | 0.726376 | 0.733707 | 0.706709 | 0.699637 | 0.715755 |
| 0.678564 | 0.653877 | 0.737626 | 0.722878 | 0.739617 | 0.743729 | 0.717421 | 0.708228 | 0.713992 |
| 0.674379 | 0.655376 | 0.735872 | 0.731173 | 0.731874 | 0.739112 | 0.725006 | 0.714885 | 0.723541 |
| 0.67533 | 0.655428 | 0.73239 | 0.728843 | 0.735482 | 0.733874 | 0.711366 | 0.71222 | 0.72254 |
| 0.67508 | 0.654774 | 0.735768 | 0.725714 | 0.720369 | 0.742946 | 0.709329 | 0.719234 | 0.716965 |
| 0.676144 | 0.650177 | 0.737101 | 0.716846 | 0.726837 | 0.731172 | 0.713704 | 0.6977 | 0.719319 |
| 0.678951 | 0.661944 | 0.746099 | 0.740333 | 0.734239 | 0.732751 | 0.719431 | 0.709522 | 0.725268 |
| 0.669675 | 0.65283 | 0.73592 | 0.724971 | 0.728948 | 0.750889 | 0.724943 | 0.715537 | 0.716184 |
| 0.666524 | 0.65 | 0.732638 | 0.72019 | 0.72938 | 0.724882 | 0.70914 | 0.707215 | 0.713821 |
| 0.674989 | 0.654706 | 0.736519 | 0.725742 | 0.732007 | 0.749737 | 0.712107 | 0.718785 | 0.73226 |
| 0.678225 | 0.654317 | 0.73991 | 0.727048 | 0.728762 | 0.740879 | 0.716727 | 0.712079 | 0.719673 |
| 0.676237 | 0.656133 | 0.743597 | 0.724797 | 0.732361 | 0.741269 | 0.720428 | 0.719547 | 0.720571 |
| 0.669283 | 0.653933 | 0.7397 | 0.725565 | 0.729846 | 0.738825 | 0.716265 | 0.721298 | 0.719118 |
| 0.670344 | 0.653423 | 0.738513 | 0.717287 | 0.733224 | 0.746526 | 0.716899 | 0.713639 | 0.719574 |
| 0.670452 | 0.651424 | 0.74112 | 0.718406 | 0.723953 | 0.736154 | 0.71458 | 0.715333 | 0.706816 |
| 0.67231 | 0.652343 | 0.741762 | 0.726478 | 0.731223 | 0.738503 | 0.721014 | 0.708626 | 0.722451 |
| 0.671811 | 0.653633 | 0.740275 | 0.728642 | 0.732793 | 0.749062 | 0.71969 | 0.716573 | 0.710896 |
| 0.674622 | 0.659402 | 0.740033 | 0.727835 | 0.743087 | 0.747889 | 0.730636 | 0.724525 | 0.729204 |
| 0.673198 | 0.657677 | 0.734725 | 0.718624 | 0.728545 | 0.732997 | 0.718226 | 0.709586 | 0.710029 |
| 0.671997 | 0.65811 | 0.741304 | 0.725163 | 0.735914 | 0.742212 | 0.714806 | 0.721269 | 0.713949 |
| 0.675535 | 0.656858 | 0.745902 | 0.733244 | 0.733611 | 0.742307 | 0.724494 | 0.715789 | 0.719158 |
| 0.677899 | 0.649427 | 0.740684 | 0.727518 | 0.731305 | 0.740435 | 0.720968 | 0.720584 | 0.719104 |
| 0.67376 | 0.660265 | 0.744742 | 0.733359 | 0.733985 | 0.745073 | 0.73304 | 0.718937 | 0.715896 |
| 0.670187 | 0.656385 | 0.738117 | 0.725897 | 0.728618 | 0.74296 | 0.72778 | 0.711604 | 0.713989 |
| 0.675412 | 0.658041 | 0.742401 | 0.727203 | 0.730061 | 0.760194 | 0.727607 | 0.723108 | 0.72563 |
| 0.671814 | 0.658272 | 0.741066 | 0.728849 | 0.732653 | 0.751958 | 0.730309 | 0.723374 | 0.727164 |
| 0.668809 | 0.653465 | 0.739496 | 0.72677 | 0.733285 | 0.748268 | 0.72619 | 0.719114 | 0.714957 |
| 0.671885 | 0.657683 | 0.744564 | 0.729131 | 0.732982 | 0.742647 | 0.726119 | 0.72 | 0.722754 |
| 0.679031 | 0.660953 | 0.741926 | 0.730828 | 0.727859 | 0.741379 | 0.723696 | 0.714166 | 0.719588 |
| 0.678953 | 0.65563 | 0.737794 | 0.733359 | 0.730081 | 0.747267 | 0.727691 | 0.713229 | 0.722535 |
| 0.681906 | 0.659269 | 0.74068 | 0.731168 | 0.732696 | 0.754927 | 0.733042 | 0.719898 | 0.72517 |
| 0.674707 | 0.656127 | 0.730428 | 0.722553 | 0.728654 | 0.739709 | 0.714383 | 0.717985 | 0.722354 |
| 0.674111 | 0.658292 | 0.741132 | 0.730311 | 0.731787 | 0.745798 | 0.726335 | 0.715695 | 0.72118 |
| 0.673123 | 0.654498 | 0.7326 | 0.72936 | 0.727355 | 0.743331 | 0.722872 | 0.711691 | 0.71155 |

| | | | | | | | | |
|---|---|---|---|---|---|---|---|---|
| 0.662114 | 0.648883 | 0.726433 | 0.73591 | 0.722509 | 0.728403 | 0.716611 | 0.721925 | 0.716847 |
| 0.719101 | 0.703388 | 0.715783 | 0.715123 | 0.719482 | 0.721857 | 0.703272 | 0.697405 | 0.708418 |
| 0.677922 | 0.65143 | 0.726065 | 0.722768 | 0.717029 | 0.735431 | 0.71966 | 0.705034 | 0.706213 |
| 0.678104 | 0.654492 | 0.74695 | 0.725404 | 0.730549 | 0.740988 | 0.721005 | 0.716042 | 0.725648 |
| 0.673094 | 0.654746 | 0.731039 | 0.721673 | 0.717626 | 0.733662 | 0.721216 | 0.707499 | 0.712112 |
| 0.669495 | 0.650247 | 0.736147 | 0.724725 | 0.723018 | 0.744093 | 0.727632 | 0.716495 | 0.714712 |
| 0.676464 | 0.649768 | 0.747033 | 0.725223 | 0.72516 | 0.742845 | 0.721652 | 0.721581 | 0.722524 |
| 0.665456 | 0.647864 | 0.738277 | 0.723728 | 0.72256 | 0.735435 | 0.727681 | 0.710924 | 0.710494 |
| 0.672982 | 0.646639 | 0.736763 | 0.725138 | 0.736756 | 0.737769 | 0.722184 | 0.709795 | 0.716264 |
| 0.683843 | 0.66446 | 0.733755 | 0.729766 | 0.737253 | 0.739987 | 0.719274 | 0.722492 | 0.72204 |
| 0.678486 | 0.660638 | 0.740393 | 0.732525 | 0.730838 | 0.747595 | 0.730883 | 0.712155 | 0.727381 |
| 0.678859 | 0.657852 | 0.736159 | 0.73094 | 0.732127 | 0.735943 | 0.723963 | 0.711331 | 0.71598 |
| 0.672994 | 0.659273 | 0.750417 | 0.728042 | 0.732033 | 0.742352 | 0.725504 | 0.712997 | 0.722766 |
| 0.675804 | 0.656872 | 0.733453 | 0.733384 | 0.735665 | 0.743041 | 0.725023 | 0.727453 | 0.722436 |
| 0.679504 | 0.656898 | 0.735188 | 0.730688 | 0.731359 | 0.73788 | 0.731054 | 0.712155 | 0.723284 |
| 0.66504 | 0.65033 | 0.734452 | 0.727601 | 0.730563 | 0.744321 | 0.719791 | 0.709732 | 0.716431 |
| 0.679523 | 0.655223 | 0.733645 | 0.727536 | 0.732065 | 0.736328 | 0.721484 | 0.715925 | 0.720812 |
| 0.675342 | 0.655907 | 0.723698 | 0.718027 | 0.72453 | 0.735116 | 0.717037 | 0.708688 | 0.710868 |
| 0.673543 | 0.656823 | 0.745198 | 0.736114 | 0.736183 | 0.752597 | 0.73363 | 0.724296 | 0.731047 |
| 0.670149 | 0.653245 | 0.738146 | 0.726547 | 0.733554 | 0.746784 | 0.717543 | 0.710451 | 0.718797 |
| 0.675699 | 0.653493 | 0.735032 | 0.730903 | 0.72972 | 0.745443 | 0.719274 | 0.721888 | 0.723983 |
| 0.66221 | 0.645107 | 0.744443 | 0.734409 | 0.735556 | 0.734432 | 0.716946 | 0.711869 | 0.726335 |
| 0.668539 | 0.649672 | 0.745316 | 0.731081 | 0.7375 | 0.740338 | 0.707556 | 0.720824 | 0.722784 |
| 0.663358 | 0.656347 | 0.742559 | 0.729497 | 0.741217 | 0.73738 | 0.714873 | 0.715093 | 0.726272 |
| 0.672955 | 0.662689 | 0.749392 | 0.743228 | 0.734557 | 0.742539 | 0.719161 | 0.720744 | 0.732227 |
| 0.669787 | 0.652654 | 0.749695 | 0.731494 | 0.744331 | 0.735865 | 0.704457 | 0.718037 | 0.732772 |
| 0.673094 | 0.653538 | 0.741573 | 0.727166 | 0.735658 | 0.737324 | 0.715888 | 0.721658 | 0.72374 |
| 0.668712 | 0.659384 | 0.7449 | 0.73121 | 0.745891 | 0.729377 | 0.722635 | 0.718394 | 0.720923 |
| 0.674685 | 0.649725 | 0.744747 | 0.728697 | 0.737936 | 0.737465 | 0.72337 | 0.715069 | 0.721208 |
| 0.742166 | 0.733674 | 0.656032 | 0.655369 | 0.660926 | 0.65728 | 0.654045 | 0.634621 | 0.64307 |
| 0.759591 | 0.742975 | 0.659959 | 0.65836 | 0.660328 | 0.680065 | 0.656091 | 0.643265 | 0.635641 |
| 0.755927 | 0.741002 | 0.659574 | 0.659432 | 0.656157 | 0.668548 | 0.645584 | 0.639967 | 0.64494 |
| 0.747962 | 0.80233 | 0.661596 | 0.657331 | 0.652925 | 0.669452 | 0.642417 | 0.639841 | 0.639528 |
| 0.761137 | 0.748116 | 0.663751 | 0.659707 | 0.652988 | 0.676459 | 0.655507 | 0.638986 | 0.63593 |
| 0.675078 | 0.651901 | 0.747554 | 0.740972 | 0.732854 | 0.749894 | 0.728631 | 0.728843 | 0.731141 |
| 0.664996 | 0.658115 | 0.746203 | 0.737473 | 0.730162 | 0.747475 | 0.724936 | 0.722143 | 0.721874 |
| 0.670998 | 0.652046 | 0.74947 | 0.739768 | 0.738436 | 0.751049 | 0.732486 | 0.721845 | 0.726893 |
| 0.672459 | 0.657514 | 0.75072 | 0.741313 | 0.744944 | 0.759115 | 0.741898 | 0.734174 | 0.735321 |
| 0.670771 | 0.648494 | 0.749962 | 0.738963 | 0.734905 | 0.753953 | 0.737907 | 0.73553 | 0.730631 |
| 0.672395 | 0.65783 | 0.749131 | 0.742272 | 0.741075 | 0.749738 | 0.734444 | 0.721515 | 0.735329 |
| 0.675016 | 0.656065 | 0.743438 | 0.737515 | 0.738226 | 0.743767 | 0.719835 | 0.715179 | 0.723943 |
| 0.667682 | 0.652138 | 0.747902 | 0.736984 | 0.736088 | 0.740927 | 0.732345 | 0.722761 | 0.735719 |
| 0.663835 | 0.652996 | 0.74415 | 0.731017 | 0.740286 | 0.749623 | 0.73376 | 0.72969 | 0.728927 |
| 0.667354 | 0.653853 | 0.740654 | 0.735939 | 0.738211 | 0.742732 | 0.738715 | 0.738018 | 0.744064 |
| 0.666812 | 0.652152 | 0.744369 | 0.743494 | 0.732446 | 0.752425 | 0.727443 | 0.719723 | 0.724572 |
| 0.66855 | 0.653426 | 0.747243 | 0.739798 | 0.744229 | 0.750317 | 0.733178 | 0.721203 | 0.726999 |

| | | | | | | | | |
|---|---|---|---|---|---|---|---|---|
| 0.676302 | 0.657275 | 0.757202 | 0.745964 | 0.738915 | 0.743337 | 0.722068 | 0.732582 | 0.743389 |
| 0.657186 | 0.653095 | 0.747764 | 0.740604 | 0.736058 | 0.741672 | 0.73125 | 0.723921 | 0.731032 |
| 0.67342 | 0.657312 | 0.753452 | 0.741482 | 0.745957 | 0.766025 | 0.73666 | 0.729469 | 0.738046 |
| 0.669262 | 0.653606 | 0.750153 | 0.732525 | 0.74093 | 0.744475 | 0.731178 | 0.714651 | 0.734639 |
| 0.669519 | 0.648017 | 0.756923 | 0.745647 | 0.739043 | 0.752233 | 0.725576 | 0.721699 | 0.731824 |
| 0.671878 | 0.65226 | 0.747599 | 0.741373 | 0.742951 | 0.75746 | 0.739272 | 0.729683 | 0.732244 |
| 0.673703 | 0.655532 | 0.755757 | 0.741527 | 0.747628 | 0.754203 | 0.727315 | 0.722957 | 0.734295 |
| 0.669236 | 0.658586 | 0.753035 | 0.744222 | 0.751384 | 0.756665 | 0.735845 | 0.727325 | 0.742355 |
| 0.673272 | 0.652418 | 0.757156 | 0.745898 | 0.740314 | 0.746234 | 0.721721 | 0.709065 | 0.735653 |
| 0.669251 | 0.654093 | 0.756369 | 0.731317 | 0.73102 | 0.739929 | 0.730501 | 0.726749 | 0.729454 |
| 0.670063 | 0.65014 | 0.751569 | 0.744801 | 0.736544 | 0.757846 | 0.729398 | 0.727821 | 0.726426 |
| 0.677106 | 0.649186 | 0.747863 | 0.740806 | 0.734025 | 0.756201 | 0.73134 | 0.73395 | 0.727273 |
| 0.665401 | 0.648803 | 0.755524 | 0.735306 | 0.733235 | 0.758146 | 0.730543 | 0.717691 | 0.723835 |
| 0.67423 | 0.65049 | 0.759777 | 0.74834 | 0.73713 | 0.744896 | 0.731497 | 0.72644 | 0.740707 |
| 0.671609 | 0.654195 | 0.735325 | 0.727608 | 0.724401 | 0.736831 | 0.715826 | 0.711299 | 0.708086 |
| 0.678517 | 0.665347 | 0.736984 | 0.731533 | 0.740029 | 0.744181 | 0.720393 | 0.723301 | 0.71382 |
| 0.672421 | 0.657993 | 0.727219 | 0.720948 | 0.73105 | 0.741882 | 0.717024 | 0.72241 | 0.709191 |
| 0.675687 | 0.659627 | 0.742452 | 0.725456 | 0.735932 | 0.743547 | 0.727315 | 0.711783 | 0.712732 |
| 0.670726 | 0.658716 | 0.733109 | 0.72759 | 0.735919 | 0.744319 | 0.724468 | 0.70906 | 0.717312 |
| 0.676484 | 0.65543 | 0.73489 | 0.723011 | 0.732611 | 0.740756 | 0.719035 | 0.704616 | 0.717411 |
| 0.678619 | 0.658557 | 0.750038 | 0.734982 | 0.741846 | 0.743557 | 0.732833 | 0.717772 | 0.716212 |
| 0.676406 | 0.658561 | 0.732471 | 0.725384 | 0.733871 | 0.733947 | 0.716964 | 0.710594 | 0.722172 |
| 0.671731 | 0.660257 | 0.740673 | 0.726181 | 0.732737 | 0.741963 | 0.725349 | 0.721523 | 0.721739 |
| 0.666296 | 0.659099 | 0.761534 | 0.746609 | 0.750884 | 0.763735 | 0.746141 | 0.742438 | 0.738513 |
| 0.672933 | 0.661667 | 0.739422 | 0.73035 | 0.731011 | 0.736466 | 0.722549 | 0.71686 | 0.719878 |
| 0.766811 | 0.766661 | 0.663289 | 0.655704 | 0.655503 | 0.664377 | 0.642337 | 0.636862 | 0.640273 |
| 0.669819 | 0.650943 | 0.742788 | 0.730902 | 0.733874 | 0.743611 | 0.731409 | 0.718952 | 0.726381 |
| 0.671268 | 0.65626 | 0.740707 | 0.72458 | 0.728762 | 0.74144 | 0.724611 | 0.713314 | 0.710179 |
| 0.676066 | 0.662238 | 0.73486 | 0.72501 | 0.733565 | 0.74203 | 0.724698 | 0.709024 | 0.72322 |
| 0.675918 | 0.653051 | 0.741834 | 0.727866 | 0.735145 | 0.753211 | 0.729748 | 0.714407 | 0.719081 |
| 0.67634 | 0.654491 | 0.745661 | 0.73157 | 0.731944 | 0.738298 | 0.726193 | 0.728377 | 0.727465 |
| 0.681698 | 0.653872 | 0.742 | 0.732353 | 0.730243 | 0.74469 | 0.729065 | 0.72055 | 0.717038 |
| 0.67066 | 0.653916 | 0.735325 | 0.730925 | 0.733927 | 0.743465 | 0.719487 | 0.711874 | 0.711929 |
| 0.66674 | 0.656125 | 0.744282 | 0.720534 | 0.730397 | 0.733546 | 0.715443 | 0.711758 | 0.720443 |
| 0.677485 | 0.652588 | 0.734725 | 0.720117 | 0.734011 | 0.738879 | 0.719606 | 0.714492 | 0.706477 |
| 0.669305 | 0.649085 | 0.737395 | 0.721315 | 0.725812 | 0.729979 | 0.713039 | 0.714245 | 0.70731 |
| 0.674786 | 0.657721 | 0.74197 | 0.736517 | 0.736679 | 0.743781 | 0.731656 | 0.716634 | 0.727382 |
| 0.66977 | 0.654941 | 0.743182 | 0.722594 | 0.731103 | 0.738931 | 0.724319 | 0.720193 | 0.718252 |
| 0.6717 | 0.66363 | 0.736 | 0.728476 | 0.736823 | 0.75216 | 0.723288 | 0.719637 | 0.719094 |
| 0.672707 | 0.662659 | 0.733803 | 0.7257 | 0.7246 | 0.741302 | 0.724452 | 0.720266 | 0.722521 |
| 0.669166 | 0.647039 | 0.73766 | 0.726765 | 0.734002 | 0.733642 | 0.715763 | 0.711544 | 0.721448 |
| 0.679034 | 0.657372 | 0.74148 | 0.729978 | 0.727572 | 0.746548 | 0.724468 | 0.720023 | 0.719479 |
| 0.670936 | 0.654145 | 0.736705 | 0.718326 | 0.73664 | 0.744568 | 0.719339 | 0.716367 | 0.717385 |
| 0.670987 | 0.654554 | 0.740382 | 0.725729 | 0.731424 | 0.741051 | 0.731034 | 0.715909 | 0.728274 |
| 0.675816 | 0.65625 | 0.746417 | 0.74012 | 0.738345 | 0.74728 | 0.736818 | 0.722317 | 0.738576 |
| 0.676977 | 0.653132 | 0.737923 | 0.737679 | 0.733152 | 0.747784 | 0.73218 | 0.719303 | 0.728062 |

| | | | | | | | | |
|---|---|---|---|---|---|---|---|---|
| 0.673791 | 0.648891 | 0.741648 | 0.737847 | 0.727716 | 0.747494 | 0.734548 | 0.731078 | 0.730324 |
| 0.679063 | 0.655352 | 0.750301 | 0.744611 | 0.737894 | 0.754156 | 0.742712 | 0.73587 | 0.744283 |
| 0.670726 | 0.65325 | 0.743157 | 0.728569 | 0.740334 | 0.750423 | 0.731685 | 0.733564 | 0.729722 |
| 0.653727 | 0.644327 | 0.715125 | 0.727603 | 0.72924 | 0.722883 | 0.732872 | 0.699462 | 0.709302 |
| 0.669241 | 0.649226 | 0.742076 | 0.73425 | 0.728823 | 0.74215 | 0.733287 | 0.723217 | 0.725987 |
| 0.672582 | 0.642276 | 0.737879 | 0.722678 | 0.727587 | 0.747894 | 0.724138 | 0.721905 | 0.715135 |
| 0.669462 | 0.655716 | 0.740097 | 0.729719 | 0.729012 | 0.75079 | 0.734168 | 0.732092 | 0.736985 |
| 0.668382 | 0.64922 | 0.739802 | 0.729183 | 0.731672 | 0.744868 | 0.735057 | 0.725206 | 0.723505 |
| 0.681454 | 0.658101 | 0.748639 | 0.731478 | 0.730457 | 0.746605 | 0.729355 | 0.724928 | 0.726703 |
| 0.670677 | 0.659431 | 0.742136 | 0.733525 | 0.733454 | 0.74692 | 0.732932 | 0.725843 | 0.725677 |
| 0.674464 | 0.654653 | 0.74159 | 0.72825 | 0.723828 | 0.746928 | 0.729648 | 0.719422 | 0.724933 |
| 0.674555 | 0.653498 | 0.744034 | 0.738142 | 0.730993 | 0.745752 | 0.729686 | 0.72393 | 0.718638 |
| 0.672 | 0.660526 | 0.739569 | 0.73054 | 0.728059 | 0.741336 | 0.726897 | 0.720057 | 0.719905 |
| 0.664539 | 0.650597 | 0.748112 | 0.737868 | 0.73373 | 0.745992 | 0.732118 | 0.721298 | 0.722977 |
| 0.675384 | 0.653852 | 0.747765 | 0.738237 | 0.734141 | 0.753733 | 0.737781 | 0.730168 | 0.729923 |
| 0.674092 | 0.653298 | 0.739786 | 0.731881 | 0.731175 | 0.747158 | 0.733241 | 0.720659 | 0.722781 |
| 0.676244 | 0.653978 | 0.750414 | 0.730189 | 0.744565 | 0.747869 | 0.742554 | 0.73195 | 0.735303 |
| 0.670188 | 0.650708 | 0.745187 | 0.735645 | 0.736909 | 0.747961 | 0.733441 | 0.726989 | 0.730126 |
| 0.675594 | 0.65152 | 0.739176 | 0.725081 | 0.730394 | 0.745127 | 0.724494 | 0.716515 | 0.726593 |
| 0.673272 | 0.645979 | 0.738987 | 0.735943 | 0.734175 | 0.745713 | 0.730743 | 0.729054 | 0.72989 |
| 0.677043 | 0.651019 | 0.748181 | 0.733897 | 0.734491 | 0.745287 | 0.740861 | 0.718218 | 0.72214 |
| 0.675359 | 0.653808 | 0.745208 | 0.737819 | 0.737473 | 0.75972 | 0.735709 | 0.732605 | 0.734976 |
| 0.696667 | 0.695423 | 0.684025 | 0.685786 | 0.674357 | 0.680277 | 0.672219 | 0.671004 | 0.668213 |
| 0.708702 | 0.72522 | 0.662591 | 0.672332 | 0.660191 | 0.648849 | 0.660543 | 0.653796 | 0.652154 |
| 1 | 0.728825 | 0.659125 | 0.66427 | 0.65313 | 0.666783 | 0.63848 | 0.642961 | 0.641158 |
| 0.728825 | 1 | 0.653537 | 0.65102 | 0.649182 | 0.650182 | 0.642084 | 0.638542 | 0.646972 |
| 0.659125 | 0.653537 | 1 | 0.754993 | 0.759758 | 0.748062 | 0.750585 | 0.747516 | 0.76163 |
| 0.66427 | 0.65102 | 0.754993 | 1 | 0.737691 | 0.76229 | 0.757392 | 0.749012 | 0.785831 |
| 0.65313 | 0.649182 | 0.759758 | 0.737691 | 1 | 0.754753 | 0.74014 | 0.754695 | 0.746181 |
| 0.666783 | 0.650182 | 0.748062 | 0.76229 | 0.754753 | 1 | 0.749904 | 0.846475 | 0.750353 |
| 0.63848 | 0.642084 | 0.750585 | 0.757392 | 0.74014 | 0.749904 | 1 | 0.742184 | 0.740757 |
| 0.642961 | 0.638542 | 0.747516 | 0.749012 | 0.754695 | 0.846475 | 0.742184 | 1 | 0.795767 |
| 0.641158 | 0.646972 | 0.76163 | 0.785831 | 0.746181 | 0.750353 | 0.740757 | 0.795767 | 1 |
| 0.733921 | 0.731495 | 0.656378 | 0.66068 | 0.658692 | 0.659039 | 0.650988 | 0.649689 | 0.649336 |
| 0.673284 | 0.651601 | 0.760079 | 0.754362 | 0.743799 | 0.762946 | 0.746657 | 0.747926 | 0.738229 |
| 0.656908 | 0.651302 | 0.753749 | 0.747644 | 0.757188 | 0.759112 | 0.740528 | 0.74742 | 0.74108 |
| 0.666009 | 0.658974 | 0.750278 | 0.755162 | 0.756879 | 0.761186 | 0.748663 | 0.743796 | 0.747586 |
| 0.646739 | 0.647305 | 0.751298 | 0.741196 | 0.738482 | 0.745407 | 0.745648 | 0.745627 | 0.752692 |
| 0.662318 | 0.650345 | 0.751506 | 0.751172 | 0.7504 | 0.759408 | 0.743172 | 0.742927 | 0.739006 |
| 0.657796 | 0.654188 | 0.739154 | 0.734255 | 0.729317 | 0.754741 | 0.728012 | 0.725787 | 0.719793 |
| 0.653453 | 0.641572 | 0.738984 | 0.730623 | 0.739644 | 0.75097 | 0.733093 | 0.724029 | 0.730839 |
| 0.650282 | 0.648608 | 0.74281 | 0.742857 | 0.73852 | 0.745688 | 0.73206 | 0.727652 | 0.732663 |
| 0.660993 | 0.649694 | 0.744308 | 0.731898 | 0.746292 | 0.746848 | 0.742938 | 0.732436 | 0.727922 |
| 0.662879 | 0.648656 | 0.742935 | 0.741841 | 0.747271 | 0.74591 | 0.754403 | 0.743559 | 0.730894 |
| 0.661041 | 0.65608 | 0.741976 | 0.745321 | 0.743679 | 0.752423 | 0.739626 | 0.751946 | 0.754793 |
| 0.634434 | 0.644155 | 0.730933 | 0.735268 | 0.736188 | 0.746261 | 0.71484 | 0.735775 | 0.729869 |

| | | | | | | | | |
|---|---|---|---|---|---|---|---|---|
| 0.671223 | 0.657833 | 0.7656 | 0.750747 | 0.754396 | 0.767905 | 0.747679 | 0.750074 | 0.760308 |
| 0.64619 | 0.639505 | 0.760815 | 0.747158 | 0.743967 | 0.762048 | 0.754819 | 0.750165 | 0.73771 |
| 0.649186 | 0.650137 | 0.746891 | 0.745734 | 0.737888 | 0.739192 | 0.740772 | 0.73747 | 0.737522 |
| 0.657134 | 0.652491 | 0.751278 | 0.749256 | 0.738722 | 0.756168 | 0.741499 | 0.736824 | 0.745246 |
| 0.647577 | 0.655127 | 0.748154 | 0.739483 | 0.752524 | 0.746751 | 0.757374 | 0.735557 | 0.753488 |
| 0.647701 | 0.661553 | 0.760731 | 0.750789 | 0.743965 | 0.747475 | 0.755422 | 0.75987 | 0.749879 |
| 0.650535 | 0.65172 | 0.745614 | 0.756365 | 0.749244 | 0.744911 | 0.755396 | 0.727727 | 0.744424 |
| 0.651446 | 0.643639 | 0.764145 | 0.748263 | 0.736959 | 0.746161 | 0.739245 | 0.747928 | 0.76589 |
| 0.657934 | 0.641885 | 0.752209 | 0.743792 | 0.742989 | 0.746399 | 0.746536 | 0.741099 | 0.734651 |
| 0.669033 | 0.659359 | 0.759442 | 0.738594 | 0.748461 | 0.758591 | 0.727294 | 0.730259 | 0.727735 |
| 0.669143 | 0.660572 | 0.757648 | 0.736194 | 0.739349 | 0.753333 | 0.7223 | 0.738074 | 0.725875 |
| 0.6674 | 0.661089 | 0.754278 | 0.740115 | 0.739965 | 0.744359 | 0.72932 | 0.72457 | 0.743887 |
| 0.656799 | 0.661398 | 0.743035 | 0.737043 | 0.738945 | 0.735214 | 0.726479 | 0.714286 | 0.72405 |
| 0.664935 | 0.657451 | 0.760987 | 0.754378 | 0.738594 | 0.74773 | 0.736068 | 0.734796 | 0.738355 |
| 0.655355 | 0.654012 | 0.753995 | 0.737148 | 0.744921 | 0.746839 | 0.73668 | 0.736793 | 0.726716 |

| | plot 22 | | | | | | | plot |
|---|---|---|---|---|---|---|---|---|
| Wild_plot_ | Wild_plot_ | Wild_plot_ | Wild_plot_ | Wild_plot_ | Wild_plot_ | Wild_plot_ | Wild_plot_ | Wild_plot_ |
| 0.722929 | 0.698673 | 0.674874 | 0.663464 | 0.653929 | 0.673183 | 0.666956 | 0.656331 | 0.656891 |
| 0.724215 | 0.707002 | 0.679079 | 0.674849 | 0.661799 | 0.681699 | 0.676199 | 0.668732 | 0.666063 |
| 0.724068 | 0.697979 | 0.668005 | 0.664069 | 0.653832 | 0.671499 | 0.662611 | 0.651102 | 0.657586 |
| 0.7299 | 0.705707 | 0.681864 | 0.668835 | 0.662473 | 0.676246 | 0.672397 | 0.661236 | 0.669384 |
| 0.726422 | 0.70253 | 0.672452 | 0.670754 | 0.651159 | 0.67421 | 0.662541 | 0.655361 | 0.660468 |
| 0.723012 | 0.706405 | 0.681405 | 0.669169 | 0.656189 | 0.683382 | 0.673472 | 0.663413 | 0.668814 |
| 0.659893 | 0.760836 | 0.738908 | 0.731458 | 0.723239 | 0.735202 | 0.724481 | 0.726969 | 0.724213 |
| 0.655662 | 0.760058 | 0.74246 | 0.722563 | 0.716596 | 0.736493 | 0.724129 | 0.729054 | 0.72962 |
| 0.727894 | 0.701937 | 0.678354 | 0.665585 | 0.652182 | 0.677525 | 0.672784 | 0.665693 | 0.667472 |
| 0.728619 | 0.701252 | 0.673435 | 0.668881 | 0.652158 | 0.674242 | 0.665387 | 0.656771 | 0.659424 |
| 0.719501 | 0.702016 | 0.67601 | 0.671176 | 0.656949 | 0.677525 | 0.671013 | 0.661508 | 0.664361 |
| 0.731213 | 0.697387 | 0.669727 | 0.662328 | 0.647373 | 0.66341 | 0.654111 | 0.649402 | 0.646089 |
| 0.73206 | 0.708511 | 0.682356 | 0.674385 | 0.657348 | 0.682148 | 0.670053 | 0.6623 | 0.667624 |
| 0.734562 | 0.717002 | 0.688515 | 0.678338 | 0.667985 | 0.691976 | 0.680174 | 0.676421 | 0.68057 |
| 0.718761 | 0.726686 | 0.704383 | 0.702018 | 0.679245 | 0.700304 | 0.697645 | 0.689331 | 0.690859 |
| 0.739866 | 0.707453 | 0.680894 | 0.674873 | 0.657285 | 0.684628 | 0.675217 | 0.667537 | 0.670955 |
| 0.72449 | 0.709935 | 0.674752 | 0.675269 | 0.654764 | 0.679792 | 0.674384 | 0.657673 | 0.668585 |
| 0.699928 | 0.73638 | 0.711326 | 0.704231 | 0.68267 | 0.709285 | 0.704246 | 0.69829 | 0.695606 |
| 0.733445 | 0.697141 | 0.675395 | 0.662848 | 0.64178 | 0.669589 | 0.664516 | 0.652955 | 0.656113 |
| 0.730891 | 0.705063 | 0.680018 | 0.668321 | 0.653412 | 0.675073 | 0.670015 | 0.657729 | 0.664498 |
| 0.738936 | 0.702855 | 0.671964 | 0.66306 | 0.648118 | 0.672365 | 0.666956 | 0.659283 | 0.662174 |
| 0.740748 | 0.705293 | 0.678962 | 0.670916 | 0.654993 | 0.678584 | 0.666295 | 0.657911 | 0.662236 |
| 0.739026 | 0.708711 | 0.684898 | 0.682014 | 0.660953 | 0.679326 | 0.675133 | 0.663909 | 0.669746 |
| 0.744117 | 0.705083 | 0.674798 | 0.672427 | 0.654496 | 0.678483 | 0.670764 | 0.660838 | 0.659873 |
| 0.732898 | 0.704671 | 0.68037 | 0.67318 | 0.660804 | 0.681283 | 0.67127 | 0.663842 | 0.662887 |
| 0.736725 | 0.703538 | 0.676313 | 0.668959 | 0.649503 | 0.676217 | 0.670523 | 0.661092 | 0.665165 |
| 0.722048 | 0.699384 | 0.674592 | 0.670367 | 0.64643 | 0.670902 | 0.661901 | 0.661434 | 0.659732 |
| 0.730621 | 0.702745 | 0.673561 | 0.671258 | 0.650853 | 0.679152 | 0.668842 | 0.664673 | 0.663423 |
| 0.733837 | 0.707795 | 0.682018 | 0.675503 | 0.662903 | 0.683125 | 0.677524 | 0.666769 | 0.672221 |
| 0.719402 | 0.703955 | 0.676102 | 0.669975 | 0.651933 | 0.679538 | 0.672258 | 0.662087 | 0.664143 |
| 0.733037 | 0.702709 | 0.675059 | 0.67258 | 0.648783 | 0.676649 | 0.666255 | 0.654521 | 0.666515 |
| 0.729535 | 0.701747 | 0.672003 | 0.666319 | 0.650446 | 0.671491 | 0.663224 | 0.660294 | 0.659271 |
| 0.733715 | 0.7012 | 0.672916 | 0.669695 | 0.648057 | 0.672245 | 0.668719 | 0.656925 | 0.66164 |
| 0.730246 | 0.705082 | 0.673292 | 0.667313 | 0.65296 | 0.675416 | 0.669747 | 0.662389 | 0.662562 |
| 0.732001 | 0.699593 | 0.67169 | 0.66671 | 0.647101 | 0.668277 | 0.663494 | 0.653564 | 0.656851 |
| 0.739831 | 0.703481 | 0.672538 | 0.669324 | 0.65355 | 0.674673 | 0.667949 | 0.663473 | 0.662491 |
| 0.726722 | 0.706038 | 0.67891 | 0.673459 | 0.653307 | 0.678085 | 0.67317 | 0.664324 | 0.668274 |
| 0.71744 | 0.705302 | 0.67623 | 0.67311 | 0.657519 | 0.680853 | 0.667497 | 0.66646 | 0.667377 |
| 0.726188 | 0.708955 | 0.679502 | 0.675257 | 0.659692 | 0.68468 | 0.676379 | 0.668714 | 0.671536 |

| | | | | | | | | |
|---|---|---|---|---|---|---|---|---|
| 0.704562 | 0.742139 | 0.715406 | 0.70742 | 0.695817 | 0.713571 | 0.711064 | 0.70295 | 0.705371 |
| 0.719076 | 0.696178 | 0.666592 | 0.659061 | 0.641503 | 0.665587 | 0.657716 | 0.64629 | 0.650388 |
| 0.730504 | 0.699593 | 0.667414 | 0.66658 | 0.648673 | 0.670267 | 0.660971 | 0.65659 | 0.654302 |
| 0.728091 | 0.711034 | 0.677624 | 0.676195 | 0.659339 | 0.681706 | 0.672999 | 0.66442 | 0.664761 |
| 0.734875 | 0.702824 | 0.678543 | 0.670875 | 0.651275 | 0.671299 | 0.66317 | 0.662669 | 0.663199 |
| 0.739017 | 0.708487 | 0.675879 | 0.66449 | 0.651534 | 0.6752 | 0.670149 | 0.659552 | 0.660636 |
| 0.729695 | 0.700091 | 0.671277 | 0.662693 | 0.645042 | 0.672273 | 0.660736 | 0.655567 | 0.656428 |
| 0.734477 | 0.705185 | 0.680208 | 0.671294 | 0.656407 | 0.675715 | 0.671308 | 0.66309 | 0.668532 |
| 0.722925 | 0.706386 | 0.678543 | 0.673459 | 0.657606 | 0.681642 | 0.669444 | 0.659719 | 0.661531 |
| 0.726401 | 0.704872 | 0.683676 | 0.669389 | 0.656479 | 0.682839 | 0.673013 | 0.676135 | 0.670023 |
| 0.729379 | 0.696061 | 0.669184 | 0.661192 | 0.639161 | 0.670401 | 0.658214 | 0.650916 | 0.651945 |
| 0.730784 | 0.697315 | 0.670234 | 0.661751 | 0.636622 | 0.66825 | 0.657914 | 0.650146 | 0.64772 |
| 0.740343 | 0.701759 | 0.675105 | 0.668907 | 0.649855 | 0.670474 | 0.663386 | 0.656352 | 0.6543 |
| 0.728455 | 0.697973 | 0.674242 | 0.662842 | 0.648774 | 0.675516 | 0.667331 | 0.657077 | 0.657151 |
| 0.724022 | 0.698616 | 0.671979 | 0.662855 | 0.652529 | 0.677112 | 0.664753 | 0.657396 | 0.658043 |
| 0.725546 | 0.698145 | 0.67599 | 0.667108 | 0.646697 | 0.673891 | 0.66312 | 0.657928 | 0.661402 |
| 0.725487 | 0.698713 | 0.669244 | 0.663015 | 0.644894 | 0.666818 | 0.659848 | 0.650558 | 0.654438 |
| 0.728271 | 0.70499 | 0.67982 | 0.666493 | 0.646074 | 0.676946 | 0.668196 | 0.660169 | 0.66305 |
| 0.735092 | 0.705153 | 0.673937 | 0.671013 | 0.651864 | 0.678326 | 0.670847 | 0.663638 | 0.663803 |
| 0.727087 | 0.701114 | 0.673064 | 0.666754 | 0.64833 | 0.667641 | 0.66236 | 0.654018 | 0.651187 |
| 0.730123 | 0.700434 | 0.669334 | 0.665049 | 0.648619 | 0.664972 | 0.659582 | 0.649371 | 0.654675 |
| 0.717283 | 0.69844 | 0.674185 | 0.664321 | 0.643138 | 0.67235 | 0.661145 | 0.652388 | 0.660257 |
| 0.729969 | 0.698364 | 0.671419 | 0.663871 | 0.652652 | 0.66957 | 0.666175 | 0.6547 | 0.661213 |
| 0.737632 | 0.707863 | 0.68726 | 0.678132 | 0.662657 | 0.684828 | 0.677659 | 0.671495 | 0.6708 |
| 0.714624 | 0.716874 | 0.691243 | 0.683709 | 0.666904 | 0.694209 | 0.682427 | 0.678368 | 0.674436 |
| 0.658593 | 0.759762 | 0.743082 | 0.72385 | 0.719985 | 0.736479 | 0.7254 | 0.730874 | 0.730485 |
| 0.654963 | 0.764497 | 0.744021 | 0.73784 | 0.720569 | 0.747706 | 0.72882 | 0.72447 | 0.731023 |
| 0.653542 | 0.766898 | 0.743569 | 0.737193 | 0.723186 | 0.743433 | 0.73273 | 0.725345 | 0.731623 |
| 0.731797 | 0.697143 | 0.670428 | 0.665243 | 0.647622 | 0.672764 | 0.666171 | 0.664402 | 0.665412 |
| 0.735503 | 0.699709 | 0.678939 | 0.669506 | 0.654392 | 0.673453 | 0.663511 | 0.661567 | 0.658441 |
| 0.717224 | 0.705341 | 0.680451 | 0.67395 | 0.657559 | 0.683207 | 0.668971 | 0.661812 | 0.668281 |
| 0.720322 | 0.707736 | 0.682007 | 0.680654 | 0.662972 | 0.687415 | 0.675615 | 0.669684 | 0.674604 |
| 0.73269 | 0.701701 | 0.674717 | 0.670419 | 0.65433 | 0.674384 | 0.670039 | 0.658567 | 0.664492 |
| 0.744147 | 0.703629 | 0.672627 | 0.674227 | 0.654969 | 0.673107 | 0.662185 | 0.656154 | 0.662027 |
| 0.736085 | 0.701548 | 0.676361 | 0.67048 | 0.655926 | 0.673896 | 0.670771 | 0.662751 | 0.66783 |
| 0.733918 | 0.704747 | 0.675445 | 0.668757 | 0.653105 | 0.673231 | 0.668155 | 0.659196 | 0.663645 |
| 0.740262 | 0.704423 | 0.681166 | 0.670047 | 0.655831 | 0.677728 | 0.67087 | 0.659643 | 0.665367 |
| 0.759751 | 0.720064 | 0.691907 | 0.691423 | 0.671372 | 0.697395 | 0.687594 | 0.675282 | 0.684995 |
| 0.732303 | 0.697788 | 0.669273 | 0.67021 | 0.646146 | 0.668068 | 0.667629 | 0.656555 | 0.663362 |
| 0.744894 | 0.702774 | 0.675219 | 0.666581 | 0.649151 | 0.674793 | 0.667731 | 0.660562 | 0.660304 |
| 0.732642 | 0.69874 | 0.67195 | 0.665108 | 0.650089 | 0.67301 | 0.667822 | 0.6556 | 0.660006 |
| 0.757121 | 0.719653 | 0.688227 | 0.687111 | 0.668305 | 0.693044 | 0.682434 | 0.674534 | 0.681853 |
| 0.742847 | 0.703288 | 0.678346 | 0.670496 | 0.658589 | 0.678624 | 0.674534 | 0.661449 | 0.664904 |
| 0.732179 | 0.700215 | 0.666407 | 0.66658 | 0.649494 | 0.672149 | 0.658133 | 0.653729 | 0.657045 |
| 0.761034 | 0.700836 | 0.674865 | 0.666537 | 0.651466 | 0.675764 | 0.668402 | 0.660197 | 0.664858 |
| 0.73258 | 0.702995 | 0.672433 | 0.670657 | 0.654059 | 0.677714 | 0.668524 | 0.656398 | 0.663101 |

| | | | | | | | | |
|---|---|---|---|---|---|---|---|---|
| 0.733506 | 0.702128 | 0.678363 | 0.671063 | 0.65607 | 0.673559 | 0.668609 | 0.6556 | 0.660938 |
| 0.746471 | 0.703028 | 0.67874 | 0.67208 | 0.659825 | 0.67655 | 0.665562 | 0.660968 | 0.665565 |
| 0.746842 | 0.704918 | 0.679275 | 0.67048 | 0.655989 | 0.676249 | 0.668887 | 0.658983 | 0.663159 |
| 0.753561 | 0.711096 | 0.688095 | 0.68167 | 0.663677 | 0.688664 | 0.683462 | 0.67466 | 0.678175 |
| 0.771659 | 0.709219 | 0.687549 | 0.677003 | 0.664407 | 0.687296 | 0.679896 | 0.669517 | 0.67615 |
| 0.734792 | 0.702048 | 0.679426 | 0.671199 | 0.657215 | 0.678616 | 0.667535 | 0.663586 | 0.658786 |
| 0.740488 | 0.70404 | 0.67709 | 0.672427 | 0.651811 | 0.679775 | 0.671157 | 0.663456 | 0.668119 |
| 0.752012 | 0.704658 | 0.68108 | 0.671384 | 0.659946 | 0.680667 | 0.67352 | 0.666511 | 0.666515 |
| 0.741863 | 0.702845 | 0.676321 | 0.669027 | 0.647372 | 0.675807 | 0.666294 | 0.661477 | 0.669105 |
| 0.735409 | 0.69941 | 0.673297 | 0.666884 | 0.646357 | 0.669058 | 0.663741 | 0.657709 | 0.657725 |
| 0.730769 | 0.703343 | 0.674111 | 0.666108 | 0.652888 | 0.675757 | 0.665645 | 0.658274 | 0.658871 |
| 0.744569 | 0.705758 | 0.676314 | 0.677645 | 0.661488 | 0.675335 | 0.669782 | 0.659522 | 0.666011 |
| 0.740146 | 0.704803 | 0.674785 | 0.671125 | 0.653098 | 0.674276 | 0.667414 | 0.662388 | 0.665599 |
| 0.733172 | 0.702251 | 0.677518 | 0.669027 | 0.652111 | 0.673458 | 0.661759 | 0.656597 | 0.660466 |
| 0.728537 | 0.701076 | 0.678462 | 0.668836 | 0.663152 | 0.674884 | 0.669481 | 0.662619 | 0.663101 |
| 0.737321 | 0.702826 | 0.676939 | 0.663944 | 0.657336 | 0.673659 | 0.665722 | 0.65573 | 0.664863 |
| 0.736557 | 0.70294 | 0.67728 | 0.668019 | 0.65529 | 0.680549 | 0.669661 | 0.658856 | 0.664945 |
| 0.735126 | 0.703824 | 0.67586 | 0.6726 | 0.658962 | 0.675824 | 0.668105 | 0.661656 | 0.666767 |
| 0.739251 | 0.705936 | 0.677173 | 0.668621 | 0.658891 | 0.680452 | 0.666378 | 0.662088 | 0.665812 |
| 0.72914 | 0.704344 | 0.681448 | 0.672194 | 0.66558 | 0.678802 | 0.672813 | 0.666151 | 0.668644 |
| 0.778452 | 0.705838 | 0.68367 | 0.671087 | 0.65601 | 0.682328 | 0.670629 | 0.664252 | 0.669177 |
| 0.722547 | 0.701329 | 0.669264 | 0.663018 | 0.647965 | 0.673672 | 0.665152 | 0.655466 | 0.661428 |
| 0.732591 | 0.701703 | 0.679986 | 0.666493 | 0.661453 | 0.678713 | 0.672768 | 0.66333 | 0.661698 |
| 0.748374 | 0.703414 | 0.678308 | 0.67157 | 0.659331 | 0.681117 | 0.669256 | 0.663809 | 0.667674 |
| 0.739881 | 0.705586 | 0.68033 | 0.671579 | 0.65666 | 0.68034 | 0.674065 | 0.660983 | 0.671714 |
| 0.735006 | 0.70439 | 0.682547 | 0.674156 | 0.653314 | 0.680264 | 0.671053 | 0.663621 | 0.67082 |
| 0.736065 | 0.704395 | 0.677758 | 0.671877 | 0.649735 | 0.678755 | 0.669824 | 0.660268 | 0.666567 |
| 0.728848 | 0.702031 | 0.676623 | 0.669861 | 0.654031 | 0.6795 | 0.669463 | 0.655765 | 0.663008 |
| 0.717292 | 0.702501 | 0.676151 | 0.671581 | 0.653536 | 0.67655 | 0.668597 | 0.65898 | 0.663462 |
| 0.739163 | 0.70614 | 0.683084 | 0.675715 | 0.652151 | 0.682832 | 0.677254 | 0.669651 | 0.6727 |
| 0.768812 | 0.707695 | 0.682474 | 0.67594 | 0.661276 | 0.682121 | 0.674557 | 0.665391 | 0.670934 |
| 0.734611 | 0.701832 | 0.676667 | 0.67499 | 0.656769 | 0.676827 | 0.674453 | 0.665043 | 0.662291 |
| 0.742396 | 0.707543 | 0.682004 | 0.674352 | 0.654832 | 0.678888 | 0.677411 | 0.668064 | 0.673144 |
| 0.72965 | 0.710238 | 0.678907 | 0.676667 | 0.658353 | 0.684355 | 0.674809 | 0.6626 | 0.666464 |
| 0.732928 | 0.705996 | 0.674975 | 0.67324 | 0.654155 | 0.677372 | 0.669581 | 0.664417 | 0.664211 |
| 0.729755 | 0.706728 | 0.679638 | 0.673657 | 0.661145 | 0.678044 | 0.66708 | 0.660409 | 0.665408 |
| 0.734547 | 0.699381 | 0.670822 | 0.666368 | 0.645551 | 0.674929 | 0.67127 | 0.654914 | 0.661844 |
| 0.744217 | 0.682174 | 0.671129 | 0.66597 | 0.65118 | 0.671846 | 0.667023 | 0.656738 | 0.666027 |
| 0.753948 | 0.686845 | 0.676246 | 0.668345 | 0.656916 | 0.67378 | 0.673687 | 0.668048 | 0.665476 |
| 0.723421 | 0.681045 | 0.666625 | 0.65938 | 0.655553 | 0.669754 | 0.66298 | 0.659019 | 0.65868 |
| 0.734835 | 0.680312 | 0.668348 | 0.671747 | 0.654538 | 0.668319 | 0.666905 | 0.651389 | 0.659028 |
| 0.730353 | 0.663503 | 0.656861 | 0.65907 | 0.64558 | 0.653311 | 0.647921 | 0.641049 | 0.651261 |
| 0.740346 | 0.666344 | 0.66019 | 0.66734 | 0.660501 | 0.661688 | 0.658335 | 0.647461 | 0.652044 |
| 0.736754 | 0.669059 | 0.662354 | 0.660552 | 0.649482 | 0.658934 | 0.655838 | 0.651078 | 0.650757 |
| 0.738656 | 0.665225 | 0.658145 | 0.659146 | 0.656501 | 0.662574 | 0.659258 | 0.657273 | 0.654806 |
| 0.74502 | 0.663192 | 0.666907 | 0.662575 | 0.648638 | 0.662106 | 0.661817 | 0.651329 | 0.66123 |
| 0.734476 | 0.669335 | 0.669427 | 0.673518 | 0.656899 | 0.671371 | 0.668931 | 0.659351 | 0.662274 |

| | | | | | | | | |
|---|---|---|---|---|---|---|---|---|
| 0.752056 | 0.671469 | 0.66486 | 0.658901 | 0.655101 | 0.667476 | 0.665348 | 0.661775 | 0.658758 |
| 0.731627 | 0.667703 | 0.669998 | 0.670052 | 0.653589 | 0.668995 | 0.667362 | 0.662954 | 0.665469 |
| 0.744831 | 0.666312 | 0.664583 | 0.663497 | 0.662315 | 0.67035 | 0.663252 | 0.662191 | 0.663361 |
| 0.728722 | 0.659348 | 0.660146 | 0.662415 | 0.64488 | 0.661027 | 0.660196 | 0.647979 | 0.654871 |
| 0.744636 | 0.667481 | 0.662393 | 0.665267 | 0.652862 | 0.658145 | 0.653608 | 0.652796 | 0.653275 |
| 0.767189 | 0.66484 | 0.6688 | 0.661612 | 0.650929 | 0.670132 | 0.66451 | 0.653461 | 0.660179 |
| 0.737697 | 0.664595 | 0.668429 | 0.666372 | 0.657246 | 0.666774 | 0.666382 | 0.662124 | 0.655112 |
| 0.743246 | 0.659474 | 0.65628 | 0.662985 | 0.648649 | 0.658796 | 0.657435 | 0.64836 | 0.659167 |
| 0.729335 | 0.667402 | 0.661724 | 0.656807 | 0.658326 | 0.661629 | 0.658164 | 0.650458 | 0.661248 |
| 0.731185 | 0.66285 | 0.655822 | 0.655535 | 0.652132 | 0.664432 | 0.65339 | 0.655727 | 0.649189 |
| 0.753927 | 0.655585 | 0.661155 | 0.658005 | 0.649334 | 0.654808 | 0.656524 | 0.649875 | 0.650442 |
| 0.714857 | 0.693116 | 0.663721 | 0.664395 | 0.645778 | 0.663284 | 0.659572 | 0.652436 | 0.65546 |
| 0.726537 | 0.703823 | 0.67957 | 0.669778 | 0.655068 | 0.672316 | 0.669793 | 0.660665 | 0.664911 |
| 0.723666 | 0.685219 | 0.674944 | 0.674847 | 0.660259 | 0.682119 | 0.671431 | 0.666149 | 0.667854 |
| 0.735936 | 0.711098 | 0.685402 | 0.6783 | 0.662938 | 0.690948 | 0.67639 | 0.669723 | 0.670089 |
| 0.732285 | 0.700706 | 0.672882 | 0.665668 | 0.66094 | 0.674385 | 0.667657 | 0.65414 | 0.666013 |
| 0.727572 | 0.702776 | 0.68282 | 0.672425 | 0.656 | 0.678599 | 0.670436 | 0.664988 | 0.66587 |
| 0.724386 | 0.709028 | 0.686319 | 0.675304 | 0.663667 | 0.688039 | 0.677311 | 0.67275 | 0.678185 |
| 0.732811 | 0.691634 | 0.667734 | 0.655045 | 0.643973 | 0.664898 | 0.65453 | 0.654184 | 0.65332 |
| 0.728699 | 0.7165 | 0.685014 | 0.679049 | 0.660151 | 0.687164 | 0.676608 | 0.669148 | 0.671542 |
| 0.744637 | 0.688294 | 0.682885 | 0.683121 | 0.668581 | 0.685141 | 0.679795 | 0.673282 | 0.675054 |
| 0.727021 | 0.707792 | 0.683919 | 0.674879 | 0.658862 | 0.683228 | 0.670504 | 0.666 | 0.665811 |
| 0.723231 | 0.708684 | 0.682685 | 0.673195 | 0.665108 | 0.686777 | 0.675679 | 0.668825 | 0.671308 |
| 0.720996 | 0.681099 | 0.676623 | 0.674616 | 0.670705 | 0.676035 | 0.669832 | 0.669449 | 0.66447 |
| 0.704541 | 0.677094 | 0.660275 | 0.650221 | 0.63575 | 0.658086 | 0.655764 | 0.639717 | 0.647796 |
| 0.738847 | 0.696743 | 0.688387 | 0.685813 | 0.675064 | 0.687126 | 0.68977 | 0.682685 | 0.681988 |
| 0.713856 | 0.703331 | 0.671289 | 0.66463 | 0.6501 | 0.670921 | 0.663729 | 0.656495 | 0.661536 |
| 0.723708 | 0.711876 | 0.681828 | 0.675521 | 0.661651 | 0.68235 | 0.671147 | 0.664018 | 0.667068 |
| 0.706208 | 0.685501 | 0.65251 | 0.644641 | 0.636018 | 0.656322 | 0.648792 | 0.638556 | 0.641636 |
| 0.732127 | 0.71632 | 0.69211 | 0.68323 | 0.667732 | 0.692316 | 0.684295 | 0.683677 | 0.679581 |
| 0.730414 | 0.717033 | 0.690439 | 0.684756 | 0.670165 | 0.692153 | 0.683149 | 0.676632 | 0.679575 |
| 0.710709 | 0.694288 | 0.671014 | 0.660336 | 0.649378 | 0.667375 | 0.662591 | 0.651336 | 0.662451 |
| 0.714024 | 0.690915 | 0.667858 | 0.658399 | 0.64519 | 0.670282 | 0.658725 | 0.644518 | 0.654951 |
| 0.713582 | 0.71859 | 0.695696 | 0.6883 | 0.670839 | 0.695598 | 0.683372 | 0.678089 | 0.681241 |
| 0.747696 | 0.717191 | 0.689889 | 0.68176 | 0.662963 | 0.69442 | 0.688046 | 0.673479 | 0.676996 |
| 0.719832 | 0.720298 | 0.69604 | 0.685737 | 0.673788 | 0.696613 | 0.682824 | 0.681356 | 0.678073 |
| 0.68935 | 0.669849 | 0.647921 | 0.638586 | 0.622531 | 0.648905 | 0.639223 | 0.638266 | 0.634253 |
| 0.709019 | 0.71538 | 0.690766 | 0.681599 | 0.675729 | 0.686125 | 0.683063 | 0.674354 | 0.680539 |
| 0.672225 | 0.673789 | 0.640134 | 0.63591 | 0.621354 | 0.646486 | 0.635265 | 0.62637 | 0.631192 |
| 0.671011 | 0.664104 | 0.636121 | 0.624106 | 0.61438 | 0.634201 | 0.62547 | 0.618222 | 0.617024 |
| 0.723696 | 0.71592 | 0.693733 | 0.681205 | 0.66839 | 0.68611 | 0.685484 | 0.674977 | 0.68052 |
| 0.678274 | 0.666508 | 0.635194 | 0.62761 | 0.618687 | 0.636335 | 0.633283 | 0.623969 | 0.626314 |
| 0.66379 | 0.656925 | 0.631077 | 0.618838 | 0.605091 | 0.624736 | 0.62388 | 0.612107 | 0.616464 |
| 0.656271 | 0.654048 | 0.627185 | 0.614597 | 0.60446 | 0.622107 | 0.619661 | 0.605672 | 0.613234 |
| 0.719285 | 0.72725 | 0.699842 | 0.694658 | 0.679296 | 0.700146 | 0.695847 | 0.690326 | 0.687614 |
| 0.651492 | 0.656297 | 0.627196 | 0.616302 | 0.604146 | 0.625645 | 0.623906 | 0.611167 | 0.618285 |
| 0.724012 | 0.719863 | 0.69299 | 0.688232 | 0.674236 | 0.694692 | 0.689271 | 0.684194 | 0.684848 |

| | | | | | | | | |
|---|---|---|---|---|---|---|---|---|
| 0.719455 | 0.724731 | 0.697612 | 0.693614 | 0.67439 | 0.699052 | 0.690132 | 0.686034 | 0.685495 |
| 0.715371 | 0.720217 | 0.692969 | 0.685688 | 0.670777 | 0.694763 | 0.68302 | 0.679048 | 0.68445 |
| 0.707185 | 0.738063 | 0.714707 | 0.710371 | 0.699134 | 0.719134 | 0.710855 | 0.701624 | 0.706758 |
| 0.642353 | 0.644907 | 0.614057 | 0.60622 | 0.591508 | 0.614914 | 0.608015 | 0.601075 | 0.601581 |
| 0.654372 | 0.649126 | 0.624877 | 0.617125 | 0.605679 | 0.622881 | 0.619359 | 0.608466 | 0.617191 |
| 0.724132 | 0.729873 | 0.703329 | 0.694693 | 0.675184 | 0.703973 | 0.697878 | 0.690602 | 0.695407 |
| 0.639658 | 0.64976 | 0.623564 | 0.61444 | 0.598257 | 0.623356 | 0.613722 | 0.600509 | 0.612333 |
| 0.68256 | 0.72553 | 0.704965 | 0.700533 | 0.687923 | 0.700711 | 0.695316 | 0.691995 | 0.694735 |
| 0.712157 | 0.728325 | 0.702638 | 0.695292 | 0.680755 | 0.705006 | 0.700966 | 0.692224 | 0.695798 |
| 0.670539 | 0.726962 | 0.704169 | 0.700416 | 0.683892 | 0.704443 | 0.692796 | 0.693986 | 0.69435 |
| 0.698494 | 0.727465 | 0.704958 | 0.694105 | 0.682141 | 0.702591 | 0.695465 | 0.688314 | 0.695833 |
| 0.705426 | 0.732484 | 0.706667 | 0.756757 | 0.79661 | 0.755245 | 0.774648 | 0.766423 | 0.761905 |
| 0.648774 | 0.745429 | 0.752346 | 0.751465 | 0.745963 | 0.741146 | 0.736854 | 0.737533 | 0.747276 |
| 0.647234 | 0.743777 | 0.738478 | 0.745154 | 0.749213 | 0.740308 | 0.730649 | 0.725789 | 0.72872 |
| 0.646109 | 0.754118 | 0.75234 | 0.758756 | 0.745986 | 0.748758 | 0.737673 | 0.731952 | 0.743024 |
| 0.654339 | 0.757931 | 0.755398 | 0.745892 | 0.749777 | 0.747294 | 0.743337 | 0.734372 | 0.736418 |
| 0.583743 | 0.59322 | 0.581774 | 0.57351 | 0.563216 | 0.584015 | 0.577008 | 0.567972 | 0.574612 |
| 0.647348 | 0.752169 | 0.753739 | 0.756053 | 0.743319 | 0.749811 | 0.737825 | 0.73309 | 0.739778 |
| 0.691891 | 0.730443 | 0.709256 | 0.703876 | 0.68699 | 0.711658 | 0.701236 | 0.700141 | 0.694811 |
| 0.686163 | 0.734826 | 0.706406 | 0.705645 | 0.683537 | 0.707495 | 0.706883 | 0.702498 | 0.703472 |
| 0.693375 | 0.733965 | 0.711761 | 0.711466 | 0.687968 | 0.708852 | 0.703313 | 0.697319 | 0.702657 |
| 0.548029 | 0.591094 | 0.553585 | 0.541543 | 0.529511 | 0.554186 | 0.547453 | 0.53863 | 0.539671 |
| 0.526757 | 0.57809 | 0.542567 | 0.52798 | 0.515307 | 0.545256 | 0.533477 | 0.525281 | 0.531638 |
| 0.654067 | 0.755411 | 0.737064 | 0.735053 | 0.717728 | 0.727676 | 0.72125 | 0.721725 | 0.722335 |
| 0.654506 | 0.754531 | 0.735548 | 0.722823 | 0.708994 | 0.729472 | 0.725565 | 0.7251 | 0.720563 |
| 0.641236 | 0.753393 | 0.731101 | 0.723148 | 0.717662 | 0.728516 | 0.717341 | 0.720595 | 0.710825 |
| 0.645298 | 0.761146 | 0.735695 | 0.735207 | 0.718738 | 0.735373 | 0.72544 | 0.716754 | 0.718556 |
| 0.658969 | 0.758683 | 0.73331 | 0.729475 | 0.717138 | 0.73863 | 0.723347 | 0.723316 | 0.725109 |
| 0.659372 | 0.759575 | 0.746601 | 0.737298 | 0.73469 | 0.738723 | 0.732214 | 0.732417 | 0.731868 |
| 0.658946 | 0.758128 | 0.739785 | 0.734131 | 0.71809 | 0.739965 | 0.73313 | 0.729572 | 0.729341 |
| 0.659112 | 0.758392 | 0.740541 | 0.735951 | 0.723364 | 0.736312 | 0.729963 | 0.720588 | 0.734375 |
| 0.665299 | 0.76532 | 0.746146 | 0.742965 | 0.721657 | 0.740732 | 0.735031 | 0.726595 | 0.731206 |
| 0.661881 | 0.757628 | 0.736511 | 0.728094 | 0.714233 | 0.733746 | 0.726748 | 0.721082 | 0.724359 |
| 0.660876 | 0.754899 | 0.729298 | 0.732071 | 0.709254 | 0.728575 | 0.729212 | 0.72 | 0.727216 |
| 0.662491 | 0.762118 | 0.7415 | 0.733008 | 0.719657 | 0.741066 | 0.735839 | 0.725693 | 0.733103 |
| 0.650802 | 0.756016 | 0.72915 | 0.733523 | 0.709487 | 0.738584 | 0.727168 | 0.722054 | 0.724653 |
| 0.658266 | 0.753627 | 0.734279 | 0.733072 | 0.716261 | 0.738571 | 0.729726 | 0.726925 | 0.729831 |
| 0.65611 | 0.758668 | 0.740922 | 0.730541 | 0.725093 | 0.738471 | 0.727996 | 0.719549 | 0.730589 |
| 0.653116 | 0.758393 | 0.738264 | 0.724759 | 0.717549 | 0.731044 | 0.726485 | 0.72255 | 0.731067 |
| 0.660283 | 0.75309 | 0.728425 | 0.726051 | 0.713653 | 0.73235 | 0.728898 | 0.722933 | 0.724676 |
| 0.655504 | 0.745541 | 0.72684 | 0.742698 | 0.732194 | 0.737024 | 0.724056 | 0.724942 | 0.734769 |
| 0.660126 | 0.736166 | 0.734439 | 0.745132 | 0.751238 | 0.745335 | 0.734342 | 0.731384 | 0.737869 |
| 0.662706 | 0.760003 | 0.737536 | 0.732553 | 0.711802 | 0.733943 | 0.730917 | 0.719163 | 0.722257 |
| 0.665712 | 0.765979 | 0.744897 | 0.742202 | 0.719298 | 0.74443 | 0.735765 | 0.727693 | 0.732084 |
| 0.650769 | 0.714231 | 0.715648 | 0.72015 | 0.7122 | 0.715356 | 0.723409 | 0.711754 | 0.721756 |
| 0.657843 | 0.762659 | 0.743226 | 0.733181 | 0.71837 | 0.73911 | 0.73217 | 0.72358 | 0.722811 |
| 0.647667 | 0.752944 | 0.731606 | 0.726595 | 0.705524 | 0.731177 | 0.731836 | 0.719226 | 0.723218 |

| | | | | | | | | |
|---|---|---|---|---|---|---|---|---|
| 0.662698 | 0.758452 | 0.739439 | 0.733171 | 0.714496 | 0.728874 | 0.731177 | 0.727157 | 0.724887 |
| 0.653995 | 0.76127 | 0.73997 | 0.731498 | 0.720832 | 0.73405 | 0.733503 | 0.725249 | 0.726313 |
| 0.657777 | 0.76188 | 0.737813 | 0.738436 | 0.718954 | 0.740902 | 0.73779 | 0.726996 | 0.73327 |
| 0.655075 | 0.761942 | 0.740444 | 0.733649 | 0.718281 | 0.732946 | 0.729255 | 0.72908 | 0.724063 |
| 0.668267 | 0.763034 | 0.738949 | 0.744557 | 0.724755 | 0.7444 | 0.739003 | 0.733704 | 0.734691 |
| 0.661328 | 0.75503 | 0.737619 | 0.734884 | 0.717491 | 0.735746 | 0.729681 | 0.72359 | 0.719017 |
| 0.665305 | 0.757449 | 0.736927 | 0.732741 | 0.720044 | 0.732463 | 0.727041 | 0.730654 | 0.720415 |
| 0.648038 | 0.749848 | 0.729532 | 0.723605 | 0.709098 | 0.729002 | 0.724474 | 0.706454 | 0.716793 |
| 0.650213 | 0.754396 | 0.735748 | 0.730692 | 0.708241 | 0.736536 | 0.730962 | 0.723624 | 0.722144 |
| 0.654046 | 0.75435 | 0.733677 | 0.725645 | 0.703348 | 0.728326 | 0.726284 | 0.713898 | 0.714286 |
| 0.66096 | 0.763232 | 0.737661 | 0.728166 | 0.718067 | 0.736114 | 0.726532 | 0.72603 | 0.72313 |
| 0.653846 | 0.756231 | 0.736352 | 0.732619 | 0.721154 | 0.726887 | 0.727273 | 0.718464 | 0.724346 |
| 0.647597 | 0.750869 | 0.723064 | 0.720071 | 0.707976 | 0.725228 | 0.721426 | 0.714172 | 0.716418 |
| 0.65245 | 0.753008 | 0.729865 | 0.728821 | 0.709471 | 0.731315 | 0.718363 | 0.720495 | 0.713698 |
| 0.662405 | 0.763765 | 0.734218 | 0.732984 | 0.713104 | 0.738893 | 0.727701 | 0.72398 | 0.724047 |
| 0.655822 | 0.761349 | 0.74177 | 0.738005 | 0.726651 | 0.737999 | 0.731368 | 0.727591 | 0.73004 |
| 0.660724 | 0.755028 | 0.741436 | 0.733006 | 0.718523 | 0.739862 | 0.732284 | 0.728264 | 0.728838 |
| 0.651507 | 0.748948 | 0.73345 | 0.726654 | 0.711517 | 0.727124 | 0.714213 | 0.715083 | 0.721586 |
| 0.653481 | 0.753472 | 0.734566 | 0.729369 | 0.716809 | 0.73481 | 0.722364 | 0.723581 | 0.719187 |
| 0.654229 | 0.755557 | 0.735671 | 0.730161 | 0.723042 | 0.736541 | 0.725819 | 0.71875 | 0.726209 |
| 0.646315 | 0.752118 | 0.727101 | 0.720985 | 0.711078 | 0.730255 | 0.714785 | 0.715207 | 0.713205 |
| 0.651908 | 0.752948 | 0.730616 | 0.728721 | 0.714847 | 0.733255 | 0.723696 | 0.717374 | 0.719767 |
| 0.655491 | 0.762381 | 0.736392 | 0.737288 | 0.723954 | 0.738949 | 0.728404 | 0.725512 | 0.728342 |
| 0.646787 | 0.757602 | 0.737111 | 0.73346 | 0.716803 | 0.730291 | 0.719974 | 0.722682 | 0.718998 |
| 0.650135 | 0.758235 | 0.733614 | 0.723627 | 0.717002 | 0.733193 | 0.716357 | 0.718929 | 0.717676 |
| 0.651055 | 0.754501 | 0.735129 | 0.729495 | 0.715642 | 0.734452 | 0.727706 | 0.726356 | 0.721145 |
| 0.646667 | 0.747542 | 0.722756 | 0.721538 | 0.706257 | 0.724862 | 0.714267 | 0.713324 | 0.718021 |
| 0.650822 | 0.754771 | 0.728776 | 0.729527 | 0.714179 | 0.729365 | 0.714249 | 0.724077 | 0.71201 |
| 0.646975 | 0.753548 | 0.730079 | 0.724133 | 0.711528 | 0.731753 | 0.719173 | 0.721576 | 0.714397 |
| 0.654337 | 0.756267 | 0.737033 | 0.735698 | 0.713092 | 0.731616 | 0.725042 | 0.719217 | 0.719285 |
| 0.656484 | 0.756408 | 0.736114 | 0.729792 | 0.717896 | 0.735915 | 0.728273 | 0.724769 | 0.726021 |
| 0.661112 | 0.755723 | 0.733536 | 0.723372 | 0.720917 | 0.733373 | 0.725198 | 0.72175 | 0.724209 |
| 0.667321 | 0.761018 | 0.742025 | 0.730749 | 0.72211 | 0.734948 | 0.730979 | 0.729157 | 0.732423 |
| 0.663002 | 0.76461 | 0.745878 | 0.741544 | 0.724581 | 0.746656 | 0.73461 | 0.729084 | 0.733877 |
| 0.647048 | 0.750043 | 0.72996 | 0.724902 | 0.711078 | 0.728175 | 0.719954 | 0.724972 | 0.716698 |
| 0.661558 | 0.752042 | 0.733934 | 0.727581 | 0.720112 | 0.741278 | 0.728248 | 0.724552 | 0.730606 |
| 0.6485 | 0.752921 | 0.734233 | 0.723947 | 0.718901 | 0.732547 | 0.724872 | 0.721418 | 0.721937 |
| 0.653764 | 0.752528 | 0.735006 | 0.730421 | 0.714867 | 0.734734 | 0.725297 | 0.72356 | 0.724509 |
| 0.649186 | 0.748471 | 0.724748 | 0.71754 | 0.706832 | 0.726317 | 0.710968 | 0.707567 | 0.713835 |
| 0.650313 | 0.749911 | 0.728073 | 0.727521 | 0.716462 | 0.733136 | 0.727743 | 0.721027 | 0.721183 |
| 0.644065 | 0.743281 | 0.72145 | 0.715466 | 0.699397 | 0.721845 | 0.715109 | 0.714704 | 0.719528 |
| 0.651589 | 0.749007 | 0.732391 | 0.731761 | 0.712786 | 0.72736 | 0.721521 | 0.720273 | 0.726143 |
| 0.647999 | 0.741889 | 0.7233 | 0.713832 | 0.709358 | 0.719342 | 0.715699 | 0.706446 | 0.716793 |
| 0.644071 | 0.742642 | 0.724729 | 0.724768 | 0.711391 | 0.730128 | 0.717137 | 0.714451 | 0.717687 |
| 0.656545 | 0.753523 | 0.733905 | 0.727849 | 0.726703 | 0.735035 | 0.728069 | 0.724384 | 0.723299 |
| 0.650588 | 0.743379 | 0.725297 | 0.714494 | 0.710327 | 0.725067 | 0.710211 | 0.713624 | 0.719709 |

| | | | | | | | | |
|---|---|---|---|---|---|---|---|---|
| 0.659079 | 0.755999 | 0.733909 | 0.72891 | 0.704815 | 0.735425 | 0.728278 | 0.725832 | 0.726062 |
| 0.659494 | 0.760135 | 0.74285 | 0.733569 | 0.723034 | 0.740628 | 0.735788 | 0.738191 | 0.739453 |
| 0.65219 | 0.756025 | 0.733396 | 0.732563 | 0.710877 | 0.735131 | 0.725884 | 0.728097 | 0.725681 |
| 0.656395 | 0.759561 | 0.736313 | 0.729111 | 0.7186 | 0.73454 | 0.7216 | 0.719174 | 0.72346 |
| 0.667563 | 0.762839 | 0.752671 | 0.741062 | 0.724907 | 0.740322 | 0.735759 | 0.731419 | 0.735221 |
| 0.658359 | 0.757159 | 0.744154 | 0.738719 | 0.721214 | 0.745236 | 0.735042 | 0.728497 | 0.73723 |
| 0.651657 | 0.75124 | 0.739177 | 0.728834 | 0.71857 | 0.735936 | 0.731291 | 0.726411 | 0.726292 |
| 0.653442 | 0.757508 | 0.735342 | 0.733433 | 0.7228 | 0.732303 | 0.728298 | 0.723996 | 0.725246 |
| 0.662818 | 0.757816 | 0.737512 | 0.729117 | 0.721442 | 0.73886 | 0.728672 | 0.72484 | 0.730274 |
| 0.652716 | 0.754613 | 0.732543 | 0.724231 | 0.712435 | 0.725029 | 0.726876 | 0.722178 | 0.722526 |
| 0.655111 | 0.756699 | 0.734576 | 0.732661 | 0.719582 | 0.73187 | 0.727319 | 0.719931 | 0.717468 |
| 0.658865 | 0.757817 | 0.737232 | 0.734699 | 0.723735 | 0.740361 | 0.737579 | 0.732188 | 0.729785 |
| 0.655356 | 0.76497 | 0.744387 | 0.730028 | 0.717806 | 0.734511 | 0.728041 | 0.729618 | 0.722845 |
| 0.657041 | 0.761076 | 0.738353 | 0.732676 | 0.715257 | 0.740806 | 0.731056 | 0.731322 | 0.727287 |
| 0.657666 | 0.763624 | 0.743348 | 0.74207 | 0.718785 | 0.739418 | 0.734949 | 0.733037 | 0.731886 |
| 0.655401 | 0.763142 | 0.740922 | 0.735262 | 0.724125 | 0.739817 | 0.730008 | 0.72624 | 0.726603 |
| 0.661522 | 0.763255 | 0.74111 | 0.734057 | 0.725263 | 0.737413 | 0.724408 | 0.727344 | 0.726233 |
| 0.64936 | 0.758007 | 0.742738 | 0.739838 | 0.724157 | 0.73907 | 0.725572 | 0.725157 | 0.722171 |
| 0.659605 | 0.765616 | 0.745533 | 0.735005 | 0.715683 | 0.741838 | 0.734972 | 0.735322 | 0.730069 |
| 0.650709 | 0.763521 | 0.744281 | 0.73481 | 0.72246 | 0.740517 | 0.738434 | 0.733078 | 0.735837 |
| 0.658506 | 0.757693 | 0.731036 | 0.731967 | 0.720643 | 0.738001 | 0.723647 | 0.726479 | 0.72818 |
| 0.65615 | 0.760903 | 0.738092 | 0.732467 | 0.723377 | 0.737106 | 0.738951 | 0.736255 | 0.732988 |
| 0.655063 | 0.759365 | 0.733402 | 0.7351 | 0.727374 | 0.735737 | 0.728811 | 0.732593 | 0.721009 |
| 0.654855 | 0.763399 | 0.744224 | 0.742009 | 0.727222 | 0.745822 | 0.734866 | 0.735219 | 0.729373 |
| 0.657259 | 0.762018 | 0.737024 | 0.734503 | 0.728653 | 0.742405 | 0.732646 | 0.732103 | 0.729839 |
| 0.659975 | 0.758544 | 0.746078 | 0.733965 | 0.726698 | 0.744306 | 0.728731 | 0.730506 | 0.724828 |
| 0.657123 | 0.763473 | 0.734852 | 0.730372 | 0.72181 | 0.740288 | 0.731536 | 0.727504 | 0.728732 |
| 0.654953 | 0.763512 | 0.74457 | 0.723798 | 0.731784 | 0.740248 | 0.737178 | 0.734442 | 0.729684 |
| 0.660515 | 0.767147 | 0.745467 | 0.737835 | 0.719993 | 0.748306 | 0.735313 | 0.732242 | 0.734424 |
| 0.664332 | 0.765933 | 0.751808 | 0.741173 | 0.72306 | 0.746626 | 0.735565 | 0.738633 | 0.7317 |
| 0.655958 | 0.755847 | 0.736626 | 0.736936 | 0.724883 | 0.742171 | 0.72938 | 0.729639 | 0.722561 |
| 0.659936 | 0.764471 | 0.74122 | 0.740766 | 0.730381 | 0.75227 | 0.738368 | 0.734335 | 0.727798 |
| 0.660613 | 0.766345 | 0.746093 | 0.742194 | 0.724465 | 0.747009 | 0.732203 | 0.73408 | 0.731493 |
| 0.659548 | 0.763545 | 0.744117 | 0.739386 | 0.724028 | 0.742844 | 0.728133 | 0.73199 | 0.725971 |
| 0.664336 | 0.762831 | 0.74435 | 0.744416 | 0.727765 | 0.749466 | 0.729674 | 0.736944 | 0.731108 |
| 0.662403 | 0.764669 | 0.746036 | 0.734452 | 0.728249 | 0.752161 | 0.736849 | 0.735366 | 0.729219 |
| 0.664456 | 0.765712 | 0.745812 | 0.734318 | 0.727138 | 0.749737 | 0.740812 | 0.737794 | 0.73569 |
| 0.65996 | 0.765943 | 0.745621 | 0.742217 | 0.728499 | 0.750148 | 0.730507 | 0.73597 | 0.732067 |
| 0.66087 | 0.766066 | 0.747113 | 0.733154 | 0.7175 | 0.741337 | 0.733095 | 0.73363 | 0.732394 |
| 0.662312 | 0.762485 | 0.746642 | 0.736162 | 0.732022 | 0.746131 | 0.728962 | 0.729673 | 0.733449 |
| 0.663186 | 0.76641 | 0.750787 | 0.732519 | 0.728471 | 0.743846 | 0.731122 | 0.733567 | 0.731278 |
| 0.662778 | 0.7705 | 0.744132 | 0.738395 | 0.725831 | 0.745741 | 0.735808 | 0.730831 | 0.737024 |
| 0.658228 | 0.772458 | 0.748052 | 0.740911 | 0.718762 | 0.742638 | 0.737279 | 0.732973 | 0.740286 |
| 0.662845 | 0.760325 | 0.744076 | 0.741255 | 0.721129 | 0.748434 | 0.734385 | 0.729466 | 0.729925 |
| 0.658506 | 0.764563 | 0.743238 | 0.739644 | 0.723906 | 0.747499 | 0.732212 | 0.729459 | 0.728742 |
| 0.64992 | 0.763261 | 0.743744 | 0.733623 | 0.720075 | 0.744633 | 0.728516 | 0.72473 | 0.7269 |

| | | | | | | | | |
|---|---|---|---|---|---|---|---|---|
| 0.662802 | 0.750024 | 0.740671 | 0.726671 | 0.725195 | 0.740989 | 0.729204 | 0.728822 | 0.727092 |
| 0.691042 | 0.743625 | 0.723409 | 0.714608 | 0.6992 | 0.726696 | 0.717358 | 0.711027 | 0.71356 |
| 0.653317 | 0.765501 | 0.731619 | 0.733897 | 0.709767 | 0.739485 | 0.730309 | 0.727097 | 0.725959 |
| 0.656438 | 0.763243 | 0.740201 | 0.74452 | 0.730534 | 0.740792 | 0.734856 | 0.731773 | 0.732504 |
| 0.65708 | 0.761968 | 0.730928 | 0.735927 | 0.708738 | 0.738975 | 0.729232 | 0.718659 | 0.721835 |
| 0.660633 | 0.768312 | 0.738748 | 0.737481 | 0.722473 | 0.745721 | 0.732866 | 0.726498 | 0.721029 |
| 0.655228 | 0.7628 | 0.748504 | 0.746304 | 0.724484 | 0.742457 | 0.740971 | 0.739074 | 0.731058 |
| 0.659124 | 0.756234 | 0.737135 | 0.73543 | 0.71163 | 0.734037 | 0.72673 | 0.72772 | 0.720378 |
| 0.644951 | 0.767328 | 0.74598 | 0.745813 | 0.728842 | 0.743868 | 0.733103 | 0.731327 | 0.72757 |
| 0.665286 | 0.761917 | 0.74368 | 0.738238 | 0.722295 | 0.743536 | 0.738651 | 0.734828 | 0.73066 |
| 0.658212 | 0.768023 | 0.743748 | 0.739566 | 0.725683 | 0.744621 | 0.729899 | 0.733504 | 0.724836 |
| 0.664393 | 0.760732 | 0.738309 | 0.735698 | 0.724022 | 0.743272 | 0.73729 | 0.7368 | 0.734034 |
| 0.66405 | 0.768499 | 0.746706 | 0.739824 | 0.733271 | 0.751345 | 0.734529 | 0.7317 | 0.735952 |
| 0.660106 | 0.764394 | 0.743215 | 0.734247 | 0.729775 | 0.74497 | 0.734785 | 0.727646 | 0.736505 |
| 0.658537 | 0.763888 | 0.739845 | 0.73984 | 0.728678 | 0.749793 | 0.731538 | 0.733355 | 0.731746 |
| 0.654846 | 0.765847 | 0.741018 | 0.735077 | 0.720666 | 0.743729 | 0.733147 | 0.725173 | 0.731228 |
| 0.66074 | 0.764335 | 0.739201 | 0.732926 | 0.720692 | 0.737659 | 0.740238 | 0.731453 | 0.736892 |
| 0.657782 | 0.764711 | 0.739574 | 0.737563 | 0.721373 | 0.735075 | 0.734407 | 0.731398 | 0.734402 |
| 0.660119 | 0.761544 | 0.754316 | 0.742786 | 0.731703 | 0.752485 | 0.736711 | 0.736102 | 0.735158 |
| 0.65471 | 0.760978 | 0.742032 | 0.738909 | 0.726828 | 0.743946 | 0.739966 | 0.732221 | 0.73295 |
| 0.662121 | 0.766 | 0.746106 | 0.744345 | 0.726664 | 0.744459 | 0.733651 | 0.735915 | 0.731566 |
| 0.642082 | 0.752802 | 0.738669 | 0.729561 | 0.725785 | 0.74306 | 0.726675 | 0.726381 | 0.721531 |
| 0.653402 | 0.754482 | 0.742637 | 0.737006 | 0.725219 | 0.743875 | 0.733798 | 0.727666 | 0.729353 |
| 0.654885 | 0.764966 | 0.746494 | 0.739655 | 0.728186 | 0.750471 | 0.734941 | 0.727258 | 0.730129 |
| 0.659173 | 0.767591 | 0.748734 | 0.747509 | 0.735355 | 0.748283 | 0.734974 | 0.734586 | 0.735971 |
| 0.653564 | 0.762667 | 0.743659 | 0.732798 | 0.723396 | 0.742016 | 0.731571 | 0.726836 | 0.734007 |
| 0.653498 | 0.762697 | 0.747246 | 0.739586 | 0.728032 | 0.744844 | 0.7282 | 0.734482 | 0.725087 |
| 0.660384 | 0.765887 | 0.754753 | 0.748029 | 0.728778 | 0.748775 | 0.733229 | 0.739832 | 0.732775 |
| 0.654691 | 0.766454 | 0.75012 | 0.736326 | 0.734771 | 0.745793 | 0.729865 | 0.730826 | 0.735922 |
| 0.736204 | 0.68944 | 0.672268 | 0.667748 | 0.650066 | 0.669239 | 0.664826 | 0.660458 | 0.652508 |
| 0.740011 | 0.700572 | 0.674427 | 0.665446 | 0.650458 | 0.670993 | 0.665627 | 0.655803 | 0.666768 |
| 0.725134 | 0.699043 | 0.66881 | 0.665709 | 0.649409 | 0.668741 | 0.661296 | 0.649801 | 0.655193 |
| 0.743105 | 0.692343 | 0.668642 | 0.664802 | 0.651141 | 0.670735 | 0.66497 | 0.651029 | 0.658323 |
| 0.733616 | 0.697981 | 0.67117 | 0.663392 | 0.645881 | 0.671894 | 0.668227 | 0.658816 | 0.661119 |
| 0.648915 | 0.767326 | 0.75 | 0.743969 | 0.73872 | 0.751824 | 0.733169 | 0.730312 | 0.729956 |
| 0.651621 | 0.769318 | 0.750235 | 0.747912 | 0.731657 | 0.748907 | 0.736406 | 0.728059 | 0.731988 |
| 0.652151 | 0.771361 | 0.751934 | 0.746598 | 0.73316 | 0.752466 | 0.733752 | 0.731591 | 0.738488 |
| 0.653221 | 0.770707 | 0.754029 | 0.751538 | 0.741911 | 0.755254 | 0.735665 | 0.733643 | 0.737157 |
| 0.645726 | 0.769284 | 0.745274 | 0.743024 | 0.737647 | 0.749148 | 0.732719 | 0.734272 | 0.734953 |
| 0.660246 | 0.770438 | 0.754717 | 0.752452 | 0.743912 | 0.758495 | 0.739192 | 0.735819 | 0.739943 |
| 0.65602 | 0.76832 | 0.753225 | 0.744164 | 0.733271 | 0.750532 | 0.733867 | 0.733002 | 0.729899 |
| 0.649822 | 0.769366 | 0.751123 | 0.739692 | 0.734694 | 0.754032 | 0.733497 | 0.73175 | 0.737413 |
| 0.646201 | 0.76653 | 0.750121 | 0.743371 | 0.736058 | 0.746441 | 0.728782 | 0.727065 | 0.729764 |
| 0.652841 | 0.762365 | 0.747417 | 0.747786 | 0.732318 | 0.743408 | 0.731041 | 0.728934 | 0.725932 |
| 0.655091 | 0.767919 | 0.748091 | 0.741891 | 0.732536 | 0.757601 | 0.735256 | 0.728731 | 0.737271 |
| 0.653535 | 0.774559 | 0.751419 | 0.74425 | 0.738109 | 0.757953 | 0.738058 | 0.739208 | 0.742161 |

| | | | | | | | | |
|---|---|---|---|---|---|---|---|---|
| 0.664309 | 0.777168 | 0.759671 | 0.746533 | 0.737318 | 0.753713 | 0.745343 | 0.735992 | 0.741864 |
| 0.65508 | 0.777467 | 0.757526 | 0.748631 | 0.740057 | 0.758261 | 0.739192 | 0.735422 | 0.743715 |
| 0.658884 | 0.770478 | 0.766438 | 0.751637 | 0.740299 | 0.751708 | 0.741644 | 0.740948 | 0.737893 |
| 0.650116 | 0.772131 | 0.746795 | 0.744561 | 0.727662 | 0.751449 | 0.729298 | 0.738011 | 0.725991 |
| 0.651529 | 0.769869 | 0.754116 | 0.752215 | 0.734648 | 0.753765 | 0.741952 | 0.74227 | 0.737337 |
| 0.65375 | 0.76934 | 0.754071 | 0.747634 | 0.741102 | 0.75656 | 0.737497 | 0.738877 | 0.73831 |
| 0.659624 | 0.776506 | 0.750917 | 0.748596 | 0.739621 | 0.757129 | 0.742422 | 0.740962 | 0.74517 |
| 0.658726 | 0.776149 | 0.753936 | 0.752286 | 0.752055 | 0.748984 | 0.73595 | 0.740656 | 0.737647 |
| 0.658651 | 0.77134 | 0.749615 | 0.744672 | 0.727835 | 0.751455 | 0.742786 | 0.742645 | 0.7407 |
| 0.651046 | 0.77297 | 0.752565 | 0.749658 | 0.735835 | 0.753519 | 0.737555 | 0.737383 | 0.736701 |
| 0.653522 | 0.769995 | 0.752961 | 0.746097 | 0.728611 | 0.751424 | 0.735522 | 0.739501 | 0.734019 |
| 0.654011 | 0.767964 | 0.753927 | 0.745342 | 0.729377 | 0.752469 | 0.734876 | 0.738739 | 0.734326 |
| 0.654245 | 0.765813 | 0.754223 | 0.745775 | 0.728814 | 0.74768 | 0.741508 | 0.7403 | 0.734347 |
| 0.657841 | 0.769442 | 0.761988 | 0.75025 | 0.741085 | 0.754178 | 0.735433 | 0.73255 | 0.735973 |
| 0.658335 | 0.76065 | 0.738587 | 0.733879 | 0.726828 | 0.73952 | 0.727026 | 0.720923 | 0.730216 |
| 0.659274 | 0.76467 | 0.749765 | 0.746246 | 0.729078 | 0.7505 | 0.74053 | 0.732545 | 0.733271 |
| 0.656559 | 0.759251 | 0.739699 | 0.731825 | 0.724766 | 0.737818 | 0.728709 | 0.724084 | 0.723562 |
| 0.665169 | 0.764188 | 0.743406 | 0.738823 | 0.729252 | 0.741637 | 0.729949 | 0.723913 | 0.736111 |
| 0.661578 | 0.761926 | 0.747937 | 0.74 | 0.733763 | 0.745905 | 0.729733 | 0.728009 | 0.733684 |
| 0.657443 | 0.764118 | 0.744602 | 0.742636 | 0.728657 | 0.740931 | 0.732767 | 0.727757 | 0.72904 |
| 0.657364 | 0.762953 | 0.744455 | 0.741678 | 0.720674 | 0.747036 | 0.730473 | 0.730352 | 0.734191 |
| 0.661591 | 0.761764 | 0.741175 | 0.739089 | 0.716939 | 0.741512 | 0.73131 | 0.728341 | 0.723296 |
| 0.659398 | 0.76048 | 0.745949 | 0.739323 | 0.725155 | 0.742433 | 0.736986 | 0.733624 | 0.731178 |
| 0.658426 | 0.777778 | 0.759927 | 0.762753 | 0.747848 | 0.757009 | 0.739569 | 0.740844 | 0.744715 |
| 0.66642 | 0.759842 | 0.749848 | 0.743139 | 0.730217 | 0.747314 | 0.741432 | 0.727958 | 0.734928 |
| 0.743729 | 0.695736 | 0.670276 | 0.664268 | 0.648102 | 0.669688 | 0.660354 | 0.658635 | 0.656335 |
| 0.66129 | 0.761386 | 0.742562 | 0.738115 | 0.727611 | 0.745518 | 0.740303 | 0.736355 | 0.732584 |
| 0.656677 | 0.767296 | 0.743294 | 0.740409 | 0.727442 | 0.743011 | 0.733701 | 0.722002 | 0.727117 |
| 0.662225 | 0.769378 | 0.751684 | 0.739627 | 0.728379 | 0.745603 | 0.733428 | 0.729063 | 0.733827 |
| 0.654877 | 0.767488 | 0.744561 | 0.739378 | 0.731163 | 0.751146 | 0.732846 | 0.736909 | 0.725074 |
| 0.657985 | 0.762178 | 0.742624 | 0.744228 | 0.725446 | 0.747707 | 0.738391 | 0.734228 | 0.73852 |
| 0.657205 | 0.762378 | 0.749702 | 0.738738 | 0.731096 | 0.744382 | 0.734491 | 0.729639 | 0.727344 |
| 0.653715 | 0.763139 | 0.740924 | 0.737143 | 0.72809 | 0.744389 | 0.732928 | 0.733301 | 0.729101 |
| 0.655729 | 0.762552 | 0.737373 | 0.734806 | 0.726262 | 0.747989 | 0.732472 | 0.738374 | 0.730739 |
| 0.656108 | 0.75721 | 0.738408 | 0.730859 | 0.722254 | 0.743043 | 0.732612 | 0.725255 | 0.731121 |
| 0.651829 | 0.758976 | 0.737195 | 0.735153 | 0.72454 | 0.739993 | 0.728864 | 0.721023 | 0.725738 |
| 0.660252 | 0.7622 | 0.746868 | 0.74274 | 0.731739 | 0.745352 | 0.738565 | 0.736317 | 0.740694 |
| 0.660205 | 0.75974 | 0.74953 | 0.74374 | 0.726126 | 0.733628 | 0.736414 | 0.73001 | 0.72953 |
| 0.653324 | 0.766347 | 0.740042 | 0.742904 | 0.727103 | 0.748411 | 0.735446 | 0.736131 | 0.737899 |
| 0.659066 | 0.766667 | 0.743376 | 0.735326 | 0.726651 | 0.744038 | 0.736268 | 0.734558 | 0.734173 |
| 0.650116 | 0.760733 | 0.746196 | 0.735101 | 0.721785 | 0.741989 | 0.729186 | 0.726192 | 0.731433 |
| 0.658594 | 0.767951 | 0.742453 | 0.734287 | 0.729436 | 0.744092 | 0.736365 | 0.730308 | 0.738941 |
| 0.663291 | 0.7564 | 0.74718 | 0.745001 | 0.731233 | 0.738194 | 0.736393 | 0.732007 | 0.725269 |
| 0.654376 | 0.759457 | 0.743841 | 0.740816 | 0.720684 | 0.747239 | 0.735142 | 0.736234 | 0.732547 |
| 0.660057 | 0.771263 | 0.753954 | 0.746053 | 0.73948 | 0.757165 | 0.74569 | 0.734824 | 0.747951 |
| 0.661439 | 0.76753 | 0.746659 | 0.740821 | 0.731173 | 0.753893 | 0.741548 | 0.730197 | 0.74029 |

| | | | | | | | | |
|---|---|---|---|---|---|---|---|---|
| 0.651887 | 0.76718 | 0.749796 | 0.740144 | 0.727425 | 0.751407 | 0.740484 | 0.733917 | 0.741768 |
| 0.653907 | 0.773111 | 0.758621 | 0.752101 | 0.73691 | 0.760563 | 0.744201 | 0.746034 | 0.742172 |
| 0.651631 | 0.764588 | 0.747559 | 0.740336 | 0.730588 | 0.75341 | 0.739762 | 0.735701 | 0.73893 |
| 0.651515 | 0.735049 | 0.723608 | 0.723338 | 0.721339 | 0.734581 | 0.722682 | 0.720326 | 0.720098 |
| 0.656616 | 0.769065 | 0.74582 | 0.74712 | 0.730406 | 0.750817 | 0.739142 | 0.735731 | 0.738229 |
| 0.657005 | 0.764711 | 0.74783 | 0.738254 | 0.727751 | 0.748999 | 0.745144 | 0.736435 | 0.737739 |
| 0.65949 | 0.769562 | 0.754202 | 0.751165 | 0.729957 | 0.755208 | 0.737187 | 0.735831 | 0.740519 |
| 0.65334 | 0.767558 | 0.745482 | 0.745215 | 0.723125 | 0.746439 | 0.739208 | 0.737629 | 0.741141 |
| 0.663558 | 0.77626 | 0.756766 | 0.74908 | 0.738911 | 0.757084 | 0.740273 | 0.73872 | 0.74185 |
| 0.66009 | 0.774732 | 0.750556 | 0.745427 | 0.734194 | 0.755071 | 0.733974 | 0.736302 | 0.74342 |
| 0.653832 | 0.769184 | 0.753916 | 0.742764 | 0.732133 | 0.745499 | 0.734145 | 0.731408 | 0.736233 |
| 0.658069 | 0.771089 | 0.754677 | 0.748419 | 0.734498 | 0.755676 | 0.745992 | 0.739419 | 0.742161 |
| 0.660698 | 0.768057 | 0.746626 | 0.748029 | 0.739783 | 0.746985 | 0.731748 | 0.732918 | 0.73224 |
| 0.654971 | 0.773854 | 0.746451 | 0.743086 | 0.735701 | 0.749883 | 0.737629 | 0.728862 | 0.744369 |
| 0.655368 | 0.770212 | 0.755683 | 0.748944 | 0.739869 | 0.750439 | 0.73913 | 0.738894 | 0.739638 |
| 0.651345 | 0.765106 | 0.742052 | 0.738355 | 0.726884 | 0.750587 | 0.736136 | 0.732545 | 0.739329 |
| 0.657927 | 0.771659 | 0.757389 | 0.748175 | 0.737729 | 0.757679 | 0.747619 | 0.740764 | 0.750942 |
| 0.655313 | 0.772114 | 0.753792 | 0.746744 | 0.736576 | 0.750993 | 0.736815 | 0.738444 | 0.738144 |
| 0.656016 | 0.764205 | 0.742555 | 0.733008 | 0.728172 | 0.74918 | 0.7371 | 0.724984 | 0.737057 |
| 0.648601 | 0.767886 | 0.749971 | 0.747747 | 0.735694 | 0.750645 | 0.738515 | 0.735506 | 0.736619 |
| 0.64947 | 0.773895 | 0.75475 | 0.741244 | 0.738821 | 0.752374 | 0.744042 | 0.743812 | 0.742947 |
| 0.663254 | 0.768363 | 0.751839 | 0.74635 | 0.731207 | 0.754156 | 0.743702 | 0.743169 | 0.745771 |
| 0.687713 | 0.683884 | 0.678593 | 0.678395 | 0.683903 | 0.687462 | 0.685023 | 0.680923 | 0.688583 |
| 0.71135 | 0.66643 | 0.659596 | 0.663859 | 0.655924 | 0.662062 | 0.654484 | 0.657674 | 0.657372 |
| 0.733921 | 0.673284 | 0.656908 | 0.666009 | 0.646739 | 0.662318 | 0.657796 | 0.653453 | 0.650282 |
| 0.731495 | 0.651601 | 0.651302 | 0.658974 | 0.647305 | 0.650345 | 0.654188 | 0.641572 | 0.648608 |
| 0.656378 | 0.760079 | 0.753749 | 0.750278 | 0.751298 | 0.751506 | 0.739154 | 0.738984 | 0.74281 |
| 0.66068 | 0.754362 | 0.747644 | 0.755162 | 0.741196 | 0.751172 | 0.734255 | 0.730623 | 0.742857 |
| 0.658692 | 0.743799 | 0.757188 | 0.756879 | 0.738482 | 0.7504 | 0.729317 | 0.739644 | 0.73852 |
| 0.659039 | 0.762946 | 0.759112 | 0.761186 | 0.745407 | 0.759408 | 0.754741 | 0.75097 | 0.745688 |
| 0.650988 | 0.746657 | 0.740528 | 0.748663 | 0.745648 | 0.743172 | 0.728012 | 0.733093 | 0.73206 |
| 0.649689 | 0.747926 | 0.74742 | 0.743796 | 0.745627 | 0.742927 | 0.725787 | 0.724029 | 0.727652 |
| 0.649336 | 0.738229 | 0.74108 | 0.747586 | 0.752692 | 0.739006 | 0.719793 | 0.730839 | 0.732663 |
| 1 | 0.656417 | 0.658692 | 0.651878 | 0.654583 | 0.661319 | 0.660427 | 0.659495 | 0.659796 |
| 0.656417 | 1 | 0.765827 | 0.772783 | 0.753476 | 0.757404 | 0.746433 | 0.745839 | 0.748503 |
| 0.658692 | 0.765827 | 1 | 0.768215 | 0.760847 | 0.755632 | 0.749409 | 0.747944 | 0.750264 |
| 0.651878 | 0.772783 | 0.768215 | 1 | 0.760941 | 0.757449 | 0.750203 | 0.753542 | 0.74241 |
| 0.654583 | 0.753476 | 0.760847 | 0.760941 | 1 | 0.750102 | 0.737372 | 0.745122 | 0.738804 |
| 0.661319 | 0.757404 | 0.755632 | 0.757449 | 0.750102 | 1 | 0.765824 | 0.769996 | 0.754252 |
| 0.660427 | 0.746433 | 0.749409 | 0.750203 | 0.737372 | 0.765824 | 1 | 0.847716 | 0.838747 |
| 0.659495 | 0.745839 | 0.747944 | 0.753542 | 0.745122 | 0.769996 | 0.847716 | 1 | 0.739357 |
| 0.659796 | 0.748503 | 0.750264 | 0.74241 | 0.738804 | 0.754252 | 0.838747 | 0.739357 | 1 |
| 0.659564 | 0.748681 | 0.74798 | 0.742058 | 0.74103 | 0.756713 | 0.749741 | 0.760692 | 0.741051 |
| 0.658681 | 0.74516 | 0.741189 | 0.741305 | 0.748722 | 0.747912 | 0.736388 | 0.73172 | 0.747233 |
| 0.668357 | 0.745459 | 0.747177 | 0.756354 | 0.764643 | 0.777592 | 0.757273 | 0.773798 | 0.760188 |
| 0.654032 | 0.739767 | 0.737335 | 0.74113 | 0.743088 | 0.738856 | 0.768759 | 0.770292 | 0.745238 |

| | | | | | | | | |
|---|---|---|---|---|---|---|---|---|
| 0.665197 | 0.775193 | 0.768049 | 0.759994 | 0.751585 | 0.762316 | 0.751976 | 0.749005 | 0.747541 |
| 0.653503 | 0.759206 | 0.760447 | 0.752308 | 0.755451 | 0.755353 | 0.746339 | 0.744089 | 0.743494 |
| 0.659 | 0.760407 | 0.746611 | 0.748423 | 0.733364 | 0.747401 | 0.73405 | 0.738804 | 0.740998 |
| 0.663705 | 0.748421 | 0.748884 | 0.754957 | 0.755703 | 0.752455 | 0.736728 | 0.738298 | 0.74749 |
| 0.662637 | 0.763711 | 0.768771 | 0.758637 | 0.76905 | 0.75493 | 0.734301 | 0.745332 | 0.74057 |
| 0.669861 | 0.748749 | 0.759711 | 0.751135 | 0.760559 | 0.743493 | 0.740954 | 0.746784 | 0.743457 |
| 0.653405 | 0.741262 | 0.745134 | 0.754026 | 0.755333 | 0.747774 | 0.729183 | 0.742035 | 0.736842 |
| 0.655357 | 0.745165 | 0.752335 | 0.753086 | 0.758229 | 0.751433 | 0.738205 | 0.744548 | 0.729674 |
| 0.651906 | 0.734746 | 0.752502 | 0.750639 | 0.747896 | 0.744895 | 0.738205 | 0.727462 | 0.740471 |
| 0.662514 | 0.768916 | 0.754976 | 0.751862 | 0.741109 | 0.755517 | 0.743576 | 0.740307 | 0.738365 |
| 0.66469 | 0.76297 | 0.756015 | 0.74784 | 0.742609 | 0.749781 | 0.741643 | 0.738413 | 0.734046 |
| 0.670103 | 0.754268 | 0.752982 | 0.742615 | 0.741582 | 0.759467 | 0.739792 | 0.743095 | 0.743038 |
| 0.663978 | 0.752263 | 0.744078 | 0.742183 | 0.730367 | 0.747245 | 0.733383 | 0.73688 | 0.736767 |
| 0.65812 | 0.760735 | 0.746892 | 0.761347 | 0.736786 | 0.758455 | 0.740856 | 0.743734 | 0.741198 |
| 0.659236 | 0.754188 | 0.746552 | 0.74972 | 0.738837 | 0.754532 | 0.743252 | 0.740399 | 0.738381 |

| t 23 | | | | | | | | plot 24 |
|---|---|---|---|---|---|---|---|---|
| Wild_plot_ | Wild_plot_ | Wild_plot_ | Wild_plot_ | Wild_plot_ | Wild_plot_ | Wild_plot_ | Wild_plot_ | Wild_plot_ |
| 0.655442 | 0.655153 | 0.653799 | 0.646665 | 0.685917 | 0.652572 | 0.658273 | 0.659211 | 0.65345 |
| 0.662009 | 0.660302 | 0.654949 | 0.651065 | 0.696809 | 0.662091 | 0.662453 | 0.668853 | 0.659961 |
| 0.649371 | 0.653073 | 0.649028 | 0.64181 | 0.686123 | 0.654435 | 0.656317 | 0.660667 | 0.650137 |
| 0.661386 | 0.664377 | 0.662759 | 0.647507 | 0.691689 | 0.663781 | 0.671362 | 0.664847 | 0.658141 |
| 0.652498 | 0.661162 | 0.662952 | 0.633417 | 0.685058 | 0.651994 | 0.662534 | 0.660985 | 0.660359 |
| 0.667071 | 0.670588 | 0.665925 | 0.646966 | 0.695868 | 0.667117 | 0.67066 | 0.668039 | 0.664846 |
| 0.724838 | 0.72704 | 0.721282 | 0.716368 | 0.750843 | 0.727935 | 0.728469 | 0.735083 | 0.736391 |
| 0.720511 | 0.715208 | 0.71052 | 0.712927 | 0.753136 | 0.723762 | 0.719141 | 0.730108 | 0.725082 |
| 0.661711 | 0.667813 | 0.658591 | 0.652951 | 0.691738 | 0.662464 | 0.670893 | 0.664697 | 0.65835 |
| 0.65806 | 0.656041 | 0.660284 | 0.653303 | 0.68813 | 0.654146 | 0.66285 | 0.66287 | 0.665674 |
| 0.657789 | 0.662162 | 0.659349 | 0.645707 | 0.691169 | 0.658681 | 0.66426 | 0.665051 | 0.65736 |
| 0.64658 | 0.650625 | 0.650174 | 0.634695 | 0.681101 | 0.644949 | 0.650423 | 0.647369 | 0.656268 |
| 0.660864 | 0.658754 | 0.661783 | 0.641635 | 0.696815 | 0.670558 | 0.66546 | 0.665143 | 0.66588 |
| 0.66916 | 0.669275 | 0.674286 | 0.648119 | 0.699548 | 0.671133 | 0.677541 | 0.680918 | 0.675212 |
| 0.680744 | 0.689205 | 0.683791 | 0.676631 | 0.71699 | 0.689683 | 0.691007 | 0.686587 | 0.680487 |
| 0.662615 | 0.660975 | 0.658483 | 0.660462 | 0.6966 | 0.666802 | 0.669617 | 0.668885 | 0.665818 |
| 0.658056 | 0.659078 | 0.656475 | 0.640215 | 0.693399 | 0.660421 | 0.666375 | 0.661391 | 0.659207 |
| 0.695978 | 0.692718 | 0.685636 | 0.669398 | 0.720115 | 0.696005 | 0.696149 | 0.695864 | 0.691766 |
| 0.650321 | 0.652562 | 0.656335 | 0.639208 | 0.685153 | 0.652827 | 0.660717 | 0.657421 | 0.649892 |
| 0.658952 | 0.658857 | 0.664188 | 0.651714 | 0.691871 | 0.661645 | 0.669288 | 0.66411 | 0.660106 |
| 0.653813 | 0.651901 | 0.65555 | 0.634463 | 0.690604 | 0.656575 | 0.664339 | 0.657189 | 0.653975 |
| 0.656864 | 0.660486 | 0.651358 | 0.647473 | 0.692138 | 0.661823 | 0.661556 | 0.662983 | 0.65942 |
| 0.665273 | 0.664131 | 0.667992 | 0.64665 | 0.696964 | 0.667614 | 0.669658 | 0.664654 | 0.664374 |
| 0.656038 | 0.658 | 0.653061 | 0.64832 | 0.692751 | 0.660382 | 0.666377 | 0.66213 | 0.65978 |
| 0.656761 | 0.658448 | 0.65377 | 0.641551 | 0.693383 | 0.660249 | 0.666909 | 0.669087 | 0.663583 |
| 0.654721 | 0.657495 | 0.658651 | 0.646928 | 0.69051 | 0.65793 | 0.664692 | 0.667307 | 0.657199 |
| 0.656315 | 0.654455 | 0.654865 | 0.639785 | 0.682619 | 0.651975 | 0.659015 | 0.661934 | 0.65524 |
| 0.650567 | 0.656185 | 0.659459 | 0.634192 | 0.693273 | 0.656024 | 0.665901 | 0.661667 | 0.65214 |
| 0.666376 | 0.667751 | 0.659458 | 0.648113 | 0.696271 | 0.666622 | 0.671499 | 0.671209 | 0.664118 |
| 0.657886 | 0.662263 | 0.662648 | 0.651598 | 0.692122 | 0.660503 | 0.668031 | 0.664982 | 0.668918 |
| 0.653012 | 0.660377 | 0.657895 | 0.639029 | 0.692352 | 0.659517 | 0.66363 | 0.660027 | 0.661768 |
| 0.647858 | 0.657825 | 0.642355 | 0.636819 | 0.684005 | 0.658901 | 0.658996 | 0.659518 | 0.649273 |
| 0.656228 | 0.655478 | 0.652961 | 0.645694 | 0.685872 | 0.65776 | 0.663828 | 0.66134 | 0.658101 |
| 0.657296 | 0.65473 | 0.652547 | 0.6399 | 0.688334 | 0.658667 | 0.662744 | 0.665394 | 0.65401 |
| 0.650805 | 0.65089 | 0.650198 | 0.642926 | 0.684554 | 0.654609 | 0.663759 | 0.659138 | 0.649065 |
| 0.655809 | 0.656304 | 0.658215 | 0.64736 | 0.690323 | 0.662551 | 0.6656 | 0.660894 | 0.664423 |
| 0.662094 | 0.662945 | 0.664183 | 0.650829 | 0.692792 | 0.664149 | 0.670406 | 0.666227 | 0.656274 |
| 0.658793 | 0.663456 | 0.655026 | 0.646188 | 0.695246 | 0.658271 | 0.671788 | 0.666667 | 0.664878 |
| 0.659434 | 0.66584 | 0.66067 | 0.639126 | 0.697636 | 0.662935 | 0.665223 | 0.669821 | 0.663143 |

| | | | | | | | | |
|---|---|---|---|---|---|---|---|---|
| 0.69884 | 0.700712 | 0.705436 | 0.678526 | 0.726033 | 0.704427 | 0.707253 | 0.703421 | 0.704315 |
| 0.650474 | 0.649331 | 0.649901 | 0.629782 | 0.681398 | 0.64687 | 0.653301 | 0.654996 | 0.645444 |
| 0.653266 | 0.652215 | 0.659796 | 0.639516 | 0.685684 | 0.648794 | 0.655098 | 0.66224 | 0.653808 |
| 0.664873 | 0.670292 | 0.66131 | 0.652615 | 0.697332 | 0.662095 | 0.668401 | 0.668676 | 0.670789 |
| 0.651283 | 0.658049 | 0.652475 | 0.639339 | 0.690104 | 0.654685 | 0.659206 | 0.666716 | 0.654004 |
| 0.659589 | 0.663482 | 0.654046 | 0.647811 | 0.694182 | 0.663268 | 0.663242 | 0.663473 | 0.661411 |
| 0.655018 | 0.64835 | 0.647189 | 0.63375 | 0.684998 | 0.655786 | 0.658091 | 0.65569 | 0.652986 |
| 0.655908 | 0.666284 | 0.659664 | 0.650783 | 0.696228 | 0.664948 | 0.662619 | 0.663951 | 0.658024 |
| 0.660658 | 0.666731 | 0.658234 | 0.654405 | 0.69416 | 0.663353 | 0.674535 | 0.667757 | 0.66166 |
| 0.655647 | 0.666161 | 0.668567 | 0.653274 | 0.693207 | 0.668112 | 0.671009 | 0.673545 | 0.663861 |
| 0.646977 | 0.652839 | 0.646562 | 0.637909 | 0.686485 | 0.648051 | 0.657409 | 0.659182 | 0.653459 |
| 0.649065 | 0.658621 | 0.649344 | 0.635959 | 0.687928 | 0.64931 | 0.655422 | 0.658136 | 0.645772 |
| 0.651542 | 0.65663 | 0.648272 | 0.638181 | 0.687198 | 0.659414 | 0.656831 | 0.661504 | 0.656027 |
| 0.653799 | 0.657925 | 0.648823 | 0.637876 | 0.685317 | 0.65333 | 0.661822 | 0.658374 | 0.650294 |
| 0.647469 | 0.655491 | 0.647192 | 0.640351 | 0.68756 | 0.653778 | 0.653337 | 0.659257 | 0.64794 |
| 0.651534 | 0.656728 | 0.656949 | 0.634374 | 0.688472 | 0.652364 | 0.656616 | 0.659047 | 0.65509 |
| 0.649842 | 0.651261 | 0.646708 | 0.636511 | 0.68262 | 0.650826 | 0.65429 | 0.660544 | 0.646896 |
| 0.657163 | 0.65798 | 0.654074 | 0.642789 | 0.690577 | 0.660664 | 0.658537 | 0.663155 | 0.66088 |
| 0.65681 | 0.655896 | 0.658391 | 0.65619 | 0.69396 | 0.661978 | 0.662866 | 0.663366 | 0.663237 |
| 0.651963 | 0.654088 | 0.657292 | 0.63486 | 0.685249 | 0.653603 | 0.656544 | 0.660296 | 0.656657 |
| 0.645724 | 0.647227 | 0.641975 | 0.63648 | 0.685321 | 0.650434 | 0.655585 | 0.655551 | 0.651034 |
| 0.657312 | 0.653715 | 0.647232 | 0.636827 | 0.685231 | 0.652637 | 0.659242 | 0.661993 | 0.650508 |
| 0.646928 | 0.658657 | 0.64524 | 0.628241 | 0.692764 | 0.656006 | 0.660133 | 0.662269 | 0.650291 |
| 0.663123 | 0.666348 | 0.666998 | 0.647397 | 0.701224 | 0.66961 | 0.675983 | 0.678497 | 0.667059 |
| 0.665514 | 0.67804 | 0.673827 | 0.654188 | 0.708538 | 0.677778 | 0.680318 | 0.677443 | 0.667383 |
| 0.719863 | 0.715938 | 0.711982 | 0.712969 | 0.753819 | 0.724268 | 0.718881 | 0.72973 | 0.725655 |
| 0.721467 | 0.729081 | 0.724209 | 0.713815 | 0.757836 | 0.737772 | 0.733802 | 0.726994 | 0.743368 |
| 0.72357 | 0.729687 | 0.724254 | 0.713194 | 0.757174 | 0.733314 | 0.729277 | 0.727766 | 0.739681 |
| 0.654791 | 0.648445 | 0.654215 | 0.643401 | 0.691024 | 0.658086 | 0.659841 | 0.66269 | 0.658732 |
| 0.651082 | 0.654413 | 0.655104 | 0.642426 | 0.688544 | 0.660194 | 0.661955 | 0.661747 | 0.653633 |
| 0.671459 | 0.666476 | 0.664505 | 0.647208 | 0.693542 | 0.665357 | 0.665025 | 0.663961 | 0.663601 |
| 0.665741 | 0.67072 | 0.671797 | 0.662388 | 0.694493 | 0.665854 | 0.669171 | 0.67095 | 0.662639 |
| 0.656114 | 0.65662 | 0.660777 | 0.650983 | 0.690276 | 0.657694 | 0.661898 | 0.66163 | 0.65214 |
| 0.656146 | 0.656339 | 0.657856 | 0.635182 | 0.689011 | 0.658048 | 0.661402 | 0.661488 | 0.660444 |
| 0.655538 | 0.65946 | 0.660415 | 0.652951 | 0.694176 | 0.659929 | 0.663718 | 0.665924 | 0.661955 |
| 0.658625 | 0.660601 | 0.658193 | 0.643634 | 0.692552 | 0.657969 | 0.663813 | 0.664146 | 0.660067 |
| 0.653336 | 0.661027 | 0.657524 | 0.649714 | 0.692028 | 0.662712 | 0.668547 | 0.663719 | 0.661773 |
| 0.683903 | 0.677209 | 0.676889 | 0.661146 | 0.704837 | 0.680977 | 0.684087 | 0.686587 | 0.680586 |
| 0.650504 | 0.657626 | 0.65985 | 0.644095 | 0.687972 | 0.658931 | 0.6615 | 0.65922 | 0.654261 |
| 0.654111 | 0.662802 | 0.653317 | 0.642274 | 0.690784 | 0.658032 | 0.656857 | 0.659907 | 0.654273 |
| 0.650577 | 0.656422 | 0.650821 | 0.637805 | 0.686047 | 0.651592 | 0.661193 | 0.65742 | 0.657126 |
| 0.676958 | 0.675953 | 0.678074 | 0.662472 | 0.707502 | 0.677769 | 0.685347 | 0.681825 | 0.674409 |
| 0.66025 | 0.658718 | 0.663027 | 0.648502 | 0.694198 | 0.661131 | 0.666715 | 0.669082 | 0.665029 |
| 0.649543 | 0.651838 | 0.658344 | 0.633365 | 0.686093 | 0.651755 | 0.660622 | 0.659154 | 0.650572 |
| 0.662026 | 0.666092 | 0.653304 | 0.643901 | 0.690123 | 0.660306 | 0.662614 | 0.667259 | 0.656631 |
| 0.655758 | 0.6626 | 0.659896 | 0.645151 | 0.691758 | 0.663058 | 0.663475 | 0.665532 | 0.657751 |

| | | | | | | | | |
|---|---|---|---|---|---|---|---|---|
| 0.652513 | 0.658245 | 0.660237 | 0.64148 | 0.689301 | 0.659643 | 0.658149 | 0.662531 | 0.662615 |
| 0.661352 | 0.660156 | 0.660428 | 0.646066 | 0.689797 | 0.655848 | 0.663215 | 0.66848 | 0.658038 |
| 0.651239 | 0.662819 | 0.65694 | 0.645437 | 0.691967 | 0.664818 | 0.666234 | 0.662702 | 0.66359 |
| 0.668819 | 0.673959 | 0.666667 | 0.654633 | 0.700203 | 0.673925 | 0.675908 | 0.678291 | 0.677765 |
| 0.659272 | 0.670353 | 0.665437 | 0.655227 | 0.699596 | 0.666846 | 0.671751 | 0.668184 | 0.663933 |
| 0.653287 | 0.664188 | 0.660359 | 0.638871 | 0.692694 | 0.652847 | 0.658864 | 0.666667 | 0.66123 |
| 0.653356 | 0.664948 | 0.660912 | 0.648313 | 0.694057 | 0.667655 | 0.6697 | 0.665737 | 0.658183 |
| 0.660024 | 0.663845 | 0.663922 | 0.653995 | 0.693065 | 0.661192 | 0.668612 | 0.667062 | 0.6641 |
| 0.657433 | 0.655324 | 0.65156 | 0.637349 | 0.691062 | 0.653596 | 0.655392 | 0.660881 | 0.65174 |
| 0.650433 | 0.659372 | 0.658379 | 0.63798 | 0.687284 | 0.653359 | 0.663126 | 0.666271 | 0.660299 |
| 0.654699 | 0.659943 | 0.653245 | 0.653456 | 0.689223 | 0.655393 | 0.661233 | 0.656714 | 0.661865 |
| 0.660325 | 0.664168 | 0.664343 | 0.646419 | 0.693858 | 0.662524 | 0.665793 | 0.671798 | 0.664554 |
| 0.656709 | 0.66129 | 0.659639 | 0.645927 | 0.687561 | 0.655813 | 0.663553 | 0.667759 | 0.660102 |
| 0.655232 | 0.665899 | 0.66034 | 0.644809 | 0.688771 | 0.658331 | 0.661545 | 0.662522 | 0.656238 |
| 0.653488 | 0.661447 | 0.661881 | 0.644041 | 0.691067 | 0.655545 | 0.661274 | 0.663902 | 0.656232 |
| 0.652835 | 0.654872 | 0.656583 | 0.650032 | 0.684778 | 0.656086 | 0.660758 | 0.661767 | 0.656593 |
| 0.657043 | 0.65694 | 0.654414 | 0.640845 | 0.692048 | 0.660454 | 0.670025 | 0.65931 | 0.653362 |
| 0.657548 | 0.657596 | 0.661051 | 0.648811 | 0.695778 | 0.659131 | 0.661392 | 0.662162 | 0.660885 |
| 0.656347 | 0.660819 | 0.662556 | 0.653048 | 0.689956 | 0.662836 | 0.669022 | 0.670016 | 0.654078 |
| 0.65687 | 0.661127 | 0.660345 | 0.648095 | 0.692174 | 0.661825 | 0.665696 | 0.667608 | 0.661037 |
| 0.659327 | 0.66102 | 0.664356 | 0.648967 | 0.692953 | 0.66306 | 0.666086 | 0.669978 | 0.657967 |
| 0.655285 | 0.657731 | 0.658907 | 0.642386 | 0.687757 | 0.654611 | 0.662984 | 0.657306 | 0.65866 |
| 0.656093 | 0.666032 | 0.654245 | 0.645008 | 0.693249 | 0.660194 | 0.669241 | 0.668239 | 0.663069 |
| 0.660024 | 0.662859 | 0.662713 | 0.641677 | 0.694735 | 0.661484 | 0.665849 | 0.664501 | 0.658704 |
| 0.654366 | 0.663029 | 0.652024 | 0.652077 | 0.695707 | 0.662517 | 0.66565 | 0.668993 | 0.660841 |
| 0.658797 | 0.667238 | 0.662891 | 0.652809 | 0.693108 | 0.665406 | 0.665853 | 0.665934 | 0.658778 |
| 0.658688 | 0.66654 | 0.655762 | 0.654649 | 0.69194 | 0.659286 | 0.666762 | 0.663639 | 0.6591 |
| 0.655554 | 0.659655 | 0.654909 | 0.634965 | 0.687548 | 0.656078 | 0.662496 | 0.660006 | 0.651836 |
| 0.658961 | 0.657959 | 0.660228 | 0.649396 | 0.69157 | 0.65477 | 0.66388 | 0.666422 | 0.660196 |
| 0.660584 | 0.670636 | 0.660833 | 0.653405 | 0.694989 | 0.661787 | 0.670057 | 0.669732 | 0.667708 |
| 0.659154 | 0.667048 | 0.664602 | 0.655921 | 0.695526 | 0.664909 | 0.671165 | 0.663883 | 0.665817 |
| 0.652716 | 0.65399 | 0.65803 | 0.645446 | 0.689721 | 0.661221 | 0.667525 | 0.663501 | 0.659319 |
| 0.660098 | 0.665391 | 0.670175 | 0.648267 | 0.696095 | 0.661284 | 0.667541 | 0.668506 | 0.66266 |
| 0.66273 | 0.672881 | 0.663857 | 0.653761 | 0.698163 | 0.663366 | 0.668511 | 0.667111 | 0.670408 |
| 0.661049 | 0.65668 | 0.66155 | 0.646004 | 0.690797 | 0.6554 | 0.665754 | 0.668586 | 0.658527 |
| 0.656071 | 0.662154 | 0.662722 | 0.641521 | 0.694404 | 0.659951 | 0.663471 | 0.666815 | 0.663462 |
| 0.651911 | 0.657923 | 0.655672 | 0.633393 | 0.684583 | 0.654655 | 0.653119 | 0.654036 | 0.657435 |
| 0.658338 | 0.662394 | 0.657948 | 0.651862 | 0.680119 | 0.659222 | 0.664412 | 0.656684 | 0.662869 |
| 0.661039 | 0.665792 | 0.654545 | 0.648063 | 0.68465 | 0.66205 | 0.66439 | 0.668315 | 0.662738 |
| 0.648464 | 0.653501 | 0.657463 | 0.646174 | 0.676273 | 0.656025 | 0.65458 | 0.658292 | 0.653322 |
| 0.651221 | 0.660202 | 0.656596 | 0.646858 | 0.676749 | 0.655756 | 0.649608 | 0.656261 | 0.660101 |
| 0.643914 | 0.649502 | 0.654706 | 0.637306 | 0.66633 | 0.646658 | 0.649932 | 0.650087 | 0.647449 |
| 0.65476 | 0.659464 | 0.665106 | 0.643869 | 0.665565 | 0.649439 | 0.656271 | 0.657316 | 0.659598 |
| 0.647932 | 0.663232 | 0.646512 | 0.642208 | 0.672923 | 0.646013 | 0.655918 | 0.6539 | 0.655632 |
| 0.657377 | 0.65857 | 0.668936 | 0.646685 | 0.669228 | 0.655509 | 0.653894 | 0.659711 | 0.662156 |
| 0.648945 | 0.659398 | 0.662878 | 0.649635 | 0.673024 | 0.653377 | 0.658934 | 0.658875 | 0.662742 |
| 0.657842 | 0.664139 | 0.659676 | 0.655755 | 0.674721 | 0.658799 | 0.655471 | 0.657343 | 0.662629 |

| | | | | | | | | |
|---|---|---|---|---|---|---|---|---|
| 0.654294 | 0.657679 | 0.651233 | 0.636301 | 0.670522 | 0.653846 | 0.653528 | 0.654797 | 0.653292 |
| 0.658372 | 0.662952 | 0.668903 | 0.658098 | 0.678072 | 0.655966 | 0.663164 | 0.665861 | 0.662818 |
| 0.656666 | 0.662526 | 0.662665 | 0.645138 | 0.66948 | 0.64715 | 0.661329 | 0.666855 | 0.656821 |
| 0.650523 | 0.650033 | 0.655868 | 0.64623 | 0.670222 | 0.645721 | 0.654604 | 0.65656 | 0.660041 |
| 0.65235 | 0.659271 | 0.658659 | 0.641875 | 0.670032 | 0.651635 | 0.656041 | 0.663077 | 0.66493 |
| 0.647625 | 0.658788 | 0.665003 | 0.649142 | 0.666283 | 0.655439 | 0.665349 | 0.659904 | 0.664433 |
| 0.653435 | 0.665484 | 0.661169 | 0.651481 | 0.666914 | 0.653127 | 0.657845 | 0.660552 | 0.654906 |
| 0.647605 | 0.65132 | 0.661189 | 0.639823 | 0.667104 | 0.6525 | 0.658868 | 0.656774 | 0.656295 |
| 0.658839 | 0.661337 | 0.665664 | 0.649888 | 0.66758 | 0.655299 | 0.661378 | 0.656969 | 0.657638 |
| 0.651519 | 0.657308 | 0.658331 | 0.647736 | 0.662463 | 0.648225 | 0.650192 | 0.657764 | 0.648813 |
| 0.650348 | 0.653948 | 0.665455 | 0.6423 | 0.662594 | 0.652597 | 0.652541 | 0.651002 | 0.660862 |
| 0.650652 | 0.650038 | 0.653285 | 0.635076 | 0.68003 | 0.646184 | 0.654049 | 0.651594 | 0.656699 |
| 0.658068 | 0.658486 | 0.666584 | 0.636221 | 0.692814 | 0.659508 | 0.661397 | 0.658901 | 0.652503 |
| 0.664768 | 0.676127 | 0.673138 | 0.656448 | 0.682734 | 0.665742 | 0.670588 | 0.670359 | 0.668158 |
| 0.663582 | 0.660259 | 0.664688 | 0.645767 | 0.698769 | 0.668027 | 0.670741 | 0.674985 | 0.66575 |
| 0.657366 | 0.660252 | 0.663087 | 0.641141 | 0.685701 | 0.658269 | 0.660712 | 0.654032 | 0.654542 |
| 0.66435 | 0.664649 | 0.657521 | 0.653055 | 0.691391 | 0.66104 | 0.668158 | 0.667694 | 0.662257 |
| 0.666085 | 0.666349 | 0.67156 | 0.652146 | 0.701995 | 0.667977 | 0.677065 | 0.667355 | 0.673977 |
| 0.641074 | 0.648613 | 0.641231 | 0.626431 | 0.680324 | 0.647495 | 0.65022 | 0.653256 | 0.642625 |
| 0.663979 | 0.665663 | 0.673035 | 0.657008 | 0.7002 | 0.671858 | 0.674322 | 0.675573 | 0.668414 |
| 0.670277 | 0.671327 | 0.67694 | 0.66328 | 0.69066 | 0.672769 | 0.676851 | 0.679737 | 0.673201 |
| 0.660259 | 0.668259 | 0.660429 | 0.645933 | 0.696654 | 0.671392 | 0.672719 | 0.670028 | 0.663327 |
| 0.670056 | 0.664826 | 0.663843 | 0.655084 | 0.700968 | 0.665807 | 0.674829 | 0.670658 | 0.667785 |
| 0.666471 | 0.660941 | 0.665037 | 0.654985 | 0.685483 | 0.663309 | 0.660726 | 0.66501 | 0.669711 |
| 0.640714 | 0.641921 | 0.645327 | 0.630449 | 0.670093 | 0.643098 | 0.648947 | 0.647094 | 0.634627 |
| 0.673427 | 0.681808 | 0.684456 | 0.671005 | 0.691963 | 0.668243 | 0.678655 | 0.678414 | 0.677674 |
| 0.64841 | 0.656971 | 0.655998 | 0.643768 | 0.688414 | 0.662099 | 0.661114 | 0.664041 | 0.659009 |
| 0.667304 | 0.662811 | 0.662142 | 0.643084 | 0.698819 | 0.665584 | 0.671488 | 0.668675 | 0.669557 |
| 0.635024 | 0.635311 | 0.643062 | 0.624769 | 0.66586 | 0.644903 | 0.646695 | 0.642816 | 0.637911 |
| 0.67728 | 0.675671 | 0.671337 | 0.659292 | 0.706939 | 0.678125 | 0.673484 | 0.677282 | 0.669835 |
| 0.67583 | 0.676705 | 0.668302 | 0.6541 | 0.70902 | 0.6779 | 0.67549 | 0.680558 | 0.670739 |
| 0.651963 | 0.653046 | 0.651611 | 0.644916 | 0.68 | 0.653392 | 0.655212 | 0.658476 | 0.651862 |
| 0.643237 | 0.652182 | 0.646026 | 0.639406 | 0.680137 | 0.648353 | 0.651894 | 0.655611 | 0.647127 |
| 0.671878 | 0.680598 | 0.677524 | 0.660618 | 0.7051 | 0.678892 | 0.679174 | 0.684601 | 0.67679 |
| 0.669374 | 0.671881 | 0.674063 | 0.65869 | 0.702087 | 0.677051 | 0.673571 | 0.675537 | 0.676328 |
| 0.672683 | 0.671267 | 0.683114 | 0.663593 | 0.706844 | 0.682404 | 0.68243 | 0.684857 | 0.675855 |
| 0.629717 | 0.627261 | 0.634597 | 0.623805 | 0.659758 | 0.632541 | 0.636211 | 0.631333 | 0.62603 |
| 0.667359 | 0.670465 | 0.669537 | 0.652201 | 0.703636 | 0.672516 | 0.68074 | 0.676835 | 0.67287 |
| 0.619574 | 0.626987 | 0.628154 | 0.611922 | 0.657293 | 0.630311 | 0.628228 | 0.631171 | 0.624069 |
| 0.611558 | 0.613735 | 0.614162 | 0.593928 | 0.648012 | 0.62135 | 0.615984 | 0.617385 | 0.617631 |
| 0.675178 | 0.675409 | 0.672597 | 0.661563 | 0.702876 | 0.674027 | 0.677601 | 0.677253 | 0.674696 |
| 0.615862 | 0.624865 | 0.62254 | 0.60524 | 0.649022 | 0.624393 | 0.618491 | 0.622691 | 0.625023 |
| 0.604082 | 0.60952 | 0.605611 | 0.598442 | 0.642683 | 0.613345 | 0.610346 | 0.616918 | 0.610194 |
| 0.60058 | 0.610953 | 0.601145 | 0.599402 | 0.639068 | 0.609599 | 0.612899 | 0.609558 | 0.603661 |
| 0.676328 | 0.682452 | 0.68789 | 0.675048 | 0.711113 | 0.68456 | 0.683511 | 0.684086 | 0.683448 |
| 0.60342 | 0.60813 | 0.60383 | 0.590329 | 0.641561 | 0.608426 | 0.609895 | 0.613313 | 0.603932 |
| 0.676757 | 0.685115 | 0.67911 | 0.665609 | 0.706182 | 0.676753 | 0.680626 | 0.683673 | 0.676373 |

| | | | | | | | | |
|---|---|---|---|---|---|---|---|---|
| 0.688241 | 0.682085 | 0.675037 | 0.661071 | 0.710292 | 0.684503 | 0.68559 | 0.687039 | 0.69337 |
| 0.673856 | 0.673913 | 0.683042 | 0.667199 | 0.707957 | 0.676784 | 0.682705 | 0.68253 | 0.680709 |
| 0.704774 | 0.694439 | 0.700673 | 0.678208 | 0.725446 | 0.706108 | 0.703022 | 0.705725 | 0.705615 |
| 0.590641 | 0.595719 | 0.592288 | 0.580334 | 0.628744 | 0.597984 | 0.597817 | 0.600924 | 0.591804 |
| 0.604578 | 0.606548 | 0.615312 | 0.597332 | 0.632546 | 0.608729 | 0.606711 | 0.615221 | 0.610647 |
| 0.684137 | 0.692634 | 0.687924 | 0.670294 | 0.712137 | 0.68814 | 0.691299 | 0.689423 | 0.68446 |
| 0.598756 | 0.607778 | 0.598353 | 0.596028 | 0.63742 | 0.609726 | 0.61019 | 0.60972 | 0.603451 |
| 0.686229 | 0.689202 | 0.691061 | 0.667313 | 0.713868 | 0.687845 | 0.693989 | 0.694063 | 0.685555 |
| 0.675882 | 0.692441 | 0.691884 | 0.678846 | 0.716873 | 0.689095 | 0.692466 | 0.68789 | 0.687177 |
| 0.682701 | 0.685312 | 0.687643 | 0.665688 | 0.715551 | 0.68743 | 0.68737 | 0.694313 | 0.686594 |
| 0.683931 | 0.692864 | 0.689655 | 0.673821 | 0.717387 | 0.692957 | 0.693763 | 0.691043 | 0.685954 |
| 0.75969 | 0.768657 | 0.760331 | 0.747664 | 0.777778 | 0.783784 | 0.81746 | 0.736486 | 0.795455 |
| 0.743191 | 0.748422 | 0.744358 | 0.730337 | 0.759732 | 0.744325 | 0.741654 | 0.744551 | 0.754354 |
| 0.73252 | 0.737134 | 0.743722 | 0.726559 | 0.750041 | 0.744396 | 0.735367 | 0.741686 | 0.743946 |
| 0.741053 | 0.747012 | 0.741505 | 0.738516 | 0.760914 | 0.750808 | 0.751869 | 0.760959 | 0.745237 |
| 0.734498 | 0.741399 | 0.74933 | 0.724314 | 0.757304 | 0.746514 | 0.742958 | 0.748404 | 0.745779 |
| 0.567263 | 0.568945 | 0.561525 | 0.552137 | 0.590395 | 0.570049 | 0.565682 | 0.570029 | 0.561235 |
| 0.734005 | 0.738381 | 0.745396 | 0.736383 | 0.767555 | 0.750861 | 0.750228 | 0.75657 | 0.751992 |
| 0.693369 | 0.699162 | 0.68994 | 0.678053 | 0.716624 | 0.694494 | 0.69672 | 0.695423 | 0.695251 |
| 0.694075 | 0.695005 | 0.696738 | 0.680274 | 0.720292 | 0.697067 | 0.698082 | 0.698665 | 0.694652 |
| 0.690434 | 0.695151 | 0.698875 | 0.680208 | 0.72161 | 0.697833 | 0.703158 | 0.700258 | 0.684672 |
| 0.531573 | 0.533898 | 0.526983 | 0.520531 | 0.563602 | 0.535851 | 0.536535 | 0.543796 | 0.530926 |
| 0.517477 | 0.521875 | 0.507501 | 0.504698 | 0.554257 | 0.525108 | 0.524707 | 0.525546 | 0.518567 |
| 0.724582 | 0.707279 | 0.719938 | 0.711303 | 0.743239 | 0.727157 | 0.725265 | 0.72834 | 0.725349 |
| 0.729159 | 0.720388 | 0.716261 | 0.706155 | 0.745145 | 0.728984 | 0.727506 | 0.727217 | 0.727254 |
| 0.711007 | 0.704375 | 0.720942 | 0.701894 | 0.740143 | 0.717606 | 0.717473 | 0.714439 | 0.714402 |
| 0.71951 | 0.717015 | 0.715057 | 0.700654 | 0.750025 | 0.727963 | 0.721799 | 0.722138 | 0.720439 |
| 0.721161 | 0.713386 | 0.720235 | 0.714804 | 0.747032 | 0.72596 | 0.728237 | 0.722146 | 0.729741 |
| 0.734924 | 0.719402 | 0.73537 | 0.727545 | 0.75035 | 0.731019 | 0.731571 | 0.728874 | 0.723174 |
| 0.722737 | 0.724037 | 0.729481 | 0.706942 | 0.751185 | 0.731075 | 0.727747 | 0.730594 | 0.725 |
| 0.724488 | 0.71865 | 0.718419 | 0.697429 | 0.745662 | 0.72285 | 0.723225 | 0.729509 | 0.726213 |
| 0.727338 | 0.724618 | 0.727881 | 0.727303 | 0.749459 | 0.719833 | 0.729528 | 0.735249 | 0.726478 |
| 0.723324 | 0.720542 | 0.714395 | 0.694652 | 0.744528 | 0.713966 | 0.721493 | 0.723776 | 0.7164 |
| 0.725148 | 0.706534 | 0.712652 | 0.70026 | 0.742331 | 0.716989 | 0.714899 | 0.711936 | 0.708543 |
| 0.730205 | 0.715907 | 0.728208 | 0.710958 | 0.750124 | 0.722682 | 0.728347 | 0.728289 | 0.725932 |
| 0.717246 | 0.708698 | 0.710211 | 0.708706 | 0.743138 | 0.72507 | 0.726766 | 0.727273 | 0.715096 |
| 0.722842 | 0.716974 | 0.716463 | 0.715702 | 0.744455 | 0.718829 | 0.722057 | 0.726464 | 0.721311 |
| 0.723715 | 0.720134 | 0.720082 | 0.708017 | 0.749205 | 0.726699 | 0.72715 | 0.734154 | 0.720798 |
| 0.716209 | 0.718349 | 0.720978 | 0.700197 | 0.747453 | 0.715899 | 0.728386 | 0.717674 | 0.722845 |
| 0.72043 | 0.711332 | 0.72553 | 0.705999 | 0.74428 | 0.715753 | 0.720608 | 0.720463 | 0.712249 |
| 0.731388 | 0.732047 | 0.736509 | 0.73102 | 0.739687 | 0.730655 | 0.727594 | 0.732203 | 0.729927 |
| 0.739084 | 0.743335 | 0.743553 | 0.73182 | 0.744403 | 0.731149 | 0.739995 | 0.739287 | 0.737881 |
| 0.727126 | 0.727344 | 0.727179 | 0.715374 | 0.742056 | 0.724605 | 0.730327 | 0.728624 | 0.734506 |
| 0.738247 | 0.731187 | 0.737196 | 0.719391 | 0.755909 | 0.729263 | 0.734356 | 0.728302 | 0.731249 |
| 0.714324 | 0.714737 | 0.72244 | 0.699914 | 0.721949 | 0.712783 | 0.714012 | 0.710255 | 0.719344 |
| 0.725995 | 0.725186 | 0.716871 | 0.716694 | 0.750544 | 0.726674 | 0.729932 | 0.725209 | 0.725872 |
| 0.718727 | 0.712968 | 0.720072 | 0.702488 | 0.741875 | 0.71769 | 0.725019 | 0.71997 | 0.717489 |

| | | | | | | | | |
|---|---|---|---|---|---|---|---|---|
| 0.722582 | 0.715259 | 0.720285 | 0.703594 | 0.747748 | 0.720324 | 0.729369 | 0.72439 | 0.725244 |
| 0.725641 | 0.714119 | 0.720783 | 0.711118 | 0.746211 | 0.724321 | 0.725409 | 0.72154 | 0.72731 |
| 0.724232 | 0.725377 | 0.727131 | 0.71438 | 0.747618 | 0.731069 | 0.72268 | 0.726205 | 0.728239 |
| 0.724802 | 0.71967 | 0.723758 | 0.716239 | 0.748591 | 0.720859 | 0.727668 | 0.722603 | 0.726072 |
| 0.731461 | 0.731717 | 0.727089 | 0.715824 | 0.755885 | 0.728518 | 0.73492 | 0.729833 | 0.735857 |
| 0.729842 | 0.72549 | 0.716578 | 0.706981 | 0.74826 | 0.719085 | 0.726335 | 0.729326 | 0.729035 |
| 0.723866 | 0.718939 | 0.717057 | 0.706326 | 0.749778 | 0.722624 | 0.725534 | 0.718096 | 0.730612 |
| 0.723264 | 0.715936 | 0.715325 | 0.702496 | 0.736763 | 0.71265 | 0.718451 | 0.717758 | 0.715907 |
| 0.713873 | 0.716833 | 0.710761 | 0.700751 | 0.748688 | 0.723461 | 0.726551 | 0.716366 | 0.72264 |
| 0.71535 | 0.709754 | 0.71355 | 0.70363 | 0.738001 | 0.713906 | 0.722054 | 0.721023 | 0.713672 |
| 0.715746 | 0.719253 | 0.722265 | 0.710692 | 0.754199 | 0.727681 | 0.727749 | 0.724117 | 0.722718 |
| 0.723474 | 0.719305 | 0.718361 | 0.703125 | 0.743077 | 0.720176 | 0.723133 | 0.722171 | 0.730491 |
| 0.712471 | 0.706185 | 0.70976 | 0.691196 | 0.736154 | 0.71207 | 0.721776 | 0.714858 | 0.709902 |
| 0.725667 | 0.7204 | 0.72403 | 0.704447 | 0.741853 | 0.710736 | 0.720519 | 0.718132 | 0.722384 |
| 0.726765 | 0.717849 | 0.719294 | 0.703486 | 0.751381 | 0.729479 | 0.72257 | 0.727134 | 0.726152 |
| 0.727584 | 0.72904 | 0.722591 | 0.716123 | 0.752277 | 0.730748 | 0.728103 | 0.731752 | 0.733823 |
| 0.73161 | 0.723211 | 0.722107 | 0.716617 | 0.745055 | 0.727916 | 0.730485 | 0.730745 | 0.727851 |
| 0.719486 | 0.720939 | 0.713623 | 0.700925 | 0.742817 | 0.725429 | 0.718773 | 0.72194 | 0.718858 |
| 0.724493 | 0.71623 | 0.72175 | 0.704463 | 0.744101 | 0.724463 | 0.71985 | 0.723557 | 0.71847 |
| 0.720872 | 0.717828 | 0.719528 | 0.700426 | 0.744983 | 0.73241 | 0.722582 | 0.717381 | 0.724356 |
| 0.716396 | 0.709207 | 0.713619 | 0.702513 | 0.737598 | 0.723319 | 0.716157 | 0.714086 | 0.716124 |
| 0.722638 | 0.709091 | 0.714763 | 0.703704 | 0.743703 | 0.723302 | 0.716012 | 0.715473 | 0.720066 |
| 0.724335 | 0.719922 | 0.722322 | 0.713399 | 0.747225 | 0.731995 | 0.729948 | 0.723217 | 0.728545 |
| 0.719305 | 0.723107 | 0.713555 | 0.713481 | 0.748221 | 0.728771 | 0.722189 | 0.72567 | 0.723089 |
| 0.720783 | 0.712451 | 0.72 | 0.710943 | 0.741085 | 0.72523 | 0.721004 | 0.721811 | 0.715514 |
| 0.719899 | 0.711804 | 0.715865 | 0.714666 | 0.740042 | 0.720721 | 0.728346 | 0.718923 | 0.724343 |
| 0.711664 | 0.711477 | 0.708301 | 0.705803 | 0.736974 | 0.716847 | 0.716781 | 0.717706 | 0.707272 |
| 0.715998 | 0.711283 | 0.718799 | 0.713194 | 0.74011 | 0.71594 | 0.716139 | 0.71768 | 0.719869 |
| 0.717093 | 0.717736 | 0.715203 | 0.703569 | 0.739814 | 0.721542 | 0.721754 | 0.71573 | 0.726563 |
| 0.719409 | 0.725153 | 0.714322 | 0.707631 | 0.744746 | 0.721948 | 0.721798 | 0.72269 | 0.720277 |
| 0.723746 | 0.723216 | 0.726432 | 0.713485 | 0.749157 | 0.727427 | 0.727136 | 0.725358 | 0.722392 |
| 0.723307 | 0.716081 | 0.720229 | 0.712601 | 0.742517 | 0.727376 | 0.726885 | 0.724786 | 0.72637 |
| 0.727289 | 0.721994 | 0.73074 | 0.719987 | 0.752985 | 0.73522 | 0.734221 | 0.729322 | 0.732549 |
| 0.734597 | 0.727578 | 0.731308 | 0.714761 | 0.755273 | 0.738082 | 0.724071 | 0.737352 | 0.728461 |
| 0.717972 | 0.709449 | 0.708611 | 0.698008 | 0.740328 | 0.719177 | 0.717964 | 0.720376 | 0.726019 |
| 0.719978 | 0.719181 | 0.72066 | 0.713386 | 0.743495 | 0.725697 | 0.725889 | 0.71989 | 0.720016 |
| 0.721288 | 0.719589 | 0.709801 | 0.711836 | 0.739587 | 0.717169 | 0.721199 | 0.724238 | 0.720342 |
| 0.714642 | 0.713306 | 0.715162 | 0.707309 | 0.742533 | 0.719714 | 0.723121 | 0.718023 | 0.720978 |
| 0.70856 | 0.708135 | 0.712536 | 0.700134 | 0.736879 | 0.72049 | 0.716545 | 0.711289 | 0.709598 |
| 0.72199 | 0.712404 | 0.718061 | 0.707575 | 0.746054 | 0.724294 | 0.723175 | 0.719339 | 0.724759 |
| 0.709017 | 0.71013 | 0.704995 | 0.697962 | 0.733077 | 0.715984 | 0.714329 | 0.706534 | 0.718002 |
| 0.720206 | 0.71875 | 0.716476 | 0.715292 | 0.738061 | 0.72129 | 0.723571 | 0.713841 | 0.719305 |
| 0.711631 | 0.69982 | 0.702022 | 0.697235 | 0.728497 | 0.715442 | 0.713293 | 0.713658 | 0.711194 |
| 0.721796 | 0.714836 | 0.709763 | 0.699088 | 0.738639 | 0.719321 | 0.713952 | 0.716358 | 0.710758 |
| 0.718927 | 0.71026 | 0.720864 | 0.707415 | 0.744374 | 0.729486 | 0.723339 | 0.726833 | 0.727725 |
| 0.710486 | 0.710521 | 0.699755 | 0.709587 | 0.733555 | 0.720374 | 0.711377 | 0.718429 | 0.711986 |

| | | | | | | | | |
|---|---|---|---|---|---|---|---|---|
| 0.720448 | 0.716315 | 0.726067 | 0.708223 | 0.741299 | 0.724771 | 0.726436 | 0.719932 | 0.728471 |
| 0.734586 | 0.727417 | 0.733663 | 0.727303 | 0.750425 | 0.73136 | 0.729876 | 0.73281 | 0.73024 |
| 0.719758 | 0.722991 | 0.718303 | 0.713901 | 0.748114 | 0.725846 | 0.720884 | 0.721266 | 0.724683 |
| 0.728749 | 0.725598 | 0.725354 | 0.714617 | 0.74958 | 0.726369 | 0.723657 | 0.727813 | 0.722177 |
| 0.739336 | 0.739156 | 0.733126 | 0.7229 | 0.75895 | 0.738129 | 0.737705 | 0.738903 | 0.733647 |
| 0.737604 | 0.729187 | 0.731105 | 0.723566 | 0.755283 | 0.731186 | 0.735263 | 0.734609 | 0.72934 |
| 0.732107 | 0.726145 | 0.720197 | 0.716522 | 0.746576 | 0.72739 | 0.720054 | 0.723893 | 0.723954 |
| 0.722553 | 0.725277 | 0.735036 | 0.707591 | 0.747573 | 0.723581 | 0.724891 | 0.73225 | 0.719772 |
| 0.734587 | 0.73188 | 0.716649 | 0.709656 | 0.751399 | 0.730819 | 0.732941 | 0.730181 | 0.732861 |
| 0.71637 | 0.727002 | 0.713079 | 0.707171 | 0.745016 | 0.723177 | 0.716843 | 0.72281 | 0.717338 |
| 0.727534 | 0.720749 | 0.72002 | 0.700913 | 0.742574 | 0.72782 | 0.720781 | 0.72364 | 0.727328 |
| 0.733829 | 0.7262 | 0.721441 | 0.716189 | 0.74758 | 0.728614 | 0.722803 | 0.72791 | 0.726384 |
| 0.72765 | 0.730018 | 0.722493 | 0.706211 | 0.749677 | 0.726774 | 0.720719 | 0.72698 | 0.728886 |
| 0.729208 | 0.732716 | 0.729521 | 0.720992 | 0.751865 | 0.726085 | 0.729774 | 0.727717 | 0.729774 |
| 0.734088 | 0.729825 | 0.72848 | 0.713485 | 0.755086 | 0.733847 | 0.73112 | 0.735353 | 0.734531 |
| 0.723627 | 0.73177 | 0.726732 | 0.714618 | 0.747545 | 0.722543 | 0.719615 | 0.731244 | 0.732923 |
| 0.724273 | 0.726651 | 0.720978 | 0.716975 | 0.753279 | 0.730769 | 0.730671 | 0.736666 | 0.736574 |
| 0.721096 | 0.72777 | 0.715949 | 0.706037 | 0.755468 | 0.730834 | 0.72703 | 0.730939 | 0.731495 |
| 0.732476 | 0.73869 | 0.737471 | 0.713858 | 0.753057 | 0.732506 | 0.729429 | 0.730551 | 0.73749 |
| 0.728375 | 0.727255 | 0.737442 | 0.719008 | 0.749061 | 0.739364 | 0.729336 | 0.736543 | 0.738414 |
| 0.723549 | 0.729767 | 0.714249 | 0.709287 | 0.745053 | 0.723238 | 0.719168 | 0.718278 | 0.727808 |
| 0.729837 | 0.729474 | 0.729786 | 0.717898 | 0.750868 | 0.729939 | 0.722164 | 0.729071 | 0.729675 |
| 0.728202 | 0.723367 | 0.723812 | 0.720938 | 0.750295 | 0.72529 | 0.724683 | 0.726511 | 0.724791 |
| 0.72742 | 0.72875 | 0.726553 | 0.715364 | 0.752494 | 0.737777 | 0.726082 | 0.73775 | 0.723099 |
| 0.732377 | 0.730596 | 0.733025 | 0.714052 | 0.753982 | 0.735459 | 0.730518 | 0.732837 | 0.730512 |
| 0.733321 | 0.727903 | 0.731289 | 0.720588 | 0.755264 | 0.734552 | 0.724283 | 0.727538 | 0.739489 |
| 0.726945 | 0.722691 | 0.722821 | 0.708815 | 0.753635 | 0.736233 | 0.734194 | 0.734907 | 0.723469 |
| 0.724181 | 0.723375 | 0.721532 | 0.702826 | 0.748107 | 0.735348 | 0.726719 | 0.726624 | 0.724235 |
| 0.734844 | 0.733557 | 0.726224 | 0.707597 | 0.762193 | 0.739707 | 0.728019 | 0.737575 | 0.737983 |
| 0.731925 | 0.736327 | 0.732453 | 0.725249 | 0.762127 | 0.740455 | 0.731343 | 0.748199 | 0.740037 |
| 0.730692 | 0.727038 | 0.724023 | 0.709464 | 0.750701 | 0.733646 | 0.725585 | 0.726448 | 0.730422 |
| 0.730602 | 0.733728 | 0.726197 | 0.719973 | 0.753487 | 0.733841 | 0.726591 | 0.726161 | 0.725228 |
| 0.732211 | 0.737773 | 0.730233 | 0.717219 | 0.757612 | 0.740457 | 0.73085 | 0.732853 | 0.737316 |
| 0.724932 | 0.731809 | 0.722752 | 0.721909 | 0.755663 | 0.731902 | 0.734091 | 0.732623 | 0.7331 |
| 0.728048 | 0.736257 | 0.7307 | 0.720544 | 0.755351 | 0.740071 | 0.730101 | 0.737021 | 0.74023 |
| 0.737199 | 0.726748 | 0.730381 | 0.718469 | 0.751834 | 0.738054 | 0.734351 | 0.731164 | 0.730469 |
| 0.733515 | 0.730301 | 0.73581 | 0.715471 | 0.759182 | 0.736768 | 0.738074 | 0.733927 | 0.725538 |
| 0.734426 | 0.734059 | 0.727626 | 0.725025 | 0.751378 | 0.739397 | 0.730932 | 0.741345 | 0.74113 |
| 0.737607 | 0.731467 | 0.721156 | 0.70918 | 0.75144 | 0.740589 | 0.72821 | 0.728868 | 0.73527 |
| 0.730972 | 0.725946 | 0.729366 | 0.715367 | 0.7498 | 0.733475 | 0.733364 | 0.736899 | 0.73614 |
| 0.726376 | 0.726614 | 0.734747 | 0.719697 | 0.757006 | 0.735253 | 0.732047 | 0.734227 | 0.736469 |
| 0.73887 | 0.732291 | 0.728686 | 0.725876 | 0.75358 | 0.739229 | 0.730042 | 0.736404 | 0.739847 |
| 0.740338 | 0.728487 | 0.733093 | 0.729215 | 0.754767 | 0.74117 | 0.735484 | 0.737773 | 0.740132 |
| 0.729412 | 0.725559 | 0.720183 | 0.723369 | 0.752483 | 0.73269 | 0.733434 | 0.734955 | 0.727698 |
| 0.725777 | 0.726233 | 0.732759 | 0.714004 | 0.751843 | 0.7382 | 0.732528 | 0.736793 | 0.734167 |
| 0.725394 | 0.727488 | 0.729982 | 0.709964 | 0.749124 | 0.737404 | 0.72844 | 0.737127 | 0.730605 |

| | | | | | | | | |
|---|---|---|---|---|---|---|---|---|
| 0.722382 | 0.731907 | 0.725173 | 0.723865 | 0.750083 | 0.725288 | 0.722576 | 0.729637 | 0.729318 |
| 0.71089 | 0.708922 | 0.701312 | 0.703211 | 0.734503 | 0.711916 | 0.707989 | 0.714847 | 0.705425 |
| 0.733859 | 0.732072 | 0.719104 | 0.709529 | 0.756122 | 0.728099 | 0.723893 | 0.721456 | 0.731974 |
| 0.73013 | 0.731534 | 0.728325 | 0.727394 | 0.754947 | 0.732723 | 0.733562 | 0.73602 | 0.737249 |
| 0.725854 | 0.728763 | 0.723735 | 0.713289 | 0.747326 | 0.724996 | 0.72256 | 0.721324 | 0.724803 |
| 0.723763 | 0.720428 | 0.721164 | 0.725385 | 0.750526 | 0.72628 | 0.72916 | 0.726108 | 0.726019 |
| 0.733198 | 0.739596 | 0.732002 | 0.714716 | 0.759024 | 0.740656 | 0.73976 | 0.735621 | 0.737435 |
| 0.731205 | 0.735401 | 0.730373 | 0.722846 | 0.755267 | 0.730775 | 0.726625 | 0.721786 | 0.727102 |
| 0.734262 | 0.732844 | 0.735499 | 0.715838 | 0.755507 | 0.730634 | 0.727953 | 0.729738 | 0.728752 |
| 0.727339 | 0.728653 | 0.72884 | 0.72393 | 0.756188 | 0.729432 | 0.733424 | 0.729846 | 0.736463 |
| 0.734046 | 0.73033 | 0.723481 | 0.717168 | 0.749333 | 0.731972 | 0.732951 | 0.733129 | 0.74081 |
| 0.730294 | 0.72609 | 0.722882 | 0.718315 | 0.754919 | 0.732693 | 0.730966 | 0.729925 | 0.722517 |
| 0.733478 | 0.734868 | 0.727577 | 0.712541 | 0.754879 | 0.741296 | 0.734305 | 0.73893 | 0.73889 |
| 0.724539 | 0.730187 | 0.735104 | 0.716127 | 0.750271 | 0.729412 | 0.729185 | 0.728532 | 0.722662 |
| 0.736535 | 0.733704 | 0.729424 | 0.723496 | 0.752187 | 0.731069 | 0.728289 | 0.731978 | 0.725331 |
| 0.734977 | 0.73358 | 0.728965 | 0.719057 | 0.754309 | 0.7285 | 0.729111 | 0.729697 | 0.72632 |
| 0.724475 | 0.730335 | 0.726848 | 0.723645 | 0.752537 | 0.730197 | 0.726848 | 0.72751 | 0.731396 |
| 0.720318 | 0.729072 | 0.727884 | 0.71755 | 0.753223 | 0.728412 | 0.722929 | 0.727511 | 0.728615 |
| 0.742455 | 0.737001 | 0.739472 | 0.72551 | 0.753236 | 0.739118 | 0.73964 | 0.743727 | 0.752599 |
| 0.735776 | 0.728603 | 0.741765 | 0.720993 | 0.75144 | 0.729803 | 0.728733 | 0.725829 | 0.732354 |
| 0.732668 | 0.730067 | 0.735614 | 0.727426 | 0.751472 | 0.7304 | 0.732562 | 0.737459 | 0.728163 |
| 0.729132 | 0.728548 | 0.721709 | 0.714286 | 0.752089 | 0.726252 | 0.721459 | 0.728968 | 0.726741 |
| 0.728644 | 0.723661 | 0.729129 | 0.713573 | 0.753901 | 0.726189 | 0.729352 | 0.728668 | 0.726382 |
| 0.72798 | 0.725281 | 0.724994 | 0.71556 | 0.756835 | 0.726966 | 0.735582 | 0.730587 | 0.732409 |
| 0.738377 | 0.735883 | 0.729102 | 0.719855 | 0.761052 | 0.740202 | 0.73574 | 0.737095 | 0.731722 |
| 0.730629 | 0.729237 | 0.726102 | 0.716617 | 0.754174 | 0.729237 | 0.730223 | 0.727259 | 0.731533 |
| 0.731203 | 0.731678 | 0.720615 | 0.72585 | 0.750433 | 0.727534 | 0.734775 | 0.730926 | 0.732614 |
| 0.736081 | 0.733693 | 0.729568 | 0.723411 | 0.755104 | 0.733438 | 0.738706 | 0.734992 | 0.740367 |
| 0.739703 | 0.732114 | 0.731433 | 0.718521 | 0.758763 | 0.731373 | 0.732821 | 0.729942 | 0.733762 |
| 0.649483 | 0.658586 | 0.652731 | 0.64191 | 0.680482 | 0.653907 | 0.654568 | 0.656077 | 0.652401 |
| 0.652947 | 0.657834 | 0.653357 | 0.640764 | 0.687784 | 0.653763 | 0.660125 | 0.655249 | 0.651789 |
| 0.653338 | 0.651752 | 0.651616 | 0.635407 | 0.685012 | 0.652156 | 0.655641 | 0.654283 | 0.646325 |
| 0.649211 | 0.657459 | 0.65736 | 0.646445 | 0.683055 | 0.64827 | 0.657414 | 0.66017 | 0.662942 |
| 0.646584 | 0.659328 | 0.653429 | 0.640227 | 0.683965 | 0.658058 | 0.6616 | 0.660756 | 0.650301 |
| 0.72898 | 0.730769 | 0.731612 | 0.727364 | 0.761987 | 0.739734 | 0.733596 | 0.74193 | 0.733636 |
| 0.729254 | 0.734625 | 0.734476 | 0.72076 | 0.764468 | 0.736015 | 0.734858 | 0.738088 | 0.736008 |
| 0.732095 | 0.746448 | 0.73189 | 0.718472 | 0.763673 | 0.742514 | 0.747991 | 0.747677 | 0.737544 |
| 0.73469 | 0.735973 | 0.742976 | 0.726818 | 0.76666 | 0.743044 | 0.73747 | 0.747946 | 0.737712 |
| 0.731933 | 0.729142 | 0.73702 | 0.725705 | 0.766898 | 0.743448 | 0.735117 | 0.74209 | 0.740412 |
| 0.746589 | 0.73826 | 0.738581 | 0.740566 | 0.764929 | 0.744657 | 0.746226 | 0.745595 | 0.744301 |
| 0.73523 | 0.734017 | 0.741708 | 0.724826 | 0.760528 | 0.740308 | 0.738648 | 0.736801 | 0.745391 |
| 0.732847 | 0.742412 | 0.732935 | 0.709911 | 0.760351 | 0.737493 | 0.734922 | 0.732628 | 0.735018 |
| 0.725501 | 0.726942 | 0.72948 | 0.718443 | 0.756564 | 0.742511 | 0.740873 | 0.735632 | 0.738958 |
| 0.726515 | 0.734623 | 0.728733 | 0.726004 | 0.760833 | 0.74404 | 0.740527 | 0.738501 | 0.744924 |
| 0.731519 | 0.734223 | 0.737856 | 0.714858 | 0.759498 | 0.741141 | 0.731532 | 0.737312 | 0.738259 |
| 0.739849 | 0.744873 | 0.734391 | 0.725008 | 0.763264 | 0.744831 | 0.742438 | 0.745409 | 0.731918 |

| | | | | | | | | |
|---|---|---|---|---|---|---|---|---|
| 0.736437 | 0.732667 | 0.747042 | 0.721317 | 0.769998 | 0.746626 | 0.748467 | 0.751252 | 0.745956 |
| 0.743091 | 0.736097 | 0.737984 | 0.724184 | 0.766294 | 0.756115 | 0.747907 | 0.749186 | 0.748292 |
| 0.737584 | 0.741942 | 0.734552 | 0.730196 | 0.766216 | 0.753313 | 0.758074 | 0.74454 | 0.743742 |
| 0.723868 | 0.722906 | 0.728029 | 0.716413 | 0.764937 | 0.743155 | 0.747903 | 0.742964 | 0.735409 |
| 0.732883 | 0.72884 | 0.738164 | 0.713671 | 0.762528 | 0.744146 | 0.743895 | 0.754019 | 0.748705 |
| 0.736525 | 0.734238 | 0.734921 | 0.719934 | 0.771937 | 0.745232 | 0.754486 | 0.756693 | 0.747729 |
| 0.739967 | 0.729633 | 0.738238 | 0.727909 | 0.767837 | 0.747965 | 0.743543 | 0.752619 | 0.744268 |
| 0.745996 | 0.745955 | 0.744764 | 0.724681 | 0.769663 | 0.748629 | 0.752776 | 0.756271 | 0.752194 |
| 0.735558 | 0.734942 | 0.732709 | 0.720294 | 0.766024 | 0.748069 | 0.746104 | 0.750117 | 0.743117 |
| 0.741282 | 0.73577 | 0.735957 | 0.721279 | 0.762153 | 0.747143 | 0.753015 | 0.748092 | 0.749845 |
| 0.733503 | 0.732178 | 0.724929 | 0.722018 | 0.762638 | 0.748429 | 0.750229 | 0.747318 | 0.739905 |
| 0.739442 | 0.729708 | 0.729183 | 0.725312 | 0.757799 | 0.745816 | 0.744271 | 0.745318 | 0.738996 |
| 0.732169 | 0.731559 | 0.730515 | 0.721047 | 0.76084 | 0.743534 | 0.745517 | 0.747607 | 0.73809 |
| 0.745676 | 0.735143 | 0.737503 | 0.728043 | 0.765707 | 0.745269 | 0.743252 | 0.745627 | 0.744324 |
| 0.721728 | 0.731887 | 0.728725 | 0.726875 | 0.749471 | 0.7315 | 0.728388 | 0.726909 | 0.726843 |
| 0.727734 | 0.741242 | 0.740078 | 0.720715 | 0.761991 | 0.736479 | 0.731301 | 0.734308 | 0.734423 |
| 0.721406 | 0.732066 | 0.733008 | 0.713303 | 0.751846 | 0.730475 | 0.722972 | 0.729693 | 0.726522 |
| 0.731938 | 0.72645 | 0.729716 | 0.72123 | 0.752041 | 0.735793 | 0.732477 | 0.734722 | 0.733894 |
| 0.73168 | 0.738177 | 0.733753 | 0.72027 | 0.752986 | 0.738448 | 0.72954 | 0.737312 | 0.736853 |
| 0.738307 | 0.733387 | 0.724895 | 0.721489 | 0.749516 | 0.734688 | 0.728229 | 0.72186 | 0.734369 |
| 0.726974 | 0.737286 | 0.729737 | 0.713856 | 0.756247 | 0.737378 | 0.739377 | 0.730481 | 0.730205 |
| 0.725253 | 0.729384 | 0.725552 | 0.719639 | 0.746963 | 0.728478 | 0.718154 | 0.726298 | 0.725669 |
| 0.735912 | 0.728417 | 0.737559 | 0.718112 | 0.759763 | 0.734991 | 0.73247 | 0.738293 | 0.743302 |
| 0.746038 | 0.74578 | 0.746959 | 0.731121 | 0.771486 | 0.758939 | 0.756719 | 0.761512 | 0.757001 |
| 0.739106 | 0.730675 | 0.737975 | 0.727459 | 0.761134 | 0.734004 | 0.733023 | 0.739234 | 0.74338 |
| 0.657904 | 0.656691 | 0.65568 | 0.638995 | 0.68263 | 0.649939 | 0.657172 | 0.658569 | 0.650235 |
| 0.732091 | 0.736582 | 0.728979 | 0.720676 | 0.754843 | 0.73181 | 0.73003 | 0.740889 | 0.728703 |
| 0.732594 | 0.72738 | 0.729161 | 0.720266 | 0.756294 | 0.735257 | 0.731891 | 0.738326 | 0.731712 |
| 0.732302 | 0.727976 | 0.727249 | 0.724545 | 0.75564 | 0.738288 | 0.734252 | 0.731836 | 0.739992 |
| 0.727322 | 0.731881 | 0.727273 | 0.724024 | 0.755529 | 0.737407 | 0.732831 | 0.739747 | 0.73276 |
| 0.732944 | 0.733174 | 0.734076 | 0.723504 | 0.758055 | 0.741165 | 0.732999 | 0.740347 | 0.732606 |
| 0.730896 | 0.734213 | 0.729919 | 0.72035 | 0.755852 | 0.737973 | 0.735901 | 0.74292 | 0.734339 |
| 0.726828 | 0.730174 | 0.726522 | 0.712261 | 0.750552 | 0.733087 | 0.731461 | 0.727608 | 0.729963 |
| 0.731023 | 0.734269 | 0.732157 | 0.724126 | 0.754001 | 0.737863 | 0.732388 | 0.736569 | 0.726731 |
| 0.726201 | 0.728978 | 0.730021 | 0.721373 | 0.751084 | 0.725809 | 0.72698 | 0.733386 | 0.731057 |
| 0.727339 | 0.727345 | 0.722165 | 0.721643 | 0.753824 | 0.73235 | 0.724408 | 0.732653 | 0.730864 |
| 0.736861 | 0.733797 | 0.734161 | 0.718553 | 0.756154 | 0.739328 | 0.735201 | 0.741561 | 0.738611 |
| 0.732175 | 0.72645 | 0.730077 | 0.71438 | 0.754625 | 0.732458 | 0.732124 | 0.730722 | 0.733238 |
| 0.729514 | 0.729458 | 0.730819 | 0.72849 | 0.755792 | 0.739285 | 0.726916 | 0.735248 | 0.734391 |
| 0.72619 | 0.737427 | 0.728725 | 0.720514 | 0.757342 | 0.735231 | 0.732565 | 0.730572 | 0.727666 |
| 0.727918 | 0.726157 | 0.735522 | 0.71178 | 0.749822 | 0.726511 | 0.728703 | 0.729726 | 0.736418 |
| 0.734175 | 0.728407 | 0.73015 | 0.724786 | 0.756261 | 0.734044 | 0.730093 | 0.728565 | 0.738822 |
| 0.730713 | 0.727998 | 0.725019 | 0.723397 | 0.750994 | 0.731397 | 0.727314 | 0.732804 | 0.73658 |
| 0.73739 | 0.737229 | 0.738269 | 0.715474 | 0.752432 | 0.73842 | 0.734978 | 0.732286 | 0.73432 |
| 0.7423 | 0.742527 | 0.737115 | 0.722001 | 0.760416 | 0.743251 | 0.743971 | 0.745697 | 0.748031 |
| 0.734023 | 0.739633 | 0.736883 | 0.715143 | 0.758376 | 0.74025 | 0.736866 | 0.743848 | 0.737914 |

| | | | | | | | | |
|---|---|---|---|---|---|---|---|---|
| 0.741788 | 0.734019 | 0.745511 | 0.722259 | 0.753425 | 0.734821 | 0.729925 | 0.734466 | 0.732665 |
| 0.743571 | 0.746195 | 0.744252 | 0.72392 | 0.765491 | 0.748513 | 0.747582 | 0.75 | 0.746807 |
| 0.745231 | 0.73778 | 0.740412 | 0.72849 | 0.754159 | 0.74348 | 0.733283 | 0.732223 | 0.737005 |
| 0.739989 | 0.72515 | 0.732075 | 0.716312 | 0.743699 | 0.727525 | 0.71898 | 0.727549 | 0.711895 |
| 0.736405 | 0.739925 | 0.725852 | 0.719351 | 0.759322 | 0.737308 | 0.730109 | 0.728795 | 0.727899 |
| 0.725515 | 0.737525 | 0.732152 | 0.726138 | 0.757057 | 0.737509 | 0.73913 | 0.735735 | 0.726095 |
| 0.737663 | 0.740637 | 0.739402 | 0.739336 | 0.767037 | 0.743812 | 0.742948 | 0.735692 | 0.741536 |
| 0.733008 | 0.736423 | 0.743849 | 0.720452 | 0.756286 | 0.739492 | 0.73769 | 0.736389 | 0.739522 |
| 0.73947 | 0.733834 | 0.73869 | 0.723482 | 0.760255 | 0.744212 | 0.745948 | 0.74587 | 0.744056 |
| 0.737396 | 0.731326 | 0.737115 | 0.733378 | 0.763431 | 0.737587 | 0.741429 | 0.73745 | 0.74002 |
| 0.731324 | 0.728636 | 0.73071 | 0.722792 | 0.758024 | 0.735079 | 0.732913 | 0.743614 | 0.741497 |
| 0.729538 | 0.737577 | 0.741952 | 0.729711 | 0.762926 | 0.737953 | 0.739988 | 0.739528 | 0.741016 |
| 0.73429 | 0.734443 | 0.729479 | 0.716324 | 0.755611 | 0.74146 | 0.734949 | 0.73646 | 0.736336 |
| 0.739217 | 0.735782 | 0.737899 | 0.720159 | 0.763425 | 0.737525 | 0.73826 | 0.740638 | 0.731837 |
| 0.741912 | 0.741993 | 0.736856 | 0.725588 | 0.761297 | 0.74083 | 0.748333 | 0.741414 | 0.736648 |
| 0.734395 | 0.729687 | 0.732903 | 0.721946 | 0.756209 | 0.735718 | 0.731546 | 0.734691 | 0.72255 |
| 0.742485 | 0.751344 | 0.738163 | 0.736194 | 0.762907 | 0.743434 | 0.740602 | 0.747038 | 0.73232 |
| 0.750046 | 0.745234 | 0.73885 | 0.718263 | 0.766296 | 0.741876 | 0.742379 | 0.739877 | 0.740809 |
| 0.727207 | 0.735137 | 0.735034 | 0.718677 | 0.750273 | 0.73551 | 0.726619 | 0.741906 | 0.729945 |
| 0.736041 | 0.73512 | 0.73885 | 0.724801 | 0.755891 | 0.737191 | 0.736621 | 0.737335 | 0.730675 |
| 0.74 | 0.743396 | 0.740337 | 0.725242 | 0.760655 | 0.744239 | 0.739483 | 0.734591 | 0.739765 |
| 0.738773 | 0.748443 | 0.752806 | 0.735166 | 0.758923 | 0.741665 | 0.744034 | 0.745477 | 0.736356 |
| 0.682523 | 0.675862 | 0.689381 | 0.691645 | 0.684072 | 0.674754 | 0.688234 | 0.680484 | 0.684994 |
| 0.675949 | 0.653007 | 0.672485 | 0.660099 | 0.665377 | 0.655828 | 0.657013 | 0.662311 | 0.676305 |
| 0.660993 | 0.662879 | 0.661041 | 0.634434 | 0.671223 | 0.64619 | 0.649186 | 0.657134 | 0.647577 |
| 0.649694 | 0.648656 | 0.65608 | 0.644155 | 0.657833 | 0.639505 | 0.650137 | 0.652491 | 0.655127 |
| 0.744308 | 0.742935 | 0.741976 | 0.730933 | 0.7656 | 0.760815 | 0.746891 | 0.751278 | 0.748154 |
| 0.731898 | 0.741841 | 0.745321 | 0.735268 | 0.750747 | 0.747158 | 0.745734 | 0.749256 | 0.739483 |
| 0.746292 | 0.747271 | 0.743679 | 0.736188 | 0.754396 | 0.743967 | 0.737888 | 0.738722 | 0.752524 |
| 0.746848 | 0.74591 | 0.752423 | 0.746261 | 0.767905 | 0.762048 | 0.739192 | 0.756168 | 0.746751 |
| 0.742938 | 0.754403 | 0.739626 | 0.71484 | 0.747679 | 0.754819 | 0.740772 | 0.741499 | 0.757374 |
| 0.732436 | 0.743559 | 0.751946 | 0.735775 | 0.750074 | 0.750165 | 0.73747 | 0.736824 | 0.735557 |
| 0.727922 | 0.730894 | 0.754793 | 0.729869 | 0.760308 | 0.73771 | 0.737522 | 0.745246 | 0.753488 |
| 0.659564 | 0.658681 | 0.668357 | 0.654032 | 0.665197 | 0.653503 | 0.659 | 0.663705 | 0.662637 |
| 0.748681 | 0.74516 | 0.745459 | 0.739767 | 0.775193 | 0.759206 | 0.760407 | 0.748421 | 0.763711 |
| 0.74798 | 0.741189 | 0.747177 | 0.737335 | 0.768049 | 0.760447 | 0.746611 | 0.748884 | 0.768771 |
| 0.742058 | 0.741305 | 0.756354 | 0.74113 | 0.759994 | 0.752308 | 0.748423 | 0.754957 | 0.758637 |
| 0.74103 | 0.748722 | 0.764643 | 0.743088 | 0.751585 | 0.755451 | 0.733364 | 0.755703 | 0.76905 |
| 0.756713 | 0.747912 | 0.777592 | 0.738856 | 0.762316 | 0.755353 | 0.747401 | 0.752455 | 0.75493 |
| 0.749741 | 0.736388 | 0.757273 | 0.768759 | 0.751976 | 0.746339 | 0.73405 | 0.736728 | 0.734301 |
| 0.760692 | 0.73172 | 0.773798 | 0.770292 | 0.749005 | 0.744089 | 0.738804 | 0.738298 | 0.745332 |
| 0.741051 | 0.747233 | 0.760188 | 0.745238 | 0.747541 | 0.743494 | 0.740998 | 0.74749 | 0.74057 |
| 1 | 0.758583 | 0.758081 | 0.739697 | 0.740097 | 0.7409 | 0.739503 | 0.733425 | 0.743454 |
| 0.758583 | 1 | 0.75009 | 0.746584 | 0.749442 | 0.741204 | 0.741485 | 0.744209 | 0.7472 |
| 0.758081 | 0.75009 | 1 | 0.823688 | 0.742273 | 0.749481 | 0.747424 | 0.743856 | 0.757497 |
| 0.739697 | 0.746584 | 0.823688 | 1 | 0.735155 | 0.742452 | 0.748926 | 0.731895 | 0.741618 |

| | | | | | | | | |
|---|---|---|---|---|---|---|---|---|
| 0.740097 | 0.749442 | 0.742273 | 0.735155 | 1 | 0.778407 | 0.765337 | 0.760418 | 0.769784 |
| 0.7409 | 0.741204 | 0.749481 | 0.742452 | 0.778407 | 1 | 0.764594 | 0.756543 | 0.75629 |
| 0.739503 | 0.741485 | 0.747424 | 0.748926 | 0.765337 | 0.764594 | 1 | 0.749599 | 0.754028 |
| 0.733425 | 0.744209 | 0.743856 | 0.731895 | 0.760418 | 0.756543 | 0.749599 | 1 | 0.750445 |
| 0.743454 | 0.7472 | 0.757497 | 0.741618 | 0.769784 | 0.75629 | 0.754028 | 0.750445 | 1 |
| 0.746662 | 0.74814 | 0.767591 | 0.752425 | 0.77344 | 0.764415 | 0.761163 | 0.755562 | 0.780762 |
| 0.741728 | 0.753238 | 0.761188 | 0.732794 | 0.78946 | 0.775389 | 0.759847 | 0.75612 | 0.764228 |
| 0.742065 | 0.738086 | 0.773053 | 0.761287 | 0.774449 | 0.754291 | 0.773484 | 0.746729 | 0.76953 |
| 0.736783 | 0.751719 | 0.75442 | 0.733796 | 0.757001 | 0.752465 | 0.750428 | 0.752249 | 0.755638 |
| 0.741714 | 0.736288 | 0.742872 | 0.715709 | 0.758864 | 0.748794 | 0.736427 | 0.745305 | 0.750367 |
| 0.742596 | 0.736906 | 0.735578 | 0.721493 | 0.76451 | 0.747443 | 0.741642 | 0.738985 | 0.747786 |
| 0.749612 | 0.742475 | 0.740831 | 0.731308 | 0.759881 | 0.745131 | 0.746367 | 0.738523 | 0.747303 |
| 0.740458 | 0.740982 | 0.738602 | 0.731586 | 0.749034 | 0.736495 | 0.73645 | 0.738579 | 0.741533 |
| 0.740526 | 0.746457 | 0.74641 | 0.733236 | 0.760249 | 0.742318 | 0.737936 | 0.751515 | 0.743572 |
| 0.743073 | 0.737352 | 0.740304 | 0.730057 | 0.75426 | 0.741687 | 0.730489 | 0.739463 | 0.739711 |

| | | | | | | plot 25 | | |
|---|---|---|---|---|---|---|---|---|
| Wild_plot_ | Wild_plot_ | Wild_plot_ | Wild_plot_ | Wild_plot_ | Wild_plot_ | Wild_plot_ | Wild_plot_ | Wild_plot_ |
| 0.65873 | 0.646552 | 0.645425 | 0.641791 | 0.686674 | 0.68705 | 0.686272 | 0.669625 | 0.676783 |
| 0.669915 | 0.661546 | 0.659434 | 0.650659 | 0.694474 | 0.69158 | 0.689214 | 0.680254 | 0.689262 |
| 0.653975 | 0.645229 | 0.638769 | 0.644228 | 0.685262 | 0.682771 | 0.676214 | 0.674874 | 0.674503 |
| 0.664734 | 0.656762 | 0.648325 | 0.651283 | 0.692919 | 0.692465 | 0.6855 | 0.676936 | 0.681872 |
| 0.660757 | 0.652408 | 0.644648 | 0.645214 | 0.68986 | 0.686271 | 0.682177 | 0.677132 | 0.680836 |
| 0.666906 | 0.657276 | 0.657267 | 0.658696 | 0.697388 | 0.695825 | 0.689387 | 0.680327 | 0.68452 |
| 0.722469 | 0.718426 | 0.723068 | 0.713926 | 0.75541 | 0.750452 | 0.745892 | 0.737905 | 0.747741 |
| 0.72112 | 0.719572 | 0.713083 | 0.717525 | 0.754092 | 0.749496 | 0.747892 | 0.738512 | 0.73991 |
| 0.658396 | 0.652204 | 0.649605 | 0.651972 | 0.691045 | 0.690978 | 0.685114 | 0.677612 | 0.684335 |
| 0.655537 | 0.647209 | 0.647331 | 0.645998 | 0.686226 | 0.686752 | 0.682436 | 0.675932 | 0.681226 |
| 0.659661 | 0.656856 | 0.646615 | 0.652399 | 0.692797 | 0.688684 | 0.681949 | 0.677246 | 0.681523 |
| 0.655572 | 0.647183 | 0.640443 | 0.636242 | 0.688227 | 0.685009 | 0.675918 | 0.670256 | 0.671418 |
| 0.671973 | 0.666197 | 0.65159 | 0.654947 | 0.696759 | 0.692405 | 0.685358 | 0.68449 | 0.688152 |
| 0.682018 | 0.664401 | 0.663366 | 0.660336 | 0.70413 | 0.702224 | 0.695335 | 0.687125 | 0.695621 |
| 0.689571 | 0.679027 | 0.676164 | 0.677784 | 0.720839 | 0.713281 | 0.71056 | 0.703865 | 0.702409 |
| 0.67157 | 0.660576 | 0.653919 | 0.656532 | 0.69545 | 0.69104 | 0.690942 | 0.679112 | 0.687951 |
| 0.669494 | 0.651462 | 0.651436 | 0.646755 | 0.69649 | 0.69258 | 0.688491 | 0.679495 | 0.685986 |
| 0.700652 | 0.68235 | 0.684584 | 0.681203 | 0.725302 | 0.721702 | 0.715522 | 0.711231 | 0.709967 |
| 0.655785 | 0.641457 | 0.643226 | 0.647752 | 0.685048 | 0.684539 | 0.680645 | 0.677673 | 0.670977 |
| 0.664906 | 0.65309 | 0.643516 | 0.649699 | 0.689225 | 0.688072 | 0.682859 | 0.676412 | 0.682583 |
| 0.657742 | 0.645576 | 0.643678 | 0.642157 | 0.684602 | 0.686493 | 0.68101 | 0.671993 | 0.678398 |
| 0.664399 | 0.652409 | 0.648033 | 0.654947 | 0.688447 | 0.688555 | 0.684453 | 0.676919 | 0.677982 |
| 0.670926 | 0.657063 | 0.655809 | 0.652657 | 0.697476 | 0.695054 | 0.691921 | 0.677548 | 0.686701 |
| 0.658877 | 0.647733 | 0.647059 | 0.644336 | 0.693994 | 0.693021 | 0.685163 | 0.681818 | 0.67974 |
| 0.6687 | 0.663534 | 0.653878 | 0.652519 | 0.695532 | 0.688398 | 0.689035 | 0.681867 | 0.685989 |
| 0.662506 | 0.64993 | 0.64398 | 0.650068 | 0.691437 | 0.686628 | 0.680697 | 0.675443 | 0.679498 |
| 0.65006 | 0.649193 | 0.649032 | 0.646272 | 0.687831 | 0.686486 | 0.680557 | 0.674623 | 0.674317 |
| 0.662919 | 0.655452 | 0.647807 | 0.648714 | 0.691851 | 0.687488 | 0.687713 | 0.680486 | 0.684725 |
| 0.672543 | 0.667136 | 0.662657 | 0.662195 | 0.697407 | 0.696294 | 0.69078 | 0.67909 | 0.689139 |
| 0.663366 | 0.656243 | 0.655941 | 0.651756 | 0.695267 | 0.692541 | 0.685185 | 0.680406 | 0.681091 |
| 0.654031 | 0.656769 | 0.642896 | 0.650527 | 0.693257 | 0.689733 | 0.684948 | 0.67656 | 0.680708 |
| 0.66001 | 0.652768 | 0.640785 | 0.644961 | 0.685378 | 0.685507 | 0.676902 | 0.673731 | 0.675565 |
| 0.65359 | 0.648324 | 0.648355 | 0.647792 | 0.694035 | 0.685918 | 0.683426 | 0.674779 | 0.681033 |
| 0.665462 | 0.663097 | 0.646361 | 0.652901 | 0.695188 | 0.692367 | 0.690664 | 0.677855 | 0.681333 |
| 0.657768 | 0.649287 | 0.641396 | 0.641275 | 0.687004 | 0.684062 | 0.68 | 0.674754 | 0.677808 |
| 0.664491 | 0.648465 | 0.654585 | 0.654375 | 0.693662 | 0.688489 | 0.684244 | 0.678588 | 0.678318 |
| 0.668589 | 0.655981 | 0.657794 | 0.644875 | 0.695039 | 0.691243 | 0.686387 | 0.680496 | 0.68559 |
| 0.663833 | 0.661332 | 0.656741 | 0.658794 | 0.694051 | 0.689367 | 0.683182 | 0.675219 | 0.682517 |
| 0.666827 | 0.664637 | 0.644951 | 0.652517 | 0.700123 | 0.692935 | 0.69041 | 0.679322 | 0.686585 |

| | | | | | | | | |
|---|---|---|---|---|---|---|---|---|
| 0.703713 | 0.689103 | 0.691314 | 0.686135 | 0.731962 | 0.725228 | 0.726584 | 0.716402 | 0.716014 |
| 0.652538 | 0.643817 | 0.636217 | 0.643379 | 0.680612 | 0.679389 | 0.672008 | 0.663765 | 0.665227 |
| 0.660972 | 0.648364 | 0.639318 | 0.646418 | 0.682863 | 0.680882 | 0.676558 | 0.671787 | 0.675386 |
| 0.671822 | 0.658231 | 0.660593 | 0.659771 | 0.694502 | 0.692501 | 0.690919 | 0.679118 | 0.686597 |
| 0.65961 | 0.654206 | 0.644192 | 0.654277 | 0.687818 | 0.688149 | 0.678781 | 0.675252 | 0.675948 |
| 0.665381 | 0.655026 | 0.653184 | 0.653286 | 0.690099 | 0.693876 | 0.686037 | 0.681385 | 0.687619 |
| 0.658355 | 0.64665 | 0.643109 | 0.646788 | 0.688296 | 0.681823 | 0.681426 | 0.67062 | 0.676088 |
| 0.661928 | 0.655261 | 0.644155 | 0.652399 | 0.691686 | 0.69212 | 0.68455 | 0.675942 | 0.681 |
| 0.666667 | 0.657634 | 0.650041 | 0.65 | 0.693092 | 0.691704 | 0.6906 | 0.680265 | 0.688134 |
| 0.664661 | 0.657123 | 0.646025 | 0.656011 | 0.695068 | 0.695077 | 0.691739 | 0.685734 | 0.686903 |
| 0.659559 | 0.641421 | 0.644142 | 0.640885 | 0.688029 | 0.68539 | 0.677623 | 0.669487 | 0.674882 |
| 0.658182 | 0.65113 | 0.643013 | 0.643798 | 0.687453 | 0.682799 | 0.676552 | 0.668584 | 0.67505 |
| 0.665527 | 0.646503 | 0.643094 | 0.645954 | 0.690612 | 0.68739 | 0.679449 | 0.671689 | 0.675464 |
| 0.6627 | 0.644398 | 0.641975 | 0.645981 | 0.688462 | 0.683067 | 0.679352 | 0.673642 | 0.675272 |
| 0.655742 | 0.64645 | 0.630494 | 0.640969 | 0.689401 | 0.687666 | 0.680505 | 0.674941 | 0.681726 |
| 0.663812 | 0.645085 | 0.63863 | 0.639353 | 0.686342 | 0.684195 | 0.677918 | 0.676551 | 0.677924 |
| 0.657108 | 0.638713 | 0.63609 | 0.641992 | 0.685215 | 0.686348 | 0.673941 | 0.671445 | 0.673277 |
| 0.66287 | 0.651786 | 0.638738 | 0.658739 | 0.695462 | 0.693615 | 0.68124 | 0.679354 | 0.681276 |
| 0.667392 | 0.657746 | 0.64958 | 0.650698 | 0.6983 | 0.692338 | 0.689899 | 0.680724 | 0.683807 |
| 0.661434 | 0.647677 | 0.64828 | 0.643948 | 0.687298 | 0.68457 | 0.678387 | 0.672509 | 0.674793 |
| 0.657607 | 0.644684 | 0.635194 | 0.644365 | 0.68528 | 0.683845 | 0.678318 | 0.67122 | 0.675544 |
| 0.65961 | 0.649895 | 0.636091 | 0.636758 | 0.685939 | 0.683437 | 0.676219 | 0.673072 | 0.677251 |
| 0.650651 | 0.652507 | 0.641033 | 0.639106 | 0.690353 | 0.68715 | 0.685748 | 0.672708 | 0.677537 |
| 0.668438 | 0.663545 | 0.650815 | 0.65521 | 0.70146 | 0.698534 | 0.690002 | 0.681712 | 0.690993 |
| 0.671898 | 0.674071 | 0.662677 | 0.671331 | 0.705654 | 0.704883 | 0.695136 | 0.692407 | 0.69833 |
| 0.722126 | 0.721395 | 0.711722 | 0.720236 | 0.75431 | 0.749823 | 0.748441 | 0.739003 | 0.742885 |
| 0.728177 | 0.727977 | 0.727787 | 0.719495 | 0.752036 | 0.75329 | 0.746873 | 0.738075 | 0.739474 |
| 0.727683 | 0.723107 | 0.718322 | 0.717466 | 0.753826 | 0.758023 | 0.749972 | 0.738807 | 0.747778 |
| 0.659172 | 0.652529 | 0.640389 | 0.650273 | 0.68658 | 0.683485 | 0.6813 | 0.676172 | 0.679316 |
| 0.659927 | 0.645101 | 0.649003 | 0.648796 | 0.686203 | 0.686327 | 0.680758 | 0.676502 | 0.677052 |
| 0.670843 | 0.65924 | 0.655941 | 0.661017 | 0.694627 | 0.691585 | 0.687547 | 0.681577 | 0.683682 |
| 0.666347 | 0.66146 | 0.656242 | 0.663838 | 0.697291 | 0.694325 | 0.690809 | 0.687214 | 0.685876 |
| 0.660895 | 0.64791 | 0.646915 | 0.639935 | 0.689345 | 0.688976 | 0.677984 | 0.675711 | 0.678712 |
| 0.666425 | 0.650059 | 0.65708 | 0.653195 | 0.691076 | 0.690977 | 0.683459 | 0.676474 | 0.68343 |
| 0.669409 | 0.652629 | 0.650514 | 0.647746 | 0.689797 | 0.690647 | 0.682832 | 0.680033 | 0.680993 |
| 0.666667 | 0.649045 | 0.653028 | 0.64778 | 0.691563 | 0.689837 | 0.683584 | 0.674792 | 0.682135 |
| 0.665308 | 0.651201 | 0.648113 | 0.648143 | 0.691619 | 0.686483 | 0.682812 | 0.676781 | 0.680399 |
| 0.683771 | 0.674496 | 0.672869 | 0.67053 | 0.70884 | 0.707046 | 0.699535 | 0.693778 | 0.700441 |
| 0.668615 | 0.646356 | 0.641648 | 0.638108 | 0.683147 | 0.682382 | 0.677321 | 0.672357 | 0.673752 |
| 0.663797 | 0.652981 | 0.641178 | 0.64484 | 0.691997 | 0.687874 | 0.683397 | 0.675978 | 0.680445 |
| 0.663872 | 0.648316 | 0.649418 | 0.64353 | 0.686619 | 0.684598 | 0.677683 | 0.668569 | 0.676619 |
| 0.6836 | 0.678081 | 0.66713 | 0.666115 | 0.707039 | 0.705546 | 0.700814 | 0.696676 | 0.701244 |
| 0.669919 | 0.654554 | 0.655549 | 0.651972 | 0.693777 | 0.690145 | 0.687822 | 0.680216 | 0.681556 |
| 0.654941 | 0.650783 | 0.643338 | 0.645946 | 0.686169 | 0.684704 | 0.682186 | 0.669225 | 0.679039 |
| 0.661982 | 0.652661 | 0.642005 | 0.647107 | 0.693708 | 0.689803 | 0.687446 | 0.679428 | 0.681527 |
| 0.667469 | 0.653156 | 0.652316 | 0.646454 | 0.692059 | 0.689739 | 0.683483 | 0.675228 | 0.68623 |

| | | | | | | | | |
|---|---|---|---|---|---|---|---|---|
| 0.665053 | 0.652804 | 0.647202 | 0.654432 | 0.692928 | 0.689221 | 0.67896 | 0.673893 | 0.680823 |
| 0.663072 | 0.653541 | 0.643784 | 0.642973 | 0.691473 | 0.687579 | 0.687113 | 0.681792 | 0.682947 |
| 0.665867 | 0.654779 | 0.652304 | 0.652517 | 0.693841 | 0.688967 | 0.686314 | 0.674907 | 0.67792 |
| 0.674407 | 0.668938 | 0.659265 | 0.663734 | 0.701786 | 0.699155 | 0.694928 | 0.687759 | 0.694097 |
| 0.670907 | 0.662679 | 0.6567 | 0.662288 | 0.699991 | 0.695678 | 0.691214 | 0.679459 | 0.689407 |
| 0.668765 | 0.649217 | 0.650493 | 0.650954 | 0.694122 | 0.689528 | 0.6867 | 0.677203 | 0.684086 |
| 0.666106 | 0.663008 | 0.654704 | 0.656761 | 0.69658 | 0.691749 | 0.685586 | 0.681845 | 0.685617 |
| 0.667881 | 0.664455 | 0.654121 | 0.658651 | 0.696746 | 0.692905 | 0.687843 | 0.68112 | 0.686405 |
| 0.653883 | 0.65073 | 0.643267 | 0.644608 | 0.694505 | 0.687445 | 0.680913 | 0.680846 | 0.682234 |
| 0.658249 | 0.654669 | 0.640608 | 0.650206 | 0.687447 | 0.684154 | 0.682514 | 0.6728 | 0.676332 |
| 0.66223 | 0.652224 | 0.646174 | 0.638628 | 0.688895 | 0.685912 | 0.683312 | 0.676905 | 0.679131 |
| 0.659096 | 0.657512 | 0.654974 | 0.65375 | 0.690144 | 0.687555 | 0.683518 | 0.679953 | 0.682301 |
| 0.664239 | 0.65966 | 0.640638 | 0.642234 | 0.690958 | 0.685541 | 0.682627 | 0.673695 | 0.684236 |
| 0.654132 | 0.655514 | 0.649699 | 0.65374 | 0.691242 | 0.688252 | 0.684921 | 0.674682 | 0.683407 |
| 0.664823 | 0.657605 | 0.650668 | 0.645788 | 0.692447 | 0.6856 | 0.682199 | 0.677358 | 0.684473 |
| 0.662572 | 0.653185 | 0.65023 | 0.640706 | 0.68696 | 0.684516 | 0.680767 | 0.669107 | 0.675349 |
| 0.66335 | 0.647695 | 0.654172 | 0.644542 | 0.692652 | 0.688874 | 0.682254 | 0.675873 | 0.677717 |
| 0.665065 | 0.658151 | 0.648568 | 0.646216 | 0.690995 | 0.687852 | 0.681605 | 0.669993 | 0.68185 |
| 0.663938 | 0.658048 | 0.651068 | 0.655323 | 0.695805 | 0.69156 | 0.685916 | 0.676226 | 0.68589 |
| 0.662715 | 0.661786 | 0.649575 | 0.648893 | 0.690025 | 0.689952 | 0.684742 | 0.673515 | 0.681538 |
| 0.660183 | 0.658697 | 0.649375 | 0.65851 | 0.690725 | 0.691623 | 0.687821 | 0.679709 | 0.684279 |
| 0.659929 | 0.648755 | 0.644885 | 0.645963 | 0.687435 | 0.683263 | 0.681084 | 0.6741 | 0.681375 |
| 0.666187 | 0.656594 | 0.653898 | 0.650815 | 0.693266 | 0.688232 | 0.684272 | 0.676914 | 0.681154 |
| 0.665388 | 0.652002 | 0.646964 | 0.656516 | 0.69321 | 0.691471 | 0.683633 | 0.678434 | 0.681818 |
| 0.660101 | 0.65049 | 0.644898 | 0.648501 | 0.689865 | 0.689193 | 0.685813 | 0.67673 | 0.68034 |
| 0.661972 | 0.659986 | 0.64891 | 0.652068 | 0.692829 | 0.693494 | 0.687819 | 0.680518 | 0.685926 |
| 0.661667 | 0.654114 | 0.6428 | 0.643836 | 0.691402 | 0.686029 | 0.680229 | 0.676196 | 0.679841 |
| 0.656175 | 0.651738 | 0.635756 | 0.652293 | 0.689337 | 0.686537 | 0.685791 | 0.678647 | 0.680924 |
| 0.665785 | 0.65072 | 0.644769 | 0.651156 | 0.692315 | 0.689357 | 0.686486 | 0.676374 | 0.685087 |
| 0.668982 | 0.665256 | 0.654694 | 0.649714 | 0.695095 | 0.692504 | 0.687783 | 0.684198 | 0.680725 |
| 0.667623 | 0.660417 | 0.65698 | 0.655958 | 0.694791 | 0.695754 | 0.689559 | 0.677484 | 0.685205 |
| 0.659816 | 0.64653 | 0.652244 | 0.647027 | 0.69225 | 0.690855 | 0.685289 | 0.675272 | 0.681117 |
| 0.670634 | 0.65991 | 0.656336 | 0.651429 | 0.694774 | 0.695336 | 0.688487 | 0.686084 | 0.683573 |
| 0.674763 | 0.660448 | 0.6614 | 0.661366 | 0.695484 | 0.693111 | 0.691089 | 0.678525 | 0.686606 |
| 0.662605 | 0.65739 | 0.646851 | 0.649132 | 0.694775 | 0.687811 | 0.682745 | 0.677268 | 0.684839 |
| 0.664181 | 0.651174 | 0.653261 | 0.653616 | 0.692388 | 0.688891 | 0.685432 | 0.676391 | 0.679185 |
| 0.652486 | 0.648816 | 0.634561 | 0.63967 | 0.688371 | 0.684057 | 0.678383 | 0.665361 | 0.677757 |
| 0.660891 | 0.653413 | 0.64958 | 0.651195 | 0.684551 | 0.68192 | 0.683617 | 0.678018 | 0.675966 |
| 0.6718 | 0.654594 | 0.651814 | 0.653846 | 0.684811 | 0.681176 | 0.681294 | 0.679165 | 0.677765 |
| 0.659327 | 0.65179 | 0.639739 | 0.651057 | 0.675305 | 0.674677 | 0.673368 | 0.671706 | 0.67615 |
| 0.665787 | 0.645955 | 0.650468 | 0.646077 | 0.676963 | 0.679133 | 0.670946 | 0.668791 | 0.670254 |
| 0.649435 | 0.650802 | 0.646135 | 0.639493 | 0.668366 | 0.669968 | 0.662791 | 0.659311 | 0.660995 |
| 0.665423 | 0.657158 | 0.652044 | 0.646172 | 0.668705 | 0.665336 | 0.665987 | 0.668497 | 0.665922 |
| 0.662816 | 0.654431 | 0.648215 | 0.64018 | 0.67051 | 0.668622 | 0.664704 | 0.667397 | 0.666865 |
| 0.665638 | 0.664151 | 0.649985 | 0.65599 | 0.670506 | 0.667437 | 0.669581 | 0.666211 | 0.66615 |
| 0.671081 | 0.654654 | 0.654639 | 0.645604 | 0.667664 | 0.668078 | 0.670362 | 0.667896 | 0.664849 |
| 0.667939 | 0.66112 | 0.655947 | 0.646356 | 0.671325 | 0.672991 | 0.672459 | 0.672439 | 0.676689 |

| | | | | | | | | |
|---|---|---|---|---|---|---|---|---|
| 0.660317 | 0.659802 | 0.646578 | 0.645347 | 0.673921 | 0.673772 | 0.673594 | 0.669833 | 0.670835 |
| 0.671578 | 0.655247 | 0.660731 | 0.651814 | 0.673221 | 0.675676 | 0.672786 | 0.672116 | 0.67506 |
| 0.661936 | 0.65625 | 0.656003 | 0.653375 | 0.668297 | 0.673379 | 0.67187 | 0.66719 | 0.668651 |
| 0.664638 | 0.658367 | 0.648988 | 0.646178 | 0.669185 | 0.665599 | 0.664419 | 0.665744 | 0.662658 |
| 0.660197 | 0.654983 | 0.651354 | 0.642879 | 0.666132 | 0.671246 | 0.670103 | 0.668612 | 0.663459 |
| 0.665708 | 0.65935 | 0.658902 | 0.647905 | 0.668006 | 0.66773 | 0.67089 | 0.668761 | 0.667048 |
| 0.674766 | 0.651768 | 0.661679 | 0.653608 | 0.666075 | 0.669257 | 0.66906 | 0.669906 | 0.667621 |
| 0.664366 | 0.661952 | 0.654896 | 0.649481 | 0.66111 | 0.665008 | 0.668213 | 0.667098 | 0.667278 |
| 0.658508 | 0.654416 | 0.641753 | 0.647527 | 0.668086 | 0.672096 | 0.669416 | 0.665161 | 0.666082 |
| 0.671633 | 0.65604 | 0.644166 | 0.647018 | 0.662381 | 0.665105 | 0.664273 | 0.663042 | 0.661871 |
| 0.67287 | 0.66399 | 0.65 | 0.641962 | 0.659817 | 0.656979 | 0.668004 | 0.658657 | 0.665889 |
| 0.653381 | 0.644806 | 0.631607 | 0.647233 | 0.684151 | 0.67913 | 0.67514 | 0.669013 | 0.673322 |
| 0.663165 | 0.661744 | 0.648612 | 0.657909 | 0.690767 | 0.688249 | 0.681176 | 0.673946 | 0.681262 |
| 0.671518 | 0.672654 | 0.656747 | 0.655723 | 0.6829 | 0.685512 | 0.684429 | 0.676919 | 0.685888 |
| 0.667475 | 0.657371 | 0.652304 | 0.660695 | 0.700594 | 0.699348 | 0.694657 | 0.68299 | 0.686636 |
| 0.663046 | 0.65717 | 0.651182 | 0.648671 | 0.686032 | 0.684794 | 0.681299 | 0.676284 | 0.680646 |
| 0.668588 | 0.658554 | 0.651081 | 0.652906 | 0.691721 | 0.689268 | 0.685532 | 0.685329 | 0.68929 |
| 0.678832 | 0.663925 | 0.66201 | 0.659592 | 0.702099 | 0.694995 | 0.69064 | 0.684849 | 0.688638 |
| 0.648833 | 0.645095 | 0.635977 | 0.638808 | 0.683458 | 0.679579 | 0.671952 | 0.665328 | 0.668365 |
| 0.6673 | 0.668988 | 0.663784 | 0.659085 | 0.701598 | 0.703119 | 0.69478 | 0.686359 | 0.696713 |
| 0.679827 | 0.675439 | 0.67337 | 0.667269 | 0.697152 | 0.692328 | 0.687329 | 0.681024 | 0.689732 |
| 0.664402 | 0.658973 | 0.659757 | 0.655476 | 0.694334 | 0.694077 | 0.689729 | 0.682616 | 0.684173 |
| 0.672714 | 0.658365 | 0.662188 | 0.65202 | 0.701028 | 0.695512 | 0.691497 | 0.686254 | 0.691138 |
| 0.672961 | 0.660852 | 0.655255 | 0.652644 | 0.681382 | 0.682159 | 0.679614 | 0.679256 | 0.684142 |
| 0.645246 | 0.641294 | 0.635113 | 0.62961 | 0.668788 | 0.667558 | 0.66587 | 0.660257 | 0.66361 |
| 0.689861 | 0.677675 | 0.674065 | 0.664815 | 0.690152 | 0.691836 | 0.688489 | 0.68656 | 0.689061 |
| 0.654954 | 0.653435 | 0.637721 | 0.641651 | 0.692233 | 0.687824 | 0.678914 | 0.674814 | 0.676921 |
| 0.672539 | 0.658576 | 0.65648 | 0.653867 | 0.698748 | 0.696152 | 0.693609 | 0.683742 | 0.69427 |
| 0.638683 | 0.636797 | 0.627925 | 0.633669 | 0.668501 | 0.665725 | 0.662858 | 0.653246 | 0.660835 |
| 0.675077 | 0.669844 | 0.662261 | 0.661334 | 0.708518 | 0.706316 | 0.701494 | 0.689583 | 0.694575 |
| 0.676197 | 0.666589 | 0.667208 | 0.662524 | 0.706065 | 0.704532 | 0.698376 | 0.694419 | 0.696487 |
| 0.659118 | 0.646596 | 0.639547 | 0.6503 | 0.678944 | 0.683613 | 0.679316 | 0.670781 | 0.669477 |
| 0.653541 | 0.644869 | 0.638739 | 0.638949 | 0.679813 | 0.682541 | 0.67815 | 0.670542 | 0.673515 |
| 0.669863 | 0.676595 | 0.665393 | 0.668949 | 0.708262 | 0.706418 | 0.702407 | 0.693921 | 0.700154 |
| 0.665013 | 0.670613 | 0.654877 | 0.657174 | 0.703274 | 0.702677 | 0.70243 | 0.687442 | 0.693162 |
| 0.676323 | 0.674637 | 0.666398 | 0.670492 | 0.709607 | 0.705635 | 0.701211 | 0.691687 | 0.695878 |
| 0.636555 | 0.629087 | 0.622537 | 0.619147 | 0.660284 | 0.656476 | 0.653656 | 0.646482 | 0.650279 |
| 0.680774 | 0.673791 | 0.66821 | 0.664313 | 0.704623 | 0.701618 | 0.694852 | 0.689308 | 0.690684 |
| 0.629536 | 0.628016 | 0.617854 | 0.615204 | 0.657275 | 0.656495 | 0.653811 | 0.644611 | 0.65 |
| 0.621005 | 0.605572 | 0.606452 | 0.610925 | 0.648294 | 0.643969 | 0.64603 | 0.631136 | 0.634111 |
| 0.679182 | 0.671352 | 0.669473 | 0.661321 | 0.701424 | 0.702682 | 0.697793 | 0.690426 | 0.688735 |
| 0.629689 | 0.611465 | 0.604204 | 0.609107 | 0.65281 | 0.649866 | 0.647834 | 0.632797 | 0.638586 |
| 0.609569 | 0.601184 | 0.598985 | 0.608953 | 0.639886 | 0.640235 | 0.63459 | 0.623904 | 0.629605 |
| 0.608861 | 0.602003 | 0.605216 | 0.607106 | 0.640524 | 0.636055 | 0.63298 | 0.627276 | 0.626968 |
| 0.685569 | 0.677049 | 0.668287 | 0.675683 | 0.716996 | 0.709997 | 0.706625 | 0.699251 | 0.701651 |
| 0.606719 | 0.602716 | 0.598267 | 0.596227 | 0.641459 | 0.637575 | 0.635587 | 0.623195 | 0.627072 |
| 0.68149 | 0.669618 | 0.668838 | 0.669471 | 0.706268 | 0.705423 | 0.702173 | 0.692533 | 0.695069 |

| | | | | | | | | |
|---|---|---|---|---|---|---|---|---|
| 0.685728 | 0.678755 | 0.677508 | 0.675734 | 0.712105 | 0.713143 | 0.707257 | 0.700247 | 0.699176 |
| 0.673661 | 0.664104 | 0.670688 | 0.667485 | 0.707275 | 0.708459 | 0.706444 | 0.694591 | 0.69433 |
| 0.69935 | 0.695519 | 0.689101 | 0.696895 | 0.729973 | 0.727721 | 0.723491 | 0.717209 | 0.71809 |
| 0.597035 | 0.592239 | 0.583105 | 0.590152 | 0.626829 | 0.62937 | 0.620312 | 0.613158 | 0.615893 |
| 0.612053 | 0.604181 | 0.596469 | 0.597269 | 0.637859 | 0.633219 | 0.630117 | 0.622548 | 0.622436 |
| 0.688867 | 0.675441 | 0.679537 | 0.672527 | 0.711551 | 0.712042 | 0.708975 | 0.697792 | 0.699384 |
| 0.60887 | 0.601471 | 0.59109 | 0.592768 | 0.637835 | 0.636446 | 0.635592 | 0.624959 | 0.630068 |
| 0.690406 | 0.678275 | 0.675235 | 0.67438 | 0.716244 | 0.713875 | 0.706491 | 0.701675 | 0.701809 |
| 0.695175 | 0.678098 | 0.679484 | 0.676334 | 0.717706 | 0.715215 | 0.713189 | 0.702604 | 0.71073 |
| 0.68784 | 0.677296 | 0.67652 | 0.675676 | 0.71991 | 0.715599 | 0.709277 | 0.696817 | 0.707959 |
| 0.695299 | 0.676204 | 0.680548 | 0.671107 | 0.719392 | 0.71532 | 0.716923 | 0.702945 | 0.702836 |
| 0.821705 | 0.785047 | 0.806452 | 0.783019 | 0.727891 | 0.728477 | 0.794521 | 0.751773 | 0.776119 |
| 0.743382 | 0.757895 | 0.749104 | 0.753545 | 0.742857 | 0.73706 | 0.748009 | 0.745871 | 0.745309 |
| 0.747669 | 0.74488 | 0.753363 | 0.730115 | 0.740548 | 0.737387 | 0.743643 | 0.733417 | 0.738249 |
| 0.74977 | 0.749102 | 0.752206 | 0.742282 | 0.752766 | 0.750079 | 0.75323 | 0.743911 | 0.754178 |
| 0.75205 | 0.744935 | 0.739879 | 0.749181 | 0.74692 | 0.74278 | 0.73889 | 0.737731 | 0.744144 |
| 0.572433 | 0.57039 | 0.56351 | 0.560102 | 0.590532 | 0.591352 | 0.589063 | 0.584845 | 0.582531 |
| 0.751332 | 0.758936 | 0.750871 | 0.744031 | 0.749658 | 0.740997 | 0.741973 | 0.74434 | 0.742983 |
| 0.691819 | 0.679962 | 0.684867 | 0.680407 | 0.723182 | 0.721764 | 0.71903 | 0.711651 | 0.707953 |
| 0.696582 | 0.687817 | 0.686512 | 0.683049 | 0.722937 | 0.716702 | 0.718302 | 0.705485 | 0.707966 |
| 0.698499 | 0.686303 | 0.68605 | 0.682601 | 0.727414 | 0.720434 | 0.717699 | 0.70896 | 0.711487 |
| 0.541619 | 0.521903 | 0.516425 | 0.51718 | 0.568434 | 0.567793 | 0.564672 | 0.550632 | 0.555132 |
| 0.519244 | 0.511325 | 0.502845 | 0.509723 | 0.557712 | 0.554252 | 0.550904 | 0.538592 | 0.542445 |
| 0.725697 | 0.712524 | 0.721036 | 0.721096 | 0.754704 | 0.74972 | 0.740258 | 0.734788 | 0.735045 |
| 0.730507 | 0.706464 | 0.714043 | 0.712657 | 0.755481 | 0.752579 | 0.744127 | 0.737277 | 0.738177 |
| 0.710912 | 0.710134 | 0.705882 | 0.705269 | 0.74279 | 0.741374 | 0.738362 | 0.725488 | 0.730826 |
| 0.715318 | 0.718826 | 0.709442 | 0.713761 | 0.755963 | 0.751664 | 0.740737 | 0.732697 | 0.743747 |
| 0.715633 | 0.712938 | 0.714847 | 0.716098 | 0.75493 | 0.751612 | 0.744235 | 0.733325 | 0.737861 |
| 0.73573 | 0.72038 | 0.726292 | 0.720215 | 0.755657 | 0.752236 | 0.746831 | 0.741071 | 0.74963 |
| 0.719057 | 0.71408 | 0.719978 | 0.711926 | 0.754745 | 0.753096 | 0.748401 | 0.739838 | 0.741681 |
| 0.72658 | 0.721444 | 0.711642 | 0.722936 | 0.748534 | 0.744416 | 0.735832 | 0.730836 | 0.73797 |
| 0.715483 | 0.721077 | 0.706588 | 0.719085 | 0.756823 | 0.755018 | 0.747573 | 0.742556 | 0.747652 |
| 0.706863 | 0.701607 | 0.705193 | 0.700334 | 0.750025 | 0.741845 | 0.734873 | 0.729101 | 0.736816 |
| 0.718076 | 0.703178 | 0.696523 | 0.703003 | 0.744413 | 0.74129 | 0.738429 | 0.728921 | 0.737059 |
| 0.723821 | 0.717756 | 0.710092 | 0.709218 | 0.757864 | 0.752697 | 0.743905 | 0.736727 | 0.739104 |
| 0.714712 | 0.706689 | 0.707399 | 0.710497 | 0.749803 | 0.742918 | 0.735803 | 0.731492 | 0.740327 |
| 0.715171 | 0.712052 | 0.710681 | 0.71076 | 0.748161 | 0.747154 | 0.739529 | 0.732415 | 0.733512 |
| 0.715451 | 0.713834 | 0.71043 | 0.71614 | 0.75539 | 0.749523 | 0.742178 | 0.732664 | 0.740763 |
| 0.720371 | 0.718086 | 0.70729 | 0.717324 | 0.747464 | 0.748824 | 0.737606 | 0.731075 | 0.739302 |
| 0.710882 | 0.710089 | 0.70845 | 0.708602 | 0.746179 | 0.747216 | 0.740435 | 0.731593 | 0.736356 |
| 0.741967 | 0.732154 | 0.72779 | 0.733976 | 0.776586 | 0.781417 | 0.782137 | 0.795135 | 0.800207 |
| 0.737491 | 0.742657 | 0.735878 | 0.741119 | 0.760747 | 0.752489 | 0.759894 | 0.74979 | 0.752321 |
| 0.716007 | 0.715665 | 0.708567 | 0.712528 | 0.752244 | 0.743439 | 0.73622 | 0.732716 | 0.734432 |
| 0.727861 | 0.715876 | 0.71364 | 0.726604 | 0.759366 | 0.756819 | 0.74906 | 0.741104 | 0.746378 |
| 0.714627 | 0.712187 | 0.713313 | 0.713789 | 0.719493 | 0.71319 | 0.715886 | 0.719585 | 0.718249 |
| 0.721703 | 0.710173 | 0.711317 | 0.710003 | 0.75498 | 0.753779 | 0.743194 | 0.731122 | 0.7432 |
| 0.71238 | 0.711333 | 0.699637 | 0.70906 | 0.748144 | 0.741224 | 0.730533 | 0.727284 | 0.73672 |

| | | | | | | | | |
|---|---|---|---|---|---|---|---|---|
| 0.713792 | 0.711971 | 0.70392 | 0.715695 | 0.748952 | 0.747616 | 0.740037 | 0.7383 | 0.742815 |
| 0.715929 | 0.716112 | 0.713367 | 0.713408 | 0.750947 | 0.747376 | 0.738845 | 0.738204 | 0.738084 |
| 0.712732 | 0.717211 | 0.708555 | 0.710259 | 0.752145 | 0.746285 | 0.74205 | 0.733893 | 0.74192 |
| 0.715307 | 0.71321 | 0.708928 | 0.719989 | 0.753053 | 0.75098 | 0.743771 | 0.736836 | 0.741111 |
| 0.72562 | 0.718245 | 0.719089 | 0.722844 | 0.758819 | 0.752757 | 0.7544 | 0.742351 | 0.749878 |
| 0.716123 | 0.710875 | 0.698124 | 0.705422 | 0.756232 | 0.747462 | 0.743434 | 0.738326 | 0.740475 |
| 0.711515 | 0.703721 | 0.70302 | 0.711297 | 0.7564 | 0.753542 | 0.745126 | 0.739749 | 0.737417 |
| 0.701238 | 0.699682 | 0.696721 | 0.697055 | 0.749181 | 0.742172 | 0.730748 | 0.715276 | 0.725403 |
| 0.705103 | 0.707892 | 0.705669 | 0.701725 | 0.75122 | 0.745238 | 0.736076 | 0.730128 | 0.735957 |
| 0.713076 | 0.706227 | 0.701606 | 0.709375 | 0.748011 | 0.738302 | 0.733754 | 0.727273 | 0.730445 |
| 0.717154 | 0.713107 | 0.709024 | 0.707017 | 0.751658 | 0.750503 | 0.742212 | 0.737128 | 0.739703 |
| 0.718977 | 0.704074 | 0.705615 | 0.712034 | 0.753831 | 0.747768 | 0.739057 | 0.733989 | 0.740129 |
| 0.705664 | 0.704236 | 0.698225 | 0.69797 | 0.744034 | 0.738821 | 0.728855 | 0.723633 | 0.727929 |
| 0.710725 | 0.710278 | 0.699517 | 0.708417 | 0.750354 | 0.742366 | 0.735576 | 0.728667 | 0.730726 |
| 0.708354 | 0.709515 | 0.707774 | 0.71581 | 0.75175 | 0.746704 | 0.741791 | 0.734328 | 0.7395 |
| 0.727295 | 0.729406 | 0.725634 | 0.716923 | 0.76022 | 0.751257 | 0.746007 | 0.740126 | 0.749267 |
| 0.726861 | 0.718591 | 0.723217 | 0.715909 | 0.756453 | 0.753475 | 0.742458 | 0.73615 | 0.7423 |
| 0.716328 | 0.70804 | 0.707096 | 0.707145 | 0.749336 | 0.743178 | 0.734642 | 0.731425 | 0.740481 |
| 0.713117 | 0.713664 | 0.711917 | 0.719457 | 0.752448 | 0.751665 | 0.743089 | 0.73343 | 0.743489 |
| 0.713143 | 0.718735 | 0.713962 | 0.717905 | 0.753716 | 0.750759 | 0.743145 | 0.731974 | 0.738774 |
| 0.71134 | 0.698872 | 0.698833 | 0.708536 | 0.744431 | 0.738563 | 0.73683 | 0.728612 | 0.733845 |
| 0.713102 | 0.712299 | 0.713838 | 0.723465 | 0.755465 | 0.74954 | 0.744416 | 0.731213 | 0.740461 |
| 0.720701 | 0.722397 | 0.717697 | 0.724118 | 0.759026 | 0.754734 | 0.748787 | 0.741456 | 0.745585 |
| 0.720808 | 0.71584 | 0.710393 | 0.72068 | 0.748289 | 0.745458 | 0.735926 | 0.731341 | 0.741392 |
| 0.718384 | 0.709935 | 0.713154 | 0.710741 | 0.752455 | 0.747966 | 0.742633 | 0.735161 | 0.739978 |
| 0.713858 | 0.709215 | 0.714854 | 0.724335 | 0.752689 | 0.746003 | 0.738971 | 0.7315 | 0.74136 |
| 0.709903 | 0.705031 | 0.70151 | 0.704881 | 0.746989 | 0.740268 | 0.733452 | 0.725101 | 0.734941 |
| 0.711649 | 0.704838 | 0.701467 | 0.714123 | 0.746455 | 0.744533 | 0.737635 | 0.726884 | 0.735567 |
| 0.727889 | 0.70493 | 0.714205 | 0.716719 | 0.753275 | 0.746159 | 0.742623 | 0.733966 | 0.739488 |
| 0.714891 | 0.711959 | 0.711788 | 0.7176 | 0.751475 | 0.751185 | 0.737984 | 0.736554 | 0.739024 |
| 0.72002 | 0.715433 | 0.712475 | 0.712598 | 0.753138 | 0.749344 | 0.741339 | 0.738512 | 0.744488 |
| 0.722573 | 0.714462 | 0.712238 | 0.71502 | 0.749454 | 0.747092 | 0.741885 | 0.736268 | 0.74274 |
| 0.731518 | 0.714286 | 0.727067 | 0.721519 | 0.759187 | 0.753152 | 0.744973 | 0.739188 | 0.747079 |
| 0.723058 | 0.720127 | 0.713881 | 0.726781 | 0.757271 | 0.752273 | 0.746374 | 0.737757 | 0.746775 |
| 0.727617 | 0.701424 | 0.706633 | 0.708558 | 0.750911 | 0.746627 | 0.73845 | 0.730432 | 0.734411 |
| 0.727432 | 0.706453 | 0.720397 | 0.714042 | 0.756349 | 0.754177 | 0.747597 | 0.735065 | 0.745342 |
| 0.719721 | 0.702872 | 0.721591 | 0.712398 | 0.749852 | 0.742326 | 0.738066 | 0.727482 | 0.736887 |
| 0.713547 | 0.713214 | 0.71296 | 0.713124 | 0.744934 | 0.747264 | 0.743896 | 0.731403 | 0.737713 |
| 0.714898 | 0.704545 | 0.699035 | 0.705158 | 0.743084 | 0.73882 | 0.730184 | 0.724016 | 0.732726 |
| 0.720248 | 0.707973 | 0.718274 | 0.716714 | 0.746215 | 0.744872 | 0.740328 | 0.726816 | 0.732781 |
| 0.710392 | 0.702483 | 0.697395 | 0.699138 | 0.741147 | 0.733724 | 0.730308 | 0.721198 | 0.726661 |
| 0.714214 | 0.712844 | 0.714902 | 0.721868 | 0.750637 | 0.744854 | 0.734745 | 0.73109 | 0.732449 |
| 0.71275 | 0.704138 | 0.702054 | 0.706223 | 0.739736 | 0.732819 | 0.727314 | 0.721162 | 0.73187 |
| 0.712055 | 0.700197 | 0.711914 | 0.717003 | 0.743919 | 0.739855 | 0.734794 | 0.727112 | 0.735548 |
| 0.715795 | 0.716356 | 0.716867 | 0.718885 | 0.746962 | 0.749896 | 0.739333 | 0.7334 | 0.736604 |
| 0.707555 | 0.70793 | 0.703804 | 0.708333 | 0.740137 | 0.733131 | 0.7256 | 0.721656 | 0.735473 |

| | | | | | | | | |
|---|---|---|---|---|---|---|---|---|
| 0.71738 | 0.709356 | 0.709586 | 0.714693 | 0.757429 | 0.749596 | 0.740918 | 0.733228 | 0.738057 |
| 0.721637 | 0.722398 | 0.7246 | 0.72273 | 0.752912 | 0.749036 | 0.74986 | 0.741067 | 0.742706 |
| 0.718797 | 0.715027 | 0.714616 | 0.715234 | 0.751043 | 0.746409 | 0.741617 | 0.732694 | 0.734578 |
| 0.718189 | 0.709554 | 0.716167 | 0.717861 | 0.746383 | 0.743536 | 0.740044 | 0.7255 | 0.737862 |
| 0.733584 | 0.725547 | 0.721102 | 0.733238 | 0.761679 | 0.761306 | 0.75478 | 0.743892 | 0.750704 |
| 0.72462 | 0.722601 | 0.718254 | 0.726016 | 0.755503 | 0.753947 | 0.747894 | 0.742538 | 0.74388 |
| 0.709718 | 0.717088 | 0.708381 | 0.712984 | 0.75267 | 0.752953 | 0.739581 | 0.734165 | 0.736554 |
| 0.714712 | 0.721816 | 0.717477 | 0.718065 | 0.752029 | 0.748368 | 0.741277 | 0.741658 | 0.741541 |
| 0.724682 | 0.714286 | 0.721644 | 0.727588 | 0.74922 | 0.75 | 0.742363 | 0.743638 | 0.740471 |
| 0.706743 | 0.714911 | 0.704757 | 0.717324 | 0.743838 | 0.736837 | 0.732703 | 0.729658 | 0.733957 |
| 0.711898 | 0.715868 | 0.71341 | 0.718093 | 0.752231 | 0.750499 | 0.742321 | 0.733782 | 0.741001 |
| 0.718031 | 0.715494 | 0.710386 | 0.721781 | 0.752831 | 0.755591 | 0.744109 | 0.733967 | 0.742777 |
| 0.720278 | 0.713698 | 0.723621 | 0.721907 | 0.752344 | 0.74902 | 0.741505 | 0.73368 | 0.740429 |
| 0.72371 | 0.716872 | 0.714005 | 0.724548 | 0.74656 | 0.743585 | 0.74185 | 0.741826 | 0.736682 |
| 0.717702 | 0.724079 | 0.715527 | 0.717483 | 0.756786 | 0.751203 | 0.745904 | 0.73871 | 0.752256 |
| 0.714572 | 0.718162 | 0.720884 | 0.721888 | 0.746199 | 0.745548 | 0.739943 | 0.736855 | 0.7408 |
| 0.72418 | 0.718915 | 0.719989 | 0.717863 | 0.756449 | 0.752254 | 0.746952 | 0.737803 | 0.747476 |
| 0.726185 | 0.7212 | 0.719136 | 0.723374 | 0.753877 | 0.751794 | 0.751016 | 0.744781 | 0.737857 |
| 0.730063 | 0.720885 | 0.719184 | 0.721953 | 0.753882 | 0.752486 | 0.747673 | 0.741272 | 0.743219 |
| 0.730567 | 0.724539 | 0.731241 | 0.722504 | 0.748713 | 0.752742 | 0.746662 | 0.741081 | 0.741444 |
| 0.717182 | 0.711803 | 0.717876 | 0.715491 | 0.748218 | 0.749095 | 0.738082 | 0.73533 | 0.738877 |
| 0.720268 | 0.720292 | 0.71213 | 0.717495 | 0.752402 | 0.744751 | 0.745746 | 0.736919 | 0.738347 |
| 0.72112 | 0.718171 | 0.715941 | 0.728454 | 0.749781 | 0.747376 | 0.745351 | 0.735283 | 0.744067 |
| 0.720873 | 0.727273 | 0.719358 | 0.722767 | 0.755363 | 0.75358 | 0.748433 | 0.73833 | 0.745024 |
| 0.727655 | 0.724105 | 0.725169 | 0.727579 | 0.757994 | 0.753191 | 0.746 | 0.734091 | 0.742258 |
| 0.728432 | 0.720107 | 0.726439 | 0.721056 | 0.757171 | 0.750781 | 0.746227 | 0.733196 | 0.744328 |
| 0.718835 | 0.71619 | 0.715254 | 0.722144 | 0.757211 | 0.753665 | 0.743491 | 0.735052 | 0.742861 |
| 0.735228 | 0.72301 | 0.722222 | 0.716505 | 0.756062 | 0.747279 | 0.744007 | 0.743376 | 0.741638 |
| 0.731354 | 0.725724 | 0.734112 | 0.723529 | 0.759871 | 0.754389 | 0.751656 | 0.739487 | 0.747297 |
| 0.740329 | 0.737102 | 0.731693 | 0.736345 | 0.763456 | 0.757134 | 0.752678 | 0.74766 | 0.751772 |
| 0.725101 | 0.719581 | 0.714813 | 0.726059 | 0.750074 | 0.746175 | 0.741447 | 0.736617 | 0.739467 |
| 0.725223 | 0.719912 | 0.717637 | 0.723332 | 0.755377 | 0.754629 | 0.749917 | 0.739896 | 0.746874 |
| 0.725378 | 0.728867 | 0.722854 | 0.730769 | 0.758117 | 0.757228 | 0.752689 | 0.744254 | 0.745579 |
| 0.724888 | 0.725514 | 0.721541 | 0.727456 | 0.755773 | 0.749718 | 0.748959 | 0.738971 | 0.737063 |
| 0.733683 | 0.732222 | 0.728199 | 0.72843 | 0.756565 | 0.748775 | 0.752727 | 0.747505 | 0.744212 |
| 0.744221 | 0.726675 | 0.727376 | 0.729923 | 0.757849 | 0.751575 | 0.745216 | 0.742315 | 0.745191 |
| 0.724095 | 0.720603 | 0.727143 | 0.727169 | 0.760897 | 0.753117 | 0.747562 | 0.744048 | 0.747862 |
| 0.727543 | 0.731863 | 0.728437 | 0.720812 | 0.761687 | 0.757752 | 0.749104 | 0.746541 | 0.748611 |
| 0.724112 | 0.725176 | 0.717883 | 0.720123 | 0.759339 | 0.753648 | 0.746049 | 0.74349 | 0.74223 |
| 0.736261 | 0.717291 | 0.719943 | 0.728673 | 0.75384 | 0.753768 | 0.750391 | 0.735812 | 0.744243 |
| 0.733201 | 0.719383 | 0.724544 | 0.720695 | 0.756433 | 0.753036 | 0.753314 | 0.741877 | 0.742455 |
| 0.723372 | 0.728096 | 0.724148 | 0.728208 | 0.765164 | 0.759179 | 0.752763 | 0.740615 | 0.749172 |
| 0.734184 | 0.72871 | 0.728103 | 0.735277 | 0.761942 | 0.756654 | 0.754481 | 0.753899 | 0.752869 |
| 0.725726 | 0.704442 | 0.725406 | 0.724737 | 0.759431 | 0.751018 | 0.750729 | 0.744326 | 0.745627 |
| 0.724431 | 0.716164 | 0.723242 | 0.735849 | 0.756413 | 0.751848 | 0.749361 | 0.744348 | 0.749908 |
| 0.720988 | 0.716618 | 0.717137 | 0.728522 | 0.754887 | 0.750994 | 0.747762 | 0.739301 | 0.745086 |

| | | | | | | | | |
|---|---|---|---|---|---|---|---|---|
| 0.719162 | 0.732326 | 0.719065 | 0.722895 | 0.745934 | 0.744946 | 0.740285 | 0.738154 | 0.735306 |
| 0.709229 | 0.704518 | 0.702583 | 0.70253 | 0.735735 | 0.733407 | 0.728651 | 0.72103 | 0.724363 |
| 0.716915 | 0.727004 | 0.713839 | 0.715503 | 0.752979 | 0.747528 | 0.740749 | 0.737429 | 0.741747 |
| 0.732232 | 0.734949 | 0.724711 | 0.729538 | 0.763553 | 0.757506 | 0.754557 | 0.742325 | 0.750061 |
| 0.713534 | 0.718404 | 0.713474 | 0.719523 | 0.753146 | 0.748939 | 0.742421 | 0.736614 | 0.74014 |
| 0.719166 | 0.714951 | 0.715466 | 0.721009 | 0.752202 | 0.748781 | 0.744539 | 0.736977 | 0.74013 |
| 0.732466 | 0.739021 | 0.736305 | 0.723521 | 0.753594 | 0.754835 | 0.745726 | 0.741292 | 0.748397 |
| 0.718916 | 0.720676 | 0.71531 | 0.71601 | 0.750619 | 0.749022 | 0.743002 | 0.73539 | 0.742642 |
| 0.731781 | 0.721189 | 0.718936 | 0.72114 | 0.756667 | 0.753907 | 0.748317 | 0.741289 | 0.746846 |
| 0.730721 | 0.720488 | 0.719613 | 0.732133 | 0.761919 | 0.755011 | 0.748138 | 0.75083 | 0.74917 |
| 0.733034 | 0.727096 | 0.724246 | 0.729714 | 0.760776 | 0.757686 | 0.752543 | 0.743867 | 0.750581 |
| 0.730846 | 0.722101 | 0.727195 | 0.733466 | 0.754795 | 0.752551 | 0.749334 | 0.743149 | 0.743455 |
| 0.721522 | 0.723643 | 0.715011 | 0.729606 | 0.767307 | 0.756918 | 0.755612 | 0.742542 | 0.747646 |
| 0.727386 | 0.722894 | 0.716288 | 0.732614 | 0.762021 | 0.756558 | 0.74725 | 0.746336 | 0.749511 |
| 0.72614 | 0.720448 | 0.721088 | 0.730923 | 0.755108 | 0.752626 | 0.746187 | 0.743527 | 0.746512 |
| 0.718497 | 0.72261 | 0.725049 | 0.724341 | 0.754444 | 0.753806 | 0.747317 | 0.741468 | 0.743964 |
| 0.72842 | 0.726626 | 0.714692 | 0.726545 | 0.749388 | 0.749495 | 0.745539 | 0.740426 | 0.747692 |
| 0.723104 | 0.715639 | 0.712472 | 0.723471 | 0.748309 | 0.749495 | 0.743846 | 0.741833 | 0.741647 |
| 0.737069 | 0.734472 | 0.73047 | 0.733007 | 0.758134 | 0.756665 | 0.754617 | 0.751006 | 0.754259 |
| 0.722152 | 0.720491 | 0.717795 | 0.736295 | 0.751645 | 0.747763 | 0.750171 | 0.747817 | 0.745886 |
| 0.718415 | 0.724933 | 0.712913 | 0.729668 | 0.753298 | 0.751945 | 0.748664 | 0.741979 | 0.746531 |
| 0.715441 | 0.717656 | 0.709164 | 0.723157 | 0.748092 | 0.747589 | 0.746153 | 0.733226 | 0.742341 |
| 0.724009 | 0.7238 | 0.71696 | 0.722994 | 0.757799 | 0.755819 | 0.740303 | 0.740709 | 0.745595 |
| 0.725915 | 0.728669 | 0.715459 | 0.725529 | 0.756451 | 0.754283 | 0.744622 | 0.743934 | 0.751354 |
| 0.736802 | 0.735417 | 0.72996 | 0.736317 | 0.761696 | 0.761088 | 0.755851 | 0.747483 | 0.751706 |
| 0.719564 | 0.723581 | 0.718652 | 0.726627 | 0.753967 | 0.753931 | 0.744593 | 0.74328 | 0.749351 |
| 0.735347 | 0.721728 | 0.724667 | 0.732501 | 0.753574 | 0.749432 | 0.748413 | 0.74086 | 0.746278 |
| 0.735968 | 0.724663 | 0.720534 | 0.732117 | 0.755504 | 0.755271 | 0.750311 | 0.744923 | 0.748251 |
| 0.727067 | 0.723441 | 0.721526 | 0.731326 | 0.754073 | 0.756397 | 0.748573 | 0.742775 | 0.745093 |
| 0.667688 | 0.649472 | 0.63803 | 0.643367 | 0.683667 | 0.682132 | 0.675692 | 0.673488 | 0.67914 |
| 0.661825 | 0.648585 | 0.645908 | 0.651374 | 0.687231 | 0.685378 | 0.678875 | 0.676551 | 0.673802 |
| 0.658478 | 0.648105 | 0.637524 | 0.640893 | 0.682425 | 0.681849 | 0.677354 | 0.675106 | 0.677886 |
| 0.662594 | 0.645764 | 0.646266 | 0.641504 | 0.686343 | 0.681414 | 0.677011 | 0.674991 | 0.676463 |
| 0.658152 | 0.651244 | 0.641067 | 0.641525 | 0.685663 | 0.686894 | 0.680387 | 0.672114 | 0.675717 |
| 0.730187 | 0.740292 | 0.726961 | 0.743162 | 0.761603 | 0.75482 | 0.751384 | 0.737508 | 0.745113 |
| 0.733082 | 0.730882 | 0.723902 | 0.7402 | 0.760008 | 0.752624 | 0.75462 | 0.741447 | 0.740309 |
| 0.731169 | 0.7295 | 0.729676 | 0.741046 | 0.757354 | 0.755971 | 0.752481 | 0.742669 | 0.743401 |
| 0.734985 | 0.731078 | 0.738129 | 0.735973 | 0.7653 | 0.756281 | 0.759655 | 0.742707 | 0.751815 |
| 0.731871 | 0.732535 | 0.735908 | 0.737532 | 0.762854 | 0.758292 | 0.751753 | 0.741065 | 0.74785 |
| 0.7335 | 0.736881 | 0.731437 | 0.736423 | 0.763168 | 0.757981 | 0.755709 | 0.748512 | 0.753234 |
| 0.725461 | 0.737217 | 0.727662 | 0.734607 | 0.761393 | 0.758829 | 0.755807 | 0.748346 | 0.754804 |
| 0.731059 | 0.721614 | 0.725457 | 0.731812 | 0.754372 | 0.754994 | 0.748465 | 0.744121 | 0.744704 |
| 0.729957 | 0.733881 | 0.727273 | 0.732009 | 0.754584 | 0.750498 | 0.746472 | 0.741094 | 0.747113 |
| 0.728619 | 0.736341 | 0.727246 | 0.738925 | 0.757045 | 0.750583 | 0.749653 | 0.740101 | 0.749648 |
| 0.726334 | 0.735009 | 0.723178 | 0.732818 | 0.751906 | 0.749227 | 0.754 | 0.736855 | 0.740695 |
| 0.731837 | 0.739763 | 0.726184 | 0.740006 | 0.757922 | 0.753598 | 0.751399 | 0.747966 | 0.74758 |

| | | | | | | | | |
|---|---|---|---|---|---|---|---|---|
| 0.748473 | 0.7438 | 0.736071 | 0.738395 | 0.767153 | 0.770118 | 0.764094 | 0.751999 | 0.760588 |
| 0.741159 | 0.744094 | 0.734326 | 0.733485 | 0.766581 | 0.760867 | 0.760789 | 0.749816 | 0.75314 |
| 0.746737 | 0.748217 | 0.742498 | 0.754501 | 0.766325 | 0.760645 | 0.759536 | 0.750917 | 0.760261 |
| 0.74625 | 0.731994 | 0.738484 | 0.741245 | 0.760773 | 0.755235 | 0.758949 | 0.750092 | 0.750737 |
| 0.744456 | 0.737749 | 0.739966 | 0.743014 | 0.765799 | 0.765339 | 0.757032 | 0.755949 | 0.753598 |
| 0.74584 | 0.753652 | 0.750573 | 0.749281 | 0.765713 | 0.761791 | 0.754877 | 0.745721 | 0.753471 |
| 0.749809 | 0.737596 | 0.7409 | 0.748489 | 0.770639 | 0.768008 | 0.764939 | 0.756556 | 0.762196 |
| 0.753582 | 0.745568 | 0.741344 | 0.735547 | 0.760112 | 0.758244 | 0.759973 | 0.754275 | 0.758893 |
| 0.732915 | 0.742454 | 0.728988 | 0.736662 | 0.760648 | 0.759206 | 0.754609 | 0.745926 | 0.755769 |
| 0.741522 | 0.744958 | 0.74568 | 0.742407 | 0.763837 | 0.761129 | 0.753244 | 0.744331 | 0.751539 |
| 0.743927 | 0.737568 | 0.734116 | 0.746169 | 0.758138 | 0.762691 | 0.757831 | 0.752797 | 0.752627 |
| 0.735481 | 0.736558 | 0.728543 | 0.745172 | 0.760707 | 0.758638 | 0.759649 | 0.753548 | 0.752795 |
| 0.742666 | 0.737767 | 0.739194 | 0.746073 | 0.756057 | 0.756585 | 0.750232 | 0.745379 | 0.749616 |
| 0.750065 | 0.741724 | 0.743787 | 0.731083 | 0.766184 | 0.766844 | 0.76729 | 0.751543 | 0.759199 |
| 0.723565 | 0.721408 | 0.714853 | 0.721573 | 0.751592 | 0.744768 | 0.745287 | 0.736784 | 0.743822 |
| 0.73243 | 0.727096 | 0.721497 | 0.725834 | 0.759283 | 0.759421 | 0.755494 | 0.749817 | 0.745802 |
| 0.721352 | 0.721555 | 0.718662 | 0.718976 | 0.743899 | 0.744554 | 0.745467 | 0.73744 | 0.735645 |
| 0.733833 | 0.724062 | 0.717617 | 0.723235 | 0.760556 | 0.760329 | 0.753844 | 0.750214 | 0.745026 |
| 0.718719 | 0.718164 | 0.707289 | 0.723307 | 0.756245 | 0.748865 | 0.74697 | 0.74371 | 0.740213 |
| 0.721746 | 0.719882 | 0.712493 | 0.714326 | 0.749249 | 0.752769 | 0.739391 | 0.734867 | 0.736178 |
| 0.729892 | 0.724602 | 0.720216 | 0.729576 | 0.76164 | 0.755465 | 0.749662 | 0.750839 | 0.746438 |
| 0.722827 | 0.717509 | 0.707499 | 0.722397 | 0.754529 | 0.751913 | 0.737571 | 0.737502 | 0.743292 |
| 0.73473 | 0.725649 | 0.722397 | 0.726204 | 0.753287 | 0.751598 | 0.752601 | 0.745509 | 0.743574 |
| 0.767028 | 0.75944 | 0.756339 | 0.751092 | 0.76644 | 0.764944 | 0.760598 | 0.756944 | 0.758529 |
| 0.740072 | 0.722608 | 0.726824 | 0.735028 | 0.757996 | 0.756728 | 0.758313 | 0.752756 | 0.751082 |
| 0.660382 | 0.642191 | 0.641334 | 0.642896 | 0.687243 | 0.685591 | 0.680336 | 0.67351 | 0.677213 |
| 0.731726 | 0.727162 | 0.717311 | 0.720943 | 0.755168 | 0.748897 | 0.751162 | 0.741826 | 0.745241 |
| 0.740276 | 0.728134 | 0.724109 | 0.737158 | 0.75814 | 0.752542 | 0.751219 | 0.740082 | 0.741581 |
| 0.728608 | 0.725533 | 0.730991 | 0.729529 | 0.754998 | 0.755946 | 0.753836 | 0.739734 | 0.748401 |
| 0.738346 | 0.721792 | 0.728702 | 0.726602 | 0.759002 | 0.757551 | 0.755082 | 0.742624 | 0.742046 |
| 0.737417 | 0.721243 | 0.727794 | 0.725949 | 0.753803 | 0.748579 | 0.744874 | 0.742458 | 0.745527 |
| 0.737468 | 0.721937 | 0.727743 | 0.736311 | 0.759237 | 0.754359 | 0.751669 | 0.745782 | 0.744881 |
| 0.722042 | 0.718849 | 0.717804 | 0.724711 | 0.753096 | 0.750025 | 0.745609 | 0.737549 | 0.738122 |
| 0.728737 | 0.725553 | 0.723598 | 0.728392 | 0.755046 | 0.750494 | 0.74776 | 0.741892 | 0.742814 |
| 0.739086 | 0.724752 | 0.717241 | 0.726173 | 0.748204 | 0.741442 | 0.744438 | 0.737803 | 0.734109 |
| 0.725872 | 0.722562 | 0.723979 | 0.718803 | 0.754458 | 0.750564 | 0.745852 | 0.734214 | 0.740332 |
| 0.731677 | 0.723238 | 0.725719 | 0.729553 | 0.759844 | 0.753039 | 0.74578 | 0.74164 | 0.744649 |
| 0.72673 | 0.710488 | 0.721781 | 0.722946 | 0.757773 | 0.755686 | 0.746088 | 0.745663 | 0.748558 |
| 0.726841 | 0.729498 | 0.720467 | 0.726136 | 0.757956 | 0.752942 | 0.748245 | 0.741211 | 0.745209 |
| 0.71997 | 0.715898 | 0.716486 | 0.71754 | 0.757242 | 0.758876 | 0.745467 | 0.740888 | 0.744931 |
| 0.731242 | 0.722615 | 0.717854 | 0.716266 | 0.75891 | 0.759913 | 0.744558 | 0.735594 | 0.743871 |
| 0.732517 | 0.720247 | 0.714988 | 0.726984 | 0.761065 | 0.756135 | 0.750565 | 0.743984 | 0.748754 |
| 0.729382 | 0.719482 | 0.717137 | 0.729071 | 0.753572 | 0.748508 | 0.748555 | 0.742475 | 0.744596 |
| 0.726276 | 0.723617 | 0.723392 | 0.726129 | 0.753925 | 0.752067 | 0.746509 | 0.733543 | 0.74856 |
| 0.730836 | 0.741197 | 0.737625 | 0.733447 | 0.766363 | 0.760903 | 0.755284 | 0.748746 | 0.751976 |
| 0.731126 | 0.72587 | 0.731476 | 0.730539 | 0.76153 | 0.757919 | 0.753195 | 0.748773 | 0.749142 |

| | | | | | | | | |
|---|---|---|---|---|---|---|---|---|
| 0.732546 | 0.729432 | 0.728465 | 0.732026 | 0.755682 | 0.753476 | 0.747035 | 0.738385 | 0.755082 |
| 0.737092 | 0.737098 | 0.737397 | 0.740311 | 0.766903 | 0.763812 | 0.7596 | 0.75417 | 0.758591 |
| 0.727318 | 0.724054 | 0.721414 | 0.727868 | 0.75597 | 0.750507 | 0.752182 | 0.735161 | 0.752659 |
| 0.721118 | 0.720028 | 0.720285 | 0.708562 | 0.734435 | 0.724565 | 0.725506 | 0.707795 | 0.723719 |
| 0.734958 | 0.725448 | 0.719461 | 0.724662 | 0.760297 | 0.756426 | 0.746308 | 0.740597 | 0.745722 |
| 0.725044 | 0.727965 | 0.73142 | 0.721841 | 0.75732 | 0.752144 | 0.754388 | 0.742644 | 0.743835 |
| 0.737001 | 0.731563 | 0.7374 | 0.738506 | 0.760393 | 0.755997 | 0.752949 | 0.74458 | 0.747266 |
| 0.732036 | 0.729106 | 0.722977 | 0.733504 | 0.764217 | 0.755652 | 0.749807 | 0.742421 | 0.749237 |
| 0.732899 | 0.737555 | 0.727713 | 0.741434 | 0.76146 | 0.759184 | 0.755365 | 0.749299 | 0.750092 |
| 0.731512 | 0.719815 | 0.725825 | 0.734 | 0.763971 | 0.758254 | 0.754082 | 0.743163 | 0.751628 |
| 0.728979 | 0.721105 | 0.723103 | 0.722473 | 0.760094 | 0.755296 | 0.752499 | 0.738614 | 0.746698 |
| 0.739219 | 0.724087 | 0.726231 | 0.739193 | 0.761341 | 0.753753 | 0.75265 | 0.74641 | 0.749106 |
| 0.732317 | 0.719055 | 0.732877 | 0.729131 | 0.766104 | 0.759445 | 0.75294 | 0.744523 | 0.751623 |
| 0.73755 | 0.735101 | 0.728919 | 0.740582 | 0.768868 | 0.762922 | 0.752772 | 0.745241 | 0.749695 |
| 0.739391 | 0.737717 | 0.731659 | 0.733069 | 0.76226 | 0.759289 | 0.75058 | 0.740893 | 0.746883 |
| 0.725935 | 0.726074 | 0.722285 | 0.719977 | 0.761184 | 0.754873 | 0.743399 | 0.742282 | 0.74767 |
| 0.735411 | 0.739152 | 0.730845 | 0.731326 | 0.762189 | 0.759068 | 0.756017 | 0.751758 | 0.748298 |
| 0.736038 | 0.73878 | 0.731874 | 0.737489 | 0.764862 | 0.760924 | 0.746334 | 0.748755 | 0.749573 |
| 0.724807 | 0.729412 | 0.721939 | 0.72771 | 0.760778 | 0.761639 | 0.749668 | 0.745771 | 0.748045 |
| 0.734679 | 0.727893 | 0.730051 | 0.731458 | 0.761198 | 0.757405 | 0.748502 | 0.746379 | 0.753025 |
| 0.740787 | 0.735807 | 0.737939 | 0.739565 | 0.768852 | 0.761765 | 0.755032 | 0.751004 | 0.754775 |
| 0.743635 | 0.732515 | 0.732744 | 0.734127 | 0.763047 | 0.760417 | 0.75425 | 0.747867 | 0.751751 |
| 0.685042 | 0.676355 | 0.692564 | 0.664766 | 0.685875 | 0.68559 | 0.685558 | 0.678968 | 0.688268 |
| 0.679662 | 0.654369 | 0.666089 | 0.655496 | 0.664562 | 0.664312 | 0.666833 | 0.664275 | 0.657596 |
| 0.647701 | 0.650535 | 0.651446 | 0.657934 | 0.669033 | 0.669143 | 0.6674 | 0.656799 | 0.664935 |
| 0.661553 | 0.65172 | 0.643639 | 0.641885 | 0.659359 | 0.660572 | 0.661089 | 0.661398 | 0.657451 |
| 0.760731 | 0.745614 | 0.764145 | 0.752209 | 0.759442 | 0.757648 | 0.754278 | 0.743035 | 0.760987 |
| 0.750789 | 0.756365 | 0.748263 | 0.743792 | 0.738594 | 0.736194 | 0.740115 | 0.737043 | 0.754378 |
| 0.743965 | 0.749244 | 0.736959 | 0.742989 | 0.748461 | 0.739349 | 0.739965 | 0.738945 | 0.738594 |
| 0.747475 | 0.744911 | 0.746161 | 0.746399 | 0.758591 | 0.753333 | 0.744359 | 0.735214 | 0.74773 |
| 0.755422 | 0.755396 | 0.739245 | 0.746536 | 0.727294 | 0.7223 | 0.72932 | 0.726479 | 0.736068 |
| 0.75987 | 0.727727 | 0.747928 | 0.741099 | 0.730259 | 0.738074 | 0.72457 | 0.714286 | 0.734796 |
| 0.749879 | 0.744424 | 0.76589 | 0.734651 | 0.727735 | 0.725875 | 0.743887 | 0.72405 | 0.738355 |
| 0.669861 | 0.653405 | 0.655357 | 0.651906 | 0.662514 | 0.66469 | 0.670103 | 0.663978 | 0.65812 |
| 0.748749 | 0.741262 | 0.745165 | 0.734746 | 0.768916 | 0.76297 | 0.754268 | 0.752263 | 0.760735 |
| 0.759711 | 0.745134 | 0.752335 | 0.752502 | 0.754976 | 0.756015 | 0.752982 | 0.744078 | 0.746892 |
| 0.751135 | 0.754026 | 0.753086 | 0.750639 | 0.751862 | 0.74784 | 0.742615 | 0.742183 | 0.761347 |
| 0.760559 | 0.755333 | 0.758229 | 0.747896 | 0.741109 | 0.742609 | 0.741582 | 0.730367 | 0.736786 |
| 0.743493 | 0.747774 | 0.751433 | 0.744895 | 0.755517 | 0.749781 | 0.759467 | 0.747245 | 0.758455 |
| 0.740954 | 0.729183 | 0.738205 | 0.738205 | 0.743576 | 0.741643 | 0.739792 | 0.733383 | 0.740856 |
| 0.746784 | 0.742035 | 0.744548 | 0.727462 | 0.740307 | 0.738413 | 0.743095 | 0.73688 | 0.743734 |
| 0.743457 | 0.736842 | 0.729674 | 0.740471 | 0.738365 | 0.734046 | 0.743038 | 0.736767 | 0.741198 |
| 0.746662 | 0.741728 | 0.742065 | 0.736783 | 0.741714 | 0.742596 | 0.749612 | 0.740458 | 0.740526 |
| 0.74814 | 0.753238 | 0.738086 | 0.751719 | 0.736288 | 0.736906 | 0.742475 | 0.740982 | 0.746457 |
| 0.767591 | 0.761188 | 0.773053 | 0.75442 | 0.742872 | 0.735578 | 0.740831 | 0.738602 | 0.74641 |
| 0.752425 | 0.732794 | 0.761287 | 0.733796 | 0.715709 | 0.721493 | 0.731308 | 0.731586 | 0.733236 |

| | | | | | | | | |
|---|---|---|---|---|---|---|---|---|
| 0.77344 | 0.78946 | 0.774449 | 0.757001 | 0.758864 | 0.76451 | 0.759881 | 0.749034 | 0.760249 |
| 0.764415 | 0.775389 | 0.754291 | 0.752465 | 0.748794 | 0.747443 | 0.745131 | 0.736495 | 0.742318 |
| 0.761163 | 0.759847 | 0.773484 | 0.750428 | 0.736427 | 0.741642 | 0.746367 | 0.73645 | 0.737936 |
| 0.755562 | 0.75612 | 0.746729 | 0.752249 | 0.745305 | 0.738985 | 0.738523 | 0.738579 | 0.751515 |
| 0.780762 | 0.764228 | 0.76953 | 0.755638 | 0.750367 | 0.747786 | 0.747303 | 0.741533 | 0.743572 |
| 1 | 0.768852 | 0.869811 | 0.766619 | 0.745685 | 0.741886 | 0.751713 | 0.744274 | 0.74869 |
| 0.768852 | 1 | 0.764938 | 0.74965 | 0.744267 | 0.737351 | 0.735444 | 0.732383 | 0.749116 |
| 0.869811 | 0.764938 | 1 | 0.765617 | 0.745958 | 0.738879 | 0.750149 | 0.736969 | 0.746386 |
| 0.766619 | 0.74965 | 0.765617 | 1 | 0.728547 | 0.73586 | 0.734102 | 0.740043 | 0.738913 |
| 0.745685 | 0.744267 | 0.745958 | 0.728547 | 1 | 0.871898 | 0.802451 | 0.801722 | 0.811949 |
| 0.741886 | 0.737351 | 0.738879 | 0.73586 | 0.871898 | 1 | 0.807607 | 0.801811 | 0.808557 |
| 0.751713 | 0.735444 | 0.750149 | 0.734102 | 0.802451 | 0.807607 | 1 | 0.806501 | 0.794661 |
| 0.744274 | 0.732383 | 0.736969 | 0.740043 | 0.801722 | 0.801811 | 0.806501 | 1 | 0.810311 |
| 0.74869 | 0.749116 | 0.746386 | 0.738913 | 0.811949 | 0.808557 | 0.794661 | 0.810311 | 1 |
| 0.756382 | 0.734076 | 0.744379 | 0.746223 | 0.818533 | 0.806538 | 0.803844 | 0.801023 | 0.861028 |

| |
|---|
| Wild_plot_25_2230 |
| 0.67197528 |
| 0.67806098 |
| 0.66670585 |
| 0.67331288 |
| 0.66949552 |
| 0.68205554 |
| 0.73984039 |
| 0.73979341 |
| 0.67869907 |
| 0.6732521 |
| 0.67020273 |
| 0.66270168 |
| 0.67760275 |
| 0.68279953 |
| 0.70103707 |
| 0.68189953 |
| 0.67448785 |
| 0.70226843 |
| 0.66548589 |
| 0.67139815 |
| 0.6696207 |
| 0.67155183 |
| 0.67828768 |
| 0.67413201 |
| 0.67940021 |
| 0.67160668 |
| 0.66899739 |
| 0.669848 |
| 0.67897424 |
| 0.67775518 |
| 0.67466445 |
| 0.66572803 |
| 0.66737139 |
| 0.67360376 |
| 0.66987294 |
| 0.67162757 |
| 0.67836946 |
| 0.67936088 |
| 0.68439843 |

| |
|---|
| 0.71715109 |
| 0.65776841 |
| 0.66266337 |
| 0.67667845 |
| 0.66981913 |
| 0.67603286 |
| 0.66898339 |
| 0.67741935 |
| 0.67697266 |
| 0.67901965 |
| 0.66568012 |
| 0.66327737 |
| 0.67033493 |
| 0.66769596 |
| 0.66940476 |
| 0.67111271 |
| 0.66399522 |
| 0.67183615 |
| 0.68245823 |
| 0.66789712 |
| 0.66849054 |
| 0.66891972 |
| 0.66654921 |
| 0.68252081 |
| 0.69289461 |
| 0.74065392 |
| 0.74162562 |
| 0.7417235 |
| 0.67211174 |
| 0.67242004 |
| 0.68173685 |
| 0.67980941 |
| 0.66976855 |
| 0.67003928 |
| 0.67019757 |
| 0.66922351 |
| 0.67024287 |
| 0.68922156 |
| 0.66558792 |
| 0.67096472 |
| 0.66772375 |
| 0.68963861 |
| 0.67469165 |
| 0.67063539 |
| 0.6785757 |
| 0.67279194 |

| |
|---|
| 0.66965867 |
| 0.67426749 |
| 0.67048541 |
| 0.68615237 |
| 0.6852424 |
| 0.67013355 |
| 0.67683937 |
| 0.67376055 |
| 0.6749226 |
| 0.66694342 |
| 0.670895 |
| 0.67414122 |
| 0.67303787 |
| 0.67312059 |
| 0.6714472 |
| 0.66839806 |
| 0.67303787 |
| 0.66996583 |
| 0.67532314 |
| 0.67162226 |
| 0.67233841 |
| 0.66873973 |
| 0.67041154 |
| 0.67401296 |
| 0.67443243 |
| 0.68057366 |
| 0.67483218 |
| 0.67333886 |
| 0.67587999 |
| 0.67433818 |
| 0.67598957 |
| 0.67193537 |
| 0.67937836 |
| 0.67694883 |
| 0.67431084 |
| 0.67109319 |
| 0.66625817 |
| 0.67782263 |
| 0.6690095 |
| 0.66719347 |
| 0.66462481 |
| 0.6538185 |
| 0.65812592 |
| 0.6613357 |
| 0.67038069 |
| 0.66482202 |
| 0.66850417 |

| |
|---|
| 0.66755595 |
| 0.66374589 |
| 0.66597741 |
| 0.66203993 |
| 0.65925362 |
| 0.65758755 |
| 0.66438242 |
| 0.65634777 |
| 0.66415869 |
| 0.66163566 |
| 0.66097419 |
| 0.66697903 |
| 0.67591827 |
| 0.67766746 |
| 0.68175829 |
| 0.67001774 |
| 0.67596313 |
| 0.682165 |
| 0.65790692 |
| 0.68321374 |
| 0.68230622 |
| 0.68057881 |
| 0.68684211 |
| 0.67336826 |
| 0.65574163 |
| 0.68040603 |
| 0.66904474 |
| 0.68094447 |
| 0.65166094 |
| 0.69061947 |
| 0.68964706 |
| 0.66717846 |
| 0.6651791 |
| 0.69273009 |
| 0.68660568 |
| 0.69372256 |
| 0.6486054 |
| 0.685284 |
| 0.64365989 |
| 0.6291064 |
| 0.68622811 |
| 0.63223279 |
| 0.62430447 |
| 0.6223511 |
| 0.69728577 |
| 0.62526433 |
| 0.689178 |

| |
|---|
| 0.69150821 |
| 0.69215268 |
| 0.71130206 |
| 0.61393721 |
| 0.6209622 |
| 0.69473183 |
| 0.61983743 |
| 0.69470255 |
| 0.7044093 |
| 0.70263994 |
| 0.70218678 |
| 0.79285714 |
| 0.74593863 |
| 0.74209468 |
| 0.74393483 |
| 0.74268628 |
| 0.58120119 |
| 0.74806508 |
| 0.70432895 |
| 0.70251855 |
| 0.70814039 |
| 0.55483798 |
| 0.54058722 |
| 0.73094503 |
| 0.73513846 |
| 0.72504615 |
| 0.73852025 |
| 0.73877752 |
| 0.73863636 |
| 0.73756098 |
| 0.73017274 |
| 0.74233129 |
| 0.73583282 |
| 0.73156162 |
| 0.73872319 |
| 0.73145528 |
| 0.73400343 |
| 0.73415083 |
| 0.7329778 |
| 0.72969017 |
| 0.78144371 |
| 0.75353622 |
| 0.73404125 |
| 0.7397428 |
| 0.72135025 |
| 0.74028529 |
| 0.73293222 |

| |
|---|
| 0.7377892 |
| 0.73459196 |
| 0.7379242 |
| 0.73975637 |
| 0.74173075 |
| 0.73983839 |
| 0.73783685 |
| 0.72434308 |
| 0.73200538 |
| 0.73351478 |
| 0.73977147 |
| 0.73490059 |
| 0.7281049 |
| 0.73140133 |
| 0.73968914 |
| 0.73778701 |
| 0.73952732 |
| 0.72777778 |
| 0.73780788 |
| 0.7369782 |
| 0.72940589 |
| 0.73391304 |
| 0.7352691 |
| 0.72796185 |
| 0.73546152 |
| 0.73389977 |
| 0.72113479 |
| 0.72640575 |
| 0.73229218 |
| 0.73372854 |
| 0.73644653 |
| 0.73464069 |
| 0.73731088 |
| 0.73837998 |
| 0.73310268 |
| 0.74074534 |
| 0.72715021 |
| 0.73415536 |
| 0.72356008 |
| 0.72928589 |
| 0.72422032 |
| 0.72757098 |
| 0.7206718 |
| 0.72325911 |
| 0.73387199 |
| 0.7207256 |

| |
|---|
| 0.73753248 |
| 0.74292453 |
| 0.73606414 |
| 0.73611451 |
| 0.74482504 |
| 0.73816224 |
| 0.73270794 |
| 0.73264781 |
| 0.73877917 |
| 0.72968328 |
| 0.7343712 |
| 0.73923709 |
| 0.74002954 |
| 0.74467036 |
| 0.74307768 |
| 0.73585603 |
| 0.73915163 |
| 0.73625304 |
| 0.74217016 |
| 0.74 |
| 0.73951949 |
| 0.73830858 |
| 0.74085403 |
| 0.7447668 |
| 0.74149246 |
| 0.73860145 |
| 0.74311024 |
| 0.74143836 |
| 0.74213297 |
| 0.75235464 |
| 0.74191944 |
| 0.74274423 |
| 0.748205 |
| 0.73794393 |
| 0.74508829 |
| 0.74656585 |
| 0.74318684 |
| 0.74726096 |
| 0.74495571 |
| 0.74346587 |
| 0.73557341 |
| 0.74696857 |
| 0.74802859 |
| 0.74242424 |
| 0.74311927 |
| 0.73923267 |

| |
|---|
| 0.72998805 |
| 0.72150668 |
| 0.73441195 |
| 0.74609807 |
| 0.73647799 |
| 0.73563645 |
| 0.74340194 |
| 0.74275827 |
| 0.74375999 |
| 0.74714782 |
| 0.74475179 |
| 0.73530135 |
| 0.74701023 |
| 0.74242981 |
| 0.73870529 |
| 0.73926231 |
| 0.74563251 |
| 0.73967197 |
| 0.74579082 |
| 0.74326205 |
| 0.74317817 |
| 0.73965321 |
| 0.74142187 |
| 0.74388133 |
| 0.7495005 |
| 0.74287137 |
| 0.74339193 |
| 0.74554356 |
| 0.74264706 |
| 0.67369775 |
| 0.66805374 |
| 0.6690827 |
| 0.67187303 |
| 0.66635382 |
| 0.7437211 |
| 0.74878626 |
| 0.74539839 |
| 0.75433813 |
| 0.74758723 |
| 0.7520436 |
| 0.74525846 |
| 0.74272986 |
| 0.73895378 |
| 0.75371879 |
| 0.73675 |
| 0.74928098 |

| |
|---|
| 0.75739719 |
| 0.74264888 |
| 0.75339926 |
| 0.7464299 |
| 0.75056208 |
| 0.74834271 |
| 0.76552854 |
| 0.75427996 |
| 0.75084724 |
| 0.73989868 |
| 0.74617775 |
| 0.74943876 |
| 0.74294148 |
| 0.74490596 |
| 0.7301705 |
| 0.74672273 |
| 0.7323654 |
| 0.74265251 |
| 0.74001763 |
| 0.7427099 |
| 0.75056661 |
| 0.72979673 |
| 0.74368956 |
| 0.75545907 |
| 0.74798103 |
| 0.66817209 |
| 0.7377451 |
| 0.74509804 |
| 0.74137503 |
| 0.74610438 |
| 0.74041594 |
| 0.74814255 |
| 0.73979402 |
| 0.74327906 |
| 0.73666288 |
| 0.73771308 |
| 0.74460925 |
| 0.74659063 |
| 0.73961424 |
| 0.73802958 |
| 0.7432585 |
| 0.74707437 |
| 0.74305468 |
| 0.74102849 |
| 0.75018587 |
| 0.74582767 |

| |
|---|
| 0.74596327 |
| 0.75307579 |
| 0.74487641 |
| 0.72469983 |
| 0.74271012 |
| 0.74190748 |
| 0.74180429 |
| 0.74499939 |
| 0.74777778 |
| 0.73937085 |
| 0.73677767 |
| 0.74621682 |
| 0.7470798 |
| 0.74824399 |
| 0.74428326 |
| 0.74219039 |
| 0.74769571 |
| 0.749168 |
| 0.74539156 |
| 0.74562485 |
| 0.74941394 |
| 0.74950593 |
| 0.68214506 |
| 0.65932642 |
| 0.65535525 |
| 0.6540116 |
| 0.75399473 |
| 0.73714777 |
| 0.74492099 |
| 0.74683858 |
| 0.73668033 |
| 0.73679275 |
| 0.72671608 |
| 0.65923567 |
| 0.75418786 |
| 0.74655172 |
| 0.7497196 |
| 0.73883741 |
| 0.75453237 |
| 0.74325153 |
| 0.74039882 |
| 0.73838054 |
| 0.74307253 |
| 0.73735152 |
| 0.74030354 |
| 0.73005698 |

| |
|---|
| 0.75426044 |
| 0.74168714 |
| 0.73048907 |
| 0.73946292 |
| 0.73971054 |
| 0.7563818 |
| 0.73407643 |
| 0.74437943 |
| 0.74622263 |
| 0.81853282 |
| 0.80653841 |
| 0.80384398 |
| 0.80102348 |
| 0.86102767 |
| 1 |
